## Supplementary material for "Genomic diversity of 39 samples of *Pyropia* species grown in Japan": S1 Fig

>P_haitanensis_NC_021189
tatatgatcaataattaaagaatctttttttgtaagaattcctgaaatatacaaaactttt
tatttaatatagaaaaagtaaattagaataagttgaacatatttt-ttctactttaaacc
ttttgattgag------attctaaatagaaataaaaatgttgacatagtttatatctata
ggaataaaaatgaaacttgcagtttatggtaaaggcggtataggtaaatctacaactagt
tgtaatatttctgtagctctttcaaaaagaggcaaaagagttctacaaattggctgtgat
cctaagcatgatagtacatttaccctaacagggtttttaatcccaacaattatcgacacg
ttacaatctaaagattaccattatgaagatgtttggcctgaagatgttatctataaagga
tatggaggtgtagactgtgttgaagctggcggacctccggctggtgctggttgtggaggc
tatgttgtgggtgagacagttaaattattaaaagaacttaatgcttttgatgaatatgat
atcattttatttgatgttttaggtgacgttgtatgtggaggctttgcagctccattaaat
tatgcagattattgcttgattattacagataatgggttcgatgctttgtttgcagccaat
agaattgctgcttcagtacgagaaaaagctagaacacattccttgcgacttgctggactg
gttggtaatagaacagataaaagagatctaatcgataagtatatagattgtgttccaatg
cctgtattagaagtattacccttaattgaagatattcgtgtgtctcgagtaaaaggtaaa
actttatttgaaatgtctgaaactgataaagatttagcatatgtgtgtgattactatcta
aatatcgcagaccaattaattactaggccagaaggcgtagtccctaaagagtctccagac
agagagttgttcagtcttttatctgattttttatttaaatcctaaatctaatgtaggacaa
gctgtcgcagatcaagaagaattagatttaatgatagtgtaaataatattttaagcataa
cataaaaaggaataagataatgtctacagttcaatcagatgctcttacttttgaatgtga
aacagggaattatcatactttttgtccaattagttgtgtttcttggttatatcaaaaaat
tgaagatagtttttttcttagttattggaactaaaacttgtggttatttttttgcaaaacgc
aatgggggtaatgattttcgctgagcctagatatgctatggcagagttagaagaaggaga
tatctcagctaagcttaatgattatggtgaacttcgtagattgtgcttgcaaattaagaa
agatagaaatcctagtgttattttttggattggaacatgcacaacagaaattataaaaat
ggaccttgaaggaattgctccaaaattagaagcggaaattcgtgttcctattgtagttgc
tagagctaatggcttagattatgcttttacacaaggtgaagacacggttttagcagctat
ggctcagcgttgtccttcaaatttaaataatcaagtaggcagtaatagtcgcgaaatttc
ttcccatatccccttagttttatttggatctctgcccaacccagtggtaacgcagttaac
tatggaattaaaaaaacaaggtattcatgtctctggctggttaccatctaagaggtatac
tgaattaccggttatacaagaaggctattatgttgcaggagttaatccatttcttagtcg
aacagctactacattaatgagacgaagaaaaacaaaattaattggtgcgccatttccaat
aggcccagatggtactagagcctggattgaaaaaatttgctcagtattaaatattcaacc
tattggactagcagatagagagcaagaaatctgggattctttagaagattatattgcttt
aatacgaggcaagtccgtttttttttatgggagataatttattagaggtctccctagcaag
atttcttactcggtgtggaatgactgtgtatgaaatcggtattccatacatggataaacg

ctatcaagcagcagaattagctttattgaaagccacatgtgacgaaatgaatgtcatgat
gccaacgattgtagaaaaacctgataactataatcaagtagatcgaattcgtgatttgaa
accagacttagttattactggtatggctcatgcaaatcctttagaagctagaggcattaa
tacaaaatggtctgtggagtttacctttgcacaaattcatggttttacaaatgcaagaga
tattttagaactagttacaagaccattgcgacgcaatcttagtttgtcagaattaggctg
ggatgtctatagcaagcaaagttagtattagt-------tgcgacttacatggcataaaa
tccttgtaagtct--tctatattatgctaaagaagcatttcttgcccgcccatcttctgc
ccaagcttgtaaattttctatttgttctttatctgtaaatgctaaaggtacaaattgctt
tatagctattttaatatcttctgtattaaattctctctgttcactaaaagctgtatgcat
actttctataatcgcttgctcgatttcagcacctgagaatttattacataataaacttaa
ttgtttaatatcatattcttgccatgattttggtctaacttttgataaatgtatcctaaa
aattgattctctttccttgcgattaggtaagtctaagaaaaaaatctcatcaaaccgtcc
tttccgtagcatttctgaaggtaaactttgaattttattagctgttgcgacgacaaatac
gggagcttttttttctgaaagccaagttataaatgttccaaatactcttgcactagtacc
actatctccttggctatacaagccagaaaaagccttatctatttcgtcaatccataatat
acaaggcgataatccttcagcaatatttatcatatccctcattctggattctgactcgcc
aactagtccaccaaataattttccaatatcaagacgtagtaacggcaaagtccaatcctt
ggcaatagctttagctgttaaagattttcctgttccttgtataccaactagcagtaaacc
tttaggggaaggaataccataattcaaactttgtttcgaaaaagagcgagatctttttttg
tagccattgctttagtacatctaatccaccaatatctttattcactttattgtaaggata
aaattctagtagacacgtttggttgataatttgtctttttttcttcaataataatcggtaa
gctgcgagaatcgagttgactatattgtgcaataatttttgtaatcactttgcggattct
atctattgatagaccttggcaggatttggtcatattattaactagttctgaatctagatc
aagatttacagctttacttaatcttgtgatttctttttttgatttctaaaagacttggtaa
aggtaaatccattattgtcacaatatctcttaaagcaaaaggtatatttattttacaaga
aataataattatatgtttagattgtgctttgattactttagctaaattgcgaagtttcct
agttaggactatttcatttaaaaaagagtcgaagtcttttaaaatgaaaatattcaaaga
ttcattatctaagttttcaataaactctaatgctagtaaaggattccgttttgcataacc
attatcgttagggttgctagtatatccatctacaaaatcccagcaataaacttgttgatt
gtcagagcagtgtaatttatttttttataatatattcgagtctttcttcttctctagtatt
tattacaataataggatgtcgagactttaatagtaatcgtaaatcttgtgtgaaattcat
gtcagtatttgtccccatcagacttttgtacttcaccaatta----aagactaaataaga
ttatctaacgttgcatggatatattcaaaagtc------atgtatctattttctaatcta
ttatgatttcagacttaactaaccataattttttttcttcctttgcgtcgtctactattt
aaaatagagcgtccgcatggtgttttcattctagcccggaaaccagaaactttaattttt
tttctttttgatccttgtaatgttctcttagtcattatgtctcctgtattgtaaaagatt

aaatattattttatcatcttttatatatcttgattatatagatctgtattaatatcaaag
tt-aaacgcaataatttatattaatcaagaattatttatataatattaagaattaagtat
----------tagctttatgatattagctctgcctattttttatttaagtatcctgacag
cgtttttgctaattttgaattggttagtttttcagcagctaaaaacaattttgtcattag
aatctcaatttaagtattttattgaaaaaagtcaaaatgatatattggcagtagaagaaa
gctttgcttttgcaaaggtttgtgtagcaaagaagtgcttttctagagcgattatcgaag
gtcaattggcaatgaaaaagagctcagaattaaacgagaaagagagcaatataataattg
ccaatttgcataacatgattggttttatttattctgaagcacagcaactaaatattgcaa
aaaacttttatcagcaagccttacaaatagaccctaattatgtcacagctttaaataatt
tagctaaaatttacgaagacaccaaggattttaaaaaagcagaggcactatataataagg
tattaactcttaacttagacaataaaatcgctaatagaaggaaggaatttataacaaaga
tca---------gaaatacttaatattatttataatcgggatagcaggatttgaacctgc
gacatcctgctcccaaagcaggcgcgctaccaaactgcgctatatcccgttagacataat
tagtactatataccttttctacaattatgtctactatttcagtgttttatctaggatacc
aaaacatgcctttaactccatcaggatctttaatgaatcctaatttttttgtaaaaactaa
ctacatcaggttctgcaaataatgtaatagtgctaatttctgcttgtcttaattgtttta
ttaattggtgtatcacgactttccctaaacctaaaccttgaaaatctggatgaataacaa
catcccagatagtagcattgaatccattatctgatgttgctctcgcaaatcctacgagtt
tagtatttgtatctttcttttgtattaaagaaatgataatcgaactattttttaatgcga
tttttactttttttagaggtcttttaacccatcctaccgaatcacacagctgttctaatt
cgtataagttaatatttttagtgctactgagatagatatctttgaattcggctctattac
aagttttatctaaaacaacaagtttttttgaaatttttttcagagttacttttgatatgtg
agttttgaaaaaaatttttccagaagatcataatgattatgatagaat----taaatata
tatatattatgatattgccttaatgaacaacatattaatatatgtccagtttttaatatg
gagttattaagaaagaaaattgtttgaaggtatagaaataaaaattttttgtaaaatcttt
caataaaagtatataaagagctcttttttaagattaacatctattagtattaatacagtct
cgcaattgagatgaaacaacatattattctatatctaatgccttttaaaggagttatttg
tgaaaaaatctatgtttctaacctgcctacttgctttgttattgcttagtaacccgataa
tagctaatgcagaagtggctgggctagtgccttgtaaagattcggcggcattcaataagc
gtatggttaatagtgttaaaaaacttaaagctagattagctaaatatgatgcagatactc
cgccagcgttagctttaaataagcaaatagaaaaaactaaaactagatttgctacttatg
gtagagcaggcctattgtgtggtacagatggtttacctcatttaatctctgacggtcgat
ggagtagagccggagactttgtttttccaggactgctatttttatacattacaggctgga
ttggttgggtaggtcgaggatatcttctatccgttgctaaaactagcaagcccactgaaa
aagaaattatttagatgtgccattagctattaaattcatgacatctggattcgcatggc
cgttagctgcttggcaagaatttagtagtgggcaactagttgcctccaatgacgatatta

ctgtttcaccccgttagcaaaaaatttatatgaataataattttaccaaatacttatcaa
cagcacctgtagttggcgtgttatggatgacgttcacagctggttttattatagaattaa
accggtttttcccagatgttttatatttttatttgtaaataaataa--aatctcataaa-
---aaccacacaaatatagtaaaacactattatttgtgtggttttt----atatagaata
taactattta-tttttacggaaattctttctttcgactattttaatattagattttacat
aataaaaaa-acagtaatattatgagtttagtaagccaaattatagttaatgctgatgat
gaattaagatatccaacaatcggagaattacagtcaattcaagactacttaagtacagga
agtaatagaattagaatcgcaactattattcgggataaagaaaaagagattatacaaaaa
gcaagtaaacaaatttttcagttacatccagaatatatagctccaggaggtaatgctgca
ggttctagaaaaagatcattatgcctacgtgattatggctggtacttaagattaatcact
tatggagtattagctggtgataaagattctatagaaacgattggtatcatcggtgttaga
gaaatgtataattctttaggtgttcctattattggaatgttagacgcaattcaatgtcta
aaagaagcttctctagaaatgcttggtcaagatgatatcaaaattattgcgccttatttt
gactacataattcgaggcatgtcataaatagtaaattcaattaattagttgaataaaagc
ttgtataatgttatgattagttcaagcccgaaagcatataattgtaaagactaaatcttg
tcagaaccggaaggtagcagcaataatgtttaattacaacagttatga-ttttcgggtgt
ttactatttgtactcaatagtctctcattctaattatgatgctatccataaatt----aa
taatatataatccataaaacctttcagtaaaaataa-ttaataatttgatattaatacgg
aaaacatctatactttttttataaaacttctttatatctcatctaaacaatggaaactta
acttaatgggtgttcatattttatcaacaggttcgtctgtcccaaatttttctgtagaga
ataaacaatttgaagatatgatagaaacttctgactattggattttaacaagaacaggaa
taaaaaaaagacatctagctccttcttctacttctttaactaagttagctgcagaagctg
ctaataatgctttatctgcagctaatttaaaacctaccgatattaatttaattatactag
ccacatctacgcctgatgatttatttggtagtgcaagccagctacaagcagaaatcggtg
caactgcatctgttgcttttgatattacagctgcttgctccgggttcattgttgcattag
taacagcagctcagtttatccagactggttcatatgacaatatcttagtcgttggagcag
atacaatgtcgagatggattaattggtcagatagaactacctgtatcttatttggcgatg
gtgcaggagcagtagtagtgggacaaagccttaaaaatagtattttaggttttaagttat
gtacagatggccagctaaatagtcatttgcaattgatgaatacaccaaaaaataatcaaa
aatttggatttatagatattcctcatggaaattataattctataacaatgaatggtaaag
aagtatacaaatttgctgtatttcaagttccaatggtaattagacaatgtttagataatt
taaatatatcaatagatgaagtagattggtttatattgcatcaggcaaatactagaattt
tagaagcaattgcgagtagattgtcaatacctttttctaaaatgattactaatttagagc
actatggtaatacatctgcagcatcaattcctttagcgttagatgaagccattaaatcca
ataaaattaagccaggtcaaaccattgttttgtcaggttttggagctggtctaacttggg
gtgcgattgtcttgaagtggtaat---tataatattgcggatgacgagaatcgaactcgt

aaagctttcgctacacacccctcaagcgtgcgtgtataccaatttcaccacatccgcact
ttttatcacgaata-ataaaaatttataatatttatttaaatattaatataa--tgtgta
tatatatatacattcttgtcagaaactctcttatattttattttaagtgataaaattgag
t----ttagtatagatgataaaatttatttacaatcataaaaataaataaatatataa
ggaaaataaaatatgacaccgtccttatcaagctttttaaacagtctaattcttggggcc
gtaattgtagttgtccccataactttagcacttttatttgttagtcaaaaagacagaaca
attcgatcctaacaaagactttgaaacatatggagaagcaatttgagattaataataaac
tatttttcaaacggatatgctctccatatatttctgtatt--ttggaataggttgttt-t
ctttactttgcaatagcctgcacagtgta-----------ttttgttctcatttactctt
tta----taacgatacgtatgtcagg-ttaataatag--tttttttattctgtatctttc
atttaatattaaagtattttatatacatattgttatgttcaagtaataaagaaaagctca
tgtctttgtccaattggcctcttaaaaaagaaaactccgaagcatatactagcaagtata
caaagcaaatcacaatccctgatggattgtggataaaatgcttcgattgcggtctgttaa
tgtattctaaagttctaaaaaggaatttaaaagtttgtcctcaatgtagctatcattttc
aagcttctagcagtgaaagaattgatcagttgatagacaaaggtagctggcgacctatgg
atattcatttaatttctacagatccattaggtttcaaagatcaaaagctttatagtcaaa
ggttaaaagatactgctcttaagactggcttacaagatgcagtccagacgggtattggta
ctatgcaaggtagacaagtctgcttaggtattatggacttccgatttatgggaggaagta
tgggatctgttgtaggcgaaaaattaacaaggctactggaaagagcaactcaagaaaaat
tacctgcaattatactttgtgcatcgggcggcgcaagaatgcaagaaggcatgttaagtt
taatgcagatggcaaaaatttcttctgcgttggaaatgcataaaaaagcaaatctcttgt
atatatctgttttaacatctccaactacaggaggcgtcacagctagctttgctatgctcg
gagatctaattattgcagaaccaaaagctcttattgcatttgctggtagacgagttatag
aacaaactattaaagaagacttgccagataattttcaaagttcagaatatctatttgagc
atggttttctagatttaattgtaccaagaactcagcttagatcaaagctaatacaaattt
tatcattccatactcatactaagtaagtagtttgacattgatcataatattgaaatgcta
gtaaaaattaagatattaagaaattttcactaaagatatttgttgactaactagaattac
aattactattgtagaaaaatttgtcttaattctatttttttcagtcttttaactaattg-
tttttacgactttaaataattcttagtttaatacaaatttataaagagaata-----cat
actaagctaaatatagtaatattatgttttcctataatactgacctatcaaatttcgagg
aattttatgcttaaaagatcttcttggcttgctactttattggggctactaactgtagtt
tctataagtacgcatgcaaatgccatagagctagacgaggcaacaagaactgtcccatta
gaatcttctggaagaactgtagttcttacgccagaacaagttaagcgaggcaagcgatta
ttcaataattcttgtgctatttgtcataatggtggtattacaaaaacaaatccgaatatt
ggacttgacccagagtctttaggcttagcaacgccacaaagagatacaatcgaggggtta
gttgactatatgaaagatccaacaagctatgatggcgcagagtcaattgcagaactgcat

ccaagtattaaaagtgccgaaattttcccaaaaatgcgtaatttaacggatgaagattta
tttacaattgctggtcatatcttgctccaacctaaaattgtttccgaaaaatggggcgga
ggaaaaatttactattaaaggttcaaacacaataatcctacaagtactagtttatgca-t
ttattgcccaagtatgattattttgttagatcatgatatatattaaaggtggagtttgct
aatagtaagtta-tttttgcctaaacttgcttacattattcgtttagttattaaaaagga
gaagttaaattgaaaaagacgctttcagttcttttcactgtatttagtttttttgtgata
ggttttgcacaaattgcttttgctgcagatctagataatggagaaaaagttttttctgct
aattgtgcggcatgtcatgcgggtggtaacaatgctatcatgccagataaaactttaaaa
aaagatgtgcttgaagctaacagtatgaatagtattgatgctatcacgtatcaagtgaaa
aacggtaaaaatgctatgcctgcttttggaggtagactagttgatgaagatattgaagat
gcagcaaattatgtattatctcaatctgaaaaaggttggtaattatacttaattttgcac
agtattaaaggatagacaatctatttaattattcattagattgtctattctttgtttatg
---ttattatatatagatatttatacaatactga-attatgatgaaaagaatacctgcaa
ttcttgtactagaagatggtacttattacaaaggatggtcatttaagcaaagaaatccag
caattacaattggagaagtggtttttaatactggaatgacaggttatcaagaaatcatta
cagatcctagctacttccagcaaattgtcaccttcacatacccagaaattggtaatacag
gtattaacagccaagatgttgaatctcataatattagtattaaaggcctggtggcaaaaa
atatttgtaaaatttcaagtaactggagagagcaagaatctttagttaaatatttaagcc
gttatgatattcctttcatctttggactagatactcgatccttaactcaatatttacgtc
aatttggcacaatgaatggttgtatttctactaataatttaaatcatagttatttaaagc
aaagaatttctgaagttcctggaatgcaaggcttagatttaattccatatgtaacaacag
acagcatttatccttgggacgaaaaaagtttgcctatctggtatttaaacgataaaagaa
gagtgcaattagatacccagttgaaagttgttgttatagattttggagttaaactaaaca
tactgagacgactagctacacttggatgtcagataactgttgtgcctgcacaaacatctg
tagaagatattttagctaataaacctgatggtatattactttctaatggcccaggtgatc
catctgtagtgtattatggaattaagacagtgaaaaatttattagatcataatgtaccta
ttttcggtatttgtatgggacatcaaattttaaatttggcccttgacgctaaaactttta
aactcaaatttggtcatagaggtattaatcatccatctggattggaacagcaagttgaaa
taactagtcaaaaccatggtttcgcagtcgatttacaatctgtcttcaagtcatctgtac
aagttactcatttaatttaaatgatactactgttgctggaataggccatagtcaaaatc
catatttctctgtacaatatcatccagaatcgagcccgggccctcatgatgctgactatt
tattcgaaaattttgtagaaataatgataaaatctaagcgtaaagttgc-----------
ttaataatcttcccatgctttatgattaaataaagctgataatacttgtactccccgtaa
ttgattgaagagaggtaaatgcccctttggaccataagaattccattgaaattcatcagg
gtatctacaaggaatatgatcaacttcccacccaatcttttgccagaattttccccaatc
ttttccaacactaagccatatttgcctctgaataaaaagcccaaatttacctttttgagtg

tgtatgccacaatttatcaatagtctgaagatcttgggcaggtatttttttatatctgt
aaaatatagccagttgcgattttgtgcattcacaccagctaattgaataagtttttgttg
agttaattgatctgcctcttgcaaatcttgttgtgttagcaacacttgtaaatctttata
attcatgtgtttcgcagaacgtaatgggacaataccctcgggacataagtcagtagtaaa
cctaatgatttcttgatttttactattcaataacttgttatagatcaaaccatcgacgca
attacttttataattgggcctactaatcctttcaagaaatagatccgctaaactttttaa
ttgcatagagtcattattatgtatagtctcaataatatcaagttgttttttga---tatt
atttgaactagtattttcacttaattctaaaaactgaa------------attgcatttg
attt-ttacattaatttaaatagtg----attatttcttaaaaagaatttga-ttttttt
gaaaattagaagaacgacgaattatttctagaagtgttttcataatttgattgaatatgt
ttctccccgcataaatgataatttgaccaattaaaaaaattgcactttttattttaggta
caaagaagtcttgtacctcaagtagaaatgtaattactatttgcattgatgataatgtat
ataattcatgtgatctacaagcatatatatactgacaatttaaaccttcctgataaaacc
cccatactttgtaacgactttcgtaaattgctcgcggtctctcaatataatttttaatta
aagtattccaaactaaattatttttaatttttctagttttctatcagataaaaatttga
cattacaaagatagcttgcatttgtttcatgtactgtatcaaaattttgaaccaacatat
gtgcaataatattacttaactgaattaaatagttctcaagaaaaatttcaacttgtttaa
taggtacatttgaagtaaataaattgaaagtcttattcacaccttgagaggatccaaaaa
gtagctgaataattagaatttgtagtaacatcttatagtcatatattaggtagtttaagt
cattaatatcttggcaaatatcattatagtctagttcgtataacctacaaaatttttga
cacatctatgaaataaatcaattaaaatattttcagttagaatcgtaatatcatttaaac
ctaaatttgaattataaatttctaacaggatttttctaattctgttaatattatttta
ataagtttctttttacttcagttcgaaaaacatctaaaattaatacttcctgagaacaat
tagttaattttctattaatttttatagaagttctaactaatagctctgcaacctcttgat
tgagtatgtgtccctgtgagctcggccaataattattcacggtatatcatagtaaggaat
atttaag-----tttatacgataacttttaatacactgaaaagcaagtttatcactgta
tagccattag-------cttttacagatcttaatagttatttttcaaataaagttatatg
atattagcgaccaagctaattttattaaaatttataattaataatgacaatttattactt
tgcacttgccagtcaaaactttctcttagtacaagagcccttggaagaagtttttagaga
gagagttaactattatcagtcaaataataaagcaatcgatttttggctagtgccaaatcc
ttctttttttagaaaaaccagaaatgatttcgtttaaaaatcttgtacctaatgatgctgt
agccataatctctactaacccagtatttattaattggttaaagttaagaattggttatgt
ctgtgttggacaatttgaagatgatttacaactttctgaagagtctttaaacattactgt
tgcaccagaaaagatttaatcatacttttattgccactaggaat-aattat-----ttaa
acggcaaatataaatt-caaatccttgtttttttctatagtttatcgcttagtattttatt
ctgagatggagtgacaaacagtgtaagtatttcttgactaaatgtttcagttgctttgga

tttatacctattagggttgactataatagacagcattcgcttaatagtgacattttcaat
ctgagcccaatgtacaatccctagctctaactcctttgcaatagctgaaacagaaacaaa
agcagctcctaaacctgactgcacggcatttttatagcttctatagaatttaattccat
ctctattttaaagcgactgctatcaataccatgttgactgagcactttatctatgacttt
tctgattgttgattgagtgtctaatgcaataaatttaagtctatacaggtcttctttttg
gatgtcttctaatttagaaaacggatgcgatgttggaagtattagtgctaattcatcttc
ggcataagatgtaacttgcaaaatatcttgtaattcagtaggtacttcccctccaataat
tgctaggtcaacttgaccattagctacgctccaggaaatgagtctagtggaatgtacttg
tagttgaacagctacttgtggatatctttgcctgaatagtcctattaatcttggcatcaa
atatgtcccagttgtctgacttgctccaataattaatgtaccaccttggaggttttgtaa
gtcatcaagagctcggcatgtttcttcacataaagctaaaattcttcctccataccgtaa
aagtaaacttcccgcctcagttaaagttgctttttattacctctctcaaagagggaaac
atttaattggcgctctaaattttgaatttgtaaactaatagctggctgagaaacatataa
gctattagctgccttttaaaactgccttctttggcaattgcttttaatattcttaattg
atccaatgtaaatggaagatctgtcattaaagaattgtagtataatg---------tata
ttatttaatattgatcaatttatgcattcgccaagttaaatatttgctattacgcttcac
tgctttgaaaataataaaaagtataatgttaaatacactgtattagattttaatatctgt
ttaatgttaaatttatataataaatatcg------taattttaattataagatataggag
acaacaatatggactctagacttttaattgtattaataccagttttagcagctgcatctt
gggctatttataatattggtagagtcgcgctacaacaatttaagaaaatgacatcttaat
gatgatgagaatata-atcttaaaattttatgagatagagaactttaaaatctctatctc
tattgtatatcttaataaaaaaagtaagta--acccttgtacttactgttattacctatt
aagatgatgtac-agtacatatatttataataagtttaagctaacttatt-----tttta
taattaactaaattaataacacaattatggctgttccaaagaaaagaacatctaaagcta
aaaaaaatgcacgtaaagcaaattggaaaaatcaagcaagagtcgaagcacaaaaagctt
tgtctttagcaaaatcagtattaactagaaaatcgaatggatttgtttataatccttctg
aaacagcagatacctatactaactaagagtcttt-ttactatcaagctagccaagtatca
aaaaatctattttaagtaatttttatatat----taaacatttggaaatcattctatttg
ataataaaataaatagcagattaaacaaaattacggcaacaaattatgcagcaaagctag
atgaaactaatgaaatttggttatttaattgcatcgaaaatatccagcacactttttga
aaagtcaattaaaatcaagtcatattactaaaattttcgtttctggtactagctttgaat
atgtagcaggtttacctggactattatccagcttaacactaagtggtagaataaatccca
ttagcatatatggcccaaagtctttggaatggtacctgaaagcttctaccaaatattctc
aaactaatttttcttttcctattaattttcataatgtgaaatatggagaacaagttatta
atcaattctatacagtagtttgtttaccgttgagccaggacaatttactgtatggattta
ctatcttaaaaaaagaaaagccaggagcatttaaattagcaaaagctaaaaccttaaata

ttactcaaggtcctatatatggcaaattaaaagaaagaaataattttctaagtcccgatg
gttattatctatgtggcaaagatttctcgttaagtacaacaatgggacacaaaatatctg
ttccactattagtcagatattctagaattatttcagagatgcattggttttgctcttatt
ctattcgatgtaacacttatctacatcagcaaaccataaaacatttgccaaacaatattt
taaccgatactatgaaatctcaaatatctcaagataatagttgcattgaataataattat
tcttgatatataatagtctcgacaatataaataacatacaagagcgaatttcttg-----
----ttatatattaaatttataattatttatgacatacgcaattattgaagcaagtggca
aacaactttgggtagaggaaggacgctattatgatttaaatcgcatacctttgatccag
gccaatcaattatactgggtaaagttttattgttaaataaaaatggacaagtgactttag
gacatccttgcatagcaggagtaacaataaaagctacaataatgaggcatttacgaggaa
aaaaaataactgtttttaaaatgaaacccaagaaaaaaatgagattaaaaaaaggacatc
gacaagaattgactcgtttaatgattaattcaatagtaaactaggtctagatacagttgt
aactgagattgcttcaa-aagattttgaatatttaactacaatacatataatggcacata
aaaaaggtagtggtagcacaagaaatggtagagactctaattctaaacgcttaggagtga
aaaaatatggaagcgaacaagtaagagcaggtaatattttagttagacaacgaggcacga
aagtcaagccaggtcaaaatgttgggagaggaaacgatgacactttattttcgctgatta
atggctttgtatttttcgaaaagtctaaccaaaagcaaaaaacaattagtgtttatgctg
caaaaaaatagccaaaataattggtgcaagatgacctgaat----attttatttcactt
acactggcaccataaacttgatttattaagaaatggagttccttaatgaaccaatgaatt
ttattaaagatttgctttgaaatgtaatgacaaacactattgtaatttcctgtctacaca
atatggctgctattctatattgtggtcagatacagaaactagtagtagcaaatgctcatt
atcaagtaagtgatatttacctaggttgtgtagataaaatcttctcaggaataaatgcag
catttattgacctgggaaaaaatgagtacagtggattcatccatattagtgatactggtc
cgcttaaaaagaaatattatgttaatagtattactaacattttaacaattcgacaaaaaa
ttttagtacaaattactaaagaaccaactttaaataaaggaccaaggcttactgctaata
ttacattatcgggtcgatatattgtactaatgccttttagtcaagcaatttgcatatctc
gaaaaatacatgatgaagatgaacgtcattatttgaaagctttagctattctaattaaac
cagcaacaatgggcttattgttccgaccttctgctgtaggcattgatgaagaaattatac
taagcgaactaaaaaatttaaaagagcaatggaactttgttcaaaaatctgcaatcaaca
gctatgcacctggtctcctatataaggatgaggatattattaaaaaggtaattagagatt
tttatagtaataatacaaataatatcgtaattgattcaaacttgggattaaaacaattga
gttattatatccatacttggcactgcaataattctagtacagctccttcaattaagcttt
atagtaacaatcaatgtatactagatgcttttggtattaatcaagcaatttctagagcgc
taattccaaaagttgatcttatacttggaggctatatgtttattgaaactttagaagctt
ttacaattattgatgtaaattccggatcttttaataactctactagtgcgcgagaaacag
ttttaaaaacaaattgttctgcagcaacagaaatagcttaccagctgcaaattagaaata

ttgcaggtgtaattataattgattttattgatatggaatcacagagagatcaattacaat
tattagagcacttcaataaagagctagcacttgatgatgcaaagcctcaaattgtacagt
tatctgaattaggcctcgttgagttaactagaagaaggcagggcaaaagtttatatgagt
taattagtagtgacttcaattactttcatctccttatgcaatcagagcagactaagtctc
ttgaaagaatagatcaaaaaaagcaaatcttccagactcttaatagatcatggctatctg
tagaaattaatactattaataaagtatttttcaaaaatcaaattttatatacttacca
acttttatttagcacgtgatttatatgtaattaacagtaatctcacttataaacaaaatt
gtttattcactactagattgaaacttatctactctcaagagtataataaaatcttgcctt
gtactggttatagttgtctcatcagttctaatatagactataatcaaaagttccttttgt
gaactgtcggtttaaattgt----------------------------------------
--------------------taaaaaagaagccctctcttgtatgtgagagggcttc-tt
ttttagcttactaactattctagtgaattaattagaaattaaccattgacagctggagct
gctagagctactggcaaggattcaccagaagctagatctagagggaagttatgagcatta
cgttcatgcattacttccatacctaggttagcacggttgatgatatcagcccatgtattg
attacacgaccttggctatcaacaacagactggttaaagttgaaaccattcaagttgaat
gccattgtgcttacagataaagctgttagccagataccaactacaggccatagacctaag
aagaagtgtagagaacgagagttgttgaaactagcgtattggaagattaaacgaccgaag
tagccatgagctgcaacgatgttgtaagtttcttcttcttgtccgaatttgtaaccatag
ttagcagattcattttcgcttgtttcacgaattaagctagatgtaactagggatccgtgc
atagcactgaacagagaaccaccgaatacaccagcaacacctaattggtggaatggatgc
attagaatgttgtgctcagcttggaacacaagcatgaagttaaatgttccggagatacct
agaggcattccatcagagaaactaccttggccaattgggtatactaggaatactgctgct
gctgctgctactggagcagtaaaagcaacggagatccatggacgcatacctaggcggtag
cttagttcccactcacgaccaatgtagcaagctacgccagttaggaaatgaagaacaact
agttggtaaggaccaccgttgtataaccattcgtctaaagaagcagcttcccaaattggg
tagaagtgaataccgatagctgcagaacttggaataacagcaccagaaatgatgttattt
ccgtatagaagggaaccagcaactggttcacgaattccatcaatgtctactggaggtgca
gctacgaatgcaatgatgaatacagatgtggcagttaataaagttggaatcattagaaca
ccaaaccaaccaatgtataggcggttttcagtactagtaatccaagagcagaaacgttcc
cacaagctagcgctttcgcgtctttgtaaagtagcagtcataatttttatcagtttta
aggatttaactaagatgcttcccaaacaatattttgtctgttaaatatattattacttaa
gatcgttttaaatgagaaca--tttttaaaaaaca-taattaaatttaatctcgataact
aagaaaatataattattgagatattattatataattgaa----------aaataaattt
tt-act-------------ttttgcagaattagtatgattttaagggtataataatagta
atgcatactaatgctgcttattttgatttcaacagtttacataatcgagctttacttaaa
tacgtatgtcacactttacaaaaattcagacgactatacaagacttaaatctgttaaagc

atgctttaacagatttaggtttaatttggtctatgaattctagtcatatacaagttgatg
aaaatagtcagcacaaagtagatatcttgatcagacaagataatttatcacatattggct
ttatctggaatgataataagtaccacttagttgctgacttgcaactttgggaacaacctt
ggtctttagaagtattttagataaattatcgcagaagtatgcttaccattctattatcc
aagaaacgaaaaaacaaggctttgaaaaaatggagcaaatctcgcaaaacgatggatcaa
ttaagttagttgtgcaacgttggagttgttaattaatatagaaatgcgggcggagagact
tgaactctcacgagattatctcactagaacctaaatctagcgcgtctaccaattccacca
cgcccgcactgctgtagt--aataatactatatcacaattcatgaataaaaacaaactca
-agaaataaattagctaccaagtttaaatgcatcaacaaatagaccatatgttataccaa
ttttgaagttattaattaaattgattttattttaatttttttgattagagtaaacga
tcttactagttaactctgttgctgcaactattagactagctgcaatgattccccagtctc
ctgtttgcccaggaatcgtagataatcctgttgaaataaagaagcctaataacaagctga
ttaaaccagttgttaactcacttaaagaatagtacagtttatcatttaagttttctacta
aatatgcaaaaaaggtagatagtcgagttttaatcataatatttaataacggagttttaa
aaggataataa-----------agattaatgtttttagatccttttgaaacccgtcaata
caaaaattgctttttg-----------tttaaagatcttgctcacttaaagcattaataa
gatgttgaaatggttcttgtataatgtctttacgtttaattgattgtaaatcaagttctt
cttctattatttcttgcaataattgtatgcttcttaccgttggagcaacaggcacagaca
atgaattataagtatctttgaggccgtttaatacacgctgatctagaatattagtgctac
cagagactaatgcataactagcatagcgaaggtagtattctatatctcttagacatgcag
catatcttctagttgtataagaatttcctccaggtcttaataattcgggttgctcttcgt
ataattgagcagcagcctcttttaatatatttgtagcctgattattgataatttcagcta
tttttattctatccaatccacttgaaaagaaagattctatttgtccgacagctgctttat
ctaaataacgaccagtaagatcgtagcgatttaatattgctgttatagcatcttgcataa
attaattctcctac-ttgctatcagagtggtattatgaatgactagtat-----tgtaa-
----------tataatcatcact-------------------------------------
-----------gtttgtataaatagagtatatgtagtttttaattactttttttataaaa
attaagaaataaagattttt-taataggtttacaattttcttaatataactcaggttct
caaccgagataagttttgtaacacat--aagctaaaataaagatgaatgagtaatcttga
------------------------------------------------------------
------------------------------------------------------------
------------------------------------------------------------
------------------------------------------------------------
------------------------------------------------------------
------------------------------------------------------------
------------------------------------------------------------

```
------------------------------------------------------------
------------------------------------------------------------
------------------------------------------------------------
--------------------------ataagaataatatgcatcaa--------------
------------------------------------------------------------
------------------------------------------------------------
------------------------------------------------------------
------------------------------------------------------------
------------------------------------------------------------
------------------------------------------------------------
------------------------------------------------------------
------------------------------------------------------------
------------------------------------------------------------
------------------------------------------------------------
------------------------------------------------------------
------------------------------------------------------------
------------------------------------------------------------
------------------------------------------------------------
------------------------------------------------------------
------------------------------------------------------------
------------------------------------------------------------
------------------------------------------------------------
------------------------------------------------------------
------------------------------------------------------------
------------------------------------------------------------
------------------------------------------------------------
-----------------agtatagatagtcttttactccagcatcaaaaaagatatttaa
cacatattgaaaaaattgattatgtaagtaaaaatctaaatttagataaaattagctatt
ttataattacaaccaataagaataacaaagatatccttgtagaacaattttgttataatg
cagatggttatacttattcaatattatttaaaggtcaaacagctaaacacctttgttaca
tagttcttcagcttaaacaattaaatataagaatatctacagctcatgctctatatttag
gacgagaactaatgaagtcggaattaactcttatattagatcaacaatatatccaagact
aaa--cttcatagatattaattatatttgtgctaatatagtattaataagggcatgtaac
tcagtggatagagtatcagattccggttctgatggccgtgggttcgaatccctccctgcc
cggcgagtatttagaattct-gtggcaagatatgt-------tagtatctaatttaataa
tattaaaaatattggggctgccaggtttctacattataaaaacaagaaaatatgaaagtt
```

aaacaagctctttgaaagagcctctagtaaaattaataaatgcagaaaacaatattgttt
ctttttctcgaaaattagctgttgcttaaatactgttaattttggcattcgacgtgtta
agctcttattcgcggcgaatattctgttagagttgctcttagtataagaaaagtttcaaa
atctatttatatttatgtttttacctatttttatttaaagtaaggttagta-ttttcaa
tttttataatggacgcgggttcgaatcccgccagctccatcaatttaact---------t
tcataattaactaatccaatctttttaagaaccagaaatataattggagcaaatatggat
tttataaaaatcacagaacaagcttttaagcaaattgtagtcttgaaaaatgattacaat
aatgatgtacatttaagaataggtgtcagacaaggtggctgttctggcatgtcatactca
atgaattttgaagatgtagctaacttacaagagaccgatgaaaaactgattttagatgac
ttttctgttgtttgtgatcccaaaagtcttttatatctttatggattgtcattagatttt
agcaacgacttaataggaggagggttccaattctccaacccaaatgccagccagacttgt
ggttgtggtaaatctttctcaggctaaa-------gctactttgttttatattttgtatt
atacgtgttactcttataatcagccattataaatacaaaagtatttttattgaaagctct
ttaaactagtcaaagccttatgaaaactactttaaagaggttttatt-tattatggcagc
ttaaataagtatttatcatcaattgatgaaagt---------acaaaaattcactccctt
gttattgcctaattttaattagttaatttttttattgatacatgctaatattt-attagcc
ctgatttattaggaatttataatttctttttttataaaagagcttgtcagtttctagcta-
----------------------aaatagattatatatttt-attcatttataaaaactc
aacatgaatcttgctcaagttacttgcaactacttcaatacgaaaactctcaataagttt
tattgcaatacggaatatgtttcctatggtaattttagtcatctcaatttatcaaagata
aatttgccatcttctgccaaaataattgcatttggactccctcaatttgattttattcct
acaaagattaaaaactctatccgagaagcagtggtaatatctaccatcgatgacagtaat
atttttgctgattcaactaataattggataaaaaaacataaagataaaaatcaccactct
cttttctataaaaaaatttttaaagctttactgcatcaaaaaataaaagtcgtgacagat
tcagc------taaaaattctaaaatagataaagagaagggcttgattttgagtcaaaaa
tatatattaaagaacagttggaagccatccataatgtcttcttactttgaaaacaaaaaa
aataatgaggttgctaatgtatctaaaaaatttttaatcgatcagctatctgcatcacct
gtgtttgtagtaaaaaaatggttttaatgaaattattttaggccacccagcattccgtata
aagagaggtgtaattagtaattacactcactcgttttcaaatttattaaagcaatctcag
ccaacttatcctatatctactggcttattctttttttcatccagatgacgcgtttgaattt
aaaaactttataaagtcggttaatccgttggctagtgagcatatggaaataaatgtagaa
cctgttggcctacactttgcttataaaatgaacagaaatgtatcttctgatactcaattt
ttattcattccagattttaaagaggtaggcgatttattatttaaatataaaaaagataag
cacttaatctttcacaaggatcagtgtcatggtaaaaattttttttcaggggcaacctata
tatattattcagcctactattttcaaagatcgaagcggaaacctaaatactattcaattt
actggactaaatgataatagagaagttatttttactagtcttgaagctgctaataaatca

tggacaaattttataaaaaacaattcacaattaaaatcaattagaaaaccgactttatta
gtctataatctagaaagtttttaaaagatcaagaacgattgaataaacaagattttcaa
aaatttgttgtgattactaataaaaaagcttatcttgctacaaaagagttggtagctgta
cccgattctaatagcgttttaaagcatttaaaattaaatatgaagcctaaactcttcttt
ataaaattatgggtaagacggctattttccaccttaacttatgaat-aaatgatgtt-gg
atttattttcttgaatagtattcaataactagtaattcatttagttgtaaagcaacccat
tctctgtcaataactccattgacttttcctgatagattagatttatttaattctaagtgg
ctaggaatgttagctaatccagggaacgctaaatagttttccactagtcttttagatgtt
tcccgagattttacactaattaattctcctggcttacattgataactacatatagatact
acttgtccattaatgcaaatgtgaccatgatttactagctgtcttgcagcaggaattgta
ggcgccattccaagtctaaaaacagtattatcaagtctcatttctagaagctgtaataaa
atttgacctgtagatccttggagtttttagcggccttaatatacttaaagagttgtttt
tcacttaatccataattaaagcgcaattttttgcttttcttctaatctgacagcatattca
gaaggttttctagatttttgtccgtgctcccctggtggataagattgtttaattgttttt
ctacttagcccaggcaaatcacctagtcgacgggaaatgcgtactcgtggtcctctgtat
ctagacatattttataaattttccgaatattaat-gattaag----------taatgagt
agctatttttaacacaaaaagagtattaaaacaaagc-ttttgctaaaaaattttcatg
aaaaatatag--------------------tgctttac-tttttaagcgggtagcgggaa
tcgaacccgcatcattagcttggaaggctaaggttttaccactaaactatacccgcaata
ttcaataaactgtaacaggcttaatataacataactatcttattcttagatattattaaa
attgacaagaag--caaattatttatgctccttctaaaactacatcaagaaaacgagcaa
tctctgcagcttgctcttcaatttctgaaagcaataatggttgacccactctagttaatg
gaattactttttttatcttttgtgcataagtaaatttctcgtcgtggattcaaaccttctt
taatatctattttttattgacttaatatcttttatattgaattgaagacaaatttgtctat
tttttcctgggaaacctaaacgaaaaattttaacagtaccttcatctttattaaactcat
tatatccagcacctatgtcccaaataatagttaaccataaaaataaactaagaaagattc
cgatgctgccatagaatgtcataacaataccttgcggaataaatattaattctgttgaag
tcgtaaaaggcaataaattaatttgaaagtaacttgataatcctgcaagaagaaatcctg
aagctccaacaaagattatggtagcccaccagtaattactaagtcttcttgaacctaaaa
ttagatcttttctaacagagtctatagataatgtttttttcataatttgtt-ttttaaaa
caata-tactaatttgacatataaaaattttgacaatttttttataagcgttttagattt
attaaatgagtttgatagcttttaatcctaaaaataatcctgttgctaaaccaaaaaaag
ctgcaagaaatattatatatcctagaaaaactgaca---tatta----ttattattattg
ttacaatagtgtttacaaaaatagttgaaatctattactaaagtattatattacgttaaa
tgcttcgctgtctacatgaaaaaactttttttcagtttttagcgtcgtatttatattagaa
ataatattgtaataattttattaaacttctggagaacagaatcttatggcagaatttatt

aagccttataatgacgatccttttgtaggaaatctatctacaccagttagtacatcgagt
tttagtaaaggattattaggaaatttaccagcttatcgtcgaggtctatctccacttcta
cgaggcctggagataggaatggcacatggatacttttaattggtccttttgataagtta
ggtcctctacgaggtacagatgtagctttacttgctggattttatcttcagttggcttg
attattattctcactacatgtttatctatgtatggaaacgtatccttttctagaacagat
tcaaaagatccattgcaaacagcggaaggttgggggcagtttactgctggattcttagtt
ggagcagtaggtggctcaggatttgcttacttattgctagctaatattcctgtactccaa
actgcaggtttaagcttgttttcttaagctagtaaggggacttgaacccgtaacctactg
attacaaatcagttgctctaccaattgagctatactagcgctcagctaattgtaacatat
aaagatatttgttgcaatctggtattttcagatgcaacaaatatctttttttatttagaca
gaactgttaaattaatatatgtatgatatgttaattagaatcagagctatctgactcagc
agaatattcttgatcacagtttaaataataacaaattgtttgatctagacatgttgcgga
ccaaccaacaggtgtttctcctgataattcttgcatatctccagaattgtaaaaatcatc
acttggcttattactatttgcttgcataaatatcactcccaaaacactctgaacttaatg
tgaagtaattgaacttctcaataatatattaacacagaatataaaatatggtgcttattc
accaattgtaacaatattatgaaatctacaagttctttgtaagtgttttaatagcttttg
ctgaaacaagcattttaatccatttattcttctctcttgaccatactttttttctgttgca
aattaacttttgtaattttttttgttcgtttgtgagaatgagagactgcgtacccattat
tcgctactttccctgtaagctgacatttttttgacataacgtt------attaacttata
tatttatttaaagtacttgccaaagttgactttggtacagcaccaattaccgtgtcaact
ctttctcccgctttaaaaatcattaaagtaggtatacttctaataccatattcagctgca
attgtaggattatcgtctgtgtttatttttactaccttaatagatgactcatattcttct
gcaatctcatccactacaggagagaccattctacaaggaccacaccacggcgcccaaaaa
tctacgagtacaggtaagttattattaataacttcttgtttgaaagaggcatctgtaact
tgagatactgacatatttttttaacctttaatatatgttccttgctaagcaaatcttatca
caaatttcaagaaaaatcttctatagtgaattactattagtgttgaattaa---ttt-tt
tactagacattaaaaaaacttattacctatataaataaagtggtggccgagtgatatctt
gatagtgataatttttctattaaggttgaacactagaaatcatttttcaatcatatattt
ttttgaaaa--acttctttgtgaccatgaaagc-taagtatattacaatatattttatta
tttgtaacttatttcttataatggtataaacaacgcaaaagatactgccttataatcaag
gaggaatacatgtctcaatccgtagaatcacggactaggattaaaagcgaacgttacgaa
tctggagtaatcccttacgctaaaatgggttactgggatgctgactatgtaattaaagaa
acagatgttttagctctgttcagaatcactcctcagccgggtgttgaccctattgaagca
tctgctgcaattgccggtgaatcttcaacagctacttggacagttgtgtggactgacttg
ctaacagcttgcgacttatatagagcaaaagcatatagagtagatccagttccaaacgtt
gcagatcaatactttgcctatattgcttatgatattgatctatttgaagaaggttctatt

gcaaacttaactgcttcaatcattggtaacgtttttggatttaaagctgttaaagctctt
cgtctagaagatatgcgtatgccggtagcttacctaaaaactttccaaggtcctgcaact
ggattaattgtagaacgtgagcgaatggataaatttggtagacctttcttgggtgcaaca
gttaagcctaaactaggtctgtctggtaaaaactatggaagagtagtatacgaaggttta
aaaggtggtcttgacttccttaaagacgatgaaaatattaattctcaaccatttatgcgt
tggaaagaaagatttttatattccatggaaggtgtaaataaagcatctgctgctgctggc
gaaattaaaggccattaccttaatgtaacagctgcgaccatggaagatatgtatgaaaga
gcagaattctccaaagaggttggtagtatcatttgtatgattgaccttgtaattggctat
actgctattcaaaccatggcaatatgggctcgtaagcatgatatgattttacatttacat
agagcaggtaactctacttactctcgtcaaaaaaatcatggtatgaacttccgagttatt
tgcaaatggatgcgaatggcaggtgttgaccatattcacgcaggtacagttgtaggtaaa
cttgaaggagatcctttaatgattaaaggcttctacaacactttacttgaaagtgaaaca
gatatcaatctacctcaaggtctattctttgctcaaaactgggcatctctacgtaaagtt
gtaccagtagcatctggtggtattcatgctggtcaaatgcaccaacttcttgattactta
ggcgatgatgtagttcttcagtttggtggtggtacaattggacatcctgatggtatccaa
gcaggtgcgactgcaaatagagtagcattagagtccatggttatggcaagaaatgaaggt
cgtaattatgtagcagaaggtccacaaatcttaagagacgctgctaaaacttgtggacct
ctacaaacagctttagacctatggaaagatattagttttaactatacttccacagacaca
gctgatttcgttgagactccaacagcaaacatctagtttaatgactacttgctaatgctt
aatttggcaaattgtaagtagaattgacttataacaataaggagcatagaatagtgagag
taacacaagggaccttttccttccttccagacctaactgatgaacaaattaataagcaac
ttgcttatatcgtttctaaagggttttcagcaaatgttgagtatactgacgatcctcatc
caagaaactcatattgggaattatggggattacctttatttgatgtaaaagatgcatctg
ctgttatgtatgagattagttcatgcagaaaagcaaaacctaattattatattaaagtta
atgcttttgataatactcgaggtattgaaagttgtgtactatcttttattgtaaatagac
ctattaacgagccagggttcttattacaacgccaagactttgaaggtagaactatgaaat
atagtttacatagttatgctactgaaaagcctgaaggagctagatattaatttcaattaa
tatctaaaacttatgagttagttacccctctttaatatcttaattactaaattaattttg
attttaaagagggtaaatttcgaaataaattctagttaaattagaattaacaatccaaa
atacacagaaataatgcaatcacaagacataatctccaacgatactcttgttaacttaca
agaagaatatgatagaacacaaatccaggaagttcttaatgaattaaatcaagaacttat
aggattagttcctgtaaaaactagaattcgcgaaattgcagctttattactgattgatag
attgcgcagaaaactagaactagtatctggtaatccaggattacatatgtcatttacagg
tagtcctggaactggcaaaacaacagtagccatgaaaatggcagatattttacatagact
tggctatataaaaaaaggacatttattgacggttacacgagatgatcttgtaggccaata
tattggccatactgctcccaaaactaaagaagttctaaagcaggccatgggaggagtttt

gtttattgatgaagcctattatttatataaagcagataatgaaagagattatggctctga
agcaattgaaattttactacaagtaatggaaaatcagcgaaatgacttggttgtcatctt
tgccgggtacaaagatagaatggaaaaattctatgaatcaaatccgggactttcttctag
agtagctaatcatgtcgactttccagattatacttcggacgagttattacaaatagctaa
aatgatgatagaagaacaacaatattgttttacagaagaggcagataaaactctgctaga
atatactgagcgaaggatgaaacagccatatttgctaatgcgagaagtattcgtaatgc
tattgatagagctagaatgaggcaagccaatagaatttttgcaagtggagaaaaagtttt
gacaaaagctgatttagtaacaattgaagcagaggacatcttgaaaagtagattattttc
attgcccaattcttaataattaagtagcttaatacaacgagatatcataaaaaaactttc
accataagttgcaaatttctttaaaaacctatactattaacatagtaaaggtg-tggcgg
catagccaagtggtaaggcagaggattgcaaatcctttatcccccagttcaaatctgggt
gccgcctagtactaaaaa--gggggtgtggtggaatggtagacacaacagacttaaaatc
tgttga-ttttaataatcgtgagggttcaagtccctccacccccatgtattaacttataa
aaaaaca---atgtgtatttgtattaattgtaaccatattactgaatgtgatacttatca
tttaatagaatctcagcacaaacagctgcatcttactgaaaccccagctttcatacctaa
atgccctgtagtttgtgttaatatatctagtaatagcagccatactcaaatcgattggga
tttagtggaatgcttatcatttgtggaaaaaccaaatagttggaatttaggctgtgatta
gttgtaaaaagaattatgtggtatacgcaataactctatcttttaaatactctgtattaa
ttttagatgttcttaccaacaatatttttcctgttcttgctatctctattataccaaatt
tcgttaataattgttcgatggcaacaatctttcctggatcaccagtaacctctataatta
aaagatcttccgcgatatctactatcttagctctaaaaattttcacaatttctaaagctt
ctgttctggtctgagaattaatcctaatcttaattaacattaattctctttcaactgaag
gaatatttgttacatcttgcacgttcagaatatttactagtttatataattgtttagtga
gttgttcaatagttctattgtctccctgaactaccatcgtaattctagaaactccgattt
gttctgctggtccaactgctaaacttgcaatattgaagcctcgacgtgcaaatagaccag
atattcttgaaagtactcctgcttcatcttgaactaaaactgataaggtatgtttcatga
aattttaatgttatactgcttgatctattacgtatattctatgaaaattacagaaaaaat
gctagttactatataactaaattattccgactaaacaaaatcccaccatcagacggcata
tctattttgtaatgttatac-----------tatttaatgatattctaattctaaataga
atgctaaatttttttaaaatttgttaggaacttgtgctagaaaaatacaaaaacagtaaga
aactctctcgagatctaccccaaatcaacgatcgtattagatttccaaaagttcgagtaa
ttgacgatgaaggtgaacaattaggtatttttgtacctgaagaagctatccaattagctg
cccaacaaggattagatttagttgttgttagtgacaagtctgatccaccagtttgccgaa
tattggactatggtaaatataagtttacacaagaaaaaagagctagagaagccaaaaaaa
aacaacataacagtagtgttaaagaagtcaagatgagatacaagatagaagagcatgatt
ataaagttagaattaatcaagcatctaagttcatccaagcaggagataaagtaaaagtca

ctataacattccgtggtcgcgaaattcagcattctaatttagctatagatttgttaaaca
aaatggcaagcgacttaacttcgatatctgaaattcagcaagctccatcacgagatggaa
gaaatgttataatgcttttatctcccaagaaagcgagtt-aaatgattataatttaaatg
catctggctggattcgaaccagcgacgtctctttcgaaatggcggattatgagtccgctg
ccttcggcccctcggccacagatgcatcacttaaagtttataactcataaagtatataaa
aatcaatgcttgtttctttttcatgacaccta-aaaaaattattattcattcatattatg
ataggtcgcaactgctgaaggggaaacttggtttagatacctaaatatccagtatttaaa
tactgtgtccagtataactggaaaagttgaaataaatagaaaaataaagtcacgactttc
aggtagtcctaaatgccttaaaattacttcaataataacttcccagccatgaggagaatg
aaaaccaacaaacatgtcagtaaaaagaataattagaaaagctttcgcagtatcactaag
gccataaataatttcatttaaaaaagatttaacaattgcaatttgcctttgtccagttat
cattaataagataaaaacaaatattgatacaatatcggacaaaatatttttacagcgtt
tgcactttcatttgcataatattctcctaattctctagccttcagctgtactctttttc
tataatttcataagatatatcttctgatgggttcaataaaacttcaaagtgaatttttc
ttcaaacctttgtagttcagcgaaagctctctcttcttgagatgaatttaaaaaaatttt
aggttgttcctggttccataaataatcaatacaaggaccaaacacaaaaaaattttgaagc
ttggttcactaatactggagagataaataataataaaatatattttacagaagtaatagt
ttgatgcctagatattctaaattcttctatggcttcagattctccattcggatctaactc
ttttcgaaacttttcaaaagtattagtaatagacctaggtattggcccaactttttcaaa
ggcaaattgattatttcgttttaaattccaatatttcatttttattgctgctaaaattt
ctattaatcaataattgatcacgtattcactataaatattaattctagaactaaacatat
aaaacaatagttatgaataataaaatacaattaaactttcattatgcattattactatct
tttactacagcaatgtatggatataaatcttatcgtgcaactatgaatacactgttaatc
aaatgtgaagatagccattcattattgtctggtaaacatactgcctttattcctataatt
atattaaaaaaaaatctttaaactccctaaaatcagaacctcacaatactgagttgtgtc
ttaatggaatgggaaatagaaaattcattgacaaaattgcaagtttgctaagccgtttag
aaaaacagatgcactatactgaacatcttaatcaaataaaaactagtgggtacttttctt
tactttatattaacagtaaagcaatctatacaaaaaaaattagtaatatttttctattac
ctaatgatgttttaaaaagaatttatattcataataagaaaaaaaagcttatacctcata
cattcttaataaaattatttcaaaaacaaataggttacccgaaaaattttgcgaacttaa
acctagccttatcaaaaattatgcaatggtatgctgatagaggatatcaatggtctttag
tggagatcaagcaggctgctgatccatcgtatctcattctgaatatccatgaaggattag
ttaagacaattacaacagagtattatactttgtcttataaaaaattacgcagtatatcaa
gtacggaatcaatagagaagtatcttggagtaacagtaggacttcctttaaatattaatg
ttttgcaaaaaaaaattaactatttaaaagataatcaattagttggcaacattatttata
gtattgagagatcgagcaataactctatgaatttagatatcaaatttcaaatacaagaac

taaaagataaagatttactagtacttgtagaaagatcatctaaaacatctcaaattattt
ctcacgtatatgacttaatagatttagaatgc---agttcctctaattggctagcagctt
ccaatataatgtcactatcgactagcaaaatacaagcatgctattttgatagtaaaatcg
attctgaatacgtctatacagatgctgttaatttaataaagcggttaacttattccattt
ctcatcaaaaaacactaataattgacgattttaatttacaaaatttactaagctggacaa
aaagaaatactattggattccaattgtatttacggaatttggatcatgccaaatcttttt
gtctattcagtctaaaatgcatccagagtggacttaatataaaaatttcgtatctaaatc
cttcattaagcattagtcgtaatttgagttttcaaattgcagtacaaataattaaacaat
atcaaacaagtaaacaatcagctctatctt------------------------------
------tgtttttaaaaagggagatttagcacaattcattttgaaagtatatgttcat
atcattttacatcttatttttcgatatctgaaaagatattactttctcgagttgtgtata
tagattccatattttacaattctgaaaatataggtctttataataataataaaaatcaca
ttataagtagctatgatatctttagacacaataagaaattattatatcaagaatttctga
ccttattgttaagtatacgttatcaaaatttcaattatttagactggcctttaaaaggct
atttatttgaaatgaagtctttatattttaccccatttcagaaaagcaatttttttgaacc
actcttgtcttgacaactacaaaaacctatttttttcataaaatttacttaaaacaaatct
ctcattttagtttaccactgtatttcagagcccatcttaaccatattttagttaatacta
ttaaatgtcagtcgaatttaaatatggaaactatatatctattattaagcgattatccta
ttaaatacactctgcttaagtctattgttactttttctataaaagttaggatggaatatt
taatacctataagccggagcatcagagtatctttgttttataattatctagactgttttt
taatgagacaatctaaacacttagcatatatactggaagatctgtcaggccataaacaat
tccaaagtttttttattggaaaaattttcatggggatttggaattcagcttaaattaccaa
ttaagcagatgccacccctttcaattgaatatactgtaaacagtggtcgccacttttgtc
tttacctgcatatttattatcaaagataattattattaaaatgaaaaatttaatttcttt
ttttgaccgaagcaaaggaaaatggatttcacagcgaacaacctatgaactacgtaataa
gaatatgagttcagtacaatctcaaatgcgaatacaatcaggtaatttgttatcaggatc
tacgcgacttgcatcattgaattggggagaaatttttagacaggttaaccataattctta
ttctcaagatacaagcaaatataatggccagtttaacttagaatttagtaatcatctaaa
taatcgtaggctattgacatcatgcacagtaacagatcctagtctagttaattttaaaac
tcgatatggtagtacaactatagatgaaacatattggtttgcaacaagtaatttacgtct
aagtactagcattattaaaaaatttgatacttgtgttgcagtatctttttgttcagaaat
caaagtttaa-attctgtataaaaacaactaatagctaaaa-agttattagttgctttta
tatatctaatttttgttttagagagtatataaaatattatattattctaaatctttacgg
ttaggatttctagaaggatcattagataagaatccaaaaacaaacagagaaataaagaaa
atgacagtagtataaacaaatattttcaaagtaaacatatttttatccttaaaagctatt
agtgtg----atatttatatatc-acacataattttatatcacgatataagttctggcgc

tattttataaacaaatcaatatctttatttttttagtaatgatttatacttcaatatgcgt
gatgcactaaatactaaatctttttgtacacgcttctcactatttttttaatattttgtga
tttctatgaagtttaccttcaattactacataatctaatttcttcagcaatttaaaagtt
tctaaagacttttttattccaaataattaagttaattaccgctaatctttttctttgtaaa
agcctaactttcaacttaattatttggtgtctactatcactaattttaatactcttacac
ctcaaaatttgaactagtaaggtacagctattcat-attaat-ttaa------tcttgta
actaatctattatgacagataa-atgaaaaaatcaaaatgaaatttcattatcttgttga
ccttgttcataactaatatcttttaatacttttaatattttcccgaaatactcttgcaaa
tatttttctaacgatattacttctgagctatcaatgcttaaaatactataaacttcattc
atcgaagcagtaaattgctccccactagttaaaatctcggcaaatgctaaacgatcagaa
atattccatgtccattgaagggttttagtgatttttcttaaagctgttaatagtccaatt
ggaatttgtgatatttgtgtttttttgcccagataacttttcacataatttaataatttca
gctgatgtccaggccgtgttaccaactaatggcagcactctgttttctgtagaaggcact
ccaagacttttttataactagctttgccgcatcctgagtatcaatatatgcaataggtgta
gactcccctgtaacccatactgattttttatccaaaataggaattgcatattggttaatt
aatccttgaaaaaaccctcccaaagaaaataccgtgtactttatatcagatttttgaaga
aagtctactacttgtgattttaagttcatcaatggaacagctgggtacttatctgcattc
aatattgaaaagaaaataaatctttcaactttagcagctttagcggcctcaattaaggca
attttaccatctaaatctatttttgctgcattgtaaggatctgtaggacgagatgtagac
gcatctatgactgctgttactccacaaaaagattgcaaaatactttcaggtaattttaaa
tcaccatatacaagttctgcgccccattctttgagaaaagctgactttcgtaagtttctc
accatgcatttcacattgtagccttcatcaagagcacgccttacaatttgacgtcctaga
gttcccgtcgctccaattactagaagagtcatattaattaaaaataagtatgtataaaga
cattggtttggtcgacagcagagattaaattaggtagagagggaatcgaaccctcatgac
cgaagtcgtcacattttgagtgtgatgcgtataccaatttcgccatctaccctattatag
accaactactacaagtaatttcgtctttagcgtaaaaataatactaagtttcaatagata
tgtcaacttttgagtttatattaataaaaaaccggagtatcttatgtccacttttgaact
aattccttatagattatatttaagtcaatctagtacgtggatgcataggatgaaagcaga
aattaaaatatatatattaacgatattatggatttcaatttttattttctcttactgtaa
actatgcattgttgccctaagtttaattgttcttagccttactgtaaagaataaacaaaa
cattatccagagacatttattgcgaactttagttatgacaatggtaactaccttcttatc
attcagcatagctactaattacaaacaatattctcaaccaaaacaatcacaagatttatc
ttactataatacagctaaaaattcttacatctatcaaaatatattgctacctaatagtcg
taagatcacaacaaaacaactgaggtgcactttaaagcctgcactctatttttttattac
tatatattcaattaaactagtcatgataacgacttccccagaaattttagtgattactgc
ttatagatctaggatctttaatacaatattaaatcgtgaattattatttactttcttatt

gtcatcacatattgttaccaatataattagtaaattagataaacttgttcaagtagctaa
cttaagaggtagcttaaatttatataaatcgtccacaagacttttatatttctttttt
gatgtttcaaatctttttttagagattattaaggaatcaaaagaaatatctcaagctct
ttatactagaaatcttaatcaagaaaacagtaatttttaaaaatatatagagccagatc
gggtattagtgattggttaaatgcaattattgggaccttatattttattattttaggatt
agtataaca-aatctaatcatccttctgactctggtaatacgaatagtaatagacttaag
tcgtggtgtaaatatgaaaaat-tcagatattatataaataattaaatgttttattatgg
tcaatactcaaaatcaaatttcacaaacttcagagcttgattatatagttaatcaacctt
ataaatatggatttaaaacttctgttgaatcagaacaatttcctagaggaatagatgaaa
atatagtaagattaatttctaagaaaaaaaatgagcctgaatatttattaaactttaggc
ttaaagcatacaaaaaaatggaaaaaaatgagtagtccatcatgggccaatctcaagaatc
ctaatatagatttaacagtattatttattacgctgttccgaagttaaagaaggaattaa
acagcttagatgaggttgacccagaaattcttgatacttttaataaattgggcatctctc
tcaatgaacaaaaacgaatttcaaatgtagctgttgacgctgtttttgacagtgtttcta
ttgcaactacttttaagaaagaattatctgcagctggtgttatattctgctcgatttccg
aagctatacgagattatccgaatttaattcataagtatttaggtactgttgtgcccgctg
gggacaattactttgctgcattaaattctgcagtatttagtgatggttctttttgctata
ttcctcctaatactgtttgtcctctagaattatcaacttatttcgtattaataacgaag
aatctggtcaatttgaaagaacactcattgtagctgatcgtggtagtaaagtaagctatt
tagaaggttgtactgcccctcaatttgatacaaaccaattgcatgcagcaattgtagaat
tagtagctcttgatggcgctgaaattaaatactctactgtacaaaattggtatgctggca
atcaagagggcaaaggtggtatatacaatttcgttactaagcgaggcttatgctctggca
ataattctaaaatttcatggacacaagtagaaactgggtcagcaattacctggaaatacc
ccagttgtattttagctggccatagttctcaaggagaattttattctgtagcgttaacaa
ataattatcaagaagctgatactgggactaaaatgattcatattggtaataatactaaaa
gtagaatcatctctaaagggatttctgctggtaaatcaaaaaacagttacagaggactag
taaaaattggcgctcaatcatttaattctcgcaattattctcaatgtgattctttattaa
taggtcaatcgtctcaagctaatacttttccttatattcaagtacagaatccgacatcga
aagtagaacatgaagcatctacttcaaaaattagcgaagatcagatttttttactttttac
aaagaggtatcagtttagaagaatctgtagctcttatgatcagcgggttttgtaaagatg
tattcaatgaacttcctatggagtttgcagctgaagccgatcgtttactgagtttaaaat
tagaaggaactgtaggatgagtcaaactattttagaaattaaagatttgtatgcgtctat
tggtgagacaacaattttaaaggggtaaacttatctatacaagaaggcgaaatccatgc
gattatgggacctaatggttcgggtaaaagtacactatcaaaagtaattgcagggcatcc
agcttactctttaatcagtggagacatcttatttttggaaaaagtattcttgaaatgga
gccagatgagagagcaagagctggtatttttcttgcttttcagtatcctgttgaaattcc

aggagtaagcaatgccgatttttacgcattgcattaaatgctcgtagaaaatttcaagg
attatcagaatttagccccttagaatttttcaattaataaatgaaaaataaaccttgt
tggcatgcaagaaagtttcttgacaagaaatgtcaatgaaggattttctggcggagaaaa
aaagcgtaacgaaattcttcaaatggctttgctagatagtaacttatctattttagatga
aacagattctggattagatatagatgcacttagagtagtagcaaaaggaattaatacttt
agcaaaatcaacaaattcaattattttaattactcattatcaaagattattggattatat
tattccagattttgttcatattatgagtaacggaaaaattgtcaaaactggcaatcttac
tcttgcacaagatctagaaaaaaatgggtatgactggattgtacagagttaaaaatttgt
taaataaggcaatactgagactaaagtatcttctttagtctcaatattattgatcaacag
aaag-----ttatttaggctttactaaaaccttgcttaacatcttcaatggctcttttta
acagttcttcactagatgcgtctaatttttactgctacgaatactttcaccaaattctg
gtttagaattctttaaatcttcccgtaattctttaataaattcagccacttttgatactt
caatatcatctaaataaccgttaattccagtataaataatagctgtttgttcttcaactg
gaataggagaattttgagcctgcttcaaaatttctctaagacgttgacctcttgcaagct
gattttgagttgctttgtctaaatcagatgcgaactgagaaaaagcttctaattcagcaa
attgtgccaactccaattttaatttgcctgccacttgtttcatcgcttttatttgagcag
cagatcctactctagagacagaaataccgacattaatagctggtctaatacctgagttaa
ataaatcacccgataaaaaaatttgaccatctgtaattgaaattacattcgttggaatat
atgcagacacatctcccgcttgagtttcaataataggaagagcagtcatactaccgcctc
ctagttctgaattcaatttagctgctctttctagcaatctagaatgtaaatagaatacat
cacctggatatgcttcacgtccaggcggtcttcttagtaaaagagacatttgacgatagg
cttgggcttgctttgttaaatcatcataaataaccagggttgctttacctttatacataa
aatattcggctaaagctgcccccgtataaggagcaatatattgtaaagttgcaggactat
ctgcattagctgcgactataatagtataatcaagagctcctttatcttgtagtgaagata
caacttgtgccactgaagatgccttttgtcctatcgcaacatatacacaaacgacatctt
gcccttttgattaataattgtatccagcgcaacagctgttttacctgtttgacggtcac
caataattaattctcgttgacctcgaccaattgggatcatagaatcaatagctgtgattc
ctgtttgcattggttcacaaactgattgacgtccaattataccaggagccatagactcaa
ttaatcttgttgaattacttgcaggttcacctttatcatctattggacgagctaagggat
caactactctacctaaaaaggcgtcaccaactggaatttgagcaattctaccagtccctt
ttacagaactaccctctagaatgtctctaccgtctcccattagtactacacccacattgt
cactttccaaattcaaggcaacaccaatggttttatcttcaaattcaagcaattcaccag
ccataacttcatcgagcccgtacactcgagcaataccatcaccaacttgtaaaactgttc
ctatattggctacttctatatcttgatcatacttttcaatttgttgacgaataatactac
ttatttcatcaggtctaatgtttaccatgtttttaatattctttttattaatgggaaatt
ctaaagtggtaattactcagattaatactatctg--tttttactgtattttaagttgct

actacatcaaggtgtgaagccatctgccttagttgccctctaatactagtatcaatgact
ttcgatccaatctgaatagtgaaaccaccaattaactctggatcgacagaaacaattagt
ttaacttctttagcctgggtcataaccttaatttttctgttagcagattttcttggtca
gagtttaaagcaatagaagtattaatattggcaattgttagagattccatttgatatgca
agttctaaatacttactagctatagcatccaacattgcaattcttttacgatcaactaaa
atcattagaaatgatagtgtattttcagtaatttgatcaccgaaagtggcaataatgact
tgctttttgcttctgcagtttttaatggattagct---aaaaaatttttaagcttgtca
gattgcaataaaatattttgtattaactttatatctatactcacctttctgtaactttt
tttgttttagctaattcaagaagagctacagcataaggctgcgcgatttttataacaacg
ttattactgctcatagttggtctcctaacttagcaatattattatcaatgatacgtagtt
gcatttcagagctcatttgattttctagctgcaaagtaactcgtttaagagcaagaaatg
ttatttgctgctggatttgccttcgaatttgtttttcagctgtttcaatatttgacttgc
ctgtcatagcaagtctttcaatatctaattgaccttgagctaatatagagcttctaactt
ttcctgccgttagctgagcttctttttgatttgatctataataatttgagtctgagcaa
gttgtttttctgattctgacaatctagcactagcttgctctagtctttcttctgattctt
gtatagctgctaagacttttatctgtcttgcatttaaactagatccaagaaattgtttca
aaacataaataagaccaaataaaagcaacaatatgttaataacatttgcttcaaaaatat
cagaattaaaaccgaatttatgttcactactatgttctgataaaatagtaattatttgtg
gtatttttactatattattcatttgaaattct----------tactattttaatatttta
cttaggaatcatgattttaaagattgactactcaatagtttagctttaatttggtcgctt
aaagtatccacctgatcttccaaagttttaagagcctcttctttttgaatatttagctgt
tttgatgcttcagcaatcaattttcagcattcaattgagctttttaatatcttcagag
actatactttgcgcttctttttgagaagcagctatctttaattgtgcattacgacgagct
tcagataaatcttgctcatattttgcagctaactcatctgctttaacaagcatagaagat
gctgtagtcaaagttgtgcggatatattcatcacgttcatctagtactctagttactggt
ttataaaaaatagtattcagtaaaaccattaacgtaagaaactgtaatgccattaaaggc
aaagtaccattgaagtcaaataagccaccttcaatttcttccgctaatagaaatggtaaa
taaatcatgttactaattatgttgatagttaataatttggtttcaattaagaaaattaac
tttatagtgcgcttagtcttattagactaagcgcaaatgattattttaaccaacgtatgg
attagcaaataatagagacagcgcaacaactagaccataaattgttaaagattccataaa
tgctaaactcagaagcagcgtgcctctaatttttccctctacttctggttgtctagcaat
accttccacagcattagctgctgcactaccttgaccaataccaggacctattgcagcaag
acctaccgcaagaccggcagcaataacagatgcagctgaaacgattgaatccataataaa
ttttgatttttttgttatataagtagaaaattgacaaatttcggataatacttaaaatcta
accaatgaaatattggagattcaaatat-----tgtaattatagtatactactcttctcc
atgaccttccattgcttccccaatataagctgcggataaggtagaaaaaattagtgcttg

gattgaactagcgaataagcctagtatcataactggcaacggtatcaaaattggaattaa
cagagtaaatacagaaacaactaattcatctgctaagacatttccaaatagacgaaaact
taaagataatggtttcgtaaagtcttctaaaatattgataggcaataatactggagtagg
ttgaatgtacctagcaaagtaacctaaaccttttttacttaagcctgcataaaagtaagc
tagagaagttagtaaagataaagctacagttgtattaatatcatttgttggagcagctag
ctcaccttccggaagttgaattaacttccaagggatcaatgcacctgcccaattacaacc
tagaatgaataaaaacagggtagcaatatatggaacccaagggcgatattcatgctctcc
gatctggtttttagcaatatcttgtaagaattctaatataaactccatgaaattttgaaa
tttttcaggaattcgctgaagatttctagtacctaggaaagatatagttaatagtgtagc
aattactaaccaagatacaataaaaacctgaccatgtaattgtagactacctattttcca
atatagatgtttcccgacttctactgctgacaagctagtatatggattaaaatctatgag
actgttttgatacataataaatatttcttttaaatagtacaagaatcaaaaaataagttt
tttgaagaattaatatgcaattagtttatttgcacgattagattttttaataaattaaaag
caaaaataaattctaatagagatagctatctttagtagtgattatttttttgtaaaaaaca
attattgctacctataatgcttttatgtttaattatatacccatatatgaattatttaat
atgattataggcatttcgctacaaactttattgttttatccatttt----aaaaattctt
ctaatttacttattatttaaaatatctgcaacttcatccgcaaagttagctttcatattt
tcttcgccttctcccaatacaaatctaacaaatcttctaattttaatattttcgcctaaa
acggaaatattttgattaattaaatcttgaatactaatatcttgattgcgaatgaacatt
tgatctaataacgaaagttctttcagcctctttttcattcgcccttcgataattgctcct
attctatcagctggcttgtttttcaaatcatctttaccagcttcaactctttttttctgaa
ctgataacttcatctggtatatgctgtgttgatacatatgccacatttggcgatgctgct
atttgcatagcgatgtctttagctaatttttgaaattctggacgtcgggccacaaaatca
gtttcacagttcacttcaactaatacaccgattcttcctcctgtgtgaatatagctttct
aatagaccttcaagtgcagtacgattagatttttttgttagctgaagctaatcctttttgt
cttaatgattctaaagcttttttcttcattaccattactagcttctaaagctttttttgcag
tccatcatacctgccccagtcttatcacgtaaggcttttacagtttgagctgaaatttgt
aatgtcatatattttcaaaaatttttaagtatttaaattaactaaaacaaattattataa
ttcactttccacagagcgagctacttcctgtacatctgtttgcccatatttaccatcata
aattgcatctgctatcttaccgacaatgagtttaattgatctaatagcatcgtcgttagc
tggaattggaatattaataatttcgggactgcagttcgtgtcaagcatgcaaatagttgg
gatacctaatttcaaacattcttgtatagctgttgtttctctcttttgatctacaactac
aacaatatcaggcaaacgagtcatattttttaataccatttaaatgtttacgaagtttatc
taattctctccgtagaacagcagcttctttttttaggcagttgatcaataataccactttt
gtcttgctcttctaactgtctaaggcggctgactcgggatttgattgtcacccaattagt
taacattcctcctaaccatctttggttaacataataagaatcacatctttgtgcttcctg

agcaactattcctgcggcttgtctttttgttcctaaaaataaaactttcttgccatcgga
cgaagcctttttataaattcacaagcttctgtaagtaattgagcagtctgcactagatc
aataatatgtatgccatttctttctgtataaatatacggaaacattttaggattccatct
gcgagcttgatgtccaaaatgcacgcctgcttctaataactctgctaaggtaacgacagc
cataggattttaata-ttaaataatgtaatttttcgggaaatccctttttc----aaac
taattcaactcatttaacttattgcaattccaataaaccaataataaattaacatatttt
tacagtcgattaag-aa--aaaattgttttactttattcaatctacacttttattgctaa
agtagttacgcgcggttctgtcgtctaaaataatatcatctaaatcatctctcacagacg
acacactactttcgctactgatatctggattccaatctttttttctaacgacgctctat
tatgattgtcatacgtattaaaaccagtgccagctggtataagtctaccgataataacat
tttctttcaaacctcttaaccaatccagtttaccagatatagcagcttcagttaatactt
tcgtcgtttcttgaaaacttgctgcagaaataaaactttctgtattgagagatgcttgcg
taatacctaacagcacgggtcgataggaagcattcagcttatttcttaaagtcatagctt
tattggtttgctcaatcttttgcaactctacaagttccccaggcaaatagccggtttctt
caccattctctattttcactttagaagtcatctgtcggactatgacttctacatgtttat
cagaaatattaactccttgagattgataaactaattgtacttctttaacaagtaataact
gtatttcttgaaaacttaatctagctgcttcatataaagctagtccgcattcaatatata
ggttgaagcttgcatcaagaatatcatgaggatttaataatacatctgtttttttttcgcg
cttcaagaatttcttcaattcttggaaggccttgtataatatcaccagtttttgctctgt
caaatacgagaatagctagtgtttctcctcttctaattaatgcattattatcaacatgta
aaattgccccgctagaaactaggtaaggccgagcaattcttaatgttattgattgagaag
acacgtgagtaatttgtccagagtctaaagaaaccatattttcagcaatcaaatcaccgc
aacgaatccaatcaccaacacatactttagttttagctttttctatattaaaagttttct
tgtcagaagatgttccaattagaattcgtctactagtattcttttcagaataaatttctt
cgaccataccttcactcattgcaataatttctgtactagcaacaaccgtcaatggcttga
tatgttcaccatttttgacaataattctagtttgactttgttgtttttctgatttattga
catcaatacttttaattgataatgtttcaaaagttgaaagcctcaaatcaaaataattag
tatcactttctttattagaaaactcaattgaggatattaaactatcatctgtatgctcaa
tttcagctacaagattggttgtaattaaatgaactccatctatcgatttcaccctttctc
cgtcacgaaaaggtgtccttttaactagctttaatttagtttggctcggtttttgtaatg
ccaaattatcaattaagctagattttatttccggtattgaataaacaataacaggtctaa
tgaggatataaggcatttgctcattttctatatactcccaataaactaacttatctgttg
aaatgttgttatgcaatgtttctccaggtcttaaaaacccccctactttttttcattgttac
tatgtaattcgtttagtttatatatagatccaggcttaataatgatttcacgaactattc
catccttctgaattatctcaacaattccagagcttttagaaaaaatatttttgactagct
cagttccactttctataatatccccattataaactaataatagtgatgcatctttattaa

tttcgtgtgtttcctctgagatccataatatatagccaggactcaaaatttcataagcat
ctttttctgaacccactttctttttgatacattcaagtctaaatatttaataataccac
cacttgtagttttgtaagtatcagatattagctcagcgactgtatagccatcgtaaa---
tttttgattggggacagctttaaaaataaatttctgtttttttctgtttctaaaatgt
aagattctttttattaattaaatctgtataaatataacaaccgggaattacaatagact
cagtaataacttgtatattagtaatgctgctattagcattctgctgtatcctgacttccc
cagcatagtgatttatcaactctgtctgcgcaagtattgtgccagctttagttgtatcct
ctttattaacaatcacattagccgagtcagatatattataaacttctccagagagaaccc
aaatcaacccacctgttttagtgattcgtgtagtatattgattattatctgtttcttcaa
ctgttaagttagagaaacagattttaccagaaaggtctgacactacatgtttttgagccc
tttctgtcatcagtctatttctgcgaggtgattcagcaatcacctgatcttttgatactg
attgcccctcatgtagcagaagagtggttcctttacttaagttaattattgacttttgat
ttcttctagatcgaattgtaacttgggttggattctttgtaattaaggcttgttctccat
gccttgtacgaacatctttatatgattcaatatctagatctgttagctgaccatctatgg
gtgcgtaaatttgctctgccaattcaccagtaaatacacccccgtatgaaatgttctca
ttgttaactgggtgccaggctccccaatagattgagcagcaataatgccaacagcttctc
ctaaatcaactaagcggccatgagcgagattccatccgtaacaatactgacaaacagaac
ttctcgaattacatgtaacaggtgatctcaccaaaactttttt---gatgccagttttag
ttatctctttggctaatttagggctaatatcttgctttgtgtgagcaatcaaaacatttg
tatctggatggaacacactctctgctaatactcggccagacaaagcttgctctagattga
ttagtatttttgtgtatctactaagtcttctagtattatacctttatttgttccacagt
ctacttctctaattataacgtcttgagaaacgtcaacaagtcgacgcgttaagtagcccg
aatctgctgttctcaaagcagtatcaactaatcctttcttgctccgtatgatgatataa
agtagtccgtaacagttaatccttctctaaaattacttgatattggaaggtcaataattt
gtccttgcggatcagccataagccctctcataccaactaattgtctaacttgagaaatat
tgcctctagctccagagaaagccatcatataaacagcatttaaagggtctgtttctttaa
agtacttaattacttcctgttttaatgattcgcttgcattgttccatgtgtcaataactt
tttgaaatcgttcgaccgctgtaatttcccctcggcggtacttattttcggtagctttaa
tttcttcaattgtacctaataaaagactttgtttgctcggaggaatccttaaatcttcca
aactcaaagagattcctgcctgcgtagcgtaatgaaatccaagatctttaagtttatcag
ccatattagctgctcgtgctataccgtaattacgaaaagcccaaacaattaaatttttaa
gttcgtttttatcaataaccctatttgaaaaacttggctgcgcaagacttcttttactat
ccactcagtgttctccatattaattatttatt----taactacgccactaaagactcttg
aataattttgttgaagattattcggccagctgttgttcgaatatattgcacaattctctg
ctgttcagcatcctcttttataatatggttattataaaattttgttctactattatcact
gtgcttttcgattttaattagctcagtagtttggtcaccatctacaaatccatcaaaacg

agcccatatataagagtgcaaatcaactttcttttgctcataagccatcactacatcttc
taagctagcaaaatattgattcgctcccttctgttcagaaggattattagccgtcaaata
ataacaacccagaaccatatcttgactaggcatgataattggctgacccgtagctggaga
aagaaaattatgaggagccaacattagcaatctagcttcggcttgtgcttctaaagatag
aggaacatgaacagccatttgatcaccatcaaaatcagcgttaaatgcaggacagactaa
gggatgcagcttaatagctcttccttctacaaggattggttcgaatgcttgaatacctaa
tctgtgtagtgttggagctctatttaaaagtacagggtgaccttgaataacttcatttaa
cacattccaaatagaaggttcatttttttgaatcatttttttgcagctttaatattatt
cactaagccctgtaagattaatctatgaattacaaagggttgaaataattcaagagccat
ttctcgcggcaatccacattgatgaagttttaaatgagggcctacgacaattacggatct
tccagaataatccactcttttacctaataagttttgtctaaagcgtccttgtttcccttc
tataatatctgataaggattttagcggtctattattcgcacctacaacagttctgcctct
acgtccattatccatcagagaatcaacagcctcttgtagcatcctttttttcatttctaat
aatgatttctggagctaaaattgacttcagacgcgataatctattgtttctgttaatgat
tctacgataaaactcatttaaatctgccgttgcgaatctccctccatccaattggaccat
aggtcgtaaatctggaggtataacagggattactgtaaataccatccaagagggatctgc
gcctgttgctataaagttctcaatcaatcttaagcgtttcatttttttattaaattttaa
tgaaggtgtcttaaaactttttggtggatttgttgcttcagatcgtaaagtttcagcaat
gtattctaaatctaaatctttcaatagtttttgaatagcttctgcgcctatcccaacttc
tacttgattattttcattttgattttggtagatttcctcttcaaggctcttccattcata
gccttcgagcagttgcttatattttaaattgatatttgtattagattgtgtaactacata
agagtgaaaataaacgatttttttctacctctttaacttttaaatctaatgctaaagcgat
ataacttgtacttccttttaaataccaaacatgagttactggtgatgctaattcaatata
agccatccgatgtcttctcactcgagattctgttacttctactccacatcgctcgcagac
tatacctttatagcgaaagcgtttatatttaccacagtgacattcccaatccttaactgg
gccaaaaatttttttcgcagaataaaccatccatttcaggttttaaagttctataattaat
tgtttctggctttgtaatttcaccaacaatttgtccgttgggtaaacttctttcacccca
atgtcgaattttctcgggagaagctaaactaatctttacatagtcaaaatgttgctcaaa
ctgtgtcataaatttgaataccttgaaattaaataattaaccaacttatttacatatgcc
atattcgaaaatgca----ttaataaagaaattgttcaaaatcatcaactggtggagtat
cataactagtgcgatctgctcgattctctttagtgtcagacatcaaatcgacttcaactg
tacgtctttgaccgtcttcaaatagtttcaatttatgtactgcaatgtctaatcctaatg
actgaagctctcgcattaagaccttaaaagattctggggtcccaggcttaggtataggct
ttcctttcactattgcatttaatgcttcatttctagcttgcatgtcatccgactttacag
ttagtaattcttgtagagtatatgctgctccaaaggcttctaaagcccatacttccattt
cgcctaatctttgacctccatgctgggctctacctcctaatggttgctgtgtaactaaag

agtaaggaccagtagatctcgcatgaatcttatcatctacaaggtgaacaagcttaagca
tataagctctaccaattgtaacaggattatcaaaagattctccagttctgccatcgaaga
cttgcattttaccaggatgcagatcattaaaaagccatttattattagtgagtaatgatg
cttcttttaatttctattaacaagtgctcttgaggcttctgctccatacatctcgtcaa
aaggtattattttaaatcgtttacctaaatagccacctgctagacctagtaagcactcaa
atacttggccaacattcattctggaaggtactcctaaaggatttaaaacaatatctacgg
gggtaccatcagataagtaaggcatgtcttgtttaggtaagattctagaaataatcccct
tattaccatgacgaccagccattttatcgcccacttgaatttttctcttctgggcaacat
aaacgcgaatcatagcattcgtaccaggagggagttcatctcctttttgcctagtaaata
ctctaacattaacgacccttcctttagccgcattaggcaatcttaaagatgtatctctaa
catctctagccttttctccaaaaatagcccttagtaatttaccttcaggtaattgatcag
cttcgcctttaggagtaattttccctactaagatatctcctgcttcaacccaagagccgc
ctaccacaattccatttcgatctaaatcttttaaagagttatcacttacatttgggatct
ctctagtaatttcttctggtcctaatttagtttggcgacattctacttcatacttctcaa
tatgaatagaagtgtataaatcatcataaactagcctttcactaattaagaatgcatctt
cgtagttataaccttcccaaggcatataagcaacgagaatattccttcctagagcaattt
ctcctccatcagtggatgcaccatcagctaaagtttgaccgacaactattttctctccca
cccaaacaattggacgctgattaatacaagtatcttggttagagcgataatatttttca
aacgataatggactgttctaccattattatcttgaatacctattttattagcagaaacat
aattaacataacctgcagttctacttataacaaccatgccagagtctcgcgcaattttag
tctcaagacctgttccgatgattggcttttctggatataataatggcacagcttgtctct
gcatattagaccccattaaggctctatttgcatcatcatgctctaaaaaaggaattaatg
atgttgcagctgaaatgacttgaataggtgaaattgcaatgtaatctacttggctcggtg
ttgttgtaataaattcttgacgataacgtacaggaataatatctccttcaatataatttt
gagaattaactttaacatctcctggagctactcgaaaatcatcttcctcatctgctgtta
agtagatgggactctgttgataaattacttgaccttcatttaccggataaaggggggttt
caataaagccaaaaatattaactctagcacaggttgccagtgagccaattaaacccgcgt
tcggaccttctggtgtttcaattgggcagatccttccatagtgacttggatgtagatctc
gcacagcaaatccagctctatctttattaaaacctcctggacctaaagcactaattcttc
ttttatgggttagttctgcaactggatttgtttgatccataaattgagaaagctgactag
atccaaaaaattctcttactgaggcaattaacggcttaggattaattaggttagataagc
ttaatgagtctatatcacatatcatcattctttctctaataatacgttctaagcgattaa
gacctactctaaattgattttgcagcaactctccaactgaacggacccttcgattaccaa
ggtgatctatatcatcaaggttacctgaatttttatctttaatattaattaaataatcaa
tcgaggataagatatcttgaggagacaacactcgaaaattcttaggtatattaagaccta
atttttgttaattttatatcgaccaacttctcctaagtcatatctctttggatcaaaaa

agcgagaataaagcatctgcttagcaacagcaactgttgctggctcattaggtctaagct
tagagtagactataagcaatgcttcttcatctgttacttcttcgacattgtttttccaa
tttctttagctaattctttaattgaatataattgagatgcagagatgagaaaagcatatt
tactcagccccttttcaatctcgtctttattaaggccaatagcacggagaaaaatataag
catttactttatgggtcttatctattcttatccaaatttctcctttagggtcaatttcaa
atttcaaccaggagcctcgattagaaatcagactagcactgtaaatctgcttgccatttt
tatctatctcttgtttatagtaaattcctgggctacgaattatttgattaataataactc
tttctgtgccagatacaatgaaagtccctctgttagtcattattggcaagtctccaatga
agaccaatctcttcttgtatttttatttttatttccttctcaacactaagctgtaaat
gtgttgatgataaatctagtgacttgatactttttccatgatctttagaaggtagatcaa
tgtcttttcttgtcaacttcgcagggacatatatttgagcactataggttctatctctat
tttttgcctgtctaacactgtaacgagggaattttatttatactcattaccaaacaatt
gcagttcaagccgactagtaggatctaatatatttggaaagaattcaagtacttcagtca
taccttctaatagaaaccatttaaaactggctctttgaatttcaactaaatctggaagaa
gtttattttttaagcttatacgttgaaccatagatctcctttataacaaattctgcaatc
gaagagtttaa-tactatcaacagagtacttataggtaatacctgagtatagttcactac
attatttcttctatagacatgactagaacaaagtattttattcaacaaaaaagattttta
aaattt---ttcacaatcaatatagatataaaaatattacagctattaaaaatag-----
----------------gaagacgaaattaatacttgtctgtctgtcaaatagttgacatc
aagatagacaagtattatttatttaatcaattctg-tttttttgagcgcaagataatgct
ctcgcgagtctagctttttttctggcaccagtattaggatgaaatgctcctttttttatt
gctttatcaatcttgctatatacttgagaaatacttaattgcacatcgttaagattagca
ctttctagattatcaatatttagcaggcatcttttagttaaagtttttacaaccgattta
tatttacggttaattagacgatttctttccgaagttttaatgcgtttaatcgcagaaagg
ttcttagccacagtaaatccaataaaactttttataaaatatttttagaaacgaaaataaa
caaattagagatctattcaaaaagggaatatgcgccgtattataacattaaagaaaaaca
ataggcaatagttcactgagggctcttgttagtacaaactaaatattgtcgttgcataga
a-tttaaaaaatagtattaaat-----taaactaaccagacatatagatatatatgaaaa
aaattgaagctattattagaccttttaaacttaatgaggtaaaacttgccttagtcaggg
aaggtattggagggatgactgttgttaaggtctctggttttggaaggcaaaaaggacaaa
cagaaagatataaaggatctgaatattctatcgatattattgataaaataaaaatcgaaa
tcattattagcgatgataaagtagataaaatcacagagacaattattaaagcctcgaaaa
caggagaaattggagatggaaaaatattcattagtagcgtcgaaaaagtcataagaatta
gaactaatgatttagattccgaagctttataaaactcacactttttgagacttgatact
tcttcacgaaacatgagataataaacgtttattgtataataataactaaggtataataat
ataatggctaaaagcaaaggtgcccgcatcgtaataactttagagtgctctaataaaact

ggagattttagcaagaagagaaaacctggcgtttttcgatatacaactactaaaaatcga
cgaaatacaccaaacagaattgaattaaaaaaattctgtcctaattgtaatcagcattgt
ctattcaaagagattaagtaattattaattttactaaaat--aatcatattatggccata
tatagaaaaagaatatctccaattaagcccaaagagtctgtagattacaaagatattgac
ttgctaagaaaattcattacagagcaaggtaaaatactacctaagcgatccactggattg
acttcaaaacaacaaaaaaaacttactaaagctattaaacaagccagaatactttcccta
ctaccgtttttaaataaagattaaatatatagaaatatgctttagattgtttcctattaa
catt-------tccatgaatttatttactttaattaccgaatatactgagc---------
----------tactataaagcagctatttactactaattttgttcattttataattaaga
aaaa--atctaacttgttgttaaaatagattt-----agacttataatcgaacctctttt
aatgaattatattacatagtttatttaaaagaggagtcagttttatttgagttaataaat
agtacttttgtattctttagaccagtataaagtatttaagataaagattttttgtttaggt
acaagatcaaacgaatgaatcttatctccgggttgccataattgaaattcagaaatgaaa
attccacattcgttaccagcttggatttcttcaacatcttctctaacccgttttaaagat
tctatctttccctcataaataacttcatcttcccgaatgactttaatccaagagtttttc
aataatttattattcgtgactctacatcctgctatttttctattagccaaagaaaataca
gtgcttacttctgcttctcctactggtacttctgaatattccggatcaagtagatctttc
attcttcctttaatatcttcaatcaatgcataaataatttgataattttcgactagtata
ttaaattttgctgcagcttgcttcgctccaggagcaaaattagtattaaagccaatgaga
ttagcatttgtcgtcgacgctaattcaacatctgtagcagtaatttcgccaggcataatt
gaaacgacatttatttgtactttgctctgaggaaattgtgataaagaatctaggattgct
tctgtagagccttgattatctgttttaataattaaagaaatttgcttgcttagatcttta
gaattagtagttctcaatatatctaacgtaatacgactatttagtgctttctgcttctgt
gcgattaatgaatttttttgaagtattttcaagagctttaagctttgcttctttatcattt
ttcactactaaagctatctctcctatgacaggaacagccgataatccccaaatctcaaca
acagatgatggtattgctgaattaatctttttctgttcattattaatgatagctcgaatc
tttgcataagctgatccaataaccatattgtcgctcgtatgcagcgtaccattttggatc
aaaagtgttgctactggcccatgggatttgtctaagtgagcctcaactataacaccttgt
gcaggctgtgttggatcagcctgtaagttttctaactcagctaataaggtaatggtttct
aataatttatcaatattgtagccggtaacagaactaataggaattacaggaacttgtcca
ccaagtgtctcagacatcacattgtgttttaataagtcctgctcaataatatttgtattt
gtgaagtctttatctatcttagatattgctaccacaaaaggaacatcagctttttgaata
tgatgaatagcctctacggtttgaggtttaataccatcgtctgcagctataataataata
gctacatctgttaatttagctcctctagatctcatacttgtaaaagcttcgtgccctgga
gtatctaaaaagataatttttttgtttattagctttattgatatattcaacttcataagct
gcaatagtttgtgtaattcctccaatttctttattagcattgttagaattatgaatgtaa

tctaataaggttgttttaccatgatctacatgccccataactgtaacaatcggaggtctt
ttaacatagttaccaactttatggagagagctagaagtatctaaattagcaggatacaag
ttattattttcttttgtgtttgagtcaactcctataccaaaattatttgctactgatgct
attattgaagcatctattgtttcattaatagtaactggtattcctttaaggaataaatat
tttatcacatctgtttcttgaacacaaattagtttagataactcttgaatagtaagagga
ttggtaatactaatagattctggtggagaattagataaactgctggtttggtcttggctt
tgagttaatgatatgtttttttgtcttatagacttttttagtattagtaaatttttttata
acttgagcttttggtttaggtggacgcattagagaaatagccaagtctcctgcagtctgc
ggaatcatagaattagaacctctaaagttctct---tcctcatcatcaatatgtattttg
gtctttattttt-tttcgttgtctgttctta--tttttttttattatctgctagatcatga
accttattaaaatttttgtttttttttatctagtttgggtggagaatttaattctaaatgc
ggttcactactagctgtatgcgtatcagattgatcaagatctaagggtaaaacactgtct
ttatgggctgattctaatctaattttatatataatttggggatttttttaaatctataatt
ttttcagaagc---tatatttggactggatgcaggggatctattttcgaaattttgatta
tttaaaaacatgaatattgattttatgatctaatagtaaattttaaatttcaatacatag
ctcaaagtaaagaaaaatcagatatct------ataatactaatat---agaaaattaat
ctttgttgtattagatttaatataacactaaagtcttttgttagttatacaagtttaaaa
ctttt-tgtgatgataaacaacaacttctgctttattgagatacaatcaatttatttaga
tataactgtttttcaa-aggtttttaaa----ttgtctaatattgtgttcgttatgaatt
tttattaaagtttgaaaa-ctataaaatagagtcataatcattaataatctccacgccac
ttatgccacgatctcaaaaaaatgataattttattgacaaaacgtttacagtattagctg
atattgtattaaaaatacttcccacaagcaaagaagaaaaagaagctttttcttactaca
gagacggaatgtctgcacaatctgaaggcgagtatgcagaagctttagaaaactattacg
aagctttaaaactagaagaagatccatatgatagaagttatattttatataatataggtc
ttatttatgctagtaatggtgaatatgtaaaagctttagaatactatcatcaaggattag
aactaaactttaagttacctcaagcactcaacaatatagccgtaatataccattatcaag
gagttcaagcggtcgaagataaagatacagaattatctaaattaatgtttgacaaggccg
ctcagtattggcaacaagcgattaagttagcacctgacaattatattgaagctcagaatt
ggttaaaaacaacaggaaggatgagaaatatacaaggatattaaaatatttaggatataa
t---tatattaattaatgaagcag----aattagagatgcttttaccctataaagcattt
gtagtat--------ataatatactaattac------tgtataaaaatcaacataataaa
cgaaattaaatctattgactatatttatgaatacaaataaacataatggttagtaaaaca
aaaagcatcggatctgtcactcaaatcattggaccagttttagacatagcatttcctaac
ggacaacttccaaaagtattcaatgccctcaaagtagaaagctcagagggaactattact
tgtgaagttcaacaacttttaggtgataacaaagtaagagctgtatctatgagttcaact
gaaggactacaaagaggagttcaagttcttgatacaggatctcctatatccgttccagta

ggtgcagatactcttgggcgcatttttaacgttttaggtgaacctgtagacaatttagga
ccagtaaattctgaaagtactttacctattcatagaccagcacctgcttttactaagtta
gaaacaaagccaaatatttttgaaacaggtattaaggtagtcgatttattagctccttat
agaagaggaggcaaaattggtttatttggggtgccggagtaggtaagacagtattaatt
atggaactgattaataacatcgctaaagctcatggcggtgtatctgtatttggaggtgtt
ggcgaaagaactagagagggtaatgacctctatatggaaatgaaagagtctaaagtaatt
gatgcagacaatttgaaagagtctaaagtcgcactcgtatatggtcaaatgaatgaacct
ccaggggcccgtatgcgtgttggcttgactgcattaacaatggcagaatactttagagat
attaataaacaagacgttctattgtttattgataatatttttcggtttgtccaagcagga
tcagaagtatctgctctactaggacgtatgccatctgctgttggttaccaaccaactttg
gcaaccgagatgggtgcattacaagaaagaattacttcaacaacagaaggttcaattacg
tctattcaagctgtttatgtaccagctgacgatttaacagatccagctcctgcaaccaca
tttgcacatttagatgcaacaacagtgctttcacggaatttagcagcaaaaggaatctat
cccgcagttgatccactagattcaacttcaacaatgctacagcctggaattgttggaact
gagcattattctacagctcaagaggtaaaatccactcttcaaagatataaagagttacaa
gatattatcgccattctaggtcttgacgaactctctgaagaagataggcaaactgtatcc
agagctagaaaaattgaaagatttttatctcagcctttcttcgttgctgaagtatttact
ggctcacctggaaaatatgtatctttagaggatgcaattaaaggttttcagatgattta
aaaggtaaattagatgacttacctgaacaagcattttatctagtaggtgatattgacgaa
gcgatacagaaagcggatagcatgaaagattaatataattaacaatgactttaaatataa
gaattattgcacctgatcggactgtttgggatgcagaagcacaagaaattattttaccaa
gtagcactggacagcttggtattttaacaggacatgcgcctctacttacagctttagata
taggagttatgagagttagagtagataaagaatggatgccgattgttttgatgggtggat
ttgcagaaatagagaacaatcagttaactattcttgtaaatggtgcagaagaagctactc
aaattgacttatcagaagcagaaaaaaatttagaaactgcaactcaattattaaatgatg
cctcttctaataaagagaaaatagaggcaacacaaaaaattagaaaagctagggctagag
tacaagctgcaacagcagctattgtataataacatttt------gctaaaaatataaaac
cttagtaattattactaag-------gttttatgttttatttatgtatttatatacaact
tgagtaagcttagcttaaaccagaacaaatataatcaaaatataaacccatttctttgcc
agcatctggtccaactaaactaatagtaacttctttcatagctaaaatcgcttgaatggt
tgcaccaataggaacacctagagaattataggtttcttttagaccatttaatactctttc
ttccaaaatagatggatctccagctaacattccataagtagcatagcgtaagtaataatc
taaatctcgaatacatgcagcataacgtcgggttgtatacatattacctcctggacgagt
aatatccgagtaaagtaaagattttgctacagattctttaataattgttgctgcattagc
tgcaatagtagcggctgctcttactctaagctcacctgtctgaaaatagcctcttaattt
ttcaacagagctatcatctaaatatttaccttgaacgtccgctgcattaataacagaagt

aattgcgtcttgcataacttgtaaaaattccttaattttaaatatctaatactaaagggg
tatataacaaaataactgaacttttttataaataccagtactactgcatggcacccaaagt
ataatcaaagtaaaaaccagcttctgctgaatcttcaccagcaagcagtgagcaagctac
acttttcatacatttcacgccttcagcgactcccgaaattggtgttcctaaagaattata
catttctttaacaccgactagtccaatttcttcaataggcgttacgtctccagcaactat
tccgtaagttactaggcgaagataataatcaaggtctcgtaaacaggtagctgtcatctc
ttctccataagcgtttccaccaggagaaactacatcaggtctttttttgaaatagctgttg
accgccttgttttacaatacgctcacgattatctgttaaaatttgagctattcttagacg
acgttgtcctgataagacaaaacttttaattctgtctaattcaccaggacttaaatatct
tgcttctgcatctgcatttacaattgactttgtaacaatactcatggataaaactcctgt
aatgcttt--taaaaaagcttaatactgattatttatgatataactcagaaagtagtacg
ccttttaaaaaaattcatagctatagcacgttggtgatatatgaagtaggacaaatactt
ggataggtactaaaacaagaaagttacttatcgtaccttattttattgattgccgagtac
tttttttaaaactaggtactacaattgatatatcttgcttagtaaggcgattatataatgt
ttctgtgttagggaaattagctgctggcaaagttggaaatcttctatacggaacagtata
tttaccatatactgaattatactcagtactttcaactaaagtactaataaatgcagacaa
tccttttgaagctaaaatttggttaaagaatctaatttctgcctgattattaggagctcg
tcctaagatatgcttcgtacctaattctataacttttgtatttggatatggctgataaaa
ctctttaccatataattcagataatgctaatttttctactaattctttaacattaatttc
cctgctcaaaaaggatgcttcaatatctaaaaactctcctccaatactaaaagaattaag
atctctttcaaaaatctggcgatatgcggctcttaaagcttgctctaatacttctttact
actattgtcgtttacttcaaatataatagattgatctcttaaagagctaactccttgaga
aatacgagactggatatcattgccgcttcttatttcttttacagcgcctaattcaacaaa
tctagaagacttgctagtaatgactttcttaaatctttggtcaataatacctggccttaa
gcttcttagagcgataccagctggtgtactatatctttcataaggaacagtactatcacc
aaaagtttctatatactctgagctatcaactattgcatctatcacttgatagtagccctg
cttatatgcaatatcaaaatatttattaatttcttggcggccatatgtaggacgacccaa
taaacggttatggatatactcaatagctttacagatatataatggctcccagtaaagtga
tctaaatatacttgacttcgctaattgtctaacaaattcacgcactgtaatttgaccatc
ttttaactgactttctattggcttaagaagtaatttttcttcttgatatacttctctgcc
aaaaactcttaaataagcaacttttgtaacggcatcgactgaattttcaaaattttcaga
agagtcagcttttacaagaggattggataatttaaaaatttttggacccaaagatcctgc
agacttaggtcttacctgcggattactgatctggttatagattcctgggccacatctaac
cagaattctcctagtatctttaccaaaaagagcttgtcgtttcctaggatccttattttc
tttaggaaagattgctccaaattgaatcgacagtggatcattgcctattccatatggatg
ctgatcaggtaatgcttgcttgtaatcactaaatagggttataaactgaggcaccttacg

aaaaggagcactataattaagaagatctatttgaggtccccaattacgacactcttgagg
ctcttcacctaaatttcggaaataaggtacagtttcttctccaaaataatctgtatattc
agcagaatttaaaattgcatttactagcccacttaaaccagtagctgatagaatagcaaa
atatttttggaattcctctaaagagctgggccctctaccaaggaaatgtctaaatgctaa
ttctaatgctcggctatttacaaaaggctcataaaattgctttctatagatacttgaagt
tcctaaagaacggataaattccttaatggagatttggccatttttacttgcgactctaa
gttagataaagataaatcatatgctttagcaatatctctctcaaagatttgtctgtagca
cgctttaactacagtatttttttcatctgcagataaactaggcttcataacaaaacgagg
cgtagaaactcctgcttgtgcataaatctgaggaaggcgcaaaccttgtaaatccccaga
tgttcttttccttaatttatcagtcaaagatggagcttcaaattcagaaattacaacatt
aaaatactctttgaccagatcttgacctttaatatcttcttcaaagattaaaagagctgt
tcgtctcatctctctgagggcaacaattgctgccgcgcttgagcaagcattatcaattaa
ttctcttaatcctctaatattaaccgataaaatattgggatcacctgatacaattgcata
tgttaaataccgtaaaaaccagtctaaatctcgcagagattttctcattcttgtagtccc
atatctcaagacatttatgggtttgaatcctggaggagtagctcctccagcattaaataa
agaacgaaaactctgcccaaaatctccttgaatattgccagatagctcattaattttgtc
ttgagaactttgatctccagcaataataacagctgcttgtggacgttctaaataagaaat
tgcagacccgcctacaaatattttgtctgcagctcgagctactagaatattggcatttt
agttaaaatatctgcaacttctaatctttttgtccagaatttagaaaagaaactagttg
atttaattctcctagctgtaaaaatcggtcttgctgctctgcctgagtaatagttaaaat
tgaggcagtccgataaagctgggggcgtgctaatgggcttccgccacttgctttgatact
cattactttatttatctccttaacttatattacttgctagctaattcaaatatcacttaa
tatttctaactatttaaagatactacgcaagcacatttgttttctatattattaggtact
agtagttatagtacatagagtcacacaactaacttctataaatacaaattttgtaatact
tctattaacatttatcaaaagtaaatataattgataactcttttatattaaaatgcaatt
attaagagagtgatgctaaataaagtataattagtaagttttatagatagtaaaatt-ca
agatttattctacaaacattaatca------------cttgactgtttgacaattatacg
attgttcttgcttttaattggtttagactagtttcacatttttctaacgtaaacaatat
taataagcaatttgtatgaatttaaaaccacagaataacctattaactaacggaaatata
aaccaacaagatgtacctgtaaatgatatgccaatgtcaattactgagcatttagaggaa
ttaagacaaaggactttatttgtatttttattctttttgcttgctgcaactttgagtttc
acacagatcaaaataattgttgccatattgcaagctccagctattggtattaaattttta
caattagcgccaggagaatattttttttcttctattaaggttgcaatatattcaggaatt
gttgcaacaagtccattcggggtataccaagttatcttgtatatactacctggactaact
ggcagagagcgaaaaattattttgcctatcctaattagttcagtcttactttttataata
ggcggtatatttgcttactttattttagcaccagctgctttaacatttttgatcagctat

ggatctgacattgtcgaaccactatggtcttttgaacaatacttcgactttattttacta
cttttatttagcacaggattagctttcgaaattcctatcatacagttactactaggtgtt
tcaggaacactatcttccaatcagatgattcgagcttggaggtatattattattatatcg
acaattgctggagccattttaactccctcaactgatccagttacacaaatcatcatgtct
tctgctgtcttagtgctttattttagtggaattgtcatattgttacttttaaaaaagtaa
atgtagt--------aaactttcatttcatattatacttttcttgtg-----ctaagcaa
tggagttt--atatagcttttagttcttaacttagatgaatattgaagta--------tt
aagctcttttaaatatttaatttagtcttataatttaaccaatatg--------------
------------------------------------------------------------
------------------------------------------------------------
------------------------------------------------------------
------------------------------------------------------------
------------------------------------------------------------
------------------------------------------------------------
--------------------------------------------------tatatattgg
c---ttttttcatcaattatttgcactttacatttctgtacta----ttctatgatgctt
gggactctttatgtaatctgtttactgaat----gacgttgaagaaaattcagcataaat
tttgttaatttataccaatataaaaagtattttattgtgagttttatttatataaaggct
tattcattatatatttataacagtaagtctaattaatagtttttttatgcttaattaaagt
aataactcttaatccttttggcagaaaagtctgtcttaatattttatc--aaaaaaacag
aaaaaatgaagctaaatagtctaattaacttaattcaaaagtctatatgttcttgtacat
ttttattaattattttaaatattatttgtgtcgctcctaattctagtaatgcatttccaa
tttatgcacaacaagcttacgaaagtccaagagaagcaactggtcgaatagtatgtgcca
attgtcatcttgctcaaaagccagtagaaatagaagcacctcaatcggtattacctaata
ctgtttttgaaactattgtaaaaattccatatgacagtagtgctaaacagattttaggta
atggaagtaaaggagggttaaatgtcggagctgttgtcatattgcctgaaggatttaagc
tggctcctgctaatagattatctacagagttaaaggaaaaaactaaaaatctatatattc
aaccatatagcgctaaacaagataacattttagtaattggacctattcccggcgacaaga
atagagaaatagtttttccaatactctctcctgatcctgcaaaagataaaaaagctcatt
tttttaagtacccaatatatgttggtggaaatagaggtagggggcaactttatcctacag
gagataaaagcaataacaatgttgttacagctttaagtagtggcaaagttaataagattg
aggttcttgagaaaggggatttgtagttcatgtgactaatagtaacaatatagaatcaa
ctcaaaagatttccccaggtcttgaacttaaggttaaagaaggtgatacagttcaattag
atcaagctttaaatagtgacccgaatgttggtggtttcggccaaaatgaaacagaaatag
ttctacaaagtccaaacagaattaagggcatgattgttttctttttttgctagtgttttag
ctcaaattttctttgtattaaagaaaaaacaattcgaaaaagttcaggcagcagaaatga

atttctaaagcaagataaaactggtacttactataattaaagtg-aaaacactttattca
aagaatgtaccagttttcatgctatataaccgttgcttttgaagacattataatgtcttt
aacagagtctacaatttgactaggctgaattacagtagcctgttctaagctaccattata
tggtgttggtatatcctgcgaagatagtcttactacagggacatctaattcatcaaaaag
gtgttcattgatttgtgcaattaactctgctccaattccagcagttttcatgcattcttc
tacaattaaaactttgtgagttttcttgactgacacagatatagattctatatctagagg
ttttaaagatataaggtctatcacttctggatcgtaaccttcttttagcaaagctggtaa
tgcttgaatcacatgatgcctcattcgagagtaagttaaaatagtaatatcttttcctgt
acgcacaaattcagctttattaagaggtaatagatattcttcttcaggaatttcttcttg
taaattgtaaagcaaaacatgttcaaagaagacaactggattattatctcgaattgcaga
tttcaataatccttttgcattatatggagtagagcaagcaactatttttaaacctggaat
agcttgaaaataagcttctaatctctgagaatgttctgcgcctaattgtctaccaactcc
tcccggtcccctaatgactaaaggtaatgtaaagtttcctccggaggtatagcgcaacat
gcctgcattattagaaatttgattaaatgctaacaataaaaagctcatattcataccttc
aacaattggtcttaaccctgttatagctgcaccaatcgccataccagtaaaactattctc
tgctattggtgtatctaaaactcttaaatccccatacttactatgcaaatctttagttac
cttataagatccgccgtagtgacctacatcttcgcctattacacatacagtcgaatcttt
tgccatttcctcatctgttgccgctcttaatgcgtcaaacataaagattttactcatttt
tgtatttaatttttataatttcactactaatctaaattaaaagttaagatattaaatttt
c-tcacgtatctgaaaaaagatatcgtttaagttctgagatattaggctctggactggaa
atagcaaattgcacagattgttctaaatctttttaacagcagtttgaatctcgttgagt
tcacttatgtcggcgatcttattatctaaaatatatttttaagtttttgattggatct
cttgcaatccaagcttctttttcttgtcttgatcttagttcatcaggatctgcaagagag
tgaccacggaatctatatgttaaagcttctattaatgttgggccttcaccatgtctggct
ctctgaactgcttgaatcgctgcttgccttacagctagtacatccattccatcaacttcc
ataccaggaagaccaaaagcttctgccttttatgtatttccggcatggaagaagatcgg
tgatgtgccatacctatagcccactgattattttcaacaataaatataataggtaacttc
caaagaactgccatatttaaacattcgaaaaattgaccattattagtagtaccatcacca
aaaaagcaagctgtaactctcaaatcttcagtttcttttagtacttgctgacgataaata
ctttggaaagctgcacctgttgcgacaggtataccttctgcaataaaagcaaacccacct
aaaaaattatgaggagcagaaaaaatatgcattgaacctcctctgcctttgctacaacca
gtttcttttccaaatagctcggccattacatttttcgagggaacacctttacttaaagcg
tgtacatgatctcgataagtgctacatacataatctgtagaattgagaagcttaattaca
cctgtagacacagcttcttgaccattataaagatgcacaaaaccaaacatctttcctttg
taatacatctgagcacatatgtcttcaaaatttctgcctaacaacatatcttcatacaag
actaataaagtcttttt------attaagaacagttgaattatagttagttagcggcaat

tgaactttcttaggataactcataatttataaaggaacctct----------tattatac
ataggaattttgaagtaaaagttatcaatagaaaacacacagataaaaatttataaatta
ttatatgc-ttttactaaagacttcaatagttatttgcaactagttcagtatatcaaatt
aattttgcttatactacagatgtctaaagcttagatgaaaagaaa-tagttaaa------
------------------------ataaataaaaataaacagaattattagtgagaaaat
gattattattaatagataattacttaatatttctaaatattttcaaattaatggctatat
catcatctttattttaccctgtcgagaaagagttatacagtctagaaacaaatttgcaag
ctgtggctgggacacgtcatccaattttatacgcagcggcgaaacatctgttcgaagctg
gaggaaaacgagttagaccagctctcgttcttttagtggctaaagcaacatctgagaaac
aagatattaatactggacagagaaggttagcagaaattactgaaattatacatactgcaa
gtttagtacatgacgatattattgacgaatgtacaacgcggagaggagtcaagactgtgc
acaatttgtttaatactaaaattgctgtattagctggagatttttttatttgcacaatctt
cttggtatttagctaatattgaaaatttagcagtagtgaaagctatttctaaagtaatta
cagattttgcagaaggcgaaattaggcaaggattagttcactttgaccctagtatttcaa
tagatgcttacattgaaaagtcattttataagactgcttcacttattgctgctagctgtc
gaggagcagctatgctgaatgattctaatactcaagttaatcatgatctatatctttacg
gtaaacatatgggcttagcatttcaaattatggacgatgttttagatataattggttcta
ctaaaagcttagggaaacctgctggcgcagatctagccaatggtaatttaacatctccta
ttctattttcacttactcaagaaccagatttaaacaatcttattcaaagagaattttgta
ataggacagatgtagcctcagcgttatttcttataaaaaaaaagcggaggaattacaaagg
ctcaagacttggccaaagaacaggtacaggcggcactttgctgtctccagtttttaccaa
aatctgcacctgtatctagcttaaaagaattgacacatttcataatcacaagattgtcat
agaacacttgctaaaattaaaatgctattaactaatttttttactatctctacaatagtct
gaaaagtttggatatcattacttgctaactgtgctagcattttacgatttaatgcaatat
tctctttttttaacgcactaataaaagtactgtacttcatgccttgactatgagctgcag
catttattctagttatccaaagacgacgaaaatctctcttttttcctttttcgaccaacat
aagaatagcgaagagatttgagaacttgctgtttagctgttcggaataaacaattatgtg
cacctttaaagcctttagccagcttaaatatttttgcccgtcttttttttgcaacattac
ctcttttgactctactcataaaatattac-ttacaattaaattattaacagcttcctaga
ttcaaatgcttatcg----attataaataaggtaaatttacagcgatattcttaatatct
cgtgtgtcaactaagcaggtacaagaaagatttctttttttgcttcgatgatttttttttgt
aataaatgacttttagaagctttatgtcgcagaaattttcctgatgaagacactttgaat
ctttttgctatcgcttttgatgtttttaacttaggcatattgtaaacttttaatgattaa
caaaagtatgaaaagtctgaatttatcaaacttagacttttcaatacttaggataatctt
aacacaaattattttacttactatagaggttaacttaaacaatttatatatatataatagca
aataattaatccgtagctactaaagttgatttatatttagctttaaatttttgtatagta

ttagcaattagcattgttattattttaagggtacttgagtttaattcttcgaacattttc
atatttaatgtgaatgatatattcgcttctgacacaatattttgcatttgatcatccgat
aaaggaatcatatccaatgcagctctatacttatctttaaacactttgtcatcttgtatt
tggtcaaagtcataaaacctagtacctctttcatctgataaattcatcgcacctctagcg
attttctttaagatttggcctccagataaatcacctaagtaacgcgtataagcatgagca
actaataattccggctgtttacagcctatactatgaattctatcaacatagattattgtt
gcaggagatggtttaataagatttaaccaatctgctccataataataacttaaatcttct
gacagacttatttttctattgagttctgtaaaatatataggtttaatagctgggtgatct
ttattagaaaacagttcttcttctatagcacaataaacgaaataaagattagcaaccaat
ttacgatatgatttcttgtctacgacacctcccaaaaaagatttcacaaaactaacattt
tctgccatactatgagacttagtagtgccttctcttaattcatttgctaaagtattaacc
ataatatcaatcttccatttta-aaagataacaaattaaac---ataaacagttatctat
atattta------agttgtactaattaagctgctgcaaaataatttttgattttattgg
atctggattcattgttttatcacctggcttccaattagctggacaaacttcatctggatg
agattgtacatattgaattgcttgcagaactctcaaagtttcttctacactcctaccaaa
ctcgagattgttaattgtagaatactgaataattcctttaggatctataataaatagtcc
tcttagcgctacaccatcactgttcaaaacattataagcagcactaatctctttttttaa
gtctgatactaatggataggcaagatctcctaatccaccggattctctgtctgtctgtaa
ccaagcaagatgagaatactcgctatcaaccgaaacacctagaacttctgtgttaagctc
agagaaagcatcatatttatcactaaatgctgtaatttctgtagggcaaacaaatgtaaa
atctaaagggtaaaagaataagataatatacttatttttaaatctgataattttaacgt
tttaaattcttggtcataaacagctgtagctgaaaaatcaggagctagctgacctactcg
aagacaatttggtccagaaatcattaatttttcccagaagaattgatatagttatgttat
ttaataaataattatataacaaaatttagtatatatgaaaatgctatcagtcttttaaaa
attac--------tcaatggcttccaaaatactattttatagcggggaatggatttgaa
ccattgaccttcgggttatgagcccgacgagctaccagactgctctaccccgcggatatt
taatatcatattagtccatctaaatagataaaagcaaatatattatataattcaaaagta
aaagaataagagcacaggagcactaattattaggcttaaaaatgacaaagtatttttat
tttattgagtagaggaaagacttcatataagccgatgaatcgatctttaagaagaacttc
cgatggcttcacccaaatttgaccgtcataccacccagactcttcatagaataccgtagc
actcattaaacgttttactacgtatgaccatcctaaatataatctgattagaataaatcc
agttattaaactcgaggtaataaattctgagaagacaaattttaaaggcagcttggtaat
tggaaaaattgaaagcaatactggactaactaatagaccattaagtagtagtgtaacagt
gattttcctgttatatgattgactacttaaggttggccagcagaagaaccaggaatttt
taaagaagtatactcattaacaggttgctgctcttttggtacaggacactgagtgttata
taaattcattgaaatttaaaattgttatactagaaattgtttttttatctgctttagat-

-ttaggga--------gttaattgattgcaattataatagttagtatcagaaatacgacg
ataaacaaccaagtaactttgtttaatgtgttttcagtactacgagtattgctaaaaaac
tgattctgagctccagcacttcccagaccttcagatttgggattatgtatcagaatagta
aaaaccaatataatagttgatacataccaaaagaattttaaaatttgttccattgtgtaa
aaattaataaaaaatagcacaaagacaatcgtctaaaaaatattgattgtcttattatat
taagaaagatttattctccctgcacttttaaaagtaaaaagcctattgctaaccctacta
aaactaagactgaacttaatacagcactttctacaatttctccgaccatt-gtgtttttc
cttattattttattatta---tcattttgtaaaaaatttgatacctattaaa-ctaaaat
ccatttcttccccaaaccaccatcgcaatagagaaagtaaacattgccattaaagctgcc
catcctaaacttaaaatatccataatttataatattc--ctgaaaataaggtctcttata
cttaaactataatattcttaatattaaaaatatcattatataataagctaattataacag
agtactacttaattcgtatttttagtgtatttcaagaagattgac-tttttctaaaatac
ttaacaagtattaaattgtaaaaaatttctgcaat----ttttgcattattatacctatt
aatttggaatgtaaatatcaatttaaaatcgttagtaatttgatttttttcaattgtaca
att---aacatatgtaaaaacataaagattcatagcagctaagttga-----tggtctac
tataatttatcaat--aaaaaggaagttagtatttaaattatgcaaacaactataagtaa
tggaaaaacgtctagcaaggaaactctactaacccctaggttctatacgactgattttga
agaaatggctaatatggatatttcagaaaatcaagaagatttctttgctattttagaaga
atttcgagctgattataatagtgagcactttattcgagatgaagaatttaatcaatcttg
gtccaatttagaagataagactaaatctttatttattgaatttctagaaagatcttgcac
tgcagaattttcaggtttttttactatacaaagaattatcacgaaaattaaaagatagtaa
tccaattattgctgaatgtttttttactaatgtctagagatgaagcccggcatgcaggttt
tttgaataaagctataggagatttcaatttatctttagatttaggctttctaacaaagag
tagaaaatatactttcttctctcctaagtttatttttatgcaacttatctttcagaaaa
aataggatactggagatatataactatttaccgtcatcttgagcagcaccctgagcatcg
catctatccgatttttagattctttgaaaattggtgccaagatgaaaatcgtcatggaga
ctttttttgccgcattactcaaatcacaaccacatttttttaaacgactggaaagctaaaat
gtggtgcagatttttttctattaagtgtatttgcaacaatgtatttaaatgattttcagag
aatggattttttataatgcaataggtctagattctagacaatatgatatgcaagtaatacg
aaaaactaatgaaagtgctgctagagttttccctgtcgctttagatgtagataacccaaa
attcttcaaatatttagatacttgcgcatgtgacaatagggctctaatagatatcgataa
acaaaatttcccattgttaattaaagctattataaaaaatgcctctctattactccctaat
tattagtttgctaaaaatatatttaattaaaccgatcgattcaaagaccgtctggaatac
agttcgataatcaaatcacataaagatatattttcataatacacactatctcattattga
tataaatcaaaatgaaaagctcacttct-----gtcaggaaggagcttttattttattca
caaataaacaatatggttctatagatt-tagaactaagaacttactttttcttttgcttc

atacataacttctaaggtgattagatttttagaacaatcgcgtgcaaatttttctgtgtt
tctttttactttacctctaacaaaacctggaattttacttaattctttctgtgcttcttg
tgaccagttaatactatcaacagcagagatccctacgcttaatgcttccgttgtatcatg
accaccaaatatttctaataagtgatcttccatacccaatgtgaatgagttatagactaa
atcagcaatttgatttgtaccttcatagccaaggaatggacgataactcaaaggaaaatt
ttgtatatgtactggtgaagaaataacaccacaaggtatatttaaacgtttaccaatatg
tctttccatctgtgttccgaaaattgctgcaggctctgttttagcaattaaatctccaat
taaaccatggtcatctgaaataattacttcatcgcagaattcttgaacttgttctttaaa
ccaatcttcatcatatttgcaataagtgccacaccaagtaacatgtatacccatttcctg
gtgtaaaattcgagttattgctgcagcatgagttgcatctccgaatacaatagctttttt
accagttaaattttggcagtcaattgatctagaaaaccaggcagattgcgagataaatct
agtttgttcatctatataattttcgtagtctacaatggctcctaatgtagttaataactg
ttctattgatcggatacatgcagctgtttgtacaatacccataggggtagtatcaacata
aggcatattaaattccttctctaaataacgagctgtcattagtccggtctctctatatgg
gacaaaattaaaccatgcagatggtaaatttttttagatcctgaaccgacgcattttcggg
aataatttgattaattttaatatctaaatcttgaaataatctttttaattctgaaatatc
atgctgattatgaaatcctaaacttacagctccgataatatttacagatggtttttctgt
cttttgagttaaaacctgattattagatttagctttttccatgtaaaaagtaacaatctg
ttccagagttctatcaccagcttgaagttcatttaccctataatgattaacatcagctaa
taaaacatcagcttctgtttctatagaggctctgctaacaaaattttgtaaatcttcttg
caatatactagaagtacaagttggtgttaaaataactaaatctgggttttcttctctatc
ttttctagtaatgttttcaacaactttttcttgtgaaccacgtgctagtacatgtctatc
tacaacacttgctgtaacaggcgtaaaatctctgtctctttcaagcattgaacgcataac
attaaaatagtcatccccgagaggcgcatgcataattgcatgaactttttttgaaagaact
agcaattctaagagttccaatatgagcagggcctgcatacatccaataagctaatttcat
attaaattttcctatgatttatttagacatagtaatacttttaaagccaaataataaata
aaattcgaacatctaatttaactccttaatgatttttttcatg-tttttagatagagcgta
acattatatcaagtcaaaatt-------------agagaaaaatgactaatgaataaagc
ttgaaagtaaagatttcttgcagttatagtttttctttaagtaaacatgtaaaaattgta
ctatatataaatgtactatatataaatttttttttactgaatattggtaatgccagataca
attaacttaaatatgccttccccaacgtttggtggaagcactggcggttggttaagagcc
gcagaagtagaagaaaagtatgcaataacatggacaggcaaaaatgaaagtaagtttgaa
atgccgaccggtggtacagcaactatgagaaatggagagaacttactttatttagctaaa
aaagagcagtgtttagctttaggtactcaattaaaaagtaaatttaaaatatctgactat
aaaatttacagagtatttccaaacggagaagtacaatacttacatccgaaagatggtgtt
tttcctgaaaaagttaatgctggaagagctagcgttaacagcgttgatcattctattggg

cgaaatgttaatcctgtagatgtcaaatttacaaataaggcaacttacgattagttgata
aaaaaacctagacataaaataacttttatatagtctaggtttttgataaagtatactaag
agtatggaaaggtggccgagtggttgaaggcgtctgatttgaaatcagttgaactt--ta
tagttccgtgggttcgaatcccaccctttcctttataagact------------ataact
taattattactata----aattatacgctaatagtattcagtttagacttaatgaaatct
acaacttgcgataaattagaaatttttcggcatcttcatctggtatttctataccaaat
ttttcctctattgccattactaactcaactgtatcgagagaatctgctccaagatcactt
gaaaaatttgcttccctggtcactatttttttcaattcctaactgttctgctacaata
tcttgaacttttcaaaaatttcattatcttgcataatatcttgtttcctataaggtgtt
ttcaaaaattttctggctcatattattgtctaactctaattaattagtaattttgagttg
aggtccaaatagttctacctatatgatatactattattctggataaatacgaagtttgcg
gtatggctcttctccaaaactacgagctctcaactggtagttatcaactaaattatgctg
cattttgcgaatataggctgaacgaggtatcaactgaactgtcgatttttcgttcaagat
aatagtttcaatagccagtttagcttcttgaagggcttgaatctcatcaaatttttatt
ttcacaaagttctacccaattaagatcagaagaagtatgaatatttaatatttttctcag
tgcacgagtaatttgtgggactgtactattctgaattgtatatataataatttgttttga
ttttgcaatttgtcttaattttgtattttgctttacttgattacgcaatgctaaaatagc
atctgacttttctatctctttagttaaaataattggtaaatctaaagaagaaataactga
cgcaatatgctgccaacttaaagaataagcatataaatactgattcggcatttctaaggg
caaagatgggttattaatatttataggagcattaatagtagttgacaataaagatgtatc
acgattttgtaatttattcaaatcggatggcttatttattaattctctatactgcaaaac
agcatgtgacggttgtacactaattggtttacggaaactagaaatatttgacggcaaaac
ttctaaagactgtgacggatagcatttaattaaaatgcggccattagcttgaatttgtct
tttttgtacaaaaggttgatggccctgtaaaatttgatctatagtttctttcactttatt
atggactatccaaacagtgcgttcatggatttcaattgcaatttgaaaagcgggcgcggc
ttttctttctaatatgcttttctgagtacctcttctttttgcttcatcatcacctaatgt
gacatattgaatacctccaattaagtctgacagtgtaggatttttaattaaactttctaa
atgatttccatgtgctgtacctactaattgcactcctctttctgcgatagttcgagcagc
caatgcttctagctctgtaccaatttcatctataataatgacttcaggcatatgattttc
taccgcttctatcattacctgatgctgtaaatcaggccttgcaacttgcattctccgagc
tctacctatagctggatgtggaatgtcaccatcgccagctatttcattagaagtatctat
aataacaactctttttttccatttcatctgctaagacacgagcaatttctcgaacagctgt
tgttttaccaactccaggttttcctaaaagtaaaattgaatcaccttgttctagtaaatc
cctgataatactaatggtgccaaaaactgctcgtccaacacgacaagttaaaccaataat
acttccttctctattacgcatggaactaatacgatgcagggtcttttctataccagctcg
attatctccgctaaaattaccaactttttttacacaataatctaagtcttgccaactaat

agatctttgagataagtattctggcctatctggaaatcttgcttctggacggcgacctaa
atccatcactacttctattaagtgatttctattagggtgttgtttcaggggctctttgac
aaagttcggtaagatttctagtaatttatctaagtcatctgcaataagcatgattgaata
tataataattaactaatcagatagaaaaattaagattctaaagattacaatacatattaa
agaatattgtgctcgtaatcaattttatacttaaattcaataagaaatta-cagatggta
aacaatagattaaactatgtacagtaaataaattggtcaaaattaagagtcatgttaata
tgtttattggacagaaagttcgtattaagtatagtaagcaaaaagttgctaatgacatag
cagataaagtcggtaatcgcggaattattagaggcataaaatttatcaacagtcaatgtg
ttactattattgttgaatttgaaaatcacactagactttggatgtttaaggaggagctaa
tctatttaaatgaat------ttaaacaataagataagattaattt----tattatatat
aaatttttacgagtcgttcaaattatacattttcaaggagtattgttaattgtgcaaatt
actattaaaaaattacaagacttactttcatcagtacaaagaaagaagatacagacattc
aaactgaaacaggataattttgaacttctattaaataaaacttacaaaaaaactaatcca
gaaatattaccttcacagaact---ctgtattacaacacaatcaagctaacataattaag
tcaacacctaataacaatggtccttctgttagtaataataacgttacagaatatgccact
atcgtctctccgatggtggggacattctatcactcacctgctcctggtgaaaaaattttt
gtgcaagttggggatgaagtcaaattcaatcaaacagtttgtattattgaagcaatgaag
ttgatgaatgaaattgaagctgaaattgaaggaaaaattatagaaattctagttaaagat
ggcgatatcgtagattgtggtcaagccttaatgaaagttgaaacataatgccaatatttc
cattttacatattgttaacagaaattatttagatagaaaaattagcttataatataatta
tgaaaacttatttcgtaaaggcac-aaaaagttagtatataaaattatttactttatctg
ccaaacatagaaataagcgttttgattccttgtctcaagagaggagaatctcaatgacaa
ttagctcaaaagagcaagagacaaagaaggtaaaaatctcggttgataaaaatcccgtag
atacttctttcgaaaaatgggcccaaccaggtcatttttctcgtacactagcaaaaggac
caaaaactactacttggatttggaatcttcatgctgatgctcacgacttcgatagtcaaa
ccagttctttagaagaagtttcccgtaaaattttcagcgcccattttggacagctttctg
taatatttttatggcttagtggaatgtattttcacggagcacgtttctctaactatgttg
cttggttaagtaatccaacaggtatcaaaccgagcgcacaagttgtttggcctatagttg
gccaagagattttaaatggagatgtaggtggtggctttcaaggcgtgcaagttacatcag
gatggttccagctttggagagcatcaggaattactacagaatttcaactttactgtactg
ctatcggtggattaggtatggctgctttaatgttgtttgcaggatggttccattaccata
aagcagctccaaagctagagtggttccaaaatgttgaatcaatgatgaatcaccacttag
ctggtcttttaggcttaggctgcttaggatgggcaggccatcagatccatttatctttac
ctattaataaacttttagactctggtgtatctcctcaagagattccattgccacatgaat
ttttaattaatagagagcttatggctcagctttacccaagtttcagtaaagggttagttc
catttttcactttaaattgggctgaatattctgactttttgactttcaaggaggtctca

accctgttactggcggtctatggttgagtgatactgctcatcatcatctggcattagctg
ttttatttcttgctgcaggccacatgtatagaaccaattggggtattggacatagtatga
aagaaattttagaagctcacaaaggtccttttactggtaacggtcatgaaggtctctatg
agattcttacaacttcttggcacgcgcagcttgctattaatttagctatgatgggatcat
taagtataatagtggcacatcatatgtatgctatgcctccatatccatatattgctactg
attatccaactcaattatcactattcacccaccatatgtggattggaggattttgtattg
ttggcgcaggagcacatgcttctatctttatggtaagagattataatcctgctgaaaatt
ataataacctcttagatagaattattagacatcgcgatgctattgtctctcatttaaatt
gggtatgcatatttcttggattccattcatttggtttatatatccacaatgatactatga
gagcacttggaagatcccaagacatgttttcggatacagctatacaattgcaacctattt
tcgctcaatgggttcaaagtatacatactctagctcctggtaatacagctcctaatgcat
tagcaacagctagttatgcatttggaggagatgtcgtttctgttggaaataaagtcgcaa
tgatgcctatttctctaggaacagctgattttctggttcaccacatacatgcatttacta
tccacgtaactgtacttatcttagtgaaaggattccttttctctagaaactctcgactaa
ttcctgataaggcaaatcttggctttagatttccatgtgatggacctggcagaggcggta
cttgccaagtgtctggctgggatcatgttttccttggagtattttggatgtacaactctc
tatctgtagcaattttttcactttagttggaaaatgcaatcagatgtttggggtagcgtgt
ctccatccggaaatgtttctcatatcactggtggtaattttgcgcaaagtgcaattacga
tcaatggatggttaagagatttcttatgggctcaagcatctcaggttattcaatcatatg
gttccgcactatctgcatatggactaattttcttggcagcacattttgtgtgggcattta
gcttaatgtttttatttagtggcagaggttattggcaagaacttatagagtcaattgtat
gggcacataataaaataaaagttgctcctgcaatacaaccaagagctttaagtattactc
aaggaagagcagtcggtgttgcacactacttattaggtggaattggtacaacttgggcat
tctttttagcaagaattatttcagtaggctaatagtgaaaataggataaaaa-aacaatt
atggcaacaaaatttcctaagtttagccaagctttatcacaagatcctacaactcgaagg
atttggtatggtattgctacggcgcatgactttgaaagtcatgatggaatgacagaagaa
aatttatatcagaagatatttgcttcacactttgggcatttagcaatcatcttttttatgg
acatcgggcaacttattccatgtagcttggcaaggtaactttgaacagtgggtattaaat
cccttaaaagttaaaccaattgctcacgcaatttgggatccacactttggcaaccagct
ttaaaggcttttagtaaaggtggatcagcatatccagtaaacatagcatactccggagta
tatcactggtggtacacaattggaatgagaagcaatcaagacctgtattctggggcttta
ttcttgctcgttttatcagctttacttttatttggtgggtggctacatctacaaccaaaa
ttcaagcctggtttatcttggtttaaaaataatgaatcaagattaaatcatcatttgtct
ggattattcggagttagctctttagcttggacaggccacctagtacatgttgctatacca
gaagcaagaggtcaacatgtaggatgggataattttacaacagtattgcctcatccagct
ggtttacaaccatttttttagcggtaattggagtgtatatgctcaaagcccagataccgct

caacatatattcggaactaatgaaggggcaggtacagccattttaaccttcttaggtgga
tttcaccctcaaagtcaatctttatggctaactgacatggctcatcaccatttagctatt
gctgtagtcttcattgttgctggacatatgtatagaactaattggggaattggacataat
ttaaaagatattttagatgctcatagaccgcctagtggcagattaggagctggacataaa
ggactatttgataccatcactaattctttacatatacaattaggattagcattagcctca
cttggcgtaattacttctttagtagctcagcatatgtacgctatgcctccatatgctttc
atggctaaagattttaccacacaagcatctttatacactcatcatcaatatattgctgga
tttttaatggtgggagcatttgctcatggtgcaatattctttgttcgagattatgatcca
gaacaaaataagaataatgttttagctcgcatgctagaacataaagaagctatcatttct
catttaagttgggtaactctgtttttagggtttcacaccttaggtctttatgtccataat
gacacaatgattgcttttggtactccagaaaaacaaattttaattgagccagtatttgct
cagtggattcaagcttcttccgggaaagcactttacggttttgatgtgttactatcatct
tctagcaatattgcaacacaagctggtagcaatatttggttaccaggttggctagaggca
attaatagcggaaagaattcattgttcttaacaatcggtcctggtgatttttttagttcat
catgctattgcattaggattacatactactaccttaattttagtgaaaggcgctttagat
gcacgaggatctaagctgatgcctgacaaaaaagatttcggttatagttttccttgcgat
ggccctggtagaggcggtacctgtgatatatctgcatgggatgcattttacttagctgta
ttttggatgttaaataccataggatgggtaacattttattggcattggaaacatattact
atttggcaaggcaatgcaactcaatttaatgagtcttctacatatctaatgggatggttt
agagattacttatggttaaattcttctccattaattaatggttacaatccatatggtatg
aacaatttatcagtatgggcatggatgtttctatttggacatttagtgtgggcaacagga
tttatgttcttaatctcttggcgtggttattggcaagagttaatagagactctagcatgg
gcacatgagcgtacacctttagcgaacttaattcgttggaaagataaacctgttgcgtta
tcaattgttcaagccaggctagtaggtttagcacatt-tttctgtaggatatgtattaac
ttatgcagcttttgttttagcctcaacagcaggcaaatttggttaaagtaga-----att
taattaattgcaagaaaaaaaagtcagtcatgata-gtttcgtgactgacttttttttct
tgacaatagttaattattaaagcttaacttcaatatctacaccagaaggtagatttaact
tcattaatgcatctattgtttgtgaagaaggttgatgaatatcaataatccttctatgag
atctggtttcaaaatgttctcttgagtctttatctacgtgaggagaacgtaagacacagt
aaattcgcctttttgtaggtaaagggattggtcctactgcaatagcattagttctagatg
ccgtatcgactattttatcacaagatgtattaagaatactcgaatcgtatgctttaagct
taatcctaatttttgtctgctgaataagtgtcatattgattaaatcttattatttaagaa
ttttagagacaacacctgcgcctacagtacggccaccttctctaatagcaaaacgcatac
cttgttcaattgcaattgcattaattaattcagcagtcatcttaattctatcaccaggca
taaccatttcagcatcagtaccatcatcagcagtaaactgattaatagttccagttacat
cagttgttctaacataaaactgaggtctatagcctggaaaaaatggagtatgtcttccac

cttcttctttagttaaaatatagacttcagcttcaaactgagtgtgaggagtaattgtac
caggtttagctagcaccattcctctttcaatatctttttctgaacacctctcaaaagaa
ttccaatattatcacctgccagaccttcttctagcgtcttttggaacatttctaatccag
taatagttgtggtccgagtctctctcaagccaacaatttcaattgtatcaccaactttta
taattcctctttcgatcctaccagtagcaacagttccacgtcctgtaatagaaaaaacat
cttctacagccattaagaaagttttgtcgacatctctttctggtgttggaatataagtat
caactgcctccataagtgaaaaaatcttatcaacccatttatcttcacccgctttaatag
tcgggttatttgtgactgcttctaaggctaataaagctgagccggcaacaaaaggaatat
catctccaggaaagtcatactggcttaacaactctctaccttctaattcaactaattcta
acaactcttcatcatccacttgatcttctttattcaagaataccactaatgtaggaacac
caacttgttttgctagtaagatatgttcacgagtttgtggcattggaccatctgctgcag
agactactaaaatagctccatccatttgagctgcacccgtaatcatatttttacatagt
cagcatgacctggacaatccacatgagcatagtgacggttatctgtttcatattctacat
gagcagtattaatagtaatacctctagctttttcttctggcgcagcatcaatttcatcaa
actttttgctgaggtggaacctacagttgataaagtagctgaaattgcagctgttaaag
ttgttttaccatggtcaacatggccaattgttccgatgttaacatgtggtttttttacgtt
caaatttagatcgagccatatttctattctccttatagtaaaagttgctttttggttac
ttttaacgataaatcttataaattgaaagacttattatttaatatagataaggtaaaata
agatatagttaatttaatatctatagtgtgcaaacgctttatttgcttccgccatcctat
gtgtttcttctctttttctaatggaactgcctgtttcattagcggcatccatgatttcgt
tggccaattttattgccatacttttaccagatctttctcgagaaaatctagtaatccatc
ttaatgctaaatttgtacctctatatgctcttacctctattggaacttgataagtagatc
cgcctactcttctggctttcacttccactaaaggagtgatatttcgaatagctttttcta
gaatatttaaagggtctgcttctgttctatctttaactatatctaaagcttgatatataa
ttctttgtgctaaagtttttttaccacttttcaaaatacgaacagttagcatactaacca
gtctacttttatataaaggatctggtgatgcgaaccttttttttagctgtattacgacggg
acataatattttaaatttgttattgtgataatttgtt-taatagaagatatttaagattt
aggctttttttgtcccatatttagatcgacttttacaacgatcttttactccagcagcatc
taatgtgccacgaactacatgatatcgtactccaggtaaatctttaattctgccccctct
aattagcactactgaatgctcttgaatattatgaccaacaccaggaatataagctgtaac
ttcaaatcctgatgtcaatctcacacgtgcaacttttcttaaagcggaatttggtttctt
gggagtcgtcgtatatactcttgtacagacacctcttctttgagggcaactttgaagagc
tggagattttgtttttttatgtatttttcgtctttcagatctaacaagttgttgaattgt
tggcataataaaattaattttaagtctttaatagatattattcgaacttgctgcgcatga
attaattagcctttatttatcttgtatttacgcataaatctttctactcgcccttctgta
tcaataattctttgtgaaccagtaaaaaatgggtgatttcctgaccaaatatccacatgt

agctcaggctttgttgagccaattgtcataattaattggccgtcacaataaacttttgct
tctgggtaccaactaggatgtatgttatcttttgccattgcttctataattgtataattc
tatcaataataaatattaacgcttagaaaattgaggagcttttcttgctttcttcaaacc
atatttttttctctcttttactcgaggatctcttgttaaatacccttcagatttaagtgc
gtttctattttctggattaattgagcacaatgctcttgcgacacctaaacgaattgcatc
tgcttgtcctgttaatcctcctcctctagcatttacatgtatatcatactgatttaataa
tcctagaacttgtaacggtgcatacgaaactcttaagtaattaggactaaactgaagata
tgactcaccaggtataccgtttataattaaattgcctgaacctgggaccagtcttacttg
tgcaacagaacacttccggcgacctgttccagaatagattgcacgagttttaattaattc
tgtcgacataatattccttattaactaaataaatatacattctttatactatatactctt
gcggcttttgtgcgatatgtggatgaagtggacctgcatagactttaagcttcgtaaata
attttcttcctaatggacctttaggaagcatacctttaaccgatttctcaataattctat
taggcaatcttctctgtagctgctcaaaggtttctacttttagtccaccaggctgcccag
aatgtcttctatataatttctgattagttttatttccgcttacatatatttgagaagaat
taatcacaattacataatctccagtatctagataaggcgtataggaaggcttatttttac
cccttaatatactagaaatatgagtcgatatcctaccaagtgtctgattttagcatcta
taatataccaatttggattagtatttaatgatggtacttgcgttttgttcatgaaaagat
taatacttttagttaacaataagagagtaggtaaatgctgcacatataatgtcgatacaa
ataatattt-tgtaatatta-------------ttgataaagatagctttcatgacagtt
acttaaattgtttgttagcaagaaaacttatagttatcaataagtgataata-ttgtggc
aatataaaataggttattcactttttttcttttggcaaacttatacctaattttttttgt-
agagcatatataacttcttcagctgatttttggccaaagtttttaatttctagtagttct
tcttgagagtaatctagtaaatctgcaatagaatgaatttgagctcgttttaagcaatta
tatgctcgaacagatagctggagttcttctattaaaacttgactaatttttttatcttct
ttgctacgataattatctgctgatttaaagtctaaatttctcagtgagcagaaaaggtta
gttaaaactgtagcaccttggcttatagcttcttggggagagatactcccatttgtccat
atttgtaaaattaacctgtcttttatgctattagtacctatacgtacttcttctacttta
taattgactttattaactggcataaaaactgagtctacttgcagaaaatctacagataat
tcatctactgctttttctgctaaacgatatccacaactttttttcaattttaaactccatt
tcgaatattgtattattgcaaatagttgcaatatactgcctaggatctactacttctata
tcagaagatagttcaaataatccagctgtaactatagctggcccctggactctaattctg
ccaatttgagattctttattatggcttttaaatactacttcttttagattcaatagtatt
tctagcacatcttctcttaccccaggaattgtagagaactcatggtttactcctgcaatg
cgcacagcgactatagcggtaccttcaagatctgataatattgatcttcttaaagcattg
cctaatgtaatgccttgtccttggtttaatgtttctattacaaaactaccgtactgtcca
cgtgctccatctgttcttgattctacacattcaatttgaaattgagccacctaagcaagc

tcc-ttaaaaataatcaatggttagtaactctgcctgcgataagaaagaatctatacacg
gcgtttcttggggggacggcagccattatgaggtacaggagtaatatcttttattaaagt
aatctctaatcctgcagcttgtaatgctcgaattgctgtttcgcgacctgctcctggacc
atttactaaaacttctgtttgacgcagaccttgatccattgcctgtttagcagctttttc
agctgcagtttgagctgcaaaaggtgtacctttttagctcccttgaatccactagcacc
agaggatgaccaagataacgtttcacctttaaattagtaattgtaacaatcgtgttatt
aaatgtagatttaatatgtgtaacaccattaactgcattacgtctattttttttttactcc
agatttttttaattgtctagccatcgtgtttcgtctattatgtaattatttaaaatatta
tttccttggagctttttttcttacctgctactgttttttttacctcctcttcttgttcttgc
attagttcttgttctttggcctcttaaaggaagaccaagacgatgccttctacctctata
agtactgatctccataagtcttttaatactcatagattcgaaacgtttgaggtcgccttc
aatctgataactagattctaaaatttctcttatactaacaacctgttgatcatttaaatc
ttgacacttaatatcagcatcgacgtttgtttttttcaatatttcttttgagcgagatag
tccaataccataaatatatgttaaagctatttctattctttattccttggaagatctac
tccagcaattctagccacgttgtgctttctccaataaataatatttac-gtttacaacag
aagattaca-gaattagccttgtctttgcttatgtttaggattagtacaaattaccatga
cttttctgtgacgacgaattattctacatttttcacacattttgcgaacagaaggacgaa
ctttcatattaa----ctctatatactaggaatattactg---------tattaattatt
tacatgtgaaatgtcaataatatgtttgattactttaatttctatagctttcacttcgtc
atactatcatatttttttgatatgacatatgtttgaatttgcttagctgtgtctattgcc
actccaactaaaataagtaaagatgtcgctccaagacctctcaagttttgtatctgagta
acctttttctattataaacggaatcagcgctactgtaaataaaaacgaagctccgagaaat
gttagtctatttaatataacttgtaaataatcaatggtcgcttgacctggacgaatattc
ggaatacttgcacccatttttttttaaatttattgcaatgtcttctggattcattactatc
gatgtataaaaataactaaaaaaaagaattaatgcacaataaagaatgaggtatagcgaa
ccgttaggacaaaatagatatagaatttggagtaacgttttattttgaattatttgagtt
aaataagatggaagagccatagatgcagatgcaaaaacaataggcataactccaccttga
tttaatttcaaaggcaagtaactgttcggatccaaaattgaagattttcctagttgtctt
gctgaaataattttaattcttcttgttccttcttgtacgcaaattgtaattataatcatc
aataagaagatagcaatgaacaatccaaacttaagccccgcattgctatagtttgtatca
aaaaatgactgtgtaaaattttttggtaatcctgatacaatattttgaaaaataagcaaa
gaggctccattcccaattccttttttctgtaattaactcagataaccacataatgatcatg
gagcctgccgttaaagctaaaacagactcacaaacaaatgtaaaattccagttaaataca
tacggctttacccatatagagattgctccagattgcaaagtggcccatccaagagacaaa
tatcgggtaatttgagttatttttttggcgtcctagctcaccttcttctttctgtaatttt
tctaaatcaggaataattttagtgagtagctgcattacaattgaagaattaatataagga

acaatacctaaggcaaagatccctattgttgaaaatcctcctccagaaaaaatattcaaa
aaatttactaaagtattttttctacgcttgcataaaaagcatcatggtctatacctgga
acaggtataaaaattcctaagcgcgctaagactaataaaaaaagagtaaatataatgcga
tttcttaagtcactttttggctcataaataaatt-----ttaaaaaagacaccga--gt
tagttaagaaccctattccaatacttcgacaaatttaaaatagggcgctggcaacaagct
atagaagtaatggca-taatatcttatgttgaataaagactttctatagaaactcctcga
tctttagctgcttcgctaaaagttcttaattgagttaggccatttaaaacagctcttgca
ttatttagtgtattattagatcctaactgtttagctagaatattttgtacacctgataat
tctaatactgtccttacagagccaccagcaattacaccagaaccaggtgcagaaggcctt
agtataactttggctgcaccagaaattccattaataggatgaggtatagaattagatttt
gtaagtggcactgtaactaaatgtttttagcatcagttactcctttttaactgctcca
attacatcactagcttttcctactccaacaccaacttggccctgctcattaccaacaatg
agaataactcgaaagcttagtttcttgcctccttttacaactttagtgactcttttaact
tgtacaactctttcttcccaaccactatctttatctttgcctttactctgtttttacga
ttagccattttaagcgttattcctaa----ttaatta-tgactttaaaaacccatacctg
cttccttagcagcttcagctaaggctttaactcttccatgatataatttccctcctctat
caaagacaacattcgtaataccttctttaatagacttgtgagcgagctgttgaccaacta
cgcgagaagtgtcacaatttgagcctattttatcaaattcttgattatttaaattaacag
atgatgttgagactaatgtgattccttttgtatcatcaattatttgtgcatatatatgtt
tattggatctaaatacgcatagacgcggcctacttgaagttccctgaactttttgcgaa
ctcttttatgcttatgaattctagtctgcttagtatttattttcattattatttaccttt
tccagcttttccagcttttcttctgacaaattcaccttgatatctgattcccttcccctt
gtaaggttcgggaggtctagtagagcggatagttgaagcaacctgacctacaacttcttt
atctatacctgaaacagtaatattggtattattttcaactgtaatttgaatattttcagg
aggttttattttgactacatggctataaccaacgcttaatattaaatcttgattatctat
ctgtgaacgataacctacaccttgtatttgtagtttcttagtaaacccattagaaacacc
ttcaatcatgttactaataagagttctggataatccatgtaattgattagccatttttgt
ctgtttacttgtcctaacagaaattttatcattaagtatctctaagttaataccagccgg
cagagttctagatagagtacccttaggtcctttaaccgtgatggtatgaccatcaaactg
agtactaagattcgccggcaataaaattatttttttccaatacgagacat-ataccacc
tataaaataattaccaaatatagcataatatttcaccaccaagaccatcatgacgagctt
gtctatctgtcataacacctttagaagtagaaataagagcgattcccaaacccccaagaa
ctctaggtagttctttatggtttgcatatactcttaatccaggcttactaattcttttca
gagcagtaataactggttgacgatttttaccgttatattttagagaaatcattaaatgca
tatctatgccttcacccatttgttcaaaattttgaacaaatccttcttcttttagtactg
ttgccatattgcatgtcattttcgttgctgggacttgcacaatttgatgtcttgctaagt

ttgcattacgaatacgtgtcagcatatcggcgatcgtatcgttgaccacc-agatcctcc
ttacgataaattgaattatttaatatgtttttaagattccttaaaaggcataccaagctt
tttcagcaaagctagcccttcttgatctgttttggcggtagttacaattgatatatcaga
gccgcgaatttgatcaatattatcgtagtctatttctggaaaaattaactgttcgcgcaa
acctaaattatagttacctttgccatcaaaactttttggactaattcctctgaagtctct
aattcttggcagggttaatttgattagcttctctaaaaaagagaacatcttgtcttttcg
cagatgtactacaatgccaataggaacttcttctcgaattttaaacccagcaatggattt
tttagcttttgtgacaatcggtttttgtccagtgattaatgttaattcttgaatactact
ttcaagagctttagcattctgagaagcttcacctagcccacggttaatagtgattttcgt
aacacgaggaacctcatgtacatttttatattggaattcctcttttaaagcttgaataac
agttgttttatatttttcttttaatcccattactattatgtttt-acaaattatttaata
agttcacccgtctttttcagttttcgga-ttttttggcctttatcgttaatgattactga
agatctactagcaatattattctgttcactaaatagcatgacattagaactatgtattgg
agcttcaaatttaataatttctccggtctcaccttcttgttgaggtttttttatgttttac
ttttaaattaattcctttaacaaccactttacttgttttgtggatgattgcaataattgc
gcctatttttgttttatcacttccagaaataacttggactaagtcaccctttttcaattt
aatttttgtattaactttagtttttttgaaagaacttctcatcatattacctccggcgca
agagagactattttagaaaaatttttatctctcaactctcgagctataggtccaaatact
cttgtaccacgaggattattatcttgattaataataacggctgcattatctccaaatcta
atactcataccatcagttcttcttaaagctttgcgagttcttactactacagctctgaca
acatcagatcttttaactggcatattaggagatgcatctttaactactccaataatgaca
tcacctatcgatgcataagaaggattactagtacctaatactcgaatacacattattttt
ttagcaccactattatccgcaacattaagatagctttgagtctgtatcatacttttattc
tatataagattaattatcaaaagacttagataagatattaaccatcgtccaacactttgt
acgacttaaaggccgtgtttctcgtattgtaactacatctcctggtgcgcattcattatt
ctcatcatgtactttatattttttttgtgcggatcatagtttttgcatatttctatgagc
aattctattttctacagctactactatagttttattcattttattactgacaactttacc
agttgtttcttttaaaggcatagtttttatc--ttatatatatatacagtgaagttaaat
tataaaataattatgtggctggcttactccgagacttttcaacagttaaaagttgagcta
atctatgcttggaatgcttaaataagtgaggctgaaaatcttgtcttgtagctcttttca
gccttaagtcaaatagcactcttttaattacaagtatttcttcagctaaagaagaattat
ccatttttgtaacatctgatattttagggaaggtcatactttaagattctattgtatttc
gaactataaatttggtctttattggtaatttataagaagctagtttcatagcttgctgag
cagtttttttgtggcacgccagtaatttcaaatagaatgtgaccaggcctaataactgcga
cccaatattccggagctcctttaccagatcccattcgggtttctgcaggacgcgctgtta
caggtttgtcaggaaataccctaatccagagcttacctcctcgcctaacatatctcgtaa

tagttctgcgagtagcctctatttgtctagatgttaaccatactggttctatagcttgca
atgcataatcaccgaatgcgattgtattacctttactagcagaacctttcattctgcctc
tgtgttgttttctaaactttgttttcttgggacttagcataaaaattaaacaatttaaat
aattaagaagctgttagatcagaagtttcctgattatatacagattctgtgcttttcgat
tctggcaaaatttcgcctttaaagagccaaactttaactccaagtacaccatatgtagtg
tgagcttgtcgatgacaataatcaatatcagctcttaaagtttgtaaaggaacacgacct
tctctaacccattcgctcctagcaatctctgcgccattaagacgcccagaaacttgtatt
ttaacgccttgtgtatttgctctttgcgtcctttgaacagcttgtctcacagctcgacgg
aaggcaactctttttctaattgctgagtaataaattctgcgactaaggttgcttcagaa
tcgggatctgcaatctctaaaacattgactctgatttgtttactcgaatctagaattgaa
gataacgattttcgtagagactcaattcctgccccagactttccaagcacaatgccaggt
cgagctgttgctattaaaatttcgacttgatcaactttacgattgatttcgattttagca
atgctagcattactaagtttagagtgtataaacgaacgaattttatgatcttcttgtaaa
agaactggatagtctttagaattggcgaaccatgaagaacgatgtttttgggtaatgcct
attcggaaacccaaaggatgaatttttttgacccacagtatccttttctataaattaaaag
ttataattactaattgactttgaatac----aagactacaattctgagacacataatgta
atgtgacaagttggtttatgtatagggaatgctcttccttgtgctcttggctgaaatctt
tttagcgtagggcctttatctgcaaaagccttactaataaataactcacttttgttcagt
ccatcattatgctcagcattggcagcagcagactctaaaatctgctttatataagaacac
actcgatatggcataaattccagaataattaatgcttcttgatatttcctacctctaatt
tgatctaatacacgacgaactttatgtggagaaagacgaatatatttccctactgcttta
gtttcttttacgtttttttgtaatactcataattatagtttaacgacgcgcttttctatca
ccttttatgtgagtacggaaagttcttgttggaacaaactctcctaatttgtgacctacc
atttgatctgatacaaagacaggaaaatgttgtttaccattataaacagctattgtatgc
cccaccatatctggaataatcgtagatgctcttgaccaagtttttaaaacttctttttttc
ccaacagtatttaatgcttctatacgtttaagaagactcacatcaataaaagggccttta
tgtatagatcttgacataataataaattattagattcttaataaaataaattctatttac
gg-gacgcaaaatgtatggggtgctatacttattgggattccgtgttttaacacctaatg
ctggtttgccccaaggtgttaccggacgagctctcccaattggagatttaccttcaccac
caccatgtgggtggtctactggattcattacgacacctctaacagtaggtcttttaccta
gccaacgatttctaccagctttaccaagagtaatattactagcgtcaatattaccaactt
gccctatagtagcatagcattcttttcgaatcatacgaacttcgcttgagggcagtttca
ccgtgacaaaagagccttctttggctactatttgagcataagttcctgctgcacgaacaa
tttgccctccgcaagaaggtcttaattctatattgtgaacagctgtgcctagaggaatag
ttgataaaggtaatgcattgccaacttcaataggcgcagtgggaccggacagcaccattg
ctccaacacgcaatgatcgaggatgtaaaatgtaccttttttctccgtcaagataatgca

ataatgctattctagcatttctgttcggatcatattcaatagaggcaactttagcaacta
cgttgtgcctgtttcttttaaaatcaattaatctatactgctgtttatgtccaccacctt
tatgacgacacgtaataacacctctattgttgcgacctttgcaaaaatgatgtttaacta
ttaaggattttctggcttatcagtagtaatctctgaaaaagtagaaactgttctatttc
ttgttccaggcgtataggcccggtataaacgaattgccatataggaaaatttaaaaagag
taaatatttta------ctgtgtaatacaattaagtttctgggaataagttaatagaatc
ttctggcgcgagtgttacgattgctctcttatagtgaggccgtttacctacaaatctacc
aatactcctttttttcttaggtggatggcaagtgttaacacctgtaacttgaacattgaa
aatatactgtatcgcagctttaatattaattttagttgctttgggatcaacagcaaaaca
gtactggttttcttctaataattttgtagttttatcagtaatgattgggtatttaactaa
atctaataagcctcttgaatcaatgctatccattatatacctcttgtattttagataaag
catcaactgtaataataattttatgcgcagctaaaagggccatgatattcaaagtgtcag
ctgaaataatttctacattatttaaattgcgaatggaaagatagacattcggatcttttt
tatccactacaactaaaactttttttgtttaaatcaagattccatcggtgaatagcttcca
taaacaatttagtttttggctgctgaaaataactattaaagttttctacaattaacgtat
tgactgatttattatttaacgttgtccttaaagctaattgtctttctttttttattcattt
ttttagtaaaactacgaggcttgggtccaaagattacaccaccacctctccataaaggtg
agcgtatagaaccggctcttgctctaccagttcctttttggcgccatggtttgcggcctc
cacctcggacttcacttcttgttttggtattagcagaaccttgtcgtttttcattccttt
gtcttactaaagctctatgaactaagtacataccagagtcttgactcactttaagattca
aatctgcattaccacttatttcacctttccaattatagacttgataatttaattgtgttt
taactgtcattagatatatcgcgtgaaagaatattgttatttactaatttttactaaatt
accaggcttacctgggacagctccttttacaatcaaaaaatcattttctgagttaatact
tacaatttgcaaatttttttatcgtaactttttttattaccaagttggccagccatattttt
tcctgggtaaaccctcccaggcgtcgtaccagctccaatagaaccaggttgcctatggtt
tttcgagccatgagacattggacctctactaaaatgatgtctcttttgatagcctgaaaa
gcctttacctatgcttttagaagaaacattgattgtctggcctactttaaacatatctgt
agacaagatttgactaacttcaaagtcatcagtagattttactaagtactcacgcaaata
tttcagtgggggagcttgtgacttctttaaatgtcctaataatggtttatttaacttttg
ttcaacaacttgcttgtacccaacttgaatagcattatagccatccttaggtaaagattt
aatttgagtaacaatacacggcccaacctgaattacagtaactggaattgacaagccagc
ttcatcaaagaattgggtcatacctactttagtaccgagtataccaacagacactagatt
tctcctttttgagagatttatactattatcgtacatgaaatttttacttcaaattgcctg
aag--taaactgaactaagagatgtctgagattaatcatgacaacataaagagaaaaaac
tttacttattcaatctaaaatagaaatagtttttttgtccagttacattatgtatagatta
aaggcaaacttattatattatttatacactacg---------------------------

----tgttcaaaataatatgaaccagaagacatgaattcttttag-----atttgtacaa
cattacattattagtactgtctaaaatata----gatagaatattgaatttctacctctc
tagaa-------aaactaaaattgtgttcggtaactacaccttttatttacatatttat
tgtgcagtatatatgcatatctaatacagcacaggtaaaaatatagc----tgtatagcg
aatatacttacaagaattagagaggtattaatgggtaaagttgttggaattgacttagga
acaaccaattctgtgattgctgttatggaaggaggtaaacctactgtaataccaaacgca
gagggttttagaaccactgcttctgttgttgcatatactaaaagtggagataaacttgtg
ggacaaattgctaggaggcaagctgttattaacccagagaatactttctactcagtcaaa
agatttatagggcgtaaacaaaatgaaatttctcaagaaattagacagacatcatataat
gttaagactagtggatccagcataaaaattgcctgtcctgcacttgataaagattttgca
cccgaagagatttccgctcaagtactcagaaaacttgttgaagatgctagcacatactta
ggcgagactgttacacaagcagttataaccgtacccgcttactttaatgactcacaaaga
caagctacgaaagatgccggtaaaatagccggtttagatgtactaagaattatcaatgaa
cctactgcagcttccttatcttatggactagacaaacaaaataatgaaacaattctagta
tttgaccttggaggcggaacatttgatgtgtccgttttagaagttggagatggagtattt
gaagtactttcaacttctggagatacacatttaggtggagatgactttgaccaacaaatt
gtggaatggttaatcaaagatttcaagcaaaatgaaggtattgatcttgcaaaagataga
caagcacttcaaagattaaccgaagctgcagaaaaagcaaaaatagaactatcgaattta
actcaaacagaaattaatcttccatttattactgctacacaagatggtcccaagcactta
gaaaaaactgtaactaggggtaaatttgaagaactttgctctaatctaatagacaaatgc
agtattcctgtaaataatgctctaaaagatgcaaaattagaagcttccagtattgatgaa
gtagttttagtaggtggatctacaagaattcctgctatacaacagatggttaaaagatta
ataggtaaagatcccaatcaaagtgtaaacccagatgaagttgttgctattggagcagct
gtacaagctggcgttttagctggcgaagtcaaagatattctattattagatgtgacacca
ttgtctttaggagtagaaactttaggtggggtaatgacgaagattataccaagaaacact
actattccgacaaaaaaatcagaagtattttctacagctgtagacaatcaaccaaatgta
gaaatccaagtgcttcaaggtgaaagagagttgacaaaagacaacaaaagcttaggaaca
ttccgattagatggtattatgcctgcacctagaggcgttcctcaaattgaagttacattt
gatattgatgctaatggaattttgtctgttaaagcaaaagagcaagctacaggcaaagaa
caatcaattactatatctggagcttctactttacctaaagatgatgttgaaagaatggta
aaagaagctgaagaaaatttcgatacagaccaaaaaagaagaaagaatattgatactaag
aatcaagcagaatctttatgttaccaggctgaaaaacaaattaaagaatttgaagataaa
atcagccaagatttgaaaacaaagatagaggaactaattacagagcttagatctagttta
gagaaagaagaatatgaaaatattgaatctatctctcaacgactacagaactcattaatg
gatattggaaaaatggctgctcaggctgaaagtaaaaatacaaacacaaaagatgatggc
actgtaattgacacagatttctctgaagctaaataaaaaatt-agcgggtaacgcgattc

gaacgcgcgacatcaaccttggcaaggttgcgctctaccactgagctatacccgcacaga
atatattattacaaacatggtaacatttgtcaatctgt--------agttatat---agt
tgagctattaatcattaaactatagattacaaacttgttaaaaacaaaaaaactattatc
gtttttaaa-aaaagaagtacttatataag-tatataagtacttcttttattcaaaaa-
-c-aaaatataattagatatcagtt-------ttaaactactaaggaattaaccaagcta
gtaacaattactagtcctgtccataagcctgcccctgtatagattaacccttttgattgt
tcccattgaccgggtgatgctagaacaacagggacgcctactactaacacagtcgataat
acaattaataatagcactaataattgaattgcaataatcatttattctattctcctaaaa
aattcaaatgtgaataatattgtatataatacaatgaagctttataccttcatcaattta
tctatgaagaaataatttaaaattattgttttctcaataaatattaaattaagtattaaa
atctttattgtaaacaataacactatctttttttaa-atagataaaaaatttaaagaaag
tctttttaaaaaaaatttagaataaggtgatatgctgtataaaaagctaagtattatta
gttattagagaaattttaaaataagaggtaaaacatatgaattcagccctttttttagcg
aaactaccggaagcttatgcagttttcaaaccgatcattgatattttacctgtaattcct
gtatttttctacttttagcttttgtttggcaagcagcaattggttttagataaccttga
agctataacaaaaatagcaaatataaaaagtaaaatttttgcgctctaatcttatatct
tacacaataataactattatggttgaacccttactatcaggaattgttcttggccttatt
cctatcactttatttggattattagtagcagcttatctacaatatcaacgtggaaatcaa
ttaggactataaagcctaggttatagatatttataaaagctgtaataattatggtttta
taaatatctacaaaccagctaattataaatacagaaaaaatttattgaaaacagattga
taattggcaatactgattttatagtaagatagcgaatctttcataagataataactattt
taaagttatttgaacccattaaactttctatatgacaagctgcttctttctatacaccaa
cccttcttgaattggatttgctttatttaaaaaagctaatacttgtgaaagatctttaac
actggcttcgctacggaatggattctctaatttactatactcgggtctcttaataagaac
ttttattacagccattagcgaaacatcatttagtgcaatcaatctactggcttcaagaga
tttctcatcaactataggtggatcaaacatctcattaaacttttcggtgtttaaactagc
gaaaatatttatttgactattgttatttttttacaatcttcatcatcatcgttagcagct
tttaaaatcacaacgccctctggattagcgctggttaatctataattagtacgcgtatca
tcgctagttattatcgaaaatacttgtggatctctttcgacataaagccaagacttttc
aaaggatatggctccttgtatagcacaagtgtctagaaaacctacagtaagattgtagtt
acgacccattatacaagcacaagcccacacacctagatttataaacgactgttggcttcg
agcattttacaagaaattgctttcagtcgccgatccatagtaattacaccttccgcata
tttagtgctgacagtagcttataaaacctcatctttagcaaacccttgtacaattctaat
ataatcggataccaagaagtgatctattatatcaacatgcaaagtagggtaatacggatg
cgggtctataggaaagggatggatttttaattctagtaagtagccaaacataataaactt
ggattcaaataaaggtaattttatattaaagtatgatatagatctctttcatatattcta

gacatttgagtgtaagacaatcatttctttactatttataatcttgatttgtgatgaaat
agcaaaattcttatcgccataagcgctcttatatagaaatgtctgagttatgaaatctac
atcgactaataaagaaaaatatatttatttattattttcaatattatcaatgaaaatact
tttagtataaatatactgatttttttctagtaaaattgtttaccagctcgacttagtaac
tccaggtaaaagacattcatgagacatttctcgaaaaacatggcgagataaaccaaaatc
tctataataccctctacttctaccagttaaccagcatctgtttctacctctaacggctgc
gctgtttctaggcatttcttgaagtctttgtcttagttccattttgtcagaaaagcttat
agctttttttagttgatctttaatagccagtctttttaaagagtattttttttaagctt
ttccctttttaagctctctttggatcatatttttttttgccatattataa-aacttactta
aaaaattacttaatcaatatactagctaaaaaatctctgaactgcaacatttatatggtc
aaaaacatatagcataaa-------------atagatattttatgaattatact---act
ttatgctattgaaagaaatttatttaataaagttcttgttcaacatgagttgaaatagtg
caatcagatgtaggataagcaatacatgttaaaacataacctgctaaaagctgatcatca
tctaagaatgattgatcacgttggtctacaccccccttcagtaactttacctgcgcaagtt
gagcaagctccagctctacaggaatacggaagttcaacaccttcttcttctgctgcatct
aggatgtatgtatcgtccgcacagttaaatgtaacatcaatattttcatcttcacatact
aaatgaattttataatctgcca--ttttttttactccagaataaagatatgcttgaaactt
aataa--taaaac-aatttaaaggacttgcccaaa-attgtatattagtttcattatgca
taatagactgcaaaaaaaagaaaataatatactattgtaagaaattaagctacatttcata
ctaaatagaactattgcaggttcaaagtatatatataaattcatatagttaaaattattt
ttatcatttaaataactaaacttaatactct-agacctttagattaactgca------ca
gctaatcttttacaataaacaaataagtaattatgacttcaatgataaatcaacaaatag
aactaaagcctattcttcgatttcagataccacatttatattcgagttttcttatacagg
aaatcaaatctttaacaaaaagactctttttgcaagtttggagaagacctgcgacattga
tggcaggtattattcagcctttattatggctaattctattcggtgggctttttttataatg
ctcccgtgaatttatttactattaacacaagctataactgttttttaagctctggaatta
taattttcacatcgtttactggtgcacttaattcaggtcttcctttaatgtttgatcgag
aattcgggtttttaaatcgcttattaacggctcctttaatatctagaacttctattattt
tatcttctgctacttttatgacttgtattagtttaatacaagttatattcatagttacag
cctcattatttatgggaaattctcccttaaacagtgatagtgcgatagtttttggactta
taattctattagtaactgtgggagttacaatacttagtttagctttatcttttactctgc
ctggtcatattgaactattagcacttatttagtaatcaatttgcctttttttattttcta
gtacggctctggcccctctatatttcatgccaccttggctccagttaattgctagtctaa
atcctttaagctatgcaatagaaggtacaagatatttatattccagcatacattggaatt
ttacagagtctattataaaaattagttggggagatgtttgtttagggcaaattattgcat
tattagttattctagatataattgcagcctaccttgtctctaagatactaaaagctaaac

ttaatta-----------------aaattttaataaaaaatttattaaatttattttaga
aagattttttactatataatactagtagtagtatggataattgaaatagaaatagtttttc
aaaaccaaagctattatcaacaatgataaacatttgtaagaaagtcaacaaagtatgttc
ttattcataggaggcatgtagtcaatgggactaccatggtaccgtgtacacacggttgtt
ttaaatgatcctggacgcttaattgcagtccatctaatgcataccgcacttgtagcaggt
tgggcaggatctatggcattatacgaattagctgtatttgatccttcagatccagtatta
aatccaatgtggagacaaggaatgtttgttatgccattcatggcaaggcttggcgtaaca
gattcttggggaggatggagtattactggagaaagtgtttccaatcctggattatggagc
tttgaaggagtagctttaactcatatagttctttctggaatgctttttctagctgctatt
tggcattgggtttattgggatttagaattatttcgagatccaagaactggagaaccagct
ttagacttgccaaaaattttcggaattcatctattgctatcaagtttactttgttttgga
tttggtgcttttcatgtaactggattattcggaccaggaatgtgggtatctgatggttac
ggagtaacaggtaaagtactgccagtggctccggcgtggggaccagaaggatttaaccca
tttaatcctggaggagttgcgtcacaccacattgctgcaggtactgtaggtatattagct
ggtgtttttcatttaactgtgagaccgccgcaaagactataccgcgctctaagaatgggt
aatatcgaaacagtattatcaagtagtatctctgctgtttttttctcagcttttgttact
tgtggaacaatgtggtacggctctgcaactacacctattgagttatttggtccaactcga
tatcaatgggatagtggatattttcagcaagaaattgaaaaacgagtagagaacgctatt
gctgatggagctgctcctagcgaagcatggtcaagaattccagataagttagccttctat
gattatattggtaataatccagcaaaaggtggtttattcagagctggccctatgaataaa
ggagacggagttgctgaagcatggcttggacatcctgtatttcaagataaagaaggaaga
gaactgagtgtccgcagaatgcctgctttcttcgaaacttttccggtcattttagtcgat
aaagacggtatcatacgagctgacattccatttaggagagcagaatctaagtacagtata
gaacaagtaggtgtaacagctagtttttatggcggaaaattaaatggccaagtatttaat
gatgctcctagcgttaaaaaatatgcaagaaaagcccaattaggggaggtatttgaattt
gaccgaactacactagaatcagatggtgtattcagaagtagtccaagaggctggtttaca
tttggtcatgctaatttcgctttaatatttttctttggacatctttggcatggttcaaga
actatatttagagatgtgtttgcaggaattggagccgaagtaactgaacaagtagagttt
ggagctttccaaaaattaggcgacagaagcagtaagaagcaaggagctgtataatgct-a
agccaaatacagtctttatc---aaatatttccaatattaaaatagttcaggaaaagttt
attgtctaatagttactaattaggagaacctatggaagccttggtttatgtatttttatt
gacaggaacactaatggttatattctttgcgatctttttttagagaacctccaagaatagc
taaatagctttattaagtttaaccttacaataatagaaccactttccaacttattagtgg
agagtggttctattgtaggttacagtatattattcttcatgctcttcaaaaggatctctg
agatcttttgaagcaggaccaaatgccgtatatattgaatagcctgtaatacctaaaagc
aaacttgaaataaaaatactaagaactgttgcagtttccataattcaatcagtattaaga

tgaactttctctataatgatattatcaatattaaaacatttaattattaaatttatcaaa
atatattatggcacttagaactagacttggagaaattttaagacccttaaattcagagta
tggaaaagttgctccaggttggggaacaactcctatcatggggattttcatgttactatt
tttcttatttttgttgatcattttgcaaatctacaattcatcactagtactagaaaatgt
agatgtagattgggctactttaggtag----ctaattatacttttttctttaa--------
tttaataacattaaagataat----------------aagcgatcgctattataacataa
tagcgatcgcttaatttctactttactatata--caaaagtaattattttcctaagtcaa
ttaattcgctatttgcaaagctattagtattgacattattataattaactttttcaaatc
ttactagtactggatacttgataccacttttatctattgctgcaacagttccaagttctt
gataccagtatgattcctttctcaatatttttacttttgagccacgttccatatttttag
tcctaagatagattattaaagattagtataatagataaagaaataaaaat-aaaaaaata
cataaaattagtaccgacaat---------taaatagtaaaaggagagagagggattcga
accctcgataactacagttatgacaattttcgaaattgttgcaataaacctctctgccat
ctctcctaaattaaaactcttttccccgaattagataagaataacatagaactaatgta
gttgcctagaaaattaaaactatgttaaatataaattataatctttctatagatctcttt
acaaaatatgtacacagattctatatcagcttatttgaagatttttaatcttatacaat
ttttactaaataaataaactctctcataacatgaaactatcttggaaaacacttcttctt
tggtctttaccaatatttgtcgttgggttctttttctggcaaggtttttaggtccgact
actacagatgttggtagtaatatcgcaagttctagaatgacatatggaagattcttagaa
tatttagatatgggttgggttaaacgagttgacctatatgaaaataatcacacagcaatc
gtcgaagctgttggaccagaattaggcaacagggttcaacgaattcgagttgagctgcca
gcaagtgcaccagaattaattacaaagctacgcaaagctaatgttgatttagatgctcat
cctcctaaaagcacaagtgcagtatggggactattaggtaatttactatttcctctattg
ttagttggaggcttagcttttctatttagaagatctaatagtgctagtggcgggcctggt
caagcaatgtcatttggtaaatcaaaagctttatttcaaatggaagccaaaacaggagta
gtatttaatgatgttgctggagttgaagaagcaaaagaagagtttcaagaagtagtaact
tttttaaaacaacctgaatcatttactgctgttggtgccaaaataccaaaaggcgtttta
ttagttggtcctcctggtacaggaaaaacattattagcaaaggcaattgctggtgaagca
agcgttcctttttttagtatttcaggctcagaatttgtggaaatgtttgttggagttggt
gcttctcgcgtaagagacctattcaaaaaagcaaaagacaatgcaccttgcattgttttt
attgatgaaattgatgctgtcggaagacaaagaggaacaggtgtcggaggtggaaatgat
gaaagggaacaaacattaaatcagctattaactgaaatggatggctttgaaggaatact
ggtgttattgtaattgctgcaactaacagagctgacattctagattctgcattattacgg
ccaggaagatttgacaggcaagtgtctgtggatgttcctgattttaaaggtagattagca
attcttgaggttcatgctaaaaataaaaaaatggagcctaaagtatctttagaaactatt
gcaagacgaactccaggcttttcgggagctgatttagctaacttattaaatgaagctgct

attttaacagctcgaagaagaaagaatgcaatgactatgtcagaaattgatacatcaatt
gatagagtggtagctggaatggaaggcacacctttaattgatagtaaaagtaaaagatta
attgcataccatgaggtaggccatgcaataataggaagcttattagaacatcatgatcct
gttcaaaaagtcaccttaataccaagggggcaagctagaggcttaacatggtttactcct
agcgatgatcaaagtttaatctcaaggtcgcaaatactagcacgtatcgttggtgctctt
ggtggaagagctgcagaagagattatttcggagatgcagaagtaaccactggtgctagt
aatgatttgcagcaagttacatcaatggctagacaaatggtaactcgattcgggatgtct
aaaattggcccattatctcttgaaagtcaaggtggagaccctttcttgggtagaggaatg
ggaggaggctcagaatactcagatgaggttgcaactaatattgataaacaagtacgagaa
attgtgagcgaatgttataagcaggctaaagaaattattgcagataatcgagtagttatg
gatagattagttgatctactaattgaaaaagaaacaattgaaggcaatgaatttagagat
attgttaaagaatacactgcaattccagagaaaaattactataaatctcaattttaa---
---------------atgatacgggactgacgggattcgaacccgcaacttccgccgtga
cagggcggtgctctaaccagttgaactacagtcccaataaaggg---tacagacagataa
tctcataattagcttgtgattgtcaaattatactgtaaaacatgaaaagta---------
--------tcctcaataatattttcatgtttttattataaagccaattgattaatgca
taaggagagagagggattcgaaccctcggtacgaagttaatcgtacagtagattagcaat
ctaccgctttcgaccactcagccacctctccaaagagtcttacgtactcaatatatcaaa
tattattaaattttaacagctgtgcataaatatttatggagataagcggattcgaaccgc
tgaccctctcaatgccattgagatgctctaccaactgagctataacccc---tttcttaa
ggctattaagaaatggctcaagcgggatttgaacctgcgaccttgggcttatgaatcccc
tgctctaaccactgagctactgagccataataaaaaccattgataaatactataccataa
tttatcttaaaatataaatcaataaaaaagtctttactacaagtacatagatcttatatt
g----------------tccagatt----------------------aacatttcaaaga
tattaaattaccctctacgatatactctcaattcttttcactttat-tggcagtatttca
atattcagcttgtaa---------------------------------------------
--------------------aataagt---------------------------------
----------------------------------------------------aatttttt
g-----------------------------------------------------------
------------------------------------------------------------
------------------------------------------------------------
------------------------------------------------------------
------------------------------------------------------------
------------------------------------------------------------
-------------------------------caagaatatatagaa--------------
----------------------------------------------tcttgctacatatt

------------------------------------------------------------
------------------tacattattgttaataatt------------------------
------------------------------------------------------------
------------------------------------------------------------
---agcctttgctaaacta-----------------------------------------
------------------------------------------------------------
------taaatctatgaat-----------------------------------------
------------------------------------------------------------
------------------------------------------------------------
------------------------------------------------------------
-------------------------------------tttagtagatt------------
-----agactaaataattatgtcttttaaattaggtaaaagtaaagaaagtatgtgagca
aaatatcttaa-----------------ttttt---------------------------
------------------------------------------------------------
------------------------------------------------------------
-------------------------------------ttagaaatattttattttcaaca
tttt-tttataatagagcagttaaaaactgaataaatagctaatatc-----ttatacaa
ccaacatcgagccaatgccttgatcagttaatatttctagcaagagagcgtgatcaattc
taccatccaaaatatgagctgaagctactccttgggctaaagagcgaatgcagcaattaa
ctttggggatcatacctccggagatcacctcggtctgagttaagtctctagcttgctgta
tatttaagtgactaatgagtgttgtaggatctgaagagttacgtaaaataccaggagtat
ctgttaacaaaattagtttttccgcattaagggctgctgctacttctcccgctacagtat
cagcattaatattgtatgactgaccttgtttatctgccgcaacactagcaattactggta
tgtagttattatgaattaagatctttaatagcttagtatcaacattctgtacttctccaa
caaagccaagattcgctttaccgctcggtctaggagtaataagtaaaccatctttccctg
acaaacctacacttttacctccttgcttattaatcgttgctacaagatccttattgactc
gtccaactaataccatttccacaatatccatagtaggctgatctgtcaccctaacaccgt
tctcaaattgtgataaaatttttagacgatctaaccacaaattaatttcaggacctccac
catgaactaaaataggacgcaaccctataaaagaaagaaaaacgacatcactgattactt
gatcttttaatttttggtttttcatagctgctcccccatattttattacaataattcttg
aggaaaattgttgaatatatggtagggcctcacttaagactttcactctttctgaattag
tcagcatgaataaaatagaattataatatataaatttatttgcaggacttatcttaattt
aatagatactagtaagtactttaaaacttttagaaaataaacatattattaatag-----
----ttaaaagtataaggatgataattatgagtaatttttggaataatatattaaaattt
ccacgatttttagttagtgtcattttcggattagttttaataattattagtccttttttc
gtgttatttaaaaaaccattgacaagtttttttttcatcatatcattaaccggtttaatc

gcaattttagcggcaataa-tacagaaaatgctcaatatcgaatgttgttgagagaatta
atttaatttaattaaagtttttgaacatatataatattatattatcattaactgtgataa
tttaaatttcttttctat--aaaaagtcatacgtattaaatattatttgatagatattat
gttattaaaaccaatagttagctctgaaaaaactggtgcttttgccctttagatagtat
tgttagacatggtgttaagcatatatttggttatccaggcggagctattcttcctatcta
tgatgaactttatgcttgggaagagtcttctttaattaaacatattcttgttcgacatga
gcagggagctgctcacgctgcagattcttattccagatcaacaggagaagttggagtttg
ttttgctacatctggtccaggagcaactaatcttgtttcaggtatagctaccgcgcatat
tgattctgtacctatactagctgttactggtcaagttggaagagcgtttataggtacaga
tgcgttccaggaagtagatatctttggaattactctccctattgtcaagcattcttatgt
agttcgagatcccagagatatgtctaggattgttgcagaatcattctttatttgtaaaca
tggtcgaccaggccctgttttgattgatgtacctaaagatgtaggattagagaagtttaa
ttatttttctattgatccaggcaaagtcaatatcccaggttgtagaccagttactacctt
gaagtctagacaaatacttatggcagctaaaatgatacaacaatctagtcaacctttgtt
gtatataggaggaggagctataatctctgattcacatcaaattattaaagaacttgttga
cttgtataaaatacctgtcactactactttaatggggaaagggattttaatgaagacag
tgaatattgtttgggaatgttgggtatgcatggtaccgcatacgctaattttgcagttag
tgaatgtgatctattaattgcattaggagctagatttgatgatagggtgacagggaagtt
agacgagtttgcttgtaacgcgcaagtaattcatgtagatattgatcctgctgaagtcgg
aaagaatagaattcctcaggttgctatcgttggtgatgttacagaagttgttactgcaat
attaaatctactaaaaacttctttccctccttatccggagcagattgtatcttggcaaga
gcgaattaatcgttggcgcgaacagtatccattactggttcccaaaaaatcaacaagtat
ttcgcctcaagaaatacttgttgcagctaataaactagcccaaaatgcttattttactac
agatgttggtcagcatcaaatgtggtctgctcaattttaaaagttaaagctaaacactg
gctttcaagtgccggtttaggtacaatgggatatggattacctgcagcaattggtgctca
agtagcacatcctaatgatgtagttatttgtattagtggtgattctagctttcaaatgaa
tatgcaagaactaggaactatcgcacaatatcagttacctgtcaaaattattattattaa
taatcgttggcaaggaatggttagacaatggcaacaagcttttatggcgaaagatactc
tcattcccgaatgatagaaggagcaccaaatttccaaaagcttgcagaagcctttggtat
taaggcttttactattaatagtaggcaaagtatgcaatcgtccttagaagctgctatcga
gcatcatggtccagttttgctagactgtcaagttacagaaaatgaaaattgctatccaat
ggttgctccgggtaaaagtaactcacaaatgataggcattgccaaacctcaaagaggtac
tgcctcaaattatattaatgcaaatatctagattag------------------------
------------------------------------------------------------
------------------------------------------------------------
------------------------------------------------------------

---------------aaggtctatattccctcaaaaacaaagtg-gattttc----tttt
atgagtacatatttctaatagtatatacaatttttttaaaagatagcctttgacgagaatt
gaactcgtgaccttccccttaccaaggggatgctctacctctgagccacaaaggc-tttt
tattatattgggccgggttggatttgaaccaacgtaggcaaagccagcggatttacagtc
cgcccccattaaccactcgggcaccgaccctaattagtttttattcaaaacataacttaa
acagtatcataatttagttgcttaaaacaagtcttttcataaataaaact-attaagtga
tggacatagctggactcgaaccagcgactttcacgatgtcgacgtgacactctaaccagc
tgagttacatgtccaaaaccttaataattattataacactatttattgattttataagta
tctatttttaattaaagatttgctttaaacgacctgctttaccaagaagatctcttagat
aatatagtttagctctccgtactttagctcttctaataactttaataaatgttattcgtg
gagaattgagaaaaaaaacgcgctcaactccaataccttgaaatgagcatcttaaagtta
agctagtatttaagctatttttttcttcttctagccaagattactccttcgcataattgtt
ctctcgttttattaccctccttaaccatcaatccaagttgtattgtatcgccaactctaa
tctcaggtaagtcttgtttaataaaagaagtctccacacttttcatgagcttacttattt
ttgtactatttgctttcatatctagt-ttagccttataatg-ttttttgcaatattatac
attaaaacaataaatatgaaaattattagtagataagcagagtagtatctcagaatcttt
gtattatttattgctagactagatttacttcattttgtgttatcaatgtatttgcttaac
tatcaaaaaattaattcgctaattatagaataatgaccaacttatatttcta----atag
aaatacaaaattcttttatattaaat--taacctttagatatttaattcatt--------
------------------------------------------------------------
------------------------------------------------------------
------------------------------------------------------------
------------------------------------------------------------
------------------------------------------------------------
------------------------------------------------------------
------------------------------------------------------------
------------------------------------------------------------
------------------------------------------------------------
------------------------------------------------------------
------------------------------------------------------------
------------------------------------------------------------
------------------------------------------------------------
------------------------------------------------------------
------------------------------------------------------------
------------------------------------------------------------
------------------------------------------------------------

```
------------------------------------------------------------
------------------------------------------------------------
------------------------------------------------------------
------------------------------------------------------------
------------------------------------------------------------
------------------------------------------------------------
------------------------------------------------------------
------------------------------------------------------------
------------------------------------------------------------
------------------------------------------------------------
------------------------------------------------------------
------------------------------------------------------------
------------------------------------------------------------
------------------------------------------------------------
------------------------------------------------------------
------------------------------------------------------------
------------------------------------------------------------
------------------------------------------------------------
------------------------------------------------------------
------------------------------------------------------------
------------------------------------------------------------
---------aattat---------------------------------------------
------------------------------------------------------------
------------------------------------------------------------
------------------------------------------------------------
------------------------------------------------------------
------------------------------------------------------------
------------------------------------------------------------
------------------------------------------------------------
------------------------------------------------------------
------------------------------------------------------------
------------------------------------------------------------
------------------------------------------------------------
------------------------------------------------------------
------------------------------------------------------------
------------------------------------------------------------
```

------------------------------------------------------------
------------------------------------------------------------
------------------------------------------------------------
------------------------------------------------------------
------------------------------------------------------------
------------------------------------------------------------
------tacccgtaatttgttatgatctaagtatctcacccaattatttttttattgatg
tta------------ctatatttaaatacttatttttttgctaaagaattttacacataa
aacagagtaatcaaaacaacaagattaaggcttaagtatttattagttttgtgttataat
catttactatcaaaggtagtggcataacataaaacataaacaatgtttgaacgctttact
gaaaaagctataaaagtaatcatgttagctcaggaagaagctcgacgcttagggcacaac
tttgtcggaactgagcaaatattattaggattagtaggtgaaggtaccggggttgcagcc
caagtattaaaatctatgaacgtgaatctaaaagatgcaagagtcgaagttgaaaaaatt
ataggaagaggatctggttttgtagcggttgaaattcctttcactcctcgagcaaaaaga
gtactagaattgtctttagaggaagcacgtcaacttggccataattatattggtactgaa
catttgctaatgggcttagtcagagaaggggagggagttgcagcgagagtcttagaaaat
ttagcagttgatgtttcttcaattagagcagaagttatacaaatgctgggagaaaatgca
gaagctaatgtaagtggaagtaatactacacaggctagaagcaaaacacctacattagaa
gagttcggctctaatttaactcaaatggctatggaaggtggtttagatcctgtagttgga
agacaaaaagagatagaacgtgttattcaaattctagggagaaggactaaaaacaatcct
gttctaattggggaaccaggtgtaggtaaaacagcaattgcagaaggcttagctcaaaga
atagctaatagagatgttccttctatattagaagataaattagttattactctagatgtt
ggcttattagttgctggcactaaatatagaggtgaatttgaggaaagactaaaacgtatt
atggacgaaatcaaatcggctgataatgtaatactagttattgatgaagtccatacatta
atcggggctggggctgcagagggcgcaatagatgcagccaatttactaaaacctgctttg
gcacgaggagaattgcaatgtataggtgcaacaactttagaagaatatagaaaacatata
gaaaaagacccagcactagagagaagatttcagcctgttgtagttggagagcctagtgtt
gaagaaacaattgaaatcttatttggtcttcgagaccgttatgaaaaacatcatcaatta
acaatgtctgatggcgctttggcggcagctgcgaaatatgctaatcaatatatttctgat
cgattcttgccagataaagcaattgatttaattgatgaagctggttctagagttcgttta
ctaaattctcaactacctccagctgcaagagaattggataaagagttaagagctgtatta
aaaacaaaagatgaagctattagagctcaaaaatatgaaacggcagaacaatatagagca
agagaaatggaaattaaagctcaaattgcagcaattgctcaaagcaaaaaaaatgagccc
gatttaaacttagaagatcctgttgtaacagaagatgatattgctgaaattgttgctgct
tggactggtataccagtaactaagctgactaaaagtgagtcagaaaaattaatgcacatg
gaagaaactttacatgggcgtatcattggccaagacgaggcagttgtagctgtatctaga

gctataagacgagcaagagtaggtctgaaaaatccgaatagaccaattgcaagtttcatt
ttttctggaccaacaggtgtaggtaaaacagaattgacaaaagctttggcttcttatttt
tttgggtctgaagcttctatgatacgtttagatatgtcagaatatatggaaagacatact
gtatctaaattaattggttctcctcccggctatgtaggttatagtgagggtggttatcta
acagaagcagtaagaaaaaaaccatatactgtaattctttttgatgaaattgaaaaggct
catcctgatattttaatcttcttcttcaaattttagaagatggcagattaacagatgcc
aaaggtcgaactattgattttaagaatactcttctaataatgacttccaatattggatct
aaagtcattgaaaaaggaggaggtagtttaggattcgagttgtcagaagaccaaacagaa
tctcaatacactagagtcagatctttagtaaatgaagaattaaaacaatactttagacca
gagtttctaaaccgattagatgaaattatcgtatttcgtcagcttaccaaagatgaagtt
agagaaattgcagaactgatgttgaatgaagtttttgccagaattaagcaacaggatatt
caactaaatgtaacagaacgttttaaacaacgactagtagaagaaggctataatccaagt
tacggagctcgaccacttagacgagcagtgatgagattattagaagatagtttagcggaa
gaagttttatcaggtaaaattaaagctggcgatagtgctgtagtagatgtcactagcgaa
ggagaagtaacagttttactaggtgaaaaattagaactgctaacgtaaa------aaata
atttattaatctaaaagcattgaatataattt--attcaatgctttt--gaagaaaatta
acaagatgggttgctataattgacaacctctgatattctttatatgtatata---aagcc
gggatagctcagttggtagagcagtggattgaaaatcctcgtgtcaccagttcaaatctg
gttcttggcat-tttaaatatttgaaaagtttgctagatgagcatcaaattttagttgaa
acatacctgttggtccatttctatgttttgctattataatttctgtcaagtctcctccta
cagcttcttgagtataataactctctctatatagcattataactaaatctgcatcttgct
ctatagaattgtgtacgataaagttgttcgcgataaaattatgtaagagttgacattcta
aatcaaatacttgttgcaagctaataaggcgaacatgtttaagattctcaaaaattaaaa
ataaagatggacaagaatttaagagagtttttctttctttaatggcgtt------tattt
gaatagcaagcatattattgtgattaatttcatcgcatcttttccatccttgtgttgtta
atagcttgtgattgcttgttaactttatatacttgccagattgtgtaataatttcataca
cagtttttttacctgtcttggaaatattagctttactgtatctaaataagtttgtttttt
gtgcagtaaagctattaacttca-atatgcttatgatgaaatttaaatagaatttg--ag
tctgactatgtaacggtaaatgaagatgactaaatttcgagatacatccactttctctta
gatcagataataaaggtcttttatcgctccgagtttcaagattcctattcagttgagata
atactagtataggtaaatttaagtcttttgctaatatttttaatgatctagtaattagag
atagttcttgagatctattatcagattgtctgctatcttgtaataattgaagatagtcta
taataattagttctatatttttcccttgtaattttaagaattgtactttcgtttttatag
tgtccgtagaaatttttgcgctatcatcaatgtaaagattcagatctgcaagagctttgc
tttttcgaataacatactgccattcattattattaagttggccagattgaatttttttgac
tatttaagtgacattcttgagctaaaatcctccttaggagttgttctatagacatttcta

agctaaataaaataacgaacgatttctctttgctcattatatatctagctacattaatag
caaatgctgttttccccattgaaggtcttcctgcaagaacaatcaaatctgacttttgaa
aaccttgtgtaatagaatctaattctgtaaaaccggaaagtacgctactatttatactaa
ttttttttgttttgtctaaatgaacaagcagattagcaagaatggtagctagtgtttgtg
tatctttctcttcaaggatttcgtaagcttttgttaattggctaacaatagaggttatat
tattctgtctaacaggctggtttgaacagctaattaaaaataatgaatctccagatatta
aaagtaatcttttaatataattatctataataactgacgaatattcatctaaaacatcag
ataaaggtgcttgttctattagattaataacttcgtctagttcattcaagtgtctaatta
tttttctgtttttaagttggctacacaatttcctatagttactctttttatatgattaa
cattttctaggacagctctgtaaagtaatgaattcgatgtaaaataaaaaaaatcaggag
agagtttttctactgattttagtagagagactgattgtttagttaatattgtactaatta
atattttttcagctaaaatattatgaggaggtaaatatttatga------atactgatta
gttcctcc-ttgtaacattattctaata--------agactaattaattactagcttaat
ttgactctggcagaacttgtagttggatgttggcagttacctggtttaatagcttaacct
ctatattataaatgccaactgttttaatctcgggtaaaaggatattttgtttttcaatat
ctatatttgtagcattttttaatagtttgtgaaatttctttttctgtgacgcttccaaaaa
tattttgcccatcccctgtttttttagtcacgctgaacttctgtatttcctctagaagtt
gctgaattttttttgcatcttctttagctattttaaatttttcttcttttatggctgcgt
ataattttagcttttttaaaataccgtttgttgcgactattgccattttatttgggatta
aaaagttccttgcataaccagcagcaacttttacaagatcattacttttgccaagttttt
gaatattctcgtttagcacaacttttataactttttttacccatatttattctgaattcta
aaaa----atatatataattttattatacttgttgtacccagatttt----aataggaag
tatttacttgttgacatagcatctccgtgaaactgataatgaattttcatctttattctg
gtataataatttatacaacgtttt-----attaggagaggtggccgagtggttgaaggcg
cagcattggaaatgctgtttaggagtaatcttaacgagggttcgaatccctctctctccg
tataa--aaattatgataatctttatatagtccagtcaaaacttccaataatgtttctcc
atgcatttaaaccacctttgactctcacaatattgagcttttgagctattaataacttag
aagcaaatagagatctagaatctaagctacagtatacaaagcaaattttatctttacaat
ttatattagagtaatgtattgttttgagttgttttaaaggtatatttaatgagtgaatta
agtgatttttttttatattcttcttgatttctgacatcgattaaaatatattttaggtctt
tgcgatttaaaagtttttgtaattgaacaacattgatttctcttataataagattaggtt
gatcatagctgtatctgttccagtattttttaga-------ggataatacaaatttagta
tttaaaattttaaacttactaaatgataaagttaaagcgttataagttaatatagtaccg
cttaatatagacttgtaaccaagaataattttaatagcctcagttgcttgaagtgtacca
ataatacctggtaataaacctaaaacaccagaattgctgcatgtatctatttcactattc
ctatttacgactttattgtggaagtctcgataacttggacctccttgatagttaaataca

ctaacttgtccttcaaattgaaaaatagccccataaatatgtattttatttaattctagg
caagtatcatttagcagatatcgagtacgaaaattgtctgatccatctataataatatcg
tagtttttaattatatcaaatgagttgttataggttagtcttgcttcaaaagcttttact
gtacattttggattcaaatctaatatttttttttagctatttctacttttgagtatcct
atatcgtgaactgtatatataatttgcctttgtaaattagaaagatcaataatatcgtcg
tccataataccgatgcttccaactccagcagctgcaaggtaaattattgcaggagagcca
agcccaccagcaccaataaatagcaccttagctttttttaatctttcttgcccctctaat
tgaatctgtggtaacactaaatgtttagaatatcttgtatattcttctaaagagtatttt
atattatctgctttaaagttcagcat-aaaatttatatgttaattaataacaacta----
--gaaactaattgataagcttattgagatgtttttgtaaactaggttcttctttatcta
aaatagctccttgcgtaggacagacctgaatacaaatactgcagtcaatacaggcagaaa
agtcaatccagtaccaattggtattaatactattttgtccttgtccttggtgaatacaag
ctactggacaagcagatacacattcggcaactccaatgcatttttcggttacaattgtat
gagacatgttatttaattaat-aaaagataaactatttacatctagtagttttattgtat
actaaattgacatacgcta-ccaagccagtaattaaaataatctacttttatagtttact
gatttgatctgcgcctttagttcagctggtagaacgcaggtctccaaaacctgatgtcga
gggttcaagtccttcagggcgcgcttagtttttgttatatttgcatagtagcattttacc
tttttgcaaacatcaaaatgatatgatataaaacatcatattctatatatttgatgtttt
atgtgccagtactg-ttttttttatatgttattattggtaagttaattatattaattctat
ttttataatcaataaaaaatagctgaaattaagtgcagattcacttagtga-----tttt
atatatataactaaaaaaattaaaaactatggctaaaaaagttaccggaattgttaagtt
ggcattaaatgcaggtaaagctactcctgcaccacctgtcgggcctgctttaggtcagca
tggtgtaaatattgtcatgttctgtaaagagtataatgcgcgtactgcagataaatcagg
actagtaattcctgtagaaatttcaatatatgaagatagaagtttcacttttatacttaa
aactcctcctgcttctgtactaattgccaaagctgctggcttaaataaaggttcaggcga
accaaacacaaaaaaaaataggtagtataacgtataagcagttagaatctattgcagaaac
taagctaccggatttaaatactaacaatgtacctcaagctatgaaaatagttgaaggtac
agctaaaaatatgggaatcctaattaaagattaaaaaatttcaattataattatta-tt-
-tttatgaaaaaaatttcacgtcgacttaaaatattaaaatctcaagttgagcctaaact
ttataatattgatgaagcaatatcgatgttaaaggcaacatcaaatgctaaatttaagga
aactgcagaagctcatatttgtttagggctaaatcctaagtatgcagatcagcagcttag
agcaacagtcatattaccaaaaggaacaggtaaatcaataaaagtagcagttattgccaa
aggagaaaaattagcagaagcaattagtgcaggagctgatgtaagtggttcagaagaact
catcgatgaaatattgaaaggtagattagattttgacaaattaatagcaacgccagatgt
tatgcctctaatagcgaagcttggccgtgttttaggacctagaggattaatgccttctcc
caaagccggcaccgttaccatagaagtaggcaaggctgtgaatgaatttaaaggcgggaa

agtcgagtatcgagttgatagaactggcataattcatgtaccttttggaaagtccagttt
tcctcaagaagatctagtcttaaatctgcaaacaattaaagaatcaattgatagaaataa
gccttctggtgcaaaagggaagtactggaaaactttttttcttatctagtaccatgggacc
atccatccaaatcgatattaatagtcttctatgaaaaatatttatttaattacataactt
gaataatatattatttcaaatta-taattggctagcaatgttaatagaataatttgctta
ttataccttatctgcaccaaaaatttaatatatctcttactaattatgagtacaaaagt
tgaaaatattttagaagaactaaaatctttaaatcttctagaagctgctgaactagtaaa
acagatagaagaaacattcgatgttgatgcgtctgctgctacaggaggcatgatgatgtc
tgcaccagctgtatcaccagccgcatctgaagttgaggagaaaacagaatttgatgttgt
tttagaagaagtacctgcacctaaaaagatttctgttctaaaagttgttcgctctttaac
cggcttaggcttgaaagaagcaaaagatcttgttgaatccacgcctaaagttctaaaaga
aggtgcatctaaagatgattcagaagctatgaaaaaacagttagaagatgctggcgcaac
agttgttattaagtaataaa-t--agtagtgcttttgtttagaagtaaacaaaagcac-t
ttttatttaatagaattaattagactagaaaagtccaagtgtaattgcattattaatagg
catagtcgcaccaatacctaaccaaacacttacaactgtaccaattaagaaaacagttgt
tgcgattggtcgtctaaaaggattttgaaacttattgacattttcaatgaatggaactgt
aagtaaccctgctggtacagctgccatacttagaacacccaatagcttattaggaattac
tcttaacagattaaatgttggaaagaaataccattctggtaaaatttctaatggagtagc
aaatggattagacttttcacctatagatgaaggctctaaaattgctaacccaatactgca
tgcaaaagttccaataattacaactggaaagacatataatagatcatttggccaagctgg
ttctccataataattatggcccattccttttgctaatttagctcttagttttggatctgt
taaatcaggtttttttaagaattgacatgttatttttttgtgtttaatataaaattgagtct
aattactttatcaaagtgtagttatacattgttataatggtccggaaattccttgtttgc
gtatcattaaaaaatgcataagcatgaaaatagcagttagtaatggaagtacaaaagtat
gtaaactgtagaatcttgttaaagtaccttgtcctacactaactcctcctcttaataatt
ctacaatacttcctccaataactggaacagcatctggaacgcctgtaacaatttttacag
cccaatatccaatttgatcccatggtaaagaataacctgttaccccaaatgaaacagtaa
gtactcctaaaattacacctgtcacccatgtcaattctctaggtttttttaaaaccacccg
ttaaatagacacgaaatacatgtaaaatcatcatcaagaccatcatactagctgaccatc
tatgaattgatctaataagccacccgaagttcacatcagtcataatatattctaccgatg
tgaaagcttctgcaactgtaggtctatagtagaatgtcattgcaaatccagttgcaactt
gaattagaaaagatacaaatacaatcccgcctaagcaataaaaaatattaacatgaggtg
gtacatatttgctagaaatatcatcagcaattgcttgaatttctaatctttcttcaaacc
agtcataaattttactcataaaacagcttcaaaaagc-tcttcaatata-tttttgcggt
taagctacttttgattcaatag------ttattgaacttatatt-atttattatctaca
cttattaattattataacatctttctatctattt--------------------aaaatt

ctctgactcaataaaacaggactatgaacgacgatcttttttttctgtatagaaatcaat
ttagtctttagcaattcagataaaatccttgttacacttactctactactccctataatt
tgcgctaacattttatgcgtcatagcaaaattaagcataataccattattactctgagtc
ccattttgctctgctagtaataataataagcttacaagtctacttgtaatacttttgtgt
gacatgatttctacaaaatgataagtttttctgagcaacatattaaatgatctataaaa
aataaattaaatagtgggtagttactacacgctcttataatagtgttatagttaatacaa
gcaatttttgttttacctattgcttctgcttcatagtagaagttatcattagctaactct
aattgtccgaatgtatctccattagttaataaattgagtgttattttatgtgtgctttta
aagacttttgtaataattaaagatcctattaatataatgtatagtctagagctgctatca
aatagaataatatcattaggctgtagtagaaaaatttgataactaatttattactatta
aaaaatagcaaccatggattataagagggaaaattatttgaactactttcagaaaatgtt
gagatagatttttttgatttaaatgattttagtcataattcatactttat-a------
---aaaaatactctaataat-taagaactgaaattaaatgtcatacaagttgatgctagt
tgagaatgacattgtattatcaaaagccattcaagaatacttaatagatcaaggatttaa
tgtttacattgccagtaatggattagaagcatttaatttagctaatcaatataatttcaa
tttgataatttctgatataatgatgccattagttaatggctatgaattattagaaaaact
tcaaaatactaaggcgctatcaaagataccagttattttttgacagccaaaggaatgac
taaagatagaataaaaggttatgacatgggatgttacggttacttatctaaaccatttga
cccagaagagttagtttcacttattaaaaacttaattgctagagacatatcgcaacatac
atctacagaaaaatgtgacatctctaagcagtcaattaaccaaaaaatacatttgactcc
tagagaagcaagtgttttagaccttgtgattgacggactaactaacaaagaaatatctac
aatattagatactagtattagaaatgtagaaaaatatgttagtcgacttttacaaaaaac
taatatgaagaatagaactttactagttaaatattctataaataataaattactagataa
tgtaactaatcaaagggcgaatgacgggactcgaacccgcgaatgatggagccacaaccc
attgccttaaccccttggctacactcgccatattataaatatcactataacttgtttgtt
gatatatcgtctactctgtaagtcatttatttatctaacatattatctaaatttaatctt
ccatgacgtataataatattagtattcaaattaatggagagccgtttaattgctcaaagc
ctatttccttgcaattttattaaattatcttgattttaattctgagcatatagcagttg
agctaaacgacactttgttacctgagcctctttttcactcaacttacttaaatgatcaag
ataaactagaggttattactattgtaggtggtggataaatttcattaactaa--------
---------ctattttaaatgcttcattgataaggataaacgtccctgcttcttgtccac
atgaataatgacagctttaatagtatcgcctattttaaattgtgacgatatctgatctaa
ttgttttatgttaatttcagagatatgtacaagacctttaagattgccaacttttataaa
taatccgtaaggtgtaatctgatttataattccttcaataatattaccaacaattaaatt
tgatgaggcttgagcgattaaagctcttctgtggcttagtattagattattagatctttc
ttcaacattgagtagctttaacttgataaattcattattagccttatcactttttttggaa

attcccgagatgagaatttggtacaaatccagatataccttcaaggctaatgatcatccc
tcctttattaaatcctttattcgaacattaagtaaagaatcttcagctaataattgtcg
aattcttttccatgctcttatatattcaaggcgccgaattgataaaattaattgttttga
tcgaatattataatctagtaaaaagaattctcttgtatcattaatgtttaaagaagtgaa
attatttaaatcttgattacttgacacctcttgaataggtaagtatgcagatattggtgt
tcctatatcaaccaaaacaccatttaactcaaagctaaatattgtaccagcgacaatatc
gccaagatttaaatcgtatttgtatttttgcaaaacagctgcaaaatttctgtgagtaaa
tccttcattattttttgtcatattgaatagtctgaattaaagattgattttgtacaaaat
attaagttgaataaacttctgtcgtattatatgaaaatttttaaattcgttctgtggtta
ttcctgatctaaacgtattagaatatacggaagacataaaaagtaaacgtagatagctac
agatttaaccaatatacagtctgaggaaagtccgggctccataaatacaatttatgctgg
agagatcccagtgtaggcaattatgaggatagtaccacagaaataaaccgccagaataaa
a----tatactagctggtaagggtgcaaaggcaagttaaaagcttaccaaaaatactgta
aagtatttgttaggtaaacccctaaaatggagcaaagctgctaaacaaatatttgtgtat
attttatatagtttagtttaaaatactgcatgaagttattggtaacaataactctagagg
aatagctatcctttacaatatata-tctattatattgaaagaacagaacccggcttatgt
agtacttttatgtcttacattattctggctattgccgaatctctaagtttaagtgttgt
atgatttttcttgtagctcatgttgtttatcatctatctctaaatttgttaaagttttt
tgactagagcgatatgtaaatctcaaacctatactttttttctgttctccaagggattta
tttgtatattgatcaaataaccttatgctttctaaatctttatcattaaactgatctagt
aattttaataagtaattaaattccatctttttaggtattattaatgaaagatctcttgtg
atagatggatatttagagtaaggtcgaatttgataatttaaataattaaattgattatga
gaatcttctaaaatgtttaaattaatttcaagaacaaaaagttttgcagtaagacctaat
ttactataggttacttgattgagttcactgaatatgccaatattagtattattataaatt
aatgttgcagatctatccttttgaatgaaattcactctactgtcaattatttctggtttt
gtccattcaattttctatttatttttgaaaaaaatttctacaattcctttagcttca
taccaatttaaagagtggactggttgtgaccatgtagatctaatatttaaattaccacct
aatatcatggctagattagttgtttcgacaattttgttgtgaacaggattaaaaactgtt
cctatttcaaaactgtctacagtttgattactttgcttaagattataaagattagattca
attaatccctcgagcaaactatagcgtaaattagagtaatcttggataagaggattattt
agactaacacctcctagagatttcactaaagaataatgaactaattcagttaaccctaaa
tttcttaaaatatttctacttttatctatgaattttctcttttgtgataatttttacta
aattgaatacttggcactgagcttttaaatttatgatagccataaattcttgcaatctct
tcaataatatctatttctcgaaaaacatcttgtcttcgataatttggaacactaatttt
atactttcttgatttttatgaataatagcaaaatttagagaatgaagaatattttgaatt
tcttgaaagtacaaaaaacgtacctgattattgttactgattggtcctaaaatatcttga

acttttttaattgataaatctatatttagagtatcattcattgttttctctgttgagaat
gtctgcctaatattcccgccagctaagtctattattaaagagatagtttcaagataagca
ttttcccagttgtctgtatttagtcctcgttcctgcctaactgaactttctgtcctaatg
ttaagatctcgggaagactgtctaactatagattgtttaaatatagcagattctacaaat
atagatgttgtattagaatccgtatcaaagttgttattggatcctatgccagccacacta
actacatttgtattcacttgagtcactaaaatcttcttatttaactcaatattttcatta
tttaacttaattttagtactttttccaacattaaagttactttgaattgttatagtatct
tcagg---------agtaattttattaaaatcaataatattaataggttgtccccatttt
agcataatataattgcttatatctactaacaaatttctatgtgtaaagccagaggacttt
aaacgatttttagccaatttggagaatctataattttaatttcatcgataattgcagaa
aaatagttattacaatgaatcaagctatgattattgagtatatt-ttgttgtctagaaga
gtatttattactaaattgatttatagtttttaacattaaagatccagttaaagcggataa
ctctcttgaaagacctatcatgcttaatacatccgatctattagctgttgatgatatatc
taatacatgatcaatctgattgcctactgtaatagattcaatagcttctacttcaaaacc
tgcctgtgttaattgactagttaaatgaactagatctattgtttctatattagcaagttc
ttttaaccaatttaaagaaactttcataatgaattgccatttatcttatataattattta
ttttacttgtgattaaaaactaatttaaaaaattcatcaaattaatataagttatttgat
gaatttgcttatacgatatattcttgtgctatattactttcaggtcgtagtaattaagaa
ttagcttttgtaaaagtcagtccattatcttgagcatacctttccataaatctcataaat
ctatcccaatcatcaggacttttaatcacatgtacagcttctattgcttgtggtttacca
ttaataaacttagcattgacatctcttgtaataatttgtccctctgtgtccattagatac
atgcctgttatttcacctttatctgccatactagcatctaaaattttgggatttgtaaat
ctgaaagtagctgttccggtactaccatctctagatcttgttaagcgaacatctggcacg
acttcttcattgatgccttgaataaattgaattgttgccatatcaatttt-tttatttgt
atataatgtatattggataaaactgatttctaattctttaatgcttgttgaaaaactgtt
ttaaaagcttatagtagcagcagatcaaattttgtgtactatataaaactgtaacttttc
tttttttg----------------------------------------------------
------------------------------------------------------------
------------------------------------------------------------
------------------------------------------------------------
------------------------------------------------------------
----------------------atctgttcaaagaactaacttattataagctaatcata
ctaatattagccatataatgttctgaagt-gaaattaa-gggttgtagctcagatggata
gagcaagcgcctcctaagcgctaggccagcggttcaagtccgctcaaccctgttaataaa
caaaatt---ttgctgaattctacaaataaattagttgaaaaaagaggaaaaaaacaaaa
cagc-----aaatagattctgagtaacactttttattgaa-----aaaaatgatcttaac

tttgataatagaagaattttgtgaaaaaatatttaaaatttttgatctctatattcagaaa
aagtttttttaaaatataaactgaaa-----taaaagtcttgagtaacaataggaattaa
attatttaatttttttgaaattgatctaaaaaactttatatgaagagaatagttttttttct
ctttctctattcagtatttgcctccataattattacttcattcattcacaattaataaaa
tggattttg---------------------------------------------------
------------------------------------------------------------
-----------------------------ctttaatgaatgtttatgttatttgattata
ttccatactatataagagtttttctcaacttgtgagagtcaatttgtagttttaatttta
taatatactttatctttggagattgatctacattcaccttcacacatctcttttattgcc
tattaggagaaactatatactgaaatttaacaataattgtgaaaatcgtaggttccttca
acatattattatcctttcatttttgtctccactgctgatcaaagcaacagtgattagact
tattcatttagctggttctagtgagagaagttttatacaagccg--gacacttatttaaa
t---gtgttattaccaaagctgttaaaataatagccattaagattcaatgacttttttagt
taatagtttataatcgccatgtttgatttagtactgatatatagttacaagtagttctac
tattgataaaattgacacttttgtgactgtataagctttcgatagaaaaactatgatatt
attcataatgaaaaagcgagtattgaagtttgttgctttcataacaaaaaatatgtttag
tcaatgatgaattttgctagcgatccgagtagtaaatctgtcacatgtaaagatgaaaat
tatatatttgagaatactctctatatctgtatatagcggttggcataacttatcagattt
gatttgcagaatcagtatactaagagatcgcaaaatggttattctaagaaaattttgtct
cagtaatttcacttctaatagatacttgttagtaataaacttttaattcttttatgaaaa
aacttcataatatcaatatttattagaactggggtgggaggattcgaacctgcgaatggc
ggagtcaaagtccgctgccttaccacttggctacaccccaacttatagcaaatataatat
cagtcacataccccgccgtgtcaatagattattcgcattatttaataacttttttagattt
ttgtaaaatagaatcttttttttcaaataagatgtactatgttcaaattgcattatgctc
atgtcttcttgttgttgattggacatccattttatggtcactgttccttgttcaacttct
tgattgcctataataatacaagctacagctcttttcttattagcttgtttgatttgttta
ccaaagttacttgagctaacatctatttctattgtaaaaagtttttgatgtaaaaagcgc
attattttcatgccaacttcattagctttttttaccttgtgtagcaatataaaaatcaaca
tatttacttggtaactctatattctctttcgcaagaagtagtaaacgttctaatcctatt
gcacatccgacagctggtgttgctacgcctcctaattgcttcactaagctatcataccttt
cctccaccacaaactgtatcttgaccttttgatgttaatgtcttaatttcaaaagcagta
tcattataatagtctaatcctctaactaacttgtcatttatgttataaggaatatcaagt
aattttaaataattacaaacagcatcaaaatgtctttgtgacgaaatacttaaaaaatta
gaaatcttgggagcctcagctaatatttttttgtgtatttatattttttgaatctagaatc
ctaataggattactaattagtctcttttgtgaatcaggatctaagtcgtcatagtactgt
ttcagatattcttgcaattttgcttgataaatactacgatcttctgcttttccgattgaa

ttaagatcaagctttaagtctttgagattaagattattaaatatacttatagctaaatgt
attacttcactatctgctcttgcatctaggctgccaagaaattcaataccaagttggtga
aattgtctttgtcgtccactttgtggcctttcatacctgaacattggcccgctataccat
aatctttgtaaactatgatgatagttcattttattttcgataaaagctcttacaatgcct
gcagtgccttcaggtcttaatgtaatatctctattgcttcgatcatgaaatctgtacatt
tctttatttacaatatctgtatctcttccaatacctctatcatataaatcgctactttca
aaaataggtgttctaatttcttgatagtttgcacattctaataagttggaaattttgtta
tgtataaattgccagtattgaagctcttcgggcagaatatcttttgttcctctaatagct
tgaatttttgccatgataattttcggatcaccttcttaatgtgtttgcattaaatagcag
ttgtagactgttattcagtcatgtattatacattag--aacatgttgatg-tattttatg
aaatacgggcaaggagggattcgaaccccgacaccatggttcgtagccatgtgctctaa
tccactgagctacaagcccacttaacaagctatcttaatatatcattctttttcaatgta
tacaactatattttgtct------atttatatttgaaagttgtctgaaagt-aaaaaat
gt----atttatctttaacaattagagttagtcatgtataattttccatttattaaaact
ttgcaca--aatatcgacaaacttaattatgtaattatcttatactgataatttcagaag
aa-cttgtgcttttacttcttgatcga------ctagtatctaattcatctatattccaa
at-aaaaaatcttatctaaaatgagtaagcaaattctatatcaagatgatgcacgtaaag
cattagaaaaaggcatggacattttaacagaagctgtttctgttaccttgggacctaaag
gaagaaacgtcgttttagaaaaaaaatttggggctcctcaaattattaacgatggagtaa
caatcgcaaaagaaattagcctagaaaatcatattgaaaatactggagttgcacttattc
gacaagctgcatctaaaacaaatgatgtcgctggtgatggaacaacaacagctacagtac
tggcatcagcaattgttaaacaaggcatgagaaacgttgctgcagggtcaaatcctatgg
ctataaaaaaaggcatagagaaagcaacaaattttgtagtgagtaaaatagctgaatatg
ctaaaccagtagaagataaaaaagctattatacaagttgcttcgctatcatcgggcaatg
atatagaagtaggtaaaatgatagcgaatgctatagataaagtgggaagggaaggtgtaa
tttctctagaagaaggcaagtcaactaatacgattcttgaaattacagaaggcatgcagt
tcgaaaaaggctttatttctccttattttgttacagataccgagcgaatggaagtccttc
aagaaaatccatttattctctttacagacaagaaaatcactttagtgcagcaagaacttg
ttccactccttgagcaaatagcaaaaacatctcgtcctttattaataatagctgaagata
tagagaaagaagcattagctacgattgtagttaacaaattaagaggcattttaaatgttg
ttgcggttagggctccggggtttggtgatagaagaaaatctttgcttgaagatatgagta
ttttaactaatggacaagtaatcactgaggatgcaggtttatcgcttgatacagttcaat
tagatatgttagggaaagctcgcagagttattgtaactaaagactcaactacaattattg
cagatggtcatgaagttaaagttaaatcaagatgtgaacaaatcaagcggcaaatagaga
cgagcgattctctttatgaaagggaaaaattacaagaacgattagcaaagctttctggtg
gcgtagctgttataaaagttggtgcagctacagaaacagagatgaaagataaaaaattaa

ggctggaagatgccattaatgcaacaaaagctgctattgaagaaggaattgtcccaggag
ggggagctactaatgttcacatttctagcgagttatttacatgggctaaaaacaatttag
ttgaagatgaattaatcggagctctaatagttgaaagagcagttacttatcctttaagac
gtatagcttttaatgcgggagataacggagcagtagtagtagaaaaagttaaaagtaacg
attttcacataggctatgacgcagcaaatggcacaattgtaaatatgtacgatgcaggta
ttatagatcctgcaaaagttgctcgatcagctttacaaaatgcagcttctattgcagcga
tggttttaaccacagaatgtattgtagttgataaagccaatgactaagtta---------
-----tagataatattaaacaaagaataatttccatagctagaaattattctttgtttca
attaagctaattaattgatcaaatgtataattttataataaagccgaagtttaaatgta
cctctttgctcaactactaagtagagaagatacaagctcgtaattcctaatttatcaatt
aaagtattattatcatatatggattttcaagagatctaaaaaatgacattttcgtaaaa
taatatcgaaatcttcttttgtagttatttgataaatcattactgtatgtagattgatct
ccaattgcgtttaaaacttttaaaataatgtttttaagtctctatctgactttatctta
gaatcggtaatcaattgatttaattgaatatcaaaactaagattaagaatattacttgtt
acggatgttcgattttgaatatctatagatttgacagccaagaagattaaacttttattg
ttcattgtaaaatgtaagacaattgatattgtattctttaaatacta-atttatcttct
tgtgcgtttatagttttggagaaaaataattaatattatta--------tattaattact
gcccgaaaaaataaattctggaaatctagcttgagcctttgatagtgagtgtttttacc
ttcttcttgccagtaattaataacttctgttgtcttatttagtaattcaattctgtctct
ttcagcaatccatggtttggattctagctcggcttttaattcttcccaagcatcattcct
aggccagaaaaaataagaagtaagtggacttgtaccatttccaacaatttggtcaatagc
aatagcaatattattttccaaccaaagtacttttagtgtaaattttgacaagtgattatc
tccattgtaaatagcaaacatatgtatatacagtaaagtactacattagttattttttt
agcaacaagctgagctttatttaaaatattctgtaccgttctagtagtttgtgcaccttg
tgctaatctctgtaaaattttgaaatatcaatacgagtttcattagttataggattata
gaatcccaattcttctatagccttgccatctcgtttgtttttgctatccattacaacaat
tctgtagcttggctgtttttttcttccatatctctttaatcttaacttgaccatcttagt
cccctcgcttattttttaatatctttattaaattatatcttacgatattatgcaataat
tcgaacaattaagcagactcaagcctgacattattaaatattaccattaaacactggaga
aaaataatacctagcataggtgacatgtccatcccgaacattgggggaatactgcctcta
aagagctttaaatatggatcggtaattcgatttaatgaacagaatggttcattataccag
ttcacagttggaaaccatgccaatgataatttaagtaaaattaaaattaaataaatctca
gaaaaattagctattgatccaagtaataaatttaatgtaccaggaagagtattcatacgt
attttataattattatatagttcattatactaatagtttgagtatatattataagctctg
taaataaat-tagctaaacaaaaaacttattcttaaaaatatcttctcttgaacaatttt
cttcattgatagcactgttaactttcgttgtataaatttctaatttagaccaattatttc

ctagctgatttagttgagatgttctgcattctaaacatgctatttttatattatttaaaa
tagctcctttttttttctaaatcttttaagactggtgtaataattgcttcatctaaaaata
aatccaagatgagaatttttgtaaatgaatttaattgagctggaacattattaacaatgt
tgttgctattaactagagctatattagctgtcggaagtaaagctcttgctccctctgcta
gtataaatccataaggcataataatgacaataatatacttataatgactgttaataagat
ttacattgatattagtttttaattcaatagtttctgttactaaccattctctcattattt
cataagtaatccatttacctagttctgagcaagcagttcttaaaattgtacctggattat
tgctattttcgagaatcccagcccaatgttgtattaaagggtgggaagctatattaattt
tgagttgcatgttttgctaaatatatttaacatttaatttattaattgtcgctatagttt
atgttcactgagtattattaaaatagcat----atatattaaagaaagcaattat-aaaa
aatgaaaaaaactctttggctttggggttcacagatagtgctgaaacttggaatggtag
atttgcgatgattggttttatagcagttattattattgaagtagttacaggacaaggact
attatatctaataggtgttatatcataataaaaaatcaaaaaagttcttgtaataaatta
caagaac-tttttga-ctgaatgtaaatctaatcgagtggtctcatagatagcactggtt
cattctctctgttaatacctttttcaaatccagcagctgcagctcttgctctaccagcat
gccaaagatgtccaataaataggaagaatcctaagaagaaatgagatgttgttaaccaag
atctaggagatacatagttaactgagttaatttctgtagctacaccgccaacggagttca
atgaacccagtggggcatgagtcatatattctgctgctcgtctttcttgccaaggctgaa
tatcatttttaattttattcaaatcaagaccatttggtcctctaagaggttcaacccaag
gagctcttaagtcccaaaacctcatagtttcaccaccaaaaataatctctccactaggag
atctcattagatattttcctaagccagtaggcccttgagaagatgcaacattagcaccta
atctttgatctctaactaagaaagtgaaagcttgagcttgtgaagcttcaggaccagtag
gtccataaaactcacttggataagctgtattattataccaaacaaaattagaagctgtaa
gacccatgattgataaagcacctaagctgtaagataagtatgcttcaccagaccatacaa
aagctcttctagcccaagcaaaaggttttgtaagtatgtgccaaattcctccagcgatgc
aaataatgcctatccaaacatgaccaccaacaaggtcttccatgttatttacactaacga
tccagccatcaccgccaaatggagattttaagacgtacccgaaaataactaaaggattaa
gagtaggattgctaacaaatctaacatcaccgccacctggagcccaagtatcatataccc
caccaatgaacagagctttgataactagtaaaaaagctccaatacctagtaaaactaaat
gtataccaagtatagttgtcattttgtttttatctctccaatcatatccgaaaaaaggaa
aagattcttcaagagtatcgggtccgattagtgaatgataaagaccaccaaatccaagaa
cggctgaagaaattaaatgaactacacctactacaaagtatggatatgtattaaaaattt
cgccgcctggacctactccccagcctagtgtagctaagtgtggaattaaaataaatcctt
gttcatacagaggtttttctggcacaaagtgagctacctcaaaaagagtcatagcaccag
tccaaaagaccattataccggcatgagcaacatgagcgccaagcaatttgccagaaacgt
taattaagcgtgcattgccagaccaccaggcaaatccggtagattcaatgtctctaccgc

caacaccaacattcgtattaaagggcgtttccacgtggtaaaacctcctcagggaatata
aagttttcatgaggttgatcctgagcagccatccaagagcgaataccttcgtttaatagg
atgttcttagtataaaaagtttcaaactcaggatcttctgcagctcttagttcttgagaa
acaaaatcgtaagctcttaagttaagagctagtccaacaattccaaatgcacttgtccac
aatccagttactggtacaaatagcatgaaaaaatgtagccaacgtttattggaaaaagct
acaccaaaaatctgtgaccagaatctgttcgctgttaccattgaataagtttcttcagac
tgtgtaggcgtaaatgcccggaaagtatctgcagcatcaccatcttcaaataatgtattt
tgcacggtcgcaccatgaatagcacatagtaaagcaccacctaagatcccggcaacaccc
atcatgtggaatggattcaaagtccaattatgaaatccctgtaagaatagcagaaatcta
aaaattgcagctactccaagacttggagcaaagaaccagcttgcttgacccaaagggtac
attaagaacacagatacaaatactgcaattggtccagaaaaagcaatagcattgtatggt
cttagaccaactaacctagcaatttcaaattgtcttaagcaaaatccaattaatccaaac
gatccatgtaaagcaatgaatgcccataggccgccaatttggcaccagcgagtaaaatct
ccttgagcttcaggtccccaaaggaaaagaagggaatgacccatgctgtttgctggggta
gaaacagcagcagtcaaaaagttacatccttctaggtatgaacttgctagtccatgagta
taccaagaagtcacaaaagtagttccagttagccagccgcctacagcaaggtaagcgcaa
ggaaacagaagtagtccagaccaacccacgaatacgaatcggtctctttttagccaatcg
tctacaagatcaaaccaccacgagtttttcttgtccaattgctatggtcataattta
atctccagagcaaatttaataagtaatttgttcaaataaaca---ctttagatctaatct
aaaattttacttttctttacggtcaatataattatacgtcaaaatcaacaaaaattcacg
tttttaataaca-tttttttagctcgttaagttaactatctaagaat-------------
taactgtttttaaattaataaaaattataacttgttcatttctataattctctgaaata
gatatcctgttcctcttgctgttaaaatcaagtctggattacttggatcatcttctagtt
tagctcttagccgtgagatatgcacatcaacaactctagtgtctacatgtcgttctggtg
tatatccccagacctcttgtaagatagatgctctagaaaaaggttcgccagccttactca
ctaaaagttcaagaagactaaactccattcctgtcaaacgaactctctcattatttttat
aaacttgtcttttattcgtgtctatctttaaaaatccaatactaattatgccagaatttg
cgacgccaagattagttgttattttatcagctcttctaagaacagatcgaatacgagctt
ccagctctttaggagaaaatggtttaactacataatcatcagcacctatttctagtccag
taatcctgtcgcacacttcgcctaaagctgttaacattattataggaacatcagattctt
ttcgtagttcttgacaaactccgtaaccatcgagtttaggcatcatcacatctaaaacga
caaggcttggatattcttgtctgaacactattaaagcttcctctccgtttgaagctgtta
tgacttcgtaacctattattgttaatcttgtttctaaaattcttcttatgctggcctcat
cgtcaacaacaagaatttttttctttctggttctccaa-tttcttatttactcctataagt
caacaagcataactactataatcttaaatttgtacctgtttgagtattctaatttgctgt
aaaa-----cttatttattataaattttataagtt-----tagattaattggtatcagt-

-cattatatcttgaaatatctgtttgtcatttatctc---------tattttaatgcata
agttgattcaagttttcagaattcagtatataaata-ctttttagacaagtctttgacat
tgtattttctgaataaaaaaggggttgacaaaccggctgatagaaagtattctacattga
tataaa----ttactgatcataaacaaacagtaa-----attttttagagtagcataag
tactttaat
>Pyr_45
tatatgatcaataattaaagaatctttttgtaagaattcctgaaatatacaaaactttt
tatttaatatagaaaaagtaaattagaataagttgaacatatttt-ttctactttaaacc
ttttgattgag------attctaaatagaaataaaaatgttgacatagtttatatctata
ggaataaaaatgaaacttgcagtttatggtaaaggcggtataggtaaatctacaactagt
tgtaatatttctgtagctctttcaaaaagaggcaaaagagttctacaaattggctgtgat
cctaagcatgatagtacatttaccctaacagggtttttaatcccaacaattatcgacacg
ttacaatctaaagattaccattatgaagatgtttggcctgaagatgttatctataaagga
tatggaggtgtagactgtgttgaagctggcggacctccggctggtgctggttgtggaggc
tatgttgtgggtgagacagttaaattattaaaagaacttaatgcttttgatgaatatgat
atcattttatttgatgttttaggtgacgttgtatgtggaggctttgcagctccattaaat
tatgcagattattgcttgattattacagataatgggttcgatgctttgtttgcagccaat
agaattgctgcttcagtacgagaaaaagctagaacacattccctgcgacttgctggactg
gttggtaatagaacagataaaagagatctaatcgataagtatatagattgtgttccaatg
cctgtattagaagtattacccttaattgaagatattcgtgtgtctcgagtaaaaggtaaa
actttatttgaaatgtctgaaactgataaagatttagcatatgtgtgtgattactatcta
aatatcgcagaccaattaattactaggccagaaggcgtagtccctaaagagtctccagac
agagagttgttcagtcttttatctgattttatttaaatcctaaatctaatgtaggacaa
gctgtcgcagatcaagaagaattagatttaatgatagtgtaaataatattttaagcataa
cataaaaaggaataagataatgtctacagttcaatcagatgctcttacttttgaatgtga
aacagggaattatcatacttttgtccaattagttgtgtttcttggttatatcaaaaaat
tgaagatagtttttcttagttattggaactaaaacttgtggttatttttttgcaaaacgc
aatgggggtaatgattttcgctgagcctagatatgctatggcagagttagaagaaggaga
tatctcagctaagcttaatgattatggtgaacttcgtagattgtgcttgcaaattaagaa
agatagaaatcctagtgttattttttggattggaacatgcacaacagaaattataaaaat
ggaccttgaaggaattgctccaaaattagaagcggaaattcgtgttcctattgtagttgc
tagagctaatggcttagattatgcttttacacaaggtgaagacacggttttagcagctat
ggctcagcgttgtccttcaaatttaaataatcaagtaggcagtaatagtcgcgaaatttc
ttcccatatccccttagttttatttggatctctgcccaacccagtggtaacgcagttaac
tatggaattaaaaaaacaaggtattcatgtctctggctggttaccatctaagaggtatac
tgaattaccggttatacaagaaggctattatgttgcaggagttaatccatttcttagtcg

aacagctactacattaatgagacgaagaaaaacaaaattaattggtgcgccatttccaat
aggcccagatggtactagagcctggattgaaaaaatttgctcagtattaaatattcaacc
tattggactagcagatagagagcaagaaatctgggattctttagaagattatattgcttt
aatacgaggcaagtccgtttttttatgggagataatttattagaggtctccctagcaag
atttcttactcggtgtggaatgactgtgtatgaaatcggtattccatacatggataaacg
ctatcaagcagcagaattagctttattgaaagccacatgtgacgaaatgaatgtcatgat
gccaacgattgtagaaaaacctgataactataatcaagtagatcgaattcgtgatttgaa
accagacttagttattactggtatggctcatgcaaatcctttagaagctagaggcattaa
tacaaaatggtctgtggagtttacctttgcacaaattcatggttttacaaatgcaagaga
tattttagaactagttacaagaccattgcgacgcaatcttagtttgtcagaattaggctg
ggatgtctatagcaagcaaagttagtattagt-------tgcgacttacatggcataaaa
tccttgtaagtct--tctatattatgctaaagaagcatttcttgcccgcccatcttctgc
ccaagcttgtaaattttctatttgttctttatctgtaaatgctaaaggtacaaattgctt
tatagctattttaatatcttctgtattaaattctctctgttcactaaaagctgtatgcat
actttctataatcgcttgctcgatttcagcacctgagaatttattacataataaacttaa
ttgtttaatatcatattcttgccatgattttggtctaacttttgataaatgtatcctaaa
aattgattctctttccttgcgattaggtaagtctaagaaaaaaatctcatcaaaccgtcc
tttccgtagcatttctgaaggtaaactttgaattttattagctgttgcgacgacaaatac
gggagctttttttctgaaagccaagttataaatgttccaaatactcttgcactagtacc
actatctccttggctatacaagccagaaaaagccttatctatttcgtcaatccataatat
acaaggcgataatccttcagcaatatttatcatatccctcattctggattctgactcgcc
aactagtccaccaaataattttccaatatcaagacgtagtaacggcaaagtccaatcctt
ggcaatagctttagctgttaaagattttcctgttccttgtataccaactagcagtaaacc
tttaggggaaggaataccataattcaaactttgtttcgaaaaagagcgagatctttttg
tagccattgctttagtacatctaatccaccaatatctttattcactttattgtaaggata
aaattctagtagacacgtttggttgataatttgtctttttttcttcaataataatcggtaa
gctgcgagaatcgagttgactatattgtgcaataatttttgtaatcactttgcggattct
atctattgatagaccttggcaggatttggtcatattattaactagttctgaatctagatc
aagatttacagctttacttaatcttgtgatttctttttttgatttctaaaagacttggtaa
aggtaaatccattattgtcacaatatctcttaaagcaaaaggtatatttattttacaaga
aataataattatatgtttagattgtgctttgattactttagctaaattgcgaagtttcct
agttaggactatttcatttaaaaaagagtcgaagtcttttaaaatgaaaatattcaaaga
ttcattatctaagttttcaataaactctaatgctagtaaaggattccgttttgcataacc
attatcgttagggttgctagtatatccatctacaaaatcccagcaataaacttgttgatt
gtcagagcagtgtaatttatttttataatatattcgagtctttcttcttctctagtatt
tattacaataataggatgtcgagactttaatagtaatcgtaaatcttgtgtgaaattcat

gtcagtatttgtccccatcagacttttgtacttcaccaatta----aagactaaataaga
ttatctaacgttgcatggatatattcaaaagtc------atgtatctattttctaatcta
ttatgatttcagacttaactaaccataatttttttcttcctttgcgtcgtctactattt
aaaatagagcgtccgcatggtgttttcattctagcccggaaaccagaaactttaatttt
tttctttttgatccttgtaatgttctcttagtcattatgtctcctgtattgtaaaagatt
aaatattattttatcatcttttatatatcttgattatatagatctgtattaatatcaaag
tt-aaacgcaataatttatattaatcaagaattatttatataatattaagaattaagtat
----------tagctttatgatattagctctgcctatttttatttaagtatcctgacag
cgtttttgctaattttgaattggttagtttttcagcagctaaaaacaattttgtcattag
aatctcaatttaagtattttattgaaaaaagtcaaaatgatatattggcagtagaagaaa
gctttgcttttgcaaaggtttgtgtagcaaagaagtgcttttctagagcgattatcgaag
gtcaattggcaatgaaaaagagctcagaattaaacgagaaagagagcaatataataattg
ccaatttgcataacatgattggttttatttattctgaagcacagcaactaaatattgcaa
aaaacttttatcagcaagccttacaaatagaccctaattatgtcacagctttaaataatt
tagctaaaatttacgaagacaccaaggattttaaaaaagcagaggcactatataataagg
tattaactcttaacttagacaataaaatcgctaatagaaggaaggaatttataacaaaga
tca---------gaaatacttaatattatttataatcgggatagcaggatttgaacctgc
gacatcctgctcccaaagcaggcgcgctaccaaactgcgctatatcccgttagacataat
tagtactatataccttttctacaattatgtctactatttcagtgttttatctaggatacc
aaaacatgcctttaactccatcaggatctttaatgaatcctaatttttttgtaaaaactaa
ctacatcaggttctgcaaataatgtaatagtgctaatttctgcttgtcttaattgtttta
ttaattggtgtatcacgactttccctaaacctaaaccttgaaaatctggatgaataacaa
catcccagatagtagcattgaatccattatctgatgttgctctcgcaaatcctacgagtt
tagtatttgtatctttcttttgtattaaagaaatgataatcgaactatttttttaatgcga
tttttacttttttttagaggtcttttaacccatcctaccgaatcacacagctgttctaatt
cgtataagttaatatttttagtgctactgagatagatatctttgaattcggctctattac
aagttttatctaaaacaacaagtttttttgaaattttttttcagagttacttttgatatgtg
agttttgaaaaaaatttttccagaagatcataatgattatgatagaat----taaatata
tatatattatgatattgccttaatgaacaacatattaatatatgtccagtttttaatatg
gagttattaagaaagaaaattgtttgaaggtatagaaataaaaatttttgtaaaatcttt
caataaaagtatataaagagctctttttaagattaacatctattagtatcaatacagtct
cgcaattgagatgaaacaacatattattctatatctaatgccttttaaaggagttatttg
tgaaaaaatctatgtttctaacctgcctacttgctttgttattacttagtaacccgataa
tagctaatgcagaagtggctgggctagtgccttgtaaagattcggcggcattcaataagc
gtatggttaatagtgttaaaaaacttaaagctagattagctaaatatgatgcagatactc
cgccagcgttagctttaaataagcaatagaaaaaactaaaactagatttgctacttatg

gtagagcaggcctattgtgtggtacagatggtttacctcatttaatctctgacggtcgat
ggagtagagccggagactttgtttttccaggactgctatttttatacattacaggctgga
ttggttgggtaggtcgaggatatcttctatccgttgctaaaactagcaagcccactgaaa
aagaaattatttagatgtgccattagctattaaattcatgacatctggattcgcatggc
cgttagctgcttggcaagaatttagtagtgggcaactagttgcctccaatgacgatatta
ctgtttcaccccgttagcaaaaaatttatatgaataataattttaccaaatacttatcaa
cagcacctgtagttggcgtgttatggatgacgttcacagctggttttattatagaattaa
accggtttttcccagatgttttatatttttatttgtaaataaataa--aatctcataaa-
---aaccacacaaatatagtaaaacactattatttgtgtggttttt----atatagaata
taactattta-tttttacggaaattctttctttcgactattttaatattagattttacat
aataaaaaa-acagtaatattatgagtttagtaagccaaattatagttaatgctgatgat
gaattaagatatccaacaatcggagaattacagtcaattcaagactacttaagtacagga
agtaatagaattagaatcgcaactattattcgggataaagaaaaagagattatacaaaaa
gcaagtaaacaaatttttcagttacatccagaatatatagctccaggaggtaatgctgca
ggttctagaaaaagatcattatgcctacgtgattatggctggtacttaagattaatcact
tatggagtattagctggtgataaagattctatagaaacgattggtatcatcggtgttaga
gaaatgtataattctttaggtgttcctattattggaatgttagacgcaattcaatgtcta
aaagaagcttctctagaaatgcttggtcaagatgatatcaaaattattgcgccttatttt
gactacataattcgaggcatgtcataaatagtaaattcaattaattagttgaataaaagc
ttgtataatgttatgattagttcaagcccgaaagcatataattgtaaagactaaatcttg
tcagaaccggaaggtagcagcaataatgtttaattacaacagttatga-ttttcgggtgt
ttactatttgtactcaatagtctctcattctaattatgatgctatccataaatt----aa
taatatataatccataaaacctttcagtaaaaataa-ttaataatttgatattaatacgg
aaaacatctatactttttttataaaacttctttatatctcatctaaacaatggaaactta
acttaatgggtgttcatattttatcaacaggttcgtctgtcccaaatttttctgtagaga
ataaacaatttgaagatatgatagaaacttctgactattggattttaacaagaacaggaa
taaaaaaaagacatctagctccttcttctacttctttaactaagttagctgcagaagctg
ctaataatgctttatctgcagctaatttaaaacctaccgatattaatttaattatactag
ccacatctacgcctgatgatttatttggtagtgcaagccagctacaagcagaaatcggtg
caactgcatctgttgcttttgatattacagctgcttgctccgggttcattgttgcattag
taacagcagctcagtttatccagactggttcatatgacaatatcttagtcgttggagcag
atacaatgtcgagatggattaattggtcagatagaactacctgtatcttatttggcgatg
gtgcaggagcagtagtagtgggacaaagccttaaaaatagtattttaggttttaagttat
gtacagatggccagctaaatagtcatttgcaattgatgaatacaccaaaaaataatcaaa
aatttggatttatagatattcctcatggaaattataattctataacaatgaatggtaaag
aagtatacaaatttgctgtatttcaagttccaatggtaattagacaatgtttagataatt

taaatatatcaatagatgaagtagattggtttatattgcatcaggcaaatactagaattt
tagaagcaattgcgagtagattgtcaataccttttctaaaatgattactaatttagagc
actatggtaatacatctgcagcatcaattcctttagcgttagatgaagccattaaatcca
ataaaattaagccaggtcaaaccattgttttgtcaggttttggagctggtctaacttggg
gtgcgattgtcttgaagtggtaat---tataatattgcggatgacgagaatcgaactcgt
aaagctttcgctacacacccctcaagcgtgcgtgtataccaatttcaccacatccgcact
ttttatcacgaata-ataaaaatttataatatttatttaaatattaatataa--tgtgta
tatatatatacattcttgtcagaaactctcttatattttattttaagtgataaaattgag
t----ttagtatagatgataaaattttattttacaatcataaaaataaataaatatataa
ggaaaataaaatatgacaccgtccttatcaagctttttaaacagtctaattcttggggcc
gtaattgtagttgtccccataactttagcacttttatttgttagtcaaaaagacagaaca
attcgatcctaacaaagactttgaaacatatggagaagcaatttgagattaataataaac
tatttttcaaacggatatgctctccatatatttctgtatt--ttggaataggttgttt-t
ctttactttgcaatagcctgcacagtgta-----------ttttgttctcatttactctt
tta----taacgatacgtatgtcagg-ttaataatag--tttttttattctgtatctttc
atttaatattaaagtattttatatacatattgttatgttcaagtaataaagaaaagctca
tgtctttgtccaattggcctcttaaaaaagaaaactccgaagcatatactagcaagtata
caaagcaaatcacaatccctgatggattgtggataaaatgcttcgattgcggtctgttaa
tgtattctaaagttctaaaaaggaatttaaaagtttgtcctcaatgtagctatcattttc
aagcttctagcagtgaaagaattgatcagttgatagacaaaggtagctggcgacctatgg
atattcatttaatttctacagatccattaggtttcaaagatcaaaagctttatagtcaaa
ggttaaaagatactgctcttaagactggcttacaagatgcagtccagacgggtattggta
ctatgcaaggtagacaagtctgcttaggtattatggacttccgatttatgggaggaagta
tgggatctgttgtaggcgaaaaattaacaaggctactggaaagagcaactcaagaaaaat
tacctgcaattatactttgtgcatcgggcggcgcaagaatgcaagaaggcatgttaagtt
taatgcagatggcaaaaatttcttctgcgttggaaatgcataaaaaagcaaatctcttgt
atatatctgttttaacatctccaactacaggaggcgtcacagctagctttgctatgctcg
gagatctaattattgcagaaccaaaagctcttattgcatttgctggtagacgagttatag
aacaaactattaaagaagacttgccagataattttcaaagttcagaatatctatttgagc
atggttttctagatttaattgtaccaagaactcagcttagatcaaagctaatacaaattt
tatcattccatactcatactaagtaagtagtttgacattgatcataatattgaaatgcta
gtaaaaattaagatattaagaaattttcactaaagatatttgttgactaactagaattac
aattactattgtagaaaaatttgtcttaattctatttttttcagtcttttaactaattgt
tttttacgactttaaataattcttagtttaatacaaatttataaagagaata-----cat
actaagctaaatatagtaatattatgttttcctataatactgacctatcaaatttcgagg
aattttatgcttaaaagatcttcttggcttgctactttattggggctactaactgtagtt

tctataagtacgcatgcaaatgccatagagctagacgaggcaacaagaactgtcccatta
gaatcttctggaagaactgtagttcttacgccagaacaagttaagcgaggcaagcgatta
ttcaataattcttgtgctatttgtcataatggtggtattacaaaaacaaatccgaatatt
ggacttgacccagagtctttaggcttagcaacgccacaaagagatacaatcgaggggtta
gttgactatatgaaagatccaacaagctatgatggcgcagagtcaattgcagaactgcat
ccaagtattaaaagtgccgaaattttcccaaaaatgcgtaatttaacggatgaagattta
tttacaattgctggtcatatcttgctccaacctaaaattgtttccgaaaaatggggcgga
ggaaaaatttactattaaaggttcaaacacaataatcctacaagtactagtttatgca-t
ttattgcccaagtatgattattttgttagatcatgatatatattaaaggtggagtttgct
aatagtaagtta-tttttgcctaaacttgcttacattattcgtttagttattaaaaagga
gaagttaaattgaaaaagacgctttcagttcttttcactgtatttagtttttttgtgata
ggttttgcacaaattgcttttgctgcagatctagataatggagaaaaagtttttttctgct
aattgtgcggcatgtcatgcgggtggtaacaatgctatcatgccagataaaactttaaaa
aaagatgtgcttgaagctaacagtatgaatagtattgatgctatcacgtatcaagtgaaa
aacggtaaaaatgctatgcctgcttttggaggtagactagttgatgaagatattgaagat
gcagcaaattatgtattatctcaatctgaaaaaggttggtaattatacttaattttgcac
agtattaaaggatagacaatctatttaattattcattagattgtctattctttatttatg
---ttattatatatagatatttatacaatactga-attatgatgaaaagaatacctgcaa
ttcttgtactagaagatggtacttattacaaaggatggtcatttaagcaaagaaatccag
caattacaattggagaagtggtttttaatactggaatgacaggttatcaagaaattatta
cagatcctagctacttccagcaaattgtcaccttcacatacccagaaattggtaatacag
gtattaacagccaagatgttgaatctcataatattagtattaaaggcctggtggcaaaaa
atatttgtaaaatttcaagtaactggagagagcaagaatctttagttaaatatttaagcc
gttatgatattcctttcatctttggactagatactcgatccttaactcaatatttacgtc
aatttggcacaatgaatggttgtatttctactaataatttaaatcatagttatttaaagc
aaagaatttctgaagttcctggaatgcaaggcttagatttaattccatatgtaacaacag
acagcatttatccttgggacgaaaaaagtttgcctatctggtatttaaacgataaaagaa
gagtgcaattagatacccagttgaaagttgttgttatagattttggagttaaactaaaca
tactgagacgactagctacacttggatgtcagataactgttgtgcctgcacaaacatctg
tagaagatattttagctaataaacctgatggtatattactttctaatggcccaggtgatc
catctgtagtgtattatggaattaagacagtgaaaaaatttattagatcataatgtaccta
ttttcggtatttgtatgggacatcaaattttaaatttggcccttgacgctaaaactttta
aactcaaatttggtcatagaggtattaatcatccatctggattggaacagcaagttgaaa
taactagtcaaaaccatggtttcgcagtcgatttacaatctgtcttcaagtcatctgtac
aagttactcattttaatttaaatgatactactgttgctggaataggccatagtcaaaatc
catatttctctgtacaatatcatccagaatcgagcccgggccctcatgatgctgactatt

tattcgaaaattttgtagaaataatgataaaatctaagcgtaaagttgc-----------
ttaataatcttcccatgctttatgattaaataaagctgataatacttgtactccccgtaa
ttgattgaagagaggtaaatgcccctttggaccataagaattccattgaaattcatcagg
gtatctacaaggaatatgatcaacttcccacccaatcttttgccagaattttccccaatc
ttttccaacactaagccatatttgcctctgaataaaaagcccaaatttacctttgagtg
tgtatgccacaatttatcaatagtctgaagatcttgggcaggtattttttttatatctgt
aaaatatagccagttgcgattttgtgcattcacaccagctaattgaataagtttttgttg
agttaattgatctgcctcttgcaaatcttgttgtgttagcaacacttgtaaatctttata
attcatgtgtttcgcagaacgtaatgggacaataccctcgggacataagtcagtagtaaa
cctaatgatttcttgattttactattcaataacttgttatagatcaaaccatcgacgca
attacttttataattgggcctactaatcctttcaagaaatagatccgctaaactttttaa
ttgcatagagtcattattatgtatagtctcaataatatcaagttgtttttttga---tatt
atttgaactagtattttcacttaattctaaaaactgaa------------attgcatttg
attt-ttacattaatttaaatagtg----attatttcttaaaaagaatttga-ttttttt
gaaaattagaagaacgacgaattatttctagaagtgttttcataatttgattgaatatgt
ttctccccgcataaatgataatttgaccaattaaaaaaattgcacttttattttaggta
caaagaagtcttgtacctcaagtagaaatgtaattactatttgcattgatgataatgtat
ataattcatgtgatctacaagcatatatatactgacaatttaaaccttcctgataaaacc
cccatactttgtaacgactttcgtaaattgctcgcggtctctcaatataatttttaatta
aagtattccaaactaaattatttttaatttttctagttttctatcagataaaaaatttga
cattacaaagatagcttgcatttgtttcatgtactgtatcaaaattttgaaccaacatat
gtgcaataatattacttaactgaattaaatagttctcaagaaaaatttcaacttgtttaa
taggtacatttgaagtaaataaattgaaagtcttattcacaccttgagaggatccaaaaa
gtagctgaataattagaatttgtagtaacatcttatagtcatatattaggtagtttaagt
cattaatatcttggcaaatatcattatagtctagttcgtataacctacaaaatttttga
cacatctatgaaataaatcaattaaaatattttcagttagaatcgtaatatcatttaaac
ctaaatttgaattataaatttctaacaggatttttctaattctgttaatattattttta
ataagtttctttttacttcagttcgaaaaacatctaaaattaatacttcctgagaacaat
tagttaatttctattaattttatagaagttctaactaatagctctgcaacctcttgat
tgagcatgtgtccctgtgagctcggccaataattattcacggtatatcatagtaaggaat
atttaag-----tttatacgataacttttaatacactgaaaagcaagtttatcactgta
tagccattag-------cttttacagatcttaatagttatttttcaaataaagttatatg
atattagcgaccaagctaattttattaaaatttataattaataatgacaatttattactt
tgcacttgccagtcaaaactttctcttagtacaagagcccttggaagaagtttttagaga
gagagttaactattatcagtcaaataataaagcaatcgatttttggctagtgccaaatcc
ttctttttagaaaaaccagaaatgatttcgtttaaaaatcttgtacctaatgatgctgt

agccataatctctactaacccagtatttattaattggttaaagttaagaattggttatgt
ctgtgttggacaatttgaagatgatttacaactttctgaagagtctttaaacattactgt
tgcaccagaaaagatttaatcatactttattgccactaggaat-aattat-----ttaa
acggcaaatataaatt-caaatccttgtttttctatagtttatcgcttagtattttatt
ctgagatggagtgacaaacagtgtaagtatttcttgactaaatgtttcagttgctttgga
tttatacctattagggttgactataatagacagcattcgcttaatagtgacattttcaat
ctgagcccaatgtacaatccctagctctaactcctttgcaatagctgaaacagaaacaaa
agcagctcctaaacctgactgcacggcatttttatagcttctatagaatttaattccat
ctctatttaaagcgactgctatcaataccatgttgactgagcactttatctatgacttt
tctgattgttgattgagtgtctaatgcaataaatttaagtctatacaggtcttcttttg
gatgtcttctaatttagaaaacggatgcgatgttggaagtattagtgctaattcatcttc
ggcataagatgtaacttgcaaaatatcttgtaattcagtaggtacttcccctccaataat
tgctaggtcaacttgaccattagctacgctccaggaaatgagtctagtggaatgtacttg
tagttgaacagctacttgtggatatctttgcctgaatagtcctattaatcttggcatcaa
atatgtcccagttgtctgacttgctccaataattaatgtaccaccttggaggttttgtaa
gtcatcaagagctcggcatgtttcttcacataaagctaaaattcttcctccataccgtaa
aagtaaacttcccgcctcagttaaagttgctttttattacctctctcaaagagggaaac
atttaattggcgctctaaattttgaatttgtaaactaatagctggctgagaaacatataa
gctattagctgcctttttaaaactgccttctttggcaattgcttttaatattcttaattg
atccaatgtaaatggaagatctgtcattaaagaattgtagtataatg---------tata
ttatttaatattgatcaatttatgcattcgccaagttaaatatttgctattacgcttcac
tgctttgaaaataataaaaagtataatgttaaatacactgtattagattttaatatctgt
ttaatgttaaatttatataataaatatcg------taattttaattataagatataggag
acaacaatatggactctagacttttaattgtattaataccagttttagcagctgcatctt
gggctatttataatattggtagagtcgcgctacaacaatttaagaaaatgacatcttaat
gatgatgagaatata-atcttaaaattttatgagatagagaactttaaaatctctatctc
tattgtatatcttaataaaaaaagtaagta--acccttgtacttactgttattacctatt
aagatgatgtac-agtacatatatttataataagtttaagctaacttatt-----tttta
taattaactaaattaataacacaattatggctgttccaaagaaaagaacatctaaagcta
aaaaaaatgcacgtaaagcaaattggaaaaatcaagcaagagtcgaagcacaaaaagctt
tgtctttagcaaaatcagtattaactagaaaatcgaatggatttgtttataatccttctg
aaacagcagatacctatactaactaagagtcttt-ttactatcaagctagccaagtatca
aaaaatctattttaagtaatttttatatat----taaacatttggaaatcattctatttg
ataataaaataaatagcagattaaacaaaattacggcaacaaattatgcagcaaagctag
atgaaactaatgaaatttggttatttaattgcatcgaaaatatccagcacactttttga
aaagtcaattaaaatcaagtcatattactaaaattttcgtttctggtactagctttgaat

atgtagcaggtttacctggactattatccagcttaacactaagtggtagaataaatccca
ttagcatatatggcccaaagtctttggaatggtacctgaaagcttctaccaaatattctc
aaactaattttcttttcctattaattttataatgtgaaatatggagaacaagttatta
atcaattctatacagtagtttgtttaccgttgagccaggacaatttactgtatggattta
ctatcttaaaaaaagaaaagccaggagcatttaaattagcaaaagctaaaaccttaaata
ttactcaaggtcctatatatggcaaattaaaagaaagaaataattttctaagtcccgatg
gttattatctatgtggcaaagatttctcgttaagtacaacaatgggacacaaaatatctg
ttccactattagtcagatattctagaattatttcagagatgcattggttttgctcttatt
ctattcgatgtaacacttatctacatcagcaaaccataaaacatttgccaaacaatattt
taaccgatactatgaaatctcaaatatctcaagataatagttgcattgaataataattat
tcttgatatataatagtctcgacaatataaataacatacaagagcgaatttcttg-----
----ttatatattaaatttataattatttatgacatacgcaattattgaagcaagtggca
aacaactttgggtagaggaaggacgctattatgatttaaatcgcataccttttgatccag
gccaatcaattatactgggtaaagttttattgttaaataaaaaatggacaagtgactttag
gacatccttgcatagcaggagtaacaataaaagctacaataatgaggcatttacgaggaa
aaaaaataactgtttttaaaatgaaacccaagaaaaaaatgagattaaaaaaaggacatc
gacaagaattgactcgtttaatgattaattcaatagtaaactaggtctagatacagttgt
aactgagattgcttcaa-aagattttgaatatttaactacaatacatataatggcacata
aaaaaggtagtggtagcacaagaaatggtagagactctaattctaaacgcttaggagtga
aaaaatatggaagcgaacaagtaagagcaggtaatattttagttagacaacgaggcacga
aagtcaagccaggtcaaaatgttgggagaggaaacgatgacactttattttcgctgatta
atggctttgtatttttcgaaaagtctaaccaaaagcaaaaaacaattagtgtttatgctg
caaaaaaatagccaaaataattggtgcaagatgacctgaat----attttatttcactt
acactggcaccataaacttgatttattaagaaatggagttccttaatgaaccaatgaatt
ttattaaagatttgctttgaaatgtaatgacaaacactattgtaatttcctgtctacaca
atatggctgctattctatattgtggtcagatacagaaactagtagtagcaaatgctcatt
atcaagtaagtgatatttacctaggttgtgtagataaaatcttctcaggaataaatgcag
catttattgacctgggaaaaaatgagtacagtggattcatccatattagtgatactggtc
cgcttaaaaagaaatattatgttaatagtattactaacattttaacaattcgacaaaaaa
ttttagtacaaattactaaagaaccaactttaaataaaggaccaaggcttactgctaata
ttacattatcgggtcgatatattgtactaatgccttttagtcaagcaatttgcatatctc
gaaaaatacatgatgaagatgaacgtcattatttgaaagctttagctattctaattaaac
cagcaacaatgggcttattgttccgaccttctgctgtaggcattgatgaagaaattatac
taagcgaactaaaaaatttaaaagagcaatggaactttgttcaaaaatctgcaatcaaca
gctatgcacctggtctcctatataaggatgaggatattattaaaaaggtaattagagatt
tttatagtaataatacaaataatatcgtaattgattcaaacttgggattaaaacaattga

gttattatatccatacttggcactgcaataattctagtacagctccttcaattaagcttt
atagtaacaatcaatgtatactagatgcttttggtattaatcaagcaatttctagagcgc
taattccaaaagttgatcttatacttggaggctatatgtttattgaaactttagaagctt
ttacaattattgatgtaaattccggatcttttaataactctactagtgcgcgagaaacag
ttttaaaaacaaattgttctgcagcaacagaaatagcttaccagctgcaaattagaaata
ttgcaggtgtaattataattgattttattgatatggaatcacagagagatcaattacaat
tattagagcacttcaataaagagctagcacttgatgatgcaaagcctcaaattgtacagt
tatctgaattaggcctcgttgagttaactagaagaaggcagggcaaaagtttatatgagt
taattagtagtgacttcaattactttcatctccttatgcaatcagagcagactaagtctc
ttgaaagaatagatcaaaaaaagcaaatcttccagactcttaatagatcatggctatctg
tagaaattaatactattaataaagtatttttcaaaaatcaaatttttatatacttacca
acttttatttagcacgtgatttatatgtaattaacagtaatctcacttataaacaaaatt
gtttattcactactagattgaaacttatctactctcaagagtataataaaatcttgcctt
gtactggttatagttgtctcatcagttctaatatagactataatcaaaagttccttttgt
gaactgtcggtttaaattgt----------------------------------------
--------------------taaaaaagaagccctctcttgtatgtgagagggcttc-tt
ttttagcttactaactattctagtgaattaattagaaattaaccattgacagctggagct
gctagagctactggcaaggattcaccagaagctagatctagagggaagttatgagcatta
cgttcatgcattacttccatacctaggttagcacggttgatgatatcagcccatgtattg
attacacgaccttggctatcaacaacagactggttaaagttgaaaccattcaagttgaat
gccattgtgcttacagataaagctgttagccagataccaactacaggccatagacctaag
aagaagtgtagagaacgagagttgttgaaactagcgtattggaagattaaacgaccgaag
tagccatgagctgcaacgatgttgtaagtttcttcttcttgtccgaatttgtaaccatag
ttagcagattcattttcgcttgtttcacgaattaagctagatgtaactagggatccgtgc
atagcactgaacagagaaccaccgaatacaccagcaacacctaattggtggaatggatgc
attagaatgttgtgctcagcttggaacacaagcatgaagttaaatgttccggagatacct
agaggcattccatcagagaaactaccttggccaattgggtatactaggaatactgctgct
gctgctgctactggagcagtaaaagcaacggagatccatggacgcatacctaggcggtag
cttagttcccactcacgaccaatgtagcaagctacgccagttaggaaatgaagaacaact
agttggtaaggaccaccgttgtataaccattcgtctaaagaagcagcttcccaaattggg
tagaagtgaataccgatagctgcagaacttggaataacagcaccagaaatgatgttattt
ccgtatagaagggaaccagcaactggttcacgaattccatcaatgtctactggaggtgca
gctacgaatgcaatgatgaatacagatgtggcagttaataaagttggaatcattagaaca
ccaaaccaaccaatgtataggcggttttcagtactagtaatccaagagcagaaacgttcc
cacaagctagcgctttcgcgtctttgtaaagtagcagtcataatttttatcagtttta
aggatttaactaagatgcttcccaaacaatattttgtctgttaaatatattattacttaa

gatcgttttaaatgagaaca--tttttaaaaaaca-taattaaatttaatctcgataact
aagaaaatataattattgagatattattatataattgaa----------aaaataaattt
tt-act-------------ttttgcagaattagtatgattttaagggtataataatagta
atgcatactaatgctgcttattttgatttcaacagtttacataatcgagctttacttaaa
tacgtatgtcacactttacaaaaattcagacgactatacaagacttaaatctgttaaagc
atgctttaacagatttaggtttaatttggtctatgaattctagtcatatacaagttgatg
aaaatagtcagcacaaagtagatatcttgatcagacaagataatttatcacatattggct
ttatctggaatgataataagtaccacttagttgctgacttgcaactttgggaacaacctt
ggtctttagaagtatttttagataaattatcgcagaagtatgcttaccattctattatcc
aagaaacgaaaaaacaaggctttgaaaaaatggagcaaatctcgcaaaacgatggatcaa
ttaagttagttgtgcaacgttggagttgttaattaatatagaaatgcgggcggagagact
tgaactctcacgagattatctcactagaacctaaatctagcgcgtctaccaattccacca
cgcccgcactgctgtagt--aataatactatatcacaattcatgaataaaaacaaactca
-agaaataaattagctaccaagtttaaatgcatcaacaaatagaccatatgttataccaa
ttttgaagttattaattaaattgatttttatttttaattttttttgattagagtaaacga
tcttactagttaactctgttgctgcaactattagactagctgcaatgattccccagtctc
ctgtttgcccaggaatcgtagataatcctgttgaaataaagaagcctaataacaagctga
ttaaaccagttgttaactcacttaaagaatagtacagtttatcatttaagttttctacta
aatatgcaaaaaaggtagatagtcgagttttaatcataatatttaataacggagttttaa
aaggataataa-----------agattaatgtttttagatccttttgaaacccgtcaata
caaaaattgctttttg-----------tttaaagatcttgctcacttaaagcattaataa
gatgttgaaatggttcttgtataatgtctttacgtttaattgattgtaaatcaagttctt
cttctattatttcttgcaataattgtatgcttcttaccgttggagcaacaggcacagaca
atgaattataagtatctttgaggccgtttaatacacgctgatctagaatattagtgctac
cagagactaatgcataactagcatagcgaaggtagtattctatatctcttagacatgcag
catatcttctagttgtataagaatttcctccaggtcttaataattcgggttgctcttcgt
ataattgagcagcagcctcttttaatatatttgtagcctgattattgataatttcagcta
tttttattctatccaatccacttgaaaagaaagattctatttgtccgacagctgctttat
ctaaataacgaccagtaagatcgtagcgatttaatattgctgttatagcatcttgcataa
attaattctcctac-ttgctatcagagtggtattatgaatgactagtat-----tgtaa-
----------tataatcatcact-------------------------------------
-----------gtttgtataaatagagtatatgtagtttttaattactttttttataaaa
attaagaaataaagatttttt-taataggtttacaatttcttaatataactcaggttct
caaccgagataagttttgtaacacat--aagctaaaataaagatgaatgagtaatcttga
------------------------------------------------------------
------------------------------------------------------------

------------------------------------------------------------
------------------------------------------------------------
------------------------------------------------------------
------------------------------------------------------------
------------------------------------------------------------
------------------------------------------------------------
------------------------------------------------------------
------------------------------------------------------------
--------------------------ataagaataatatgcatcaa--------------
------------------------------------------------------------
------------------------------------------------------------
------------------------------------------------------------
------------------------------------------------------------
------------------------------------------------------------
------------------------------------------------------------
------------------------------------------------------------
------------------------------------------------------------
------------------------------------------------------------
------------------------------------------------------------
------------------------------------------------------------
------------------------------------------------------------
------------------------------------------------------------
------------------------------------------------------------
------------------------------------------------------------
------------------------------------------------------------
------------------------------------------------------------
------------------------------------------------------------
------------------------------------------------------------
------------------------------------------------------------
------------------------------------------------------------
------------------------------------------------------------
-----------------agtatagatagtcttttactccagcatcaaaaaagatatttaa
cacatattgaaaaaattgattatgtaagtaaaaatctaaatttagataaaattagctatt
ttataattacaaccaataagaataacaaagatatccttgtagaacaattttgttataatg
cagatggttatacttattcaatattatttaaaggtcaaacagctaaacacctttgttaca
tagttcttcagcttaaacaattaaatataagaatatctacagctcatgctctatatttag

gacgagaactaatgaagtcggaattaactcttatattagatcaacaatatatccaagact
aaa--cttcatagatattaattatatttgtgctaatatagtattaataagggcatgtaac
tcagtggatagagtatcagattccggttctgatggccgtgggttcgaatccctccctgcc
cggcgagtatttagaattct-gtggcaagatatgt-------tagtatctaatttaataa
tattaaaaatattggggctgccaggtttctacattataaaaacaagaaaatatgaaagtt
aaacaagctctttgaaagagcctctagtaaaattaataaatgcagaaaacaatattgttt
ctttttctcgaaaattagctgttgcttaaatactgttaatttttggcattcgacgtgtta
agctcttattcgcggcgaatattctgttagagttgctcttagtataagaaaagtttcaaa
atctatttatatttatgtttttttacctatttttatttaaagtaaggttagta-ttttcaa
tttttataatggacgcgggttcgaatcccgccagctccatcaatttaact---------t
tcataattaactaatccaatctttttaagaaccagaaatataattggagcaaatatggat
tttataaaaatcacagaacaagcttttaagcaaattgtagtcttgaaaaatgattacaat
aatgatgtacatttaagaataggtgtcagacaaggtggctgttctggcatgtcatactca
atgaattttgaagatgtagctaacttacaagagaccgatgaaaaactgatttagatgac
ttttctgttgtttgtgatcccaaaagtcttttatatctttatggattgtcattagatttt
agcaacgacttaataggaggagggttccaattctccaacccaaatgccagccagacttgt
ggttgtggtaaatctttctcaggctaaa-------gctactttgttttatattttgtatt
atacgtgttactcttataatcagccattataaatacaaaagtatttttattgaaagctct
ttaaactagtcaaagccttatgaaaactactttaaagaggttttatt-tattatggcagc
ttaaataagtatttatcatcaattgatgaaagt---------acaaaaattcactccctt
gttattgcctaattttaattagttaatttttattgatacatgctaatattt-attagcc
ctgatttattaggaatttataatttctttttttataaaagagcttgtcagtttctagcta-
----------------------aaatagattatatatttt-attcattttataaaaactc
aacatgaatcttgctcaagttacttgcaactacttcaatacgaaaactctcaataagttt
tattgcaatacggaatatgtttcctatggtaattttagtcatctcaatttatcaaagata
aatttgccatcttctgccaaaataattgcatttggactccctcaatttgattttattcct
acaaagattaaaaactctatccgagaagcagtggtaatatctaccatcgatgacagtaat
atttttgctgattcaactaataattggataaaaaaacataaagataaaaatcaccactct
cttttctataaaaaaattttaaagctttactgcatcaaaaaataaaagtcgtgacagat
tcagc------taaaaattctaaaatagataaagagaagggcttgattttgagtcaaaaa
tatatattaaagaacagttggaagccatccataatgtcttcttactttgaaaacaaaaaa
aataatgaggttgctaatgtatctaaaaaatttttaatcgatcagctatctgcatcacct
gtgtttgtagtaaaaaatggttttaatgaaattattttaggccacccagcattccgtata
aagagaggtgtaattagtaattacactcactcgttttcaaatttattaaagcaatctcag
ccaacttatcctatatctactggcttattctttttttcatccagatgacgcgtttgaattt
aaaaactttataaagtcggttaatccgttggctagtgagcatatggaaataaatgtagaa

cctgttggcctacactttgcttataaaatgaacagaaatgtatcttctgatactcaattt
ttattcattccagattttaaagaggtaggcgatttattatttaaatataaaaaagataag
cacttaatctttcacaaggatcagtgtcatggtaaaaatttttttcaggggcaacctata
tatattattcagcctactattttcaaagatcgaagcggaaacctaaatactattcaattt
actggactaaatgataatagagaagttatttttactagtcttgaagctgctaataaatca
tggacaaattttataaaaaacaattcacaattaaaatcaattagaaaaccgactttatta
gtctataatctagaaagtttttttaaaagatcaagaacgattgaataaacaagattttcaa
aaatttgttgtgattactaataaaaaagcttatcttgctacaaaagagttggtagctgta
cccgattctaatagcgttttaaagcatttaaaattaaatatgaagcctaaactcttcttt
ataaaattatgggtaagacggctattttccaccttaacttatgaat-aaatgatgtt-gg
atttattttcttgaatagtattcaataactagtaattcatttagttgtaaagcaacccat
tctctgtcaataactccattgacttttcctgatagattagatttatttaattctaagtgg
ctaggaatgttagctaatccagggaacgctaaatagttttccactagtcttttagatgtt
tcccgagattttacactaattaattctcctggcttacattgataactacatatagatact
acttgtccattaatgcaaatgtgaccatgatttactagctgtcttgcagcaggaattgta
ggcgccattccaagtctaaaaacagtattatcaagtctcatttctagaagctgtaataaa
atttgacctgtagatccttggagtttttttagcggccttaatatacttaaagagttgtttt
tcacttaatccataattaaagcgcaattttttgcttttcttctaatctgacagcatattca
gaaggttttctagatttttgtccgtgctcccctggtggataagattgtttaattgttttt
ctacttagcccaggcaaatcacctagtcgacgggaaatgcgtactcgtggtcctctgtat
ctagacatattttataaattttccgaatattaat-gattaag----------taatgagt
agctatttttaacacaaaaagagtattaaaacaaagc-ttttgctaaaaaattttcatg
aaaaatatag--------------------tgctttac-tttttaagcgggtagcgggaa
tcgaacccgcatcattagcttggaaggctaaggttttaccactaaactatacccgcaata
ttcaataaactgtaacaggcttaatataacataactatcttattcttagatattattaaa
attgacaagaag--caaattatttatgctccttctaaaactacatcaagaaaacgagcaa
tctctgcagcttgctcttcaatttctgaaagcaataatggttgacccactctagttaatg
gaattactttttatcttttgtgcataagtaaatttctcgtcgtggattcaaaccttctt
taatatctattttattgacttaatatcttttatattgaattgaagacaaatttgtctat
tttttcctgggaaacctaaacgaaaaattttaacagtaccttcatctttattaaactcat
tatatccagcacctatgtcccaaataatagttaaccataaaaataaactaagaaagattc
cgatgctgccatagaatgtcataacaataccttgcggaataaatattaattctgttgaag
tcgtaaaaggcaataaattaatttgaaagtaacttgataatcctgcaagaagaaatcctg
aagctccaacaaagattatggtagcccaccagtaattactaagtcttcttgaacctaaaa
ttagatcttttctaacagagtctatagataatgtttttttcataatttgtt-ttttaaaa
caata-tactaatttgacatataaaaattttgacaatttttttataagcgttttagattt

attaaatgagtttgatagcttttaatcctaaaaataatcctgttgctaaaccaaaaaaag
ctgcaagaaatattatatatcctagaaaaactgaca---tatta----ttattattattg
ttacaatagtgtttacaaaaatagttgaaatctattactaaagtattatattacgttaaa
tgcttcgctgtctacatgaaaaaactttttcagttttagcgtcgtatttatattagaa
ataatattgtaataatttattaaacttctggagaacagaatcttatggcagaatttatt
aagccttataatgacgatccttttgtaggaaatctatctacaccagttagtacatcgagt
tttagtaaaggattattaggaaatttaccagcttatcgtcgaggtctatctccacttcta
cgaggcctggagataggaatggcacatggatacttttaattggtcctttgataagtta
ggtcctctacgaggtacagatgtagctttacttgctggattttatcttcagttggcttg
attattattctcactacatgtttatctatgtatggaaacgtatccttttctagaacagat
tcaaaagatccattgcaaacagcggaaggttgggggcagtttactgctggattcttagtt
ggagcagtaggtggctcaggatttgcttacttattgctagctaatattcctgtactccaa
actgcaggtttaagcttgttttcttaagctagtaaggggacttgaacccgtaacctactg
attacaaatcagttgctctaccaattgagctatactagcgctcagctaattgtaacatat
aaagatatttgttgcaatctggtattttcagatgcaacaaatatctttttttatttagaca
gaactgttaaattaatatatgtatgatatgttaattagaatcagagctatctgactcagc
agaatattcttgatcacagtttaaataataacaaattgtttgatctagacatgttgcgga
ccaaccaacaggtgtttctcctgataattcttgcatatctccagaattgtaaaaatcatc
acttggcttattactatttgcttgcataaatatcactcccaaaacactctgaacttaatg
tgaagtaattgaacttctcaataatatattaacacagaatataaaatatggtgcttattc
accaattgtaacaatattatgaaatctacaagttctttgtaagtgttttaatagcttttg
ctgaaacaagcattttaatccatttattcttctctcttgaccatactttttctgttgca
aattaacttttgtaatttttttgttcgtttgtgagaatgagagactgcgtacccattat
tcgctactttccctgtaagctgacatttttttgacataacgtt------attaacttata
tatttatttaaagtacttgccaaagttgactttggtacagcaccaattaccgtgtcaact
ctttctcccgctttaaaaatcattaaagtaggtatacttctaataccatattcagctgca
attgtaggattatcgtctgtgtttatttttactaccttaatagatgactcatattcttct
gcaatctcatccactacaggagagaccattctacaaggaccacaccacggcgcccaaaaa
tctacgagtacaggtaagttattattaataacttcttgtttgaaagaggcatctgtaact
tgagatactgacatatttttttaacctttaatatatgttccttgctaagcaaatcttatca
caaatttcaagaaaaaatcttctatagtgaattactattagtgttgaattaa---ttt-tt
tactagacattaaaaaaacttattacctatataaataaagtggtggccgagtgatatctt
gatagtgataatttttctattaaggttgaacactagaaatcatttttcaatcatatattt
ttttgaaaa--acttctttgtgaccatgaaagc-taagtatattacaatatattttatta
tttgtaacttatttcttataatggtataaacaacgcaaaagatactgccttataatcaag
gaggaatacatgtctcaatccgtagaatcacggactaggattaaaagcgaacgttacgaa

tctggagtaatcccttacgctaaaatgggttactgggatgctgactatgtaattaaagaa
acagatgttttagctctgttcagaatcactcctcagccgggtgttgaccctattgaagca
tctgctgcaattgccggtgaatcttcaacagctacttggacagttgtgtggactgacttg
ctaacagcttgcgacttatatagagcaaaagcatatagagtagatccagttccaaacgtt
gcagatcaatactttgcctatattgcttatgatattgatctatttgaagaaggttctatt
gcaaacttaactgcttcaatcattggtaacgtttttggatttaaagctgttaaagctctt
cgtctagaagatatgcgtatgccggtagcttacctaaaaactttccaaggtcctgcaact
ggattaattgtagaacgtgagcgaatggataaatttggtagacctttcttgggtgcaaca
gttaagcctaaactaggtctgtctggtaaaaactatggaagagtagtatacgaaggttta
aaaggtggtcttgacttccttaaagacgatgaaaatattaattctcaaccatttatgcgt
tggaaagaaagattttatattccatggaaggtgtaaataaagcatctgctgctgctggc
gaaattaaaggccattaccttaatgtaacagctgcgaccatggaagatatgtatgaaaga
gcagaattctccaaagaggttggtagtatcatttgtatgattgaccttgtaattggctat
actgctattcaaaccatggcaatatgggctcgtaagcatgatatgattttacatttacat
agagcaggtaactctacttactctcgtcaaaaaaatcatggtatgaacttccgagttatt
tgcaaatggatgcgaatggcaggtgttgaccatattcacgcaggtacagttgtaggtaaa
cttgaaggagatcctttaatgattaaaggcttctacaacactttacttgaaagtgaaaca
gatatcaatctacctcaaggtctattctttgctcaaaactgggcatctctacgtaaagtt
gtaccagtagcatctggtggtattcatgctggtcaaatgcaccaacttcttgattactta
ggcgatgatgtagttcttcagtttggtggtggtacaattggacatcctgatggtatccaa
gcaggtgcgactgcaaatagagtagcattagagtccatggttatggcaagaaatgaaggt
cgtaattatgtagcagaaggtccacaaatcttaagagacgctgctaaaacttgtggacct
ctacaaacagctttagacctatggaaagatattagttttaactatacttccacagacaca
gctgatttcgttgagactccaacagcaaacatctagtttaatgactacttgctaatgctt
aatttggcaaattgtaagtagaattgacttataacaataaggagcatagaatagtgagag
taacacaagggaccttttccttccttccagacctaactgatgaacaaattaataagcaac
ttgcttatatcgtttctaaagggttttcagcaaatgttgagtatactgacgatcctcatc
caagaaactcatattgggaattatggggattacctttatttgatgtaaaagatgcatctg
ctgttatgtatgagattagttcatgcagaaaagcaaaacctaattattatattaaagtta
atgcttttgataatactcgaggtattgaaagttgtgtactatcttttattgtaaatagac
ctattaacgagccagggttcttattacaacgccaagactttgaaggtagaactatgaaat
atagtttacatagttatgctactgaaaagcctgaaggagctagatattaatttcaattaa
tatctaaaacttatgagttagttacccctctttaatatcttaattactaaattaattttg
atttaaagaggggtaaatttcgaaataaattctagttaaattagaattaacaatccaaa
atacacagaaataatgcaatcacaagacataatctccaacgatactcttgttaacttaca
agaagaatatgatagaacacaaatccaggaagttcttaatgaattaaatcaagaacttat

aggattagttcctgtaaaaactagaattcgcgaaattgcagctttattactgattgatag
attgcgcagaaaactagaactagtatctggtaatccaggattacatatgtcatttacagg
tagtcctggaactggcaaaacaacagtagccatgaaaatggcagatattttacatagact
tggctatataaaaaaggacatttattgacggttacacgagatgatcttgtaggccaata
tattggccatactgctcccaaaactaaagaagttctaaagcaggccatgggaggagtttt
gtttattgatgaagcctattatttatataaagcagataatgaaagagattatggctctga
agcaattgaaattttactacaagtaatggaaaatcagcgaaatgacttggttgtcatctt
tgccgggtacaaagatagaatggaaaaattctatgaatcaaatccgggactttcttctag
agtagctaatcatgtcgactttccagattatacttcggacgagttattacaaatagctaa
aatgatgatagaagaacaacaatattgttttacagaagaggcagataaaactctgctaga
atatactgagcgaaggatgaaacagccatattttgctaatgcgagaagtattcgtaatgc
tattgatagagctagaatgaggcaagccaatagaattttgcaagtggagaaaaagtttt
gacaaaagctgatttagtaacaattgaagcagaggacatcttgaaaagtagattattttc
attgcccaattcttaataattaagtagcttaatacaacgagatatcataaaaaaactttc
accataagttgcaaatttctttaaaaacctatactattaacatagtaaaggtg-tggcgg
catagccaagtggtaaggcagaggattgcaaatcctttatccccagttcaaatctgggt
gccgcctagtactaaaaa--gggggtgtggtggaatggtagacacaacagacttaaaatc
tgttga-ttttaataatcgtgagggttcaagtccctccacccccatgtattaacttataa
aaaaaca---atgtgtatttgtattaattgtaaccatattactgaatgtgatacttatca
tttaatagaatctcagcacaaacagctgcatcttactgaaaccccagctttcatacctaa
atgccctgtagtttgtgttaatatatctagtaatagcagccatactcaaatcgattggga
tttagtggaatgcttatcatttgtggaaaaaccaaatagttggaatttaggctgtgatta
gttgtaaaaagaattatgtggtatacgcaataactctatcttttaaatactctgtattaa
ttttagatgttcttaccaacaatatttttcctgttcttgctatctctattataccaaatt
tcgttaataattgttcgatggcaacaatctttcctggatcaccagtaacctctataatta
aaagatcttccgcgatatctactatcttagctctaaaaattttcacaatttctaaagctt
ctgttctggtctgagaattaatcctaatcttaattaacattaattctctttcaactgaag
gaatatttgttacatcttgcacgttcagaatatttactagtttatataattgtttagtga
gttgttcaatagttctattgtctccctgaactaccatcgtaattctagaaactccgattt
gttctgctggtccaactgctaaacttgcaatattgaagcctcgacgtgcaaatagaccag
atattcttgaaagtactcctgcttcatcttgaactaaaactgataaggtatgtttcatga
aattttaatgttatactgcttgatctattacgtatattctatgaaaattacagaaaaaat
gctagttactatataactaaattattccgactaaacaaaatcccaccatcagacggcata
tctattttgtaatgttatac-----------tatttaatgatattctaattctaaataga
atgctaaatttttttaaaatttgttaggaacttgtgctagaaaaatacaaaaacagtaaga
aactctctcgagatctaccccaaatcaacgatcgtattagatttccaaaagttcgagtaa

ttgacgatgaaggtgaacaattaggtatttttgtacctgaagaagctatccaattagctg
cccaacaaggattagatttagttgttgttagtgacaagtctgatccaccagtttgccgaa
tattggactatggtaaatataagtttacacaagaaaaaagagctagagaagccaaaaaaa
aacaacataacagtagtgttaaagaagtcaagatgagatacaagatagaagagcatgatt
ataaagttagaattaatcaagcatctaagttcatccaagcaggagataaagtaaaagtca
ctataacattccgtggtcgcgaaattcagcattctaatttagctatagatttgttaaaca
aaatggcaagcgacttaacttcgatatctgaaattcagcaagctccatcacgagatggaa
gaaatgttataatgcttttatctcccaagaaagcgagtt-aaatgattataatttaaatg
catctggctggattcgaaccagcgacgtctctttcgaaatggcggattatgagtccgctg
ccttcggcccctcggccacagatgcatcacttaaagtttataactcataaagtatataaa
aatcaatgcttgtttctttttcatgacaccta-aaaaaattattattcattcatattatg
ataggtcgcaactgctgaaggggaaacttggtttagatacctaaatatccagtatttaaa
tactgtgtccagtataactggaaaagttgaaataaatagaaaaataaagtcacgactttc
aggtagtcctaaatgccttaaaattacttcaataataacttcccagccatgaggagaatg
aaaaccaacaaacatgtcagtaaaaagaataattagaaaagctttcgcagtatcactaag
gccataaataatttcatttaaaaaagatttaacaattgcaatttgcctttgtccagttat
cattaataagataaaaacaaatattgatacaatatcggacaaaatatttttttacagcgtt
tgcactttcatttgcataatattctcctaattctctagccttcagctgtactcttttttc
tataatttcataagatatatcttctgatgggttcaataagacttcaaagtgaatttttttc
ttcaaacctttgtagttcagcgaaagctctctcttcttgagatgaatttaaaaaaatttt
aggttgttcctggttccataaataatcaatacaaggaccaaacacaaaaaatttttgaagc
ttggttcactaatactggagagataaataataataaaatatattttacagaagtaatagt
ttgatgcctagatattctaaattcttctatggcttcagattctccattcggatctaactc
ttttcgaaacttttcaaaagtattagtaatagacctaggtattggcccaactttttcaaa
ggcaaattgattatttcgttttaaattccaatatttcatttttttattgctgctaaaattt
ctattaatcaataattgatcacgtattcactataaatattaattctagaactaaacatat
aaaacaatagttatgaataataaaatacaattaaactttcattatgcattattactatct
tttactacagcaatgtatggatataaatcttatcgtgcaactatgaatacactgttaatc
aaatgtgaagatagccattcattattgtctggtaaacatactgcctttattcctataatt
atattaaaaaaaaatctttaaactccctaaaatcagaacctcacaatactgagttgtgtc
ttaatggaatgggaaatagaaaattcattgacaaaattgcaagtttgctaagccgtttag
aaaaacagatgcactatactgaacatcttaatcaaataaaaactagtgggtacttttctt
tactttatattaacagtaaagcaatctatacaaaaaaaaattagtaatatttttctattac
ctaatgatgttttaaaaagaatttatattcataataagaaaaaaaagcttatacctcata
cattcttaataaaattatttcaaaaacaaataggttacccgaaaaattttgcgaacttaa
acctagccttatcaaaaattatgcaatggtatgctgatagaggatatcaatggtctttag

tggagatcaagcaggctgctgatccatcgtatctcattctgaatatccatgaaggattag
ttaagacaattacaacagagtattatactttgtcttataaaaaattacgcagtatatcaa
gtacggaatcaatagagaagtatcttggagtaacagtaggacttcctttaaatattaatg
ttttgcaaaaaaaaattaactatttaaaagataatcaattagttggcaacattatttata
gtattgagagatcgagcaataactctatgaatttagatatcaaatttcaaatacaagaac
taaaagataaagatttactagtacttgtagaaagatcatctaaaacatctcaaattattt
ctcacgtatatgacttaatagatttagaatgc---agttcctctaattggctagcagctt
ccaatataatgtcactatcgactagcaaaatacaagcatgctattttgatagtaaaatcg
attctgaatacgtctatacagatgctgttaatttaataaagcggttaacttattccattt
ctcatcaaaaaacactaataattgacgattttaatttacaaaatttactaagctggacaa
aaagaaatactattggattccaattgtatttacggaatttggatcatgccaaatcttttt
gtctattcagtctaaaatgcatccagagtggacttaatataaaaatttcgtatctaaatc
cttcattaagcattagtcgtaatttgagttttcaaattgcagtacaaataattaaacaat
atcaaacaagtaaacaatcagctctatctt------------------------------
------tgtttttaaaaagggagatttagcacaattcatttttgaaagtatatgttcat
atcattttacatcttatttttcgatatctgaaaagatattactttctcgagttgtgtata
tagattccatattttacaattctgaaaatataggtctttataataataataaaaatcaca
ttataagtagctatgatatctttagacacaataagaaattattatatcaagaatttctga
ccttattgttaagtatacgttatcaaaatttcaattatttagactggcctttaaaaggct
atttatttgaaatgaagtctttatattttaccccatttcagaaaagcaatttttttgaacc
actcttgtcttgacaactacaaaaacctatttttttcataaaatttacttaaaacaaatct
ctcattttagtttaccactgtatttcagagcccatcttaaccatattttagttaatacta
ttaaatgtcagtcgaatttaaatatggaaactatatatctattattaagcgattatccta
ttaaatacactctgcttaagtctattgttactttttctataaaagttaggatggaatatt
taatacctataagccggagcatcagagtatctttgttttataattatctagactgttttt
taatgagacaatctaaacacttagcatatatactggaagatctgtcaggccataaacaat
tccaaagtttttttattggaaaaattttcatggggatttggaattcagcttaaattaccaa
ttaagcagatgccacccctttcaattgaatatactgtaaacagtggtcgccacttttgtc
tttacctgcatatttattatcaaagataattattattaaaatgaaaaatttaatttcttt
ttttgaccgaagcaaaggaaaatggatttcacagcgaacaacctatgaactacgtaataa
gaatatgagttcagtacaatctcaaatgcgaatacaatcaggtaatttgttatcaggatc
tacgcgacttgcatcattgaattggggagaaatttttagacaggttaaccataattctta
ttctcaagatacaagcaaatataatggccagtttaacttagaatttagtaatcatctaaa
taatcgtaggctattgacatcatgcacagtaacagatcctagtctagttaattttaaaac
tcgatatggtagtacaactatagatgaaacatattggtttgcaacaagtaatttacgtct
aagtactagcattattaaaaaatttgatacttgtgttgcagtatctttttgttcagaaat

caaagtttaa-attctgtataaaaacaactaatagctaaaa-agttattagttgctttta
tatatctaatttttgttttagagagtatataaaatattatattattctaaatctttacgg
ttaggatttctagaaggatcattagataagaatccaaaaacaaacagagaaataaagaaa
atgacagtagtataaacaaatattttcaaagtaaacatatttttatccttaaaagctatt
agtgtg----atatttatatatc-acacataattttatatcacgatataagttctggcgc
tattttataaacaaatcaatatctttatttttagtaatgatttatacttcaatatgcgt
gatgcactaaatactaaatctttttgtacacgcttctcactatttttaatattttgtga
tttctatgaagtttaccttcaattactacataatctaatttcttcagcaatttaaaagtt
tctaaagactttttattccaaataattaagttaattaccgctaatctttttctttgtaaa
agcctaactttcaacttaattatttggtgtctactatcactaattttaatactcttacac
ctcaaaatttgaactagtaaggtacagctattcat-attaat-ttaa------tcttgta
actaatctattatgacagataa-atgaaaaaatcaaaatgaaatttcattatcttgttga
ccttgttcataactaatatcttttaatacttttaatattttcccgaaatactcttgcaaa
tatttttctaacgatattacttctgagctatcaatgcttaaaatactataaacttcattc
atcgaagcagtaaattgctccccactagttaaaatctcggcaaatgctaaacgatcagaa
atattccatgtccattgaagggttttagtgatttttcttaaagctgttaatagtccaatt
ggaatttgtgatatttgtgtttttttgcccagataacttttcacataatttaataatttca
gctgatgtccaggccgtgttaccaactaatggcagcactctgttttctgtagaaggcact
ccaagactttttataactagctttgccgcatcctgagtatcaatatatgcaataggtgta
gactcccctgtaacccatactgatttttttatccaaaataggaattgcatattggttaatt
aatccttgaaaaaaccctcccaaagaaaataccgtgtactttatatcagattttttgaaga
aagtctactacttgtgattttaagttcatcaatggaacagctgggtacttatctgcattc
aatattgaaaagaaaataaatctttcaactttagcagctttagcggcctcaattaaggca
attttaccatctaaatctatttttgctgcattgtaaggatctgtaggacgagatgtagac
gcatctatgactgctgttactccacaaaaagattgcaaaatactttcaggtaattttaaa
tcaccatatacaagttctgcgccccattctttgagaaaagctgactttcgtaagtttctc
accatgcatttcacattgtagccttcatcaagagcacgccttacaatttgacgtcctaga
gttcccgtcgctccaattactagaagagtcatattaattaaaaataagtatgtataaaga
cattggtttggtcgacagcagagattaaattaggtagagagggaatcgaaccctcatgac
cgaagtcgtcacattttgagtgtgatgcgtataccaatttcgccatctaccctattatag
accaactactacaagtaatttcgtctttagcgtaaaaataatactaagtttcaatagata
tgtcaacttttgagtttatattaataaaaaaccggagtatcttatgtccacttttgaact
aattccttatagattatatttaagtcaatctagtacgtggatgcataggatgaaagcaga
aattaaaatatatatattaacgatattatggatttcaattttttattttctcttactgtaa
actatgcattgttgccctaagtttaattgttcttagccttactgtaaagaataaacaaaa
cattatccagagacatttattgcgaactttagttatgacaatggtaactaccttcttatc

attcagcatagctactaattacaaacaatattctcaaccaaaacaatcacaagatttatc
ttactataatacagctaaaaattcttacatctatcaaaatatattgctacctaatagtcg
taagatcacaacaaaacaactgaggtgcactttaaagcctgcactctattttttttattac
tatatattcaattaaactagtcatgataacgacttccccagaaattttagtgattactgc
ttatagatctaggatctttaatacaatattaaatcgtgaattattatttactttcttatt
gtcatcacatattgttaccaatataattagtaaattagataaacttgttcaagtagctaa
cttaagaggtagcttaaatttatataaatcgtccacaagacttttttatattttcttttttt
gatgtttcaaatcttttttttagagattattaaggaatcaaaagaaatatctcaagctct
ttatactagaaatcttaatcaagaaaacagtaatttttttaaaaatatatagagccagatc
gggtattagtgattggttaaatgcaattattgggaccttatattttattattttaggatt
agtataaca-aatctaatcatccttctgactctggtaatacgaatagtaatagacttaag
tcgtggtgtaaatatgaaaaat-tcagatattatataaataattaaatgttttattatgg
tcaatactcaaaatcaaatttcacaaacttcagagcttgattatatagttaatcaacctt
ataaatatggatttaaaacttctgttgaatcagaacaatttcctagaggaatagatgaaa
atatagtaagattaatttctaagaaaaaaaatgagcctgaatatttattaaactttaggc
ttaaagcatacaaaaaatggaaaaaaatgagtagtccatcatgggccaatctcaagaatc
ctaatatagattttaacagtattatttattacgctgttccgaagttaaagaaggaattaa
acagcttagatgaggttgacccagaaattcttgatacttttaataaattgggcatctctc
tcaatgaacaaaaacgaatttcaaatgtagctgttgacgctgtttttgacagtgtttcta
ttgcaactacttttaagaaagaattatctgcagctggtgttatattctgctcgatttccg
aagctatacgagattatccgaatttaattcataagtatttaggtactgttgtgcccgctg
gggacaattactttgctgcattaaattctgcagtatttagtgatggttctttttgctata
ttcctcctaatactgtttgtcctctagaattatcaacttattttcgtattaataacgaag
aatctggtcaatttgaaagaacactcattgtagctgatcgtggtagtaaagtaagctatt
tagaaggttgtactgcccctcaatttgatacaaaccaattgcatgcagcaattgtagaat
tagtagctcttgatggcgctgaaattaaatactctactgtacaaaattggtatgctggca
atcaagagggcaaaggtggtatatacaatttcgttactaagcgaggcttatgctctggca
ataattctaaaatttcatggacacaagtagaaactgggtcagcaattacctggaaatacc
ccagttgtattttagctggccatagttctcaaggagaattttattctgtagcgttaacaa
ataattatcaagaagctgatactgggactaaaatgattcatattggtaataatactaaaa
gtagaatcatctctaaagggatttctgctggtaaatcaaaaaacagttacagaggactag
taaaaattggcgctcaatcatttaattctcgcaattattctcaatgtgattctttattaa
taggtcaatcgtctcaagctaatacttttccttatattcaagtacagaatccgacatcga
aagtagaacatgaagcatctacttcaaaaattagcgaagatcagatttttttactttttac
aaagaggtatcagtttagaagaatctgtagctcttatgatcagcgggttttgtaaagatg
tattcaatgaacttcctatggagtttgcagctgaagccgatcgtttactgagtttaaaat

tagaaggaactgtaggatgagtcaaactattttagaaattaaagatttgtatgcgtctat
tggtgagacaacaattttaaaggggtaaacttatctatacaagaaggcgaaatccatgc
gattatgggacctaatggttcgggtaaaagtacactatcaaaagtaattgcagggcatcc
agcttactctttaatcagtggagacatcttatttttggaaaaagtattcttgaaatgga
gccagatgagagagcaagagctggtatttttcttgcttttcagtatcctgttgaaattcc
aggagtaagcaatgccgatttttttacgcattgcattaaatgctcgtagaaaatttcaagg
attatcagaatttagccccttagaatttttcaattaataaatgaaaaaataaaccttgt
tggcatgcaagaaagtttcttgacaagaaatgtcaatgaaggattttctggcggagaaaa
aaagcgtaacgaaattcttcaaatggctttgctagatagtaacttatctattttagatga
aacagattctggattagatatagatgcacttagagtagtagcaaaaggaattaatacttt
agcaaaatcaacaaattcaattattttaattactcattatcaaagattattggattatat
tattccagattttgttcatattatgagtaacggaaaaattgtcaaaactggcaatcttac
tcttgcacaagatctagaaaaaaaatgggtatgactggattgtacagagttaaaaatttgt
taaataaggcaatactgagactaaagtatcttctttagtctcaatattattgatcaacag
aaag-----ttatttaggctttactaaaaccttgcttaacatcttcaatggctcttttta
acagttcttcactagatgcgtctaatttttttactgctacgaatactttcaccaaattctg
gtttagaattctttaaatcttcccgtaattctttaataaattcagccacttttgatactt
caatatcatctaaataaccgttaattccagtataaataatagctgtttgttcttcaactg
gaataggagaattttgagcctgcttcaaaatttctctaagacgttgacctcttgcaagct
gattttgagttgctttgtctaaatcagatgcgaactgagaaaaagcttctaattcagcaa
attgtgccaactccaattttaatttgcctgccacttgtttcatcgcttttatttgagcag
cagatcctactctagagacagaaataccgacattaatagctggtctaatacctgagttaa
ataaatcacccgataaaaaaaatttgaccatctgtaattgaaattacattcgttggaatat
atgcagacacatctcccgcttgagtttcaataataggaagagcagtcatactaccgcctc
ctagttctgaattcaatttagctgctctttctagcaatctagaatgtaaatagaatacat
cacctggatatgcttcacgtccaggcggtcttcttagtaaaagagacatttgacgatagg
cttgggcttgctttgttaaatcatcataaataaccagggttgctttacctttatacataa
aatattcggctaaagctgcccccgtataaggagcaatatattgtaaagttgcaggactat
ctgcattagctgcgactataatagtataatcaagagctcctttatcttgtagtgaagata
caacttgtgccactgaagatgccttttgtcctatcgcaacatatacacaaacgacatctt
gcccttttttgattaataattgtatccagcgcaacagctgttttacctgtttgacggtcac
caataattaattctcgttgacctcgaccaattgggatcatagaatcaatagctgtgattc
ctgtttgcattggttcacaaactgattgacgtccaattataccaggagccatagactcaa
ttaatcttgttgaattacttgcaggttcacctttatcatctattggacgagctaagggat
caactactctacctaaaaaggcatcaccaactggaatttgagcaattctaccagtccctt
ttacagaactaccctctagaatgtctctaccgtctcccattagtactacacccacattgt

cactttccaaattcaaggcaacaccaatggttttatcttcaaattcaagcaattcaccag
ccataacttcatcgagcccgtacactcgagcaataccatcaccaacttgtaaaactgttc
ctatattggctacttctatatcttgatcatacttttcaatttgttgacgaataatactac
ttatttcatcaggtctaatgtttaccatgttttaatattctttttattaatgggaaatt
ctaaagtggtaattactcagattaatactatctg--tttttactgtattttaagttgct
actacatcaaggtgtgaagccatctgccttagttgccctctaatactagtatcaatgact
ttcgatccaatctgaatagtgaaaccaccaattaactctggatcgacagaaacaattagt
ttaacttctttagcctgggtcataaccttaatttttctgttagcagattttcttggtca
gagtttaaagcaatagaagtattaatattggcaattgttagagattccatttgatatgca
agttctaaatacttactagctatagcatccaacattgcaattcttttacgatcaactaaa
atcattagaaatgatagtgtattttcagtaatttgatcaccgaaagtggcaataatgact
tgctttttgcttctgcagtttttaatggattagct---aaaaaattttaagcttgtca
gattgcaataaaatattttgtattaactttatatctatactcaccttttctgtaactttt
tttgttttagctaattcaagaagagctacagcataaggctgcgcgattttataacaacg
ttattactgctcatagttggtctcctaacttagcaatattattatcaatgatacgtagtt
gcatttcagagctcatttgattttctagctgcaaagtaactcgtttaagagcaagaaatg
ttatttgctgctggatttgccttcgaatttgtttttcagctgtttcaatatttgacttgc
ctgtcatagcaagtctttcaatatctaattgaccttgagctaatatagagcttctaactt
ttcctgccgttagctgagcttctttttgatttgatctataataatttgagtctgagcaa
gttgtttttctgattctgacaatctagcactagcttgctctagtctttcttctgattctt
gtatagctgctaagacttttatctgtcttgcatttaaactagatccaagaaattgtttca
aaacataaataagaccaaataaaagcaacaatatgttaataacatttgcttcaaaaatat
cagaattaaaaccgaatttatgttcactactatgttctgataaaatagtaattatttgtg
gtatttttactatattattcatttgaaattct----------tactattttaatatttta
cttaggaatcatgattttaaagattgactactcaatagtttagctttaatttggtcgctt
aaagtatccacctgatcttccaaagttttaagagcctcttctttttgaatatttagctgt
tttgatgcttcagcaatcaatttttcagcattcaattgagctttttttaatatcttcagag
actatactttgcgcttctttttgagaagcagctatctttaattgtgcattacgacgagct
tcagataaatcttgctcatattttgcagctaactcatctgctttaacaagcatagaagat
gctgtagtcaaagttgtgcggatatattcatcacgttcatctagtactctagttactggt
ttataaaaaatagtattcagtaaaaccattaacgtaagaaactgtaatgccattaaaggc
aaagtaccattgaagtcaaataagccaccttcaatttcttccgctaatagaaatggtaaa
taaatcatgttactaattatgttgatagttaataatttggtttcaattaagaaaattaac
tttatagtgcgcttagtcttattagactaagcgcaaatgattattttaaccaacgtatgg
attagcaaataatagagacagcgcaacaactagaccataaattgttaaagattccataaa
tgctaaactcagaagcagcgtgcctctaatttttccctctacttctggttgtctagcaat

accttccacagcattagctgctgcactaccttgaccaataccaggacctattgcagcaag
acctaccgcaagaccggcagcaataacagatgcagctgaaacgattgaatccataataaa
ttttgatttttttgttatataagtagaaaattgacaaatttcggataatacttaaaatcta
accaatgaaatattggagattcaaatat-----tgtaattatagtatactactcttctcc
atgaccttccattgcttccccaatataagctgcggataaggtagaaaaaattagtgcttg
gattgaactagcgaataagcctagtatcataactggcaacggtatcaaaattggaattaa
cagagtaaatacagaaacaactaattcatctgctaagacatttccaaatagacgaaaact
taaagataatggtttcgtaaagtcttctaaaatattgataggcaataatactggagtagg
ttgaatgtacctagcaaagtaacctaaaccttttttacttaagcctgcataaaagtaagc
tagagaagttagtaaagataaagctacagttgtattaatatcatttgttggagcagctag
ctcaccttccggaagttgaattaacttccaagggatcaatgcacctgcccaattacaacc
tagaatgaataaaaacagggtagcaatatatggaacccaagggcgatattcatgctctcc
gatctggtttttagcaatatcttgtaagaattctaatataaactccatgaaattttgaaa
tttttcaggaattcgctgaagatttctagtacctaggaaagatatagttaatagtgtagc
aattactaaccaagatacaataaaaacctgaccatgtaattgtagactacctattttcca
atatagatgtttcccgacttctactgctgacaagctagtatatggattaaaatctatgag
actgttttgatacataataaatatttcttttaaatagtacaagaatcaaaaaataagttt
tttgaagaattaatatgcaattagtttatttgcacgattagatttttaataaattaaaag
caaaaataaattctaatagagatagctatctttagtagtgattatttttttgtaaaaaaca
attattgctacctataatgcttttatgtttaattatatacccatatatgaattatttaat
atgattataggcatttcgctacaaactttattgttttatccatttt----aaaaattctt
ctaatttacttattatttaaaatatctgcaacttcatccgcaaagttagctttcatattt
tcttcgccttctcccaatacaaatctaacaaatcttctaattttaatattttcgcctaaa
acggaaatattttgattaattaaatcttgaatactaatatcttgattgcgaatgaacatt
tgatctaataacgaaagttctttcagcctctttttcattcgcccttcgataattgctcct
attctatcagctggcttgtttttcaaatcatctttaccagcttcaactctttttttctgaa
ctgataacttcatctggtatatgctgtgttgatacatatgccacatttggcgatgctgct
atttgcatagcgatgtctttagctaattttgaaattctggacgtcgggccacaaaatca
gtttcacagttcacttcaactaatacaccgattcttcctcctgtgtgaatatagctttct
aatagaccttcaagtgcagtacgattagatttttttgttagctgaagctaatccttttttgt
cttaatgattctaaagctttttcttcattaccattactagcttctaaagctttttttgcag
tccatcatacctgccccagtcttatcacgtaaggcttttacagtttgagctgaaatttgt
aatgtcatatattttcaaaaatttttaagtatttaaattaactaaaacaaattattataa
ttcactttccacagagcgagctacttcctgtacatctgtttgcccatatttaccatcata
aattgcatctgctatcttaccgacaattagtttaattgatctaatagcatcgtcgttagc
tggaattggaatattaataatttcgggactgcagttcgtgtcaagcatgcaaatagttgg

gataccctaatttcaaacattcttgtatagctgttgtttctctcttttgatctacaactac
aacaatatcaggcaaacgagtcatattttaataccatttaaatgtttacgaagtttatc
taattctctccgtagaacagcagcttctttttaggcagttgatcaataataccactttt
gtcttgctcttctaactgtctaaggcggctgactcgggatttgattgtcacccaattagt
taacattcctcctaaccatctttggttaacataataagaatcacatctttgtgcttcctg
agcaactattcctgcggcttgtctttttgttcctaaaaataaaactttcttgccatcgga
cgaagcctttttataaattcacaagcttctgtaagtaattgagcagtctgcactagatc
aataatatgtatgccatttctttctgtataaatatacggaaacattttaggattccatct
gcgagcttgatgtccaaaatgcacgcctgcttctaataactctgctaaggtaacgacagc
cataggattttaata-ttaaataatgtaatttttcgggaaatccctttttc----aaac
taattcaactcatttaacttattgcaattccaataaaccaataataaattaacatatttt
tacagtcgattaag-aa--aaaattgttttactttattcaatctacacttttattgctaa
agtagttacgcgcggttctgtcgtctaaaataatatcatctaaatcatctctcacagacg
acacactactttcgctactgatatctggattccaatctttttttctaacgacgctctat
tatgattgtcatacgtattaaaaccagtgccagctggtataagtctaccgataataacat
tttctttcaaacctcttaaccaatccagtttaccagatatagcagcttcagttaatactt
tcgtcgtttcttgaaaacttgctgcagaaataaaactttctgtattgagagatgcttgcg
taatacctaacagcacgggtcgataggaagcattcagcttatttcttaaagtcatagctt
tattggtttgctcaatcttttgcaactctacaagttccccaggcaaatagccggtttctt
caccattctctattttcactttagaagtcatctgtcggactatgacttctacatgtttat
cagaaatattaactccttgagattgataaactaattgtacttctttaacaagtaataact
gtatttcttgaaaacttaatctagctgcttcatataaagctagtccgcattcaatatata
ggttgaagcttgcatcaagaatatcatgaggatttaataatacatctgtttttttttcgcg
cttcaagaatttcttcaattcttggaaggccttgtataatatcaccagtttttgctctgt
caaatacgagaatagctagtgtttctcctcttctaattaatgcattattatcaacatgta
aaattgccccgctagaaactaggtaaggccgagcaattcttaatgttattgattgagaag
acacgtgagtaatttgtccagagtctaaagaaaccatattttcagcaatcaaatcaccgc
aacgaatccaatcaccaacacatactttagttttagctttttctatattaaaagttttct
tgtcagaagatgttccaattagaattcgtctactagtattcttttcagaataaatttctt
cgaccataccttcactcattgcaataatttctgtactagcaacaaccgtcaatggcttga
tatgttcaccatttttgacaataattctagtttgactttgttgtttttctgatttattga
catcaatacttttaattgataatgtttcaaaagttgaaagcctcaaatcaaaataattag
tatcactttctttattagaaaactcaattgaggatattaaactatcatctgtatgctcaa
tttcagctacaagattggttgtaattaaatgaactccatctatcgatttcaccctttctc
cgtcacgaaaaggtgtccttttaactagctttaatttagtttggctcggtttttgtaatg
ccaaattatcaattaagctagattttatttccggtattgaataaacaataacaggtctaa

tgaggatataaggcatttgctcattttctatatactcccaataaactaacttatctgttg
aaatgttgttatgcaatgtttctccaggtcttaaaaacccccctactttttttcattgttac
tatgtaattcgtttagtttatatatagatccaggcttaataatgatttcacgaactattc
catccttctgaattatctcaacaattccagagcttttagaaaaaatatttttgactagct
cagttccactttctataatatccccattataaactaataatagtgatgcatctttattaa
tttcgtgtgtttcctctgagatccataatatatagccaggactcaaaatttcataagcat
ctttttctgaacccactttctttttttgatacattcaagtctaaatatttaataataccac
cacttgtagttttgtaagtatcagatattagctcagcgactgtatagccatcgtaaa---
ttttttgattggggacagctttaaaaataaatttctgtttttttttctgtttctaaaatgt
aagattcttttttattaattaaatctgtataaatataacaaccgggaattacaatagact
cagtaataacttgtatattagtaatgctgctattagcattctgctgtatcctgacttccc
cagcatagtgatttatcaactctgtctgcgcaagtattgtgccagctttagttgtatcct
ctttattaacaatcacattagccgagtcagatatattataaacttctccagagagaaccc
aaatcaacccacctgttttagtgattcgtgtagtatattgattattatctgtttcttcaa
ctgttaagttagagaaacagattttaccagaaaggtctgacactacatgtttttgagccc
tttctgtcatcagtctatttctgcgaggtgattcagcaatcacctgatcttttgatactg
attgaccctcatgtagcagaagagtggttcctttacttaagttaattattgacttttgat
ttcttctagatcgaattgtaacttgggttggattctttgtaattaaggcttgttctccat
gccttgtacgaacatctttatatgattcaatatctagatctgttagctgaccatctatgg
gtgcgtaaatttgctctgccaattcaccagtaaatacacccccccgtatgaaatgttctca
ttgttaactgggtgccaggctccccaatagattgagcagcaataatgccaacagcttctc
ctaaatcaactaagcggccatgagcgagattccatccgtaacaatactgacaaacagaac
ttctcgaattacatgtaacaggtgatctcaccaaaactttttt---gatgccagttttag
ttatctctttggctaatttagggctaatatcttgctttgtgtgagcaatcaaaacatttg
tatctggatggaacacactctctgctaatactcggccagacaaagcttgctctagattga
ttagtatttttttgtgtatctactaagtcttctagtattatacctttatttgttccacagt
ctacttctctaattataacgtcttgagaaacgtcaacaagtcgacgcgttaagtagcccg
aatctgctgttctcaaagcagtatcaactaatcctttttcttgctccgtatgatgatataa
agtagtccgtaacagttaatccttctctaaaattacttgatattggaaggtcaataattt
gtccttgcggatcagccataagccctctcataccaactaattgtctaacttgagaaatat
tgcctctagctccagagaaagccatcatataaacagcatttaaagggtctgtttctttaa
agtacttaattacttcctgttttaatgattcgcttgcattgttccatgtgtcaataactt
tttgaaatcgttcgaccgctgtaatttcccctcggcggtacttattttcggtagctttaa
tttcttcaattgtacctaataaaagactttgtttgctcggaggaatccttaaatcttcca
aactcaaagagattcctgcctgcgtagcgtaatgaaatccaagatctttaagtttatcag
ccatattagctgctcgtgctataccgtaattacgaaaagcccaaacaattaaatttttaa

gttcgttttatcaataaccctatttgaaaaacttggctgcgcaagacttcttttactat
ccactcagtgttctccatattaattatttatt----taactacgccactaaagactcttg
aataatttgttgaagattattcggccagctgttgttcgaatatattgcacaattctctg
ctgttcagcatcctcttttataatatggttattataaaattttgttctactattatcact
gtgcttttcgattttaattagctcagtagtttggtcaccatctacaaatccatcaaaacg
agcccatatataagagtgcaaatcaactttcttttgctcataagccatcactacatcttc
taagctagcaaaatattgattcgctcccttctgttcagaaggattattagccgtcaaata
ataacaacccagaaccatatcttgactaggcatgataattggctgacccgtagctggaga
aagaaaattatgaggagccaacattagcaatctagcttcggcttgtgcttctaaagatag
aggaacatgaacagccatttgatcaccatcaaaatcagcgttaaatgcaggacagactaa
gggatgcagcttaatagctcttccttctacaaggattggttcgaatgcttgaatacctaa
tctgtgtagtgttggagctctatttaaaagtacagggtgaccttgaataacttcatttaa
cacattccaaatagaaggttcatttttttgaatcatttttttttgcagctttaatattatt
cactaagccctgtaagattaatctatgaattacaaagggttgaaataattcaagagccat
ttctcgcggcaatccacattgatgaagttttaaatgagggcctacgacaattacggatct
tccagaataatccactcttttacctaataagttttgtctaaagcgtccttgtttcccttc
tataatatctgataaggattttagcggtctattattcgcacctacaacagttctgcctct
acgtccattatccatcagagaatcaacagcctcttgtagcatcctttttttcatttctaat
aatgatttctggagctaaaattgacttcagacgcgataatctattgtttctgttaatgat
tctacgataaaactcatttaaatctgccgttgcgaatctccctccatccaattggaccat
aggtcgtaaatctggaggtataacagggattactgtaaataccatccaagaggatctgc
gcctgttgctataaagttctcaatcaatcttaagcgtttcatttttttattaaattttaa
tgaaggtgtcttaaaactttttggtggatttgttgcttcagatcgtaaagtttcagcaat
gtattctaaatctaaatctttcaatagtttttgaatagcttctgcgcctatcccaacttc
tacttgattattttcattttgattttggtagatttcctcttcaaggctcttccattcata
gccttcgagcagttgcttatattttaaattgatatttgtattagattgtgtaactacata
agagtgaaaataaacgatttttttctacctctttaacttttaaatctaatgctaaagcgat
ataacttgtacttccttttaaataccaaacatgagttactggtgatgctaattcaatata
agccatccgatgtcttctcactcgagattctgttacttctactccacatcgctcgcagac
tatacctttatagcgaaagcgtttatatttaccacagtgacattcccaatccttaactgg
gccaaaaatttttttcgcagaataaaccatccatttcaggttttaaagttctataattaat
tgtttctggctttgtaatttcaccaacaatttgtccgttgggtaaacttctttcacccca
atgtcgaattttctcgggagaagctaaactaatctttacatagtcaaaatgttgctcaaa
ctgtgtcataaatttgaataccttgaaattaaataattaaccaacttatttacatatgcc
atattcgaaaatgca----ttaataaagaaattgttcaaaatcatcaactggtggagtat
cataactagtgcgatctgctcgattctctttagtgtcagacatcaaatcgacttcaactg

tacgtctttgaccgtcttcaaatagtttcaatttatgtactgcaatgtctaatcctaatg
actgaagctctcgcattaagaccttaaaagattctggggtcccaggcttaggtataggct
ttcctttcactattgcatttaatgcttcatttctagcttgcatgtcatccgactttacag
ttagtaattcttgtagagtatatgctgctccaaaggcttctaaagcccatacttccattt
cgcctaatctttgacctccatgctgggctctacctcctaatggttgctgtgtaactaaag
agtaaggaccagtagatctcgcatgaatcttatcatctacaaggtgaacaagcttaagca
tataagctctaccaattgtaacaggattatcaaaagattctccagttctgccatcgaaga
cttgcattttaccaggatgcagatcattaaaaagccatttattattagtgagtaatgatg
cttcttttaattttctattaacaagtgctcttgaggcttctgctccatacatctcgtcaa
aaggtattattttaaatcgtttacctaaatagccacctgctagacctagtaagcactcaa
atacttggccaacattcattctggaaggtactcctaaaggatttaaaacaatatctacgg
gggtaccatcagataagtaaggcatgtcttgtttaggtaagattctagaaataatcccct
tattaccatgacgaccagccattttatcgcccacttgaattttctcttctgggcaacat
aaacgcgaatcatagcattcgtaccaggagggagttcatctccttttgcctagtaaata
ctctaacattaacgacccttcctttagccgcattaggcaatcttaaagatgtatctctaa
catctctagccttttctccaaaaatagcccttagtaatttaccttcaggtaattgatcag
cttcgcctttaggagtaattttccctactaagatatctcctgcttcaacccaagagccac
ctaccacaattccatttcgatctaaatcttttaaagagttatcacttacatttgggatct
ctctagtaatttcttctggtcctaatttagtttggcgacattctacttcatacttctcaa
tatgaatagaagtgtataaatcatcataaactagcctttcactaattaagaatgcatctt
cgtagttataaccttcccaaggcatataagcaacgagaatattccttcctagagcaattt
ctcctccatcagtggatgcaccatcagctaaagtttgaccgacaactattttctctccca
cccaaacaattggacgctgattaatacaagtatcttggttagagcgataatattttttca
aacgataatggactgttctaccattattatcttgaatacctattttattagcagaaacat
aattaacataacctgcagttctacttataacaaccatgccagagtctcgcgcaattttag
tctcaagacctgttccgatgattggcttttctggatataataatggcacagcttgtctct
gcatattagaccccattaaggctctatttgcatcatcatgctctaaaaaaggaattaatg
atgttgcagctgaaatgacttgaataggtgaaattgcaatgtaatctacttggctcggtg
ttgttgtaataaattcttgacgataacgtacaggaataatatctccttcaatataatttt
gagaattaactttaacatctcctggagctactcgaaaatcatcttcctcatctgctgtta
agtagatgggactctgttgataaattacttgaccttcatttaccggataaaaggggggttt
caataaagccaaaaatattaactctagcacaggttgccagtgagccaattaaacccgcgt
tcggaccttctggtgtttcaattgggcagatccttccatagtgacttggatgtagatctc
gcacagcaaatccagctctatctttattaaaacctcctggacctaaagcactaattcttc
ttttatgggttagttctgcaactggatttgtttgatccataaattgagaaagctgactag
atccaaaaaattctcttactgaggcaattaacggcttaggattaattaggttagataagc

ttaatgagtctatatcacatatcatcattctttctctaataatacgttctaagcgattaa
gacctactctaaattgattttgcagcaactctccaactgaacggacccttcgattaccaa
ggtgatctatatcatcaaggttacctgaattttttatctttaatattaattaaataatcaa
tcgaggataagatatcttgaggagacaacactcgaaaattcttaggtatattaagaccta
attttttgttaattttatatcgaccaacttctcctaagtcatatctctttggatcaaaaa
agcgagaataaagcatctgcttagcaacagcaactgttgctggctcattaggtctaagct
tagagtagactataagcaatgcttcttcatctgttacttcttcgacattgtttttttccaa
tttctttagctaattctttaattgaatataattgagatgcagagatgagaaaagcatatt
tactcagccccttttcaatctcgtctttattaaggccaatagcacggagaaaaatataag
catttactttatgggtcttatctattcttatccaaatttctcctttagggtcaatttcaa
atttcaaccaggagcctcgattagaaatcagactagcactgtaaatctgcttgccatttt
tatctatctcttgtttatagtaaattcctgggctacgaattatttgattaataataactc
tttctgtgccagatacaatgaaagtccctctgttagtcattattggcaagtctccaatga
agaccaatctcttcttgtatttttttatttttttatttccttctcaacactaagctgtaaat
gtgttgatgataaatctagtgacttgatacttttttccatgatctttagaaggtagatcaa
tgtcttttcttgtcaacttcgcagggacatatatttgagcactataggttctatctctat
tttttgcctgtctaacactgtaacgagggaattttattttatactcattaccaaacaatt
gcagttcaagccgactagtaggatctaatatatttggaaagaattcaagtacttcagtca
taccttctaatagaaaccatttaaaactggctctttgaatttcaactaaatctggaagaa
gtttatttttttaagcttatacgttgaaccatagatctcctttataacaaattctgcaatc
gaagagtttaa-tactatcaacagagtacttataggtaatacctgagtatagttcactac
attatttcttctatagacatgactagaacaaagtattttattcaacaaaaaagattttta
aaattt---ttcacaatcaatatagatataaaaatattacagctattaaaaatag-----
----------------gaagacgaaattaatacttgtctgtctgtcaaatagttgacatc
aagatagacaagtattatttatttaatcaattctgttttttttgagcgcaagataatgct
ctcgcgagtctagctttttttctggcaccagtattaggatgaaatgctcctttttttatt
gctttatcaatcttgctatatacttgagaaatacttaattgcacatcgttaagattagca
ctttctagattatcaatatttagcaggcatcttttagttaaagtttttacaaccgattta
tatttacggttaattagacgatttctttccgaagttttaatgcgtttaatcgcagaaagg
ttcttagccacagtaaatccaataaaacttttataaaatatttttagaaacgaaaataca
caaattagagatctattcaaaaagggaatatgcgccgtattataacattaaagaaaaaca
ataggcaatagttcactgagggctcttgttagtacaaactaaatattgtcgttgcataga
a-tttaaaaaatagtattaaat-----taaactaaccagacatatagatatatatgaaaa
aaattgaagctattattagacctttttaaacttaatgaggtaaaacttgccttagtcaggg
aaggtattggagggatgactgttgttaaggtctctggttttggaaggcaaaaaggacaaa
cagaaagatataaaggatctgaatattctatcgatattattgataaaataaaaatcgaaa

tcattattagcgatgataaagtagataaaatcacagagacaattattaaagcctcgaaaa
caggagaaattggagatggaaaaatattcattagtagcgtcgaaaaagtcataagaatta
gaactaatgatttagattccgaagctttataaaactcacactttttttgagacttgatact
tcttcacgaaacatgagataataaacgtttattgtataataataactaaggtataataat
ataatggctaaaagcaaaggtgcccgcatcgtaataactttagagtgctctaataaaact
ggagatttagcaagaagagaaaacctggcgtttttcgatatacaactactaaaaatcga
cgaaatacaccaaacagaattgaattaaaaaaattctgtcctaattgtaatcagcattgt
ctattcaaagagattaagtaattattaattttactaaaat--aatcatattatggccata
tatagaaaaagaatatctccaattaagcccaaagagtctgtagattacaaagatattgac
ttgctaagaaaattcattacagagcaaggtaaaatactacctaagcgatccactggattg
acttcaaaacaacaaaaaaaacttactaaagctattaaacaagccagaatactttcccta
ctaccgtttttaaataaagattaaatatatacaaatatgctttagattgtttcctattaa
catt-------tccatgaatttatttactttaattaccgaatatactgagc---------
----------tactataaagcagctatttactactaattttgttcattttataattaaga
aaaa--atctaacttgttgttaaaatagattt-----agacttataatcgaacctctttt
aatgaattatattacatagtttatttaaaagaggagtcagttttatttgagttaataaat
agtacttttgtattctttagaccagtataaagtatttaagataaagatttttgtttaggt
acaagatcaaacgaatgaatcttatctccgggttgccataattgaaattcagaaatgaaa
attccacattcgttaccagcttggatttcttcaacatcttctctaacccgttttaaagat
tctatctttccctcataaataacttcatcttcccgaatgactttaatccaagagtttttc
aataatttattattcgtgactctacatcctgctatttttctattagccaaagaaaataca
gtgcttacttctgcttctcctactggtacttctgaatattccggatcaagtagatctttc
attcttcctttaatatcttcaatcaatgcataaataatttgataattttcgactagtata
ttaaattttgctgcagcttgcttcgctccaggagcaaaattagtattaaagccaatgaga
ttagcatttgtcgtcgacgctaattcaacatctgtagcagtaatttcgccaggcataatt
gaaacgacatttaattgtactttgctctgaggaaattgtgataaagaatctaggattgct
tctgtagagccttgattatctgttttaataattaaagaaatttgcttgcttagatcttta
gaattagtagttctcaatatatctaacgtaatacgactatttagtgctttctgcttctgt
gcgattaatgaatttttttgaagtattttcaagagctttaagctttgcttctttatcattt
ttcactactaaagctatctctcctatgacaggaacagccgataatccccaaatctcaaca
acagatgatggtattgctgaattaatctttttctgttcattattaatgatagctcgaatc
tttgcataagctgatccaataaccatattgtcgctcgtatgcagcgtaccattttggatc
aaaagtgttgctactggcccatgggatttgtctaagtgagcctcaactataacaccttgt
gcaggctgtgttggatcagcctgtaagttttctaactcagctaataaggtaatggtttct
aataatttatcaatattgtagccggtaacagaactaataggaattacaggaacttgtcca
ccaagtgtctcagacatcacattgtgttttaataagtcctgctcaataatatttgtattt

gtgaagtctttatctatcttagatattgctaccacaaaaggaacatcagctttttgaata
tgatgaatagcctctacggtttgaggtttaataccatcgtctgcagctataataataata
gctacatctgttaatttagctcctctagatctcatacttgtaaaagcttcgtgccctgga
gtatctaaaaagataatttttgtttattagctttattgatatattcaacttcataagct
gcaatagtttgtgtaattcctccaatttctttattagcattgttagaattatgaatgtaa
tctaataaggttgttttaccatgatctacatgccccataactgtaacaatcggaggtctt
ttaacatagttaccaactttatggagagagctagaagtatctaaattagcaggatacaag
ttattatttcttttgtgtttgagtcaactcctataccaaaattatttgctactgatgct
attattgaagcatctattgtttcattaatagtaactggtattcctttaaggaataaatat
tttatcacatctgtttcttgaacacaaattagtttagataactcttgaatagtaagagga
ttggtaatactaatagattctggtggagaattagataaactgctggtttggtcttggctt
tgagttaatgatatgtttttttgtcttatagacttttagtattagtaaatttttttata
acttgagcttttggtttaggtggacgcattagagaaatagccaagtctcctgcagtctgc
ggaatcatagaattagaacctctaaagttctct---tcctcatcatcaatatgtattttg
gtctttattttt-tttcgttgtctgttctta--tttttttattatctgctagatcatga
accttattaaaattttgttttttttatctagtttgggtggagaatttaattctaaatgc
ggttcactactagctgtatgcgtatcagattgatcaagatctaagggtaaaacactgtct
ttatgggctgattctaatctaattttatatataatttggggatttttaaatctataatt
ttttcagaagc---tatatttggactggatgcaggggatctattttcgaaattttgatta
tttaaaaacatgaatattgattttatgatctaatagtaaattttaaatttcaatacatag
ctcaaagtaaagaaaaatcagatatct------ataatactaatat---agaaaattaat
ctttgttgtattagatttaatataacactaaagtcttttgttagttatacaagtttaaaa
ctttt-tgtgatgataaacaacaacttctgctttattgagatacaatcaatttatttaga
tataactgtttttcaa-aggtttttaaa----ttgtctaatattgtgttcgttatgaatt
tttattaaagtttgaaaa-ctataaaatagagtcataatcattaataatctccacgccac
ttatgccacgatctcaaaaaaatgataatttattgacaaaacgtttacagtattagctg
atattgtattaaaaatacttcccacaagcaaagaagaaaaagaagcttttttcttactaca
gagacggaatgtctgcacaatctgaaggcgagtatgcagaagctttagaaaactattacg
aagctttaaaactagaagaagatccatatgatagaagttatattttatataatataggtc
ttatttatgctagtaatggtgaatatgtaaaagctttagaatactatcatcaaggattag
aactaaactttaagttacctcaagcactcaacaatatagccgtaatataccattatcaag
gagttcaagcggtcgaagataaagatacagaattatctaaattaatgtttgacaaggccg
ctcagtattggcaacaagcgattaagttagcacctgacaattatattgaagctcagaatt
ggttaaaaacaacaggaaggatgagaaatatacaaggatattaaaatatttaggatataa
t---tatattaattaatgaagcag----aattagagatgcttttaccctataaagcattt
gtagtat--------ataatatactaattac------tgtataaaaatcaacataataaa

cgaaattaaatctattgactatatttatgaatacaaataaacataatggttagtaaaaca
aaaagcatcggatctgtcactcaaatcattggaccagttttagacatagcatttcctaac
ggacaacttccaaaagtattcaatgccctcaaagtagaaagctcagagggaactattact
tgtgaagttcaacaacttttaggtgataacaaagtaagagctgtatctatgagttcaact
gaaggactacaaagaggagttcaagttcttgatacaggatctcctatatccgttccagta
ggtgcagatactcttgggcgcatttttaacgttttaggtgaacctgtagacaatttagga
ccagtaaattctgaaagtactttacctattcatagaccagcacctgcttttactaagtta
gaaacaaagccaaatattttgaaacaggtattaaggtagtcgatttattagctccttat
agaagaggaggcaaaattggtttatttggggtgccggagtaggtaagacagtattaatt
atggaactgattaataacatcgctaaagctcatggcggtgtatctgtatttggaggtgtt
ggcgaaagaactagagagggtaatgacctctatatggaaatgaaagagtctaaagtaatt
gatgcagacaatttgaaagagtctaaagtcgcactcgtatatggtcaaatgaatgaacct
ccaggggcccgtatgcgtgttggcttgactgcattaacaatggcagaatactttagagat
attaataaacaagacgttctattgtttattgataatatttttcggtttgtccaagcagga
tcagaagtatctgctctactaggacgtatgccatctgctgttggttaccaaccaactttg
gcaaccgagatgggtgcattacaagaaagaattacttcaacaacagaaggttcaattacg
tctattcaagctgtttatgtaccagctgacgatttaacagatccagctcctgcaaccaca
tttgcacatttagatgcaacaacagtgctttcacggaatttagcagcaaaaggaatctat
cccgcagttgatccactagattcaacttcaacaatgctacagcctggaattgttggaact
gagcattattctacagctcaagaggtaaaatccactcttcaaagatataaagagttacaa
gatattatcgccattctaggtcttgacgaactctctgaagaagataggcaaactgtatcc
agagctagaaaaattgaaagatttttatctcagcctttcttcgttgctgaagtatttact
ggctcacctggaaaatatgtatctttagaggatgcaattaaaggttttcagatgatttta
aaaggtaaattagatgacttacctgaacaagcattttatctagtaggtgatattgacgaa
gcgatacagaaagcggatagcatgaaagattaatataattaacaatgactttaaatataa
gaattattgcacctgatcggactgtttgggatgcagaagcacaagaaattatttaccaa
gtagcactggacagcttggtattttaacaggacatgcgcctctacttacagctttagata
taggagttatgagagttagagtagataaagaatggatgccgattgttttgatgggtggat
ttgcagaaatagagaacaatcagttaactattcttgtaaatggtgcagaagaagctactc
aaattgacttatcagaagcagaaaaaaatttagaaactgcaactcaattattaaatgatg
cctcttctaataaagagaaaatagaggcaacacaaaaaattagaaaagctagggctagag
tacaagctgcaacagcagctattgtataataacatttt------gctaaaaatataaaac
cttagtaattattactaag-------gttttatgttttatttatgtatttatatacaact
tgagtaagcttagcttaaaccagaacaaatataatcaaaatataaacccatttctttgcc
agcatctggtccaactaaactaatagtaacttctttcatagctaaaatcgcttgaatggt
tgcaccaataggaacacctagagaattataggtttcttttagaccatttaatactctttc

ttccaaaatagatggatctccagctaacattccataagtagcatagcgtaagtaataatc
taaatctcgaatacatgcagcataacgtcgggttgtatacatattacctcctggacgagt
aatatccgagtaaagtaaagattttgctacagattctttaataattgttgctgcattagc
tgcaatagtagcggctgctcttactctaagctcacctgtctgaaaatagcctcttaattt
ttcaacagagctatcatctaaatatttaccttgaacgtccgctgcattaataacagaagt
aattgcgtcttgcataacttgtaaaaattccttaattttaaatatctaatactaaagggg
tatataacaaaataactgaacttttataaataccagtactactgcatggcacccaaagt
ataatcaaagtaaaaaccagcttctgctgaatcttcaccagcaagcagtgagcaagctac
acttttcatacatttcacgccttcagcgactcccgaaattggtgttcctaaagaattata
catttctttaacaccgactagtccaatttcttcaataggcgttacgtctccagcaactat
tccgtaagttactaggcgaagataataatcaaggtctcgtaaacaggtagctgtcatctc
ttctccataagcgtttccaccaggagaaactacatcaggtctttttgaaatagctgttg
accgccttgttttacaatacgctcacgattatctgttaaaatttgagctattcttagacg
acgttgtcctgataagacaaaacttttaattctgtctaattcaccaggacttaaatatct
tgcttctgcatctgcatttacaattgactttgtaacaatactcatggataaaactcctgt
aatgcttt--taaaaaagcttaatactgattatttatgatataactcagaaagtagtacg
ccttttaaaaaaattcatagctatagcacgttggtgatatatgaagtaggacaaatactt
ggataggtactaaaacaagaaagttacttatcgtaccttattttattgattgccgagtac
ttttttaaaactaggtactacaattgatatatcttgcttagtaaggcgattatataatgt
ttctgtgttagggaaattagctgctggcaaagttggaaatcttctatacggaacagtata
tttaccatatactgaattatactcagtactttcaactaaagtactaataaatgcagacaa
tccttttgaagctaaaatttggttaaagaatctaatttctgcctgattattaggagctcg
tcctaagatatgcttcgtacctaattctataacttttgtatttggatatggctgataaaa
ctctttaccatataattcagataatgctaatttttctactaattctttaacattaatttc
cctgctcaaaaaggatgcttcaatatctaaaaactctcctccaatactaaaagaattaag
atctctttcaaaaatctggcgatatgcggctcttaaagcttgctctaatacttctttact
actattgtcgtttacttcaaatataatagattgatctcttaaagagctaactccttgaga
aatacgagactggatatcattgccgcttcttatttcttttacagcgcctaattcaacaaa
tctagaagacttgctagtaatgactttcttaaatctttggtcaatgatacctggccttaa
gcttcttagagcgataccagctggtgtactatatctttcataaggaacagtactatcacc
aaaagtttctatatactctgagctatcaactattgcatctatcacttgatagtagccctg
cttatatgcaatatcaaaatatttattaatttcttggcggccatatgtaggacgacccaa
taaacggttatggatatactcaatagctttacagatatataatggctcccagtaaagtga
tctaaatatacttgacttcgctaattgtctaacaaattcacgcactgtaatttgaccatc
ttttaactgactttctattggcttaagaagtaatttttcttcttgatatacttctctgcc
aaaaactcttaaataagcaacttttgtaacggcatcgactgaattttcaaaattttcaga

agagtcagcttttacaagaggattggataatttaaaaatttttggacccaaagatcctgc
agacttaggtcttacctgcggattactgatctggttatagattcctgggccacatctaac
cagaattctcctagtatctttaccaaaaagagcttgtcgtttcctaggatccttattttc
tttaggaaagattgctccaaattgaatcgacagtggatcattgcctattccatatggatg
ctgatcaggtaatgcttgcttgtaatcactaaatagggttataaactgaggcaccttacg
aaaaggagcactataattaagaagatctatttgaggtccccaattacgacactcttgagg
ctcttcacctaaatttcggaaataaggtacagtttcttctccaaaataatctgtatattc
agcagaatttaaaattgcatttactagcccacttaaaccagtagctgatagaatagcaaa
atatttttggaattcctctaaagagctgggccctctaccaaggaaatgtctaaatgctaa
ttctaatgctcggctatttacaaaaggctcataaaattgctttctatagatacttgaagt
tcctaaagaacggataaattccttaatggagatttggccatttttttacttgcgactctaa
gttagataaagataaatcatatgctttagcaatatctctctcaaagatttgtctgtagca
cgctttaactacagtatttttttcatctgcagataaactaggcttcataacaaaacgagg
cgtagaaactcctgcttgtgcataaatctgaggaaggcgcaaaccttgtaaatccccaga
tgttcttttccttaatttatcagtcaaagatggagcttcaaattcagaaattacaacatt
aaaatactctttgaccagatcttgacctttaatatcttcttcaaagattaaaagagctgt
tcgtctcatctctctgagggcaacaattgctgccgcgcttgagcaagcattatcaattaa
ttctcttaatcctctaatattaaccgataaaatattgggatcacctgatacaattgcata
tgttaaataccgtaaaaaccagtctaaatctcgcagagattttctcattcttgtagtccc
atatctcaagacatttatgggtttgaatcctggaggagtagctcctccagcattaaataa
agaacgaaaactctgcccaaaatctccttgaatattgccagatagctcattaattttgtc
ttgagaactttgatctccagcaataataacagctgcttgtggacgttctaaataagaaat
tgcagacccgcctacaaatattttgtctgcagctcgagctactagaatattggcattttt
agttaaaatatctgcaacttctaatctttttttgtccagaatttagaaaagaaactagttg
atttaattctcctagctgtaaaaatcggtcttgctgctctgcctgagtaatagttaaaat
tgaggcagtccgataaagctgggggcgtgctaatgggcttccgccacttgctttgatact
cattactttatttatctccttaacttatattacttgctagctaattcaaatatcacttaa
tatttctaactatttaaagatactacgcaagcacatttgttttctatattattaggtact
agtagttatagtacatagagtcacacaactaacttctataaatacaaattttgtaatact
tctattaacatttatcaaaagtaaatataattgataactcttttatattaaaatgcaatt
attaagagagtgatgctaaataaagtataattagtaagttttatagatagtaaaatt-ca
agatttattctacaaacattaatca------------cttgactgtttgacaattatacg
attgttcttgcttttaattggtttagactagtttcacatttttttctaacgtaaacaatat
taataagcaatttgtatgaatttaaaaccacagaataacctattaactaacggaaatata
aaccaacaagatgtacctgtaaatgatatgccaatgtcaattactgagcatttagaggaa
ttaagacaaaggactttatttgtatttttattctttttgcttgctgcaactttgagtttc

acacagatcaaaataattgttgccatattgcaagctccagctattggtattaaattttta
caattagcgccaggagaatatttttttttcttctattaaggttgcaatatattcaggaatt
gttgcaacaagtccattcggggtataccaagttatcttgtatatactacctggactaact
ggcagagagcgaaaaattatttgcctatcctaattagttcagtcttacttttataata
ggcggtatatttgcttactttattttagcaccagctgctttaacattttgatcagctat
ggatctgacattgtcgaaccactgtggtcttttgaacaatacttcgactttattttacta
cttttatttagcacaggattagctttcgaaattcctatcatacagttactactaggtgtt
tcaggaacactatcttccaatcagatgattcgagcttggaggtatattattattatatcg
acaattgctggagccattttaactccctcaactgatccagttacacaaatcatcatgtct
tctgctgtcttagtgctttattttagtggaattgtcatattgttacttttaaaaaagtaa
atgtagt--------aaactttcatttcatattatacttttcttgtg-----ctaagcaa
tggagttt--atatagcttttagttcttaacttagatgaatattgaagta--------tt
aagctcttttaaatatttaatttagtcttataatttaaccaatatg--------------
------------------------------------------------------------
------------------------------------------------------------
------------------------------------------------------------
------------------------------------------------------------
------------------------------------------------------------
------------------------------------------------------------
--------------------------------------------------tatatattgg
c---ttttttcatcaattatttgcactttacatttctgtacta----ttctatgatgctt
gggactctttatgtaatctgtttactgaat----gacgttgaagaaaattcagcataaat
tttgttaatttataccaatataaaaagtattttattgtgagttttatttatataaaggct
tattcattatatatttataacagtaagtctaattaatagtttttttatgcttaattaaagt
aataactcttaatccttttggcagaaaagtctgtcttaatattttatc--aaaaaaacag
aaaaaatgaagctaaatagtctaattaacttaattcaaaagtctatatgttcttgtacat
ttttattaattattttaaatattatttgtgtcgctcctaattctagtaatgcatttccaa
tttatgcacaacaagcttacgaaagtccaagagaagcaactggtcgaatagtatgtgcca
attgtcatcttgctcaaaagccagtagaaatagaagcacctcaatcggtattacctaata
ctgtttttgaaactattgtaaaaattccatatgacagtagtgctaaacagattttaggta
atggaagtaaaggagggttaaatgtcggagctgttgtcatattgcctgaaggatttaagc
tggctcctgctaatagattatctacagagttaaaggaaaaactaaaaatctatatattc
aaccatatagcgctaaacaagataacattttagtaattggacctattcccggcgacaaga
atagagaaatagtttttccaatactctctcctgatcctgcaaaagataaaaaagctcatt
tttttaagtacccaatatatgttggtggaaatagaggtagggggcaactttatcctacag
gagataaaagcaataacaatgttgttacagctttaagtagtggcaaagttaataagattg

aggttcttgagaaaggggatttgtagttcatgtgactaatagtaacaatatagaatcaa
ctcaaaagatttccccaggtcttgaacttaaggttaaagaaggtgatacagttcaattag
atcaagctttaaatagtgacccgaatgttggtggtttcggccaaaatgaaacagaaatag
ttctacaaagtccaaacagaattaagggcatgattgttttctttttgctagtgttttag
ctcaaattttctttgtattaaagaaaaaacaattcgaaaaagttcaggcagcagaaatga
atttctaaagcaagataaaactggtacttactataattaaagtg-aaaacactttattca
aagaatgtaccagttttcatgctatataaccgttgcttttgaagacattataatgtcttt
aacagagtctacaatttgactaggctgaattacagtagcctgttctaagctaccattata
tggtgttggtatatcctgcgaagatagtcttactacagggacatctaattcatcaaaaag
gtgttcattgatttgtgcaattaactctgctccaattccagcagttttcatgcattcttc
tacaattaaaactttgtgagttttcttgactgacacagatatagattctatatctagagg
ttttaaagatataaggtctatcacttctggatcgtaaccttcttttagcaaagctggtaa
tgcttgaatcacatgatgcctcattcgagagtaagttaaaatagtaatatcttttcctgt
acgcacaaattcagctttattaagaggtaatagatattcttcttcaggaatttcttcttg
taaattgtaaagcaaaacatgttcaaagaagacaactggattattatctcgaattgcaga
tttcaataatccttttgcattatatggagtagagcaagcaactatttttaaacctggaat
agcttgaaaataagcttctaatctctgagaatgttctgcgcctaattgtctaccaactcc
tcccggtcccctaatgactaaaggtaatgtaaagtttcctccggaggtatagcgcaacat
gcctgcattattagaaatttgattaaatgctaacaataaaaagctcatattcataccttc
aacaattggtcttaaccctgttatagctgcaccaatcgccataccagtaaaactattctc
tgctattggtgtatctaaaactcttaaatccccatacttactatgcaaatctttagttac
cttataagatccgccgtagtgacctacatcttcgcctattacacatacagtcgaatcttt
tgccatttcctcatctgttgccgctcttaatgcgtcaaacataaagatttactcatttt
tgtatttaattttataatttcactactaatctaaattaaaagttaagatattaaatttt
c-tcacgtatctgaaaaaagatattgtttaagttctgagatattaggctctggactggaa
atagcaaattgcacagattgttctaaatctttttaacagcagtttgaatctcgttgagt
tcacttatgtcggcgatcttattatctaaaatatattttttaagtttttgattggatct
cttgcaatccaagcttctttttcttgtcttgatcttagttcatcaggatctgcaagagag
tgaccacggaatctatatgttaaagcttctattaatgttgggccttcaccatgtctggct
ctctgaactgcttgaatcgctgcttgccttacagctagtacatccattccatcaacttcc
ataccaggaagaccaaaagcttctgcctttttatgtatttccggcatggaagaagatcgg
tgatgtgccatacctatagcccactgattattttcaacaataaatataataggtaacttc
caaagaactgccatatttaaacattcgaaaaattgaccattattagtagtaccatcacca
aaaaagcaagctgtaactctcaaatcttcagtttcttttagtacttgctgacgataaata
ctttggaaagctgcacctgttgcgacaggtataccttctgcaataaaagcaaacccacct
aaaaaattatgaggagcagaaaaaatatgcattgaacctcctctgcctttgctacaacca

gtttcttttccaaatagctcggccattacatttttcgagggaacacctttacttaaagcg
tgtacatgatctcgataagtgctacatacataatctgtagaattgagaagcttaattaca
cctgtagacacagcttcttgaccattataaagatgcacaaaaccaaacatctttcctttg
taatacatctgagcacatatgtcttcaaaatttctgcctaacaacatatcttcatacaag
actaataaagtcttttt------attaagaacagttgaattatagttagttagcggcaat
tgaactttcttaggataactcataatttataaaggaacctct----------tattatac
ataggaattttgaagtaaaagttatcaatagaaaacacacagataaaaatttataaatta
ttatatgc-ttttactaaagacttcaatagttatttgcaactagttcagtatatcaaatt
aattttgcttatactacagatgtctaaagcttagatgaaaagaaa-tagttaaa------
------------------------ataaataaaaataaacagaattattagtgagaaaat
gattattattaatagataattacttaatatttctaaatattttcaaattaatggctatat
catcatctttattttaccctgtcgagaaagagttatacagtctagaaacaaatttgcaag
ctgtggctgggacacgtcatccaattttatacgcagcggcgaaacatctgttcgaagctg
gaggaaaacgagttagaccagctctcgttcttttagtggctaaagcaacatctgagaaac
aagatattaatactggacagagaaggttagcagaaattactgaaattatacatactgcaa
gtttagtacatgacgatattattgacgaatgtacaacgcggagaggagtcaagactgtgc
acaatttgtttaatactaaaattgctgtattagctggagatttttttatttgcacaatctt
cttggtatttagctaatattgaaaatttagcagtagtgaaagctatttctaaagtaatta
cagattttgcagaaggcgaaattaggcaaggattagttcactttgaccctagtatttcaa
tagatgcttacattgaaaagtcattttataagactgcttcacttattgctgctagctgtc
gaggagcagctatgctgaatgattctaatactcaagttaatcatgatctatatctttacg
gtaaacatatgggcttagcatttcaaattatggacgatgttttagatataattggttcta
ctaaaagcttagggaaacctgctggcgcagatctagccaatggtaatttaacatctccta
ttctattttcacttactcaagaaccagatttaaacaatcttattcaaagagaattttgta
ataggacagatgtagcctcagcgttatttcttataaaaaaaagcggaggaattacaaagg
ctcaagacttggccaaagaacaggtacaggcggcactttgctgtctccagtttttaccaa
aatctgcacctgtatctagcttaaaagaattgacacatttcataatcacaagattgtcat
agaacacttgctaaaattaaaatgctattaactaatttttttactatctctacaatagtct
gaaaagtttggatatcattacttgctaactgtgctagcattttacgatttaatgcaatat
tctctttttttaacgcactaataaaagtactgtacttcatgccttgactatgagctgcag
catttattctagttatccaaagacgacgaaaatctctctttttcctttttcgaccaacat
aagaatagcgaagagatttgagaacttgctgtttagctgttcggaataaacaattatgtg
cacctttaaagcctttagccagcttaaatatttttgcccgtcttttttttgcaacattac
ctcttttgactctactcataaaatattac-ttacaattaaattattaacagcttcctaga
ttcaaatgcttatcg----attataaataaggtaaatttacagcgatattcttaatatct
cgtgtgtcaactaagcaggtacaagaaagattcctttttttgcttcgatgatttttttttgt

aataaatgacttttagaagctttatgtcgcagaaattttcctgatgaagacactttgaat
ctttttgctatcgcttttgatgtttttaacttaggcatattgtaaacttttaatgattaa
caaaagtatgaaaagtctgaatttatcaaacttagacttttcaatacttaggataatctt
aacacaaattattttacttactatagaggttaacttaaacaatttatatatataatagca
aataattaatccgtagctactaaagttgatttatatttagctttaaatttttgtatagta
ttagcaattagcattgttattattttaagggtacttgagtttaattcttcgaacattttc
atatttaatgtgaatgatatattcgcttctgacacaatattttgcatttgatcatccgat
aaaggaatcatatccaatgcagctctatacttatctttaaacactttgtcatcttgtatt
tggtcaaagtcataaaacctagtacctctttcatctgataaattcatcgcacctctagcg
atttctttaagatttggcctccagataaatcacctaagtaacgcgtataagcatgagca
actaataattccggctgtttacagcctatactatgaattctatcaacatagattattgtt
gcaggagatggtttaataagatttaaccaatctgctccataataataacttaaatcttct
gacagacttattttttctattgagttctgtaaaatatataggtttaatagctgggtgatct
ttattagaaaacagttcttcttctatagcacaataaacgaaataaagattagcaaccaat
ttacgatatgatttcttgtctacgacacctcccaaaaaagatttcacaaaactaacattt
tctgccatactatgagacttagtagtgccttctcttaattcatttgctaaagtattaacc
ataatatcaatcttccatttta-aaagataacaaattaaac---ataaacagttatctat
atattta------agttgtactaattaagctgctgcaaaataatttttttgattttattgg
atctggattcattgttttatcacctggcttccaattagctggacaaacttcatctggatg
agattgtacatattgaattgcttgcagaactctcaaagtttcttctacactcctaccaaa
ctcgagattgttaattgtagaatactgaataattcctttaggatctataataaatagtcc
tcttagcgctacaccatcactgttcaaaacattataagcagcactaatctctttttttaa
gtctgatactaatggataggcaagatctcctaatccaccggattctctgtctgtctgtaa
ccaagcaagatgagaatactcgctatcaaccgaaacacctagaacttctgtgttaagctc
agagaaagcatcatatttatcactaaatgctgtaatttctgtagggcaaacaaatgtaaa
atctaaagggtaaaagaataagataatatacttatttttaaatctgataattttaacgt
tttaaattcttggtcataaacagctgtagctgaaaaatcaggagctagctgacctactcg
aagacaatttggtccagaaatcattaatttttcccagaagaattgatatagttatgttat
ttaataaataattatataacaaaatttagtatatatgaaaatgctatcagtcttttaaaa
attac--------tcaatggcttccaaaatactatttttatagcggggaatggatttgaa
ccattgaccttcgggttatgagcccgacgagctaccagactgctctaccccgcggatatt
taatatcatattagtccatctaaatagataaaagcaaatatattatataattcaaaagta
aaagaataagagcacaggagcactaattattaggcttaaaaatgacaaagtatttttat
tttattgagtagaggaaagacttcatataagccgatgaatcgatctttaagaagaacttc
cgatggcttcacccaaatttgaccgtcataccacccagactcttcatagaataccgtagc
actcattaaacgttttactacgtatgaccatcctaaatataatctgattagaataaatcc

agttattaaactcgaggtaataaattctgagaagacaaattttaaaggcagcttggtaat
tggaaaaattgaaagcaatactggactaactaatagaccattaagtagtagtgtaacagt
gattttcctgttatatgattgactacttaaggttggccagcagaagaaccaggaatttt
taaagaagtatactcattaacaggttgctgctcttttggtacaggacactgagtgttata
taaattcattgaaatttaaaattgttatactagaaattgtttttttatctgctttagat-
-ttaggga--------gttaattgattgcaattataatagttagtatcagaaatacgacg
ataaacaaccaagtaactttgtttaatgtgttttcagtactacgagtattgctaaaaaac
tgattctgagctccagcacttcccagaccttcagatttgggattatgtatcagaatagta
aaaaccaatataatagttgatacataccaaaagaattttaaatttgttccattgtgtaa
aaattaataaaaaatagcacaaagacaatcgtctaaaaaatattgattgtcttattatat
taagaaagatttattctccctgcacttttaaaagtaaaaagcctattgctaaccctacta
aaactaagactgaacttaatacagcactttctacaatttctccgaccatt-gtgtttttc
cttattattttattatta---tcattttgtaaaaaatttgatacctattaaa-ctaaaat
ccatttcttccccaaaccaccatcgcaatagagaaagtaaacattgccattaaagctgcc
catcctaaacttaaaatatccataatttataatattc--ctgaaaataaggtctcttata
cttaaactataatattcttaatattaaaaatatcattatataataagctaattataacag
agtactacttaattcgtatttttagtgtatttcaagaagattgac-tttttctaaaatac
ttaacaagtattaaattgtaaaaaatttctgcaat----ttttgcattattatacctatt
aatttggaatgtaaatatcaatttaaaatcgttagtaattttgatttttttcaattgtaca
att---aacatatgtaaaaacataaagattcatagcagctaagttga-----tggtctac
tataatttatcaat--aaaaaggaagttagtatttaaattatgcaaacaactataagtaa
tggaaaaacgtctagcaaggaaactctactaacccctaggttctatacgactgattttga
agaaatggctaatatggatatttcagaaaatcaagaagatttctttgctattttagaaga
atttcgagctgattataatagtgagcactttattcgagatgaagaatttaatcaatcttg
gtccaatttagaagataagactaaatctttatttattgaatttctagaaagatcttgcac
tgcagaattttcaggtttttttactatacaaagaattatcacgaaaattaaaagatagtaa
tccaattattgctgaatgtttttttactaatgtctagagatgaagcccggcatgcaggttt
tttgaataaagctataggagatttcaatttatctttagatttaggctttctaacaaagag
tagaaaatatactttcttctctcctaagtttattttttatgcaacttatctttcagaaaa
aataggatactggagatatataactatttaccgtcatcttgagcagcaccctgagcatcg
catctatccgatttttagattctttgaaaattggtgccaagatgaaaatcgtcatggaga
ctttttttgccgcattactcaaatcacaaccacatttttttaaacgactggaaagctaaaat
gtggtgcagatttttttctattaagtgtatttgcaacaatgtatttaaatgattttcagag
aatggatttttataatgcaataggtctagattctagacaatatgatatgcaagtaatacg
aaaaactaatgaaagtgctgctagagttttccctgtcgctttagatgtagataacccaaa
attcttcaaatatttagatacttgcgcatgtgacaatagggctctaatagatatcgataa

acaaaatttcccattgttaattaaagctattataaaaatgcctctctattactccctaat
tattagtttgctaaaaatatatttaattaaaccgatcgattcaaagaccgtctggaatac
agttcgataatcaaatcacataaagatatatttcataatacacactatctcattattga
tataaatcaaaatgaaaagctcacttct-----gtcaggaaggagcttttattttattca
caaataaacaatatggttctatagatt-tagaactaagaacttactttttcttttgcttc
atacataacttctaaggtgattagattttagaacaatcgcgtgcaaatttttctgtgtt
tctttttactttacctctaacaaaacctggaattttacttaattctttctgtgcttcttg
tgaccagttaatactatcaacagcagagatccctacgcttaatgcttccgttgtatcatg
accaccaaatatttctaataagtgatcttccatacccaatgtgaatgagttatagactaa
atcagcaatttgatttgtaccttcatagccaaggaatggacgataactcaaaggaaaatt
ttgtatatgtactggtgaagaaataacaccacaaggtatatttaaacgtttaccaatatg
tctttccatctgtgttccgaaaattgctgcaggctctgttttagcaattaaatctccaat
taaaccatggtcatctgaaataattacttcatcgcagaattcttgaacttgttctttaaa
ccaatcttcatcatatttgcaataagtgccacaccaagtaacatgtatacccatttcctg
gtgtaaaattcgagttattgctgcagcatgagttgcatctccgaatacaatagctttttt
accagttaaattttggcagtcaattgatctagaaaaccaggcagattgcgagataaatct
agtttgttcatctatataattttcgtagtctacaatggctcctaatgtagttaataactg
ttctattgatcggatacatgcagctgtttgtacaatacccatagggggtagtatcaacata
aggcatattaaattccttctctaaataacgagctgtcattagtccggtctctctatatgg
gacaaaattaaaccatgcagatggtaaatttttagatcctgaaccgacgcattttcggg
aataatttgattaattttaatatctaaatcttgaaataatctttttaattctgaaatatc
atgctgattatgaaatcctaaacttacagctccgataatatttacagatggtttttctgt
cttttgagttaaaacctgattattagatttagctttttccatgtaaaaagtaacaatctg
ttccagagttctatcaccagcttgaagttcatttaccctataatgattaacatcagctaa
taaaacatcagcttctgtttctatagaggctctgctaacaaaatttgtaaatcttcttg
caatatactagaagtacaagttggtgttaaaataactaaatctgggttttcttctctatc
ttttctagtaatgttttcaacaactttttcttgtgaaccacgtgctagtacatgtctatc
tacaacacttgctgtaacaggcgtaaaatctctgtctctttcaagcattgaacgcataac
attaaaatagtcatccccgagaggcgcatgcataattgcatgaactttttttgaaagaact
agcaattctaagagttccaatatgagcagggcctgcatacatccaataagctaatttcat
attaaattttcctatgatttatttagacatagtaatacttttaaagccaaataataaata
aaattcgaacatctaatttaactccttaatgatttttttcatg-tttttagatagagcgta
acattatatcaagtcaaaatt-------------agagaaaaatgactaatgaataaagc
ttgaaagtaaagatttcttgcagttatagtttttctttaagtaaacatgtaaaaattgta
ctatatataaatgtactatatataaattttttttactgaatattggtaatgccagataca
attaacttaaatatgccttccccaacgtttggtggaagcactggcggttggttaagagcc

gcagaagtagaagaaaagtatgcaataacatggacaggcaaaaatgaaagtaagtttgaa
atgccgaccggtggtacagcaactatgagaaatggagagaacttactttatttagctaaa
aaagagcagtgtttagctttaggtactcaattaaaaagtaaatttaaaatatctgactat
aaaatttacagagtatttccaaacggagaagtacaatacttacatccgaaagatggtgtt
tttcctgaaaaagttaatgctggaagagctagcgttaacagcgttgatcattctattggg
cgaaatgttaatcctgtagatgtcaaatttacaaataaggcaacttacgattagttgata
aaaaaacctagacataaaataacttttatatagtctaggtttttgataaagtatactaag
agtatggaaaggtggccgagtggttgaaggcgtctgatttgaaatcagttgaactt--ta
tagttccgtgggttcgaatcccaccctttcctttataagact------------ataact
taattattactata----aattatacgctaatagtattcagtttagacttaatgaaatct
acaacttgcgataaattagaaatttttcggcatcttcatctggtatttctataccaaat
ttttcctctattgccattactaactcaactgtatcgagagaatctgctccaagatcactt
gaaaaatttgcttccctggtcactattttttttcaattcctaactgttctgctacaata
tcttgaactttttcaaaaatttcattatcttgcataatatcttgtttcctataaggtgtt
ttcaaaaattttctggctcatattattgtctaactctaattaattagtaattttgagttg
aggtccaaatagttctacctatatgatatactattattctggataaatacgaagtttgcg
gtatggctcttctccaaaactacgagctctcaactggtagttatcaactaaattatgctg
cattttgcgaatataggctgaacgaggtatcaactgaactgtcgatttttcgttcaagat
aatagtttcaatagccagtttagcttcttgaagggcttgaatctcatcaaatttttttatt
ttcacaaagttctacccaattaagatcagaagaagtatgaatatttaatatttttctcag
tgcacgagtaatttgtgggactgtactattctgaattgtatatataataatttgttttga
ttttgcaatttgtcttaattttgtattttgctttacttgattacgcaatgctaaaatagc
atctgacttttctatctctttagttaaaataattggtaaatctaaagaagaaataactga
cgcaatatgctgccaacttaaagaataagcatataaatactgattcggcatttctaaggg
caaagatgggttattaatatttataggagcattaatagtagttgacaataaagatgtatc
acgattttgtaatttattcaaatcggatggcttatttattaattctctatactgcaaaac
agcatgtgacggttgtacactaattggtttacggaaactagaaatatttgacggcaaaac
ttctaaagactgtgacggatagcatttaattaaaatgcggccattagcttgaatttgtct
tttttgtacaaaaggttgatggccctgtaaaatttgatctatagtttctttcactttatt
atggactatccaaacagtgcgttcatggatttcaattgcaatttgaaaagcgggcgcggc
ttttctttctaatatgcttttctgagtacctcttctttttgcttcatcatcacctaatgt
gacatattgaatacctccaattaagtctgacagtgtaggattttttaattaaactttctaa
atgatttccatgtgctgtacctactaattgcactcctctttctgcgatagttcgagcagc
caatgcttctagctctgtaccaatttcatctataataatgacttcaggcatatgattttc
taccgcttctatcattacctgatgctgtaaatcaggccttgcaacttgcattctccgagc
tctacctatagctggatgtggaatgtcaccatcgccagctatttcattagaagtatctat

aataacaactctttttttccatttcatctgctaagacacgagcaatttctcgaacagctgt
tgttttaccaactccaggttttcctaaaagtaaaattgaatcaccttgttctagtaaatc
cctgataatactaatggtgccaaaaactgctcgtccaacacgacaagttaaaccaataat
acttccttctctattacgcatggaactaatacgatgcagggtcttttctataccagctcg
attatctccgctaaaattaccaactttttttacacaataatctaagtcttgccaactaat
agatctttgagataagtattctggcctatctggaaatcttgcttctggacggcgacctaa
atccatcactacttctattaagtgatttctattagggtgttgtttcaggggctctttgac
aaagttcggtaagatttctagtaatttatctaagtcatctgcaataagcatgattgaata
tataataattaactaatcagatagaaaaattaagattctaaagattacaatacatattaa
agaatattgtgctcgtaatcaattttatacttaaattcaataagaaatta-cagatggta
aacaatagattaaactatgtacagtaaataaattggtcaaaattaagagtcatgttaata
tgtttattggacagaaagttcgtattaagtatagtaagcaaaaagttgctaatgacatag
cagataaagtcggtaatcgcggaattattagaggcataaaatttatcaacagtcaatgtg
ttactattattgttgaatttgaaaatcacactagactttggatgtttaaggaggagctaa
tctatttaaatgaat------ttaaacaataagataagattaattt----tattatatat
aaatttttacgagtcgttcaaattatacattttcaaggagtattgttaattgtgcaaatt
actattaaaaaattacaagacttactttcatcagtacaaagaaagaagatacagacattc
aaactgaaacaggataattttgaacttctattaaataaaacttacaaaaaaactaatcca
gaaatattaccttcacagaact---ctgtattacaacacaatcaagctaacataattaag
tcaacacctaataacaatggtccttctgttagtaataataacgttacagaatatgccact
atcgtctctccgatggtggggacattctatcactcacctgctcctggtgaaaaaattttt
gtgcaagttggggatgaagtcaaattcaatcaaacagtttgtattattgaagcaatgaag
ttgatgaatgaaattgaagctgaaattgaaggaaaaattatagaaattctagttaaagat
ggcgatatcgtagattgtggtcaagccttaatgaaagttgaaacataatgccaatatttc
cattttacatattgttaacagaaattatttagatagaaaaattagcttataatataatta
tgaaaacttatttcgtaaaggcac-aaaaagttagtatataaaattatttactttatctg
ccaaacatagaaataagcgttttgattccttgtctcaagagaggagaatctcaatgacaa
ttagctcaaaagagcaagagacaaagaaggtaaaaatctcggttgataaaaatcccgtag
atacttctttcgaaaaatgggcccaaccaggtcatttttctcgtacactagcaaaaggac
caaaaactactacttggatttggaatcttcatgctgatgctcacgacttcgatagtcaaa
ccagttctttagaagaagtttcccgtaaaattttcagcgcccattttggacagctttctg
taatatttttatggcttagtggaatgtattttcacggagcacgtttctctaactatgttg
cttggttaagtaatccaacaggtatcaaaccgagcgcacaagttgtttggcctatagttg
gccaagagattttaaatggagatgtaggtggtggctttcaaggcgtgcaagttacatcag
gatggttccagctttggagagcatcaggaattactacagaatttcaactttactgtactg
ctatcggtggattaggtatggctgctttaatgttgtttgcaggatggttccattaccata

aagcagctccaaagctagagtggttccaaaatgttgaatcaatgatgaatcaccacttag
ctggtcttttaggcttaggctgcttaggatgggcaggccatcagatccatttatctttac
ctattaataaacttttagactctggtgtatctcctcaagagattccattgccacatgaat
ttttaattaatagagagcttatggctcagctttacccaagtttcagtaaagggttagttc
catttttcactttaaattgggctgaatattctgactttttgactttcaagggaggtctca
accctgttactggcggtctatggttgagtgatactgctcatcatcatctggcattagctg
ttttatttcttgctgcaggccacatgtatagaaccaattggggtattggacatagtatga
aagaaattttagaagctcacaaaggtcctttttactggtaacggtcatgaaggtctctatg
agattcttacaacttcttggcacgcgcagcttgctattaatttagctatgatgggatcat
taagtataatagtggcacatcatatgtatgctatgcctccatatccatatattgctactg
attatccaactcaattatcactattcacccaccatatgtggattggaggattttgtattg
ttggcgcaggagcacatgcttctatctttatggtaagagattataatcctgctgaaaatt
ataataacctcttagatagaattattagacatcgcgatgctattgtctctcatttaaatt
gggtatgcatatttcttggattccattcatttggtttatatatccacaatgatactatga
gagcacttggaagatcccaagacatgttttcggatacagctatacaattgcaacctattt
tcgctcaatgggttcaaagtatacatactctagctcctggtaatacagctcctaatgcat
tagcaacagctagttatgcatttggaggagatgtcgtttctgttggaaataaagtcgcaa
tgatgcctatttctctaggaacagctgattttctggttcaccacatacatgcatttacta
tccacgtaactgtacttatcttagtgaaaggattccttttctctagaaactctcgactaa
ttcctgataaggcaaatcttggctttagatttccatgtgatggacctggcagaggcggta
cttgccaagtgtctggctgggatcatgttttccttggagtattttggatgtacaactctc
tatctgtagcaatttttcactttagttggaaaatgcaatcagatgtttggggtagcgtgt
ctccatccggaaatgtttctcatatcactggtggtaattttgcgcaaagtgcaattacga
tcaatggatggttaagagatttcttatgggctcaagcatctcaggttattcaatcatatg
gttccgcactatctgcatatggactaattttcttggcagcacattttgtgtgggcattta
gcttaatgttttttatttagtggcagaggttattggcaagaacttatagagtcaattgtat
gggcacataataaaataaaagttgctcctgcaatacaaccaagagctttaagtattactc
aaggaagagcagtcggtgttgcacactacttattaggtggaattggtacaacttgggcat
tctttttagcaagaattatttcagtaggctaatagtgaaaataggataaaaa-aacaatt
atggcaacaaaatttcctaagtttagccaagctttatcacaagatcctacaactcgaagg
atttggtatggtattgctacggcgcatgactttgaaagtcatgatggaatgacagaagaa
aatttatatcagaagatatttgcttcacactttgggcatttagcaatcatctttttatgg
acatcgggcaacttattccatgtagcttggcaaggtaactttgaacagtgggtattaaat
cccttaaaagttaaaccaattgctcacgcaatttgggatccacactttgggcaaccagct
ttaaaggcttttagtaaaggtggatcagcatatccagtaaacatagcatactccggagta
tatcactggtggtacacaattggaatgagaagcaatcaagacctgtattctggggcttta

ttcttgctcgttttatcagctttactttatttggtgggtggctacatctacaaccaaaa
ttcaagcctggtttatcttggtttaaaaataatgaatcaagattaaatcatcatttgtct
ggattattcggagttagctctttagcttggacaggccacctagtacatgttgctatacca
gaagcaagaggtcaacatgtaggatgggataatttacaacagtattgcctcatccagct
ggtttacaaccatttttagcggtaattggagtgtatatgctcaaagcccagataccgct
caacatatattcggaactaatgaaggggcaggtacagccattttaaccttcttaggtgga
tttcaccctcaaagtcaatctttatggctaactgacatggctcatcaccatttagctatt
gctgtagtcttcattgttgctggacatatgtatagaactaattggggaattggacataat
ttaaaagatattttagatgctcatagaccgcctagtggcagattaggagctggacataaa
ggactatttgataccatcactaattctttacatatacaattaggattagcattagcctca
cttggcgtaattacttctttagtagctcagcatatgtacgctatgcctccatatgctttc
atggctaaagatttaccacacaagcatctttatacactcatcatcaatatattgctgga
tttttaatggtgggagcatttgctcatggtgcaatattctttgttcgagattatgatcca
gaacaaaataagaataatgttttagctcgcatgctagaacataaagaagctatcatttct
catttaagttgggtaactctgtttttagggtttcacaccttaggtctttatgtccataat
gacacaatgattgcttttggtactccagaaaaacaaattttaattgagccagtatttgct
cagtggattcaagcttcttccgggaaagcactttacggttttgatgtgttactatcatct
tctagcaatattgcaacacaagctggtagcaatatttggttaccaggttggctagaggca
attaatagcggaaagaattcattgttcttaacaatcggtcctggtgatttttttagttcat
catgctattgcattaggattacatactactaccttaattttagtgaaaggcgctttagat
gcacgaggatctaagctgatgcctgacaaaaaagatttcggttatagttttccttgcgat
ggccctggtagaggcggtacctgtgatatatctgcatgggatgcattttacttagctgta
ttttggatgttaaataccataggatgggtaacattttattggcattggaaacatattact
atttggcaaggcaatgcaactcaatttaatgagtcttctacatatctaatgggatggttt
agagattacttatggttaaattcttctccattaattaatggttacaatccatatggtatg
aacaatttatcagtatgggcatggatgtttctatttggacatttagtgtgggcaacagga
tttatgttcttaatctcttggcgtggttattggcaagagttaatagagactctagcatgg
gcacatgagcgtacacctttagcgaacttaattcgttggaaagataaacctgttgcgtta
tcaattgttcaagccaggctagtaggtttagcacatt-tttctgtaggatatgtattaac
ttatgcagcttttgttttagcctcaacagcaggcaaatttggttaaagtaga-----att
taattaattgcaagaaaaaaaagtcagtcatgata-gtttcgtgactgactttttttct
tgacaatagttaattattaaagcttaacttcaatatctacaccagaaggtagatttaact
tcattaatgcatctattgtttgtgaagaaggttgatgaatatcaataatccttctatgag
atctggtttcaaaatgttctcttgagtctttatctacgtgaggagaacgtaagacacagt
aaattcgcctttttgtaggtaaagggattggtcctactgcaatagcattagttctagatg
ccgtatcgactattttatcacaagatgtattaagaatactcgaatcgtatgctttaagct

taatcctaatttttgtctgctgaataagtgtcatattgattaaatcttattatttaagaa
ttttagagacaacacctgcgcctacagtacggccaccttctctaatagcaaaacgcatac
cttgttcaattgcaattgcattaattaattcagcagtcatcttaattctatcaccaggca
taaccatttcagcatcagtaccatcatcagcagtaaactgattaatagttccagttacat
cagttgttctaacataaaactgaggtctatagcctggaaaaaatggagtatgtcttccac
cttcttctttagttaaaatatagacttcagcttcaaactgagtgtgaggagtaattgtac
caggtttagctagcaccattcctctttcaatatctttttctgaacacctctcaaaagaa
ttccaatattatcacctgccagaccttcttctagcgtcttttggaacatttctaatccag
taatagttgtggtccgagtctctctcaagccaacaatttcaattgtatcaccaactttta
taattcctctttcgatcctaccagtagcaacagttccacgtcctgtaatagaaaaaacat
cttctacagccattaagaaagttttgtcgacatctctttctggtgttggaatataagtat
caactgcctccataagtgaaaaaatcttatcaacccatttatcttcacccgctttaatag
tcgggttatttgtgactgcttctaaggctaataaagctgagccggcaacaaaaggaatat
catctccaggaaagtcatactggcttaacaactctctaccttctaattcaactaattcta
acaactcttcatcatccacttgatcttctttattcaagaataccactaatgtaggaacac
caacttgttttgctagtaagatatgttcacgagtttgtggcattggaccatctgctgcag
agactactaaaatagctccatccatttgagctgcacccgtaatcatatttttttacatagt
cagcatgacctggacaatccacatgagcatagtgacggttatctgtttcatattctacat
gagcagtattaatagtaatacctctagctttttcttctggcgcagcatcaatttcatcaa
actttttgctgaggtggaacctacagttgataaagtagctgaaattgcagctgttaaag
ttgttttaccatggtcaacatggccaattgttccgatgttaacatgtggtttttttacgtt
caaatttagatcgagccatattttctattctccttatagtaaaagttgctttttggttac
ttttaacgataaatcttataaattgaaagacttattatttaatatagataaggtaaaata
agatatagttaatttaatatctatagtgtgcaaacgctttatttgcttccgccatcctat
gtgtttcttctctttttctaatggaactgcctgtttcattagcggcatccatgatttcgt
tggccaattttattgccatacttttaccagatctttctcgagaaaatctagtaatccatc
ttaatgctaaatttgtacctctatatgctcttacctctattggaacttgataagtagatc
cgcctactcttctggctttcacttccactaaaggagtgatatttcgaatagctttttcta
gaatatttaaagggtctgcttctgttctatctttaactatatctaaagcttgatatataa
ttctttgtgctaaagtttttttaccacttttcaaaatacgaacagttagcatactaacca
gtctacttttatataaaggatctggtgatgcgaacctttttttagctgtattacgacggg
acataatattttaaatttgttattgtgataatttgtt-taatagaagatatttaagattt
aggctttttttgtcccatatttagatcgacttttacaacgatcttttactccagcagcatc
taatgtgccacgaactacatgatatcgtactccaggtaaatctttaattctgccccctct
aattagcactactgaatgctcttgaatattatgaccaacaccaggaatataagctgtaac
ttcaaatcctgatgtcaatctcacacgtgcaacttttcttaaagcggaatttggtttctt

gggagtcgtcgtatatactcttgtacagacacctcttctttgagggcaactttgaagagc
tggagattttgtttttttatgtatttttcgtctttcagatctaacaagttgttgaattgt
tggcataataaaattaattttaagtctttaatagatattattcgaacttgctgcgcatga
attaattagcctttatttatcttgtatttacgcataaatctttctactcgcccttctgta
tcaataattctttgtgaaccagtaaaaaatgggtgatttcctgaccaaatatccacatgt
agctcaggctttgttgagccaattgtcataattaattggccgtcacaataaacttttgct
tctgggtaccaactaggatgtatgttatcttttgccattgcttctataattgtataattc
tatcaataataaatattaacgcttagaaaattgaggagcttttcttgctttcttcaaacc
atatttttttctctcttttactcgaggatctcttgttaaatacccttcagatttaagtgc
gtttctattttctggattaattgagcacaatgctcttgcgacacctaaacgaattgcatc
tgcttgtcctgttaatcctcctcctctagcatttacatgtatatcatactgatttaataa
tcctagaacttgtaacggtgcatacgaaactcttaagtaattaggactaaactgaagata
tgactcaccaggtataccgtttataattaaattgcctgaacctgggaccagtcttacttg
tgcaacagaacacttccggcgacctgttccagaatagattgcacgagttttaattaattc
tgtcgacataatattccttattaactaaataaatatacattctttatactatatactctt
gcggcttttgtgcgatatgtggatgaagtggacctgcatagactttaagcttcgtaaata
attttcttcctaatggacctttaggaagcatacctttaaccgatttctcaataattctat
taggcaatcttctctgtagctgctcaaaggtttctacttttagtccaccaggctgcccag
aatgtcttctatataatttctgattagttttatttccgcttacatatatttgagaagaat
taatcacaattacataatctccagtatctagataaggcgtataggaaggcttatttttac
cccttaatatactagaaatatgagtcgatatcctaccaagtgtctgattttttagcatcta
taatataccaatttggattagtatttaatgatggtacttgcgttttgttcatgaaaagat
taatacttttagttaacaataagagagtaggtaaatgctgcacatataatgtcgatacaa
ataatattt-tgtaatatta-------------ttgataaagatagctttcatgacagtt
acttaaattgtttgttagcaagaaaacttatagttatcaataagtgataata-ttgtggc
aatataaaataggttactcactttttttcttttggcaaacttatacctaattttttttgt-
agagcatatataacttcttcagctgattttttggccaaagtttttaatttctagtagttct
tcttgagagtaatctagtaaatctgcaatagaatgaatttgagctcgttttaagcaatta
tatgctcgaacagatagctggagttcttctattaaaacttgactaatttttttatcttct
ttgctacgataattatctgctgatttaaagtctaaatttctcagtgagcagaaaaggtta
gttaaaactgtagcaccttggcttatagcttcttggggagagatactcccatttgtccat
atttgtaaaattaacctgtcttttatgctattagtacctatacgtacttcttctacttta
taattgactttattaactggcataaaaactgagtctacttgcagaaaatctacagataat
tcatctactgctttttctgctaaacgatatccacaactttttttcaattttaaactccatt
tcgaatattgtattattgcaaatagttgcaatatactgcctaggatctactacttctata
tcagaagatagttcaaataatccagctgtaactatagctggcccctggactctaattctg

ccaatttgagattctttattatggcttttaaatactacttcttttagattcaatagtatt
tctagcacatcttctcttaccccaggaattgtagagaactcatggtttactcctgcaatg
cgcacagcgactatagcggtaccttcaagatctgataatattgatcttcttaaagcattg
cctaatgtaatgccttgtccttggtttaatgtttctattacaaaactaccgtactgtcca
cgtgctccatctgttcttgattctacacattcaatttgaaattgagccacctaagcaagc
tcc-ttaaaaataatcaatggttagtaactctgcctgcgataagaaagaatctatacacg
gcgtttcttgggggggacggcagccattatgaggtacaggagtaatatcttttattaaagt
aatctctaatcctgcagcttgtaatgctcgaattgctgtttcgcgacctgctcctggacc
atttactaaaacttctgtttgacgcagaccttgatccattgcctgtttagcagctttttc
agctgcagtttgagctgcaaaaggtgtacctttttagctcccttgaatccactagcacc
agaggatgaccaagataacgtttcaccttttaaattagtaattgtaacaatcgtgttatt
aaatgtagatttaatatgtgtaacaccattaactgcattacgtctattttttttactcc
agattttttaattgtctagccatcgtgtttcgtctattatgtaattatttaaaatatta
tttccttggagctttttcttacctgctactgtttttttacctcctcttcttgttcttgc
attagttcttgttctttggcctcttaaaggaagaccaagacgatgccttctacctctata
agtactgatctccataagtcttttaatactcatagattcgaaacgtttgaggtcgccttc
aatctgataactagattctaaaatttctcttatactaacaacctgttgatcatttaaatc
ttgacacttaatatcagcatcgacgtttgttttttcaatatttcttttgagcgagatag
tccaataccataaatatatgttaaagctatttctattcttttattccttggaagatctac
tccagcaattctagccacgttgtgctttctccaataaataatatttac-gtttacaacag
aagattaca-gaattagccttgtctttgcttatgtttaggattagtacaaattaccatga
cttttctgtgacgacgaattattctacatttttcacacattttgcgaacagaaggacgaa
ctttcatattaa----ctctatatactaggaatattactg---------tattaattatt
tacatgtgaaatgtcaataatatgtttgattactttaatttctatagctttcacttcgtc
atactatcatattttttgatatgacatatgtttgaatttgcttagctgtgtctattgcc
actccaactaaaataagtaaagatgtcgctccaagacctctcaagttttgtatctgagta
accttttctattataaacggaatcagcgctactgtaaataaaaacgaagctccgagaaat
gttagtctatttaatataacttgtaaataatcaatggtcgcttgacctggacgaatattc
ggaatacttgcacccatttttttaaatttattgcaatgtcttctggattcattactatc
gatgtataaaaataactaaaaaaaagaattaatgcacaataaagaatgaggtatagcgaa
ccgttaggacaaaatagatatagaatttggagtaacgttttattttgaattatttgagtt
aaataagatggaagagccatagatgcagatgcaaaaacaataggcataactccaccttga
tttaatttcaaaggcaagtaactgttcggatccaaaattgaagattttcctagttgtctt
gctgaaataattttaattcttcttgttccttcttgtacgcaaattgtaattataatcatc
aataagaagatagcaatgaacaatccaaacttaagccccgcattgctatagtttgtatca
aaaaatgactgtgtaaaatttttttggtaatcctgatacaatattttgaaaaataagcaaa

gaggctccattcccaattcctttttctgtaattaactcagataaccacataatgatcatg
gagcctgccgttaaagctaaaacagactcacaaacaaatgtaaaattccagttaaataca
tacggctttacccatatagagattgctccagattgcaaagtggcccatccaagagacaaa
tatcgggtaatttgagttatttttggcgtcctagctcaccttcttctttctgtaatttt
tctaaatcaggaataattttagtgagtagctgcattacaattgaagaattaatataagga
acaatacctaaggcaaagatccctattgttgaaaatcctcctccagaaaaaatattcaaa
aaatttactaaagtatttttttctacgcttgcataaaaagcatcatggtctatacctgga
acaggtataaaaattcctaagcgcgctaagactaataaaaaaagagtaaatataatgcga
tttcttaagtcactttttttggctcataaataaatt-----ttaaaaaagacaccga--gt
tagttaagaaccctattccaatacttcgacaaatttaaaatagggcgctggcaacaagct
atagaagtaatggca-taatatcttatgttgaataaagactttctatagaaactcctcga
tctttagctgcttcgctaaaagttcttaattgagttaggccatttaaaacagctcttgca
ttatttagtgtattattagatcctaactgtttagctagaatattttgtacacctgataat
tctaatactgtccttacagagccaccagcaattacaccagaaccaggtgcagaaggcctt
agtataactttggctgcaccagaaattccattaataggatgaggtatagaattagatttt
gtaagtggcactgtaactaaatgtttttttagcatcagttactcctttttttaactgctcca
attacatcactagcttttcctactccaacaccaacttggccctgctcattaccaacaatg
agaataactcgaaagcttagtttcttgcctccttttacaactttagtgactcttttaact
tgtacaactctttcttcccaaccactatctttatctttgcctttactctgtttttttacga
ttagccattttaagcgttattcctaa----ttaatta-tgactttaaaaacccatacctg
cttccttagcagcttcagctaaggctttaactcttccatgatataatttccctcctctat
caaagacaacattcgtaataccttctttaatagacttgtgagcgagctgttgaccaacta
cgcgagaagtgtcacaatttgagcctattttatcaaattcttgattatttaaattaacag
atgatgttgagactaatgtgattccttttgtatcatcaattatttgtgcatatatatgtt
tattggatctaaatacgcatagacgcggcctacttgaagttccctgaactttttttgcgaa
ctcttttatgcttatgaattctagtctgcttagtatttattttcattattatttaccttt
tccagcttttccagcttttcttctgacaaattcaccttgatatctgattcccttcccctt
gtaaggttcgggaggtctagtagagcggatagttgaagcaacctgacctacaacttcttt
atctatacctgaaacagtaatattggtattattttcaactgtaatttgaatattttcagg
aggttttattttgactacatggctataaccaacgcttaatattaaatcttgattatctat
ctgtgaacgataacctacaccttgtatttgtagtttcttagtaaacccattagaaacacc
ttcaatcatgttactaataagagttctggataatccatgtaattgattagccatttttgt
ctgtttacttgtcctaacagaaattttatcattaagtatctctaagttaataccagccgg
cagagttctagatagagtacccttaggtcctttaaccgtgatggtatgaccatcaaactg
agtactaagattcgccggcaataaaattattttttttccaatacgagacat-ataccacc
tataaaataattaccaaatatagcataatatttcaccaccaagaccatcatgacgagctt

gtctatctgtcataacacctttagaagtagaaataagagcgattcccaaacccccaagaa
ctctaggtagttctttatggtttgcatatactcttaatccaggcttactaattcttttca
gagcagtaataactggttgacgattttttaccgttatattttagagaaatcattaaatgca
tatctatgccttcacccatttgttcaaaattttgaacaaatccttcttcttttagtactg
ttgccatattgcatgtcattttcgttgctgggacttgcacaatttgatgtcttgctaagt
ttgcattacgaatacgtgtcagcatatcggcgatcgtatcgttgaccacc-agatcctcc
ttacgataaattgaattatttaatatgtttttaagattccttaaaaggcataccaagctt
tttcagcaaagctagcccttcttgatctgttttggcggtagttacaattgatatatcaga
gccgcgaatttgatcaatattatcgtagtctatttctggaaaaattaactgttcgcgcaa
acctaaattatagttacctttgccatcaaaactttttggactaattcctctgaagtctct
aattcttggcagggttaatttgattagcttctctaaaaaagagaacatcttgtcttttcg
cagatgtactacaatgccaataggaacttcttctcgaattttaaacccagcaatggattt
tttagcttttgtgacaatcggtttttgtccagtgattaatgttaattcttgaatactact
ttcaagagctttagcattctgagaagcttcacctagcccacggttaatagtgattttcgt
aacacgaggaacctcatgtacatttttatattggaattcctcttttaaagcttgaataac
agttgttttatatttttcttttaatcccattactattatgtttt-acaaattatttaata
agttcacccgtctttttcagttttcgga-ttttttggcctttatcgttaatgattactga
agatctactagcaatattattctgttcactaaatagcatgacattagaactatgtattgg
agcttcaaatttaataatttctccggtctcaccttcttgttgaggtttttttatgttttac
ttttaaattaattcctttaacaaccactttacttgttttgtggatgattgcaataattgc
gcctatttttgttttatcacttccagaaataacttggactaagtcacccttttttcaattt
aatttttgtattaactttagtttttttgaaagaacttctcatcatattacctccggcgca
agagagactattttagaaaaatttttatctctcaactctcgagctataggtccaaatact
cttgtaccacgaggattattatcttgattaataataacggctgcattatctccaaatcta
atactcataccatcagttcttcttaaagctttgcgagttcttactactacagctctgaca
acatcagatcttttaactggcatattaggagatgcatctttaactactccaataatgaca
tcacctatcgatgcataagaaggattactagtacctaatactcgaatacacattattttt
ttagcgccactattatccgcaacattaagatagctttgagtctgtatcatacttttattc
tatataagattaattatcaaaagacttagataagatattaaccatcgtccaacactttgt
acgacttaaaggccgtgtttctcgtattgtaactacatctcctggtgcgcattcattatt
ctcatcatgtactttatatttttttgtgcggatcatagtttttgcatattttctatgagc
aattctattttctacagctactactatagttttattcattttattactgacaactttacc
agttgtttcttttaaaggcatagtttttatc--ttatatatatatacagtgaagttaaat
tataaaataattatgtggctggcttactccgagacttttcaacagttaaaagttgagcta
atctatgcttggaatgcttaaataagtgaggctgaaaatcttgtcttgtagctcttttca
gccttaagtcaaatagcactcttttaattacaagtatttcttcagctaaagaagaattat

ccatttttgtaacatctgatattttagggaaggtcatactttaagattctattgtatttc

gaactataaatttggtctttattggtaatttataagaagctagtttcatagcttgctgag

cagtttttgtggcacgccagtaatttcaaatagaatgtgaccaggcctaataactgcga

cccaatattccggagctcctttaccagatcccattcgggtttctgcaggacgcgctgtta

caggtttgtcaggaaataccctaatccagagcttacctcctcgcctaacatatctcgtaa

tagttctgcgagtagcctctatttgtctagatgttaaccatactggttctatagcttgca

atgcataatcaccgaatgcgattgtattacctttactagcagaacctttcattctgcctc

tgtgttgttttctaaactttgttttcttgggacttagcataaaaattaaacaatttaaat

aattaagaagctgttagatcagaagtttcctgattatatacagattctgtgcttttcgat

tctggcaaaatttcgcctttaaagagccaaactttaactccaagtacaccatatgtagtg

tgagcttgtcgatgacaataatcaatatcagctcttaaagtttgtaaaggaacacgacct

tctctaacccattcgctcctagcaatctctgcgccattaagacgcccagaaacttgtatt

ttaacgccttgtgtatttgctctttgcgtcctttgaacagcttgtctcacagctcgacgg

aaggcaactctttttttctaattgctgagtaataaattctgcgactaaggttgcttcagaa

tcgggatctgcaatctctaaaacattgactctgatttgtttactcgaatctagaattgaa

gataacgattttcgtagagactcaattcctgccccagactttccaagcacaatgccaggt

cgagctgttgctattaaaatttcgacttgatcaactttacgattgatttcgattttagca

atgctagcattactaagtttagagtgtataaacgaacgaattttatgatcttcttgtaaa

agaactggatagtctttagaattggcgaaccatgaagaacgatgtttttgggtaatgcct

attcggaaacccaaaggatgaatttttttgacccacagtatccttttctataaattaaaag

ttataattactaattgactttgaatac----aagactacaattctgagacacataatgta

atgtgacaagttggtttatgtatagggaatgctcttccttgtgctcttggctgaaatctt

tttagcgtagggcctttatctgcaaaagccttactaataaataactcacttttgttcagt

ccatcattatgctcagcattggcagcagcagactctaaaatctgctttatataagaacac

actcgatatggcataaattccagaataattaatgcttcttgatatttcctacctctaatt

tgatctaatacacgacgaactttatgtggagaaagacgaatatatttccctactgcttta

gtttcttttacgtttttttgtaatactcataattatagtttaacgacgcgcttttctatca

ccttttatgtgagtacggaaagttcttgttggaacaaactctcctaatttgtgacctacc

atttgatctgatacaaagacaggaaaatgttgtttaccattataaacagctattgtatgc

cccaccatatctggaataatcgtagatgctcttgaccaagtttttaaaacttcttttttc

ccaacagtatttaatgcttctatacgtttaagaagactcacatcaataaaagggccttta

tgtatagatcttgacataataataaattattagattcttaataaaataaattctatttac

ggcgacgcaaaatgtatggggtgctatacttattgggattccgtgttttaacacctaatg

ctggtttgccccaaggtgttaccggacgagctctcccaattggagatttaccttcaccac

caccatgtgggtggtctactggattcattacgacacctctaacagtaggtcttttaccta

gccaacgatttctaccagctttaccaagagtaatattactagcgtcaatattaccaactt

gccctatagtagcatagcattcttttcgaatcatacgaacttcgcttgagggcagtttca
ccgtgacaaaagagccttctttggctactatttgagcataagttcctgctgcacgaacaa
tttgccctccgcaagaaggtcttaattctatattgtgaacagctgtgcctagaggaatag
ttgataaaggtaatgcattgccaacttcaataggcgcagtgggaccggacagcaccattg
ctccaacacgcaatgatcgaggatgtaaaatgtacctttttttctccgtcaagataatgca
ataatgctattctagcatttctgttcggatcatattcaatagaggcaactttagcaacta
cgttgtgcctgtttcttttaaaatcaattaatctatactgctgtttatgtccaccacctt
tatgacgacacgtaataacacctctattgttgcgacctttgcaaaaatgatgtttaacta
ttaaggatttttctggcttatcagtagtaatctctgaaaaagtagaaactgttctatttc
ttgttccaggcgtataggcccggtataaacgaattgccatataggaaaatttaaaaagag
taaatatttta------ctgtgtaatacaattaagtttctgggaataagttaatagaatc
ttctggcgcgagtgttacgattgctctcttatagtgaggccgtttacctacaaatctacc
aatactcctttttttcttaggtggatggcaagtgttaacacctgtaacttgaacattgaa
aatatactgtatcgcagctttaatattaattttagttgctttgggatcaacagcaaaaca
gtactggttttcttctaataattttgtagttttatcagtaatgattgggtatttaactaa
atctaataagcctcttgaatcaatgctatccattatatacctcttgtattttagataaag
catcaactgtaataataattttatgcgcagctaaaagggccatgatattcaaagtgtcag
ctgaaataatttctacattatttaaattgcgaatggaaagatagacattcggatcttttt
tatccactacaactaaaactttttttgtttaaatcaagattccatcggtgaatagcttcca
taaacaatttagtttttggctgctgaaaataactattaaagttttctacaattaacgtat
tgactgatttattatttaacgttgtccttaaagctaattgtctttctttttttattcattt
ttttagtaaaactacgaggcttgggtccaaagattacaccaccacctctccataaaggtg
agcgtatagaaccggctcttgctctaccagttcctttttggcgccatggtttgcggcctc
cacctcggacttcacttcttgttttggtattagcagaaccttgtcgtttttcattccttt
gtcttactaaagctctatgaactaagtacataccagagtcttgactcactttaagattca
aatctgcattaccacttatttcacctttccaattatagacttgataatttaattgtgttt
taactgtcattagatatatcgcgtgaaagaatattgttatttactaatttttactaaatt
accaggcttacctgggacagctccttttacaatcaacaaatcattttctgagttaatact
tataatttgcaaatttttttatcgtaactttttttattaccaagttggccagccatattttt
tcctgggtaaaccctcccaggcgtcgtaccagctccaatagaaccaggttgcctatggtt
tttcgagccatgagacattggacctctactaaaatgatgtctcttttgatagcctgaaaa
gcctttacctatgcttttagaagaaacattgattgtctggcctactttaaacatatctgt
agacaagatttgactaacttcaaagtcatcagtagattttactaagtactcacgcaaata
tttcagtgggggagcttgtgacttctttaaatgtcctaataatggtttatttaacttttg
ttcaacaacttgcttgtacccaacttgaatagcattatagccatccttaggtaaagattt
aatttgagtaacaatacacggcccaacctgaattacagtaactggaattgacaagccagc

ttcatcaaagaattgggtcatacctactttagtaccgagtataccaacagacactagatt
tctccttttttgagagatttatactattatcgtacatgaaatttttacttcaaattgcctg
aag--taaactgaactaagagatgtctgagattaatcatgacaacataaagagaaaaaac
tttacttattcaatctaaaatagaaatagtttttgtccagttacattatgtatagatta
aaggcaaacttattatattatttatacactacg---------------------------
----tgttcaaaataatatgaaccagaagacatgaattcttttag-----atttgtacaa
cattacattattagtactgtctaaaatata----gatagaatattgaatttctacctctc
tagaa-------aaactaaaattgtgttcggtaactacaccttttatttacatatttat
tgtgcagtatatatgcatatctaatacagcacaggtaaaaatatagc----tgtatagcg
aatatacttacaagaattagagaggtattaatgggtaaagttgttggaattgacttagga
acaaccaattctgtgattgctgttatggaaggaggtaaacctactgtaataccaaacgca
gagggttttagaaccactgcttctgttgttgcatatactaaaagtggagataaacttgtg
ggacaaattgctaggaggcaagctgttattaacccagagaatactttctactcagtcaaa
agatttatagggcgtaaacaaaatgaaatttctcaagaaattagacagacatcatataat
gttaagactagtggatccagcataaaaattgcctgtcctgcacttgataaagattttgca
cccgaagagatttccgctcaagtactcagaaaacttgttgaagatgctagcacatactta
ggcgagactgttacacaagcagttataaccgtacccgcttactttaatgactcacaaaga
caagctacgaaagatgccggtaaaatagccggtttagatgtactaagaattattaatgaa
cctactgcagcttccttatcttatggactagacaaacaaaataatgaaacaattctagta
tttgaccttggaggcggaacatttgatgtgtccgttttagaagttggagatggagtattt
gaagtactttcaacttctggagatacacatttaggtggagatgactttgaccaacaaatt
gtggaatggttaatcaaagatttcaagcaaaatgaaggtattgatcttgcaaaagataga
caagcacttcaaagattaaccgaagctgcagaaaaagcaaaaatagaactatcgaattta
actcaaacagaaattaatcttccatttattactgctacacaagatggtcccaagcactta
gaaaaaactgtaactaggggtaaatttgaagaactttgctctaatctaatagacaaatgc
agtattcctgtaaataatgctctaaaagatgcaaaattagaagcttccagtattgatgaa
gtagttttagtaggtggatctacaagaattcctgctatacaacagatggttaaaagatta
ataggtaaagatcccaatcaaagtgtaaacccagatgaagttgttgctattggagcagct
gtacaagctggcgttttagctggcgaagtcaaagatattctattattagatgtgacacca
ttgtctttaggagtagaaactttaggtggggtaatgacgaagattataccaagaaacact
actattccgacaaaaaaatcagaagtattttctacagctgtagacaatcaaccaaatgta
gaaatccaagtgcttcaaggtgaaagagagttgacaaaagacaacaaaagcttaggaaca
ttccgattagatggtattatgcctgcacctagaggcgttcctcaaattgaagttacattt
gatattgatgctaatggaattttgtctgttaaagcaaaagagaaagctacaggcaaagaa
caatcaattactatatctggagcttctactttacctaaagatgatgttgaaagaatggta
aaagaagctgaagaaaatttcgatacagaccaaaaaagaagaaagaatattgatactaag

aatcaagcagaatctttatgttaccaggctgaaaaacaaattaaagaatttgaagataaa
atcagccaagatttgaaaacaaagatagaggaactaattacagagcttagatctagttta
gagaaagaagaatatgaaaatattgaatctatctctcaacgactacagaactcattaatg
gatattggaaaaatggctgctcaggctgaaagtaaaaatacaaacacaaaagatgatggc
actgtaattgacacagatttctctgaagctaaataaaaaatt-agcgggtaacgcgattc
gaacgcgcgacatcaaccttggcaaggttgcgctctaccactgagctatacccgcacaga
atatattattacaaacatggtaacatttgtcaatctgt--------agttatat---agt
tgagctattaatcattaaactatagattacaaacttgttaaaaacaaaaaaactattatc
gttttttaaa-aaaagaagtacttatataag-tatataagtacttcttttattcaaaaa-
-c-aaaatataattagatatcagtt-------ttaaactactaaggaattaaccaagcta
gtaacaattactagtcctgtccataagcctgcccctgtatagattaaccccttttgattgt
tcccattgaccgggtgatgctagaacaacagggacgcctactactaacacagtcgataat
acaattaataatagcactaataattgaattgcaataatcatttattctattctcctaaaa
aattcaaatgtgaataatattgtatataatacaatgaagctttataccttcatcaattta
tctatgaagaaataatttaaaattattgttttctcaataaatattaaattaagtattaaa
atctttattgtaaacaataacactatcttttttaa-atagataaaaaatttaaagaaag
tctttttaaaaaaaatttagaataaggtgatatgctgtataaaaagctaagtattatta
gttattagagaaattttaaaataagaggtaaaacatatgaattcagccctttttttagcg
aaactaccggaagcttatgcagttttcaaaccgatcattgatattttacctgtaattcct
gtattttttctacttttagcttttgtttggcaagcagcaattggttttagataaccttga
agctataacaaaaaatagcaaatataaaaagtaaaatttttttgcgctctaatcttatatct
tacacaataataactattatggttgaacccttactatcaggaattgttcttggccttatt
cctatcactttatttggattattagtagcagcttatctacaatatcaacgtggaaatcaa
ttaggactataaagcctaggttatagatatttataaaagctgtaataattatggtttta
taaatatctacaaaccagctaattataaatacagaaaaaatttattgaaaacagattga
taattggcaatactgattttatagtaagatagcgaatctttcataagataataactattt
taaagttatttgaacccattaaactttctatatgacaagctgcttctttctatacaccaa
cccttcttgaattggatttgctttatttaaaaaagctaatacttgtgaaagatctttaac
actggcttcgctacggaatggattctctaatttactatactcgggtctcttaataagaac
ttttattacagccattagcgaaacatcatttagtgcaatcaatctactggcttcaagaga
tttctcatcaactataggtggatcaaacatctcattaaacttttcggtgtttaaactagc
gaaaatatttatttgactattgttatttttttacaatcttcatcatcatcgttagcagct
tttaaaatcacaacgccctctggattagcgctggttaatctataattagtacgcgtatca
tcgctagttattatcgaaaatacttgtggatctctttcgacataaagccaagactttttc
aaaggatatggctccttgtatagcacaagtgtctagaaaacctacagtaagattgtagtt
acgacccattatacaagcacaagcccccacacctagatttataaacgactgttggcttcg

agcatttttacaagaaattgctttcagtcgccgatccatagtaattacaccttccgcata
tttagtgctgacagtagcttataaaacctcatctttagcaaacccttgtacaattctaat
ataatcggataccaagaagtgatctattatatcaacatgcaaagtagggtaatacggatg
cggttctataggaaagggatggattttaattctagtaagtagccaaacataataaactt
ggattcaaataaaggtaattttatattaaagtatgatatagatctctttcatatattcta
gacatttgagtgtaagacaatcatttctttactatttataatcttgatttgtgatgaaat
agcaaaattcttatcgccataagcgctcttatatagaaatgtctgagttatgaaatctac
atcgactaataaagaaaaatatatttatttattattttcaatattatcaatgaaaatact
tttagtataaatatactgatttttttctagtaaaattgtttaccagctcgacttagtaac
tccaggtaaaagacattcatgagacatttctcgaaaaacatggcgagataaaccaaaatc
tctataataccctctacttctaccagttaaccagcatctgtttctacctctaacggctgc
gctgtttctaggcatttcttgaagtctttgtcttagttccattttgtcagaaaagcttat
agctttttttagttgatctttaatagccagtctttttaaagagtattttttttttaagctt
ttccctttttaagctctctttggatcatattttttttttgccatattataa-aacttactta
aaaaattacttaatcaatatactagctaaaaaatctctgaactgcaacatttatatggtc
aaaaacatatagcataaa-------------atagatattttatgaattatact---act
ttatgctattgaaagaaatttatttaataaagttcttgttcaacatgagttgaaatagtg
caatcagatgtaggataagcaatacatgttaaaacataacctgctaaaagctgatcatca
tctaagaatgattgatcacgttggtctacacccccttcagtaactttacctgcgcaagtt
gagcaagctccagctctacaggaatacggaagttcaacaccttcttcttctgctgcatct
aggatgtatgtatcgtccgcacagttaaatgtaacatcaatattttcatcttcacatact
aaatgaattttataatctgcca--ttttttttactccagaataaagatatgcttgaaactt
aataa--taaaac-aatttaaaggacttgcccaaa-attgtatattagtttcattatgca
taatagactgcaaaaaaagaaaataatatactattgtaagaaattaagctacatttcata
ctaaatagaactattgcaggttcaaagtatatatataaattcatatagttaaaattattt
ttatcatttaaataactaaacttaatactct-agacctttagattaactgca------ca
gctaatcttttacaataaacaaataagtaattatgacttcaatgataaatcaacaaatag
aactaaagcctattcttcgatttcagataccacatttatattcgagttttcttatacagg
aaatcaaatctttaacaaaaagactctttttgcaagtttggagaagacctgcgacattga
tggcaggtattattcagcctttattatggctaattctattcggtgggctttttttataatg
ctcccgtgaatttatttactattaacacaagctataactgttttttaagctctggaatta
taattttcacatcgtttactggtgcacttaattcaggtcttcctttaatgtttgatcgag
aattcgggtttttaaatcgcttattaacggctcctttaatatctagaacttctattattt
tatcttctgctacttttatgacttgtattagtttaatacaagttatattcatagttacag
cctcattatttatgggaaattctcccttaaacagtgatagtgcgatagtttttggactta
taattctattagtaactgtgggagttacaatacttagtttagctttatcttttactctgc

ctggtcatattgaactattagcacttattttagtaatcaatttgcctttttttattttcta
gtacggctctggcccctctatatttcatgccaccttggctccagttaattgctagtctaa
atcctttaagctatgcaatagaaggtacaagatatttatattccagcatacattggaatt
ttacagagtctattataaaaattagttggggagatgtttgtttagggcaaattattgcat
tattagttattctagatataattgcagcctaccttgtctctaagatactaaaagctaaac
ttaatta-----------------aaattttaataaaaaatttattaaatttattttaga
aagattttactatataatactagtagtagtatggataattgaaatagaaatagtttttc
aaaaccaaagctattatcaacaatgataaacatttgtaagaaagtcaacaaagtatgttc
ttattcataggaggcatgtagtcaatgggactaccatggtaccgtgtacacacggttgtt
ttaaatgatcctggacgcttaattgcagtccatctaatgcataccgcacttgtagcaggt
tgggcaggatctatggcattatacgaattagctgtatttgatccttcagatccagtatta
aatccaatgtggagacaaggaatgtttgttatgccattcatggcaaggcttggcgtaaca
gattcttggggaggatggagtattactggagaaagtgtttccaatcctggattatggagc
tttgaaggagtagctttaactcatatagttctttctggaatgctttttctagctgctatt
tggcattgggtttattgggatttagaattatttcgagatccaagaactggagaaccagct
ttagacttgccaaaaattttcggaattcatctattgctatcaagtttactttgttttgga
tttggtgcttttcatgtaactggattattcggaccaggaatgtgggtatctgatggttac
ggagtaacaggtaaagtactgccagtggctccggcgtggggaccagaaggatttaaccca
tttaatcctggaggagttgcgtcacaccacattgctgcaggtactgtaggtatattagct
ggtgtttttcatttaactgtgagaccgccgcaaagactataccgcgctctaagaatgggt
aatatcgaaacagtattatcaagtagtatctctgctgtttttttctcagcttttgttact
tgtggaacaatgtggtacggctctgcaactacacctattgagttatttggtccaactcga
tatcaatgggatagtggatattttcagcaagaaattgaaaaacgagtagagaacgctatt
gctgatggagctgctcctagcgaagcatggtcaagaattccagataagttagccttctat
gattatattggtaataatccagcaaaaggtggtttattcagagctggccctatgaataaa
ggagacggagttgctgaagcatggcttggacatcctgtatttcaagataaagaaggaaga
gaactgagtgtccgcagaatgcctgctttcttcgaaacttttccggtcattttagtcgat
aaagacggtatcatacgagctgacattccatttaggagagcagaatctaagtacagtata
gaacaagtaggtgtaacagctagtttttatggcggaaaattaaatggccaagtatttaat
gatgctcctagcgttaaaaaatatgcaagaaaagcccaattaggggaggtatttgaattt
gaccgaactacactagaatcagatggtgtattcagaagtagtccaagaggctggtttaca
tttggtcatgctaatttcgctttaatatttttctttggacatctttggcatggttcaaga
actatatttagagatgtgtttgcaggaattggagccgaagtaactgaacaagtagagttt
ggagctttccaaaaattaggcgacagaagcagtaagaagcaaggagctgtataatgct-a
agccaaatacagtctttatc---aaatatttccaatattaaaatagttcaggaaaagttt
attgtctaatagttactaattaggagaacctatggaagccttggtttatgtatttttatt

gacaggaacactaatggttatattctttgcgatctttttagagaacctccaagaatagc
taaatagctttattaagtttaaccttacaataatagaaccactttccaacttattagtgg
agagtggttctattgtaggttacagtatattattcttcatgctcttcaaaaggatctctg
agatcttttgaagcaggaccaaatgccgtatatattgaatagcctgtaatacctaaaagc
aaacttgaaataaaaatactaagaactgttgcagtttccataattcaatcagtattaaga
tgaactttctctataatgatattatcaatattaaaacatttaattattaaatttatcaaa
atatattatggcacttagaactagacttggagaaattttaagacccttaaattcagagta
tggaaaagttgctccaggttggggaacaactcctatcatggggattttcatgttactatt
tttcttatttttgttgatcattttgcaaatctacaattcatcactagtactagaaaatgt
agatgtagattgggctactttaggtag----ctaattatacttttttctttaa--------
tttaataacattaaagataat----------------aagcgatcgctattataacataa
tagcgatcgcttaatttctactttactatata--caaaagtaattattttcctaagtcaa
ttaattcgctatttgcaaagctattagtattgacattattataattaacttttttcaaatc
ttactagtactggatacttgataccactttatctattgctgcaacagttccaagttctt
gataccagtatgattcctttctcaatatttttacttttgagccacgttccatatttttag
tcctaagatagattattaaagattagtataatagataaagaaataaaaat-aaaaaaata
cataaaattagtaccgacaat---------taaatagtaaaaggagagagagaggattcga
accctcgataactacagttatgacaattttcgaaattgttgcaataaacctctctgccat
ctctcctaaattaaaactctttttccccgaattagataagaataacatagaactaatgta
gttgcctagaaaattaaaactatgttaaatataaattataatctttctatagatctcttt
acaaaatatgtacacagattctatatcagcttatttgaagatttttttaatcttatacaat
ttttactaaataaataaactctctcataacatgaaactatcttggaaaacacttcttctt
tggtctttaccaatatttgtcgttgggttctttttctggcaaggtttttttaggtccgact
actacagatgttggtagtaatatcgcaagttctagaatgacatatggaagattcttagaa
tatttagatatgggttgggttaaacgagttgacctatatgaaaataatcacacagcaatc
gtcgaagctgttggaccagaattaggcaacagggttcaacgaattcgagttgagctgcca
gcaagtgcaccagaattaattacaaagctacgcaaagctaatgttgatttagatgctcat
cctcctaaaagcacaagtgcagtatggggactattaggtaatttactatttcctctattg
ttagttggaggcttagcttttctatttagaagatctaatagtgctagtggcgggcctggt
caagcaatgtcatttggtaaatcaaaagctttatttcaaatggaagccaaaacaggagta
gtatttaatgatgttgctggagttgaagaagcaaaagaagagtttcaagaagtagtaact
tttttaaaacaacctgaatcatttactgctgttggtgccaaaataccaaaaggcgtttta
ttagttggtcctcctggtacaggaaaaacattattagcaaaggcaattgctggtgaagca
agcgttccttttttttagtatttcaggctcagaatttgtggaaatgtttgttggagttggt
gcttctcgcgtaagagacctattcaaaaaagcaaaagacaatgcaccttgcattgttttt
attgatgaaattgatgctgtcggaagacaaagaggaacaggtgtcggaggtggaaatgat

gaaagggaacaaacattaaatcagctattaactgaaatggatggctttgaagggaatact
ggtgttattgtaattgctgcaactaacagagctgacattctagattctgcattattacgg
ccaggaagatttgacaggcaagtgtctgtggatgttcctgattttaaaggtagattagca
attcttgaggttcatgctaaaaataaaaaaatggagcctaaagtatctttagaaactatt
gcaagacgaactccaggcttttcgggagctgatttagctaacttattaaatgaagctgct
atttaacagctcgaagaagaaagaatgcaatgactatgtcagaaattgatacatcaatt
gatagagtggtagctggaatggaaggcacacctttaattgatagtaaaagtaaaagatta
attgcataccatgaggtaggccatgcaataataggaagcttattagaacatcatgatcct
gttcaaaaagtcaccttaataccaagggggcaagctagaggcttaacatggtttactcct
agcgatgatcaaagtttaatctcaaggtcgcaaatactagcacgtatcgttggtgctctt
ggtggaagagctgcagaagagattattttcggagatgcagaagtaaccactggtgctagt
aatgatttgcagcaagttacatcaatggctagacaaatggtaactcgattcgggatgtct
aaaattggcccattatctcttgaaagtcaaggtggagaccctttcttgggtagaggaatg
ggaggaggctcagaatactcagatgaggttgcaactaatattgataaacaagtacgagaa
attgtgagcgaatgttataagcaggctaaagaaattattgcagataatcgagtagttatg
gatagattagttgatctactaattgaaaaagaaacaattgaaggcaatgaatttagagat
attgttaaagaatacactgcaattccagagaaaaattactataaatctcaattttaa---
---------------atgatacgggactgacgggattcgaacccgcaacttccgccgtga
cagggcggtgctctaaccagttgaactacagtcccaataaaggg---tacagacagataa
tctcataattagcttgtgattgtcaaattatactgtaaaacatgaaaagta---------
---------tcctcaataatatttttcatgttttttattataaagccaattgattaatgca
taaggagagagagggattcgaaccctcggtacgaagttaatcgtacagtagattagcaat
ctaccgctttcgaccactcagccacctctccaaagagtcttacgtactcaatatatcaaa
tattattaaattttaacagctgtgcataaatatttatggagataagcggattcgaaccgc
tgaccctctcaatgccattgagatgctctaccaactgagctataacccc---tttcttaa
ggctattaagaaatggctcaagcgggatttgaacctgcgaccttgggcttatgaatcccc
tgctctaaccactgagctactgagccataataaaaaccattgataaatactataccataa
tttatcttaaaatataaatcaataaaaaagtctttactacaagtacatagatcttatatt
g----------------tccagatt----------------------aacatttcaaaga
tattaaattaccctctacgatatactctcaattcttttcactttat-tggcagtatttca
atattcagcttgtaa---------------------------------------------
--------------------aataagt---------------------------------
---------------------------------------------------aattttttt
g-----------------------------------------------------------
------------------------------------------------------------
------------------------------------------------------------

------------------------------------------------------------
------------------------------------------------------------
------------------------------------------------------------
-------------------------------caagaatatatagaa--------------
----------------------------------------------tcttgctacatatt
------------------------------------------------------------
------------------tacattattgttaataatt-----------------------
------------------------------------------------------------
------------------------------------------------------------
---agcctttgctaaacta-----------------------------------------
------------------------------------------------------------
------taaatctatgaat-----------------------------------------
------------------------------------------------------------
------------------------------------------------------------
------------------------------------------------------------
-------------------------------------tttagtagatt------------
-----agactaaataattatgtcttttaaattaggtaaaagtaaagaaagtatgtgagca
aaatatcttaa-----------------ttttt---------------------------
------------------------------------------------------------
------------------------------------------------------------
-------------------------------------ttagaaatattttattttcaaca
tttt-tttataatagagcagttaaaaactgaataaatagctaatatc-----ttatacaa
ccaacatcgagccaatgccttgatcagttaatatttctagcaagagagcgtgatcaattc
taccatccaaaatatgagctgaagctactccttgggctaaagagcgaatgcagcaattaa
ctttggggatcatacctccggagatcacctcggtctgagttaagtctctagcttgctgta
tatttaagtgactaatgagtgttgtaggatctgaagagttacgtaaaataccaggagtat
ctgttaacaaaattagtttttccgcattaagggctgctgctacttctcccgctacagtat
cagcattaatattgtatgactgaccttgtttatctgccgcaacactagcaattactggta
tgtagttattatgaattaagatctttaatagcttagtatcaacattctgtacttctccaa
caaagccaagattcgctttaccgctcggtctaggagtaataagtaaaccatctttccctg
acaaacctacacttttacctccttgcttattaatcgttgctacaagatccttattgactc
gtccaactaataccatttccacaatatccatagtaggctgatctgtcaccctaacaccgt
tctcaaattgtgataaaatttttagacgatctaaccacaaattaatttcaggacctccac
catgaactaaaataggacgcaaccctataaaagaaagaaaaacgacatcactgattactt
gatcttttaatttttggtttttcatagctgctcccccatattttattacaataattcttg
aggaaaattgttgaatatatggtagggcctcacttaagactttcactctttctgaattag

tcagcatgaataaaatagaattataatatataaatttatttgcaggacttatcttaattt
aatagatactagtaagtactttaaaacttttagaaaataaacatattattaatag-----
----ttaaaagtataaggatgataattatgagtaattttggaataatatattaaaattt
ccacgattttagttagtgtcattttcggattagttttaataattattagtcctttttc
gtgttatttaaaaaaccattgacaagttttttttcatcatatcattaaccggtttaatc
gcaattttagcggcaataa-tacagaaaatgctcaatatcgaatgttgttgagagaatta
atttaatttaattaaagtttttgaacatatataatattatattatcattaactgtgataa
tttaaatttcttttctat--aaaaagtcatacgtattaaatattatttgatagatattat
gttattaaaaccaatagttagctctgaaaaaactggtgcttttgcccttttagatagtat
tgttagacatggtgttaagcatatatttggttatccaggcggagctattcttcctatcta
tgatgaactttatgcttgggaagagtcttctttaattaaacatattcttgttcgacatga
gcagggagctgctcacgctgcagattcttattccagatcaacaggagaagttggagtttg
ttttgctacatctggtccaggagcaactaatcttgtttcaggtatagctaccgcgcatat
tgattctgtacctatactagctgttactggtcaagttggaagagcgtttataggtacaga
tgcgttccaggaagtagatatctttggaattactctccctattgtcaagcattcttatgt
agttcgagatcccagagatatgtctaggattgttgcagaatcattctttatttgtaaaca
tggtcgaccaggccctgttttgattgatgtacctaaagatgtaggattagagaagtttaa
ttatttttctattgatccaggcaaagtcaatatcccaggttgtagaccagttactacctt
gaagtctagacaaatacttatggcagctaaaatgatacaacaatctagtcaacctttgtt
gtatataggaggaggagctataatctctgattcacatcaaattattaaagaacttgttga
cttgtataaaatacctgtcactactactttaatggggaaagggattttaatgaagacag
tgaatattgtttgggaatgttgggtatgcatggtaccgcatacgctaattttgcagttag
tgaatgtgatctattaattgcattaggagctagatttgatgatagggtgacagggaagtt
agacgagtttgcttgtaacgcgcaagtaattcatgtagatattgatcctgctgaagtcgg
aaagaatagaattcctcaggttgctatcgttggtgatgttacagaagttgttactgcaat
attaaatctactaaaaacttctttccctccttatccggagcagattgtatcttggcaaga
gcgaattaatcgttggcgcgaacagtatccattactggttcccaaaaaatcaacaagtat
ttcgcctcaagaaatacttgttgcagctaataaactagcccaaaatgcttattttactac
agatgttggtcagcatcaaatgtggtctgctcaattttttaaaagttaaagctaaacactg
gctttcaagtgccggtttaggtacaatgggatatggattacctgcagcaattggtgctca
agtagcacatcctaatgatgtagttatttgtattagtggtgattctagctttcaaatgaa
tatgcaagaactaggaactatcgcacaatatcagttacctgtcaaaattattattattaa
taatcgttggcaaggaatggttagacaatggcaacaagctttttatggcgaaagatactc
tcattcccgaatgatagaaggagcaccaaatttccaaaagcttgcagaagcctttggtat
taaggcttttactattaatagtaggcaaagtatgcaatcgtccttagaagctgctatcga
gcatcatggtccagttttgctagactgtcaagttacagaaaatgaaaattgctatccaat

ggttgctccgggtaaaagtaactcacaaatgataggcattgccaaacctcaaagaggtac
tgcctcaaattatattaatgcaaatatctagattag------------------------
------------------------------------------------------------
------------------------------------------------------------
------------------------------------------------------------
---------------aaggtctatattccctcaaaaacaaagtg-gattttc----tttt
atgagtacatatttctaatagtatatacaatttttaaaagatagcctttgacgagaatt
gaactcgtgaccttccccttaccaaggggatgctctacctctgagccacaaaggc-tttt
tattatattgggccgggttggatttgaaccaacgtaggcaaagccagcggatttacagtc
cgcccccattaaccactcgggcaccgaccctaattagtttttattcaaaacataacttaa
acagtatcataatttagttgcttaaaacaagtcttttcataaataaaact-attaagtga
tggacatagctggactcgaaccagcgactttcacgatgtcgacgtgacactctaaccagc
tgagttacatgtccaaaaccttaataattattataacactatttattgattttataagta
tctatttttaattaaagatttgctttaaacgacctgctttaccaagaagatctcttagat
aatatagtttagctctccgtactttagctcttctaataactttaataaatgttattcgtg
gagaattgagaaaaaaaacgcgctcaactccaataccttgaaatgagcatcttaaagtta
agctagtatttaagctatttttcttcttctagccaagattactccttcgcataattgtt
ctctcgttttattaccctccttaaccatcaatccaagttgtattgtatcgccaactctaa
tctcaggtaagtcttgtttaataaaagaagtctccacacttttcatgagcttacttattt
ttgtactatttgctttcatatctagt-ttagctttataatg-ttttttgcaatattatac
attaaaacaataaatatgaaaattattagtagataagcagagtagtatctcagaatcttt
gtattatttattgctagactagatttacttcattttgtgttatcaatgtatttgcttaac
tatcaaaaaattaattcgctaattatagaataatgaccaacttatatttcta----atag
aaatacaaaattcttttatattaaat--taacctttagatatttaattcattgaaaagac
attaggatcaccgtccgaatacatacctgaaagttttaaatttggtatggcaaacgcaca
gatgaacatagaacagctcacaaaacgtttatataatttagtcatatatgtattaataaa
gatgcatcgctattaccggtgctcgatagactaatttgctacgaggctttgaggtcttta
acttagttttattagacttaccacttttcatcagtttagttcccttgccatgatcaacaa
cgcgcgaagtgtgtacatatgaaacactaactggtggcattgctgactgaaatttgtgag
cttttaaaaagacttgttcacaatcagaaaatgcaagtaaaacttcatcatcaacatcga
ttgagctagaagtatttatagatgtatcgccggacactagctcgttagagaacacgttat
cgggtattccgtaaatattaattgtggctggactactaagtcctataaaacgcaatgata
cttttagattatgtataaatcctgtagtagagtcatttagtccacgtgtatttggtattg
gcaacattaacaaattattatgactattcttaggataaagtatactaataaatttccatt
ttataataacagactgcaacctacgtataggcatcaaaaatttaaacaatagatcgttta
agaaagtttggacggatagacctgtaaaaactgattgcttgctattattcttaaaaacta

aaaccagcgagtaatctgtaaccgaatcaacaaaaccataaacaagtttaattttcacat
atttagtaggcgacatatatgtataagcaagctgatcaaacatcgtgtaatttaagaatg
atttatagacgaatgaattgtatttattaatctggcatatttcgaggtggtaagcgcaaa
cgacgatctgaagatgggacatcaatcgtaacaagattttgattgagcgcagcagcctca
cgtgccaacctatatctataattacaatagcgaataaatctgtagactagataaataata
aaaacaatgagtagtgaaaatccaaagtgtacaactaaacgaatgtatcggcataactga
taataggtcacttcattaagcgctatactaaaaactttaccggtcattccttcgcaatta
tcgtaaagtgttttcccatcaatcaaaccaaaagtcagatcaaaaggtttcttcatttca
ggcgtaatacgatccacagctggattcctaggaggaacaccttttaagagatcaataacc
ccattagtagtattgctaacaaattcatacgtatttgcagcaggattcgcagaagaggta
acagcaccatcgttcatcatagtttcatacaattgtgaaatcgtataagcacagaccggg
cgttctacctccttcgataagagtgtacctgctttggtaccaccttcgagaacaatttt
cgggcatcggtacgagttatagtcggattggcttttaggaaagctgtaactgcttttcgc
ttggtataagtatctgttgcgagtccagcaactatttgggctagaggtaacaccgtagct
ttaatagggaccggtataagaaatgatgctacagattttagacgtttatgttgatctggc
ataagagtacccttgattttaggtataaaaattgcggggacaaattatttacaaaatcac
ctattttcatattttgtcccagtagccttactaagcaacgcccgtcaaatgacgtaaaac
aggtgtattttaggggattaagccccaacccggcaaaatttgctaaattttcaattaaaa
tctcctttcaaaacatttggtttttatattctattactataacttcatttttgggttttc
ttaaggcctaaccgcgacttatttatattcactattcaaacttacttcaattcatttgtt
agtgttacactcgagccgtcattagcgatgtttcgctagtgtgtgtgaattattgcatca
gtgtttataccttagattgctattgtgttttactagttcactagttgacatgcgattgt
aaacgtgttatcatgtcttatttcctttattgttagaacataatatgagcataaatttag
cactgcaaaatcaaccagttcgtggtaggcgccttttgcatgatcattcatatcgccaac
gcaaagctgcggcaggacaatgtcaattaagtgtctgggtacctaaaaatcactacgatg
gtttaagagtcttattgacagatcatgatgtattatctaaagttctatcttattactata
gtattcaatcgaatccaccacacattaagcgaaatattcgtacaacatcacctaaagtta
ttgaaagaacactcggtagttaccagtctagcaagattagtcgccatactaatgattatc
aaataagtgatatgtctattaaaccagaagaaaaatttgataatgagacgaatttcataa
cggaagaattagctgataacaagttaaataaattatcagaagaatttgatgcaaagctta
taaatctatctacaaatattgatagtagatttaatcttttatctgaacaatttaccgcgt
taaacactagcgttttaacgctaacttcaaaagttgcggagatttcctcacaacaagttg
tgctaagacttgattgtgagaagaattttaacaagctaaatgataaaataaatgaacaat
cttttacgggtgaaactggtatcgatatgtttgataatacgaaaactcattcaaataaca
atgattcaacagaattagattctgatttagaaggatatgatgtgtgtgaattccctattg
tggataaatacgatctaatagatctttcaaaaatctcaggcaatacgttgcgagcgtata

atataagtcgatttacagatgcaagtaaatctactcagacaaagtttttagcagattata
agttggcatgtttttgtaaacatattatggagaaaatggttgcaaacgaagagccaacac
cctcattaagagcgttggctcgtgttgcaacatctaaaggctatggatatataaatccca
agactggaaagcccagtcttatttataaaagtagagttaaaagagttatgcaatctttca
atattccgtttaatacttaatagccaacagtttgcattccaaatttattatgcgttgttt
ataatttatataatgtgttctttatatctttttctaaatctacaactaatttattcctag
gatcttcacagaattgtctatgaatatgagctctttcaacttttgttagtgttgatctat
aggaacgtttaccagctatgatagccatgccaatcactaaaatagctctttgcaaatcaa
gattatttcctgtttgaaagaacatttcatagccttgcttatatttactgtataattgag
cttcctcccttttccaagaggattcctgagtttcgctttcaaactgaaactgtgataatg
ataattctcgaagctttttaacagattgtatcgattgtccaggatgtatacagttacgat
tgtatttgccaaggttttgtagatactgtttagtttttaatattcgactttcgacaatcg
gtttgattcgaggtatgtcagtaggattttcgttaatggctttaaataaattttgtaccg
gattgtaatgaactaatttattatcactagttattacataatcttgatttttgtatcctc
tcatttttattgaagggactatatttttatgtcgagtttgataatttgccttaccatgtg
ttgccaattcggtgaatatagcttcgacttcgacatcaaatagttttataatcgttagat
ctggtaacacgattcttaaatgctcggtaatcatttttgagattgttgaattatccagat
caataaatcttaatttattagtatcgccatcataaattaaacgggttttataatcaatgt
ctcccaaaggttttagagatatttgtctggtttttctactgtgttcaagatcatataaac
ttacggcagaaagtttgcgatctttttttcgaatgtacaatttttttcaaggtctaatgaca
cactatcagttatagtttgtgtaccattcagggttttctcaagataccaagccatagctc
tggctctcgccgtaacattgttcccgacaacggctttactaaccggaaattttggtgaaa
ccaatataccgtaaatcgtgttgattaatagtttaattaacttattcattgaactagtgt
ccggattatctttaggatacttttctcttaaatcaatcagacgtcctagtatcaatttat
caatatttacactaaaccaattatgatcttgtttattagatgttataacttcaaaattct
ctttgtcttgtttcaatttccaagtgttaacacctgtatgtagtttcaccttttcttcat
agtccgaaaaattattaactcttgcagattttggataaaatacacccgatataactttta
agcggtctaaatattcttttcgctgagctggtttacaaaccttatccaaccactcaagat
cggcgtatgtaaaaggagctaaatgaacttctcgactataatacttagtagattcatctc
taaatacattgtctgaatcttgataatcagtatcactcaattctaaatctgttaggtttt
taggcggataccaagacataaaccagtcatttttatgttctaataaatatcctgggtctg
ttgaaacacgaaaataccaaagaccatctactaactcatgtccaatatctgctaaagttt
ctctaagagttttatactgattgttctgagcagatttcttatattctataagtatcggtc
ggccaataggaaattcttgtacttttaaacaattaccataacagcttttgatatcaaccc
tggcaaaaatacctcgggacgctgtgtgcaatggtttattattaaagcagcgtccgccat
tagtttttgaaagataaatcccagtttgatttttatttgcaaacaggcttgtagtagaac

ctttcttgatatgtgacatcattgtacgcgaactaattttaagttcattttgcaatgcaa
ctgtaaataattcagctacggtcgtaccaatggtcaaccctgtacgcttatataacttgc
caaaacccaactccttataaatctttttaataagatgttcttgatttaatagcgcttcat
agacaagtagatcgcttgtagcgtattttataaaacgatttggatctgttattagtacat
ctaacatgttactcttttcagatactgttataagttctttatctataatagatgtacccg
tgttggctaagaagcttcgtaaagatatagctccttggatagcgcaggtatctgtgaatc
ctagctttatattgtactccttgccaaatatacgtattgcaataggtatatctatatatg
gaatacaccttctagaactaactctagtatatgctttaaacgcctttccattaatatac
aaccatgattctcgattttgttcgcagttgcttctaataatctttccttaacacttccgc
taacaatacgataacaatcagccaataagaaatgtccgtatatatctatatatatagttt
gattatctgactgtggtttaacagtttctgctgtataatttaatttgtttaaatagtcta
atactatacccgtaatttgttatgatctaagtatctcacccaattattttttttattgatg
tta--------------ctatatttaaatacttattttcgctaaagaattttacacataa
aacagagtaatcaaaacaacaagattaaggcttaagtatttattagttttgtgttataat
catttactatcaaaggtagtggcataacataaaacataaacaatgtttgaacgctttact
gaaaaagctataaaagtaatcatgttagctcaggaagaagctcgacgcttagggcacaac
tttgtcggaactgagcaaatattattaggattagtaggtgaaggtaccggggttgcagcc
caagtattaaaatctatgaacgtgaatctaaaagatgcaagagtcgaagttgaaaaaatt
ataggaagaggatctggttttgtagcggttgaaattcctttcactcctcgagcaaaaaga
gtactagaattgtctttagaggaagcacgtcaacttggccataattatattggtactgaa
catttgctaatgggcttagtcagagaaggggagggagttgcagcgagagtcttagaaaat
ttagcagttgatgtttcttcaattagagcagaagttatacaaatgctgggagaaaatgca
gaagctaatgtaagtggaagtaatactacacaggctagaagcaaaacacctacattagaa
gagttcggctctaatttaactcaaatggctatggaaggtggtttagatcctgtagttgga
agacaaaaagagatagaacgtgttattcaaattctagggagaaggactaaaaacaatcct
gttctaattggggaaccaggtgtaggtaaaacagcaattgcagaaggcttagctcaaaga
atagctaatagagatgttccttctatattagaagataaattagttattactctagatgtt
ggcttattagttgctggcactaaatatagaggtgaatttgaggaaagactaaaacgtatt
atggacgaaatcaaatcggctgataatgtaatactagttattgatgaagtccatacatta
atcggggctggggctgcagagggcgcaatagatgcagccaatttactaaaacctgctttg
gcacgaggagagaattgcaatgtataggtgcaacaactttagaagaatatagaaaacatata
gaaaaagacccagcactagagagaagatttcagcctgttgtagttggagagcctagtgtt
gaagaaacaattgaaatcttatttggtcttcgagaccgttatgaaaaacatcatcaatta
acaatgtctgatggcgctttggcggcagctgcgaaatatgctaatcaatatatttctgat
cgattcttgccagataaagcaattgatttaattgatgaagctggttctagagttcgttta
ctaaattctcaactacctccagctgcaagagaattggataaagagttaagagctgtatta

aaaacaaaagatgaagctattagagctcaaaaatatgaaacggcagaacaatatagagca
agagaaatggaaattaaagctcaaattgcagcaattgctcaaagcaaaaaaaatgagccc
gatttaaacttagaagatcctgttgtaacagaagatgatattgctgaaattgttgctgct
tggactggtataccagtaactaagctgactaaaagtgagtcagaaaaattaatgcacatg
gaagaaactttacatgggcgtatcattggccaagacgaggcagttgtagctgtatctaga
gctataagacgagcaagagtaggtctgaaaaatccgaatagaccaattgcaagtttcatt
ttttctggaccaacaggtgtaggtaaaacagaattgacaaaagctttggcttcttatttt
tttgggtctgaagcttctatgatacgtttagatatgtcagaatatatggaaagacatact
gtatctaaattaattggttctcctcccggctatgtaggttatagtgagggtggttatcta
acagaagcagtaagaaaaaaaccatatactgtaattctttttgatgaaattgaaaaggct
catcctgatatttttaatcttcttcttcaaattttagaagatggcagattaacagatgcc
aaaggtcgaactattgattttaagaatactcttctaataatgacttccaatattggatct
aaagtcattgaaaaaggaggaggtagtttaggattcgagttgtcagaagaccaaacagaa
tctcaatacactagagtcagatctttagtaaatgaagaattaaaacaatactttagacca
gagtttctaaaccgattagatgaaattatcgtatttcgtcagcttaccaaagatgaagtt
agagaaattgcagaactgatgttgaatgaagtttttgccagaattaagcaacaggatatt
caactaaatgtaacagaacgttttaaacaacgactagtagaagaaggctataatccaagt
tacggagctcgaccacttagacgagcagtgatgagattattagaagatagtttagcggaa
gaagttttatcaggtaaaattaaagctggcgatagtgctgtagtagatgtcactagcgaa
ggagaagtaacagttttactaggtgaaaaattagaactgctaacgtaaa------aaata
atttattaatctaaaagcattgaatataattt--attcaatgcttttt--gaagaaaatta
acaagatgggttgctataattgacaacctctgatattctttatatgtatata---aagcc
gggatagctcagttggtagagcagtggattgaaaatcctcgtgtcaccagttcaaatctg
gttcttggcat-tttaaatatttgaaaagtttgctagatgagcatcaaattttagttgaa
acatacctgttggtccatttctatgttttgctattataatttctgtcaagtctcctccta
cagcttcttgagtataataactctctctatatagcattataactaaatctgcatcttgct
ctatagaattgtgtacgataaagttgttcgcgataaaattatgtaagagttgacattcta
aatcaaatacttgttgcaagctaataaggcgaacatgtttaagattctcaaaaattaaaa
ataaagatggacaagaatttaagagagtttttctttctttaatggcgtt------tattt
gaatagcaagcatattattgtgattaatttcatcgcatcttttccatccttgtgttgtta
atagcttgtgattgcttgttaactttatatacttgccagattgtgtaataatttcataca
cagtttttttacctgtcttggaaatattagctttactgtatctaaataagtttgtttttt
gtgcagtaaagctattaacttca-atatgcttatgatgaaatttaaatagaatttg--ag
tctgactatgtaacggtaaatgaagatgactaaatttcgagatacatccactttctctta
gatcagataataaaggtcttttatcgctccgagtttcaagattcctattcagttgagata
atactagtataggtaaatttaagtcttttgctaatatttttaatgatctagtaattagag

atagttcttgagatctattatcagattgtctgctatcttgtaataattgaagatagtcta
taataattagttctatattttccсttgtaattttaagaattgtactttcgttttatag
tgtccgtagaaattttgcgctatcatcaatgtaaagattcagatctgcaagagctttgc
tttttcgaataacatactgccattcattattattaagttggccagattgaatttttgac
tatttaagtgacattcttgagctaaaatcctccttaggagttgttctatagacatttcta
agctaaataaaataacgaacgatttctctttgctcattatatatctagctacattaatag
caaatgctgttttccccattgaaggtcttcctgcaagaacaatcaaatctgacttttgaa
aaccttgtgtaatagaatctaattctgtaaaaccggaaagtacgctactatttatactaa
ttttttttgttttgtctaaatgaacaagcagattagcaagaatggtagctagtgtttgtg
tatctttctcttcaaggatttcgtaagcttttgttaattggctaacaatagaggttatat
tattctgtctaacaggctggtttgaacagctaattaaaaataatgaatctccagatatta
aaagtaatcttttaatataattatctataataactgacgaatattcatctaaaacatcag
ataaaggtgcttgttctattagattaataacttcgtctagttcattcaagtgtctaatta
ttttttctgtttttaagttggctacacaatttcctatagttactcttttatatgattaa
cattttctaggacagctctgtaaagtaatgaattcgatgtaaaataaaaaaaatcaggag
agagtttttctactgattttagtagagagactgattgtttagttaatattgtactaatta
atattttttcagctaaaatattatgaggaggtaaatatttatga------atactgatta
gttcctcc-ttgtaacattattctaata--------agactaattaattactagcttaat
ttgactctggcagaacttgtagttggatgttggcagttacctggtttaatagcttaacct
ctatattataaatgccaactgttttaatctcgggtaaaaggatattttgtttttcaatat
ctatatttgtagcatttttaatagtttgtgaaatttctttttctgtgacgcttccaaaaa
tattttgcccatcccctgtttttttagtcacgctgaacttctgtatttcctctagaagtt
gctgaatttttttgcatcttctttagctatttaaatttttcttcttttatggctgcgt
ataattttagctttttaaaataccgtttgttgcgactattgccattttatttgggatta
aaaagttccttgcataaccagcagcaacttttacaagatcattacttttgccaagttttt
gaatattctcgtttagcacaacttttataacttttttacccatatttattctgaattcta
aaaa----atatatataattttattatacttgttgtacccagatttt----aataggaag
tatttacttgttgacatagcatctccgtgaaactgataatgaattttcatctttattctg
gtataataatttatacaacgtttt-----attaggagaggtggccgagtggttgaaggcg
cagcattggaaatgctgtttaggagtaatcttaacgagggttcgaatccctctctctccg
tataa--aaattatgataatctttatatagtccagtcaaaacttccaataatgtttctcc
atgcatttaaaccacctttgactctcacaatattgagcttttgagctattaataacttag
aagcaaatagagatctagaatctaagctacagtatacaaagcaaattttatctttacaat
ttatattagagtaatgtattgttttgagttgttttaaaggtatatttaatgagtgaatta
agtgatttttttgatattcttcttgatttctgacatcgattaaaatatattttaggtctt
tgcgatttaaaagtttttgtaattgaacaacattgatttctcttataataagattaggtt

gatcatagctgtatctgttccagtatttttaga-------ggataatacaaatttagta
tttaaaattttaaacttactaaatgataaagttaaagcgttataagttaatatagtaccg
cttaatatagacttgtaaccaagaataattttaatagcctcagttgcttgaagtgtacca
ataatacctggtaataaacctaaaacaccagaattgctgcatgtatctatttcactattc
ctatttacgactttattgtggaagtctcgataacttggacctccttgatagttaaataca
ctaacttgtccttcaaattgaaaaatagccccataaatatgtattttatttaattctagg
caagtatcatttagcagatatcgagtacgaaaattgtctgatccatctataataatatcg
tagtttttaattatatcaaatgagttgttataggttagtcttgcttcaaaagcttttact
gtacattttggattcaaatctaatatttttttttagctatttctacttttgagtatcct
atatcgtgaactgtatatataatttgcctttgtaaattagaaagatcaataatatcgtcg
tccataataccgatgcttccaactccagcagctgcaaggtaaattattgcaggagagcca
agcccaccagcaccaataaatagcaccttagctttttttaatctttcttgcccctctaat
tgaatctgtggtaacactaaatgtttagaatatcttgtatattcttctaaagagtatttt
atattatctgctttaaagttcagcat-aaaatttatatgttaattaataacaacta----
--gaaactaattgataagcttattgagatgtttttgtaaactaggttcttctttatcta
aaatagctccttgcgtaggacagacctgaatacaaatactgcagtcaatacaggcagaaa
agtcaatccagtaccaattggtattaatactattttgtccttgtccttggtgaatacaag
ctactggacaagcagatacacattcggcaactccaatgcatttttcggttacaattgtat
gagacatgttatttaattaat-aaaagataaactatttacatctagtagttttattgtat
actaaattgacatacgcta-ccaagccagtaattaaaataatctacttttatagtttact
gatttgatctgcgcctttagttcagctggtagaacgcaggtctccaaaacctgatgtcga
gggttcaagtccttcagggcgcgcttagtttttgttatatttgcatagtagcattttacc
tttttgcaaacatcaaaatgatatgatataaaacatcatattctatatatttgatgtttt
atgtgccagtactg-tttttttatatgttattattggtaagttaattatattaattctat
ttttataatcaataaaaaatagctgaaattaagtgcagattcacttagtga-----tttt
atatatataactaaaaaaattaaaaactatggctaaaaaagttaccggaattgttaagtt
ggcattaaatgcaggtaaagctactcctgcaccacctgtcgggcctgctttaggtcagca
tggtgtaaatattgtcatgttctgtaaagagtataatgcgcgtactgcagataaatcagg
actagtaattcctgtagaaatttcaatatatgaagatagaagtttcacttttatacttaa
aactcctcctgcttctgtactaattgccaaagctgctggcttaaataaaggttcaggcga
accaaacacaaaaaaaaataggtagtataacgtataagcagttagaatctattgcagaaac
taagctaccggatttaaatactaacaatgtacctcaagctatgaaaatagttgaaggtac
agctaaaaatatgggaatcctaattaaagattaaaaaatttcaattataattatta-tt-
-tttatgaaaaaaatttcacgtcgacttaaaatattaaaatctcaagttgagcctaaact
ttataatattgatgaagcaatatcgatgttaaaggcaacatcaaatgctaaatttaagga
aactgcagaagctcatatttgtttagggctaaatcctaagtatgcagatcagcagcttag

agcaacagtcatattaccaaaaggaacaggtaaatcaataaaagtagcagttattgccaa
aggagaaaaattagcagaagcaattagtgcaggagctgatgtaagtggttcagaagaact
catcgatgaaatattgaaaggtagattagattttgacaaattaatagcaacgccagatgt
tatgcctctaatagcgaagcttggccgtgttttaggacctagaggattaatgccttctcc
caaagccggcaccgttaccatagaagtaggcaaggctgtgaatgaatttaaaggcgggaa
agtcgagtatcgagttgatagaactggcataattcatgtaccttttggaaagtccagttt
tcctcaagaagatctagtcttaaatctgcaaacaattaaagaatcaattgatagaaataa
gccttctggtgcaaaagggaagtactggaaaactttttcttatctagtaccatgggacc
atccatccaaatcgatattaatagtcttctatgaaaaatatttatttaattacataactt
gaataatatattatttcaaatta-taattggctagcaatgttaatagaataatttgctta
ttataccttatctgcaccaaaaattttaatatatctcttactaattatgagtacaaaagt
tgaaaatattttagaagaactaaaatctttaaatcttctagaagctgctgaactagtaaa
acagatagaagaaacattcgatgttgatgcgtctgctgctacaggaggcatgatgatgtc
tgcaccagctgtatcaccagccgcatctgaagttgaggagaaaacagaatttgatgttgt
tttagaagaagtacctgcacctaaaaagatttctgttctaaaagttgttcgctctttaac
cggcttaggcttgaaagaagcaaaagatcttgttgaatccacgcctaaagttctaaaaga
aggtgcatctaaagatgattcagaagctatgaaaaaacagttagaagatgctggcgcaac
agttgttattaagtaataaa-t--agtagtgcttttgtttagaagtaaacaaaagcac-t
ttttatttaatagaattaattagactagaaaagtccaagtgtaattgcattattaatagg
catagtcgcaccaatacctaaccaaacacttacaactgtaccaattaagaaaacagttgt
tgcgattggtcgtctaaaaggattttgaaacttattgacattttcaatgaatggaactgt
aagtaaccctgctggtacagctgccatacttagaacacccaatagcttattaggaattac
tcttaacagattaaatgttggaaagaaataccattctggtaaaatttctaatggagtagc
aaatggattagacttttcacctatagatgaaggctctaaaattgctaacccaatactgca
tgcaaaagttccaataattacaactggaaagacatataatagatcatttggccaagctgg
ttctccataataattatggcccattcctttgctaatttagctcttagttttggatctgt
taaatcaggtttttttaagaattgacatgttattttttgtgtttaatataaaattgagtct
aattactttatcaaagtgtagttatacattgttataatggtccggaaattccttgtttgc
gtatcattaaaaaatgcataagcatgaaaatagcagttagtaatggaagtacaaaagtat
gtaaactgtagaatcttgttaaagtaccttgtcctacactaactcctcctcttaataatt
ctacaatacttcctccaataactggaacagcatctggaacgcctgtaacaatttttacag
cccaatatccaatttgatcccatggtaaagaataacctgttaccccaaatgaaacagtaa
gtactcctaaaattacacctgtcacccatgtcaattctctaggtttttttaaaaccacccg
ttaaatagacacgaaatacatgtaaaatcatcatcaagaccatcatactagctgaccatc
tatgaattgatctaataagccacccgaagttcacatcagtcataatatattctaccgatg
tgaaagcttctgcaactgtaggtctatagtagaatgtcattgcaaatccagttgcaactt

gaattagaaaagatacaaatacaatcccgcctaagcaataaaaaatattaacatgaggtg
gtacatatttgctagaaatatcatcagcaattgcttgaatttctaatctttcttcaaacc
agtcataaattttactcataaaacagcttcaaaaagc-tcttcaatata-tttttgcggt
taagctacttttgattcaatag------ttattgaacttatatt-attttattatctaca
cttattaattattataacatctttctatctattt--------------------aaaatt
ctctgactcaataaaacaggactatgaacgacgatctttttttctgtatagaaatcaat
ttagtctttagcaattcagataaaatccttgttacacttactctactactccctataatt
tgcgctaacattttatgcgtcatagcaaaattaagcataataccattattactctgagtc
ccattttgctctgctagtaataataataagcttacaagtctacttgtaatacttttgtgt
gacatgatttctacaaaatgataagtttttctgagcaacatattaaatgatctataaaa
aataaattaaatagtgggtagttactacacgctcttataatagtgttatagttaatacaa
gcaattttgttttacctattgcttctgcttcatagtagaagttatcattagctaactct
aattgtccgaatgtatctccattagttaataaattgagtgttattttatgtgtgctttta
aagacttttgtaataattaaagatcctattaatataatgtatagtctagagctgctatca
aatagaataatatcattaggctgtagtagaaaaatttgataactaattttattactatta
aaaaatagcaaccatggattataagagggaaaattatttgaactactttcagaaaatgtt
gagatagatttttttgatttaaatgatttttagtcataattcatactttat-a------
---aaaaatactctaataat-taagaactgaaattaaatgtcatacaagttgatgctagt
tgagaatgacattgtattatcaaaagccattcaagaatacttaatagatcaaggatttaa
tgtttacattgccagtaatggattagaagcatttaatttagctaatcaatataatttcaa
tttgataatttctgatataatgatgccattagttaatggctatgaattattagaaaaact
tcaaaatactaaggcgctatcaaagataccagttatttttttgacagccaaaggaatgac
taaagatagaataaaaggttatgacatgggatgttacggttacttatctaaaccatttga
cccagaagagttagtttcacttattaaaaacttaattgctagagacatatcgcaacatac
atctacagaaaaatgtgacatctctaagcagtcaattaaccaaaaaatacatttgactcc
tagagaagcaagtgttttagaccttgtgattgacggactaactaacaaagaaatatctac
aatattagatactagtattagaaatgtagaaaaatatgttagtcgacttttacaaaaaac
taatatgaagaatagaactttactagttaaatattctataaataataaattactagataa
tgtaactaatcaaagggcgaatgacgggactcgaacccgcgaatgatggagccacaaccc
attgccttaaccccttggctacactcgccatattataaatatcactataacttgtttgtt
gatatatcgtctactctgtaagtcatttatttatctaacatattatctaaatttaatctt
ccatgacgtataataatattagtattcaaattaatggagagccgtttaattgctcaaagc
ctatttccttgcaatttttattaaattatcttgattttaattctgagcatatagcagttg
agctaaacgacactttgttacctgagcctctttttcactcaacttacttaaatgatcaag
ataaactagaggttattactattgtaggtggtggataaatttcattaactaa--------
---------ctattttaaatgcttcattgataaggataaacgtccctgcttcttgtccac

atgaataatgacagctttaatagtatcgcctattttaaattgtgacgatatctgatctaa
ttgttttatgttaatttcagagatatgtacaagacctttaagattgccaacttttataaa
taatccgtaaggtgtaatctgatttataattccttcaataatattaccaacaattaaatt
tgatgaggcttgagcgattaaagctcttctgtggcttagtattagattattagatctttc
ttcaacattgagtagctttaacttgataaattcattattagccttatcactttttttggaa
attcccgagatgagaatttggtacaaatccagatataccttcaaggctaatgatcatccc
tcctttattaaatccttttattcgaacattaagtaaagaatcttcagctaataattgtcg
aattcttttccatgctcttatatattcaaggcgccgaattgataaaattaattgttttga
tcgaatattataatctagtaaaaagaattctcttgtatcattaatgtttaaagaagtgaa
attatttaaatcttgattacttgacacctcttgaataggtaagtatgcagatattggtgt
tcctatatcaaccaaaacaccatttaactcaaagctaaatattgtaccagcgacaatatc
gccaagatttaaatcgtatttgtatttttgtaaaacagctgcaaaatttctgtgagtaaa
tccttcattatttttttgtcatattgaatagtctgaattaaagattgattttgtacaaaat
attaagttgaataaacttctgtcgtattatatgaaaatttttaaattcgttctgtggtta
ttcctgatctaaacgtattagaatatacggaagacataaaaagtaaacgtagatagctac
agatttaaccaatatacagtctgaggaaagtccgggctccataaatacaatttatgctgg
agagatcccagtgtaggcaattatgaggatagtaccacagaaataaaccgccagaataaa
a----tatactagctggtaagggtgcaaaggcaagttaaaagcttaccaaaaatactgta
aagtatttgttaggtaaacccctaaaatggagcaaagctgctaaacaaatatttgtgtat
attttatatagtttagtttaaaatactgcatgaagttattggtaacaataactctagagg
aatagctatcctttacaatatata-tctattatattgaaagaacagaacccggcttatgt
agtacttttttatgtcttacattattctggctattgccgaatctctaagtttaagtgttgt
atgatttttttcttgtagctcatgttgtttatcatctatctctaaatttgttaaagttttt
tgactagagcgatatgtaaatctcaaacctatactttttttttctgttctccaagggattta
tttgtatattgatcaaataaccttatgctttctaaatctttatcattaaactgatctagt
aattttaataagtaattaaattccatctttttaggtattattaatgaaagatctcttgtg
atagatggatatttagagtaaggtcgaatttgataatttaaataattaaattgattatga
gaatcttctaaaatgtttaaattaatttcaagaacaaaaagttttgcagtaagacctaat
ttactataggttacttgattgagttcactgaatatgccaatattagtattattataaatt
aatgttgcagatctatccttttgaatgaaattcactctactgtcaattatttctggtttt
gtccattcaatttttctatttatttttttgaaaaaaatttctacaattcctttagcttca
taccaatttaaagagtggactggttgtgaccatgtagatctaatatttaaattaccacct
aatatcatggctagattagttgtttcgacaattttgttgtgaacaggattaaaaactgtt
cctatttcaaaactgtctacagtttgattactttgcttaagattataaagattagattca
attaatccctcgagcaaactatagcgtaaattagagtaatcttggataagaggattattt
agactaacacctcctagagatttcactaaagaataatgaactaattcagttaaccctaaa

tttcttaaaatatttctacttttatctatgaattttctcttttgtgataatttttacta

aattgaatacttggcactgagcttttaaatttatgatagccataaattcttgcaatctct

tcaataatatctatttctcgaaaaacatcttgtcttcgataatttggaacactaatttt

atactttcttgattttatgaataatagtaaaatttagagaatgaagaatattttgaatt

tcttgaaagtacaaaaaacgtacctgattattgttactgattggtcctaaaatatcttga

actttttaattgataaatctatatttagagtatcattcattgttttctctgttgagaat

gtctgcctaatattcccgccagctaagtctattattaaagagatagtttcaagataagca

ttttcccagttgtctgtatttagtcctcgttcctgcctaactgaactttctgtcctaatg

ttaagatctcgggaagactgtctaactatagattgtttaaatatagcagattctacaaat

atagatgttgtattagaatccgtatcaaagttgttattggatcctatgccagccacacta

actacatttgtattcacttgagtcactaaaatcttcttatttaactcaatattttcatta

tttaacttaattttagtactttttccaacattaaagttactttgaattgttatagtatct

tcagg---------agtaattttattaaaatcaataatattaataggttgtccccatttt

agcataatataattgcttatatctactaacaaatttctatgtgtaaagccagaggacttt

aaacgatttttagccaatttggagaatctataattttaatttcatcgataattgcagaa

aaatagttattacaatgaatcaagctatgattattgagtatatt-ttgttgtctagaaga

gtatttattactaaattgatttatagtttttaacattaaagatccagttaaagcggataa

ctctcttgaaagacctatcatgcttaatacatccgatctattagctgttgatgatatatc

taatacatgatcaatctgattgcctactgtaatagattcaatagcttctacttcaaaacc

tgcctgtgttaattgactagttaaatgaactagatctattgtttctatattagcaagttc

ttttaaccaatttaaagaaactttcataatgaattgccatttatcttatataattattta

ttttacttgtgattaaaaactaatttaaaaaattcatcaaattaatataagttatttgat

gaatttgcttatacgatatattcttgtgctatattactttcaggtcgtagtaattaagaa

ttagcttttgtaaaagtcagtccattatcttgagcatacctttccataaatctcataaat

ctatcccaatcatcaggacttttaatcacatgtacagcttctattgcttgtggtttacca

ttaataaacttagcattgacatctcttgtaataatttgtccctctgtgtccattagatac

atgcctgttatttcacctttatctgccatactagcatctaaaattttgggatttgtaaat

ctgaaagtagctgttccggtactaccatctctagatcttgttaagcgaacatctggcacg

acttcttcattgatgccttgaataaattgaattgttgccatatcaatttt-tttatttgt

atataatgtatattggataaaactgatttctaattctttaatgcttgttgaaaaactgtt

ttaaaagcttatagtagcagcagatcaaattttgtgtactatataaaactgtaactttc

tttttttg----------------------------------------------------

------------------------------------------------------------

------------------------------------------------------------

------------------------------------------------------------

------------------------------------------------------------

----------------------atctgttcaaagaactaacttattataagctaatcata
ctaatattagccatataatgttctgaagt-gaaattaa-gggttgtagctcagatggata
gagcaagcgcctcctaagcgctaggccagcggttcaagtccgctcaaccctgttaataaa
caaaatt---ttgctgaattctacaaataaattagttgaaaaaagaggaaaaaaacaaaa
cagc-----aaatagattctgagtaacacttttattgaa-----aaaaatgatcttaac
tttgataatagaagaattttgtgaaaaaatatttaaaatttgatctctatattcagaaa
aagtttttttaaaatataaactgaaa-----taaaagtcttgagtaacaataggaattaa
attatttaatttttgaaattgatctaaaaaactttatatgaagagaatagtttttttct
ctttctctattcagtatttgcctccataattattacttcattcattcacaattaataaaa
tggattttg---------------------------------------------------
------------------------------------------------------------
-----------------------------ctttaatgaatgtttatgttatttgattata
ttccatactatataagagtttttctcaacttgtgagagtcaatttgtagttttaattta
taatatactttatctttggagattgatctacattcaccttcacacatctcttttattgcc
tattaggagaaactatatactgaaatttaacaataattgtgaaaatcgtaggttccttca
acatattattatcctttcatttttgtctccactgctgatcaaagcaacagtgattagact
tattcatttagctggttctagtgagagaagttttatacaagccg--gacacttatttaaa
t---gtgttattaccaaagctgttaaaataatagccattaagattcaataacttttttagt
taatagtttataatcgccatgtttgatttagtactgatatatagttacaagtagttctac
tattgataaaattgacacttttgtgactgtataagctttcgatagaaaaactatgatatt
attcataatgaaaaagcgagtattgaagtttgttgctttcataacaaaaaatatgtttag
tcaatgatgaattttgctagcgatccgagtagtaaatctgtcacatgtaaagatgaaaat
tatatatttgagaatactctctatatctgtatatagcggttggcataacttatcagattt
gatttgcagaatcagtatactaagagatcgcaaaatggttattctaagaaaattttgtct
cagtaatttcacttctaatagatacttgttagtaataaacttttaattcttttatgaaaa
aacttcataatatcaatatttattagaactggggtgggaggattcgaacctgcgaatggc
ggagtcaaagtccgctgccttaccacttggctacaccccaacttatagcaaatataatat
cagtcacataccccgccgtgtcaatagattattcgcattatttaataacttttttagattt
ttgtaaaatagaatcttttttttcaaataagatgtactatgttcaaattgcattatgctc
atgtcttcttgttgttgattggacatccattttatggtcactgttccttgttcaacttct
tgattgcctataataatacaagctacagctcttttcttattagcttgtttgatttgttta
ccaaagttacttgagctaacatctatttctattgtaaaaagtttttgatgtaaaaagcgc
attattttcatgccaacttcattagctttttaccttgtgtagcaatataaaaatcaaca
tatttacttggtaactctatattctctttcgcaagaagtagtaaacgttctaatcctatt
gcacatccgacagctggtgttgctacgcctcctaattgcttcactaagctatcataccttt
cctccaccacaaactgtatcttgaccttttgatgttaatgtcttaatttcaaaagcagta

tcattataatagtctaatcctctaactaacttgtcatttatgttataaggaatatcaagt
aattttaaataattacaaacagcatcaaaatgtctttgtgacgaaatacttaaaaaatta
gaaatcttgggagcctcagctaatattttttgtgtatttatatttttttgaatctagaatc
ctaataggattactaattagtctcttttgtgaatcaggatctaagtcgtcatagtactgt
ttcagatattcttgcaattttgcttgataaatactacgatcttctgcttttccgattgaa
ttaagatcaagctttaagtctttgagattaagattattaaatatacttatagctaaatgt
attacttcactatctgctcttgcatctaggctgccaagaaattcaataccaagttggtga
aattgtctttgtcgtccactttgtggcctttcatacctgaacattggcccgctataccat
aatctttgtaaactatgatgatagttcattttattttcgataaaagctcttacaatgcct
gcagtgccttcaggtcttaatgtaatatctctattgcttcgatcatgaaatctgtacatt
tctttatttacaatatctgtatctcttccaatacctctatcatataaatcgctactttca
aaaataggtgttctaatttcttgatagtttgcacattctaataagttggaaattttgtta
tgtataaattgccagtattgaagctcttcgggcagaatatcttttgttcctctaatagct
tgaatttttgccatgataattttcggatcaccttcttaatgtgtttgcattaaatagcag
ttgtagactgttattcagtcatgtattatacattag--aacatgttgatg-tattttatg
aaatacgggcaaggagggattcgaacccccgacaccatggttcgtagccatgtgctctaa
tccactgagctacaagcccacttaacaagctatcttaatatatcattctttttcaatgta
tacaactatattttgtct------atttatatttgaaagttgtctgaaagt-aaaaaat
gt----atttatctttaacaattagagttagtcatgtataattttccatttattaaaact
ttgcaca--aatatcgacaaacttaattatgtaattatcttatactgataatttcagaag
aa-cttgtgcttttacttcttgatcga------ctagtatctaattcatctatattccaa
at-aaaaaatcttatctaaaatgagtaagcaaattctatatcaagatgatgcacgtaaag
cattagaaaaaggcatggacattttaacagaagctgtttctgttaccttgggacctaaag
gaagaaacgtcgttttagaaaaaaaatttggggctcctcaaattattaacgatggagtaa
caatcgcaaaagaaattagcctagaaaatcatattgaaaatactggagttgcacttattc
gacaagctgcatctaaaacaaatgatgtcgctggtgatggaacaacaacagctacagtac
tggcatcagcaattgttaaacaaggcatgagaaacgttgctgcagggtcaaatcctatgg
ctataaaaaaaggcatagagaaagcaacaaattttgtagtgagtaaaatagctgaatatg
ctaaaccagtagaagataaaaaagctattatacaagttgcttcgctatcatcgggcaatg
atatagaagtaggtaaaatgatagcgaatgctatagataaagtgggaagggaaggtgtaa
tttctctagaagaaggcaagtcaactaatacgattcttgaaattacagaaggcatgcagt
tcgaaaaaggctttatttctccttattttgttacagataccgagcgaatggaagtccttc
aagaaaatccatttattctctttacagacaagaaaatcactttagtgcagcaagaacttg
ttccactccttgagcaaatagcaaaaacatctcgtcctttattaataatagctgaagata
tagagaaagaagcattagctacgattgtagttaacaaattaagaggcattttaaatgttg
ttgcggttagggctccggggtttggtgatagaagaaaatctttgcttgaagatatgagta

ttttaactaatggacaagtaatcactgaggatgcaggtttatcgcttgatacagttcaat
tagatatgttagggaaagctcgcagagttattgtaactaaagactcaactacaattattg
cagatggtcatgaagttaaagttaaatcaagatgtgaacaaatcaagcggcaaatagaga
cgagcgattctctttatgaaagggaaaaattacaagaacgattagcaaagctttctggtg
gcgtagctgttataaaagttggtgcagctacagaaacagagatgaaagataaaaaattaa
ggctggaagatgccattaatgcaacaaaagctgctattgaagaaggaattgtcccaggag
gggagctactaatgttcacatttctagcgagttatttacatgggctaaaaacaatttag
ttgaagatgaattaatcggagctctaatagttgaaagagcagttacttatcctttaagac
gtatagcttttaatgcgggagataacggagcagtagtagtagaaaaagttaaaagtaacg
attttcacataggctatgacgcagcaaatggcacaattgtaaatatgtacgatgcaggta
ttatagatcctgcaaaagttgctcgatcagctttacaaaatgcagcttctattgcagcga
tggttttaaccacagaatgtattgtagttgataaagccaatgactaagtta---------
-----tagataatattaaacaaagaataatttccatagctagaaattattctttgtttca
attaagctaattaattgatcaaatgtataatttttataataaagccgaagtttaaatgta
cctctttgctcaactactaagtagagaagatacaagctcgtaattcctaatttatcaatt
aaagtattattatcatatatggattttcaagagatctaaaaaatgacattttcgtaaaa
taatatcgaaatcttcttttgtagttatttgataaatcattactgtatgtagattgatct
ccaattgcgtttaaaacttttaaaataatgtttttttaagtctctatctgactttatctta
gaatcggtaatcaattgatttaattgaatatcaaaactaagattaagaatattacttgtt
acggatgttcgattttgaatatctatagatttgacagccaagaagattaaacttttattg
ttcattgtaaaatgtaagacaattgatattgtattctttaaatacta-attttatcttct
tgtgcgtttatagttttggagaaaaataattaatattatta--------tattaattact
gcccgaaaaaataaattctggaaatctagcttgagcctttgatagtgagtgtttttttacc
ttcttcttgccagtaattaataacttctgttgtcttatttagtaattcaattctgtctct
ttcagcaatccatggtttggattctagctcggcttttaattcttcccaagcatcattcct
aggccagaaaaaataagaagtaagtggacttgtaccatttccaacaatttggtcaatagc
aatagcaatattattttccaaccaaagtacttttagtgtaaattttgacaagtgattatc
tccattgtaaatagcaaacatatgtatatacagtaaagtactacattagttattttttt
agcaacaagctgagctttatttaaaatattctgtaccgttctagtagtttgtgcaccttg
tgctaatctctgtaaaatttttgaaatatcaatacgagtttcattagttataggattata
gaatcccaattcttctatagccttgccatctcgtttgtttttgctatccattacaacaat
tctgtagcttggctgtttttttcttccatatctctttaatcttaacttgaccatcttagt
cccctcgcttattttttaatatctttattaaattatatcttacgatattatgcaataat
tcgaacaattaagcagactcaagcctgacattattaaatattaccattaaacactggaga
aaaataatacctagcataggtgacatgtccatcccgaacattgggggaatactgcctcta
aagagctttaaatatggatcggtaattcgatttaatgaacagaatggttcattataccag

ttcacagttggaaaccatgccaatgataatttaagtaaaattaaaattaaataaatctca
gaaaaattagctattgatccaagtaataaatttaatgtaccaggaagagtattcatacgt
attttataattattatatagttcattatactaatagtttgagtatatattataagctctg
taaataaat-tagctaaacaaaaaacttattcttaaaatatcttctcttgaacaatttt
cttcattgatagcactgttaactttcgttgtataaatttctaatttagaccaattatttc
ctagctgatttagttgagatgttctgcattctaaacatgctatttttatattatttaaaa
tagctccttttttttctaaatcttttaagactggtgtaataattgcttcatctaaaaata
aatccaagatgagaatttttgtaaatgaatttaattgagctggaacattattaacaatgt
tgttgctattaactagagctatattagctgtcggaagtaaagctcttgctccctctgcta
gtataaatccataaggcataataatgacaataatatacttataatgactgttaataagat
ttacattgatattagtttttaattcaatagtttctgttactaaccattctctcattattt
cataagtaatccatttacctagttctgagcaagcagttcttaaaattgtacctggattat
tgctattttcgagaatcccagcccaatgttgtattaaagggtgggaagctatattaattt
tgagttgcatgttttgctaaatatatttaacatttaatttattaattgtcgctatagttt
atgttcactgagtattattaaaatagcat----atatattaaagaaagcaattat-aaaa
aatgaaaaaaactctttggctttggggttcacagatagtgctgaaacttggaatggtag
atttgcgatgattggttttatagcagttattattattgaagtagttacaggacaaggact
attatatctaataggtgttatatcataataaaaaatcaaaaaagttcttgtaataaatta
caagaactttttga-ctgaatgtaaatctaatcgagtggtctcatagatagcactggtt
cattctctctgttaatacctttttcaaatccagcagctgcagctcttgctctaccagcat
gccaaagatgtccaataaataggaagaatcctaagaagaaatgagatgttgttaaccaag
atctaggagatacatagttaactgagttaatttctgtagctacaccgccaacggagttca
atgaacccagtggggcatgagtcatatattctgctgctcgtctttcttgccaaggctgaa
tatcattttaattttattcaaatcaagaccatttggtcctctaagaggttcaacccaag
gagctcttaagtcccaaaacctcatagtttcaccaccaaaaataatctctccactaggag
atctcattagatattttcctaagccagtaggcccttgagaagatgcaacattagcaccta
atctttgatctctaactaagaaagtgaaagcttgagcttgtgaagcttcaggaccagtag
gtccataaaactcacttggataagctgtattattataccaaacaaaattagaagctgtaa
gacccatgattgataaagcacctaagctgtaagataagtatgcttcaccagaccatacaa
aagctcttctagcccaagcaaaaggttttgtaagtatgtgccaaattcctccagcgatgc
aaataatgcctatccaaacatgaccaccaacaaggtcttccatgttatttacactaacga
tccagccatcaccgccaaatggagattttaagacgtacccgaaaataactaaaggattaa
gagtaggattgctaacaaatctaacatcaccgccacctggagcccaagtatcatataccc
caccaatgaacagagctttgataactagtaaaaaagctccaatacctagtaaaactaaat
gtataccaagtatagttgtcattttgtttttatctctccaatcatatccgaaaaaaggaa
aagattcttcaagagtatcgggtccgattagtgaatgataaagaccaccaaatccaagaa

cggctgaagaaattaaatgaactacacctactacaaagtatggatatgtattaaaaattt
cgccgcctggacctactccccagcctagtgtagctaagtgtggaattaaaataaatcctt
gttcatacagaggtttttctggcacaaagtgagctacctcaaaaagagtcatagcaccag
tccaaaagaccattataccggcatgagcaacatgagcgccaagcaatttgccagaaacgt
taattaagcgtgcattgccagaccaccaggcaaatccggtagattcaatgtctctaccgc
caacaccaacattcgtattaaagggcgtttccacgtggtaaaacctcctcagggaatata
aagttttcatgaggttgatcctgagcagccatccaagagcgaataccttcgtttaatagg
atgttcttagtataaaaagtttcaaactcaggatcttctgcagctcttagttcttgagaa
acaaaatcgtaagctcttaagttaagagctagtccaacaattccaaatgcacttgtccac
aatccagttactggtacaaatagcatgaaaaaatgtagccaacgtttattggaaaaagct
acaccaaaaatctgtgaccagaatctgttcgctgttaccattgaataagtttcttcagac
tgtgtaggcgtaaatgcccggaaagtatctgcagcatcaccatcttcaaataatgtattt
tgcacggtcgcaccatgaatagcacatagtaaagcaccacctaagatcccggcaacaccc
atcatgtggaatggattcaaagtccaattatgaaatccctgtaagaatagcagaaatcta
aaaattgcagctactccaagacttggagcaaagaaccagcttgcttgacccaaagggtac
attaagaacacagatacaaatactgcaattggtccagaaaaagcaatagcattgtatggt
cttagaccaactaacctagcaatttcaaattgtcttaagcaaaatccaattaatccaaac
gatccatgtaaagcaatgaatgcccataggccgccaatttggcaccagcgagtaaaatct
ccttgagcttcaggtccccaaaggaaaagaagggaatgacccatgctgtttgctggggta
gaaacagcagcagtcaaaaagttacatccttctaggtatgaacttgctagtccatgagta
taccaagaagtcacaaaagtagttccagttagccagccgcctacagcaaggtaagcgcaa
ggaaacagaagtagtccagaccaacccacgaatacgaatcggtctctttttagccaatcg
tctacaagatcaaaaccaccacgagtttttttcttgtccaattgctatggtcataattta
atctccagagcaaatttaataagtaatttgttcaaataaaca---ctttagatctaatct
aaaattttactttttctttacggtcaatataattatacgtcaaaatcaacaaaaattcacg
tttttaataaca-tttttttagctcgttaagttaactatctaagaat-------------
taactgtttttttaaattaataaaaattataacttgttcatttctataattctctgaaata
gatatcctgttcctcttgctgttaaaatcaagtctggattacttggatcatcttctagtt
tagctcttagccgtgagatatgcacatcaacaactctagtgtctacatgtcgttctggtg
tatatccccagacctcttgtaagatagatgctctagaaaaaggttcgccagccttactca
ctaaaagttcaagaagactaaactccattcctgtcaaacgaactctctcattatttttat
aaacttgtcttttattcgtgtctatctttaaaaatccaatactaattatgccagaatttg
cgacgccaagattagttgttattttatcagctcttctaagaacagatcgaatacgagctt
ccagctctttaggagaaaatggtttaactacataatcatcagcacctatttctagtccag
taatcctgtcgcacacttcgcctaaagctgttaacattattataggaacatcagattctt
ttcgtagttcttgacaaactccgtaaccatcgagtttaggcatcatcacatctaaaacga

caaggcttggatattcttgtctgaacactattaaagcttcctctccgtttgaagctgtta
tgacttcgtaacctattattgttaatcttgtttctaaaattcttcttatgctggcctcat
cgtcaacaacaagaatttttctttctggttctccaa-tttcttatttactcctataagt
caacaagcataactactataatcttaaatttgtacctgtttgagtattctaatttgctgt
aaaa-----cttatttattataaattttataagtt-----tagattaattggtatcagt-
-cattatatcttgaaatatctgtttgtcatttatctc---------tattttaatgcata
agttgattcaagttttcagaattcagtatataaata-ctttttagacaagtctttgacat
tgtattttctgaataaaaaaggggttgacaaaccggctgatagaaagtattctacattga
tataaa----ttactgatcataaacaaacagtaa-----atttttttagagtagcataag
tactttaat
>Pyr_35
tatatgattaataattgcaacatcttttatgtaagaattcctaaaatatacaaaactttt
tatttaatatataaaaagtaaattagaataagttgagcatattttgttctactttagagc
ttttgattgag------attctaaatagaaacaaagatgtttacatagtttatatctata
ggaataaaaatgaaacttgcagtttatggtaaaggtggtataggcaaatctacaactagt
tgtaatatttctgtagctctttcaaaaagaggcaaaaaagttctacaaattggctgtgat
cctaagcatgatagtacatttactctaacagggtttttaatcccaacaattatcgacacg
ttacaatctaaagattaccactacgaggatgtttggcctgaagatgtgatttataaagga
tatggaggtgtagactgtgttgaagctggcgggcctcccgctggtgctggttgtggaggc
tatgttgtaggtgaaacagttaaactcttaaaagaacttaatgcttttgatgaatatgat
attattttatttgatgttttaggtgatgttgtatgtggaggctttgcagctccattaaac
tatgcagattattgtctgattattacagataatgggttcgatgctttatttgcagccaat
agaattgctgcttcagtacgagaaaaagctagaacacattctttgcgacttgctggatta
gttggtaatagaacagataaaagagatctaattgataagtatatagattgtgttccaatg
cctgtattagaagttttaccattgattgaagatattcgtgtgtcccgagtaaaaggtaaa
actttatttgaaatgtctgaaactgaccaagatttagcgtatgtatgtgattactatcta
aatattgcagaccaattaattactaggccagaaggcgtagtccctaaagagtctcctgac
agagagctgtttagtcttttatctgatttttacttaaatcctaaatctaatgtaggacaa
actgtagtagatcaagaagaattagatttaatgatagtgtaaataatattttaagcataa
gataaaaaggaataagacaatgtctaaaattcaatcagacgctcttacttttgaatgtga
aacaggaaactatcatacttttttgtccaattagttgtgtttcttggttatatcaaaaaat
tgaagatagtttttttcttagttataggaacgaaaacttgtggttattttttgcaaaacgc
aatgggggtaatgattttcgctgagcctaggtatgctatggcagaattagaagaaggaga
catttcagctaagctcaatgactatggtgaacttcgtagattgtgcttgcaaattaagaa
agatagaaatcccagcgttatttttggattggaacatgtacaacagaaattataaaaat
ggaccttgaaggaattgctccaaaattagaagcagaaattcgcgttcctattgtagttgc

tagagctaatggtttagattatgcttttacacaaggtgaagacactgttttagcagctat
ggctcagcgttgtccttcaaatttaaatatgcaagcaggtagtaatagtcatgaaaattc
ttctcatattcctttagttttatttggctctcttcctgacccagtggtaactcaactaac
tatggaattaaaaaaacaaggtattcatgtttctggctggttgccatctaagaggtatac
tgaattaccggttatacaagaaggctattatgttgctggagtcaatccatttcttagtcg
aacagctactacattaatgagacgaagaaaaacaaaattaattggtgcaccatttccaat
aggcccagatggtactagagcctggattgaaaaaatttgctcagtattaagtattcaacc
tgttgggctagcagatagagagcaagaaatctgggactctttagaagattatattacttt
aatacgaggcaagtctgtctttttttatgggagataatttattagaagtctccttagcaag
atttcttactcggtgtggaatgactgtatatgaaattggtattccatacatggataaacg
ttatcaagcagcagaattagctttattaaaaaccacatgtgataaaatgaaggtcatgat
gccaacaattgtagaaaagccagataactataatcaagtagatcgaattcgtgatttaaa
gcctgatttagttattactggtatggctcatgcaaatcctttagaagctagaggaattaa
tacaaaatggtctgtggagtttacctttgcgcaaattcatggttttacgaatgcaagaga
tattttagaactagttacaagaccattgcgacgcaatcttagtttatcagaattaggctg
ggacgtttatagtaagcaaaggtagtattaatt---aattgcgacttacatggtataaaa
tccttgtaagtct--tatatattatgctaaagaagcatttcttgctcggccatcttctgc
ccaagcttgtaaattctctacttgttctttatctgtaaatgctaaaggtataaactgctt
tatagctattttaatatcttctgtattaaattctctttgctcactaaaagctgtatgcat
actttctataattgcttgctcaatttcagccccagagaatttattacataataaacttaa
ttgcttaatatcatattcttgccatgattttggtcgaacttttgataaatggatcctaaa
aattgactctctttccttgcgattaggtaaatctaagaaaaaaatttcatcaaaccttcc
ctttcttagcatttcagaaggtaaactttgaattttattagctgttgcgacaacaaatac
gggagtctttttttctgagagccaagttatgaatgttccgaatactcttgcactagtacc
actatcaccttggctatacaagccagaaaaagctttatctatttcgtcaatccataatat
acaaggcgataatccttctgcaatgtttatcatttctctcattttttgattccgactcgcc
aactaatcctccgaataattttccaatgtcaagacgtagtaaaggtaaagtccaatcgtt
tgcaatggctttagcggttaaagatttgcctgtgccttgtataccaactagtagtaaccc
tttaggtgaaggaataccataatttaaactttgttttgaaaaagagcgagaccttttttg
tagccattgctttagtacatctaatccaccaatatctctatttactttattatagggata
aaattctagtagacgtgtttggttgataatttgtctttttttcttcaataataatcggtaa
gctgcgagaatctatttgattatattgtgcaataattttggtaattactttgcggatttt
atctattgatagaccttggcaagattttgtcatattattaactagttctgaatcaagatc
aagatttaccgctttacttaatcttgtgatttctttttttgatttctagaagacttggtaa
aggcaaatctattattgttacaatatctcttaacgcaaaaggtatatttattttacaaga
aataataattatgtgtttagcttgtgtttttattacttttgccaaattgcgaactttcct

agttagaactatttcatttaagaaagaatcgaaatcttttaaaataaaaatattcaaaga
ttcactatctaaattttcaataaactctaatgctagcaaaggattccgttttgcataacc
attatcgttagggttactagtgtatccatctacaaaatcccagcaataaacttgctgatt
gtcagaacaatgtaatttatttttataatatattctagtctttcttcttctctagtatt
tattacaataacaggatatcgagactttaatagtaatcgtaaatcttgcgtgaaattcat
gtcaatattttccccattagactttatacttcaccaatta----aagactaaataaga
ttatctaatattgcatgtatatattcaaaagcc------atgtatctattttctaatcta
ttgtgatttccaacctaactagtcatgatttttttcttcctttgcgccgtctactattt
aaaatagagcgtccacatggtgttttcattctggcgcggaaaccggaaactctgattttt
tttcttttcgatccctgtagtgttctcttagtcattatgtctcctgtgttgtaaaagatt
aaatattattttatcatcttttatatgtcttgataatatatctctgtattaatatcaaag
tt-aaatgcaataatttatattaatcaagaattatttatataatattaagaattaattat
----------tagctttatgatattagctttgcctatttttttatttaagtatcctgacaa
tgtttttgctaattttgaattggttaatctttcagcagctaaaaataatttgtcattag
aatcacaatttaaatattttattgaaaaaagccaaaatgatatattatcagtagaagaaa
gctttgcctttgcaaaagcttgtgtagcaaagaagtgcttttctagagctattatcgaag
gccaattagcgataaaaaagagcttacaattaaatgataaagagggcagtataataattg
ctaatttgcataatatgcttggttttatttattctgaagcaaaacaactaaatatcgcaa
aaagcttttaccagcaagctttacaaataaaccctaattatgttgtagctttaaataatt
tagctaaagtttacgaagataccaaagatttgacaaaagcagaggccatatatgataagg
tattaactctcaacttagataataaaatagctaatagaaggaaaaactatataacaaaaa
aca---------gaaatgcttaatgttatttacaatcgggatagcaggatttgaacctgc
gacatcctgctcccaaagcaggcgcgctaccaaactgcgctatatcccgttagacataac
tagtactatatacctttatgtaattatgtctaccgtttcagtgttttatctaggatacc
aaaacatgcctttaactccatctggatctttaatgaatcctaatttttttataaaaactaa
ctacatcaggttctgcaaataatgtaatagtgctaatttctgcttgtctcaattgtttta
ttaattgatgtataactacttttcctaaacctacgccctgaaaatcaggatggataacaa
catcccagatagtagcattgaatccattatctgatgttgctctagcaaatcccactagtc
gagtattcgcatctttttttgtattaaagaaatgataatcgaactatttttaatgcga
tttttaccttttttagaggtcttttaacccatcctaccgaatcacaaagctgctccaatt
cgtataagttaatatttttagtgctactcagatagatatctttaaattcggctctattac
aagttttgtctaaaacgacaagtttttgaaattttttcagaattacttttgatatgtg
agttttgaaaaaaatttttccagaagatcataatgattatgatagaattaaataaatata
tatatattatgatattgccttaataaacaacatattaatatatgtccagtttttaatacg
aagctattaagaaagaagattgttttaagatatagaaataaaaatttttgtaaaatcttt
taagaaaagtatataaagagctctttttaagattaacatctattagtataaataaagtct

tgcaattgagatgagacaacatattattctatatctaatgccttttaaaggagttatttg
tgaaaaaatccatgtttctaacctgcctacttgctttattactgcttagcaacccgataa
tagctaatgcagaagtagctgggctagtgccttgtaaagattcggcggcattcaataagc
gtatggttaatagtgttaaaaaacttaaagctagattagctaaatacgatgcagatactc
ccccagcgttagctttaaataagcaaatagaaaaaactaaaacaagatttgctacttatg
gtagagcaggtctattatgtggcacagatggcttacctcatttaatttctgacggtcgat
ggagtagagccggagacttcgtttttcctggattactattcttatatattacaggctgga
ttggttgggtaggtagaggatatctttatctgttgctaaaactagcaagcctactgaga
aagaaattattttagatgtgccattagctattaaatttatgacatctggattcgcatggc
cattagctgcttggcaagaatttagtagtgggcaactgattgcctctaatgatgatatta
ctgtttcaccccgttagtaaaaaatttatatgaataataattttaccaaatatttatcaa
cagcacctgtagttggtgtgttatggatgacatttacagcaggttttattatagaattaa
accgtttttcccagatgttttatatttttatttataaataaataa--aatcttataaa-
--gaaccacacaaatatagtaaaacactattatttgtgtggttctt----atataaaaca
tcattatttatttttactgattttttttctttcgactattttaatattagattttatat
aataaaaaa-acatcaatattatgagtttagtaagccaaattatagttaatgctgatgat
gaattaagatatccaacaatcggagaattgcagtcaattcaagactatttaagtacaggc
agtaataggattagaattgcaactattattagggataaagaacaagaaattgtacaaaga
gcgagtaaacaaatttttcagctacatccagaatatatagctccaggaggtaatgctgca
ggttccagaaaaagatcattatgcttgcgtgattacggttggtacttaagactaattacg
tatggagtattagctggtgataaagattctatagaaacgattggtatcataggtgttaga
gaaatgtataattctttaggtgttcctataattggaatgttagatgcaattcaatgttta
aaagaagcttctctagagatgcttggtcaagatgacattagaattatttctccttatttt
gattacataattcgaggtatgtcataaatagtaaattgaattatttggttgaataaaagc
ctatacaatgctataatcagtttaagcccgaaagcatataattgtaaaaactaaatcttg
tcagaaccggaaggtagcagcaataatgtttaattacaaaagttatga-ttttcgggtgt
ttactatttgtacttaatagtctctaattctaattatgatattatccataaatc----aa
taatagataactcatgaaacctttcagtaaaaataa-ttaataatttgatattaatacgg
aaaccacctatac-tttttataaaatttctttatatctcatctaaacaatggaaactta
acttaatgggtgttcatattttatcaacaggttcgtctgtcccaaattttctgtagaga
atcaacaatttgaagatatgatagagacttctgaccattggatttcaacaagaacaggaa
taaaaaaaagacatctagctccttcttctacttctttaactaagttagctgcagaagccg
ctaataacgccttatctgcagctaatttaaaacctactgatattaatttaattatactag
ctacatctacgcctgatgatttatttggtagtgcaagccagctacaagcagaaataggtg
caaccgcatctgttgcttttgatattacagccgcctgctcaggattcattgtggcattag
taacagcagctcagtttattgagactggttcatatgacaatattttagttgttggagcag

atataatgtcgagatggattaattggtcagataggactacctgtattttatttggtgatg
gtgcaggagcagtagtggtgggacaaagcttaaaaaacagtattttaggttttaagttat
gtacagatggccagctaaatagtcatttgcaattgatgaatactccagaaaataatcaaa
aatttggattcatagatattcctcatggaaattataattctataacaatgaatggaaaag
aagtatacaaatttgctgtatttcaagttccaatagtaattagacaatgtttagataatt
taaatatttcaatagatgaagtggattggtttatattacatcaggcaaatactagaattt
tagaagcaattgcgagtagattgtcaatacctttttctaagatgattactaatttagaga
actatggtaatacatctgcagcatcaattcctttagcgttagatgaagctattaaatcta
ataaaattaaaccgggtcaaactattgttttgtcaggtttcggagctggtttaacttggg
gcgcgattgtcttgaagtggtaat---tagaatattgcggatgacgagactcgaactcgt
acagctttcgctacacacccctcaagcgtgcgtgtataccaatttcaccacatccgcatt
ttttattatgaata-ataaaaatttatcacatttatttaaatattaatataa--tgtgta
tatatatatacattcttgtcagaaactctcttatattttcttttaagtgataaaattgag
t----ttagtatagataataaaatttttatttacaatcataa----aaataaatatataa
ggaaaataaaatatgacaccgtccttatcaagctttttaaatagtctaattcttggggca
gtaattgtagttgttcccataacgttagcacttttatttgttagtcaaaaagacagaaca
attcgatcctaacaaggacttggaaacatatggagaagcaatttaagattaataataaac
tattttttaaatgtatatgctctccatatatttctacatt--ttgaaagaatttttgc-t
ttttactttgcaatagccttcacagtgtattctgtt-tttttttgtcctcacttactctt
tta----taaagatacaaatgtcagg-ttgataatagt-tttttttattctgtatctttc
atttaatatcaaagtattttatatacatattattatgttcaattaataaagaaaagctca
tgtctttgtctaattggcctctcaaaaaagaaaactccgaagcatatactactaagaatt
caaagcaaatcacaattcctgatggattgtggataaaatgctttgattgcggtctgttaa
tgtattctaaagttttaaaacgaaatttaaaagtttgtcctcaatgtagctatcattttc
aagcttctagcaatgaaagaattgatcaattaatagacagaggtagctggcgacctatgg
atgttcatttaatttctacagatccattaggttttaaagatcaaaagctttatagtcaaa
ggttaaaagatacagctctcaagactgggttacaagatgcagtccagacgggcattggta
atatgcaaggtagacaagtctgtttaggcattatggacttccggtttatgggaggaagta
tgggatctgtcgtaggcgaaaaattaacaaggctactggaaaaagcaactcaagaaaaat
tacctgcaattatactctgtgcatctggcggagcaagaatgcaggaaggcatgctaagtt
taatgcagatggcgaaaatttcttctgcactagaaatgcataaaaaaagcaaatctcttgt
atatctctgttttaacatctccaacgacagggggtgttacggctagctttgctatgcttg
gagatctaattattgcagaaccaaaagctcttattgcatttgctggcagacgagttatag
agcaaactattaaagaagacttgccagacaattttcaaagttcagaatatctatttgagc
atggtttcctagatttaattgtaccaagaactcagcttagatcaaagctaatacaaattt
tatcattacacaaccaaagtaagtaattagtgtcacattgatcgtaacattaaagtgcga

atataaattaagatattaagaaattttcactaaagatatttgttgactaactagaattac
aattactattgtagaaaaatttgtcttaattcta-tttttttcagtcttttaactaattg-
tttctatgactttaaataattgttagtttaatacaaatttataaagagaata-----cat
actaagctaaatatagtaatattatgtttccctataatactgacctatcaaatttcgagg
aattttatgcttaaaagatcttcttggcttgctactatattgggactactaactgtagtt
tctctaagtacgcatgcaaatgccatagagctagacgaggcaacaagaactgtcccatta
gaatcttctggaagaactgtagttcttacgccagaacaagttaagcgaggcaagcgatta
ttcaataattcttgtgctatttgtcataatggtggaatcacaaaaacaaatccaaatatt
ggacttgacccagaatctttaggcttagcaacgccacaaagagatacaattgaaggttta
gttgattatatgaaagatccaacaagttatgatggcgcagagtcaattgcagaattacac
ccaagtattaaaagtgccgaaatttttcccaaaatgcgtaatttaacggatgaagattta
tttacaattgctggtcatatcttgcttcaacctaaaattgtttccgaaaaatggggagga
ggaaaaatttactattaagggttcaaacgcaatagtcttataagtactagtttatgta-t
ttatttcctaagtgtgattattttgttagatcgtgatatatattaaagatggagtttgtt
gatagtaagttatttttttgcctaaacttacttgcattattcgtctagttattaaaaagga
gaagttaaattgaaaaagacgctttcagttcttttcactgtgtttagtttttttatgata
ggttttgcacaaattgtttttgctgcagatctagataatggagaaaaagtttttttctgct
aattgcgcggcatgtcatgcgggtggtaataatgctatcatgccagataaaactttaaaa
aaagatgtacttgaagctaatagtatgaatagtattgatgctatcacttaccaagtgaaa
aatggcaaaaatgctatgcctgcttttggaggaagactagttgatgaagatattgaagat
gcagcaaattatgtattatctcaatctgaaaaaggttggtaattatacttgattttgctc
aatattaaaggatagacaatctatttagttgttcattagattgtctattctttgtttctg
---ttattatatatagatatttacacaatactga-attatgatgaaaagaataccagcaa
ttctcgtactagaagatggtacttattataaggggtggtcatttaagcaaagaaagccag
taatgacaattggagaagtagtttttaatactggaatgacaggatatcaagaaataatta
cagatcctagctacttccagcaaattgtcactttcacatacccagaaattggtaatacag
gtattaataatcaagatgttgaatctcataatattagcattaaaggtcttgttgcgaaaa
atatttgtaaaatttcaagtagctggagggagcaagaatctttagtcaaatatttaagcc
gcaatgatattcctttcattttttggaatagatactcgatctgtaactcaatatttgcgtc
aatttggcacaatgaatggctgtatttctactaataatttaaatcatagttatttaaagc
aaagactttctcaagttcctggaatgcagggttcagatttaattccatatgtaacaacac
aaagcatatatccttgggatgaaaaaagtttgcccagctggtatttaactgataaaagta
gagtcgcactcgatacccaattgaaagtaattgttatagattttggagttaaactcaaca
tactaagacgattagctacactgggatgtcagataactgttgtgcctgcgcaaacatctg
tagaagatattttagctaataaacctgacggtatcctactttctaatggtccaggagatc
catctgtagtgtattatggaattaaaacagtaaaaaatttgttaaatcacaatgtaccta

tttttggtatttgtatgggacatcaaattttaaatttggcccttaacgctaaaactttta
aacttaaatttggtcatagaggtattaaccatccatctggattggaacagcaagttgaaa
taactagtcaaaaccacggtttcgcagttgacttgcaatctgtcttcaaatcatctgtac
aagttactcatttaatttaaatgataccactgttgctggaataggtcatagtcaaagtc
catatttctctgtacaatatcatccagaatcgagtccaggccctcatgatgccgattatt
tatttgaaaattttgtagaaataatgataaaatctaagcgtaaagttaa-----------
ttaataatcttcccatgcttcatgattaaatagcgctgataatacttgtactcctcgtaa
ttgattgaagagaggtagatgccctcttggaccataagaattccattgaaattcatcagg
atatctacaaggaatattatcaacttcccacccaatcttttgccagaacttttcccaatc
tttgccgacactaagccatatttgcctttgaataaaaagcccaaatttacctttttgagta
tgtatgccataatttatcaatagtctgaagatcttgagcaggtatttttttatatctgt
aaaatataaccaattgcgattttgtgcattcacgccagctaattgaataagcttttgttg
agttaattgatctgctttgagcaaatcttgttgtgttagcaactcttgtaaatctttgta
attcatatgtttagcagaatttaatggaacaataccctcgggacacaagctagtagtcaa
tctgacgatttcttgatttttactatttaataacttgttgtagattaagccatcgacgca
attacttttataatcgggtctattaatcctctcaaaaaatagatcagctaaacctttttaa
ttgtataaagtcattattatgtatagtctcaataatatcaagttgtcttttga---tatt
atttgacctagtactcttacttagttctaaaagctgaactcgaatttgattttgcatttg
attt-ttaaattaacttaaatagta----attagttcttaaagagaatttgattttttt
gaaaattagaagaacgacgaattatttctaaaagtgttttcataatttgattgaatatgt
tttgtcccgtataaatgataatttgaccaattagaaagactgtgctttttatttttaggaa
caaagaagtcttgtaactcaagtagaaatgtaatcaatatttgtattgatgataatgtat
ataattcattcgatctgcaagcatatatatactgacaatttaaaccttcctggtaaaacc
cccatactttgtaacgactttcataaattgctcgaggtctttcaatataattcttaatta
aagtattccaaattaaattatttttaatttttctagttttctatcagataaaaatttga
cattgcaaagatagctcgcatttgtttcatgtactgtatcaaaattttgaactaacatat
gtgcaataatattgcttaactgaattaaatagttttcaagaaaaacctcaacttgtttaa
ttggtacatttgaagtaaatagattaaaagtcttatttacacccttagaggatccaaaaa
gtagctgaataattagaatttgtagtaacatcctatagtcgtatattaggtagtttaaat
cattaatatcttggcaaatattattatagttaagttcatacaaactacaaaatcttttga
cacatctatgaaacaaatcaattaaaatattctcagttagagtcgtaatctcatttaaac
ctagatttgaattataaatttctaaaaggattttctctaactctgacaatattattttta
ataagtttctttttacttcagttcgaaaaacatctaaaattaacacttcctgagaacaat
tagttaatttctattaatttttatagaagttctaactaatagttctgcgacttcttgat
tgagtgtctgtccctgcgagctcggccaataattattcacgttatgtcatagtaagaaat
atttaaattaattttatacgataacttttttaatacaccgaaaagcaagtttattactgca

tagctattag-------gttttacagatcttaatagttatttttcaaataaaggtatatg
atattagcgactaagctaattttattgaaattta----taataatgacaatttactactt
tgcacttgccagtcaaaactttcttttagtacaagagcccttggaagaagtttttagaga
gagagttaattattaccagtcaaacaataaagtaatcgatttttggctaataccaaatcc
ttctttcttagaaaagccagaaatgatttcgtttaaagatcttgtacctaatgatgctgt
agctatagtctctactaacccagtatttattaattggttaaagcttagaattggttatgt
ctgtattggacaattcgaagatgatctacagctgtccgaagaatctttgaacataactgt
tacacaggaaaaaatttaatcatgcttctattaccactagtaac-aaata------ttaa
acggcaaatataatttacaaacacttatttttcttatagtttatcgcttattattttatg
ttgagatggagtgacaaacagtgtcagtatttcttgactaaatgtttcagttgctttgga
tttatatctgttggggttgactataatagacagcattcgttttatagtgacattttcaat
ctgagcccaatgtacaatcccaagttctaactccttggcaatagctgaaacagaaacaaa
agccgctcccaaacctgattgcactgcatttttttatagcttctatagaatttaattccat
ttctattttgaaacgactgctatcaataccatgttgacttagcactttatctatgacttt
tctgattgttgactgagtatctaatgcaataaatttaagtctgtatagatcttctttttg
tatgtcttctagtttagaaaatggatgtgatgttggaagtattagtgctaattcatcttc
agcataagatgtaacttgaagaacatcttgcaattcagtaggtacttcccctccaataat
tgctagatcaacttggccattagctacgctccaagaaatgagcctagtggaatgtacttg
taattgaacagctacttgcggatatctttgcctaaatagtcctattaatcttggcatcaa
atatgtcccagttgtttgactagctccaataattaatgtgccaccttgtaagttctgtag
atcatcaagagctcgacatgtttcttcacataaagctaaaattctacctccatatcgtaa
aagtagatttcctgcctcagttaaagttgctttttttattgcctctttcaaagagggaaac
atttaattggcgctctaaattttgaatttgtaaactaatagctggctgagaaacatataa
gctattagctgccttttttaaaactgccttctttggcaattgcttttaatattcttaattg
atccaatgtaaatggaagatctgtcattaaagaattgtagtataatg---------tata
ttatttagtattgatcaatttatgcattcgccaagttaaatatttgctattacgcttcac
tattttgaaaataataaaaagtataatgttaaatacactgtattagattttaacatctgt
ttgatgttaaatttatacaatagatatcgtaaatttaattttaactataaaatataggag
aaaac---atggactctagacttttaattgtattaataccagttttagcagctgcatctt
gggcagtttataatattggtagagttgcactgcaacaatttaggaaaatgacatcttagt
gatgctcagactata-atttaaaattttatgagatagagaacttgaagatctctatctc
tattgtataccttaataaaaaaagtcagta--actcttgtacttactgttattacctatt
aagatgatgtac-agtaaatacattcctaataagtttaagctaacttatt-----tttta
taattaattaaattaataacacaattatggctgttccaaagaaaagaacatctaaagcta
aaaaaaatgcacgtaaagcaaattggaaaaaccaagcaaaaggcgaagcacaaaaagcct
tgtctttagcaaaatcagtattaactagaaaatctaatggatttgtttataatctttctg

agacagcagataattctactgactaggagtattt-ctactattaagctaactaagtatcc
aaaaaaacattttaagtaatttttatatat----taaacacttggaaatcattctatttg
ataataaaataaatagcagattaaacaaaattacagcaacaaattatgcagcaaaactag
atcaaactagtgagatctggttatttaattgtatcgaaaatattcaacatactttttga
aaagtcaattaaaatcgaatcatattactaaaattttcatttctggtactagttttgaat
atgcggcaggtttacctggactattatctagcttaacactaagtggtagaataaatccaa
ttagtatatacggcccaaagtctctgaaacggtatcttcaagcttgtaccaaatattctc
aaactaatttttcttttcctattaattttcataatgtaaaatatggagaacaagttgtcg
atgaactgtatacagtagtttgtttacctctaagtaagaaaagtttactgtatggattca
ctatcttaaaaaaagaaaagcagggagcatttaaagtagcaaaagctaaaatcttaaaca
ttactcaaggtcctatatatggcaaattaaaagaaagagataattttctaagtccagatg
gttattatttatgtggtaatgatttctcgtcaagcacaacaatgggacataaaatatctc
ttccactattagtcaggtattctagaattatttcagagatgcattggttttgctcttatc
ctattagatttaacacgtattcacatcagcaagccataaaatatttgccgcacaatattt
taactgatattatgaaatcccaaatatctcaagataatagtttcgttgaataatacttat
tcttgatatataatagtctcggcaatctaaataacatacaagaaagaatctcttg-----
----ttctatattaaatttataatcacttatgacgtacgcaattattgaagcaagtggca
aacaactttgggtagaggaaggacgctattatgatttaaatcgtatacctgttgatccag
gccaatcgattatactgggtaaagttttattgttaaacaagaatggagaagtgactttag
gacgtccgtgtatagcgggagttaccataaaagccacagtaatgaggcatttaagaggaa
aaaaattaactgtttttaaaatgaaaccaaagaaaaaaatgagattgaaaaaaggccatc
gacaagaattaactcgtttaatgattgattcaataataaactagg------tacagttga
gactaagatttttttcaacaagagtttgaacatttaactaaaatatatataatggcacata
aaaaaggtagtggtagcacaagaaatggtagagactctaattctaagcgcttgggagtga
aaaaatatggcggcgagcaagtaatagcaggtaatattttaattagacaacgtggaacta
aagttaagccgggccaaaatgttggtaaaggaaaagatgacactttgttttcactgatta
acggcttcgtgtttttttgaaaagtctaaccaaaagcaaaaaaacaattagtgtttattctg
caaaaaaatagccaaaataattggtgcaagctgacttgaat----atttttatttcactt
acactagcaccataaacttgatttattaagaaatggagttccttaatgaaccaatgaatt
ttattaaagatttgctttaaaatgtaatgacaaacactattgtaatttcctgtctacaca
atatggctgctattctatattgtggtcagatacagaaactagtagtagcaaatgctcatt
atcaagtaagtgatatttacctaggttgtgtagataaaatcttctcaggaataaatgcag
catttattgacctgggaaagaatgagtacagtggcttcatccatattagtgacactggtc
cgcttaaaaagaaatattatgttaatagtattactaacattttaacaattcgacaaaaaa
ttttagtacaaattactaaagagccaactttaaataaaggaccaaggctcactgctaata
ttacattgtcaggccgatatattgtactaatgccttttagccaagcaatttgcatatctc

gaaaaatacatgatgaagatgaacgtcattatttaaagtctttagctatcctaatcaagc
cagcaacaatgggcttactatttagaccttctgctgtaggtattgatgaagaaattatat
taagtgaactaaaaaatttaaaagagcaatggaactttgttcaaaaatctgcaatcaata
gctatgcacctgtccttctatataaagatgaagatattattaaaaaggtaattagagatt
tttataataataatacaaataatattgtaattgattcaaatttaggattaaaacaattga
attattatatccatacctggcattgtaataattcaagtacagctccttcaattaagcttt
atagtaataatcaatgtatattagatgcttttggtattaatcaagcaatttctcgagctc
taattccaaaagttgatcttatacttggaggctatatgtttattgaaacattagaagctt
ttacaattattgatgtcaattctggatcttttaataactctactagtgcgcgagaaacag
tcttaaaaacaaattgttctgcagcaacagaaatagcttaccagttacaaattagaaata
ttgcgggtgtaattataattgattttattgatatggaatcacagagagatcaattgcagt
tattagagcactttaataaagagttagcacttgatgatgcaaaacctcaaattgtacagt
tatctgaattgggtttagttgaattaactagaagaagacaaggcaaaagcttatatgagt
taattagtagtgactttaattactttcatttctttatccaagcagagcgaattaagtctt
ttaaaagaatagatcacaaaaagcaaacacttcagactcttaatagatcatggttatcta
cagaaattaatactattaataaagtatttttcaaaaatccaatttttatatactggcta
acttttacttggcgcgcgatttacatatagttgatagtaatcttacttataagcaaaatc
atttattaactactagattgaaactaatttactctcaagattataacaaaatcttgcctt
ttactatccatagccatctaatcagctcgagtatagactataatcagaagtttttgttgt
aaaatatttgtttaaattgt----------------------------------------
--------------------taaaaaagaagccctctcttttataagagagggcttc-tt
ttttagcttactaactattctagctaattaattagaaattaaccgttaacagctggagct
gttagagctactggtaaagattcgccagaagctagatctagagggaagttatgagcatta
cgttcatgcattacttccatacctaggttagcacggttgatgatatcagcccatgtattg
attacacgaccttggctatcaacaacggattggttaaagttaaaaccatttaagttgaat
gccattgtgcttacagataaagctgttagccagataccaactacaggccatagacctaag
aagaaatgtagagaacgagagttgttgaaactagcgtattggaagattaaacgaccgaag
tagccatgagctgcaacgatgttataagtttcttcttcttgtccgaatttgtaaccgtag
ttagcagattcgttttcgcttgtttcacgaattaagctagatgtaactagggatccgtgc
atagcactgaacagagaaccaccgaatacaccagcaacacctaattggtggaatgggtgc
attaaaatgttgtgctcagcttggaacacaagcatgaagttaaatgttccggagatacct
agaggcattccatcagagaaactaccttggccaattgggtatactaggaatactgctgct
gctgctgctactggagcagtaaaagcaacggaaatccatggacgcatacctaggcggtag
cttagttcccactcacgaccaatatagcaagctacgccagttaggaaatgaagaacaact
agttggtaaggaccaccgttgtataaccattcgtctaaagaagcagcttcccaaattggg
tagaagtgaataccgatagctgcagaacttggaataacagcaccagaaatgatgttgttt

ccgtatagaagggaaccagcaactggttcacgaattccatcaatgtctactggaggtgca
gctacgaatgcaatgatgaatacagatgtggcagttaataaagttggaatcattagcaca
ccgaaccaaccaatgtataggcgattttcagtactagtaatccaagagcagaaacgttcc
cacaagctagcgctttcgcgtctttgtaaagtagcagtcataatttttatcaattttta
aggatttaactaagatgcttcccaaacaatattttgtctgttagatatattattacttag
aatcttgtaaaatgagaacgt-tttttcaaaaata-taattaaatttagtctcaataact
a--aaaatgtacttattaagctactattatatacttgaa----------gaaataaattt
ttcact-------------ttttctaaaattagtataattttaaggatataataatagta
atgcatactaatgctgcttattttgatttcaacagtttacataattgagccttacttaaa
tacgcatgtcacactttacaaaaattcagacgacaatacaagacttaaatctgttaaagc
atgctttaacagatttaggtttgacttggtgtatgaattctagtcatatacaagttggtg
aaaatagtcaacacaaagtagatatattgattaaacaggataatctatcgcatattggct
ttatctggaatgataataaatataacctagttgctgatttacaactttgggaacaacctt
ggtctttagaagtctttttagataaattgtcccaaaaatatgcttaccattctattatta
aagaaacaaagaagcaaggttttgaaaaaatggagcaaacctcgcaaaacgatggatcaa
ttaagttagttgtgcaacgttggagctgttaattactatataaatgcgggcggagagact
tgaactctcacgagattacctcactagaacctaaatctagcgcgtctaccaattccacca
cgcccgcattgacatgac--aatagcattgtatcatagttaatctgaaaaaacaaactaa
-aaagataaactagctgccaagtttaaatgcatcaacaaataggccgtatgttataccaa
tcttgaagttattaattaaattaatctttatatttaatttttttcgattagagtaaacga
ttttactagttaactctgttgctgcaactattaaactggctgcaatgattccccaatctc
ctgtttgcccaggaattgtagataatcctgttgaaataaagaagcctaataacagactga
ttaagccagttgttaactcatttagagaatagtaaagtttatcacttaagtttttttactg
aatacgcaagaaaggtagataatctagttttaatcataatatctaataacggagttttaa
aaggataacaa-----------agatgaatgtttttaaatccttttgaaacccgtcaata
caaaaattacttttttg-----------tttaaaggtcttgttcactcaaagcattaatca
gatgttgaaatggttcttgtataatgtctttacgtttaattgactgtaagtctagttctt
cttctattatttcttgcaataattgtatgcttcttactgttggagcaataggtacagata
atgaattgtaagtatctttgagcccgtttaatacacgctgatctagaatgttggtgctac
cagagactagtgcataactagcataacgaagatagtattctatgtctcttaaacacgcgg
catatcttcttgtggtataagaatttcctccaggtcttaataattcgggttgctcttcgt
acaattgagcggcagcctctttcaagatgttcgtagcttgattattaataatttcagcta
tttttattctatctagcccacttgagaagaaagactccatttgtccgacagctgctttat
ctagataacgaccagtaagatcgtaacgatttaatattgctgttatcgcatcttgcataa
attaattctcctac-ttgctatcagagcggtattatgaatgacttatat-----tataat
catgaatgtttataatatttatttaaaatctattctggattgatcagtttgcaattaaaa

acagatatagattttatataaatagagtatacgtagtttttaattaattttttgataaaa
attaagaaatacagattttttt-taataggtttgtaattttcttaatataactcaggttct
cgaccgaaataagttttgtaacacat--aagctagaacaaagatgagtaggtagtatt--
------------------------------------------------------------
------------------------------------------------------------
------------------------------------------------------------
------------------------------------------------------------
------------------------------------------------------------
------------------------------------------------------------
------------------------------------------------------------
------------------------------------------------------------
------------------------------------------------------------
------------------------------------------------------------
---------------------------taataataatatgcatcaa--------------
------------------------------------------------------------
------------------------------------------------------------
------------------------------------------------------------
------------------------------------------------------------
------------------------------------------------------------
------------------------------------------------------------
------------------------------------------------------------
------------------------------------------------------------
------------------------------------------------------------
------------------------------------------------------------
------------------------------------------------------------
------------------------------------------------------------
------------------------------------------------------------
------------------------------------------------------------
------------------------------------------------------------
------------------------------------------------------------
------------------------------------------------------------
------------------------------------------------------------
------------------------------------------------------------
------------------------------------------------------------
------------------------------------------------------------
------------------------------------------------------------

-----------------agtataaataatctcttaatacagcatcaaaaaaggtatttaa
cacatattgaaaaatttgattatttaaataaaaacttaaatttagataagattagctatt
ttataattacaactaataaaaatgataaatacatccttgtagaacaattctgttataatg
cagacggttctgtttattcaatattatttaaaggtcagacagctaaacacctttgctata
taattcttcagcttaaacaattaaatataagaatatcttcagctcatgctttatatttag
gaagagaactaatgaagtctgaattaactcttgtattagatcaacaatatatccaagact
aaa--ctgcatagatattaattgtatttgtgctagtataatattactaagggcatgtaac
tcagtggatagagtatcagattccggttctgatagccgtgggttcgaatccctccttgcc
cgaagagtatttagaatact-gtaacaagatacat-------tgggatcttgtttagtaa
tagtaaacatgttggggctgcaaggtttctacattatgaaaataagaaaatatgaaagta
agacaagctcttt-agagagtttctagtaaaattaataaatgcagaaaacaatattgttt
ctttttctcgaaaattagctattgcttaaatactgttaatttttgacattcgaagcgtta
aactcttatgcgtgacgaatattctgttagagttgctcttagtataagaaaagttataaa
atttatttatatttatgtttttttacctatttttatttaaagtaaggttagta-ttttcaa
tttttataatggacgtgggttcgagtcccaccagctccatcaatttaact---------t
tcataattaactaatccaatctttttaataaccagaaatataattggagcaaatatggat
ttcataaaaattacagaacaagcttttaagcaaattacagtcctgaaaagcaattatgat
caggatgtacatttaagaataggtgtaagacaaggaggctgctctggtatgtcatactca
atgaattttgaagatgtagttaatctaaaagagactgatgaaaaacttattttagatgac
ttttctgttgtttgtgatcccaaaagccttttgtatctttacggattatcattggatttt
agcaacgatttaataggaggagggtttcaattttccaacccaaatgctagccagacttgt
ggttgcggtaaatctttctctgggtaaac-attgtgctattt--ttttatattttgtact
gtacgtgttactcttgtaattag-catcataaatacacaaatatctttattgaaagctct
ttaaactagtcaaagacttataaaaactactttaaggagactttatt-tattcttgtagc
ttaagtaaatatttatgattaaatgatgaaagt---------acgaaaattcactttatc
attattgtctaattttaattagttaa-tttttattgatatatgctaatcttt-attagac
tcgatttattgaaaatttatag-ttcttttttctaagaaagtctgtaaatttctagcta-
----------------------aaataaattatatatttt-attcattttataaaaaatc
aacatgaatcttgctcaagttacttgcgagtactctagtacaaaaattcttaataagttt
tattgcaatacggaatgtgtttcccctggtgattttagccacctcaaattat--------
-------catcctctgcaaaaataattgcttttggaattcctcaatttgattttatccct
acaaaaattaaaaactctattagagaagcagtagtcatatctactgtcgatgatagtaat
atttttgctgattcaactaataattggataaaaaaacataaagataaaaatcaccattct
cttttctataaaaaaatttttaaagctttactaaatcaaaaaatcaaagtagttatagat
cgagc------taataatactacaatagataaagagaagaattcaattatgagtcaaaaa
tatgtttggggtaatagttggaagccatcaataatatcatcttacttcgaaaacaaaaaa

aatcatgaagtttctaatgcatctaaaaagtttctaattgatcagttgtcggcatcacct
gtctttgtagtaaaaaatggttttaatgaaattattttaggccacccagcattccgtatt
aagagaggaagcattagcaatcaaactcactcgtttttaaatttattaaagcaatctcag
caaaattatcctatatctactggtttattctttttacatccagatgacgcatttgaattt
aaaaactttataaaatctgttaatccgttggctagtcagcagatggagataaatgtagaa
cctgtcggccttcactttgcttataaaatgagcagaaatctatcttctgatactcaattt
ttgttcattccagattttaaagaggtaggcgatttattatttaaacatcaaggcaataag
aacttaatctttcacaaggatcagcgtcacggcaaacatttttttcagggacaacctata
tatattattcaacctactacttttaaagatcgaagtggcaagctaaatactattcaattt
actggactaagtgataacagagaagttgttttcactagtcttgaagccgctaataaatca
tggacaaattttataaaaaataattcacaattaaaatcaatcagtaagccgactttatta
gtctataatctagaaagtttttaaaagatcaagaacgattgaataaacaaaattttcaa
aaattcgttgtagttactaataaaaaagcttatcttactacaagagagttaataactgca
ccagattctaatagcgttttaaagcatttaaaattaaatatgaagcctaaactcttcttt
gtgaaattatgggtaagacggctattttccaccttgacttatgaat-aaatgacgtt-gt
atttattttcttgaatagtattcgataactaatagttcatttagttgtaaagcaacccat
tctctgtcaataactccattgactttccctgataaattagatttattcaattctaagtgg
ctaggaatattagctagtccaggaaacgctaaatagttttctactagtcttttagatgtt
tgctgatcttttacactaattaattctcctggtttacattgataactgcatatagatact
acttgttcgttaatacaaatgtgaccatgattcactagctgtcttgcagcaggaatggtg
ggtgccattccaagtctaaaaacagtattgtcaagtctcatttctaagagctgtaataaa
atttgacctgtagatccttgaagtttttagcagccttaacatacttaaatagttgtttt
tcacttagtccataattaaagcgtaacttctgtttttcttctaatcttacggcatactca
gaaggttttctagattttgtccatgttcccctggtggataagaccttttaattgctttt
ctgctaagtccaggtaaatcgcctagtcgacgggaaatgcgcactcgtggtcctctatat
ctagacatattttataaattctccgaatattaat-gattaag----------taatgagt
agctatttttaacacaaaaagagtattaaaaccaagc-ttttactaaaaaatctttata
caaaatagag---------------------tgctttactttttaagcgggtagcggaa
tcgaacccgcatcattagcttggaaggctaaggttttaccactaaactatacccgcaatg
ttcaataaattgtgacgcgcttaatataacataattatcttatgtttagatattatctaa
tctagaaaagaggtgaaattatttatgctccttctaaaactacatcaagaaagcgagcaa
tctctgcagcctgctcttcaacttctgaaagtaataatggctgacccactctagttaacg
gaattactcttttatcttttgtgcataaataaatttctcgtcgtggatttaaaccttctt
taatatctattcttattgatttaatttcttttatattcaactgaagacaaatttgtctat
ttttgcctggaaaacctaaccgaaaaattttaacagtaccttcatctttattaaattcat
tgtacccagcacctatgtcccaaataatagttagccacaaaaataaactaagaaagattc

cgatgctgccatagaatgtcataacaataccttgcggaataaatactaattctgttgaag
ttgtaaaaggcaataaatcaatttgaaagtaacttgataatcccgcaaggagaaaccctg
aagctccaataaagattattgtagcccaccaataattactaaatcttcgggaacccaaaa
ttaaatcttttctaacagagtttatagataatgctttttcataatttgttactttaaaa
caata-tact---------------aattttgacaatttttt--tcagtgttttagattt
gttaaatgagtttgatagcttttaatcctaaaaataatcctgttgctaaaccaaaaaaag
cagcaagaaatattgtgtaccctagaaaaactgacatattatta----ttattattattg
ttacaatagtgtttacaaaaatagttgaaatctattactagagtattatattacgttaaa
tgcttcgctgtctacatgaaaaaactttttttcagtttttagcattgtatttacattagaa
ataatattgtaataatttattaaacttctggagaacagaatcttatggcagaatttatc
aagccttataatgatgacccctttgtaggaaatttatctacaccaattagtacatcaagc
tttagtaaaggactattagggaatctaccagcttatcgtcgaggcctatctccacttcta
cgaggactagaaatcggaatggcacacggatacttttttaattggtccttttgataagtta
ggtcctttgcgaggtacagatgtagccttactagcaggattttttatcttcagttggctta
attattattttgactacatgcttatctatgtacggaaacgtatcttttactagaacagat
tcaaaagatccactgcaaacagctgaaggttggggtcagtttactgctggattcttagtt
ggagcagtaggcggctcaggatttgcttacttgttgctagctaatattcctgtacttcaa
acggcaggtttgagcttattttcttaagctagtaaggggacttgaacccgtaacctactg
attacaaatcagttgctctaccaattgagctatactagcacctaattaattataacatat
aaagatatttgttgcaatctggtattttcagatgcaacaaatatctttttttatttagaca
gagctgttaaattagtatatctataatatgttaattagaatctgaactatctgactcagc
agaatattcttgatcacagtttaaataataacaaattgtttgatctagacatgttgcgga
ccaaccaacaggtgtttctcctgataattcttgcatatctccagaactataaaaatcatc
actcggcttattactatttgcttgcataaatatcactcccaaaacactcttaacttaatt
tgaagtaatttaacttctcaataatatattaacatataatacaaaacatggtgcttattc
attaattgtaacaataatatgaaatttacaaggtctgagtaagtgttttaatggcttttg
ttgaaacaagcattttaatccatttattcttctctcttgaccatactttttttatgttgta
aattaacttttttgtaattttttttgttcgcttgtgagaatgagaaactgcgtacccattat
tagctactttccctgtaagctgacatttttttcgacataatttt------attaacttata
tatttatttaaagtacttgctagagttgactttggcacggcaccaattaccgtatcaact
ctttctccagctttaaaaatcattaaagtaggtatgcttctaataccatattctgctgca
attgttggattatcgtccgtgtttatttttactaccttaatagatgactcgtattcttct
gcaatttcatcaactacaggagaaaccattctacaaggaccacaccatggcgcccaaaaa
tctacgagtacaggtaagttattattaataacttcttgtttgaaagaggcatctgtaact
tgagatactgacatattttttaacctttaatatatgttccttgctaagcaaatcttatca
caaattttaggaaaaatcttctatagttaattactattagtgttgaattaattgttt-tt

aattagacattaaaaaaacttattacctatataaataaagtggtggccgagtgatatctt
gatagtgatagtttttctattaaggttgaacaatagaaataatttttcaagcatata-tt
ttttgaaaa--acttctttgtgaccatgaaagcttaattatattacaatatattttatta
tttgtaacttagttctgataatggtataaacaacgcaaaagatactgccttataatcaag
gaggaatacatgtctcaatccgtagaatcacggactaggattaaaagcgaacgttacgaa
tctggagtaatcccttacgctaaaatgggttactgggatgctgactatgtaattaaagaa
acagatattttagctctgtttagaattactcctcagccgggtgttgaccctattgaagca
tctgctgcaattgcaggtgaatcttcaacagctacttggacagttgtatggactgatttg
ctaacagcttgcgatttatatagagcaaaagcatatagagtagatccagttccaaacgtt
gcagatcaatactttgcttatattgcttatgatattgatttatttgaagaaggttctatt
gcaaacttaactgcttcaattattggtaacgtttttggatttaaagctgttaaagctctt
cgtctagaagatatgcgtatgccagtagcttacttaaaaactttccaaggtcctgcaact
ggactaattgtagagcgtgagcgtatggataaatttggcaggcctttcttgggtgcgaca
gttaagcctaaactaggtttgtctggtaaaaattatggaagagtagtatacgaaggtcta
aaaggtggtcttgatttccttaaagatgatgaaaatattaactcacaaccatttatgcgt
tggagagaaagattttatattccatggaaggtgtaaataaagcatctgcttcggctggt
gaaattaaaggtcattaccttaacgtaacagctgcgactatggaagatatgtatgaaaga
gcagaattctccaaagaagtcggcagtatcatttgcatgattgaccttgtaattggttat
actgctattcaaagcatggcgatatgggctcgtaagcatgacatgattttacatttacat
agagcaggtaactctacttactctcgtcaaaaaaatcatggtatgaacttccgagttatt
tgcaaatggatgcgtatggcaggtgttgaccatatccatgcaggtacggttgtaggtaag
cttgaaggagatcctttaatgattaaaggtttctacaatactttacttgaaagtgaaaca
gatatcaatttacctcaaggtctattctttgctcaaaactgggcatctctacgtaaagtt
gtgccagtagcatctggtggtattcatgctggtcaaatgcaccagcttcttgattactta
ggtgacgatgtagttcttcagtttggtggtggtacaattggacatcctgatggtatccaa
gcaggcgcgactgcaaatagagtagccttagagtccatggttatggcaagaaatgaaggt
cgtaactatgtagcagaaggtcctcaaatcttaagagacgctgctaaaacttgtggacct
ctacaaacagctttagatttatggaaagatattagttttaactatacttccacagataca
gctgatttcgttgagactccaacagcaaacatctagtttaatgactacttgctaatgctt
gatttagcaaattgtaagtagaattaacttataataataaggagcatagaatagtgagac
taacacaagggactttttccttccttccagatctaactgatgagcaaattaataaacagc
ttacttacatcgtttctaaaggtttatcagcaaacgttgagtacactgatgatcctcatc
caagaaattcctattgggaattgtggggtttacctttatttgatgtaaaagatgcatctg
ctgttatgtatgagattagctcatgcagaaaagcaaaacctaattattatgttaaagtta
atgcttttgataatactcgaggtattgaaagttgtgtaatgtcttttatcgtaaatagac
ctgctaatgagccaggattcttattacaacgccaagactttgaaggtagaactatgaaat

atagtcttcatagctatgctactgaaaagcctgaaggagctagatattaattttaattaa
tagctaaaacctattaactggttacccctctttgatatcttaattactaaattaattttg
attttaaggaggggcaaatttcgaaataaattctagttaaattagaatttacaattaaaa
atacacagaaataatgcaatcacaagatataatctccaacgatactcttgttaacttaca
agaagaatatgatagaacacaaatccaagaagttcttaacgaattaaatcaagaacttat
aggattagttcctgtaaaaactagaattcgcgaaattgcagctctattactaattgatag
actacgtagacaactagaactggtatctggtaatccagggttacatatgtcatttacagg
tagcccagggactggcaaaacaacagtggccatgaaaatggcggatattttgcatagact
tggctatataaaaaaagggcatttattaacagttacacgagacgatcttgtggggcaata
tattggtcatactgctcccaaaactaaagaagttctaaagcaggcaatgggaggagttct
gtttattgatgaagcttattatttatataaggcagacaatgaaagagattatggctctga
agcaatcgaaattttactacaagtaatggaaaatcaacgaaatgacttagttgtcatttt
tgctggttacaaagacagaatggaaaaattctatgaatcaaatccaggactttcttctag
agtagctaatcatgtcgatttcccagattacacttctgatgagctattgcaaatagctaa
aatgatgatagaggaacagcaatattgttttacggaagaggctgataaaactcttttaga
atatactgagcgaagaatgaaacagccatattttgctaatgcgaggagtattcgcaatgc
tattgatagagctagaatgaggcaagccaatagaatttttgcaagtggagaaaaagtttt
gacaaaagctgatttagtaacaattgaagcagaggatatcttgaaaagtagattattttc
attacctaattcttagtgattaagtagtttgataaaactgtatattatgaaaaaactttt
ataataagttgcaaatttctttgaaaacctatactattaatgtattataggtg-tggcgg
catagccaagtggtaaggcagaggattgcaaatccttcatcccccagttcaaatctgggt
gccgcctagtactaaaaa--gggggtgtggtggaatggtagacacaacagacttaaaatc
tgttga-ttttaataatcgtgagggttcaagtccctccacccccacatattaacttataa
caaaaaacagatgtgtatttgtattaattgtaaccatattactgaatgtgatacttatca
tttaatagaatcccagcacaaacaacttcatctaactgagactccatcattcatacctaa
atgccctgttgtttgtgttaatatatctaacaatagtagcaatatccaaattgattggga
tttagtcgaatgcttatcatttgtggagaaaccgaatagttggaatttaggctgtgatta
gttgtgcaaaaaattatgtggtgtacgcaataactctatcttttagatactctgtattaa
ttttagatgttcttactaacagaattttccctgttcttgctatctctataataccgaatt
tagttaataattgttcaatggcaacaatctttcctggatcgccagtaacctctataatta
aaaagttttccgcgacatctactatcttagctctaaaaattttcacaatttctaaagctt
ctgttctggtctgagaattaattctaatcttaattaacattaattctctttcaactgaag
gaatattggttacgtcttgtacgttaagaatatttactaatttatataattgtttggtga
gttgttcaatagttctgttgtctccctgaactaccatcgtaattctagaaactccgattt
gttctgctggtccaactgctaaacttgcaatattaaagcctctgcgggcaaatagaccag
atattcttgaaagtactcccgcttcatcttgaactaaaactgataaggtgtgtttcatga

aattttaatattatactatttgatctattaaatatattctatgaaaattatagaaaaaat
actagttaatatatagcgaaattattcaaactaaacaaaatcctattatcagatagcata
ttttttttgtaatgttatactattaatctagtatttaatgatattctaattctaaataga
atgctaaatttttaaaatttgttaggaacttgtgttagaaaaatacaaaaacagtaaaa
aactctctcgagatctaccccaaatcaacgatcgtattagatttccaaaagtccgagtaa
ttgacgatgaaggtgaacagttaggtatttttgtacctgaagaagctatgcaattagctg
cccaacaagggttagacttagttgttgttagtgataagtctgatccaccagtttgcagaa
tattggattatggtaaatataagtttacacaggaaaaaagagccagagaagctaagaaaa
aacaacataatagtagtgtaaaagaagtaaagatgcgatataagatagaagagcatgatt
ataaagttagaattaatcaagcatctaagtttatccaagcaggagataaagtaaaagtaa
ctataacatttcgtggtcgcgaaattcagcattctaatttagctatagatttgttaaaca
aaatggcaagcgacttgacttcgatatctgaaattcagcaagctccatcacgagatggaa
gaaacgttataatgcttttatctcccaagaaagtgagct-aaatgatttaatttaaatg
catctggctggattcgaaccagcgacgtctctttcgaaatggcggattatgagtccgctg
ccttcggcccctcggccacagatgcatcacttaaagtttataactcataaagtatct-aa
aatcaatatctatttcttgttcatgacatcta-aaaaaattattactcattcatattgtg
atatgtcgcaactgctgacggagaaacttggttcaagtagcgaaatatccaatatttaaa
tattgtgtctagtatgactggaaaagttgaaataaatagaaaaataaagtcacgactttc
aggtaatcctagatgccttaaaattacttcaataataacttcccagccatgaggagaatg
aaaaccaacaaacatatcggtgaaaagaatgattagaaaagcttttgcagtgtcactaag
gccataaataatttcatttaaaaaagatttgacaattgcaatttgtctttgtccagttat
cattaataatataaaaacaaatattgatacaatatcggacaaaatgtttttcacagcatt
cgcactttcatttgcatagtattctcctagttctctggccttcagctgcactctttttc
tataatttcataagatatatcttccgatgggttcaataagacttcaaagtgaattttctc
ttcaaacctttgtagttcagcaaaagctctctcttcttgagatgaatttaaaaaaatttt
aggttgttcttggttccataaataatcaatacaaggaccaaacacaaaaaactttgaagc
ttgattaactaggactggagagataaataataataaaatatattttacggaagtaatagt
ttgatgtctagatattctaaactcttctatggcttcggattctccatttggatctaactc
tttgcgaaacttttcaaaagtattagtaattgacctaggtattggtccaactttttcaaa
ggcaaattgattatttcgttttaaattccaatatttcatttttattgctgctaaaattt
ctattaatcaataattggtcacgtattcactataaatattaattctagaactaaacatat
aaaacaatagttatgaataataaaatacaattaagctttaattacgcattgttactatct
cttgctacaacaatatatggatgtagatcttatcgtgaaactatgagtacactattaata
aaatatgaagatagctatccattattgtctgatagacatactgcctttattcctataatt
gtattaaaaaaaaatctttaaactccctaaaatcagaacctcacaatactaagttgtgtc
ttaatggaatgggaaatagaaaatttattgataaaattgcaaatttgctaagccgtttag

aaaaacagatacactatactgaacatattaatcaaataaaaactagtgggtacttttctt
tactttatgttaacagtaaagcaatgtatacaaaagaaattagcaatatctttatattgc
caaatgatgttctaaaaagaatttatattcagaataaacaaaaaaagcttatacctgata
cattcttaataaaattatttcaaaaacaagtaggttacccgagaaattttgcaaacttaa
acctagccttgtcaaaaattatgcaatggtattctgatagaggatatcaatggtctttag
tggaaattaaacaagctactgacccatctcatctgatccttaatatccatgaaggattag
ttaaaacaattacaacagagtattatactttgtctcgtaaaaaatcacctagcatctcta
gtacagaattaatagagaagtatcttggagtaaaagtcggacttcctttaaatattaatt
ttttacaaaaaaaaattaactatttaaaagataatcaattagttggaaacattatttata
gtattgagcgatcagccaataactctatgaatctagatattaaatttcaaatacaagaac
taaaagataaagatttactagtacttgtagaaagatcatccaaaacttctcaaattatct
ctcacgcatatgacttactaggtgtagaacgcaggagttctcctaattgcctggcagatt
ccacaataatgtcattatcgactattaaattacaagcatgctattttaattataaaattg
attttgaacagaaatatatagatgctattaatctaatcaaactgttaacttattccattt
ttcgtcaaaaaacactaataagtgatagttttaatttacagactttagtcagctggataa
aaagaaatactattggatttcaactgtatttacgaaatttggatcgtgcaaaatcttttt
gtctattcagtctgaaatgtattcagagtggactgaatataaagatttcatatctaaatc
cttcattaatagttaatcgaaatttgagttttcagcttgcaatacaaataattaagcaat
atcacacgaatcaacaatcagttccatctt------------------------------
------tgtttttagcacaaggagatctggtacaattaatttttgaaagtatatgtacat
atcattttacatcttgtttttcgatatctgaaaaaatattactttctcgagttgtttata
tagattccatattttacaattctgaaaatttaagtctttgtaacaagaataaaaatcaat
ttgtaaatagctgtgatatcttgagacaaaataaaaaattattgcagcaaaaatttctta
ctttattgttaagcatccattatcaaaattttaattatttggattggcctttaaaaggcc
atttatttgaaatgaggtttttatactttaccccatttcagaacagcaatttttttaaact
actcttttcttgataactacaaaaacctgtttttcataaaatttatttaaaacaaatct
ctcactttaatttaccattgtgttttagagaccatcttaaccatattttagttagtacta
ttcaatgtcagtcaaatttagatatggagactgtttatctattgttaagcaattatcctc
ttgaatatactctatttaagtctattgttactttttctcttaaagttaggatggaatatt
taatacctataagcaggagtattagagtatctttgttttataattatctagattgttttt
tcataaaacaatctaaaaaagtagcatatgcgctccaagatctatcgagcagtaaacaaa
ttcgaagtttttattggaaaaattttcatggggatttggaattcagctaagattaccaa
ttaaacagatgccacccctttcaattgaatacactttaaacagtggtaggcacttttgta
tttatctacatatttactattaaagatggctagtattaaaatgaaaaatttaatttcttt
ttttaaccgaagcaaaggaaaatggatttcacaaagaacaacttatgaactatataataa
gaatatgagttcagtacaatctcaaatgcaaataaagtcaaatgaattattgttagaatc

tgcaggacttgcgtcactaaattggggagagatttctagacaagttaaccataattctta
ttctcaaaatacaatcagaaataataataagtttaacttagaatttagtaatcagctaaa
taatcgtaagctattgacattatgcacggtaacagatcctagtctaattaattttaaaac
tcgatatggtagtactactgtggatgaaacatattggtttgcaacaagtaaattgcgcct
aagtaccagtattattaaaaaatttgatacttgtattgcagtatctttttgttcagaaat
caaagtttaa-attctgtataaaagcaactaatagctgaaa-agttattagttgcttta
tggatctaagttttgttttatagagtatataaaataatatattattctaaatctttacgg
ttaggatttctagaaggatcattagataaaaatccaaaaacaaacagagaaataaagaaa
atcacagtagtataaacaaatattttcaaagtaaacatatttacatccttaaaagctatt
agtgtg----atatttatatatc-acacataattttatatcacgatatgagtctcggtgc
tattttataaacaaatcaatatctttatttttagcaatgatttatactttaatatgcgt
gatacactaaatattaaatctttttgtatacgcttcttactattttaaatactttatta
tttctatggagtttaccttcaattattacgtagtctagtttcttcagtaatttgaaaatc
tctaaagatttttattccaaataattaagttgattgtagctaatcttttcctttgtaaa
agcctgacttccaacttaatgattttgtttctattatcactaattttaatacttttacac
cttaaaatttgaactagcaatgtacagctattcat-atttgtattaa------tctcata
actaatctattatgccagataa-atgaaaaaattaaaatgagatctcgttatcttgttga
acttgttcataactaatatcttttaaaacctttaatattttcccgaaatactcttgcaaa
tatttttctagcgatattacttctgagctatcaatgcctaaaatactgtaaacttcattc
atggaagcagtaaattgctctccactagttaaaatctctgcaaatgctagacgatcagaa
atattccatgtccattgaagggttttagtaattttccttaaagctgttaataatccaatt
ggaatttgtgatatttgtgtttttttgtccagataacttttcacataatttaataatttcc
gctgatgtccaagctgtgttaccaactagtggcagtattctattttctgtggaaggcact
ccaagacttttataactaattttgctgcatcctgagtatcaatatatgcaatcggcgta
gactctcctgtaacccatactgatttcttatccaaaataggaattgcatattggttaatt
aatccctgaaaaaatcctcccagagaaaataccgtgtactttatattagattttcgaaga
aaatctactacttgcgacttcaagttcattaatggaacagctgggtatttatctgcattt
aatattgaaaagaaaataaatctttcaacttcagcagctttggctgcctcaattaaggca
attttaccatctaaatctatttttgctgcattataaggatctgtaggacgagatgtagaa
gcatctatgactgctgttactccacaaaaagattgcaaaatactttcaggtaattttaaa
tcaccataaacaagttctgccccccattctttgagaaaagctgactttcgtaaatttctt
accatgcattttacattatagccttcatctagagcacgccttacaatttgacgtcctaga
gtccctgttgctccaattactagaagagtcatattaattaaaaataagtatgtataaaga
cattgatttcatcgtcgacggagagtacattaggtagagagggaatcgaaccctcatgac
cgaagtcgtcacattttgagtgtgatgcgtataccaatttcgccatctaccctattatag
accaactactacaagtaatttcgtctttagcgtaaaaataatactaagttttaatagata

tgtcaacttttttagtctatattaacaaaaaacaggagtatcttatgtctacttttgaatt
aattccttatagattatatttaagtcaatctagtacatggatgcatatgatgaaagcaga
aattaaaatctatatattaacactattatggatttcagttttcatttttttcttactttaa
actatttattattgcgctaagtttaattatgattagcctgactataagaaataaacaaaa
tattatacaaaaacatttcttgcaaaccttaatcatgacagtggtaactaccttcttgtc
atttagcttgactactagttacaaacaatattcgcaactaaaacaatcacaaaacttatc
ttacaataataaagctaaagatccttacaattatcaaaatatacaaatatctagtggtct
taagattcgaaaaaaacaattaagatgcagtttaaagcctgcactctattttttttattac
tatatattcaatgaaactagtcatgataacgacttccccagaaattttagtgattactgc
ttatagatctaggatcttcagtgtgctatttaataatgaattattatttaccttcttatt
gtcgtcacatattgttactaatgtaattagtaaattaaataaaatcattcaagtagctag
tttaagaggtagcttaaatttatataaatcatccacaaaacttttaatatttttcttttttt
aatatttcaagtcttttttttttagagatcattaaggaatcaaaagaaatatctcaagctct
ttatactagaaatcttaatcaagaaaacagtaattttttaaaaatatatagggccaaatc
taatattaatgattgtttaattgcgattattggaaccttatatttcattattttaggagt
agcataaca-aaccgaattatctttccgactctagtaatacgaatagtaatagacttaag
tagtagttagaatatgtaaaat-tcaaatattatataaataattaaatgttttattatgg
tcaatactcaaaatccaattgcacaaacttcagagcttgattatattgtcaatcaacctt
acaaatacgggtttaaaacttctgttgaatcagaacaatttcctagaggcattaatgaaa
atatcgtaagattaatttctaagaagaaaaatgagcccgagtatttattaaacttcaggc
ttaaagcatacaaaaaatggaaaaaaatgagcagtccatcatgggctcatcttaagcatc
cgaatatagacttcaacaatattatttattatgctgttccgaaattaaagaaagaactaa
atagtttagatgaagttgatccagaaattcttgacacttttaacaaattgggtatctctc
tcaatgaacaaaaacgaatttccaatgtagccgttgacgctgtttttgatagtgtttcta
ttgcaactactttttaagaaagaattatctgaagctggtgttatattctgttcaatttctg
aagctatacgggattatcccaatttaattaataagtatttgggtactgttgtaccagctg
gggacaattattttgctgcattaaattctgcagtatttagtgatggctctttttgctata
ttcctcctgatacagtttgtcctctagaattatcaacttattttcgcattaataacgaag
aatctggtcaatttgaaagaacattaattgtagctgatcgtggtagtaaagtgagttatt
tagaaggttgtaccgcgcctcaatttgatacaaaccaattgcatgcagcaattgtagaat
tagtagctcttgatggtgccgaaattaaatactctaccgtacaaaattggtatgctggta
atcaagaaggcgaaggtggcatatataattttgttactaaaagaggcttatgctctggca
ataactctaaaatttcatggacacaagtagaaactgggtcagcaattacctggaaatacc
ctagctgtattttagctggccatagttctcaaggagaattttattctgtagctttaacaa
ataattatcaagaagctgatacaggtactaagatgattcacattggtaataatactaaaa
gtagaatcatctctaaggggatttctgctggtaaatcgaaaaacagctatagaggactag

ttaaaattggccctcagtcatttaattctcgtaattactctcaatgtgattctttattaa
taggtcaatcgtctcaggctaatacttttccttatattcaagtacagaatccaacatcaa
aagtagaacatgaagcatctacttcaaaaattagcgaagatcagattttttattttttac
aaagaggtattagtttagaagaatctgtagctcttatgatcagcgggttttgcaaagatg
tatttaatgaacttcctatggagtttgcagctgaagccgatcgtttacttagtttaaaat
tagaaggaactgtaggatgagtcaaaccattttagaaattaaagatttatatgcgtctat
tggagaaacagtaattttaaagggtgtaaacttatctatacaagaaggcgaaatccatgc
aattatgggacctaatggttccggtaaaagtaccctatccaaagtaattgcagggcatcc
agcatactctttaataagtggagacatcttattttttgggaaaagtattcttgcaatgga
gccagatgagagagcaagagctggtatctttcttgcttttcagtatcccgtcgaaattcc
aggagtcagtaatgcagatttttttacgaattgcattaaacgctcgtagaaaattcaaagg
attatcagaatttagccccctagaatttttcaattaataaatgaaaagataaagcttgt
tggcatgcaagaaagtttcttgacaagaaatgttaatgaaggattttctggcggagagaa
aaagcgtaacgaaattcttcaaatggctttgctagatagcaacttatctattctagatga
aacagattctgggttagatatagatgcacttagagtagtcgcaaaaggcattaatacttt
agcaaactcaacaaattcaattatttaattactcattatcaaagattattggattatat
cattccagattttgttcatattatgagtaacggcaaaattgtaaaaaccggcaatctcac
tcttgcaaaagatctggaaaagcatgggtatgactggattgtacagagctaaatatttgt
taaatacaaaaatactgagactaaagtattttctttagtctcaatattttgaccaacaa
aaag-----ttatttaggctttatcgaaaccttgtttaacatcttcaattgcttttttta
gcagttcttcactagatgcgtctaatttttagtgctacgaatactttccccaaattctg
gtttagaattctttaaatcttctctcaattctttgatgaactcagccacttttgacacct
caatgtcatctaaataaccgttaattccagtataaataatagctgtttgttcttcaactg
gaataggagagttctgagcttgctttaaaatttctctaagacgttgacctcgtgccagtt
gattttgagttgctttgtctaaatcagatgcgaattgagaaaaagcttctaattcagcaa
attgtgctaactccaattttaatttgcctgccacttgtttcatcgcttttatttgagcag
cagatcctactctagagacagaaataccgacgttaatagcgggcctaatacctgagttaa
acaaatcgcccgatagaaaaatttgaccatccgtaattgaaattacatttgttggtatat
atgcggatacatccccggcttgagtttcaataattggaagggcagtcatactaccgcctc
ctaattctgagttcaatttagctgctctttctagcaatctagagtgtaaataaaatacat
cacctggatatgcttcacgcccaggaggtcttctcagtagaagagacatttgacgatagg
cttgcgcttgctttgttaaatcatcataaataactagtgtcgctttacctttatacataa
aatactctgctaaagctgcacctgtataaggagcaatatattgcaacgttgctggactat
ctgcattagctgcaactataatagtataatcaagagctcctttatcttgtagtgaagata
caacttgagccactgacgatgctttttgtcctattgcaacatatacacaaacgacatctt
gccctttctggttaatgattgtatccagcgcaacagctgttttacctgtttgacggtcac

caataattaattctctttgacctcgaccaattgggatcatagaatcaatagctgtgattc
ctgtttgcataggttcacaaactgattgacgtccaattatgccaggggccatagactcaa
ttaatcttgttgcgttacttgcaggttcacctttgtcatctattggacgagctaagggat
caaccactctacctaaaaaagcatcgccaactggaatttgagcaattctaccagttcctt
ttacagagctgccctctaaaatatctctaccatctcccatcagtacaacacccacgttat
cactttctaaatttaaggcgacaccaatggttttgtcttcaaattcaagcaattcaccag
ccataacttcatcaagcccatatactcgagcaataccatcaccaacttgtaaaactgttc
ctatattggctacttctacatcttgatcatacttttcaatttgttgacgaataatactac
ttatttcatcaggtctaatgtttaccatgtttttaatattctttttattaatgggaaatt
ctaaagtggtaattactcagattaatactatctg--tcttttactgcattttaagttact
actacattaaggtgtgaagccatctgccttagttgtcctctgatactagtatcaataact
ttcgatccaatctgaatagtgaaaccaccaattagctcgggatcgacagagacaattagt
ttgacttctttagactgggtcatcaccttgatttttctgtcagcagatttcttgatca
gagcttaaagcaatagaagtattaatattagcaattgttagagattccatttgatatgca
agttcgagatacttactagctatagcatctaacattgcaattcttttgcgatcaactaaa
atcagtaaaaatgataatgtattttcgctaatttgatcactgaaagtggcaataatgact
tgctttttgcttctgcagtttttaatggattagct---aaaaagtttttcagcttgtca
gattgcaataaaatattttgtattgactgtatatcttgactcaccttttctgtaactttt
tttattttagctaactcaagaagagctacggcataaggctgagctattttcataacaatg
ttattactgctcatagttgatctcctaacttagcaatattattatcagtgatacgtagct
gcatttcagagcttatttgattttctagctgcacagtaactcgtttaagagcaagaaatg
ttatttgctgctggatttgcctgcgaatttgtttttcagctgtttcgatatttgacttgc
ctgtcatagccagtctttcaatatctaattgaccttgagctaatatagaacttctaactt
ttcctgctgtcagctgagcctctttttgatttgatctataataatttgagtctgagcaa
gttgcttttctgattctgataatctagcacttgcttgctctagtctttcttctgattctt
gtatagctgctaagacttttatctgtcttgcatttaaactagatccaagaaattgtttca
gcacgtaaataagaccaaataaaagcaacaagatattgataacattcgcttcaaaaatat
cagaattaaagccgaatttatgttcactactatgttctgataaaatagtaattatttgtg
gtatgtttactatattattcatttagaattct----------tactattttaatatttta
cttaggagtcatgattttaaagattgactacttaatagtttggctttaatttggtcgctt
aaagtatccacttgatcttccaaagttttaagagcttcttctttttgagtgtttagctgt
tttgatgcttcagcaattaatttttcagcattcaattgagctttttttaatatcttcagag
actatattttgcgcttctttttgagaagaagctatctttagttgtgcattacgacgagct
tcggataaatcttgttcatattttgcagctaactcgtctgctttaacaagcatagaagat
gcagtagttaaagttgtgcgaatatattcatcacgttcatctagtactctagttactggt
ttataaaaaatagtatttaataaaaccattaaagtaagaaactgtaatgccattaaaggc

aaagtaccattgaagtcaaataagccccccttcaatttcttccgctaatagaaatggtaaa
taaatcattgtactaattatgtttacagttaataatttggtctcaattaagaaaattaac
tttatagtgcgcttagtctgattagactaagcgcaaatgattattttaaccaacgtatgg
attagcaaataatagagacagtgcaacaactagaccataaattgttaaagattccataaa
tgctaaacttagaagtaatgtgcctctaatttttccttctacttctggttgtctagcaat
accttccacagcattagcggctgcactaccttgaccaataccaggacctattgcagcaag
acctacagcaagaccggcagcgataacggatgcagctgaaacgattgaatccataatgaa
ttttgatttttttgttatataagtagaaaattgacaaatttcggataatatttaaaatcta
aacaatagaatatcgtagattcaaatat-----tgtaattatagtatactactcttctcc
atgaccttccattgcttccccaatataagctgcagacaaagtggaaaaaaattagtgcttg
gattgaactagcaaataatcctaataccataactggcaacggtatcaaaattggaattag
caaagtaaatacagcgacaactaattcatctgctaagacatttccgaatagacgaaaact
taaagataatggcttagtaaaatcttctaaaatattgataggtaataatactggagtagg
ttgaatatacctggcaaagtaacctaacccctttttttacttaagcctgcataaaagtaagc
taaagaagttagtaaagataatgctacagttgtattaatatcatttgttggagcagctag
ctcaccttctggaagttgaattaacttccaaggaattaatgcgcctgcccaattacaacc
tagaatgaataaaaatagagtagcaatatatggaacccaaggacgatattcatgctctcc
aatctgattttagcaatgtcttgtaagaattctaatataaactccataaaattttgaaa
tttttcaggaattctttgaagattcctggtacccaggaaagatattgttaatagtgtagc
aattactaaccaagacacaataaaaacttgaccatgtaattgtaaactacctattttcca
atatagatgttttccgacttctactgctgacaagctagtatatggattaaaatctatgag
actgttttgatacataataaatatttcttttaaatagtgcaagaatcaaaaaataagttt
tttgaagaattaatatgcaattagtttattggcaagattaggtttttaataaattaagag
caaaaataaattctaatagagatgggtatctttagtagtgattatttttttgtaaaaaaca
attattgctacctataatgcttttatgtttaattatatacccatatattaattatttaat
atgattataagtatttcgctacaaactttattgttttatctattct----aggtgttatt
ctaactcatttcttatttaaaatatctgcaacttcatcggcaaaattagctctggtattt
tcttctcctccacctaatacgaatctaacaaatcttctgattttatattctcacctaaa
acggaaatattttgattgattaaatcttcaatactaatatcttgattgcgaatgaacatt
tggtctaataatgaaagttctttcagcctctttttcattcgcccttcaataattccgtct
attctatcggcgggcttgtttttcaagtcatctttaccagcttcaactctttttttctaaa
ctgataacttcagctggtatgtgctgtgttgatacgtattccacatttggcgatgctgct
atttgcatagcaatatctttagctaatttttgaaattctggacggcgtgccacaaaatcc
gtttcacagtttacttcaactaatacacctattcttcctccggtgtgaatatagctttct
aatagaccttcaattgcagttcgactagatttcttattagctgaagctaacccttttttgt
cttaatgattctaaagctttttcttcattaccattattagcttctaaagctttttttacag

tccatcatgcctgcaccagttttatcacgtaaggctttcacagtttgagcagaaatttgt
agtgtcatatattttcaaaaatttttaagtatttaaattaactaaaacaaattattataa
tttactttcaacagagctagctatttcttctacatccgtctgcccatatttgccatcata
aattgcgtctgctatttaccaacaattagtttaattgatctaatagcatcatcattagc
tggaattggaatattaataatttcgggactgcagttagtgtcaagcatgcaaatagttgg
gatacccaatttcaaacactcttgtatagctgttgtttctcgcttttgatctataactac
aacgacatcaggcagacgagtcatatttttaataccgtttaaatgtttacgaagtttatc
taactctctccgtagaacagcagcttctttttaggtagttgatcaataataccacttgt
atcttgctcttctaactgtctaagacgactgactcgagatttgattgttacccaattagt
taacattcctcctaaccatctttggttaatgtaatacgaatcacatctttgtgcttcttg
ggcaactattcctgccgcttgtctttttgttcctaaaaataaaactttcttaccatcaga
cgatgcctttttataaactcacaagcttctgtcagtaattgagcagtctgcactagatc
aataatatgtataccatttctttctgtatagatatacggaaacattttagggttccatct
acgagcttgatgtccaaaatgcacgcctgcttctaataattctgctaaggtaacaacagc
cataagattattaata-ttaaataatgtgattattcgggaaatcccttttttt----aaac
tgat-----tcacttaacttattgcaattctaatcaaccgataataaattaacacacttt
tacagtagattaagaaa--aaaatctttttactttatttaatctacacttttattactaa
aatagttacgcgcggttctgtcgtctaaaataatatcgtctaaatcatctctaacggatg
acacactactcccgctactggtatctgaattccaggctttttttatctaacgacactttat
tatgattgtcgtacgtattaaagccagtaccagctggtataagtctaccgataataacat
tttctttcaaacctctcaaccaatccaatttaccagatatagcagcttcagttaatactt
ttgtggtttcttggaaacttgctgcagaaataaaactttctgtattgagagatgcttgcg
taatacctaataacacggggcgatatgaggcattcaacttgtttcttagtgtcatagctt
tatttgtttgttcaatcttttgcaactctacaagttctccaggtaaatagccggtttctt
caccattttctattttcactttagaggtcatctgtcggactataacttctacatgtttat
cagaaatattaactccttgggattgataaactaattgtacttctttaacaagtagtaact
gtatctcttgaaaacttaatctagctgcttcatatagagccagtccgcattcaatatata
gattgaaactcgcatcaagaatatcatgaggatttaataacacatccgtctttttttcgtg
cttcaagaatttcttcaattcttggaaggccttgtataatatcaccagtttttgctctat
caaatacgagaatagctaatgtttctcctcttctaattaatgcattattgtcaacatgta
gaattgcgccgttagaaactaaatagggcctagcaattcttaatgtaattgattgagaag
atacttgaataatttgtccagagtctaaagaaaccatattttcagtaatagaatcgccgc
aacgaatccaatcgccaacatttattttagttttggcttcttctatattaaaaactttt
tatcagaagatgttgcaactagaatccgtctactagtattcttttcagaataaatttctt
caacgacaccttcactaattgcaataatttctgtgcttgcaacaaccgttaatggtttaa
tatattgaccatttttaacaataattctagtttgactttgttgtttttctgatttattga

catcaatacttttaattgatagtgtctcaaaagttgaaagcctcaagtcaaaataattat
taccgctttctttagcagagaattcaattgaagatattaaactatcatctgcatgctcaa
tttcagctacaagattggttgtaactaaatgaactccatctatcgatttcaccctttctc
cgtcacgaaaagctgttcttttaactagctttaatttagtttgactaggtttttgtgatg
ccaaattatcaattaagctagattttgtttccggtattgaataaacaataacaggtctaa
taaggatataaggcatttgctcattttctatatattcccaataaactaacttatctgtgg
aaatattattatgcaatgtttctccaggtcttaaaaacccccctactttttttcattattac
tatacacttcatttagtttatagatagatcctggcttaataatgatttcacgaactattc
catccttctggattatctcaacaattcctgagcttttagaaaaaacatttttgaccaact
cagttccactttctataatatccccattataaactagcaaaagtgatgcatctttattaa
tctcatgtgtttcttctgagatccataatatgtagccagggctcaaaatttcataagcat
ctttctctgaacccactttctttttttgaaacatttaaatctaaatatttaatgatgccac
cacttgtagttttataagtatcagatattagctcagcaactgtatcaccgtcgtgaa---
tttttttgatttggaacagctttgaaaataaatttttgtttttttttctgtttctaaaatat
aagattcttttttattaattaaatctgtgtaaatataacaacctggaattataatagact
ctgtaataacttgtatattagtaatgctgctattagcatttttttgtatcctgacttccc
ccgcatagtgatttattaactctgtctgcgcgagtattgtgccagctgtaattttatctt
ctttatcaacaatcacgttggccgagtcagatatactataaacttctcctgagagaaccc
aaatcaatccacccgttttagtaattcgtgtagtatattgattattatctgtttcttcaa
ctgttaagttagagaaacagattttaccagacaggtcggatactacatgtttttgagccc
tttctgtcatcagtctatttctacgaggtgattcagcaatcacctgatcttttgatacta
attggccatcacgcaataagagagtagttcctttacttaagttaattattgacttctgat
ttcttctagatcgaattgtaacttgcgttggatttttttgtaattaaggcttgctctccat
gccttgtgcgaacatttatatatgattcaatatctagatccgttagctggccatctatgg
gagcataaatttgttctgctaattcaccagtaaatacacccccagtgtggaacgttctca
ttgttaactgagtgccgggttccccaatagattgagcagcaataatgcccacagcttctc
ccaaatcaactaagcgaccatgagcgagattccatccgtaacaatactgacaaacagaac
ttctcgaattacatgtcacaggtgatctaaccagaactttttt---aataccagctctag
ttatctccctggctaatttagggctaatatcttgctttgtatgagcaatcaaaacattcg
tatctggatgaaacacattttctgctaatactcgaccagacaaagcttgttctaaattga
ttaatgtttttttgtgtgtctactaaatcttctagtattatacctttatttgttctacagt
ctacttctctaatgataacgtcttgagaaacgtcaacaagtcgacgcgttaagtagccgg
aatctgctgttctcaaagcagtatcaactaatcctttttcttgctccgtacgatgatataa
aatagtctgtaacagttaatccttctctaaaattacttgatattggtaagtcaataattt
gtccttgcggatcagccataagccctctcataccaaccagttgtctgacttgagaaatat
tgcctctagctccagagaaagccatcatataaacagcatttaaggatctgtctctttaa

agtacttaattacttcttgttttaatgattcgcttgcattgttccatgtatcaataactt
tttgaaatcgttcgactgctgtaatttcccctcggcggtatttattttcggtagctttaa
tttcttcaattgtgcccaataaaagactttgtttgctaggaggaatccttaaatcttcca
gacttaacgagattcctgcctgcgtagcgtaatggaatccaagatctttaagcttatcag
ccatattagctgctcgtgctataccatagttacgaaaagcccaaacaattaaattcttaa
gttcattttatcaataaccttatttgaaaaacttggctgtgcaagacttcttctattat
ccactcaatcttctccatattaattatgtatt----taactacgccactaaagattcttg
aataattttgttaaagattatccgaccagctgttgttcgaatatattgcacaattctatg
ttggtcagcatcttcttttatgatgtgattattaaaaaattttgtgctactattatcact
gtgtttttcaattttaattggctcagtagtttgatcaccatctacaaatccatcaaaacg
agcccatatataagaatgcaagtcaactttcttctgctcataagccatcactacatcttc
taagctagcaaaatattgatttgccccctgctgttgacacggattattagcagtcaaata
gtaacaacccagaaccatatcttgactaggcatgataattggctgacctgtagctggaga
aagaaaattatgaggagctaacattagcaatctagcttcggcttgtgcttctagagatag
aggaacatgcacggccatttgatctccatcaaaatcagcattaaatgcaggacaaactaa
gggatgaagcttaatagctcttccttctacaagaataggttcaaatgcttgaatacctaa
tctgtgtaatgtcggagctctattcaacagtacagggtgaccttgaataacttcatttaa
aacattccaaatagaaggttcattttctgaatcatttttttagctgctttaatattatt
aactaagccctgtaagattaatctgtggatcacaaagggttgaaataactctagagccat
ttctcgtggcaatccacattgatgaagttttaaatgaggacctacaacaattacagatct
tcccgagtaatctactcttttgcctaataagttttgtctaaagcgtccttgtttaccttc
tataatatctgataaggattttaaaggcctattattcgcacccacaacagtcctacctct
acgtccattatccatcagagagtcaacagcctcttgtagcattcttttctcgtttctaat
aataatttccggagccaaaattgatttcagacgtgataatctattatttctattaatgat
tcgacgataaaactcatttaaatctgccgttgcaaatctgcctccatccaattggaccat
aggtcgtaaatctggaggtataacagggattactgtaaataccatccaagaaggatctgc
acctgttgctataaaattctcaatcaatcttaagcgtttcatttttttattaaattttaa
agaaggtgtcttgaaacttttggtggatttgttgcttcagatcgtaaattttctgcaat
gtattctaaatctaaatctttcaacaatttctgaatagcttctgcacctattccaacttc
tacttgattactttcattttgattttggtaaatttcctcttcaagacttttccattcata
accttctagcagctgcttgtattttaaattgatatttgtattagattgtgtaactacata
ggagtgaaaatagacaatttttttctacttcttttacttttaaatctaatgctaaagcgat
atagcttgtacttccctttaaataccaaacatgagttactggtgatgctaattcaatata
agccatccgatgtcttcgcactcgagattctgttacttctactccacaccgttcacagac
tataccttttatagcgaaagcgtttatatttaccacaatgacattcccaatctttaactgg
gccaaaaatttttcgcagaataaaccatccatttctggttttaaagttctataattaat

tgtttctggttttgtaatctcaccaacaatttgtccattgggcaaacttctttcacccca
ttgacgaatttttttcgggagaagctagactaatctttacatagtcaaaatattgttcaaa
ctgtgtcataaatttgaataccttaaaattaagtaattaaccaaattatttacatatgcc
atattcaaaaatgaa----ttaataaagaaattgttcaaaatcatctactgggggagtat
cataattgctccgatctgctcgattgtctttagagtcagacatcaaatcgacttcgactg
tccgtctttgaccatcttcaaatagtttcaatttatgtactgcaatatctaatcctagtg
actgaagctctcgcattaaaaccttaaaagattcgggggtccctggcttagggataggct
ttcctttaactattgcattcaatgcttcatttctagcttgcatatcatctgactttacag
ttaataattcttgaagagtatatgctgctccaaaagcttctaaagcccatacttccattt
ctcccaatctttgacctccatgctgagctctacctcctaatggttgctgcgtaactaaag
agtaaggaccagtggatctcgcatgaatcttatcatctacaaggtgaacaagtttaagca
tataagctctaccaattgtaacaggattatcaaaacactctcctgttctgccgtcgaaga
cttgcattttaccaggatgctgatcattaaaaagccatttattattagtcagtaatgaag
cttcttttaactttctattcacaagtgctcttgaggcttccgctccatacatctcgtcaa
aaggtattattttaaatcgtttacctaaatatccacctgctagtcctaataagcactcaa
atacttggccgacattcattctagaaggaacacctaaaggatttaaaacaatatctacag
gggtaccatcagataggtaaggcatatcttgtttaggtagaattctagaaataattccct
tattgccatgacgaccagccattttatcgcctacttgaattttcctcttttgcgcaacgt
aaacacgaatcatcgcatttgtaccaggaggaagttcatctcctttttgcctagtaaata
ccctaacattaaccacccttcctttagccgcgttaggtaatcttagcgatgtatctctga
catctctagccttttctccaaaaatagcccttaataatttaccttcaggtaattgatctg
cttcgcctttaggagtaatttttcctactaaaacatctcctgcttcaacccaagaaccac
ctaccacgattccatttctatctaaatcttttaaagagttatcacttacattagggattt
ccctagtgatttcttctggtcctaatttagtttggcgacattctacttcatacttttcga
tatgaatagaagtgtataaatcatcataaactagcctttcactaattaaaaatgcatctt
cgtagttgtaaccttcccaaggcatgtaagcaacgaggacatttcttcctagagcaattt
ccccgccatcagtagatgcaccatcagctaaagtttggccaacaactatttttttctccca
cccacacgattggacgctgattaatacaagtatcttggttagagcgataatatttttttca
aacgataatggactgttctaccattattatcttggatgcctattttattagcagaaacat
aattaacataacctgcagttctactgataacaaccatcccagagtctcgagcaattttag
tctcaaggcctgttccaatgattggcttttctggatataacaatggtacagcctgtcttt
gcatattagaccccattaaggccctatttgcatcatcatgctccaaaaaaaggaattaatg
atgttgcagctgaaatgacttgaataggtgaaattgcaatataatctacttgacttggtg
ttgttgtaataaattcttgacggtaacgtacaggaataatatcgccttcaatataatttt
gagaattaactttaacatcgcctggagcaactctaaaatcatcttcctcatctgctgtta
aatacataggactatcttgataaattacttgaccttcgtgtaccagataaaaaggagttt

cgataaagccaaaaatattaactctggcacaggttgccagcgagccaattaaacctgcgt
ttggaccttcaggtgtttcgattgggcagatccttccatagtgacttgggtgcagatcgc
gaacagcaaatcctgctctatctttattaaagcctcctggacctagagcactaattcttc
gtttatgggttaattctgcaactggatttgtttgatccataaattgagaaagttgactag
atccaaaaaattctcttactgatgcaattaacggcttgggattaattagattagataaac
tcagcgagtctatatcgcatatcatcattctctctctaataatacgttctaaacgattga
gacctactctaaactgattttgcaataactctccaactgaacgaacccgtcgattgccaa
gatgatctatatcatcaaggttacctgagtttttatctttaatattaattaaataatcaa
tcgacgataaaatatcttgaggggagagcacacgaaaagttttaggtatattaagaccta
atttttgttaatcttatgtcgacccacttctcctaagtcatatctctttgggtcgaaaa
agcgagaataaagcatttgcttagcaacagcaactgttgctggctcattcggtctcagtt
tagagtagactataagcagtgcttcttcatctgttacttcttcgatattatttttccaa
tttctttagctaattctttaactgaatatggttgggatgcagagataagaaaggcgtatt
tgctcagtcctttttcaatctcgtctttattaaggccaatagcgcgaagaaaaatataag
catttactttatgagtcttatctattcttatccaaatttctcctttaggatcaatttcaa
attttaaccaagagcctcgattagaaatcaggctagcactgtaaatttgcttgccatttt
tatctatctcttgtttatagtaaattcctgggctgcggattatttgattaataataactc
tttctgtaccagatacaataaaagttcctctgttagtcattattggcaagtctccaatga
agactaatcttttcttgtatttttttgtttttatttgcttttcagtactaagctgtaaat
gtgttgatgataaatctagtgatttgatatttttccatgatctttagaaggtaaatcaa
tatcttttctcgtcaactttgcaggaacgtatatttgggcactataagttttatctctat
tttttgcctgtctaacactgtaacgaggaaattttattttatactcattaccaaatagtt
gcagttcgagtcgattagtaggatctgatatattcggaaagaattcaagtacttcagtca
taccttctaatagaaaccatttaaaactggctctctgaatttcgactaaatctggaagaa
gtttatttttaagcttatacgttgaaccatagatctcctttataacaaattttgaaatt
gaagagtttaa-tactatcaatagagtacttgcacgtcatacctgagcatagttcactac
attctttcttctctagacacaactagaacactatattttattcaacaaaaaagattttta
aaattt---tt-----acaatatagatataaaagtccttcggctatcaaaaaccgttgaa
gttattaaaa-acacaaaagacgaaattaatacttgtccgtctgtcaagtatttgacatc
aagatagacaagtattatttatttaatcaattctg-tttttttgagcgtaagataatgct
ctcgcaagtctagctttttttctagcgccagtattagggtggaatgctcctttttttatt
gctttatcaatcttgctatatacttgagaaacacttgattgtacatcattaagattagaa
ctttctagattatcaatatttagcaggcatcttttagttaaagtttttacaaccgattta
tacttacggttaattagacgatttctttccgaagttttaatgcgtttaatcgcagaaagg
ttcttagccacagtaaatccaataaaacttttataaaatatttttagaaacgaaaataca
caaatttaagatctattcaaaaaaggaatatgcgccgtattataacattaaagaaaaaca

ataggcaatagttcactaagagctcttgttagtacaaaataaatattgtcattgcacaaa
a-tttacaaaatagtattaaat-----taaactaaccagacatataaatatatatgaaaa
aaattgaagctattattagaccttttaaacttaatgaagtaaaacttgccttagttaagg
aaggtattgggggatgactgttgttaatgtctctggttttggaagacaaaaaggccaaa
cagaaaggtataaaggatctgaatattctattgatattattgataaaataaaaatcgaaa
tcattattagcgatgacaaagtagataaaatcacagagacaattattaaagcttcgaaaa
caggagaaattggagatggaaaaatatttattagtagcatcgaaaaagtcataagaatta
gaactaatgatttggattccgaagctttataaaatttacactttttgagacttgatact
ttttcatgaaacatgagataataagcgtttattgtataaaaacaactaaggtataata--
ataatggctaaaagcaaaggtgcccgcatcgtaataactttagagtgctctgataaagct
ggcgattttagtcataaaaggaaacctggcgttttccgatatacaactactaaaaatcga
cgaaacacaccaagtagaattgaattaagaaaattctgccctaattgtaatcagcattgt
ctattcaaagagattaaatagttattaatttattaaaat--aatcatattatggccgtg
tatagaaaaagaatatctccaattaagcccacagagtctgtagattacaaagatattgac
ttgctaaggaaattcattacagagcaaggcaaaatactaccaaaaagatccaccggatta
acttcgaaacaacaaaaaaaacttactaaagcgattaaacaagccaggatactttctcta
ttgcctttttaaataaagattaaatatataccattatgcttcatattgtttcctattaa
tatt-------tttatggatttatttactgtaagtatcgaatatactgggt---------
----------tgcgacaaaaaagctgtttattaccaattttgttcattttataattaata
aaaa--atccaacttactattaaactagattc-----agacttatagcagtacctctttt
aatgaattatattgtacaattcatttaaaagaggagttcattttattctagttaataaat
aatcctcctgtctttttatgcctatacacagttttaaaataaagatttttgtttaggt
ataagatcaaatgaatgaatcctatcaccggattgccataattgaaattcagagataaaa
attccacattcattaccagcttggatttcttcaacatcttctctaacccgctttaaagac
tctatttttccttcataaataacttcatgttccctaataactttaatccaagaattttc
agtaatttattattagtgactctacatcctgctatttttctattagccaaagaaaagaca
gtgcttacttcggcttctcctactggcacttctgaatattcagggtcaagtagctcttcc
attcttcctgtaacatcttcaatcaatgcataaataatttgataattttcaactagtata
ttcaattttgctgcagcttgctttgctccaggagcaaaattagtattgaaaccaagaaga
ttagcatttgttgtcgatgctaattcaacatctgtagcagtaatttcgccaggcataatt
aaaacaacgtttaattgcaccttactttggggaaattgtgataaagaatctaagattgct
tctgtagagccttgattatctgttttaataattaaagaaatttgcttacttaaatcttta
gaattagtggctcccaaggtgtctaacgtaatacgactattaagtgctttctgtttctgg
acgattagtgaatttttttgaagtattttcaagagctttaagctttgcttctttatcattc
ttaactactagagctatctctcctgtgacaggaacagccgacagtccccaaatttcaaca
acagatgatggtgttgctgaattaatctttttttgttgattattaataatagctcgaatc

tttgcgtaagctgatccaataaccatattatcgctcgtatgcagtgtaccattttggatt
aaaagggttgctactggcccatgagatttgtctaagtgagcctcgactataacacctttc
gcaggctgtgttggatcagcctgcaagttttctaactcagctaataaggtaatggtttct
aataatttatcaatattgtagccagtaatagaactgataggaattacaggaacttgtcca
ccaagtgtctcagacatcacattgtgttttaataagtcctgttcgataatatctgtgttt
gcagagtctttatctatcttagatattgctaccacaaaaggaacattagcttttttgaata
tgctgaatagcctctacagtctgaggtttaatgccatcgtctgcagctataataataata
gctacatctgttaaattagctcctctagatctcatacttgtaaaagcttcatgacctgga
gtgtctaaaaaaataatttttttgcttatgctcattattgatatactcaacttcataagct
gcaatagtttgtgtaattcctccgatttctttattggcgttgttagagttatgaatataa
tctagtaaggttgttttgccatggtctacatgccccattactgtaacaatcggaggtctc
ctaatatagttgcaagcttcattaagagagctagaagtatctaaattggcagaatataag
ttattattttcttttatttctgagtcaactgatataccaaaattattcgctactgatgat
attattgaagtatctattgtttgattaatagtaactgatattccttgaaggaataaatac
tttataatatctgtttcttgaacacaaattagtttagataactcttgaatagtaagagga
ttagtaataataatagattctggtggagaatcagatacactgatggcttgattttggctt
tgagttaatgatattttttttttgttttgtagacttttttagtattagtaattttttttata
acttgagcttgtggtttaggtggacgcattagagagatagctaaatctcctgcagtctgc
ggaatactagaattagcacctctaaaattctct---tcctcatcatcaatatgtattttg
gtctttattttc-tttcgttgtctgtttttg--tttttttttattatctgctaaatcatga
accttattgaaattttttattttttttatctagtttaggtggagaattcaactctaaatga
ggttcactactcgctgtatgggtatctgatttatcaagatctaagtgcaacaaattatct
gtatgggctgattctaatctaattttatatataatttggggactttttaaatctataatt
ttttcagaaga---tatatttggactaggtgcaagagatctattttcgaaattttgatta
tttaaaaacatgactgttgagtttatgatctaatagtaaattttcaatttcaatacatag
cttaaagtaaagaagaatcacatatct------ataatactaatat---agaaaattaat
ttttgttgtattagatttaatataacactaaagtcttttgttagttatacaagtttaaaa
ctttt-tataatgataaacactacattctagtttattgatatacaatctatttatttaaa
tagtactgtttttcaa-aagtttttaaa----ttatctaatattgtgttcgttatgaatt
tttattaaagtttgaaag-ctatacaattaagtaataatcattaataatctccacgccac
ttatgccacgatctcaaaaaaatgataattttattgacaaaacgtttacagtattagctg
atattgtattaaaaatacttcctacaagcaaagaagaaaaagaagcctttttcttactaca
gagacggcatgtctgcacaatctgaaggagagtatgctgaagctctagaaaactattatg
aagctttaaaactagaagaagatccatatgatagaagttatattttatataatataggtc
tcatttatgctagtaatggtgaatatgtaaaagctttagaatattatcatcaaggattag
aactaaactttaagttgcctcaagctctcaacaatatagccgtaatataccattatcaag

gagttcaagcagttgaagataaagatacagagctatctaaattaatgtttgacaaggctg
ctcagtattggcagcaagccattaagttagcaccggataattacattgaagctcagaatt
ggttaaaaacaacaggaaggatgagaaatatacaaggatattaaaatatttaggatataa
t---tatattaattaatgactcag----aattaaggactcttttactctatagcacattt
gtattat--------ataatatactaattgc------tgtataagaatcaacgtaataaa
cgaaattaaatctattgactatatttatgaatacaaataagcataatggttagtaaaaca
aaaagtactggatctgttactcaaatcattggaccagttttagatattgcatttcctaat
ggacaacttccaaaagtattcaatgcactcaaagtagaaagctcagaaggaactattact
tgtgaagttcaacaacttttaggtgacaacaaagtacgagctgtgtctatgagctcaact
gaaggactacaaagaggagtgaaagttcttgataccggatctcctatagccgttccagta
ggcgccgatactcttgggcgtatttttaacgttttaggggaacctgtagacaatttagga
ccagtcaattctgaaagtactttacctattcatagacccgcaccagcttttactaagtta
gaaacaaagccaaatattttcgaaacaggcattaaggtcgtcgacttattggctccttat
agaagaggaggaaaaattggcttatttggaggtgctggagtaggcaagactgtattaatt
atggaactgattaataacattgctaaagctcatggtggtgtatctgtatttggtggtgtt
ggcgagagaactagagaaggaaatgatctttatatggagatgaaagagtcgaaagtaatt
aatgcagataatttgaaagagtcaaaagtggcactcgtatatggtcaaatgaatgaacct
ccaggagcacgtatgcgtgttggcttaactgcattaacaatggcagaatactttagagat
attaataaacaagacgttctattgtttattgataatatttttcgatttgtccaagcagga
tcagaagtatctgctctactaggacgtatgccatctgctgttggttaccaaccaactttg
gcaaccgaaatgggtgcattacaagaaagaattacttcaacaacagaaggatcgattaca
tctattcaagctgtttatgtaccagctgatgatttaacagatccagctcctgcaactaca
tttgcacatttagatgcaacaacagtgctttcacgaaatttagcagcaaagggcatttat
cccgcagttgatccgttagattcaacttcaacaatgctacagcctggaattgttggcact
gagcattattctacagctcaagaggtaaaatcaactcttcaacgatataaagaattacaa
gatattattgccattctaggtcttgacgaactttctgaagaagataggcaaactgtatcc
agagctagaaaaattgaaagatttttatctcaacctttcttcgttgctgaagtatttact
ggatcacctggaaaatatgtatctttagaagatgcaattaaaggttttcaaatgatttta
aaaggcaaattggatgacttacctgaacaagcattttatctagtaggtgatattgatgaa
gcgatacagaaagcggatagcatgaaagattaacataattaacaatgactttaaatataa
gaattattgcacctgaccggactgtttgggatgcagaggcacaagaaattattttaccca
gtagcactggacaacttggtattttaacaggccatgcccctctacttactgctttagata
taggagttatgagagttagagtagataaggaatggatgccgattgttttgatgggtggat
tcgcagaaatagaaaacaatcagttaactattcttgtaaatggcgcagaagaagctaatc
aaattgatttatcagaagcagaaaaaaatttagaaaatgcaactcaattattaagtgatg
cctcttctaataaagaaaaaaatagaggcaacacaaaaaaattcgaaaagctagagctagag

tacaggctgcaacagcagctattgtataa-aacatttt------gctgaaaatataaaac
cttagtaataactactaag--------------gttttatttgtgtatttataggcaact
taagtaaggttagcttaaaccagaacaaatataatcaaaatataaacccatctctttacc
cgcatctggtccaactaaactaatagtaacttctttcatagctaaaattgcttggattgt
tgcaccaatagggacacctaaagaattatatgtctcttttagaccatttaatactctttc
ttccaagatagatggatctccagctaacatcccataagtagcatagcgtaaataataatc
taaatcgcgaatgcatgcagcataacgtcgagttgtgtacatattacctcctgggcgggt
aatatccgagtaaagtaaagattttgctacagattctttaataattgttgctgcattagc
tgcgatagtagcagcagctcttactcttagttcacctgtctgaaaatagcctcttaattt
ttcaacagagctatcatctaagtatttcccttgaacatcggctgcattaataacagaagt
aattgcgtcttgcataaattgtaaaaattccttaattttaaatatttaatactaaagggg
tatataacaaaataactgaacttttatatataccagtattactgcatagcacccaaagt
ataatcaaagtaaaaaccagcttctgctgaatcttcaccagcaagtaatgagcaagcttc
acttttcatacatttgacaccttcagcgactcccgaaattggtgttcctaaagaattata
catttctttaacgccgactagtccaatttcttcaataggcgtcacatctccagcaactat
tccataagttactagacgaagataataatcaaggtctcttaaacaggtagctgtcatctc
ttctccataagcatttccaccgggagagacgacatcaggtctttttttgaaacagctgttg
accgccctgctttacaatacgctcacgattatctgttaaaatttgagctattcttaaacg
acgttgtccagataagacaaaacttttaattctatctaattcaccaggacttaaatatct
tgcttctgcatctgcatttacaattgacttggtaacaatactcatggataaaactcctgt
aatgcttt--ttaaaaagcttaatactaattatttatgatataactcagaaagtagtgcg
cattttagaaaaatttatagctacagcacgttggtgatgtgtaaagcagcacaaatactt
atgtaagtactaaaacaagaaagttgcatgtcgtaccttattttattggttaccaagtac
ttttttaaaactaggtaccacaattgatacatcttgtttagtaaggcgattatataatgt
ttccgtattcgggaaattagctgccggcaaagttggaaatcttctataaggaacagtata
tttgccatatactgaattatactcagtactttcaactaaattactaatgaatgcagataa
tccttttgaagctaaaatttggttaaagaatctaatttctgcctgattattaggagctcg
tcctaagatatgcttcgtacctagttctataacttttgtatttggatatggctgataaaa
ttctttaccatataattcagataatgctaatttttctactaattctttaacattgatttt
cttattcaagaaggatgcttcaatatctaaaaactctcctccgatactaaaagaatttag
atctctttcaaaaatctggcgatatgtcgctcttaaagcttgctctaatacttcttgatt
actattattgtttacttcaaatataatagattgatctcttaaagagctaactccttgcga
gatacgagactggatgtcattgctacttcttatttcttttacagtacctagttcaacaaa
tctagcagacttgctagtaatgactttttaaatctctggtcaataatgcctggtcttaa
gcttcttagagcaataccggctggtgtactatatctttcataaggcacagtactatcacc
aaaagtttctatatactcagaactatcaattattgcatctatcacttgatagtaaccctg

tttatatgcaatatcaaaatacttattaatttcttgacggccatatgtaggacgacccaa
taagcggttgtgaatatactcaatagctttgcagatatataatggctcccagtaaagtga
tctaaatatactcgatttggctaattgtctaacaaattcacgaactgtaatttgaccatc
ttttagttgactttctattggcttaagaagtagtttttcttcttgatacacgtctctacc
aaagactcttaaataagcaactttcgtaacaacattgactgaattttcaaaattttcaga
agagtcagcttttacgagagtgttggataatttaaaaattttggggcccaaagatcctgc
agacttaggtcttacctgtggattactaatctggttgtagattccggggccacgcctaac
taaaattcttctagtatctttgccaaaaagagcttgtctttttctaggatccttattttc
tttaggaaaaattgctccaaattgaatcgacaatggatcattgcctattccatatggatg
ctgatcaggtaatgcttgtttgtaatcactaaataatgttataaactgaggtaccttacg
aaaaggagcactatagttaagaagatctatttgcggcccccaattacgacactcttgagg
ctcttcgcctaaattccggaagtaaggtacagtttcttctccaaaataatctgtatattc
agaagaatttaaaattgcatttactagtccacttaaaccagtagctgacagaatagcaaa
atatttttgaaattcctccaaagagctaggccctctaccaagaaaatgtctaaatgctaa
ttctagtgctcggctatttacaaaaggctcatagaattgctttctataaatacttgaggt
tcctaacgaacggacaaattctttaatggaaatttgaccatttttacttgcgactctaa
gttagataaagacaagtcatatgctttggcaatatctctttcaaagatttgtctataaca
tgctttcactacggtattttttcgtctgcagacaaactaggcttcataacaaaacgagg
cgtagaaactcctgcctgtgcgtaaatttgaggaaggcgcaaaccttgtaaatctccaga
tgttctttttcttaacttatcggtcaaagatggagcttcaaattcagaaattacaacatt
aaaatactctttgactaaatcttgacctttaatatcttcttcaaatatcaatagagctgt
tcgtctcatctctctaagggccacaattgctgctgcgcttgagcaagcattatcaattaa
ttctcttagtcctctaatattaacagataaaatattgggatcacctgatacaattgcata
tgttaagtaccgtaaaaaccagtctaaatctcttaaagattttctcattcttgtagtacc
gtatctcaagacatttataggtttaaatccaggaggggtagctcctccagcattaaataa
agaacgaaaactctgcccaaaatctccttgaatatttccagatagctcattaattttgtc
ttgagaactttgatctcctgcaatcataacagctgcctgtggacgctctaaataagaaat
tgcagacccacctacaaatattttatctgcagctcgagctactagaatattggcattttt
agttaaaatgtctgcaacttctaatctttttttgcccagaatttagaaaagaaactagttg
atttaattctcccagctgtaaaaatcggtcttgctgctctgcctgagtaatagttaaaat
tgaggcagtccgataaagctgggggcgtgctaatgggcttccgccacttgctttaatact
cattactttatttatctccttaactgatattacttgctagcgaattcaaatatcacctga
tatttctaactatttaaagatactatgcaagtacatttgttttctatattattaggtact
agtagttatagtacatagagtcgcgcaactaacttctataaatacaaattttgtaatact
tctattaacatttatcaaaagtgaatataattgatgactcttttatattaaaatgcaatt
agtgagagattaattctaaataaagtatacttagtaagttttatagatagtaaaatt-ca

agacttattctacaaacattaatca------------cttgactgtttgacaattatatg
attgttcttgcttttaattggtttagactagtttcacatttttctaacgtaaacaatat
caataagcaatttgtatgaatttagaaccacagaataacctattaactaacaacaatata
aaccaacaagatatacccgaaaatgatataccaatgtcaattacggaacatttagaggaa
ttgagacaaagaactttatttgtattttattctttttgcttgctgcaactttaagtttc
acacagattaaaataattgtcgctatactacaagctccagctattggtattaaattttta
caattagcgccaggggaatatttttttcgtctattaaagttgcaatatattctggaatt
gttgcaacagctccgttcggtttataccaagttatcttatatatactacctggactaact
ggcagagagcgaaaaataattttacctatactaattagttcagtcttccttttgtaaca
ggcggtatatttgcttactttatttggcaccagctgctttaacctttttgatcagctat
ggatctgacattgtagaaccattgtggtcttttgaacaatatttggactttattttacta
cttttatttagcacaggattagctttcgagatccctatcatacaattgttactaggtgtt
tcaggtgcattatcttccaatcagatgatccgagcttggaggtatattattattatatcg
acaatagctggagccattttaactccttcaactgatccagttactcaaattatcatgtct
tctgctgttctagtgctttattttagtggaatcgttatattactacttttaaaaaagtaa
atatagt--------aagctttatttcatactatatttttcttatc-----ctaaaaaa
tggagtct--atataacttgtggttcttaacttagattaatattgaagcattcggctttt
agactctttaaaaatatttagtcttattatggaactaatataatatg-------------
------------------------------------------------------------
------------------------------------------------------------
------------------------------------------------------------
------------------------------------------------------------
------------------------------------------------------------
------------------------------------------------------------
--------------------------------------------------tatatatcga
t---tttttcatcaattatttgtgattttcatttttgtacta----ctttatgattctt
gcaactatttatgtaatctatttactgaat----aataataaagaaaattcagcgtaaat
tatgataatctataccaatataaaaagtattttattgtgagttttatttatataaaggct
tattcattatatatttataacagtaagtctaattaatagtttttcatgcttaattaaagt
aataactcttaatccttttggcaaaacagttcgtcttaatattttatc---aaaaaacag
aaaaaatgaagctaaatagtctaattaacttaattcaaaagtctatatgttcttgtacac
ttttattaattattttaaacattatttgtgtcgcacctaattctagtagtgcatttccaa
tctatgcacaacaagcttacgaaagtccaagagaagcaactggtagaatagtatgtgcca
attgtcatcttgctcaaaagccagtagaaatagaagcgcctcaagcggtactacctaata
ctgtttttgaaactgttgtaaaaattccatatgacagtagtgctaagcaaattttaggta
atggaagtaaaggagggttaaatgtcggagctgttgtaatattacctgaaggatttaagt

tggctcctgctaatagattatctacagagttaaaggaaaaaaccaaaaatctatacattc
aaccatatagctctaaacaaaataatattttagtaatcggacctattcctggtgacaaga
atagagaaatagtttttccaatactctctcctgatcctgcaaaagataaaaaagctcatt
tttttaagtacccaatatatgttggtggaaatagaggtcggggtcaactttatcctacag
gagataagagcaataacaatgttgttgcagctttaagtagtggtaaagttaataagattg
aggtccttgagaaaggtggatttatagttcatgtgactaacagtaacaatgtagaatcaa
ctcaaaagatttcccaaggtcttgaacttaaggttaaagaaggtgatacagttcaattag
atcaggctttaaatagtgatcctaatgtcggtggtttcggtcaaaatgaaacagaaatag
ttctacaaagtccgaacagaattaaggggatgattgttttctttttttgctagtgttttag
ctcaaatttttctttgtattaaagaaaaaacaattcgaaaaagttcaggcagcagaaatga
atttttaacgcaagataaaactggtacttgctataattaaagtg-taaacactttattca
aagaatgtaccagtttttatattatataaccattgtttttgaagacagtataatattttt
aacagattctataatttgactaggttgaattacagtagcttgttctaagctaccattata
tggcgttggtatatcctgcgaagatagtcttactacaggaacatctaaatcgtcaaaaag
atgttcattgatttgtgcaattagctccgctccaattccagctgttttcatacattcttc
tacaatcaaaactttgtgagttttcttgactgagacagatatagactctatgtctagagg
ttttaaagatataagatctataacttctggatcgtaaccttcttctagcaaagctggtag
tgcttgaataacatgatgacgcattctagagtaagttagaatagtaatatcttttcctat
acgcacaaattcagctttgttaagaggcaataggtattcttctttaggaatttcttcttg
taaattgtaaagtaaaacatgctcaaagaagacaactggattattatctcgaattgcaga
tttcagtaatccttttgcattatatggagtagagcaagcaactatttttaaacctggaat
agcttgaaaataagcttctaatctctgagaatgttctgcgcctaattgtcgaccaactcc
tcccggtcccctaatgactaaaggcaacgtaaagtttcctccggaagtatagcgcaacat
gcctgcattattagaaatttggttaaatgctaataacagaaagctcatattcatgccttc
aacaatcggtcttaaccctgttatggctgcaccaatcgccatacccgtaaaactattctc
tgctattggtgtatctagaactcttaaatctccatacttgctgtgcaaatctttagtcac
cttataagagccaccatagtgacctacatcttctcctatcacacacacagtcgaatcgtt
ttccatttcctcatctgttgccgctcttagtgcgtcaaacataaagattttactcatttt
tatatttagtttttatggtttaactactactcagaagtaaaagttaagatatccgatttt
cttcaggtatctgaaaaaagatatcgtttaagttctgagatattaggctctggactggaa
atagcaaatttcacagattgctctaaatctttttttaacagaagtttgaatttcattgagt
tcatttatgtcggcaatcttattatctaaaatatatttttttaagtttttttgattggatct
cttgcaatccaagcttctttttcttgtcttgatcttaattcatcaggatctgcaagagaa
tgaccgcggaatctatatgttaaagcttctattaatgtcgggccttcaccctgtctagct
ctctgaactgcttgaatcgctgcttgccttacagctagtacatccattccatcaacttca
ataccaggaagaccaaaagcttctgccttttatgtatttctggtattgaggaagatcgg

tggtgtgccatacctatagcccactgattattttcaacaataaatataataggtaacttc
caaagaactgccatatttaaacattcaaaaaattgaccgttattagtagtcccatcaccg
aaaaagcaagctgtaactctcaaatcttcagtttcttttaacacttgctgacgataaata
ctttggaaagctgcacctgtcgcaacaggtatgccttctgcaataaaagcaaatccacct
aaaaaattatgaggtgcagaaaaaatatgcattgaacctcctctgcctttgctacaacca
gtttcttttccaaatagctcagccattacattttagatggaacgcctttacttaaagca
tgtacatgatctctataagtactacagacataatctgtagaattaagaaccttaattaca
cctgtagaaacagcttcttgaccattgtaaagatgcacaaagccaaacatctttcctttg
taatacatttgagcacacatatcttcaaaatttctacctaacaacatatcttcatataag
actaataaagtattctt------attaagaccagttgaattatagttagttagcggcaat
tgaactttcttaggataactcataatttataaagaaacctct----------tattatac
ataggaattttgaagtaaaagttatcaatagaaaacacacagataaaaattcataaatta
ttatatgcttttattaaagactttaataattatttgcaactagttcagtatatcaaatt
aattttgcttatactacaggtgtctaaaacttagattacaaggga-tagtggaaaagtgc
ttaaaagaatag-taaaaacaaatagaaatagaaataaacagaattatgcgtaagaaaat
gactattattaatagataattacttaatatttctaaatattttcaaattaatggctatat
catcatctttattttaccctgttgagaaagaattgtacagtttggaaacaaatctaaaag
ctgtagccggaacacgtcatccaattctatacgcagcagcaaaacatctgttcgaagctg
gaggaaaacgagttagaccagctctcgttttttagtggctaaagcaacatcagagaaac
aagatattaatactggacagagaaggttagcagaaattaccgaaattatacataccgcaa
gtttagtacatgacgatattattgacgaatgtacaacgcgtagaggagtcaagactgtgc
ataatttatttaatacgaagattgctgtattagctggagatttttatttgcacaatctt
cttggtatttagctaatattgaaaatttagcagtcgtaaaagctatttctaaagtaatta
cagattttgcagaaggcgagatcaggcagggcttagttcattttgaccctagtatttcaa
tagatgcttacattgaaaaatcattttataagactgcttcactcattgctgctagctgtc
gaggagcagctatgctgaatggttctagtactcaggttaataatgatttatatctttacg
gtaaacatatgggcctagcatttcaaattatggatgacgttttagatataactggttcta
ctaaaagcttggggaaacctgccggtgcagatctaactaatggtaatttaacatctccta
ttctgttttcacttactaaagaattagatttaaaccgccttattcaaagagaattttgta
acagtacagatgtagcctcagcattatttcttattaaaaaaagcggagggattacaaaag
ctcaagacttggctaaagaacaggtacaggcagcactctactgcctaaagtttttaccaa
aatctgcacctgtatctagcttaaaagaattgacacatttcataatcacaagattgtcat
agaatacttgctaaaattaaaatgctattaactaatttttcactgtttctaaaatagtct
gaaaagtttgaatatcattacttgctaactgtgctagcattttacgatttaatgcaatat
tctctttttcaatgcactaataaaagtactatacttcataccttgactatgagctgcgg
catttattctggttatccaaaggcgacgaaaatctctctttttcctttttcgaccaacat

aagaatatctcagcgctttaagaacttgctgtttagctgttcgaaataaacacttatgtg
ccccttaaagcctttagctagcttaaatatttcgctcgtcttttttttgcaacattac
ctcttttaactctactcataaaatattac-ttaaaattaatttattaccggcttcctcag
tttgaattgttatcg----attataaataaggcaaatttatagcgatatttttaatatct
cttgaatcagctgaacatgtagaagaaagatgtcttctttgctttgacgacttcttttgt
aataaatgactttagaagctttatgtcgcaaaaatttcccgatgaagacactttaaat
ctttttgctatcgcttttgatgtttttaacttaggcatattataaacttttaatgattaa
caaaagtatgaaaagtctaaagatattaaacttagacttttcaatacttaagataatctt
aacacaaattattttacttactatagagatttactgaaataatttatgtatataacagta
agtaattaatcagtagctgctaaagttgatttatatttagctttaaatttttgtatagta
ttagcaattagcattgttattattttaagagtacttgaatttaattcttcgaacattctc
atatttaatgtgaatgaaatatttgcttctgatacaatattttgtatttgctcctccgat
aaaggaatcatattcagtgaagctctatacttagctttaaagactttgtcatcttgtatt
tggtcaaagtcataaaacctagtacctctttcatcggataaattcatcgcacttctagcg
attttttttaagatttggcctccagataaatcacctaaatagcgggtataagcatgagca
actaataattctggctgtttacagcctatactatgaattctatcaacatatattcttgtt
gcaggagatggtttaataagattcaaccaatctgctccataataataacttaaatcttct
gacagacttattttctattaagttctgtgaaatatataggttggatagctgggtgatct
ttattagaaaatagttcttcttctatagcgcaataaacgaaataaagattagcaactaat
ttacgatatgatttcttatctacgacacctcccaaaaaagatttcacaaaactaacattt
tctgccatactatgagacttagtagtgccttctcttaattcattcgctaaagtattaacc
ataatatcaatcttccatttta-aaagataaccgattaaat---ataaaaagttatctat
atattta------cgttttactaattaagctgctgcaaaataatttttgattttattgg
gtctggattcatcgttttatcaccaggcttccaattagctggacacacttcatctggatg
agattgtacatattgaattgcttgtaggactctcaaagtttcttctacactcctaccaaa
ctcgagattgttaattgtagaatactgaataattcctttgggatctataataaatagtcc
tcttaacgctataccatcactgtttaaaacattataagcagcactaatctcttttttttaa
gtctgatactagtggatatgcaagatctcctaagccaccagattctctatctgtttgtaa
ccaggcaagatgagaatactcgctatccaccgaaacacctagaacttctgtgttgagctc
ggaaaaagcatcatacttatcactaaatgctgtaatttctgtaggacaaacaaatgtaaa
atctaaaggataaaaaaataagataatgtacttatttttaaatctgataattttaaagt
tttaaattcttggtcataaacagctgtagctgaaaaatcaggagctagctgacctactcg
aagacaatttggtccagaaatcattaatttttcccagaagaattgatataattgtgttat
ttaataaataattatataacaaaatttagtatatattaaaatgctatcagtctttttttaa
attac--------tcaatgactctacgaatactatttttatagcggggaatggatttgaa
ccattgaccttcgggttatgagcccgacgagctaccagactgctctaccccgcggataat

taaaatcatattaatccatctaaatagataaaagcaaatatattataagattcaaaaata
aaagaataaaagcacaggagcactaattaataagctcaaaaatgataaagtatttttat
tttatttaacagagggaaaacttcatataagccgataaagcgatctttaagaagaatttc
cgatggctttacccaaatttgaccgtcataccaaccagattcttcatagaataccgtagc
actcattaaacgttttactacgtatgaccatcctaaatatagtctgattaaaataaatcc
agttattaaacttgaggtaataaattctgaaaaaacaaatttcagaggcagcttggtaat
tggaaaaattgaaagcaatactggactaactaataaaccattaagtagcagtgtaaccgt
aatttttctgttgtataattgactgcttagagttggccaacaaaaaaaccaagaatttct
tagagaagtgtattcgttaacaggttgctgctctttgggtacaggacactgagtattata
taaattcattgaaatttaaaattattatactagaaattgtttttttatctgatctaaatg
attagcga--------gttaattaattgcaattatagtagttagtatcagaaataaaact
agaaataaccaagtaactttatttaatgcattttcagtactacgagtattgctaaaaaat
tgattctgagctccgacacttcctagaccttcagatttgggattatgtatcaaaatagta
aaaactaatataattgttactgcataccacaagaatttcaaaatttgttccattatatac
aagttaataaaaaatagcacaaagacaatcgtctaaaagatattgattgtcttatcatat
tgaaaaatatttattcaccctgtacttttaacagtaaaaatcctattgccaaccctacta
aaactaaaactgaacttaagacagcactttctacaatttctccgaccatt-atatttttc
cttattattttattatta---taattttataaaaaagttgatacctattaaa-ctagaag
ccgtttcttccccaaactaccattgcaatagaaaaggtaaacattgccattagagctgcc
catcctaaacttaaaatatccataatttataatattc--ctgaaaataaggtctcttgta
cttaaactataatattcttaatattaaaaatatcattatataataagctaattataacag
agtactactgaattagtacttttagtgcatttaaagaagattgaa-tttttgcgaagtac
ttagcaagtattaatttgtaaaaaatttctgccat----ttttgcattattatacctatt
aaattgaaatgtaagtattactcaaaaatcgttagtaattttgatttttttcaattgtacc
att---aacatatgtaaaaacataaagattcatataagctaagttaa-----tgttctac
tataatttattaat---aaaaggaagctggtattaaaatcatgcaaacaactataagtaa
tgggaaaacgtctagcaaagaaactctactaacacctcgattttatacgactgattttga
agaaatggctaatatggatatttcagaaaatcaacaagatttctttgctattttagaaga
atttcgagctgattataatagtgaacattttattcgagacgaagaatttaatcagtcttg
gtccagcttagaagataagactaaatctttatttattgaatttctagaaagatcttgcac
cgcagaattttcaggtttttttgctatacaaagaattatcacgaaaattaaaagatagtaa
tccgattattgccgaatgctttttgctaatgtctagagatgaagcacgccatgcaggttt
tttgaataaagctataggagatttcaatttatctttagatttaggctttctaacgaagag
tcgaaaatataccttcttttctcctaagtttattttttatgcaacttatctttcagaaaa
aataggatactggagatacataactatttatcgtcatcttgagcagcatcctgaacatcg
catctatccaatctttagattctttgaaaattggtgccaagatgaaaatcgtcatggaga

ctttttttgccgcattacttaaatcgcaaccacatttttttgaacgactggaaagctaaaat
gtggtgcagattttttctattaagtgtatttgcaacaatgtatctaaatgattttcaaag
aattgatttttataatgcaataggcctagattctagacagtatgatatgcaagtaatacg
aaaaactaatgaaagtgctgctagagtttttcctgttgctttagatgtagataatccaaa
gttctttaaatatttagatacttgcgcatgtgacaatagagctctaatagatatcgataa
gcaaaatttcccattattaattaaagctattataaaaactcctctatattgctccctaac
tattagtttactaaaaatatatttgattaaaccgattgattcgaaaacagtgtggaatac
agttcgataatcaaataacataagaataaatcttaatgacgcaggctatctagttattaa
tataacgcagactgaaaagctcacttct-----gccaagaaggagctt---ttttattaa
caaatagacaatatggttctataaatt-tagaactaggaacttactttttcttttgcttc
atacataacttctagggtgattagatttttggaacaatcacgtgcaaatttttctgtatt
tctttttactttacctctaacaaaacccggaatttactcaattctttctgtgcttcttg
tgaccagttaatactatcaacagcagagattcctacacttaatgcttctgttgtatcatg
accaccaaatatttctaataagtgatcttccatgcccaacgtgaatgagttgtacactaa
atctgcaatttgatttgtaccttcatagccaaggaatggacgataacttaagggaaaatt
ttgtatatgtactggtgaagaaataaccccacaaggtatatttaatcgtttaccaatatg
tctttccatctgtgttccaaaaaattgctgcaggttctgttttagcaattaaatctccaat
taaaccatgatcatcggaaataattacttcatcacagaattcttggacttgttctttaaa
ccactcttcgtcatatttgcaataagtgccacaccaagtaacatgtatgcccatttcttg
atgtaaaattcgagttattgctgcagcatgagttgcatctccgaatacaatagcctttttt
gccagttagattttggcagtcaattgatctagaaaaccaagcagattgagagataaatct
agtttgttcatctatatacttttcatagtctacaatagctcctaatgtagttaataactt
ttgtattgagcggatacatgctgctgtttgcacaattcccataggtgtagtatcaacata
aggcatgttaaattctttttctaaataacgagctgtcattagcccggtctctctataggg
aacaaaattaaaccatgcagatggtaaatttttttaaatcttgaacggatgcattttcggg
aataatttgattaatcttaatatctagatcttgaaatagtctttttaattctgaaatatc
atgctgattatgaaatcctaaacttacagctccaataatattaacagatggggtttctgt
ctttcgaattaaaactttattgttagattgcgctttttccatataaaaagtaacaatttg
ttccaaagttctatcaccagcttgaagttcgtttactctgtaatgattaacgtcagctaa
taaaacatcagcttctgtttctatagaggctctgctaacaaaatttgtaaatcttcttg
taagatactagaagtacatgttggcgtcaaaataactaagtctgggttttcttctctgtc
ttttctagtaatattctcaacaactttttcttgcgaaccacgtgctagtacatgtctatc
tacaacactcgctgtaacaggcgtaaagtctctgtctcgttcaagcattgaacgcatgac
attaaaatagtcgtctccgagaggtgcatgcataattgcatgaactttttttgaaagagct
agcaattctaagagttccaatatgggcagggcctgcatacatccaataagctaatttcat
attaaattttcctatgatttatttagacatagtaatacttttaaagccaaataataaata

aaagtcaaacatctaatttaactctttaataatttttttcatg-tttttagataaagatta
acattacattgtagtcaaatc-------------agaaataaatgagtaatgaataaagc
ttgaaagtaaagatttcttgcagttatagtttttctttaagtaaacatgtcaaaattgta
ctatatataaatgtactatatataaacttttttttactgaatattggtaatgccagataca
attaacttaaatatgccttccccaacgtttggcggaagcactggcggttggttaagagcc
gcagaagtagaagaaaagtatgcaataacatggaccggcaaaaatgacagtaagtttgaa
atgccaaccggtggcacagcaactatgagaaatggggagaacttgctttatttagctaaa
aaagagcagtgtttagctttaggtactcaattaaaaagtaaatttaaaatatctgactat
aaaatttacagagtgttcccgaacggagaagtgcaatatttacatcctaaagatggtgtt
tttcctgaaaaagttaatgctggaagagctagtgttaacagcgttgatcattctattggg
aaaaatgttaatcctgtagatgtcaaatttacaaataaggcaacttacgattagtcaatc
aaaaaacctagacataaaattactttatatagtctaggtttttgatagagtatactagt
attatggaaaggtggccgagtggttgaaggcgtctgatttgaaatcagttgaactg--ta
tcgttccgtgggttcgaatcccaccctttcctttataaagct------------ataat-
--------------------ttatacactaatagtattaagtttagacttaataaaatct
acaacttgcgataaattagaaatttttcggcgtcttcatctggtatttctataccgaat
ttttcctctattgccattactaactctactgtatctaaagaatctgcgccaaggtcgctt
gaaaaatttgcttctctagttacgatttttctttcgattcctaactgctctgctacaata
tcttgaactttttcaaaaatttcattatcttgcataatatttgttttcctctaaggtgtt
atcaaaaatttttcggctcatattattatctaactctaatcaattagtaattttgagtta
agatctaaatatttctacctatatgatatactattattctggataaatacgaagtttgcg
atatggctcttctccaaaactacgagctctcaactggtagttatcaactaaattatgttg
catttttcgaatataagctgaacgaggtatcaactgaactgttgaattttcgttcaagat
aatagtttcaatagccaatttagcttcttgaagggcttgaatttcatcaaatttttttatt
tttacaaagttctacccaattaagatcagaagaagtatgaatatttaatattttttctcag
tgcacgagtaatttgtgggactgtactattctgaattgtatatataataatttgttttga
ttttgcaatttgtcttaattttgtattttgctttacttgattacgcaatgctagaatagc
atctgacttttctatctctttagttaaaataattggtaaatctaaagaagaaataactga
tgcaatatgatgccaactcaaagaataagcatataaatactgattaggcacttctaatgg
------tggtgtattaatatttataggagcgttaatagtagttgacaataaagatgtatc
acgattttgtaatttattcaaatcggatgacttatttttttaattctctgtactgcagaac
aggatgtgcctgttgtatattaattggttggcggaagctagaaatatttgatggcaaaac
ttctaaagattgtgatggatagcatttaattaaaatgcggccattagcttgaatttgtct
tttttgtacaaaaggttgatggccctgtaaaatttgatctatagtttctcttacttgatt
atggactatccaaacagttcgttcatgaatttcaattgcaatttgaaaagcaggcgcggc
ttttctttctagtatgcttttctgagtacctcttctttttgcttcatcatcacctaatgt

aacatattgaattcctccaattaagtctgctaatgtaggatttttaattaaactttctaa
atgatttccatgtgctgtacctactaattggactcctctttctgctatagttcgagcggc
taatgcttccagctctgtaccaatttcatctataataatcacttcgggcatatgattttc
taccgcttctatcattacctgatgctgtaaatccggccttgcaacttgcattctcctggc
tctacctatagctggatgtggaatatcaccatcaccagctatttcattagaagtatctat
aatgacaactctttttttccatttcatctgctaagacacgagcaatttctcgaacagctgt
tgttttaccaactccaggttttcctaaaagtaaaattgaatcaccttgttctagtaaatc
tctaataatactaattgtaccgaaaactgctcgtccaacacgacaggttaaaccaataat
acttccttctctattacgcatggaactaatacgatgcaaggtttttttctatgccagctcg
attatctccgctaaaattaccaacttttttttacacagtagtctaagtcttgccaactaat
agatctttgagataagtattctggcttatctggaaatcttgcttccggacggcgacctaa
atccatcactacttctattaaatgatttctactagggtgctgtttaaggggctctttgac
aaagttcggtaagatttctagcaacttatctaagtcatctgcaataagcatgattgaata
tataataattaattaatcagatagaaaaattaagattctaaagattacaatacatattaa
agaatattatgctcgtaatcaattttatacttaaattaaataagaaattg-cagatggaa
aagactagattaaaccgtatacagtaaatagattggtcaaaattaatagtcatgttaata
tgtttattggacagaaagttcgtattaagtatagcaagcaaaaagttgctaatgacatag
cagataaggttggtgatcgaggaattattagaggcgtaaagtttatcaatagtcaatgtg
ttactgttattgttgaatttgacaatcatactagactgtggatgtttagagaggagctaa
tttgtttaaatgaat------ttaaaaaatgatataagattaattt----tattatatag
aaattttcatgagtcgttcaaattatacattttaaaggagtattgttaattgtgcaaatt
actattaaaaaattacaagacttactttcatcagtacaaagaaaaaagatacagacattt
aaactaaaacaggatgattttgaacttctattaaataaaacttataagaaaactaattca
gaaatattacctttacaaaact---ctgtattacagcatagtcaagctgcaataattaaa
tcaacatctaataacactagtcgatctattagtaacaatgatgttacagaatatgccact
atcgtctctccaatggtgggaacattttatcactcacctgctcctggtgagaaaattttt
gtgcaagttggagatgaagtcaaattcaatcaaacagtttgtattattgaagcaatgaag
ttgatgaatgaaatagaagctgaagttgaagggaaaattatcgaaattcttgttaaagat
ggcgatatcgtcgattgtggtcaagccttaatgaaggttgaaacatgatgacaatctttc
cattttacatattgttaacagaaattatttagatagaaaaattagtttatactatagtta
tgaaaacttatttcgtaaagacac-ataaagttattatgtaaaataatttactttatctg
ccaaacatagaaataagcgttttgattccttgtctcaagagaggagaatctcaatgacaa
ttagctcaaaagagcaagagacaaagaaggtaaaaatctcggttgataaaaatcccgtag
atacttctttcgaaaaatgggcccaaccaggtcatttttctcgtacactagcaaaaggac
caaaaactactacttggatttggaatcttcatgctgatgctcacgacttcgatagtcaaa
ccagttctttagaagaagtttctcgtaaaatttttagcgcccattttggacaactttctg

taatatttttatggcttagtggaatgtattttcacggagcacgcttctctaactatgttg
cttggttaagtaatccaacaggtattaaaccaagcgctcaagttgtttggcctatagttg
gccaagagatttaaatggtgatgtcggtggtggctttcaaggcgtgcaagttacatcag
gctggttccagctatggagagcatccggaattactacggaattccaactttactgtactg
ctattggcggattagctatggctgctttaatgttgtttgcaggatggtttcattaccaca
aagcagctccaaagttagagtggtttcaaaatgttgaatcaatgatgaatcaccacttag
ctggtcttttagggttaggctgcttaggctgggcaggccatcaaatccatttatctttac
caattaataaactcttagactccggtgtatctccgcaagagattccactgccacatgaat
ttttaattaatagagagctgatggctcagctttacccaagttttagtaaaggattagttc
cgtttttcactttaaactgggctgaatattctgactttctaactttcaaaggagggctta
accctgtcactggcggtttatggctaagtgatactgctcatcatcatctggcattagctg
ttctatttcttgctgcaggccacatgtatagaaccaattggggtattgggcacagtatga
aagaaattttagaagctcacaaaggtccttttactggtaacggtcatgaggggctctatg
aaattcttacaacttcttggcacgcgcaacttgctattaatttagctatgatgggatcat
taagcatcatagtggcacatcatatgtatgctatgcctccatatccatatattgctactg
attacccaactcaattatcactatttactcaccatatgtggattggaggattttgtgtcg
ttggcgcaggagcacatgcttctatatttatggtaagagattataatcctgctgaaaatt
ataataatttgttggatagagttattaggcatcgcgatgctattgtctctcatttaaatt
gggtatgcatatttcttgggttccattcatttggtttatatatccataacgatactatga
gagcactaggaagatcccaagacatgttctcagatacagctatacaattgcaacctattt
tcgctcaatgggtacaaagtatacatactttagctcctggtaatacagctcccaatgcgt
tagcaacagctagttatgcattcggaggagatgtcgtttctgttggaaataaagttgcaa
tgatgccgatttctttaggaacagctgattttctagttcaccatatacatgcatttacta
ttcacgtaactgtacttatcttagttaaaggtttccttttctcaagaaactctcgactaa
ttcctgataaggcaaatcttggctttagatttccatgtgatggccctggtagaggcggta
cttgccaagtatctggctgggatcatgttttccttggactattctggatgtataattctc
tatctgtagcaattttttcactttagttggaaaatgcaatcagatgtttggggtagcgtgt
ctccatctggaaatgtttcccacatcactggtggtaattttgcgcaaagcgcaattacaa
tcaatggatggttaagagatttcttatgggctcaagcatctcaggttattcagtcatatg
gttcggcactatctgcatatggactaattttcttggcagcacattttgtatgggcattta
gtttgatgttcttatttagtggtaggggttattggcaagaacttatagagtcaatcgtat
gggcacataacaaaataaaagtcgctcctgcaattcaaccaagagccttaagcattactc
aaggaagagcagtcggtgttgcacactacttactaggtggaattggtacaacttgggcat
tctttttagcaagaattatttcagtaggctaatagtgaaaataggataaaaaaaacaatt
atggcaacaaaatttcctaagtttagccaagctttatcacaagatcctacaactcgcagg
atttggtatggtattgctacggcgcatgactttgaaagtcatgatggaatgacagaagaa

aatttatatcaaaagatatttgcttcgcactttggacatctagcaatcatctttttatgg
acatccggcaatttatttcacgtcgcttggcagggcaactttgaacagtgggtgttaaat
cctttaaaagttaagccaattgctcacgcaatttgggatccacactttggacaaccagct
ttaaaagcttttagtaaaggtgggtcagcatatccagtaaatatcgcatactccggcgta
tatcactggtggtatacaattggaatgagaagcaatcaagacctatattctggggcttta
ttcttacttgttttatcagctttacttctatttggtgggtggctacatctacaacccaaa
tttaagcctggtttgtcttggtttaagaacaatgaatcaagattaaatcatcatttatct
ggattgtttggcgttagctctttagcttggacaggtcacttagtacacgttgctatacca
gaagcaagaggacaacatgtaggatgggataattttacaacagtattacctcatccagct
ggtctacaaccatttttttagcggcaactggagtgtgtatgctcaaagcccagatacagct
caacacgtattcggcactaatgagggtgcaggtacagctattttaactttttttaggtgga
tttcaccctcaaagtcagtctttatggctaactgacatggcccatcaccatttagctatt
gctgtagtcttcattgttgctggacatatgtatagaactaattggggggattggacataat
ttaaaagatattttagatgctcatagaccgcctagtggcagattaggagctggacataaa
ggactatttgataccattactaattctttacatatacaactgggattagcattagcttca
cttggcgtaattacttctttagtagctcaacatatgtatgctatgcctccatatgctttc
atggctaaagattttactacgcaagcatctttgtacactcatcatcaatatattgctgga
tttttaatggtaggagcatttgctcatggtgcaatatttttttgttcgagattatgatcca
gaacaaaataagaataatgttttagctcgcatgctagaacataaagaagctatcatttct
catttaagttgggtaactctatttttagggtttcacacattaggtctttatgtacacaat
gacactatgattgcttttggtacaccagaaaaacaaattttaattgagccagtatttgct
caatggattcaagcttcttcaggaaaagcactttacggttttgatgtattactatcatct
tctagcaatattgcaacacaagctggtagtaacatttggctaccaggttggctagaggca
attaatagcggaaagaattcattgttcttaacaatcggtcctggtgatttttttagttcat
catgctattgcattaggcttacatacaactaccttaattttggtgaaaggcgctttagat
gcgagaggatctaaacttatgcctgacaaaaaagatttcggctatagtttcccttgtgat
ggtcctggtagaggcggtacctgtgatatatctgcatgggatgcattttacttggctgtc
ttttggatgctaaataccataggctgggtaacattttattggcattggaaacatattact
atttggcaaggtaatgcaactcaatttaatgagtcttctacctatctaatgggatggttt
agagattacttatggttaaattcttctccattaattaatggttacaatccatatggtatg
aacaatttatcagtatggtcatggatgttcctatttggacatttagtatgggcaacagga
tttatgttcttaatctcttggcgtggttattggcaagagttaatagaaactctggcatgg
gcacatgagcgtacacctttagcaaacttaattcgttggaaagataaacctgtggcatta
tcaattgttcaggccagattagtaggtttagcacatt-tttcagtaggatatgtattaac
ttatgcagcttttgtattagcttcaacagcaggcaaatttggttaaggtaaa-----att
taattaattgca--ataaaaaagtcagtcatgaat-ctatcatgactgacttttttttat

tgacaatagttacttattaaagtttaacttcaatatctataccagaaggtagatttaact
tcattaatgcatctattgtttgagaagaaggttgatgaatatcgataatccttctgtgag
atcttatttcaaaatgttctcttgagtctttatctacatgaggagaacgtaagacgcagt
agattcttctttttgtaggtaaagggatcggccctactgcaatagcgttagttctagatg
ctgtgtcgactattttatcacaagatgcattaaggatactcgaattatatgctttaagct
taatcctaatttttgtttgctgagtaagtgtcatattgagtaaatcttattatttaagaa
ttttggagacaacacctgcacctacagtgcggccaccttctctaatagcaaaacgcatac
cttgctcaattgcaattgcattaattaattcagcagtcattttaattctatcaccaggca
tcaccatttcagcatcggtaccatcatcagcagtaaactgattaatagtacccgttacat
cagttgttctaacataaaactgaggtctgtagcctggaaaaaatggagtatgtcttccac
cttcttctttagttaaaatataaacttcagcttcaaactgagtgtgaggtgtaattgtcc
caggtttagctaataccattcctctttcaatgtctttttctgaacgcctcgcaaaagaa
ttccaatattatcacctgctagaccttcttctagcgtcttttggaacatttctaatccag
taatagttgttgtccgagtctctctcaagccaacaatttcaattgtatcaccgactttta
taattcctctttcgatcctaccagtagcaacagttccacgtcctgtaatagaaaagacat
cttctacagccattaagaaagttttgtcgacatctctttcaggtgttggaatgtaagtat
caactgcgtccataagtgaaaaaatcttatcaacccatttatcttccccctgcttaatag
tcgggttctttgtaactgcttctagggctaacaaagcagagccagcaacaaaaggaatgt
catctccaggaaagtcatattggcttaacagttctctaccttctaattcaactaattcta
acaactcttcatcatctacttgatcttctttatttaagaatactactaatgtaggaacac
caacttgttttgctaacaagatatgttcacgagtttgtggcattggaccatctgctgcag
aaacaactaaaatagctccatccatttgagcagcacccgtaatcatatttttacatagt
cagcatgacctggacaatccacatgagcatagtgacggttatctgtttcatattccacat
gagcagtattaatagtaatacctctagctttttcttctggcgcagcatcaatttcatcaa
actttttgctgaggtggaacctaaagttgataaagtcgctgagattgcagctgttagag
ttgttttaccatggtcaacgtgaccaattgttccgatattaacatgtggtttttacgtt
caaatttagatcgagccatatttttgctttccttatagtacaagttgcttaatggttac
ttttaacgatagatcttacagattgaaagatttattatttgatatagacagtg-----ca
agggatggttaatctaatatcgatagtgtgcaaacgctttatttgcttccgccatcctat
gtgtttcttctctttttctaatagagctacctgtttcattggctgcatccataatttcgt
tagccaattttattgccatacttttaccagatctttctcgagaaaatctagtaatccatc
ttaatgctaaatttgtacctctatatgctcttacttctattggtacttgataagtagagc
ctcctacccttctggctttcacttccactaaaggagtgatatttcgaatagctttttcta
aaatatttaaaggatctaattcagtcctatctttaactatatctaaagcttgatatataa
ttctttgtgccaaagtttttttaccactttttaaaatacgaacagttagcatactaacaa
gtctgcttttatatagagggtctggtgacgcgaacctttttttagctgtattacgacgag

acataatatttgaaatttgttattgtgataatttgtt-taatagaagatatttaagattt
aggctttttgtaccatatttagatcgacttttacaacgatccttaactccagcagcatc
taatgtaccacgcactacatgatatcgtactccaggtaaatctttaattctccctcccct
aattagcactactgaatgctcttgaatattgtgaccaacaccaggaatataagctgtaac
ttcaaatcctgatgtcaatctcacacgtgcaactttcttaaagcagaattgggtttctt
aggagtcgtagtatatactcttgtgcagacacctcttctttgaggacaactttgaagagc
tggagattttgtttttttatgtattttacgtctttcagatctaacaagttgttgaattgt
tggcataataaaattgatttaagtctttaatagatattatttgaacttgctacacgtga
attaattagtctttatttatcttgtatttacgcataaatctttctactcgaccttctgtg
tcaataattctttgtgaaccagtaaagaacggatgatttcctgaccagatatccacatgt
aactcaggctttgttgaaccaattgtcataattaattggccatcacaatagacttttgct
tctgggtaccaatttggatgtatgttatttttgccattgcttatataattg--------
tatcaataataaatattaacgcttagaaaattgaggagcttttcttgctttctttaaacc
atattttttctctcctttactcgtggatctcttgttaaataccccttcagatttaagtgc
gtttctattctctggattaattgagcacaatgctcttgcgacacctaaacgaattgcatc
tgcttgccccgttaatcctcctcctctagcatttacatggatatcatactgatttaataa
tcctagaacctgcaatggtgcatgcgatactcttaagtaattaggactaaattgaagata
tgactcaccaggtataccgtttataattagattacctgaacctgggactaatcttacctg
tgcaacagaacacttgcggcgacctgttccagaatagattgcacgagttttaattaattc
tgtcgacataatattcctttataactaaataaatatacattctttatactatatactctt
gtggcttttgcgcgatatgtgggtgaattggaccagcataaactttaagcttcgtaaata
atttcttcctaatggacctttaggaagcatgcctttaaccgatttttcaataattctat
taggcaatcttgtctggagctgctcaaatgtttctacttttagtccaccaggctgtccag
aatgtcttctatatagtttttgattagttttattgccgcttatatgtacttgagaagagt
taatcacaattacataatctccagtatctagataaggcgtataagaaggcttatttttac
cccttaagatattagaaatatgagtagatatcctacctaatgtctgattttagcgtcta
taatataccaatttggattagtatttaatgatggtgcttgcgttttgttcatgaaaagat
taatactttagttggcaataagagaataggtaaatgctgtacatgtaatttcgatacaa
ataatacttgtaaaatatta-------------ttgacaaagagaactgtcgtgacagtt
atttaaattatttgttagcaagaaaacttctaattatcagtaatttataata-ttgtggt
aatataaaacaggttaatcactttttttcttttggcaaacttatacctaatttttttttgt-
agggcatatattacttcttcagctgatttttggccaaagtttttaatttctagcagttct
tcttgagagtaatctagtaaatctgcaatagaatgaatttgagctcgctttaagcaattg
tatgctctaacagacaactgcagttcttcgattaaaacttgactaatttttttatcttct
ttactacgatagttatctgctgatttaaagtccaaatttcttagtgagcagaaaaggttc
gttaaaactgtggcaccttggcttatagcttcttggggagaaatactcccatttgtccat

atttgtaaaattaaccgatcttttatactattagtacctatacgtacttcttctacttta
taattgactttattaactggcataaagactgaatctacttgcaagaaatctaaagataat
tcatctactgctttttctgctaaacgatatccacagttttttttcaattttaaactccatt
tcgaatattgtattattgcaaatagttgcaatatactgcctaggatctactacttctata
tcagaagataattcaaacaatccagctgtaacgatagctggtccctggactctaattcta
ccaatttgagattcttgattatgacttttaaatactacttcttttaggttcaataatatt
tccaaaacatcttctcttactccaggaattgtagaaaactcatggtttactcctgcaatg
cgcacagcaactatagcagtaccttcaagatctgataatattgatcttcttaaagcatta
cctaatgtaatgccttgtccttggtttaatgcttctattacaaaactaccgtactgtcca
cgtgctccatctgttcttgattctacacattcaatttgaaattgagccacctaagcaagc
tcc-ttaaaaataatcaatgattagtaattttgtcgacaataagaaagaatctatacacg
gcgtttcttaggaggacggcatccattgtgaggcacaggagtaatatcctttattaaagt
aatctctaaacctgcggcttgtaatgctcgaattgctgtttctcgacctgcacccggacc
atttactaaaacttcagtttgacgcataccttggtccattgcctgtttagcagcttttttc
agctgcagtttgagctgcaaaaggtgtaccttttttagctcctttgaatccactagaacc
agaggatgaccaagataatgtttcaccttttaaattagtaatagtaacaatcgtgttatt
aaatgtagatttaatatgtgtaacaccattaactgcattgcgtttgttctttcttactcc
agattttttaattgtctagccatcttgtttcgtctattatataattagttacaatatta
tttccttggagctttttcttgcctgctactgttttttttacctcctcttcgtgttctcgc
attagttcttgttctttggcctcttaaaggaagaccaagacgatgccttctacctctata
agtactaatttccataagtcttttaatactcatcgattcaaaacgtttgagatctccttc
aatctgataactagattctaaaatttctcttatgctaacaacctgttgatcatttagatc
ttgacacttagtatcagcatcaatgttggtttttttttaatatttcttttgaacgagatag
cccaatgccataaatatatgttaaagctatctctattcttttgtttttttggaagatctac
tccagcaattcgagccacgttgtgctttctccaataaataatgtttac-gtttatgacag
aagattaca-ggattagccttgtctttgcttgtgtttaggattagtgcaaattaccatga
cttttctgtgacgacgaattattctacatttttcacacattttgcggacagaaggacgaa
ctttcatattaa----ctctatatactaggaatattactg---------tattaattatt
tacatcaaaaatgttaataatatatttgattactttaattgttatagctttcacttcgtc
atactatcatattttttttgatatgacatatgtttgaatttgcttagccgtgtctattgcg
actccaactaaaataagtaaagatgtcgctccaagacctcttaagttttggatctgagta
accttttctattataaacggaatcagcgctactgtaaataagaacgaggctccgagaaat
gtcagtctatttaatataacttgtaaataatcaatagtcgcttgacctggacgaatattc
ggaatactggcacccatttttttttaaatttattgcgatgtcttctggattcattactatt
gatgtataaaaaatagctaaaaaaaaagaattaacgcacagtaaagaataaggtacagcgaa
ccgttgggacaaaatagatatagaatttggagtattgttttattttgaattatttgagtt

aaataagatggtagagccatagatgcagatgcaaaaacaataggcataactccaccttga
tttaatttcaaaggcaagtaactgttcggatccagaattgaggattttcctaactgcctt
gctgaaataattttaattcttcttgttccttcttgtacacaaattgtaattacaatcatc
aataaaaatataacaatgaacaatccaaacttaagacccggattactatagtttgcatca
aaaaatgactgtgtaaagtttttttggtagtcctgatacaatattttggaaaataagcaaa
gaggctccatttccgattcctttttctgtaattaactcagataaccacataatgatcatg
gaacctgccgttaaagccaaaacagactcacaaacaaatgtaaagttccaattaaataca
tacggctttacccatatagatattgctccagattgtaaagtagcccatccaagagacaaa
tatcgggtaatttgagtgatttttttggcgtcccagctcaccttcttctttctgtaatttt
tctaaatcaggaataattttcgtgagtagttgcattacaattgaagaattaatataagga
acaatacccaaagcaaaaatccctattgttgagaatcctcctccagaaaagatattcagg
aaatttactaaagtgttcttttctacgcttgcataaaaagcatcatggtctatacctgga
acaggtataaaaatgcctaagcgcgctaagactaatagaaaaagagtaaatataatgcga
tttcttaaatcacttttttgtctcataaataaatta---aaaaaaaaagacactga--gt
tagataaaaaccctattccaatacttgtataagtttaaaatagggaaatgg---caagct
atagaagtagcgcta-taatatcttatgtcgaataaagactttctataggaactcctcga
tctttagctgcttcgctaaaagttcttaattgagttaggccgtttaaaacagctctggca
ttgttgagtgtattattagatcccaactgtttagctagaatgttctgtacgcctgataat
tctaatactgtccttacagaacctccagcaattacaccagaaccaggcgcggaaggcctt
agaataacttttgctgcaccagaaattccgttaataggatgaggtatagaattaaatttt
gtaagtggcactgtaactaaatgtttttttggcatcagttactcctttttttaactgctcca
attacatcactagcttttcctactccgacaccaacttgcccctgctcattaccaacaata
agaataactcgaaagcttagcttcttgcctccttttacaactttagtaactcttttaact
tgtacaactctttcttcccaaccactatctttatctttgcctttactctgtttttttacga
ttagccattttaagcactattcctaa----ttaatta-tgactttaaaaacccatacctg
cttccttagcagcttcagctaaggctttaactcttccatgatataatttcctcctctgt
caaagacaacatttgtaatgccttctttcatagacttttcagctagctgtttaccaacta
ctcgagaagtgtcacaatttgagcctattttatcaagttcttgattatttaaattaacag
atgacgttgcgactaatgtaattccttgtgtatcatcaattatttgtgcatatatgtgtt
tattagatctaaatatacatagacgcggcctacttgaagttccttgaacttttttttcgaa
ctcttttatgcttatgaattctagtctgcttagtatttattttcattattatttacccttt
tccggcttttccggcttttcttctgacaaattcgccttggtatctaattcctttaccctt
ataaggttccggaggtctagtggatcgaatagttgaagcaacttgcccaacaacttcttt
gtctatacctgaaacagtaatatttgtgttattttcaactgtaattttaatattttcggg
aggttgtattttgactacatggctatagccaacgcttaatattaaatcttgattatctat
ttgtgagcgataacctacaccttgtatttgtagcttcttagtaaacccattagaaacacc

ttcaatcatattactaataagagttctggataatccatgtaattgactagccatttttgt
ctgcccacttgtcttgacagaaatttatcattaaatatctctaagttaataccatccgg
caaagttcgcgatagagtacccttcggtcctttcaccgtgatggtacggccatcaaactc
agtactaagatttgtcggcaataaaattattttttttccaatacgagacat-attccacc
tataaaatagttaccaaatatagcataatatttcaccaccaagaccatcatgacgggctt
gtctatctgtcataacacctttagaagtagaaataagagcgattcctaaacccccaagaa
ctctaggtagttccttgtgatttgcatagactcttaatccaggcttactaattcttttca
gagcagtaataactggttgacgatttttaccgttatactttagagaaatcattaaatgtg
tctctatgccttcacccatttgttcaaaattttgaacaaatccttcttcttttagtactg
ttgccatattgcatgtcattttcgttgctggaacttgcacaatttgatgtcttgctaagt
ttgcgttacgaatacgtgttagcatatcggcgatcgtatcgttgaccacc-agatcctcc
ttacgataaactgaattatttaatatttttttcaagattccttaaaaggcataccaagctt
ttttaacaaagctaacccttcttgatctgttttggctgtagttacaattgatatatcaag
acctcgaatttgatcaatattgtcgtagtctatttctggaaaaattaattgttcgcgcaa
acctaaattataattacctttgccatcaaagctttttggactaattcctctgaaatctct
aattcttggcaaggttaaattgattagcttctctaaaaaagaatacatcttatcttttcg
tagatgtactacaattccaataggcacttcttctcgaattttaaaaccagcaatagattt
tttagcttttgtaacaattggtttttgtccagttattagtgttaattctttaatactact
ttccaaagctttagcattttgagaagcttcacctaaaccacgattaatagtgattttcgt
aaaacgaggaacttcatgtacattttttatattggaattcctcttttaaagcttgagtaac
agttgttttatattttttcttttaatcctattaccattatctttt-acaaattatttaata
agttcacctgtctttttcaatttccgga-tttttgacctttatcgttaattatgactga
aaatctactggcaatattattttgttcactaaacaacatgacattagaactatgtattgg
agcttcaaatttgataacttctcctgtctcaccttcttgctgaggtttttttatgttttac
ttttagattaattcctttaacaattactttacttgtcttatggatgattgcaataatttc
gcctgtcttggttttatcgcttccagaaataacttgaactaaatcacccttttttcaattt
aatttttgcattaactttagtttttttcaaagaacttttcattatattacctccggtgca
agagagactattttagaaaaatttttatctctcaactctcgagctataggtccaaatact
cttgtgccgcggggattattatcctgattaataataacagctgcattatctccgaatcta
atactcataccgtcagttcttctcaaggctttgcgagttcttactactacagctctaaca
acatcagatcttttaactggcatattaggtgaagcatctttaactactccgatgatgaca
tcacctatagatgcataagcaggattactagtacctaatactctaatacacattattttt
ttagcgccactattatccgcaacattaagatagctttgagtctgtatcatacttttattc
tatataagattaattatcaaaagacttagataagatattaaccatcgtccaacacttcgt
gcgacttaaaggccgtgtttcttgtattgtaactatatctcctgctgcgcattcgttatt
ttcatcatgtactttatatttttttgtgcggatcatggttttcgcatattttctatgagc

aattctattttctacagctactactatagttttattcattttattacttacaactttacc
agttgtttcttttaaaggcatagtttttatc--ttttatatat----aatgaaattaagt
tataaagtaattatatagctggtttatttcgagacttttcaatagttaaaagttgagcta
atctatgcttagaatgtttgaataagtgaggctgaaaatcttgtcttgtagctcttttca
gtcttaactcaaaaagcgttcttttaattacaagtatttcttcagctaaggaagaatcat
ccatttgtgtaacatctgatattttagggaaagtcatgctttaagattctattatatttc
gaactataaatttggtcttcattggtaatttataagaagctagtttcatagcttgctgag
cagttctttgtggcacaccagtaatttcaaataaaatgtgaccaggcctaataactgcga
cccaatattctggagctcctttaccagatcccattctagtttctgcaggacgcgccgtta
caggtttgtctggaaacaccctaatccatagcttacctcctcgtcttacatatctagtga
tagttcttctagtagcctctatttgtctcgaggttaaccatactggttctatagcttgca
atgcataatcaccgaatgcaattgtattgcctttactagcagaacctttcattctgcctc
tgtgttgttttctgaactttgttttcttggggcttagcataagaattaaacaatttaaat
aattaagaagctgttggatcagaagtttcctgattatatacagattctgtgctctttgat
tctggcaaaatttctcctttaaagagccacactttaactccaagtactccatatgtagtg
tgagcttgtcgatgacaataatcaatgtcagctcttaaagtttgtaaagggacgcggcct
tctctaacccattcacttctagcaatctctgcgccattaagacgtccagacacttgtatt
ttaacaccttgtgtatttgctctttgtgccctttgaacagcctgtctcacagctcgacga
aaggcaactctttttttctaactgctgagtaataaattctgcgactaaggttgcttcagaa
tctggatctgcaatttctacaacattgactctaatttgtttgctcggatctaaaattgaa
gataataattttcttaaagattcaattcctgctccagattttccaagtacaataccaggt
cgagctgttgctattaaaatttcgacttgatcaactttacgattaatttcgattttagca
atgctagcattactaagtttagaatatataaacgaacgaattttatgatcttcttctaaa
agaactggatagtctttagagttggcgaaccatgaagaacgatgtttttgggtaatgcct
attcggaaacccaaaggatgaatttttttgacccacagtatccttttctataaattaaaag
ttgtgattactaattgacttgaaatac----aagactacaactctgagacacataacgta
atgtgacaagttggtttatgtataggaaatgctcttccttgtgctcttggttgaaatctt
tttagcgtagggcctttatctgcaaaagccttgctaataaataactgacttttattcagg
ccatcattatgctcagcatttgcagcagcagactctaaaatttgctttatataagagcac
actcgatatggcataaattctaaaataattaatgcttcttgatatttcctgcctctaatt
tgatctaatacacgacggactttatgtggagaaagacgaatgtatttccctactgcttta
gtttcttttacgtttttttgtaatgctcataattatagtttaacgacgggcttttctatca
ccttttacgtgggtacggaaagttcttgttggaacaaactcccctaatttatggcctacc
atttggtcagatacaaagacaggaaaatgttgtttaccattgtaaacagctattgtatgc
ccaaccatatctggaataatagtagatgctctcgaccaggtttttaaaacttctttttttc
cccaaagtatttaatgcttctatgcggttaagaagactcacatcaataaaagggccttta

tgtatagatcttgacataataataaattattagattctaaatgaaataaattctatttac
ggcgacgcaaaacgtatgggttgctatatttattgggatttctcgttttaacacccaatg
ctggtttgccccaaggtgttactggacgagctctcccaattggagatttaccttcaccac
caccatgtgggtgatctactggattcattacaacacctctaactgtaggtcttttaccta
accaacgatttctaccagctttaccaagagtgatattactagcatcaatattgccaactt
gcccaatagtagcatagcattcttttctaatcatacgaacttcgcttgagggtagtttca
cagtgacaaaagagccttccttggctactatttgagcataagttcctgctgcacgaacaa
tttgtcctccgcaagaaggtcgtaattctatattgtgaacggcggttcctagaggaatac
tggataaaggtaatgcatttccaacttcaataggcgcagtgggaccagaaagcactattg
ctccaacacgcaatgatcgagggtgcaaaatataccttttttctccatcaagataatgca
ataatgctattctagcatttctattagggtcatattcaatagaggccactttagcaacta
tattatgcctgtttcttttaaaatcaattaatctatactgctgcttatgtccgccacctt
tatgacgacatgtaataacacctctattgttgcgacctttgcaaaaatgatgtttaacta
ttaaagatttctctggcttatcagtagtaatctctgaaaaagtagaaactgttctatttc
ttgtcccaggcgtataggcccggtataaacgaattgccatataaggaaatttaaaaagag
taaatatgtga------ctgtgtaatacgattaagtttctggaaataaattaatagaatc
ttctggcacaagtgttacgattgctctcttgtaatgaggtcttctacctacaaatctacc
aatacttcttttttcttaggtggatggcaagtgttaacacctttgacttgaacattaaa
aatatactgtatggcagctttaatattaattttggttgctttgggatcaacagcaaaaca
gtactgattttcttctaataattttgttgttttatcagtaatgattggatatttaactaa
atccaataaacctcttgaatcaatgctatccattatatacctcttgtattttagataaag
catcaactgtaataatgattttatgcgcagctaaaagggccatgatattcaaagtatcag
ctgaaataatttctacattatgtaaattgcgaatggaaagataaacattcggatcttttt
tatccaccacaactagaactttttatttaaatcaagattccatcggtgaatagcttcca
taaacagtttagtctttggctgctgaaagtagctattaaagttttctacaatcaaagtat
taactgatttattatttaatgccgtccttaaagctaattgtctttctttttgttcattt
ttttagtaaaactgcgaggctttggtccaaatattactccaccgcctctccataagggtg
agcggatagaaccagctcttgctctacctgttccttttggcgccatggtttgcggcctc
caccctgaacttcacttcttgttttagtattagcagaaccttgccgtttttcgctacttt
gttttactaaagctctatgaactaaatacataccagagtcttggcttaccttaagattta
aatctgcgctaccacttatttgaccttcccaattatagacttgataatttaattgtgttt
taactgtcatgagatatatcgagtgaaagaatattgttatttactaattttactaaagt
tccaggtttacctgggacagctccttttacaatcaacaaatcattctttgagttaatact
tacaatttgcaaatttttaccgtaactttttattaccaagttgaccagccatattttt
tcctgggtagaccctcccaggcgttgtcccagcaccaatagaaccgggctgcctatgatt
tttcgaaccatgagacattggacctctactaaaatgatgcctcttttgatagcctgaaaa

gcctttacctatgcttttagaagaaacatcgatttttgacccactttaaacatatctgt
ggacaaaatttgactgacttcaaagtcatcagtagatttcactaagtactcactcaaata
ttttagtgggggagcttgtgatttttttaaatgtcctaatagtggtttatttaactttg
ttcaacaacttgcttataaccaatttgaatagcgttgtagccatcgcgaggtaaggcttt
aatttgagtaacaacacatggcccaacttgaattacagtaactggaattgacaagccatc
ttcatcaaagaattgagtcatacctactttagtaccgagtataccaacagacactagatt
tctcctttttgagagatttatactatcatcatacataaaa-ttttactccaaattgcttg
aag--taaaccaaactaagagatgcatgacattaatcatggcaacataaagag-aaaaac
tttacttattcaatctaaaatagaaacagtttttgtccagttacattatatataaatta
aaagtaagcttattatattatttacacactac----------------------------
-------------taatatgaagaacaaggaatgtcttcttttag-----ttttacacaa
cattgcattattagtactgtctcaaatataaatgaataaaatattaaaatttcacctctc
tagagaaactacaaactacaattgtgttcggtaactacaccttttatttacatatttat
tgtgcagtatatatgcatatatagtacagcgcacgtaaaaatatagc----tgtataaca
aatatacttataagaattaaagaggtattaatgggtaaagttgttggaatcgacttagga
acaaccaattcagtgattgctgttatggagggaggtaaaccgactgtaataccaaacgca
gaaggttttagaactactgcttctgttgttgcatatactaaaagtggagataaacttgtg
ggacaaattgctagaaggcaggctgttattaacccagagaatactttctactcagtcaaa
aggtttataggacgtaaacaaaatgagatttcccaagaaattagacagacatcatataat
gttaagactagtggatcaagcataaaaattgcctgtcctgcacttaataaagattttgca
cccgaagaaatttctgctcaagtactgagaaaacttgttgaagatgctagtacatactta
ggcgagactgttacacaagcagttattaccgtacctgcatactttaatgactcacaaaga
caagctaccaaagatgcaggtaaaatagccgggttagatgtactaaggattattaacgaa
cctactgcagcctccttgtcttatggactagacaaacaaaataatgaaacaattttagta
tttgatcttggaggaggaacattcgacgtgtccgttttagaagttggagatggagtattt
gaagtgctttcaacttctggagatacgcatttaggtggagatgactttgaccaacaaatt
gtggaatggctaataaaagattttaaacaaaatgaaggtattgatcttggtaaagataga
caagcacttcaaagattaaccgaggctgcagaaaaagcaaaaatagaactatcaaattta
actcaaacagaaattaatcttccatttattactgctacacaagatggtccaaaacatttg
gaaaaaactgtaactaggggggaaatttgaagaactttgctcgaatctaatagacaaatgc
agtattcctgtaaataatgctctaaaagatgcaaaactagaagcttccagtattgatgaa
gttgttttagtaggtggatctacaagaattccagctatacaacaaatggtcaaaagatta
ataggtaaagatcccaatcaaagtgtaaacccagatgaagtggttgctattggagcagct
gtacaagctggtgttttagctggtgaagttaaagatattctactattagatgtgacaccg
ttatctttaggagtagaaactttaggtggggtgatgacgaagattataccaagaaatact
actattccgacaaaaaaatcagaagtgttttctacagctgtagacaatcaaccaaatgta

gaaatccaagtacttcaaggcgaaagagagttaaccaaagacaacaaaagcttaggaacg
tttcgactagatggtattatgcctgcacctagaggcgttcctcagattgaagttacattt
gatattgatgctaatggaattttatctgttaaagcaaaagaaaaagctacaggcaaagaa
cagtcaattactatatctggagcttctactttacctaaagatgatgttgatagaatggta
aaagaagctgaagagaattttgatacggaccaaaaaagaagaaaaaatattgatattaag
aatcaagcagaatctttatcttaccaggctgaaaaacaaattaaggaatttgaagataaa
attagccaagatttaaaaacaaagatagaagggctaattatagaccttagagctagttta
gagaaagaagaatatgagaaaattgaatctatctcccaacaactacaaaactcgttaatg
gatattggcaaaatggctgctcagtctgaaagtaaaaatacaaacacaaaagatgatgac
actgtaattgacacagatttctctgaagctaaataagaaatt-agcgggtaacgcgattc
gaacgcgcgacatcaaccttggcaaggttgcgctctaccactgagctatacccgcataga
gtgtattattacaaatatggtaacatttgtcaatctgt--------aattatat---agt
tgagttactaattcttaaactataaattaaacacctggtaaaaatacaaa----attttt
gtttttgaat-aaaagaagtacttatataac-gatataagtacttcttttattcaaaag-
-caaaaatataattaaatatcagtg-------ttaaactactaaggaatttactaagcta
gtaacaattactaatccagtccataggcctgctcctgtgtagattaacccttttgattgt
tcccattgaccaggtgatgctagaacaacaggtacgcctactactaacacggtcgacaga
acaattaataatagcactaataattgaatcgcaataatcatttattatattctcctgaaa
aattcaaatgtgaataatattgtatataatacaatgaagctttataccttcattaattta
tctatgaagaaataatttaaaattattgttttctcaataaatattaaattaagtattaaa
atctttattgtaaacaataacactatctttttttaa-atagataaaaaatttaaagaaag
tctttttaaaaaaaatttaggataaggtgatatgctgtataaaaagttaagtattatta
gttattagagaaatttcaaaataagaggtaaaccatatgaactcagccctttttttagca
aaactacccgaagcttatgcagttttcaaaccaatcattgatattttgcctgtaattcct
gtgttctttctacttctagcttttgtttggcaagcagcaatcggttttagataactttta
agttatgacaaaaatagcaaatataaagagtgaaatttttttaagctctaatcgtatatct
tacacaataataactattatggttgaaccattactatcaggaattgttcttggccttatt
cctatcactttattcggattattagtagcagcttatctacaatatcaacgtggaaatcaa
ttaggactttaaagcctaaattatagatatttataaaaactataattattgtagtcttta
taaatatctacaaaccagctaattataaatacagaaaaaatttattgaaaccagattaa
taattgacaaaattgatatgatagtataa-------------------------------
------------------------------------------------------------
------------------------------------------------------------
------------------------------------------------------------
------------------------------------------------------------
------------------------------------------------------------

------------------------------------------------------------
------------------------------------------------------------
------------------------------------------------------------
------------------------------------------------------------
------------------------------------------------------------
------------------------------------------------------------
------------------------------------------------------------
------------------------------------------------------------
------------------------------------------------------------
------------------------------------------------------------
------------------------------------------------------------
------------------------------------------------------------
------------------------------------------------------------
----------------------------taataaaaattttaccaactcgacttagtaac
tccaggtaaaagacattcatgagacatttctcgaaacacatggcgagataacccaaaatc
tctgtaataccccctacttctaccagttaaccagcacctatttctacctctaacggctgc
gctgtttctaggcatttcttgaagtctttgtcttagttccattttgtcagaaaagcttat
agcttctttcagctgatctttaatagcgagtcttttcaagtagtattttttttcaagctt
ttctcttttaagctctctttggatcatatttttttttgccatattgtaa-aatttacttt
aaaaattatttaatcaatatactagctaaaaaatctctgaactgcaacatttatataatc
aaaaacatatagcataaagcaatgccataatataaatatttaattaatggtact---act
ttatgctattgacccaaatttatttaataaagttcttgctcaacatgagttgaaatagtg
caatcagattgaggataagcaatacatgttaaaacataacctgctaacatctgatcatca
tctaagaatgattgatcagattggtctacaagaccttccgtaactttgcctgcgcaagtt
gaacaagctccagctctacaggaatatggaagttcaataccttcttcttctgctgcatct
aggatatatgtatcatctgcacatttaaatgtaacatcaatgttttcgtcttcgcatact
aaatgaattttatactctgcca---tttttttactccataataaagatacacttgaaactt
aataaagcaaaat-aatctaaaggatttacccaaagattgtatattagtttcattatgca
taatagactgcaaaaaaaggaaataatagactattttgagaaatcaggctacatttcata
ctaaatagaactatagtaggttcaaagtatatatataaattcatatagttaaaattattt
ttatcatttaaataactaaacttaacaatct-aaacttttagattaatttac------ta
tttaatcttttacaataagcaaataagtaattatgacttcaatgatgaatcaacaaatag
agctgaagcctattcttcgatttcagataccatatttatattcaggcttccttatacagg
aaattaaggctttaacgaccagactttttctgcaagtttggagaagacctgctacattga
tggcaggtattattcagcctttattatggttaattttatttgggggtcttttttataatg
ctccagtgaatttatttactattaacacgagctataactgtttcttaagctctggaatta

taattttcacatcgtttactggtgcactcaattcaggccttcctttaatgtttgacagag
aattcggattttaaatcgattattaacagctcctttagtgtctagaacttctattattt
tatcttctgctacttttatgacttgtattagcttaatacaagttatatttatagttacag
cttcattatttatgggaaattctcctttaaacagtgatagcgctataattttggattta
taattctattagtaactgtgggagttacaatgcttagtttagctttatctttactctgc
ctggtcatattgaactattagcacttattttagtaattaatttgccttttctattttcta
gtacggctttggctcctctgtatttcatgccaccttggctgcagttgattgctagtctga
atcctttaagttacgcaatagaaggtacaagatatttatattccagcatacattggaact
ttacagagtgcattatcaaaattagttggggagatgtttgtttaggacagattatttcac
tattagttattttagatgtaattgcagcctatgttgtttctaagatattaaaagctaaac
ttaatta-----------------aaattttaataaaaaatttattaaatttattttaga
aagattttactatataatactattagtagtatggataattgaaataggaatagtttttc
aaaaccaaagctattatcaacaatgataaacatttgtaagaaagtcaacaaagtatgttc
ttattcataggaggcatgtagtcaatgggactaccatggtatcgtgtacacacggttgtt
ttaaatgatcctggacgcttaattgcagtacatctaatgcatactgcacttgtagcaggt
tgggcaggatctatggcattatacgaattggctgtatttgatccttcagatccagtatta
aatccaatgtggagacaaggaatgtttgttatgccattcatggcaaggcttggcgtaaca
gattcttggggaggatggagtattactggagaaagcgtctccaatcctggattgtggagt
tttgaaggagtagctttaacccatatagttctttctggaatgctttttctggctgctatt
tggcattgggtttattgggatttagaattatttcgagatccaagaactggcgaaccagct
ttagacttgccaaaaatttttgggattcatctattgctatcaagtttactgtgctttgga
ttcggtgcttttcatgtaactggattatttggaccaggaatgtgggtatctgatggttac
ggagtaacaggtaaagtgctgccagtggctccggcgtggggaccagaaggatttaacccg
tttaatcccggaggagttgcatcacaccacattgctgcaggtactgtaggtatattagct
ggtgtttttcatttaactgttagaccaccgcaaagattataccgcgctctaagaatgggt
aatatcgaaactgtattatcgagtagtatctcggctgtttttttctcagcttttgttact
tgcggaacaatgtggtatggctctgcaactacacctgtcgagttatttggtccaactaga
tatcaatgggatagtggatattttcaacaagaaattgaaaaacgagtcgaaaacgctatt
gctgatggggctgctcctagcgaagcatggtcaagaattccagataagttagctttctat
gactatattggtaacaatccagcaaaaggtggtttattcagagctggccctatgaataaa
ggagacggagttgctgaagcatggcttggacatcctgtatttcaggataaagaagggaga
gaactaagtgttcgcagaatgcctgctttcttcgaaacgtttcccgtcattttagttgat
aaagatggcatcatacgagctgacattccatttaggagagcagagtctaagtacagtatc
gaacaagtgggtgtaaccgctagtttttatggcggcaaattaaatggccaagtatttaac
gatgctcctagcgttaagaagtatgcaaggaaagctcaattaggtgaagttttcgaattt
gaccgaactacactagaatcagacggtgtattcagaagtagtccaagaggctggtttaca

tttggtcatgctaatttcgctttaatctttttctttggacatctttggcatggttcacga
actatctttagagatgtattcgcaggaatcggagctgaagtaactgaacaagtagaattt
ggagctttccaaaaattaggcgacagaagcagtaaaaagcaaggcgctgtataagact-a
catcaaacacagtccttatt---caatatttccagtattaaaataggtcaaaaaaagttt
attatctaatagttactaattaggagaacatatggaagccttggtttatgtatttttatt
gacaggaacactaatggttatattcttcgcgatctttttagagaacctccaagaatagc
taaataactttattaggtttccccttacaataatagaaccactttccagcttattagtgg
aaagtggttctattgtaagttacagtatattattcttcatgctcttcaaaaggatctcta
agatcttttgaagcaggaccaaatgccgtatatattgaataacctgtaatacctaaaagc
aaactcgaaatgaaaatactaagaactgttgcagtttccataattcaatcagtattaagg
tgaactttttctataatgatattatcaatattaaaacatttaattattaaatttatcaaa
atatattatggcacttagaactagacttggagagatcttaagacctttaaattcagagta
tggaaaagttgctccaggttggggaacgactcctatcatggggattttatgttactatt
tttcttatttttgttgatcatttacaaatctacaattcatcactagtactagagaatgt
agatgtagattgggctactttaggtagctaactaattacactgttgctttaa--------
tttaataacattaaaaataat----------------aagcgatcactattatgtcataa
tagcgatcgctcaaattgtgttttactatata--caaaagtaattattttcctaagtcaa
ttaattcgctatctgcaaaactattcgtattaacattattataattaactttttcaaatc
ttactagtactggatacttgataccgcttttatccattgctgcaactgtcccaacttctt
gataccagtatgattcctttctcaatatttttacttttgagcctcgttccatatttttaa
tcctaagataggttattaaagatcagtataatagataaagaaataaaattaaaaaaaata
catgaaattagtaccgacaat---------ttaatcataaaaggagagagagaggattcga
accctcgataactacagttatgacaattttcgaaattgttgcaataaaccactctgccat
ctctcctaaactaaaactcttttttcggaattagataagaataacatagaacgaaggta
gttgcctagaaaattaaaactatgttaaatataaattataatctttctatagatctcttt
acaaaatatgtacacagattctatatcagcttatttcaagatttttaatcttacacaat
ttttactaaataaataaactctctcataatatgaaactatcttggaaaacacttcttctt
tggtctttaccaatatttgtcgttggtttctttttctggcaaggtttttaggtccgact
actacagatgttggtagtaatattgcaagttctagaatgacatacggaagattcttagaa
tatttggatatgggttgggttaaacgagttgacctatacgagaataaccacacagcaatc
gtagaagctgttggtccggaattaggcaatagagttcaacgaattcgagttgagctgcca
gcgagtgcaccggagttaattaccaaactacggaaggctaatgttgatttagatgctcat
cctcctaaaagtacaagtgctgtatggggactattaggtaatttattatttccgctcttg
ttggttggaggcttagcttttctatttagaagatctaatagtgctagtggtgggcctggt
caagcaatgtcatttggtaaatcgaaagctctgtttcagatggaagccaaaacaggagtc
gtatttaatgatgttgctggagttgaagaggccaaagaagagtttcaagaagtagtaact

tttttaaaacaacctgaatcatttactgctgttggtgcaaaaataccaaaaggtgttttg
ttagttggtcctcctggtacaggaaaaacgttattagcaaaagcaattgctggtgaagca
agtgttcctttttttagtatctcaggctcagaatttgtagaaatgtttgttggggttggt
gcttctcgcgtaagagacctgttcaaaaaagcaaaagacaatgcaccttgcattgttttt
attgatgaaattgatgctgtgggcagacaaagaggaacaggtgtcggaggtggaaatgat
gaaagagagcaaacattaaatcagctattaactgaaatggatggctttgaaggaatact
ggtgttattgtaattgctgctactaatagagctgatattctagattctgcattattaagg
ccaggaagatttgacaggcaagtatctgtggatgttcctgactttaaaggtagattagca
attcttgaggtgcatgctaaaaataaaaaaatggaacctaaagtatctttagaaactatt
gcaagacgaactccaggcttttcgggagctgatttagctaatttattaaatgaagctgct
attttaactgctcgaagaagaaaggatgccatgaccatgtcagaaattgatacatcaatt
gatagagtggtagctggaatggaaggcacgcctttaattgatagtaaaagtaaaagatta
attgcatatcatgaggtaggccacgcaataataggaagcttattagagcatcatgatcct
gttcaaaaagtcaccttaataccaagggggcaagctagaggcttaacatggtttactcct
agcgatgatcaaagtttaatctcaaggtcacaaatactagcacgtatcgtcggtgctctt
ggtggaagagctgcagaagaaattattttcggagatgcagaagttactactggtgcgagt
aatgatttacagcaagttacatcaatggctagacaaatggtaactcgattcggatgtct
aaaattggcccgttatctcttgaaagtcaaggtggagaccctttcttaggtagaggaatg
ggaggaggttcagaatactcagatgaagtcgcaactaatattgataaacaagtacgagaa
attgtcagtgaatgctataaacaggctaaagaaattgttgcagataatcgagtagttatg
gatagattagttgatctactaattgaaaaagaaacaatcgaaggcaatgaatttagggat
attgttaaagaatacactgctattccagaaaaaaattattataaatctcaattttaa---
---------------atagtacgggactgacgggattcgaacccgcaacttccgccgtga
cagggcggtgctctaaccagttgaactacagtccc-accaagac---tataaataaataa
tctcacaattagcttgtaattgtcaaattatactattaaacatgagaagta---------
-----------ttattaatacttttcatgttctttattacagagtcaattaatccattta
caaggagagagagggattcgaaccctcggtacgaaattaatcgtacagcagattagcaat
ctgccgctttcgaccactcagccacctctccaaaaatttttatatactgaatatagcaaa
tattattaagttttaacagatatgcataaacatttatggagctaagcggattcgaaccgc
tgaccctctcaatgccattgagatgctctaccaactgagctataacccc---ttttttga
ggctattaagaaatggctcaagcgggatttgaacctgcgaccttgggcttatgaatcccc
tgctctaaccactgagctactgagccataatcaaaactattactaaatactataccatag
attactttaaaatataaatcaatgaaaaaatatttactacaagtgcctcgattttatatt
g----------------accaatt------------------------agcatttaaaga
tatccaaat-ttttttacgatatgctatcagttcttttcatttaat-taggagtatttca
atgttcaacttccaaagactttgtatcttgttatatatttaatagaattacactctttt

ttctgacataaactattaataataagtttcttattttgataaaaaatcttgttttctccc
agaaaaacaagattttgggtgttttaacgctaactgacgccagagaaaaaaatttttt
gccgctgcgtggcggggctttggtaacattaagccgcccccgcgcctgtaacaaatttat
tttgcaacatttttaacgctacaaacgccacactataaaaagttttcaaaaattacaag
tataagctcttatttagacttaattattccaaaaagaagggtcaaaaaattcgattaaa
tctgttaatagaatttttgtgaaaacctttaatatcatatgtttcaactatattatcc
gccaactttctctctctaatttgattcaacgctattagcaaagagctcaaataagcagaa
ttaattctaggcatagttctttgccttgtcagtttaacaattgcagatgaagttttattc
aacacctcactcaactctctactactgatatcaagattgtacataacatcttgaagcttc
actcctaaagctctaaatacggcattcttatctagttcaactttggttttggtacatttt
ttaggatacgttagaaataatttttagcatttattctaaattttttcttttgtgacgga
tttcttttttttctgacataaactattaataataagttttctattttgataaaaaattct
gtttttctgggaaaaaacagaattttggcattttagcgccagctgacgccggaaaaaa
aatttttttcgccgctgcgcggcggggctttggtagcattaagccccgcccacgctcgta
acaagtttttttctgaaatattttttaacgccacgaacactacactatgaaaagttttcaa
aaattacaagtataggctcttatttagacttaattatctgaaaagaaaggctcaaaaat
ttctattaaatctattgatagaatttttttgtgaaaactttttaatatcatacgtttcatc
catattactcttcaacttcttttatataactggattcaacgctattagccaagagctcaa
ataagcagaattaattctaggcatagtttttagccttttcagcttaacaattgcagataa
agttttatttaaagcctcacctagctctttactactaatatcaagattgtacataatatc
ttgaagcttccaaagcacctgaacagctgcttcaaccttcaatgaatccttattcctgtc
tattaagattgaataattatgtctctttaattagccaaaagcaaaggaaatatctaagta
aaatatcttac-----------------tttcttataaagac-----------aattta
tttaacagcatatt----------gtacaagcaaaattacatagtcaatatatatttctg
aaatcaagatgttgcaagtcttatttttatatatattaaatagacttaagggttatgaa
ttctgtaattaaaagacatttattgacttcg-----cttagtaatatttgattcttagca
ttgt-ttaaatacagattagttagaaactgaataaatagataatatc-----ttatacaa
ccagcatcgagccaatgccttgatcagttaatatttctagcaagagagcatgatcaattc
taccatctaaaatatgcgctgaagctactccttgagctaaagaacgaatacagcagttaa
cttttgggatcattccccgtaaatcacttctgtctgggttaagtctctagcttgctgta
tattcaagtgactaattagtgttgtaggatctgaagaattacgtaaaatgccaggagtat
ctgttagtaaaattaatttttcagcattaagggctgctgctatttctcccgccacagtat
cagcattaatattgtatgactgaccttgtttatccgctgcaacactagcaattacgggta
tatagttattattaattaagatcttcaatagcttagtatcaacattttgcacttctccaa
caaagccaagatttgctttaccacttggtctcggagtaatgagtaacccatctttaccag
acaagcctacacttttacctccttgcttattaatcgttgctacaagatctttattgactc

gtccaactaataccatttctacaatatccatagtaggctgatctgtcaccccttacaccat
tatcaaattgtggtaaaattttagacgatctagccagaaattaatttctggacctccac
catgaactaaaataggacgcaacccgataaaagaaagaaaaaccagatcactgattactt
gatcttttaattttgattttcatagctgctcccccgtatttatcacaataattcttg
aagaaaattgttgaatatacggtaaggcctcacttaatactttcactctttctgaattag
tcagcatgaataaaatagaattataatatgtaaatttatttgcagagcttatattaattt
aacagatactaacaagctattcaaaactttcagagaataaacctattattaataattaaa
atatttaaaaatataaggataataattatgagtaattttggaataatatattaaaattt
ccacgatttttagttagtgtcatttttggattagttttaataattattagtcctttttttc
gtgttatttaaaaaaccaatgacaagtttttttttcatcatatcattagccggtttaatc
gcagttttagcggtaataa-taaaaaaaaatgctcaatatcgaatgttgttgagaggatta
atttaatttaattaaaatttttgaacatatataatattatattatcattaactatgataa
tttaagtttcttttctata-aaaaaatcctacatattaaatattacttgatagatattat
gtcattaaaacaaatagttggttctgaaaaaactggtgcatttgctcttcttgatagtat
tgttagacatgggattaaacatatatttggttacccaggcggagctatccttcctatata
tgatgaactttatgtttgggaagaatcttccctaattaaacatattcttgttcgacatga
gcaaggagctgctcatgctgcagattcttattctagatcaacaggacaagttggtgtttg
ttttgctacatcaggtccgggagcaaccaatcttgtttctggtatagctacggcacatat
tgactctgtacccatactagctgtcactggtcaagttgggagagcatttataggtacaga
tgcctttcaagaagtagatattttttggtattactcttcctattgtgaagcattcttatgt
agttcgtgatcctagagatatggctagaattgttgcggaagcatttttttatttgcaaaca
tggtcgaccaggacctgttttgattgatgtccctaaagatgtaggcctagaaaagtttaa
ttatttttctattgatccagggaaagtgaatattccaggttgtagaccgattactacttt
gaagtctagacaaatacttatggcagctaagatgatacaacaatctagtcaacctttgct
gtacataggaggaggagcgataatctctgactctcaccaaattattaaagaacttgttga
tttatataaaatacctgttactactactctaatgggtaaaggtattttttaatgaagatag
tgaatattgcttgggaatgttgggtatgcatggtactgcatatgctaattttgcagttag
cgaatgtgacctcttaattgcattaggggctagatttgatgatagagtaacaggaaaact
agacgagtttgcttgcaatgctcaagtaattcatgtagatattgatcctgccgaagtcgg
aaagaatagaattcctcaagttgctattgttggtgatgttacagaagttgttactgcaat
cttaaatttactaaaaacttcttttcctccttatccggagcagattgtatcttggcaaga
gcgaattaaccgttggcgccaacagtatccattgctggttccaaaaaaaatcaacaagtat
ttcacctcaagaaatacttgttgcagctaataaattagctcaaaatgcttattttactac
agatgttggtcagcatcaaatgtggtctgctcaatttttaaaagtgaaggctaagcactg
gctttcaagtgctgggttaggtacgatgggatatggattgcctgcagcaattggtgctca
agtagcacatcctaatgatgtagttatttgtattagtggtgattctagttttcaaatgaa

tatgcaagaactaggaactattgcgcaatatcagttacctgtaaaaattattattattaa
taatcgttggcaaggaatggttagacaatggcagcaggctttttatggtgaaagatactc
tcattcacgcatgacagaaggagccccaaattttcaaaagcttgcagaagctttcgggat
taaagcttttactattaatagtaggcaaaatatgcaatcttcattagaagctgctattga
gcatcatggtccagttttgctagactgtcaagttacagaaaatgaaaattgctacccaat
ggttgctccaggtaaaagtaatgcacaaatgataggtattgctaaacctcaaagaggtac
tgcctcaaactatgttaatacaaatatctaaataga------------------------
------------------------------------------------------------
------------------------------------------------------------
------------------------------------------------------------
---------------aagattcgtattttctcaaaaatataatgaaaatctt----tttt
ctgagtatatatttctaacattatat-taaatttttagacgatagcctttgacgagaatt
gaactcgtgaccttccccttaccaaggggatgctctacctctgagccacaaaggcttttt
tgttatattgggccgggttggatttgaaccaacgtaggcgaagccagcggatttacagtc
cgcccccattaaccactcgggcaccgaccctaattaatttttatgtaaaatataacttaa
acaatatcataagttagttgcttaaaacaagtcttttcataaataaaact-agtaagtaa
tggacatagctggactcgaaccagcgactttcacgatgtcgacgtgacactctaaccagc
tgagttacatgtccaaaaccttaataattattataacactatttattgattgtataagta
tctatttctaattaaagatttgttttaagcgacctgctttaccgagaagatctcttagat
aatatagcttagctctgcgtactttagctcttctaataactttaacaaatgttatgcgtg
gtgaattaagaaaaaaaacgcgctcaactccaataccttgaaatgagcatcttaaagtta
aactagtatttaagctatttttttcttcttctggccaagattactccttcgcataattgtt
cccgggtttttttaccctctttaaccatcaatccaagttgtactgtgtcgccaactctaa
tttcaggcacgtcttgtttaataaaaggattctctacacttttcatgagcttacttatct
tggtattatttgctttcatatatagcattagcctcataatg-ttttttgcaatattatac
attaaaacaatcaatatgaaaattattagtaaataagcaaaataacatcgagatagtctt
-----------------------------------------------------------t
catctaaaaacc------------------ttataagcaactgttcattaca----gtgc
aaatctgggaaatttttaaaacaaatagtataaagtggcagtctaactcact--------
------------------------------------------------------------
------------------------------------------------------------
------------------------------------------------------------
------------------------------------------------------------
------------------------------------------------------------
------------------------------------------------------------
------------------------------------------------------------

-------------------------------agtatcacgtccttt------------tt
ctttta------------------------------------------------------
-------------------------------------------actcatagtattttatt
ttatcgcgac--------------------------------------------------
------------------------------------------------------------
------------------------------------------------------------
------------------------------------------------------------
------------------------------------------------------------
--------------taaaacttcagttgta------------------------------
------------------------------------------------------------
------------------------------------------------------------
--------------tttgagagttatattcaacactttgtt-------------------
------------------------------------------------------------
------------------------------------------------------------
------------------------------------------------------------
---------------------------taaaatttagcattatataa-------------
---tttaatttttttagtaaaagaatgagcattttg------------------------
------------------------------------------------------------
--------------------caattcggcaact---------------------------
------------------------------------------------------------
------------------------------------------------------------
------------------------------------------------------------
------------------------------------------------------------
--------------------ttgctatctttcacactcctgacttcatttt---------
------------------------------------------------------------
------------------------------cttcaataaagta-----------------
-----------------------------tttactatataactaaatga-----------
------------------------------------------------------------
-----------------------------------------ggtcat-------------
-----------------------------------------catagaaatcacta-----
--------------------------------------------------------gatt
ctataagattgtattcatcatatatt-------------gtatcgtgttacctg------
-------------tgatagtcatgtatct-------------------------------
--atttgtagagcgatcataacactcaaggaataacttcgttgcaggataaattt-----
------------------------------------------------------------
------------------------------------------------------------

-----actagctctctagcggtg-----------------------ttcacagtaagcaa
tgata------------------------------cttgccaataagat------gtaag
cctttactactaatgctcctttcgacatatttgtc-------------------------
------------------------------------------------------------
-------atgaaatttacttattatttccgaaattttagg--------------------
------------------------------------------------------------
------------------------------------------------------------
------------------------------------------------------------
-----------------------------------------agaatcatataat------
--------tttattaattaatag-------------------------------------
--------------gtgctctgtatttc--------------------------------
------------------------------------------------------------
----------------------------gacaatcactaaaac-----------------
------------------------------------------------------------
----------------------------------------------------------ta
ataatttctca-------------------------------------------------
----------------------------------------------------------tg
agttaactccatctatatcaataaaa--------------------------aagtacgg
tgttctagtgaa------------------------------------------------
------------------------------------------------------------
------------------------------------------------------------
------------------------------------------------------------
-gacaaatcctaatctaataataaca----------------------------------
------------------------------------------------------------
------------------------------------------------------------
------------------------------------------------------------
----------------------------------tcttctagctgcttatttaaatgaa-
------------------------------------------------------------
------------------------------------------------------------
------------------------------------------------------------
---------------------------------------------------------cat
agtccgacaagttaa------------atttagtacaaaa--------------------
------------------------------------------------------------
------------------------------------------------------------
------------------------------------------------------------
------------------------------------------------------------

------------------------------------------------------------
------------------------------------------------------------
------------------------------------------------------------
------------------------------------------------------------
------------------------------------------------------------
---tttcgttatataaaattat--------------------------------------
------------------------------------------------------------
------------------------------------------------------------
------------------------------------------------------------
--------------ttttcacata------------------------------------
------------------------------------------------------------
------------------------------------------------------------
------------------------------------------------------------
------------------------------------------------------------
---------------------------------------------------------ttc
gatggacgaattat----------------------------------------------
------gaactgtaatttactataatttaattcttttgtcctatta-ttgattggcactg
tta-------------ctatctttaaatatttattctccactcaagaattgtatacatac
aactgaatgatcaaaacaataaaattaaggcttaagtattgattagttttgtgttataat
tagttactatcaaaggtagtggcataacacaaagcataaacaatgtttgaacgctttact
gaaaaagctataaaagtaataatgctagctcaggaagaagctagacgcttaggacataac
tttgtcggaactgagcaaatactattaggattagtaggtgaaggtactgggattgcagcc
caagtcttaaaatctatgaacgtaaatttaaaagatgcaagagttgaagtagaaaaaatt
ataggaagaggatctggttttgtagcggttgaaattcctttcactcctcgagcaaaaaga
gtactagaattatctttagaagaagcacgtcaacttggccataattatattggtaccgaa
catttgttaatgggtttagtcagagaaggagaaggagttgcagcgagagtcttagaaaat
ttagcagttgatgtttcttcgattagggcagaagttatacaaatgctaggagaaaatgca
gaagctaacgtaagtggaagtaatactacacaggctagaagcaagacacctacattagaa
gagtttggctctaatctaactcagatggctatggaaggtggcttagatcctgtagttgga
agacaaaaagagatagaacgtgttattcaaattttagggcgaagaactaaaaacaatcct
gttctaattggcgaaccaggtgtaggtaaaacagcaattgcagaaggtttagctcaaaga
atagctaatagagatgttccttctatattagaagacaaattagttattactctcgatgtt
ggtttattagttgctggcactaaatatagaggtgaatttgaggaaagactaaaacgtatt
atggatgaaatcaaatcagctgataatgtaatactagttattgatgaagtccatacatta
atcggagctggggctgcagagggagcaatagatgcagccaatctactgaagcctgcttta
gcacgaggagaattgcagtgtataggcgcaacaaccttggaagaatataggaagcatata

gaaaaagatccagcattagagcgaagatttcagccggttgtagttggagagccaagtgtt
gaagaaacaatcgaaatcttatttggtcttcgagaccgttatgaaaaacatcatcaatta
acaatgtctgacggcgctttggctgctgctgctaaatatgctaaccaatatatttctgat
cgatttttgccagataaagcaattgatttaattgatgaagctggttctagggttcgttta
ctaaattctcaattacctccagctgcaagagaattagataaagagttaagagctgtattg
aaaacaaaagatgaagctattagggctcaaaagtatgaaacagcagagcaatatagagca
agagaaatggaaattaaagctcaaattgcagcaattgctcaaagcaaaaaaaaatgagcct
gatttaaacttagaagatcctgttgtgaccgaagatgatattgctgaaattgttgctgct
tggactggtataccagtaactaagttaactaaaagtgagtcagaaaaaattaatgcacatg
gaagaaactttacatgggcggatcattggccaagatgaagcagttgtagctgtatctaga
gctataagacgagcaagagtaggtctaaaaaatcctaatagaccaattgcaagcttcatt
ttctctggaccaacaggtgtaggtaaaacagaattaacaaaagctttggcttcttatttt
tttgggtctgaagcttctatgatacgtttagatatgtcagaatatatggaaagacatact
gtatctaaactaattggttctcctcccggctatgtaggttatagtgaaggtgggtatctg
acagaagcagtaagaaaaaaaccatatactgtaattctttttgatgaaattgaaaaagct
catcctgatattttaatcttcttcttcaaattttagaagatggcagattaacagatgca
aaaggtcgaactattgattttaagaataccttctaataatgacttccaatattggatct
aaagtcattgaaaaaggaggaggtagtctaggattcgagttatcagaagatcaaacagaa
tctcaatatactagagtcaggtctttagtaaatgaagaattaaaacaatactttagacca
gagtttttaaaccgcttagatgaaattatcgtatttcgtcagcttactaaagatgaagtg
agagaaattgcagaactgatgttgaatgaagtttttgccagaattaagcaacaagatatt
cagctaaatgtaacagaacgttttaagcagcgactagtagaagaaggttacaatccaagt
tatggagctcgaccacttagacgagcagtgatgaggttattagaagatagtttagccgaa
gaagttttgtcaggtaaaattaaagctggtgatagtgctgtagtagatgttactaatgaa
ggagaggtgaccgttttattaggtgaaaaactagaactactaacttaaa------aaata
atttgttaatcggaaagcattgaataaactat--attcaatgctttt-agaagaaaatta
ataagatgggttgctataattgacaacctctgatattctttatatgtatata---aagcc
gggatagctcagttggtagagcagtggattgaaaatcctcgtgtcaccagttcaaatctg
gttcttggcat-tttaaacgtttgaaaagtttgctagatcagcatcaaattttagttgga
atatacctgtagggccatttctatgttttgctattataatttctgtcagatctcctccta
cagtttcttgagtgtagtaactttctctatacagcattataaccaaatctgcatcttgct
ctatagaattatgtactataaagttactcgcgataaaattacgtaagtgctgacagtcta
aatcgaatactgtttgtaagttaataacacttatgtgtttaagattttcaaaaactaaaa
ataaagaaggaaaagaattgcagatagactttttttctttaatagtatt------cattt
gaatagcgagcatattatcctgatcaatttcatcacatcttttccatccttgcgttgtta
atagcttgtgattgcttgttaaatgtatatactttccagattgagtaataatttcatata

tagtctttttttcctgtttttgaaatactagctttactagttttaaataagtttgattttt
gtacagtaaagctagtaacttca-atacgcttatgatgaaaattcaatagaatttg--ag
tttgaccatgtaacggtacatgaagatgactaaattttgatatacacccactttctctta
aatcagacaataaaggtcttttatcacttcgagtttcaagattcctattcagttgagata
atactagtataggtaaatttaagtcttttgctaatatttttagtgatcgagtaattaatg
atagttcttgagatctattatcagattgcctgctatcttgtaataattgaagatagtcta
taataattagttctatatttttttccttgtaattttaagaattgtacttttgtttttatag
tgtccgtagaaattttttgcactatcatcaatgtaaagattgagatctgcaagagctttac
ttctttcaataacatattgccattcatcattattaagttgacctgattgaatttttttgac
cgtttaagtgacattcttgagctaaaattcttcttagtagttgttctatcgacatttcta
agctaaataaaataacgaacgattcctctttgttaattatatatctagctacattaatag
caaatgccgttttgcccattgaaggtcttcctgcaagaataattaaatcggacttttgaa
aaccttgtgtaatagaatccagttctgtaaaaccggaaagtacgctactatttatactaa
tttttttgttgtgtctaaatgaacaagtagattagcaagaattgtagctagtgtctgag
tatctttctcttcaaggatttcgtaagcttttgttaattgactaacaatagaggttatat
tattttgcttaacaggctggtttgaacagctaattaaaaataatgaatctccagatatta
gaagtaatcttttaatataattgtcgataatgattgatgaatattcatctacgatatcag
ataaaggtgcttgttctattagactaagaacttcgtttagttcactcaagtgcctgagta
ttttctctgtttttaagttaactaaaaaatttcctatacttatcttttttgtatgattga
tattttctagggcagctttgtaaagtaatgaatttgatgtaaaataaaaaaaaatcaggag
agagcttatctactgattttagtaaagagattgattgcttagttaatattgtagtaatta
atatttttcagctaaaatattatgaggaggcaaatatttgtaa------atactgataa
gttcctcc-ttgtaacattattctaaca--------atactaattaattactaacttaat
ttgattctggaagaacttgtagttgaacgttggcagttacctgacttaaaagtttaatct
ctatattataaattccaactgtgttaatctcaggtaaaaagatattttgtttttcaatgt
ctatatttgtagcatttttaataatttgtgcaatctctttttctgtgacactaccaaaaa
tatgttgcccatctcctgtttttttacttacgctgaacttttgtatttcttctagaagtt
gttgaatttttttttgcatcttccttagctattttttaattttttcttcctttatggctgcgt
aaaatttttgctgttttaaaataccatttgttgcgattatcgccattttatttggaatta
aaaaatttcttgcataaccagcagcaacttttacaagatcattacttttaccaagttttt
gaatattctcttttaatacaacttttataactttttttacccatctttactctgaattcta
aaaa----atctatataatttattatacctgttgtgcccatattttgacagacagaaag
tattcacttgttgccataatatctgtgtaaagcggataatgaattttcatctttattctg
gtataataagtcgtacaacattat-----gttaggagaggtggccgagtggttgaaggcg
cagcattggaaatgctgtttaggagcaatcttaacgagggttcgaatccctctctctccg
tacaa--aaatgattataatacttatatagtccagtcaaaatttccaataatgtttctcc

atgcatttaaaccaccttgtactctcacaatatttagtttttgagctattagcaactgag
aagcaaataaagatctagaatctaagctgcagtacacaaagcaaactttattttctaaat
ttatattagagtaatgtatttttttgagttgttctaaaggtatatttaatgcgtgagtta
agtgatttttttatattcgtcttgatttctgacatcgattaaaatatatttaggtctt
tgcgatttaaaagtttttgcaattgcacaacattaatctctcttataataagattaagtt
tatcaaagctgtatttgttccagtgttgtttttt-------agataatacaaatttagta
tttaaaattttaaacttgctaaatgataaagttaaggcgttgtaggttaatatcatacca
cttaatatacacttatagccaagaataattttaatagcttcagttgcttgaagtgtacca
ataatacctggcaatagtcctaaaacaccagaattgctacatgtatctatttcactatct
tcatttttttactttattgtaaaagtctcgataacttggacctccttgatagttaaataca
ctaatttgtccttcaaattgaaaaatagctccataaatatgtattttattcaactcaagg
caagtgtcattcagcaggtatcgagtgttgaaattgtctgatccatctataataatatcg
tatttttttaacgatgtcaattgagttttgagaggttagtcttgttgcaaaaacttttact
ttacatttcggatttaaatctaata-tttttttttagctatttccacttttgaatatcct
atatcgtgaattgtatatataatctgtctctgtaaattagaaagatcaataatgtcgtcg
tctataataccgatgctgccaactccagcagcagcaaggtaaattatgccaggagaaccc
aggcctccggcaccaacgaataacaccttcgcttttttttaatctttcttgtccctctaat
tgaacttgtggtaacaataagtgtttagaatatcttgtatattcgtctaaagagtatttt
atattctctactttaaagttcagcataaaaatttataagttaattaataccgacta----
--gaaactaattgataagcttattaagatgacttttgtaaactaggttcttctgtatcca
aaatagcgccttgtgtaggacagacttgaatacaaatactacagtcaatacaggcagaaa
agtcaatccagtaccaattagtgttaatactatttttttccttgtccctggtgaatacaag
ctactggacaagcggctacacattcagcaactccaatgcatttttcggttacaattgtat
gagacatgttatttaattaat-aaaagataaactgtttacatctagtagttttattgtat
actagattggcatatgctatccaagtcaataattaaagtaatctacttttatagcttact
aatttgatctgcgcctttagttcagttggtagaacgcaggtctccaaaacctgatgtcga
gggttcaagtccttcagggcgcgcttagttgtttttacagtctcatagtagcattttatc
tttttgcaagcatcaaaataatatgatataaaacatcatattcaatatattggatatttt
atgtgctaggactg-ttttttttatatgttattattggtaagttgattatattaattccat
ttgtataatcaat-aaaaataactgaaattaaatgcagatgtactcaatagtc-tttttt
atatataaaacttgaaaaattaaaaactatggctaaaaaagttaccggaattgttaaatt
ggcattaaacgcagggaaagctactcctgcaccacctgtagggcctgctttaggtcagca
tggcgtaaatattgttatgttctgtaaagagtataatgcgcgtactgcagataaatcagg
actggtaattcctgtcgaaatttcaatatatgaagacagaagctttacttttatacttaa
aactccgccggcctctgtactaattgctaaagctgctggcttaaataaaggttcgggagc
accaaacacaaaaaaagtaggtagtataacatataagcagttagaagctattgcagaaac

gaaactacctgatttaaatactaacaatataccccaagctatgaaaatagttggaggtac
agctaaaaatatgggaattctaattaaagattaaaaaatttcaattatatttatta-tt-
-tttatgaaaaaaatttcacgccgacttaaaatattaaaatctcaagttgaacctaaact
tcatactattgatgaagcaatatctatagtaaaagcaacatcaaatgctaaatttaaaga
aactgcagaagcacatatttgtttaggtttaaaccctaaatacgcagatcagcagcttag
agcaacagtcatattaccaaaaggaacaggaaaattaataaaagtagcagttattgctaa
aggagaaaaattaacagaagcaattagtgcaggagctgatgtaagtggttcagaagacct
aatcgatgaaatactaaaaggtagattagattttgataaattaatagcaactccagacgt
tatgcctctgatagcaaagcttggccgtgttttaggacccagaggattaatgccttcccc
taaagccggcaccgttaccacagaagtgggcaaggctgtaaatgaatttaaaggcggaaa
agtcgaatatcgagttgatagaactggcataattcatgtaccttttggaaagtcaagttt
tcccaaagaagatttagtcttaaatcttcaaacaattaaagaatcaattgataggaataa
gccttctggtgcaaaaggaaagtactggaaaactttttttcttatctagtactatgggacc
gtctatccaaatcgatattaatagtcttttatgaaaact-tttgtgtaattacataactt
gaataatatattatttcaaatta-taatttattagcaatgttaataaaatagtctgctta
ttatatcttatctgcgctaaaaattttaatatatatcttactaattatgagtacaaaggt
tgaaaatatcttagaagaactaaaatctttaaatcttctagaagctgctgaactagtaaa
acaaatagaaaaaacattcgatgttgatgcatctgcttcttcaggaggtatgatgatgtc
tgcgccagctgtatcaccagctgcatctgaagtcgaagagaaaacggaatttgatgttgt
tctagaagaagtgcctgcacctaaaaagatttctgtcctaaaagttgttcgttctctgac
cggcttaggcttgaaagaagcaaaagatcttgttgaatccacgcctaaagttctgaaaga
aggtgcgtctaaagatgactcagaagctatgaaaaaacagttagaagatgctggtgcaac
ggttgttgttaagtaataca-t--aaaagtgcttttgtttgtaaataaacaaaagcac-t
ttttatttaatagaacaaattagactagaaaagcccaagtgtaattgcattattaatagg
catagttgcaccaatacctaaccaaatactgacaactgtgccaattaagaaaacagttgt
tgcgattggtcttctaaaaggattttgaaacttgttgacgttttcaataaatggaacagt
aagcaatcctgctggtacagctgccatacttagaacccctaatagcttattcggaatcac
tcttaacagattaaatgttggaaagaaataccattctggtaaaatttccaatggagtagc
aaatggattagacttttcacctatagacgaaggctctaaaattgctagtccaatactaca
tgcaaaagttccaataattacaactggaaagacgtataataggtcatttggccaagctgg
ttcaccataataattatgacccattccttttgctaatttagctcttaattttggatctgt
taaatcaggtttttttaagaattgacatgttattttttgtgtttaatataaaattgagttt
aattacttgatcaaagtgtaactatacattgttacagtggtccggaaattccttgcttgc
gtatcattaaaaaatgcataagcatgaaaactgcagttagtaatggaagtacaaaagtat
gtaaactgtagaatcttgttaaagttccttgtcctacactgactcctcctcttaataatt
ctacgatgctccctccgacaactggaacagcatctggaacacctgtaacaatttttacag

cccaatatccaatttgatcccatggtaaagaataacctgttacaccaaatgaaacagtaa
gtactcctaaaattacacctgttacccatgtcaattctctaggctttttaaagccacctg
ttaaatagacacggaatacatgtaaaatcatcatcaagaccatcatactagctgaccatc
tatgaattgatctaataagccatccaaagtttacatcagtcataatatactctactgacg
tgaaagcttctgcaactgtaggtctatagtagaatgtcattgcaaatccagttgcaactt
gaattagaaaagatacaaatacaatcccgcctaagcaataaaaaatattaacatgaggtg
gtacatatttgctagaaatatcatcggcaattgcttgaatttctagtctttcttcaaacc
agtcataaattttactcataaaatagcttcaaaaagctttttaatata-tttttgcggt
taagctac-tttgctttagtag------ttattgaacttatatt-attttattgtccata
cttactaattattataacacctttttatttattt--------------------aaaatt
ctttgactcaataaaacaggactatgaactacgatctttttttttctgcatagagattaat
ttggttttcagtaatccagataaaattcttgtcacacttactctactacttcctataatt
tgtgccaacattttgtgtgtcatagcaaaattaaggataataccattattactttgcgtt
ccattttgctctgctaataataataataagcttacaagtctgcttgtaatacttttatgt
gaaatgatttctgtaaaatgataagctttttctgagcaatatattaaatgatttacaaaa
aacaaattaaatggcgggtagttactgcatgctcttataatagtattatagttaatacaa
gcaattttagttttacctattgcttctgcttcatagtaaaagttgtcattagctaattct
aattgtccgaatgtatctccagtagttaataaattaagtgttattttatgggtactttta
aagacttttgtaataattaaagaacctattaatataatatatagtctagaactgctatca
aacaaaataatatcattgggcttaagtaaaaaaacttgataactaattttattactatta
aaaaatagtagccatggattataagaaaaaaatttccccgaactactttctgaaaatatt
gaaatagatttttttggatttaaatgatttttagtcataattactactttataa------
---aaaaatactctaataat-taataacggaaattaaatgtcatacaagttgatgctcgt
tgagaatgacattgtattatcaaaagccattcaagaatacttaacagatcaaggatttaa
cgtttacactgccaaaaatggattagaagcatttgagttagctaatcaatataatttcaa
tttgataatttctgatataatgatgccactagttaatggttatgaattattagaaaaact
tcaaaatactaagtctttatcaaaaataccagttatttttttgacagctaaaggaatgac
taaagatagaataaaaggttatgacatgggatgttatggttatttgtctaaaccatttga
cccagaagagctagtttcacttataaaaaatttaattgctagagatgtattgcaaaatag
atcgatagaaaaaggtaacatcccaaagcattcaattaacaataaaatacatttaacgcc
tagagaagcaagtattttagaccttgtgattgatggactaactaataaagaaatatctat
catattagatactagtatcagaaatgtagaaaaatatgttagtcgacttttacaaaaaac
taatatgaagaatagaaccttactagtgaaatattctataaataataatctaatagataa
ttcagttaatcaaagggcgaatgacgggactcgaacccgcgaatgatggagccacaaccc
attgccttaaccccttggccacactcgccatatcataaatatcactataacttgtttgct
gacatatcgtctactctctaagtggtttatttatctaacatagtatctaaatttaatcct

ttatgatgcataataatattagtattcaaattaatggagagccatttaattgctcagagc
ctatctcattgcaattttattaaattatcttgatttaattctgggcgtgtagcagttg
agctaaacgatactttattacctgagtctcttttcactcaacttacttgaatgatcaag
ataaactagaagttattactattgtcggcggtggataaattctatcaactaa--------
---------ttatttagatgcttcattgatagagataaacgcccttgtttcttatccac
atgaataataacagctttaatagtatcgcctatttaaattgtgatgatatctggtctaa
ttgttttatattaatttcagagatatgcacaagccctttaagattgccaactcttataaa
taatccgtaaggtgtaatctgatttataattccttcgataatattgccaacaattaaatt
tgatgaggcttgggctattaaagctcttctgtggcttagtattaaattattagatttctc
ttcaacattgagcagctttagcttgataaattcattattagcttgatcatttttttgaaa
attaccaagatgagaattgggcacaaatcctgatataccttcaaggctaatgatcatccc
tcctttattaaatcctttgattctaacattaagtaaagaatcttcagctaataattgtct
gattcttttccaagctcttatatattcgagacgtcgaattgataaaattaattgctttga
ttgaatattataatctagtaaaaaaaattctcttgtatcattaatgtttaaagaagtaaa
attatttaaatcttgattacttgagacctcttgaatcggtaagtacgccgatattggtgt
tcctatatcaaccaacacgccatttagctcaaagctaaatatagtgccagctacaatatc
gccaagatttaaatcatacttatatttttgcaaaacagcagcaaaatttctgtgtgtaaa
tccttcattatttttgtcatattgaatagtctgaattaaagattaattttgtacagcat
attaagttaaataaacttttattgcacgatatgaaaatttttttaatttgctctgttgtca
ttccttatttaaacgtattagaatatacggaagacatgaaaagtaaacgtagatagctac
agatttaa-taatacacagtctgaggaaagtccgggctccataaatacaatttatgctgg
agaaagcccagtgtaagaaattatgaggatagtaccacagaaataaaccaccagcgtaag
g----tatattagctggtaagggtgcaaaggcaagttaaaagcttaccaaaaatactgta
aagtatttgttaggtaaacccctaaaatggagcaaagctactaaacaaatatttgtgtat
attttgtatagtttagtttaaaatactgcatgaagttattggtaacaataactctagagg
aatagctatcctttacaatatatc-tatattatattgaaagaacagaacccggcttatgt
agtactttttatgtcttatatttctctggttattgccgaatctctaaatttaagtgttgt
ataatttttttttgtagctcatgttgtttattatctatatctaaattagttaaagttttt
tgattagagcgatatgtaaatctcaaacctatactttttttctgttctccaagggattta
tttatatattgatcaaataaccttgtgctttctaaatctttatcattaaacttatctaat
aattttaataagtaattaacttccatatttttaggtattattaatgaaagatcccttata
atagatggatatttggagtaaggtcgaatttgataatttaaataattaaattgattatca
gaatcttctaaaatgcttaaattaatttcaagaacaaaaatttttgtagtaacacccaaa
ttactataagtcaccttgttgagttcactgaatatgccaatattagtattattataaagt
aatgttgcagatctattcctatgaattaaatttactctactatcaaccacttcttgtttt
ttccattcaatttgtctattcatttttttgaaaaaaattctccacgattcctttagcctca

taccaattcaacgagtgggctggttgtgaccatgtagatctaatatctaaattaccacct
aatatcatagctagattagttgtttcgacaatttcgtcttcaacagtattaaaaactgtt
cctatttcaaaactatctatagtttgattactttgtttaagattataaagattagattca
attaatccttcgagtaaactatagcgtaaattagaatggtcttggataagaggattattg
agactaatctctccgagggatttaactaaagaataatgaactaactcagttaaccctaaa
tttcttaaaatatttctactttatctataaatttctttttgcgatactttttacta
aattgaacggttggcactacgcttttaaatttatgatagccataaattcgtgcaatttct
tcaattatatctatttctcgaaaaacatcttgtcttctataatttggaacactaattctt
atattttcttgatctttatgaataatagtaaaatttagagagtaaagaatatcttcaatt
tcttgaaagtataagaaacgcacctgaccattattattgattggccctaaaatatcttga
actttttaagtgatacatctatatttagaatctttttattgttttctctgtggagaat
gtttctcgaatgttcccaccagctaagtctattatcaaagacatagtctcaagataagca
tttttccaattatctgtatttagtcctcgttcttgcctaactgaactttctgtcctgata
ttaagatctcgggaagattgtctcactagagactgtttaaatatagcagattctataaat
atagacgttgtagtagaacttgtatcaaagatgttgttggaccctacaccagctacgcta
attacatctgcatttatttgagtcactaaaatatttttatttagcttaatattttcatta
tttaactgaattttagtatttttccatcataaaagttgctttgaattgttatgttatgt
tcagg---------agtaagtttattaaaatcaataatattaataggttgtccccatttc
agcataatataattgcttatatccactaataaatttttattcgtaaagccgcaggattgt
aggcgatttttagccaatttggagaatctgtaatttcaattttatcaataattgcagaa
aaatagttattacaatgaatcaagctatgattggtaagtatatt-ttgtttttagagaa
gtgcttaccactaggttgatttatagtttgcaacattgaagatccagttatagccaagac
ttctcttgaaagacctatcatgcttaatacgtctgatctattagctgttgatgagatatc
taatatatgatcaatttgattatctactataatagattcaacagcctctacttcaaaacc
tgcctgcgtcaattgattagttaaattatctagatctattgtttctagattagcaagttc
tttcaaccaatttaaagaaactttcataattaattgccatctatcttatatatatattta
-tttacttgtgactaaaaactaatttaaaaaattcatcaaattaatataagttatttgat
gaatttgcttatacaatatattattgtgctatcttacttttggatggtagtaattaagaa
ctagcttttgtaaaagttagcccgttatcttgagcatacctttccataaatctcataaat
ctatcccaatcatcaggacttttaattacatgtacagcttctattgcttgcggtttacca
ttaataaacttagcattaacatctctggtaataatttgtccctctgtgtccattagatac
atacctgttatttcacctttgtctgccatactagcatctaaaattttgggatttgtaaat
ctgaaagtagcggttcccgtactaccatctctagatcttgttaaacgaacatctggcaca
acttcttcgttgatgccttgaataaattgaattgttgccatatcaatttt-tttatttgt
atataatgtatattggataaaactgatttctaattctttaatgcttgttgaaaaactgtt
ttaaaagcttatagtagcagtagatcaaattttatgtattatataaaactgtaactttc

-----ttg----------------------------------------------------
------------------------------------------------------------
------------------------------------------------------------
------------------------------------------------------------
------------------------------------------------------------
----------------------atctgttcaaagaactaacttattataagctaatcata
ctaatattagccatataatgttctgaagtagaaattag-gggttgtagctcagatggata
gagcaagcgcctcctaagcgctaggccagtggttcaagtccgctcaaccctgttaataaa
caaagtt---ttgctaaattttaccgataaatcagttgaaaaaagattaagagcacaaaa
cagg-----aaacagattctaaatatcacttttttattgaaa---gaaaagcgatcttaac
tctgacaatagaagaattttttg-aaaaatatactaagtttcaatgtttatatttagaag
aaa-tttttaaaaacgtaaacagaaa-----ta----ttttgagtaatactgaaaattaa
attatttaatttcttgaaattaatttaaaaactttatatgaaatgaactatctttt---
--ctctttattcagtatttgtctacacaattatta----attaattcaaaattaatcaaa
tggcttttgggttgaagatgaatttttaaactatagcaatagatcagtaactcgattatt
attctatatagattctgtttagaatttagtttgctctttaaaaaagaagcacttattatt
tgctagatagaaattcctaaagataatatttacaatgaatgtccgtgttatttaactaca
ttccatactttatagtaaattttgccaatttacgagag-------------ttaattcta
caacat-cttggttcctggaagttgaattttat---ctttttacatttctcttttcgtc
tattacgagaaatcatatatctaaatttaatgataatagtaagaatcatggatttcttaa
ctatattactatccttttacctttattcctattgctaatcacagcagcag----------
---ttatttagtcaattctaatgctagaaatttaatacaagccagcgataattctttaaa
tttgttgttgttatcgaagttgttaacataataaccattaagattcaatgatttgttagt
tgatattctgcatctgcaattcttgacataatactgttagatagttgtaagtagttttat
tattgatataattgatattatcttgattgtatattaatttgata----------------
---------gaaaaaacgagtatttaagtttcctgctttcatgtcaaaaaatatgtttag
ttaacgatgaattttactagcaagccgattagtaactttatcatatgtaaaaataaagtt
tgtatattctagaatattctcta------tatattgcggcttgcaaaacttctcagagtt
gatttacagagccagtgggctaaaag-taacaaaatggtaattccacgaaaattttgtac
cagtgatatgacttctaataaacacttaagagtaaacaattcttaatttttgtattaaga
agcttcatagtattattaattgttagaactggggtgggaggattcgaacctgcgaatggc
ggagtcaaagtccgctgccttaccacttggctacaccccaacttatagaaactataatag
cagtcacataccctaccatgtcaagagactattctccttatttaataactttttagactt
ttgtaaaatagaattttctttttcaaataagatgtactatttttaaactgtatcattctc
atgtcttcttgcttctgattcgacatccatttgatggtaactgtttcttgttcaacttct
ttattgcctataataatacaagctacagctttttttcttattggcttgtttgatttgttta

ctaaagttacttgagctaatatctatctctatcgtaaaaaattgttgacgtaaaaatcgc
attattctcataccaacctcattagctttttaccctgtgtagcaatgtaaaaatcaata
tacttatttggtaactctatttttctttagcaaggagtagtaatcgctctaatcctatt
gcacatccaacagctggtgttgctgcgcctcctaactgctgcactaggctatcatatctt
cctccaccacacactgtatcttgaccttttgatgttaatgtcttaatttcaaaggcagta
tcattataatagtctaatcctctaactaacttatcatttatgttataaggaatatcaagt
aaattcaaataatgacaaacatcatcaaaatatttttgtgacgaaagacttaaaaaatca
gaaatcttgggagcctcagttagtatttttgtgtattagtatttttgaatctagaatc
ctaataggattactaattagtctcttctgtgaatcaggatctaattcatcatagtactga
ttaagatattcttgtaattttgcttgataaatactacgatcttctaccttgccaattgaa
ttaagatcaagcttcaagtctgtgagattaagactcttaaatatacttgtagccaaatgt
attacttcactatctgctcttgcatctaggctaccaagaaattcaataccaagttggtga
aattgtcgttgtcgtccgctttgtggtctttcatatctaaacattggcccgctataccat
aatctctgtaacctgtgatgatagctcatttatttcaatgaaagctcttacaatgctt
gcagtaccttcaggtctcaatgtaatatctctattgctgcgatcgttaaatctgtacatt
tctttatttacaatatccgtatctctaccaatacttctatcatataaatcgctgttttca
aaaataggtgttctgatttcttgatagtttgcacattctaataagttggaaattttgttg
tgtatgaactgccaatattgaagctcttcgggcagaatgtcttttgttcctctaatagct
tgaatttttgccatgataattttcggatcaccttcttaatgtgtctgcactaaatagtag
ttgtagactattattcagtcatgtattatacattag--aacatgttgatg-tattttata
aaatacgggcaaggagggattcgaaccccccgacaccatggttcgtagccatgtgctctaa
tccactgagctacaagcccacttaacaagttacctaactatatcactgttctttcatgta
tacaactatattttgtct------atttatatttgaaagttttgtcaaagtaaaaaaat
tt----atttatatttaatagttaaagttagtcatttataattcctcatttattacaact
ttgtaca--aatatcggcaaacttaattatgaaattatcttatactgataatttcctcat
aa-ctcgtgcttttactttttgatcga------ctagtatctaattaatgtatatttcaa
at-aaaaaatcttacctaaaatgagcaagcaaattctatatcaagatgatgcacgtaaag
cattagaaaaaggcatggacattttaaccgaagctgtttctgttacattaggacctaagg
gaagaaatgttgtcttagaaaaaaaatttggagctcctcaaattattaacgatggtgtaa
ccatcgcaaaagaaattagtctagaaaatcatattgaaaatactggagttgcacttattc
gacaagctgcatctaaaacaaatgacgttgctggtgatggaacaacaacagctacagtac
tggcatcagcaattgtcaaacaaggcatgagaaacgttgctgcaggttcaaatcctatgg
ctattaaaaaaggcatagaaaaagcaacaaattttgtagtgagcaaaatagctgaatatg
ctaaaccagtagaagataaaaaagcaattatacaagttgcttctctatcatcaggcaatg
atatggaagtagggaaaatgatagcgaatgctatagataaagtcggccgagaaggcgtaa
tttctttagaagaaggcaagtcaactaatacggttcttgaaattacagaaggaatgcagt

ttgaaaaaggctttatttctccgtactttgtgaccgacaccgagcgaatggaagtccttc
aagaaaatccatttattctatttacagacaagaaaattactttagtgcagcaagaacttg
ttccacttcttgaacaaatagcaaaaacatctcgccctttattaataatagctgaagata
tagagaaagaagcattagccacgattgtagttaacaaattaagaggtattttaaatgttg
tcgcggttagggctcctggatttggtgacagaagaaaatctttacttgaagatatgagta
ttttaactaatgggcaagtcgtcactgaagatgcaggtttatcactggatacagttcaac
tagatatgctaggaaaagctcgcagagttattataactaaagattcaaccacaattattg
cagatggccatgaaattaaagtaaaatcaagatgtgaacaaattaagcgacaaatagaga
caagcgattctctttatgaaagggaaaaattgcaagaacgattagcaaagctttctggtg
gtgtagctgttataaaagttggtgcagctactgaaacagagatgaaagataaaaaattaa
ggttagaagatgcaattaatgcaacaaaagctgctattgaagaaggaattgttccaggag
ggggagctactaatgtccacatctctagcgagttatttacatgggcgaaaaacaatttag
ttgaagacgaattaatcggagctctaatagttgaaagagcaattacttatcctttaagac
gtatagcttttaatgcaggagataatggagcagtagcggtggagaaagttaaaagtaatg
attttcatataggctataatgccgcaaatggcgcaattgtaaacatgtatgatgcaggta
ttatagatcctgcaaaagttgctagatcagctctgcaaaatgcagcctctattgcagcaa
tggttttaactacagaatgtattgttgttgataaagctaatgactaagttt---------
-----taaataatattaaacaaagaataatttctatctataggcattattctttgttcta
attaagttaattaattgagcaaatgtataatttttataataaagccaaagcttaaatgtg
cctctttgctcaactattaagtagaggagatacaagcctgtaattcctaaattatcaatt
aaagcagtattattatctatcgatttctcaaaagatctaaaaaatgacattttttgtaaaa
tagtatcggaatcttcttttgtagttatttgataaatcattactacatttagattgatct
gcaactgcatttaaaacttttaaaataatgctttttaagtctttatctgactgtatctta
gaatcgctaataaattgagttaattgaatatcaaaactacaattaagaaaattactggcg
acagatgtctgattttgaatatctatagatttgatagccaagaagattaaatttttattg
ttcattgtaaaatgtaagacaattaatgttgtattctttaaatacta-atttatcttct
tgtgtatttatagttttggataaaaataattaatattatta--------tattaattact
tcccgaaaaaataaattcaggaaatctagcctgagcctttgatagtgagtgtttttacc
ttcttcttgccagtaattaataacttctgttgttttatttaatagctcaatcctatctct
ttcagcaatccatggcttagattctagttcggcttttaattcttcccaagcatcatttct
tggccagaaaaaataagaagtaagtggacttgtaccatttccaacaatttgatcaattgc
aatagcaatattgttttctaaccaaagtacctttagtgtaaattttgacaagtgattatc
cccattataaatagtaaatatatgtatatacagtaaaatgctgtattagttgttctcttt
agcaacaatctgagctttatttaaaatattttgtaccgttttagtagcttgtgcaccttg
tgctaatctctgtaaaatttttgaaatgtcaatacgagtttcattagtgattggattata
aaatcctaattcttctatagctttgccatctcgtttatttctgctatccattacgacaat

tctgtagcttggttgtttttttcttccatatctttttaatctcaacttaaccatcttagt
caccccgcttattttttt--atatctttattaaattatatcttacgatattatgcaataat
tcgaacaattaagcagactcaagcctgacattattaaatattaccattaaacactggaga
aaaataatacctagcataggtgacatgtccattccaaacattggaggaatactgcctcta
aagagctttaaatatggatcagtaattcggtttaaggaacagaatggttcattataccag
ttcacagttggaaaccatgccaatgacaatttaagtaaaattaaaattaaataaatctca
gaaaaattagctattgatccaagtaataaatttaatgtaccaggaagagtattcatacgt
atttttataattattatatagttcattatactaatagtttgagtatatatcataagctcta
taaataaat-tagctaaacaaaaaacttattcttaaaaatatcttctcttgaacaatctt
cttcattgatagtactgttaactttttgttgtataaacttctaatttagaccaattatgtc
ctaactgatttagctgagatgttccgcattctaaacatgctatttttacattatttaaaa
taacaccttttttttactaaatcttttaaaactggtgtaataattgtttcatctaaaaata
aatctaagattagaacttttgtaaatgaatttaactgatctgggacattattagtaatat
tattgccattaactaaagctatactagctgtaggaagcaaagctcgcgctccttctgcta
gtataaatccataaggcataataatgacaataatatacttataatgactgttaataagat
ttatattgatattagtttttaattcaatagtttctgttactaaccattctctcattattt
cataagtaatccacttacctaattctgagcaagcagtccttaaaatcgtacttggattat
tgttattttcgagaataccagcccaatgttgtattagagggtgcgaagctatattaattt
tgagttgcatgttttgttaaatatatttaacatttaatttgttaattgtgtctttggttt
ttgttcactgagtattattaaaatagcat----atatattaaagaaaacaattat-ataa
aatgaaaaaaaactctttggctttggggtttcacagatagtgctgaaacttggaatggtag
atttgcaatgattggttttatagcagttatctttatcgaaatattgacgggacaaggact
attatatttaataggtgttatctaataataaaaaatcaaaaaagttcttgtaatacatta
caagaactttttttga-atgaatataaatctaatcgagcggtctcatagatagtactggtt
cattttctctattaatacctttttcaaatccagcagcagcagctcttgctctaccagcat
gccaaagatgtccaataaataggaaaaatcctaagaagaaatgagatgttgttaaccaag
atctaggagatacatagttaactgagttaatttctgtagctacaccgccaacagagttca
aagaacctagtggcgcatgagtcatatattctgctgcacgtctttcttgccaaggttgaa
tatcattttttaattttattcaaatcaagaccatttggtcctctaagaggttcaacccaag
gtgcacgcaagtcccaaaatctcatagtttcaccaccaaaaataatctctccactaggag
atctcattaagtatttgcccaagccagtaggtccttgagaagacgcaacattagcaccta
atctttgatctctaactaagaaagtaaaagcttgagcttgtgaagcttcaggaccagtag
gtccataaaattcacttggataagctgtattattgtaccaaacaaaattagaagctgtaa
ggcccatgattgataaagcacccaaactgtaagataagtatgcttcaccagaccatacaa
aagctcttctagcccaagcaaaaggttttgtaagtatgtgccaaattcctccagcaatac
aaataatacctatccaaacatgaccaccaacaaggtcttccatgttatttacactaacga

tccatccatcgccgccaaatggagatttcaagacgtatccgaaaataactaaaggattaa
gagtaggattgctgacaaatctaacatctccaccacctggagcccaagtatcatacactc
caccaatgaacagagctttaatcactaataagaaagctccaatgcctagtaacactaagt
gtataccaagtatggttgtcattttatttttatctctccaatcatatccaaagaaaggga
aagattcttcaagagtatcaggtccgattagtgaatgataaagaccaccaaatccaagaa
cggctgaagaaattaaatgcactacacctactacaaagtatgggtatgtattaaaaattt
cgccgcctggacccacaccccagcctagtgtagctaagtgcggaattagaataaatcctt
gctcatacagaggcttttctggtacaaagtgagctacctcaaaaagagtcatagcaccag
tccaaaataccattatacctgcatgagcaacatgagcgccaagtaatttgccagaaacgt
taattaaacgtgcattgccagaccaccaggcaaatccggtagattcaatgtctctaccgc
caacaccaacatttgtattaaagggcgtttccacgtggtaaaacctcctcaggaatata
aagttttcatgaggttgatcctgagcagccatccaagagcgaataccttcatttagcagg
atattcttagtatagaaagtttcaaactcagggtcttctgcagctctcagttcttgagaa
acaaaatcgtaagctcttaagttaagagctagtccaacaattccaaacgcacttgtccac
aatccagttacaggtacaaatagcatgaaaaaatgtagccagcgtttattagaaaaagct
acaccaaaaatttgtgaccagaatctgttcgctgttaccattgaataagtttcttcagac
tgtgtaggtgtaaatgcacgaaaagtatctgcagcatcaccatcttcaaacagtgtattt
tgtacggtcgcaccatgaatagcgcataataaagcaccacctaagattccggcaacaccc
atcatgtggaatggatttagcgtccaattatgaaatccctgtaagaatagcaaaaatcta
aaaattgcagctacgccaagacttggagcaaaaaaccaacttgcctgacccaaagggtac
attaagaaaacagatacaaatactgcaatcggtccagaaaaagcaatagcattgtatggt
ctaagaccaactaatctagcaatttcaaattgtcttaagcaaaatccaattaatccaaaa
gatccatgtaaagcaataaatgcccataagccgccaatttggcaccagcgagtaaaatct
ccttgagcttcaggtccccaaaggaaaagaagggaatgccccatgctgtttgctggcgta
gaaacagcagcagtcaagaagttgcatccttctaggtatgaactagctagtccatgagta
taccaagaagttacaaaagtagtcccagttagccagccgcctacagcaaggtaagcgcaa
ggaaacagaagtagtccagaccaacccacgaatacgaatcggtctcttttttagccaatcg
tctacaagatcaaaaccgccacgagtttttttcttgtccaattgctatggtcataattta
atctccagagcaaatttaataagtaatttgtttaaataaaca---ctttagatctagtct
aaaatttacttttctttacggtcaatataattataagtcaaaatcaacaaaaattcacg
tttttaacaata-tttttttagctcgttaagttaattattttaaagt-------------
taactattttttttaattaatcaaaatcatagtttgttcatttctataattctttgaaata
aatatcctgttcctcttgctgttaaaattaagtctggattacttggatcatcttctagtt
tagctcttaggcgtgaaatgtgcacgtctacaactcttgtgtctacgtgtcgttctggcg
tatatccccaaacctcttgtaaaatagatgctctagaaaaaggttcgccagccttactca
ctaaaagttcaagaagactgaactccattcctgttaagcgaactcgttcattgtttttgt

aaacttgtcttttattggtatctatctttaaaaatccaatactaattatgccagaatttg
ggactccaagattagctgttattttatcagcccttctaagaacagatcgaatgcgagctt
ctagctctttaggagaaaatggtttaactacataatcatcagcacctatttctagtccag
taattctgtcacacacttcgcctaaagctgttaacattattataggaacatcagattctt
tccgtagttcctggcaaactccgtaaccatcgagtttaggcatcattacatctaaaacga
caaggctaggatattcttttcgaaatactattaaagcttcctctccgtttgaagctgtta
tgacttcgtaacctattattgttaatcttgtttctaaaattcttcttatgctggcttcat
cgtcaacaacaagaatttttttcttttctggttctccaa-tttcttatttactcctataagt
caataagcacaactactataatcttaaatttgtaactgtttgagtatcttaatttgctgc
aaaa-----cttatttactataaattgtataattc-----tatattaattggtatcagt-
-cattatacctgcaaatagccgtttgtcagttatgtc---------tattttaatgcata
agttaattcaagttttcagaattcagtatataaata-ctttttagacaagccttttacat
tctattttctaaataaaaaagggggttgacaaactagctgatagaaagtattctacattaa
tataaataatttactgattataaa-gagcagtaa-----atttctttagagtattacaat
tgctttaat
>Pyr_19
tatatgata---aattaagacgtctcttatgtgagaatttctaaaatatacaaaactttt
tatttaatatagaaaaactaaattaaaataagttgaacataccttgctctaatctaaacc
tcttaattgag------attctaattagagataaaagtatttactta--ttgtatctata
ggaataaaaatgaaacttgcagtttatggtaaaggtggtatcggtaaatctacaactagt
tgcaatatttctgtagcactttcaaaaagaggtaaaaaagttctacaaattggctgcgac
cctaaacatgacagtacgtttacactaacgggatttttaatcccaacaattatcgatact
cttcaatccaaggattaccactatgaggacgtttggcctgaagatgtaatctataaagga
tatggcggtgtagactgtgttgaagctggtggaccgccggctggcgctggttgcggaggc
tacgttgtaggtgaaacagttaaactttaaaagaattaaatgcttttgatgaatatgat
attatcttatttgatgttttaggtgatgttgtatgtggaggttttgcagctccattaaat
tatgcagactactgcttaatcattacagacaacggatttgatgctttatttgcagctaat
agaatagcagcttcagtacgagaaaaagctagaacgcactctctgagattagctggactc
gttggtaatagaacagataaaagagatctaattgataaatatatagattgtgttccaatg
ccagttttagaagtcttgcccttgattgaagatattagagtgtccagagtaaaaggtaaa
actttatttgaaatggcagaaattgataaggatttagcatatgtatgcgattactatttg
aatattgcagatcagttgattacaaggccagaaggtgtagttcctaaagaatctcccgat
agagaattatttagtcttttatctgatttttacttaaatcctaaatcaaatgtaggacaa
gacaaagtagatcaagaagaattagatttgatgatagtgtaaataatattttcagaataa
tataacaaggaataagataatgtctacagctcaatcagatgctcttacttttgaatgtga
aacaggtaattatcatactttttgtcccatcagttgtgtttcttggttatatcaaaaaat

tgaagacagtttttcttagttataggaactaaaacttgtggctatttcttgcaaaatgc
aatgggagtaatgatttttgctgaaccgagatatgccatggcagaattagaagaaggaga
tatttcagctaagctaaatgattatggtgaactacgtagattatgcttacaaataaaaaa
agatagaaatcctagtgttatattttggattggaacatgtacaacagaaattataaaaat
ggatctggaaggaattgctccaaaattggaagcagaaattcgtgtccctatagtagttgc
tagagctaatggtttagattatgcctttactcaaggtgaagatacagttctagcagctat
ggctcaaaggtgtccattaaatttaaataatcaatcagataatacgagtcttaaaatttc
tcctcatattcctttggttttatttggatctcttccggattcagttgtaacccagttaac
tatggaattaaaaaaacaaggtatttttgtttctggttggttgccatcaaagaggtatac
tgaattaccggttataaaagaaggatattatgttgcgggagttaatccatttcttagtcg
cacagctactacactaatgagacgccgaaaaacgaaactaattggtgcaccatttccaat
aggtccagatggtactcgggcctggattgaaaaaatctgttcagtaatgaatgtaaagcc
tattggcttagaagacagagaaaaagaaatttgggcttctttagaagactatatttcttt
aatacgaggtaaatcagtttttttatgggtgataatttactagaagtttctcttgcaag
atttttaactagatgtggaatgaccgtatatgaaattggtatcccttatatggataaacg
ctatcaagcagcagagctagctttattaaaagctacatgtgataaaatgaacgttatgat
gccaacaattgtagaaaaaccagataattataaccaagtagaccgaatccgtgatctgaa
acccgacctagttatcactggtatggctcatgcaaatcctttagaagccagaggtattaa
tacaaaatggtcagtagaatttacatttgcacaaattcatggctttaccaatgcaagaga
tattcttgaactcgtcacaagaccattacgacgaaatcttagtttatcagaattaggctg
ggatgtgtatagcaagcaaagctagtatgcttc----tttatgacttacatggtataaaa
gccttgtaagtct--tttattttatgtcaaagaagcatttcttgctctgccatcaccagc
ccaagcttgtaaactttctacttgttctttatctgtaaatgctaaaggcacaaattgttt
gatagctattttaatatcttcagtactaaattctctctcttcactaaacgctgtatgcat
actttcaataattgcttgctcaatttcagcgccagaaaatttattacataataaactcaa
ttcatgaatatcatattcttgccacgatcttggtctaacttttgataaatgtatcttaaa
aattgactctctctcttgacgattaggtaaatctaagaaaaaaatttcatcgaacctgcc
tttccttaacatttcggaaggtaaactctgaattttattagcagttgcgacaacaaatac
tggagctttttctctgatagccaggtaataaatgtgccaaacactcttgcactagtacc
actatcaccttggctatataaaccagaaaaagctttatctatttcatcaatccataatat
gcaaggcgataatccttcagagatatttatcatttctctcattttggattccgattcacc
aactagtccaccaaacagttttcccatgtcaagacgtagcagaggcaacatccaatcgct
tgcaatagctttcgcggttaaagatttacccgtcccctgtataccaactagcaataaacc
tttgggcgaaggaatgccgtaatttaaactttgcttcgaaaatgagcgagaccttttttg
tagccattgttttaatatatctaagccaccaatatctctatttactgtattataaggata
aaattctaatagacacgtttgattaataatttgcctttttttcttcaataataataggtaa

gctccgagagtctatctgattatattgtgcaataattttagtaatcactttgcggattct
atctatcgataatccttggcaggatttgttatattattaacgagctctgaattcaaatt
aagattcaaagttttactcaatctcctgatctctttttgatttctagaagactaggtaa
gggtaaatctattactgttatgatatcgcttagagcaaaaggtatattgattttgcagga
aataataacaatgtgcttagattgcgttttgattattttagctaaattgcgaagtttcct
aattaatactatttcatttaggaaagagtcaaaatcttttaaaagaaaaagatttaaaga
ttcattatttaagttctcaataaattctaatgctaataaaggatttctttttgcataacc
attatcgctaggattgttagtgtatccatctacaaagtcccagcaatacacttgttgatt
atttgaacagtttaagctgtgttttataatgtactctagtctatcttcttctctagtatt
tactacaataataggatatcgagattttaaaagtaatcgtaaatcttgagtgaaattcat
gtcaatattttttccatcagacttttgtacttcaccaattg----aagactaaataaga
ttatctaatattgcttggatatattcaaaagtc------atctctctattttctaaatta
ttgctattattaacttaacttaccatgattttttttcttccttttcttcgtcgactattt
aatatagagcgtccacatggtgttttcattcttgctctgaaaccagaaactctaatttttt
tttcttttagatccttgtagtgttcttttagtcatgatgtttcctctgttgtaaaagatt
aaatactattttatcatcttttatatatctttatgatatagatatatattaatatcaaag
tt-aacagcaataatttatattaatcaagaactatttatataatattaaccattaaatat
----------taactttatgatattggctttgcccatttttttatttaagtattttgacca
tctttttgctaatcttaaactggttaattttccagcagctaaaaacgattttgtcattag
aatctcaatttaagtattttgttgataaaagccaaaatggtaaattagaacccgacgaaa
gttttgcttttgcaaaagtatgtgtagctaaaaaattttttttccaaagctattattgaag
gtcaattagctttgaaaaaaatatcgagaccttaataccttagataataatatagtaattg
ccaatttatataatatgctaggttttatttattttgaagcaggccaaataagttttgcca
agaacttttatgagcaagccttacaaataaaccctaattatgttgtagctttaaataacc
tggcaaaaatttatgaagaagtaaaagatttaaaaaagctgaatctctatatgataaag
tgttaattcttaacttaaataataaaactgctaatagaaggaaaaatttatagcaaaga
cta---------aaaatgtctaatgcggtttgtaatcgggatagcaggatttgaacctgc
gacatcctgctcccaaagcaggcgcgctaccaaactgcgctatatcccgtaagacataat
tagtactatatacctttatctaattatgtctactctttctgtattttatctaggatacc
aaaacatgcctttaactccatctggatccttaataaatcctaatttttttataaaaactaa
caacatcaggctctgcaaatagtgtaatagtgctaatttctgcttgtctcaattgctgta
ttaattgatgtatgactactttccctaaacctaaaccttggaaatctggatgaataacta
cgtcccagatagttgcattaaatccgttgtctgatgttgctctagcaaatcctacaagtt
tactatttgcatcttttttttgtattaaagaaataataatagaactattttttaatgcga
tctttactttttttagaggtcttttaacccatccgactgaatcacagagctgctccaatt
cgtataaattaatatttttattgctactcagatagatatctttaaattcgattctatcgc

aagttttatctaaaacaataagtttttgaaatttttttcagagctagcattgatatcgg
agttttgaaaaaaatttttccagaagatcataatgattatgatataatta--aataaata
tatatattacaatattgctttaacaaacaatatattaatatatgctcaaatttttacatg
aatctattaagatagaagattgttaaaatatagagaaataaaaattttgtaaaatcttt
caataaaatcatataaagagtgcttttaagactaacatctattattataaataaagtct
cgcaattgagatgaaataacacattattctatgtctaatgccttttagaggagttatttg
tgaaaaaatctatgtttctaaccggtttacttgctttactaattatgagcaacccaataa
tagctaacgcagaagtagctggattggtaccatgcaaagattctgcggcattcaacaagc
gtatggtaaatagtgttaaaaaacttcaagccagattatctaaatatgatgcagataccc
caccagcgttagctttaaataagcaaatagagaaaactaaaactagatttgcaacttatg
gtcgagcaggtttgttgtgtggtactgatggattgccacatttaatttctgatggtagat
ggagtagagccggggactttgtttttccaggactattattcttgtatattacaggatgga
ttggctgggtaggtagaggatatcttttatctgttgctaagactagtaagccaacagaaa
aggaaatcattttagatgtgccattagctatcaaatttatgtcatccggatttgcatggc
cgctagcggcttggcaagaatttagtagtggacaattaattgcttccaatgacgatatta
ctgtttcaccccgttagtaaaaaatttatatgaataataattttaccaaatacttatcaa
cagcacctgtaattggtgtattgtggatgacgtttacagccggatttattatagaattga
accgctttttcccagatgtcttatacttttactt----ataaataa--gatcttataaa-
--aaactacaccaatatagagaaatactattatttgtgtagtttttaa-tatatataaca
taattattta-tttttaaaaatatttatcttttcgattattttaagattagatcttacat
aataaaata-atatcaatattatgagcttagtaacccaaattatagttaatgctgatgat
gaattaagatatcctacaattggagaattgcagtcaattcaagactacttaactacaggg
agtaatagaattagaattgctactattattagagataaagaaaaggagattatacagaaa
gctagtaaacaaatttttcagttacatccagaatatatagctccaggtggtaatgcagca
ggttctaggaaaagatcgttatgcttacgtgattatggttggtatctcagactaattaca
tatggcgtattagctggggataaagattctatagaaacaattggtattataggtgtaaga
gaaatgtacaattctttaggcgtacctataattggaatgttagatgcgatccaatgttta
aaggaagcatctttagaactacttggtcaagatgatattagaatcattgctccttatttt
gattatataattcgtggaatgtcataaatcgtagcttcaattaattagttgaataaaagc
ttttataatgttataattacaataagcccgaaagcatataattgtaaaaactaaatcttg
tcagaaccggaaggtagcagcaataatgtttaaatacaaaagttatga-ttttcgggtgt
ttactgttttcatctaatagtctcgacttttaattagtatgctatgtataaatctgtata
tgatatgtaattcatcaaacctttcagcaaaaacaatttaataatttgatattaatacgt
aaaacacctatac-ttttttataaaatttctttatatctcatctaaacaatggaaactta
acttaatgggtgttcatattttatcaacaggttcgtctgtcccaaatttttctgtagaga
atcaacaatttgaagatatgatcgaaacttccgaccattggatttcaacaagaacaggaa

taaaaaaaaagacatctagctccatcttctatttctttaactaaattagcagcagaagctg
ccaacaaagccttatatgcagctaacttaaaacctactgagatcagtttaattattttag
ctacgtctacgcctgatgatttatttggtagtgctagtcagttgcaagcagaaataggtg
caacaacatcggtagcttttgatattacggctgcctgctccggtttcattgttgctttag
taacagcagctcagttcattcaaactggttcctatgacaatattttagttgttggagcag
atacaatgtctagatggatcaattggtcagatagaactacctgtattttatttggtgatg
gtgctggagcagtagtattggggcaaagccgcaaaaaatagtattttaggctttaagttat
gtacagacggtcagctaaacagtcatttacaattaatgaataaacctgtaaataatcaaa
agtttggtgttacagaaattcctcatggaaactataattccataacaatgaatggcaagg
aagtttacaagttcgctgtatttcaagttccaacagtaattagacaatgtttgaataatt
taaatatttcaatagatgaagttgattggtttatattgcatcaagcaaacactagaatta
tagaggcaattgcgagcagattatcagtaccttttctaaaatgattacgaacctagagc
attatggaaatacatctgcagcgtcaattcctttagcgttagatgaagctattcaatcca
ataaaattcaaccaggccaaattattgttttatctggttttggagccggcttgacttggg
gagcaattgtcttgaagtggtgat---ttatatattgcggatgacgagactcgaactcgt
aaagctttcgctacacacccctcaagcgtgcgtgtataccaatttcaccacatccgcatt
ttttatcatgaata-ttaaaaatttataatatttatctaaatattcatataatgtgtgta
tatatatatacattcttgtcagaaactctcttatatttatttaagtgataaaattgag
t----ttagtatagatagtaaaattttattttaaaatcataa----aaataaatatataa
ggaaactaaaatatgacaccgtccttatcaagtttttgaatagtcttattcttggggca
gtaattgtagtcgttcccataactttggctcttttatttgttagtcaaaaagacaggaca
attcggtcctaaaaataacttagaaagatatggagaagcaatttaagatgaataatgaaa
tatttttcataataatacgctctccatacattgctatact--ttagaagaa--------t
ttttattttgtaacatcctgtaaagtttattccgctctttcttttcct-atttactctc
tta----taatgatacaaatgtcagg-ttaataatag--ttttttatttttttatctttc
atttatatcaaagtattttatatacgtattattatgttcaattaataaagaaaagctca
tgtctttgtctaattggcctctcaaaaaagaaaattctgaagcatataatattaagaatt
caaaacaaatcacaattcctgatggtttatgggtaaaatgcttcgactgtggtctattaa
tgtattctaaagtactaaagcgaaatttaaaagtttgccctcaatgtagttaccattttc
aagcttctagtaacgaaagaattgatcaattgatagaccaaggtagttggcaaccaatgg
atgttcacttgatctctacagatccattaggttttaaagatcaaaagctttatagtcaaa
ggttaaaagatactgctctgaagactggcctgcaagatgcagttcaaacaggtactggta
ctatgcaaggtaaaaaagtatgcttaggtatcatggattttagattcatgggggaagca
tgggatctgttgtaggcgaaaaactaacaagactactagaaaaagcaactcaagataagt
tgcctgcaattatactttgtgcatcaggcggagctagaatgcaagaaggtatgttgagct
taatgcaaatggcaaaaatttcttctgctctagaaatgcataaaaaagaaaatttactat

atatatctgttttaacctctcctacaacaggggggtgtaacagccagttttgctatgcttg
gagacttaattattgcagagccgaaagctcttattgcatttgctggtagacgagttatag
aacaaacaatcaaagaagacttgccagataattttcaaagttcagaatatttatttgaac
atggtttcctagatttaattgtacccagaactcagcttagatcaaagttaatccaaattt
tgtccttacataatcatagtaagtaattaat----tattgactataatataaaaatgcta
gtaaaaattaagatattaagaaattttcactaaagatatttgtagactaactagaattac
aattactattgtagaaaaatttgtcttaatgtca-t-ttttcagtctttgaacaaattg-
tttttaggactttaaataattgttagtttaatacaaatttataaagaaaata-----cat
actaagctaaatatagtaatattatgttttcctataatactgacctatcaaatttcgagg
aaatttatgcttaaaagatcttcttggcttgcggctttattgggactattaacagtggtt
tctacaagtacgcatacatatgccatagagttagacgaggcaacaagaactgttccatta
gaatcttctggcagaactgtaattcttacaccagaacaagttaaaagaggtaagcgatta
tttaataattcttgtgctatttgccataatggtggtatcacaaaaacaaatccgaatatt
ggacttgacccagaatctttaggattagctacgccacaaagagataccattgagggacta
gttgactatatgaaagatccgactagttatgatggtgcagagtcaatcgcagaattgcat
ccaagtattaaaagtgctgaaattttcctaaaatgcgtaatctaaccgatgaagacccta
ttcacaattgctggtcatatcttgctccaacctaaaattgtttctgaaaagtggggcgga
ggaaaaatttattattagaacttcaaagaactaaatcttgtgaatactagtttatgta-t
ttgttgcttaagtgtgattattttgttagatcatgatatatattaaagatggagtttgtt
gaaagcaagttatttcttagccgaacttgcttacactattcgtctagttattaaaaagga
gacgttaaattgaaaaagaagttttcagttcttttcactgtttttagtttttttgtaata
ggtttcgcacaaattgcttttgctgcagatctagataatggagaaaaagtttttttctgct
aattgtgcagcatgtcatgctggcggtaataacgctattatgccagataaaaccttaaaa
aaagatgtacttgaagctaatagtatgaatactattgatgctattacttatcaagtacaa
aatggtaaaaatgccatgcctgctttcggaggtagactggttgatgaagatattgaagat
gcagcaaattatgtattatctcaatctgaaaaaggttggtaattatacttgattttatcc
tgtattaaagaatagacaatctatttagttattcattagattgtctattctttgtttatg
---ctattatatagagatatttacacaatatt----ttatgatgaaaagaatacctgcaa
ttcttgtactagaagacggtacttattataaaggatggtcattccagcaagataaacaag
agattactattggtgaagtagtttttaatactggaatgacaggatatcaagaaataatca
cagatcctagttacttccatcaaattgtcgcttttacttaccccgaaattgggaatacag
gtattaataatcaagatattgaatctcacagtattagtattaaaggacttattgcaaaaa
acatttgtaaaatttcaagcagctggagagagcagcaatctttatttaagtatttaagta
gtaataatattccttttattttttggaatagatacaaggtctttaacccaatacttgcgtc
aatttggtacaatgaacggttgtatctctactgataatctaaaccatagttacttaaaac
agaaaatttgcgagattccaagtatgcaaggtttagatttaatcccgcatgtaactacaa

gaaatgtttacccctgggatgaaaaaagtttgccgaattggtatttgacagataacatga
gagtgcaccgagttattcagttgaaagttattgttatagattttggagttaaactaaata
tactcagaagactagctacacttggatgtcagataacggttgtgcctgcccacactcctt
taaaagatattttggcttaccaacctgatggcatattactctctaatggtccaggagatc
catctgcagtacattacgggatccagacagttacaaacttactagattacaatgtgccta
tatttgggatttgtatggggcatcaaatttaaatttagctcttaaagctaaaactttca
aacttaaatttggtcatagaggtattaaccatccatccggattgaaccagcaagttgaaa
taactagtcaaaaccatggctttgcagttgacttgacttcggtttttgaatctcctgtaa
gagtgactcattttaatctaaatgacactactattgcaggaactggacataatcaaagtc
cttatttttctgtgcaatatcatccagaatcgagcccaggccctcatgatgctgattatc
tattcgaaagttttatagaaataatgacaaagtccaagaagaaagttag-----------
ttagtaattttcccatgctttatgagtgaataaagcggatagtacttgtactcctcgtaa
ttgattgaagagaggtaaatgccctttggggccatgagaattccattgaaattcttcagg
atatctgcaaggaattctatcaacttcccagccgatttttgccagaatttttcccaatc
ttttccaacactaagccatacttgtctttgaacaaataacccaaatttaccctttgagtg
tgtatgccataaattatcaatagtttgtaaatcttgagcaggtatttttttatatctgt
aaaatatagccaattacgagtttgtgcatttacaccagctagttgaataagtttttgttg
agttagctgatctgctttgagcaaatcccgatgagtcagtaacacttgtaaatctttgta
attcatctgttgagcagagcgcaatggaacaatgccatcgggaaataagttagaggcaaa
tttcactatttcttgattcttactgttgagtaacttttcataaattaaaccgtcaacaca
attacttttatagttaggcccagtaatcctctcaaacaatagatctgctaaatcttttaa
ctcgatagagtcgttgctatttatattctcaataatttcaagttgttgcttga---catt
attagacttagtatccttattcaattctaaaagctgggctcggatttgatttggcatctg
ttttgtttaattagttttaatagtcataaataaggtcttatagagaattagattgttttt
tgaaattagaagagcgccgcagtatttctaaagaagttcgcataatttgattcaataagt
tttgtcctgcataaattataatttgaccaattaaaaaaattgtgtttttatttttaggaa
caaaaaaatcttgtacctcaagcagaaaagtaattagtatttgcattgatgacaacatct
gtaattcattcgatctgcaagcatatatatactgacagtttaagccttcttgataaaatc
cccatactttgtaacgactttcgtaaattgctcgtggtcgctcaagatagtttttaatta
tagtattccaaattaggttattctttaatttttctaatttctatcagatagaaatttta
cattacagagataacttgcatttgtctcatttacagtattgaaattttgaactaacatat
gagcaataatgttgcttagctgaattaaataattttcaagaaaaatttcgacctgcttaa
ctggcatattgtaagtaaagagattaaaagttttattcacagtttgggatgacccaaaaa
gtagttgaataattagaatttgtagtagcatcttataatcgttcagtaggtagttcaaat
cattaatatcttggtaaatatcaatatagtccagttcatataacttacaaaatcttttga
cgcatctatggaataaatcaattaaaatattttcggtcaagtttgtaatatcatttacat

ctaaatttgaattataaatttctaaaagtatcttttccagttcagctaatataattttta
ataagttccttttacttcggtgcgaaaaacatctaaaattaacacttcctgagaacaat
tagttaatctcttattaatctttatagaagttctaactaacagctccgctacctcctgat
tgagtgttggtccctgagagcttggccaataattattcacgttataccatagtaagaaat
attgaatttaatgttatacgataacttttaatatactgaaaagcaagttagttactcta
tagcaattag-------gttttacagatcttaatacttatttatcaaataaagttatatt
atagtagcgactgagctaattttattaaaattta----taataatgacaacttattactt
tgcccttgcgagtcaaaatttctattagtacaagaaccactagaagaagtctttagaga
gagagttaactactatcagtcaaataataaagcaattgattttggctaataccaaaccc
ttctttttagagaaaccagaaatgatttcatttaagaatcttgtacctaaagaggctgt
agctataatctctactaatccaatatttattaattggttaaagctaagaattggctatat
ttgtattggacaatttgaagataacctacaactttctgaagaatcgttaaatattactgt
tttaaaagacaaaatttaacaatgcttttattataactagtgtt-aattattttggttaa
gttatagttatagtcttttagtattta-gatgcttatagtttatcacttagtattttatg
ctgagatggagtgacaaatagtgtaagtatttcttggctaaatgtttcagttgccttaga
tttatatctatttggattaactataatagatagcattcgttttatagtaacattttcaat
ttgagcccaatgtacaattccaagttccaattccttagcaatagctgaaacggaaacaaa
ggcagctcctagccctgattgcacagcgtttttgatagcttctatagaattcaattccat
ctctattttaaagcgactgctgtcaataccatgttgactgagtactttatctatgacttt
tctaattgttgattgagtatctaacgcaataaatctaagcctgtataagtcttcttttg
aatgtctcctagtttagaaaaaggatgtgattttggtaatatcagtgctaattcgtcttc
tgcataagaagtaacttgcaaaacatcttgtaattcagtaggcacttctcctccaataat
tgctaaatcaacttggccattagctacgctccatgaaataagtctagttgaatgtacttg
caattgaacagccacttgtggatatctttgcctaaatagtccgattaatcttggcatcaa
atatgtcccagttgtctggctagccccaataattaatgtgccaccttgtaagttttgtaa
gtcgtcaagagctcgacaagtttcttcgcagagggctaaaattctgcccccgtatcgtaa
aagaagacttcctgcctcagttaaagttgccttcttattacctctttcaaatagggaaac
atttaattggcgctctaaattttgaatttgcaaactaatagccggctgagaaacatacaa
gctattagctgctttttaaaactgccctctttggcaattgcttttaatattcttaactg
atccaatgtaaatggaaggtctgtcattaaagaattgtagtataatg---------tata
ttatttagtattgatcaatgtatgcatttgccaaccaaaatatttgttattacgcttcat
tatttttaaaataataaaaagtataatgttaaataagctgtattaaaatttaatatctgt
ttgatattaaatttatactaaagatatcgtaaatttaattctaactataaaatataggag
acaat---atggactctagacttttagttgtactaataccagttttagcagcagcatctt
gggcagtttacaatattggtagagttgcattacagcaatttagaaaaatgacatcttagt
tttgcttagaatata-attttagactttatgagatagaggactttaaaatccctatctc

tattgtatatcttaataaaaaaat-caggc--ccccttgtacttactgttattacacatt
aagattattgat-agtaccta----gataataagtttaagctaacttatttttattttta
taattaattaaattaataacacaattatggctgttccaaagaaaagaacatctaaagcta
aaaaaaatgcacgcaaagcaaattggaaaaatcaagcaaaaactgaagcacaaaaagctt
tatctttagcaaaatcagtattaactagaaaatccaatggatttatttataatctaactg
aagcgtcagatacttttagtgattagaagtgttt-ttactacaagtctgaaaaagtatta
cgcaacaagttctaagtaatttttatctat----taatcatttgaaaatcattctatttg
ataataaaataaatagcagattaaataaaattacagtaacaaattatgcagcaaagctag
atcaaactagtgaaatttggttgttcaattgtattgaaaatattcagcatattttttaa
aaagtcaattaaaatcaagtcatattactaaaattttatttctggtactagctttgaat
ataccgcaggtctaccaggactattatccagcttaacactaagtggtagactacatccta
ttagtatatatagtcctcagtctctcaaaaagtatcttgaagtatgtaccaaatattccc
aaactaatttttcttttcctattaattttcataatttacaatacggagaacaagttgtta
gccaattttatacagtaatttgtttaccgttgagtaaaaagagcttgctgtatggattta
ttattttaaaaaaagaaaagcatggagtatttaaattagcacaagctaaaaccttgaata
ttcttcaaggacctatatatggaaaactcaaagaaaaagataatttttaagtccagatg
gttactatctatccggccaagacttctcttctaatacaataatgggacataaaatatcgc
ttcctttattagtcagatattctagaattatttctgagatgcattggttttgctcttatc
ctattagattaaatacttattcgcatcagcaagtagtcaagtgtttaccacataatgttt
taactgatattatgaagtcgcaaatatatcaagataacagttttgttgaataatacttat
tcttgatatataatggtctcggtaatccaaataacatactagaaataatttcttgaattt
atttgttcatattaaatttacaatcatttatgacatacgcaattattgaagcaagtggca
cccagctttggatagaagaaggccgctactatgatttaaatcatatacctgttgatccag
gtcagtcgattatattaggaaaagtcttattattaaataaaaaatgggcaggttactttag
gacgtccttgtatagaaggtgttactataaaagctacagtaatgaggcacttgcgaggaa
agaaaataactgttttcaagatgaaaccaaagaagaaaatgagattaaaaaaaggtcatc
gacaagaattgactcgtttaatgatcgattctataacctcttaga------taaaatcta
gactcatattttt--aa-aaactttttaatacttaactaaaatagatataatggcacata
aaaaaggtagtggtagtacgagaaatggcagagactctaatgccaagcgtttaggtgtta
aaaaatatggtggagagcaagtaacagcaggtaatattttaatcagacaacggggaacta
aagttaagcctggccaaaatgttggaaaaggaaaagatgatacattattttcgctaatcg
atggtttcgtgctgtttgaaaagtcaaatcaaaagcaaaaaacaattagtgtttattctg
ctaagaaatagttaaaacaattggtgcaagataacttgagt----gtatttattgcactt
acagtggcaccatatatttgatttattaagaaatggagttccttaatgaaccaatgaatt
ttattaaagatttgctgtgaaatttaatgacaaacactattgtaatttcctgtctacaca
atatggctgctattctatattgtggtcagatacagaaactagtagtagcaaatgctcatt

atcaagtaagtgatatctacctaggttgtgttgataaaatcttctcaggaataaatgcgg
catttattaacctaggaaagaatgagtacagtggttttatacatatcagtgatacgggtc
cgcttaaaaagaaatattatgtcaataatattactaacattttaacaatacgacaaaaaa
ttttagtacaaatcattaaagagccaactttaaataaaggtccaaggctcactgccaata
ttacattatcaggtcgatatattgtactaatgccttttagtcaatcaatctgcatatctc
gaaaaatacatgatgaagatgagcgtcattatttgaaatctttagctattttaattaaac
cagcaacaatgggcttgctatttagaccttctgctgtaggcgtcgatgaggaaattatat
taagcgaattaaaaaatttaaaagaacaatggaactttgttcaaaaatctgcaattaata
gtcattcacctgttcttctatataaagatgaagatattgttaaaaaggtaatccgagatt
tttataataataatacaaacaatatagtaattgattcaaacctgggattaaaacaattaa
attattatatccacacttggcactgtaataactctagtacagttcccaagatgaagcttt
atagtaataatcaatgtatactagatgcttttggtatcaatcaggcaatttccagagctc
ttattccaaaagttgatcttatacttggtggctatatgtttattgaaactttagaagctt
ttactattattgacgtcaattctggatctttcaataattctactagtgcacgagaaacag
ttttaaaaaccaattgttctgcagcaacagaaatagcttatcagttacaaattagaaata
ttactggtgtaattataatcgacttcattgatatggaatcacaaagagatcaattgcagt
tattagaacactttaataaagagctggcacttgatgatgctaaaccacaaattgtacagt
tatctgaattaggtctagttgaattgactagaagaagacagggcaaaagtttatacgagc
taattagtagtgattctaattacttttatttttcacacaatcagagagatcccagtctc
ttaagagatttaatgatagacaacagaaacagcagacttttaatagatcttggctatcta
tagagattaatactattaacaaggtctttttcaaaagtcaaatttgtgcagacctgcta
acttttacctaattcgtaatctctatatagttaaaagtagtattacttataaacaaaatt
atttattaactcacagatcaaaacttatttattctcaagaatacaacaaagtattaccaa
gca------gttattatctagctagtctcaacaggaatagtaatcaggagttcctatctt
aagatttttatatcaactgtt---------------------------------------
-------------------gaaaaaaagaagctctctcttatataagagagagcttcttt
ttttagcttactaacta-tctagttagttaattaaaaattaaccgtaaacagctggagct
gttagagctactggtaaagattcaccagaagctagatctagagggaagttgtgagcgtta
cgttcgtgcattacttccatacctaggttagcacggttgatgatatcagcccatgtgtta
attacacgaccttggctatcaacaacagattggttaaagttaaaaccattcaagttaaat
gccattgtgcttacagataaagctgttagccagataccaactacaggccatagacctaag
aagaaatgtagagaacgagagttgttgaaactagcatattggaagattaaacgaccgaag
tagccatgagctgcaacgatgttataagtttcttcttcttgtccgaatttgtaaccatag
ttagcagattcgttttcgcttgtttcacgaattaagctagatgtaactagggatccgtgc
atagcactgaacaaagaaccaccaaacacaccagcaacacctagttggtggaatgggtgc
attaaaatgttgtgctcagcttggaatacaagcatgaagttaaatgttccggagatacct

agaggcattccatcagagaaactaccttggccaattgggtatactaggaatactgctgct
gctgctgctactggagcagtaaaagcaacggaaatccatggacgcatacctaggcggtag
cttagttcccactcacgaccaatgtagcaagctacgccagttagaaaatgaagaacaact
aattggtaaggaccaccgttgtatagccattcgtctaaagaagcagcttcccagattggg
tagaagtgaataccgatagctgcagaacttggaataacagcaccagaaataatgttgttt
ccgtatagaagggaaccagcaactggctcacgaattccatcaatgtctactggaggtgca
gctacgaatgcaatgatgaatacagatgtggcagttaatagagttggaatcattaataca
ccaaaccaaccaatgtataggcggttttcagtactagtaatccaagagcagaaacgttcc
cacaagctagcgctttcgcgtctttgtaaagtagcagtcataatttttttatcaattttta
aggagttaacaaagatgcttcccaaacaataatttgtctgttatggaaattattacttaa
ggccttttcaaatgagaacattttaaaaaaaaaca-tagcaaaatctaatctcagtaact
aa-aaatt-taattgttgatatattgcttaatacttcaa-----------taataaattt
tt-gct-------------ttttacaaaattaacaccattttagggatataataataata
atgcatattaatgctgcttattttgattttaatagtttatataattttgccttacttaca
tacgcatgtcacactttacaaaaattcagacgaccatacaagatttaaatctgttaaagc
atgctttaacagatttgggtttgacttggcaaatgaattctagtcatataaaagttggtg
aaaataatcagcataaggtagatatcttaattaaacaagataatttgtcacatattggct
ttatttggaatgataataaataccacttagttgccgacttgcaactttgggaacaaccct
ggtctctagaagtatttttggataaattatctcagaagtatgcttaccactctattatcg
aagaaacaaagaagcaaggctttgaaaaaaatgcagcagatttcacaaaaagatggatcaa
ttaagttaattgtgcaacgctggaattattaactaaaatataaatgcgggcggagagact
tgaactctcacgagattatctcactagaacctaaatctagcgcgtctaccaattccacca
cgcccgcactgttaaaac--aatattactttatcataatctatactaaaacacaaactaa
-tagtataaactagcttccaagcttaaaagcgtctacaaataaaccatatgttataccaa
ttttgaagttgttgaataaattaatttttatattgaattgcttttggctagagtagacta
ttttactagtcaattcagttgctgcaactattagactggctgcaataatacccccaatctc
ctgtttgtccaggaattgtagataaacctgttgaaataaaaaaatcctaacaataaactaa
ttaagccagttgttaactcgctcaaagaataataaagcttattatttaagtttttttatta
agtaagcaaaaaaagttgaaagtctagttttaatcatagtatttagtaacggagttttaa
aaggatgacaagtattaacag-tcattaatattgtcgtatccttttaaaactcgttaaga
caaaaagctgctcttgt---------ttttaaaggtcttgttcgcttaaagcattaataa
gatgctgaaacggctcttgtataatatctttgcgtttaattgattgcaaatctaattctt
cttctatgatttcttgtaataactgtatgcttcttactgttggagcaacaggtacagata
gtgaattgtaagtatcttttagcccatttaatacacgctgatcgagaatatttgtactac
cagaaaccaaggcataactagtataccgtaggtaatattctatatctcttagacaagctg
catatcttcttgttgtataagaatttcctccaggccttaataattcaggctgttcttcgt

```
acaactgagctgcggcctcttttaaaatattggtagcttgatcattaataatttcagcta
tttttattcgatctagtccacttgaaaagaaagattctagttgtccgacagctgctttat
ctagataacgaccagtaagatcgtaacgatttaatattgctgttatagcatcttgcataa
attaattctcctac-ttgctatcagagcggtattataaatgactaatattgtaatataat
cattagtgtttatgaaatttatttaaaacttattctttattgcttaatgtgcaattaaaa
actcgcatagattttacatatatagagtatacgtagtttttaatta-ttttttttataaaa
attaagaaataaagctttttt-taagaggttcgtgaatttcttaatacaatctaagcttt
ccgcccaaataagttttgtaaagtac--------aaaaaaagattattcaataatagtaa
------------------------------------------------------------
------------------------------------------------------------
------------------------------------------------------------
------------------------------------------------------------
------------------------------------------------------------
------------------------------------------------------------
------------------------------------------------------------
------------------------------------------------------------
------------------------------------------------------------
------------------------------------------------------------
--------------------------acaatattaatatgtatcaa--------------
------------------------------------------------------------
------------------------------------------------------------
------------------------------------------------------------
------------------------------------------------------------
------------------------------------------------------------
------------------------------------------------------------
------------------------------------------------------------
------------------------------------------------------------
------------------------------------------------------------
------------------------------------------------------------
------------------------------------------------------------
------------------------------------------------------------
------------------------------------------------------------
------------------------------------------------------------
------------------------------------------------------------
------------------------------------------------------------
------------------------------------------------------------
```

------------------------------------------------------------
------------------------------------------------------------
------------------------------------------------------------
------------------------------------------------------------
------------------------------------------------------------
-----------------agtataaataatctttactccagcatcaaaaaaaatatttaa
catatgttgatacaattgattattcaagtgaaaacttagatttagaccaatttagctatt
ttatcattacaattaataaagctgataaatgtattcttatagagcaattttgttataatg
ctgaaggtcatatttactcaatattcttcaaaggtccaacagctaaacacctttcttcca
taatttgtcattttcaacaattaaatacaataatatccacagctcatgcaatatatttag
gaagagagctaatgaagtcagaattagctctcgtgctagatcaacaatatattcaagatt
aaatttttttaagtattaattatatttatgctagtataatattggtaagggcatgtaac
tcagtggatagagtatcagattccgattctgatggccgtgggttcgaatcccgccctgcc
c---gagtattcagattact-acactaaaacatct-------taatagctagtttgataa
tagtatgggcattggggctgcaaggtttctacattatgaaaagaagaaaatata--aaaa
aaacaagctctccaaaagagcttttagtacaattaataaatgcagaaaacaatattgttt
ctttttctcgaaaattagctcttgcttaaatactgttaattttggtattcgacatgtta
aactcttattcatggcgaatattctgtcagagtcgctcttagtttaagaaaagtcag-aa
aaatatttatctttacgtttt-tacctatttttatttaaataaaggttagta-ttttcaa
cttttatagtggacgtgggttcgagtcccaccagctccatcaatttgatt---------t
gtataattagctagttcaacctttttaataaccagatatataattggaggaaatatggat
tttataacaattactaagcccgctttaaagcaaattgcaatcttgaaaaatgatcatgaa
aatgatgtacatctaagaataggtgttagacaaggcggatgttcaggtatgtcatactca
atgaattttgaacatgttgacaatttaaaagatacagatgaacggcttcttctagataat
ttttctgttgtctgcgaccctaaaagtcttctttatctttatggcttatcattagatttt
agctcggaattaataggaggaggatttcaatttctaacccgaatgctagccaaacttgt
ggctgtggtaaatctttttcaggctaaca-ctctaacaggtttttttatatattttgtatt
acacatagtattttagcaagtcg-tttaatagagacaagcaatgctctgttaaaaac---
----------------------------taatttcaagagttttttattgtattatt-----
----ttatatatttattatcaactacttgaaaatggattagttaagcaatctaatttgtt
atcagtatcaaactttaagtgattaactttttattgatatatgctaatcttt-ataagcc
taaacgtatctgtacattacaattttttgaaataaaaagagtcttaacataactgtcaa-
----------------------gaagaaattata----------tattttataaatactc
aacatgaatctgtcacaacttattgttaattcctctatgatgaaaattcctaataacttt
tattgcaatacgaaatttatttcttctgagaatttaagtcacttacaattatcaaaaaca
aatttattatctcctacaaaaataattgcttttggactacctaattttgatttcatctct

acacagattacaagttctattagggaatcagttgttgtatctagtattgatgataataat
ctttttgctgccccaactaatgattggataaaaaaacacaaggataaaaatcatcattct
ttgttttataagaaaatctttaaagctttgctaaaccgtaaaatcagagttgttgatgag
ccatt------ttataactctgaaataaataaagaaaagagcataatgacaagtcaaaga
tacgtctggggtaaaagctggaagccttcaattattttatcttgcttaaaaaacaaaaaa
aatagcaaaatttctgacgcatcgaaaaaatttttaactgaacagttatctgcatcacca
gtatttgtggtaaaaaatggttttaatgaaattattttaggtcacccattatctcgtgtt
aaaacaggaagagttaataatctgacgcatgcattttctaatttattaaatcagtctaat
gaaacttatcctatatctaccggtttattctttttttcatccagatgatgcatttgaattc
aaagactttataatatcggttaatccactggcggccaaacatatggagataagcgtagaa
cccgttggtttacactttgcttataaaatgaatagagatagatcgtcagatactcaattt
tcgtttattccagatttaaagaagtaggagatttattattcaaatatagaaaagatcat
catttagtttttcataagaatcagcattatggcaaagatttttttcaggggcaaccaata
tatatgattcagccgatcacttttaaagatcgatccggaaagctgaatattatcaaattt
actgggctaaatgacacaagagaaattatttttactaatcttgaagctgctaatagatca
tggacacattttataaagaggaatccacaattaaaatcagttaaaaatcctactttatta
gtttataatctagaaagtttttttaaaagatcaagaacgattggataatcaagatttacac
aaatttgttgtgatcactaataaatcatcttatcttaccacaaaagagttaatagcttta
accgattctaatagctttttttaagcacttaaaattaaatataaagcctaaactcttttttt
gtgaaactatggggaagacggttattttccaccttaacttatgaat-aaatgctatt-gc
ctttatttcctcgaataatattctataactagtaattcattgagttgtaaagcgacccat
tccctatcaatgactccattgactttccctgataaattagatttatttaattctaagtgg
ctaggaatattagctaatccagggaaagctaagtaattttcgactagttttcgagatgct
tcttgatttttcacactaattgattctcctggtttacattgataactacagatagatact
acttgtccattaatacaaatatgaccatggttcactagttgtctagcggcaggaattgta
ggagccataccaagtctaaaaacagtattgtctagtctcatctctagtagttgtaacaaa
atttgacctgtcgacccttgtaatttttttagctgcttttacgtacttaaataattgtttc
tcacttaatccataattaaatcgcaacttctgtttttcctctaatcttacagcatactca
gaaggttttctagacttttgtccatgttctccaggtggataagatcgcttaattgctttt
ctactgagtccgggtaaatcacctaatctacgggaaatgcgtactcgtggccctctgtat
ctagacatattatttaaattctcctaatgtaaatggattaag----------taatgagt
agctatttttttaacacaaaaagagcattaaaacaaagc-ttttgctaaaaaatctttata
agaaatataggcttggacgaaa---cacactatgatgc-tttttaagcgggtagcgggaa
tcgaacccgcatcattagcttggaaggctaaggttttaccactaaactatacccgcaata
ttaaacagattgtcactaactcaatataacatagttatgttaagttaagataatcttaaa
tttaatagaaat-tcaaattatttacgctccttctaaaactacatcaagaaaacgagcaa

tttcggcagcttgttcttcaacttctgataataataatggttgaccaaccctagttaatg
gaattactcttttatcttttgtacataaataaatttcacgtcttggattgagcccctctt
taatatctatttttattgatttgatttcttttatactaaactgaagacagatttttctat
tctttccagggaaaccaagccgaaaaatttttacaatacctttatctttgttgaattcat
tatatccagcccctatattccaaataatagttagccatagaaacaaactcaggaagattc
ctatactgccatagaatgtcattacaataccttgtggaataaataccaaatcggttgaag
ttgtaaagggtaacaaatcaacttgaaaataacttgatagtccagcaagaagaaagccca
aagccccaataaatattatagtagcccaccagtaattactcagtcttcgtgaacctaaaa
ttaaatcttttctaactgagtgtatagataatgtttttttcataatttgttcttttaaaa
cagta-tactaatgcaaaatggaaaaattttgacaattttt--attagcgctttagattt
tttaaataagcttgattgcttttagccctaaaaatagtcctgttgctaacccaaaaaaag
cagcaagaaatattgtatatcctaaaaaaactgaca---tatta----ttattattattg
ttacaatagtgtttacaaaaatagtggaaatctattactagggtattatattacgttaaa
tgccttgctgtctacatgaaaaaactttttcagtttttagcattgtgttcatattagaa
ataatattctagtcaatttattaaacttctggagaacagaatcttatgtcagaatttatc
aagccttataacgatgatccttttgtaggcaatttgtccacgccagttagtacatcaagt
tttagtaaaggacttctaggaaatctaccagcttaccgtcgaggtttatctccgcttctt
agaggattagagataggaatggcacatggatacttttaattggaccttttgataaattg
ggccctttacgaggtacagatgtagcattactagcaggatttctatcttcggtcggtctc
attattattctcaccacatgtttatccatgtatggtaatgtatcttttactagagcagac
tcgaaagatccattacaaacttctgaaggctggggacaattcactgcaggatttctagtt
ggagcagtaggtggttcaggatttgcttatttattgctagctaatatacctgtattacag
actgcaggtcttagtttattttcttaagctagtaaggggacttgaacccgtaacctactg
attacaaatcagttgctctaccaattgagctatactagcacctttttcatcataacatat
aaagatatttgttgcaatctgttattttcagatgtaacaaatatctttttttatttaaaca
gaactgttatattaacac-------gtatattaattagaatctgaattatctgactctgc
agaatattcttgatcacagtttagataataacaaattgtttgatctagacaagttgcaga
ccaaccaacaggtgtttcacctgataattcttgcatatctccagaattgtaaaaatcatc
atttggcttattactgtttgcttgcataaatatcactcccaaaacactctgaacttaatg
tgaagtaatttaacttctctataatatattaacacaaaatattaaatatggtgattattc
tttaattgtaaagatgcggctaactttataaagtttgagtgagagtcttgatagcttttg
ttgaaataaacattttaatccatttattttgctctctcgaccatactttttttatgttgta
aattaactttttgtagtttttttgttcgtttatgagaatgagatactgcatatccattat
tagctactttccccgtaagctgacactttttttgacatgatttt-------ttaacttata
tatttatttagagtacttgctaaagttgactttggcacagctccaatcactgtgtcaact
ctttctcctgctttaaagatcattaaagtaggtatacttctaatgccatattcagctgca

atagtaggattttcgtctgtgtttatttttactaccttaatagatgattcatattcttcc
gcaatttcatcaactacaggagagaccattctacaaggaccacaccatgacgcccaaaaa
tctacgagtacaggtaagttattgttaataacttcttgtttgaaagaggcatctgtaact
tgagatactgacatattctttaacctttaatatatgttccttgctaagcaaatcttatca
caaatttcaagaaaaatcttctatagtgaattactattagtgttgaattaatttttaaa
aattagacattaaaaaaacttattacctatataaataaagtggtggccgagtgatatctt
gatagtgatagtttttatattaaggttgaacactagaaataacttttcaagcaaatttt
atttgaaaa--acttctctgtgaccataaaaacttaattatattacaatatattttatta
tttgtaacttagttctgataatggtataaacaacgcaaaagatactgccttataatcaag
gaggaatacatgtctcaatccgtagaatcacggactaggattaaaagcgaacgttacgaa
tctggagtaatcccttacgctaaaatgggctactgggatgctgactatgtaattaaagaa
acagatattttagctcttttcagaatcactcctcaaccaggtgttgacccgattgaagca
tctgctgcaattgcaggtgaatcttcaacagctacttggacagttgtatggactgattta
ttaacagcttgtgacttatacagagcaaaagcatatcgagtagatccagttccaaacgtg
gcagatcaatattttgcttacatagcttatgatattgatttatttgaagaaggttccatt
gcgaacttaactgcttcaattattggtaatgtttttggttttaaagctgttaaagctctt
cgtttggaagacatgcgcatgccagtagcttatctaaaaacgttccaaggtcctgcaact
ggattgattgtagaacgtgagcgtatggataagttcggtagacctttcttaggtgctaca
gttaagcctaaactaggtttatctggcaaaaactacggaagagttgtatacgaaggctta
aaaggaggtcttgatttcctgaaagatgatgagaatattaactcacaaccatttatgcgt
tggagagaaagatttttatattctatggaaggtgtaaataaagcatctgcttctgctggc
gaaattaaaggtcattaccttaacgtaacagctgcgacaatggaagatatgtatgagaga
gccgaattctctaaagaggttggtagtatcatttgtatgattgaccttgtgattggttat
actgcgattcaaagtatggcaatttgggctcgtaaacatgacatgattttacatttacat
agagctggtaactcaacttactctcgtcaaaaaaatcatggtatgaacttccgagttatt
tgcaaatggatgcgtatggctggtgttgaccatattcacgcaggtacagttgtaggtaag
ctggaaggagatcctttaatgattaaaggcttctacaatactctacttgaaagcgaaaca
gatataaacctacctcaaggtctgttctttgctcaaaattgggcttccctacgtaaagtt
gtaccagtagcatctggtggtattcatgctggtcaaatgcaccaacttcttgattactta
ggtgatgatgtggttcttcagtttggtggtggtacaattggacatcctgatggtatccaa
gcaggtgcaactgctaacagagtagcactagagtccatggttatggcaagaaatgaaggc
cgtaactatgtagcagaaggtccacaaatcttgagggacgctgctaaacttgtgggcct
ctacaaacagctttagatttatggaaagatattagttttaactatacttccacagataca
gctgatttcgttgagactccaacagcaaacatctagtttaatgacgacttactaatactt
taaacagtcaattgtaagtagaattaacttataacaataaggagcacagaatagtgagac
taacacaagggactttttccttccttccagatttaactgatgagcaaattaataaacagc

tttcttatatcgtttctaaaggcttatcagcaaacgttgagtatactgacgatcctcatc
caagaaactcctattgggaactgtggggtttacctttatttgatgtaaaagatgcttctg
ccgttatgtatgaaattagctcatgcagaaaagcaaaacctaattattatgttaaagtta
acgcttttgataatactagaggtattgaaagttgtgtaatgtctttcattgtaaatagac
ctgctaatgaaccaggattcttattacaacgccaagacttcgaaggtagaactatgaagt
acagtcttcatagctatgctactgaaaagcctgaaggagctag------atattaattaa
gaattaatattggctaatta--ctcccctctttaaaatcacaattgctaagttaattatg
attttaaagaggggtaatttcccaagtaaattttgactaaa-tagattggataattaaaa
atacacagaaataatgcaatcacaggatataatttccaacgatactcttgttaatttaca
agaagaatatgatagaacacaaatccaagaagttttaaatgagttaaatcaagaacttat
aggattagtacctgtaaagaccagaattcgcgaaattgctgcgctattattgattgacag
attacgtagaaaactagaactagtttctggtaatccaggattacacatgtcatttacagg
tagtccaggaactggtaaaactacagttgctatgaaaatggcggatattttgcatagact
tggatatataaaaaaaggacatttgttaacagttacaagagatgatcttgtaggtcaata
tattggacatactgcacctaaaaccaaggaagttcttaagcaagcaatgggcggagtttt
atttattgacgaagcttactatctatataaagcagataatgaaagagactatggctctga
agcaattgagattttattacaagtaatggaaaaccaaagaaatgacttagttgttatctt
tgctggatataaagatagaatggaaaaattctacgaatccaatccaggactctcttctag
agtagctaatcatgtagactttccagattatacttcagatgaattattacaaatagctaa
aatgatgatagaagaacagcagtactgttttacagaagaagcagacaaaactcttttaga
gtatactgagcgaagaatgaaacagccttattttgctaatgcaagaagcattcgcaatgc
tattgacagggctagaatgagacaagccaataggatttttgccagtggagaaaaagtatt
aacaaaagctgatttagtaacgattgaagcagaagatatcttgaaaagtagattattttc
attacctgatgcttaatattcacgtgatttaatgaaactatatattattaaaaag-cttc
ataatcagttgcaaatttttttgaaaaaccccttattatcatcttaatgaaggtg-tggcgg
catagccaagtggtaaggcagaggattgcaaatccttcatcccccagttcaaatctgggt
gccgcctagtactaaaaa--gggggtgtggtggaatggtagacacaacagacttaaaatc
tgttgatttttagtaatcgtgagggttcaagtccctccacccccatatattaacttataa
ataaaaagagatgtgtatttgtattaattgcaaccatattacaaaatgtaagacctatca
cttaatagaatctcaacacaagcagcctcatattactagaagtccattgtttatacctaa
atatcctgtagttcatgttaatatatctaacaatggtacctataatcaaattgattggga
tttagtggagtgtttatcatttgtagaaaaaccgaatagttggaatttagacgctaatta
gttacaagaaaaattatgtactataagcaacaactctatttttttagatactcggtattga
ttttggatgttctaaccaataagattttacctgttcttgctatctctataatgccaaatt
tagttagtaattgttcaatagcaacaatctttcctggatctccagtaacttctataatta
aaagatcttctgcaatatctactatcttagctctaaaaatttttacaatttctaaagctt

ctgttctagtctgagaattgatcttaatcttaattaacattagttctctttcaactgaag
gaatatttgttacatcttgaacattaaggatatttactaacttatataattgtttagtaa
gttgttcaatagttctattgtctccttgaactaccatcgtgattctagagaccccaattt
gctctgctggtccgactgctaaacttgcgatattaaagcctctccgggcaaatagaccag
atattcttgacagtactccggcttcatcttgaactaaaactgataaggtgtgtttcatga
aaaattaattttaatctatttgatctattaagtatattctaagaaaattacagaaaaaat
actagttagtctacaactaaattatctaaactaaacaaaatcctactatcagataatata
tctaccttgtaatgctatactattaacttagtatttaatgatattctaattgcaaataga
atgct-aatttttaaaatttgttaggaacttgtgctagaaaaatacaaaaacagtaaaa
aactctctcgagatttacctcaaatcaatgatcgcatcagatttccgaaagtccgagtaa
ttgatgacgaaggtgaacaactaggtatttttgtgcctgaagaagctatacaattagctg
tccaacaaggtttagacttagttgttgttagtgataaatcggacccgccagtatgccgaa
tattagactatggtaaatataagtttacacaagaaaaaagagctagagaagctaagaaaa
agcaacataacagtagtattaaagaagtgaaaatgcgatacaagatagaagagcatgatt
ataaagttagaataaatcaggcatctaaatttattcaagcaggagacaaagtaaaagcaa
ctatcacatttcgggggcgtgaaatccagcactctaatttagctatagatttattgaata
aaatggcaagcgatctaaatacaatagctgaaattcagcaagctccatcaagagatggca
gaaatgttataatgctcttatctcccaaaaaagttggctaaactaattttatttaaatg
catctggctggattcgaaccagcgacgtctctttcgaaatggcggattatgagtccgctg
ccttcggcccctcggccacagatgcatcacttaaagtttataactcataaagcatataaa
aatcaatgtttagctatttttaataaagtctataagagtctattattcattcatattgtg
atatgtagcaactgctgaaggggaaacttgatttaagtatcgaaatatccagtatttaaa
aattgtgtcgagtatgactggaaaagttgaaataaataaaaaaataaaatctcgactttc
tggaagtcctaaatgtcttaaaattacttcaataataacttcccatccatgaggagaatg
aaaacccacaaacatatcagtaaaaagaatgattaaaaaagctttagctgtatcactaag
accataaataatttcatttaaaaaagatttgacgactgcaatttgtctttgtccagttat
cattagtaatataaagacaagtattgatacaagatcggataaaatatttttaacagcatt
tgcactttcgtttgcataatattcgcccaattctctagccttcagctgtactcttttttc
aatgatttcgtaagatatatcttccgaaggattcagaagaacttcaaagtgaattttctc
ttcaaatctttgtaattcagcaaaagctctttcctcttgagaggaatttaaaaaaatttt
aggctgttcttggttccataaataatcaatacaaggtccaaatacaaaaaactttgaagc
ctgattaacgagtactggagagatgaataatagtaaaatatattttacggaagtaatagt
ttggtgcctagatattctaaactcctctatagcttcagattcgccattaggatctaactc
tttacgaaacttttcaaaagtattagtaattgacctaggtattggcccaaccttttcaaa
ggcgaattgattatttcgttttaaattccaatatttcatttttattgctgctaaaattt
ctattaatcaataattgatcacgtattcactataaatattaattctagaactaaacatat

aaaacaatatttatgaaaaataaaatacaattaaaaattaattatgcattagttctatct
gtggctacagcaatgtatggatataaatcttatcagccaactataaatacactcttaatt
aaagatgaaaaaaactggccttgtttctctaaaaggcatactgcctttattcctataatt
gtattaaaaaaaaatcatcaaaatccctgaactcagaacctagaaatagtaagttatatt
ttaatggaattggaagtagaaaatttatcgagagaattgttaatctacttgaatatttag
aaaaacaggcaaactatactgaatatctgaaccaaattaaaactagtggttatttttctt
tactctaccaggacagcaatacaatgtataagaaaaatattagtaatatttttttattac
ctaatgatattctaaaaagaatctatatttgcaataagaaaaaaaagcttataccacatg
cattcttaataaatttatttcggaaacagataggctatccaaaaagttttgctaattta
attgggccttctccaaaattatgaaatggtattatgatagaggataccaatggtctctag
ttgaagttaaacaagcgtctgatgcatcttctattgtaattgatattcatgaaggggtag
tgaagacaattataacagaatactacactttatcttataaaagactctcaggtatctcat
acatagagtcaatagagcagtatcttggagtaagagtcggagctccattaaatataattg
atttacaaaaaaagattagttacttaaaagataaccaactagttggcgacattatttata
gtattgaacgatcaaataataactcaatgagtttggatgtcaaatttcagatacaagaat
taaaagataaagagataatagtgcttgcagaaagttcttcta------------ttatct
cgcatgcatgtgatttactgaatc---aatataga---------aataggctagtagctt
ctaatataatttcactatcaactaataaattacacgcatgctattttaattataaactcg
actatcaatataagtatataaacacctctaatctcgtacaattgttaacttactctattt
cttgtaggaagacactacttagccattattttaatttacaaactttaataagctggacaa
aaaaaaatactattggatttcagctatatttacgaaatctaagctttggaaaagcttttt
gtgtgttcagtatgaaatttatcaaaaatggacttaatataagaattttatatataaatc
cttcattgatagttgatcagaattttatctttcaatttgctatacaaataattaagcaat
atcatactgcta------aaccaccagcct------------------------------
------tatttttaagcaatctagatcttgaacaatatgttgctgaaagtctattaatgt
accatttacatcttgcttctcaatatctgaaaaaatattattatctcggattatgcata
cagattctttatttttcaactctgagaatctccattttcattttgatgatagaaatcacg
cgatcaacagttatgataattttaaacaaaatacaaaaatatttttatcaagagtttctct
ctttattatttagtttacgttatcaaaattttaattatttaggttggccttcaaaaggcc
attttttgaaataaaatctttatacttggctccatttcaaaagagtg------------
---atttttctgatagtcgtaagaccctattcttccacaaaatgtcgttaaagcaagtat
ctaattttaatttaccaatatcttttagaagccactttaaccatattttagttagtacta
ttaaatgccagtcaaatttgaatatgagaactgtatctttgctattaattgattcccctg
ctgaatatatgctatacaaatctattcttaatttctctattaaagttaggatgcagtatt
tcatacctatgagcaacagtataagagtatctctattctataattatttagattgttttt
taataagatcatctcagagtttcattcacatatggcaagatttaagaactcttaaaccaa

ttccaaatttttggttaaaaaaattttcatatggagttggaattcagcttaaattaccaa
ttaaacagatgccacctctttcaattgaatatactataactagtagtcgctacttttgta
tttaccttcgtacttattatcaacgataactattactaaaatgaaaaatttaatttcttt
ttttgatcgcagcaaagggaagtggatttcacaaagaacaacttatgaattatctaataa
aaacatgagttcagcacagtctcagatgataatgaaaataggcaaatcattgtcaggatc
tataatacttgcatcgttaaattggggagacatttatagacaagttgcccattacgctaa
aagtcatagccggagtgaatataataataagtttaacttacaatttaataatcaactaaa
taaccataaattattgacgctgtgcatagtgacagatcctagtctaatcagttttaaaac
tcgctatggaagtactacaatagatgaaacatattggtttgcaacaaataatttacgtct
aagtactagtattgttaaacgattcaatacttgtgtggcagtatctttttgttcagaaat
taaagtttaa-actctatatagaagcaactaataaccttta-ggctattagttgcttcta
tatatctaaattttctattaaaaattat----aatactatattattctaagtctttacga
ttaggattcctagaaggatcattagagaggaagccaaagacaaacaaagaaataaagaaa
ataacggtagtatatacaaagattttcaaagtaaacatattttcatccttaaaagctatt
agtgcg----atatttacatatctccacataattttatatcatgatatgagttccagtac
tattttataaataaatcaatatctttatttttttagtaatgatttatatttgaatatgcgt
gatgtactaaatactaaatcttttccgatttgttttgtcttgtttgtaaatattttgtta
tttctatggagtttaccttcaattatcacatagtctaacttctttagctgtctaaaagta
tatgaagatttttttgttccaaatagttagattgattataactaatcgttttctttttaac
agtcttgcttttaacttaattatttggcttttattttcactaattctgatactcttacaa
ctaagaatttggactagcagagtacagctattcataatttataacaa------tttttta
aagaatctactaaaatagatgattttacaaaactaaaatgatatttcattatcttgttga
ccttgctcataagtaatgtctttcaatacttttaatatttttccaaagtactcttgcagg
tatttttctaaagatattacttccaacctgtcaattccaagaatactgtaaacttcatcc
atcggagctgtaaattgatctccgctagttaaaatttcagcaaatgctaaacgatcagaa
atattccatgtccattgaagagttttagtaatttttcttaaggcttttaataatccaatc
ggaatttgtgagatctgtgtttttttgcccagataatttttcacataatttaataatttca
gctgatgtccaagcggtattaccaactagcggcaggattctattttctgtggaaggtact
ccaagacttttttataactaacttggctgcatcctgagtatcaatgtatgcaatgggcgta
gattctcctgtaacccagactgacttcttgtctaaaataggaatggcatactggttgatt
aaaccttgaaaaaatcctcctaaagaaaacacagtatatttcacattggacttctgaagg
aaatctactacctgagattttaagctcattaatggaacatctgggtatttctctgaattt
agtattgaaaagaaaataaatctttcaactttagccgctttagctgcttcaattaatgca
atttttccatctaaatctatttttttctgcattgtaaggatcagtaggacgagatgtggaa
gcatctatgactgctgttactccacaaaaagattgcaaaatactttcgggtaattttaaa
tcaccatatataagttctgctccccactctttgagaaaagcagactttcttaaatttctt

accatgcattttacattgtagccttcatctaaagctcgtcttacaatttgacgtcctaaa
gttccggttgctccaattactaaaagagtcatattagttaaaaataagtatgtataaaga
cattgatttggttgacggcagagagtaaattaggtagagagggaatcgaaccctcatgac
cgaagtcgtcacattttgagtgtgatgcgtataccaatttcgccatctaccctattatag
accaactactacaagtaatttagtctttagcgtaaagataatactaagtttcaatagata
tgtcaacttttaaatttatattaataaaaaaacggagtctcttatgaccacttttgaatt
aattccttatagattgtatttaagtcaatctaacacatggatgcatagaataaaagcaga
aattaagatatatatagtaatgcttttatggatttcaattttacttttcttactttaa
actatgtattattgctttaagcttaattgcaataagttttactataagaagtaaacagaa
tattattcaacaacatttgttacagacgttattaatgacgttcttaactactgttttgtc
ttttagtgtggctattaattataaacaatatgcagaacaagaacaatctcagtatttatc
tgattccaagaaatataaaaattctaggagctactattatatacaaatagctaacgatca
aaggatcaaaagacaatacttaattactactttaaagccttccttgtactttttttattac
tatatattctattaaactagttatgataacaacctctccagaagttttagtaattactat
ttatagatctagtataataaataaaatatttaaaaatgaattgctgtttatcttccttct
ttcgtcacatattgttactagcattattaatagagttgataaagttattcaagtaactag
cttaagaggaagcttgagtttatataattcatttacaaggcttttaatgttttctttgtt
aatatttcaagtcttctttttagagattatccgggaatcaaaagagatagctcaagctct
ttatactagaaatctcaatcaagaaaacaataatttttttgaaaatatatacagtaaaatc
tcactttagtgatcggctaaatataattattagcactttgtactttattattttagctat
agcgtaataaaatccacttattcttctgactctagtaatacgaagagtaataaaattagt
tagtagtttaaatagatagaat-taacatattatataaataattaaatgtttcattatgg
tcaatactcaaaatcaaatttcacaaacttcagaacttgattatatagttaatcaacctt
ataaatatggctttaaaacttctgttgaatctgagcaatttcctaggggaataagtgaag
atattgtccgcttgatctcaaagaaaaaagatgagcctgaatatctgttgaattttaggc
ttagagcatataaaaaatggaaaaagatgagtaatccgtcatgggctcatattaagcatc
cgaatatagactttaataatattatttattatgctgttcctaaattaaaaaaagaattga
aaagtctagatgaggttgacccagaaattcttgacacttttaataagttaggtatatctt
taaatgagcaaaagcgaatttccaatgtagccgttgatgctgtttttgatagtgtatcta
ttgcaactacttttaagaaagaactgtctgaagctggtgttatattctgttcaatttctg
aagctataagggattatccagaattaattaaaaaatacttaggtactgttgtcccagctg
gtgataattattttgctgcattgaactctgcggtatttagtgatggctctttttgctata
tccccccctaatacagtttgccctttagaattatcaacttactttcgtattaacaacgaag
aatctgggcaatttgaaaggacactaattattgctgatcgtggcagtaaagtaagttatc
ttgaaggttgtaccgctcctcaatttgacacaaatcaattacatgcagcgattgtagaat
taatagctcttgaaggagccgaaattaaatattctacagtacagaattggtatgctggta

acaaagaaggtaaaggcggtatatacaattttgttactaaacgaggcttctgctcgggta
ataattcaaaaatttcatggactcaagtagaaactggatctgcaattacctggaaatatc
caagctgtattttagctggtgataattctcagggagaattttactctgtagctttaacaa
ataattatcaagaggctgatacagggactaaaatgatccatattggcaacaatacaaaaa
gtagaattatttctaaaggtatttctgcgggtagatcgaaaaacagttatcgaggcctag
taaaaattggccctcagtcatttaattcacgtaattattctcaatgcgattctttattaa
ttggtcaatcatctcaagctaatacatttccttatattcaagtacagaatccaacatcaa
aagtagaacatgaagcatccacttcaaaaattagcgaggaccaaatttttatttttttac
aaagagggatcaatttagaagaatctattgctcttatgatcagtggtttttgtaaagatg
tctttaatgaattgcctatggaatttgctactgaagctgatcgtttactgagtttaaaat
tagagggaactgtaggatgagccagattattttagagatcaaagatctatatgcttctgt
tggtaaaacaacaattttaaaaggggtaaacttatctattcgagccggcgagatacatgc
gattatgggtcctaacggttcaggtaaaagtacattatcaaaagtaattgcaggacatcc
agcatattcattaataagcggagacattttatttttttggacacagtattcttgaaatgga
gccagacgagagggcaagagctggtatttttttagcttttcagtatcctgttgaaattcc
gggagtcagtaattctgatttcctaagaattgcattaaatgctcgcagaaagtttcaagg
attatcggagtttagccccttagaatttttttcaactaataatagaaaaagtagatcttgt
tggcatgcaagaaagtttttttgacaagaaatgttaatgaaggattttctggtggagaaaa
aaaacgtaacgaaattcttcaaatggctttactagatagcaacttatctattttagacga
aacagattctggactagatattgatgcattacgagtagttgcaaaaggaattaacacttt
agctaaatcaacaaattcaattattttgattactcactatcaaagattactagattatat
tattccagattttgttcatattatgagtaacggaaaaattgtaaaaactggtaatgttac
tcttgcccaagatctagaaaagcacggatatgactggattacgcaaacatagctatttag
tag-------aatattgagactaaagaatattctttagtctcaatatt-tttgcaaatac
-aagtt-aattatttaagctttgtcaaaaccttgcttaacatcttcaattgctcttttta
gcaattcttcgctagctggttctaatttttttagtactacgaacattttcaccaaattctg
gtttagaatttttttaaatcttctcgaagttctctaataaattcagccacttttgaaactt
caatatcatctagatagccattaattccagtataaataatagcagtctgttcttcaactg
gaataggagaattttgagcttgcttcaaaatttctcgaagacgttgacctcttgctagtt
gattttgggttgctttatctagatcagatgcaaactgagaaaaagcttctaattcagcaa
attgtgctaactctagttttaatttgcctgctacttgtttcatggcttttatttgcgcag
cagagcctactctggagactgagataccaacattaatagctggtctaatccctgagttaa
ataagtctccagataaaaatatttgtccatctgtaattgaaattacatttgttggtatat
atgcagatacgtccccagcttgagtttcaatgataggaagagctgtcatactacctccac
ctaattcagcattcaatttagctgctctctctagcaatctagagtgtaaatagaatacat
caccaggataggcttcccgtcccggaggtcttctaagcaaaagagacatctgacgataag

cttgagcttgcttagttaaatcatcataaataactaatgttgctttacctttgtacataa
aatattcagctaaagccgcacctgtataaggagcaatatattgtaaagtcgcaggactgt
ctgcattagctgcaactataatagtataatcgagagctcctttatcctgtagtgaagata
ctacttgagctactgaagatgccttttgtcctatcgcaacatatacacaaacaacatctt
gacctttttgattaataatcgtgtccaatgcaacagctgttttacctgtttgacggtcac
caataattaattctcgctgacctctaccaattgggatcatagagtcaatagcagttatcc
ctgtttgcataggctcacaaactgattgccttccaattataccaggggccattgactcaa
ttagccttgttccattactcgcaggttcacctttatcatctattggccgagctaatggat
caacaactctacctaaaaaagcatctccgaccggaatttgagcaatgcgaccagtacctt
ttacagaactgccctctaaaatatccctaccatctcccatgagtactacaccaacgttat
cactttctaaatttaaagcaactccaatagttttatcttcaaattcaagtaattcaccag
ccataacttcatcaagtccatatactcgagcaataccatcaccaacttgcaaaactgttc
ctatattggctacttctacatcctgatcatacttttcaatttgttgacgaataatactac
ttatttcatcaggtctaatatttaccatattttaatattattttattaattggaaatt
ctaaagcggtaattagtcagactaatgctatctgtctttttttactttattttaagttgct
acaacatcaaggtgtgaagccatctgtcttagctggcctctaatactagtatcaataact
ttcgatccaatctgaatagtaaagccaccaattagctcgggatcgactgaaataattagt
ttaacttctttagctttagtcataaccttgatttttttctgttaataaagtttcctgatca
gggcttaaagcaatagaagtattaatattcgcaattgttaaagattccatttggtacgcc
agttccaaatatttgctagctatagcatctaacattccaattcgcttgcgatcaactaaa
atcattagaaatgataatgtattttcgctaatttgatcaccaacagttgcaataattact
tctttttttgactctacagtttttagtggattagct---aaaaaatttttagcttgtca
gattgcgctaaaatattttgtattgactgaatatcttggctcaccttttcagtaactttt
ttagttttagctaaatcaagaagagctactgcataaggctgagctatttttataacaaca
ttgttactgctcatagttgatctcctaacttagcaatattattatcaattatacgtagtt
gcatttcggagctcatttgattttccagctgcaaagtaactcttttaagagcaagaaatg
ttatttgctgctggatttgccttcggatctgttgttcagctgtttctatatttgatttac
ctgttatagcaagtctttcaatgtccaattggccttgagctaatatggaacttctaactt
ttccagctgttagctgagcttctttcttaatttgatctataataatttgagtttgagcaa
gttgtttttctgattctgacaatctagcacttgcttgctctagtctttcttcggattctt
ggatagctgctaaaacttttatctgtcttgcacttaaacttgatccgagaaattgcttca
aaacataaataagaccaaataaaagcaacaatatattaataacatttgcttcaaaaatat
cagaattaaagccgaatttgtgttcactactatgttctgataaaatagtaattatttgtg
gtatttttactatattattcatttaaaattca----------tactattttaatatttta
ctctggaatcatgattttaaagattgaccacttaatagtttagctttaatttgatcgctc
aaagtatcaacctgattttctaaagttttaagagcttcttctttttgaatatttaactgt

ttcgatgcttcagcaatcagtttctctgcgttcaattgagctttttttatatcttctgag
actatattttgagcttctttttgagaagaggctatttttaattgtgcattgcggcgtgct
tcagatagatcttcttcatattttgcagctagctcatctgctttaacaagcatagacgat
gcagtagtcaatgttgttcgaatatattcgtctcgttcatctagtactttagttacaggt
ttgtaaaaaatattatttagtaaaaccattaaagtaagaaactgtaatgccattaaaggc
aaagtaccattgaagtcaaataagccaccttcaatttcttcggctaatagaaatggtaaa
tcaatcattttactaattatgtttatagttaataatttggtttcaattaagaaaaataac
tttatagtgcgcttagtctcatcagactaagcgcaaatgattattttaaccaacgtatgg
attagcgaacaataaagacagcgcaacaaccaggccgtaaattgttaaagattccataaa
tgctaaacttagaagaagtgtacctctaatctttccttctacttctggctgtctagcaat
accttcgacagcattagctgctgcactaccttgaccaatgcctgggcctattgccgcaag
acctacagcaagaccagcagcgataacggatgcggctgaaacgattgaatccataatgaa
ttttaatttttttgttatataagtagaaaattgacaaatttcgtataatacttaaaatcta
tataattaagtataatagattctaatat-----tgtaattatagtatattactcttctcc
atgaccttccattgcttctccaatataagctgcagacagagtagaaaaaaattagtgcttg
tattgaactcgcaaataatccgagtatcataactggtaatggtattaaaattggaattag
cagagtaaatacagaaacaactaattcatctgctaagacgtttccaaacaatcgaaaact
taaagatagtggtttcgtaaaatcttctagaatattaattggtagtaatactggagttgg
ctgaatatatctagcaaagtaacctaacccttttttacttaaacctgcgtaaaaataagc
tacagatgtgagtagagacaaggccacagttgtattaatgtcatttgttggggcagctag
ctcaccttcaggaaggtgaattaacttccaaggaattaatgcacctgcccagttacagcc
tagaataaatagaaataaagttgcaatgtaaggcacccacgggcgatattcatgctcacc
aatctgatttttttgcaatatcttgcagaaattctaatataaattccataaaattctgaaa
tttttcaggaattctttgtagatttctagtacctagaaatgataaagttaaaagtgtagc
aattactagccaagagacaatgaaaacttgcccatgcaattgtaggctacctattttcca
atatagatgttttccaacttctactgctgataagctagtatatgggttaagatctatgag
actgttttgatacataataaatatttcttttaaatagtgcaagaatcaaaaaaataag-tt
tttgaagaattaacatgccattaatttattcttaaaattagattattaataaattaagag
caaaaataaattctaatagagatagctatctttagtagtgattatttttttgtaaaaaaca
attattgctacctataatgcttttacgtttaattatatacccatatattaattatttaat
atgattataggtattttgctacaaactttattgttttatccattct----agagattgtt
atagtttatttttttatttaacatacctgcgacttcatctgcaaaattaactttcaaattt
tcttctccccctcccagaagaaatcttacaaatcttctaattttaatattttcacctaga
acagcaatattttggtttattaaatcttcaatactaatatcttgattgcgaataaacatt
tgatctaataatgaaagctccttaagtctctttttcattcgtccttcaatgattgtttcg
attctatcaactggtttgtttttcaagtcatctttaccagcttcaattctttgttccaaa

tcaataatttcactaggtatatcctctgttgatacgtattccacattaggcgatgccgct
atttgcatagcaatatctttagctaacttttggaactctggacgtcgtgctacaaaatct
gtttcacaattgacttcaactaatacacctattcttccgcctgtatgaatataactctct
agtaaaccttcaatagcggtacgattagatttttgttagctgaagctagccctttttgt
cttaatgattctaaagctttttcttcattgccattattagcttgtaaagctttcttgcag
tccatcatacctgctccagtcttatcacgtaaggcttttacgatttgagcagaaatttgt
agtgtcatatgctttcaaaaatttttaagtatttaaatt----------aattattataa
ttcactttgagtaggatgagttatttcttctaaatctgtttgcccgtatttgccatcata
aattgcatctgctattttacctacaattagtttaatggacctaatggcatcatcattagc
tgggattggaatattaataatttctggactacagttagtatcaagtatacaaatagttgg
aatacctaattttaaacactcttgtatagctgttgtttctctcttttgatctacaactac
aacaatatcaggtaaacgagtcatattttaataccatttagatgtttgcgaagtttatc
taattctcttcgtaaaacagcagcttctttttcggtaactgatcaatcataccactttt
atcttgttcttctagctgtctcagacgactaactcgagatttaatcgttacccaattagt
taacatcccccaagccatctctggttaacataataagaatcacagcgttgtgcttcctg
tgccactataccagctgcttgtctttttgttcctagaaataaaactttttaccatccga
cgaagcctgttttataaattcacaagcttctgttaataattgtgcagtttggactagatc
aataatatgtatgccatttctttctgtataaatatagggaaacattttaggattccatct
acgtgcttgatgtccaaaatgaacacctgcttctagtaattctgctaatgtaacaatagc
cataagattattaatatttaaataatgtaattattcgggaaatcccttttttttagaaac
tgat-----tcaactaacttatttgaattttagtaaactaataataaattaacacacttt
taaagcagatcaataaaatagatcctgaagaattcat----tctacacttttattgctaa
aataattacgcgcggttctgtcgtctaaaatgatatcatctaaatcatctcgtacagatg
aaatagtactttcgtcatcagtatttggagttaaattctttttctctaacgatgagctat
tatagttatcatacatattgaaagcagtaccagctggtataagtctacctataataacat
tttctttcagacctcttaaccaatctaatttaccagatatagcagcttcagttaatactt
ttgtagtttcttgaaaacttgctgcagaaataaagctttctgtatttaaggaagcttgag
taatacctaatagtacaggcctatatgatgcattaatcttatgtctcaaagtcatggctt
tattagtctgttcaatcttttgtaactctacaagctctccaggtaaatagcctgtttctt
caccattttctattttcactttagaagtcatctgccgtactattacttctacatgcttat
cagaaatattgactccttgagattgatatactaattgtacttctttaacaagtagtaact
gaatctcttgaaagcttaatcgagcagcttcatataaggctaatccacattcaatataca
aattaaaacttgcgtctaggatatcatgaggattcaacaatacgtctgttttctttcttg
cttcaagaatttcttcaattcttggaagaccttgtataatgtcacctgtttttgctctat
caaaaaccaacacagctaatgtttctccccttctaattaatgcattgttattaacatgta
aaatggcaccattagaaaccaagtaaggtctagcaattcttaatgtaactgatctagaag

atatttcaataatttgtccagagtctaaagaagtcatattttcagtaataaaatcaccgc
agcgaatccaatcacccacacttactttaactattgattttcctatattaaaggtctttt
tatcagaagatgtcgcaatcaaaatgcgcctactagtattcttttcagaataaatttcct
caacagtaccttcactcattgcaataatctcagtactagcaacaactgtaaatggcttga
tatattcaccatttttacaataattctagtttgactttgttgcttctctgatttattta
catcgatacttttaattgataatgtttcaaaagttgacagccttaagtcaaaataattat
tgccactttctttagcagaaaactcaattgaggatattaaagtatcatctcgatgctcaa
tttcagccacaagattagttgttactaaatgaactccttctatagatttcactctttctc
cgtcccgaaaaggagttcttttaactagctttagtttagtttgacttggtttctgtgaaa
ctagattctcaattagactggatttgtttctggtattgaataaacaataacaggtctaa
taagaatataaggcatttgctcgttttctatatactcccaataaactaacttatctgtag
aaatattattatgaagtgtttctccaggtcttaaaaaacctctactttatcattattac
tatacacctcactaagtttatagatagatccaggcttaataataatttctcgaactattc
catccttttggattatttcaacaatacctgagcttttagaaaaaatattttaactagct
ctgttccactttctataacatctccattatgcactagtagtagtgatgaatctttattaa
tttcgtgagtttcttctgaaatccataagatatatcccggacttaaaatttcgtaagcat
ctttatctgaacctgttttctttttgatacatttaagtctaaatacttaataatacccc
cacttgccgttttgtacgtatcagatattagttctgcgactgtgtaaccatcttgaa---
ttttttggtttgggatagctttgaaaagaaatttttgtttttttctgtttctaaaatat
acgattctttttgtggattacatctgtataaatataacaacctggaattacaatagatt
cagtaataacttgtatattagtaatactgctattagtattctggtgtatccttacttctc
cagcataatgatttattaattctgtctgcgcaagtatcgtaccagctttagttttatctt
ctttatgaacaataacattagcggaatcagatatactataaacttcacctgagagcaccc
agatcaagcctccagttttagtaattcttgtagtatattgattattatctgtttcttcaa
ctgttaagttggagaaacatattttgccagagagatctgaaactacatgtttttgagctc
tttctgtcatgagtcgatttctacggggtgattcagcaatcacttgatccttcgatacta
attcaccatcacatattaaaagagtagtatctttacttaagttaattattgacttttgat
ttttcttagatctgattgtaacttgggttggctttttttgttattaatgcttgctcaccat
gtctagtacgaacatctgtatacgattcaatatctagatctatcagttgaccatctatag
gtgagtaaatttgttctgctaattcaccggtaaatactcctccagtatgaaaggtcctca
ttgtcaattgagtgccaggctcaccaatagattgagccgcaataataccaacagcctctc
ctaaatcaactaagcgaccatgtgcaagattccatccgtaacaatactgacaaacagaac
ttctagagttacatgtaacaggagatcttactaaaactttttt---gataccagctttag
ttatctcttttgctaacttaggactgacatcttggttcgtatgagcaattaaacaatttg
tctctggatggaatacattttctgctaatactcggccagataaagcttgctctagattaa
ttaatattttctgcgtatctactaaatcttctagtatgatgcctttattggtcttacagt

ctacttctctaattattacatcttgagaaacatcaacaagtcgacgcgtcaaatagccgg
aatctgctgttcttaaagcagtatcaactaatcctttccttgctccgtatgatgatataa
aataatcagtaacagttagtccttctctaaaattactcgatattggaagatcaataattt
gtccctgtggatcagccataagccccctcatccccactaactgtcttacttgagaaatat
tacctctagcacctgagaacgccatcatataaacagcatttagaggatctgtctctttaa
agtatttaataacttcttgtttcaaggattcactcgcattattccacgtatcaataactt
tttgaaatcgttcaactgctgtaatctcgcctcggcgatatttgttttcagtggccttaa
tatcttcaatcgtgcctaataagagactttgcttgctgggaggaattcttaaatcttcta
aacttaaagaaattcctgcttgggtcgcataatgaaatccaagatctttaagcttgtcgg
ccatattggctgctcgggctataccgtaattacgaaaagcccagacaattaaatttttta
gttcgtttttatcaataaccttatttgaaaaacttggctgtgcaagacttcgtctattat
ccactaaatcttctccatattaaatatttgtt--attaactatgctactaaagactcttg
aataattttattaaagattatacggccagctgttgttctaatatattgcacaattctctg
attttcagcatcttcttttatgatatggttattaaaaaattttgttgtactcttgtcaga
atgtttttcaatcttaattggatcatgaggttgatcaccatctactaatccatcaaaacg
agcccatatataagaatgtaaatcaagtttcttttgctcataagccataactacatcttc
taagcttgcaaaatactgattagatccctgctgctgagatggattattagctgtcaaata
atagcaacctaaaaccatgtcttggcttggcataataattggttgacctgttgctggaga
gagaaaattatgagggctaacattagcaatctagcttcagcttgagcttctaaagacaa
aggaacatgaacagccatttgatctccgtcaaaatcagcgttaaacgctgggcatactaa
tggatgcagcttaatagctcttccttctacaagaataggttcaaatgcttgaatacctaa
tctgtgtagcgtgggagctctattcaatagtactgggtgcccttgaataacttcatttaa
aacattccaaatagaagattcatttttttgaatcatctttttggcagctttaatattgtt
aactaatccctgcaaaattagtctgtggataacaaaaggctgaaataactccaaagccat
ttctcgtggcaatccacattgatgtagttttaaatgaggacctacaactattacagatct
accagaataatctactcttttacccaacaagttttgtctaaaacgaccttgttttccctc
tataatatctgataaagattttaatggtctgttatttgcacctacaacagttctacctct
acgaccattatccatgagagagtcaacggcctcttgcagcattctttttttcatttctgat
aataatctccggagctaaaattgactttagacgagataatctattatttctattaataat
tcgacgataaaactcatttaaatctgccgtcgcaaatctacctccatctaattggaccat
aggccgtaagtcgggaggtatcacaggaataactgtgaataccatccaagaaggatctgc
accggttgctatgaaattttcaatcaatcttaagcgtttcatttttttattaaattttaa
tgaaggtgtcttaaaactttttggtggatttgtagcttcaaaccttaaggtttcagctat
gtcctctaaatctaaatctttcaacagtttttgaatagcttctgcacctattcctacttc
tacttggttattctcatcttgattttggtagatttcctcttcaaggctcttccattcgta
accttcgagaagctgcttgtattttaaattaatatctgtattagattgcgtaactacata

agaatgaaaataaacaatttttctacttctttgacttttaagtctaatgctaaagcaat
atagcttgtacttccttttaagtaccaaacatgagttactggtgaggaaagttcaatgta
agccatccgatgtcttctcacgcgtgattctgtaacttctacgccacatcgttcacaaac
tatgcctttgtagcgaaaacgtttatatttaccacagtgacattcccaatccttaactgg
gccaaaaattttctcacagaataaaccatccatttctggctttaaagttctgtaattaat
tgtttcaggctttgtgatttcaccaacaatttgtccgttaggcaaacttctttcgcccca
ctgtcgaatttttttcgggagaagctaaactaatttttacgtagtcaaaatattgctcaaa
ctttgtcataaatttgaataccttaaaatcaaataattagccaatttatttacatgcttc
atattcaaaaatgaatatcttaataaaggaattgttcaaaatcgtctactggaggagtat
cataattagaacgatctgctcggttatctttagaatccgacatcaagtcaacttcaactg
ttcgcctttggccatcttcaaatagtttcaatttatgtactgcaatgtctaaacctaatg
attgcagctctcgcattaagactttaaaagattccggtgttccaggcttcggtataggct
tccctttaactattgcgtttaatgcttcatttctagcttgcatatcatccgactttacag
ttaataattcttgcaaagtatatgctgcaccaaaagcttctaaagcccacacttccattt
cacctaatctttgacctccatgctgtgctctacctcccaatggctgctgcgtaaccagag
agtagggaccagtggatcttgcgtggatttatcatcaacaagatgaacaagcttaagca
tataagctctaccaactgtaacaggattatcgaaaggctctcctgttctaccatcaaaaa
cttgcattttacctggatgctgatcattaaaaagccatttattgctagtaatcaatgagg
cttcttttaattttctatttacaagtgctcttgatgcttctgctccatacatctcatcaa
aaggtattattttaaatctttttcctaaatagccacctgctaaacccagtaagcactcaa
atacttgaccaacattcattctagaagggacacctaaaggatttaaaacaatgtctactg
gtgttccatctgataaataaggcatatcttgtttaggtaagatcctagaaatgatacctt
tattgccgtgacgaccagccattttatcacccacttggatttttctcttctgcgcaacat
agacgcgaatcatagcatttgttccgggaggcagctcatctcctttctgcctagtaaaga
ccctaaccttaactactctaccttttgcagcattaggaagccgtaaagatgtatctctca
catcccgagccttttctccgaaaatagctcgcaacaatttaccttctggtaattgatcag
cttcacctttaggagtaatttttcctactaaaatatctccagcttcaacccaagaaccac
cgacaacaattccatttctatctaaatccttcagagagttatcgctgacgttaggaatct
ctctagtgatttcttctggtcctaatttagtttgccggcattctacttcatacttttcga
tatgaatagaagtatacaagtcatcataaactaatctttcgctaattaaaaacgcatctt
cgtagttataaccttcccaaggcatataagcaacaagaatatttctccctaaagcgattt
ctcctccgtctgtagatgcgccatcagctaaagtttggccaacaactatttttttctccta
cccaaacaattggacgctgattaatacaagtatcttggttagaccgataatattttttta
agcgataatgaactgttctaccattagtatcctgaatgcctattttattagcagaaacat
aatttacataaccagatgttctgctaataacaaccataccagagtctcgagctattttag
tttcaagaccagtaccaattattggcttttctggatataataatggaacagcttgccttt

gcatattagatcccattagagctctattcgcatcgtcatgttccaaaaaaggtattaatg
atgttgcagctgaaattacttgaataggtgaaattgcaatataatctacttgggttggcg
ttgttgtaataaactcttgacgataacgtacaggaataatatctccttcaatgtaatatt
gtttactaactttaacatctcctggagctactcgaaaatcatcttcctcatctgctgtta
gatagacaggactattgtgataaatcacttgaccttggtgtaccggataaaatggggttt
ctatgaaaccaaaaacattgacccgcgcacaagttgctagtgaaccaattaaacctgcgt
tcggaccttccggagtttcaattgggcaaattctaccatagtgactaggatgtaaatcac
gaactgcaaaacctgccctgtctttattaaaacctcctggacctaaggcactaattcttc
tcttgtgagttagttctgcaactggattagtttggtccataaattgagaaagctgactag
atccaaagaattctctcactgacgcaattaatggcttgggattaattagatttgataaac
tcaatgaatctatatcacatatcatcattctttctctaataatgcgttctaaacgattga
gacctactctgaattggttctgcagcaattctccaactgaacgaactcttctattaccaa
gatgatctatatcatcaagattacctgagtttttatccttaatattaattagataatcaa
ttgaggataagatatcttgaggagacaatactcggaacgttttaggtatattgagaccta
gtttttattaattttatatctaccgacttcacctaagtcatatcttttaggatcaaaaa
aacgagagtaaagcatctgcttagcaacagcaactgttgctggttcattaggtctaagct
tagaataaactatgagaagggcttcctcatcagttacttcttcaatattatttttccaa
tttctttagctaattctttaactgaatatatttgggatgcagaaataagaaaagcatatt
tactaagtcctttttcaatttcatctttattgagcccaatagctcgtagaaaaatataag
catttactttgtgggttttatctattcttatccaaatttctcccttaggatcaatttcga
attttaaccaagaacctctattagaaattaggcttgcactataaatctgcttaccatttt
tgtctatctcttgtttataatagattcctggactacggattatttggttaataataactc
tttctgtaccagatacaataaaagttcctctatttgtcattattggcaagtctccaataa
aaaccaatctctttttgtacttttttatttttatctgcttttcagcactgagctgtagat
gggttgatgataaatctaatgatttaatattttttcctgatctttagaaggtagatcaa
tatcttttcttgttaactttacaggaacatatatttgagcactataagttttatccctat
tttttgcctgtctgacactataacgaggaaattttatcttatattcattaccaaataatt
gtaattcgagtcgactagtagggtctgatatcttcggaaaaaattcaagaacttccgtca
aaccttctaataaaaaccatttaaaactggccctctgaatttcaactaaatctggaagaa
gtttatttttaagcttatacgttggaccatagatctcctttataacaaattttgaaatc
gaagagtttaa-tactattaatagagtactggtagataatgtataagcatattttgctac
attatttcttctctagacataactagaacataatattttattcaa--aaaaagattttaa
aaatt----ttcac--acactatatatataagcactttgctgcta--aaagaaagttcaa
gctatataa-------aaaaacgaaattaatacttgtctgtctgtcaagtaattgacatc
aagatagacaagtattatttatttaatcaattctg-tttttttgagcaaaagacaatgct
ctcgcaagtctagctttttttctagcgcctgtattagggtgaaatgccccttttttttaca

gctttgtcaatcttgctatatacttgagaaatacttgactgcacatcattgaaattatca
ctttctaaattatcaatatttagaaggcatcttttagttaaagttttgacaaccgattta
tacttacggttaatgagacgatttctttccgaagttttaatgcgtttaatcgcagaaagg
ttcttagccacagtaaatccaataaaactttataaaatatttttagaaacgaaaataca
caaatctgagatctattcaaaaagggaatatgcggcgtattataacattaaagaaaaaca
ataggcaatagttcaccaagagctcttgttattacacaatagagattgtcattgcgaaaa
actttgtaagatagtatttaattagaataaactaactagacatataaatatatatgaaaa
aaattgaagctattattagaccttttaagcttaatgaagtaaaatttgcgctagtcaagg
aaggtattggaggaatgactgttatcaaagtttctggatttgggagacagaagggtcaaa
ctgaaagatataaaggatctgaatactctattgatatcattgataaaataaaaattgaaa
ttattattaacgatgataaagtagagaaaattgtagagacgattattaaggcttctaaaa
caggggaaattggagatgggaaaatatttattagtagtattgaaagagtaataagaatca
ggactaatgacttagattttgaagctttatagtatttttacttt-tttagacttgatact
tcttcactaaacatgagataataagcgcttattgtatacttataactaaggtgtaata--
ctaatggctaaaagtaaaggtgcacggatcgtaataactttagagtgctctgataaagct
ggagattttgctcaaaaaaggaaacctggcgtttttcgatatacaactactaaaaataga
cgaaatacaccaaatagaattgaattaaacaagttttgtcctaattgcaatcaacattgt
atcttcaaagaaattaaatagttatttatcttattaaaat--aatcatattatggctgta
tatagaaaaagaatatctccaattaaaccaacagaggctgttgactacaaagatattgac
ttgctaagaaaatttattacagaacagggcaaaatactgcctaaacggtcaaccggattg
acttcgaagcagcaaaaaaaacttactaaagcgattaaacaagctagaatcctttcttta
ttaccttttttaaataaagattaa--atatatggcctaacttcacatttttttcatctaaa
tattagaaacttctatagatttgtttagcaaaatgatctaacatactgaat---------
----------tagtactaaaa-tctaatcattataagttttgttcattttataattaata
atagcactataattta---ttaagctagattc-----agacataaagtgcaacctctttt
aattgattatattatatagtctatttaaaagaggaaatttttttgcttc----------t
agcaattcaagattctttcagctttaacgatatatttaaaacaaagatttttgtttaggt
atcagatcaaatgagtgaattttatctcccgactgccataattgaaattcagagatgaaa
attccacattcgttaccagcttgaatttcttcgacatcttctctaacgcgttttaaagat
tcaatttttccttgataaataacttttttcttcccggatcactttaatccaagagtttttt
aatagcttattattgatcactcgacatcctgctatttttctatttgctaacgagaatact
gtgcttacttctgcttctcccaccggtacttctgagtattcaggatcaagtaagtcttcc
atccttcttttaatgtcttcaattaaggcataaatgatttgatagttttctattaatata
ttcaactttgccgatgcttgctttgttcctggagcaaagttagtattgaacccaatgaga
atagaatttgtggttgaagcaagttcaacatctgtagcagtaatctctccaggcatgatt
gacacgacattgagttgaactttactctgaggaaattgagataaagaatctaggattgct

tctgtagagccttgattatctgttttaataactaaagaaatttgcttactaatatcttta
gaattagtgtttttttaaagtatctaatgtaattcgactattaagtgctctttgtttctgt
ataattgatgagtctttagaggtattttcaatcgctttaagtttggcttctttatcactt
ttaactgctaaagctatttctcctgtagctggtacagatgacaatccccaaatttcaacg
acggatgagggtattgctaaattaatttctctttagcgttgctaattatagctcgaatc
tttgcgtaagctgatccaataaccaaattatcactaatattcaatgtaccattttgtatt
aatagagttgctactggaccatgcgatttatctaagtgggcttctattataataccttgt
gcaggctgtgttgggtcagctttcaaattctctaattcagctaacaaggtaatagtttct
aataatttatcaacattttgaccagtaagagaactaataggaattataggcacttgtcca
ccgagtttctcagacatgacattgtgttttaataaatcttgctcaataatatctatattc
gaaccagctttatctatcttagatattgctacgacaaaagggacattcgctttttgaata
tgatggatagcctctatggtttgaggtttaactccgtcatcagcagctataataataata
gcgacatctgttaaattagctcctctggacctcatacttgtgaaagcttcatggcctgga
gtatccaagaaaacaatttttttgtttattagctttcttaatatactcaacttcataagct
gcaatggcttgagtgattcctccaatttctttattagcattattagattttcgaatataa
tctaatagcgttgttttgccatgatctacatgtcccatcactgtgacaattggaggtctt
ttaatataattgccagtttcataaagaacattagagttatctaaattactaggaagtcca
ttattatttcctctacatttgattctactgctatcccaaaattatcagctactgatgat
attattgaggcgtctatggtttgattcatagtaactgatattcttttaagaaataaatat
tttataatatctgtttcttgaacacaaattagtttagataattcttgaatagttagagga
ttagtaatactaatagactctggcggagaatttgattgactgcttgttgaattttgacca
ggagttaatgctactttttttctgtcttatagatttttttattattagttagtttttttgcg
acttccacttttggcttaggtggacgcattaaagagatagctaaatctcctgcagtctgc
gcaacattagaattagagtctctaaagttgtca---tcatcatcgtcaatatgtattttc
gtttttagtttt-ttgcgctgtctattctta--tttttttactatctactaagtcgtga
atttattaaaattttttactttttttatctagtttgggtggagaacttaattctagatgc
tgttcaccacc---tgtatgtgattctgatttatcaaggtctaaatgtaacagatcatca
ttattaactgactctaatctaattttatagataatctgagggtttttcaaatctataatt
gtttcagaagagttgatatttgagctagatcgagaggatctattttcaaagttttgatta
tttaaaaacataaatgttggttttattatctaatagtaactttctaattttaatacagag
cttaatataaacaaaaatcgaatatctgtataaataatacaaatgt---agagaattaat
ctctgttgtattagatttaatataacactaaagtcgtttgttcgttatacaagtttaaaa
ctttt-tatgatgataaacactagattctagtttattggtatacaatctatttatctaaa
taatactgtttttgtc-gggcgtttaaa----ttatctaatgttatattcattatgaaat
tctattgaaatttaaaaa-ctcaacaattaagtaagaatcgttaatcacctccacgccac
ttatgccacgatctcaaaaaaaatgataattttattgacaaaacttttacagtattagctg

atattgtattaaaaatacttcctacaagtaaagaggaaaaggaagctttttcttactatc
gagatggaatgtctgcacaatctgaaggtgagtatgcagaagctttagaaaattattatg
aggctttaaaactagaagaagatccatatgatagaagctatattttatacaatatagggc
ttatttatgctagtaatggagagtatgtaaaagctttagaatactaccatcaaggattag
aactaaattttaaattacctcaggcccttaataatatagccgtgatatatcattaccaag
gagttcaagcaattgaggataaagatacagagctatctaaattaatgtttgacaaggcag
ctcaatattggcaacaggctattaagttagctcctgataattatattgaagctcaaaatt
ggctgaaaacaacaggacgaatgaggaatatacaaggatattaatatatttgagatataa
taaagatatcaattaataaattaaaattagttaaagaaactgatactatatagagtactt
-tgatat--------atgttatactagttga------tctctaagacttagctcaataac
cgaaattaaatctattgactatattaatgaatacaaataagcataatggttagtacaaca
aaaagtaccggatctgttactcaaattatcggaccagttttagatattgcatttcctaac
ggacagcttccgaaagtattcaatgcactcaaagtacaaagctcagaagcaactattact
tgtgaagtacaacaacttttgggtgacaacaaggtacgagctgtttctatgagttccact
gaaggactacaaagaggggtcgaagttattgatactggatcccctatatctgttcctgta
ggtacagatactcttggacgtattttttaatgttttaggtgaacctgtagataatttgggt
cccgttaattctgagagtactttacctatccatcgaccagcacctgcttttactaagtta
gagacaaaaccaaatatttttgaaacaggtattaaagttgtcgatttacttgctccctat
agaagaggtgggaaaattggtttatttggaggtgctggagtaggtaaaactgtattaatt
atggaactaattaataacatcgctaaagctcatggtggagtatctgtatttggaggcgtt
ggtgaaagaacaagagaaggaaacgacctatatatggaaatgaaggagtctaaagtaatt
aatgcagataacctgaaagaatctaaagtagcattagtatatggtcaaatgaatgaacct
cctggagcgcgtatgcgcgttggcttaactgcattgacaatggcagagtactttagagat
attaacaaacaagacgttctattgtttattgataatattttcggtttgtacaagctgga
tcagaagtgtcagctctactaggccgtatgccatctgctgtgggttaccagccaactcta
gcaactgaaatgggagcactgcaagaaagaattacttcaacaacagaaggatcaattaca
tctattcaagctgtatatgtaccggctgatgatttaacagacccagctccagccactaca
tttgcacatttagatgcaacaacagtactttcgaggaacttggcagcaaaaggaatttat
cctgcagttgatccactggattcaacatcaacaatgctacagcctggaattgttggaact
gagcattattctactgctcaagaggtaaaatcaactttacaaagatataaagaactgcaa
gatattattgctattcttggtcttgatgaactttcagaggaagatagacagactgtctca
agagcgagaaaaattgaaagatttttatctcaaccttttttcgtggcggaagtgtttact
ggatcacctgggaaatatgtatctttggaagatgcaattaaagggtttcaaatgatctta
aaaggcaatttagacgacttgcctgagcaggcattttatctagtaggtgatatagatgaa
gctatacaaaaagctgacagcatgaaagattaatataattaacgatgactttaaatataa
gaattattgctcctgatcggactgtttgggatgcagaagcacaagaaattattttaccaa

gtagtacaggacaacttggtattttaacaggccatgcacctttgcttacagctttagata
ttggagttatgagagtaagagtagataaagaatggatgccgattgttttgctgggcgggt
ttgcagaaatagagaacaatcaattaactattctagtcaatggggcagaagaagctagtc
aaattgatttatcagaagcagaaaaaaatttagacactgcaactcaacttttaagtgatg
cctcatctaataaagaaaaaatagaagcaacacaaaaaatacgaaaagctagagctcgag
tacaagctgcaacagctgcaactccataa-aagattcc------tcttaaaacataaaac
tttagtaattattactaaa-------gttttatgttttatttgtatggttataggcaact
taaataaacttagcttagacctgaacaaatataatcaaaatataatcccatctctttacc
tgcatctggtccaactaaactaatagtaacttctttcatagctaaaatagcttgaatcgt
tgcaccaataggaacacctagggaattgtatgtttcttttagaccatttaatactctttc
ttcaagaatagatggatctccagctagcattccataagtagcatagcgtaagtagtagtc
taaatcacgaatgcatgcagcataacgtcgagttgtatacatgttaccacctgggcgtgt
aatatccgaataaagtagagattttgctacggattctttaataattgttgctgcattagc
tgcaatagtagcagcagctcttactcgtagctcacctgtttgaaaatagcctcttaattt
ttcaacagagctatcatctaaatatttaccttggacatcagctgcattaataacagaagt
aattgcgtcttgcataactttcacaaattccttaattttaaatatctaatacta-ggggg
tatataacaaaattactagatcctttagcgatatcagtattactgcatagcacccaaagt
ataatcaaagtaaaaaccagcttctgctgaatcttcacctgcaagcagtgagcaagctac
acttttcatgcattttacgccttcagcaactcctgaaattggtgttcctaaagaattata
catttctttaacacccactaatccaatttcttcaataggagttacatcgccagcaactat
tccgtaagttactaagcgaagataatagtcaagatctcgtagacaagtagcagtcatctc
ttcgccgtaagcatttccaccaggagaaactacatcaggtctttttgaaacagctgttg
gccgccttgctttacaatacgctcacgattatctgttaaaatttgagctattcttaaacg
acgttgtccagataaaacaaaacttttaattctatctaattccccaggacttaaatatct
tgcttctgcatctgcatttacaattgactttgtaacaatactcatggataaaactcctgt
aatgcttt-aaaaaaaagcttaatactaattatttatgatataactcagaaagtaatgta
ctttttgag-aattttataaccggaacacgttcatgatatacgaggcaggataggtactg
aggtaagtactaaaacgagaaagttatatgtaggaacttctgttattgatttcctagtac
tttttaaaacttggtaccacaattgatatatcctgcttagtcaaacgattataaagtgt
ttgtgtattaggaaagttagctgctggtaaagttggaaatcttctgtatggaacagtatc
tttaccatataccgcattgtattcatcactttcaactaaattacttataaatgcagataa
gccttttgaagctaaaatctgattcaagaatctaatttctgcctgattattaggagctcg
tcctaagatatgcttagtacctaattctataacttttgtatttggatatggctgataaaa
ttctttaccatatagttcagatagcgccaatttttctacgagttctttgacacagatttg
tctattcaagaaagctgactcaatatccaagaattcaccgccgatactaaaagagttaag
atctctttcgaaaatttgacgatatgctgctcttaaagcttgttctaacatttctttgtt

actatcactattcacttcaaagacaatagattgatctcttaaagaagttactccttgtga
aatacgagacttaatatcattactactcctcatttctttgacagcacctaattcaacaaa
cctagcagatttactagcaataactttttttgaatctctggtcaataattcctggtcttaa
gcttcttaaagcaacaccagctggcgtgctatatctttcatacggaacagtattatctcc
aaaagtttctgtgtactcagagctatcaagtattgcgtctattacttggtaatagccttg
cttatatgcaatatcaaaatatttattaatttcttgccgaccatatgtagggcgacccag
taagcggttatgaatatattcaatagctttacaaatatataatggctcccaataaagaga
tctaaatatacttgattttgctaactgtctaacaaattcgcgaactgtaatttgattgtc
ttttagctgactttctattggcttgagaattagtttttcttcttgatacacttctctacc
aaaaactcttaaataagcaacttttgtgacaacttctactgaattttcaaaattttggga
tgaatctgactttataagagcactggataatttaaagatttttggtcccagggatcctgc
tgacttaggtcttacctgtggattactaatctgattgtagattcctggcccgcgtctaac
tagtatccttcttgtatctttaccaaaaagggcttgtctttttcttggatctttattttc
tttagggaaaatagctccaaattgaattgataatggatcattacctatgccatatggatg
ttgatcaggtaatgattgcttataatcactaaataaagttataaattgaggaaccttacg
aaaaggcgcactataactgagaagatctatttgaggcccccaattacggcactcttgggg
ctcttcacctaaatttctaaagtaaggcacagtctcttcgccgaaataatctgtatactc
agaagaatttaaaattgcatttactaaaccacttaaaccagtagatgataaaatagcaaa
atatttttgaaactcttctagagagctaggacctctacccaagaaatgtctaaatgctaa
ttctaatgctcgactattgacaaaaggttcataaaattgctttctataaatacttgaagt
tcctaacgaacgaataaattccttaattgaaatttggccatttttttacttgtgactccaa
attcgataaagagaggtcatatgctttcgcaatatctctttcaaaaatttgtctataaca
tgctttcacaacagtatttttctcatcagcagataaactaggcttcataacaaagcgagg
tgttgacactcctgcctgtacataagtctgaggaagacgtaaaccttgtaaatctccaga
tattctttttcttaatttatcagtcaaagatggagcttcaaattcagaaatcacaacatt
aaaatactctttaactaaatcttgaccttttatatcttcctcaaagattagtaaggctgt
ccgtctcatctctctaagagccacaattgctgctgcacttgagcaagcattatcaattag
ctctctcaatcctcgtatattaacagataaaatattagggtcgcctgatacaattgcata
tgtcaaataccgtaaaaaccagtctaaatctcgaagagattttctcattcttgtggtgcc
gtatctcaagacgtttataggtttaaatcctggcggagtagctcccccagcattaaataa
agatcggaaactttggccaaaatctccttggatatttcctgataactcattaattttatc
ttgagaactttgatcacctgcgataataacagctgcttgaggacgttctaaataagaaat
tgcagacccacctacgaatattttatctgctgctcgtgctacgagaatatttgcattttt
agttaaaatatctgcaacttccagtctttttttgtccagaatttaaaaaagcaactaattg
attcaattcaccaagctgtaaaaatcggtcttgttgctctgcctgagtaatagttaaaat
tgaggcagtccgataaagctgggggcgtgctaatgggcttccgccacttgctttgatact

cattactttatttatctccttaactgatattgcttgctagcgaatacaaatatgacttga
tatttctaaatatttaaagatactatgcaagcaca-ctgttttctatattattaggcact
agtacttatagtacaaagagtcttacaactaacttctataaatacaaattttgtaatact
tctattaacatttatcaaaaatatatttaagtgctcactcttttatattaaaattcaatt
agtgagagattgacgtgaaataaagtatactaagtaagttttatagataggaaaattaaa
agatttattctacaaacattaatca------------cttgaatgtttagcaattataca
attgttattgcttttaattggtttagactagtttt-tattttttctaacataaacaatat
taatgagcaatttgtatgaatttaaagccacaaaacaacctattaactaacgaaaatata
aatcacatagatatacccgaaaatgatataccaatgtcaattacggaacatttagaagaa
ttaagacaacgaactttatttgtgtttttattctttttatttgctacaactataagtttc
acacaattaagataattgttgcaatactgcaagctcctgcttttggtattaaatttttta
caattggcaccaggagaatattttttttcatctattaaagttgcaatatattgcgggatt
gttgcaacaactccgtttgccgtttatcaagttatattatacatactcccaggactaact
cgaaaagaaagaaaaattatttacctctattaattagttctgtattgcttttttatcaca
ggtgggatatttgcttactttgttctcgcaccagcagctttaacatttttaattagctat
gggtctgatattgtagaaccactatggtctttcgaacaatactttgactttattctatta
cttttacttagtacaggattagcatttgagataccagtcatacaattattgctcggtgtt
tctgggacattttcttctaatcaaatgatacgagcttggagatatattattattatagca
acaattgctggagctattttgaccccttcaactgatcccgttacacaattaataatgtct
tcagctgttttattactttattttggtggaattgttatattattagtcttaaaaaagtag
agtaaat---------attttcta-ttaatccaatatatttgttatc-----tgggcaac
taatctataaaaacaatttct------------------agatttaagtatcaggcttct
aaatatttcaa---------gtctctttttatcatttaaataaaatc-------------
------------------------------------------------------------
------------------------------------------------------------
------------------------------------------------------------
------------------------------------------------------------
------------------------------------------------------------
------------------------------------------------------------
--------------------------------------------------gatacattgg
cttgtttttccatcaattatttgtattgtgcac-tttttagcactatctttattattctt
agaattatttccttaatctatttattaaat----agatatgaagagatttcagtattgat
tatggtaatctatatcaacgtaaaaagtattttcttatgagttttatttatataaaggct
tattcattatatagttataacagtaagtctaattaatag-ttttcatgcttaattaaagt
aataactcttaattcttttggcagaaaaatctgtcttaatattttatc--aaaaaaatag
-aaaaatgaagctaaatagtctaattaacttaattcaaaagtctatatattcttgtacac

ttttattaattattttaaatattatttgtgttgcacctaattctagtaatgcatttccaa
tttatgcgcagcaagcttatgaaagtccaagagaagcaactggtaggatagtatgtgcca
attgtcatcttgctcaaaagcctgtggagatagaagcgcctcaagcggtactacctaata
ctgttttcgagactgttgtgaagattccatatgacagcaatgctaaacagattttaggta
atggcagtaaaggaggcttaaatgttggagcggttgtaatattacctgaaggatttaagt
tagctcctgctaatagattatctacagagttaaaagaaaagactaaaaatctttacattc
agccgtacagtgctaaacaagataatattttagtaattggacctatttctggtgataaaa
atagagaaatagttttcccaatactatctcccgatcctgcaaaagataaaaaagctcatt
ttttcaagtatccaatatatgttggcggaaatagaggacgaggccagatttatccaacgg
gtgataaaagcaacaataatattgtctctgctttaagcagtggtaaaattaataaaattg
aatcactagacaagggtggatttataatacatgtgactaacagtagcaatgtagagtcaa
cacaaaagatttcacctggtctcgaacttaaagtgaaagaaggggatacaattcagcttg
atcaagctttgaatagtgatccaaatgtaggtggttttggccagaacgaaacagaaatag
ttttacagagtccaaatagaattaagggcatgattgttttctttttcgctagtgttctag
ctcaaattttcttcgtattaaagaaaaaacaatttgaaaaagttcaagcagctgaaatga
atttttaaaggaatataaaactggcacttactataaataaagta-taaacactttattca
caggatgtaccagtctttatgttatgtaatactcgctttgacggaggttatgatactttt
aacagaatctacaatttggctaggctgaattactgtagcctgctctaagcttccattata
tggtgtgggtagatcctgcgaagatagcctcacgacaggagcatctaattcatcaaaaag
atattcattaatttgtgcaattagctctgctccaattccggctgttttcatacactcttc
tacaattaaaactttgtgagttttctttactgagatagatatagagtctatatccagagg
ttttaaagatataagatctataacttctggatcataaccttcttttaataaggccggtaa
tgcttggataacatgatgcctcattctggaataggttaaaatcgtaatatcttttccttt
tcgtacaaattcagctttattaagaggtaagaaatattcttcttgaggaatctcttcttg
taaattgtaaagtagaacgtgctcaaagaatacaactggattattgtctcgaattgcaga
tttcagtaatccttttgcgttataaggagtagaacaagcaactatttttaagccggggat
ggcttggaaataagcttccagtctctgagaatgctctgcacctaactgtctacccacacc
tcccggtcctctaataactaaaggcaacgtaaaattacctccagaagtataacgtaacat
tcctgcattattagaaatttgattgaatgctaataataaaaaactcatattcataccttc
aacaattggtcttagccctgttatagctgcaccaattgccataccagtaaagctattttc
tgctattggtgtatcaaggactcttaaatccccatatttactatgcaaatctttagttac
cttataagatccaccatagtgaccaacatcttcgcctattacgcagacagttggatcttt
ttccatttcttcatctgttgccgctcgtaaagcgtcaaacataaagattttactcatttt
aatatttgatttttatgatttgaccactgttttaaagtaacagttaagactccatattct
atttagttatctgcaaaaagatatcgtttcaattctgaaatgttaggttctggactggaa
atagcaaatttcacggcttgttccaattctgttttaacagcagcctgaatcccattaagt

tcacctatatcagcaatttcattatctagaatatatttttttgagttttttgataggatct
cttgccacccaagcctctttttcttgtcttgatcttagttcatcaggatctgccagagag
tgaccacgaaatctatatgttaatgcttctattaaggttggaccatcaccttggcgagct
ctttgaactgcttgttttgcagcttgtcttacagctagcacatccattccatcaacttca
atcccaggaagcccaaaagcttccgctttttttatgtatttcaggtattgaagaagaccgg
tgatgtgccatacctatagcccactgattatttcaacaacaaatataataggtagtttc
cagagaactgccatattcaaacattcaaaaaaattgcccattgttggtagtcccgtcacca
aaaaagcaagccgtgactcttaaatcgtctgtttctttaagtacttgctgacggtagata
ctttgaaaagctgcccctgttgcaactggtataccttctgcaataaaagcaaagccaccc
aaaaaattgtgaggcgcagaaaaaatatgcatcgaacctcctctgcctttactacaacca
gtctcttttccaaacaactcagccattacatttttagatgggacgcctttacttaaagcg
tgtacatggtctcgataggtactgcaaacataatcagtcggattcagaagtttaattaca
cctgtagaaacagcttcttgaccattataaagatgaacaaaaccaaacatctttccttta
taatacatctgagcacacatatcttcaaaatttctgcctaacaacatatcttcatataaa
actaataaattactcttattaagattaagacctgttgaattataattagtcagcggcaat
tgaactttcttaggataactcataatttgtaaagaaacctct----------tattgtaa
ataggaattttgaagaaaaagctatcaatagaaaacatacagataaaaacttataaatta
ttgtatgtttttactaaagactttaataattatttgacactagttcagtatatcaaatt
aattttgcttatactacagatgtctgaaacttagacaaaaaagtagtaa-aaaaaattta
ttaaaagaatagttaaaaacagatataaatagaagtaaacagaattatt-gaaaaaaaat
gattattattaattaa----tacttaatatttctaaatattttacaattaatggctatgt
caccatctttattttatcctgttgaacaagaactatgtagtcttgaaaaaaatctgaaag
ctgttgccgggactcgtcatccaattttatatgccgcagcaaagcatctattcgatgctg
gaggaaaacgagttagaccagctcttgtgtttttagtggctaaagcaacctctgagaagc
aagatataaatactggacaaaaaaggctagcagaaattactgaaattatacatactgcta
gtttggtacatgatgatattatcgacgagtgcacaacacgtagaggagtcaaaactgtac
ataatttattcaatactaagattgctgtgctagcaggagatttttttatttgcacagtctt
cttggtatttagctaatattgaaaatttagcagtagtcaaagctatttctaaagtcatta
ccgactttgccgaaggagaaattaggcaaggcttagttcatttcgatcctaatatttcaa
tagatgcttacattgagaaatcattttacaagactgcttcactcattgctgctagttgtc
gcggtgcagctatgcttaatggttgcaatcatcaaataaataatgatctttatctttacg
gtaaacatatgggattagcatttcaaattatggacgatgttttagatataactggttcta
ctcaaagcttagggaaacctgctggcgctgatctaataaatgggaatttaacctctccta
tcctttttttcacttactcaagaaacaagtttaaatgatcttattgatagagagttctgta
atagtacagacgtagcctcagcattatttcttataaaaagaagcggaggaattacaaaag
ctaaagatttagctaaagaacaggtgcaggcggcactttcctgccttcagtttttaccac

aatctacacctgtatctagtttaaaagaattaacacatttcataatcacaagattgtcgt
agaggacttgctaaaattaaaatgttattaactaatttttttactgtttctaaaatagtct
gaaaagtttgaatatcattactagctagctgcgctagcattttacgatttaacgcaatat
tctcttttttaatgcactaataaaagtactataattcatgccttgattgtgagcagcag
catttattctagttatccaaagacgacgaaaatctctctttttccttttgcgaccaacat
aagaatatcgaagagctttaaggacttgctgcttagctgttcgaaataaacatttatgtg
cacctttaaagcctttagctagcttaaagattttagctcgtcttttttttgcaacgttac
ctcttttaactctactcataaaatattac-ttaaattttaattattaccagcttactaaa
ttcgtatactctttg----attataaataaggtaattttattgcaatatttttaatatcc
tttaagtcaaccaaacaagtagaagagagatgtcttctttgttttgatgactttttttgt
aataaatgacttttagaagctttatgtcgaagaaattttccggatgaagagactttaaat
ctttttgctattgcttttgatgtttttaacttaggcatattataaacttttaatgattaa
caaaagcatgaaaagtctgaggatattaatcttagacctttcaatacttagaataatctt
aacacaatttattttacttactacataggttaactgaaacaatttatatatataatacta
atcagttaatcaaccgcagctaaagtggatttgtattttgctttgaacttttgtatagta
ttagcaattagcattgttataattttaagactacttgaatttaattcttcgaacattttc
atatttaatgtaaatgaaatattagcttctgagacaatattttgaatttgaacatcagac
aaaggaatcatatctaatgcagctctatattgatctttaaataatttatcgtcttttatt
tggtcaaaatcgtaaaattttgttcctcctgaatctgataaattcatagcacctctagct
atttttttaaaatttggccgccagaaagatcacccaaatagcgcgtataagcatgagcg
actaataattcaggttgtttatgtcctataatgtgaattctatcaacatagattttttgta
gcaggagatggttcaataaaatctgaccaatctgatccgtaataataatttaaatcttca
gatagactctctttcctattgagctctgtaaaatatataggtttaatagctggatgatct
ttattagaaaataactcttcctctattgcacaatagacaaagtatagatttgcaaccaat
ttacgatatgacttcttatctacaactcctcccaaaaaagatttaacaaaactaacattt
tctgccatactatgggacttagtagtaccttctcttagttcattcgctaaagtattaatc
ataatatcaatattccattttt-aaagacaacaaattaaca---ataaatagttatccgt
atatttatacttgagttgtgctaactaggctgctgcaaaataatttttttgattttattgg
atctggattcattgttttgtcaccaggcttccaattagctggacaaacttcatctggatg
agattgtacatattgaattgcttgtaacacccttaaagtttcttctacactcctgccaaa
ctctaaattattgattgtagagtactgaataattcctttcgggtctataataaatagtcc
tcttagagctacaccatcactattcaaaacattataagccgcactgatctctttttttaa
gtctgatactaatggatatgaaagatctcctaatccacccgattctctgtcagtctgtaa
ccaggcaagatgagaatactcactatcaaccgagactccaagaacttctgtattaagttc
ggaaaaagcattatacttatcgctaaatgctgtaatttctgtagggcatacaaatgtaaa
atctaaaggataaaagaataagacaatatacttgttcttgaaatcagataattttaacgt

tttaaattcttggtcataaacagctgtagctgaaaaatcaggagctagctgacctactcg
aagacaatttggtccagaaatcattaatttttcccagaagaattgatataattttgttat
ttaataaataattatataacaaaatttagtatatattaaaatactaccagtctttttaa
attac--------taaatgatataatcaatactatctttatagcggggaatggatttgaa
ccattgaccttcgggttatgagcccgacgagctaccagactgctctaccccgcggataac
tagaattatattagtctagatatatatataaaagcaaatgcataatagattttagaagta
aaaaaatagaagaacaggtccactgatcattaaacttagaaacgataaagtatttttat
cttatttaaaagaggaaagacttcatataaaccaataaatctatcttttaaaagaatttc
ggacggctttacccaaatttgaccatcataccagcctgattcttcataaaatactgtagc
actcattaatcgttttactacgtatgaccagcctaaatataatctgattaaaataaatcc
tgtcattaggctggaagtaataaattctgaaaagaaaaatttcagaggtagtttattgat
tggaaaaattgataataatataggactaattaataaacaattaattagtagtgcaatagt
tatttttttattataagatctgcgacttaaagttggccagcaaaagaaccaagaatttt
caaagaagtatattcatgaacaggttgctgctcctttgggacaggacattgagtattata
taaattcattgacattaaaaagtattatactaaagac--tttctttatagtatctaatat
aggtacgacattaaatattaactaactgcaattatagtagtgagtaaaagaaataaaact
aaaaatagccaagtaattttatttaaagtactctcagtactacgagtattactaaagaac
tgattctgagcaccgacacttcctaagccttcagatttaggattatgtattaaaatagtg
aagattaatattattgttgatgaataccaaaaaaattttaaaatttgttccattatgtaa
gagttaat-aaaaatagcacaaagacaatcgtctaaaaggtattgattgtcttatcatat
taaaaagtatttattctccctgtacttttaaaagtaaaaatcctattgccagccctacca
aaactaggactgaacttaatacggcactttctacaatttctccgaccatt-ttatttttc
cttattactttattattg---tcattcttttgacaagattgtattgagctaa-ctaaaat
ccgtttctaccccaaactaccattgcaattgaaaatgtaaacattgccattagagctgcc
catcccaaacttaaaatatccataatttataatactc--ctaaaaataagatctcttta
cttaaactataatattcttaatattaaaaatatcattatataataagctaattataacag
agtactacttaattcgtatttttggtgtatttaaagaagattgaatttttttcgaggtac
tggataagtatttcatcctaagaaccatctttaac----ttttgatttattatactgctt
aaatctaaatata--tagcactcaagaatcattagtaattatgatgtttttaattatacg
attaacaacatatgtcaaaatagaagaattcatagcgactcagttaa-----tatgctaa
tataatttatcaat--aaaaaggaagttagtatttcaattatgcaaacaactataaataa
tgggcaaccgtctagtaaagaaactttactaacacctagattctacacgactgactttga
agagatggctaatatggatatttccggcaatcaagaagatttttttggctatcctcgaaga
atttcgagctgactataatagtgaacactttattagagatgaagagtttaatcaatcttg
gtctaatttagaacataaaactaaatccttatttattgagtttttagaaagatcttgtac
cgcagaattttcaggtttttttactatataaagaattatccagaaaattaaaagacagaaa

tccggttatagctgagtgctttttattgatgtctagagatgaagctaggcatgctggttt
tttaaataaagctatcggagacttcaatttatctttagatttgggattttaacgaagag
tcgcaagtatactttttctctcctaaatttatttttatgcaacctatctttctgaaaa
aattggatactggagatacataactatttaccgccatcttgaacaatatccagaacaccg
tatttatccaatctttagatttttgaaaattggtgtcaggatgaaaatcgtcatggtga
ttttttgctgctttgcttaaatctcaacctcatttttaaatgactggaaagcaaaaat
gtggtgcaggtttttctattaagtgtatttgcaactatgtacttgaacgactttcaaag
aattgattttataatgccataggcctagactctagacagtatgatatgcaagtaatacg
caaaaccaacgaaagtgccgctagagttttccctgttgctttagacgtggacaatccaaa
atttttcaaatatttagatacttgtgcatgtgataatagagctctgatagatattgacaa
taacaattctccattattcattaaatctattgtaaagatacctctatatttttccttatt
tacaaatttactgaagatatatttgattaagccaatagactcaaaaacagtatggaatac
agttcgatagttaaatacaagggaatactttttattataaaaagctatctaattattaa
tacaagat------aaaagcttccttct-----cgatggaaggagcttttattttgtcaa
aaa-tatataatgggattatatagatc-tagaactaagaacttactttttcttttgcttc
atacataacctctaaagtaattatatttttagaacaatcacgggcaaatttctctgtatt
tctttttactttacctctaacaaacccaggaattttatttaattctttctgtgcttcttc
tgaccaattaatactatcaacagcagaaattcctacacttaatgcttctgttgtgtcatg
accaccaaatatttccagcaaatgatcttccataccaagtgtaaatgaattatatactaa
gtcagcaatttgatttgttccctcataaccaagaaatggacggtaacttaaaggaaagtt
ctggatatgcactggtgaagaaataacgccacaagggatatttaaacgcttaccaatatg
tctttccatctgtgttccaaaaattgctgcaggttcagttttagcaattaaatctccaat
taaaccatgatcatctgaaacaattacttcatcgcaaaattcttgaacttgatctttaaa
ccactcttcatcatatttgcaataagttccacaccaagcgacatttatacccatttcctg
gtgtaaaattcgagttattgctgcagcatgagttgcatctccaaatacaatagccttttt
acctgttaaattctgacaatcaattgatctggaaaaccatgctgattgtgaaataaatcg
ggtttgttcatctatgtatttttcgtaatctacagcagctcccgatgcatttaataattg
ctgtattgatcggatgcatgcagctgtttgaacaatacccataggtgtaatgtcaacata
aggcatattaaactctttctctaaatatcgagctgtcattagtccagtttctctatatgg
aataaaattaaaccatgcagatggtaatttttttaaatcttgaactgatgcattttctgg
aataatctggtttatttggatatctagatcttgaaataacctttttaattctgctatatc
atgctgattgtgaaatcctaagcttactgctccaataatattaacagatggagttttcgt
cttttgagttaaaacttgattattagatttagctttttccatatagaaagcgacaatttg
ttctaacgttctatcacctgcttgcagttcatttactctataatgattaacatcagctaa
taaaacatctgcttctgtttctatagatgctctgctaacaaagttttgcaaatcctcttg
caaaatactcgaagtgcaagttggtgttaaaataactaaatcagggctttcttctctgtc

ttttctagtgatattttctacaactttttcttgcgaaccacgggctaacacatgtctatc
cacaacacttgctgtaacaggagtaaaatctctgtctctttcaagcattgaacgcattac
attaaaataatcgtctccgagaggcgcatgcataattgcatgaactttttttaaaagaact
ggcaattcttaaagttccaatatgagcagggcctgcatacatccaataagctaatttcat
attaaattttcctatgatttatttagacatagtaatacttttaaagctaactaacaaaca
ttagaaaaacatctaatttaactccttgataatttttgtaatatttttttagatgaagctta
tcattataaatatttaaaaaatataattcagaatagaaattaataactgttagatcaagc
ttgaaagtaaagatttcttgcagttatagttttttctttaagtaaacatgtaaaaaattgta
ctatatataaatgtactatatataaactttttttactgaatattggtaatgccagataca
attaacttaaatatgccttccccaacgtttggtggaagcactggcggttggttaagagcc
gcagaagtagaagaaaagtatgcgataacatggacaggtaaaaaatgaaagtaagtttgaa
atgccaaccggtggtaccgcaactatgcgaaatggagaaaacttactttatttagctaag
aaagaacaatgtttagctttaggtactcaactaaaaagtaaatttaaaatatctgactat
aagatttacagagtttttcctaacggagaagtgcaatatttgcatcctaaagatggagtc
ttcccagaaaaagttaatgctggaagagctagcgtcaacagtgttgatcattccattgga
caaaatattaatcctgtagatgtcaaattcatgaataaagcaacttacgattagtctctc
-aaaaacctagacataaaataacttttatatagtctaggtttttgataaggtatactgat
agtatggaagggtggccgagtggttgaaggcgtctgatttgaaatcagttgaacta--tc
tggttccgtgggttcgaatcccacccttccgttataaaact------------atagtt
tgcttctcaaggcaagctagctatacactgatagtattaagtttagacttaataaagtct
acaacttgtgacaggttagaaatttttttcggcatcctcatctggtatctctatactaaat
ttttcttctatggccattactagctcaactgtgtctaaagaatcggctcccaggtcactt
gaaaaatttgcttccctggtcactatttttttttcaattcctagctgttctgctacaata
tcttgaacttttttcaaagatttcattatcttgcataatatattatttcctctaaggtgtt
atcaaaaatttttttggcttatattattacccaactctaacaaattagtaatttttagtcg
agattaaaatatttatacctatatgatatactattattctggataaatacgaagtttacg
atatggctcttctccgaaactacgagctctcaactggtagttgtcaactagattatgctg
catttttcggatataggctgaccgaggtgtaagctgaactattgaactttcattcaagat
aatagtttcaacagccaatttagcttcttgaagagcctggatctcatcaaactttttatt
tttacagagttctacccaattaagaccagatgaagtatgaatgtttaatatttttctgag
tgcacaagtaatttgtggaaccgtactgttctgaattgtatatataataatttgttttga
ttttgcaatttgtctaagttttgtattttgctttacttgattacgtagcgctagaatagc
atcagacttttctatctccttggttaaaataattggtaaatctagtgaagaaataactga
tgtaatatgctgccaactcaaagaataagcatataagtactgatgaggtatttctaattg
caaagattgtttattaatattaataggggcattgaaagtagttgacaataaagatgtatc
acgattttgtgatttattcaaatctaaggacttatttcttaattctctatattgtaaaac

aggagacgcctgctgtgtgttaattgatttacgactattagagatgtttacaggtaagac
ttctacagattgcgatggataacatttaattaaaattctaccattagcttgaatttgtct
tttttgcacaaaaggttgatggccttgcaaaatttgatctattgtctctttaactttatt
atgaactatccaaacatttcgctcatgtatttcgatcgcaatctgaaaagctggcgatgc
ttttctttctagtatactcttctgcgtacctctacgtttagcttcatcgtcacccaatgt
tacatattgaattccgccgattaggtccgctaatgttggattttttaattaaactttctaa
atgatttccatgcgctgtacctactaattggactcctctttctgctatagttcgagcagc
taacgcttctagctccgttccaatttcatctataataataacctcgggcatgtgattctc
aacagcttctatcattacttgatgttgcaaatcgggccttgcgacttgcattcttctagc
tctaccaatagccgggtgaggaatatctccgtcaccagctatttcattcgaagtatctat
aataactactctttttttccatttcatcagctaaaacacgagccatttctcgaactgcggt
tgttttaccaactcctggttttcctaagagtaaaattgaatcacctcgctctagtaaatc
tcgaataatactaattgtaccaaaaactgctctgccaacacgacaggttaagccgataac
actaccctctctattacgcagtgaactaatacgatgcaatgtttttttcgataccagctcg
attatcgccgctaaaattaccaatttttttttacacagtaatctaagtcttgccaactaat
agatctttgagataaatattctggattatctggaaatctggcttctggacgtcgacctaa
atccataacaacttctattaaattatttctattgggatgctgttttaaaggttctttgac
aaaatttggcaaaatttctagcaacttatctaagtcatctgcaataagcatgattgagta
tataataattaattaatcaaatagaagattcaagattctaaagattacaataaatattaa
agaatattataattggaatcaattttatatttaaattgaacaggaattta-tagatgtaa
aagaactgattaaactttacgcaataaatcgattggtcaaaattaatagtcgtattaaca
tgtttattggacaaaaagttcgtattaaatatagcaaacaaaaagttgctagtgatatag
cagataaagttggtgaattaggagttattaagggaataaagtttattaatagtcaatgtg
ttactattattgttgagtttgacaatcatactaggctttggatgtttagagaggaattaa
tctgtctaaatgaat------taa-------aaacataatttattttatttattatacag
aatttcttatgagtcgttaacataataaattttaaaggagtacttttaattgtgcaaatt
actattaaaaaattacaagacttactttcgtctgtacaaagaaaaaagatccagacatta
aaactaaaacaagataagtttgaacttctattaaataagacctataaaaaggtcaatcaa
gaaataataccttcgcagaaatctgctgtattacaaaatagtccatctacgacaattaaa
tcaataaatagtacaaaaaaatcttctgttgtcaatgaagaccgtacagaatatgccact
attgtttctccaatggttggaacgttttatcattcacctgctcctggtgaaaaaattttt
gtacgagttggcgatgaggtcaaattcaatcaaacagtttgtattattgaggcaatgaag
ttaatgaacgaaatcgaagcggaaattgaaggaaaaattatagaaattcttgttaaagat
ggtgatatagtagattgcgggcaagctttaatgaaggttgaaatataatgctaattttcc
cattttacatattgttaacagaaattatttagatagaaaaattagcttatactatagtta
tgaaaacttatttcgcaaaggcacaaaaaaactaatatataaaattattt-ttttatctg

ccaaaaatagaagtaagcgttttgattccttgtctcaagagaggagaatctcaatggcaa
ttagctcaaaagagcaagagacaaagaaggtaaaaatctcggttgataaaaatcccgtag
atacttctttcgaaaaatgggcccaaccaggccatttttctcgtacactagcaaaaggac
caaaaactactacttggatttggaatcttcatgctgatgctcacgacttcgatagtcaaa
ccagttctttagaagaagtttcacgtaagattttcagtgcacattttgggcaactgtctg
taatatttttatggcttagtggaatgtattttcacggagcccgcttctctaactatgttg
cttggttaagtaatccaacaggtattaagccaagtgcacaggttgtttggcctatagttg
gacaagaaattttaaatggcgatgtaggtggtggctttcaaggagtacaagttacatctg
gatggttccaactgtggagagcatcaggaattactacagaatttcagctttactgtactg
ctattggcggattagctatggctgctttaatgctgtttgcaggatggtttcattatcata
aagcagctccaaagttagaatggtttcaaaatgttgagtcaatgatgaatcaccatttag
ctgggcttttaggcttaggctgtttaggctgggcaggtcatcaaatccatttgtctttgc
ctattaataagctactagattctggtgtgtctccgcaagaaatcccactacctcatgaat
ttttaattaatagagagcttatggctcagctgtatccaagttttagtaaaggattagttc
cattctttactttaaattgggctgaatattccgactttttaacttttaaaggaggtttaa
atcctgttactggaggtttatggctaagtgatactgctcatcatcatctagcattagctg
ttctatttcttgctgcaggtcatatgtatagaaccaattggggtattggacatagcatga
aagaaattctagaagctcataaaggacctttaccggcaacggtcacgaaggtctatatg
aaattcttacaacttcttggcatgcacagcttgcaattaatttagctatgatgggatctt
taagcataattgtagcacatcacatgtacgctatgcctccttatccatatattgctactg
attacccgactcaattatcgcttttcactcatcatatgtggattggaggattttgtattg
ttggagcaggagcgcatgcttctatatttatggtaagggattataatcctgcagaaaatt
ataacaatcttttagatagagttattaggcatcgagatgctattgtttctcatctaaatt
gggtatgtatatttcttggattccattcatttggtttatacattcacaatgatactatgc
gggcacttggaagatctcaagatatgttctctgatacagctatacagttacaacctattt
ttgctcaatgggtacaaagtatacacactttagctcctggaaatacagctccaaatgcgt
tagcaacagctagttatgcatttggaggagatgttgtttctgttggtaacaaagttgcaa
tgatgcctatttctttaggtactgcagatttttagttcaccatatacatgcatttacta
ttcatgtaactgttttaattttagttaaaggattccttttctcgagaaactctagactaa
ttcctgataaggccaatcttggcttcagatttccatgtgatggacctggtagaggtggta
cttgccaagtatctggctgggatcatgtttttcttggcttattctggatgtacaattctc
tgtctgtagcaatttttcactttagttggaaaatgcaatccgatgtttggggtagtgtat
ctccgtccggaaatgtttctcatattactggtggtaattttgcacaaagtgcaattacaa
tcaatggatggttaagagatttcctttgggctcaagcatctcaggttattcaatcatacg
gttctgcgctatctgcatatggattaatttcttagcagcacattttgtatgggcattca
gtttgatgttcttatttagtggtagaggttattggcaagagcttatagaatcaatcgtat

gggcgcataataaaataaaagttgctcctgcaattcaaccaagagctttaagtattactc
aaggtagagcagtcggtgttgcacactacttattaggtggaattggtacaacttgggcat
tctttttagcaagaattatttcagtaggctaatagtgaaaataggat-aaaaaaacaatt
atggcaacaaaatttcctaagtttagccaagctttatcacaagatcctacaactagaagg
atttggtatggtattgctacggcacatgactttgaaagtcatgatggaatgacagaagaa
aatttatatcagaagatatttgcttcgcactttggacatctagcaattatcttcttatgg
acatctggtaatttattccatgtagcttggcaaggcaactttgaacagtgggtattaaat
cctttgaaagttaaaccaattgctcatgcaatttgggatccacattttggacaaccagct
ttaaaagcttttagtaaaggggggctcagcttatccagtaaatatagcatattctggcgta
tatcactggtggtacactattggtatgagaagcaatcaagacctatattctggggcttta
ttcttactagttttatcagctttactcttatttggagggtggctacatctacaaccaaaa
ttcaagcctggtttatcctggtttaaaaataacgaatcaagattaaatcatcatttatct
ggattatttggggttagttctttagcctggacaggtcatttagtacatgttgctatacct
gaggcaagaggacaacatgtaggatgggataattttacaacagtattacctcatccagcc
ggtttacaaccatttttcagtggtaattggagtgtatatgctcaaaatccagatacagct
caacatttattcggaactaatgaaggtgcgggtacagcaattttaacatttctgggggga
ttccatcctcaaagtcagtctttatggctaactgacatggctcatcaccatttggctatt
gcagtagtattcattgttgctggacatatgtataggactaattggggggattgggcacaat
ctaaaagatattttagatgctcatagaccacctagtggtagattaggagctggacataaa
gggttatttgatactattactaattctttacacatgcagttaggattagcattagcttcc
ctaggtgtaattacttcgttggtagctcaacatatgtatgctatgcctccatatgctttc
atggctaaagatttcacaactcaagcgtctttgtacacacatcatcaatatattgctggg
tttctaatggttggagcttttgctcatggggcaatattctttgttcgagactatgaccct
gaacagaataaagataatgttttagctcgcatgctagaacataaagaagctatcatttct
catctaagttgggtaactctattttttagggtttcatacattaggcctttatgttcacaac
gacacaatgattgcttttggaactcctgaaaaacaaattttaattgagccggtatttgct
caatggattcaagcttcttcaggaaaagcactttacggttttgatgtgttactatcatcc
tctactaatgtcgcaacacaagctggtagcaatatttggctgccaggctggttagaagca
attaatagcggaaaaaattcattgtttttaacaattggtcctggtgacttcttagttcat
catgcaattgcattgggattacatactactacgttaattttagttaaaggtgctttagat
gcgagaggctctaaacttatgccggacaaaaaagactttggttatagttttccttgcgat
ggacctggcagaggtggcacctgtgatatatctgcatgggatgcgttctatttagctgta
ttttggatgctgaatacgataggttgggtaacattttattggcattggaaacatattaca
atatggcaaggcaatgcaactcaattcaatgagtcttcaacttatctaatgggatggttt
agagattacttatggctaaattcttctccattaattaatggttataatccatatggcatg
aataatttatcagtatggtcatggatgttcttatttgggcatttagtatgggcaacagga

tttatgttcttgatctcctggcgtggctattggcaagagttaattgaaactctagcctgg
gcgcatgaacgtacacctttagcaaacttgattcgttggaaagataaacctgttgcgcta
tcaattgtacaggcaagattagtaggtttagcacatt-tttctgtaggatatgtattaac
ttacgcagcttttgtattagcttcaacagcaggcaaatttggttagactaaa-----ttc
aaattagctaa-----aaaaaagtcagtcataata-gttttatgactgactttttggat
ttaaaacaaaaacttactaaagtttcacctcaatatctacaccggaaggcaaatttaact
tcattaatgcatctatcgtttgagaagaaggttgatgaatatcaataattcttctatgag
atcttatttcaaaatgttctcttgagtctttatctacatgtggagaacgtaagacacaat
aaattcttcttcttgtaggtaaaggaattggccctactgcaacagcattagttctagatg
cggtatctagtattttgtcacacgacgtattaagtatactagaattatatgctttcagtt
taattctgatttttgtctgctgagtaattgtcatgttaaaaagatcttattatttaagaa
ttttagagacaacacctgcacctacggtacggccaccttctctaatagcgaaacgcatac
cttgctcaattgcaattgcattaattaattcagcagtcatcttaattctgtcaccaggca
tgaccatttccgcatcagtaccatcatcagcagtaaactgattaatagtaccagttacat
cagttgttctaacataaaactgaggtctatatcctgggaaaaatggagtatgtcttccgc
cctcctctttagttagaatataaacttctgcttcaaattgagtatgaggtgtaattgtac
caggtttagctaataccatacctctttcaatatctttttttctgcacgcctcttaaaagaa
ttccaatattgtcacccgctagaccttcttctagcgtttttgaaacatttctaatccag
taatagtcgttgtacgagtttctcgtaaacctacaatttcaattgtatcaccaactttaa
taatgcctctttcaattctaccagtggcaacagttccacgtcctgtgatagaaaaaacat
cttctacagccattaagaaagttttatcgacatctctctccggtgttggaatatacgtgt
caactgcttccataagtgaaaaaatcttgtcaacccatttatcttcaccttgcttaatag
ctgggtttttcgtcacagcttctaatgctaataaggcagaacctgcaacaaaaggaatat
catctccaggaaagtcgtattgacttaataattcccttccttctaattctactaattcta
gtagctcttcgtcatctacttgatcttccttatttaggaatactactaatgtgggaacac
ctacttgttttgctaataaaatatgttcacgagtttgtggcattggaccatctgccgcag
atacaactaaaatagctccatccatctgagcagcacctgtaatcatatttttttacgtaat
cggcatggcctggacaatctacgtgagcataatgacgattatctgtttcgtattcaacat
gagcagtattaatagtaatacctctagctttttcttctggagcggcatcaatttcatcaa
atttttttgctgcagtagaacctaaagttgataaagtcgcagagatcgcagctgttaaag
ttgtcttaccatgatcaacatgaccaattgttccgatattgacatgaggtttttttacgtt
caaatttagatcgagccatgctttttgttttccttatagtaaaagttgctttatagttac
ttttaacgataaatcttgtgaattgaaagatttattatttaatacagacaaga-----ta
aaatataactaatctaatatcgatagtgagcaaacgctttatttgcttctgccattctat
gtgtctcctctctctttctaatagaatttcctgtttcattagctgcatccataatttcgt
tagctaattttatagacatacttttaccagatctttctctagaaaatttagtgatccatc

ttaacgctaaattcgtacctctataagctcgtacttcaataggaacttggtaagtagaac
caccaactcttctagcttttacttctactagaggagtaatatttcggatagctttttcta
aaatatttaaaggatctgattctgttctctctttaacgatatctaaagcctgatatataa
ttctttgagctaaagtttttttaccacttttcaaaatacgaacagttaacatacttacga
gtctgcttttatataaaggatcaggtgatgcgaacctttttttagctgtattacgacgag
acatagtatttaaaattggttattgtaataa-tgttc-tattagaagttatttaagattt
aggctttttttgtcccatatttagatcggcttttacgacgatcctttactcctgcagcatc
taaagtaccacgtactacatgatatcggactccaggcaaatctttaattcggccacccct
aatcagtaccacagaatgctcttgaatattatgacctacgccaggaatataagctgtaac
ttcaaatccagatgttaatctaacgcgcgctacttttcttaaagcagagttcggtttttt
aggagtcgtagtatatactcttgtacaaacacctcttctttgagggcaactttgaagagc
tggagattttgtttttttatgtatttttcgtctttcggatctaacaagttgttgaatggt
tggcataataaaattgatttaaagtctttaatagatactatttgaacctgctttagatga
cttggctagtttttatttatcttatacttacgcataaatctttctactctaccttctgta
tcaataattctctgtgaaccagtaaagaatggatggtttcctgaccagatatctacgtgt
agttctggctttgtggacccaatcgtcataattaactggccgtcacaataaacttttgct
tctggataccaatttggatgtatattatcttttgccatggtttctataattatataaatg
tatcaataataactattaacgcttagaaaattgaggagcctttcttgctttctttaaacc
atattttttctctcttttactcttggatctcttgttaaataaccctcagacttgagtgt
cgttctattttctggattaattgagcacaatgctcttgcaacacctaaacgaattgcatc
ggcttgccctgtcaaacctcctcctctagcattcacatgtatatcatactgatttaatag
tcctaaaacttgtaatggtgcatatgaaactcttaagtaattaggactaaattgaagata
agactctcctggtataccgtttataattaaattacctgatcctgggactagtcttacttg
tgcaacagaacacttccggcgacctgttccagaatagattgcacgagttttaattaattc
tgtcgacataacattccttgataactaaataaatatacattctttatactatatactctt
gtggcttttgcgcgacatgtgggtgaattgggccagaatatactttaagcttcgtaaata
gttttctgcctaatggacctttaggaagcatacctttaaccgatttttcaataattctgt
taggtaatcttgtctgaagctgataaaatgtttcaacttttaatccgccaggttgtccag
aatgtcttctgtatagtttttgattcgttttattgccacttacagatacgtgagcagagt
taataacaattacataatctccagtgtctagatagggtgtataagaaggcttgtttttgc
ctcttaagatattagaaatatgagtagatattctaccaagtgtctgatttttagcgtcta
taacataccaatgagaattagtatttaatgagggtgattgcgttttattcatgaaaagat
taatacttttggttgacaataagagaatatgtaaatggtttacattcaatttctatacaa
gtaatattt-aaaaatatta-------------ttaacaaagagaacttttctaacagtt
atttcaagtgtttgtcagcaagaaaatttctagttctcaataacttatactatttttaat
aatataaaatatattaatcactttttttcttttggcaaacttatacctagttttttttgc-

aaggcatatataacttcttcagctgattttggccaaaattttaatttctagcagttct
tcttgagagtaatctagtaaatctgcaatagaatggatttgagctcgctttaagcagtta
taagccctaacagataactgcagttcttctattaaaacttgactaatttttttatcttct
ttactacgataattatctgctgatttaaagtctaaatttcttagtgaacaaaaaagatta
gttaaaactgtagctccttgacttattgcttcttgaggagagatgctcccatttgtccag
atttgtataatcaatctatctttatgctgttgctaccaatacggacttcttctacttta
taattaactttattaactggcataaaaacagagtctacttgcaaaaaatccacagataat
tcatctactgcttttcagctaagcgatacccgcaatttttttcaattttaaactccatt
tcaaatattgtattattgcaaatagttgcaatatactgtctaggatctactacctctata
tcagaagataattcaaataggccggctgtaactatagctggcccttgaactctaattcga
ccaatttgagattctttgttataacttttaaatactacttcttttagattaagtaatatt
tctaacacatcttctcttaccccagggattgtagaaaactcatggttcactccagcaatg
cgtacagcaactatagcagtaccttcaagatctgacaatattgatcttcttaaagcatta
cctaatgtaataccttgtccttgatttaatggttctattacaaaactaccgtactgccca
cgcgccccatgtgttcttgactctacgcattcaatttgaaattgagccacctaagaaagc
tcc-tttaaaataatcagtgattagtaagcctctctaaaataagaaggaatttatactcg
gcgtttcttaggaggacgacatccattatgaggtacaggagtaatatcttttattagagt
aatctctaatcctgcagcctgcaaagctctaattgctgtttctcgacccgctcctggacc
atttactaaaacttcggtttggcgcataccttgatccatagcttgtttagctgctttttc
agctgcggtttgagctgcaaaaggtgttcctttttagctcccttaaatccacttgcacc
agaggatgaccatgataatgtctctcctttcaaattagtaatagtaacaattgtattatt
gaatgtagatttaatatgtgtaacaccgttaactgcattacgtttagtttttcttgctcc
ggatttttttatttgtctagccatcgtgtctcgtctattatatagttaattagaagatta
ttttcttggagctttttcttacctgctactgtttttttacctcctctgcgtgttctagc
attagttctagttctttgtcctctcaaaggaagaccaagacggtgtcttctacctctata
agtactaatttccataagtctcttaatgctcatagactcaaaacgtttgagatccccttc
aatctgatagttagactcaagaatttctcttatactgacaacttgttgatcatttaaatc
ttgacacttaatatcaacatctatctttgtttttctaatatttctttcgaacgcgatag
tccaataccataaatatatgttaaagctatctctattcttttgtttcttggaagatctac
tccagcaattctggccactttgtattgtctccaataaataatatgtat-ttttataacgg
tagtttata-agcttagccttgtctctgcttatgtttaggattagtgcaaattaccatca
cctttctgtgacgccgaattattctacatttttcacacatttttcgaacagaaggacgaa
ctttcatattaa----ttctatatactaggatattactatattaactctattaattatc
gacattaaaaacgtcaataatatattgtattactttaactattttagattttacttggtc
atactatcatattttttttgatatgacgtatgtctgaatttgcttagctgtgtctattgcg
actccaactaaaataagtaaagaagtagccccaagacctcttaagttttggatctgagta

acctttctattataaacggaatcagcgctactgtaaataaaaatgaggctcctaggaat
gtcagtctatttaatataacttgtaaataatcaatagtcgcttgacctggacgaatatta
ggaatgcttgcacccatttttttagatttgtagcaatatcttctggattcattactatt
gatgtataaaaatagctaaagaaaaggatcaaagcacaataaagaagaaggtatagtgaa
ccattaggacaaaacaagtataggatttggagtaatgttttattttgaataatttgggtt
agataagatgggagggccatagatgcagatgcaaaaacaataggcataactccaccttga
tttaatttcaagggcaagtaactatttggatccaaaattgaagattttcctagctgtctt
gctgaaataattttaattcttcttgttccttcttgtacacaaattgtaattattatcatt
aataaaaagatcgctataaataatccaaacttgagacttgcattactataattagcatca
aaaaatgactgtgtaaaattctttggtagtcctgacacaatgttttgaaaaataagcaaa
gaggctccattaccaattcctttttctgtaattaactctgataaccacataataatcata
gagccagctgttaaagctaaaacagactcgcagacaaattcgacattccaattaaaaaca
tatggttttacccatatagagattgctccagattgtaaagtcgcccaacctagagctaag
taccttgtaatttgagttatttttggcggcccaattcaccctcttccttctgtaatttt
tccagattggggacaattttcgtaagtagctgcattacaattgaagagttaatataagga
acaatacctaacgcaaaaatccctattgttgaaaaacctcctccagaaaaaatatttaga
aaatttactaaagtatttttttctacacttgcataaaaggcgtcatgatctatacctgga
acaggtataaatattcctaaacgtgctaaaactaatagaaaaagggtaaagataatacga
tttctcaggtcactttttttggctcataaataaattttaaatgaaaagagatactttt--tt
tatgtaaaaaccctattccaagacttatataagtttaaaatagggaagtaa---aatatt
atag------taata-taatattctatgtagaatacaagctttctataggcactcctcga
tctttagctgcttcactaaaagttcttaattgagttaagccatttagaacagctcgcgcg
ttatttaatgtattattagaacctaattgcttagctaatatattctgtacacccgacaat
tctaggactgtccttacagaacctccggcaattacacctgaaccaggtgcagagggcctt
aatataactttcgcggcaccagaaattccattaataggatgaggtatagagttcgatttt
gtcagtggaactgtaactagatgtttttttagcatctgttactcctttttttactgcacca
attacatcgcttgcttttcccacgccgacaccgacttggccttgctcgttaccaacaaca
agaataactcgaaagcttagtttttttacctccttttacaactttagtaactcttttaact
tgtacgactctttcttcccagccactatctttatcttttcctttgctctgtttttttacga
ttggccattttaagaattattcctaattagttaatta-cgcttctagaaacccatgcctg
cttccttagcagcttcagccaaggccttaactcttccatgatataactttccacctctat
cgaagacaacatttttaatgccttctttcatagactgttctgctaactgtttaccaacta
cgcgagaagtatcacagtttggtcttatattatcggattctttactatttaaattaacag
atgatgtagctactaatgtaataccttgtgtatcatcaattatttgtgcgtatatatgtt
tgttagatctaaatacacacaggcgtggcctacttgaggttccttgaactttttttcgaa
ctcttttatgcttatgaattctagtttgtttagtgtttagtttcattattatttaccttt

tccagctttcccagcttttcttctaacaaattcaccttgatatctaattccttttccttt
ataaggctcaggaggcctaatagaacgaatagttgaagcaacctgaccgacaacttcttt
atctatgccagagacagtaatatttgtattattttcaactttaatttcaatattagcagg
aggtttaattttcactacatgactatagccaacatttaagattaaatcttggttatcaat
ttgagaacgatagcctacaccttgtatttgcagtttttttaaaaaatccattagaaacacc
ttcaatcatattactaataagagttctggatagcccatgtaactgactagccatttttgt
ttgtccactcgtcttgacagcaatcgtgtcattaagtatttctaaattaataccagctgg
taaagttctggatagggtacctttaggtcctgtcacagtgatagtctgaccatcaaactg
agtactaagatttgtcggcaataaaattattttttttccaatacgagacat-attccacc
tataaaatagttaccaaatatagcataatatttcaccaccaagaccatcatgacgagctt
gtctatctgtcataacacctctagaagtagaaataagagcgatgcctaaacctccaagaa
ctctgggtagctctttgtgatttgcgtagactcttagtccgggtttactaattcttttca
gagcagtaataactggttgacgattttttaccattatacttaagagaaatcattaaattag
tctctataccttcacctatttgttcgaaattttgaacaaatccttcttcttttagtactg
ttgccatattgcatgtcattttcgttgctggaacttgcacaatttgatgtcttgctaagt
ttgcgttacgaatacgtgtcagcatatcggcgatcgtatcgttgaccaccttgatcctcc
ttgcga-----tgaattatttaaaatattttcaagattccctgaaaggcatccctagctt
ttttaacaaagctagaccttcttgatctgttttagctgtagtgacaattgatatatctag
accacgaatttgatcaatattatcatagtctatttctggaaagatcaactgttcacgtaa
acctaaattgtaattacctttgccgtcaaaactttttaggactaattcctctgaagtctct
aattcttggtaatgttaaattaattagtttctctaaaaaagaatacatcttgtcttttct
tagatgcacaacaattccaataggaacctcttctcgaattttaaatcctgcaatagattt
tttagcttttgtgacaatcggtttttgtccagttattaatgttaattcttgaatactact
ttcaagagccttagcattctgagaagcttcacctaagccacggttaatagtaattttagt
aaacctaggaacttcatgtacatttttgtactgaaattcatcttttaaagattgagtaac
agttgttttatattttctttttaatcctattgccatggtatttttatcaattatttaata
agttcgccagtctttttcagttttcgaa-ttttttggcctttatcattaattattactga
ggatcgactagcaatattattttgctcactgaataacataacattagaggtatgtattgg
agcttcaaatttgataatttctcccgtttctccttcttgttgaggtttttttatgcttcac
tttgagattaattcctttaacaatcaccttacttgttttatgaatgattgcaataatttc
acctgtttttgttttatcacttccagaaataacttgaactaagtcccctttttttaattt
aatttttgtattattttttgattgttttagaagaacttttcattatactacctccggtgca
agagacactattttagaaaaatttttatctcttaactctcgggctataggtccgaagact
cttgtgccacgtggattattatcttgattaatgataactgccgcattatctccaaatcta
atgctcataccgtcagttctcccttaaagctttacgagttcttactacaacagctctgaca
acatcagatcgtttaactggcatattgggagacgcatctttgaccactccaataataaca

tccccaatagatgcataagaaggattactagtgcctaatactctaatacacattattttt
ttagcaccactattatctgcaacattaagatagctttgagtctgtatcatacttttattc
tgtataaaattaattatcaaaagacttagataagatattaaccattgtccagcactttgt
gcgacttaaaggtcttgtctcttgtattgtaacaatatcaccgattgtgcattcattatt
ttcatcatgcgctttatatttctttgtccggatcatagtttttgcgtactttctatgaga
gattctattttctacagctactactatagttttattcattttatcgcttactactttacc
tgttgtttcttttaaaggcatagtctttatattttttatatat---------aattaaat
tttaagattattgcacggttgacttattacgagacttttcaacagttaaaagttgagcta
atctatgcttagaatgtttaaatagatgaggctgaaaatcttgccttgtggctcttttta
gccttaaatcaaaaagctctcttttttattacaaggatttcttcagctaaagaagaagagt
ccatgtttgtaacatctgatattttagggaaagtcatgttttaggattctgttgtattcc
gaactataaatttggtcttgattggtaatttataagaagctaatttcatagcttgttgag
cagtttttttgtggtacacctgtaatttcaaatagaatatgtccaggcttaataactgcaa
cccaatattctggagctcctttacctgaccccatgcgagtttcagctggacgggcagtga
ctggtttatctggaaatactctaatccataatttgccacctcttctaacatatctagtaa
tagttcttcgagtagcttctatttgtctagaagttaaccaaactggctctgtcgcttgta
acgcataatcaccgaacgcaattgtgttgcctttactagcagaacctttcattctacctc
tgtgttgttttctaaattttgttttcttggggcttagcataaaaattaaataataataat
aattaagaagctgttgaatcaggagtttcctgattataagcagattctagacttttcgat
tctggcaaaatttctcctttaaaaagccaaactttttactccgagcacaccatacgtagtg
tgagcttggcgatgacaataatcaatatcagctcttaaagtttgtaaaggtacacgacct
tctctaacccattcacttctagctatctctgcaccattaagccgaccagatacttgaatt
ttaacaccctgtgtatttgctctttgagctctttgaactgcttgtctaacagctctacga
aaggcaactctttttttctagctgttgagtaataaattctgctactaaggttgcttcagaa
tctggatctgcaatctctataacattgactctgagttgtttgctagggcctagaattaag
gacaacgattttcttaaagactcaattccagctccagattttccaagtacaattccaggt
cgagctgttgctatgagaatttctacttgatcaactttacgattgatttcaattttagca
atactagcattactaagttttgaatgtataaatgagcgaattttatgatcttcttgtaaa
agaactgggtagtctttagagttagcaaaccatgaagaacgatgtttttgggtaatgcct
atgcgaaaacccaaaggatgaatttttttgacccacagtatccttgattataaattaaaag
ctataatgactaattgacgttaag---------agactacagttctgagacacctaatgta
atatgacaagttggcttatgtattggaaaggctcttccttgggctctaggctgaaatctt
tttaaagtcgggcctttgtctgcaaaggctttactaacaaataattggtttttatttaat
ccatcattatgctcagcattcgctgcagcagattccaggatttgctttatatgagaacag
actcggtatggcataaattccagaataattaatgcttcttggtattttctacctctaatt
tggtctaatacacggcggactttatgcggagaaagacgaatgtatttccctactgcttta

gtttcttttacgtttttttgtaatactcatatttatagtttaacgacgggcttttctgtca
ccttttatgtgagtgcggaaagttcttgttgggacaaactctcctaatttatgtcctacc
atttggtctgacacaaagacaggaaaatgttgtttgccattataaacagctattgtatgc
cctaccatatcgggaataatagtagatgctctagaccaagtttttaggacttctttttt
cctgaaatatttaatgcttctatacgtttaagaagactaacatcaataaaagggccttta
tgtatagatcttgacataataataaatcattagattttgataaaataaattctatttac
ggcgacgtaagacgtacgggttactatatttatttggatttcgtgttttaacacccaatg
caggtttaccccaaggtgttacaggacgagctcttccaattggagatttgccttcaccac
caccatgtggatggtctacaggattcataacaacacctctaacagtaggtcttttaccta
accaacgatttctaccagctttaccaagagtaatattactagcatcaatattaccaactt
gtccaatagtagcatagcattcttttcgaatcattcgaacttcacttgaaggcaatttta
cagtaacaaaagcaccttcttttgctactatttgagcataagttcctgctgcacgaacaa
tctgtcctccgcacgaaggtcttagttctatattatgtactgctgttcctaaaggaatac
tagataaaggcaaagcatttccaacttcgataggggcagtagaaccagaaagaaccatag
atcctacacttagagatcgagggtgcagaatgtatcttttctccccatcaagatagtgta
ataatgcaattcttgcatttctattcggatcatattcaatagaagcaactttagcagcta
tattatgcctatttcttttaaagtcaattaatctgtactgctgcttatgtccaccacctt
tatgacgacaggtaatgacacctctattatttcgacctttgcagaaatggtgtttaacta
ttaatgatttttctggcttatcagtagtaatctctgaaaaggtagaaactgttctatttc
ttgttcctggtgtgtaggcacgatataaacgaattgccatataagaagatttgagaagag
taaatgttgaa------ctatgtaatatatttaagtttctggaaacaaattgatagaatc
ttttgaggcaagtgtcacaattgctttcttgtaatgtggtcgcttgcctacgaatctgcc
gatgctccttttttttcttaggaggatggcaagtgttaacacctgtaacttgcacgttgaa
aatatactgtatagcagccttaatattaattttgttgctttgggatcaacagcaaaaca
atactgatttcttctaataatttagttgtcttatcagtaatgatgggatatttaactag
atctaataaacctcttgaatcaatgctatccattatatacctcttgtattttagataaag
catcaactgtaataataattttatgtgcagctaacaaagccataatatttagtgtatcag
ctgaaataagttctacgttgtgtagattgcgaatagaaagataaacgttcggatcttttt
tgtctactatgactaagactttttgtttaaatcaagattccatcgatgtatagcttcca
taaatagtttagtcctgggctgatgaaaataactgttaaaattttctacaatcaatgtat
taactgatttgttatttaatgctgtccttaatgctaattgtctttctttttattcattt
ttttggtaaaactacgaggttttggcccaaatattacaccaccgcctctccatagaggtg
aacggatagaacctgctcttgctcgaccagtcccttttgacgccatggtttacgacctc
cacctcgaacttcacttcttgttttagtattcgcagaaccctgacgcttttcattacttt
gctttactaacgctctatggactaagtacatccctgaatcttgactaaccttaagattta
aatcagcgttgccacttacttgaccttcccaattatagacttgataatttaattgtgtgt

taactgtcatgaaatatatcgtatgaaagaatattgctacttactaattttactaaagc
gccaggtttaccaggaacagctcccttaacgatcaacaaatcattctctgagttaatact
tacaatttgcaaatttttattgtaactttttattaccaagttggccagccatgttttt
acctggataaactcttccaggcgttgtaccagctccaatagaaccaggttgtctatgatt
tttcgaaccatgtgacattgggcctctactaaaatgatgcctcttttggtaaccggagaa
acctttacctacacttctagaagagacattaatcttttggcctacttgaaatagatctgt
agatagtatttggctaacttcgaaatcatcagtagatttcacttcgtattcacgtaagta
ttttagtgggggagcctgtgatttttttaagtgtcccaacaatggtttatttaactttttg
ttcagcaacttgcttataaccgacttgaatagcattatagccatcagtagatatagcttt
aatttgagtaataatacatggcccaacttgaattacagtaactggaattgataaaccagc
ttcatcaaagaattgggtcatacctactttagtaccgagtataccaacagacactagatt
tctccttttttgagagatttatactatcattgtacataaaaattctacttcatattgactg
aag--taaattaaactgactgatgcatgatattattcatgacaacataaaggtttaaaac
tttacctatttaatctaaaatagaaacagctttttgtccagttacagcgtatataaatta
gaagcatgcttattaaattttttatagactact-gaaatttcactgatattgatattcgt
aacttacttgcagtaat--------atcaacatgtataagtttaattgcctagactagaa
ccagctattattagaaatgattcaagtatagataaaagcgcatttgaacttttatctatt
gacaaatatagtagaatatattgaggttcggtaactacaccttttatttttcacatttat
tgtgca--gtatatatacatctaatgcagtaaacataagaatataac----tgtatagca
aatgtaattacaaaaattagagaggtattaatgggcaaagtcgttggaattgatcttgga
acaacgaattctgtaattgctgttatggaaggaggtaaacctaccgtcataccgaatgca
gaaggttttagaactacagcttctgttgttgcatatactaaaagtggagataaacttgtc
ggacaaattgccaggaggcaagctgttattaacccagaaaacactttctactctgtcaaa
agattcataggacgtaaacagaatgaaatttcgcaagagattcggcaaacatcatataat
gttaaaactagtggatcaagcataaaaattgcttgccctgcactgaataaagattttgct
ccagaagaaatttcagctcaagtactgagaaaacttgttgaagatgctagtacctactta
ggtgagactgttacacaagccgtaataactgtaccggcttactttaacgattcacaaaga
caagctactaaagacgcaggtaaaatagcaggcttagatgtattgagaattattaatgag
cctactgcagcttctttgtcttatggactagacaaacaaaataatgaaacaatactagta
tttgaccttggcgggggcacatttgatgtatctgtattagaagttggagatggagtattt
gaagtactctcaacttctggagatacacatttaggtggagatgactttgatcagcagatt
gtagaatggctaatcaaagattttaaacaaaatgaaggtattgatcttggtaaagataga
caagcacttcaaagattaaccgaagctgcagaaaaagcaaagatagaactatcaaattta
actcaaacagagattaatcttccttttattactgctacacaagatggtccaaagcattta
gaaaagactgtaactagaggtaagtttgaagaactttgttcaaatttaatagacaaatgt
agtattcctgtaaacaatgctctgaaagatgcaaaactagaagcttccagtattgatgaa

gttgttttagttggtggatctacaaggattccagccatacagcaaatggttaaaagatta
attggtaaagatccaaaccagagcgtcaatccagatgaagttgttgctattggtgcagcc
gtacaagctggagttttagcaggcgaagtcaaagatattctattactagatgtgacgccg
ttatctttaggagtggaaactttaggtggcgtgatgacaaagattataccaagaaacact
actattcctacaaaaaaatcagaagtattttctacagctgtagataatcaaccaaatgtg
gaaattcaagtacttcaaggcgaaagagaactaacaaaagacaataagagcttagggacg
ttccgattagatggcattatgcctgcacctagggggttcctcaaattgaagttacgttt
gacattgacgctaacggaattttatctgtaaaagccaaagaaaaggctactggtaaagag
caatcaattactatatctggagcatcaactctacctaaagatgatgttgaaagaatggta
aaagaagccgaagaaaatttcgatacagaccaaaaaagaagaaagaacattgacacaaag
aatcaagcagagtccctatgttaccaagctgaaaagcaagttaaagagtttgaagacaaa
attagccaagatttaaaaataaaaatagaggagctaattacagagcttagatctagtcta
gaaaaagaagaatatgacaatatcgaatccatttctcagcaattacaaaattctctgatg
gacattggaaaaaatgctgctcagactgaaagtaaagatgcaaaagcaaaggatgacaac
accgtgattgacactgatttctctgaagctaagt-aaaagta-agcgggtaacgcgattc
gaacgcgcgacatcaaccttggcaaggttgcgctctaccactgagctatacccgcatatg
ttgtattattacaaaaattaagatatttgtcaatctgtta------atctatac---atc
tagactattaatttccgaatcacatattagctaaatgttgtaaatataacaattgctata
actttttaa-------aagtacttatatcat-tatataagtactttttatgtt-------
-caaaaatacaaataagtattatag-------ctaaactactaaggaattaactaagcta
gtaacaattactaatcctgtccataagccagctcccgtatatattaatccttttgattgc
tcccattgacctggggatgctaaaacaacaggtacacctactactaacacggtcgacaat
gcaattaataatagcactaataattggatcgcaataatcatttaatatgttctcctgaaa
aatttaaatgttaataatactgtatataatacaatgaagcttttgaccttcagcaattta
tctatgaagaaataatttaaaattattgttttctaaataaaccttaaattaagtattaaa
atctttattgtaaataataatactatatttttttta---tagataaaaaatttaaagaaag
tctttttaaaaaaaaattaagataagatgatatgctgtataaaaagctaagtattatta
attattagagaaatttaaa-ataagaggtaaaacatatgaactcagccctttttttggca
aaactaccagaagcttatgccgttttaaaccgatcattgatatcttacccgtaattcct
gtattttttcttcttttagcttttgtttggcaagcagcaattggttttagataaaattta
ggttataaaaagaataacaaatatacaaagtgacatttttttgagctctaatcttatatct
tacacaataataactattatggttgaacccttactatcaggaattgttcttggacttatt
cctattactttattcggattattagtagcagcttatttacaatatcaacggggcaaccaa
ttaggactgtaaacgctaagtcatagatatttataaaaactacaataattatagttttta
taaatatctacaaataagctaattttaaatacaaagataatattattgaaactagagtga
taacgaaaaagatttatataa---taa---------------------------------

------------------------------------------------------------
------------------------------------------------------------
------------------------------------------------------------
------------------------------------------------------------
------------------------------------------------------------
------------------------------------------------------------
------------------------------------------------------------
------------------------------------------------------------
------------------------------------------------------------
------------------------------------------------------------
------------------------------------------------------------
------------------------------------------------------------
------------------------------------------------------------
------------------------------------------------------------
------------------------------------------------------------
------------------------------------------------------------
------------------------------------------------------------
------------------------------------------------------------
---------------------------caaactaattgtttaccagctcgacttagtaac
tcctggtaaaagacattcatgagacatttccctaaaaacatggcgagataatccaaaatc
tctataataccctctacttctcccggttaaccaacatctatttctccctctaacaggggc
gctgtttctaggcatttcttgaagtttttgtcttaaatccattttgtcagaaaagctgcc
agcttctttcagctgatctttaatagccaatctctttaagtaatatttttttgcaagttt
cgatctcttaatctctctttggatcatattctttttagccatgctgtag-aactctcttt
taaaattatttattcagtatattagctaaaaaagctcctatctgcaacttttatctattt
aaaaataaatagcataaagtagtaaaatgcaattgatatttaatcaacgatacc---act
ttatgctattatcaaaaatttatttaataaagttcttgttcaacatgagttgaaatagtg
caatcagatgaagggtatgcaatacaggttaacacataaccagctaaaagttgatcatca
tctaagaatgattgatcagcttgatctacagttccttctgtaactttacctgcacaagtt
gaacaagctccagctctacaagagtaaggcagttcaattccttcttcttctgctgcatct
aaaatatatgtatcctctgcacaattaaatgtaacatcaattccttcttcttcacatagt
aaatgaattttataatcagccatttctttttactccataataaagatacacttgaaactt
aataaagcaaaat-aatttaaagaacttaccaaaaaattatatactagtttcattatgca
tgatagactggaaaaaaagaaaataatatactattctgaaaaatcaggctacatttccta
ctaaatagaactatagtaggtccaaagtatatacataaattcatatagttaaaattattt
ttatcatttaaatagctgaacttaatagta---------taggttgatcatc------aa

gctagtcttctgc---caataaataattaattatgatttcaatgcttaatcaacaagtag
aacttaagcctattcttaaattccagataacacaaatatattcacattctcttattcagg
aaattaaagcattaataaccagacttgttttgcaagtttggaggaggccggctacattga
tggcaggtattatgcaacctttgttatggttaattctatttggtgggcttttctacaatg
ctcctataaatttgtttactatcaatacaaactataattgtttttttgagctctgggatta
taattttcacatcttttactggagctttaaattcaggtcttccattaatgtttgatagag
aatttggattttaaatagattattaacggctccgttagtatcgagggcttctatcattt
tatcttctgctacttttatgacttgtattagtttaatacaagttatatttatagttacag
cttctcttttatgggaaactcacctctaaatagcgatagtgctatgattttggactta
taatcctattagtgactgtaggagttacaatgcttagtttagctttatctttcactctgc
caggtcatattgagctattagcatttattttagtagttaacttacccttttattttcta
gtacggctttagctcctttatatttcatgccaccatggcttcagttaattgcaagtctca
atccattgagttatgccatagaaggtacaagatatttatattcaagcgtaaactggaatt
ttacagagtgtgtgattaagattagctggggagatatttgtttagggcagattattgttt
tattaatcgctttagatataatggcagcttatgttgtgtctaagatattaaaagctaaac
ttaatta------------------aaatttaataaaaaatttattaaatttatttaaca
aagattttactatataatactaatagtagtatggataattgaaatagaaatagtttttc
aaaaccaaagctattatcaacaatgataaacatttgtaagaaagtcaacaaagtatgttc
ttattcataggaggcatgtagtcaatgggactaccatggtaccgtgtacacacggttgtt
ttaaatgatcctggacgcttaattgcagtccacctaatgcatactgcacttgtagcaggt
tgggcagggtctatggcattatacgaactagctgtatttgatccttcagatccagtacta
aatccaatgtggcgacaaggcatgtttgttatgccatttatggctagactgggcgtaaca
gattcatggggaggatggagtataacaggagaaagcgtatccaatcctggactgtggagt
tttgaaggtgtcgcgttaactcatatagtcctttctggcatgcttttcctagctgctatt
tggcattgggtttattgggatttagaattatttagagatccacgaactggcgaaccagct
ttagatttaccgaaaatttttggaattcatttattgctatcaagtctactttgtttcgga
tttggagctttccatgtgactggacttttggaccaggaatgtgggtatcagatgggtat
ggagtaaccggaaaggtgctaccagtagctccagcatggggaccagaaggattcaatccg
tttaatcctggaggagttgcatctcaccacattgccgcaggtactgtaggtatattagct
ggtgttttccatttaactgttagaccaccacaaagactgtatcgtgctctaagaatgggt
aatattgaaactgtattatcaagtagtatctctgctgtttttttctcagcttttgtgact
tgtggtacgatgtggtacggctctgcaactacgcctattgaattatttggtccaactaga
tatcagtgggatagtggatattttcagcaagaaattgagaaacgagtagaaaatgctatt
gctgatggtgctgcacctagcgaagcatggtcaaggattcctgacaagttggcattctat
gactatattggtaataatccagcaaaaggaggattattcagagcaggccctatgaataaa
ggcgacggagttgcagaagcatggcttggacatcctgtattccaggataaagaaggaaga

gagcttagtgttcgcagaatgcctgctttttcgaaacttttcctgtaattttagttgat
aaagatggtattatacgagctgacattccgtttagaagagcagagtccaagtatagtatt
gaacaagtaggtgtaacagctagtttttatggtggcaaattaaatggccaagttttcaac
gatgctcctagcgttaaaaaatatgcaaggaaagctcaattaggtgaggtatttgaattt
gatcgaactacattagaatcagatggagtatttaggagtagtcctagaggctggtttaca
ttcggtcatgctaatttcgctttaatttcttctttggacatctttggcatggttcaaga
actatcttccgggatgtatttgcgggaatcggagctgaagttactgaacaagttgaattt
ggagctttccaaaaattaggagatagaagtagtaaaaaacaaggagctgtataagaccaa
agtcaaagacagtatttgtc---aaatatttccaatattaaaatagttaagacaaagttt
attatctaatagttactaatcaggagaatttatggaagccttagtctatgtatttttact
aacaggaacgctgatggttatattctttgcgatctttttagagaacctccaagaatagc
aaagtaacttgattccgtttaaacttataataatagaaccactttccaacttattagtgg
aaagtggttctattgtagattacattgtattattcttcatgctcttcaaaaggatctcga
aggtcttttgaagcaggaccaaatgctgtgtatatcgagtaacctgtaatacctaaaagc
agactcgaaataaaaatactaagaactgttgcagtttccataattcaagcagtattaaga
tgaactttttctataatgatattatcaatattaaaacatttaattattaaaattatcaaa
atatactatggcacttagaactagacttggagaaattttaagacccttaaattcagagta
tggaaaagttgctccaggttggggcacaactcctattatgggaattttcatgttactatt
tttcttgtttttattaattattttacaaatctataattcttcattagtattagaaaatgt
agatgtagattgggctactttaggtag----ctaattagataagttctttga--------
ttaaacaatattaaagataac----------------aagcgatcgctattagattatag
tagcgatcgcttaaaaaatgttttactgtatatccagagaaaattatttacctaaattaa
ttaactcgctatctgcaaaactatttgtattaacattattatagttaactttttcaaacc
ttactaatactggatatttgataccacttttatcaattgctgcaacagtaccaatttcct
gataccagtatgattcctttctcaatatttttacttttgagcctcgttccatatttttag
tccttagatagattattaaagattattatattgaagaattttaaaaaactagacaaaata
tataaaagtactaccgacaattgtaatacctaactaataaaaggagagagagggattcga
accctcgataacaaaagttatgacaattttcgaaattgttgcaataaaccgctctgccat
ctctcctaaattaaaatactcttttagaaaattagacaagaataacatagaactaaggta
gttgcctagaaaattaaaaatatgttaaatataaattataatctttctatagatatattt
acaaaatatgtacatagaatctatatcagcttatttcaagatttttttaatcttatacaat
ttttactaaataaataaattctctcataatatgaaactatcttggaaaacacttcttctt
tggtctttaccaatatttgtcgttggtttctttttctggcaaggtttttttaggtccaact
actacagatgttggcagtaatatcgcaagttctagaatgacatatggacgatttttagaa
tatttagatatgggttgggtgaaacgggttgacctctatgaaaataatcatacagcaatt
gtagaagctgttggtccagaattaggaaatagagttcaacgaattcgagttgaactgcca

gcaagtgcgccagaattaattacaaaattacgcaaagccaatgttgatctagatgctcat
ccccctaaaagtacaagtgcagtatggggactattaggcaatttactatttcctttacta
ttagttggcggattagctttcttatttagaagatctaataatgctagtggtgggcctggt
caagcaatgtcatttggcaaatcgaaagctttgtttcaaatggaggctaaaacaggagtg
gtatttaatgatgtagctggagttgaagaggcaaaggaggaatttcaagaagtagtaaca
tttttaaaacagcctgaatcatttactgctgttggtgcaaaaataccaaaaggcgtttta
ttagttggtcctcctggcacaggcaaaacattactagcaaaagctattgctggcgaggca
agtgttcctttttttagtatctcaggctcagaatttgtagaaatgtttgttggtgttggc
gcttctcgtgtgagagacctattcaagaaagcaaaagacaatgcgccttgtatcgttttt
attgatgaaattgatgctgttggtagacaaagaggaacaggcgttggaggtggtaatgat
gaaagggaacaaacattaaatcagctattaactgaaatggatggctttgaaggaaatact
ggtgttattgtaattgccgctactaacagagctgatattttggattctgcattattaaga
cctggaagatttgatagacaagtttctgtagatgtaccagattttaaaggcaggttagca
attcttgaagttcatgctaaaaataagaaaatggaacctaaagtatctttagaaactatt
gccagaagaactcccggcttttcaggagctgatttagctaacttactaaatgaggctgct
atcttaacggctcgacgaagaaaaaatgcaatgactatgtctgaaattgatacatcaatt
gatcgagtagtagctgggatggaaggcactcctttaattgatagtaaaagcaagagatta
attgcatatcacgaagtgggtcacgcaataataggcagtttattagagcatcatgatcct
gtgcaaaaagttacattaataccaagagggcaagcaagaggcttaacttggtttactcct
agtgatgatcaaagtctaatatcaagatcccaaatactagcccgtatcgtaggtgctctt
ggtggtagagctgcagaagaaatcattttcggcgacgcagaagttaccacgggtgcaagt
aatgatttgcagcaagtaacatcaatggccagacaaatggtcactcgatttggcatgtct
aaaattggacctttatctcttgaaagtcaaggaggagacccatttttaggtagaggcatg
ggaggaggctcagaatattcagatgaagttgcaactaatattgataaacaagtaagggaa
attgtaagtgaatgctatgcgcaagctaaaaacattattatagataatcgagtagtgatg
gatagattagttgatttactaattgagaaagaaacaattgaaggcaatgaatttagagac
atcgttaaggaatatacagcaattcctgaaaaaaattactacatatcacaattttaa---
---------------attagacgggactgacgggattcgaacccgcaacttccgccgtga
cagggcggtgctctaaccagttgaactacagtccc-aaaaagataaatacaaatagataa
tctcatagttagcttaaacttgtcaaattagaatattaagtatgaagagta---------
--------tttgtaatagtactcttcatatttaata-tagtagcgaaaatatttcttta
gaaggagagagagggattcgaaccctcggtacgaaattaatcgtacagcagattagcaat
ctgccgctttcgaccactcagccacctctccatga---tctatatataaaatatatcaaa
tatacttatatttaataggtgtgcataaatatttatggagctaagcggattcgaaccgc
tgaccctctcaatgccattgagatgctctaccaactgagctataaccc----ctttatga
ggatattaagaaatggctcaagcgggatttgaacctgcgaccttgggcttatgaatcccc

tgctctaaccactgagctactgagccatagtataaaatattattaattattataccataa
attatgctcaaatataaatcaataaaagaataattactatca-ttcactactttta----
-----------------tacaagttgtaaatattcggacaagatttcaaaattaaaaatg
tctttaactttttgtttga-------attatttgtttacgaaa-----------atattg
gtatttaaaattaac---------------------------------------------
--------------------aataaatttg------------------------------
---------------------------------------------------tctttattt
g-----------------------------------------------------------
------------------------------------------------------------
------------------------------------------------------------
------------------------------------------------------------
------------------------------------------------------------
-------tacaca-----------------------------------------------
-------------------------------caaaaatattggaaa--------------
------------------------ttttcatcaatt--------aattatactaattagt
------------------------------------------------------------
------------------tagactataaccgacgtac-----------------------
------------------------------------------------------------
------------------------------------------------------------
---aagaattatttaacta-----------------------------------------
------------------------------------------------------------
---cgttaaat-------------------------------------------------
------------------------------------------------------------
------------------------------------------------------------
------------------------------------------------------------
-------------------------------------tgtgatgcatt------------
-----gggcttgatcaaaacttctctaaaatcaacaaaatctagt---------------
-----------------------------------------------------aattcaa
gtaaatcaaatact----------acagacgcttaaaaaaataata--------------
caattagaatcgtaaacatttagtttttga-ttatgttaaataaatttgattatcataga
tttaatatttaaaagtgttctattacttgta-----ttgagtaactgt-aatatttaatg
tcttctttttttagataataatcataatataagctaaaaaatattttt---tattaaacaa
ctagcatagagccaataccttgatcagttaatatttctagcaagagagcatgatctattc
taccatctagaatatgcgctgaagctacaccctgagctaaagaacgaatacaacaattca
ctttaggaatcatgccaccagaaattacttctgtttgagttaaatctctagcttcctgta
tattcagatgactaattaacgttgaaggatctgaagaattacgtaaaataccaggcgtat

ctgtgagtagaattaatttttcggcattaagagcggctgctatttctcctgcgacagtat
cagcattaatattatacgattggccttgcttgtctgctgcaacactagctataacaggta
tataattattattaattaatatcttcagtaatttagtatcaatattttgtacttctccaa
caaagccaaggtttgcttttccatttggtcttggagtaataagtaatccatctttccccg
acaaacctacactttttaccaccttgtttattaatagttgcgacaagatctttgttgactc
ttccaactaaaaccatttccactatatccatagtaggttgatctgttactctaacgccgt
catcgaattttggtaatattttaaggcgatctaaccaaaaattaatttctggtcctccac
catgaactaaaatgggacgtagtcctataaaggacagaaacacaagatcactaatcactt
gatctttttgtttttgatttttcatagccgatcccccatattttataacaataattctgg
aagaaaattgctgaatataaggcagagcttcacttaagactcttactctttctgaattag
tcaacataaataaagtaggattatgatatataaaattatttgcacaacttgtgttaattt
aaaaagtaccagtaggtacttc-aaaacttgaaaaaatagacatgttactaaaaa-----
----ttaaaactataaggattataattatgaataaattttgggataatgttttaagattt
ccacgatttttagtcagcgtcattttaggattagttttgataattattagtcctttttttc
gtgttattaaaaaaaccattgacaagtttttttttcatcatatcattgacgggattaatc
acagttttagcgataataa-tacaaaaaatgctaaatatcgaatgttgttgagagaaata
aataaatttaattaaaatttttgaacatatataatattatattatcatttaatatgataa
tct-attttcttttccat---aaaacttataagtattaaatattatttgataaatattat
gccactaaaacaaatagttagttctgaaaaaaactggtgcctttgcacttctggatagtat
tgttaggcacggagttaaacatatatttggttatccaggtggagccattcttcctatcta
tgatgagctttacgcttgggaggaagcctatctaattaaacacattcttgttcgccatga
gcaaggcgctgctcatgctgcagattcttattctaggtcaacgggagaggttggagtatg
ctttgccacttctggtccaggggcaactaatcttgtctcgggtatagctacagcacacat
tgattctgtacctatattagctataacaggtcaagttggaagagcttttattggcacaga
tgcttttcaagaagtagatatttttgggattacacttcctattgtgaaacattcatatgt
agtccgtgatcctagagacctgtctagaattgttgcggaagcatttttttatttgtaaaca
tggtagaccaggtccagtattaattgatgttcctaaagatgtaggattagagaagtttaa
ttatttttctgttgaacccggaaaggttaatattcctggctgtaggccaattaccagcct
aaagtcaagacaaatccttatggcagctaaaatgatacagcaatccagccaaccattgtt
gtatattggtggaggagctataatctctgattcacatcaaattattaaagaacttgttga
tttttataaaatacctgttactactactttgatggggaaggggatttttaatgaggacag
tgactattgtctaggaatgttaggtatgcatggtactgcgtatgctaattttgcagttag
tgagtgtgatcttttaattgctttaggagctagatttgatgatagagttactggaaaatt
agatgaatttgcttgtaatgcccaagtgattcacgtagatattgatcctgctgaagtagg
aaaaaataggattcctcaagtcgctattgtcggtgacgtagcagaagttgttagtgcaat
attaaatttattaaagacttcttttccccttatccagatcagattacatcttggcaaga

aagaattaatcgttggcgtcaacagtatcctttactggttcctagaaaatcaacaagcat
ttcacctcaagagattcttgttgcaactaataaattagcccaaaatgcttattttactac
agatgttggccagcatcaaatgtggtcagctcaatttctgaaagtaaaagctaagcattg
gctttcaagtgctggattaggcacgatgggttatggtttacctgcagcaattggtgctca
agtagcacatcctaatgacgtagttatttgtattagtggtgattctagttttcaaatgaa
tatgcaagagttaggaactatcgcgcaataccagctaccggttaaaattattattattaa
taatcgatggcaagggatggttagacaatggcaacaggctttctatggtgaaagatattc
acactcaagaatgacagaaggagcacctgattttcaaaagcttgcagaagcttttggtat
taaagcttttactattaataataggcaaaatatgcaatctgctttacaagctgctattga
tcatcctggtccagttttattagattgccaagttacagaaaatgaaaactgttatccaat
ggttgctcctggaaaaagtaatgcacaaatgataggtatagctaaaccgcagagaggaac
tgcttccaactatattaataacagcgttggaagtaa------------------------
------------------------------------------------------------
------------------------------------------------------------
------------------------------------------------------------
---------------atagttagtattttctt-aaaacagaata-aagtaaatgaagtat
ctaaatgtaattttcttatactatta-aaaactatgtaagaaaagcctttgacgagaatt
gaactcgtgaccttccccttaccaaggggatgctctacctctgagccacaaaggc-tttt
tactatattgggccgggttggatttgaaccaacgtaggcgaagccagcggatttacagtc
cgcccccattaaccactcgggcaccgaccctagttaaaccttatttaagctatcatatac
aagatatcataattatgttgctaaaaacaagtacttttataaataaaactgaataagtaa
tggacatagctggacttgaaccagcgactttcacgatgtcgacgtgacactctaaccaac
tgagttacatgtccaaaactttaatcattattataacactgtttagtgattttataagta
tttatttctaattgaagatttgttttaatcggccggctttaccgagcagatctcttagat
aatacagtttagctctacgtattttagctcttctaataactctgacaaaagttattcgtg
gtgaattaagaaaaaatactctttcaactccaataccttgaaatgagcatcttaaagtta
aactagtatttaagctttttttcttcttctggctaatactactccttcacataattgtt
cacgagttttactaccttccttaaccattagtcccagttgtattgtatcgccaactttaa
tttcagatacttctgttttgacaagaggtatctcaacacttttcattagctgacttagct
ttgtattatttgctttcatatataggattagtcttgtgaca-tgttttgaaatattatac
attaaaaaaacaaatatgaaaaattttagcgaataagcatattagtatttataaaacttt
tttttattt---------------------------------------------------
--------------------------------------------cgattata--------
----------------catatccatt------------atattcaact------------
------------------------------------------------------------
------------------------------------------------------------

------------------------------------------------------------
------------------------------------------------------------
------------------------------------------------------------
------------------------------------------------------------
------------------------------------------------------------
------------------------------------------------------------
------------------------------------------------------------
------------------------------------------------------------
------------------------------------------------------------
------------------------------------------------------------
------------------------------------------------------------
------------------------------------------------------------
------------------------------------------------------------
------------------------------------------------------------
------------------------------------------------------------
------------------------------------------------------------
------------------------------------------------------------
------------------------------------------------------------
------------------------------------------------------------
------------------------------------------------------------
------------------------------------------------------------
------------------------------------------------------------
------------------------------------------------------------
------------------------------------------------------------
------------------------------------------------------------
------------------------------------------------------------
------------------------------------------------------------
------------------------------------------------------------
------------------------------------------------------------
------------------------------------------------------------
------------------------------------------------------------
------------------------------------------------------------
------------------------------------------------------------
------------------------------------------------------------
------------------------------------------------------------
------------------------------------------------------------

------------------------------------------------------------
------------------------------------------------------------
------------------------------------------------------------
------------------------------------------------------------
------------------------------------------------------------
------------------------------------------------------------
------------------------------------------------------------
------------------------------------------------------------
------------------------------------------------------------
----------acattcttccatagtttc--------------------------------
------------------------------------------------------------
------------------------------------------------------------
------------------------------------------------------------
------------------------------------------------------------
--------tttgataatcagtaa-------------------------------------
------------------------------------------------------------
------------------------------------------------------------
------------------------------------------------------------
------------------------------------------------------------
------------------------------------------------------------
------------------------------------------------------------
------------------------------------------------------------
------------------------------------------------------------
------------------------------------------------------------
------------------------------------------------------------
------------------------------------------------------------
------------------------------------------------------------
------------------------------------------------------------
------------------------------------------------------------
------------------------------------------------------------
------------------------------------------------------------
------------------------------------------------------------
------------------------------------------------------------
------------------------------------------------------------
------------------------------------------------------------
------------------------------------------------------------

---------------------------atcta----------------------------
------------------------------------------------------------
------------------------------------------------------------
------------------------------------------------------------
------------------------------------------------------------
------------------------------------------------------------
------------------------------------------------------------
------------------------------------------------------------
------------------------------------------------------------
------------------------------------------------------------
------------------------------------------------------------
------------------------------------------------------------
------------------------------------------------------------
------------------------------------------------------------
------------------------------------------------------------
------------------------------------------------------------
------------------------------------------------------------
------------------------------------------------------------
------------------------------------------------------------
------------------------------------------------------------
------------------------------------------------------------
------------------------------------------------------------
----gatgatcacatctcattttcaaacatggcttatgtatgaaaaagttctata---aa
aatctaatgactaaaataataaaattaaggcttaagtatttcttagttttgtgttataat
tagttactatcaaaggtagtggcataacacgaagcataaa-aatgttcgaacgctttact
gaaaaagctataaaagtcataatgctagcacaggaagaagctagacgcttaggacataat
tttgtcggaactgagcagatattattaggattagttggcgaaggtactggaatcgcagct
caagtcttgaaatcgatgaatgtgaatctaaaagatgcaagagttgaagttgaaaaaatt
ataggaagagggtctggctttgtagcggttgaaattcctttcactcctcgagcaaaaaga
gtactagaattatccttagaggaagcacgccaactaggccataattatattggtacagaa
cacttgctaatgggcttagtccgagaaggagaaggagttgcggcaagagttttagaaaat
ttggcagttgatgtttcttcaattagagctgaagttatacaaatgcttggagaaaatgcg
gaagccaatgtaagtggaagcaatactacgcaggctagaagtaaaacaccaacactagaa
gagtttggatctaacttaactcaaatggctattgaaggtggtttagatcctgtagtcgga
agacaaaaggaaatagaacgagttatccaaatcttaggtagaagaactaaaaataatccc
gtcttaattggggagcctggtgtaggtaagacagcgattgcggaaggattagctcaaaga

attgctaatagagacgtaccttctattttagaagataaattagttattactcttgatgtt
ggtttattagtagctggaactaaatatagaggtgaatttgaagaaagactaaaacgtatt
atggatgagattaaatcagctgataatgtaatattagtgattgatgaagttcatacattg
attggagctggtgctgcagaaggagcaatagatgcagctaatctgcttaagccagcttta
gcaaggggagaattgcaatgtataggtgcaacaactttagaagaatatagaaaacatata
gaaaaagatccagcattagaaagaagatttcaaccagttgtagttggagagccaagtgtt
gaagaaacaattgaaattttatttggtcttagagaccgttatgaaaagcaccatcaatta
acaatgtcagatggagctttggctgcagctgctaaatatgctaatcagtatatttctgac
cgcttttttgccagataaagcaattgacttaattgatgaggctggttctagagttcgttta
ctaaattctcaattacctccggctgctagagaattagataaagagttaagagctgtatta
aaaacaaaagatgaagctattagagctcaaaaatatgaaacagcagagcagtatagagca
agagaaatggaaattaaagctcaaattgcagcaattgctcaaagtaaaaagaatgagcct
gatttaaatttagaagatcctgttgttacagaagatgatattgctgaaattgttgctgca
tggactggtataccagtaactaagcttactaaaagtgagtcagaaaaattaatgcacatg
gaagaaactttgcatggacgcattattggtcaagatgaagcggttgtagctgtctctaga
gcaataagacgcgcaagagtaggtctaaaaaatcctaacagaccaattgcaagctttatt
ttttccggaccgacgggtgttggaaaaacagaattaacaaaagctttagcttcttatttc
ttcggttcagaagcttctatgatacggctagatatgtcagaatacatggaaagacacact
gtatctaaactaattggttctcccccaggatatgtaggttatagtgaaggtggttatcta
acagaagcggtaagaaaaaaaccatatactgttattttatttgacgaaattgaaaaagct
catccggatattttttaatctacttcttcaaattttagaagatggccgactaacagatgca
aaaggtagaactattgatttcaagaatactcttttgattatgacttctaacattggatct
aaagttattgaaaaaggaggaggaggtttaggctttgaattatcagaagatcaaacagaa
tcccaatatactagagtacgatctttagtaaatgaagaattgaaacaatactttagacca
gagtttctaaatcgattagatgaaattattgtatttcgtcagcttactaaagatgaagtt
agagaaattgcagaattaatgctcaatgaagtctttgcgagaattaagcagcaagatatt
caattaaatgtaacagaacgatttaagcaacgattagtagaagaaggatataacccaagc
tatggagctagaccacttagacgagcagtaatgaggcttttagaagatagtttggctgaa
gaagttttatctggtaaaattaaagctggtgacagtgcagtagtagatgttactaatgaa
ggtgaagttacagttttattaggtgaaaaattagaactgttaacataaa------aaaca
atttattaat--taaagcattgaagaggttat--attcaatgctat---gaaaaaaaata
ataagatgggttgctataattgacaacctctgatactctgtatttgtatata---aagcc
gggatagctcagttggtagagcagtggattgaaaatcctcgtgtcaccagttcaaatctg
gttcttggcat-tttaagcattagaaaattttgctatctgagcattaaattttaattgaa
atgtccctgtgggaccattcctatgttttgctattataacttctgtgaagtctctatttg
tagtttcttgtgtatagtaactttctctatatagcattattactaaatctgcatcttgtt

ctatagaattatgaacaataaaattatttgaaataaaattgcacaaaggtttacattcta
aatcaaacactatttgtaaacttataatgtttattttttgtatattttcaaaagtcacag
atgaaa------aagaatgaaaaatatttttttgttcttcaaagttttt------tattt
gaatagcaatcatatcattttgattaatttcatcacatcttttccacccttgtgttgtga
atagtttatggttacttgttaattgtatatatttgccagctccagtaataatttttatata
cagtttttttttcctgtttttgaaatattagatttaattgctagaaataactgtagtttgt
gcgcagtaaagttaacgacttca-atctttttataatagcaatcaaataatatttg--ag
gttgattatataatgatgtctgaagataattgaatttagatatacacccactttctctta
aatcggataacaaaggccttttgtcacttcgagtttcaaggtttctatttagttgagata
atactaagataggtaagcttaaatctttgctaatattttaagtgatcgagtaattaatg
atagttcttgagatctattgtcagattttctactctcctgtagtaattgaagataatcta
taataattaattctatatttttttccttgcaatttaaagaatttcacttttgtctttatag
tttcagtagaaatcttagcgctatcatcaatataaaggttaaggttagcaagatttttac
tttttttgaataacatattgccagtcatcattattaagctgaccagattgaatttttttggc
cgtttaaattacactcttgagctaaaattcttcttagtagttgctctgtcgacatctcca
agctaaataaaataacataagatttttttctgattgatgacatatctagttatattaatag
caaaagcagttttccccattgaaggtcttcctgcaaggataattaaatctgacttttgaa
acccttgtgttatagagtcgagttctgtaaaaccagaaagtatactactatttatactaa
tctttttcctggtatccaaatggactaatagacttgcaagaattgtagctaatgtttgcg
tatctttttcgtcaagtatttcataagcttctgttaattgactaacaatagaagctatat
tttttttgttcaataagttgttttgagcagctaatcaaacataatgaatctccacatgcta
aaagtaatcttttaatgtagttatctataacgaccgctgaatattcgcttatagcatcag
ataggggagcttgttctattagactaaaaacttcatttagctcatccaagtcttgaatca
ttttttcatttttttaagttaatgaaaaaatttcttacatttttttgtctttgcatggttaa
tattttcgagtgctgctctataaagtagtgaagttgatgtaaaataaaaaaaaatcaggag
agatttttttctactgattttagtaatgatatcgattgctgagttaatattatactaatta
atattttttcagctaaaatattatgaggaggtagatatttataa------atacttatta
attcctcc-ttgtaacatcactcttatggtattagtgtattagttaactattgatttaat
tggattctggaagaacttgcaatttaatatttgcagttacttgatgcaatagcttaattt
ctacatcgtaaatgccaattgttttcacctctggtaaagagatactttgtttttcaatgt
ctatgttggtagtgtttttaataatttgtgaaatttctttttctgtgacactaccaaaaa
tattttggccatctcctgtttttttactaacgctaaacttttgtatttcttctagaagtt
gcttaacctttttagcatcttctttagctatttttaatttttcttctcttatagctgcat
gaaatctttgttgttttaaaataccatttgtggcaactaccgccattttattcgggatta
aaaaatttcttgcataaccagaagctactttgataaggtcattactcttgccaagctttt
gaatattctcttttagcacaacatttacaactttttttactcatctt-------aattcta

aaaa----atctatattatcctattatacctgttgtatacagatttt----aataggaat
tatttatttgttgagataatattcttgtaaaaacaaaggtgatttttattttt-ttctg
atataattattttcacaataattc-----attaggagaggtggccgagtggttgaaggcg
cagcattggaaatgctgtttaggagcaatcttaacgagggttcgaatccctctctctccg
tgcaa--aaaatataaaaagttttatagagtccaatctaaatctccaataacattcttcc
aagcatttaatccaccccgaactctaacaatatttagtttttgagctattaaaaattcag
aagcaaatattgatcgagaatctaagctacagtaaacaaaacaaattttatcttgtaaat
tgatattagagtaattcatttcttttatttttgtaaaggtagatttaatgaatgtatta
aatgacttttatgatactcttcatgatttcttacatctattaaaatatattgtggattcc
tactaattaaaaacctttgtaattgaataacactaatttctcgtacaaatgtactaaatt
tactgccataatattcgttccaatgcttttgtgt-------ggataatacaaatttagta
tttataatcttaaatttcttaaatgatgaagttaatgcgttgtacgttaatatagtccca
cttaatatagacttataacccagaacgattttgacagcttcagtggcttgaagtgtacca
ataatacctggtaatagacctaaaactccggaattactacaagtatctcttgcactctct
ttattttcaatttcactgtagaagtctcgataaacaggtccaccttgataattaaataca
ctaacttgtccttcaaattgaaaaatggcaccataaatatgtattttatttaactcaagg
caagtattattcaagacatatcgagtatcgaagttatcagatccatctataataatatca
tattctctaataatgtcgattgaattctcgtagcttagtcttgtgttgaacacagttact
atacactgtggatttaaatctaata-tttttttttagctatttctacttttgaatatcct
atatcatgcatcgtgtataaaatttgcctctgtaaattagaaagatcaactatgtcatca
tccataatgccaatgctcccaattcccacagcagcaaggtaaataattcctggagatcct
agcccaccagcaccaatgaataaaacttttgctttttttaatctttcttgtccttctaat
tgaatttgtggcaatactaaatgcttagaatatcttgtatattcctctaaggagtatttt
gtattgtctgttttaaaatttagcataaaaatttatagtttatttaatagtactta----
--aaagctagctgataagtttattaaaatgacttttgtaaacttggttcttctctatcta
aaatagctccttgtgtaggacagacctgaacacaaatactacaatcaatacaggcagaaa
agtcaatccaataccaattagtattactattgtttttcccctttccttgatgaatacaag
ctactgggcaagcttccacgcattcagcaactccaatacatttttctgttacaattgtat
gagacatattatttaattaat-aaaagataagctatttacatctagtagttttgttgtat
actaaatttacatatgctatccagtccaatatttaaggtagtctatttttacagtttact
aatttgatatgcgcctttagttcagttggtagaacgcaggtctccaaaacctgatgtcga
gggttcaagtccttcagggcgcgtttagtttttatcgtaattcaataatatcattttatc
tttttgcaaacatcaaaatgatatgatataaaagatcatattttatacattgaatattct
atgtactagtactg-ttttttaatatgttattattgataagttaattatactaattctat
ttttgtaataagt-aaaaacagttctaattaagtatgaacgtatttaatagtc-ttttgt
atatac--aacttaaaaaattttaaactatggctaaaaaagttactggaattgttaaact

agcattaaatgcaggtaaagctactcctgcgccacctgtcggacctgctttaggacagca
tggcgtgaatattgttatgttctgcaaagagtacaatgcacgtaccgctgagaagtcagg
actagtgattcctgtagagatctcaatatatgaagatagaagttttacttttatacttaa
aactcctcctgcttctgtactaattgctaaagctgctggactaaataaaggttcaggtga
accaaataccaaaaaagtaggcagcataacaaataaacagttagaatctattgcagaaac
aaaactgcccgatttgaacactaacaatatacctcaagctatgaaaatagttggaggtac
ggctaaaaatatgggaattctgattaaagatt-aaaagtttcaattatatttatta-ta-
-tttatgaaaaaaatttcacgtcgacttaaaacattacaatcaaaagttgagcctaaact
ttacactattaatgaagcagtatctatattaaaagaaacatcaaatgctaaatttaagga
aactgcagaagctcatattgctctaggtttaaatcctaaatatgcagaccagcaactcag
agcaacagttatattacctaaaggcacaggcaaattgataaaagtagcagttattactaa
gggagaaaaattaacagaagcaattagtgcgggagctgatgttagtggttccgaagaact
aattgatgaaatactgaaaggtagattagattttgataagctaatagcaacaccagatgt
tatgcctttaatagcaaagttgggaagagtattaggacctcgagggttaatgccttctcc
taaagcaggcacggtaacattgaacgtagcaaaagccgtaaatgaatttaaaggggtaa
agttgaatacagagttgatagaacaggtataattcatgtgccttttggaaaatctagttt
ttcacaagaagatctagttttgaacctccaaacaattaaagaatcgattgatagaaataa
gccttctggagcaaaagggaagtactggaaaactttttcttatctagcaccatggggcc
atctgttcaaatcgatatcactagccttctatgaaaaatttttgtatagttatagaactt
gaataatctattatttcaaatta-taatttgctagcattattaacgtaataatttgctta
ctatatcttatctgcattaaaaacttt-ttatatatcttattaattatgagtacaaaggt
tgaaaatatcttggaagagttaaaatctttaaaccttctagaggctgctgaactagttaa
acaaatagaagaaacgtttgatgttgatgcatctgcggcttcaggaggtatgatgatggc
cgcaccaacttcagcaccagcatctgctgaggttgaagagaagacagaatttgatgttgt
cctagaggaagtcccagcccctaaaaagatttctgtattaaaagttgttcgctctctaac
tggtttgggtttaaaagaagcaaaagatcttgttgaatctacgccaaaagtcttaaaaga
agctgcatctaaagacgatgcagaaactatgaaaaaacagctagaagatgctggtgcgac
agttattgttaagtaataa-----agaggcgcttctgtttatacttaaacagaagcgcct
ctttatttattataggcaattatactagaaaagtccaagtgtaattgcattgttgatagg
catagttgcaccaatacctagccaaatacttacaactgtaccaattaaaaaaacagtggt
tgcaattggtcttctaaatggattttgaaatttgttaacattttcaatgaatggaacagt
aagtaaaccagctggtacagcagccatacttagaacacctaatagtttattaggaattac
tcttaatagattaaatgttggaaagaagtaccattctggtaaaatctctaatggagttgc
aaatggattcgacttctcacctatagaagatggttctaaaattgcgagtccaatactaca
ggcaaaagtgccaataattacaactggaaagacatataatagatcatttggccaagctgg
ctctccgtaataattatggcccattccttttgttaatttagctctcagctttggatctgt

taaatctggtttttttaagaattgacatatta-tttttttgtttaatataaagattagtct
aatcactgttttaaagcgtaattttcaactattataatggtccagaaattccttgtttac
gtatcattaaaaaatgcataagcatgaaaacggcagttagtagcggaagtacaaaagtgt
gtaaactatagaacctagttaaagttccttggcctacactaactcctcctcttaacagtt
ctactatactcgctccaacaactggaacagcatcaggaacaccagtaacaatttttacag
cccagtatccgatttgatcccacggcaaagaataacctgttaccccaaaagaaacagtaa
gcactcctaaaattacacctgttacccatgtcaattctctaggtttttttgaaacctcctg
ttaaataaacccggaatacatgtagaatcatcatcaagaccatcatgctcgctgaccatc
tatgaattgatctaataagccatccaaagtttacatcagtcatgatgtactctactgatg
taaaagcttctgcaactgtaggtctataatagaatgtcattgcaaatccagttgcaactt
gaattagaaaagatacaaatacaataccccctaagcaataaaaaatattaacatgaggtg
gtacatacttactagaaatatcgtcagcaattgcttgaatttctaatctctcttcaaacc
agtcataaattttactcataaaatagcttcaaaaagctcttctaatata--ttttgcgtt
taagctactattaattcaatag------ttattgaacttatattgtttttataatcaata
cctattaaatattataacatctttttattgattt--------------------aaaatt
ctctgactcagtaaaacagggctatgcactataattctttttttttgtatagagattaat
ttcgtttttagtaatttagctaaaattcttgttacactgactctattactacctataatc
tgtgctaaaattttatgtgtaatagtaaagttaagcataataccattattaacttgagtt
ccattttgctcagataataataataataagctcacaagtctactggttatacttttatgt
gaaataacttctataaaatgataagcttttgccgagcaaaatattaaatgatttacaaaa
aatagattgaatggagcgcagttactacatgcttttataatagtagtatagttaatacag
gcaacttctgttttatctattgcttctgcttcgtagtaaaaattatcgtcgaccaattct
atttgtccaaatgtatctcctgtagttaataagttaagagttactttatgtgtatttctc
aggactttgtaataattaaagatcctatcagtataatgtataatctagagctgctatta
aacaaaagagtatcattttgtctaagtgaaaaaatttgataattaattttattacgatta
aaaaatagtagccatggattgcaagaaaaaacatcttctgaactactttcagagaatatt
ggaacaaaattttcttgatttaaacaatttttagtcataattactactttataa------
---aaaaatattctaataat-taagaactgaaattaaatgttatataaattgatgttagt
tgagaatgacattgcattatcaaaggccattcaagaatacttaatagatcaaggcttcaa
tgtttatattgccaataatggattagaagcactgaatctagcttatcaatataatttcaa
tttgataatttctgatataatgatgcccctagttaatggctatgaactattagcggaact
tcaaaaaaacaaagcgttatccaaaatacctgttattttttttgacagctaaaggaatgac
gaaagacagaataaaaggttacgacatagggtgctatggttatttatctaaaccatttga
ccctaatgagttactctcaattattaacaacttaatcgctagagatgccttaaaagaagc
ttctttacaaaactcggaaatatcaaaccaggaattaaatcataaaatacgtttgacgcc
tagagaaaaaagtattttagatcttgttgttgatggacttactaataaagaaatatctac

tatattgaatactagtgttagaaatgtagaaaaatatgttagtcgactcttacacaaaac
taatatgaagaataggactttgctagttaagtattctataaataataatttattaaataa
taagatcaatgaaagggcgaatgacgggactcgaacccgcgaatgatggagccacaaccc
attgccttaaccccttggccacactcgccatattacatatatcactatagctcgtttttt
gatatatcgtctagttttcgagttatatatttatataaaacaatatttactttgaatttt
-tatgatacatagcaatattagtattcaaattaatggagaaccatttaattgctcaaagc
ccatctcattgcaatttttattaaattatcttgactttaattccgagcgtgtagcagtcg
agctaaatgacattctgttaccagagactctgtttcactcaacttatttgaatgatcaag
ataagctagaggttattaccattgtaggtggtggataatctatttttacat---------
---agttctctatttcaaatgtttcatagatagagataatcttccttgtttttatctac
atgaataataacagctttaatagtatcacctattttaaattgtgatgatatctgctctag
ttctttgatattaatctcagaaatatggacaagacctttgagatttcctacctttataaa
taacccgtaaggtgtaatctgatttattatgccttcaataatattaccaacaattaaatt
agatgatgcttgagatattagagctcttctatgactgagtattaaattattagatttctc
ttcaacattgagtaattttagcttaataaatttattattaaattgctcactttttttgaaa
attaccaagatgagaattaggtacaaatccagatataccttcaaggttaattatcattcc
tcctttattgaatccttttatcattacattaagtaaagagtcttcagctaatagttgcct
aattcttttccatgctcttatatattcaaggcgacgaattgataagattaattgtttcga
ctgaatattataatctaataaaaaaaattctcttgtatcattaatatttaaagaagtgaa
attatttaaatcttgattactagatacttcttgaataggcaagtatgcagatattggtgt
tcctatatctactaataccccatttaattcaaagctaaatatagttccagctacaatatc
tccaagattcaaatcatacttgtatttttgtaaaacagctgcaaaatttctatgagtaaa
tccgtcattatttttttgtcatattgaatagtatgaattaaagagtaactttgtacaatat
attaaggtaaacaaatgtttgtcgaattgtatg-aagaattttaatttattcggttgtaa
ttaaacatataaacatattaaaatatttaggagacataaaaagtagacgtagatagctgc
agatttaa-taatatataatctgaggaaagtccgggctccacaaatacaatttatgctgg
agaaatcccagtgtaggtaactgcgaggatagtaccacagaaaaaaaccgccagaataag
t----tatattagctggtaagggtgcaaaggtaagttaaaagcttaccaaaagtactgca
aagtatttgttaggtaaaccccctaaaatggagcaaagctactaaacaaatatttgtgtat
attttatctagtttagttgaaaatactgcatgaagttattagtaacaataactcgagagg
aatagctatcctttacaatatatc-tctattatattgaaagaacagaacccggcttatgt
aatacttattatgtcttgtattattatgtctattgtcgaatctctaaatttaactgtttg
acaatttttttctgtagctgatgttgtttataatctatttctgaattggttaaagttttc
tcattagatcgatatgtaaatcttagacctatactttttttcccttttccaattgattta
tctatatattgatcaaataatgttgtactttctaaatctttatcataaaattgatccaat
gattttaataagtatttaatttccatattcttaggtattattaaagagagatctcttatg

atcgatggatacttagagtatggtttaatctgataatttaaataattgaattcattatga
caatcttctaaaatgtttaagttaacttcaagaacaaaaagttttgtgttaagtcctaat
tcactatattttgcttgattaagttcactgaatatgccaataacgttattattatagatt
agtgtggcaaatttcccttttgaatgaactttattctaccatcgactatattttttttt
gtccactcaatttgtctattaatcttttgaaaaaaattttctacgatccctttagcttca
taccaatttaaagaatgcgcaggatgtgaccatgtagatcggatatctagattaccaccc
aatatgatagctagattagtagtttccataattttgttttgatcattatgaaatactgtt
cctatctcaaagctatctatagtttggttgctttttttaatattataaagatttgattca
attaatccttccagcaaactgctgcgtaaattagaataatcttgaataagaggattgttt
aaagttatatttcctttggactttattaaagaataatgaactaattcagttagtcctaga
tttcttaaaatacttctacttttatctatgaattgtctctttgtcgataactttttgttg
aattgaatatttggtactgagctcctgaatttatgataaccataaattctggcaatttct
tcaataatatctatttctcgaaaaacatcttgttgtctataatttggaacagtaacttct
atattttccttattttgataaataagatcaaaatttaaagagtggagaatattttgaatt
tcttcaaagaataaaaaacgtgtttgaccattatatataattggtcctaaaatgtcatga
actttttaattgataaattaatattaagagcattatcagttgtttttctcgatagaaa
gtctctctaacatttccgcatgttaaatctgtgattaaagaaagagcttcaaaatgtgca
tttttccaattatctacatttagcccacgttcttgtctaattgaactttctgttcgaata
ttaagaactcgagaagactttctaaccacagattgtttaaatatagcagactctacaagt
atcgatgttgtattataatctgtatcaagattagagttgcttccaataccagctatacta
attatatttgtatttacttgagtaaccaaaatatttttgtttaacccaatattttcattg
tttaacttaatctgatcattattttcaataggaaaatcgcttcgaattgttagactattt
tcatgattcatattattaattttattcaaatcaacaacattaatgggctgtccccatttc
aacataatataattgcttatatctgttaacaaatttctgtgtgtaaacccagaagagagc
aatcgatttttagccaatctggagaatctttaattcttatttcgtctataatggcagca
aaataattgtcgcaatttaataaattgtgatcacttattatagt-ctccttttagagaa
atatctattattgggttgatatatagctgataacattgaggaccctgttaaagccgagac
ttctcttgatagaccaatcatgcttaacacgtctgacctgttagctgttgatgatatatc
taatacatgatcaatctcacctccaattgtaatagattcaatagcttcaacttcaaaacc
tgcttgtgttaattgattagttaaattatctatatctattgttttgatattaacaagttc
tttcaaccaatttaaagaaactttcataattaattgtcatctatcttatagaagtattta
ttttacttatgattaaaaatcaatttaaaaaattcatcaaattaatataagttatttgat
gaatttgcttatacaatatattattgtgatatcttacttttatatggcacttattaagaa
ttagcttttgtaaaagttagcccattatcttgggcatatctctccataaatctcatgaat
ctatcccaatcatcaggacttttaataacatggacagcttctattgcttgtggcttacca
ttaataaacttggcattaacatctcttgtaataatttgtccctctgtgtccatcagatac

atacctgtgatttcacctttatctgccatactagcatctaaaattttgggatttgtaaac
cgaaaagttgctgttcctgtactaccatctcgagaccttgttaaacgtacatctggtaca
acttcttcattgataccttgaataaattgaattgttgccatatcaatcttctttatttat
atataatttatattggataaaactgatttctagttcttgaatgcttgttgagaaactgtt
ttaaaagcttatagtagcagtagatcaaattttatgtattatataaaactgtaacttttc
-ttttttg----------------------------------------------------
------------------------------------------------------------
------------------------------------------------------------
------------------------------------------------------------
------------------------------------------------------------
----------------------atctgttcaaagaattaacttattataagctaatcata
cgaatattagccatataatgtttgtaagt-gaaattaaggggttgtagctcagaaggata
gagcaagcgcctcctaagcgctaggtcagcggttcaagtccgctcaaccctgttaacaag
ttaagtagtattatttaacttgttctgcaaatcagtattgatttaagttaaagttcttga
tagctaaaacaaaagaatctgagaaatatttttattta----------aattatcttaac
tctgaatatctgataattattaa-aaagttct----------------------------
----ctttttaagacacaaggtgatgaagataagatatttaaactaataatggaaattaa
gtttttgaagttatagaagccaat--ataatactttctattaacaga-----------cg
cttgcttctattcctatttgtttacaaga--aatt----atgaatatcaaaata------
--attttagctcca-------------------------------------ctgat----
------------------------------------------------------------
-----------------------------tactaataaata-------------------
------------------tttatttcagtctagacaag------atgaatcttcatggta
taagag------------------------------------------------------
------------------------------------------------------------
------------------------------------------------------------
------------------------------------------------------------
------------------------------------------------------------
--------------agctattctctgtgtcatacttatataaaatcaggat-------ac
tgtt--------------------------------------------------------
------------------------------------------------------------
------------------------------------------------------------
------------------------------------------------------------
------------------------------------------------------------
------------------------------------------------------------
---------------gtcattattataactggggtgggaggattcgaacctgcgaatggc

ggagtcaaagtccgctgccttaccacttggctacaccccaacatgtgaatattataatag
cagtcgtataccctgccatgtcaatagactgttgtctttatttaataattttttgctgctc
ttgtagaataaaatctttttttcaaataagatacgctatgtttaagttgaacgactgtt
atgttttcttgcttttgactaattagccacttgatagtaattattccttctgcaatttcg
ttacttcctataataatacaagctacagctcttttttatttgcttgcttaatctgttta
ctaaaattacttgagctaacatctatttctattgtgaaaaactgctgatgtaaaaaacgc
attacttgcataccagtttcgttagctttttcaccttgtgtagcaatataaaaatcgata
gatttattaggcaagtttatattatcctttgcaagcagtaataggcgttctaatcctatt
gcacatccaacagctggtgttgctttgccacctaattggtgaactaagctgtcatatctt
ccaccaccgcatatagtatcttgcccttttgatgtcaatgtcttaatttcgaaagcagta
tcattataataatctaatccgcgaactaacttattatttatattgtagggaatatcaagt
aattttagataattgcaaacgtcatcaaagtgtttttgtgattccaaacttaaaaaatca
gaaatttttggagcctctgttagtatttttttgtgtattagcatcttttgaatctaaaatt
ctaattggattactagttaatctttttttgtgagtcagtgtctaaatcatcatggtacttt
gtaagataatctagtaatttaacttgataaatactacgatcctctacttttccaattgaa
ttcagatcaagttttaagttatgcagattgagattgttaaagatgctcatagctaaatgt
ataacttcactatctgctcttgcgtctaggctaccaataaactcaatgccaagttggtga
aattgtcgttgtcgtccactttgcggtctttcgtacctgaacattggaccgctataccat
aatctttgtaaactatgttggtaactcattttattttcaataaaagatctcacaatcccg
gcagtaccttctggccttaaggtaatatctctattgctgcgatcatgaaaccggtacatt
tctttatttacaatatcagtattttcaccaatactcctatcatataattcactattttca
aaaataggtgttctaatttcttggtaatttgcacattctaataatttggaaattttatta
tgtataaactgccaatattgaatttcatccggaagaatatcttttgtccctctaatagcc
tgaatttttgccatgataatttttttgattaccttcttaatatttctgcattaaatagcag
ttgtagactggtattcagtcatgtattatacattag--aacatactggtg-tattttata
aaatacgggcaaggagggattcgaaccccccgacaccatggttcgtagccatgtgctctaa
tccactgagctacaagcccacttaataagtacgttaactatatcattattctctaatgta
tacaactatattcttttt------atcttacatttgaaatttttgtaaaagtgaaaaatt
at----gtttgtttttacaattggaactattcatatataatgcttaatttattataact
ttgtaaa--aatatctataagtttaattgtatgattctactatatt-gtaatcttgtcat
aattttgcgctcttatttttcaattag------ctagcatctaaataacttatatttcaa
ataaaaaaaatcttatctaaaatgagtaaacaaattctatatcaggatgatgcaagaaaag
cactagaaaaaggcatggatattttaacagaagctgtttctgtgactttaggaccaaaag
gaagaaatgttgtcttggaaaaaaaaatttggtgcccctcaaattattaatgatggtgtaa
cgattgcaaaagaaattagtttagaaaatcatattgaaaatactggagtcgcactgatta
gacaagcagcatctaaaacaaatgatgtagccggtgatggtacaacaacagctacagtac

ttgcttcagcaattgttaagcaaggcatgagaaatgttgcagcaggttcaaatcctatgg
ctattaaaaaaggtattgaaaaagcaacaaattttgttgtcagtaaaatagctgaatatg
ctaagccagtagaagatacaaaagctattatacaagttgcttctttgtcatcaggaaatg
atatagaggtaggtaaaatgatagcgaatgctatagaaaaagttggaagagaaggagtta
tttctttagaagaaggaaaatcaactaatacgattcttgagatcacagaaggaatgcagt
ttgaaaaaggctttatttctccttatttcgttacagatacagaacgaatggaagttcttc
aagaaaacccttttattttatttacagacaaaaaaattactttggtacagcaagaacttg
ttccattgcttgagcaaattgcaaaaacgtctcggcctttattaataatagctgaagaca
tcgagaaagaagcattagccacaattgtagttaataaattaagagggattttgaatgttg
tagcggttagagctcctggatttggtgatagaagaaaatctttacttgaagatatgagta
tcttaacgaatggacaagtaattactgaagatgcaggtttatcacttgatacagttcaat
tagatatgttaggaaaagcccgaagagttattgttactaaagattcaacaacaataattg
cagatggtcatgaaattaaagttaaatcaagatgtgagcaaattaagcggcaaatagaga
cgagtgactctttatatgaaagagaaaaattgcaagaacgattagcaaagctttctggag
gagttgccgtcattaaagttggtgcagctacagaaacagagatgaaagataaaaaactaa
gactagaagatgcaattaatgcaacaaaagcagcaattgaagaaggtattgtaccaggag
gaggcgctactaatgtccatatctctagtgaattatttacatgggctaaaaacaatttaa
ctgaagatgaattaattggtgctttaatagttgaacgagctgtgacctatcctttaagac
gaatagcttttaatgcaggtgacaatggagcagtaatagtagaaaaagttaaaagtcatg
atttccatataggctacgatgcagcaaacagtaatattgtaaatatgtatgatagaggta
ttatagatccagcgaaagtagctagatcagctttgcaaaacgcagcttctattgcagcaa
tggttttaactactgaatgtattgtagttgataaagttgatgattgatttata------a
gataatagataatctcaaacaaagaataatttctaattatagaaattattctttgtttca
attaagctaattaattgatcaaatgtataattttataataaaaccaaagtttaaacaca
tcttctttttcaactattaagtagagaagatataagcccgtaatacctaaattatcaatt
aaaatattattttcatatatagattttctaaagatctaaaaaatgacattttttgaaaaa
taataccgaaatcttcttttgtagttatttgctaactcattactgcctaaagattgattt
ccaattgaattcagaacctttaacagaacactttttaaatctttatctgactttatttta
gaatcaataataatttgattcaattgcatgtcaaaactaagattaaaaatactacttgtt
aaagatatttgattttgaatatctagagattttatagccaagaagattaaacttttattg
ttcattgtaaaatataagacaattaatgatgtattctttaaatacta-aatttatcttct
tgtgcttctatcgtttttttgt-aaaataattaatgctttaatttttttagttttagttact
acccgaaaaaataaattctggaaatctagcttgagctttagctagagagtgtttttttcc
ttcttcttgccagtaattgataacttctgttgttttgtttaataactcaattctatctct
ttcagcaatccacggcttagattctagttctgcttttaattcttcccaagcatcatttct
tggccagaaaaaataagaagtaagtggactagtgccattcccaacaatttgatcaatagc

aatagcaatattattttctaaccaaagtacttttagtgtaaattttgacaagtttttatc
tccattattaatattaagtgttgg----tacagtagaatactgtactagttattttttttc
agcaagaatttgagctttgtttaaaatattttgtactgttttagtagcttgtgcaccttg
agacaatcgttgcaaaatttttgaaatgtcaacgcgagtttcattagttataggattata
aaatcctaattcctctatagctttaccatcgcgtttatttctgctatccattacaacaat
tctatagcttggttgtttttttcttccgtatcgttttaaccttaacttgaccatattatt
cacctcgcttattttttt--atatctttattaaattatatcttacgacattatgcaataat
tcaaacaattaagcagattcaagtctaacgttattaaatattaccattaaacattgtaga
aaaataatacctaacataggtgacatgtccatgccaaacataggaggaatgcttcctctg
aatagttttaaatatggatcagtaattctatttaatgaacagaatggctcattgtaccaa
tttacagttggaaaccatgccaatgataatttaagtaaaattaaaattaaataaatctca
gaaaaattagctattgatccaagtaataaatttaatgtaccaggaagagtattcatacgt
attttataattattatatacttcattatactaatagtttgattatatattataatctctg
taaataaat-cagctaaacaaaaaacttatccttaaagctatcttctcgcgagcattttt
ctttatctacattagtatttacttttgtcgtataaatttctagtttagaccaattttgtc
ctaacttatttaattgagatgttccacattctaaacatgctatttttatattgctcaaaa
cagcaccttttttgtataaaatcattcaataatggagtaatagttgcttcatccaaaaata
aatctaagacaaggatttttgtaaatgaatttaattgagctgggacattgttaataatat
tattataattaactaaaactatattagctgtaggaagtaaagctcgtgctccttccgcta
atataaagccataaggcataataattacaataatatatttataatgactgttaataagac
ttatagtggtattagcttttaattcgattgtttctgttactaaccactctctcattattt
cgtaagtgatccacttacctagttctgagcaagcagttcttaaaattgtacctggattat
tgttgttttcgagaacccctgaccaatgttgtattaatggatgggatgctatattaattt
ggagttgcatgttttgttaaatatatttaacatttaatgtgttgattgaaacttcaattt
ctatttgttgagtattattaaaatatcat----atatattaaagaaaataattat-aaaa
aatgaaaaaaagtctttggttgtggggttttacagatagtgctgaaacttggaatggtag
attcgcaatgattggttttatatcagttatttttatcgaagtagttacagggcaaggatt
attatatttaataggtatgatgtcataataaaaatccaaaaaagttcttgtaatgtttta
caagaactttttttgg-atttttataaatctaatctagcggtctcatagatagaacaggtt
cattctctctattgataccttttttcaaatccagcagcagcagctcttgctctgcctgcat
gccaaagatgtccaataaataggaagaatcctaggaagaaatgagatgttgttaaccaag
atctaggagatacatagttaactgaattaatttctgtagctacaccaccaacagaattca
aagaacctagtggtgcgtgagtcatgtactctgcagcacgcctttcctgccacggttgaa
tatcgttcttgattttattcaaatcaagaccatttggtcctctaagaggttcaacccaag
gagctcgcaagtcccaaaatctcatggtttcaccaccaaaaataatttctccactaggag
atctcattaggtatttacctaatccagtaggtccttgagaagacgcaacattggcaccta

gtctttgatctctaactaagaaagtaaaagcttgagcttgtgaagcttcaggaccagtag
gtccatagaattcactaggataagctgtgttgttataccaaacaaaattagaagctgtaa
gacccatgattgataaagcacctagactataagacaagtaagcctcaccagaccatacaa
aagctcttctagcccaagcaaaaggttttgttagtatatgccaaattccgccagcaatac
aaataatgccaatccagacatgaccacctacaagatcttccatgttatttacacttacga
tccatccatcaccgccaaacggagactttaaaacataaccaaagataactaggggattaa
gtgtaggattgctaacaaatctaacatctccaccacctggagcccaagtgtcgtataccc
caccaacaaataaagctttaataactaatagaaaagcgccaatgcctagtaatactaaat
gtataccaagtattgttgtcattttgtttttatctcgccaatcgtatccgaagaaaggaa
atgattcttcaagagtatcagggccaatcagagaatgataaagaccaccaaagccaagaa
cagctgaagagattaaatgcactacgcctactacaaagtatgggtatgtattgaaaattt
cgccgcctggaccaacaccccagcctagtgtagctaagtgcggaattagaataaaccctt
gttcgtacaaaggcttctcgggaacaaagtgcgctacctcaaaaagagtcatggcaccag
tccaaaagaccattatacctgcatgagcaacgtgagcaccaagtaatttgccagaaacgt
taattaagcgtgcattgccagaccaccaggcaaacccggtagactcaatgtctctaccgc
caacaccaacagttgtattaaagggcgtttccacgtggtaaaacctcctcagggaatata
aagttttcatgaggttgatcctgagcagccatccaagagcgaataccttcgttcaataga
atgtttttagtataaaaagtttcaaactcaggatcttccgcagctctcaactcttgagaa
acaaaatcgtaagctcttagatttagagctagcccaacaattccaaatgcgcttgtccat
aatccagttactggcacaaataacatgaaaaaatgtagccaacgtttattagaaaaagca
acaccaaaaatctgtgaccagaatctgttagctgtcaccatcgaataagtttcttcagat
tgtgttggagtaaatgcgcgaaaagtatctgcagcatccccatcttcgaataaggtattc
tgtacagttgcaccgtgaatagcacatagtagagcaccacctaagattcctgcaacgccc
atcatatggaacggattcagagtccagttatgaaatccttgcaaaaatagcaaaaatcta
aaaattgcagcaactccaaggcttggggcaaagaaccagcttgcttggcctagagggtac
attagaaagacggagacgaatactgcaattggcccagaaaacgcaatagcgttgtatggt
ctaagaccaactagcctagcaatttcaaattgtcttaaacagaatccaattagtccaaat
gatccatgtaaagctataaatgcccacagaccaccaatttggcaccagcgagtaaaatct
ccttgagcttcaggtccccaaaggaaaagaagtgaatgtcccatgctgtttgctggcgtg
gaaacagccgcagttaaaaaattgcatccttctagatatgaactggctagcccatgagta
taccaagaagtaacaaaagtagttccagttagccagccacctacagcaaggtaagcgcaa
ggaaacagaagtagtccagaccaacctacgaatacaaatcggtctctttttagccaatcg
tctacaagatcaaaaccaccacgagtttttttcttgtccaattgctatggtcataattta
atctccagagcaaatttaataagtaatttgtttaaataaaca---ctttagatatattct
aaaattttacttttctttacggtcaatataattataagtcaaaactaaccaaaattcacg
tttttaataaca-ttttttagctcattaagttaattttttttaatatatgaacataaata

gagttattgttt-aactcaaagtactcataatttattcatttctataatcctttgaaaga
gatatcctgttcctcgtgctgttaaaatcaagtcgggactacttggatcatcttctagtt
tagctctcaatcgtgaaatatgaacatcaacaactctcgtgtcaacatgtcgttcgggag
tatagccccagacttcttgcaaaatagaagctctagaaaaaggttctccagccttactaa
ctaaaagctcgagaagactgaattccatacctgttaagcgaactctttcattatttttat
aaacttgtcttttattggtatctattttagaaatccaatactaataatcccagagtttg
ggactccaagactagtggtgattttatcagctcttctaagaacagaacgaatgcgagcct
ctagttccttaggagaaaatggtttaacgacatagtcatcagcacctatttctaatccgg
taattctatcgcatacttcacctaacgctgttagcattattataggaacatctgattctt
ttcgaagttcttgacaaacaccgtaaccatctaatttaggcatcataacatctaaaacaa
caaggctaggatactcttttctaaacactagtaaagcttcttctccatttgaagcagtta
taacttcgtaacctattattgttagtcttgtttctaaaatccttcttatgctggcctcat
catcaacaacgagaatcttttctttctggttctccaatttttttatttactcctataaat
cactgataatgattactataaccttaaattttagctctatgagtatcttcatttctttt
agaacttagtttagttattataaattctgcaagtctagtttgcattaattagtaccagt-
-aaatatttttgt-----------tttaagttatgac---------tattttaatccata
agttgactctagtttgcagaattcaatatatacata-ttttctagccaagtaatttgtat
tctattttctaaataaaaaagggggttgacaaactagttgatagaaagtattctacattac
tataag----ttaattctaaaaaagaaattctaa-----atcttctttcggagtaataaa
tactttaat
>Pyr_27
tatatgata---aattaagacgtctcttatgtgagaatttctaaaatatacaaaactttt
tatttaatatagaaaaactaaattaaaataagttgaacataccttgctctaatctaaacc
tcttaattgag------attctaattagagataaaagtatttactta--ttgtatctata
ggaataaaaatgaaacttgcagtttatggtaaaggtggtatcggtaaatctacaactagt
tgcaatatttctgtagcactttcaaaaagaggtaaaaaagttctacaaattggctgcgac
cctaaacatgacagtacgtttacactaacgggattttaatcccaacaattatcgatact
cttcaatccaaggattaccactatgaggacgtttggcctgaagatgtaatctataaagga
tatggcggtgtagactgtgttgaagctggtggaccgccggctggcgctggttgcggaggc
tacgttgtaggtgaaacagttaaactttaaaagaattaaatgcttttgatgaatatgat
attatcttatttgatgttttaggtgatgttgtatgtggaggttttgcagctccattaaat
tatgcagactactgcttaatcattacagacaacggatttgatgctttatttgcagctaat
agaatagcagcttcagtacgagaaaaagctagaacgcactctctgagattagctggactc
gttggtaatagaacagataaaagagatctaattgataaatatatagattgtgttccaatg
ccagttttagaagtcttgcccttgattgaagatattagagtgtccagagtaaaaggtaaa
actttatttgaaatggcagaaattgataaggatttagcatatgtatgcgattactatttg

aatattgcagatcagttgattacaaggccagaaggtgtagttcctaaagaatctcccgat
agagaattatttagtctgttatctgattttacttaaatcctaaatcaaatgtaggacaa
gacaaagtagatcaagaagaattagatttgatgatagtgtaaataatattttcagaataa
tataacaaggaataagataatgtctacagctcaatcagatgctcttacttttgaatgtga
aacaggtaattatcatacttttgtcccatcagttgtgtttcttggttatatcaaaaaat
tgaagacagtttttcttagttataggaactaaaacttgtggctatttcttgcaaaatgc
aatgggagtaatgatttttgctgaaccgagatatgccatggcagaattagaagaaggaga
tatttcagctaagctaaatgattatggtgaactacgtagattatgcttacaaataaaaaa
agatagaaatcctagtgttatattttggattggaacatgtacaacagaaattataaaaat
ggatctggaaggaattgctccaaaattggaagcagaaattcgtgtccctatagtagttgc
tagagctaatggtttagattatgcctttactcaaggtgaagatacagttctagcagctat
ggctcaaaggtgtccattaaatttaaataatcaatcagataatacgagtcttaaaatttc
tcctcatattcctttggttttatttggatctcttccggattcagttgtaacccagttaac
tatggaattaaaaaaacaaggtatttttgtttctggttggttgccatcaaagaggtatac
tgaattaccggttataaaagaaggatattatgttgcgggagttaatccatttcttagtcg
cacagctactacactaatgagacgccgaaaaacgaaactaattggtgcaccatttccaat
aggtccagatggtactcgggcctggattgaaaaaatctgttcagtaatgaatgtaaagcc
tattggcttagaagacagagaaaaagaaatttgggcttctttagaagactatatttcttt
aatacgaggtaaatcagtttttttatgggtgataatttactagaagtttctcttgcaag
atttttaactagatgtggaatgaccgtatatgaaattggtatcccttatatggataaacg
ctatcaagcagcagagctagctttattaaaagctacatgtgataaaatgaacgttatgat
gccaacaattgtagaaaaaccagataattataaccaagtagaccgaatccgtgatctgaa
acccgacctagttatcactggtatggctcatgcaaatcctttagaagccagaggtattaa
tacaaaatggtcagtagaatttacatttgcacaaattcatggctttaccaatgcaagaga
tattcttgaactcgtcacaagaccattacgacgaaatcttagtttatcagaattaggctg
ggatgtgtatagcaagcaaagctagtatgcttc----tttatgacttacatggtataaaa
gccttgtaagtct--tttattttatgtcaaagaagcatttcttgctctgccatcaccagc
ccaagcttgtaaactttctacttgttctttatctgtaaatgctaaaggcacaaattgttt
gatagctattttaatatcttcagtactaaattctctctcttcactaaacgctgtatgcat
actttcaataattgcttgctcaatttcagcgccagaaaatttattacataataaactcaa
ttcatgaatatcatattcttgccacgatcttggtctaacttttgataaatgtatcttaaa
aattgactctctctcttgacgattaggtaaatctaagaaaaaaatttcatcgaacctgcc
tttccttaacatttcggaaggtaaactctgaattttattagcagttgcgacaacaaatac
tggagctttttctctgatagccaggtaataaatgtgccaaacactcttgcactagtacc
actatcaccttggctatataaaccagaaaaagctttatctatttcatcaatccataatat
gcaaggcgataatccttcagagatatttatcatttctctcattttggattccgattcacc

aactagtccaccaaacagttttcccatgtcaagacgtagcagaggcaacatccaatcgct
tgcaatagctttcgcggttaaagatttacccgtcccctgtataccaactagcaataaacc
tttgggcgaaggaatgccgtaatttaaactttgcttcgaaaatgagcgagaccttttttg
tagccattgttttaatatatctaagccaccaatatctctatttactgtattataaggata
aaattctaatagacacgtttgattaataatttgcctttttttcttcaataataataggtaa
gctccgagagtctatctgattatattgtgcaataattttagtaatcactttgcggattct
atctatcgataatccttggcaggattttgttatattattaacgagctctgaattcaaatt
aagattcaaagttttactcaatctcctgatctctttttttgatttctagaagactaggtaa
gggtaaatctattactgttatgatatcgcttagagcaaaaggtatattgattttgcagga
aataataacaatgtgcttagattgcgttttgattattttagctaaattgcgaagtttcct
aattaatactatttcatttaggaaagagtcaaaatcttttaaaagaaaaagatttaaaga
ttcattatttaagttctcaataaattctaatgctaataaaggatttctttttgcataacc
attatcgctaggattgttagtgtatccatctacaaagtcccagcaatacacttgttgatt
atttgaacagtttaagctgtgttttataatgtactctagtctatcttcttctctagtatt
tactacaataataggatatcgagattttaaaagtaatcgtaaatcttgagtgaaattcat
gtcaatattttttttccatcagacttttgtacttcaccaattg----aagactaaataaga
ttatctaatattgcttggatatattcaaaagtc------atctctctattttctaaatta
ttgctattattaacttaacttaccatgattttttttcttccttttcttcgtcgactattt
aatatagagcgtccacatggtgttttcattcttgctctgaaaccagaaactctaattttt
tttcttttagatccttgtagtgttcttttagtcatgatgtttcctctgttgtaaaagatt
aaatactattttatcatcttttatatatctttatgatatagatatatattaatatcaaag
tt-aacagcaataatttatattaatcaagaactatttatataatattaaccattaaatat
----------taactttatgatattggctttgcccatttttttatttaagtattttgacca
tctttttgctaatcttaaactggttaattttccagcagctaaaaacgattttgtcattag
aatctcaatttaagtattttgttgataaaagccaaaatggtaaattagaacccgacgaaa
gttttgcttttgcaaaagtatgtgtagctaaaaaatttttttccaaagctattattgaag
gtcaattagctttgaaaaaatatcgagaccttaataccttagataataatatagtaattg
ccaatttatataatatgctaggttttatttattttgaagcaggccaaataagttttgcaa
agaacttttatgagcaagccttacaaataaaccctaattatgttgtagctttaaataacc
tggcaaaaatttatgaagaagtaaaagattttaaaaaagctgaatctctatatgataaag
tgttaattcttaacttaaataataaaactgctaatagaaggaaaaattttatagcaaaga
cta---------aaaatgtctaatgcggtttgtaatcgggatagcaggatttgaacctgc
gacatcctgctcccaaagcaggcgcgctaccaaactgcgctatatcccgtaagacataat
tagtactatataccttttatctaattatgtctactctttctgtattttatctaggatacc
aaaacatgcctttaactccatctggatccttaataaatcctaatttttttataaaaactaa
caacatcaggctctgcaaatagtgtaatagtgctaatttctgcttgtctcaattgctgta

ttaattgctgtatgactactttccctaaacctaaaccttggaaatctggatgaataacta
cgtcccagatagttgcattaaatccgttgtctgatgttgctctagcaaatcctacaagtt
tactatttgcatctttttttgtattaaagaaataataatagaactatttttaatgcga
tctttactttttttagaggtcttttaacccatccgactgaatcacagagctgctccaatt
cgtataaattaatatttttattgctactcagatagatatctttaaattcgattctatcgc
aagttttatctaaaacaataagtttttgaaattttttcagagctagcattgatatcgg
agttttgaaaaaaatttttccagaagatcataatgattatgatataatta--aataaata
tatatattacaatattgctttaacaaacaatatattaatatatgctcaaatttttacatg
aatctattaagatagaagattgttaaaatatagagaaataaaaattttgtaaaatcttt
caataaaatcatataaagagtgctttttaagactaacatctattattataaataaagtct
cgcaattgagatgaaataacacattattctatgtctaatgccttttagaggagttatttg
tgaaaaaatctatgtttctaaccggtttacttgctttactaattatgagcaacccaataa
tagctaacgcagaagtagctggattggtaccatgcaaagattctgcggcattcaacaagc
gtatggtaaatagtgttaaaaaacttcaagccagattatctaaatatgatgcagataccc
caccagcgttagctttaaataagcaaatagagaaaactaaaactagatttgcaacttatg
gtcgagcaggtttgttgtgtggtactgatggattgccacatttaatttctgatggtagat
ggagtagagccggggactttgtttttccaggactattattcttgtatattacaggatgga
ttggctgggtaggtagaggatatcttttatctgttgctaagactagtaagccaacagaaa
aggaaatcattttagatgtgccattagctatcaaatttatgtcatccggatttgcatggc
cgctagcggcttggcaagaatttagtagtggacaattaattgcttccaatgacgatatta
ctgtttcaccccgttagtaaaaaatttatatgaataataattttaccaaatacttatcaa
cagcacctgtaattggtgtattgtggatgacgtttacagccggatttattatagaattga
accgctttttcccagatgtcttatacttttactt----ataaataa--gatcttataaa-
--aaactacaccaatatagagaaatactattatttgtgtagtttttaa-tatatataaca
taattattta-tttttaaaaatatttatcttttcgattattttaagattagatcttacat
aataaaata-atatcaatattatgagcttagtaacccaaattatagttaatgctgatgat
gaattaagatatcctacaattggagaattgcagtcaattcaagactacttaactacaggg
agtaatagaattagaattgctactattattagagataaagaaaaggagattatacagaaa
gctagtaaacaaatttttcagttacatccagaatatatagctccaggtggtaatgcagca
ggttctaggaaaagatcgttatgcttacgtgattatggttggtatctcagactaattaca
tatggcgtattagctggggataaagattctatagaaacaattggtattataggtgtaaga
gaaatgtacaattctttaggcgtacctataattggaatgttagatgcgatccaatgttta
aaggaagcatctttagaactacttggtcaagatgatattagaatcattgctccttatttt
gattatataattcgtggaatgtcataaatcgtagcttcaattaattagttgaataaaagc
ttttataatgttataattacaataagcccgaaagcatataattgtaaaaactaaatcttg
tcagaaccggaaggtagcagcaataatgtttaaatacaaaagttatga-ttttcgggtgt

ttactgttttcatctaatagtctcgacttttaattagtatgctatgtataaatctgtata
tgatatgtaattcatcaaacctttcagcaaaaacaatttaataatttgatattaatacgt
aaaacacctatac-tttttataaaatttctttatatctcatctaaacaatggaaactta
acttaatgggtgttcatattttatcaacaggttcgtctgtcccaaatttttctgtagaga
atcaacaatttgaagatatgatcgaaacttccgaccattggatttcaacaagaacaggaa
taaaaaaaagacatctagctccatcttctatttctttaactaaattagcagcagaagctg
ccaacaaagccttatatgcagctaacttaaaacctactgagatcagtttaattattttag
ctacgtctacgcctgatgatttatttggtagtgctagtcagttgcaagcagaaataggtg
caacaacatcggtagcttttgatattacggctgcctgctccggtttcattgttgctttag
taacagcagctcagttcattcaaactggttcctatgacaatattttagttgttggagcag
atacaatgtctagatggatcaattggtcagatagaactacctgtattttatttggtgatg
gtgctggagcagtagtattggggcaaagccgcaaaaatagtattttaggctttaagttat
gtacagacggtcagctaaacagtcatttacaattaatgaataaacctgtaaataatcaaa
agtttggtgttacagaaattcctcatggaaactataattccataacaatgaatggcaagg
aagtttacaagttcgctgtatttcaagttccaacagtaattagacaatgtttgaataatt
taaatatttcaatagatgaagttgattggtttatattgcatcaagcaaacactagaatta
tagaggcaattgcgagcagattatcagtaccttttctaaaatgattacgaacctagagc
attatggaaatacatctgcagcgtcaattcctttagcgttagatgaagctattcaatcca
ataaaattcaaccaggccaaattattgttttatctggttttggagccggcttgacttggg
gagcaattgtcttgaagtggtgat---ttatatattgcggatgacgagactcgaactcgt
aaagctttcgctacacacccctcaagcgtgcgtgtataccaatttcaccacatccgcatt
ttttatcatgaata-ttaaaaatttataatatttatctaaatattcatataatgtgtgta
tatatatatacattcttgtcagaaactctcttatattttattttaagtgataaaattgag
t----ttagtatagatagtaaaattttattttaaaatcataa----aaataaatatataa
ggaaactaaaatatgacaccgtccttatcaagtttttgaatagtcttattcttggggca
gtaattgtagtcgttcccataactttggctcttttatttgttagtcaaaaagacaggaca
attcggtcctaaaaataacttagaaagatatggagaagcaatttaagatgaataatgaaa
tatttttcataataatacgctctccatacattgctatact--ttagaagaa--------t
ttttattttgtaacatcctgtaaagtttattccgctctttctttttcct-atttactctc
tta----taatgatacaaatgtcagg-ttaataatag--tttttttatttttttatctttc
attttatatcaaagtattttatatacgtattattatgttcaattaataaagaaaagctca
tgtctttgtctaattggcctctcaaaaaagaaaattctgaagcatataacattaagaatt
caaaacaaatcacaattcctgatggtttatgggtaaaatgcttcgactgtggtctattaa
tgtattctaaagtactaaagcgaaatttaaaagtttgccctcaatgtagttaccattttc
aagcttctagtaacgaaagaattgatcaattgatagaccaaggtagttggcaaccaatgg
atgttcacttgatctctacagatccattaggttttaaagatcaaaagctttatagtcaaa

ggttaaaagatactgctctgaagactggcctgcaagatgcagttcaaacaggtactggta
ctatgcaaggtaaaaaagtatgcttaggtatcatggattttagattcatggggggaagca
tgggatctgttgtaggcgaaaaactaacaagactactagaaaaagcaactcaagataagt
tgcctgcaattatactttgtgcatcaggcggagctagaatgcaagaaggtatgttgagct
taatgcaaatggcaaaaatttcttctgctctagaaatgcataaaaaagaaaatttactat
atatatctgttttaacctctcctacaacaggggggtgtaacagccagttttgctatgcttg
gagacttaattattgcagagccgaaagctcttattgcatttgctggtagacgagttatag
aacaaacaatcaaagaagacttgccagataattttcaaagttcagaatatttatttgaac
atggtttcctagatttaattgtacccagaactcagcttagatcaaagttaatccaaattt
tgtccttacataatcatagtaagtaattaat----tattgactataatataaaaatgcta
gtaaaaattaagatattaagaaattttcactaaagatatttgtagactaactagaattac
aattactattgtagaaaaatttgtcttaatgtca-t-ttttcagtctttgaacaaattg-
tttttaggactttaaataattgttagtttaatacaaatttataaagaaaata-----cat
actaagctaaatatagtaatattatgttttcctataatactgacctatcaaatttcgagg
aaatttatgcttaaaagatcttcttggcttgcggctttattgggactattaacagtggtt
tctacaagtacgcatacatatgccatagagttagacgaggcaacaagaactgttccatta
gaatcttctggcagaactgtaattcttacaccagaacaagttaaaagaggtaagcgatta
tttaataattcttgtgctatttgccataatggtggtatcacaaaaacaaatccgaatatt
ggacttgacccagaatctttaggattagctacgccacaaagagataccattgagggacta
gttgactatatgaaagatccgactagttatgatggtgcagagtcaatcgcagaattgcat
ccaagtattaaaagtgctgaaatttttcctaaaatgcgtaatctaaccgatgaagaccta
ttcacaattgctggtcatatcttgctccaacctaaaattgtttctgaaaagtggggcgga
ggaaaaatttattattagaacttcaaagaactaaatcttgtgaatactagtttatgta-t
ttgttgcttaagtgtgattattttgttagatcatgatatatattaaagatggagtttgtt
gaaagcaagttatttcttagccgaacttgcttacactattcgtctagttattaaaaagga
gacgttaaattgaaaaagaagttttcagttcttttcactgtttttagttttttttgtaata
ggtttcgcacaaattgcttttgctgcagatctagataatggagaaaaagtttttttctgct
aattgtgcagcatgtcatgctggcggtaataacgctattatgccagataaaaccttaaaa
aaagatgtacttgaagctaatagtatgaatactattgatgctattacttatcaagtacaa
aatggtaaaaatgccatgcctgctttcggaggtagactggttgatgaagatattgaagat
gcagcaaattatgtattatctcaatctgaaaaaggttggtaattatacttgattttatcc
tgtattaaagaatagacaatctatttagttattcattagattgtctattctttgtttatg
---ctattatatagagatatttacacaatatt----ttatgatgaaaagaatacctgcaa
ttcttgtactagaagacggtacttattataaaggatggtcattccagcaagataaacaag
agattactattggtgaagtggtttttaatactggaatgacaggatatcaagaaataatca
cagatcctagttacttccatcaaattgtcgcttttacttaccccgaaattgggaatacag

gtattaataatcaagatattgaatctcacagtattagtattaaaggacttattgcaaaaa
acatttgtaaaatttcaagcagctggagagagcagcaatctttatttaagtatttaagta
gtaataatattccttttatttttggaatagatacaaggtctttaacccaatacttgcgtc
aatttggtacaatgaacggttgtatctctactgataatctaaaccatagttacttaaaac
agaaaatttgcgagattccaagtatgcaaggtttagatttaatcccgcatgtaactacaa
gaaatgtttacccctgggatgaaaaaagtttgccgaattggtatttgacagataacatga
gagtgcaccgagttattcagttgaaagttattgttatagattttggagttaaactaaata
tactcagaagactagctacacttggatgtcagataacggttgtgcctgcccacactcctt
taaaagatattttggcttaccaacctgatggcatattactctctaatggtccaggagatc
catctgcagtacattacgggatccagacagttacaaacttactagattacaatgtgccta
tatttgggatttgtatggggcatcaaattttaaatttagctcttaaagctaaaactttca
aacttaaatttggtcatagaggtattaaccatccatccggattgaaccagcaagttgaaa
taactagtcaaaaccatggctttgcagttgacttgacttcggtttttgaatctcctgtaa
gagtgactcatttaatctaaatgacactactattgcaggaactggacataatcaaagtc
cttatttttctgtgcaatatcatccagaatcgagcccaggccctcatgatgctgattatc
tattcgaaagttttatagaaataatgacaaagtccaagaagaaagttag-----------
ttagtaattttcccatgctttatgagtgaataaagcggatagtacttgtactcctcgtaa
ttgattgaagagaggtaaatgccctttggggccatgagaattccattgaaattcttcagg
atatctgcaaggaattctatcaacttcccagccgatttttttgccagaatttttcccaatc
ttttccaacactaagccatacttgtctttgaacaaataacccaaatttaccctttgagtg
tgtatgccataaattatcaatagtttgtaaatcttgagcaggtattttttttatatctgt
aaaatatagccaattacgagtttgtgcatttacaccagctagttgaataagtttttgttg
agttagctgatctgctttgagcaaatcccgatgagtcagtaacacttgtaaatctttgta
attcatctgttgagcagagcgcaatggaacaatgccatcgggaaataagttagaggcaaa
tttcactatttcttgattcttactgttgagtaacttttcataaattaaaccgtcaacaca
attacttttatagttaggcccagtaatcctctcaaacaatagatctgctaaatcttttaa
ctcgatagagtcgttgctatttatattctcaataatttcaagttgttgcttga---catt
attagacttagtatccttattcaattctaaaagctgggctcggatttgatttggcatctg
ttttgtttaattagttttaatagtcataaataaggtcttatagagaattagattgttttt
tgaaattagaagagcgccgcagtatttctaaagaagttcgcataatttgattcaataagt
tttgtcctgcataaattataatttgaccaattaaaaaaattgtgtttttttatttaggaa
caaaaaaatcttgtacctcaagcagaaaagtaattagtatttgcattgatgacaacatct
gtaattcattcgatctgcaagcatatatatactgacagtttaagccttcttgataaaatc
cccatactttgtaacgactttcgtaaattgctcgtggtcgctcaagatagtttttaatta
tagtattccaaattaggttattctttaatttttctaatttctatcagatagaaatttta
cattacagagataacttgcatttgtctcatttacagtattgaaattttgaactaacatat

gagcaataatgttgcttagctgaattaaataattttcaagaaaaatttcgacctgcttaa
ctggcatattgtaagtaaagagattaaaagttttattcacagtttgggatgacccaaaaa
gtagttgaataattagaatttgtagtagcatcttataatcgttcagtaggtagttcaaat
cattaatatcttggtaaatatcaatatagtccagttcatataacttacaaaatcttttga
cgcatctatggaataaatcaattaaaatattttcggtcaagtttgtaatatcatttacat
ctaaatttgaattataaatttctaaaagtatcttttctagttcagctaatataattttta
ataagttccttttttacttcggtgcgaaaaacatctaaaattaacacttcctgagaacaat
tagttaatctcttattaatctttatagaagttctaactaacagctccgctacctcctgat
tgagtgttggtccctgagagcttggccaataattattcacgttataccatagtaagaaat
attgaatttaatgttatacgataacttttttaatatactgaaaagcaagttagttactcta
tagcaattag-------gttttacagatcttaatacttatttatcaaataaagttatatt
atagtagcgactgagctaattttattaaaattta----taataatgacaacttattactt
tgcccttgcgagtcaaaattttctattagtacaagaaccactagaagaagtctttagaga
gagagttaactactatcagtcaaataataaagcaattgatttttggctaataccaaaccc
ttctttttagagaaaccagaaatgatttcatttaagaatcttgtacctaaagaggctgt
agctataatctctactaatccaatatttattaattggttaaagctaagaattggctatat
ttgtattggacaatttgaagataacctacaactttctgaagaatcgttaaatattactgt
tttaaaagacaaaatttaacaatgcttttattataactagtgtt-aattattttggttaa
gttatagttatagtcttttagtattta-gatgcttatagtttatcacttagtattttatg
ctgagatggagtgacaaatagtgtaagtatttcttggctaaatgtttcagttgccttaga
tttatatctatttggattaactataatagatagcattcgttttatagtaacattttcaat
ttgagcccaatgtacaattccaagttccaattccttagcaatagctgaaacggaaacaaa
ggcagctcctagccctgattgcacagcgtttttgatagcttctatagaattcaattccat
ctctattttaaagcgactgctgtcaataccatgttgactgagtactttatctatgacttt
tctaattgttgattgagtatctaacgcaataaatctaagcctgtataagtcttcttttttg
aatgtctcctagtttagaaaaaggatgtgattttggtaatatcagtgctaattcgtcttc
tgcataagaagtaacttgcaaaacatcttgtaattcagtaggcacttctcctccaataat
tgctaaatcaacttggccattagctacgctccatgaaataagtctagttgaatgtacttg
caattgaacagccacttgtggatatctttgcctaaatagtccgattaatcttggcatcaa
atatgtcccagttgtctggctagccccaataattaatgtgccaccttgtaagttttgtaa
gtcgtcaagagctcgacaagtttcttcgcagagggctaaaattctgcccccgtatcgtaa
aagaagacttcctgcctcagttaaagttgccttcttattacctctttcaaatagggaaac
atttaattggcgctctaaattttgaatttgcaaactaatagccggctgagaaacatacaa
gctattagctgctttttaaaactgccctctttggcaattgcttttaatattcttaactg
atccaatgtaaatggaaggtctgtcattaaagaattgtagtataatg---------tata
ttatttagtattgatcaatgtatgcatttgccaaccaaaatatttgttattacgcttcat

tatttttaaaataataaaaagtataatgttaaataagctgtattaaaatttaatatctgt
ttgatattaaatttatactaaagatatcgtaaatttaattctaactataaaatataggag
acaat---atggactctagacttttagttgtactaataccagttttagcagcagcatctt
gggcagtttacaatattggtagagttgcattacagcaatttagaaaaatgacatcttagt
tttgcttagaatata-atttttagactttatgagatagaggactttaaaatccctatctc
tattgtatatcttaataaaaaaat-caggc--ccccttgtacttactgttattacacatt
aagattattgat-agtaccta----gataataagtttaagctaacttattttttatttta
taattaattaaattaataacacaattatggctgttccaaagaaaagaacatctaaagcta
aaaaaaatgcacgcaaagcaaattggaaaaatcaagcaaaaactgaagcacaaaaagctt
tatctttagcaaaatcagtattaactagaaaatccaatggatttatttataatctaactg
aagcgtcagatacttttagtgattagaagtgttt-ttactacaagtctgaaaaagtatta
cgcaacaagttctaagtaatttttatctat----taatcatttgaaaatcattctatttg
ataataaaataaatagcagattaaataaaattacagtaacaaattatgcagcaaagctag
atcaaactagtgaaatttggttgttcaattgtattgaaaatattcagcatattttttaa
aaagtcaattaaaatcaagtcatattactaaaatttttatttctggtactagctttgaat
ataccgcaggtctaccaggactattatccagcttaacactaagtggtagactacatccta
ttagtatatatagtcctcagtctctcaaaaagtatcttgaagtatgtaccaaatattccc
aaactaatttttcttttcctattaatttcataatttacaatacggagaacaagttgtta
gccaattttatacagtaatttgtttaccgttgagtaaaaagagcttgctgtatggattta
ttattttaaaaaaagaaaagcatggagtatttaaattagcacaagctaaaaccttgaata
ttcttcaaggacctatatatggaaaactcaaagaaaaagataatttttaagtccagatg
gttactatctatccggccaagacttctcttctaatacaataatgggacataaaatatcgc
ttcctttattagtcagatattctagaattatttctgagatgcattggttttgctcttatc
ctattagattaaatacttattcgcatcagcaagtagtcaagtgtttaccacataatgttt
taactgatattatgaagtcgcaaatatatcaagataacagttttgttgaataatacttat
tcttgatatataatggtctcggtaatccaaataacatactagaaataatttcttgaattt
atttgttcatattaaatttacaatcatttatgacatacgcaattattgaagcaagtggca
cccagctttggatagaagaaggccgctactatgatttaaatcatatacctgttgatccag
gtcagtcgattatattaggaaaagtcttattattaaataaaaaatgggcaggttactttag
gacgtccttgtatagaaggtgttactataaaagctacagtaatgaggcacttgcgaggaa
agaaaataactgttttcaagatgaaaccaaagaagaaaatgagattaaaaaaaggtcatc
gacaagaattgactcgtttaatgatcgattctataacctcttaga------taaaatcta
gactcatattttt--aa-aaacttttaatacttaactaaaatagatataatggcacata
aaaaaggtagtggtagtacgagaaatggcagagactctaatgccaagcgtttaggtgtta
aaaaatatggtggagagcaagtaacagcaggtaatattttaatcagacaacggggaacta
aagttaagcctggccaaaatgttggaaaaggaaaagatgatacattattttcgctaatcg

atggtttcgtgctgtttgaaaagtcaaatcaaaagcaaaaaacaattagtgtttattctg
ctaagaaatagttaaaacaattggtgcaagataacttgagt----gtatttattgcactt
acagtggcaccatatatttgatttattaagaaatggagttccttaatgaaccaatgaatt
ttattaaagatttgctgtgaaatttaatgacaaacactattgtaatttcctgtctacaca
atatggctgctattctatattgtggtcagatacagaaactagtagtagcaaatgctcatt
atcaagtaagtgatatctacctaggttgtgttgataaaatcttctcaggaataaatgcgg
catttattaacctaggaaagaatgagtacagtggttttatacatatcagtgatacgggtc
cgcttaaaaagaaatattatgtcaataatattactaacattttaacaatacgacaaaaaa
ttttagtacaaatcattaaagagccaactttaaataaaggtccaaggctcactgccaata
ttacattatcaggtcgatatattgtactaatgccttttagtcaatcaatctgcatatctc
gaaaaatacatgatgaagatgagcgtcattatttgaaatctttagctattttaattaaac
cagcaacaatgggcttgctatttagaccttctgctgtaggcgtcgatgaggaaattatat
taagcgaattaaaaaatttaaaagaacaatggaactttgttcaaaaatctgcaattaata
gtcattcacctgttcttctatataaagatgaagatattgttaaaaaggtaatccgagatt
tttataataataatacaaacaatatagtaattgattcaaacctgggattaaaacaattaa
attattatatccacacttggcactgtaataactctagtacagttcccaagatgaagcttt
atagtaataatcaatgtatactagatgcttttggtatcaatcaggcaatttccagagctc
ttattccaaaagttgatcttatacttggtggctatatgtttattgaaactttagaagctt
ttactattattgacgtcaattctggatctttcaataattctactagtgcacgagaaacag
ttttaaaaaccaattgttctgcagcaacagaaatagcttatcagttacaaattagaaata
ttactggtgtaattataatcgacttcattgatatggaatcacaaagagatcaattgcagt
tattagaacactttaataaagagctggcacttgatgatgctaaaccacaaattgtacagt
tatctgaattaggtctagttgaattgactagaagaagacagggcaaaagtttatacgagc
taattagtagtgattctaattactttttatttttcacacaatcagagagatcccagtctc
ttaagagatttaatgatagacaacagaaacagcagacttttaatagatcttggctatcta
tagagattaatactattaacaaggtctttttcaaaagtcaaatttgtgcagacctgcta
acttttacctaattcgtaatctctatatagttaaaagtagtattacttataaacaaaatt
atttattaactcacagatcaaaacttatttattctcaagaatacaacaaagtattaccaa
gca------gttattatctagctagtctcaacaggaatagtaatcaggagttcctatctt
aagatttttatatcaactgtt---------------------------------------
-------------------gaaaaaaagaagctctctcttatataagagagagcttcttt
ttttagcttactaacta-tctagttagttaattaaaaattaaccgtaaacagctggagct
gttagagctactggtaaagattcaccagaagctagatctagagggaagttgtgagcgtta
cgttcgtgcattacttccatacctaggttagcacggttgatgatatcagcccatgtgtta
attacacgaccttggctatcaacaacagattggttaaagttaaaaccattcaagttaaat
gccattgtgcttacagataaagctgttagccagataccaactacaggccatagacctaag

aagaaatgtagagaacgagagttgttgaaactagcatattggaagattaaacgaccgaag
tagccatgagctgcaacgatgttataagtttcttcttcttgtccgaatttgtaaccatag
ttagcagattcgttttcgcttgtttcacgaattaagctagatgtaactagggatccgtgc
atagcactgaacaaagaaccaccaaacacaccagcaacacctagttggtggaatgggtgc
attaaaatgttgtgctcagcttggaatacaagcatgaagttaaatgtcccggagatacct
agaggcattccatcagagaaactaccttggccaattgggtatactaggaatactgctgct
gctgctgctactggagcagtaaaagcaacggaaatccatggacgcatacctaggcggtag
cttagttcccactcacgaccaatgtagcaagctacgccagttagaaaatgaagaacaact
aattggtaaggaccaccgttgtatagccattcgtctaaagaagcagcttcccagattggg
tagaagtgaataccgatagctgcagaacttggaataacagcaccagaaataatgttgttt
ccgtatagaagggaaccagcaactggctcacgaattccatcaatgtctactggaggtgca
gctacgaatgcaatgatgaatacagatgtggcagttaatagagttggaatcattaataca
ccaaaccaaccaatgtataggcggttttcagtactagtaatccaagagcagaaacgttcc
cacaagctagcgctttcgcgtctttgtaaagtagcagtcataatttttatcaattttta
aggagttaacaaagatgcttcccaaacaataatttgtctgttatggaaattattacttaa
ggccttttcaaatgagaacattttaaaaaaaaaca-tagcaaaatctaatctcagtaact
aa-aaatt-taattgttgatatattgcttaatacttcaa-----------taataaattt
tt-gct-------------ttttacaaaattaacaccatttagggat---ataataata
atgcatattaatgctgcttattttgattttaatagtttatataattttgccttacttaca
tacgcatgtcacactttacaaaaattcagacgaccatacaagatttaaatctgttaaagc
atgctttaacagatttgggtttgacttggcaaatgaattctagtcatataaaagttggtg
aaaataatcagcataaggtagatatcttaattaaacaagataatttgtcacatattggct
ttatttggaatgataataaataccacttagttgccgacttgcaactttgggaacaaccct
ggtctctagaagtattttggataaattatctcagaagtatgcttaccactctattatcg
aagaaacaaagaagcaaggctttgaaaaaatgcagcagatttcacaaaaagatggatcaa
ttaagttaattgtgcaacgctggaattattaactaaaatataaatgcgggcggagagact
tgaactctcacgagattatctcactagaacctaaatctagcgcgtctaccaattccacca
cgcccgcactgttaaaac--aatattactttatcataatctatactaaaacacaaactaa
-tagtataaactagcttccaagcttaaaagcgtctacaaataaaccatatgttataccaa
ttttgaagttgttgaataaattaatttttatattgaattgcttttggctagagtagacta
ttttactagtcaattcagttgctgcaactattagactggctgcaataataccccaatctc
ctgtttgtccaggaattgtagataaacctgttgaaataaaaaatcctaacaataaactaa
ttaagccagttgttaactcgctcaaagaataataaagcttattatttaagtttttttatta
agtaagcaaaaaaagttgaaagtctagttttaatcatagtatttagtaacggagttttaa
aaggatgacaagtattaacag-tcattaatattgtcgtatccttttaaaactcgttaaga
caaaaagctgctcttgt---------ttttaaaggtcttgttcgcttaaagcattaataa

gatgctgaaacggctcttgtataatatctttgcgtttaattgattgcaaatctaattctt
cttctatgatttcttgtaataactgtatgcttcttactgttggagcaacaggtacagata
gtgaattgtaagtatcttttagcccatttaatacacgctgatcgagaatatttgtactac
cagaaaccaaggcataactagtataccgtaggtaatattctatatctcttagacaagctg
catatcttcttgttgtataagaatttcctccaggccttaataattcaggctgttcttcgt
acaactgagctgcggcctcttttaaaatattggtagcttgatcattaataatttcagcta
tttttattcgatctagtccacttgaaaagaaagattctagttgtccgacagctgctttat
ctagataacgaccagtaagatcgtaacgatttaatattgctgttatagcatcttgcataa
attaattctcctac-ttgctatcagagcggtattataaatgactaatattgtaatataat
cattagtgtttatgaaatttatttaaaacttattctttattgcttaatgtgcaattaaaa
actcgcatagattttacatatatagagtatacgtagtttttaatta-tttttttataaaa
attaagaaataaagctttttt-taagaggttcgtgaatttcttaatacaatctaagcttt
ccgcccaaataagttttgtaaagtac--------aaaaaaagattattcaataatagtaa
------------------------------------------------------------
------------------------------------------------------------
------------------------------------------------------------
------------------------------------------------------------
------------------------------------------------------------
------------------------------------------------------------
------------------------------------------------------------
------------------------------------------------------------
------------------------------------------------------------
------------------------------------------------------------
--------------------------acaatattaatatgtatcaa--------------
------------------------------------------------------------
------------------------------------------------------------
------------------------------------------------------------
------------------------------------------------------------
------------------------------------------------------------
------------------------------------------------------------
------------------------------------------------------------
------------------------------------------------------------
------------------------------------------------------------
------------------------------------------------------------
------------------------------------------------------------
------------------------------------------------------------

------------------------------------------------------------
------------------------------------------------------------
------------------------------------------------------------
------------------------------------------------------------
------------------------------------------------------------
------------------------------------------------------------
------------------------------------------------------------
------------------------------------------------------------
------------------------------------------------------------
------------------------------------------------------------
-----------------agtataaataatcttttactccagcatcaaaaaaaatatttaa
catatgttgatacaattgattattcaagtgaaaacttagatttagaccaatttagctatt
ttatcattacaattaataaagctgataaatgtattcttatagagcaattttgttataatg
ctgaaggtcatatttactcaatattcttcaaaggtccaacagctaaacacctttcttcca
taatttgtcattttcaacaattaaatacaataatatccacagctcatgcaatatatttag
gaagagagctaatgaagtcagaattagctctcgtgctagatcaacaatatattcaagatt
aaatttttttaagtattaattatatttatgctagtataatattggtaagggcatgtaac
tcagtggatagagtatcagattccgattctgatggccgtgggttcgaatcccgccctgcc
c---gagtattcagattact-acactaaaacatct-------taatagctagtttgataa
tagtatgggcattggggctgcaaggtttctacattatgaaaagaagaaaatata--aaaa
aaacaagctctccaaaagagcttttagtacaattaataaatgcagaaaacaatattgttt
ctttttctcgaaaattagctcttgcttaaatactgttaattttggtattcgacatgtta
aactcttattcatggcgaatattctgtcagagtcgctcttagtttaagaaaagtcag-aa
aaatatttatctttacgtttt-tacctatttttatttaaataaaggttagta-ttttcaa
cttttatagtggacgtgggttcgagtcccaccagctccatcaatttgatt---------t
gtataattagctagttcaaccttttaataaccagatatataattggaggaaatatggat
tttataacaattactaagcccgctttaaagcaaattgcaatcttgaaaaatgatcatgaa
aatgatgtacatctaagaataggtgttagacaaggcggatgttcaggtatgtcatactca
atgaattttgaacatgttgacaatttaaaagatacagatgaacggcttcttctagataat
ttttctgttgtctgcgaccctaaaagtcttctttatctttatggcttatcattagatttt
agctcggaattaataggaggaggatttcaattttctaacccgaatgctagccaaacttgt
ggctgtggtaaatctttttcaggctaaca-ctctaacaggtttttatatattttgtatt
acacatagtattttagcaagtcg-tttaatagagacaagcaatgctctgttaaaaac---
---------------------------taatttcaagagtttttattgtattatt-----
----ttatatatttattatcaactacttgaaaatggattagttaagcaatctaatttgtt
atcagtatcaaactctaagtgattaactttttattgatatatgctaatcttt-ataagcc

taaacgtatctgtacattacaatttttgacataaaaagagtcttaacataactgtcaa-
----------------------gaagaaattata----------tattttataaatactc
aacatgaatctgtcacaacttattgttaattcctctatgatgaaaattcctaataacttt
tattgcaatacgaaatttatttcttctgagaatttaagtcacttacaattatcaaaaaca
aatttattatctcctacaaaaataattgcttttggactacctaattttgatttcatctct
acacagattacaagttctattagggaatcagttgttgtatctagtattgatgataataat
ctttttgctgccccaactaatgattggataaaaaaacacaaggataaaaatcatcattct
ttgttttataagaaaatctttaaagctttgctaaaccgtaaaatcagagttgttgatgag
ccatt------ttataactctgaaataaataaagaaaagagcataatgacaagtcaaaga
tacgtctggggtaaaagctggaagccttcaattattttatcttgcttaaaaaacaaaaaa
aatagcaaaatttctgacgcatcgaaaaaatttttaactgaacagttatctgcatcacca
gtatttgtggtaaaaaatggttttaatgaaattattttaggtcacccattatctcgtgtt
aaaacaggaagagttaataatctgacgcatgcattttctaatttattaaatcagtctaat
gaaacttatcctatatctaccggtttattctttttcatccagatgatgcatttgaattc
aaagactttataatatcggttaatccactggcggccaaacatatggagataagcgtagaa
cccgttggtttacactttgcttataaaatgaatagagatagatcgtcagatactcaattt
tcgtttattccagattttaaagaagtaggagatttattattcaaatatagaaaagatcat
catttagtttttcataagaatcagcattatggcaaagatttttttcaggggcaaccaata
tatatgattcagccgatcacttttaaagatcgatccggaaagctgaatattatcaaattt
actgggctaaatgacacaagagaaattatttttactaatcttgaagctgctaatagatca
tggacacattttataaagaggaatccacaattaaaatcagttaaaaatcctactttatta
gtttataatctagaaagtttttaaaagatcaagaacgattggataatcaagatttacac
aaatttgttgtgatcactaataaatcatcttatcttaccacaaaagagttaatagcttta
accgattctaatagctttttaagcacttaaaattaaatataaagcctaaactcttttt
gtgaaactatggggaagacggttattttccaccttaacttatgaat-aaatgctatt-gc
ctttatttcctcgaataatattctataactagtaattcattgagttgtaaagcgacccat
tccctatcaatgactccattgactttccctgataaattagatttatttaattctaagtgg
ctaggaatattagctaatccagggaaagctaagtaattttcgactagttttcgagatgct
tcttgatttttcacactaattgattctcctggtttacattgataactacagatagatact
acttgtccattaatacaaatatgaccatggttcactagttgtctagcggcaggaattgta
ggagccataccaagtctaaaaacagtattgtctagtctcatctctagtagttgtaacaaa
atttgacctgtcgacccttgtaatttttagctgcttttacgtacttaaataattgtttc
tcacttaatccataattaaatcgcaacttctgtttttcctctaatcttacagcatactca
gaaggttttctagacttttgtccatgttctccaggtggataagatcgcttaattgctttt
ctactgagtccgggtaaatcacctaatctacgggaaatgcgtactcgtggccctctgtat
ctagacatattatttaaattctcctaatgtaaatggattaag----------taatgagt

agctatttttaacacaaaaagagcattaaaacaaagc-ttttgctaaaaaatctttata
agaaatataggcttggacgaaa---cacactatgatgc-tttttaagcgggtagcgggaa
tcgaacccgcatcattagcttggaaggctaaggttttaccactaaactatacccgcaata
ttaaacagattgtcactaactcaatataacatagttatgttaagttaagataatcttaaa
tttaatagaaat-tcaaattatttacgctccttctaaaactacatcaagaaaacgagcaa
tttcggcagcttgttcttcaacttctgataataataatggttgaccaaccctagttaatg
gaattactcttttatcttttgtacataaataaatttcacgtcttggattgagcccctctt
taatatctattttattgatttgatttctttatactaaactgaagacagattttctat
tctttccagggaaacctagccgaaaaatttttacaataccttatctttgttgaattcat
tatatccagcccctatattccaaataatagttagccatagaaacaaactcaggaagattc
ctatactgccatagaatgtcattacaataccttgtggaataaataccaaatcggttgaag
ttgtaaagggtaacaaatcaacttgaaaataacttgatagtccagcaagaagaaagccca
aagccccaataaatattatagtagcccaccagtaattactcagtcttcgtgaacctaaaa
ttaaatcttttctaactgagtgtatagataatgtttttttcataatttgttcttttaaaa
cagta-tactaatgcaaaatggaaaaattttgacaattttt--attagcgctttagattt
tttaaataagcttgattgcttttagccctaaaaatagtcctgttgctaacccaaaaaaag
cagcaagaaatattgtatatcctaaaaaaactgaca---tatta----ttattattattg
ttacaatagtgtttacaaaaatagtggaaatctattactagggtattatattacgttaaa
tgccttgctgtctacatgaaaaaactttttcagtttttagcattgtgttcatattagaa
ataatattctagtcaatttattaaacttctggagaacagaatcttatgtcagaatttatc
aagccttataacgatgatccttttgtaggcaatttgtccacgccagttagtacatcaagt
tttagtaaaggacttctaggaaatctaccagcttaccgtcgaggtttatctccgcttctt
agaggattagagataggaatggcacatggatacttttttaattggaccttttgataaattg
ggccctttacgaggtacagatgtagcattactagcaggatttctatcttcggtcggtctc
attattattctcaccacatgtttatccatgtatggtaatgtatcttttactagagcagac
tcgaaagatccattacaaacttctgaaggctggggacaattcactgcaggatttctagtt
ggagcagtaggtggttcaggatttgcttatttattgctagctaatatacctgtattacag
actgcaggtcttagtttattttcttaagctagtaaggggacttgaacccgtaacctactg
attacaaatcagttgctctaccaattgagctatactagcaccttttttcatcataacatat
aaagatatttgttgcaatctgttattttcagatgtaacaaatatctttttttatttaaaca
gaactgttatattaacac-------gtatattaattagaatctgaattatctgactctgc
agaatattcttgatcacagtttagataataacaaattgtttgatctagacaagttgcaga
ccaaccaacaggtgtttcacctgataattcttgcatatctccagaattgtaaaaatcatc
atttggcttattactgtttgcttgcataaatatcactcccaaaacactctgaacttaatg
tgaagtaatttaacttctctataatatattaacacaaaatattaaatatggtgattattc
tttaattgtaaagatgcggctaactttataaagtttgagtgagagtcttgatagcttttg

ttgaaataaacattttaatccatttattttgctctctcgaccatactttttttatgttgta
aattaactttttgtagtttttttgttcgtttatgagaatgagatactgcatatccattat
tagctactttccccgtaagctgacactttttttgacatgatttt-------ttaacttata
tatttatttagagtacttgctaaagttgactttggcacagctccaatcactgtgtcaact
ctttctcctgctttaaagatcattaaagtaggtatacttctaatgccatattcagctgca
atagtaggattttcgtctgtgtttattttttactaccttaatagatgattcatattcttcc
gcaatttcatcaactacaggagagaccattctacaaggaccacaccatgacgcccaaaaa
tctacgagtacaggtaagttattgttaataacttcttgtttgaaagaggcatctgtaact
tgagatactgacatattctttaacctttaatatatgttccttgctaagcaaatcttatca
caaatttcaagaaaaatcttctatagtgaattactattagtgttgaattaatttttttaaa
aattagacattaaaaaaacttattacctatataaataaagtggtggccgagtgatatctt
gatagtgatagttttttatattaaggttgaacactagaaataacttttcaagcaaattttt
atttgaaaa--acttctctgtgaccataaaaacttaattatattacaatatattttatta
tttgtaacttagttctgataatggtataaacaacgcaaaagatactgccttataatcaag
gaggaatacatgtctcaatccgtagaatcacggactaggattaaaagcgaacgttacgaa
tctggagtaatcccttacgctaaaatgggctactgggatgctgactatgtaattaaagaa
acagatattttagctcttttcagaatcactcctcaaccaggtgttgacccgattgaagca
tctgctgcaattgcaggtgaatcttcaacagctacttggacagttgtatggactgattta
ttaacagcttgtgacttatacagagcaaaagcatatcgagtagatccagttccaaacgtg
gcagatcaatattttgcttacatagcttatgatattgatttatttgaagaaggttccatt
gcgaacttaactgcttcaattattggtaatgtttttggttttaaagctgttaaagctctt
cgtttggaagacatgcgcatgccagtagcttatctaaaaacgttccaaggtcctgcaact
ggattgattgtagaacgtgagcgtatggataagttcggtagacctttcttaggtgctaca
gttaagcctaaactaggtttatctggcaaaaactacggaagagttgtatacgaaggctta
aaaggaggtcttgatttcctgaaagatgatgagaatattaactcacaaccatttatgcgt
tggagagaaagatttttatattctatggaaggtgtaaataaagcatctgcttctgctggc
gaaattaaaggtcattaccttaacgtaacagctgcgacaatggaagatatgtatgagaga
gccgaattctctaaagaggttggtagtatcatttgtatgattgaccttgtgattggttat
actgcgattcaaagtatggcaatttgggctcgtaaacatgacatgattttacatttacat
agagctggtaactcaacttactctcgtcaaaaaaatcatggtatgaacttccgagttatt
tgcaaatggatgcgtatggctggtgttgaccatattcacgcaggtacagttgtaggtaag
ctggaaggagatcctttaatgattaaaggcttctacaatactctacttgaaagcgaaaca
gatataaacctacctcaaggtctgttctttgctcaaaattgggcttccctacgtaaagtt
gtaccagtagcatctggtggtattcatgctggtcaaatgcaccaacttcttgattactta
ggtgatgatgtggttcttcagtttggtggtggtacaattggacatcctgatggtatccaa
gcaggtgcaactgctaacagagtagcactagagtccatggttatggcaagaaatgaaggc

cgtaactatgtagcagaaggtccacaaatcttgagggacgctgctaaaacttgtgggcct
ctacaaacagctttagatttatggaaagatattagttttaactatacttccacagataca
gctgatttcgttgagactccaacagcaaacatctagtttaatgacgacttactaatactt
taaacagtcaattgtaagtagaattaacttataacaataaggagcacagaatagtgagac
taacacaagggacttttccttccttccagatttaactgatgagcaaattaataaacagc
tttcttatatcgtttctaaaggcttatcagcaaacgttgagtatactgacgatcctcatc
caagaaactcctattgggaactgtggggtttacctttatttgatgtaaaagatgcttctg
ccgttatgtacgaaattagctcatgcagaaaagcaaaacctaattattatgttaaagtta
acgcttttgataatactagaggtattgaaagttgtgtaatgtctttcattgtaaatagac
ctgctaatgaaccaggattcttattacaacgccaagacttcgaaggtagaactatgaagt
acagtcttcatagctatgctactgaaaagcctgaaggagctag------atattaattaa
gaattaatattggctaatta--ctcccctctttaaaatcacaattactaagttaattatg
attttaaagaggggtaatttcccaagtaaattttgactaaa-tagattggataattaaaa
atacacagaaataatgcaatcacaggatataatttccaacgatactcttgttaatttaca
agaagaatatgatagaacacaaatccaagaagttttaaatgagttaaatcaagaacttat
aggattagtacctgtaaagaccagaattcgcgaaattgctgcgctattattgattgacag
attacgtagaaaactagaactagtttctggtaatccaggattacacatgtcatttacagg
tagtccaggaactggtaaaactacagttgctatgaaaatggcggatattttgcatagact
tggatatataaaaaaaggacatttgttaacagttacaagagatgatcttgtaggtcaata
tattggacatactgcacctaaaaccaaggaagttcttaagcaagcaatgggcggagtttt
atttattgacgaagcttactatctatataaagcagataatgaaagagactatggctctga
agcaattgagattttattacaagtaatggaaaaccaaagaaatgacttagttgttatctt
tgctggatataaagatagaatggaaaaattctacgaatccaatccaggactctcttctag
agtagctaatcatgtagactttccagattatacttcagatgaattattacaaatagctaa
aatgatgatagaagaacagcagtactgttttacagaagaagcagacaaaactcttttaga
gtatactgagcgaagaatgaaacagccttatttgctaatgcaagaagcattcgcaatgc
tattgacagggctagaatgagacaagccaataggatttttgccagtggagaaaaagtatt
aacaaaagctgatttagtaacgattgaagcagaagatatcttgaaaagtagattattttc
attacctgatgcttaatattcacgtgatttaatgaaactatatattattaaaaag-cttc
ataatcagttgcaaattttttgaaaaacccttattatcatcttaatgaaggtg-tggcgg
catagccaagtggtaaggcagaggattgcaaatccttcatcccccagttcaaatctgggt
gccgcctagtactaaaaa--gggggtgtggtggaatggtagacacaacagacttaaaatc
tgttgattttagtaatcgtgagggttcaagtccctccacccccatatattaacttataa
ataaaaagagatgtgtatttgtattaattgcaaccatattacaaaatgtaagacctatca
cttaatagaatctcaacacaagcagcctcatattactagaagtccattgtttatacctaa
atatcctgtagttcatgttaatatatctaacaatggtacctataatcaaattgattggga

tttagtggagtgtttatcatttgtagaaaaaccgaatagttggaatttagacgctaatta
gttacaagaaaaattatgtactataagcaacaactctatttttttagatactcggtattga
ttttggatgttctaaccaataagattttacctgttcttgctatctctataatgccaaatt
tagttagtaattgttcaatagcaacaatctttcctggatctccagtaacttctataatta
aaagatcttctgcaatatctactatcttagctctaaaaatttttacaatttctaaagctt
ctgttctagtctgagaattgatcttaatcttaattaacattagttctctttcaactgaag
gaatatttgttacatcttgaacattaaggatatttactaacttatataattgtttagtaa
gttgttcaatagttctattgtctccttgaactaccatcgtgattctagagaccccaattt
gctctgctggtccgactgctaaacttgcgatattaaagcctctccgggcaaatagaccag
atattcttgacagtactccggcttcatcttgaactaaaactgataaggtgtgtttcatga
aaaattaattttaatctatttgatctattaagtatattctaagaaaattacagaaaaaat
actagttagtctacaactaaattatctaaactaaacaaaatcctactatcagataatata
tctaccttgtaatgctatactattaacttagtatttaatgatattctaattgcaaataga
atgct-aatttttaaaatttgttaggaacttgtgctagaaaaatacaaaaacagtaaaa
aactctctcgagatttacctcaaatcaatgatcgcatcagatttccgaaagtccgagtaa
ttgatgacgaaggtgaacaactaggtatttttgtgcctgaagaagctatacaattagctg
tccaacaaggtttagacttagttgttgttagtgataaatcggacccgccagtatgccgaa
tattagactatggtaaatataagtttacacaagaaaaaagagctagagaagctaagaaaa
agcaacataacagtagtattaaagaagtgaaaatgcgatacaagatagaagagcatgatt
ataaagttagaataaatcaggcatctaaatttattcaagcaggagacaaagtaaaagcaa
ctatcacatttcgggggcgtgaaatccagcactctaatttagctatagatttattgaata
aaatggcaagcgatctaaatacaatagctgaaattcagcaagctccatcaagagatggca
gaaatgttataatgctcttatctcccaaaaaagttggctaaactaatttttatttaaatg
catctggctggattcgaaccagcgacgtctctttcgaaatggcggattatgagtccgctg
ccttcggcccctcggccacagatgcatcacttaaagtttataactcataaagcatataaa
aatcaatgtttagctatttttaataaagtctataagagtctattattcattcatattgtg
atatgtagcaactgctgaaggggaaacttgatttaagtatcgaaatatccagtatttaaa
aattgtgtcgagtatgactggaaaagttgaaataaataaaaaaataaaatctcgactttc
tggaagtcctaaatgtcttaaaattacttcaataataacttcccatccatgaggagaatg
aaaacccacaaacatatcagtaaaaagaatgattaaaaaagctttagctgtatcactaag
accataaataatttcatttaaaaaagatttgacgactgcaatttgtctttgtccagttat
cattagtaatataaagacaagtattgatacaagatcggataaaatatttttaacagcatt
tgcactttcgtttgcataatattcgcccaattctctagccttcagctgtactctttttc
aatgatttcgtaagatatatcttccgaaggattcagaagaacttcaaagtgaattttctc
ttcaaatctttgtaattcagcaaaagctctttcctcttgagaggaatttaaaaaaatttt
aggctgttcttggttccataaataatcaatacaaggtccaaatacaaaaaactttgaagc

ctgattaacgagtactggagagatgaataatagtaaaatatattttacggaagtaatagt
ttggtgcctagatattctaaactcctctatagcttcagattcgccattaggatctaactc
tttacgaaacttttcaaaagtattagtaattgacctaggtattggcccaaccttttcaaa
ggcgaattgattatttcgttttaaattccaatatttcatttttattgctgctaaaattt
ctattaatcaataattgatcacgtattcactataaatattaattctagaactaaacatat
aaaacaatatttatgaaaaataaaatacaattaaaaattaattatgcattagttctatct
gtggctacagcaatgtatggatataaatcttatcagccaactataaatacactcttaatt
aaagatgaaaaaaactggccttgtttctctaaaaggcatactgcctttattcctataatt
gtattaaaaaaaaatcatcaaaatccctgaactcagaacctagaaatagtaagttatatt
ttaatggaattggaagtagaaaatttatcgagagaattgttaatctacttgaatatttag
aaaaacaggcaaactatactgaatatctgaaccaaattaaaactagtggttatttttctt
tactctaccaggacagcaatacaatgtataagaaaaatattagtaatattttttattac
ctaatgatattctaaaaagaatctatatttgcaataagaaaaaaaaagcttataccacatg
cattcttaataaatttatttcggaaacagataggctatccaaaaagttttgctaatttta
attgggccttctccaaaattatgaaatggtattatgatagaggataccaatggtctctag
ttgaagttaaacaagcgtctgatgcatcttctattgtaattgatattcatgaaggggtag
tgaagacaattataacagaatactacactttatcttataaaagactctcaggtatctcat
acatagagtcaatagagcagtatcttggagtaagagtcggagctccattaaatataattg
atttacaaaaaaagattagttacttaaaagataaccaactagttggcgacattatttata
gtattgaacgatcaaataataactcaatgagtttggatgtcaaatttcagatacaagaat
taaaagataaagagataatagtgcttgcagaaagttcttcta------------ttatct
cgcatgcatgtgatttactgaatc---aatataga---------aataggctagtagctt
ctaatataatttcactatcaactaataaattacacgcatgctattttaattataaactcg
actatcaatataagtatataaacacctctaatctcgtacaattgttaacttactctattt
cttgtaggaagacactacttagccattattttaatttacaaactttaataagctggacaa
aaaaaaatactattggatttcagctatatttacgaaatctaagctttggaaaagcttttt
gtgtgttcagtatgaaatttatcaaaaatggacttaatataagaattttatatataaatc
cttcattgatagttgatcagaattttatctttcaatttgctatacaaataattaagcaat
atcatactgcta------aaccaccagcct------------------------------
------tatttttaagcaatctagatcttgaacaatatgttgctgaaagtctattaatgt
accattttacatcttgcttctcaatatctgaaaaaatattattatctcggattatgcata
cagattctttatttttcaactctgagaatctccattttcattttgatgatagaaatcacg
cgatcaacagttatgataattttaaacaaaatacaaaaatattttatcaagagtttctct
ctttattatttagtttacgttatcaaaattttaattatttaggttggccttcaaaaggcc
attttttgaaataaaatctttatacttggctccatttcaaaagagtg------------
---atttttctgatagtcgtaagaccctattcttccacaaaatgtcgttaaagcaagtat

ctaattttaatttaccaatatcttttagaagccactttaaccatattttagttagtacta
ttaaatgccagtcaaatttgaatatgagaactgtatctttgctattaattgattcccctg
ctgaatatatgctatacaaatctattcttaatttctctattaaagttaggatgcagtatt
tcatacctatgagcaacagtataagagtatctctattctataattatttagattgttttt
taataagatcatctcagagtttcattcacatatggcaagatttaagaactcttaaaccaa
ttccacattttggttaaaaaaattttcatatggagttggaattcagcttaaattaccaa
ttaaacagatgccacctctttcaattgaatatactataactagtagtcgctacttttgta
tttaccttcgtacttattatcaacgataactattactaaaatgaaaaatttaatttcttt
ttttgatcgcagcaaagggaagtggatttcacaaagaacaacttatgaattatctaataa
aaacatgagttcagcacagtctcagatgataatgaaaataggcaaatcattgtcaggatc
tataatacttgcatcgttaaattggggagacatttatagacaagttgcccattacgctaa
aagtcatagccggagtgaatataataataagtttaacttacaatttaataatcaactaaa
taaccataaattattgacgctgtgcatagtgacagatcctagtctaatcagttttaaaac
tcgctatggaagtactacaatagatgaaacatattggtttgcaacaaataatttacgtct
aagtactagtattgttaaacgattcaatacttgtgtggcagtatctttttgttcagaaat
taaagtttaa-actctatatagaagcaactaataaccttta-ggctattagttgcttcta
tatatctaaattttctattaaaaattat----aatactatattattctaagtctttacga
ttaggattcctagaaggatcattagagaggaagccaaagacaaacaaagaaataaagaaa
ataacggtagtatatacaaagattttcaaagtaaacatattttcatccttaaaagctatt
agtgcg----atatttacatatctccacataattttatatcatgatatgagttccagtac
tattttataaataaatcaatatctttattttttagtaatgatttatatttgaatatgcgt
gatgtactaaatactaaatcttttccgatttgttttgtcttgtttgtaaatattttgtta
tttctatggagtttaccttcaattatcacatagtctaacttctttagctgtctaaaagta
tatgaagatttttttgttccaaatagttagattgattataactaatcgttttctttttaac
agtcttgcttttaacttaattatttggctttttattttcactaattctgatactcttacaa
ctaagaatttggactagcagagtacagctattcataatttataacaa------tttttta
aagaatctactaaaatagatgattttacaaaactaaaatgatatttcattatcttgttga
ccttgctcataagtaatgtctttcaatacttttaatatttttccaaagtactcttgcagg
tatttttctaaagatattacttccaacctgtcaattccaagaatactgtaaacttcatcc
atcggagctgtaaattgatctccgctagttaaaatttcagcaaatgctaaacgatcagaa
atattccatgtccattgaagagttttagtaatttttcttaaggcttttaataatccaatc
ggaatttgtgagatctgtgtttttttgcccagataatttttcacataatttaataatttca
gctgatgtccaagcggtattaccaactagcggcaggattctattttctgtggaaggtact
ccaagacttttttataactaacttggctgcatcctgagtatcaatgtatgcaatgggcgta
gattctcctgtaacccagactgacttcttgtctaaaataggaatggcatactggttgatt
aaaccttgaaaaaatcctcctaaagaaaacacagtatatttcacattggacttctgaagg

aaatctactacctgagattttaagctcattaatggaacatctgggtatttctctgaattt
agtattgaaaagaaaataaatctttcaactttagccgctttagctgcttcaattaatgca
atttttccatctaaatctatttttttctgcattgtaaggatcagtaggacgagatgtggaa
gcatctatgactgctgttactccacaaaaagattgcaaaatactttcgggtaattttaaa
tcaccatatataagttctgctccccactctttgagaaaagcagactttcttaaatttctt
accatgcattttacattgtagccttcatctaaagctcgtcttacaatttgacgtcctaaa
gttccggttgctccaattactaaaagagtcatattagttaaaaataagtatgtataaaga
cattgatttggttgacagcagagagtaaattaggtagagagggaatcgaaccctcatgac
cgaagtcgtcacattttgagtgtgatgcgtataccaatttcgccatctaccctattatag
accaactactacaagtaatttagtctttagcgtaaagataatactaagtttcaatagata
tgtcaacttttaaatttatattaataaaaaaacggagtctcttatgaccacttttgaatt
aattccttatagattgtatttaagtcaatctaacacatggatgcatagaataaaagcaga
aattaagatatatatagtaatgcttttatggatttcaatttttactttttcttactttaa
actatgtattattgctttaagcttaattgcaataagttttactataagaagtaaacagaa
tattattcaacaacatttgttacagacgttattaatgacgttcttaactactgttttgtc
ttttagtgtggctattaattataaacaatatgcagaacaagaacaatctcagtatttatc
tgattccaagaaatataaaaattctaggagctactattatatacaaatagctaacgatca
aaggatcaaaagacaatacttaattactactttaaagccttccttgtactttttttattac
tatatattctattaaactagttatgataacaacctctccagaagttttagtaattactat
ttatagatctagtataataaataaaatatttaaaaatgaattgctgtttatcttccttct
ttcgtcacatattgttactagcattattaatagagttgataaagttattcaagtaactag
cttaagaggaagcttgagtttatataattcatttacaaggcttttaatgttttctttgtt
aatatttcaagtcttctttttagagattatccgggaatcaaaagagatagctcaagctct
ttatactagaaatctcaatcaagaaaacaataatttttttgaaaatatatacagtagaatc
tcactttagtgatcggctaaatataattattagcactttgtactttattattttagctat
agcgtaataaaatccacttattcttctgactctagtaatacgaagagtaataaaattagt
tagtagtttaaatagatagaat-taacatattatataaataattaaatgtttcattatgg
tcaatactcaaaatcaaatttcacaaacttcagaacttgattatatagttaatcaacctt
ataaatatggctttaaaacttctgttgaatctgagcaatttcctaggggaataagtgaag
atattgtccgcttgatctcaaagaaaaaagatgagcctgaatatctgttgaattttaggc
ttagagcatataaaaaatggaaaaagatgagtaatccgtcatgggctcatattaagcatc
cgaatatagactttaataatattatttattatgctgttcctaaattaaaaaaagaattga
aaagtctagatgaggttgacccagaaattcttgacacttttaataagttaggtatatctt
taaatgagcaaaagcgaatttccaatgtagccgttgatgctgtttttgatagtgtatcta
ttgcaactacttttaagaaagaactgtctgaagctggtgttatattctgttcaatttctg
aagctataagggattatccagaattaattaaaaaatacttaggtactgttgtcccagctg

gtgataattattttgctgcattgaactctgcggtatttagtgatggctctttttgctata
tcccccctaatacagtttgccctttagaattatcaacttactttcgtattaacaacgaag
aatctgggcaatttgaaaggacactaattattgctgatcgtggcagtaaagtaagttatc
ttgaaggttgtaccgctcctcaatttgacacaaatcaattacatgcagcgattgtagaat
taatagctcttgaaggagccgaaattaaatattctacagtacagaattggtatgctggta
acaaagaaggtaaaggcggtatatacaattttgttactaaacgaggcttctgctcgggta
ataattcaaaaatttcatggactcaagtagaaactggatctgcaattacctggaaatatc
caagctgtatttttagctggtgataattctcagggagaattttactctgtagctttaacaa
ataattatcaagaggctgatacagggactaaaatgatccatattggcaacaatacaaaaa
gtagaattatttctaaaggtatttctgcgggtagatcgaaaaacagttatcgaggcctag
taaaaattggccctcagtcatttaattcacgtaattattctcaatgcgattctttattaa
ttggtcaatcatctcaagctaatacatttccttatattcaagtacagaatccaacatcaa
aagtagaacatgaagcatccacttcaaaaattagcgaggaccaaatttttttatttttac
aaagagggatcaatttagaagaatctattgctcttatgatcagtggtttttgtaaagatg
tctttaatgaattgcctatggaatttgctactgaagctgatcgtttactgagtttaaaat
tagagggaactgtaggatgagccagattattttagagatcaaagatctatatgcttctgt
tggtaaaacaacaattttaaaaggggtaaacttatctattcgagccggcgagatacatgc
gattatgggtcctaacggttcaggtaaaagtacattatcaaaagtaattgcaggacatcc
agcatattcattaataagcggagacattttatttttggacacagtattcttgaaatgga
gccagacgagagggcaagagctggtatttttttagcttttcagtatcctgttgaaattcc
gggagtcagtaattctgatttcctaagaattgcattaaatgctcgcagaaagtttcaagg
attatcggagtttagccccttagaatttttcaactaataatagaaaaagtagatcttgt
tggcatgcaagaaagtttttttgacaagaaatgttaatgaaggattttctggtggagaaaa
aaaacgtaacgaaattcttcaaatggctttactagatagcaacttatctattttagacga
aacagattctggactagatattgatgcattacgagtagttgcaaaaggaattaacacttt
agctaaatcaacaaattcaattattttgattactcactatcaaagattactagattatat
tattccagattttgttcatattatgagtaacggaaaaattgtaaaaactggtaatgttac
tcttgcccaagatctagaaaagcacggatatgactggattacgcaaacatagctatttag
tag-------aatattgagactaaagaatattctttagtctcaatatt-tttgcaaatac
-aagtt-aattatttaagctttgtcaaaaccttgcttaacatcttcaattgctcttttta
gcaattcttcgctagctggttctaatttttagtactacgaacattttcaccaaattctg
gtttagaatttttaaatcttctcgaagttctctaataaattcagccacttttgaaactt
caatatcatctagatagccattaattccagtataaataatagcagtctgttcttcaactg
gaataggagaattttgagcttgcttcaaaatttctcgaagacgttgacctcttgctagtt
gattttgggttgctttatctagatcagatgcaaactgagaaaaagcttctaattcagcaa
attgtgctaactctagttttaatttgcctgctacttgtttcatggcttttatttgcgcag

cagagcctactctggagactgagataccaacattaatagctggtctaatccctgagttaa
ataagtctccagataaaaatatttgtccatctgtaattgaaattacatttgttggtatat
atgcagatacgtccccagcttgagtttcaatgataggaagagctgtcatactacctccac
ctaattcagcattcaatttagctgctctctctagcaatctagagtgtaaatagaatacat
caccaggataggcttcccgtcccggaggtcttctaagcaaaagagacatctgacgataag
cttgagcttgcttagttaaatcatcataaataactaatgttgctttacctttgtacataa
aatattcagctaaagccgcacctgtataaggagcaatatattgtaaagtcgcaggactgt
ctgcattagctgcaactataatagtataatcgagagctcctttatcctgtagtgaagata
ctacttgagctactgaagatgccttttgtcctatcgcaacatatacacaaacaacatctt
gacctttttgattaataatcgtgtccaatgcaacagctgttttacctgtttgacggtcac
caataattaattctcgctgacctctaccaattgggatcatagagtcaatagcagttatcc
ctgtttgcataggctcacaaactgattgccttccaattataccaggggccattgactcaa
ttagccttgttccattactcgcaggttcacctttatcatctattggccgagctaatggat
caacaactctacctaaaaaagcatctccgaccggaatttgagcaatgcgaccagtacctt
ttacagaactgccctctaaaatatccctaccatctcccatgagtactacaccaacgttat
cactttctaaatttaaagcaactccaatagttttatcttcaaattcaagtaattcaccag
ccataacttcatcaagtccatatactcgagcaataccatcaccaacttgcaaaactgttc
ctatattggctacttctacatcctgatcatacttttcaatttgttgacgaataatactac
ttatttcatcaggtctaatatttaccatatttttaatattatttttattaattggaaatt
ctaaagcggtaattagtcagactaatgctatctgtctttttttactttattttaagttgct
acaacatcaaggtgtgaagccatctgtcttagctggcctctaatactagtatcaataact
ttcgatccaatctgaatagtaaagccaccaattagctcgggatcgactgaaataattagt
ttaacttctttagctttagtcataaccttgatttttttctgttaataaagtttcctgatca
gggcttaaagcaatagaagtattaatattcgcaattgttaaagattccatttggtacgcc
agttccaaatatttgctagctatagcatctaacattccaattcgcttgcgatcaactaaa
atcattagaaatgataatgtattttcgctaatttgatcaccaacagttgcaataattact
tcttttttttgactctacagtttttagtggattagct---aaaaaaatttttttagcttgtca
gattgcgctaaaatattttgtattgactgaatatcttggctcaccttttcagtaactttt
ttagttttagctaaatcaagaagagctactgcataaggctgagctatttttataacaaca
ttgttactgctcatagttgatctcctaacttagcaatattattatcaattatacgtagtt
gcatttcggagctcatttgattttccagctgcaaagtaactcttttaagagcaagaaatg
ttatttgctgctggatttgccttcggatctgttgttcagctgtttctatatttgatttac
ctgttatagcaagtctttcaatgtccaattggccttgagctaatatggaacttctaactt
ttccagctgttagctgagcttctttcttaatttgatctataataatttgagtttgagcaa
gttgtttttctgattctgacaatctagcacttgcttgctctagtctttcttcggattctt
ggatagctgctaaaacttttatctgtcttgcacttaaacttgatccgagaaattgcttca

aaacataaataagaccaaataaaagcaacaatatattaataacatttgcttcaaaaatat
cagaattaaagccgaatttgtgttcactactatgttctgataaaatagtaattatttgtg
gtatttttactatattattcatttaaaattca----------tactattttaatatttta
ctctggaatcatgatttaaagattgaccacttaatagtttagctttaatttgatcgctc
aaagtatcaacctgattttctaaagttttaagagcttcttcttttgaatatttaactgt
ttggatgcttcagcaatcagtttctctgcgttcaattgagctttttttatatcttctgag
actatattttgagcttctttttgagaagaggctatttttaattgtgcattgcggcgtgct
tcagatagatcttcttcatattttgcagctagctcatctgctttaacaagcatagacgat
gcagtagtcaatgttgttcgaatatattcgtctcgttcatctagtactttagttacaggt
ttgtaaaaaatattatttagtaaaaccattaaagtaagaaactgtaatgccattaaaggc
aaagtaccattgaagtcaaataagccaccttcaatttcttcggctaatagaaatggtaaa
tcaatcattttactaattatgtttatagttaataatttggtttcaattaagaaaaataac
tttatagtgcgcttagtctcatcagactaagcgcaaatgattattttaaccaacgtatgg
attagcgaacaataaagacagcgcaacaaccaggccgtaaattgttaaagattccataaa
tgctaaacttagaagaagtgtacctctaatctttccttctacttctggctgtctagcaat
accttcgacagcattagctgctgcactaccttgaccaatgcctgggcctattgccgcaag
acctacagcaagaccagcagcgataacggatgcagctgaaacgattgaatccataatgaa
ttttaatttttttgttatataagtagaaaattgacaaatttcgtataatacttaaaatcta
tataattaagtataatagattctaatat-----tgtaattatagtatattactcttctcc
atgaccttccattgcttctccaatataagctgcagacagagtagaaaaaattagtgcttg
tattgaactcgcaaataatccgagtatcataactggtaatggtattaaaattggaattag
cagagtaaatacagaaacaactaattcatctgctaagacgtttccaaacaatcgaaaact
taaagatagtggtttcgtaaaatcttctagaatattaattggtagtaatactggagttgg
ctgaatatatctagcaaagtaacctaacccttttttacttaaacctgcgtaaaaataagc
tacagatgtgagtagagacaaggccacagttgtattaatgtcatttgttggggcagctag
ctcaccttcaggaaggtgaattaacttccaaggaattaatgcacctgcccagttacaacc
tagaataaatagaaataaagttgcaatgtaaggcacccacgggcgatattcatgctcacc
aatctgatttttttgcaatatcttgcagaaattctaatataaattccataaaattctgaaa
tttttcaggaattctttgtagatttctagtacctagaaatgataaagttaaaagtgtagc
aattactagccaagagacaatgaaaacttgcccatgcaattgtaggctacctattttcca
atatagatgttttccaacttctactgctgataagctagtatatgggttaagatctatgag
actgttttgatacataataaatatttcttttaaatagtgcaagaatcaaaaaaataag-tt
tttgaagaattaacatgccattaatttattcttaaaattagattattaataaattaagag
caaaaataaattctaatagagatagctatctttagtagtgattatttttttgtaaaaaaca
attattgctacctataatgcttttacgtttaattatatacccatatattaattatttaat
atgattataggtattttgctacaaactttattgttttatccattct----agagattgtt

atagtttattttttatttaacatacctgcgacttcatctgcaaaattaactttcaaattt
tcttctccccctcccagaagaaatcttacaaatcttctaattttaatattttcacctaga
acagcaatattttggtttattaaatcttcaatactaatatcttgattgcgaataaacatt
tgatctaataatgaaagctccttaagtctcttttcattcgtccttcaatgattgtttcg
attctatcaactggtttgtttttcaagtcatctttaccagcttcaattctttgttccaaa
tcaataatttcactaggtatatcctctgttgatacgtattccacattaggcgatgccgct
atttgcatagcaatatctttagctaacttttggaactctggacgtcgtgctacaaaatct
gtttcacaattgacttcaactaatacacctattcttccgcctgtatgaatataactctct
agtaaaccttcaatagcggtacgattagatttttgttagctgaagctagccctttttgt
cttaatgattctaaagctttttcttcattgccattattagcttgtaaagctttcttgcag
tccatcatacctgctccagtcttatcacgtaaggcttttacgatttgagcagaaatttgt
agtgtcatatgctttcaaaaattttaagtatttaaatt----------aattattataa
ttcactttgagtaggatgagttatttcttctaaatctgtttgcccgtatttgccatcata
aattgcatctgctatttacctacaattagtttaatggacctaatggcatcatcattagc
tgggattggaatattaataatttctggactacagttagtatcaagtatacaaatagttgg
aatacctaattttaaacactcttgtatagctgttgtttctctcttttgatctacaactac
aacaatatcaggtaaacgagtcatattttaataccatttagatgtttgcgaagtttatc
taattctcttcgtaaaacagcagcttctttttcggtaactgatcaatcataccactttt
atcttgttcttctagctgtctcagacgactaactcgagatttaatcgttacccaattagt
taacatcccccaagccatctctggttaacataataagaatcacagcgttgtgcttcctg
tgccactataccagctgcttgtcttttgttcctagaaataaaactttttaccatccga
cgaagcctgttttataaattcacaagcttctgttaataattgtgcagtttggactagatc
aataatatgtatgccatttctttctgtataaatatagggaaacattttaggattccatct
acgtgcttgatgtccaaaatgaacacctgcttctagtaattctgctaatgtaacaatagc
cataagattattaatatttaaataatgtaattattcgggaaatccctttttttagaaac
tgat-----tcaactaacttatttgaattttagtaaactaataataaattaacacacttt
taaagcagatcaataaaatagatcctgaagaattcat----tctacacttttattgctaa
aataattacgcgcggttctgtcgtctaaaatgatatcatctaaatcatctcgtacagatg
aaatagtactttcgtcatcagtatttggagttaaattctttttctctaacgatgagctat
tatagttatcatacatattgaaaccagtaccagctggtataagtctacctataataacat
tttctttcagacctcttaaccaatctaatttaccagatatagcagcttcagttaatactt
ttgtagtttcttgaaaacttgctgcagaaataaagctttctgtatttaaggaagcttgag
taatacctaatagtacaggcctatatgatgcattaatcttatgtctcaaagtcatggctt
tattagtctgttcaatcttttgtaactctacaagctctccaggtaaatagcctgtttctt
caccattttctattttcactttagaagtcatctgccgtactattacttctacatgcttat
cagaaatattgactccttgagattgatatactaattgtacttctttaacaagtagtaact

gaatctcttgaaagcttaatcgagcagcttcatataaggctaatccacattcaatataca
aattaaaacttgcgtctaggatatcatgaggattcaacaatacgtctgttttctttcttg
cttcaagaatttcttcaattcttggaagaccttgtataatgtcacctgtttttgctctat
caaaaaccaacacagctaatgtttctccccttctaattaatgcattgttattaacatgta
aaatggcaccattagaaaccaagtaaggtctagcaattcttaatgtaactgatctagaag
atatttcaataatttgtccagagtctaaagaagtcatattttcagtaataaaatcaccgc
agcgaatccaatcacccacacttactttaactattgattttcctatattaaaggtctttt
tatcagaagatgtcgcaatcaaaatgcgcctactagtattcttttcagaataaatttcct
caacagtaccttcactcattgcaataatctcagtactagcaacaactgtaaatggcttga
tatattcaccatttttacaataattctagtttgactttgttgcttctctgatttattta
catcgatacttttaattgataatgtttcaaaagttgacagccttaagtcaaaataattat
tgccactttctttagcagaaaactcaattgaggatattaaagtatcatctcgatgctcaa
tttcagccacaagattagttgttactaaatgaactccttctatagatttcactctttctc
cgtcccgaaaaggagttcttttaactagctttagtttagtttgacttggtttctgtgaaa
ctagattctcaattagactggattttgtttctggtattgaataaacaataacaggtctaa
taagaatataaggcatttgctcgttttctatatactcccaataaactaacttatctgtag
aaatattattatgaagtgtttctccaggtcttaaaaaacctctacttttatcattattac
tatacacctcactaagtttatagatagatccaggcttaataataatttctcgaactattc
catccttttggattatttcaacaatacctgagcttttagaaaaaatatttttaactagct
ctgttccactttctataacatctccattatgcactagtagtagtgatgaatctttattaa
tttcgtgagtttcttctgaaatccataagatatatcccggacttaaaatttcgtaagcat
ctttatctgaacctgttttctttttgatacatttaagtctaaatacttaataataccc c
cacttgccgttttgtacgtatcagatattagttctgcgactgtgtaaccatcttgaa---
ttttttggtttgggatagctttgaaaagaaatttttgtttttttttctgtttctaaaatat
acgattctttttgtggattacatctgtataaatataacaacctggaattacaatagatt
cagtaataacttgtatattagtaatactgctattagtattctggtgtatccttacttctc
cagcataatgatttattaattctgtctgcgcaagtatcgtaccagctttagttttatctt
ctttatgaacaataacattagcggaatcagatatactataaacttcacctgagagcaccc
agatcaagcctccagttttagtaattcttgtagtatattgattattatctgtttcttcaa
ctgttaagttggagaaacatattttgccagagagatctgaaactacatgtttttgagctc
tttctgtcatgagtcgatttctacggggtgattcagcaatcacttgatccttcgatacta
attcaccatcacatattaaaagagtagtatctttacttaagttaattattgacttttgat
ttttcttagatctgattgtaacttgggttggctttttttgttattaatgcttgctcaccat
gtctagtacgaacatctgtatacgattcaatatctagatctatcagttgaccatctatag
gtgagtaaatttgttctgctaattcaccggtaaatactcctccagtatgaaaggtcctca
ttgtcaattgagtgccaggctcaccaatagattgagccgcaataataccaacagcctctc

ctaaatcaactaagcgaccatgtgcaagattccatccgtaacaatactgacaaacagaac
ttctagagttacatgtaacaggagatcttactaaaactttttt---gataccagctttag
ttatctcttttgctaacttaggactgacatcttggttcgtatgagcaattaaacaatttg
tctctggatggaatacattttctgctaatactcggccagataaagcttgctctagattaa
ttaatattttctgcgtatctactaaatcttctagtatgatgcctttattggtcttacagt
ctacttctctaattattacatcttgagaaacatcaacaagtcgacgcgtcaaatagccgg
aatctgctgttcttaaagcagtatcaactaatcctttccttgctccgtatgatgatataa
aataatcagtaacagttagtccttctctaaaattactcgatattggaagatcaataattt
gtccctgtggatcagccataagccccctcatccccactaactgtcttacttgagaaatat
tacctctagcacctgagaacgccatcatataaacagcatttagaggatctgtctctttaa
agtatttaataacttcttgtttcaaggattcactcgcattattccacgtatcaataactt
tttgaaatcgttcaactgctgtaatctcgcctcggcgatatttgttttcagtggccttaa
tatcttcaatcgtgcctaataagagactttgcttgctgggaggaattcttaaatcttcta
aacttaaagaaattcctgcttgggtcgcataatgaaatccaagatctttaagcttgtcgg
ccatattggctgctcgggctataccgtaattacgaaaagcccagacaattaaattttta
gttcgtttttatcaataaccttatttgaaaaacttggctgtgcaagacttcgtctattat
ccactaaatcttctccatattaaatatttgtt--attaactatgctactaaagactcttg
aataattttattaaagattatacggccagctgttgttctaatatattgcacaattctctg
attttcagcatcttcttttatgatatggttattaaaaaattttgttgtactcttgtcaga
atgtttttcaatcttaattggatcatgaggttgatcaccatctactaatccatcaaaacg
agcccatatataagaatgtaaatcaagtttcttttgctcataagccataactacatcttc
taagcttgcaaaatactgattagatccctgctgctgagatggattattagctgtcaaata
atagcaacctaaaaccatgtcttggcttggcataataattggttgacctgttgctggaga
gagaaaattatgaggggctaacattagcaatctagcttcagcttgagcttctaaagacaa
aggaacatgaacagccatttgatctccgtcaaaatcagcgttaaacgctgggcatactaa
tggatgcagcttaatagctcttccttctacaagaataggttcaaatgcttgaatacctaa
tctgtgtagcgtgggagctctattcaatagtactgggtgcccttgaataacttcatttaa
aacattccaaatagaagattcatttttttgaatcatctttttggcagctttaatattgtt
aactaatccctgcaaaattagtctgtggataacaaaaggctgaaataactccaaagccat
ttctcgtggcaatccacattgatgtagttttaaatgaggacctacaactattacagatct
accggaataatctactcttttacccaacaagttttgtctaaaacgaccttgttttccctc
tataatatctgataaagatttaatggtctgttatttgcacctacaacagttctacctct
acgaccattatccatgagagagtcaacggcctcttgcagcattctttttttcatttctgat
aataatctccggagctaaaattgactttagacgagataatctattatttctattaataat
tcgacgataaaactcatttaaatctgccgtcgcaaatctacctccatctaattggaccat
aggccgtaagtcgggaggtatcacaggaataactgtgaataccatccaagaaggatctgc

accggttgctatgaaattttcaatcaatcttaagcgtttcatttttttattaaattttaa
tgaaggtgtcttaaaactttttggtggatttgtagcttcaaaccttaaggtttcagctat
gtcctctaaatctaaatctttcaacagtttttgaatagcttctgcacctattcctacttc
tacttggttattctcatcttgattttggtagatttcctcttcaaggctcttccattcgta
accttcgagaagctgcttgtattttaaattaatatctgtattagattgcgtaactacata
agaatgaaaataaacaatttttttctacttctttgacttttaagtctaatgctaaagcaat
atagcttgtacttccttttaagtaccaaacatgagttactggtgaggaaagttcaatgta
agccatccgatgtcttctcacgcgtgattctgtaacttctacgccacatcgttcacaaac
tatgcctttgtagcgaaaacgtttatatttaccacagtgacattcccaatccttaactgg
gccaaaaattttctcacagaataaaccatccatttctggctttaaagttctgtaattaat
tgtttcaggctttgtgatttcaccaacaatttgtccgttaggcaaacttctttcgcccca
ctgtcgaatttttttcgggagaagctaaactaatttttacgtagtcaaaatattgctcaaa
ctttgtcataaatttgaataccttaaaatcaaataattagccaatttatttacatgcttc
atattcaaaaatgaatatcttaataaaggaattgttcaaaatcgtctactggaggagtat
cataattagaacgatctgctcggttatctttagaatccgacatcaagtcaacttcaactg
ttcgcctttggccatcttcaaatagtttcaatttatgtactgcaatgtctaaacctaatg
attgcagctctcgcattaagactttaaaagattccggtgttccaggcttcggtataggct
tccctttaactattgcgtttaatgcttcatttctagcttgcatatcatccgactttacag
ttaataattcttgcaaagtatatgctgcaccaaaagcttctaaagcccacacttccattt
cacctaatctttgacctccatgctgtgctctacctcccaatggctgctgcgtaaccagag
agtagggaccagtggatcttgcgtggattttatcatcaacaagatgaacaagcttaagca
tataagctctaccaactgtaacaggattatcgaaaggctctcctgttctaccatcaaaaa
cttgcattttacctggatgctgatcattaaaaagccatttattgctagtaatcaatgatg
cttcttttaatttctatttacaagtgctcttgatgcttctgctccatacatctcatcaa
aaggtattattttaaatctttttcctaaatagccacctgctaaacccagtaagcactcaa
atacttgaccaacattcattctagaagggacacctaaaggatttaaaacaatgtctactg
gtgttccatctgataaataaggcatatcttgtttaggtaagatcctagaaatgatacctt
tattgccgtgacgaccagccattttatcacccacttggatttttctcttctgcgcaacat
agacgcgaatcatagcatttgttccgggaggcagctcatctcctttctgcctagtaaaga
ccctaaccttaactactctaccttttgcagcattaggaagccgtaaagatgtatctctca
catcccgagccttttctccgaaaatagctcgcaacaatttaccttctggtaattgatcag
cttcacctttaggagtaatttttcctactaaaatatctccagcttcaacccaagaaccac
cgacaacaattccatttctatctaaatccttcagagagttatcgctgacgttaggaatct
ctctagtgatttcttctggtcctaatttagtttgccggcattctacttcatacttttcga
tatgaatagaagtatacaagtcatcataaactaatctttcgctaattaaaaacgcatctt
cgtagttataaccttcccaaggcatataagcaacaagaatatttctccctaaagcgattt

ctcctccgtctgtagatgcgccatcagctaaagtttggccaacaactatttttctccta
cccaaacaattggacgctgattaatacaagtatcttggttagaccgataatatttttta
agcgataatgaactgttctaccattagtatcctgaatgcctattttattagcagaaacat
aatttacataaccagatgttctgctaataacaaccataccagagtctcgagctattttag
tttcaagaccagtaccaattattggcttttctggatataataatggaacagcttgccttt
gcatattagatcccattagagctctattcgcatcgtcatgttccaaaaaaggtattaatg
atgttgcagctgaaattacttgaataggtgaaattgcaatataatctacttgggttggcg
ttgttgtaataaactcttgacgataacgtacaggaataatatctccttcaatgtaatatt
gtttactaactttaacatctcctggagctactcgaaaatcatcttcctcatctgctgtta
gatagacaggactattgtgataaatcacttgaccttggtgtaccggataaaatggggttt
ctatgaaaccaaaaacattgacccgcgcacaagttgctagtgaaccaattaaacctgcgt
tcggaccttccggagtttcaattgggcaaattctaccatagtgactaggatgtaaatcac
gaactgcaaaacctgccctgtctttattaaaacctcctggacctaaggcactaattcttc
tcttgtgagttagttctgcaactggattagtttggtccataaattgagaaagctgactag
atccaaagaattctctcactgacgcaattaatggcttgggattaattagatttgataaac
tcaatgaatctatatcacatatcatcattctttctctaataatgcgttctaaacgattga
gacctactctgaattggttctgcagcaattctccaactgaacgaactcttctattaccaa
gatgatctatatcatcaagattacctgagtttttatccttaatattgattagataatcaa
ttgaggataagatatcttgaggagacaatactcggaacgttttaggtatattgagaccta
gtttttattaatttatatctaccgacttcacctaagtcatatcttttaggatcaaaaa
aacgagagtaaagcatctgcttagcaacagcaactgttgctggttcattaggtctaagct
tagaataaactatgagaagggcttcctcatcagttacttcttcaatattatttttccaa
tttctttagctaattctttaactgaatatatttgggatgcagaaataagaaaagcatatt
tactaagtcctttttcaatttcatctttattgagcccaatagctcgtagaaaaatataag
catttactttgtgggttttatctattcttatccaaatttctcccttaggatcaatttcga
attttaaccaagaacctctattagaaattaggcttgcactataaatctgcttaccatttt
tgtctatctcttgtttataatagattcctggactacggattatttggttaataataactc
tttctgtaccagatacaataaaagttcctctatttgtcattattggcaagtctccaataa
aaaccaatctctttttgtacttttttatttttatctgcttttcagcactgagctgtagat
gggttgatgataaatctaatgatttaatattttttcctggtctttagaaggtagatcaa
tatcttttcttgttaactttacaggaacatatatttgagcactataagttttatccctat
tttttgcctgtctgacactataacgaggaaattttatcttatattcattaccaaataatt
gtaattcgagtcgactagtagggtctgatatcttcggaaaaaaattcaagaacttccgtca
aaccttctaataaaaaccatttaaaactggccctctgaatttcaactaaatctggaagaa
gtttattttttaagcttatacgttggaccatagatctcctttataacaaattttgaaatc
gaagagtttaa-tactattaatagagtactggtagataatgtataagcatattttgctac

attatttcttctctagacataactagaacataatattttattcaa--aaaaagattttaa
aaatt----ttcac--acactatatatataagcactttgctgcta--aaagaaagttcaa
gctatataa-------aaaaacgaaattaatacttgtctgtctgtcaagtaattgacatc
aagatagacaagtattatttatttaatcaattctg-tttttttgagcaaaagacaatgct
ctcgcaagtctagctttttttctagcgcctgtattagggtgaaatgcccctttttttaca
gctttgtcaatcttgctatatacttgagaaatacttgactgcacatcattgaaattatca
ctttctaaattatcaatatttagaaggcatcttttagttaaagttttgacaaccgattta
tacttacggttaatgagacgatttctttccgaagttttaatgcgtttaatcgcagaaagg
ttcttagccacagtaaatccaataaaacttttataaaatatttttagaaacgaaaataca
caaatctgagatctattcaaaaagggaatatgcggcgtattataacattaaagaaaaaca
ataggcaatagttcaccaagagctcttgttattacacaatagagattgtcattgcgaaaa
actttgtaagatagtatttaattagaataaactaactagacatataaatatatatgaaaa
aaattgaagctattattagaccttttaagcttaatgaagtaaaatttgcgctagtcaagg
aaggtattggaggaatgactgttatcaaagtttctggatttgggagacagaagggtcaaa
ctgaaagatataaaggatctgaatactctattgatatcattgataaaataaaaattgaaa
ttattattaacgatgataaagtagagaaaattgtagagacgattattaaggcttctaaaa
caggggaaattggagatgggaaaatatttattagtagtattgaaagagtaataagaatca
ggactaatgacttagattttgaagctttatagtatttttacttt-tttagacttgatact
tcttcactaaacatgagataataagcgcttattgtatacttataactaaggtgtaata--
ctaatggctaaaagtaaaggtgcacggatcgtaataactttagagtgctctgataaagct
ggagattttgctcaaaaaaggaaacctggcgtttttcgatatacaactactaaaaataga
cgaaatacaccaaatagaattgaattaaacaagttttgtcctaattgcaatcaacattgt
atcttcaaagaaattaaatagttatttatcttattaaaat--aatcatattatggctgta
tatagaaaaagaatatctccaattaaaccaacagaggctgttgactacaaagatattgac
ttgctaagaaaatttattacagaacagggcaaaatactgcctaaacggtcaaccggattg
acttcgaagcagcaaaaaaaacttactaaagcgattaaacaagctagaatcctttcttta
ttacctttttttaaataaagattaa--atatatggcctaacttcacatttttttcatctaaa
tattagaaacttctatagatttgtttagcaaaatgatctaacatactgaat---------
----------tagtactaaaa-tctaatcattataagttttgttcattttataattaata
atagcactataattta---ttaagctagattc-----agacataaagtgcaacctctttt
aattgattatattatatagtctatttaaaagaggaaatttttttgcttc----------t
agcaattcaagattctttcagctttaacgatatatttaaaacaaagatttttgtttaggt
atcagatcaaatgagtgaattttatctcccgactgccataattgaaattcagagatgaaa
attccacattcgttaccagcttgaatttcttcgacatcttctctaacgcgttttaaagat
tcaatttttccttgataaataactttttcttcccggatcactttaatccaagagtttttt
aatagcttattattgatcactcgacatcctgctatttttctatttgctaacgagaatact

gtgcttacttctgcttctcccaccggtacttctgagtattcaggatcaagtaagtcttcc
atccttcttttaatgtcttcaattaaggcataaatgatttgatagttttctattaatata
ttcaactttgccgatgcttgctttgttcctggagcaaagttggtattgaacccaatgaga
atagaatttgtggttgaagcaagttcaacatctgtagcagtaatctctccaggcatgatt
gacacgacattgagttgaactttactctgaggaaattgagataaagaatctaggattgct
tctgtagagccttgattatctgttttaataactaaagaaatttgcttactaatatcttta
gaattagtgtttttaaagtatctaatgtaattcgactattaagtgctctttgtttctgt
ataattgatgagtctttagaggtattttcaatcgctttaagtttggcttctttatcactt
ttaactgctaaagctatttctcctgtagctggtacagatgacaatccccaaatttcaacg
acggatgagggtattgctaaattaatttctctttagcgttgctaattatagctcgaatc
tttgcgtaagctgatccaataaccaaattatcactaatattcaatgtaccattttgtatt
aatagagttgctactggaccatgcgatttatctaagtgggcttctattataataccttgt
gcaggctgtgttgggtcagctttcaaattctctaattcagctaacaaggtaatagtttct
aataatttatcaacatttgaccagtaagagaactaataggaattataggcacttgtcca
ccgagtttctcagacatgacattgtgttttaataaatcttgctcaataatatctatattc
gaaccagctttatctatcttagatattgctacgacaaaagggacattcgcttttttgaata
tgatggatagcctctatggtttgaggtttaactccgtcatcagcagctataataataata
gcgacatctgttaaattagctcctctggacctcatacttgtgaaagcttcatggcctgga
gtatccaagaaaacaatttttttgtttattagctttcttaatatactcaacttcataagct
gcaatggcttgagtgattcctccaatttctttattagcattattagattttcgaatataa
tctaatagcgttgttttgccatgatctacatgtcccatcactgtgacaattggaggtctt
ttaatataattgccagtttcataaagaacattagagttatctaaattactaggaagtcca
ttattattttcctctacatttgattctactgctatcccaaaattatcagctactgatgat
attattgaggcgtctatggtttgattcatagtaactgatattcttttaagaaataaatat
tttataatatctgtttcttgaacacaaattagtttagataattcttgaatagttagagga
ttagtaatactaatagactctggcggagaatttgattgactgcttgttgaattttgacca
ggagttaatgctactttttttctgtcttatagatttttttattattagttagttttttttgcg
acttccacttttggcttaggtggacgcattaaagagatagctaaatctcctgcagtctgc
gcaacattagaattagagtctctaaagttgtca---tcatcatcgtcaatatgtattttc
gtttttagtttt-ttgcgctgtctattctta--tttttttttactatctactaagtcgtga
attttattaaaatttttactttttttatctagtttgggtggagaacttaattctagatgc
tgttcaccacc---tgtatgtgattctgatttatcaaggtctaaatgtaacagatcatca
ttattaactgactctaatctaattttatagataatctgagggtttttcaaatctataatt
gtttcagaagagttgatatttgagctagatcgagaggatctattttcaaagttttgatta
tttaaaaacataaatgttggttttattatctaatagtaactttctaattttaatacagag
cttaatataaacaaaaatcgaatatctgtataaataatacaaatgt---agagaattaat

ctctgttgtattagatttaatataacactaaagtcgtttgttcgttatacaagtttaaaa
cttttt-tatgatgataaacactagattctagtttattggtatacaatctatttatctaaa
taatactgtttttgtc-gggcgtttaaa----ttatctaatgttatattcattatgaaat
tctattgaaatttaaaaa-ctcaacaattaagtaagaatcgttaatcacctccacgccac
ttatgccacgatctcaaaaaaatgataattttattgacaaaacttttacagtattagctg
atattgtattaaaaatacttcctacaagtaaagaggaaaaggaagcttttttcttactatc
gagatggaatgtctgcacaatctgaaggtgagtatgcagaagctttagaaaattattatg
aggctttaaaactagaagaagatccatatgatagaagctatattttatacaatatagggc
ttatttatgctagtaatggagagtatgtaaaagctttagaatactaccatcaaggattag
aactaaattttaaattacctcaggcccttaataatatagccgtgatatatcattaccaag
gagttcaagcaattgaggataaagatacagagctatctaaattaatgtttgacaaggcag
ctcaatattggcaacaggctattaagttagctcctgataattatattgaagctcaaaatt
ggctgaaaacaacaggacgaatgaggaatatacaaggatattaatatatttgagatataa
taaagatatcaattaataaattaaaattagttaaagaaactgatactatatagagtactt
-tgatat--------atgttatactagttga------tctctaagacttagctcaataac
cgaaattaaatctattgactatattaatgaatacaaataagcataatggttagtacaaca
aaaagtaccggatctgttactcaaattatcggaccagttttagatattgcatttcctaac
ggacagcttccgaaagtattcaatgcactcaaagtacaaagctcagaagcaactattact
tgtgaagtacaacaactgttgggtgacaacaaggtacgagctgtttctatgagttccact
gaaggactacaaagaggggtcgaagttattgatactggatcccctatatctgttcctgta
ggtacagatactcttggacgtatttttaatgttttaggtgaacctgtagataatttgggt
cccgttaattctgagagtactttacctatccatcgaccagcacctgcttttactaagtta
gagacaaaaccaaatatttttgaaacaggtattaaagttgtcgatttacttgctccctat
agaagaggtgggaaaattggtttatttggaggtgctggagtaggtaaaactgtattaatt
atggaactaattaataacatcgctaaagctcatggtggagtatctgtatttggaggcgtt
ggtgaaagaacaagagaaggaaacgacctatatatggaaatgaaggagtctaaagtaatt
aatgcagataacctgaaagaatctaaagtagcattagtatatggtcaaatgaatgaacct
cctggagcgcgtatgcgcgttggcttaactgcattgacaatggcagagtactttagagat
attaacaaacaagacgttctattgtttattgataatatttttcggtttgtacaagctgga
tcagaagtgtcagctctactaggccgtatgccatctgctgtgggttaccagccaactcta
gcaactgaaatgggagcactgcaagaaagaattacttcaacaacagaaggatcaattaca
tctattcaagctgtatatgtaccggctgatgatttaacagacccagctccagccactaca
tttgcacatttagatgcaacaacagtactttcgaggaacttggcagcaaaaggaatttat
cctgcagttgatccactggattcaacatcaacaatgctacagcctggaattgttggaact
gagcattattctactgctcaagaggtaaaatcaactttacaaagatataaagaactgcaa
gatattattgctattcttggtcttgatgaactttcagaggaagatagacagactgtctca

agagcgagaaaaattgaaagatttttatctcaaccttttttcgtggcggaagtgtttact
ggatcacctgggaaatatgtatctttggaagatgcaattaaagggtttcaaatgatctta
aaaggcaatttagacgacttgcctgagcaggcattttatctagtaggtgatatagatgaa
gctatacaaaaagctgacagcatgaaagattaatataattaacgatgactttaaatataa
gaattattgctcctgatcggactgtttgggatgcagaagcacaagaaattattttaccaa
gtagtacaggacaacttggtattttaacaggccatgcacctttgcttacagctttagata
ttggagttatgagagtaagagtagataaagaatggatgccgattgttttgctgggcgggt
ttgcagaaatagagaacaatcaattaactattctagtcaatggggcagaagaagctagtc
aaattgatttatcagaagcagaaaaaaatttagacactgcaactcaacttttaagtgatg
cctcatctaataaagaaaaaatagaagcaacacaaaaaatacgaaaagctagagctcgag
tacaagctgcaacagctgcaactccataa-aagattcc------tcttaaaacataaaac
tttagtaattattactaaa-------gttttatgttttatttgtatggttataggcaact
taaataaacttagcttagacctgaacaaatataatcaaaatataatcccatctctttacc
tgcatctggtccaactaaactaatagtaacttctttcatagctaaaatagcttgaatcgt
tgcaccaataggaacacctagggaattgtatgtttcttttagaccatttaatactctttc
ttcaagaatagatggatctccagctagcattccataagtagcatagcgtaagtagtagtc
taaatcacgaatgcatgcagcataacgtcgagttgtatacatgttaccacctgggcgtgt
aatatccgaataaagtagagattttgctacggattctttaataattgttgctgcattagc
tgcaatagtagcagcagctcttactcgtagctcacctgtttgaaaatagcctcttaattt
ttcaacagagctatcatctaaatatttaccttggacatcagctgcattaataacagaagt
aattgcgtcttgcataactttcacaaattccttaattttaaatatctaatacta-ggggg
tatataacaaaattactagatcctttagcgatatcagtattactgcatagcacccaaagt
ataatcaaagtaaaaaccagcttctgctgaatcttcacctgcaagcagtgagcaagctac
acttttcatgcattttacgccttcagcaactcctgaaattggtgttcctaaagaattata
catttctttaacacccactaatccaatttcttcaataggagttacatcgccagcaactat
tccgtaagttactaagcgaagataatagtcaagatctcgtagacaagtagcagtcatctc
ttcgccgtaagcatttccaccaggagaaactacatcaggtctttttttgaaacagttgttg
gccgccttgctttacaatacgctcacgattatctgttaaaatttgagctattcttaaacg
acgttgtccagataaaacaaaacttttaattctatctaattccccaggacttaaatatct
tgcttctgcatctgcatttacaattgactttgtaacaatactcatggataaaactcctgt
aatgcttt-aaaaaaaagcttaatactaattatttatgatataactcagaaagtaatgta
ctttttgag-aattttataaccgaaacacgttcatgatatacgaggcaggataggtactg
aggtaagtactaaaacgagaaagttatatgtaggaacttctgttattgatttcctagtac
tttttaaaacttggtaccacaattgatatatcctgcttagtcaaacgattataaagtgt
ttctgtattaggaaagttagctgctggtaaagttggaaatcttctgtatggaacagtatc
tttaccatataccgcattgtattcatcactttcaactaaattacttataaatgcagataa

gccttttgaagctaaaatctgattcaagaatctaatttctgcctgattattaggagctcg
tcctaagatatgcttagtacctaattctataacttttgtatttggatatggctgataaaa
ttctttaccatatagttcagatagcgccaatttttctacgagttctttgacacagatttg
tctattcaagaaagctgactcaatatccaagaattcaccgccgatactaaaagagttaag
atctctttcgaaaatttgacgatatgctgctcttaaagcttgttctaacatttctttgtt
actatcactattcacttcaaagacaatagattgatctcttaaagaagttactccttgtga
aatacgagacttaatatcattactactcctcatttctttgacagcacctaattcaacaaa
cctagcagatttactagcaataactttttgaatctctggtcaataattcctggtcttaa
gcttcttaaagcaacaccagctggcgtactatatctttcatacggaacagtattatctcc
aaaagtttctgtgtactcagagctatcaagtattgcgtctattacttggtaatagccttg
cttatatgcaatatcaaaatatttattaatttcttgccgaccatatgtagggcgacccag
taagcggttatgaatatattcaatagctttacaaatatataatggctcccaataaagaga
tctaaatatacttgattttgctaactgtctaacaaattcgcgaactgtaatttgattgtc
ttttagctgactttctattggcttgagaattagtttttcttcttgatacacttctctacc
aaaaactcttaaataagcaacttttgtgacaacttctactgaattttcaaaattttggga
tgaatctgactttataagagcactggataatttaaagattttggtcccagggatcctgc
tgacttaggtcttacctgtggattactaatctgattgtagattcctggcccgcgtctaac
tagtatccttcttgtatctttaccaaaaagggcttgtctttttcttggatctttattttc
tttagggaaaatagctccaaattgaattgataatggatcattacctatgccatatggatg
ttgatcaggtaatgattgcttataatcactaaataaagttataaattgaggaaccttacg
aaaaggcgcactataactgagaagatctatttgaggcccccaattacggcactcttgggg
ctcttcacctaaatttctaaagtaaggcacagtctcttcgccgaaataatctgtatactc
agaagaatttaaaattgcatttactaaaccacttaaaccagtagatgataaaatagcaaa
atatttttgaaactcttctagagagctaggacctctacccaagaaatgtctaaatgctaa
ttctaatgctcgactattgacaaaaggttcataaaattgctttctataaatacttgaagt
tcctaacgaacgaataaattccttaattgaaatttggccatttttttacttgtgactccaa
attcgataaagagaggtcatatgctttcgcaatatctctttcaaaaatttgtctataaca
tgctttcacaacagtatttttctcatcagcagataaactaggcttcataacaaagcgagg
tgttgacactcctgcctgtacataagtctgaggaagacgtaaaccttgtaaatctccaga
tattctttttcttaatttatcagtcaaagatggagcttcaaattcagaaatcacaacatt
aaaatactctttaactaaatcttgaccttttatatcttcctcaaagattagtaaggctgt
ccgtctcatctctctaagagccacaattgctgctgcacttgagcaagcattatcaattag
ctctctcaatcctcgtatattaacagataaaatattagggtcgcctgatacaattgcata
tgtcaaataccgtaaaaaccagtctaaatctcgaagagattttctcattcttgtggtgcc
gtatctcaagacgtttataggtttaaatcctggcggagtagctcccccagcattaaataa
agatcggaaactttggccaaaatctccttggatatttcctgataactcattaattttatc

ttgagaactttgatcacctgcgataataacagctgcttgaggacgttctaaataagaaat
tgcagacccacctacgaatattttatctgctgctcgtgctacgagaatatttgcattttt
agttaaaatatctgcaacttccagtctttttgtccagaatttaaaaaagcaactaattg
attcaattcaccaagctgtaaaaatcggtcttgttgctctgcctgagtaatagttaaaat
tgaggcagtccgataaagctgggggcgtgctaatgggcttccgccacttgctttgatact
cattactttatttatctccttaactgatattgcttgctagcgaatacaaatatgacttga
tatttctaaatatttaaagatactatgcaagcaca-ctgttttctatattattaggtact
agtacttatagtacaaagagtcttacaactaacttctataaatacaaattttgtaatact
tctattaacatttatcaaaaatatatttaagtgctcactcttttatattaaaattcaatt
agtgagagattgacgtgaaataaagtatactaagtaagttttatagataggaaaattaaa
agatttattctacaaacattaatca------------cttgaatgtttagcaattataca
attgttattgcttttaattggtttagactagtttt-tatttttctaacataaacaatat
taatgagcaatttgtatgaatttaaagccacaaaacaacctattaactaacgaaaatata
aatcacatagatatacccgaaaatgatataccaatgtcaattacggaacatttagaagaa
ttaagacaacgaactttatttgtgtttttattctttttatttgctacaactataagtttc
acacaaattaagataattgttgcaatactgcaagctcctgcttttggtattaaattttta
caattggcaccaggagaatattttttttcatctattaaagttgcaatatattgcgggatt
gttgcaacaactccgtttgccgtttatcaagttatattatacatactcccaggactaact
cgaaaagaaagaaaaattattttacctctattaattagttctgtattgctttttatcaca
ggtgggatatttgcttactttgttctcgcaccagcagctttaacattttttaattagctat
gggtctgatattgtagaaccactatggtctttcgaacaatactttgactttattctatta
cttttacttagtacaggattagcatttgagataccagtcatacaattattgctcggtgtt
tctgggacattttcttctaatcaaatgatacgagcttggagatatattattattatagca
acaattgctggagctattttgaccccttcaactgatcccgttacacaattaataatgtct
tcagctgttttattactttattttggtggaattgttatattattagtcttaaaaaagtag
agtaaat---------attttcta-ttaatccaatatatttgttatc-----tgggcaac
taatctataaaaacaatttct------------------agatttaagtatcaggcttct
aaatatttcaa---------gtctctttttatcatttaaataaaatc-------------
------------------------------------------------------------
------------------------------------------------------------
------------------------------------------------------------
------------------------------------------------------------
------------------------------------------------------------
------------------------------------------------------------
--------------------------------------------------gatacattgg
cttgtttttccatcaattatttgtattgtgcac-tttttagcactatctttattattctt

agaattatttccttaatctatttattaaat----agatatgaagagatttcagtattgat
tatggtaatctatatcaacgtaaaaagtattttcttatgagttttatttatataaaggct
tattcattatatagttataacagtaagtctaattaatag-ttttcatgcttaattaaagt
aataactcttaattcttttggcagaaaaatctgtcttaatattttatc--aaaaaaatag
-aaaaatgaagctaaatagtctaattaacttaattcaaaagtctatatattcttgtacac
ttttattaattattttaaatattatttgtgttgcacctaattctagtaatgcatttccaa
tttatgcgcagcaagcttatgaaagtccaagagaagcaactggtaggatagtatgtgcca
attgtcatcttgctcaaaagcctgtggagatagaagcgcctcaagcggtactacctaata
ctgttttcgagactgttgtgaagattccatatgacagcaatgctaaacagattttaggta
atggcagtaaaggaggcttaaatgttggagcggttgtaatattacctgaaggatttaagt
tagctcctgctaatagattatctacagagttaaaagaaaagactaaaaatctttacattc
agccgtacagtgctaaacaagataatattttagtaattggacctatttctggtgataaaa
atagagaaatagttttcccaatactatctcccgatcctgcaaaagataaaaaagctcatt
ttttcaagtatccaatatatgttggcggaaatagaggacgaggccagatttatccaacgg
gtgataaaagcaacaataatattgtctctgctttaagcagtggtaaaattaataaaattg
aatcactagacaagggtggatttataatacatgtgactaacagtagcaatgtagagtcaa
cacaaaagatttcacctggtctcgaacttaaagtgaaagaaggggatacaattcagcttg
atcaagctttgaatagtgatccaaatgtaggtggttttggccagaacgaaacagaaatag
ttttacagagtccaaatagaattaagggcatgattgttttctttttcgctagtgttctag
ctcaaatttcttcgtattaaagaaaaaacaatttgaaaaagttcaagcagctgaaatga
atttttaaaggaatataaaactggcacttactataaataaagta-taaacactttattca
caggatgtaccagtctttatgttatgtaatactcgctttgacggaggttatgatactttt
aacagaatctacaatttggctaggctgaattactgtagcctgctctaagcttccattata
tggtgtgggtagatcctgcgaagatagcctcacgacaggagcatctaattcatcaaaaag
atattcattaatttgtgcaattagctctgctccaattccggctgttttcatacactcttc
tacaattaaaactttgtgagttttctttactgagatagatatagagtctatatccagagg
ttttaaagatataagatctataacttctggatcataaccttcttttaataaggccggtaa
tgcttggataacatgatgcctcattctggaataggttaaaatcgtaatatcttttccttt
tcgtacaaattcagctttattaagaggtaagaaatattcttcttgaggaatctcttcttg
taaattgtaaagtagaacgtgctcaaagaatacaactggattattgtctcgaattgcaga
tttcagtaatccttttgcgttataaggagtagaacaagcaactatttttaagccggggat
ggcttggaaataagcttccagtctctgagaatgctctgcacctaactgtctacccacacc
tcccggtcctctaataactaaaggcaacgtaaaattacctccagaagtataacgtaacat
tcctgcattattagaaatttgattgaatgctaataataaaaaactcatattcataccttc
aacaattggtcttagccctgttatagctgcaccaattgccataccagtaaagctattttc
tgctattggtgtatcaaggactcttaaatccccatatttactatgcaaatctttagttac

cttataagatccaccatagtgaccaacatcttcgcctattacgcagacagttggatcttt
ttccatttcttcatctgttgccgctcgtaaagcgtcaaacataaagatttactcatttt
aatatttgatttttatgatttgaccactgttttaaagtaacagttaagactccatattct
atttagttatctgcaaaaagatatcgtttcaattctgaaatgttaggttctggactggaa
atagcaaatttcacggcttgttccaattctgttttaacagcagcctgaatcccattaagt
tcacctatatcagcaatttcattatctagaatatatttttgagtttttgataggatct
cttgccacccaagcctctttttcttgtcttgatcttagttcatcaggatctgccagagag
tgaccacgaaatctatatgttaatgcttctattaaggttggaccatcaccttggcgagct
ctttgaactgcttgttttgcagcttgtcttacagctagcacatccattccatcaacttca
atcccaggaagcccaaaagcttccgctttttatgtatttcaggtattgaagaagaccgg
tgatgtgccatacctatagcccactgattattttcaacaacaaatataataggtagtttc
cagagaactgccatattcaaacattcaaaaaattgcccattgttggtagtcccgtcacca
aaaaagcaagccgtgactcttaaatcgtctgtttctttaagtacttgctgacggtagata
ctttgaaaagctgcccctgttgcaactggtataccttctgcaataaaagcaaagccaccc
aaaaaattgtgaggcgcagaaaaaatatgcatcgaacctcctctgcctttactacaacca
gtctcttttccaaacaactcagccattacattttagatgggacgcctttacttaaagcg
tgtacatggtctcgataggtactgcaaacataatcagtcggattcagaagtttaattaca
cctgtagaaacagcttcttgaccattataaagatgaacaaaaccaaacatctttccttta
taatacatctgagcacacatatcttcaaaatttctgcctaacaacatatcttcatataaa
actaataaattactcttattaagattaagacctgttgaattataattagtcagcggcaat
tgaactttcttaggataactcataatttgtaaagaaacctct----------tattgtaa
ataggaattttgaagaaaaagctatcaatagaaaacatacagataaaaacttataaatta
ttgtatgtttttactaaagactttaataattatttgacactagttcagtatatcaaatt
aattttgcttatactacagatgtctgaaacttagacaaaaaagtagtaa-aaaaaattta
ttaaaagaatagttaaaaacagatataaatagaagtaaacagaattatt-gaaaaaaaat
gattattattaattaa----tacttaatatttctaaatattttacaattaatggctatgt
caccatctttattttatcctgttgaacaagaactatgtagtcttgaaaaaaatctgaaag
ctgttgccgggactcgtcatccaattttatatgccgcagcaaagcatctattcgatgctg
gaggaaaacgagttagaccagctcttgtgtttttagtggctaaagcaacctctgagaagc
aagatataaatactggacaaaaaaggctagcagaaattactgaaattatacatactgcta
gtttggtacatgatgatattatcgacgagtgcacaacacgtagaggagtcaaaactgtac
ataatttattcaatactaagattgctgtgctagcaggagatttttttatttgcacagtctt
cttggtatttagctaatattgaaaatttagcagtagtcaaagctatttctaaagtcatta
ccgactttgccgaaggagaaattaggcaaggcttagttcatttcgatcctaatatttcaa
tagatgcttacattgagaaatcattttacaagactgcttcactcattgctgctagttgtc
gcggtgcagctatgcttaatggttgcaatcatcaaataaataatgatctttatctttacg

gtaaacatatgggattagcatttcaaattatggacgatgttttagatataactggttcta
ctcaaagcttagggaaacctgctggcgctgatctaataaatgggaatttaacctctccta
tcctttttcacttactcaagaaacaagtttaaatgatcttattgatagagagttctgta
atagtacagacgtagcctcagcattatttcttataaaaagaagcggaggaattacaaaag
ctaaagatttagctaaagaacaggtgcaggcggcactttcctgccttcagtttttaccac
aatctacacctgtatctagtttaaaagaattaacacatttcataatcacaagattgtcgt
agaggacttgctaaaattaaaatgttattaactaatttttttactgtttctaaaatagtct
gaaaagtttgaatatcattactagctagctgcgctagcattttacgatttaacgcaatat
tctcttttttaatgcactaataaaagtactataattcatgccttgattgtgagcagcag
catttattctagttatccaaagacgacgaaaatctctcttttccttttgcgaccaacat
aagaatatcgaagagctttaaggacttgctgcttagctgttcgaaataaacatttatgtg
cacctttaaagcctttagctagcttaaagatttagctcgtcttttttttgcaacgttac
ctcttttaactctactcataaaatattac-ttaaattttaattattaccagcttactaaa
ttcgtatactctttg----attataaataaggtaattttattgcaatatttttaatatcc
tttaagtcaaccaaacaagtagaagagagatgtcttctttgttttgatgactttttttgt
aataaatgactttagaagctttatgtcgaagaaattttccggatgaagagactttaaat
ctttttgctattgcttttgatgtttttaacttaggcatattataaacttttaatgattaa
caaaagcatgaaaagtctgaggatattaatcttagacctttcaatacttagaataatctt
aacacaatttattttacttactacataggttaactgaaacaatttatatatataatacta
atcagttaatcaaccgcagctaaagtggatttgtattttgctttgaacttttgtatagta
ttagcaattagcattgttataattttaagactacttgaatttaattcttcgaacattttc
atatttaatgtaaatgaaatattagcttctgagacaatattttgaatttgaacatcagac
aaaggaatcatatctaatgcagctctatattgatctttaaataatttatcgtcttttatt
tggtcaaaatcgtaaaattttgttcctcctgaatctgataaattcatagcacctctagct
atttttttttaaaatttggccgccagaaagatcacccaaatagcgcgtataagcatgagcg
actaataattcaggttgtttatgtcctataatgtgaattctatcaacatagattttttgta
gcaggagatggttcaataaaatctgaccaatctgatccgtaataataatttaaatcttca
gatagactctctttcctattgagctctgtaaaatatataggtttaatagctggatgatct
ttattagaaaataactcttcctctattgcacaatagacaaagtatagatttgcaaccaat
ttacgatatgacttcttatctacaactcctcccaaaaaaagatttaacaaaactaacattt
tctgccatactatgggacttagtagtaccttctcttagttcattcgctaaagtattaatc
ataatatcaatattccattttt-aaagacaacaaattaaca---ataaatagttatccgt
atatttatacttgagttgtgctaactaggctgctgcaaaataattttttgattttattgg
atctggattcattgttttgtcaccaggcttccaattagctggacaaacttcatctggatg
agattgtacatattgaattgcttgtaacacccttaaagtttcttctacactcctgccaaa
ctctaaattattgattgtagagtactgaataattcctttcgggtctataataaatagtcc

tcttagagctacaccatcactattcaaaacattataagccgcactgatctcttttttttaa
gtctgatactaatggatatgaaagatctcctaatccacccgattctctgtcagtctgtaa
ccaggcaagatgagaatactcgctatcaaccgagactccaagaacttctgtattaagttc
ggaaaaagcattatacttatcgctaaatgctgtaatttctgtagggcatacaaatgtaaa
atctaaaggataaaagaataagacaatatacttgttcttgaaatcagataattttaacgt
tttaaattcttggtcataaacagctgtagctgaaaaatcaggagctagctgacctactcg
aagacaatttggtccagaaatcattaatttttcccagaagaattgatataattttgttat
ttaataaataattatataacaaaatttagtatatattaaaatactaccagtcttttttaa
attac--------taaatgatataatcaatactatctttatagcggggaatggatttgaa
ccattgaccttcgggttatgagcccgacgagctaccagactgctctaccccgcggataac
tagaattatattagtctagatatatatataaaagcaaatgcataatagattttagaagta
aaaaaatagaagaacaggtccactgatcattaaacttagaaacgataaagtatttttttat
cttatttaaaagaggaaagacttcatataaaccaataaatctatcttttaaaagaatttc
ggacggctttacccaaatttgaccatcataccagcctgattcttcataaaatactgtagc
actcattaatcgttttactacgtatgaccagcctaaatataatctgattaaaataaatcc
tgtcattaggctggaagtaataaattctgaaaagaaaaatttcagaggtagtttattgat
tggaaaaattgataataatataggactaattaataaacaattaattagtagtgcaatagt
tatttttttattataagatctgcgacttaaagttggccaacaaaagaaccaagaattttt
caaagaagtatattcatgaacaggttgctgctcctttgggacaggacattgagtattata
taaattcattgacattaaaaagtattatactaaagac--tttctttatagtatctaatat
aggtacgacattaaatattaactaactgcaattatagtagtgagtaaaagaaataaaact
aaaaatagccaagtaattttatttaaagtactctcagtactacgagtattactaaagaac
tgattctgagcaccgacacttcctaagccttcagatttaggattatgtattaaaatagtg
aagattaatattattgttgatgaataccaaaaaaattttaaaatttgttccattatgtaa
gagttaat-aaaaatagcacaaagacaatcgtctaaaaggtattgattgtcttatcatat
taaaaagtatttattctccctgtacttttaaaagtaaaaatcctattgccagccctacca
aaactaggactgaacttaatacggcactttctacaatttctccgaccatt-ttatttttc
cttattactttattattg---tcattcttttgacaagattgtattgagctaa-ctaaaat
ccgtttctaccccaaactaccattgcaattgaaaatgtaaacattgccattagagctgcc
catcccaaacttaaaatatccataatttataatactc--ctaaaaataagatctcttta
cttaaactataatattcttaatattaaaaatatcattatataataagctaattataacag
agtactacttaattcgtatttttggtgtatttaaagaagattgaatttttttcgaggtac
tggataagtatttcatcctaagaaccatctttaac----ttttgatttattatactgctt
aaatctaaatata--tagcactcaagaatcattagtaattatgatgtttttaattatacg
attaacaacatatgtcaaaatagaagaattcatagcgactcagttaa-----tatgctaa
tataatttatcaat--aaaaaggaagttagtatttcaattatgcaaacaactataaataa

tgggcaaccgtctagtaaagaaactttactaacacctagattctacacgactgactttga
agagatggctaatatggatatttccggcaatcaagaagatttttttggctatcctcgaaga
atttcgagctgactataatagtgaacactttattagagatgaagagtttaatcaatcttg
gtctaatttagaacataaaactaaatccttatttattgagtttttagaaagatcttgtac
cgcagaattttcaggtttttttactatataaagaattatccagaaaattaaaagacagaaa
tccggttatagctgagtgcttttttattgatgtctagagatgaagctaggcatgctggttt
tttaaataaagctatcggagacttcaatttatctttagatttgggattttttaacgaagag
tcgcaagtatactttttttctctcctaaatttatttttttatgcaacctatctttctgaaaa
aattggatactggagatacataactatttaccgccatcttgaacaatatccagaacaccg
tatttatccaatctttagatttttttgaaaattggtgtcaggatgaaaatcgtcatggtga
ttttttttgctgctttgcttaaatctcaacctcattttttaaatgactggaaagcaaaaat
gtggtgcaggtttttttctattaagtgtatttgcaactatgtacttgaacgactttcaaag
aattgatttttataatgccataggcctagactctagacagtatgatatgcaagtaatacg
caaaaccaacgaaagtgccgctagagttttccctgttgctttagacgtggacaatccaaa
atttttcaaatatttagatacttgtgcatgtgataatagagctctgatagatattgacaa
taacaattctccattattcattaaatctattgtaaagatacctctatatttttccttatt
tacaaatttactgaagatatatttgattaagccaatagactcaaaaacagtatggaatac
agttcgatagttaaatacaagggaatactttttttattataaaaagctatctaattattaa
tacaagat------aaaagcttccttct-----cgatggaaggagcttttattttgtcaa
aaa-tatataatgggattatatagatc-tagaactaagaacttacttttttcttttgcttc
atacataacctctaaagtaattatattttttagaacaatcacgggcaaatttctctgtatt
tctttttactttacctctaacaaacccaggaatttttatttaattctttctgtgcttcttc
tgaccaattaatactatcaacagcagaaattcctacacttaatgcttctgttgtgtcatg
accaccaaatatttccagcaaatgatcttccataccaagtgtaaatgaattatatactaa
gtcagcaatttgatttgttccctcataaccaagaaatggacggtaacttaaaggaaagtt
ctggatatgcactggtgaagaaataacgccacaagggatatttaaacgcttaccaatatg
tctttccatctgtgttccaaaaaattgctgcaggttcagttttagcaattaaatctccaat
taaaccatgatcatctgaaacaattacttcatcgcaaaattcttgaacttgatctttaaa
ccactcttcatcatatttgcaataagttccacaccaagcgacatttatacccatttcctg
gtgtaaaattcgagttattgctgcagcatgagttgcatctccaaatacaatagcctttttt
acctgttaaattctgacaatcaattgatctggaaaaccatgctgattgtgaaataaatcg
ggtttgttcatctatgtatttttttcgtaatctacagcagctcccgatgcatttaataattg
ctgtattgatcggatgcatgcagctgtttgaacaatacccataggtgtaatgtcaacata
aggcatattaaactctttctctaaatatcgagctgtcattagtccagtttctctatatgg
aataaaattaaaccatgcagatggtaatttttttttaaatcttgaactgatgcattttctgg
aataatctggtttatttggatatctagatcttgaaataaccttttttaattctgctatatc

atgctgattgtgaaatcctaagcttactgctccaataatattaacagatggagttttcgt
cttttgagttaaaacttgattattagatttagctttttccatatagaaagcgacaatttg
ttctaacgttctatcacctgcttgcagttcatttactctataatgattaacatcagctaa
taaaacatctgcttctgtttctatagatgctctgctaacaaagttttgcaaatcctcttg
caaaatactcgaagtgcaagttggtgttaaaataactaaatcagggctttcttctctgtc
ttttctagtgatattttctacaactttttcttgcgaaccacgggctaacacatgtctatc
cacaacacttgctgtaacaggagtaaaatctctgtctctttcaagcattgaacgcattac
attaaaataatcgtctccgagaggcgcatgcataattgcatgaacttttttaaaagaact
ggcaattcttaaagttccaatatgagcagggcctgcatacatccaataagctaatttcat
attaaattttcctatgatttatttagacatagtaatactttttaaagctaactaacaaaca
ttagaaaaacatctaatttaactccttgataattttgtaatattttttagatgaagctta
tcattataaatatttaaaaaatataattcagaatagaaattaataactgttagatcaagc
ttgaaagtaaagatttcttgcagttatagtttttctttaagtaaacatgtaaaaaattgta
ctatatataaatgtactatatataaactttttttactgaatattggtaatgccagataca
attaacttaaatatgccttccccaacgtttggtggaagcactggcggttggttaagagcc
gcagaagtagaagaaaagtatgcgataacatggacaggtaaaaatgaaagtaagtttgaa
atgccaaccggtggtaccgcaactatgcgaaatggagaaaacttactttatttagctaag
aaagaacaatgtttagctttaggtactcaactaaaaagtaaatttaaaatatctgactat
aagatttacagagtttttcctaacggagaagtgcaatatttgcatcctaaagatggagtc
ttcccagaaaaagttaatgctggaagagctagcgtcaacagtgttgatcattccattgga
caaaatattaatcctgtagatgtcaaattcatgaataaagcaacttacgattagtctctc
-aaaaacctagacataaaataacttttatatagtctaggtttttgataaggtatactgat
agtatggaagggtggccgagtggttgaaggcgtctgatttgaaatcagttgaacta--tc
tggttccgtgggttcgaatcccacccttttccgttataaaact------------atagtt
tgcttctcaaggcaagctagctatacactgatagtattaagtttagacttaataaagtct
acaacttgtgacaggttagaaatttttttcggcatcctcatctggtatctctatactaaat
ttttcttctatggccattactagctcaactgtgtctaaagaatcggctcccaggtcactt
gaaaaatttgcttccctggtcactattttttttttcaattcctagttgttctgctacaata
tcttgaacttttttcaaagatttcattatcttgcataatatattatttcctctaaggtgtt
atcaaaaatttttttggcttatattattacccaactctaacaaattagtaatttttagtcg
agattaaaatatttatacctatatgatatactattattctggataaatacgaagtttacg
atatggctcttctccgaaactacgagctctcaactggtagttgtcaactagattatgctg
catttttcggatataggctgaccgaggtgtaagctgaactattgaactttcattcaagat
aatagtttcaacagccaatttagcttcttgaagagcctggatctcatcaaactttttatt
tttacagagttctacccaattaagaccagatgaagtatgaatgtttaatatttttctgag
tgcacgagtaatttgtggaaccgtactgttctgaattgtatatataataatttgttttga

ttttgcaatttgtctaagttttgtattttgctttacttgattacgtagcgctagaatagc
atcagacttttctatctccttggttaaaataattggtaaatctagtgaagaaataactga
tgtaatatgctgccaactcaaagaataagcatataagtactgatgaggtatttctaattg
caaagattgtttattaatattaataggggcattgaaagtagttgacaataaagatgtatc
acgattttgtgatttattcaaatctaaggacttatttcttaattctctatattgtaaaac
aggagacgcctgctgtgtgttaattgatttacgactattagagatgtttacaggtaagac
ttctacagattgcgatggataacatttaattaaaattctaccattagcttgaatttgtct
tttttgcacaaaaggttgatggccttgcaaaatttgatctattgtctctttaactttatt
atgaactatccaaacatttcgctcatgtatttcgatcgcaatctgaaaagctggcgatgc
ttttctttctagtatactcttctgcgtacctctacgtttagcttcatcgtcacccaatgt
tacatattgaattccgccgattaggtccgctaatgttggatttttaattaaactttctaa
atgatttccatgcgctgtacctactaattggactcctctttctgctatagttcgagctgc
taacgcttctagctccgttccaatttcatctataataataacctcgggcatgtgattctc
aacagcttctatcattacttgatgttgcaaatcgggccttgcgacttgcattcttctagc
tctaccaatagccgggtgaggaatatctccgtcaccagctatttcattcgaagtatctat
aataactactctttttttccatttcatcagctaaaacacgagccatttctcgaactgcggt
tgttttaccaactcctggttttcctaagagtaaaattgaatcacctcgctctagtaaatc
tcgaataatactaattgtaccaaaaactgctctgccaacacgacaggttaagccgataac
actaccctctctattacgcagtgaactaatacgatgcaatgtttttttcgataccagctcg
attatcgccgctaaaattaccaatttttttttacacagtaatctaagtcttgccaactaat
agatctttgagataaatattctggattatctggaaatctggcttctggacgtcgacctaa
atccataacaacttctattaaattatttctattgggatgctgttttaaaggttctttgac
aaaatttggcaaaatttctagcaacttatctaagtcatctgcaataagcatgattgagta
tataataattaattaatcaaatagaagattcaagattctaaagattacaatacatattaa
agaatattataattggaatcaattttatatttaaattgaacaggaattta-tagatgtaa
aagaactgattaaactttacgcaataaatcgattggtcaaaattaatagtcgtattaaca
tgtttattggacaaaaagttcgtattaaatatagcaaacaaaaagttgctagtgatatag
cagataaagttggtgaattaggagttattaagggaataaagtttattaatagtcaatgtg
ttactattattgttgagtttgacaatcatactaggctttggatgtttagagaggaattaa
tctgtctaaatgaat------taa-------aaacataatttattttatttattatacag
aatttcttatgagtcgttaacataataaattttaaaggagtacttttaattgtgcaaatt
actattaaaaaattacaagacttactttcgtctgtacaaagaaaaaagatccagacatta
aaactaaaacaagataagtttgaacttctattaaataagacctataaaaaggtcaatcaa
gaaataataccttcgcagaaatctgctgtattacaaaatagtccatctacgacaattaaa
tcaataaatagtacaaaaaaatcttctgttgtcaatgaagaccgtacagaatatgccact
attgtttctccaatggttggaacgttttatcattcacctgctcctggtgaaaaaatttttt

gtacgagttggcgatgaggtcaaattcaatcaaacagtttgtattattgaggcaatgaag
ttaatgaacgaaatcgaagcggaaattgaaggaaaaattatagaaattcttgttaaagat
ggtgatatagtagattgcgggcaagctttaatgaaggttgaaatataatgctaattttcc
cattttacatattgttaacagaaattatttagatagaaaaattagcttatactatagtta
tgaaaacttatttcgcaaaggcacaaaaaaactaatatataaaattattt-ttttatctg
ccaaaaatagaagtaagcgttttgattccttgtctcaagagaggagaatctcaatggcaa
ttagctcaaaagagcaagagacaaagaaggtaaaaatctcggttgataaaaatcccgtag
atacttctttcgaaaaatgggcccaaccaggccatttttctcgtacactagcaaaaggac
caaaaactactacttggatttggaatcttcatgctgatgctcacgacttcgatagtcaaa
ccagttctttagaagaagtttcacgtaagattttcagtgcacattttgggcaactgtctg
taatatttttatggcttagtggaatgtattttcacggagcccgcttctctaactatgttg
cttggttaagtaatccaacaggtattaagccaagtgcacaggttgtttggcctatagttg
gacaagaaattttaaatggcgatgtaggtggtggctttcaaggagtacaagttacatctg
gatggttccaactgtggagagcatcaggaattactacagaatttcagctttactgtactg
ctattggcggattagctatggctgctttaatgctgtttgcaggatggtttcattatcata
aagcagctccaaagttagaatggtttcaaaatgttgagtcaatgatgaatcaccatttag
ctgggcttttaggcttaggctgtttaggctgggcaggtcatcaaatccatttgtctttgc
ctattaataagctactagattctggtgtgtctccgcaagaaatcccactacctcatgaat
ttttaattaatagagagcttatggctcagctgtatccaagttttagtaaaggattagttc
cattctttactttaaattgggctgaatattccgacttttttaacttttaaaggaggtttaa
atcctgttactggaggtttatggctaagtgatactgctcatcatcatctagcattagctg
ttctatttcttgctgcaggtcatatgtatagaaccaattggggtattggacatagcatga
aagaaattctagaagctcataaaggaccttttaccggcaacggtcacgaaggtctatatg
aaattcttacaacttcttggcatgcacagcttgcaattaatttagctatgatgggatctt
taagcataattgtagcacatcacatgtacgctatgcctccttatccatatattgctactg
attacccgactcaattatcgcttttcactcatcatatgtggattggaggattttgtattg
ttggagcaggagcgcatgcttctatatttatggtaagggattataatcctgcagaaaatt
ataacaatcttttagatagagttattaggcatcgagatgctattgtttctcatctaaatt
gggtatgtatatttcttggattccattcatttggtttatacattcacaatgatactatgc
gggcacttggaagatctcaagatatgttctctgatacagctatacagttacaacctattt
ttgctcaatgggtacaaagtatacacactttagctcctggaaatacagctccaaatgcgt
tagcaacagctagttatgcatttggaggagatgttgtttctgttggtaacaaagttgcaa
tgatgcctatttctttaggtactgcagatttttttagttcaccatatacatgcatttacta
ttcatgtaactgttttaattttagttaaaggattccttttctcgagaaactctagactaa
ttcctgataaggccaatcttggcttcagatttccatgtgatggacctggtagaggtggta
cttgccaagtatctggctgggatcatgtttttcttggcttattctggatgtacaattctc

tgtctgtagcaatttttcactttagttggaaaatgcaatccgatgtttggggtagtgtat
ctccgtccggaaatgtttctcatattactggtggtaattttgcacaaagtgcaattacaa
tcaatggatggttaagagatttcctttgggctcaagcatctcaggttattcaatcatacg
gttctgcgctatctgcatatggattaattttcttagcagcacattttgtatgggcattca
gtttgatgttcttatttagtggtagaggttattggcaagagcttatagaatcaatcgtat
gggcgcataataaaataaaagttgctcctgcaattcaaccaagagctttaagtattactc
aaggtagagcagtcggtgttgcacactacttattaggtggaattggtacaacttgggcat
tctttttagcaagaattatttcagtaggctaatagtgaaaataggat-aaaaaaacaatt
atggcaacaaaatttcctaagtttagccaagctttatcacaagatcctacaactagaagg
atttggtatggtattgctacggcacatgactttgaaagtcatgatggaatgacagaagaa
aatttatatcagaagatatttgcttcgcactttggacatctagcaattatcttcttatgg
acatctggtaatttattccatgtagcttggcaaggcaactttgaacagtgggtattaaat
cctttgaaagttaaaccaattgctcatgcaatttgggatccacattttggacaaccagct
ttaaaagcttttagtaaaggggctcagcttatccagtaaatatagcatattctggcgta
tatcactggtggtacactattggtatgagaagcaatcaagacctatattctggggcttta
ttcttactagttttatcagctttactcttatttggagggtggctacatctacaaccaaaa
ttcaagcctggtttatcctggtttaaaaataacgaatcaagattaaatcatcatttatct
ggattatttggggttagttctttagcctggacaggtcatttagtacatgttgctatacct
gaggcaagaggacaacatgtaggatgggataattttacaacagtattacctcatccagcc
ggtttacaaccatttttcagtggtaattggagtgtatatgctcaaaatccagatacagct
caacatttattcggaactaatgaaggtgcgggtacagcaattttaacatttctgggggga
ttccatcctcaaagtcagtctttatggctaactgacatggctcatcaccatttggctatt
gcagtagtattcattgttgctggacatatgtataggactaattggggggattgggcacaat
ctaaaagatattttagatgctcatagaccacctagtggtagattaggagctggacataaa
gggttatttgatactattactaattctttacacatgcagttaggattagcattagcttcc
ctaggtgtaattacttcgttggtagctcaacatatgtatgctatgcctccatatgctttc
atggctaaagatttcacaactcaagcgtctttgtacacacatcatcaatatattgctggg
tttctaatggttggagcttttgctcatggggcaatattctttgttcgagactatgaccct
gaacagaataaagataatgttttagctcgcatgctagaacataaagaagctatcatttct
catctaagttgggtaactctatttttagggtttcatacattaggcctttatgttcacaac
gacacaatgattgcttttggaactcctgaaaaacaaattttaattgagccggtatttgct
caatggattcaagcttcttcaggaaaagcactttacggttttgatgtgttactatcatcc
tctactaatgtcgcaacacaagctggtagcaatatttggctgccaggctggttagaagca
attaatagcggaaaaaattcattgtttttaacaattggtcctggtgacttcttagttcat
catgcaattgcattgggattacatactactacgttaattttagttaaaggtgctttagat
gcgagaggctctaaacttatgccggacaaaaaagactttggttatagttttccttgcgat

ggacctggcagaggtggcacctgtgatatatctgcatgggatgcgttctatttagctgta
ttttggatgctgaatacgataggttgggtaacattttattggcattggaaacatattaca
atatggcaaggcaatgcaactcaattcaatgagtcttcaacttatctaatgggatggttt
agagattacttatggctaaattcttctccattaattaatggttataatccatatggcatg
aataatttatcagtatggtcatggatgttcttatttgggcatttagtatgggcaacagga
tttatgttcttgatctcctggcgtggctattggcaagagttaattgaaactctagcctgg
gcgcatgaacgtacacctttagcaaacttgattcgttggaaagataaacctgttgcgcta
tcaattgtacaggcaagattagtaggtttagcacatt-tttctgtaggatatgtattaac
ttacgcagcttttgtattagcttcaacagcaggcaaatttggttagactaaa-----ttc
aaattagctaa-----aaaaaagtcagtcataata-gttttatgactgactttttggat
ttaaaacaaaaacttactaaagtttcacctcaatatctacaccggaaggcaaatttaact
tcattaatgcatctatcgtttgagaagaaggttgatgaatatcaataattcttctatgag
atcttatttcaaaatgttctcttgagtctttatctacatgtggagaacgtaagacacaat
aaattcttcttcttgtaggtaaaggaattggccctactgcaacagcattagttctagatg
cggtatctagtattttgtcacacgacgtattaagtatactagaattatatgctttcagtt
taattctgatttttgtctgctgagtaattgtcatgttaaaaagatcttattatttaagaa
ttttagagacaacacctgcacctacggtacggccaccttctctaatagcgaaacgcatac
cttgctcaattgcaattgcattaattaattcagcagtcatcttaattctgtcaccaggca
tgaccatttccgcatcagtaccatcatcagcagtaaactgattaatagtaccagttacat
cagttgttctaacataaaactgaggtctatatcctgggaaaaatggagtatgtcttccgc
cctcctctttagttagaatataaacttctgcttcaaattgagtatgaggtgtaattgtac
caggtttagctaataccatacctctttcaatatctttttttctgcacgcctcttaaaagaa
ttccaatattgtcacccgctagaccttcttctagcgtttttttgaaacatttctaatccag
taatagtcgttgtacgagtttctcgtaaacctacaatttcaattgtatcaccaactttaa
taatgcctctttcaattctaccagtggcaacagttccacgtcctgtgatagaaaaaacat
cttctacagccattaagaaagttttatcgacatctctctccggtgttggaatatacgtgt
caactgcttccataagtgaaaaaatcttgtcaacccatttatcttcaccttgcttaatag
ctgggtttttcgtcacagcttctaatgctaataaggcagaacctgcaacaaaaggaatat
catctccaggaaagtcgtattgacttaataattcccttccttctaattctactaattcta
gtagctcttcgtcatctacttgatcttccttatttaggaatactactaatgtgggaacac
ctacttgttttgctaataaaatatgttcacgagtttgtggcattggaccatctgccgcag
atacaactaaaatagctccatccatctgagcagcacctgtaatcatatttttttacgtaat
cggcatggcctggacaatctacgtgagcataatgacgattatctgtttcgtattcaacat
gagcagtattaatagtaatacctctagctttttcttctggagcggcatcaatttcatcaa
atttttttgctgcagtagaacctaaagttgataaagtcgcagagatcgcagctgttaaag
ttgtcttaccatgatcaacatgaccaattgttccgatattgacatgaggtttttttacgtt

caaatttagatcgagccatgctttttgttttccttatagtaaaagttgctttatagttac
ttttaacgataaatcttgtgaattgaaagatttattatttaatacagacaaga-----ta
aaatataactaatctaatatcgatagtgagcaaacgctttatttgcttctgccattctat
gtgtatcctctctctttctaatagaatttcctgtttcattagctgcatccataatttcgt
tagctaattttatagacatacttttaccagatctttctctagaaaatttagtgatccatc
ttaacgctaaattcgtacctctataagctcgtacttcaataggaacttggtaagtagaac
caccaactcttctagcttttacttctactagaggagtaatatttcggatagctttttcta
aaatatttaaaggatctgattctgttctctctttaacgatatctaaagcctgatatataa
ttctttgagctaaagtttttttaccacttttcaaaatacgaacagttaacatacttacga
gtctgcttttatataaaggatcaggtgatgcgaacctttttttagctgtattacgacgag
acatagtatttaaaattggttattgtaataa-tgttc-tattagaagttatttaagattt
aggctttttttgtcccatatttagatcggcttttacgacgatcctttactcctgcagcatc
taaagtaccacgtactacatgatatcggactccaggcaaatctttaattcggccacccct
aatcagtaccacagaatgctcttgaatattatgacctacgccaggaatataagctgtaac
ttcaaatccagatgttaatctaacgcgcgctacttttcttaaagcagagttcggtttttt
aggagtcgtagtatatactcttgtacaaacacctcttctttgagggcaactttgaagagc
tggagattttgtttttttatgtatttttcgtctttcggatctaacaagttgttgaatggt
tggcataataaaattgatttaaagtctttaatagatactatttgaacctgctttagatga
cttggctagtttttatttatcttatacttacgcataaatctttctactctaccttctgta
tcaataattctctgtgaaccagtaaagaatggatggtttcctgaccagatatctacgtgt
agttctggctttgtggacccaatcgtcataattaactggccgtcacaataaactttttgct
tctggataccaatttggatgtatattatcttttgccatggtttctataattatataaatg
tatcaataataactattaacgcttagaaaattgaggagcctttcttgctttctttaaacc
atatttttttctctcttttactcttggatctcttgttaaataaccctcagacttgagtgt
cgttctattttctggattaattgagcacaatgctcttgcaacacctaaacgaattgcatc
ggcttgccctgtcaaacctcctcctctagcattcacatgtatatcatactgatttaatag
tcctaaaacttgtaatggtgcatatgaaactcttaagtaattaggactaaattgaagata
agactctcctggtataccgtttataattaaattacctgatcctgggactagtcttacttg
tgcaacagaacacttccggcgacctgttccagaatagattgcacgagttttaattaattc
tgtcgacataacattccttgataactaaataaatatacattctttatactatatactctt
gtggcttttgcgcgacatgtgggtgaattgggccagaatatactttaagcttcgtaaata
gttttctgcctaatggacctttaggaagcatacctttaaccgatttttcaataattctgt
taggtaatcttgtctgaagctgataaaatgtttcaacttttaatccgccaggttgtccag
aatgtcttctgtatagtttttgattcgttttattgccacttacagatacgtgagcagagt
taataacaattacataatctccagtgtctagatagggtgtataagaaggcttgtttttgc
ctcttaagatattagaaatatgagtagatattctaccaagtgtctgatttttagcgtcta

taacataccaatgagaattagtatttaatgagggtgattgcgttttattcatgaaaagat
taatacttttggttgacaataagagaatatgtaaatggtttacattcaatttctatacaa
gtaatattt-aaaaatatta-------------ttaacaaagagaacttttctaacagtt
atttcaagtgtttgtcagcaagaaaatttctagttctcaataacttatactatttttaat
aatataaaatatattaatcactttttcttttggcaaacttatacctagttttttttgc-
aaggcatatataacttcttcagctgattttggccaaaattttaatttctagcagttct
tcttgagagtaatctagtaaatctgcaatagaatggatttgagctcgctttaagcagtta
taagccctaacagataactgcagttcttctattaaaacttgactaatttttttatcttct
ttactacgataattatctgctgatttaaagtctaaatttcttagtgaacaaaaaagatta
gttaaaactgtagctccttgacttattgcttcttgaggagagatgctcccatttgtccag
atttgtataatcaatctatcttttatgctgttgctaccaatacggacttcttctacttta
taattaactttattaactggcataaaaacagagtctacttgcaaaaaatccacagataat
tcatctactgcttttcagctaagcgatacccgcaattttttttcaatttaaactccatt
tcaaatattgtattattgcaaatagttgcaatatactgtctaggatctactacctctata
tcagaagataattcaaataggccggctgtaactatagctggcccttgaactctaattcga
ccaatttgagattctttgttataacttttaaatactacttcttttagattaagtaatatt
tctaacacatcttctcttaccccagggattgtagaaaactcatggttcactccagcaatg
cgtacagcaactatagcagtaccttcaagatctgacaatattgatcttcttaaagcatta
cctaatgtaataccttgtccttgatttaatggttctattacaaaactaccgtactgccca
cgcgccccatgtgttcttgactctacgcattcaatttgaaattgagccacctaagaaagc
tcc-tttaaaataatcagtgattagtaagcctctctaaaataagaaggaatttatactcg
gcgtttcttaggaggacgacatccattatgaggtacaggagtaatatcttttattagagt
aatctctaatcctgcagcctgcaaagctctaattgctgtttctcgacccgctcctggacc
atttactaaaacttcggtttggcgcataccttgatccatagcttgtttagctgcttttttc
agctgcggtttgagctgcaaaaggtgttcctttttttagctcccttaaatccacttgcacc
agaggatgaccatgataatgtctctcctttcaaattagtaatagtaacaattgtattatt
gaatgtagatttaatatgtgtaacaccgttaactgcattacgtttagtttttcttgctcc
ggatttttttatttgtctagccatcgtgtctcgtctattatatagttaattagaagatta
ttttcttggagctttttcttacctgctactgttttttacctcctctgcgtgttctagc
attagttctagttctttgtcctctcaaaggaagaccaagacggtgtcttctacctctata
agtactaatttccataagtctcttaatgctcatagactcaaaacgtttgagatccccttc
aatctgatagttagactcaagaatttctcttatactgacaacttgttgatcatttaaatc
ttgacacttaatatcaacatctatctttgtttttctaatatttctttcgaacgagatag
tccaataccataaatatatgttaaagctatctctattcttttgtttcttggaagatctac
tccagcaattctggccactttgtattgtctccaataaataatatgtat-ttttataacgg
tagtttata-agcttagccttgtctctgcttatgtttaggattagtgcaaattaccatca

cctttctgtgacgccgaattattctacatttttcacacatttttcgaacagaaggacgaa
ctttcatattaa----ttctatatactaggatattactatattaactctattaattatc
gacattaaaaacgtcaataatatattgtattactttaactattttagattttacttggtc
atactatcatatttttttgatatgacgtatgtctgaatttgcttagctgtgtctattgcg
actccaactaaaataagtaaagaagtagccccaagacctcttaagttttggatctgagta
accttttctattataaacggaatcagcgctactgtaaataaaaatgaggctcctaggaat
gtcagtctatttaatataacttgtaaataatcaatagtcgcttgacctggacgaatatta
ggaatgcttgcacccatttttttagatttgtagcaatatcttctggattcattactatt
gatgtataaaaatagctaaagaaaaggatcaaagcacaataaagaagaaggtatagtgaa
ccattaggacaaaacaagtataggatttggagtaatgttttattttgaataatttgggtt
agataagatgggagggccatagatgcagatgcaaaaacaataggcataactccaccttga
tttaatttcaagggcaagtaactatttggatccaaaattgaagattttcctagctgtctt
gctgaaataatttaattcttcttgttccttcttgtacacaaattgtaattattatcatt
aataaaaagatcgctataaataatccaaacttgagacttgcattactataattagcatca
aaaaatgactgtgtaaaattctttggtagtcctgacacaatgttttgaaaaataagcaaa
gaggctccattaccaattccttttctgtaattaactctgataaccacataataatcata
gagccagctgttaaagctaaaacagactcgcagacaaattcgacattccaattaaaaaca
tatggttttacccatatagagattgctccagattgtaaagtcgcccaacctagagctaag
taccttgtaatttgagttattttttggcggcccaattcaccctcttccttctgtaatttt
tccagattggggacaattttcgtaagtagctgcattacaattgaagagttaatataagga
acaatacctaacgcaaaaatccctattgttgaaaaacctcctccagaaaaaatatttaga
aaatttactaaagtatttttttctacacttgcataaaaggcgtcatgatctatacctgga
acaggtataaatattcctaaacgtgctaaaactaatagaaaaagggtaaagataatacga
tttctcaggtcactttttttggctcataaataaattttaaatgaaaagagatactttt--tt
tatgtaaaaaccctattccaagacttatataagtttaaaatagggaagtaa---aatatt
atag------taata-taatattctatgtagaatacaagctttctataggcactcctcga
tctttagctgcttcactaaaagttcttaattgagttaagccatttagaacagctcgcgcg
ttatttaatgtattattagaacctaattgcttagctaatatattctgtacacccgacaat
tctaggactgtccttacagaacctccggcaattacacctgaaccaggtgcagagggcctt
aatataactttcgcggcaccagaaattccattaataggatgaggtatagagttcgatttt
gtcagtggaactgtaactagatgtttttttagcatctgttactcctttttttactgcacca
attacatcgcttgcttttcccacgccgacaccgacttggccttgctcgttaccaacaaca
agaataactcgaaagcttagtttttttacctccttttacaactttagtaactcttttaact
tgtacgactctttcttcccagccactatctttatcttttcctttgctctgtttttttacga
ttggccattttaagaattattcctaattagttaatta-cgcttctagaaacccatgcctg
cttccttagcagcttcagccaaggccttaactcttccatgatataactttccacctctat

cgaagacaacatttttaatgccttctttcatagactgttctgctaactgtttaccaacta
cgcgagaagtatcacagtttggtcttatattatcggattctttactatttaaattaacag
atgatgtagctactaatgtaataccttgtgtatcatcaattatttgtgcgtatatatgtt
tgttagatctaaatacacacaggcgtggcctacttgaggttccttgaactttttttcgaa
ctcttttatgcttatgaattctagtttgtttagtgtttagtttcattattatttaccttt
tccagctttcccagcttttcttctaacaaattcaccttgatatctaattccttttccttt
ataaggctcaggaggcctaatagaacgaatagttgaagcaacctgaccgacaacttcttt
atctatgccagagacagtaatatttgtattattttcaactttaatttcaatattagcagg
aggtttaattttcactacatgactatagccaacatttaagattaaatcttggttatcaat
ttgagaacgatagcctacaccttgtatttgcagtttttttaaaaaatccattagaaacacc
ttcaatcatattactaataagagttctggatagcccatgtaactgactagccatttttgt
ttgtccactcgtcttgacagcaatcgtgtcattaagtatttctaaattaataccagctgg
taaagttctggatagggtacctttaggtcctgtcacagtgatagtctgaccatcaaactg
agtactaagatttgtcggcaataaaattatttttttttccaatacgagacat-attccacc
tataaaatagttaccaaatatagcataatatttcaccaccaagaccatcatgacgagctt
gtctatctgtcataacacctctagaagtagaaataagagcgatgcctaaacctccaagaa
ctctgggtagctctttgtgatttgcgtagactcttagtccgggtttactaattcttttca
gagcagtaataactggttgacgattttaccattatacttaagagaaatcattaaattag
tctctataccttcacctatttgttcgaaattttgaacaaatccttcttcttttagtactg
ttgccatattgcatgtcattttcgttgctggaacttgcacaatttgatgtcttgctaagt
ttgcgttacgaatacgtgtcagcatatcggcgatcgtatcgttgaccaccttgatcctcc
ttgcga-----tgaattatttaaaatattttcaagattccctgaaaggcatccctagctt
ttttaacaaagctagaccttcttgatctgttttagctgtagtgacaattgatatatctag
accacgaatttgatcaatattatcatagtctatttctggaaagatcaactgttcacgtaa
acctaaattgtaattacctttgccgtcaaaacttttaggactaattcctctgaagtctct
aattcttggtaatgttaaattaattagtttctctaaaaaagaatacatcttgtcttttct
tagatgcacaacaattccaataggaacctcttctcgaattttaaatcctgcaatagattt
tttagcttttgtgacaatcggtttttgtccagttattaatgttaattcttgaatactact
ttcaagagccttagcattctgagaagcttcacctaagccacggttaatagtaattttagt
aaacctaggaacttcatgtacatttttgtactgaaattcatcttttaaagattgagtaac
agttgttttatatttttcttttaatcctattgccatggtatttttatcaattatttaata
agttcgccagtctttttcagttttcgaa-ttttttggcctttatcattaattattactga
ggatcgactagcaatattattttgctcactgaataacataacattagaggtatgtattgg
agcttcaaatttgataatttctcccgtttctccttcttgttgaggtttttttatgcttcac
tttgagattaattcctttaacaatcaccttacttgttttatgcatgattgcaataatttc
acctgtttttgttttatcacttccagaaataacttgaactaagtcccctttttttaattt

aatttttgtattattttttgattgttttagaagaacttttcattatactacctccggtgca
agagacactatttttagaaaaattttttatctcttaactctcgggctataggtccgaagact
cttgtgccacgtggattattatcttgattaatgataactgccgcattatctccaaatcta
atgctcataccgtcagttctccttaaagctttacgagttcttactacaacagctctgaca
acatcagatcgtttaactggcatattgggagacgcatctttgaccactccaataataaca
tccccaatagatgcataagaaggattactagtgcctaatactctaatacacattattttt
ttagcaccactattatctgcaacattaagatagctttgagtctgtatcatacttttattc
tgtataaaattaattatcaaaagacttagataagatattaaccattgtccagcactttgt
gcgacttaaaggtcttgtctcttgtattgtaacaatatcaccgattgtgcattcattatt
ttcatcatgcgctttatatttctttgtccggatcatagtttttgcgtactttctatgaga
gattctattttctacagctactactatagttttattcattttatcgcttactactttacc
tgttgtttcttttaaaggcatagtctttatattttttatatat---------aattaaat
tttaagattattgcacggttgacttattacgagacttttcaacagttaaaagttgagcta
atctatgcttagaatgtttaaatagatgaggctgaaaatcttgccttgtggctcttttta
gccttaaatcaaaaagctctctttttattacaaggatttcttcagctaaagaagaagagt
ccatgtttgtaacatctgatattttagggaaagtcatgttttaggattctgttgtattcc
gaactataaatttggtcttgattggtaatttataagaagctaatttcatagcttgttgag
cagtttttgtggtacacctgtaatttcaaatagaatatgtccaggcttaataactgcaa
cccaatattctggagctcctttacctgaccccatgcgagtttcagctggacgggcagtga
ctggtttatctggaaatactctaatccataatttgccacctcttctaacatatctagtaa
tagttcttcgagtagcttctatttgtctagaagttaaccaaactggctctgtcgcttgta
acgcataatcaccgaacgcaattgtgttgcctttactagcagaacctttcattctacctc
tgtgttgttttctaaattttgttttcttggggcttagcataaaaattaaataataataat
aattaagaagctgttgaatcaggagtttcctgattataagcagattctagacttttcgat
tctggcaaaatttctcctttaaaaagccaaacttttactccgagcacaccatacgtagtg
tgagcttggcgatgacaataatcaatatcagctcttaaagtttgtaaaggtacacgacct
tctctaacccattcacttctagctatctctgcaccattaagccgaccagatacttgaatt
ttaacaccctgtgtatttgctctttgagctctttgaactgcttgtctaacagctctacga
aaggcaactctttttttctagctgttgagtaataaattctgctactaaggttgcttcagaa
tctggatctgcaatctctataacattgactctgagttgtttgctagggcctagaattaag
gacaacgattttcttaaagactcaattccagctccagattttccaagtacaattccaggt
cgagctgttgctatgagaatttctacttgatcaactttacgattgatttcaattttagca
atactagcattactaagttttgaatgtataaatgagcgaattttatgatcttcttgtaaa
agaactgggtagtctttagagttagcaaaccatgaagaacgatgtttttgggtaatgcct
atgcgaaaacccaaaggatgaatttttttgacccacagtatccttgattataaattaaaag
ctataatgactaattgacgttaag--------agactacagttctgagacacctaatgta

atatgacaagttggcttatgtattggaaaggctcttccttgggctctaggctgaaatctt
tttaaagtcgggcctttgtctgcaaaggctttactaacaaataattggttttatttaat
ccatcattatgctcagcattcgctgcagcagattccaggatttgctttatatgagaacag
actcggtatggcataaattccagaataattaatgcttcttggtattttctacctctaatt
tggtctaatacacggcggactttatgcggagaaagacgaatgtatttccctactgcttta
gtttcttttacgtttttgtaatactcatatttatagtttaacgacgggcttttctgtca
ccttttatgtgagtgcggaaagttcttgttgggacaaactctcctaatttatgtcctacc
atttggtctgacacaaagacaggaaaatgttgtttgccattataaacagctattgtatgc
cctaccatatcgggaataatagtagatgctctagaccaagtttttaggacttcttttttt
cctgaaatatttaatgcttctatacgtttaagaagactaacatcaataaaagggccttta
tgtatagatcttgacataataataaatcattagatttttgataaaataaattctatttac
ggcgacgtaagacgtacgggttactatatttatttggatttcgtgttttaacacccaatg
caggtttaccccaaggtgttacaggacgagctcttccaattggagatttgccttcaccac
caccatgtggatggtctacaggattcataacaacacctctaacagtaggtcttttaccta
accaacgatttctaccagctttaccaagagtaatattactagcatcaatattaccaactt
gtccaatagtagcatagcattcttttcgaatcattcgaacttcacttgaaggcaatttta
cagtaacaaaagcaccttcttttgctactatttgagcataagttcctgctgcacgaacaa
tctgtcctccgcacgaaggtcttagttctatattatgtactgctgttcctaaaggaatac
tagataaaggcaaagcatttccaacttcgataggggcagtagaaccagaaagaaccatag
atcctacacttagagatcgagggtgcagaatgtatcttttctccccatcaagatagtgta
ataatgcaattcttgcatttctattcggatcatattcaatagaagcaactttagcagcta
tattatgcctatttcttttaaagtcaattaatctgtactgctgcttatgtccaccacctt
tatgacgacaggtaatgacacctctattatttcgacctttgcagaaatggtgtttaacta
ttaatgatttttctggcttatcagtagtaatctctgaaaaggtagaaactgttctatttc
ttgttcctggtgtgtaggcacgatataaacgaattgccatataagaagatttgagaagag
taaatgttgaa------ctatgtaatatatttaagtttctggaaacaaattgatagaatc
ttttgaggcaagtgtcacaattgctttcttgtaatgtggtcgcttgcctacgaatctgcc
gatgctcctttttttcttaggaggatggcaagtgttaacacctgtaacttgcacgttgaa
aatatactgtatagcagccttaatattaatttttgttgctttgggatcaacagcaaaaca
atactgattttcttctaataatttagttgtcttatcagtaatgatgggatatttaactaa
atctaataaacctcttgaatcaatgctatccattatatacctcttgtattttagataaag
catcaactgtaataataattttatgtgcagctaacaaagccataatatttagtgtatcag
ctgaaataagttctacgttgtgtagattgcgaatagaaagataaacgttcggatcttttt
tgtctactatgactaagactttttttgtttaaatcaagattccatcgatgtatagcttcca
taaatagtttagtcctgggctgatgaaaataactgttaaaattttctacaatcaatgtat
taactgatttgttatttaatgctgtccttaatgctaattgtctttctttttttattcattt

ttttggtaaaactacgaggttttggcccaaatattacaccaccgcctctccatagaggtg
aacggatagaacctgctcttgctcgaccagtcccttttgacgccatggtttacgacctc
cacctcgaacttcacttcttgttttagtattcgcagaaccctgacgcttttcattacttt
gctttactaacgctctatggactaagtacatccctgaatcttgactaaccttaagattta
aatcagcgttgccacttacttgaccttcccaattatagacttgataatttaattgtgtgt
taactgtcatgaaatatatcgtatgaaagaatattgctacttactaattttttactaaagc
gccaggtttaccaggaacagctcccttaacgatcaacaaatcattctctgagttaatact
tacaatttgcaaatttttttattgtaactttttattaccaagttggccagccatgttttt
acctggataaactcttccaggcgttgtaccagctccaatagaaccaggttgtctatgatt
tttcgaaccatgtgacattgggcctctactaaaatgatgcctcttttggtaaccggagaa
acctttacctacacttctagaagagacattaatcttttggcctacttgaaatagatctgt
agatagtatttggctaacttcgaaatcatcagtagatttcacttcgtattcacgtaagta
ttttagtgggggagcctgtgatttttttaagtgtcccaacaatggtttatttaacttttg
ttcagcaacttgcttataaccgacttgaatagcattatagccatcagtagatatagcttt
aatttgagtaataatacatggcccaacttgaattacagtaactggaattgataaaccagc
ttcatcaaagaattgggtcatacctactttagtaccgagtataccaacagacactagatt
tctcctttttgagagatttatactatcattgtacataaaaattctacttcatattgactg
aag--taaattaaactgactgatgcatgatattattcatgacaacataaaggtttaaaac
tttacctatttaatctaaaatagaaacagctttttgtccagttacagcgtatataaatta
gaagcatgcttattaaatttttttatagactact-gaaatttcactgatattgatattcgt
aacttacttgcagtaat--------atcaacatgtataagtttaattgcctagactagaa
ccagctattattagaaatgattcaagtatagataaaagcgcatttgaacttttatctatt
gacaaatatagtagaatatattgaggttcggtaactacaccttttttattttcacatttat
tgtgca--gtatatatacatctaatgcagtaaacataggaatataac----tgtatagca
aatgtaattacaaaaattagagaggtattaatgggcaaagtcgttggaattgatcttgga
acaacgaattctgtaattgctgttatggaaggaggtaaacctaccgtcataccgaatgca
gaaggttttagaactacagcttctgttgttgcatatactaaaagtggagataaacttgtc
ggacaaattgccaggaggcaagctgttattaacccagaaaacactttctactctgtcaaa
agattcataggacgtaaacagaatgaaatttcgcaagagattcggcaaacatcatataat
gttaaaactagtggatcaagcataaaaaattgcttgccctgcactgaataaagattttgct
ccagaagaaatttcagctcaagtactgagaaaacttgttgaagatgctagtacctactta
ggtgagactgttacacaagccgtaataactgtaccggcttactttaacgattcacaaaga
caagctactaaagacgcaggtaaaatagcaggcttagatgtattgagaattattaatgag
cctactgcagcttctttgtcttatggactagacaaacaaaataatgaaacaatactagta
tttgaccttggcgggggcacatttgatgtatctgtattagaagttggagatggagtattt
gaagtactctcaacttctggagatacacatttaggtggagatgactttgatcagcagatt

gtagaatggctaatcaaagatttaaacaaaatgaaggtattgatcttggtaaagataga
caagcacttcaaagattaaccgaagctgcagaaaaagcaaagatagaactatcaaattta
actcaaacagagattaatcttccttttattactgctacacaagatggtccaaagcattta
gaaaagactgtaactagaggtaagtttgaagaactttgttcaaatttaatagacaaatgt
agtattcctgtaaacaatgctctgaaagatgcaaaactagaagcttccagtattgatgaa
gttgttttagttggtggatctacaaggattccagccatacagcaaatggttaaaagatta
attggtaaagatccaaaccagagcgtcaatccagatgaagttgttgctattggtgcagcc
gtacaagctggagttttagcaggcgaagtcaaagatattctattactagatgtgacgccg
ttatctttaggagtggaaactttaggtggcgtgatgacaaagattataccaagaaacact
actattcctacaaaaaaatcagaagtattttctacagctgtagataatcaaccaaatgtg
gaaattcaagtacttcaaggcgaaagagaactaacaaaagacaataagagcttagggacg
ttccgattagatggcattatgcctgcacctaggggggttcctcaaattgaagttacgttt
gacattgacgctaacggaattttatctgtaaaagccaaagaaaaggctactggtaaagag
caatcaattactatatctggagcatcaactctacctaaagatgatgttgaaagaatggta
aaagaagccgaagaaaatttcgatacagaccaaaaaagaagaaagaacattgacacaaag
aatcaagcagagtccctatgttaccaagctgaaaagcaagttaaagagtttgaagacaaa
attagccaagatttaaaaataaaaatagaggagctaattacagagcttagatctagtcta
gaaaaagaagaatatgacaatatcgaatccatttctcagcaattacaaaattctctgatg
gacattggaaaaaatgctgctcagactgaaagtaaagatgcaaaagcaaaggatgacaac
accgtgattgacactgatttctctgaagctaagt-aaaagta-agcgggtaacgcgattc
gaacgcgcgacatcaaccttggcaaggttgcgctctaccactgagctatacccgcatatg
ttgtattattacaaaaattaagatatttgtcaatctgtta------atctatac---atc
tagactattaatttccgaatcacatattagctaaatgttgtaaatataacaattgctata
actttttaa-------aagtacttatatcat-tatataagtacttttttatgtt-------
-caaaaatacaaataagtattatag-------ctaaactactaaggaattaactaagcta
gtaacaattactaatcctgtccataagccagctcccgtatatattaatccttttgattgc
tcccattgacctggggatgctaaaacaacaggtacacctactactaacacggtcgacaat
gcaattaataatagcactaataattggatcgcaataatcatttaatatgttctcctgaaa
aatttaaatgttaataatactgtatataatacaatgaagcttttgaccttcagcaattta
tctatgaagaaataatttaaaattattgttttataaataaaccttaaattaagtattaaa
atctttattgtaaataataatactatattttttta---tagataaaaaatttaaagaaag
tctttttttaaaaaaaaattaagataagatgatatgctgtataaaaagctaagtattatta
attattagagaaatttaaa-ataagaggtaaaacatatgaactcagccctttttttggca
aaactaccagaagcttatgccgtttttaaaccgatcattgatatcttacccgtaattcct
gtattttttcttcttttagcttttgtttggcaagcagcaattggttttagataaaattta
ggttataaaaagaataacaaatatacagagtgacatttttttgagctctaatcttatatct

tacacaataataactattatggttgaacccttactatcaggaattgttcttggacttatt
cctattactttattcggattattagtagcagcttatttacaatatcaacggggcaaccaa
ttaggactgtaaacgctaaatcatagatatttataaaaactacaataattatagtttttta
taaatatctacaaataagctaatttaaatacaaagataatattattgaaactagagtga
taacgaaaaagatttatataa---taa---------------------------------
------------------------------------------------------------
------------------------------------------------------------
------------------------------------------------------------
------------------------------------------------------------
------------------------------------------------------------
------------------------------------------------------------
------------------------------------------------------------
------------------------------------------------------------
------------------------------------------------------------
------------------------------------------------------------
------------------------------------------------------------
------------------------------------------------------------
------------------------------------------------------------
------------------------------------------------------------
------------------------------------------------------------
------------------------------------------------------------
------------------------------------------------------------
---------------------------caaactaattgtttaccagctcgacttagtaac
tcctggtaaaagacattcatgagacatttccctaaaaacatggcgagataatccaaaatc
tctataataccctctacttctcccggttaaccaacatctatttctccctctaacaggggc
gctgtttctaggcatttcttgaagtttttgtcttaaatccattttgtcagaaaagctgcc
agcttctttcagctgatctttaatagccaatctctttaagtaatattttttgcaagttt
cgatctcttaatctctctttggatcatattctttttagccatgctgtag-aactctcttt
taaaattatttattcagtatattagctaaaaaagctcctatctgcaacttttatctattt
aaaaataaatagcataaagtagtaaaatgcaattgatatttaatcaacgatacc---act
ttatgctattatcaaaaatttatttaataaagttcttgttcaacatgagttgaaatagtg
caatcagatgaagggtatgcaatacaggttaacacataaccagctaaaagttgatcatca
tctaagaatgattgatcagcttgatctacagttccttctgtaactttacctgcacaagtt
gaacaagctccagctctacaagagtaaggcagttcaattccttcttcttctgctgcatct
aaaatatatgtatcctctgcacaattaaatgtaacatcaattccttcttcttcacatagt

aaatgaattttataatcagccatttctttttactccataataaagatacacttgaaactt
aataaagcaaaat-aatttaaagaacttaccaaaaaattatatactagtttcattatgca
tgatagactggaaaaaaagaaaataatatactattctgaaaaatcaggctacatttccta
ctaaatagaactatagtaggtccaaagtatatacataaattcatatagttaaaattattt
ttatcatttaaatagctgaacttaatagta---------taggttgatcatc------aa
gctagtcttctgc---caataaataattaattatgatttcaatgcttaatcaacaagtag
aacttaagcctattcttaaattccagataacacaaatatattcacattctcttattcagg
aaattaaagcattaataaccagacttgttttgcaagtttggaggaggccggctacattga
tggcaggtattatgcaacctttgttatggttaattctatttggtgggcttttctacaatg
ctcctataaatttgtttactatcaatacaaactataattgtttttttgagctctgggatta
taattttcacatcttttactggagctttaaattcaggtcttccattaatgtttgatagag
aatttggatttttaaatagattattaacggctccgttagtatcgagggcttctatcattt
tatcttctgctacttttatgacttgtattagtttaatacaagttatatttatagttacag
cttctcttttttatgggaaactcacctctaaatagcgatagtgctatgatttttggactta
taatcctattagtaactgtaggagttacaatgcttagtttagctttatctttcactctgc
caggtcatattgagctattagcatttattttagtagttaacttacccttttattttcta
gtacggctttagctcctttatatttcatgccaccatggcttcagttaattgcaagtctca
atccattgagttatgccatagaaggtacaagatatttatattcaagcgtaaactggaatt
ttacagagtgtgtgattaagattagctggggagatatttgtttagggcagattattgttt
tattaatcgctttagatataatggcagcttatgttgtgtctaagatattaaaagctaaac
ttaatta-----------------aaattttaataaaaaatttattaaatttatttaaca
aagatttttactatataatactaatagtagtatggataattgaaatagaaatagtttttc
aaaaccaaagctattatcaacaatgataaacatttgtaagaaagtcaacaaagtatgttc
ttattcataggaggcatgtagtcaatgggactaccatggtaccgtgtacacacggttgtt
ttaaatgatcctggacgcttaattgcagtccacctaatgcatactgcacttgtagcaggt
tgggcagggtctatggcattatacgaactagctgtatttgatccttcagatccagtacta
aatccaatgtggcgacaaggcatgtttgttatgccatttatggctagactgggcgtaaca
gattcatggggaggatggagtataacaggagaaagcgtatccaatcctggactgtggagt
tttgaaggtgtcgcgttaactcatatagtcctttctggcatgcttttcctagctgctatt
tggcattgggtttattgggatttagaattatttagagatccacgaactggcgaaccagct
ttagatttaccgaaaatttttggaattcatttattgctatcaagtctactttgtttcgga
tttggagctttccatgtgactggacttttggaccaggaatgtgggtatcagatgggtat
ggagtaaccggaaaggtgctaccagtagctccagcatggggaccagaaggattcaatccg
tttaatcctggaggagttgcatctcaccacattgccgcaggtactgtaggtatattagct
ggtgttttccatttaactgttagaccaccacaaagactgtatcgtgctctaagaatgggt
aatattgaaactgtattatcaagtagtatctctgctgtttttttctcagcttttgtgact

tgtggtacgatgtggtacggctctgcaactacgcctattgaattatttggtccaactaga
tatcagtgggatagtggatattttcagcaagaaattgagaaacgagtagaaaatgctatt
gctgatggtgctgcacctagcgaagcatggtcaaggattcctgacaagttggcattctat
gactatattggtaataatccagcaaaaggaggattattcagagcaggccctatgaataaa
ggcgacggagttgcagaagcatggcttggacatcctgtattccaggataaagaaggaaga
gagcttagtgttcgcagaatgcctgctttttcgaaacttttcctgtaattttagttgat
aaagatggtattatacgagctgacattccgtttagaagagcagagtccaagtatagtatt
gaacaagtaggtgtaacagctagtttttatggtggcaaattaaatggccaagttttcaac
gatgctcctagcgttaaaaaatatgcaaggaaagctcaattaggtgaggtatttgaattt
gatcgaactacattagaatcagatggagtatttaggagtagtcctagaggctggtttaca
ttcggtcatgctaatttcgctttaattttcttctttggacatctttggcatggttcaaga
actatcttccgggatgtatttgcgggaatcggagctgaagttactgaacaagttgaattt
ggagctttccaaaaattaggagatagaagtagtaaaaaacaaggagctgtataagaccaa
agtcaaagacagtatttgtc---aaatatttccaatattaaaatagttaagacaaagttt
attatctaatagttactaatcaggagaatttatggaagccttagtctatgtatttttact
aacaggaacgctgatggttatattctttgcgatctttttagagaacctccaagaatagc
aaagtaacttgattccgtttaaacttataataatagaaccactttccaacttattagtgg
aaagtggttctattgtagattacattgtattattcttcatgctcttcaaaaggatctcga
aggtcttttgaagcaggaccaaatgctgtgtatatcgagtaacctgtaatacctaaaagc
agactcgaaataaaaatactaagaactgttgcagtttccataattcaagcagtattaaga
tgaactttttctataatgatattatcaatattaaaacatttaattattaaaattatcaaa
atatactatggcacttagaactagacttggagaaattttaagacccttaaattcagagta
tggaaaagttgctccaggttggggcacaactcctattatgggaattttcatgttactatt
tttcttgttttattaattattttacaaatctataattcttcattagtattagaaaatgt
agatgtagattgggctactttaggtag----ctaattagataagttctttga--------
ttaaacaatattaaagataac----------------aagcgatcgctattagattatag
tagcgatcgcttaaaaaatgttttactgtatatccagagaaaattatttacctaaattaa
ttaactcgctatctgcaaaactatttgtattaacattattatagttaactttttcaaacc
ttactaatactggatatttgataccacttttatcaattgctgcaacagtaccaatttcct
gataccagtatgattcctttctcaatatttttacttttgagcctcgttccatatttttag
tccttagatagattattaaagattattatattgaagaattttaaaaaactagacaaaata
tataaaagtactaccgacaattgtaatacctaactaataaaaggagagagagagggattcga
accctcgataacaaaagttatgacaattttcgaaattgttgcaataaaccgctctgccat
ctctcctaaattaaaatactcttttagaaaattagacaagaataacatagaactaaggta
gttgcctagaaaattaaaaatatgttaaatataaattataatctttctatagatatattt
acaaaatatgtacatagaatctatatcagcttatttcaagatttttaatcttatacaat

ttttactaaataaataaattctctcataatatgaaactatcttggaaaacacttcttctt
tggtctttaccaatatttgtcgttggtttctttttctggcaaggtttttaggtccaact
actacagatgttggcagtaatatcgcaagttctagaatgacatatggacgatttttagaa
tatttagatatgggttgggtgaaacgggttgacctctatgaaaataatcatacagcaatt
gtagaagctgttggtccagaattaggaaatagagttcaacgaattcgagttgaactgcca
gcaagtgcgccagaattaattacaaaattacgcaaagccaatgttgatctagatgctcat
ccccctaaaagtacaagtgcagtatggggactattaggcaatttactatttcctttacta
ttagttggcggattagctttcttatttagaagatctaataatgctagtggtgggcctggt
caagcaatgtcatttggcaaatcgaaagctttgtttcaaatggaggctaaaacaggagtg
gtatttaatgatgtagctggagttgaagaggcaaaggaggaatttcaagaagtagtaaca
tttttaaaacagcctgaatcatttactgctgttggtgcaaaaataccaaaaggcgtttta
ttagttggtcctcctggcacaggcaaaacattactagcaaaagctattgctggcgaggca
agtgttccttttttagtatctcaggctcagaatttgtagaaatgtttgttggtgttggc
gcttctcgtgtgagagacctattcaagaaagcaaaagacaatgcgccttgtatcgttttt
attgatgaaattgatgctgttggtagacaaagaggaacaggcgttggaggtggtaatgat
gaaagggaacaaacattaaatcagctattaactgaaatggatggctttgaaggaaatact
ggtgttattgtaattgccgctactaacagagctgatattttggattctgcattattaaga
cctggaagatttgatagacaagtttctgtagatgtaccagattttaaaggcaggttagca
attcttgaagttcatgctaaaaataagaaaatggaacctaaagtatctttagaaactatt
gccagaagaactcccggcttttcaggagctgatttagctaacttactaaatgaggctgct
atcttaacggctcgacgaagaaaaaatgcaatgactatgtctgaaattgatacatcaatt
gatcgagtagtagctgggatggaaggcactcctttaattgatagtaaaagcaagagatta
attgcatatcacgaagtgggtcacgcaataataggcagtttattagagcatcatgatcct
gtgcaaaaagttacattaataccaagagggcaagcaagaggcttaacttggtttactcct
agtgatgatcaaagtctaatatcaagatcccaaatactagcccgtatcgtaggtgctctt
ggtggtagagctgcagaagaaatcattttcggcgacgcagaagttaccacgggtgcaagt
aatgatttgcagcaagtaacatcaatggccagacaaatggtcactcgatttggcatgtct
aaaattggacctttatctcttgaaagtcaaggaggagacccattttttaggtagaggcatg
ggaggaggctcagaatattcagatgaagttgcaactaatattgataaacaagtaagggaa
attgtaagtgaatgctatgcgcaagctaaaaacattattatagataatcgagtagtgatg
gatagattagttgatttactaattgagaaagaaacaattgaaggcaatgaatttagagac
atcgttaaggaatatacagcaattcctgaaaaaaattactacatatcacaattttaa---
---------------atgagacgggactgacgggattcgaacccgcaacttccgccgtga
cagggcggtgctctaaccagttgaactacagtccc-aaaaagataaatacaaatagataa
tctcatagttagcttaaacttgtcaaattagaatattaagtatgaagagta---------
--------tttgtaatagtactcttcatatttaata-tagtagcgaaaatatttcttta

gaaggagagagagggattcgaaccctcggtacgaaattaatcgtacagcagattagcaat
ctgccgctttcgaccactcagccacctctccatga---tctatatataaaatatatcaaa
tatacttatattttaataggtgtgcataaatatttatggagctaagcggattcgaaccgc
tgaccctctcaatgccattgagatgctctaccaactgagctataaccc----ctttatga
ggatattaagaaatggctcaagcgggatttgaacctgcgaccttgggcttatgaatcccc
tgctctaaccactgagctactgagccatagtataaaatattattaattattataccataa
attatgctcaaatataaatcaataaaagaataattactatca-ttcactactttta----
------------------tacaagttgtaaatattcggacaagatttcaaaattaaaatg
tctttaactttttgtttga-------attatttgtttacgaaa-----------atattg
gtatttaaaattaac---------------------------------------------
--------------------aataaatttg------------------------------
---------------------------------------------------tctttattt
a-----------------------------------------------------------
------------------------------------------------------------
------------------------------------------------------------
------------------------------------------------------------
------------------------------------------------------------
-------tacaca-----------------------------------------------
-------------------------------caaaaatattggaaa--------------
------------------------ttttcatcaatt--------aattatactaattagt
------------------------------------------------------------
------------------tagactataaccgacgtac-----------------------
------------------------------------------------------------
------------------------------------------------------------
---aagaattatttaacta-----------------------------------------
------------------------------------------------------------
---cgttaaat-------------------------------------------------
------------------------------------------------------------
------------------------------------------------------------
------------------------------------------------------------
-------------------------------------tgtgatgcatt------------
-----gggcttgatcaaaacttctctaaaatcaacaaaatctagt---------------
-----------------------------------------------------aattcaa
gtaaatcaaatact----------acagacgcttaaaaaaataata--------------
caattagaatcgtaaacatttagtttttga-ttatgttaaataaatttgattatcataga
tttaatatttaaaagtgttctattacttgta-----ttgagtaactgt-aatatttaatg

tcttctttttagataataatcataatataagctaaaaaatattttt---tattaaacaa
ctagcatagagccaataccttgatcagttaatatttctagcaagagagcatgatctattc
taccatctagaatatgcgctgaagctacaccctgagctaaagaacgaatacaacaattca
ctttaggaatcatgccaccagaaattacttctgtttgagttaaatctctagcttcctgta
tattcagatgactaattaacgttgaaggatctgaagaattacgtaaaataccaggcgtat
ctgtgagtagaattaatttttcggcattaagagcggctgctatttctcctgcgacagtat
cagcattaatattatacgattggccttgcttgtctgctgcaacactagctataacaggta
tataattattattaattaatatcttcagtaatttagtatcaatattttgtacttctccaa
caaagccaaggtttgcttttccatttggtcttggagtaataagtaatccatctttccccg
acaaacctacacttttaccaccttgtttattaatagttgcgacaagatctttgttgactc
ttccaactaaaaccatttccactatatccatagtaggttgatctgttactctaacgccgt
catcgaattttggtaatattttaaggcgatctaaccaaaaattaatttctggtcctccac
catgaactaaaatgggacgtagtcctataaaggacagaaacacaagatcactaatcactt
gatctttttgtttttgatttttcatagccgatcccccatattttataacaataattctgg
aagaaaattgctgaatataaggcagagcttcacttaagactcttactctttctgaattag
tcaacataaataaagtaggattatgatatataaaattatttgcacaacttgtgttaattt
aaaaagtaccagtaggtacttc-aaaacttgaaaaaatagacatgttactaaaaa-----
----ttaaaactataaggattataattatgaataaattttgggataatgttttaagattt
ccacgattttagtcagcgtcattttaggattagttttgataattattagtccttttttc
gtgttattaaaaaaaccattgacaagtttttttttcatcatatcattgacgggattaatc
acagttttagcgataataa-tacaaaaaatgctaaatatcgaatgttgttgagagaaata
aataaatttaattaaaatttttgaacatatataatattatattatcatttaatatgataa
tct-attttcttttccat---aaaacttataagtattaaatattatttgataaatattat
gccactaaaacaaatagttagttctgaaaaaactggtgcctttgcacttctggatagtat
tgttaggcacggagttaaacatatatttggttatccaggtggagccattcttcctatcta
tgatgagctttacgcttgggaggaagcctatctaattaaacacattcttgttcgccatga
gcaaggcgctgctcatgctgcagattcttattctaggtcaacgggagaggttggagtatg
ctttgccacttctggtccaggggcaactaatcttgtctcgggtatagctacagcacacat
tgattctgtacctatattagctataacaggtcaagttggaagagcttttattggcacaga
tgcttttcaagaagtagatatttttgggattacacttcctattgtgaaacattcatatgt
agtccgtgatcctagagacctgtctagaattgttgcggaagcatttttttatttgtaaaca
tggtagaccaggtccagtattaattgatgttcctaaagatgtaggattagagaagtttaa
ttatttttctgttgaacccggaaaggttaatattcctggctgtaggccaattaccagcct
aaagtcaagacaaatccttatggcagctaaaatgatacagcaatccagccaaccattgtt
gtatattggtggaggagctataatctctgattcacatcaaattattaaagaacttgttga
tttctataaaatacctgttactactactttgatggggaaggggattttaatgaggacag

tgactattgtctaggaatgttaggtatgcatggtactgcgtatgctaattttgcagttag
tgagtgtgatcttttaattgctttaggagctagatttgatgatagagttactggaaaatt
agatgaatttgcttgtaatgcccaagtgattcacgtagatattgatcctgctgaagtagg
aaaaaataggattcctcaagtcgctattgtcggtgacgtagcagaagttgttagtgcaat
attaaatttattaaagacttcttttccccttatccagatcagattacatcttggcaaga
aagaattaatcgttggcgtcaacagtatcctttactggttcctagaaaatcaacaagcat
ttcacctcaagagattcttgttgcaactaataaattagcccaaaatgcttattttactac
agatgttggccagcatcaaatgtggtcagctcaatttctgaaagtaaaagctaagcattg
gctttcaagtgctggattaggcacgatgggttatggtttacctgcagcaattggtgctca
agtagcacatcctaatgacgtagttatttgtattagtggtgattctagttttcaaatgaa
tatgcaagagttaggaactatcgcgcaataccagctaccggttaaaattattattattaa
taatcgatggcaagggatggttagacaatggcaacaggctttctatggtgaaagatattc
acactcaagaatgacagaaggagcacctgattttcaaaagcttgcagaagcttttggtat
taaagcttttactattaataataggcaaaatatgcaatctgctttacaagctgctattga
tcatcctggtccagttttattagattgccaagttacagaaaatgaaaactgttatccaat
ggttgctcctggaaaaagtaatgcacaaatgataggtatagctaaaccgcagagaggaac
tgcttccaactatattaataacagcgttggaagtaa------------------------
------------------------------------------------------------
------------------------------------------------------------
------------------------------------------------------------
---------------atagttagtattttctt-aaaacagaata-aagtaaatgaagtat
ctaaatgtaattttcttatactatta-aaaactatgtaagaaaagcctttgacgagaatt
gaactcgtgaccttccccttaccaaggggatgctctacctctgagccacaaaggc-tttt
tactatattgggccgggttggatttgaaccaacgtaggcgaagccagcggatttacagtc
cgcccccattaaccactcgggcaccgaccctagttaaaccttatttaagctatcatatac
aagatatcataattatgtttctaaaaacaagtacttttataaataaaactgaataagtaa
tggacatagctggacttgaaccagcgactttcacgatgtcgacgtgacactctaaccaac
tgagttacatgtccaaaactttaatcattattataacactgtttagtgattttataagta
tttatttctaattgaagatttgttttaatcggccggctttaccgagcagatctcttagat
aatacagtttagctctacgtattttagctcttctaataactctgacaaaagttattcgtg
gtgaattaagaaaaaatactctttcaactccaataccttgaaatgagcatcttaaagtta
aactagtatttaagctttttttttcttcttctggctaatactactccttcacataattgtt
cacgagttttactaccttccttaaccattagtcccagttgtattgtatcgccaactttaa
tttcagatacttctgttttgacaagaggtatctcaacacttttcattagctgacttagct
ttgtattatttgctttcatatataggattagtcttgtgaca-tgttttgaaatattatac
attaaaaaaacaaatatgaaaaattttagcgaataagcatattagtatttataaaacttt

tttttattt---------------------------------------------------
--------------------------------------------cgattata--------
----------------catatccatt------------atattcaact------------
------------------------------------------------------------
------------------------------------------------------------
------------------------------------------------------------
------------------------------------------------------------
------------------------------------------------------------
------------------------------------------------------------
------------------------------------------------------------
------------------------------------------------------------
------------------------------------------------------------
------------------------------------------------------------
------------------------------------------------------------
------------------------------------------------------------
------------------------------------------------------------
------------------------------------------------------------
------------------------------------------------------------
------------------------------------------------------------
------------------------------------------------------------
------------------------------------------------------------
------------------------------------------------------------
------------------------------------------------------------
------------------------------------------------------------
------------------------------------------------------------
------------------------------------------------------------
------------------------------------------------------------
------------------------------------------------------------
------------------------------------------------------------
------------------------------------------------------------
------------------------------------------------------------
------------------------------------------------------------
------------------------------------------------------------
------------------------------------------------------------
------------------------------------------------------------
------------------------------------------------------------

------------------------------------------------------------
------------------------------------------------------------
------------------------------------------------------------
------------------------------------------------------------
------------------------------------------------------------
------------------------------------------------------------
------------------------------------------------------------
------------------------------------------------------------
------------------------------------------------------------
------------------------------------------------------------
------------------------------------------------------------
------------------------------------------------------------
------------------------------------------------------------
------------------------------------------------------------
----------acattcttccatagtttc--------------------------------
------------------------------------------------------------
------------------------------------------------------------
------------------------------------------------------------
------------------------------------------------------------
--------tttgataatcagtaa-------------------------------------
------------------------------------------------------------
------------------------------------------------------------
------------------------------------------------------------
------------------------------------------------------------
------------------------------------------------------------
------------------------------------------------------------
------------------------------------------------------------
------------------------------------------------------------
------------------------------------------------------------
------------------------------------------------------------
------------------------------------------------------------
------------------------------------------------------------
------------------------------------------------------------
------------------------------------------------------------
------------------------------------------------------------
------------------------------------------------------------

------------------------------------------------------------
------------------------------------------------------------
------------------------------------------------------------
------------------------------------------------------------
------------------------------------------------------------
---------------------------atcta----------------------------
------------------------------------------------------------
------------------------------------------------------------
------------------------------------------------------------
------------------------------------------------------------
------------------------------------------------------------
------------------------------------------------------------
------------------------------------------------------------
------------------------------------------------------------
------------------------------------------------------------
------------------------------------------------------------
------------------------------------------------------------
------------------------------------------------------------
------------------------------------------------------------
------------------------------------------------------------
------------------------------------------------------------
------------------------------------------------------------
------------------------------------------------------------
------------------------------------------------------------
------------------------------------------------------------
------------------------------------------------------------
------------------------------------------------------------
----gatgatcacatctcattttcaaacatggcttatgtatgaaaaagttctata---aa
aatctaatgactaaaataataaaattaaggcttaagtatttcttagttttgtgttataat
tagttactatcaaaggtagtggcataacacgaagcataaa-aatgttcgaacgctttact
gaaaaagctataaaagtcataatgctagcacaggaagaagctagacgcttaggacataat
tttgtcggaactgagcagatattattaggattagttggcgaaggtactggaatcgcagct
caagtcttgaaatcgatgaatgtgaatctaaaagatgcaagagttgaagttgaaaaaatt
ataggaagagggtctggctttgtagcggttgaaattcctttcactcctcgagcaaaaaga
gtactagaattatccttagaggaagcacgccaactaggccataattatattggtacagaa
cacttgctaatgggcttagtccgagaaggagaaggagttgcggcaagagttttagaaaat

ttggcagttgatgtttcttcaattagagctgaagttatacaaatgcttggagaaaatgcg
gaagccaatgtaagtggaagcaatactacgcaggctagaagtaaaacaccaacactagaa
gagtttggatctaacttaactcaaatggctattgaaggtggtttagatcctgtagtcgga
agacaaaaggaaatagaacgagttatccaaatcttaggtagaagaactaaaaataatccc
gtcttaattggggagcctggtgtaggtaagacagcgattgcggaaggattagctcaaaga
attgctaatagagacgtaccttctattttagaagataaattagttattactcttgatgtt
ggtttattagtagctggaactaaatatagaggtgaatttgaagaaagactaaaacgtatt
atggatgagattaaatcagctgataatgtaatattagtgattgatgaagttcatacattg
attggagctggtgctgcagaaggagcaatagatgccgctaatctgcttaagccagcttta
gcaaggggagaattgcaatgtataggtgcaacaactttagaagaatatagaaaacatata
gaaaaagatccagcattagaaagaagatttcaaccagttgtagttggagagccaagtgtt
gaagaaacaattgaaattttatttggtcttagagaccgttatgaaaagcaccatcaatta
acaatgtcagatggagctttggctgcagctgctaaatatgctaatcagtatatttctgac
cgctttttgccagataaagcaattgacttaattgatgaggctggttctagagtccgttta
ctaaattctcaattacctccggctgctagagaattagataaagagttaagagctgtatta
aaaacaaaagatgaagctattagagctcaaaaatatgaaacagcagagcagtatagagca
agagaaatggaaattaaagctcaaattgcagcaattgctcaaagtaaaaagaatgagcct
gatttaaatttagaagatcctgttgttacagaagatgatattgctgaaattgttgctgca
tggactggtataccagtaactaagcttactaaaagtgagtcagaaaaattaatgcacatg
gaagaaactttgcatggacgcattattggtcaagatgaagcggttgtagctgtctctaga
gcaataagacgcgcaagagtaggtctaaaaaatcctaacagaccaattgcaagctttatt
ttttccggaccgacgggtgttggaaaaacagaattaacaaaagctttagcttcttatttc
ttcggttcagaagcttctatgatacggctagatatgtcagaatacatggaaagacacact
gtatctaaactaattggttctcccccaggatatgtaggttatagtgaaggtggttatcta
acagaagcggtaagaaaaaaaccatatactgttattttatttgacgaaattgaaaaagct
catccggatattttaatctacttcttcaaattttagaagatggccgactaacagatgca
aaaggtagaactattgatttcaagaatactcttttgattatgacttctaacattggatct
aaagttattgaaaaaggaggaggaggtttaggctttgaattatcagaagatcaaacagaa
tcccaatatactagagtacgatctttagtaaatgaagaattgaaacaatactttagacca
gagtttctaaatcgattagatgaaattattgtatttcgtcagcttactaaagatgaagtt
agagaaattgcagaattaatgctcaatgaagtctttgcgagaattaagcagcaagatatt
caattaaatgtaacagaacgatttaagcaacgattagtagaagaaggatataacccaagc
tatggagctagaccacttagacgagcagtaatgaggcttttagaagatagtttggctgaa
gaagttttatctggtaaaattaaagctggtgacagtgcagtagtagatgttactaatgaa
ggtgaagttacagttttattaggtgaaaaattagaactgttaacataaa------aaaca
atttattaat--taaagcattgaagaggttat--attcaatgctat---gaaaaaaata

ataagatgggttgctataattgacaacctctgatactctgtatttgtatata---aagcc
gggatagctcagttggtagagcagtggattgaaaatcctcgtgtcaccagttcaaatctg
gttcttggcat-tttaagcattagaaaattttgctatctgagcattaaattttaattgaa
atgtccctgtgggaccattcctatgttttgctattataacttctgtgaagtctctacttg
tagtttcttgtgtatagtaactttctctatatagcattattactaaatctgcatcttgtt
ctatagaattatgaacaataaaattatttgaaataaaattgcacaaaggtttacattcta
aatcaaacactatttgtaaacttataatgtttatttttttgtatattttcaaaagtcacag
atgaaa------aagaatgaaaaatattttttgttcttcaaagttttt------tattt
gaatagcaatcatatcattttgattaatttcatcacatcttttccacccttgtgttgtga
atagtttatggttacttgttaattgtatatatttgccagctccagtaataatttatata
cagttttttttcctgtttttgaaatattagatttaattgctagaaataactgtagtttgt
gcgcagtaaagttaacgacttca-atctttttataatagcaatcaaataatatttg--ag
gttgattatataatgatgtctgaagataattgaatttagatatacacccactttctctta
aatcggataacaaaggccttttgtcacttcgagtttcaaggtttctatttagttgagata
atactaagataggtaagcttaaatcttttgctaatattttaagtgatcgagtaattaatg
atagttcttgagatctattgtcagattttctactctcctgtagtaattgaagataatcta
taataattaattctatattttttccttgcaatttaaagaatttcacttttgtctttatag
tttcagtagaaatcttagcgctatcatcaatataaaggttaaggttagcaagatttttac
tttttgaataacatattgccagtcatcattattaagctgaccagattgaatttttttggc
cgtttaaattacactcttgagctaaaattcttcttagtagttgctctgtcgacatctcca
agctaaataaaataacataagatttttttctgattgatgacatatctagttatattaatag
caaaagcagttttccccattgaaggtcttcctgcaaggataattaaatctgacttttgaa
acccttgtgttatagagtcgagttctgtaaaaccagaaagtatactattatttatactaa
tctttttcctggtatccaaatggactaatagacttgcaagaattgtagctaatgtttgcg
tatctttttcgtcaagtatttcataagcttctgttaattgactaacaatagaagctatat
ttttttgttcaataagttgttttgagcagctaatcaaacataatgaatctccacatgcta
aaagtaatcttttaatgtagttatctataacgaccgctgaatattcgcttatagcatcag
ataggggagcttgttctattagactaaaaacttcatttagctcatccaagtcttgaatca
ttttttcatttttttaagttaatgaaaaaatttcttacatttttttgtctttgcatggttaa
tattttcgagtgctgctctataaagtagtgaagttgatgtaaaataaaaaaaaatcaggag
agatttttttctactgattttagtaatgatatcgattgctgagttaatattatactaatta
atattttttcagctaaaatattatgaggaggtagatatttataa------atacttatta
attcctcc-ttgtaacatcactcttatggtattagtgtattagttaactattgatttaat
tggattctggaagaacttgcaatttaatatttgcagttacttgatgcaatagcttaattt
ctacatcgtaaatgccaattgttttcacctctggtaaagagatactttgtttttcaatgt
ctatgttggtagtgtttttaataatttgtgaaatttctttttctgtgacactaccaaaaa

tattttggccatctcctgtttttttactaacgctaaactttttgtatttcttctagaagtt
gcttaaccttttttagcatcttctttagctattttttaattttttcttctcttatagctgcat
gaaatctttgttgttttaaaataccatttgtggcaactaccgccatttttattcgggatta
aaaaatttcttgcataaccagaagctactttgataaggtcattactcttgccaagctttt
gaatattctcttttagcacaacatttacaactttttttactcatctt-------aattcta
aaaa----atctatattatcctattatacctgttgtatacagatttt----aataggaat
tatttatttgttgagataatattcttgtaaaaacaaaggtgatttttttatttt-ttctg
atataattattttcacaataattc-----attaggagaggtggccgagtggttgaaggcg
cagcattggaaatgctgtttaggagcaatcttaacgagggttcgaatccctctctctccg
tgcaa--aaaatataaaaagtttatagagtccaatctaaatctccaataacattcttcc
aagcatttaatccaccccgaactctaacaatatttagtttttgagctattaaaaattcag
aagcaaatattgatcgagaatctaagctacagtaaacaaaacaaattttatcttgtaaat
tgatattagagtaattcatttcttttattttttgtaaaggtagatttaatgaatgtatta
aatgacttttatgatactcttcatgatttcttacatctattaaaatatattgtggattcc
tactaattaaaaacctttgtaattgaataacactaatttctcgtacaaatgtactaaatt
tactgccataatattcgttccaatgcttttttgt-------ggataatacaaatttagta
tttataatcttaaatttcttaaatgatgaagttaatgcgttgtacgttaatatagtccca
cttaatatagacttataacccagaacgattttgacagcttcagtggcttgaagtgtacca
ataatacctggtaatagacctaaaactccggaattactacaagtatctcttgcactctct
ttattttcaatttcactgtagaagtctcgataaacaggtccaccttgataattaaataca
ctaacttgtccttcaaattgaaaaatggcaccataaatatgtattttatttaactcaagg
caagtattattcaagacatatcgagtatcgaagttatcagatccatctataataatatca
tattctctaataatgtcgattgaattctcgtagcttagtcttgtgttgaacacagttact
atacactgtggatttaaatctaata-tttttttttagctatttctactttttgaatatcct
atatcatgcatcgtgtataaaatttgcctctgtaaattagaaagatcaactatgtcatca
tccataatgccaatgctcccaattcccgcagcagcaaggtaaataattcctggagatcct
agcccaccagcaccaatgaataaaacttttgctttttttaatctttcttgtccttctaat
tgaatttgtggcaatactaaatgcttagaatatcttgtatattcctctaaggagtatttt
gtattgtctgttttaaaatttagcataaaaatttatagtttatttaatagtactta----
--aaagctagctgataagtttattaaaatgacttttgtaaacttggttcttctctatcta
aaatagctccttgtgtaggacagacctgaacacaaatactacaatcaatacaggcagaaa
agtcaatccaataccaattagtattactattgtttttcccctttccttgatgaatacaag
ctactgggcaagcttccacgcattcagcaactccaatacatttttctgttacaattgtat
gagacatattatttaattaat-aaaagataagctatttacatctagtagttttgttgtat
actaaatttacatatgctatccagtccaatatttaaggtagtctatttttacagtttact
aatttgatatgcgcctttagttcagttggtagaacgcaggtctccaaaacctgatgtcga

gggttcaagtccttcagggcgcgtttagtttttatcgtaattcaataatatcattttatc
tttttgcaaacatcaaaatgatatgatataaaagatcatattttatacattgaatattct
atgtactagtactg-tttttaatatgttattattgataagttaattatactaattctat
ttttgtaataagt-aaaaacagttctaattaagtatgaacgtatttaatagtc-ttttgt
atatac--aacttaaaaaattttaaactatggctaaaaaagttactggaattgttaaact
agcattaaatgcaggtaaagctactcctgcgccacctgtcggacctgctttaggacagca
tggcgtgaatattgttatgttctgcaaagagtacaatgcacgtaccgctgagaagtcagg
actagtgattcctgtagagatctcaatatatgaagatagaagttttacttttatacttaa
aactcctcctgcttctgtactaattgctaaagctgctggactaaataaaggttcaggtga
accaaataccaaaaaagtaggcagcataacaaataaacagttagaatctattgcagaaac
aaaactgcccgatttgaacactaacaatatacctcaagctatgaaaatagttggaggtac
ggctaaaaatatgggaattctgattaaagatt-aaaagtttcaattatatttatta-ta-
-tttatgaaaaaaatttcacgtcgacttaaaacattaaaatcaaaagttgagcctaaact
ttacactattaatgaagcagtatctatattaaaagaaacatcaaatgctaaatttaagga
aactgcagaagctcatattgctctaggtttaaatcctaaatatgcagaccagcaactcag
agcaacagttatattacctaaaggcacaggcaaattgataaaagtagcagttattactaa
gggagaaaaattaacagaagcaattagtgcgggagctgatgttagtggttccgaagaact
aattgatgaaatactgaaaggtagattagattttgataagctaatagcaacaccagatgt
tatgcctttaatagcaaagttgggaagagtattaggacctcgagggttaatgccttctcc
taaagcaggcacggtaacattgaacgtagcaaaagccgtaaatgaatttaaaggggggtaa
agttgaatacagagttgatagaacaggtataattcatgtgccttttggaaaatctagttt
ttcacaagaagatctagttttgaacctccaaacaattaaagaatcgattgatagaaataa
gccttctggagcaaaagggaagtactggaaaactttttcttatctagcaccatggggcc
atctgttcaaatcgatatcactagccttctatgaaaaattttgtatagttatagaactt
gaataatctattatttcaaatta-taatttgctagcattattaacgtaataatttgctta
ctatatcttatctgcattaaaaacttt-ttatatatcttattaattatgagtacaaaggt
tgaaaatatcttggaagagttaaaatctttaaaccttctagaggctgctgaactagttaa
acaaatagaagaaacgtttgatgttgatgcatctgcggcttcaggaggtatgatgatggc
cgcaccaacttcagcaccagcatctgctgaggttgaagagaagacagaatttgatgttgt
cctagaggaagtcccagcccctaaaaagatttctgtattaaaagttgttcgctctctaac
tggtttgggtttaaaagaagcaaaagatcttgttgaatctacgccaaaagtcttaaaaga
agctgcatctaaagacgatgcagaaactatgaaaaaacagctagaagatgctggtgcgac
agttattgttaagtaataa-----agaggcgcttctgtttatacttaaacagaagcgcct
ctttatttattataggcaattatactagaaaagtccaagtgtaattgcattgttgatagg
catagttgcaccaatacctagccaaatacttacaactgtaccaattaaaaaaacagtggt
tgcaattggtcttctaaatggattttgaaatttgttaacattttcaatgaatggaacagt

aagtaaaccagctggtacagcagccatacttagaacacctaatagtttattaggaattac
tcttaatagattaaatgttggaaagaagtaccattctggtaaaatctctaatggagttgc
aaatggattcgacttctcacctatagaagatggttctaaaattgcgagtccaatactaca
ggcaaaagtgccaataattacaactggaaagacatataatagatcatttggccaagctgg
ctctccgtaataattatggcccattccttttgctaatttagctctcagctttggatctgt
taaatctggtttttttaagaattgacatatta-tttttttgtttaatataaagattagtct
aatcactgttttaaagcgtaattttcaactattataatggtccagaaattccttgtttac
gtatcattaaaaaatgcataagcatgaaaacggcagttagtagcggaagtacaaaagtgt
gtaaactatagaacctagttaaagttccttggcctacactaactcctcctcttaacagtt
ctactatactcgctccaacaactggaacagcatcaggaacaccagtaacaatttttacag
cccagtatccgatttgatcccacggcaaagaataacctgttaccccaaaagaaacagtaa
gcactcctaaaattacacctgttacccatgtcaattctctaggtttttttgaaacctcctg
ttaaataaacccggaatacatgtagaatcatcatcaagaccatcatgctcgctgaccatc
tatgaattgatctaataagccatccaaagtttacatcagtcatgatgtactctactgatg
taaaagcttctgcaactgtaggtctataatagaatgtcattgcaaatccagttgcaactt
gaattagaaaagatacaaatacaataccccctaagcaataaaaaatattaacatgaggtg
gtacatacttactagaaatatcgtcagcaattgcttgaatttctaatctctcttcaaacc
agtcataaattttactcataaaatagcttcaaaaagctcttctaatata--ttttgcgtt
taagctactattaattcaatag------ttattgaacttatattatttttataatcaata
cctattaaatattataacatctttttattgattt--------------------aaaatt
ctctgactcagtaaaacagggctatgcactataattcttttttttttgtatagagattaat
ttcgtttttagtaatttagctaaaattcttgttacactgactctattactacctataatc
tgtgctaaaattttatgtgtaatagtaaagttaagcataataccattattaacttgagtt
ccattttgctcagataataataataataagctcacaagtctactggttatacttttatgt
gaaataacttctataaaatgataagcttttgccgagcaaaatattaaatgatttacaaaa
aatagattgaatggagcgcagttactacatgcttttataatagtagtatagttaatacag
gcaacttctgttttatctattgcttctgcttcgtagtaaaaattatcgtcgaccaattct
atttgtccaaatgtatttcctgtagttaataagttaagagttactttatgtgtatttctc
aggacttttgtaataattaaagatcctatcagtataatgtataatctagagctgctatta
aacaaaagagtatcattttgtctaagtgaaaaaatttgataattaatttattacgatta
aaaaatagtagccatggattgcaagaaaaaacatcttctgaactactttcagagaatatt
ggaacaaaattttcttgatttaaacaattttagtcataattactactttataa------
---aaaaatattctaataat-taagaactgaaattaaatgttatataaattgatgttagt
tgagaatgacattgcattatcaaaggccattcaagaatacttaatagatcaaggcttcaa
tgtttatattgccaataatggattagaagcactgaatctagcttatcaatataatttcaa
tttgataatttctgatataatgatgcccctagttaatggctatgaactattagcggaact

tcaaaaaaacaaagcgttatccaaaatacctgttattttttgacagctaaaggaatgac
gaaagacagaataaaaggttacgacatagggtgctatggttatttatctaaaccatttga
ccctaatgagttactctcaattattaacaacttaatcgctagagatgccttaaaagaagc
ttctttacaaaactcggaaatatcaaaccaggaattaaatcataaaatacgtttgacgcc
tagagaaaaaagtattttagatcttgttgttgatggacttactaataaagaaatatctac
tatattgaatactagtgttagaaatgtagaaaaatatgttagtcgactcttacacaaaac
taatatgaagaataggactttgctagttaagtattctataaataataatttattaaataa
taagatcaatgaaagggcgaatgacgggactcgaacccgcgaatgatggagccacaaccc
attgccttaaccccttggccacactcgccatattacatatatcactatagctcgtttttt
gatatatcgtctagttttcgagttatatatttatataaaacaatatttactttgaatttt
-tatgatacatagcaatattagtattcaaattaatggagaaccatttaattgctcaaagc
ccatctcattgcaattttttattaaattatcttgactttaattccgagcgtgtagcagtcg
agctaaatgacattctgttaccagagactctgtttcactcaacttatttgaatgatcaag
ataagctagaggttattaccattgtaggtggtggataatctatttttacat---------
---agttctctatttcaaatgtttcatagatagagataatcttccttgtttttatctac
atgaataataacagctttaatagtatcacctatttaaattgtgatgatatctgctctag
ttctttgatattaatctcagaaatatggacaagacctttgagatttcctacctttataaa
taacccgtaaggtgtaatctgatttattatgccttcaataatattaccaacaattaaatt
agatgatgcttgagatattagagctcttctatgactgagtattaaattattagatttctc
ttcaacattgagtaattttagcttaataaatttattattaaattgctcactttttttgaaa
attaccaagatgagaattaggtacaaatccagatataccttcaaggttaattatcattcc
tcctttattgaatccttttatcattacattaagtaaagagtcttcagctaatagttgcct
aattcttttccatgctcttatatattcaaggcgacgaattgataagattaattgtttcga
ctgaatattataatctaataaaaaaaattctcttgtatcattaatatttaaagaagtgaa
attatttaaatcttgattactagatacttcttgaataggcaagtatgcagatattggtgt
tcctatatctactaataccccatttaattcaaagctaaatatagttccagctacaatatc
tccaagattcaaatcatacttgtatttttgtaaaacagctgcaaaatttctatgagtaaa
tccgtcattatttttttgtcatattgaatagtatgaattaaagagtaactttgtacaatat
attaaggtaaacaaatgtttgtcgaattgtatg-aagaattttaatttattcggttgtaa
ttaaacatataaacatattaaaatatttaggagacataaaaagtagacgtagatagctgc
agatttaa-taatatataatctgaggaaagtccgggctccacaaatacaatttatgctgg
agaaatcccagtgtaggtaactgcgaggatagtaccacagaaaaaaaccgccagaataag
t----tatattagctggtaagggtgcaaaggtaagttaaaagcttaccaaaagtactgca
aagtatttgttaggtaaaccccctaaaatggagcaaagctactaaacaaatatttgtgtat
attttatctagtttagttgaaaatactgcatgaagttattagtaacaataactcgagagg
aatagctatcctttacaatatatc-tctattatattgaaagaacagaacccggcttatgt

aatacttattatgtcttgtattattatgtctattgtcgaatctctaaatttaactgtttg
acaatttttttctgtagctgatgttgtttataatctatttctgaattggttaaagttttc
tcattagatcgatatgtaaatcttagacctatactttttttccctttccaattgattta
tctatatattgatcaaataatgttgtactttctaaatctttatcataaaattgatccaat
gattttaataagtatttaatttccatattcttaggtattattaaagagagatctcttatg
atcgatggatacttagagtatggtttaatctgataatttaaataattgaattcattatga
caatcttctaaaatgtttaagttaacttcaagaacaaaaagttttgtgttaagtcctaat
tcactatattttgcttgattaagttcactgaatatgccaataacgttattattatagatt
agtgtggcaaatttccctttttgaatgaactttattctaccatcgactatattttttttt
gtccactcaatttgtctattaatcttttgaaaaaaatttctacgatccctttagcttca
taccaatttaaagaatgcgcaggatgtgaccatgtagatcggatatctagattaccaccc
aatatgatagctagattagtagtttccataattttgttttgatcattatgaaatactgtt
cctatctcaaagctatctatagtttggttgctttttttaatattataaagatttgattca
attaatccttccagcaaactgctgcgtaaattagaataatcttgaataagaggattgttt
aaagttatatttcctttggactttattaaagaataatgaactaattcagttagtcctaga
tttcttaaaatacttctacttttatctatgaattgtctctttgtcgataacttttttgttg
aattgaatatttggtactgagctcctgaatttatgataaccataaattctggcaatttct
tcaataatatctatttctcgaaaaacatcttgtcgtctataatttggaacagtaacttct
atattttccttattttgataaataagatcaaaatttaaagagtggagaatattttgaatt
tcttcaaagaataaaaaacgtgtttgaccattatatataattggtcctaaaatgtcatga
actttttttaattgataaattaatattaagagcattatcagttgtttttttctcgatagaaa
gtctctctaacatttccgcatgttaaatctgtgattaaagaaagagcttcaaaatgtgca
tttttccaattatctacatttagcccacgttcttgtctaattgaactttctgttcgaata
ttaagaactcgagaagactttctaaccacagattgtttaaatatagcagactctacaagt
atcgatgttgtattataatctgtatcaagattagagttgcttccaataccagctatacta
attatatttgtatttacttgagtaaccaaaatatttttgtttaacccaatattttcattg
tttaacttaatctgatcattattttcaataggaaaatcgcttcgaattgttagactattt
tcatgattcatattattaattttattcaaatcaacaacattaatgggctgtccccatttc
aacataatataattgcttatatctgttaacaaatttctgtgtgtaaacccagaagagagc
aatcgatttttttagccaatctggagaatctttaattcttatttcgtctataatggcagca
aaataattgtcgcaatttaataaattgtgatcacttattatagt-ctcctttttagagaa
atatctattattgggttgatatatagctgataacattgaggaccctgttaaagccgagac
ttctcttgatagaccaatcatgcttaacacgtctgacctgttagctgttgatgatatatc
taatacatgatcaatctcacctccaattgtaatagattcaatagcttcaacttcaaaacc
tgcttgtgttaattgattagttaaattatctatatctattgttttgatattaacaagttc
tttcaaccaatttaaagaaactttcataattaattgtcatctatcttatagaagtattta

ttttacttatgattaaaaatcaatttaaaaaattcatcaaattaatataagttatttgat
gaatttgcttatacaatatattattgtgatatcttacttttatatggcacttattaagaa
ttagcttttgtaaaagttagcccattatcttgggcatatctctccataaatctcatgaat
ctatcccaatcatcaggacttttaataacatggacagcttctattgcttgtggcttacca
ttaataaacttggcattaacatctcttgtaataatttgtccctctgtgtccatcagatac
atacctgtgatttcacctttatctgccatactagcatctaaaattttgggatttgtaaac
cgaaaagttgctgttcctgtactaccatctcgagaccttgttaaacgtacatctggtaca
acttcttcattgataccttgaataaattgaattgttgccatatcaatcttctttatttat
atataatttatattggataaaactgatttctagttcttgaatgcttgttgagaaactgtt
ttaaaagcttatagtagcagtagatcaaattttatgtattatataaaactgtaacttttc
-ttttttg-----------------------------------------------------
------------------------------------------------------------
------------------------------------------------------------
------------------------------------------------------------
------------------------------------------------------------
----------------------atctgttcaaagaattaacttattataagctaatcata
cgaatattagccatataatgtttgtaagt-gaaattaaggggttgtagctcagaaggata
gagcaagcgcctcctaagcgctaggtcagcggttcaagtccgctcaaccctgttaacaag
ttaagtagtattatttaacttgttctgcaaatcagtattgatttaagttaaagttcttga
tagctaaaacaaaagaatctgagaaatatttttattta----------aattatcttaac
tctgaatatctgataattattaa-aaagttct----------------------------
----ctttttaagacacaaggtgatg-----aagatatttaaactaataatggaaattaa
gtttttgaagttatagaagccaat--ataatactttctattaacaga-----------cg
cttgcttctattcctatttgtttacaaga--aatt----atgaatatcaaaata------
--attttagctcca-------------------------------------ctgat----
------------------------------------------------------------
-----------------------------tactaataaata-------------------
------------------tttatttcagtctagacaag------atgaatcttcatggta
taagag------------------------------------------------------
------------------------------------------------------------
------------------------------------------------------------
------------------------------------------------------------
------------------------------------------------------------
--------------atgctattctctgtgtcatacttatataaaatcaggat-------ac
tgtt--------------------------------------------------------
------------------------------------------------------------

------------------------------------------------------------
------------------------------------------------------------
------------------------------------------------------------
------------------------------------------------------------
---------------gtcattattataactggggtgggaggattcgaacctgcgaatggc
ggagtcaaagtccgctgccttaccacttggctacaccccaacatgtgaatattataatag
cagtcgtataccctgccatgtcaatagactgttgtctttatttaataatttttgctgctc
ttgtagaataaaatctttttttttcaaataagatacgctatgtttaagttgaacgactgtt
atgttttcttgcttttgactaattagccacttgatagtaattattccttctgcaatttcg
ttacttcctataataatacaagctacagctctttttttatttgcttgcttaatctgttta
ctaaaattacttgagctaacatctatttctattgtgaaaaactgctgatgtaaaaaacgc
attacttgcataccagtttcgttagctttttcaccttgtgtagcaatataaaaatcgata
gatttattaggcaagtttatattatcctttgcaagcagtaataggcgttctaatcctatt
gcacatccaacagctggtgttgctttgccacctaattggtgaactaagctgtcatatctt
ccaccaccgcatatagtatcttgcccttttgatgtcaatgtcttaatttcgaaagcagta
tcattataataatctaatccgcgaactaacttattatttatattgtagggaatatcaagt
aattttagataattgcaaacgtcatcaaagtgtttttgtgattccaaacttaaaaaatca
gaaattttggagcctctgttagtatttttttgtgtattagcatcttttgaatctaaaatt
ctaattggattactagttaatctttttttgtgagtcagtgtctaaatcatcatggtacttt
gtaagataatctagtaatttaacttgataaatactacgatcctctacttttccaattgaa
ttcagatcaagttttaagttatgcagattgagattgttaaagatgctcatagctaaatgt
ataacttcactatctgctcttgcgtctaggctaccaataaactcaatgccaagttggtga
aattgtcgttgtcgtccactttgcggtctttcgtacctgaacattggaccgctataccat
aatctttgtaaactatgttggtaactcattttattttcaataaaagatctcacaatcccg
gcagtaccttctggccttaaggtaatatctctattgctgcgatcatgaaaccggtacatt
tctttatttacaatatcagtattttcaccaatactcctatcatataattcactattttca
aaaataggtgttctaatttcttggtaatttgcacattctaataatttggaaattttatta
tgtataaactgccaatattgaatttcatccggaagaatatcttttgtccctctaatagcc
tgaattttgccatgataatttttttgattaccttcttaatatttctgcattaaatagcag
ttgtagactggtattcagtcatgtattatacattag--aacatactggtg-tattttata
aaatacgggcaaggagggattcgaacccccgacaccatggttcgtagccatgtgctctaa
tccactgagctacaagcccacttaataagtacgttaactatatcattattctctaatgta
tacaactatattcttttt------atcttacatttgaaattttgtaaaagtgaaaaatt
at----gtttgtttttacaattggaactattcatatataatgcttaatttattataact
ttgtaaa--aatatctataagtttaattgtatgattctactatatt-gtaatcttgtcat
aattttgcgctcttatttttcaattag------ctagcatctaaataacttatatttcaa

ataaaaaaatcttatctaaaatgagtaaacaaattctatatcaggatgatgcaagaaaag
cactagaaaaaggcatggatattttaacagaagctgtttctgtgactttaggaccaaaag
gaagaaatgttgtcttggaaaaaaaatttggtgcccctcaaattattaatgatggtgtaa
cgattgcaaaagaaattagtttagaaaatcatattgaaaatactggagtcgcactgatta
gacaagcagcatctaaaacaaatgatgtagccggtgatggtacaacaacagctacagtac
ttgcttcagcaattgttaagcaaggcatgagaaatgttgcagcaggttcaaatcctatgg
ctattaaaaaaggtattgaaaaagcaacaaattttgttgtcagtaaaatagctgaatatg
ctaagccagtagaagatacaaaagctattatacaagttgcttctttgtcatcaggaaatg
atatagaggtaggtaaaatgatagcgaatgctatagaaaaagttggaagagaaggagtta
tttctttagaagaaggaaaatcaactaatacgattcttgagatcacagaaggaatgcagt
ttgaaaaaggctttatttctccttatttcgttacagatacagaacgaatggaagttcttc
aagaaaacccttttattttatttacagacaaaaaaattactttggtacagcaagaacttg
ttccattgcttgagcaaattgcaaaaacgtctcggcctttattaataatagctgaagaca
tcgagaaagaagcattagccacaattgtagttaataaattaagagggattttgaatgttg
tagcggttagagctcctggatttggtgatagaagaaaatctttacttgaagatatgagta
tcttaacgaatggacaagtaattactgaagatgcaggtttatcacttgatacagttcaat
tagatatgttaggaaaagcccgaagagttattgttactaaagactcaacaacaataattg
cagatggtcatgaaattaaagttaaatcaagatgtgagcaaattaagcggcaaatagaga
cgagtgactctttatatgaaagagaaaaattgcaagaacgattagcaaagctttctggag
gagttgccgtcattaaagttggtgcagctacagaaacagagatgaaagataaaaaactaa
gactagaagatgcaattaatgcaacaaaagcagcaattgaagaaggtattgtaccaggag
gaggagctactaatgtccatatctctagtgaattatttacatgggctaaaaacaatttaa
ctgaagatgaattaattggtgctttaatagttgaacgagctgtgacctatcctttaagac
gaatagcttttaatgcaggtgacaatggagcagtaatagtagaaaaagttaaaagtcatg
atttccatataggctacgatgcagcaaacagtaatattgtaaatatgtatgatagaggta
ttatagatccagcgaaagtagctagatcagctttgcaaaacgcagcttctattgcagcaa
tggttttaactactgaatgtattgtagttgataaagttgatgattgatttata------a
gataatagataatctcaaacaaagaataatttctaattatagaaattattctttgtttca
attaagctaattaattgatcaaatgtataatttttataataaaaccaaagtttaaacaca
tcttctttttcaactattaagtagagaagatataagcccgtaatacctaaattatcaatt
aaaatattattttcatatatagatttttctaaagatctaaaaaatgacattttttgaaaaa
taataccgaaatcttcttttgtagttatttgctaactcattactgcctaaagattgattt
ccaattgaattcagaacctttaacagaacactttttaaatctttatctgactttatttta
gaatcaataataatttgattcaattgcatgtcaaaactaagattaaaaatactacttgtt
aaagatatttgattttgaatatctagagattttatagccaagaagattaaacttttattg
ttcattgtaaaatataagacaattaatgatgtattctttaaatacta-aatttatcttct

tgtgcttctatcgttttttgt-aaaataattaatgctttaatttttttagttttagttact
acccgaaaaaataaattctggaaatctagcttgagctttagctagagagtgtttttttcc
ttcttcttgccagtaattgataacttctgttgttttgtttaataactcaattctatctct
ttcagcaatccacggcttagattctagttctgcttttaattcttcccaagcatcatttct
tggccagaaaaaataagaagtaagtggactagtgccattcccaacaatttgatcaatagc
aatagcaatattattttctaaccaaagtactttagtgtaaattttgacaagtttttatc
tccattattaatattaagtgttgg----tacagtagaatactgtactagttattttttttc
agcaagaatttgagctttgtttaaaatattttgtactgttttagtagcttgtgcaccttg
agacaatcgttgcaaaatttttgaaatgtcaacgcgagtttcattagttataggattata
aaatcctaattcctctatagctttaccatcgcgtttatttctgctatccattacaacaat
tctatagcttggttgtttttttcttccgtatcgttttaaccttaacttgaccatattatt
cacctcgcttatttttt--atatctttattaaattatatcttacgacattatgcaataat
tcaaacaattaagcagattcaagtctaacgttattaaatattaccattaaacattgtaga
aaaataatacctaacataggtgacatgtccatgccaaacataggaggaatgcttcctctg
aatagttttaaatatggatcagtaattctatttaatgaacagaatggctcattgtaccaa
tttacagttggaaaccatgccaatgataatttaagtaaaattaaaattaaataaatctca
gaaaaattagctattgatccaagtaataaatttaatgtaccaggaagagtattcatacgt
atttataattattatatacttcattatactaatagtttgattatatattataatctctg
taaataaat-cagctaaacaaaaaacttatccttaaagctatcttctcgcgagcattttt
ctttatctacattagtatttacttttgtcgtataaatttctagtttagaccaattttgtc
ctaacttatttaattgagatgttccacattctaaacatgctatttttatattgctcaaaa
cagcacctttttgtataaaatcattcaataatggcgtaatagttgcttcatccaaaaata
aatctaagacaaggatttttgtaaatgaatttaattgagctgggacattgttaataatat
tattataattaactaaaactatattagctgtaggaagtaaagctcgtgctccttccgcta
atataaagccataaggcataataattacaataatatatttataatgactgttaataagac
ttatagtggtattagcttttaattcgattgtttctgttactaaccactctctcattattt
cgtaagtgatccacttacctagttctgagcaagcagttcttaaaattgtacctggattat
tgttgttttcgagaacccctgaccaatgttgtattaatggatgggatgctatattaattt
ggagttgcatgttttgttaaatatatttaacatttaatgtgttgattgaaacttcaattt
ctatttgttgagtattattaaaatatcat----atatattaaagaaaataattat-aaaa
aatgaaaaaaagtctttggttgtggggttttacagatagtgctgaaacttggaatggtag
attcgcaatgattggttttatatcagttatttttatcgaagtagttacagggcaaggatt
attatatttaataggtatgatgtcataataaaaatccaaaaaagttcttgtaatgtttta
caagaactttttttgg-atttttataaatctaatctagcggtctcatagatagaacaggtt
cattctctctattgataccttttttcaaatccagcagcagcagctcttgctctgcctgcat
gccaaagatgtccaataaataggaagaatcctaggaagaaatgagatgttgttaaccaag

atctaggagatacatagttaactgaattaatttctgtagctacaccaccaacagaattca
aagaacctagtggtgcgtgagtcatgtactctgcagcacgcctttcctgccacggttgaa
tatcgttcttgattttattcaaatcaagaccatttggtcctctaagaggttcaacccaag
gagctcgcaagtcccaaaatctcatggtttcaccaccaaaaataatttctccactaggag
atctcattaggtatttacctaatccagtaggtccttgagaagacgcaacattggcaccta
gtctttgatctctaactaagaaagtaaaagcttgagcttgtgaagcttcaggaccagtag
gtccatagaattcactaggataagctgtgttgttataccaaacaaaattagaagctgtaa
gacccatgattgataaagcacctagactataagacaagtaagcctcaccagaccatacaa
aagctcttctagcccaagcaaaaggttttgttagtatatgccaaattccgccagcaatac
aaataatgccaatccagacatgaccacctacaagatcttccatgttatttacacttacga
tccatccatcaccgccaaacggagactttaaaacataaccaaagataactaggggattaa
gtgtaggattgctaacaaatctaacatctccaccacctggagcccaagtgtcgtataccc
caccaacaaataaagctttaataactaatagaaaagcgccaatgcctagtaatactaaat
gtataccaagtattgttgtcattttgttttatctcgccaatcgtatccgaagaaaggaa
atgattcttcaagagtatcagggccaatcagagaatgataaagaccaccaaagccaagaa
cagctgaagagattaaatgcactacgcctactacaaagtatgggtatgtattgaaaattt
cgccgcctggaccaacaccccagcctagtgtagctaagtgcggaattagaataaaccctt
gttcgtacaaaggcttctcgggaacaaagtgcgctacctcaaaaagagtcatggcaccag
tccaaaagaccattatacctgcatgagcaacgtgagcaccaagtaatttgccagaaacgt
taattaagcgtgcattgccagaccaccaggcaaacccggtagactcaatgtctctaccgc
caacaccaacagttgtattaaagggcgtttccacgtggtaaaacctcctcagggaatata
aagttttcatgaggttgatcctgagcagccatccaagagcgaataccttcgttcaataga
atgtttttagtataaaaagtttcaaactcaggatcttccgcagctctcaactcttgagaa
acaaaatcgtaagctcttagatttagagctagcccaacaattccaaatgcgcttgtccat
aatccagttactggcacaaataacatgaaaaaatgtagccaacgtttattagaaaaagca
acaccaaaaatctgtgaccagaatctgttagctgtcaccatcgaataagtttcttcagat
tgtgttggagtaaatgcgcgaaaagtatctgcagcatccccatcttcgaataaggtattc
tgtacagttgcaccgtgaatagcacatagtagagcaccacctaagattcctgcaacgccc
atcatatggaacggattcagagtccagttatgaaatccttgcaaaaatagcaaaaatcta
aaaattgcagcaactccaaggcttggggcaaagaaccagcttgcttggcctagagggtac
attagaaagacggagacgaatactgcaattggcccagaaaacgcaatagcgttgtatggt
ctaagaccaactagcctagcaatttcaaattgtcttaaacagaatccaattagtccaaat
gatccatgtaaagctataaatgcccacagaccaccaatttggcaccagcgagtaaaatct
ccttgagcttcaggtccccaaaggaaaagaagtgaatgtcccatgctgtttgctggcgtg
gaaacagccgcagttaaaaaattgcatccttctagatatgaactggctagcccatgagta
taccaagaagtaacaaaagtagttccagttagccagccacctacagcaaggtaagcgcaa

ggaaacagaagtagtccagaccaacctacgaatacaaatcggtctctttttagccaatcg
tctacaagatcaaaaccaccacgagtttttcttgtccaattgctatggtcataattta
atctccagagcaaatttaataagtaatttgtttaaataaaca---ctttagatatattct
aaaatttactttctttacggtcaatataattataagtcaaaactaaccaaaattcacg
tttttaataaca-ttttttagctcattaagttaatttttttaatatatgaacataaata
gagttattgttt-aactcaaagtactcataatttattcatttctataatcctttgaaaga
gatatcctgttcctcgtgctgttaaaatcaagtcgggactacttggatcatcttctagtt
tagctctcaatcgtgaaatatgaacatcaacaactctcgtgtcaacatgtcgttcgggag
tatagccccagacttcttgcaaaatagaagctctagaaaaaggttctccagccttactaa
ctaaaagctcgagaagactgaattccatacctgttaagcgaactctttcattatttttat
aaacttgtcttttattggtatctatttttagaaatccaatactaataatcccagagtttg
ggactccaagactagtggtgattttatcagctcttctaagaacagaacgaatgcgagcct
ctagttccttaggagaaaatggtttaacgacatagtcatcagcacctatttctaatccgg
taattctatcgcatacttcacctaacgctgttagcattattataggaacatctgattctt
ttcgaagttcttgacaaacaccgtaaccatctaatttaggcatcataacatctaaaacaa
caaggctaggatactcttttctaaacactagtaaagcttcttctccatttgaagcagtta
taacttcgtaacctattattgttagtcttgtttctaaaatccttcttatgctggcctcat
catcaacaacgagaatcttttctttctggttctccaatttttttatttactcctataaat
cactgataatgattactataaccttaaattttagctctatgagtatcttcatttctttt
agaacttagtttagttattataaattctgcaagtctagtttgcattaattagtaccagt-
-aaatattttgt------------tttaagttatgac---------tattttaatccata
agttgactctagtttgcagaattcaatatatacata-ttttctagccaagtaatttgtat
tctattttctaaataaaaaaggggttgacaaactagttgatagaaagtattctacattac
tataag----ttaattctaaaaaagaaattctaa-----atcttctttcggagtaataaa
tactttaat
>Pyr_1_10_11_12_13_14_15_16_17_18_20_21_22_24_26_28_30_34_36_38_40_
41_42_P_yezoensis_MK695880
tatatgata---aattaagacatctgttatgtgagaatttctaaaatatacaaactttt
tatttaatatagaaaaagtaaattaaaataagttgaacatactttgctctaatctaaacc
tcttaattgag------attctaattagagacaaaagtatttacttattttgtatctata
ggaataaaaatgaaacttgcagtttatggtaaaggtggtataggtaaatctacaactagt
tgcaatatttctgtagcactttcaaaaagaggtaaaaaagttctacaaattggctgcgac
cctaaacatgatagtacatttacgctaacaggtttttaatcccaacaattatcgatact
cttcaatccaaggattaccactatgaggacgtttggcctgaagatgtaatctataaggga
tatggcggtgtagactgtgttgaagctggtggaccgccggctggcgctggttgcggaggc
tacgttgtaggtgaaacagttaaacttttaaaagaattaaatgcttttgatgaatacgat

attatcttatttgatgttttaggtgatgttgtatgtggaggttttgcagctccattaaat
tatgcagactactgcttaatcattacagacaatggatttgatgctttatttgcagctaat
agaatagcagcttcagtacgagaaaaagctagaacgcactctctgagattagctggactt
gttggtaatagaacagataaaagagatctaattgataaatatatagattgtgttccaatg
ccagtattagaagtcttgcccttgattgaagatattagagtgtccagagtaaaaggtaaa
actttatttgaaatggcagaaattgataaggatttagcatatgtatgcgattactatttg
aatattgcagatcagttgattacaaggccagaaggtgtagttcctaaagaatctccggat
agagaattatttagtcttttatctgatttttacttaaatcctaaatcaaaggtaggacaa
gaaaaagtagatcaagaagaattagatttgatgatagtgtaaataatattttcagcataa
tataacaaggaataagataatgtctacagctcaatcagatgctcttacttttgaatgtga
aacaggtaattatcatacttttttgtcccatcagttgcgtttcttggttatatcaaaaaat
tgaagacagtttttttcttagttatagggactaaaacttgtggttatttcttgcaaaatgc
aatgggagtaatgatttttgctgaaccgagatatgctatggcagaattagaggaaggaga
tatttcagctaagctaaatgattatggtgaactccgtagattatgcttacaaataaaaaa
agatagaaatcctagtgttatattttggattggaacatgtacaaccgaaattataaaaat
ggatctggaaggaattgctccaaaattggaagcagaaattcgtgttcctatagtagttgc
tagagctaatggtttagattatgcctttacccaaggtgaagatacagttctagcagctat
ggctcaaaggtgtccattaaatttaaagaatcaatcagataacacgagtcttaaaacttc
tcctcatattcctttagttttatttggatctcttccggatccagttgtaacccagttaac
tatggaattaaaaaaacaaggtatttttgtttctggttggttgccatcaaagaggtatac
tgaattaccggttataaaagaaggatattatgttgcgggagttaatccatttcttagtcg
cacagctactacactaatgagacgccgaaaaacaaaactaattggtgcaccatttccaat
aggtccagatggtactcgagcctggattgaaaaaatctgttcagtaatgaatgtaaagcc
tattggattagaagatagagaaaaagcaatttgggcttctttagaagactatatttcttt
aatccgaggtaaatcagtttttttttatgggtgataatttactagaagtatctcttgcaag
atttttaactagatgtggaatgactgtatatgaaattggtattccttatatggataaacg
ctatcaagcagcagagctagctttattaaaagctacatgtgataaaatgaacgttatgat
gccaacaattgtagaaaaaccagataattataatcaagtagaccgaatccgtgatctgaa
acccgatttagttatcactggtatggctcatgcaaatcctttagaagccagaggtattaa
tacaaaatggtcagtagaattcacatttgcacaaattcatggctttaccaatgcaagaga
tattcttgaactcgtcacaagaccattacggcgaaatcttagtttgtcagaattaggctg
ggatgtttatagcaagcaaagctagtattcctc----tttatgacttacatggtataaaa
gccttgtaagtct--tttattttatgttaaagaagcatttcttgctctgccatcaccagc
ccaagcttgtaaactttctacttgttctttgtctgtaaatgctaaaggcacaaattgttt
gatagctattttaatatcttcagtactaaattctctctcttcactaaacgccgtatgcat
actttcaataattgcttgctcaatttcagcgccagaaaatttattacataataaactcaa

ttcatgaatatcatatttttgccacgatcttggtctaacttttgataaatgtatcttaaa
aattgattctctctcttgacgattaggtaaatctaagaaaaaaatttcatcgaacctgcc
tttccttaacatttcggaaggtaaactttgaattttattagctgttgcgacaacaaatac
tggagctttttctctgatagccaggtaataaatgtaccaaatactctggcactagtacc
actatcaccttgactatgtaaaccagaaaaagctttatctatttcatcaatccataatat
gcaaggtgataatccttcagagatatttatcatttctctcattttgattccgattcacc
aactagtccaccaaacagttttcccatgtcaagacgtagcagaggcagcatccaatcgct
tgcaatagctttcgcggttaaggatttacccgtcccctgtataccaactagcaataaacc
tttgggcgaaggaataccgtaatttaaactttgcttcgaaaatgagcgagaccttttttg
tagccattgttttaatacatctaagccaccaatatccctatttactttactataaggata
aaattctaatagatgcgtttgattaataatttgcctttttttcttcaataataataggtaa
gctccgagagtctatctgattatattgtgcaataattttagtaatcactttgcggattct
atctatcgataatccttggcaggattttgttatattattaacgagctctgaattcaaatt
aagattcaaagctttactcaatcttatgatctctttttttgatttctagaagactaggtaa
gggcaaatctattactgttatgatatcattcagagcaaaaggtatattgattttgcaaga
aataataataatgtgcttagattgcgttttgattattttagctaaattgcgaagtttcct
aattaatactatttcgtttaggaaagagtcaaaatcttttaaaagaaaaagatttaaaga
ttcattattcaagttttcaataaattctaatgctaataaaggatttctttttgcataacc
attatcgctaggattgttagtgtatccatctacaaagtcccaggaatatacttgctgact
gtttgaacagtttaagctgtgttttataatatactctagtctatcttcttctctagtatt
tattacaataataggatatcgagattttaaaagtaatcgtaaatcttgagtgaaattcat
gtcaatatttgtttccatcagacttttgtacttcaccaattgaagaaagactaaataaga
ttatctaatattccttggatatattcaaaagtc------atctctctattttctaaatta
ttgctattactaatttaacttaccatgattttttttcttcctttcttcgtcgactattt
aatatagagcgtccacatggtgttttcattcttgctctgaaaccagaaactctaatttt
tttcttttagatccttgtagtgttctttagtcatgatgtctcctctgttgtaaaagatt
aaatactattttatcatcttttatatatctttatgatatagatatgtattaatatcaaag
tt-aacagcaataatttatattaatcaagaactatttatataatattaaacattaaatat
----------taactttatgatattggctttgcccatttttttatttaagtattttgacca
tctttttgctaatcttaaactggttaatttttcagcagctaaaaacgattttgttattag
aatctcaatttaagtattttgttgataaaagccaaaatcgtaaattagaacccgacgaaa
gttttgcttttgcaaaagtatgtgtagctaaaaaatatttttctaaagctattattgaag
gtcaattagctttaaaaaactatcgagacctgaatatcttagataataatatagtaattg
ctaatttgtataatatgctaggttttatttattttgaagcaggccaaacaagttttgcaa
agaacttttatgagcaagccttacaaataaaccctaattatgttgtagctttaaataacc
tggcaaaaatttatgaagaggtaaaagatttaaaaaaagccgaatctctatacgataaag

tgttaactcttaacttaaataataaaactgctaatagaaggaaagatttatagcaaaaa
cta---------aaaacatctaatactatttgtaatcgggatagcaggatttgaacctgc
gacatcctgctcccaaagcaggcgcgctaccaaactgcgctatatcccgtaagacataat
tattactatataccttttctctaattatgtctactctttctgtattttatctaggatacc
aaaacatgcctttaactccatctggatctttgataaatcctaatttttataaaaactaa
caacatcaggttctgcgaatagtgtaatagtgctaatttctgcctgtctcaattgctgta
ttaattggtgtatgactacttttcctaaacctaaaccttggaaatctggatgaataacta
cgtcccagatagttgcattaaatccgttgtctgatgttgctctagcaaatcctacaagtt
tactatttgcatcttttttttgtattaaagaaataataatagaactatttttaatgcga
tctttactttttttagaggtcttttaacccatccgactgaatcacaaagctgctccaatt
catataaattaatattcttattgctactcagatagatatctttaaattcgattttatcgc
aagttttatctaaaacaataagtttttgaaatttttttcagagctagcattgatattgg
agttttgaaaaaaatttttccagaagatcataatgattatgatataatta--aataaata
tatatattacaatattgctttaacaaacaatatattaatatatgctcaaatttttacatg
aatctattaagatagaagattgttaaaatatagagaaataaaaatttttgtaaaatcttt
caataaaatgatataaagagtgcttttaagactaacatctattattataaataaagtct
cgaaattgagatgaaataacacattattctatgtctaatgccttttaaaggagttatttg
tgaaaaaatctatgtgtctaacctgtttacttgctttactaattatgagcaacccaataa
tagctaacgcagaagtagctggattggtgccctgcaaagattctgcggcattcaataagc
gtatggtaaatagtgtgaaaaaacttcaagccagattagctaaatatgatgcagacacgc
caccagcgttagctttaaataagcaaatagagaaaactaaaactagatttgcaacttatg
gtcgagcaggtttgctgtgtggtactgatggattgccacatttaatttctgatggtcgat
ggagtagagccggggactttgtttttccaggactattattcttgtatattacaggatgga
ttggctgggtaggtagaggatatcttttatctgttgctaagactagtaagccaacagaaa
aggaaatcattttagatgtgccattagctatcaaatttatgtcatccggatttgcatggc
cgctagcggcttggcaagaatttagtagtggacaattaattgcttccaatgacgatatta
ctgtttcaccccgttagtaaaaaatttatatgaataataattttaccaaatacttatcaa
cagcacctgtaattggtgtattgtggatgacgtttacagccggatttattatagaattga
accgctttttcccagatgtcttatacttttactt----ataaataa--gatcgtataaa-
--aaactacataaatatagagagatactattatttgtgtagtttta---atatataaca
taattattta-tttttaaaaatgtttatcttttcgattattttaatattagatcttacat
aataaaata-atatcaatattatgagcttagtaacccaaattatagttaatgccgatgat
gaattaagatatcctacaattggagaattgcagtcaattcaagactacttaactacagga
agcaatagaattagaattgctactattattagagataaagaaaaggagattatacagaaa
gctagtaagcaaatttttcagttacatccagaatatatagctccaggtggtaatgcagca
ggttctaggaaaagatcgttatgcttacgtgattatggttggtatctcagactaattaca

tacggagtattagctggggataaagattctatagaaacaattggtattataggagtaaga
gaaatgtacaattctttaggtgtacctataattggaatgttagatgcaatccagtgttta
aaggaagcatctttagaaatgcttggtcaagatgatattagaatcatttctccttatttt
gattatataattcgtggaatgtcataaatcatagcttcaattaattagttgaataaaagc
ttttataatgttataattacaataagcccgagagcatataattgtaaaaactaaatcttg
tcagaaccggaaggtagcagcaataatgtttcaatacaaaagttatga-ttttcgggtgt
ttattgtttttatctaatagtctcgacttttaattagtatgctatgtataaatctgtata
tgacatgtaatccatcaaaccttccagcaaaaataatttaataatttgatattaatacgt
aaaacacctatac-tcttttataaaatttctttatatctcatctaaacaatggaaactta
acttaatgggtgttcatattttatcaacaggttcgtctgtcccaaattttctgtagaga
atcaacaatttgaagacatgatcgaaacttccgaccattggatttcaacaagaacaggaa
taaaaaaaagacatctagctccatcttctacttctttaactaaattagcggcagaagctg
caaacaaagccttatatgcagctaacttaaaacctactgagatcagtttaattattttag
ctacgtctacgcctgatgatttatttggtagtgctagccagttgcaagcggaaataggtg
caacaacatcagtagcttttgatattacggctgcctgctccggttttattgttgctttag
taacagcagctcagttcattcaaactggttcctatgacaatattttagttgttggagcag
acacaatgtctagatggattaattggtcagatagaactacctgtattttatttggtgatg
gtgctggagcagtagtattggggcaaagcctcaaaaatagtattttaggctttaagttat
gtacagacggtcagctaaacagtcatttacaattaatgaataaacctgtaaataatcaaa
aatttggtggtacagaaattcctcatggaaactataattccataacaatgaatggcaagg
aagtgtacaagttcgctgtatttcaagttccaacagtaattagacaatgtttgaataatt
taaacatttcaatagatgaagttgattggtttatattgcatcaagcaaacactagaatca
tagaagcaattgcgagcagattatcagtacctttttctaaaatgattacgaacttagagc
attatggaaatacatctgcagcgtcaatccctttagcgttagatgaagctattcaatcca
ataaaattcaaccaggtcaaattattgttttatctggttttggagcaggcttgacttggg
gagcaattgtcttgaagtggtgat---ttatatattgcggatgacgagactcgaactcgt
aaagctttcgctacacacccctcaagcgtgcgtgtataccaatttcaccacatccgcatt
ttttaccatgaata-ttaaaaatttataatatttatctaaatattcatataa--tgtgta
tatatatatacattcttgtcagaaactctcttatattttattttaagtgataaaattgag
t----ttagtatagatagtaaaattttattttaaaatcataa----aaataaatatataa
ggaaaataaaatatgacaccatctttatcaagtttttttgaatagtcttattcttggggca
gtaattgtagtcgttcccataactttggctcttttatttgttagtcaaaaagacaggaca
atccggtcctaaaaataacttagaaagatatggagaagcaatttaagatgaataatgaaa
tatttttcatactaataagctctccatatattgctatact--ttagaagaattttattt
ttttattttgtaacatcctgtaaagtttattccgttctttctttttccc-atttactctc
ttataattaatgatacaaatgtcagg-ttaataatag---ttttttattctgtatctttc

---------------attttatatacgtattattatgttcaattaataaagaaaagctca
tgtctttgtctaattggcctctcaaaaaagaaaattctgaagcatataatattaagaatt
caaaacaaatcacaattcctgatggtttatggataaaatgcttcgactgtggtctattaa
tgtattctaaagtactgaagagaaatttaaaagtttgccctcaatgtagttatcattttc
aagcttctagtaacgaaagaattgatcaattgatagaccaaggtagttggcaaccaatgg
atgttcacttgatctctacagatccattaggctttaaagaccaaaagctttatagtcaaa
ggttaaaagatactgctttcaagactggcctgcaagacgcagttcaaacaggtactggga
ctatgcaaggtaaaaaagtatgcttaggtattatggattttagattcatgggggaagca
tgggatctgttgtaggcgaaaaactaacaagactgctagaaaaagcaactcaagaaaagt
tgcctgcaattatactttgtgcatcaggcggagctagaatgcaagaaggtatgttgagct
taatgcaaatggcaaaaatttcttctgctctagaaatgcataaaaaagaaaatctactat
atatatctgttttaacctctcccacaacagggggtgtaacagctagttttgctatgcttg
gagacttaattattgcagagccaaaagctcttattgcatttgctggtagacgagttatag
aacaaacaatcaaagaagacttgccagataattttcaaagttcagaatatttatttgaac
atggtttcctagatttaattgtatccagaactcagcttagatcaaagttaatacaaattt
tgtccttacataatcatagtaagtaatgaattgaatattgactataatatcaaaatgcta
gtaaaaattaagatattaagaaattttcactaaagatatttgtagactaactagaattac
aattactattgtagaaaaatttgtcttaatgcca-tttttcagtcttttaacaaattg-
tttttatgactttaaataattgttagtttaatacaaatttataaagagaata-----cat
actaagctaaatatagtaatattatgttttcctataatactgacctatcaaatttcgagg
aaatttatgcttaaaagatcttcttggcttgcggctttattgggactattaacagtagtt
tctacaagtacgcatacatatgccatagagttagacgaggcaacaagaactgttccatta
gaatcttctggcagaactgtaattcttacaccagaacaagttaaaagaggcaagcgatta
tttaataattcttgtgctatttgccataatggtggtatcacgaaaacaaatccgaatatt
ggacttgacccagaatctttaggattagctacgccacaaagagataccattgaaggacta
gttgactatatgaaagatccgactagttatgatggtgcagagtcaatcgcagaattgcat
ccaagtattaaaagtgctgaaatttttcctaaaatgcgcaatctaaccgatgaagaccta
tttacaatcgcaggtcatatcttacttcaacctaaaattgtttctgaaaagtggggcgga
ggaaaaatttactattagaacttcaaagacctaaaccttgtgaatactagtttatgta-t
ttgttgcttaagtgtgattattttgttagatcatgatatatattaaagatggagtttgtt
gacagcaagttacttttacccgaacttgtttacactattcgtctagttattaaaaagga
gacgttaaattgaagaagaagctttcagttcttttcactgtttttagttttttgtaata
ggtttcgcacaaattgcttttgctgcagatctagataatggagaaaaagtttttctgct
aattgtgcagcatgtcatgctggcggtaataacgccattatgccagataaaaccttaaaa
aaagatgtacttgaagctaatagtatgaatactattgatgctattacttatcaagtacaa
aatggtaaaaatgccatgcctgctttcggaggtagactggttgatgaagatattgaagat

gcagcaaattatgtattatctcaatctgaaaaaggttggtaattatacttgattttatcc
tgtattaaagaatagacaatctatttagttgttcattagattgtctattctttgtttatg
---ctattatataaagatatttacacaatatt----ttatgatgaaaagaatacccgcaa
ttcttgtactagaagacggtgcatattataaaggatggtcattccagcaagataaacaag
agattactattggtgaagtagtttttaatactggaatgacaggatatcaagaaataatca
cagatcctagttacttccatcaaattgtcgcttttacctacccggaaattgggaatacag
gtattaataatcaagatattgaatctcacagtattagtattaaaggacttattgcaaaaa
atatttgtaaaatttcaagcagctggagagagcagcaatctttagttaagtatttaagta
gtaataatattccttttattttcggaatagatacaaggtctttaacccaatacttgcgtc
aatttggtacaatgaacggttgtatctctactgataatttaaatcatagttacttaaaac
agaaaatttgtgagattccaagtatgcaaggtttagatttaatcccgcatgtaactacaa
gaaatgtttacccctgggatgaaaaaagttttccgaattggtatttgacagataacatta
gagtacaccgagttattcagttgaaagttattgttatagattttggagttaaactaaata
tactcagaagactagctacacttggatgtcagataacggttgtgcctgcccacactcctt
taaaagatattttggcttaccagcctgatggtatattactctctaatggtccaggagatc
catcagcagtacattacggcatccagacagttacaaatttactagattacaatgtgccta
tatttgggatttgtatggggcatcaaattttaaatttagctcttaaagctaaactttca
aacttaaatttggccatagaggtattaaccatccatccggattgaaccagcaagttgaaa
taactagtcaaaatcatggctttgcagttgaattgacttcggtttttgaatctcctgtaa
gagtgactcattttaatctaaatgacactactattgcaggaactggacataatcaaagtc
cttatttttctgtgcaatatcatccagaatcgagcccaggccctcatgatgctgattatc
tattcgaaagttttatagaaataatgacaaagtccaagaataaagttag-----------
ttagtaattttcccatgctttatgagtgaataaagcggatagtacttgtactcctcgtaa
ttgattaaagagaggcaaatgtcctctggggccatgagaattccattgaaattcctcagg
atatctgcaaggaattctatcaacttcccagccgatttttgccagaattttccccaatc
ttttccaacactaagccatatttgtctttgaacaaatagcccaaatttaccttttgagtg
tgtatgccataacttatcaatagtttgtaaatcttgagcaggtatttttttatatctgt
aaaatatagccaattacgagtttgtgcatttacaccagctagttgaataagcttttgttg
agttagctgatctgctttgagcaaatcccgatgagtcagtaacatttgtaaatctttgta
attcatctgttgagcagagcgcaatggaacaatgccatcgggacataagttagaagcaaa
tttcactatttcttgatttttactgttaagtaacttttcataaattaaaccgtcaacaca
attactttatagttaggcccagtaatcctctcaaaaaatagatctgctaaatcttttaa
ctcgatagagtcgttgctatttatattctcaataatttcaagttgttgcttga---catt
attagacttagtattcttattcaattctaaaagctgggctcggatttgatttggcatctg
atttgtttaattagttttaatagtc----ataaggtcttagagagaattagattgttttt
tgaaattagaagagcgccgcagtatttctaaagaagttcgcataatttgattcaataagt

tttgtcctgcataaattataatttgaccaattaaaaaaattgtgctttttattttaggaa
caaaaaaatcttgtacctcaagcagaaaagtaattattatttgcattggtgacaacatct
gtaattcattcgatctgcaagcatatatatactgacagtttaagccttcttgataaaatc
cccatactttgtaacgactttcgtaaattgctcgtggtcgctcaagatagtttttaatta
tagtattccaaattaggttattctttaatttttctaattttctatcagatagaaatttta
cattacagagataacttgcatttgtctcatttacagtattaaaattttgaactaacatat
gagcaataatgttgcttagctgaattaaataattttcaagaaaaatttcgacctgtttaa
ctggcatattgtaagtaaagagattaaaagttttattcacagtttgagatgacccaaaaa
gtagctgaataattaaaatttgtagtagcattttatagtcgttcagtagatagttcaaat
catgaatatcttggtaaatgtcaatacagtccagttcatataacttacaaaatcttttga
tacacctatggaataaatcaattaaaatattttcggtcaagtttgtaatatcatttacat
ctaaatttgaattataaatttctaaaagtatcttttccagttcagctaatatgattttta
ataagtttctttttacttcagtgcgaaaaacatctaaaattaacacttcctgagagcaat
tagttaatctcttattaatctttatagaagttctaactaatagctctgctacctcttgat
tgagtgttggtccctgagagcttggccaataattattcacgttataccatagtaagaaat
attgaatttaatgttatacgataacttttttaatatactgaaaagcaagttaattactcta
tagcaactag-------tttttacagatcttaatagttatttatcaaataaagttatatg
atagtagcgattgagctaattttattaaaattta----taataatgacaacttattactt
cgcccttgcgagtcaaaattttctattagtacaagaaccactagaagaagtttttagaga
gagagttaactactatcagtcaaataataaagcaattgatttttggctaataccaaaccc
ttcttttctagagaaaccagaaatgatttcatttaaaaaccttgtacctaaagacgctgt
agctataatctctactaatccaatatttattaattggttaaagctaagaattggctatat
ctgtattggacaatttgaagataacctacaactttctgaagaatcgttaaatattactgt
tttaacagacaaaatttaacaatgcttttattatgactagtgttaaattattttggttaa
gttataattatagtcttttagtattta-gactcttatagtttatcacttagtattttatg
ctgagatggagtgacaaatagtgtaagtatttcttggctaaacgtttcagttgccttaga
tttatatctatttggattaactataatggatagcattcgttttatagtaacattttcaat
ttgagcccaatgtacaattccaagttccaattccttagcaatagctgaaacggaaacaaa
ggcagctcctagccctgattgcacagcgtttttgatagcttctatagaattcaattccat
ctctatttttaaagcgactgctgtcaataccatgttgactgagtactttatctatcacttt
tctaattgttgattgagtatctaatgcaataaatctaagcctgtataagtcttctttttg
aatgtctcctagtttagaaaaaggatgtgatttgggtaatataagtgctaattcgtcttc
cgcataagaagtaacctgcaaaacatcttgcaattcagtaggcacttctcctccaataat
tgctaaatcaacttgaccattagctacgctccatgaaataagtctagttgaatgtacttg
caattgaacagccacttgtggatatctttgcctaaatagtccgattaatcttggcatcaa
atatgtcccagttgtctggctagctccaataattaatgtgccaccttgtaagttttgtaa

gtcgtcaagagctcgacaagtttcttcgcagagagctaaaattctgcccccgtatcgtaa
aagaagacttcctgcctcagttaaagttgccttcttattacctctttcaaatagggaaac
atttaattggcgctctaaattttgaatttgcaaactaatagctggctgagaaacatacaa
gctattagctgctttttttaaaactgccctctttggcaattgcttttaatattcttaactg
atccaatgtaaatggaaggtctgtcattaaagaattgtagtataatg---------tata
ttatttagtattgatcaatgtatgcatttgccaactaaaatatttgtcattacgcttcat
tattttcaaaataataaaaagtataatgttaaataagctgtattaaaatttagtatctgt
ttgatattaaatttatacaagagatatcgtgaatttaattctaactataaaatataggag
acaat---atggactctagacttttagttgtactaataccagttttagcagcagcatctt
gggcagtttacaatattggtagagttgcactacagcaatttagaaaaatgacatcttagt
tttgcttagaatata-atttttagactttatgagatagaggactttaaaatccctatctc
tattgtatatcttaataaaaaaag-aaagc--ccccttgtacttactgttattacacatt
aatattattggt-ggtaccta----gataataagtttaagctaacttatttttttattta
taattaattaaattaataacacaattatggctgttccaaagaaaagaacatctaaagcta
aaaaaaatgcacgcaaagcaaattggaaaaatcaagcaaaaactgaagctcaaagagctt
tgtctttagcaaaatcagtattaactagaaaatccaatggatttatttataatctaactg
aagcatcagatacttttagcgattagaagtgttt-ttactacaagtctgaaaaagtatta
cgcaacaagttctcagtaatttttatctat----taatcatttgaaaatcattctatttg
ataataaaataaatagcagattaaataaaatcacattaacaaattatgcagcaaagctag
atcaaactagtgaaatttggttgttcaattgtattgaaaatattcagcatattttttta
aaagtcaattaaaatcaagtcatattactaaaatttttatttctggtactagctttgaat
ataccgcaggtctaccaggattattatccagcttaacactaagtggtagactacatccta
ttagtatatatagtccccagtctctcaaaaagtatcttgaagcatgtaccaaatattccc
aaactaatttttcttttcctattaattttcataatttacaatacggaggacaagttgtta
accaattttatacagtaatttgtttaccgttgagcaaaaagagcctgctgtatggattta
ttattctaaaaaaagaaaagcagggagtatttaatttagcacaagctaaaaccttgaata
ttcttcaaggacctatatatggaaaactcaaagaaaaggataatttttaagtccagatg
gttactatctatccggccaagacttctcttctaatacaataatgggacataaaatatcgc
ttccgttattagtcagatattctagaattatttctgagatgcattggttttgctcttatc
ctattagattaaatacttattcgcatcagcaaggagtaaagtgtttaccacataatgttt
taactgatattatgaaatcccaaatatatcaagataacagttttgttgaataatacttat
tcttgatatataatggtctcggtaatctaaataaaatactagaaataatttcttaaattt
atttgtctatattaaatttacaatcatttatgacatacgcaattattgaagcaagtggca
cgcagctttggatagaagaaggccgctactatgatttaaatcatatacctgttgatccag
gtcagtcgattatattaggaaaagtcttattattgaataaaaatggggaggttactttag
gacgcccttgtatagaaggggttacgataaaggctacagtaatgaggcacttgcgaggaa

agaagataactgttttcaaaatgaaaccaaagaagaaaatgagattaaaaaaaggtcatc
gacaagaattgactcgtttaatgatcgattctataacatcttaga------taaaatcta
gactcatattttt--aa-aaactttttaatatttaactaaaatagatataatggcacata
aaaaaggtagtggtagtacaagaaatggcagagactctaattccaagcgtttaggtgtta
aaaaatatggtggagagcaagtaacagcaggtaatattttaatcagacaacggggaacta
aagttaagcctggccaaaatgttggaaaaggaaaagatgatacattgttttctctaattg
atggtttcgtgctgtttgaaaagtcaaatcaaaagcaaaaaacaattagtgtttattctg
ctaagaaatagttaaaacaattggtgcaagataacttgagt----gtatttatttcactt
acagtggcaccatatatttgatttattaagaaatggagttccttaatgaaccaatgaatt
ttattaaagatttgctgtgaaatttaatgacaaacactattgtaatttcctgtctacaca
atatggctgctattctatattgtggtcagatacagaaactagtagtagcaaatgctcatt
atcaagtaagtgatatctacctaggttgtgttgataaaatcttctcaggaataaatgcgg
catttattaacctaggaaagaatgagtacagtggttttatacatatcagtgataccggtc
cgcttaaaaagaaatattatgtcaataatattactaacattttaacaatacggcaaaaaa
ttttagtacaaatcattaaagagccaactttaaataaaggtccaaggctcactgccaata
ttacattatcaggtcgatatattgtattaatgccttttagtcaatcaatctgtatatctc
gaaaaatatatgatgaagatgagcgtcattatttgaaatctttagctattttaattaaac
cggcaacaatgggcttgctatttagaccttctgctgtaggtgtcgatgaggaaataatat
taagcgaattaaaaaatctaaaagaacaatggaactttgttcaaaaatctgcaattaata
gttattcacctgttcttctatataaagatgaagatattgttaaaaaagtaatccgagatt
tttataataataatacaaacaatatagtaattgattcaaacctgggattaaaacaattaa
attattatatccacacttggcactgtaataactctagcacagttcctaagattaagcttt
atagtaataatcaatgtatactagatgcttttggtatcaatcaggcaatttccagagctc
tcattccaaaagttgatcttatacttggtggctatatgtttattgaaactttagaagctt
ttactattattgacgttaattctggatcttttaataattccactagtgcacgagaaacag
ttttaaaaaccaactgttctgcagcaacagaaatagcttatcagttacaaattagaaata
ttactggtgtaattataatcgacttcattgatatggaatcacaaagagatcaattgcaat
tattagaacactttaataaagagctatcacttgatgatgctaaaccacaaattgtacagt
tatctgaattaggtttagttgaattgactagaagaagacagggcaaaagtttgtacgagt
taattagtagtgattctaattactttatttttcacacaatcagagagatctcagtctc
ttaagagattcgatgatagacagcagaaacaacagatttttaataaatcttggctatctg
cagagattaatactattaacaaggtctttttcaaaagtcaaatttgtgcagacctgcta
acttttacctaattcgtaatctctatatagttaaaagtagtattacttataaacaaaatt
atttattaactcacagatcaaaacttatttattctaaagaatacagcaaagtattaccaa
gca------gttattatctagctagtctcaataagaatagtaatcaagagttcctatctt
aagattgttatatcaactgtt---------------------------------------

-------------------gaaaaaaagaagctctctcttatataagagagagcttc-tt
ttatagcttactaacta-tctagttagttaattaaaaattaaccgttaacagctggagct
gttagagctactggtaaagattcaccagaagctagatctagagggaagttgtgagcgtta
cgttcgtgcattacttccatacctaggttagcacggttgatgatatcagcccatgtgtta
attacacgaccttggctatcaacaacagattggttaaagttaaaaccattcaagttgaat
gccattgtgcttacagataaagctgttagccagataccaactacaggccatagacctaag
aagaaatgtagagaacgagagttgttgaaactagcatattggaagattaaacgaccgaag
tagccatgagctgcaacgatgttataagtttcttcttcttgtccgaatttgtaaccatag
ttagcagattcgttttcgcttgtttcacgaattaagctagatgtaactagggatccgtgc
atagcactgaacagagaaccaccaaacacaccagcaacacctagttggtggaatgggtgc
attaaaatgttgtgctcagcttggaatacaagcatgaagttaaatgttccggagatacct
agaggcattccatcagagaaactaccttggccaattgggtatactaggaatactgctgct
gctgctgctactggagcagtaaaagcaacggaaatccatggacgcatacctaggcggtag
cttagttcccactcacgaccaatgtagcaagctacgccagttaggaaatgaagaacaact
aattggtaaggaccaccgttgtatagccattcgtctaaagaagcagcttcccagattggg
tagaagtgaataccgatagctgcagaacttggaataacagcaccagaaatgatgttgttt
ccgtatagaagggaaccagcaactggctcacgaattccatcaatgtctactggaggtgca
gctacgaatgcaatgatgaatacagatgtggcagttaatagagttggaatcattaataca
ccaaaccaaccaatgtataggcggttttcagtactagtaatccaagagcagaaacgttcc
cacaagctagcgctttcgcgtctttgtaaagtagcagtcataatttttatcaattttta
aggatttaacaaagatgcttcccaaacaataagttgtctgttatgaaaattattacttaa
ggccttttcaaatgagaacatttttaaaaaaaaca-tagcaaaatctaatctcagtaact
aa-aattt-tacttgttcatatattgcttaataattcaa-----------taataaattt
tt-gct-------------ttttgcaaaattaacactattttagggat---ataataata
atgcatattaatgctgcttattttgattttaatagtttatataattttgccctacttaaa
tacgcatgtcacactttacaaaaattcagacgaccatacaagatttaaatctgttaaagc
atgctttaacagatttgggtttgatttggcaaatgaattctagtcatataaaagttggtg
aaaataatcagcataaggtagatattttaattaaacaagataacttgtcacatattggct
ttacttggaatgataatagatatcacttagttgccgacttacaactttgggaacaaccct
ggtctctagaagtatttttggataagttatctcagaagtatgcttactactctatcatcg
aagaaacaaagaagcaaggctttgaaaaaatgcagcagatttcacaaaaagatggatcaa
ttaagttaattgtgcaacgctggaattattaactaacatacaaatgcgggcggagagact
tgaactctcacgagattatctcactagaacctaaatctagcgcgtctaccaattccacca
cgcccgcactgttgagac--aatattattctatcataatctatactaaaacacaaactaa
-tagtataaactagcttccaagcttaaaagcgtctacaaatagaccgtatgttataccaa
ttttgaagttgttgaataaattaatttttatatttaattgcttgtggctagagtagacta

ttttactagtcaactcagttgctgcaactattagactggctgcaataataccccaatctc
ctgtttgtccaggaatcgtagacaaacctgttgaaataaaaaatcctagcaataaactaa
ttaagccagttgttaactcgctcaaagaataataaagcttattatttaagtttttttatta
agtaagcaaaaaaagttgaaagtctagttttaatcatagtatttagtaacggagttttaa
aaggatgacaagtattaacag-tcattaatattgtcgtatccttttaaaactcgttaata
caaacagctgttcttgt---------ttttaaaggtcttgttcgcttaaagcattaataa
gatgctgaaacggctcttgtataatatctgtgcgtttaattgattgcaaatctaattctt
cttctatgatttcttgtaataactgtatgcttcttactgttggagcaacaggtacagata
gtgaattgtaagtatcttttagcccatttaatacacgctgatcgagaatatttgtactac
cagaaaccaaggcataactagcataccgtaggtaatattctatatcccttagacaagctg
catatcttcttgttgtataagaatttcctccaggccttaataattcaggctgttcttcgt
acaactgagctgcggcctcttttaaaatattggtagcttgatcattaataatttcagcta
tttttattcgatctagtccactcgaaaagaaagattctagttgtccgacagctgtcttat
ctagataacgaccagtaagatcgtaacgatttaatattgctgttatagcatcttgcataa
attaattctcctac-ttgctatcagagcggtattataaatgactaatattgtaatataat
cattagtgtttatgaaatttatttaaaacttattctttattgcttaatgtgcaattaaaa
actcgcatagattttacacatatagagtatacgtagtttttaatta-ttttttataaaa
accaagaaataaagctttttt-taagaggttcgtgaatttgttaatataacatgagcttt
cctcctaaataagttttgtaaagtac--aaagtaaaaaaacgattagtcaataatagtaa
------------------------------------------------------------
------------------------------------------------------------
------------------------------------------------------------
------------------------------------------------------------
------------------------------------------------------------
------------------------------------------------------------
------------------------------------------------------------
------------------------------------------------------------
------------------------------------------------------------
------------------------------------------------------------
--------------------------acaatattaatatgtatcaa--------------
------------------------------------------------------------
------------------------------------------------------------
------------------------------------------------------------
------------------------------------------------------------
------------------------------------------------------------
------------------------------------------------------------

------------------------------------------------------------
------------------------------------------------------------
------------------------------------------------------------
------------------------------------------------------------
------------------------------------------------------------
------------------------------------------------------------
------------------------------------------------------------
------------------------------------------------------------
------------------------------------------------------------
------------------------------------------------------------
------------------------------------------------------------
------------------------------------------------------------
------------------------------------------------------------
------------------------------------------------------------
------------------------------------------------------------
------------------------------------------------------------
-----------------agtataaataagcttttactccagcatcaaaaaaaatatttga
gatatgttgatacaactgattattcaagtgaaaacttagatttagaccaatttagctatt
ttatcattacaattaataaaactgataaatgtattcttatagagcaattttgttataatg
ctgaaggccatatttactcaatattcttcaaaggtccaacagctaaacatctttcttcta
taatttgtcattttcaacaattaaatacaataatatctacagctcatgccatatatttag
gacgagagctaatgaagtcagaattagctctcgtgctagatcaacaatatattcaagatt
aaa--ttttgtaagtattaattatatttatgctagtataatattagtaagggcatgtaac
tcagtggatagagtatcagattccgattctgatggccgtgggttcgaatcccgccctgcc
c---gagtactcagagtact-atactaaaacatct-------taatagctagtttgataa
tagtatgggcattggggctgcaaggtttctacattatgaaaagaagaaaatatg--aaaa
aaacaagctctccaaaagagcttttagtacaatcaataaatgcagaaaacaatattgttt
ctttttctcgaaaattagcttttgcttaaacactgttaatttttggcattcgacatgtta
aactcttattcatggcgaatattctgtcagagtcgctcttagtttaagaaaagtcagaaa
aaatatttatctttacgtttt-tacctatttttatttaaataaaggttagta-ttttcaa
cttttatagtggacgtgggttcgagtcccaccagctccatcaatttgatt---------t
gtataattagctagtttaacctttttaataaccagatatataattggaggaaatatgggt
ttcataacaattactaagcccgctttaaagcaaattgcaatcttgaaaaatgatcatgaa
aatgatgtacatctaagaataggtgttagacaaggcggatgttcaggtatgtcatactca
atgaattttgaacatgttgacaaattaaaagatacagatgaacggcttcgtctagataat
ttttctgttgtctgcgaccctaaaagtcttctttatctttatggattatcattagatttt

agctcggaattaataggaggaggatttcaattttctaacccgaatgccagccaaacttgt
ggctgcggtaaatctttttcaggctgacc-ctcgaacacgtttttatatattttgtatt
atatatattattttagcaagtcg-tttattagagacaagcaatgctctgttaaaaac---
----------------------------taatttcaagagttttattgtattatt-----
----ttatatatttattatcaactacttgaaaataaattagctaagcaatctaatttgtt
atcagtaccaaactttaaatgattaacttttattgatatatgctaatcttt-atcagcc
taaacatatctgtacattacaatcttttgacataaaaagagtcttagcatacctctcaa-
----------------------aaataaattcaa----------tattttataaatactc
aacatgaatctgtcacaacttatcgttaattcttctatgatgaaaattcctaataacttt
tattgcaatataaaagctatttcttctgagaatttaagtcacttacaattatcaaaaaca
aatttattatctcctacaaaaataattgcttttggactacctaaatttgattttatttct
acacagattacaagttctattagagaaacagttgttgtgtctactattgatgataataat
ctttttgctgccccaactaatgattggataaaaaaacacaaggataaaaatcatcattct
ttgttttataaaaaagtttttaaagctttgctaaaccataaaattagagttgttgctgag
ccact------tcataactccgaaataaataaagaaaagagcataatggcaagtcaaaga
tacatctggggtaaaagctggaagccttcaattattttatcttgcttaaaaaacaaaaaa
aatagcaaaatttctgatgcatcgaaaaaatttttaactgaacagttatctgcatcacca
gtatttgtggtaaaaaatggttttaatgaaattattttaggtcacccattatctcgtgtt
aaaagaggaagagttaataatctgatgcatgcgttttctaatttattaaatcagtctaat
gcaacttatcctatatctactggtctattctttttttcatccagatgatgcatttgaattt
aaagactttataatatcggtcaatccactggcggccaaacatatggagataagcgtagag
cctgttggtctacactttgcttataaaatgaatagaaatatatcgtcagatactcaattt
tcgtttattccagattttaaagaagtaggagatttattattcaaatatagaaaaggtaat
catttagtttttcataagaatcagcattatggcaaagatttttttcaggggcaaccaata
tatatgattcagccgatcacttttaaagatcgagccggaaagctaaatattatcaaattt
acggggctaaatgacacaagagaaattatttttactaatcttgaagctgctaatagatca
tgggcacattttataaagaggaattcacaattaaaatcaattaaaaatcctactctatta
gtctataatttagaaagtttttttaaaagatcaagaacgattggataatcaagatttaaat
aagtttgtggtgatcactaataaagaatcttatcttgccacaaaagagttaatagcttta
cccgattctaatagcttttctaagcacttaaaattaaatataaagcctaaactcttttt
gtgaaactatgggtaagacggttattttccaccttaacttatgaat-aaatgctatt-gc
ctttattttctcgaataatattctataactagtaattcattgagttgtaaagcaacccat
tccctatcaatgactccattgactttccctgataaattggatttattcaattctaagtgg
ctaggaatattagctagtccagggaaagctaagtaattttctactagttttcgagatgct
tcttgatttttacactaattgattctcctggtttacattgataactacagatagatact
acttgtccattaatacaaatatgaccatggttcactagttgtctagcggcaggaattgta

ggagccataccaagtctaaaaacagtattgtctagcctcatctctagtagctgtaacaaa
atttgacctgtcgacccttgtagtttttagctgcttttacgtacttaaataactgtttc
tcacttaatccataattaaatcgcaacttctgtttttcttctaatcttacagcatactca
gaaggttttctggacttttgtccatgttctccaggtggataagatcttttaatcgctttt
ctactgagtccgggtaaatcacctaatctacgggaaatgcgtactcgtggtcctctgtat
ctagacatattatttaaattctcctaatgtaaat-gattaag----------taatgagt
agctatttttaacacaaaaagagcattaaaacaaagc-ttttgctaaaaaatatttaca
aaaaatataggtttggacaaaa---tacattactatgc-tttttaagcgggtagcgggaa
tcgaacccgcatcattagcttggaaggctaaggttttaccactaaactatacccgcaata
ttcaacagattgtcactaactcaatataacatagttatgttaagttaagataatcttaaa
tttaatagaatt-ttaaattatttatgctccttctaaaaccacatcaagaaaacgagcaa
tttcggcagcttgttcttcaacttctgataataataatggttgaccgaccctagttaatg
gaattactcttttatcttttgtacataaataaatttcacgtcttggatttagcccctctt
taatatctatttttattgatttgatttctttatattaaactgaagacagatttttctat
tctttccagggaaacctagccgaaaaatttttacaatacctttatctttgttaaattcat
tatatccagcccctatattccaaataatagttagccacagaaacaaactcagaaagactc
ctatactgccatagaatgtcattacaataccttgtggaataaataccaaatcggttgaat
ttgtaaaaggtaacaaatcaacttgaaaataacttgatagtccagcaagaagaaagccca
aagccccaataaatattatagtagcccaccagtaattactcagtcttcgtgaacctaaaa
ttaaatcttttctaactgagtgtatagataatgtttttttcataatttgttcttttaaaa
cagta-tactaatgcaaaatggaaaaattttgacaattttt--atgcgcgctttagattt
tttaaataagcttgattgcttttagtcctaaaaacagtcctgttgctaacccaaaaaaag
cagcaagaaatattatatatcctagaaaaactgaca---tattattatttattattattg
ttacaatagtgtttacaaaaatagtgaaaatctattactacggtattatattacgttaaa
tgccttgctgtctacaggaaaaaactttttcagtttttagcattgtgtttatattagaa
ataatattctagtaaatttattaaacttctggagaacagaatcttatgtcagaatttatc
aagccttataacgatgatccttttgtaggcaatttgtccacgccagttagtacgtcaagt
tttagtaaaggacttctaggaaatctaccagcttaccgtcgaggtttatctccgcttctt
agaggattagaaataggaatggcacatggatacttttaattggaccttttgataaattg
ggccctttacgaggtacagatgtagcgttactagcgggatttctatcttcggtcggcctc
attattattctcactacatgtttatccatgtatggtaatgtatcttttactagagcagat
tcgaaagatccactacaaacttctgaaggctggggacaattcactgcaggatttctagtt
ggagcagtaggtggttcaggatttgcttatttattgctagctaatatacctgtattacag
actgcaggtcttagtttattctcttaagctagtaaggggacttgaacccgtaacctactg
attacaaatcagttgctctaccaattgagctatactagcattcagttcataataacatat
aaagatatttgttgcaatctgctattttcagatgtaacaaatatctttttttatttaaaca

gaactgttatgttaacac-------gtatattaattagaatctgaattatctgactctgc
agaagattcctgatcacagtttagataataacaaattgtttgatctagacaagttgcaga
ccaaccaacaggtgtttctcctgataattcttgcatatctacagaattataaaaattatc
atttggcttattactgtttgcttgcataaatatcactcccaaaacactctgaacttaatg
tgaagtaatttaacttctcaataatatattaacacaaaatattaaatatggtgattattc
tttaattgtaaagatgctgctaactttataaagtttgagtgagagtcttgatagcttttg
ttgaaataagcattttaacccatttattttgcccgctcgaccatactttttgtgttgta
aattaactttttgtagttttttgttcgtttatgagaatgagatacggcatatccattat
tagctacttttcccgtaagctgacactttttgacataatttt-------ctaacttata
tatttatttagagtacttgctaaagttgacttgggcacagctccaatcactgtgtctact
ctttctcctgctttaaagatcattaaagtaggtatacttctaatgccatattcagccgca
atagtaggattatcgtctgtgtttatttttactaccttaatagatgattcatattcttcc
gcaatttcatcaactacaggagaaaccattctacaaggaccacaccatggcgcccaaaaa
tctacgagtacaggtaagttattgttaataacttcttgtttgaaagaggcatctgtaact
tgagatactgacatattctttaacctttaatatatgttccttgctaagcaaatcttatca
caaatttcaagaaaaatcttctatagtgaattactattagtgttgaattatttttttaaa
aattagacattaaaaaaacttattacctatataaataaagtggtggccgagtgatatctt
gatagtgataatttttatattaaggttgaacactagaaataactttcaagcaaactctt
atttgaaaa--atttctctgtgaccataaaaacttaattatattacaatatattttatta
tttgtaacttagttctgataatggtataaacaacgcaaaagatactgccttataatcaag
gaggaatacatgtctcaatccgtagaatcacggactaggattaaaagcgaacgttacgaa
tctggagtaatcccttacgctaaaatgggctactgggatgctgactatgtgattaaagaa
acagatattctagctcttttcagaatcactcctcaaccaggtgttgacccgattgaagca
tctgctgcaattgcaggtgaatcttcaacagctacttggacagttgtatggactgattta
ttaacagcttgtgacttatacagagcaaaagcatatcgagtagatccagttccaaacgtg
gcagatcaatattttgcttacatagcttatgatattgatttgtttgaagaaggttccatt
gcgaacttaactgcttcaattattggtaacgtttttgggtttaaagctgttaaagctctt
cgcttggaagatatgcgtatgccagtagcttatctaaaaacgttccaaggtcctgcaact
ggattgattgtagaacgtgagcgtatggataagttcggtagacctttcttaggtgctaca
gttaaacctaaactaggtttatctggcaaaaactacggaagagttgtatacgaaggcctg
aaaggcggtcttgatttccttaaagatgatgagaatattaactcacaaccatttatgcgt
tggagagaaagattttatattctatggaaggtgtaaataaagcatcggcttctgctggc
gaaattaaaggtcattaccttaacgtaacagccgcgacaatggaagatatgtatgagaga
gccgaattctctaaagaggttggtagtatcatttgtatgattgaccttgtgattggttat
actgcgattcaaagtatggcaatttgggctcgtaaacatgacatgattttacatttacat
agagctggtaactcaacttactctcgtcaaaaaaatcatggtatgaacttccgagttatt

tgcaaatggatgcgtatggctggtgttgaccatattcacgcaggtacagttgtaggtaag
cttgaaggagatcctttaatgattaaaggcttctacaatactctacttgaaagcgacaca
gatatcaacctacctcaaggtctgttctttgctcaaaattgggcttccctacgtaaagtt
gtaccagtagcatctggtggtattcatgctggtcaaatgcaccaacttcttgattactta
ggtgatgatgtagttcttcagtttggtggtggtacaattggacatcctgatggtatccaa
gcaggtgcaactgctaacagagtagcactagagtccatggttatggcaagaaatgaaggc
cgtaactatgtagcagaaggtccacaaatcttgagggacgctgctaaaacttgtgggcct
ctacaaacagctttagatttatggaaagatattagttttaactacacttccacagataca
gctgatttcgttgagactccaacagcaaacatctagtttaatgactacttactgatactt
taaatagttaattgtaagtgaaattaacttataacaataaggagcatagaatagtgagac
taacacaagggactttttccttccttccagatttaactgatgagcaaattaataaacagc
ttgcttatatcgtttctaaaggcttatcagcaaacgttgagtatactgacgatcctcatc
caagaaactcctattgggaactgtggggtttacctttatttgatgtaaaagatgcttctg
ctgttatgtacgaaattagctcatgcagaaaagcaaaacctaattattatgttaaagtta
acgcttttgataatactagaggtattgaaagttgtgtaatgtctttcattgtaaatagac
ctgctaatgaaccaggattcttattacaacgccaagacttcgaaggtagaactatgaagt
atagtcttcatagctatgctactgaaaagcctgaaggagctaggtattaatattaattct
taattaatattggctaatta--ttcccctcttaaaaatcataattaatttagtaattgtg
atttttaagagggatagcttcccaagtaaattttgactaaa-tagattgaataattaaaa
atacacagaaataatgcaatcacaggatataatttccaacgatactcttgttaatttaca
agaagaatatgatagaacacaaatccaagaagttttaaatgagttaaatcaagaacttat
aggattagtgcctgtaaagaccagaattcgcgaaattgctgcgctattattgattgacag
attacgcagaaaactagaactagtttctggtaatccaggattacacatgtcatttacagg
tagtccaggaactggtaaaactacagttgctatgaaaatggctgatattttgcacagact
tggatatataaaaaaagggcatttgttgacagttacaagagatgatcttgtaggtcaata
tattggacatactgcccctaaaactaaggaagttcttaaacaagcaatgggaggagtttt
atttattgacgaagcttactatctatataaagcagataatgaaagagactatggctctga
ggcaattgaaattttattacaagtaatggaaaaccaaagaaatgacttagttgttatctt
tgctggatataaagatagaatggaaaaattctacgaatccaacccaggactctcttctag
agtagctaatcatgtagacttcccagattatacttcagatgaattattacaaatagctaa
aatgatgatagaagaacagcagtactgttttacagaagaagcagataaaactcttttaga
gtataccgagcgaagaatgaaacagccttattttgctaatgcaagaagtattcgcaatgc
tattgacagggctagaatgagacaagccaataggattttttgccagtggagaaaaagtatt
aacaaaagctgatttggtaacgattgaagcagaagatatcttgaaaagtagattattttc
attacctaatgcttaatatacacgtgatttcatgaaactatatattattaaaaag-tttc
ataatcagttgcaaatttcttgaaaaacctttattattatcttaatgtaggtg-tggcgg

catagccaagtggtaaggcagaggattgcaaatccttcatcccccagttcaaatctgggt
gccgcctagtactaaaaaagggggggtgtggtggaatggtagacacaacagacttaaaatc
tgttgatttttagtaatcgtgagggttcaagtccctccacccccatatattgatttataa
ataaaaagagatgtgtatttgtattaattgcaaccatattaccaaatgtaatacctatca
cttaatagaatctcaacataaacagcctcatcttactagaagtccattgtttatacctaa
atatcctgtagttcatgttaatatatctaacaactgtacctataatcaaattgattggga
tttagtggagtgtttatcatttgtagaaaaaccgaatagttggaatttagacgctaatta
gttacaagaaaaattatgtaccataagcaacaactctatttttagatactcggtattaa
ttttggatgttctaaccagtaagattttacccgttcttgctatctctataatgccaaatt
tagttagtaattgttcaatagcaacaatctttcccggatctccagtaacctctataatta
aaagatcttctgcaatgtctactatcttagctctaaaaattcttacaatttctaaagctt
ctgttctagtctgagaattgatctgaatcttaattaacattagttctctttcaactgaag
gaatatttgttacatcttgaacgttaaggatatttactaacttatataattgtttcgtaa
gttgttcaatagttctattgtctccttgaactaccattgtgattctagagaccccaattt
gctctgctggtccaactgctaaacttgcgatattaaagcctctccgggcaaatagaccag
atattcttgacagtactccggcttcatcttgaactaaaactgataaggtgtgtttcatga
aaaattaaatttaatctatttgatctattaagtatattctaagaaaattacagaaaaaat
actagttagtctataactaaattatccaaactaaacaaaatcctactatcagataatata
tctaccttgtaatgctatactattaacttagtatttaatgatattctaattgcaaataga
atgct-aatttttttaaatttgttaggaacttgtgctagaaaaatacaaaaacagtaaaa
aactctctcgagatttacctcaaatcaatgatcgcattagatttccaaaagtccgagtaa
ttgatgacgaaggtgaacaactaggtatttttgtgcctgaagaagctatacaattagctg
tccaacaaggtttagacttagttgttgttagtgataaatcggacccgccagtatgccgaa
tattagactatggtaaatataagtttacacaagaaaaaagagctagagaagctaagaaaa
agcaacataacagtagtattaaagaagtaaaaatgcgatataagatagaagagcatgatt
ataaagttagaataaaccaggcatccaaatttattcaagcaggagataaagtaaaagcaa
ctatcacatttcgggggcgtgaaatccagcactctaatttagctatagatttattgaata
agatggcaagcgatctaaatgcaatagctgaaattcagcaagctccatcaagagatggca
gaaatgtcataatgctcttatctcccaaaaaagttagctaaactaatttttatttaaatg
catctggctggattcgaaccagcgacgtctctttcgaaatggcggattatgagtccgctg
ccttcggcccctcggccacagatgcatcacttaaagtttataactcataaagcatataaa
aatcaatgtttagctatttttaataaagtctataaaagtctattattcattcatattgtg
atatgtcgcaactgctgaaggggaaacttgatttaagtatcgaaatatccagtatttaaa
aattgtgtcgagtataactggaaaagttgaaataaataaaaaaataaaatctcgactttc
cggaagtcctaaatgtcttaaaattacttcaataataacttcccatccatgaggagaatg
aaaacccacaaacatatcagtaaaaagaatgattaaaaaagctttagctgtatcactaag

accataaataatttcatttaaaaaagatttaacgactgcaatttgtctttgtccagttat
cattagtaatataaagacaagtattgatacaagatcggataaaatattttaacagcatt
tgcactttcgtttgcataatattctcccaactctctagccttcagctgcactctttttc
aatgatttcgtaagatatatcttccgaaggattcagaagaacttcaaagtgaattttctc
ttcaaatctttgtaattcagcaaaagctctttcctcttgagaggaatttaaaaaaatttt
aggctgttcttggttccataaataatcaatacaaggtccaaatacaaaaaatttgaagc
ctgattaactagtactggagagatgaataataataaaatatattttacggaagtaatagt
ttggtgtctagatattctaaactcctctatagcttcagattcgccattaggatctaactc
tttacgaaacttttcaaaagtattagtaattgacctaggtattggcccaactttttcaaa
ggcgaattgattatttcgttttaaattccaatatttcatttttattgctgctaaaattt
ctattaatcaataattgatcacgtattcactataaatattaattctagaactaaacatat
aaaacaatatttatgaaaaataaaatacaattaaaaattaattatgcattagttctatct
gcagctacagcagtttatggatataaatcttatcagccaactataaatacactcttaatc
aaagatgaagaaaactggcctttttctctaaaaggcagactgcctttattcctataatt
gtattaaaaaaaaatcatcaaaatccctgaactcagaacctagaaatagtaagttatatc
ttaatggaattggaagtagaaaatttatcgacagaattgtcactctacttggatatttag
aaaaacaggcaaactatactgaatacttgaaccaaataaaaacaagtggttatttttctt
tagtctacctagacagtgatataatgtataagaaaaatattagtaacatttttttactac
ctaatgatattctaaaaagaatctatatttgcaataagaaaaaaaagcttataccgcatt
cattcttaataaatttatttcagaaacagataggctatccaaaaagttttgctaatttaa
attgggccttctctaaaattataaaatggtattatgatagaggataccaatggtctctag
ttgaagttaaacaagcgtctgatgcatcttctattgtaattgatattcatgaaggggtag
tgaagacaattataacagaatactacactctatcttataaaagagtctcaggtatcttat
gcgtagagtcaatagagcagtaccttggagtaagagtgggagctccactaaatataattg
atttacaaaaaaaaattacttacttaaaagataatcaactagttggcgacattatttata
gtattgaacgatcaaataatagctcaatgagtttggatatcaaatttcaaatacaagaat
taaaagataaagagataatagtgcttgcagaaagttcttcta------------ttatct
cgcatgcatgtaatttactcaatc---aatataga---------aataggctagtagctt
ctaatatagtttcactatcaactaataaattacacgcatgctattttaattataaacttg
actatcaatataagtatataaacacctctaatcttataaaattgttagcttattctattt
cttgtaggaagacactacttattcattattttaatttacaaactttaataagctgtacaa
aaaaaaatactattggatttcagctatatttacgaaatctaagttttgggaaagctttct
gtgtgcttagtatgaaatttatcaaaaatggacttaatataaaaattttatatataaatc
cttcattgatagttgatcagaattttgtatttcaatttgctatacaaatcattaagcaat
atcatactgcta------aacctccagctt------------------------------
------tatttttaacgaatctagatcttgaacaatatgttgctgaaagtttattaatgt

accattttacatcttgcttctcaatatctgaaaaaatattattatctcgaattatgcata
cagattctttatttttcaactctgaaa------attttcattttgatgatagaactaacg
cgatcaacagttatgatacttttaaacaaaatacaaaaatattttatcaagagtttcttt
ctttattattaagtttacgttatcaaaattttaattatttaggttggcctttaaaaggcc
attttttgaaataaaatctttatacttggctccatttcaaaagagtg------------
---atttttctgatagtcgtaagacgctattcttccacaaaatgtcgttaaagcaagtat
ctaattttaatttaccagtatcttttaaaagccaccttaaccatattttagttagtacta
ttaaatgtcagtcaaatttgaatatgagaactgtatctttgctattaattgattcacctg
ctgaatatctgctatacaaatctattcttaatttctctattaaagttaggatgcagtatt
tcatacctatgagcaacaatataagattatctctattttataattatttagattgttttt
taataagatcatctcagagttgcattcacatatggcaagatctaagaactcttacaccaa
ttcaaaatttttggttaaaaaaatttcatatggagctggaattcagcttaaattaccaa
ttaagcagatgccacctctttcaattgaatatactgtaactagtagtcgctacttttgta
tttaccttcgtacttattatcaacgataactattactaaaatgaaaaatttaatttcttt
ttttgatcgcagcaaagggaaatggatttcacaaagaacaacttatgaattatctaataa
aaacatgagttcggtacagtctcagatgacaatgaaaataggcaactcattgtcaggatc
tataatacttgcatcgttaaattgggggacatttatagacaagttgcccattacgctaa
aaatcacagccggagtgaatatgataataagtttaacttgcaattcggtaatcaactaaa
taatcataaattattaacgctgtgcatagtgacagatcctagtctaattagttttaaaac
tcgctatggaagtactaccatagatgaaacatattggttcgcaacaaataatttacgtct
aagtactagtattgttaaacgattcaacacttgtgtggcagtatctttttgttcagaaat
taaagtttaa-actctatatagaagcaactaatagcctttta-ggctattagttgcttcta
tgtatctaaattttgtattaaaaattat----aatactatattattctaaatctttacga
ttgggattcctggaaggatcattagagaggaacccaaagacaaacaaagaaataaagaaa
ataacggtagtatatacaaagattttcaaagtaaacatattttcatccttaaaagctatt
agtgcg----atatttacatatctccacataattttatatcatgatataagttccagcac
tattttataaataaatcaatatctttatttttttagtaatgatttatatttaaatatgcgt
gatgtactaaatactaaatcttttccgacttgttttgtcttgtttgtaaatattttgtta
tttctatggagtttaccttcaattatcacatagtctaacttctttagctgtttaaaagta
tatgaagatttttttgttccaaatagttagattgattataactaatcgttttctttttaac
agtcttgcttttaacttaattatttggcttttattttcactaattctgatactcttacaa
ctaagaatttggactagaagagtacagctattcataatttataacaa------tttttta
aaaaatctactaaaatagatgattttacagaactaaaatgatatttcattatcttgttga
ccttgctcataagtaatatctttcaatacttttaatatttttccaaaatactcttgcagg
tatttttctaaagatattacttccaacctgtcaattccaagaatactgtaaacttcatcc
atcggagctgtaaattgctctccgctacttaaaatttcggcaaatgctaaacgatcagaa

atattccatgtccattgaagagttttagtaattcttcttaaggcttttaataatccaatc
ggaatttgtgagatctgtgtttttttgcccagataatttttcacataatttaataatttca
gctgatgtccaagcggtattaccaactaacggcagggtcctattttctgtggaaggtact
ccaagacttttataactaacttggctgcatcttgagtatcaatgtatgcaatgggtgta
gattctcctgtaacccagactgatttcttgtctaaaataggaatggcatattggttgatt
aaaccttgaaaaaatcctcctaaagaaaacacagtatatttcacattggacttctgaagg
aaatctactacctgagactttaagttcatcaatggaacatctgggtatttctctgaattt
aatattgaaaagaaaataaatctttcaactttagccgctttagctgcttcaattaatgca
attttcccatctaaatctattttttctgcattataaggatcagtaggacgagatgtagaa
gcatctatgattgctgttaccccacaaaaagattgcaaaatactttcgggtaattttaaa
tcaccatatataagttctgctccccactctttgagaaaggcagactttcttaaatttctt
accatgcattttacattgtagccttcatctaaagctcgtcttacaatttgacgtcctaaa
gttccggttgctccaattactaaaagagtcatattagttaaaaataagtatgtataaaga
cattgatttggttgacggcagagagtaaattaggtagagagggaatcgaaccctcatgac
cgaagtcgtcacattttgagtgtgatgcgtataccaatttcgccatctaccctattatag
accaactactacaagtaatttagtctttagcgtaaaaataatactaagtttcaatagata
tgtcaacttttaaatttatattaataaaaaaacggagtctcttatggccacttttgaatt
aattccttatagattgtatttgagtcaatctaacacatggatccatagaataaaagcaga
aattaagatatatatagtagcgcttttatggatttcaattttatttttttcttactttaa
actatgtattattgctttaagcttaattgccataagttttactataagaagtaaacagaa
tattattcaaaaacatttgttacaaacgttattaatgacgttcttaactactgttttgtc
ttttagtgtggctattagttataaacaatatgcagaacaagaacaatcgcagtatttatc
tgattccaagaaatataaaaattccagcagctactattatatacaaatagcgaacgatca
aaggatcaaaagacaatacttaattactactttaaagccttccttgtactttttttattac
tatatattctattaaactagttatgataacaacctctccagaagttttagtaattactat
ttatagatctaggataataaataaaatatttaaaaatgaattgctatttatcttccttct
ttcgtcacatattgttactagtattattaatagaatcgataaagtgattcaagtaactag
cttaagaggaagcttgaatttatataattcattgacaaggcctttaatgttttctttgtt
aatatttcaagtcttctttttggagattattcgggagtcaaaagaaatagctcaagctct
ttatactagaaatctcaatcaagaaaacaataatttttttgaaaatatatacagtaaaatc
taactttagtgatcggctcaatataattattagcactttgtactttattattttagctct
agcgtaatagaatccacttattcttctgactctagtaatacgaagagtaataaaattagt
tagtagtttaaatagatagaat-taagatattatataaataattaaatgtttcattatgg
tcaatactcaaaatcaaatttcacaaacttcagaacttgattatatagttaatcaacctt
ataaatatggctttaaaacttctgttgaatctgagcaatttcctaggggaataagtgaag
atattgtccgcttgatctccaagaaaaaagatgagcctgaatatctattgaattttaggc

ttaaagcatataaaaaatggaaaaagatgagtagtccgtcatgggctcatattaagcatc
cgaatatagactttaatacgattatttattacgctgttcctaaattaaaaaaagaattga
aaagtctagatgaggttgatccagaaattcttgacacttttaacaagttaggtatatctt
taaatgagcaaaagcgaatttccaatgtagctgttgatgctgtttttgatagtgtatcta
ttgcaactacttttaagaaagaactgtctgaagctggtgttatattctgttcaatttctg
aagctataagggattatccagaattaataaaaaaatacttaggtactgttgtaccagctg
gtgataattattttgctgcattgaactctgcggtatttagtgatggctctttttgctata
tccctcctaatacagtttgccctttagaattatcaacttactttcgtattaacaacgaag
aatctgggcaatttgaaaggacactaattattgctgatcgtggcagtaaagtaagttatc
ttgaaggttgtactgctcctcaatttgacacaaatcaattacatgcagcgattgtagaat
tagtagctcttgaaggagccgaaattaaatattctacagtacagaattggtatgctggta
acaaagaaggtaaaggtggtatatacaatttgttactaaacgaggcttatgctcgggta
acaattcaaaaatttcatggactcaagtagaaactggatctgcaattacctggaaatatc
caagctgtattttagctggtgaaaattctcagggagaatttactctgtagctttaacaa
ataactatcaagaggctgatacagggactaaaatgatccatattggtaacaatacaaaaa
gtagaattatttctaaaggtatttctgcgggtagatcgaaaaacagttatcgaggcctag
taaaagttggccctcagtcatttaattctcgtaattattctcaatgcgattctttattaa
ttggtcaatcatctcaagctaatacatttccttatattcaagtacagaatccaacatcaa
aagtagaacatgaagcatccacttcaaaaattagcgaggaccaaatttttatttttac
aaagagggatcaatttagaagaatctattgctcttatgatcagtggtttttgtaaagatg
tctttaatgaattgcctatggaatttgctactgaagctgatcgtttactgagtttaaaat
tagagggaactgtaggatgagccagactattttagaaatcaaagatttatatgcttctgt
tggtgaaacaacaattttaaaagggtaaacttatctattcgagccggcgaaatacatgc
gattatggggcctaacggttcaggtaaaagtacattatcaaaagtaattgcaggacatcc
agcgtattcactaataagcggagatattttatttttggacaaagtattcttgaaatgga
gccagacgagagggcaaaagcaggtatttttttagcttttcagtatcctgttgaaattcc
tggagtcagtaattctgatttcttaagaattgcattaaatgctcgcagaaagtttcaagg
attatcggagtttagccccttagaatttttcaactaataacagaaaaaatagatcttgt
tggcatgcaagaaagtttttttgacaagaaatgttaatgaaggatttctggtggagaaaa
aaaacgtaacgaaattcttcaaatggctttactagatagcaaaatatctattttagatga
aacagattctggactagatattgatgcattacgagtagttgcaaaaggaattaacacttt
agctaaatcaacaaattcaattattttgattactcactatcaaagattattagattatat
tattccagattttgttcatattatgagtaacgggcaaattgtaaaaactggtagtgttac
tcttgcccaagatctagaaaagcacggatatgactggattacgcagacataactatttac
tac-------aatattgagactaaagaatattctttagtctcaatatt-tttgcaaatgc
aaagtt-aattatttaagctttatcaaaaccttgcttaacatcttcaattgctcttttta

gcaattcttcactaactggttctaatttttttagtactacgaacattttcaccaaattctg
gtttagaattctttaaatcttctcgaagttctctaataaattcagctacttttgaaactt
caatatcatctagatagccattaattccagtgtaaataatagcagtctgttcttcaactg
gaataggagaattttgggcttgcttcaaaatttctcgaagacgttgacctcttgctagtt
gattttgggttgctttatctagatcagatgcaaactgagaaaaagcttctaattcagcaa
attgtgctaactctagttttaacttgcctgccacttgtttcatggcttttatttgcgcag
cagagcctactctagagactgagatcccaacattaatagctggtctaatccctgagttaa
ataagtcaccagatagaaatatttgtccatctgtaattgaaattacatttgttggtatat
atgcagatacgtccccagcttgagtttcaataatagggagagctgtcatactacctccac
ctaattcagcattcaatttagctgctctctctagcaatctagagtgtaaatagaatacat
ccccaggataggcttctcgtcccggaggtcttctaagcaaaagagacatctgacgataag
cttgagcttgcttagttaaatcatcataaataactaatgttgctttacctttgtacataa
aatattcagctaaagccgcacctgtataaggagcaatatattgtaaagtcgcaggactat
ctgcattagctgcaactataatagtataatcaagagctcctttatcctgtagtgaagata
ctacttgggctactgaagatgccttttgtcctatcgcaacatatacacaaacaacatctt
ggccttttttgattaataattgtgtccaatgcaacagctgttttacctgtttgacggtcac
caataattaattctcgctgacctctaccaattgggatcatagagtcaatagcagttatcc
ctgtttgcataggttcacacactgattgccttccaattataccaggggccattgactcaa
ttagccttgttccattacttgcaggttcacctttatcatctattggccgagctaatggat
caacaactctacctaaaaaagcatctccgaccggaatttgagcaattcgaccagtccctt
ttacggaactgccctctaaaatatccctaccatctcccatgagtactacaccaacgttat
cactttctaaatttaaagcaaccccaatagttttatcttcaaattcaagtagttcaccag
ccatgacttcatcaagtccatatactcgagcgataccatcaccaacttgtaaaactgttc
ctatattggctacttctacatcctgatcatacttttcaatttgttgacgaataatactac
ttatttcatcaggtctaatatttaccatattttaatattattttttattaattggaaatt
ctaaagcggtaattagtcagactaatgctatctgtctttttttactttattttaagttact
acaacatcaaggtgtgaagccatttgtcttagctgtcctctaatactagtatcaatgact
tttgatccaatctgaatagtaaagccaccaattagctcgggctcgactgaaataattagt
ttaacttctttagccttagtcataaccttgatttttttctgttaatagagtttcctgatca
gggcttaaagcaatagaagtattaatatttgcaattgttaaagattccatttggtaagcc
agttccaaatacttactagctatagcatctaacattccaattcgcttgcgatcaactaaa
atcattagaaatgataatgtattttcgttaatttgatcgccaacagttgcaataattact
tctttttttgcctctacagtttttaatggattagct---aaaaaaatttttagcttgtca
gattgcgctaaaatattttgtattgactgaatatcttggctcaccttttcagtaactttt
ttagttttagctaaatcaagaagagctactgcataaggctgagctatttttataacaaca
ttgttactgctcatagttgatctcctaacttagcaatattattatcaattatacgcagtt

gcatttcggagctcatttgattttccagctgcaaagtaactcttttaagagcaagaaatg
ttatttgttgctggatttgccttcggatctgtttttcagctgtttctatatttgatttac
ctgttatggcaagtctttcaatatctaattggccttgagctaatatggaacttctaactt
ttccagctgttagctgagcttctttcttaatttgatctataatgatttgagtttgagcaa
gttgtttttctgattctgacaatctagcacttgcttgctctagtctttcttcggattctt
gtatagctgctaaaacttttatctgtcttgcatttaaacttgatccgagaaattgcttca
aaacataaataagaccaaataaaagcaacaatatattaataacatttgcttcaaaaatat
cagaattaaagccgaatttgtgttcactactatgttctgataaaatagtaattatttgtg
gtatttttactatattattcatttaaaattct----------tactattttaatatttta
ctctggaatcatgattttcaagattgactacttaataatttaactttaatttgatcgctc
aaagtgtcaacctgattttctaaagttttaagagcttcttctttttgaacatttaactgt
ttcgatgcttcagcaatcagtttctctgcattcaattgagctttttttatatcttctgag
actatattttgagcttctttttgagaagaggctattttttaattgtgcattgcggcgagct
tcagatagatcttcttcatattttgcagctagctcatctgctttaacaagcatagacgat
gcagtagtcaatgttgttcgaatatattcatctcgttcgtctagtactttagttacaggt
ttataaaaaatagtatttaataaaaccattaaagtaagaaactgtaatgccattaaaggc
aaagtaccattgaagtcaaataagccaccttcaatttcttcggctaatagaaatggtaaa
tcaatcattttactaattatgtttatagttaataattttgtttcaattaagaaaaataac
tttatagtgcgcttagtctcatcagactaagcgcaaatgattattttaaccaacgtatgg
attagcgaacaataaagacaacgcaacaactaggccgtaaattgttaaagattccataaa
tgctaaacttagaagaagtgtaccccgaatctttccttctacttctggctgtctagcaat
accttcgacagcattagctgctgcactaccttgaccaatgcctgggcctattgcagcaag
acctacagcaagaccggcagcgataacggatgcggctgaaacgattgaatccataatgaa
ttttaatttttttgttatatgagtagaaaattgacaaatttcgtataatacttaaaatcta
tataattaagtataatagattctaatat-----tgtaattatagtatattactcttctcc
atgaccttccattgcttctccaatataagctgcggacaaagtagaaaagattaatgcctg
tattgaactcgcaaataatccgagtatcataactggtaatggtattaaaattggaattag
caaagtaaatacagaaacaactaattcatctgctaagacgtttccaaataatcgaaaact
taaagatagtggtttcgtaaaatcttctagaatattgattggtagtaatactggagttgg
ctgaatatatctagcaaagtagcctaaccctttttttacttaaacctgcataaaaaataagc
taaagatgtaagtaaagacagggccacagttgtattaatgtcatttgttggggcagctag
ctcaccttcaggcaagtgaattaacttccaaggaattaatgcacctgcccagttacaacc
tagaataaatagaaataaagttgcaatgtaaggcacccaagggcgatattcatgctcacc
aatctgatttttttgcaatatcttgcagaaattctaatataaattccataaaattctgaaa
tttttcaggaattctttgtagatttctagtacctagaaatgataaagttaaaagtgtagc
aattactaaccaagagacaatgaaaacttgcccatgcaattgtaagctacctattttcca

atatagatgttttccaacttctactgctgataaactattatatgggttaaaatctatgag
actgttttgatacataataaatattt-ttttaaatagtgcaagaatcaaaaaataag-tt
tttgaagaattaatatgcaattaatttattcttaaaattagactattaataaattaaaag
caaaaataaattctaatagagatagctattttagtagtgattatttttttgtaaaaaaca
attattgctacctataatgctttttacgtttaattatatacccatatattaattatttaat
atgattataggtattttgctacaaactttattgttttatccattct----aaagattgtt
atagtttatttttttatttaaaatatctgcgacttcatctgcaaaattaactttcaaattt
tcttctccccctcccagaagaaatcttacaaatcttctaattttaatattttcacctaga
acagcaatattttggtttattaaatcttcaatactaatatcttgattgcgaataaacatt
tgatctaataatgatagctccttaagtctctttttcattcgtccttcaatgattgtttcg
attctatcaactggtttgtttttcaagtcatctttaccagcttcaactctttgttccaga
ttaataatttcactaggtatatcctgtgttgatacgtattccacattaggcgatgccgct
atttgcatagcaatatctttagctaacttttggaactctggacgtcgtgctacaaaatct
gtttcacagttgacttcaactaatacacctattcttccgccagtatgaatataactctct
agtaaaccttcaatagcggtacgattagatttcttgttagctgaagccagcccttttttgt
cttaatgattctaaagctttttcttcattgccattattagcttgtaaagctttcttgcag
tccatcatacctgctccagtcttatcacgtaaggcttttacgacttgagcagaaatttgt
agtgtcatatgctttcaaaaatttttaagtatttaaatt----------aattattataa
ttcactttgagtaggatgagttatttcttctaaatctgtttgcccgtatttaccatcata
aattgcatctgctattttacctacaattagtttaattgacctaatggcatcatcattagc
tgggattggaatattaataatttctggactacaattagtatcaagtatacaaatagttgg
aatacctaattttaaacactcttgtatagctgttgtttctctcttttgatctacaactac
aacaatatcaggtaaacgagtcatattttttaataccatttagatgtttacgaagtttatc
taattctcttcgtaaaacagcggcttctttttttgggtaactgatcaatcataccactttt
atcttgttcttctagctgtctcagacgactaactcgagacttaatcgttacccaattagt
taacattcctccaagccatctctggttaacataataagaatcacagcgttgtgcttcctg
tgccactataccagcggcttgtctttttgttcctagaaataaaactttttttaccatccga
cgaagcctgttttataagttcacaagcttctgttaataattgtgcagtttggactagatc
aataatatgtatgccatttctttctgtataaatatatggaaacattttaggattccatct
acgtgcttgatgtccaaaatgaacacctgcttctagtaattctgctaatgtaacaatagc
cataagattattaatatttaaataatgtaattattcgggaaatccc-tttgtttagaaac
tgat-----tcaactaacttatttgaattttagtaaactaataataaattaacacacttt
taaagcagatcaacaaaatagatcctgaagaattcat----tctacacttttattgctaa
aataattacgcgcggttctgtcgtctaaaatgatatcatctaaatcatctcgtacagatg
aaatagtactttcgtcattagtatttgcagttaaattctttttctctaacgatgagccat
tacagttatcatacatattgaagccggtaccagctggtataagccgacctataataacat

tttctttcagacctcttaaccaatctaatttaccagatatagcagcttcagttaatactt
ttgtagtttcttgaaaacttgctgcagaaataaagctttctgtatttaaggaagcttgag
taatacctaatagcacaggcctatatgatgcattaatcttatttctcaaggttatggctt
tattagtctgttcaatcttctgtaactctacaagctctccaggtaaatagcctgtttctt
caccattttctattttcactttagaagtcatctgccgtactattacttctacatgcttat
cagaaatattcactccttgagattgatataccaattgtacttctttaacaagtagtaact
gaatctcttgaaagcttaatcgagcagcttcatataaggctaatccacattcaatataca
aatcaaaacttgcatcgaggatatcatgaggatttaacaatacgtctgttttctttcttg
cttcaagaatttcttcaattcttggaagcccttgtataatgtcacctgtttttgctctat
caaaaaccaatatagctaatgtttctccccttctaattaatgcattgttatcaacatgta
aaatagcaccattagaaaccaagtaaggtctagcaattcttaatgtaactgatctagaag
aaatttcaataatttgtccagagtctaaagaagtcatattttcagtaataaaatcaccac
aacgaatccaatcacccacacttactttaactgttgattttcctatattaaaggtctttt
tatcagaagatgtcgcaatcaaaatgcgcctgctagtattcttttcagaataaatctcct
caacagtaccttcactcattgcaataatctcagtactagcaacaactgtaaatggcttga
tatattcaccatttttacaataattctagtttgactttgttgcttctctgatttattta
catcaatacttttaattgataatgtttcaaaagttgacagccttaagtcaaataattat
tgccactttctttagcagaaaactcaattgaggatattaaattatcatctcgatgctcaa
tttcagccacaagattagttgttactaaatgaactccttctatagatttcactctttctc
cgtcccgaaaaggagttcttttaactagctttagtttagtttgacttggtttttgtgaaa
ctagattttcaattagactggattttgtttctggtattgaataaacaataacaggtctaa
taagaatatagggcatttgctcgttttctatatactcccaataaactaacttatctgtag
aaatattattatgaagtctttctccaggtcttaaaaaacctctacttttatcattattac
tatacacctcactaagtttatagatagatccaggcttaataataatttctcgaactattc
catccttttcgattatttcaacaatacctgagcttttagaaaaaatatttttaactagct
ctgttccactttctataacatctccattatgcactagtattagtgatgaatctttattaa
tttcgtgagtttcttctgaaatccataagatatatcccggacttaaaatttcgtaggcat
ctttatctaagcctgtttttcttttttgatacatttaagtctaaatatttaataataccac
cacttgtcgttttatacgtatcagatattagttctgcgactgtgtaaccatcttgaa---
ttttttggtttggaatagctttgaaaagaaatttttgttttttttctgtttctaaaatat
aagattctttttttgtggattacatctgtataaatataacaacctggaattacaatagatt
cagtaataacttgtatattggtaatactgctattagtattctggtgtatccttacttctc
cagcataatgatttattaattctgtctgtgcaagtatagtaccagctttagttttatctt
ctttatgaacaataacattagcagaatcagatatactataaacttcacctgagagcaccc
agatcaagcctccagttttagtaattcttgtagtatattgattattatctgtttcttcaa
ctgttaaattggagaaacagattttgccagagagatctgaaactacatgtttttgagctc

tttctgtcatgagtcgatttctgcggggtgattcagcaatcacttgatccttcgatacta
attcaccatcacatattaaaagagtagtacctttacttaagttaattattgacttctgat
ttttcttagatctgattgtaacttgggttggctttttttgttattaatgcttgctcaccat
gtctggtacgaacatctgtatacgattcaatatctagatctatcaattgaccatctatag
gtgcgtagatttgttctgctaattcaccggtaaatactcctccagtatgaaaggtcctca
ttgtcaattgagtgccaggctcaccaatagattgagccgcaataataccaacagcctctc
ctaaatcaactaagcggccatgtgcaagattccatccgtaacaatactgacaaacagaac
ttctagagttacatgtaacaggagatcttactaaactttttt---gataccagctttag
ttatctcttttgctaacttaggactgacatcttggttcgtatgagcaattaaacaatttg
tctctggatggaatacatttctgctaatactcggccagataaagcttgttctagattaa
ttaatattttctgcgtatctactaaatcttctagtatgatgcctttattggtcttacagt
ctacttctctaattattacatcttgagaaacatcgacaagtcgacgcgtcaaatagccgg
aatctgctgttcttaaagcagtatcaactaatccttttcttgctccgtatgatgatataa
aataatcagtaacagttagtccttctctaaaattactcgatattggaagatcaataattt
gtccctgtggatcagccataagccctctcatgcccactaactgtcttacttgagaaatat
tacctctagcacctgagaacgccatcatataaacagcatttagaggatctgtctctttaa
agtatttaataacttcttgtttcaaggattcactcgcattattccacgtatcaataactt
tttgaaatcgttcaactgctgtaatctcacctcggcgatatttgttttcagtggccttaa
tatcttcaatcgtgcctaataaaagactttgcttactaggaggaattcttaaatcttcta
aacttaaagaaattcctgcctgggtcgcataatgaaatccaagatctttaagcttgtcgg
ccatattggctgctcgggctataccataattacgaaaagcccagacaattaaatttttta
gttcgttttatcaataaccttatttgaaaaacttggctgtgcaagacttcgtctattat
ccactaaatcttctccatattaaatgttcgtt--attaactatgctactaaagactcttg
aataattttattaaagattatacggccagctgttgttctaatatattgcacaattctctg
attttcagcgtcttcttttatgatatggttattaaaaaattttgttgtactattgtcaga
atgtttttcaatcttaatcggatcatgaggttgatcaccatctactaatccatcaaaacg
agcccatatataagaatgtaaatcaattttcttctgttcataagccataactacatcttc
taagcttgcaaaatactgattagatccctgctgctgagatggattattagctgttaaata
atagcaacctaaaaccatgtcttggcttggcataataattggttgacctgttgctggaga
gagaaaattatgagggctaacattagcaatctagcttcggcttgagcctctaaagacaa
aggaacatgaacagccatttgatctccgtcaaaatcagcgttaaacgctgggcatactaa
tggatgaagcttaatagctcttccttctacaagaataggttcaaatgcttgaatacctaa
tctgtgtagtgtgggagctctgttcaatagtactgggtgaccttgaataacttcatttaa
gacattccaaatagaagattcatttttttgaatcatctttttggcagctttaatattgtt
aactaatccctgcaaaattagtctgtgaataacaaaaggctgaaataactccaaagccat
ttctcgtggcaatccacattgatgtagttttaaatgaggacctacaacaattacagatct

accggaataatctactcttttacctagcaagttttgtctaaaacgaccttgttttccttc
tataatatctgacaaagatttaatggtctattatttgcacctacaacagttctacctct
acgaccattatccatgagagagtcaacggcctcttgcagcattctttttcatttctgat
aataatctccggagctaaaattgactttagacgagataatctattatttctattaataat
tcgacgataaaactcattcaaatctgccgtcgcaaatctacctccatctaattggaccat
aggccgtaagtctggaggtatcacaggaatgactgtgaataccatccaagaaggatctgc
acctgttgctatgaaattttcaatcaatcttaagcgtttcatttttttattgaattttaa
tgaaggtgtcttaaaactttttggtggatttgtagcttcagaccttaaggtttcagctat
gtgctctaaatctaaatctttcaacagtttctgaatagcttctgcacctattcctacttc
tacttggttattctcatcttgattttggtagatttcctcttcaagacttttccattcgta
accttcgagcagttgcttgtattttaaattaatatctgtattagattgcgtaactacata
agaatgaaaataaacaatttttctacttctttgacttttaagtctaatgctaaagcaat
atagcttgtacttccttttaagtaccaaacatgagttactggtgaggcaagttcaatata
agccatccgatgtcttctcacgcgtgattctgtaacttctacgccacatcgttcacaaac
tatacctttgtagcgaaaacgtttatatttaccacagtgacattcccaatccttaactgg
gccaaaaattttctcacagaataaaccatccatttctggctttaaagttctgtaattaat
tgtttcaggctttgtaatttcaccaacaatttgtccgttgggcaaacttctttcgcccca
ctgtcgaatttttttcgggagaagctaaactaatttttacgtagtcaaaatattgctcaaa
ctttgtcataaatttgaataccttaaaatcaaataattagccaatttatttacatgcttc
atattgaaaaatgaatatcttaataaaggaattgttcaaaatcgtctactggaggagtat
cataattagaacgatctactcggttatctttagaatccgacatcaagtcaacttcaactg
tccgtctttggccatcttcaaatagtttcaatttatgtactgcaatgtctaaacctaatg
actgcagctctcgcattaagactttaaaagattctggagttccaggcttcggtataggct
tccctttaactattgcatttaatgcttcatttctagcttgcatatcatccgactttacag
ttaataattcttgcaaagtatatgctgcaccaaaagcttctaaagcccacacttccattt
cacctaatctttgacctccatgctgtgctctacctcctaatggctgctgcgtaactagag
agtaaggaccagtggatcttgcatggattttatcatcaacaagatgaacaagcttaagca
tataagctctaccaactgtaacaggattatcaaaaggctctcctgttctaccatcaaaaa
cttgcattttacctggatgctgatcattaaaaagccatttattgctagtaatcaatgatg
cttcttttaattttctatttacaagtgctcttgatgcttctgctccatacatttcatcaa
aaggtattattttaaatctctttcctaaatagccacctgctaagcctagtaggcactcaa
atacttgaccaacattcattctagaaggtacacctaaaggatttaaaacaatgtccactg
gtgttccatctgataaataaggcatatcttgtttaggtaagatcctagaaatgatacctt
tattgccgtgacgaccagccattttatcacccacttggattttctcttctgcgcaacat
agacgcgaatcatagcatttgttccgggaggcagctcatctcctttctgcctagtaaaaa
ccctaaccttaactactctacctttttgcagcattaggaagccgtaaagatgtatctctca

catctcgagccttttctccaaaaatagctcgcaacaatttaccttctggtagttgatcag
cttcacctttaggagtaatttttcctactaaaatatctccagcttcaacccaagaaccac
cgacaacaattccatttctatctaaatccttcagagagttatcgctgacgttaggaatct
ctctagtaatttcttctggtcctaatttagtttgacggcattctacttcatacttttcga
tatggatagaagtatacaagtcatcataaactaatctttcgctaattaaaaatgcatctt
cgtagttataaccttcccaaggcatataagcaactagaatatttctccctaaagcgattt
cccctccgtctgtggatgcgccatcagctaaagtttggccaacaactatttttctccca
cccaaacaattgggcgctgattaatacaagtatcttggttagaccgataatattttttta
agcgataatgaactgttctaccactattatcctgaatgcctattttattagcagaaacat
aatttacatgaccagatgttctgctaataacaaccataccagagtctcgagctattttag
tctcaagaccagtaccaattattggcttttctggatataataatggaacagcttgccttt
gcatattagatcccattagagctctattcgcatcatcatgttctaaaaaaggtattaatg
atgttgcagctgaaattacttgaataggtgaaattgcaatataatctacttgggttgggg
ttgttgtaataaactcttgacgataacgtacaggaataatatctccttcaatgtaatgtt
gcttactaactttaacatctcctggagctactcggaaatcatcttcctcatctgctgtta
gatagacaggactattgtgataaatcacttgaccttggtttaccggataaaatggggttt
ctatgaaaccaaaaacattgacccgcgcacaggttgctagtgaaccaattaaacctgcgt
tcggaccttccggagtttcaattgggcaaattctaccatagtgactaggatgtaaatcac
gaactgcaaaacctgccctgtctttattaaaacctcctggacctaaggcactaattcttc
tcttgtgagttagttctgcaactggattagtttggtccataaattgagaaagctgactag
atccaaagaattctctcactgacgcaattaatggcttgggattaattagatttgataaac
tcaatgaatctatatcacatatcatcattctctctctaataatgcgttctaaacgattga
gacctactctgaattggttctgcagcaattctccaactgaacgaactcttctattaccaa
gatgatctatatcatcaagattacctgagtttttatccttaatattaattagataatcaa
ttgaggataagatgtcttgaggagacaatactcggaacgttttaggtatattgagaccta
gtttttattaattttatatctaccaacttcacctaagtcatatcttttaggatcaaaaa
aacgagagtaaagcatctgcttagcaacagcaactgttgctggttcattaggtctaagct
tagaataaactatgagcagggcttcctcatcagttacttcttcaatattatttttttccaa
tttctttagctaattctttaactgaatatatttgggatgcagaaataagaaaagcatatt
tactaagtcctttttcaatttcatctttattgagtccaatagctcgcagaaaaatataag
catttactttgtgggtcttatctattcttatccaaatttctcccttaggatcaatttcaa
attttaaccaagaacctctattagaaattaggcttgcactataaatctgcttaccatttt
tgtctatctcttgtttataatagattcctggactacggattatttggttaataataactc
tttctgtcccagatacaataaaagttcctctatttgtcattattggcaagtctccaataa
aaactaatctctttttgtactttttatttttttatctgcttttcagcactgagctgtagat
gggttgatgataaatctaatgatttaatatttttttcttggtttttagaaggtagatcaa

tatcttttcttgttaactttgcaggaacatatatttgagcactataagttttatccctat
tttttgcctgtctgacactataacgaggaaattttatcttgtactcattaccaaataatt
gtaattcgagtcgactagtagggtctgatatcttcggaaaaatttcaagaacttccgtca
aaccttctaataaaaaccatttaaaactggccctctgaatttcaactaaatctggaagaa
gtttatttttaagcttatacgttgaaccatagatctcctttataacaaattttgaaatc
gaagagtttaa-tactattaatagagtactggtagataatgcataagcatagtttgctac
attatttcttctctagacataactagaacatcatattttattcaa--aaaaagattttta
aaatttttcttcac--acacactatctataagtactttgctgcta--aaagatagttcaa
gctattaaa-------aaaaacgaaattaatacttgtctgtctgtcaattaattgacatc
aagatagacaagtattatttatttaatcaattctg-tttttctgagcaaaagacaatgct
ctcgcaagtctagcttttttctagcgcctgtattagggtgaaatgcccctttttttaca
gctttatcaatcttactatatacttgagaaatacttgactgcacatcattgaaattatca
ctttctaaattatcaatatttagaaggcatcttttagttaaagttttgacaaccgattta
tacttacggttaatgagacgatttctttccgaagttttaatgcgtttaatcgcagaaagg
ttcttagccacagtaaatccaataaaacttttataaaatatttttagaaacgaaaataca
caaatctgggatctattcaaaaagggaatatgcggcgtattataacattaaagaaaaaca
ataggcaatagttcaacaagagctcttgttattacacaatagagattgtcattgcgaaaa
actttgtaagatagtatttaattaaaataaactaactagacatataaatatatatgaaaa
aaattgaagctattattagacctttcaagcttaatgaagtaaaacttgctctagtcaagg
aaggtattggaggaatgacggttatcaaagtttctggatttgggagacagaagggtcaaa
ctgaaagatataaaggatccgaatactctattgatatcattgataaaataaaaattgaaa
ttattattagcgatgataaagtagagaaaattgtagagacgattattaaggcttctaaaa
caggggaaattggagatggaaaaatatttattagtagtattgaaagagtaataagaatca
ggactaatgacttaaattttgaagctttatagtatttttacttt-tttagacttgatact
tcttcactaaacatgagataataagcgtttattgtatacttataactaaggtgtaata--
ctaatggctaaaagcaaaggtgcacgaatcgtaataactttagagtgctctgataaggct
ggagagtttgctcaaaaaaggaaacctggcgtttttcgatatacaactactaaaaataga
cgaaatacaccaagtagaattgaattaaacaagttttgtcctaattgtaatcagcattgt
atcttcaaagaaattaaatagttatttatcttattaaaat--aatcatattatggctata
tatagaaaaagaatatctccaattaagccaacagatgctgttgactacaaagatattgac
ctgctaagaaaatttattacagaacaaggcaaaatactgcctaaaaggtcaactggattg
acttcgaagcagcaaaaaaaacttactaaagcgattaaacaagctagaatcctttcttta
ttacctttttaaataaagattaa--atatatagcctaacttcatatttttccatctaaa
tattaaaaatttctatagatttgtttattaaaatgatctaacatactgaat---------
----------tactaagaaga-tctactcattacaagttttgttcattttataattaata
atagcattataattta---ttaagctagattc-----agacataaag-gtaacctctttt

aattgattatattatatagtctatttaaaagaggaaattttttttgcttc----------t
agcaattcaagattctttaagctttaacaaggtatttaaaacaaagatttttgtttaggt
atcagatcaaatgaatgaatcttatctcccgactgccataattgaaattcagagatgaaa
attccacattcgttgccagcttgaatttcttcgacatcttctctaacgcgctttaaagat
tcaatttttccttgataaataactttttcttctcgaatcactttaatccaagagtttttt
aatagcttattattgatcactcgacatcctgctatttttctatttgctaacgagaatact
gtacttacttctgcttctcccaccggtacttctgaatattcaggatcaagtaagtcttcc
atccttcttttaatgtcttcaattaaggcataaatgatttgatagttttctattaatata
ttcgactttgccgatgcttgctttgttcctggagcaaagttggtattgaacccaatgaga
gtagaatttgtggttgaagctagttcaacatctgtagcagtaatttctccaggcatgatt
gagacaacattgagttgaactttactctgaggaaattgagataaagaatctaggattgct
tctgtagagccttgattatctgttttaataattaaagaaatttgcttactaatatcttta
gaattagtgtttttttaaagtatctaatgtaattcgactattaagtgctctttgtttctgt
ataattgatgagtctttagaggtattttcaatcgctttaagtttggcttctttatcactt
ttaactgctaaagctatttctcctgtagctggtacagatgacaatccccaaatttcaaca
acagatgatggtattgctaaattaattttctctttagcgttgttaattataactcgaatc
tttgcgtaagctgatccaataaccaaattatcactaatattcaatgtaccattttgtatt
aatagagttgctactggaccatgcgatttatctaaatgggcttctattataataccttgt
gcaggctgtgttggatcagctttcaaatcctctaattcagctaacaaggtaatagtttct
aataatttatcaacattttgaccagtaagagaactaataggaattataggcacttggcca
ccaagtttctccgacatgacattgtattttaataaatcttgctcaataatatctgtatta
gaaccagctttatctatcttagatattgctatgacaaaagggacattcgctttttgaata
tgattaatagcctctatggtttgaggtttaactccgtcatcagcagctataataataata
gcgacatctgttaaattagctcccctagacctcatacttgtaaaagcttcatggcctgga
gtatccaagaaaacaatttttttgtttattatctttcttaatatactcaacttcataagct
gcaatggcttgagtaattcctccaatttctttgttagcattattagattttcgaatataa
tctaatagcgttgttttgccatgatctacatgtcccataactgtgacaattggaggtctt
ttaatataattgccagtttcataaaaagcattagagttatctaaattactagaaagtcca
ttattattttcctttacatttgattctactgctataccaaaattatcagccactgatgat
attattgaggcgtctatggtttgattcatagtaactgatattcttttaagaaataaatat
tttataatatctgtttcttgaacacaaattagtttagataattcttgaatagttagagga
ttagtaatactaatagactctggcggagaattagattgaatgcttgctgaattttgacta
ggagttacggctactttttttctgtcttatagatttttttattactagtcagtttttttaca
acttccacttttggcttaggtggacgcattaaagagatagctaaatctcctgcagtctgc
gaaacattagaattagagtctctaaagttgtca---tcatcatcgtcaatatgtatttta
gtttttattttt-ttgcgctgtctattctta--tttttttttactatctactaagtcgtga

attttattaaaatttttactttttttatctagtttaggtggagaacttaattctagatgc
tgttcaccacc---tgtatgtgattctgatttatcaaggtctaaatttaacagattatca
ttattgactgactctaatctaattttatagataatctgaggatttttcaaatctacaatt
gtttcagaagaattgatatttgagctagatctagaggatctattttcaaagttttgatta
tttaaaaacataaatattggttttattatctaatagtaactttctaattttaatacagag
cttaatataaacaagaatcgaatatctgtataaataatacaaatgt---agagaattaat
ctatgttgtattagatttaatataacactaaagtcctttgttcgttatacaagtttaaaa
ctttt-tatgatgataaacactagattctagtttattggtatacaatctatttatctaaa
taatactgtttttgtcagggcttttaaa----ttatctaatgttatgttcattatgaaat
tctattaaaatttaaaaa-ctcaacaattaagtaagaatcgttaatcacctccacgccac
ttatgccacgatctcaaaaaaaatgataattttattgacaaaacttttacagtattagctg
atattgtattaaaaatacttcctacaagtaaagaggaaaaagaagctttctcttactatc
gagatggaatgtctgcacaatctgaaggtgagtatgcggaagctttagaaaattattatg
aggctttaaaactagaagaagatccatatgatagaagctatattttatataatatagggc
ttatttatgctagtaatggagagtatgtgaaagctttagaatactaccatcaaggattag
aactaaattttaaattacctcaggcccttaataatatagccgtgatataccattaccaag
gagttcaagcaattgaggataaagatacagagctatctaaattaatgtttgataaggcgg
ctcaatattggcagcaggctattaagttagctcctgataattatattgaagctcaaaatt
ggctgaaaacgacaggacgaatgaggaatatacaaggatattaatatatttaagatataa
taaagatatcaattaataaattaaaattagttaaagaaactggtactatatagagtactt
-tactatatattataatattatactagttgatctctatctctaagacttagctgaataac
cgaaattaaatctattgactatattcatgaatacaaataagcataatggttagtacaaca
aaaagtaccggatctgttactcaaattatcggaccagttttagatattgcatttcctaac
ggacagcttccgaaagtattcaacgcactcaaagtacaaagctcagaaggaactattact
tgtgaagtacaacaacttttgggtgacaacaaggtacgagctgtttctatgagttccact
gaaggactacaaagaggggtagaagttattgatactggatcccctatatctgttcctgta
ggtacagatactcttggacgtatttttaatgttttaggtgaacctgtagataatttgggt
cccgttgattctgagagtactttacctatccatcgaccagcacctgcttttactaagcta
gagacaaaaccaaatatttttgaaacaggtattaaagtcgtcgatttacttgctccttat
agaagaggtgggaaaattggtttattcggaggtgctggagtaggtaaaactgtattaatt
atggaactaattaataacatcgctaaagctcatggtggagtatctgtatttggaggcgtt
ggtgaaagaacaagagaaggaaacgacctatatatggaaatgaaagagtctaaagtaatt
gatgcagataatctgaaagaatctaaagtagcattagtatatggtcaaatgaatgaacct
cctggagcacgtatgcgcgttggcttaactgcattgacaatggcagaatactttagagat
attaacaaacaagacgttctattgtttattgataatatttttcggtttgtacaagctgga
tcagaagtatcagctctactaggccgtatgccatctgctgtgggttaccaaccaactcta

gcaactgaaatgggagcactgcaagaaagaattacttcaacaacagaaggatcaattaca
tctattcaagctgtatatgtgccggctgatgatttaacagacccagctccggccactaca
tttgcacatttagatgcaacaacagtactttcgaggaacttggcggcaaaaggaatttac
cctgcagtggatccactggattcaacatcaacaatgctacagcctggaattgttggaact
gagcattattctactgctcaagaggtaaaatcaactttacaaagatataaagaactgcaa
gatattattgctattcttggtcttgatgaactttcagaggaagacagacaaactgtatca
agagcaagaaaaattgaaagatttttatctcaacctttttcgtggcagaagtgtttact
ggatcacctgggaaatatgtatctttggaagatgcaattaaagggtttcaaatgatctta
aaaggcaatttagacgacttgcctgagcaggcattttatctagtaggtgatatagatgaa
gctatacaaaaagctgacagcatgaaagattaatataattaacgatgactttaaatataa
gaattattgctcctgatcggactgtttgggatgcagaagcacaagaaattattttaccaa
gtagtacagggcaacttggtattttaacaggccatgcacctttgcttacagctttagata
ttggagttatgagagtaagagtagataaagaatggatgccgattgttttgctgggcggtt
ttgcagaaatagagaacaatcaattaactattctggttaatggcgcagaagaagctagtc
aaattgatttatcagaagcagaaaaaaatttagacactgcaactcaactcttaagtgatg
cctcgtctaataaagaaaaaatagaagcaacacaaaaaatacgaaaagctcgagctcgag
tacaagctgcaacagcagcaacttcgtaa-aagattcc------tcttaaaacataaaac
tttagtaattattactaaa-------gttttatgttttatttgtatggttataggcagct
taaataaacttagcttagacctgaacaaatataatcaaaatataatcccatctctttacc
tgcatctggtccaactaaactaatagtaacttctttcatagctaaaatagcttgaatcgt
tgcaccaataggaacacctagagaattgtatgtttcttttagaccatttaatactctttc
ttcaaggatagatggatctccagctagcattccataagtagcatagcgtaagtagtagtc
taaatcacggatgcatgcagcataacgtcgagttgtatacatattaccacctgggcgtgt
aatatccgaataaagtagagattttgctacggattctttaataattgttgctgcattagc
tgcaatagtagcagcagctctgactcgtagctcacctgtttgaaaatagcctcttaattt
ttcaacagagctatcatctaaatatttaccttggacatcagctgcattaataacagaagt
aattgcgtcttgcataactttcacacattccttaattttaaatatctaatacta--gggg
tatataacaaaattactagatcttttagcgatatcagtattactgcatagcacccaaagt
ataatcaaagtaaaaaccagcttctgctgaatcttcacctgcaagcagtgagcaagctac
acttttcatgcattttacaccttcagcaactcctgaaattggtgttcctaaagaattata
catttctttaacacccactaatccaatttcttcaataggagttacatcgccagcaactat
tccgtaagttactaagcgaagataatagtccagatctcgtagacaagtagcagtcatctc
ttcgccataagcatttccaccaggagaaactacatcaggtctttttttgaaacagttgttg
gccgccttgctttacaatacgctcacgattatctgttaaaatttgagctattcttaaacg
acgttgtccagataaaacaaaacttttaattctatctaattccccaggacttaaatatct
tgcttctgcatctgcatttacaattgactttgtaacaatactcatggataaaactcctgt

aatgcttt--aaaaaaagcttaatactaattatttatgatataactcagaaagtaatgta
ctttttgag-aattttctaaccgaaacacgttcatgatatatgaagcaggatagatactg
aggtaagtactaaaacgagaaagttatatgtaggaacttctgttattgatttcctagtac
tttttaaaacttggtaccacaattgatatatcctgcttagtcaaacgattatagagtgt
ttctgtattaggaaagttagctgctggtaaagtcggaaatcttctgtatggaacagtatc
tttgccgtataccgcattgtattcaacactttcaactaaattacttataaatgcagataa
gccttttgaagctaaaatttgattaaagaatctaatttctgcctgattattaggagctct
tcctaagatatgcttagtacctaattctataacttttgtatttggatatggctgataaaa
ttctttaccatatagttcagatagcgccagtttttctactagttctttgacacagatttg
tctattcaagaaagctgacttaatatccaagaattcaccgccgacactaaaagagttaag
atctctttcgaaaatttgacgatatgctgctcttaaagcttgttctaacatttctttgtt
actatcactattcacttcaaagacaatagattgatctcttaaagaagttactccttgtga
aatacgagactgaatatcattactactcctcatttctttgacagtgcctaattcaacaaa
cctagcagatttactagcaataactttttgaatctctggtcaataattcctggtcttaa
gcttcttaaagcaacaccagctggagtactatatctttcataaggaacagtattatctcc
aaaagtttctgtgtactctgagctatcaattattgtgtctattacttggtaatagccttg
cttatatgcaatatcaaaatatttattaatttcttgccgaccatatgtagggcgacccag
taagcggttatgaatatattcaatagctttacaaatatataatggctcccaataaagaga
tctaaatatacttgattttgctaactgtctaacaaactcgcgaactgtaatttgattatc
ttttagctgactttctattggcttgagaattagtttttcttcttgatacacttctctgcc
aaaaactcttaaataagcaacttttgtgacaacttctactgaattttcaaaattttggga
tgaatctgactttataagagcgttggacaatttaaagattttggtcccagggatcctgc
tgacttaggtcttacctgtggattactaatctgattgtagattcctggcccacgtctaac
taatatccttcttgtatctttaccaaaaagagcttgtctttttcttggatctttattttc
tttagggaaaatagctccaaattgaattgataatggatcattacctattccatatggatg
ttgatcaggtaatgattgcttataatcactaaataaagttataaattgaggaaccttacg
aaaaggcgcactataggtgagaagatctatttgaggcccccaattacggcactcttgagg
ctcttcacctaaatttctaaagtaaggcacagtctcttcaccgaaataatctgtatactc
agaagaatttaaaagtgcattcactaaaccacttaaaccagtagatgataaaatagcaaa
atatttttgaaactcttctaaagagctaggacctctacccaagaaatgtctaaatgctaa
ttctaatgctcgactattgacaaaaggttcataaaattgctttctataaatacttgaagt
tcctaacgaacgaataaattccttaattgaaatttggccatttttttacttgcgactccaa
attcgataaagagaggtcatatgctttagcaatatctctttcaaaaatttgtctataaca
tgctttcacaacaatatttttctcatcagcagataaactaggcttcataacaaaacgagg
tgttgacactcctgcctgtacataagtctgaggaagacgcaaaccttgtaaatctccaga
tattctttttcttaatttatcagtcaaagatggagcttcaaattcagaaatcacaacatt

aaaatactctttaactaaatcttgaccttttatatcttcctcaaagattaataatgctgt
ccgtctcatctctctaagagccacaattgctgctgcacttgagcaagcgttatcaattaa
ctctctcaatcctcgtatattaacagataaaatattagggtcgcctgatacaattgcata
tgtcaaataccgtaaaaaccagtctaaatctcgaagagattttctcattcttgtagtacc
gtatctcaagacgtttataggtttaaaccctggcggagtagctcccccagcattaaataa
agatcggaaactttggccaaaatctccttggatatttcctgataactcattaattttatc
ttgagaactttgatctcctgcaataataacagctgcttgcggacgttctaaataagaaat
tgcagacccacctacgaatattttatctgctgctcgtgctaccagaatatttgcattttt
agttaaaatatctgcaacttccagtctttttgtccagaatttaaaaaagaaactaattg
atttaattcaccaagctgtaaaaatcggtcttgttgctctgcctgagtaatagttaaaat
tgaggcagtccgataaagctgggggcgtgctaatgggcttccgccacttgctttgatact
cattactttatttatctccttaactgatattgcttgctagcgaatacaaatatagcttga
tatttctaaatatttaaagatactatgcaagcaca-ctgttttctatattattaggtact
agtacttatagtacagagagtcttataactaacttctataaatacaaatttgtaatact
tctattaacatttgtcaaaaaaatatttaagtgctcactcttttatattaaaattcaatt
agtgagaggttgatgtgaaataaagtatactaagtaagttttatagataggaaaattaaa
agatttattctacaaacattaatca------------cttgaatgtttagcaattgtaca
attgttattgcttttaattggtttagactagttttatattttttctaacataaacaatat
taatgagcaatttgtatgaatttaaagccacaaaacaacctattaactaacgaaaatata
gatcatgtagatatacccgaaaatgatataccaatgtcaattacggaacatttagaagaa
ttaagacaacggactttatttgtgtttttattctttttgtttgctacaactataagtttc
acacaaattaagatcattgttgcaatactgcaagctcctgctgttggtattaagtttta
caactggcaccaggagaatatttttttcatctattaaggttgcaatatattgcgggatt
gttgcaacaactccgtttgccgtttatcaagttatattatacatactcccaggactaact
ggaaaagaaagaaaaattattttgcctctattaattagttctgtattgctttttattaca
ggtggtatatttgcttactttgttctcgcaccagcagctttaacattttaattagctat
gggtctgatattgtagaaccattatggtctttcgaacaatactttgactttattctatta
cttttacttagtacaggattagcatttgagataccaatcatacaattattgctcggtgtt
tcagggacattttcttctagtcaaatgatacgagcttggagatatattattattatagca
acaattgctggagctattctgaccccttcgactgatcccgttacacaattaataatgtct
tcagctgttttattactttattttggtggaattgttatattattagtcttaaaaaagtag
agtaaat---------attttcta-ttaatccaatatatttgttatc-----tgggcaaa
taatctat-aaaacaattgct------------------agatttaagtatcaggcttct
aaatatttcaa---------gtctcttttatcattgaaatacaatc-------------
------------------------------------------------------------
------------------------------------------------------------

------------------------------------------------------------
------------------------------------------------------------
------------------------------------------------------------
------------------------------------------------------------
--------------------------------------------------gatacattgg
cttgtttttcatcaattatttgtgttgcacac-tttttagtactctctttattgttctt
agaattatttctttaatctatttattaaat----agatatgaagagatttaagtattgat
tatggtaatctatatcaacgtaaaaagtattttattgtgagttttatttatataaaggct
tattcattatatagttataacagtaagtctaattaatag-ttttcatgcttaattaaagt
aataactcttaattcttttggcagaaaaatctgtcttaatattttatc--aaaaaaacag
-aaaaatgaagctaaatagtctaattaacttaattcaaaagtctatatattcttgtacac
ttttactaattattttaaatattatttgtgtcgcacctaattctagtaatgcatttccaa
tttatgcgcaacaagcttatgaaagtccaagagaagcaactggtaggatagtatgtgcca
attgtcatcttgctcaaaagcctgtggagatagaagcgcctcaagcagtactacctaata
ctgttttcgagactgttgtgaagattccatatgacagcaatgctaaacagattttaggta
atggcagtaaaggaggcttaaatgttggagccgttgtaatattacctgaaggatttaagt
tagctcctgttaatagattatctacagagttaaaagaaaagactagaaatctttacattc
agccgtacagtgctaaacaagacaacattttagtaattggacctatttctggtgataaaa
atagagaaatagttttcccaatactatctcccgatcctgcaaaagataaaaaagctcatt
ttttcaagtatccaatatatgttggcggaaatagaggacgaggccagatttatccaacgg
gtgacaaaagcaacaataatattgtctctgctttaagcagtggtaaaattaataaaattg
aattactagacaaaggtggatttataatacatgtgactaacagtagcaatgtggagtcaa
cacagaagatttcacctggtctcgaacttagagtaaaagaaggggatacaattcagcttg
atcaagctttgaatagtgatccaaatgtaggtggttttggtcagaacgaaacagaaatag
ttttacagagtccaaatagaattaagggcatgattgttttctttttcgctagtgttctag
ctcaaattttcttcgtattaaagaaaaaacaatttgaaaaagttcaagcagctgaaatga
atttttaaagcaatataaaactggcacttactataaagaaagta-taaacactttattca
caggatgtaccagtttttatgttatgtaataatcgctttgacggaagttatgatactttt
aacagaatctacaatctgactaggctgaattactgtagcctgctctaagcttccattata
tggtgtgggtatatcctgcgaagatagcctcacgacaggagcatctaattcatcaaaaag
atactcattaatttgtgcaattagctctgctccaattccggctgttttcatacactcttc
tacaattaaaactttgtgagttttctttactgagatagatatagagtctatatccagagg
ttttaaagatataagatctataacttctggatcataaccttcttttaataaggccggtaa
tgcttggataacatgatgcctcattctagaataggttaaaatcgtaatatcttttccttt
tcgtacaaattcaactttattgagaggtaagaaatattcttcttgaggaatctcttcttg
taaattataaagtagaacgtgctcaaagaatacaactggattattgtctcgaattgcaga

tttcagtaatccttttgcattataaggagtagaacaagcaactatttttaagccggggat
ggcttggaaataagcttccagtctctgagaatgctctgcacctaactgcctaccaactcc
tcccggtcctctaataactaaaggcaatgtaaaattacctccagaagtataacgtaacat
tcctgcattattagaaatttgattgaatgctaataataaaaaactcatgttcataccttc
aacaattggtcttagccctgttatagctgcaccaattgccataccagtaaagctattttc
tgctattggtgtatcaagaactctcaaatccccatatttgctatgcaaatctttagttac
cttataagatccaccatagtgaccaacatcttctcctattacgcagacagttggatcttt
tgccatttcctcatctgttgccgctcgtaaagcgtcaaacataaagatttactcatttt
aatatttgatttttatgatttgaccactgttttaaagtaacagttaagacaccatattct
atttagttatctgcaaaaagatatcgtttcaattctgacatgttaggttctggactggaa
atagcaaatttcacagcttgttccaattcggttttaacagcattctgaatctcattaagt
tcacctatattagcaatttcattatctagaatatatttttgagtttttgataggatct
cttgccacccaagcctctttttcttgtcttgatcttagttcatcaggatctgcgagagag
tgaccacgaaatctatatgttaatgcttctattaaggttggaccatcaccttggcgagct
ctttgaactgcttgttttgcagcttgtcttacagctagcacatccattccatcaacttca
atcccaggaagcccaaaagcttctgctttttatgtatttcaggtattgaagaagaccgg
tgatgtgccatacctatagcccattgattattttcaacaacaaatataataggtagtttc
cagagaactgccatattcagacattcaaaaaattgcccattattggtagtcccatcacca
aaaaagcaagccgtgactcttaaatcctctgtttctttaagtacttgctggcggtagata
ctttgaaaggctgcccctgttgcaaccggtataccttccgcaataaaagcaaagccacct
aaaaaattgtgaggcgcagaaaaaatatgcatcgaacctcctctgcctttactacaacca
gtctcttttccaaataactcagccatcacattttagatgggacgcctttacttaaagca
tgtacatggtctcgataggtactgcaaacataatcagttggattcagaagtttaattaca
cctgtagaaacagcttcttgaccattataaagatgaacaaaaccgaacatctttccttta
taatacatctgagcacacatatcttcaaaatttctgcctaacaacatgtcttcatataaa
actaataaattactcttattaagattaagacctgttgaattatagttagtcagcggcaat
tgaactttcttaggataactcataatttgtaaagaaacctct----------tattgtaa
ataggaattttgaagaaacagctatcaatagaaaacacacagataaaaacttataaatta
taatatgtttttactaaagactttaataattatttgaaactagttcagtatatcaaatt
aattttgcttatactacggatgtctgaaacttatataaaaaagtagt-a-aaaaagttta
ttaaaagaatagttaaaaacagatataaatagaagtaaacagaattatt-gaaaaaaaat
gattattattaattaa----tacttaatatttctaaatattttacaattaatggctatgt
caccatctttattttatcctgttgaacaagaactatgtagtcttgaaaaaaatctgaaag
ctgttgctgggactcgtcatccaattttatatgccgcagcaaagcatctattcgatgctg
gaggaaaacgagttagaccagctcttgtatttttagtggctaaagcaacctctgagaagc
aagatataaatactggacaaaaaaggctagcagaaattactgaaattatacatactgcta

gtttggtacatgatgatattattgacgagtgcacaacacgtagaggagtcaaaactgtac
ataatttattcaatactaagattgctgtgctagcaggagatttttatttgcacagtctt
cttggtatttagctaatattgaaaatttagccgtagtcaaagctatttctaaagtcatca
ccgactttgcagaaggagaaattaggcaaggcttagttcatttcgatcccagtatttcaa
tagatgcttacattgagaaatcattttacaagactgcttcactcattgctgctagttgtc
ggggtgcagctatgcttaatggttccaatcatcaaataaataatgatctttatctttacg
gtaaacatatgggattagcatttcaaattatggacgatgttctagatataactggttcta
ctaagagcttagggaaacctgctggcgctgatctaataaatggaaatttgacctctcctc
tcctttttcacttactcaagaagcaagtttaaatgatcttattgatagggagttctgta
atagtacagatatagcctcaacattatttcttataaaaagaagcgggggaattacaaaag
ctaaagatttagctaaagaacaggtgcaggcggcactttcctgccttcagtttttaccac
aatctacacctgtatctagtttaaaagaattaacacatttcataatcacaagattgtcat
aaaggacttgctaaaattaaaatgttattaactaatttttttactgtttctaaaacagtct
gaaaagtttgaatatcattactagctagctgcgctagcattttacgatttaacgcaatat
tctcttttttttaatgcactaataaaagtactataattcatgccttgattgtgagcagcag
catttattctagttatccaaagacgacgaaaatctctcttttttccttttttcgaccaacat
aagaatatcgaagagctttaaggacttgctgcttagctgttcgaaataaacatttatgtg
cacctttaaagcctttagctagcttaaagattttagctcgtctttttttttgcaacgttac
ctcttttaactctactcataaaatattac-ttaaattttaattattaccagcttactaaa
ttcttatactctttg----attataaataaggtaatttcattgcaatgtttttaatatct
tttaagtcaaccgaacaagtagaagaaagatgtcttctttgttttgatgactttttttgt
aataaatgacttttagaagctttatgtcgaagaaattttccggatgaagagactttaaat
ctttttgctattgcttttgatgtttttaacttaggcatattataaacttttaatgattaa
caaaagcatgaaaagtctgaggatatcaatcttagacctttcaatacttagaataatctt
aacacaatttattttacttactacataggttaactgaaacaatttatatatataatacta
atcagttaatcaaccatagctaaagtggatttatattttgctttgaacttttgtatagta
ttagcaattagcattgttataattttaagactgcttgaattcaattcttcgaacattttc
atatttaatgtaaatgaaatattggcttctgagacaatattttgaatttgaacatcagac
aagggaatcatatctaatgcagctctatattgatctttaaataatttatcgtcttttatt
tggtcaaaatcgtaaaatttcgttcctcctgaatctgataaattcatagcccctctagct
atttttttttaaaatttggccgccagaaaggtcacccaaatagcgcgtataagcatgagca
actaataattcaggttgtttatgtcctatagtgtgaattctatcaacataaattttgta
gcaggagatggttcaataaaatctaaccaatctgatccataataatagtttaaatcttca
gatagactcgctttcctattaagctctgtaaaatatataggtttaatagctggatgattt
ttattagaaaataactcttcctctattgcacaatagacaaagtatagatttgcaaccaat
ttacgatatgacttcttatctacaactcctcccaaaaaagatttaacaaaactaacattt

tctgccatactatgggacttagtagtaccttctcttagttcattcgctaaagtattaacc
ataatatcaatattccattttt-aaagataacaaattaaca---ataaatagttatccgt
atatttataattgagttgcgctaactaggctgctgcaaaataatttttttgattttattgg
atctggattcattgttttgtcaccaggcttccaattagctggacaaacttcatctggatg
agattgtacatattgaattgcttgtaaaaccctcaaagtttcttctacactcctgccaaa
ctctagattattgattgtagagtactgaataattcctttcgggtctataataaatagtcc
tcttagagctacaccatcactattcaaaacattataagctgcactgatctcttttttttaa
gtctgatactaatggatatgaaagatctcctaatccacctgattctctgtcagtctgtaa
ccaagcaagatgagaatactcactatcaaccgagactccaagaacttctgtattaagttc
cgaaaaagcattatacttatcgctaaatgctgtaatttctgtagggcacacaaatgtaaa
atctaaaggataaaagaataagacaatatacttattcttgagatcagataattttaacgt
tttaaattcttggtcataaacagctgtagctgaaaaatcaggagctagctgacctactcg
aagacaatttggtccagaaatcattaatttttcccagaagaattgatataattttgttat
ttaataaataattatataacaaaatttagtatatattaaaatactacaagtctttttttaa
attac--------taaatgatataatcaatactatctttatagcggggaatggatttgaa
ccattgaccttcgggttatgagcccgacgagctaccagactgctctaccccgcggataac
taaaatcatattagtctacatatatatataaaagcaaatgcataatagattttagaagta
aaaaaatagaagaactggtccactgatcattaaacttagaaacgacaaagtattttttat
cttatttaaaagagggaagacttcatataaaccaataaatctatcttttaaaagaatttc
ggacggctttacccaaatttgaccgtcataccagcctgattcttcataaaatactgtagc
actcattaatcgttttactacgtatgaccagcctaaatataatctgattaaaataaatcc
tgtcattaggctggaagtaataaattctgaaaagaaaaatttcaaaggtagtttagtgat
tggaaaaattgataataatataggactaactaataaacagttaagtagtagtgcaatagt
tattttttattataagatctacggcttaaagttggccagcaaaagaaccaagaattttt
caaagaagtatattcatgaacaggttgctgctcctttggcacaggacattgagtattata
taaattcattgacattaaaaagtattataataaagac--tttgtttatattatctaacat
aggtacgacattaaatattaactaactgcaattatagtagtgagtagaagaaataaaact
aaaaatagccaagtaattttatttaaagtattctcagtactacgagtattactaaagaac
tgattctgagcacctacacttcctaaaccttcggatttaggattgtgtattagaatagtg
aagattaatattattgtagatgagtaccaaaaaaatttaaaatttgttccattatgtaa
aagttaat-aaaaatagcacaaagacaatcgtctaaaagatattgattgtcttatcatat
taaaaagtatttattcaccctgtacttttaaaagtaaaaatcctattgccagccctacca
aaactaggactgaacttaatacagcactttctacaatttctccgaccatt-ttatttttc
cttattattttattattg---tcattcttttgacaagattatattgagctaa-ctaaaat
ccgtttctaccccaaactaccattgcgattgaaaatgtaaacattgccattagagctgcc
catcccaaacttaaaatatccataatttataatattc--ctaaaaataagatctcttta

cttaaactataatattcttaatattaaaaatatcattatataataagctaattataacag
agtactacttaattcgtatttttagtgtatttaaagaagattgaa-ttttttcgaggtac
tggataagtatttcatcctaagaaccatagtcaac----ttttgatttattataatgctt
aaatctaaatata--tatcactcaagaatcattagtaattatgatgtttttaattatacg
att---aacatatttcaaaatagaagaattcatagcgactaagttaa-----tatgctaa
tataatttatcaat--aaaaaggaagttagtatttcaattatgcaaacaactataaataa
tgggcaaacgtctagtaaagaaactttactaacacctagattctatacgactgactttga
agagatggctaatatggatatttccggcaatcaagaagatttttttggctatcctcgaaga
atttcgagctgactataatagtgaacattttattagagacgaagagtttaatcaatcttg
gtctaatttagaacataaaactaaatccttatttattgagtttttagaaagatcttgtac
cgcagaattttcaggtttttttactatataaggaattatccagaaaattaaaagacagaaa
tccggttatagctgagtgctttttattaatgtctagagatgaagctaggcatgccggttt
tttaaataaagctatcggggacttcaatttatctttagatttaggatttttaacgaagag
tcgcaagtatactttttttctcacctaaatttatttttatgcaacctatctttctgaaaa
aattggatactggagatacataactatttaccgccatcttgaacaacatccagaacaccg
tatttatccaatctttagatttttttgaaaattggtgtcaggatgaaaatcgtcacggaga
tttttttgctgctttgctcaaatcccaacctcatttttttaaatgactggaaagcaaaaat
gtggtgcagatttttttctattaagtgtatttgcaacaatgtacttaaacgactttcaaag
aattgattttttataatgccataggcctagactctagacagtatgatatgcaagtaatacg
aaaaactaacgaaagtgccgctagagttttcccggttgctttagacgtggacaatccaaa
atttttcaaatatttagatacttgtgcatgtgataatagagctctgatcgatattgataa
taacaattctccattattcattaaatctatcgtaaagatacctctatatttttccttatt
tgcaaatttactaaagatatatttgattaagccaatagactcaaaaacagtatggaatac
agttcgatagttaaatacaagggaatgctttttttcttataaaaagctatctagtgattaa
tataaaat------aaaagcttccttct-----cgatggaaggagcttttattttgtcaa
ga---atataatgggattatatagatc-tagaactaagaacttactttttcttttgcttc
atacataacctctaaagtaattagatttttagaacaatcacgggcaaatttctctgtatt
tctttttactttacctctaacaaacccaggaattttatttaattctttctgtgcttcttc
tgaccaattaatactatcaacagcagaaattcctatacttaatgcttctgttgtatcatg
accaccaaatatttctagcaaatgatcttccataccaagtgtaaacgaattatatactaa
atcagcaatttgatttgttccctcataaccaagaaatggacggtaacttaaaggaaagtt
ttgtatatgcactggtgaagaaataacgccacaaggaatatttagacgcttaccaatatg
tctttccatctgtgttccaaaaattgctgcaggttcagttttagcaattaaatctccaat
taaaccatgatcatctgaaacgattacttcatcgcagaattcttgaacttgatctttaaa
ccactcttcatcatatttgcaataagttccacaccaggcgacatgtatacccatttcctg
gtgtaaaattcgagttattgctgcagcatgagttgcatctccaaatacaatagccttttt

acctgttaaattctgacaatcaattgatcttgaaaaccatgcagattgtgaaataaatcg
ggtttgctcgtctatgtatttttcgtagtctacagcagctcccaatgcatttactaattg
ctgtattgatcggatgcatgcagctgtttgaacaatacccataggtgtaatgtcaacata
aggcatattaaattctttctctaaatatcgagctgtcattagtccagtttctctatatgg
aataaaattaaaccatgcagagggtaatttttttaaatcttgaactgatgcattttctgg
aataatttggtttatttggatatctagatcttgaaataacctttttaattctgctatatc
atgttgattatgaaatcctaagcttactgctccaataatattaacagatggagttttagt
cttttgagttaaaacttgattattagatttagctttttccatatagaaagtgacaatttg
ttctaatgttctatcacctgcttgcagttcatttactctataatgattaacatcagctaa
taaaacatctgcttctgtttctatagatgctctgctaacaaagttttgcaaatcttcttg
caaaatactcgaagtacaagttggtgttagaataactaaatcagggctttcttctctgtc
ttttctagtaatattttctacaactttttcttgagaaccacgggctaacacatgtctatc
tacaacacttgctgtaacaggagtaaaatctctgtctctttcaagcattgaacgcattac
attaaaataatcgtctccgagaggcgcatgcataattgcatgaactttttttaaaagaact
ggcaattcttaaagttccaatatgagcagggcctgcatacatccaataagctaatttcat
attaaattttcctatgatttatttagacatagtaatactttttaaagctaactaacaaaca
ttagccaaacatctaatttaactccttgacaattttgtaatatttttttaaatgaagctta
tcattataaatatttaaaaaatataattcagaataga----aataactgttagatcaagc
ttgaaagtaaagatttcttgcagttatagtttttctttaagtaaacatgtaaaaattgta
ctatatataaatgtactatatataaacttttttttactgaatattggtaatgccagataca
attaacttaaatatgccttccccaacgtttggtggaagcactggcggttggttaagagcc
gcagaagtagaagaaaagtatgcgataacatggacaggcaaaaatgaaagtaagtttgaa
atgccaactggtggtaccgcaactatgcgaaatggagaaaacttactctatttagctaag
aaagaacaatgtttagctttaggtactcaactaaaaagtaaatttaaaatatctgactat
aagatttacagagtttttcctaacggagaagtgcaatatttgcatcctaaagatggagtc
tttccagaaaaagttaatactgggagagctagcgtcaacagtgttgatcattctattgga
caaaatattaatcctgtagatgtcaaattcatgaataaagcaacttacgattagtctctc
-aaaaacctagacataaaataactttatatagtctaggtttttgataatgtatactgat
agtatggaagggtggccgagtggttgaaggcgtctgatttgaaatcagttgaacta--tc
tggttccgtgggttcgaatcccaccctttccgttataaaact------------atagtt
ggcttctcaaggcaagctagctatacactgatagtattaagtttagacttaataaagtct
acaacttgtgacaggttagaaatttttttcggcatcctcatctggtatctctatactaaat
ttttcttctatggccattaccagctcaactgtatctagagaatcggctcccaggtcactt
gaaaaatttgcttccctagtcactatttttttttcaattcctagctgttctgctacaata
tcttgaactttttcaaagatttcattatcttgcataatatattatttcctctaaggtgtt
atcaaaaatttttcggcttatattattatccaactctaaaaaattagtaatttttagtcg

agattaaaatatttatacctatatgatatactattattctggataaatacgaagtttacg
atatggctcttctccaaaactacgagctctcaactggtagttgtcaactagattatgctg
catttttcgtatataggctgatcgaggtgtaagctgaactattgaactttcattcaagat
aatagtttcaatagccaatttagcttcttgaagagcctggatctcatcaaaatttttatt
tttacagagttctacccaattaagaccagatgaagtatgaatgtttaatatttttctgag
tgcacgagtaatttgtggaaccgtactattctgaattgtatatataataatttgttttga
ttttgcaatttgtctaagttttgtattttgctttacttgattacgtagcgctagaatagc
atcagacttttctatctccttggttaaaataattggtagatctagtgaagaaataactga
tgtaatatgctgccaactcaaagaataagcatataagtactgatgaggtatttccaattg
caaagattgtttattaatattaataggggcattgaaagtagttgacaataaagatgtatc
acgattttgtgatttattcaaatctaaggacttatttcttaattctctatattgtaaaac
cggaggcgcctgctgtgcgttaattgattttcgagcattagagatgtttacaggtaagac
ttctatagattgcgatggataacatttaattaaaattctaccattagcttgaatttgtct
tttttgcacaaaaggttgatggccttgcaaaatttgatctattgtctctttaactttatt
atgaactatccaaacatttcgctcgtgtatttcgatcgcaatctgaaaagcaggctcagc
ttttctttctagtatactcttttgcgtacctctacgtttagcttcatcgtcacctaatgt
tacatattgaattccaccaattaggtccgccaatgttggattttttaattaaactttctaa
atgatttccatgcgctgtacctactaattgaactcctctttctgctatagttcgagcagc
taacgcttctaattccgtgccaatttcatctataataataacctcgggcatgtgattttc
aacagcttctatcattacttgatgttgcaaatcgggccttgcgacttgcattcttctagc
tctaccaatagctggatgaggaatatctccgtcaccagctatttcatttgaagtatctat
aataaccactctttttttccatttcgtcagctaaaacacgggccatttctcgaactgcggt
tgttttaccaactcctggttttcctaagagtaaaattgaatcaccttgctgtagtaaatc
tcgaataatactaattgtaccaaagactgctctgccgacacgacaggttaagccaataac
actaccttctctattacgcagtgaactaatacgatgcaaggtttttttcgataccagctcg
attatcgccgctaaaattaccaactttttttacacaataatctaagtcttgccaactaat
agatctttgagataaatattctgggttatctggaaatctagcttctggacgacgacctaa
atccataacgacttctattaaattatttctattaggatgctgttgtaaaggttctttgac
aaaatttggcaaaatttctagcaacttatctaagtcatctgcaataagcatgattgagta
tataataattaattaatcaaatagaagattcaagattctaaagattacaatacatattaa
agaatattatgattggaatcaattttatacttaaattgaacaggaagtta-tagatgtaa
aagaactgattaaactttacgcaataaatcgattggtcaaaattaatagtcgtgttaaca
tgtttattggacaaaaagttcgtattaaatatagcaaacaaaaagttgctagtgatatag
cagataaagttggtgaattaggagttattaagggaataaagtttattaatagtcaatgtg
ttactattattgttgagtttgacaatcacactaggctttggatgtttagagaggaattaa
tctgtctaaatgaat------taaaaacatgaaacataatttattttatttattatacaa

aatttcttatgagtcgttaaaataataaattttaaaggagtacttttaattgtgcaaatt
actattaaaaaattacaagacttactttcgtctgtacaaagaaaaaagatccagacatta
aaactaaaacaaggtaagtttgaacttctattaaataagacctataaaaaagtcaatcaa
gaaattataccttcgcagaaatctgctgtattacaaaatagtccatctacgataattaaa
tcaataaataatacaaaaaaatcttctgttgtcaatgaagaccgtacagaatatgccact
attgtttctccaatggttggaacgttttatcattcacctgctcctggtgaaaaaattttt
gtacaagttggcgatgaggtcaaattcaatcaaacagtctgtattattgaagcaatgaag
ttaatgaacgaaatcgaagcagaaattgaaggcaagattatagaaattcttgttaaagat
ggtgatatagtagattgcgggcaagccttaatgaaggttgaaacataatgctaatttttc
cattttacatattgttaacagaaattatttagatagaaaaattagcttatactatagtta
tgaaaacttatttcgcaaaggcac-taaaaactaatatataaaattatttactttatctg
ccaaaaatagaaataagcgttttgattccttgtctcaagagaggagaatctcaatggcaa
ttagctcaaaagagcaagagacaaagaaggtaaaaatctcggttgataaaaatcccgtag
atacttctttcgaaaagtgggcccaaccaggccatttttctcgtacactagcaaaaggac
caaaaactactacttggatttggaatcttcatgctgatgctcacgacttcgatagtcaaa
ccagttctttagaagaagtttcacgtaagattttcagtgcacattttgggcagctgtctg
taatatttttatggcttagtggaatgtattttcacggagcccgcttctctaattatgttg
cttggttaagtaatccaacaggtattaagccaagtgcgcaggttgtttggcctatagttg
ggcaagaaattttaaatggcgatgtaggtggtggctttcaaggagtacaagttacatctg
gatggttccaactgtggagagcatcaggaattactacagaatttcagctttactgtactg
ctattggcggattagctatggctgctttaatgctgtttgcaggatggtttcattatcata
aagctgctccaaagttagaatggtttcaaaatgttgaatcaatgatgaatcaccatttag
ctgggcttttaggcttaggctgtttaggctggacaggccatcaaatccatttgtctttgc
ctattaataagctactagattctggcgtgtctccgcaagaaattccactacctcatgagt
ttttaattaatagagagcttatggctcagctgtatccaagttttagtaaaggattagttc
cattctttactttaaattgggctgaatattccgacttttttaacttttaaaggaggtttaa
accctgttactggaggtttatggctaagtgatactgctcatcatcatctagcattagctg
ttctatttcttgctgcaggtcatatgtatagaaccaattggggtattggacatagcatga
aagaaattctagaagctcacaaaggaccttttaccggcaacggtcacgaaggtctatatg
aaattcttacaacttcttggcatgcacagcttgcaattaatttagctatgatgggatctt
taagcatcattgtagcacatcacatgtatgcaatgcctccttatccatatattgctactg
attacccaactcagttatcgctcttcactcatcatatgtggattggaggattttgtattg
ttggagcaggagcgcatgcttctatatttatggtaagagattataaccctgcagaaaatt
ataacaatcttttagatagagtcattaggcatcgagatgctattgtttctcatctaaatt
gggtatgtatatttcttggattccattcatttggtttatacattcacaatgatactatgc
gggcacttggaagatctcaagatatgttctctgatacagctatacagttacaacctattt

ttgctcaatgggtacaaagtatacacactttagctcctggaaatacagctccaaatgcat
tagcaacagctagttatgcatttggaggagatattgtttctgttggtaacaaagttgcaa
tgatgcctatttctttaggtactgcagatttttagttcaccatatacatgcatttacta
ttcatgtaactgttttaattttagttaaagggttccttttctcaagaaactctagactaa
ttcctgacaaggccaatcttggcttcaggtttccatgtgatggacctggtagaggtggta
cttgccaagtgtctggctgggatcatgtttttcttggcttattctggatgtacaattctc
tgtctgtagcaatttttcactttagttggaaaatgcaatcagatgtttggggtagtgtat
ctccgtctggaaatgtttctcatattactggcggtaattttgcacagagtgcaattacaa
tcaatggatggctgagagatttcctttgggctcaagcatctcaagttattcaatcatacg
gttctgcgctatctgcatatggattaatttcttagcagcacattttgtatgggcattca
gtttgatgttcttatttagtggtagaggttattggcaagagcttatagaatcaatcgtat
gggcgcacaataagataaaagttgctcctgcaattcaaccaagagctttaagtattactc
aaggtagagcagtcggtgttgcacactacttattaggtggaattggtacaacttgggcat
tctttttagcgagaattatttcagtaggctaatagtgaaaataggataaaaaaaacaatt
atggcaacaaaatttcctaagtttagccaagctttatcacaagatcctacaactagaagg
atttggtatggtattgctacggcacatgactttgaaagtcatgatggaatgacagaagaa
aatttatatcaaaagatattcgcttcgcactttggacatctagcaattatcttcttatgg
acatctggtaatttattccatgtagcttggcaaggcaactttgaacagtgggtattaaat
cctttgaaagttaaaccaattgctcatgcaatttgggatccgcattttggacaacctgct
ttgaaagcttttagtaaaggtggctcagcttatccagtaaatatagcatattctggcgta
taccactggtggtatactattggtatgagaagcaatcaagacctgtattctggggcttta
ttcttactagttttatcagctttactcttatttggagggtggctacatctacaaccaaaa
ttcaagcctggtttatcatggtttaaaaataacgaatcaagattaaatcatcatttatct
ggattatttggggttagttctttagcctggacaggtcatttagtacatgttgctatacct
gaggcaagaggacaacatgtaggatgggataattttacaaccgtattacctcatccagcc
ggtttacagccgtttttcagtggtaattggagtgtatatgctcaaaatccagatacagct
caacatttattcggaactaatgaaggtgcgggtacagcaattctgacatttctaggagga
tttcatcctcaaagtcagtctttgtggctaactgatatggctcatcaccatttggctatt
gcagtagtattcattgttgctggacatatgtataggactaattggggaattgggcacaat
ctaaaagatattttagatgctcatagaccacctagtggtagattaggagctggacataaa
gggctatttgatactattactaattctttacacatacagttgggattagcattggcttcc
ctaggtgtaattacttcgttggtagctcaacatatgtatgctatgcctccatatgctttc
atggctaaagatttcacaactcaagcatctttgtacacacatcatcaatatattgctggg
tttctaatggttggagcttttgctcatggggcaatattctttgttcgagactatgaccct
gaacagaataaagataatgttttggctcgtatgctagaacataaagaagctatcatttct
catttaagttgggtaactctatttttagggtttcatacattaggcctttatgttcacaat

gatacaatgattgcttttggaactcctgaaaaacaaattctaattgagccggtatttgct
caatggattcaagcctcttcagggaaagcactttatgggtttgatgtgttactatcatcc
tctactaatatcgcaacacaagctggtagcaatatttggctgccaggctggttagaagcg
attaatagcggaaaaaattcattgtttttaacaattggtcctggtgacttcttagttcat
catgcaattgcattgggattacatactactacgttaattttagttaaaggtgctttagat
gcaagaggctctaaacttatgccggacaaaaaagactttggatatagttttccttgcgat
ggacctggcagaggcggcacctgtgatatatctgcatgggatgcgttctatttagctgta
ttttggatgctaaatacaataggttgggtaacattttattggcattggaaacatattaca
atatggcaaggcaatgcaactcaattcaatgagtcttcaacttatctaatgggatggttt
agagattacttatggctaaattcttctccattaattaatggttataatccatatggcatg
aataatttatcagtatggtcatggatgttcctatttggacatttagtatgggcaacagga
tttatgttcttgatctcctggcgtggctattggcaagagttaattgaaactctagcatgg
gcgcatgaacgtacacctttagcaaacttgattcgttggaaagataaacctgttgcatta
tcaattgtacaggcaagattagtaggtttagcacatt-tttctgtaggatatgtattaac
ttacgcagcttttgtattagcttcaacagcaggcaaatttggttagactaaa-----ttc
gaattagctaa-----aaaaaagtcagtcataata-gttttatgactgactttttagat
ttaaaacaaaaacttactaaagtttcacctcaatatctacaccagaaggcaaatttaact
tcattaatgcatctatcgtttgagaagaaggttgatgaatatcaataattcttctgtgag
atcttatttcaaagtgttctcttgagtctttatctacatgtggagaacgtaaaacacaat
aaattcttctttttgtcggtaaaggaattggccctactgcaacggcattagttctagatg
cggtatctagtattttgtcacacgacgtattaagtataatagaattatatgctttcagtt
taattctgatttttgtctgctgagtaattgtcatgttaaaaagatcttattatttaagaa
ttttagagacaacacctgcgcctacggtacggccaccttctctaatagcgaaacgcatac
cttgctcaattgcaattgcattaattaattcagcagtcatcttaattctgtcaccaggca
taaccatttctgcatcagtaccatcatcagcagtaaactgattaatagtaccagttacat
cagttgttctaacataaaactgaggtctatatcctggaaaaaatggagtatgtcttccgc
cctcctctttagttagaatataaacttctgcttcaaattgagtatgaggtgtaattgtac
caggtttagctaataccatacctctttcaatatctttttttctgcacacctcttaaaagaa
ttccaatattatcacccgctagaccttcttctaacgtttttttgaaacatttctaatccag
taatagtcgttgtacgagtttctcgtaaacctacaatttcaattgtgtcaccaactttaa
taatgcctctttcaattctaccagtggcaacagttccacgtcctgtaatagaaaaaacat
cttctacagccattaagaaagttttatcgacatctctctccggcgttggaatatatgtgt
caactgcttccataagtgaaaaaatcttgtcaacccatttatcttcaccttgcttaatag
ctgggtttttcgtcacagcttctaatgccaataaggcagaacctgcaacaaaaggaatat
catctccaggaaagtcgtattgacttaataattcccttccttctaattctactaattcta
gtagctcttcgtcatctacttgatcttccttatttaggaatactactaatgtaggaacac

ctacttgttttgctaataaaatatgttcacgagtttgtggcattggaccgtctgccgcag
atacaactaaaatagctccatccatctgagcagcacccgtaatcatatttttttacgtagt
cggcatggcctggacaatctacgtgagcataatgacgattatctgtttcgtattcaacat
gagcagtattaatagtaatacctctagctttttcttctggagcggcatcaatttcatcaa
atttttttcgctgcagtagaccctaaagttgataaagtcgcagagatcgctgctgttaaag
ttgtcttaccatgatcaacatgaccaattgtgccaatattgacatgaggtttttttacgtt
caaatttagatcgagccatgctttttgttttccttatagtaaaagttgctttatagttac
ttttaacgataaatcttgtggattgaaagatttattatttaatacagacaaga-----ta
caatataactaatctaatatcgatagtgagcaaacgctttatttgcttctgccattctat
gtgtatcctctctctttctaatagaatttcctgtttcattagctgcatccataatttcat
tagctaatttcatagacatacttttaccagatctgtctctagagaatttagtaatccatc
ttagtgctaaattcgtacctctataagctcgtacttcaataggaacttggtaagtagaac
caccaactcttcttgcttttacttctactagaggagtaatatttcggatggcttttttcta
aaatatttaaaggatctgattctgttctctctttaacgatatctaaagcctgatatataa
ttctttgagataaagtttttttaccacttttcaaaatacgaacagttaacatacttacga
gtctgcttttatataaaggatcaggtgatgcaaaccttttttttagctgtattacgacgag
acatagtatttaaaattggttattgtaataa-tgttc-tattagcaggtatttaagattt
aggcttttttgtcccatatttagatcggcttttacgacgatcttttactcctgcagcatc
taaagtaccacgtactacatgatatcggactccaggcaaatctttaattcggccgcccct
aatcagtaccacagaatgctcttgaatattatgacctacgccaggaatataagctgtaac
ttcaaatccagatgttaatctaacacgcgctacttttcttaaagcagagttcggtttttt
aggagttgtagtatatactcttgtacaaacacctcttctttgagggcaactttgaagagc
tggagattttgtttttttatgtatttttcgtctttcggatctaacaagttgttgaattgt
tggcataataaaattgatttaaagtctttaatagatactatttgaacttgctttacatga
attggctagtttttatttatcttatacttacgcataaatctttctactctaccttctgta
tcaataattctctgtgaaccagtaaagaatggatggtttcctgaccagatatctacgtgt
agttctggctttgtggacccaatcgtcataattaactggccgtcacagtaaacttttgct
tctggataccaatttggatgtatattatcttttgccattgtttttataattatataaatg
tatcaataataactattaacgcttagaaaattgaggagcctttctggctttctttaaacc
atattttttttctctcttttactcttggatctcttgttaaatagccttcagacttaagtgt
cgttctattttctggattaattgagcacaatgctcttgcaacacctaaacgaattgcatc
agcttgccctgtcaaacctcctcctctagcatttacatggatatcatactggtttagtag
tcctaaaacttgtaatggtgcatatgaaactcttaagtaattaggactaaattgaagata
agactctcctggtataccgtttataattaaattacctgaccctgggactagtcttacttg
tgcaacagaacacttccggcgacctgttccagaatagattgcgcgagttttaattaattc
tgtcgacataacattccttgataactaaataaatatacattctttatactatatactctt

gtggcttttgcgcgacatgtgggtgaattgggccagaatacactttaagcttcgtaaata
gttttctgcctaatggacctttaggaagcatacctttaaccgatttttcaataattctgt
taggtaatcttgtctgaagctgatcaaatgtttcaacttttaatccgccaggctgtccag
aatgtcttctgtatagtttttgattcgttttatttccacttacagatacgtgagcagaat
taataacgattacataatctccagtatctagatagggtgtataagaaggcttgttttac
ctcttaagatattagaaatatgagtagatattctaccaagtgtctgattttagcgtcta
taacataccaatgagaattagtatttaatgagggtgattgcgttttattcatgaaaagat
taatacttttggttgacaataagagaatatgtaaatggtttacattcaatttctatacaa
gtagtattt-aaaaatatta--------------ttaacaaagagaactttctaacagtt
atttaaattgtttgtcagcaagaaaatttctagttatcaataacttatactatttttaat
aatataaaatatattaatcactttttttcttttggcaaacttatacctagttttttttgc-
aaggcatatataacttcttcagctgatttctggccaaaatttttaatttctagcagttct
tcttgagagtaatctagtaaatctgcaatagaatggatttgagctcgctttaagcagtta
taagccctaacagataactgcagttcttctattaaaacttgactaatttttttatcttct
ttgctacgataattatctgctgatttaaagtctaaatttcttagtgaacaaaaaagatta
gttaaaactgtagctccttgacttattgcttcttgaggagagatgctcccatttgtccag
atttgtataatcaatctatcttttatgctgttgctaccaatacggacttcttctacttta
taattaactttattaactggcataaaaacagagtctacttgcaaaaaatccacagataat
tcatctacagctttttcagctaagcgatacccgcaatttttttcaattttaaactccatt
tcaaatattgtattattgcaaatagttgcaatatactgtctaggatctactacctctata
tcagaagataattcaaataggccggctgtaactatagctggcccttgaactctaattcga
ccaatttgagattctttgttataacttttaaatactacttcttttagattaagtaatatt
tctaacacatcttctcttaccccaggaattgtagaaaactcatggttcactccagcaatc
cgtacagcaactatagcagtaccttcaagatctgacaatattgatcttctcaaagcatta
cctaatgtaataccttgtccttgatttaatggttctattacaaaactaccgtactgccca
cgcgccccatctgttcttgactctatgcattcaatttgaaattgagccacctaagaaagc
tcc-tttaaaataatcagtgattagtaagtctctctaaaacaagaaggaatttatactcg
gcgtttcttaggagggcgacatccattatgaggtacaggagtaatatcttttattagagt
aatctctaatcctgcagcctgcaaagctctaattgctgtttctcgacccgctcctggacc
atttactagaacttcagtttggcgcataccttgatccatagcttgtctagctgctttttc
agctgctgtttgagctgcaaaaggtgttcctttttttagctcccttaaatccacttgcacc
agaggatgaccatgataatgtttctccttttaaattagtaatagtaacaattgtattatt
gaatgtagatttaatatgtgtaataccgttaactgcattacgtttagtttttcttgctcc
ggatttttttatttgtctagccatcgtgtctcgtctattatatagttaattaaaagatta
ttttcttggagctttttttcttacctgctactgtttttttacctcctctgcgtgttctagc
attagttctagttctttgtcctctcaaaggaagaccaagacggtgtcttctacctctata

agtactaatttccataagtctcttaatgctcatagactcaaaacgtttgagatccccttc
aatctgataattagactcaagaatttctcttatactgacaacttgttgatcatttaaatc
ttgacacttaatatcagcgtctatgtttgtttttctaatatttctttcgagcgagatag
tccaataccataaatatatgttaaagctatctctattcttttgtttcttggaagatctac
tccagcaattctggccactttgtcttgtctccaataaataatatgtat-ttttataacgg
ttgtttata-agcttagccttgtctctgcttatgtttaggattagtgcaaattaccatca
cctttctgtgacgccgaattattctacatttttcacacatttttcgaacagaaggacgaa
ctttcatattaa----ctctatatactaggaatattactatattaactctattaattatc
gacattaaaaatgtcaacaatatattgtactactttaactgttgtagattttacttcgtc
atactatcatatttttttgatatgacgtatgtctgaatttgcttagctgtgtctattgcg
actccaactaaaataagtaaagacgtagccccgagacctcttaagttttggatctgagta
accttttctattataaacggaatcagcgctactgtaaataaaaacgaggctcctaggaat
gtcagtctatttaatatgacttgtaaataatcaatagtcgcttgacctggacgaatatta
ggaatgcttgcacccatttttttaaatttatagcaatatcttctggattcattactatt
gatgtataaaaatagctaaagaaaaggatcaaagcacaataaagaagaaggtatagtgaa
ccattaggacaaaacaggtataggatttggagtaatgttttattttgaataatttgggtt
agataagatgggagggccatagatgcagatgcaaaaacaataggcataactccaccttga
tttaatttcaagggcaagtaactatttggatccaaaattgaagattttcctagctgtctt
gctgaaataattttaattcttcttgttccttcttgtacacaaattgtaattattatcatt
aataaaaagatcgctataaataatccgaacttaagacttgcattactataactagcatca
aaaaatgactgtgtaaaattctttggtagtcctgacacaatgttttgaaaaataagtaaa
gaggctccattaccaattcccttttccgtaattaactctgataaccacataataatcata
gaacctgctgttaaagctaaaacagactcgcagacaaatgcaaaattccaattaaaaaca
tatggttttacccatatagagattgctccagattgtaaagtcgcccaacctagagctaag
taccttgtaatttgagttatttttggcggcccaattcaccttcttccttctgtaatttt
tccagattggggacaattttcgtaagtagctgcattacaattgaagagttaatataagga
acaatacctaacgcaaaaatccctattgttgaaaaacctcctccagaaaaaatattcaga
aaatttactaaagtatttttttctacacttgcataaaaggcgtcatgatctatacctgga
acaggtataaatattcctaaacgtgctaaaactaatagaaagagagtaaagataatacga
tttcttagatcactttttggctcataaataaattttaaatgaaaagagatactttt--tt
tatgtaaaaaccctattccaagacttgtataagtttaaaatagggaagtaa---aatatt
atagtaatattaata-taatattctatgtagaatacaggctttctattggaactcctcga
tctttagctgcttcactaaaagttcttaattgagttaagccattgagaacagctcgcgcg
ttatttaatgtattattagaacctaattgcttagctaatatattctgtacacccgacaat
tcgaggactgtccttacagaacctccggcaattacacctgagccaggtgcagagggcctt
aatataactttcgcggcaccagaaattccgttaataggatgaggtatagaattcgatttt

gtcagtggaactgtaactagatgtttttttagcatctgttactccttttttttactgcacca
attacatcgcttgcttttcccacgccgacaccaacttggccttgctcattaccaataaca
agaataactcgaaagcttagtttttttacctccttttacaactttagtaactcttttaact
tgtacgactctttcttcccagccactatctttatcttttcctttgctctgtttttttacga
ttggccattttaagaattattcctaattagttaatta-cgattctagaaacccatgcctg
cttccttagcagcttcagccaaggccttaactcttccatgatataactttcctcctctat
cgaagacgacatttttaatgccttctttcatagactgttctgctaactgtttaccaacta
cgcgagaagtatcacagtttggtcttatgttatcagattctttattatttaaattaacag
atgatgtagctactaatgtaataccttgtgtgtcatcaattatttgtgcgtatatatgtt
tattagatctaaatacacacagacgaggcctacttgaggttccttgaactttttttccgaa
ctcttttatgcttatgaattctagtttgtttagtgtttagtttcattattatttaccttt
tccagctttcccagcttttcttctaacaaattcaccttgatatctaattccttttccttt
ataaggctcaggaggcctaatagaacgaatagttgaagcaacctgaccgacaacttcttt
atctatgccagagacagtaatatttgtgttgttttcaactttaatttcaatattggcagg
aggtttaattttcactacatgactatagccaacacttaagattaaatcttggttatcaat
ttgagaacgatagcctacaccttgtatttgcagtttttttaaaaaatccattagaaacacc
ttcaatcatattactaataagagttctggatagcccatgtaactgactagccatttttgt
ttgtccactcgtcttgacagcaattgtgtcgttaagtatttctaaattaataccagctgg
taaagttctggatagggtacctttaggtcctgtcacggtgatagtctgaccatcgaactg
agtactaagatttgtcggcaataaaattattttttttccaatacgagacat-attccacc
tataaaatagttaccaaatatagcataatatttcaccaccaagaccatcatgacgagctt
gtctatctgtcataacgcctctagaagtagaaataagagcgatgcctaaacctccaagaa
ctctgggcaactctttatgatttgcgtagactcttagtccgggtttactaattcttttca
gagcagtaataactggttgacgatttttaccattatacttaagagaaatcattaaatgag
tctctataccttcacccatttgttcgaaattttgaacaaatccttcttctttcagtactg
ttgccatattgcatgtcattttcgttgctggaacttgcacaatttgatgtcttgctaagt
ttgcgttacgaatacgtgtcagcatatcggcgatcgtatcgttgaccaccttgatcctcc
ttgcga-----tgaattatttgaaatattttcaagattccctgaaaggcatccctagctt
ttttaacaaagctagaccttcttgatctgttttagctgtagtgacaattgatatatctaa
accacgaatttgatcaatattatcatagtctatttctggaaagatcaactgttcacgtaa
acctaaattgtaattacctttgccgtcaaaacttctaggactaattcctctgaagtctct
aattcttggtaatgttaaattaattaatttctctaaaaaagaatacatcttgtcttttct
tagatgcacaacaattccgataggaacctcttctcgaattttaaatcctgcaatagattt
tttagcttttgtgacaattggtttttgtccagttattaatgtcaattcttgaatactact
ttcaagagccttagcattctgagaagcttcacctaagccacggttaatagtaattttagt
aaacctaggaacttcatgtacatttttgtactgaaattcatcttttaaagattgagtaac

agttgttttatatttttcttttaatcctattgccattatattttcatcaattatttaata
agttcgccagtctttttcagttttcgaa-ttttttggcctttatcattaattattactga
ggatcgactagcaatattattttgttcactaaataacataacgttagaggtatgtattgg
agcttcaaatttgataatttctcccgtttctccttcttgttgaggtttttttatgcttcac
tttgagattaattcctttaacaatcaccttatttgttttataaataattgcaataatttc
acctgtttttgttttatcacttccagaaataacttgaactaagtcccctttttttaattt
aatttttgtattatttttggttgttttagaaggacctttcattatactacctccggtgca
agagaaactatttagaaaaattttatctcttaactctcgggctataggtccgaagact
cttgtgccacgtggattattatcttgattaatgataactgccgcattatctccaaatcta
atgctcataccgtcagttctccgtaaagctttacgagttcttactacaacggctctgaca
acatcagatcgtttaactggcatattgggagacgcatctttgactactccaataataaca
tctccaatagatgcataagaaggattactagtgcctagtactctaatacacattattttt
ctagcaccactattatctgcaacattaagatagctttgagtctgtatcatacttttattc
tgtataagattaattatcaaaagacttagataagatattaaccattgtccaacactttgt
gcgacttaaaggtcgtgtctcttgtattgtaacaatatcgccgattgtgcattcattatt
ttcatcatgcgctttatatttctttgtccggatcatagtttttgcgtactttctatgaga
gattctattttctacagctactactatagttttattcattttatcgcttactactttacc
tgttgtttcttttaaaggcatagtctttatatttttttatatat---------aattaaat
tttaagattattacatggttgacttagtacgagacttttcaacagttaaaagttgggcta
atctatgcttagaatgtttaaatagatgaggctggaaatcttgccttgtggctcttttta
gccttaaatcaaaaagctctcttttttattacaaggatttcttcagctaaagaagaagagt
ccaggtttgtaacatctgatattttagggaaagtcatattttaggattctgttgtattcc
gaactataaatttagtcttgattggtaatttataagaagctaatttcatagcttgttgag
cagtttttttgtggtacacctgtaatttcaaatagaatatgtccaggcttaataactgcaa
cccaatattctggagctcctttacctgaccccatgcgagtttcagctggacggcagtga
ctggtttatctggaaatactctaatccatagtttaccacctcttctcacatatctagtaa
tagttcttcgagtagcttctatttgtctagaagttaaccaaactggctctgtcgcttgta
atgcataatcaccgaacgcaattgtgttacctttgctagcagaacctttcattctacctc
tatgttgttttctaaattttgttttcttggggcttagcataaaaattaaaaaataataat
aattaagaagctgttgaatcaggagtttcctgattataagcagattctagacttttcgat
tctggcaaaatttctcctttaaaaagccaaactttttactccgagcacaccatacgtagtg
tgagcttggcgatgacaataatcaatatcggctcttaaagtttgtaaaggtacacgacct
tctctaacccattcacttctagctatctctgcaccattaagccgaccagatacttgaatt
ttaacaccctgtgtatttgctctttgagctctttgaactgcttgtctgacagctctacga
aaggccactctttttttctagctgttgagtaataaattctgctactaaggttgcttcagaa
tctggatctgcaatctctacaacatttactctgagttgtttgctagggtctaggattaag

gacagcgattttcttaaagactcaattcccgctccagattttccaagtacgattcctggt
cgagctgttgctatgagaatttctacttgatcaactttacgattaatttcaattttagca
atactagcattactaagttttgaatgtataaatgagcgaattttatgatcttcttgtaaa
agaactgggtagtctttagagttagcaaaccatgaagaacgatgtttttgggtaatgcct
atgcgaaagcccaaaggatgaatttttttgacccacagtatccttgattataagttaaaag
ctataatgactaattgacgttaaa--------agactacagttctgaaacacctaatgta
atatgacaagttggtttatgtattggaaaggctcttccttgtgctctaggctgaaatctt
tttaaagtcgggcctttgtctgcaaaggctttactaacaaacaattggtttttatttaat
ccatcattatgctcagcattcgctgcagcagattccaggatttgctttatatgagaacag
actcggtatggcataaattccagaataattaatgcttcttggtattttctacctctaatt
tggtctaatacacgacggactttatgcggagaaagacgaatgtatttccctactgcttta
gtttcttttacgtttttttgtaatactcatagttatagtttaacgacgggcttttctgtca
ccttttacgtgagtgcggaaagttcttgttggaacaaactctcctaatttatgtcctacc
atttggtctgacacaaagacaggaaaatgttgtttaccattataaacagctattgtatgc
cctaccatatcgggaataatagtagatgctctcgaccaagtttttaggacttctttttt
cctgaaatatttaatgcttctatacgtttaagaagactaacatcaataaaagggccttta
tgtatagatcttgacataataataaatcattagattcttaataaaataaattctatttac
ggcgacgtaagacgtacgggttactatatttatttggatttcgtgttttaacacccaatg
caggtttaccccaaggtgttacaggacgagctcgtccaattggagatttgccttcaccac
caccatgtggatggtctacaggattcatgacaacacctctaacagtaggtcttttaccta
accaacgatttctaccagctttaccaagagtaatattactagcatcaatattaccaactt
gtccaatagtagcgtagcattcttttcgaatcattcgaacttcgcttgaaggcaattta
cagtaacaaaagtaccttcttttgctactatttgagcataagttcctgctgcacgaacaa
tctgtcctccgcacgaaggtcttaattctatattatgtactgctgttcctaaaggaatac
tagataaaggcaaagcatttccaacttcgataggggcagtaggaccagaaagaaccatag
atcctacactgagagatcgagggtgtagaatatatcttttctcaccatcaagatagtgta
ataatgcaattcttgcatttctattcggatcatattcaatagaagcaactttagcaacta
tattatgtctatttcttttaaagtcaattaatctgtactgctgcttatgtccaccacctt
tatgacgacaggtaatgacacctctattatttcgacctttgcagaaatggtgtttaacta
ttaatgatttttctggcttatcagtagtaatctctgaaaaggtagaaaccgttctatttc
ttgtccctggtgtgtaggcacgatataaacgaattgccatataagaagatttgagaagag
taaatgttgaa------ctatgtgatatatttaagtttctggaaacaaattaatagaatc
ttctgaggcaagtgttacaattgctttcttgtaatgtggtcgcttccctacgaatctacc
gatgctcctttttttcttaggaggatgacaagtgttaacacctgtaacttgcacgttgaa
aatatactgtatagcagccttaatattaatttttgttgctttgggatcaacggcaaaaca
atactgattctcttctaataatttagttgtcttatcagtaatgatgggatatttaaccag

atctaataaacctcttgaatcaatgctatccattatatacctcttgtattttagataaag
catcaactgtaataataattttatgtgcagctaacaaagccataatatttagtgtatcag
ctgaaataagttctacgttgtgtagattgcgaatagaaagataaacattcggatcttttt
tgtctactatgactaaaactttttatttaaatcaagattccatcgatgtatagcttcca
taaatagtttagtcttgggctgatgaaaataactgttaaaattttctacaatcagtgtat
taactgatttgttatttaatgctgtccttaatgctaattgtctttctttttattcattt
ttttggtaaaactacgaggttttggcccaaatattacaccaccacctctccatagaggtg
aacggatagaacctgctcttgctcgaccagtccctttttgacgccatggtttacgacctc
cacctcgaacttcacttcttgttttagtattagcagaaccctggcgcttttcattacttt
gctttactaaagctctatggactaagtacatcccagaatcttgactgaccttaagattta
aatcagcgttgccacttacttgaccttcccaattatagacttgataatttaattgcgtgt
taactgtcatgaaatatatcgtatgaaagaatattgctacttactaatttttaccaaagc
gccaggtttaccaggaacagctcccttaacgatcaacaaatcattctctgagttaatact
tacaatttgcaaatttttttattgtaactttttttattaccaagttggccagccatgtttttt
acctggataaactcttccaggcgttgtaccggctccaatagagccaggttgcctatgatt
tttcgaaccatgtgacattgggcctctactaaaatgatgcctcttttggtaaccggagaa
acctttacctacacttctagaagagacattaatcttttggcctacttgaaatagatctgt
agatagtatttggctaacttcgaaatcatcagtagatttcatttcgtattcacgtaaata
ttttagtgggggagcctgtgattttttttaagtgtcccaacaatggtttatttaatttttg
ttcagcaacttgcttgtaaccgacttgaatagcattgtagccatcagtagatacagcttt
aatttgagtaataacacatggcccaacttgaattacagtaactggaattgataaaccagc
ttcatcaaagaattgggtcatacctactttagtaccgagtataccaacagacactagatt
tctcctttttgagagatttatactatcattgtacataaaaattctacttcatagttactg
aag--taaattaaactaacagatgcatgatattattcatgacaacataaaggttaaaaac
tttacctatttaatctaaaatagaaacagctttttgtctagttacagcgtatataaatta
gaagtatgcttattaaatttttttataaactact-tagatttaactgatattgatatccgt
aacttacttgcagtaatatc----aatcaacatgtataagtttaattgcctaaactagaa
ccagctattattagaaataattcaagtatagataaaagcgtatttgaacttttatatatt
gacaaatataatggaatatattgaggttcggtaactacacctttttatttttacatttat
tgtgca--gtatatatacatctaatgcagtaaacataagaatataac----tgtatagca
aatgtaattacacaaattagagaggtattaatgggcaaagtcgttggaattgatcttgga
acaacgaattctgtaattgctgttatggaaggaggtaaacctaccgtcataccgaatgca
gaaggttttagaactacagcttctgttgttgcatatactaaaagtggagataaacttgta
ggacaaattgccaggaggcaagctgttattaacccagaaaacactttctactctgtcaaa
agattcataggacgtaaacagaatgaaatttcgcaagagattcggcaaacatcatataat
gttaaaactagtggatcaagcataaaaattgcttgccctgcactgaataaagattttgct

ccagaagaaatttcagctcaagtactgagaaaacttgttgaagatgctagtacgtactta
ggtgagactgttacacaagcagtaataactgtacctgcttactttaacgattcacaaaga
caagctactaaagacgcaggtaaaatagcaggcttagatgtattgagaattattaatgag
cctaccgcagcttctttgtcttatggactagacaagcaaaataatgaaacaatactagta
tttgaccttggtgggggcacatttgatgtatctgtattagaagttggagatggagtattt
gaagtactctcaacttctggagatacacatttaggcggagatgactttgatcagcagatt
gtagaatggctaatcaaagattttaaacagaatgaaggtattgatcttggtaaagataga
caagcacttcagagattgaccgaggctgcagaaaaagcaaagatagagctgtcaaattta
actcaaacagagatcaatcttccttttattactgccacacaagatggtccaaaacattta
gaaaaaactgtaactagagggaagtttgaagaactttgttcaaatttaatagataaatgt
agtatccctgtaaataatgctctgaaagatgcaaaactagaagcttccagtattgatgaa
gttgttttagttggtggatctacaaggattccagccatacagcaaatggttaaaagatta
attggtaaagatccaaaccaaagtgtcaatccagatgaagttgttgctattggtgcagcc
gtacaagctggagttttagcaggcgaagtcaaagatattctattactagatgtgacgccg
ttatctttaggagtggaaactctgggtggcgtgatgacaaagattataccaagaaatact
actattcctacaaaaaaatcagaagtattttctacagctgtagataatcaaccaaatgtg
gaaattcaagtacttcaaggcgaaagagaactaacaaaagacaataaaagcttaggacg
ttccgattagatggcattatgcctgcacctagaggagttcctcaaattgaagttaccttt
gatattgacgctaacggaattttatctgtaaaagccaaagaaaaggctactggtaaagag
caatcaattactatatctggagcatcaactttacctaaagatgatgttgaaagaatggta
aaagaagctgaagaaaatttcgatacagaccaaaaaagaagaaagaatattgacacaaag
aatcaagcagagtccctatgttaccaagctgaaaagcaagttaaagagtttgaagacaaa
attagccaagatttaaaaataaaaatagaagagctaattacagagcttagatctagtcta
gagaaagaagaatatgacaatattgaatctatttctcagcaattacaaaatgctctgatg
gacattggaaaaaatgctgctcagactgaaagtaaagatacaaaagcgaaggatgacgac
actgtgattgacactgatttctctgaagctaagt-aaaagta-agcgggtaacgcgattc
gaacgcgcgacatcaaccttggcaaggttgcgctctaccactgagctatacccgcataga
ttgtattattacaaaaattaagatatttgtcaatctgtta------atctatac---atc
tagactattaattttggaattacatattagctaactgttgtaaatataacaattgctata
attttttaa-------aagtacttatatcat-tatataagtacttttatatt-------
-caaaaatacaaataagtattatag-------ctaaactactaaggaattaactaagcta
gtaacaattactaatcctgtccataagccggctcctgtatatattaatcctttgattgc
tcccattggccaggggatgctaaaacaacaggtacgcctactactaacacagtcgataat
gcaattaataatagcactaataattggatcgcaataatcatttaatatgttctcctgaaa
aattcaaatgttaataatactgtatataatacaatgaagcttttgaccttcagcaattta
tctatgaagaaataatttaaaattattgttttctaaataaaccttaaattaagtattaaa

atctttattgtaaacaataatactatattttttta--gtagataaaaaatttaaagaaag
tctttttaaaaaaaaattaagataaggtgatatgctgtataaaaagctaagtattatta
attattagagaaatttaaa-ataagaggtaaaacatatgaactcagcccttttttttagca
aaactaccagaagcttatgccattttttaaaccgatcattgatatcttacccgtaattcct
gtatttttcttcttttagcttttgtttggcaagcagcaattggttttagataacattta
ggttataaaaagaataacaaatatacagagtgccaattttgagctctaatcttatatct
tacacaataataactattatggttgaacccttactatcaggaattgttcttggacttatt
cctattactttattcggattattagtagcagcttatttacaatatcaacggggcaaccaa
ttaggactgtaaagcctaaattatagatatttataaaaactacaataattatagtttta
taaatatctacaaataagctaatttaaatacagaaataatactattgaaactagagtga
taacaaaaaaaatttatgtaa---taa---------------------------------
------------------------------------------------------------
------------------------------------------------------------
------------------------------------------------------------
------------------------------------------------------------
------------------------------------------------------------
------------------------------------------------------------
------------------------------------------------------------
------------------------------------------------------------
------------------------------------------------------------
------------------------------------------------------------
------------------------------------------------------------
------------------------------------------------------------
------------------------------------------------------------
------------------------------------------------------------
------------------------------------------------------------
------------------------------------------------------------
------------------------------------------------------------
------------------------------------------------------------
---------------------------caagctaattgtttaccagcttgacttagtaac
tcctggtaaaagacattcatgagacatttctcgaaaaacatggcgagataatccaaaatc
tcgataataacctctacttctacctgttaaccaacatctatttctccctctaacagcggc
gctgtttctaggcatttcttggagtttttgtcttaaatccattttgtcagaaaagctgcc
agcttctttcagttgatctttaatagccagtctttttaaataatattttttgcaagttt
cgatctcttaatctctctttggatcatattctttttagccatgctgtaa-aactctcttt
taaaattatttattcaatatattagctaaaaagctcctgcctgcaacttttatatattt

aaaaataaatagcataaagtagtgaaatgcaattgatatttaatcaacgatacc---act
ttatgctattatcaaaaatttatttagtaaagttcttgttcaacatgagttgaaatagtg
caatcagatgaaggatatgcaatacaggttaaaacatagccagctaaaagttgatcatca
tctaagaatgattgatcagcttgatctacagttccttctgtaactttacctgcacaagtt
gaacaagctccagctctacaagagtaaggaagttcaattccttcttcttctgctgcatct
agaatatatgtatcctccgcacaattaaatgtgacatcaattccttcttcttcacatagt
aaatgaattttataatcagccatttcttcttactccttaataaagatacacttgaaactt
aataaagcaaaat-aatttaaagaacttaccaaaaaattatatactagtttcattatgca
tgatagactggaaaaaaagaaaatagtatactattctgaaaaatcaagctacatttccta
ctaaatagaactatagtaggtccaaagtatatacataaattcatatagttaaaattactt
ttatcatttaaatagctgaacttgatagta---------tagattgatcatc------aa
gctagtcttctgc---caataaataattaattatgatttcaatgcttaatcaacaagtag
aacttaagcctattcttaaatttcagataacacaaatatattcgcattctcttattcagg
aaattaaagccttaataactagacttgttttgcaagtttggagaaggccggctacattga
tggcaggtattatccaacctttattatggttaattctatttggtgggcttttctacaatg
ctcctataaatttgttcactattaatacaagctataattgtttttttgagctctgggatta
taatttttacctcttttactggagctttgaattcaggtcttccattaatgtttgatagag
aatttggattttaaatagattattaacggctcctttagtatcgaggacttctatcattt
tatcttctgctacttttatgacttgtattagtttaatacaagttgtatttatagttacag
cttctcttttatgggaaactcacctctaaatagcgatagtactatgattttggactta
tgattctattagtgactgtaggagttacaatgcttagtttagctttatctttcactctgc
caggtcatattgagctattagcatttattttagtagttaacttgcccttttttatttcta
gtacggctttagctcctttatatttcatgccgccatggcttcagttaattgcaagtctca
atccattgagttatgcaatagaaggtacaagatacttatattcaagcgtaaactggaatt
ttacagagtgtgtgattaagattagctggggagatatttgtttagggcaaattattattt
tattaatcgctttagatataatggcagcttatcttgtgtctaatatattaaaagctaaac
ttaatta------------------aaattttaataaaaaatttattaaatttattttaca
aagatttttactatataatactaatagtagtatggataattgaaatagaaatagtttttc
aaaaccaaagctattatcaacaatgataaacatttgtaagaaagtcaacaaagtatgttc
ttattcataggaggcatgtagtcaatgggactaccatggtaccgtgtacacacggttgtt
ttaaatgatcctggacggttaattgcagtccacctaatgcatactgcacttgtagcgggt
tgggcaggatctatggcattatacgaactagctgtatttgatccttcagatccagtatta
aatccaatgtggcgacaaggcatgtttgttatgccatttatggctagactgggcgtaaca
gattcatggggaggatggagtataacaggagagagcgtatccaatcctggactgtggagt
tttgaaggtgtagccttaactcatatagtcctttctggcatgctttttctagctgctatt
tggcattgggtttattgggatttggaattatttagagatccacgaactggcgagccagcc

ttagatttaccgaaaatttttggaattcatttattgctatcaagtctactttgtttcgga
tttggagctttccatgtaactggacttttggaccaggaatgtgggtatcagatgggtac
ggagtaaccggaaaggtattaccagtagctccagcatggggaccagaaggattcaacccg
tttaatcctggaggggttgcatctcaccacattgccgcaggtactgtaggtatattagct
ggtgttttccatttaactgttagaccaccacaaagactgtatcgtgctctaagaatgggt
aatattgaaactgtattatcaagtagtatctctgctgtttttttctcagcttttgtgact
tgtggtacgatgtggtacggttctgcaactacgcctattgaattatttggtccaactaga
tatcagtgggatagtggatattttcagcaagaaattgagaaacgagtagaaaatgctatt
gctgatggtgctgcacctagcgaagcatggtcaagaattcctgacaagttggcattctat
gactatattggtaataatccagcaaaaggaggattattcagagcgggccctatgaataaa
ggcgacggagttgctgaagcatggcttggacatcctgtattccaagataaagaaggaaga
gagcttagtgttcgcagaatgcctgctttttttcgaaacttttcctgtaattttagttgat
aaagatggtattatacgagctgacattccatttagaagagcagagtctaagtatagtatt
gaacaagtaggtgtaacagctagtttttatggtggcaaattaaatggccaagttttcaac
gatgctcctagcgttaaaaaatatgcaaggaaagctcaattaggtgaggtatttgaattt
gatcgaactacattagaatcagatggagtgtttagaagtagtcctagaggctggtttaca
ttcggtcatgctaatttcgctttaatttcttctttggacatctttggcatggttcaaga
actatcttccgggatgtgtttgcgggaatcggagctgaagttactgaacaagttgaattt
ggagctttccaaaaattaggagatagaagtagtaaaaaacaaggagctgtataagaccaa
agtcaaagatagtctttatc---aaatatttctaatattaaaatagttaagacaaagttt
attatctaatagttactaatcaggagaatttatggaagccttagtctatgtatttttact
aacaggaacgctaatggttatattctttgcgatctttttagagaacctccaagaatagc
aaagtaacttgattccgtttaaacttataataatagaaccactttccaacttattagtgg
aaagtggttttattgtagattacattgtattattcttcatgctcttcaaaaggatctcgg
agatcttttgaagcaggaccaaatgctgtgtatatcgagtaacctgtaatacctaaaagc
agactcgaaataaaaatactaagaactgttgcagtttccataatttaagcagtattaaga
tgaacttttttctataatgatattatcaatattaaaacatttaattattaaaattatcaaa
atatactatggcacttagaactagacttggagaaattttaagacccttaaattcagagta
tggaaaagttgctccaggttggggcacaactcctattatgggaattttcatgttactatt
tttcttgtttttattaatcattttacaaatctataattcttcattagtattagaaaatgt
agatgtagattgggctactttaggtag----ctaactagataagttctttga--------
ttacacaatattaaagacaac----------------aagcgatcgctattagattatag
tagcgatcgctttaaaagtattttactatatatccagagaaaattatttacctaaattaa
ttaactcggtatctgcaaaactatttgtgttaacattattatagttaactttttcaaacc
ttactaatactggatatttgataccacttttatcaattgctgcaacagtaccaatttcct
gataccagtacgattcttttctcaatatttttacttttgagcctctttccatatttttag

tcctcagatagattattaaagattattatattaaaaaattaaaaaaaactaaacaaaata
tataaaaatactaccgacaattgtaatatataactaataaaaggagagagagggattcga
accctcgataacaaaagttatgacaattttcgaaattgttgcaataaaccactctgccat
ctctcctgaattaaaatactgttttaaaaaattagacaagaataacatagaactaaggta
gttgcctagaaaattaaaactatgttaaatataaattataatctttctatagatatattt
aaaaaatatgtacatagaatctatatcagcttacttcaagatttttaatcttatacaat
ttttactaaataaataaattctctcataatatgaaactatcttggaaaacacttcttctt
tggtctttaccaatatttgtaattggtttctttttctggcaaggtttttaggtccaact
actacagatgttggcagtaatatcgcaagttctagaatgacatatggacgatttttagaa
tatttagatatgggttgggtgaaacgggttgacctctatgaaaataatcatacagcaatt
gtagaagctgttgggccagaattaggaaacagggttcaacgaattcgagttgaactgcca
gcaagtgcgccagaattaattacaaaattacgcaaagccaatgttgatctagatgctcac
ccccctaaaagtacaagtgcagtatggggactattaggcaatttactatttcctttacta
ttagttggcgggttagctttcttatttagaagatctaataatgctagtggtggacctggt
caagcaatgtcatttggcaaatcgaaagctttatttcaaatggaggctaaaacgggagta
gtatttaatgatgtagctggagttgaagaggcaaaggaagaatttcaagaagtggtaaca
tttttaaaacagcctgaatcatttactgctgttggtgcaaaaataccaaaaggcgtttta
ttagttggacctcctggcacaggcaaaacattactagcaaaagctattgctggcgaggct
agtgttcctttttttagtatctcaggctcagaatttgtagaaatgtttgttggtgttggc
gcttctcgtgtgagagacctattcaagaaagcaaaagacaatgcgccttgtatcgttttt
attgatgaaattgatgctgttggtagacaacgaggaacaggtgttggaggtggtaatgat
gaaagggaacaaacattaaatcaactattgactgaaatggatggctttgaaggaaatact
ggtgttattgtaattgccgctactaacagagctgatattttagattctgcattattaaga
cctggaagatttgatagacaagtttctgtagatgtaccagattttaaaggcaggttagca
attcttgaagttcatgctaaaaataagaaaatggaacctaaagtatctttagaaaccatt
gccagaagaactcccggcttttcaggagctgatttagctaacttactaaatgaggctgct
atcttaacggctcgacgaagaaaaaatgcaatgactatgtctgaaattgatacatcaatt
gatcgagtagtagctgggatggaaggcactcctttaattgacagtaaaagcaaaagatta
attgcgtatcacgaagtgggtcacgcaataataggcagtttattagagcatcatgatcct
gtgcaaaaagttacattaataccaagagggcaagcaagaggcttaacttggtttactcct
agtgatgatcaaagtctaatatcaagatctcaaatactagcccgtatcgtaggtgctctt
ggtggcagagctgcagaagaaatcattttcggtgacgcagaagttactactggtgcaagt
aatgatttacagcaagtaacatcaatggcaagacaaatggtcactcgatttggaatgtct
aaaattggacctttatctcttgaaagccaaggaggagacccatttttaggtagaggcatg
ggaggaggctcagaatattcagatgaagttgcaactaatattgataagcaagtaagggaa
attgtcagtgaatgctatgcacaagctaaacacattattatagataatcgagtagtgata

gatagattagttgatttactaattgaaaaagaaacaattgaaggcaatgaatttagagac
atcgttaaggaatacactgcaattcctgaaaaaaattactacatatcacaattttaa---
---------------attaaacgggactgacgggattcgaacccgcaacttccgccgtga
cagggcggtgctctaaccagttgaactacagtccc-aaaaagatgaatacccagagataa
tctcatagttagcttaaacttgtcaaattagaatattaaatatgaagagta---------
--------tttgtaataatactcttcatatttgata-tagtagctaaaatatttctttta
gaaggagagagagggattcgaaccctcggtacgaagttaatcgtacagcagattagcaat
ctgccgctttcgaccactcagccacctctccatga---tatatattatatatatatcaaa
tatacttatattttaataggtgtgcataaatatttatggagctaagcggattcgaaccgc
tgaccctctcaatgccattgagatgctctaccaactgagctataaccc----ctttggta
agatattaagaaatggctcaagcgggatttgaacctgcgaccttgggcttatgaatcccc
tgctctaaccactgagctactgagccatagtataaaatactattaaatattataccataa
attatggtcaaatataaatcaataaaagaataattactatca-tttactacttttg----
-----------------tgcaagttgtaaatattcgaataaaatttcaaaattaaaaatg
tctttaactttttgttaga-------atta-ttgattacgaaa-----------atattg
atatttagaattaac---------------------------------------------
--------------------aataaatttg------------------------------
---------------------------------------------------tctttattt
g-----------------------------------------------------------
------------------------------------------------------------
------------------------------------------------------------
------------------------------------------------------------
------------------------------------------------------------
-------tagaca-----------------------------------------------
-------------------------------caaaaatattggaaa--------------
------------------------ttttcatcaat-----------------taattagt
------------------------------------------------------------
------------------taaactataaccgacgtac-----------------------
------------------------------------------------------------
------------------------------------------------------------
---aagaattattgaacta-----------------------------------------
------------------------------------------------------------
---cgttaaat-------------------------------------------------
------------------------------------------------------------
------------------------------------------------------------
------------------------------------------------------------

-----------------------------------tgtgatgcatt------------
-----gggcttgatcaaaacttctctaaaatcaatagaatatagt---------------
-----------------------------------------------------aattcaa
gtaaatcaaatactaacaaatgtgacaaacgcttaaaaaaataata--------------
caattagaatcgtaaaaatttggtttttga-ctgtgttaaataaatttgattatcataga
tttaatatttaaaagtgttctattacttgta-----ttgagtaattgt-aatatttaata
tcttctcttttagataataatcataatataagctaaaaaatagtttt---tattaaacaa
ctagcatagagccaataccttgatcagttaatatttctagcaagagagcatgatctattc
taccatctaaaatatgcgctgaagctacaccctgagctaaagaacgaatacaacaattca
ctttaggaatcatgccaccagaaattactgctgtttgagttaaatctctagcttcctgta
tattcagatgactaattaatgttgaaggatctgaagaattacgtaaaatgccgggcgtat
ctgtgagtagaattagtttttcggcattaagagcggctgctatttctcctgcgacagtat
cagcattaatattatacgattggccttccttgtctgctgcaacactagctataacaggta
tataattattattaattaatatcttcagtaatttagtatcaatattttgtacttctccaa
caaagccaaggtttgcttttccatttggtctcggagtaataagtaatccatcttttcccg
acaaacctacacttttaccaccttgtttattaatagttgcgacaagatctttattgactc
ttccaactaaaaccatttccactatatccatggtaggttgatctgttactctaacgccat
catcgaattttggtaatattttgaggcgatctaaccaaaaattaatttctggtcctccac
catgaactaaaatgggacgtagtcctataaaagacagaaaaacaagatcactaatcactt
gatcttttagtttctgattttcatagccgctcccccatatttataacaataattctgg
aagaaaattgctgaatataaggcagagcttcacttaagacttttactctttctgaattag
tcaacataaataaagtaggattgtgatatataaaattatatacacagcttgtgttaattt
aaaagataccaatgggt---------acttgaaaaaatagaaatattactaacag-----
----ttaaaactatagggattataattatgaataaattttgggataatgtattaagattt
ccacgattttagtcagcgtcattttaggattaattttgataattattagtcctttttc
gtgttattaaaaaaaccactgacaagtttttttttcatcatatcattggccggattaatc
acagttttggcaataataa-tacaaaaaatgataaatatcgaatgttgttgagagaaata
aataaatttaattaaaattttgaacatatataatattatattatcattaaatatgataa
ttt-atttccttttctat---aaaacccataaatattaaatattatttgataaatattat
gccactaaaacaaagagttagctctgaaaagaccggtgccttcgcacttttggatagtat
tgttaggcacggagttaaacatatatttggttatccaggtggagctattcttcctattta
tgatgagctttatgcttgggaagaagcctctctaattaaacatatccttgttcgtcatga
gcaaggcgctgctcatgctgcagattcttattctagatcaacaggagaggttggagtatg
cttcgctacttctggcccaggagcaactaatcttgtctcaggtatagctacagcacatat
tgattctgtacctatattagctataacaggtcaagttgggagagcttttattggtacaga
tgctttccaagaagtagatatttttgggattacacttcctattgtaaaacattcatatgt

agttcgtgaccctagagacatgtctagaattgttgcggaagcatttttttatttgtaaaca
cggtagaccaggtccagtattaattgatgttcctaaagatgtaggattagagaagtttaa
ttatttttctgttgagcccggaaaagttaatattcctggctgtaggccaattaccagcct
aaagtcaagacaaatccttatggcagctaaaatgatacagcaatctagccagccattgtt
gtatattggtggaggagccataatctctgattcacatcaaattattaaagaacttgttga
tttttataaaatacctgttactactactttgatggggaaggggattttttaatgaggatag
cgactattgtctagggatgttaggtatgcatggtactgcgtatgctaattttgcagttag
tgagtgcgatcttttaattgctttaggagctagatttgatgatagagttactggaaaatt
agatgaatttgcttgtaatgcccaagtgattcacgtagatattgatcctgctgaagtagg
aaaaaataggattcctcaagttgctattgtcggtgacgtagcagaagttgttagtgaaat
attgaatttattaaagacttctttccccccttatccagagcagattatatcttggcaaga
aagaattaatcgttggcgtcaacagtatcctttactggttcctagaaaatcaacaagcat
ttcacctcaagagattcttgttgcaactaataaattagcccaaaatgcttattttactac
agatgttggccagcatcaaatgtggtcagctcaatttctgaaagtaaaagctaagcattg
gctttcaagtgctggattaggcacgatgggttatggtttacctgcagcaattggcgctca
agtagcacatccaaatgacgtagtcatttgtattagtggtgattctagttttcaaatgaa
tatgcaagagttaggaactatcgcgcaataccagttaccagttaaaattattattattaa
taatcgatggcaagggatggttagacaatggcaacaagctttctatggtgaaaggtattc
acactcaagaatgacagaaggagcacctgattttcaaaagcttgcagaagcttttggtat
taaagcttttactattaataataggcaaaatatgcaatctgctttacaagttgctattga
ttatcctggtccagttttattagattgccaagttacagaaaatgaaaactgttatccaat
ggttgctcctggaaaaagtaatgcacaaatgataggtatcgctaaaccgcagagaggtac
tgcttccaactatattaataatagtgtttaaagtaa------------------------
------------------------------------------------------------
------------------------------------------------------------
------------------------------------------------------------
---------------atagttagtattttta-aacaaagaata-aagtaca--------
-taaatgtaatttttttatactatta-aaaactatgtaagaacagcctttgacgagaatt
gaactcgtgaccttccccttaccaaggggatgctctacctctgagccacaaaggc-tttt
tactatattgggccgggttggatttgaaccaacgtaggcgaagccagcggatttacagtc
cgcccccattaaccactcgggcaccgaccctaattaaaccttattcaagatatgacctat
aagatatcataactatgttgctaaaaacaagtactttgataaataaaactgaataagtaa
tggacatagctggacttgaaccagcgactttcacgatgtcgacgtgacactctaaccaac
tgagttacatgtccaaaactttaattattattataacactgtttagtgattttataagta
tctatttctaattgaagatttgttttaatcgtccggctttaccgagcagatctcttagat
aatacagtttagctctacgtattttagctcttctaatcactttgacaaaagttattcgcg

gtgaattgagaaaaaatactctttcaactccaataccttgaaatgagcatcttaaagtta

aactagtatttaagcttttttttcttcttctcgctaatactactccttcacataattgtt

cccgagttttactaccttccttaaccattagtccaagttgtattgtatcgccaactttaa

tttcaggtacttctgttttgataaaaggtatctcaacacttttcattagctgacttagct

ttgtactatttgctttcatatataggattagtcttgtgaca-tgttttgaaatattatac

attaaaaaaacaaatatgaaaaattttagcgaataagcatactagtatttataaaac-tc

tttttatct---------------------------------------------------

--------------------------------------------ccattata--------

----------------cagatccatt------------atattcaact------------

------------------------------------------------------------

------------------------------------------------------------

------------------------------------------------------------

------------------------------------------------------------

------------------------------------------------------------

------------------------------------------------------------

------------------------------------------------------------

------------------------------------------------------------

------------------------------------------------------------

------------------------------------------------------------

------------------------------------------------------------

------------------------------------------------------------

------------------------------------------------------------

------------------------------------------------------------

------------------------------------------------------------

------------------------------------------------------------

------------------------------------------------------------

------------------------------------------------------------

------------------------------------------------------------

------------------------------------------------------------

------------------------------------------------------------

------------------------------------------------------------

------------------------------------------------------------

------------------------------------------------------------

------------------------------------------------------------

------------------------------------------------------------

------------------------------------------------------------

------------------------------------------------------------
------------------------------------------------------------
------------------------------------------------------------
------------------------------------------------------------
------------------------------------------------------------
------------------------------------------------------------
------------------------------------------------------------
------------------------------------------------------------
------------------------------------------------------------
------------------------------------------------------------
------------------------------------------------------------
------------------------------------------------------------
------------------------------------------------------------
------------------------------------------------------------
------------------------------------------------------------
------------------------------------------------------------
------------------------------------------------------------
------------------------------------------------------------
------------------------------------------------------------
------------------------------------------------------------
----------agattcttttatagtttc--------------------------------
------------------------------------------------------------
------------------------------------------------------------
------------------------------------------------------------
------------------------------------------------------------
--------tttgataatcagtac-------------------------------------
------------------------------------------------------------
------------------------------------------------------------
------------------------------------------------------------
------------------------------------------------------------
------------------------------------------------------------
------------------------------------------------------------
------------------------------------------------------------
------------------------------------------------------------
------------------------------------------------------------
------------------------------------------------------------

------------------------------------------------------------
------------------------------------------------------------
------------------------------------------------------------
------------------------------------------------------------
------------------------------------------------------------
------------------------------------------------------------
------------------------------------------------------------
------------------------------------------------------------
------------------------------------------------------------
------------------------------------------------------------
------------------------------------------------------------
---------------------------atcta----------------------------
------------------------------------------------------------
------------------------------------------------------------
------------------------------------------------------------
------------------------------------------------------------
------------------------------------------------------------
------------------------------------------------------------
------------------------------------------------------------
------------------------------------------------------------
------------------------------------------------------------
------------------------------------------------------------
------------------------------------------------------------
------------------------------------------------------------
------------------------------------------------------------
------------------------------------------------------------
------------------------------------------------------------
------------------------------------------------------------
------------------------------------------------------------
------------------------------------------------------------
------------------------------------------------------------
------------------------------------------------------------
------------------------------------------------------------
----gatgatcaaatattattttcaaacatggcttatgtatgaaaaagttctata---aa
aacctaatgac--------taaaattagggcttaagtatttcttagttttgtgttataat
tagttactatcaaaggtagtggcataacacgaagcatcaa-aatgttcgaacgctttact

gaaaaagctataaaagtcataatgctagcacaggaagaagctagacgcttaggacataat
tttgtcggaactgagcagatactattaggattagttggtgaaggtactggaatcgcagct
caagttttgaaatcgatgaatgtgaatctaaaagatgcaagagttgaagtcgaaaaaatt
ataggaagagggtctggttttgtagcggttgaaattcctttcactcctcgagcaaaaaga
gtactagaattatctttagaggaagcacgtcaactaggccataactatattggcacagaa
cacttgctaatgggcttagtccgagaaggagaaggagttgcggcaagagttttagaaaat
ctggcagtcgatgtttcttcaattagagctgaagttatacaaatgctcggagaaaatgcg
gaagccaatgtaagtggaagcaatactacgcaagctagaagtaaaacaccaacattagaa
gagttcggatctaacttaactcaaatggctatggaaggtggtttagatcctgtagtcgga
agacaaaaggaaatagaacgagttattcaaatcttaggtagaagaactaaaaataatcct
gtcttaattggggagcctggtgtaggtaagacagcgattgcggaaggattagctcaaaga
attgctaatagagatgtaccttctatttagaagataaattagttattactcttgatgta
ggtctattagtagccggaactaaatatagaggtgaatttgaagaaaggctaaaacgtatt
atggatgagattaaatcagctgataatgtaatattagtgattgatgaagttcatacattg
attggggctggtgctgcagaaggagcaatagatgcagctaatctgcttaagccagcttta
gcaaggggagaattgcaatgtataggtgcaacaactttagaagaatatagaaaacatata
gaaaaagatccagcattagaaagaagatttcaaccagttgtagttggagagccaagtgtt
gaagaaacaattgaaattttgtttggtcttagagaccgttatgaaaagcaccatcaatta
acaatgtcagatggagctttggctgcagctgctaaatacgctaatcagtatatttctgac
cgatttttgccagataaagcaattgatttaattgatgaagctggttctagagttcgttta
ctaaattctcaattacctcctgctgccagagaattagataaagagttaagagctgtatta
aaaacaaaagatgaagctattagagctcaaaaatatgaaacagcagagcagtatagagca
agagaaatggaaattaaagctcaaattgcagcaattgctcaaagtaaaaagaatgagcct
gatttaaatttagaagatcctgttgttacagaagatgatattgctgaaattgttgctgca
tggactggtataccagtaactaagcttactaaaagtgagtcagaaaaaattaatgcacatg
gaagaaactttgcatggacgtattattggtcaagacgaagcggttgtagctgtctctaga
gcgataagacgcgcaagagtaggtctaaaaaatcctaacagaccaattgcaagctttatt
ttttccggaccgacgggtgtaggaaaaacagaattaacaaaagctttggcttcttatttc
tttggttcagaagcttctatgatacggctagatatgtcagaatacatggaaagacacact
gtatctaaactaattggttctcctccaggatatgtgggttatagtgaaggtggttatcta
acagaagcggtaagaaaaaaaccatatactgtcatcttatttgacgaaattgaaaaagct
catccggatatttttaatctacttcttcaaattttagaagatggccgactaacagatgca
aaaggtagaactattgattttaagaatactcttttaattatgacttctaatattggatct
aaagttattgaaaaaggaggaggtagtttaggctttgaattatcagaagatcaaacagaa
tcccaatatactagagtacgatctttagtaaatgaagaactgaaacaatactttagacca
gagtttctaaatcgattagatgaaattattgtatttcgtcagcttactaaagatgaagtt

agagaaattgcagaattaatgcttaatgaagtctttgcgagaattaagcagcaagatatt
caattaaatgtaacagaacgatttaagcaacgattagtagaagaaggatataatccaagc
tatggagctagaccacttagacgagcagtaatgaggcttttagaagatagtctggctgaa
gaagttttatctggtaaaattaaagctggtgatagtgcagtagtagatgttactaatgaa
ggagaagttacagttttattaggtgaaaaattagaactgttaacataaa------aaaca
atttattaat--taaagcattgaagaggttat--attcaatgctat---gaaaaaaatta
ataagatgggttgctataattgacaacctctgatactctttatttgtatata---aagcc
gggatagctcagttggtagagcagtggattgaaaatcctcgtgtcaccagttcaaatctg
gttcttggcat-tttaagcatcagaaaattttgctatctgagcattaaattttaattgaa
atgttcctgtcggaccattcctgtgttttgctattataacttctgtgaagtctctacttc
tagtttcttgtgtatagtaactttctctatatagcattattactaaatctgcatcttgtt
ctatagaattatgaacaataaagttgtttgaaataaaattgcataaaggtttacattcta
aatcaaacactatttgtaaacttgtaacgtttatttttttgtatactttcaaaagtcaaag
atgaaa------aagaattaaaaacatttttttgttcttcaaagttttt------tattt
gaatagcaatcatgtcattttgattaattttatcacatcttttccatccttgtgttgtga
gtagtttatggttacttgttaattgtatatatttcccagcttcggtaataattttatata
cagtttttttccctgttttcgaaatactacatttaattgctagaaataactgttgttttt
gcgcactaaagttaatgacttca-atctttttataatagcaattaaataatatttg--aa
gttgattatataatgatgtctgaagataattgaatttggatatacacccactttctctta
aatcggataacaaaggccttttgtcactccgagtttcaaggtttctatttagttgagata
atgccaagataggcaagcttaaatcttttgctaatattttaagtgatcgagtaattaatg
atagttcttgagatctattgtcagattgtctactctcctgcagcaattgaagataatcta
taataattaattctatatttttttccttgtaatttaaagaattttacttttgtctttatag
tgtcagtagaaatcttagcgctatcatcaatataaagattaaggttagcaagagttttac
ttttttgaataacatattgccattcatcattattaagctgaccagattgaatttttttggc
cgtttaaattacactcttgagctaaaattcttcttagtagttgctctgtagacatctcta
agctaaataaaataacataagatcttttctgattaattacatatctagttatattaatag
caaaagcagttttccccattgaaggtcttcctgcaagaataattaaatctgacttttgaa
acccttgtgttatagagtcgagttctgtaaaaccagaaagtatactactatttatactaa
tctttttccttgtatccaaatggactaataggtttgcaagaattgtagctaatgtttgcg
tatctttttcgtcaagtatttcataagcttttgttaattgactaacaatagaagttatgt
tttttttgttcaataagttgttttgagcagctaattaaacataatgaatctccacatgcta
aaagtaatcttttaatgtagttatctataacgactgctgaatattcgcttatagtatcag
ataggggagcttgttctattagactaaaaacttcatttagctcattcaaatcttgaatca
ttttttcatttttttaagttaatgaaaaaaatttcttacgctttttttgtctttgcatggttaa
cattttcgagtgctgctctataaagtaatgaagttgatgtaaaataaaaaaaaatcaggag

agatttttctactgattttagtaatgatatcgattgctgagttaatattatactaatta
atatttttcagctaaaatattatgaggaggtagatatttataa------atacttatta
cttcctcc-ttgtaacattactcttgtg--------gtattagttaactattgatttaat
ttgattctggaagaacttgcaatttaatatttgcagttacttgatgtaatagcttaattt
ctacatcgtaaatgccaattgttttcacgtctggtaaagagatactttgtttttcaatgt
ctatgttagtagtgtttttaataatttgtgaaatttctttttctgtgacactaccaaaaa
tattgtggccatctcctgtttttttactaacgctaaacctttgtatttcttctagaagtt
gcttgactttttagcattttctttagctgtttttaattttttcttctcttatagctgcat
aaaatttttgttgttttaaaataccatttgtggcaactgccgccattttattcgggatta
aaaaatttcttgcataaccagaagctactttgataacgtcattactcttgccaagttttt
gaatattctcttttagcacaacatttataactttttttactcatctt-------aattcta
aaaa----atctgtattatcctattatacctgttgtatacagatttt----aatagtaag
tatttatttgttgatgtaatattcttgtaaaa--aaaggtaatttttttatttttt-ttctg
atataatcactttcacaataattc-----attaggagaggtggccgagtggttgaaggcg
cagcattggaaatgctgtttaggagcaatcttaacgagggttcgaatccctctctctccg
tacaa--aaaatataaaaagttttatagagtccaatctaaatctccaataatattcttcc
aagcatttaatccaccccgaactctaacaatatttagtttttgagctattaaaaacttag
aagcaaatattgatctagaatctaagctacagtaaacaaaacaaattttatcttgtaaat
tgatatgagagtaattcattcctttttatttttttgtaaaggtagatttaatgaatgtatta
aatgacttttatgatattcttcatgatttcttacatctattaaaatatattgtggattcc
tactaattaaaaacttttgtaattgaataacactaatttctcgtacaaatgtactagatt
tactgccataatatttgttccaatgctttttttgt-------ggataatacaaatttagta
tttataatcttaaatttgttaaatgatgaagttaatgagttgtacgttaatatagttcca
cttaatatagacttataacccagaacgattttgacagcttcagttgcttgaagtgtacca
acaatacccggtaatagacctaaaactccggaattactacaagtatctcttgcgctctct
ttattttcagtttcactgtagaagtctcgataaacaggtccaccttgataattaaataca
ctaacttgtccttcaaattgaaaaaatggcaccataaatatgtattttatttaactctagg
caagtatcattcaagagatatcgagtatcgaagttatcagatccatctataataatatca
tattgtctaataatgtcgattgaattctcatagcttagtcttgttttgaacacagttact
atacactgtggatttaaatctaata-tttttttttagctatttctacttttgaatatcct
atatcatgcatcgtgtataaaatttgcctctgtaaattagagagatcaattatgtcatcg
tctataattccaatgctgccaattcctgcagcagcaaggtaaatgattcctggagatcct
agcccaccagccccaatgaataaaacttttgcttcttttaatctttcttgtccttctaat
tgaatttgtggcaatactaaatgcttagaatatcttgtatattcttctagtgagtatttt
gtattttctgttttaaaatttagcataaaaatttatagtttatttaatagcactta----
--aaagctagctgataagtttattaaaatgacttttgtaaacttggttcttctctatcta

aaatagctccttgtgtaggacagacctgaacacaaatactacaatcaatacaggcagaaa
agtcaatccaataccaattagtgttattattgtttttccctttccttgatgaatacaag
ctacggggcaagcttccacgcattcagaaactccaatacatttttctgttacaattgtat
gagacatattatttaattaat-aaaagataagctatttacatctagtagttttgttgtat
actaaatttacatatgctatccaggccaatatttaaggtagtctatttctacagtttact
aatttgatatgcgcctttagttcagttggtagaacgcaggtctccaaaacctgatgtcga
gggttcaagtccttcagggcgcgtttag-ttttatcgtaattcaataatatcattttatc
tttttgcaaacatcaaaatgatatgatataaaagatcatattttatacattgaatattct
atgtactagtactg-ttttttaatatgttattattgataagttaattatactaattctat
ttttgtaataagt-aaaaacagttctaattaagtatgcacgtatttaatagtc-ttttgt
atatat--aacttgaaaaaatttaaactatggctaaaaaagttactggaattgttaaact
agcattaaatgcaggtaaagctactcctgcgccacctgtcggacctgctttaggacagca
cggcgtgaatattgttatgttctgcaaagagtacaatgcacgtactgctgacaagtcagg
actagtaatccctgtagagatctcaatatatgaagatagaagttttactttatacttaa
aactcctcctgcttctgtactaatcgctaaagctgctggactaaataaaggttcgggtga
accaaataccaaaaaagtaggcagcataacaaataaacagttagaatctattgcagaaac
aaaacttcccgatttgaacactaacaatatacctcaagctatgaaaatagttggaggtac
ggctaaaaatatgggaattctaattaaagatt-aaaagtttcaattatatttacta-tt-
-tttatgaaaaaattttcacgtcgacttacaacattaaaatcaaaagttgagcctaaact
ttacactattaatgaagcagtatctatattaaaagcaacgtcaaatgctaaatttaagga
aactgcagaagctcatattgctctaggtttaaatcctaaatatgcagaccagcaactcag
agcaacagttatattaccgaaaggtacaggtaaattgataaaagtagcagttattgctaa
gggagaaaaattaacagaagcaattagtgcgggagctgatgtcagtggctccgaagaact
aattgatgaaatactgaaaggtagattagattttgataagctgatagcaacaccagatgt
tatgcctttaatagcaaagctgggacgagtattaggccctcgagggttaatgccttctcc
taaagcaggcacggtaacattggacgtagcaaaagccgtgaatgaatttaaaggggtaa
agttgaatacagagttgatagaacaggtataattcatgtaccttttggaaaatctagttt
ttcacaagaagatctagttttgaacctccaaacaattaaagaatcgattgatagaaacaa
gccttctggagcaaaagggaagtactggaaaactttttttcttatctagcaccatggggcc
atctattcaaattgacatcactagccttctatgaaaaattttgtataattatagaactt
gaataatctattatttcaaatta-taatttgctagcattattaacgtaacaacttgctta
ctatatcttatctgcattaaaaacttt-ttatatatcttattaattatgagtacaaaggt
tgaaaatatcttggaagagctaaaatctttaaaccttctagaggctgctgaactagttaa
acaaatagaagaaacgtttgatgttgatgcatctgcggcttcaggaggtatgatgatggc
cacaccaacttcagcaccagcgtctgctgaggttgaagagaagacagaatttgatgttgt
cctagaggaagtcccagcacctaaaaagatttctgtattaaaagttgttcgctctctgac

tggtttgggtttaaaagaagcaaaagatcttgttgaatctactccaaaagtcttaaaaga
aggtgcttctaaagatgatgcagaaactatgaaaaaacagctagaagatgctggcgcaac
agttattgttaagtaatga-----agaagcgcttctgtttaagtataaacagaagtgcct
ctttatttattataagcaattatactagaaaagtccaagtgtaattgcattgttgatagg
catagttgcaccaatacctagccaaatacttacaacggtaccaattaagaaaacagtggt
tgcaattggtcttctaaatggattttgaaatttgttaacattttcaatgaatggaacagt
aagtaaaccagctggtacagcagccatacttagaacacctaatagtttattaggaatcac
tcttaatagattaaatgttggaaagaagtaccattctggtaaaatctctaatggagtcgc
aaatggattagacttttcacctatagaagatggttctaaaattgcgagtccaatactgca
ggcaaaagtgccaataattacaactggaaagacgtataatagatcatttggccaagctgg
ctctccgtaataattatggcccattccttttgctaatttagctctcagcttcggatctgt
taaatctggtttttaagaattgacatattg-tttttgtgtttaatataaagattagtct
aattactgttttaaagtgtaattttcaactattataatggtccagaaattccttgtttac
gtatcattaaaaaatgcataagcatgaaaacggcagttagtagcggaagtacaaaagtgt
gtaaactatagaacctagttaaagttccttggcctacactaactcctcctcttaacaatt
ctactatactcgctccgacaactggaacagcatcagggacacctgtaacaatttttacag
cccagtatccgatttgatcccatggcaaagaataacctgttaccccaaaagaaacagtaa
gcactcctaaaattacacctgttacccatgtcaattctctaggtttttgaaacctcctg
ttaaataaacacggaatacatgtagaatcatcatcaagaccatcatgctcgctgaccatc
tatgaattgatctaataagccatccaaagtttacatcagtcataatgtactctactgatg
taaaagcttctgcaactgtaggtctataatagaatgtcattgcaaatccagttgcaactt
gaattagaaaagatacaaatacaatacccccctaagcaataaaaaatattaacatgaggtg
gtacatacttactagaaatatcgtcagcaattgcttgaatttctaatctctcttcaaacc
agtcataaattttactcataaaatagcttcaaaaagctcttctaatata--ttttgcgtt
taagctactattaattcaatag------ttattgaacttatatt-tttttataatcaata
cttattaactattataacatcttttattgattt---------------------aaaatt
ctctgacttagtaaaacagggctatgtactataattcttttttttgtatagagattaat
ttcgtttttagtaaattagctaaaattcttgttacactgactctattactacctataatc
tgtgctaaaattttatgtgtaatagtaaagttaagcataataccattattaacttgagtt
ccattttgctcagctaataataataataagctcgccagtctactggttatacttttatgt
gaaatgatttctataaaatgataagcttttgccgagcaaaatactaaatgatttacaaaa
aataaattgaatggagcgcagttactacatgcttttataatagtagtatagttaatacag
gcaacttctgttttatctattgcttctgcttcgtagtaaaaattatcgtcgaccaattct
atttgtccaaatgtatctcctgtagttaataagttaagagttactttatgtgtatttctc
aggacttttgtaataattaaagatcctatcaatataatgtataatctagagctgctatta
aacaaaagagtatcatttttcttaagtgaaaaaatttgataattaattttattacgatta

aaaaatagtagccatggattgcaagaaaaaaaatcacctgaactattttcagagaatatt
ggaatacaattttcttgatttaaacaatttttagtcataattactactttataa------
---aaaaatattctaataat-taagaactgaaattaaatgtcatataaattgatgttagt
tgagaatgacattgtattatcaaaggccattcaagaatacttaatagatcaaggcttcaa
tgtttatattgccaataatggattagaagcactgaatctagcttatcaatataatttcga
cttgataatttctgatataatgatgccgctagttaatggctatgaactattagcaaaact
taaaaaaaacaaagcattatccaaaatacctgttatctttttgacagctaaaggaatgac
taaagacagaataaaaggctacgacatggggtgctatggttatttatctaaaccatttga
tcctaatgagttgctttcaattattaacaacttaatcgctagagatgtcttaaaagaagc
ttctttacaaaactcggcaacatcaaaccagcaattaaatcataaaatacgtttgacacc
tagagaaaaaagtattttagatcttgttgttgatggacttactaataaagaaatatctac
tatactgaatactagtgttagaaatgtagaaaaatatgttagtcgactcttacacaaaac
taatatgaagaataggactttgctagtcaagtattctataaataataatctattaaataa
tgagatcaatgaaagggcgaatgacgggactcgaacccgcgaatgatggagccacaaccc
attgccttaaccccttggccacactcgccatattacatacaccactatagcttgtttttt
gatatatcgtctagttttcgagttatatatttatctaaaacaatatttactttgaatttt
-tatgatacatagcaatattagtattcaaattaatggggaaccatttaattgctcaaagc
ccatctcattgcaatttttattaaattatcttgattttaattccgagcgtgtagcagtcg
agctaaatgacattcttttaccagagactctgtttcactcaacttatttgaatgatcaag
ataagctagaggttattaccattgtaggtggtggataatctatttgtacatac-------
-acagttttctatctcaaatgtttcatagatagagataatcttccttgtttttatctac
atgaataataacagctttaatagtgtcacctattttaaattgtgatgatatctgctctaa
ttctttgatattaatctcagaaatatggacaagacctttgagatttcctacctttataaa
taatccgtaaggtgtaatctgatttattatgccttcaataatattaccaacaattaaatt
agatgatgcttgagatattagagctcttctatgactcagtattaaattattagatttctc
ttcgacattgagtaattttagcttaataaatttattattaaattgctcactttttgaaa
attaccgagatgagaattaggtacaaatccagatataccttcaaggttaattatcattcc
tcccttattgaatccttttatcattacattaagtaaagagtcttcagctaatagttgcct
aattcttttccatgctcttatatattcaaggcgacgaattgataatattaattgtttcga
ctgaatattataatctaataaaaaaaattctcttgtgtcattaatgtttaaagaagtgaa
attatttaaatcttgattactagatacttcttgaataggcaagtatgcagatattggtgt
tcctatatctactaataccccatttaattcaaagctaaatatagttccagctacaatatc
tccaagattcaaatcatacttgtatttttgtaaaacagctgcaaaatttctatgagtaaa
tccttcattatttttttgtcatattgaatagtatgaattaaagagtaactttgtacaatat
attaaggtaaataaatgtttgtcgaattgtatgaaaaaattttaatttgttcggttgtaa
ttaagtatataaacatattaaaatatctaggagacatgaaaagtagacgtagatagctgc

agatttaa-tactatataatctgaggaaagtccgggctccacaaatacaatttatgctgg
agaaatcccagtgtaggtaactgcgaggatagtaccacagaaaaaaaccgccagaataag
t----tatattagctggtaagggtgcaaaggcaagttaaaagcttaccaaaagtactgca
aagtatttgttaggtaaacccctaaaatggagcaaagctactaaacaaatatttgtgtat
attttatctagtttagttgaaaatactgcatgaagttattagtaacaataactcgagagg
aatagctatcctttacaatatatc-tcgattatattgaaagaacagaacccggcttatgt
agtacttttatgtcttgtattattatgtctattgtcgaatctttaaatttaactgtttg
acaatttttttttgtagctgatgttgtttataatctatttctgaattggttaaagttttt
tcattagatcgatatgtaaatcttagacctatactttttttctcttttccaattgattca
tctatatattgatcaaataatgttgtactttctaaatctttatcataaaattgatctaat
aattttaataagtatttaatttccatattcttaggtattattaaagagagatctcttatg
atcgatggatacttagagtatggttgaatctgataacttaaataattgaattcattatga
caatcttctaaaatgtttaagttaacttcaagaaaaaaaagttttgtattgagtcctaat
tcactatatgttgcttgattaagttcactgaatatgccaataataatattgttatagatt
agtgtggcaaattttcctttttgaatgaactttattctaccatcgactatatcttttttt
gtccactcaatttgtctattaatcttttgaaaaaaatttctacgatccctttagcttca
taccaatttaaagaatgcgcagggtgtgaccatgtagatcggatatctagattaccaccc
aatatgatagctagattagtagtttctacaattttgttttgatcattatgaaatactgtt
cctatctcaaagctatctatagtttggttgctttgtttaatattataaagatttgattca
attaatccttccagcaaactgccgcgtaaattagaataatcttgaataagaggattgttt
aaagctatatttcctttggactttattaaagaataatgaactaattcagttaatcctaga
tttcttaaaatacttctacttttgtctatgaattttctctttgtggataaccttttgttg
aattgaatatttggtactgagctcctgaatttatgataaccataaattctagcaatttct
tcaataacatctatttctcgaaaaacatcttctcgtctataatttggaacagtaacttct
atattttccttattttgataaataagatcaaaatttaaagagtgaagaatattttgaatt
tcttcaaagaataaaaaacgtgtttgaccattatacataattggtcctaaaatgtcatga
actttttaattgataaattaatattaagagcatgatcagttattttttctcgacagaaa
gtatctctaatatttccgcatgttaaatctgtgattaaagcaagagcttcaaaatgtgca
tttttccaattatctacatttagcccacgttcttgtctaattgaactttctgttcgaata
ttaagaactcgagaagactttctaactacagattgtttaaatatagcagactctacaagt
atcgattttgtattataatctgtatcaaaatcagagttgcttccaataccagctatacta
gttacatttgtatttacttgagtaactaaaatatttttgtttaactcaatattttcattg
tttaacttaatctgatcattacttccaagaggaaaattgcttcgaattgttagactattt
ttatgattcatattattaattttattcaaatcaataatattaataggctgtccccatttc
aacataatataattgcttatatctgttaacaaatttctgtgtgtaaacccagaagagagc
aaccgatttttagccaatctggagaatctttaactcttatttcgtctataatggcagca

aaataattgtcacaatttaataaatcgtgattactgattatagt-ctcctttttagagaa
atatctattattgggttgatgtatagctgataacattgaggaccctgttaaagccgagac
ttctcttgatagaccaatcatgcttaacacgtctgacctgttagctgttgatgatatatc
taatatatgatcaatctcacctccaattataatagattcaatagcttcaacttcaaaacc
tgcttgtgttagttgattagttaaattgtctatatctattgttttgatattaccaagttc
tttcaaccaatttaaagaaactttcataattaattgtcatctatcttatagaagtattta
ttttacttatgattaaaaatcaatttaaaaaattcatcaaattaatataagttatttgat
gaatttgcttatacaatatattattgtgatatcttacttttatatggcagttattaagaa
ctagcttttgtaaaagttagcccattatcttgggcatatctttccataaatctcatgaat
ctatcccaatcatcaggacttttaataacatgtacagcttctattgcttgtggtttacca
ttaataaacttggcattaacatctcttgtaataatttgtccctctgtgtccatcaggtac
atacctgtgatttcacctttgtctgccatactagcatctaaaattttgggatttgtaaac
cgaaaagttgctgttcctgtactaccatctcgagaccttgttaaacgtacatctggtaca
acttcttcattgataccttgaataaattgaattgttgccatatcaatcttctttatttgt
atataatttatattggataaaactgatttctagttcttgaatgcttgttgagaaactgtt
ttaaaagcttatagtatcagtagatcaaattttatgtattatataaaactgtaactttc
-ttttttg----------------------------------------------------
------------------------------------------------------------
------------------------------------------------------------
------------------------------------------------------------
------------------------------------------------------------
----------------------atctgttcaaagaactaacttattataagctaatcata
cgaatattagccatataatgttcgtaagt-gaaattaaggggttgtagctcagaaggata
gagcaagcgcctcctaagcgctaggtcagcggttcaagtccgctcaaccctgttaacgaa
ttaagtagtattatttaacttgttctgcaaatcagtattgatctaagttaaagtacttga
cagctaaaacaaaagaatctgagaaatattttcatttaaagaacgaacaattatcttaac
tctaaatatctgataattagtaa-aaaattct----------------------------
----ctttttaaaacacaagatgatg-----aagatatttaaactaataattgaaattaa
gtttttgaagttatagaagccaat--ataatactttctattaacaga-----------tg
cttgcttctattactatctgtttacaaga--aatt----atgagtatcaaaata------
--attttagctcca-------------------------------------ctgac----
------------------------------------------------------------
-----------------------------tactaataaata-------------------
------------------tttattttagtctagacaag------atgaatcttcatggta
taacag------------------------------------------------------
------------------------------------------------------------

------------------------------------------------------------
------------------------------------------------------------
------------------------------------------------------------
--------------agctattctttgtgtcatacttatataaaattaggat-------ac
tgtt--------------------------------------------------------
------------------------------------------------------------
------------------------------------------------------------
------------------------------------------------------------
------------------------------------------------------------
------------------------------------------------------------
---------------ttcattaccataactggggtgggaggattcgaacctgcgaatggc
ggagtcaaagtccgctgccttaccacttggctacaccccaacatgtgaacattataatag
cagtcgtataccctgccatgtcaatagactattgtctttatttaataatttttgctgctc
ttgtagaataaaatcttttttttcaaataagatacgctatgtttaagttgcacgactgtt
atgttttcttgcttttgactgaacagccacttgatagtaattattccttctgcaatttcg
ttacttcctataataatacaagctacagctctttttttatttgcttgcttaatctgctta
ctaaaattacttgagctaacatcgatttctattttaaaaaactgctgatgtaaaaaacgc
attatttgcataccagtttcgttagcttttgtaccttgtgtagcaatataaaaatcgata
gatttattaggcaagtctatattatcttttgcaatcagtaataggcgttctaatcctatt
gcacatccaacagctggtgttgctttgccacctaattggtgaactaagctgtcatatctt
ccaccaccacatatagtatcttgcccttttgatgtcaatgtcttaatttcgaaagcagta
tcattataataatctaatcctcgaactaacttattatttatattgtagggaatattaagt
aattttagataattgcaaacgtcaccaaagtgttgttgtgattccaaacttaaaaaatca
gaaatttttggagcctctgtcagtatttttgtgtattagaatcttttgaatctaaaatt
ctaattggattactagttaatctttttgtgagtcagtgtctaaatcatcatggtacttt
gtaagataatctcgtaatttaacttgataaatactacgatcctctacttttccaattgaa
ttcagatcaagtttcaagttgtgcagattgagattgttaaatatgctcatagctaaatgt
ataacttcactatctgctcttgcatctaagctaccaataaactcaatgccaagttggtga
aattgtcgttgtcgtccactttgtggtctttcgtacctgaacattggaccgctataccat
aatctttgtaaactatggtggtaactcatttatttcaataaaagatctcacaataccg
gcagtaccttctggccttaaggtaatatctctattgctgcgatcatgaaaccgatacatt
tctttatttacaatatcagtatcttcaccaatacccctatcatataattcactattttca
aaaataggtgttctaatttcttggtaatttgcacattctaataatttggaaattttatta
tgtataaactgccaatattgaatttcatcgggaagaatatcttttgtccctctaatagcc
tgaatttttgccatgataatttttgattaccttcttaatatgtctgcattaaatagcag
ttgtagactgttattcagtcatgtattatacattag--aacatactgatg-taatttata

aagtacgggcaaggagggattcgaacccccgacaccatggttcgtagccatgtgctctaa
tccactgagctacaagcccacttaataagtaccttaactatatcattgttcactaatgta
tacaactatattcttttt------atcttacatgtgaaatttttgtaaaagtaaaaaatt
at----gtttgtttttacaattggaactattcatatataatgcttaatttattataact
ttgtaaa--aatatctataagtttaattgtatgattctattatattggtaatcttgttat
aattttgcgctcttatctttcaattgg------ctagcatctaattaacttatatttcaa
ataaaaaaatcttatctaaaatgagtaaacaaattctatatcaggatgatgccagaaaag
cattagaaaaaggcatggatattttaacagaagctgtttctgtgactttaggaccaaaag
gaagaaatgttgtcttagaaaaaaaaattcggtgcccctcaaattattaatgatggtgtaa
cgattgcaaaggaaattagtttagaaaatcatattgaaaataccggagtcgcactgatta
gacaagcagcatctaaaacaaatgatgtagctggtgatggtacaacaacagctacagtac
tggcttcagcaattgttaagcaaggaatgagaaatgttgcagcaggttcgaatccgatgg
ctattaaaaaaggtattgaaaaagcaacaaattttgttgttagtaaaatagctgagtatg
ctaagccagtagaagatacaaaagctattatacaagttgcttctttgtcatcaggaaatg
atatagaggtaggtaaaatgatagcgaatgctatagaaaaagttggcagagaaggagtta
tttctttagaagaagggaaatcaactaatacgattcttgagatcacagaaggaatgcagt
ttgaaaaaggctttatttctccttatttcgttacagatacagaacgaatggaagttcttc
aagaaaatcctttttatttatttacagacaaaaaaattactttggtgcagcaagaacttg
tgccattgcttgagcaaattgcaaaaacatctcggcctttattaataatagctgaagaca
tcgagaaagaagcattagccacaattgtagttaataaattaagagggattttgaatgttg
tagcggttagagctcctggattcggtgatagaagaaaatctttacttgaagatatgagta
tcttaacgaacggacaagtaattactgaagatgcaggtttatcacttgatacagttcaat
tagatatgttaggaaaagcccgaagagttattgttactaaagactcgacaacaataattg
cagatggtcatgaaattaaagttaaatcaagatgtgagcaaattaagcggcaaatagaga
cgagtgactctttatatgaaagagaaaaattgcaagaacgattagcaaagctttctggag
gagttgccgtcattaaagttggtgcagctacagaaacagagatgaaagataaaaaactaa
gactagaagatgcaattaatgcaacaaaagcagcaattgaagaaggtattgtaccaggag
gaggagctactaatgtccatatctctagtgaattatttacatgggctaaaaacaatttag
ttgaagatgaattaattggtgctttaatagttgaacgagctgtgacctatccgttaagac
gaatagcttttaatgcaggtgataatggagcagtaatagtggaaaaagttaaaagtcacg
attttcatataggctatgatgcagcaaacggtaatattgtaaatatgtacgatagaggta
ttatagatccagcgaaagtagctagatcagctttgcaaaacgcagcttctattgcagcaa
tggttttaactactgaatgtattgtagttgataaagtcgatgattgagctata------a
gataatagataatctcaaacaaagaataatttctaattatagaaattattctttgtttca
attaatctaattaattgatcaaatgtataatttttataataaagccaaagtttaaacacg
tcttctttttcaactattaagtagagaagatataagcccgtaatacctaaattatcaatt

aaaatattattctcatatatagatttttctaaagatctaaaaaatgacatttttgaaaaa
taataccgaaatcttctcttatagttatttgctaaatcattactgtctaaagattgattt
ccaattgaattcagaacctttaacagaatactttttaaatctttatctgactttatttga
gaatcaataataatttgattcaattgtatgtcaaaactaagattaaaaatattacttgtt
aaagatatttgattttgaatatctagagattttatagccaagaagattaaacttttattg
ttcattgtaaaatataagacaattgatgatgtattctttaaatacta-aatttatcttct
tgtgcttctatcgttttttgt-aaaataattaatgttttaatttttttagttttagttact
acccgaaaaaataaattctggaaatctagcttgagctttagctagagagtgtttttttccc
ttcttcttgccagtaattgataacttctgttgttttgtttaataactcaattctgtctct
ttcagcaatccacggcttggattctagttctgcttttaattcttcccaagcatcattcct
tggccagaaaaaataagaagtaagtggactagtgccatttccaacaatctgatcaatagc
aatagcaatattattttctaaccaaagtacttttagtgtaaattttgacaagtttttatc
tccattattaatattaagtattgg----tacagtagaatactgtactagttatttttttc
agcaacaatttgagctttgtttaaaatgtttttttaccgtttagtagcttgtgcaccttg
agataatcgttgcaaaatttttgaaatgttaacacgagtttcattatttataggattata
aaatcctaattcttctatagctttaccatcgcgtttatttctgctatccattacaacaat
tctatagcttggttgttttttttcttccgtatcgctttaaccttaacttgaccatattagt
cacctcgcttattttttt--atatctttattaaattatatcttacgacattatgcaataat
tcaaacaattaagctgattcaagtctaacattattaaatattaccattaaacattgtaga
aaaataatacctaacataggtgacatgtccattccaaacatgggaggaatgcttcctctg
aatagttttaagtatggatcagtaattctatttaatgaacagaatggttcattgtaccaa
tttacagttggaaaccatgccaatgataatttaagtaaaattaaaattaaataaatctca
gaaaaattagctatcgatccaagcaataaatttaatgtaccaggaagagtgttcatacgt
attttataattattatatacttcgttatactaatagtttgattatatattataatcgatg
taaataaat-cagctaaacaaaaaacttatccttaaaatcatcttctcttgagcactttt
ctttatctacattagtatttacttttgtcgtataaatttccagtttagaccaaatctgtc
ctaactgatttaattgagatgttccacattctaaacaggctatttttatattgctcaaaa
cagcacctttttttataaaatcgttcaataatggcgtaatagttgcttcatccaaaaata
aatctaagacaagtatttttgtaaatgaatttaattgatctgggacattgttaataatat
cattataattaactaaaactatattagctgtaggaagtaaagatcgtgctccttccgcta
agataaagccataaggcataataattacaataatatatttataatgactgctaataagac
ttatagtggtattagcttttaattcgattgtttctgttactagccactctctcattattt
cgtaagtaatccatttgcctagttctgagcaagcagttcttaaaatcgtacctggattat
tgtcgtttttgagaatccctgaccaatgttgtattaatggatgggatgctatattaattt
ggagttgcatgttttgttaaatatatttaacatttaatttgttgattgaatcttcaattc
ctatttgttgagtattattaaaatagcat----atatattaaagaaaataattat-aaaa

aatgaaaaaaagtctttggctgtggggttttacagatagtgctgaaacttggaatggtag
atttgcaatgattggttttatatcagttatttttatcgaagtagttacaggacaaggatt
attatatttaataggtatgatgtcataataaaaatccaaaaaagttcttgtaatttttta
taagaactttttgg-attttataaatctaatctagcggtctcatagatagaacaggtt
cattctctctattgatacctttttcaaatccagcagcagcagctcttgctctgcctgcat
gccaaagatgtccaataaataggaagaatcctaggaagaaatgagatgttgttaaccaag
atctaggagatacatagttaactgaattaatttctgtagctacaccaccaacggaattca
aagaacctagtggtgcgtgagtcatatattctgcggcacgcctttcctgccacggttgaa
tatcgttcttgattttatttaaatcaagaccatttggtcctctaagaggctcaacccaag
gagctcgcaagtcccaaaatctcatggtttcaccaccaaaaataatttctccactaggag
atctcattaggtatttacctaatccagtaggtccttgagaagatgcaacattcgcaccta
gtctttgatctctaactaagaaagtaaaagcttgagcttgtgaagcttcaggaccagttg
gtccatagaattcactaggataagctgtgttgttataccaaacaaaattagaagccgtaa
gccccatgattgataaagcacctagactataagataagtaagcctcaccagaccatacaa
aagctcttctagcccaagcaaaaggttttgttagtatatgccagattccgccagcaatac
aaataatgccaatccagacatgaccacctacaagatcttccatgttatttacacttacga
tccatccatcaccgccaaacggagactttaagacataaccaaagataactaggggattaa
gtgtaggattgctaacaaatctaacatctccaccacctggagcccaagtgtcgtataccc
caccaataaataaagctttaataactaacagaaaagcgccaatgcctaataacactaagt
gtataccaagtattgttgtcattttgtttttatcccgccaatcgtatccgaagaaaggaa
atgattcttcaagagtatcagggccaatcagagaatgataaagacctccaaagccaagaa
cagctgaagaaattaaatgcactacgcctactacaaagtatgggtatgtattgaaaatct
cgccgcctggaccaacaccccagcctagtgtagctaagtgcggaattagaataaaccctt
gttcgtacaaaggcttctcgggaacaaaatgtgctacctcaaaaagagtcatggcaccag
tccaaaagaccattatacctgcatgagcaacatgagcaccaagcaatttgccagaaacgt
taattaagcgtgcattgccagaccaccaggcaaacccggtagactcaatgtctctaccgc
caacaccaacagttgtattaaagggcgtttccacgtggtaaaacctcctcagggaatata
aagttttcatgaggttgatcttgagcagccatccaagagcgaataccttcgttcaatagg
atattttagtataaaaagtttcaaactcaggatcttccgcagctctcaactcttgagaa
acaaaatcgtaagctcttaaatttagagctagtccaacaattccaaatgcgcttgtccat
aatccagttactggtacaaataacatgaaaaaatgtagccaacgtttattagaaaaagca
acaccaaaaatctgtgaccagaatctattagctgttaccatcgaataagtttcttcagat
tgtgttggagtaaatgcacgaaaagtatctgcagcatcaccatcttcgaataaggtattc
tgtacagttgcaccgtgaatagcacatagtaaagcaccacctaagattcctgcaacgccc
atcatatggaacggattcagggtccagttatgaaatccttgcaaaaatagcaaaaatcta
aaaattgcagcaactccaaggcttggggcaaagaaccagctcgcttggcctagagggtac

attagaaagacggaaacgaatactgcaattggtccagaaaatgcaatagcgttgtatggt
ctaagaccaactagcctagcaatttcaaattgtcttaaacagaatccaattagtccaaat
gatccatgtaaagctataaatgcccacagaccaccaatttggcaccagcgagtaaaatct
ccttgagcttcaggtccccaaaggaaaagaagtgaatgtcccatgctgtttgctggggta
gaaacagccgcagttaaaaaattgcatccttctagatatgaactggctagcccatgagta
taccaagaagtaacaaaagtagtcccagttagccagccacctacagcaaggtaagcgcaa
ggaaacagaagtagtccagaccaacctacgaatacaaatcggtctctttttagccaatcg
tctacaagatcaaaaccaccacgagtttttcttgtccaattgctatggtcataatttta
atctccagagcaaatttaataagtaatttgtttaaataaaca---ctttagatatactct
aaaattttacttttctttacggtcaatataattataagtcaaaagtaaccaaaattcacg
cttttaataaca-tttttttagctcattaagttaa-ttttttaatatataaacagtaata
gagttattgttt-aacttacaatactcacaatttattcatttctataatcctttgaaaaa
gatatcctgttcctcgtgctgttaaaatcaagtcgggattacttggatcatcttctagtt
tagctctcaatcgtgaaatatgaacatcaacaactctcgtgtcaacatgccgttcgggag
tatagccccaaacttcttgcaagatagaagctctagaaaaaggttctccagccttactaa
ctaaaagctcgagaagactgaattccatacctgttaatcgaactctttcattatttttat
aaacttgtcttttgttagtatctattttaaaaatccaatactaataatcccagagtttg
ggactccaagactagtggtgattttatcagctcttcgaagaacagaacgaatgcgagctt
ctagctccttaggagaaaatggtttaacgacatagtcatcagcacctatttctaatccgg
taattctatcgcagacttcacctaacgctgttagcattataataggaacatctgattctt
ttcgaagttcttgacaaacaccgtaaccatctagtttaggcatcataacatctaaaacaa
caaggctaggatactcttttctaaacacgagtaaagcttcttctccatttgaagcagtga
taacttcataacctattattgttagtcttgtttctaaaattcttcttatgctggcctcat
cgtcaacaacgagaatcttttctttctggttctccaa-tttcttatttactcctgtaact
caccgatcatgattaccataaccttaaattttagcgctatgagtatctttattcacttt
agaa-----tttagttattataaattttgcaagtctagtttacattaattagtaccagt-
-caatatttttgt-----------tttaagttatgac---------tattttaatccata
agttgactctagtttgcagaattcaatatatacata-ttttctacccaagtaaattgtat
tctattttctaaataaaaaaggggttgacaaaccagttgatagaaagtattctacattac
tataag----ttaattccaacaaagaagttttaa-----atcttctttcggagtaatgaa
tactttaat
>Pyr_23
tatatgata---aattaagacatctgttatgtgagaatttctaaaatatacaaaactttt
tatttaatatagaaaaagtaaattaaaataagttgaacatactttgctctaatctaaacc
tcttaattgag------attctaattagagacaaaagtatttacttattttgtatctata
ggaataaaaatgaaacttgcagtttatggtaaaggtggtataggtaaatctacaactagt

tgcaatatttctgtagcactttcaaaaagaggtaaaaaagttctacaaattggctgcgac
cctaaacatgatagtacatttacgctaacaggtttttaatcccaacaattatcgatact
cttcaatccaaggattaccactatgaggacgtttggcctgaagatgtaatctataaggga
tatggcggtgtagactgtgttgaagctggtggaccgccggctggcgctggttgcggaggc
tacgttgtaggtgaaacagttaaacttttaaaagaattaaatgcttttgatgaatacgat
attatcttatttgatgttttaggtgatgttgtatgtggaggttttgcagctccattaaat
tatgcagactactgcttaatcattacagacaatggatttgatgctttatttgcagctaat
agaatagcagcttcagtacgagaaaaagctagaacgcactctctgagattagctggactt
gttggtaatagaacagataaaagagatctaattgataaatatatagattgtgttccaatg
ccagtattagaagtcttgcccttgattgaagatattagagtgtccagagtaaaaggtaaa
actttatttgaaatggcagaaattgataaggatttagcatatgtatgcgattactatttg
aatattgcagatcagttgattacaaggccagaaggtgtagttcctaaagaatctccggat
agagaattatttagtcttttatctgattttacttaaatcctaaatcaaaggtaggacaa
gaaaaagtagatcaagaagaattagatttgatgatagtgtaaataatattttcagcataa
tataacaaggaataagataatgtctacagctcaatcagatgctcttacttttgaatgtga
aacaggtaattatcatactttttgtcccatcagttgcgtttcttggttatatcaaaaaat
tgaagacagtttttttcttagttatagggactaaaacttgtggttatttcttgcaaaatgc
aatgggagtaatgatttttgctgaaccgagatatgctatggcagaattagaggaaggaga
tatttcagctaagctaaatgattatggtgaactccgtagattatgcttacaaataaaaaa
agatagaaatcctagtgttatattttggattggaacatgtacaaccgaaattataaaaat
ggatctggaaggaattgctccaaaattggaagcagaaattcgtgttcctatagtagttgc
tagagctaatggtttagattatgcctttacccaaggtgaagatacagttctagcagctat
ggctcaaaggtgtccattaaatttaaagaatcaatcagataacacgagtcttaaaacttc
tcctcatattcctttagttttatttggatctcttccggatccagttgtaacccagttaac
tatggaattaaaaaaacaaggtatttttgtttctggttggttgccatcaaagaggtatac
tgaattaccggttataaaagaaggatattatgttgcgggagttaatccatttcttagtcg
cacagctactacactaatgagacgccgaaaaacaaaactaattggtgcaccatttccaat
aggtccagatggtactcgagcctggattgaaaaaatctgttcagtaatgaatgtaaagcc
tattggattagaagatagagaaaaagcaatttgggcttctttagaagactatatttcttt
aatccgaggtaaatcagtttttttttatgggtgataatttactagaagtatctcttgcaag
atttttaactagatgtggaatgactgtatatgaaattggtattccttatatggataaacg
ctatcaagcagcagagctagctttattaaaagctacatgtgataaaatgaacgttatgat
gccaacaattgtagaaaaaccagataattataatcaagtagaccgaatccgtgatctgaa
acccgatttagttatcactggtatggctcatgcaaatcctttagaagccagaggtattaa
tacaaaatggtcagtagaattcacatttgcacaaattcatggctttaccaatgcaagaga
tattcttgaactcgtcacaagaccattacggcgaaatcttagtttgtcagaattaggctg

ggatgtttatagcaagcaaagctagtattcctc----tttatgacttacatggtataaaa
gccttgtaagtct--tttattttatgttaaagaagcatttcttgctctgccatcaccagc
ccaagcttgtaaactttctacttgttctttgtctgtaaatgctaaaggcacaaattgttt
gatagctattttaatatcttcagtactaaattctctctcttcactaaacgccgtatgcat
actttcaataattgcttgctcaatttcagcgccagaaaatttattacataataaactcaa
ttcatgaatatcatatttttgccacgatcttggtctaacttttgataaatgtatcttaaa
aattgattctctctcttgacgattaggtaaatctaagaaaaaaatttcatcgaacctgcc
tttccttaacatttcggaaggtaaactttgaattttattagctgttgcgacaacaaatac
tggagctttttctctgatagccaggtaataaatgtaccaaatactctggcactagtacc
actatcaccttgactatgtaaaccagaaaaagctttatctatttcatcaatccataatat
gcaaggtgataatccttcagagatatttatcatttctctcatttttgattccgattcacc
aactagtccaccaaacagttttcccatgtcaagacgtagcagaggcagcatccaatcgct
tgcaatagctttcgcggttaaggatttacccgtcccctgtataccaactagcaataaacc
tttgggcgaaggaataccgtaatttaaactttgcttcgaaaatgagcgagaccttttttg
tagccattgttttaatacatctaagccaccaatatccctatttactttactataaggata
aaattctaatagatgcgtttgattaataatttgcctttttttcttcaataataataggtaa
gctccgagagtctatctgattatattgtgcaataattttagtaatcactttgcggattct
atctatcgataatccttggcaggattttgttatattattaacgagctctgaattcaaatt
aagattcaaagctttactcaatcttatgatctctttttttgatttctagaagactaggtaa
gggcaaatctattactgttatgatatcattcagagcaaaaggtatattgattttgcaaga
aataataataatgtgcttagattgcgttttgattattttagctaaattgcgaagtttcct
aattaatactatttcgtttaggaaagagtcaaaatcttttaaaagaaaaagatttaaaga
ttcattattcaagttttcaataaattctaatgctaataaaggatttctttttgcataacc
attatcgctaggattgttagtgtatccatctacaaagtcccaggaatatacttgctgact
gtttgaacagtttaagctgtgttttataatatactctagtctatcttcttctctagtatt
tattacaataataggatatcgagattttaaaagtaatcgtaaatcttgagtgaaattcat
gtcaatatttgtttccatcagacttttgtacttcaccaattgaagaaagactaaataaga
ttatctaatattccttggatatattcaaaagtc------atctctctattttctaaatta
ttgctattactaatttaacttaccatgatttttttcttccttttcttcgtcgactattt
aatatagagcgtccacatggtgttttcattcttgctctgaaaccagaaactctaatttt
tttcttttagatccttgtagtgttcttttagtcatgatgtctcctctgttgtaaaagatt
aaatactattttatcatcttttatatatctttatgatatagatatgtattaatatcaaag
tt-aacagcaataatttatattaatcaagaactatttatataatattaaacattaaatat
----------taactttatgatattggctttgcccatttttttatttaagtattttgacca
tctttttgctaatcttaaactggttaatttttcagcagctaaaaacgattttgttattag
aatctcaatttaagtattttgttgataaaagccaaaatcgtaaattagaacccgacgaaa

gttttgcttttgcaaaagtatgtgtagctaaaaaatatttttctaaagctattattgaag
gtcaattagctttaaaaaactatcgagacctgaatatcttagataataatatagtaattg
ctaatttgtataatatgctaggttttatttattttgaagcaggccaaacaagttttgcaa
agaacttttatgagcaagccttacaaataaaccctaattatgttgtagctttaaataacc
tggcaaaaatttatgaagaggtaaaagatttaaaaaaagccgaatctctatacgataaag
tgttaactcttaacttaaataataaaactgctaatagaaggaaagatttatagcaaaaa
cta---------aaaacatctaatactatttgtaatcgggatagcaggatttgaacctgc
gacatcctgctcccaaagcaggcgcgctaccaaactgcgctatatcccgtaagacataat
tattactatatacctttctctaattatgtctactctttctgtattttatctaggatacc
aaaacatgcctttaactccatctggatctttgataaatcctaatttttataaaaactaa
caacatcaggttctgcgaatagtgtaatagtgctaatttctgcctgtctcaattgctgta
ttaattggtgtatgactacttttcctaaacctaaccttggaaatctggatgaataacta
cgtcccagatagttgcattaaatccgttgtctgatgttgctctagcaaatcctacaagtt
tactatttgcatcttttttttgtattaaagaaataataatagaactatttttttaatgcga
tctttactttttttagaggtcttttaacccatccgactgaatcacaaagctgctccaatt
catataaattaatattcttattgctactcagatagatatctttaaattcgattttatcgc
aagttttatctaaaacaataagtttttttgaaattttttttcagagctagcattgatattgg
agttttgaaaaaaatttttccagaagatcataatgattatgatataatta--aataaata
tatatattacaatattgctttaacaaacaatatattaatatatgctcaaatttttacatg
aatctattaagatagaagattgttaaaatatagagaaataaaaatttttgtaaaatcttt
caataaaatgatataaagagtgcttttttaagactaacatctattattataaataaagtct
cgaaattgagatgaaataacacattattctatgtctaatgccttttaaaggagttatttg
tgaaaaaatctatgtgtctaacctgtttacttgctttactaattatgagcaacccaataa
tagctaacgcagaagtagctggattggtgccctgcaaagattctgcggcattcaataagc
gtatggtaaatagtgtgaaaaaacttcaagccagattagctaaatatgatgcagacacgc
caccagcgttagctttaaataagcaaatagagaaaactaaaactagatttgcaacttatg
gtcgagcaggtttgctgtgtggtactgatggattgccacatttaatttctgatggtcgat
ggagtagagccggggactttgtttttccaggactattattcttgtatattacaggatgga
ttggctgggtaggtagaggatatcttttatctgttgctaagactagtaagccaacagaaa
aggaaatcattttagatgtgccattagctatcaaatttatgtcatccggatttgcatggc
cgctagcggcttggcaagaatttagtagtggacaattaattgcttccaatgacgatatta
ctgtttcaccccgttagtaaaaaatttatatgaataataattttaccaaatacttatcaa
cagcacctgtaattggtgtattgtggatgacgtttacagccggatttattatagaattga
accgctttttcccagatgtcttatacttttactt----ataaataa--gatcgtataaa-
--aaactacataaatatagagagatactattatttgtgtagtttttta---atatataaca
taattattta-tttttaaaaatgtttatcttttcgattattttaatattagatcttacat

aataaaata-atatcaatattatgagcttagtaacccaaattatagttaatgccgatgat
gaattaagatatcctacaattggagaattgcagtcaattcaagactacttaactacagga
agcaatagaattagaattgctactattattagagataaagaaaaggagattatacagaaa
gctagtaagcaaatttttcagttacatccagaatatatagctccaggtggtaatgcagca
ggttctaggaaaagatcgttatgcttacgtgattatggttggtatctcagactaattaca
tacggagtattagctggggataaagattctatagaaacaattggtattataggagtaaga
gaaatgtacaattctttaggtgtacctataattggaatgttagatgcaatccagtgttta
aaggaagcatctttagaaatgcttggtcaagatgatattagaatcatttctccttatttt
gattatataattcgtggaatgtcataaatcatagcttcaattaattagttgaataaaagc
ttttataatgttataattacaataagcccgagagcatataattgtaaaaactaaatcttg
tcagaaccggaaggtagcagcaataatgtttcaatacaaaagttatga-ttttcgggtgt
ttattgtttttatctaatagtctcgacttttaattagtatgctatgtataaatctgtata
tgacatgtaatccatcaaaccttccagcaaaaataatttaataatttgatattaatacgt
aaaacacctatac-tcttttataaaatttctttatatctcatctaaacaatggaaactta
acttaatgggtgttcatattttatcaacaggttcgtctgtcccaaatttttctgtagaga
atcaacaatttgaagacatgatcgaaacttccgaccattggatttcaacaagaacaggaa
taaaaaaaagacatctagctccatcttctacttctttaactaaattagcggcagaagctg
caaacaaagccttatatgcagctaacttaaaacctactgagatcagtttaattattttag
ctacgtctacgcctgatgatttatttggtagtgctagccagttgcaagcggaaataggtg
caacaacatcagtagcttttgatattacggctgcctgctccggttttattgttgctttag
taacagcagctcagttcattcaaactggttcctatgacaatattttagttgttggagcag
acacaatgtctagatggattaattggtcagatagaactacctgtattttatttggtgatg
gtgctggagcagtagtattggggcaaagcctcaaaaatagtattttaggctttaagttat
gtacagacggtcagctaaacagtcatttacaattaatgaataaacctgtaaataatcaaa
aatttggtggtacagaaattcctcatggaaactataattccataacaatgaatggcaagg
aagtgtacaagttcgctgtatttcaagttccaacagtaattagacaatgtttgaataatt
taaacatttcaatagatgaagttgattggtttatattgcatcaagcaaacactagaatca
tagaagcaattgcgagcagattatcagtaccttttctaaaatgattacgaacttagagc
attatggaaatacatctgcagcgtcaatccctttagcgttagatgaagctattcaatcca
ataaaattcaaccaggtcaaattattgttttatctggttttggagcaggcttgacttggg
gagcaattgtcttgaagtggtgat---ttatatattgcggatgacgagactcgaactcgt
aaagctttcgctacacacccctcaagcgtgcgtgtataccaatttcaccacatccgcatt
ttttaccatgaata-ttaaaaatttataatatttatctaaatattcatataa--tgtgta
tatatatatacattcttgtcagaaactctcttatattttattttaagtgataaaattgag
t----ttagtatagatagtaaaattttattttaaaatcataa----aaataaatatataa
ggaaaataaaatatgacaccatctttatcaagtttttttgaatagtcttattcttggggca

gtaattgtagtcgttcccataactttggctcttttatttgttagtcaaaaagacaggaca
atccggtcctaaaaataacttagaaagatatggagaagcaatttaagatgaataatgaaa
tatttttcatactaataagctctccatatattgctatact--ttagaagaatttttattt
ttttattttgtaacatcctgtaaagtttattccgttctttctttttccc-atttactctc
ttataattaatgatacaaatgtcagg-ttaataatag---ttttttattctgtatctttc
---------------attttatatacgtattattatgttcaattaataaagaaaagctca
tgtctttgtctaattggcctctcaaaaaagaaaattctgaagcatataatattaagaatt
caaaacaaatcacaattcctgatggtttatggataaaatgcttcgactgtggtctattaa
tgtattctaaagtactgaagagaaatttaaaagtttgccctcaatgtagttatcattttc
aagcttctagtaacgaaagaattgatcaattgatagaccaaggtagttggcaaccaatgg
atgttcacttgatctctacagatccattaggctttaaagaccaaaagctttatagtcaaa
ggttaaaagatactgctttcaagactggcctgcaagacgcagttcaaacaggtactggga
ctatgcaaggtaaaaaagtatgcttaggtattatggattttagattcatgggggaagca
tgggatctgttgtaggcgaaaaactaacaagactgctagaaaaagcaactcaagaaaagt
tgcctgcaattatactttgtgcatcaggcggagctagaatgcaagaaggtatgttgagct
taatgcaaatggcaaaaatttcttctgctctagaaatgcataaaaaagaaaatctactat
atatatctgttttaacctctcccacaacaggggggtgtaacagctagttttgctatgcttg
gagacttaattattgcagagccaaaagctcttattgcatttgctggtagacgagttatag
aacaaacaatcaaagaagacttgccagataattttcaaagttcagaatatttatttgaac
atggtttcctagatttaattgtatccagaactcagcttagatcaaagttaatacaaattt
tgtccttacataatcatagtaagtaatgaattgaatattgactataatatcaaaatgcta
gtaaaaattaagatattaagaaattttcactaaagatatttgtagactaactagaattac
aattactattgtagaaaaatttgtcttaatgcta-ttttttcagtcttttaacaaattg-
tttttatgactttaaataattgttagtttaatacaaatttataaagagaata-----cat
actaagctaaatatagtaatattatgttttcctataatactgacctatcaaatttcgagg
aaatttatgcttaaaagatcttcttggcttgcggctttattgggactattaacagtagtt
tctacaagtacgcatacatatgccatagagttagacgaggcaacaagaactgttccatta
gaatcttctggcagaactgtaattcttacaccagaacaagttaaaagaggcaagcgatta
tttaataattcttgtgctatttgccataatggtggtatcacgaaaacaaatccgaatatt
ggacttgacccagaatctttaggattagctacgccacaaagagataccattgaaggacta
gttgactatatgaaagatccgactagttatgatggtgcagagtcaatcgcagaattgcat
ccaagtattaaaagtgctgaaatttttcctaaaatgcgcaatctaaccgatgaagaccta
tttacaatcgcaggtcatatcttacttcaacctaaaattgtttctgaaaagtggggcgga
ggaaaaatttactattagaacttcaaagacctaaaccttgtgaatactagtttatgta-t
ttgttgcttaagtgtgattattttgttagatcatgatatatattaaagatggagtttgtt
gacagcaagttactttttacccgaacttgtttacactattcgtctagttattaaaaagga

gacgttaaattgaagaagaagctttcagttcttttcactgtttttagtttttttgtaata
ggtttcgcacaaattgcttttgctgcagatctagataatggagaaaaagtttttttctgct
aattgtgcagcatgtcatgctggcggtaataacgccattatgccagataaaaccttaaaa
aaagatgtacttgaagctaatagtatgaatactattgatgctattacttatcaagtacaa
aatggtaaaaatgccatgcctgctttcggaggtagactggttgatgaagatattgaagat
gcagcaaattatgtattatctcaatctgaaaaaggttggtaattatacttgattttatcc
tgtattaaagaatagacaatctatttagttgttcattagattgtctattctttgtttatg
---ctattatataaagatatttacacaatatt----ttatgatgaaaagaatacccgcaa
ttcttgtactagaagacggtgcatattataaaggatggtcattccagcaagataaacaag
agattactattggtgaagtagtttttaatactggaatgacaggatatcaagaaataatca
cagatcctagttacttccatcaaattgtcgcttttacctacccggaaattgggaatacag
gtattaataatcaagatattgaatctcacagtattagtattaaaggacttattgcaaaaa
atatttgtaaaatttcaagcagctggagagagcagcaatctttagttaagtatttaagta
gtaataatattccttttattttcggaatagatacaaggtctttaacccaatacttgcgtc
aatttggtacaatgaacggttgtatctctactgataatttaaatcatagttacttaaaac
agaaaatttgtgagattccaagtatgcaaggtttagatttaatcccgcatgtaactacaa
gaaatgtttacccctgggatgaaaaaagttttccgaattggtatttgacagataacatta
gagtacaccgagttattcagttgaaagttattgttatagatttggagttaaactaaata
tactcagaagactagctacacttggatgtcagataacggttgtgcctgcccacactcctt
taaaagatattttggcttaccagcctgatggtatattactctctaatggtccaggagatc
catcagcagtacattacggcatccagacagttacaaatttactagattacaatgtgccta
tatttgggatttgtatggggcatcaaattttaaatttagctcttaaagctaaaactttca
aacttaaatttggccatagaggtattaaccatccatccggattgaaccagcaagttgaaa
taactagtcaaaatcatggctttgcagttgaattgacttcggtttttgaatctcctgtaa
gagtgactcattttaatctaaatgacactactattgcaggaactggacataatcaaagtc
cttatttttctgtgcaatatcatccagaatcgagcccaggccctcatgatgctgattatc
tattcgaaagttttatagaaataatgacaaagtccaagaataaagttag-----------
ttagtaattttcccatgctttatgagtgaataaagcggatagtacttgtactcctcgtaa
ttgattaaagagaggcaaatgtcctctggggccatgagaattccattgaaattcctcagg
atatctgcaaggaattctatcaacttcccagccgatttttgccagaattttccccaatc
ttttccaacactaagccatatttgtctttgaacaaatagcccaaatttaccttttgagtg
tgtatgccataacttatcaatagtttgtaaatcttgagcaggtatttttttatatctgt
aaaatatagccaattacgagtttgtgcatttacaccagctagttgaataagcttttgttg
agttagctgatctgctttgagcaaatcccgatgagtcagtaacatttgtaaatctttgta
attcatctgttgagcagagcgcaatggaacaatgccatcgggacataagttagaagcaaa
tttcactatttcttgatttttactgttaagtaacttttcataaattaaaccgtcaacaca

attacttttatagttaggcccagtaatcctctcaaaaaatagatctgctaaatcttttaa
ctcgatagagtcgttgctatttatattctcaataatttcaagttgttgcttga---catt
attagacttagtattcttattcaattctaaaagctgggctcggatttgatttggcatctg
atttgtttaattagttttaatagtc----ataaggtcttagagagaattagattgttttt
tgaaattagaagagcgccgcagtatttctaaagaagttcgcataatttgattcaataagt
tttgtcctgcataaattataatttgaccaattaaaaaaattgtgctttttattttaggaa
caaaaaaatcttgtacctcaagcagaaaagtaattattatttgcattggtgacaacatct
gtaattcattcgatctgcaagcatatatatactgacagtttaagccttcttgataaaatc
cccatactttgtaacgactttcgtaaattgctcgtggtcgctcaagatagtttttaatta
tagtattccaaattaggttattctttaatttttctaatttctatcagatagaaattta
cattacagagataacttgcatttgtctcatttacagtattaaaattttgaactaacatat
gagcaataatgttgcttagctgaattaaataattttcaagaaaaatttcgacctgtttaa
ctggcatattgtaagtaaagagattaaaagttttattcacagtttgagatgacccaaaaa
gtagctgaataattaaaatttgtagtagcattttatagtcgttcagtagatagttcaaat
catgaatatcttggtaaatgtcaatacagtccagttcatataacttacaaaatcttttga
tacacctatggaataaatcaattaaaatattttcggtcaagtttgtaatatcatttacat
ctaaatttgaattataaatttctaaaagtatcttttccagttcagctaatatgattttta
ataagtttctttttacttcagtgcgaaaaacatctaaaattaacacttcctgagagcaat
tagttaatctcttattaatctttatagaagttctaactaatagctctgctacctcttgat
tgagtgttggtccctgagagcttggccaataattattcacgttataccatagtaagaaat
attgaatttaatgttatacgataactttttaatatactgaaaagcaagttaattactcta
tagcaactag-------tttttacagatcttaatagttatttatcaaataaagttatatg
atagtagcgattgagctaattttattaaattta----taataatgacaacttattactt
cgcccttgcgagtcaaaatttctattagtacaagaaccactagaagaagtttttagaga
gagagttaactactatcagtcaaataataaagcaattgatttttggctaataccaaaccc
ttcttttctagagaaaccagaaatgatttcatttaaaaaccttgtacctaaagacgctgt
agctataatctctactaatccaatatttattaattggttaaagctaagaattggctatat
ctgtattggacaatttgaagataacctacaactttctgaagaatcgttaaatattactgt
tttaacagacaaaatttaacaatgcttttattatgactagtgttaaattattttggttaa
gttataattatagtcttttagtattta-gactcttatagtttatcacttagtattttatg
ctgagatggagtgacaaatagtgtaagtatttcttggctaaacgtttcagttgccttaga
tttatatctatttggattaactataatggatagcattcgttttatagtaacattttcaat
ttgagcccaatgtacaattccaagttccaattccttagcaatagctgaaacggaaacaaa
ggcagctcctagccctgattgcacagcgtttttgatagcttctatagaattcaattccat
ctctattttaaagcgactgctgtcaataccatgttgactgagtactttatctatcacttt
tctaattgttgattgagtatctaatgcaataaatctaagcctgtataagtcttcttttg

aatgtctcctagtttagaaaaaggatgtgatttgggtaatataagtgctaattcgtcttc
cgcataagaagtaacctgcaaaacatcttgcaattcagtaggcacttctcctccaataat
tgctaaatcaacttgaccattagctacgctccatgaaataagtctagttgaatgtacttg
caattgaacagccacttgtggatatctttgcctaaatagtccgattaatcttggcatcaa
atatgtcccagttgtctggctagctccaataattaatgtgccaccttgtaagttttgtaa
gtcgtcaagagctcgacaagtttcttcgcagagagctaaaattctgcccccgtatcgtaa
aagaagacttcctgcctcagttaaagttgccttcttattacctctttcaaatagggaaac
atttaattggcgctctaaattttgaatttgcaaactaatagctggctgagaaacatacaa
gctattagctgctttttttaaaactgccctctttggcaattgcttttaatattcttaactg
atccaatgtaaatggaaggtctgtcattaaagaattgtagtataatg---------tata
ttatttagtattgatcaatgtatgcatttgccaactaaaatatttgtcattacgcttcat
tattttcaaaataataaaaagtataatgttaaataagctgtattaaaatttagtatctgt
ttgatattaaatttatacaagagatatcgtgaatttaattctaactataaaatataggag
acaat---atggactctagacttttagttgtactaataccagttttagcagcagcatctt
gggcagtttacaatattggtagagttgcactacagcaatttagaaaaatgacatcttagt
tttgcttagaatata-atttttagactttatgagatagaggactttaaaatccctatctc
tattgtatatcttaataaaaaaag-aaagc--ccccttgtacttactgttattacacatt
aatattattggt-ggtaccta----gataataagtttaagctaacttatttttttattta
taattaattaaattaataacacaattatggctgttccaaagaaaagaacatctaaagcta
aaaaaaatgcacgcaaagcaaattggaaaaatcaagcaaaaactgaagctcaaagagctt
tgtctttagcaaaatcagtattaactagaaaatccaatggatttatttataatctaactg
aagcatcagatacttttagcgattagaagtgttt-ttactacaagtctgaaaaagtatta
cgcaacaagttctcagtaatttttatctat----taatcatttgaaaatcattctatttg
ataataaaataaatagcagattaaataaaatcacattaacaaattatgcagcaaagctag
atcaaactagtgaaatttggttgttcaattgtattgaaaatattcagcatattttttta
aaagtcaattaaaatcaagtcatattactaaaattttatttctggtactagctttgaat
ataccgcaggtctaccaggattattatccagcttaacactaagtggtagactacatccta
ttagtatatatagtccccagtctctcaaaaagtatcttgaagcatgtaccaaatattccc
aaactaatttttcttttcctattaattttcataatttacaatacggaggacaagttgtta
accaattttatacagtaatttgtttaccgttgagcaaaaagagcctgctgtatggattta
ttattctaaaaaaagaaaagcagggagtatttaatttagcacaagctaaaaccttgaata
ttcttcaaggacctatatatggaaaactcaaagaaaaggataatttttttaagtccagatg
gttactatctatccggccaagacttctcttctaatacaataatgggacataaaatatcgc
ttccgttattagtcagatattctagaattatttctgagatgcattggttttgctcttatc
ctattagattaaatacttattcgcatcagcaaggagtaaagtgtttaccacataatgttt
taactgatattatgaaatcccaaatatatcaagataacagttttgttgaataatacttat

tcttgatatataatggtctcggtaatctaaataaaatactagaaataatttcttaaattt
atttgtctatattaaatttacaatcatttatgacatacgcaattattgaagcaagtggca
cgcagctttggatagaagaaggccgctactatgatttaaatcatatacctgttgatccag
gtcagtcgattatattaggaaaagtcttattattgaataaaaatggggaggttactttag
gacgcccttgtatagaaggggttacgataaaggctacagtaatgaggcacttgcgaggaa
agaagataactgttttcaaaatgaaaccaaagaagaaaatgagattaaaaaaaggtcatc
gacaagaattgactcgtttaatgatcgattctataacatcttaga------taaaatcta
gactcatattttt--aa-aaacttttaatatttaactaaaatagatataatggcacata
aaaaaggtagtggtagtacaagaaatggcagagactctaattccaagcgtttaggtgtta
aaaaatatggtggagagcaagtaacagcaggtaatattttaatcagacaacggggaacta
aagttaagcctggccaaaatgttggaaaaggaaaagatgatacattgttttctctaattg
atggtttcgtgctgtttgaaaagtcaaatcaaaagcaaaaaacaattagtgtttattctg
ctaagaaatagttaaaacaattggtgcaagataacttgagt----gtatttatttcactt
acagtggcaccatatatttgatttattaagaaatggagttccttaatgaaccaatgaatt
ttattaaagatttgctgtgaaatttaatgacaaacactattgtaatttcctgtctacaca
atatggctgctattctatattgtggtcagatacagaaactagtagtagcaaatgctcatt
atcaagtaagtgatatctacctaggttgtgttgataaaatcttctcaggaataaatgcgg
catttattaacctaggaaagaatgagtacagtggttttatacatatcagtgataccggtc
cgcttaaaaagaaatattatgtcaataatattactaacattttaacaatacggcaaaaaa
ttttagtacaaatcattaaagagccaactttaaataaaggtccaaggctcactgccaata
ttacattatcaggtcgatatattgtattaatgccttttagtcaatcaatctgtatatctc
gaaaaatatatgatgaagatgagcgtcattatttgaaatctttagctattttaattaaac
cggcaacaatgggcttgctatttagaccttctgctgtaggtgtcgatgaggaaataatat
taagcgaattaaaaaatctaaaagaacaatggaactttgttcaaaaatctgcaattaata
gttattcacctgttcttctatataaagatgaagatattgttaaaaaagtaatccgagatt
tttataataataatacaaacaatatagtaattgattcaaacctgggattaaaacaattaa
attattatatccacacttggcactgtaataactctagcacagttcctaagattaagcttt
atagtaataatcaatgtatactagatgcttttggtatcaatcaggcaatttccagagctc
tcattccaaaagttgatcttatacttggtggctatatgtttattgaaactttagaagctt
ttactattattgacgttaattctggatcttttaataattccactagtgcacgagaaacag
ttttaaaaaccaactgttctgcagcaacagaaatagcttatcagttacaaattagaaata
ttactggtgtaattataatcgacttcattgatatggaatcacaaagagatcaattgcaat
tattagaacactttaataaagagctatcacttgatgatgctaaaccacaaattgtacagt
tatctgaattaggtttagttgaattgactagaagaagacagggcaaaagtttgtacgagt
taattagtagtgattctaattacttttattttttcacacaatcagagagatctcagtctc
ttaagagattcgatgatagacagcagaaacaacagatttttaataaatcttggctatctg

cagagattaatactattaacaaggtctttttcaaaagtcaaatttgtgcagacctgcta
acttttacctaattcgtaatctctatatagttaaaagtagtattacttataaacaaaatt
atttattaactcacagatcaaaacttatttattctaaagaatacagcaaagtattaccaa
gca------gttattatctagctagtctcaataagaatagtaatcaagagttcctatctt
aagattgttatatcaactgtt---------------------------------------
-------------------gaaaaaaagaagctctctcttatataagagagagcttc-tt
ttatagcttactaacta-tctagttagttaattaaaaattaaccgttaacagctggagct
gttagagctactggtaaagattcaccagaagctagatctagagggaagttgtgagcgtta
cgttcgtgcattacttccatacctaggttagcacggttgatgatatcagcccatgtgtta
attacacgaccttggctatcaacaacagattggttaaagttaaaaccattcaagttgaat
gccattgtgcttacagataaagctgttagccagataccaactacaggccatagacctaag
aagaaatgtagagaacgagagttgttgaaactagcatattggaagattaaacgaccgaag
tagccatgagctgcaacgatgttataagtttcttcttcttgtccgaatttgtaaccatag
ttagcagattcgttttcgcttgtttcacgaattaagctagatgtaactagggatccgtgc
atagcactgaacagagaaccaccaaacacaccagcaacacctagttggtggaatgggtgc
attaaaatgttgtgctcagcttggaatacaagcatgaagttaaatgttccggagatacct
agaggcattccatcagagaaactaccttggccaattgggtatactaggaatactgctgct
gctgctgctactggagcagtaaaagcaacggaaatccatggacgcatacctaggcggtag
cttagttcccactcacgaccaatgtagcaagctacgccagttaggaaatgaagaacaact
aattggtaaggaccaccgttgtatagccattcgtctaaagaagcagcttcccagattggg
tagaagtgaataccgatagctgcagaacttggaataacagcaccagaaatgatgttgttt
ccgtatagaagggaaccagcaactggctcacgaattccatcaatgtctactggaggtgca
gctacgaatgcaatgatgaatacagatgtggcagttaatagagttggaatcattaataca
ccaaaccaaccaatgtataggcggttttcagtactagtaatccaagagcagaaacgttcc
cacaagctagcgctttcgcgtctttgtaaagtagcagtcataatttttttatcaattttta
aggatttaacaaagatgcttcccaaacaataagttgtctgttatgaaaattattacttaa
ggccttttcaaatgagaacatttttaaaaaaaaca-tagcaaaatctaatctcagtaact
aa-aattt-tacttgttcatatattgcttaataattcaa-----------taataaattt
tt-gct-------------ttttgcaaaattaacactattttagggat---ataataata
atgcatattaatgctgcttattttgattttaatagtttatataattttgccctacttaaa
tacgcatgtcacactttacaaaaattcagacgaccatacaagatttaaatctgttaaagc
atgctttaacagatttgggtttgatttggcaaatgaattctagtcatataaaagttggtg
aaaataatcagcataaggtagatattttaattaaacaagataacttgtcacatattggct
ttacttggaatgataatagatatcacttagttgccgacttacaactttgggaacaaccct
ggtctctagaagtatttttggataagttatctcagaagtatgcttactactctatcatcg
aagaaacaaagaagcaaggctttgaaaaaaatgcagcagatttcacaaaaagatggatcaa

ttaagttaattgtgcaacgctggaattattaactaacatacaaatgcgggcggagagact
tgaactctcacgagattatctcactagaacctaaatctagcgcgtctaccaattccacca
cgcccgcactgttgagac--aatattattctatcataatctatactaaaacacaaactaa
-tagtataaactagcttccaagcttaaaagcgtctacaaatagaccgtatgttataccaa
ttttgaagttgttgaataaattaattttatatttaattgcttgtggctagagtagacta
ttttactagtcaactcagttgctgcaactattagactggctgcaataataccccaatctc
ctgtttgtccaggaatcgtagacaaacctgttgaaataaaaaatcctagcaataaactaa
ttaagccagttgttaactcgctcaaagaataataaagcttattatttaagtttttatta
agtaagcaaaaaaagttgaaagtctagttttaatcatagtatttagtaacggagttttaa
aaggatgacaagtattaacag-tcattaatattgtcgtatccttttaaaactcgttaata
caaacagctgttcttgt---------ttttaaaggtcttgttcgcttaaagcattaataa
gatgctgaaacggctcttgtataatatctgtgcgtttaattgattgcaaatctaattctt
cttctatgatttcttgtaataactgtatgcttcttactgttggagcaacaggtacagata
gtgaattgtaagtatcttttagcccatttaatacacgctgatcgagaatatttgtactac
cagaaaccaaggcataactagcataccgtaggtaatattctatatcccttagacaagctg
catatcttcttgttgtataagaatttcctccaggccttaataattcaggctgttcttcgt
acaactgagctgcggcctcttttaaaatattggtagcttgatcattaataatttcagcta
tttttattcgatctagtccactcgaaaagaaagattctagttgtccgacagctgtcttat
ctagataacgaccagtaagatcgtaacgatttaatattgctgttatagcatcttgcataa
attaattctcctac-ttgctatcagagcggtattataaatgactaatattgtaatataat
cattagtgtttatgaaatttatttaaaacttattctttattgcttaatgtgcaattaaaa
actcgcatagattttacacatatagagtatacgtagtttttaatta-tttttttataaaa
accaagaaataaagctttttt-taagaggttcgtgaatttgttaatataacatgagcttt
cctcctaaataagttttgtaaagtac--aaagtaaaaaaacgattagtcaataatagtaa
------------------------------------------------------------
------------------------------------------------------------
------------------------------------------------------------
------------------------------------------------------------
------------------------------------------------------------
------------------------------------------------------------
------------------------------------------------------------
------------------------------------------------------------
------------------------------------------------------------
------------------------------------------------------------
--------------------------acaatattaatatgtatcaa--------------
------------------------------------------------------------

------------------------------------------------------------
------------------------------------------------------------
------------------------------------------------------------
------------------------------------------------------------
------------------------------------------------------------
------------------------------------------------------------
------------------------------------------------------------
------------------------------------------------------------
------------------------------------------------------------
------------------------------------------------------------
------------------------------------------------------------
------------------------------------------------------------
------------------------------------------------------------
------------------------------------------------------------
------------------------------------------------------------
------------------------------------------------------------
------------------------------------------------------------
------------------------------------------------------------
------------------------------------------------------------
------------------------------------------------------------
------------------------------------------------------------
-----------------agtataaataagcttttactccagcatcaaaaaaaatatttga
gatatgttgatacaactgattattcaagtgaaaacttagatttagaccaatttagctatt
ttatcattacaattaataaaactgataaatgtattcttatagagcaattttgttataatg
ctgaaggccatatttactcaatattcttcaaaggtccaacagctaaacatctttcttcta
taatttgtcattttcaacaattaaatacaataatatctacagctcatgccatatatttag
gacgagagctaatgaagtcagaattagctctcgtgctagatcaacaatatattcaagatt
aaa--ttttgtaagtattaattatatttatgctagtataatattagtaagggcatgtaac
tcagtggatagagtatcagattccgattctgatggccgtgggttcgaatcccgccctgcc
c---gagtactcagagtact-atactaaaacatct-------taatagctagtttgataa
tagtatgggcattggggctgcaaggtttctacattatgaaaagaagaaaatatg--aaaa
aaacaagctctccaaaagagcttttagtacaatcaataaatgcagaaaacaatattgttt
ctttttctcgaaaattagcttttgcttaaacactgttaatttttggcattcgacatgtta
aactcttattcatggcgaatattctgtcagagtcgctcttagtttaagaaaagtcagaaa
aaatatttatctttacgtttt-tacctatttttatttaaataaaggttagta-ttttcaa
cttttatagtggacgtgggttcgagtcccaccagctccatcaatttgatt---------t

gtataattagctagtttaaccttttaataaccagatatataattggaggaaatatgggt
ttcataacaattactaagcccgctttaaagcaaattgcaatcttgaaaaatgatcatgaa
aatgatgtacatctaagaataggtgttagacaaggcggatgttcaggtatgtcatactca
atgaattttgaacatgttgacaaattaaaagatacagatgaacggcttcgtctagataat
ttttctgttgtctgcgaccctaaaagtcttctttatctttatggattatcattagatttt
agctcggaattaataggaggaggatttcaatttctaacccgaatgccagccaaacttgt
ggctgcggtaaatctttttcaggctgacc-ctcgaacacgtttttatatattttgtatt
atatatattattttagcaagtcg-tttattagagacaagcaatgctctgttaaaaac---
----------------------------taatttcaagagttttattgtattatt-----
----ttatatatttattatcaactacttgaaaataaattagctaagcaatctaatttgtt
atcagtaccaaactttaaatgattaacttttattgatatatgctaatcttt-atcagcc
taaacatatctgtacattacaatcttttgacataaaaagagtcttagcatacctctcaa-
----------------------aaataaattcaa----------tattttataaatactc
aacatgaatctgtcacaacttatcgttaattcttctatgatgaaaattcctaataacttt
tattgcaatataaaagctatttcttctgagaatttaagtcacttacaattatcaaaaaca
aatttattatctcctacaaaaataattgcttttggactacctaaatttgattttatttct
acacagattacaagttctattagagaaacagttgttgtgtctactattgatgataataat
ctttttgctgccccaactaatgattggataaaaaaacacaaggataaaaatcatcattct
ttgttttataaaaaagtttttaaagctttgctaaaccataaaattagagttgttgctgag
ccact------tcataactccgaaataaataaagaaaagagcataatggcaagtcaaaga
tacatctggggtaaaagctggaagccttcaattattttatcttgcttaaaaaacaaaaaa
aatagcaaaatttctgatgcatcgaaaaaatttttaactgaacagttatctgcatcacca
gtatttgtggtaaaaaatggttttaatgaaattattttaggtcacccattatctcgtgtt
aaaagaggaagagttaataatctgatgcatgcgttttctaatttattaaatcagtctaat
gcaacttatcctatatctactggtctattctttttttcatccagatgatgcatttgaattt
aaagactttataatatcggtcaatccactggcggccaaacatatggagataagcgtagag
cctgttggtctacactttgcttataaaatgaatagaaatatatcgtcagatactcaattt
tcgtttattccagattttaaagaagtaggagatttattattcaaatatagaaaaggtaat
catttagtttttcataagaatcagcattatggcaaagatttttttcaggggcaaccaata
tatatgattcagccgatcacttttaaagatcgagccggaaagctaaatattatcaaattt
acggggctaaatgacacaagagaaattatttttactaatcttgaagctgctaatagatca
tgggcacattttataaagaggaattcacaattaaaatcaattaaaaatcctactctatta
gtctataatttagaaagtttttttaaaagatcaagaacgattggataatcaagatttaaat
aagtttgtggtgatcactaataaagaatcttatcttgccacaaaagagttaatagcttta
cccgattctaatagcttttctaagcacttaaaattaaatataaagcctaaactctttttt
gtgaaactatgggtaagacggttattttccaccttaacttatgaat-aaatgctatt-gc

ctttattttctcgaataatattctataactagtaattcattgagttgtaaagcaacccat
tccctatcaatgactccattgactttccctgataaattggatttattcaattctaagtgg
ctaggaatattagctagtccagggaaagctaagtaattttctactagttttcgagatgct
tcttgatttttacactaattgattctcctggtttacattgataactacagatagatact
acttgtccattaatacaaatatgaccatggttcactagttgtctagcggcaggaattgta
ggagccataccaagtctaaaaacagtattgtctagcctcatctctagtagctgtaacaaa
atttgacctgtcgacccttgtagtttttagctgcttttacgtacttaaataactgtttc
tcacttaatccataattaaatcgcaacttctgtttttcttctaatcttacagcatactca
gaaggttttctggacttttgtccatgttctccaggtggataagatcttttaatcgctttt
ctactgagtccgggtaaatcacctaatctacgggaaatgcgtactcgtggtcctctgtat
ctagacatattatttaaattctcctaatgtaaat-gattaag----------taatgagt
agctatttttaacacaaaaagagcattaaaacaaagc-ttttgctaaaaaatatttaca
aaaaatataggtttggacaaaa---tacattactatgc-tttttaagcgggtagcgggaa
tcgaacccgcatcattagcttggaaggctaaggttttaccactaaactatacccgcaata
ttcaacagattgtcactaactcaatataacatagttatgttaagttaagataatcttaaa
tttaatagaatt-ttaaattatttatgctccttctaaaaccacatcaagaaaacgagcaa
tttcggcagcttgttcttcaacttctgataataataatggttgaccgaccctagttaatg
gaattactcttttatcttttgtacataaataaatttcacgtcttggatttagcccctctt
taatatctatttttattgatttgatttcttttatattaaactgaagacagatttttctat
tctttccagggaaacctagccgaaaaattttacaatacctttatctttgttaaattcat
tatatccagcccctatattccaaataatagttagccacagaaacaaactcagaaagactc
ctatactgccatagaatgtcattacaataccttgtggaataaataccaaatcggttgaat
ttgtaaaaggtaacaaatcaacttgaaaataacttgatagtccagcaagaagaaagccca
aagccccaataaatattatagtagcccaccagtaattactcagtcttcgtgaacctaaaa
ttaaatcttttctaactgagtgtatagataatgtttttttcataatttgttcttttaaaa
cagta-tactaatgcaaaatggaaaaattttgacaatttt--atgcgcgctttagattt
tttaaataagcttgattgcttttagtcctaaaaacagtcctgttgctaacccaaaaaaag
cagcaagaaatattatatatcctagaaaaactgaca---tattattatttattattattg
ttacaatagtgtttacaaaaatagtgaaaatctattactacggtattatattacgttaaa
tgccttgctgtctacaggaaaaaactttttcagtttttagcattgtgtttatattagaa
ataatattctagtaaatttattaaacttctggagaacagaatcttatgtcagaatttatc
aagccttataacgatgatccttttgtaggcaatttgtccacgccagttagtacgtcaagt
tttagtaaaggacttctaggaaatctaccagcttaccgtcgaggtttatctccgcttctt
agaggattagaaataggaatggcacatggatacttttaattggaccttttgataaattg
ggcccttacgaggtacagatgtagcgttactagcgggatttctatcttcggtcggcctc
attattattctcactacatgtttatccatgtatggtaatgtatcttttactagagcagat

tcgaaagatccactacaaacttctgaaggctggggacaattcactgcaggatttctagtt
ggagcagtaggtggttcaggatttgcttatttattgctagctaatatacctgtattacag
actgcaggtcttagtttattctcttaagctagtaaggggacttgaacccgtaacctactg
attacaaatcagttgctctaccaattgagctatactagcattcagttcataataacatat
aaagatatttgttgcaatctgctattttcagatgtaacaaatatctttttttatttaaaca
gaactgttatgttaacac-------gtatattaattagaatctgaattatctgactctgc
agaagattcctgatcacagtttagataataacaaattgtttgatctagacaagttgcaga
ccaaccaacaggtgtttctcctgataattcttgcatatctacagaattataaaaattatc
atttggcttattactgtttgcttgcataaatatcactcccaaaacactctgaacttaatg
tgaagtaatttaacttctcaataatatattaacacaaaatattaaatatggtgattattc
tttaattgtaaagatgctgctaactttataaagtttgagtgagagtcttgatagcttttg
ttgaaataagcattttaacccatttatttgcccgctcgaccatactttttgtgttgta
aattaactttttgtagtttttttgttcgtttatgagaatgagatacggcatatccattat
tagctacttttcccgtaagctgacactttttgacataatttt-------ctaacttata
tatttatttagagtacttgctaaagttgacttgggcacagctccaatcactgtgtctact
ctttctcctgctttaaagatcattaaagtaggtatacttctaatgccatattcagccgca
atagtaggattatcgtctgtgtttatttttactaccttaatagatgattcatattcttcc
gcaatttcatcaactacaggagaaaccattctacaaggaccacaccatggcgcccaaaaa
tctacgagtacaggtaagttattgttaataacttcttgtttgaaagaggcatctgtaact
tgagatactgacatattctttaacctttaatatatgttccttgctaagcaaatcttatca
caaatttcaagaaaaatcttctatagtgaattactattagtgttgaattattttttaaa
aattagacattaaaaaaacttattacctatataaataaagtggtggccgagtgatatctt
gatagtgataatttttatattaaggttgaacactagaaataacttttcaagcaaactctt
atttgaaaa--atttctctgtgaccataaaaacttaattatattacaatatattttatta
tttgtaacttagttctgataatggtataaacaacgcaaaagatactgccttataatcaag
gaggaatacatgtctcaatccgtagaatcacggactaggattaaaagcgaacgttacgaa
tctggagtaatcccttacgctaaaatgggctactgggatgctgactatgtgattaaagaa
acagatattctagctcttttcagaatcactcctcaaccaggtgttgacccgattgaagca
tctgctgcaattgcaggtgaatcttcaacagctacttggacagttgtatggactgattta
ttaacagcttgtgacttatacagagcaaaagcatatcgagtagatccagttccaaacgtg
gcagatcaatattttgcttacatagcttatgatattgatttgtttgaagaaggttccatt
gcgaacttaactgcttcaattattggtaacgtttttgggtttaaagctgttaaagctctt
cgcttggaagatatgcgtatgccagtagcttatctaaaaacgttccaaggtcctgcaact
ggattgattgtagaacgtgagcgtatggataagttcggtagacctttcttaggtgctaca
gttaaacctaaactaggtttatctggcaaaaactacggaagagttgtatacgaaggcctg
aaaggcggtcttgatttccttaaagatgatgagaatattaactcacaaccatttatgcgt

tggagagaaagatttttatattctatggaaggtgtaaataaagcatcggcttctgctggc
gaaattaaaggtcattaccttaacgtaacagccgcgacaatggaagatatgtatgagaga
gccgaattctctaaagaggttggtagtatcatttgtatgattgaccttgtgattggttat
actgcgattcaaagtatggcaatttgggctcgtaaacatgacatgattttacatttacat
agagctggtaactcaacttactctcgtcaaaaaaatcatggtatgaacttccgagttatt
tgcaaatggatgcgtatggctggtgttgaccatattcacgcaggtacagttgtaggtaag
cttgaaggagatcctttaatgattaaaggcttctacaatactctacttgaaagcgacaca
gatatcaacctacctcaaggtctgttctttgctcaaaattgggcttccctacgtaaagtt
gtaccagtagcatctggtggtattcatgctggtcaaatgcaccaacttcttgattactta
ggtgatgatgtagttcttcagtttggtggtggtacaattggacatcctgatggtatccaa
gcaggtgcaactgctaacagagtagcactagagtccatggttatggcaagaaatgaaggc
cgtaactatgtagcagaaggtccacaaatcttgagggacgctgctaaacttgtgggcct
ctacaaacagctttagatttatggaaagatattagttttaactacacttccacagataca
gctgatttcgttgagactccaacagcaaacatctagtttaatgactacttactgatactt
taaatagttaattgtaagtgaaattaacttataacaataaggagcatagaatagtgagac
taacacaagggactttttccttccttccagatttaactgatgagcaaattaataaacagc
ttgcttatatcgtttctaaaggcttatcagcaaacgttgagtatactgacgatcctcatc
caagaaactcctattgggaactgtggggtttacctttatttgatgtaaaagatgcttctg
ctgttatgtacgaaattagctcatgcagaaaagcaaaacctaattattatgttaaagtta
acgcttttgataatactagaggtattgaaagttgtgtaatgtctttcattgtaaatagac
ctgctaatgaaccaggattcttattacaacgccaagacttcgaaggtagaactatgaagt
atagtcttcatagctatgctactgaaaagcctgaaggagctaggtattaatattaattct
taattaatattggctaatta--ttcccctcttaaaaatcataattaatttagtaattgtg
atttttaagagggatagcttcccaagtaaattttgactaaa-tagattgaataattaaaa
atacacagaaataatgcaatcacaggatataatttccaacgatactcttgttaatttaca
agaagaatatgatagaacacaaatccaagaagttttaaatgagttaaatcaagaacttat
aggattagtgcctgtaaagaccagaattcgcgaaattgctgcgctattattgattgacag
attacgcagaaaactagaactagtttctggtaatccaggattacacatgtcatttacagg
tagtccaggaactggtaaaactacagttgctatgaaaatggctgatattttgcacagact
tggatatataaaaaaagggcatttgttgacagttacaagagatgatcttgtaggtcaata
tattggacatactgcccctaaaactaaggaagttcttaaacaagcaatgggaggagtttt
atttattgacgaagcttactatctatataaagcagataatgaaagagactatggctctga
ggcaattgaaattttattacaagtaatggaaaaccaaagaaatgacttagttgttatctt
tgctggatataaagatagaatggaaaaattctacgaatccaacccaggactctcttctag
agtagctaatcatgtagacttcccagattatacttcagatgaattattacaaatagctaa
aatgatgatagaagaacagcagtactgttttacagaagaagcagataaaactcttttaga

gtataccgagcgaagaatgaaacagccttattttgctaatgcaagaagtattcgcaatgc
tattgacagggctagaatgagacaagccaataggattttgccagtggagaaaaagtatt
aacaaaagctgatttggtaacgattgaagcagaagatatcttgaaaagtagattattttc
attacctaatgcttaatatacacgtgatttcatgaaactatatattattaaaaag-tttc
ataatcagttgcaaatttcttgaaaaacctttattattatcttaatgtaggtg-tggcgg
catagccaagtggtaaggcagaggattgcaaatccttcatcccccagttcaaatctgggt
gccgcctagtactaaaaaagggggtgtggtggaatggtagacacaacagacttaaaatc
tgttgatttttagtaatcgtgagggttcaagtccctccacccccatatattgatttataa
ataaaaagagatgtgtatttgtattaattgcaaccatattaccaaatgtaatacctatca
cttaatagaatctcaacataaacagcctcatcttactagaagtccattgtttatacctaa
atatcctgtagttcatgttaatatatctaacaactgtacctataatcaaattgattggga
tttagtggagtgtttatcatttgtagaaaaaccgaatagttggaatttagacgctaatta
gttacaagaaaaattatgtaccataagcaacaactctatttttttagatactcggtattaa
ttttggatgttctaaccagtaagattttacccgttcttgctatctctataatgccaaatt
tagttagtaattgttcaatagcaacaatctttcccggatctccagtaacctctataatta
aaagatcttctgcaatgtctactatcttagctctaaaaattcttacaatttctaaagctt
ctgttctagtctgagaattgatctgaatcttaattaacattagttctctttcaactgaag
gaatatttgttacatcttgaacgttaaggatatttactaacttatataattgtttcgtaa
gttgttcaatagttctattgtctccttgaactaccattgtgattctagagaccccaattt
gctctgctggtccaactgctaaacttgcgatattaaagcctctccgggcaaatagaccag
atattcttgacagtactccggcttcatcttgaactaaaactgataaggtgtgtttcatga
aaaattaaatttaatctatttgatctattaagtatattctaagaaaattacagaaaaaat
actagttagtctataactaaattatccaaactaaacaaaatcctactatcagataatata
tctaccttgtaatgctatactattaacttagtatttaatgatattctaattgcaaataga
atgct-aatttttttaaatttgttaggaacttgtgctagaaaaatacaaaaacagtaaaa
aactctctcgagatttacctcaaatcaatgatcgcattagatttccaaaagtccgagtaa
ttgatgacgaaggtgaacaactaggtatttttgtgcctgaagaagctatacaattagctg
tccaacaaggtttagacttagttgttgttagtgataaatcggacccgccagtatgccgaa
tattagactatggtaaatataagtttacacaagaaaaaagagctagagaagctaagaaaa
agcaacataacagtagtattaaagaagtaaaaatgcgatataagatagaagagcatgatt
ataaagttagaataaaccaggcatccaaatttattcaagcaggagataaagtaaaagcaa
ctatcacatttcgggggcgtgaaatccagcactctaatttagctatagatttattgaata
agatggcaagcgatctaaatgcaatagctgaaattcagcaagctccatcaagagatggca
gaaatgtcataatgctcttatctcccaaaaaagttagctaaactaatttttatttaaatg
catctggctggattcgaaccagcgacgtctctttcgaaatggcggattatgagtccgctg
ccttcggcccctcggccacagatgcatcacttaaagtttataactcataaagcatataaa

aatcaatgtttagctatttttaataaagtctataaaagtctattattcattcatattgtg
atatgtcgcaactgctgaaggggaaacttgatttaagtatcgaaatatccagtatttaaa
aattgtgtcgagtataactggaaaagttgaaataaataaaaaaataaaatctcgactttc
cggaagtcctaaatgtcttaaaattacttcaataataacttcccatccatgaggagaatg
aaaacccacaaacatatcagtaaaaagaatgattaaaaaagctttagctgtatcactaag
accataaataatttcatttaaaaaagatttaacgactgcaatttgtctttgtccagttat
cattagtaatataaagacaagtattgatacaagatcggataaaatatttttaacagcatt
tgcactttcgtttgcataatattctcccaactctctagccttcagctgcactctttttc
aatgatttcgtaagatatatcttccgaaggattcagaagaacttcaaagtgaattttctc
ttcaaatctttgtaattcagcaaaagctctttcctcttgagaggaatttaaaaaaatttt
aggctgttcttggttccataaataatcaatacaaggtccaaatacaaaaaattttgaagc
ctgattaactagtactggagagatgaataataataaaatatattttacggaagtaatagt
ttggtgtctagatattctaaactcctctatagcttcagattcgccattaggatctaactc
tttacgaaacttttcaaaagtattagtaattgacctaggtattggcccaactttttcaaa
ggcgaattgattatttcgttttaaattccaatatttcatttttattgctgctaaaattt
ctattaatcaataattgatcacgtattcactataaatattaattctagaactaaacatat
aaaacaatatttatgaaaaataaaatacaattaaaaattaattatgcattagttctatct
gcagctacagcagtttatggatataaatcttatcagccaactataaatacactcttaatc
aaagatgaagaaaactggcctttttctctaaaaggcagactgcctttattcctataatt
gtattaaaaaaaaatcatcaaaatccctgaactcagaacctagaaatagtaagttatatc
ttaatggaattggaagtagaaaatttatcgacagaattgtcactctacttggatatttag
aaaaacaggcaaactatactgaatacttgaaccaaataaaaacaagtggttatttttctt
tagtctacctagacagtgatataatgtataagaaaaatattagtaacatttttttactac
ctaatgatattctaaaaagaatctatatttgcaataagaaaaaaaagcttataccgcatt
cattcttaataaatttatttcagaaacagataggctatccaaaaagttttgctaatttaa
attgggccttctctaaaattataaaatggtattatgatagaggataccaatggtctctag
ttgaagttaaacaagcgtctgatgcatcttctattgtaattgatattcatgaaggggtag
tgaagacaattataacagaatactacactctatcttataaaagagtctcaggtatcttat
gcgtagagtcaatagagcagtaccttggagtaagagtgggagctccactaaatataattg
atttacaaaaaaaaattacttacttaaaagataatcaactagttggcgacattatttata
gtattgaacgatcaaataatagctcaatgagtttggatatcaaatttcaaatacaagaat
taaaagataaagagataatagtgcttgcagaaagttcttcta------------ttatct
cgcatgcatgtaatttactcaatc---aatataga---------aataggctagtagctt
ctaatatagtttcactatcaactaataaattacacgcatgctattttaattataaacttg
actatcaatataagtatataaacacctctaatcttataaaattgttagcttattctattt
cttgtaggaagacactacttattcattattttaatttacaaactttaataagctgtacaa

aaaaaaatactattggatttcagctatatttacgaaatctaagttttgggaaagctttct
gtgtgcttagtatgaaatttatcaaaaatggacttaatataaaaatttatatataaatc
cttcattgatagttgatcagaattttgtatttcaatttgctatacaaatcattaagcaat
atcatactgcta------aacctccagctt------------------------------
------tatttttaacgaatctagatcttgaacaatatgttgctgaaagtttattaatgt
accattttacatcttgcttctcaatatctgaaaaaatattattatctcgaattatgcata
cagattctttatttttcaactctgaaa------attttcattttgatgatagaactaacg
cgatcaacagttatgatactttttaaacaaaatacaaaaatattttatcaagagtttcttt
ctttattattaagtttacgttatcaaaattttaattatttaggttggcctttaaaaggcc
attttttgaaataaaatctttatacttggctccatttcaaaagagtg-------------
---attttctgatagtcgtaagacgctattcttccacaaaatgtcgttaaagcaagtat
ctaattttaatttaccagtatcttttaaaagccaccttaaccatattttagttagtacta
ttaaatgtcagtcaaatttgaatatgagaactgtatctttgctattaattgattcacctg
ctgaatatctgctatacaaatctattcttaatttctctattaaagttaggatgcagtatt
tcatacctatgagcaacaatataagattatctctattttataattatttagattgttttt
taataagatcatctcagagttgcattcacatatggcaagatctaagaactcttacaccaa
ttcaaaatttttggttaaaaaaattttcatatggagctggaattcagcttaaattaccaa
ttaagcagatgccacctctttcaattgaatatactgtaactagtagtcgctacttttgta
tttaccttcgtacttattatcaacgataactattactaaaatgaaaaatttaatttcttt
ttttgatcgcagcaaagggaaatggatttcacaaagaacaacttatgaattatctaataa
aaacatgagttcggtacagtctcagatgacaatgaaaataggcaactcattgtcaggatc
tataatacttgcatcgttaaattgggggacatttatagacaagttgcccattacgctaa
aaatcacagccggagtgaatatgataataagtttaacttgcaattcggtaatcaactaaa
taatcataaattattaacgctgtgcatagtgacagatcctagtctaattagttttaaaac
tcgctatggaagtactaccatagatgaaacatattggttcgcaacaaataatttacgtct
aagtactagtattgttaaacgattcaacacttgtgtggcagtatctttttgttcagaaat
taaagtttaa-actctatatagaagcaactaatagccttta-ggctattagttgcttcta
tgtatctaaattttgtattaaaaattat----aatactatattattctaaatctttacga
ttgggattcctggaaggatcattagagaggaacccaaagacaaacaaagaaataaagaaa
ataacggtagtatatacaaagattttcaaagtaaacatattttcatccttaaaagctatt
agtgcg----atatttacatatctccacataattttatatcatgatataagttccagcac
tattttataaataaatcaatatctttattttttagtaatgatttatatttaaatatgcgt
gatgtactaaatactaaatcttttccgacttgttttgtcttgtttgtaaatattttgtta
tttctatggagtttaccttcaattatcacatagtctaacttctttagctgtttaaaagta
tatgaagatttttttgttccaaatagttagattgattataactaatcgttttctttttaac
agtcttgcttttaacttaattatttggcttttattttcactaattctgatactcttacaa

ctaagaatttggactagaagagtacagctattcataatttataacaa------tttttta
aaaaatctactaaaatagatgattttacagaactaaaatgatatttcattatcttgttga
ccttgctcataagtaatatctttcaatacttttaatatttttccaaaatactcttgcagg
tatttttctaaagatattacttccaacctgtcaattccaagaatactgtaaacttcatcc
atcggagctgtaaattgctctccgctacttaaaatttcggcaaatgctaaacgatcagaa
atattccatgtccattgaagagttttagtaattcttcttaaggcttttaataatccaatc
ggaatttgtgagatctgtgtttttttgcccagataatttttcacataatttaataatttca
gctgatgtccaagcggtattaccaactaacggcagggtcctattttctgtggaaggtact
ccaagacttttttataactaacttggctgcatcttgagtatcaatgtatgcaatgggtgta
gattctcctgtaacccagactgatttcttgtctaaaataggaatggcatattggttgatt
aaaccttgaaaaaatcctcctaaagaaaacacagtatatttcacattggacttctgaagg
aaatctactacctgagactttaagttcatcaatggaacatctgggtatttctctgaattt
aatattgaaaagaaaataaatctttcaactttagccgctttagctgcttcaattaatgca
atttcccatctaaatctatttttttctgcattataaggatcagtaggacgagatgtagaa
gcatctatgattgctgttaccccacaaaaagattgcaaaatactttcgggtaattttaaa
tcaccatatataagttctgctccccactctttgagaaaggcagactttcttaaatttctt
accatgcattttacattgtagccttcatctaaagctcgtcttacaatttgacgtcctaaa
gttccggttgctccaattactaaaagagtcatattagttaaaaataagtatgtataaaga
cattgatttggttgacggcagagagtaaattaggtagagagggaatcgaaccctcatgac
cgaagtcgtcacattttgagtgtgatgcgtataccaatttcgccatctaccctattatag
accaactactacaagtaatttagtctttagcgtaaaaataatactaagtttcaatagata
tgtcaacttttaaatttatattaataaaaaaacggagtctcttatggccacttttgaatt
aattccttatagattgtatttgagtcaatctaacacatggatccatagaataaaagcaga
aattaagatatatatagtagcgcttttatggatttcaattttttattttttcttactttaa
actatgtattattgctttaagcttaattgccataagttttactataagaagtaaacagaa
tattattcaaaaacatttgttacaaacgttattaatgacgttcttaactactgttttgtc
ttttagtgtggctattagttataaacaatatgcagaacaagaacaatcgcagtatttatc
tgattccaagaaatataaaaattccagcagctactattatatacaaatagcgaacgatca
aaggatcaaaagacaatacttaattactactttaaagccttccttgtactttttttattac
tatatattctattaaactagttatgataacaacctctccagaagttttagtaattactat
ttatagatctaggataataaataaaatatttaaaaatgaattgctatttatcttccttct
ttcgtcacatattgttactagtattattaatagaatcgataaagtgattcaagtaactag
cttaagaggaagcttgaatttatataattcattgacaaggcctttaatgttttctttgtt
aatatttcaagtcttctttttggagattattcgggagtcaaaagaaatagctcaagctct
ttatactagaaatctcaatcaagaaaacaataatttttttgaaaatatatacagtaaaatc
taactttagtgatcggctcaatataattattagcactttgtactttattattttagctct

agcgtaatagaatccacttattcttctgactctagtaatacgaagagtaataaaattagt
tagtagtttaaatagatagaat-taagatattatataaataattaaatgtttcattatgg
tcaatactcaaaatcaaatttcacaaacttcagaacttgattatatagttaatcaacctt
ataaatatggctttaaaacttctgttgaatctgagcaatttcctaggggaataagtgaag
atattgtccgcttgatctccaagaaaaaagatgagcctgaatatctattgaattttaggc
ttaaagcatataaaaaatggaaaaagatgagtagtccgtcatgggctcatattaagcatc
cgaatatagactttaatacgattatttattacgctgttcctaaattaaaaaaagaattga
aaagtctagatgaggttgatccagaaattcttgacacttttaacaagttaggtatatctt
taaatgagcaaaagcgaatttccaatgtagctgttgatgctgtttttgatagtgtatcta
ttgcaactacttttaagaaagaactgtctgaagctggtgttatattctgttcaatttctg
aagctataagggattatccagaattaataaaaaaatacttaggtactgttgtaccagctg
gtgataattattttgctgcattgaactctgcggtatttagtgatggctctttttgctata
tccctcctaatacagtttgccctttagaattatcaacttactttcgtattaacaacgaag
aatctgggcaatttgaaaggacactaattattgctgatcgtggcagtaaagtaagttatc
ttgaaggttgtactgctcctcaatttgacacaaatcaattacatgcagcgattgtagaat
tagtagctcttgaaggagccgaaattaaatattctacagtacagaattggtatgctggta
acaaagaaggtaaaggtggtatatacaattttgttactaaacgaggcttatgctcgggta
acaattcaaaaatttcatggactcaagtagaaactggatctgcaattacctggaaatatc
caagctgtattttagctggtgaaaattctcagggagaattttactctgtagctttaacaa
ataactatcaagaggctgatacagggactaaaatgatccatattggtaacaatacaaaaa
gtagaattatttctaaaggtatttctgcgggtagatcgaaaaacagttatcgaggcctag
taaaagttggccctcagtcatttaattctcgtaattattctcaatgcgattctttattaa
ttggtcaatcatctcaagctaatacatttccttatattcaagtacagaatccaacatcaa
aagtagaacatgaagcatccacttcaaaaattagcgaggaccaaatttttatttttttac
aaagagggatcaatttagaagaatctattgctcttatgatcagtggtttttgtaaagatg
tctttaatgaattgcctatggaatttgctactgaagctgatcgtttactgagtttaaaat
tagagggaactgtaggatgagccagactattttagaaatcaaagatttatatgcttctgt
tggtgaaacaacaattttaaaaggggtaaacttatctattcgagccggcgaaatacatgc
gattatggggcctaacggttcaggtaaaagtacattatcaaaagtaattgcaggacatcc
agcgtattcactaataagcggagatattttatttttttggacaaagtattcttgaaatgga
gccagacgagagggcaaaagcaggtatttttttagcttttcagtatcctgttgaaattcc
tggagtcagtaattctgatttcttaagaattgcattaaatgctcgcagaaagtttcaagg
attatcggagtttagccccttagaatttttttcaactaataacagaaaaaatagatcttgt
tggcatgcaagaaagtttttttgacaagaaatgttaatgaaggattttctggtggagaaaa
aaaacgtaacgaaattcttcaaatggctttactagatagcaaaatatctattttagatga
aacagattctggactagatattgatgcattacgagtagttgcaaaaggaattaacacttt

agctaaatcaacaaattcaattattttgattactcactatcaaagattattagattatat
tattccagattttgttcatattatgagtaacgggcaaattgtaaaaactggtagtgttac
tcttgcccaagatctagaaaagcacggatatgactggattacgcagacataactatttac
tac-------aatattgagactaaagaatattctttagtctcaatatt-tttgcaaatgc
aaagtt-aattatttaagctttatcaaaaccttgcttaacatcttcaattgctcttttta
gcaattcttcactaactggttctaatttttagtactacgaacattttcaccaaattctg
gtttagaattctttaaatcttctcgaagttctctaataaattcagctacttttgaaactt
caatatcatctagatagccattaattccagtgtaaataatagcagtctgttcttcaactg
gaataggagaattttgggcttgcttcaaaatttctcgaagacgttgacctcttgctagtt
gattttgggttgctttatctagatcagatgcaaactgagaaaaagcttctaattcagcaa
attgtgctaactctagttttaacttgcctgccacttgtttcatggcttttatttgcgcag
cagagcctactctagagactgagatcccaacattaatagctggtctaatccctgagttaa
ataagtcaccagatagaaatatttgtccatctgtaattgaaattacatttgttggtatat
atgcagatacgtccccagcttgagtttcaataatagggagagctgtcatactacctccac
ctaattcagcattcaatttagctgctctctctagcaatctagagtgtaaatagaatacat
ccccaggataggcttctcgtcccggaggtcttctaagcaaaagagacatctgacgataag
cttgagcttgcttagttaaatcatcataaataactaatgttgctttacctttgtacataa
aatattcagctaaagccgcacctgtataaggagcaatatattgtaaagtcgcaggactat
ctgcattagctgcaactataatagtataatcaagagctcctttatcctgtagtgaagata
ctacttgggctactgaagatgccttttgtcctatcgcaacatatacacaaacaacatctt
ggcctttttgattaataattgtgtccaatgcaacagctgttttacctgtttgacggtcac
caataattaattctcgctgacctctaccaattgggatcatagagtcaatagcagttatcc
ctgtttgcataggttcacacactgattgccttccaattataccaggggccattgactcaa
ttagccttgttccattacttgcaggttcacctttatcatctattggccgagctaatggat
caacaactctacctaaaaaagcatctccgaccggaatttgagcaattcgaccagtccctt
ttacggaactgccctctaaaatatccctaccatctcccatgagtactacaccaacgttat
cactttctaaatttaaagcaaccccaatagttttatcttcaaattcaagtagttcaccag
ccatgacttcatcaagtccatatactcgagcgataccatcaccaacttgtaaaactgttc
ctatattggctacttctacatcctgatcatacttttcaatttgttgacgaataatactac
ttatttcatcaggtctaatatttaccatattttaatattatttttattaattggaaatt
ctaaagcggtaattagtcagactaatgctatctgtctttttttactttattttaagttact
acaacatcaaggtgtgaagccatttgtcttagctgtcctctaatactagtatcaatgact
tttgatccaatctgaatagtaaagccaccaattagctcgggctcgactgaaataattagt
ttaacttctttagccttagtcataaccttgatttttttctgttaatagagtttcctgatca
gggcttaaagcaatagaagtattaatatttgcaattgttaaagattccatttggtaagcc
agttccaaatacttactagctatagcatctaacattccaattcgcttgcgatcaactaaa

atcattagaaatgataatgtattttcgttaatttgatcgccaacagttgcaataattact
tcttttttgcctctacagtttttaatggattagct---aaaaaaattttagcttgtca
gattgcgctaaaatattttgtattgactgaatatcttggctcaccttttcagtaactttt
ttagttttagctaaatcaagaagagctactgcataaggctgagctattttataacaaca
ttgttactgctcatagttgatctcctaacttagcaatattattatcaattatacgcagtt
gcatttcggagctcatttgattttccagctgcaaagtaactcttttaagagcaagaaatg
ttatttgttgctggatttgccttcggatctgtttttcagctgtttctatatttgatttac
ctgttatggcaagtctttcaatatctaattggccttgagctaatatggaacttctaactt
ttccagctgttagctgagcttctttcttaatttgatctataatgatttgagtttgagcaa
gttgtttttctgattctgacaatctagcacttgcttgctctagtctttcttcggattctt
gtatagctgctaaaacttttatctgtcttgcatttaaacttgatccgagaaattgcttca
aaacataaataagaccaaataaaagcaacaatatattaataacatttgcttcaaaaatat
cagaattaaagccgaatttgtgttcactactatgttctgataaaatagtaattatttgtg
gtatttttactatattattcatttaaaattct----------tactattttaatatttta
ctctggaatcatgattttcaagattgactacttaataatttaactttaatttgatcgctc
aaagtgtcaacctgatttctaaagttttaagagcttcttcttttgaacatttaactgt
ttcgatgcttcagcaatcagtttctctgcattcaattgagctttttttatatcttctgag
actatatttgagcttctttttgagaagaggctattttaattgtgcattgcggcgagct
tcagatagatcttcttcatattttgcagctagctcatctgctttaacaagcatagacgat
gcagtagtcaatgttgttcgaatatattcatctcgttcgtctagtactttagttacaggt
ttataaaaaatagtatttaataaaaccattaaagtaagaaactgtaatgccattaaaggc
aaagtaccattgaagtcaaataagccaccttcaatttcttcggctaatagaaatggtaaa
tcaatcattttactaattatgtttatagttaataattttgtttcaattaagaaaaataac
tttatagtgcgcttagtctcatcagactaagcgcaaatgattattttaaccaacgtatgg
attagcgaacaataaagacaacgcaacaactaggccgtaaattgttaaagattccataaa
tgctaaacttagaagaagtgtaccccgaatctttccttctacttctggctgtctagcaat
accttcgacagcattagctgctgcactaccttgaccaatgcctgggcctattgcagcaag
acctacagcaagaccggcagcgataacggatgcggctgaaacgattgaatccataatgaa
ttttaatttttttgttatatgagtagaaaattgacaaatttcgtataatacttaaaatcta
tataattaagtataatagattctaatat-----tgtaattatagtatattactcttctcc
atgaccttccattgcttctccaatataagctgcggacaaagtagaaaagattaatgcctg
tattgaactcgcaaataatccgagtatcataactggtaatggtattaaaattggaattag
caaagtaaatacagaaacaactaattcatctgctaagacgtttccaaataatcgaaaact
taaagatagtggtttcgtaaaatcttctagaatattgattggtagtaatactggagttgg
ctgaatatatctagcaaagtagcctaaccctttttacttaaacctgcataaaaataagc
taaagatgtaagtaaagacagggccacagttgtattaatgtcatttgttggggcagctag

ctcaccttcaggcaagtgaattaacttccaaggaattaatgcacctgcccagttacaacc
tagaataaatagaaataaagttgcaatgtaaggcacccaagggcgatattcatgctcacc
aatctgatttttgcaatatcttgcagaaattctaatataaattccataaaattctgaaa
tttttcaggaattctttgtagatttctagtacctagaaatgataaagttaaaagtgtagc
aattactaaccaagagacaatgaaaacttgcccatgcaattgtaagctacctattttcca
atatagatgttttccaacttctactgctgataaactattatatgggttaaaatctatgag
actgttttgatacataataaatattt-ttttaaatagtgcaagaatcaaaaaataag-tt
tttgaagaattaatatgcaattaatttattcttaaaattagactattaataaattaaaag
caaaaataaattctaatagagatagctatttttagtagtgattatttttttgtaaaaaaca
attattgctacctataatgcttttacgtttaattatatacccatatattaattatttaat
atgattataggtattttgctacaaactttattgttttatccattct----aaagattgtt
atagtttatttttttatttaaaatatctgcgacttcatctgcaaaattaactttcaaattt
tcttctcccccctcccagaagaaatcttacaaatcttctaattttaatattttcacctaga
acagcaatattttggtttattaaatcttcaatactaatatcttgattgcgaataaacatt
tgatctaataatgatagctccttaagtctctttttcattcgtccttcaatgattgtttcg
attctatcaactggtttgtttttcaagtcatctttaccagcttcaactctttgttccaga
ttaataatttcactaggtatatcctgtgttgatacgtattccacattaggcgatgccgct
atttgcatagcaatatctttagctaacttttggaactctggacgtcgtgctacaaaatct
gtttcacagttgacttcaactaatacacctattcttccgccagtatgaatataactctct
agtaaaccttcaatagcggtacgattagatttcttgttagctgaagccagccctttttgt
cttaatgattctaaagctttttcttcattgccattattagcttgtaaagctttcttgcag
tccatcatacctgctccagtcttatcacgtaaggcttttacgacttgagcagaaatttgt
agtgtcatatgctttcaaaaatttttaagtatttaaatt----------aattattataa
ttcactttgagtaggatgagttatttcttctaaatctgtttgcccgtatttaccatcata
aattgcatctgctattttacctacaattagtttaattgacctaatggcatcatcattagc
tgggattggaatattaataatttctggactacaattagtatcaagtatacaaatagttgg
aatacctaattttaaacactcttgtatagctgttgtttctctcttttgatctacaactac
aacaatatcaggtaaacgagtcatatttttaataccatttagatgtttacgaagtttatc
taattctcttcgtaaaacagcggcttctttttttgggtaactgatcaatcataccactttt
atcttgttcttctagctgtctcagacgactaactcgagacttaatcgttacccaattagt
taacattcctccaagccatctctggttaacataataagaatcacagcgttgtgcttcctg
tgccactataccagcggcttgtctttttgttcctagaaataaaactttttttaccatccga
cgaagcctgttttataagttcacaagcttctgttaataattgtgcagtttggactagatc
aataatatgtatgccatttctttctgtataaatatatggaaacattttaggattccatct
acgtgcttgatgtccaaaatgaacacctgcttctagtaattctgctaatgtaacaatagc
cataagattattaatatttaaataatgtaattattcgggaaatccc-tttgtttagaaac

tgat-----tcaactaacttatttgaattttagtaaactaataataaattaacacacttt
taaagcagatcaacaaaatagatcctgaagaattcat----tctacacttttattgctaa
aataattacgcgcggttctgtcgtctaaaatgatatcatctaaatcatctcgtacagatg
aaatagtactttcgtcattagtatttgcagttaaattcttttctctaacgatgagccat
tacagttatcatacatattgaagccggtaccagctggtataagccgacctataataacat
tttctttcagacctcttaaccaatctaatttaccagatatagcagcttcagttaatactt
ttgtagtttcttgaaaacttgctgcagaaataaagctttctgtatttaaggaagcttgag
taatacctaatagcacaggcctatatgatgcattaatcttatttctcaaggttatggctt
tattagtctgttcaatcttctgtaactctacaagctctccaggtaaatagcctgtttctt
caccattttctattttcactttagaagtcatctgccgtactattacttctacatgcttat
cagaaatattcactccttgagattgatataccaattgtacttctttaacaagtagtaact
gaatctcttgaaagcttaatcgagcagcttcatataaggctaatccacattcaatataca
aatcaaaacttgcatcgaggatatcatgaggatttaacaatacgtctgttttctttcttg
cttcaagaatttcttcaattcttggaagcccttgtataatgtcacctgtttttgctctat
caaaaaccaatatagctaatgtttctccccttctaattaatgcattgttatcaacatgta
aaatagcaccattagaaaccaagtaaggtctagcaattcttaatgtaactgatctagaag
aaatttcaataatttgtccagagtctaaagaagtcatattttcagtaataaaatcaccac
aacgaatccaatcacccacacttactttaactgttgattttcctatattaaaggtctttt
tatcagaagatgtcgcaatcaaaatgcgcctgctagtattcttttcagaataaatctcct
caacagtaccttcactcattgcaataatctcagtactagcaacaactgtaaatggcttga
tatattcaccatttttacaataattctagtttgactttgttgcttctctgatttattta
catcaatacttttaattgataatgtttcaaaagttgacagccttaagtcaaaataattat
tgccactttctttagcagaaaactcaattgaggatattaaattatcatctcgatgctcaa
tttcagccacaagattagttgttactaaatgaactccttctatagatttcactctttctc
cgtcccgaaaaggagttcttttaactagctttagtttagtttgacttggtttttgtgaaa
ctagattttcaattagactggattttgtttctggtattgaataaacaataacaggtctaa
taagaatatagggcatttgctcgttttctatatactcccaataaactaacttatctgtag
aaatattattatgaagtctttctccaggtcttaaaaaacctctactttatcattattac
tatacacctcactaagtttatagatagatccaggcttaataataatttctcgaactattc
catccttttcgattatttcaacaatacctgagcttttagaaaaaatatttttaactagct
ctgttccactttctataacatctccattatgcactagtattagtgatgaatctttattaa
tttcgtgagtttcttctgaaatccataagatatatcccggacttaaaatttcgtaggcat
ctttatctaagcctgttttcttttttgatacatttaagtctaaatatttaataataccac
cacttgtcgttttatacgtatcagatattagttctgcgactgtgtaaccatcttgaa---
ttttttggtttggaatagctttgaaaagaaatttttgtttttttttctgtttctaaaatat
aagattctttttgtggattacatctgtataaatataacaacctggaattacaatagatt

cagtaataacttgtatattggtaatactgctattagtattctggtgtatccttacttctc
cagcataatgatttattaattctgtctgtgcaagtatagtaccagctttagttttatctt
ctttatgaacaataacattagcagaatcagatatactataaacttcacctgagagcaccc
agatcaagcctccagttttagtaattcttgtagtatattgattattatctgtttcttcaa
ctgttaaattggagaaacagattttgccagagagatctgaaactacatgtttttgagctc
tttctgtcatgagtcgatttctgcggggtgattcagcaatcacttgatccttcgatacta
attcaccatcacatattaaaagagtagtacctttacttaagttaattattgacttctgat
ttttcttagatctgattgtaacttgggttggctttttttgttattaatgcttgctcaccat
gtctggtacgaacatctgtatacgattcaatatctagatctatcaattgaccatctatag
gtgcgtagatttgttctgctaattcaccggtaaatactcctccagtatgaaaggtcctca
ttgtcaattgagtgccaggctcaccaatagattgagccgcaataataccaacagcctctc
ctaaatcaactaagcggccatgtgcaagattccatccgtaacaatactgacaaacagaac
ttctagagttacatgtaacaggagatcttactaaaactttttt---gataccagctttag
ttatctcttttgctaacttaggactgacatcttggttcgtatgagcaattaaacaatttg
tctctggatggaatacattttctgctaatactcggccagataaagcttgttctagattaa
ttaatattttctgcgtatctactaaatcttctagtatgatgcctttattggtcttacagt
ctacttctctaattattacatcttgagaaacatcgacaagtcgacgcgtcaaatagccgg
aatctgctgttcttaaagcagtatcaactaatccttttcttgctccgtatgatgatataa
aataatcagtaacagttagtccttctctaaaattactcgatattggaagatcaataattt
gtccctgtggatcagccataagccctctcatgcccactaactgtcttacttgagaaatat
tacctctagcacctgagaacgccatcatataaacagcatttagaggatctgtctctttaa
agtatttaataacttcttgtttcaaggattcactcgcattattccacgtatcaataactt
tttgaaatcgttcaactgctgtaatctcacctcggcgatatttgttttcagtggccttaa
tatcttcaatcgtgcctaataaaagactttgcttactaggaggaattcttaaatcttcta
aacttaaagaaattcctgcctgggtcgcataatgaaatccaagatctttaagcttgtcgg
ccatattggctgctcgggctataccataattacgaaaagcccagacaattaaatttttta
gttcgtttttatcaataaccttatttgaaaaacttggctgtgcaagacttcgtctattat
ccactaaatcttctccatattaaatgttcgtt--attaactatgctactaaagactcttg
aataattttattaaagattatacggccagctgttgttctaatatattgcacaattctctg
attttcagcgtcttcttttatgatatggttattaaaaaattttgttgtactattgtcaga
atgtttttcaatcttaatcggatcatgaggttgatcaccatctactaatccatcaaaacg
agcccatatataagaatgtaaatcaattttcttctgttcataagccataactacatcttc
taagcttgcaaaatactgattagatccctgctgctgagatggattattagctgttaaata
atagcaacctaaaaccatgtcttggcttggcataataattggttgacctgttgctggaga
gagaaaattatgaggggctaacattagcaatctagcttcggcttgagcctctaaagacaa
aggaacatgaacagccatttgatctccgtcaaaatcagcgttaaacgctgggcatactaa

tggatgaagcttaatagctcttccttctacaagaataggttcaaatgcttgaatacctaa
tctgtgtagtgtgggagctctgttcaatagtactgggtgaccttgaataacttcatttaa
gacattccaaatagaagattcatttttttgaatcatctttttggcagctttaatattgtt
aactaatccctgcaaaattagtctgtgaataacaaaaggctgaaataactccaaagccat
ttctcgtggcaatccacattgatgtagttttaaatgaggacctacaacaattacagatct
accggaataatctactcttttacctagcaagttttgtctaaaacgaccttgttttccttc
tataatatctgacaaagattttaatggtctattatttgcacctacaacagttctacctct
acgaccattatccatgagagagtcaacggcctcttgcagcattctttttttcatttctgat
aataatctccggagctaaaattgactttagacgagataatctattatttctattaataat
tcgacgataaaactcattcaaatctgccgtcgcaaatctacctccatctaattggaccat
aggccgtaagtctggaggtatcacaggaatgactgtgaataccatccaagaaggatctgc
acctgttgctatgaaattttcaatcaatcttaagcgtttcatttttttattgaattttaa
tgaaggtgtcttaaaactttttggtggatttgtagcttcagaccttaaggtttcagctat
gtgctctaaatctaaatctttcaacagtttctgaatagcttctgcacctattcctacttc
tacttggttattctcatcttgattttggtagatttcctcttcaagacttttccattcgta
accttcgagcagttgcttgtattttaaattaatatctgtattagattgcgtaactacata
agaatgaaaataaacaatttttttctacttctttgacttttaagtctaatgctaaagcaat
atagcttgtacttccttttaagtaccaaacatgagttactggtgaggcaagttcaatata
agccatccgatgtcttctcacgcgtgattctgtaacttctacgccacatcgttcacaaac
tatacctttgtagcgaaaacgtttatatttaccacagtgacattcccaatccttaactgg
gccaaaaattttctcacagaataaaccatccatttctggctttaaagttctgtaattaat
tgtttcaggctttgtaatttcaccaacaatttgtccgttgggcaaacttctttcgcccca
ctgtcgaatttttttcgggagaagctaaactaatttttacgtagtcaaaatattgctcaaa
ctttgtcataaatttgaataccttaaaatcaaataattagccaatttatttacatgcttc
atattgaaaaatgaatatcttaataaaggaattgttcaaaatcgtctactggaggagtat
cataattagaacgatctactcggttatctttagaatccgacatcaagtcaacttcaactg
tccgtctttggccatcttcaaatagtttcaatttatgtactgcaatgtctaaacctaatg
actgcagctctcgcattaagactttaaaagattctggagttccaggcttcggtataggct
tccctttaactattgcatttaatgcttcatttctagcttgcatatcatccgactttacag
ttaataattcttgcaaagtatatgctgcaccaaaagcttctaaagcccacacttccattt
cacctaatctttgacctccatgctgtgctctacctcctaatggctgctgcgtaactagag
agtaaggaccagtggatcttgcatggattttatcatcaacaagatgaacaagcttaagca
tataagctctaccaactgtaacaggattatcaaaaggctctcctgttctaccatcaaaaa
cttgcattttacctggatgctgatcattaaaaagccatttattgctagtaatcaatgatg
cttcttttaattttctatttacaagtgctcttgatgcttctgctccatacatttcatcaa
aaggtattattttaaatctctttcctaaatagccacctgctaagcctagtaggcactcaa

atacttgaccaacattcattctagaaggtacacctaaaggatttaaaacaatgtccactg
gtgttccatctgataaataaggcatatcttgtttaggtaagatcctagaaatgatacctt
tattgccgtgacgaccagccattttatcacccacttggatttttctcttctgcgcaacat
agacgcgaatcatagcatttgttccgggaggcagctcatctcctttctgcctagtaaaaa
ccctaaccttaactactctaccttttgcagcattaggaagccgtaaagatgtatctctca
catctcgagccttttctccaaaaatagctcgcaacaatttaccttctggtagttgatcag
cttcacctttaggagtaatttttcctactaaaatatctccagcttcaacccaagaaccac
cgacaacaattccatttctatctaaatccttcagagagttatcgctgacgttaggaatct
ctctagtaatttcttctggtcctaatttagtttgacggcattctacttcatacttttcga
tatggatagaagtatacaagtcatcataaactaatctttcgctaattaaaaatgcatctt
cgtagttataaccttcccaaggcatataagcaactagaatatttctccctaaagcgattt
cccctccgtctgtggatgcgccatcagctaaagtttggccaacaactatttttctccca
cccaaacaattgggcgctgattaatacaagtatcttggttagaccgataatattttttta
agcgataatgaactgttctaccactattatcctgaatgcctattttattagcagaaacat
aatttacatgaccagatgttctgctaataacaaccataccagagtctcgagctattttag
tctcaagaccagtaccaattattggcttttctggatataataatggaacagcttgccttt
gcatattagatcccattagagctctattcgcatcatcatgttctaaaaaaggtattaatg
atgttgcagctgaaattacttgaataggtgaaattgcaatataatctacttgggttgggg
ttgttgtaataaactcttgacgataacgtacaggaataatatctccttcaatgtaatgtt
gcttactaactttaacatctcctggagctactcggaaatcatcttcctcatctgctgtta
gatagacaggactattgtgataaatcacttgaccttggtttaccggataaaatggggttt
ctatgaaaccaaaaacattgacccgcgcacaggttgctagtgaaccaattaaacctgcgt
tcggaccttccggagtttcaattgggcaaattctaccatagtgactaggatgtaaatcac
gaactgcaaaacctgccctgtctttattaaaacctcctggacctaaggcactaattcttc
tcttgtgagttagttctgcaactggattagtttggtccataaattgagaaagctgactag
atccaaagaattctctcactgacgcaattaatggcttgggattaattagatttgataaac
tcaatgaatctatatcacatatcatcattctctctctaataatgcgttctaaacgattga
gacctactctgaattggttctgcagcaattctccaactgaacgaactcttctattaccaa
gatgatctatatcatcaagattacctgagtttttatccttaatattaattagataatcaa
ttgaggataagatgtcttgaggagacaatactcggaacgttttaggtatattgagaccta
gtttttattaatttatatctaccaacttcacctaagtcatatcttttaggatcaaaaa
aacgagagtaaagcatctgcttagcaacagcaactgttgctggttcattaggtctaagct
tagaataaactatgagcagggcttcctcatcagttacttcttcaatattatttttccaa
tttctttagctaattctttaactgaatatatttgggatgcagaaataagaaaagcatatt
tactaagtccttttttcaatttcatctttattgagtccaatagctcgcagaaaaatataag
catttactttgtgggtcttatctattcttatccaaatttctcccttaggatcaatttcaa

attttaaccaagaacctctattagaaattaggcttgcactataaatctgcttaccatttt
tgtctatctcttgtttataatagattcctggactacggattatttggttaataataactc
tttctgtcccagatacaataaaagttcctctatttgtcattattggcaagtctccaataa
aaactaatctctttttgtacttttttatttttatctgcttttcagcactgagctgtagat
gggttgatgataaatctaatgatttaatattttttcttggtttttagaaggtagatcaa
tatcttttcttgttaactttgcaggaacatatatttgagcactataagttttatccctat
tttttgcctgtctgacactataacgaggaaattttatcttgtactcattaccaaataatt
gtaattcgagtcgactagtagggtctgatatcttcggaaaaatttcaagaacttccgtca
aaccttctaataaaaaccatttaaaactggccctctgaatttcaactaaatctggaagaa
gtttattttttaagcttatacgttgaaccatagatctcctttataacaaattttgaaatc
gaagagtttaa-tactattaatagagtactggtagataatgcataagcatagtttgctac
attatttcttctctagacataactagaacatcatattttattcaa--aaaaagattttta
aaatttttcttcac--acacactatctataagtactttgctgcta--aaagatagttcaa
gctattaaa-------aaaaacgaaattaatacttgtctgtctgtcaattaattgacatc
aagatagacaagtattatttatttaatcaattctg-tttttctgagcaaaagacaatgct
ctcgcaagtctagcttttttttctagcgcctgtattagggtgaaatgcccctttttttaca
gctttatcaatcttactatatacttgagaaatacttgactgcacatcattgaaattatca
ctttctaaattatcaatatttagaaggcatcttttagttaaagttttgacaaccgattta
tacttacggttaatgagacgatttctttccgaagttttaatgcgtttaatcgcagaaagg
ttcttagccacagtaaatccaataaaactttataaaatatttttagaaacgaaaataca
caaatctgggatctattcaaaaagggaatatgcggcgtattataacattaaagaaaaaca
ataggcaatagttcaacaagagctcttgttattacacaatagagattgtcattgcgaaaa
actttgtaagatagtatttaattaaaataaactaactagacatataaatatatatgaaaa
aaattgaagctattattagacctttcaagcttaatgaagtaaaacttgctctagtcaagg
aaggtattggaggaatgacggttatcaaagtttctggatttgggagacagaagggtcaaa
ctgaaagatataaaggatccgaatactctattgatatcattgataaaataaaaattgaaa
ttattattagcgatgataaagtagagaaaattgtagagacgattattaaggcttctaaaa
caggggaaattggagatggaaaaatatttattagtagtattgaaagagtaataagaatca
ggactaatgacttaaattttgaagctttatagtatttttacttt-tttagacttgatact
tcttcactaaacatgagataataagcgtttattgtatacttataactaaggtgtaata--
ctaatggctaaaagcaaaggtgcacgaatcgtaataactttagagtgctctgataaggct
ggagagtttgctcaaaaaaggaaacctggcgtttttcgatatacaactactaaaaataga
cgaaatacaccaagtagaattgaattaaacaagttttgtcctaattgtaatcagcattgt
atcttcaaagaaattaaatagttatttatcttattaaaat--aatcatattatggctata
tatagaaaaagaatatctccaattaagccaacagatgctgttgactacaaagatattgac
ctgctaagaaaatttattacagaacaaggcaaaatactgcctaaaaggtcaactggattg

acttcgaagcagcaaaaaaaacttactaaagcgattaaacaagctagaatcctttcttta
ttaccttttttaaataaagattaa--atatatagcctaacttcatatttttccatctaaa
tattaaaaatttctatagatttgtttattaaaatgatctaacatactgaat---------
----------tactaagaaga-tctactcattacaagttttgttcattttataattaata
atagcattataattta---ttaagctagattc-----agacataaag-gtaacctctttt
aattgattatattatatagtctatttaaaagaggaaatttttttgcttc----------t
agcaattcaagattctttaagctttaacaaggtatttaaaacaaagatttttgtttaggt
atcagatcaaatgaatgaatcttatctcccgactgccataattgaaattcagagatgaaa
attccacattcgttgccagcttgaatttcttcgacatcttctctaacgcgctttaaagat
tcaatttttccttgataaataacttttttcttctcgaatcactttaatccaagagtttttt
aatagcttattattgatcactcgacatcctgctatttttctatttgctaacgagaatact
gtacttacttctgcttctcccaccggtacttctgaatattcaggatcaagtaagtcttcc
atccttcttttaatgtcttcaattaaggcataaatgatttgatagttttctattaatata
ttcgactttgccgatgcttgctttgttcctggagcaaagttggtattgaacccaatgaga
gtagaatttgtggttgaagctagttcaacatctgtagcagtaatttctccaggcatgatt
gagacaacattgagttgaactttactctgaggaaattgagataaagaatctaggattgct
tctgtagagccttgattatctgttttaataattaaagaaatttgcttactaatatcttta
gaattagtgtttttttaaagtatctaatgtaattcgactattaagtgctctttgtttctgt
ataattgatgagtctttagaggtattttcaatcgctttaagtttggcttctttatcactt
ttaactgctaaagctatttctcctgtagctggtacagatgacaatccccaaatttcaaca
acagatgatggtattgctaaattaatttctctttagcgttgttaattataactcgaatc
tttgcgtaagctgatccaataaccaaattatcactaatattcaatgtaccattttgtatt
aatagagttgctactggaccatgcgatttatctaaatgggcttctattataataccttgt
gcaggctgtgttggatcagctttcaaatcctctaattcagctaacaaggtaatagtttct
aataatttatcaacattttgaccagtaagagaactaataggaattataggcacttggcca
ccaagtttctccgacatgacattgtattttaataaatcttgctcaataatatctgtatta
gaaccagctttatctatcttagatattgctatgacaaaagggacattcgctttttgaata
tgattaatagcctctatggtttgaggtttaactccgtcatcagcagctataataataata
gcgacatctgttaaattagctccccctagacctcatacttgtaaaagcttcatggcctgga
gtatccaagaaaacaatttttttgtttattatctttcttaatatactcaacttcataagct
gcaatggcttgagtaattcctccaatttctttgttagcattattagattttcgaatataa
tctaatagcgttgttttgccatgatctacatgtcccataactgtgacaattggaggtctt
ttaatataattgccagtttcataaaaagcattagagttatctaaattactagaaagtcca
ttattattttcctttacatttgattctactgctataccaaaattatcagccactgatgat
attattgaggcgtctatggtttgattcatagtaactgatattcttttaagaaataaatat
tttataatatctgtttcttgaacacaaattagtttagataattcttgaatagttagagga

ttagtaatactaatagactctggcggagaattagattgaatgcttgctgaattttgacta
ggagttacggctactttttctgtcttatagatttttattactagtcagtttttttaca
acttccacttttggcttaggtggacgcattaaagagatagctaaatctcctgcagtctgc
gaaacattagaattagagtctctaaagttgtca---tcatcatcgtcaatatgtatttta
gtttttattttt-ttgcgctgtctattctta--ttttttttactatctactaagtcgtga
atttattaaaatttttactttttttatctagtttaggtggagaacttaattctagatgc
tgttcaccacc---tgtatgtgattctgatttatcaaggtctaaatttaacagattatca
ttattgactgactctaatctaattttatagataatctgaggatttttcaaatctacaatt
gtttcagaagaattgatatttgagctagatctagaggatctattttcaaagttttgatta
tttaaaaacataaatattggttttattatctaatagtaactttctaattttaatacagag
cttaatataaacaagaatcgaatatctgtataaataatacaaatgt---agagaattaat
ctatgttgtattagatttaatataacactaaagtcctttgttcgttatacaagtttaaaa
ctttt-tatgatgataaacactagattctagtttattggtatacaatctatttatctaaa
taatactgtttttgtcagggcttttaaa----ttatctaatgttatgttcattatgaaat
tctattaaaatttaaaaa-ctcaacaattaagtaagaatcgttaatcacctccacgccac
ttatgccacgatctcaaaaaaatgataattttattgacaaaacttttacagtattagctg
atattgtattaaaaatacttcctacaagtaaagaggaaaaagaagctttctcttactatc
gagatggaatgtctgcacaatctgaaggtgagtatgcggaagctttagaaaattattatg
aggctttaaaactagaagaagatccatatgatagaagctatattttatataatatagggc
ttatttatgctagtaatggagagtatgtgaaagctttagaatactaccatcaaggattag
aactaaattttaaattacctcaggcccttaataatatagccgtgatataccattaccaag
gagttcaagcaattgaggataaagatacagagctatctaaattaatgtttgataaggcgg
ctcaatattggcagcaggctattaagttagctcctgataattatattgaagctcaaaatt
ggctgaaaacgacaggacgaatgaggaatatacaaggatattaatatatttaagatataa
taaagatatcaattaataaattaaaattagttaaagaaactggtactatatagagtactt
-tactatatattataatattatactagttgatctctatctctaagacttagctgaataac
cgaaattaaatctattgactatattcatgaatacaaataagcataatggttagtacaaca
aaaagtaccggatctgttactcaaattatcggaccagttttagatattgcatttcctaac
ggacagcttccgaaagtattcaacgcactcaaagtacaaagctcagaaggaactattact
tgtgaagtacaacaacttttgggtgacaacaaggtacgagctgtttctatgagttccact
gaaggactacaaagaggggtagaagttattgatactggatcccctatatctgttcctgta
ggtacagatactcttggacgtatttttaatgttttaggtgaacctgtagataatttgggt
cccgttgattctgagagtactttacctatccatcgaccagcacctgcttttactaagcta
gagacaaaaccaaatatttttgaaacaggtattaaagtcgtcgatttacttgctccttat
agaagaggtgggaaaattggtttattcggaggtgctggagtaggtaaaactgtattaatt
atggaactaattaataacatcgctaaagctcatggtggagtatctgtatttggaggcgtt

ggtgaaagaacaagagaaggaaacgacctatatatggaaatgaaagagtctaaagtaatt
gatgcagataatctgaaagaatctaaagtagcattagtatatggtcaaatgaatgaacct
cctggagcacgtatgcgcgttggcttaactgcattgacaatggcagaatactttagagat
attaacaaacaagacgttctattgtttattgataatatttttcggtttgtacaagctgga
tcagaagtatcagctctactaggccgtatgccatctgctgtgggttaccaaccaactcta
gcaactgaaatgggagcactgcaagaaagaattacttcaacaacagaaggatcaattaca
tctattcaagctgtatatgtgccggctgatgatttaacagacccagctccggccactaca
tttgcacatttagatgcaacaacagtactttcgaggaacttggcggcaaaaggaatttac
cctgcagtggatccactggattcaacatcaacaatgctacagcctggaattgttggaact
gagcattattctactgctcaagaggtaaaatcaactttacaaagatataaagaactgcaa
gatattattgctattcttggtcttgatgaactttcagaggaagacagacaaactgtatca
agagcaagaaaaattgaaagatttttatctcaacctttttcgtggcagaagtgtttact
ggatcacctgggaaatatgtatctttggaagatgcaattaaagggtttcaaatgatctta
aaaggcaatttagacgacttgcctgagcaggcattttatctagtaggtgatatagatgaa
gctatacaaaaagctgacagcatgaaagattaatataattaacgatgactttaaatataa
gaattattgctcctgatcggactgtttgggatgcagaagcacaagaaattattttaccaa
gtagtacagggcaacttggtattttaacaggccatgcacctttgcttacagctttagata
ttggagttatgagagtaagagtagataaagaatggatgccgattgttttgctgggcggtt
ttgcagaaatagagaacaatcaattaactattctggttaatggcgcagaagaagctagtc
aaattgatttatcagaagcagaaaaaaatttagacactgcaactcaactcttaagtgatg
cctcgtctaataaagaaaaaatagaagcaacacaaaaaatacgaaaagctcgagctcgag
tacaagctgcaacagcagcaacttcgtaa-aagattcc------tcttaaaacataaaac
tttagtaattattactaaa-------gttttatgttttatttgtatggttataggcagct
taaataaacttagcttagacctgaacaaatataatcaaaatataatcccatctctttacc
tgcatctggtccaactaaactaatagtaacttctttcatagctaaaatagcttgaatcgt
tgcaccaataggaacacctagagaattgtatgtttcttttagaccatttaatactctttc
ttcaaggatagatggatctccagctagcattccataagtagcatagcgtaagtagtagtc
taaatcacggatgcatgcagcataacgtcgagttgtatacatattaccacctgggcgtgt
aatatccgaataaagtagagattttgctacggattctttaataattgttgctgcattagc
tgcaatagtagcagcagctctgactcgtagctcacctgtttgaaaatagcctcttaattt
ttcaacagagctatcatctaaatatttaccttggacatcagctgcattaataacagaagt
aattgcgtcttgcataactttcacacattccttaattttaaatatctaatacta--gggg
tatataacaaaattactagatcttttagcgatatcagtattactgcatagcacccaaagt
ataatcaaagtaaaaaccagcttctgctgaatcttcacctgcaagcagtgagcaagctac
acttttcatgcattttacaccttcagcaactcctgaaattggtgttcctaaagaattata
catttctttaacacccactaatccaatttcttcaataggagttacatcgccagcaactat

tccgtaagttactaagcgaagataatagtccagatctcgtagacaagtagcagtcatctc
ttcgccataagcatttccaccaggagaaactacatcaggtctttttttgaaacagttgttg
gccgccttgctttacaatacgctcacgattatctgttaaaatttgagctattcttaaacg
acgttgtccagataaaacaaaactttttaattctatctaattccccaggacttaaatatct
tgcttctgcatctgcatttacaattgactttgtaacaatactcatggataaaactcctgt
aatgcttt--aaaaaaagcttaatactaattatttatgatataactcagaaagtaatgta
ctttttgag-aattttctaaccgaaacacgttcatgatatatgaagcaggatagatactg
aggtaagtactaaaacgagaaagttatatgtaggaacttctgttattgatttcctagtac
tttttaaaacttggtaccacaattgatatatcctgcttagtcaaacgattatagagtgt
ttctgtattaggaaagttagctgctggtaaagtcggaaatcttctgtatggaacagtatc
tttgccgtataccgcattgtattcaacactttcaactaaattacttataaatgcagataa
gccttttgaagctaaaatttgattaaagaatctaatttctgcctgattattaggagctct
tcctaagatatgcttagtacctaattctataactttgtatttggatatggctgataaaa
ttctttaccatatagttcagatagcgccagtttttctactagttctttgacacagatttg
tctattcaagaaagctgacttaatatccaagaattcaccgccgacactaaaagagttaag
atctctttcgaaaatttgacgatatgctgctcttaaagcttgttctaacatttctttgtt
actatcactattcacttcaaagacaatagattgatctcttaaagaagttactccttgtga
aatacgagactgaatatcattactactcctcatttctttgacagtgcctaattcaacaaa
cctagcagatttactagcaataactttttgaatctctggtcaataattcctggtcttaa
gcttcttaaagcaacaccagctggagtactatatctttcataaggaacagtattatctcc
aaaagtttctgtgtactctgagctatcaattattgtgtctattacttggtaatagccttg
cttatatgcaatatcaaaatatttattaatttcttgccgaccatatgtagggcgacccag
taagcggttatgaatatattcaatagctttacaaatatataatggctcccaataaagaga
tctaaatatacttgattttgctaactgtctaacaaactcgcgaactgtaatttgattatc
ttttagctgactttctattggcttgagaattagtttttcttcttgatacacttctctgcc
aaaaactcttaaataagcaacttttgtgacaacttctactgaattttcaaaattttggga
tgaatctgactttataagagcgttggacaatttaaagatttttggtcccagggatcctgc
tgacttaggtcttacctgtggattactaatctgattgtagattcctggcccacgtctaac
taatatccttcttgtatctttaccaaaaagagcttgtctttttcttggatctttattttc
tttagggaaaatagctccaaattgaattgataatggatcattacctattccatatggatg
ttgatcaggtaatgattgcttataatcactaaataaagttataaattgaggaaccttacg
aaaaggcgcactataggtgagaagatctatttgaggcccccaattacggcactcttgagg
ctcttcacctaaatttctaaagtaaggcacagtctcttcaccgaaataatctgtatactc
agaagaatttaaaagtgcattcactaaaccacttaaaccagtagatgataaaatagcaaa
atatttttgaaactcttctaaagagctaggacctctacccaagaaatgtctaaatgctaa
ttctaatgctcgactattgacaaaaggttcataaaattgctttctataaatacttgaagt

tcctaacgaacgaataaattccttaattgaaatttggccatttttacttgcgactccaa
attcgataaagagaggtcatatgctttagcaatatctctttcaaaaatttgtctataaca
tgctttcacaacaatatttttctcatcagcagataaactaggcttcataacaaaacgagg
tgttgacactcctgcctgtacataagtctgaggaagacgcaaaccttgtaaatctccaga
tattctttttcttaatttatcagtcaaagatggagcttcaaattcagaaatcacaacatt
aaaatactctttaactaaatcttgacctttatatcttcctcaaagattaataatgctgt
ccgtctcatctctctaagagccacaattgctgctgcacttgagcaagcgttatcaattaa
ctctctcaatcctcgtatattaacagataaaatattagggtcgcctgatacaattgcata
tgtcaaataccgtaaaaaccagtctaaatctcgaagagattttctcattcttgtagtacc
gtatctcaagacgtttataggtttaaaccctggcggagtagctcccccagcattaaataa
agatcggaaactttggccaaaatctccttggatatttcctgataactcattaattttatc
ttgagaactttgatctcctgcaataataacagctgcttgcggacgttctaaataagaaat
tgcagacccacctacgaatattttatctgctgctcgtgctaccagaatatttgcattttt
agttaaaatatctgcaacttccagtctttttttgtccagaatttaaaaaagaaactaattg
atttaattcaccaagctgtaaaaatcggtcttgttgctctgcctgagtaatagttaaaat
tgaggcagtccgataaagctgggggcgtgctaatgggcttccgccacttgctttgatact
cattactttatttatctccttaactgatattgcttgctagcgaatacaaatatagcttga
tatttctaaatatttaaagatactatgcaagcaca-ctgttttctatattattaggtact
agtacttatagtacagagagtcttataactaacttctataaatacaaattttgtaatact
tctattaacatttgtcaaaaaaatatttaagtgctcactcttttatattaaaattcaatt
agtgagaggttgatgtgaaataaagtatactaagtaagttttatagataggaaaattaaa
agatttattctacaaacattaatca------------cttgaatgtttagcaattgtaca
attgttattgcttttaattggtttagactagttttatatttttctaacataaacaatat
taatgagcaatttgtatgaatttaaagccacaaaacaacctattaactaacgaaaatata
gatcatgtagatatacccgaaaatgatataccaatgtcaattacggaacatttagaagaa
ttaagacaacggactttatttgtgtttttattctttttgtttgctacaactataagtttc
acacaaattaagatcattgttgcaatactgcaagctcctgctgttggtattaagtttta
caactggcaccaggagaatatttttttcatctattaaggttgcaatatattgcggatt
gttgcaacaactccgtttgccgtttatcaagttatattatacatactcccaggactaact
ggaaaagaaagaaaaattattttgcctctattaattagttctgtattgctttttattaca
ggtggtatatttgcttactttgttctcgcaccagcagctttaacatttttaattagctat
gggtctgatattgtagaaccattatggtctttcgaacaatactttgactttattctatta
cttttacttagtacaggattagcatttgagataccaatcatacaattattgctcggtgtt
tcagggacatttcttctagtcaaatgatacgagcttggagatatattattattatagca
acaattgctggagctattctgaccccttcgactgatcccgttacacaattaataatgtct
tcagctgttttattactttattttggtggaattgttatattattagtcttaaaaaagtag

agtaaat---------attttcta-ttaatccaatatatttgttatc-----tggcaaa
taatctat-aaaacaattgct------------------agatttaagtatcaggcttct
aaatatttcaa---------gtctcttttatcattgaaatacaatc--------------
------------------------------------------------------------
------------------------------------------------------------
------------------------------------------------------------
------------------------------------------------------------
------------------------------------------------------------
------------------------------------------------------------
--------------------------------------------------gatacattgg
cttgtttttcatcaattatttgtgttgcacac-tttttagtactctctttattgttctt
agaattatttctttaatctatttattaaat----agatatgaagagatttaagtattgat
tatggtaatctatatcaacgtaaaaagtattttattgtgagttttatttatataaaggct
tattcattatatagttataacagtaagtctaattaatag-ttttcatgcttaattaaagt
aataactcttaattcttttggcagaaaaatctgtcttaatattttatc--aaaaaaacag
-aaaaatgaagctaaatagtctaattaacttaattcaaaagtctatatattcttgtacac
ttttactaattattttaaatattatttgtgtcgcacctaattctagtaatgcatttccaa
tttatgcgcaacaagcttatgaaagtccaagagaagcaactggtaggatagtatgtgcca
attgtcatcttgctcaaaagcctgtggagatagaagcgcctcaagcagtactacctaata
ctgttttcgagactgttgtgaagattccatatgacagcaatgctaaacagattttaggta
atggcagtaaaggaggcttaaatgttggagccgttgtaatattacctgaaggatttaagt
tagctcctgttaatagattatctacagagttaaaagaaaagactagaaatctttacattc
agccgtacagtgctaaacaagacaacattttagtaattggacctatttctggtgataaaa
atagagaaatagttttcccaatactatctcccgatcctgcaaaagataaaaaagctcatt
ttttcaagtatccaatatatgttggcggaaatagaggacgaggccagatttatccaacgg
gtgacaaaagcaacaataatattgtctctgctttaagcagtggtaaaattaataaaattg
aattactagacaaaggtggatttataatacatgtgactaacagtagcaatgtggagtcaa
cacagaagatttcacctggtctcgaacttagagtaaaagaaggggatacaattcagcttg
atcaagctttgaatagtgatccaaatgtaggtggttttggtcagaacgaaacagaaatag
ttttacagagtccaaatagaattaagggcatgattgttttctttttcgctagtgttctag
ctcaaattttcttcgtattaaagaaaaaacaatttgaaaaagttcaagcagctgaaatga
atttttaaagcaatataaaactggcacttactataaagaaagta-taaacactttattca
caggatgtaccagtttttatgttatgtaataatcgctttgacggaagttatgatactttt
aacagaatctacaatctgactaggctgaattactgtagcctgctctaagcttccattata
tggtgtgggtatatcctgcgaagatagcctcacgacaggagcatctaattcatcaaaaag
atactcattaatttgtgcaattagctctgctccaattccggctgttttcatacactcttc

tacaattaaaactttgtgagttttctttactgagatagatatagagtctatatccagagg
ttttaaagatataagatctataacttctggatcataaccttcttttaataaggccggtaa
tgcttggataacatgatgcctcattctagaataggttaaaatcgtaatatcttttccttt
tcgtacaaattcaactttattgagaggtaagaaatattcttcttgaggaatctcttcttg
taaattataaagtagaacgtgctcaaagaatacaactggattattgtctcgaattgcaga
tttcagtaatccttttgcattataaggagtagaacaagcaactatttttaagccggggat
ggcttggaaataagcttccagtctctgagaatgctctgcacctaactgcctaccaactcc
tcccggtcctctaataactaaaggcaatgtaaaattacctccagaagtataacgtaacat
tcctgcattattagaaatttgattgaatgctaataataaaaaactcatgttcataccttc
aacaattggtcttagccctgttatagctgcaccaattgccataccagtaaagctattttc
tgctattggtgtatcaagaactctcaaatccccatatttgctatgcaaatctttagttac
cttataagatccaccatagtgaccaacatcttctcctattacgcagacagttggatcttt
tgccatttcctcatctgttgccgctcgtaaagcgtcaaacataaagattttactcatttt
aatatttgatttttatgatttgaccactgttttaaagtaacagttaagacaccatattct
atttagttatctgcaaaaagatatcgtttcaattctgacatgttaggttctggactggaa
atagcaaatttcacagcttgttccaattcggttttaacagcattctgaatctcattaagt
tcacctatattagcaatttcattatctagaatatatttttttgagtttttttgataggatct
cttgccacccaagcctctttttcttgtcttgatcttagttcatcaggatctgcgagagag
tgaccacgaaatctatatgttaatgcttctattaaggttggaccatcaccttggcgagct
ctttgaactgcttgttttgcagcttgtcttacagctagcacatccattccatcaacttca
atcccaggaagcccaaaagcttctgctttttttatgtatttcaggtattgaagaagaccgg
tgatgtgccatacctatagcccattgattattttcaacaacaaatataataggtagtttc
cagagaactgccatattcagacattcaaaaaattgcccattattggtagtcccatcacca
aaaaagcaagccgtgactcttaaatcctctgtttctttaagtacttgctggcggtagata
ctttgaaaggctgcccctgttgcaaccggtataccttccgcaataaaagcaaagccacct
aaaaaattgtgaggcgcagaaaaaatatgcatcgaacctcctctgcctttactacaacca
gtctcttttccaaataactcagccatcacatttttagatgggacgcctttacttaaagca
tgtacatggtctcgataggtactgcaaacataatcagttggattcagaagtttaattaca
cctgtagaaacagcttcttgaccattataaagatgaacaaaaccgaacatctttccttta
taatacatctgagcacacatatcttcaaaatttctgcctaacaacatgtcttcatataaa
actaataaattactcttattaagattaagacctgttgaattatagttagtcagcggcaat
tgaactttcttaggataactcataatttgtaaagaaacctct----------tattgtaa
ataggaattttgaagaaacagctatcaatagaaaacacacagataaaaacttataaatta
taatatgtttttactaaagactttaataattatttgaaactagttcagtatatcaaatt
aattttgcttatactacggatgtctgaaacttatataaaaaagtagt-a-aaaaagttta
ttaaaagaatagttaaaaacagatataaatagaagtaaacagaattatt-gaaaaaaat

gattattattaattaa----tacttaatatttctaaatattttacaattaatggctatgt
caccatctttattttatcctgttgaacaagaactatgtagtcttgaaaaaaatctgaaag
ctgttgctgggactcgtcatccaattttatatgccgcagcaaagcatctattcgatgctg
gaggaaaacgagttagaccagctcttgtattttagtggctaaagcaacctctgagaagc
aagatataaatactggacaaaaaaggctagcagaaattactgaaattatacatactgcta
gtttggtacatgatgatattattgacgagtgcacaacacgtagaggagtcaaaactgtac
ataatttattcaatactaagattgctgtgctagcaggagatttttttatttgcacagtctt
cttggtatttagctaatattgaaaatttagccgtagtcaaagctatttctaaagtcatca
ccgactttgcagaaggagaaattaggcaaggcttagttcatttcgatcccagtatttcaa
tagatgcttacattgagaaatcattttacaagactgcttcactcattgctgctagttgtc
ggggtgcagctatgcttaatggttccaatcatcaaataaataatgatctttatctttacg
gtaaacatatgggattagcatttcaaattatggacgatgttctagatataactggttcta
ctaagagcttagggaaacctgctggcgctgatctaataaatggaaatttgacctctcctc
tcctttttcacttactcaagaagcaagtttaaatgatcttattgatagggagttctgta
atagtacagatatagcctcaacattatttcttataaaaagaagcgggggaattacaaaag
ctaaagatttagctaaagaacaggtgcaggcggcactttcctgccttcagtttttaccac
aatctacacctgtatctagtttaaaagaattaacacatttcataatcacaagattgtcat
aaaggacttgctaaaattaaaatgttattaactaatttttttactgtttctaaaacagtct
gaaaagtttgaatatcattactagctagctgcgctagcattttacgatttaacgcaatat
tctcttttttttaatgcactaataaaagtactataattcatgccttgattgtgagcagcag
catttattctagttatccaaagacgacgaaaatctctctttttcctttttcgaccaacat
aagaatatcgaagagctttaaggacttgctgcttagctgttcgaaataaacatttatgtg
cacctttaaagcctttagctagcttaaagattttagctcgtcttttttttgcaacgttac
ctcttttaactctactcataaaatattac-ttaaattttaattattaccagcttactaaa
ttcttatactctttg----attataaataaggtaatttcattgcaatgtttttaatatct
tttaagtcaaccgaacaagtagaagaaagatgtcttctttgttttgatgactttttttgt
aataaatgacttttagaagctttatgtcgaagaaattttccggatgaagagactttaaat
ctttttgctattgcttttgatgtttttaacttaggcatattataaacttttaatgattaa
caaaagcatgaaaagtctgaggatatcaatcttagacctttcaatacttagaataatctt
aacacaatttattttacttactacataggttaactgaaacaatttatatatataatacta
atcagttaatcaaccatagctaaagtggatttatattttgctttgaacttttgtatagta
ttagcaattagcattgttataattttaagactgcttgaattcaattcttcgaacattttc
atatttaatgtaaatgaaatattggcttctgagacaatattttgaatttgaacatcagac
aagggaatcatatctaatgcagctctatattgatctttaaataatttatcgtcttttatt
tggtcaaaatcgtaaaatttcgttcctcctgaatctgataaattcatagcccctctagct
atttttttaaaatttggccgccagaaaggtcacccaaatagcgcgtataagcatgagca

actaataattcaggttgtttatgtcctatagtgtgaattctatcaacataaattttgta
gcaggagatggttcaataaaatctaaccaatctgatccataataatagtttaaatcttca
gatagactcgctttcctattaagctctgtaaaatatataggtttaatagctggatgattt
ttattagaaaataactcttcctctattgcacaatagacaaagtatagatttgcaaccaat
ttacgatatgacttcttatctacaactcctcccaaaaaagatttaacaaaactaacattt
tctgccatactatgggacttagtagtaccttctcttagttcattcgctaaagtattaacc
ataatatcaatattccattttt-aaagataacaaattaaca---ataaatagttatccgt
atatttataattgagttgcgctaactaggctgctgcaaaataatttttgattttattgg
atctggattcattgttttgtcaccaggcttccaattagctggacaaacttcatctggatg
agattgtacatattgaattgcttgtaaaaccctcaaagtttcttctacactcctgccaaa
ctctagattattgattgtagagtactgaataattcctttcgggtctataataaatagtcc
tcttagagctacaccatcactattcaaaacattataagctgcactgatctcttttttaa
gtctgatactaatggatatgaaagatctcctaatccacctgattctctgtcagtctgtaa
ccaagcaagatgagaatactcactatcaaccgagactccaagaacttctgtattaagttc
cgaaaaagcattatacttatcgctaaatgctgtaatttctgtagggcacacaaatgtaaa
atctaaaggataaaagaataagacaatatacttattcttgagatcagataattttaacgt
tttaaattcttggtcataaacagctgtagctgaaaaatcaggagctagctgacctactcg
aagacaatttggtccagaaatcattaatttttcccagaagaattgatataattttgttat
ttaataaataattatataacaaaatttagtatatattaaaatactacaagtctttttaa
attac--------taaatgatataatcaatactatctttatagcggggaatggatttgaa
ccattgaccttcgggttatgagcccgacgagctaccagactgctctaccccgcggataac
taaaatcatattagtctacatatatatataaaagcaaatgcataatagattttagaagta
aaaaaatagaagaactggtccactgatcattaaacttagaaacgacaaagtatttttat
cttatttaaaagagggaagacttcatataaaccaataaatctatcttttaaaagaatttc
ggacggctttacccaaatttgaccgtcataccagcctgattcttcataaaatactgtagc
actcattaatcgttttactacgtatgaccagcctaaatataatctgattaaaataaatcc
tgtcattaggctggaagtaataaattctgaaaagaaaaatttcaaaggtagtttagtgat
tggaaaaattgataataatataggactaactaataaacagttaagtagtagtgcaatagt
tattttttattataagatctacggcttaaagttggccagcaaaagaaccaagaattttt
caaagaagtatattcatgaacaggttgctgctcctttggcacaggacattgagtattata
taaattcattgacattaaaaagtattataataaagac--tttgtttatattatctaacat
aggtacgacattaaatattaactaactgcaattatagtagtgagtagaagaaataaaact
aaaaatagccaagtaattttatttaaagtattctcagtactacgagtattactaaagaac
tgattctgagcacctacacttcctaaaccttcggatttaggattgtgtattagaatagtg
aagattaatattattgtagatgagtaccaaaaaaattttaaaatttgttccattatgtaa
aagttaat-aaaaatagcacaaagacaatcgtctaaaagatattgattgtcttatcatat

taaaaagtatttattcaccctgtacttttaaaagtaaaaatcctattgccagccctacca
aaactaggactgaacttaatacagcactttctacaatttctccgaccatt-ttattttc
cttattattttattattg---tcattcttttgacaagattatattgagctaa-ctaaaat
ccgtttctaccccaaactaccattgcgattgaaaatgtaaacattgccattagagctgcc
catcccaaacttaaaatatccataatttataatattc--ctaaaaataagatctcttta
cttaaactataatattcttaatattaaaaatatcattatataataagctaattataacag
agtactacttaattcgtatttttagtgtatttaaagaagattgaa-tttttcgaggtac
tggataagtatttcatcctaagaaccatagtcaac----ttttgatttattataatgctt
aaatctaaatata--tatcactcaagaatcattagtaattatgatgtttttaattatacg
att---aacatatttcaaaatagaagaattcatagcgactaagttaa-----tatgctaa
tataatttatcaat--aaaaaggaagttagtatttcaattatgcaaacaactataaataa
tgggcaaacgtctagtaaagaaactttactaacacctagattctatacgactgactttga
agagatggctaatatggatatttccggcaatcaagaagatttttggctatcctcgaaga
atttcgagctgactataatagtgaacatttattagagacgaagagtttaatcaatcttg
gtctaatttagaacataaaactaaatccttatttattgagtttttagaaagatcttgtac
cgcagaattttcaggtttttactatataaggaattatccagaaaattaaaagacagaaa
tccggttatagctgagtgctttttattaatgtctagagatgaagctaggcatgccggttt
tttaaataaagctatcggggacttcaatttatctttagatttaggattttaacgaagag
tcgcaagtatactttttctcacctaaatttatttttatgcaacctatctttctgaaaa
aattggatactggagatacataactatttaccgccatcttgaacaacatccagaacaccg
tatttatccaatctttagatttttgaaaattggtgtcaggatgaaaatcgtcacggaga
tttttttgctgctttgctcaaatcccaacctcatttttaaatgactggaaagcaaaaat
gtggtgcagatttttctattaagtgtatttgcaacaatgtacttaaacgactttcaaag
aattgattttataatgccataggcctagactctagacagtatgatatgcaagtaatacg
aaaaactaacgaaagtgccgctagagttttcccggttgctttagacgtggacaatccaaa
atttttcaaatatttagatacttgtgcatgtgataatagagctctgatcgatattgataa
taacaattctccattattcattaaatctatcgtaaagatacctctatatttttccttatt
tgcaaatttactaaagatatatttgattaagccaatagactcaaaaacagtatggaatac
agttcgatagttaaatacaagggaatgctttttttcttataaaaagctatctagtgattaa
tataaaat------aaaagcttccttct-----cgatggaaggagcttttattttgtcaa
ga---atataatgggattatatagatc-tagaactaagaacttactttttcttttgcttc
atacataacctctaaagtaattagattttagaacaatcacgggcaaatttctctgtatt
tctttttactttacctctaacaaacccaggaattttatttaattctttctgtgcttcttc
tgaccaattaatactatcaacagcagaaattcctatacttaatgcttctgttgtatcatg
accaccaaatatttctagcaaatgatcttccataccaagtgtaaacgaattatatactaa
atcagcaatttgatttgttccctcataaccaagaaatggacggtaacttaaaggaaagtt

ttgtatatgcactggtgaagaaataacgccacaaggaatatttagacgcttaccaatatg
tctttccatctgtgttccaaaaattgctgcaggttcagttttagcaattaaatctccaat
taaaccatgatcatctgaaacgattacttcatcgcagaattcttgaacttgatctttaaa
ccactcttcatcatatttgcaataagttccacaccaggcgacatgtatacccatttcctg
gtgtaaaattcgagttattgctgcagcatgagttgcatctccaaatacaatagccttttt
acctgttaaattctgacaatcaattgatcttgaaaaccatgcagattgtgaaataaatcg
ggtttgctcgtctatgtatttttcgtagtctacagcagctcccaatgcatttactaattg
ctgtattgatcggatgcatgcagctgtttgaacaatacccataggtgtaatgtcaacata
aggcatattaaattctttctctaaatatcgagctgtcattagtccagtttctctatatgg
aataaaattaaaccatgcagagggtaatttttttaaatcttgaactgatgcattttctgg
aataatttggtttatttggatatctagatcttgaaataaccttttttaattctgctatatc
atgttgattatgaaatcctaagcttactgctccaataatattaacagatggagttttagt
cttttgagttaaaacttgattattagatttagctttttccatatagaaagtgacaatttg
ttctaatgttctatcacctgcttgcagttcatttactctataatgattaacatcagctaa
taaaacatctgcttctgtttctatagatgctctgctaacaaagttttgcaaatcttcttg
caaaatactcgaagtacaagttggtgttagaataactaaatcagggctttcttctctgtc
ttttctagtaatattttctacaactttttcttgagaaccacgggctaacacatgtctatc
tacaacacttgctgtaacaggagtaaaatctctgtctctttcaagcattgaacgcattac
attaaaataatcgtctccgagaggcgcatgcataattgcatgaactttttttaaaagaact
ggcaattcttaaagttccaatatgagcagggcctgcatacatccaataagctaatttcat
attaaattttcctatgatttatttagacatagtaatactttttaaagctaactaacaaaca
ttagccaaacatctaatttaactccttgacaattttgtaatattttttaaatgaagctta
tcattataaatatttaaaaaatataattcagaataga----aataactgttagatcaagc
ttgaaagtaaagatttcttgcagttatagtttttctttaagtaaacatgtaaaaattgta
ctatatataaatgtactatatataaacttttttttactgaatattggtaatgccagataca
attaacttaaatatgccttccccaacgtttggtggaagcactggcggttggttaagagcc
gcagaagtagaagaaaagtatgcgataacatggacaggcaaaaaatgaaagtaagtttgaa
atgccaactggtggtaccgcaactatgcgaaatggagaaaacttactctatttagctaag
aaagaacaatgtttagctttaggtactcaactaaaaagtaaatttaaaatatctgactat
aagatttacagagtttttcctaacggagaagtgcaatatttgcatcctaaagatggagtc
tttccagaaaaagttaatactgggagagctagcgtcaacagtgttgatcattctattgga
caaaatattaatcctgtagatgtcaaattcatgaataaagcaacttacgattagtctctc
-aaaaacctagacataaaataacttttatatagtctaggtttttgataatgtatactgat
agtatggaagggtggccgagtggttgaaggcgtctgatttgaaatcagttgaacta--tc
tggttccgtgggttcgaatcccaccctttccgttataaaact------------atagtt
ggcttctcaaggcaagctagctatacactgatagtattaagtttagacttaataaagtct

acaacttgtgacaggttagaaatttttttcggcatcctcatctggtatctctatactaaat
ttttcttctatggccattaccagctcaactgtatctagagaatcggctcccaggtcactt
gaaaaatttgcttccctagtcactattttttttttcaattcctagctgttctgctacaata
tcttgaacttttttcaaagatttcattatcttgcataatatattatttcctctaaggtgtt
atcaaaaatttttcggcttatattattatccaactctaaaaaattagtaattttttagtcg
agattaaaatatttatacctatatgatatactattattctggataaatacgaagtttacg
atatggctcttctccaaaactacgagctctcaactggtagttgtcaactagattatgctg
catttttcgtatataggctgatcgaggtgtaagctgaactattgaactttcattcaagat
aatagtttcaatagccaatttagcttcttgaagagcctggatctcatcaaaatttttatt
tttacagagttctacccaattaagaccagatgaagtatgaatgtttaatatttttctgag
tgcacgagtaatttgtggaaccgtactattctgaattgtatatataataatttgttttga
ttttgcaatttgtctaagttttgtattttgctttacttgattacgtagcgctagaatagc
atcagactttttctatctccttggttaaaataattggtagatctagtgaagaaataactga
tgtaatatgctgccaactcaaagaataagcatataagtactgatgaggtatttccaattg
caaagattgtttattaatattaatagggggcattgaaagtagttgacaataaagatgtatc
acgattttgtgatttattcaaatctaaggacttatttcttaattctctatattgtaaaac
cggaggcgcctgctgtgcgttaattgattttcgagcattagagatgtttacaggtaagac
ttctatagattgcgatggataacatttaattaaaattctaccattagcttgaatttgtct
tttttgcacaaaaggttgatggccttgcaaaatttgatctattgtctctttaactttatt
atgaactatccaaacatttcgctcgtgtatttcgatcgcaatctgaaaagcaggctcagc
ttttctttctagtatactcttttgcgtacctctacgtttagcttcatcgtcacctaatgt
tacatattgaattccaccaattaggtccgccaatgttggattttttaattaaactttctaa
atgatttccatgcgctgtacctactaattgaactcctctttctgctatagttcgagcagc
taacgcttctaattccgtgccaatttcatctataataataacctcgggcatgtgattttc
aacagcttctatcattacttgatgttgcaaatcgggccttgcgacttgcattcttctagc
tctaccaatagctggatgaggaatatctccgtcaccagctatttcatttgaagtatctat
aataaccactcttttttttccatttcgtcagctaaaacacgggccatttctcgaactgcggt
tgttttaccaactcctggtttttcctaagagtaaaattgaatcaccttgctgtagtaaatc
tcgaataatactaattgtaccaaagactgctctgccgacacgacaggttaagccaataac
actaccttctctattacgcagtgaactaatacgatgcaaggtttttttcgataccagctcg
attatcgccgctaaaattaccaactttttttacacaataatctaagtcttgccaactaat
agatctttgagataaatattctgggttatctggaaatctagcttctggacgacgacctaa
atccataacgacttctattaaattatttctattaggatgctgttgtaaaggttctttgac
aaaatttggcaaaatttctagcaacttatctaagtcatctgcaataagcatgattgagta
tataataattaattaatcaaatagaagattcaagattctaaagattacaatacatattaa
agaatattatgattggaatcaattttatacttaaattgaacaggaagtta-tagatgtaa

aagaactgattaaactttacgcaataaatcgattggtcaaaattaatagtcgtgttaaca
tgtttattggacaaaaagttcgtattaaatatagcaaacaaaaagttgctagtgatatag
cagataaagttggtgaattaggagttattaagggaataaagtttattaatagtcaatgtg
ttactattattgttgagtttgacaatcacactaggctttggatgtttagagaggaattaa
tctgtctaaatgaat------taaaaacatgaaacataatttattttatttattatacaa
aatttcttatgagtcgttaaaataataaattttaaaggagtacttttaattgtgcaaatt
actattaaaaaattacaagacttactttcgtctgtacaaagaaaaaagatccagacatta
aaactaaaacaaggtaagtttgaacttctattaaataagacctataaaaaagtcaatcaa
gaaattataccttcgcagaaatctgctgtattacaaaatagtccatctacgataattaaa
tcaataaataatacaaaaaaatcttctgttgtcaatgaagaccgtacagaatatgccact
attgtttctccaatggttggaacgttttatcattcacctgctcctggtgaaaaaatttt
gtacaagttggcgatgaggtcaaattcaatcaaacagtctgtattattgaagcaatgaag
ttaatgaacgaaatcgaagcagaaattgaaggcaagattatagaaattcttgttaaagat
ggtgatatagtagattgcgggcaagccttaatgaaggttgaaacataatgctaattttc
cattttacatattgttaacagaaattatttagatagaaaaattagcttatactatagtta
tgaaaacttatttcgcaaaggcac-taaaaactaatatataaaattatttactttatctg
ccaaaaatagaaataagcgttttgattccttgtctcaagagaggagaatctcaatggcaa
ttagctcaaaagagcaagagacaaagaaggtaaaaatctcggttgataaaaatcccgtag
atacttctttcgaaaagtgggcccaaccaggccatttttctcgtacactagcaaaaggac
caaaaactactacttggatttggaatcttcatgctgatgctcacgacttcgatagtcaaa
ccagttctttagaagaagtttcacgtaagattttcagtgcacattttgggcagctgtctg
taatatttttatggcttagtggaatgtattttcacggagcccgcttctctaattatgttg
cttggttaagtaatccaacaggtattaagccaagtgcgcaggttgtttggcctatagttg
ggcaagaaattttaaatggcgatgtaggtggtggctttcaaggagtacaagttacatctg
gatggttccaactgtggagagcatcaggaattactacagaatttcagctttactgtactg
ctattggcggattagctatggctgctttaatgctgtttgcaggatggtttcattatcata
aagctgctccaaagttagaatggtttcaaaatgttgaatcaatgatgaatcaccatttag
ctgggcttttaggcttaggctgtttaggctggacaggccatcaaatccatttgtctttgc
ctattaataagctactagattctggcgtgtctccgcaagaaattccactacctcatgagt
ttttaattaatagagagcttatggctcagctgtatccaagttttagtaaaggattagttc
cattctttactttaaattgggctgaatattccgactttttaacttttaaaggaggtttaa
accctgttactggaggtttatggctaagtgatactgctcatcatcatctagcattagctg
ttctatttcttgctgcaggtcatatgtatagaaccaattggggtattggacatagcatga
aagaaattctagaagctcacaaaggaccttttaccggcaacggtcacgaaggtctatatg
aaattcttacaacttcttggcatgcacagcttgcaattaatttagctatgatgggatctt
taagcatcattgtagcacatcacatgtatgcaatgcctccttatccatatattgctactg

attacccaactcagttatcgctcttcactcatcatatgtggattggaggattttgtattg
ttggagcaggagcgcatgcttctatatttatggtaagagattataaccctgcagaaaatt
ataacaatcttttagatagagtcattaggcatcgagatgctattgtttctcatctaaatt
gggtatgtatatttcttggattccattcatttggtttatacattcacaatgatactatgc
gggcacttggaagatctcaagatatgttctctgatacagctatacagttacaacctattt
ttgctcaatgggtacaaagtatacacactttagctcctggaaatacagctccaaatgcat
tagcaacagctagttatgcatttggaggagatattgtttctgttggtaacaaagttgcaa
tgatgcctatttctttaggtactgcagatttttagttcaccatatacatgcatttacta
ttcatgtaactgttttaattttagttaaagggttccttttctcaagaaactctagactaa
ttcctgacaaggccaatcttggcttcaggtttccatgtgatggacctggtagaggtggta
cttgccaagtgtctggctgggatcatgtttttcttggcttattctggatgtacaattctc
tgtctgtagcaatttttcactttagttggaaaatgcaatcagatgtttggggtagtgtat
ctccgtctggaaatgtttctcatattactggcggtaattttgcacagagtgcaattacaa
tcaatggatggctgagagatttcctttgggctcaagcatctcaagttattcaatcatacg
gttctgcgctatctgcatatggattaattttcttagcagcacattttgtatgggcattca
gtttgatgttcttatttagtggtagaggttattggcaagagcttatagaatcaatcgtat
gggcgcacaataagataaaagttgctcctgcaattcaaccaagagctttaagtattactc
aaggtagagcagtcggtgttgcacactacttattaggtggaattggtacaacttgggcat
tctttttagcgagaattatttcagtaggctaatagtgaaaataggataaaaaaaacaatt
atggcaacaaaatttcctaagtttagccaagctttatcacaagatcctacaactagaagg
atttggtatggtattgctacggcacatgactttgaaagtcatgatggaatgacagaagaa
aatttatatcaaaagatattcgcttcgcactttggacatctagcaattatcttcttatgg
acatctggtaatttattccatgtagcttggcaaggcaactttgaacagtgggtattaaat
cctttgaaagttaaaccaattgctcatgcaatttgggatccgcattttggacaacctgct
ttgaaagcttttagtaaaggtggctcagcttatccagtaaatatagcatattctggcgta
taccactggtggtatactattggtatgagaagcaatcaagacctgtattctggggcttta
ttcttactagttttatcagctttactcttatttggagggtggctacatctacaaccaaaa
ttcaagcctggtttatcatggtttaaaaataacgaatcaagattaaatcatcatttatct
ggattatttggggttagttctttagcctggacaggtcatttagtacatgttgctatacct
gaggcaagaggacaacatgtaggatgggataattttacaaccgtattacctcatccagcc
ggtttacagccgtttttcagtggtaattggagtgtatatgctcaaaatccagatacagct
caacatttattcggaactaatgaaggtgcgggtacagcaattctgacatttctaggagga
tttcatcctcaaagtcagtctttgtggctaactgatatggctcatcaccatttggctatt
gcagtagtattcattgttgctggacatatgtataggactaattggggaattgggcacaat
ctaaaagatattttagatgctcatagaccacctagtggtagattaggagctggacataaa
gggctatttgatactattactaattctttacacatacagttgggattagcattggcttcc

ctaggtgtaattacttcgttggtagctcaacatatgtatgctatgcctccatatgctttc
atggctaaagatttcacaactcaagcatctttgtacacacatcatcaatatattgctggg
tttctaatggttggagcttttgctcatggggcaatattctttgttcgagactatgaccct
gaacagaataaagataatgttttggctcgtatgctagaacataaagaagctatcatttct
catttaagttgggtaactctattttagggtttcatacattaggcctttatgttcacaat
gatacaatgattgcttttggaactcctgaaaaacaaattctaattgagccggtatttgct
caatggattcaagcctcttcagggaaagcactttatgggtttgatgtgttactatcatcc
tctactaatatcgcaacacaagctggtagcaatatttggctgccaggctggttagaagcg
attaatagcggaaaaaattcattgtttttaacaattggtcctggtgacttcttagttcat
catgcaattgcattgggattacatactactacgttaattttagttaaaggtgctttagat
gcaagaggctctaaacttatgccggacaaaaaagactttggatatagttttccttgcgat
ggacctggcagaggcggcacctgtgatatatctgcatgggatgcgttctatttagctgta
ttttggatgctaaatacaataggttgggtaacattttattggcattggaaacatattaca
atatggcaaggcaatgcaactcaattcaatgagtcttcaacttatctaatgggatggttt
agagattacttatggctaaattcttctccattaattaatggttataatccatatggcatg
aataatttatcagtatggtcatggatgttcctatttggacatttagtatgggcaacagga
tttatgttcttgatctcctggcgtggctattggcaagagttaattgaaactctagcatgg
gcgcatgaacgtacacctttagcaaacttgattcgttggaaagataaacctgttgcatta
tcaattgtacaggcaagattagtaggtttagcacatt-tttctgtaggatatgtattaac
ttacgcagcttttgtattagcttcaacagcaggcaaatttggttagactaaa-----ttc
gaattagctaa-----aaaaaagtcagtcataata-gttttatgactgactttttagat
ttaaaacaaaaacttactaaagtttcacctcaatatctacaccagaaggcaaatttaact
tcattaatgcatctatcgtttgagaagaaggttgatgaatatcaataattcttctgtgag
atcttatttcaaagtgttctcttgagtctttatctacatgtggagaacgtaaaacacaat
aaattcttctttttgtcggtaaaggaattggccctactgcaacggcattagttctagatg
cggtatctagtattttgtcacacgacgtattaagtataatagaattatatgctttcagtt
taattctgatttttgtctgctgagtaattgtcatgttaaaaagatcttattatttaagaa
ttttagagacaacacctgcgcctacggtacggccaccttctctaatagcgaaacgcatac
cttgctcaattgcaattgcattaattaattcagcagtcatcttaattctgtcaccaggca
taaccatttctgcatcagtaccatcatcagcagtaaactgattaatagtaccagttacat
cagttgttctaacataaaactgaggtctatatcctggaaaaaatggagtatgtcttccgc
cctcctctttagttagaatataaacttctgcttcaaattgagtatgaggtgtaattgtac
caggtttagctaataccatacctctttcaatatctttttctgcacacctcttaaaagaa
ttccaatattatcacccgctagaccttcttctaacgtttttgaaacatttctaatccag
taatagtcgttgtacgagtttctcgtaaacctacaatttcaattgtgtcaccaactttaa
taatgcctctttcaattctaccagtggcaacagttccacgtcctgtaatagaaaaaacat

cttctacagccattaagaaagttttatcgacatctctctccggcgttggaatatatgtgt
caactgcttccataagtgaaaaaatcttgtcaacccatttatcttcaccttgcttaatag
ctgggtttttcgtcacagcttctaatgccaataaggcagaacctgcaacaaaaggaatat
catctccaggaaagtcgtattgacttaataattcccttccttctaattctactaattcta
gtagctcttcgtcatctacttgatcttccttatttaggaatactactaatgtaggaacac
ctacttgttttgctaataaaatatgttcacgagtttgtggcattggaccgtctgccgcag
atacaactaaaatagctccatccatctgagcagcacccgtaatcatatttttttacgtagt
cggcatggcctggacaatctacgtgagcataatgacgattatctgtttcgtattcaacat
gagcagtattaatagtaatacctctagctttttcttctggagcggcatcaatttcatcaa
atttttttcgctgcagtagaccctaaagttgataaagtcgcagagatcgctgctgttaaag
ttgtcttaccatgatcaacatgaccaattgtgccaatattgacatgaggtttttttacgtt
caaatttagatcgagccatgctttttgttttccttatagtaaaagttgctttatagttac
ttttaacgataaatcttgtggattgaaagatttattatttaatacagacaaga-----ta
caatataactaatctaatatcgatagtgagcaaacgctttatttgcttctgccattctat
gtgtatcctctctctttctaatagaatttcctgtttcattagctgcatccataatttcat
tagctaatttcatagacatactttttaccagatctgtctctagagaatttagtaatccatc
ttagtgctaaattcgtacctctataagctcgtacttcaataggaacttggtaagtagaac
caccaactcttcttgcttttacttctactagaggagtaatatttcggatggctttttcta
aaatatttaaaggatctgattctgttctctctttaacgatatctaaagcctgatatataa
ttctttgagataaagttttttttaccacttttcaaaatacgaacagttaacatacttacga
gtctgcttttatataaaggatcaggtgatgcaaacctttttttagctgtattacgacgag
acatagtatttaaaattggttattgtaataa-tgttc-tattagcaggtatttaagattt
aggctttttttgtcccatatttagatcggcttttacgacgatcttttactcctgcagcatc
taaagtaccacgtactacatgatatcggactccaggcaaatctttaattcggccgcccct
aatcagtaccacagaatgctcttgaatattatgacctacgccaggaatataagctgtaac
ttcaaatccagatgttaatctaacacgcgctactttttcttaaagcagagttcggtttttt
aggagttgtagtatatactcttgtacaaacacctcttctttgagggcaactttgaagagc
tggagattttgtttttttatgtatttttcgtctttcggatctaacaagttgttgaattgt
tggcataataaaattgatttaaagtctttaatagatactatttgaacttgctttacatga
attggctagtttttatttatcttatacttacgcataaatctttctactctaccttctgta
tcaataattctctgtgaaccagtaaagaatggatggtttcctgaccagatatctacgtgt
agttctggctttgtggacccaatcgtcataattaactggccgtcacagtaaacttttgct
tctggataccaatttggatgtatattatcttttgccattgtttttataattatataaatg
tatcaataataactattaacgcttagaaaattgaggagcctttctggctttctttaaacc
atatttttttctctcttttactcttggatctcttgttaaatagccttcagacttaagtgt
cgttctattttctggattaattgagcacaatgctcttgcaacacctaaacgaattgcatc

agcttgccctgtcaaacctcctcctctagcatttacatggatatcatactggtttagtag
tcctaaaacttgtaatggtgcatatgaaactcttaagtaattaggactaaattgaagata
agactctcctggtataccgtttataattaaattacctgaccctgggactagtcttacttg
tgcaacagaacacttccggcgacctgttccagaatagattgcgcgagttttaattaattc
tgtcgacataacattccttgataactaaataaatatacattctttatactatatactctt
gtggcttttgcgcgacatgtgggtgaattgggccagaatacactttaagcttcgtaaata
gttttctgcctaatggacctttaggaagcatacctttaaccgatttttcaataattctgt
taggtaatcttgtctgaagctgatcaaatgtttcaacttttaatccgccaggctgtccag
aatgtcttctgtatagtttttgattcgttttatttccacttacagatacgtgagcagaat
taataacgattacataatctccagtatctagatagggtgtataagaaggcttgtttttac
ctcttaagatattagaaatatgagtagatattctaccaagtgtctgatttttagcgtcta
taacataccaatgagaattagtatttaatgagggtgattgcgttttattcatgaaaagat
taatacttttggttgacaataagagaatatgtaaatggtttacattcaatttctatacaa
gtagtattt-aaaaatatta-------------ttaacaaagagaacttttctaacagtt
atttaaattgtttgtcagcaagaaaatttctagttatcaataacttatactatttttaat
aatataaaatatattaatcactttttcttttggcaaacttatacctagtttttttgc-
aaggcatatataacttcttcagctgatttctggccaaaattttaatttctagcagttct
tcttgagagtaatctagtaaatctgcaatagaatggatttgagctcgctttaagcagtta
taagccctaacagataactgcagttcttctattaaaacttgactaattttttatcttct
ttgctacgataattatctgctgatttaaagtctaaatttcttagtgaacaaaaaagatta
gttaaaactgtagctccttgacttattgcttcttgaggagagatgctcccatttgtccag
atttgtataatcaatctatcttttatgctgttgctaccaatacggacttcttctacttta
taattaactttattaactggcataaaaacagagtctacttgcaaaaaatccacagataat
tcatctacagctttttcagctaagcgatacccgcaatttttttcaattttaaactccatt
tcaaatattgtattattgcaaatagttgcaatatactgtctaggatctactacctctata
tcagaagataattcaaataggccggctgtaactatagctggcccttgaactctaattcga
ccaatttgagattctttgttataacttttaaatactacttcttttagattaagtaatatt
tctaacacatcttctcttaccccaggaattgtagaaaactcatggttcactccagcaatc
cgtacagcaactatagcagtaccttcaagatctgacaatattgatcttctcaaagcatta
cctaatgtaataccttgtccttgatttaatggttctattacaaaactaccgtactgccca
cgcgccccatctgttcttgactctatgcattcaatttgaaattgagccacctaagaaagc
tcc-tttaaaataatcagtgattagtaagtctctctaaaacaagaaggaatttatactcg
gcgtttcttaggagggcgacatccattatgaggtacaggagtaatatcttttattagagt
aatctctaatcctgcagcctgcaaagctctaattgctgtttctcgacccgctcctggacc
atttactagaacttcagtttggcgcataccttgatccatagcttgtctagctgctttttc
agctgctgtttgagctgcaaaaggtgttcctttttttagctcccttaaatccacttgcacc

agaggatgaccatgataatgtttctccttttaaattagtaatagtaacaattgtattatt
gaatgtagatttaatatgtgtaataccgttaactgcattacgtttagtttttcttgctcc
ggatttttttatttgtctagccatcgtgtctcgtctattatatagttaattaaaagatta
ttttcttggagctttttcttacctgctactgtttttttacctcctctgcgtgttctagc
attagttctagttctttgtcctctcaaaggaagaccaagacggtgtcttctacctctata
agtactaatttccataagtctcttaatgctcatagactcaaaacgtttgagatccccttc
aatctgataattagactcaagaatttctcttatactgacaacttgttgatcatttaaatc
ttgacacttaatatcagcgtctatgtttgtttttctaatatttctttcgagcgagatag
tccaataccataaatatatgttaaagctatctctattcttttgtttcttggaagatctac
tccagcaattctggccactttgtcttgtctccaataaataatatgtat-ttttataacgg
ttgtttata-agcttagccttgtctctgcttatgtttaggattagtgcaaattaccatca
cctttctgtgacgccgaattattctacatttttcacacatttttcgaacagaaggacgaa
ctttcatattaa----ctctatatactaggaatattactatattaactctattaattatc
gacattaaaaatgtcaacaatatattgtactactttaactgttgtagatttacttcgtc
atactatcatattttttgatatgacgtatgtctgaatttgcttagctgtgtctattgcg
actccaactaaaataagtaaagacgtagccccgagacctcttaagttttggatctgagta
accttttctattataaacggaatcagcgctactgtaaataaaaacgaggctcctaggaat
gtcagtctatttaatatgacttgtaaataatcaatagtcgcttgacctggacgaatatta
ggaatgcttgcacccatttttttaaatttatagcaatatcttctggattcattactatt
gatgtataaaaatagctaaagaaaaggatcaaagcacaataaagaagaaggtatagtgaa
ccattaggacaaaacaggtataggatttggagtaatgttttattttgaataatttgggtt
agataagatgggagggccatagatgcagatgcaaaaacaataggcataactccaccttga
tttaatttcaagggcaagtaactatttggatccaaaattgaagattttcctagctgtctt
gctgaaataattttaattcttcttgttccttcttgtacacaaattgtaattattatcatt
aataaaaagatcgctataaataatccgaacttaagacttgcattactataactagcatca
aaaaatgactgtgtaaaattctttggtagtcctgacacaatgttttgaaaaataagtaaa
gaggctccattaccaattcccttttccgtaattaactctgataaccacataataatcata
gaacctgctgttaaagctaaaacagactcgcagacaaatgcaaaattccaattaaaaaca
tatggttttacccatatagagattgctccagattgtaaagtcgcccaacctagagctaag
taccttgtaatttgagttatttttggcggcccaattcaccttcttccttctgtaatttt
tccagattggggacaattttcgtaagtagctgcattacaattgaagagttaatataagga
acaatacctaacgcaaaaatccctattgttgaaaaacctcctccagaaaaaatattcaga
aaatttactaaagtattttttctacacttgcataaaaggcgtcatgatctatacctgga
acaggtataaatattcctaaacgtgctaaaactaatagaaagagagtaaagataatacga
tttcttagatcactttttggctcataaataaattttaaatgaaaagagatacttt--tt
tatgtaaaaaccctattccaagacttgtataagtttaaaatagggaagtaa---aatatt

atagtaatattaata-taatattctatgtagaatacaggctttctattggaactcctcga
tctttagctgcttcactaaaagttcttaattgagttaagccattgagaacagctcgcgcg
ttatttaatgtattattagaacctaattgcttagctaatatattctgtacacccgacaat
tcgaggactgtccttacagaacctccggcaattacacctgagccaggtgcagagggcctt
aatataactttcgcggcaccagaaattccgttaataggatgaggtatagaattcgatttt
gtcagtggaactgtaactagatgtttttagcatctgttactcctttttttactgcacca
attacatcgcttgcttttcccacgccgacaccaacttggccttgctcattaccaataaca
agaataactcgaaagcttagtttttacctccttttacaactttagtaactcttttaact
tgtacgactctttcttcccagccactatctttatcttttcctttgctctgtttttacga
ttggccattttaagaattattcctaattagttaatta-cgattctagaaacccatgcctg
cttccttagcagcttcagccaaggccttaactcttccatgatataactttcctcctctat
cgaagacgacattttaatgccttctttcatagactgttctgctaactgtttaccaacta
cgcgagaagtatcacagtttggtcttatgttatcagattctttattatttaaattaacag
atgatgtagctactaatgtaataccttgtgtgtcatcaattatttgtgcgtatatatgtt
tattagatctaaatacacacagacgaggcctacttgaggttccttgaactttttccgaa
ctcttttatgcttatgaattctagtttgtttagtgtttagtttcattattatttaccttt
tccagctttcccagcttttcttctaacaaattcaccttgatatctaattccttttccttt
ataaggctcaggaggcctaatagaacgaatagttgaagcaacctgaccgacaacttcttt
atctatgccagagacagtaatatttgtgttgttttcaactttaatttcaatattggcagg
aggtttaattttcactacatgactatagccaacacttaagattaaatcttggttatcaat
ttgagaacgatagcctacaccttgtatttgcagtttttaaaaaatccattagaaacacc
ttcaatcatattactaataagagttctggatagcccatgtaactgactagccattttgt
ttgtccactcgtcttgacagcaattgtgtcgttaagtatttctaaattaataccagctgg
taaagttctggatagggtacctttaggtcctgtcacggtgatagtctgaccatcgaactg
agtactaagatttgtcggcaataaaattatttttttccaatacgagacat-attccacc
tataaaatagttaccaaatatagcataatatttcaccaccaagaccatcatgacgagctt
gtctatctgtcataacgcctctagaagtagaaataagagcgatgcctaaacctccaagaa
ctctgggcaactctttatgatttgcgtagactcttagtccgggtttactaattcttttca
gagcagtaataactggttgacgatttttaccattatacttaagagaaatcattaaatgag
tctctataccttcacccatttgttcgaaattttgaacaaatccttcttctttcagtactg
ttgccatattgcatgtcattttcgttgctggaacttgcacaatttgatgtcttgctaagt
ttgcgttacgaatacgtgtcagcatatcggcgatcgtatcgttgaccaccttgatcctcc
ttgcga-----tgaattatttgaaatattttcaagattccctgaaaggcatccctagctt
ttttaacaaagctagaccttcttgatctgttttagctgtagtgacaattgatatatctaa
accacgaatttgatcaatattatcatagtctatttctggaaagatcaactgttcacgtaa
acctaaattgtaattacctttgccgtcaaaacttctaggactaattcctctgaagtctct

aattcttggtaatgttaaattaattaatttctctaaaaaagaatacatcttgtcttttct
tagatgcacaacaattccgataggaacctcttctcgaattttaaatcctgcaatagattt
tttagcttttgtgacaattggtttttgtccagttattaatgtcaattcttgaatactact
ttcaagagccttagcattctgagaagcttcacctaagccacggttaatagtaattttagt
aaacctaggaacttcatgtacatttttgtactgaaattcatcttttaaagattgagtaac
agttgttttatatttttcttttaatcctattgccattatatttcatcaattatttaata
agttcgccagtctttttcagttttcgaa-ttttttggcctttatcattaattattactga
ggatcgactagcaatattatttgttcactaaataacataacgttagaggtatgtattgg
agcttcaaatttgataatttctcccgtttctccttcttgttgaggtttttttatgcttcac
tttgagattaattcctttaacaatcaccttatttgttttataaataattgcaataatttc
acctgtttttgttttatcacttccagaaataacttgaactaagtcccctttttttaattt
aatttttgtattattttttggttgttttagaaggacctttcattatactacctccggtgca
agagaaactatttagaaaaattttatctcttaactctcgggctataggtccgaagact
cttgtgccacgtggattattatcttgattaatgataactgccgcattatctccaaatcta
atgctcataccgtcagttctccgtaaagctttacgagttcttactacaacggctctgaca
acatcagatcgtttaactggcatattgggagacgcatctttgactactccaataataaca
tctccaatagatgcataagaaggattactagtgcctagtactctaatacacattattttt
ctagcaccactattatctgcaacattaagatagctttgagtctgtatcatacttttattc
tgtataagattaattatcaaaagacttagataagatattaaccattgtccaacactttgt
gcgacttaaaggtcgtgtctcttgtattgtaacaatatcgccgattgtgcattcattatt
ttcatcatgcgctttatatttctttgtccggatcatagtttttgcgtactttctatgaga
gattctattttctacagctactactatagttttattcatttttatcgcttactactttacc
tgttgtttcttttaaaggcatagtctttatattttttatatat---------aattaaat
tttaagattattacatggttgacttagtacgagacttttcaacagttaaaagttgggcta
atctatgcttagaatgtttaaatagatgaggctggaaatcttgccttgtggctcttttta
gccttaaatcaaaaagctctcttttttattacaaggatttcttcagctaaagaagaagagt
ccaggtttgtaacatctgatattttagggaaagtcatattttaggattctgttgtattcc
gaactataaatttagtcttgattggtaatttataagaagctaatttcatagcttgttgag
cagtttttttgtggtacacctgtaatttcaaatagaatatgtccaggcttaataactgcaa
cccaatattctggagctcctttacctgaccccatgcgagtttcagctggacgggcagtga
ctggtttatctggaaatactctaatccatagtttaccacctcttctcacatatctagtaa
tagttcttcgagtagcttctatttgtctagaagttaaccaaactggctctgtcgcttgta
atgcataatcaccgaacgcaattgtgttacctttgctagcagaacctttcattctacctc
tatgttgttttctaaattttgttttcttggggcttagcataaaaattaaaaaataataat
aattaagaagctgttgaatcaggagtttcctgattataagcagattctagacttttcgat
tctggcaaaatttctcctttaaaaagccaaacttttactccgagcacaccatacgtagtg

tgagcttggcgatgacaataatcaatatcggctcttaaagtttgtaaaggtacacgacct
tctctaacccattcacttctagctatctctgcaccattaagccgaccagatacttgaatt
ttaacaccctgtgtatttgctctttgagctctttgaactgcttgtctgacagctctacga
aaggccactctttttctagctgttgagtaataaattctgctactaaggttgcttcagaa
tctggatctgcaatctctacaacatttactctgagttgtttgctagggtctaggattaag
gacagcgattttcttaaagactcaattcccgctccagattttccaagtacgattcctggt
cgagctgttgctatgagaatttctacttgatcaactttacgattaatttcaattttagca
atactagcattactaagttttgaatgtataaatgagcgaattttatgatcttcttgtaaa
agaactgggtagtctttagagttagcaaaccatgaagaacgatgtttttgggtaatgcct
atgcgaaagcccaaaggatgaatttttttgacccacagtatccttgattataagttaaaag
ctataatgactaattgacgttaaa--------agactacagttctgaaacacctaatgta
atatgacaagttggtttatgtattggaaaggctcttccttgtgctctaggctgaaatctt
tttaaagtcgggcctttgtctgcaaaggctttactaacaaacaattggtttttatttaat
ccatcattatgctcagcattcgctgcagcagattccaggatttgctttatatgagaacag
actcggtatggcataaattccagaataattaatgcttcttggtattttctacctctaatt
tggtctaatacacgacggactttatgcggagaaagacgaatgtatttccctactgcttta
gtttcttttacgtttttttgtaatactcatagttatagtttaacgacgggcttttctgtca
ccttttacgtgagtgcggaaagttcttgttggaacaaactctcctaatttatgtcctacc
atttggtctgacacaaagacaggaaaatgttgtttaccattataaacagctattgtatgc
cctaccatatcgggaataatagtagatgctctcgaccaagtttttaggacttctttttttt
cctgaaatatttaatgcttctatacgtttaagaagactaacatcaataaaagggccttta
tgtatagatcttgacataataataaatcattagattcttaataaaataaattctatttac
ggcgacgtaagacgtacgggttactatatttatttggatttcgtgttttaacacccaatg
caggtttaccccaaggtgttacaggacgagctcgtccaattggagatttgccttcaccac
caccatgtggatggtctacaggattcatgacaacacctctaacagtaggtcttttacctta
accaacgatttctaccagctttaccaagagtaatattactagcatcaatattaccaactt
gtccaatagtagcgtagcattcttttcgaatcattcgaacttcgcttgaaggcaatttta
cagtaacaaaagtaccttcttttgctactatttgagcataagttcctgctgcacgaacaa
tctgtcctccgcacgaaggtcttaattctatattatgtactgctgttcctaaaggaatac
tagataaaggcaaagcatttccaacttcgataggggcagtaggaccagaaagaaccatag
atcctacactgagagatcgagggtgtagaatatatcttttctcaccatcaagatagtgta
ataatgcaattcttgcatttctattcggatcatattcaatagaagcaactttagcaacta
tattatgtctatttcttttaaagtcaattaatctgtactgctgcttatgtccaccacctt
tatgacgacaggtaatgacacctctattatttcgacctttgcagaaatggtgtttaacta
ttaatgatttttctggcttatcagtagtaatctctgaaaaggtagaaaccgttctatttc
ttgtccctggtgtgtaggcacgatataaacgaattgccatataagaagatttgagaagag

taaatgttgaa------ctatgtgatatatttaagtttctggaaacaaattaatagaatc
ttctgaggcaagtgttacaattgctttcttgtaatgtggtcgcttccctacgaatctacc
gatgctcctttttttcttaggaggatgacaagtgttaacacctgtaacttgcacgttgaa
aatatactgtatagcagccttaatattaattttgttgctttgggatcaacggcaaaaca
atactgattctcttctaataatttagttgtcttatcagtaatgatgggatatttaaccag
atctaataaacctcttgaatcaatgctatccattatatacctcttgtattttagataaag
catcaactgtaataataattttatgtgcagctaacaaagccataatatttagtgtatcag
ctgaaataagttctacgttgtgtagattgcgaatagaaagataaacattcggatcttttt
tgtctactatgactaaaactttttatttaaatcaagattccatcgatgtatagcttcca
taaatagtttagtcttgggctgatgaaaataactgttaaaattttctacaatcagtgtat
taactgatttgttatttaatgctgtccttaatgctaattgtctttctttttttattcattt
ttttggtaaaactacgaggttttggcccaaatattacaccaccacctctccatagaggtg
aacggatagaacctgctcttgctcgaccagtccctttttgacgccatggtttacgacctc
cacctcgaacttcacttcttgttttagtattagcagaaccctggcgcttttcattacttt
gctttactaaagctctatggactaagtacatcccagaatcttgactgaccttaagattta
aatcagcgttgccacttacttgaccttcccaattatagacttgataatttaattgcgtgt
taactgtcatgaaatatatcgtatgaaagaatattgctacttactaattttaccaaagc
gccaggtttaccaggaacagctcccttaacgatcaacaaatcattctctgagttaatact
tacaatttgcaaatttttattgtaactttttattaccaagttggccagccatgttttt
acctggataaactcttccaggcgttgtaccggctccaatagagccaggttgcctatgatt
tttcgaaccatgtgacattgggcctctactaaaatgatgcctcttttggtaaccggagaa
acctttacctacacttctagaagagacattaatcttttggcctacttgaaatagatctgt
agatagtatttggctaacttcgaaatcatcagtagatttcatttcgtattcacgtaaata
ttttagtgggggagcctgtgatttttttaagtgtcccaacaatggtttatttaattttttg
ttcagcaacttgcttgtaaccgacttgaatagcattgtagccatcagtagatacagcttt
aatttgagtaataacacatggcccaacttgaattacagtaactggaattgataaaccagc
ttcatcaaagaattgggtcatacctactttagtaccgagtataccaacagacactagatt
tctcctttttgagagatttatactatcattgtacataaaaattctacttcatagttactg
aag--taaattaaactaacagatgcatgatattattcatgacaacataaaggttaaaaac
tttacctatttaatctaaaatagaaacagctttttgtctagttacagcgtatataaatta
gaagtatgcttattaaatttttttataaactact-tagatttaactgatattgatatccgt
aacttacttgcagtaatatc----aatcaacatgtataagtttaattgcctaaactagaa
ccagctattattagaaataattcaagtatagataaaagcgtatttgaacttttatatatt
gacaaatataatggaatatattgaggttcggtaactacaccttttttattttttacatttat
tgtgca--gtatatatacatctaatgcagtaaacataagaatataac----tgtatagca
aatgtaattacacaaattagagaggtattaatgggcaaagtcgttggaattgatcttgga

acaacgaattctgtaattgctgttatggaaggaggtaaacctaccgtcataccgaatgca
gaaggttttagaactacagcttctgttgttgcatatactaaaagtggagataaacttgta
ggacaaattgccaggaggcaagctgttattaacccagaaaacactttctactctgtcaaa
agattcataggacgtaaacagaatgaaatttcgcaagagattcggcaaacatcatataat
gttaaaactagtggatcaagcataaaaattgcttgccctgcactgaataaagattttgct
ccagaagaaatttcagctcaagtactgagaaaacttgttgaagatgctagtacgtactta
ggtgagactgttacacaagcagtaataactgtacctgcttactttaacgattcacaaaga
caagctactaaagacgcaggtaaaatagcaggcttagatgtattgagaattattaatgag
cctaccgcagcttctttgtcttatggactagacaagcaaaataatgaaacaatactagta
tttgaccttggtgggggcacatttgatgtatctgtattagaagttggagatggagtattt
gaagtactctcaacttctggagatacacatttaggcggagatgactttgatcagcagatt
gtagaatggctaatcaaagattttaaacagaatgaaggtattgatcttggtaaagataga
caagcacttcagagattgaccgaggctgcagaaaaagcaaagatagagctgtcaaattta
actcaaacagagatcaatcttccttttattactgccacacaagatggtccaaaacattta
gaaaaaactgtaactagagggaagtttgaagaactttgttcaaatttaatagataaatgt
agtatccctgtaaataatgctctgaaagatgcaaaactagaagcttccagtattgatgaa
gttgttttagttggtggatctacaaggattccagccatacagcaaatggttaaaagatta
attggtaaagatccaaaccaaagtgtcaatccagatgaagttgttgctattggtgcagcc
gtacaagctggagttttagcaggcgaagtcaaagatattctattactagatgtgacgccg
ttatctttaggagtggaaactctgggtggcgtgatgacaaagattataccaagaaatact
actattcctacaaaaaaatcagaagtattttctacagctgtagataatcaaccaaatgtg
gaaattcaagtacttcaaggcgaaagagaactaacaaaagacaataaaagcttagggacg
ttccgattagatggcattatgcctgcacctagaggagttcctcaaattgaagttaccttt
gatattgacgctaacggaattttatctgtaaaagccaaagaaaaggctactggtaaagag
caatcaattactatatctggagcatcaactttacctaaagatgatgttgaaagaatggta
aaagaagctgaagaaaatttcgatacagaccaaaaaagaagaaagaatattgacacaaag
aatcaagcagagtccctatgttaccaagctgaaaagcaagttaaagagtttgaagacaaa
attagccaagatttaaaaataaaaatagaagagctaattacagagcttagatctagtcta
gagaaagaagaatatgacaatattgaatctatttctcagcaattacaaaatgctctgatg
gacattggaaaaaatgctgctcagactgaaagtaaagatacaaaagcgaaggatgacgac
actgtgattgacactgatttctctgaagctaagt-aaaagta-agcgggtaacgcgattc
gaacgcgcgacatcaaccttggcaaggttgcgctctaccactgagctatacccgcataga
ttgtattattacaaaaattaagatatttgtcaatctgtta------atctatac---atc
tagactattaattttggaattacatattagctaactgttgtaaatataacaattgctata
attttttaa-------aagtacttatatcat-tatataagtacttttttatatt-------
-caaaaatacaaataagtattatag-------ctaaactactaaggaattaactaagcta

gtaacaattactaatcctgtccataagccggctcctgtatatattaatccttttgattgc
tcccattggccaggggatgctaaaacaacaggtacgcctactactaacacagtcgataat
gcaattaataatagcactaataattggatcgcaataatcatttaatatgttctcctgaaa
aattcaaatgttaataatactgtatataatacaatgaagcttttgaccttcagcaattta
tctatgaagaaataatttaaaattattgttttctaaataaaccttaaattaagtattaaa
atctttattgtaaacaataatactatatttttttta--gtagataaaaaatttaaagaaag
tcttttttaaaaaaaaattaagataaggtgatatgctgtataaaaagctaagtattatta
attattagagaaatttaaa-ataagaggtaaaacatatgaactcagccctttttttagca
aaactaccagaagcttatgccatttttaaaccgatcattgatatcttacccgtaattcct
gtattttttcttcttttagcttttgtttggcaagcagcaattggttttagataacattta
ggttataaaaagaataacaaatatacagagtgccaatttttgagctctaatcttatatct
tacacaataataactattatggttgaacccttactatcaggaattgttcttggacttatt
cctattactttattcggattattagtagcagcttatttacaatatcaacggggcaaccaa
ttaggactgtaaagcctaaattatagatatttataaaaactacaataattatagttttta
taaatatctacaaataagctaattttaaatacagaaataatactattgaaactagagtga
taacaaaaaaaatttatgtaa---taa---------------------------------
------------------------------------------------------------
------------------------------------------------------------
------------------------------------------------------------
------------------------------------------------------------
------------------------------------------------------------
------------------------------------------------------------
------------------------------------------------------------
------------------------------------------------------------
------------------------------------------------------------
------------------------------------------------------------
------------------------------------------------------------
------------------------------------------------------------
------------------------------------------------------------
------------------------------------------------------------
------------------------------------------------------------
------------------------------------------------------------
------------------------------------------------------------
------------------------------------------------------------
---------------------------caagctaattgtttaccagcttgacttagtaac
tcctggtaaaagacattcatgagacatttctcgaaaaacatggcgagataatccaaaatc

tcgataataacctctacttctacctgttaaccaacatctatttctccctctaacagcggc
gctgtttctaggcatttcttggagtttttgtcttaaatccattttgtcagaaaagctgcc
agcttctttcagttgatctttaatagccagtctttttaaataatatttttttgcaagttt
cgatctcttaatctctctttggatcatattctttttagccatgctgtaa-aactctcttt
taaaattatttattcaatatattagctaaaaaagctcctgcctgcaacttttatatattt
aaaaataaatagcataaagtagtgaaatgcaattgatatttaatcaacgatacc---act
ttatgctattatcaaaaatttatttagtaaagttcttgttcaacatgagttgaaatagtg
caatcagatgaaggatatgcaatacaggttaaaacatagccagctaaaagttgatcatca
tctaagaatgattgatcagcttgatctacagttccttctgtaactttacctgcacaagtt
gaacaagctccagctctacaagagtaaggaagttcaattccttcttcttctgctgcatct
agaatatatgtatcctccgcacaattaaatgtgacatcaattccttcttcttcacatagt
aaatgaattttataatcagccatttcttcttactccttaataaagatacacttgaaactt
aataaagcaaaat-aatttaaagaacttaccaaaaaattatatactagtttcattatgca
tgatagactggaaaaaaagaaaatagtatactattctgaaaaatcaagctacatttccta
ctaaatagaactatagtaggtccaaagtatatacataaattcatatagttaaaattactt
ttatcatttaaatagctgaacttgatagta---------tagattgatcatc------aa
gctagtcttctgc---caataaataattaattatgatttcaatgcttaatcaacaagtag
aacttaagcctattcttaaatttcagataacacaaatatattcgcattctcttattcagg
aaattaaagccttaataactagacttgttttgcaagtttggagaaggccggctacattga
tggcaggtattatccaacctttattatggttaattctatttggtgggcttttctacaatg
ctcctataaatttgttcactattaatacaagctataattgtttttttgagctctgggatta
taatttttacctcttttactggagctttgaattcaggtcttccattaatgtttgatagag
aatttggatttttaaatagattattaacggctcctttagtatcgaggacttctatcattt
tatcttctgctacttttatgacttgtattagtttaatacaagttgtatttatagttacag
cttctcttttttatgggaaactcacctctaaatagcgatagtactatgatttttggactta
tgattctattagtgactgtaggagttacaatgcttagtttagctttatctttcactctgc
caggtcatattgagctattagcatttattttagtagttaacttgcccttttttattttcta
gtacggctttagctcctttatatttcatgccgccatggcttcagttaattgcaagtctca
atccattgagttatgcaatagaaggtacaagatacttatattcaagcgtaaactggaatt
ttacagagtgtgtgattaagattagctggggagatatttgtttagggcaaattattattt
tattaatcgctttagatataatggcagcttatcttgtgtctaatatattaaaagctaaac
ttaatta------------------aaatttaataaaaaatttattaaatttattttaca
aagatttttactatataatactaatagtagtatggataattgaaatagaaatagtttttc
aaaaccaaagctattatcaacaatgataaacatttgtaagaaagtcaacaaagtatgttc
ttattcataggaggcatgtagtcaatgggactaccatggtaccgtgtacacacggttgtt
ttaaatgatcctggacggttaattgcagtccacctaatgcatactgcacttgtagcgggt

tgggcaggatctatggcattatacgaactagctgtatttgatccttcagatccagtatta
aatccaatgtggcgacaaggcatgtttgttatgccatttatggctagactgggcgtaaca
gattcatggggaggatggagtataacaggagagagcgtatccaatcctggactgtggagt
tttgaaggtgtagccttaactcatatagtcctttctggcatgctttttctagctgctatt
tggcattgggtttattgggatttggaattatttagagatccacgaactggcgagccagcc
ttagatttaccgaaaatttttggaattcatttattgctatcaagtctactttgtttcgga
tttggagctttccatgtaactggacttttggaccaggaatgtgggtatcagatgggtac
ggagtaaccggaaaggtattaccagtagctccagcatggggaccagaaggattcaacccg
tttaatcctggaggggttgcatctcaccacattgccgcaggtactgtaggtatattagct
ggtgttttccatttaactgttagaccaccacaaagactgtatcgtgctctaagaatgggt
aatattgaaactgtattatcaagtagtatctctgctgtttttttctcagcttttgtgact
tgtggtacgatgtggtacggttctgcaactacgcctattgaattatttggtccaactaga
tatcagtgggatagtggatattttcagcaagaaattgagaaacgagtagaaaatgctatt
gctgatggtgctgcacctagcgaagcatggtcaagaattcctgacaagttggcattctat
gactatattggtaataatccagcaaaaggaggattattcagagcgggccctatgaataaa
ggcgacggagttgctgaagcatggcttggacatcctgtattccaagataaagaaggaaga
gagcttagtgttcgcagaatgcctgctttttcgaaacttttcctgtaattttagttgat
aaagatggtattatacgagctgacattccatttagaagagcagagtctaagtatagtatt
gaacaagtaggtgtaacagctagtttttatggtggcaaattaaatggccaagttttcaac
gatgctcctagcgttaaaaaatatgcaaggaaagctcaattaggtgaggtatttgaattt
gatcgaactacattagaatcagatggagtgtttagaagtagtcctagaggctggtttaca
ttcggtcatgctaatttcgctttaatttcttctttggacatctttggcatggttcaaga
actatcttccgggatgtgtttgcgggaatcggagctgaagttactgaacaagttgaattt
ggagctttccaaaaattaggagatagaagtagtaaaaaacaaggagctgtataagaccaa
agtcaaagatagtctttatc---aaatatttctaatattaaaatagttaagacaaagttt
attatctaatagttactaatcaggagaatttatggaagccttagtctatgtatttttact
aacaggaacgctaatggttatattctttgcgatctttttagagaacctccaagaatagc
aaagtaacttgattccgtttaaacttataataatagaaccactttccaacttattagtgg
aaagtggttttattgtagattacattgtattattcttcatgctcttcaaaaggatctcgg
agatcttttgaagcaggaccaaatgctgtgtatatcgagtaacctgtaatacctaaaagc
agactcgaaataaaaatactaagaactgttgcagtttccataatttaagcagtattaaga
tgaactttttctataatgatattatcaatattaaaacatttaattattaaaattatcaaa
atatactatggcacttagaactagacttggagaaattttaagacccttaaattcagagta
tggaaaagttgctccaggttggggcacaactcctattatgggaattttcatgttactatt
tttcttgtttttattaatcattttacaaatctataattcttcattagtattagaaaatgt
agatgtagattgggctactttaggtag----ctaactagataagttctttga--------

ttacacaatattaaagacaac----------------aagcgatcgctattagattatag
tagcgatcgctttaaaagtattttactatatatccagagaaaattatttacctaaattaa
ttaactcggtatctgcaaaactatttgtgttaacattattatagttaactttttcaaacc
ttactaatactggatatttgataccactttatcaattgctgcaacagtaccaatttcct
gataccagtacgattcttttctcaatatttttacttttgagcctctttccatatttttag
tcctcagatagattattaaagattattatattaaaaaattaaaaaaaactaaacaaaata
tataaaaatactaccgacaattgtaatatataactaataaaaggagagagagggattcga
accctcgataacaaaagttatgacaattttcgaaattgttgcaataaaccactctgccat
ctctcctgaattaaaatactgttttaaaaaattagacaagaataacatagaactaaggta
gttgcctagaaaattaaaactatgttaaatataaattataatctttctatagatatattt
aaaaaatatgtacatagaatctatatcagcttacttcaagatttttaatcttatacaat
ttttactaaataaataaattctctcataatatgaaactatcttggaaaacacttcttctt
tggtctttaccaatatttgtaattggtttctttttctggcaaggtttttaggtccaact
actacagatgttggcagtaatatcgcaagttctagaatgacatatggacgatttttagaa
tatttagatatgggttgggtgaaacgggttgacctctatgaaaataatcatacagcaatt
gtagaagctgttgggccagaattaggaaacagggttcaacgaattcgagttgaactgcca
gcaagtgcgccagaattaattacaaaattacgcaaagccaatgttgatctagatgctcac
ccccctaaaagtacaagtgcagtatggggactattaggcaatttactatttcctttacta
ttagttggcgggttagctttcttatttagaagatctaataatgctagtggtggacctggt
caagcaatgtcatttggcaaatcgaaagctttatttcaaatggaggctaaaacgggagta
gtatttaatgatgtagctggagttgaagaggcaaaggaagaatttcaagaagtggtaaca
tttttaaaacagcctgaatcatttactgctgttggtgcaaaaataccaaaaggcgtttta
ttagttggacctcctggcacaggcaaaacattactagcaaaagctattgctggcgaggct
agtgttccttttttagtatctcaggctcagaatttgtagaaatgtttgttggtgttggc
gcttctcgtgtgagagacctattcaagaaagcaaaagacaatgcgccttgtatcgttttt
attgatgaaattgatgctgttggtagacaacgaggaacaggtgttggaggtggtaatgat
gaaagggaacaaacattaaatcaactattgactgaaatggatggctttgaaggaaatact
ggtgttattgtaattgccgctactaacagagctgatattttagattctgcattattaaga
cctggaagatttgatagacaagtttctgtagatgtaccagattttaaaggcaggttagca
attcttgaagttcatgctaaaaataagaaaatggaacctaaagtatctttagaaaccatt
gccagaagaactcccggcttttcaggagctgatttagctaacttactaaatgaggctgct
atcttaacggctcgacgaagaaaaaaatgcaatgactatgtctgaaattgatacatcaatt
gatcgagtagtagctgggatggaaggcactcctttaattgacagtaaaagcaaaagatta
attgcgtatcacgaagtgggtcacgcaataataggcagtttattagagcatcatgatcct
gtgcaaaaagttacattaataccaagagggcaagcaagaggcttaacttggtttactcct
agtgatgatcaaagtctaatatcaagatctcaaatactagcccgtatcgtaggtgctctt

ggtggcagagctgcagaagaaatcattttcggtgacgcagaagttactactggtgcaagt
aatgatttacagcaagtaacatcaatggcaagacaaatggtcactcgatttggaatgtct
aaaattggacctttatctcttgaaagccaaggaggagacccatttttaggtagaggcatg
ggaggaggctcagaatattcagatgaagttgcaactaatattgataagcaagtaagggaa
attgtcagtgaatgctatgcacaagctaaacacattattatagataatcgagtagtgata
gatagattagttgatttactaattgaaaaagaaacaattgaaggcaatgaatttagagac
atcgttaaggaatacactgcaattcctgaaaaaaattactacatatcacaattttaa---
---------------attaaacgggactgacgggattcgaacccgcaacttccgccgtga
cagggcggtgctctaaccagttgaactacagtccc-aaaaagatgaatacccagagataa
tctcatagttagcttaaacttgtcaaattagaatattaaatatgaagagta---------
--------tttgtaataatactcttcatatttgata-tagtagctaaaatatttctttta
gaaggagagagagggattcgaaccctcggtacgaagttaatcgtacagcagattagcaat
ctgccgctttcgaccactcagccacctctccatga---tatatattatatatatatcaaa
tatacttatattttaataggtgtgcataaatatttatggagctaagcggattcgaaccgc
tgaccctctcaatgccattgagatgctctaccaactgagctataaccc----ctttggta
agatattaagaaatggctcaagcgggatttgaacctgcgaccttgggcttatgaatcccc
tgctctaaccactgagctactgagccatagtataaaatactattaaatattataccataa
attatggtcaaatataaatcaataaaagaataattactatca-tttactacttttg----
-----------------tgcaagttgtaaatattcgaataaaatttcaaaattaaaaatg
tctttaactttttgttaga-------atta-ttgattacgaaa-----------atattg
atatttagaattaac---------------------------------------------
--------------------aataaatttg------------------------------
---------------------------------------------------tctttattt
g-----------------------------------------------------------
------------------------------------------------------------
------------------------------------------------------------
------------------------------------------------------------
------------------------------------------------------------
-------tagaca-----------------------------------------------
-------------------------------caaaaatattggaaa--------------
------------------------ttttcatcaat-----------------taattagt
------------------------------------------------------------
------------------taaactataaccgacgtac-----------------------
------------------------------------------------------------
------------------------------------------------------------
---aagaattattgaacta-----------------------------------------

------------------------------------------------------------
---cgttaaat-------------------------------------------------
------------------------------------------------------------
------------------------------------------------------------
------------------------------------------------------------
-------------------------------------tgtgatgcatt------------
-----gggcttgatcaaaacttctctaaaatcaatagaatatagt---------------
-----------------------------------------------------aattcaa
gtaaatcaaatactaacaaatgtgacaaacgcttaaaaaaataata--------------
caattagaatcgtaaaaatttggtttttga-ctgtgttaaataaatttgattatcataga
tttaatatttaaaagtgttctattacttgta-----ttgagtaattgt-aatatttaata
tcttctcttttagataataatcataatataagctaaaaaatagtttt---tattaaacaa
ctagcatagagccaataccttgatcagttaatatttctagcaagagagcatgatctattc
taccatctaaaatatgcgctgaagctacaccctgagctaaagaacgaatacaacaattca
ctttaggaatcatgccaccagaaattactgctgtttgagttaaatctctagcttcctgta
tattcagatgactaattaatgttgaaggatctgaagaattacgtaaaatgccgggcgtat
ctgtgagtagaattagtttttcggcattaagagcggctgctatttctcctgcgacagtat
cagcattaatattatacgattggccttccttgtctgctgcaacactagctataacaggta
tataattattattaattaatatcttcagtaatttagtatcaatattttgtacttctccaa
caaagccaaggtttgcttttccatttggtctcggagtaataagtaatccatcttttcccg
acaaacctacactttttaccaccttgtttattaatagttgcgacaagatctttattgactc
ttccaactaaaaccatttccactatatccatggtaggttgatctgttactctaacgccat
catcgaattttggtaatattttgaggcgatctaaccaaaaattaatttctggtcctccac
catgaactaaaatgggacgtagtcctataaaagacagaaaaacaagatcactaatcactt
gatcttttagtttctgatttttcatagccgctcccccatattttataacaataattctgg
aagaaaattgctgaatataaggcagagcttcacttaagactttactctttctgaattag
tcaacataaataaagtaggattgtgatatataaaattatatacacagcttgtgttaattt
aaaagataccaatgggt---------acttgaaaaaatagaaatattactaacag-----
----ttaaaactatagggattataattatgaataaattttgggataatgtattaagattt
ccacgatttttagtcagcgtcattttaggattaattttgataattattagtcctttttc
gtgttattaaaaaaaccactgacaagtttttttttcatcatatcattggccggattaatc
acagttttggcaataataa-tacaaaaaatgataaatatcgaatgttgttgagagaaata
aataaatttaattaaaatttttgaacatatataatattatattatcattaaatatgataa
ttt-atttcctttctat---aaaacccataaatattaaatattatttgataaatattat
gccactaaaacaaagagttagctctgaaaagaccggtgccttcgcacttttggatagtat
tgttaggcacggagttaaacatatatttggttatccaggtggagctattcttcctattta

```
tgatgagctttatgcttgggaagaagcctctctaattaaacatatccttgttcgtcatga
gcaaggcgctgctcatgctgcagattcttattctagatcaacaggagaggttggagtatg
cttcgctacttctggcccaggagcaactaatcttgtctcaggtatagctacagcacatat
tgattctgtacctatattagctataacaggtcaagttgggagagctttattggtacaga
tgctttccaagaagtagatatttttgggattacacttcctattgtaaaacattcatatgt
agttcgtgaccctagagacatgtctagaattgttgcggaagcatttttttatttgtaaaca
cggtagaccaggtccagtattaattgatgttcctaaagatgtaggattagagaagtttaa
ttatttttctgttgagcccggaaaagttaatattcctggctgtaggccaattaccagcct
aaagtcaagacaaatccttatggcagctaaaatgatacagcaatctagccagccattgtt
gtatattggtggaggagccataatctctgattcacatcaaattattaaagaacttgttga
tttttataaaatacctgttactactactttgatggggaaggggatttttaatgaggatag
cgactattgtctagggatgttaggtatgcatggtactgcgtatgctaattttgcagttag
tgagtgcgatcttttaattgctttaggagctagatttgatgatagagttactggaaaatt
agatgaatttgcttgtaatgcccaagtgattcacgtagatattgatcctgctgaagtagg
aaaaaataggattcctcaagttgctattgtcggtgacgtagcagaagttgttagtgaaat
attgaatttattaaagacttctttccccccttatccagagcagattatatcttggcaaga
aagaattaatcgttggcgtcaacagtatcctttactggttcctagaaaatcaacaagcat
ttcacctcaagagattcttgttgcaactaataaattagcccaaaatgcttattttactac
agatgttggccagcatcaaatgtggtcagctcaatttctgaaagtaaaagctaagcattg
gctttcaagtgctggattaggcacgatgggttatggtttacctgcagcaattggcgctca
agtagcacatccaaatgacgtagtcatttgtattagtggtgattctagttttcaaatgaa
tatgcaagagttaggaactatcgcgcaataccagttaccagttaaaattattattattaa
taatcgatggcaagggatggttagacaatggcaacaagctttctatggtgaaaggtattc
acactcaagaatgacagaaggagcacctgattttcaaaagcttgcagaagcttttggtat
taaagcttttactattaataataggcaaaatatgcaatctgctttacaagttgctattga
ttatcctggtccagttttattagattgccaagttacagaaaatgaaaactgttatccaat
ggttgctcctggaaaaagtaatgcacaaatgataggtatcgctaaaccgcagagaggtac
tgcttccaactatattaataatagtgtttaaagtaa------------------------
------------------------------------------------------------
------------------------------------------------------------
------------------------------------------------------------
---------------atagttagtatttttta-aacaaagaata-aagtaca--------
-taaatgtaatttttttatactatta-aaaactatgtaagaacagcctttgacgagaatt
gaactcgtgaccttccccttaccaaggggatgctctacctctgagccacaaaggc-tttt
tactatattgggccgggttggatttgaaccaacgtaggcgaagccagcggatttacagtc
cgcccccattaaccactcgggcaccgaccctaattaaaccttattcaagatatgacctat
```

```
aagatatcataactatgttgctaaaaacaagtactttgataaataaaactgaataagtaa
tggacatagctggacttgaaccagcgactttcacgatgtcgacgtgacactctaaccaac
tgagttacatgtccaaaactttaattattattataacactgtttagtgattttataagta
tctatttctaattgaagatttgttttaatcgtccggctttaccgagcagatctcttagat
aatacagtttagctctacgtattttagctcttctaatcactttgacaaaagttattcgcg
gtgaattgagaaaaaatactctttcaactccaataccttgaaatgagcatcttaaagtta
aactagtatttaagcttttttttcttcttctcgctaatactactccttcacataattgtt
cccgagttttactaccttccttaaccattagtccaagttgtattgtatcgccaactttaa
tttcaggtacttctgttttgataaaaggtatctcaacacttttcattagctgacttagct
ttgtactatttgctttcatatataggattagtcttgtgaca-tgttttgaaatattatac
attaaaaaaacaaatatgaaaaattttagcgaataagcatactagtatttataaaac-tc
tttttatct---------------------------------------------------
--------------------------------------------ccattata--------
----------------cagatccatt------------atattcaact------------
------------------------------------------------------------
------------------------------------------------------------
------------------------------------------------------------
------------------------------------------------------------
------------------------------------------------------------
------------------------------------------------------------
------------------------------------------------------------
------------------------------------------------------------
------------------------------------------------------------
------------------------------------------------------------
------------------------------------------------------------
------------------------------------------------------------
------------------------------------------------------------
------------------------------------------------------------
------------------------------------------------------------
------------------------------------------------------------
------------------------------------------------------------
------------------------------------------------------------
------------------------------------------------------------
------------------------------------------------------------
------------------------------------------------------------
------------------------------------------------------------
```

------------------------------------------------------------
------------------------------------------------------------
------------------------------------------------------------
------------------------------------------------------------
------------------------------------------------------------
------------------------------------------------------------
------------------------------------------------------------
------------------------------------------------------------
------------------------------------------------------------
------------------------------------------------------------
------------------------------------------------------------
------------------------------------------------------------
------------------------------------------------------------
------------------------------------------------------------
------------------------------------------------------------
------------------------------------------------------------
------------------------------------------------------------
------------------------------------------------------------
------------------------------------------------------------
------------------------------------------------------------
------------------------------------------------------------
------------------------------------------------------------
------------------------------------------------------------
------------------------------------------------------------
------------------------------------------------------------
----------agattcttttatagtttc--------------------------------
------------------------------------------------------------
------------------------------------------------------------
------------------------------------------------------------
------------------------------------------------------------
--------tttgataatcagtac-------------------------------------
------------------------------------------------------------
------------------------------------------------------------
------------------------------------------------------------
------------------------------------------------------------
------------------------------------------------------------

```
------------------------------------------------------------
------------------------------------------------------------
------------------------------------------------------------
------------------------------------------------------------
------------------------------------------------------------
------------------------------------------------------------
------------------------------------------------------------
------------------------------------------------------------
------------------------------------------------------------
------------------------------------------------------------
------------------------------------------------------------
------------------------------------------------------------
------------------------------------------------------------
------------------------------------------------------------
------------------------------------------------------------
------------------------------------------------------------
---------------------------atcta----------------------------
------------------------------------------------------------
------------------------------------------------------------
------------------------------------------------------------
------------------------------------------------------------
------------------------------------------------------------
------------------------------------------------------------
------------------------------------------------------------
------------------------------------------------------------
------------------------------------------------------------
------------------------------------------------------------
------------------------------------------------------------
------------------------------------------------------------
------------------------------------------------------------
------------------------------------------------------------
------------------------------------------------------------
------------------------------------------------------------
------------------------------------------------------------
------------------------------------------------------------
------------------------------------------------------------
```

------------------------------------------------------------
------------------------------------------------------------
----gatgatcaaatattattttcaaacatggcttatgtatgaaaaagttctata---aa
aacctaatgac--------taaaattagggcttaagtatttcttagttttgtgttataat
tagttactatcaaaggtagtggcataacacgaagcatcaa-aatgttcgaacgctttact
gaaaaagctataaaagtcataatgctagcacaggaagaagctagacgcttaggacataat
tttgtcggaactgagcagatactattaggattagttggtgaaggtactggaatcgcagct
caagttttgaaatcgatgaatgtgaatctaaaagatgcaagagttgaagtcgaaaaaatt
ataggaagagggtctggttttgtagcggttgaaattcctttcactcctcgagcaaaaaga
gtactagaattatctttagaggaagcacgtcaactaggccataactatattggcacagaa
cacttgctaatgggcttagtccgagaaggagaaggagttgcggcaagagttttagaaaat
ctggcagtcgatgtttcttcaattagagctgaagttatacaaatgctcggagaaaatgcg
gaagccaatgtaagtggaagcaatactacgcaagctagaagtaaaacaccaacattagaa
gagttcggatctaacttaactcaaatggctatggaaggtggtttagatcctgtagtcgga
agacaaaaggaaatagaacgagttattcaaatcttaggtagaagaactaaaaataatcct
gtcttaattggggagcctggtgtaggtaagacagcgattgcggaaggattagctcaaaga
attgctaatagagatgtaccttctattttagaagataaattagttattactcttgatgta
ggtctattagtagccggaactaaatatagaggtgaatttgaagaaaggctaaaacgtatt
atggatgagattaaatcagctgataatgtaatattagtgattgatgaagttcatacattg
attggggctggtgctgcagaaggagcaatagatgcagctaatctgcttaagccagcttta
gcaaggggagaattgcaatgtataggtgcaacaactttagaagaatatagaaaacatata
gaaaaagatccagcattagaaagaagatttcaaccagttgtagttggagagccaagtgtt
gaagaaacaattgaaattttgtttggtcttagagaccgttatgaaaagcaccatcaatta
acaatgtcagatggagctttggctgcagctgctaaatacgctaatcagtatatttctgac
cgattttttgccagataaagcaattgatttaattgatgaagctggttctagagttcgttta
ctaaattctcaattacctcctgctgccagagaattagataaagagttaagagctgtatta
aaaacaaaagatgaagctattagagctcaaaaatatgaaacagcagagcagtatagagca
agagaaatggaaattaaagctcaaattgcagcaattgctcaaagtaaaaagaatgagcct
gatttaaatttagaagatcctgttgttacagaagatgatattgctgaaattgttgctgca
tggactggtataccagtaactaagcttactaaaagtgagtcagaaaaattaatgcacatg
gaagaaactttgcatggacgtattattggtcaagacgaagcggttgtagctgtctctaga
gcgataagacgcgcaagagtaggtctaaaaaatcctaacagaccaattgcaagctttatt
ttttccggaccgacgggtgtaggaaaaacagaattaacaaaagctttggcttcttatttc
tttggttcagaagcttctatgatacggctagatatgtcagaatacatggaaagacacact
gtatctaaactaattggttctcctccaggatatgtgggttatagtgaaggtggttatcta
acagaagcggtaagaaaaaaaccatatactgtcatcttatttgacgaaattgaaaaagct

catccggatatttttaatctacttcttcaaattttagaagatggccgactaacagatgca
aaaggtagaactattgattttaagaatactcttttaattatgacttctaatattggatct
aaagttattgaaaaaggaggaggtagtttaggctttgaattatcagaagatcaaacagaa
tcccaatatactagagtacgatctttagtaaatgaagaactgaaacaatactttagacca
gagtttctaaatcgattagatgaaattattgtatttcgtcagcttactaaagatgaagtt
agagaaattgcagaattaatgcttaatgaagtctttgcgagaattaagcagcaagatatt
caattaaatgtaacagaacgatttaagcaacgattagtagaagaaggatataatccaagc
tatggagctagaccacttagacgagcagtaatgaggcttttagaagatagtctggctgaa
gaagttttatctggtaaaattaaagctggtgatagtgcagtagtagatgttactaatgaa
ggagaagttacagttttattaggtgaaaaattagaactgttaacataaa------aaaca
atttattaat--taaagcattgaagaggttat--attcaatgctat---gaaaaaaatta
ataagatgggttgctataattgacaacctctgatactctttatttgtatata---aagcc
gggatagctcagttggtagagcagtggattgaaaatcctcgtgtcaccagttcaaatctg
gttcttggcat-tttaagcatcagaaaatttgctatctgagcattaaattttaattgaa
atgttcctgtcggaccattcctgtgttttgctattataacttctgtgaagtctctacttc
tagtttcttgtgtatagtaactttctctatatagcattattactaaatctgcatcttgtt
ctatagaattatgaacaataaagttgtttgaaataaaattgcataaaggtttacattcta
aatcaaacactatttgtaaacttgtaacgtttatttttttgtatactttcaaaagtcaaag
atgaaa------aagaattaaaaacatttttttgttcttcaaagttttt------tattt
gaatagcaatcatgtcattttgattaattttatcacatcttttccatccttgtgttgtga
gtagtttatggttacttgttaattgtatatatttcccagcttcggtaataattttatata
cagtttttttccctgttttcgaaatactacatttaattgctagaaataactgttgttttt
gcgcactaaagttaatgacttca-atctttttataatagcaattaaataatatttg--aa
gttgattatataatgatgtctgaagataattgaatttggatatacacccactttctctta
aatcggataacaaaggccttttgtcactccgagtttcaaggtttctatttagttgagata
atgccaagataggcaagcttaaatcttttgctaatattttaagtgatcgagtaattaatg
atagttcttgagatctattgtcagattgtctactctcctgcagcaattgaagataatcta
taataattaattctatatttttttccttgtaatttaaagaatttttacttttgtctttatag
tgtcagtagaaatcttagcgctatcatcaatataaagattaaggttagcaagagttttac
ttttttgaataacatattgccattcatcattattaagctgaccagattgaatttttttggc
cgtttaaattacactcttgagctaaaattcttcttagtagttgctctgtagacatctcta
agctaaataaaataacataagatcttttctgattaattacatatctagttatattaatag
caaaagcagttttccccattgaaggtcttcctgcaagaataattaaatctgacttttgaa
acccttgtgttatagagtcgagttctgtaaaaccagaaagtatactactatttatactaa
tctttttccttgtatccaaatggactaataggtttgcaagaattgtagctaatgtttgcg
tatctttttcgtcaagtatttcataagcttttgttaattgactaacaatagaagttatgt

ttttttgttcaataagttgttttgagcagctaattaaacataatgaatctccacatgcta
aaagtaatcttttaatgtagttatctataacgactgctgaatattcgcttatagtatcag
ataggggagcttgttctattagactaaaaacttcatttagctcattcaaatcttgaatca
tttttcatttttaagttaatgaaaaaatttcttacgcttttgtctttgcatggttaa
cattttcgagtgctgctctataaagtaatgaagttgatgtaaaataaaaaaaatcaggag
agatttttctactgattttagtaatgatatcgattgctgagttaatattatactaatta
atatttttcagctaaaatattatgaggaggtagatatttataa------atacttatta
cttcctcc-ttgtaacattactcttgtg--------gtattagttaactattgatttaat
ttgattctggaagaacttgcaatttaatatttgcagttacttgatgtaatagcttaattt
ctacatcgtaaatgccaattgttttcacgtctggtaaagagatactttgtttttcaatgt
ctatgttagtagtgtttttaataatttgtgaaatttctttttctgtgacactaccaaaaa
tattgtggccatctcctgtttttttactaacgctaaacctttgtatttcttctagaagtt
gcttgactttttagcatttctttagctgtttttaatttttcttctcttatagctgcat
aaaattttgttgttttaaaataccatttgtggcaactgccgccattttattcgggatta
aaaaatttcttgcataaccagaagctactttgataacgtcattactcttgccaagttttt
gaatattctcttttagcacaacatttataactttttactcatctt-------aattcta
aaaa----atctgtattatcctattatacctgttgtatacagatttt----aatagtaag
tatttatttgttgatgtaatattcttgtaaaa--aaaggtaatttttattttt-ttctg
atataatcactttcacaataattc-----attaggagaggtggccgagtggttgaaggcg
cagcattggaaatgctgtttaggagcaatcttaacgagggttcgaatccctctctctccg
tacaa--aaaatataaaaagttttatagagtccaatctaaatctccaataatattcttcc
aagcatttaatccaccccgaactctaacaatatttagtttttgagctattaaaaacttag
aagcaaatattgatctagaatctaagctacagtaaacaaaacaaattttatcttgtaaat
tgatatgagagtaattcattcctttatttttgtaaaggtagatttaatgaatgtatta
aatgacttttatgatattcttcatgatttcttacatctattaaaatatattgtggattcc
tactaattaaaactttgtaattgaataacactaatttctcgtacaaatgtactagatt
tactgccataatatttgttccaatgctttttgt-------ggataatacaaatttagta
tttataatcttaaatttgttaaatgatgaagttaatgagttgtacgttaatatagttcca
cttaatatagacttataacccagaacgattttgacagcttcagttgcttgaagtgtacca
acaatacccggtaatagacctaaaactccggaattactacaagtatctcttgcgctctct
ttattttcagtttcactgtagaagtctcgataaacaggtccaccttgataattaaataca
ctaacttgtccttcaaattgaaaaatggcaccataaatatgtattttatttaactctagg
caagtatcattcaagagatatcgagtatcgaagttatcagatccatctataataatatca
tattgtctaataatgtcgattgaattctcatagcttagtcttgttttgaacacagttact
atacactgtggatttaaatctaata-tttttttttagctatttctacttttgaatatcct
atatcatgcatcgtgtataaaatttgcctctgtaaattagagagatcaattatgtcatcg

tctataattccaatgctgccaattcctgcagcagcaaggtaaatgattcctggagatcct
agcccaccagccccaatgaataaaacttttgcttcttttaatctttcttgtccttctaat
tgaatttgtggcaatactaaatgcttagaatatcttgtatattcttctagtgagtatttt
gtattttctgttttaaaatttagcataaaaatttatagtttatttaatagcactta----
--aaagctagctgataagtttattaaaatgacttttgtaaacttggttcttctctatcta
aaatagctccttgtgtaggacagacctgaacacaaatactacaatcaatacaggcagaaa
agtcaatccaataccaattagtgttattattgtttttccctttccttgatgaatacaag
ctacggggcaagcttccacgcattcagaaactccaatacattttctgttacaattgtat
gagacatattatttaattaat-aaaagataagctatttacatctagtagttttgttgtat
actaaatttacatatgctatccaggccaatatttaaggtagtctatttctacagtttact
aatttgatatgcgcctttagttcagttggtagaacgcaggtctccaaaacctgatgtcga
gggttcaagtccttcagggcgcgtttag-ttttatcgtaattcaataatatcattttatc
tttttgcaaacatcaaaatgatatgatataaaagatcatattttatacattgaatattct
atgtactagtactg-tttttaatatgttattattgataagttaattatactaattctat
ttttgtaataagt-aaaaacagttctaattaagtatgcacgtatttaatagtc-ttttgt
atatat--aacttgaaaaaatttaaactatggctaaaaaagttactggaattgttaaact
agcattaaatgcaggtaaagctactcctgcgccacctgtcggacctgctttaggacagca
cggcgtgaatattgttatgttctgcaaagagtacaatgcacgtactgctgacaagtcagg
actagtaatccctgtagagatctcaatatatgaagatagaagttttacttttatacttaa
aactcctcctgcttctgtactaatcgctaaagctgctggactaaataaaggttcgggtga
accaaataccaaaaaagtaggcagcataacaaataaacagttagaatctattgcagaaac
aaaacttcccgatttgaacactaacaatatacctcaagctatgaaaatagttggaggtac
ggctaaaaatatgggaattctaattaaagatt-aaaagtttcaattatatttacta-tt-
-tttatgaaaaaattttcacgtcgacttacaacattaaaatcaaaagttgagcctaaact
ttacactattaatgaagcagtatctatattaaaagcaacgtcaaatgctaaatttaagga
aactgcagaagctcatattgctctaggtttaaatcctaaatatgcagaccagcaactcag
agcaacagttatattaccgaaaggtacaggtaaattgataaaagtagcagttattgctaa
gggagaaaaattaacagaagcaattagtgcgggagctgatgtcagtggctccgaagaact
aattgatgaaatactgaaaggtagattagattttgataagctgatagcaacaccagatgt
tatgcctttaatagcaaagctgggacgagtattaggccctcgagggttaatgccttctcc
taaagcaggcacggtaacattggacgtagcaaaagccgtgaatgaatttaaaggggtaa
agttgaatacagagttgatagaacaggtataattcatgtaccttttggaaaatctagttt
ttcacaagaagatctagttttgaacctccaaacaattaaagaatcgattgatagaaacaa
gccttctggagcaaaagggaagtactggaaaactttttcttatctagcaccatggggcc
atctattcaaattgacatcactagccttctatgaaaaattttgtataattatagaactt
gaataatctattatttcaaatta-taatttgctagcattattaacgtaacaacttgctta

ctatatcttatctgcattaaaaactttt-ttatatatcttattaattatgagtacaaaggt
tgaaaatatcttggaagagctaaaatctttaaaccttctagaggctgctgaactagttaa
acaaatagaagaaacgtttgatgttgatgcatctgcggcttcaggaggtatgatgatggc
cacaccaacttcagcaccagcgtctgctgaggttgaagagaagacagaatttgatgttgt
cctagaggaagtcccagcacctaaaaagatttctgtattaaaagttgttcgctctctgac
tggtttgggtttaaaagaagcaaaagatcttgttgaatctactccaaaagtcttaaaaga
aggtgcttctaaagatgatgcagaaactatgaaaaaacagctagaagatgctggcgcaac
agttattgttaagtaatga-----agaagcgcttctgtttaagtataaacagaagtgcct
ctttatttattataagcaattatactagaaaagtccaagtgtaattgcattgttgatagg
catagttgcaccaatacctagccaaatacttacaacggtaccaattaagaaaacagtggt
tgcaattggtcttctaaatggattttgaaatttgttaacattttcaatgaatggaacagt
aagtaaaccagctggtacagcagccatacttagaacacctaatagtttattaggaatcac
tcttaatagattaaatgttggaaagaagtaccattctggtaaaatctctaatggagtcgc
aaatggattagacttttcacctatagaagatggttctaaaattgcgagtccaatactgca
ggcaaaagtgccaataattacaactggaaagacgtataatagatcatttggccaagctgg
ctctccgtaataattatggcccattccttttgctaatttagctctcagcttcggatctgt
taaatctggtttttttaagaattgacatattg-tttttgtgtttaatataaagattagtct
aattactgttttaaagtgtaattttcaactattataatggtccagaaattccttgtttac
gtatcattaaaaaatgcataagcatgaaaacggcagttagtagcggaagtacaaaagtgt
gtaaactatagaacctagttaaagttccttggcctacactaactcctcctcttaacaatt
ctactatactcgctccgacaactggaacagcatcagggacacctgtaacaatttttacag
cccagtatccgatttgatcccatggcaaagaataacctgttaccccaaaagaaacagtaa
gcactcctaaaattacacctgttacccatgtcaattctctaggtttttttgaaacctcctg
ttaaataaacacggaatacatgtagaatcatcatcaagaccatcatgctcgctgaccatc
tatgaattgatctaataagccatccaaagtttacatcagtcataatgtactctactgatg
taaaagcttctgcaactgtaggtctataatagaatgtcattgcaaatccagttgcaactt
gaattagaaaagatacaaatacaatacccccctaagcaataaaaaatattaacatgaggtg
gtacatacttactagaaatatcgtcagcaattgcttgaatttctaatctctcttcaaacc
agtcataaattttactcataaaatagcttcaaaaagctcttctaatata--ttttgcgtt
taagctactattaattcaatag------ttattgaacttatatt-tttttataatcaata
cttattaactattataacatctttttattgattt--------------------aaaatt
ctctgacttagtaaaacagggctatgtactataattcttttttttttgtatagagattaat
ttcgtttttagtaaattagctaaaattcttgttacactgactctattactacctataatc
tgtgctaaaattttatgtgtaatagtaaagttaagcataataccattattaacttgagtt
ccattttgctcagctaataataataataagctcgccagtctactggttatacttttatgt
gaaatgatttctataaaatgataagcttttgccgagcaaaatactaaatgatttacaaaa

aataaattgaatggagcgcagttactacatgcttttataatagtagtatagttaatacag
gcaacttctgttttatctattgcttctgcttcgtagtaaaaattatcgtcgaccaattct
atttgtccaaatgtatctcctgtagttaataagttaagagttactttatgtgtatttctc
aggacttttgtaataattaaagatcctatcaatataatgtataatctagagctgctatta
aacaaaagagtatcatttttcttaagtgaaaaaatttgataattaatttttattacgatta
aaaaatagtagccatggattgcaagaaaaaaaatcacctgaactattttcagagaatatt
ggaatacaattttcttgatttaaacaatttttagtcataattactactttataa------
---aaaaatattctaataat-taagaactgaaattaaatgtcatataaattgatgttagt
tgagaatgacattgtattatcaaaggccattcaagaatacttaatagatcaaggcttcaa
tgtttatattgccaataatggattagaagcactgaatctagcttatcaatataatttcga
cttgataatttctgatataatgatgccgctagttaatggctatgaactattagcaaaact
taaaaaaaacaaagcattatccaaaatacctgttatctttttgacagctaaaggaatgac
taaagacagaataaaaggctacgacatggggtgctatggttatttatctaaaccatttga
tcctaatgagttgctttcaattattaacaacttaatcgctagagatgtcttaaaagaagc
ttctttacaaaactcggcaacatcaaaccagcaattaaatcataaaatacgtttgacacc
tagagaaaaaagtatttagatcttgttgttgatggacttactaataaagaaatatctac
tatactgaatactagtgttagaaatgtagaaaaatatgttagtcgactcttacacaaaac
taatatgaagaataggactttgctagtcaagtattctataaataataatctattaaataa
tgagatcaatgaaagggcgaatgacgggactcgaacccgcgaatgatggagccacaaccc
attgccttaaccccttggccacactcgccatattacatacaccactatagcttgtttttt
gatatatcgtctagttttcgagttatatatttatctaaaacaatatttactttgaatttt
-tatgatacatagcaatattagtattcaaattaatggggaaccatttaattgctcaaagc
ccatctcattgcaatttttattaaattatcttgattttaattccgagcgtgtagcagtcg
agctaaatgacattcttttaccagagactctgtttcactcaacttatttgaatgatcaag
ataagctagaggttattaccattgtaggtggtggataatctatttgtacatac-------
-acagttttctatctcaaatgtttcatagatagagataatcttccttgtttttatctac
atgaataataacagctttaatagtgtcacctattttaaattgtgatgatatctgctctaa
ttctttgatattaatctcagaaatatggacaagacctttgagatttcctacctttataaa
taatccgtaaggtgtaatctgatttattatgccttcaataatattaccaacaattaaatt
agatgatgcttgagatattagagctcttctatgactcagtattaaattattagatttctc
ttcgacattgagtaattttagcttaataaatttattattaaattgctcactttttgaaa
attaccgagatgagaattaggtacaaatccagatataccttcaaggttaattatcattcc
tcccttattgaatccttttatcattacattaagtaaagagtcttcagctaatagttgcct
aattcttttccatgctcttatatattcaaggcgacgaattgataatattaattgtttcga
ctgaatattataatctaataaaaaaaattctcttgtgtcattaatgtttaaagaagtgaa
attatttaaatcttgattactagatacttcttgaataggcaagtatgcagatattggtgt

tcctatatctactaataccccatttaattcaaagctaaatatagttccagctacaatatc
tccaagattcaaatcatacttgtatttttgtaaaacagctgcaaaatttctatgagtaaa
tccttcattatttttttgtcatattgaatagtatgaattaaagagtaactttgtacaatat
attaaggtaaataaatgtttgtcgaattgtatgaaaaaattttaatttgttcggttgtaa
ttaagtatataaacatattaaaatatctaggagacatgaaaagtagacgtagatagctgc
agatttaa-tactatataatctgaggaaagtccgggctccacaaatacaatttatgctgg
agaaatcccagtgtaggtaactgcgaggatagtaccacagaaaaaaaccgccagaataag
t----tatattagctggtaagggtgcaaaggcaagttaaaagcttaccaaaagtactgca
aagtatttgttaggtaaacccctaaaatggagcaaagctactaaacaaatatttgtgtat
attttatctagtttagttgaaaatactgcatgaagttattagtaacaataactcgagagg
aatagctatcctttacaatatatc-tcgattatattgaaagaacagaacccggcttatgt
agtactttttatgtcttgtattattatgtctattgtcgaatctttaaatttaactgtttg
acaatttttttttgtagctgatgttgtttataatctatttctgaattggttaaagttttt
tcattagatcgatatgtaaatcttagacctatactttttttctcttttccaattgattca
tctatatattgatcaaataatgttgtactttctaaatctttatcataaaattgatctaat
aattttaataagtatttaatttccatattcttaggtattattaaagagagatctcttatg
atcgatggatacttagagtatggttgaatctgataacttaaataattgaattcattatga
caatcttctaaaatgtttaagttaacttcaagaaaaaaaagttttgtattgagtcctaat
tcactatatgttgcttgattaagttcactgaatatgccaataataatattgttatagatt
agtgtggcaaattttcctttttgaatgaactttattctaccatcgactatatcttttttt
gtccactcaatttgtctattaatcttttgaaaaaaattttctacgatccctttagcttca
taccaatttaaagaatgcgcagggtgtgaccatgtagatcggatatctagattaccaccc
aatatgatagctagattagtagtttctacaattttgttttgatcattatgaaatactgtt
cctatctcaaagctatctatagtttggttgctttgtttaatattataaagatttgattca
attaatccttccagcaaactgccgcgtaaattagaataatcttgaataagaggattgttt
aaagctatatttcctttggactttattaaagaataatgaactaattcagttaatcctaga
tttcttaaaatacttctacttttgtctatgaattttctctttgtggataaccttttgttg
aattgaatatttggtactgagctcctgaatttatgataaccataaattctagcaatttct
tcaataacatctatttctcgaaaaacatcttctcgtctataatttggaacagtaacttct
atattttccttattttgataaataagatcaaaatttaaagagtgaagaatattttgaatt
tcttcaaagaataaaaaacgtgtttgaccattatacataattggtcctaaaatgtcatga
actttttaattgataaattaatattaagagcatgatcagttatttttctcgacagaaa
gtatctctaatatttccgcatgttaaatctgtgattaaagcaagagcttcaaaatgtgca
tttttccaattatctacatttagcccacgttcttgtctaattgaactttctgttcgaata
ttaagaactcgagaagactttctaactacagattgtttaaatatagcagactctacaagt
atcgattttgtattataatctgtatcaaaatcagagttgcttccaataccagctatacta

```
gttacatttgtatttacttgagtaactaaaatatttttgtttaactcaatattttcattg
tttaacttaatctgatcattacttccaagaggaaaattgcttcgaattgttagactattt
ttatgattcatattattaattttattcaaatcaataatattaataggctgtccccatttc
aacataatataattgcttatatctgttaacaaatttctgtgtgtaaacccagaagagagc
aaccgatttttagccaatctggagaatctttaactcttatttcgtctataatggcagca
aaataattgtcacaatttaataaatcgtgattactgattatagt-ctcctttttagagaa
atatctattattgggttgatgtatagctgataacattgaggaccctgttaaagccgagac
ttctcttgatagaccaatcatgcttaacacgtctgacctgttagctgttgatgatatatc
taatatatgatcaatctcacctccaattataatagattcaatagcttcaacttcaaaacc
tgcttgtgttagttgattagttaaattgtctatatctattgttttgatattaccaagttc
tttcaaccaatttaaagaaactttcataattaattgtcatctatcttatagaagtattta
ttttacttatgattaaaaatcaatttaaaaaattcatcaaattaatataagttatttgat
gaatttgcttatacaatatattattgtgatatcttacttttatatggcagttattaagaa
ctagcttttgtaaaagttagcccattatcttgggcatatctttccataaatctcatgaat
ctatcccaatcatcaggacttttaataacatgtacagcttctattgcttgtggtttacca
ttaataaacttggcattaacatctcttgtaataatttgtccctctgtgtccatcaggtac
atacctgtgatttcacctttgtctgccatactagcatctaaaattttgggatttgtaaac
cgaaaagttgctgttcctgtactaccatctcgagaccttgttaaacgtacatctggtaca
acttcttcattgataccttgaataaattgaattgttgccatatcaatcttctttatttgt
atataatttatattggataaaactgatttctagttcttgaatgcttgttgagaaactgtt
ttaaaagcttatagtatcagtagatcaaattttatgtattatataaaactgtaacttttc
-tttttg-----------------------------------------------------
------------------------------------------------------------
------------------------------------------------------------
------------------------------------------------------------
------------------------------------------------------------
----------------------atctgttcaaagaactaacttattataagctaatcata
cgaatattagccatataatgttcgtaagt-gaaattaaggggttgtagctcagaaggata
gagcaagcgcctcctaagcgctaggtcagcggttcaagtccgctcaaccctgttaacgaa
ttaagtagtattatttaacttgttctgcaaatcagtattgatctaagttaaagtacttga
cagctaaaacaaaagaatctgagaaatattttcatttaaagaacgaacaattatcttaac
tctaaatatctgataattagtaa-aaaattct----------------------------
----ctttttaaaacacaagatgatg-----aagatatttaaactaataattgaaattaa
gtttttgaagttatagaagccaat--ataatactttctattaacaga-----------tg
cttgcttctattactatctgtttacaaga--aatt----atgagtatcaaaata------
--attttagctcca-------------------------------------ctgac----
```

------------------------------------------------------------
-----------------------------tactaataaata-------------------
------------------tttattttagtctagacaag------atgaatcttcatggta
taacag------------------------------------------------------
------------------------------------------------------------
------------------------------------------------------------
------------------------------------------------------------
------------------------------------------------------------
--------------agctattctttgtgtcatacttatataaaattaggat-------ac
tgtt--------------------------------------------------------
------------------------------------------------------------
------------------------------------------------------------
------------------------------------------------------------
------------------------------------------------------------
------------------------------------------------------------
---------------ttcattaccataactggggtgggaggattcgaacctgcgaatggc
ggagtcaaagtccgctgccttaccacttggctacaccccaacatgtgaacattataatag
cagtcgtataccctgccatgtcaatagactattgtctttatttaataatttttgctgctc
ttgtagaataaaatcttttttttcaaataagatacgctatgtttaagttgcacgactgtt
atgttttcttgcttttgactgaacagccacttgatagtaattattccttctgcaatttcg
ttacttcctataataatacaagctacagctctttttttatttgcttgcttaatctgctta
ctaaaattacttgagctaacatcgatttctattttaaaaaactgctgatgtaaaaaacgc
attatttgcataccagtttcgttagcttttgtaccttgtgtagcaatataaaaatcgata
gatttattaggcaagtctatattatcttttgcaatcagtaataggcgttctaatcctatt
gcacatccaacagctggtgttgctttgccacctaattggtgaactaagctgtcatatctt
ccaccaccacatatagtatcttgcccttttgatgtcaatgtcttaatttcgaaagcagta
tcattataataatctaatcctcgaactaacttattatttatattgtagggaatattaagt
aattttagataattgcaaacgtcaccaaagtgttgttgtgattccaaacttaaaaaatca
gaaatttttggagcctctgtcagtatttttgtgtattagaatcttttgaatctaaaatt
ctaattggattactagttaatctttttgtgagtcagtgtctaaatcatcatggtacttt
gtaagataatctcgtaatttaacttgataaatactacgatcctctacttttccaattgaa
ttcagatcaagtttcaagttgtgcagattgagattgttaaatatgctcatagctaaatgt
ataacttcactatctgctcttgcatctaagctaccaataaactcaatgccaagttggtga
aattgtcgttgtcgtccactttgtggtctttcgtacctgaacattggaccgctataccat
aatctttgtaaactatggtggtaactcattttattttcaataaaagatctcacaataccg
gcagtaccttctggccttaaggtaatatctctattgctgcgatcatgaaaccgatacatt

tctttatttacaatatcagtatcttcaccaatacccctatcatataattcactattttca
aaaataggtgttctaatttcttggtaatttgcacattctaataatttggaaattttatta
tgtataaactgccaatattgaatttcatcgggaagaatatcttttgtccctctaatagcc
tgaattttgccatgataatttttgattaccttcttaatatgtctgcattaaatagcag
ttgtagactgttattcagtcatgtattatacattag--aacatactgatg-taatttata
aagtacgggcaaggagggattcgaacccccgacaccatggttcgtagccatgtgctctaa
tccactgagctacaagcccacttaataagtaccttaactatatcattgttcactaatgta
tacaactatattcttttt------atcttacatgtgaaattttttgtaaaagtaaaaaatt
at----gtttgtttttacaattggaactattcatatataatgcttaatttattataact
ttgtaaa--aatatctataagtttaattgtatgattctattatattggtaatcttgttat
aattttgcgctcttatctttcaattgg------ctagcatctaattaacttatatttcaa
ataaaaaaatcttatctaaaatgagtaaacaaattctatatcaggatgatgccagaaaag
cattagaaaaaggcatggatattttaacagaagctgtttctgtgactttaggaccaaaag
gaagaaatgttgtcttagaaaaaaaattcggtgcccctcaaattattaatgatggtgtaa
cgattgcaaaggaaattagtttagaaaatcatattgaaaataccggagtcgcactgatta
gacaagcagcatctaaaacaaatgatgtagctggtgatggtacaacaacagctacagtac
tggcttcagcaattgttaagcaaggaatgagaaatgttgcagcaggttcgaatccgatgg
ctattaaaaaaggtattgaaaaagcaacaaattttgttgttagtaaaatagctgagtatg
ctaagccagtagaagatacaaaagctattatacaagttgcttctttgtcatcaggaaatg
atatagaggtaggtaaaatgatagcgaatgctatagaaaaagttggcagagaaggagtta
tttctttagaagaagggaaatcaactaatacgattcttgagatcacagaaggaatgcagt
ttgaaaaaggctttatttctccttatttcgttacagatacagaacgaatggaagttcttc
aagaaaatccttttattttatttacagacaaaaaaattactttggtgcagcaagaacttg
tgccattgcttgagcaaattgcaaaaacatctcggcctttattaataatagctgaagaca
tcgagaaagaagcattagccacaattgtagttaataaattaagagggattttgaatgttg
tagcggttagagctcctggattcggtgatagaagaaaatctttacttgaagatatgagta
tcttaacgaacggacaagtaattactgaagatgcaggtttatcacttgatacagttcaat
tagatatgttaggaaaagcccgaagagttattgttactaaagactcgacaacaataattg
cagatggtcatgaaattaaagttaaatcaagatgtgagcaaattaagcggcaaatagaga
cgagtgactctttatatgaaagagaaaaattgcaagaacgattagcaaagctttctggag
gagttgccgtcattaaagttggtgcagctacagaaacagagatgaaagataaaaaactaa
gactagaagatgcaattaatgcaacaaaagcagcaattgaagaaggtattgtaccaggag
gaggagctactaatgtccatatctctagtgaattatttacatgggctaaaaacaatttag
ttgaagatgaattaattggtgctttaatagttgaacgagctgtgacctatccgttaagac
gaatagcttttaatgcaggtgataatggagcagtaatagtggaaaaagttaaaagtcacg
attttcatataggctatgatgcagcaaacggtaatattgtaaatatgtacgatagaggta

ttatagatccagcgaaagtagctagatcagctttgcaaaacgcagcttctattgcagcaa
tggttttaactactgaatgtattgtagttgataaagtcgatgattgagctata------a
gataatagataatctcaaacaaagaataatttctaattatagaaattattctttgtttca
attaatctaattaattgatcaaatgtataattttttataataaagccaaagtttaaacacg
tcttcttttcaactattaagtagagaagatataagcccgtaatacctaaattatcaatt
aaaatattattctcatatatagattttctaaagatctaaaaaatgacattttgaaaaa
taataccgaaatcttctcttatagttatttgctaaatcattactgtctaaagattgattt
ccaattgaattcagaacctttaacagaatacttttaaatctttatctgactttatttga
gaatcaataataatttgattcaattgtatgtcaaaactaagattaaaaatattacttgtt
aaagatatttgattttgaatatctagagattttatagccaagaagattaaacttttattg
ttcattgtaaaatataagacaattgatgatgtattctttaaatacta-aatttatcttct
tgtgcttctatcgtttttttgt-aaaataattaatgttttaatttttttagttttagttact
acccgaaaaaataaattctggaaatctagcttgagctttagctagagagtgtttttttccc
ttcttcttgccagtaattgataacttctgttgttttgtttaataactcaattctgtctct
ttcagcaatccacggcttggattctagttctgcttttaattcttcccaagcatcattcct
tggccagaaaaaataagaagtaagtggactagtgccatttccaacaatctgatcaatagc
aatagcaatattattttctaaccaaagtacttttagtgtaaattttgacaagtttttatc
tccattattaatattaagtattgg----tacagtagaatactgtactagttattttttttc
agcaacaatttgagctttgtttaaaatgtttttttaccgttttagtagcttgtgcaccttg
agataatcgttgcaaaattttttgaaatgttaacacgagtttcattatttataggattata
aaatcctaattcttctatagctttaccatcgcgtttatttctgctatccattacaacaat
tctatagcttggttgtttttttcttccgtatcgctttaaccttaacttgaccatattagt
cacctcgcttattttttt--atatctttattaaattatatcttacgacattatgcaataat
tcaaacaattaagctgattcaagtctaacattattaaatattaccattaaacattgtaga
aaaataatacctaacataggtgacatgtccattccaaacatgggaggaatgcttcctctg
aatagttttaagtatggatcagtaattctatttaatgaacagaatggttcattgtaccaa
tttacagttggaaaccatgccaatgataatttaagtaaaattaaaattaaataaatctca
gaaaaattagctatcgatccaagcaataaatttaatgtaccaggaagagtgttcatacgt
attttataattattatatacttcgttatactaatagtttgattatatattataatcgatg
taaataaat-cagctaaacaaaaaacttatccttaaaatcatcttctcttgagcactttt
ctttatctacattagtatttacttttgtcgtataaatttccagtttagaccaaatctgtc
ctaactgatttaattgagatgttccacattctaaacaggctatttttatattgctcaaaa
cagcacctttttttataaaatcgttcaataatggcgtaatagttgcttcatccaaaaata
aatctaagacaagtatttttgtaaatgaatttaattgatctgggacattgttaataatat
cattataattaactaaaactatattagctgtaggaagtaaagatcgtgctccttccgcta
agataaagccataaggcataataattacaataatatatttataatgactgctaataagac

ttatagtggtattagcttttaattcgattgtttctgttactagccactctctcattattt
cgtaagtaatccatttgcctagttctgagcaagcagttcttaaaatcgtacctggattat
tgtcgtttttgagaatccctgaccaatgttgtattaatggatgggatgctatattaattt
ggagttgcatgttttgttaaatatatttaacatttaatttgttgattgaatcttcaattc
ctatttgttgagtattattaaaatagcat----atatattaaagaaaataattat-aaaa
aatgaaaaaaagtctttggctgtggggttttacagatagtgctgaaacttggaatggtag
atttgcaatgattggttttatatcagttatttttatcgaagtagttacaggacaaggatt
attatatttaataggtatgatgtcataataaaaatccaaaaaagttcttgtaattttta
taagaactttttgg-atttttataaatctaatctagcggtctcatagatagaacaggtt
cattctctctattgataccttttcaaatccagcagcagcagctcttgctctgcctgcat
gccaaagatgtccaataaataggaagaatcctaggaagaaatgagatgttgttaaccaag
atctaggagatacatagttaactgaattaatttctgtagctacaccaccaacggaattca
aagaacctagtggtgcgtgagtcatatattctgcggcacgcctttcctgccacggttgaa
tatcgttcttgatttatttaaatcaagaccatttggtcctctaagaggctcaacccaag
gagctcgcaagtcccaaaatctcatggtttcaccaccaaaaataatttctccactaggag
atctcattaggtatttacctaatccagtaggtccttgagaagatgcaacattcgcaccta
gtctttgatctctaactaagaaagtaaaagcttgagcttgtgaagcttcaggaccagttg
gtccatagaattcactaggataagctgtgttgttataccaaacaaaattagaagccgtaa
gccccatgattgataaagcacctagactataagataagtaagcctcaccagaccatacaa
aagctcttctagcccaagcaaaaggttttgttagtatatgccagattccgccagcaatac
aaataatgccaatccagacatgaccacctacaagatcttccatgttatttacacttacga
tccatccatcaccgccaaacggagactttaagacataaccaaagataactaggggattaa
gtgtaggattgctaacaaatctaacatctccaccacctggagcccaagtgtcgtataccc
caccaataaataaagctttaataactaacagaaaagcgccaatgcctaataacactaagt
gtataccaagtattgttgtcattttgtttttatcccgccaatcgtatccgaagaaaggaa
atgattcttcaagagtatcagggccaatcagagaatgataaagacctccaaagccaagaa
cagctgaagaaattaaatgcactacgcctactacaaagtatgggtatgtattgaaaatct
cgccgcctggaccaacaccccagcctagtgtagctaagtgcggaattagaataaaccctt
gttcgtacaaaggcttctcgggaacaaaatgtgctacctcaaaaagagtcatggcaccag
tccaaaagaccattatacctgcatgagcaacatgagcaccaagcaatttgccagaaacgt
taattaagcgtgcattgccagaccaccaggcaaacccggtagactcaatgtctctaccgc
caacaccaacagttgtattaaagggcgtttccacgtggtaaaacctcctcagggaatata
aagttttcatgaggttgatcttgagcagccatccaagagcgaataccttcgttcaatagg
atatttttagtataaaaagtttcaaactcaggatcttccgcagctctcaactcttgagaa
acaaaatcgtaagctcttaaatttagagctagtccaacaattccaaatgcgcttgtccat
aatccagttactggtacaaataacatgaaaaaatgtagccaacgtttattagaaaaagca

acaccaaaaatctgtgaccagaatctattagctgttaccatcgaataagtttcttcagat
tgtgttggagtaaatgcacgaaaagtatctgcagcatcaccatcttcgaataaggtattc
tgtacagttgcaccgtgaatagcacatagtaaagcaccacctaagattcctgcaacgccc
atcatatggaacggattcagggtccagttatgaaatccttgcaaaaatagcaaaaatcta
aaaattgcagcaactccaaggcttggggcaaagaaccagctcgcttggcctagagggtac
attagaaagacggaaacgaatactgcaattggtccagaaaatgcaatagcgttgtatggt
ctaagaccaactagcctagcaatttcaaattgtcttaaacagaatccaattagtccaaat
gatccatgtaaagctataaatgcccacagaccaccaatttggcaccagcgagtaaaatct
ccttgagcttcaggtccccaaaggaaaagaagtgaatgtcccatgctgtttgctggggta
gaaacagccgcagttaaaaaattgcatccttctagatatgaactggctagcccatgagta
taccaagaagtaacaaaagtagtcccagttagccagccacctacagcaaggtaagcgcaa
ggaaacagaagtagtccagaccaacctacgaatacaaatcggtctctttttagccaatcg
tctacaagatcaaaaccaccacgagtttttcttgtccaattgctatggtcataatttta
atctccagagcaaatttaataagtaatttgtttaaataaaca---ctttagatatactct
aaaattttacttttctttacggtcaatataattataagtcaaaagtaaccaaaattcacg
cttttaataaca-ttttttagctcattaagttaa-ttttttaatatataaacagtaata
gagttattgttt-aacttacaatactcacaatttattcatttctataatcctttgaaaaa
gatatcctgttcctcgtgctgttaaaatcaagtcgggattacttggatcatcttctagtt
tagctctcaatcgtgaaatatgaacatcaacaactctcgtgtcaacatgccgttcgggag
tatagccccaaacttcttgcaagatagaagctctagaaaaaggttctccagccttactaa
ctaaaagctcgagaagactgaattccatacctgttaatcgaactctttcattatttttat
aaacttgtcttttgttagtatctatttttaaaaatccaatactaataatcccagagtttg
ggactccaagactagtggtgattttatcagctcttcgaagaacagaacgaatgcgagctt
ctagctccttaggagaaaatggtttaacgacatagtcatcagcacctatttctaatccgg
taattctatcgcagacttcacctaacgctgttagcattataataggaacatctgattctt
ttcgaagttcttgacaaacaccgtaaccatctagtttaggcatcataacatctaaaacaa
caaggctaggatactcttttctaaacacgagtaaagcttcttctccatttgaagcagtga
taacttcataacctattattgttagtcttgtttctaaaattcttcttatgctggcctcat
cgtcaacaacgagaatcttttctttctggttctccaa-tttcttatttactcctgtaact
caccgatcatgattaccataaccttaaattttagcgctatgagtatctttattcacttt
agaa-----tttagttattataaattttgcaagtctagtttacattaattagtaccagt-
-caatatttttgt-----------tttaagttatgac---------tattttaatccata
agttgactctagtttgcagaattcaatatatacata-ttttctacccaagtaaattgtat
tctattttctaaataaaaaaggggttgacaaaccagttgatagaaagtattctacattac
tataag----ttaattccaacaaagaagttttaa-----atcttctttcggagtaatgaa
tactttaat

>Pyr_25_39
tatatgata---aattaagacatctgttatgtgagaatttctaaaatatacaaaactttt
tatttaatatagaaaaagtaaattaaaataagttgaacatactttgctctaatctaaacc
tcttaattgag------attctaattagagacaaaagtatttacttattttgtatctata
ggaataaaaatgaaacttgcagtttatggtaaaggtggtataggtaaatctacaactagt
tgcaatatttctgtagcactttcaaaaagaggtaaaaaagttctacaaattggctgcgac
cctaaacatgatagtacatttacgctaacaggtttttttaatcccaacaattatcgatact
cttcaatccaaggattaccactatgaggacgtttggcctgaagatgtaatctataaggga
tatggcggtgtagactgtgttgaagctggtggaccgccggctggcgctggttgcggaggc
tacgttgtaggtgaaacagttaaacttttaaaagaattaaatgcttttgatgaatacgat
attatcttatttgatgttttaggtgatgttgtatgtggaggttttgcagctccattaaat
tatgcagactactgcttaatcattacagacaatggatttgatgctttatttgcagctaat
agaatagcagcttcagtacgagaaaaagctagaacgcactctctgagattagctggactt
gttggtaatagaacagataaaagagatctaattgataaatatatagattgtgttccaatg
ccagtattagaagtcttgcccttgattgaagatattagagtgtccagagtaaaaggtaaa
actttatttgaaatggcagaaattgataaggatttagcatatgtatgcgattactatttg
aatattgcagatcagttgattacaaggccagaaggtgtagttcctaaagaatctccggat
agagaattatttagtcttttatctgattttacttaaatcctaaatcaaaggtaggacaa
gaaaaagtagatcaagaagaattagatttgatgatagtgtaaataatattttcagcataa
tataacaaggaataagataatgtctacagctcaatcagatgctcttacttttgaatgtga
aacaggtaattatcatactttttgtcccatcagttgcgtttcttggttatatcaaaaaat
tgaagacagtttttttcttagttatagggactaaaacttgtggttatttcttgcaaaatgc
aatgggagtaatgatttttgctgaaccgagatatgctatggcagaattagaggaaggaga
tatttcagctaagctaaatgattatggtgaactccgtagattatgcttacaaataaaaaa
agatagaaatcctagtgttatattttggattggaacatgtacaaccgaaattataaaaat
ggatctggaaggaattgctccaaaattggaagcagaaattcgtgttcctatagtagttgc
tagagctaatggtttagattatgcctttacccaaggtgaagatacagttctagcagctat
ggctcaaaggtgtccattaaatttaaagaatcaatcagataacacgagtcttaaaacttc
tcctcatattcctttagttttatttggatctcttccggatccagttgtaacccagttaac
tatggaattaaaaaaacaaggtatttttgtttctggttggttgccatcaaagaggtatac
tgaattaccggttataaaagaaggatattatgttgcgggagttaatccatttcttagtcg
cacagctactacactaatgagacgccgaaaaacaaaactaattggtgcaccatttccaat
aggtccagatggtactcgagcctggattgaaaaaatctgttcagtaatgaatgtaaagcc
tattggattagaagatagagaaaaagcaatttgggcttctttagaagactatatttcttt
aatccgaggtaaatcagtttttttttatgggtgataatttactagaagtatctcttgcaag
atttttaactagatgtggaatgactgtatatgaaattggtattccttatatggataaacg

ctatcaagcagcagagctagctttattaaaagctacatgtgataaaatgaacgttatgat
gccaacaattgtagaaaaaccagataattataatcaagtagaccgaatccgtgatctgaa
acccgatttagttatcactggtatggctcatgcaaatcctttagaagccagaggtattaa
tacaaaatggtcagtagaattcacatttgcacaaattcatggctttaccaatgcaagaga
tattcttgaactcgtcacaagaccattacggcgaaatcttagtttgtcagaattaggctg
ggatgtttatagcaagcaaagctagtattcctc----tttatgacttacatggtataaaa
gccttgtaagtct--tttattttatgttaaagaagcatttcttgctctgccatcaccagc
ccaagcttgtaaactttctacttgttctttgtctgtaaatgctaaaggcacaaattgttt
gatagctattttaatatcttcagtactaaattctctctcttcactaaacgccgtatgcat
actttcaataattgcttgctcaatttcagcgccagaaaatttattacataataaactcaa
ttcatgaatatcatatttttgccacgatcttggtctaacttttgataaatgtatcttaaa
aattgattctctctcttgacgattaggtaaatctaagaaaaaaatttcatcgaacctgcc
tttccttaacatttcggaaggtaaactttgaattttattagctgttgcgacaacaaatac
tggagctttttctctgatagccaggtaataaatgtaccaaatactctggcactagtacc
actatcaccttgactatgtaaaccagaaaaagctttatctatttcatcaatccataatat
gcaaggtgataatccttcagagatatttatcatttctctcattttgattccgattcacc
aactagtccaccaaacagttttcccatgtcaagacgtagcagaggcagcatccaatcgct
tgcaatagctttcgcggttaaggatttacccgtcccctgtataccaactagcaataaacc
tttgggcgaaggaataccgtaatttaaactttgcttcgaaaatgagcgagaccttttttg
tagccattgttttaatacatctaagccaccaatatccctatttactttactataaggata
aaattctaatagatgcgtttgattaataatttgcctttttttcttcaataataataggtaa
gctccgagagtctatctgattatattgtgcaataattttagtaatcactttgcggattct
atctatcgataatccttggcaggattttgttatattattaacgagctctgaattcaaatt
aagattcaaagctttactcaatcttatgatctctttttgatttctagaagactaggtaa
gggcaaatctattactgttatgatatcattcagagcaaaaggtatattgattttgcaaga
aataataataatgtgcttagattgcgttttgattattttagctaaattgcgaagtttcct
aattaatactatttcgtttaggaaagagtcaaaatcttttaaaagaaaaagatttaaaga
ttcattattcaagttttcaataaattctaatgctaataaaggatttctttttgcataacc
attatcgctaggattgttagtgtatccatctacaaagtcccaggaatatacttgctgact
gtttgaacagtttaagctgtgttttataatatactctagtctatcttcttctctagtatt
tattacaataataggatatcgagattttaaaagtaatcgtaaatcttgagtgaaattcat
gtcaatatttgtttccatcagacttttgtacttcaccaattgaagaaagactaaataaga
ttatctaatattccttggatatattcaaaagtc------atctctctattttctaaatta
ttgctattactaatttaacttaccatgatttttttcttccttttcttcgtcgactattt
aatatagagcgtccacatggtgttttcattcttgctctgaaaccagaaactctaatttt
tttcttttagatccttgtagtgttcttttagtcatgatgtctcctctgttgtaaaagatt

aaatactattttatcatcttttatatatctttatgatatagatatgtattaatatcaaag
tt-aacagcaataatttatattaatcaagaactatttatataatattaaacattaaatat
----------taactttatgatattggctttgcccatttttttatttaagtattttgacca
tctttttgctaatcttaaactggttaattttcagcagctaaaaacgattttgttattag
aatctcaatttaagtattttgttgataaaagccaaaatcgtaaattagaacccgacgaaa
gttttgcttttgcaaaagtatgtgtagctaaaaaatatttttctaaagctattattgaag
gtcaattagctttaaaaaactatcgagacctgaatatcttagataataatatagtaattg
ctaatttgtataatatgctaggttttatttattttgaagcaggccaaacaagttttgcaa
agaacttttatgagcaagccttacaaataaaccctaattatgttgtagctttaaataacc
tggcaaaaatttatgaagaggtaaaagatttaaaaaaagccgaatctctatacgataaag
tgttaactcttaacttaaataataaaactgctaatagaaggaaagattttatagcaaaaa
cta---------aaaacatctaatactatttgtaatcgggatagcaggatttgaacctgc
gacatcctgctcccaaagcaggcgcgctaccaaactgcgctatatcccgtaagacataat
tattactatatacctttttctctaattatgtctactctttctgtattttatctaggatacc
aaaacatgcctttaactccatctggatctttgataaatcctaatttttttataaaaactaa
caacatcaggttctgcgaatagtgtaatagtgctaatttctgcctgtctcaattgctgta
ttaattggtgtatgactacttttcctaaacctaaaccttggaaatctggatgaataacta
cgtcccagatagttgcattaaatccgttgtctgatgttgctctagcaaatcctacaagtt
tactatttgcatcttttttttgtattaaagaaataataatagaactatttttttaatgcga
tctttactttttttagaggtcttttaacccatccgactgaatcacaaagctgctccaatt
catataaattaatattcttattgctactcagatagatatctttaaattcgattttatcgc
aagttttatctaaaacaataagtttttgaaatttttttcagagctagcattgatattgg
agttttgaaaaaaatttttccagaagatcataatgattatgatataatta--aataaata
tatatattacaatattgctttaacaaacaatatattaatatatgctcaaatttttacatg
aatctattaagatagaagattgttaaaatatagagaaataaaaattttttgtaaaatcttt
caataaaatgatataaagagtgcttttttaagactaacatctattattataaataaagtct
cgaaattgagatgaaataacacattattctatgtctaatgccttttaaaggagttatttg
tgaaaaaatctatgtgtctaacctgtttacttgctttactaattatgagcaacccaataa
tagctaacgcagaagtagctggattggtgccctgcaaagattctgcggcattcaataagc
gtatggtaaatagtgtgaaaaaacttcaagccagattagctaaatatgatgcagacacgc
caccagcgttagctttaaataagcaaatagagaaaactaaaactagatttgcaacttatg
gtcgagcaggtttgctgtgtggtactgatggattgccacatttaatttctgatggtcgat
ggagtagagccggggactttgtttttccaggactattattcttgtatattacaggatgga
ttggctgggtaggtagaggatatcttttatctgttgctaagactagtaagccaacagaaa
aggaaatcattttagatgtgccattagctatcaaatttatgtcatccggatttgcatggc
cgctagcggcttggcaagaatttagtagtggacaattaattgcttccaatgacgatatta

ctgtttcaccccgttagtaaaaaatttatatgaataataattttaccaaatacttatcaa
cagcacctgtaattggtgtattgtggatgacgtttacagccggatttattatagaattga
accgctttttcccagatgtcttatacttttactt----ataaataa--gatcgtataaa-
--aaactacataaatatagagagatactattatttgtgtagttttta---atatataaca
taattattta-tttttaaaaatgtttatcttttcgattattttaatattagatcttacat
aataaaata-atatcaatattatgagcttagtaacccaaattatagttaatgccgatgat
gaattaagatatcctacaattggagaattgcagtcaattcaagactacttaactacagga
agcaatagaattagaattgctactattattagagataaagaaaaggagattatacagaaa
gctagtaagcaaatttttcagttacatccagaatatatagctccaggtggtaatgcagca
ggttctaggaaaagatcgttatgcttacgtgattatggttggtatctcagactaattaca
tacggagtattagctggggataaagattctatagaaacaattggtattataggagtaaga
gaaatgtacaattctttaggtgtacctataattggaatgttagatgcaatccagtgttta
aaggaagcatctttagaaatgcttggtcaagatgatattagaatcatttctccttatttt
gattatataattcgtggaatgtcataaatcatagcttcaattaattagttgaataaaagc
ttttataatgttataattacaataagcccgagagcatataattgtaaaaactaaatcttg
tcagaaccggaaggtagcagcaataatgtttcaatacaaaagttatga-ttttcgggtgt
ttattgtttttatctaatagtctcgacttttaattagtatgctatgtataaatctgtata
tgacatgtaatccatcaaaccttccagcaaaaataatttaataatttgatattaatacgt
aaaacacctatac-tcttttataaaatttctttatatctcatctaaacaatggaaactta
acttaatgggtgttcatattttatcaacaggttcgtctgtcccaaatttttctgtagaga
atcaacaatttgaagacatgatcgaaacttccgaccattggatttcaacaagaacaggaa
taaaaaaaagacatctagctccatcttctacttctttaactaaattagcggcagaagctg
caaacaaagccttatatgcagctaacttaaaacctactgagatcagtttaattattttag
ctacgtctacgcctgatgatttatttggtagtgctagccagttgcaagcggaaataggtg
caacaacatcagtagcttttgatattacggctgcctgctccggttttattgttgctttag
taacagcagctcagttcattcaaactggttcctatgacaatattttagttgttggagcag
acacaatgtctagatggattaattggtcagatagaactacctgtattttatttggtgatg
gtgctggagcagtagtattggggcaaagcctcaaaaatagtattttaggctttaagttat
gtacagacggtcagctaaacagtcatttacaattaatgaataaacctgtaaataatcaaa
aatttggtggtacagaaattcctcatggaaactataattccataacaatgaatggcaagg
aagtgtacaagttcgctgtatttcaagttccaacagtaattagacaatgtttgaataatt
taaacatttcaatagatgaagttgattggtttatattgcatcaagcaaacactagaatca
tagaagcaattgcgagcagattatcagtacctttttctaaaatgattacgaacttagagc
attatggaaatacatctgcagcgtcaatccctttagcgttagatgaagctattcaatcca
ataaaattcaaccaggtcaaattattgttttatctggttttggagcaggcttgacttggg
gagcaattgtcttgaagtggtgat---ttatatattgcggatgacgagactcgaactcgt

aaagctttcgctacacacccctcaagcgtgcgtgtataccaatttcaccacatccgcatt
ttttaccatgaata-ttaaaaatttataatatttatctaaatattcatataa--tgtgta
tatatatatacattcttgtcagaaactctcttatattttattttaagtgataaaattgag
t----ttagtatagatagtaaaatttttatttaaaatcataa----aaataaatatataa
ggaaaataaaatatgacaccatctttatcaagtttttttgaatagtcttattcttggggca
gtaattgtagtcgttcccataactttggctcttttatttgttagtcaaaaagacaggaca
atccggtcctaaaaataacttagaaagatatggagaagcaatttaagatgaataatgaaa
tatttttcatactaataagctctccatatattgctatact--ttagaagaatttttattt
ttttattttgtaacatcctgtaaagtttattccgttctttctttttccc-atttactctc
ttataattaatgatacaaatgtcagg-ttaataatag---tttttttattctgtatctttc
---------------atttatatacgtattattatgttcaattaataaagaaaagctca
tgtctttgtctaattggcctctcaaaaaagaaaattctgaagcatataatattaagaatt
caaaacaaatcacaattcctgatggtttatggataaaatgcttcgactgtggtctattaa
tgtattctaaagtactgaagagaaatttaaaagtttgccctcaatgtagttatcattttc
aagcttctagtaacgaaagaattgatcaattgatagaccaaggtagttggcaaccaatgg
atgttcacttgatctctacagatccattaggctttaaagaccaaaagctttatagtcaaa
ggttaaaagatactgctttcaagactggcctgcaagacgcagttcaaacaggtactggga
ctatgcaaggtaaaaaagtatgcttaggtattatggattttagattcatggggggaagca
tgggatctgttgtaggcgaaaaactaacaagactgctagaaaaagcaactcaagaaaagt
tgcctgcaattatactttgtgcatcaggcggagctagaatgcaagaaggtatgttgagct
taatgcaaatggcaaaaatttcttctgctctagaaatgcataaaaaagaaaatctactat
atatatctgttttaacctctcccacaacaggggggtgtaacagctagttttgctatgcttg
gagacttaattattgcagagccaaaagctcttattgcatttgctggtagacgagttatag
aacaaacaatcaaagaagacttgccagataattttcaaagttcagaatatttatttgaac
atggtttcctagatttaattgtatccagaactcagcttagatcaaagttaatacaaattt
tgtccttacataatcatagtaagtaatgaattgaatattgactataatatcaaaatgcta
gtaaaaattaagatattaagaaattttcactaaagatatttgtagactaactagaattac
aattactattgtagaaaaatttgtcttaatgcca-ttttttcagtcttttaacaaattg-
tttttatgactttaaataattgttagtttaatacaaatttataaagagaata-----cat
actaagctaaatatagtaatattatgttttcctataatactgacctatcaaatttcgagg
aaatttatgcttaaaagatcttcttggcttgcggctttattgggactattaacagtagtt
tctacaagtacgcatacatatgccatagagttagacgaggcaacaagaactgttccatta
gaatcttctggcagaactgtaattcttacaccagaacaagttaaaagaggcaagcgatta
tttaataattcttgtgctatttgccataatggtggtatcacgaaaacaaatccgaatatt
ggacttgacccagaatctttaggattagctacgccacaaagagataccattgaaggacta
gttgactatatgaaagatccgactagttatgatggtgcagagtcaatcgcagaattgcat

ccaagtattaaaagtgctgaaatttttcctaaaatgcgcaatctaaccgatgaagaccta
tttacaatcgcaggtcatatcttacttcaacctaaaattgtttctgaaaagtggggcgga
ggaaaaatttactattagaacttcaaagacctaaaccttgtgaatactagtttatgta-t
ttgttgcttaagtgtgattattttgttagatcatgatatatattaaagatggagtttgtt
gacagcaagttacttttacccgaacttgtttacactattcgtctagttattaaaaagga
gacgttaaattgaagaagaagctttcagttcttttcactgtttttagtttttttgtaata
ggtttcgcacaaattgcttttgctgcagatctagataatggagaaaaagtttttttctgct
aattgtgcagcatgtcatgctggcggtaataacgccattatgccagataaaaccttaaaa
aaagatgtacttgaagctaatagtatgaatactattgatgctattacttatcaagtacaa
aatggtaaaaatgccatgcctgctttcggaggtagactggttgatgaagatattgaagat
gcagcaaattatgtattatctcaatctgaaaaaggttggtaattatacttgattttatcc
tgtattaaagaatagacaatctatttagttgttcattagattgtctattctttgtttatg
---ctattatataaagatatttacacaatatt----ttatgatgaaaagaatacccgcaa
ttcttgtactagaagacggtgcatattataaaggatggtcattccagcaagataaacaag
agattactattggtgaagtagtttttaatactggaatgacaggatatcaagaaataatca
cagatcctagttacttccatcaaattgtcgcttttacctacccggaaattgggaatacag
gtattaataatcaagatattgaatctcacagtattagtattaaaggacttattgcaaaaa
atatttgtaaaatttcaagcagctggagagagcagcaatctttagttaagtatttaagta
gtaataatattccttttattttcggaatagatacaaggtctttaacccaatacttgcgtc
aatttggtacaatgaacggttgtatctctactgataatttaaatcatagttacttaaaac
agaaaatttgtgagattccaagtatgcaaggtttagatttaatcccgcatgtaactacaa
gaaatgtttacccctgggatgaaaaaagttttccgaattggtatttgacagataacatta
gagtacaccgagttattcagttgaaagttattgttatagattttggagttaaactaaata
tactcagaagactagctacacttggatgtcagataacggttgtgcctgcccacactcctt
taaaagatattttggcttaccagcctgatggtatattactctctaatggtccaggagatc
catcagcagtacattacggcatccagacagttacaaatttactagattacaatgtgccta
tatttgggatttgtatggggcatcaaattttaaatttagctcttaaagctaaaactttca
aacttaaatttggccatagaggtattaaccatccatccggattgaaccagcaagttgaaa
taactagtcaaaatcatggctttgcagttgaattgacttcggtttttgaatctcctgtaa
gagtgactcatttaatctaaatgacactactattgcaggaactggacataatcaaagtc
cttatttttctgtgcaatatcatccagaatcgagcccaggccctcatgatgctgattatc
tattcgaaagttttatagaaataatgacaaagtccaagaataaagttag-----------
ttagtaattttcccatgctttatgagtgaataaagcggatagtacttgtactcctcgtaa
ttgattaaagagaggcaaatgtcctctggggccatgagaattccattgaaattcctcagg
atatctgcaaggaattctatcaacttcccagccgatttttttgccagaattttccccaatc
ttttccaacactaagccatatttgtctttgaacaaatagcccaaatttaccttttgagtg

tgtatgccataacttatcaatagtttgtaaatcttgagcaggtattttttttatatctgt
aaaatatagccaattacgagtttgtgcatttacaccagctagttgaataagcttttgttg
agttagctgatctgctttgagcaaatcccgatgagtcagtaacatttgtaaatctttgta
attcatctgttgagcagagcgcaatggaacaatgccatcgggacataagttagaagcaaa
tttcactatttcttgatttttactgttaagtaacttttcataaattaaaccgtcaacaca
attacttttatagttaggcccagtaatcctctcaaaaaatagatctgctaaatcttttaa
ctcgatagagtcgttgctatttatattctcaataatttcaagttgttgcttga---catt
attagacttagtattcttattcaattctaaaagctgggctcggatttgatttggcatctg
atttgtttaattagttttaatagtc----ataaggtcttagagagaattagattgttttt
tgaaattagaagagcgccgcagtatttctaaagaagttcgcataatttgattcaataagt
tttgtcctgcataaattataatttgaccaattaaaaaaattgtgctttttattttaggaa
caaaaaaatcttgtacctcaagcagaaaagtaattattatttgcattggtgacaacatct
gtaattcattcgatctgcaagcatatatatactgacagtttaagccttcttgataaaatc
cccatactttgtaacgactttcgtaaattgctcgtggtcgctcaagatagtttttaatta
tagtattccaaattaggttattctttaatttttctaattttctatcagatagaaatttta
cattacagagataacttgcatttgtctcatttacagtattaaaattttgaactaacatat
gagcaataatgttgcttagctgaattaaataattttcaagaaaaatttcgacctgtttaa
ctggcatattgtaagtaaagagattaaaagtttattcacagtttgagatgacccaaaaa
gtagctgaataattaaaatttgtagtagcattttatagtcgttcagtagatagttcaaat
catgaatatcttggtaaatgtcaatacagtccagttcatataacttacaaaatcttttga
tacacctatggaataaatcaattaaaatattttcggtcaagtttgtaatatcatttacat
ctaaatttgaattataaatttctaaaagtatcttttccagttcagctaatatgattttta
ataagtttctttttacttcagtgcgaaaaacatctaaaattaacacttcctgagagcaat
tagttaatctcttattaatctttatagaagttctaactaatagctctgctacctcttgat
tgagtgttggtccctgagagcttggccaataattattcacgttataccatagtaagaaat
attgaatttaatgttatacgataacttttaatatactgaaaagcaagttaattactcta
tagcaactag-------tttttacagatcttaatagttatttatcaaataaagttatatg
atagtagcgattgagctaattttattaaaattta----taataatgacaacttattactt
cgcccttgcgagtcaaaattttctattagtacaagaaccactagaagaagtttttagaga
gagagttaactactatcagtcaaataataaagcaattgatttttggctaataccaaaccc
ttcttttctagagaaaccagaaatgatttcatttaaaaaccttgtacctaaagacgctgt
agctataatctctactaatccaatatttattaattggttaaagctaagaattggctatat
ctgtattggacaatttgaagataacctacaactttctgaagaatcgttaaatattactgt
tttaacagacaaaatttaacaatgcttttattatgactagtgttaaattattttggttaa
gttataattatagtcttttagtattta-gactcttatagtttatcacttagtattttatg
ctgagatggagtgacaaatagtgtaagtatttcttggctaaacgtttcagttgccttaga

tttatatctatttggattaactataatggatagcattcgttttatagtaacattttcaat
ttgagcccaatgtacaattccaagttccaattccttagcaatagctgaaacggaaacaaa
ggcagctcctagccctgattgcacagcgtttttgatagcttctatagaattcaattccat
ctctatttaaagcgactgctgtcaataccatgttgactgagtactttatctatcacttt
tctaattgttgattgagtatctaatgcaataaatctaagcctgtataagtcttcttttg
aatgtctcctagtttagaaaaaggatgtgatttgggtaatataagtgctaattcgtcttc
cgcataagaagtaacctgcaaaacatcttgcaattcagtaggcacttctcctccaataat
tgctaaatcaacttgaccattagctacgctccatgaaataagtctagttgaatgtacttg
caattgaacagccacttgtggatatctttgcctaaatagtccgattaatcttggcatcaa
atatgtcccagttgtctggctagctccaataattaatgtgccaccttgtaagttttgtaa
gtcgtcaagagctcgacaagtttcttcgcagagagctaaaattctgcccccgtatcgtaa
aagaagacttcctgcctcagttaaagttgccttcttattacctctttcaaatagggaaac
atttaattggcgctctaaattttgaatttgcaaactaatagctggctgagaaacatacaa
gctattagctgctttttaaaactgccctctttggcaattgcttttaatattcttaactg
atccaatgtaaatggaaggtctgtcattaaagaattgtagtataatg---------tata
ttatttagtattgatcaatgtatgcatttgccaactaaaatatttgtcattacgcttcat
tattttcaaaataataaaaagtataatgttaaataagctgtattaaaatttagtatctgt
ttgatattaaatttatacaagagatatcgtgaatttaattctaactataaaatataggag
acaat---atggactctagacttttagttgtactaataccagttttagcagcagcatctt
gggcagtttacaatattggtagagttgcactacagcaatttagaaaaatgacatcttagt
tttgcttagaatata-atttttagactttatgagatagaggactttaaaatccctatctc
tattgtatatcttaataaaaaaag-aaagc--ccccttgtacttactgttattacacatt
aatattattggt-ggtaccta----gataataagtttaagctaacttatttttatttta
taattaattaaattaataacacaattatggctgttccaaagaaaagaacatctaaagcta
aaaaaaatgcacgcaaagcaaattggaaaaatcaagcaaaaactgaagctcaaagagctt
tgtctttagcaaaatcagtattaactagaaaatccaatggatttatttataatctaactg
aagcatcagatacttttagcgattagaagtgttt-ttactacaagtctgaaaaagtatta
cgcaacaagttctcagtaatttttatctat----taatcatttgaaaatcattctatttg
ataataaaataaatagcagattaaataaaatcacattaacaaattatgcagcaaagctag
atcaaactagtgaaatttggttgttcaattgtattgaaaatattcagcatatttttttta
aaagtcaattaaaatcaagtcatattactaaaattttttatttctggtactagctttgaat
ataccgcaggtctaccaggattattatccagcttaacactaagtggtagactacatccta
ttagtatatatagtccccagtctctcaaaaagtatcttgaagcatgtaccaaatattccc
aaactaatttttcttttcctattaattttcataatttacaatacggaggacaagttgtta
accaattttatacagtaatttgtttaccgttgagcaaaaagagcctgctgtatggattta
ttattctaaaaaaagaaaagcagggagtatttaatttagcacaagctaaaaccttgaata

ttcttcaaggacctatatatggaaaactcaaagaaaaggataatttttttaagtccagatg
gttactatctatccggccaagacttctcttctaatacaataatgggacataaaatatcgc
ttccgttattagtcagatattctagaattatttctgagatgcattggttttgctcttatc
ctattagattaaatacttattcgcatcagcaaggagtaaagtgtttaccacataatgttt
taactgatattatgaaatcccaaatatatcaagataacagttttgttgaataatacttat
tcttgatatataatggtctcggtaatctaaataaaatactagaaataatttcttaaattt
atttgtctatattaaatttacaatcatttatgacatacgcaattattgaagcaagtggca
cgcagctttggatagaagaaggccgctactatgatttaaatcatatacctgttgatccag
gtcagtcgattatattaggaaaagtcttattattgaataaaaaatggggaggttactttag
gacgcccttgtatagaaggggttacgataaaggctacagtaatgaggcacttgcgaggaa
agaagataactgttttcaaaatgaaaccaaagaagaaaatgagattaaaaaaaggtcatc
gacaagaattgactcgtttaatgatcgattctataacatcttaga------taaaatcta
gactcatattttt--aa-aaactttttaatatttaactaaaatagatataatggcacata
aaaaaggtagtggtagtacaagaaatggcagagactctaattccaagcgtttaggtgtta
aaaaatatggtggagagcaagtaacagcaggtaatattttaatcagacaacggggaacta
aagttaagcctggccaaaatgttggaaaaggaaaagatgatacattgttttctctaattg
atggtttcgtgctgtttgaaaagtcaaatcaaaagcaaaaaacaattagtgtttattctg
ctaagaaatagttaaaacaattggtgcaagataacttgagt----gtatttatttcactt
acagtggcaccatatatttgatttattaagaaatggagttccttaatgaaccaatgaatt
ttattaaagatttgctgtgaaatttaatgacaaacactattgtaatttcctgtctacaca
atatggctgctattctatattgtggtcagatacagaaactagtagtagcaaatgctcatt
atcaagtaagtgatatctacctaggttgtgttgataaaatcttctcaggaataaatgcgg
catttattaacctaggaaagaatgagtacagtggttttatacatatcagtgataccggtc
cgcttaaaaagaaatattatgtcaataatattactaacattttaacaatacggcaaaaaa
ttttagtacaaatcattaaagagccaactttaaataaaggtccaaggctcactgccaata
ttacattatcaggtcgatatattgtattaatgccttttagtcaatcaatctgtatatctc
gaaaaatatatgatgaagatgagcgtcattatttgaaatctttagctattttaattaaac
cggcaacaatgggcttgctatttagaccttctgctgtaggtgtcgatgaggaaataatat
taagcgaattaaaaaatctaaaagaacaatggaactttgttcaaaaatctgcaattaata
gttattcacctgttcttctatataaagatgaagatattgttaaaaaagtaatccgagatt
tttataataataatacaaacaatatagtaattgattcaaacctgggattaaaacaattaa
attattatatccacacttggcactgtaataactctagcacagttcctaagattaagcttt
atagtaataatcaatgtatactagatgcttttggtatcaatcaggcaatttccagagctc
tcattccaaaagttgatcttatacttggtggctatatgtttattgaaactttagaagctt
ttactattattgacgttaattctggatcttttaataattccactagtgcacgagaaacag
ttttaaaaaccaactgttctgcagcaacagaaatagcttatcagttacaaattagaaata

ttactggtgtaattataatcgacttcattgatatggaatcacaaagagatcaattgcaat
tattagaacactttaataaagagctatcacttgatgatgctaaaccacaaattgtacagt
tatctgaattaggtttagttgaattgactagaagaagacagggcaaaagtttgtacgagt
taattagtagtgattctaattactttatttttcacacaatcagagagatctcagtctc
ttaagagattcgatgatagacagcagaaacaacagattttaataaatcttggctatctg
cagagattaatactattaacaaggtctttttcaaaagtcaaatttgtgcagacctgcta
acttttacctaattcgtaatctctatatagttaaaagtagtattacttataaacaaaatt
atttattaactcacagatcaaaacttatttattctaaagaatacagcaaagtattaccaa
gca------gttattatctagctagtctcaataagaatagtaatcaagagttcctatctt
aagattgttatatcaactgtt---------------------------------------
-------------------gaaaaaaagaagctctctcttatataagagagagcttc-tt
ttatagcttactaacta-tctagttagttaattaaaaattaaccgttaacagctggagct
gttagagctactggtaaagattcaccagaagctagatctagagggaagttgtgagcgtta
cgttcgtgcattacttccatacctaggttagcacggttgatgatatcagcccatgtgtta
attacacgaccttggctatcaacaacagattggttaaagttaaaaccattcaagttgaat
gccattgtgcttacagataaagctgttagccagataccaactacaggccatagacctaag
aagaaatgtagagaacgagagttgttgaaactagcatattggaagattaaacgaccgaag
tagccatgagctgcaacgatgttataagtttcttcttcttgtccgaatttgtaaccatag
ttagcagattcgttttcgcttgtttcacgaattaagctagatgtaactagggatccgtgc
atagcactgaacagagaaccaccaaacacaccagcaacacctagttggtggaatgggtgc
attaaaatgttgtgctcagcttggaatacaagcatgaagttaaatgttccggagatacct
agaggcattccatcagagaaactaccttggccaattgggtatactaggaatactgctgct
gctgctgctactggagcagtaaaagcaacggaaatccatggacgcatacctaggcggtag
cttagttcccactcacgaccaatgtagcaagctacgccagttaggaaatgaagaacaact
aattggtaaggaccaccgttgtatagccattcgtctaaagaagcagcttcccagattggg
tagaagtgaataccgatagctgcagaacttggaataacagcaccagaaatgatgttgttt
ccgtatagaagggaaccagcaactggctcacgaattccatcaatgtctactggaggtgca
gctacgaatgcaatgatgaatacagatgtggcagttaatagagttggaatcattaataca
ccaaaccaaccaatgtataggcggttttcagtactagtaatccaagagcagaaacgttcc
cacaagctagcgctttcgcgtctttgtaaagtagcagtcataatttttatcaatttta
aggatttaacaaagatgcttcccaaacaataagttgtctgttatgaaaattattacttaa
ggccttttcaaatgagaacattttaaaaaaaaca-tagcaaaatctaatctcagtaact
aa-aattt-tacttgttcatatattgcttaataattcaa-----------taataaattt
tt-gct-------------ttttgcaaaattaacactattttagggat---ataataata
atgcatattaatgctgcttattttgattttaatagtttatataattttgccctacttaaa
tacgcatgtcacactttacaaaaattcagacgaccatacaagatttaaatctgttaaagc

atgctttaacagatttgggtttgatttggcaaatgaattctagtcatataaaagttggtg
aaaataatcagcataaggtagatattttaattaaacaagataacttgtcacatattggct
ttacttggaatgataatagatatcacttagttgccgacttacaactttgggaacaaccct
ggtctctagaagtatttttggataagttatctcagaagtatgcttactactctatcatcg
aagaaacaaagaagcaaggctttgaaaaaatgcagcagatttcacaaaaagatggatcaa
ttaagttaattgtgcaacgctggaattattaactaacatacaaatgcgggcggagagact
tgaactctcacgagattatctcactagaacctaaatctagcgcgtctaccaattccacca
cgcccgcactgttgagac--aatattattctatcataatctatactaaaacacaaactaa
-tagtataaactagcttccaagcttaaaagcgtctacaaatagaccgtatgttataccaa
ttttgaagttgttgaataaattaatttttatatttaattgcttgtggctagagtagacta
ttttactagtcaactcagttgctgcaactattagactggctgcaataataccccaatctc
ctgtttgtccaggaatcgtagacaaacctgttgaaataaaaaatcctagcaataaactaa
ttaagccagttgttaactcgctcaaagaataataaagcttattatttaagtttttttatta
agtaagcaaaaaaagttgaaagtctagttttaatcatagtatttagtaacggagttttaa
aaggatgacaagtattaacag-tcattaatattgtcgtatccttttaaaactcgttaata
caaacagctgttcttgt---------ttttaaaggtcttgttcgcttaaagcattaataa
gatgctgaaacggctcttgtataatatctgtgcgtttaattgattgcaaatctaattctt
cttctatgatttcttgtaataactgtatgcttcttactgttggagcaacaggtacagata
gtgaattgtaagtatcttttagcccatttaatacacgctgatcgagaatatttgtactac
cagaaaccaaggcataactagcataccgtaggtaatattctatatcccttagacaagctg
catatcttcttgttgtataagaatttcctccaggccttaataattcaggctgttcttcgt
acaactgagctgcggcctcttttaaaatattggtagcttgatcattaataatttcagcta
tttttattcgatctagtccactcgaaaagaaagattctagttgtccgacagctgtcttat
ctagataacgaccagtaagatcgtaacgatttaatattgctgttatagcatcttgcataa
attaattctcctac-ttgctatcagagcggtattataaatgactaatattgtaatataat
cattagtgtttatgaaatttatttaaaacttattctttattgcttaatgtgcaattaaaa
actcgcatagattttacacatatagagtatacgtagtttttaatta-tttttttataaaa
accaagaaataaagctttttt-taagaggttcgtgaatttgttaatataacatgagcttt
cctcctaaataagttttgtaaagtac--aaagtaaaaaaacgattagtcaataatagtaa
------------------------------------------------------------
------------------------------------------------------------
------------------------------------------------------------
------------------------------------------------------------
------------------------------------------------------------
------------------------------------------------------------
------------------------------------------------------------

------------------------------------------------------------
------------------------------------------------------------
------------------------------------------------------------
--------------------------acaatattaatatgtatcaa--------------
------------------------------------------------------------
------------------------------------------------------------
------------------------------------------------------------
------------------------------------------------------------
------------------------------------------------------------
------------------------------------------------------------
------------------------------------------------------------
------------------------------------------------------------
------------------------------------------------------------
------------------------------------------------------------
------------------------------------------------------------
------------------------------------------------------------
------------------------------------------------------------
------------------------------------------------------------
------------------------------------------------------------
------------------------------------------------------------
------------------------------------------------------------
------------------------------------------------------------
------------------------------------------------------------
------------------------------------------------------------
------------------------------------------------------------
------------------------------------------------------------
-----------------agtataaataagcttttactccagcatcaaaaaaaatatttga
gatatgttgatacaactgattattcaagtgaaaacttagatttagaccaatttagctatt
ttatcattacaattaataaaactgataaatgtattcttatagagcaattttgttataatg
ctgaaggccatatttactcaatattcttcaaaggtccaacagctaaacatctttcttcta
taatttgtcattttcaacaattaaatacaataatatctacagctcatgccatatatttag
gacgagagctaatgaagtcagaattagctctcgtgctagatcaacaatatattcaagatt
aaa--ttttgtaagtattaattatatttatgctagtataatattagtaagggcatgtaac
tcagtggatagagtatcagattccgattctgatggccgtgggttcgaatcccgccctgcc
c---gagtactcagagtact-atactaaaacatct-------taatagctagtttgataa
tagtatgggcattggggctgcaaggtttctacattatgaaaagaagaaaatatg--aaaa

aaacaagctctccaaaagagcttttagtacaatcaataaatgcagaaaacaatattgttt
ctttttctcgaaaattagcttttgcttaaacactgttaattttggcattcgacatgtta
aactcttattcatggcgaatattctgtcagagtcgctcttagtttaagaaaagtcagaaa
aaatatttatctttacgtttt-tacctattttatttaaataaaggttagta-ttttcaa
cttttatagtggacgtgggttcgagtcccaccagctccatcaatttgatt---------t
gtataattagctagtttaaccttttaataaccagatatataattggaggaaatatgggt
ttcataacaattactaagcccgctttaaagcaaattgcaatcttgaaaaatgatcatgaa
aatgatgtacatctaagaataggtgttagacaaggcggatgttcaggtatgtcatactca
atgaattttgaacatgttgacaaattaaaagatacagatgaacggcttcgtctagataat
ttttctgttgtctgcgaccctaaaagtcttctttatctttatggattatcattagatttt
agctcggaattaataggaggaggatttcaattttctaacccgaatgccagccaaacttgt
ggctgcggtaaatctttttcaggctgacc-ctcgaacacgtttttatatattttgtatt
atatatattattttagcaagtcg-tttattagagacaagcaatgctctgttaaaaac---
---------------------------taatttcaagagttttattgtattatt-----
----ttatatatttattatcaactacttgaaaataaattagctaagcaatctaatttgtt
atcagtaccaaactttaaatgattaacttttattgatatatgctaatcttt-atcagcc
taaacatatctgtacattacaatcttttgacataaaaagagtcttagcatacctctcaa-
----------------------aaataaattcaa----------tattttataaatactc
aacatgaatctgtcacaacttatcgttaattcttctatgatgaaaattcctaataacttt
tattgcaatataaaagctatttcttctgagaatttaagtcacttacaattatcaaaaaca
aatttattatctcctacaaaaataattgcttttggactacctaaatttgattttatttct
acacagattacaagttctattagagaaacagttgttgtgtctactattgatgataataat
ctttttgctgccccaactaatgattggataaaaaaacacaaggataaaaatcatcattct
ttgttttataaaaaagtttttaaagctttgctaaaccataaaattagagttgttgctgag
ccact------tcataactccgaaataaataaagaaaagagcataatggcaagtcaaaga
tacatctggggtaaaagctggaagccttcaattattttatcttgcttaaaaaacaaaaaa
aatagcaaaatttctgatgcatcgaaaaaatttttaactgaacagttatctgcatcacca
gtatttgtggtaaaaaatggttttaatgaaattattttaggtcacccattatctcgtgtt
aaaagaggaagagttaataatctgatgcatgcgttttctaatttattaaatcagtctaat
gcaacttatcctatatctactggtctattctttttcatccagatgatgcatttgaattt
aaagactttataatatcggtcaatccactggcggccaaacatatggagataagcgtagag
cctgttggtctacactttgcttataaaatgaatagaaatatatcgtcagatactcaattt
tcgtttattccagattttaaagaagtaggagatttattattcaaatatagaaaaggtaat
catttagtttttcataagaatcagcattatggcaaagatttttttcaggggcaaccaata
tatatgattcagccgatcacttttaaagatcgagccggaaagctaaatattatcaaattt
acggggctaaatgacacaagagaaattatttttactaatcttgaagctgctaatagatca

tgggcacattttataaagaggaattcacaattaaaatcaattaaaaatcctactctatta
gtctataatttagaaagtttttaaaagatcaagaacgattggataatcaagatttaaat
aagtttgtggtgatcactaataaagaatcttatcttgccacaaaagagttaatagcttta
cccgattctaatagcttttctaagcacttaaaattaaatataaagcctaaactcttttt
gtgaaactatgggtaagacggttattttccaccttaacttatgaat-aaatgctatt-gc
ctttattttctcgaataatattctataactagtaattcattgagttgtaaagcaacccat
tccctatcaatgactccattgactttccctgataaattggatttattcaattctaagtgg
ctaggaatattagctagtccagggaaagctaagtaattttctactagttttcgagatgct
tcttgatttttacactaattgattctcctggtttacattgataactacagatagatact
acttgtccattaatacaaatatgaccatggttcactagttgtctagcggcaggaattgta
ggagccataccaagtctaaaaacagtattgtctagcctcatctctagtagctgtaacaaa
atttgacctgtcgacccttgtagtttttagctgcttttacgtacttaaataactgtttc
tcacttaatccataattaaatcgcaacttctgtttttcttctaatcttacagcatactca
gaaggttttctggacttttgtccatgttctccaggtggataagatcttttaatcgctttt
ctactgagtccgggtaaatcacctaatctacgggaaatgcgtactcgtggtcctctgtat
ctagacatattatttaaattctcctaatgtaaat-gattaag----------taatgagt
agctatttttaacacaaaaagagcattaaaacaaagc-ttttgctaaaaaatatttaca
aaaaatataggtttggacaaaa---tacattactatgc-tttttaagcgggtagcgggaa
tcgaacccgcatcattagcttggaaggctaaggttttaccactaaactatacccgcaata
ttcaacagattgtcactaactcaatataacatagttatgttaagttaagataatcttaaa
tttaatagaatt-ttaaattatttatgctccttctaaaaccacatcaagaaaacgagcaa
tttcggcagcttgttcttcaacttctgataataataatggttgaccgaccctagttaatg
gaattactcttttatcttttgtacataaataaatttcacgtcttggatttagcccctctt
taatatctatttttattgatttgatttcttttatattaaactgaagacagatttttctat
tctttccagggaaacctagccgaaaaatttttacaatacctttatctttgttaaattcat
tatatccagcccctatattccaaataatagttagccacagaaacaaactcagaaagactc
ctatactgccatagaatgtcattacaataccttgtggaataaataccaaatcggttgaat
ttgtaaaaggtaacaaatcaacttgaaaataacttgatagtccagcaagaagaaagccca
aagccccaataaatattatagtagcccaccagtaattactcagtcttcgtgaacctaaaa
ttaaatcttttctaactgagtgtatagataatgtttttttcataatttgttcttttaaaa
cagta-tactaatgcaaaatggaaaaattttgacaattttt--atgcgcgctttagattt
tttaaataagcttgattgcttttagtcctaaaaacagtcctgttgctaacccaaaaaaag
cagcaagaaatattatatatcctagaaaaactgaca---tattattatttattattattg
ttacaatagtgtttacaaaaatagtgaaaatctattactacggtattatattacgttaaa
tgccttgctgtctacaggaaaaaactttttcagtttttagcattgtgtttatattagaa
ataatattctagtaaatttattaaacttctggagaacagaatcttatgtcagaatttatc

aagccttataacgatgatccttttgtaggcaatttgtccacgccagttagtacgtcaagt
tttagtaaaggacttctaggaaatctaccagcttaccgtcgaggtttatctccgcttctt
agaggattagaaataggaatggcacatggatacttttaattggaccttttgataaattg
ggccctttacgaggtacagatgtagcgttactagcgggatttctatcttcggtcggcctc
attattattctcactacatgtttatccatgtatggtaatgtatcttttactagagcagat
tcgaaagatccactacaaacttctgaaggctggggacaattcactgcaggatttctagtt
ggagcagtaggtggttcaggatttgcttatttattgctagctaatatacctgtattacag
actgcaggtcttagtttattctcttaagctagtaaggggacttgaacccgtaacctactg
attacaaatcagttgctctaccaattgagctatactagcattcagttcataataacatat
aaagatatttgttgcaatctgctattttcagatgtaacaaatatctttttatttaaaca
gaactgttatgttaacac-------gtatattaattagaatctgaattatctgactctgc
agaagattcctgatcacagtttagataataacaaattgtttgatctagacaagttgcaga
ccaaccaacaggtgtttctcctgataattcttgcatatctacagaattataaaaattatc
atttggcttattactgtttgcttgcataaatatcactcccaaaacactctgaacttaatg
tgaagtaatttaacttctcaataatatattaacacaaaatattaaatatggtgattattc
tttaattgtaaagatgctgctaactttataaagtttgagtgagagtcttgatagcttttg
ttgaaataagcattttaacccatttattttgcccgctcgaccatactttttgtgttgta
aattaacttttgtagtttttttgttcgtttatgagaatgagatacggcatatccattat
tagctacttttcccgtaagctgacactttttgacataatttt-------ctaacttata
tatttatttagagtacttgctaaagttgacttgggcacagctccaatcactgtgtctact
ctttctcctgctttaaagatcattaaagtaggtatacttctaatgccatattcagccgca
atagtaggattatcgtctgtgtttatttttactaccttaatagatgattcatattcttcc
gcaatttcatcaactacaggagaaaccattctacaaggaccacaccatggcgcccaaaaa
tctacgagtacaggtaagttattgttaataacttcttgtttgaaagaggcatctgtaact
tgagatactgacatattctttaacctttaatatatgttccttgctaagcaaatcttatca
caaatttcaagaaaaatcttctatagtgaattactattagtgttgaattattttttaaa
aattagacattaaaaaaacttattacctatataaataaagtggtggccgagtgatatctt
gatagtgataattttatattaaggttgaacactagaaataacttttcaagcaaactctt
atttgaaaa--atttctctgtgaccataaaaacttaattatattacaatatattttatta
tttgtaacttagttctgataatggtataaacaacgcaaaagatactgccttataatcaag
gaggaatacatgtctcaatccgtagaatcacggactaggattaaaagcgaacgttacgaa
tctggagtaatcccttacgctaaaatgggctactgggatgctgactatgtgattaaagaa
acagatattctagctcttttcagaatcactcctcaaccaggtgttgacccgattgaagca
tctgctgcaattgcaggtgaatcttcaacagctacttggacagttgtatggactgattta
ttaacagcttgtgacttatacagagcaaaagcatatcgagtagatccagttccaaacgtg
gcagatcaatattttgcttacatagcttatgatattgatttgtttgaagaaggttccatt

gcgaacttaactgcttcaattattggtaacgtttttgggtttaaagctgttaaagctctt
cgcttggaagatatgcgtatgccagtagcttatctaaaaacgttccaaggtcctgcaact
ggattgattgtagaacgtgagcgtatggataagttcggtagacctttcttaggtgctaca
gttaaacctaaactaggtttatctggcaaaaactacggaagagttgtatacgaaggcctg
aaaggcggtcttgatttccttaaagatgatgagaatattaactcacaaccatttatgcgt
tggagagaaagatttttatattctatggaaggtgtaaataaagcatcggcttctgctggc
gaaattaaaggtcattaccttaacgtaacagccgcgacaatggaagatatgtatgagaga
gccgaattctctaaagaggttggtagtatcatttgtatgattgaccttgtgattggttat
actgcgattcaaagtatggcaatttgggctcgtaaacatgacatgattttacatttacat
agagctggtaactcaacttactctcgtcaaaaaaatcatggtatgaacttccgagttatt
tgcaaatggatgcgtatggctggtgttgaccatattcacgcaggtacagttgtaggtaag
cttgaaggagatcctttaatgattaaaggcttctacaatactctacttgaaagcgacaca
gatatcaacctacctcaaggtctgttctttgctcaaaattgggcttccctacgtaaagtt
gtaccagtagcatctggtggtattcatgctggtcaaatgcaccaacttcttgattactta
ggtgatgatgtagttcttcagtttggtggtggtacaattggacatcctgatggtatccaa
gcaggtgcaactgctaacagagtagcactagagtccatggttatggcaagaaatgaaggc
cgtaactatgtagcagaaggtccacaaatcttgagggacgctgctaaaacttgtgggcct
ctacaaacagctttagatttatggaaagatattagttttaactacacttccacagataca
gctgatttcgttgagactccaacagcaaacatctagtttaatgactacttactgatactt
taaatagttaattgtaagtgaaattaacttataacaataaggagcatagaatagtgagac
taacacaagggactttttccttccttccagatttaactgatgagcaaattaataaacagc
ttgcttatatcgtttctaaaggcttatcagcaaacgttgagtatactgacgatcctcatc
caagaaactcctattgggaactgtggggtttacctttatttgatgtaaaagatgcttctg
ctgttatgtacgaaattagctcatgcagaaaagcaaaacctaattattatgttaaagtta
acgcttttgataatactagaggtattgaaagttgtgtaatgtctttcattgtaaatagac
ctgctaatgaaccaggattcttattacaacgccaagacttcgaaggtagaactatgaagt
atagtcttcatagctatgctactgaaaagcctgaaggagctaggtattaatattaattct
taattaatattggctaatta--ttcccctcttaaaaatcataattaatttagtaattgtg
atttttaagagggatagcttcccaagtaaattttgactaaa-tagattgaataattaaaa
atacacagaaataatgcaatcacaggatataatttccaacgatactcttgttaatttaca
agaagaatatgatagaacacaaatccaagaagttttaaatgagttaaatcaagaacttat
aggattagtgcctgtaaagaccagaattcgcgaaattgctgcgctattattgattgacag
attacgcagaaaactagaactagtttctggtaatccaggattacacatgtcatttacagg
tagtccaggaactggtaaaactacagttgctatgaaaatggctgatattttgcacagact
tggatatataaaaaaagggcatttgttgacagttacaagagatgatcttgtaggtcaata
tattggacatactgcccctaaaactaaggaagttcttaaacaagcaatgggaggagtttt

atttattgacgaagcttactatctatataaagcagataatgaaagagactatggctctga
ggcaattgaaattttattacaagtaatggaaaaccaaagaaatgacttagttgttatctt
tgctggatataaagatagaatggaaaaattctacgaatccaacccaggactctcttctag
agtagctaatcatgtagacttcccagattatacttcagatgaattattacaaatagctaa
aatgatgatagaagaacagcagtactgttttacagaagaagcagataaaactcttttaga
gtataccgagcgaagaatgaaacagccttattttgctaatgcaagaagtattcgcaatgc
tattgacagggctagaatgagacaagccaataggattttttgccagtggagaaaaagtatt
aacaaaagctgatttggtaacgattgaagcagaagatatcttgaaaagtagattattttc
attacctaatgcttaatatacacgtgatttcatgaaactatatattattaaaaag-tttc
ataatcagttgcaaatttcttgaaaaacctttattattatcttaatgtaggtg-tggcgg
catagccaagtggtaaggcagaggattgcaaatccttcatcccccagttcaaatctgggt
gccgcctagtactaaaaaagggggtgtggtggaatggtagacacaacagacttaaaatc
tgttgattttagtaatcgtgagggttcaagtccctccacccccatatattgatttataa
ataaaaagagatgtgtatttgtattaattgcaaccatattaccaaatgtaatacctatca
cttaatagaatctcaacataaacagcctcatcttactagaagtccattgtttatacctaa
atatcctgtagttcatgttaatatatctaacaactgtacctataatcaaattgattggga
tttagtggagtgtttatcatttgtagaaaaaccgaatagttggaatttagacgctaatta
gttacaagaaaaattatgtaccataagcaacaactctatttttttagatactcggtattaa
ttttggatgttctaaccagtaagattttacccgttcttgctatctctataatgccaaatt
tagttagtaattgttcaatagcaacaatctttcccggatctccagtaacctctataatta
aaagatcttctgcaatgtctactatcttagctctaaaaattcttacaatttctaaagctt
ctgttctagtctgagaattgatctgaatcttaattaacattagttctctttcaactgaag
gaatatttgttacatcttgaacgttaaggatatttactaacttatataattgtttcgtaa
gttgttcaatagttctattgtctccttgaactaccattgtgattctagagaccccaattt
gctctgctggtccaactgctaaacttgcgatattaaagcctctccgggcaaatagaccag
atattcttgacagtactccggcttcatcttgaactaaaactgataaggtgtgtttcatga
aaaattaaatttaatctatttgatctattaagtatattctaagaaaattacagaaaaaat
actagttagtctataactaaattatccaaactaaacaaaatcctactatcagataatata
tctaccttgtaatgctatactattaacttagtatttaatgatattctaattgcaaataga
atgct-aatttttttaaatttgttaggaacttgtgctagaaaaatacaaaaacagtaaaa
aactctctcgagatttacctcaaatcaatgatcgcattagatttccaaaagtccgagtaa
ttgatgacgaaggtgaacaactaggtatttttgtgcctgaagaagctatacaattagctg
tccaacaaggtttagacttagttgttgttagtgataaatcggacccgccagtatgccgaa
tattagactatggtaaatataagtttacacaagaaaaaagagctagagaagctaagaaaa
agcaacataacagtagtattaaagaagtaaaaatgcgatataagatagaagagcatgatt
ataaagttagaataaaccaggcatccaaatttattcaagcaggagataaagtaaaagcaa

ctatcacatttcgggggcgtgaaatccagcactctaatttagctatagatttattgaata
agatggcaagcgatctaaatgcaatagctgaaattcagcaagctccatcaagagatggca
gaaatgtcataatgctcttatctcccaaaaaagttagctaaactaattttttatttaaatg
catctggctggattcgaaccagcgacgtctctttcgaaatggcggattatgagtccgctg
ccttcggcccctcggccacagatgcatcacttaaagtttataactcataaagcatataaa
aatcaatgtttagctattttttaataaagtctataaaagtctattattcattcatattgtg
atatgtcgcaactgctgaaggggaaacttgatttaagtatcgaaatatccagtatttaaa
aattgtgtcgagtataactggaaaagttgaaataaataaaaaaataaaatctcgactttc
cggaagtcctaaatgtcttaaaattacttcaataataacttcccatccatgaggagaatg
aaaacccacaaacatatcagtaaaaagaatgattaaaaaagctttagctgtatcactaag
accataaataatttcatttaaaaaagatttaacgactgcaatttgtctttgtccagttat
cattagtaatataaagacaagtattgatacaagatcggataaaatatttttaacagcatt
tgcactttcgtttgcataatattctcccaactctctagccttcagctgcactctttttttc
aatgatttcgtaagatatatcttccgaaggattcagaagaacttcaaagtgaatttttctc
ttcaaatctttgtaattcagcaaaagctctttcctcttgagaggaatttaaaaaaatttt
aggctgttcttggttccataaataatcaatacaaggtccaaatacaaaaaaattttgaagc
ctgattaactagtactggagagatgaataataataaaatatattttacggaagtaatagt
ttggtgtctagatattctaaactcctctatagcttcagattcgccattaggatctaactc
tttacgaaacttttcaaaagtattagtaattgacctaggtattggcccaactttttcaaa
ggcgaattgattatttcgttttaaattccaatatttcatttttttattgctgctaaaattt
ctattaatcaataattgatcacgtattcactataaatattaattctagaactaaacatat
aaaacaatatttatgaaaaataaaatacaattaaaaattaattatgcattagttctatct
gcagctacagcagtttatggatataaatcttatcagccaactataaatacactcttaatc
aaagatgaagaaaactggcctttttttctctaaaaggcagactgcctttattcctataatt
gtattaaaaaaaaatcatcaaaatccctgaactcagaacctagaaatagtaagttatatc
ttaatggaattggaagtagaaaatttatcgacagaattgtcactctacttggatatttag
aaaaacaggcaaactatactgaatacttgaaccaaataaaaacaagtggttatttttctt
tagtctacctagacagtgatataatgtataagaaaaatattagtaacatttttttactac
ctaatgatattctaaaaagaatctatatttgcaataagaaaaaaaagcttataccgcatt
cattcttaataaatttatttcagaaacagataggctatccaaaaagttttgctaatttaa
attgggccttctctaaaattataaaatggtattatgatagaggataccaatggtctctag
ttgaagttaaacaagcgtctgatgcatcttctattgtaattgatattcatgaaggggtag
tgaagacaattataacagaatactacactctatcttataaaagagtctcaggtatcttat
gcgtagagtcaatagagcagtaccttggagtaagagtgggagctccactaaatataattg
atttacaaaaaaaaaattacttacttaaaagataatcaactagttggcgacattatttata
gtattgaacgatcaaataatagctcaatgagtttggatatcaaatttcaaatacaagaat

taaaagataaagagataatagtgcttgcagaaagttcttcta------------ttatct
cgcatgcatgtaatttactcaatc---aatataga---------aataggctagtagctt
ctaatatagtttcactatcaactaataaattacacgcatgctattttaattataaacttg
actatcaatataagtatataaacacctctaatcttataaaattgttagcttattctattt
cttgtaggaagacactacttattcattattttaatttacaaactttaataagctgtacaa
aaaaaaatactattggatttcagctatatttacgaaatctaagttttgggaaagctttct
gtgtgcttagtatgaaatttatcaaaaatggacttaatataaaaatttatatataaatc
cttcattgatagttgatcagaattttgtatttcaatttgctatacaaatcattaagcaat
atcatactgcta------aacctccagctt------------------------------
------tatttttaacgaatctagatcttgaacaatatgttgctgaaagtttattaatgt
accattttacatcttgcttctcaatatctgaaaaaatattattatctcgaattatgcata
cagattctttatttttcaactctgaaa------attttcattttgatgatagaactaacg
cgatcaacagttatgatacttttaaacaaaatacaaaaatatttatcaagagtttcttt
ctttattattaagtttacgttatcaaaatttaattatttaggttggcctttaaaaggcc
attttttgaaataaaatctttatacttggctccatttcaaaagagtg------------
---atttttctgatagtcgtaagacgctattcttccacaaaatgtcgttaaagcaagtat
ctaattttaatttaccagtatcttttaaaagccaccttaaccatattttagttagtacta
ttaaatgtcagtcaaatttgaatatgagaactgtatctttgctattaattgattcacctg
ctgaatatctgctatacaaatctattcttaatttctctattaaagttaggatgcagtatt
tcatacctatgagcaacaatataagattatctctattttataattatttagattgttttt
taataagatcatctcagagttgcattcacatatggcaagatctaagaactcttacaccaa
ttcaaaatttttggttaaaaaaatttcatatggagctggaattcagcttaaattaccaa
ttaagcagatgccacctctttcaattgaatatactgtaactagtagtcgctacttttgta
tttaccttcgtacttattatcaacgataactattactaaaatgaaaaatttaatttcttt
ttttgatcgcagcaaagggaaatggatttcacaaagaacaacttatgaattatctaataa
aaacatgagttcggtacagtctcagatgacaatgaaaataggcaactcattgtcaggatc
tataatacttgcatcgttaaattgggggacatttatagacaagttgcccattacgctaa
aaatcacagccggagtgaatatgataataagtttaacttgcaattcggtaatcaactaaa
taatcataaattattaacgctgtgcatagtgacagatcctagtctaattagttttaaaac
tcgctatggaagtactaccatagatgaaacatattggttcgcaacaaataatttacgtct
aagtactagtattgttaaacgattcaacacttgtgtggcagtatctttttgttcagaaat
taaagtttaa-actctatatagaagcaactaatagcctttta-ggctattagttgcttcta
tgtatctaaattttgtattaaaaattat----aatactatattattctaaatctttacga
ttgggattcctggaaggatcattagagaggaacccaaagacaaacaaagaaataaagaaa
ataacggtagtatatacaaagattttcaaagtaaacatattttcatccttaaaagctatt
agtgcg----atatttacatatctccacataattttatatcatgatataagttccagcac

tattttataaataaatcaatatctttatttttagtaatgatttatatttaaatatgcgt

gatgtactaaatactaaatcttttccgacttgttttgtcttgtttgtaaatattttgtta

tttctatggagtttaccttcaattatcacatagtctaacttctttagctgtttaaaagta

tatgaagatttttgttccaaatagttagattgattataactaatcgttttctttttaac

agtcttgcttttaacttaattatttggcttttattttcactaattctgatactcttacaa

ctaagaatttggactagaagagtacagctattcataatttataacaa------tttttta

aaaaatctactaaaatagatgattttacagaactaaaatgatatttcattatcttgttga

ccttgctcataagtaatatctttcaatacttttaatatttttccaaaatactcttgcagg

tatttttctaaagatattacttccaacctgtcaattccaagaatactgtaaacttcatcc

atcggagctgtaaattgctctccgctacttaaaatttcggcaaatgctaaacgatcagaa

atattccatgtccattgaagagttttagtaattcttcttaaggcttttaataatccaatc

ggaatttgtgagatctgtgtttttttgcccagataatttttcacataatttaataatttca

gctgatgtccaagcggtattaccaactaacggcagggtcctattttctgtggaaggtact

ccaagacttttataactaacttggctgcatcttgagtatcaatgtatgcaatgggtgta

gattctcctgtaacccagactgatttcttgtctaaaataggaatggcatattggttgatt

aaaccttgaaaaaatcctcctaaagaaaacacagtatatttcacattggacttctgaagg

aaatctactacctgagactttaagttcatcaatggaacatctgggtatttctctgaattt

aatattgaaaagaaaataaatctttcaactttagccgctttagctgcttcaattaatgca

attttcccatctaaatctatttttttctgcattataaggatcagtaggacgagatgtagaa

gcatctatgattgctgttaccccacaaaaagattgcaaaatactttcgggtaattttaaa

tcaccatatataagttctgctccccactctttgagaaaggcagactttcttaaatttctt

accatgcattttacattgtagccttcatctaaagctcgtcttacaatttgacgtcctaaa

gttccggttgctccaattactaaaagagtcatattagttaaaaataagtatgtataaaga

cattgatttggttgacggcagagagtaaattaggtagagagggaatcgaaccctcatgac

cgaagtcgtcacattttgagtgtgatgcgtataccaatttcgccatctaccctattatag

accaactactacaagtaatttagtctttagcgtaaaaataatactaagtttcaatagata

tgtcaacttttaaatttatattaataaaaaaacggagtctcttatggccacttttgaatt

aattccttatagattgtatttgagtcaatctaacacatggatccatagaataaaagcaga

aattaagatatatatagtagcgcttttatggatttcaatttttatttttttcttactttaa

actatgtattattgctttaagcttaattgccataagttttactataagaagtaaacagaa

tattattcaaaaacatttgttacaaacgttattaatgacgttcttaactactgttttgtc

ttttagtgtggctattagttataaacaatatgcagaacaagaacaatcgcagtatttatc

tgattccaagaaatataaaaattccagcagctactattatatacaaatagcgaacgatca

aaggatcaaaagacaatacttaattactactttaaagccttccttgtactttttttattac

tatatattctattaaactagttatgataacaacctctccagaagttttagtaattactat

ttatagatctaggataataaataaaatatttaaaaatgaattgctatttatcttccttct

ttcgtcacatattgttactagtattattaatagaatcgataaagtgattcaagtaactag
cttaagaggaagcttgaatttatataattcattgacaaggcctttaatgttttctttgtt
aatatttcaagtcttctttttggagattattcgggagtcaaaagaaatagctcaagctct
ttatactagaaatctcaatcaagaaaacaataatttttgaaaatatatacagtaaaatc
taactttagtgatcggctcaatataattattagcactttgtactttattattttagctct
agcgtaatagaatccacttattcttctgactctagtaatacgaagagtaataaaattagt
tagtagtttaaatagatagaat-taagatattatataaataattaaatgtttcattatgg
tcaatactcaaaatcaaatttcacaaacttcagaacttgattatatagttaatcaacctt
ataaatatggctttaaaacttctgttgaatctgagcaatttcctaggggaataagtgaag
atattgtccgcttgatctccaagaaaaaagatgagcctgaatatctattgaattttaggc
ttaaagcatataaaaaatggaaaaagatgagtagtccgtcatgggctcatattaagcatc
cgaatatagactttaatacgattatttattacgctgttcctaaattaaaaaaagaattga
aaagtctagatgaggttgatccagaaattcttgacacttttaacaagttaggtatatctt
taaatgagcaaaagcgaatttccaatgtagctgttgatgctgtttttgatagtgtatcta
ttgcaactacttttaagaaagaactgtctgaagctggtgttatattctgttcaatttctg
aagctataagggattatccagaattaataaaaaaatacttaggtactgttgtaccagctg
gtgataattattttgctgcattgaactctgcggtatttagtgatggctctttttgctata
tccctcctaatacagtttgccctttagaattatcaacttactttcgtattaacaacgaag
aatctgggcaatttgaaaggacactaattattgctgatcgtggcagtaaagtaagttatc
ttgaaggttgtactgctcctcaatttgacacaaatcaattacatgcagcgattgtagaat
tagtagctcttgaaggagccgaaattaaatattctacagtacagaattggtatgctggta
acaaagaaggtaaaggtggtatatacaattttgttactaaacgaggcttatgctcgggta
acaattcaaaaatttcatggactcaagtagaaactggatctgcaattacctggaaatatc
caagctgtattttagctggtgaaaattctcagggagaattttactctgtagctttaacaa
ataactatcaagaggctgatacagggactaaaatgatccatattggtaacaatacaaaaa
gtagaattatttctaaaggtatttctgcgggtagatcgaaaaacagttatcgaggcctag
taaaagttggccctcagtcatttaattctcgtaattattctcaatgcgattctttattaa
ttggtcaatcatctcaagctaatacatttccttatattcaagtacagaatccaacatcaa
aagtagaacatgaagcatccacttcaaaaattagcgaggaccaaatttttattttttac
aaagagggatcaatttagaagaatctattgctcttatgatcagtggtttttgtaaagatg
tctttaatgaattgcctatggaatttgctactgaagctgatcgtttactgagtttaaaat
tagagggaactgtaggatgagccagactattttagaaatcaaagatttatatgcttctgt
tggtgaaacaacaattttaaaaggggtaaacttatctattcgagccggcgaaatacatgc
gattatggggcctaacggttcaggtaaaagtacattatcaaaagtaattgcaggacatcc
agcgtattcactaataagcggagatattttatttttggacaaagtattcttgaaatgga
gccagacgagagggcaaaagcaggtatttttttagcttttcagtatcctgttgaaattcc

tggagtcagtaattctgatttcttaagaattgcattaaatgctcgcagaaagtttcaagg
attatcggagtttagccccttagaatttttcaactaataacagaaaaaatagatcttgt
tggcatgcaagaaagtttttgacaagaaatgttaatgaaggattttctggtggagaaaa
aaaacgtaacgaaattcttcaaatggctttactagatagcaaaatatctatttagatga
aacagattctggactagatattgatgcattacgagtagttgcaaaaggaattaacacttt
agctaaatcaacaaattcaattattttgattactcactatcaaagattattagattatat
tattccagattttgttcatattatgagtaacgggcaaattgtaaaaactggtagtgttac
tcttgcccaagatctagaaaagcacggatatgactggattacgcagacataactatttac
tac-------aatattgagactaaagaatattctttagtctcaatatt-tttgcaaatgc
aaagtt-aattatttaagctttatcaaaaccttgcttaacatcttcaattgctcttttta
gcaattcttcactaactggttctaatttttagtactacgaacattttcaccaaattctg
gtttagaattctttaaatcttctcgaagttctctaataaattcagctacttttgaaactt
caatatcatctagatagccattaattccagtgtaaataatagcagtctgttcttcaactg
gaataggagaattttgggcttgcttcaaaatttctcgaagacgttgacctcttgctagtt
gattttgggttgctttatctagatcagatgcaaactgagaaaaagcttctaattcagcaa
attgtgctaactctagttttaacttgcctgccacttgtttcatggcttttatttgcgcag
cagagcctactctagagactgagatcccaacattaatagctggtctaatccctgagttaa
ataagtcaccagatagaaatatttgtccatctgtaattgaaattacatttgttggtatat
atgcagatacgtccccagcttgagtttcaataatagggagagctgtcatactacctccac
ctaattcagcattcaatttagctgctctctctagcaatctagagtgtaaatagaatacat
ccccaggataggcttctcgtcccggaggtcttctaagcaaaagagacatctgacgataag
cttgagcttgcttagttaaatcatcataaataactaatgttgctttacctttgtacataa
aatattcagctaaagccgcacctgtataaggagcaatatattgtaaagtcgcaggactat
ctgcattagctgcaactataatagtataatcaagagctcctttatcctgtagtgaagata
ctacttgggctactgaagatgccttttgtcctatcgcaacatatacacaaacaacatctt
ggcctttttgattaataattgtgtccaatgcaacagctgttttacctgtttgacggtcac
caataattaattctcgctgacctctaccaattgggatcatagagtcaatagcagttatcc
ctgtttgcataggttcacacactgattgccttccaattataccaggggccattgactcaa
ttagccttgttccattacttgcaggttcacctttatcatctattggccgagctaatggat
caacaactctacctaaaaaagcatctccgaccggaatttgagcaattcgaccagtccctt
ttacggaactgccctctaaaatatccctaccatctcccatgagtactacaccaacgttat
cactttctaaatttaaagcaaccccaatagttttatcttcaaattcaagtagttcaccag
ccatgacttcatcaagtccatatactcgagcgataccatcaccaacttgtaaaactgttc
ctatattggctacttctacatcctgatcatacttttcaatttgttgacgaataatactac
ttatttcatcaggtctaatatttaccatatttttaatattatttttattaattggaaatt
ctaaagcggtaattagtcagactaatgctatctgtctttttttactttattttaagttact

acaacatcaaggtgtgaagccatttgtcttagctgtcctctaatactagtatcaatgact
tttgatccaatctgaatagtaaagccaccaattagctcgggctcgactgaaataattagt
ttaacttctttagccttagtcataaccttgatttttctgttaatagagtttcctgatca
gggcttaaagcaatagaagtattaatatttgcaattgttaaagattccatttggtaagcc
agttccaaatacttactagctatagcatctaacattccaattcgcttgcgatcaactaaa
atcattagaaatgataatgtattttcgttaatttgatcgccaacagttgcaataattact
tctttttttgcctctacagtttttaatggattagct---aaaaaaatttttagcttgtca
gattgcgctaaaatattttgtattgactgaatatcttggctcaccttttcagtaactttt
ttagttttagctaaatcaagaagagctactgcataaggctgagctatttttataacaaca
ttgttactgctcatagttgatctcctaacttagcaatattattatcaattatacgcagtt
gcatttcggagctcatttgattttccagctgcaaagtaactcttttaagagcaagaaatg
ttatttgttgctggatttgccttcggatctgtttttcagctgtttctatatttgatttac
ctgttatggcaagtctttcaatatctaattggccttgagctaatatggaacttctaactt
ttccagctgttagctgagcttctttcttaatttgatctataatgatttgagtttgagcaa
gttgtttttctgattctgacaatctagcacttgcttgctctagtctttcttcggattctt
gtatagctgctaaaacttttatctgtcttgcatttaaacttgatccgagaaattgcttca
aaacataaataagaccaaataaaagcaacaatatattaataacatttgcttcaaaaatat
cagaattaaagccgaatttgtgttcactactatgttctgataaaatagtaattatttgtg
gtatttttactatattattcatttaaaattct----------tactattttaatatttta
ctctggaatcatgattttcaagattgactacttaataatttaactttaatttgatcgctc
aaagtgtcaacctgattttctaaagttttaagagcttcttctttttgaacatttaactgt
ttcgatgcttcagcaatcagtttctctgcattcaattgagctttttttatatcttctgag
actatattttgagcttctttttgagaagaggctatttttaattgtgcattgcggcgagct
tcagatagatcttcttcatattttgcagctagctcatctgctttaacaagcatagacgat
gcagtagtcaatgttgttcgaatatattcatctcgttcgtctagtactttagttacaggt
ttataaaaaatagtatttaataaaaccattaaagtaagaaactgtaatgccattaaaggc
aaagtaccattgaagtcaaataagccaccttcaatttcttcggctaatagaaatggtaaa
tcaatcattttactaattatgtttatagttaataattttgtttcaattaagaaaaataac
tttatagtgcgcttagtctcatcagactaagcgcaaatgattattttaaccaacgtatgg
attagcgaacaataaagacaacgcaacaactaggccgtaaattgttaaagattccataaa
tgctaaacttagaagaagtgtaccccgaatctttccttctacttctggctgtctagcaat
accttcgacagcattagctgctgcactaccttgaccaatgcctgggcctattgcagcaag
acctacagcaagaccggcagcgataacggatgcggctgaaacgattgaatccataatgaa
ttttaatttttttgttatatgagtagaaaattgacaaatttcgtataatacttaaaatcta
tataattaagtataatagattctaatat-----tgtaattatagtatattactcttctcc
atgaccttccattgcttctccaatataagctgcggacaaagtagaaaagattaatgcctg

tattgaactcgcaaataatccgagtatcataactggtaatggtattaaaattggaattag
caaagtaaatacagaaacaactaattcatctgctaagacgtttccaaataatcgaaaact
taaagatagtggtttcgtaaaatcttctagaatattgattggtagtaatactggagttgg
ctgaatatatctagcaaagtagcctaacccttttttacttaaacctgcataaaaataagc
taaagatgtaagtaaagacagggccacagttgtattaatgtcatttgttggggcagctag
ctcaccttcaggcaagtgaattaacttccaaggaattaatgcacctgcccagttacaacc
tagaataaatagaaataaagttgcaatgtaaggcacccaagggcgatattcatgctcacc
aatctgatttttgcaatatcttgcagaaattctaatataaattccataaaattctgaaa
tttttcaggaattctttgtagatttctagtacctagaaatgataaagttaaaagtgtagc
aattactaaccaagagacaatgaaaacttgcccatgcaattgtaagctacctattttcca
atatagatgttttccaacttctactgctgataaactattatatgggttaaaatctatgag
actgttttgatacataataaatattt-ttttaaatagtgcaagaatcaaaaaataag-tt
tttgaagaattaatatgcaattaatttattcttaaaattagactattaataaattaaaag
caaaaataaattctaatagagatagctattttagtagtgattatttttttgtaaaaaaca
attattgctacctataatgcttttacgtttaattatatacccatatattaattatttaat
atgattataggtattttgctacaaactttattgttttatccattct----aaagattgtt
atagtttatttttatttaaaatatctgcgacttcatctgcaaaattaactttcaaattt
tcttctcccccctcccagaagaaatcttacaaatcttctaattttaatattttcacctaga
acagcaatattttggtttattaaatcttcaatactaatatcttgattgcgaataaacatt
tgatctaataatgatagctccttaagtctcttttttcattcgtccttcaatgattgtttcg
attctatcaactggtttgtttttcaagtcatctttaccagcttcaactctttgttccaga
ttaataatttcactaggtatatcctgtgttgatacgtattccacattaggcgatgccgct
atttgcatagcaatatctttagctaacttttggaactctggacgtcgtgctacaaaatct
gtttcacagttgacttcaactaatacacctattcttccgccagtatgaatataactctct
agtaaaccttcaatagcggtacgattagatttcttgttagctgaagccagccctttttgt
cttaatgattctaaagctttttcttcattgccattattagcttgtaaagctttcttgcag
tccatcatacctgctccagtcttatcacgtaaggcttttacgacttgagcagaaatttgt
agtgtcatatgctttcaaaaatttttaagtatttaaatt----------aattattataa
ttcactttgagtaggatgagttatttcttctaaatctgtttgcccgtatttaccatcata
aattgcatctgctattttacctacaattagtttaattgacctaatggcatcatcattagc
tgggattggaatattaataatttctggactacaattagtatcaagtatacaaatagttgg
aatacctaattttaaacactcttgtatagctgttgtttctctcttttgatctacaactac
aacaatatcaggtaaacgagtcatatttttaataccatttagatgtttacgaagtttatc
taattctcttcgtaaaacagcggcttctttttttgggtaactgatcaatcataccactttt
atcttgttcttctagctgtctcagacgactaactcgagacttaatcgttacccaattagt
taacattcctccaagccatctctggttaacataataagaatcacagcgttgtgcttcctg

tgccactataccagcggcttgtctttttgttcctagaaataaaactttttaccatccga
cgaagcctgttttataagttcacaagcttctgttaataattgtgcagtttggactagatc
aataatatgtatgccatttctttctgtataaatatatggaaacattttaggattccatct
acgtgcttgatgtccaaaatgaacacctgcttctagtaattctgctaatgtaacaatagc
cataagattattaatatttaaataatgtaattattcgggaaatccc-tttgtttagaaac
tgat-----tcaactaacttatttgaattttagtaaactaataataaattaacacacttt
taaagcagatcaacaaaatagatcctgaagaattcat----tctacacttttattgctaa
aataattacgcgcggttctgtcgtctaaaatgatatcatctaaatcatctcgtacagatg
aaatagtactttcgtcattagtatttgcagttaaattctttttctctaacgatgagccat
tacagttatcatacatattgaagccggtaccagctggtataagccgacctataataacat
tttctttcagacctcttaaccaatctaatttaccagatatagcagcttcagttaatactt
ttgtagtttcttgaaaacttgctgcagaaataaagctttctgtatttaaggaagcttgag
taatacctaatagcacaggcctatatgatgcattaatcttatttctcaaggttatggctt
tattagtctgttcaatcttctgtaactctacaagctctccaggtaaatagcctgtttctt
caccattttctattttcactttagaagtcatctgccgtactattacttctacatgcttat
cagaaatattcactccttgagattgatataccaattgtacttctttaacaagtagtaact
gaatctcttgaaagcttaatcgagcagcttcatataaggctaatccacattcaatataca
aatcaaaacttgcatcgaggatatcatgaggatttaacaatacgtctgttttctttcttg
cttcaagaatttcttcaattcttggaagcccttgtataatgtcacctgtttttgctctat
caaaaaccaatatagctaatgtttctccccttctaattaatgcattgttatcaacatgta
aaatagcaccattagaaaccaagtaaggtctagcaattcttaatgtaactgatctagaag
aaatttcaataatttgtccagagtctaaagaagtcatattttcagtaataaaatcaccac
aacgaatccaatcacccacacttactttaactgttgattttcctatattaaaggtctttt
tatcagaagatgtcgcaatcaaaatgcgcctgctagtattcttttcagaataaatctcct
caacagtaccttcactcattgcaataatctcagtactagcaacaactgtaaatggcttga
tatattcaccatttttttacaataattctagtttgactttgttgcttctctgatttattta
catcaatacttttaattgataatgtttcaaaagttgacagccttaagtcaaaataattat
tgccactttctttagcagaaaactcaattgaggatattaaattatcatctcgatgctcaa
tttcagccacaagattagttgttactaaatgaactccttctatagatttcactctttctc
cgtcccgaaaaggagttcttttaactagctttagtttagtttgacttggtttttgtgaaa
ctagattttcaattagactggattttgtttctggtattgaataaacaataacaggtctaa
taagaatatagggcatttgctcgttttctatatactcccaataaactaacttatctgtag
aaatattattatgaagtctttctccaggtcttaaaaaacctctacttttatcattattac
tatacacctcactaagtttatagatagatccaggcttaataataatttctcgaactattc
catccttttcgattatttcaacaatacctgagcttttagaaaaaatatttttaactagct
ctgttccactttctataacatctccattatgcactagtattagtgatgaatctttattaa

tttcgtgagtttcttctgaaatccataagatatatcccggacttaaaatttcgtaggcat
ctttatctaagcctgttttctttttgatacatttaagtctaaatatttaataataccac
cacttgtcgttttatacgtatcagatattagttctgcgactgtgtaaccatcttgaa---
tttttggtttggaatagctttgaaaagaaattttgttttttttctgtttctaaaatat
aagattctttttgtggattacatctgtataaatataacaacctggaattacaatagatt
cagtaataacttgtatattggtaatactgctattagtattctggtgtatccttacttctc
cagcataatgatttattaattctgtctgtgcaagtatagtaccagctttagttttatctt
ctttatgaacaataacattagcagaatcagatatactataaacttcacctgagagcaccc
agatcaagcctccagttttagtaattcttgtagtatattgattattatctgtttcttcaa
ctgttaaattggagaaacagattttgccagagagatctgaaactacatgtttttgagctc
tttctgtcatgagtcgatttctgcggggtgattcagcaatcacttgatccttcgatacta
attcaccatcacatattaaaagagtagtacctttacttaagttaattattgacttctgat
ttttcttagatctgattgtaacttgggttggctttttttgttattaatgcttgctcaccat
gtctggtacgaacatctgtatacgattcaatatctagatctatcaattgaccatctatag
gtgcgtagatttgttctgctaattcaccggtaaatactcctccagtatgaaaggtcctca
ttgtcaattgagtgccaggctcaccaatagattgagccgcaataataccaacagcctctc
ctaaatcaactaagcggccatgtgcaagattccatccgtaacaatactgacaaacagaac
ttctagagttacatgtaacaggagatcttactaaaactttttt---gataccagctttag
ttatctcttttgctaacttaggactgacatcttggttcgtatgagcaattaaacaatttg
tctctggatggaatacattttctgctaatactcggccagataaagcttgttctagattaa
ttaatattttctgcgtatctactaaatcttctagtatgatgcctttattggtcttacagt
ctacttctctaattattacatcttgagaaacatcgacaagtcgacgcgtcaaatagccgg
aatctgctgttcttaaagcagtatcaactaatccttttcttgctccgtatgatgatataa
aataatcagtaacagttagtccttctctaaaattactcgatattggaagatcaataattt
gtccctgtggatcagccataagccctctcatgcccactaactgtcttacttgagaaatat
tacctctagcacctgagaacgccatcatataaacagcatttagaggatctgtctctttaa
agtatttaataacttcttgtttcaaggattcactcgcattattccacgtatcaataactt
tttgaaatcgttcaactgctgtaatctcacctcggcgatatttgttttcagtggccttaa
tatcttcaatcgtgcctaataaaagactttgcttactaggaggaattcttaaatcttcta
aacttaaagaaattcctgcctgggtcgcataatgaaatccaagatctttaagcttgtcgg
ccatattggctgctcgggctataccataattacgaaaagcccagacaattaaatttttta
gttcgttttttatcaataaccttatttgaaaaacttggctgtgcaagacttcgtctattat
ccactaaatcttctccatattaaatgttcgtt--attaactatgctactaaagactcttg
aataattttattaaagattatacggccagctgttgttctaatatattgcacaattctctg
attttcagcgtcttcttttatgatatggttattaaaaaattttgttgtactattgtcaga
atgtttttcaatcttaatcggatcatgaggttgatcaccatctactaatccatcaaaacg

agcccatatataagaatgtaaatcaattttcttctgttcataagccataactacatcttc
taagcttgcaaaatactgattagatccctgctgctgagatggattattagctgttaaata
atagcaacctaaaaccatgtcttggcttggcataataattggttgacctgttgctggaga
gagaaaattatgaggggctaacattagcaatctagcttcggcttgagcctctaaagacaa
aggaacatgaacagccatttgatctccgtcaaaatcagcgttaaacgctgggcatactaa
tggatgaagcttaatagctcttccttctacaagaataggttcaaatgcttgaatacctaa
tctgtgtagtgtgggagctctgttcaatagtactgggtgaccttgaataacttcatttaa
gacattccaaatagaagattcatttttttgaatcatctttttggcagctttaatattgtt
aactaatccctgcaaaattagtctgtgaataacaaaaggctgaaataactccaaagccat
ttctcgtggcaatccacattgatgtagttttaaatgaggacctacaacaattacagatct
accggaataatctactcttttacctagcaagttttgtctaaaacgaccttgttttccttc
tataatatctgacaaagattttaatggtctattatttgcacctacaacagttctacctct
acgaccattatccatgagagagtcaacggcctcttgcagcattctttttttcatttctgat
aataatctccggagctaaaattgactttagacgagataatctattatttctattaataat
tcgacgataaaactcattcaaatctgccgtcgcaaatctacctccatctaattggaccat
aggccgtaagtctggaggtatcacaggaatgactgtgaataccatccaagaaggatctgc
acctgttgctatgaaattttcaatcaatcttaagcgtttcatttttttattgaattttaa
tgaaggtgtcttaaaactttttggtggatttgtagcttcagaccttaaggtttcagctat
gtgctctaaatctaaatctttcaacagtttctgaatagcttctgcacctattcctacttc
tacttggttattctcatcttgattttggtagatttcctcttcaagacttttccattcgta
accttcgagcagttgcttgtattttaaattaatatctgtattagattgcgtaactacata
agaatgaaaataaacaatttttctacttctttgacttttaagtctaatgctaaagcaat
atagcttgtacttccttttaagtaccaaacatgagttactggtgaggcaagttcaatata
agccatccgatgtcttctcacgcgtgattctgtaacttctacgccacatcgttcacaaac
tatacctttgtagcgaaaacgtttatatttaccacagtgacattcccaatccttaactgg
gccaaaaattttctcacagaataaaccatccatttctggctttaaagttctgtaattaat
tgtttcaggctttgtaatttcaccaacaatttgtccgttgggcaaacttctttcgcccca
ctgtcgaatttttttcgggagaagctaaactaatttttacgtagtcaaaatattgctcaaa
ctttgtcataaatttgaataccttaaaatcaaataattagccaatttatttacatgcttc
atattgaaaaatgaatatcttaataaaggaattgttcaaaatcgtctactggaggagtat
cataattagaacgatctactcggttatctttagaatccgacatcaagtcaacttcaactg
tccgtctttggccatcttcaaatagtttcaatttatgtactgcaatgtctaaacctaatg
actgcagctctcgcattaagactttaaaagattctggagttccaggcttcggtataggct
tccctttaactattgcatttaatgcttcatttctagcttgcatatcatccgactttacag
ttaataattcttgcaaagtatatgctgcaccaaaagcttctaaagcccacacttccattt
cacctaatctttgacctccatgctgtgctctacctcctaatggctgctgcgtaactagag

agtaaggaccagtggatcttgcatggattttatcatcaacaagatgaacaagcttaagca
tataagctctaccaactgtaacaggattatcaaaaggctctcctgttctaccatcaaaaa
cttgcattttacctggatgctgatcattaaaaagccatttattgctagtaatcaatgatg
cttcttttaatttctatttacaagtgctcttgatgcttctgctccatacatttcatcaa
aaggtattattttaaatctctttcctaaatagccacctgctaagcctagtaggcactcaa
atacttgaccaacattcattctagaaggtacacctaaaggatttaaaacaatgtccactg
gtgttccatctgataaataaggcatatcttgtttaggtaagatcctagaaatgatacctt
tattgccgtgacgaccagccattttatcacccacttggatttttctcttctgcgcaacat
agacgcgaatcatagcatttgttccgggaggcagctcatctcctttctgcctagtaaaaa
ccctaaccttaactactctaccttttgcagcattaggaagccgtaaagatgtatctctca
catctcgagccttttctccaaaaatagctcgcaacaatttaccttctggtagttgatcag
cttcacctttaggagtaatttttcctactaaaatatctccagcttcaacccaagaaccac
cgacaacaattccatttctatctaaatccttcagagagttatcgctgacgttaggaatct
ctctagtaatttcttctggtcctaatttagtttgacggcattctacttcatacttttcga
tatggatagaagtatacaagtcatcataaactaatctttcgctaattaaaaatgcatctt
cgtagttataaccttcccaaggcatataagcaactagaatatttctccctaaagcgattt
cccctccgtctgtggatgcgccatcagctaaagtttggccaacaactatttttttctccca
cccaaacaattgggcgctgattaatacaagtatcttggttagaccgataatattttttta
agcgataatgaactgttctaccactattatcctgaatgcctattttattagcagaaacat
aatttacatgaccagatgttctgctaataacaaccataccagagtctcgagctattttag
tctcaagaccagtaccaattattggcttttctggatataataatggaacagcttgccttt
gcatattagatcccattagagctctattcgcatcatcatgttctaaaaaaggtattaatg
atgttgcagctgaaattacttgaataggtgaaattgcaatataatctacttgggttgggg
ttgttgtaataaactcttgacgataacgtacaggaataatatctccttcaatgtaatgtt
gcttactaactttaacatctcctggagctactcggaaatcatcttcctcatctgctgtta
gatagacaggactattgtgataaatcacttgaccttggtttaccggataaaatggggttt
ctatgaaaccaaaaacattgacccgcgcacaggttgctagtgaaccaattaaacctgcgt
tcggaccttccggagtttcaattgggcaaattctaccatagtgactaggatgtaaatcac
gaactgcaaaacctgccctgtctttattaaaacctcctggacctaaggcactaattcttc
tcttgtgagttagttctgcaactggattagtttggtccataaattgagaaagctgactag
atccaaagaattctctcactgacgcaattaatggcttgggattaattagatttgataaac
tcaatgaatctatatcacatatcatcattctctctctaataatgcgttctaaacgattga
gacctactctgaattggttctgcagcaattctccaactgaacgaactcttctattaccaa
gatgatctatatcatcaagattacctgagtttttatccttaatattaattagataatcaa
ttgaggataagatgtcttgaggagacaatactcggaacgttttaggtatattgagaccta
gtttttattaattttatatctaccaacttcacctaagtcatatcttttaggatcaaaaa

aacgagagtaaagcatctgcttagcaacagcaactgttgctggttcattaggtctaagct
tagaataaactatgagcagggcttcctcatcagttacttcttcaatattatttttccaa
tttctttagctaattctttaactgaatatatttgggatgcagaaataagaaaagcatatt
tactaagtcctttttcaatttcatctttattgagtccaatagctcgcagaaaaatataag
catttactttgtgggtcttatctattcttatccaaatttctcccttaggatcaatttcaa
attttaaccaagaacctctattagaaattaggcttgcactataaatctgcttaccatttt
tgtctatctcttgtttataatagattcctggactacggattatttggttaataataactc
tttctgtcccagatacaataaaagttcctctatttgtcattattggcaagtctccaataa
aaactaatctctttttgtactttttatttttttatctgcttttcagcactgagctgtagat
gggttgatgataaatctaatgatttaatattttttttcttggtttttagaaggtagatcaa
tatcttttcttgttaactttgcaggaacatatatttgagcactataagttttatccctat
tttttgcctgtctgacactataacgaggaaattttatcttgtactcattaccaaataatt
gtaattcgagtcgactagtagggtctgatatcttcggaaaaatttcaagaacttccgtca
aaccttctaataaaaaccatttaaaactggccctctgaatttcaactaaatctggaagaa
gtttatttttttaagcttatacgttgaaccatagatctcctttataacaaattttgaaatc
gaagagtttaa-tactattaatagagtactggtagataatgcataagcatagtttgctac
attatttcttctctagacataactagaacatcatattttattcaa--aaaaagattttta
aaatttttcttcac--acacactatctataagtactttgctgcta--aaagatagttcaa
gctattaaa-------aaaaacgaaattaatacttgtctgtctgtcaattaattgacatc
aagatagacaagtattatttatttaatcaattctg-tttttctgagcaaaagacaatgct
ctcgcaagtctagctttttttctagcgcctgtattagggtgaaatgcccctttttttaca
gctttatcaatcttactatatacttgagaaatacttgactgcacatcattgaaattatca
ctttctaaattatcaatatttagaaggcatcttttagttaaagttttgacaaccgattta
tacttacggttaatgagacgatttctttccgaagttttaatgcgtttaatcgcagaaagg
ttcttagccacagtaaatccaataaaactttttataaaatatttttagaaacgaaaataca
caaatctgggatctattcaaaaagggaatatgcggcgtattataacattaaagaaaaaca
ataggcaatagttcaacaagagctcttgttattacacaatagagattgtcattgcgaaaa
actttgtaagatagtatttaattaaaataaactaactagacatataaatatatatgaaaa
aaattgaagctattattagacctttcaagcttaatgaagtaaaacttgctctagtcaagg
aaggtattggaggaatgacggttatcaaagtttctggatttgggagacagaagggtcaaa
ctgaaagatataaaggatccgaatactctattgatatcattgataaaataaaaaattgaaa
ttattattagcgatgataaagtagagaaaattgtagagacgattattaaggcttctaaaa
caggggaaattggagatggaaaaatatttattagtagtattgaaagagtaataagaatca
ggactaatgacttaaattttgaagctttatagtatttttacttt-tttagacttgatact
tcttcactaaacatgagataataagcgtttattgtatacttataactaaggtgtaata--
ctaatggctaaaagcaaaggtgcacgaatcgtaataactttagagtgctctgataaggct

ggagagtttgctcaaaaaaggaaacctggcgtttttcgatatacaactactaaaaataga
cgaaatacaccaagtagaattgaattaaacaagttttgtcctaattgtaatcagcattgt
atcttcaaagaaattaaatagttatttatcttattaaaat--aatcatattatggctata
tatagaaaaagaatatctccaattaagccaacagatgctgttgactacaaagatattgac
ctgctaagaaaatttattacagaacaaggcaaaatactgcctaaaaggtcaactggattg
acttcgaagcagcaaaaaaaacttactaaagcgattaaacaagctagaatcctttcttta
ttaccttttttaaataaagattaa--atatatagcctaacttcatatttttccatctaaa
tattaaaaatttctatagatttgtttattaaaatgatctaacatactgaat---------
----------tactaagaaga-tctactcattacaagttttgttcattttataattaata
atagcattataattta---ttaagctagattc-----agacataaag-gtaacctctttt
aattgattatattatatagtctatttaaaagaggaaatttttttgcttc----------t
agcaattcaagattctttaagctttaacaaggtatttaaaacaaagattttttgtttaggt
atcagatcaaatgaatgaatcttatctcccgactgccataattgaaattcagagatgaaa
attccacattcgttgccagcttgaatttcttcgacatcttctctaacgcgctttaaagat
tcaatttttccttgataaataactttttcttctcgaatcactttaatccaagagtttttt
aatagcttattattgatcactcgacatcctgctatttttctatttgctaacgagaatact
gtacttacttctgcttctcccaccggtacttctgaatattcaggatcaagtaagtcttcc
atccttcttttaatgtcttcaattaaggcataaatgatttgatagtttctattaatata
ttcgactttgccgatgcttgctttgttcctggagcaaagttggtattgaacccaatgaga
gtagaatttgtggttgaagctagttcaacatctgtagcagtaatttctccaggcatgatt
gagacaacattgagttgaactttactctgaggaaattgagataaagaatctaggattgct
tctgtagagccttgattatctgttttaataattaaagaaatttgcttactaatatcttta
gaattagtgtttttttaaagtatctaatgtaattcgactattaagtgctctttgtttctgt
ataattgatgagtctttagaggtattttcaatcgctttaagtttggcttctttatcactt
ttaactgctaaagctatttctcctgtagctggtacagatgacaatccccaaatttcaaca
acagatgatggtattgctaaattaattttctctttagcgttgttaattataactcgaatc
tttgcgtaagctgatccaataaccaaattatcactaatattcaatgtaccattttgtatt
aatagagttgctactggaccatgcgatttatctaaatgggcttctattataataccttgt
gcaggctgtgttggatcagctttcaaatcctctaattcagctaacaaggtaatagtttct
aataatttatcaacattttgaccagtaagagaactaataggaattataggcacttggcca
ccaagtttctccgacatgacattgtattttaataaatcttgctcaataatatctgtatta
gaaccagctttatctatcttagatattgctatgacaaaagggacattcgctttttgaata
tgattaatagcctctatggtttgaggtttaactccgtcatcagcagctataataataata
gcgacatctgttaaattagctccccctagacctcatacttgtaaaagcttcatggcctgga
gtatccaagaaaacaatttttttgtttattatctttcttaatatactcaacttcataagct
gcaatggcttgagtaattcctccaatttctttgttagcattattagattttcgaatataa

tctaatagcgttgttttgccatgatctacatgtcccataactgtgacaattggaggtctt
ttaatataattgccagtttcataaaaagcattagagttatctaaattactagaaagtcca
ttattattttcctttacatttgattctactgctataccaaaattatcagccactgatgat
attattgaggcgtctatggtttgattcatagtaactgatattcttttaagaaataaatat
tttataatatctgtttcttgaacacaaattagtttagataattcttgaatagttagagga
ttagtaatactaatagactctggcggagaattagattgaatgcttgctgaattttgacta
ggagttacggctactttttctgtcttatagatttttattactagtcagtttttttaca
acttccacttttggcttaggtggacgcattaaagagatagctaaatctcctgcagtctgc
gaaacattagaattagagtctctaaagttgtca---tcatcatcgtcaatatgtatttta
gtttttattttt-ttgcgctgtctattctta--tttttttactatctactaagtcgtga
attttattaaaatttttactttttttatctagtttaggtggagaacttaattctagatgc
tgttcaccacc---tgtatgtgattctgatttatcaaggtctaaatttaacagattatca
ttattgactgactctaatctaattttatagataatctgaggatttttcaaatctacaatt
gtttcagaagaattgatatttgagctagatctagaggatctattttcaaagttttgatta
tttaaaaacataaatattggttttattatctaatagtaactttctaattttaatacagag
cttaatataaacaagaatcgaatatctgtataaataatacaaatgt---agagaattaat
ctatgttgtattagatttaatataacactaaagtcctttgttcgttatacaagtttaaaa
ctttt-tatgatgataaacactagattctagtttattggtatacaatctatttatctaaa
taatactgtttttgtcagggcttttaaa----ttatctaatgttatgttcattatgaaat
tctattaaaatttaaaaa-ctcaacaattaagtaagaatcgttaatcacctccacgccac
ttatgccacgatctcaaaaaaatgataattttattgacaaaacttttacagtattagctg
atattgtattaaaaatacttcctacaagtaaagaggaaaaagaagctttctcttactatc
gagatggaatgtctgcacaatctgaaggtgagtatgcggaagctttagaaaattattatg
aggctttaaaactagaagaagatccatatgatagaagctatattttatataatatagggc
ttatttatgctagtaatggagagtatgtgaaagctttagaatactaccatcaaggattag
aactaaattttaaattacctcaggcccttaataatatagccgtgatataccattaccaag
gagttcaagcaattgaggataaagatacagagctatctaaattaatgtttgataaggcgg
ctcaatattggcagcaggctattaagttagctcctgataattatattgaagctcaaaatt
ggctgaaaacgacaggacgaatgaggaatatacaaggatattaatatatttaagatataa
taaagatatcaattaataaattaaaattagttaaagaaactggtactatatagagtactt
-tactatatattataatattatactagttgatctctatctctaagacttagctgaataac
cgaaattaaatctattgactatattcatgaatacaaataagcataatggttagtacaaca
aaaagtaccggatctgttactcaaattatcggaccagttttagatattgcatttcctaac
ggacagcttccgaaagtattcaacgcactcaaagtacaaagctcagaaggaactattact
tgtgaagtacaacaacttttgggtgacaacaaggtacgagctgtttctatgagttccact
gaaggactacaaagaggggtagaagttattgatactggatcccctatatctgttcctgta

ggtacagatactcttggacgtatttttaatgttttaggtgaacctgtagataatttgggt
cccgttgattctgagagtactttacctatccatcgaccagcacctgcttttactaagcta
gagacaaaaccaaatatttttgaaacaggtattaaagtcgtcgatttacttgctccttat
agaagaggtgggaaaattggtttattcggaggtgctggagtaggtaaaactgtattaatt
atggaactaattaataacatcgctaaagctcatggtggagtatctgtatttggaggcgtt
ggtgaaagaacaagagaaggaaacgacctatatatggaaatgaaagagtctaaagtaatt
gatgcagataatctgaaagaatctaaagtagcattagtatatggtcaaatgaatgaacct
cctggagcacgtatgcgcgttggcttaactgcattgacaatggcagaatactttagagat
attaacaaacaagacgttctattgtttattgataatatttttcggtttgtacaagctgga
tcagaagtatcagctctactaggccgtatgccatctgctgtgggttaccaaccaactcta
gcaactgaaatgggagcactgcaagaaagaattacttcaacaacagaaggatcaattaca
tctattcaagctgtatatgtgccggctgatgatttaacagacccagctccggccactaca
tttgcacatttagatgcaacaacagtactttcgaggaacttggcggcaaaaggaatttac
cctgcagtggatccactggattcaacatcaacaatgctacagcctggaattgttggaact
gagcattattctactgctcaagaggtaaaatcaactttacaaagatataaagaactgcaa
gatattattgctattcttggtcttgatgaactttcagaggaagacagacaaactgtatca
agagcaagaaaaattgaaagatttttatctcaacctttttttcgtggcagaagtgtttact
ggatcacctgggaaatatgtatctttggaagatgcaattaaagggtttcaaatgatctta
aaaggcaatttagacgacttgcctgagcaggcattttatctagtaggtgatatagatgaa
gctatacaaaaagctgacagcatgaaagattaatataattaacgatgactttaaatataa
gaattattgctcctgatcggactgtttgggatgcagaagcacaagaaattattttaccaa
gtagtacagggcaacttggtattttaacaggccatgcacctttgcttacagctttagata
ttggagttatgagagtaagagtagataaagaatggatgccgattgttttgctgggcggtt
ttgcagaaatagagaacaatcaattaactattctggttaatggcgcagaagaagctagtc
aaattgatttatcagaagcagaaaaaaatttagacactgcaactcaactcttaagtgatg
cctcgtctaataaagaaaaaatagaagcaacacaaaaaatacgaaaagctcgagctcgag
tacaagctgcaacagcagcaacttcgtaa-aagattcc------tcttaaaacataaaac
tttagtaattattactaaa-------gttttatgttttatttgtatggttataggcagct
taaataaacttagcttagacctgaacaaatataatcaaaatataatcccatctctttacc
tgcatctggtccaactaaactaatagtaacttctttcatagctaaaatagcttgaatcgt
tgcaccaataggaacacctagagaattgtatgtttcttttagaccatttaatactctttc
ttcaaggatagatggatctccagctagcattccataagtagcatagcgtaagtagtagtc
taaatcacggatgcatgcagcataacgtcgagttgtatacatattaccacctgggcgtgt
aatatccgaataaagtagagattttgctacggattctttaataattgttgctgcattagc
tgcaatagtagcagcagctctgactcgtagctcacctgtttgaaaatagcctcttaattt
ttcaacagagctatcatctaaatatttaccttggacatcagctgcattaataacagaagt

aattgcgtcttgcataactttcacacattccttaattttaaatatctaatacta--gggg
tatataacaaaattactagatcttttagcgatatcagtattactgcatagcacccaaagt
ataatcaaagtaaaaaccagcttctgctgaatcttcacctgcaagcagtgagcaagctac
acttttcatgcattttacaccttcagcaactcctgaaattggtgttcctaaagaattata
catttctttaacacccactaatccaatttcttcaataggagttacatcgccagcaactat
tccgtaagttactaagcgaagataatagtccagatctcgtagacaagtagcagtcatctc
ttcgccataagcatttccaccaggagaaactacatcaggtctttttgaaacagttgttg
gccgccttgctttacaatacgctcacgattatctgttaaaatttgagctattcttaaacg
acgttgtccagataaaacaaaactttaattctatctaattccccaggacttaaatatct
tgcttctgcatctgcatttacaattgactttgtaacaatactcatggataaaactcctgt
aatgcttt--aaaaaaagcttaatactaattatttatgatataactcagaaagtaatgta
ctttttgag-aattttctaaccgaaacacgttcatgatatatgaagcaggatagatactg
aggtaagtactaaaacgagaaagttatatgtaggaacttctgttattgatttcctagtac
tttttaaaacttggtaccacaattgatatatcctgcttagtcaaacgattatagagtgt
ttctgtattaggaaagttagctgctggtaaagtcggaaatcttctgtatggaacagtatc
tttgccgtataccgcattgtattcaacactttcaactaaattacttataaatgcagataa
gccttttgaagctaaaatttgattaaagaatctaatttctgcctgattattaggagctct
tcctaagatatgcttagtacctaattctataactttgtatttggatatggctgataaaa
ttctttaccatatagttcagatagcgccagtttttctactagttctttgacacagatttg
tctattcaagaaagctgacttaatatccaagaattcaccgccgacactaaaagagttaag
atctctttcgaaaatttgacgatatgctgctcttaaagcttgttctaacatttctttgtt
actatcactattcacttcaaagacaatagattgatctcttaaagaagttactccttgtga
aatacgagactgaatatcattactactcctcatttctttgacagtgcctaattcaacaaa
cctagcagatttactagcaataactttttgaatctctggtcaataattcctggtcttaa
gcttcttaaagcaacaccagctggagtactatatctttcataaggaacagtattatctcc
aaaagtttctgtgtactctgagctatcaattattgtgtctattacttggtaatagccttg
cttatatgcaatatcaaaatatttattaatttcttgccgaccatatgtagggcgacccag
taagcggttatgaatatattcaatagctttacaaatatataatggctcccaataaagaga
tctaaatatacttgattttgctaactgtctaacaaactcgcgaactgtaatttgattatc
ttttagctgactttctattggcttgagaattagtttttcttcttgatacacttctctgcc
aaaaactcttaaataagcaacttttgtgacaacttctactgaattttcaaaattttggga
tgaatctgactttataagagcgttggacaatttaaagatttttggtcccagggatcctgc
tgacttaggtcttacctgtggattactaatctgattgtagattcctggcccacgtctaac
taatatccttcttgtatctttaccaaaaagagcttgtctttttcttggatctttattttc
tttagggaaaatagctccaaattgaattgataatggatcattacctattccatatggatg
ttgatcaggtaatgattgcttataatcactaaataaagttataaattgaggaaccttacg

aaaaggcgcactataggtgagaagatctatttgaggcccccaattacggcactcttgagg
ctcttcacctaaatttctaaagtaaggcacagtctcttcaccgaaataatctgtatactc
agaagaatttaaaagtgcattcactaaaccacttaaaccagtagatgataaaatagcaaa
atatttttgaaactcttctaaagagctaggacctctacccaagaaatgtctaaatgctaa
ttctaatgctcgactattgacaaaaggttcataaaattgctttctataaatacttgaagt
tcctaacgaacgaataaattccttaattgaaatttggccatttttacttgcgactccaa
attcgataaagagaggtcatatgctttagcaatatctctttcaaaaatttgtctataaca
tgctttcacaacaatatttttctcatcagcagataaactaggcttcataacaaaacgagg
tgttgacactcctgcctgtacataagtctgaggaagacgcaaaccttgtaaatctccaga
tattctttttcttaatttatcagtcaaagatggagcttcaaattcagaaatcacaacatt
aaaatactctttaactaaatcttgaccttttatatcttcctcaaagattaataatgctgt
ccgtctcatctctctaagagccacaattgctgctgcacttgagcaagcgttatcaattaa
ctctctcaatcctcgtatattaacagataaaatattagggtcgcctgatacaattgcata
tgtcaaataccgtaaaaaccagtctaaatctcgaagagattttctcattcttgtagtacc
gtatctcaagacgtttataggtttaaaccctggcggagtagctcccccagcattaaataa
agatcggaaactttggccaaaatctccttggatatttcctgataactcattaattttatc
ttgagaactttgatctcctgcaataataacagctgcttgcggacgttctaaataagaaat
tgcagacccacctacgaatatttatctgctgctcgtgctaccagaatatttgcattttt
agttaaaatatctgcaacttccagtctttttgtccagaatttaaaaaagaaactaattg
atttaattcaccaagctgtaaaaatcggtcttgttgctctgcctgagtaatagttaaaat
tgaggcagtccgataaagctgggggcgtgctaatgggcttccgccacttgctttgatact
cattactttatttatctccttaactgatattgcttgctagcgaatacaaatatagcttga
tatttctaaatatttaaagatactatgcaagcaca-ctgttttctatattattaggtact
agtacttatagtacagagagtcttataactaacttctataaatacaaattttgtaatact
tctattaacatttgtcaaaaaaatatttaagtgctcactcttttatattaaaattcaatt
agtgagaggttgatgtgaaataaagtatactaagtaagttttatagataggaaaattaaa
agatttattctacaaacattaatca------------cttgaatgtttagcaattgtaca
attgttattgcttttaattggtttagactagttttatatttttctaacataaacaatat
taatgagcaatttgtatgaatttaaagccacaaaacaacctattaactaacgaaaatata
gatcatgtagatatacccgaaaatgatataccaatgtcaattacggaacatttagaagaa
ttaagacaacggactttatttgtgtttttattctttttgtttgctacaactataagtttc
acacaaattaagatcattgttgcaatactgcaagctcctgctgttggtattaagttttta
caactggcaccaggagaatattttttttcatctattaaggttgcaatatattgcgggatt
gttgcaacaactccgtttgccgtttatcaagttatattatacatactcccaggactaact
ggaaaagaaagaaaaattattttgcctctattaattagttctgtattgctttttattaca
ggtggtatatttgcttactttgttctcgcaccagcagctttaacatttttaattagctat

gggtctgatattgtagaaccattatggtctttcgaacaatactttgactttattctatta
cttttacttagtacaggattagcatttgagataccaatcatacaattattgctcggtgtt
tcagggacattttcttctagtcaaatgatacgagcttggagatatattattattatagca
acaattgctggagctattctgaccccttcgactgatcccgttacacaattaataatgtct
tcagctgttttattactttattttggtggaattgttatattattagtcttaaaaaagtag
agtaaat---------atttcta-ttaatccaatatatttgttatc-----tggcaaa
taatctat-aaaacaattgct------------------agatttaagtatcaggcttct
aaatatttcaa---------gtctcttttatcattgaaatacaatc-------------
------------------------------------------------------------
------------------------------------------------------------
------------------------------------------------------------
------------------------------------------------------------
------------------------------------------------------------
------------------------------------------------------------
--------------------------------------------------gatacattgg
cttgtttttcatcaattatttgtgttgcacac-tttttagtactctctttattgttctt
agaattatttctttaatctatttattaaat----agatatgaagagatttaagtattgat
tatggtaatctatatcaacgtaaaaagtattttattgtgagttttatttatataaaggct
tattcattatatagttataacagtaagtctaattaatag-ttttcatgcttaattaaagt
aataactcttaattcttttggcagaaaaatctgtcttaatattttatc--aaaaaaacag
-aaaaatgaagctaaatagtctaattaacttaattcaaaagtctatatattcttgtacac
ttttactaattattttaaatattatttgtgtcgcacctaattctagtaatgcatttccaa
tttatgcgcaacaagcttatgaaagtccaagagaagcaactggtaggatagtatgtgcca
attgtcatcttgctcaaaagcctgtggagatagaagcgcctcaagcagtactacctaata
ctgttttcgagactgttgtgaagattccatatgacagcaatgctaaacagattttaggta
atggcagtaaaggaggcttaaatgttggagccgttgtaatattacctgaaggatttaagt
tagctcctgttaatagattatctacagagttaaaagaaaagactagaaatctttacattc
agccgtacagtgctaaacaagacaacattttagtaattggacctatttctggtgataaaa
atagagaaatagttttcccaatactatctcccgatcctgcaaaagataaaaaagctcatt
ttttcaagtatccaatatatgttggcggaaatagaggacgaggccagatttatccaacgg
gtgacaaaagcaacaataatattgtctctgctttaagcagtggtaaaattaataaaattg
aattactagacaaaggtggatttataatacatgtgactaacagtagcaatgtggagtcaa
cacagaagatttcacctggtctcgaacttagagtaaaagaaggggatacaattcagcttg
atcaagctttgaatagtgatccaaatgtaggtggttttggtcagaacgaaacagaaatag
ttttacagagtccaaatagaattaagggcatgattgttttctttttcgctagtgttctag
ctcaaattttcttcgtattaaagaaaaaacaatttgaaaaagttcaagcagctgaaatga

atttttaaagcaatataaaactggcacttactataaagaaagta-taaacactttattca
caggatgtaccagtttttatgttatgtaataatcgctttgacggaagttatgatactttt
aacagaatctacaatctgactaggctgaattactgtagcctgctctaagcttccattata
tggtgtgggtatatcctgcgaagatagcctcacgacaggagcatctaattcatcaaaaag
atactcattaatttgtgcaattagctctgctccaattccggctgttttcatacactcttc
tacaattaaaactttgtgagttttctttactgagatagatatagagtctatatccagagg
ttttaaagatataagatctataacttctggatcataaccttcttttaataaggccggtaa
tgcttggataacatgatgcctcattctagaataggttaaaatcgtaatatcttttccttt
tcgtacaaattcaactttattgagaggtaagaaatattcttcttgaggaatctcttcttg
taaattataaagtagaacgtgctcaaagaatacaactggattattgtctcgaattgcaga
tttcagtaatccttttgcattataaggagtagaacaagcaactatttttaagccggggat
ggcttggaaataagcttccagtctctgagaatgctctgcacctaactgcctaccaactcc
tcccggtcctctaataactaaaggcaatgtaaaattacctccagaagtataacgtaacat
tcctgcattattagaaatttgattgaatgctaataataaaaaactcatgttcataccttc
aacaattggtcttagccctgttatagctgcaccaattgccataccagtaaagctattttc
tgctattggtgtatcaagaactctcaaatccccatatttgctatgcaaatctttagttac
cttataagatccaccatagtgaccaacatcttctcctattacgcagacagttggatcttt
tgccatttcctcatctgttgccgctcgtaaagcgtcaaacataaagattttactcatttt
aatatttgatttttatgatttgaccactgttttaaagtaacagttaagacaccatattct
atttagttatctgcaaaaagatatcgtttcaattctgacatgttaggttctggactggaa
atagcaaatttcacagcttgttccaattcggttttaacagcattctgaatctcattaagt
tcacctatattagcaatttcattatctagaatatatttttttgagtttttttgataggatct
cttgccacccaagcctctttttcttgtcttgatcttagttcatcaggatctgcgagagag
tgaccacgaaatctatatgttaatgcttctattaaggttggaccatcaccttggcgagct
ctttgaactgcttgttttgcagcttgtcttacagctagcacatccattccatcaacttca
atcccaggaagcccaaaagcttctgctttttttatgtatttcaggtattgaagaagaccgg
tgatgtgccatacctatagcccattgattattttcaacaacaaatataataggtagtttc
cagagaactgccatattcagacattcaaaaaattgcccattattggtagtcccatcacca
aaaaagcaagccgtgactcttaaatcctctgtttctttaagtacttgctggcggtagata
ctttgaaaggctgcccctgttgcaaccggtataccttccgcaataaaagcaaagccacct
aaaaaattgtgaggcgcagaaaaaatatgcatcgaacctcctctgcctttactacaacca
gtctcttttccaaataactcagccatcacatttttagatgggacgcctttacttaaagca
tgtacatggtctcgataggtactgcaaacataatcagttggattcagaagtttaattaca
cctgtagaaacagcttcttgaccattataaagatgaacaaaaccgaacatctttccttta
taatacatctgagcacacatatcttcaaaatttctgcctaacaacatgtcttcatataaa
actaataaattactcttattaagattaagacctgttgaattatagttagtcagcggcaat

tgaactttcttaggataactcataatttgtaaagaaacctct----------tattgtaa
ataggaattttgaagaaacagctatcaatagaaaacacacagataaaaacttataaatta
taatatgtttttactaaagactttaataattatttgaaactagttcagtatatcaaatt
aattttgcttatactacggatgtctgaaacttatataaaaaagtagt-a-aaaaagttta
ttaaaagaatagttaaaaacagatataaatagaagtaaacagaattatt-gaaaaaaaat
gattattattaattaa----tacttaatatttctaaatattttacaattaatggctatgt
caccatctttattttatcctgttgaacaagaactatgtagtcttgaaaaaaatctgaaag
ctgttgctgggactcgtcatccaattttatatgccgcagcaaagcatctattcgatgctg
gaggaaaacgagttagaccagctcttgtatttttagtggctaaagcaacctctgagaagc
aagatataaatactggacaaaaaaggctagcagaaattactgaaattatacatactgcta
gtttggtacatgatgatattattgacgagtgcacaacacgtagaggagtcaaaactgtac
ataatttattcaatactaagattgctgtgctagcaggagatttttttatttgcacagtctt
cttggtatttagctaatattgaaaatttagccgtagtcaaagctatttctaaagtcatca
ccgactttgcagaaggagaaattaggcaaggcttagttcatttcgatcccagtatttcaa
tagatgcttacattgagaaatcattttacaagactgcttcactcattgctgctagttgtc
ggggtgcagctatgcttaatggttccaatcatcaaataaataatgatctttatctttacg
gtaaacatatgggattagcatttcaaattatggacgatgttctagatataactggttcta
ctaagagcttagggaaacctgctggcgctgatctaataaatggaaatttgacctctcctc
tcctttttttcacttactcaagaagcaagtttaaatgatcttattgatagggagttctgta
atagtacagatatagcctcaacattatttcttataaaaagaagcgggggaattacaaaag
ctaaagatttagctaaagaacaggtgcaggcggcactttcctgccttcagtttttaccac
aatctacacctgtatctagtttaaaagaattaacacatttcataatcacaagattgtcat
aaaggacttgctaaaattaaaatgttattaactaatttttttactgtttctaaaacagtct
gaaaagtttgaatatcattactagctagctgcgctagcattttacgatttaacgcaatat
tctcttttttttaatgcactaataaaagtactataattcatgccttgattgtgagcagcag
catttattctagttatccaaagacgacgaaaatctctctttttcctttttcgaccaacat
aagaatatcgaagagctttaaggacttgctgcttagctgttcgaaataaacatttatgtg
cacctttaaagcctttagctagcttaaagattttagctcgtctttttttttgcaacgttac
ctcttttaactctactcataaaatattac-ttaaattttaattattaccagcttactaaa
ttcttatactctttg----attataaataaggtaatttcattgcaatgtttttaatatct
tttaagtcaaccgaacaagtagaagaaagatgtcttctttgttttgatgactttttttgt
aataaatgacttttagaagctttatgtcgaagaaattttccggatgaagagactttaaat
ctttttgctattgcttttgatgtttttaacttaggcatattataaacttttaatgattaa
caaaagcatgaaaagtctgaggatatcaatcttagacctttcaatacttagaataatctt
aacacaatttattttacttactacataggttaactgaaacaatttatatatataatacta
atcagttaatcaaccatagctaaagtggatttatattttgctttgaacttttgtatagta

ttagcaattagcattgttataattttaagactgcttgaattcaattcttcgaacattttc
atatttaatgtaaatgaaatattggcttctgagacaatattttgaatttgaacatcagac
aagggaatcatatctaatgcagctctatattgatctttaaataatttatcgtcttttatt
tggtcaaaatcgtaaaatttcgttcctcctgaatctgataaattcatagcccctctagct
atttttttaaaatttggccgccagaaaggtcacccaaatagcgcgtataagcatgagca
actaataattcaggttgtttatgtcctatagtgtgaattctatcaacataaatttttgta
gcaggagatggttcaataaaatctaaccaatctgatccataataatagtttaaatcttca
gatagactcgctttcctattaagctctgtaaaatatataggtttaatagctggatgattt
ttattagaaaataactcttcctctattgcacaatagacaaagtatagatttgcaaccaat
ttacgatatgacttcttatctacaactcctcccaaaaaagatttaacaaaactaacattt
tctgccatactatgggacttagtagtaccttctcttagttcattcgctaaagtattaacc
ataatatcaatattccattttt-aaagataacaaattaaca---ataaatagttatccgt
atatttataattgagttgcgctaactaggctgctgcaaaataatttttgattttattgg
atctggattcattgttttgtcaccaggcttccaattagctggacaaacttcatctggatg
agattgtacatattgaattgcttgtaaaaccctcaaagtttcttctacactcctgccaaa
ctctagattattgattgtagagtactgaataattcctttcgggtctataataaatagtcc
tcttagagctacaccatcactattcaaaacattataagctgcactgatctcttttttaa
gtctgatactaatggatatgaaagatctcctaatccacctgattctctgtcagtctgtaa
ccaagcaagatgagaatactcactatcaaccgagactccaagaacttctgtattaagttc
cgaaaaagcattatacttatcgctaaatgctgtaatttctgtagggcacacaaatgtaaa
atctaaaggataaaagaataagacaatatacttattcttgagatcagataattttaacgt
tttaaattcttggtcataaacagctgtagctgaaaaatcaggagctagctgacctactcg
aagacaatttggtccagaaatcattaattttcccagaagaattgatataattttgttat
ttaataaataattatataacaaaatttagtatatattaaaatactacaagtctttttaa
attac--------taaatgatataatcaatactatctttatagcggggaatggatttgaa
ccattgaccttcgggttatgagcccgacgagctaccagactgctctaccccgcggataac
taaaatcatattagtctacatatatatataaaagcaaatgcataatagattttagaagta
aaaaaatagaagaactggtccactgatcattaaacttagaaacgacaaagtatttttat
cttatttaaaagagggaagacttcatataaaccaataaatctatcttttaaaagaatttc
ggacggctttacccaaatttgaccgtcataccagcctgattcttcataaaatactgtagc
actcattaatcgttttactacgtatgaccagcctaaatataatctgattaaaataaatcc
tgtcattaggctggaagtaataaattctgaaaagaaaaatttcaaaggtagtttagtgat
tggaaaaattgataataatataggactaactaataaacagttaagtagtagtgcaatagt
tattttttattataagatctacggcttaaagttggccagcaaaagaaccaagaatttt
caaagaagtatattcatgaacaggttgctgctcctttggcacaggacattgagtattata
taaattcattgacattaaaaagtattataataaagac--tttgtttatattatctaacat

aggtacgacattaaatattaactaactgcaattatagtagtgagtagaagaaataaaact
aaaaatagccaagtaattttatttaaagtattctcagtactacgagtattactaaagaac
tgattctgagcacctacacttcctaaaccttcggatttaggattgtgtattagaatagtg
aagattaatattattgtagatgagtaccaaaaaaattttaaaatttgttccattatgtaa
aagttaat-aaaaatagcacaaagacaatcgtctaaaagatattgattgtcttatcatat
taaaaagtatttattcaccctgtacttttaaaagtaaaaatcctattgccagccctacca
aaactaggactgaacttaatacagcactttctacaatttctccgaccatt-ttatttttc
cttattattttattattg---tcattcttttgacaagattatattgagctaa-ctaaaat
ccgtttctaccccaaactaccattgcgattgaaaatgtaaacattgccattagagctgcc
catcccaaacttaaaatatccataatttataatattc--ctaaaaataagatctctttta
cttaaactataatattcttaatattaaaaatatcattatataataagctaattataacag
agtactacttaattcgtatttttagtgtatttaaagaagattgaa-tttttcgaggtac
tggataagtatttcatcctaagaaccatagtcaac----ttttgatttattataatgctt
aaatctaaatata--tatcactcaagaatcattagtaattatgatgtttttaattatacg
att---aacatatttcaaaatagaagaattcatagcgactaagttaa-----tatgctaa
tataatttatcaat--aaaaaggaagttagtatttcaattatgcaaacaactataaataa
tgggcaaacgtctagtaaagaaactttactaacacctagattctatacgactgactttga
agagatggctaatatggatatttccggcaatcaagaagatttttttggctatcctcgaaga
atttcgagctgactataatagtgaacattttattagagacgaagagtttaatcaatcttg
gtctaatttagaacataaaactaaatccttatttattgagtttttagaaagatcttgtac
cgcagaattttcaggtttttttactatataaggaattatccagaaaattaaaagacagaaa
tccggttatagctgagtgcttttttattaatgtctagagatgaagctaggcatgccggttt
tttaaataaagctatcggggacttcaatttatctttagatttaggatttttaacgaagag
tcgcaagtatactttttctcacctaaatttattttttatgcaacctatctttctgaaaa
aattggatactggagatacataactatttaccgccatcttgaacaacatccagaacaccg
tatttatccaatctttagatttttttgaaaattggtgtcaggatgaaaatcgtcacggaga
tttttttgctgctttgctcaaatcccaacctcattttttaaatgactggaaagcaaaaat
gtggtgcagatttttttctattaagtgtatttgcaacaatgtacttaaacgactttcaaag
aattgattttataatgccataggcctagactctagacagtatgatatgcaagtaatacg
aaaaactaacgaaagtgccgctagagttttcccggttgctttagacgtggacaatccaaa
atttttcaaatatttagatacttgtgcatgtgataatagagctctgatcgatattgataa
taacaattctccattattcattaaatctatcgtaaagatacctctatatttttccttatt
tgcaaatttactaaagatatatttgattaagccaatagactcaaaaacagtatggaatac
agttcgatagttaaatacaagggaatgctttttttcttataaaaagctatctagtgattaa
tataaaat------aaaagcttccttct-----cgatggaaggagcttttattttgtcaa
ga---atataatgggattatatagatc-tagaactaagaacttacttttttcttttgcttc

atacataacctctaaagtaattagatttttagaacaatcacgggcaaatttctctgtatt
tctttttactttacctctaacaaacccaggaatttatttaattctttctgtgcttcttc
tgaccaattaatactatcaacagcagaaattcctatacttaatgcttctgttgtatcatg
accaccaaatatttctagcaaatgatcttccataccaagtgtaaacgaattatatactaa
atcagcaatttgatttgttccctcataaccaagaaatggacggtaacttaaaggaaagtt
ttgtatatgcactggtgaagaaataacgccacaaggaatatttagacgcttaccaatatg
tctttccatctgtgttccaaaaattgctgcaggttcagttttagcaattaaatctccaat
taaaccatgatcatctgaaacgattacttcatcgcagaattcttgaacttgatctttaaa
ccactcttcatcatatttgcaataagttccacaccaggcgacatgtatacccatttcctg
gtgtaaaattcgagttattgctgcagcatgagttgcatctccaaatacaatagccttttt
acctgttaaattctgacaatcaattgatcttgaaaaccatgcagattgtgaaataaatcg
ggtttgctcgtctatgtatttttcgtagtctacagcagctcccaatgcatttactaattg
ctgtattgatcggatgcatgcagctgtttgaacaatacccataggtgtaatgtcaacata
aggcatattaaattctttctctaaatatcgagctgtcattagtccagtttctctatatgg
aataaaattaaaccatgcagagggtaatttttttaaatcttgaactgatgcattttctgg
aataatttggtttatttggatatctagatcttgaaataacctttttaattctgctatatc
atgttgattatgaaatcctaagcttactgctccaataatattaacagatggagttttagt
cttttgagttaaaacttgattattagatttagctttttccatatagaaagtgacaatttg
ttctaatgttctatcacctgcttgcagttcatttactctataatgattaacatcagctaa
taaaacatctgcttctgtttctatagatgctctgctaacaaagttttgcaaatcttcttg
caaaatactcgaagtacaagttggtgttagaataactaaatcagggctttcttctctgtc
ttttctagtaatattttctacaactttttcttgagaaccacgggctaacacatgtctatc
tacaacacttgctgtaacaggagtaaaatctctgtctctttcaagcattgaacgcattac
attaaaataatcgtctccgagaggcgcatgcataattgcatgaactttttttaaaagaact
ggcaattcttaaagttccaatatgagcagggcctgcatacatccaataagctaatttcat
attaaattttcctatgatttatttagacatagtaatacttttaaagctaactaacaaaca
ttagccaaacatctaatttaactccttgacaattttgtaatattttttaaatgaagctta
tcattataaatatttaaaaaatataattcagaataga----aataactgttagatcaagc
ttgaaagtaaagatttcttgcagttatagtttttctttaagtaaacatgtaaaaattgta
ctatatataaatgtactatatataaactttttttactgaatattggtaatgccagataca
attaacttaaatatgccttccccaacgtttggtggaagcactggcggttggttaagagcc
gcagaagtagaagaaaagtatgcgataacatggacaggcaaaaatgaaagtaagtttgaa
atgccaactggtggtaccgcaactatgcgaaatggagaaaacttactctatttagctaag
aaagaacaatgtttagctttaggtactcaactaaaaagtaaatttaaaatatctgactat
aagatttacagagtttttcctaacggagaagtgcaatatttgcatcctaaagatggagtc
tttccagaaaaagttaatactgggagagctagcgtcaacagtgttgatcattctattgga

caaaatattaatcctgtagatgtcaaattcatgaataaagcaacttacgattagtctctc
-aaaaacctagacataaaataacttttatatagtctaggtttttgataatgtatactgat
agtatggaagggtggccgagtggttgaaggcgtctgatttgaaatcagttgaacta--tc
tggttccgtgggttcgaatcccaccctttccgttataaaact------------atagtt
ggcttctcaaggcaagctagctatacactgatagtattaagtttagacttaataaagtct
acaacttgtgacaggttagaaatttttcggcatcctcatctggtatctctatactaaat
ttttcttctatggccattaccagctcaactgtatctagagaatcggctcccaggtcactt
gaaaaatttgcttccctagtcactatttttttttcaattcctagctgttctgctacaata
tcttgaactttttcaaagatttcattatcttgcataatatattatttcctctaaggtgtt
atcaaaaatttttcggcttatattattatccaactctaaaaaattagtaatttttagtcg
agattaaaatatttatacctatatgatatactattattctggataaatacgaagtttacg
atatggctcttctccaaaactacgagctctcaactggtagttgtcaactagattatgctg
catttttcgtatataggctgatcgaggtgtaagctgaactattgaactttcattcaagat
aatagtttcaatagccaatttagcttcttgaagagcctggatctcatcaaaatttttatt
tttacagagttctacccaattaagaccagatgaagtatgaatgtttaatatttttctgag
tgcacgagtaatttgtggaaccgtactattctgaattgtatatataataatttgttttga
ttttgcaatttgtctaagttttgtattttgctttacttgattacgtagcgctagaatagc
atcagactttctatctccttggttaaaataattggtagatctagtgaagaaataactga
tgtaatatgctgccaactcaaagaataagcatataagtactgatgaggtatttccaattg
caaagattgtttattaatattaataggggcattgaaagtagttgacaataaagatgtatc
acgattttgtgatttattcaaatctaaggacttatttcttaattctctatattgtaaaac
cggaggcgcctgctgtgcgttaattgattttcgagcattagagatgtttacaggtaagac
ttctatagattgcgatggataacatttaattaaaattctaccattagcttgaatttgtct
tttttgcacaaaaggttgatggccttgcaaaatttgatctattgtctctttaactttatt
atgaactatccaaacatttcgctcgtgtatttcgatcgcaatctgaaaagcaggctcagc
ttttctttctagtatactcttttgcgtacctctacgtttagcttcatcgtcacctaatgt
tacatattgaattccaccaattaggtccgccaatgttggatttttaattaaactttctaa
atgatttccatgcgctgtacctactaattgaactcctctttctgctatagttcgagcagc
taacgcttctaattccgtgccaatttcatctataataataacctcgggcatgtgattttc
aacagcttctatcattacttgatgttgcaaatcgggccttgcgacttgcattcttctagc
tctaccaatagctggatgaggaatatctccgtcaccagctatttcatttgaagtatctat
aataaccactctttttttccatttcgtcagctaaaacacgggccatttctcgaactgcggt
tgttttaccaactcctggttttcctaagagtaaaattgaatcaccttgctgtagtaaatc
tcgaataatactaattgtaccaaagactgctctgccgacacgacaggttaagccaataac
actaccttctctattacgcagtgaactaatacgatgcaaggtttttttcgataccagctcg
attatcgccgctaaaattaccaactttttttacacaataatctaagtcttgccaactaat

agatctttgagataaatattctgggttatctggaaatctagcttctggacgacgacctaa
atccataacgacttctattaaattatttctattaggatgctgttgtaaaggttctttgac
aaaatttggcaaaatttctagcaacttatctaagtcatctgcaataagcatgattgagta
tataataattaattaatcaaatagaagattcaagattctaaagattacaatacatattaa
agaatattatgattggaatcaattttatacttaaattgaacaggaagtta-tagatgtaa
aagaactgattaaactttacgcaataaatcgattggtcaaaattaatagtcatgttaaca
tgtttattggacaaaaagttcgtattaaatatagcaaacaaaaagttgctagtgatatag
cagataaagttggtgaattaggagttattaagggaataaagtttattaatagtcaatgtg
ttactattattgttgagtttgacaatcacactaggctttggatgtttagagaggaattaa
tctgtctaaatgaat------taaaaacatgaaacataatttatttatttattatacaa
aatttcttatgagtcgttaaaataataaattttaaaggagtacttttaattgtgcaaatt
actattaaaaaattacaagacttactttcgtctgtacaaagaaaaaagatccagacatta
aaactaaaacaaggtaagtttgaacttctattaaataagacctataaaaaagtcaatcaa
gaaattataccttcgcagaaatctgctgtattacaaaatagtccatctacgataattaaa
tcaataaataatacaaaaaaatcttctgttgtcaatgaagaccgtacagaatatgccact
attgtttctccaatggttggaacgttttatcattcacctgctcctggtgaaaaaatttt
gtacaagttggcgatgaggtcaaattcaatcaaacagtctgtattattgaagcaatgaag
ttaatgaacgaaatcgaagcagaaattgaaggcaagattatagaaattcttgttaaagat
ggtgatatagtagattgcgggcaagccttaatgaaggttgaaacataatgctaattttc
cattttacatattgttaacagaaattatttagatagaaaaattagcttatactatagtta
tgaaaacttatttcgcaaaggcac-taaaaactaatatataaaattatttactttatctg
ccaaaaatagaaataagcgttttgattccttgtctcaagagaggagaatctcaatggcaa
ttagctcaaaagagcaagagacaaagaaggtaaaaatctcggttgataaaaatcccgtag
atacttctttcgaaaagtgggcccaaccaggccatttttctcgtacactagcaaaaggac
caaaaactactacttggatttggaatcttcatgctgatgctcacgacttcgatagtcaaa
ccagttctttagaagaagtttcacgtaagattttcagtgcacattttgggcagctgtctg
taatatttttatggcttagtggaatgtattttcacggagcccgcttctctaattatgttg
cttggttaagtaatccaacaggtattaagccaagtgcgcaggttgtttggcctatagttg
ggcaagaaattttaaatggcgatgtaggtggtggctttcaaggagtacaagttacatctg
gatggttccaactgtggagagcatcaggaattactacagaatttcagctttactgtactg
ctattggcggattagctatggctgctttaatgctgtttgcaggatggtttcattatcata
aagctgctccaaagttagaatggtttcaaaatgttgaatcaatgatgaatcaccatttag
ctgggcttttaggcttaggctgtttaggctggacaggccatcaaatccatttgtctttgc
ctattaataagctactagattctggcgtgtctccgcaagaaattccactacctcatgagt
ttttaattaatagagagcttatggctcagctgtatccaagttttagtaaaggattagttc
cattctttactttaaattgggctgaatattccgacttttaacttttaaaggaggtttaa

accctgttactggaggtttatggctaagtgatactgctcatcatcatctagcattagctg
ttctatttcttgctgcaggtcatatgtatagaaccaattggggtattggacatagcatga
aagaaattctagaagctcacaaaggaccttttaccggcaacggtcacgaaggtctatatg
aaattcttacaacttcttggcatgcacagcttgcaattaatttagctatgatgggatctt
taagcatcattgtagcacatcacatgtatgcaatgcctccttatccatatattgctactg
attacccaactcagttatcgctcttcactcatcatatgtggattggaggattttgtattg
ttggagcaggagcgcatgcttctatatttatggtaagagattataaccctgcagaaaatt
ataacaatcttttagatagagtcattaggcatcgagatgctattgtttctcatctaaatt
gggtatgtatatttcttggattccattcatttggtttatacattcacaatgatactatgc
gggcacttggaagatctcaagatatgttctctgatacagctatacagttacaacctattt
ttgctcaatgggtacaaagtatacacactttagctcctggaaatacagctccaaatgcat
tagcaacagctagttatgcatttggaggagatattgtttctgttggtaacaaagttgcaa
tgatgcctatttctttaggtactgcagatttttagttcaccatatacatgcatttacta
ttcatgtaactgttttaattttagttaaagggttccttttctcaagaaactctagactaa
ttcctgacaaggccaatcttggcttcaggtttccatgtgatggacctggtagaggtggta
cttgccaagtgtctggctgggatcatgtttttcttggcttattctggatgtacaattctc
tgtctgtagcaatttttcactttagttggaaaatgcaatcagatgtttggggtagtgtat
ctccgtctggaaatgtttctcatattactggcggtaattttgcacagagtgcaattacaa
tcaatggatggctgagagatttcctttgggctcaagcatctcaagttattcaatcatacg
gttctgcgctatctgcatatggattaatttcttagcagcacattttgtatgggcattca
gtttgatgttcttatttagtggtagaggttattggcaagagcttatagaatcaatcgtat
gggcgcacaataagataaaagttgctcctgcaattcaaccaagagctttaagtattactc
aaggtagagcagtcggtgttgcacactacttattaggtggaattggtacaacttgggcat
tctttttagcgagaattatttcagtaggctaatagtgaaaataggataaaaaaaacaatt
atggcaacaaaatttcctaagtttagccaagctttatcacaagatcctacaactagaagg
atttggtatggtattgctacggcacatgactttgaaagtcatgatggaatgacagaagaa
aatttatatcaaaagatattcgcttcgcactttggacatctagcaattatcttcttatgg
acatctggtaatttattccatgtagcttggcaaggcaactttgaacagtgggtattaaat
cctttgaaagttaaaccaattgctcatgcaatttgggatccgcattttggacaacctgct
ttgaaagcttttagtaaaggtggctcagcttatccagtaaatatagcatattctggcgta
taccactggtggtatactattggtatgagaagcaatcaagacctgtattctggggcttta
ttcttactagttttatcagctttactcttatttggagggtggctacatctacaaccaaaa
ttcaagcctggtttatcatggtttaaaaataacgaatcaagattaaatcatcatttatct
ggattatttggggttagttctttagcctggacaggtcatttagtacatgttgctatacct
gaggcaagaggacaacatgtaggatgggataattttacaaccgtattacctcatccagcc
ggtttacagccgtttttcagtggtaattggagtgtatatgctcaaaatccagatacagct

caacatttattcggaactaatgaaggtgcgggtacagcaattctgacatttctaggagga
tttcatcctcaaagtcagtctttgtggctaactgatatggctcatcaccatttggctatt
gcagtagtattcattgttgctggacatatgtataggactaattggggaattgggcacaat
ctaaaagatatttagatgctcatagaccacctagtggtagattaggagctggacataaa
gggctatttgatactattactaattctttacacatacagttgggattagcattggcttcc
ctaggtgtaattacttcgttggtagctcaacatatgtatgctatgcctccatatgctttc
atggctaaagatttcacaactcaagcatctttgtacacacatcatcaatatattgctggg
tttctaatggttggagcttttgctcatggggcaatattctttgttcgagactatgaccct
gaacagaataaagataatgttttggctcgtatgctagaacataaagaagctatcatttct
catttaagttgggtaactctattttagggtttcatacattaggcctttatgttcacaat
gatacaatgattgcttttggaactcctgaaaaacaaattctaattgagccggtatttgct
caatggattcaagcctcttcagggaaagcactttatgggtttgatgtgttactatcatcc
tctactaatatcgcaacacaagctggtagcaatatttggctgccaggctggttagaagcg
attaatagcggaaaaaattcattgttttaacaattggtcctggtgacttcttagttcat
catgcaattgcattgggattacatactactacgttaattttagttaaaggtgctttagat
gcaagaggctctaaacttatgccggacaaaaaagactttggatatagttttccttgcgat
ggacctggcagaggcggcacctgtgatatatctgcatgggatgcgttctatttagctgta
ttttggatgctaaatacaataggttgggtaacattttattggcattggaaacatattaca
atatggcaaggcaatgcaactcaattcaatgagtcttcaacttatctaatgggatggttt
agagattacttatggctaaattcttctccattaattaatggttataatccatatggcatg
aataatttatcagtatggtcatggatgttcctatttggacatttagtatgggcaacagga
tttatgttcttgatctcctggcgtggctattggcaagagttaattgaaactctagcatgg
gcgcatgaacgtacacctttagcaaacttgattcgttggaaagataaacctgttgcatta
tcaattgtacaggcaagattagtaggtttagcacatt-tttctgtaggatatgtattaac
ttacgcagcttttgtattagcttcaacagcaggcaaatttggttagactaaa-----ttc
gaattagctaa-----aaaaaagtcagtcataata-gttttatgactgactttttagat
ttaaaacaaaaacttactaaagtttcacctcaatatctacaccagaaggcaaatttaact
tcattaatgcatctatcgtttgagaagaaggttgatgaatatcaataattcttctgtgag
atcttatttcaaagtgttctcttgagtctttatctacatgtggagaacgtaaaacacaat
aaattcttctttttgtcggtaaaggaattggccctactgcaacggcattagttctagatg
cggtatctagtattttgtcacacgacgtattaagtataatagaattatatgctttcagtt
taattctgatttttgtctgctgagtaattgtcatgttaaaaagatcttattatttaagaa
ttttagagacaacacctgcgcctacggtacggccaccttctctaatagcgaaacgcatac
cttgctcaattgcaattgcattaattaattcagcagtcatcttaattctgtcaccaggca
taaccatttctgcatcagtaccatcatcagcagtaaactgattaatagtaccagttacat
cagttgttctaacataaaactgaggtctatatcctggaaaaaatggagtatgtcttccgc

cctcctctttagttagaatataaacttctgcttcaaattgagtatgaggtgtaattgtac
caggtttagctaataccatacctctttcaatatctttttctgcacacctcttaaaagaa
ttccaatattatcacccgctagaccttcttctaacgtttttgaaacatttctaatccag
taatagtcgttgtacgagtttctcgtaaacctacaatttcaattgtgtcaccaactttaa
taatgcctctttcaattctaccagtggcaacagttccacgtcctgtaatagaaaaaacat
cttctacagccattaagaaagttttatcgacatctctctccggcgttggaatatatgtgt
caactgcttccataagtgaaaaaatcttgtcaacccatttatcttcaccttgcttaatag
ctgggtttttcgtcacagcttctaatgccaataaggcagaacctgcaacaaaaggaatat
catctccaggaaagtcgtattgacttaataattcccttccttctaattctactaattcta
gtagctcttcgtcatctacttgatcttccttatttaggaatactactaatgtaggaacac
ctacttgttttgctaataaaatatgttcacgagtttgtggcattggaccgtctgccgcag
atacaactaaaatagctccatccatctgagcagcacccgtaatcatatttttacgtagt
cggcatggcctggacaatctacgtgagcataatgacgattatctgtttcgtattcaacat
gagcagtattaatagtaatacctctagctttttcttctggagcggcatcaatttcatcaa
atttttcgctgcagtagaccctaaagttgataaagtcgcagagatcgctgctgttaaag
ttgtcttaccatgatcaacatgaccaattgtgccaatattgacatgaggtttttacgtt
caaatttagatcgagccatgcttttgttttccttatagtaaaagttgctttatagttac
ttttaacgataaatcttgtggattgaaagatttattatttaatacagacaaga-----ta
caatataactaatctaatatcgatagtgagcaaacgctttatttgcttctgccattctat
gtgtatcctctctctttctaatagaatttcctgtttcattagctgcatccataatttcat
tagctaatttcatagacatacttttaccagatctgtctctagagaatttagtaatccatc
ttagtgctaaattcgtacctctataagctcgtacttcaataggaacttggtaagtagaac
caccaactcttcttgcttttacttctactagaggagtaatatttcggatggctttttcta
aaatatttaaaggatctgattctgttctctctttaacgatatctaaagcctgatatataa
ttctttgagataaagtttttttaccacttttcaaaatacgaacagttaacatacttacga
gtctgcttttatataaaggatcaggtgatgcaaacctttttttagctgtattacgacgag
acatagtatttaaaattggttattgtaataa-tgttc-tattagcaggtatttaagattt
aggctttttttgtcccatatttagatcggcttttacgacgatcttttactcctgcagcatc
taaagtaccacgtactacatgatatcggactccaggcaaatctttaattcggccgcccct
aatcagtaccacagaatgctcttgaatattatgacctacgccaggaatataagctgtaac
ttcaaatccagatgttaatctaacacgcgctacttttcttaaagcagagttcggtttttt
aggagttgtagtatatactcttgtacaaacacctcttctttgagggcaactttgaagagc
tggagattttgtttttttatgtatttttcgtctttcggatctaacaagttgttgaattgt
tggcataataaaattgatttaaagtctttaatagatactatttgaacttgctttacatga
attggctagtttttatttatcttatacttacgcataaatctttctactctaccttctgta
tcaataattctctgtgaaccagtaaagaatggatggtttcctgaccagatatctacgtgt

agttctggctttgtggacccaatcgtcataattaactggccgtcacagtaaacttttgct
tctggataccaatttggatgtatattatcttttgccattgtttttataattatataaatg
tatcaataataactattaacgcttagaaaattgaggagcctttctggctttctttaaacc
atatttttttctctcttttactcttggatctcttgttaaatagccttcagacttaagtgt
cgttctattttctggattaattgagcacaatgctcttgcaacacctaaacgaattgcatc
agcttgccctgtcaaacctcctcctctagcatttacatggatatcatactggtttagtag
tcctaaaacttgtaatggtgcatatgaaactcttaagtaattaggactaaattgaagata
agactctcctggtataccgtttataattaaattacctgaccctgggactagtcttacttg
tgcaacagaacacttccggcgacctgttccagaatagattgcgcgagttttaattaattc
tgtcgacataacattccttgataactaaataaatatacattctttatactatatactctt
gtggcttttgcgcgacatgtgggtgaattgggccagaatacactttaagcttcgtaaata
gttttctgcctaatggacctttaggaagcatacctttaaccgatttttcaataattctgt
taggtaatcttgtctgaagctgatcaaatgtttcaacttttaatccgccaggctgtccag
aatgtcttctgtatagtttttgattcgttttatttccacttacagatacgtgagcagaat
taataacgattacataatctccagtatctagatagggtgtataagaaggcttgtttttac
ctcttaagatattagaaatatgagtagatattctaccaagtgtctgattttagcgtcta
taacataccaatgagaattagtatttaatgagggtgattgcgttttattcatgaaaagat
taatacttttggttgacaataagagaatatgtaaatggtttacattcaatttctatacaa
gtagtattt-aaaaatatta--------------ttaacaaagagaactttctaacagtt
atttaaattgtttgtcagcaagaaaatttctagttatcaataacttatactatttttaat
aatataaaatatattaatcactttttttcttttggcaaacttatacctagttttttttgc-
aaggcatatataacttcttcagctgatttctggccaaaatttttaatttctagcagttct
tcttgagagtaatctagtaaatctgcaatagaatggatttgagctcgctttaagcagtta
taagccctaacagataactgcagttcttctattaaaacttgactaatttttttatcttct
ttgctacgataattatctgctgatttaaagtctaaatttcttagtgaacaaaaaagatta
gttaaaactgtagctccttgacttattgcttcttgaggagagatgctcccatttgtccag
atttgtataatcaatctatcttttatgctgttgctaccaatacggacttcttctacttta
taattaactttattaactggcataaaaacagagtctacttgcaaaaaatccacagataat
tcatctacagctttttcagctaagcgatacccgcaatttttttcaattttaaactccatt
tcaaatattgtattattgcaaatagttgcaatatactgtctaggatctactacctctata
tcagaagataattcaaataggccggctgtaactatagctggcccttgaactctaattcga
ccaatttgagattctttgttataacttttaaatactacttcttttagattaagtaatatt
tctaacacatcttctcttaccccaggaattgtagaaaactcatggttcactccagcaatc
cgtacagcaactatagcagtaccttcaagatctgacaatattgatcttctcaaagcatta
cctaatgtaataccttgtccttgatttaatggttctattacaaaactaccgtactgccca
cgcgccccatctgttcttgactctatgcattcaatttgaaattgagccacctaagaaagc

tcc-tttaaaataatcagtgattagtaagtctctctaaaacaagaaggaatttatactcg
gcgtttcttaggagggcgacatccattatgaggtacaggagtaatatcttttattagagt
aatctctaatcctgcagcctgcaaagctctaattgctgtttctcgacccgctcctggacc
atttactagaacttcagtttggcgcataccttgatccatagcttgtctagctgctttttc
agctgctgtttgagctgcaaaaggtgttcctttttagctcccttaaatccacttgcacc
agaggatgaccatgataatgtttctcctttaaattagtaatagtaacaattgtattatt
gaatgtagatttaatatgtgtaataccgttaactgcattacgtttagtttttcttgctcc
ggattttttatttgtctagccatcgtgtctcgtctattatatagttaattaaaagatta
ttttcttggagctttttcttacctgctactgttttttacctcctctgcgtgttctagc
attagttctagttctttgtcctctcaaaggaagaccaagacggtgtcttctacctctata
agtactaatttccataagtctcttaatgctcatagactcaaaacgtttgagatccccttc
aatctgataattagactcaagaatttctcttatactgacaacttgttgatcatttaaatc
ttgacacttaatatcagcgtctatgtttgtttttctaatatttctttcgagcgagatag
tccaataccataaatatatgttaaagctatctctattcttttgtttcttggaagatctac
tccagcaattctggccactttgtcttgtctccaataaataatatgtat-ttttataacgg
ttgtttata-agcttagccttgtctctgcttatgtttaggattagtgcaaattaccatca
cctttctgtgacgccgaattattctacatttttcacacatttttcgaacagaaggacgaa
ctttcatattaa----ctctatatactaggaatattactatattaactctattaattatc
gacattaaaaatgtcaacaatatattgtactactttaactgttgtagattttacttcgtc
atactatcatattttttgatatgacgtatgtctgaatttgcttagctgtgtctattgcg
actccaactaaaataagtaaagacgtagccccgagacctcttaagttttggatctgagta
accttttctattataaacggaatcagcgctactgtaaataaaaacgaggctcctaggaat
gtcagtctatttaatatgacttgtaaataatcaatagtcgcttgacctggacgaatatta
ggaatgcttgcacccatttttttaaatttatagcaatatcttctggattcattactatt
gatgtataaaaatagctaaagaaaaggatcaaagcacaataaagaagaaggtatagtgaa
ccattaggacaaaacaggtataggatttggagtaatgttttattttgaataatttgggtt
agataagatgggagggccatagatgcagatgcaaaaacaataggcataactccaccttga
tttaatttcaagggcaagtaactatttggatccaaaattgaagattttcctagctgtctt
gctgaaataatttaattcttcttgttccttcttgtacacaaattgtaattattatcatt
aataaaaagatcgctataaataatccgaacttaagacttgcattactataactagcatca
aaaaatgactgtgtaaaattctttggtagtcctgacacaatgttttgaaaaataagtaaa
gaggctccattaccaattcccttttccgtaattaactctgataaccacataataatcata
gaacctgctgttaaagctaaaacagactcgcagacaaatgcaaaattccaattaaaaaca
tatggttttacccatatagagattgctccagattgtaaagtcgcccaacctagagctaag
taccttgtaatttgagttatttttggcggcccaattcaccttcttccttctgtaatttt
tccagattggggacaattttcgtaagtagctgcattacaattgaagagttaatataagga

acaatacctaacgcaaaaatccctattgttgaaaaacctcctccagaaaaaatattcaga
aaatttactaaagtatttttttctacacttgcataaaaggcgtcatgatctatacctgga
acaggtataaatattcctaaacgtgctaaaactaatagaaagagagtaaagataatacga
tttcttagatcactttttttggctcataaataaattttaaatgaaaagagatactttt--tt
tatgtaaaaaccctattccaagacttgtataagtttaaaatagggaagtaa---aatatt
atagtaatattaata-taatattctatgtagaatacaggctttctattggaactcctcga
tctttagctgcttcactaaaagttcttaattgagttaagccattgagaacagctcgcgcg
ttatttaatgtattattagaacctaattgcttagctaatatattctgtacacccgacaat
tcgaggactgtccttacagaacctccggcaattacacctgagccaggtgcagagggcctt
aatataactttcgcggcaccagaaattccgttaataggatgaggtatagaattcgatttt
gtcagtggaactgtaactagatgtttttttagcatctgttactcctttttttactgcacca
attacatcgcttgcttttcccacgccgacaccaacttggccttgctcattaccaataaca
agaataactcgaaagcttagtttttttacctccttttacaactttagtaactcttttaact
tgtacgactctttcttcccagccactatctttatcttttcctttgctctgtttttttacga
ttggccattttaagaattattcctaattagttaatta-cgattctagaaacccatgcctg
cttccttagcagcttcagccaaggccttaactcttccatgatataactttcctcctctat
cgaagacgacatttttaatgccttctttcatagactgttctgctaactgtttaccaacta
cgcgagaagtatcacagtttggtcttatgttatcagattctttattatttaaattaacag
atgatgtagctactaatgtaataccttgtgtgtcatcaattatttgtgcgtatatatgtt
tattagatctaaatacacacagacgaggcctacttgaggttccttgaactttttttccgaa
ctcttttatgcttatgaattctagtttgtttagtgtttagtttcattattatttaccttt
tccagctttcccagcttttcttctaacaaattcaccttgatatctaattccttttccttt
ataaggctcaggaggcctaatagaacgaatagttgaagcaacctgaccgacaacttcttt
atctatgccagagacagtaatatttgtgttgttttcaactttaatttcaatattggcagg
aggtttaattttcactacatgactatagccaacacttaagattaaatcttggttatcaat
ttgagaacgatagcctacaccttgtatttgcagtttttttaaaaaatccattagaaacacc
ttcaatcatattactaataagagttctggatagcccatgtaactgactagccatttttgt
ttgtccactcgtcttgacagcaattgtgtcgttaagtatttctaaattaataccagctgg
taaagttctggatagggtacctttaggtcctgtcacggtgatagtctgaccatcgaactg
agtactaagatttgtcggcaataaaattatttttttttccaatacgagacat-attccacc
tataaaatagttaccaaatatagcataatatttcaccaccaagaccatcatgacgagctt
gtctatctgtcataacgcctctagaagtagaaataagagcgatgcctaaacctccaagaa
ctctgggcaactctttatgatttgcgtagactcttagtccgggtttactaattcttttca
gagcagtaataactggttgacgatttttaccattatacttaagagaaatcattaaatgag
tctctataccttcacccatttgttcgaaattttgaacaaatccttcttctttcagtactg
ttgccatattgcatgtcattttcgttgctggaacttgcacaatttgatgtcttgctaagt

ttgcgttacgaatacgtgtcagcatatcggcgatcgtatcgttgaccaccttgatcctcc
ttgcga-----tgaattatttgaaatattttcaagattccctgaaaggcatccctagctt
ttttaacaaagctagaccttcttgatctgttttagctgtagtgacaattgatatatctaa
accacgaatttgatcaatattatcatagtctatttctggaaagatcaactgttcacgtaa
acctaaattgtaattacctttgccgtcaaaacttctaggactaattcctctgaagtctct
aattcttggtaatgttaaattaattaatttctctaaaaaagaatacatcttgtcttttct
tagatgcacaacaattccgataggaacctcttctcgaattttaaatcctgcaatagattt
tttagcttttgtgacaattggtttttgtccagttattaatgtcaattcttgaatactact
ttcaagagccttagcattctgagaagcttcacctaagccacggttaatagtaattttagt
aaacctaggaacttcatgtacatttttgtactgaaattcatcttttaaagattgagtaac
agttgttttatatttttcttttaatcctattgccattatattttcatcaattatttaata
agttcgccagtctttttcagttttcgaa-tttttggcctttatcattaattattactga
ggatcgactagcaatattattttgttcactaaataacataacgttagaggtatgtattgg
agcttcaaatttgataatttctcccgtttctccttcttgttgaggtttttatgcttcac
tttgagattaattcctttaacaatcaccttatttgttttataaataattgcaataatttc
acctgtttttgttttatcacttccagaaataacttgaactaagtcccctttttttaattt
aatttttgtattattttttggttgttttagaaggacctttcattatactacctccggtgca
agagaaactattttagaaaaatttttatctcttaactctcgggctataggtccgaagact
cttgtgccacgtggattattatcttgattaatgataactgccgcattatctccaaatcta
atgctcataccgtcagttctccgtaaagctttacgagttcttactacaacggctctgaca
acatcagatcgtttaactggcatattgggagacgcatctttgactactccaataataaca
tctccaatagatgcataagaaggattactagtgcctagtactctaatacacattattttt
ctagcaccactattatctgcaacattaagatagctttgagtctgtatcatacttttattc
tgtataagattaattatcaaaagacttagataagatattaaccattgtccaacactttgt
gcgacttaaaggtcgtgtctcttgtattgtaacaatatcgccgattgtgcattcattatt
ttcatcatgcgctttatatttctttgtccggatcatagtttttgcgtactttctatgaga
gattctattttctacagctactactatagttttattcattttatcgcttactactttacc
tgttgtttcttttaaaggcatagtctttatatttttttatatat---------aattaaat
tttaagattattacatggttgacttagtacgagacttttcaacagttaaaagttgggcta
atctatgcttagaatgtttaaatagatgaggctggaaatcttgccttgtggctcttttta
gccttaaatcaaaaagctctctttttattacaaggatttcttcagctaaagaagaagagt
ccaggtttgtaacatctgatattttagggaaagtcatattttaggattctgttgtattcc
gaactataaatttagtcttgattggtaatttataagaagctaatttcatagcttgttgag
cagtttttttgtggtacacctgtaatttcaaatagaatatgtccaggcttaataactgcaa
cccaatattctggagctcctttacctgaccccatgcgagtttcagctggacgggcagtga
ctggtttatctggaaatactctaatccatagtttaccacctcttctcacatatctagtaa

tagttcttcgagtagcttctatttgtctagaagttaaccaaactggctctgtcgcttgta
atgcataatcaccgaacgcaattgtgttacctttgctagcagaacctttcattctacctc
tatgttgttttctaaattttgttttcttggggcttagcataaaaattaaaaaataataat
aattaagaagctgttgaatcaggagtttcctgattataagcagattctagacttttcgat
tctggcaaaatttctcctttaaaaagccaaacttttactccgagcacaccatacgtagtg
tgagcttggcgatgacaataatcaatatcggctcttaaagtttgtaaaggtacacgacct
tctctaacccattcacttctagctatctctgcaccattaagccgaccagatacttgaatt
ttaacaccctgtgtatttgctctttgagctctttgaactgcttgtctgacagctctacga
aaggccactctttttctagctgttgagtaataaattctgctactaaggttgcttcagaa
tctggatctgcaatctctacaacatttactctgagttgtttgctagggtctaggattaag
gacagcgattttcttaaagactcaattcccgctccagattttccaagtacgattcctggt
cgagctgttgctatgagaatttctacttgatcaactttacgattaatttcaattttagca
atactagcattactaagttttgaatgtataaatgagcgaattttatgatcttcttgtaaa
agaactgggtagtctttagagttagcaaaccatgaagaacgatgtttttgggtaatgcct
atgcgaaagcccaaaggatgaatttttttgacccacagtatccttgattataagttaaaag
ctataatgactaattgacgttaaa--------agactacagttctgaaacacctaatgta
atatgacaagttggtttatgtattggaaaggctcttccttgtgctctaggctgaaatctt
tttaaagtcgggcctttgtctgcaaaggctttactaacaaacaattggttttatttaat
ccatcattatgctcagcattcgctgcagcagattccaggatttgctttatatgagaacag
actcggtatggcataaattccagaataattaatgcttcttggtattttctacctctaatt
tggtctaatacacgacggactttatgcggagaaagacgaatgtatttccctactgcttta
gtttcttttacgtttttttgtaatactcatagttatagtttaacgacgggcttttctgtca
ccttttacgtgagtgcggaaagttcttgttggaacaaactctcctaatttatgtcctacc
atttggtctgacacaaagacaggaaaatgttgtttaccattataaacagctattgtatgc
cctaccatatcgggaataatagtagatgctctcgaccaagtttttaggacttctttttttt
cctgaaatatttaatgcttctatacgtttaagaagactaacatcaataaaagggccttta
tgtatagatcttgacataataataaatcattagattcttaataaaataaattctatttac
ggcgacgtaagacgtacgggttactatatttatttggatttcgtgttttaacacccaatg
caggtttaccccaaggtgttacaggacgagctcgtccaattggagatttgccttcaccac
caccatgtggatggtctacaggattcatgacaacacctctaacagtaggtcttttaccta
accaacgatttctaccagctttaccaagagtaatattactagcatcaatattaccaactt
gtccaatagtagcgtagcattcttttcgaatcattcgaacttcgcttgaaggcaattta
cagtaacaaaagtaccttcttttgctactatttgagcataagttcctgctgcacgaacaa
tctgtcctccgcacgaaggtcttaattctatattatgtactgctgttcctaaaggaatac
tagataaaggcaaagcatttccaacttcgataggggcagtaggaccagaaagaaccatag
atcctacactgagagatcgagggtgtagaatatatcttttctcaccatcaagatagtgta

ataatgcaattcttgcatttctattcggatcatattcaatagaagcaactttagcaacta
tattatgtctatttcttttaaagtcaattaatctgtactgctgcttatgtccaccacctt
tatgacgacaggtaatgacacctctattatttcgacctttgcagaaatggtgtttaacta
ttaatgattttctggcttatcagtagtaatctctgaaaaggtagaaaccgttctatttc
ttgtccctggtgtgtaggcacgatataaacgaattgccatataagaagatttgagaagag
taaatgttgaa------ctatgtgatatatttaagtttctggaaacaaattaatagaatc
ttctgaggcaagtgttacaattgctttcttgtaatgtggtcgcttccctacgaatctacc
gatgctcctttttttcttaggaggatgacaagtgttaacacctgtaacttgcacgttgaa
aatatactgtatagcagccttaatattaattttgttgctttgggatcaacggcaaaaca
atactgattctcttctaataatttagttgtcttatcagtaatgatgggatatttaaccag
atctaataaacctcttgaatcaatgctatccattatatacctcttgtattttagataaag
catcaactgtaataataattttatgtgcagctaacaaagccataatatttagtgtatcag
ctgaaataagttctacgttgtgtagattgcgaatagaaagataaacattcggatcttttt
tgtctactatgactaaaactttttatttaaatcaagattccatcgatgtatagcttcca
taaatagtttagtcttgggctgatgaaaataactgttaaaattttctacaatcagtgtat
taactgatttgttatttaatgctgtccttaatgctaattgtctttctttttattcattt
ttttggtaaaactacgaggttttggcccaaatattacaccaccacctctccatagaggtg
aacggatagaacctgctcttgctcgaccagtccctttttgacgccatggtttacgacctc
cacctcgaacttcacttcttgttttagtattagcagaaccctggcgcttttcattacttt
gctttactaaagctctatggactaagtacatcccagaatcttgactgaccttaagattta
aatcagcgttgccacttacttgaccttcccaattatagacttgataatttaattgcgtgt
taactgtcatgaaatatatcgtatgaaagaatattgctacttactaattttaccaaagc
gccaggtttaccaggaacagctcccttaacgatcaacaaatcattctctgagttaatact
tacaatttgcaaatttttattgtaactttttattaccaagttggccagccatgttttt
acctggataaactcttccaggcgttgtaccggctccaatagagccaggttgcctatgatt
tttcgaaccatgtgacattgggcctctactaaaatgatgcctcttttggtaaccggagaa
acctttacctacacttctagaagagacattaatcttttggcctacttgaaatagatctgt
agatagtatttggctaacttcgaaatcatcagtagatttcatttcgtattcacgtaaata
ttttagtgggggagcctgtgatttttttaagtgtcccaacaatggtttatttaattttg
ttcagcaacttgcttgtaaccgacttgaatagcattgtagccatcagtagatacagcttt
aatttgagtaataacacatggcccaacttgaattacagtaactggaattgataaaccagc
ttcatcaaagaattgggtcatacctactttagtaccgagtataccaacagacactagatt
tctccttttgagagatttatactatcattgtacataaaaattctacttcatagttactg
aag--taaattaaactaacagatgcatgatattattcatgacaacataaaggttaaaaac
tttacctatttaatctaaaatagaaacagctttttgtctagttacagcgtatataaatta
gaagtatgcttattaaatttttataaactact-tagatttaactgatattgatatccgt

aacttacttgcagtaatatc----aatcaacatgtataagtttaattgcctaaactagaa
ccagctattattagaaataattcaagtatagataaaagcgtatttgaacttttatatatt
gacaaatataatggaatatattgaggttcggtaactacaccttttttatttttacatttat
tgtgca--gtatatatacatctaatgcagtaaacataagaatataac----tgtatagca
aatgtaattacacaaattagagaggtattaatgggcaaagtcgttggaattgatcttgga
acaacgaattctgtaattgctgttatggaaggaggtaaacctaccgtcataccgaatgca
gaaggttttagaactacagcttctgttgttgcatatactaaaagtggagataaacttgta
ggacaaattgccaggaggcaagctgttattaacccagaaaacactttctactctgtcaaa
agattcataggacgtaaacagaatgaaatttcgcaagagattcggcaaacatcatataat
gttaaaactagtggatcaagcataaaaattgcttgccctgcactgaataaagattttgct
ccagaagaaatttcagctcaagtactgagaaaacttgttgaagatgctagtacgtactta
ggtgagactgttacacaagcagtaataactgtacctgcttactttaacgattcacaaaga
caagctactaaagacgcaggtaaaatagcaggcttagatgtattgagaattattaatgag
cctaccgcagcttctttgtcttatggactagacaagcaaaataatgaaacaatactagta
tttgaccttggtgggggcacatttgatgtatctgtattagaagttggagatggagtattt
gaagtactctcaacttctggagatacacatttaggcggagatgactttgatcagcagatt
gtagaatggctaatcaaagattttaaacagaatgaaggtattgatcttggtaaagataga
caagcacttcagagattgaccgaggctgcagaaaaagcaaagatagagctgtcaaattta
actcaaacagagatcaatcttccttttattactgccacacaagatggtccaaaacattta
gaaaaaactgtaactagagggaagtttgaagaactttgttcaaatttaatagataaatgt
agtatccctgtaaataatgctctgaaagatgcaaaactagaagcttccagtattgatgaa
gttgttttagttggtggatctacaaggattccagccatacagcaaatggttaaaagatta
attggtaaagatccaaaccaaagtgtcaatccagatgaagttgttgctattggtgcagcc
gtacaagctggagttttagcaggcgaagtcaaagatattctattactagatgtgacgccg
ttatctttaggagtggaaactctgggtggcgtgatgacaaagattataccaagaaatact
actattcctacaaaaaaatcagaagtattttctacagctgtagataatcaaccaaatgtg
gaaattcaagtacttcaaggcgaaagagaactaacaaaagacaataaaagcttagggacg
ttccgattagatggcattatgcctgcacctagaggagttcctcaaattgaagttaccttt
gatattgacgctaacggaattttatctgtaaaagccaaagaaaaggctactggtaaagag
caatcaattactatatctggagcatcaactttacctaaagatgatgttgaaagaatggta
aaagaagctgaagaaaatttcgatacagaccaaaaaagaagaaagaatattgacacaaag
aatcaagcagagtccctatgttaccaagctgaaaagcaagttaaagagtttgaagacaaa
attagccaagatttaaaaataaaaatagaagagctaattacagagcttagatctagtcta
gagaaagaagaatatgacaatattgaatctatttctcagcaattacaaaatgctctgatg
gacattggaaaaaatgctgctcagactgaaagtaaagatacaaaagcgaaggatgacgac
actgtgattgacactgatttctctgaagctaagt-aaaagta-agcgggtaacgcgattc

gaacgcgcgacatcaaccttggcaaggttgcgctctaccactgagctatacccgcataga
ttgtattattacaaaaattaagatatttgtcaatctgtta------atctatac---atc
tagactattaattttggaattacatattagctaactgttgtaaatataacaattgctata
attttttaa-------aagtacttatatcat-tatataagtactttttatatt-------
-caaaaatacaaataagtattatag-------ctaaactactaaggaattaactaagcta
gtaacaattactaatcctgtccataagccggctcctgtatatattaatccttttgattgc
tcccattggccaggggatgctaaaacaacaggtacgcctactactaacacagtcgataat
gcaattaataatagcactaataattggatcgcaataatcatttaatatgttctcctgaaa
aattcaaatgttaataatactgtatataatacaatgaagcttttgaccttcagcaattta
tctatgaagaaataatttaaaattattgttttctaaataaaccttaaattaagtattaaa
atctttattgtaaacaataatactatattttttta--gtagataaaaaatttaaagaaag
tcttttttaaaaaaaaattaagataaggtgatatgctgtataaaaagctaagtattatta
attattagagaaatttaaa-ataagaggtaaaacatatgaactcagccctttttttagca
aaactaccagaagcttatgccattttaaaccgatcattgatatcttacccgtaattcct
gtatttttcttcttttagcttttgtttggcaagcagcaattggttttagataacattta
ggttataaaaagaataacaaatatacagagtgccaattttgagctctaatcttatatct
tacacaataataactattatggttgaacccttactatcaggaattgttcttggacttatt
cctattactttattcggattattagtagcagcttatttacaatatcaacggggcaaccaa
ttaggactgtaaagcctaaattatagatatttataaaaactacaataattatagtttta
taaatatctacaaataagctaattttaaatacagaaataatactattgaaactagagtga
taacaaaaaaaatttatgtaa---taa---------------------------------
------------------------------------------------------------
------------------------------------------------------------
------------------------------------------------------------
------------------------------------------------------------
------------------------------------------------------------
------------------------------------------------------------
------------------------------------------------------------
------------------------------------------------------------
------------------------------------------------------------
------------------------------------------------------------
------------------------------------------------------------
------------------------------------------------------------
------------------------------------------------------------
------------------------------------------------------------
------------------------------------------------------------

------------------------------------------------------------
------------------------------------------------------------
------------------------------------------------------------
---------------------------caagctaattgtttaccagcttgacttagtaac
tcctggtaaaagacattcatgagacatttctcgaaaaacatggcgagataatccaaaatc
tcgataataacctctacttctacctgttaaccaacatctatttctccctctaacagcggc
gctgtttctaggcatttcttggagtttttgtcttaaatccattttgtcagaaaagctgcc
agcttctttcagttgatctttaatagccagtctttttaaataatatttttttgcaagttt
cgatctcttaatctctctttggatcatattctttttagccatgctgtaa-aactctcttt
taaaattatttattcaatatattagctaaaaaagctcctgcctgcaacttttatatattt
aaaaataaatagcataaagtagtgaaatgcaattgatatttaatcaacgatacc---act
ttatgctattatcaaaaatttatttagtaaagttcttgttcaacatgagttgaaatagtg
caatcagatgaaggatatgcaatacaggttaaaacatagccagctaaaagttgatcatca
tctaagaatgattgatcagcttgatctacagttccttctgtaactttacctgcacaagtt
gaacaagctccagctctacaagagtaaggaagttcaattccttcttcttctgctgcatct
agaatatatgtatcctccgcacaattaaatgtgacatcaattccttcttcttcacatagt
aaatgaattttataatcagccatttcttcttactccttaataaagatacacttgaaactt
aataaagcaaaat-aatttaaagaacttaccaaaaaattatatactagtttcattatgca
tgatagactggaaaaaaagaaaatagtatactattctgaaaaatcaagctacatttccta
ctaaatagaactatagtaggtccaaagtatatacataaattcatatagttaaaattactt
ttatcatttaaatagctgaacttgatagta---------tagattgatcatc------aa
gctagtcttctgc---caataaataattaattatgatttcaatgcttaatcaacaagtag
aacttaagcctattcttaaatttcagataacacaaatatattcgcattctcttattcagg
aaattaaagccttaataactagacttgttttgcaagtttggagaaggccggctacattga
tggcaggtattatccaacctttattatggttaattctatttggtgggcttttctacaatg
ctcctataaatttgttcactattaatacaagctataattgtttttgagctctgggatta
taatttttacctcttttactggagctttgaattcaggtcttccattaatgtttgatagag
aatttggatttttaaatagattattaacggctcctttagtatcgaggacttctatcattt
tatcttctgctacttttatgacttgtattagtttaatacaagttgtatttatagttacag
cttctcttttttatgggaaactcacctctaaatagcgatagtactatgatttttggactta
tgattctattagtgactgtaggagttacaatgcttagtttagctttatctttcactctgc
caggtcatattgagctattagcatttatttagtagttaacttgcccttttttattttcta
gtacggctttagctcctttatatttcatgccgccatggcttcagttaattgcaagtctca
atccattgagttatgcaatagaaggtacaagatacttatattcaagcgtaaactggaatt
ttacagagtgtgtgattaagattagctggggagatatttgtttagggcaaattattattt
tattaatcgctttagatataatggcagcttatcttgtgtctaatatattaaaagctaaac

ttaatta-----------------aaattttaataaaaaatttattaaatttattttaca
aagattttttactatataatactaatagtagtatggataattgaaatagaaatagtttttc
aaaaccaaagctattatcaacaatgataaacatttgtaagaaagtcaacaaagtatgttc
ttattcataggaggcatgtagtcaatgggactaccatggtaccgtgtacacacggttgtt
ttaaatgatcctggacggttaattgcagtccacctaatgcatactgcacttgtagcgggt
tgggcaggatctatggcattatacgaactagctgtatttgatccttcagatccagtatta
aatccaatgtggcgacaaggcatgtttgttatgccatttatggctagactgggcgtaaca
gattcatggggaggatggagtataacaggagagagcgtatccaatcctggactgtggagt
tttgaaggtgtagccttaactcatatagtcctttctggcatgctttttctagctgctatt
tggcattgggtttattgggatttggaattatttagagatccacgaactggcgagccagcc
ttagatttaccgaaaatttttggaattcatttattgctatcaagtctactttgtttcgga
tttggagctttccatgtaactggacttttggaccaggaatgtgggtatcagatgggtac
ggagtaaccggaaaggtattaccagtagctccagcatggggaccagaaggattcaacccg
tttaatcctggaggggttgcatctcaccacattgccgcaggtactgtaggtatattagct
ggtgttttccatttaactgttagaccaccacaaagactgtatcgtgctctaagaatgggt
aatattgaaactgtattatcaagtagtatctctgctgtttttttctcagcttttgtgact
tgtggtacgatgtggtacggttctgcaactacgcctattgaattatttggtccaactaga
tatcagtgggatagtggatattttcagcaagaaattgagaaacgagtagaaaatgctatt
gctgatggtgctgcacctagcgaagcatggtcaagaattcctgacaagttggcattctat
gactatattggtaataatccagcaaaaggaggattattcagagcgggccctatgaataa
ggcgacggagttgctgaagcatggcttggacatcctgtattccaagataaagaaggaaga
gagcttagtgttcgcagaatgcctgctttttcgaaacttttcctgtaattttagttgat
aaagatggtattatacgagctgacattccatttagaagagcagagtctaagtatagtatt
gaacaagtaggtgtaacagctagtttttatggtggcaaattaaatggccaagttttcaac
gatgctcctagcgttaaaaaatatgcaaggaaagctcaattaggtgaggtatttgaattt
gatcgaactacattagaatcagatggagtgtttagaagtagtcctagaggctggtttaca
ttcggtcatgctaatttcgctttaattttcttctttggacatctttggcatggttcaaga
actatcttccgggatgtgtttgcgggaatcggagctgaagttactgaacaagttgaattt
ggagctttccaaaaattaggagatagaagtagtaaaaaacaaggagctgtataagaccaa
agtcaaagatagtctttatc---aaatatttctaatattaaaatagttaagacaaagttt
attatctaatagttactaatcaggagaatttatggaagccttagtctatgtatttttact
aacaggaacgctaatggttatattctttgcgatctttttagagaacctccaagaatagc
aaagtaacttgattccgtttaaacttataataatagaaccactttccaacttattagtgg
aaagtggttttattgtagattacattgtattattcttcatgctcttcaaaaggatctcgg
agatcttttgaagcaggaccaaatgctgtgtatatcgagtaacctgtaatacctaaaagc
agactcgaaataaaaatactaagaactgttgcagtttccataatttaagcagtattaaga

tgaactttttctataatgatattatcaatattaaaacatttaattattaaaattatcaaa
atatactatggcacttagaactagacttggagaaattttaagacccttaaattcagagta
tggaaaagttgctccaggttggggcacaactcctattatgggaattttcatgttactatt
tttcttgttttattaatcattttacaaatctataattcttcattagtattagaaaatgt
agatgtagattgggctactttaggtag----ctaactagataagttctttga--------
ttacacaatattaaagacaac----------------aagcgatcgctattagattatag
tagcgatcgctttaaaagtattttactatatatccagagaaaattatttacctaaattaa
ttaactcggtatctgcaaaactatttgtgttaacattattatagttaactttttcaaacc
ttactaatactggatatttgataccacttttatcaattgctgcaacagtaccaatttcct
gataccagtacgattcttttctcaatatttttacttttgagcctctttccatatttttag
tcctcagatagattattaaagattattatattaaaaaattaaaaaaaactaaacaaaata
tataaaaatactaccgacaattgtaatatataactaataaaaggagagagagggattcga
accctcgataacaaaagttatgacaattttcgaaattgttgcaataaaccactctgccat
ctctcctgaattaaaatactgttttaaaaaattagacaagaataacatagaactaaggta
gttgcctagaaaattaaaactatgttaaatataaattataatctttctatagatatattt
aaaaaatatgtacatagaatctatatcagcttacttcaagatttttaatcttatacaat
ttttactaaataaataaattctctcataatatgaaactatcttggaaaacacttcttctt
tggtctttaccaatatttgtaattggtttctttttctggcaaggtttttaggtccaact
actacagatgttggcagtaatatcgcaagttctagaatgacatatggacgatttttagaa
tatttagatatgggttgggtgaaacgggttgacctctatgaaaataatcatacagcaatt
gtagaagctgttgggccagaattaggaaacagggttcaacgaattcgagttgaactgcca
gcaagtgcgccagaattaattacaaaattacgcaaagccaatgttgatctagatgctcac
ccccctaaaagtacaagtgcagtatggggactattaggcaatttactatttcctttacta
ttagttggcgggttagctttcttatttagaagatctaataatgctagtggtggacctggt
caagcaatgtcatttggcaaatcgaaagctttatttcaaatggaggctaaaacgggagta
gtatttaatgatgtagctggagttgaagaggcaaaggaagaatttcaagaagtggtaaca
tttttaaaacagcctgaatcatttactgctgttggtgcaaaaataccaaaaggcgtttta
ttagttggacctcctggcacaggcaaaacattactagcaaaagctattgctggcgaggct
agtgttcctttttttagtatctcaggctcagaatttgtagaaatgtttgttggtgttggc
gcttctcgtgtgagagacctattcaagaaagcaaaagacaatgcgccttgtatcgttttt
attgatgaaattgatgctgttggtagacaacgaggaacaggtgttggaggtggtaatgat
gaaagggaacaaacattaaatcaactattgactgaaatggatggctttgaaggaaatact
ggtgttattgtaattgccgctactaacagagctgatattttagattctgcattattaaga
cctggaagatttgatagacaagtttctgtagatgtaccagattttaaaggcaggttagca
attcttgaagttcatgctaaaaataagaaaatggaacctaaagtatctttagaaaccatt
gccagaagaactcccggcttttcaggagctgatttagctaacttactaaatgaggctgct

atcttaacggctcgacgaagaaaaaatgcaatgactatgtctgaaattgatacatcaatt
gatcgagtagtagctgggatggaaggcactcctttaattgacagtaaaagcaaaagatta
attgcgtatcacgaagtgggtcacgcaataataggcagtttattagagcatcatgatcct
gtgcaaaaagttacattaataccaagagggcaagcaagaggcttaacttggtttactcct
agtgatgatcaaagtctaatatcaagatctcaaatactagcccgtatcgtaggtgctctt
ggtggcagagctgcagaagaaatcattttcggtgacgcagaagttactactggtgcaagt
aatgatttacagcaagtaacatcaatggcaagacaaatggtcactcgatttggaatgtct
aaaattggacctttatctcttgaaagccaaggaggagacccatttttaggtagaggcatg
ggaggaggctcagaatattcagatgaagttgcaactaatattgataagcaagtaagggaa
attgtcagtgaatgctatgcacaagctaaacacattattatagataatcgagtagtgata
gatagattagttgatttactaattgaaaaagaaacaattgaaggcaatgaatttagagac
atcgttaaggaatacactgcaattcctgaaaaaaattactacatatcacaattttaa---
---------------attaaacggactgacgggattcgaacccgcaacttccgccgtga
cagggcggtgctctaaccagttgaactacagtccc-aaaaagatgaatacccagagataa
tctcatagttagcttaaacttgtcaaattagaatattaaatatgaagagta---------
--------tttgtaataatactcttcatatttgata-tagtagctaaaatatttcttta
gaaggagagagagggattcgaaccctcggtacgaagttaatcgtacagcagattagcaat
ctgccgctttcgaccactcagccacctctccatga---tatatattatatatatatcaaa
tatacttatattttaataggtgtgcataaatatttatggagctaagcggattcgaaccgc
tgaccctctcaatgccattgagatgctctaccaactgagctataaccc----ctttggta
agatattaagaaatggctcaagcgggatttgaacctgcgaccttgggcttatgaatcccc
tgctctaaccactgagctactgagccatagtataaaatactattaaatattataccataa
attatggtcaaatataaatcaataaaagaataattactatca-tttactacttttg----
-----------------tgcaagttgtaaatattcgaataaaatttcaaattaaaatg
tctttaacttttgttaga-------atta-ttgattacgaaa-----------atattg
atatttagaattaac---------------------------------------------
--------------------aataaatttg------------------------------
---------------------------------------------------tctttattt
g-----------------------------------------------------------
------------------------------------------------------------
------------------------------------------------------------
------------------------------------------------------------
------------------------------------------------------------
-------tagaca-----------------------------------------------
-------------------------------caaaaatattggaaa--------------
------------------------ttttcatcaat-----------------taattagt

------------------------------------------------------------
------------------taaactataaccgacgtac-----------------------
------------------------------------------------------------
------------------------------------------------------------
---aagaattattgaacta-----------------------------------------
------------------------------------------------------------
---cgttaaat-------------------------------------------------
------------------------------------------------------------
------------------------------------------------------------
------------------------------------------------------------
-------------------------------------tgtgatgcatt------------
-----gggcttgatcaaaacttctctaaaatcaatagaatatagt---------------
-----------------------------------------------------aattcaa
gtaaatcaaatactaacaaatgtgacaaacgcttaaaaaaataata--------------
caattagaatcgtaaaaatttggtttttga-ctgtgttaaataaatttgattatcataga
tttaatatttaaaagtgttctattacttgta-----ttgagtaattgt-aatatttaata
tcttctcttttagataataatcataatataagctaaaaaatagtttt---tattaaacaa
ctagcatagagccaataccttgatcagttaatatttctagcaagagagcatgatctattc
taccatctaaaatatgcgctgaagctacaccctgagctaaagaacgaatacaacaattca
ctttaggaatcatgccaccagaaattactgctgtttgagttaaatctctagcttcctgta
tattcagatgactaattaatgttgaaggatctgaagaattacgtaaaatgccgggcgtat
ctgtgagtagaattagtttttcggcattaagagcggctgctatttctcctgcgacagtat
cagcattaatattatacgattggccttccttgtctgctgcaacactagctataacaggta
tataattattattaattaatatcttcagtaatttagtatcaatattttgtacttctccaa
caaagccaaggtttgcttttccatttggtctcggagtaataagtaatccatcttttcccg
acaaacctacacttttaccaccttgtttattaatagttgcgacaagatctttattgactc
ttccaactaaaaccatttccactatatccatggtaggttgatctgttactctaacgccat
catcgaattttggtaatattttgaggcgatctaaccaaaaattaatttctggtcctccac
catgaactaaaatgggacgtagtcctataaaagacagaaaaacaagatcactaatcactt
gatcttttagtttctgatttttcatagccgctcccccatatttataacaataattctgg
aagaaaattgctgaatataaggcagagcttcacttaagacttttactctttctgaattag
tcaacataaataaagtaggattgtgatatataaaattatatacacagcttgtgttaattt
aaaagataccaatgggt---------acttgaaaaaatagaaatattactaacag-----
----ttaaactatagggattataattatgaataaattttgggataatgtattaagattt
ccacgatttttagtcagcgtcattttaggattaattttgataattattagtcctttttc
gtgttattaaaaaaaccactgacaagtttttttttcatcatatcattggccggattaatc

acagttttggcaataataa-tacaaaaaatgataaatatcgaatgttgttgagagaaata
aataaatttaattaaaatttttgaacatatataatattatattatcattaaatatgataa
ttt-atttccttttctat---aaaacccataaatattaaatattatttgataaatattat
gccactaaaacaaagagttagctctgaaaagaccggtgccttcgcactttggatagtat
tgttaggcacggagttaaacatatatttggttatccaggtggagctattcttcctattta
tgatgagctttatgcttgggaagaagcctctctaattaaacatatccttgttcgtcatga
gcaaggcgctgctcatgctgcagattcttattctagatcaacaggagaggttggagtatg
cttcgctacttctggcccaggagcaactaatcttgtctcaggtatagctacagcacatat
tgattctgtacctatattagctataacaggtcaagttgggagagcttttattggtacaga
tgctttccaagaagtagatatttttgggattacacttcctattgtaaaacattcatatgt
agttcgtgaccctagagacatgtctagaattgttgcggaagcatttttttatttgtaaaca
cggtagaccaggtccagtattaattgatgttcctaaagatgtaggattagagaagtttaa
ttatttttctgttgagcccggaaaagttaatattcctggctgtaggccaattaccagcct
aaagtcaagacaaatccttatggcagctaaaatgatacagcaatctagccagccattgtt
gtatattggtggaggagccataatctctgattcacatcaaattattaaagaacttgttga
tttttataaaatacctgttactactactttgatggggaaggggatttttaatgaggatag
cgactattgtctagggatgttaggtatgcatggtactgcgtatgctaattttgcagttag
tgagtgcgatcttttaattgctttaggagctagatttgatgatagagttactggaaaatt
agatgaatttgcttgtaatgcccaagtgattcacgtagatattgatcctgctgaagtagg
aaaaaataggattcctcaagttgctattgtcggtgacgtagcagaagttgttagtgaaat
attgaatttattaaagacttctttccccccttatccagagcagattatatcttggcaaga
aagaattaatcgttggcgtcaacagtatcctttactggttcctagaaaatcaacaagcat
ttcacctcaagagattcttgttgcaactaataaattagcccaaaatgcttattttactac
agatgttggccagcatcaaatgtggtcagctcaatttctgaaagtaaaagctaagcattg
gctttcaagtgctggattaggcacgatgggttatggtttacctgcagcaattggcgctca
agtagcacatccaaatgacgtagtcatttgtattagtggtgattctagttttcaaatgaa
tatgcaagagttaggaactatcgcgcaataccagttaccagttaaaattattattattaa
taatcgatggcaagggatggttagacaatggcaacaagctttctatggtgaaaggtattc
acactcaagaatgacagaaggagcacctgattttcaaaagcttgcagaagcttttggtat
taaagcttttactattaataataggcaaaatatgcaatctgctttacaagttgctattga
ttatcctggtccagttttattagattgccaagttacagaaaatgaaaactgttatccaat
ggttgctcctggaaaaagtaatgcacaaatgataggtatcgctaaaccgcagagaggtac
tgcttccaactatattaataatagtgtttaaagtaa------------------------
------------------------------------------------------------
------------------------------------------------------------
------------------------------------------------------------

---------------atagttagtatttttta-aacaaagaata-aagtaca--------
-taaatgtaatttttttatactatta-aaaactatgtaagaacagcctttgacgagaatt
gaactcgtgaccttccccttaccaaggggatgctctacctctgagccacaaaggc-tttt
tactatattgggccgggttggatttgaaccaacgtaggcgaagccagcggatttacagtc
cgcccccattaaccactcgggcaccgaccctaattaaaccttattcaagatatgacctat
aagatatcataactatgttgctaaaaacaagtactttgataaataaaactgaataagtaa
tggacatagctggacttgaaccagcgactttcacgatgtcgacgtgacactctaaccaac
tgagttacatgtccaaaactttaattattattataacactgtttagtgattttataagta
tctatttctaattgaagatttgttttaatcgtccggctttaccgagcagatctcttagat
aatacagtttagctctacgtattttagctcttctaatcactttgacaaaagttattcgcg
gtgaattgagaaaaaatactctttcaactccaataccttgaaatgagcatcttaaagtta
aactagtatttaagcttttttttcttcttctcgctaatactactccttcacataattgtt
cccgagttttactaccttccttaaccattagtccaagttgtattgtatcgccaactttaa
tttcaggtacttctgttttgataaaaggtatctcaacacttttcattagctgacttagct
ttgtactatttgctttcatatataggattagtcttgtgaca-tgttttgaaatattatac
attaaaaaaacaaatatgaaaaattttagcgaataagcatactagtatttataaaac-tc
tttttatct---------------------------------------------------
--------------------------------------------ccattata--------
----------------cagatccatt------------atattcaact------------
------------------------------------------------------------
------------------------------------------------------------
------------------------------------------------------------
------------------------------------------------------------
------------------------------------------------------------
------------------------------------------------------------
------------------------------------------------------------
------------------------------------------------------------
------------------------------------------------------------
------------------------------------------------------------
------------------------------------------------------------
------------------------------------------------------------
------------------------------------------------------------
------------------------------------------------------------
------------------------------------------------------------
------------------------------------------------------------
------------------------------------------------------------

```
------------------------------------------------------------
------------------------------------------------------------
------------------------------------------------------------
------------------------------------------------------------
------------------------------------------------------------
------------------------------------------------------------
------------------------------------------------------------
------------------------------------------------------------
------------------------------------------------------------
------------------------------------------------------------
------------------------------------------------------------
------------------------------------------------------------
------------------------------------------------------------
------------------------------------------------------------
------------------------------------------------------------
------------------------------------------------------------
------------------------------------------------------------
------------------------------------------------------------
------------------------------------------------------------
------------------------------------------------------------
------------------------------------------------------------
------------------------------------------------------------
------------------------------------------------------------
------------------------------------------------------------
------------------------------------------------------------
------------------------------------------------------------
------------------------------------------------------------
------------------------------------------------------------
------------------------------------------------------------
------------------------------------------------------------
----------agattcttttatagtttc--------------------------------
------------------------------------------------------------
------------------------------------------------------------
------------------------------------------------------------
------------------------------------------------------------
--------tttgataatcagtac-------------------------------------
```

```
------------------------------------------------------------
------------------------------------------------------------
------------------------------------------------------------
------------------------------------------------------------
------------------------------------------------------------
------------------------------------------------------------
------------------------------------------------------------
------------------------------------------------------------
------------------------------------------------------------
------------------------------------------------------------
------------------------------------------------------------
------------------------------------------------------------
------------------------------------------------------------
------------------------------------------------------------
------------------------------------------------------------
------------------------------------------------------------
------------------------------------------------------------
------------------------------------------------------------
------------------------------------------------------------
------------------------------------------------------------
------------------------------------------------------------
---------------------------atcta----------------------------
------------------------------------------------------------
------------------------------------------------------------
------------------------------------------------------------
------------------------------------------------------------
------------------------------------------------------------
------------------------------------------------------------
------------------------------------------------------------
------------------------------------------------------------
------------------------------------------------------------
------------------------------------------------------------
------------------------------------------------------------
------------------------------------------------------------
------------------------------------------------------------
------------------------------------------------------------
```

------------------------------------------------------------
------------------------------------------------------------
------------------------------------------------------------
------------------------------------------------------------
------------------------------------------------------------
------------------------------------------------------------
------------------------------------------------------------
----gatgatcaaatattattttcaaacatggcttatgtatgaaaaagttctata---aa
aacctaatgac--------taaaattagggcttaagtatttcttagttttgtgttataat
tagttactatcaaaggtagtggcataacacgaagcatcaa-aatgttcgaacgctttact
gaaaaagctataaaagtcataatgctagcacaggaagaagctagacgcttaggacataat
tttgtcggaactgagcagatactattaggattagttggtgaaggtactggaatcgcagct
caagttttgaaatcgatgaatgtgaatctaaaagatgcaagagttgaagtcgaaaaaatt
ataggaagagggtctggttttgtagcggttgaaattcctttcactcctcgagcaaaaaga
gtactagaattatctttagaggaagcacgtcaactaggccataactatattggcacagaa
cacttgctaatgggcttagtccgagaaggagaaggagttgcggcaagagttttagaaaat
ctggcagtcgatgtttcttcaattagagctgaagttatacaaatgctcggagaaaatgcg
gaagccaatgtaagtggaagcaatactacgcaagctagaagtaaaacaccaacattagaa
gagttcggatctaacttaactcaaatggctatggaaggtggtttagatcctgtagtcgga
agacaaaaggaaatagaacgagttattcaaatcttaggtagaagaactaaaaataatcct
gtcttaattggggagcctggtgtaggtaagacagcgattgcggaaggattagctcaaaga
attgctaatagagatgtaccttctattttagaagataaattagttattactcttgatgta
ggtctattagtagccggaactaaatatagaggtgaatttgaagaaaggctaaaacgtatt
atggatgagattaaatcagctgataatgtaatattagtgattgatgaagttcatacattg
attggggctggtgctgcagaaggagcaatagatgcagctaatctgcttaagccagcttta
gcaaggggagaattgcaatgtataggtgcaacaactttagaagaatatagaaaacatata
gaaaaagatccagcattagaaagaagatttcaaccagttgtagttggagagccaagtgtt
gaagaaacaattgaaattttgtttggtcttagagaccgttatgaaaagcaccatcaatta
acaatgtcagatggagctttggctgcagctgctaaatacgctaatcagtatatttctgac
cgatttttgccagataaagcaattgatttaattgatgaagctggttctagagttcgttta
ctaaattctcaattacctcctgctgccagagaattagataaagagttaagagctgtatta
aaaacaaaagatgaagctattagagctcaaaaatatgaaacagcagagcagtatagagca
agagaaatggaaattaaagctcaaattgcagcaattgctcaaagtaaaaagaatgagcct
gatttaaatttagaagatcctgttgttacagaagatgatattgctgaaattgttgctgca
tggactggtataccagtaactaagcttactaaaagtgagtcagaaaaattaatgcacatg
gaagaaactttgcatggacgtattattggtcaagacgaagcggttgtagctgtctctaga

gcgataagacgcgcaagagtaggtctaaaaaatcctaacagaccaattgcaagctttatt
ttttccggaccgacgggtgtaggaaaaacagaattaacaaaagctttggcttcttatttc
tttggttcagaagcttctatgatacggctagatatgtcagaatacatggaaagacacact
gtatctaaactaattggttctcctccaggatatgtgggttatagtgaaggtggttatcta
acagaagcggtaagaaaaaaaccatatactgtcatcttatttgacgaaattgaaaaagct
catccggatatttttaatctacttcttcaaattttagaagatggccgactaacagatgca
aaaggtagaactattgattttaagaatactcttttaattatgacttctaatattggatct
aaagttattgaaaaaggaggaggtagtttaggctttgaattatcagaagatcaaacagaa
tcccaatatactagagtacgatctttagtaaatgaagaactgaaacaatactttagacca
gagtttctaaatcgattagatgaaattattgtatttcgtcagcttactaaagatgaagtt
agagaaattgcagaattaatgcttaatgaagtctttgcgagaattaagcagcaagatatt
caattaaatgtaacagaacgatttaagcaacgattagtagaagaaggatataatccaagc
tatggagctagaccacttagacgagcagtaatgaggcttttagaagatagtctggctgaa
gaagttttatctggtaaaattaaagctggtgatagtgcagtagtagatgttactaatgaa
ggagaagttacagttttattaggtgaaaaattagaactgttaacataaa------aaaca
atttattaat--taaagcattgaagaggttat--attcaatgctat---gaaaaaaatta
ataagatgggttgctataattgacaacctctgatactctttatttgtatata---aagcc
gggatagctcagttggtagagcagtggattgaaaatcctcgtgtcaccagttcaaatctg
gttcttggcat-tttaagcatcagaaaattttgctatctgagcattaaattttaattgaa
atgttcctgtcggaccattcctgtgttttgctattataacttctgtgaagtctctacttc
tagtttcttgtgtatagtaactttctctatatagcattattactaaatctgcatcttgtt
ctatagaattatgaacaataaagttgtttgaaataaaattgcataaaggtttacattcta
aatcaaacactatttgtaaacttgtaacgtttatttttttgtatactttcaaaagtcaaag
atgaaa------aagaattaaaaacatttttttgttcttcaaagttttt------tattt
gaatagcaatcatgtcattttgattaattttatcacatcttttccatccttgtgttgtga
gtagtttatggttacttgttaattgtatatatttcccagcttcggtaataattttatata
cagtttttttccctgttttcgaaatactacatttaattgctagaaataactgttgttttt
gcgcactaaagttaatgacttca-atcttttttataatagcaattaaataatatttg--aa
gttgattatataatgatgtctgaagataattgaatttggatatacacccactttctctta
aatcggataacaaaggccttttgtcactccgagtttcaaggtttctatttagttgagata
atgccaagataggcaagcttaaatcttttgctaatattttaagtgatcgagtaattaatg
atagttcttgagatctattgtcagattgtctactctcctgcagcaattgaagataatcta
taataattaattctatatttttttccttgtaatttaaagaattttacttttgtctttatag
tgtcagtagaaatcttagcgctatcatcaatataaagattaaggttagcaagagttttac
ttttttgaataacatattgccattcatcattattaagctgaccagattgaatttttttggc
cgtttaaattacactcttgagctaaaattcttcttagtagttgctctgtagacatctcta

agctaaataaaataacataagatcttttctgattaattacatatctagttatattaatag
caaaagcagttttccccattgaaggtcttcctgcaagaataattaaatctgacttttgaa
acccttgtgttatagagtcgagttctgtaaaaccagaaagtatactactatttatactaa
tctttttccttgtatccaaatggactaataggtttgcaagaattgtagctaatgtttgcg
tatctttttcgtcaagtatttcataagcttttgttaattgactaacaatagaagttatgt
ttttttgttcaataagttgttttgagcagctaattaaacataatgaatctccacatgcta
aaagtaatcttttaatgtagttatctataacgactgctgaatattcgcttatagtatcag
ataggggagcttgttctattagactaaaaacttcatttagctcattcaaatcttgaatca
ttttttcatttttaagttaatgaaaaaatttcttacgctttttgtctttgcatggttaa
cattttcgagtgctgctctataaagtaatgaagttgatgtaaaataaaaaaaatcaggag
agatttttttctactgattttagtaatgatatcgattgctgagttaatattatactaatta
atattttttcagctaaaatattatgaggaggtagatatttataa------atacttatta
cttcctcc-ttgtaacattactcttgtg--------gtattagttaactattgatttaat
ttgattctggaagaacttgcaatttaatatttgcagttacttgatgtaatagcttaattt
ctacatcgtaaatgccaattgttttcacgtctggtaaagagatactttgtttttcaatgt
ctatgttagtagtgtttttaataatttgtgaaatttctttttctgtgacactaccaaaaa
tattgtggccatctcctgtttttttactaacgctaaacctttgtatttcttctagaagtt
gcttgactttttttagcatttctttagctgtttttaatttttcttctcttatagctgcat
aaaattttgttgttttaaaataccatttgtggcaactgccgccattttattcgggatta
aaaaatttcttgcataaccagaagctactttgataacgtcattactcttgccaagttttt
gaatattctcttttagcacaacatttataacttttttactcatctt-------aattcta
aaaa----atctgtattatcctattatacctgttgtatacagatttt----aatagtaag
tatttatttgttgatgtaatattcttgtaaaa--aaaggtaatttttttattttt-ttctg
atataatcactttcacaataattc-----attaggagaggtggccgagtggttgaaggcg
cagcattggaaatgctgtttaggagcaatcttaacgagggttcgaatccctctctctccg
tacaa--aaaatataaaaagttttatagagtccaatctaaatctccaataatattcttcc
aagcatttaatccaccccgaactctaacaatatttagtttttgagctattaaaaacttag
aagcaaatattgatctagaatctaagctacagtaaacaaaacaaattttatcttgtaaat
tgatatgagagtaattcattcctttttatttttttgtaaaggtagatttaatgaatgtatta
aatgacttttatgatattcttcatgatttcttacatctattaaaatatattgtggattcc
tactaattaaaaacttttgtaattgaataacactaatttctcgtacaaatgtactagatt
tactgccataatatttgttccaatgctttttttgt-------ggataatacaaatttagta
tttataatcttaaatttgttaaatgatgaagttaatgagttgtacgttaatatagttcca
cttaatatagacttataacccagaacgattttgacagcttcagttgcttgaagtgtacca
acaatacccggtaatagacctaaaactccggaattactacaagtatctcttgcgctctct
ttattttcagtttcactgtagaagtctcgataaacaggtccaccttgataattaaataca

ctaacttgtccttcaaattgaaaaatggcaccataaatatgtattttatttaactctagg
caagtatcattcaagagatatcgagtatcgaagttatcagatccatctataataatatca
tattgtctaataatgtcgattgaattctcatagcttagtcttgttttgaacacagttact
atacactgtggatttaaatctaata-tttttttttagctatttctactttgaatatcct
atatcatgcatcgtgtataaaatttgcctctgtaaattagagagatcaattatgtcatcg
tctataattccaatgctgccaattcctgcagcagcaaggtaaatgattcctggagatcct
agcccaccagccccaatgaataaaacttttgcttcttttaatctttcttgtccttctaat
tgaatttgtggcaatactaaatgcttagaatatcttgtatattcttctagtgagtatttt
gtattttctgttttaaaatttagcataaaaatttatagtttatttaatagcactta----
--aaagctagctgataagtttattaaaatgacttttgtaaacttggttcttctctatcta
aaatagctccttgtgtaggacagacctgaacacaaatactacaatcaatacaggcagaaa
agtcaatccaataccaattagtgttattattgtttttttccctttccttgatgaatacaag
ctacggggcaagcttccacgcattcagaaactccaatacatttttctgttacaattgtat
gagacatattatttaattaat-aaaagataagctatttacatctagtagttttgttgtat
actaaatttacatatgctatccaggccaatatttaaggtagtctatttctacagtttact
aatttgatatgcgcctttagttcagttggtagaacgcaggtctccaaaacctgatgtcga
gggttcaagtccttcagggcgcgtttag-ttttatcgtaattcaataatatcattttatc
tttttgcaaacatcaaaatgatatgatataaaagatcatattttatacattgaatattct
atgtactagtactg-ttttttaatatgttattattgataagttaattatactaattctat
ttttgtaataagt-aaaaacagttctaattaagtatgcacgtatttaatagtc-ttttgt
atatat--aacttgaaaaaatttaaactatggctaaaaaagttactggaattgttaaact
agcattaaatgcaggtaaagctactcctgcgccacctgtcggacctgctttaggacagca
cggcgtgaatattgttatgttctgcaaagagtacaatgcacgtactgctgacaagtcagg
actagtaatccctgtagagatctcaatatatgaagatagaagttttacttttatacttaa
aactcctcctgcttctgtactaatcgctaaagctgctggactaaataaaggttcgggtga
accaaataccaaaaaagtaggcagcataacaaataaacagttagaatctattgcagaaac
aaaacttcccgatttgaacactaacaatatacctcaagctatgaaaatagttggaggtac
ggctaaaaatatgggaattctaattaaagatt-aaaagtttcaattatatttacta-tt-
-tttatgaaaaaattttcacgtcgacttacaacattaaaatcaaaagttgagcctaaact
ttacactattaatgaagcagtatctatattaaaagcaacgtcaaatgctaaatttaagga
aactgcagaagctcatattgctctaggtttaaatcctaaatatgcagaccagcaactcag
agcaacagttatattaccgaaaggtacaggtaaattgataaaagtagcagttattgctaa
gggagaaaaattaacagaagcaattagtgcgggagctgatgtcagtggctccgaagaact
aattgatgaaatactgaaaggtagattagattttgataagctgatagcaacaccagatgt
tatgcctttaatagcaaagctgggacgagtattaggccctcgagggttaatgccttctcc
taaagcaggcacggtaacattggacgtagcaaaagccgtgaatgaatttaaaggggtaa

agttgaatacagagttgatagaacaggtataattcatgtaccttttggaaaatctagttt
ttcacaagaagatctagttttgaacctccaaacaattaaagaatcgattgatagaaacaa
gccttctggagcaaaagggaagtactggaaaactttttttcttatctagcaccatggggcc
atctattcaaattgacatcactagccttctatgaaaaattttgtataattatagaactt
gaataatctattatttcaaatta-taatttgctagcattattaacgtaacaacttgctta
ctatatcttatctgcattaaaaacttt-ttatatatcttattaattatgagtacaaaggt
tgaaaatatcttggaagagctaaaatctttaaaccttctagaggctgctgaactagttaa
acaaatagaagaaacgtttgatgttgatgcatctgcggcttcaggaggtatgatgatggc
cacaccaacttcagcaccagcgtctgctgaggttgaagagaagacagaatttgatgttgt
cctagaggaagtcccagcacctaaaaagatttctgtattaaaagttgttcgctctctgac
tggtttgggtttaaaagaagcaaaagatcttgttgaatctactccaaaagtcttaaaaga
aggtgcttctaaagatgatgcagaaactatgaaaaaacagctagaagatgctggcgcaac
agttattgttaagtaatga-----agaagcgcttctgtttaagtataaacagaagtgcct
ctttatttattataagcaattatactagaaaagtccaagtgtaattgcattgttgatagg
catagttgcaccaatacctagccaaatacttacaacggtaccaattaagaaaacagtggt
tgcaattggtcttctaaatggattttgaaatttgttaacattttcaatgaatggaacagt
aagtaaaccagctggtacagcagccatacttagaacacctaatagtttattaggaatcac
tcttaatagattaaatgttggaaagaagtaccattctggtaaaatctctaatggagtcgc
aaatggattagacttttcacctatagaagatggttctaaaattgcgagtccaatactgca
ggcaaaagtgccaataattacaactggaaagacgtataatagatcatttggccaagctgg
ctctccgtaataattatggcccattccttttgctaatttagctctcagcttcggatctgt
taaatctggtttttttaagaattgacatattg-tttttgtgtttaatataaagattagtct
aattactgttttaaagtgtaattttcaactattataatggtccagaaattccttgtttac
gtatcattaaaaaatgcataagcatgaaaacggcagttagtagcggaagtacaaaagtgt
gtaaactatagaacctagttaaagttccttggcctacactaactcctcctcttaacaatt
ctactatactcgctccgacaactggaacagcatcagggacacctgtaacaatttttacag
cccagtatccgatttgatcccatggcaaagaataacctgttaccccaaaagaaacagtaa
gcactcctaaaattacacctgttacccatgtcaattctctaggtttttttgaaacctcctg
ttaaataaacacggaatacatgtagaatcatcatcaagaccatcatgctcgctgaccatc
tatgaattgatctaataagccatccaaagtttacatcagtcataatgtactctactgatg
taaaagcttctgcaactgtaggtctataatagaatgtcattgcaaatccagttgcaactt
gaattagaaaagatacaaatacaatacccccctaagcaataaaaaatattaacatgaggtg
gtacatacttactagaaatatcgtcagcaattgcttgaatttctaatctctcttcaaacc
agtcataaattttactcataaaatagcttcaaaaagctcttctaatata--ttttgcgtt
taagctactattaattcaatag------ttattgaacttatatt-tttttataatcaata
cttattaactattataacatctttttattgattt--------------------aaaatt

ctctgacttagtaaaacagggctatgtactataattcttttttttgtatagagattaat
ttcgttttagtaaattagctaaaattcttgttacactgactctattactacctataatc
tgtgctaaaattttatgtgtaatagtaaagttaagcataataccattattaacttgagtt
ccattttgctcagctaataataataataagctcgccagtctactggttatacttttatgt
gaaatgatttctataaaatgataagcttttgccgagcaaaatactaaatgatttacaaaa
aataaattgaatggagcgcagttactacatgcttttataatagtagtatagttaatacag
gcaacttctgttttatctattgcttctgcttcgtagtaaaaattatcgtcgaccaattct
atttgtccaaatgtatctcctgtagttaataagttaagagttactttatgtgtatttctc
aggacttttgtaataattaaagatcctatcaatataatgtataatctagagctgctatta
aacaaaagagtatcattttcttaagtgaaaaaatttgataattaatttattacgatta
aaaaatagtagccatggattgcaagaaaaaaaatcacctgaactattttcagagaatatt
ggaatacaattttcttgatttaaacaattttagtcataattactactttataa------
---aaaaatattctaataat-taagaactgaaattaaatgtcatataaattgatgttagt
tgagaatgacattgtattatcaaaggccattcaagaatacttaatagatcaaggcttcaa
tgtttatattgccaataatggattagaagcactgaatctagcttatcaatataatttcga
cttgataatttctgatataatgatgccgctagttaatggctatgaactattagcaaaact
taaaaaaaacaaagcattatccaaaatacctgttatctttttgacagctaaaggaatgac
taaagacagaataaaaggctacgacatggggtgctatggttatttatctaaaccatttga
tcctaatgagttgctttcaattattaacaacttaatcgctagagatgtcttaaaagaagc
ttctttacaaaactcggcaacatcaaaccagcaattaaatcataaaatacgtttgacacc
tagagaaaaaagtattttagatcttgttgttgatggacttactaataaagaaatatctac
tatactgaatactagtgttagaaatgtagaaaaatatgttagtcgactcttacacaaaac
taatatgaagaataggactttgctagtcaagtattctataaataataatctattaaataa
tgagatcaatgaaagggcgaatgacgggactcgaacccgcgaatgatggagccacaaccc
attgccttaaccccttggccacactcgccatattacatacaccactatagcttgtttttt
gatatatcgtctagttttcgagttatatatttatctaaaacaatatttactttgaatttt
-tatgatacatagcaatattagtattcaaattaatggggaaccatttaattgctcaaagc
ccatctcattgcaattttattaaattatcttgattttaattccgagcgtgtagcagtcg
agctaaatgacattcttttaccagagactctgtttcactcaacttatttgaatgatcaag
ataagctagaggttattaccattgtaggtggtggataatctatttgtacatac-------
-acagttttctatctcaaatgtttcatagatagagataatcttccttgtttttatctac
atgaataataacagctttaatagtgtcacctattttaaattgtgatgatatctgctctaa
ttctttgatattaatctcagaaatatggacaagacctttgagatttcctacctttataaa
taatccgtaaggtgtaatctgatttattatgccttcaataatattaccaacaattaaatt
agatgatgcttgagatattagagctcttctatgactcagtattaaattattagatttctc
ttcgacattgagtaattttagcttaataaatttattattaaattgctcactttttttgaaa

attaccgagatgagaattaggtacaaatccagatataccttcaaggttaattatcattcc
tcccttattgaatcctttatcattacattaagtaaagagtcttcagctaatagttgcct
aattcttttccatgctcttatatattcaaggcgacgaattgataatattaattgtttcga
ctgaatattataatctaataaaaaaaattctcttgtgtcattaatgtttaaagaagtgaa
attatttaaatcttgattactagatacttcttgaataggcaagtatgcagatattggtgt
tcctatatctactaataccccatttaattcaaagctaaatatagttccagctacaatatc
tccaagattcaaatcatacttgtatttttgtaaaacagctgcaaaatttctatgagtaaa
tccttcattattttttgtcatattgaatagtatgaattaaagagtaactttgtacaatat
attaaggtaaataaatgtttgtcgaattgtatgaaaaaattttaatttgttcggttgtaa
ttaagtatataaacatattaaaatatctaggagacatgaaaagtagacgtagatagctgc
agatttaa-tactatataatctgaggaaagtccgggctccacaaatacaatttatgctgg
agaaatcccagtgtaggtaactgcgaggatagtaccacagaaaaaaaccgccagaataag
t----tatattagctggtaagggtgcaaaggcaagttaaaagcttaccaaaagtactgca
aagtatttgttaggtaaacccctaaaatggagcaaagctactaaacaaatatttgtgtat
attttatctagtttagttgaaaatactgcatgaagttattagtaacaataactcgagagg
aatagctatcctttacaatatatc-tcgattatattgaaagaacagaacccggcttatgt
agtacttttatgtcttgtattattatgtctattgtcgaatctttaaatttaactgtttg
acaattttttttgtagctgatgttgtttataatctatttctgaattggttaaagttttt
tcattagatcgatatgtaaatcttagacctatactttttttctcttttccaattgattca
tctatatattgatcaaataatgttgtactttctaaatctttatcataaaattgatctaat
aattttaataagtatttaatttccatattcttaggtattattaaagagagatctcttatg
atcgatggatacttagagtatggttgaatctgataacttaaataattgaattcattatga
caatcttctaaaatgtttaagttaacttcaagaaaaaaaaagttttgtattgagtcctaat
tcactatatgttgcttgattaagttcactgaatatgccaataataatattgttatagatt
agtgtggcaaattttcctttttgaatgaactttattctaccatcgactatatcttttttt
gtccactcaatttgtctattaatcttttgaaaaaaattttctacgatccctttagcttca
taccaatttaaagaatgcgcagggtgtgaccatgtagatcggatatctagattaccaccc
aatatgatagctagattagtagtttctacaattttgttttgatcattatgaaatactgtt
cctatctcaaagctatctatagtttggttgctttgtttaatattataaagatttgattca
attaatccttccagcaaactgccgcgtaaattagaataatcttgaataagaggattgttt
aaagctatatttcctttggactttattaaagaataatgaactaattcagttaatcctaga
tttcttaaaatacttctacttttgtctatgaattttctctttgtggataaccttttgttg
aattgaatatttggtactgagctcctgaatttatgataaccataaattctagcaatttct
tcaataacatctatttctcgaaaaacatcttctcgtctataatttggaacagtaacttct
atattttccttattttgataaataagatcaaaatttaaagagtgaagaatattttgaatt
tcttcaaagaataaaaaacgtgtttgaccattatacataattggtcctaaaatgtcatga

actttttttaattgataaattaatattaagagcatgatcagttattttttctcgacagaaa
gtatctctaatatttccgcatgttaaatctgtgattaaagcaagagcttcaaaatgtgca
tttttccaattatctacatttagcccacgttcttgtctaattgaactttctgttcgaata
ttaagaactcgagaagactttctaactacagattgtttaaatatagcagactctacaagt
atcgattttgtattataatctgtatcaaaatcagagttgcttccaataccagctatacta
gttacatttgtatttacttgagtaactaaaatatttttgtttaactcaatattttcattg
tttaacttaatctgatcattacttccaagaggaaaattgcttcgaattgttagactattt
ttatgattcatattattaatttttattcaaatcaataatattaataggctgtccccatttc
aacataatataattgcttatatctgttaacaaatttctgtgtgtaaacccagaagagagc
aaccgatttttttagccaatctggagaatctttaactcttatttcgtctataatggcagca
aaataattgtcacaatttaataaatcgtgattactgattatagt-ctccttttttagagaa
atatctattattgggttgatgtatagctgataacattgaggaccctgttaaagccgagac
ttctcttgatagaccaatcatgcttaacacgtctgacctgttagctgttgatgatatatc
taatatatgatcaatctcacctccaattataatagattcaatagcttcaacttcaaaacc
tgcttgtgttagttgattagttaaattgtctatatctattgttttgatattaccaagttc
tttcaaccaatttaaagaaactttcataattaattgtcatctatcttatagaagtattta
ttttacttatgattaaaaatcaatttaaaaaattcatcaaattaatataagttatttgat
gaatttgcttatacaatatattattgtgatatcttactttttatatggcagttattaagaa
ctagcttttgtaaaagttagcccattatcttgggcatatctttccataaatctcatgaat
ctatcccaatcatcaggacttttaataacatgtacagcttctattgcttgtggtttacca
ttaataaacttggcattaacatctcttgtaataatttgtccctctgtgtccatcaggtac
atacctgtgatttcacctttgtctgccatactagcatctaaaatttttgggatttgtaaac
cgaaaagttgctgttcctgtactaccatctcgagaccttgttaaacgtacatctggtaca
acttcttcattgataccttgaataaattgaattgttgccatatcaatcttctttatttgt
atataatttatattggataaaactgatttctagttcttgaatgcttgttgagaaactgtt
ttaaaagcttatagtatcagtagatcaaattttatgtattatataaaactgtaacttttc
-ttttttg-----------------------------------------------------
------------------------------------------------------------
------------------------------------------------------------
------------------------------------------------------------
------------------------------------------------------------
----------------------atctgttcaaagaactaacttattataagctaatcata
cgaatattagccatataatgttcgtaagt-gaaattaaggggttgtagctcagaaggata
gagcaagcgcctcctaagcgctaggtcagcggttcaagtccgctcaaccctgttaacgaa
ttaagtagtattatttaacttgttctgcaaatcagtattgatctaagttaaagtacttga
cagctaaaacaaaagaatctgagaaatattttcatttaaagaacgaacaattatcttaac

tctaaatatctgataattagtaa-aaaattct----------------------------
----cttttaaaacacaagatgatg-----aagatatttaaactaataattgaaattaa
gtttttgaagttatagaagccaat--ataatactttctattaacaga-----------tg
cttgcttctattactatctgtttacaaga--aatt----atgagtatcaaaata------
--atttagctcca--------------------------------------ctgac----
------------------------------------------------------------
-----------------------------tactaataaata-------------------
------------------tttattttagtctagacaag------atgaatcttcatggta
taacag------------------------------------------------------
------------------------------------------------------------
------------------------------------------------------------
------------------------------------------------------------
------------------------------------------------------------
--------------agctattctttgtgtcatacttatataaaattaggat-------ac
tgtt--------------------------------------------------------
------------------------------------------------------------
------------------------------------------------------------
------------------------------------------------------------
------------------------------------------------------------
------------------------------------------------------------
---------------ttcattaccataactggggtgggaggattcgaacctgcgaatggc
ggagtcaaagtccgctgccttaccacttggctacaccccaacatgtgaacattataatag
cagtcgtataccctgccatgtcaatagactattgtctttatttaataatttttgctgctc
ttgtagaataaaatctttttttcaaataagatacgctatgtttaagttgcacgactgtt
atgttttcttgcttttgactgaacagccacttgatagtaattattccttctgcaatttcg
ttacttcctataataatacaagctacagctctttttttatttgcttgcttaatctgctta
ctaaaattacttgagctaacatcgatttctattttaaaaaactgctgatgtaaaaaacgc
attatttgcataccagtttcgttagcttttgtaccttgtgtagcaatataaaaatcgata
gatttattaggcaagtctatattatcttttgcaatcagtaataggcgttctaatcctatt
gcacatccaacagctggtgttgctttgccacctaattggtgaactaagctgtcatatctt
ccaccaccacatatagtatcttgcccttttgatgtcaatgtcttaatttcgaaagcagta
tcattataataatctaatcctcgaactaacttattatttatattgtagggaatattaagt
aattttagataattgcaaacgtcaccaaagtgttgttgtgattccaaacttaaaaaatca
gaaatttttggagcctctgtcagtatttttttgtgtattagaatcttttgaatctaaaatt
ctaattggattactagttaatctttttttgtgagtcagtgtctaaatcatcatggtacttt
gtaagataatctcgtaatttaacttgataaatactacgatcctctacttttccaattgaa

ttcagatcaagtttcaagttgtgcagattgagattgttaaatatgctcatagctaaatgt
ataacttcactatctgctcttgcatctaagctaccaataaactcaatgccaagttggtga
aattgtcgttgtcgtccactttgtggtctttcgtacctgaacattggaccgctataccat
aatctttgtaaactatggtggtaactcattttattttcaataaaagatctcacaataccg
gcagtaccttctggccttaaggtaatatctctattgctgcgatcatgaaaccgatacatt
tctttatttacaatatcagtatcttcaccaatacccctatcatataattcactattttca
aaaataggtgttctaatttcttggtaatttgcacattctaataatttggaaattttatta
tgtataaactgccaatattgaatttcatcgggaagaatatcttttgtccctctaatagcc
tgaatttttgccatgataatttttgattaccttcttaatatgtctgcattaaatagcag
ttgtagactgttattcagtcatgtattatacattag--aacatactgatg-taatttata
aagtacgggcaaggagggattcgaacccccgacaccatggttcgtagccatgtgctctaa
tccactgagctacaagcccacttaataagtaccttaactatatcattgttcactaatgta
tacaactatattctttt------atcttacatgtgaaattttgtaaaagtaaaaaatt
at----gtttgtttttacaattggaactattcatatataatgcttaatttattataact
ttgtaaa--aatatctataagtttaattgtatgattctattatattggtaatcttgttat
aattttgcgctcttatctttcaattgg------ctagcatctaattaacttatatttcaa
ataaaaaaatcttatctaaaatgagtaaacaaattctatatcaggatgatgccagaaaag
cattagaaaaaggcatggatattttaacagaagctgtttctgtgactttaggaccaaaag
gaagaaatgttgtcttagaaaaaaaattcggtgcccctcaaattattaatgatggtgtaa
cgattgcaaaggaaattagtttagaaaatcatattgaaaataccggagtcgcactgatta
gacaagcagcatctaaaacaaatgatgtagctggtgatggtacaacaacagctacagtac
tggcttcagcaattgttaagcaaggaatgagaaatgttgcagcaggttcgaatccgatgg
ctattaaaaaaggtattgaaaaagcaacaaattttgttgttagtaaaatagctgagtatg
ctaagccagtagaagatacaaaagctattatacaagttgcttctttgtcatcaggaaatg
atatagaggtaggtaaaatgatagcgaatgctatagaaaaagttggcagagaaggagtta
tttctttagaagaagggaaatcaactaatacgattcttgagatcacagaaggaatgcagt
ttgaaaaaggctttatttctccttatttcgttacagatacagaacgaatggaagttcttc
aagaaaatccttttattttatttacagacaaaaaaattactttggtgcagcaagaacttg
tgccattgcttgagcaaattgcaaaaacatctcggcctttattaataatagctgaagaca
tcgagaaagaagcattagccacaattgtagttaataaattaagagggattttgaatgttg
tagcggttagagctcctggattcggtgatagaagaaaatctttacttgaagatatgagta
tcttaacgaacggacaagtaattactgaagatgcaggtttatcacttgatacagttcaat
tagatatgttaggaaaagcccgaagagttattgttactaaagactcgacaacaataattg
cagatggtcatgaaattaaagttaaatcaagatgtgagcaaattaagcggcaaatagaga
cgagtgactctttatatgaaagagaaaaattgcaagaacgattagcaaagctttctggag
gagttgccgtcattaaagttggtgcagctacagaaacagagatgaaagataaaaaactaa

gactagaagatgcaattaatgcaacaaaagcagcaattgaagaaggtattgtaccaggag
gaggagctactaatgtccatatctctagtgaattatttacatgggctaaaaacaatttag
ttgaagatgaattaattggtgctttaatagttgaacgagctgtgacctatccgttaagac
gaatagcttttaatgcaggtgataatggagcagtaatagtggaaaaagttaaaagtcacg
attttcatataggctatgatgcagcaaacggtaatattgtaaatatgtacgatagaggta
ttatagatccagcgaaagtagctagatcagctttgcaaaacgcagcttctattgcagcaa
tggttttaactactgaatgtattgtagttgataaagtcgatgattgagctata------a
gataatagataatctcaaacaaagaataatttctaattatagaaattattctttgtttca
attaatctaattaattgatcaaatgtataatttttataataaagccaaagtttaaacacg
tcttctttttcaactattaagtagagaagatataagcccgtaatacctaaattatcaatt
aaaatattattctcatatatagattttctaaagatctaaaaaatgacatttttgaaaaa
taataccgaaatcttctcttatagttatttgctaaatcattactgtctaaagattgattt
ccaattgaattcagaacctttaacagaatactttttaaatctttatctgactttatttga
gaatcaataataatttgattcaattgtatgtcaaaactaagattaaaaatattacttgtt
aaagatatttgattttgaatatctagagattttatagccaagaagattaaacttttattg
ttcattgtaaaatataagacaattgatgatgtattctttaaatacta-aatttatcttct
tgtgcttctatcgtttttgt-aaaataattaatgttttaatttttagttttagttact
acccgaaaaaataaattctggaaatctagcttgagctttagctagagagtgtttttccc
ttcttcttgccagtaattgataacttctgttgttttgtttaataactcaattctgtctct
ttcagcaatccacggcttggattctagttctgcttttaattcttcccaagcatcattcct
tggccagaaaaaataagaagtaagtggactagtgccatttccaacaatctgatcaatagc
aatagcaatattattttctaaccaaagtactttagtgtaaattttgacaagtttttatc
tccattattaatattaagtattgg----tacagtagaatactgtactagttattttttc
agcaacaatttgagctttgtttaaaatgtttttaccgttttagtagcttgtgcaccttg
agataatcgttgcaaaatttttgaaatgttaacacgagtttcattatttataggattata
aaatcctaattcttctatagctttaccatcgcgtttatttctgctatccattacaacaat
tctatagcttggttgtttttttcttccgtatcgctttaaccttaacttgaccatattagt
cacctcgcttatttttt--atatctttattaaattatatcttacgacattatgcaataat
tcaaacaattaagctgattcaagtctaacattattaaatattaccattaaacattgtaga
aaaataatacctaacataggtgacatgtccattccaaacatgggaggaatgcttcctctg
aatagttttaagtatggatcagtaattctatttaatgaacagaatggttcattgtaccaa
tttacagttggaaaccatgccaatgataatttaagtaaaattaaaattaaataaatctca
gaaaaattagctatcgatccaagcaataaatttaatgtaccaggaagagtgttcatacgt
attttataattattatatacttcgttatactaatagtttgattatatattataatcgatg
taaataaat-cagctaaacaaaaaacttatccttaaaatcatcttctcttgagcactttt
ctttatctacattagtatttacttttgtcgtataaatttccagtttagaccaaatctgtc

ctaactgatttaattgagatgttccacattctaaacaggctatttttatattgctcaaaa
cagcaccttttttttataaaatcgttcaataatggcgtaatagttgcttcatccaaaaata
aatctaagacaagtatttttgtaaatgaatttaattgatctgggacattgttaataatat
cattataattaactaaaactatattagctgtaggaagtaaagatcgtgctccttccgcta
agataaagccataaggcataataattacaataatatatttataatgactgctaataagac
ttatagtggtattagcttttaattcgattgtttctgttactagccactctctcattattt
cgtaagtaatccatttgcctagttctgagcaagcagttcttaaaatcgtacctggattat
tgtcgtttttgagaatccctgaccaatgttgtattaatggatgggatgctatattaattt
ggagttgcatgttttgttaaatatatttaacatttaatttgttgattgaatcttcaattc
ctatttgttgagtattattaaaatagcat----atatattaaagaaaataattat-aaaa
aatgaaaaaaagtctttggctgtggggttttacagatagtgctgaaacttggaatggtag
atttgcaatgattggttttatatcagttatttttatcgaagtagttacaggacaaggatt
attatatttaataggtatgatgtcataataaaaatccaaaaaagttcttgtaattttta
taagaactttttttgg-atttttataaatctaatctagcggtctcatagatagaacaggtt
cattctctctattgataccttttttcaaatccagcagcagcagctcttgctctgcctgcat
gccaaagatgtccaataaataggaagaatcctaggaagaaatgagatgttgttaaccaag
atctaggagatacatagttaactgaattaatttctgtagctacaccaccaacggaattca
aagaacctagtggtgcgtgagtcatatattctgcggcacgcctttcctgccacggttgaa
tatcgttcttgatttttatttaaatcaagaccatttggtcctctaagaggctcaacccaag
gagctcgcaagtcccaaaatctcatggtttcaccaccaaaaaataatttctccactaggag
atctcattaggtatttacctaatccagtaggtccttgagaagatgcaacattcgcaccta
gtctttgatctctaactaagaaagtaaaagcttgagcttgtgaagcttcaggaccagttg
gtccatagaattcactaggataagctgtgttgttataccaaacaaaattagaagccgtaa
gccccatgattgataaagcacctagactataagataagtaagcctcaccagaccatacaa
aagctcttctagcccaagcaaaaggttttgttagtatatgccagattccgccagcaatac
aaataatgccaatccagacatgaccacctacaagatcttccatgttatttacacttacga
tccatccatcaccgccaaacggagactttaagacataaccaaagataactaggggattaa
gtgtaggattgctaacaaatctaacatctccaccacctggagcccaagtgtcgtataccc
caccaataaataaagctttaataactaacagaaaagcgccaatgcctaataacactaagt
gtataccaagtattgttgtcattttgttttatcccgccaatcgtatccgaagaaaggaa
atgattcttcaagagtatcagggccaatcagagaatgataaagacctccaaagccaagaa
cagctgaagaaattaaatgcactacgcctactacaaagtatgggtatgtattgaaaatct
cgccgcctggaccaacaccccagcctagtgtagctaagtgcggaattagaataaaccctt
gttcgtacaaaggcttctcgggaacaaaatgtgctacctcaaaaagagtcatggcaccag
tccaaaagaccattatacctgcatgagcaacatgagcaccaagcaatttgccagaaacgt
taattaagcgtgcattgccagaccaccaggcaaacccggtagactcaatgtctctaccgc

caacaccaacagttgtattaaagggcgtttccacgtggtaaaacctcctcagggaatata
aagttttcatgaggttgatcttgagcagccatccaagagcgaataccttcgttcaatagg
atatttttagtataaaaagtttcaaactcaggatcttccgcagctctcaactcttgagaa
acaaaatcgtaagctcttaaatttagagctagtccaacaattccaaatgcgcttgtccat
aatccagttactggtacaaataacatgaaaaaatgtagccaacgtttattagaaaaagca
acaccaaaaatctgtgaccagaatctattagctgttaccatcgaataagtttcttcagat
tgtgttggagtaaatgcacgaaaagtatctgcagcatcaccatcttcgaataaggtattc
tgtacagttgcaccgtgaatagcacatagtaaagcaccacctaagattcctgcaacgccc
atcatatggaacggattcagggtccagttatgaaatccttgcaaaaatagcaaaaatcta
aaaattgcagcaactccaaggcttggggcaaagaaccagctcgcttggcctagagggtac
attagaaagacggaaacgaatactgcaattggtccagaaaatgcaatagcgttgtatggt
ctaagaccaactagcctagcaatttcaaattgtcttaaacagaatccaattagtccaaat
gatccatgtaaagctataaatgcccacagaccaccaatttggcaccagcgagtaaaatct
ccttgagcttcaggtccccaaaggaaaagaagtgaatgtcccatgctgtttgctggggta
gaaacagccgcagttaaaaaattgcatccttctagatatgaactggctagcccatgagta
taccaagaagtaacaaaagtagtcccagttagccagccacctacagcaaggtaagcgcaa
ggaaacagaagtagtccagaccaacctacgaatacaaatcggtctctttttagccaatcg
tctacaagatcaaaaccaccacgagtttttcttgtccaattgctatggtcataattta
atctccagagcaaatttaataagtaatttgtttaaataaaca---ctttagatatactct
aaaattttactttctttacggtcaatataattataagtcaaaagtaaccaaaattcacg
cttttaataaca-tttttttagctcattaagttaa-ttttttaatatataaacagtaata
gagttattgttt-aacttacaatactcacaatttattcatttctataatcctttgaaaaa
gatatcctgttcctcgtgctgttaaaatcaagtcgggattacttggatcatcttctagtt
tagctctcaatcgtgaaatatgaacatcaacaactctcgtgtcaacatgccgttcgggag
tatagccccaaacttcttgcaagatagaagctctagaaaaaggttctccagccttactaa
ctaaaagctcgagaagactgaattccatacctgttaatcgaactctttcattatttttat
aaacttgtcttttgttagtatctatttttaaaaatccaatactaataatcccagagtttg
ggactccaagactagtggtgattttatcagctcttcgaagaacagaacgaatgcgagctt
ctagctccttaggagaaaatggtttaacgacatagtcatcagcacctatttctaatccgg
taattctatcgcagacttcacctaacgctgttagcattataataggaacatctgattctt
ttcgaagttcttgacaaacaccgtaaccatctagtttaggcatcataacatctaaaacaa
caaggctaggatactcttttctaaacacgagtaaagcttcttctccatttgaagcagtga
taacttcataacctattattgttagtcttgtttctaaaattcttcttatgctggcctcat
cgtcaacaacgagaatcttttctttctggttctccaa-tttcttatttactcctgtaact
caccgatcatgattaccataaccttaaatttttagcgctatgagtatctttattcacttt
agaa-----tttagttattataaattttgcaagtctagtttacattaattagtaccagt-

-caatattttgt-----------tttaagttatgac---------tattttaatccata
agttgactctagtttgcagaattcaatatatacata-ttttctacccaagtaaattgtat
tctattttctaaataaaaaagggttgacaaaccagttgatagaaagtattctacattac
tataag----ttaattccaacaaagaagttttaa-----atcttctttcggagtaatgaa
tactttaat
>Pyr_29_33
tatatgata---aattaagacatctgttatgtgagaatttctaaaatatacaaaactttt
tatttaatatagaaaaagtaaattaaaataagttgaacatactttgctctaatctaaacc
tcttaattgag------attctaattagagacaaaagtatttacttattttgtatctata
ggaataaaaatgaaacttgcagtttatggtaaaggtggtataggtaaatctacaactagt
tgcaatatttctgtagcactttcaaaaagaggtaaaaaagttctacaaattggctgcgac
cctaaacatgatagtacatttacgctaacaggtttttaatcccaacaattatcgatact
cttcaatccaaggattaccactatgaggacgtttggcctgaagatgtaatctataaggga
tatggcggtgtagactgtgttgaagctggtggaccgccggctggcgctggttgcggaggc
tacgttgtaggtgaaacagttaaacttttaaaagaattaaatgcttttgatgaatacgat
attatcttatttgatgttttaggtgatgttgtatgtggaggttttgcagctccattaaat
tatgcagactactgcttaatcattacagacaatggatttgatgctttatttgcagctaat
agaatagcagcttcagtacgagaaaaagctagaacgcactctctgagattagctggactt
gttggtaatagaacagataaaagagatctaattgataaatatatagattgtgttccaatg
ccagtattagaagtcttgcccttgattgaagatattagagtgtccagagtaaaaggtaaa
actttatttgaaatggcagaaattgataaggatttagcatatgtatgcgattactatttg
aatattgcagatcagttgattacaaggccagaaggtgtagttcctaaagaatctccggat
agagaattatttagtcttttatctgatttttacttaaatcctaaatcaaaggtaggacaa
gaaaaagtagatcaagaagaattagatttgatgatagtgtaaataatattttcagcataa
tataacaaggaataagataatgtctacagctcaatcagatgctcttacttttgaatgtga
aacaggtaattatcatacttttttgtcccatcagttgcgtttcttggttatatcaaaaaat
tgaagacagtttttttcttagttatagggactaaaacttgtggttatttcttgcaaaatgc
aatgggagtaatgatttttgctgaaccgagatatgctatggcagaattagaggaaggaga
tatttcagctaagctaaatgattatggtgaactccgtagattatgcttacaaataaaaaa
agatagaaatcctagtgttatattttggattggaacatgtacaaccgaaattataaaaat
ggatctggaaggaattgctccaaaattggaagcagaaattcgtgttcctatagtagttgc
tagagctaatggtttagattatgcctttacccaaggtgaagatacagttctagcagctat
ggctcaaaggtgtccattaaatttaaagaatcaatcagataacacgagtcttaaacttc
tcctcatattcctttagttttatttggatctcttccggatccagttgtaacccagttaac
tatggaattaaaaaaacaaggtatttttgtttctggttggttgccatcaaagaggtatac
tgaattaccggttataaaagaaggatattatgttgcgggagttaatccatttcttagtcg

cacagctactacactaatgagacgccgaaaaacaaaactaattggtgcaccatttccaat
aggtccagatggtactcgagcctggattgaaaaaatctgttcagtaatgaatgtaaagcc
tattggattagaagatagagaaaaagcaatttgggcttctttagaagactatatttcttt
aatccgaggtaaatcagtttttttatgggtgataatttactagaagtatctcttgcaag
atttttaactagatgtggaatgactgtatatgaaattggtattccttatatggataaacg
ctatcaagcagcagagctagctttattaaaagctacatgtgataaaatgaacgttatgat
gccaacaattgtagaaaaaccagataattataatcaagtagaccgaatccgtgatctgaa
acccgatttagttatcactggtatggctcatgcaaatcctttagaagccagaggtattaa
tacaaaatggtcagtagaattcacatttgcacaaattcatggctttaccaatgcaagaga
tattcttgaactcgtcacaagaccattacggcgaaatcttagtttgtcagaattaggctg
ggatgtttatagcaagcaaagctagtattcctc----tttatgacttacatggtataaaa
gccttgtaagtct--tttattttatgttaaagaagcatttcttgctctgccatcaccagc
ccaagcttgtaaactttctacttgttctttgtctgtaaatgctaaaggcacaaattgttt
gatagctattttaatatcttcagtactaaattctctctcttcactaaacgccgtatgcat
actttcaataattgcttgctcaatttcagcgccagaaaatttattacataataaactcaa
ttcatgaatatcatatttttgccacgatcttggtctaacttttgataaatgtatcttaaa
aattgattctctctcttgacgattaggtaaatctaagaaaaaaatttcatcgaacctgcc
tttccttaacatttcggaaggtaaactttgaattttattagctgttgcgacaacaaatac
tggagctttttctctgatagccaggtaataaatgtaccaaatactctggcactagtacc
actatcaccttgactatgtaaaccagaaaaagctttatctatttcatcaatccataatat
gcaaggtgataatccttcagagatatttatcatttctctcattttttgattccgattcacc
aactagtccaccaaacagttttcccatgtcaagacgtagcagaggcagcatccaatcgct
tgcaatagctttcgcggttaaggatttacccgtcccctgtataccaactagcaataaacc
tttgggcgaaggaataccgtaatttaaactttgcttcgaaaatgagcgagaccttttttg
tagccattgttttaatacatctaagccaccaatatccctatttactttactataaggata
aaattctaatagatgcgtttgattaataatttgcctttttttcttcaataataataggtaa
gctccgagagtctatctgattatattgtgcaataattttagtaatcactttgcggattct
atctatcgataatccttggcaggattttgttatattattaacgagctctgaattcaaatt
aagattcaaagctttactcaatcttatgatctctttttttgatttctagaagactaggtaa
gggcaaatctattactgttatgatatcattcagagcaaaaggtatattgattttgcaaga
aataataataatgtgcttagattgcgttttgattattttagctaaattgcgaagtttcct
aattaatactatttcgtttaggaaagagtcaaaatcttttaaaagaaaaagatttaaaga
ttcattattcaagttttcaataaattctaatgctaataaaggatttctttttgcataacc
attatcgctaggattgttagtgtatccatctacaaagtcccaggaatatacttgctgact
gtttgaacagtttaagctgtgttttataatatactctagtctatcttcttctctagtatt
tattacaataataggatatcgagattttaaaagtaatcgtaaatcttgagtgaaattcat

gtcaatatttgtttccatcagacttttgtacttcaccaattgaagaaagactaaataaga
ttatctaatattccttggatatattcaaaagtc------atctctctatttctaaatta
ttgctattactaatttaacttaccatgattttttttcttccttttcttcgtcgactattt
aatatagagcgtccacatggtgttttcattcttgctctgaaaccagaaactctaatttt
tttcttttagatccttgtagtgttcttttagtcatgatgtctcctctgttgtaaaagatt
aaatactattttatcatcttttatatatctttatgatatagatatgtattaatatcaaag
tt-aacagcaataatttatattaatcaagaactatttatataatattaaacattaaatat
----------taactttatgatattggctttgcccatttttttatttaagtattttgacca
tctttttgctaatcttaaactggttaatttttcagcagctaaaaacgatttgttattag
aatctcaatttaagtattttgttgataaaagccaaaatcgtaaattagaacccgacgaaa
gttttgcttttgcaaaagtatgtgtagctaaaaaatatttttctaaagctattattgaag
gtcaattagctttaaaaaactatcgagacctgaatatcttagataataatatagtaattg
ctaatttgtataatatgctaggttttatttattttgaagcaggccaaacaagttttgcaa
agaacttttatgagcaagccttacaaataaaccctaattatgttgtagctttaaataacc
tggcaaaaatttatgaagaggtaaaagatttaaaaaaagccgaatctctatacgataaag
tgttaactcttaacttaaataataaaactgctaatagaaggaaagatttatagcaaaaa
cta---------aaaacatctaatactatttgtaatcgggatagcaggatttgaacctgc
gacatcctgctcccaaagcaggcgcgctaccaaactgcgctatatcccgtaagacataat
tattactatataccttttctctaattatgtctactctttctgtattttatctaggatacc
aaaacatgcctttaactccatctggatctttgataaatcctaatttttttataaaaactaa
caacatcaggttctgcgaatagtgtaatagtgctaatttctgcctgtctcaattgctgta
ttaattggtgtatgactactttcctaaacctaaaccttggaaatctggatgaataacta
cgtcccagatagttgcattaaatccgttgtctgatgttgctctagcaaatcctacaagtt
tactatttgcatctttttttgtattaaagaaataataatagaactatttttaatgcga
tctttactttttttagaggtcttttaacccatccgactgaatcacaaagctgctccaatt
catataaattaatattcttattgctactcagatagatatctttaaattcgattttatcgc
aagttttatctaaaacaataagtttttgaaatttttttcagagctagcattgatattgg
agttttgaaaaaaatttttccagaagatcataatgattatgatataatta--aataaata
tatatattacaatattgctttaacaaacaatatattaatatatgctcaaatttttacatg
aatctattaagatagaagattgttaaaatatagagaaataaaaatttttgtaaaatcttt
caataaaatgatataaagagtgctttttaagactaacatctattattataaataaagtct
cgaaattgagatgaaataacacattattctatgtctaatgccttttaaaggagttatttg
tgaaaaaatctatgtgtctaacctgtttacttgctttactaattatgagcaacccaataa
tagctaacgcagaagtagctggattggtgccctgcaaagattctgcggcattcaataagc
gtatggtaaatagtgtgaaaaaacttcaagccagattagctaaatatgatgcagacacgc
caccagcgttagctttaaataagcaaatagagaaaactaaaactagatttgcaacttatg

gtcgagcaggtttgctgtgtggtactgatggattgccacatttaatttctgatggtcgat
ggagtagagccggggactttgtttttccaggactattattcttgtatattacaggatgga
ttggctgggtaggtagaggatatcttttatctgttgctaagactagtaagccaacagaaa
aggaaatcatttagatgtgccattagctatcaaatttatgtcatccggatttgcatggc
cgctagcggcttggcaagaatttagtagtggacaattaattgcttccaatgacgatatta
ctgtttcaccccgttagtaaaaaatttatatgaataataatttaccaaatacttatcaa
cagcacctgtaattggtgtattgtggatgacgtttacagccggatttattatagaattga
accgctttttcccagatgtcttatacttttactt----ataaataa--gatcgtataaa-
--aaactacataaatatagagagatactattatttgtgtagtttta---atatataaca
taattattta-tttttaaaaatgtttatcttttcgattattttaatattagatcttacat
aataaaata-atatcaatattatgagcttagtaacccaaattatagttaatgccgatgat
gaattaagatatcctacaattggagaattgcagtcaattcaagactacttaactacagga
agcaatagaattagaattgctactattattagagataaagaaaaggagattatacagaaa
gctagtaagcaaatttttcagttacatccagaatatatagctccaggtggtaatgcagca
ggttctaggaaaagatcgttatgcttacgtgattatggttggtatctcagactaattaca
tacggagtattagctggggataaagattctatagaaacaattggtattataggagtaaga
gaaatgtacaattctttaggtgtacctataattggaatgttagatgcaatccagtgttta
aaggaagcatctttagaaatgcttggtcaagatgatattagaatcatttctccttatttt
gattatataattcgtggaatgtcataaatcatagcttcaattaattagttgaataaaagc
ttttataatgttataattacaataagcccgagagcatataattgtaaaaactaaatcttg
tcagaaccggaaggtagcagcaataatgtttcaatacaaaagttatga-ttttcgggtgt
ttattgtttttatctaatagtctcgacttttaattagtatgctatgtataaatctgtata
tgacatgtaatccatcaaaccttccagcaaaaataatttaataatttgatattaatacgt
aaaacacctatac-tcttttataaaatttctttatatctcatctaaacaatggaaactta
acttaatgggtgttcatattttatcaacaggttcgtctgtcccaaatttttctgtagaga
atcaacaatttgaagacatgatcgaaacttccgaccattggatttcaacaagaacaggaa
taaaaaaaagacatctagctccatcttctacttctttaactaaattagcggcagaagctg
caaacaaagccttatatgcagctaacttaaaacctactgagatcagtttaattattttag
ctacgtctacgcctgatgatttatttggtagtgctagccagttgcaagcggaaataggtg
caacaacatcagtagcttttgatattacggctgcctgctccggttttattgttgctttag
taacagcagctcagttcattcaaactggttcctatgacaatattttagttgttggagcag
acacaatgtctagatggattaattggtcagatagaactacctgtattttatttggtgatg
gtgctggagcagtagtattggggcaaagcctcaaaaatagtattttaggctttaagttat
gtacagacggtcagctaaacagtcatttacaattaatgaataaacctgtaaataatcaaa
aatttggtggtacagaaattcctcatggaaactataattccataacaatgaatggcaagg
aagtgtacaagttcgctgtatttcaagttccaacagtaattagacaatgtttgaataatt

taaacatttcaatagatgaagttgattggtttatattgcatcaagcaaacactagaatca
tagaagcaattgcgagcagattatcagtacctttttctaaaatgattacgaacttagagc
attatggaaatacatctgcagcgtcaatccctttagcgttagatgaagctattcaatcca
ataaaattcaaccaggtcaaattattgttttatctggttttggagcaggcttgacttggg
gagcaattgtcttgaagtggtgat---ttatatattgcggatgacgagactcgaactcgt
aaagctttcgctacacacccctcaagcgtgcgtgtataccaatttcaccacatccgcatt
ttttaccatgaata-ttaaaaatttataatatttatctaaatattcatataa--tgtgta
tatatatatacattcttgtcagaaactctcttatatttatttaagtgataaaattgag
t----ttagtatagatagtaaaattttatttaaaatcataa----aaataaatatataa
ggaaaataaaatatgacaccatctttatcaagtttttgaatagtcttattcttggggca
gtaattgtagtcgttcccataactttggctcttttatttgttagtcaaaaagacaggaca
atccggtcctaaaaataacttagaaagatatggagaagcaatttaagatgaataatgaaa
tatttttcatactaataagctctccatatattgctatact--ttagaagaatttttattt
ttttatttgtaacatcctgtaaagtttattccgttctttctttttccc-atttactctc
ttataattaatgatacaaatgtcagg-ttaataatag---ttttttattctgtatctttc
---------------atttatatacgtattattatgttcaattaataaagaaaagctca
tgtctttgtctaattggcctctcaaaaaagaaaattctgaagcatataatattaagaatt
caaaacaaatcacaattcctgatggtttatggataaaatgcttcgactgtggtctattaa
tgtattctaaagtactgaagagaaatttaaaagtttgccctcaatgtagttatcattttc
aagcttctagtaacgaaagaattgatcaattgatagaccaaggtagttggcaaccaatgg
atgttcacttgatctctacagatccattaggctttaaagaccaaaagctttatagtcaaa
ggttaaaagatactgctttcaagactggcctgcaagacgcagttcaaacaggtactggga
ctatgcaaggtaaaaaagtatgcttaggtattatggattttagattcatgggggggaagca
tgggatctgttgtaggcgaaaaactaacaagactgctagaaaaagcaactcaagaaaagt
tgcctgcaattatactttgtgcatcaggcggagctagaatgcaagaaggtatgttgagct
taatgcaaatggcaaaaatttcttctgctctagaaatgcataaaaaagaaaatctactat
atatatctgttttaacctctcccacaacaggggggtgtaacagctagttttgctatgcttg
gagacttaattattgcagagccaaaagctcttattgcatttgctggtagacgagttatag
aacaaacaatcaaagaagacttgccagataattttcaaagttcagaatatttatttgaac
atggtttcctagatttaattgtatccagaactcagcttagatcaaagttaatacaaattt
tgtccttacataatcatagtaagtaatgaattgaatattgactataatatcaaaatgcta
gtaaaaattaagatattaagaaattttcactaaagatatttgtagactaactagaattac
aattactattgtagaaaaatttgtcttaatgcca-ttttttcagtcttttaacaaattg-
tttttatgactttaaataattgttagtttaatacaaatttataaagagaata-----cat
actaagctaaatatagtaatattatgttttcctataatactgacctatcaaatttcgagg
aaatttatgcttaaaagatcttcttggcttgcggctttattgggactattaacagtagtt

tctacaagtacgcatacatatgccatagagttagacgaggcaacaagaactgttccatta
gaatcttctggcagaactgtaattcttacaccagaacaagttaaaagaggcaagcgatta
tttaataattcttgtgctatttgccataatggtggtatcacgaaaacaaatccgaatatt
ggacttgacccagaatctttaggattagctacgccacaaagagataccattgaaggacta
gttgactatatgaaagatccgactagttatgatggtgcagagtcaatcgcagaattgcat
ccaagtattaaaagtgctgaaatttttcctaaaatgcgcaatctaaccgatgaagaccta
tttacaatcgcaggtcatatcttacttcaacctaaaattgtttctgaaaagtggggcgga
ggaaaaatttactattagaacttcaaagacctaaaccttgtgaatactagtttatgta-t
ttgttgcttaagtgtgattattttgttagatcatgatatatattaaagatggagtttgtt
gacagcaagttacttttacccgaacttgtttacactattcgtctagttattaaaaagga
gacgttaaattgaagaagaagctttcagttcttttcactgtttttagtttttttgtaata
ggtttcgcacaaattgcttttgctgcagatctagataatggagaaaaagtttttttctgct
aattgtgcagcatgtcatgctggcggtaataacgccattatgccagataaaaccttaaaa
aaagatgtacttgaagctaatagtatgaatactattgatgctattacttatcaagtacaa
aatggtaaaaatgccatgcctgctttcggaggtagactggttgatgaagatattgaagat
gcagcaaattatgtattatctcaatctgaaaaaggttggtaattatacttgattttatcc
tgtattaaagaatagacaatctatttagttgttcattagattgtctattctttgtttatg
---ctattatataaagatatttacacaatatt----ttatgatgaaaagaatacccgcaa
ttcttgtactagaagacggtgcatattataaaggatggtcattccagcaagataaacaag
agattactattggtgaagtagtttttaatactggaatgacaggatatcaagaaataatca
cagatcctagttacttccatcaaattgtcgctttttacctacccggaaattgggaatacag
gtattaataatcaagatattgaatctcacagtattagtattaaaggacttattgcaaaaa
atatttgtaaaatttcaagcagctggagagagcagcaatctttagttaagtatttaagta
gtaataatattccttttatttcggaatagatacaaggtctttaacccaatacttgcgtc
aatttggtacaatgaacggttgtatctctactgataatttaaatcatagttacttaaaac
agaaaatttgtgagattccaagtatgcaaggtttagatttaatcccgcatgtaactacaa
gaaatgtttacccctgggatgaaaaaagttttccgaattggtatttgacagataacatta
gagtacaccgagttattcagttgaaagttattgttatagattttggagttaaactaaata
tactcagaagactagctacacttggatgtcagataacggttgtgcctgcccacactcctt
taaaagatattttggcttaccagcctgatggtatattactctctaatggtccaggagatc
catcagcagtacattacggcatccagacagttacaaatttactagattacaatgtgccta
tatttgggatttgtatggggcatcaaattttaaatttagctcttaaagctaaaactttca
aacttaaatttggccatagaggtattaaccatccatccggattgaaccagcaagttgaaa
taactagtcaaaatcatggctttgcagttgaattgacttcggtttttgaatctcctgtaa
gagtgactcattttaatctaaatgacactactattgcaggaactggacataatcaaagtc
cttatttttctgtgcaatatcatccagaatcgagcccaggccctcatgatgctgattatc

tattcgaaaattttatagaaataatgacaaagtccaagaataaagttag-----------
ttagtaattttcccatgctttatgagtgaataaagcggatagtacttgtactcctcgtaa
ttgattaaagagaggcaaatgtcctctggggccatgagaattccattgaaattcctcagg
atatctgcaaggaattctatcaacttcccagccgatttttgccagaattttccccaatc
ttttccaacactaagccatatttgtctttgaacaaatagcccaaatttacctttgagtg
tgtatgccataacttatcaatagtttgtaaatcttgagcaggtattttttttatatctgt
aaaatatagccaattacgagtttgtgcatttacaccagctagttgaataagcttttgttg
agttagctgatctgctttgagcaaatcccgatgagtcagtaacatttgtaaatctttgta
attcatctgttgagcagagcgcaatggaacaatgccatcgggacataagttagaagcaaa
tttcactatttcttgattttactgttaagtaacttttcataaattaaaccgtcaacaca
attacttttatagttaggcccagtaatcctctcaaaaaatagatctgctaaatcttttaa
ctcgatagagtcgttgctatttatattctcaataatttcaagttgttgcttga---catt
attagacttagtattcttattcaattctaaaagctgggctcggatttgatttggcatctg
atttgtttaattagttttaatagtc----ataaggtcttagagagaattagattgttttt
tgaaattagaagagcgccgcagtatttctaaagaagttcgcataatttgattcaataagt
tttgtcctgcataaattataatttgaccaattaaaaaaattgtgcttttttatttttaggaa
caaaaaaatcttgtacctcaagcagaaaagtaattattatttgcattggtgacaacatct
gtaattcattcgatctgcaagcatatatatactgacagtttaagccttcttgataaaatc
cccatactttgtaacgactttcgtaaattgctcgtggtcgctcaagatagtttttaatta
tagtattccaaattaggttattctttaatttttctaatttctatcagatagaaatttta
cattacagagataacttgcatttgtctcatttacagtattaaaattttgaactaacatat
gagcaataatgttgcttagctgaattaaataattttcaagaaaaatttcgacctgtttaa
ctggcatattgtaagtaaagagattaaaagttttattcacagtttgagatgacccaaaaa
gtagctgaataattaaaatttgtagtagcattttatagtcgttcagtagatagttcaaat
catgaatatcttggtaaatgtcaatacagtccagttcatataacttacaaaatcttttga
tacacctatggaataaatcaattaaaatattttcggtcaagtttgtaatatcatttacat
ctaaatttgaattataaatttctaaaagtatcttttccagttcagctaatatgattttta
ataagtttctttttacttcagtgcgaaaaacatctaaaattaacacttcctgagagcaat
tagttaatctcttattaatctttatagaagttctaactaatagctctgctacctcttgat
tgagtgttggtccctgagagcttggccaataattattcacgttataccatagtaagaaat
attgaatttaatgttatacgataacttttttaatatactgaaaagcaagttaattactcta
tagcaactag-------tttttacagatcttaatagttatttatcaaataaagttatatg
atagtagcgattgagctaattttattaaaattta----taataatgacaacttattactt
cgcccttgcgagtcaaaattttctattagtacaagaaccactagaagaagtttttagaga
gagagttaactactatcagtcaaataataaagcaattgatttttggctaataccaaaccc
ttcttttctagagaaaccagaaatgatttcatttaaaaaccttgtacctaaagacgctgt

agctataatctctactaatccaatatttattaattggttaaagctaagaattggctatat
ctgtattggacaatttgaagataacctacaactttctgaagaatcgttaaatattactgt
tttaacagacaaaatttaacaatgcttttattatgactagtgttaaattattttggttaa
gttataattatagtcttttagtattta-gactcttatagtttatcacttagtattttatg
ctgagatggagtgacaaatagtgtaagtatttcttggctaaacgtttcagttgccttaga
tttatatctatttggattaactataatggatagcattcgttttatagtaacattttcaat
ttgagcccaatgtacaattccaagttccaattccttagcaatagctgaaacggaaacaaa
ggcagctcctagccctgattgcacagcgtttttgatagcttctatagaattcaattccat
ctctatttaaagcgactgctgtcaataccatgttgactgagtactttatctatcacttt
tctaattgttgattgagtatctaatgcaataaatctaagcctgtataagtcttcttttg
aatgtctcctagtttagaaaaaggatgtgatttgggtaatataagtgctaattcgtcttc
cgcataagaagtaacctgcaaaacatcttgcaattcagtaggcacttctcctccaataat
tgctaaatcaacttgaccattagctacgctccatgaaataagtctagttgaatgtacttg
caattgaacagccacttgtggatatctttgcctaaatagtccgattaatcttggcatcaa
atatgtcccagttgtctggctagctccaataattaatgtgccaccttgtaagttttgtaa
gtcgtcaagagctcgacaagtttcttcgcagagagctaaaattctgcccccgtatcgtaa
aagaagacttcctgcctcagttaaagttgccttcttattacctctttcaaatagggaaac
atttaattggcgctctaaattttgaatttgcaaactaatagctggctgagaaacatacaa
gctattagctgctttttaaaactgccctctttggcaattgcttttaatattcttaactg
atccaatgtaaatggaaggtctgtcattaaagaattgtagtataatg---------tata
ttatttagtattgatcaatgtatgcatttgccaactaaaatatttgtcattacgcttcat
tattttcaaaataataaaaagtataatgttaaataagctgtattaaaatttagtatctgt
ttgatattaaatttatacaagagatatcgtgaatttaattctaactataaaatataggag
acaat---atggactctagacttttagttgtactaataccagttttagcagcagcatctt
gggcagtttacaatattggtagagttgcactacagcaatttagaaaaatgacatcttagt
tttgcttagaatata-atttttagactttatgagatagaggactttaaaatccctatctc
tattgtatatcttaataaaaaaag-aaagc--ccccttgtacttactgttattacacatt
aatattattggt-ggtaccta----gataataagtttaagctaacttatttttatttta
taattaattaaattaataacacaattatggctgttccaaagaaaagaacatctaaagcta
aaaaaaatgcacgcaaagcaaattggaaaaatcaagcaaaaactgaagctcaaagagctt
tgtctttagcaaaatcagtattaactagaaaatccaatggatttatttataatctaactg
aagcatcagatacttttagcgattagaagtgttt-ttactacaagtctgaaaaagtatta
cgcaacaagttctcagtaatttttatctat----taatcatttgaaaatcattctatttg
ataataaaataaatagcagattaaataaaatcacattaacaaattatgcagcaaagctag
atcaaactagtgaaatttggttgttcaattgtattgaaaatattcagcatattttttta
aaagtcaattaaaatcaagtcatattactaaaatttttatttctggtactagctttgaat

ataccgcaggtctaccaggattattatccagcttaacactaagtggtagactacatccta
ttagtatatatagtccccagtctctcaaaaagtatcttgaagcatgtaccaaatattccc
aaactaatttttcttttcctattaattttcataatttacaatacggaggacaagttgtta
accaatttatacagtaatttgtttaccgttgagcaaaaagagcctgctgtatggattta
ttattctaaaaaaagaaaagcagggagtatttaatttagcacaagctaaaaccttgaata
ttcttcaaggacctatatatggaaaactcaaagaaaaggataatttttaagtccagatg
gttactatctatccggccaagacttctcttctaatacaataatgggacataaaatatcgc
ttccgttattagtcagatattctagaattatttctgagatgcattggttttgctcttatc
ctattagattaaatacttattcgcatcagcaaggagtaaagtgtttaccacataatgttt
taactgatattatgaaatcccaaatatatcaagataacagttttgttgaataatacttat
tcttgatatataatggtctcggtaatctaaataaaatactagaaataatttcttaaattt
atttgtctatattaaatttacaatcatttatgacatacgcaattattgaagcaagtggca
cgcagctttggatagaagaaggccgctactatgatttaaatcatatacctgttgatccag
gtcagtcgattatattaggaaaagtcttattattgaataaaaaatggggaggttactttag
gacgcccttgtatagaaggggttacgataaaggctacagtaatgaggcacttgcgaggaa
agaagataactgttttcaaaatgaaaccaaagaagaaaatgagattaaaaaaaggtcatc
gacaagaattgactcgtttaatgatcgattctataacatcttaga------taaaatcta
gactcatatttttt--aa-aaacttttaatatttaactaaaatagatataatggcacata
aaaaaggtagtggtagtacaagaaatggcagagactctaattccaagcgtttaggtgtta
aaaaatatggtggagagcaagtaacagcaggtaatattttaatcagacaacggggaacta
aagttaagcctggccaaaatgttggaaaaggaaaagatgatacattgttttctctaattg
atggtttcgtgctgtttgaaaagtcaaatcaaaagcaaaaaaacaattagtgtttattctg
ctaagaaatagttaaaacaattggtgcaagataacttgagt----gtatttatttcactt
acagtggcaccatatatttgatttattaagaaatggagttccttaatgaaccaatgaatt
ttattaaagatttgctgtgaaatttaatgacaaacactattgtaatttcctgtctacaca
atatggctgctattctatattgtggtcagatacagaaactagtagtagcaaatgctcatt
atcaagtaagtgatatctacctaggttgtgttgataaaatcttctcaggaataaatgcgg
catttattaacctaggaaagaatgagtacagtggttttatacatatcagtgataccggtc
cgcttaaaaagaaatattatgtcaataatattactaacattttaacaatacggcaaaaaa
ttttagtacaaatcattaaagagccaactttaaataaaggtccaaggctcactgccaata
ttacattatcaggtcgatatattgtattaatgccttttagtcaatcaatctgtatatctc
gaaaaatatatgatgaagatgagcgtcattatttgaaatctttagctattttaattaaac
cggcaacaatgggcttgctatttagaccttctgctgtaggtgtcgatgaggaaataatat
taagcgaattaaaaaatctaaaagaacaatggaactttgttcaaaaatctgcaattaata
gttattcacctgttcttctatataaagatgaagatattgttaaaaaagtaatccgagatt
tttataataataatacaaacaatatagtaattgattcaaacctgggattaaaacaattaa

attattatatccacacttggcactgtaataactctagcacagttcctaagattaagcttt
atagtaataatcaatgtatactagatgcttttggtatcaatcaggcaatttccagagctc
tcattccaaaagttgatcttatacttggtggctatatgtttattgaaactttagaagctt
ttactattattgacgttaattctggatcttttaataattccactagtgcacgagaaacag
ttttaaaaaccaactgttctgcagcaacagaaatagcttatcagttacaaattagaaata
ttactggtgtaattataatcgacttcattgatatggaatcacaaagagatcaattgcaat
tattagaacactttaataaagagctatcacttgatgatgctaaaccacaaattgtacagt
tatctgaattaggtttagttgaattgactagaagaagacagggcaaaagtttgtacgagt
taattagtagtgattctaattactttatttttcacacaatcagagagatctcagtctc
ttaagagattcgatgatagacagcagaaacaacagattttaataaatcttggctatctg
cagagattaatactattaacaaggtctttttcaaaagtcaaatttgtgcagacctgcta
acttttacctaattcgtaatctctatatagttaaaagtagtattacttataaacaaaatt
atttattaactcacagatcaaaacttatttattctaaagaatacagcaaagtattaccaa
gca------gttattatctagctagtctcaataagaatagtaatcaagagttcctatctt
aagattgttatatcaactgtt---------------------------------------
-------------------gaaaaaaagaagctctctcttatataagagagagcttc-tt
ttatagcttactaacta-tctagttagttaattaaaaattaaccgttaacagctggagct
gttagagctactggtaaagattcaccagaagctagatctagagggaagttgtgagcgtta
cgttcgtgcattacttccatacctaggttagcacggttgatgatatcagcccatgtgtta
attacacgaccttggctatcaacaacagattggttaaagttaaaaccattcaagttgaat
gccattgtgcttacagataaagctgttagccagataccaactacaggccatagacctaag
aagaaatgtagagaacgagagttgttgaaactagcatattggaagattaaacgaccgaag
tagccatgagctgcaacgatgttataagtttcttcttcttgtccgaatttgtaaccatag
ttagcagattcgttttcgcttgtttcacgaattaagctagatgtaactagggatccgtgc
atagcactgaacagagaaccaccaaacacaccagcaacacctagttggtggaatgggtgc
attaaaatgttgtgctcagcttggaatacaagcatgaagttaaatgttccggagatacct
agaggcattccatcagagaaactaccttggccaattgggtatactaggaatactgctgct
gctgctgctactggagcagtaaaagcaacggaaatccatggacgcatacctaggcggtag
cttagttcccactcacgaccaatgtagcaagctacgccagttaggaaatgaagaacaact
aattggtaaggaccaccgttgtatagccattcgtctaaagaagcagcttcccagattggg
tagaagtgaataccgatagctgcagaacttggaataacagcaccagaaatgatgttgttt
ccgtatagaagggaaccagcaactggctcacgaattccatcaatgtctactggaggtgca
gctacgaatgcaatgatgaatacagatgtggcagttaatagagttggaatcattaataca
ccaaaccaaccaatgtataggcggttttcagtactagtaatccaagagcagaaacgttcc
cacaagctagcgctttcgcgtctttgtaaagtagcagtcataatttttttatcaatttta
aggatttaacaaagatgcttcccaaacaataagttgtctgttatgaaaattattacttaa

ggccttttcaaatgagaacatttttaaaaaaaaca-tagcaaaatctaatctcagtaact
aa-aattt-tacttgttcatatattgcttaataattcaa-----------taataaattt
tt-gct-------------ttttgcaaaattaacactattttagggat---ataataata
atgcatattaatgctgcttattttgattttaatagtttatataattttgccctacttaaa
tacgcatgtcacactttacaaaaattcagacgaccatacaagatttaaatctgttaaagc
atgctttaacagatttgggtttgatttggcaaatgaattctagtcatataaaagttggtg
aaaataatcagcataaggtagatattttaattaaacaagataacttgtcacatattggct
ttacttggaatgataatagatatcacttagttgccgacttacaactttgggaacaaccct
ggtctctagaagtatttttggataagttatctcagaagtatgcttactactctatcatcg
aagaaacaaagaagcaaggctttgaaaaaatgcagcagatttcacaaaaagatggatcaa
ttaagttaattgtgcaacgctggaattattaactaacatacaaatgcgggcggagagact
tgaactctcacgagattatctcactagaacctaaatctagcgcgtctaccaattccacca
cgcccgcactgttgagac--aatattattctatcataatctatactaaaacacaaactaa
-tagtataaactagcttccaagcttaaaagcgtctacaaatagaccgtatgttataccaa
ttttgaagttgttgaataaattaatttttatatttaattgcttgtggctagagtagacta
ttttactagtcaactcagttgctgcaactattagactggctgcaataataccccaatctc
ctgtttgtccaggaatcgtagacaaacctgttgaaataaaaaatcctagcaataaactaa
ttaagccagttgttaactcgctcaaagaataataaagcttattatttaagtttttatta
agtaagcaaaaaaagttgaaagtctagttttaatcatagtatttagtaacggagttttaa
aaggatgacaagtattaacag-tcattaatattgtcgtatccttttaaaactcgttaata
caaacagctgttcttgt---------ttttaaaggtcttgttcgcttaaagcattaataa
gatgctgaaacggctcttgtataatatctgtgcgtttaattgattgcaaatctaattctt
cttctatgatttcttgtaataactgtatgcttcttactgttggagcaacaggtacagata
gtgaattgtaagtatcttttagcccatttaatacacgctgatcgagaatatttgtactac
cagaaaccaaggcataactagcataccgtaggtaatattctatatcccttagacaagctg
catatcttcttgttgtataagaatttcctccaggccttaataattcaggctgttcttcgt
acaactgagctgcggcctcttttaaaatattggtagcttgatcattaataatttcagcta
tttttattcgatctagtccactcgaaaagaaagattctagttgtccgacagctgtcttat
ctagataacgaccagtaagatcgtaacgatttaatattgctgttatagcatcttgcataa
attaattctcctac-ttgctatcagagcggtattataaatgactaatattgtaatataat
cattagtgtttatgaaatttatttaaaacttattctttattgcttaatgtgcaattaaaa
actcgcatagattttacacatatagagtatacgtagtttttaatta-tttttttataaaa
accaagaaataaagctttttt-taagaggttcgtgaatttgttaatataacatgagcttt
cctcctaaataagttttgtaaagtac--aaagtaaaaaaacgattagtcaataatagtaa
------------------------------------------------------------
------------------------------------------------------------

------------------------------------------------------------
------------------------------------------------------------
------------------------------------------------------------
------------------------------------------------------------
------------------------------------------------------------
------------------------------------------------------------
------------------------------------------------------------
------------------------------------------------------------
--------------------------acaatattaatatgtatcaa--------------
------------------------------------------------------------
------------------------------------------------------------
------------------------------------------------------------
------------------------------------------------------------
------------------------------------------------------------
------------------------------------------------------------
------------------------------------------------------------
------------------------------------------------------------
------------------------------------------------------------
------------------------------------------------------------
------------------------------------------------------------
------------------------------------------------------------
------------------------------------------------------------
------------------------------------------------------------
------------------------------------------------------------
------------------------------------------------------------
------------------------------------------------------------
------------------------------------------------------------
------------------------------------------------------------
------------------------------------------------------------
------------------------------------------------------------
------------------------------------------------------------
-----------------agtataaataagcttttactccagcatcaaaaaaaatatttga
gatatgttgatacaactgattattcaagtgaaaacttagatttagaccaatttagctatt
ttatcattacaattaataaaactgataaatgtattcttatagagcaattttgttataatg
ctgaaggccatatttactcaatattcttcaaaggtccaacagctaaacatctttcttcta
taatttgtcattttcaacaattaaatacaataatatctacagctcatgccatatatttag

gacgagagctaatgaagtcagaattagctctcgtgctagatcaacaatatattcaagatt
aaa--ttttgtaagtattaattatatttatgctagtataatattagtaagggcatgtaac
tcagtggatagagtatcagattccgattctgatggccgtgggttcgaatcccgccctgcc
c---gagtactcagagtact-atactaaaacatct-------taatagctagtttgataa
tagtatgggcattggggctgcaaggtttctacattatgaaaagaagaaaatatg--aaaa
aaacaagctctccaaaagagcttttagtacaatcaataaatgcagaaaacaatattgttt
ctttttctcgaaaattagcttttgcttaaacactgttaatttttggcattcgacatgtta
aactcttattcatggcgaatattctgtcagagtcgctcttagtttaagaaaagtcagaaa
aaatatttatctttacgtttt-tacctatttttatttaaataaaggttagta-ttttcaa
cttttatagtggacgtgggttcgagtcccaccagctccatcaatttgatt---------t
gtataattagctagtttaacctttttaataaccagatatataattggaggaaatatgggt
ttcataacaattactaagcccgctttaaagcaaattgcaatcttgaaaaatgatcatgaa
aatgatgtacatctaagaataggtgttagacaaggcggatgttcaggtatgtcatactca
atgaattttgaacatgttgacaaattaaaagatacagatgaacggcttcgtctagataat
ttttctgttgtctgcgaccctaaaagtcttctttatctttatggattatcattagatttt
agctcggaattaataggaggaggatttcaattttctaacccgaatgccagccaaacttgt
ggctgcggtaaatctttttcaggctgacc-ctcgaacacgtttttttatatatttttgtatt
atatatattattttagcaagtcg-tttattagagacaagcaatgctctgttaaaaac---
---------------------------taatttcaagagtttttattgtattatt-----
----ttatatatttattatcaactacttgaaaataaattagctaagcaatctaatttgtt
atcagtaccaaactttaaatgattaactttttattgatatatgctaatcttt-atcagcc
taaacatatctgtacattacaatcttttgacataaaaagagtcttagcatacctctcaa-
----------------------aaataaattcaa----------tattttataaatactc
aacatgaatctgtcacaacttatcgttaattcttctatgatgaaaattcctaataacttt
tattgcaatataaaagctatttcttctgagaatttaagtcacttacaattatcaaaaaca
aatttattatctcctacaaaaataattgcttttggactacctaaatttgattttatttct
acacagattacaagttctattagagaaacagttgttgtgtctactattgatgataataat
ctttttgctgccccaactaatgattggataaaaaaacacaaggataaaaatcatcattct
ttgttttataaaaaagtttttaaagctttgctaaaccataaaattagagttgttgctgag
ccact------tcataactccgaaataaataaagaaaagagcataatggcaagtcaaaga
tacatctggggtaaaagctggaagccttcaattattttatcttgcttaaaaaacaaaaaa
aatagcaaaatttctgatgcatcgaaaaaatttttaactgaacagttatctgcatcacca
gtatttgtggtaaaaaatggttttaatgaaattattttaggtcacccattatctcgtgtt
aaaagaggaagagttaataatctgatgcatgcgttttctaatttattaaatcagtctaat
gcaacttatcctatatctactggtctattctttttttcatccagatgatgcatttgaattt
aaagactttataatatcggtcaatccactggcggccaaacatatggagataagcgtagag

cctgttggtctacactttgcttataaaatgaatagaaatatatcgtcagatactcaattt
tcgtttattccagattttaaagaagtaggagatttattattcaaatatagaaaaggtaat
catttagtttttcataagaatcagcattatggcaaagatttttttcaggggcaaccaata
tatatgattcagccgatcacttttaaagatcgagccggaaagctaaatattatcaaattt
acggggctaaatgacacaagagaaattatttttactaatcttgaagctgctaatagatca
tgggcacattttataaagaggaattcacaattaaaatcaattaaaaatcctactctatta
gtctataatttagaaagtttttttaaaagatcaagaacgattggataatcaagatttaaat
aagtttgtggtgatcactaataaagaatcttatcttgccacaaaagagttaatagcttta
cccgattctaatagcttttctaagcacttaaaattaaatataaagcctaaactcttttttt
gtgaaactatgggtaagacggttattttccaccttaacttatgaat-aaatgctatt-gc
ctttattttctcgaataatattctataactagtaattcattgagttgtaaagcaacccat
tccctatcaatgactccattgactttccctgataaattggatttattcaattctaagtgg
ctaggaatattagctagtccagggaaagctaagtaattttctactagttttcgagatgct
tcttgatttttacactaattgattctcctggtttacattgataactacagatagatact
acttgtccattaatacaaatatgaccatggttcactagttgtctagcggcaggaattgta
ggagccataccaagtctaaaaacagtattgtctagcctcatctctagtagctgtaacaaa
atttgacctgtcgacccttgtagtttttagctgcttttacgtacttaaataactgtttc
tcacttaatccataattaaatcgcaacttctgtttttcttctaatcttacagcatactca
gaaggttttctggacttttgtccatgttctccaggtggataagatcttttaatcgctttt
ctactgagtccgggtaaatcacctaatctacgggaaatgcgtactcgtggtcctctgtat
ctagacatattatttaaattctcctaatgtaaat-gattaag----------taatgagt
agctatttttttaacacaaaaagagcattaaaacaaagc-ttttgctaaaaaatatttaca
aaaaatataggtttggacaaaa---tacattactatgc-tttttaagcgggtagcgggaa
tcgaacccgcatcattagcttggaaggctaaggttttaccactaaactatacccgcaata
ttcaacagattgtcactaactcaatataacatagttatgttaagttaagataatcttaaa
tttaatagaatt-ttaaattatttatgctccttctaaaaccacatcaagaaaacgagcaa
tttcggcagcttgttcttcaacttctgataataataatggttgaccgaccctagttaatg
gaattactcttttatcttttgtacataaataaatttcacgtcttggatttagcccctctt
taatatctatttttattgatttgatttcttttatattaaactgaagacagatttttctat
tctttccagggaaacctagccgaaaaatttttacaatacctttatctttgttaaattcat
tatatccagcccctatattccaaataatagttagccacagaaacaaactcagaaagactc
ctatactgccatagaatgtcattacaataccttgtggaataaataccaaatcggttgaat
ttgtaaaaggtaacaaatcaacttgaaaataacttgatagtccagcaagaagaaagccca
aagccccaataaatattatagtagcccaccagtaattactcagtcttcgtgaacctaaaa
ttaaatcttttctaactgagtgtatagataatgtttttttcataatttgttcttttaaaa
cagta-tactaatgcaaaatggaaaaattttgacaatttt--atgcgcgctttagattt

tttaaataagcttgattgcttttagtcctaaaaacagtcctgttgctaacccaaaaaaag
cagcaagaaatattatatatcctagaaaaactgaca---tattattatttattattattg
ttacaatagtgtttacaaaaatagtgaaaatctattactacggtattatattacgttaaa
tgccttgctgtctacaggaaaaaactttttcagttttagcattgtgtttatattagaa
ataatattctagtaaatttattaaacttctggagaacagaatcttatgtcagaatttatc
aagccttataacgatgatccttttgtaggcaatttgtccacgccagttagtacgtcaagt
tttagtaaaggacttctaggaaatctaccagcttaccgtcgaggtttatctccgcttctt
agaggattagaaataggaatggcacatggatacttttaattggaccttttgataaattg
ggccctttacgaggtacagatgtagcgttactagcgggatttctatcttcggtcggcctc
attattattctcactacatgtttatccatgtatggtaatgtatcttttactagagcagat
tcgaaagatccactacaaacttctgaaggctggggacaattcactgcaggatttctagtt
ggagcagtaggtggttcaggatttgcttatttattgctagctaatatacctgtattacag
actgcaggtcttagtttattctcttaagctagtaaggggacttgaacccgtaacctactg
attacaaatcagttgctctaccaattgagctatactagcattcagttcataataacatat
aaagatatttgttgcaatctgctattttcagatgtaacaaatatctttttttatttaaaca
gaactgttatgttaacac-------gtatattaattagaatctgaattatctgactctgc
agaagattcctgatcacagtttagataataacaaattgtttgatctagacaagttgcaga
ccaaccaacaggtgtttctcctgataattcttgcatatctacagaattataaaaattatc
atttggcttattactgtttgcttgcataaatatcactcccaaaacactctgaacttaatg
tgaagtaatttaacttctcaataatatattaacacaaaatattaaatatggtgattattc
tttaattgtaaagatgctgctaactttataaagtttgagtgagagtcttgatagcttttg
ttgaaataagcattttaacccatttattttgcccgctcgaccatactttttgtgttgta
aattaactttttgtagtttttttgttcgtttatgagaatgagatacggcatatccattat
tagctacttttcccgtaagctgacactttttgacataatttt-------ctaacttata
tatttatttagagtacttgctaaagttgacttgggcacagctccaatcactgtgtctact
ctttctcctgctttaaagatcattaaagtaggtatacttctaatgccatattcagccgca
atagtaggattatcgtctgtgtttatttttactaccttaatagatgattcatattcttcc
gcaatttcatcaactacaggagaaaccattctacaaggaccacaccatggcgcccaaaaa
tctacgagtacaggtaagttattgttaataacttcttgtttgaaagaggcatctgtaact
tgagatactgacatattctttaacctttaatatatgttccttgctaagcaaatcttatca
caaatttcaagaaaaaatcttctatagtgaattactattagtgttgaattattttttaaa
aattagacattaaaaaaacttattacctatataaataaagtggtggccgagtgatatctt
gatagtgataatttttatattaaggttgaacactagaaataacttttcaagcaaactctt
atttgaaaa--atttctctgtgaccataaaaacttaattatattacaatatattttatta
tttgtaacttagttctgataatggtataaacaacgcaaaagatactgccttataatcaag
gaggaatacatgtctcaatccgtagaatcacggactaggattaaaagcgaacgttacgaa

tctggagtaatcccttacgctaaaatgggctactgggatgctgactatgtgattaaagaa
acagatattctagctcttttcagaatcactcctcaaccaggtgttgacccgattgaagca
tctgctgcaattgcaggtgaatcttcaacagctacttggacagttgtatggactgattta
ttaacagcttgtgacttatacagagcaaaagcatatcgagtagatccagttccaaacgtg
gcagatcaatattttgcttacatagcttatgatattgatttgtttgaagaaggttccatt
gcgaacttaactgcttcaattattggtaacgtttttgggtttaaagctgttaaagctctt
cgcttggaagatatgcgtatgccagtagcttatctaaaaacgttccaaggtcctgcaact
ggattgattgtagaacgtgagcgtatggataagttcggtagacctttcttaggtgctaca
gttaaacctaaactaggtttatctggcaaaaactacggaagagttgtatacgaaggcctg
aaaggcggtcttgatttccttaaagatgatgagaatattaactcacaaccatttatgcgt
tggagagaaagattttatattctatggaaggtgtaaataaagcatcggcttctgctggc
gaaattaaaggtcattaccttaacgtaacagccgcgacaatggaagatatgtatgagaga
gccgaattctctaaagaggttggtagtatcatttgtatgattgaccttgtgattggttat
actgcgattcaaagtatggcaatttgggctcgtaaacatgacatgattttacatttacat
agagctggtaactcaacttactctcgtcaaaaaaatcatggtatgaacttccgagttatt
tgcaaatggatgcgtatggctggtgttgaccatattcacgcaggtacagttgtaggtaag
cttgaaggagatcctttaatgattaaaggcttctacaatactctacttgaaagcgacaca
gatatcaacctacctcaaggtctgttctttgctcaaaattgggcttccctacgtaaagtt
gtaccagtagcatctggtggtattcatgctggtcaaatgcaccaacttcttgattactta
ggtgatgatgtagttcttcagtttggtggtggtacaattggacatcctgatggtatccaa
gcaggtgcaactgctaacagagtagcactagagtccatggttatggcaagaaatgaaggc
cgtaactatgtagcagaaggtccacaaatcttgagggacgctgctaaacttgtgggcct
ctacaaacagctttagatttatggaaagatattagttttaactacacttccacagataca
gctgatttcgttgagactccaacagcaaacatctagtttaatgactacttactgatactt
taaatagttaattgtaagtgaaattaacttataacaataaggagcatagaatagtgagac
taacacaagggacttttccttccttccagatttaactgatgagcaaattaataaacagc
ttgcttatatcgtttctaaaggcttatcagcaaacgttgagtatactgacgatcctcatc
caagaaactcctattgggaactgtggggtttacctttatttgatgtaaaagatgcttctg
ctgttatgtacgaaattagctcatgcagaaaagcaaaacctaattattatgttaaagtta
acgcttttgataatactagaggtattgaaagttgtgtaatgtctttcattgtaaatagac
ctgctaatgaaccaggattcttattacaacgccaagacttcgaaggtagaactatgaagt
atagtcttcatagctatgctactgaaaagcctgaaggagctaggtattaatattaattct
taattaatattggctaatta--ttcccctcttaaaaatcataattaatttagtaattgtg
atttttaagagggatagcttcccaagtaaattttgactaaa-tagattgaataattaaaa
atacacagaaataatgcaatcacaggatataatttccaacgatactcttgttaatttaca
agaagaatatgatagaacacaaatccaagaagttttaaatgagttaaatcaagaacttat

aggattagtgcctgtaaagaccagaattcgcgaaattgctgcgctattattgattgacag
attacgcagaaaactagaactagtttctggtaatccaggattacacatgtcatttacagg
tagtccaggaactggtaaaactacagttgctatgaaaatggctgatattttgcacagact
tggatatataaaaaaagggcatttgttgacagttacaagagatgatcttgtaggtcaata
tattggacatactgccccctaaaactaaggaagttcttaaacaagcaatgggaggagtttt
atttattgacgaagcttactatctatataaagcagataatgaaagagactatggctctga
ggcaattgaaattttattacaagtaatggaaaaccaaagaaatgacttagttgttatctt
tgctggatataaagatagaatggaaaaattctacgaatccaacccaggactctcttctag
agtagctaatcatgtagacttcccagattatacttcagatgaattattacaaatagctaa
aatgatgatagaagaacagcagtactgttttacagaagaagcagataaaactcttttaga
gtataccgagcgaagaatgaaacagccttattttgctaatgcaagaagtattcgcaatgc
tattgacagggctagaatgagacaagccaataggattttgccagtggagaaaaagtatt
aacaaaagctgatttggtaacgattgaagcagaagatatcttgaaaagtagattattttc
attacctaatgcttaatatacacgtgatttcatgaaactatatattattaaaaag-tttc
ataatcagttgcaaatttcttgaaaaacctttattattatcttaatgtaggtg-tggcgg
catagccaagtggtaaggcagaggattgcaaatccttcatcccccagttcaaatctgggt
gccgcctagtactaaaaaaggggggtgtggtggaatggtagacacaacagacttaaaatc
tgttgatttttagtaatcgtgagggttcaagtccctccacccccatatattgatttataa
ataaaaagagatgtgtatttgtattaattgcaaccatattaccaaatgtaatacctatca
cttaatagaatctcaacataaacagcctcatcttactagaagtccattgtttatacctaa
atatcctgtagttcatgttaatatatctaacaactgtacctataatcaaattgattggga
tttagtggagtgtttatcatttgtagaaaaaccgaatagttggaatttagacgctaatta
gttacaagaaaaattatgtaccataagcaacaactctatttttttagatactcggtattaa
ttttggatgttctaaccagtaagattttacccgttcttgctatctctataatgccaaatt
tagttagtaattgttcaatagcaacaatctttcccggatctccagtaacctctataatta
aaagatcttctgcaatgtctactatcttagctctaaaaattcttacaatttctaaagctt
ctgttctagtctgagaattgatctgaatcttaattaacattagttctctttcaactgaag
gaatatttgttacatcttgaacgttaaggatatttactaacttatataattgtttcgtaa
gttgttcaatagttctattgtctccttgaactaccattgtgattctagagaccccaattt
gctctgctggtccaactgctaaacttgcgatattaaagcctctccgggcaaatagaccag
atattcttgacagtactccggcttcatcttgaactaaaactgataaggtgtgtttcatga
aaaattaaatttaatctatttgatctattaagtatattctaagaaaattacagaaaaaat
actagttagtctataactaaattatccaaactaaacaaaatcctactatcagataatata
tctaccttgtaatgctatactattaacttagtatttaatgatattctaattgcaaataga
atgct-aatttttttaaatttgttaggaacttgtgctagaaaaatacaaaaacagtaaaa
aactctctcgagatttacctcaaatcaatgatcgcattagatttccaaaagtccgagtaa

ttgatgacgaaggtgaacaactaggtatttttgtgcctgaagaagctatacaattagctg
tccaacaaggtttagacttagttgttgttagtgataaatcggacccgccagtatgccgaa
tattagactatggtaaatataagtttacacaagaaaaaagagctagagaagctaagaaaa
agcaacataacagtagtattaaagaagtaaaaatgcgatataagatagaagagcatgatt
ataaagttagaataaaccaggcatccaaatttattcaagcaggagataaagtaaaagcaa
ctatcacatttcgggggcgtgaaatccagcactctaatttagctatagatttattgaata
agatggcaagcgatctaaatgcaatagctgaaattcagcaagctccatcaagagatggca
gaaatgtcataatgctcttatctcccaaaaaagttagctaaactaattttttatttaaatg
catctggctggattcgaaccagcgacgtctctttcgaaatggcggattatgagtccgctg
ccttcggcccctcggccacagatgcatcacttaaagtttataactcataaagcatataaa
aatcaatgtttagctatttttaataaagtctataaaagtctattattcattcatattgtg
atatgtcgcaactgctgaaggggaaacttgatttaagtatcgaaatatccagtatttaaa
aattgtgtcgagtataactggaaaagttgaaataaataaaaaaataaaatctcgactttc
cggaagtcctaaatgtcttaaaattacttcaataataacttcccatccatgaggagaatg
aaaacccacaaacatatcagtaaaaagaatgattaaaaaagctttagctgtatcactaag
accataaataatttcatttaaaaaagatttaacgactgcaatttgtctttgtccagttat
cattagtaatataaagacaagtattgatacaagatcggataaaatatttttaacagcatt
tgcactttcgtttgcataatattctcccaactctctagccttcagctgcactctttttc
aatgatttcgtaagatatatcttccgaaggattcagaagaacttcaaagtgaattttctc
ttcaaatctttgtaattcagcaaaagctctttcctcttgagaggaatttaaaaaaatttt
aggctgttcttggttccataaataatcaatacaaggtccaaatacaaaaaattttgaagc
ctgattaactagtactggagagatgaataataataaaatatattttacggaagtaatagt
ttggtgtctagatattctaaactcctctatagcttcagattcgccattaggatctaactc
tttacgaaacttttcaaaagtattagtaattgacctaggtattggcccaactttttcaaa
ggcgaattgattatttcgttttaaattccaatatttcatttttttattgctgctaaaattt
ctattaatcaataattgatcacgtattcactataaatattaattctagaactaaacatat
aaaacaatatttatgaaaaataaaatacaattaaaaattaattatgcattagttctatct
gcagctacagcagtttatggatataaatcttatcagccaactataaatacactcttaatc
aaagatgaagaaaactggcctttttctctaaaaggcagactgcctttattcctataatt
gtattaaaaaaaaatcatcaaaatccctgaactcagaacctagaaatagtaagttatatc
ttaatggaattggaagtagaaaatttatcgacagaattgtcactctacttggatatttag
aaaaacaggcaaactatactgaatacttgaaccaaataaaaacaagtggttatttttctt
tagtctacctagacagtgatataatgtataagaaaaatattagtaacatttttttactac
ctaatgatattctaaaaagaatctatatttgcaataagaaaaaaaagcttataccgcatt
cattcttaataaatttatttcagaaacagataggctatccaaaaagttttgctaatttaa
attgggccttctctaaaattataaaatggtattatgatagaggataccaatggtctctag

ttgaagttaaacaagcgtctgatgcatcttctattgtaattgatattcatgaaggggtag
tgaagacaattataacagaatactacactctatcttataaaagagtctcaggtatcttat
gcgtagagtcaatagagcagtaccttggagtaagagtgggagctccactaaatataattg
atttacaaaaaaaaattacttacttaaaagataatcaactagttggcgacattatttata
gtattgaacgatcaaataatagctcaatgagtttggatatcaaatttcaaatacaagaat
taaaagataaagagataatagtgcttgcagaaagttcttcta------------ttatct
cgcatgcatgtaatttactcaatc---aatataga---------aataggctagtagctt
ctaatatagtttcactatcaactaataaattacacgcatgctattttaattataaacttg
actatcaatataagtatataaacacctctaatcttataaaattgttagcttattctattt
cttgtaggaagacactacttattcattatttaatttacaaactttaataagctgtacaa
aaaaaaatactattggatttcagctatatttacgaaatctaagttttgggaaagctttct
gtgtgcttagtatgaaatttatcaaaaatggacttaatataaaaatttatatataaatc
cttcattgatagttgatcagaattttgtatttcaatttgctatacaaatcattaagcaat
atcatactgcta------aacctccagctt------------------------------
------tatttttaacgaatctagatcttgaacaatatgttgctgaaagtttattaatgt
accattttacatcttgcttctcaatatctgaaaaaatattattatctcgaattatgcata
cagattctttatttttcaactctgaaa------attttcattttgatgatagaactaacg
cgatcaacagttatgatactttaaacaaaatacaaaaatatttatcaagagtttcttt
ctttattattaagtttacgttatcaaaattttaattatttaggttggcctttaaaaggcc
atttttttgaaataaaatctttatacttggctccatttcaaaagagtg------------
---atttttctgatagtcgtaagacgctattcttccacaaaatgtcgttaaagcaagtat
ctaattttaatttaccagtatcttttaaaagccaccttaaccatattttagttagtacta
ttaaatgtcagtcaaatttgaatatgagaactgtatctttgctattaattgattcacctg
ctgaatatctgctatacaaatctattcttaatttctctattaaagttaggatgcagtatt
tcatacctatgagcaacaatataagattatctctattttataattatttagattgttttt
taataagatcatctcagagttgcattcacatatggcaagatctaagaactcttacaccaa
ttcaaaatttttggttaaaaaaattttcatatggagctggaattcagcttaaattaccaa
ttaagcagatgccacctctttcaattgaatatactgtaactagtagtcgctacttttgta
tttaccttcgtacttattatcaacgataactattactaaaatgaaaaatttaatttcttt
ttttgatcgcagcaaagggaaatggatttcacaaagaacaacttatgaattatctaataa
aaacatgagttcggtacagtctcagatgacaatgaaaataggcaactcattgtcaggatc
tataatacttgcatcgttaaattgggggacatttatagacaagttgcccattacgctaa
aaatcacagccggagtgaatatgataataagtttaacttgcaattcggtaatcaactaaa
taatcataaattattaacgctgtgcatagtgacagatcctagtctaattagttttaaaac
tcgctatggaagtactaccatagatgaaacatattggttcgcaacaaataatttacgtct
aagtactagtattgttaaacgattcaacacttgtgtggcagtatctttttgttcagaaat

taaagtttaa-actctatatagaagcaactaatagccttta-ggctattagttgcttcta
tgtatctaaattttgtattaaaaattat----aatactatattattctaaatctttacga
ttgggattcctggaaggatcattagagaggaacccaaagacaaacaaagaaataaagaaa
ataacggtagtatatacaaagattttcaaagtaaacatatttcatccttaaaagctatt
agtgcg----atatttacatatctccacataattttatatcatgatataagttccagcac
tattttataaataaatcaatatctttatttttagtaatgatttatatttaaatatgcgt
gatgtactaaatactaaatcttttccgacttgttttgtcttgtttgtaaatattttgtta
tttctatggagtttaccttcaattatcacatagtctaacttctttagctgtttaaaagta
tatgaagatttttgttccaaatagttagattgattataactaatcgttttctttttaac
agtcttgcttttaacttaattatttggctttttatttcactaattctgatactcttacaa
ctaagaatttggactagaagagtacagctattcataatttataacaa------tttttta
aaaaatctactaaaatagatgatttacagaactaaaatgatatttcattatcttgttga
ccttgctcataagtaatatctttcaatacttttaatatttttccaaaatactcttgcagg
tatttttctaaagatattacttccaacctgtcaattccaagaatactgtaaacttcatcc
atcggagctgtaaattgctctccgctacttaaaatttcggcaaatgctaaacgatcagaa
atattccatgtccattgaagagttttagtaattcttcttaaggcttttaataatccaatc
ggaatttgtgagatctgtgtttttttgcccagataatttttcacataatttaataatttca
gctgatgtccaagcggtattaccaactaacggcagggtcctattttctgtggaaggtact
ccaagactttttataactaacttggctgcatcttgagtatcaatgtatgcaatgggtgta
gattctcctgtaacccagactgatttcttgtctaaaataggaatggcatattggttgatt
aaaccttgaaaaaatcctcctaaagaaaacacagtatatttcacattggacttctgaagg
aaatctactacctgagactttaagttcatcaatggaacatctgggtatttctctgaattt
aatattgaaaagaaaataaatctttcaactttagccgctttagctgcttcaattaatgca
atttttcccatctaaatctatttttttctgcattataaggatcagtaggacgagatgtagaa
gcatctatgattgctgttaccccacaaaaagattgcaaaatactttcgggtaattttaaa
tcaccatatataagttctgctccccactctttgagaaaggcagactttcttaaatttctt
accatgcattttacattgtagccttcatctaaagctcgtcttacaatttgacgtcctaaa
gttccggttgctccaattactaaaagagtcatattagttaaaaataagtatgtataaaga
cattgatttggttgacggcagagagtaaattaggtagagagggaatcgaaccctcatgac
cgaagtcgtcacattttgagtgtgatgcgtataccaatttcgccatctaccctattatag
accaactactacaagtaatttagtctttagcgtaaaaataatactaagtttcaatagata
tgtcaacttttaaatttatattaataaaaaaacggagtctcttatggccacttttgaatt
aattccttatagattgtatttgagtcaatctaacacatggatccatagaataaaagcaga
aattaagatatatatagtagcgcttttatggatttcaattttatttttttcttactttaa
actatgtattattgctttaagcttaattgccataagttttactataagaagtaaacagaa
tattattcaaaaacatttgttacaaacgttattaatgacgttcttaactactgttttgtc

ttttagtgtggctattagttataaacaatatgcagaacaagaacaatcgcagtatttatc
tgattccaagaaatataaaaattccagcagctactattatatacaaatagcgaacgatca
aaggatcaaaagacaatacttaattactactttaaagccttccttgtactttttttattac
tatatattctattaaactagttatgataacaacctctccagaagttttagtaattactat
ttatagatctaggataataaataaaatatttaaaaatgaattgctatttatcttccttct
ttcgtcacatattgttactagtattattaatagaatcgataaagtgattcaagtaactag
cttaagaggaagcttgaatttatataattcattgacaaggcctttaatgttttctttgtt
aatatttcaagtcttctttttggagattattcgggagtcaaaagaaatagctcaagctct
ttatactagaaatctcaatcaagaaaacaataatttttttgaaaatatatacagtaaaatc
taactttagtgatcggctcaatataattattagcactttgtactttattattttagctct
agcgtaatagaatccacttattcttctgactctagtaatacgaagagtaataaaattagt
tagtagtttaaatagatagaat-taagatattatataaataattaaatgtttcattatgg
tcaatactcaaaatcaaatttcacaaacttcagaacttgattatatagttaatcaacctt
ataaatatggctttaaaacttctgttgaatctgagcaatttcctaggggaataagtgaag
atattgtccgcttgatctccaagaaaaaagatgagcctgaatatctattgaattttaggc
ttaaagcatataaaaaatggaaaaagatgagtagtccgtcatgggctcatattaagcatc
cgaatatagactttaatacgattatttattacgctgttcctaaattaaaaaaagaattga
aaagtctagatgaggttgatccagaaattcttgacacttttaacaagttaggtatatctt
taaatgagcaaaagcgaatttccaatgtagctgttgatgctgtttttgatagtgtatcta
ttgcaactacttttaagaaagaactgtctgaagctggtgttatattctgttcaatttctg
aagctataagggattatccagaattaataaaaaaatacttaggtactgttgtaccagctg
gtgataattattttgctgcattgaactctgcggtatttagtgatggctctttttgctata
tccctcctaatacagtttgccctttagaattatcaacttactttcgtattaacaacgaag
aatctgggcaatttgaaaggacactaattattgctgatcgtggcagtaaagtaagttatc
ttgaaggttgtactgctcctcaatttgacacaaatcaattacatgcagcgattgtagaat
tagtagctcttgaaggagccgaaattaaatattctacagtacagaattggtatgctggta
acaaagaaggtaaaggtggtatatacaattttgttactaaacgaggcttatgctcgggta
acaattcaaaaatttcatggactcaagtagaaactggatctgcaattacctggaaatatc
caagctgtattttagctggtgaaaattctcagggagaattttactctgtagctttaacaa
ataactatcaagaggctgatacagggactaaaatgatccatattggtaacaatacaaaaa
gtagaattatttctaaaggtatttctgcgggtagatcgaaaaacagttatcgaggcctag
taaaagttggccctcagtcatttaattctcgtaattattctcaatgcgattctttattaa
ttggtcaatcatctcaagctaatacatttccttatattcaagtacagaatccaacatcaa
aagtagaacatgaagcatccacttcaaaaattagcgaggaccaaatttttatttttttac
aaagagggatcaatttagaagaatctattgctcttatgatcagtggtttttgtaaagatg
tctttaatgaattgcctatggaatttgctactgaagctgatcgtttactgagtttaaaat

tagagggaactgtaggatgagccagactattttagaaatcaaagatttatatgcttctgt
tggtgaaacaacaattttaaaaggggtaaacttatctattcgagccggcgaaatacatgc
gattatggggcctaacggttcaggtaaaagtacattatcaaaagtaattgcaggacatcc
agcgtattcactaataagcggagatattttatttttggacaaagtattcttgaaatgga
gccagacgagagggcaaaagcaggtatttttttagcttttcagtatcctgttgaaattcc
tggagtcagtaattctgatttcttaagaattgcattaaatgctcgcagaaagtttcaagg
attatcggagtttagccccttagaatttttcaactaataacagaaaaaatagatcttgt
tggcatgcaagaaagtttttttgacaagaaatgttaatgaaggattttctggtggagaaaa
aaaacgtaacgaaattcttcaaatggctttactagatagcaaaatatctattttagatga
aacagattctggactagatattgatgcattacgagtagttgcaaaaggaattaacacttt
agctaaatcaacaaattcaattattttgattactcactatcaaagattattagattatat
tattccagattttgttcatattatgagtaacgggcaaattgtaaaaactggtagtgttac
tcttgcccaagatctagaaaagcacggatatgactggattacgcagacataactatttac
tac-------aatattgagactaaagaatattctttagtctcaatatt-tttgcaaatgc
aaagtt-aattatttaagctttatcaaaaccttgcttaacatcttcaattgctcttttta
gcaattcttcactaactggttctaatttttagtactacgaacattttcaccaaattctg
gtttagaattctttaaatcttctcgaagttctctaataaattcagctacttttgaaactt
caatatcatctagatagccattaattccagtgtaaataatagcagtctgttcttcaactg
gaataggagaattttgggcttgcttcaaaatttctcgaagacgttgacctcttgctagtt
gattttgggttgctttatctagatcagatgcaaactgagaaaaagcttctaattcagcaa
attgtgctaactctagttttaacttgcctgccacttgtttcatggcttttatttgcgcag
cagagcctactctagagactgagatcccaacattaatagctggtctaatccctgagttaa
ataagtcaccagatagaaatatttgtccatctgtaattgaaattacatttgttggtatat
atgcagatacgtccccagcttgagtttcaataatagggagagctgtcatactacctccac
ctaattcagcattcaatttagctgctctctctagcaatctagagtgtaaatagaatacat
ccccaggataggcttctcgtcccggaggtcttctaagcaaaagagacatctgacgataag
cttgagcttgcttagttaaatcatcataaataactaatgttgctttacctttgtacataa
aatattcagctaaagccgcacctgtataaggagcaatatattgtaaagtcgcaggactat
ctgcattagctgcaactataatagtataatcaagagctcctttatcctgtagtgaagata
ctacttgggctactgaagatgccttttgtcctatcgcaacatatacacaaacaacatctt
ggccttttgattaataattgtgtccaatgcaacagctgttttacctgtttgacggtcac
caataattaattctcgctgacctctaccaattgggatcatagagtcaatagcagttatcc
ctgtttgcataggttcacacactgattgccttccaattataccaggggccattgactcaa
ttagccttgttccattacttgcaggttcacctttatcatctattggccgagctaatggat
caacaactctacctaaaaaagcatctccgaccggaatttgagcaattcgaccagtccctt
ttacggaactgccctctaaaatatccctaccatctcccatgagtactacaccaacgttat

cactttctaaatttaaagcaaccccaatagttttatcttcaaattcaagtagttcaccag
ccatgacttcatcaagtccatatactcgagcgataccatcaccaacttgtaaaactgttc
ctatattggctacttctacatcctgatcatacttttcaatttgttgacgaataatactac
ttatttcatcaggtctaatatttaccatattttaatattatttttattaattggaaatt
ctaaagcggtaattagtcagactaatgctatctgtctttttttactttattttaagttact
acaacatcaaggtgtgaagccatttgtcttagctgtcctctaatactagtatcaatgact
tttgatccaatctgaatagtaaagccaccaattagctcgggctcgactgaaataattagt
ttaacttctttagccttagtcataaccttgatttttttctgttaatagagtttcctgatca
gggcttaaagcaatagaagtattaatatttgcaattgttaaagattccatttggtaagcc
agttccaaatacttactagctatagcatctaacattccaattcgcttgcgatcaactaaa
atcattagaaatgataatgtattttcgttaatttgatcgccaacagttgcaataattact
tcttttttttgcctctacagtttttaatggattagct---aaaaaaattttttagcttgtca
gattgcgctaaaatattttgtattgactgaatatcttggctcacctttttcagtaactttt
ttagttttagctaaatcaagaagagctactgcataaggctgagctatttttataacaaca
ttgttactgctcatagttgatctcctaacttagcaatattattatcaattatacgcagtt
gcatttcggagctcatttgattttccagctgcaaagtaactcttttaagagcaagaaatg
ttatttgttgctggatttgccttcggatctgtttttcagctgtttctatatttgatttac
ctgttatggcaagtctttcaatatctaattggccttgagctaatatggaacttctaactt
ttccagctgttagctgagcttctttcttaatttgatctataatgatttgagtttgagcaa
gttgtttttctgattctgacaatctagcacttgcttgctctagtctttcttcggattctt
gtatagctgctaaaacttttatctgtcttgcatttaaacttgatccgagaaattgcttca
aaacataaataagaccaaataaaagcaacaatatattaataacatttgcttcaaaaatat
cagaattaaagccgaatttgtgttcactactatgttctgataaaatagtaattatttgtg
gtatttttactatattattcatttaaaattct----------tactattttaatatttta
ctctggaatcatgattttcaagattgactacttaataatttaactttaatttgatcgctc
aaagtgtcaacctgattttctaaagttttaagagcttcttctttttgaacatttaactgt
ttcgatgcttcagcaatcagtttctctgcattcaattgagctttttttatatcttctgag
actatattttgagcttctttttgagaagaggctatttttaattgtgcattgcggcgagct
tcagatagatcttcttcatattttgcagctagctcatctgctttaacaagcatagacgat
gcagtagtcaatgttgttcgaatatattcatctcgttcgtctagtactttagttacaggt
ttataaaaaatagtatttaataaaaccattaaagtaagaaactgtaatgccattaaaggc
aaagtaccattgaagtcaaataagccaccttcaatttcttcggctaatagaaatggtaaa
tcaatcattttactaattatgtttatagttaataattttgtttcaattaagaaaaataac
tttatagtgcgcttagtctcatcagactaagcgcaaatgattattttaaccaacgtatgg
attagcgaacaataaagacaacgcaacaactaggccgtaaattgttaaagattccataaa
tgctaaacttagaagaagtgtaccccgaatctttccttctacttctggctgtctagcaat

accttcgacagcattagctgctgcactaccttgaccaatgcctgggcctattgcagcaag
acctacagcaagaccggcagcgataacggatgcggctgaaacgattgaatccataatgaa
ttttaatttttttgttatatgagtagaaaattgacaaatttcgtataatacttaaaatcta
tataattaagtataatagattctaatat-----tgtaattatagtatattactcttctcc
atgaccttccattgcttctccaatataagctgcggacaaagtagaaaagattaatgcctg
tattgaactcgcaaataatccgagtatcataactggtaatggtattaaaattggaattag
caaagtaaatacagaaacaactaattcatctgctaagacgtttccaaataatcgaaaact
taaagatagtggtttcgtaaaatcttctagaatattgattggtagtaatactggagttgg
ctgaatatatctagcaaagtagcctaacccttttttacttaaacctgcataaaaataagc
taaagatgtaagtaaagacagggccacagttgtattaatgtcatttgttggggcagctag
ctcaccttcaggcaagtgaattaacttccaaggaattaatgcacctgcccagttacaacc
tagaataaatagaaataaagttgcaatgtaaggcacccaagggcgatattcatgctcacc
aatctgatttttttgcaatatcttgcagaaattctaatataaattccataaaattctgaaa
tttttcaggaattctttgtagatttctagtacctagaaatgataaagttaaaagtgtagc
aattactaaccaagagacaatgaaaacttgcccatgcaattgtaagctacctattttcca
atatagatgttttccaacttctactgctgataaactattatatgggttaaaatctatgag
actgttttgatacataataaatatttt-ttttaaatagtgcaagaatcaaaaaaataag-tt
tttgaagaattaatatgcaattaatttattcttaaaattagactattaataaattaaaag
caaaaataaattctaatagagatagctatttttagtagtgattatttttttgtaaaaaaca
attattgctacctataatgcttttacgtttaattatatacccatatattaattatttaat
atgattataggtattttgctacaaactttattgttttatccattct----aaagattgtt
atagtttattttttatttaaaatatctgcgacttcatctgcaaaattaactttcaaattt
tcttctccccctcccagaagaaatcttacaaatcttctaattttaatattttcacctaga
acagcaatattttggtttattaaatcttcaatactaatatcttgattgcgaataaacatt
tgatctaataatgatagctccttaagtctctttttcattcgtccttcaatgattgtttcg
attctatcaactggtttgtttttcaagtcatctttaccagcttcaactctttgttccaga
ttaataatttcactaggtatatcctgtgttgatacgtattccacattaggcgatgccgct
atttgcatagcaatatctttagctaacttttggaactctggacgtcgtgctacaaaatct
gtttcacagttgacttcaactaatacacctattcttccgccagtatgaatataactctct
agtaaaccttcaatagcggtacgattagatttcttgttagctgaagccagcccttttttgt
cttaatgattctaaagctttttcttcattgccattattagcttgtaaagctttcttgcag
tccatcatacctgctccagtcttatcacgtaaggcttttacgacttgagcagaaatttgt
agtgtcatatgctttcaaaaatttttaagtatttaaatt----------aattattataa
ttcactttgagtaggatgagttatttcttctaaatctgtttgcccgtatttaccatcata
aattgcatctgctattttacctacaattagtttaattgacctaatggcatcatcattagc
tgggattggaatattaataatttctggactacaattagtatcaagtatacaaatagttgg

aatacctaattttaaacactcttgtatagctgttgtttctctcttttgatctacaactac
aacaatatcaggtaaacgagtcatattttttaataccatttagatgtttacgaagtttatc
taattctcttcgtaaaacagcggcttcttttttgggtaactgatcaatcataccactttt
atcttgttcttctagctgtctcagacgactaactcgagacttaatcgttacccaattagt
taacattcctccaagccatctctggttaacataataagaatcacagcgttgtgcttcctg
tgccactataccagcggcttgtcttttttgttcctagaaataaaactttttttaccatccga
cgaagcctgttttataagttcacaagcttctgttaataattgtgcagtttggactagatc
aataatatgtatgccatttctttctgtataaatatatggaaacattttaggattccatct
acgtgcttgatgtccaaaatgaacacctgcttctagtaattctgctaatgtaacaatagc
cataagattattaatatttaaataatgtaattattcgggaaatccc-tttgtttagaaac
tgat-----tcaactaacttatttgaattttagtaaactaataataaattaacacacttt
taaagcagatcaacaaaatagatcctgaagaattcat----tctacactttttattgctaa
aataattacgcgcggttctgtcgtctaaaatgatatcatctaaatcatctcgtacagatg
aaatagtactttcgtcattagtatttgcagttaaattctttttctctaacgatgagccat
tacagttatcatacatattgaagccggtaccagctggtataagccgacctataataacat
tttctttcagacctcttaaccaatctaatttaccagatatagcagcttcagttaatactt
ttgtagtttcttgaaaacttgctgcagaaataaagctttctgtatttaaggaagcttgag
taatacctaatagcacaggcctatatgatgcattaatcttatttctcaaggttatggctt
tattagtctgttcaatcttctgtaactctacaagctctccaggtaaatagcctgtttctt
caccattttctattttcactttagaagtcatctgccgtactattacttctacatgcttat
cagaaatattcactccttgagattgatataccaattgtacttctttaacaagtagtaact
gaatctcttgaaagcttaatcgagcagcttcatataaggctaatccacattcaatataca
aatcaaaacttgcatcgaggatatcatgaggatttaacaatacgtctgttttctttcttg
cttcaagaatttcttcaattcttggaagcccttgtataatgtcacctgtttttgctctat
caaaaaccaatatagctaatgtttctccccttctaattaatgcattgttatcaacatgta
aaatagcaccattagaaaccaagtaaggtctagcaattcttaatgtaactgatctagaag
aaatttcaataatttgtccagagtctaaagaagtcatattttcagtaataaaatcaccac
aacgaatccaatcacccacacttactttaactgttgattttcctatattaaaggtctttt
tatcagaagatgtcgcaatcaaaatgcgcctgctagtattcttttcagaataaatctcct
caacagtaccttcactcattgcaataatctcagtactagcaacaactgtaaatggcttga
tatattcaccatttttttacaataattctagtttgactttgttgcttctctgatttattta
catcaatacttttaattgataatgtttcaaaagttgacagccttaagtcaaaataattat
tgccactttctttagcagaaaactcaattgaggatattaaattatcatctcgatgctcaa
tttcagccacaagattagttgttactaaatgaactccttctatagatttcactctttctc
cgtcccgaaaaggagttcttttaactagctttagtttagtttgacttggtttttgtgaaa
ctagattttcaattagactggattttgtttctggtattgaataaacaataacaggtctaa

taagaatatagggcatttgctcgttttctatatactcccaataaactaacttatctgtag
aaatattattatgaagtctttctccaggtcttaaaaaacctctactttatcattattac
tatacacctcactaagtttatagatagatccaggcttaataataatttctcgaactattc
catccttttcgattatttcaacaatacctgagcttttagaaaaaatattttaactagct
ctgttccactttctataacatctccattatgcactagtattagtgatgaatctttattaa
tttcgtgagtttcttctgaaatccataagatatatcccggacttaaaatttcgtaggcat
ctttatctaagcctgttttctttttgatacatttaagtctaaatatttaataataccac
cacttgtcgttttatacgtatcagatattagttctgcgactgtgtaaccatcttgaa---
ttttttggtttggaatagctttgaaaagaaattttgtttttttctgtttctaaaatat
aagattctttttgtggattacatctgtataaatataacaacctggaattacaatagatt
cagtaataacttgtatattggtaatactgctattagtattctggtgtatccttacttctc
cagcataatgatttattaattctgtctgtgcaagtatagtaccagctttagttttatctt
ctttatgaacaataacattagcagaatcagatatactataaacttcacctgagagcaccc
agatcaagcctccagttttagtaattcttgtagtatattgattattatctgtttcttcaa
ctgttaaattggagaaacagattttgccagagagatctgaaactacatgtttttgagctc
tttctgtcatgagtcgatttctgcggggtgattcagcaatcacttgatccttcgatacta
attcaccatcacatattaaaagagtagtacctttacttaagttaattattgacttctgat
ttttcttagatctgattgtaacttgggttggctttttgttattaatgcttgctcaccat
gtctggtacgaacatctgtatacgattcaatatctagatctatcaattgaccatctatag
gtgcgtagatttgttctgctaattcaccggtaaatactcctccagtatgaaaggtcctca
ttgtcaattgagtgccaggctcaccaatagattgagccgcaataataccaacagcctctc
ctaaatcaactaagcggccatgtgcaagattccatccgtaacaatactgacaaacagaac
ttctagagttacatgtaacaggagatcttactaaaactttttt---gataccagctttag
ttatctcttttgctaacttaggactgacatcttggttcgtatgagcaattaaacaatttg
tctctggatggaatacatttctgctaatactcggccagataaagcttgttctagattaa
ttaatattttctgcgtatctactaaatcttctagtatgatgcctttattggtcttacagt
ctacttctctaattattacatcttgagaaacatcgacaagtcgacgcgtcaaatagccgg
aatctgctgttcttaaagcagtatcaactaatccttttcttgctccgtatgatgatataa
aataatcagtaacagttagtccttctctaaaattactcgatattggaagatcaataattt
gtccctgtggatcagccataagccctctcatgcccactaactgtcttacttgagaaatat
tacctctagcacctgagaacgccatcatataaacagcatttagaggatctgtctctttaa
agtatttaataacttcttgtttcaaggattcactcgcattattccacgtatcaataactt
tttgaaatcgttcaactgctgtaatctcacctcggcgatatttgttttcagtggccttaa
tatcttcaatcgtgcctaataaaagactttgcttactaggaggaattcttaaatcttcta
aacttaaagaaattcctgcctgggtcgcataatgaaatccaagatctttaagcttgtcgg
ccatattggctgctcgggctataccataattacgaaaagcccagacaattaaattttta

gttcgtttttatcaataaccttatttgaaaaacttggctgtgcaagacttcgtctattat
ccactaaatcttctccatattaaatgttcgtt--attaactatgctactaaagactcttg
aataattttattaaagattatacggccagctgttgttctaatatattgcacaattctctg
attttcagcgtcttcttttatgatatggttattaaaaaatttgttgtactattgtcaga
atgtttttcaatcttaatcggatcatgaggttgatcaccatctactaatccatcaaaacg
agcccatatataagaatgtaaatcaattttcttctgttcataagccataactacatcttc
taagcttgcaaaatactgattagatccctgctgctgagatggattattagctgttaaata
atagcaacctaaaaccatgtcttggcttggcataataattggttgacctgttgctggaga
gagaaaattatgaggggctaacattagcaatctagcttcggcttgagcctctaaagacaa
aggaacatgaacagccatttgatctccgtcaaaatcagcgttaaacgctgggcatactaa
tggatgaagcttaatagctcttccttctacaagaataggttcaaatgcttgaatacctaa
tctgtgtagtgtgggagctctgttcaatagtactgggtgaccttgaataacttcatttaa
gacattccaaatagaagattcatttttttgaatcatctttttggcagctttaatattgtt
aactaatccctgcaaaattagtctgtgaataacaaaaggctgaaataactccaaagccat
ttctcgtggcaatccacattgatgtagttttaaatgaggacctacaacaattacagatct
accggaataatctactcttttacctagcaagttttgtctaaaacgaccttgttttccttc
tataatatctgacaaagattttaatggtctattatttgcacctacaacagttctacctct
acgaccattatccatgagagagtcaacggcctcttgcagcattctttttttcatttctgat
aataatctccggagctaaaattgactttagacgagataatctattatttctattaataat
tcgacgataaaactcattcaaatctgccgtcgcaaatctacctccatctaattggaccat
aggccgtaagtctggaggtatcacaggaatgactgtgaataccatccaagaaggatctgc
acctgttgctatgaaattttcaatcaatcttaagcgtttcatttttttattgaattttaa
tgaaggtgtcttaaaactttttggtggatttgtagcttcagaccttaaggtttcagctat
gtgctctaaatctaaatctttcaacagtttctgaatagcttctgcacctattcctacttc
tacttggttattctcatcttgattttggtagatttcctcttcaagacttttccattcgta
accttcgagcagttgcttgtattttaaattaatatctgtattagattgcgtaactacata
agaatgaaaataaacaatttttttctacttctttgacttttaagtctaatgctaaagcaat
atagcttgtacttccttttaagtaccaaacatgagttactggtgaggcaagttcaatata
agccatccgatgtcttctcacgcgtgattctgtaacttctacgccacatcgttcacaaac
tatacctttgtagcgaaaacgtttatatttaccacagtgacattcccaatccttaactgg
gccaaaaattttctcacagaataaaccatccatttctggctttaaagttctgtaattaat
tgtttcaggctttgtaatttcaccaacaatttgtccgttgggcaaacttctttcgcccca
ctgtcgaatttttttcgggagaagctaaactaatttttacgtagtcaaaatattgctcaaa
ctttgtcataaatttgaataccttaaaatcaaataattagccaatttatttacatgcttc
atattgaaaaatgaatatcttaataaaggaattgttcaaaatcgtctactggaggagtat
cataattagaacgatctactcggttatctttagaatccgacatcaagtcaacttcaactg

tccgtctttggccatcttcaaatagtttcaatttatgtactgcaatgtctaaacctaatg
actgcagctctcgcattaagactttaaaagattctggagttccaggcttcggtataggct
tccctttaactattgcatttaatgcttcatttctagcttgcatatcatccgactttacag
ttaataattcttgcaaagtatatgctgcaccaaaagcttctaaagcccacacttccattt
cacctaatctttgacctccatgctgtgctctacctcctaatggctgctgcgtaactagag
agtaaggaccagtggatcttgcatggatttatcatcaacaagatgaacaagcttaagca
tataagctctaccaactgtaacaggattatcaaaaggctctcctgttctaccatcaaaaa
cttgcattttacctggatgctgatcattaaaaagccatttattgctagtaatcaatgatg
cttcttttaatttctatttacaagtgctcttgatgcttctgctccatacatttcatcaa
aaggtattattttaaatctctttcctaaatagccacctgctaagcctagtaggcactcaa
atacttgaccaacattcattctagaaggtacacctaaaggatttaaaacaatgtccactg
gtgttccatctgataaataaggcatatcttgtttaggtaagatcctagaaatgatacctt
tattgccgtgacgaccagccattttatcacccacttggattttctcttctgcgcaacat
agacgcgaatcatagcatttgttccgggaggcagctcatctcctttctgcctagtaaaaa
ccctaaccttaactactctaccttttgcagcattaggaagccgtaaagatgtatctctca
catctcgagccttttctccaaaaatagctcgcaacaatttaccttctggtagttgatcag
cttcacctttaggagtaatttttcctactaaaatatctccagcttcaacccaagaaccac
cgacaacaattccatttctatctaaatccttcagagagttatcgctgacgttaggaatct
ctctagtaatttcttctggtcctaatttagtttgacggcattctacttcatacttttcga
tatggatagaagtatacaagtcatcataaactaatctttcgctaattaaaaatgcatctt
cgtagttataaccttcccaaggcatataagcaactagaatatttctccctaaagcgattt
cccctccgtctgtggatgcgccatcagctaaagtttggccaacaactatttttttctccca
cccaaacaattgggcgctgattaatacaagtatcttggttagaccgataatattttttta
agcgataatgaactgttctaccactattatcctgaatgcctattttattagcagaaacat
aatttacatgaccagatgttctgctaataacaaccataccagagtctcgagctattttag
tctcaagaccagtaccaattattggcttttctggatataataatggaacagcttgccttt
gcatattagatcccattagagctctattcgcatcatcatgttctaaaaaaggtattaatg
atgttgcagctgaaattacttgaataggtgaaattgcaatataatctacttgggttgggg
ttgttgtaataaactcttgacgataacgtacaggaataatatctccttcaatgtaatgtt
gcttactaactttaacatctcctggagctactcggaaatcatcttcctcatctgctgtta
gatagacaggactattgtgataaatcacttgaccttggtttaccggataaaatggggttt
ctatgaaaccaaaaacattgacccgcgcacaggttgctagtgaaccaattaaacctgcgt
tcggaccttccggagtttcaattgggcaaattctaccatagtgactaggatgtaaatcac
gaactgcaaaacctgccctgtctttattaaaacctcctggacctaaggcactaattcttc
tcttgtgagttagttctgcaactggattagtttggtccataaattgagaaagctgactag
atccaaagaattctctcactgacgcaattaatggcttgggattaattagatttgataaac

tcaatgaatctatatcacatatcatcattctctctctaataatgcgttctaaacgattga
gacctactctgaattggttctgcagcaattctccaactgaacgaactcttctattaccaa
gatgatctatatcatcaagattacctgagtttttatccttaatattaattagataatcaa
ttgaggataagatgtcttgaggagacaatactcggaacgttttaggtatattgagaccta
gtttttttattaatttatatctaccaacttcacctaagtcatatcttttaggatcaaaaa
aacgagagtaaagcatctgcttagcaacagcaactgttgctggttcattaggtctaagct
tagaataaactatgagcagggcttcctcatcagttacttcttcaatattatttttttccaa
tttctttagctaattctttaactgaatatatttgggatgcagaaataagaaaagcatatt
tactaagtcctttttcaatttcatctttattgagtccaatagctcgcagaaaaatataag
catttactttgtgggtcttatctattcttatccaaatttctcccttaggatcaatttcaa
attttaaccaagaacctctattagaaattaggcttgcactataaatctgcttaccatttt
tgtctatctcttgtttataatagattcctggactacggattatttggttaataataactc
tttctgtcccagatacaataaaagttcctctatttgtcattattggcaagtctccaataa
aaactaatctctttttgtacttttttatttttatctgcttttcagcactgagctgtagat
gggttgatgataaatctaatgatttaatatttttttcttggtttttagaaggtagatcaa
tatcttttcttgttaactttgcaggaacatatatttgagcactataagttttatccctat
tttttgcctgtctgacactataacgaggaaattttatcttgtactcattaccaaataatt
gtaattcgagtcgactagtagggtctgatatcttcggaaaaatttcaagaacttccgtca
aaccttctaataaaaaccatttaaaactggccctctgaatttcaactaaatctggaagaa
gtttattttttaagcttatacgttgaaccatagatctcctttataacaaattttgaaatc
gaagagtttaa-tactattaatagagtactggtagataatgcataagcatagtttgctac
attatttcttctctagacataactagaacatcatattttattcaa--aaaaagattttta
aaatttttcttcac--acacactatctataagtactttgctgcta--aaagatagttcaa
gctattaaa-------aaaaacgaaattaatacttgtctgtctgtcaattaattgacatc
aagatagacaagtattatttatttaatcaattctg-tttttctgagcaaaagacaatgct
ctcgcaagtctagctttttttctagcgcctgtattagggtgaaatgcccctttttttaca
gctttatcaatcttactatatacttgagaaatacttgactgcacatcattgaaattatca
ctttctaaattatcaatatttagaaggcatcttttagttaaagttttgacaaccgattta
tacttacggttaatgagacgatttctttccgaagttttaatgcgtttaatcgcagaaagg
ttcttagccacagtaaatccaataaaacttttataaaatatttttagaaacgaaaataca
caaatctgggatctattcaaaaagggaatatgcggcgtattataacattaaagaaaaaca
ataggcaatagttcaacaagagctcttgttattacacaatagagattgtcattgcgaaaa
actttgtaagatagtatttaattaaaataaactaactagacatataaatatatatgaaaa
aaattgaagctattattagacctttcaagcttaatgaagtaaaacttgctctagtcaagg
aaggtattggaggaatgacggttatcaaagtttctggatttgggagacagaagggtcaaa
ctgaaagatataaaggatccgaatactctattgatatcattgataaaataaaaattgaaa

ttattattagcgatgataaagtagagaaaattgtagagacgattattaaggcttctaaaa
caggggaaattggagatggaaaaatatttattagtagtattgaaagagtaataagaatca
ggactaatgacttaaattttgaagctttatagtatttttacttt-tttagacttgatact
tcttcactaaacatgagataataagcgtttattgtatacttataactaaggtgtaata--
ctaatggctaaaagcaaaggtgcacgaatcgtaataactttagagtgctctgataaggct
ggagagtttgctcaaaaaaggaaacctggcgtttttcgatatacaactactaaaaataga
cgaaatacaccaagtagaattgaattaaacaagttttgtcctaattgtaatcagcattgt
atcttcaaagaaattaaatagttatttatcttattaaaat--aatcatattatggctata
tatagaaaaagaatatctccaattaagccaacagatgctgttgactacaaagatattgac
ctgctaagaaaatttattacagaacaaggcaaaatactgcctaaaaggtcaactggattg
acttcgaagcagcaaaaaaaacttactaaagcgattaaacaagctagaatcctttcttta
ttaccttttttaaataaagattaa--atatatagcctaacttcatattttccatctaaa
tattaaaaatttctatagatttgtttattaaaatgatctaacatactgaat---------
----------tactaagaaga-tctactcattacaagttttgttcattttataattaata
atagcattataattta---ttaagctagattc-----agacataaag-gtaacctctttt
aattgattatattatatagtctatttaaaagaggaaatttttttgcttc----------t
agcaattcaagattctttaagctttaacaaggtatttaaaacaaagattttttgtttaggt
atcagatcaaatgaatgaatcttatctcccgactgccataattgaaattcagagatgaaa
attccacattcgttgccagcttgaatttcttcgacatcttctctaacgcgctttaaagat
tcaatttttccttgataaataactttttcttctcgaatcactttaatccaagagtttttt
aatagcttattattgatcactcgacatcctgctatttttctatttgctaacgagaatact
gtacttacttctgcttctcccaccggtacttctgaatattcaggatcaagtaagtcttcc
atccttcttttaatgtcttcaattaaggcataaatgatttgatagttttctattaatata
ttcgactttgccgatgcttgctttgttcctggagcaaagttggtattgaacccaatgaga
gtagaatttgtggttgaagctagttcaacatctgtagcagtaatttctccaggcatgatt
gagacaacattgagttgaactttactctgaggaaattgagataaagaatctaggattgct
tctgtagagccttgattatctgttttaataattaaagaaatttgcttactaatatcttta
gaattagtgtttttttaaagtatctaatgtaattcgactattaagtgctctttgtttctgt
ataattgatgagtctttagaggtattttcaatcgctttaagtttggcttctttatcactt
ttaactgctaaagctatttctcctgtagctggtacagatgacaatccccaaatttcaaca
acagatgatggtattgctaaattaattttctctttagcgttgttaattataactcgaatc
tttgcgtaagctgatccaataaccaaattatcactaatattcaatgtaccattttgtatt
aatagagttgctactggaccatgcgatttatctaaatgggcttctattatataataccttgt
gcaggctgtgttggatcagctttcaaatcctctaattcagctaacaaggtaatagtttct
aataatttatcaacattttgaccagtaagagaactaataggaattataggcacttggcca
ccaagtttctccgacatgacattgtattttaataaatcttgctcaataatatctgtatta

gaaccagctttatctatcttagatattgctatgacaaaagggacattcgctttttgaata
tgattaatagcctctatggtttgaggtttaactccgtcatcagcagctataataataata
gcgacatctgttaaattagctcccctagacctcatacttgtaaaagcttcatggcctgga
gtatccaagaaaacaatttttgtttattatctttcttaatatactcaacttcataagct
gcaatggcttgagtaattcctccaatttctttgttagcattattagattttcgaatataa
tctaatagcgttgttttgccatgatctacatgtcccataactgtgacaattggaggtctt
ttaatataattgccagtttcataaaaagcattagagttatctaaattactagaaagtcca
ttattattttcctttacatttgattctactgctataccaaaattatcagccactgatgat
attattgaggcgtctatggtttgattcatagtaactgatattcttttaagaaataaatat
tttataatatctgtttcttgaacacaaattagtttagataattcttgaatagttagagga
ttagtaatactaatagactctggcggagaattagattgaatgcttgctgaattttgacta
ggagttacggctactttttctgtcttatagatttttattactagtcagtttttttaca
acttccacttttggcttaggtggacgcattaaagagatagctaaatctcctgcagtctgc
gaaacattagaattagagtctctaaagttgtca---tcatcatcgtcaatatgtatttta
gtttttattttt-ttgcgctgtctattctta--tttttttactatctactaagtcgtga
atttattaaaatttttactttttttatctagtttaggtggagaacttaattctagatgc
tgttcaccacc---tgtatgtgattctgatttatcaaggtctaaatttaacagattatca
ttattgactgactctaatctaattttatagataatctgaggattttcaaatctacaatt
gtttcagaagaattgatatttgagctagatctagaggatctattttcaaagttttgatta
tttaaaaacataaatattggttttattatctaatagtaactttctaattttaatacagag
cttaatataaacaagaatcgaatatctgtataaataatacaaatgt---agagaattaat
ctatgttgtattagatttaatataacactaaagtcctttgttcgttatacaagtttaaaa
ctttt-tatgatgataaacactagattctagtttattggtatacaatctatttatctaaa
taatactgtttttgtcagggcttttaaa----ttatctaatgttatgttcattatgaaat
tctattaaaatttaaaaa-ctcaacaattaagtaagaatcgttaatcacctccacgccac
ttatgccacgatctcaaaaaaatgataatttttattgacaaaacttttacagtattagctg
atattgtattaaaaatacttcctacaagtaaagaggaaaaagaagctttctcttactatc
gagatggaatgtctgcacaatctgaaggtgagtatgcggaagctttagaaaattattatg
aggctttaaaactagaagaagatccatatgatagaagctatattttatataatatagggc
ttatttatgctagtaatggagagtatgtgaaagctttagaatactaccatcaaggattag
aactaaattttaaattacctcaggcccttaataatatagccgtgatataccattaccaag
gagttcaagcaattgaggataaagatacagagctatctaaattaatgtttgataaggcgg
ctcaatattggcagcaggctattaagttagctcctgataattatattgaagctcaaaatt
ggctgaaaacgacaggacgaatgaggaatatacaaggatattaatatatttaagatataa
taaagatatcaattaataaattaaaattagttaaagaaactggtactatatagagtactt
-tactatatattataatattatactagttgatctctatctctaagacttagctgaataac

cgaaattaaatctattgactatattcatgaatacaaataagcataatggttagtacaaca
aaaagtaccggatctgttactcaaattatcggaccagttttagatattgcatttcctaac
ggacagcttccgaaagtattcaacgcactcaaagtacaaagctcagaaggaactattact
tgtgaagtacaacaacttttgggtgacaacaaggtacgagctgtttctatgagttccact
gaaggactacaaagaggggtagaagttattgatactggatcccctatatctgttcctgta
ggtacagatactcttggacgtatttttaatgttttaggtgaacctgtagataatttgggt
cccgttgattctgagagtactttacctatccatcgaccagcacctgcttttactaagcta
gagacaaaaccaaatattttgaaacaggtattaaagtcgtcgatttacttgctccttat
agaagaggtgggaaaattggtttattcggaggtgctggagtaggtaaaactgtattaatt
atggaactaattaataacatcgctaaagctcatggtggagtatctgtatttggaggcgtt
ggtgaaagaacaagagaaggaaacgacctatatatggaaatgaaagagtctaaagtaatt
gatgcagataatctgaaagaatctaaagtagcattagtatatggtcaaatgaatgaacct
cctggagcacgtatgcgcgttggcttaactgcattgacaatggcagaatactttagagat
attaacaaacaagacgttctattgtttattgataatatttttcggtttgtacaagctgga
tcagaagtatcagctctactaggccgtatgccatctgctgtgggttaccaaccaactcta
gcaactgaaatgggagcactgcaagaaagaattacttcaacaacagaaggatcaattaca
tctattcaagctgtatatgtgccggctgatgatttaacagacccagctccggccactaca
tttgcacatttagatgcaacaacagtactttcgaggaacttggcggcaaaaggaatttac
cctgcagtggatccactggattcaacatcaacaatgctacagcctggaattgttggaact
gagcattattctactgctcaagaggtaaaatcaactttacaaagatataaagaactgcaa
gatattattgctattcttggtcttgatgaactttcagaggaagacagacaaactgtatca
agagcaagaaaaattgaaagatttttatctcaacctttttcgtggcagaagtgtttact
ggatcacctgggaaatatgtatctttggaagatgcaattaaagggtttcaaatgatctta
aaaggcaatttagacgacttgcctgagcaggcattttatctagtaggtgatatagatgaa
gctatacaaaaagctgacagcatgaaagattaatataattaacgatgactttaaatataa
gaattattgctcctgatcggactgtttgggatgcagaagcacaagaaattatttaccaa
gtagtacagggcaacttggtattttaacaggccatgcacctttgcttacagctttagata
ttggagttatgagagtaagagtagataaagaatggatgccgattgttttgctgggcggtt
ttgcagaaatagagaacaatcaattaactattctggttaatggcgcagaagaagctagtc
aaattgatttatcagaagcagaaaaaaatttagacactgcaactcaactcttaagtgatg
cctcgtctaataaagaaaaaatagaagcaacacaaaaaatacgaaaagctcgagctcgag
tacaagctgcaacagcagcaacttcgtaa-aagattcc------tcttaaaacataaaac
tttagtaattattactaaa-------gttttatgttttatttgtatggttataggcagct
taaataaacttagcttagacctgaacaaatataatcaaaatataatcccatctctttacc
tgcatctggtccaactaaactaatagtaacttctttcatagctaaaatagcttgaatcgt
tgcaccaataggaacacctagagaattgtatgtttcttttagaccatttaatactctttc

ttcaaggatagatggatctccagctagcattccataagtagcatagcgtaagtagtagtc
taaatcacggatgcatgcagcataacgtcgagttgtatacatattaccacctgggcgtgt
aatatccgaataaagtagagattttgctacggattctttaataattgttgctgcattagc
tgcaatagtagcagcagctctgactcgtagctcacctgtttgaaaatagcctcttaattt
ttcaacagagctatcatctaaatatttaccttggacatcagctgcattaataacagaagt
aattgcgtcttgcataactttcacacattccttaattttaaatatctaatacta--gggg
tatataacaaaattactagatcttttagcgatatcagtattactgcatagcacccaaagt
ataatcaaagtaaaaaccagcttctgctgaatcttcacctgcaagcagtgagcaagctac
acttttcatgcattttacaccttcagcaactcctgaaattggtgttcctaaagaattata
catttctttaacacccactaatccaatttcttcaataggagttacatcgccagcaactat
tccgtaagttactaagcgaagataatagtccagatctcgtagacaagtagcagtcatctc
ttcgccataagcatttccaccaggagaaactacatcaggtctttttgaaacagttgttg
gccgccttgctttacaatacgctcacgattatctgttaaaatttgagctattcttaaacg
acgttgtccagataaaacaaaactttaattctatctaattccccaggacttaaatatct
tgcttctgcatctgcatttacaattgactttgtaacaatactcatggataaaactcctgt
aatgcttt--aaaaaaagcttaatactaattatttatgatataactcagaaagtaatgta
ctttttgag-aattttctaaccgaaacacgttcatgatatatgaagcaggatagatactg
aggtaagtactaaaacgagaaagttatatgtaggaacttctgttattgatttcctagtac
ttttttaaaacttggtaccacaattgatatatcctgcttagtcaaacgattatagagtgt
ttctgtattaggaaagttagctgctggtaaagtcggaaatcttctgtatggaacagtatc
tttgccgtataccgcattgtattcaacactttcaactaaattacttataaatgcagataa
gccttttgaagctaaaatttgattaaagaatctaatttctgcctgattattaggagctct
tcctaagatatgcttagtacctaattctataacttttgtatttggatatggctgataaaa
ttctttaccatatagttcagatagcgccagtttttctactagttctttgacacagatttg
tctattcaagaaagctgacttaatatccaagaattcaccgccgacactaaaagagttaag
atctctttcgaaaatttgacgatatgctgctcttaaagcttgttctaacatttctttgtt
actatcactattcacttcaaagacaatagattgatctcttaaagaagttactccttgtga
aatacgagactgaatatcattactactcctcatttctttgacagtgcctaattcaacaaa
cctagcagatttactagcaataactttttgaatctctggtcaataattcctggtcttaa
gcttcttaaagcaacaccagctggagtactatatctttcataaggaacagtattatctcc
aaaagtttctgtgtactctgagctatcaattattgtgtctattacttggtaatagccttg
cttatatgcaatatcaaaatatttattaatttcttgccgaccatatgtagggcgacccag
taagcggttatgaatatattcaatagctttacaaatatataatggctcccaataaagaga
tctaaatatacttgattttgctaactgtctaacaaactcgcgaactgtaatttgattatc
ttttagctgactttctattggcttgagaattagtttttcttcttgatacacttctctgcc
aaaaactcttaaataagcaacttttgtgacaacttctactgaattttcaaaattttggga

tgaatctgactttataagagcgttggacaatttaaagatttttggtcccagggatcctgc
tgacttaggtcttacctgtggattactaatctgattgtagattcctggcccacgtctaac
taatatccttcttgtatctttaccaaaaagagcttgtctttttcttggatctttattttc
tttagggaaaatagctccaaattgaattgataatggatcattacctattccatatggatg
ttgatcaggtaatgattgcttataatcactaaataaagttataaattgaggaaccttacg
aaaaggcgcactataggtgagaagatctatttgaggcccccaattacggcactcttgagg
ctcttcacctaaatttctaaagtaaggcacagtctcttcaccgaaataatctgtatactc
agaagaatttaaaagtgcattcactaaaccacttaaaccagtagatgataaaatagcaaa
atatttttgaaactcttctaaagagctaggacctctacccaagaaatgtctaaatgctaa
ttctaatgctcgactattgacaaaaggttcataaaattgctttctataaatacttgaagt
tcctaacgaacgaataaattccttaattgaaatttggccatttttttacttgcgactccaa
attcgataaagagaggtcatatgctttagcaatatctctttcaaaaatttgtctataaca
tgctttcacaacaatatttttctcatcagcagataaactaggcttcataacaaaacgagg
tgttgacactcctgcctgtacataagtctgaggaagacgcaaaccttgtaaatctccaga
tattctttttcttaatttatcagtcaaagatggagcttcaaattcagaaatcacaacatt
aaaatactctttaactaaatcttgaccttttatatcttcctcaaagattaataatgctgt
ccgtctcatctctctaagagccacaattgctgctgcacttgagcaagcgttatcaattaa
ctctctcaatcctcgtatattaacagataaaatattagggtcgcctgatacaattgcata
tgtcaaataccgtaaaaaccagtctaaatctcgaagagattttctcattcttgtagtacc
gtatctcaagacgtttataggtttaaaccctggcggagtagctcccccagcattaaataa
agatcggaaactttggccaaaatctccttggatatttcctgataactcattaattttatc
ttgagaactttgatctcctgcaataataacagctgcttgcggacgttctaaataagaaat
tgcagacccacctacgaatattttatctgctgctcgtgctaccagaatatttgcattttt
agttaaaatatctgcaacttccagtctttttttgtccagaatttaaaaaagaaactaattg
atttaattcaccaagctgtaaaaatcggtcttgttgctctgcctgagtaatagttaaaat
tgaggcagtccgataaagctgggggcgtgctaatgggcttccgccacttgctttgatact
cattactttatttatctccttaactgatattgcttgctagcgaatacaaatatagcttga
tatttctaaatatttaaagatactatgcaagcaca-ctgttttctatattattaggtact
agtacttatagtacagagagtcttataactaacttctataaatacaaattttgtaatact
tctattaacatttgtcaaaaaaatatttaagtgctcactcttttatattaaaattcaatt
agtgagaggttgatgtgaaataaagtatactaagtaagttttatagataggaaaattaaa
agatttattctacaaacattaatca------------cttgaatgtttagcaattgtaca
attgttattgcttttaattggtttagactagttttatatttttttctaacataaacaatat
taatgagcaatttgtatgaatttaaagccacaaaacaacctattaactaacgaaaatata
gatcatgtagatatacccgaaaatgatataccaatgtcaattacggaacatttagaagaa
ttaagacaacggactttatttgtgtttttattctttttgtttgctacaactataagtttc

acacaaattaagatcattgttgcaatactgcaagctcctgctgttggtattaagtttta
caactggcaccaggagaatatttttttcatctattaaggttgcaatatattgcgggatt
gttgcaacaactccgtttgccgtttatcaagttatattatacatactcccaggactaact
ggaaaagaaagaaaaattattttgcctctattaattagttctgtattgcttttattaca
ggtggtatatttgcttactttgttctcgcaccagcagctttaacattttaattagctat
gggtctgatattgtagaaccattatggtctttcgaacaatactttgactttattctatta
cttttacttagtacaggattagcatttgagataccaatcatacaattattgctcggtgtt
tcagggacattttcttctagtcaaatgatacgagcttggagatatattattattatagca
acaattgctggagctattctgaccccttcgactgatcccgttacacaattaataatgtct
tcagctgttttattactttattttggtggaattgttatattattagtcttaaaaaagtag
agtaaat---------attttcta-ttaatccaatatatttgttatc-----tgggcaaa
taatctat-aaaacaattgct------------------agatttaagtatcaggcttct
aaatatttcaa---------gtctctttttatcattgaaatacaatc-------------
------------------------------------------------------------
------------------------------------------------------------
------------------------------------------------------------
------------------------------------------------------------
------------------------------------------------------------
------------------------------------------------------------
--------------------------------------------------gatacattgg
cttgtttttcatcaattatttgtgttgcacac-tttttagtactctctttattgttctt
agaattatttctttaatctatttattaaat----agatatgaagagatttaagtattgat
tatggtaatctatatcaacgtaaaaagtattttattgtgagttttatttatataaaggct
tattcattatatagttataacagtaagtctaattaatag-ttttcatgcttaattaaagt
aataactcttaattcttttggcagaaaaatctgtcttaatattttatc--aaaaaaacag
-aaaaatgaagctaaatagtctaattaacttaattcaaaagtctatatattcttgtacac
ttttactaattattttaaatattatttgtgtcgcacctaattctagtaatgcatttccaa
tttatgcgcaacaagcttatgaaagtccaagagaagcaactggtaggatagtatgtgcca
attgtcatcttgctcaaaagcctgtggagatagaagcgcctcaagcagtactacctaata
ctgttttcgagactgttgtgaagattccatatgacagcaatgctaaacagattttaggta
atggcagtaaaggaggcttaaatgttggagccgttgtaatattacctgaaggatttaagt
tagctcctgttaatagattatctacagagttaaaagaaaagactagaaatctttacattc
agccgtacagtgctaaacaagacaacattttagtaattggacctatttctggtgataaaa
atagagaaatagttttcccaatactatctcccgatcctgcaaaagataaaaaagctcatt
ttttcaagtatccaatatatgttggcggaaatagaggacgaggccagatttatccaacgg
gtgacaaaagcaacaataatattgtctctgctttaagcagtggtaaaattaataaaattg

aattactagacaaaggtggatttataatacatgtgactaacagtagcaatgtggagtcaa
cacagaagatttcacctggtctcgaacttagagtaaaagaaggggatacaattcagcttg
atcaagctttgaatagtgatccaaatgtaggtggttttggtcagaacgaaacagaaatag
ttttacagagtccaaatagaattaagggcatgattgttttctttttcgctagtgttctag
ctcaaatttcttcgtattaaagaaaaaacaatttgaaaaagttcaagcagctgaaatga
atttttaaagcaatataaaactggcacttactataaagaaagta-taaacactttattca
caggatgtaccagtttttatgttatgtaataatcgctttgacggaagttatgatactttt
aacagaatctacaatctgactaggctgaattactgtagcctgctctaagcttccattata
tggtgtgggtatatcctgcgaagatagcctcacgacaggagcatctaattcatcaaaaag
atactcattaatttgtgcaattagctctgctccaattccggctgttttcatacactcttc
tacaattaaaactttgtgagttttctttactgagatagatatagagtctatatccagagg
ttttaaagatataagatctataacttctggatcataaccttcttttaataaggccggtaa
tgcttggataacatgatgcctcattctagaataggttaaaatcgtaatatcttttccttt
tcgtacaaattcaactttattgagaggtaagaaatattcttcttgaggaatctcttcttg
taaattataaagtagaacgtgctcaaagaatacaactggattattgtctcgaattgcaga
tttcagtaatccttttgcattataaggagtagaacaagcaactatttttaagccggggat
ggcttggaaataagcttccagtctctgagaatgctctgcacctaactgcctaccaactcc
tcccggtcctctaataactaaaggcaatgtaaaattacctccagaagtataacgtaacat
tcctgcattattagaaatttgattgaatgctaataataaaaaactcatgttcataccttc
aacaattggtcttagccctgttatagctgcaccaattgccataccagtaaagctattttc
tgctattggtgtatcaagaactctcaaatccccatatttgctatgcaaatctttagttac
cttataagatccaccatagtgaccaacatcttctcctattacgcagacagttggatcttt
tgccatttcctcatctgttgccgctcgtaaagcgtcaaacataaagatttactcatttt
aatatttgatttttatgatttgaccactgttttaaagtaacagttaagacaccatattct
atttagttatctgcaaaaagatatcgtttcaattctgacatgttaggttctggactggaa
atagcaaatttcacagcttgttccaattcggttttaacagcattctgaatctcattaagt
tcacctatattagcaatttcattatctagaatatatttttttgagtttttttgataggatct
cttgccacccaagcctctttttcttgtcttgatcttagttcatcaggatctgcgagagag
tgaccacgaaatctatatgttaatgcttctattaaggttggaccatcaccttggcgagct
ctttgaactgcttgttttgcagcttgtcttacagctagcacatccattccatcaacttca
atcccaggaagcccaaaagcttctgctttttttatgtatttcaggtattgaagaagaccgg
tgatgtgccatacctatagcccattgattattttcaacaacaaatataataggtagtttc
cagagaactgccatattcagacattcaaaaaattgcccattattggtagtcccatcacca
aaaaagcaagccgtgactcttaaatcctctgtttctttaagtacttgctggcggtagata
ctttgaaaggctgcccctgttgcaaccggtataccttccgcaataaaagcaaagccacct
aaaaaattgtgaggcgcagaaaaaatatgcatcgaacctcctctgcctttactacaacca

gtctcttttccaaataactcagccatcacatttttagatgggacgcctttacttaaagca
tgtacatggtctcgataggtactgcaaacataatcagttggattcagaagtttaattaca
cctgtagaaacagcttcttgaccattataaagatgaacaaaaccgaacatctttccttta
taatacatctgagcacacatatcttcaaaatttctgcctaacaacatgtcttcatataaa
actaataaattactcttattaagattaagacctgttgaattatagttagtcagcggcaat
tgaactttcttaggataactcataatttgtaaagaaacctct----------tattgtaa
ataggaattttgaagaaacagctatcaatagaaaacacacagataaaaacttataaatta
taatatgtttttactaaagactttaataattatttgaaactagttcagtatatcaaatt
aattttgcttatactacggatgtctgaaacttatataaaaaagtagt-a-aaaaagttta
ttaaaagaatagttaaaaacagatataaatagaagtaaacagaattatt-gaaaaaaaat
gattattattaattaa----tacttaatatttctaaatattttacaattaatggctatgt
caccatctttattttatcctgttgaacaagaactatgtagtcttgaaaaaaatctgaaag
ctgttgctgggactcgtcatccaattttatatgccgcagcaaagcatctattcgatgctg
gaggaaaacgagttagaccagctcttgtatttttagtggctaaagcaacctctgagaagc
aagatataaatactggacaaaaaaggctagcagaaattactgaaattatacatactgcta
gtttggtacatgatgatattattgacgagtgcacaacacgtagaggagtcaaaactgtac
ataatttattcaatactaagattgctgtgctagcaggagatttttttatttgcacagtctt
cttggtatttagctaatattgaaaatttagccgtagtcaaagctatttctaaagtcatca
ccgactttgcagaaggagaaattaggcaaggcttagttcatttcgatcccagtatttcaa
tagatgcttacattgagaaatcattttacaagactgcttcactcattgctgctagttgtc
ggggtgcagctatgcttaatggttccaatcatcaaataaataatgatctttatctttacg
gtaaacatatgggattagcatttcaaattatggacgatgttctagatataactggttcta
ctaagagcttagggaaacctgctggcgctgatctaataaatggaaatttgacctctcctc
tcctttttcacttactcaagaagcaagtttaaatgatcttattgatagggagttctgta
atagtacagatatagcctcaacattatttcttataaaaagaagcggggaattacaaaag
ctaaagatttagctaaagaacaggtgcaggcggcactttcctgccttcagtttttaccac
aatctacacctgtatctagtttaaaagaattaacacatttcataatcacaagattgtcat
aaaggacttgctaaaattaaaatgttattaactaatttttttactgtttctaaaacagtct
gaaaagtttgaatatcattactagctagctgcgctagcattttacgatttaacgcaatat
tctctttttttaatgcactaataaaagtactataattcatgccttgattgtgagcagcag
catttattctagttatccaaagacgacgaaaatctctctttttcctttttcgaccaacat
aagaatatcgaagagctttaaggacttgctgcttagctgttcgaaataaacatttatgtg
cacctttaaagcctttagctagcttaaagattttagctcgtcttttttttgcaacgttac
ctcttttaactctactcataaaatattac-ttaaattttaattattaccagcttactaaa
ttcttatactctttg----attataaataaggtaatttcattgcaatgtttttaatatct
tttaagtcaaccgaacaagtagaagaaagatgtcttctttgttttgatgactttttttgt

aataaatgacttttagaagctttatgtcgaagaaattttccggatgaagagactttaaat
ctttttgctattgcttttgatgtttttaacttaggcatattataaacttttaatgattaa
caaaagcatgaaaagtctgaggatatcaatcttagacctttcaatacttagaataatctt
aacacaatttatttacttactacataggttaactgaaacaatttatatatataatacta
atcagttaatcaaccatagctaaagtggatttatatttttgctttgaacttttgtatagta
ttagcaattagcattgttataattttaagactgcttgaattcaattcttcgaacattttc
atatttaatgtaaatgaaatattggcttctgagacaatattttgaatttgaacatcagac
aagggaatcatatctaatgcagctctatattgatctttaaataatttatcgtcttttatt
tggtcaaaatcgtaaaatttcgttcctcctgaatctgataaattcatagcccctctagct
atttttttaaaatttggccgccagaaaggtcacccaaatagcgcgtataagcatgagca
actaataattcaggttgtttatgtcctatagtgtgaattctatcaacataaatttttgta
gcaggagatggttcaataaaatctaaccaatctgatccataataatagtttaaatcttca
gatagactcgctttcctattaagctctgtaaaatatataggtttaatagctggatgattt
ttattagaaaataactcttcctctattgcacaatagacaaagtatagatttgcaaccaat
ttacgatatgacttcttatctacaactcctcccaaaaaagatttaacaaaactaacattt
tctgccatactatgggacttagtagtaccttctcttagttcattcgctaaagtattaacc
ataatatcaatattccattttt-aaagataacaaattaaca---ataaatagttatccgt
atatttataattgagttgcgctaactaggctgctgcaaaataatttttttgattttattgg
atctggattcattgttttgtcaccaggcttccaattagctggacaaacttcatctggatg
agattgtacatattgaattgcttgtaaaaccctcaaagtttcttctacactcctgccaaa
ctctagattattgattgtagagtactgaataattcctttcgggtctataataaatagtcc
tcttagagctacaccatcactattcaaaacattataagctgcactgatctctttttttaa
gtctgatactaatggatatgaaagatctcctaatccacctgattctctgtcagtctgtaa
ccaagcaagatgagaatactcactatcaaccgagactccaagaacttctgtattaagttc
cgaaaaagcattatacttatcgctaaatgctgtaatttctgtagggcacacaaatgtaaa
atctaaaggataaaagaataagacaatatacttattcttgagatcagataattttaacgt
tttaaattcttggtcataaacagctgtagctgaaaaatcaggagctagctgacctactcg
aagacaatttggtccagaaatcattaatttttcccagaagaattgatataattttgttat
ttaataaataattatataacaaaatttagtatatattaaaatactacaagtctttttaa
attac--------taaatgatataatcaatactatctttatagcggggaatggatttgaa
ccattgaccttcgggttatgagcccgacgagctaccagactgctctaccccgcggataac
taaaatcatattagtctacatatatatataaaagcaaatgcataatagatttagaagta
aaaaaatagaagaactggtccactgatcattaaacttagaaacgacaaagtatttttat
cttatttaaaagagggaagacttcatataaaccaataaatctatcttttaaaagaatttc
ggacggctttacccaaatttgaccgtcataccagcctgattcttcataaaatactgtagc
actcattaatcgttttactacgtatgaccagcctaaatataatctgattaaaataaatcc

tgtcattaggctggaagtaataaattctgaaaagaaaaatttcaaaggtagtttagtgat
tggaaaaattgataataatataggactaactaataaacagttaagtagtagtgcaatagt
tattttttattataagatctacggcttaaagttggccagcaaaagaaccaagaattttt
caaagaagtatattcatgaacaggttgctgctcctttggcacaggacattgagtattata
taaattcattgacattaaaaagtattataataaagac--tttgtttatattatctaacat
aggtacgacattaaatattaactaactgcaattatagtagtgagtagaagaaataaaact
aaaaatagccaagtaattttatttaaagtattctcagtactacgagtattactaaagaac
tgattctgagcacctacacttcctaaaccttcggatttaggattgtgtattagaatagtg
aagattaatattattgtagatgagtaccaaaaaaattttaaaatttgttccattatgtaa
aagttaat-aaaaatagcacaaagacaatcgtctaaaagatattgattgtcttatcatat
taaaaagtatttattcaccctgtacttttaaaagtaaaaatcctattgccagccctacca
aaactaggactgaacttaatacagcactttctacaatttctccgaccatt-ttatttttc
cttattattttattattg---tcattcttttgacaagattatattgagctaa-ctaaaat
ccgtttctaccccaaactaccattgcgattgaaaatgtaaacattgccattagagctgcc
catcccaaacttaaaatatccataatttataatattc--ctaaaaataagatctcttta
cttaaactataatattcttaatattaaaaatatcattatataataagctaattataacag
agtactacttaattcgtatttttagtgtatttaaagaagattgaa-ttttttcgaggtac
tggataagtatttcatcctaagaaccatagtcaac----ttttgatttattataatgctt
aaatctaaatata--tatcactcaagaatcattagtaattatgatgtttttaattatacg
att---aacatatttcaaaatagaagaattcatagcgactaagttaa-----tatgctaa
tataatttatcaat--aaaaaggaagttagtatttcaattatgcaaacaactataaataa
tgggcaaacgtctagtaaagaaactttactaacacctagattctatacgactgactttga
agagatggctaatatggatatttccggcaatcaagaagatttttttggctatcctcgaaga
atttcgagctgactataatagtgaacattttattagagacgaagagtttaatcaatcttg
gtctaatttagaacataaaactaaatccttatttattgagtttttagaaagatcttgtac
cgcagaattttcaggtttttactatataaggaattatccagaaaattaaaagacagaaa
tccggttatagctgagtgctttttattaatgtctagagatgaagctaggcatgccggttt
tttaaataaagctatcggggacttcaatttatctttagatttaggatttttaacgaagag
tcgcaagtatactttttttctcacctaaatttattttttatgcaacctatctttctgaaaa
aattggatactggagatacataactatttaccgccatcttgaacaacatccagaacaccg
tatttatccaatctttagatttttttgaaaattggtgtcaggatgaaaatcgtcacggaga
tttttttgctgctttgctcaaatcccaacctcatttttttaaatgactggaaagcaaaaat
gtggtgcagatttttttctattaagtgtatttgcaacaatgtacttaaacgactttcaaag
aattgatttttataatgccataggcctagactctagacagtatgatatgcaagtaatacg
aaaaactaacgaaagtgccgctagagttttcccggttgctttagacgtggacaatccaaa
atttttcaaatatttagatacttgtgcatgtgataatagagctctgatcgatattgataa

taacaattctccattattcattaaatctatcgtaaagatacctctatatttttccttatt
tgcaaatttactaaagatatatttgattaagccaatagactcaaaaacagtatggaatac
agttcgatagttaaatacaagggaatgctttttttcttataaaaagctatctagtgattaa
tataaaat------aaaagcttccttct-----cgatggaaggagcttttattttgtcaa
ga---atataatgggattatatagatc-tagaactaagaacttactttttcttttgcttc
atacataacctctaaagtaattagattttagaacaatcacgggcaaatttctctgtatt
tctttttactttacctctaacaaacccaggaattttatttaattctttctgtgcttcttc
tgaccaattaatactatcaacagcagaaattcctatacttaatgcttctgttgtatcatg
accaccaaatatttctagcaaatgatcttccataccaagtgtaaacgaattatatactaa
atcagcaatttgatttgttccctcataaccaagaaatggacggtaacttaaaggaaagtt
ttgtatatgcactggtgaagaaataacgccacaaggaatatttagacgcttaccaatatg
tctttccatctgtgttccaaaaattgctgcaggttcagttttagcaattaaatctccaat
taaaccatgatcatctgaaacgattacttcatcgcagaattcttgaacttgatctttaaa
ccactcttcatcatatttgcaataagttccacaccaggcgacatgtatacccatttcctg
gtgtaaaattcgagttattgctgcagcatgagttgcatctccaaatacaatagccttttt
acctgttaaattctgacaatcaattgatcttgaaaaccatgcagattgtgaaataaatcg
ggtttgctcgtctatgtatttttcgtagtctacagcagctcccaatgcatttactaattg
ctgtattgatcggatgcatgcagctgtttgaacaatacccataggtgtaatgtcaacata
aggcatattaaattctttctctaaatatcgagctgtcattagtccagtttctctatatgg
aataaaattaaaccatgcagagggtaatttttttaaatcttgaactgatgcattttctgg
aataatttggtttatttggatatctagatcttgaaataacctttttaattctgctatatc
atgttgattatgaaatcctaagcttactgctccaataatattaacagatggagttttagt
cttttgagttaaaacttgattattagatttagctttttccatatagaaagtgacaatttg
ttctaatgttctatcacctgcttgcagttcatttactctataatgattaacatcagctaa
taaaacatctgcttctgtttctatagatgctctgctaacaaagttttgcaaatcttcttg
caaaatactcgaagtacaagttggtgttagaataactaaatcagggctttcttctctgtc
ttttctagtaatattttctacaactttttcttgagaaccacgggctaacacatgtctatc
tacaacacttgctgtaacaggagtaaaatctctgtctctttcaagcattgaacgcattac
attaaaataatcgtctccgagaggcgcatgcataattgcatgaactttttttaaaagaact
ggcaattcttaaagttccaatatgagcagggcctgcatacatccaataagctaatttcat
attaaattttcctatgatttatttagacatagtaatacttttaaagctaactaacaaaca
ttagccaaacatctaatttaactccttgacaattttgtaatattttttaaatgaagctta
tcattataaatatttaaaaaatataattcagaataga----aataactgttagatcaagc
ttgaaagtaaagatttcttgcagttatagtttttctttaagtaaacatgtaaaaattgta
ctatatataaatgtactatatataaactttttttactgaatattggtaatgccagataca
attaacttaaatatgccttccccaacgtttggtggaagcactggcggttggttaagagcc

gcagaagtagaagaaaagtatgcgataacatggacaggcaaaaatgaaagtaagtttgaa
atgccaactggtggtaccgcaactatgcgaaatggagaaaacttactctatttagctaag
aaagaacaatgtttagctttaggtactcaactaaaaagtaaatttaaaatatctgactat
aagatttacagagtttttcctaacggagaagtgcaatatttgcatcctaaagatggagtc
tttccagaaaaagttaatactgggagagctagcgtcaacagtgttgatcattctattgga
caaaatattaatcctgtagatgtcaaattcatgaataaagcaacttacgattagtctctc
-aaaaacctagacataaaataacttttatatagtctaggtttttgataatgtatactgat
agtatggaagggtggccgagtggttgaaggcgtctgatttgaaatcagttgaacta--tc
tggttccgtgggttcgaatcccaccctttccgttataaaact------------atagtt
ggcttctcaaggcaagctagctatacactgatagtattaagtttagacttaataaagtct
acaacttgtgacaggttagaaatttttcggcatcctcatctggtatctctatactaaat
ttttcttctatggccattaccagctcaactgtatctagagaatcggctcccaggtcactt
gaaaaatttgcttccctagtcactatttttttttcaattcctagctgttctgctacaata
tcttgaactttttcaaagatttcattatcttgcataatatattatttcctctaaggtgtt
atcaaaaatttttcggcttatattattatccaactctaaaaaattagtaatttttagtcg
agattaaaatatttatacctatatgatatactattattctggataaatacgaagtttacg
atatggctcttctccaaaactacgagctctcaactggtagttgtcaactagattatgctg
catttttcgtatataggctgatcgaggtgtaagctgaactattgaactttcattcaagat
aatagtttcaatagccaatttagcttcttgaagagcctggatctcatcaaaatttttatt
tttacagagttctacccaattaagaccagatgaagtatgaatgtttaatatttttctgag
tgcacgagtaatttgtggaaccgtactattctgaattgtatatataataatttgttttga
ttttgcaatttgtctaagttttgtattttgctttacttgattacgtagcgctagaatagc
atcagacttttctatctccttggttaaaataattggtagatctagtgaagaaataactga
tgtaatatgctgccaactcaaagaataagcatataagtactgatgaggtatttccaattg
caaagattgtttattaatattaatagggcattgaaagtagttgacaataaagatgtatc
acgattttgtgatttattcaaatctaaggacttatttcttaattctctatattgtaaaac
cggaggcgcctgctgtgcgttaattgattttcgagcattagagatgtttacaggtaagac
ttctatagattgcgatggataacatttaattaaaattctaccattagcttgaatttgtct
tttttgcacaaaaggttgatggccttgcaaaatttgatctattgtctctttaactttatt
atgaactatccaaacatttcgctcgtgtatttcgatcgcaatctgaaaagcaggctcagc
ttttctttctagtatactcttttgcgtacctctacgtttagcttcatcgtcacctaatgt
tacatattgaattccaccaattaggtccgccaatgttggattttaattaaactttctaa
atgatttccatgcgctgtacctactaattgaactcctctttctgctatagttcgagcagc
taacgcttctaattccgtgccaatttcatctataataataacctcgggcatgtgattttc
aacagcttctatcattacttgatgttgcaaatcgggccttgcgacttgcattcttctagc
tctaccaatagctggatgaggaatatctccgtcaccagctatttcatttgaagtatctat

aataaccactctttttttccatttcgtcagctaaaacacgggccatttctcgaactgcggt
tgttttaccaactcctggttttcctaagagtaaaattgaatcaccttgctgtagtaaatc
tcgaataatactaattgtaccaaagactgctctgccgacacgacaggttaagccaataac
actaccttctctattacgcagtgaactaatacgatgcaaggtttttttcgataccagctcg
attatcgccgctaaaattaccaacttttttttacacaataatctaagtcttgccaactaat
agatctttgagataaatattctgggttatctggaaatctagcttctggacgacgacctaa
atccataacgacttctattaaattatttctattaggatgctgttgtaaaggttctttgac
aaaatttggcaaaatttctagcaacttatctaagtcatctgcaataagcatgattgagta
tataataattaattaatcaaatagaagattcaagattctaaagattacaatacatattaa
agaatattatgattggaatcaattttatacttaaattgaacaggaagtta-tagatgtaa
aagaactgattaaactttacgcaataaatcgattggtcaaaattaatagtcgtgttaaca
tgtttattggacaaaaagttcgtattaaatatagcaaacaaaaagttgctagtgatatag
cagataaagttggtgaattaggagttattaagggaataaagtttattaatagtcaatgtg
ttactattattgttgagtttgacaatcacactaggctttggatgtttagagaggaattaa
tctgtctaaatgaat------taaaaacatgaaacataatttattttatttattatacaa
aatttcttatgagtcgttaaaataataaattttaaaggagtacttttaattgtgcaaatt
actattaaaaaattacaagacttactttcgtctgtacaaagaaaaaaagatccagacatta
aaactaaaacaaggtaagtttgaacttctattaaataagacctataaaaaagtcaatcaa
gaaattataccttcgcagaaatctgctgtattacaaaatagtccatctacgataattaaa
tcaataaataatacaaaaaaatcttctgttgtcaatgaagaccgtacagaatatgccact
attgtttctccaatggttggaacgttttatcattcacctgctcctggtgaaaaaatttt
gtacaagttggcgatgaggtcaaattcaatcaaacagtctgtattattgaagcaatgaag
ttaatgaacgaaatcgaagcagaaattgaaggcaagattatagaaattcttgttaaagat
ggtgatatagtagattgcgggcaagccttaatgaaggttgaaacataatgctaattttc
catttacatattgttaacagaaattatttagatagaaaaattagcttatactatagtta
tgaaaacttatttcgcaaaggcac-taaaaactaatatataaaattatttactttatctg
ccaaaaatagaaataagcgttttgattccttgtctcaagagaggagaatctcaatggcaa
ttagctcaaaagagcaagagacaaagaaggtaaaaatctcggttgataaaaatcccgtag
atacttctttcgaaaagtgggcccaaccaggccatttttctcgtacactagcaaaaggac
caaaaactactacttggatttggaatcttcatgctgatgctcacgacttcgatagtcaaa
ccagttctttagaagaagtttcacgtaagattttcagtgcacattttgggcagctgtctg
taatatttttatggcttagtggaatgtattttcacggagcccgcttctctaattatgttg
cttggttaagtaatccaacaggtattaagccaagtgcgcaggttgtttggcctatagttg
ggcaagaaattttaaatggcgatgtaggtggtggctttcaaggagtacaagttacatctg
gatggttccaactgtggagagcatcaggaattactacagaatttcagctttactgtactg
ctattggcggattagctatggctgctttaatgctgtttgcaggatggtttcattatcata

aagctgctccaaagttagaatggtttcaaaatgttgaatcaatgatgaatcaccatttag
ctgggcttttaggcttaggctgtttaggctggacaggccatcaaatccatttgtctttgc
ctattaataagctactagattctggcgtgtctccgcaagaaattccactacctcatgagt
ttttaattaatagagagcttatggctcagctgtatccaagttttagtaaaggattagttc
cattctttactttaaattgggctgaatattccgactttttaacttttaaaggaggtttaa
accctgttactggaggtttatggctaagtgatactgctcatcatcatctagcattagctg
ttctatttcttgctgcaggtcatatgtatagaaccaattggggtattggacatagcatga
aagaaattctagaagctcacaaaggacctttaccggcaacggtcacgaaggtctatatg
aaattcttacaacttcttggcatgcacagcttgcaattaatttagctatgatgggatctt
taagcatcattgtagcacatcacatgtatgcaatgcctccttatccatatattgctactg
attacccaactcagttatcgctcttcactcatcatatgtggattggaggattttgtattg
ttggagcaggagcgcatgcttctatatttatggtaagagattataaccctgcagaaaatt
ataacaatcttttagatagagtcattaggcatcgagatgctattgtttctcatctaaatt
gggtatgtatatttcttggattccattcatttggtttatacattcacaatgatactatgc
gggcacttggaagatctcaagatatgttctctgatacagctatacagttacaacctattt
ttgctcaatgggtacaaagtatacacactttagctcctggaaatacagctccaaatgcat
tagcaacagctagttatgcatttggaggagatattgtttctgttggtaacaaagttgcaa
tgatgcctatttctttaggtactgcagatttttagttcaccatatacatgcatttacta
ttcatgtaactgttttaattttagttaaagggttccttttctcaagaaactctagactaa
ttcctgacaaggccaatcttggcttcaggtttccatgtgatggacctggtagaggtggta
cttgccaagtgtctggctgggatcatgtttttcttggcttattctggatgtacaattctc
tgtctgtagcaatttttcactttagttggaaaatgcaatcagatgtttggggtagtgtat
ctccgtctggaaatgtttctcatattactggcggtaattttgcacagagtgcaattacaa
tcaatggatggctgagagatttcctttgggctcaagcatctcaagttattcaatcatacg
gttctgcgctatctgcatatggattaattttcttagcagcacattttgtatgggcattca
gtttgatgttcttatttagtggtagaggttattggcaagagcttatagaatcaatcgtat
gggcgcacaataagataaaagttgctcctgcaattcaaccaagagctttaagtattactc
aaggtagagcagtcggtgttgcacactacttattaggtggaattggtacaacttgggcat
tctttttagcgagaattatttcagtaggctaatagtgaaaataggataaaaaaaacaatt
atggcaacaaaatttcctaagtttagccaagctttatcacaagatcctacaactagaagg
atttggtatggtattgctacggcacatgactttgaaagtcatgatggaatgacagaagaa
aatttatatcaaaagatattcgcttcgcactttggacatctagcaattatcttcttatgg
acatctggtaatttattccatgtagcttggcaaggcaactttgaacagtgggtattaaat
cctttgaaagttaaaccaattgctcatgcaatttgggatccgcattttggacaacctgct
ttgaaagcttttagtaaaggtggctcagcttatccagtaaatatagcatattctggcgta
taccactggtggtatactattggtatgagaagcaatcaagacctgtattctggggcttta

ttcttactagttttatcagctttactcttatttggagggtggctacatctacaaccaaaa
ttcaagcctggtttatcatggtttaaaaataacgaatcaagattaaatcatcatttatct
ggattatttggggttagttctttagcctggacaggtcatttagtacatgttgctatacct
gaggcaagaggacaacatgtaggatgggataatttacaaccgtattacctcatccagcc
ggtttacagccgtttttcagtggtaattggagtgtatatgctcaaaatccagatacagct
caacatttattcggaactaatgaaggtgcgggtacagcaattctgacatttctaggagga
tttcatcctcaaagtcagtctttgtggctaactgatatggctcatcaccatttggctatt
gcagtagtattcattgttgctggacatatgtataggactaattggggaattgggcacaat
ctaaaagatattttagatgctcatagaccacctagtggtagattaggagctggacataaa
gggctatttgatactattactaattctttacacatacagttgggattagcattggcttcc
ctaggtgtaattacttcgttggtagctcaacatatgtatgctatgcctccatatgctttc
atggctaaagatttcacaactcaagcatctttgtacacacatcatcaatatattgctggg
tttctaatggttggagcttttgctcatggggcaatattctttgttcgagactatgaccct
gaacagaataaagataatgttttggctcgtatgctagaacataaagaagctatcatttct
catttaagttgggtaactctatttttagggtttcatacattaggcctttatgttcacaat
gatacaatgattgcttttggaactcctgaaaaacaaattctaattgagccggtatttgct
caatggattcaagcctcttcagggaaagcactttatgggtttgatgtgttactatcatcc
tctactaatatcgcaacacaagctggtagcaatatttggctgccaggctggttagaagcg
attaatagcggaaaaaattcattgtttttaacaattggtcctggtgacttcttagttcat
catgcaattgcattgggattacatactactacgttaattttagttaaaggtgctttagat
gcaagaggctctaaacttatgccggacaaaaaagactttggatatagttttccttgcgat
ggacctggcagaggcggcacctgtgatatatctgcatgggatgcgttctatttagctgta
ttttggatgctaaatacaataggttgggtaacattttattggcattggaaacatattaca
atatggcaaggcaatgcaactcaattcaatgagtcttcaacttatctaatgggatggttt
agagattacttatggctaaattcttctccattaattaatggttataatccatatggcatg
aataatttatcagtatggtcatggatgttcctatttggacatttagtatgggcaacagga
tttatgttcttgatctcctggcgtggctattggcaagagttaattgaaactctagcatgg
gcgcatgaacgtacacctttagcaaacttgattcgttggaaagataaacctgttgcatta
tcaattgtacaggcaagattagtaggtttagcacatt-tttctgtaggatatgtattaac
ttacgcagcttttgtattagcttcaacagcaggcaaatttggttagactaaa-----ttc
gaattagctaa-----aaaaaagtcagtcataata-gttttatgactgactttttagat
ttaaaacaaaaacttactaaagtttcacctcaatatctacaccagaaggcaaatttaact
tcattaatgcatctatcgtttgagaagaaggttgatgaatatcaataattcttctgtgag
atcttatttcaaagtgttctcttgagtctttatctacatgtggagaacgtaaaacacaat
aaattcttctttttgtcggtaaaggaattggccctactgcaacggcattagttctagatg
cggtatctagtattttgtcacacgacgtattaagtataatagaattatatgctttcagtt

taattctgatttttgtctgctgagtaattgtcatgttaaaaagatcttattatttaagaa
ttttagagacaacacctgcgcctacggtacggccaccttctctaatagcgaaacgcatac
cttgctcaattgcaattgcattaattaattcagcagtcatcttaattctgtcaccaggca
taaccatttctgcatcagtaccatcatcagcagtaaactgattaatagtaccagttacat
cagttgttctaacataaaactgaggtctatatcctggaaaaaatggagtatgtcttccgc
cctcctctttagttagaatataaacttctgcttcaaattgagtatgaggtgtaattgtac
caggtttagctaataccatacctctttcaatatctttttctgcacacctcttaaaagaa
ttccaatattatcacccgctagaccttcttctaacgtttttgaaacatttctaatccag
taatagtcgttgtacgagtttctcgtaaacctacaatttcaattgtgtcaccaactttaa
taatgcctctttcaattctaccagtggcaacagttccacgtcctgtaatagaaaaaacat
cttctacagccattaagaaagttttatcgacatctctctccggcgttggaatatatgtgt
caactgcttccataagtgaaaaaatcttgtcaacccatttatcttcaccttgcttaatag
ctgggtttttcgtcacagcttctaatgccaataaggcagaacctgcaacaaaaggaatat
catctccaggaaagtcgtattgacttaataattcccttccttctaattctactaattcta
gtagctcttcgtcatctacttgatcttccttatttaggaatactactaatgtaggaacac
ctacttgttttgctaataaaatatgttcacgagtttgtggcattggaccgtctgccgcag
atacaactaaaatagctccatccatctgagcagcacccgtaatcatatttttttacgtagt
cggcatggcctggacaatctacgtgagcataatgacgattatctgtttcgtattcaacat
gagcagtattaatagtaatacctctagctttttcttctggagcggcatcaatttcatcaa
atttttcgctgcagtagaccctaaagttgataaagtcgcagagatcgctgctgttaaag
ttgtcttaccatgatcaacatgaccaattgtgccaatattgacatgaggtttttttacgtt
caaatttagatcgagccatgctttttgttttccttatagtaaaagttgctttatagttac
ttttaacgataaatcttgtggattgaaagatttattatttaatacagacaaga-----ta
caatataactaatctaatatcgatagtgagcaaacgctttatttgcttctgccattctat
gtgtatcctctctctttctaatagaatttcctgtttcattagctgcatccataatttcat
tagctaatttcatagacatacttttaccagatctgtctctagagaatttagtaatccatc
ttagtgctaaattcgtacctctataagctcgtacttcaataggaacttggtaagtagaac
caccaactcttcttgcttttacttctactagaggagtaatatttcggatggctttttcta
aaatatttaaaggatctgattctgttctctctttaacgatatctaaagcctgatatataa
ttctttgagataaagtttttttaccacttttcaaaatacgaacagttaacatacttacga
gtctgcttttatataaaggatcaggtgatgcaaacctttttttagctgtattacgacgag
acatagtatttaaaattggttattgtaataa-tgttc-tattagcaggtatttaagattt
aggctttttttgtcccatatttagatcggcttttacgacgatcttttactcctgcagcatc
taaagtaccacgtactacatgatatcggactccaggcaaatctttaattcggccgcccct
aatcagtaccacagaatgctcttgaatattatgacctacgccaggaatataagctgtaac
ttcaaatccagatgttaatctaacacgcgctacttttcttaaagcagagttcggtttttt

aggagttgtagtatatactcttgtacaaacacctcttctttgagggcaactttgaagagc
tggagattttgttttttatgtatttttcgtctttcggatctaacaagttgttgaattgt
tggcataataaaattgatttaaagtctttaatagatactatttgaacttgctttacatga
attggctagttttatttatcttatacttacgcataaatctttctactctaccttctgta
tcaataattctctgtgaaccagtaaagaatggatggtttcctgaccagatatctacgtgt
agttctggctttgtggacccaatcgtcataattaactggccgtcacagtaaacttttgct
tctggataccaatttggatgtatattatcttttgccattgtttttataattatataaatg
tatcaataataactattaacgcttagaaaattgaggagcctttctggctttctttaaacc
atatttttttctctctttactcttggatctcttgttaaatagccttcagacttaagtgt
cgttctattttctggattaattgagcacaatgctcttgcaacacctaaacgaattgcatc
agcttgccctgtcaaacctcctcctctagcatttacatggatatcatactggtttagtag
tcctaaaacttgtaatggtgcatatgaaactcttaagtaattaggactaaattgaagata
agactctcctggtataccgtttataattaaattacctgaccctgggactagtcttacttg
tgcaacagaacacttccggcgacctgttccagaatagattgcgcgagttttaattaattc
tgtcgacataacattccttgataactaaataaatatacattctttatactatatactctt
gtggcttttgcgcgacatgtgggtgaattgggccagaatacactttaagcttcgtaaata
gttttctgcctaatggacctttaggaagcatacctttaaccgatttttcaataattctgt
taggtaatcttgtctgaagctgatcaaatgtttcaacttttaatccgccaggctgtccag
aatgtcttctgtatagtttttgattcgttttatttccacttacagatacgtgagcagaat
taataacgattacataatctccagtatctagatagggtgtataagaaggcttgtttttac
ctcttaagatattagaaatatgagtagatattctaccaagtgtctgattttagcgtcta
taacataccaatgagaattagtatttaatgagggtgattgcgttttattcatgaaaagat
taatacttttggttgacaataagagaatatgtaaatggtttacattcaatttctatacaa
gtagtattt-aaaaatatta-------------ttaacaaagagaactttctaacagtt
atttaaattgtttgtcagcaagaaaatttctagttatcaataacttatactatttttaat
aatataaaatatattaatcactttttcttttggcaaacttatacctagttttttttgc-
aaggcatatataacttcttcagctgatttctggccaaaatttttaatttctagcagttct
tcttgagagtaatctagtaaatctgcaatagaatggatttgagctcgctttaagcagtta
taagccctaacagataactgcagttcttctattaaaacttgactaatttttttatcttct
ttgctacgataattatctgctgatttaaagtctaaatttcttagtgaacaaaaaagatta
gttaaaactgtagctccttgacttattgcttcttgaggagagatgctcccatttgtccag
atttgtataatcaatctatcttttatgctgttgctaccaatacggacttcttctacttta
taattaactttattaactggcataaaaacagagtctacttgcaaaaaatccacagataat
tcatctacagctttttcagctaagcgatacccgcaatttttttcaattttaaactccatt
tcaaatattgtattattgcaaatagttgcaatatactgtctaggatctactacctctata
tcagaagataattcaaataggccggctgtaactatagctggcccttgaactctaattcga

ccaatttgagattctttgttataacttttaaatactacttcttttagattaagtaatatt
tctaacacatcttctcttaccccaggaattgtagaaaactcatggttcactccagcaatc
cgtacagcaactatagcagtaccttcaagatctgacaatattgatcttctcaaagcatta
cctaatgtaataccttgtccttgatttaatggttctattacaaaactaccgtactgccca
cgcgccccatctgttcttgactctatgcattcaatttgaaattgagccacctaagaaagc
tcc-tttaaaataatcagtgattagtaagtctctctaaaacaagaaggaatttatactcg
gcgtttcttaggagggcgacatccattatgaggtacaggagtaatatcttttattagagt
aatctctaatcctgcagcctgcaaagctctaattgctgtttctcgacccgctcctggacc
atttactagaacttcagtttggcgcataccttgatccatagcttgtctagctgctttttc
agctgctgtttgagctgcaaaaggtgttcctttttttagctcccttaaatccacttgcacc
agaggatgaccatgataatgtttctccttttaaattagtaatagtaacaattgtattatt
gaatgtagatttaatatgtgtaataccgttaactgcattacgtttagtttttcttgctcc
ggatttttttatttgtctagccatcgtgtctcgtctattatatagttaattaaaagatta
ttttcttggagctttttttcttacctgctactgttttttttacctcctctgcgtgttctagc
attagttctagttctttgtcctctcaaaggaagaccaagacggtgtcttctacctctata
agtactaatttccataagtctcttaatgctcatagactcaaaacgtttgagatccccttc
aatctgataattagactcaagaatttctcttatactgacaacttgttgatcatttaaatc
ttgacacttaatatcagcgtctatgtttgtttttctaatatttctttcgagcgagatag
tccaataccataaatatatgttaaagctatctctattcttttgtttcttggaagatctac
tccagcaattctggccactttgtcttgtctccaataaataatatgtat-ttttataacgg
ttgtttata-agcttagccttgtctctgcttatgtttaggattagtgcaaattaccatca
cctttctgtgacgccgaattattctacatttttcacacatttttcgaacagaaggacgaa
ctttcatattaa----ctctatatactaggaatattactatattaactctattaattatc
gacattaaaaatgtcaacaatatattgtactactttaactgttgtagattttacttcgtc
atactatcatattttttttgatatgacgtatgtctgaatttgcttagctgtgtctattgcg
actccaactaaaataagtaaagacgtagccccgagacctcttaagttttggatctgagta
acctttttctattataaacggaatcagcgctactgtaaataaaaacgaggctcctaggaat
gtcagtctatttaatatgacttgtaaataatcaatagtcgcttgacctggacgaatatta
ggaatgcttgcacccatttttttttaaatttatagcaatatcttctggattcattactatt
gatgtataaaaatagctaaagaaaaggatcaaagcacaataaagaagaaggtatagtgaa
ccattaggacaaaacaggtataggatttggagtaatgttttattttgaataatttgggtt
agataagatgggagggccatagatgcagatgcaaaaacaataggcataactccaccttga
tttaatttcaagggcaagtaactatttggatccaaaattgaagattttcctagctgtctt
gctgaaataattttaattcttcttgttccttcttgtacacaaattgtaattattatcatt
aataaaaagatcgctataaataatccgaacttaagacttgcattactataactagcatca
aaaaatgactgtgtaaaattctttggtagtcctgacacaatgttttgaaaaataagtaaa

gaggctccattaccaattcccttttccgtaattaactctgataaccacataataatcata
gaacctgctgttaaagctaaaacagactcgcagacaaatgcaaaattccaattaaaaaca
tatggttttacccatatagagattgctccagattgtaaagtcgcccaacctagagctaag
taccttgtaatttgagttattttttggcggcccaattcaccttcttccttctgtaatttt
tccagattggggacaattttcgtaagtagctgcattacaattgaagagttaatataagga
acaatacctaacgcaaaaatccctattgttgaaaaacctcctccagaaaaaatattcaga
aaatttactaaagtatttttttctacacttgcataaaaggcgtcatgatctatacctgga
acaggtataaatattcctaaacgtgctaaaactaatagaaagagagtaaagataatacga
tttcttagatcactttttttggctcataaataaatttaaatgaaaagagatactttt--tt
tatgtaaaaaccctattccaagacttgtataagtttaaaatagggaagtaa---aatatt
atagtaatattaata-taatattctatgtagaatacaggctttctattggaactcctcga
tctttagctgcttcactaaaagttcttaattgagttaagccattgagaacagctcgcgcg
ttatttaatgtattattagaacctaattgcttagctaatatattctgtacacccgacaat
tcgaggactgtccttacagaacctccggcaattacacctgagccaggtgcagagggcctt
aatataactttcgcggcaccagaaattccgttaataggatgaggtatagaattcgatttt
gtcagtggaactgtaactagatgtttttttagcatctgttactccttttttttactgcacca
attacatcgcttgcttttcccacgccgacaccaacttggccttgctcattaccaataaca
agaataactcgaaagcttagtttttttacctccttttacaactttagtaactcttttaact
tgtacgactctttcttcccagccactatctttatcttttcctttgctctgtttttttacga
ttggccattttaagaattattcctaattagttaatta-cgattctagaaacccatgcctg
cttccttagcagcttcagccaaggccttaactcttccatgatataactttcctcctctat
cgaagacgacattttttaatgccttctttcatagactgttctgctaactgtttaccaacta
cgcgagaagtatcacagtttggtcttatgttatcagattctttattatttaaattaacag
atgatgtagctactaatgtaataccttgtgtgtcatcaattatttgtgcgtatatatgtt
tattagatctaaatacacacagacgaggcctacttgaggttccttgaactttttttccgaa
ctcttttatgcttatgaattctagtttgtttagtgtttagtttcattattatttaccttt
tccagctttcccagcttttcttctaacaaattcaccttgatatctaattccttttccttt
ataaggctcaggaggcctaatagaacgaatagttgaagcaacctgaccgacaacttcttt
atctatgccagagacagtaatatttgtgttgttttcaactttaatttcaatattggcagg
aggtttaattttcactacatgactatagccaacacttaagattaaatcttggttatcaat
ttgagaacgatagcctacaccttgtatttgcagtttttttaaaaaaatccattagaaacacc
ttcaatcatattactaataagagttctggatagcccatgtaactgactagccatttttgt
ttgtccactcgtcttgacagcaattgtgtcgttaagtatttctaaattaataccagctgg
taaagttctggatagggtacctttaggtcctgtcacggtgatagtctgaccatcgaactg
agtactaagatttgtcggcaataaaattattttttttccaatacgagacat-attccacc
tataaaatagttaccaaatatagcataatatttcaccaccaagaccatcatgacgagctt

gtctatctgtcataacgcctctagaagtagaaataagagcgatgcctaaacctccaagaa
ctctgggcaactctttatgatttgcgtagactcttagtccgggtttactaattcttttca
gagcagtaataactggttgacgattttaccattatacttaagagaaatcattaaatgag
tctctataccttcacccatttgttcgaaattttgaacaaatccttcttctttcagtactg
ttgccatattgcatgtcattttcgttgctggaacttgcacaatttgatgtcttgctaagt
ttgcgttacgaatacgtgtcagcatatcggcgatcgtatcgttgaccaccttgatcctcc
ttgcga-----tgaattatttgaaatattttcaagattccctgaaaggcatccctagctt
ttttaacaaagctagaccttcttgatctgttttagctgtagtgacaattgatatatctaa
accacgaatttgatcaatattatcatagtctatttctggaaagatcaactgttcacgtaa
acctaaattgtaattacctttgccgtcaaaacttctaggactaattcctctgaagtctct
aattcttggtaatgttaaattaattaatttctctaaaaaagaatacatcttgtcttttct
tagatgcacaacaattccgataggaacctcttctcgaattttaaatcctgcaatagattt
tttagcttttgtgacaattggtttttgtccagttattaatgtcaattcttgaatactact
ttcaagagccttagcattctgagaagcttcacctaagccacggttaatagtaattttagt
aaacctaggaacttcatgtacatttttgtactgaaattcatcttttaaagattgagtaac
agttgttttatattttctttttaatcctattgccattatattttcatcaattatttaata
agttcgccagtctttttcagttttcgaa-ttttttggcctttatcattaattattactga
ggatcgactagcaatattattttgttcactaaataacataacgttagaggtatgtattgg
agcttcaaatttgataatttctcccgtttctccttcttgttgaggtttttttatgcttcac
tttgagattaattcctttaacaatcaccttatttgttttataaataattgcaataatttc
acctgtttttgttttatcacttccagaaataacttgaactaagtccccttttttttaattt
aatttttgtattatttttggttgttttagaaggacctttcattatactacctccggtgca
agagaaactattttagaaaaatttttatctcttaactctcgggctataggtccgaagact
cttgtgccacgtggattattatcttgattaatgataactgccgcattatctccaaatcta
atgctcataccgtcagttctccgtaaagctttacgagttcttactacaacggctctgaca
acatcagatcgtttaactggcatattgggagacgcatctttgactactccaataataaca
tctccaatagatgcataagaaggattactagtgcctagtactctaatacacattattttt
ctagcaccactattatctgcaacattaagatagctttgagtctgtatcatacttttattc
tgtataagattaattatcaaaagacttagataagatattaaccattgtccaacactttgt
gcgacttaaaggtcgtgtctcttgtattgtaacaatatcgccgattgtgcattcattatt
ttcatcatgcgctttatatttctttgtccggatcatagtttttgcgtactttctatgaga
gattctattttctacagctactactatagttttattcattttatcgcttactactttacc
tgttgtttcttttaaaggcatagtctttatatttttatatat---------aattaaat
tttaagattattacatggttgacttagtacgagacttttcaacagttaaaagttgggcta
atctatgcttagaatgtttaaatagatgaggctggaaatcttgccttgtggctcttttta
gccttaaatcaaaaagctctctttttattacaaggatttcttcagctaaagaagaagagt

ccaggtttgtaacatctgatattttagggaaagtcatattttaggattctgttgtattcc
gaactataaatttagtcttgattggtaatttataagaagctaatttcatagcttgttgag
cagtttttttgtggtacacctgtaatttcaaatagaatatgtccaggcttaataactgcaa
cccaatattctggagctcctttacctgaccccatgcgagtttcagctggacgggcagtga
ctggtttatctggaaatactctaatccatagtttaccacctcttctcacatatctagtaa
tagttcttcgagtagcttctatttgtctagaagttaaccaaactggctctgtcgcttgta
atgcataatcaccgaacgcaattgtgttacctttgctagcagaacctttcattctacctc
tatgttgttttctaaattttgttttcttggggcttagcataaaaattaaaaaataataat
aattaagaagctgttgaatcaggagtttcctgattataagcagattctagacttttcgat
tctggcaaaatttctcctttaaaaagccaaactttactccgagcacaccatacgtagtg
tgagcttggcgatgacaataatcaatatcggctcttaaagtttgtaaaggtacacgacct
tctctaacccattcacttctagctatctctgcaccattaagccgaccagatacttgaatt
ttaacaccctgtgtatttgctctttgagctctttgaactgcttgtctgacagctctacga
aaggccactctttttctagctgttgagtaataaattctgctactaaggttgcttcagaa
tctggatctgcaatctctacaacatttactctgagttgtttgctagggtctaggattaag
gacagcgattttcttaaagactcaattcccgctccagattttccaagtacgattcctggt
cgagctgttgctatgagaatttctacttgatcaactttacgattaatttcaattttagca
atactagcattactaagttttgaatgtataaatgagcgaattttatgatcttcttgtaaa
agaactgggtagtctttagagttagcaaaccatgaagaacgatgtttttgggtaatgcct
atgcgaaagcccaaaggatgaatttttttgacccacagtatccttgattataagttaaaag
ctataatgactaattgacgttaaa--------agactacagttctgaaacacctaatgta
atatgacaagttggtttatgtattggaaaggctcttccttgtgctctaggctgaaatctt
tttaaagtcgggcctttgtctgcaaaggctttactaacaaacaattggtttttatttaat
ccatcattatgctcagcattcgctgcagcagattccaggatttgctttatatgagaacag
actcggtatggcataaattccagaataattaatgcttcttggtattttctacctctaatt
tggtctaatacacgacggactttatgcggagaaagacgaatgtatttccctactgcttta
gtttcttttacgtttttttgtaatactcatagttatagtttaacgacgggcttttctgtca
ccttttacgtgagtgcggaaagttcttgttggaacaaactctcctaatttatgtcctacc
atttggtctgacacaaagacaggaaaatgttgtttaccattataaacagctattgtatgc
cctaccatatcgggaataatagtagatgctctcgaccaagtttttaggacttcttttttt
cctgaaatatttaatgcttctatacgtttaagaagactaacatcaataaaagggccttta
tgtatagatcttgacataataataaatcattagattcttaataaaataaattctatttac
ggcgacgtaagacgtacgggttactatatttatttggatttcgtgttttaacacccaatg
caggtttaccccaaggtgttacaggacgagctcgtccaattggagatttgccttcaccac
caccatgtggatggtctacaggattcatgacaacacctctaacagtaggtcttttaccta
accaacgatttctaccagctttaccaagagtaatattactagcatcaatattaccaactt

gtccaatagtagcgtagcattcttttcgaatcattcgaacttcgcttgaaggcaattta
cagtaacaaaagtaccttcttttgctactatttgagcataagttcctgctgcacgaacaa
tctgtcctccgcacgaaggtcttaattctatattatgtactgctgttcctaaaggaatac
tagataaaggcaaagcatttccaacttcgataggggcagtaggaccagaaagaaccatag
atcctacactgagagatcgagggtgtagaatatatcttttctcaccatcaagatagtgta
ataatgcaattcttgcatttctattcggatcatattcaatagaagcaactttagcaacta
tattatgtctatttcttttaaagtcaattaatctgtactgctgcttatgtccaccacctt
tatgacgacaggtaatgacacctctattatttcgacctttgcagaaatggtgtttaacta
ttaatgatttttctggcttatcagtagtaatctctgaaaaggtagaaaccgttctatttc
ttgtccctggtgtgtaggcacgatataaacgaattgccatataagaagatttgagaagag
taaatgttgaa------ctatgtgatatatttaagtttctggaaacaaattaatagaatc
ttctgaggcaagtgttacaattgctttcttgtaatgtggtcgcttccctacgaatctacc
gatgctcctttttttcttaggaggatgacaagtgttaacacctgtaacttgcacgttgaa
aatatactgtatagcagccttaatattaatttttgttgctttgggatcaacggcaaaaca
atactgattctcttctaataatttagttgtcttatcagtaatgatgggatatttaaccag
atctaataaacctcttgaatcaatgctatccattatatacctcttgtattttagataaag
catcaactgtaataataattttatgtgcagctaacaaagccataatatttagtgtatcag
ctgaaataagttctacgttgtgtagattgcgaatagaaagataaacattcggatcttttt
tgtctactatgactaaaactttttatttaaatcaagattccatcgatgtatagcttcca
taaatagtttagtcttgggctgatgaaaataactgttaaaattttctacaatcagtgtat
taactgatttgttatttaatgctgtccttaatgctaattgtctttctttttattcattt
ttttggtaaaactacgaggttttggcccaaatattacaccaccacctctccatagaggtg
aacggatagaacctgctcttgctcgaccagtccctttttgacgccatggtttacgacctc
cacctcgaacttcacttcttgttttagtattagcagaaccctggcgcttttcattacttt
gctttactaaagctctatggactaagtacatcccagaatcttgactgaccttaagattta
aatcagcgttgccacttacttgaccttcccaattatagacttgataatttaattgcgtgt
taactgtcatgaaatatatcgtatgaaagaatattgctacttactaattttaccaaagc
gccaggtttaccaggaacagctcccttaacgatcaacaaatcattctctgagttaatact
tacaatttgcaatttttttattgtaactttttttattaccaagttggccagccatgttttt
acctggataaactcttccaggcgttgtaccggctccaatagagccaggttgcctatgatt
tttcgaaccatgtgacattgggcctctactaaaatgatgcctcttttggtaaccggagaa
acctttacctacacttctagaagagacattaatcttttggcctacttgaaatagatctgt
agatagtatttggctaacttcgaaatcatcagtagatttcatttcgtattcacgtaaata
ttttagtgggggagcctgtgattttttttaagtgtcccaacaatggtttatttaattttg
ttcagcaacttgcttgtaaccgacttgaatagcattgtagccatcagtagatacagcttt
aatttgagtaataacacatggcccaacttgaattacagtaactggaattgataaaccagc

ttcatcaaagaattgggtcatacctactttagtaccgagtataccaacagacactagatt
tctcctttttgagagatttatactatcattgtacataaaaattctacttcatagttactg
aag--taaattaaactaacagatgcatgatattattcatgacaacataaaggttaaaaac
tttacctatttaatctaaaatagaaacagcttttttgtctagttacagcgtatataaatta
gaagtatgcttattaaatttttttataaactact-tagatttaactgatattgatatccgt
aacttacttgcagtaatatc----aatcaacatgtataagtttaattgcctaaactagaa
ccagctattattagaaataattcaagtatagataaaagcgtatttgaacttttatatatt
gacaaatataatggaatatattgaggttcggtaactacaccttttttattttttacatttat
tgtgca--gtatatatacatctaatgcagtaaacataagaatataac----tgtatagca
aatgtaattacacaaattagagaggtattaatgggcaaagtcgttggaattgatcttgga
acaacgaattctgtaattgctgttatggaaggaggtaaacctaccgtcataccgaatgca
gaaggttttagaactacagcttctgttgttgcatatactaaaagtggagataaacttgta
ggacaaattgccaggaggcaagctgttattaacccagaaaacactttctactctgtcaaa
agattcataggacgtaaacagaatgaaatttcgcaagagattcggcaaacatcatataat
gttaaaactagtggatcaagcataaaaattgcttgccctgcactgaataaagattttgct
ccagaagaaatttcagctcaagtactgagaaaacttgttgaagatgctagtacgtactta
ggtgagactgttacacaagcagtaataactgtacctgcttactttaacgattcacaaaga
caagctactaaagacgcaggtaaaatagcaggcttagatgtattgagaattattaatgag
cctaccgcagcttctttgtcttatggactagacaagcaaaataatgaaacaatactagta
tttgaccttggtgggggcacatttgatgtatctgtattagaagttggagatggagtattt
gaagtactctcaacttctggagatacacatttaggcggagatgactttgatcagcagatt
gtagaatggctaatcaaagatttttaaacagaatgaaggtattgatcttggtaaagataga
caagcacttcagagattgaccgaggctgcagaaaaagcaaagatagagctgtcaaattta
actcaaacagagatcaatcttccttttattactgccacacaagatggtccaaaacattta
gaaaaaactgtaactagagggaagtttgaagaactttgttcaaatttaatagataaatgt
agtatccctgtaaataatgctctgaaagatgcaaaactagaagcttccagtattgatgaa
gttgttttagttggtggatctacaaggattccagccatacagcaaatggttaaaagatta
attggtaaagatccaaaccaaagtgtcaatccagatgaagttgttgctattggtgcagcc
gtacaagctggagttttagcaggcgaagtcaaagatattctattactagatgtgacgccg
ttatctttaggagtggaaactctgggtggcgtgatgacaaagattataccaagaaatact
actattcctacaaaaaaatcagaagtattttctacagctgtagataatcaaccaaatgtg
gaaattcaagtacttcaaggcgaaagagaactaacaaaagacaataaaagcttagggacg
ttccgattagatggcattatgcctgcacctagaggagttcctcaaattgaagttaccttt
gatattgacgctaacggaattttatctgtaaaagccaaagaaaaggctactggtaaagag
caatcaattactatatctggagcatcaactttacctaaagatgatgttgaaagaatggta
aaagaagctgaagaaaatttcgatacagaccaaaaaagaagaaagaatattgacacaaag

aatcaagcagagtccctatgttaccaagctgaaaagcaagttaaagagtttgaagacaaa
attagccaagatttaaaaataaaaatagaagagctaattacagagcttagatctagtcta
gagaaagaagaatatgacaatattgaatctatttctcagcaattacaaaatgctctgatg
gacattggaaaaaatgctgctcagactgaaagtaaagatacaaaagcgaaggatgacgac
actgtgattgacactgatttctctgaagctaagt-aaaagta-agcgggtaacgcgattc
gaacgcgcgacatcaaccttggcaaggttgcgctctaccactgagctatacccgcataga
ttgtattattacaaaaattaagatatttgtcaatctgtta------atctatac---atc
tagactattaattttggaattacatattagctaactgttgtaaatataacaattgctata
atttttaa-------aagtacttatatcat-tatataagtacttttttatatt-------
-caaaaatacaaataagtattatag-------ctaaactactaaggaattaactaagcta
gtaacaattactaatcctgtccataagccggctcctgtatatattaatccttttgattgc
tcccattggccaggggatgctaaaacaacaggtacgcctactactaacacagtcgataat
gcaattaataatagcactaataattggatcgcaataatcatttaatatgttctcctgaaa
aattcaaatgttaataatactgtatataatacaatgaagcttttgaccttcagcaattta
tctatgaagaaataatttaaaattattgttttctaaataaaccttaaattaagtattaaa
atctttattgtaaacaataatactatatttttttta--gtagataaaaaatttaaagaaag
tcttttttaaaaaaaaattaagataaggtgatatgctgtataaaaagctaagtattatta
attattagagaaatttaaa-ataagaggtaaaacatatgaactcagcccttttttttagca
aaactaccagaagcttatgccatttttaaaccgatcattgatatcttacccgtaattcct
gtattttttcttcttttagcttttgtttggcaagcagcaattggttttagataacattta
ggttataaaaagaataacaaatatacagagtgccaatttttgagctctaatcttatatct
tacacaataataactattatggttgaacccttactatcaggaattgttcttggacttatt
cctattactttattcggattattagtagcagcttatttacaatatcaacggggcaaccaa
ttaggactgtaaagcctaaattatagatatttataaaaactacaataattatagttttta
taaatatctacaaataagctaattttaaatacagaaataatactattgaaactagagtga
taacaaaaaaaatttatgtaa---taa---------------------------------
------------------------------------------------------------
------------------------------------------------------------
------------------------------------------------------------
------------------------------------------------------------
------------------------------------------------------------
------------------------------------------------------------
------------------------------------------------------------
------------------------------------------------------------
------------------------------------------------------------
------------------------------------------------------------

------------------------------------------------------------
------------------------------------------------------------
------------------------------------------------------------
------------------------------------------------------------
------------------------------------------------------------
------------------------------------------------------------
------------------------------------------------------------
------------------------------------------------------------
---------------------------caagctaattgtttaccagcttgacttagtaac
tcctggtaaaagacattcatgagacatttctcgaaaaacatggcgagataatccaaaatc
tcgataataacctctacttctacctgttaaccaacatctatttctccctctaacagcggc
gctgtttctaggcatttcttggagtttttgtcttaaatccatttgtcagaaaagctgcc
agcttctttcagttgatctttaatagccagtctttttaaataatattttttgcaagttt
cgatctcttaatctctctttggatcatattctttttagccatgctgtaa-aactctcttt
taaaattatttattcaatatattagctaaaaaagctcctgcctgcaacttttatatattt
aaaaataaatagcataaagtagtgaaatgcaattgatatttaatcaacgatacc---act
ttatgctattatcaaaaatttatttagtaaagttcttgttcaacatgagttgaaatagtg
caatcagatgaaggatatgcaatacaggttaaaacatagccagctaaaagttgatcatca
tctaagaatgattgatcagcttgatctacagttccttctgtaactttacctgcacaagtt
gaacaagctccagctctacaagagtaaggaagttcaattccttcttcttctgctgcatct
agaatatatgtatcctccgcacaattaaatgtgacatcaattccttcttcttcacatagt
aaatgaattttataatcagccatttcttcttactccttaataaagatacacttgaaactt
aataaagcaaaat-aatttaaagaacttaccaaaaaattatatactagtttcattatgca
tgatagactggaaaaaaagaaaatagtatactattctgaaaaatcaagctacatttccta
ctaaatagaactatagtaggtccaaagtatatacataaattcatatagttaaaattactt
ttatcatttaaatagctgaacttgatagta---------tagattgatcatc------aa
gctagtcttctgc---caataaataattaattatgatttcaatgcttaatcaacaagtag
aacttaagcctattcttaaatttcagataacacaaatatattcgcattctcttattcagg
aaattaaagccttaataactagacttgttttgcaagtttggagaaggccggctacattga
tggcaggtattatccaacctttattatggttaattctatttggtgggcttttctacaatg
ctcctataaatttgttcactattaatacaagctataattgtttttttgagctctgggatta
taatttttacctcttttactggagctttgaattcaggtcttccattaatgtttgatagag
aatttggattttaaatagattattaacggctcctttagtatcgaggacttctatcattt
tatcttctgctacttttatgacttgtattagtttaatacaagttgtatttatagttacag
cttctctttttatgggaaactcacctctaaatagcgatagtactatgatttttggactta
tgattctattagtgactgtaggagttacaatgcttagtttagctttatctttcactctgc

caggtcatattgagctattagcatttattttagtagttaacttgcccttttttatttcta
gtacggctttagctcctttatatttcatgccgccatggcttcagttaattgcaagtctca
atccattgagttatgcaatagaaggtacaagatacttatattcaagcgtaaactggaatt
ttacagagtgtgtgattaagattagctggggagatatttgtttagggcaaattattattt
tattaatcgctttagatataatggcagcttatcttgtgtctaatatattaaaagctaaac
ttaatta-----------------aaatttaataaaaaatttattaaatttattttaca
aagattttttactatataatactaatagtagtatggataattgaaatagaaatagtttttc
aaaaccaaagctattatcaacaatgataaacatttgtaagaaagtcaacaaagtatgttc
ttattcataggaggcatgtagtcaatgggactaccatggtaccgtgtacacacggttgtt
ttaaatgatcctggacggttaattgcagtccacctaatgcatactgcacttgtagcgggt
tgggcaggatctatggcattatacgaactagctgtatttgatccttcagatccagtatta
aatccaatgtggcgacaaggcatgtttgttatgccatttatggctagactgggcgtaaca
gattcatggggaggatggagtataacaggagagagcgtatccaatcctggactgtggagt
tttgaaggtgtagccttaactcatatagtcctttctggcatgctttttctagctgctatt
tggcattgggtttattgggatttggaattatttagagatccacgaactggcgagccagcc
ttagatttaccgaaaattttggaattcatttattgctatcaagtctactttgtttcgga
tttggagctttccatgtaactggacttttggaccaggaatgtgggtatcagatgggtac
ggagtaaccggaaaggtattaccagtagctccagcatggggaccagaaggattcaacccg
tttaatcctggaggggttgcatctcaccacattgccgcaggtactgtaggtatattagct
ggtgttttccatttaactgttagaccaccacaaagactgtatcgtgctctaagaatgggt
aatattgaaactgtattatcaagtagtatctctgctgttttttttctcagcttttgtgact
tgtggtacgatgtggtacggttctgcaactacgcctattgaattatttggtccaactaga
tatcagtgggatagtggatattttcagcaagaaattgagaaacgagtagaaaatgctatt
gctgatggtgctgcacctagcgaagcatggtcaagaattcctgacaagttggcattctat
gactatattggtaataatccagcaaaaggaggattattcagagcgggccctatgaataaa
ggcgacggagttgctgaagcatggcttggacatcctgtattccaagataaagaaggaaga
gagcttagtgttcgcagaatgcctgctttttttcgaaacttttcctgtaattttagttgat
aaagatggtattatacgagctgacattccatttagaagagcagagtctaagtatagtatt
gaacaagtaggtgtaacagctagtttttatggtggcaaattaaatggccaagttttcaac
gatgctcctagcgttaaaaaatatgcaaggaaagctcaattaggtgaggtatttgaattt
gatcgaactacattagaatcagatggagtgtttagaagtagtcctagaggctggtttaca
ttcggtcatgctaatttcgctttaattttcttctttggacatctttggcatggttcaaga
actatcttccgggatgtgtttgcgggaatcggagctgaagttactgaacaagttgaattt
ggagctttccaaaaattaggagatagaagtagtaaaaaacaaggagctgtataagaccaa
agtcaaagatagtctttatc---aaatatttctaatattaaaatagttaagacaaagttt
attatctaatagttactaatcaggagaatttatggaagccttagtctatgtattttttact

aacaggaacgctaatggttatattctttgcgatctttttagagaacctccaagaatagc
aaagtaacttgattccgtttaaacttataataatagaaccactttccaacttattagtgg
aaagtggttttattgtagattacattgtattattcttcatgctcttcaaaaggatctcgg
agatcttttgaagcaggaccaaatgctgtgtatatcgagtaacctgtaatacctaaaagc
agactcgaaataaaaatactaagaactgttgcagtttccataatttaagcagtattaaga
tgaactttttctataatgatattatcaatattaaaacatttaattattaaaattatcaaa
atatactatggcacttagaactagacttggagaaattttaagacccttaaattcagagta
tggaaaagttgctccaggttggggcacaactcctattatgggaattttcatgttactatt
tttcttgtttttattaatcatttacaaatctataattcttcattagtattagaaaatgt
agatgtagattgggctactttaggtag----ctaactagataagttctttga--------
ttacacaatattaaagacaac----------------aagcgatcgctattagattatag
tagcgatcgctttaaaagtattttactatatatccagagaaaattatttacctaaattaa
ttaactcggtatctgcaaaactatttgtgttaacattattatagttaactttttcaaacc
ttactaatactggatatttgataccactttatcaattgctgcaacagtaccaatttcct
gataccagtacgattcttttctcaatatttttacttttgagcctctttccatatttttag
tcctcagatagattattaaagattattatattaaaaaattaaaaaaaactaaacaaaata
tataaaaatactaccgacaattgtaatatataactaataaaaggagagagagaggattcga
accctcgataacaaaagttatgacaattttcgaaattgttgcaataaaccactctgccat
ctctcctgaattaaaatactgttttaaaaaattagacaagaataacatagaactaaggta
gttgcctagaaaattaaaactatgttaaatataaattataatctttctatagatatattt
aaaaaatatgtacatagaatctatatcagcttacttcaagatttttaatcttatacaat
ttttactaaataaataaattctctcataatatgaaactatcttggaaaacacttcttctt
tggtctttaccaatatttgtaattggtttctttttctggcaaggtttttaggtccaact
actacagatgttggcagtaatatcgcaagttctagaatgacatatggacgatttttagaa
tatttagatatgggttgggtgaaacgggttgacctctatgaaaataatcatacagcaatt
gtagaagctgttgggccagaattaggaaacagggttcaacgaattcgagttgaactgcca
gcaagtgcgccagaattaattacaaaattacgcaaagccaatgttgatctagatgctcac
ccccctaaaagtacaagtgcagtatggggactattaggcaatttactatttcctttacta
ttagttggcgggttagctttcttatttagaagatctaataatgctagtggtggacctggt
caagcaatgtcatttggcaaatcgaaagctttatttcaaatggaggctaaaacgggagta
gtatttaatgatgtagctggagttgaagaggcaaaggaagaatttcaagaagtggtaaca
tttttaaaacagcctgaatcatttactgctgttggtgcaaaaataccaaaaggcgtttta
ttagttggacctcctggcacaggcaaaacattactagcaaaagctattgctggcgaggct
agtgttcctttttttagtatctcaggctcagaatttgtagaaatgtttgttggtgttggc
gcttctcgtgtgagagacctattcaagaaagcaaaagacaatgcgccttgtatcgttttt
attgatgaaattgatgctgttggtagacaacgaggaacaggtgttggaggtggtaatgat

gaaagggaacaaacattaaatcaactattgactgaaatggatggctttgaaggaaatact
ggtgttattgtaattgccgctactaacagagctgatatttagattctgcattattaaga
cctggaagatttgatagacaagtttctgtagatgtaccagattttaaaggcaggttagca
attcttgaagttcatgctaaaaataagaaaatggaacctaaagtatctttagaaaccatt
gccagaagaactcccggcttttcaggagctgatttagctaacttactaaatgaggctgct
atcttaacggctcgacgaagaaaaaatgcaatgactatgtctgaaattgatacatcaatt
gatcgagtagtagctgggatggaaggcactcctttaattgacagtaaaagcaaaagatta
attgcgtatcacgaagtgggtcacgcaataataggcagtttattagagcatcatgatcct
gtgcaaaaagttacattaataccaagagggcaagcaagaggcttaacttggtttactcct
agtgatgatcaaagtctaatatcaagatctcaaatactagcccgtatcgtaggtgctctt
ggtggcagagctgcagaagaaatcattttcggtgacgcagaagttactactggtgcaagt
aatgatttacagcaagtaacatcaatggcaagacaaatggtcactcgatttggaatgtct
aaaattggacctttatctcttgaaagccaaggaggagacccattttttaggtagaggcatg
ggaggaggctcagaatattcagatgaagttgcaactaatattgataagcaagtaagggaa
attgtcagtgaatgctatgcacaagctaaacacattattatagataatcgagtagtgata
gatagattagttgatttactaattgaaaaagaaacaattgaaggcaatgaatttagagac
atcgttaaggaatacactgcaattcctgaaaaaaattactacatatcacaattttaa---
---------------attaaacgggactgacgggattcgaacccgcaacttccgccgtga
cagggcggtgctctaaccagttgaactacagtccc-aaaaagatgaatacccagagataa
tctcatagttagcttaaacttgtcaaattagaatattaaatatgaagagta---------
--------tttgtaataatactcttcatatttgata-tagtagctaaaatatttctttta
gaaggagagagagggattcgaaccctcggtacgaagttaatcgtacagcagattagcaat
ctgccgctttcgaccactcagccacctctccatga---tatatattatatatatatcaaa
tatacttatatttaataggtgtgcataaatatttatggagctaagcggattcgaaccgc
tgaccctctcaatgccattgagatgctctaccaactgagctataaccc----ctttggta
agatattaagaaatggctcaagcgggatttgaacctgcgaccttgggcttatgaatcccc
tgctctaaccactgagctactgagccatagtataaaatactattaaatattataccataa
attatggtcaaatataaatcaataaaagaataattactatca-tttactacttttg----
-----------------tgcaagttgtaaatattcgaataaaatttcaaaattaaaaatg
tctttaactttttgttaga-------atta-ttgattacgaaa-----------atattg
atatttagaattaac---------------------------------------------
--------------------aataaattttg-----------------------------
-----------------------------------------------------tctttattt
g-----------------------------------------------------------
------------------------------------------------------------
------------------------------------------------------------

------------------------------------------------------------
------------------------------------------------------------
-------tagaca-----------------------------------------------
-------------------------------caaaaatattggaaa--------------
------------------------ttttcatcaat-----------------taattagt
------------------------------------------------------------
------------------taaactataaccgacgtac-----------------------
------------------------------------------------------------
------------------------------------------------------------
---aagaattattgaacta-----------------------------------------
------------------------------------------------------------
---cgttaaat-------------------------------------------------
------------------------------------------------------------
------------------------------------------------------------
------------------------------------------------------------
-------------------------------------tgtgatgcatt------------
-----gggcttgatcaaaacttctctaaaatcaatagaatatagt---------------
-----------------------------------------------------aattcaa
gtaaatcaaatactaacaaatgtgacaaacgcttaaaaaaataata--------------
caattagaatcgtaaaaatttggtttttga-ctgtgttaaataaatttgattatcataga
tttaatatttaaaagtgttctattacttgta-----ttgagtaattgt-aatatttaata
tcttctcttttagataataatcataatataagctaaaaaatagtttt---tattaaacaa
ctagcatagagccaataccttgatcagttaatatttctagcaagagagcatgatctattc
taccatctaaaatatgcgctgaagctacaccctgagctaaagaacgaatacaacaattca
ctttaggaatcatgccaccagaaattactgctgtttgagttaaatctctagcttcctgta
tattcagatgactaattaatgttgaaggatctgaagaattacgtaaaatgccgggcgtat
ctgtgagtagaattagtttttcggcattaagagcggctgctatttctcctgcgacagtat
cagcattaatattatacgattggccttccttgtctgctgcaacactagctataacaggta
tataattattattaattaatatcttcagtaatttagtatcaatattttgtacttctccaa
caaagccaaggtttgcttttccatttggtctcggagtaataagtaatccatcttttcccg
acaaacctacacttttaccaccttgtttattaatagttgcgacaagatctttattgactc
ttccaactaaaaccatttccactatatccatggtaggttgatctgttactctaacgccat
catcgaattttggtaatattttgaggcgatctaaccaaaaattaatttctggtcctccac
catgaactaaaatgggacgtagtcctataaaagacagaaaaacaagatcactaatcactt
gatcttttagtttctgatttttcatagccgctcccccatattttataacaataattctgg
aagaaaattgctgaatataaggcagagcttcacttaagacttttactctttctgaattag

tcaacataaataaagtaggattgtgatatataaaattatatacacagcttgtgttaattt
aaaagataccaatgggt---------acttgaaaaaatagaaatattactaacag-----
----ttaaaactatagggattataattatgaataaattttgggataatgtattaagattt
ccacgattttagtcagcgtcattttaggattaattttgataattattagtcctttttc
gtgttattaaaaaaaccactgacaagtttttttttcatcatatcattggccggattaatc
acagttttggcaataataa-tacaaaaaatgataaatatcgaatgttgttgagagaaata
aataaatttaattaaaatttttgaacatatataatattatattatcattaaatatgataa
ttt-atttcctttctat---aaaacccataaatattaaatattatttgataaatattat
gccactaaaacaaagagttagctctgaaaagaccggtgccttcgcacttttggatagtat
tgttaggcacggagttaaacatatatttggttatccaggtggagctattcttcctattta
tgatgagctttatgcttgggaagaagcctctctaattaaacatatccttgttcgtcatga
gcaaggcgctgctcatgctgcagattcttattctagatcaacaggagaggttggagtatg
cttcgctacttctggcccaggagcaactaatcttgtctcaggtatagctacagcacatat
tgattctgtacctatattagctataacaggtcaagttgggagagcttttattggtacaga
tgctttccaagaagtagatatttttgggattacacttcctattgtaaaacattcatatgt
agttcgtgaccctagagacatgtctagaattgttgcggaagcatttttatttgtaaaca
cggtagaccaggtccagtattaattgatgttcctaaagatgtaggattagagaagtttaa
ttatttttctgttgagcccggaaaagttaatattcctggctgtaggccaattaccagcct
aaagtcaagacaaatccttatggcagctaaaatgatacagcaatctagccagccattgtt
gtatattggtggaggagccataatctctgattcacatcaaattattaaagaacttgttga
tttttataaaatacctgttactactactttgatggggaaggggattttaatgaggatag
cgactattgtctagggatgttaggtatgcatggtactgcgtatgctaattttgcagttag
tgagtgcgatcttttaattgctttaggagctagatttgatgatagagttactggaaaatt
agatgaatttgcttgtaatgcccaagtgattcacgtagatattgatcctgctgaagtagg
aaaaaataggattcctcaagttgctattgtcggtgacgtagcagaagttgttagtgaaat
attgaatttattaaagacttctttccccccttatccagagcagattatatcttggcaaga
aagaattaatcgttggcgtcaacagtatcctttactggttcctagaaaatcaacaagcat
ttcacctcaagagattcttgttgcaactaataaattagcccaaaatgcttattttactac
agatgttggccagcatcaaatgtggtcagctcaatttctgaaagtaaaagctaagcattg
gctttcaagtgctggattaggcacgatgggttatggtttacctgcagcaattggcgctca
agtagcacatccaaatgacgtagtcatttgtattagtggtgattctagttttcaaatgaa
tatgcaagagttaggaactatcgcgcaataccagttaccagttaaaattattattattaa
taatcgatggcaagggatggttagacaatggcaacaagctttctatggtgaaaggtattc
acactcaagaatgacagaaggagcacctgattttcaaaagcttgcagaagcttttggtat
taaagcttttactattaataataggcaaaatatgcaatctgctttacaagttgctattga
ttatcctggtccagttttattagattgccaagttacagaaaatgaaaactgttatccaat

ggttgctcctggaaaaagtaatgcacaaatgataggtatcgctaaaccgcagagaggtac
tgcttccaactatattaataatagtgtttaaagtaa------------------------
------------------------------------------------------------
------------------------------------------------------------
------------------------------------------------------------
---------------atagttagtatttttta-aacaaagaata-aagtaca--------
-taaatgtaatttttttatactatta-aaaactatgtaagaacagcctttgacgagaatt
gaactcgtgaccttccccttaccaaggggatgctctacctctgagccacaaaggc-tttt
tactatattgggccgggttggatttgaaccaacgtaggcgaagccagcggatttacagtc
cgcccccattaaccactcgggcaccgaccctaattaaaccttattcaagatatgacctat
aagatatcataactatgttgctaaaaacaagtactttgataaataaaactgaataagtaa
tggacatagctggacttgaaccagcgactttcacgatgtcgacgtgacactctaaccaac
tgagttacatgtccaaaactttaattattattataacactgtttagtgattttataagta
tctatttctaattgaagatttgttttaatcgtccggctttaccgagcagatctcttagat
aatacagtttagctctacgtattttagctcttctaatcactttgacaaaagttattcgcg
gtgaattgagaaaaaatactctttcaactccaataccttgaaatgagcatcttaaagtta
aactagtatttaagcttttttttcttcttctcgctaatactactccttcacataattgtt
cccgagttttactaccttccttaaccattagtccaagttgtattgtatcgccaactttaa
tttcaggtacttctgttttgataaaaggtatctcaacacttttcattagctgacttagct
ttgtactatttgctttcatatataggattagtcttgtgaca-tgttttgaaatattatac
attaaaaaaacaaatatgaaaaattttagcgaataagcatactagtatttataaaac-tc
tttttatct---------------------------------------------------
--------------------------------------------ccattata--------
----------------cagatccatt------------atattcaact------------
------------------------------------------------------------
------------------------------------------------------------
------------------------------------------------------------
------------------------------------------------------------
------------------------------------------------------------
------------------------------------------------------------
------------------------------------------------------------
------------------------------------------------------------
------------------------------------------------------------
------------------------------------------------------------
------------------------------------------------------------
------------------------------------------------------------

------------------------------------------------------------
------------------------------------------------------------
------------------------------------------------------------
------------------------------------------------------------
------------------------------------------------------------
------------------------------------------------------------
------------------------------------------------------------
------------------------------------------------------------
------------------------------------------------------------
------------------------------------------------------------
------------------------------------------------------------
------------------------------------------------------------
------------------------------------------------------------
------------------------------------------------------------
------------------------------------------------------------
------------------------------------------------------------
------------------------------------------------------------
------------------------------------------------------------
------------------------------------------------------------
------------------------------------------------------------
------------------------------------------------------------
------------------------------------------------------------
------------------------------------------------------------
------------------------------------------------------------
------------------------------------------------------------
------------------------------------------------------------
------------------------------------------------------------
------------------------------------------------------------
------------------------------------------------------------
------------------------------------------------------------
------------------------------------------------------------
------------------------------------------------------------
------------------------------------------------------------
------------------------------------------------------------
------------------------------------------------------------
----------agattcttttatagtttc--------------------------------

------------------------------------------------------------
------------------------------------------------------------
------------------------------------------------------------
------------------------------------------------------------
--------tttgataatcagtac-------------------------------------
------------------------------------------------------------
------------------------------------------------------------
------------------------------------------------------------
------------------------------------------------------------
------------------------------------------------------------
------------------------------------------------------------
------------------------------------------------------------
------------------------------------------------------------
------------------------------------------------------------
------------------------------------------------------------
------------------------------------------------------------
------------------------------------------------------------
------------------------------------------------------------
------------------------------------------------------------
------------------------------------------------------------
------------------------------------------------------------
------------------------------------------------------------
------------------------------------------------------------
------------------------------------------------------------
------------------------------------------------------------
------------------------------------------------------------
---------------------------atcta----------------------------
------------------------------------------------------------
------------------------------------------------------------
------------------------------------------------------------
------------------------------------------------------------
------------------------------------------------------------
------------------------------------------------------------
------------------------------------------------------------
------------------------------------------------------------
------------------------------------------------------------

------------------------------------------------------------
------------------------------------------------------------
------------------------------------------------------------
------------------------------------------------------------
------------------------------------------------------------
------------------------------------------------------------
------------------------------------------------------------
------------------------------------------------------------
------------------------------------------------------------
------------------------------------------------------------
------------------------------------------------------------
------------------------------------------------------------
----gatgatcaaatattattttcaaacatggcttatgtatgaaaaagttctata---aa
aacctaatgac--------taaaattagggcttaagtatttcttagttttgtgttataat
tagttactatcaaaggtagtggcataacacgaagcatcaa-aatgttcgaacgctttact
gaaaaagctataaaagtcataatgctagcacaggaagaagctagacgcttaggacataat
tttgtcggaactgagcagatactattaggattagttggtgaaggtactggaatcgcagct
caagttttgaaatcgatgaatgtgaatctaaaagatgcaagagttgaagtcgaaaaaatt
ataggaagagggtctggttttgtagcggttgaaattcctttcactcctcgagcaaaaaga
gtactagaattatctttagaggaagcacgtcaactaggccataactatattggcacagaa
cacttgctaatgggcttagtccgagaaggagaaggagttgcggcaagagttttagaaaat
ctggcagtcgatgtttcttcaattagagctgaagttatacaaatgctcggagaaaatgcg
gaagccaatgtaagtggaagcaatactacgcaagctagaagtaaaacaccaacattagaa
gagttcggatctaacttaactcaaatggctatggaaggtggtttagatcctgtagtcgga
agacaaaaggaaatagaacgagttattcaaatcttaggtagaagaactaaaaataatcct
gtcttaattggggagcctggtgtaggtaagacagcgattgcggaaggattagctcaaaga
attgctaatagagatgtaccttctatttagaagataaattagttattactcttgatgta
ggtctattagtagccggaactaaatatagaggtgaatttgaagaaaggctaaaacgtatt
atggatgagattaaatcagctgataatgtaatattagtgattgatgaagttcatacattg
attggggctggtgctgcagaaggagcaatagatgcagctaatctgcttaagccagcttta
gcaaggggagaattgcaatgtataggtgcaacaactttagaagaatatagaaaacatata
gaaaaagatccagcattagaaagaagatttcaaccagttgtagttggagagccaagtgtt
gaagaaacaattgaaattttgtttggtcttagagaccgttatgaaaagcaccatcaatta
acaatgtcagatggagctttggctgcagctgctaaatacgctaatcagtatatttctgac
cgatttttgccagataaagcaattgatttaattgatgaagctggttctagagttcgttta
ctaaattctcaattacctcctgctgccagagaattagataaagagttaagagctgtatta

aaaacaaaagatgaagctattagagctcaaaaatatgaaacagcagagcagtatagagca
agagaaatggaaattaaagctcaaattgcagcaattgctcaaagtaaaaagaatgagcct
gatttaaatttagaagatcctgttgttacagaagatgatattgctgaaattgttgctgca
tggactggtataccagtaactaagcttactaaaagtgagtcagaaaaattaatgcacatg
gaagaaactttgcatggacgtattattggtcaagacgaagcggttgtagctgtctctaga
gcgataagacgcgcaagagtaggtctaaaaaatcctaacagaccaattgcaagctttatt
ttttccggaccgacgggtgtaggaaaaacagaattaacaaaagctttggcttcttatttc
tttggttcagaagcttctatgatacggctagatatgtcagaatacatggaaagacacact
gtatctaaactaattggttctcctccaggatatgtgggttatagtgaaggtggttatcta
acagaagcggtaagaaaaaaaccatatactgtcatcttatttgacgaaattgaaaaagct
catccggatattttaatctacttcttcaaattttagaagatggccgactaacagatgca
aaaggtagaactattgattttaagaatactcttttaattatgacttctaatattggatct
aaagttattgaaaaaggaggaggtagtttaggctttgaattatcagaagatcaaacagaa
tcccaatatactagagtacgatctttagtaaatgaagaactgaaacaatactttagacca
gagtttctaaatcgattagatgaaattattgtatttcgtcagcttactaaagatgaagtt
agagaaattgcagaattaatgcttaatgaagtctttgcgagaattaagcagcaagatatt
caattaaatgtaacagaacgatttaagcaacgattagtagaagaaggatataatccaagc
tatggagctagaccacttagacgagcagtaatgaggcttttagaagatagtctggctgaa
gaagttttatctggtaaaattaaagctggtgatagtgcagtagtagatgttactaatgaa
ggagaagttacagttttattaggtgaaaaattagaactgttaacataaa------aaaca
atttattaat--taaagcattgaagaggttat--attcaatgctat---gaaaaaaatta
ataagatgggttgctataattgacaacctctgatactctttatttgtatata---aagcc
gggatagctcagttggtagagcagtggattgaaaatcctcgtgtcaccagttcaaatctg
gttcttggcat-tttaagcatcagaaaattttgctatctgagcattaaattttaattgaa
atgttcctgtcggaccattcctgtgttttgctattataacttctgtgaagtctctacttc
tagtttcttgtgtatagtaactttctctatatagcattattactaaatctgcatcttgtt
ctatagaattatgaacaataaagttgtttgaaataaaattgcataaaggtttacattcta
aatcaaacactatttgtaaacttgtaacgtttatttttttgtatactttcaaaagtcaaag
atgaaa------aagaattaaaaacatttttttgttcttcaaagttttt------tattt
gaatagcaatcatgtcattttgattaattttatcacatcttttccatccttgtgttgtga
gtagtttatggttacttgttaattgtatatatttcccagcttcggtaataattttatata
cagtttttttccctgttttcgaaatactacatttaattgctagaaataactgttgttttt
gcgcactaaagttaatgacttca-atctttttataatagcaattaaataatatttg--aa
gttgattatataatgatgtctgaagataattgaatttggatatacacccactttctctta
aatcggataacaaaggccttttgtcactccgagtttcaaggtttctatttagttgagata
atgccaagataggcaagcttaaatcttttgctaatattttaagtgatcgagtaattaatg

atagttcttgagatctattgtcagattgtctactctcctgcagcaattgaagataatcta
taataattaattctatatttttccttgtaatttaaagaatttactttgtctttatag
tgtcagtagaaatcttagcgctatcatcaatataaagattaaggttagcaagagttttac
tttttgaataacatattgccattcatcattattaagctgaccagattgaatttttggc
cgtttaaattacactcttgagctaaaattcttcttagtagttgctctgtagacatctcta
agctaaataaaataacataagatcttttctgattaattacatatctagttatattaatag
caaaagcagttttccccattgaaggtcttcctgcaagaataattaaatctgacttttgaa
acccttgtgttatagagtcgagttctgtaaaaccagaaagtatactactatttatactaa
tctttttccttgtatccaaatggactaataggtttgcaagaattgtagctaatgtttgcg
tatctttttcgtcaagtatttcataagcttttgttaattgactaacaatagaagttatgt
ttttttgttcaataagttgttttgagcagctaattaaacataatgaatctccacatgcta
aaagtaatcttttaatgtagttatctataacgactgctgaatattcgcttatagtatcag
ataggggagcttgttctattagactaaaaacttcatttagctcattcaaatcttgaatca
ttttttcatttttaagttaatgaaaaaatttcttacgctttttgtctttgcatggttaa
cattttcgagtgctgctctataaagtaatgaagttgatgtaaaataaaaaaaatcaggag
agattttttctactgattttagtaatgatatcgattgctgagttaatattatactaatta
atattttttcagctaaaatattatgaggaggtagatatttataa------atacttatta
cttcctcc-ttgtaacattactcttgtg--------gtattagttaactattgatttaat
ttgattctggaagaacttgcaatttaatatttgcagttacttgatgtaatagcttaattt
ctacatcgtaaatgccaattgttttcacgtctggtaaagagatactttgtttttcaatgt
ctatgttagtagtgtttttaataatttgtgaaatttctttttctgtgacactaccaaaaa
tattgtggccatctcctgtttttttactaacgctaaacctttgtatttcttctagaagtt
gcttgactttttagcattttctttagctgtttttaatttttcttctcttatagctgcat
aaaatttttgttgttttaaaataccatttgtggcaactgccgccattttattcgggatta
aaaaatttcttgcataaccagaagctactttgataacgtcattactcttgccaagttttt
gaatattctctttagcacaacatttataactttttttactcatctt-------aattcta
aaaa----atctgtattatcctattatacctgttgtatacagatttt----aatagtaag
tatttatttgttgatgtaatattcttgtaaaa--aaaggtaatttttttatttttt-ttctg
atataatcactttcacaataattc-----attaggagaggtggccgagtggttgaaggcg
cagcattggaaatgctgtttaggagcaatcttaacgagggttcgaatccctctctctccg
tacaa--aaaatataaaaagttttatagagtccaatctaaatctccaataatattcttcc
aagcatttaatccaccccgaactctaacaatatttagtttttgagctattaaaaacttag
aagcaaatattgatctagaatctaagctacagtaaacaaaacaaattttatcttgtaaat
tgatatgagagtaattcattcctttttattttttgtaaaggtagatttaatgaatgtatta
aatgacttttatgatattcttcatgatttcttacatctattaaaatatattgtggattcc
tactaattaaaaacttttgtaattgaataacactaatttctcgtacaaatgtactagatt

tactgccataatatttgttccaatgctttttgt-------ggataatacaaatttagta
tttataatcttaaatttgttaaatgatgaagttaatgagttgtacgttaatatagttcca
cttaatatagacttataacccagaacgattttgacagcttcagttgcttgaagtgtacca
acaatacccggtaatagacctaaaactccggaattactacaagtatctcttgcgctctct
ttattttcagtttcactgtagaagtctcgataaacaggtccaccttgataattaaataca
ctaacttgtccttcaaattgaaaaatggcaccataaatatgtattttatttaactctagg
caagtatcattcaagagatatcgagtatcgaagttatcagatccatctataataatatca
tattgtctaataatgtcgattgaattctcatagcttagtcttgttttgaacacagttact
atacactgtggatttaaatctaata-tttttttttagctatttctacttttgaatatcct
atatcatgcatcgtgtataaaatttgcctctgtaaattagagagatcaattatgtcatcg
tctataattccaatgctgccaattcctgcagcagcaaggtaaatgattcctggagatcct
agcccaccagccccaatgaataaaacttttgcttcttttaatctttcttgtccttctaat
tgaatttgtggcaatactaaatgcttagaatatcttgtatattcttctagtgagtatttt
gtattttctgttttaaaatttagcataaaaatttatagtttatttaatagcactta----
--aaagctagctgataagtttattaaaatgacttttgtaaacttggttcttctctatcta
aaatagctccttgtgtaggacagacctgaacacaaatactacaatcaatacaggcagaaa
agtcaatccaataccaattagtgttattattgtttttccctttccttgatgaatacaag
ctacggggcaagcttccacgcattcagaaactccaatacatttttctgttacaattgtat
gagacatattatttaattaat-aaaagataagctatttacatctagtagttttgttgtat
actaaatttacatatgctatccaggccaatatttaaggtagtctatttctacagtttact
aatttgatatgcgcctttagttcagttggtagaacgcaggtctccaaaacctgatgtcga
gggttcaagtccttcagggcgcgtttag-ttttatcgtaattcaataatatcattttatc
tttttgcaaacatcaaaatgatatgatataaaagatcatattttatacattgaatattct
atgtactagtactg-ttttttaatatgttattattgataagttaattatactaattctat
ttttgtaataagt-aaaaacagttctaattaagtatgcacgtatttaatagtc-ttttgt
atatat--aacttgaaaaaatttaaactatggctaaaaaagttactggaattgttaaact
agcattaaatgcaggtaaagctactcctgcgccacctgtcggacctgctttaggacagca
cggcgtgaatattgttatgttctgcaaagagtacaatgcacgtactgctgacaagtcagg
actagtaatccctgtagagatctcaatatatgaagatagaagttttacttttatacttaa
aactcctcctgcttctgtactaatcgctaaagctgctggactaaataaaggttcgggtga
accaaataccaaaaaaagtaggcagcataacaaataaacagttagaatctattgcagaaac
aaaacttcccgatttgaacactaacaatatacctcaagctatgaaaatagttggaggtac
ggctaaaaatatgggaattctaattaaagatt-aaaagtttcaattatatttacta-tt-
-tttatgaaaaaattttcacgtcgacttacaacattaaaatcaaaagttgagcctaaact
ttacactattaatgaagcagtatctatattaaaagcaacgtcaaatgctaaatttaagga
aactgcagaagctcatattgctctaggtttaaatcctaaatatgcagaccagcaactcag

agcaacagttatattaccgaaaggtacaggtaaattgataaaagtagcagttattgctaa
gggagaaaaattaacagaagcaattagtgcgggagctgatgtcagtggctccgaagaact
aattgatgaaatactgaaaggtagattagattttgataagctgatagcaacaccagatgt
tatgcctttaatagcaaagctgggacgagtattaggccctcgagggttaatgccttctcc
taaagcaggcacggtaacattggacgtagcaaaagccgtgaatgaatttaaaggggggtaa
agttgaatacagagttgatagaacaggtataattcatgtaccttttggaaaatctagttt
ttcacaagaagatctagttttgaacctccaaacaattaaagaatcgattgatagaaacaa
gccttctggagcaaaagggaagtactggaaaactttttcttatctagcaccatggggcc
atctattcaaattgacatcactagccttctatgaaaaatttttgtataattatagaactt
gaataatctattatttcaaatta-taatttgctagcattattaacgtaacaacttgctta
ctatatcttatctgcattaaaaacttt-ttatatatcttattaattatgagtacaaaggt
tgaaaatatcttggaagagctaaaatctttaaaccttctagaggctgctgaactagttaa
acaaatagaagaaacgtttgatgttgatgcatctgcggcttcaggaggtatgatgatggc
cacaccaacttcagcaccagcgtctgctgaggttgaagagaagacagaatttgatgttgt
cctagaggaagtcccagcacctaaaaagatttctgtattaaaagttgttcgctctctgac
tggtttgggtttaaaagaagcaaaagatcttgttgaatctactccaaaagtcttaaaaga
aggtgcttctaaagatgatgcagaaactatgaaaaaacagctagaagatgctggcgcaac
agttattgttaagtaatga-----agaagcgcttctgtttaagtataaacagaagtgcct
ctttatttattataagcaattatactagaaaagtccaagtgtaattgcattgttgatagg
catagttgcaccaatacctagccaaatacttacaacggtaccaattaagaaaacagtggt
tgcaattggtcttctaaatggattttgaaatttgttaacattttcaatgaatggaacagt
aagtaaaccagctggtacagcagccatacttagaacacctaatagtttattaggaatcac
tcttaatagattaaatgttggaaagaagtaccattctggtaaaatctctaatggagtcgc
aaatggattagacttttcacctatagaagatggttctaaaattgcgagtccaatactgca
ggcaaaagtgccaataattacaactggaaagacgtataatagatcatttggccaagctgg
ctctccgtaataattatggcccattcctttgctaatttagctctcagcttcggatctgt
taaatctggtttttaagaattgacatattg-tttttgtgtttaatataaagattagtct
aattactgttttaaagtgtaattttcaactattataatggtccagaaattccttgtttac
gtatcattaaaaaatgcataagcatgaaaacggcagttagtagcggaagtacaaaagtgt
gtaaactatagaacctagttaaagttccttggcctacactaactcctcctcttaacaatt
ctactatactcgctccgacaactggaacagcatcagggacacctgtaacaatttttacag
cccagtatccgatttgatcccatggcaaagaataacctgttaccccaaaagaaacagtaa
gcactcctaaaattacacctgttacccatgtcaattctctaggtttttgaaacctcctg
ttaaataaacacggaatacatgtagaatcatcatcaagaccatcatgctcgctgaccatc
tatgaattgatctaataagccatccaaagtttacatcagtcataatgtactctactgatg
taaaagcttctgcaactgtaggtctataatagaatgtcattgcaaatccagttgcaactt

gaattagaaaagatacaaatacaataccccctaagcaataaaaaatattaacatgaggtg
gtacatacttactagaaatatcgtcagcaattgcttgaatttctaatctctcttcaaacc
agtcataaattttactcataaaatagcttcaaaaagctcttctaatata--ttttgcgtt
taagctactattaattcaatag------ttattgaacttatatt-tttttataatcaata
cttattaactattataacatcttttattgattt---------------------aaaatt
ctctgacttagtaaaacagggctatgtactataattctttttttttgtatagagattaat
ttcgtttttagtaaattagctaaaattcttgttacactgactctattactacctataatc
tgtgctaaaattttatgtgtaatagtaaagttaagcataataccattattaacttgagtt
ccattttgctcagctaataataataataagctcgccagtctactggttatacttttatgt
gaaatgatttctataaaatgataagcttttgccgagcaaaatactaaatgatttacaaaa
aataaattgaatggagcgcagttactacatgcttttataatagtagtatagttaatacag
gcaacttctgttttatctattgcttctgcttcgtagtaaaaattatcgtcgaccaattct
atttgtccaaatgtatctcctgtagttaataagttaagagttactttatgtgtatttctc
aggacttttgtaataattaaagatcctatcaatataatgtataatctagagctgctatta
aacaaaagagtatcatttttcttaagtgaaaaaatttgataattaattttattacgatta
aaaaatagtagccatggattgcaagaaaaaaaatcacctgaactattttcagagaatatt
ggaatacaattttcttgatttaaacaatttttagtcataattactactttataa------
---aaaaatattctaataat-taagaactgaaattaaatgtcatataaattgatgttagt
tgagaatgacattgtattatcaaaggccattcaagaatacttaatagatcaaggcttcaa
tgtttatattgccaataatggattagaagcactgaatctagcttatcaatataatttcga
cttgataatttctgatataatgatgccgctagttaatggctatgaactattagcaaaact
taaaaaaaacaaagcattatccaaaatacctgttatctttttgacagctaaaggaatgac
taaagacagaataaaaggctacgacatggggtgctatggttatttatctaaaccatttga
tcctaatgagttgctttcaattattaacaacttaatcgctagagatgtcttaaaagaagc
ttctttacaaaactcggcaacatcaaaccagcaattaaatcataaaatacgtttgacacc
tagagaaaaaagtattttagatcttgttgttgatggacttactaataaagaaatatctac
tatactgaatactagtgttagaaatgtagaaaaatatgttagtcgactcttacacaaaac
taatatgaagaataggactttgctagtcaagtattctataaataataatctattaaataa
tgagatcaatgaaagggcgaatgacgggactcgaacccgcgaatgatggagccacaaccc
attgccttaaccccttggccacactcgccatattacatacaccactatagcttgtttttt
gatatatcgtctagttttcgagttatatatttatctaaaacaatatttactttgaatttt
-tatgatacatagcaatattagtattcaaattaatggggaaccatttaattgctcaaagc
ccatctcattgcaatttttattaaattatcttgattttaattccgagcgtgtagcagtcg
agctaaatgacattcttttaccagagactctgtttcactcaacttatttgaatgatcaag
ataagctagaggttattaccattgtaggtggtggataatctatttgtacatac-------
-acagttttctatctcaaatgtttcatagatagagataatcttccttgtttttatctac

atgaataataacagctttaatagtgtcacctattttaaattgtgatgatatctgctctaa
ttctttgatattaatctcagaaatatggacaagacctttgagatttcctacctttataaa
taatccgtaaggtgtaatctgatttattatgccttcaataatattaccaacaattaaatt
agatgatgcttgagatattagagctcttctatgactcagtattaaattattagatttctc
ttcgacattgagtaattttagcttaataaatttattattaaattgctcactttttttgaaa
attaccgagatgagaattaggtacaaatccagatataccttcaaggttaattatcattcc
tcccttattgaatccttttatcattacattaagtaaagagtcttcagctaatagttgcct
aattcttttccatgctcttatatattcaaggcgacgaattgataatattaattgtttcga
ctgaatattataatctaataaaaaaaattctcttgtgtcattaatgtttaaagaagtgaa
attatttaaatcttgattactagatacttcttgaataggcaagtatgcagatattggtgt
tcctatatctactaataccccatttaattcaaagctaaatatagttccagctacaatatc
tccaagattcaaatcatacttgtattttttgtaaaacagctgcaaaatttctatgagtaaa
tccttcattatttttttgtcatattgaatagtatgaattaaagagtaactttgtacaatat
attaaggtaaataaatgtttgtcgaattgtatgaaaaaattttaatttgttcggttgtaa
ttaagtatataaacatattaaaatatctaggagacatgaaaagtagacgtagatagctgc
agatttaa-tactatataatctgaggaaagtccgggctccacaaatacaatttatgctgg
agaaatcccagtgtaggtaactgcgaggatagtaccacagaaaaaaaaccgccagaataag
t----tatattagctggtaagggtgcaaaggcaagttaaaagcttaccaaaagtactgca
aagtatttgttaggtaaacccctaaaatggagcaaagctactaaacaaatatttgtgtat
attttatctagtttagttgaaaatactgcatgaagttattagtaacaataactcgagagg
aatagctatcctttacaatatatc-tcgattatattgaaagaacagaacccggcttatgt
agtacttttatgtcttgtattattatgtctattgtcgaatctttaaatttaactgtttg
acaattttttttttgtagctgatgttgtttataatctatttctgaattggttaaagttttt
tcattagatcgatatgtaaatcttagacctatacttttttttctcttttccaattgattca
tctatatattgatcaaataatgttgtactttctaaatctttatcataaaattgatctaat
aattttaataagtatttaatttccatattcttaggtattattaaagagagatctcttatg
atcgatggatacttagagtatggttgaatctgataacttaaataattgaattcattatga
caatcttctaaaatgtttaagttaacttcaagaaaaaaaaagttttgtattgagtcctaat
tcactatatgttgcttgattaagttcactgaatatgccaataataatattgttatagatt
agtgtggcaaattttcctttttgaatgaactttattctaccatcgactatatcttttttt
gtccactcaatttgtctattaatcttttgaaaaaaattttctacgatccctttagcttca
taccaatttaaagaatgcgcagggtgtgaccatgtagatcggatatctagattaccaccc
aatatgatagctagattagtagtttctacaattttgttttgatcattatgaaatactgtt
cctatctcaaagctatctatagtttggttgctttgtttaatattataaagatttgattca
attaatccttccagcaaactgccgcgtaaattagaataatcttgaataagaggattgttt
aaagctatatttcctttggactttattaaagaataatgaactaattcagttaatcctaga

tttcttaaaatacttctacttttgtctatgaattttctctttgtggataaccttttgttg
aattgaatatttggtactgagctcctgaatttatgataaccataaattctagcaatttct
tcaataacatctatttctcgaaaaacatcttctcgtctataatttggaacagtaacttct
atattttccttattttgataaataagatcaaaatttaaagagtgaagaatattttgaatt
tcttcaaagaataaaaaacgtgtttgaccattatacataattggtcctaaaatgtcatga
actttttaattgataaattaatattaagagcatgatcagttatttttctcgacagaaa
gtatctctaatatttccgcatgttaaatctgtgattaaagcaagagcttcaaaatgtgca
tttttccaattatctacatttagcccacgttcttgtctaattgaactttctgttcgaata
ttaagaactcgagaagactttctaactacagattgtttaaatatagcagactctacaagt
atcgattttgtattataatctgtatcaaaatcagagttgcttccaataccagctatacta
gttacatttgtatttacttgagtaactaaaatatttttgtttaactcaatattttcattg
tttaacttaatctgatcattacttccaagaggaaaattgcttcgaattgttagactattt
ttatgattcatattattaattttattcaaatcaataatattaataggctgtccccatttc
aacataatataattgcttatatctgttaacaaatttctgtgtgtaaacccagaagagagc
aaccgatttttttagccaatctggagaatctttaactcttatttcgtctataatggcagca
aaataattgtcacaatttaataaatcgtgattactgattatagt-ctcctttttagagaa
atatctattattgggttgatgtatagctgataacattgaggaccctgttaaagccgagac
ttctcttgatagaccaatcatgcttaacacgtctgacctgttagctgttgatgatatatc
taatatatgatcaatctcacctccaattataatagattcaatagcttcaacttcaaaacc
tgcttgtgttagttgattagttaaattgtctatatctattgttttgatattaccaagttc
tttcaaccaatttaaagaaactttcataattaattgtcatctatcttatagaagtattta
ttttacttatgattaaaaatcaatttaaaaaattcatcaaattaatataagttatttgat
gaatttgcttatacaatatattattgtgatatcttacttttatatggcagttattaagaa
ctagcttttgtaaaagttagcccattatcttgggcatatctttccataaatctcatgaat
ctatcccaatcatcaggacttttaataacatgtacagcttctattgcttgtggtttacca
ttaataaacttggcattaacatctcttgtaataatttgtccctctgtgtccatcaggtac
atacctgtgatttcacctttgtctgccatactagcatctaaaattttgggatttgtaaac
cgaaaagttgctgttcctgtactaccatctcgagaccttgttaaacgtacatctggtaca
acttcttcattgataccttgaataaattgaattgttgccatatcaatcttctttatttgt
atataatttatattggataaaactgatttctagttcttgaatgcttgttgagaaactgtt
ttaaaagcttatagtatcagtagatcaaattttatgtattatataaaactgtaactttc
-ttttttg----------------------------------------------------
------------------------------------------------------------
------------------------------------------------------------
------------------------------------------------------------
------------------------------------------------------------

----------------------atctgttcaaagaactaacttattataagctaatcata
cgaatattagccatataatgttcgtaagt-gaaattaaggggttgtagctcagaaggata
gagcaagcgcctcctaagcgctaggtcagcggttcaagtccgctcaaccctgttaacgaa
ttaagtagtattatttaacttgttctgcaaatcagtattgatctaagttaaagtacttga
cagctaaaacaaaagaatctgagaaatattttcatttaaagaacgaacaattatcttaac
tctaaatatctgataattagtaa-aaaattct----------------------------
----ctttttaaaacacaagatgatg-----aagatatttaaactaataattgaaattaa
gtttttgaagttatagaagccaat--ataatactttctattaacaga-----------tg
cttgcttctattactatctgtttacaaga--aatt----atgagtatcaaaata------
--attttagctcca-------------------------------------ctgac----
------------------------------------------------------------
-----------------------------tactaataaata-------------------
------------------tttattttagtctagacaag------atgaatcttcatggta
taacag------------------------------------------------------
------------------------------------------------------------
------------------------------------------------------------
------------------------------------------------------------
------------------------------------------------------------
--------------agctattctttgtgtcatacttatataaaattaggat-------ac
tgtt--------------------------------------------------------
------------------------------------------------------------
------------------------------------------------------------
------------------------------------------------------------
------------------------------------------------------------
------------------------------------------------------------
---------------ttcattaccataactggggtgggaggattcgaacctgcgaatggc
ggagtcaaagtccgctgccttaccacttggctacaccccaacatgtgaacattataatag
cagtcgtataccctgccatgtcaatagactattgtctttatttaataattttgctgctc
ttgtagaataaaatcttttttttcaaataagatacgctatgtttaagttgcacgactgtt
atgttttcttgcttttgactgaacagccacttgatagtaattattccttctgcaatttcg
ttacttcctataataatacaagctacagctctttttttatttgcttgcttaatctgctta
ctaaaattacttgagctaacatcgatttctattttaaaaaactgctgatgtaaaaaacgc
attatttgcataccagtttcgttagcttttgtaccttgtgtagcaatataaaaatcgata
gatttattaggcaagtctatattatcttttgcaatcagtaataggcgttctaatcctatt
gcacatccaacagctggtgttgctttgccacctaattggtgaactaagctgtcatatctt
ccaccaccacatatagtatcttgcccttttgatgtcaatgtcttaatttcgaaagcagta

tcattataataatctaatcctcgaactaacttattatttatattgtagggaatattaagt
aattttagataattgcaaacgtcaccaaagtgttgttgtgattccaaacttaaaaaatca
gaaattttggagcctctgtcagtatttttgtgtattagaatcttttgaatctaaaatt
ctaattggattactagttaatctttttgtgagtcagtgtctaaatcatcatggtacttt
gtaagataatctcgtaatttaacttgataaatactacgatcctctacttttccaattgaa
ttcagatcaagtttcaagttgtgcagattgagattgttaaatatgctcatagctaaatgt
ataacttcactatctgctcttgcatctaagctaccaataaactcaatgccaagttggtga
aattgtcgttgtcgtccactttgtggtctttcgtacctgaacattggaccgctataccat
aatctttgtaaactatggtggtaactcattttattttcaataaaagatctcacaataccg
gcagtaccttctggccttaaggtaatatctctattgctgcgatcatgaaaccgatacatt
tctttatttacaatatcagtatcttcaccaataccccctatcatataattcactattttca
aaaataggtgttctaatttcttggtaatttgcacattctaataatttggaaattttatta
tgtataaactgccaatattgaatttcatcgggaagaatatcttttgtccctctaatagcc
tgaatttttgccatgataatttttgattaccttcttaatatgtctgcattaaatagcag
ttgtagactgttattcagtcatgtattatacattag--aacatactgatg-taatttata
aagtacgggcaaggagggattcgaacccccgacaccatggttcgtagccatgtgctctaa
tccactgagctacaagcccacttaataagtaccttaactatatcattgttcactaatgta
tacaactatattcttttt------atcttacatgtgaaattttgtaaaagtaaaaaatt
at----gtttgtttttacaattggaactattcatatataatgcttaatttattataact
ttgtaaa--aatatctataagtttaattgtatgattctattatattggtaatcttgttat
aattttgcgctcttatctttcaattgg------ctagcatctaattaacttatatttcaa
ataaaaaaatcttatctaaaatgagtaaacaaattctatatcaggatgatgccagaaaag
cattagaaaaaggcatggatattttaacagaagctgtttctgtgactttaggaccaaaag
gaagaaatgttgtcttagaaaaaaaattcggtgcccctcaaattattaatgatggtgtaa
cgattgcaaaggaaattagtttagaaaatcatattgaaaataccggagtcgcactgatta
gacaagcagcatctaaaacaaatgatgtagctggtgatggtacaacaacagctacagtac
tggcttcagcaattgttaagcaaggaatgagaaatgttgcagcaggttcgaatccgatgg
ctattaaaaaaggtattgaaaaagcaacaaattttgttgttagtaaaatagctgagtatg
ctaagccagtagaagatacaaaagctattatacaagttgcttctttgtcatcaggaaatg
atatagaggtaggtaaaatgatagcgaatgctatagaaaaagttggcagagaaggagtta
tttctttagaagaagggaaatcaactaatacgattcttgagatcacagaaggaatgcagt
ttgaaaaaggctttatttctccttatttcgttacagatacagaacgaatggaagttcttc
aagaaaatccttttattttatttacagacaaaaaaattactttggtgcagcaagaacttg
tgccattgcttgagcaaattgcaaaaacatctcggcctttattaataatagctgaagaca
tcgagaaagaagcattagccacaattgtagttaataaattaagagggattttgaatgttg
tagcggttagagctcctggattcggtgatagaagaaaatctttacttgaagatatgagta

tcttaacgaacggacaagtaattactgaagatgcaggtttatcacttgatacagttcaat
tagatatgttaggaaaagcccgaagagttattgttactaaagactcgacaacaataattg
cagatggtcatgaaattaaagttaaatcaagatgtgagcaaattaagcggcaaatagaga
cgagtgactctttatatgaaagagaaaaattgcaagaacgattagcaaagctttctggag
gagttgccgtcattaaagttggtgcagctacagaaacagagatgaaagataaaaaactaa
gactagaagatgcaattaatgcaacaaaagcagcaattgaagaaggtattgtaccaggag
gaggagctactaatgtccatatctctagtgaattatttacatgggctaaaaacaatttag
ttgaagatgaattaattggtgctttaatagttgaacgagctgtgacctatccgttaagac
gaatagcttttaatgcaggtgataatggagcagtaatagtggaaaaagttaaaagtcacg
attttcatataggctatgatgcagcaaacggtaatattgtaaatatgtacgatagaggta
ttatagatccagcgaaagtagctagatcagctttgcaaaacgcagcttctattgcagcaa
tggttttaactactgaatgtattgtagttgataaagtcgatgattgagctata------a
gataatagataatctcaaacaaagaataatttctaattatagaaattattctttgtttca
attaatctaattaattgatcaaatgtataatttttataataaagccaaagtttaaacacg
tcttctttttcaactattaagtagagaagatataagcccgtaatacctaaattatcaatt
aaaatattattctcatatatagatttttctaaagatctaaaaaatgacattttgaaaaa
taataccgaaatcttctcttatagttatttgctaaatcattactgtctaaagattgattt
ccaattgaattcagaacctttaacagaatacttttaaatctttatctgactttatttga
gaatcaataataatttgattcaattgtatgtcaaaactaagattaaaaatattacttgtt
aaagatatttgattttgaatatctagagattttatagccaagaagattaaacttttattg
ttcattgtaaaatataagacaattgatgatgtattctttaaatacta-aatttatcttct
tgtgcttctatcgtttttttgt-aaaataattaatgttttaatttttttagttttagttact
acccgaaaaaataaattctggaaatctagcttgagctttagctagagagtgtttttttccc
ttcttcttgccagtaattgataacttctgttgttttgtttaataactcaattctgtctct
ttcagcaatccacggcttggattctagttctgcttttaattcttcccaagcatcattcct
tggccagaaaaaataagaagtaagtggactagtgccatttccaacaatctgatcaatagc
aatagcaatattattttctaaccaaagtactttagtgtaaattttgacaagtttttatc
tccattattaatattaagtattgg----tacagtagaatactgtactagttatttttttc
agcaacaatttgagctttgtttaaaatgtttttttaccgttttagtagcttgtgcaccttg
agataatcgttgcaaaatttttgaaatgttaacacgagtttcattatttataggattata
aaatcctaattcttctatagctttaccatcgcgtttatttctgctatccattacaacaat
tctatagcttggttgtttttttcttccgtatcgctttaaccttaacttgaccatattagt
cacctcgcttatttttt--atatctttattaaattatatcttacgacattatgcaataat
tcaaacaattaagctgattcaagtctaacattattaaatattaccattaaacattgtaga
aaaataatacctaacataggtgacatgtccattccaaacatgggaggaatgcttcctctg
aatagttttaagtatggatcagtaattctatttaatgaacagaatggttcattgtaccaa

tttacagttggaaaccatgccaatgataatttaagtaaaattaaaattaaataaatctca
gaaaaattagctatcgatccaagcaataaatttaatgtaccaggaagagtgttcatacgt
attttataattattatatacttcgttatactaatagtttgattatatattataatcgatg
taaataaat-cagctaaacaaaaaacttatccttaaaatcatcttctcttgagcactttt
ctttatctacattagtatttacttttgtcgtataaatttccagtttagaccaaatctgtc
ctaactgatttaattgagatgttccacattctaaacaggctatttttatattgctcaaaa
cagcaccttttttttataaaatcgttcaataatggcgtaatagttgcttcatccaaaaata
aatctaagacaagtatttttgtaaatgaatttaattgatctgggacattgttaataatat
cattataattaactaaaactatattagctgtaggaagtaaagatcgtgctccttccgcta
agataaagccataaggcataataattacaataatatatttataatgactgctaataagac
ttatagtggtattagcttttaattcgattgtttctgttactagccactctctcattattt
cgtaagtaatccatttgcctagttctgagcaagcagttcttaaaatcgtacctggattat
tgtcgtttttgagaatccctgaccaatgttgtattaatggatgggatgctatattaattt
ggagttgcatgttttgttaaatatatttaacatttaatttgttgattgaatcttcaattc
ctatttgttgagtattattaaaatagcat----atatattaaagaaaataattat-aaaa
aatgaaaaaaagtctttggctgtggggttttacagatagtgctgaaacttggaatggtag
atttgcaatgattggttttatatcagttatttttatcgaagtagttacaggacaaggatt
attatatttaataggtatgatgtcataataaaaatccaaaaaagttcttgtaattttta
taagaactttttttgg-atttttataaatctaatctagcggtctcatagatagaacaggtt
cattctctctattgataccttttttcaaatccagcagcagcagctcttgctctgcctgcat
gccaaagatgtccaataaataggaagaatcctaggaagaaatgagatgttgttaaccaag
atctaggagatacatagttaactgaattaatttctgtagctacaccaccaacggaattca
aagaacctagtggtgcgtgagtcatatattctgcggcacgcctttcctgccacggttgaa
tatcgttcttgattttatttaaatcaagaccatttggtcctctaagaggctcaacccaag
gagctcgcaagtcccaaaatctcatggtttcaccaccaaaaataatttctccactaggag
atctcattaggtatttacctaatccagtaggtccttgagaagatgcaacattcgcaccta
gtctttgatctctaactaagaaagtaaaagcttgagcttgtgaagcttcaggaccagttg
gtccatagaattcactaggataagctgtgttgttataccaaacaaaattagaagccgtaa
gccccatgattgataaagcacctagactataagataagtaagcctcaccagaccatacaa
aagctcttctagcccaagcaaaaggttttgttagtatatgccagattccgccagcaatac
aaataatgccaatccagacatgaccacctacaagatcttccatgttatttacacttacga
tccatccatcaccgccaaacggagactttaagacataaccaaagataactaggggattaa
gtgtaggattgctaacaaatctaacatctccaccacctggagcccaagtgtcgtataccc
caccaataaataaagctttaataactaacagaaaagcgccaatgcctaataacactaagt
gtataccaagtattgttgtcattttgtttttatcccgccaatcgtatccgaagaaaggaa
atgattcttcaagagtatcagggccaatcagagaatgataaagacctccaaagccaagaa

cagctgaagaaattaaatgcactacgcctactacaaagtatgggtatgtattgaaaatct
cgccgcctggaccaacaccccagcctagtgtagctaagtgcggaattagaataaaccctt
gttcgtacaaaggcttctcgggaacaaaatgtgctacctcaaaaagagtcatggcaccag
tccaaaagaccattatacctgcatgagcaacatgagcaccaagcaatttgccagaaacgt
taattaagcgtgcattgccagaccaccaggcaaacccggtagactcaatgtctctaccgc
caacaccaacagttgtattaaagggcgtttccacgtggtaaaacctcctcagggaatata
aagttttcatgaggttgatcttgagcagccatccaagagcgaataccttcgttcaatagg
atatttttagtataaaaagtttcaaactcaggatcttccgcagctctcaactcttgagaa
acaaaatcgtaagctcttaaatttagagctagtccaacaattccaaatgcgcttgtccat
aatccagttactggtacaaataacatgaaaaaatgtagccaacgtttattagaaaaagca
acaccaaaaatctgtgaccagaatctattagctgttaccatcgaataagtttcttcagat
tgtgttggagtaaatgcacgaaaagtatctgcagcatcaccatcttcgaataaggtattc
tgtacagttgcaccgtgaatagcacatagtaaagcaccacctaagattcctgcaacgccc
atcatatggaacggattcagggtccagttatgaaatccttgcaaaaatagcaaaaatcta
aaaattgcagcaactccaaggcttggggcaaagaaccagctcgcttggcctagagggtac
attagaaagacggaaacgaatactgcaattggtccagaaaatgcaatagcgttgtatggt
ctaagaccaactagcctagcaatttcaaattgtcttaaacagaatccaattagtccaaat
gatccatgtaaagctataaatgcccacagaccaccaatttggcaccagcgagtaaaatct
ccttgagcttcaggtccccaaaggaaaagaagtgaatgtcccatgctgtttgctggggta
gaaacagccgcagttaaaaaattgcatccttctagatatgaactggctagcccatgagta
taccaagaagtaacaaaagtagtcccagttagccagccacctacagcaaggtaagcgcaa
ggaaacagaagtagtccagaccaacctacgaatacaaatcggtctctttttagccaatcg
tctacaagatcaaaaccaccacgagtttttttcttgtccaattgctatggtcataattta
atctccagagcaaatttaataagtaatttgtttaaataaaca---ctttagatatactct
aaaattttactttctttacggtcaatataattataagtcaaaagtaaccaaaattcacg
cttttaataaca-tttttttagctcattaagttaa-ttttttaatatataaacagtaata
gagttattgttt-aacttacaatactcacaatttattcatttctataatcctttgaaaaa
gatatcctgttcctcgtgctgttaaaatcaagtcgggattacttggatcatcttctagtt
tagctctcaatcgtgaaatatgaacatcaacaactctcgtgtcaacatgccgttcgggag
tatagccccaaacttcttgcaagatagaagctctagaaaaaggttctccagccttactaa
ctaaaagctcgagaagactgaattccatacctgttaatcgaactctttcattatttttat
aaacttgtcttttgttagtatctatttttaaaaatccaatactaataatcccagagtttg
ggactccaagactagtggtgattttatcagctcttcgaagaacagaacgaatgcgagctt
ctagctccttaggagaaaatggtttaacgacatagtcatcagcacctatttctaatccgg
taattctatcgcagacttcacctaacgctgttagcattataataggaacatctgattctt
ttcgaagttcttgacaaacaccgtaaccatctagtttaggcatcataacatctaaaacaa

caaggctaggatactcttttctaaacacgagtaaagcttcttctccatttgaagcagtga
taacttcataacctattattgttagtcttgtttctaaaattcttcttatgctggcctcat
cgtcaacaacgagaatcttttctttctggttctccaa-tttcttatttactcctgtaact
caccgatcatgattaccataaccttaaattttagcgctatgagtatctttattcacttt
agaa-----tttagttattataaattttgcaagtctagtttacattaattagtaccagt-
-caatattttgt-----------tttaagttatgac---------tattttaatccata
agttgactctagtttgcagaattcaatatatacata-ttttctacccaagtaaattgtat
tctattttctaaataaaaaagggttgacaaaccagttgatagaaagtattctacattac
tataag----ttaattccaacaaagaagttttaa-----atcttctttcggagtaatgaa
tactttaat
>Pyr_2_3_4_7_8_9
tatatgata---aattaagacatctgttatgtgagaatttctaaaatatacaaaactttt
tatttaatatagaaaaagtaaattaaaataagttgaacatactttgctctaatctaaacc
tcttaattgag------attctaattagagacaaaagtatttacttattttgtatctata
ggaataaaaatgaaacttgcagtttatggtaaaggtggtataggtaaatctacaactagt
tgcaatatttctgtagcactttcaaaaagaggtaaaaaagttctacaaattggctgcgac
cctaaacatgatagtacatttacgctaacaggtttttaatcccaacaattatcgatact
cttcaatccaaggattaccactatgaggacgtttggcctgaagatgtaatctataaggga
tatggcggtgtagactgtgttgaagctggtggaccgccggctggcgctggttgcggaggc
tacgttgtaggtgaaacagttaaacttttaaaagaattaaatgcttttgatgaatacgat
attatcttatttgatgttttaggtgatgttgtatgtggaggttttgcagctccattaaat
tatgcagactactgcttaatcattacagacaatggatttgatgctttatttgcagctaat
agaatagcagcttcagtacgagaaaaagctagaacgcactctctgagattagctggactt
gttggtaatagaacagataaaagagatctaattgataaatatatagattgtgttccaatg
ccagtattagaagtcttgcccttgattgaagatattagagtgtccagagtaaaaggtaaa
actttatttgaaatggcagaaattgataaggatttagcatatgtatgcgattactatttg
aatattgcagatcagttgattacaaggccagaaggtgtagttcctaaagaatctccggat
agagaattatttagtcttttatctgatttttacttaaatcctaaatcaaaggtaggacaa
gaaaaagtagatcaagaagaattagatttgatgatagtgtaaataatattttcagcataa
tataacaaggaataagataatgtctacagctcaatcagatgctcttacttttgaatgtga
aacaggtaattatcatactttttgtcccatcagttgcgtttcttggttatatcaaaaaat
tgaagacagtttttttcttagttatagggactaaaacttgtggttatttcttgcaaaatgc
aatgggagtaatgatttttgctgaaccgagatatgctatggcagaattagaggaaggaga
tatttcagctaagctaaatgattatggtgaactccgtagattatgcttacaaataaaaaa
agatagaaatcctagtgttatattttggattggaacatgtacaaccgaaattataaaaat
ggatctggaaggaattgctccaaaattggaagcagaaattcgtgttcctatagtagttgc

tagagctaatggtttagattatgcctttacccaaggtgaagatacagttctagcagctat
ggctcaaaggtgtccattaaatttaaagaatcaatcagataacacgagtcttaaaacttc
tcctcatattcctttagttttatttggatctcttccggatccagttgtaacccagttaac
tatggaattaaaaaaacaaggtatttttgtttctggttggttgccatcaaagaggtatac
tgaattaccggttataaaagaaggatattatgttgcgggagttaatccatttcttagtcg
cacagctactacactaatgagacgccgaaaaacaaaactaattggtgcaccatttccaat
aggtccagatggtactcgagcctggattgaaaaaatctgttcagtaatgaatgtaaagcc
tattggattagaagatagagaaaaagcaatttgggcttctttagaagactatatttcttt
aatccgaggtaaatcagtttttttatgggtgataatttactagaagtatctcttgcaag
atttttaactagatgtggaatgactgtatatgaaattggtattccttatatggataaacg
ctatcaagcagcagagctagctttattaaaagctacatgtgataaaatgaacgttatgat
gccaacaattgtagaaaaaccagataattataatcaagtagaccgaatccgtgatctgaa
acccgatttagttatcactggtatggctcatgcaaatcctttagaagccagaggtattaa
tacaaaatggtcagtagaattcacatttgcacaaattcatggctttaccaatgcaagaga
tattcttgaactcgtcacaagaccattacggcgaaatcttagtttgtcagaattaggctg
ggatgtttatagcaagcaaagctagtattcctc----tttatgacttacatggtataaaa
gccttgtaagtct--tttattttatgttaaagaagcatttcttgctctgccatcaccagc
ccaagcttgtaaactttctacttgttctttgtctgtaaatgctaaaggcacaaattgttt
gatagctattttaatatcttcagtactaaattctctctcttcactaaacgccgtatgcat
actttcaataattgcttgctcaatttcagcgccagaaaatttattacataataaactcaa
ttcatgaatatcatatttttgccacgatcttggtctaacttttgataaatgtatcttaaa
aattgattctctctcttgacgattaggtaaatctaagaaaaaaatttcatcgaacctgcc
tttccttaacatttcggaaggtaaactttgaattttattagctgttgcgacaacaaatac
tggagctttttctctgatagccaggtaataaatgtaccaaatactctggcactagtacc
actatcaccttgactatgtaaaccagaaaaagctttatctatttcatcaatccataatat
gcaaggtgataatccttcagagatatttatcatttctctcattttgattccgattcacc
aactagtccaccaaacagttttcccatgtcaagacgtagcagaggcagcatccaatcgct
tgcaatagctttcgcggttaaggatttacccgtcccctgtataccaactagcaataaacc
tttgggcgaaggaataccgtaatttaaactttgcttcgaaaatgagcgagaccttttttg
tagccattgttttaatacatctaagccaccaatatccctatttactttactataaggata
aaattctaatagatgcgtttgattaataatttgccttttttcttcaataataataggtaa
gctccgagagtctatctgattatattgtgcaataattttagtaatcactttgcggattct
atctatcgataatccttggcaggattttgttatattattaacgagctctgaattcaaatt
aagattcaaagctttactcaatcttatgatctctttttgatttctagaagactaggtaa
gggcaaatctattactgttatgatatcattcagagcaaaaggtatattgattttgcaaga
aataataataatgtgcttagattgcgttttgattattttagctaaattgcgaagtttcct

aattaatactatttcgtttaggaaagagtcaaaatcttttaaaagaaaaagatttaaaga
ttcattattcaagttttcaataaattctaatgctaataaaggatttctttttgcataacc
attatcgctaggattgttagtgtatccatctacaaagtcccaggaatatacttgctgact
gtttgaacagtttaagctgtgttttataatatactctagtctatcttcttctctagtatt
tattacaataataggatatcgagattttaaaagtaatcgtaaatcttgagtgaaattcat
gtcaatatttgtttccatcagacttttgtacttcaccaattgaagaaagactaaataaga
ttatctaatattccttggatatattcaaaagtc------atctctctattttctaaatta
ttgctattactaatttaacttaccatgatttttttcttcctttcttcgtcgactattt
aatatagagcgtccacatggtgttttcattcttgctctgaaaccagaaactctaattttt
tttcttttagatccttgtagtgttctttagtcatgatgtctcctctgttgtaaaagatt
aaatactattttatcatcttttatatatctttatgatatagatatgtattaatatcaaag
tt-aacagcaataatttatattaatcaagaactatttatataatattaaacattaaatat
----------taactttatgatattggctttgcccatttttatttaagtattttgacca
tctttttgctaatcttaaactggttaatttttcagcagctaaaaacgatttgttattag
aatctcaatttaagtattttgttgataaaagccaaaatcgtaaattagaacccgacgaaa
gttttgcttttgcaaaagtatgtgtagctaaaaaatatttttctaaagctattattgaag
gtcaattagctttaaaaaactatcgagacctgaatatcttagataataatatagtaattg
ctaatttgtataatatgctaggttttatttattttgaagcaggccaaacaagttttgcaa
agaacttttatgagcaagccttacaaataaaccctaattatgttgtagctttaaataacc
tggcaaaaatttatgaagaggtaaaagatttaaaaaaagccgaatctctatacgataaag
tgttaactcttaacttaaataataaaactgctaatagaaggaaagatttatagcaaaaa
cta---------aaaacatctaatactatttgtaatcgggatagcaggatttgaacctgc
gacatcctgctcccaaagcaggcgcgctaccaaactgcgctatatcccgtaagacataat
tattactatataccttttctctaattatgtctactctttctgtattttatctaggatacc
aaaacatgcctttaactccatctggatctttgataaatcctaatttttataaaaactaa
caacatcaggttctgcgaatagtgtaatagtgctaatttctgcctgtctcaattgctgta
ttaattggtgtatgactacttttcctaaacctaaaccttggaaatctggatgaataacta
cgtcccagatagttgcattaaatccgttgtctgatgttgctctagcaaatcctacaagtt
tactatttgcatcttttttttgtattaaagaaataataatagaactatttttaatgcga
tctttactttttttagaggtcttttaacccatccgactgaatcacaaagctgctccaatt
catataaattaatattcttattgctactcagatagatatctttaaattcgattttatcgc
aagttttatctaaaacaataagtttttgaaatttttttcagagctagcattgatattgg
agttttgaaaaaaatttttccagaagatcataatgattatgatataatta--aataaata
tatatattacaatattgctttaacaaacaatatattaatatatgctcaaatttttacatg
aatctattaagatagaagattgttaaaatatagagaaataaaaatttttgtaaaatcttt
caataaaatgatataaagagtgctttttaagactaacatctattattataaataaagtct

cgaaattgagatgaaataacacattattctatgtctaatgccttttaaaggagttatttg
tgaaaaaatctatgtgtctaacctgtttacttgctttactaattatgagcaacccaataa
tagctaacgcagaagtagctggattggtgccctgcaaagattctgcggcattcaataagc
gtatggtaaatagtgtgaaaaaacttcaagccagattagctaaatatgatgcagacacgc
caccagcgttagctttaaataagcaaatagagaaaactaaaactagatttgcaacttatg
gtcgagcaggtttgctgtgtggtactgatggattgccacatttaatttctgatggtcgat
ggagtagagccggggactttgtttttccaggactattattcttgtatattacaggatgga
ttggctgggtaggtagaggatatcttttatctgttgctaagactagtaagccaacagaaa
aggaaatcattttagatgtgccattagctatcaaatttatgtcatccggatttgcatggc
cgctagcggcttggcaagaatttagtagtggacaattaattgcttccaatgacgatatta
ctgtttcaccccgttagtaaaaaatttatatgaataataattttaccaaatacttatcaa
cagcacctgtaattggtgtattgtggatgacgtttacagccggatttattatagaattga
accgctttttcccagatgtcttatacttttactt----ataaataa--gatcgtataaa-
--aaactacataaatatagagagatactattatttgtgtagtttta---atatataaca
taattattta-tttttaaaaatgtttatcttttcgattattttaatattagatcttacat
aataaaata-atatcaatattatgagcttagtaacccaaattatagttaatgccgatgat
gaattaagatatcctacaattggagaattgcagtcaattcaagactacttaactacagga
agcaatagaattagaattgctactattattagagataaagaaaaggagattatacagaaa
gctagtaagcaaatttttcagttacatccagaatatatagctccaggtggtaatgcagca
ggttctaggaaaagatcgttatgcttacgtgattatggttggtatctcagactaattaca
tacggagtattagctggggataaagattctatagaaacaattggtattataggagtaaga
gaaatgtacaattctttaggtgtacctataattggaatgttagatgcaatccagtgttta
aaggaagcatctttagaaatgcttggtcaagatgatattagaatcatttctccttatttt
gattatataattcgtggaatgtcataaatcatagcttcaattaattagttgaataaaagc
ttttataatgttataattacaataagcccgagagcatataattgtaaaaactaaatcttg
tcagaaccggaaggtagcagcaataatgtttcaatacaaaagttatga-ttttcgggtgt
ttattgtttttatctaatagtctcgacttttaattagtatgctatgtataaatctgtata
tgacatgtaatccatcaaaccttccagcaaaaataatttaataatttgatattaatacgt
aaaacacctatac-tcttttataaaatttctttatatctcatctaaacaatggaaactta
acttaatgggtgttcatattttatcaacaggttcgtctgtcccaaatttttctgtagaga
atcaacaatttgaagacatgatcgaaacttccgaccattggatttcaacaagaacaggaa
taaaaaaaagacatctagctccatcttctacttctttaactaaattagcggcagaagctg
caaacaaagccttatatgcagctaacttaaaacctactgagatcagtttaattattttag
ctacgtctacgcctgatgatttatttggtagtgctagccagttgcaagcggaaataggtg
caacaacatcagtagcttttgatattacggctgcctgctccggttttattgttgctttag
taacagcagctcagttcattcaaactggttcctatgacaatattttagttgttggagcag

acacaatgtctagatggattaattggtcagatagaactacctgtattttatttggtgatg
gtgctggagcagtagtattggggcaaagcctcaaaaatagtattttaggctttaagttat
gtacagacggtcagctaaacagtcatttacaattaatgaataaacctgtaaataatcaaa
aatttggtggtacagaaattcctcatggaaactataattccataacaatgaatggcaagg
aagtgtacaagttcgctgtatttcaagttccaacagtaattagacaatgtttgaataatt
taaacatttcaatagatgaagttgattggtttatattgcatcaagcaaacactagaatca
tagaagcaattgcgagcagattatcagtacctttttctaaaatgattacgaacttagagc
attatggaaatacatctgcagcgtcaatccctttagcgttagatgaagctattcaatcca
ataaaattcaaccaggtcaaattattgttttatctggttttggagcaggcttgacttggg
gagcaattgtcttgaagtggtgat---ttatatattgcggatgacgagactcgaactcgt
aaagctttcgctacacacccctcaagcgtgcgtgtataccaatttcaccacatccgcatt
ttttaccatgaata-ttaaaaatttataatatttatctaaatattcatataa--tgtgta
tatatatatacattcttgtcagaaactctcttatattttattttaagtgataaaattgag
t----ttagtatagatagtaaaattttattttaaaatcataa----aaataaatatataa
ggaaaataaaatatgacaccatctttatcaagtttttttgaatagtcttattcttggggca
gtaattgtagtcgttcccataactttggctcttttatttgttagtcaaaaagacaggaca
atccggtcctaaaaataacttagaaagatatggagaagcaatttaagatgaataatgaaa
tatttttcatactaataagctctccatatattgctatact--ttagaagaatttttattt
ttttattttgtaacatcctgtaaagtttattccgttctttctttttccc-atttactctc
ttataattaatgatacaaatgtcagg-ttaataatag---tttttttattctgtatctttc
---------------attttatatacgtattattatgttcaattaataaagaaaagctca
tgtctttgtctaattggcctctcaaaaaagaaaattctgaagcatataatattaagaatt
caaaacaaatcacaattcctgatggtttatggataaaatgcttcgactgtggtctattaa
tgtattctaaagtactgaagagaaatttaaaagtttgccctcaatgtagttatcattttc
aagcttctagtaacgaaagaattgatcaattgatagaccaaggtagttggcaaccaatgg
atgttcacttgatctctacagatccattaggctttaaagaccaaaagctttatagtcaaa
ggttaaaagatactgctttcaagactggcctgcaagacgcagttcaaacaggtactggga
ctatgcaaggtaaaaaagtatgcttaggtattatggattttagattcatgggggaagca
tgggatctgttgtaggcgaaaaactaacaagactgctagaaaaagcaactcaagaaaagt
tgcctgcaattatactttgtgcatcaggcggagctagaatgcaagaaggtatgttgagct
taatgcaaatggcaaaaatttcttctgctctagaaatgcataaaaaagaaaatctactat
atatatctgttttaacctctcccacaacaggggggtgtaacagctagttttgctatgcttg
gagacttaattattgcagagccaaaagctcttattgcatttgctggtagacgagttatag
aacaaacaatcaaagaagacttgccagataattttcaaagttcagaatatttatttgaac
atggtttcctagatttaattgtatccagaactcagcttagatcaaagttaatacaaattt
tgtccttacataatcatagtaagtaatgaattgaatattgactataatatcaaaatgcta

gtaaaaattaagatattaagaaattttcactaaagatatttgtagactaactagaattac
aattactattgtagaaaaatttgtcttaatgcca-ttttttcagtcttttaacaaattg-
tttttatgactttaaataattgttagtttaatacaaatttataaagagaata-----cat
actaagctaaatatagtaatattatgttttcctataatactgacctatcaaatttcgagg
aaatttatgcttaaaagatcttcttggcttgcggctttattgggactattaacagtagtt
tctacaagtacgcatacatatgccatagagttagacgaggcaacaagaactgttccatta
gaatcttctggcagaactgtaattcttacaccagaacaagttaaaagaggcaagcgatta
tttaataattcttgtgctatttgccataatggtggtatcacgaaaacaaatccgaatatt
ggacttgacccagaatctttaggattagctacgccacaaagagataccattgaaggacta
gttgactatatgaaagatccgactagttatgatggtgcagagtcaatcgcagaattgcat
ccaagtattaaaagtgctgaaatttttcctaaaatgcgcaatctaaccgatgaagaccta
tttacaatcgcaggtcatatcttacttcaacctaaaattgtttctgaaaagtggggcgga
ggaaaaatttactattagaacttcaaagacctaaaccttgtgaatactagtttatgta-t
ttgttgcttaagtgtgattattttgttagatcatgatatatattaaagatggagtttgtt
gacagcaagttactttttacccgaacttgtttacactattcgtctagttattaaaaagga
gacgttaaattgaagaagaagctttcagttcttttcactgtttttagtttttttgtaata
ggtttcgcacaaattgcttttgctgcagatctagataatggagaaaaagtttttttctgct
aattgtgcagcatgtcatgctggcggtaataacgccattatgccagataaaaccttaaaa
aaagatgtacttgaagctaatagtatgaatactattgatgctattacttatcaagtacaa
aatggtaaaaatgccatgcctgctttcggaggtagactggttgatgaagatattgaagat
gcagcaaattatgtattatctcaatctgaaaaaggttggtaattatacttgattttatcc
tgtattaaagaatagacaatctatttagttgttcattagattgtctattctttgtttatg
---ctattatataaagatatttacacaatatt----ttatgatgaaaagaatacccgcaa
ttcttgtactagaagacggtgcatattataaaggatggtcattccagcaagataaacaag
agattactattggtgaagtagtttttaatactggaatgacaggatatcaagaaataatca
cagatcctagttacttccatcaaattgtcgcttttacctacccggaaattgggaatacag
gtattaataatcaagatattgaatctcacagtattagtattaaaggacttattgcaaaaa
atatttgtaaaatttcaagcagctggagagagcagcaatctttagttaagtatttaagta
gtaataatattccttttattttcggaatagatacaaggtctttaacccaatacttgcgtc
aatttggtacaatgaacggttgtatctctactgataatttaaatcatagttacttaaaac
agaaaatttgtgagattccaagtatgcaaggtttagatttaatcccgcatgtaactacaa
gaaatgtttacccctgggatgaaaaaagttttccgaattggtatttgacagataacatta
gagtacaccgagttattcagttgaaagttattgttatagattttggagttaaactaaata
tactcagaagactagctacacttggatgtcagataacggttgtgcctgcccacactcctt
taaaagatattttggcttaccagcctgatggtatattactctctaatggtccaggagatc
catcagcagtacattacggcatccagacagttacaaatttactagattacaatgtgccta

tatttgggatttgtatggggcatcaaattttaaatttagctcttaaagctaaaactttca
aacttaaatttggccatagaggtattaaccatccatccggattgaaccagcaagttgaaa
taactagtcaaaatcatggctttgcagttgaattgacttcggtttttgaatctcctgtaa
gagtgactcatttaatctaaatgacactactattgcaggaactggacataatcaaagtc
cttatttttctgtgcaatatcatccagaatcgagcccaggccctcatgatgctgattatc
tattcgaaagttttatagaaataatgacaaagtccaagaataaagttag-----------
ttagtaattttcccatgctttatgagtgaataaagcggatagtacttgtactcctcgtaa
ttgattaaagagaggcaaatgtcctctggggccatgagaattccattgaaattcctcagg
atatctgcaaggaattctatcaacttcccagccgatttttgccagaattttccccaatc
ttttccaacactaagccatatttgtctttgaacaaatagcccaaatttacctttgagtg
tgtatgccataacttatcaatagtttgtaaatcttgagcaggtatttttttatatctgt
aaaatatagccaattacgagtttgtgcatttacaccagctagttgaataagcttttgttg
agttagctgatctgctttgagcaaatcccgatgagtcagtaacatttgtaaatctttgta
attcatctgttgagcagagcgcaatggaacaatgccatcgggacataagttagaagcaaa
tttcactatttcttgatttttactgttaagtaacttttcataaattaaaccgtcaacaca
attacttttatagttaggcccagtaatcctctcaaaaaatagatctgctaaatcttttaa
ctcgatagagtcgttgctatttatattctcaataatttcaagttgttgcttga---catt
attagacttagtattcttattcaattctaaaagctgggctcggatttgatttggcatctg
atttgtttaattagttttaatagtc----ataaggtcttagagagaattagattgttttt
tgaaattagaagagcgccgcagtatttctaaagaagttcgcataatttgattcaataagt
tttgtcctgcataaattataatttgaccaattaaaaaaattgtgctttttatttttaggaa
caaaaaaatcttgtacctcaagcagaaaagtaattattatttgcattggtgacaacatct
gtaattcattcgatctgcaagcatatatatactgacagtttaagccttcttgataaaatc
cccatactttgtaacgactttcgtaaattgctcgtggtcgctcaagatagtttttaatta
tagtattccaaattaggttattctttaatttttctaatttctatcagatagaaattttta
cattacagagataacttgcatttgtctcatttacagtattaaaattttgaactaacatat
gagcaataatgttgcttagctgaattaaataattttcaagaaaaatttcgacctgtttaa
ctggcatattgtaagtaaagagattaaaagttttattcacagtttgagatgacccaaaaa
gtagctgaataattaaaatttgtagtagcattttatagtcgttcagtagatagttcaaat
catgaatatcttggtaaatgtcaatacagtccagttcatataacttacaaaatcttttga
tacacctatggaataaatcaattaaaatattttcggtcaagtttgtaatatcatttacat
ctaaatttgaattataaatttctaaaagtatcttttccagttcagctaatatgattttta
ataagtttctttttacttcagtgcgaaaaacatctaaaattaacacttcctgagagcaat
tagttaatctcttattaatctttatagaagttctaactaatagctctgctacctcttgat
tgagtgttggtccctgagagcttggccaataattattcacgttataccatagtaagaaat
attgaatttaatgttatacgataactttttaatatactgaaaagcaagttaattactcta

tagcaactag-------tttttacagatcttaatagttatttatcaaataaagttatatg
atagtagcgattgagctaattttattaaaattta----taataatgacaacttattactt
cgcccttgcgagtcaaaattttctattagtacaagaaccactagaagaagtttttagaga
gagagttaactactatcagtcaaataataaagcaattgattttggctaataccaaaccc
ttcttttctagagaaaccagaaatgatttcatttaaaaaccttgtacctaaagacgctgt
agctataatctctactaatccaatatttattaattggttaaagctaagaattggctatat
ctgtattggacaatttgaagataacctacaactttctgaagaatcgttaaatattactgt
tttaacagacaaaatttaacaatgcttttattatgactagtgttaaattattttggttaa
gttataattatagtcttttagtattta-gactcttatagtttatcacttagtattttatg
ctgagatggagtgacaaatagtgtaagtatttcttggctaaacgtttcagttgccttaga
tttatatctatttggattaactataatggatagcattcgttttatagtaacattttcaat
ttgagcccaatgtacaattccaagttccaattccttagcaatagctgaaacggaaacaaa
ggcagctcctagccctgattgcacagcgtttttgatagcttctatagaattcaattccat
ctctatttaaagcgactgctgtcaataccatgttgactgagtactttatctatcacttt
tctaattgttgattgagtatctaatgcaataaatctaagcctgtataagtcttctttttg
aatgtctcctagtttagaaaaaggatgtgatttgggtaatataagtgctaattcgtcttc
cgcataagaagtaacctgcaaaacatcttgcaattcagtaggcacttctcctccaataat
tgctaaatcaacttgaccattagctacgctccatgaaataagtctagttgaatgtacttg
caattgaacagccacttgtggatatctttgcctaaatagtccgattaatcttggcatcaa
atatgtcccagttgtctggctagctccaataattaatgtgccaccttgtaagttttgtaa
gtcgtcaagagctcgacaagtttcttcgcagagagctaaaattctgcccccgtatcgtaa
aagaagacttcctgcctcagttaaagttgccttcttattacctctttcaaatagggaaac
atttaattggcgctctaaattttgaatttgcaaactaatagctggctgagaaacatacaa
gctattagctgctttttttaaaactgccctctttggcaattgcttttaatattcttaactg
atccaatgtaaatggaaggtctgtcattaaagaattgtagtataatg---------tata
ttatttagtattgatcaatgtatgcatttgccaactaaaatatttgtcattacgcttcat
tattttcaaaataataaaaagtataatgttaaataagctgtattaaaatttagtatctgt
ttgatattaaatttatacaagagatatcgtgaatttaattctaactataaaatataggag
acaat---atggactctagacttttagttgtactaataccagttttagcagcagcatctt
gggcagtttacaatattggtagagttgcactacagcaatttagaaaaatgacatcttagt
tttgcttagaatata-atttttagactttatgagatagaggactttaaaatccctatctc
tattgtatatcttaataaaaaaag-aaagc--ccccttgtacttactgttattacacatt
aatattattggt-ggtaccta----gataataagtttaagctaacttattttttattta
taattaattaaattaataacacaattatggctgttccaaagaaaagaacatctaaagcta
aaaaaaatgcacgcaaagcaaattggaaaaatcaagcaaaaactgaagctcaaagagctt
tgtctttagcaaaatcagtattaactagaaaatccaatggatttatttataatctaactg

aagcatcagatacttttagcgattagaagtgttt-ttactacaagtctgaaaaagtatta
cgcaacaagttctcagtaatttttatctat----taatcatttgaaaatcattctatttg
ataataaaataaatagcagattaaataaaatcacattaacaaattatgcagcaaagctag
atcaaactagtgaaatttggttgttcaattgtattgaaaatattcagcatattttttta
aaagtcaattaaaatcaagtcatattactaaaattttatttctggtactagctttgaat
ataccgcaggtctaccaggattattatccagcttaacactaagtggtagactacatccta
ttagtatatatagtccccagtctctcaaaaagtatcttgaagcatgtaccaaatattccc
aaactaatttttcttttcctattaattttcataatttacaatacggaggacaagttgtta
accaatttatacagtaatttgtttaccgttgagcaaaaagagcctgctgtatggattta
ttattctaaaaaaagaaaagcagggagtatttaatttagcacaagctaaaaccttgaata
ttcttcaaggacctatatatggaaaactcaaagaaaaggataatttttaagtccagatg
gttactatctatccggccaagacttctcttctaatacaataatgggacataaaatatcgc
ttccgttattagtcagatattctagaattatttctgagatgcattggttttgctcttatc
ctattagattaaatacttattcgcatcagcaaggagtaaagtgtttaccacataatgttt
taactgatattatgaaatcccaaatatatcaagataacagttttgttgaataatacttat
tcttgatatataatggtctcggtaatctaaataaaatactagaaataatttcttaaattt
atttgtctatattaaatttacaatcatttatgacatacgcaattattgaagcaagtggca
cgcagctttggatagaagaaggccgctactatgatttaaatcatatacctgttgatccag
gtcagtcgattatattaggaaaagtcttattattgaataaaaaatggggaggttactttag
gacgcccttgtatagaaggggttacgataaaggctacagtaatgaggcacttgcgaggaa
agaagataactgttttcaaaatgaaaccaaagaagaaaatgagattaaaaaaaggtcatc
gacaagaattgactcgtttaatgatcgattctataacatcttaga------taaaatcta
gactcatattttt--aa-aaacttttaatatttaactaaaatagatataatggcacata
aaaaaggtagtggtagtacaagaaatggcagagactctaattccaagcgtttaggtgtta
aaaaatatggtggagagcaagtaacagcaggtaatattttaatcagacaacggggaacta
aagttaagcctggccaaaatgttggaaaaggaaaagatgatacattgttttctctaattg
atggtttcgtgctgtttgaaaagtcaaatcaaaagcaaaaaacaattagtgtttattctg
ctaagaaatagttaaaacaattggtgcaagataacttgagt----gtatttatttcactt
acagtggcaccatatatttgatttattaagaaatggagttccttaatgaaccaatgaatt
ttattaaagatttgctgtgaaatttaatgacaaacactattgtaatttcctgtctacaca
atatggctgctattctatattgtggtcagatacagaaactagtagtagcaaatgctcatt
atcaagtaagtgatatctacctaggttgtgttgataaaatcttctcaggaataaatgcgg
catttattaacctaggaaagaatgagtacagtggttttatacatatcagtgataccggtc
cgcttaaaaagaaatattatgtcaataatattactaacattttaacaatacggcaaaaaa
ttttagtacaaatcattaaagagccaactttaaataaaggtccaaggctcactgccaata
ttacattatcaggtcgatatattgtattaatgccttttagtcaatcaatctgtatatctc

gaaaaatatatgatgaagatgagcgtcattatttgaaatctttagctattttaattaaac
cggcaacaatgggcttgctatttagaccttctgctgtaggtgtcgatgaggaaataatat
taagcgaattaaaaaatctaaaagaacaatggaactttgttcaaaaatctgcaattaata
gttattcacctgttcttctatataaagatgaagatattgttaaaaaagtaatccgagatt
tttataataataatacaaacaatatagtaattgattcaaacctgggattaaaacaattaa
attattatatccacacttggcactgtaataactctagcacagttcctaagattaagcttt
atagtaataatcaatgtatactagatgcttttggtatcaatcaggcaatttccagagctc
tcattccaaaagttgatcttatacttggtggctatatgtttattgaaactttagaagctt
ttactattattgacgttaattctggatcttttaataattccactagtgcacgagaaacag
ttttaaaaaccaactgttctgcagcaacagaaatagcttatcagttacaaattagaaata
ttactggtgtaattataatcgacttcattgatatggaatcacaaagagatcaattgcaat
tattagaacactttaataaagagctatcacttgatgatgctaaaccacaaattgtacagt
tatctgaattaggtttagttgaattgactagaagaagacagggcaaaagtttgtacgagt
taattagtagtgattctaattactttatttttcacacaatcagagagatctcagtctc
ttaagagattcgatgatagacagcagaaacaacagatttttaataaatcttggctatctg
cagagattaatactattaacaaggtctttttcaaaagtcaaatttgtgcagacctgcta
acttttacctaattcgtaatctctatatagttaaaagtagtattacttataaacaaaatt
atttattaactcacagatcaaaacttatttattctaaagaatacagcaaagtattaccaa
gca------gttattatctagctagtctcaataagaatagtaatcaagagttcctatctt
aagattgttatatcaactgtt---------------------------------------
-------------------gaaaaaaagaagctctctcttatataagagagagcttc-tt
ttatagcttactaacta-tctagttagttaattaaaaattaaccgttaacagctggagct
gttagagctactggtaaagattcaccagaagctagatctagagggaagttgtgagcgtta
cgttcgtgcattacttccatacctaggttagcacggttgatgatatcagcccatgtgtta
attacacgaccttggctatcaacaacagattggttaaagttaaaaccattcaagttgaat
gccattgtgcttacagataaagctgttagccagataccaactacaggccatagacctaag
aagaaatgtagagaacgagagttgttgaaactagcatattggaagattaaacgaccgaag
tagccatgagctgcaacgatgttataagtttcttcttcttgtccgaatttgtaaccatag
ttagcagattcgttttcgcttgtttcacgaattaagctagatgtaactagggatccgtgc
atagcactgaacagagaaccaccaaacacaccagcaacacctagttggtggaatgggtgc
attaaaatgttgtgctcagcttggaatacaagcatgaagttaaatgttccggagatacct
agaggcattccatcagagaaactaccttggccaattgggtatactaggaatactgctgct
gctgctgctactggagcagtaaaagcaacggaaatccatggacgcatacctaggcggtag
cttagttcccactcacgaccaatgtagcaagctacgccagttaggaaatgaagaacaact
aattggtaaggaccaccgttgtatagccattcgtctaaagaagcagcttcccagattggg
tagaagtgaataccgatagctgcagaacttggaataacagcaccagaaatgatgttgttt

ccgtatagaagggaaccagcaactggctcacgaattccatcaatgtctactggaggtgca
gctacgaatgcaatgatgaatacagatgtggcagttaatagagttggaatcattaataca
ccaaaccaaccaatgtataggcggttttcagtactagtaatccaagagcagaaacgttcc
cacaagctagcgctttcgcgtctttgtaaagtagcagtcataatttttatcaattttta
aggatttaacaaagatgcttcccaaacaataagttgtctgttatgaaaattattacttaa
ggccttttcaaatgagaacattttaaaaaaaaca-tagcaaaatctaatctcagtaact
aa-aattt-tacttgttcatatattgcttaataattcaa-----------taataaattt
tt-gct-------------ttttgcaaaattaacactattttagggat---ataataata
atgcatattaatgctgcttattttgattttaatagtttatataattttgccctacttaaa
tacgcatgtcacactttacaaaaattcagacgaccatacaagatttaaatctgttaaagc
atgctttaacagatttgggtttgatttggcaaatgaattctagtcatataaaagttggtg
aaaataatcagcataaggtagatattttaattaaacaagataacttgtcacatattggct
ttacttggaatgataatagatatcacttagttgccgacttacaactttgggaacaaccct
ggtctctagaagtatttttggataagttatctcagaagtatgcttactactctatcatcg
aagaaacaaagaagcaaggctttgaaaaaaatgcagcagatttcacaaaaagatggatcaa
ttaagttaattgtgcaacgctggaattattaactaacatacaaatgcgggcggagagact
tgaactctcacgagattatctcactagaacctaaatctagcgcgtctaccaattccacca
cgcccgcactgttgagac--aatattattctatcataatctatactaaaacacaaactaa
-tagtataaactagcttccaagcttaaaagcgtctacaaatagaccgtatgttataccaa
ttttgaagttgttgaataaattaattttatatttaattgcttgtggctagagtagacta
ttttactagtcaactcagttgctgcaactattagactggctgcaataataccccaatctc
ctgtttgtccaggaatcgtagacaaacctgttgaaataaaaaatcctagcaataaactaa
ttaagccagttgttaactcgctcaaagaataataaagcttattatttaagtttttttatta
agtaagcaaaaaaagttgaaagtctagttttaatcatagtatttagtaacggagttttaa
aaggatgacaagtattaacag-tcattaatattgtcgtatccttttaaaactcgttaata
caaacagctgttcttgt---------ttttaaaggtcttgttcgcttaaagcattaataa
gatgctgaaacggctcttgtataatatctgtgcgtttaattgattgcaaatctaattctt
cttctatgatttcttgtaataactgtatgcttcttactgttggagcaacaggtacagata
gtgaattgtaagtatcttttagcccatttaatacacgctgatcgagaatatttgtactac
cagaaaccaaggcataactagcataccgtaggtaatattctatatcccttagacaagctg
catatcttcttgttgtataagaatttcctccaggccttaataattcaggctgttcttcgt
acaactgagctgcggcctcttttaaaatattggtagcttgatcattaataatttcagcta
tttttattcgatctagtccactcgaaaagaaagattctagttgtccgacagctgtcttat
ctagataacgaccagtaagatcgtaacgatttaatattgctgttatagcatcttgcataa
attaattctcctac-ttgctatcagagcggtattataaatgactaatattgtaatataat
cattagtgtttatgaaatttatttaaaacttattctttattgcttaatgtgcaattaaaa

actcgcatagattttacacatatagagtatacgtagtttttaatta-tttttttataaaa
accaagaaataaagctttttt-taagaggttcgtgaatttgttaatataacatgagcttt
cctcctaaataagttttgtaaagtac--aaagtaaaaaaacgattagtcaataatagtaa
------------------------------------------------------------
------------------------------------------------------------
------------------------------------------------------------
------------------------------------------------------------
------------------------------------------------------------
------------------------------------------------------------
------------------------------------------------------------
------------------------------------------------------------
------------------------------------------------------------
------------------------------------------------------------
--------------------------acaatattaatatgtatcaa--------------
------------------------------------------------------------
------------------------------------------------------------
------------------------------------------------------------
------------------------------------------------------------
------------------------------------------------------------
------------------------------------------------------------
------------------------------------------------------------
------------------------------------------------------------
------------------------------------------------------------
------------------------------------------------------------
------------------------------------------------------------
------------------------------------------------------------
------------------------------------------------------------
------------------------------------------------------------
------------------------------------------------------------
------------------------------------------------------------
------------------------------------------------------------
------------------------------------------------------------
------------------------------------------------------------
------------------------------------------------------------
------------------------------------------------------------
------------------------------------------------------------

-----------------agtataaataagcttttactccagcatcaaaaaaaatatttga
gatatgttgatacaactgattattcaagtgaaaacttagatttagaccaatttagctatt
ttatcattacaattaataaaactgataaatgtattcttatagagcaattttgttataatg
ctgaaggccatatttactcaatattcttcaaaggtccaacagctaaacatctttcttcta
taatttgtcattttcaacaattaaatacaataatatctacagctcatgccatatatttag
gacgagagctaatgaagtcagaattagctctcgtgctagatcaacaatatattcaagatt
aaa--ttttgtaagtattaattatatttatgctagtataatattagtaagggcatgtaac
tcagtggatagagtatcagattccgattctgatggccgtgggttcgaatcccgccctgcc
c---gagtactcagagtact-atactaaaacatct-------taatagctagtttgataa
tagtatgggcattggggctgcaaggtttctacattatgaaaagaagaaaatatg--aaaa
aaacaagctctccaaaagagcttttagtacaatcaataaatgcagaaaacaatattgttt
ctttttctcgaaaattagcttttgcttaaacactgttaatttttggcattcgacatgtta
aactcttattcatggcgaatattctgtcagagtcgctcttagtttaagaaaagtcagaaa
aaatatttatctttacgtttt-tacctatttttatttaaataaaggttagta-ttttcaa
cttttatagtggacgtgggttcgagtcccaccagctccatcaatttgatt---------t
gtataattagctagtttaaccttttaataaccagatatataattggaggaaatatgggt
ttcataacaattactaagcccgctttaaagcaaattgcaatcttgaaaaatgatcatgaa
aatgatgtacatctaagaataggtgttagacaaggcggatgttcaggtatgtcatactca
atgaattttgaacatgttgacaaattaaaagatacagatgaacggcttcgtctagataat
ttttctgttgtctgcgaccctaaaagtcttctttatctttatggattatcattagatttt
agctcggaattaataggaggaggatttcaattttctaacccgaatgccagccaaacttgt
ggctgcggtaaatctttttcaggctgacc-ctcgaacacgtttttatatattttgtatt
atatatattattttagcaagtcg-tttattagagacaagcaatgctctgttaaaaac---
---------------------------taatttcaagagtttttattgtattatt-----
----ttatatatttattatcaactacttgaaaataaattagctaagcaatctaatttgtt
atcagtaccaaactttaaatgattaacttttattgatatatgctaatcttt-atcagcc
taaacatatctgtacattacaatcttttgacataaaaagagtcttagcatacctctcaa-
----------------------aaataaattcaa----------tattttataaatactc
aacatgaatctgtcacaacttatcgttaattcttctatgatgaaaattcctaataacttt
tattgcaatataaaagctatttcttctgagaatttaagtcacttacaattatcaaaaaca
aatttattatctcctacaaaaataattgcttttggactacctaaatttgattttatttct
acacagattacaagttctattagagaaacagttgttgtgtctactattgatgataataat
ctttttgctgccccaactaatgattggataaaaaaacacaaggataaaaatcatcattct
ttgttttataaaaaagtttttaaagctttgctaaaccataaaattagagttgttgctgag
ccact------tcataactccgaaataaataaagaaaagagcataatggcaagtcaaaga
tacatctggggtaaaagctggaagccttcaattattttatcttgcttaaaaaacaaaaaa

aatagcaaaatttctgatgcatcgaaaaaatttttaactgaacagttatctgcatcacca
gtatttgtggtaaaaaatggttttaatgaaattattttaggtcacccattatctcgtgtt
aaaagaggaagagttaataatctgatgcatgcgttttctaatttattaaatcagtctaat
gcaacttatcctatatctactggtctattctttttcatccagatgatgcatttgaattt
aaagactttataatatcggtcaatccactggcggccaaacatatggagataagcgtagag
cctgttggtctacactttgcttataaaatgaatagaaatatatcgtcagatactcaattt
tcgtttattccagattttaaagaagtaggagatttattattcaaatatagaaaaggtaat
catttagtttttcataagaatcagcattatggcaaagatttttttcaggggcaaccaata
tatatgattcagccgatcacttttaaagatcgagccggaaagctaaatattatcaaattt
acggggctaaatgacacaagagaaattattttactaatcttgaagctgctaatagatca
tgggcacattttataaagaggaattcacaattaaaatcaattaaaaatcctactctatta
gtctataatttagaaagtttttttaaaagatcaagaacgattggataatcaagatttaaat
aagtttgtggtgatcactaataaagaatcttatcttgccacaaaagagttaatagcttta
cccgattctaatagcttttctaagcacttaaaattaaatataaagcctaaactctttttt
gtgaaactatgggtaagacggttattttccaccttaacttatgaat-aaatgctatt-gc
ctttattttctcgaataatattctataactagtaattcattgagttgtaaagcaacccat
tccctatcaatgactccattgactttccctgataaattggatttattcaattctaagtgg
ctaggaatattagctagtccagggaaagctaagtaattttctactagttttcgagatgct
tcttgatttttacactaattgattctcctggtttacattgataactacagatagatact
acttgtccattaatacaaatatgaccatggttcactagttgtctagcggcaggaattgta
ggagccataccaagtctaaaaacagtattgtctagcctcatctctagtagctgtaacaaa
atttgacctgtcgacccttgtagtttttagctgcttttacgtacttaaataactgtttc
tcacttaatccataattaaatcgcaacttctgtttttcttctaatcttacagcatactca
gaaggttttctggacttttgtccatgttctccaggtggataagatcttttaatcgctttt
ctactgagtccgggtaaatcacctaatctacgggaaatgcgtactcgtggtcctctgtat
ctagacatattatttaaattctcctaatgtaaat-gattaag----------taatgagt
agctatttttaacacaaaaagagcattaaaacaaagc-ttttgctaaaaaatatttaca
aaaaatataggtttggacaaaa---tacattactatgc-tttttaagcgggtagcgggaa
tcgaacccgcatcattagcttggaaggctaaggttttaccactaaactatacccgcaata
ttcaacagattgtcactaactcaatataacatagttatgttaagttaagataatcttaaa
tttaatagaatt-ttaaattatttatgctccttctaaaaccacatcaagaaaacgagcaa
tttcggcagcttgttcttcaacttctgataataataatggttgaccgaccctagttaatg
gaattactcttttatcttttgtacataaataaatttcacgtcttggatttagcccctctt
taatatctatttttattgatttgatttcttttatattaaactgaagacagatttttctat
tctttccagggaaacctagccgaaaaatttttacaatacctttatctttgttaaattcat
tatatccagcccctatattccaaataatagttagccacagaaacaaactcagaaagactc

ctatactgccatagaatgtcattacaataccttgtggaataaataccaaatcggttgaat
ttgtaaaaggtaacaaatcaacttgaaaataacttgatagtccagcaagaagaaagccca
aagccccaataaatattatagtagcccaccagtaattactcagtcttcgtgaacctaaaa
ttaaatcttttctaactgagtgtatagataatgtttttttcataatttgttcttttaaaa
cagta-tactaatgcaaaatggaaaaattttgacaattttt--atgcgcgctttagattt
tttaaataagcttgattgcttttagtcctaaaaacagtcctgttgctaacccaaaaaaag
cagcaagaaatattatatatcctagaaaaactgaca---tattattatttattattattg
ttacaatagtgtttacaaaaatagtgaaaatctattactacggtattatattacgttaaa
tgccttgctgtctacaggaaaaaactttttttcagtttttagcattgtgtttatattagaa
ataatattctagtaaatttattaaacttctggagaacagaatcttatgtcagaatttatc
aagccttataacgatgatccttttgtaggcaatttgtccacgccagttagtacgtcaagt
tttagtaaaggacttctaggaaatctaccagcttaccgtcgaggtttatctccgcttctt
agaggattagaaataggaatggcacatggatactttttaattggaccttttgataaattg
ggccctttacgaggtacagatgtagcgttactagcgggatttctatcttcggtcggcctc
attattattctcactacatgtttatccatgtatggtaatgtatcttttactagagcagat
tcgaaagatccactacaaacttctgaaggctggggacaattcactgcaggatttctagtt
ggagcagtaggtggttcaggatttgcttatttattgctagctaatatacctgtattacag
actgcaggtcttagtttattctcttaagctagtaaggggacttgaacccgtaacctactg
attacaaatcagttgctctaccaattgagctatactagcattcagttcataataacatat
aaagatatttgttgcaatctgctattttcagatgtaacaaatatctttttttatttaaaca
gaactgttatgttaacac-------gtatattaattagaatctgaattatctgactctgc
agaagattcctgatcacagtttagataataacaaattgtttgatctagacaagttgcaga
ccaaccaacaggtgtttctcctgataattcttgcatatctacagaattataaaaattatc
atttggcttattactgtttgcttgcataaatatcactcccaaaacactctgaacttaatg
tgaagtaatttaacttctcaataatatattaacacaaaatattaaatatggtgattattc
tttaattgtaaagatgctgctaactttataaagtttgagtgagagtcttgatagcttttg
ttgaaataagcattttaacccatttattttgcccgctcgaccatactttttttgtgttgta
aattaactttttgtagtttttttgttcgtttatgagaatgagatacggcatatccattat
tagctacttttcccgtaagctgacactttttttgacataatttt-------ctaacttata
tatttatttagagtacttgctaaagttgacttgggcacagctccaatcactgtgtctact
ctttctcctgctttaaagatcattaaagtaggtatacttctaatgccatattcagccgca
atagtaggattatcgtctgtgtttatttttactaccttaatagatgattcatattcttcc
gcaatttcatcaactacaggagaaaccattctacaaggaccacaccatggcgcccaaaaa
tctacgagtacaggtaagttattgttaataacttcttgtttgaaagaggcatctgtaact
tgagatactgacatattctttaacctttaatatatgttccttgctaagcaaatcttatca
caaatttcaagaaaaatcttctatagtgaattactattagtgttgaattatttttttaaa

aattagacattaaaaaaacttattacctatataaataaagtggtggccgagtgatatctt
gatagtgataatttttatattaaggttgaacactagaaataacttttcaagcaaactctt
atttgaaaa--atttctctgtgaccataaaaacttaattatattacaatatattttatta
tttgtaacttagttctgataatggtataaacaacgcaaaagatactgccttataatcaag
gaggaatacatgtctcaatccgtagaatcacggactaggattaaaagcgaacgttacgaa
tctggagtaatcccttacgctaaaatgggctactgggatgctgactatgtgattaaagaa
acagatattctagctcttttcagaatcactcctcaaccaggtgttgacccgattgaagca
tctgctgcaattgcaggtgaatcttcaacagctacttggacagttgtatggactgattta
ttaacagcttgtgacttatacagagcaaaagcatatcgagtagatccagttccaaacgtg
gcagatcaatattttgcttacatagcttatgatattgatttgtttgaagaaggttccatt
gcgaacttaactgcttcaattattggtaacgtttttgggtttaaagctgttaaagctctt
cgcttggaagatatgcgtatgccagtagcttatctaaaaacgttccaaggtcctgcaact
ggattgattgtagaacgtgagcgtatggataagttcggtagacctttcttaggtgctaca
gttaaacctaaactaggtttatctggcaaaaactacggaagagttgtatacgaaggcctg
aaaggcggtcttgatttccttaaagatgatgagaatattaactcacaaccatttatgcgt
tggagagaaagattttatattctatggaaggtgtaaataaagcatcggcttctgctggc
gaaattaaaggtcattaccttaacgtaacagccgcgacaatggaagatatgtatgagaga
gccgaattctctaaagaggttggtagtatcatttgtatgattgaccttgtgattggttat
actgcgattcaaagtatggcaatttgggctcgtaaacatgacatgattttacatttacat
agagctggtaactcaacttactctcgtcaaaaaaatcatggtatgaacttccgagttatt
tgcaaatggatgcgtatggctggtgttgaccatattcacgcaggtacagttgtaggtaag
cttgaaggagatcctttaatgattaaaggcttctacaatactctacttgaaagcgacaca
gatatcaacctacctcaaggtctgttctttgctcaaaattgggcttccctacgtaaagtt
gtaccagtagcatctggtggtattcatgctggtcaaatgcaccaacttcttgattactta
ggtgatgatgtagttcttcagtttggtggtggtacaattggacatcctgatggtatccaa
gcaggtgcaactgctaacagagtagcactagagtccatggttatggcaagaaatgaaggc
cgtaactatgtagcagaaggtccacaaatcttgagggacgctgctaaaacttgtgggcct
ctacaaacagctttagatttatggaaagatattagttttaactacacttccacagataca
gctgatttcgttgagactccaacagcaaacatctagtttaatgactacttactgatactt
taaatagttaattgtaagtgaaattaacttataacaataaggagcatagaatagtgagac
taacacaagggactttttccttccttccagatttaactgatgagcaaattaataaacagc
ttgcttatatcgtttctaaaggcttatcagcaaacgttgagtatactgacgatcctcatc
caagaaactcctattgggaactgtggggtttacctttatttgatgtaaaagatgcttctg
ctgttatgtacgaaattagctcatgcagaaaagcaaaacctaattattatgttaaagtta
acgcttttgataatactagaggtattgaaagttgtgtaatgtctttcattgtaaatagac
ctgctaatgaaccaggattcttattacaacgccaagacttcgaaggtagaactatgaagt

atagtcttcatagctatgctactgaaaagcctgaaggagctaggtattaatattaattct
taattaatattggctaatta--ttcccctcttaaaaatcataattaatttagtaattgtg
atttttaagagggatagcttcccaagtaaattttgactaaa-tagattgaataattaaaa
atacacagaaataatgcaatcacaggatataatttccaacgatactcttgttaatttaca
agaagaatatgatagaacacaaatccaagaagttttaaatgagttaaatcaagaacttat
aggattagtgcctgtaaagaccagaattcgcgaaattgctgcgctattattgattgacag
attacgcagaaaactagaactagtttctggtaatccaggattacacatgtcatttacagg
tagtccaggaactggtaaaactacagttgctatgaaaatggctgatattttgcacagact
tggatatataaaaaaagggcatttgttgacagttacaagagatgatcttgtaggtcaata
tattggacatactgccccctaaaactaaggaagttcttaaacaagcaatgggaggagtttt
atttattgacgaagcttactatctatataaagcagataatgaaagagactatggctctga
ggcaattgaaattttattacaagtaatggaaaaccaaagaaatgacttagttgttatctt
tgctggatataaagatagaatggaaaaattctacgaatccaacccaggactctcttctag
agtagctaatcatgtagacttcccagattatacttcagatgaattattacaaatagctaa
aatgatgatagaagaacagcagtactgttttacagaagaagcagataaaactcttttaga
gtataccgagcgaagaatgaaacagccttattttgctaatgcaagaagtattcgcaatgc
tattgacagggctagaatgagacaagccaataggatttttgccagtggagaaaaagtatt
aacaaaagctgatttggtaacgattgaagcagaagatatcttgaaaagtagattattttc
attacctaatgcttaatatacacgtgatttcatgaaactatatattattaaaaag-tttc
ataatcagttgcaaatttcttgaaaaacctttattattatcttaatgtaggtg-tggcgg
catagccaagtggtaaggcagaggattgcaaatccttcatcccccagttcaaatctgggt
gccgcctagtactaaaaaagggggtgtggtggaatggtagacacaacagacttaaaatc
tgttgatttttagtaatcgtgagggttcaagtccctccacccccatatattgatttataa
ataaaaagagatgtgtatttgtattaattgcaaccatattaccaaatgtaatacctatca
cttaatagaatctcaacataaacagcctcatcttactagaagtccattgtttatacctaa
atatcctgtagttcatgttaatatatctaacaactgtacctataatcaaattgattggga
tttagtggagtgtttatcatttgtagaaaaaccgaatagttggaatttagacgctaatta
gttacaagaaaaattatgtaccataagcaacaactctatttttagatactcggtattaa
ttttggatgttctaaccagtaagattttacccgttcttgctatctctataatgccaaatt
tagttagtaattgttcaatagcaacaatctttcccggatctccagtaacctctataatta
aaagatcttctgcaatgtctactatcttagctctaaaaattcttacaatttctaaagctt
ctgttctagtctgagaattgatctgaatcttaattaacattagttctctttcaactgaag
gaatatttgttacatcttgaacgttaaggatatttactaacttatataattgtttcgtaa
gttgttcaatagttctattgtctccttgaactaccattgtgattctagagaccccaattt
gctctgctggtccaactgctaaacttgcgatattaaagcctctccgggcaaatagaccag
atattcttgacagtactccggcttcatcttgaactaaaactgataaggtgtgtttcatga

aaaattaaatttaatctatttgatctattaagtatattctaagaaaattacagaaaaat
actagttagtctataactaaattatccaaactaaacaaaatcctactatcagataatata
tctaccttgtaatgctatactattaacttagtatttaatgatattctaattgcaaataga
atgct-aattttttaaatttgttaggaacttgtgctagaaaaatacaaaaacagtaaaa
aactctctcgagatttacctcaaatcaatgatcgcattagatttccaaaagtccgagtaa
ttgatgacgaaggtgaacaactaggtatttttgtgcctgaagaagctatacaattagctg
tccaacaaggtttagacttagttgttgttagtgataaatcggacccgccagtatgccgaa
tattagactatggtaaatataagtttacacaagaaaaaagagctagagaagctaagaaaa
agcaacataacagtagtattaaagaagtaaaaatgcgatataagatagaagagcatgatt
ataaagttagaataaaccaggcatccaaatttattcaagcaggagataaagtaaaagcaa
ctatcacatttcgggggcgtgaaatccagcactctaatttagctatagatttattgaata
agatggcaagcgatctaaatgcaatagctgaaattcagcaagctccatcaagagatggca
gaaatgtcataatgctcttatctcccaaaaaagttagctaaactaatttttatttaaatg
catctggctggattcgaaccagcgacgtctctttcgaaatggcggattatgagtccgctg
ccttcggcccctcggccacagatgcatcacttaaagtttataactcataaagcatataaa
aatcaatgtttagctatttttaataaagtctataaaagtctattattcattcatattgtg
atatgtcgcaactgctgaaggggaaacttgatttaagtatcgaaatatccagtatttaaa
aattgtgtcgagtataactggaaaagttgaaataaataaaaaaataaaatctcgactttc
cggaagtcctaaatgtcttaaaattacttcaataataacttcccatccatgaggagaatg
aaaacccacaaacatatcagtaaaaagaatgattaaaaaagctttagctgtatcactaag
accataaataatttcatttaaaaaagatttaacgactgcaatttgtctttgtccagttat
cattagtaatataaagacaagtattgatacaagatcggataaaatatttttaacagcatt
tgcactttcgtttgcataatattctcccaactctctagccttcagctgcactctttttc
aatgatttcgtaagatatatcttccgaaggattcagaagaacttcaaagtgaattttctc
ttcaaatctttgtaattcagcaaaagctctttcctcttgagaggaatttaaaaaaatttt
aggctgttcttggttccataaataatcaatacaaggtccaaatacaaaaaattttgaagc
ctgattaactagtactggagagatgaataataataaaatatattttacggaagtaatagt
ttggtgtctagatattctaaactcctctatagcttcagattcgccattaggatctaactc
tttacgaaacttttcaaaagtattagtaattgacctaggtattggcccaactttttcaaa
ggcgaattgattatttcgttttaaattccaatatttcatttttattgctgctaaaattt
ctattaatcaataattgatcacgtattcactataaatattaattctagaactaaacatat
aaaacaatatttatgaaaaataaaatacaattaaaaattaattatgcattagttctatct
gcagctacagcagtttatggatataaatcttatcagccaactataaatacactcttaatc
aaagatgaagaaaactggcctttttctctaaaaggcagactgcctttattcctataatt
gtattaaaaaaaaatcatcaaaatccctgaactcagaacctagaaatagtaagttatatc
ttaatggaattggaagtagaaaatttatcgacagaattgtcactctacttggatatttag

aaaaacaggcaaactatactgaatacttgaaccaaataaaaacaagtggttatttttctt
tagtctacctagacagtgatataatgtataagaaaaatattagtaacatttttttactac
ctaatgatattctaaaaagaatctatatttgcaataagaaaaaaaagcttataccgcatt
cattcttaataaatttatttcagaaacagataggctatccaaaaagttttgctaatttaa
attgggccttctctaaaattataaaatggtattatgatagaggataccaatggtctctag
ttgaagttaaacaagcgtctgatgcatcttctattgtaattgatattcatgaaggggtag
tgaagacaattataacagaatactacactctatcttataaaagagtctcaggtatcttat
gcgtagagtcaatagagcagtaccttggagtaagagtgggagctccactaaatataattg
atttacaaaaaaaaattacttacttaaaagataatcaactagttggcgacattatttata
gtattgaacgatcaaataatagctcaatgagtttggatatcaaatttcaaatacaagaat
taaaagataaagagataatagtgcttgcagaaagttcttcta------------ttatct
cgcatgcatgtaatttactcaatc---aatataga---------aataggctagtagctt
ctaatatagtttcactatcaactaataaattacacgcatgctattttaattataaacttg
actatcaatataagtatataaacacctctaatcttataaaattgttagcttattctattt
cttgtaggaagacactacttattcattattttaatttacaaactttaataagctgtacaa
aaaaaaatactattggatttcagctatatttacgaaatctaagttttgggaaagctttct
gtgtgcttagtatgaaatttatcaaaaatggacttaatataaaaattttatatataaatc
cttcattgatagttgatcagaattttgtatttcaatttgctatacaaatcattaagcaat
atcatactgcta------aacctccagctt------------------------------
------tatttttaacgaatctagatcttgaacaatatgttgctgaaagtttattaatgt
accattttacatcttgcttctcaatatctgaaaaaatattattatctcgaattatgcata
cagattctttatttttcaactctgaaa------attttcatttgatgatagaactaacg
cgatcaacagttatgatacttttaaacaaaatacaaaaatattttatcaagagtttcttt
ctttattattaagtttacgttatcaaaattttaattatttaggttggcctttaaaaggcc
atttttttgaaataaaatctttatacttggctccatttcaaaagagtg-------------
---attttctgatagtcgtaagacgctattcttccacaaaatgtcgttaaagcaagtat
ctaattttaatttaccagtatcttttaaaagccaccttaaccatattttagttagtacta
ttaaatgtcagtcaaatttgaatatgagaactgtatctttgctattaattgattcacctg
ctgaatatctgctatacaaatctattcttaatttctctattaaagttaggatgcagtatt
tcatacctatgagcaacaatataagattatctctattttataattatttagattgttttt
taataagatcatctcagagttgcattcacatatggcaagatctaagaactcttacaccaa
ttcaaaattttggttaaaaaaattttcatatggagctggaattcagcttaaattaccaa
ttaagcagatgccacctctttcaattgaatatactgtaactagtagtcgctactttgta
tttaccttcgtacttattatcaacgataactattactaaaatgaaaaatttaatttcttt
ttttgatcgcagcaaagggaaatggatttcacaaagaacaacttatgaattatctaataa
aaacatgagttcggtacagtctcagatgacaatgaaaataggcaactcattgtcaggatc

tataatacttgcatcgttaaattgggggacatttatagacaagttgcccattacgctaa
aaatcacagccggagtgaatatgataataagtttaacttgcaattcggtaatcaactaaa
taatcataaattattaacgctgtgcatagtgacagatcctagtctaattagttttaaaac
tcgctatggaagtactaccatagatgaaacatattggttcgcaacaaataatttacgtct
aagtactagtattgttaaacgattcaacacttgtgtggcagtatctttttgttcagaaat
taaagtttaa-actctatatagaagcaactaatagcctta-ggctattagttgcttcta
tgtatctaaattttgtattaaaaattat----aatactatattattctaaatctttacga
ttgggattcctggaaggatcattagagaggaacccaaagacaaacaaagaaataaagaaa
ataacggtagtatatacaaagattttcaaagtaaacatattttcatccttaaaagctatt
agtgcg----atatttacatatctccacataattttatatcatgatataagttccagcac
tattttataaataaatcaatatctttattttttagtaatgatttatatttaaatatgcgt
gatgtactaaatactaaatcttttccgacttgttttgtcttgtttgtaaatattttgtta
tttctatggagtttaccttcaattatcacatagtctaacttctttagctgtttaaaagta
tatgaagatttttgttccaaatagttagattgattataactaatcgttttctttttaac
agtcttgcttttaacttaattatttggcttttattttcactaattctgatactcttacaa
ctaagaatttggactagaagagtacagctattcataatttataacaa------tttttta
aaaaatctactaaaatagatgattttacagaactaaaatgatatttcattatcttgttga
ccttgctcataagtaatatctttcaatacttttaatatttttccaaaatactcttgcagg
tatttttctaaagatattacttccaacctgtcaattccaagaatactgtaaacttcatcc
atcggagctgtaaattgctctccgctacttaaaatttcggcaaatgctaaacgatcagaa
atattccatgtccattgaagagttttagtaattcttcttaaggcttttaataatccaatc
ggaatttgtgagatctgtgtttttttgcccagataatttttcacataatttaataatttca
gctgatgtccaagcggtattaccaactaacggcagggtcctattttctgtggaaggtact
ccaagacttttataactaacttggctgcatcttgagtatcaatgtatgcaatgggtgta
gattctcctgtaacccagactgatttcttgtctaaaataggaatggcatattggttgatt
aaaccttgaaaaaatcctcctaaagaaaacacagtatatttcacattggacttctgaagg
aaatctactacctgagactttaagttcatcaatggaacatctgggtatttctctgaattt
aatattgaaaagaaaataaatctttcaactttagccgctttagctgcttcaattaatgca
attttcccatctaaatctatttttttctgcattataaggatcagtaggacgagatgtagaa
gcatctatgattgctgttaccccacaaaaagattgcaaaatactttcgggtaattttaaa
tcaccatatataagttctgctccccactctttgagaaaggcagactttcttaaatttctt
accatgcattttacattgtagccttcatctaaagctcgtcttacaatttgacgtcctaaa
gttccggttgctccaattactaaaagagtcatattagttaaaaataagtatgtataaaga
cattgatttggttgacggcagagagtaaattaggtagagagggaatcgaaccctcatgac
cgaagtcgtcacattttgagtgtgatgcgtataccaatttcgccatctaccctattatag
accaactactacaagtaatttagtctttagcgtaaaaataatactaagtttcaatagata

tgtcaacttttaaatttatattaataaaaaaacggagtctcttatggccacttttgaatt
aattccttatagattgtatttgagtcaatctaacacatggatccatagaataaaagcaga
aattaagatatatatagtagcgcttttatggatttcaattttattttttcttactttaa
actatgtattattgctttaagcttaattgccataagttttactataagaagtaaacagaa
tattattcaaaaacatttgttacaaacgttattaatgacgttcttaactactgttttgtc
ttttagtgtggctattagttataaacaatatgcagaacaagaacaatcgcagtatttatc
tgattccaagaaatataaaaattccagcagctactattatatacaaatagcgaacgatca
aaggatcaaaagacaatacttaattactactttaaagccttccttgtactttttttattac
tatatattctattaaactagttatgataacaacctctccagaagttttagtaattactat
ttatagatctaggataataaataaaatatttaaaaatgaattgctatttatcttccttct
ttcgtcacatattgttactagtattattaatagaatcgataaagtgattcaagtaactag
cttaagaggaagcttgaatttatataattcattgacaaggcctttaatgttttctttgtt
aatatttcaagtcttctttttggagattattcgggagtcaaaagaaatagctcaagctct
ttatactagaaatctcaatcaagaaaacaataatttttttgaaaatatatacagtaaaatc
taactttagtgatcggctcaatataattattagcactttgtactttattattttagctct
agcgtaatagaatccacttattcttctgactctagtaatacgaagagtaataaaattagt
tagtagtttaaatagatagaat-taagatattatataaataattaaatgtttcattatgg
tcaatactcaaaatcaaatttcacaaacttcagaacttgattatatagttaatcaacctt
ataaatatggctttaaaacttctgttgaatctgagcaatttcctaggggaataagtgaag
atattgtccgcttgatctccaagaaaaaagatgagcctgaatatctattgaattttaggc
ttaaagcatataaaaaatggaaaaagatgagtagtccgtcatgggctcatattaagcatc
cgaatatagactttaatacgattatttattacgctgttcctaaattaaaaaaagaattga
aaagtctagatgaggttgatccagaaattcttgacacttttaacaagttaggtatatctt
taaatgagcaaaagcgaatttccaatgtagctgttgatgctgtttttgatagtgtatcta
ttgcaactacttttaagaaagaactgtctgaagctggtgttatattctgttcaatttctg
aagctataagggattatccagaattaataaaaaaatacttaggtactgttgtaccagctg
gtgataattattttgctgcattgaactctgcggtatttagtgatggctctttttgctata
tccctcctaatacagtttgccctttagaattatcaacttactttcgtattaacaacgaag
aatctgggcaatttgaaaggacactaattattgctgatcgtggcagtaaagtaagttatc
ttgaaggttgtactgctcctcaatttgacacaaatcaattacatgcagcgattgtagaat
tagtagctcttgaaggagccgaaattaaatattctacagtacagaattggtatgctggta
acaaagaaggtaaaggtggtatatacaattttgttactaaacgaggcttatgctcgggta
acaattcaaaaatttcatggactcaagtagaaactggatctgcaattacctggaaatatc
caagctgtattttagctggtgaaaattctcagggagaattttactctgtagctttaacaa
ataactatcaagaggctgatacagggactaaaatgatccatattggtaacaatacaaaaa
gtagaattatttctaaaggtatttctgcgggtagatcgaaaaacagttatcgaggcctag

taaaagttggccctcagtcatttaattctcgtaattattctcaatgcgattctttattaa
ttggtcaatcatctcaagctaatacatttccttatattcaagtacagaatccaacatcaa
aagtagaacatgaagcatccacttcaaaaattagcgaggaccaaatttttatttttac
aaagagggatcaatttagaagaatctattgctcttatgatcagtggtttttgtaaagatg
tctttaatgaattgcctatggaatttgctactgaagctgatcgtttactgagtttaaaat
tagagggaactgtaggatgagccagactattttagaaatcaaagatttatatgcttctgt
tggtgaaacaacaattttaaaaggggtaaacttatctattcgagccggcgaaatacatgc
gattatggggcctaacggttcaggtaaaagtacattatcaaaagtaattgcaggacatcc
agcgtattcactaataagcggagatattttatttttggacaaagtattcttgaaatgga
gccagacgagagggcaaaagcaggtattttttagcttttcagtatcctgttgaaattcc
tggagtcagtaattctgatttcttaagaattgcattaaatgctcgcagaaagtttcaagg
attatcggagtttagccccttagaatttttcaactaataacagaaaaaatagatcttgt
tggcatgcaagaaagtttttgacaagaaatgttaatgaaggattttctggtggagaaaa
aaaacgtaacgaaattcttcaaatggctttactagatagcaaaatatctattttagatga
aacagattctggactagatattgatgcattacgagtagttgcaaaaggaattaacacttt
agctaaatcaacaaattcaattattttgattactcactatcaaagattattagattatat
tattccagattttgttcatattatgagtaacgggcaaattgtaaaaactggtagtgttac
tcttgcccaagatctagaaaagcacggatatgactggattacgcagacataactatttac
tac-------aatattgagactaaagaatattctttagtctcaatatt-tttgcaaatgc
aaagtt-aattatttaagctttatcaaaaccttgcttaacatcttcaattgctcttttta
gcaattcttcactaactggttctaatttttagtactacgaacattttcaccaaattctg
gtttagaattctttaaatcttctcgaagttctctaataaattcagctactttgaaactt
caatatcatctagatagccattaattccagtgtaaataatagcagtctgttcttcaactg
gaataggagaattttgggcttgcttcaaaatttctcgaagacgttgacctcttgctagtt
gattttgggttgctttatctagatcagatgcaaactgagaaaaagcttctaattcagcaa
attgtgctaactctagttttaacttgcctgccacttgtttcatggcttttatttgcgcag
cagagcctactctagagactgagatcccaacattaatagctggtctaatccctgagttaa
ataagtcaccagatagaaatatttgtccatctgtaattgaaattacatttgttggtatat
atgcagatacgtccccagcttgagtttcaataatagggagagctgtcatactacctccac
ctaattcagcattcaatttagctgctctctctagcaatctagagtgtaaatagaatacat
ccccaggataggcttctcgtcccggaggtcttctaagcaaaagagacatctgacgataag
cttgagcttgcttagttaaatcatcataaataactaatgttgctttacctttgtacataa
aatattcagctaaagccgcacctgtataaggagcaatatattgtaaagtcgcaggactat
ctgcattagctgcaactataatagtataatcaagagctcctttatcctgtagtgaagata
ctacttgggctactgaagatgccttttgtcctatcgcaacatatacacaaacaacatctt
ggccttttttgattaataattgtgtccaatgcaacagctgttttacctgtttgacggtcac

caataattaattctcgctgacctctaccaattgggatcatagagtcaatagcagttatcc
ctgtttgcataggttcacacactgattgccttccaattataccaggggccattgactcaa
ttagccttgttccattacttgcaggttcacctttatcatctattggccgagctaatggat
caacaactctacctaaaaaagcatctccgaccggaatttgagcaattcgaccagtccctt
ttacggaactgccctctaaaatatccctaccatctcccatgagtactacaccaacgttat
cactttctaaatttaaagcaaccccaatagttttatcttcaaattcaagtagttcaccag
ccatgacttcatcaagtccatatactcgagcgataccatcaccaacttgtaaaactgttc
ctatattggctacttctacatcctgatcatacttttcaatttgttgacgaataatactac
ttatttcatcaggtctaatatttaccatattttaatattatttttattaattggaaatt
ctaaagcggtaattagtcagactaatgctatctgtctttttttactttattttaagttact
acaacatcaaggtgtgaagccatttgtcttagctgtcctctaatactagtatcaatgact
tttgatccaatctgaatagtaaagccaccaattagctcgggctcgactgaaataattagt
ttaacttctttagccttagtcataaccttgatttttctgttaatagagtttcctgatca
gggcttaaagcaatagaagtattaatatttgcaattgttaaagattccattggtaagcc
agttccaaatacttactagctatagcatctaacattccaattcgcttgcgatcaactaaa
atcattagaaatgataatgtattttcgttaatttgatcgccaacagttgcaataattact
tcttttttgcctctacagtttttaatggattagct---aaaaaaattttagcttgtca
gattgcgctaaaatattttgtattgactgaatatcttggctcaccttttcagtaactttt
ttagttttagctaaatcaagaagagctactgcataaggctgagctatttttataacaaca
ttgttactgctcatagttgatctcctaacttagcaatattattatcaattatacgcagtt
gcatttcggagctcatttgattttccagctgcaaagtaactcttttaagagcaagaaatg
ttatttgttgctggatttgccttcggatctgtttttcagctgtttctatatttgatttac
ctgttatggcaagtctttcaatatctaattggccttgagctaatatggaacttctaactt
ttccagctgttagctgagcttctttcttaatttgatctataatgatttgagtttgagcaa
gttgtttttctgattctgacaatctagcacttgcttgctctagtctttcttcggattctt
gtatagctgctaaaacttttatctgtcttgcatttaaacttgatccgagaaattgcttca
aaacataaataagaccaaataaaagcaacaatatattaataacatttgcttcaaaaatat
cagaattaaagccgaatttgtgttcactactatgttctgataaaatagtaattatttgtg
gtatttttactatattattcatttaaaattct----------tactatttaatatttta
ctctggaatcatgattttcaagattgactacttaataatttaactttaatttgatcgctc
aaagtgtcaacctgattttctaaagttttaagagcttcttctttttgaacatttaactgt
ttcgatgcttcagcaatcagtttctctgcattcaattgagctttttttatatcttctgag
actatattttgagcttctttttgagaagaggctatttttaattgtgcattgcggcgagct
tcagatagatcttcttcatatttgcagctagctcatctgctttaacaagcatagacgat
gcagtagtcaatgttgttcgaatatattcatctcgttcgtctagtactttagttacaggt
ttataaaaaatagtatttaataaaaccattaaagtaagaaactgtaatgccattaaaggc

aaagtaccattgaagtcaaataagccaccttcaatttcttcggctaatagaaatggtaaa
tcaatcattttactaattatgtttatagttaataattttgtttcaattaagaaaaataac
tttatagtgcgcttagtctcatcagactaagcgcaaatgattattttaaccaacgtatgg
attagcgaacaataaagacaacgcaacaactaggccgtaaattgttaaagattccataaa
tgctaaacttagaagaagtgtaccccgaatctttccttctacttctggctgtctagcaat
accttcgacagcattagctgctgcactaccttgaccaatgcctgggcctattgcagcaag
acctacagcaagaccggcagcgataacggatgcggctgaaacgattgaatccataatgaa
ttttaatttttgttatatgagtagaaaattgacaaatttcgtataatacttaaaatcta
tataattaagtataatagattctaatat-----tgtaattatagtatattactcttctcc
atgaccttccattgcttctccaatataagctgcggacaaagtagaaaagattaatgcctg
tattgaactcgcaaataatccgagtatcataactggtaatggtattaaaattggaattag
caaagtaaatacagaaacaactaattcatctgctaagacgtttccaaataatcgaaaact
taaagatagtggtttcgtaaaatcttctagaatattgattggtagtaatactggagttgg
ctgaatatatctagcaaagtagcctaacccttttttacttaaacctgcataaaaataagc
taaagatgtaagtaaagacagggccacagttgtattaatgtcatttgttggggcagctag
ctcaccttcaggcaagtgaattaacttccaaggaattaatgcacctgcccagttacaacc
tagaataaatagaaataaagttgcaatgtaaggcacccaagggcgatattcatgctcacc
aatctgatttttttgcaatatcttgcagaaattctaatataaattccataaaattctgaaa
tttttcaggaattctttgtagatttctagtacctagaaatgataaagttaaaagtgtagc
aattactaaccaagagacaatgaaaacttgcccatgcaattgtaagctacctattttcca
atatagatgttttccaacttctactgctgataaactattatatgggttaaaatctatgag
actgttttgatacataataaatattt-ttttaaatagtgcaagaatcaaaaaataag-tt
tttgaagaattaatatgcaattaatttattcttaaaattagactattaataaattaaaag
caaaaataaattctaatagagatagctatttttagtagtgattattttttgtaaaaaaca
attattgctacctataatgcttttacgtttaattatatacccatatattaattatttaat
atgattataggtattttgctacaaactttattgttttatccattct----aaagattgtt
atagtttatttttatttaaaatatctgcgacttcatctgcaaaattaactttcaaattt
tcttctccccctcccagaagaaatcttacaaatcttctaatttaatattttcacctaga
acagcaatattttggtttattaaatcttcaatactaatatcttgattgcgaataaacatt
tgatctaataatgatagctccttaagtctctttttcattcgtccttcaatgattgtttcg
attctatcaactggtttgtttttcaagtcatctttaccagcttcaactctttgttccaga
ttaataatttcactaggtatatcctgtgttgatacgtattccacattaggcgatgccgct
atttgcatagcaatatctttagctaacttttggaactctggacgtcgtgctacaaaatct
gtttcacagttgacttcaactaatacacctattcttccgccagtatgaatataactctct
agtaaaccttcaatagcggtacgattagatttcttgttagctgaagccagccctttttgt
cttaatgattctaaagctttttcttcattgccattattagcttgtaaagctttcttgcag

tccatcatacctgctccagtcttatcacgtaaggcttttacgacttgagcagaaatttgt
agtgtcatatgctttcaaaaatttttaagtatttaaatt----------aattattataa
ttcactttgagtaggatgagttatttcttctaaatctgtttgcccgtatttaccatcata
aattgcatctgctattttacctacaattagtttaattgacctaatggcatcatcattagc
tgggattggaatattaataatttctggactacaattagtatcaagtatacaaatagttgg
aatacctaattttaaacactcttgtatagctgttgtttctctcttttgatctacaactac
aacaatatcaggtaaacgagtcatattttttaataccatttagatgtttacgaagtttatc
taattctcttcgtaaaacagcggcttctttttttgggtaactgatcaatcataccactttt
atcttgttcttctagctgtctcagacgactaactcgagacttaatcgttacccaattagt
taacattcctccaagccatctctggttaacataataagaatcacagcgttgtgcttcctg
tgccactataccagcggcttgtctttttgttcctagaaataaaactttttttaccatccga
cgaagcctgttttataagttcacaagcttctgttaataattgtgcagtttggactagatc
aataatatgtatgccatttctttctgtataaatatatggaaacattttaggattccatct
acgtgcttgatgtccaaaatgaacacctgcttctagtaattctgctaatgtaacaatagc
cataagattattaatatttaaataatgtaattattcgggaaatccc-tttgtttagaaac
tgat-----tcaactaacttatttgaattttagtaaactaataataaattaacacacttt
taaagcagatcaacaaaatagatcctgaagaattcat----tctacacttttattgctaa
aataattacgcgcggttctgtcgtctaaaatgatatcatctaaatcatctcgtacagatg
aaatagtactttcgtcattagtatttgcagttaaattctttttctctaacgatgagccat
tacagttatcatacatattgaagccggtaccagctggtataagccgacctataataacat
tttctttcagacctcttaaccaatctaatttaccagatatagcagcttcagttaatactt
ttgtagtttcttgaaaacttgctgcagaaataaagctttctgtatttaaggaagcttgag
taatacctaatagcacaggcctatatgatgcattaatcttatttctcaaggttatggctt
tattagtctgttcaatcttctgtaactctacaagctctccaggtaaatagcctgtttctt
caccattttctattttcactttagaagtcatctgccgtactattacttctacatgcttat
cagaaatattcactccttgagattgatataccaattgtacttctttaacaagtagtaact
gaatctcttgaaagcttaatcgagcagcttcatataaggctaatccacattcaatataca
aatcaaaacttgcatcgaggatatcatgaggatttaacaatacgtctgttttctttcttg
cttcaagaatttcttcaattcttggaagcccttgtataatgtcacctgtttttgctctat
caaaaaccaatatagctaatgtttctccccttctaattaatgcattgttatcaacatgta
aaatagcaccattagaaaccaagtaaggtctagcaattcttaatgtaactgatctagaag
aaatttcaataatttgtccagagtctaaagaagtcatattttcagtaataaaatcaccac
aacgaatccaatcacccacacttactttaactgttgattttcctatattaaaggtctttt
tatcagaagatgtcgcaatcaaaatgcgcctgctagtattcttttcagaataaatctcct
caacagtaccttcactcattgcaataatctcagtactagcaacaactgtaaatggcttga
tatattcaccatttttttacaataattctagtttgactttgttgcttctctgatttattta

catcaatacttttaattgataatgtttcaaaagttgacagccttaagtcaaaataattat
tgccactttctttagcagaaaactcaattgaggatattaaattatcatctcgatgctcaa
tttcagccacaagattagttgttactaaatgaactccttctatagatttcactctttctc
cgtcccgaaaaggagttcttttaactagctttagtttagtttgacttggtttttgtgaaa
ctagattttcaattagactggattttgtttctggtattgaataaacaataacaggtctaa
taagaatatagggcatttgctcgttttctatatactcccaataaactaacttatctgtag
aaatattattatgaagtctttctccaggtcttaaaaaacctctactttatcattattac
tatacacctcactaagtttatagatagatccaggcttaataataatttctcgaactattc
catccttttcgattatttcaacaatacctgagcttttagaaaaaatatttttaactagct
ctgttccactttctataacatctccattatgcactagtattagtgatgaatctttattaa
tttcgtgagtttcttctgaaatccataagatatatcccggacttaaaatttcgtaggcat
ctttatctaagcctgtttttctttttgatacatttaagtctaaatatttaataataccac
cacttgtcgttttatacgtatcagatattagttctgcgactgtgtaaccatcttgaa---
ttttttggtttggaatagctttgaaaagaaattttttgtttttttctgtttctaaaatat
aagattctttttttgtggattacatctgtataaatataacaacctggaattacaatagatt
cagtaataacttgtatattggtaatactgctattagtattctggtgtatccttacttctc
cagcataatgatttattaattctgtctgtgcaagtatagtaccagctttagttttatctt
ctttatgaacaataacattagcagaatcagatatactataaacttcacctgagagcaccc
agatcaagcctccagttttagtaattcttgtagtatattgattattatctgtttcttcaa
ctgttaaattggagaaacagattttgccagagagatctgaaactacatgtttttgagctc
tttctgtcatgagtcgatttctgcggggtgattcagcaatcacttgatccttcgatacta
attcaccatcacatattaaaagagtagtacctttacttaagttaattattgacttctgat
ttttcttagatctgattgtaacttgggttggctttttttgttattaatgcttgctcaccat
gtctggtacgaacatctgtatacgattcaatatctagatctatcaattgaccatctatag
gtgcgtagatttgttctgctaattcaccggtaaatactcctccagtatgaaaggtcctca
ttgtcaattgagtgccaggctcaccaatagattgagccgcaataataccaacagcctctc
ctaaatcaactaagcggccatgtgcaagattccatccgtaacaatactgacaaacagaac
ttctagagttacatgtaacaggagatcttactaaaacttttttt---gataccagctttag
ttatctcttttgctaacttaggactgacatcttggttcgtatgagcaattaaacaatttg
tctctggatggaatacatttctgctaatactcggccagataaagcttgttctagattaa
ttaatattttctgcgtatctactaaatcttctagtatgatgcctttattggtcttacagt
ctacttctctaattattacatcttgagaaacatcgacaagtcgacgcgtcaaatagccgg
aatctgctgttcttaaagcagtatcaactaatccttttcttgctccgtatgatgatataa
aataatcagtaacagttagtccttctctaaaattactcgatattggaagatcaataattt
gtccctgtggatcagccataagccctctcatgcccactaactgtcttacttgagaaatat
tacctctagcacctgagaacgccatcatataaacagcatttagaggatctgtctctttaa

agtatttaataacttcttgtttcaaggattcactcgcattattccacgtatcaataactt
tttgaaatcgttcaactgctgtaatctcacctcggcgatatttgttttcagtggccttaa
tatcttcaatcgtgcctaataaaagactttgcttactaggaggaattcttaaatcttcta
aacttaaagaaattcctgcctgggtcgcataatgaaatccaagatctttaagcttgtcgg
ccatattggctgctcgggctataccataattacgaaaagcccagacaattaaatttttta
gttcgttttatcaataaccttatttgaaaaacttggctgtgcaagacttcgtctattat
ccactaaatcttctccatattaaatgttcgtt--attaactatgctactaaagactcttg
aataatttattaaagattatacggccagctgttgttctaatatattgcacaattctctg
attttcagcgtcttcttttatgatatggttattaaaaaattttgttgtactattgtcaga
atgtttttcaatcttaatcggatcatgaggttgatcaccatctactaatccatcaaaacg
agcccatatataagaatgtaaatcaattttcttctgttcataagccataactacatcttc
taagcttgcaaaatactgattagatccctgctgctgagatggattattagctgttaaata
atagcaacctaaaaccatgtcttggcttggcataataattggttgacctgttgctggaga
gagaaaattatgaggggctaacattagcaatctagcttcggcttgagcctctaaagacaa
aggaacatgaacagccatttgatctccgtcaaaatcagcgttaaacgctgggcatactaa
tggatgaagcttaatagctcttccttctacaagaataggttcaaatgcttgaatacctaa
tctgtgtagtgtgggagctctgttcaatagtactgggtgaccttgaataacttcatttaa
gacattccaaatagaagattcatttttttgaatcatctttttggcagctttaatattgtt
aactaatccctgcaaaattagtctgtgaataacaaaaggctgaaataactccaaagccat
ttctcgtggcaatccacattgatgtagttttaaatgaggacctacaacaattacagatct
accggaataatctactcttttacctagcaagttttgtctaaaacgaccttgttttccttc
tataatatctgacaaagattttaatggtctattatttgcacctacaacagttctacctct
acgaccattatccatgagagagtcaacggcctcttgcagcattctttttttcatttctgat
aataatctccggagctaaaattgactttagacgagataatctattatttctattaataat
tcgacgataaaactcattcaaatctgccgtcgcaaatctacctccatctaattggaccat
aggccgtaagtctggaggtatcacaggaatgactgtgaataccatccaagaaggatctgc
acctgttgctatgaaattttcaatcaatcttaagcgtttcatttttttattgaattttaa
tgaaggtgtcttaaaactttttggtggatttgtagcttcagaccttaaggtttcagctat
gtgctctaaatctaaatctttcaacagtttctgaatagcttctgcacctattcctacttc
tacttggttattctcatcttgattttggtagatttcctcttcaagacttttccattcgta
accttcgagcagttgcttgtattttaaattaatatctgtattagattgcgtaactacata
agaatgaaaataaacaatttttttctacttctttgacttttaagtctaatgctaaagcaat
atagcttgtacttccttttaagtaccaaacatgagttactggtgaggcaagttcaatata
agccatccgatgtcttctcacgcgtgattctgtaacttctacgccacatcgttcacaaac
tatacctttgtagcgaaaacgtttatatttaccacagtgacattcccaatccttaactgg
gccaaaaattttctcacagaataaaccatccatttctggctttaaagttctgtaattaat

tgtttcaggctttgtaatttcaccaacaatttgtccgttgggcaaacttctttcgcccca
ctgtcgaatttttttcgggagaagctaaactaattttttacgtagtcaaaatattgctcaaa
ctttgtcataaatttgaataccttaaaatcaaataattagccaatttatttacatgcttc
atattgaaaaatgaatatcttaataaaggaattgttcaaaatcgtctactggaggagtat
cataattagaacgatctactcggttatctttagaatccgacatcaagtcaacttcaactg
tccgtctttggccatcttcaaatagtttcaatttatgtactgcaatgtctaaacctaatg
actgcagctctcgcattaagactttaaaagattctggagttccaggcttcggtataggct
tccctttaactattgcatttaatgcttcatttctagcttgcatatcatccgactttacag
ttaataattcttgcaaagtatatgctgcaccaaaagcttctaaagcccacacttccattt
cacctaatctttgacctccatgctgtgctctacctcctaatggctgctgcgtaactagag
agtaaggaccagtggatcttgcatggattttatcatcaacaagatgaacaagcttaagca
tataagctctaccaactgtaacaggattatcaaaaggctctcctgttctaccatcaaaaa
cttgcattttacctggatgctgatcattaaaaagccatttattgctagtaatcaatgatg
cttcttttaatttctatttacaagtgctcttgatgcttctgctccatacatttcatcaa
aaggtattattttaaatctctttcctaaatagccacctgctaagcctagtaggcactcaa
atacttgaccaacattcattctagaaggtacacctaaaggatttaaaacaatgtccactg
gtgttccatctgataaataaggcatatcttgtttaggtaagatcctagaaatgatacctt
tattgccgtgacgaccagccattttatcacccacttggatttttctcttctgcgcaacat
agacgcgaatcatagcatttgttccgggaggcagctcatctcctttctgcctagtaaaaa
ccctaaccttaactactctaccttttgcagcattaggaagccgtaaagatgtatctctca
catctcgagccttttctccaaaaatagctcgcaacaatttaccttctggtagttgatcag
cttcacctttaggagtaattttttcctactaaaatatctccagcttcaacccaagaaccac
cgacaacaattccatttctatctaaatccttcagagagttatcgctgacgttaggaatct
ctctagtaatttcttctggtcctaatttagtttgacggcattctactgcatacttttcga
tatggatagaagtatacaagtcatcataaactaatctttcgctaattaaaaatgcatctt
cgtagttataaccttcccaaggcatataagcaactagaatatttctccctaaagcgattt
cccctccgtctgtggatgcgccatcagctaaagtttggccaacaactatttttttctccca
cccaaacaattgggcgctgattaatacaagtatcttggttagaccgataatatttttttta
agcgataatgaactgttctaccactattatcctgaatgcctattttattagcagaaacat
aatttacatgaccagatgttctgctaataacaaccataccagagtctcgagctattttag
tctcaagaccagtaccaattattggcttttctggatataataatggaacagcttgccttt
gcatattagatcccattagagctctattcgcatcatcatgttctaaaaaaggtattaatg
atgttgcagctgaaattacttgaataggtgaaattgcaatataatctacttgggttgggg
ttgttgtaataaactcttgacgataacgtacaggaataatatctccttcaatgtaatgtt
gcttactaactttaacatctcctggagctactcggaaatcatcttcctcatctgctgtta
gatagacaggactattgtgataaatcacttgaccttggtttaccggataaaatggggttt

ctatgaaaccaaaaacattgacccgcgcacaggttgctagtgaaccaattaaacctgcgt
tcggaccttccggagtttcaattgggcaaattctaccatagtgactaggatgtaaatcac
gaactgcaaaacctgccctgtctttattaaaacctcctggacctaaggcactaattcttc
tcttgtgagttagttctgcaactggattagtttggtccataaattgagaaagctgactag
atccaaagaattctctcactgacgcaattaatggcttgggattaattagatttgataaac
tcaatgaatctatatcacatatcatcattctctctctaataatgcgttctaaacgattga
gacctactctgaattggttctgcagcaattctccaactgaacgaactcttctattaccaa
gatgatctatatcatcaagattacctgagttttttatccttaatattaattagataatcaa
ttgaggataagatgtcttgaggagacaatactcggaacgttttaggtatattgagaccta
gtttttattaatttatatctaccaacttcacctaagtcatatcttttaggatcaaaaa
aacgagagtaaagcatctgcttagcaacagcaactgttgctggttcattaggtctaagct
tagaataaactatgagcagggcttcctcatcagttacttcttcaatattatttttttccaa
tttctttagctaattctttaactgaatatatttgggatgcagaaataagaaaagcatatt
tactaagtcctttttcaatttcatctttattgagtccaatagctcgcagaaaaatataag
catttactttgtgggtcttatctattcttatccaaatttctcccttaggatcaatttcaa
attttaaccaagaacctctattagaaattaggcttgcactataaatctgcttaccatttt
tgtctatctcttgtttataatagattcctggactacggattatttggttaataataactc
tttctgtcccagatacaataaaagttcctctatttgtcattattggcaagtctccaataa
aaactaatctctttttgtacttttttattttttatctgcttttcagcactgagctgtagat
gggttgatgataaatctaatgatttaatattttttttcttggtttttagaaggtagatcaa
tatcttttcttgttaactttgcaggaacatatatttgagcactataagttttatccctat
tttttgcctgtctgacactataacgaggaaattttatcttgtactcattaccaaataatt
gtaattcgagtcgactagtagggtctgatatcttcggaaaaaatttcaagaacttccgtca
aaccttctaataaaaaccatttaaaactggccctctgaatttcaactaaatctggaagaa
gtttattttttaagcttatacgttgaaccatagatctcctttataacaaattttgaaatc
gaagagtttaa-tactattaatagagtactggtagataatgcataagcatagtttgctac
attatttcttctctagacataactagaacatcatattttattcaa--aaaaagattttta
aaatttttcttcac--acacactatctataagtactttgctgcta--aaagatagttcaa
gctattaaa-------aaaaacgaaattaatacttgtctgtctgtcaattaattgacatc
aagatagacaagtattatttatttaatcaattctg-tttttctgagcaaaagacaatgct
ctcgcaagtctagcttttttttctagcgcctgtattagggtgaaatgccccttttttttaca
gctttatcaatcttactatatacttgagaaatacttgactgcacatcattgaaattatca
ctttctaaattatcaatatttagaaggcatcttttagttaaagttttgacaaccgattta
tacttacggttaatgagacgatttctttccgaagttttaatgcgtttaatcgcagaaagg
ttcttagccacagtaaatccaataaaacttttataaaatatttttagaaacgaaaataca
caaatctgggatctattcaaaaagggaatatgcggcgtattataacattaaagaaaaaca

ataggcaatagttcaacaagagctcttgttattacacaatagagattgtcattgcgaaaa
actttgtaagatagtatttaattaaaataaactaactagacatataaatatatatgaaaa
aaattgaagctattattagacctttcaagcttaatgaagtaaaacttgctctagtcaagg
aaggtattggaggaatgacggttatcaaagtttctggatttgggagacagaagggtcaaa
ctgaaagatataaaggatccgaatactctattgatatcattgataaaataaaaattgaaa
ttattattagcgatgataaagtagagaaaattgtagagacgattattaaggcttctaaaa
caggggaaattggagatggaaaaatatttattagtagtattgaaagagtaataagaatca
ggactaatgacttaaattttgaagctttatagtatttttacttt-tttagacttgatact
tcttcactaaacatgagataataagcgtttattgtatacttataactaaggtgtaata--
ctaatggctaaaagcaaaggtgcacgaatcgtaataactttagagtgctctgataaggct
ggagagtttgctcaaaaaaggaaacctggcgtttttcgatatacaactactaaaaataga
cgaaatacaccaagtagaattgaattaaacaagttttgtcctaattgtaatcagcattgt
atcttcaaagaaattaaatagttatttatcttattaaaat--aatcatattatggctata
tatagaaaaagaatatctccaattaagccaacagatgctgttgactacaaagatattgac
ctgctaagaaaatttattacagaacaaggcaaaatactgcctaaaaggtcaactggattg
acttcgaagcagcaaaaaaaacttactaaagcgattaaacaagctagaatcctttcttta
ttaccttttttaaataaagattaa--atatatagcctaacttcatatttttccatctaaa
tattaaaaatttctatagatttgtttattaaaatgatctaacatactgaat---------
----------tactaagaaga-tctactcattacaagttttgttcattttataattaata
atagcattataattta---ttaagctagattc-----agacataaag-gtaacctctttt
aattgattatattatatagtctatttaaaagaggaaattttttttgcttc----------t
agcaattcaagattctttaagctttaacaaggtatttaaaacaaagatttttgtttaggt
atcagatcaaatgaatgaatcttatctcccgactgccataattgaaattcagagatgaaa
attccacattcgttgccagcttgaatttcttcgacatcttctctaacgcgctttaaagat
tcaatttttccttgataaataacttttttcttctcgaatcactttaatccaagagtttttt
aatagcttattattgatcactcgacatcctgctatttttctatttgctaacgagaatact
gtacttacttctgcttctcccaccggtacttctgaatattcaggatcaagtaagtcttcc
atccttcttttaatgtcttcaattaaggcataaatgatttgatagttttctattaatata
ttcgactttgccgatgcttgctttgttcctggagcaaagttggtattgaacccaatgaga
gtagaatttgtggttgaagctagttcaacatctgtagcagtaatttctccaggcatgatt
gagacaacattgagttgaactttactctgaggaaattgagataaagaatctaggattgct
tctgtagagccttgattatctgttttaataattaaagaaatttgcttactaatatcttta
gaattagtgtttttttaaagtatctaatgtaattcgactattaagtgctctttgtttctgt
ataattgatgagtctttagaggtattttcaatcgctttaagtttggcttctttatcactt
ttaactgctaaagctatttctcctgtagctggtacagatgacaatccccaaatttcaaca
acagatgatggtattgctaaattaattttctctttagcgttgttaattataactcgaatc

tttgcgtaagctgatccaataaccaaattatcactaatattcaatgtaccattttgtatt
aatagagttgctactggaccatgcgatttatctaaatgggcttctattataataccttgt
gcaggctgtgttggatcagctttcaaatcctctaattcagctaacaaggtaatagtttct
aataatttatcaacattttgaccagtaagagaactaataggaattataggcacttggcca
ccaagtttctccgacatgacattgtattttaataaatcttgctcaataatatctgtatta
gaaccagctttatctatcttagatattgctatgacaaaagggacattcgctttttgaata
tgattaatagcctctatggtttgaggtttaactccgtcatcagcagctataataataata
gcgacatctgttaaattagctcccctagacctcatacttgtaaaagcttcatggcctgga
gtatccaagaaaacaatttttttgtttattatctttcttaatatactcaacttcataagct
gcaatggcttgagtaattcctccaatttctttgttagcattattagattttcgaatataa
tctaatagcgttgttttgccatgatctacatgtcccataactgtgacaattggaggtctt
ttaatataattgccagtttcataaaaagcattagagttatctaaattactagaaagtcca
ttattattttcctttacatttgattctactgctataccaaaattatcagccactgatgat
attattgaggcgtctatggtttgattcatagtaactgatattcttttaagaaataaatat
tttataatatctgtttcttgaacacaaattagtttagataattcttgaatagttagagga
ttagtaatactaatagactctggcggagaattagattgaatgcttgctgaattttgacta
ggagttacggctactttttttctgtcttatagattttttattactagtcagtttttttaca
acttccacttttggcttaggtggacgcattaaagagatagctaaatctcctgcagtctgc
gaaacattagaattagagtctctaaagttgtca---tcatcatcgtcaatatgtatttta
gtttttattttt-ttgcgctgtctattctta--ttttttttactatctactaagtcgtga
attttattaaaattttttactttttttatctagtttaggtggagaacttaattctagatgc
tgttcaccacc---tgtatgtgattctgatttatcaaggtctaaatttaacagattatca
ttattgactgactctaatctaattttatagataatctgaggatttttcaaatctacaatt
gtttcagaagaattgatatttgagctagatctagaggatctattttcaaagttttgatta
tttaaaaacataaatattggttttattatctaatagtaactttctaattttaatacagag
cttaatataaacaagaatcgaatatctgtataaataatacaaatgt---agagaattaat
ctatgttgtattagatttaatataacactaaagtcctttgttcgttatacaagtttaaaa
ctttt-tatgatgataaacactagattctagtttattggtatacaatctatttatctaaa
taatactgtttttgtcagggcttttaaa----ttatctaatgttatgttcattatgaaat
tctattaaaatttaaaaa-ctcaacaattaagtaagaatcgttaatcacctccacgccac
ttatgccacgatctcaaaaaaaatgataattttattgacaaaacttttacagtattagctg
atattgtattaaaaatacttcctacaagtaaagaggaaaaagaagctttctcttactatc
gagatggaatgtctgcacaatctgaaggtgagtatgcggaagctttagaaaattattatg
aggctttaaaactagaagaagatccatatgatagaagctatattttatataatatagggc
ttatttatgctagtaatggagagtatgtgaaagctttagaatactaccatcaaggattag
aactaaattttaaattacctcaggcccttaataatatagccgtgatataccattaccaag

gagttcaagcaattgaggataaagatacagagctatctaaattaatgtttgataaggcgg
ctcaatattggcagcaggctattaagttagctcctgataattatattgaagctcaaaatt
ggctgaaaacgacaggacgaatgaggaatatacaaggatattaatatatttaagatataa
taaagatatcaattaataaattaaaattagttaaagaaactggtactatatagagtactt
-tactatatattataatattatactagttgatctctatctctaagacttagctgaataac
cgaaattaaatctattgactatattcatgaatacaaataagcataatggttagtacaaca
aaaagtaccggatctgttactcaaattatcggaccagttttagatattgcatttcctaac
ggacagcttccgaaagtattcaacgcactcaaagtacaaagctcagaaggaactattact
tgtgaagtacaacaacttttgggtgacaacaaggtacgagctgtttctatgagttccact
gaaggactacaaagaggggtagaagttattgatactggatcccctatatctgttcctgta
ggtacagatactcttggacgtatttttaatgttttaggtgaacctgtagataatttgggt
cccgttgattctgagagtactttacctatccatcgaccagcacctgcttttactaagcta
gagacaaaaccaaatatttttgaaacaggtattaaagtcgtcgatttacttgctccttat
agaagaggtgggaaaattggtttattcggaggtgctggagtaggtaaaactgtattaatt
atggaactaattaataacatcgctaaagctcatggtggagtatctgtatttggaggcgtt
ggtgaaagaacaagagaaggaaacgacctatatatggaaatgaaagagtctaaagtaatt
gatgcagataatctgaaagaatctaaagtagcattagtatatggtcaaatgaatgaacct
cctggagcacgtatgcgcgttggcttaactgcattgacaatggcagaatactttagagat
attaacaaacaagacgttctattgtttattgataatatttttcggtttgtacaagctgga
tcagaagtatcagctctactaggccgtatgccatctgctgtgggttaccaaccaactcta
gcaactgaaatgggagcactgcaagaaagaattacttcaacaacagaaggatcaattaca
tctattcaagctgtatatgtgccggctgatgatttaacagacccagctccggccactaca
tttgcacatttagatgcaacaacagtactttcgaggaacttggcggcaaaaggaatttac
cctgcagtggatccactggattcaacatcaacaatgctacagcctggaattgttggaact
gagcattattctactgctcaagaggtaaaatcaactttacaaagatataaagaactgcaa
gatattattgctattcttggtcttgatgaactttcagaggaagacagacaaactgtatca
agagcaagaaaaattgaaagatttttatctcaacctttttcgtggcagaagtgtttact
ggatcacctgggaaatatgtatctttggaagatgcaattaaagggtttcaaatgatctta
aaaggcaatttagacgacttgcctgagcaggcattttatctagtaggtgatatagatgaa
gctatacaaaaagctgacagcatgaaagattaatataattaacgatgactttaaatataa
gaattattgctcctgatcggactgtttgggatgcagaagcacaagaaattattttaccaa
gtagtacagggcaacttggtattttaacaggccatgcacctttgcttacagctttagata
ttggagttatgagagtaagagtagataaagaatggatgccgattgttttgctgggcggtt
ttgcagaaatagagaacaatcaattaactattctggttaatggcgcagaagaagctagtc
aaattgatttatcagaagcagaaaaaaatttagacactgcaactcaactcttaagtgatg
cctcgtctaataaagaaaaaatagaagcaacacaaaaaatacgaaaagctcgagctcgag

tacaagctgcaacagcagcaacttcgtaa-aagattcc------tcttaaaacataaaac
tttagtaattattactaaa-------gttttatgttttatttgtatggttataggcagct
taaataaacttagcttagacctgaacaaatataatcaaaatataatcccatctctttacc
tgcatctggtccaactaaactaatagtaacttctttcatagctaaaatagcttgaatcgt
tgcaccaataggaacacctagagaattgtatgtttcttttagaccatttaatactctttc
ttcaaggatagatggatctccagctagcattccataagtagcatagcgtaagtagtagtc
taaatcacggatgcatgcagcataacgtcgagttgtatacatattaccacctgggcgtgt
aatatccgaataaagtagagattttgctacggattctttaataattgttgctgcattagc
tgcaatagtagcagcagctctgactcgtagctcacctgtttgaaaatagcctcttaattt
ttcaacagagctatcatctaaatatttaccttggacatcagctgcattaataacagaagt
aattgcgtcttgcataactttcacacattccttaattttaaatatctaatacta--gggg
tatataacaaaattactagatcttttagcgatatcagtattactgcatagcacccaaagt
ataatcaaagtaaaaaccagcttctgctgaatcttcacctgcaagcagtgagcaagctac
acttttcatgcattttacaccttcagcaactcctgaaattggtgttcctaaagaattata
catttctttaacacccactaatccaatttcttcaataggagttacatcgccagcaactat
tccgtaagttactaagcgaagataatagtccagatctcgtagacaagtagcagtcatctc
ttcgccataagcatttccaccaggagaaactacatcaggtctttttttgaaacagttgttg
gccgccttgctttacaatacgctcacgattatctgttaaaatttgagctattcttaaacg
acgttgtccagataaaacaaaacttttaattctatctaattccccaggacttaaatatct
tgcttctgcatctgcatttacaattgactttgtaacaatactcatggataaaactcctgt
aatgcttt--aaaaaaagcttaatactaattatttatgatataactcagaaagtaatgta
ctttttgag-aattttctaaccgaaacacgttcatgatatatgaagcaggatagatactg
aggtaagtactaaaacgagaaagttatatgtaggaacttctgttattgatttcctagtac
ttttttaaaacttggtaccacaattgatatatcctgcttagtcaaacgattatagagtgt
ttctgtattaggaaagttagctgctggtaaagtcggaaatcttctgtatggaacagtatc
tttgccgtataccgcattgtattcaacactttcaactaaattacttataaatgcagataa
gccttttgaagctaaaatttgattaaagaatctaatttctgcctgattattaggagctct
tcctaagatatgcttagtacctaattctataacttttgtatttggatatggctgataaaa
ttctttaccatatagttcagatagcgccagtttttctactagttctttgacacagatttg
tctattcaagaaagctgacttaatatccaagaattcaccgccgacactaaaagagttaag
atctctttcgaaaatttgacgatatgctgctcttaaagcttgttctaacatttctttgtt
actatcactattcacttcaaagacaatagattgatctcttaaagaagttactccttgtga
aatacgagactgaatatcattactactcctcatttctttgacagtgcctaattcaacaaa
cctagcagatttactagcaataactttttttgaatctctggtcaataattcctggtcttaa
gcttcttaaagcaacaccagctggagtactatatctttcataaggaacagtattatctcc
aaaagtttctgtgtactctgagctatcaattattgtgtctattacttggtaatagccttg

cttatatgcaatatcaaaatatttattaatttcttgccgaccatatgtagggcgacccag
taagcggttatgaatatattcaatagctttacaaatatataatggctcccaataaagaga
tctaaatatacttgattttgctaactgtctaacaaactcgcgaactgtaatttgattatc
ttttagctgactttctattggcttgagaattagtttttcttcttgatacacttctctgcc
aaaaactcttaaataagcaacttttgtgacaacttctactgaattttcaaaattttggga
tgaatctgactttataagagcgttggacaatttaaagattttggtcccagggatcctgc
tgacttaggtcttacctgtggattactaatctgattgtagattcctggcccacgtctaac
taatatccttcttgtatctttaccaaaaagagcttgtctttttcttggatctttattttc
tttagggaaaatagctccaaattgaattgataatggatcattacctattccatatggatg
ttgatcaggtaatgattgcttataatcactaaataaagttataaattgaggaaccttacg
aaaaggcgcactataggtgagaagatctatttgaggcccccaattacggcactcttgagg
ctcttcacctaaatttctaaagtaaggcacagtctcttcaccgaaataatctgtatactc
agaagaatttaaaagtgcattcactaaaccacttaaaccagtagatgataaaatagcaaa
atatttttgaaactcttctaaagagctaggacctctacccaagaaatgtctaaatgctaa
ttctaatgctcgactattgacaaaaggttcataaaattgctttctataaatacttgaagt
tcctaacgaacgaataaattccttaattgaaatttggccatttttttacttgcgactccaa
attcgataaagagaggtcatatgctttagcaatatctctttcaaaaatttgtctataaca
tgctttcacaacaatatttttctcatcagcagataaactaggcttcataacaaaacgagg
tgttgacactcctgcctgtacataagtctgaggaagacgcaaaccttgtaaatctccaga
tattctttttcttaatttatcagtcaaagatggagcttcaaattcagaaatcacaacatt
aaaatactctttaactaaatcttgaccttttatatcttcctcaaagattaataatgctgt
ccgtctcatctctctaagagccacaattgctgctgcacttgagcaagcgttatcaattaa
ctctctcaatcctcgtatattaacagataaaatattagggtcgcctgatacaattgcata
tgtcaaataccgtaaaaaccagtctaaatctcgaagagattttctcattcttgtagtacc
gtatctcaagacgtttataggtttaaaccctggcggagtagctcccccagcattaaataa
agatcggaaactttggccaaaatctccttggatatttcctgataactcattaattttatc
ttgagaactttgatctcctgcaataataacagctgcttgcggacgttctaaataagaaat
tgcagacccacctacgaatattttatctgctgctcgtgctaccagaatatttgcattttt
agttaaaatatctgcaacttccagtctttttttgtccagaatttaaaaaagaaactaattg
atttaattcaccaagctgtaaaaatcggtcttgttgctctgcctgagtaatagttaaaat
tgaggcagtccgataaagctgggggcgtgctaatgggcttccgccacttgctttgatact
cattactttatttatctccttaactgatattgcttgctagcgaatacaaatatagcttga
tatttctaaatatttaaagatactatgcaagcaca-ctgttttctatattattaggtact
agtacttatagtacagagagtcttataactaacttctataaatacaaatttgtaatact
tctattaacatttgtcaaaaaaatatttaagtgctcactcttttatattaaaattcaatt
agtgagaggttgatgtgaaataaagtatactaagtaagttttatagataggaaaattaaa

agatttattctacaaacattaatca------------cttgaatgtttagcaattgtaca
attgttattgcttttaattggtttagactagttttatatttttttctaacataaacaatat
taatgagcaatttgtatgaatttaaagccacaaaacaacctattaactaacgaaaatata
gatcatgtagatatacccgaaaatgatataccaatgtcaattacggaacatttagaagaa
ttaagacaacggactttatttgtgtttttattctttttgtttgctacaactataagtttc
acacaaattaagatcattgttgcaatactgcaagctcctgctgttggtattaagtttta
caactggcaccaggagaatattttttttcatctattaaggttgcaatatattgcgggatt
gttgcaacaactccgtttgccgtttatcaagttatattatacatactcccaggactaact
ggaaaagaaagaaaaattattttgcctctattaattagttctgtattgctttttattaca
ggtggtatatttgcttactttgttctcgcaccagcagctttaacatttttaattagctat
gggtctgatattgtagaaccattatggtctttcgaacaatactttgactttattctatta
cttttacttagtacaggattagcatttgagataccaatcatacaattattgctcggtgtt
tcagggacatttcttctagtcaaatgatacgagcttggagatatattattattatagca
acaattgctggagctattctgacccccttcgactgatcccgttacacaattaataatgtct
tcagctgttttattactttattttggtggaattgttatattattagtcttaaaaaagtag
agtaaat---------attttcta-ttaatccaatatatttgttatc-----tgggcaaa
taatctat-aaaacaattgct------------------agatttaagtatcaggcttct
aaatatttcaa---------gtctcttttatcattgaaatacaatc--------------
------------------------------------------------------------
------------------------------------------------------------
------------------------------------------------------------
------------------------------------------------------------
------------------------------------------------------------
------------------------------------------------------------
--------------------------------------------------gatacattgg
cttgtttttcatcaattatttgtgttgcacac-tttttagtactctctttattgttctt
agaattatttctttaatctatttattaaat----agatatgaagagatttaagtattgat
tatggtaatctatatcaacgtaaaaagtattttattgtgagttttatttatataaaggct
tattcattatatagttataacagtaagtctaattaatag-ttttcatgcttaattaaagt
aataactcttaattcttttggcagaaaaatctgtcttaatattttatc--aaaaaaacag
-aaaaatgaagctaaatagtctaattaacttaattcaaaagtctatatattcttgtacac
ttttactaattattttaaatattatttgtgtcgcacctaattctagtaatgcatttccaa
tttatgcgcaacaagcttatgaaagtccaagagaagcaactggtaggatagtatgtgcca
attgtcatcttgctcaaaagcctgtggagatagaagcgcctcaagcagtactacctaata
ctgttttcgagactgttgtgaagattccatatgacagcaatgctaaacagattttaggta
atggcagtaaaggaggcttaaatgttggagccgttgtaatattacctgaaggatttaagt

tagctcctgttaatagattatctacagagttaaaagaaaagactagaaatctttacattc
agccgtacagtgctaaacaagacaacattttagtaattggacctatttctggtgataaaa
atagagaaatagttttcccaatactatctcccgatcctgcaaaagataaaaaagctcatt
ttttcaagtatccaatatatgttggcggaaatagaggacgaggccagatttatccaacgg
gtgacaaaagcaacaataatattgtctctgctttaagcagtggtaaaattaataaaattg
aattactagacaaaggtggatttataatacatgtgactaacagtagcaatgtggagtcaa
cacagaagatttcacctggtctcgaacttagagtaaaagaaggggatacaattcagcttg
atcaagctttgaatagtgatccaaatgtaggtggttttggtcagaacgaaacagaaatag
ttttacagagtccaaatagaattaagggcatgattgttttctttttcgctagtgttctag
ctcaaattttcttcgtattaaagaaaaaacaatttgaaaaagttcaagcagctgaaatga
atttttaaagcaatataaaactggcacttactataaagaaagta-taaacactttattca
caggatgtaccagtttttatgttatgtaataatcgctttgacggaagttatgatactttt
aacagaatctacaatctgactaggctgaattactgtagcctgctctaagcttccattata
tggtgtgggtatatcctgcgaagatagcctcacgacaggagcatctaattcatcaaaaag
atactcattaatttgtgcaattagctctgctccaattccggctgttttcatacactcttc
tacaattaaaactttgtgagttttctttactgagatagatatagagtctatatccagagg
ttttaaagatataagatctataacttctggatcataaccttcttttaataaggccggtaa
tgcttggataacatgatgcctcattctagaataggttaaaatcgtaatatcttttccttt
tcgtacaaattcaactttattgagaggtaagaaatattcttcttgaggaatctcttcttg
taaattataaagtagaacgtgctcaaagaatacaactggattattgtctcgaattgcaga
tttcagtaatccttttgcattataaggagtagaacaagcaactattttttaagccggggat
ggcttggaaataagcttccagtctctgagaatgctctgcacctaactgcctaccaactcc
tcccggtcctctaataactaaaggcaatgtaaaattacctccagaagtataacgtaacat
tcctgcattattagaaatttgattgaatgctaataataaaaaactcatgttcataccttc
aacaattggtcttagccctgttatagctgcaccaattgccataccagtaaagctattttc
tgctattggtgtatcaagaactctcaaatccccatatttgctatgcaaatctttagttac
cttataagatccaccatagtgaccaacatcttctcctattacgcagacagttggatcttt
tgccatttcctcatctgttgccgctcgtaaagcgtcaaacataaagattttactcatttt
aatatttgatttttatgatttgaccactgttttaaagtaacagttaagacaccatattct
atttagttatctgcaaaaagatatcgtttcaattctgacatgttaggttctggactggaa
atagcaaatttcacagcttgttccaattcggttttaacagcattctgaatctcattaagt
tcacctatattagcaatttcattatctagaatatatttttttgagtttttttgataggatct
cttgccacccaagcctctttttcttgtcttgatcttagttcatcaggatctgcgagagag
tgaccacgaaatctatatgttaatgcttctattaaggttggaccatcaccttggcgagct
ctttgaactgcttgttttgcagcttgtcttacagctagcacatccattccatcaacttca
atcccaggaagcccaaaagcttctgctttttatgtatttcaggtattgaagaagaccgg

tgatgtgccatacctatagcccattgattattttcaacaacaaatataataggtagtttc
cagagaactgccatattcagacattcaaaaaattgcccattattggtagtcccatcacca
aaaaagcaagccgtgactcttaaatcctctgtttctttaagtacttgctggcggtagata
ctttgaaaggctgcccctgttgcaaccggtataccttccgcaataaaagcaaagccacct
aaaaaattgtgaggcgcagaaaaaatatgcatcgaacctcctctgcctttactacaacca
gtctcttttccaaataactcagccatcacattttagatgggacgcctttacttaaagca
tgtacatggtctcgataggtactgcaaacataatcagttggattcagaagtttaattaca
cctgtagaaacagcttcttgaccattataaagatgaacaaaaccgaacatctttccttta
taatacatctgagcacacatatcttcaaaatttctgcctaacaacatgtcttcatataaa
actaataaattactcttattaagattaagacctgttgaattatagttagtcagcggcaat
tgaactttcttaggataactcataatttgtaaagaaacctct----------tattgtaa
ataggaattttgaagaaacagctatcaatagaaaacacacagataaaaacttataaatta
taatatgtttttactaaagactttaataattatttgaaactagttcagtatatcaaatt
aattttgcttatactacggatgtctgaaacttatataaaaaagtagt-a-aaaaagttta
ttaaaagaatagttaaaaacagatataaatagaagtaaacagaattatt-gaaaaaaaat
gattattattaattaa----tacttaatatttctaaatattttacaattaatggctatgt
caccatctttattttatcctgttgaacaagaactatgtagtcttgaaaaaaatctgaaag
ctgttgctgggactcgtcatccaattttatatgccgcagcaaagcatctattcgatgctg
gaggaaaacgagttagaccagctcttgtatttttagtggctaaagcaacctctgagaagc
aagatataaatactggacaaaaaaggctagcagaaattactgaaattatacatactgcta
gtttggtacatgatgatattattgacgagtgcacaacacgtagaggagtcaaaactgtac
ataatttattcaatactaagattgctgtgctagcaggagatttttatttgcacagtctt
cttggtatttagctaatattgaaaatttagccgtagtcaaagctatttctaaagtcatca
ccgactttgcagaaggagaaattaggcaaggcttagttcatttcgatcccagtatttcaa
tagatgcttacattgagaaatcattttacaagactgcttcactcattgctgctagttgtc
ggggtgcagctatgcttaatggttccaatcatcaaataaataatgatctttatctttacg
gtaaacatatgggattagcatttcaaattatggacgatgttctagatataactggttcta
ctaagagcttagggaaacctgctggcgctgatctaataaatggaaatttgacctctcctc
tcctttttcacttactcaagaagcaagtttaaatgatcttattgatagggagttctgta
atagtacagatatagcctcaacattatttcttataaaaagaagcgggggaattacaaaag
ctaaagatttagctaaagaacaggtgcaggcggcactttcctgccttcagtttttaccac
aatctacacctgtatctagtttaaaagaattaacacatttcataatcacaagattgtcat
aaaggacttgctaaaattaaaatgttattaactaatttttttactgtttctaaaacagtct
gaaaagtttgaatatcattactagctagctgcgctagcattttacgatttaacgcaatat
tctcttttttttaatgcactaataaaagtactataattcatgccttgattgtgagcagcag
catttattctagttatccaaagacgacgaaaatctctctttttcctttttcgaccaacat

aagaatatcgaagagctttaaggacttgctgcttagctgttcgaaataaacatttatgtg
cacctttaaagcctttagctagcttaaagatttagctcgtctttttttgcaacgttac
ctcttttaactctactcataaaatattac-ttaaattttaattattaccagcttactaaa
ttcttatactctttg----attataaataaggtaatttcattgcaatgtttttaatatct
tttaagtcaaccgaacaagtagaagaaagatgtcttctttgttttgatgactttttttgt
aataaatgactttagaagctttatgtcgaagaaatttccggatgaagagactttaaat
ctttttgctattgcttttgatgtttttaacttaggcatattataaacttttaatgattaa
caaaagcatgaaaagtctgaggatatcaatcttagacctttcaatacttagaataatctt
aacacaatttattttacttactacataggttaactgaaacaatttatatatataatacta
atcagttaatcaaccatagctaaagtggatttatatttgctttgaactttgtatagta
ttagcaattagcattgttataattttaagactgcttgaattcaattcttcgaacattttc
atatttaatgtaaatgaaatattggcttctgagacaatatttgaatttgaacatcagac
aagggaatcatatctaatgcagctctatattgatctttaaataatttatcgtcttttatt
tggtcaaaatcgtaaaatttcgttcctcctgaatctgataaattcatagcccctctagct
atttttttaaaatttggccgccagaaaggtcacccaaatagcgcgtataagcatgagca
actaataattcaggttgtttatgtcctatagtgtgaattctatcaacataaatttttgta
gcaggagatggttcaataaaatctaaccaatctgatccataataatagtttaaatcttca
gatagactcgctttcctattaagctctgtaaaatatataggtttaatagctggatgattt
ttattagaaaataactcttcctctattgcacaatagacaaagtatagatttgcaaccaat
ttacgatatgacttcttatctacaactcctcccaaaaaagatttaacaaaactaacattt
tctgccatactatgggacttagtagtaccttctcttagttcattcgctaaagtattaacc
ataatatcaatattccattttt-aaagataacaaattaaca---ataaatagttatccgt
atatttataattgagttgcgctaactaggctgctgcaaaataatttttttgattttattgg
atctggattcattgttttgtcaccaggcttccaattagctggacaaacttcatctggatg
agattgtacatattgaattgcttgtaaaaccctcaaagtttcttctacactcctgccaaa
ctctagattattgattgtagagtactgaataattcctttcgggtctataataaatagtcc
tcttagagctacaccatcactattcaaaacattataagctgcactgatctctttttttaa
gtctgatactaatggatatgaaagatctcctaatccacctgattctctgtcagtctgtaa
ccaagcaagatgagaatactcactatcaaccgagactccaagaacttctgtattaagttc
cgaaaaagcattatacttatcgctaaatgctgtaatttctgtagggcacacaaatgtaaa
atctaaaggataaaagaataagacaatatacttattcttgagatcagataattttaacgt
tttaaattcttggtcataaacagctgtagctgaaaaatcaggagctagctgacctactcg
aagacaatttggtccagaaatcattaatttttcccagaagaattgatataattttgttat
ttaataaataattatataacaaaatttagtatatattaaaatactacaagtctttttaa
attac--------taaatgatataatcaatactatctttatagcggggaatggatttgaa
ccattgaccttcgggttatgagcccgacgagctaccagactgctctaccccgcggataac

taaaatcatattagtctacatatatatataaaagcaaatgcataatagattttagaagta
aaaaaatagaagaactggtccactgatcattaaacttagaaacgacaaagtatttttat
cttatttaaaagagggaagacttcatataaaccaataaatctatcttttaaaagaatttc
ggacggctttacccaaatttgaccgtcataccagcctgattcttcataaaatactgtagc
actcattaatcgttttactacgtatgaccagcctaaatataatctgattaaaataaatcc
tgtcattaggctggaagtaataaattctgaaaagaaaaatttcaaaggtagtttagtgat
tggaaaaattgataataatataggactaactaataaacagttaagtagtagtgcaatagt
tattttttattataagatctacggcttaaagttggccagcaaaagaaccaagaattttt
caaagaagtatattcatgaacaggttgctgctcctttggcacaggacattgagtattata
taaattcattgacattaaaaagtattataataaagac--tttgtttatattatctaacat
aggtacgacattaaatattaactaactgcaattatagtagtgagtagaagaaataaaact
aaaaatagccaagtaattttatttaaagtattctcagtactacgagtattactaaagaac
tgattctgagcacctacacttcctaaaccttcggatttaggattgtgtattagaatagtg
aagattaatattattgtagatgagtaccaaaaaaatttaaaatttgttccattatgtaa
aagttaat-aaaaatagcacaaagacaatcgtctaaaagatattgattgtcttatcatat
taaaaagtatttattcaccctgtacttttaaaagtaaaaatcctattgccagccctacca
aaactaggactgaacttaatacagcactttctacaatttctccgaccatt-ttatttttc
cttattattttattattg---tcattcttttgacaagattatattgagctaa-ctaaaat
ccgtttctaccccaaactaccattgcgattgaaaatgtaaacattgccattagagctgcc
catcccaaacttaaaatatccataatttataatattc--ctaaaaataagatctcttta
cttaaactataatattcttaatattaaaaatatcattatataataagctaattataacag
agtactacttaattcgtattttagtgtatttaaagaagattgaa-tttttcgaggtac
tggataagtatttcatcctaagaaccatagtcaac----ttttgatttattataatgctt
aaatctaaatata--tatcactcaagaatcattagtaattatgatgtttttaattatacg
att---aacatatttcaaaatagaagaattcatagcgactaagttaa-----tatgctaa
tataatttatcaat--aaaaaggaagttagtatttcaattatgcaaacaactataaataa
tgggcaaacgtctagtaaagaaactttactaacacctagattctatacgactgactttga
agagatggctaatatggatatttccggcaatcaagaagatttttttggctatcctcgaaga
atttcgagctgactataatagtgaacattttattagagacgaagagtttaatcaatcttg
gtctaatttagaacataaaactaaatccttatttattgagtttttagaaagatcttgtac
cgcagaattttcaggtttttactatataaggaattatccagaaaattaaaagacagaaa
tccggttatagctgagtgcttttttattaatgtctagagatgaagctaggcatgccggttt
tttaaataaagctatcggggacttcaatttatctttagatttaggatttttaacgaagag
tcgcaagtatactttttttctcacctaaatttatttttatgcaacctatctttctgaaaa
aattggatactggagatacataactatttaccgccatcttgaacaacatccagaacaccg
tatttatccaatctttagatttttttgaaaattggtgtcaggatgaaaatcgtcacggaga

tttttttgctgctttgctcaaatcccaacctcatttttttaaatgactggaaagcaaaaat
gtggtgcagatttttctattaagtgtatttgcaacaatgtacttaaacgactttcaaag
aattgatttttataatgccataggcctagactctagacagtatgatatgcaagtaatacg
aaaaactaacgaaagtgccgctagagttttcccggttgctttagacgtggacaatccaaa
atttttcaaatatttagatacttgtgcatgtgataatagagctctgatcgatattgataa
taacaattctccattattcattaaatctatcgtaaagatacctctatatttttccttatt
tgcaaatttactaaagatatatttgattaagccaatagactcaaaaacagtatggaatac
agttcgatagttaaatacaagggaatgctttttcttataaaaagctatctagtgattaa
tataaaat------aaaagcttccttct-----cgatggaaggagcttttatttttgtcaa
ga---atataatgggattatatagatc-tagaactaagaacttacttttcttttgcttc
atacataacctctaaagtaattagatttttagaacaatcacgggcaaatttctctgtatt
tctttttactttacctctaacaaacccaggaatttatttaattctttctgtgcttcttc
tgaccaattaatactatcaacagcagaaattcctatacttaatgcttctgttgtatcatg
accaccaaatatttctagcaaatgatcttccataccaagtgtaaacgaattatatactaa
atcagcaatttgatttgttccctcataaccaagaaatggacggtaacttaaaggaaagtt
ttgtatatgcactggtgaagaaataacgccacaaggaatatttagacgcttaccaatatg
tctttccatctgtgttccaaaaattgctgcaggttcagttttagcaattaaatctccaat
taaaccatgatcatctgaaacgattacttcatcgcagaattcttgaacttgatctttaaa
ccactcttcatcatatttgcaataagttccacaccaggcgacatgtatacccatttcctg
gtgtaaaattcgagttattgctgcagcatgagttgcatctccaaatacaatagccttttt
acctgttaaattctgacaatcaattgatcttgaaaaccatgcagattgtgaaataaatcg
ggtttgctcgtctatgtatttttcgtagtctacagcagctcccaatgcatttactaattg
ctgtattgatcggatgcatgcagctgtttgaacaatacccataggtgtaatgtcaacata
aggcatattaaattctttctctaaatatcgagctgtcattagtccagtttctctatatgg
aataaaaattaaaccatgcagagggtaattttttttaaatcttgaactgatgcattttctgg
aataatttggtttatttggatatctagatcttgaaataaccttttttaattctgctatatc
atgttgattatgaaatcctaagcttactgctccaataatattaacagatggagttttagt
cttttgagttaaaacttgattattagatttagctttttccatatagaaagtgacaatttg
ttctaatgttctatcacctgcttgcagttcatttactctataatgattaacatcagctaa
taaaacatctgcttctgtttctatagatgctctgctaacaaagttttgcaaatcttcttg
caaaatactcgaagtacaagttggtgttagaataactaaatcagggctttcttctctgtc
ttttctagtaatattttctacaacttttttcttgagaaccacgggctaacacatgtctatc
tacaacacttgctgtaacaggagtaaaatctctgtctctttcaagcattgaacgcattac
attaaaataatcgtctccgagaggcgcatgcataattgcatgaactttttttaaaagaact
ggcaattcttaaagttccaatatgagcagggcctgcatacatccaataagctaatttcat
attaaattttcctatgatttatttagacatagtaatacttttaaagctaactaacaaaca

ttagccaaacatctaatttaactccttgacaattttgtaatattttttaaatgaagctta
tcattataaatatttaaaaaatataattcagaataga----aataactgttagatcaagc
ttgaaagtaaagatttcttgcagttatagtttttctttaagtaaacatgtaaaaattgta
ctatatataaatgtactatatataaactttttttactgaatattggtaatgccagataca
attaacttaaatatgccttccccaacgtttggtggaagcactggcggttggttaagagcc
gcagaagtagaagaaaagtatgcgataacatggacaggcaaaaatgaaagtaagtttgaa
atgccaactggtggtaccgcaactatgcgaaatggagaaaacttactctatttagctaag
aaagaacaatgtttagctttaggtactcaactaaaaagtaaatttaaaatatctgactat
aagatttacagagtttttcctaacggagaagtgcaatatttgcatcctaaagatggagtc
tttccagaaaaagttaatactgggagagctagcgtcaacagtgttgatcattctattgga
caaaatattaatcctgtagatgtcaaattcatgaataaagcaacttacgattagtctctc
-aaaaacctagacataaaataactttatatagtctaggtttttgataatgtatactgat
agtatggaagggtggccgagtggttgaaggcgtctgatttgaaatcagttgaacta--tc
tggttccgtgggttcgaatcccaccctttccgttataaaact------------atagtt
ggcttctcaaggcaagctagctatacactgatagtattaagtttagacttaataaagtct
acaacttgtgacaggttagaaatttttcggcatcctcatctggtatctctatactaaat
ttttcttctatggccattaccagctcaactgtatctagagaatcggctcccaggtcactt
gaaaaatttgcttccctagtcactattttttttcaattcctagctgttctgctacaata
tcttgaactttttcaaagatttcattatcttgcataatatattatttcctctaaggtgtt
atcaaaaatttttcggcttatattattatccaactctaaaaaattagtaatttttagtcg
agattaaaatatttatacctatatgatatactattattctggataaatacgaagtttacg
atatggctcttctccaaaactacgagctctcaactggtagttgtcaactagattatgctg
catttttcgtatataggctgatcgaggtgtaagctgaactattgaactttcattcaagat
aatagtttcaatagccaatttagcttcttgaagagcctggatctcatcaaaatttttatt
tttacagagttctacccaattaagaccagatgaagtatgaatgtttaatatttttctgag
tgcacgagtaatttgtggaaccgtactattctgaattgtatatataataatttgttttga
ttttgcaatttgtctaagttttgtattttgctttacttgattacgtagcgctagaatagc
atcagacttttctatctccttggttaaaataattggtagatctagtgaagaaataactga
tgtaatatgctgccaactcaaagaataagcatataagtactgatgaggtatttccaattg
caaagattgtttattaatattaataggggcattgaaagtagttgacaataaagatgtatc
acgattttgtgatttattcaaatctaaggacttatttcttaattctctatattgtaaaac
cggaggcgcctgctgtgcgttaattgattttcgagcattagagatgtttacaggtaagac
ttctatagattgcgatggataacatttaattaaaattctaccattagcttgaatttgtct
tttttgcacaaaaggttgatggccttgcaaaatttgatctattgtctctttaactttatt
atgaactatccaaacatttcgctcgtgtatttcgatcgcaatctgaaaagcaggctcagc
ttttctttctagtatactcttttgcgtacctctacgtttagcttcatcgtcacctaatgt

tacatattgaattccaccaattaggtccgccaatgttggatttttaattaaactttctaa
atgatttccatgcgctgtacctactaattgaactcctctttctgctatagttcgagcagc
taacgcttctaattccgtgccaatttcatctataataataacctcgggcatgtgattttc
aacagcttctatcattacttgatgttgcaaatcgggccttgcgacttgcattcttctagc
tctaccaatagctggatgaggaatatctccgtcaccagctatttcatttgaagtatctat
aataaccactctttttccatttcgtcagctaaaacacgggccatttctcgaactgcggt
tgttttaccaactcctggttttcctaagagtaaaattgaatcaccttgctgtagtaaatc
tcgaataatactaattgtaccaaagactgctctgccgacacgacaggttaagccaataac
actaccttctctattacgcagtgaactaatacgatgcaaggtttttcgataccagctcg
attatcgccgctaaaattaccaactttttttacacaataatctaagtcttgccaactaat
agatctttgagataaatattctgggttatctggaaatctagcttctggacgacgacctaa
atccataacgacttctattaaattatttctattaggatgctgttgtaaaggttctttgac
aaaatttggcaaaatttctagcaacttatctaagtcatctgcaataagcatgattgagta
tataataattaattaatcaaatagaagattcaagattctaaagattacaatacatattaa
agaatattatgattggaatcaattttatacttaaattgaacaggaagtta-tagatgtaa
aagaactgattaaactttacgcaataaatcgattggtcaaaattaatagtcgtgttaaca
tgtttattggacaaaaagttcgtattaaatatagcaaacaaaaagttgctagtgatatag
cagataaagttggtgaattaggagttattaagggaataaagtttattaatagtcaatgtg
ttactattattgttgagtttgacaatcacactaggctttggatgtttagagaggaattaa
tctgtctaaatgaat------taaaaacatgaaacataatttatttatttattatacaa
aatttcttatgagtcgttaaaataataaattttaaaggagtacttttaattgtgcaaatt
actattaaaaaattacaagacttactttcgtctgtacaaagaaaaaagatccagacatta
aaactaaaacaaggtaagtttgaacttctattaaataagacctataaaaaagtcaatcaa
gaaattataccttcgcagaaatctgctgtattacaaaatagtccatctacgataattaaa
tcaataaataatacaaaaaaatcttctgttgtcaatgaagaccgtacagaatatgccact
attgtttctccaatggttggaacgttttatcattcacctgctcctggtgaaaaaatttt
gtacaagttggcgatgaggtcaaattcaatcaaacagtctgtattattgaagcaatgaag
ttaatgaacgaaatcgaagcagaaattgaaggcaagattatagaaattcttgttaaagat
ggtgatatagtagattgcgggcaagccttaatgaaggttgaaacataatgctaattttc
cattttacatattgttaacagaaattatttagatagaaaaattagcttatactatagtta
tgaaaacttatttcgcaaaggcac-taaaaactaatatataaaattatttactttatctg
ccaaaaatagaaataagcgttttgattccttgtctcaagagaggagaatctcaatggcaa
ttagctcaaaagagcaagagacaaagaaggtaaaaatctcggttgataaaaatcccgtag
atacttctttcgaaaagtgggcccaaccaggccatttttctcgtacactagcaaaaggac
caaaaactactacttggatttggaatcttcatgctgatgctcacgacttcgatagtcaaa
ccagttctttagaagaagtttcacgtaagattttcagtgcacattttgggcagctgtctg

taatatttttatggcttagtggaatgtattttcacggagcccgcttctctaattatgttg
cttggttaagtaatccaacaggtattaagccaagtgcgcaggttgtttggcctatagttg
ggcaagaaattttaaatggcgatgtaggtggtggctttcaaggagtacaagttacatctg
gatggttccaactgtggagagcatcaggaattactacagaatttcagctttactgtactg
ctattggcggattagctatggctgctttaatgctgtttgcaggatggtttcattatcata
aagctgctccaaagttagaatggtttcaaaatgttgaatcaatgatgaatcaccatttag
ctgggcttttaggcttaggctgtttaggctggacaggccatcaaatccatttgtctttgc
ctattaataagctactagattctggcgtgtctccgcaagaaattccactacctcatgagt
ttttaattaatagagagcttatggctcagctgtatccaagttttagtaaaggattagttc
cattctttactttaaattgggctgaatattccgactttttaacttttaaaggaggtttaa
accctgttactggaggtttatggctaagtgatactgctcatcatcatctagcattagctg
ttctatttcttgctgcaggtcatatgtatagaaccaattggggtattggacatagcatga
aagaaattctagaagctcacaaaggaccttttaccggcaacggtcacgaaggtctatatg
aaattcttacaacttcttggcatgcacagcttgcaattaatttagctatgatgggatctt
taagcatcattgtagcacatcacatgtatgcaatgcctccttatccatatattgctactg
attacccaactcagttatcgctcttcactcatcatatgtggattggaggattttgtattg
ttggagcaggagcgcatgcttctatatttatggtaagagattataaccctgcagaaaatt
ataacaatcttttagatagagtcattaggcatcgagatgctattgtttctcatctaaatt
gggtatgtatatttcttggattccattcatttggtttatacattcacaatgatactatgc
gggcacttggaagatctcaagatatgttctctgatacagctatacagttacaacctattt
ttgctcaatgggtacaaagtatacacactttagctcctggaaatacagctccaaatgcat
tagcaacagctagttatgcatttggaggagatattgtttctgttggtaacaaagttgcaa
tgatgcctatttctttaggtactgcagatttttagttcaccatatacatgcatttacta
ttcatgtaactgttttaattttagttaaagggttccttttctcaagaaactctagactaa
ttcctgacaaggccaatcttggcttcaggtttccatgtgatggacctggtagaggtggta
cttgccaagtgtctggctgggatcatgtttttcttggcttattctggatgtacaattctc
tgtctgtagcaatttttcactttagttggaaaatgcaatcagatgtttggggtagtgtat
ctccgtctggaaatgtttctcatattactggcggtaattttgcacagagtgcaattacaa
tcaatggatggctgagagatttcctttgggctcaagcatctcaagttattcaatcatacg
gttctgcgctatctgcatatggattaatttcttagcagcacatttgtatgggcattca
gtttgatgttcttatttagtggtagaggttattggcaagagcttatagaatcaatcgtat
gggcgcacaataagataaaagttgctcctgcaattcaaccaagagctttaagtattactc
aaggtagagcagtcggtgttgcacactacttattaggtggaattggtacaacttgggcat
tctttttagcgagaattatttcagtaggctaatagtgaaaataggataaaaaaaacaatt
atggcaacaaaatttcctaagtttagccaagctttatcacaagatcctacaactagaagg
atttggtatggtattgctacggcacatgactttgaaagtcatgatggaatgacagaagaa

aatttatatcaaaagatattcgcttcgcactttggacatctagcaattatcttcttatgg
acatctggtaatttattccatgtagcttggcaaggcaactttgaacagtgggtattaaat
cctttgaaagttaaaccaattgctcatgcaatttgggatccgcattttggacaacctgct
ttgaaagcttttagtaaaggtggctcagcttatccagtaaatatagcatattctggcgta
taccactggtggtatactattggtatgagaagcaatcaagacctgtattctggggcttta
ttcttactagttttatcagctttactcttatttggagggtggctacatctacaaccaaaa
ttcaagcctggtttatcatggtttaaaaataacgaatcaagattaaatcatcatttatct
ggattatttggggttagttctttagcctggacaggtcatttagtacatgttgctatacct
gaggcaagaggacaacatgtaggatgggataattttacaaccgtattacctcatccagcc
ggtttacagccgtttttcagtggtaattggagtgtatatgctcaaaatccagatacagct
caacatttattcggaactaatgaaggtgcgggtacagcaattctgacatttctaggagga
tttcatcctcaaagtcagtctttgtggctaactgatatggctcatcaccatttggctatt
gcagtagtattcattgttgctggacatatgtataggactaattggggaattgggcacaat
ctaaaagatatttagatgctcatagaccacctagtggtagattaggagctggacataaa
gggctatttgatactattactaattctttacacatacagttgggattagcattggcttcc
ctaggtgtaattacttcgttggtagctcaacatatgtatgctatgcctccatatgctttc
atggctaaagatttcacaactcaagcatctttgtacacacatcatcaatatattgctggg
tttctaatggttggagcttttgctcatggggcaatattctttgttcgagactatgaccct
gaacagaataaagataatgttttggctcgtatgctagaacataaagaagctatcatttct
catttaagttgggtaactctatttttagggtttcatacattaggcctttatgttcacaat
gatacaatgattgcttttggaactcctgaaaaacaaattctaattgagccggtatttgct
caatggattcaagcctcttcagggaaagcactttatgggtttgatgtgttactatcatcc
tctactaatatcgcaacacaagctggtagcaatatttggctgccaggctggttagaagcg
attaatagcggaaaaaattcattgtttttaacaattggtcctggtgacttcttagttcat
catgcaattgcattgggattacatactactacgttaattttagttaaaggtgctttagat
gcaagaggctctaaacttatgccggacaaaaaagactttggatatagttttccttgcgat
ggacctggcagaggcggcacctgtgatatatctgcatgggatgcgttctatttagctgta
ttttggatgctaaatacaataggttgggtaacattttattggcattggaaacatattaca
atatggcaaggcaatgcaactcaattcaatgagtcttcaacttatctaatgggatggttt
agagattacttatggctaaattcttctccattaattaatggttataatccatatggcatg
aataatttatcagtatggtcatggatgttcctatttggacatttagtatgggcaacagga
tttatgttcttgatctcctggcgtggctattggcaagagttaattgaaactctagcatgg
gcgcatgaacgtacacctttagcaaacttgattcgttggaaagataaacctgttgcatta
tcaattgtacaggcaagattagtaggtttagcacatt-tttctgtaggatatgtattaac
ttacgcagcttttgtattagcttcaacagcaggcaaatttggttagactaaa-----ttc
gaattagctaa-----aaaaaagtcagtcataata-gttttatgactgactttttagat

ttaaaacaaaaacttactaaagtttcacctcaatatctacaccagaaggcaaatttaact
tcattaatgcatctatcgtttgagaagaaggttgatgaatatcaataattcttctgtgag
atcttatttcaaagtgttctcttgagtctttatctacatgtggagaacgtaaaacacaat
aaattcttctttttgtcggtaaaggaattggccctactgcaacggcattagttctagatg
cggtatctagtattttgtcacacgacgtattaagtataatagaattatatgctttcagtt
taattctgatttttgtctgctgagtaattgtcatgttaaaaagatcttattatttaagaa
ttttagagacaacacctgcgcctacggtacggccaccttctctaatagcgaaacgcatac
cttgctcaattgcaattgcattaattaattcagcagtcatcttaattctgtcaccaggca
taaccatttctgcatcagtaccatcatcagcagtaaactgattaatagtaccagttacat
cagttgttctaacataaaactgaggtctatatcctggaaaaaatggagtatgtcttccgc
cctcctctttagttagaatataaacttctgcttcaaattgagtatgaggtgtaattgtac
caggtttagctaataccatacctctttcaatatctttttctgcacacctcttaaaagaa
ttccaatattatcacccgctagaccttcttctaacgtttttttgaaacatttctaatccag
taatagtcgttgtacgagtttctcgtaaacctacaatttcaattgtgtcaccaactttaa
taatgcctctttcaattctaccagtggcaacagttccacgtcctgtaatagaaaaaacat
cttctacagccattaagaaagttttatcgacatctctctccggcgttggaatatatgtgt
caactgcttccataagtgaaaaaatcttgtcaacccatttatcttcaccttgcttaatag
ctgggtttttcgtcacagcttctaatgccaataaggcagaacctgcaacaaaaggaatat
catctccaggaaagtcgtattgacttaataattcccttccttctaattctactaattcta
gtagctcttcgtcatctacttgatcttccttatttaggaatactactaatgtaggaacac
ctacttgttttgctaataaaatatgttcacgagtttgtggcattggaccgtctgccgcag
atacaactaaaatagctccatccatctgagcagcacccgtaatcatatttttttacgtagt
cggcatggcctggacaatctacgtgagcataatgacgattatctgtttcgtattcaacat
gagcagtattaatagtaatacctctagctttttcttctggagcggcatcaatttcatcaa
atttttttcgctgcagtagaccctaaagttgataaagtcgcagagatcgctgctgttaaag
ttgtcttaccatgatcaacatgaccaattgtgccaatattgacatgaggtttttttacgtt
caaatttagatcgagccatgctttttgttttccttatagtaaaagttgctttatagttac
ttttaacgataaatcttgtggattgaaagatttattatttaatacagacaaga-----ta
caatataactaatctaatatcgatagtgagcaaacgctttatttgcttctgccattctat
gtgtatcctctctctttctaatagaatttcctgtttcattagctgcatccataatttcat
tagctaatttcatagacatacttttaccagatctgtctctagagaatttagtaatccatc
ttagtgctaaattcgtacctctataagctcgtacttcaataggaacttggtaagtagaac
caccaactcttcttgcttttacttctactagaggagtaatatttcggatggctttttcta
aaatatttaaaggatctgattctgttctctctttaacgatatctaaagcctgatatataa
ttctttgagataaagtttttttaccacttttcaaaatacgaacagttaacatacttacga
gtctgcttttatataaaggatcaggtgatgcaaaccttttttttagctgtattacgacgag

acatagtatttaaaattggttattgtaataa-tgttc-tattagcaggtatttaagattt
aggctttttgtcccatatttagatcggcttttacgacgatcttttactcctgcagcatc
taaagtaccacgtactacatgatatcggactccaggcaaatctttaattcggccgcccct
aatcagtaccacagaatgctcttgaatattatgacctacgccaggaatataagctgtaac
ttcaaatccagatgttaatctaacacgcgctactttcttaaagcagagttcggtttttt
aggagttgtagtatatactcttgtacaaacacctcttctttgagggcaactttgaagagc
tggagattttgtttttttatgtatttttcgtctttcggatctaacaagttgttgaattgt
tggcataataaaattgatttaaagtctttaatagatactatttgaacttgctttacatga
attggctagtttttatttatcttatacttacgcataaatctttctactctaccttctgta
tcaataattctctgtgaaccagtaaagaatggatggtttcctgaccagatatctacgtgt
agttctggctttgtggacccaatcgtcataattaactggccgtcacagtaaacttttgct
tctggataccaatttggatgtatattatcttttgccattgtttttataattatataaatg
tatcaataataactattaacgcttagaaaattgaggagcctttctggctttctttaaacc
atatttttttctctctttactcttggatctcttgttaaatagccttcagacttaagtgt
cgttctattttctggattaattgagcacaatgctcttgcaacacctaaacgaattgcatc
agcttgccctgtcaaacctcctcctctagcatttacatggatatcatactggtttagtag
tcctaaaacttgtaatggtgcatatgaaactcttaagtaattaggactaaattgaagata
agactctcctggtataccgtttataattaaattacctgaccctgggactagtcttacttg
tgcaacagaacacttccggcgacctgttccagaatagattgcgcgagttttaattaattc
tgtcgacataacattccttgataactaaataaatatacattctttatactatatactctt
gtggcttttgcgcgacatgtgggtgaattgggccagaatacactttaagcttcgtaaata
gttttctgcctaatggacctttaggaagcatacctttaaccgatttttcaataattctgt
taggtaatcttgtctgaagctgatcaaatgtttcaacttttaatccgccaggctgtccag
aatgtcttctgtatagtttttgattcgttttatttccacttacagatacgtgagcagaat
taataacgattacataatctccagtatctagatagggtgtataagaaggcttgtttttac
ctcttaagatattagaaatatgagtagatattctaccaagtgtctgattttagcgtcta
taacataccaatgagaattagtatttaatgagggtgattgcgttttattcatgaaaagat
taatacttttggttgacaataagagaatatgtaaatggtttacattcaatttctatacaa
gtagtattt-aaaaatatta-------------ttaacaaagagaacttttctaacagtt
atttaaattgtttgtcagcaagaaaatttctagttatcaataacttatactatttttaat
aatataaaatatattaatcactttttttcttttggcaaacttatacctagttttttttgc-
aaggcatatataacttcttcagctgatttctggccaaaattttaatttctagcagttct
tcttgagagtaatctagtaaatctgcaatagaatggatttgagctcgctttaagcagtta
taagccctaacagataactgcagttcttctattaaaacttgactaatttttttatcttct
ttgctacgataattatctgctgatttaaagtctaaatttcttagtgaacaaaaaagatta
gttaaaactgtagctccttgacttattgcttcttgaggagagatgctcccatttgtccag

atttgtataatcaatctatcttttatgctgttgctaccaatacggacttcttctacttta
taattaactttattaactggcataaaaacagagtctacttgcaaaaaatccacagataat
tcatctacagctttttcagctaagcgatacccgcaatttttttcaattttaaactccatt
tcaaatattgtattattgcaaatagttgcaatatactgtctaggatctactacctctata
tcagaagataattcaaataggccggctgtaactatagctggcccttgaactctaattcga
ccaatttgagattctttgttataacttttaaatactacttcttttagattaagtaatatt
tctaacacatcttctcttaccccaggaattgtagaaaactcatggttcactccagcaatc
cgtacagcaactatagcagtaccttcaagatctgacaatattgatcttctcaaagcatta
cctaatgtaataccttgtccttgatttaatggttctattacaaaactaccgtactgccca
cgcgccccatctgttcttgactctatgcattcaatttgaaattgagccacctaagaaagc
tcc-tttaaaataatcagtgattagtaagtctctctaaaacaagaaggaatttatactcg
gcgtttcttaggagggcgacatccattatgaggtacaggagtaatatcttttattagagt
aatctctaatcctgcagcctgcaaagctctaattgctgtttctcgacccgctcctggacc
atttactagaacttcagtttggcgcataccttgatccatagcttgtctagctgctttttc
agctgctgtttgagctgcaaaaggtgttcctttttagctcccttaaatccacttgcacc
agaggatgaccatgataatgtttctccttttaaattagtaatagtaacaattgtattatt
gaatgtagatttaatatgtgtaataccgttaactgcattacgtttagtttttcttgctcc
ggatttttttatttgtctagccatcgtgtctcgtctattatatagttaattaaaagatta
ttttcttggagctttttttcttacctgctactgtttttttacctcctctgcgtgttctagc
attagttctagttctttgtcctctcaaaggaagaccaagacggtgtcttctacctctata
agtactaatttccataagtctcttaatgctcatagactcaaaacgtttgagatccccttc
aatctgataattagactcaagaatttctcttatactgacaacttgttgatcatttaaatc
ttgacacttaatatcagcgtctatgtttgtttttctaatatttctttcgagcgagatag
tccaataccataaatatatgttaaagctatctctattcttttgtttcttggaagatctac
tccagcaattctggccactttgtcttgtctccaataaataatatgtat-ttttataacgg
ttgtttata-agcttagccttgtctctgcttatgtttaggattagtgcaaattaccatca
cctttctgtgacgccgaattattctacatttttcacacatttttcgaacagaaggacgaa
ctttcatattaa----ctctatatactaggaatattactatattaactctattaattatc
gacattaaaaatgtcaacaatatattgtactactttaactgttgtagattttacttcgtc
atactatcatatttttttgatatgacgtatgtctgaatttgcttagctgtgtctattgcg
actccaactaaaataagtaaagacgtagccccgagacctcttaagttttggatctgagta
accttttctattataaacggaatcagcgctactgtaaataaaaacgaggctcctaggaat
gtcagtctatttaatatgacttgtaaataatcaatagtcgcttgacctggacgaatatta
ggaatgcttgcacccatttttttttaaatttatagcaatatcttctggattcattactatt
gatgtataaaaatagctaaagaaaaggatcaaagcacaataaagaagaaggtatagtgaa
ccattaggacaaaacaggtataggatttggagtaatgttttattttgaataatttgggtt

agataagatgggagggccatagatgcagatgcaaaaacaataggcataactccaccttga
tttaatttcaagggcaagtaactatttggatccaaaattgaagattttcctagctgtctt
gctgaaataattttaattcttcttgttccttcttgtacacaaattgtaattattatcatt
aataaaaagatcgctataaataatccgaacttaagacttgcattactataactagcatca
aaaaatgactgtgtaaaattctttggtagtcctgacacaatgttttgaaaaataagtaaa
gaggctccattaccaattccctttccgtaattaactctgataaccacataataatcata
gaacctgctgttaaagctaaaacagactcgcagacaaatgcaaaattccaattaaaaaca
tatggttttacccatatagagattgctccagattgtaaagtcgcccaacctagagctaag
taccttgtaatttgagttatttttggcggcccaattcaccttcttccttctgtaatttt
tccagattggggacaattttcgtaagtagctgcattacaattgaagagttaatataagga
acaatacctaacgcaaaaatccctattgttgaaaaacctcctccagaaaaaatattcaga
aaatttactaaagtatttttttctacacttgcataaaaggcgtcatgatctatacctgga
acaggtataaatattcctaaacgtgctaaaactaatagaaagagagtaaagataatacga
tttcttagatcactttttggctcataaataaattttaaatgaaaagagatacttt--tt
tatgtaaaaaccctattccaagacttgtataagtttaaaatagggaagtaa---aatatt
atagtaatattaata-taatattctatgtagaatacaggctttctattggaactcctcga
tctttagctgcttcactaaaagttcttaattgagttaagccattgagaacagctcgcgcg
ttatttaatgtattattagaacctaattgcttagctaatatattctgtacacccgacaat
tcgaggactgtccttacagaacctccggcaattacacctgagccaggtgcagagggcctt
aatataactttcgcggcaccagaaattccgttaataggatgaggtatagaattcgatttt
gtcagtggaactgtaactagatgtttttagcatctgttactcctttttttactgcacca
attacatcgcttgcttttcccacgccgacaccaacttggccttgctcattaccaataaca
agaataactcgaaagcttagtttttacctccttttacaactttagtaactcttttaact
tgtacgactctttcttcccagccactatctttatcttttcctttgctctgtttttacga
ttggccattttaagaattattcctaattagttaatta-cgattctagaaacccatgcctg
cttccttagcagcttcagccaaggccttaactcttccatgatataactttcctcctctat
cgaagacgacatttttaatgccttctttcatagactgttctgctaactgtttaccaacta
cgcgagaagtatcacagtttggtcttatgttatcagattctttattatttaaattaacag
atgatgtagctactaatgtaataccttgtgtgtcatcaattatttgtgcgtatatatgtt
tattagatctaaatacacacagacgaggcctacttgaggttccttgaactttttccgaa
ctcttttatgcttatgaattctagtttgtttagtgtttagtttcattattatttaccttt
tccagctttcccagcttttcttctaacaaattcaccttgatatctaattccttttccttt
ataaggctcaggaggcctaatagaacgaatagttgaagcaacctgaccgacaacttcttt
atctatgccagagacagtaatatttgtgttgttttcaactttaatttcaatattggcagg
aggtttaattttcactacatgactatagccaacacttaagattaaatcttggttatcaat
ttgagaacgatagcctacaccttgtatttgcagtttttaaaaaatccattagaaacacc

ttcaatcatattactaataagagttctggatagcccatgtaactgactagccatttttgt
ttgtccactcgtcttgacagcaattgtgtcgttaagtatttctaaattaataccagctgg
taaagttctggatagggtacctttaggtcctgtcacggtgatagtctgaccatcgaactg
agtactaagatttgtcggcaataaaattatttttttccaatacgagacat-attccacc
tataaaatagttaccaaatatagcataatatttcaccaccaagaccatcatgacgagctt
gtctatctgtcataacgcctctagaagtagaaataagagcgatgcctaaacctccaagaa
ctctgggcaactctttatgatttgcgtagactcttagtccgggtttactaattcttttca
gagcagtaataactggttgacgatttttaccattatacttaagagaaatcattaaatgag
tctctataccttcacccatttgttcgaaattttgaacaaatccttcttctttcagtactg
ttgccatattgcatgtcattttcgttgctggaacttgcacaatttgatgtcttgctaagt
ttgcgttacgaatacgtgtcagcatatcggcgatcgtatcgttgaccaccttgatcctcc
ttgcga-----tgaattatttgaaatattttcaagattccctgaaaggcatccctagctt
ttttaacaaagctagaccttcttgatctgttttagctgtagtgacaattgatatatctaa
accacgaatttgatcaatattatcatagtctatttctggaaagatcaactgttcacgtaa
acctaaattgtaattacctttgccgtcaaaacttctaggactaattcctctgaagtctct
aattcttggtaatgttaaattaattaatttctctaaaaaagaatacatcttgtcttttct
tagatgcacaacaattccgataggaacctcttctcgaattttaaatcctgcaatagattt
tttagcttttgtgacaattggtttttgtccagttattaatgtcaattcttgaatactact
ttcaagagccttagcattctgagaagcttcacctaagccacggttaatagtaattttagt
aaacctaggaacttcatgtacatttttgtactgaaattcatcttttaaagattgagtaac
agttgttttatatttttcttttaatcctattgccattatattttcatcaattatttaata
agttcgccagtctttttcagttttcgaa-tttttggcctttatcattaattattactga
ggatcgactagcaatattattttgttcactaaataacataacgttagaggtatgtattgg
agcttcaaatttgataatttctcccgtttctccttcttgttgaggtttttatgcttcac
tttgagattaattcctttaacaatcaccttatttgttttataaataattgcaataatttc
acctgttttgttttatcacttccagaaataacttgaactaagtccccttttttaattt
aatttttgtattatttttggttgttttagaaggacctttcattatactacctccggtgca
agagaaactattttagaaaaatttttatctcttaactctcgggctataggtccgaagact
cttgtgccacgtggattattatcttgattaatgataactgccgcattatctccaaatcta
atgctcataccgtcagttctccgtaaagctttacgagttcttactacaacggctctgaca
acatcagatcgtttaactggcatattgggagacgcatctttgactactccaataataaca
tctccaatagatgcataagaaggattactagtgcctagtactctaatacacattattttt
ctagcaccactattatctgcaacattaagatagctttgagtctgtatcatacttttattc
tgtataagattaattatcaaaagacttagataagatattaaccattgtccaacactttgt
gcgacttaaaggtcgtgtctcttgtattgtaacaatatcgccgattgtgcattcattatt
ttcatcatgcgctttatatttctttgtccggatcatagtttttgcgtactttctatgaga

gattctattttctacagctactactatagttttattcattttatcgcttactactttacc
tgttgtttcttttaaaggcatagtctttatattttttatatat---------aattaaat
tttaagattattacatggttgacttagtacgagacttttcaacagttaaaagttgggcta
atctatgcttagaatgtttaaatagatgaggctggaaatcttgccttgtggctcttttta
gccttaaatcaaaaagctctcttttttattacaaggatttcttcagctaaagaagaagagt
ccaggtttgtaacatctgatattttagggaaagtcatattttaggattctgttgtattcc
gaactataaatttagtcttgattggtaatttataagaagctaatttcatagcttgttgag
cagtttttttgtggtacacctgtaatttcaaatagaatatgtccaggcttaataactgcaa
cccaatattctggagctcctttacctgaccccatgcgagtttcagctggacgggcagtga
ctggtttatctggaaatactctaatccatagtttaccacctcttctcacatatctagtaa
tagttcttcgagtagcttctatttgtctagaagttaaccaaactggctctgtcgcttgta
atgcataatcaccgaacgcaattgtgttacctttgctagcagaacctttcattctacctc
tatgttgttttctaaattttgttttcttggggcttagcataaaaattaaaaaataataat
aattaagaagctgttgaatcaggagtttcctgattataagcagattctagacttttcgat
tctggcaaaatttctcctttaaaaagccaaacttttactccgagcacaccatacgtagtg
tgagcttggcgatgacaataatcaatatcggctcttaaagtttgtaaaggtacacgacct
tctctaacccattcacttctagctatctctgcaccattaagccgaccagatacttgaatt
ttaacaccctgtgtatttgctctttgagctctttgaactgcttgtctgacagctctacga
aaggccactctttttttctagctgttgagtaataaattctgctactaaggttgcttcagaa
tctggatctgcaatctctacaacatttactctgagttgtttgctagggtctaggattaag
gacagcgattttcttaaagactcaattcccgctccagattttccaagtacgattcctggt
cgagctgttgctatgagaatttctacttgatcaactttacgattaatttcaattttagca
atactagcattactaagttttgaatgtataaatgagcgaattttatgatcttcttgtaaa
agaactgggtagtctttagagttagcaaaccatgaagaacgatgtttttgggtaatgcct
atgcgaaagcccaaaggatgaatttttttgacccacagtatccttgattataagttaaaag
ctataatgactaattgacgttaaa--------agactacagttctgaaacacctaatgta
atatgacaagttggtttatgtattggaaaggctcttccttgtgctctaggctgaaatctt
tttaaagtcgggcctttgtctgcaaaggctttactaacaaacaattggtttttatttaat
ccatcattatgctcagcattcgctgcagcagattccaggatttgctttatatgagaacag
actcggtatggcataaattccagaataattaatgcttcttggtattttctacctctaatt
tggtctaatacacgacggactttatgcggagaaagacgaatgtatttccctactgcttta
gtttcttttacgtttttttgtaatactcatagttatagtttaacgacgggcttttctgtca
ccttttacgtgagtgcggaaagttcttgttggaacaaactctcctaatttatgtcctacc
atttggtctgacacaaagacaggaaaatgttgtttaccattataaacagctattgtatgc
cctaccatatcgggaataatagtagatgctctcgaccaagtttttaggacttctttttttt
cctgaaatatttaatgcttctatacgtttaagaagactaacatcaataaaagggccttta

tgtatagatcttgacataataataaatcattagattcttaataaaataaattctatttac
ggcgacgtaagacgtacgggttactatatttatttggatttcgtgttttaacacccaatg
caggtttaccccaaggtgttacaggacgagctcgtccaattggagatttgccttcaccac
caccatgtggatggtctacaggattcatgacaacacctctaacagtaggtcttttaccta
accaacgatttctaccagctttaccaagagtaatattactagcatcaatattaccaactt
gtccaatagtagcgtagcattcttttcgaatcattcgaacttcgcttgaaggcaatttta
cagtaacaaaagtaccttcttttgctactatttgagcataagttcctgctgcacgaacaa
tctgtcctccgcacgaaggtcttaattctatattatgtactgctgttcctaaaggaatac
tagataaaggcaaagcatttccaacttcgataggggcagtaggaccagaaagaaccatag
atcctacactgagagatcgagggtgtagaatatatcttttctcaccatcaagatagtgta
ataatgcaattcttgcatttctattcggatcatattcaatagaagcaactttagcaacta
tattatgtctatttcttttaaagtcaattaatctgtactgctgcttatgtccaccacctt
tatgacgacaggtaatgacacctctattatttcgacctttgcagaaatggtgtttaacta
ttaatgatttttctggcttatcagtagtaatctctgaaaaggtagaaaccgttctatttc
ttgtccctggtgtgtaggcacgatataaacgaattgccatataagaagatttgagaagag
taaatgttgaa------ctatgtgatatatttaagtttctggaaacaaattaatagaatc
ttctgaggcaagtgttacaattgctttcttgtaatgtggtcgcttccctacgaatctacc
gatgctcctttttttcttaggaggatgacaagtgttaacacctgtaacttgcacgttgaa
aatatactgtatagcagccttaatattaatttttgttgctttgggatcaacggcaaaaca
atactgattctcttctaataatttagttgtcttatcagtaatgatgggatatttaaccag
atctaataaacctcttgaatcaatgctatccattatatacctcttgtattttagataaag
catcaactgtaataataattttatgtgcagctaacaaagccataatatttagtgtatcag
ctgaaataagttctacgttgtgtagattgcgaatagaaagataaacattcggatcttttt
tgtctactatgactaaaactttttatttaaatcaagattccatcgatgtatagcttcca
taaatagtttagtcttgggctgatgaaaataactgttaaaattttctacaatcagtgtat
taactgatttgttatttaatgctgtccttaatgctaattgtctttctttttattcattt
ttttggtaaaactacgaggttttggcccaaatattacaccaccacctctccatagaggtg
aacggatagaacctgctcttgctcgaccagtcccttttttgacgccatggtttacgacctc
cacctcgaacttcacttcttgttttagtattagcagaaccctggcgcttttcattacttt
gctttactaaagctctatggactaagtacatcccagaatcttgactgaccttaagattta
aatcagcgttgccacttacttgaccttcccaattatagacttgataatttaattgcgtgt
taactgtcatgaaatatatcgtatgaaagaatattgctacttactaatttttaccaaagc
gccaggtttaccaggaacagctcccttaacgatcaacaaatcattctctgagttaatact
tacaatttgcaaatttttttattgtaactttttttattaccaagttggccagccatgttttt
acctggataaactcttccaggcgttgtaccggctccaatagagccaggttgcctatgatt
tttcgaaccatgtgacattgggcctctactaaaatgatgcctcttttggtaaccggagaa

acctttacctacacttctagaagagacattaatcttttggcctacttgaaatagatctgt
agatagtatttggctaacttcgaaatcatcagtagatttcatttcgtattcacgtaaata
ttttagtgggggagcctgtgatttttttaagtgtcccaacaatggtttatttaatttttg
ttcagcaacttgcttgtaaccgacttgaatagcattgtagccatcagtagatacagcttt
aatttgagtaataacacatggcccaacttgaattacagtaactggaattgataaaccagc
ttcatcaaagaattgggtcatacctactttagtaccgagtataccaacagacactagatt
tctcctttttgagagatttatactatcattgtacataaaaattctacttcatagttactg
aag--taaattaaactaacagatgcatgatattattcatgacaacataaaggttaaaaac
tttacctatttaatctaaaatagaaacagctttttgtctagttacagcgtatataaatta
gaagtatgcttattaaatttttttataaactact-tagatttaactgatattgatatccgt
aacttacttgcagtaatatc----aatcaacatgtataagtttaattgcctaaactagaa
ccagctattattagaaataattcaagtatagataaaagcgtatttgaacttttatatatt
gacaaatataatggaatatattgaggttcggtaactacaccttttttattttttacatttat
tgtgca--gtatatatacatctaatgcagtaaacataagaatataac----tgtatagca
aatgtaattacacaaattagagaggtattaatgggcaaagtcgttggaattgatcttgga
acaacgaattctgtaattgctgttatggaaggaggtaaacctaccgtcataccgaatgca
gaaggttttagaactacagcttctgttgttgcatatactaaaagtggagataaacttgta
ggacaaattgccaggaggcaagctgttattaacccagaaaacactttctactctgtcaaa
agattcataggacgtaaacagaatgaaatttcgcaagagattcggcaaacatcatataat
gttaaaactagtggatcaagcataaaaattgcttgccctgcactgaataaagattttgct
ccagaagaaatttcagctcaagtactgagaaaacttgttgaagatgctagtacgtactta
ggtgagactgttacacaagcagtaataactgtacctgcttactttaacgattcacaaaga
caagctactaaagacgcaggtaaaatagcaggcttagatgtattgagaattattaatgag
cctaccgcagcttctttgtcttatggactagacaagcaaaataatgaaacaatactagta
tttgaccttggtgggggcacatttgatgtatctgtattagaagttggagatggagtattt
gaagtactctcaacttctggagatacacatttaggcggagatgactttgatcagcagatt
gtagaatggctaatcaaagattttaaacagaatgaaggtattgatcttggtaaagataga
caagcacttcagagattgaccgaggctgcagaaaaagcaaagatagagctgtcaaattta
actcaaacagagatcaatcttcctttttattactgccacacaagatggtccaaaacattta
gaaaaaactgtaactagagggaagtttgaagaactttgttcaaatttaatagataaatgt
agtatccctgtaaataatgctctgaaagatgcaaaactagaagcttccagtattgatgaa
gttgttttagttggtggatctacaaggattccagccatacagcaaatggttaaaagatta
attggtaaagatccaaaccaaagtgtcaatccagatgaagttgttgctattggtgcagcc
gtacaagctggagttttagcaggcgaagtcaaagatattctattactagatgtgacgccg
ttatctttaggagtggaaactctgggtggcgtgatgacaaagattataccaagaaatact
actattcctacaaaaaaatcagaagtattttctacagctgtagataatcaaccaaatgtg

gaaattcaagtacttcaaggcgaaagagaactaacaaaagacaataaaagcttagggacg
ttccgattagatggcattatgcctgcacctagaggagttcctcaaattgaagttaccttt
gatattgacgctaacggaattttatctgtaaaagccaaagaaaaggctactggtaaagag
caatcaattactatatctggagcatcaactttacctaaagatgatgttgaaagaatggta
aaagaagctgaagaaaatttcgatacagaccaaaaaagaagaaagaatattgacacaaag
aatcaagcagagtccctatgttaccaagctgaaaagcaagttaaagagtttgaagacaaa
attagccaagatttaaaaataaaaatagaagagctaattacagagcttagatctagtcta
gagaaagaagaatatgacaatattgaatctatttctcagcaattacaaaatgctctgatg
gacattggaaaaaatgctgctcagactgaaagtaaagatacaaaagcgaaggatgacgac
actgtgattgacactgatttctctgaagctaagt-aaaagta-agcgggtaacgcgattc
gaacgcgcgacatcaaccttggcaaggttgcgctctaccactgagctatacccgcataga
ttgtattattacaaaaattaagatatttgtcaatctgtta------atctatac---atc
tagactattaattttggaattacatattagctaactgttgtaaatataacaattgctata
attttttaa-------aagtacttatatcat-tatataagtactttttatatt-------
-caaaaatacaaataagtattatag-------ctaaactactaaggaattaactaagcta
gtaacaattactaatcctgtccataagccggctcctgtatatattaatccttttgattgc
tcccattggccaggggatgctaaaacaacaggtacgcctactactaacacagtcgataat
gcaattaataatagcactaataattggatcgcaataatcatttaatatgttctcctgaaa
aattcaaatgttaataatactgtatataatacaatgaagcttttgaccttcagcaattta
tctatgaagaaataatttaaaattattgttttctaaataaaccttaaattaagtattaaa
atctttattgtaaacaataatactatattttttta--gtagataaaaaatttaaagaaag
tctttttaaaaaaaaattaagataaggtgatatgctgtataaaaagctaagtattatta
attattagagaaatttaaa-ataagaggtaaaacatatgaactcagcccttttttttagca
aaactaccagaagcttatgccattttaaaccgatcattgatatcttacccgtaattcct
gtatttttcttcttttagcttttgtttggcaagcagcaattggttttagataacattta
ggttataaaaagaataacaaatatacagagtgccaattttgagctctaatcttatatct
tacacaataataactattatggttgaacccttactatcaggaattgttcttggacttatt
cctattactttattcggattattagtagcagcttatttacaatatcaacggggcaaccaa
ttaggactgtaaagcctaaattatagatatttataaaaactacaataattatagttttta
taaatatctacaaataagctaattttaaatacagaaataatactattgaaactagagtga
taacaaaaaaaatttatgtaa---taa---------------------------------
------------------------------------------------------------
------------------------------------------------------------
------------------------------------------------------------
------------------------------------------------------------
------------------------------------------------------------

------------------------------------------------------------
------------------------------------------------------------
------------------------------------------------------------
------------------------------------------------------------
------------------------------------------------------------
------------------------------------------------------------
------------------------------------------------------------
------------------------------------------------------------
------------------------------------------------------------
------------------------------------------------------------
------------------------------------------------------------
------------------------------------------------------------
------------------------------------------------------------
---------------------------caagctaattgtttaccagcttgacttagtaac
tcctggtaaaagacattcatgagacatttctcgaaaaacatggcgagataatccaaaatc
tcgataataacctctacttctacctgttaaccaacatctatttctccctctaacagcggc
gctgtttctaggcatttcttggagtttttgtcttaaatccattttgtcagaaaagctgcc
agcttctttcagttgatctttaatagccagtctttttaaataatatttttttgcaagttt
cgatctcttaatctctctttggatcatattctttttagccatgctgtaa-aactctcttt
taaaattatttattcaatatattagctaaaaaagctcctgcctgcaacttttatatattt
aaaaataaatagcataaagtagtgaaatgcaattgatatttaatcaacgatacc---act
ttatgctattatcaaaaatttatttagtaaagttcttgttcaacatgagttgaaatagtg
caatcagatgaaggatatgcaatacaggttaaaacatagccagctaaaagttgatcatca
tctaagaatgattgatcagcttgatctacagttccttctgtaactttacctgcacaagtt
gaacaagctccagctctacaagagtaaggaagttcaattccttcttcttctgctgcatct
agaatatatgtatcctccgcacaattaaatgtgacatcaattccttcttcttcacatagt
aaatgaattttataatcagccatttcttcttactccttaataaagatacacttgaaactt
aataaagcaaaat-aatttaaagaacttaccaaaaaattatatactagtttcattatgca
tgatagactggaaaaaaagaaaatagtatactattctgaaaaatcaagctacatttccta
ctaaatagaactatagtaggtccaaagtatatacataaattcatatagttaaaattactt
ttatcatttaaatagctgaacttgatagta---------tagattgatcatc------aa
gctagtcttctgc---caataaataattaattatgatttcaatgcttaatcaacaagtag
aacttaagcctattcttaaatttcagataacacaaatatattcgcattctcttattcagg
aaattaaagccttaataactagacttgttttgcaagtttggagaaggccggctacattga
tggcaggtattatccaacctttattatggttaattctatttggtgggcttttctacaatg
ctcctataaatttgttcactattaatacaagctataattgtttttttgagctctggatta

taatttttacctcttttactggagctttgaattcaggtcttccattaatgtttgatagag
aatttggattttttaaatagattattaacggctcctttagtatcgaggacttctatcattt
tatcttctgctacttttatgacttgtattagtttaatacaagttgtatttatagttacag
cttctcttttttatgggaaactcacctctaaatagcgatagtactatgattttggactta
tgattctattagtgactgtaggagttacaatgcttagtttagctttatctttcactctgc
caggtcatattgagctattagcatttatttagtagttaacttgcccttttattttcta
gtacggctttagctcctttatatttcatgccgccatggcttcagttaattgcaagtctca
atccattgagttatgcaatagaaggtacaagatacttatattcaagcgtaaactggaatt
ttacagagtgtgtgattaagattagctggggagatatttgtttagggcaaattattattt
tattaatcgctttagatataatggcagcttatcttgtgtctaatatattaaaagctaaac
ttaatta-----------------aaattttaataaaaaatttattaaatttattttaca
aagattttactatataatactaatagtagtatggataattgaaatagaaatagtttttc
aaaaccaaagctattatcaacaatgataaacatttgtaagaaagtcaacaaagtatgttc
ttattcataggaggcatgtagtcaatgggactaccatggtaccgtgtacacacggttgtt
ttaaatgatcctggacggttaattgcagtccacctaatgcatactgcacttgtagcgggt
tgggcaggatctatggcattatacgaactagctgtatttgatccttcagatccagtatta
aatccaatgtggcgacaaggcatgtttgttatgccatttatggctagactgggcgtaaca
gattcatggggaggatggagtataacaggagagagcgtatccaatcctggactgtggagt
tttgaaggtgtagccttaactcatatagtcctttctggcatgctttttctagctgctatt
tggcattgggtttattgggatttggaattatttagagatccacgaactggcgagccagcc
ttagatttaccgaaaatttttggaattcatttattgctatcaagtctactttgtttcgga
tttggagctttccatgtaactggacttttggaccaggaatgtgggtatcagatgggtac
ggagtaaccggaaaggtattaccagtagctccagcatggggaccagaaggattcaacccg
tttaatcctggaggggttgcatctcaccacattgccgcaggtactgtaggtatattagct
ggtgttttccatttaactgttagaccaccacaaagactgtatcgtgctctaagaatgggt
aatattgaaactgtattatcaagtagtatctctgctgtttttttctcagcttttgtgact
tgtggtacgatgtggtacggttctgcaactacgcctattgaattatttggtccaactaga
tatcagtgggatagtggatattttcagcaagaaattgagaaacgagtagaaaatgctatt
gctgatggtgctgcacctagcgaagcatggtcaagaattcctgacaagttggcattctat
gactatattggtaataatccagcaaaaggaggattattcagagcgggccctatgaataaa
ggcgacggagttgctgaagcatggcttggacatcctgtattccaagataaagaaggaaga
gagcttagtgttcgcagaatgcctgctttttttcgaaacttttcctgtaattttagttgat
aaagatggtattatacgagctgacattccatttagaagagcagagtctaagtatagtatt
gaacaagtaggtgtaacagctagtttttatggtggcaaattaaatggccaagttttcaac
gatgctcctagcgttaaaaaatatgcaaggaaagctcaattaggtgaggtatttgaattt
gatcgaactacattagaatcagatggagtgtttagaagtagtcctagaggctggtttaca

ttcggtcatgctaatttcgctttaatttcttctttggacatctttggcatggttcaaga
actatcttccgggatgtgtttgcgggaatcggagctgaagttactgaacaagttgaattt
ggagctttccaaaaattaggagatagaagtagtaaaaaacaaggagctgtataagaccaa
agtcaaagatagtctttatc---aaatatttctaatattaaaatagttaagacaaagttt
attatctaatagttactaatcaggagaatttatggaagccttagtctatgtatttttact
aacaggaacgctaatggttatattctttgcgatctttttagagaacctccaagaatagc
aaagtaacttgattccgtttaaacttataataatagaaccactttccaacttattagtgg
aaagtggttttattgtagattacattgtattattcttcatgctcttcaaaaggatctcgg
agatcttttgaagcaggaccaaatgctgtgtatatcgagtaacctgtaatacctaaaagc
agactcgaaataaaaatactaagaactgttgcagtttccataatttaagcagtattaaga
tgaactttttctataatgatattatcaatattaaaacatttaattattaaaattatcaaa
atatactatggcacttagaactagacttggagaaattttaagacccttaaattcagagta
tggaaaagttgctccaggttggggcacaactcctattatgggaattttcatgttactatt
tttcttgttttattaatcatttacaaatctataattcttcattagtattagaaaatgt
agatgtagattgggctactttaggtag----ctaactagataagttctttga--------
ttacacaatattaaagacaac----------------aagcgatcgctattagattatag
tagcgatcgctttaaaagtatttactatatatccagagaaaattatttacctaaattaa
ttaactcggtatctgcaaaactatttgtgttaacattattatagttaactttttcaaacc
ttactaatactggatatttgataccacttttatcaattgctgcaacagtaccaatttcct
gataccagtacgattcttttctcaatatttttacttttgagcctctttccatatttttag
tcctcagatagattattaaagattattatattaaaaaattaaaaaaaaactaaacaaaata
tataaaaatactaccgacaattgtaatatataactaataaaaggagagagagaggattcga
accctcgataacaaaagttatgacaattttcgaaattgttgcaataaaccactctgccat
ctctcctgaattaaaatactgttttaaaaaattagacaagaataacatagaactaaggta
gttgcctagaaaattaaaactatgttaaatataaattataatctttctatagatatattt
aaaaaatatgtacatagaatctatatcagcttacttcaagatttttaatcttatacaat
ttttactaaataaataaattctctcataatatgaaactatcttggaaaacacttcttctt
tggtctttaccaatatttgtaattggtttctttttctggcaaggtttttaggtccaact
actacagatgttggcagtaatatcgcaagttctagaatgacatatggacgatttttagaa
tatttagatatgggttgggtgaaacgggttgacctctatgaaaataatcatacagcaatt
gtagaagctgttgggccagaattaggaaacagggttcaacgaattcgagttgaactgcca
gcaagtgcgccagaattaattacaaaattacgcaaagccaatgttgatctagatgctcac
ccccctaaaagtacaagtgcagtatggggactattaggcaatttactatttcctttacta
ttagttggcgggttagctttcttatttagaagatctaataatgctagtggtggacctggt
caagcaatgtcatttggcaaatcgaaagctttatttcaaatggaggctaaaacgggagta
gtatttaatgatgtagctggagttgaagaggcaaaggaagaatttcaagaagtggtaaca

tttttaaaacagcctgaatcatttactgctgttggtgcaaaaataccaaaaggcgtttta
ttagttggacctcctggcacaggcaaaacattactagcaaaagctattgctggcgaggct
agtgttcctttttttagtatctcaggctcagaatttgtagaaatgtttgttggtgttggc
gcttctcgtgtgagagacctattcaagaaagcaaaagacaatgcgccttgtatcgttttt
attgatgaaattgatgctgttggtagacaacgaggaacaggtgttggaggtggtaatgat
gaaagggaacaaacattaaatcaactattgactgaaatggatggctttgaaggaaatact
ggtgttattgtaattgccgctactaacagagctgatattttagattctgcattattaaga
cctggaagatttgatagacaagtttctgtagatgtaccagattttaaaggcaggttagca
attcttgaagttcatgctaaaaataagaaaatggaacctaaagtatctttagaaaccatt
gccagaagaactcccggcttttcaggagctgatttagctaacttactaaatgaggctgct
atcttaacggctcgacgaagaaaaaatgcaatgactatgtctgaaattgatacatcaatt
gatcgagtagtagctgggatggaaggcactcctttaattgacagtaaaagcaaaagatta
attgcgtatcacgaagtgggtcacgcaataataggcagtttattagagcatcatgatcct
gtgcaaaaagttacattaataccaagagggcaagcaagaggcttaacttggtttactcct
agtgatgatcaaagtctaatatcaagatctcaaatactagcccgtatcgtaggtgctctt
ggtggcagagctgcagaagaaatcattttcggtgacgcagaagttactactggtgcaagt
aatgatttacagcaagtaacatcaatggcaagacaaatggtcactcgatttggaatgtct
aaaattggacctttatctcttgaaagccaaggaggagacccatttttaggtagaggcatg
ggaggaggctcagaatattcagatgaagttgcaactaatattgataagcaagtaagggaa
attgtcagtgaatgctatgcacaagctaaacacattattatagataatcgagtagtgata
gatagattagttgatttactaattgaaaaagaaacaattgaaggcaatgaatttagagac
atcgttaaggaatacactgcaattcctgaaaaaaattactacatatcacaattttaa---
---------------attaaacgggactgacgggattcgaacccgcaacttccgccgtga
cagggcggtgctctaaccagttgaactacagtccc-aaaaagatgaatacccagagataa
tctcatagttagcttaaacttgtcaaattagaatattaaatatgaagagta---------
--------tttgtaataatactcttcatatttgata-tagtagctaaaatatttctttta
gaaggagagagagggattcgaaccctcggtacgaagttaatcgtacagcagattagcaat
ctgccgctttcgaccactcagccacctctccatga---tatatattatatatatatcaaa
tatacttatatttaataggtgtgcataaatatttatggagctaagcggattcgaaccgc
tgaccctctcaatgccattgagatgctctaccaactgagctataaccc----ctttggta
agatattaagaaatggctcaagcgggatttgaacctgcgaccttgggcttatgaatcccc
tgctctaaccactgagctactgagccatagtataaaatactattaaatattataccataa
attatggtcaaatataaatcaataaaagaataattactatca-tttactactttttg----
------------------tgcaagttgtaaatattcgaataaaatttcaaaattaaaaatg
tctttaactttttgttaga-------atta-ttgattacgaaa-----------atattg
atatttagaattaac---------------------------------------------

--------------------aataaatttg------------------------------
---------------------------------------------------tctttattt
g-----------------------------------------------------------
------------------------------------------------------------
------------------------------------------------------------
------------------------------------------------------------
------------------------------------------------------------
-------tagaca-----------------------------------------------
-------------------------------caaaaatattggaaa--------------
------------------------ttttcatcaat-----------------taattagt
------------------------------------------------------------
------------------taaactataaccgacgtac-----------------------
------------------------------------------------------------
------------------------------------------------------------
---aagaattattgaacta-----------------------------------------
------------------------------------------------------------
---cgttaaat-------------------------------------------------
------------------------------------------------------------
------------------------------------------------------------
------------------------------------------------------------
-------------------------------------tgtgatgcatt------------
-----gggcttgatcaaaacttctctaaaatcaatagaatatagt---------------
-----------------------------------------------------aattcaa
gtaaatcaaatactaacaaatgtgacaaacgcttaaaaaaataata--------------
caattagaatcgtaaaaatttggtttttga-ctgtgttaaataaatttgattatcataga
tttaatatttaaaagtgttctattacttgta-----ttgagtaattgt-aatatttaata
tcttctcttttagataataatcataatataagctaaaaaatagtttt---tattaaacaa
ctagcatagagccaataccttgatcagttaatatttctagcaagagagcatgatctattc
taccatctaaaatatgcgctgaagctacaccctgagctaaagaacgaatacaacaattca
ctttaggaatcatgccaccagaaattactgctgtttgagttaaatctctagcttcctgta
tattcagatgactaattaatgttgaaggatctgaagaattacgtaaaatgccgggcgtat
ctgtgagtagaattagtttttcggcattaagagcggctgctatttctcctgcgacagtat
cagcattaatattatacgattggccttccttgtctgctgcaacactagctataacaggta
tataattattattaattaatatcttcagtaatttagtatcaatattttgtacttctccaa
caaagccaaggtttgcttttccatttggtctcggagtaataagtaatccatcttttcccg
acaaacctacacttttaccaccttgtttattaatagttgcgacaagatctttattgactc

ttccaactaaaaccatttccactatatccatggtaggttgatctgttactctaacgccat
catcgaattttggtaatattttgaggcgatctaaccaaaaattaatttctggtcctccac
catgaactaaaatgggacgtagtcctataaaagacagaaaaacaagatcactaatcactt
gatcttttagtttctgattttcatagccgctcccccatatttataacaataattctgg
aagaaaattgctgaatataaggcagagcttcacttaagacttttactctttctgaattag
tcaacataaataaagtaggattgtgatatataaaattatatacacagcttgtgttaattt
aaaagataccaatgggt---------acttgaaaaaatagaaatattactaacag-----
----ttaaaactatagggattataattatgaataaattttgggataatgtattaagattt
ccacgattttagtcagcgtcattttaggattaattttgataattattagtcctttttc
gtgttattaaaaaaaccactgacaagtttttttttcatcatatcattggccggattaatc
acagttttggcaataataa-tacaaaaaatgataaatatcgaatgttgttgagagaaata
aataaatttaattaaaattttgaacatatataatattatattatcattaaatatgataa
ttt-atttcctttctat---aaaacccataaatattaaatattatttgataaatattat
gccactaaaacaaagagttagctctgaaaagaccggtgccttcgcacttttggatagtat
tgttaggcacggagttaaacatatatttggttatccaggtggagctattcttcctattta
tgatgagctttatgcttgggaagaagcctctctaattaaacatatccttgttcgtcatga
gcaaggcgctgctcatgctgcagattcttattctagatcaacaggagaggttggagtatg
cttcgctacttctggcccaggagcaactaatcttgtctcaggtatagctacagcacatat
tgattctgtacctatattagctataacaggtcaagttgggagagcttttattggtacaga
tgctttccaagaagtagatatttttgggattacacttcctattgtaaaacattcatatgt
agttcgtgaccctagagacatgtctagaattgttgcggaagcatttttatttgtaaaca
cggtagaccaggtccagtattaattgatgttcctaaagatgtaggattagagaagtttaa
ttatttttctgttgagcccggaaaagttaatattcctggctgtaggccaattaccagcct
aaagtcaagacaaatccttatggcagctaaaatgatacagcaatctagccagccattgtt
gtatattggtggaggagccataatctctgattcacatcaaattattaaagaacttgttga
tttttataaaatacctgttactactactttgatggggaaggggattttaatgaggatag
cgactattgtctagggatgttaggtatgcatggtactgcgtatgctaattttgcagttag
tgagtgcgatcttttaattgctttaggagctagatttgatgatagagttactggaaaatt
agatgaatttgcttgtaatgcccaagtgattcacgtagatattgatcctgctgaagtagg
aaaaaataggattcctcaagttgctattgtcggtgacgtagcagaagttgttagtgaaat
attgaatttattaaagacttctttccccccttatccagagcagattatatcttggcaaga
aagaattaatcgttggcgtcaacagtatcctttactggttcctagaaaatcaacaagcat
ttcacctcaagagattcttgttgcaactaataaattagcccaaaatgcttattttactac
agatgttggccagcatcaaatgtggtcagctcaatttctgaaagtaaaagctaagcattg
gctttcaagtgctggattaggcacgatgggttatggtttacctgcagcaattggcgctca
agtagcacatccaaatgacgtagtcatttgtattagtggtgattctagttttcaaatgaa

tatgcaagagttaggaactatcgcgcaataccagttaccagttaaaattattattattaa
taatcgatggcaagggatggttagacaatggcaacaagctttctatggtgaaaggtattc
acactcaagaatgacagaaggagcacctgattttcaaaagcttgcagaagcttttggtat
taaagcttttactattaataataggcaaaatatgcaatctgctttacaagttgctattga
ttatcctggtccagttttattagattgccaagttacagaaaatgaaaactgttatccaat
ggttgctcctggaaaaagtaatgcacaaatgataggtatcgctaaaccgcagagaggtac
tgcttccaactatattaataatagtgtttaaagtaa------------------------
------------------------------------------------------------
------------------------------------------------------------
------------------------------------------------------------
---------------atagttagtattttta-aacaaagaata-aagtaca---------
-taaatgtaatttttttatactatta-aaaactatgtaagaacagcctttgacgagaatt
gaactcgtgaccttccccttaccaaggggatgctctacctctgagccacaaaggc-tttt
tactatattgggccgggttggatttgaaccaacgtaggcgaagccagcggatttacagtc
cgcccccattaaccactcgggcaccgaccctaattaaaccttattcaagatatgacctat
aagatatcataactatgttgctaaaaacaagtactttgataaataaaactgaataagtaa
tggacatagctggacttgaaccagcgactttcacgatgtcgacgtgacactctaaccaac
tgagttacatgtccaaaactttaattattattataacactgtttagtgattttataagta
tctatttctaattgaagatttgttttaatcgtccggctttaccgagcagatctcttagat
aatacagtttagctctacgtattttagctcttctaatcactttgacaaaagttattcgcg
gtgaattgagaaaaaatactctttcaactccaataccttgaaatgagcatcttaaagtta
aactagtatttaagcttttttttcttcttctcgctaatactactccttcacataattgtt
cccgagttttactaccttccttaaccattagtccaagttgtattgtatcgccaactttaa
tttcaggtacttctgttttgataaaaggtatctcaacacttttcattagctgacttagct
ttgtactatttgctttcatatataggattagtcttgtgaca-tgttttgaaatattatac
attaaaaaaacaaatatgaaaaattttagcgaataagcatactagtatttataaaac-tc
tttttatct---------------------------------------------------
--------------------------------------------ccattata--------
----------------cagatccatt------------atattcaact------------
------------------------------------------------------------
------------------------------------------------------------
------------------------------------------------------------
------------------------------------------------------------
------------------------------------------------------------
------------------------------------------------------------
------------------------------------------------------------

------------------------------------------------------------
------------------------------------------------------------
------------------------------------------------------------
------------------------------------------------------------
----------agattcttttatagtttc--------------------------------
------------------------------------------------------------
------------------------------------------------------------
------------------------------------------------------------
------------------------------------------------------------
--------tttgataatcagtac-------------------------------------
------------------------------------------------------------
------------------------------------------------------------
------------------------------------------------------------
------------------------------------------------------------
------------------------------------------------------------
------------------------------------------------------------
------------------------------------------------------------
------------------------------------------------------------
------------------------------------------------------------
------------------------------------------------------------
------------------------------------------------------------
------------------------------------------------------------
------------------------------------------------------------
------------------------------------------------------------
------------------------------------------------------------
------------------------------------------------------------
------------------------------------------------------------
------------------------------------------------------------
------------------------------------------------------------
------------------------------------------------------------
------------------------------------------------------------
---------------------------atcta----------------------------
------------------------------------------------------------
------------------------------------------------------------
------------------------------------------------------------
------------------------------------------------------------

------------------------------------------------------------
------------------------------------------------------------
------------------------------------------------------------
------------------------------------------------------------
------------------------------------------------------------
------------------------------------------------------------
------------------------------------------------------------
------------------------------------------------------------
------------------------------------------------------------
------------------------------------------------------------
------------------------------------------------------------
------------------------------------------------------------
------------------------------------------------------------
------------------------------------------------------------
------------------------------------------------------------
------------------------------------------------------------
------------------------------------------------------------
----gatgatcaaatattattttcaaacatggcttatgtatgaaaaagttctata---aa
aacctaatgac--------taaaattagggcttaagtatttcttagttttgtgttataat
tagttactatcaaaggtagtggcataacacgaagcatcaa-aatgttcgaacgctttact
gaaaaagctataaaagtcataatgctagcacaggaagaagctagacgcttaggacataat
tttgtcggaactgagcagatactattaggattagttggtgaaggtactggaatcgcagct
caagttttgaaatcgatgaatgtgaatctaaaagatgcaagagttgaagtcgaaaaaatt
ataggaagagggtctggttttgtagcggttgaaattcctttcactcctcgagcaaaaaga
gtactagaattatctttagaggaagcacgtcaactaggccataactatattggcacagaa
cacttgctaatgggcttagtccgagaaggagaaggagttgcggcaagagttttagaaaat
ctggcagtcgatgtttcttcaattagagctgaagttatacaaatgctcggagaaaatgcg
gaagccaatgtaagtggaagcaatactacgcaagctagaagtaaaacaccaacattagaa
gagttcggatctaacttaactcaaatggctatggaaggtggtttagatcctgtagtcgga
agacaaaaggaaatagaacgagttattcaaatcttaggtagaagaactaaaaataatcct
gtcttaattggggagcctggtgtaggtaagacagcgattgcggaaggattagctcaaaga
attgctaatagagatgtaccttctattttagaagataaattagttattactcttgatgta
ggtctattagtagccggaactaaatatagaggtgaatttgaagaaaggctaaaacgtatt
atggatgagattaaatcagctgataatgtaatattagtgattgatgaagttcatacattg
attggggctggtgctgcagaaggagcaatagatgcagctaatctgcttaagccagcttta
gcaaggggagaattgcaatgtataggtgcaacaactttagaagaatatagaaaacatata

gaaaaagatccagcattagaaagaagatttcaaccagttgtagttggagagccaagtgtt
gaagaaacaattgaaattttgtttggtcttagagaccgttatgaaaagcaccatcaatta
acaatgtcagatggagctttggctgcagctgctaaatacgctaatcagtatatttctgac
cgattttttgccagataaagcaattgatttaattgatgaagctggttctagagttcgttta
ctaaattctcaattacctcctgctgccagagaattagataaagagttaagagctgtatta
aaaacaaaagatgaagctattagagctcaaaaatatgaaacagcagagcagtatagagca
agagaaatggaaattaaagctcaaattgcagcaattgctcaaagtaaaaagaatgagcct
gatttaaatttagaagatcctgttgttacagaagatgatattgctgaaattgttgctgca
tggactggtataccagtaactaagcttactaaaagtgagtcagaaaaattaatgcacatg
gaagaaactttgcatggacgtattattggtcaagacgaagcggttgtagctgtctctaga
gcgataagacgcgcaagagtaggtctaaaaaatcctaacagaccaattgcaagctttatt
ttttccggaccgacgggtgtaggaaaaacagaattaacaaaagctttggcttcttatttc
tttggttcagaagcttctatgatacggctagatatgtcagaatacatggaaagacacact
gtatctaaactaattggttctcctccaggatatgtgggttatagtgaaggtggttatcta
acagaagcggtaagaaaaaaaccatatactgtcatcttatttgacgaaattgaaaaagct
catccggatattttttaatctacttcttcaaattttagaagatggccgactaacagatgca
aaaggtagaactattgattttaagaatactcttttaattatgacttctaatattggatct
aaagttattgaaaaaggaggaggtagtttaggctttgaattatcagaagatcaaacagaa
tcccaatatactagagtacgatctttagtaaatgaagaactgaaacaatactttagacca
gagtttctaaatcgattagatgaaattattgtatttcgtcagcttactaaagatgaagtt
agagaaattgcagaattaatgcttaatgaagtctttgcgagaattaagcagcaagatatt
caattaaatgtaacagaacgatttaagcaacgattagtagaagaaggatataatccaagc
tatggagctagaccacttagacgagcagtaatgaggcttttagaagatagtctggctgaa
gaagttttatctggtaaaattaaagctggtgatagtgcagtagtagatgttactaatgaa
ggagaagttacagttttattaggtgaaaaattagaactgttaacataaa------aaaca
atttattaat--taaagcattgaagaggttat--attcaatgctat---gaaaaaaatta
ataagatgggttgctataattgacaacctctgatactctttatttgtatata---aagcc
gggatagctcagttggtagagcagtggattgaaaatcctcgtgtcaccagttcaaatctg
gttcttggcat-tttaagcatcagaaaattttgctatctgagcattaaattttaattgaa
atgttcctgtcggaccattcctgtgttttgctattataacttctgtgaagtctctacttc
tagtttcttgtgtatagtaactttctctatatagcattattactaaatctgcatcttgtt
ctatagaattatgaacaataaagttgtttgaaataaaattgcataaaggtttacattcta
aatcaaacactatttgtaaacttgtaacgtttattttttgtatactttcaaaagtcaaag
atgaaa------aagaattaaaaacatttttttgttcttcaaagttttt------tattt
gaatagcaatcatgtcattttgattaattttatcacatcttttccatccttgtgttgtga
gtagtttatggttacttgttaattgtatatatttcccagcttcggtaataattttatata

cagtttttttccctgttttcgaaatactacatttaattgctagaaataactgttgttttt
gcgcactaaagttaatgacttca-atctttttataatagcaattaaataatatttg--aa
gttgattatataatgatgtctgaagataattgaatttggatatacacccactttctctta
aatcggataacaaaggccttttgtcactccgagtttcaaggtttctatttagttgagata
atgccaagataggcaagcttaaatcttttgctaatattttaagtgatcgagtaattaatg
atagttcttgagatctattgtcagattgtctactctcctgcagcaattgaagataatcta
taataattaattctatatttttttccttgtaatttaaagaattttacttttgtctttatag
tgtcagtagaaatcttagcgctatcatcaatataaagattaaggttagcaagagttttac
ttttttgaataacatattgccattcatcattattaagctgaccagattgaatttttttggc
cgtttaaattacactcttgagctaaaattcttcttagtagttgctctgtagacatctcta
agctaaataaaataacataagatcttttctgattaattacatatctagttatattaatag
caaaagcagttttccccattgaaggtcttcctgcaagaataattaaatctgacttttgaa
acccttgtgttatagagtcgagttctgtaaaaccagaaagtatactactatttatactaa
tctttttccttgtatccaaatggactaataggtttgcaagaattgtagctaatgtttgcg
tatctttttcgtcaagtatttcataagcttttgttaattgactaacaatagaagttatgt
ttttttgttcaataagttgttttgagcagctaattaaacataatgaatctccacatgcta
aaagtaatcttttaatgtagttatctataacgactgctgaatattcgcttatagtatcag
ataggggagcttgttctattagactaaaaacttcatttagctcattcaaatcttgaatca
ttttttcatttttttaagttaatgaaaaaatttcttacgctttttgtctttgcatggttaa
cattttcgagtgctgctctataaagtaatgaagttgatgtaaaataaaaaaaatcaggag
agattttttctactgattttagtaatgatatcgattgctgagttaatattatactaatta
atattttttcagctaaaatattatgaggaggtagatatttataa------atacttatta
cttcctcc-ttgtaacattactcttgtg--------gtattagttaactattgatttaat
ttgattctggaagaacttgcaatttaatatttgcagttacttgatgtaatagcttaattt
ctacatcgtaaatgccaattgttttcacgtctggtaaagagatactttgtttttcaatgt
ctatgttagtagtgtttttaataatttgtgaaatttctttttctgtgacactaccaaaaa
tattgtggccatctcctgtttttttactaacgctaaacctttgtatttcttctagaagtt
gcttgactttttagcattttctttagctgtttttaatttttcttctcttatagctgcat
aaaatttttgttgttttaaaataccatttgtggcaactgccgccattttattcgggatta
aaaaatttcttgcataaccagaagctactttgataacgtcattactcttgccaagttttt
gaatattctcttttagcacaacatttataactttttttactcatctt-------aattcta
aaaa----atctgtattatcctattatacctgttgtatacagatttt----aatagtaag
tatttatttgttgatgtaatattcttgtaaaa--aaaggtaatttttttattttt-ttctg
atataatcactttcacaataattc-----attaggagaggtggccgagtggttgaaggcg
cagcattggaaatgctgtttaggagcaatcttaacgagggttcgaatccctctctctccg
tacaa--aaaatataaaaagttttatagagtccaatctaaatctccaatataatattcttcc

aagcatttaatccaccccgaactctaacaatatttagtttttgagctattaaaaacttag
aagcaaatattgatctagaatctaagctacagtaaacaaaacaaatttatcttgtaaat
tgatatgagagtaattcattccttttattttttgtaaaggtagatttaatgaatgtatta
aatgacttttatgatattcttcatgatttcttacatctattaaaatatattgtggattcc
tactaattaaaaacttttgtaattgaataacactaatttctcgtacaaatgtactagatt
tactgccataatatttgttccaatgctttttgt-------ggataatacaaatttagta
tttataatcttaaatttgttaaatgatgaagttaatgagttgtacgttaatatagttcca
cttaatatagacttataacccagaacgattttgacagcttcagttgcttgaagtgtacca
acaatacccggtaatagacctaaaactccggaattactacaagtatctcttgcgctctct
ttattttcagtttcactgtagaagtctcgataaacaggtccaccttgataattaaataca
ctaacttgtccttcaaattgaaaaatggcaccataaatatgtattttatttaactctagg
caagtatcattcaagagatatcgagtatcgaagttatcagatccatctataataatatca
tattgtctaataatgtcgattgaattctcatagcttagtcttgttttgaacacagttact
atacactgtggatttaaatctaata-tttttttttagctatttctacttttgaatatcct
atatcatgcatcgtgtataaaatttgcctctgtaaattagagagatcaattatgtcatcg
tctataattccaatgctgccaattcctgcagcagcaaggtaaatgattcctggagatcct
agcccaccagccccaatgaataaaacttttgcttcttttaatctttcttgtccttctaat
tgaatttgtggcaatactaaatgcttagaatatcttgtatattcttctagtgagtatttt
gtattttctgttttaaaatttagcataaaaatttatagtttatttaatagcactta----
--aaagctagctgataagtttattaaaatgacttttgtaaacttggttcttctctatcta
aaatagctccttgtgtaggacagacctgaacacaaatactacaatcaatacaggcagaaa
agtcaatccaataccaattagtgttattattgtttttccctttccttgatgaatacaag
ctacggggcaagcttccacgcattcagaaactccaatacatttttctgttacaattgtat
gagacatattatttaattaat-aaaagataagctatttacatctagtagttttgttgtat
actaaatttacatatgctatccaggccaatatttaaggtagtctatttctacagtttact
aatttgatatgcgcctttagttcagttggtagaacgcaggtctccaaaacctgatgtcga
gggttcaagtccttcagggcgcgtttag-ttttatcgtaattcaataatatcattttatc
tttttgcaaacatcaaaatgatatgatataaaagatcatatttatacattgaatattct
atgtactagtactg-ttttttaatatgttattattgataagttaattatactaattctat
ttttgtaataagt-aaaaacagttctaattaagtatgcacgtatttaatagtc-ttttgt
atatat--aacttgaaaaaatttaaactatggctaaaaaagttactggaattgttaaact
agcattaaatgcaggtaaagctactcctgcgccacctgtcggacctgctttaggacagca
cggcgtgaatattgttatgttctgcaaagagtacaatgcacgtactgctgacaagtcagg
actagtaatccctgtagagatctcaatatatgaagatagaagttttacttttatacttaa
aactcctcctgcttctgtactaatcgctaaagctgctggactaaataaaggttcgggtga
accaaataccaaaaaagtaggcagcataacaaataaacagttagaatctattgcagaaac

aaaacttcccgatttgaacactaacaatatacctcaagctatgaaaatagttggaggtac
ggctaaaaatatgggaattctaattaaagatt-aaaagtttcaattatatttacta-tt-
-tttatgaaaaaattttcacgtcgacttacaacattaaaatcaaaagttgagcctaaact
ttacactattaatgaagcagtatctatattaaaagcaacgtcaaatgctaaatttaagga
aactgcagaagctcatattgctctaggtttaaatcctaaatatgcagaccagcaactcag
agcaacagttatattaccgaaaggtacaggtaaattgataaaagtagcagttattgctaa
gggagaaaaattaacagaagcaattagtgcgggagctgatgtcagtggctccgaagaact
aattgatgaaatactgaaaggtagattagattttgataagctgatagcaacaccagatgt
tatgcctttaatagcaaagctgggacgagtattaggccctcgagggttaatgccttctcc
taaagcaggcacggtaacattggacgtagcaaaagccgtgaatgaatttaaaggggtaa
agttgaatacagagttgatagaacaggtataattcatgtaccttttggaaaatctagttt
ttcacaagaagatctagttttgaacctccaaacaattaaagaatcgattgatagaaacaa
gccttctggagcaaaagggaagtactggaaaactttttttcttatctagcaccatggggcc
atctattcaaattgacatcactagccttctatgaaaaattttgtataattatagaactt
gaataatctattatttcaaatta-taatttgctagcattattaacgtaacaacttgctta
ctatatcttatctgcattaaaaacttt-ttatatatcttattaattatgagtacaaaggt
tgaaaatatcttggaagagctaaaatctttaaaccttctagaggctgctgaactagttaa
acaaatagaagaaacgtttgatgttgatgcatctgcggcttcaggaggtatgatgatggc
cacaccaacttcagcaccagcgtctgctgaggttgaagagaagacagaatttgatgttgt
cctagaggaagtcccagcacctaaaaagatttctgtattaaaagttgttcgctctctgac
tggtttgggtttaaaagaagcaaaagatcttgttgaatctactccaaaagtcttaaaaga
aggtgcttctaaagatgatgcagaaactatgaaaaaacagctagaagatgctggcgcaac
agttattgttaagtaatga-----agaagcgcttctgtttaagtataaacagaagtgcct
ctttatttattataagcaattatactagaaaagtccaagtgtaattgcattgttgatagg
catagttgcaccaatacctagccaaatacttacaacggtaccaattaagaaaacagtggt
tgcaattggtcttctaaatggattttgaaatttgttaacattttcaatgaatggaacagt
aagtaaaccagctggtacagcagccatacttagaacacctaatagtttattaggaatcac
tcttaatagattaaatgttggaaagaagtaccattctggtaaaatctctaatggagtcgc
aaatggattagacttttcacctatagaagatggttctaaaattgcgagtccaatactgca
ggcaaaagtgccaataattacaactggaaagacgtataatagatcatttggccaagctgg
ctctccgtaataattatggcccattccttttgctaatttagctctcagcttcggatctgt
taaatctggtttttttaagaattgacatattg-tttttgtgtttaatataaagattagtct
aattactgttttaaagtgtaattttcaactattataatggtccagaaattccttgtttac
gtatcattaaaaaatgcataagcatgaaaacggcagttagtagcggaagtacaaaagtgt
gtaaactatagaacctagttaaagttccttggcctacactaactcctcctcttaacaatt
ctactatactcgctccgacaactggaacagcatcagggacacctgtaacaatttttacag

cccagtatccgatttgatcccatggcaaagaataacctgttaccccaaaagaaacagtaa
gcactcctaaaattacacctgttacccatgtcaattctctaggtttttgaaacctcctg
ttaaataaacacggaatacatgtagaatcatcatcaagaccatcatgctcgctgaccatc
tatgaattgatctaataagccatccaaagtttacatcagtcataatgtactctactgatg
taaaagcttctgcaactgtaggtctataatagaatgtcattgcaaatccagttgcaactt
gaattagaaaagatacaaatacaatacccctaagcaataaaaaatattaacatgaggtg
gtacatacttactagaaatatcgtcagcaattgcttgaatttctaatctctcttcaaacc
agtcataaattttactcataaaatagcttcaaaaagctcttctaatata--ttttgcgtt
taagctactattaattcaatag------ttattgaacttatatt-tttttataatcaata
cttattaactattataacatcttttattgattt--------------------aaaatt
ctctgacttagtaaaacagggctatgtactataattcttttttttgtatagagattaat
ttcgtttttagtaaattagctaaaattcttgttacactgactctattactacctataatc
tgtgctaaaattttatgtgtaatagtaaagttaagcataataccattattaacttgagtt
ccattttgctcagctaataataataataagctcgccagtctactggttatacttttatgt
gaaatgatttctataaaatgataagcttttgccgagcaaaatactaaatgatttacaaaa
aataaattgaatggagcgcagttactacatgcttttataatagtagtatagttaatacag
gcaacttctgttttatctattgcttctgcttcgtagtaaaaattatcgtcgaccaattct
atttgtccaaatgtatctcctgtagttaataagttaagagttactttatgtgtatttctc
aggacttttgtaataattaaagatcctatcaatataatgtataatctagagctgctatta
aacaaaagagtatcattttttcttaagtgaaaaaatttgataattaattttattacgatta
aaaaatagtagccatggattgcaagaaaaaaaatcacctgaactattttcagagaatatt
ggaatacaattttcttgatttaaacaatttttagtcataattactactttataa------
---aaaaatattctaataat-taagaactgaaattaaatgtcatataaattgatgttagt
tgagaatgacattgtattatcaaaggccattcaagaatacttaatagatcaaggcttcaa
tgtttatattgccaataatggattagaagcactgaatctagcttatcaatataatttcga
cttgataatttctgatataatgatgccgctagttaatggctatgaactattagcaaaact
taaaaaaaacaaagcattatccaaaatacctgttatctttttgacagctaaaggaatgac
taaagacagaataaaaggctacgacatggggtgctatggttatttatctaaaccatttga
tcctaatgagttgctttcaattattaacaacttaatcgctagagatgtcttaaaagaagc
ttctttacaaaactcggcaacatcaaaccagcaattaaatcataaaatacgtttgacacc
tagagaaaaaagtattttagatcttgttgttgatggacttactaataaagaaatatctac
tatactgaatactagtgttagaaatgtagaaaaatatgttagtcgactcttacacaaaac
taatatgaagaataggactttgctagtcaagtattctataaataataatctattaaataa
tgagatcaatgaaagggcgaatgacgggactcgaacccgcgaatgatggagccacaaccc
attgccttaaccccttggccacactcgccatattacatacaccactatagcttgtttttt
gatatatcgtctagttttcgagttatatatttatctaaaacaatatttactttgaatttt

-tatgatacatagcaatattagtattcaaattaatggggaaccatttaattgctcaaagc
ccatctcattgcaatttttattaaattatcttgattttaattccgagcgtgtagcagtcg
agctaaatgacattcttttaccagagactctgtttcactcaacttatttgaatgatcaag
ataagctagaggttattaccattgtaggtggtggataatctatttgtacatac-------
-acagttttctatctcaaatgtttcatagatagagataatcttccttgtttttatctac
atgaataataacagctttaatagtgtcacctatttaaattgtgatgatatctgctctaa
ttctttgatattaatctcagaaatatggacaagacctttgagatttcctacctttataaa
taatccgtaaggtgtaatctgatttattatgccttcaataatattaccaacaattaaatt
agatgatgcttgagatattagagctcttctatgactcagtattaaattattagatttctc
ttcgacattgagtaattttagcttaataaatttattattaaattgctcactttttgaaa
attaccgagatgagaattaggtacaaatccagatataccttcaaggttaattatcattcc
tcccttattgaatccttttatcattacattaagtaaagagtcttcagctaatagttgcct
aattcttttccatgctcttatatattcaaggcgacgaattgataatattaattgtttcga
ctgaatattataatctaataaaaaaaattctcttgtgtcattaatgtttaaagaagtgaa
attatttaaatcttgattactagatacttcttgaataggcaagtatgcagatattggtgt
tcctatatctactaataccccatttaattcaaagctaaatatagttccagctacaatatc
tccaagattcaaatcatacttgtatttttgtaaaacagctgcaaaatttctatgagtaaa
tccttcattatttttgtcatattgaatagtatgaattaaagagtaactttgtacaatat
attaaggtaaataaatgtttgtcgaattgtatgaaaaaatttttaatttgttcggttgtaa
ttaagtatataaacatattaaaatatctaggagacatgaaaagtagacgtagatagctgc
agatttaa-tactatataatctgaggaaagtccgggctccacaaatacaatttatgctgg
agaaatcccagtgtaggtaactgcgaggatagtaccacagaaaaaaaccgccagaataag
t----tatattagctggtaagggtgcaaaggcaagttaaaagcttaccaaaagtactgca
aagtatttgttaggtaaacccctaaaatggagcaaagctactaaacaaatatttgtgtat
attttatctagtttagttgaaaatactgcatgaagttattagtaacaataactcgagagg
aatagctatcctttacaatatatc-tcgattatattgaaagaacagaacccggcttatgt
agtactttttatgtcttgtattattatgtctattgtcgaatctttaaatttaactgtttg
acaatttttttttgtagctgatgttgtttataatctatttctgaattggttaaagttttt
tcattagatcgatatgtaaatcttagacctatacttttttttctcttttccaattgattca
tctatatattgatcaaataatgttgtactttctaaatctttatcataaaattgatctaat
aattttaataagtatttaatttccatattcttaggtattattaaagagagatctcttatg
atcgatggatacttagagtatggttgaatctgataacttaaataattgaattcattatga
caatcttctaaaatgtttaagttaacttcaagaaaaaaaagttttgtattgagtcctaat
tcactatatgttgcttgattaagttcactgaatatgccaataataatattgttatagatt
agtgtggcaaattttcctttttgaatgaactttattctaccatcgactatatcttttttt
gtccactcaatttgtctattaatcttttgaaaaaaattttctacgatccctttagcttca

taccaatttaaagaatgcgcagggtgtgaccatgtagatcggatatctagattaccaccc
aatatgatagctagattagtagtttctacaattttgttttgatcattatgaaatactgtt
cctatctcaaagctatctatagtttggttgctttgtttaatattataaagatttgattca
attaatccttccagcaaactgccgcgtaaattagaataatcttgaataagaggattgttt
aaagctatatttcctttggactttattaaagaataatgaactaattcagttaatcctaga
tttcttaaaatacttctacttttgtctatgaattttctctttgtggataaccttttgttg
aattgaatatttggtactgagctcctgaatttatgataaccataaattctagcaatttct
tcaataacatctatttctcgaaaaacatcttctcgtctataatttggaacagtaacttct
atattttccttattttgataaataagatcaaaatttaaagagtgaagaatattttgaatt
tcttcaaagaataaaaaacgtgtttgaccattatacataattggtcctaaaatgtcatga
actttttaattgataaattaatattaagagcatgatcagttatttttctcgacagaaa
gtatctctaatatttccgcatgttaaatctgtgattaaagcaagagcttcaaaatgtgca
tttttccaattatctacatttagcccacgttcttgtctaattgaactttctgttcgaata
ttaagaactcgagaagactttctaactacagattgtttaaatatagcagactctacaagt
atcgattttgtattataatctgtatcaaaatcagagttgcttccaataccagctatacta
gttacatttgtatttacttgagtaactaaaatatttttgtttaactcaatattttcattg
tttaacttaatctgatcattacttccaagaggaaaattgcttcgaattgttagactattt
ttatgattcatattattaattttattcaaatcaataatattaataggctgtccccatttc
aacataatataattgcttatatctgttaacaaatttctgtgtgtaaacccagaagagagc
aaccgatttttttagccaatctggagaatctttaactcttatttcgtctataatggcagca
aaataattgtcacaatttaataaatcgtgattactgattatagt-ctccttttttagagaa
atatctattattgggttgatgtatagctgataacattgaggaccctgttaaagccgagac
ttctcttgatagaccaatcatgcttaacacgtctgacctgttagctgttgatgatatatc
taatatatgatcaatctcacctccaattataatagattcaatagcttcaacttcaaaacc
tgcttgtgttagttgattagttaaattgtctatatctattgttttgatattaccaagttc
tttcaaccaatttaaagaaactttcataattaattgtcatctatcttatagaagtattta
ttttacttatgattaaaaatcaatttaaaaaattcatcaaattaatataagttatttgat
gaatttgcttatacaatatattattgtgatatcttacttttatatggcagttattaagaa
ctagcttttgtaaaagttagcccattatcttgggcatatctttccataaatctcatgaat
ctatcccaatcatcaggacttttaataacatgtacagcttctattgcttgtggtttacca
ttaataaacttggcattaacatctcttgtaataatttgtccctctgtgtccatcaggtac
atacctgtgatttcacctttgtctgccatactagcatctaaaattttgggatttgtaaac
cgaaaagttgctgttcctgtactaccatctcgagaccttgttaaacgtacatctggtaca
acttcttcattgataccttgaataaattgaattgttgccatatcaatcttctttatttgt
atataatttatattggataaaactgatttctagttcttgaatgcttgttgagaaactgtt
ttaaaagcttatagtatcagtagatcaaattttatgtattatataaaactgtaactttc

-ttttttg----------------------------------------------------
------------------------------------------------------------
------------------------------------------------------------
------------------------------------------------------------
------------------------------------------------------------
----------------------atctgttcaaagaactaacttattataagctaatcata
cgaatattagccatataatgttcgtaagt-gaaattaaggggttgtagctcagaaggata
gagcaagcgcctcctaagcgctaggtcagcggttcaagtccgctcaaccctgttaacgaa
ttaagtagtattatttaacttgttctgcaaatcagtattgatctaagttaaagtacttga
cagctaaaacaaaagaatctgagaaatattttcatttaaagaacgaacaattatcttaac
tctaaatatctgataattagtaa-aaaattct----------------------------
----ctttttaaaacacaagatgatg-----aagatatttaaactaataattgaaattaa
gtttttgaagttatagaagccaat--ataatactttctattaacaga-----------tg
cttgcttctattactatctgtttacaaga--aatt----atgagtatcaaaata------
--atttagctcca--------------------------------------ctgac----
------------------------------------------------------------
-----------------------------tactaataaata-------------------
------------------tttattttagtctagacaag------atgaatcttcatggta
taacag------------------------------------------------------
------------------------------------------------------------
------------------------------------------------------------
------------------------------------------------------------
------------------------------------------------------------
--------------agctattctttgtgtcatacttatataaaattaggat-------ac
tgtt--------------------------------------------------------
------------------------------------------------------------
------------------------------------------------------------
------------------------------------------------------------
------------------------------------------------------------
------------------------------------------------------------
---------------ttcattaccataactggggtgggaggattcgaacctgcgaatggc
ggagtcaaagtccgctgccttaccacttggctacaccccaacatgtgaacattataatag
cagtcgtataccctgccatgtcaatagactattgtctttatttaataatttttgctgctc
ttgtagaataaaatcttttttttcaaataagatacgctatgtttaagttgcacgactgtt
atgttttcttgcttttgactgaacagccacttgatagtaattattccttctgcaatttcg
ttacttcctataataatacaagctacagctctttttttatttgcttgcttaatctgctta

ctaaaattacttgagctaacatcgatttctattttaaaaaactgctgatgtaaaaaacgc
attatttgcataccagtttcgttagcttttgtaccttgtgtagcaatataaaaatcgata
gatttattaggcaagtctatattatcttttgcaatcagtaataggcgttctaatcctatt
gcacatccaacagctggtgttgctttgccacctaattggtgaactaagctgtcatatctt
ccaccaccacatatagtatcttgcccttttgatgtcaatgtcttaatttcgaaagcagta
tcattataataatctaatcctcgaactaacttattatttatattgtagggaatattaagt
aattttagataattgcaaacgtcaccaaagtgttgttgtgattccaaacttaaaaaatca
gaaatttttggagcctctgtcagtatttttgtgtattagaatcttttgaatctaaaatt
ctaattggattactagttaatctttttgtgagtcagtgtctaaatcatcatggtacttt
gtaagataatctcgtaatttaacttgataaatactacgatcctctacttttccaattgaa
ttcagatcaagtttcaagttgtgcagattgagattgttaaatatgctcatagctaaatgt
ataacttcactatctgctcttgcatctaagctaccaataaactcaatgccaagttggtga
aattgtcgttgtcgtccactttgtggtctttcgtacctgaacattggaccgctataccat
aatctttgtaaactatggtggtaactcattttattttcaataaaagatctcacaataccg
gcagtaccttctggccttaaggtaatatctctattgctgcgatcatgaaaccgatacatt
tctttatttacaatatcagtatcttcaccaatacccctatcatataattcactattttca
aaaataggtgttctaatttcttggtaatttgcacattctaataatttggaaattttatta
tgtataaactgccaatattgaatttcatcgggaagaatatcttttgtccctctaatagcc
tgaatttttgccatgataatttttttgattaccttcttaatatgtctgcattaaatagcag
ttgtagactgttattcagtcatgtattatacattag--aacatactgatg-taatttata
aagtacgggcaaggagggattcgaacccccgacaccatggttcgtagccatgtgctctaa
tccactgagctacaagcccacttaataagtaccttaactatatcattgttcactaatgta
tacaactatattcttttt------atcttacatgtgaaatttttgtaaaagtaaaaaatt
at----gtttgtttttacaattggaactattcatatataatgcttaatttattataact
ttgtaaa--aatatctataagtttaattgtatgattctattatattggtaatcttgttat
aattttgcgctcttatctttcaattgg------ctagcatctaattaacttatatttcaa
ataaaaaaatcttatctaaaatgagtaaacaaattctatatcaggatgatgccagaaaag
cattagaaaaaggcatggatattttaacagaagctgtttctgtgactttaggaccaaaag
gaagaaatgttgtcttagaaaaaaaattcggtgcccctcaaattattaatgatggtgtaa
cgattgcaaaggaaattagtttagaaaatcatattgaaaataccggagtcgcactgatta
gacaagcagcatctaaaacaaatgatgtagctggtgatggtacaacaacagctacagtac
tggcttcagcaattgttaagcaaggaatgagaaatgttgcagcaggttcgaatccgatgg
ctattaaaaaaggtattgaaaaagcaacaaattttgttgttagtaaaatagctgagtatg
ctaagccagtagaagatacaaaagctattatacaagttgcttctttgtcatcaggaaatg
atatagaggtaggtaaaatgatagcgaatgctatagaaaaagttggcagagaaggagtta
tttctttagaagaagggaaatcaactaatacgattcttgagatcacagaaggaatgcagt

ttgaaaaaggctttatttctccttatttcgttacagatacagaacgaatggaagttcttc
aagaaaatccttttattttatttacagacaaaaaaattactttggtgcagcaagaacttg
tgccattgcttgagcaaattgcaaaaacatctcggcctttattaataatagctgaagaca
tcgagaaagaagcattagccacaattgtagttaataaattaagagggattttgaatgttg
tagcggttagagctcctggattcggtgatagaagaaaatctttacttgaagatatgagta
tcttaacgaacggacaagtaattactgaagatgcaggtttatcacttgatacagttcaat
tagatatgttaggaaaagcccgaagagttattgttactaaagactcgacaacaataattg
cagatggtcatgaaattaaagttaaatcaagatgtgagcaaattaagcggcaaatagaga
cgagtgactctttatatgaaagagaaaaattgcaagaacgattagcaaagctttctggag
gagttgccgtcattaaagttggtgcagctacagaaacagagatgaaagataaaaaactaa
gactagaagatgcaattaatgcaacaaaagcagcaattgaagaaggtattgtaccaggag
gaggagctactaatgtccatatctctagtgaattatttacatgggctaaaaacaatttag
ttgaagatgaattaattggtgctttaatagttgaacgagctgtgacctatccgttaagac
gaatagcttttaatgcaggtgataatggagcagtaatagtggaaaaagttaaaagtcacg
attttcatataggctatgatgcagcaaacggtaatattgtaaatatgtacgatagaggta
ttatagatccagcgaaagtagctagatcagctttgcaaaacgcagcttctattgcagcaa
tggttttaactactgaatgtattgtagttgataaagtcgatgattgagctata------a
gataatagataatctcaaacaaagaataatttctaattatagaaattattctttgtttca
attaatctaattaattgatcaaatgtataattttataataaagccaaagtttaaacacg
tcttctttttcaactattaagtagagaagatataagcccgtaatacctaaattatcaatt
aaaatattattctcatatatagatttttctaaagatctaaaaaatgacattttttgaaaaa
taataccgaaatcttctcttatagttatttgctaaatcattactgtctaaagattgattt
ccaattgaattcagaacctttaacagaatactttttaaatctttatctgactttatttga
gaatcaataataatttgattcaattgtatgtcaaaactaagattaaaaatattacttgtt
aaagatatttgattttgaatatctagagattttatagccaagaagattaaacttttattg
ttcattgtaaaatataagacaattgatgatgtattctttaaatacta-aatttatcttct
tgtgcttctatcgttttttgt-aaaataattaatgttttaatttttttagttttagttact
acccgaaaaaaataaattctggaaatctagcttgagctttagctagagagtgtttttttccc
ttcttcttgccagtaattgataacttctgttgttttgtttaataactcaattctgtctct
ttcagcaatccacggcttggattctagttctgcttttaattcttcccaagcatcattcct
tggccagaaaaaataagaagtaagtggactagtgccatttccaacaatctgatcaatagc
aatagcaatattattttctaaccaaagtactttttagtgtaaattttgacaagtttttatc
tccattattaatattaagtattgg----tacagtagaatactgtactagttattttttttc
agcaacaatttgagctttgtttaaaatgtttttttaccgttttagtagcttgtgcaccttg
agataatcgttgcaaaatttttgaaatgttaacacgagtttcattatttataggattata
aaatcctaattcttctatagctttaccatcgcgtttatttctgctatccattacaacaat

tctatagcttggttgtttttttcttccgtatcgctttaaccttaacttgaccatattagt
cacctcgcttatttttt--atatctttattaaattatatcttacgacattatgcaataat
tcaaacaattaagctgattcaagtctaacattattaaatattaccattaaacattgtaga
aaaataatacctaacataggtgacatgtccattccaaacatgggaggaatgcttcctctg
aatagttttaagtatggatcagtaattctatttaatgaacagaatggttcattgtaccaa
tttacagttggaaaccatgccaatgataatttaagtaaaattaaaattaaataaatctca
gaaaaattagctatcgatccaagcaataaatttaatgtaccaggaagagtgttcatacgt
atttttataattattatatacttcgttatactaatagtttgattatatattataatcgatg
taaataaat-cagctaaacaaaaaacttatccttaaaatcatcttctcttgagcactttt
ctttatctacattagtatttacttttgtcgtataaatttccagtttagaccaaatctgtc
ctaactgatttaattgagatgttccacattctaaacaggctatttttatattgctcaaaa
cagcaccttttttataaaatcgttcaataatggcgtaatagttgcttcatccaaaaata
aatctaagacaagtatttttgtaaatgaatttaattgatctgggacattgttaataatat
cattataattaactaaaactatattagctgtaggaagtaaagatcgtgctccttccgcta
agataaagccataaggcataataattacaataatatatttataatgactgctaataagac
ttatagtggtattagcttttaattcgattgtttctgttactagccactctctcattattt
cgtaagtaatccatttgcctagttctgagcaagcagttcttaaaatcgtacctggattat
tgtcgtttttgagaatccctgaccaatgttgtattaatggatgggatgctatattaattt
ggagttgcatgttttgttaaatatatttaacatttaatttgttgattgaatcttcaattc
ctatttgttgagtattattaaaatagcat----atatattaaagaaaataattat-aaaa
aatgaaaaaaagtctttggctgtggggttttacagatagtgctgaaacttggaatggtag
atttgcaatgattggttttatatcagttatttttatcgaagtagttacaggacaaggatt
attatatttaataggtatgatgtcataataaaaatccaaaaaagttcttgtaattttta
taagaactttttgg-attttataaatctaatctagcggtctcatagatagaacaggtt
cattctctctattgatacctttttcaaatccagcagcagcagctcttgctctgcctgcat
gccaaagatgtccaataaataggaagaatcctaggaagaaatgagatgttgttaaccaag
atctaggagatacatagttaactgaattaatttctgtagctacaccaccaacggaattca
aagaacctagtggtgcgtgagtcatatattctgcggcacgcctttcctgccacggttgaa
tatcgttcttgatttatttaaatcaagaccatttggtcctctaagaggctcaacccaag
gagctcgcaagtcccaaaatctcatggtttcaccaccaaaaataatttctccactaggag
atctcattaggtatttacctaatccagtaggtccttgagaagatgcaacattcgcaccta
gtctttgatctctaactaagaaagtaaaagcttgagcttgtgaagcttcaggaccagttg
gtccatagaattcactaggataagctgtgttgttataccaaacaaaattagaagccgtaa
gccccatgattgataaagcacctagactataagataagtaagcctcaccagaccatacaa
aagctcttctagcccaagcaaaaggttttgttagtatatgccagattccgccagcaatac
aaataatgccaatccagacatgaccacctacaagatcttccatgttatttacacttacga

tccatccatcaccgccaaacggagactttaagacataaccaaagataactaggggattaa
gtgtaggattgctaacaaatctaacatctccaccacctggagcccaagtgtcgtataccc
caccaataaataaagctttaataactaacagaaaagcgccaatgcctaataacactaagt
gtataccaagtattgttgtcattttgtttttatcccgccaatcgtatccgaagaaaggaa
atgattcttcaagagtatcagggccaatcagagaatgataaagacctccaaagccaagaa
cagctgaagaaattaaatgcactacgcctactacaaagtatgggtatgtattgaaaatct
cgccgcctggaccaacaccccagcctagtgtagctaagtgcggaattagaataaaccctt
gttcgtacaaaggcttctcgggaacaaaatgtgctacctcaaaaagagtcatggcaccag
tccaaaagaccattatacctgcatgagcaacatgagcaccaagcaatttgccagaaacgt
taattaagcgtgcattgccagaccaccaggcaaacccggtagactcaatgtctctaccgc
caacaccaacagttgtattaaagggcgtttccacgtggtaaaacctcctcagggaatata
aagttttcatgaggttgatcttgagcagccatccaagagcgaataccttcgttcaatagg
atatttttagtataaaaagtttcaaactcaggatcttccgcagctctcaactcttgagaa
acaaaatcgtaagctcttaaatttagagctagtccaacaattccaaatgcgcttgtccat
aatccagttactggtacaaataacatgaaaaaatgtagccaacgtttattagaaaaagca
acaccaaaaatctgtgaccagaatctattagctgttaccatcgaataagtttcttcagat
tgtgttggagtaaatgcacgaaaagtatctgcagcatcaccatcttcgaataaggtattc
tgtacagttgcaccgtgaatagcacatagtaaagcaccacctaagattcctgcaacgccc
atcatatggaacggattcagggtccagttatgaaatccttgcaaaaatagcaaaaatcta
aaaattgcagcaactccaaggcttggggcaaagaaccagctcgcttggcctagagggtac
attagaaagacggaaacgaatactgcaattggtccagaaaatgcaatagcgttgtatggt
ctaagaccaactagcctagcaatttcaaattgtcttaaacagaatccaattagtccaaat
gatccatgtaaagctataaatgcccacagaccaccaatttggcaccagcgagtaaaatct
ccttgagcttcaggtccccaaaggaaaagaagtgaatgtcccatgctgtttgctggggta
gaaacagccgcagttaaaaaattgcatccttctagatatgaactggctagcccatgagta
taccaagaagtaacaaaagtagtcccagttagccagccacctacagcaaggtaagcgcaa
ggaaacagaagtagtccagaccaacctacgaatacaaatcggtctctttttagccaatcg
tctacaagatcaaaaccaccacgagtttttttcttgtccaattgctatggtcataattta
atctccagagcaaatttaataagtaatttgtttaaataaaca---ctttagatatactct
aaaatttttactttctttacggtcaatataattataagtcaaaagtaaccaaaattcacg
cttttaataaca-ttttttagctcattaagttaa-ttttttaatatataaacagtaata
gagttattgttt-aacttacaatactcacaatttattcatttctataatcctttgaaaaa
gatatcctgttcctcgtgctgttaaaatcaagtcgggattacttggatcatcttctagtt
tagctctcaatcgtgaaatatgaacatcaacaactctcgtgtcaacatgccgttcgggag
tatagccccaaacttcttgcaagatagaagctctagaaaaaggttctccagccttactaa
ctaaaagctcgagaagactgaattccatacctgttaatcgaactctttcattatttttat

aaacttgtcttttgttagtatctatttttaaaaatccaatactaataatcccagagtttg
ggactccaagactagtggtgattttatcagctcttcgaagaacagaacgaatgcgagctt
ctagctccttaggagaaaatggtttaacgacatagtcatcagcacctatttctaatccgg
taattctatcgcagacttcacctaacgctgttagcattataataggaacatctgattctt
ttcgaagttcttgacaaacaccgtaaccatctagtttaggcatcataacatctaaaacaa
caaggctaggatactcttttctaaacacgagtaaagcttcttctccatttgaagcagtga
taacttcataacctattattgttagtcttgtttctaaaattcttcttatgctggcctcat
cgtcaacaacgagaatcttttctttctggttctccaa-tttcttatttactcctgtaact
caccgatcatgattaccataaccttaaatttttagcgctatgagtatctttattcacttt
agaa-----tttagttattataaattttgcaagtctagtttacattaattagtaccagt-
-caatatttttgt-----------tttaagttatgac---------tattttaatccata
agttgactctagtttgcagaattcaatatatacata-ttttctacccaagtaaattgtat
tctattttctaaataaaaaaggggttgacaaaccagttgatagaaagtattctacattac
tataag----ttaattccaacaaagaagttttaa-----atcttctttcggagtaatgaa
tactttaat
>SRR9587917
tatatgata---aattaagacgtctgttatgtgagaatttctaaaatatacaaaactttt
tatttaatatagaaaaagtaaattaaaataagttgaacatactttgctctaatctaaacc
tcttaattgag------attctaattagagacaaaagtatttacttattttgtatctata
ggaataaaaatgaaacttgcagtttatggtaaaggtggtataggtaaatctacaactagt
tgcaatatttctgtagcactttcaaaaagaggtaaaaaagttctacaaattggctgcgac
cctaaacatgatagtacatttacgctaacaggtttttaatcccaacaattatcgatact
cttcaatccaaggattaccactatgaggacgtttggcctgaagatgtaatctataaggga
tatggcggtgtagactgtgttgaagctggtggaccgccggctggcgctggttgcggaggc
tacgttgtaggtgaaacagttaaacttttaaaagaattaaatgcttttgatgaatacgat
attatcttatttgatgttttaggtgatgttgtatgtggaggttttgcagctccattaaat
tatgcagactactgcttaatcattacagacaatggatttgatgctttatttgcagctaat
agaatagcagcttcagtacgagaaaaagctagaacgcactctctgagattagctggactt
gttggtaatagaacagataaaagagatctaattgataaatatatagattgtgttccaatg
ccagtattagaagtcttgcccttgattgaagatattagagtgtccagagtaaaaggtaaa
actttatttgaaatggcagaaattgataaggatttagcatatgtatgcgattactatttg
aatattgcagatcagttgattacaaggccagaaggtgtagttcctaaagaatctccggat
agagaattatttagtcttttatctgatttttacttaaatcctaaatcaaaggtaggacaa
gaaaaagtagatcaagaagaattagatttgatgatagtgtaaataatattttcagcataa
tataacaaggaataagataatgtctacagctcaatcagatgctcttacttttgaatgtga
aacaggtaattatcatactttttgtcccatcagttgcgtttcttggttatatcaaaaaat

tgaagacagtttttttcttagttatagggactaaaacttgtggttatttcttgcaaaatgc
aatgggagtaatgatttttgctgaaccgagatatgccatggcagaattagaggaaggaga
tatttcagctaagctaaatgattatggtgaactccgtagattatgcttacaaataaaaaa
agatagaaatcctagtgttatatttggattggaacatgtacaaccgaaattataaaaat
ggatctggaaggaattgctccaaaattggaagcagaaattcgtgttcctatagtagttgc
tagagctaatggtttagattatgcctttacccaaggtgaagatacagttctagcagctat
ggctcaaaggtgtccattaaatttaaagaatcaatcagataacacgagtcttaaaacttc
tcctcatattcctttagttttatttggatctcttccggatccagttgtaacccagttaac
tatggaattaaaaaaacaaggtattttttgtttctggttggttgccatcaaagaggtatac
tgaattaccggttataaaagaaggatattatgttgcgggagttaatccatttcttagtcg
cacagctactacactaatgagacgccgaaaaacaaaactaattggtgcaccatttccaat
aggtccagatggtactcgagcctggattgaaaaaatctgttcagtaatgaatgtaaagcc
tattggattagaagatagagaaaaagcaatttgggcttctttagaagactatatttcttt
aatccgaggtaaatcagtttttttatgggtgataatttactagaagtatctcttgcaag
atttttaactagatgtggaatgactgtatatgaaattggtattccttatatggataaacg
ctatcaagcagcagagctagctttattaaaagctacatgtgataaaatgaacgttatgat
gccaacaattgtagaaaaaccagataattataatcaagtagaccgaatccgtgatctgaa
acccgatttagttatcactggtatggctcatgcaaacccttttagaagccagaggtattaa
tacaaaatggtcagtagaattcacatttgcacaaattcatggctttaccaatgcaagaga
tattcttgaactcgtcacaagaccattacggcgaaatcttagtttgtcagaattaggctg
ggatgtttatagcaagcaaagctagtattcctctttatttatgacttacatggtataaaa
gccttgtaagtct--tttattttatgttaaagaagcatttcttactctgccatcaccagc
ccaagcttgtaaactttctacttgttctttgtctgtaaatgctaaaggcacaaactgttt
gatagctattttaatatcttcagtactaaattctctctcttcactaaacgccgtatgcat
actttcaataattgcttgctcaatttcagcgccagaaaatttattacataataaactcaa
ttcatgaatatcatattcttgccacgatcttggtctaacttttgataaatgtatcttaaa
aattgactctctctcttgacgattaggtaaatctaagaaaaaaatttcatcgaacctgcc
tttccttaacatttcggaaggtaaactttgaattttattagctgttgcgacaacaaatac
tggagctttttttctctgatagccaggtaataaatgtaccaaatactctggcactagtacc
actatcaccttgactatgtaaaccagaaaaagctttatctatttcatcaatccataatat
gcaaggcgataatccttcagagatatttatcatttctctcatttttgattccgattcacc
aactagtccaccaaacagttttcccatgtcaagacgtagcagaggcagcatccaatcgct
tgcaatagctttcgcggttaaggatttacccgtcccctgtataccaactagcaataaacc
tttgggcgaaggaataccgtaatttaaactttgcttcgaaaatgagcgagaccttttttg
tagccattgttttaatacatctaagccaccaatatccctatttactttactataaggata
aaattctaatagatgcgtttgattaataatttgtctttttttcttcaataataataggtaa

gctccgagagtctatctgattatattgtgcaataattttagtaatcactttgcggattct
atctatcgataatccttggcaggattttgttatattattaacgagctctgaattcaaatt
aagattcaaagctttactcaatcttatgatctctttttgatttctagaagactaggtaa
gggcaaatctattaccgttatgatatcattcagagcaaaaggtatattgattttgcaaga
aataataataatgtgcttagattgcgttttgattattttagctaaattgcgaagtttcct
aattaatactatttcgtttaggaaagagtcaaaatcttttaaaagaaaaagatttaaaga
ttcattattcaagttttcaataaattctaatgctaataaaggatttctttttgcataacc
attatcgctaggattgttagtgtatccatctacaaagtcccaggaatatacttgttgact
gtttgaacagtttaagctgtgttttataatatactctagtctatcttcttctctagtatt
tattacaataataggatatcgagattttaaaagtaatcgtaaatcttgagtgaaattcat
gtcaatatttgtttccatcagacttttgtacttcaccaattgaagaaagactaaataaga
ttatctaatattccttggatatattcaaaagtc------atctctctatttctaaatta
ttgctattactaatttaacttaccatgatttttttcttccttttcttcgtcgactattt
aatatagagcgtccacatggtgttttcattcttgctctgaaaccagaaactctaatttt
tttcttttagatccttgtagtgttcttttagtcatgatgtttcctctgttgtaaaagatt
aaatactattttatcatcttttatatatctttatgatatagatatgtattaatatcaaag
tt-aacagcaataatttatattaatcaagaattatttatataatattaaacattaaatat
----------taactttatgatattggctttgcccatttttatttaagtattttgacca
tctttttgctaatcttaaactggttaatttttcagcagctaaaaacgattttgttattag
aatctcaatttaagtattttgttgataaaagccaaaatcgtaaattagaacccgaagaaa
gttttgcttttgcaaaagtatgtgtagctaaaaaatatttttctaaagctattattgaag
gtcaattagctttaaaaaactatcgagacctgaatatcttagataataatatagtaattg
ctaatttgtataatatgctaggttttatttattttgaagcaggccaaacaagttttgcaa
agaacttttatgagcaagccttacaaataaaccctaattatgttgtagctttaaataacc
tggcaaaaatttacgaagaggtaaaagatttaaaaaaagccgaatctctatacgataaag
tgttaactcttaacttaaataataaaactgctaatagaaggaaagattttatagcaaaaa
cta---------aaaacatctaatactatttgtaatcgggatagcaggatttgaacctgc
gacatcctgctcccaaagcaggcgcgctaccaaactgcgctatatcccgtaagacataat
tattactatataccttttctctaattatgtctactctttctgtattttatctaggatacc
aaaacatgcctttaactccatctggatctttgataaatcctaatttttttataaaaactaa
caacatcaggttctgcgaatagtgtaatagtgctaatttctgcctgtctcaactgctgta
ttaattgatgtatgattacttttcctaaacctaaaccttggaaatctggatgaataacta
cgtcccagatagttgcattaaatccgttgtctgatgttgctctagcaaatcctacaagtt
tactatttgcatcttttttttgtattaaagaaataataatagaactatttttttaatgcga
tctttactttttttagaggtcttttaacccatccgactgaatcacaaagctgctccaatt
cgtataaattaatattcttattgctactcagatagatatctttaaattcgattttatcgc

aagttttatctaaaacaataagtttttgaaatttttttcagagctagcattgatatcgg
agttttgaaaaaaatttttccagaagatcataatgattatgatataatta--aataaata
tatatattacaatattgctttaacaaacaatatattaatatatgctcaaatttttacatg
aatctattaagatagaagattgttaaaatatagagaaataaaaatttttgtaaaatcttt
caataaaatgatataaagagtgctttttaagactaacatctattattataaataaagtct
cgaaattgagatgaaataacacattattctatgtctaatgccttttaaaggagttatttg
tgaaaaaatctatgtgtctaacctgtttacttgctttactaattatgagcaacccaataa
tagctaacgcagaagtagctggattggtgccctgcaaagattctgcggcattcaataagc
gtatggtaaatagtgtgaaaaaacttcaagccagattagctaaatatgatgcagacacgc
caccagcgttagctttaaataagcaaatagagaaaactaaaactagatttgcaacttatg
gtcgagcaggtttgctgtgtggtactgatggattgccacatttaatttctgatggtcgat
ggagtagagccggggactttgtttttccaggactattattcttgtatattacaggatgga
ttggctgggtaggtagaggatatcttttatctgttgctaagactagtaagccaacagaaa
aggaaatcattttagatgtgccattagctatcaaatttatgtcatccggatttgcatggc
cgctagcggcttggcaagaatttagtagtggacaattaattgcttccaatgacgatatta
ctgtttcaccccgttagtaaaaaatttatatgaataataattttaccaaatacttatcaa
cagcacctgtaattggtgtattgtggatgacgtttacagccggatttattatagaattga
accgctttttcccagatgtcttatacttttactt----ataaataa--gatcgtataaa-
--aaactacataaatatagagaaatactattatttgtgtagtttta---atatataaca
taattattta-tttttaaaaatgtttatcttttcgattattttaatattagatcttacat
aataaaata-atatcaatattatgagcttagtaacccaaattatagttaatgccgatgat
gaattaagatatcctacaattggagaattgcagtcaattcaagactacttaactacagga
agcaatagaattagaattgctactattattagagataaagaaaaggagattatacagaaa
gctagtaagcaaatttttcagttacatccagaatatatagctccaggtggtaatgcagca
ggttctaggaaaagatcgttatgcttacgtgattatggttggtatctcagactaattaca
tacggagtattagctggggataaagattctatagaaacaattggtattataggagtaaga
gaaatgtacaattctttaggtgtacctataattggaatgttagatgcaatccagtgttta
aaggaagcatctttagaaatgcttggtcaagatgatattagaatcatttctccttatttt
gattatataattcgtggaatgtcataaatcatagcttcaattaattagttgaataaaagc
ttttataatgttataattacaataagcccgaaagcatataattgtaaaaactaaatcttg
tcagaaccggaaggtagcagcaataatgtttcaatacaaaagttatga-ttttcgggtgt
ttattgtttttatctaatagtctcgacttttaattagtatgctatgtataaatctgtata
tgacatgtaatccatcaaaccttccagcaaaaataatttaataatttgatattaatacgt
aaaacacctatac-tcttttataaaatttctttatatctcatctaaacaatggaaactta
acttaatgggtgttcatattttatcaacaggttcgtctgtcccaaatttttctgtagaga
atcaacaatttgaagatatgatcgaaacttccgaccattggatttcaacaagaacaggaa

taaaaaaaagacatctagctccatcttctacttctttaactaaattagcggcagaagctg
caaacaaagccttatatgcagctaacttaaaacctactgagatcagtttaattattttag
ctacgtctacgcctgatgatttatttggtagtgctagccagttgcaagcagaaataggtg
caacaacatcggtagcttttgatattacggctgcctgctccggttttattgttgctttag
taacagcagctcagttcattcaaactggttcctatgacaatattttagttgttggagcag
acacaatgtctagatggattaattggtcagatagaactacctgtattttatttggtgatg
gtgctggagcagtagtattggggcaaagcctcaaaaatagtattttaggctttaagttat
gtacagatggtcagctaaacagtcatttacaattaatgaataaacctgtaaataatcaaa
aatttggtgttacagaaattcctcatggaaactataattccataacaatgaatggcaagg
aagtgtacaagttcgctgtatttcaagttccaacagtaattagacaatgtttgaataatt
taaacatttcaatagatgaagttgattggtttatattgcatcaagcaaacactagaatca
tagaagcaattgcgagcagattatcagtacctttttctaaaatgattacgaacttagagc
attatggaaatacatctgcagcgtcaatccctttagcgttagatgaagctattcaatcca
ataaaattcaaccaggccaaattattgttttatctggttttggagcaggcttgacttggg
gagcaattgtcttgaagtggtgat---ttatatattgcggatgacgagactcgaactcgt
aaagctttcgctacacacccctcaagcgtgcgtgtataccaatttcaccacatccgcatt
ttttaccatgaata-ttaaaaatttataatatttatctaaatattcatataa--tgtgta
tatatatatacattcttgtcagaaactctcttatattttattttaagtgataaaattgag
t----ttagtatagatagtaaaattttattttaaaatcataa----aaataaatatataa
ggaaaataaaatatgacaccatctttatcaagtttttttgaatagtcttattcttggggca
gtaattgtagtcgttcccataactttggctcttttatttgttagtcaaaaagacaggaca
atccggtcctaaaaataacttagaaagatatggagaagcaatttaaaatgaataataaaa
tatttttcatactaataagctctccatatattgctatact--ttagaagaatttttattt
ttttattttgtaacatcctgtaaagtttattccgttctttctttttccc-atttactctc
ttataattaatgatacaaatgtcagg-ttaataatag--tttttttattctgtatctttc
----------------atttatatacgtattattatgttcaattaataaagaaaagctca
tgtctttgtctaattggcctctcaaaaaagaaaattctgaagcatataatattaagaatt
caaaacaaatcacaattcctgatggtttatggataaaatgcttcgactgtggtctattaa
tgtattctaaagtactgaagagaaatttaaaagtttgccctcaatgtagttatcattttc
aagcttctagtaacgaaagaattgatcaattgatagaccaaggtagttggcaaccaatgg
atgttcacttgatctctacagatccattaggctttaaagaccaaaagctttatagtcaaa
ggttaaaagatactgctttcaagactggcctgcaagacgcagttcaaacaggtactggga
ctatgcaaggtaaaaaagtatgcttaggtattatggattttagattcatgggggaagca
tgggatctgttgtaggcgaaaaactaacaagactgctagaaaaagcaactcaagaaaagt
tgcctgcaattatactttgtgcatcaggcggagctagaatgcaagaaggtatgttgagct
taatgcaaatggcaaaaatttcttctgctctagaaatgcataaaaaagaaaatctactat

atatatctgttttaacctctcccacaacagggggtgtaacagctagttttgctatgcttg
gagacttaattattgcagagccaaaagctcttattgcatttgctggtagacgagttatag
aacaaacaatcaaagaagacttgccagataattttcaaagttcagaatatttatttgaac
atggtttcctagatttaattgtatccagaactcagcttagatcaaagttaatacaaattt
tgtccttacataatcatagtaagtaatgaattgaatattgactataatatcaaaatgcta
gtaaaaattaagatattaagaaattttcactaaagatatttgtagactaactagaattac
aattactattgtagaaaaatttgtcttaatgcca-ttttttcagtcttttaacaaattg-
tttttatgactttaaataattgttagtttaatacaaatttataaagagaata-----cat
actaagctaaatatagtaatattatgttttcctataatactgacctatcaaatttcgagg
aaatttatgcttaaaagatcttcttggcttgcggctttattgggactattaacagtagtt
tctacaagtacgcatacatatgccatagagttagacgaggcaacaagaactgttccatta
gaatcttctggcagaactgtaattcttacaccagaacaagttaaaagaggcaagcgatta
tttaataattcttgtgctatttgccataatggtggtatcacgaaaacaaatccgaatatt
ggacttgacccagaatctttaggattagctacgccacaaagagataccattgaaggacta
gttgactatatgaaagatccgactagttatgatggtgcagagtcaatcgcagaattgcat
ccaagtattaaaagtgctgaaattttcctaaaatgcgcaatctaaccgatgaagacctа
tttacaatcgcaggtcatatcttacttcaacctaaaattgtttctgaaaagtggggcgga
ggaaaaatttactattagaacttcaaagacctaaaccttgtgaatactagtttatgta-t
ttgttgcttaagtgtgattattttgttagatcatgatatatattaaagatggagtttgtt
gacagcaagttacttttttacccgaacttgtttacactattcgtctagttattaaaaagga
gacgttaaattgaagaagaagctttcagttcttttcactgtttttagttttttgtaata
ggtttcgcacaaattgcttttgctgcagatctagataatggagaaaaagtttttctgct
aattgtgcagcatgtcatgctggcggtaataacgccattatgccagataaaccttaaaa
aaagatgtacttgaagctaatagtatgaatactattgatgctattacttatcaagtacaa
aatggtaaaaatgccatgcctgctttcggaggtagactggttgatgaagatattgaagat
gcagcaaattatgtattatctcaatctgaaaaaggttggtaattatacttgattttatcc
tgtattaaagaatagacaatctatttagttgttcattagattgtctattctttgtttatg
---ctattatataaagatatttacacaatatt----ttatgatgaaaagaatacccgcaa
ttcttgtactagaagacggtgcgtattataaaggatggtcattccagcaagataaacaag
agattactattggtgaagtagtttttaatactggaatgacaggatatcaagaaataatca
cagatcctagttacttccatcaaattgtcgcttttacctacccggaaattgggaatacag
gtattaataatcaagatattgaatctcacagtattagtattaaaggacttattgcaaaaa
atatttgtaaaatttcaagcagctggagagagcagcaatctttagttaagtatttaagta
gtaataatattccttttattttcggaatagatacaaggtctttaacccaatacttgcgtc
aatttggtacaatgaacggttgtatctctactgataatttaaatcatagttacttaaaac
agaaaatttgtgagattccaagtatgcaaggtttagatttaatcccgcatgtaactacaa

gaaatgtttacccctgggatgaaaaaagttttccaaattggtatttgacagataacatta
gagtgcaccgagttattcagttgaaagttattgttatagattttggagttaaactaaata
tactcagaagactagctacacttggatgtcagataacggttgtgcctgcccacactcctt
taaaagatattttggcttaccagcctgatggtatattactctctaatggtccaggagatc
catcagcagtacattacggcatccagacagttacaaaattactagattacaatgtgccta
tatttgggatttgtatggggcatcaaattttaaatttagctcttaaagctaaaactttca
aacttaaatttggtcatagaggtattaaccatccatccggattgaaccagcaagttgaaa
taactagtcaaaatcatggctttgcagttgaattgacttcggtttttgaatctcctgtaa
gagtgactcattttaatctaaatgacactactattgcaggaactggacataatcaaagtc
cttatttttctgtgcaatatcatccagaatcgagcccaggccctcatgatgctgattatc
tattcgaaaattttatagaaataatgacaaagtccaagaataaagttag-----------
ttagtaattttcccatgctttatgagtgaataaagcggatagtacttgtactcctcgtaa
ttgattaaagagaggcaaatgtcctctggggccatgagaattccattgaaattcctcagg
atatctgcaaggaattctatcaacttcccagccgatttttttgccagaattttccccaatc
ttttccaacactaagccatatttgtctttgaacaaatagcccaaatttaccttttgagtg
tgtatgccataacttatcaatagtttgtaaatcttgagcaggtatttttttatatctgt
aaaatatagccaattacgagtttgtgcatttacaccagctagttgaataagcttttgttg
agttagctgatctgctttgagcaaatcccgatgagtcagtaacatttgtaaatctttgta
attcatctgttgagcagagcgcaatggaacaatgccatcgggacataagttagaagcaaa
tttcactatttcttgattttactgttaagtaacttttcataaattaaaccgtcaacaca
attacttttatagttaggcccagtaatcctctcaaaaaatagatctgctaaatcttttaa
ctcgatagagtctttgctatttatattctcaataatttcaagttgttgcttga---catt
attagacttagtattcttattcaattctaaaagctgggctcggatttgatttggcatctg
atttgtttaattagttttaatagtc----ataaggtcttagagagaattagattgttttt
tgaaattagaagagcgccgcagtatttctaaagaagttcgcataatttgattcaataagt
tttgtcctgcataaattataatttgaccaattaaaaaaattgtgctttttatttttaggaa
caaaaaaatcttgtacctcaagcagaaaagtaattattatttgcattggtgacaacatct
gtaattcattcgatctgcaagcatatatatactgacagtttaagccttcttgataaaatc
cccatactttgtaacgactttcgtaaattgctcgtggtcgctcaagatagtttttaatta
tagtattccaaattaggttattctttaattttctaattttctatcagatagaaatttta
cattacagagataacttgcatttgtctcatttacagtattaaaattttgaactaacatat
gagcaataatgttgcttagctgaattaaataattttcaagaaaaatttcgacctgcttaa
ctggcatattgtaagtaaagagattaaaagttttattcacagtttgagatgacccaaaaa
gtagctgaataattaaaatttgtagtagcatcttatagtcgttcagtagatagttcaaat
catgaatatcttggtaaatgtcaatacagtccagttcatataacttacaaaatcttttga
tacatctatggaataaatcaattaaaatattttcggtcaagtttgtaatatcatttacat

ctaaatttgaattataaatttctaaaagtatcttttccagttcagctaatatgattttta
ataagtttctttttacttcagtgcgaaaaacatctaaaattaacacttcctgagagcaat
tagttaatctcttattaatctttatagaagttctaactaatagctctgctacctcttgat
tgagtgttggtccctgagagcttggccaataattattcacgttataccatagtaagaaat
attgaatttaatgttatacgataactttttaatatactgaaaagcaagttaattactcta
tagcaactag-------tttttacagatcttaatagttatttatcaaataaagttatatg
atagtagcgattgagctaattttattaaaattta----taataatgacaacttattactt
cgcccttgcgagtcaaaatttctattagtacaagaaccactagaagaagtttttagaga
gagagttaactactatcagtcaaataataaagcaattgattttggctaataccaaaccc
ttcttttctagagaaaccagaaatgatttcatttaaaaaccttgtacctaaagacgctgt
agctataatctctactaatccaatatttattaattggttaaagctaagaatcggctatat
ctgtattgggcaatttgaagataacctacaactttctgaagaatcgttaaatattactgt
tttaacagacaaaatttaacaatgcttttattatgactagtgttaaattattttggttaa
gttataattatagtcttttagtattta-gactcttatagtttatcacttagtattttatg
ctgagatggagtgacaaatagtgtaagtatttcttggctaaacgtttcagttgccttaga
tttatatctatttggattaactataatggatagcattcgttttatagtaacattttcaat
ttgagcccaatgtacaattccaagttccaattccttagcaatagctgaaacggaaacaaa
ggcagctcctagccctgattgcacagcgtttttgatagcttctatagaattcaattccat
ctctattttaaagcgactgctgtcaataccatgttgactgagtactttatctatgacttt
tctaattgttgattgagtatctaacgcaataaatctaagcctgtataagtcttcttttg
aatgtctcctagtttagaaaaaggatgtgatttgggtaatataagtgctaattcgtcttc
cgcataagaagtaacctgcaaaacatcttgcaattcagtaggcacttctcctccaataat
tgctaaatcaacttgaccattagctacgctccatgaaataagtctagttgaatgtacttg
caattgaacagccacttgtggatatctttgcctaaatagtccgattaatcttggcatcaa
atatgtcccagttgtctggctagctccaataattaatgtgccaccttgtaagttttgtaa
gtcgtcaagagctcgacaagtttcttcgcagagagctaaaattctgccccatatcgtaa
aagaagacttcctgcctcagttaaagttgccttcttattacctctttcaaatagggaaac
attcaattggcgctctaaattttgaatttgcaaactaatagctggctgagaaacatacaa
gctattagctgctttttaaaactgccctctttggcaattgcttttaatattcttaactg
atccaatgtaaatggaaggtctgtcattaaagaattgtagtataatg---------tata
ttatttagtattgatcaatgtatgcatttgccaactaaaatatttgtcattacgcttcat
tattttcaaaataataaaaagtataatgttaaataagctgtattaaaatttagtatctgt
ttgatattagatttatacaagagatatcgtgaatttaattctaactataaaatatagga g
acaat---atggactctagacttttagttgtactaataccagttttagcagcagcatctt
gggcagtttacaatattggtagagttgcactacagcaatttagaaaaatgacatcttagt
tttgcttagaatata-atttttagactttatgagatagaggactttaaaatccctatctc

tattgtatatcttaataaaaaaag-aaagc--ccccttgtacttactgttattacacatt
aatattattggt-ggtaccta----gataataagtttaagctaacttatttttatttta
taattaattaaattaataacacaattatggctgttccaaagaaaagaacatctaaagcta
aaaaaaatgcacgcaaagcaaattggaaaaatcaagcaaaaactgaagctcaaagagctt
tgtctttagcaaaatcagtattaactagaaaatccaatggatttatttataatctaactg
aagcatcagatacttttagcgattagaagtgttt-ttactacaagtctgaaaaagtatta
cgcaacaagttctcagtaatttttatctat----taatcatttgaaaatcattctatttg
ataataaaataaatagcagattaaataaaatcacattaacaaattatgcagcaaagctag
atcaaactagtgaaatttggttgttcaattgtattgaaaatattcagcatatttttttta
aaagtcaattaaaatcaagtcatattactaaaattttatttctggtactagctttgaat
ataccgcaggtctaccaggattattatccagcttaacactaagtggtagactacatccta
ttagtatatatagtccccagtctctcaaaaagtatcttgaagcatgtaccaaatattccc
aaactaatttttcttttcctattaattttcataatttacaatacggaggacaagttgtta
accaatttatacagtaatttgtttaccgttgagcaaaaagagcctgctgtatggattta
ttattctaaaaaaagaaaagcagggagtatttaatttagcacaagctaaaaccttgaata
ttcttcaaggacctatatatggaaaactcaaagaaaaggataatttttaagtccagatg
gttactatctatccggccaagacttctcttctaatacaataatgggacataaaatatcgc
ttccgttattagtcagatattctagaattatttctgagatgcattggttttgctcttatc
ctattagattaaatacttattcgcatcagcaaggagcaaagtgtttaccacataatgttt
taactgatattatgaaatcccaaatatatcaagataacagttttgttgaataatacttat
tcttgatatataatggtctcggtaatctaaataaaatactagaaataatttcttaaattt
atttgtctatattaaatttacaatcatttatgacatacgcaattattgaagcaagtggca
cgcagctttggatagaagaaggccgctactatgatttaaatcatatacctgttgatccag
gtcagtcgattatattaggaaaagtcttattattgaataaaaatggggaggttactttag
gccgcccttgtatagaaggggttacgataaaggctacagtaatgaggcacttgcgaggaa
agaagataactgttttcaaaatgaaaccaaagaagaaaatgagattaaaaaaaggtcatc
gacaagaattgactcgtttaatgatcgattctataacatcttaga------taaaatcta
gactcatattttt--aa-aaacttttaatatttaactaaaatagatataatggcacata
aaaaaggtagtggtagtacaagaaatggcagagactctaattccaagcgtttaggtgtta
aaaaatatggtggagagcaagtaacagcaggtaatattttaatcagacaacggggaacta
aagttaagcctggccaaaatgttggaaaaggaaaagatgatacattgttttctctaattg
atggtttcgtgctgtttgaaaagtcaaatcaaaagcaaaaaacaattagtgtttattctg
ctaagaaatagttaaaacaattggtgcaagataacttgagt----gtatttatttcactt
acagtggcaccatatatttgatttattaagaaatggagttccttaatgaaccaatgaatt
ttattaaagatttgctgtgaaatttaatgacaaacactattgtaatttcctgtctacaca
atatggctgctattctatattgtggtcagatacagaaactagtagtagcaaatgctcatt

atcaagtaagtgatatctacctaggttgtgttgataaaatcttctcaggaataaatgcgg
catttattaacctaggaaagaatgagtacagtggttttatacatatcagtgataccggtc
cgcttaaaaagaaatattatgtcaataatattactaacattttaacaatacggcaaaaaa
ttttagtacaaattattaaagagccaactttaaataaaggtccaaggctcactgccaata
ttacattatcaggtcgatatattgtattaatgccttttagtcaatcaatctgtatatctc
gaaaaatatatgatgaagatgagcgtcattatttgaagtctttagctattttaattaaac
cggcaacaatgggcttgctatttagaccttctgctgtaggtgtcgatgaggaaataatat
taagcgaattaaaaaatctaaaagaacaatggaactttgttcaaaaatctgcaattaata
gttattcacctgttcttctatataaagatgaagatattgttaaaaaggtaatccgagatt
tttataataataatacaaacaatatagtaattgattcaaacctgggattaaaacaattaa
attattatatccacacttggcactgtaataactctagcacagttcctaagattaagcttt
atagtaataatcaatgtatactagatgcttttggtatcaatcaggcaatttccagagctc
tcattccaaaagttgatcttatacttggtggctatatgtttattgaaactttagaagctt
ttactattattgacgttaattctggatcttttaataattccactagtgcacgagaaacag
ttttaaaaaccaactgttctgcagcaacagaaatagcttatcagttacaaattagaaata
ttactggtgtaattataatcgacttcattgatatggaatcacaaagagatcaattgcaat
tattagaacactttaataaagagctatcacttgatgatgctaaaccacaaattgtacagt
tatctgaattaggtttagttgaattgactagaagaagacagggcaaaagtttgtacgagt
taattagtagtgattctaattacttttatttttcacacaatcagagagatctcagtctc
ttaagagattcgatgatagacagcagaaacaacagattttaataaatcttggctatctg
cagagattaatactattaacaaggtctttttcaaaagtccaatttgtgcagacctgcta
acttttacctaattcgtaatctctatatagttaaaagtagtattacttataaacaaaatt
atttattaactcacagatcaaaacttatttattctaaagaatacagcaaagtattaccaa
gca------gttattatctagctagtctcaataagaatagtaatcaagagttcctacctt
aagattgttatatcaactgtt---------------------------------------
-------------------gaaaaaaagaagctctctcttatataagagagagcttc-tt
ttgtagcttactaacta-tctagttagttaattaaaaattaaccgttaacagctggagct
gttagagctactggtaaagattcaccagaagctagatctagagggaagttgtgagcgtta
cgttcgtgcattacttccatacctaggttagcacggttgatgatatcagcccatgtgtta
attacacgaccttggctatcaacaacagattggttaaagttaaaaccattcaagttgaat
gccattgtgcttacagataaagctgttagccagataccaactacaggccatagacctaag
aagaaatgtagagaacgagagttgttgaaactagcatattggaagattaaacgaccgaag
tagccatgagctgcaacgatgttataagtttcttcttcttgtccgaatttgtaaccatag
ttagcagattcgttttcgcttgtttcacgaattaagctagatgtaactagggatccgtgc
atagcactgaacagagaaccaccaaacacaccagcaacacctagttggtggaatgggtgc
attaaaatgttgtgctcagcttggaatacaagcatgaagttaaatgttccggagatacct

agaggcattccatcagagaaactaccttggccaattgggtatactaggaatactgctgct
gctgctgctactggagcagtaaaagcaacggaaatccatggacgcatacctaggcggtag
cttagttcccactcacgaccaatgtagcaagctacgccagttaggaaatgaagaacaact
aattggtaaggaccaccgttgtatagccattcgtctaaagaagcagcttcccagattggg
tagaagtgaataccgatagctgcagaacttggaataacagcaccagaaatgatgttgttt
ccgtatagaagggaaccagcaactggctcacgaattccatcaatgtctactggaggtgca
gctacgaatgcaatgatgaatacagatgtggcagttaatagagttggaatcattaataca
ccaaaccaaccaatgtataggcggttttcagtactagtaatccaagagcagaaacgttcc
cacaagctagcgctttcgcgtctttgtaaagtagcagtcataatttttatcaattttta
aggatttaacaaagatgcttcccaaacaataagttgtctgttatggaaattattacttaa
ggccttttcaaatgagaacatttttaaaaaaaaca-tagtaaaatctaatctcagtaact
aa-aattt-tacttgttcatatattgcttaataattcaa-----------taataaattt
tt-gct-------------ttttgcaaaattaacactattttagggat---ataataata
atgcatattaatgctgcttattttgattttaatagtttatataattttgccctacttaaa
tacgcatgtcacactttacaaaaattcagacgaccatacaagatttaaatctgttaaagc
atgctttaacagatttgggtttgatttggcaaatgaattctagtcatataaaagttggtg
aaaataatcagcacaaggtagatattttaattaaacaagataatttgtcacatattggct
ttacttggaatgataatagatatcacttagttgccgacttacaactttgggaacaaccct
ggtctctagaagtatttttggataagttatctcagaagtatgcttactactctatcatcg
aagaaacaaagaagcaaggctttgaaaaaatgcagcagatttcacaaaaagatggatcaa
ttaagttaattgtgcaacgctggaattattaactaacatacaaatgcgggcggagagact
tgaactctcacgagattatctcactagaacctaaatctagcgcgtctaccaattccacca
cgcccgcactgttgagac--aatattactctatcataatctatactaaaacacaaactaa
-tagtataaactagcttccaagcttaaaagcgtctacaaatagaccgtatgttataccaa
ttttgaagttgttgaataaattaatttttatatttaattgcttgtggctagagtagacta
ttttactagtcaactcagttgctgcaactattagactggctgcaataataccccaatctc
ctgtttgtccaggaatcgtagacaaacctgttgaaataaaaaatcctagcaataaactaa
ttaagccagttgttaactcgctcaaagaataataaagcttattatttaagtttttttatta
agtaagcaaaaaaagttgaaagtctagttttaatcatagtatttagtaacggagttttaa
aaggatgacaagtattaacag-tcattaatattgtcgtatccttttaaaactcgttaata
caaacagctgttcttgt---------ttttaaaggtcttgttcgcttaaagcattaataa
gatgctgaaacggctcttgtataatatctgtgcgtttaattgattgcaaatctaattctt
cttctatgatttcttgtaataactgtatgcttcttactgttggagcaacaggtacagata
gtgaattgtaagtatcttttagcccatttaatacacgctgatcgagaatatttgtactac
cagaaaccaaggcataactagcataccgtaggtaatattctatatccctcagacaagctg
catatcttcttgttgtataagaatttcctccaggccttaataattcaggctgttcttcgt

acaactgagctgcggcctcttttaaaatattggtagcttgatcattaataatttcagcta

tttttattcgatctagtccactcgaaaagaaagattctagttgtccgacagctgtcttat

ctagataacgaccagtaagatcgtaacgatttaatattgctgttatagcatcttgcataa

attaattctcctac-ttgctatcagagcggtattataaatgactaatattgtaatataat

cattagtgtttatgaaatttatttaaaacttattctttattgcttaatgtgcaattaaaa

actcgcatagattttacacatatagagtatacgtagtttttaatta-tttttttataaaa

accaagaaataaagctttttt-taagaggttcgtgaatttgttaatataacatgagcttt

cctcctaaataagttttgtaaagtac--aaagtaaaaaaacgattagtcaataatagtaa

------------------------------------------------------------

------------------------------------------------------------

------------------------------------------------------------

------------------------------------------------------------

------------------------------------------------------------

------------------------------------------------------------

------------------------------------------------------------

------------------------------------------------------------

------------------------------------------------------------

------------------------------------------------------------

--------------------------acaatattaatatgtatcaa--------------

------------------------------------------------------------

------------------------------------------------------------

------------------------------------------------------------

------------------------------------------------------------

------------------------------------------------------------

------------------------------------------------------------

------------------------------------------------------------

------------------------------------------------------------

------------------------------------------------------------

------------------------------------------------------------

------------------------------------------------------------

------------------------------------------------------------

------------------------------------------------------------

------------------------------------------------------------

------------------------------------------------------------

------------------------------------------------------------

------------------------------------------------------------

------------------------------------------------------------
------------------------------------------------------------
------------------------------------------------------------
------------------------------------------------------------
------------------------------------------------------------
-----------------agtataaataagctttactccagcatcaaaaaaaatatttga
gatatgttgatacaactgattattcaagtgaaaacttagatttagaccaatttagctatt
ttatcattacaattaataaaactgataaatgtattcttatagagcaattttgttataatg
ctgaaggccatatttactcaatattcttcaaaggtccaacagctaaacatctttcttcta
taatttgtcattttcaacaattaaatacaataatatctacagctcatgccatatatttag
gacgagagctaatgaagtcagaattagctctcgtgctagatcaacaatatattcaagatt
aaa--ttttgtaagtattaattatatttatgctagtataatattagtaagggcatgtaac
tcagtggatagagtatcagattccgattctgatggccgtgggttcgaatcccgccctgcc
c---gagtattcagagtact-atactaaaacatct-------taatagctagtttgataa
tagtatgggcattggggctgcaaggtttctacattatgaaaagaagaaaatatg-aaaaa
aaacaagctctccaaaagagcttttagtacaatcaataaatgcagaaaacaatattgttt
ctttttctcgaaaattagctcttgcttaaacactgttaattttggcattcgacatgtta
aactcttattcatggcgaatattctgtcagagtcgctcttagtttaagaaaagtcagaaa
aaatatttatctttacgtttt-tacctatttttatttaaataaaggttagta-ttttcaa
cttttatagtggacgtgggttcgagtcccaccagctccatcaatttgatt---------t
gtataattagctagtttaaccttttaataaccagatatataattggaggaaatatgggt
ttcataacaattactaagcccgctttaaagcaaattgcaatcttgaaaaatgatcatgaa
aatgatgtacatctaagaataggtgttagacaaggcggatgttcaggtatgtcatactca
atgaattttgaacatgttgacaaattaaaagatacagatgaacggcttcgtctagataat
ttttctgttgtctgcgaccctaaaagtcttctttatctttatggattatcattagatttt
agctcggaattaataggaggaggatttcaatttctaacccgaatgccagccaaacttgt
ggctgcggtaaatctttttcaggctgacc-ctctaacacgtttttatatatatttgtatt
atatatattatttagcaagttg-tttattagagacaagcaatgctctgttaaaaac---
-----------------------------taatttcaagagttttattgtattatt-----
----ttatatatttattatcaactacttgaaaataaattagctaagcaatctaatttgtt
atcagtaccaaactttaaatgattaacttttattgatatatgctaatcttt-atcagcc
taaacatatctgtacattacaatcttttgacataaaaagagtcttaacatacctctcaa-
----------------------aaataaattcaa----------tattttataaatactc
aacatgaatctgtcacaacttatcgttaattcttctatgatgaaaattcctaataacttt
tattgcaatataaaagctatttcttctgagaatttaagtcacttacaattatcaaaaaca
aatttattatctcctacaaaaataattgcttttggactacctaaatttgattttatttct

acacagattacaagttctattagagaaacagttgttgtgtctactattgatgataataat
ctttttgctgccccaactaatgattggataaaaaaacacaaggataaaaatcatcattct
ttgttttataaaaaagtttttaaagctttgctaaaccataaaattagagttgttgctgag
ccatt------tcataactccgaaataaataaagaaaagagcataatggcaagtcaaaga
tacatctggggtaaaagctggaagccttcaattattttatcttgcttaaaaaacaaaaaa
aatagcaaaatttctgatgcatcgaaaaaatttttaactgaacagttatctgcatcacca
gtatttgtagtaaaaaatggttttaatgaaattattttaggtcacccattatctcgtgtt
aaaagaggaggagttaataatctgatgcatgcgttttctaatttattaaatcagtctaat
gcaacttatcctatatctactggtctattctttttttcatccagatgatgcatttgaattt
aaagactttataatatcggtcaatccactggcggccaaacatatggagataagcgtagag
cctgttggtctacactttgcttataaaatgaatagaaatatatcgtcagatactcaattt
tcgtttattccagatttaaagaagtaggagatttattattcaaatatagaaaaggtaat
catttagtttttcataagaatcagcattatggcaaagatttttttcaggggcaaccaata
tatatgattcagccgatcacttttaaagatcgagccggaaagctaaatattatcaaattt
acggggctaaatgacacaagagaaattatttttactaatcttgaagctgctaatagatca
tgggcacattttataaagaggaattcacaattaaaatcaattaaaaatcctactctatta
gtctataatttagaaagtttttaaaagatcaagaacgattggataatcaagatttaaat
aagtttgtggtgatcactaataaagaatcttatcttgccacaaaagagttaatagcttta
cccgattctaatagcttttctaagcacttaaaattaaatataaagcctaaactctttttt
gtgaaactatgggtaagacggttattttccaccttaacttatgaat-aaatgctatt-gc
ctttattttctcgaataatattctataactagtaattcattgagttgtaaagcaacccat
tccctatcaatgactccattgactttccctgataaattggatttattcaattctaagtgg
ctaggaatattagctagtccagggaaagctaagtaattttctactagttttcgagatgct
tcttgatttttttacactaattgattctcctggtttacattgataactacagatagatact
acttgtccattaatacaaatatgaccatggttcactagttgtctagcggcaggaattgta
ggagccataccaagtctaaaaacagtattgtctagcctcatctctagtagctgtaacaaa
atttgacctgtcgacccttgtagtttttttagctgcttttacgtacttaaataactgtttc
tcacttaatccataattaaatcgcaacttctgtttttcttctaatcttacagcatactca
gaaggttttctggactttttgtccatgttctccaggtggataagatcttttaatcgctttt
ctactgagtccgggtaaatcacctaatctacgggaaatgcgtactcgtggccctctgtat
ctagacatattatttaaattctcctaatgtaaat-gattaag----------taatgagt
agctattttttaacacaaaaagagcattaaaacaaagc-ttttgctaaaaaatatttaca
aaaaatatagatttggacaaaa---tacattactatgc-tttttaagcgggtagcgggaa
tcgaacccgcatcattagcttggaaggctaaggttttaccactaaactatacccgcaata
ttcaacagattgtcactaactcaatataacatagttatgttaagttaagataatcttaaa
tttaatagaatt-ttaaattatttatgctccttctaaaaccacatcaagaaaacgagcaa

tttcggcagcttgttcttcaacttctgataataataatggttgaccgaccctagtcaatg
gaattactcttttatcttttgtacataaataaatttcacgtcttggatttagcccctctt
taatatctattttattgatttgatttcttttatattaaactgaagacagatttttctat
tctttccagggaaacctagccgaaaaattttacgatacctttatctttgttaaattcat
tatatccagcccctatattccaaataatagttagccacagaaacaaactcagaaagactc
ctatactgccatagaatgtcattacaataccttgtggaataaataccaaatcggttgaat
ttgtaaaaggtaacaaatcaacttgaaaataacttgatagtccagcaagaagaaagccca
aagccccaataaatattatagtagcccaccagtaattactcagtcttcgtgaacctaaaa
ttaaatcttttctaactgagtgtatagataatgtttttttcataatttgttcttttaaaa
cagta-tactaatgcaaaatggaaaaatttgacaattttt--atgcgcgctttagattt
tttaaataagcttgattgcttttagtcctaaaaacagtcctgttgctaacccaaaaaaag
cagcaagaaatattatatatcctagaaaaactgacatattattattatttattattattg
ttacaatagtgtttacaaaaatagtgaaaatctattactacggtattatattacgttaaa
tgccttgctgtctacaggaaaaaactttttcagtttttagcattgtgtttatattagaa
ataatattctagtaaatttattaaacttctggagaacagaatcttatgtcagaatttatc
aagccttataacgatgatccttttgtaggcaatttgtccacgccagttagtacgtcaagt
tttagtaaaggacttctaggaaatctaccagcttaccgtcgaggtttatctccgcttctt
agaggattagaaataggaatggcacatggatacttttaattggaccttttgataaattg
ggccctttacgaggtacagatgtagcgttactagcgggatttctatcttcggtcggcctc
attattattctcactacatgtttatccatgtatggtaatgtatcttttactagagcagat
tcgaaagatccactacaaacttctgaaggctggggacaattcactgcaggatttctagtt
ggagcagtaggtggttcaggatttgcttatttattgctagctaatatacctgtattacag
actgcaggtcttagtttattctcttaagctagtaaggggacttgaacccgtaacctactg
attacaaatcagttgctctaccaattgagctatactagcattcagttcacaataacacat
aaagatatttgttgcaatctgctattttcagatgtaacaaatatctttttttatttaaaca
gaactgttatgttaacac-------gtatattaattagaatctgaattatctgactctgc
agaagattcctgatcacagtttagataataacaaattgtttgatctagacaagttgcaga
ccaaccaacaggtgtttctcctgataattcttgcatatctacagaattataaaaattatc
atttggcttattactgtttgcttgcataaatatcactcccaaaacactctgaacttaatg
tgaagtaatttaacttctcaataatatattaacacaaaatattaaatatggtgattattc
tttaattgtaaagatgctgctaactttataaagtttgagtgagagtcttgatagcttttg
ttgaaataagcattttaacccatttatttgccctctcgaccatactttttttgtgttgta
aattaactttttgtagtttttttgttcgtttatgagaatgagatacggcatatccattat
tagctacttttcccgtaagctgacactttttgacataatttt-------ctaacttata
tatttatttagagtacttgctaaagttgacttgggcacagctccaatcactgtgtctact
ctttctcctgctttaaagatcattaaagtaggtatacttctaatgccatattcagctgca

atagtaggattatcgtctgtgtttatttttactaccttaatagatgattcatattcttcc
gcaatttcatcaactacaggagaaaccattctacaaggaccacaccatggcgcccaaaaa
tctacgagtacaggtaagttattgttaataacttcttgtttgaaagaggcatctgtaact
tgagatactgacatattctttaacctttaatatatgttccttgctaagcaaatcttatca
caaatttcaagaaaaatcttctatagtgaattactattagtgttgaattatttttttaaa
aattagacattaaaaaaacttattacctatataaataaagtggtggccgagtgatatctt
gatagtgataatttttatattaaggttgaacactagaaataacttttcaagcaaactctt
gtttaaaaa--atttctctgtgaccataaaaacttaattatattacaatatattttatta
tttgtaacttagttctgataatggtataaacaacgcaaaagatactgccttataatcaag
gaggaatacatgtctcaatccgtagaatcacggactaggattaaaagcgaacgttacgaa
tctggagtaatcccctacgctaaaatgggctactgggatgctgactatgtgattaaagaa
acagatattctagctcttttcagaatcactcctcaaccaggtgttgacccgattgaagca
tctgctgcaattgcaggtgaatcttcaacagctacttggacagttgtatggactgattta
ttaacagcttgtgacttatacagagcaaaagcatatcgagtagatccagttccaaacgtg
gcagatcaatattttgcttacatagcttatgatattgatttgtttgaagaaggttccatt
gcgaacttaactgcttcaattattggtaacgtttttgggtttaaagctgttaaagctctt
cgcttggaagatatgcgtatgccagtagcttatctaaaaacgttccaaggtcctgcaact
ggattgattgtagaacgtgagcgtatggataagttcggtagacctttcttaggtgctaca
gttaaacctaaactaggtttatctggcaaaaactacggaagagttgtatacgaaggcctg
aaaggcggtcttgatttccttaaagatgatgagaatattaactcacaaccatttatgcgt
tggagagaaagatttttatattctatggaaggtgtaaataaagcatcggcttctgctggc
gaaattaaaggtcattaccttaacgtaacagccgcgacaatggaagatatgtatgagaga
gccgaattctctaaagaggttggtagtatcatttgtatgattgaccttgtgattggttat
actgcgattcaaagtatggcaatttgggctcgtaaacatgacatgattttacatttacat
agagctggtaactcaacttactctcgtcaaaaaaatcatggtatgaacttccgagttatt
tgcaaatggatgcgtatggctggtgttgaccatattcacgcaggtacagttgtaggtaag
cttgaaggagatcctttaatgattaaaggcttctacaatactctacttgaaagcgacaca
gatatcaacctacctcaaggtctgttctttgctcaaaattgggcttccctacgtaaagtt
gtaccagtagcatctggtggtattcatgctggtcaaatgcaccaacttcttgattactta
ggtgatgatgtagttcttcagtttggtggtggtacaattggacatcctgatggtatccaa
gcaggtgcaactgctaacagagtagcactagagtccatggttatggcaagaaatgaaggc
cgtaactatgtagcagaaggtccacaaatcttgagggacgctgctaaaacttgtgggcct
ctacaaacagctttagatttatggaaagatattagtttcaactacacttccacagataca
gctgatttcgttgagactccaacagcaaacatctagtttaatgactacttactgatactt
taaatagtcaattgtaagtggaattaacttataacaataaggagcatagaatagtgagac
taacacaagggactttttccttccttccagatttaactgatgagcaaattaataaacagc

ttgcttatatcgtttctaaaggcttatcagcaaacgttgagtatactgacgatcctcatc
caagaaactcctattgggaactgtggggtttacctttatttgatgtaaaagatgcttctg
ctgttatgtacgaaattagctcatgcagaaaagcaaaacctaattattatgttaaagtta
acgcttttgataatactagaggtattgaaagttgtgtaatgtctttcattgtaaatagac
ctgctaatgaaccaggattcttattacaacgccaagacttcgaaggtagaactatgaagt
atagtcttcatagctatgctactgaaaagcctgaaggagctaggtattaatattaattaa
gaattaatattggctaatta--ttaccctcttaaaaatcataattaatttagtaattgtg
atttttaagagggatagcttcccaagtaaattttgactaaa-tagattgaataattaaaa
atacacagaaataatgcaatcacaggatataatttccaacgatactcttgttaatttaca
agaagaatatgatagaacacaaatccaagaagttttaaatgagttaaatcaagaacttat
aggattagtgcctgtaaagaccagaattcgcgaaattgctgcgctattattgattgacag
attacgcagaaaactagaactagtttctggtaatccaggattacacatgtcatttacagg
tagtccaggaactggtaaaactacagttgctatgaaaatggctgatattttgcacagact
tggatatataaaaaaagggcatttgttgacagttacaagagatgatcttgtaggtcaata
tattggacatactgcccctaaaactaaggaagttcttaaacaagcaatgggaggagtttt
atttattgacgaagcttactatctatataaagcagataatgaaagagactatggctctga
ggcaattgaaattttattacaagtaatggaaaaccaaagaaatgacttagttgttatctt
tgctggatataaagatagaatggaaaaattctacgaatccaacccaggactctcttctag
agtagctaatcatgtagacttcccagattatacttcagatgaattattacaaatagctaa
aatgatgatagaagaacagcagtactgttttacagaagaagcagataaaactcttttaga
gtataccgagcgaagaatgaaacagccttattttgctaatgcaagaagtattcgcaatgc
tattgacagggctagaatgagacaagccaataggatttttgccagtggagaaaaagtatt
aacaaaagctgatttggtaacgattgaagcagaagatatcttgaaaagtagattattttc
attacctaatgcttaatatacacgtgatttcatgaaactatatattattaaaaag-tttc
ataatcagttgcaaatttcttgaaaaacctttattattatcttaatgcaggtg-tggcgg
catagccaagtggtaaggcagaggattgcaaatccttcatcccccagttcaaatctgggt
gccgcctagtactaaaaaagggggtgtggtggaatggtagacacaacagacttaaaatc
tgttgattttagtaatcgtgagggttcaagtccctccacccccatatattgatttataa
ataaaaagagatgtgtatttgtattaattgcaaccatattaccaaatgtaatacctatca
cttaatagaatctcaacataaacagcctcatcttactagaagtccattgtttatacctaa
atatcctgtagttcatgttaatatatctaacaactgtacctataatcaaattgattggga
tttagtggagtgtttatcatttgtagaaaaaccgaatagttggaatttagacgctaatta
gttacaagaaaaattatgtaccataagcaacaactctatttttagatactcggtattaa
ttttggatgttctaaccagtaagattttacccgttcttgctatctctataatgccaaatt
tagttagtaattgttcaatagcaacaatctttcccggatctccagtaacctctataatta
aaagatcttctgcaatgtctactatcttagctctaaaaattcttacaatttctaaagctt

ctgttctagtctgagaattgatctgaatcttaattaacattagttctctttcaactgaag
gaatatttgttacatcttgaacgttaaggatatttactaacttatataattgtttcgtaa
gttgttcaatagttctattgtctccttgaactaccattgtgattctagagaccccaattt
gctctgctggtccaactgctaaacttgcgatattaaaccctctccgggcaaatagaccag
atattcttgacagtactccggcttcatcttgaactaaaactgataaggtgtgtttcatga
aaaattaaatttaatctatttgatctgttaagtatattctaagaaaattacagaaaaaat
actagttagtctataactaaattatccaaactaaacaaaatcctactatcagataatata
tctaccttgtaatgctatactattaacttagtatttaatgatattctaattgcaaataga
atgct-aatttttaaaatttgttaggaacttgtgctagaaaaatacaaaaacagtaaaa
aactctctcgagatttacctcaaatcaatgatcgcattagatttccaaaagtccgagtaa
ttgatgacgaaggtgaacaactaggtatttttgtgcctgaagaagctatacaattagctg
tccaacaaggtttagacttagttgttgttagtgataaatcggacccgccagtatgccgaa
tattagactatggtaaatataagtttacacaagaaaaaagagctagagaagctaagaaaa
agcaacataacagtagtattaaagaagtaaaaatgcgatataagatagaagagcatgatt
ataaagttagaataaaccaggcatccaaatttattcaagcaggagataaagtaaaagcaa
ctatcacatttcgggggcgtgaaatccagcactctaatttagctatagatttattgaata
aaatggcaagcgatctaaatgcaatagctgaaattcagcaagctccatcaagagatggca
gaaatgtcataatgctcttatctcccaaaaaagttagctaaactaatttttatttaaatg
catctggctggattcgaaccagcgacgtctctttcgaaatggcggattatgagtccgctg
ccttcggcccctcggccacagatgcatcacttaaagtttatacctcataaagcatataaa
aatcaatgtttagctatttttaataaagtctataagagtctattattcattcatattgtg
atatgtcgcaactgctgaaggggaaacttgatttaagtatcgaaatatccagtatttaaa
aattgtgtcgagtataactggaaaagttgaaataaataaaaaaataaaatctcgactttc
cggaagtcctaaatgtcttaaaattacttcaataataacttcccatccatgaggagaatg
aaaacccacaaacatatcagtaaaaagaatgattaaaaaagctttagctgtatcactaag
accataaataatttcatttaaaaaagatttaacgactgcaatttgtctttgtccagttat
cattagtaatataaagacaagtattgatacaagatcggataaaatatttttaacagcatt
tgcactttcatttgcataatattctcccaactctctagccttcagctgcactctttttc
aatgatttcgtaagatatatcttccgaaggattcagaagaacttcaaagtgaattttctc
ttcaaatctttgtaattcagcaaaagctctttcctcttgagaggaatttaaaaaaatttt
aggctgttcttggttccataaataatcaatacaaggtccaaatacaaaaaaattttgaagc
ctgattaactagtactggagagatgaataataataaaatatattttacggaagtaatagt
ttggtgtctagatattctaaactcctctatagcttcagattcgccattaggatctaactc
tttacgaaacttttcaaaagtattagtaattgacctaggtattggcccaactttttcaaa
ggcgaattgattatttcgttttaaattccaatatttcatttttattgctgctaaaatct
ctattaatcaataattgatcacgtattcactataaatattaattctagaactaaacatat

aaaacaatatttatgaaaaataaaatacaattaaaaattaattatgcattagttctatct
gcagctacagcagtttatggatataaatcttatcagccaactataaatacactcttaatc
aaagatgaagaaaactggccttttttctctaaaaggcagactgcctttattcctataatt
gtattaaaaaaaaatcatcaaaatccctgaactcagaacctagaaatagtaagttatatc
ttaatggaattggaagtagaaaatttatcgacagaattgtcactctacttggatatttag
aaaaacaggcaaactatactgaatacttgaaccaaataaaaacaagtggttatttttctt
tagtctacctagacagtgatataatgtataagaaaaatattagtaacatttttttactac
ctaatgatattctaaaaagaatctatatttgcaataagaaaaaaaagcttataccgcatt
cattcttaataaatttatttcagaaacagataggctatccaaaaagttttgctaatttaa
attgggccttctccaaaattataaaatggtattatgatagaggataccaatggtctctag
ttgaagttaaacaagcgtctgatgcatcttctattgtaattgatattcatgaaggggtag
tgaagacaattataacagaatactacactctatcttataaaagagtctcaggtatcttat
gcgtagagtcaatagagcagtaccttggagtaagagtgggagctccactaaatataattg
atttacaaaaaaaaattacttacttaaaagataatcaactagttggcgacattatttata
gtattgaacgatcaaataatagctcaatgagtttggatatcaaatttcaaatacaagaat
taaaagataaagagataatagtgcttgcaaaaagttcttcta------------ttatct
cgcatgcatgtaatttacttaatc---aatataga---------aataggctagtagctt
ctaatatagtttcactatcaactaataaattacacgcatgctattttaattataaacttg
actatcaatataagtatataaacacgtctaatcttataaaattgttagcttattctattt
cttgtaggaagacactacttattcattattttaatttacaaactttaataagctatacaa
aaaaaaatactattggatttcagctatatttacgaaatctaagttttgggaaagctttct
gtgtgcttagtatgaaatttatcaaaaatggacttaatataaaaatttatatataaatc
cttcattgatagttgatcagaattttgtatttcaatttgctatacaaatcattaagcaat
atcatactgcta------aacctccagctt------------------------------
------tatttttaacgaatctagatcttgaacaatatgttgctgaaagtttattaatgt
accatttacatcttgcttctcaatatctgaaaaaatattattatctcgaattatgcata
cagattctttatttttcaactctgaaa------attttcattttgatgatagaactaacg
cgatcaacagttatgatacttttaaacaaaatacaaaaatattttatcaagagtttcttt
ctttattattaagtttacgttatcaaaatttttaattatttaggttggcctttaaaaggcc
attttttgaaataaaatctttatacttggctccatttcaaaagagtg------------
---atttttctgatagtcgtaagaccctattcttccacaaaatgtcgttaaagcaagtat
ctaattttaatttaccaatatcttttaaaagccaccttaaccatattttagttagtacta
ttaaatgtcagtcaaatttgaatatgagaactgtatctttgctattaattgattcacctg
ctgaatatatgctatacaaatctattcttaatttctctattaaagttaggatgcagtatt
tcatacctatgagcaacaatataagattatctctattttataattatttagattgttttt
taataagatcatctcagagtcgcattcatatatggcaagatctaagaactcttacaccaa

ttcaaaatttttggttaaaaaaattttcatatggagctggaattcagcttaaattaccaa
ttaagcagatgccacctctttcaattgaatatactgtaactagtagtcgctacttttgta
tttaccttcgtacttattatcaacgataactattactaaaatgaaaaatttaatttcttt
ttttgatcgcagcaaagggaaatggatttcacaaagaacaacttatgaattatctaataa
aaacatgagttcgatacagtctcagatgacaatgaaaataggcaactcattgtcaggatc
tataatacttgcatcgttaaattgggggacatttatagacaagttgcccattacgctaa
aaatcacagccggagtgaatatgataataagtttaacttgcaattcagtaatcaactaaa
taatcataaattattaacgctgtgcatagtgacagatcctagtctaattagttttaaaac
tcgctatggaagtactaccatagatgaaacatattggttcgcaacaaataatttacgtct
aagtactagtattgttaaacgattcaacacttgtgtggcagtatctttttgttcagaaat
taaagtttaa-actctatatagaagcaactaatagcctaaa-ggctattagttgcttcta
tatatctaaattttgtattaaaaattat----aatactatattattctaagtctttacga
ttgggattcctggaaggatcattagagaggaacccaaagacaaacaaagaaataaagaaa
ataacggtagtatatacaaagattttcaaagtaaacatattttcatccttaaaagctatt
agtgcg----atatttacatatctccacataattttatatcatgatataagttccagcac
tattttataaataaatcaatatctttatttttagtaatgatttatatttaaatatgcgt
gatgtactaaatactaaatcttttccgacttgttttgtcttgtttgtaaatattttgtta
tttctatggagtttaccttcaattatcacatagtctaacttctttagctgtttaaaagta
tatgaagatttttgttccaaatagttagattgattataactaattgttttctttttaac
agtcttgcttttaacttaattatttggcttttattttcactaattctgatactcttacaa
ctaagaatttggactagaagagtacagctattcataatttataacaa------tttttta
aaaaatctactaaaatagatgattttacagaactaaaatgatatttcattatcttgttga
ccttgctcataagtaatatctttcaatacttttaatatttttccaaaatactcttgcagg
tatttctctaaagatattacttccaacctgtcaattccaagaatactgtaaacttcatcc
atcggagctgtaaattgctctccgctacttaaaatttcggcaaatgctaaacgatcagaa
atattccatgtccattgaagagttttagtaattcttcttaaggcttttaataatccaatc
ggaatttgtgagatctgtgtttttttgcccagataatttttcacataatttaataatttca
gctgatgtccaagcggtattaccaactaacggcagggtcctattttctgtggaaggtact
ccaagacttttttataactaactttgctgcatcttgagtatcaatgtatgcaatgggtgta
gattctcctgtaacccagactgattttttgtctaaaataggaatggcatattggttgatt
aaaccttgaaaaaaatcctcctaaagaaaacacagtatatttcacattggacttctgaagg
aaatctactacctgagactttaagttcatcaatggaacatctgggtatttctctgaattt
aatattgaaaagaaaataaatctttcaactttagccgctttagctgcttcaattaatgca
attttcccatctaaatctattttttctgcattgtaaggatcagtaggacgagatgtagaa
gcatctatgattgctgttaccccacaaaaagattgcaaaatactttcgggtaattttaaa
tcaccatatataagttctgctccccactctttgagaaaggcagactttcttaaatttctt

accatgcattttacattgtagccttcatctaaagctcgtcttacaatttgacgtcctaaa
gttccggttgctccaattactaaaagagtcatattagttaaaaataagtatgtataaaga
cattgatttggttgacggcagagagtaaattaggtagagagggaatcgaaccctcatgac
cgaagtcgtcacattttgagtgtgatgcgtataccaatttcgccatctaccctattatag
accaactactacaagtaatttagtctttagcgtaaaaataatactaagtttcaatggata
tgtcaacttttaaatttatattaataaaaaaacggagtctcttatggccacttttgaatt
aattccttatagattgtatttgagtcaatctaacacatggatccatagaataaaagcaga
aattaagatatatatagtaacgcttttatggatttcaattttatttttttcttactttaa
actatgtattattgctttaagcttaattgccataagttttactataagaagtaaacagaa
tattatccaaaaacatttgttacaaacgttattaataacgttcttaactactgttttgtc
ttttagtgtggctattagttataaacaatatgcagaacaagaacaatcgcagtatttatc
tgattccaagaaatataaaaattccagcagctactattatatacaaatagctaacgatca
aaggatcaaaagacaatacttaattactactttaaagccttccttgtactttttttattac
tatatattctattaaactagttatgataacaacttctccagaagttttagtaattactat
ttatagatctaggataataaataaaatatttaaaaatgaattgctatttatcttccttct
ttcgtcacatattgttactagtattattaatagaatcgataaagtaattcaagtaactag
cttaagaggaagcttgaatttatataattcattgacaaggcctttaatgttttctttgtt
aatatttcaagtcttctttttggagattattcgggagtcaaaagaaatagctcaagctct
ttatactagaaatctcaatcaagaaaacaataatttttttgaaaatatatacagtaaaatc
taactttagtgatcggctcaatataattattagcactttgtactttattattttagctct
agcgtaatagaatccacttattcttctgactctagtaatacgaagagtaataaaattagt
tagtagtttaaatagatagaat-taagatattatataaataattaaatgtttcattatgg
ttaatactcaaaatcaaatttcacaaacttcagaacttgattatatagttaatcaacctt
ataaatatggctttaaaacttctgttgaatctgagcaatttcctaggggaataagtgaag
atattgtccgcttgatctccaagaaaaaagatgagcctgaatatctattgaattttaggc
ttaaagcatataaaaaatggaaaaagatgagtagtccgtcatgggctcatattaagcatc
cgaatatagactttaatacgattatttattacgctgttcctaaattaaaaaaagaattga
aaagtctagatgaggttgatccagaaattcttgacacttttaacaagttaggtatatctt
taaatgagcaaaagcgaatttccaatgtagccgttgatgctgtttttgatagtgtatcta
ttgcaactacttttaagaaagaactgtctgaagctggtgttatattctgttcaatttctg
aagctataagggattatccagaattaataaaaaaatacttaggtactgttgtaccagctg
gtgataattattttgctgcattgaactctgcggtatttagtgatggctctttttgctata
tccctcctaatacagtttgccctttagaattatcaacttactttcgtattaacaacgaag
aatctgggcaatttgaaaggacactaattattgctgatcgtggcagtaaagtaagttatc
ttgaaggttgtactgctcctcaatttgacacaaatcaattacatgcagcgattgtagaat
tagtagctcttgaaggagccgaaattaaatattctacagtacagaattggtatgctggta

acaaagaaggtaaaggtggtatatacaattttgttactaaacgaggcttatgctcgggta
acaattcaaaaatttcatggactcaagtagaaactggatctgcaattacctggaaatatc
caagctgtattttagctggtgaaaattctcagggagaattttactctgtagctttaacaa
ataactatcaagaggctgatacagggactaaaatgatccatattggtaacaatacaaaaa
gtagaattatttctaaaggtatttctgcgggtagatcgaaaaacagttatcgaggcctag
taaaagttggccctcagtcatttaattctcgtaattattctcaatgcgattctttattaa
ttggtcaatcatctcaagctaatacatttccctatattcaagtacagaatccaacatcaa
aagtagaacatgaagcatccacttcaaaaattagcgaggaccaaatttttatttttttac
aaagagggatcaatttagaagaatctattgctcttatgatcagtggtttttgtaaagatg
tctttaatgaattgcctatggaatttgctactgaagctgatcgtttactgagtttaaaat
tagagggaactgtaggatgagccagactattttagaaatcaaagatttatatgcttctgt
tggtgaaacaacaattttaaaaggggtaaacttatctattcgagccggcgaaatacatgc
gattatggggcctaacggttcaggtaaaagtacattatcaaaagtaattgcaggacatcc
agcgtattcactaataagcggagacattttatttttggacaaagtattcttgaaatgga
gccagacgagagggcaaaagcaggtatttttttagcttttcagtatcctgttgaaattcc
tggagtcagtaattctgatttcttaagaattgcattaaatgctcgcagaaagtttcaagg
attatcggagtttagccccttagaatttttcaactaataacagaaaaaatagatcttgt
tggcatgcaagaaagtttttgacaagaaatgttaatgaaggattttctggtggagaaaa
aaaacgtaacgaaattcttcaaatggctttactagatagcaaaatatctattttagatga
aacagattctggactagatattgatgcattacgagtagttgcaaaaggaattaacacttt
agctaaatcaacaaattcaattattttgattactcactatcaaagattattagattatat
tattccagattttgttcatattatgagtaacggacaaattgtaaaaactggtagtgttac
tcttgcccaagatctagaaaagcacggatatgactggattacgcagacataactatttac
tac-------aatattgagactaaagaatattctttagtctcaatatt-tttgcaaatgc
aaagtt-aattatttaagctttatcaaaaccttgcttaacatcttcaattgctcttttta
gcaattcttcactaactggttctaatttttagtactacgaacattttcaccaaattctg
gtttagaattctttaaatcttctcgaagttctctaataaattcagctacttttgaaactt
caatatcatctagatagccattaatgccagtgtaaataatagcagtctgttcttcaactg
gaataggagaattttgagcttgtttcaaaatttctcgaagacgttgacctcttgctagtt
gattttgggttgctttatctagatcagatgcaaactgagaaaaagcttctaattcagcaa
attgtgctaactctagttttaacttgcctgccacttgtttcatggcttttatttgcgcag
cagagcctactctagagactgagatcccaacattaatagctggtctaatccctgagttaa
ataagtcaccagatagaaatatttgtccatctgtaattgaaattacatttgttggtatat
atgcagatacgtccccagcttgagtttcaataatagggagagctgtcatactacctccac
ctaattcagcattcaatttagctgctctctctagcaatctagagtgtaaatagaatacat
ccccaggataggcttctcgtcccggaggtcttctaagcaaaagagacatctgacgataag

cttgagcttgcttagttaaatcatcataaataactaatgttgctttacctttgtacataa
aatattcagctaaagccgcacctgtataaggagcaatatattgtaaagtcgcaggactat
ctgcattagctgcaactataatagtataatcaagagctcctttatcctgtagtgaagata
ctacttgggctactgaagatgccttttgtcctatcgcaacatatacacaaacaacatctt
ggcctttttgattaataattgtgtccaatgcaacagctgttttacctgtttgacggtcac
caataattaattctcgctgacctctaccaattgggatcatagagtcaatagcagttatcc
ctgtttgcataggttcacaaactgattgccttccaattataccaggggccattgactcaa
ttagccttgttccattacttgcaggttcacctttatcatctattggccgagctaatggat
caacaactctacctaaaaaagcatctccgaccggaatttgagcaattcgaccagtccctt
ttacggaactgccctctaaaatatccctaccatctcccatgagtactacaccaacgttat
cactttctaaatttaaagcaaccccaatagttttatcttcaaattcaagtagttcaccag
ccatgacttcatcaagtccatatactcgagcgataccatcaccaacttgtaaaactgttc
ctatattggctacttctacatcctgatcatacttttcaatttgttgacgaataatactac
ttatttcatcaggtctaatatttaccatattttaatattattttttattaattggaaatt
ctaaagcggtaattagtcagactaatgctatctgtctttttttactttattttaagttact
acaacatcaaggtgtgaagccatttgtcttagctgtcctctaatactagtatcaatgact
tttgatccaatctgaatagtaaagccaccaattagctcgggctcgactgaaataattagt
ttaacttctttagccttagtcataaccttgatttttttctgttaatagagtttcctgatca
gggcttaaagcaatagaagtattaatatttgcaattgttaaagattccatttggtaagcc
agttccaaatacttactagctatagcatctaacattccaattcgcttgcgatcaactaaa
atcattagaaatgataatgtattttcgttaatttgatcgccaacagttgcaataattact
tcttttttttgcctctacagtttttaatggattagct---aaaaaaattttagcttgtca
gattgcgctaaaatattttgtattgactgaatatcttggctcaccttttcagtaactttt
ttagttttagctaaatcaagaagagctactgcataaggctgagctatttttataacaaca
ttgttactgctcatagttgatctcctaacttagcaatattattatcaattatacgcagtt
gcatttcggagctcatttgattttccagctgcaaagtaactcttttaagagcaagaaatg
ttatttgttgctggatttgccttcggatctgtttttcagctgtttctatatttgatttac
ctgttatggcaagtctttcaatatctaattggccttgagctaatatggaacttctaactt
ttccagctgttagctgagcttctttcttaatttgatctataatgatttgagtttgagcaa
gttgtttttctgattctgacaatctagcacttgcttgctctagtctttcttcggattctt
gtatagctgctaaaacttttatctgtcttgcatttaaacttgatccgagaaattgcttca
aaacataaataagaccaaataaaagcaacaatatattaataacatttgcttcaaaaatat
cagaattaaagccgaatttgtgttcactactatgttctgataaaatagtaattatttgtg
gtatttttactatattattcatttaaaattct----------tactattttaatatttta
ctctggaatcatgattttcaagattgactacttaataatttaactttaatttgatcgctc
aaagtgtcaacctgattttctaaagttttaagagcttcttctttttgaacatttaactgt

ttcgatgcttcagcaatcagtttctctgcattcaattgagctttttttatatcttctgag
actatattttgagcttctttttgagaagaggctatttttaattgtgcattgcggcgagct
tcagataaatcttcttcatattttgcagctagctcatctgctttaacaagcatagacgat
gcagtagtcaatgttgttcgaatatattcatctcgttcgtctagtactttagttacaggt
ttataaaaaatagtatttaataaaaccattaaagtaagaaactgtaatgccattaaaggc
aaagtaccattgaagtcaaataagccaccttcaatttcttcggctaatagaaatggtaaa
tcaatcattttactaattatgtttatagttaataattttgtttcaattaagaaaaataac
tttatagtgcgcttagtctcatcagactaagcgcaaatgattattttaaccaacgtatgg
attagcgaacaataaagacaacgcaacaactaggccgtaaattgttaaagattccataaa
tgctaaacttagaagaagtgtaccccgaatctttccttctacttctggctgtctagcaat
accttcgacagcattagctgctgcactaccttgaccaatgcctgggcctattgcagcaag
acctacagcaagaccggcagcgataacggatgcggctgaaacgattgaatccataatgaa
ttttaatttttttgttatatgagtagaaaattgacaaatttcgtataatacttaaaatcta
tataattaagtataatagattctaatat-----tgtaattatagtatattactcttctcc
atgaccttccattgcttctccaatataagctgcggacaaagtagaaaagattaatgcctg
tattgaactcgcaaataatccgagtatcataactggtaatggtattaaaattggaattag
caaagtaaatacagaaacaactaattcatctgctaagacgtttccaaataatcgaaaact
taaagatagtggtttcgtaaaatcttctagaatattgattgggagtaatactggagttgg
ctgaatatatctagcaaagtagcctaaccctttttacttaaacctgcataaaaataagc
taaagatgtaagtaaagacagggccacagttgtattaatgtcatttgttggggcagctag
ctcaccttcaggcaagtgaattaacttccaaggaattaatgcacctgcccagttacaacc
tagaataaatagaaataaagttgcaatgtaaggcacccaagggcgatattcatgctcacc
aatctgatttttttgcaatatcttgcagaaattctaatataaattccataaaattctggaa
tttttcaggaattctttgtagatttctagtacctagaaatgataaagttaaaagtgtagc
aattactaaccaagagacaatgaaaacttgcccatgcaattgtaagctacctattttcca
atatagatgttttccaacttctactgctgataaactattatatgggttaaaatctatgag
actgttttgatacataataaatattt-ttttaaatagtgcaagaatcaaaaaataag-tt
tttgaagaattaatatgcaattaatttattcttaaaattagactattaataaattaaaag
caaaaataaattctaatagagatagctatttttagtagtgattatttttttgtaaaaaaca
attattgctacctataatgcttttacgtttaattatatacccatatattaattatttaat
atgattataggtattttgctacaaactttattgttttatccattct----aaagattgtt
atagtttatttttttatttaaaatatctgcgacttcatctgcaaaattaactttcaaattt
tcttctccccctcccagaagaaatcttacaaatcttctaattttaatattttcacctaga
acagcaatattttggtttattaaatcttcaatactaatatcttgattgcggataaacatt
tgatctaataatgagagctccttaagtctctttttcattcgtccttcaatgattgtttcg
attctatcaactggtttgtttttcaagtcatctttaccagcttcaactctttgttccaga

ttaataatttcactaggtatatcctgtgttgatacgtattccacattaggcgatgccgct
atttgcatagcaatatctttagctaacttttggaactctggacgtcgtgctacaaaatct
gtttcacagttgacttcaactaatacacctattcttccgccagtatgaatataactctct
agtaaaccttcaatagcggtacgattagatttcttgttagctgaagccagcccttttgt
cttaatgattctaaagcttttcttcattgccattattagcttgtaaagctttcttgcag
tccatcatacctgctccagtcttatcacgtaaggcttttacgacttgagcagaaatttgt
agtgtcatatgctttcaaaaatttttaagtatttaaatt----------aattattataa
ttcactttgagtaggatgagttatttcttctaaatctgtttgcccgtatttaccatcata
aattgcatctgctattttacctacaattagtttaattgacctaatggcatcatcattagc
tgggattggaatattaataatttctggactacaattagtatcaagtatacaaatagttgg
aatacctaattttaaacactcttgtatagctgttgtttctctcttttgatctacaactac
aacaatatcaggtaaacgagtcatattttaataccatttagatgtttacgaagtttatc
taattctcttcgtaaaacagcagcttctttttgggtaactgatcaatcataccactttt
atcttgttcttctagctgtctcagacgattaactcgagacttaatcgttacccaattagt
taacattcctccaagccatctctggttaacataataagaatcacagcgttgtgcttcctg
tgccactataccagcggcttgtctttttgttcctagaaataaaactttttaccatccga
cgaagcctgttttataaattcacaagcttctgttaataattgtgcagtttggactagatc
aataatatgtatgccatttctttctgtataaatatatggaaacattttaggattccatct
acgtgcttgatgtccaaaatgaacacctgcttctagtaattctgctaatgtaacaatagc
cataagattattaatatttaaataatgtaattattcgggaaatccc-tttgtttagaaac
tgat-----tcaactaacttatttgaattttagtaaactaataataaattaacacacttt
taaagcagatcaacaaaatagatcctgaagaattcat----tctacacttttattgctaa
aataattacgcgcggttctgtcgtctaaaatgatatcatctaaatcatctcgtacagatg
aaatagtactttcgtcattagtatttgcagttaaattctttttctctaacgatgagccat
tacagttatcatacatattgaagccggtaccagccggtataagccgacctataataacat
tttctttcagacctcttaaccaatctaatttaccagatatagcagcttcagttaatactt
ttgtagtttcttgaaaacttgctgcagaaataaagctttctgtatttaaggaagcttgag
taatacctaatagcacaggcctatatgatgcattaatcttatttctcaaggttatggctt
tattagtctgttcaatcttctgtaactctacaagctctccaggtaaatagcctgtttctt
caccattttctattttcactttagaagtcatctgccgtactattacttctacatgcttat
cagaaatattcactccttgagattgatataccaattgtacttctttaacaagtagtaact
gaatctcttgaaagcttaatcgagcagcttcatataaggctaatccacattcaatataca
aatcaaaacttgcatcgaggatatcatgaggatttaacaatacgtctgttttctttcttg
cttcaagaatttcttcaattcttggaagcccttgtataatgtcacctgtttttgctctat
caaaaaccaatatagctaatgtttctccccttctaattaatgcattgttatcaacatgta
aaatagcaccattagaaaccaagtaaggtctagcaattcttaatgtaactgatctagaag

aaatttcaataatttgtccagagtctaaagaagtcatattttcagtaataaaatcaccac
aacgaatccaatcacccacacttactttaactattgattttcctatattaaaggtctttt
tatcagaagatgtcgcaatcaaaatgcgcctgctagtattcttttcagaataaatctcct
caacagtaccttcactcattgcaataatctcagtactagcaacaactgtaaatggcttga
tatattcaccatttttacaataattctagtttgactttgttgcttctctgatttattta
catcaatacttttaattgataatgtttcaaaagttgacagccttaagtcaaaataattat
tgccactttctttagcagaaaactcaattgaggatattaaattatcatctcgatgctcaa
tttcagccacaagattagttgttactaaatgaactccttctatagatttcactctttctc
cgtcccgaaaaggagttcttttaactagctttagtttagtttgacttggtttttgtgaaa
ctagattttcaattagactggatttgtttctggtattgaataaacaataacaggtctaa
taagaatatagggcatttgctcgttttctatatactcccaataaactaacttatctgtag
aaatattattatgaagtctttctccaggtcttaaaaaacctctactttatcattattac
tatacacctcactaagtttatagatagatccaggcttaataataatttctcgaactattc
catccttttcgattatttcaacaatacctgagcttttagaaaaaatattttaactagct
ctgttccactttctataacatctccattatgcactagtagtagtgatgaatctttattaa
tttcgtgagtttcttctgaaatccataagatatatcccggacttaaaatttcgtaggcat
ctttatctaagcctgttttctttttgatacatttaagtctaaatatttaataataccac
cacttgtcgttttatacgtatcagatattagttctgcgactgtgtaaccatcttgaa---
ttttttggtttggaatagctttgaaaagaaatttttgtttttttctgtttctaaaatat
aagattctttttgtggattacatctgtataaatataacaacctggaattacaatagatt
cagtaataacttgtatattggtaatactgctattagtattctggtgtatccttacttctc
cagcataatgatttattaattctgtctgtgcaagtatcgtaccagctttagttttatctt
ctttatgaacaataacattagcagaatcagatatactataaacttcacctgagagcaccc
agatcaagcctccagttttagtaattcttgtagtatattgattattatctgtttcttcaa
ctgttaagttggagaaacagattttgccagagagatctgagactacatgtttttgagctc
tttctgtcatgagtcgatttctgcggggtgattcagcaatcacttgatccttcgatacta
attcaccatcacatattaaaagagtagtacctttacttaagttaattattgacttctgat
ttttcttagatctgattgtaacttgggttggctttttttgttattaatgcttgctcaccat
gtctggtacgaacatctgtatacgattcaatatctagatctatcaattgaccatctatag
gtgcgtagatttgttctgctaattcaccggtaaatactcctccagtatgaaaggtcctca
ttgtcaattgagtgccaggctcaccaatagattgagccgcaataataccaacagcctctc
ctaaatcaactaagcggccatgtgcaagattccatccgtaacaatactgacaaacagaac
ttctagagttacatgtaacaggagatcttactaaaactttttt---gataccagctttag
ttatctcttttgctaacttaggactgacatcttggttcgtatgagcaattaaacaatttg
tctctggatggaatacattttctgctaatactcggccagataaagcttgttctagattaa
ttaatattttctgcgtatctactaaatcttctagtatgatgcctttattggtcttacagt

ctacttctctaattattacatcttgagaaacatcgacaagtcgacgcgtcaaatagccgg
aatctgctgttcttaaagcagtatcaactaatccttttcttgctccgtatgatgatataa
aataatcagtaacagttagtccttctctaaaattactcgatattggaagatcaataattt
gtccctgtggatcagccataagccctctcatgcccactaactgtcttacttgagaaatat
tacctctagcacctgagaacgccatcatataaacagcatttagaggatctgtctctttaa
agtatttaataacttcttgtttcaaggattcactcgcattattccacgtatcaataactt
tttgaaatcgttcaactgctgtaatctcacctcggcgatatttgttttcagtggccttaa
tatcttcaatcgtgcctaataaaagactttgcttactaggaggaattcttaaatcttcta
aacttaaagaaattcctgcctgggtcgcataatgaaatccaagatctttaagcttgtcgg
ccatattggctgctcggctataccataattacgaaaagcccagacaattaaatttttta
gttcgtttttatcaataaccttatttgaaaaacttggctgtgcaagacttcgtctattat
ccactaaatcttctccatattaaatgttcgtt--attaactatgctactaaagactcttg
aataattttattaaagattatacggccagctgttgttctaatatattgcacaattctctg
attttcagcgtcttcttttatgatatggttattaaaaaattttgttgtactattgtcaga
atgtttttcaatcttaatcggatcatgaggttgatcaccatctactaatccatcaaaacg
agcccatatataagaatgtaaatcaattttcttctgttcataagccataactacatcttc
taagcttgcaaaatactgattagatccctgctgctgagatggattattagctgttaaata
atagcaacctaaaaccatgtcttggcttggcataataattggttgacctgttgctggaga
gagaaaattatgaggggctaacattagcaatctagcttcggcttgagcctctaaagacaa
aggaacatgaacagccatttgatctccgtcaaaatcagcgttaaacgctgggcatactaa
tggatgaagcttaatagctcttccttctacaagaataggttcaaatgcttgaatacctaa
tctgtgtagtgtgggagctctgttcaatagtactgggtgaccttgaataacttcatttaa
gacattccaaatagaagattcatttttttgaatcatctttttggcagctttaatattgtt
aactaatccctgcaaaattagtctgtgaataacaaaaggctgaaataactccaaagccat
ttctcgtggcaatccacattgatgtagttttaaatgaggacctacaacaattacagatct
accggaataatctactcttttacctagcaagttttgtctaaaacgaccttgttttccttc
tataatatctgacaaagattttaatggtctattatttgcacctacaacagttctacctct
acgaccattatccatgagagagtcaacggcctcttgcagcattctttttttcatttctgat
aataatctccggagctaaaattgactttagacgagataatctattatttctattaataat
tcgacgataaaactcattcaaatctgccgtcgcaaatctacctccatctaattggaccat
aggccgtaagtctggaggtatcacaggaatgactgtgaataccatccaagaaggatctgc
acctgttgctatgaaattttcaatcaatcttaagcgtttcatttttttattgaattttaa
tgaaggtgtcttaaaactttttggtggatttgtagcttcagaccttaaggtttcagctat
gtgctctaaatctaaatctttcaacagtttctgaatagcttctgcacctattcctacttc
tacttggttattctcatcttgattttggtagatttcctcttcaagacttttccattcgta
accttcgagaagttgcttgtattttaaattaatatctgtattagattgcgtaactacata

agaatgaaaataaacaatttttctacttctttgacttttaagtctaatgctaaagcaat
atagcttgtacttccttttaagtaccaaacatgagttactggtgaggcaagttcaatata
agccatccgatgtcttctcacgcgtgattctgtaacttctacgccacatcgttcacaaac
tatacctttgtagcgaaaacgtttatatttaccacagtgacattcccaatccttaactgg
gccaaaaattttctcacagaataaaccatccatttctggctttaaagttctgtaattaat
tgtttcaggctttgtaatttcaccaacaatttgtccgttgggcaaacttctttcgcccca
ctgtcgaatttttttcgggagaagctaaactaatttttacgtagtcaaaatattgctcaaa
ctttgtcataaatttgaataccttaaaatcaaataattagccaatttatttacatgcttc
atattgaaaaatgaatatcttaataaaggaattgttcaaaatcgtctactggaggagtat
cataattagaacgatctactcggttatctttagaatccgacatcaagtcaacttcaactg
tccgtctttggccatcttcaaatagtttcaatttatgtactgcaatgtctaaacctaatg
actgcagctctcgcattaagactttaaaagattctggagttccaggcttcggtataggct
tccctttaactattgcatttaatgcttcatttctagcttgcatatcatccgactttacag
ttaataattcttgcaaagtatatgctgcaccaaaagcttctaaagcccacacttccattt
cacctaatctttgacctccatgctgtgctctacctcctaatggctgctgcgtaactagag
agtaaggaccagtggatcttgcatggatttatcatcaacaagatgaacaagcttaagca
tataagctctaccaactgtaacaggattatcaaaaggctctcctgttctaccatcaaaaa
cttgcattttacctggatgctgatcattaaaaagccatttattgctagtaatcaatgatg
cttcttttaattttctatttacaagtgctcttgatgcttctgctccatacatttcatcaa
aaggtattattttaaatctctttcctaaatagccacctgctaagcctagtaggcactcaa
atacttgaccaacattcattctagaaggtacacctaaaggatttaaaacaatgtccactg
gtgttccatctgataaataaggcatatcttgtttaggtaagatcctagaaatgatacctt
tattgccgtgacgaccagccattttatcacccacttggatttttctcttctgcgcaacat
agacgcgaatcatagcatttgttccgggaggcagctcatctcctttctgcctagtaaaaa
ccctaaccttaactactctaccttttgcagcattaggaagccgtaaagatgtatctctca
catctcgagccttttctccaaaaatagctcgcaacaatttaccttctggtagttgatcag
cttcacctttaggagtaatttttcctactaaaatatctccagcttcaacccaagaaccac
cgacaacaattccatttctatctaaatccttcagagagttatcgctgacgttaggaatct
ctctagtaatttcttctggtcctaatttagtttgacggcattctacttcatacttttcga
tatggatagaagtatacaagtcatcataaactaatctttcgctaattaaaaatgcatctt
cgtagttataaccttcccaaggcatataagcaactagaatatttctccctaaagcgattt
cccctccgtctgtggatgcgccatcagctaaagtttggccaacaactatttttttctccta
cccaaacaattgggcgctgattaatacaagtatcttggttagaccgataatattttttta
agcgataatgaactgttctaccactattatcctgaatgcctattttattagcagaaacat
aatttacatgaccagatgttctgctaataacaaccataccagagtctcgagctattttag
tctcaagaccagtaccaattattggcttttctggatataataatggaacagcttgccttt

gcatattagatcccattagagctctattcgcatcatcatgttccaaaaaaggtattaatg
atgttgcagctgaaattacttgaataggtgaaattgcaatataatctacttgggttgggg
ttgttgtaataaactcttgacgataacgtacaggaataatatctccttcaatgtaatgtt
gtttactaactttaacatctcctggagctactcggaaatcatcttcctcatctgctgtta
gatagacaggactattgtgataaatcacttgaccttggtttaccggataaaatggggttt
ctatgaaaccaaaaacattgacccgcgcacaggttgctagtgaaccaattaaacctgcgt
tcggaccttccggagtttcaattgggcaaattctaccatagtgactaggatgtaaatcac
gaactgcaaaacctgccctgtctttattaaaacctcctggacctaaggcactaattcttc
tcttgtgagttagttctgcaactggattagtttggtccataaattgagaaagctgactag
atccaaagaattctctcactgacgcaattaatggcttgggattaattagatttgataaac
tcaatgaatctatatcacatatcatcattctctctctaataatgcgttctaaacgattga
gacctactctgaattggttctgcagcaattctccaactgaacgaactcttctattaccaa
gatgatctatatcatcaagattacctgagtttttatccttaatattaattagataatcaa
ttgaggataagatgtcttgaggagacaatactcggaacgttttaggtatattgagaccta
gtttttattaattttatatctaccaacttcacccaagtcatatcttttaggatcaaaaa
aacgagagtaaagcatctgcttagcaacagcaactgttgctggttcattaggtctaagct
tagaataaactatgagcagggcttcctcatcagttacttcttcaatattatttttccaa
tttctttagctaattctttaactgaatatatttgggatgcagaaataagaaaagcatatt
tactaagtccttttcaatttcatctttattgagtccaatagctcgcagaaaaatataag
catttactttgtgggtcttatctattcttatccaaatttctcccttaggatcaatttcaa
attttaaccaagaacctctattagaaattaggcttgcactataaatctgcttaccatttt
tgtctatctcttgtttataatagattcctggactacggattatttggttaataataactc
tttccgtcccagatacaataaaagttcctctatttgtcattattggcaagtctccaataa
aaattaatctctttttgtatttttatttttatctgcttttcagcactgagctgtagat
gggttgatgataaatctaatgatttaatatttttttcctggtttttagaaggtagatcaa
tatcttttcttgttaactttgcaggaacatatatttgagcactataagttttatccctat
tttttgcctgtctgacactataacgaggaaattttatcttgtactcattaccaaataatt
gtaattcgagtcgactagtagggtctgatatcttcggaaaaatttcaagaacttccgtca
aaccttctaataaaaaccatttaaaactggccctctgaatttcaactaaatctggaagaa
gtttatttttttaagcttatacgttgaaccatagatctcctttataacaaattttgaaatc
gaagagtttaa-tactattaatagagtactggtagataatgcataagcatagtttgctac
attatttcttctctagacataactagaacatcatattttattcaa--aaaaagattttta
aaatttttcttcac--acacactatctataagtactttgctgcta--aaagatagttcaa
gctattaaa-------aaaaacgaaattaatacttgtctgtctgtcaattaattgacatc
aagatagacaagtattatttatttaatcaattctg-tttttctgagcaaaagacaatgct
ctcgcaagtctagctttttttctagcgcctgtattagggtgaaaggccccttttttttaca

gctttatcaatcttactatatacttgagaaatacttgactgcacatcattgaaattatca
ctttctaaattatcaatatttagaaggcatcttttagttaaagttttgacaaccgattta
tacttacggttaatgagacgatttctttccgaagttttaatgcgtttaatcgcagaaagg
ttcttagccacagtaaatccaataaaactttataaaatatttttagaaacgaaaataca
caaatctgggatctattcaaaaagggaatatgcggcgtattataacattaaagaaaaaca
ataggcaatagttcaccaagagctcttattattacacaatagagattgtcattgcgaaaa
actttgtaagatagtatttaattaaaataaactaactagacatataaatatatatgaaaa
aaattgaagctattattagacctttcaagcttaatgaagtaaaacttgctctagtcaagg
aaggtattggaggaatgacggttatcaaagtttctggatttgggagacagaagggtcaaa
ctgaaagatataaaggatccgaatactctattgatatcattgataaaataaaaattgaaa
ttattattagcgatgataaagtagagaaaattgtagagacgattattaaggcttctaaaa
caggggaaattggagatggaaaaatatttattagtagtattgaaagagtaataagaatca
ggactaatgacttaaattttgaagctttatagtatttttacttt-tttagacttgatact
tcttcactaaacatgagataataagcgtttattgtatacttataactaaggtgtaata--
ctaatggctaaaagcaaaggtgcacgaatcgtaataactttagagtgctctgataaagct
ggagagtttgctcaaaaaaggaaacctggcgtttttcgatatacaactactaaaaataga
cgaaatacaccaagtagaattgaattaaacaagttttgtcctaattgtaatcagcattgt
atcttcaaagaaattaaatagttatttatcttattaaaat--aatcatattatggctata
tacagaaaaagaatatctccaattaagccaacagatgctgttgactacaaagatattgac
ttgctaagaaaatttattacagaacaaggcaaaatactgcctaaaaggtcaactggattg
acttcgaagcagcaaaaaaaacttactaaagcgattaaacaagctagaatcctttcttta
ttacctttttaaataaagattaa--atatatagcctaacttcatattttccatctaaa
tattaaaaatttctatagatttgtttattaaaatgatctaacatactgaat---------
----------tactaagaaga-tctactcattacaagttttgttcattttataattaata
atagcatcataattta---ttaagctagattcagacaagacataaag-gtaacctctttt
aattgattatattatatagtctatttaaaagaggaaatttttttgcttc----------t
agcaattcaagattctttaagctttaacaaggtatttaaaacaaagatttttgtttaggt
atcagatcaaatgaatgaatcttatctcccgactgccataattgaaattcagagatgaaa
attccacattcgttgccagcttgaatttcttcgacatcttctctaacgcgctttaaagat
tcaatttttccttgataaataactttttcttctcgaatcactttaatccaagagtttttt
aatagcttattattgatcactcgacatcctgctatttttctatttgctaacgagaatact
gtacttacttctgcttctcccaccggtacttctgaatattcaggatcaagtaagtcttcc
atccttcttttaatgtcttcaattaaggcataaatgatttgatagttttctattaatata
ttcgactttgccgatgcttgctttgttcctggagcaaagttggtattgaacccaatgagg
gtagaatttgtggttgaagctagttcaacatctgtagcagtaatttctccaggcatgatt
gagacaacattgagttgaactttactctgaggaaattgagataaagaatctaggattgct

tctgtagagccttgattatctgttttaataattaaagaaatttgcttactaatatcttta
gaattagtgtttttaaagtatctaatgtaattcgactattaagtgctctttgtttctgt
ataattgatgagtctttagaggtattttcaatcgctttaagtttggcttctttatcactt
ttaactgctaaagctatttctcctgtagctggtacagatgacaatccccaaatttcaaca
acagatgatggtattgctaaattaatttctctttagcgttgttaattataactcgaatc
tttgcgtaagctgatccaataaccaaattatcactaatattcaatgtaccattttgtatt
aatagagttgctactggaccatgcgatttatctaaatgggcttctattataataccttgt
gcaggctgtgttggatcagctttcaaatcctctaattcagctaacaaggtaatagtttct
aataatttatcaacattttgaccagtaagagaactaataggaattataggcacttggcca
ccaagtttctccgacatgacattgtattttaataaatcttgctcaataatatctgtatta
aaaccagctttatctatcttagatattgctatgacaaaagggacattcgctttttgaata
tgattaatagcctctatggtttgaggtttaactccgtcatcagcagctataataataata
gcgacatctgttaaattagctcccctagacctcatacttgtaaaagcttcatggcctgga
gtatccaagaaaacaatttttttgtttattatctttcttaatatactcaacttcataagct
gcaattgcttgagtaattcctccaatttctttgttagcattattagattttcgaatataa
tctaatagcgttgttttgccatgatctacatgtcccatcactgtgacaattggaggtctt
ttaatataattgccagtttcataaagagcattagagtgatctaaattactagaaagtcca
ttattatttcctttacatttgattctactgctataccaaaattatcagccactgatgat
attattgaggcgtctatggtttgattcatagtaactgatattcttttaagaaataaatat
tttataatatctgtttcttgaacacaaattagtttagataattcttgaatagttagagga
ttagtaatactaatagactctggcggagaattagattgaatgcttgctgaattttgacta
ggagttactgctactttttttctgtcttatagatttttattactagtcagtttttttaca
acttccacttttggcttaggtggacgcattaaagagatagctaaatctcctgcagtctgc
gaaacattagaattagagtctctaaagttgtca---tcatcatcgtcaatatgtatttta
gtttttattttt-ttgcgctgtctattctta--tttttttactatctactaagtcgtga
atttattaaaattttactttttttatctagtttaggtggagaacttaattctagatgc
tgttcaccacc---tgtatgtgattctgatttatcaaggtctaaatttaacagattatca
ttattgactgactctaatctaattttatagataatctgaggattttcaaatctacaatt
gtttcagaagaattgatatttgagctagatctagaggatctattttcaaagttttgatta
tttaaaaacataaatattggtttttattatctaatagtaactttctaattttaatacagag
cttaatataaacaagaatcgaatatctgtataaataatacaaatgt---agagaattaat
ctatgttgtattagatttaatataacactaaagtcctttgttcgttatacaagtttaaaa
ctttt-tatgatgataaacactagattctagtttattggtatacaatctatttatctaaa
taatactgtttttgtcagggcttttaaa----ttatctaatgttatgttcattatgaaat
tctattaaaatttaaaaa-ctcaacaattaagtaagaatcgttaatcacctccacgccac
ttatgccacgatctcaaaaaaatgataattttattgacaaaacttttacagtattagctg

atattgtattaaaaatacttcctacaagtaaagaggaaaaagaagctttctcttactatc
gagatggaatgtctgcacaatctgaaggtgagtatgcggaagctttagaaaattattatg
aggctttaaaactagaggaagatccatatgatagaagctatattttatataatatagggc
ttatttatgctagtaatggtgagtatgtaaaagctttagaatactaccatcaaggattag
aactaaattttaaattacctcaggcccttaataatatagccgtgatataccattaccaag
gagttcaagcaattgaggataaagatacagagctatctaaattaatgtttgataaggcgg
ctcaatattggcagcaggctattaagttagctcctgataattatattgaagctcaaaatt
ggctgaaaacgacaggacgaatgaggaatatacaaggatattaatatatttaagatataa
taaagatatcaattaataaattaaaattagttaaagaaactggtactatatagagtactt
-tactatatattataatattatactagttga------tctctaagacttagctgaataaa
cgaaattaaatctattgactatattcatgaatacaaataagcataatggttagtacaaca
aaaagtaccggatctgttactcaaattattggaccagttttagatattgcatttcctaac
ggacagcttccgaaagtattcaacgcactcaaagtacaaagctcagaaggaactattact
tgtgaagtacaacaacttttgggtgacaacaaggtacgagctgtttctatgagttccact
gaaggactacaaagaggggtagaagttattgatactggatcccctatatctgttcctgta
ggtacagatactcttggacgtatttttaatgttttaggtgaacctgtagataatttgggt
cccgttgattctgagagtactttacctatccatcgaccagcacctgcttttactaagtta
gagacaaaaccaaatatttttgaaacaggtattaaagtcgtcgatttacttgctccttat
agaagaggtgggaaaattggtttattcggaggtgctggagtaggtaaaactgtattaatt
atggaactaattaataacatcgctaaagctcatggtggagtatctgtatttggaggcgtt
ggtgaaagaacaagagaaggaaacgacctctatatggaaatgaaagagtctaaagtaatt
gatgcagataatctgaaagaatctaaagtagcattagtatatggtcaaatgaatgaacct
cctggagcacgtatgcgcgttggcttaactgcattgacaatggcagaatactttagagat
attaacaaacaagacgttctattgtttattgataatattttcggtttgtacaagctgga
tcagaagtatcagctctactaggccgtatgccatctgctgtgggttaccaaccaactcta
gcaactgaaatgggagcacttcaagaaagaattacttcaacaacagaaggatcaattaca
tctattcaagctgtatatgtgccggctgatgatttaacagacccagctccggccactaca
tttgcacatttagatgcaacaacagtactttcgaggaacttggcggcaaaaggaatttac
cctgcagtggatccactggattcaacatcaacaatgctacagcctggaattgttggaact
gagcattattctactgctcaagaggtaaaatcaactttacaaagatataaagaactgcaa
gatattattgctattcttggtcttgatgaactttcagaggaagacagacaaactgtatca
agagcaagaaaaattgaaagatttttatctcaaccttttttcgtggcagaagtgtttact
ggatcacctgggaaatatgtatctttggaagatgcaattaaagggtttcaaatgatctta
aaaggcaatttagacgacttgcctgagcaggcattttatctagtaggtgatatagatgaa
gctatacaaaaagctgacagcatgaaagattaatataattaatgatgactttaaatataa
gaattattgctcctgatcggactgtttgggatgcagaagcacaagaaattattttaccaa

gtagtacagggcaacttggtattttaacaggccatgcacctttgcttacagctttagata
ttggagttatgagagtaagagtagataaagaatggatgccgattgttttgctgggcggtt
ttgcagaaatagagaacaatcaattaactattctggttaatggcgcagaagaagctagtc
aaattgatttatcagaagcagaaaaaaatttagacactgcaactcaactcttaagtgatg
cctcgtctaataaagaaaaaatagaagcaacacaaaaaatacgaaaagctcgagctcgag
tacaagctgcaacagcagcaacttcgtaa-aagattcc------tcttaaaacataaaac
tttagtaatgattactaaa-------gttttatgttttatttgtatggttataggcagct
taaataaacttagcttagacctgaacaaatataatcaaaatataatcccatctctttacc
tgcatctggtccaactaaactaatagtaacttctttcatagctaaaatagcttgaatcgt
tgcaccaataggaacacctagagaattgtatgtttcttttagaccatttaatactctttc
ttcaaggatagatggatctccagctagcattccataagtagcatagcgtaagtagtagtc
taaatcacggatgcatgcagcataacgtcgagttgtatacatattaccacctgggcgtgt
aatatccgaataaagtagagattttgctacggattctttaataattgttgctgcattagc
tgcaatagtagcagcagctctgactcgtagctcacctgtttgaaaatagcctcttaattt
ttcaacagagctatcatctaaatatttaccttggacatcagctgcattaataacagaagt
aattgcgtcttgcataactttcacacattccttaattttaaatatctaatacta--gggg
tatataacaaaattactagatcttttagcaatatcagtattactgcatagcacccaaagt
ataatcaaagtaaaaaccagcttctgctgaatcttcacctgcaagcagtgagcaagctac
acttttcatgcattttacaccttcagcaactcctgaaattggtgttcctaaagaattata
catttctttaacacccactaatccaatttcttcaataggagttacatcgccagcaactat
tccgtaagttactaagcgaagataatagtccagatctcgtagacaagtagcagtcatctc
ttcgccataagcatttccaccaggagaaactacatcaggtctttttgaaacagttgttg
gccgccttgctttacaatacgctcacgattatctgttaaaatttgagctattcttaaacg
acgttgtccagataaaacaaaacttttaattctatctaattccccaggacttaaatatct
tgcttctgcatctgcatttacaattgactttgtaacaatactcatggataaaactcctgt
aatgctttaaaaaaaaagcttaatactaattatttatgatataactcagaaagtaatgta
ctttttgag-aattttataaccgaaacacgttcatgatatatgaagcaggatagatactg
aggtaagtactaaaacgagaaagttatatgtaggaacttctgctattgatttcctagtac
tttttaaaacttggtaccacaattgatatatcctgcttagtcaaacgattatagagtgt
ttctgtattaggaaagttagctgctggtaaagtcggaaatcttctgtatggaacagtatc
tttgccatataccgcattgtattcaacactttcaactaaattacttataaatgcagataa
gccttttgaagctaaaatttgattaaagaatctaatttctgcctgattattaggagctct
tcctaagatatgcttagtacctaattctataacttttgtatttggatatggctgataaaa
ttctttaccatatagttcagatagcgccagtttttctactagttctttgacacagatttg
tctattcaagaaagctgacttaatatccaagaattcaccgccgacactaaaagagttaag
atctctttcgaaaatttgacgatatgctgctcttaaagcttgttctaacatttctttgtt

actatcactattgacttcaaagacaatagattgatctcttaaagaagttactccctgtga
aatacgagactgaatatcattactactcctcatttctttgacagtgcctaattcaacaaa
cctagcagatttactagcaataactttttttgaatctctggtcaataattcctggtcttaa
gcttcttaaagcaacaccagctggagtactatatctttcataaggaacagtattatctcc
aaaagtttctgtgtactccgagctatcaattattgtgtctattacttggtaatagccttg
cttatatgcaatatcaaaatatttattaatttcttgccgaccatatgtagggcgacccag
taagcggttatgaatatattcaatagctttacaaatatataatggctcccaataaagaga
tctaaatatacttgattttgctaactgtctaacaaactcgcgaactgtaatttgattatc
ttttagctgactttctattggcttgagaattagtttttcttcttgatacacttctctgcc
aaaaactcttaaataagcaacttttgtgacaacttctactgaattttcaaaattttggga
tgaatctgactttataagagcgttggacaatttaaagatttttggtcccagggatcctgc
tgacttaggtcttacctgtggattactaatctgattgtagattcctggcccacgtctaac
taatatccttcttgtatctttaccaaaaagagcttgtctttttcttggatctttattttc
tttagggaaaatagctccaaattgaattgataatggatcattacctattccatatggatg
ttgatcaggtaatgattgcttataatcactaaataaagttataaattgaggaaccttacg
aaaaggcgcactataggtgaaaagatctatttgaggcccccaattacggcactcttgagg
ctcttcacctaaatttctaaagtaaggcacagtctcttcaccgaaataatctgtatactc
agaagaatttaaaagtgcattcactaaaccacttaaaccagtagatgataaaatagcaaa
atatttttgaaactcttctaaagagctaggacctctacccaagaaatgtctaaatgctaa
ttctaatgctcgactattgacaaaaggttcataaaattgctttctataaatacttgaagt
tcctaacgaacgaatgaattccttaattgaaatttggccatttttttacttgcgactccaa
attcgataaagagaggtcatatgctttagcaatatctctttcaaaaatttgtctataaca
tgctttcacaacaatatttttctcatcagcagataaactaggcttcataacaaaacgagg
tgttgacactcctgcctgtacataagtttgaggaagacgcaaaccttgtaaatctccaga
tattctttttcttaatttatcagtcaaagatggagcttcaaattcagaaatcacaacatt
aaaatactctttaactaaatcttgaccttttatatcttcctcaaagattaataatgctgt
ccgtctcatctctctaagagccacaattgctgctgcacttgagcaagcgttatcaattaa
ctctctcaatcctcgtatattaacagataaaatattagggtcgcctgatacaattgcata
tgtcaaataccgtaaaaaccagtctaaatctcgaagagattttctcattcttgtagtacc
gtatctcaagacgtttataggtttaaaccctggcggagtagctcccccagcattaaataa
agatcggaaactttggccaaaatctccttggatatttcctgataactcattaattttatc
ttgagaactttgatctcctgcaataataacagctgcttgcggacgttctaaataagaaat
tgcagacccacctacgaatattttatctgctgctcgtgctaccagaatatttgcattttt
agttaaaatatctgcaacttccagtctttttttgtccagaatttaaaaaagaaactaattg
atttaattcaccaagctgtaaaaatcggtcttgttgctctgcctgagtaatagttaaaat
tgaggcagtccgataaagctgggggcgtgctaatgggcttccgccacttgctttgatact

cattactttatttatctccttaactgatattgcttgctagcgaatacaaatatagcttga
tatttctaaatatttaaagatactatgcaagcaca-ctgttttctatattattaggtact
agtacttatagtacagagagtcttataactaacttctataaatacaaattttgtaatact
tctattaacatttgtcaaaaaaatatttaagtgctcactcttttatattaaaattcaatt
agtgagagattgatgtgaaataaagtatactaagtaagttttatagataggaaaattaaa
agatttattctacaaacattaatca------------cttgaatgtttagcaattgtaca
attgttattgcttttaattggtttagactagttttatatttttttctaacataaacaatat
taatgagcaatttgtatgaatttaaagccacaaaacaacctattaactaacgaaaatata
aatcatgtagatatacccgaaaatgatataccaatgtcaattacggaacatttagaagaa
ttaagacaacggactttatttgtgtttttattctttttgtttgctacaactataagtttc
acacaaattaagatcattgttgcaatactgcaagctcctgctgttggtattaagtttta
caactggcaccaggagaatatttttttcatctattaaggttgcaatatattgcgggatt
gttgcaacaactccgtttgccgtttatcaagttatattatacatactcccaggactaact
ggaaaagaaagaaaaattattttgcctctattaattagttctgtattgcttttattaca
ggtggtatatttgcttactttgttctcgcaccagcagctttaacatttttaattagctat
gggtctgatattgtagaaccattatggtctttcgaacaatactttgactttattctatta
cttttacttagtacaggattagcatttgagataccaatcatacaattattgctcggtgtt
tcagggacatttcttctagtcaaatgatacgagcttggagatatattattattatagca
acaattgctggagctattctgaccccttcgactgatcccgttacacaattaataatgtct
tcagctgttttattactttattttggtggaattgttatattattagtcttaaaaaagtag
agtaaat---------atttcta-ttaatccaatatatttgttatc-----tggcaaa
taatctataaaaacaattgct------------------agatttaagtatcaggcttct
aaatatttcaa---------gtctcttttatcatttaaatacaatc-------------
------------------------------------------------------------
------------------------------------------------------------
------------------------------------------------------------
------------------------------------------------------------
------------------------------------------------------------
------------------------------------------------------------
--------------------------------------------------gatacattgg
cttgtttttttcatcaattatttgtgttgcacac-tttttagcactctctttattgttctt
agaattatttctttaatctatttattaaat----agatatgaagagatttaagtattgat
tatggtaatctatatcaacgtaaaaagtattttattgtgagttttatttatataaaggct
tattcattatatagttataacagtaagtctaattaatag-ttttcatgcttaattaaagt
aataactcttaattcttttggcagaaaaatctgtcttaatattttatc--aaaaaaacag
-aaaaatgaagctaaatagtctaattaacttaattcaaaagtctatatattcttgtacac

ttttactaattattttaaatattatttgtgtcgcacctaattctagtaatgcatttccaa
tttatgcgcaacaagcttatgaaagtccaagagaagcaactggtaggatagtatgtgcca
attgtcatcttgctcaaaagcctgtggagatagaagcgcctcaagcagtactacctaata
ctgttttcgagactgttgtgaagattccatatgacagcaatgctaaacagattttaggta
atggcagtaaaggaggcttaaatgttggagccgttgtaatattacctgaaggatttaagt
tagctcctgttaatagattatctacagagttaaaagaaaagactagaaatctttacattc
agccgtacagtgctaaacaagacaacattttagtaattggacctatttctggtgataaaa
atagagaaatagttttcccaatactatctcccgatcctgcaaaagataaaaaagctcatt
ttttcaagtatccaatatatgttggcggaaatagaggacgaggccagatttatccaacgg
gtgacaaaagcaacaataatattgtctctgctttaagcagtggtaaaattaataaaattg
aattactagacaaaggtggatttataatacatgtgactaacagtagcaatgtagagtcaa
cacagaagatttcacctggtctcgaacttagagtaaaagaaggggatacaattcagcttg
atcaagctttgaatagtgatccaaatgtaggtggttttggtcagaacgaaacagaaatag
ttttacagagtccaaatagaattaagggcatgattgttttctttttcgctagtgttctag
ctcaaattttcttcgtattaaagaaaaaacaatttgaaaaagttcaagcagctgaaatga
atttttaaagcaatataaaactggcacttactataaagaaagta-taaacactttattca
caggatgtaccagtttttatgttatgtaataatcgctttgacggaagttatgatactttt
aacagaatctacaatctgactaggctgaattactgtagcctgctctaagcttccattata
tggtgtgggtatatcctgcgaagatagcctcacgacaggagcatctaattcatcaaaaag
gtactcattaatttgtgcaattagctctgctccaattccggctgttttcatacactcttc
tacaattaaaactttgtgagttttctttactgagatagatatagagtctatatccagagg
ttttaaagatataagatctataacttctggatcataaccttcttttaataaggccggtaa
tgcttggataacatgatgcctcattctagaataggttaaaatcgtaatatcttttccttt
tcgtacaaattcaactttattgagaggtaagaaatattcttcttgaggaatctcttcttg
taaattataaagtagaacgtgctcaaagaatacaactggattattgtctcgaattgcaga
tttcagtaatccttttgcattataaggagtagaacaagcaactatttttaagccggggat
ggcttggaaataagcttccagtctctgagaatgctctgcacctaactgcctaccaactcc
tcccggtcctctaataactaaaggcaatgtaaaattacctccagaagtataacgtaacat
tcctgcattattagaaatttgattgaatgctaataataaaaaactcatgttcataccttc
aacaattggtcttagccctgttatagctgcaccaattgccataccagtaaagctattttc
tgctattggtgtatcaagaactcttaaatccccatatttgctatgcaaatctttagttac
cttataagatccaccatagtgaccaacatcttctcctattacgcagacagttggatcttt
tgccatttcctcatctgttgccgctcgtaaagcgtcaaacataaagatttactcatttt
aatatttgatttttatgatttgaccactgttttaaagtaacagttaagacaccatattct
atttagttatctgcaaaaagatatcgtttcaattctgacatgttaggttctggactggaa
atagcaaatttcacagcttgttccaattcggttttaacagcattctgaatctcattaagt

tcacctatattagcaatttcattatctagaatatatttttgagtttttgataggatct
cttgccacccaagcctctttttcttgtcttgatcttagttcatcaggatctgcgagagag
tgaccacgaaatctatatgttaatgcttctattaaggttggaccatcaccttggcgagct
ctttgaactgcttgttttgcagcttgtcttacagctagcacatccattccatcaacttca
atcccaggaagcccaaaagcttctgctttttatgtatttcaggtattgaagaagaccgg
tgatgtgccatacctatagcccactgattattttcaacaacaaatataataggtagtttc
cagagaactgccatattcagacattcaaaaaattgcccattattggtagtcccatcacca
aaaaagcaagccgtgactcttaaatcctctgtttctttaagtacttgctggcggtagata
ctttgaaaggctgcccctgttgcaaccggtataccttccgcaataaaagcaaagccacct
aaaaaattgtgaggcgcagaaaaaatatgcatcgaacctcctctgcctttactacaacca
gtctcttttccaaataactcagccatcacatttttagatgggacgcctttacttaaagca
tgtacatggtctcgataggtactgcaaacataatcagttggattcagaagtttaattaca
cctgtagaaacagcttcttgaccattataaagatgaacaaaaccaaacatctttccttta
taatacatctgagcacacatatcttcaaaatttctgcctaacaacatgtcttcatataaa
actaataaattactcttattaagattaagacctgttgaattatagttagtcagcggcaat
tgaactttcttaggataactcataatttgtaaagaaacctct----------tattgtaa
ataggaattttgaagaaacagctatcaatagaaaacacacagataaaaacttataaatta
taatatgtttttactaaagactttaataattatttgaaactagttcagtatatcaaatt
aattttgcttatactacggatgtctaaaacttatataaaaaagtagt-a-aaaaagttta
ttaaaagaatagttaaaaacagatataaatagaagtaaacagaattatt-gaaaaaaaat
gattattattaattaa----tacttaatatttctaaatattttacaattaatggctatgt
caccatctttattttatcctgttgaacaagaactatgtagtcttgaaaaaaatctgaaag
ctgttgctgggactcgtcatccaattttatatgccgcagcaaagcatctattcgatgctg
gaggaaaacgagttagaccagctcttgtatttttagtggctaaagcaacctctgagaagc
aagatataaatactggacaaaaaaggctagcagaaattactgaaattatacatactgcta
gtttggtacatgatgatattattgacgagtgcacaacacgtagaggagtcaaaactgtac
ataatttattcaatactaagattgctgtgctagcaggagatttttttatttgcacagtctt
cttggtatttagctaatattgaaaatttagccgtagtcaaagctatttctaaagtcatca
ccgactttgcagaaggagaaattaggcaaggcttagttcatttcgatcccagtatttcaa
tagatgcttacattgagaaatcattttacaagactgcttcactcattgctgctagttgtc
ggggtgcagctatgcttaatggttccaatcatcaaataaataatgatctttatctttacg
gtaaacatatgggattagcatttcaaattatggacgatgttctagatataactggttcta
ctaagagcttaggaaaacctgctggcgctgatctaataaatggaaatttgacctctcctc
tcctttttttcacttactcaagaagcaagtttaaatgatcttattgatagggagttctgta
atagtacagatatagcctcaacattatttcttataaaaagaagcgggggaattacaaaag
ctaaagatttagctaaagaacaggtgcaggcggcactttcctgccttcagtttttaccac

aatctacacctgtatctagtttaaaagaattaacacatttcataatcacaagattgtcat
aaaggacttgctaaaattaaaatgttattaactaatttttttactgtttctaaaacagtct
gaaaagtttgaatatcattactagctagctgcgctagcattttacgatttaacgcaatat
tctcttttttaatgcactaataaaagtactataattcatgccttgattgtgagcagcag
catttattctagttatccaaagacgacgaaaatctctcttttccttttcgaccaacat
aagaatatcgaagagctttaaggacttgctgcttagctgttcgaaataaacatttatgtg
cacctttaaagcctttagctagcttaaagatttagctcgtctttttttttgcaacgttac
ctcttttaactctactcataaaatattac-ttaaattttaattattaccagcttactaaa
ttcttatactctttg----attataaataaggtaatttcattgcaatgtttttaatatct
tttaagtcaaccgaacaagtagaagaaagatgtcttctttgttttgatgacttttttttgt
aataaatgacttttagaagctttatgtcgaagaaattttccggatgaagagactttaaat
ctttttgctattgcttttgatgtttttaacttaggcatattataaactttttaatgattaa
caaaagcatgaaaagtctgaggatatcaatcttagacctttcaatacttagaataatctt
aacacaatttattttacttactacataggttaactgaaacaatttatatatataatacta
atcagttaatcaaccatagctaaagtggatttatattttgctttgaacttttgtatagta
ttagcaattagcattgttataattttaagactgcttgaattcaattcttcgaacattttc
atatttaatgtaaatgaaatattggcttctgagacaatattttgaatttgaacatcagac
aagggaatcatatctaatgcagctctatattgatctttaaataatttatcgtcttttatt
tggtcaaaatcgtaaaatttcgttcctcctgaatctgataaattcatagcccctctagct
attttttttaaaatttggccaccagaaaggtcacccaaatagcgcgtataagcatgagca
actagtaattcaggttgtttatgtcctatagtgtgaattctatcaacataaattttttgta
gcaggagatggttcaataaaatctaaccaatctgatccataataatagtttaaatcttca
gatagactcgctttcctattaagctctgtaaaatatataggtttaatagctggatgattt
ttattagaaaataactcttcctctattgcacaatagacaaagtatagatttgcaaccaat
ttacgatatgacttcttatctacaactcctcccaaaaaaagatttaacaaaactaacattt
tctgccatactatgggacttagtagtaccttctcttagttcattcgctaaagtattaacc
ataatatcaatattccatttttt-aaagataacaaattaaca---ataaatagttatccgt
atatttataattgagttgcgctaactaggctgctgcaaaataatttttttgattttattgg
atctggattcattgttttgtcaccaggcttccaattagctggacaaacttcatctggatg
agattgtacatattgaattgcttgtaaaaccctcaaagtttcttctacactcctgccaaa
ctctagattattgattgtagagtactgaataattcctttcgggtctataataaatagtcc
tcttagagctacaccatcactattcaaaacattataagctgcactgatctcttttttttaa
gtctgatactaatggatatgaaagatctcctaatccacctgattctctgtcagtctgtaa
ccaagcaagatgagaatactcactatcaaccgagactccaagaacttctgtattaagttc
cgaaaaaagcattatacttatcgctaaatgctgtaatttctgtagggcacacaaatgtaaa
atctaaaggataaaagaataagacaatatacttattcttgagatcagataattttaacgt

tttaaattcttggtcataaacagctgtagctgaaaaatcaggagctagctgacctactcg
aagacaatttggtccagaaatcattaatttttcccagaagaattgatataattttgttat
ttaataaataattatataacaaaatttagtatatattaaaatactaccagtctttttaa
attac--------taaatgatataatcaatactatctttatagcggggaatggatttgaa
ccattgaccttcgggttatgagcccgacgagctaccagactgctctaccccgcggataac
taaaatcatattagtctacatatatatataaaagcaaatgcataatagatttagaagta
aaaaaatagaagaactggtccactgatcattaaacttagaaacgacaaagtatttttat
cttatttaaaagaggaaagacttcatataaaccaataaatctatcttttaaaagaatttc
ggacggctttacccaaatttgaccgtcataccagcctgattcttcataaaatactgtagc
actcattaatcgttttactacgtatgaccagcctaaatataatctgattaaaataaatcc
tgtcattaggctggaagtaataaattctgaaaagaaaaatttcaaaggtagtttagtgat
tggaaaaattgataataatataggactaactaataaacagttaagtagtagtgcaatagt
tattttttgttataagatctatggcttaaagttggccagcaaaagaaccaagaattttt
caaagaagtatattcatgaacaggttgctgctcctttggcacaggacattgagtattata
taaattcattgacattaaaaagtattataataaagac--tttgtttatattatctaacat
aggtacgacattaaatattaactaactgcaattatagtagtgagtagaagaaataaaact
aaaaatagccaagtaattttatttaaagtattctcagtactacgagtattactaaagaac
tgattctgagcacctacacttcctaaaccttcggatttaggattgtgtattagaatagtg
aagattaatattattgtagatgagtaccaaaaaaatttaaaatttgttccattatgtaa
aagttaat-aaaaatagcacaaagacaatcgtctaaaagatattgattgtcttatcatat
taaaaagtatttattcaccctgtacttttaaaagtaaaaatcctattgccagccctacca
aaactaggactgaacttaatacagcactttctacaatttctccgaccatt-ttattttc
cttattattttattattg---tcattcttttgacaagattatattgagctaa-ctaaaat
ccgtttctaccccaaactaccattgcaattgaaaatgtaaacattgccattagagctgcc
caccccaaacttaaaatatccataatttataatattc--ctaaaaataagatctcttta
cttaaactataatattcttaatattaaaaatatcattatataataagctaattataacag
agtactacttaattcgtatttttagtgtatttaaagaagattgaa-ttttttcgaggtac
tggataagtatttcatcctaagaaccatattcaac----ttttgatttattataatgctt
aaatctaaatata--tatcactcaagaatcattagtaattatgatgtttttaattatacg
att---aacatatttcaaaatagaagaattcatagcgactaagttaa-----tatgctaa
tataatttatcaat--aaaaaggaagttagtatttcaattatgcaaacaactataaataa
tgggcaaacgtctagtaaagaaactttactaacacctagattctacacgactgactttga
agagatggctaatatggatatttccggcaatcaagaagatttttttggctatcctcgaaga
atttcgagctgactataatagtgaacattttattagagatgaagagtttaatcaatcttg
gtctaatttagaacataaaactaaatccttatttattgagtttttagaaagatcttgtac
cgcagaattttcaggtttttactatataaggaattatccagaaaattaaaagacagaaa

tccggttatagctgagtgctttttattaatgtctagagatgaagctaggcatgccggttt
tttaaataaagctatcggggacttcaatttatctttagatttaggatttttaacgaagag
tcgcaagtatactttttctcacctaaatttatttttatgcaacctatctttctgaaaa
aattggatactggagatacataactatttaccgccatcttgaacaacatccagaacaccg
tatttatccaatctttagatttttgaaaattggtgtcaggatgaaaatcgtcacggaga
tttttttgctgctttgctcaaatcccaacctcatttttaaatgactggaaagcaaaaat
gtggtgcagatttttctattaagtgtatttgcaacaatgtacttaaacgactttcaaag
aattgattttataatgccataggcctagactctagacagtatgatatgcaagtaatacg
aaaaactaacgaaagtgccgctagagttttcccggttgctttagacgtggacaatccaaa
atttttcaaatatttagatacttgtgcatgtgataatagagctctgatcgatattgataa
taacaattctccattattcattaaatctatcgtaaagatacctctatatttttccttatt
tgcaaatttactaaagatatatttgattaagccaatagactcaaaaacagtatggaatac
agttcgatagttaaatacaagggaatgctttttcttataaaaagctatctagtgattaa
tataaaat------aaaagcttccttct-----cgatggaaggagcttttatttgtcaa
ga---atataatgggattatatagatc-tagaactaagaacttactttttcttttgcttc
atacataacctctaaagtaattagattttagaacaatcacgggcaaatttctctgtatt
tctttttactttacctctaacaaacccaggaattttatttaattctttctgtgcttcttc
tgaccaattaatactatcaacagcagaaattcctatacttaatgcttctgttgtatcatg
accaccaaatatttctagcaaatgatcttccataccaagtgtaaacgaattatatactaa
atcagcaatttgatttgttccctcataaccaagaaatggacggtaacttaaaggaaagtt
ttgtatatgcactggtgaagaaataacgccacaaggaatatttagacgcttaccaatatg
tctttccatctgtgttccaaaaattgctgcaggttcagttttagcaattaaatctccaat
taaaccatgatcatctgaaacgattacttcatcgcagaattcttgaacttgatctttaaa
ccactcttcatcatatttgcaataagttccacaccaggcgacatgtatacccatttcctg
gtgtaaaattcgagttattgctgcagcatgagttgcatctccaaatacaatagccttttt
acctgttaaattctgacaatcaattgatcttgaaaaccatgcagattgtgaaataaatcg
ggtttgctcgtctatgtatttttcgtagtctacagcagctcccaatgcatttactaattg
ctgtattgatcggatgcatgcagctgtttgaacaatacccataggtgtaatgtcaacata
aggcatattaaattctttctctaaatatcgagctgtcattagtccagtttctctatatgg
aataaaattaaaccatgcagagggtaatttttttaaatcttgaactgatgcattttctgg
aataatttggtttatttggatatctagatcttgaaataacctttttaattctgctatatc
atgttgattatgaaatcctaagcttactgctccaataatattaacagatggagttttagt
cttttgagttaaaacttgattattagatttagctttttccatatagaaagtgacaatttg
ttctaacgttctatcacctgcttgcagttcatttactctataatgattaacatcagctaa
taaaacatctgcttctgtttctatagatgctctgctaacaaagttttgcaaatcttcttg
caaaatactcgaagtacaagttggtgttagaataactaaatcagggctttcttctctgtc

ttttctagtaatattttctacaacttttttcttgagaaccacgggctaacacatgtctatc
tacaacacttgctgtaacaggagtaaaatctctgtctctttcaagcattgaacgcattac
attaaaataatcgtctccaagaggcgcatgcataattgcatgaactttttttaaaagaact
cgcaattcttaaagttccaatatgagcagggcctgcatacatccaataagctaatttcat
attaaattttcctatgatttatttagacatagtaatactttttaaagctaactaacaaaca
ttagccaaacatctaatttaactccttgacaattttgtaatattttttaaatgaagctta
tcattataaatatttaaaaaatataattcagaataga----aataactgttagatcaagc
ttgaaagtaaagatttcttgcagttatagttttctttaagtaaacatgtaaaaattgta
ctatatataaatgtactatatataaacttttttttactgaatattggtaatgccagataca
attaacttaaatatgccttccccaacgtttggtggaagcactggcggttggttaagagcc
gcagaagtagaagaaaagtatgcgataacatggacaggcaaaaatgaaagtaagtttgaa
atgccaactggtggtaccgcaactatgcgaaatggagaaaacttactctatttagctaag
aaagaacagtgtttagctttaggtactcaactaaaaagtaaatttaaaatatctgactat
aagatttacagagtttttcctaacggagaagtgcaatatttgcatcctaaagatggagtc
tttccagaaaaagttaatactgggagagctagcgtcaacagtgttgatcattctattgga
caaaatattaatcctgtagatgtcaaattcatgaataaagcaacttacgattagtctctc
-aaaaacctagacataaaataacttttatatagtctaggtttttgataatgtatactgat
agtatggaagggtggccgagtggttgaaggcgtctgatttgaaatcagttgaacta--tc
tggttccgtgggttcgaatcccaccctttccgttataaaact------------atagtt
ggcttctcaaggcaagctagctatacactgatagtattaagtttagacttaataaagtct
acaacttgtgacaggttagaaatttttttcggcatcctcatctggtatctctatactaaat
ttttcttctatggccattaccagctcaactgtatctagagaatcggctcccaggtcactt
gaaaaatttgcttccctagtcactattttttttttcaattcctagctgttctgctacaata
tcttgaacttttttcaaagatttcattatcttgcataatatattatttcctctaaggtgtt
atcaaaaatttttcggcttatattattatccaactctaaaaaattagtaatttttagtcg
agattaaaatatttatacctatatgatatactattattctggataaatacgaagtttacg
atatggctcttctccaaaactacgagctctcaactggtagttgtcaactagattatgctg
cattttttcgtatataggctgatcgaggtgtaagctgaactattgaactttcattcaagat
aatagtttcaatagccaatttagcttcttgaagagcctggatctcatcaaaatttttatt
tttacagagttctacccaattaagaccagatgaagtatgaatgtttaatatttttctgag
tgcacgagtaatttgtggaaccgtactattctgaattgtatatataataatttgttttga
ttttgcaatttgtctaagttttgtattttgctttacttgattacgtagcgctagaatagc
atcagacttttctatctccttggttaaaataattggtagatctagtgaagaaataactga
tgtaatatgctgccaactcaaagaataagcatataagtactgatgaggtatttccaattg
caaagattgtttattaatattaataggggcattgaaagtagttgacaataaagatgtatc
acgattttgtgatttattcaaatctaaggacttatttcttaattctctatattgtaaaac

cggaggcgcctgctgtgcgttaattgatttacgagcattagagatgtttacaggtaagac
ttctatagattgcgatggataacatttaattaaaattctaccattagcttgaatttgtct
tttttgcacaaaaggttgatggccttgcaaaatttgatctattgtctctttaactttatt
atgaactatccaaacatttcgctcgtgtatttcgatcgcaatctgaaaagcaggctcagc
ttttctttctagtatactcttttgcgtacctctacgtttagcttcatcgtcacctaatgt
tacatattgaattccaccaattaggtccgccaatgttggattttttaattaaactttctaa
atgatttccatgcgctgtacctactaattgaactcctctttctgctatagttcgagcagc
taacgcttctaattccgtgccaatttcatctataataataacctcgggcatgtgattttc
aacagcttctatcattacttgatgttgcaaatcgggccttgcgacttgcattcttctagc
tctaccaatagctggatgaggaatatctccgtcaccagctatttcatttgaagtatctat
aataactactctttttttccatttcgtcagctaaaacacgggccatttctcgaactgcggt
tgttttaccaactcctggttttcctaagagtaaaattgaatcaccttgctgtagtaaatc
tcgaataatactaattgtaccaaagactgctctgccgacacgacaggttaagccaataac
actaccttctctattacgcagtgaactaatacgatgcaaggtttttttcgataccagctcg
attatcgccgctaaaattaccaacttttttttacacaataatctaagtcttgccaactaat
agatctttgagataaatattctgggttatctggaaatctagcttctggacgacgacctaa
atccataacgacttctattaaattatttctattaggatgctgttgtaaaggttctttgac
aaaatttggcaaaatttctagcaacttatctaagtcatctgcaataagcatgattgagta
tataataattaattaatcaaatagaagattcaagattctaaagattacaatacatattaa
agaatattatgattggaatcaattttatacttaaattgaacaggaagtta-tagatgtaa
aagaactgattaaactttacgcaataaatcgattggtcaaaattaatagtcgtgttaaca
tgtttattggacaaaaagttcgtattaaatatagcaaacaaaaagttgctagtgatatag
cagataaagttggtgaattaggagttattaagggaataaagtttattaatagtcaatgtg
ttactattattgttgagtttgacaatcacactaggctttggatgtttagagaggaattaa
tctgtctaaatgaat------taaaaacatgaaacataatttattttatttattatacaa
aatttcttatgagtcgttaaaataataaattttaaaggagtacttttaattgtgcaaatt
actattaaaaaattacaagacttactttcgtctgtacaaagaaaaaagatccagacatta
aaactaaaacaaggtaagtttgaacttctattaaataagacctataaaaaagtcaatcaa
gaaattataccttcgcagaaatctgctgtattacaaaatagtccatctacgataattaaa
tcaataaataatacaaaaaaatcttctgttgttaatgaagaccgtacagaatatgccact
attgtttctccaatggttggaacgttttatcattcacctgctcctggtgaaaaaatttt
gtacaagttggcgatgaggtcaaattcaatcaaacagtctgtattattgaagcaatgaag
ttaatgaacgaaatcgaggcagaaattgaaggcaagattatagaaattcttgttaaagat
ggtgatatagtagattgcgggcaagccttaatgaaggttgaaacataatgctaattttc
cattttacatattgttaacagaaattatttagatagaaaaattagcttatactatagtta
tgaaaacttatttcgcaaaggcac-taaaaactaatatataaaattatttactttatctg

ccaaaaatagaaataagcgttttgattccttgtctcaagagaggagaatctcaatggcaa
ttagctcaaaagagcaagagacaaagaaggtaaaaatctcggttgataaaaatcccgtag
atacttctttcgaaaagtgggcccaaccaggccatttttctcgtacactagcaaaaggac
caaaaactactacttggatttggaatcttcatgctgatgctcacgacttcgatagtcaaa
ccagttctttagaagaagtttcacgtaagattttcagtgcacattttgggcagctgtctg
taatatttttatggcttagtggaatgtattttcacggagcccgcttctctaactatgttg
cttggttaagtaatccaacaggtattaagccaagtgcgcaggttgtttggcctatagttg
ggcaagaaattttaaatggcgatgtaggtggtggctttcaaggagtacaagttacatctg
gatggttccaactgtggagagcatcaggaattactacagaatttcagctttactgtactg
ctattggcggattagctatggctgctttaatgctgtttgcaggatggtttcattatcata
aagctgctccaaagttagaatggtttcaaaatgttgaatcaatgatgaatcaccatttag
ctgggctttttaggcttaggctgtttaggctgggcaggccatcaaatccatttgtctttgc
ctattaataagctactagattctggcgtgtctccgcaagaaattccactacctcatgagt
ttttaattaatagagagcttatggctcagctgtatccaagttttagtaaaggattagttc
cattctttactttaaattgggctgaatattccgactttttaacttttaaaggaggtttaa
accctgttactggaggtttatggctaagtgatactgctcatcatcatctagcattagctg
ttctatttcttgctgcaggtcatatgtatagaaccaattggggtattggacatagcatga
aagaaattctagaagctcacaaaggacctttaccggcaacggtcacgaaggtctatatg
aaattcttacaacttcttggcatgcacagcttgcaattaatttagctatgatgggatctt
taagcatcattgtagcacatcacatgtatgcaatgcctccttatccatatattgctactg
attacccgactcagttatcgctcttcactcatcatatgtggattggaggattttgtattg
ttggagcaggagcgcatgcttctatatttatggtaagggattataatcctgcagaaaatt
ataacaatcttttagatagagtcattaggcatcgagatgctattgtttctcatctaaatt
gggtatgtatatttcttggattccattcatttggtttatacattcacaatgatactatgc
gggcacttggaagatctcaagatatgttctctgatacagctatacagttacaacctattt
ttgctcaatgggtacaaagtatacacactttagctcctggaaatacagctccaaatgcat
tagcaacagctagttatgcatttggaggagatattgtttctgttggtaacaaagttgcaa
tgatgcctatttctttaggtactgcagatttttagttcaccatatacatgcatttacta
ttcatgtaactgttttaattttagttaaagggttccttttctcaagaaactctagactaa
ttcctgacaaggccaatcttggcttcaggtttccatgtgatggacctggtagaggtggta
cttgccaagtgtctggctgggatcatgtttttcttggcttattctggatgtacaattctc
tgtctgtagcaattttttcactttagttggaaaatgcaatcagatgtttggggtagtgtat
ctccgtctggaaatgtttctcatattactggcggtaattttgcacagagtgcaattacaa
tcaatggatggctgagagatttcctttgggctcaagcatctcaagttattcaatcatacg
gttctgcgctatctgcatatggattaattttcttagcagcacattttgtatgggcattca
gtttgatgttcttatttagtggtagaggttattggcaagagcttatagaatcaatcgtat

gggcgcataataagataaaagttgctcctgcaattcaaccaagagctttaagtattactc
aaggtagagcagtcggtgttgcacactacttattaggtggaattggtacaacttgggcat
tctttttagcgagaattatttcagtaggctaatagtgaaaataggataaaaaaaacaatt
atggcaacaaaatttcctaagtttagccaagctttatcacaagatcctacaactagaagg
atttggtatggtattgctacggcacatgactttgaaagtcatgatggaatgacagaagaa
aatttatatcaaaagatattcgcttcgcactttggacatctagcaattatcttcttatgg
acatctggtaatttattccatgtagcttggcaaggcaactttgaacagtgggtattaaat
cctttgaaagttaaaccaattgctcatgcaatttgggatccgcattttggacaacctgct
ttgaaagcttttagtaaaggtggctcagcttatccagtaaatatagcatattctggcgta
tatcactggtggtatactattggtatgagaagcaatcaagacctgtattctggggcttta
ttcttactagttttatcagctttactcttatttggagggtggctacatctacaaccaaaa
ttcaagcctggtttatcatggtttaaaaataacgaatcaagattaaatcatcatttatct
ggattatttggggttagttctttagcctggacaggtcatttagtacatgttgctatacct
gaggcaagaggacaacatgtaggatgggataattttacaaccgtattacctcatccagcc
ggtttacagccgtttttcagtggtaattggagtgtatatgctcaaaatccagatacagct
caacatttattcggaactaatgaaggtgcgggtacagcaattctgacatttctaggagga
tttcatcctcaaagtcagtctttgtggctaactgatatggctcatcaccatttggctatt
gcagtagtattcattgttgctggacatatgtataggactaattggggaattgggcacaat
ctaaaagatattttagatgctcatagaccacctagtggtagattaggagctggacataaa
gggctatttgatactattactaattctttacacatacagttgggattagcattggcttcc
ctaggtgtaattacttcgttggtagctcaacatatgtatgctatgcctccatatgctttc
atggctaaagatttcacaactcaagcatctttgtacacacatcatcaatatattgctggg
tttctaatggttggagcttttgctcatggggcaatattctttgttcgagactatgaccct
gaacagaataaagataatgttttagctcgtatgctagaacataaagaagctatcatttct
catttaagttgggtaactctattttagggtttcatacattaggcctttatgttcacaat
gacacaatgattgcttttggaactcctgaaaaacaaattctaattgagccggtatttgct
caatggattcaagcctcttcagggaaagcactttatgggtttgatgtgttactatcatcc
tctactaatatcgcaacacaagctggtagcaatatttggctgccaggctggttagaagcg
attaatagcggaaaaaattcattgtttttaacaattggtcctggtgacttcttagttcat
catgcaattgcattgggattacatactactacgttaattttagttaaaggtgctttagat
gcaagaggctctaaacttatgccggacaaaaaagactttggatatagttttccttgcgat
ggacctggcagaggcggcacctgtgatatatctgcatgggatgcgttctatttagctgta
ttttggatgctaaatacaataggttgggtaacattttattggcattggaaacatattaca
atatggcaaggcaatgcaactcaattcaatgagtcttcaacttatctaatgggatggttt
agagattacttatggctaaattcttctccattaattaatggttataatccatatggcatg
aataatttatcagtatggtcatggatgttcctatttggacatttagtatgggcaacagga

tttatgttcttgatctcctggcgtggctattggcaagagttaattgaaactctagcatgg
gcgcatgaacgtacacctttagcaaacttgattcgttggaaagataaacctgttgcatta
tcaattgtacaggcaagattagtaggtttagcacatt-tttctgtaggatatgtattaac
ttacgcagcttttgtattagcttcaacagcaggcaaatttggttagactaaa-----ttc
gaattagct-a-----aaaaaagtcagtcataata-gttttatgactgactttttagat
ttaaaacaaaaacttactaaagtttcacctcaatatctacaccagaaggcaaatttaact
tcattaatgcatctatcgtttgagaagaaggttgatgaatatcaataattcttctgtgag
atcttatttcaaagtgttctcttgagtctttatctacatgtggagaacgtaaaacacaat
aaattcttctttttgtcggtaaaggaattggccctactgcaacggcattagttctagatg
cggtatctagtattttgtcacacgacgtattaagtataatagaattatatgctttcagtt
taattctgatttttgtctgctgagtaattgtcatgttaaaaagatcttattatttaagaa
ttttagagacaacacctgcgcctacggtacggccaccttctctaatagcgaaacgcatac
cttgctcaattgcaattgcattaattaattcagcagtcatcttaattctgtcaccaggca
taaccatttctgcatcagtaccatcatcagcagtaaactgattaatagtaccagttacat
cagttgttctaacataaaactgaggtctatatcctggaaaaaatggagtatgtcttccgc
cctcctctttagttaaaatataaacttctgcttcaaattgagtatgaggtgtaattgtac
caggtttagctaataccatacctctttcaatatctttttctgcacacctcttaaaagaa
ttccaatattatcacccgctagaccttcttctaacgtttttgaaacatttctaatccag
taatagtcgttgtacgagtttctcgtaaacctacaatttcaattgtgtcaccaactttaa
taatgcctctttcaattctaccagtggcaacagttccacgtcctgtaatagaaaaaacat
cttctacagccattaagaaagttttatcgacatctctctccggcgttggaatatatgtgt
caactgcttccataagtgaaaaaatcttgtcaacccatttatcttcaccttgcttagtag
ctgggtttttcgtcacagcttctaatgccaataaggcagaacctgcaacaaaaggaatat
catctccaggaaagtcgtattgacttaataattcccttccttctaattctactaattcta
gtagctcttcgtcatctacttgatcttccttatttaggaatactactaatgtaggaacac
ctacttgttttgctaataaaatatgttcacgagtttgtggcattggaccgtctgccgcag
atacaactaaaatagctccatccatctgagcagcacccgtaatcatatttttttacgtagt
cggcatggcctggacaatctacgtgagcataatgacgattatctgtttcgtattcaacat
gagcagtattaatagtaatacctctagctttttcttctggagcggcatcaatttcatcaa
attttttcgctgcagtagaccctaaagttgataaagtcgcggagatcgctgctgttaaag
ttgtcttaccatgatcaacatgaccaattgtgccaatattgacatgaggttttttacgtt
caaatttagatcgagccatgctttttgttttccttatagtaaaagttgctttatagttac
ttttaacgataaatcttgtgaattgaaagatttattatttaatacagacaaga-----ta
caatataactaatctaatatcgatagtgagcaaacgctttatttgcttctgccattctat
gtgtatcctctctctttctaatagaatttcctgtttcattagctgcatccataatttcat
tagctaatttcatagacatacttttaccagatctgtctctagagaatttagtaatccatc

ttagtgctaaattcgtacctctataagctcgtacttcaataggaacttggtaagtagaac
caccaactcttcttgcttttacttctactagaggagtaatatttcggatggctttttcta
aaatatttaaaggatctgattctgttctctctttaacgatatctaaagcctgatatataa
ttctttgagataaagtttttttaccacttttcaaaatacgaacagttaacatacttacga
gtctgcttttatataaaggatcaggtgatgcaaaccttttttttagctgtattacgacgag
acatagtatttaaaattggttattgtaataa-tgttc-tattagcaggtatttaagattt
aggctttttgtcccatatttagatcggcttttacgacgatcttttactcctgcagcatc
taaagtaccacgtactacatgatatcggactccaggcaaatctttaattcggccgcccct
aatcagtaccacagaatgctcttgaatattatgacctacgccaggaatataagctgtaac
ttcaaatccagatgttaatctaacgcgcgctacttttcttaaagcagagttcggtttttt
aggagttgtagtatatactcttgtacaaacacctcttctttgagggcaactttgaagagc
tggagattttgtttttttatgtatttttcgtctttcggatctaacaagttgttgaattgt
tggcataataaaattgatttaaagtctttaatagatactatttgaacctgctttacatga
attggctagtttttatttatcttatacttacgcataaatctttctactctaccttctgta
tcaataattctctgtgaaccagtaaagaatggatggtttcctgaccagatatctacgtgt
agttctggctttgtggacccaatcgtcataattaactggccgtcacagtaaacttttgct
tctggataccaatttggatgtatattatcttttgccattgtttttataattatataaatg
tatcaataataactattaacgcttagaaaattgaggagcctttcttgctttctttaaacc
atatttttttctctcttttactcttggatctcttgttaaatagccttcagacttgagtgt
cgttctattttctggattaattgagcacaatgctcttgcaacacctaaacgaattgcatc
agcttgccctgtcaaacctcctcctctagcatttacatggatatcatactggtttagtag
tcctaaaacttgtaatggtgcatatgaaactcttaagtaattaggactaaattgaagata
agactctcctggtataccgtttataattaaattacctgaccctgggactagtcttacttg
tgcaacagaacacttccggcgacctgttccagaatagattgcgcgagttttaattaattc
tgtcgacataacattccttgataactaaataaatatacattctttatactatatactctt
gtggcttttgcgcgacatgtgggtgaattgggccagaatacactttaagcttcgtaaata
gttttctgcctaatggacctttaggaagcatacctttaaccgatttttcaataattctgt
taggtaatcttgtctgaagctgatcaaatgtttcaacttttaatccgccaggctgtccag
aatgtcttctgtatagtttttgattcgttttatttccacttacagatacgtgagcagaat
taataacgattacataatctccagtatctagatagggtgtataagaaggcttgtttttac
ctcttaagatattagaaatatgagtagatattctaccaagtgtctgatttttagcgtcta
taacataccaatgagaattagtatttaatgagggtgattgcgttttattcatgaaaagat
taatacttttggttgacaataagagaatatgtaaatggtttacattcaatttctatacaa
gtagtattt-aaaaatatta-------------ttaacaaagagaacttttctaacagtt
atttaaattgtttgtcagcaagaaaatttctagttatcaataacttatactatttttaat
aatataaaatatattaatcactttttttcttttggcaaacttatacctagttttttttgc-

aaggcatatataacttcttcagctgatttctggccaaaattttaatttctagcagttct
tcttgagagtaatctagtaaatctgcaatagaatggatttgagctcgctttaagcagtta
taagccctaacagataactgcagttcttctattaaaacttgactaattttttatcttct
ttgctacgataattatctgctgatttaaagtctaaatttcttagtgaacaaaaaagatta
gttaaaactgtagctccttgacttattgcttcttggggagatatgctcccatttgtccag
atttgtataatcaatctatctttatgctgttgctaccaatacggacttcttctacttta
taattaactttattaactggcataaaaacagagtctacttgcaaaaaatccacagataat
tcatctacagctttttcagctaagcgatacccgcaatttttttcaattttaaactccatt
tcaaatattgtgttattgcaaatagttgcaatatactgtctaggatctactacctctata
tcagaagataattcaaataggccggctgtaactatagctggcccttgaactctaattcga
ccaatttgagattctttgttataacttttaaatactacttcttttagattaagtaatatt
tctaacacatcttctcttaccccaggaattgtggaaaactcatggttcactccagcaatc
cgtacagcaactatagcagtaccttcaagatctgacaatattgatcttctcaaagcatta
cctaatgtaataccttgtccttgatttaatggttctattacaaaactaccgtactgccca
cgcgccccatctgttcttgactctatgcattcaatttgaaattgagccacctaagaaagc
tcc-tttaaaataatcagtgattagtaagtctctctaaaacaagaaggaatttatactcg
gcgtttcttaggagggcgacatccattatgaggtacaggagtaatatcttttattagagt
aatctctaatcctgcagcctgcaaagctctaattgctgtttctcgacccgctcctggacc
atttactagaacttcagtttggcgcataccttgatccatagcttgtctagctgcttttc
agctgctgtttgagctgcaaaaggtgttcctttttagctcccttaaatccacttgcacc
agaggatgaccatgataatgtttctccttttaaattagtaatagtaacaattgtattatt
gaatgtagatttaatatgtgtaataccgttaactgcattacgtttagtttttcttgctcc
ggatttttttatttgtctagccatcgtgtctcgtctattatatagttaattaaaagatta
ttttcttggagctttttcttacctgctactgtttttttacctcctctgcgtgttctagc
attagttctagttctttgtcctctcaaaggaagaccaagacggtgtcttctacctctata
agtactaatttccataagtctcttaatgctcatagactcaaaacgtttgagatccccttc
aatctgataattagactcaagaatttctcttatactgacaacttgttgatcatttaaatc
ttgacacttaatatcagcgtctatgtttgtttttctaatatttctttcgagcgagatag
tccaataccataaatatatgttaaagctatctctattcttttgtttcttggaagatctac
tccagcaattctggccactttgtcttgtctccaataaataatatgtat-ttttataacgg
ttatttata-agcttagccttgtctctgcttatgtttaggattagtgcaaattaccatca
cctttctgtgacgccgaattattctacatttttcacacatttttcgaacagaaggacgaa
ctttcatattaa----ctctatatactaggaatattactatattaactctattaattatc
gacattaaaaatgtcaacaatatattgtattactttaactgttgtagattttacttcgtc
atactatcatatttttttgatatgacgtatgtctgaatttgcttagctgtgtctattgcg
actccaactaaaataagtaaagacgtagccccgagacctcttaagttttggatctgagta

acctttttctattataaacggaatcagcgctactgtaaataaaaacgaggctcctaggaat
gtcagtctatttaatatgacttgtaaataatcaatagtcgcttgacctggacgaatatta
ggaatgcttgcacccattttttttaaatttatagcaatatcttctggattcattactatt
gatgtataaaaatagctaaagaaaaggatcaaagcacaataaagaagaaggtatagtgaa
ccattaggacaaaacaggtataggatttggagtaatgttttattttgaataatttgggtt
agataagatgggagggccatagatgcagatgcaaaaacaataggcataactccaccttga
tttaatttcaagggcaagtaactatttggatccaaaattgaagattttcctagctgtctt
gctgaaataatttttaattcttcttgttccttcttgtacacaaattgtaattattatcatt
aataaaaagatcgctataaataatccgaacttaagacttgcattactataactagcatca
aaaaatgactgtgtaaaattctttggtagtcctgacacaatgttttgaaaaataagtaaa
gaggctccattaccaattcccttttccgtaattaactctgataaccacataataatcata
gagcctgctgttaaagctaaaacagactcgcagacaaatgcaaaattccaattaaaaaca
tatggttttacccatatagagattgctccagattgtaaagtcgcccaacctagagctaag
taccttgtaatttgagttatttttttggcggcccaattcaccttcttccttctgtaatttt
tccagattggggacaattttcgtaagtagctgcattacaattgaagagttaatataagga
acaatacctaacgcaaaaatccctattgttgaaaaacctcctccagaaaaaatattcaga
aaatttactaaagtatttttttctacacttgcataaaaggcgtcatgatctatacctgga
acaggtataaatattcctaaacgtgctaaaactaatagaaagagagtaaagataatacga
tttcttagatcactttttttggctcataaataaattttaaatgaaaagagatactttt--tt
tatgtaaaaaccctattccaagacttgtataagtttaaaatagggaagtaa---aatatt
atagtaatattaata-taatattctatgtagaatacaggctttctattggaactcctcga
tctttagctgcttcactaaaagttcttaattgagttaagccattgagaacagctcgcgcg
ttatttaatgtattattagaacctaattgcttagctaatatattctgtacacccgacaat
tctaggactgtccttacagaacctccggcaattacacctgagccaggtgcagagggcctt
aatataactttcgcggcaccagaaattccgttaataggatgaggtatagaattcgatttt
gtcagtggaactgtaactagatgtttttttagcatctgttactcctttttttactgcacca
attacatcgcttgcttttcccacgccgacaccaacttggccttgctcattaccaataaca
agaataactcgaaagcttagtttttttacctccttttacaactttagtaactcttttaact
tgtacgactctttcttcccagccactatctttatcttttcctttgctctgtttttttacga
ttggccattttaagaattattcctaattagttaatta-cgattctagaaacccatgcctg
cttccttagcagcttcagccaaggccttaactcttccatgatataactttcctcctctat
cgaagacgacatttttaatgccttctttcatagactgttctgctaactgtttaccaacta
cgcgagaagtatcacagtttggtcttatgttatcagattctttattatttaaattaacag
atgatgtagctactaatgtaataccttgtgtgtcatcaattatttgtgcgtatatatgtt
tattagatctaaatacacacagacgaggcctacttgaggttccttgaactttttttccgaa
ctcttttatgcttatgaattctagtttgtttagtgtttagtttcattattatttaccttt

tccagctttcccagcttttcttctaacaaattcaccttgatatctaattccttttccttt
ataaggctcaggaggcctaatagaacgaatagttgaagcaacctgaccgacaacttcttt
atctatgccagagacagtaatatttgtgttgttttcaactttaatttcaatattggcagg
aggtttaattttcactacatgactatagccaacacttaagattaaatcttggttatcaat
ttgagaacgatagcctacaccttgtatttgcagtttttttaaaaaatccattagaaacacc
ttcaatcatattactaataagagttctggatagcccatgtaactgactagccatttttgt
ttgtccactcgtcttgacagcaattgtgtcgttaagtatttctaaattaataccagctgg
taaagttctggatagggtacctttaggtcctgtcacggtgatagtctgaccatcgaactg
agtactaagatttgtcggcaataaaattattttttttccaatacgagacat-attccacc
tataaaatagttaccaaatatagcataatatttcaccaccaagaccatcatgacgagctt
gtctatctgtcataacgcctctagaagtagaaataagagcgatgcctaaacctccaagaa
ctctgggcaactctttatgatttgcgtagactcttagtccgggtttactaattcttttca
gagcagtaataactggttgacgattttaccattatacttaagagaaatcattaaatgag
tctctataccttcacccatttgttcgaaattttgaacaaatccttcttcttttagtactg
ttgccatattgcatgtcattttcgttgctggaacttgcacaatttgatgtcttgctaagt
ttgcgttacgaatacgtgtcagcatatcggcgatcgtatcgttgaccaccttgatcctcc
ttgcga-----tgaattatttgaaatattttcaagattccctgaaaggcatccctagctt
ttttaacaaagctagaccttcttgatctgttttagctgtagtgacaattgatatatctag
accacgaatttgatcaatattatcatagtctatttctggaaagatcaactgttcacgtaa
acctaaattgtaattacctttgccatcaaaacttctaggactaattcctctaaagtctct
aattcttggtaatgttaaattaattaatttctctaaaaaagaatacatcttgtcttttct
tagatgcacaacaattccgataggaacctcttctcgaattttaaatcctgcaatagattt
tttagcttttgtgacaattggtttttgtccagttattaatgttaattcttgaatactact
ttcaagagccttagcattctgagaagcttcacctaagccacggttaatagtaattttagt
aaacctaggaacttcatgtacatttttgtactgaaattcatcttttaaagattgagtaac
agttgttttatattttttcttttaatcctattgccattatatttcatcaattatttaata
agttcgccagtctttttcagttttcgaa-ttttttggcctttatcattaattattactga
ggatcgactagcaatattattttgttcactaaataacataacgttagaggtatgtattgg
agcttcaaatttgataatttctcccgtttctccttcttgctgaggtttttttatgcttcac
tttgagattaattcctttaacaatcaccttatttgttttataaatgattgcaataatttc
acctgtttttgttttatcacttccagaaataacttgaactaagtcccctttttttaattt
aatttttgtattattttttggttgttttagaaggacctttcattatactacctccggtgca
agagaaactatttagaaaaatttttatctcttaactctcgggctataggtccgaagact
cttgtgccacgtggattattatcttgattaatgataactgccgcattatctccaaatcta
atgctcataccgtcagttctccgtaaagctttacgagttcttactacaacagctctgaca
acatcagatcgtttaactggcatattgggagacgcatctttgactactccaataataaca

tctccaatagatgcataagaaggattactagtgcctaatactctaatacacattattttt
ctagcaccactattatctgcaacattaagatagctttgagtctgtatcatacttttattc
tgtataagattaattatcaaaagacttagataagatattaaccattgtccaacactttgt
gcgacttaaaggtcgtgtctcttgtattgtaacaatatcgccgattgtgcattcattatt
ttcatcatgcgctttatatttctttgtccggatcatagtttttgcgtactttctatgaga
gattctatttctacagctactactatagttttattcatttatcgcttactactttacc
tgttgtttcttttaaaggcatagtctttatattttttatatat---------aattaaat
tttaagattattacatggttgacttagtacgagacttttcaacagttaaaagttgggcta
atctatgcttagaatgtttaaatagatgaggctggaaatcttgccttgtggctcttttta
gccttaaatcaaaaagctctcttttattacaaggatttcttcagctaaagaagaagagt
ccaggtttgtaacatctgatattttagggaaagtcatattttaggattctgttgtattcc
gaactataaatttagtcttgattggtaatttataagaagctaatttcatagcttgttgag
cagtttttgtggtacacctgtaatttcaaatagaatatgtccaggcttaataactgcaa
cccaatattctggagctcctttacctgaccccatgcgagtttcagctggacgggcagtga
ctggtttatctggaaatactctaatccatagtttaccacctcttctcacatatctagtaa
tagttcttcgagtagcttctatttgtctagaagttaaccaaactggctctgtcgcttgta
atgcataatcaccgaacgcaattgtgttacctttgctagcagaacctttcattctacctc
tatgttgttttctaaattttgttttcttggggcttagcataaaaattaaaaaataataat
aattaagaagctgttgaatcaggagtttcctgattataagcagattctagacttttcgat
tctggcaaaatttctcctttaaaaagccaaactttactccgagcacaccatacgtagtg
tgagcttggcgatgacaataatcaatatcggctcttaaagtttgtaaaggtacacgacct
tctctaacccattcacttctagctatctctgcaccattaagccgaccagatacttgaatt
ttaacaccctgtgtatttgctctttgagctctttgaactgcttgtctgacagctctacga
aaggccactctttttttctagctgttgagtaataaattctgctactaaggttgcttcagaa
tctggatctgcaatctctacaacatttactctgagttgtttgctagggtctaggattaag
gacagcgattttcttaaagactcaattcccgctccagattttccaagtacgattcctggt
cgagctgttgctatgagaatttctacttgatcaactttacgattaatttcaattttagca
atactagcattactaagttttgaatgtataaatgagcgaattttatgatcttcttgtaaa
agaactgggtagtctttagagttagcaaaccatgaagaacgatgtttttgggtaatgcct
atgcgaaagcccaaaggatgaatttttttgacccacagtatccttgattataagttaaaag
ctataatgactaattgacgttaaa--------agactacagttctgaaacacctaatgta
atatgacaagttggtttatgtattggaaaggctcttccttgtgctctaggctgaaatctt
tttaaagtcgggcctttgtctgcaaaggctttactaacaaacaattggtttttatttaat
ccatcattatgctcagcattcgctgcagcagattccaggatttgctttatatgagaacag
actcggtatggcataaattccagaataattaatgcttcttggtattttctacctctaatt
tggtctaatacacgacggactttatgcggagaaagacgaatgtatttccctactgcttta

gtttcttttacgttttttgtaatactcatagttatagtttaacgacgggcttttctgtca
ccttttacgtgagtgcggaaagttcttgttggaacaaactctcctaatttatgtcctacc
atttggtctgacacaaagacaggaaaatgttgtttaccattataaacagctattgtatgc
cctaccatatcgggaataatagtagatgctctcgaccaagttttaggacttctttttt
cctgaaatatttaatgcttctatacgtttaagaagactaacatcaataaaagggccttta
tgtatagatcttgacataataataaatcattagattcttaataaaataaattctatttac
ggcgacgtaagacgtacgggttactatatttatttggatttcgtgttttaacacccaatg
caggtttaccccaaggtgttacaggacgagctcgtccaattggagatttgccttcaccac
caccatgtggatggtctacaggattcatgacaacacctctaacagtaggtcttttaccta
accaacgatttctaccagctttaccaagagtaatattactagcgtcaatattaccaactt
gtccaatagtagcgtagcattcttttcgaatcattcgaacttcgcttgaaggcaatttta
cagtaacaaaagtaccttcttttgctactatttgagcataagttcctgctgcacgaacaa
tctgtcctccgcacgaaggtcttaattctatattatgtactgctgttcctaaaggaatac
tagataaaggcaaagcatttccaacttcgataggggcagtaggaccagaaagaaccatag
atcctacactgagagatcgagggtgtagaatatatcttttctcaccatcaagatagtgta
ataatgcaattcttgcatttctattcggatcatattcaatagaagcaactttagcaacta
tattatgtctatttcttttaaagtcaattaatctgtactgctgcttatgtccaccacctt
tatgacgacaggtaatgacacctctattatttcgacctttgcagaaatggtgtttaacta
ttaatgatttttctggcttatcagtagtaatctctgaaaaggtagaaaccgttctatttc
ttgtccctggtgtgtaggcacgatataaacgaattgccatataagaagatttgagaagag
taaatgttgaa------ctatgtgatatatttaagtttctggaaacaaattaatagaatc
ttctgaggcaagtgttacaattgctttcttgtaatgtggtcgcttccctacgaatctacc
gatgctcctttttttcttaggaggatgacaagtgttaacacctgtaacttgcacgttgaa
aatatactgtatagcagccttaatattaattttgttgctttgggatcaacggcaaaaca
atactgatttcttctaataatttagttgtcttatcagtaatgatgggatatttaaccag
atctaataaacctcttgaatcaatgctatccattatatacctcttgtattttagataaag
catcaactgtaataataattttatgtgcagctaacaaagccataatatttagtgtatcag
ctgaaataagttctacgttgtgtagattgcgaatagaaagataaacattcggatcttttt
tgtctacgatgactaaaactttttatttaaatcaagattccatcgatgtatagcttcca
taaatagtttagtcttgggctgatgaaaataactgttaaaattttctacaatcagtgtat
taactgatttgttatttaatgctgtccttaatgctaattgtctttctttttttattcattt
ttttggtaaaactacgaggttttggcccaaatattacaccaccacctctccatagaggtg
aacggatagaacctgctcttgctcgaccagtcccttttttgacgccatggtttacgacctc
cacctcgaacttcacttcttgttttagtattagcagaaccctggcgcttttcattacttt
gctttactaaagctctatggactaagtacatcccagaatcttgactgaccttaagattta
aatcagcgttgccacttacttgaccttcccaattatagacttgataatttaattgcgtgt

taactgtcatgaaatatatcgtatgaaagaatattgctacttactaatttttactaaagc
gccaggtttaccaggaacagctcccttaacgatcaacaaatcattctctgagttaatact
tacaatttgcaatttttttattgtaactttttttattaccaagttggccagccatgttttt
acctggataaactcttccaggcgttgtaccggctccaatagagccaggttgcctatgatt
tttcgaaccatgtgacattgggcctctactaaaatgatgcctcttttggtaaccggagaa
acctttacctacacttctagaagagacattaatcttttggcctacttgaaatagatctgt
agatagtatttggctaacttcgaaatcatcagtagatttcatttcgtattcacgtaaata
ttttagtgggggagcctgtgatttttttaagtgtcccaacaatggtttatttaattttttg
ttcagcaacttgcttgtaaccgacttgaatagcattgtagccatcagtagatacagcttt
aatttgagtaataacacatggcccaacttgaattacagtaactggaattgataaaccagc
ttcatcaaagaattgggtcatacctactttagtaccgagtataccaacagacactagatt
tctccttttttgagagatttatactatcattgtacataaaaattctacttcatagttactg
aag--taaattaaactaacagatgcatgatattattcatgacaacataaaggttaaaaac
tttacctatttaatctaaaatagaaacagctttttgtctagttacagcgtatataaatta
gaagtatgcttattaaatttttttataaactact-tagatttaactgatattgatatccgt
aacttacttgcagtaatatc----aatcaacatgtataagtttaattgcctaaactagaa
ccagctattattagaaataattcaagtatagataaaagcgtatttgaacttttatatatt
gacaaatataatagaatatattgaggttcggtaactacaccttttttattttttacatttat
tgtgca--gtatatatacatctaatgcagtaaacataagaatataac----tgtatagca
aatgtaattacacaaattagagaggtattaatgggcaaagtcgttggaattgatcttgga
acaacgaattctgtaattgctgttatggaaggaggtaaacctaccgtcataccgaatgca
gaaggttttagaactacagcttctgttgttgcatatactaaaagtggagataaacttgta
ggacaaattgccaggaggcaagctgttattaacccagaaaacactttctactctgtcaaa
agattcataggacgtaaacagaatgaaatttcgcaagagattcggcaaacatcatataat
gttaaaactagtggatcaagcataaaaattgcttgccctgcactgaataaagattttgct
ccagaagaaatttcagctcaagtactgagaaaacttgttgaagatgctagtacgtactta
ggtgagactgttacacaagcagtaataactgtaccggcttactttaacgattcacaaaga
caagctactaaagacgcaggtaaaatagcaggcttagatgtattgagaattattaatgag
cctaccgcagcttctttgtcttatggactagacaaacaaaataatgaaacaatactagta
tttgaccttggtgggggcacatttgatgtatctgtattagaagttggagatggagtattt
gaagtactctcaacttctggagatacacatttaggcggagatgactttgatcagcagatt
gtagaatggctaatcaaagattttaaacagaatgaaggtattgatcttggtaaagataga
caagcacttcagagattgaccgaggctgcagaaaaagcaaagatagaactgtcaaattta
actcaaacagagatcaatcttccttttattactgccacacaagatggtccaaaacattta
gaaaaaactgtaactagagggaagtttgaagaactttgttcaaatttaatagataaatgt
agtatccctgtaaataatgctctgaaagatgcaaaactagaagcttccagtattgatgaa

gttgttttagttggtggatctacaaggattccagccatacagcaaatggttaaaagatta
attggtaaagatccaaaccaaagtgtcaatccagatgaagttgttgctattggtgcagcc
gtacaagctggagttttagcaggcgaagtcaaagatattctattactagatgtgacgccg
ttatctttaggagtggaaactttgggtggcgtgatgacaaagattataccaagaaatact
actattcctacaaaaaaatcagaagtattttctacagctgtagataatcaaccaaatgtg
gaaattcaagtacttcaaggcgaaagagaactaacaaaagacaataaaagcttagggacg
ttccgattagatggcattatgcctgcacctagaggagttcctcaaattgaagttaccttt
gatattgacgctaacggaattttatctgtaaaagccaaagaaaaggctactggtaaagag
caatcaattactatatctggagcatcaactttacctaaagatgatgttgaaagaatggta
aaagaagctgaagaaaatttcgatacagaccaaaaaagaagaaagaatattgacacaaag
aatcaagcagagtccctatgttaccaagctgaaaagcaagttaaagagtttgaagacaaa
attagccaagatttaaaaataaaaatagaggagctaattacagagcttagatctagtcta
gagaaagaagaatatgacaatattgaatccatttctcagcaattacaaaatgctctgatg
gacattggaaaaaatgctgctcagactgaaagtaaagatacaaaagcgaaggatgacgac
actgtgattgacactgatttctctgaagctaagt-aaaagta-agcgggtaacgcgattc
gaacgcgcgacatcaaccttggcaaggttgcgctctaccactgagctatacccgcataga
ttgtattattacaaaaattaagatatttgtcaatctgtta------atctatac---atc
tagactattaattttggaattacatattagctaactgttgtaaatataacaattgctata
attttttaa-------aagtacttatatcat-tatataagtactttttatatt-------
-caaaaatacaaataagtattatag-------ctaaactactaaggagttaactaagcta
gtaacaattactaatcctgtccataagccggctcctgtatatattaatccttttgattgc
tcccattggccaggggatgctaaaacaacaggtacgcctactactaacacagtcgataat
gcaattaataatagcactaataattggatcgcaataatcatttaatatgttctcctgaaa
aattcaaatgttaataatactgtatataatataatgaagcttttgaccttcagcaattta
tctatgaagaaataatttaaaattattgttttctaaataaaccttaaattaagtattaaa
atctttattgtaaacaataatactatattttttta--gtagataaaaaatttaaagaaag
tcttttttaaaaaaaaattaagataaggtgatatgctgtataaaaagctaagtattatta
attattagagaaatttaaa-ataagaggtaaaacatatgaactcagcccttttttttagca
aaactaccagaagcttatgccatttttaaaccgatcattgatatcttacccgtaattcct
gtatttttcttcttttagcttttgtttggcaagcagcaattggttttagataacattta
ggttataaaaagaataacaaatatacagagtgccaattttttgagctctaatcttatatct
tacacaataataactattatggttgaacccttactatcaggaattgttcttggacttatt
cctattactttattcggattattagtagcagcttatttacaatatcaacggggcaaccaa
ttaggactgtaaagcctaaattatagatatttataaaaactacaataattatagtttta
taaatatctacaaataagctaattttaaatacagaaataatactattgaaactagagtga
taacaaaaaaaatttatgtaa---taa---------------------------------

------------------------------------------------------------
------------------------------------------------------------
------------------------------------------------------------
------------------------------------------------------------
------------------------------------------------------------
------------------------------------------------------------
------------------------------------------------------------
------------------------------------------------------------
------------------------------------------------------------
------------------------------------------------------------
------------------------------------------------------------
------------------------------------------------------------
------------------------------------------------------------
------------------------------------------------------------
------------------------------------------------------------
------------------------------------------------------------
------------------------------------------------------------
------------------------------------------------------------
---------------------------caagctaattgtttaccagctcgacttagtaac
tcctggtaaaagacattcatgagacatttctcgaaaaacatggcgagataatccaaaatc
tcgataataacctctacttctacctgttaaccaacatctatttctccctctaacagcggc
gctgtttctaggcatttcttggagtttttgtcttaaatccattttgtcagaaaagctgcc
agcttctttcagttgatctttaatagccagtctttttaaataatatttttttgcaagttt
cgatctcttaatctctctttggatcatattctttttagccatgctgtaa-aactctcttt
taaaattatttattcaatatattagctaaaaaagctcctgcctgcaacttttatatattt
aaaaataaatagcataaagtagtgaaatgcaattgatatttaatcaacgatacc---act
ttatgctattatcaaaaatttatttagtaaagttcttgttcaacatgagttgaaatagtg
caatcagatgaaggatatgcaatacaggttaaaacatagccagctaaaagttgatcatca
tctaagaatgattgatcagcttgatctacagttccttctgtaactttacctgcacaagtt
gaacaagctccagctctacaagagtaaggaagttcaattccttcttcttctgctgcatct
agaatatatgtatcctctgcacaattaaatgtgacatcaattccttcttcttcacatagt
aaatgaattttataatcagccatttcttcttactccataataaagatacacttgaaactt
aataaagcaaaat-aatttaaagaacttaccaaaaaattatatactagtttcattatgca
tgatagactggaaaaaaagaaaatagtatactattctgaaaaatcaagctacatttccta
ctaaatagaactatagtaggtccaaagtatatacataaattcatatagttaaaattactt
ttatcatttaaatagctgaacttgatagta---------tagattgatcatc------aa

gctagtcttctgc---caataaataattaattatgatttcaatgcttaatcaacaagtag
aacttaagcctattcttaaatttcagataacacaaatatattcgcattctcttattcagg
aaattaaagccttaataactagacttgttttgcaagtttggagaaggccagctacattga
tggcaggtattatccaacctttattatggttaattctatttggtgggcttttctacaatg
ctcctataaatttgttcactattaatacaagctataattgtttttttgagctctgggatta
taattttcacctcttttactggagctttgaattcaggtcttccattaatgtttgatagag
aatttggatttttaaatagattattaacggctcccttagtatcgaggacttctatcattt
tatcttctgctacttttatgacttgtattagtttaatacaagttgtatttatagttacag
cttctctttttatgggaaacccacctctaaatagcgatagtactatgatttttggactta
tgattctattagtgactgtaggagttacaatgcttagtttagctttatctttcactctgc
caggtcatattgagctattagcatttattttagtagttaacttgcccttttttattttcta
gtacggctttagctcctttatatttcatgccgccatggcttcagttaattgcaagtctca
atccattgagttatgcaatagaaggtacaagatacttatattcaagcgtaaactggaatt
ttacagagtgtgtgattaagattagctggggagatatttgtttagggcaaattattattt
tattaatcgctttagatataatggcagcttatcttgtgtctaatatattaaaagctaaac
ttaatta-----------------aaatttaataaaaaatttattaaatttattttaca
aagattttactatataatactaatagtagtatggataattgaaatagaaatagtttttc
aaaaccaaagctattatcaacaatgataaacatttgtaagaaagtcaacaaagtatgttc
ttattcataggaggcatgtagtcaatgggactaccatggtaccgtgtacacacggttgtt
ttaaatgatcctggacggttaattgcagtccacctaatgcatactgcacttgtagcgggt
tgggcaggatctatggcattatacgaactagctgtatttgatccttcagatccagtatta
aatccaatgtggcgacaaggcatgtttgttatgccatttatggctagactgggcgtaaca
gattcatggggaggatggagtataacaggagagagcgtatccaatcctggactgtggagt
tttgaaggtgtagccttaactcatatagtcctttctggcatgctttttctagctgctatt
tggcattgggtttattgggatttggaattatttagagatccacgaactggcgagccagcc
ttagatttaccgaaaatttttggaattcatttattgctatcaagtctactttgtttcgga
tttggagctttccatgtaactggacttttggaccaggaatgtgggtatcagatgggtac
ggagtaaccggaaaggtattaccagtagctccagcatggggaccagaaggattcaacccg
tttaatcctggaggagttgcatctcaccacattgccgcaggtactgtaggtatattagct
ggtgttttccatttaactgttagaccaccacaaagactgtatcgtgctctaagaatgggt
aatattgaaactgtattatcaagtagtatctctgctgtttttttctcagcttttgtgact
tgtggtacgatgtggtacggctctgcaactacgcctattgaattatttggtccaactaga
tatcagtgggatagtggatattttcagcaagaaattgagaaacgagtagaaaatgctatt
gctgatggtgctgcacctagcgaagcatggtcaagaattcctgacaagttggcattctat
gactatattggtaataatccagcaaaaggaggattattcagagcgggccctatgaataaa
ggcgacggagttgctgaagcatggcttggacatcctgtattccaagataaagaaggaaga

gagcttagtgttcgcagaatgcctgctttttcgaaactttcctgtaattttagttgat
aaagatggtattatacgagctgacattccatttagaagagcagagtctaagtatagtatt
gaacaagtaggtgtaacagctagtttttatggtggcaaattaaatggccaagttttcaac
gatgctcctagcgttaaaaaatatgcaaggaaagctcaattaggtgaggtatttgaattt
gatcgaactacattagaatcagatggagtgtttagaagtagtcctagaggctggtttaca
ttcggtcatgctaatttcgctttaatttcttctttggacatctttggcatggttcaaga
actatcttccgggatgtgtttgcgggaatcggagctgaagttactgaacaagttgaattt
ggagctttccaaaaattaggagatagaagtagtaaaaaacaaggagctgtataagaccaa
agtcaaagatagtctttatc---aaatatttctaatattaaaatagttaagacaaagttt
attatctaatagttactaatcaggagaatttatggaagccttagtctatgtatttttact
aacaggaacgctaatggttatattctttgcgatctttttagagaacctccaagaatagc
aaagtaacttgattccgtttaaacttataataatagaaccactttccaacttattagtgg
aaagtggttttattgtagattacattgtattattcttcatgctcttcaaaaggatctcgg
agatcttttgaagcaggaccaaatgctgtgtatatcgagtaacctgtaatacctaaaagc
agactcgaaataaaaatactaagaactgttgcagtttccataatttaagcagtattaaga
tgaactttttctataatgatattatcaatattaaaacatttaattattaaaattatcaaa
atatactatggcacttagaactagacttggagaaattttaagacctttaaattcagagta
tggaaaagttgctccaggttggggcacaactcctattatgggaattttcatgttactatt
tttcttgtttttattaatcattttacaaatctataattcttcattagtattagaaaatgt
agatgtagattgggctactttaggtag----ctaactagagaagttctttga--------
ttacacaatattaaagacaac----------------aagcgatcgctattagattatag
tagcgatcgctttaaaagtattttactatatatccagagaaaattatttacctaaattaa
ttaactcggtatctgcaaaactatttgtgttaacattattatagttaactttttcaaacc
ttactaatactggatatttgataccacttttatcaattgctgcaacagtaccaatttcct
gataccagtacgattcttttctcaatatttttacttttgagcctctttccatatttttag
tcctcagatagattattaaagattattatattaaaaaatt-taaaaaactaaacaaaata
tataaaaatactaccgacaattgtaatatataactaataaaaggagagagagagggattcga
accctcgataacaaaagttatgacaattttcgaaattgttgcaataaaccactctgccat
ctctcctgaattaaaatactgttttaaaaaattagacaagaataacatagaactaaggta
gttgcctagaaaattaaaactatgttaaatataaattataatctttctatagatatattt
aaaaaatatgtacatagaatctatatcagcttatttcaagatttttttaatcttatacaat
ttttactaaataaataaattctctcataatatgaaactatcttggaaaacacttcttctt
tggtctttaccaatatttgtaattggtttctttttctggcaaggtttttttaggtccaact
actacagatgttggcagtaatatcgcaagttctagaatgacatatggacgatttttagaa
tatttagatatgggttgggtgaaacgggttgacctctatgaaaataatcatacagcaatt
gtagaagctgttgggccagaattaggaaatagggttcaacgaattcgagttgaactgcca

gcaagtgcgccagaattaattacaaaattacgcaaagccaatgttgatctagatgctcac
ccccctaaaagtacaagtgcagtatggggactattaggcaatttactatttcctttacta
ttagttggcgggttagctttcttatttagaagatctaataatgctagtggtggacctggt
caagcaatgtcatttggcaaatcgaaagctttatttcaaatggaggctaaaacgggagta
gtatttaatgatgtagctggagttgaagaggcaaaggaagaatttcaagaagtggtaaca
tttttaaaacagcctgaatcatttactgctgttggtgcaaaaataccaaaaggcgtttta
ttagttggacctcctggcacaggcaaaacattactagcaaaagctattgctggcgaggct
agtgttcctttttttagtatctcaggctcagaatttgtagaaatgtttgttggtgttggc
gcttctcgtgtgagagacctattcaagaaagcaaaagacaatgcgccttgtatcgttttt
attgatgaaattgatgctgttggtagacaacgaggaacaggtgttggaggtggtaatgat
gaaagggaacaaacattaaatcaactattgactgaaatggatggctttgaaggaaatact
ggtgttattgtaattgccgctactaacagagctgatattttagattctgcattattaaga
cctggaagatttgatagacaagtttctgtagatgtaccagattttaaaggcaggttagca
attcttgaagttcatgctaaaaataagaaaatggaacctaaagtatctttagaaaccatt
gccagaagaactcccggcttttcaggagctgatttagctaacttactaaatgaggctgct
atcttaacggctcgacgaagaaaaaatgcaatgactatgtctgaaattgatacatcaatt
gatcgagtagtagccgggatggaaggcactcctttaattgacagtaaaagcaaaagatta
attgcgtatcacgaagtgggtcacgcaataataggcagtttattagagcatcatgatcct
gtgcaaaaagttacattaataccaagagggcaagcaagaggcttaacttggtttactcct
agtgatgatcaaagtctaatatcaagatctcaaatactagcccgtatcgtaggtgctctt
ggtggcagagctgcagaagaaatcattttcggtgacgcagaagttactactggtgcaagt
aatgatttacagcaagtaacatcaatggcaagacaaatggtcactcgatttggaatgtct
aaaattggacctttatctcttgaaagccaaggaggagacccatttttaggtagaggcatg
ggaggaggctcagaatattcagatgaagttgcaactaatattgataagcaagtaagggaa
attgtcagtgaatgctatgcacaagctaaacacattattatagataatcgagtagtgata
gatagattagttgatttactaattgaaaaagaaacaattgaaggcaatgaatttagagac
atcgttaaggaatacactgcaattcctgaaaaaaattactacatatcacaattttaa---
---------------attaaacgggactgacgggattcgaacccgcaacttccgccgtga
cagggcggtgctctaaccagttgaactacagtccc-aaaaagatgaatacccagagataa
tctcatagttagcttaaacttgtcaaattagaatattaaatatgaagagta---------
--------tttgtaataatactcttcatatttaata-tagtagctaaaatatttctttta
gaaggagagagagagggattcgaaccctcggtacgaagttaatcgtacagcagattagcaat
ctgccgctttcgaccactcagccacctctccatga---tatatattatatatatatcaaa
tatacttatatttaataggtgtgcataaatatttatggagctaagcggattcgaaccgc
tgaccctctcaatgccattgagatgctctaccaactgagctataaccc----ctttggta
agatattaagaaatggctcaagcgggatttgaacctgcgaccttgggcttatgaatcccc

```
tgctctaaccactgagctactgagccatagtataaaatattattaagtattataccataa
attatggtcaaatataaatcaataaaagaataattactatca-tttactacttttg----
-----------------tgcaagttgtaaatattcgaataaaatttcaaaattaaaaatg
tctttaactttttgttaga-------atta-ttgattacgaaa-----------gtattg
atatttagaattaac---------------------------------------------
--------------------aataaatttg------------------------------
---------------------------------------------------tctttattt
g-----------------------------------------------------------
------------------------------------------------------------
------------------------------------------------------------
------------------------------------------------------------
------------------------------------------------------------
-------tagaca-----------------------------------------------
-------------------------------caaaaatattggaaa--------------
------------------------ttttcatcaat-----------------taattagt
------------------------------------------------------------
------------------taaactataaccgacatac-----------------------
------------------------------------------------------------
------------------------------------------------------------
---aagaattattgaacta-----------------------------------------
------------------------------------------------------------
---cgttaaat-------------------------------------------------
------------------------------------------------------------
------------------------------------------------------------
------------------------------------------------------------
-------------------------------------tgtgatgcatt------------
-----gggcttgatcaaaacttctctaaaatcaatagaatatagt---------------
-----------------------------------------------------aattcaa
gtagatcaaatactaacaaatgtgacaaacgcttaaaaaaataata--------------
caattagaatcgtaaaaatttggtttttga-ctgtgttaaataaatttgattatcataga
tttaatatttaaaagtgttctattacttgta-----ttgagtaattgt-aatatttaata
tcttcttttttagataacaatcataatataagctaaaaaatagtttt---tattaaacaa
ctagcatagagccaataccttgatcagttaatatttctagcaagagagcatgatctattc
taccatctaaaatatgcgctgaagctacaccctgagccaaagaacgaatacaacaattca
ctttaggaatcatgccaccagaaattactgctgtttgagttaaatctctagcttcctgta
tattcagatgactaattaatgttgaaggatctgaagaattacgtaaaatgccgggcgtat
```

ctgtgagtagaattagtttttcggcattaagagcggctgctatttctcctgcgacagtat
cagcattaatattatacgattggccttccttgtctgctgcaacactagctataacaggta
tataattattattaattaatatcttcagtaatttagtatcaatattttgtacttctccaa
caaaaccaaggtttgcttttccatttggtctcggagtaataagtaatccatcttttcccg
acaaacctacacttttaccaccttgtttattaatagttgcgacaagatctttattgactc
ttccaactaaaaccatttccactatatccatggtaggttgatctgttactctaacgccat
catcgaattttggtaatattttgaggcgatctaaccaaaaattaatttctggtcctccac
catgaactaaaatgggacgtagtcctataaaagacagaaaaacaagatcactaatcactt
gatcttttagtttctgatttttcatagccgctccccccatattttataacaataattctgg
aagaaaattgctgaatataaggcagagcttcacttaagacttttactctttctgaattag
tcaacataaataaagtaggattgtgatatataaaattatatacacagcttgtgttaattt
aaaagataccaatgggt---------acttgaaaaaatagaaatattactaacag-----
----ttaaaactatagggattataattatgaataaattttgggataatgtattaagattt
cctcgattttagtcagcgtcattttaggattaattttgataattattagtcctttttc
gtgttattaaaaaaaccactgacaagtttttttttcatcatatcattggccggattaatc
acagttttggcaataataa-tacaaaaaatgataaatatcgaatgttgttgagagaaata
aataaatttaattaaaatttttgaacatatataatattatattatcattagatatgataa
ttt-atttcctttctat---aaaacccataaatattaaatattatttgataaatattat
gccactaaaacaaagagttagctctgaaaagaccggtgccttcgcacttttggatagtat
tgttaggcacggagttaaacatatatttggttatccaggtggagctattcttcctattta
tgatgagctttatgcttgggaagaagcctctctaattaaacatatccttgttcgtcatga
gcaaggcgctgctcatgctgcagattcttattctagatcaacaggagaggttggagtatg
cttcgctacttctggcccaggagcaactaatcttgtctcaggtatagctacagcacatat
tgattctgtacctatattagctataacaggtcaagttgggagagcttttattggtacaga
tgctttccaagaagtagatatttttggattacacttcctattgtaaaacattcatatgt
agttcgtgaccctagagacatgtctagaattgttgcggaagcatttttatttgtaaaca
cggtagaccaggtccagtattaattgatgttcctaaagatgtaggattagagaagtttaa
ttatttttctgttgagcccggaaaagttaatattcctggctgtaggccaattaccagcct
aaagtcaagacaaatccttatggcagctaaaatgatacagcaatctagccagccattgtt
gtatattggtggaggagccataatctctgattcacatcaaattattaaagaacttgttga
tttttataaaatacctgttactactactttgatggggaaggggatttttaatgaggatag
cgattattgtctagggatgttaggtatgcatggtactgcgtatgctaattttgcagttag
tgagtgcgatcttttaattgctttaggagctagatttgatgatagagttactggaaaatt
agatgaatttgcttgtaatgcacaagtgattcacgtagacattgatcctgctgaagtagg
aaaaaataggattcctcaagttgctattgtcggtgacgtagcagaagttgttagtgaaat
attgaatttattaaagacttctttccccccttatccagagcagattatatcttggcaaga

aagaattaatcgttggcgtcaacagtatcctttactggttcctagaaaatcaacaagcat
ttcacctcaagagattcttgttgcaactaataaattagcccaaaatgcttattttactac
agatgttggccagcatcaaatgtggtcagctcaatttctgaaagtaaaagctaagcattg
gctttcaagtgctggattaggcacgatgggttatggtttacctgcagcaattggcgctca
agtagcacatccaaatgacgtagtcatttgtattagtggtgattctagttttcaaatgaa
tatgcaagagttaggaactatcgcgcaataccagttaccagttaaaattattattattaa
taatcgatggcaagggatggttagacagtggcaacaagctttctatggtgaaaggtattc
acactcaagaatgacagaaggagcacctgattttcaaaagcttgcagaagcttttggtat
taaagcttttactattaataataggcaaaatatgcaatctgctttacaagttgctattga
ttatcctggtccagttttattagattgccaagttacagaaaatgaaaactgttatccaat
ggttgctcctggaaaaagtaatgcacaaatgataggtatcgctaaaccgcagagaggtac
tgcttccaactatattaataatagtgtttgaagtaa------------------------
------------------------------------------------------------
------------------------------------------------------------
------------------------------------------------------------
---------------atagttagtatttttta-aacaaagaata-aagtaca--------
-taaatgtattttttttatactatta-aaaactatgtaagaacagcctttgacgagaatt
gaactcgtgaccttccccttaccaaggggatgctctacctctgagccacaaaggc-tttt
tactatattgggccgggttggatttgaaccaacgtaggcgaagccagcggatttacagtc
cgcccccattaaccactcgggcaccgaccctaattaaaccttattcaagatatgacctat
aagatatcataactatgttgctaaaaacaagtactttgataaataaaactgaataagtaa
tggacatagctggacttgaaccagcgactttcacgatgtcgacgtgacactctaaccaac
tgagttacatgtccaaaactttaattatcattataacactgtttagtgattttataagta
tctatttctaattgaagatttgttttaatcgtccggctttaccgagcagatctcttagat
aatacagtttagctctacgtattttagctcttctaatcactttgacaaaagttattcgcg
gtgaattgagaaaaaatactctttcaactccaataccttgaaatgagcatcttaaagtta
aactagtatttaagcttttttttcttcttcttgctaatactactccttcacataattgtt
cccgagttttactaccttccttaaccattagtccaagttgtattgtatcgccaactttaa
tttcaggtacttctgttttgataaaaggtatctcaacacttttcattagctgacttagct
ttgtactatttgctttcatatataggattagtcttgtgaca-tgttttgaaatattatac
attaaaaaaacaaatatgaaaaattttagcgaataagcatattagtatttataaaac-tc
tttttatct---------------------------------------------------
--------------------------------------------ccattata--------
----------------cagatccatt------------atattcaact------------
------------------------------------------------------------
------------------------------------------------------------

------------------------------------------------------------
------------------------------------------------------------
------------------------------------------------------------
------------------------------------------------------------
------------------------------------------------------------
------------------------------------------------------------
------------------------------------------------------------
------------------------------------------------------------
------------------------------------------------------------
----------agattctttcatagtttc--------------------------------
------------------------------------------------------------
------------------------------------------------------------
------------------------------------------------------------
------------------------------------------------------------
--------tttgataatcagtac-------------------------------------
------------------------------------------------------------
------------------------------------------------------------
------------------------------------------------------------
------------------------------------------------------------
------------------------------------------------------------
------------------------------------------------------------
------------------------------------------------------------
------------------------------------------------------------
------------------------------------------------------------
------------------------------------------------------------
------------------------------------------------------------
------------------------------------------------------------
------------------------------------------------------------
------------------------------------------------------------
------------------------------------------------------------
------------------------------------------------------------
------------------------------------------------------------
------------------------------------------------------------
------------------------------------------------------------
------------------------------------------------------------
------------------------------------------------------------

---------------------------atcta----------------------------
------------------------------------------------------------
------------------------------------------------------------
------------------------------------------------------------
------------------------------------------------------------
------------------------------------------------------------
------------------------------------------------------------
------------------------------------------------------------
------------------------------------------------------------
------------------------------------------------------------
------------------------------------------------------------
------------------------------------------------------------
------------------------------------------------------------
------------------------------------------------------------
------------------------------------------------------------
------------------------------------------------------------
------------------------------------------------------------
------------------------------------------------------------
------------------------------------------------------------
------------------------------------------------------------
------------------------------------------------------------
------------------------------------------------------------
----gatgatcaaatattattttcaaacatggcttatgtatgaaaaagttctata---aa
aacctaatgac--------taaaattagggcttaagtatttcttagttttgtgttataat
tagttactatcaaaggtagtggcataacacgaagcatcaa-aatgttcgaacgctttact
gaaaaagctataaaagtcataatgctagcacaggaagaagctagacgcttaggacataat
tttgtcggaactgagcagatactattaggattagttggtgaaggtactggaatcgcagct
caagttttgaaatcgatgaatgtgaatctaaaagatgcaagagttgaagtcgaaaaaatt
ataggaagagggtctggttttgtagcggttgaaattcctttcactcctcgagcaaaaaga
gtactagaattatctttagaggaagcacgtcaactaggccataactatattggcacagaa
cacttgctaatgggcttagtccgagaaggagaaggagttgcggcaagagttttagaaaat
ttggcagtcgatgtttcttcaattagagctgaagttatacaaatgctcggagaaaatgcg
gaagccaatgtaagtggaagcaatactacgcaagctagaagtaaaacaccaacattagaa
gagttcggatctaacttaactcaaatggctatggaaggtggtttagatcctgtagtcgga
agacaaaaggaaatagaacgagttattcaaatcttaggtagaagaactaaaaataatcct
gtcttaattggggagcctggtgtaggtaagacagcgattgcggaaggattagctcaaaga

attgctaatagagatgtaccttctattttagaagataaattagttattactcttgatgtc
ggtctattagtagccggaactaaatatagaggtgaatttgaagaaagactaaaacgtatt
atggatgagattaaatcagctgataatgtaatattagtgattgatgaagttcatacattg
attggggctggtgctgcagaaggagcaatagatgcagctaatctgcttaagccagcttta
gcaaggggagaattgcaatgtataggtgcaacaactttagaagaatatagaaaacatata
gaaaaagatccagcattagaaagaagatttcaaccagttgtagttggagagccaagtgtt
gaagaaacaattgaaattttgtttggtcttagagaccgttatgaaaagcaccatcaatta
acaatgtcagatggagctttggctgcagctgctaaatacgctaatcagtatatttctgac
cgatttttgccagataaagcaattgatttaattgatgaagctggttctagagtccgttta
ctaaattctcaattacctcctgctgccagagaattagataaagagttaagagctgtatta
aaaacaaaagatgaagctattagagctcaaaaatatgaaacagcagagcagtatagagca
agagaaatggaaattaaagctcaaattgcagcaattgctcaaagtaaaaagaatgagcct
gatttaaatttagaagatcctgttgttacagaagatgatattgctgaaattgttgctgca
tggactggtataccagtaactaagcttactaaaagtgagtcagaaaaattaatgcacatg
gaagaaactttgcatggacgtattattggtcaagacgaagcggttgtagctgtctctaga
gcgataagacgcgcaagagtaggtctaaaaaatcctaacagaccaattgcaagctttatt
ttttccggaccgacgggtgtaggaaaaacagaattaacaaaagctttggcttcttatttc
tttggttcagaagcttctatgatacggctagatatgtcagaatacatggaaagacacact
gtatctaaactaattggttctcctccaggatatgtgggttatagtgaaggtggttatcta
acagaagcggtaagaaaaaaaccatatactgtcatcttatttgacgaaattgaaaaagct
catccggatatttttaatctacttcttcaaattttagaagatggccgactaacagatgca
aaaggcagaactattgattttaagaatactcttttaattatgacttctaatattggatct
aaagttattgaaaaaggaggaggtagtttaggctttgaattatcagaagatcaaacagaa
tcccaatatactagagtacgatctttagtaaatgaagaactgaaacaatactttagacca
gagtttctaaatcgattagatgaaattattgtatttcgtcagcttactaaagatgaagtt
agagaaattgcagaattaatgcttaatgaagtctttgcgagaattaagcagcaagatatt
caattaaatgtaacagaacgatttaagcaacgattagtagaagaaggatataatccaagc
tatggagctagaccacttagacgagcagtaatgaggcttttagaagatagtctggctgaa
gaagttttatctggtaaaattaaagctggtgatagtgcagtagtagatgttactaatgaa
ggagaagttacagttttattaggtgaaaaattagaactgttaacataaa------aaaca
atttattaat--taaagcattgaagaggttat--attcaatgctat---gaaaaaaatta
ataagatgggttgctataattgacaacctctgatattctttatttgtatata---aagcc
gggatagctcagttggtagagcagtggattgaaaatcctcgtgtcaccagttcaaatctg
gttcttggcat-tttaagcatcagaaaattttgctatctgagcattaaattttaattgaa
atgttcctgtgggaccattcctgtgttttgctattataacttctgtgaagtctctacttc
tagtttcttgtgtatagtaactttctctatatagcattattactaaatctgcatcttgtt

ctatagaattatgaacaataaagttatttgaaataaaattgcataaaggtttacattcta
aatcaaacactatttgtaaacttgtaacgtttatttttttgtatactttcaaaagtcaaag
atgaaa------aagaattaaaaacatttttttgttcttcaaagttttt------tattt
gaatagcaatcatgtcattttgattaattttatcacatcttttccatccttgtgttgtga
gtagtttatggttacttgttaattgtatatatttcccagcttcggtaataattttatata
cagtttttttccctgttttcgaaatactacatttaattgctagaaataactgtcgttttt
gcgcactaaagttaatgacttca-atctttttataatagcaattaaataatatttg--ag
gttgattatataatgatgtctgaagataattgaatttggatatacacccactttctctta
aatcggataacaaaggccttttgtcactccgagtttcaaggtttctatttagttgagata
atgccaagataggcaagcttaaatctttgctaatattttaagtgatcgagtaattaatg
atagttcttgagatctattgtcagattgtctactctcctgcagtaattgaagataatcta
taataattaattctatatttttccttgtaatttaaagaattttactttgtctttatag
tgtcagtagaaatcttagcgctatcatcaatataaagattaaggttagcaagagttttac
tttttttgaataacatattgccattcatcattattaagctgaccagattgaatttttttggc
cgtttaaattacactcttgagctaaaattcttcttagtagttgctctgtagacatctcta
agctaaataaaataacataagatcttttctgattaattacatatctagttatattaatag
caaaagcagttttccccattgaaggtcttcctgcaagaataattaaatctgacttttgaa
acccttgtgttatagagtcgagttctgtaaaaccagaaagtatactactatttatactaa
tctttttccttgtatccaaatggactaatagtcttgcaagaattgtagctaatgtttgcg
tatctttttcgtcaagtatttcataagcttttgttaattgactaacaatagaagttatgt
tttttgttcaataagttgttttgagtagctaattaaacataatgaatctccacatgcta
aaagtaatcttttaatgtagttatctataacgactgctgaatattcgcttatagtatcag
ataggggagcttgttctattagactaaaaacttcatttagctcattcaagtcttgaatca
tttttcatttttttaagttaatgaaaaaaatttcttacgctttttgtctttgcatggttaa
cattttcgagtgctgctctataaagtaatgaagttgatgtaaaataaaaaaaaatcaggag
agatttttttctactgattttagtaatgatatcgattgctgagttaatattatactaatta
atattttttcagctaaaatattatgaggaggtagatatttataa------atacttatta
cttcctcc-ttgtaacattactcttgtg--------gtattagttaactattgatttaat
ttgattctggaagaacttgcaatttaatatttgcagttacttgatgtaatagcttaattt
ctacatcgtaaatgccaattgttttcacgtctggtaaagagatactttgtttttcaatgt
ctatgttggtagtgtttttaataatttgtgaaatttctttttctgtgacactaccaaaaa
tattgtggccatctcctgttttttttactaacgctaaacctttgtatttcttctagaagtt
gcttgactttttttagcattttctttagctgtttttaatttttcttctcttatagctgcat
aaaatttttgttgttttaaaataccatttgtggcaactgccgccattttattcgggatta
aaaaatttcttgcataaccagaagctactttgataacgtcattactcttgccaagttttt
gaatattctcttttagcacaacatttataactttttttactcatctt-------aattcta

aaaa----atctgtattatcctattatacctgttgtatacagatttt----aataggaag
tatttatttgttgatataatattcttgtaaaa--aaaggtaatttttttattttt-ttctg
atataatcactttcacaataattc-----attaggagaggtggccgagtggttgaaggcg
cagcattggaaatgctgtttaggagcaatcttaacgagggttcgaatccctctctctccg
tacaa--aaaatataaaaagttttatagagtccaatctaaatctccaataatattcttcc
aagcatttaatccaccccgaactctaacaatatttagtttttgagctattaaaaactcag
aagcaaatattgatctagaatctaagctacagtaaacaaaacaaattttatcttgtaaat
tgatatgagagtaattcattcctttattttttgtaaaggtagatttaatgaatatatta
aatgacttttatgatattcttcatgatttcttacatctattaaaatatattgtggattcc
tactaattaaaaacttttgtaattgaataacactaatttctcgtacaaatgtattagatt
tactgccataatatttgttccaatgctttttgt-------ggataatacaaatttagta
tttataatcttaaatttgttaaatgatgaagttaatgagttgtacgttaatatagttcca
cttaatatagacttataacccagaacgattttgacagcttcagttgcttgaagtgtacca
acaatacccggtaagagacctaaaactccggaattactacaagtatctcttgcgctctct
ttattttcagtttcactgtagaagtctcgataaacaggtccaccttgataattaaataca
ctaacttgtccttcaaattgaaaaatggcaccataaatatgtattttatttaactctagg
caagtatcattcaagagatatcgagtatcgaagttatcagatccatctataataatatca
tattgtctaataatgtcgattgaattctcatagcttagtcttgttttgaacacagttact
atacactgtggatttaaatctaata-tttttttttagctatttctacttttgaatatcct
atatcatgcatcgtgtataaaatttgcctctgtaaattagagagatcaattatgtcatcg
tctataattccaatgctgccaattcctgcagcagcaaggtaaatgattcctggagatcct
agcccaccagcaccaatgaataaaacttttgcttcttttaatctttcttgtccttctaat
tgaatttgtggcaatactaaatgcttagaatatcttgtatattcttctagtgagtatttt
gtattttctgttttaaaatttagcataaaaatttatagtttatttaatagcactta----
--aaagctagctgataagtttattaaaatgacttttgtaaacttggttcttctctatcta
aaatagctccttgtgtaggacagacctgaacacaaatactacaatcaatacaggcagaaa
agtcaatccaataccaattagtgttattattgtttttcccctttccttgatgaatacaag
ctacggggcaagcttccacgcattcagcaactccaatacatttttctgttacaattgtat
gagacatattatttaattaat-aaaagataagctatttacatctagtagttttgttgtat
actaaatttacatatgctatccaggccaatatttaaggtagtctatttctacagtttact
aatttgatatgcgcctttagttcagttggtagaacgcaggtctccaaaacctgatgtcga
gggttcaagtccttcagggcgcgtttag-ttttatcgtaattcaataatatcattttatc
tttttgcaaacatcaaaatgatatgatataaaagatcatattttatacattgaatattct
atgtactagtactg-ttttttaatatgttattattgataagttaattatactaattctat
ttttgtaataagt-aaaaacagttctaattaagtatgcacgtatttaatagtc-ttttgt
atatat--aacttgaaaaaatttaaactatggctaaaaaagttactggaattgttaaact

agcattaaatgcaggtaaagctactcctgcgccacctgtcggacctgctttaggacagca
cggcgtgaatattgttatgttctgcaaagagtacaatgcacgtactgctgacaagtcagg
actagtaatccctgtagagatctcaatatatgaagatagaagttttacttttatacttaa
aactcctcctgcttctgtactaatcgctaaagctgctggactaaataaaggttcgggtga
accaaataccaaaaaagtaggcagcataacaaataaacagttagaatctattgcagaaac
aaaacttcccgatttgaacactaacaatatacctcaagctatgaaaatagttggaggtac
ggctaaaaatatgggaattctaattaaagatt-aaaagtttcaattatatttacta-tt-
-tttatgaaaaaattttcacgtcgacttacaacattaaaatcaaaagttgagcctaaact
ttacactattaatgaagcagtatctatattaaaagcaacgtcaaatgctaaatttaagga
aactgcagaagctcatattgctctaggtttaaatcctaaatatgcagaccagcaactcag
agcaacagttatattaccgaaaggtacaggtaaattgataaaagtagcagttattgctaa
gggagaaaaattaacagaagcaattagtgcgggagctgatgttagtggctccgaagaact
aattgatgaaatactgaaaggtagattagattttgataagctgatagcaacaccagatgt
tatgcctttaatagcaaagctgggacgagtattaggccctcgagggttaatgccttctcc
taaagcaggcacggtaacattggacgtagcaaaagccgtgaatgaatttaaaggggtaa
agttgaatacagagttgatagaacaggtataattcatgtaccttttggaaaatctagttt
ttcacaagaagatctagttttgaacctccaaacaattaaagaatcgattgatagaaacaa
gccttctggagcaaaagggaagtactggaaaactttttcttatctagcaccatggggcc
atctattcaaattgacatcactagccttctatgaaaaattttgtataattatagaactt
gaataatctattatttcaaatta-taatttgctagcattattaacgtaacaacttgctta
ctatatcttatctgcattaaaaacttt-ttatatatcttattaattatgagtacaaaggt
tgaaaatatcttggaagagctaaaatctttaaaccttctagaggctgctgaactagttaa
acaaatagaagaaacgtttgatgttgatgcatctgcggcttcaggaggtatgatgatggc
cgcaccaacttcagcaccagcgtctgctgaggttgaagagaagacagaatttgatgttgt
cctagaggaagtcccagcacctaaaaagatttctgtattaaaagctgttcgctctctgac
tggtttgggtttaaaagaagcaaaagatcttgttgaatctacgccaaaagtcttaaaaga
aggtgcttctaaagatgatgcagaaactatgaaaaaacagctagaagatgctggcgcaac
agttattgttaagtaatga-----agaagcgcttctgtttaagtataaacagaagtgcct
ctttatttattataagcaattatactagaaaagtccaagtgtaattgcattgttgatagg
catagttgcaccaatacctagccaaatacttacaacggtaccaattaagaaaacagtggt
tgcaattggtcttctaaatggattttgaaatttgttaacattttcaatgaatggaacagt
aagtaaaccagctggtacagcagccatacttagaacacctaatagtttattaggaatcac
tcttaatagattaaatgttggaaagaagtaccattctggtaaaatctctaatggagtcgc
aaatggattagacttttcacctatagaagatggttctaaaattgcgagtccaatactgca
ggcaaaagtgccaataattacaactggaaagacgtataatagatcatttggccaagctgg
ctctccgtaataattatggcccattccttttgctaatttagctctcagcttcggatctgt

taaatctggtttttttaagaattgacatattg-tttttgtgtttaatataaagattagtct
aattactgttttaaagtgtaattttcaactattataatggtccagaaattccttgtttac
gtatcattaaaaaatgcataagcatgaaaacggcagttagtagcggaagtacaaaagtgt
gtaaactatagaacctagttaaagttccttggcctacactgactcctcctcttaacaatt
ctactatactcgctccgacaactggaacagcatcagggacacctgtaacaatttttacag
cccagtatccgatttgatcccatggcaaagaataacctgttaccccaaaagaaacagtaa
gcactcctaaaattacacctgttacccatgtcaattctctaggtttttttgaaacctcctg
ttaaataaacacggaatacatgtagaatcatcatcaagaccatcatgctcgctgaccatc
tatgaattgatctaataagccatccaaagtttacatcagtcataatgtactctactgatg
taaaagcttctgcaactgtaggtctataatagaatgtcattgcaaatccagttgcaactt
gaattagaaaagatacaaatacaatacccccctaagcaataaaaaatattaacatgaggtg
gtacatacttactagaaatatcgtcagcaattgcttgaatttctaatctctcttcaaacc
agtcataaattttactcataaaatagcttc-aaaagctcttctaatata--ttttgcgtt
taagctactattaattcaatag------ttattgaacttatatt-tttttataatcaata
cttattaactattataacatctttttattgattt--------------------aaaatt
ctctgacttagtaaaacagggctatgtactataattcttttttttttgtatagagattaat
ttcgtttttagtaaattagctaaaattcttgttacactgactctattactacctataatc
tgtgctaaaattttatgtgtaatagtaaagttaagcataataccattattaacttgagtt
ccattttgctcagctaataataataataagctcgccagtctactggttatacttttatgt
gaaatgatttctataaaatgataagcttttgccgagcaaaatactaaatgatttacaaaa
aataaattgaatggagcgcagttactacacgcttttataatagtagtatagttaatacag
gcaacttctgttttatctattgcttctgcttcgtagtaaaaattatcgtcgaccaattct
atttgtccaaatgtatctcctgtagttaataagttaagagttactttatgtgtatttctc
aggacttttgtaataattaaagatcctatcaatataatgtataatctagagctgctatta
aacaaaagagtatcatttttcttaagtgaaaaaatttgataattaattttattacgatta
aaaaatagtagccatggattgcaagaaaaaaaatcacctgaactattttcagagaatatt
ggaatacaattttcttgatttaaacaatttttagtcataattactactttataa------
---aaaaatattctaataat-taagaactgaaattaaatgtcatataaattgatgttagt
tgagaatgacattgtattatcaaaggccattcaagaatacttaatagatcaaggcttcaa
tgtttatattgccaataatggattagaagcactgaatctagcttatcaatataatttcga
cttgataatttctgatataatgatgccgctagttaatggctatgaactattagcaaaact
taaaaaaaacaaagcattatccaaaatacctgttatctttttgacagctaaaggaatgac
taaagacagaataaaaggctacgacatggggtgctatggttatttatctaaaccatttga
tcctaatgagttgctttcaattattaacaacttaatcgctagagatgtcttaaaagaagc
ttctttacaaaacttggcaacatcaaaccagcaattaaatcataaaatacgtttgacacc
tagagaaaaaagtattttagatcttgttgttgatggacttactaataaagaaatatctac

tatactgaatactagtgttagaaatgtagaaaaatatgttagtcgactcttacacaaaac
taatatgaagaataggactttgctagtcaagtattctataaataataatctattaaataa
tgagatcaatgaaagggcgaatgacgggactcgaacccgcgaatgatggagccacaaccc
attgccttaacccccttggccacactcgccatattacacataccactatagcttgtttttt
gatatatcgtctagttttcgagttatatatttatctaaaacaatatttactttgaatttt
-tatgatacatagcaatattagtattcaaattaatggggaaccatttaattgctcaaagc
ccatctcattgcaatttttattaaattatcttgattttaattccgagcgtgtagcagtcg
agctaaatgacattctgttaccagagactctgtttcactcaacttatttgaatgatcaag
ataagctagaggttattaccattgtaggtggtggataatctatttgtacatac-------
-acagttctctatctcaaatgtttcatagatagagataatcttccttgtttttatctac
atgaataataacagctttaatagtgtcacctattttaaattgtgatgatatctgctctaa
ttctttgatattaatctcagaaatatggacaagacctttgagatttcctacctttataaa
taatccgtaaggtgtaatctgatttattatgccttcaataatattaccaacaattaaatt
agatgatgcttgagatattagagctcttctatgactcagtattaaattattagatttctc
ttcgacattgagtaattttagcttaataaatttattattaaattgctcactttttttgaaa
attaccgagatgagaattaggtacaaatccagatataccttcaaggttaattatcattcc
tcccttattgaatccttttatcattacattaagtaaagagtcttcagctaatagttgcct
aattcttttccatgctcttatatattcaaggcgacgaattgataatattaattgtttcga
ctgaatattataatctaataaaaaaaattctcttgtgtcattaatgtttaaagaagtgaa
attatttaaatcttgattactagatacttcttgaataggcaagtatgcagatattggtgt
tcctatatctactaataccccatttaattcaaagctaaatatagttccagctacaatatc
tccaagattcaaatcatacttgtatttttgtaaaacagctgcaaaatttctatgagtaaa
tccttcattattttttgtcatattgaatagtatgaattaaagagtaactttgtacaatat
attaaggtaaataaatgtttgtcgaattgtatgaaaaaattttaatttgttcggttgtaa
ttaagtatataaacatattaaaatatctaggagacatgaaaagtagacgtagatagctgc
agatttaa-tactatataatctgaggaaagtccgggctccacaaatacaatttatgctgg
agaaatcccagtgtaggtaactgcgaggatagtaccacagaaaaaaaccgccagaataag
t----tatattagctggtaagggtgcaaaggcaagttaaaagcttaccaaaagtactgca
aagtatttgttaggtaaacccctaaaatggagcaaagctactaaacaaatatttgtgtat
attttatctagtttagttgaaaatactgcatgaagttattagtaacaataactcgagagg
aatagctatcctttacaatatatc-tcgattatattgaaagaacagaacccggcttatgt
agtacttttttatgtcttgtattattatgtctattgtcgaatctttaaatttaactgtttg
acaattttttttttgtagctgatgttgtttataatctatttctgaattggttaaagttttt
tcattagatcgatatgtaaatcttagacctatactttttttctcttttccaattgattca
tctatatattgatcaaataatgttgtactttctaaatctttatcataaaattgatccaat
aattttaataagtatttaatttccatattcttaggtattattaaagagagatctcttatg

atcgatggatacttagagtatggttgaatctgataacttaaataattgaattcattatga
caatcttctaaaatgtttaagttaacttcaagaaaaaaaagttttgtgttgagtcctaat
tcactatatgttgcttgattaagttcactgaatatgccaataataatattgttatagatt
agtgtggcaaattttcctttttgaatgaactttattctaccatcgactatatcttttttt
gtccactcaatttgtctattaatcttttgaaaaaaattttctacgatccctttagcttca
taccaatttaaagaatgcgcagggtgtgaccatgtagatcggatatctaaattaccaccc
aatatgatagctagattattagtttctacaattttgttttgatcattatgaaatactgtt
cctatctcaaagctatctatagtttggttgctttgtttaatattataaagatttgattca
attaatccttccagcaaactgccgcgtaaattagaataatcttgaataagaggattgttt
aaagctatatttcctttggactttattaaagaataatgaactaattcagttaatcctaga
tttcttaaaatacttctacttttatctatgaattgtctctttgtggataactttttgttg
aattgaatatttggtactgagctcctgaatttatgataaccataaattctagcaatttct
tcaataacatctatttctcgaaaaacatcttctcgtctataatttggaacagtaacttct
atattttccttattttgataaataagatcaaaatttaaagagtgaagaatattttgaatt
tcttcaaagaataaaaaacgtgtttgaccattatacataattggtcctaaaatgtcatga
actttttaattgataaattaatattaagagcatgatcatttatttttctcgacagaaa
gtctctctaatatttccgcctgttagatctgtgattaaagcaagagcttcaaaatgtgca
tttttccaattatctacatttagcccacgttcttgtctaattgaactttctgttcgaata
ttaagaactcgagaagattttctaactacagattgtttaaatatagcagactctacaagt
atcgattttgtattataatctgtatcaaaatcagagttgcttccaataccagctatacta
gttacatttgtatttacttgagtaactaaaatatttttgtttaactcaatattttcattg
tttaacttaatctgatcattacttccaagaggaaaattgcttcgaattgttagactattt
ttatgattcatattattaattttattcaaatcaacaatattaataggctgtccccatttc
aacataatataattgcttatatctgttaacaaatttctgtgtgtaaacccagaagagagc
aaccgatttttagccaatctggagaatctttaactcttatttcgtctataatggcagca
aaataattgtcacaatttaataaatcgtgattactgattatagt-ctccttttagagaa
atatctattattgggttgatgtatagctgataacattgaggaccctgttaaagccgagac
ttctcttgatagaccaatcatgcttaacacgtctgacctgttagctgttgatgatatatc
taatatatgatcaatctcacttccaattataatagattcaatagcttcaacttcaaaacc
tgcttgtgttagttgattagttaaattgtctatatctattgttttgatattaccaagttc
tttcaaccaatttaaagaaactttcataattaattgtcatctatcttatagaagtattta
ttttacttatgattaaaaatcaatttaaaaaattcatcaaattaatataagttatttgat
gaatttgcttatacaatatattattgtgatatcttacttttatatggcagttattaagaa
ctagcttttgtaaaagttagcccattatcttgggcatatctttccataaatctcatgaat
ctatcccaatcatcaggacttttaataacatgtacagcttctattgcttgtggtttacca
ttaataaacttggcattaacatctcttgtaataatttgtccctctgtgtccatcaggtac

atacctgtgatttcacctttgtctgccatactagcatctaaaattttgggatttgtaaac
cgaaaagttgctgttcctgtactaccatctcgagaccttgttaaacgtacatctggtaca
acttcttcattgataccttgaataaattgaattgttgccatatcaatcttctttatttgt
atataatttatattggataaaactgatttctagttcttgaatgcttgttgagaaactgtt
ttaaaagcttatagtagcagtagatcaaattttatgtattatataaaactgtaacttttc
-ttttttg----------------------------------------------------
------------------------------------------------------------
------------------------------------------------------------
------------------------------------------------------------
------------------------------------------------------------
----------------------atctgttcaaagaactaacttattataagctaatcata
cgaatattagccatataatgttcgtaagt-gaaattaaggggttgtagctcagaaggata
gagcaagcgcctcctaagcgctaggtcagcggttcaagtccgctcaaccctgttaacgaa
ttaagtagtattatttaacttgttctgcaaatcagtattgatctaagttaaagtacttga
cagctaaaacaaaagaatctgagaaatattttcatttaaagaacgaacaattatcttaac
tctaaatatctgataattattaa-aaaattct----------------------------
----ctttttaaaacacaagatgatg-----aagatatttaaactaataattgaaattaa
gtttttgaagttatagaagccaat--ataatactttctattaacaga-----------tg
cttgcttctattactatctgtttacaaga--aatt----atgagtatcaaaata------
--attttagctcta-------------------------------------ctgac----
------------------------------------------------------------
-----------------------------tactaataaata-------------------
------------------tttattttagtctagacaag------atgaatcttcatggta
taacag------------------------------------------------------
------------------------------------------------------------
------------------------------------------------------------
------------------------------------------------------------
------------------------------------------------------------
--------------agctattctttgtgtcatacttatataaaattaggat-------ac
tgtt--------------------------------------------------------
------------------------------------------------------------
------------------------------------------------------------
------------------------------------------------------------
------------------------------------------------------------
------------------------------------------------------------
---------------ttcattattataactggggtgggaggattcgaacctgcgaatggc

ggagtcaaagtccgctgccttaccacttggctacaccccaacatgtgaacattataatag
cagtcgtataccctgccatgtcaatagactattgtctttatttaataattttgctgctc
ttgtagaataaaatcttttttttcaaataagatacgctatgtttaagttgcacgactgtt
atgttttcttgcttttgactgaacagccacttgatagtaattattccttctgcaatttcg
ttacttcctataataatacaagctacagctctttttttatttgcttgcttaatctgctta
ctaaaattacttgagctaacatcgatttctattttaaaaaactgctgatgtaaaaaacgc
attatttgcataccagtttcgttagcttttgtaccttgtgtagcaatataaaaatcgata
gattgattaggcaagtctatattatcttttgcaatcagtaataggcgttctaatcctatt
gcacatccaacagctggtgttgctttgccacctaattggtgaactaagctgtcatatctt
ccaccaccacatatagtatcttgcccttttgatgtcaatgtcttaatttcgaaagcagta
tcattataataatctaatcctcgaactaacttattatttatattgtagggaatattaagt
aattttagataattgcaaacgtcaccaaagtgtttttgtgattccaaacttaaaaaatca
gaaattttggagcctctgtcagtatttttgtgtattagaatcttttgaatctaaaatt
ctaattggattactagttaatctttttgtgagtcagtgtctaaatcatcatggtacttt
gtaagataatctcgtaatttaacttgataaatactacgatcctctacttttccaattgaa
ttcagatcaagtttcaagttgtgcagattgagattgttaaatatgctcatagctaaatgt
ataacttcactatctgctcttgcatctaagctaccaataaactcaatgccaagttggtga
aattgtcgttgtcgtccactttgtggtctttcgtacctgaacattggaccgctataccat
aatctttgtaaactatggtggtaactcattttattttcaataaaagatctcacaataccg
gcagtaccttctggccttaaggtaatatctctattgctgcgatcatgaaaccgatacatt
tctttatttacaatatcagtatcttcaccaatacccctatcatataattcactattttca
aaaataggtgttctaatttcttggtaatttgcacattctaataatttggaaattttatta
tgtataaactgccaatattgaatttcatcgggaagaatatcttttgtccctctaatagcc
tgaattttgccatgataatttttgattaccttcttaatatgtctgcattaaatagcag
ttgtagactgttattcagtcatgtattatacattag--aacatactgatg-taatttata
aagtacgggcaaggagggattcgaaccccgacaccatggttcgtagccatgtgctctaa
tccactgagctacaagcccacttaataagtaccttaactatatcattgttcactaatgta
tacaactatattctttt------atcttacatgtgaaatttttgtaaaagtaaaaaatt
at----gtttgtttttacaattggagctattcatatataatgcttaatttattataact
ttgtaaa--aatatctataagtttaattgtatgattctattatattggtaatcttgttat
aattttgcgctcttatttttcaattgg------ctagcatttaattaacttatatttcaa
ataaaaaaatcttatctaaaatgagtaaacaaattctatatcaggatgatgccagaaaag
cattagaaaaaggcatggatattttaacagaagctgtttctgtgactttaggaccaaaag
gaagaaatgttgtcttagaaaaaaaattcggtgcccctcaaattattaatgatggtgtaa
cgattgcaaaggaaattagtttagaaaatcatattgaaaataccggagtcgcactgatta
gacaagcagcatctaaaacaaatgatgtagctggtgatggtacaacaacagctacagtac

tggcttcagcaattgttaagcaaggaatgagaaatgttgcagcaggttcgaatccgatgg
ctattaaaaaaggtattgaaaaagcaacaaattttgttgttagtaaaatagctgagtatg
ctaagccagtagaagatacaaaagctattatacaagttgcttctttgtcatcaggaaatg
atatagaggtaggtaaaatgatagcgaatgctatagaaaaagttggcagagaaggagtta
tttctttagaagaagggaaatcaactaatacgattcttgagatcacagaaggaatgcagt
ttgaaaaaggctttatttctccttatttcgttacagatacagaacgaatggaagttcttc
aagaaaatccttttattttatttacagacaaaaaaattactttggtgcagcaagaacttg
tgccattgcttgagcaaattgcaaaaacatctcggcctttattaataatagctgaagaca
tcgagaaagaagcattagccacaattgtagttaataaattaagagggattttgaatgttg
tagcggttagagctcctggattcggtgatagaagaaaatctttacttgaagatatgagta
tcttaacgaacggacaagtaattactgaagatgcaggtttatcacttgatacagttcaat
tagatatgttaggaaaagcccgaagagttattgttactaaagactcgacaacaataattg
cagatggtcatgaaattaaagttaaatcaagatgtgagcaaattaagcggcaaatagaga
cgagtgactctttatatgaaagagaaaaattgcaagaacgattagcaaagctttctggag
gagttgccgtcattaaagttggtgcagctacagaaacagagatgaaagataaaaaactaa
gactagaagatgcaattaatgcaacaaaagcagcaattgaagaaggtattgtaccaggag
gaggagctactaatgtccatatctctagtgaattatttacatgggctaaaaacaatttag
ttgaagatgaattaattggtgctttaatagttgaacgagctgtgacctatccgttaagac
gaatagcttttaatgcaggtgataatggagcagtaatagtggaaaaagttaaaagtcacg
attttcatataggctatgatgcagcaaacggtaatattgtaaatatgtacgatagaggta
ttatagatccagcgaaagtagctagatcagctttgcaaaacgcagcttctattgcagcaa
tggttttaactactgaatgtattgtagttgataaagtcgatgattgagctata------a
gataatagataatctcaaacaaagaataatttctaattatagaaattattctttgtttca
attaatctaattaattgatcaaatgtataattttataataaagccaaagtttaaacacg
tcttctttttcaactattaagtagagaagatataagcccgtaatacctaaattatcaatt
aaaatattattttcatatatagatttttctaaagatctaaaaaatgacattttgaaaaa
taataccgaaatcttctcttatagttatttgctaaatcattactgtctaaagattgattt
ccaattgaattcagaacctttaacagaatacttttttaaatctttatctgactttatttga
gaatcaataataatttgattcaattgtatgtcaaaactaagattaaaatattacttgtt
aaagatatttgattttgaatatctagagattttatagccaagaagattaaacttttattg
ttcattgtaaaatataagacaattgatgatgtattctttaaatacta-aatttatcttct
tgtgcttctatcatttttttgt-aaaataattaatgttttaatttttttagttttagttact
acccgaaaaaataaattctggaaatctagcttgagctttagctagagagtgtttttttccc
ttcttcttgccagtaattgataacttctgttgttttgtttaataactcaattctgtctct
ttcagcaatccacggcttggattctagttctgcttttaattcttcccaagcatcattcct
tggccagaaaaaataagaagtaagtggactagtgccatttccaacaatctgatcaatagc

aatagcaatattattttctaaccaaagtacttttagtgtaaattttgacaagtttttatc
tccattattaatattaagtattgg----tacagtagaatactgtactagttatttttttc
agcaacaatttgagctttgtttaaaatgtttttttaccgttttagtagcttgtgcaccttg
agataatcgttgcaaaattttgaaatgttaacacgagtttcattatttataggattata
aaatcctaattcttctatagctttaccatcgcgtttatttctgctatccattacaacaat
tctatagcttggttgttttttttcttccgtatcgctttaaccttaacttgaccatattagt
cacctcgcttatttttt--atatctttattaaattatatcttacgacattatgcaataat
tcaaacaattaagctgattcaagtctaacattattaaatattaccattaaacattgtaga
aaaataatacctaacataggtgacatgtccattccaaacatgggaggaatgcttcctctg
aatagttttaagtatggatcagtaattctatttaatgaacagaatggttcattgtaccaa
tttacagttggaaaccatgccaatgataatttaagtaaaattaaaattaaataaatctca
gaaaaattagctatagatccaagcaataaatttaatgtaccaggaagagtgttcatacgt
attttataattattatatacttcgttatactaatagtttgattatatattataatcgatg
taaataaat-cagctaaacaaaaaacttatccttaaaatcatcttctcttgagcattttt
ttttatctacattagtatttacttttgtcgtataaatttccagtttagaccaaatatgtc
ctaactgatttaattgagatgttccacattctaaacaggctattttatattgctcaaaa
cagcacctttttttataaaatcgttcaataatggcgtaatagttgcttcatccaaaaata
aatctaagacaagtattttttgtaaatgaatttaattgatctgggacattgttaataatat
cattataattaactaaaactatattagctgtaggaagtaaagatcgtgctccttccgcta
agataaagccataaggcataataattacaataatatatttataatgactgctaataagac
ttatagtggtattagcttttaattcgattgtttctgttactagccactctctcattattt
cgtaagtaatccatttgcctagttctgagcaagcagttcttaaaatcgtacctggattat
tgtcgttttcgagaatccctgaccaatgttgtattaatggatgggatgctatattaattt
ggagttgcatgttttgttaaatatatttaacatttaatttgttgattgaatcttcaattc
ctatttgttgagtattattaaaatagcat----atatattaaagaaaataattat-aaaa
aatgaaaaaaagtctttggctgtggggttttacagatagtgctgaaacttggaatggtag
atttgcaatgattggttttatatcagttatttttatcgaagtagttacaggacaaggatt
attatatttaataggtatgatgtcataataaaaatccaaaaaagttcttgtaattttta
taagaactttttttgg-atttttataaatctaatctagcggtctcatagatagaacaggtt
cattctctctattgatacctttttcaaatccagcagcagcagctcttgctctgcctgcat
gccaaagatgtccaataaataggaagaatcctaggaagaaatgagatgttgttaaccaag
atctaggagatacatagttaactgaattaatttctgtagctacaccaccaacggaattca
aagaacctagtggtgcgtgagtcatatattctgcggcacgcctttcctgccacggttgaa
tatcgttcttgatttatttaaatcaagaccatttggtcctctaagaggctcaacccaag
gagctcgcaagtcccaaaatctcatggtttcaccaccaaaaataatttctccactaggag
atctcattaggtatttacctaatccagtaggtccttgagaagatgcaacattcgcaccta

gtctttgatctctaactaagaaagtaaaagcttgagcttgtgaagcttcaggaccagttg
gtccatagaattcactaggataagctgtgttgttataccaaacaaaattagaagccgtaa
gccccatgattgataaagcacctagactataagataagtaagcctcaccagaccatacaa
aagctcttctagcccaagcaaaaggttttgttagtatatgccagattccgccagcaatac
agataatgccaatccagacatgaccacctacaagatcttccatgttatttacacttacga
tccatccatcaccgccaaacggagactttaagacataaccaaagataactaggggattaa
gtgtaggattgctaacaaatctaacatctccaccacctggagcccaagtgtcgtataccc
caccaataaataaagctttaataactaacagaaaagcgccaatgcctaataacactaagt
gtataccaagtattgttgtcattttgtttttatcccgccaatcgtatccgaagaaaggaa
atgattcttcaagagtatcagggccaatcagagaatgataaagacctccaaagccaagaa
cagctgaagaaattaaatgcactacgcctactacaaagtatgggtatgtattgaaaatct
cgccgcctggaccaacaccccagcctagtgtagctaagtgcggaattagaataaaccctt
gttcgtacaaaggcttctcgggaacaaagtgtgctacctcaaaaagagtcatggcaccag
tccaaaagaccattatacctgcatgagcaacatgagcaccaagcaatttgccagaaacgt
taattaagcgtgcattgccagaccaccaggcaaacccggtagactcaatgtctctaccgc
caacaccaacagttgtattaaagggcgtttccacgtggtaaaacctcctcagggaatata
aagttttcatgaggttgatcttgagcagccatccaagagcgaataccttcgttcaatagg
atatttttagtataaaaagtttcaaactcaggatcttccgcagctctcaactcttgagaa
acaaaatcgtaagctcttaaatttagagctagtccaacaattccaaatgcgcttgtccat
aatccagttactggtacaaataacatgaaaaaatgtagccaacgtttattagagaaagca
acaccaaaaatctgtgaccagaatctattagctgttaccatcgaataagtttcttcagat
tgtgttggagtaaatgcacgaaaagtatctgcagcatcaccatcttcgaataaggtattc
tgtacagttgcaccgtgaatagcacatagtaaagcaccacctaagattcctgcaacgccc
atcatatggaacggattcagagtccagttatgaaatccttgcaaaaatagcaaaaatcta
aaaattgcagcaactccaaggcttggggcaaagaaccagctcgcttggcctagagggtac
attagaaagacggaaacgaatactgcaattggtccagaaaatgcaatagcgttgtatggt
ctaagaccaactagcctagcaatttcaaattgtcttaaacagaatccaattagtccaaat
gatccatgtaaagctataaatgcccacagaccaccaatttggcaccagcgagtaaaatct
ccttgagcttcaggtccccaaaggaaaagaagtgaatgtcccatgctgtttgctggggta
gaaacagccgcagttaaaaaattgcatccttctagatatgaactggctagcccatgagta
taccaagaagtaacaaaagtagtcccagttagccagccacctacagcaaggtaagcgcaa
ggaaacagaagtagtccagaccaacctacgaatacaaatcggtctctttttagccaatcg
tctacaagatcaaaaccaccacgagtttttttcttgtccaattgctatggtcataattta
atctccagagcaaatttaataagtaatttgtttaaataaaca---ctttagatatactct
aaaattttacttttctttacggtcaatataattataagtcaaaattaaccaaaattcacg
cttttaataaca-tttttttagctcattaagttaa-ttttttaatatataaacagtaata

gagttattgttt-aacttacaatactcataatttattcatttctataatcctttgaaaaa
gatatcctgttcctcgtgctgttaaaatcaagtcgggattacttggatcatcttctagtt
tagctctcaatcgtgaaatatgaacatcaacaactctcgtgtcaacatgccgttcgggag
tatagccccaaacttcttgcaagatagaagctctagaaaaaggttctccagccttactaa
ctaaaagctcgagaagactgaattccatacctgttaatcgaactctttcattatttttat
aaacttgtcttttgttagtatctattttagaaatccaatactaataatcccagagtttg
ggactccaagactagtggtgattttatcagctcttcgaagaacagaacgaatgcgagctt
ctagctccttaggagaaaatggtttaacgacatagtcatcagcacctatttctaatccgg
taattctatcgcagacttcacctaacgctgttagcattatgataggaacatctgattctt
ttcgaagttcttgacaaacaccgtaaccatctagtttaggcatcataacatctaaaacaa
caaggctaggatactcttttctaaacacgagtaaagcttcttctccatttgaagcagtga
taacttcataacctattattgttagtcttgtttctaaaattcttcttatgctggcctcat
cgtcaacaacgagaatcttttctttctggttctccaa-tttcttatttactcctataact
caccgatcatgattactataaccttaaattttagcgctatgagtatctttattcacttt
agaa-----tttagttattataaattttgcaagtctagtttacattaattagtaccagt-
-caatatttttgt-----------tttaagttatgac---------tattttaatccata
agttgactctagtttgcagaattcaatatatacata-ttttctagccaagtaaattgtat
tctattttctaaataaaaaagggttgacaaaccagttgatagaaagtattctacattac
tataag----ttaattccaacaaagaagttttaa-----atcttctttcggagtaatgaa
tactttaat
>SRR9587921
tatatgata---aattaagacgtctgttatgtgagaatttctaaaatatacaaaactttt
tatttaatatagaaaaagtaaattaaaataagttgaacatactttgctctaatctaaacc
tcttaattgag------attctaattagagacaaaagtatttacttattttgtatctata
ggaataaaaatgaaacttgcagtttatggtaaaggtggtataggtaaatctacaactagt
tgcaatatttctgtagcactttcaaaaagaggtaaaaaagttctacaaattggctgcgac
cctaaacatgatagtacatttacgctaacaggtttttaatcccaacaattatcgatact
cttcaatccaaggattaccactatgaggacgtttggcctgaagatgtaatctataaggga
tatggcggtgtagactgtgttgaagctggtggaccgccggctggcgctggttgcggaggc
tacgttgtaggtgaaacagttaaactttaaaagaattaaatgcttttgatgaatacgat
attatcttatttgatgttttaggtgatgttgtatgtggaggttttgcagctccattaaat
tatgcagactactgcttaatcattacagacaatggatttgatgctttatttgcagctaat
agaatagcagcttcagtacgagaaaaagctagaacgcactctctgagattagctggactt
gttggtaatagaacagataaaagagatctaattgataaatatatagattgtgttccaatg
ccagtattagaagtcttgcccttgattgaagatattagagtgtccagagtaaaaggtaaa
actttatttgaaatggcagaaattgataaggatttagcatatgtatgcgattactatttg

aatattgcagatcagttgattacaaggccagaaggtgtagttcctaaagaatctccggat
agagaattatttagtcttttatctgattttacttaaatcctaaatcaaaggtaggacaa
gaaaaagtagatcaagaagaattagatttgatgatagtgtaaataatattttcagcataa
tataacaaggaataagataatgtctacagctcaatcagatgctcttacttttgaatgtga
aacaggtaattatcatacttttttgtcccatcagttgcgtttcttggttatatcaaaaaat
tgaagacagtttttttcttagttatagggactaaaacttgtggttatttcttgcaaaatgc
aatgggagtaatgatttttgctgaaccgagatatgccatggcagaattagaggaaggaga
tatttcagctaagctaaatgattatggtgaactccgtagattatgcttacaaataaaaaa
agatagaaatcctagtgttatattttggattggaacatgtacaaccgaaattataaaaat
ggatctggaaggaattgctccaaaattggaagcagaaattcgtgttcctatagtagttgc
tagagctaatggtttagattatgcctttacccaaggtgaagatacagttctagcagctat
ggctcaaaggtgtccattaaatttaaagaatcaatcagataacacgagtcttaaaacttc
tcctcatattcctttagttttatttggatctcttccggatccagttgtaacccagttaac
tatggaattaaaaaaacaaggtatttttgtttctggttggttgccatcaaagaggtatac
tgaattaccggttataaaagaaggatattatgttgcgggagttaatccatttcttagtcg
cacagctactacactaatgagacgccgaaaaacaaaactaattggtgcaccatttccaat
aggtccagatggtactcgagcctggattgaaaaaatctgttcagtaatgaatgtaaagcc
tattggattagaagatagagaaaaagcaatttgggcttctttagaagactatatttcttt
aatccgaggtaaatcagtttttttttatgggtgataatttactagaagtatctcttgcaag
atttttaactagatgtggaatgactgtatatgaaattggtattccttatatggataaacg
ctatcaagcagcagagctagctttattaaaagctacatgtgataaaatgaacgttatgat
gccaacaattgtagaaaaaccagataattataatcaagtagaccgaatccgtgatctgaa
acccgatttagttatcactggtatggctcatgcaaaccctttagaagccagaggtattaa
tacaaaatggtcagtagaattcacatttgcacaaattcatggctttaccaatgcaagaga
tattcttgaactcgtcacaagaccattacggcgaaatcttagtttgtcagaattaggctg
ggatgtttatagcaagcaaagctagtattcctc----tttatgacttacatggtataaaa
gccttgtaagtct--tttattttatgttaaagaagcatttcttactctgccatcaccagc
ccaagcttgtaaactttctacttgttctttgtctgtaaatgctaaaggcacaaactgttt
gatagctattttaatatcttcagtactaaattctctctcttcactaaacgccgtatgcat
actttcaataattgcttgctcaatttcagcgccagaaaatttattacataataaactcaa
ttcatgaatatcatattcttgccacgatcttggtctaacttttgataaatgtatcttaaa
aattgactctctctcttgacgattaggtaaatctaagaaaaaaatttcatcgaacctgcc
tttccttaacatttcggaaggtaaactttgaattttattagctgttgcgacaacaaatac
tggagctttttctctgatagccaggtaataaatgtaccaaatactctggcactagtacc
actatcaccttgactatgtaaaccagaaaaagctttatctatttcatcaatccataatat
gcaaggcgataatccttcagagatatttatcatttctctcatttttgattccgattcacc

aactagtccaccaaacagttttcccatgtcaagacgtagcagaggcagcatccaatcgct
tgcaatagctttcgcggttaaggatttacccgtcccctgtataccaactagcaataaacc
tttgggcgaaggaataccgtaatttaaactttgcttcgaaaatgagcgagaccttttttg
tagccattgttttaatacatctaagccaccaatatccctatttactttactataaggata
aaattctaatagatgcgtttgattaataatttgtctttttttcttcaataataataggtaa
gctccgagagtctatctgattatattgtgcaataatttagtaatcactttgcggattct
atctatcgataatccttggcaggattttgttatattattaacgagctctgaattcaaatt
aagattcaaagctttactcaatcttatgatctctttttgatttctagaagactaggtaa
gggcaaatctattaccgttatgatatcattcagagcaaaaggtatattgattttgcaaga
aataataataatgtgcttagattgcgttttgattatttagctaaattgcgaagtttcct
aattaatactatttcgtttaggaaagagtcaaaatcttttaaaagaaaaagatttaaaga
ttcattattcaagttttcaataaattctaatgctaataaaggatttctttttgcataacc
attatcgctaggattgttagtgtatccatctacaaagtcccaggaatatacttgttgact
gtttgaacagtttaagctgtgttttataatatactctagtctatcttcttctctagtatt
tattacaataataggatatcgagattttaaaagtaatcgtaaatcttgagtgaaattcat
gtcaatatttgtttccatcagactttgtacttcaccaattgaagaaagactaaataaga
ttatctaatattccttggatatattcaaaagtc------atctctctatttctaaatta
ttgctattactaatttaacttaccatgattttttttcttccttttcttcgtcgactattt
aatatagagcgtccacatggtgttttcattcttgctctgaaaccagaaactctaatttttt
tttcttttagatccttgtagtgttcttttagtcatgatgtttcctctgttgtaaaagatt
aaatactattttatcatcttttatatatctttatgatatagatatgtattaatatcaaag
tt-aacagcaataatttatattaatcaagaattatttatataatattaaacattaaatat
----------taactttatgatattggctttgcccatttttttatttaagtattttgacca
tctttttgctaatcttaaactggttaatttttcagcagctaaaaacgattttgttattag
aatctcaatttaagtattttgttgataaaagccaaaatcgtaaattagaacccgaagaaa
gttttgcttttgcaaaagtatgtgtagctaaaaaatatttttctaaagctattattgaag
gtcaattagctttaaaaaactatcgagacctgaatatcttagataataatatagtaattg
ctaatttgtataatatgctaggttttatttattttgaagcaggccaaacaagttttgcaa
agaacttttatgagcaagccttacaaataaaccctaattatgttgtagctttaaataacc
tggcaaaaatttacgaagaggtaaaagatttaaaaaaagccgaatctctatacgataaag
tgttaactcttaacttaaataataaaactgctaatagaaggaaagattttatagcaaaaa
cta---------aaaacatctaatactatttgtaatcgggatagcaggatttgaacctgc
gacatcctgctcccaaagcaggcgcgctaccaaactgcgctatatcccgtaagacataat
tattactatatacctttctctaattatgtctactctttctgtattttatctaggatacc
aaaacatgcctttaactccatctggatctttgataaatcctaatttttttataaaaactaa
caacatcaggttctgcgaatagtgtaatagtgctaatttctgcctgtctcaactgctgta

ttaattgatgtatgattacttttcctaaacctaaaccttggaaatctggatgaataacta
cgtcccagatagttgcattaaatccgttgtctgatgttgctctagcaaatcctacaagtt
tactatttgcatcttttttttgtattaaagaaataataatagaactatttttaatgcga
tctttactttttttagaggtcttttaacccatccgactgaatcacaaagctgctccaatt
cgtataaattaatattcttattgctactcagatagatatctttaaattcgattttatcgc
aagttttatctaaaacaataagtttttgaaatttttttcagagctagcattgatatcgg
agttttgaaaaaaatttttccagaagatcataatgattatgatataatta--aataaata
tatatattacaatattgctttaacaaacaatatattaatatatgctcaaatttttacatg
aatctattaagatagaagattgttaaaatatagagaaataaaaatttttgtaaaatcttt
caataaaatgatataaagagtgctttttaagactaacatctattattataaataaagtct
cgaaattgagatgaaataacacattattctatgtctaatgccttttaaaggagttatttg
tgaaaaaatctatgtgtctaacctgtttacttgctttactaattatgagcaacccaataa
tagctaacgcagaagtagctggattggtgccctgcaaagattctgcggcattcaataagc
gtatggtaaatagtgtgaaaaaacttcaagccagattagctaaatatgatgcagacacgc
caccagcgttagctttaaataagcaaatagagaaaactaaaactagatttgcaacttatg
gtcgagcaggtttgctgtgtggtactgatggattgccacatttaatttctgatggtcgat
ggagtagagccggggactttgtttttccaggactattattcttgtatattacaggatgga
ttggctgggtaggtagaggatatcttttatctgttgctaagactagtaagccaacagaaa
aggaaatcattttagatgtgccattagctatcaaatttatgtcatccggatttgcatggc
cgctagcggcttggcaagaatttagtagtggacaattaattgcttccaatgacgatatta
ctgtttcaccccgttagtaaaaaatttatatgaataataattttaccaaatacttatcaa
cagcacctgtaattggtgtattgtggatgacgtttacagccggatttattatagaattga
accgctttttcccagatgtcttatacttttactt----ataaataa--gatcgtataaa-
--aaactacataaatatagagaaatactattatttgtgtagttttta---atatataaca
taattattta-tttttaaaaatgtttatcttttcgattattttaatattagatcttacat
aataaaata-atatcaatattatgagcttagtaacccaaattatagttaatgccgatgat
gaattaagatatcctacaattggagaattgcagtcaattcaagactacttaactacagga
agcaatagaattagaattgctactattattagagataaagaaaaggagattatacagaaa
gctagtaagcaaatttttcagttacatccagaatatatagctccaggtggtaatgcagca
ggttctaggaaaagatcgttatgcttacgtgattatggttggtatctcagactaattaca
tacggagtattagctggggataaagattctatagaaacaattggtattataggagtaaga
gaaatgtacaattctttaggtgtacctataattggaatgttagatgcaatccagtgttta
aaggaagcatctttagaaatgcttggtcaagatgatattagaatcatttctccttatttt
gattatataattcgtggaatgtcataaatcatagcttcaattaattagttgaataaaagc
ttttataatgttataattacaataagcccgaaagcatataattgtaaaaactaaatcttg
tcagaaccggaaggtagcagcaataatgtttcaatacaaaagttatga-ttttcgggtgt

ttattgtttttatctaatagtctcgacttttaattagtatgctatgtataaatctgtata
tgacatgtaatccatcaaaccttccagcaaaaataatttaataatttgatattaatacgt
aaaacacctatac-tcttttataaaatttctttatatctcatctaaacaatggaaactta
acttaatgggtgttcatattttatcaacaggttcgtctgtcccaaatttttctgtagaga
atcaacaatttgaagatatgatcgaaacttccgaccattggatttcaacaagaacaggaa
taaaaaaaagacatctagctccatcttctacttctttaactaaattagcggcagaagctg
caaacaaagccttatatgcagctaacttaaaacctactgagatcagtttaattattttag
ctacgtctacgcctgatgatttatttggtagtgctagccagttgcaagcagaaataggtg
caacaacatcggtagcttttgatattacggctgcctgctccggttttattgttgctttag
taacagcagctcagttcattcaaactggttcctatgacaatattttagttgttggagcag
acacaatgtctagatggattaattggtcagatagaactacctgtattttatttggtgatg
gtgctggagcagtagtattggggcaaagcctcaaaaatagtattttaggctttaagttat
gtacagatggtcagctaaacagtcatttacaattaatgaataaacctgtaaataatcaaa
aatttggtgttacagaaattcctcatggaaactataattccataacaatgaatggcaagg
aagtgtacaagttcgctgtatttcaagttccaacagtaattagacaatgtttgaataatt
taaacatttcaatagatgaagttgattggtttatattgcatcaagcaaacactagaatca
tagaagcaattgcgagcagattatcagtaccttttctaaaatgattacgaacttagagc
attatggaaatacatctgcagcgtcaatccctttagcgttagatgaagctattcaatcca
ataaaattcaaccaggccaaattattgttttatctggttttggagcaggcttgacttggg
gagcaattgtcttgaagtggtgat---ttatatattgcggatgacgagactcgaactcgt
aaagctttcgctacacacccctcaagcgtgcgtgtataccaatttcaccacatccgcatt
ttttaccatgaata-ttaaaaatttataatatttatctaaatattcatataa--tgtgta
tatatatatacattcttgtcagaaactctcttatattttattttaagtgataaaattgag
t----ttagtatagatagtaaaatttttatttaaaatcataa----aaataaatatataa
ggaaaataaaatatgacaccatctttatcaagtttttgaatagtcttattcttggggca
gtaattgtagtcgttcccataactttggctcttttatttgttagtcaaaaagacaggaca
atccggtcctaaaaataacttagaaagatatggagaagcaatttaaaatgaataataaaa
tatttttcatactaataagctctccatatattgctatact--ttagaagaatttttattt
ttttattttgtaacatcctgtaaagtttattccgttctttctttttccc-atttactctc
ttataattaatgatacaaatgtcagg-ttaataatag--tttttttattctgtatctttc
---------------attttatatacgtattattatgttcaattaataaagaaaagctca
tgtctttgtctaattggcctctcaaaaaagaaaattctgaagcatataatattaagaatt
caaaacaaatcacaattcctgatggtttatggataaaatgcttcgactgtggtctattaa
tgtattctaaagtactgaagagaaatttaaaagtttgccctcaatgtagttatcattttc
aagcttctagtaacgaaagaattgatcaattgatagaccaaggtagttggcaaccaatgg
atgttcacttgatctctacagatccattaggctttaaagaccaaaagctttatagtcaaa

ggttaaaagatactgctttcaagactggcctgcaagacgcagttcaaacaggtactggga
ctatgcaaggtaaaaaagtatgcttaggtattatggattttagattcatggggggaagca
tgggatctgttgtaggcgaaaaactaacaagactgctagaaaaagcaactcaagaaaagt
tgcctgcaattatactttgtgcatcaggcggagctagaatgcaagaaggtatgttgagct
taatgcaaatggcaaaaatttcttctgctctagaaatgcataaaaaagaaaatctactat
atatatctgttttaacctctcccacaacaggggggtgtaacagctagttttgctatgcttg
gagacttaattattgcagagccaaaagctcttattgcatttgctggtagacgagttatag
aacaaacaatcaaagaagacttgccagataattttcaaagttcagaatatttatttgaac
atggtttcctagatttaattgtatccagaactcagcttagatcaaagttaatacaaattt
tgtccttacataatcatagtaagtaatgaattgaatattgactataatatcaaaatgcta
gtaaaaattaagatattaagaaattttcactaaagatatttgtagactaactagaattac
aattactattgtagaaaaatttgtcttaatgcca-ttttttcagtcttttaacaaattg-
tttttatgactttaaataattgttagtttaatacaaatttataaagagaata-----cat
actaagctaaatatagtaatattatgttttcctataatactgacctatcaaatttcgagg
aaatttatgcttaaaagatcttcttggcttgcggctttattgggactattaacagtagtt
tctacaagtacgcatacatatgccatagagttagacgaggcaacaagaactgttccatta
gaatcttctggcagaactgtaattcttacaccagaacaagttaaaagaggcaagcgatta
tttaataattcttgtgctatttgccataatggtggtatcacgaaaacaaatccgaatatt
ggacttgacccagaatctttaggattagctacgccacaaagagataccattgaaggacta
gttgactatatgaaagatccgactagttatgatggtgcagagtcaatcgcagaattgcat
ccaagtattaaaagtgctgaaatttttcctaaaatgcgcaatctaaccgatgaagaccta
tttacaatcgcaggtcatatcttacttcaacctaaaattgtttctgaaaagtggggcgga
ggaaaaatttactattagaacttcaaagacctaaaccttgtgaatactagtttatgta-t
ttgttgcttaagtgtgattattttgttagatcatgatatatattaaagatggagtttgtt
gacagcaagttacttttacccgaacttgtttacactattcgtctagttattaaaaagga
gacgttaaattgaagaagaagctttcagttcttttcactgtttttagtttttttgtaata
ggtttcgcacaaattgcttttgctgcagatctagataatggagaaaaagtttttttctgct
aattgtgcagcatgtcatgctggcggtaataacgccattatgccagataaaaccttaaaa
aaagatgtacttgaagctaatagtatgaatactattgatgctattacttatcaagtacaa
aatggtaaaaatgccatgcctgctttcggaggtagactggttgatgaagatattgaagat
gcagcaaattatgtattatctcaatctgaaaaaggttggtaattatacttgattttatcc
tgtattaaagaatagacaatctatttagttgttcattagattgtctattctttgtttatg
---ctattatataaagatatttacacaatatt----ttatgatgaaaagaatacccgcaa
ttcttgtactagaagacggtgcgtattataaaggatggtcattccagcaagataaacaag
agattactattggtgaagtagtttttaatactggaatgacaggatatcaagaaataatca
cagatcctagttacttccatcaaattgtcgcttttacctacccggaaattgggaatacag

gtattaataatcaagatattgaatctcacagtattagtattaaaggacttattgcaaaaa
atatttgtaaaatttcaagcagctggagagagcagcaatctttagttacgtatttaagta
gtaataatattccttttattttcggaatagatacaaggtctttaacccaatacttgcgtc
aatttggtacaatgaacggttgtatctctactgataatttaaatcatagttacttaaaac
agaaaatttgtgagattccaagtatgcaaggtttagatttaatcccgcatgtaactacaa
gaaatgtttacccctgggatgaaaaaagttttccaaattggtatttgacagataacatta
gagtgcaccgagttattcagttgaaagttattgttatagattttggagttaaactaaata
tactcagaagactagctacacttggatgtcagataacggttgtgcctgcccacactcctt
taaaagatattttggcttaccagcctgatggtatattactctctaatggtccaggagatc
catcagcagtacattacggcatccagacagttacaaaattactagattacaatgtgccta
tatttgggatttgtatggggcatcaaattttaaatttagctcttaaagctaaaactttca
aacttaaatttggtcatagaggtattaaccatccatccggattgaaccagcaagttgaaa
taactagtcaaaatcatggctttgcagttgaattgacttcggtttttgaatctcctgtaa
gagtgactcatttaatctaaatgacactactattgcaggaactggacataatcaaagtc
cttatttttctgtgcaatatcatccagaatcgagcccaggccctcatgatgctgattatc
tattcgaaaattttatagaaataatgacaaagtccaagaataaagttag-----------
ttagtaattttcccatgctttatgagtgaataaagcggatagtacttgtactcctcgtaa
ttgattaaagagaggcaaatgtcctctggggccatgagaattccattgaaattcctcagg
atatctgcaaggaattctatcaacttcccagccgatttttttgccagaattttccccaatc
ttttccaacactaagccatatttgtctttgaacaaatagcccaaatttaccttttgagtg
tgtatgccataacttatcaatagtttgtaaatcttgagcaggtatttttttttatatctgt
aaaatatagccaattacgagtttgtgcatttacaccagctagttgaataagcttttgttg
agttagctgatctgctttgagcaaatcccgatgagtcagtaacatttgtaaatctttgta
attcatctgttgagcagagcgcaatggaacaatgccatcgggacataagttagaagcaaa
tttcactatttcttgattttttactgttaagtaacttttcataaattaaaccgtcaacaca
attacttttatagttaggcccagtaatcctctcaaaaaatagatctgctaaatcttttaa
ctcgatagagtctttgctatttatattctcaataatttcaagttgttgcttga---catt
attagacttagtattcttattcaattctaaaagctgggctcggatttgatttggcatctg
atttgtttaattagttttaatagtc----ataaggtcttagagagaattagattgttttt
tgaaattagaagagcgccgcagtatttctaaagaagttcgcataatttgattcaataagt
tttgtcctgcataaattataatttgaccaattaaaaaaattgtgctttttatttttaggaa
caaaaaaatcttgtacctcaagcagaaaagtaattattatttgcattggtgacaacatct
gtaattcattcgatctgcaagcatatatatactgacagtttaagccttcttgataaaatc
cccatactttgtaacgactttcgtaaattgctcgtggtcgctcaagatagtttttaatta
tagtattccaaattaggttattctttaatttttctaatttctatcagatagaaattttta
cattacagagataacttgcatttgtctcatttacagtattaaaattttgaactaacatat

gagcaataatgttgcttagctgaattaaataattttcaagaaaaatttcgacctgcttaa
ctggcatattgtaagtaaagagattaaaagttttattcacagtttgagatgacccaaaaa
gtagctgaataattaaaatttgtagtagcatcttatagtcgttcagtagatagttcaaat
catgaatatcttggtaaatgtcaatacagtccagttcatataacttacaaaatcttttga
tacatctatggaataaatcaattaaaatattttcggtcaagtttgtaatatcatttacat
ctaaatttgaattataaatttctaaaagtatcttttccagttcagctaatatgattttta
ataagtttctttttacttcagtgcgaaaaacatctaaaattaacacttcctgagagcaat
tagttaatctcttattaatctttatagaagttctaactaatagctctgctacctcttgat
tgagtgttggtccctgagagcttggccaataattattcacgttataccatagtaagaaat
attgaatttaatgttatacgataacttttaatatactgaaaagcaagttaattactcta
tagcaactag-------tttttacagatcttaatagttatttatcaaataaagttatatg
atagtagcgattgagctaattttattaaaattta----taataatgacaacttattactt
cgcccttgcgagtcaaaattttctattagtacaagaaccactagaagaagtttttagaga
gagagttaactactatcagtcaaataataaagcaattgatttttggctaataccaaaccc
ttcttttctagagaaaccagaaatgatttcatttaaaaaccttgtacctaaagacgctgt
agctataatctctactaatccaatatttattaattggttaaagctaagaatcggctatat
ctgtattgggcaatttgaagataacctacaactttctgaagaatcgttaaatattactgt
tttaacagacaaaatttaacaatgcttttattatgactagtgttaaattattttggttaa
gttataattatagtcttttagtattta-gactcttatagtttatcacttagtattttatg
ctgagatggagtgacaaatagtgtaagtatttcttggctaaacgtttcagttgccttaga
tttatatctatttggattaactataatggatagcattcgttttatagtaacattttcaat
ttgagcccaatgtacaattccaagttccaattccttagcaatagctgaaacggaaacaaa
ggcagctcctagccctgattgcacagcgtttttgatagcttctatagaattcaattccat
ctctatttaaagcgactgctgtcaataccatgttgactgagtactttatctatgacttt
tctaattgttgattgagtatctaacgcaataaatctaagcctgtataagtcttcttttg
aatgtctcctagtttagaaaaaggatgtgatttgggtaatataagtgctaattcgtcttc
cgcataagaagtaacctgcaaaacatcttgcaattcagtaggcacttctcctccaataat
tgctaaatcaacttgaccattagctacgctccatgaaataagtctagttgaatgtacttg
caattgaacagccacttgtggatatctttgcctaaatagtccgattaatcttggcatcaa
atatgtcccagttgtctggctagctccaataattaatgtgccaccttgtaagttttgtaa
gtcgtcaagagctcgacaagtttcttcgcagagagctaaaattctgcccccatatcgtaa
aagaagacttcctgcctcagttaaagttgccttcttattacctctttcaaatagggaaac
attcaattggcgctctaaattttgaatttgcaaactaatagctggctgagaaacatacaa
gctattagctgctttttaaaactgccctctttggcaattgcttttaatattcttaactg
atccaatgtaaatggaaggtctgtcattaaagaattgtagtataatg---------tata
ttatttagtattgatcaatgtatgcatttgccaactaaaatatttgtcattacgcttcat

tattttcaaaataataaaaagtataatgttaaataagctgtattaaaatttagtatctgt
ttgatattagatttatacaagagatatcgtgaatttaattctaactataaaatataggag
acaat---atggactctagacttttagttgtactaataccagttttagcagcagcatctt
gggcagtttacaatattggtagagttgcactacagcaatttagaaaaatgacatcttagt
tttgcttagaatata-atttttagactttatgagatagaggactttaaaatccctatctc
tattgtatatcttaataaaaaaag-aaagc--ccccttgtacttactgttattacacatt
aatattattggt-ggtaccta----gataataagtttaagctaacttatttttttattta
taattaattaaattaataacacaattatggctgttccaaagaaaagaacatctaaagcta
aaaaaaatgcacgcaaagcaaattggaaaaatcaagcaaaaactgaagctcaaagagctt
tgtctttagcaaaatcagtattaactagaaaatccaatggatttatttataatctaactg
aagcatcagatacttttagcgattagaagtgttt-ttactacaagtctgaaaaagtatta
cgcaacaagttctcagtaatttttatctag----taatcatttgaaaatcattctatttg
ataataaaataaatagcagattaaataaaatcacattaacaaattatgcagcaaagctag
atcaaactagtgaaatttggttgttcaattgtattgaaaatattcagcatattttttta
aaagtcaattaaaatcaagtcatattactaaaatttttatttctggtactagctttgaat
ataccgcaggtctaccaggattattatccagcttaacactaagtggtagactacatccta
ttagtatatatagtccccagtctctcaaaaagtatcttgaagcatgtaccaaatattccc
aaactaattttctttcctattaatttcataatttacaatacggaggacaagttgtta
accaattttatacagtaatttgtttaccgttgagcaaaaagagcctgctgtatggattta
ttattctaaaaaaagaaaagcagggagtatttaatttagcacaagctaaaaccttgaata
ttcttcaaggacctatatatggaaaactcaaagaaaaggataatttttttaagtccagatg
gttactatctatccggccaagacttctcttctaatacaataatgggacataaaatatcgc
ttccgttattagtcagatattctagaattatttctgagatgcattggttttgctcttatc
ctattagattaaatacttattcgcatcagcaaggagcaaagtgtttaccacataatgttt
taactgatattatgaaatcccaaatatatcaagataacagttttgttgaataatacttat
tcttgatatataatggtctcggtaatctaaataaaatactagaaataatttcttaaattt
atttgtctatattaaatttacaatcatttatgacatacgcaattattgaagcaagtggca
cgcagctttggatagaagaaggccgctactatgatttaaatcatatacctgttgatccag
gtcagtcgattatattaggaaaagtcttattattgaataaaaaatggggaggttactttag
gccgcccttgtatagaaggggttacgataaaggctacagtaatgaggcacttgcgaggaa
agaagataactgttttcaaaatgaaaccaaagaagaaaatgagattaaaaaaaggtcatc
gacaagaattgactcgtttaatgatcgattctataacatcttaga------taaaatcta
gactcatatttt--aa-aaacttttaatatttaactaaaatagatataatggcacata
aaaaaggtagtggtagtacaagaaatggcagagactctaattccaagcgtttaggtgtta
aaaaatatggtggagagcaagtaacagcaggtaatattttaatcagacaacggggaacta
aagttaagcctggccaaaatgttggaaaaggaaaagatgatacattgttttctctaattg

atggtttcgtgctgtttgaaaagtcaaatcaaaagcaaaaaacaattagtgtttattctg
ctaagaaatagttaaaacaattggtgcaagataacttgagt----gtatttatttcactt
acagtggcaccatatatttgatttattaagaaatggagttccttaatgaaccaatgaatt
ttattaaagatttgctgtgaaatttaatgacaaacactattgtaatttcctgtctacaca
atatggctgctattctatattgtggtcagatacagaaactagtagtagcaaatgctcatt
atcaagtaagtgatatctacctaggttgtgttgataaaatcttctcaggaataaatgcgg
catttattaacctaggaaagaatgagtacagtggttttatacatatcagtgataccggtc
cgcttaaaaagaaatattatgtcaataatattactaacattttaacaatacggcaaaaaa
ttttagtacaaattattaaagagccaactttaaataaaggtccaaggctcactgccaata
ttacattatcaggtcgatatattgtattaatgccttttagtcaatcaatctgtatatctc
gaaaaatatatgatgaagatgagcgtcattatttgaagtctttagctattttaattaaac
cggcaacaatgggcttgctatttagaccttctgctgtaggtgtcgatgaggaaataatat
taagcgaattaaaaaatctaaaagaacaatggaactttgttcaaaaatctgcaattaata
gttattcacctgttcttctatataaagatgaagatattgttaaaaaggtaatccgagatt
tttataataataatacaaacaatatagtaattgattcaaacctgggattaaaacaattaa
attattatatccacacttggcactgtaataactctagcacagttcctaagattaagcttt
atagtaataatcaatgtatactagatgcttttggtatcaatcaggcaatttccagagctc
tcattccaaaagttgatcttatacttggtggctatatgtttattgaaactttagaagctt
ttactattattgacgttaattctggatcttttaataattccactagtgcacgagaaacag
ttttaaaaaccaactgttctgcagcaacagaaatagcttatcagttacaaattagaaata
ttactggtgtaattataatcgacttcattgatatggaatcacaaagagatcaattgcaat
tattagaacactttaataaagagctatcacttgatgatgctaaaccacaaattgtacagt
tatctgaattaggtttagttgaattgactagaagaagacagggcaaaagtttgtacgagt
taattagtagtgattctaattactttttatttttcacacaatcagagagatctcagtctc
ttaagagattcgatgatagacagcagaaacaacagattttttaataaatcttggctatctg
cagagattaatactattaacaaggtctttttcaaaagtcaaatttgtgcagacctgcta
acttttacctaattcgtaatctctatatagttaaaagtagtattacttataaacaaaatt
atttattaactcacagatcaaaacttatttattctaaagaatacagcaaagtattaccaa
gca------gttattatctagctagtctcaataagaatagtaatcaagagttcctacctt
aagattgttatatcaactgtt---------------------------------------
-------------------gaaaaaaagaagctctctcttatataagagagagcttc-tt
ttgtagcttactaacta-tctagttagttaattaaaaattaaccgttaacagctggagct
gttagagctactggtaaagattcaccagaagctagatctagagggaagttgtgagcgtta
cgttcgtgcattacttccatacctaggttagcacggttgatgatatcagcccatgtgtta
attacacgaccttggctatcaacaacagattggttaaagttaaaaccattcaagttgaat
gccattgtgcttacagataaagctgttagccagataccaactacaggccatagacctaag

aagaaatgtagagaacgagagttgttgaaactagcatattggaagattaaacgaccgaag
tagccatgagctgcaacgatgttataagtttcttcttcttgtccgaatttgtaaccatag
ttagcagattcgttttcgcttgtttcacgaattaagctagatgtaactagggatccgtgc
atagcactgaacagagaaccaccaaacacaccagcaacacctagttggtggaatgggtgc
attaaaatgttgtgctcagcttggaatacaagcatgaagttaaatgttccggagatacct
agaggcattccatcagagaaactaccttggccaattgggtatactaggaatactgctgct
gctgctgctactggagcagtaaaagcaacggaaatccatggacgcatacctaggcggtag
cttagttcccactcacgaccaatgtagcaagctacgccagttaggaaatgaagaacaact
aattggtaaggaccaccgttgtatagccattcgtctaaagaagcagcttcccagattggg
tagaagtgaataccgatagctgcagaacttggaataacagcaccagaaatgatgttgttt
ccgtatagaagggaaccagcaactggctcacgaattccatcaatgtctactggaggtgca
gctacgaatgcaatgatgaatacagatgtggcagttaatagagttggaatcattaataca
ccaaaccaaccaatgtataggcggttttcagtactagtaatccaagagcagaaacgttcc
cacaagctagcgctttcgcgtctttgtaaagtagcagtcataatttttatcaatttta
aggatttaacaaagatgcttcccaaacaataagttgtctgttatggaaattattacttaa
ggccttttcaaatgagaacattttaaaaaaaaca-tagcaaaatctaatctcagtaact
aa-aattt-tacttgttcatatattgcttaataattcaa-----------taataaattt
tt-gct-------------ttttgcaaaattaacactatttagggat---ataataata
atgcatattaatgctgcttattttgattttaatagtttatataattttgccctacttaaa
tacgcatgtcacactttacaaaaattcagacgaccatacaagatttaaatctgttaaagc
atgctttaacagatttgggtttgatttggcaaatgaattctagtcatataaaagttggtg
aaaataatcagcacaaggtagatattttaattaaacaagataatttgtcacatattggct
ttacttggaatgataatagatatcacttagttgccgacttacaactttgggaacaaccct
ggtctctagaagtattttggataagttatctcagaagtatgcttactactctatcatcg
aagaaacaaagaagcaaggctttgaaaaaatgcagcagatttcacaaaaagatggatcaa
ttaagttaattgtgcaacgctggaattattaactaacatacaaatgcgggcggagagact
tgaactctcacgagattatctcactagaacctaaatctagcgcgtctaccaattccacca
cgcccgcactgttgagac--aatattactctatcataatctatactaaaacacaaactaa
-tagtataaactagcttccaagcttaaaagcgtctacaaatagaccgtatgttataccaa
ttttgaagttgttgaataaattaatttttatatttaattgcttgtggctagagtagacta
ttttactagtcaactcagttgctgcaactattagactggctgcaataataccccaatctc
ctgtttgtccaggaatcgtagacaaacctgttgaaataaaaaatcctagcaataaactaa
ttaagccagttgttaactcgctcaaagaataataaagcttattatttaagtttttatta
agtaagcaaaaaaagttgaaagtctagttttaatcatagtatttagtaacggagttttaa
aaggatgacaagtattaacag-tcattaatattgtcgtatccttttaaaactcgttaata
caaacagctgttcttgt---------ttttaaaggtcttgttcgcttaaagcattaataa

gatgctgaaacggctcttgtataatatctgcgcgtttaattgattgcaaatctaattctt
cttctatgatttcttgtaataactgtatgcttcttactgttggagcaacaggtacagata
gtgaattgtaagtatcttttagcccatttaatacacgctgatcgagaatatttgtactac
cagaaaccaaggcataactagcataccgtaggtaatattctatatcccttagacaagctg
catatcttcttgttgtataagaatttcctccaggccttaataattcaggctgttcttcgt
acaactgagctgcggcctcttttaaaatattggtagcttgatcattaataatttcagcta
tttttattcgatctagtccactcgaaaagaaagattctagttgtccgacagctgtcttat
ctagataacgaccagtaagatcgtaacgatttaatattgctgttatagcatcttgcataa
attaattctcctac-ttgctatcagagcggtattataaatgactaatattgtaatataat
cattagtgtttatgaaatttatttaaaacttattctttattgcttaatgtgcaattaaaa
actcgcatagattttacacatatagagtatacgtagtttttaatta-ttttttataaaa
accaagaaataaagctttttt-taagaggttcgtgaatttgttaatataacatgagcttt
cctcctaaataagttttgtaaagtac--aaagtaaaaaaacgattagtcaataatagtaa
------------------------------------------------------------
------------------------------------------------------------
------------------------------------------------------------
------------------------------------------------------------
------------------------------------------------------------
------------------------------------------------------------
------------------------------------------------------------
------------------------------------------------------------
------------------------------------------------------------
------------------------------------------------------------
--------------------------acaatattaatatgtatcaa--------------
------------------------------------------------------------
------------------------------------------------------------
------------------------------------------------------------
------------------------------------------------------------
------------------------------------------------------------
------------------------------------------------------------
------------------------------------------------------------
------------------------------------------------------------
------------------------------------------------------------
------------------------------------------------------------
------------------------------------------------------------
------------------------------------------------------------

------------------------------------------------------------
------------------------------------------------------------
------------------------------------------------------------
------------------------------------------------------------
------------------------------------------------------------
------------------------------------------------------------
------------------------------------------------------------
------------------------------------------------------------
------------------------------------------------------------
------------------------------------------------------------
-----------------agtataaataagcttttactccagcatcaaaaaaaatatttga
gatatgttgatacaactgattattcaagtgaaaacttagatttagaccaatttagctatt
ttatcattacaattaataaaactgataaatgtattcttatagagcaattttgttataatg
ctgaaggccatatttactcaatattcttcaaaggtccaacagctaaacatctttcttcta
taatttgtcattttcaacaattaaatacaataatatctacagctcatgccatatatttag
gacgagagctaatgaagtcagaattagctctcgtgctagatcaacaatatattcaagatt
aaa--ttttgtaagtattaattatatttatgctagtataatattagtaagggcatgtaac
tcagtggatagagtatcagattccgattctgatggccgtgggttcgaatcccgccctgcc
c---gagtattcagagtact-atactaaaacatct-------taatagctagtttgataa
tagtatgggcattggggctgcaaggtttctacattatgaaaagaagaaaatatg-aaaaa
aaacaagctctccaaaagagcttttagtacaatcaataaatgcagaaaacaatattgttt
ctttttctcggaaattagctcttgcttaaacactgttaattttggcattcgacatgtta
aactcttattcatggcgaatattctgtcagagtcgctcttagtttaagaaaagtcagaaa
aaatatttatctttacgtttt-tacctatttttatttaaataaaggttagta-ttttcaa
cttttatagtggacgtgggttcgagtcccaccagctccatcaatttgatt---------t
gtataattagctagtttaacctttttaataaccagatatataattggaggaaatatgggt
ttcataacaattactaagcccgctttaaagcaaattgcaatcttgaaaaatgatcatgaa
aatgatgtacatctaagaataggtgttagacaaggcggatgttcaggtatgtcatactca
atgaattttgaacatgttgacaaattaaaagatacagatgaacggcttcgtctagataat
ttttctgttgtctgcgaccctaaaagtcttctttatctttatggattatcattagatttt
agctcggaattaataggaggaggatttcaattttctaacccgaatgccagccaaacttgt
ggctgcggtaaatctttttcaggctgacc-ctctaacacgtttttttatatattttgtatt
atatatattattttagcaagttg-tttattagagacaagcaatgctctgttaaaac---
---------------------------taatttcaagagttttattgtattatt-----
----ttatatatttattatcaactacttgaaaataaattagctaagcaatctaatttgtt
atcagtaccaaactttaaatgattaactttttattgatatatgctaatcttt-atcagcc

taaacatatctgtacattacaatcttttgacataaaaagagtcttaacatacctctcaa-
----------------------aaataaattcaa----------tattttataaatactc
aacatgaatctgtcacaacttatcgttaattcttctatgatgaaaattcctaataacttt
tattgcaatataaaagctatttcttctgagaatttaagtcacttacaattatcaaaaaca
aatttattatctcctacaaaaataattgcttttggactacctaaatttgattttatttct
acacagattacaagttctattagagaaacagttgttgtgtctactattgatgataataat
ctttttgctgccccaactaatgattggataaaaaaacacaaggataaaaatcatcattct
ttgttttataaaaaagtttttaaagctttgctaaaccataaaattagagttgttgctgag
ccatt------tcataactccgaaataaataaagaaaagagcataatggcaagtcaaaga
tacatctggggtaaaagctggaagccttcaattattttatcttgcttaaaaaacaaaaaa
aatagcaaaatttctgatgcatcgaaaaaatttttaactgaacagttatctgcatcacca
gtatttgtagtaaaaaatggttttaatgaaattattttaggtcacccattatctcgtgtt
aaaagaggaggagttaataatctgatgcatgcgttttctaatttattaaatcagtctaat
gcaacttatcctatatctactggtctattctttttttcatccagatgatgcatttgaattt
aaagactttataatatcggtcaatccactggcggccaaacatatggagataagcgtagag
cctgttggtctacactttgcttataaaatgaatagaaatatatcgtcagatactcaattt
tcgtttattccagattttaaagaagtaggagatttattattcaaatatagaaaaggtaat
catttagtttttcataagaatcagcattatggcaaagatttttttcaggggcaaccaata
tatatgattcagccgatcacttttaaagatcgagccggaaagctaaatattatcaaattt
acggggctaaatgacacaagagaaattatttttactaatcttgaagctgctaatagatca
tgggcacattttataaagaggaattcacaattaaaatcaattaaaaatcctactctatta
gtctataatttagaaagtttttttaaaagatcaagaacgattggataatcaagatttaaat
aagtttgtggtgatcactaataaagaatcttatcttgccacaaaagagttaatagcttta
cccgattctaatagcttttctaagcacttaaaattaaatataaagcctaaactcttttttt
gtgaaactatgggtaagacggttattttccaccttaacttatgaat-aaatgctatt-gc
ctttattttctcgaataatattctataactagtaattcattgagttgtaaagcaacccat
tccctatcaatgactccattgactttccctgataaattggatttattcaattctaagtgg
ctaggaatattagctagtccagggaaagctaagtaattttctactagttttcgagatgct
tcttgatttttttacactaattgattctcctggtttacattgataactacagatagatact
acttgtccattaatacaaatatgaccatggttcactagttgtctagcggcaggaattgta
ggagccataccaagtctaaaaacagtattgtctagcctcatctctagtagctgtaacaaa
atttgacctgtcgacccttgtagtttttttagctgcttttacgtacttaaataactgtttc
tcacttaatccataattaaatcgcaacttctgtttttcttctaatcttacagcatactca
gaaggttttctggacttttgtccatgttctccaggtggataagatcttttaatcgctttt
ctactgagtccgggtaaatcacctaatctacgggaaatgcgtactcgtggccctctgtat
ctagacatattatttaaattctcctaatgtaaat-gattaag----------taatgagt

agctatttttaacacaaaaagagcattaaaacaaagc-ttttgctaaaaaatatttaca
aaaaatatagatttggacaaaa---tacattactatgc-tttttaagcgggtagcgggaa
tcgaacccgcatcattagcttggaaggctaaggttttaccactaaactatacccgcaata
ttcaacagattgtcactaactcaatataacatagttatgttaagttaagataatcttaaa
tttaatagaatt-ttaaattatttatgctccttctaaaaccacatcaagaaaacgagcaa
tttcggcagcttgttcttcaacttctgataataataatggttgaccgaccctagtcaatg
gaattactcttttatcttttgtacataaataaatttcacgtcttggatttagcccctctt
taatatctattttattgatttgatttcttttatattaaactgaagacagattttctat
tctttccagggaaacctagccgaaaaatttttacgatacctttatctttgttaaattcat
tatatccagcccctatattccaaataatagttagccacagaaacaaactcagaaagactc
ctatactgccatagaatgtcattacaataccttgtggaataaataccaaatcggttgaat
ttgtaaaaggtaacaaatcaacttgaaaataacttgatagtccagcaagaagaaagccca
aagccccaataaatattatagtagcccaccagtaattactcagtcttcgtgaacctaaaa
ttaaatcttttctaactgagtgtatagataatgttttttcataatttgttcttttaaaa
cagta-tactaatgcaaaatggaaaaattttgacaattttt--atgcgcgctttagattt
tttaaataagcttgattgcttttagtcctaaaaacagtcctgttgctaacccaaaaaaag
cagcaagaaatattatatatcctagaaaaactgacatattattattatttattattattg
ttacaatagtgtttacaaaaatagtgaaaatctattactacggtattatattacgttaaa
tgccttgctgtctacaggaaaaaactttttcagtttttagcattgtgtttatattagaa
ataatattctagtaaatttattaaacttctggagaacagaatcttatgtcagaatttatc
aagccttataacgatgatccttttgtaggcaatttgtccacgccagttagtacgtcaagt
tttagtaaaggacttctaggaaatctaccagcttaccgtcgaggtttatctccgcttctt
agaggattagaaataggaatggcacatggatactttttaattggaccttttgataaattg
ggccctttacgaggtacagatgtagcgttactagcgggatttctatcttcggtcggcctc
attattattctcactacatgtttatccatgtatggtaatgtatcttttactagagcagat
tcgaaagatccactacaaacttctgaaggctggggacaattcactgcaggatttctagtt
ggagcagtaggtggttcaggatttgcttatttattgctagctaatatacctgtattacag
actgcaggtcttagtttattctcttaagctagtaaggggacttgaacccgtaacctactg
attacaaatcagttgctctaccaattgagctatactagcattcagttcacaataacacat
aaagatatttgttgcaatctgctattttcagatgtaacaaatatctttttatttaaaca
gaactgttatgttaacac-------gtatattaattagaatctgaattatctgactctgc
agaagattcctgatcacagtttagataataacaaattgtttgatctagacaagttgcaga
ccaaccaacaggtgtttctcctgataattcttgcatatctacagaattataaaaattatc
atttgccttattactgtttgcttgcataaatatcactcccaaaacactctgaacttaatg
tgaagtaatttaacttctcaataatatattaacacaaaatattaaatatggtgattattc
tttaattgtaaagatgctgctaactttataaagtttgagtgagagtcttgatagcttttg

ttgaaataagcattttaacccatttatttgccctctcgaccatactttttttgtgttgta
aattaactttttgtagtttttttgttcgtttatgagaatgagatacggcatatccattat
tagctacttttcccgtaagctgacactttttttgacataatttt-------ctaacttata
tatttatttagagtacttgctaaagttgacttgggcacagctccaatcactgtgtctact
ctttctcctgctttaaagatcattaaagtaggtatacttctaatgccatattcagctgca
atagtaggattatcgtctgtgtttatttttactaccttaatagatgattcatattcttcc
gcaatttcatcaactacaggagaaaccattctacaaggaccacaccatggcgcccaaaaa
tctacgagtacaggtaagttattgttaataacttcttgtttgaaagaggcatctgtaact
tgagatactgacatattctttaacctttaatatatgttccttgctaagcaaatcttatca
caaatttcaagaaaaatcttctatagtgaattactattagtgttgaattatttttttaaa
aattagacattaaaaaaacttattacctatataaataaagtggtggccgagtgatatctt
gatagtgataattttttatattaaggttgaacactagaaataacttttcaagcaaactctt
gtttaaaaa--atttctctgtgaccataaaaacttaattatattacaatatattttatta
tttgtaacttagttctgataatggtataaacaacgcaaaagatactgccttataatcaag
gaggaatacatgtctcaatccgtagaatcacggactaggattaaaagcgaacgttacgaa
tctggagtaatcccctacgctaaaatgggctactgggatgctgactatgtgattaaagaa
acagatattctagctcttttcagaatcactcctcaaccaggtgttgacccgattgaagca
tctgctgcaattgcaggtgaatcttcaacagctacttggacagttgtatggactgattta
ttaacagcttgtgacttatacagagcaaaagcatatcgagtagatccagttccaaacgtg
gcagatcaatattttgcttacatagcttatgatattgatttgtttgaagaaggttccatt
gcgaacttaactgcttcaattattggtaacgtttttgggtttaaagctgttaaagctctt
cgcttggaagatatgcgtatgccagtagcttatctaaaaacgttccaaggtcctgcaact
ggattgattgtagaacgtgagcgtatggataagttcggtagacctttcttaggtgctaca
gttaaacctaaactaggtttatctggcaaaaactacggaagagttgtatacgaaggcctg
aaaggcggtcttgatttccttaaagatgatgagaatattaactcacaaccatttatgcgt
tggagagaaagattttatattctatggaaggtgtaaataaagcatcggcttctgctggc
gaaattaaaggtcattaccttaacgtaacagccgcgacaatggaagatatgtatgagaga
gccgaattctctaaagaggttggtagtatcatttgtatgattgaccttgtgattggttat
actgcgattcaaagtatggcaatttgggctcgtaaacatgacatgattttacatttacat
agagctggtaactcaacttactctcgtcaaaaaaatcatggtatgaacttccgagttatt
tgcaaatggatgcgtatggctggtgttgaccatattcacgcaggtacagttgtaggtaag
cttgaaggagatcctttaatgattaaaggcttctacaatactctacttgaaagcgacaca
gatatcaacctacctcaaggtctgttctttgctcaaaattgggcttccctacgtaaagtt
gtaccagtagcatctggtggtattcatgctggtcaaatgcaccaacttcttgattactta
ggtgatgatgtagttcttcagtttggtggtggtacaattggacatcctgatggtatccaa
gcaggtgcaactgctaacagagtagcactagagtccatggttatggcaagaaatgaaggc

cgtaactatgtagcagaaggtccacaaatcttgagggacgctgctaaaacttgtgggcct
ctacaaacagctttagatttatggaaagatattagtttcaactacacttccacagataca
gctgatttcgttgagactccaacagcaaacatctagtttaatgactacttactgatactt
taaatagtcaattgtaagtggaattaacttataacaataaggagcatagaatagtgagac
taacacaagggacttttccttccttccagatttaactgatgagcaaattaataaacagc
ttgcttatatcgtttctaaaggcttatcagcaaacgttgagtatactgacgatcctcatc
caagaaactcctattgggaactgtggggtttacctttatttgatgtaaaagatgcttctg
ctgttatgtacgaaattagctcatgcagaaaagcaaaacctaattattatgttaaagtta
acgcttttgataatactagaggtattgaaagttgtgtaatgtctttcattgtaaatagac
ctgctaatgaaccaggattcttattacaacgccaagacttcgaaggtagaactatgaagt
atagtcttcatagctatgctactgaaaagcctgaaggagctaggtattaatattaattaa
gaattaatattggctaatta--ttaccctcttaaaaatcataattaatttagtaattgtg
atttttaagagggatagcttcccaagtaaattttgactaaa-tagattgaataattaaaa
atacacagaaataatgcaatcacaggatataatttccaacgatactcttgttaatttaca
agaagaatatgatagaacacaaatccaagaagttttaaatgagttaaatcaagaacttat
aggattagtgcctgtaaagaccagaattcgcgaaattgctgcgctattattgattgacag
attacgcagaaaactagaactagtttctggtaatccaggattacacatgtcatttacagg
tagtccaggaactggtaaaactacagttgctatgaaaatggctgatattttgcacagact
tggatatataaaaaaagggcatttgttgacagttacaagagatgatcttgtaggtcaata
tattggacatactgcccctaaaactaaggaagttcttaaacaagcaatgggaggagtttt
atttattgacgaagcttactatctatataaagcagataatgaaagagactatggctctga
ggcaattgaaattttattacaagtaatggaaaaccaaagaaatgacttagttgttatctt
tgctggatataaagatagaatggaaaaattctacgaatccaacccaggactctcttctag
agtagctaatcatgtagacttcccagattatacttcagatgaattattacaaatagctaa
aatgatgatagaagaacagcagtactgttttacagaagaagcagataaaactcttttaga
gtataccgagcgaagaatgaaacagccttattttgctaatgcaagaagtattcgcaatgc
tattgacagggctagaatgagacaagccaataggatttttgccagtggagaaaaagtatt
aacaaaagctgatttggtaacgattgaagcagaagatatcttgaaaagtagattattttc
attacctaatgcttaatatacacgtgatttcatgaaactatatattattaaaaag-tttc
ataatcagttgcaaatttcttgaaaaacctttattattatcttaatgcaggtg-tggcgg
catagccaagtggtaaggcagaggattgcaaatccttcatcccccagttcaaatctgggt
gccgcctagtactaaaaaagggggggtgtggtggaatggtagacacaacagacttaaaatc
tgttgatttttagtaatcgtgagggttcaagtccctccacccccatatattgatttataa
ataaaaagagatgtgtatttgtattaattgcaaccatattaccaaatgtaatacctatca
cttaatagaatctcaacataaacagcctcatcttactagaagtccattgtttatacctaa
atatcctgtagttcatgttaatatatctaacaactgtacctataatcaaattgattggga

tttagtggagtgtttatcatttgtagaaaaaccgaatagttggaatttagacgctaatta
gttacaagaaaaattatgtaccataagcaacaactctatttttttagatactcggtattaa
ttttggatgttctaaccagtaagattttacccgttcttgctatctctataatgccaaatt
tagttagtaattgttcaatagcaacaatctttcccggatctccagtaacctctataatta
aaagatcttctgcaatgtctactatcttagctctaaaaattcttacaatttctaaagctt
ctgttctagtctgagaattgatctgaatcttaattaacattagttctctttcaactgaag
gaatatttgttacatcttgaacgttaaggatatttactaacttatataattgtttcgtaa
gttgttcaatagttctattgtctccttgaactaccattgtgattctagagaccccaattt
gctctgctggtccaactgctaaacttgcgatattaaaccctctccgggcaaatagaccag
atattcttgacagtactccggcttcatcttgaactaaaactgataaggtgtgtttcatga
aaaattaaatttaatctatttgatctgttaagtatattctaagaaaattacagaaaaaat
actagttagtctataactaaattatccaaactaaacaaaatcctactatcagataatata
tctaccttgtaatgctatactattaacttagtatttaatgatattctaattgcaaataga
atgct-aattttttaaaatttgttaggaacttgtgctagaaaaatacaaaaacagtaaaa
aactctctcgagatttacctcaaatcaatgatcgcattagatttccaaaagtccgagtaa
ttgatgacgaaggtgaacaactaggtatttttgtgcctgaagaagctatacaattagctg
tccaacaaggtttagacttagttgttgttagtgataaatcggacccgccagtatgccgaa
tattagactatggtaaatataagtttacacaagaaaaaagagctagagaagctaagaaaa
agcaacataacagtagtattaaagaagtaaaaatgcgatataagatagaagagcatgatt
ataaagttagaataaaccaggcatccaaatttattcaagcaggagataaagtaaaagcaa
ctatcacatttcgggggcgtgaaatccagcactctaatttagctatagatttattgaata
aaatggcaagcgatctaaatgcaatagctgaaattcagcaagctccatcaagagatggca
gaaatgtcataatgctcttatctcccaaaaaagttagctaaactaattttttatttaaatg
catctggctggattcgaaccagcgacgtctctttcgaaatggcggattatgagtccgctg
ccttcggcccctcggccacagatgcatcacttaaagtttatacctcataaagcatataaa
aatcaatgtttagctatttttaataaagtctataagagtctattattcattcatattgtg
atatgtcgcaactgctgaaggggaaacttgatttaagtatcgaaatatccagtatttaaa
aattgtgtcgagtataactggaaaagttgaaataaataaaaaaataaaatctcgactttc
cggaagtcctaaatgtcttaaaattacttcaataataacttcccatccatgaggagaatg
aaaacccacaaacatatcagtaaaaagaatgattaaaaaagctttagctgtatcactaag
accataaataatttcatttaaaaaagatttaacgactgcaatttgtctttgtccagttat
cattagtaatataaagacaagtattgatacaagatcggataaaatatttttaacagcatt
tgcactttcatttgcataatattctcccaactctctagccttcagctgcactctttttttc
aatgatttcgtaagatatatcttccgaaggattcagaagaacttcaaagtgaattttctc
ttcaaatctttgtaattcagcaaaagctctttcctcttgagaggaatttaaaaaaatttt
aggctgttcttggttccataaataatcaatacaaggtccaaatacaaaaaatttttgaagc

ctgattaactagtactggagagatgaataataataaaatatattttacggaagtaatagt
ttggtgtctagatattctaaactcctctatagcttcagattcgccattaggatctaactc
tttacgaaacttttcaaaagtattagtaattgacctaggtattggcccaactttttcaaa
ggcgaattgattatttcgttttaaattccaatatttcatttttattgctgctaaaatct
ctattaatcaataattgatcacgtattcactataaatattaattctagaactaaacatat
aaaacaatatttatgaaaaataaaatacaattaaaaattaattatgcattagttctatct
gcagctacagcagtttatggatataaatcttatcagccaactataaatacactcttaatc
aaagatgaagaaaactggcctttttctctaaaaggcagactgcctttattcctataatt
gtattaaaaaaaaatcatcaaaatccctgaactcagaacctagaaatagtaagttatatc
ttaatggaattggaagtagaaaatttatcgacagaattgtcactctacttggatatttag
aaaaacaggcaaactatactgaatacttgaaccaaataaaaacaagtggttatttttctt
tagtctacctagacagtgatataatgtataagaaaaatattagtaacatttttttactac
ctaatgatattctaaaaagaatctatatttgcaataagaaaaaaaagcttataccgcatt
cattcttaataaatttatttcagaaacagataggctatccaaaaagttttgctaatttaa
attgggccttctccaaaattataaaatggtattatgatagaggataccaatggtctctag
ttgaagttaaacaagcgtctgatgcatcttctattgtaattgatattcatgaaggggtag
tgaagacaattataacagaatactacactctatcttataaaagagtctcaggtatcttat
gcgtagagtcaatagagcagtaccttggagtaagagtgggagctccactaaatataattg
atttacaaaaaaaaattacttacttaaaagataatcaactagttggcgacattatttata
gtattgaacgatcaaataatagctcaatgagtttggatatcaaatttcaaatacaagaat
taaaagataaagagataatagtgcttgcaaaaagttcttcta------------ttatct
cgcatgcatgtaatttacttaatc---aatataga---------aataggctagtagctt
ctaatatagtttcactatcaactaataaattacacgcatgctattttaattataaacttg
actatcaatataagtatataaacacgtctaatcttataaaattgttagcttattctattt
cttgtaggaagacactacttattcattattttaatttacaaactttaataagctatacaa
aaaaaaatactattggatttcagctatatttacgaaatctaagttttgggaaagctttct
gtgtgcttagtatgaaatttatcaaaaatggacttaatataaaaatttttatatataaatc
cttcattgatagttgatcagaattttgtatttcaatttgctatacaaatcattaagcaat
atcatactgcta------aacctccagctt------------------------------
------tatttttaacgaatctagatcttgaacaatatgttgctgaaagtttattaatgt
accattttacatcttgcttctcaatatctgaaaaaatattattatctcgaattatgcata
cagattctttatttttcaactctgaaa------attttcattttgatgatagaactaacg
cgatcaacagttatgatacttttaaacaaaatacaaaaatatttttatcaagagtttcttt
ctttattattaagtttacgttatcaaaatttttaattatttaggttggcctttaaaaggcc
attttttgaaataaaatctttatacttggctccatttcaaaagagtg------------
---atttttctgatagtcgtaagaccctattcttccacaaaatgtcgttaaagcaagtat

ctaattttaatttaccaatatcttttaaaagccaccttaaccatattttagttagtacta
ttaaatgtcagtcaaatttgaatatgagaactgtatctttgctattaattgattcacctg
ctgaatatatgctatacaaatctattcttaatttctctattaaagttaggatgcagtatt
tcatacctatgagcaacaatataagattatctctattttataattatttagattgttttt
taataagatcatctcagagtcgcattcatatatggcaagatctaagaactcttacaccaa
ttcaaaattttggttaaaaaaattttcatatggagctggaattcagcttaaattaccaa
ttaagcagatgccacctctttcaattgaatatactgtaactagtagtcgctacttttgta
tttaccttcgtacttattatcaacgataactattactaaaatgaaaaatttaatttcttt
ttttgatcgcagcaaagggaaatggatttcacaaagaacaacttatgaattatctaataa
aaacatgagttcgatacagtctcagatgacaatgaaaataggcaactcattgtcaggatc
tataatacttgcatcgttaaattgggggacatttatagacaagttgcccattacgctaa
aaatcacagccggagtgaatatgataataagtttaacttgcaattcagtaatcaactaaa
taatcataaattattaacgctgtgcatagtgacagatcctagtctaattagttttaaaac
tcgctatggaagtactaccatagatgaaacatattggttcgcaacaaataatttacgtct
aagtactagtattgttaaacgattcaacacttgtgtggcagtatctttttgttcagaaat
taaagtttaa-actctatatagaagcaactaatagcctaaa-ggctattagttgcttcta
tatatctaaattttgtattaaaaattat----aatactatattattctaagtctttacga
ttgggattcctggaaggatcattagagaggaacccaaagacaaacaaagaaataaagaaa
ataacggtagtatatacaaagattttcaaagtaaacatattttcatccttaaaagctatt
agtgcg----atatttacatatctccacataattttatatcatgatataagttccagcac
tattttataaataaatcaatatctttatttttagtaatgatttatatttaaatatgcgt
gatgtactaaatactaaatcttttccgacttgttttgtcttgtttgtaaatattttgtta
tttctatggagtttaccttcaattatcacatagtctaacttctttagctgtttaaaagta
tatgaagatttttgttccaaatagttagattgattataactaattgttttctttttaac
agtcttgcttttaacttaattatttggcttttattttcactaattctgatactcttacaa
ctaagaatttggactagaagagtacagctattcataatttataacaa------tttttta
aaaaatctactaaaatagatgattttacagaactaaaatgatatttcattatcttgttga
ccttgctcataagtaatatctttcaatacttttaatatttttccaaaatactcttgcagg
tatttctctaaagatattacttccaacctgtcaattccaagaatactgtaaacttcatcc
atcggagctgtaaattgctctccgctacttaaaatttcggcaaatgctaaacgatcagaa
atattccatgtccattgaagagttttagtaattcttcttaaggcttttaataatccaatc
ggaatttgtgagatctgtgtttttttgcccagataatttttcacataatttaataatttca
gctgatgtccaagcggtattaccaactaacggcagggtcctattttctgtggaaggtact
ccaagacttttataactaactttgctgcatcttgagtatcaatgtatgcaatgggtgta
gattctcctgtaacccagactgatttttttgtctaaaataggaatggcatattggttgatt
aaaccttgaaaaaatcctcctaaagaaaacacagtatatttcacattggacttctgaagg

aaatctactacctgagactttaagttcatcaatggaacatctgggtatttctctgaattt
aatattgaaaagaaaataaatctttcaactttagccgctttagctgcttcaattaatgca
attttcccatctaaatctatttttttctgcattgtaaggatcagtaggacgagatgtagaa
gcatctatgattgctgttaccccacaaaaagattgcaaaatactttcgggtaattttaaa
tcaccatatataagttctgctccccactctttgagaaaggcagactttcttaaatttctt
accatgcattttacattgtagccttcatctaaagctcgtcttacaatttgacgtcctaaa
gttccggttgctccaattactaaaagagtcatattagttaaaaataagtatgtataaaga
cattgatttggttgacggcagagagtaaattaggtagagagggaatcgaaccctcatgac
cgaagtcgtcacattttgagtgtgatgcgtataccaatttcgccatctaccctattatag
accaactactacaagtaatttagtctttagcgtaaaaataatactaagtttcaatggata
tgtcaacttttaaatttatattaataaaaaaacggagtctcttatggccacttttgaatt
aattccttatagattgtatttgagtcaatctaacacatggatccatagaataaaagcaga
aattaagatatatatagtaacgcttttatggatttcaatttttatttttttcttactttaa
actatgtattattgctttaagcttaattgccataagttttactataagaagtaaacagaa
tattatccaaaaacatttgttacaaactttattaataacgttcttaactactgttttgtc
ttttagtgtggctattagttataaacaatatgcagaacaagaacaatcgcagtatttatc
tgattccaagaaatataaaaattccagcagctactattatatacaaatagctaacgatca
aaggatcaaaagacaatacttaattactactttaaagccttccttgtactttttttattac
tatatattctattaaactagttatgataacaacttctccagaagttttagtaattactat
ttatagatctaggataataaataaaatatttaaaaatgaattgctatttatcttccttct
ttcgtcacatattgttactagtattattaatagaatcgataaagtgattcaagtaactag
cttaagaggaagcttgaatttatataattcattgacaaggcctttaatgttttctttgtt
aatatttcaagtcttctttttggagattattcgggagtcaaaagaaatagctcaagctct
ttatactagaaatctcaatcaagaaaacaataatttttttgaaaatatatacagtaaaatc
taactttagtgatcggctcaatataattattagcactttgtactttattattttagctct
agcgtaatagaatccacttattcttctgactctagtaatacgaagagtaataaaattagt
tagtagtttaaatagatagaat-taagatattatataaataattaaatgtttcattatgg
ttaatactcaaaatcaaatttcacaaacttcagaacttgattatatagttaatcaacctt
ataaatatggctttaaaacttctgttgaatctgagcaatttcctaggggaataagtgaag
atattgtccgcttgatctccaagaaaaaagatgagcctgaatatctattgaattttaggc
ttaaagcatataaaaaaatggaaaaagatgagtagtccgtcatgggctcatattaagcatc
cgaatatagactttaatacgattatttattacgctgttcctaaattaaaaaaagaattga
aaagtctagatgaggttgatccagaaattcttgacacttttaacaagttaggtatatctt
taaatgagcaaaagcgaatttccaatgtagccgttgatgctgtttttgatagtgtatcta
ttgcaactacttttaagaaagaactgtctgaagctggtgttatattctgttcaatttctg
aagctataagggattatccagaattaataaaaaaatacttaggtactgttgtaccagctg

gtgataattattttgctgcattgaactctgcggtatttagtgatggctctttttgctata
tccctcctaatacagtttgccctttagaattatcaacttactttcgtattaacaacgaag
aatctgggcaatttgaaaggacactaattattgctgatcgtggcagtaaagtaagttatc
ttgaaggttgtactgctcctcaatttgacacaaatcaattacatgcagcgattgtagaat
tagtagctcttgaaggagccgaaattaaatattctacagtacagaattggtatgctggta
acaaagaaggtaaaggtggtatatacaattttgttactaaacgaggcttatgctcgggta
acaattcaaaaatttcatggactcaagtagaaactggatctgcaattacctggaaatatc
caagctgtattttagctggtgaaaattctcagggagaattttactctgtagctttaacaa
ataactatcaagaggctgatacagggactaaaatgatccatattggtaacaatacaaaaa
gtagaattatttctaaaggtatttctgcgggtagatcgaaaaacagttatcgaggcctag
taaaagttggccctcagtcatttaattctcgtaattattctcaatgcgattctttattaa
ttggtcaatcatctcaagctaatacatttccctatattcaagtacagaatccaacatcaa
aagtagaacatgaagcatccacttcaaaaattagcgaggaccaaatttttatttttttac
aaagagggatcaatttagaagaatctattgctcttatgatcagtggtttttgtaaagatg
tctttaatgaattgcctatggaatttgctactgaagctgatcgtttactgagtttaaaat
tagagggaactgtaggatgagccagactattttagaaatcaaagatttatatgcttctgt
tggtgaaacaacaattttaaaaggggtaaacttatctattcgagccggcgaaatacatgc
gattatggggcctaacggttcaggtaaaagtacattatcaaaagtaattgcaggacatcc
agcgtattcactaataagcggagacattttatttttttggacaaagtattcttgaaatgga
gccagacgagagggcaaaagcaggtatttttttagcttttcagtatcctgttgaaattcc
tggagtcagtaattctgatttcttaagaattgcattaaatgctcgcagaaagtttcaagg
attatcggagtttagccccttagaatttttttcaactaataacagaaaaaatagatcttgt
tggcatgcaagaaagtttttttgacaagaaatgttaatgaaggattttctggtggagaaaa
aaaacgtaacgaaattcttcaaatggctttactagatagcaaaatatctattttagatga
aacagattctggactagatattgatgcattacgagtagttgcaaaaggaattaacacttt
agctaaatcaacaaattcaattattttgattactcactatcaaagattattagattatat
tattccagattttgttcatattatgagtaacggacaaattgtaaaaactggtagtgttac
tcttgcccaagatctagaaaagcacggatatgactggattacgcagacataactatttac
tac-------aatattgagactaaagaatattctttagtctcaatatt-tttgcaaatgc
aaagtt-aattatttaagctttatcaaaaccttgcttaacatcttcaattgctcttttta
gcaattcttcactaactggttctaatttttttagtactacgaacattttcaccaaattctg
gtttagaattctttaaatcttctcgaagttctctaataaattcagctacttttgaaactt
caatatcatctagatagccattaatgccagtgtaaataatagcagtctgttcttcaactg
gaataggagaattttgagcttgtttcaaaatttctcgaagacgttgacctcttgctagtt
gattttgggttgctttatctagatcagatgcaaactgagaaaaagcttctaattcagcaa
attgtgctaactctagttttaacttgcctgccacttgtttcatggcttttatttgcgcag

cagagcctactctagagactgagatcccaacattaatagctggtctaatccctgagttaa
ataagtcaccagatagaaatatttgtccatctgtaattgaaattacatttgttggtatat
atgcagatacgtccccagcttgagtttcaataatagggagagctgtcatactacctccac
ctaattcagcattcaatttagctgctctctctagcaatctagagtgtaaatagaatacat
ccccaggataggcttctcgtcccggaggtcttctaagcaaaagagacatctgacgataag
cttgagcttgcttagttaaatcatcataaataactaatgttgctttacctttgtacataa
aatattcagctaaagccgcacctgtataaggagcaatatattgtaaagtcgcaggactat
ctgcattagctgcaactataatagtataatcaagagctcctttatcctgtagtgaagata
ctacttgggctactgaagatgccttttgtcctatcgcaacatatacacaaacaacatctt
ggccttttttgattaataattgtgtccaatgcaacagctgttttacctgtttgacggtcac
caataattaattctcgctgacctctaccaattgggatcatagagtcaatagcagttatcc
ctgtttgcataggttcacaaactgattgccttccaattataccaggggccattgactcaa
ttagccttgttccattacttgcaggttcacctttatcatctattggccgagctaatggat
caacaactctacctaaaaaagcatctccgaccggaatttgagcaattcgaccagtccctt
ttacggaactgccctctaaaatatccctaccatctcccatgagtactacaccaacgttat
cactttctaaatttaaagcaaccccaatagttttatcttcaaattcaagtagttcaccag
ccatgacttcatcaagtccatatactcgagcgataccatcaccaacttgtaaaactgttc
ctatattggctacttctacatcctgatcatacttttcaatttgttgacgaataatactac
ttatttcatcaggtctaatatttaccatattttaatattatttttattaattggaaatt
ctaaagcggtaattagtcagactaatgctatctgtctttttttactttattttaagttact
acaacatcaaggtgtgaagccatttgtcttagctgtcctctaatactagtatcaatgact
tttgatccaatctgaatagtaaagccaccaattagctcgggctcgactgaaataattagt
ttaacttctttagccttagtcataaccttgatttttttctgttaatagagtttcctgatca
gggcttaaagcaatagaagtattaatatttgcaattgttaaagattccatttggtaagcc
agttccaaatacttactagctatagcatctaacattccaattcgcttgcgatcaactaaa
atcattagaaatgataatgtattttcgttaatttgatcgccaacagttgcaataattact
tctttttttgcctctacagtttttaatggattagct---aaaaaaattttttagcttgtca
gattgcgctaaaatattttgtattgactgaatatcttggctcaccttttcagtaactttt
ttagttttagctaaatcaagaagagctactgcataaggctgagctatttttataacaaca
ttgttactgctcatagttgatctcctaacttagcaatattattatcaattatacgcagtt
gcatttcggagctcatttgattttccagctgcaaagtaactcttttaagagcaagaaatg
ttatttgttgctggatttgccttcggatctgtttttcagctgtttctatatttgatttac
ctgttatggcaagtctttcaatatctaattggccttgagctaatatggaacttctaactt
ttccagctgttagctgagcttctttcttaatttgatctataatgatttgagtttgagcaa
gttgtttttctgattctgacaatctagcacttgcttgctctagtctttcttcggattctt
gtatagctgctaaaacttttatctgtcttgcatttaaacttgatccgagaaattgcttca

aaacataaataagaccaaataaaagcaacaatatattaataacatttgcttcaaaaatat
cagaattaaagccgaatttgtgttcactactatgttctgataaaatagtaattatttgtg
gtatttttactatattattcatttaaaattct----------tactattttaatatttta
ctctggaatcatgattttcaagattgactacttaataatttaactttaatttgatcgctc
aaagtgtcaacctgattttctaaagttttaagagcttcttcttttgaacatttaactgt
ttcgatgcttcagcaatcagtttctctgcattcaattgagctttttttatatcttctgag
actatattttgagcttctttttgagaagaggctatttttaattgtgcattgcggcgagct
tcagataaatcttcttcatattttgcagctagctcatctgctttaacaagcatagacgat
gcagtagtcaatgttgttcgaatatattcatctcgttcgtctagtactttagttacaggt
ttataaaaaatagtatttaataaaaccattaaagtaagaaactgtaatgccattaaaggc
aaagtaccattgaagtcaaataagccaccttcaatttcttcggctaatagaaatggtaaa
tcaatcattttactaattatgtttatagttaataattttgtttcaattaagaaaaataac
tttatagtgcgcttagtctcatcagactaagcgcaaatgattattttaaccaacgtatgg
attagcgaacaataaagacaacgcaacaactaggccgtaaattgttaaagattccataaa
tgctaaacttagaagaagtgtaccccgaatctttccttctacttctggctgtctagcaat
accttcgacagcattagctgctgcactaccttgaccaatgcctgggcctattgcagcaag
acctacagcaagaccggcagcgataacggatgcggctgaaacgattgaatccataatgaa
ttttaatttttttgttatatgagtagaaaattgacaaatttcgtataatacttaaaatcta
tataattaagtataatagattctaatat-----tgtaattatagtatattactcttctcc
atgaccttccattgcttctccaatataagctgcggacaaagtagaaaagattaatgcctg
tattgaactcgcaaataatccgagtatcataactggtaatggtattaaaattggaattag
caaagtaaatacagaaacaactaattcatctgctaagacgtttccaaataatcgaaaact
taaagatagtggtttcgtaaaatcttctagaatattgattgggagtaatactggagttgg
ctgaatatatctagcaaagtagcctaacccttttttacttaaacctgcataaaaataagc
taaagatgtaagtaaagacagggccacagttgtattaatgtcatttgttggggcagctag
ctcaccttcaggcaagtgaattaacttccaaggaattaatgcacctgcccagttacaacc
tagaataaatagaaataaagttgcaatgtaaggcacccaagggcgatattcatgctcacc
aatctgatttttttgcaatatcttgcagaaattctaatataaattccataaaattctggaa
tttttcaggaattctttgtagatttctagtacctagaaatgataaagttaaaagtgtagc
aattactaaccaagagacaatgaaaacttgcccatgcaattgtaagctacctattttcca
atatagatgttttccaacttctactgctgataaactattatatgggttaaaatctatgag
actgttttgatacataataaatattt-ttttaaatagtgcaagaatcaaaaaaataag-tt
tttgaagaattaatatgcaattaatttattcttaaaattagactattaataaattaaaag
caaaaataaattctaatagagatagctatttttagtagtgattatttttttgtaaaaaaca
attattgctacctataatgcttttacgtttaattatatacccatatattaattatttaat
atgattataggtattttgctacaaactttattgttttatccattct----aaagattgtt

atagtttatttttatttaaaatatctgcgacttcatctgcaaaattaactttcaaattt
tcttctccccctcccagaagaaatcttacaaatcttctaattttaatattttcacctaga
acagcaatattttggtttattaaatcttcaatactaatatcttgattgcggataaacatt
tgatctaataatgagagctccttaagtctcttttcattcgtccttcaatgattgtttcg
attctatcaactggtttgtttttcaagtcatctttaccagcttcaactctttgttccaga
ttaataatttcactaggtatatcctgtgttgatacgtattccacattaggcgatgccgct
atttgcatagcaatatctttagctaacttttggaactctggacgtcgtgctacaaaatct
gtttcacagttgacttcaactaatacacctattcttccgccagtatgaatataactctct
agtaaaccttcaatagcggtacgattagatttcttgttagctgaagccagccctttttgt
cttaatgattctaaagcttttcttcattgccattattagcttgtaaagctttcttgcag
tccatcatacctgctccagtcttatcacgtaaggcttttacgacttgagcagaaatttgt
agtgtcatatgctttcaaaaattttaagtatttaaatt----------aattattataa
ttcactttgagtaggatgagttatttcttctaaatctgtttgcccgtatttaccatcata
aattgcatctgctatttacctacaattagtttaattgacctaatggcatcatcattagc
tgggattggaatattaataatttctggactacaattagtatcaagtatacaaatagttgg
aatacctaattttaaacactcttgtatagctgttgtttctctcttttgatctacaactac
aacaatatcaggtaaacgagtcatattttaataccatttagatgtttacgaagtttatc
taattctcttcgtaaaacagcagcttctttttgggtaactgatcaatcataccactttt
atcttgttcttctagctgtctcagacgattaactcgagacttaatcgttacccaattagt
taacattcctccaagccatctctggttaacataataagaatcacagcgttgtgcttcctg
tgccactataccagcggcttgtctttttgttcctagaaataaaactttttaccatccga
cgaagcctgttttataaattcacaagcttctgttaataattgtgcagtttggactagatc
aataatatgtatgccatttctttctgtataaatatatggaaacattttaggattccatct
acgcgcttgatgtccaaaatgaacacctgcttctagtaattctgctaatgtaacaatagc
cataagattattaatatttaaataatgtaattattcgggaaatccc-tttgtttagaaac
tgat-----tcaactaacttatttgaattttagtaaactaataataaattaacacacttt
taaagcagatcaacaaaatagatcctgaagaattcat----tctacacttttattgctaa
aataattacgcgcggttctgtcgtctaaaatgatatcatctaaatcatctcgtacagatg
aaatagtactttcgtcattagtatttgcagttaaattctttttctctaacgatgagccat
tacagttatcatacatattgaagccggtaccagccggtataagccgacctataataacat
tttctttcagacctcttaaccaatctaatttaccagatatagcagcttcagttaatactt
ttgtagtttcttgaaaacttgctgcagaaataaagctttctgtatttaaggaagcttgag
taatacctaatagcacaggcctatatgatgcattaatcttatttctcaaggttatggctt
tattagtctgttcaatcttctgtaactctacaagctctccaggtaaatagcctgtttctt
caccattttctattttcactttagaagtcatctgccgtactattacttctacatgcttat
cagaaatattcactccttgagattgatataccaattgtacttctttaacaagtagtaact

gaatctcttgaaagcttaatcgagcagcttcatataaggctaatccacattcaatataca
aatcaaaacttgcatcgaggatatcatgaggatttaacaatacgtctgttttctttcttg
cttcaagaatttcttcaattcttggaagcccttgtataatgtcacctgtttttgctctat
caaaaaccaatatagctaatgtttctccccttctaattaatgcattgttatcaacatgta
aaatagcaccattagaaaccaagtaaggtctagcaattcttaatgtaactgatctagaag
aaatttcaataatttgtccagagtctaaagaagtcatattttcagtaataaaatcaccac
aacgaatccaatcacccacacttactttaactattgattttcctatattaaaggtctttt
tatcagaagatgtcgcaatcaaaatgcgcctgctagtattcttttcagaataaatctcct
caacagtaccttcactcattgcaataatctcagtactagcaacaactgtaaatggcttga
tatattcaccatttttacaataattctagtttgactttgttgcttctctgatttattta
catcaatacttttaattgataatgtttcaaaagttgacagccttaagtcaaaataattat
tgccactttctttagcagaaaactcaattgaggatattaaattatcatctcgatgctcaa
tttcagccacaagattagttgttactaaatgaactccttctatagatttcactctttctc
cgtcccgaaaaggagttcttttaactagctttagtttagtttgacttggtttttgtgaaa
ctagattttcaattagactggattttgtttctggtattgaataaacaataacaggtctaa
taagaatatagggcatttgctcgttttctatatactcccaataaactaacttatctgtag
aaatattattatgaagtctttctccaggtcttaaaaaacctctacttttatcattattac
tatacacctcactaagtttatagatagatccaggcttaataataatttctcgaactattc
catccttttcgattatttcaacaatacctgagcttttagaaaaaatatttttaactagct
ctgttccactttctataacatctccattatgcactagtagtagtgatgaatctttattaa
tttcgtgagtttcttctgaaatccataagatatatcccggacttaaaatttcgtaggcat
ctttatctaagcctgttttctttttgatacatttaagtctaaatatttaataataccac
cacttgtcgttttatacgtatcagatattagttctgcgactgtgtaaccatcttgaa---
tttttggtttggaatagctttgaaaagaaatttttgtttttttctgtttctaaaatat
aagattctttttgtggattacatctgtataaatataacaacctggaattacaatagatt
cagtaataacttgtatattggtaatactgctattagtattctggtgtatccttacttctc
cagcataatgatttattaattctgtctgtgcaagtatcgtaccagctttagttttatctt
ctttatgaacaataacattagcagaatcagatatactataaacttcacctgagagcaccc
agatcaagcctccagttttagtaattcttgtagtatattgattattatctgtttcttcaa
ctgttaagttggagaaacagattttgccagagagatctgagactacatgtttttgagctc
tttctgtcatgagtcgatttctgcggggtgattcagcaatcacttgatccttcgatacta
attcaccatcacatattaaaagagtagtacctttacttaagttaattattgacttctgat
ttttcttagatctgattgtaacttgggttggctttttgttattaatgcttgctcaccat
gtctggtacgaacatctgtatacgattcaatatctagatctatcaattgaccatctatag
gtgcgtagatttgttctgctaattcaccggtaaatactcctccagtatgaaaggtcctca
ttgtcaattgagtgccaggctcaccaatagattgagccgcaataataccaacagcctctc

ctaaatcaactaagcggccatgtgcaagattccatccgtaacaatactgacaaacagaac
ttctagagttacatgtaacaggagatcttactaaaactttttt---gataccagctttag
ttatctcttttgctaacttaggactgacatcttggttcgtatgagcaattaaacaatttg
tctctggatggaatacattttctgctaatactcggccagataaagcttgttctagattaa
ttaatattttctgcgtatctactaaatcttctagtatgatgcctttattggtcttacagt
ctacttctctaattattacatcttgagaaacatcgacaagtcgacgcgtcaaatagccgg
aatctgctgttcttaaagcagtatcaactaatccttttcttgctccgtatgatgatataa
aataatcagtaacagttagtccttctctaaaattactcgatattggaagatcaataattt
gtccctgtggatcagccataagccctctcatgcccactaactgtcttacttgagaaatat
tacctctagcacctgagaacgccatcatataaacagcatttagaggatctgtctctttaa
agtatttaataacttcttgtttcaaggattcactcgcattattccacgtatcaataactt
tttgaaatcgttcaactgctgtaatctcacctcggcgatatttgttttcagtggccttaa
tatcttcaatcgtgcctaataaaagactttgcttactaggaggaattcttaaatcttcta
aacttaaagaaattcctgcctgggtcgcataatgaaatccaagatctttaagcttgtcgg
ccatattggctgctcgggctataccataattacgaaaagcccagacaattaaattttta
gttcgtttttatcaataaccttatttgaaaaacttggctgtgcaagacttcgtctattat
ccactaaatcttctccatattaaatgttcgtt--attaactatgctactaaagactcttg
aataatttattaaagattatacggccagctgttgttctaatatattgcacaattctctg
attttcagcgtcttcttttatgatatggttattaaaaaattttgttgtactattgtcaga
atgtttttcaatcttaatcggatcatgaggttgatcaccatctactaatccatcaaaacg
agcccatatataagaatgtaaatcaattttcttctgttcataagccataactacatcttc
taagcttgcaaaatactgattagatccctgctgctgagatggattattagctgttaaata
atagcaacctaaaaccatgtcttggcttggcataataattggttgacctgttgctggaga
gagaaaattatgaggggctaacattagcaatctagcttcggcttgagcctctaaagacaa
aggaacatgaacagccatttgatctccgtcaaaatcagcgttaaacgctgggcatactaa
tggatgaagcttaatagctcttccttctacaagaataggttcaaatgcttgaatacctaa
tctgtgtagtgtgggagctctgttcaatagtactgggtgaccttgaataacttcatttaa
gacattccaaatagaagattcattttttgaatcatctttttggcagctttaatattgtt
aactaatccctgcaaaattagtctgtgaataacaaaaggctgaaataactccaaagccat
ttctcgtggcaatccacattgatgtagttttaaatgaggacctacaacaattacagatct
accggaataatctactcttttacctagcaagttttgtctaaaacgaccttgttttccttc
tataatatctgacaaagattttaatggtctattatttgcacctacaacagttctacctct
acgaccattatccatgagagagtcaacggcctcttgcagcattctttttttcatttctgat
aataatctccggagctaaaattgactttagacgagataatctattatttctattaataat
tcgacgataaaactcattcaaatctgccgtcgcaaatctacctccatctaattggaccat
aggccgtaagtctggaggtatcacaggaatgactgtgaataccatccaagaaggatctgc

acctgttgctatgaaattttcaatcaatcttaagcgtttcatttttttattgaattttaa
tgaaggtgtcttaaaactttttggtggatttgtagcttcagaccttaaggtttcagctat
gtgctctaaatctaaatctttcaacagtttctgaatagcttctgcacctattcctacttc
tacttggttattctcatcttgattttggtagatttcctcttcaagacttttccattcgta
accttcgagaagttgcttgtattttaaattaatatctgtattagattgcgtaactacata
agaatgaaaataaacaatttttttctacttctttgacttttaagtctaatgctaaagcaat
atagcttgtacttccttttaagtaccaaacatgagttactggtgaggcaagttcaatata
agccatccgatgtcttctcacgcgtgattctgtaacttctacgccacatcgttcacaaac
tatacctttgtagcgaaaacgtttatatttaccacagtgacattcccaatccttaactgg
gccaaaaattttctcacagaataaaccatccatttctggctttaaagttctgtaattaat
tgtttcaggctttgtaatttcaccaacaatttgtccgttgggcaaacttctttcgcccca
ctgtcgaatttttttcgggagaagctaaactaatttttacgtagtcaaaatattgctcaaa
ctttgtcataaatttgaataccttaaaatcaaataattagccaatttatttacatgcttc
atattgaaaaatgaatatcttaataaaggaattgttcaaaatcgtctactggaggagtat
cataattagaacgatctactcggttatctttagaatccgacatcaagtcaacttcaactg
tccgtctttggccatcttcaaatagtttcaatttatgtactgcaatgtctaaacctaatg
actgcagctctcgcattaagactttaaaagattctggagttccaggcttcggtataggct
tccctttaactattgcatttaatgcttcatttctagcttgcatatcatccgactttacag
ttaataattcttgcaaagtatatgctgcaccaaaagcttctaaagcccacacttccattt
cacctaatctttgacctccatgctgtgctctacctcctaatggctgctgcgtaactagag
agtaaggaccagtggatcttgcatggattttatcatcaacaagatgaacaagcttaagca
tataagctctaccaactgtaacaggattatcaaaaggctctcctgttctaccatcaaaaa
cttgcattttacctggatgctgatcattaaaaagccatttattgctagtaatcaatgatg
cttcttttaattttctatttacaagtgctcttgatgcttctgctccatacatttcatcaa
aaggtattattttaaatctctttcctaaatagccacctgctaagcctagtaggcactcaa
atacttgaccaacattcattctagaaggtacacctaaaggatttaaaacaatgtccactg
gtgttccatctgataaataaggcatatcttgtttaggtaagatcctagaaatgatacctt
tattgccgtgacgaccagccattttatcacccacttggatttttctcttctgcgcaacat
agacgcgaatcatagcatttgttccgggaggcagctcatctcctttctgcctagtaaaaa
ccctaaccttaactactctaccttttgcagcattaggaagccgtaaagatgtatctctca
catctcgagccttttctccaaaaatagctcgcaacaatttaccttctggtagttgatcag
cttcacctttaggagtaatttttcctactaaaatatctccagcttcaacccaagaaccac
cgacaacaattccatttctatctaaatccttcagagagttatcgctgacgttaggaatct
ctctagtaatttcttctggtcctaatttagtttgacggcattctacttcatacttttcga
tatggatagaagtatacaagtcatcataaactaatctttcgctaattaaaaatgcatctt
cgtagttataaccttcccaaggcatataagcaactagaatatttctccctaaagcgattt

cccctccgtctgtggatgcgccatcagctaaagtttggccaacaactatttttctccta
cccaaacaattgggcgctgattaatacaagtatcttggttagaccgataatattttttta
agcgataatgaactgttctaccactattatcctgaatgcctattttattagcagaaacat
aatttacatgaccagatgttctgctaataacaaccataccagagtctcgagctattttag
tctcaagaccagtaccaattattggcttttctggatataataatggaacagcttgccttt
gcatattagatcccattagagctctattcgcatcatcatgttccaaaaaaggtattaatg
atgttgcagctgaaattacttgaataggtgaaattgcaatataatctacttgggttgggg
ttgttgtaataaactcttgacgataacgtacaggaataatatctccttcaatgtaatgtt
gtttactaactttaacatctcctggagctactcggaaatcatcttcctcatctgctgtta
gatagacaggactattgtgataaatcacttgaccttggtttaccggataaaatggggttt
ctatgaaaccaaaaacattgacccgcgcacaggttgctagtgaaccaattaaacctgcgt
tcggaccttccggagtttcaattgggcaaattctaccatagtgactaggatgtaaatcac
gaactgcaaaacctgccctgtctttattaaaacctcctggacctaaggcactaattcttc
tcttgtgagttagttctgcaactggattagtttggtccataaattgagaaagctgactag
atccaaagaattctctcactgacgcaattaatggcttgggattaattagatttgataaac
tcaatgaatctatatcacatatcatcattctctctctaataatgcgttctaaacgattga
gacctactctgaattggttctgcagcaattctccaactgaacgaactcttctattaccaa
gatgatctatatcatcaagattacctgagtttttatccttaatattaattagataatcaa
ttgaggataagatgtcttgaggagacaatactcggaacgttttaggtatattgagaccta
gtttttattaattttatatctaccaacttcacccaagtcatatcttttaggatcaaaaa
aacgagagtaaagcatctgcttagcaacagcaactgttgctggttcattaggtctaagct
tagaataaactatgagcagggcttcctcatcagttacttcttcaatattatttttccaa
tttctttagctaattctttaactgaatatatttgggatgcagaaataagaaaagcatatt
tactaagtccttttcaatttcatctttattgagtccaatagctcgcagaaaaatataag
catttactttgtgggtcttatctattcttatccaaatttctcccttaggatcaatttcaa
attttaaccaagaacctctattagaaattaggcttgcactataaatctgcttaccatttt
tgtctatctcttgtttataatagattcctggactacggattatttggttaataataactc
tttccgtcccagatacaataaaagttcctctatttgtcattattggcaagtctccaataa
aaattaatctctttttgtatttttttatttttatctgcttttcagcactgagctgtagat
gggttgatgataaatctaatgatttaatattttttcctggtttttagaaggtagatcaa
tatcttttcttgttaactttgcaggaacatatatttgagcactataagttttatccctat
tttttgcctgtctgacactataacgaggaaattttatcttgtactcattaccaaataatt
gtaattcgagtcgactagtagggtctgatatcttcggaaaaatttcaagaacttccgtca
aaccttctaataaaaaccatttaaaactggccctctgaatttcaactaaatctggaagaa
gtttatttttttaagcttatacgttgaaccatagatctcctttataacaaattttgaaatc
gaagagtttaa-tactattaatagagtactggtagataatgcataagcatagtttgctac

attatttcttctctagacataactagaacatcatattttattcaa--aaaaagattttta
aaatttttcttcac--acacactatctataagtactttgctgcta--aaagatagttcaa
gctattaaa-------aaaaacgaaattaatacttgtctgtctgtcaattaattgacatc
aagatagacaagtattatttatttaatcaattctg-tttttctgagcaaaagacaatgct
ctcgcgagtctagctttttttctagcgcctgtattagggtgaaaggcccctttttttaca
gctttatcaatcttactatatacttgagaaatacttgactgcacatcattgaaattatca
ctttctaaattatcaatatttagaagacatcttttagttaaagttttgacaaccgattta
tacttacggttaatgagacgatttctttccgaagttttaatgcgtttaatcgcagaaagg
ttcttagccacagtaaatccaataaaacttttataaaatatttttagaaacgaaaataca
caaatctgggatctattcaaaaagggaatatgcggcgtattataacattaaagaaaaaca
ataggcaatagttcaccaagagctcttattattacacaatagagattgtcattgcgaaaa
actttgtaagatagtatttaattaaaataaactaactagacatataaatatatatgaaaa
aaattgaagctattattagacctttcaagcttaatgaagtaaaacttgctctagtcaagg
aaggtattggaggaatgacggttatcaaagtttctggatttgggagacagaagggtcaaa
ctgaaagatataaaggatccgaatactctattgatatcattgataaaataaaaattgaaa
ttattattagcgatgataaagtagagaaaattgtagagacgattattaaggcttctaaaa
caggggaaattggagatggaaaaatatttattagtagtattgaaagagtaataagaatca
ggactaatgacttaaattttgaagctttatagtatttttacttt-tttagacttgatact
tcttcactaaacatgagataataagcgtttattgtatacttataactaaggtgtaata--
ctaatggctaaaagcaaaggtgcacgaatcgtaataactttagagtgctctgataaagct
ggagagtttgctcaaaaaaggaaacctggcgtttttcgatatacaactactaaaaataga
cgaaatacaccaagtagaattgaattaaacaagttttgtcctaattgtaatcagcattgt
atcttcaaagaaattaaatagttatttatcttattaaaat--aatcatattatggctata
tacagaaaaagaatatctccaattaagccaacagatgctgttgactacaaagatattgac
ttgctaagaaaatttattacagaacaaggcaaaatactgcctaaaaggtcaactggattg
acttcgaagcagcaaaaaaaacttactaaagcgattaaacaagctagaatcctttcttta
ttaccttttttaaataaagattaa--atatatagcctaacttcatattttttccatctaaa
tattaaaaatttctatagatttgtttattaaaatgatctaacatactgaat---------
----------tactaagaaga-tctactcattacaagttttgttcattttataattaata
atagcatcataattta---ttaagctagattcagacaagacataaag-gtaacctctttt
aattgattatattatatagtctatttaaaagaggaaattttttttgcttc----------t
agcaattcaagattctttaagctttaacaaggtatttaaaacaaagatttttgtttaggt
atcagatcaaatgaatgaatcttatctcccgactgccataattgaaattcagagatgaaa
attccacattcgttgccagcttgaatttcttcgacatcttctctaacgcgctttaaagat
tcaatttttccttgataaataactttttcttctcgaatcactttaatccaagagtttttt
aatagcttattattgatcactcgacatcctgctatttttctatttgctaacgagaatact

gtacttacttctgcttctcccaccggtacttctgaatattcaggatcaagtaagtcttcc
atccttcttttaatgtcttcaattaaggcataaatgatttgatagtttctattaatata
ttcgactttgccgatgcttgctttgttcctggagcaaagttggtattgaacccaatgagg
gtagaatttgtggttgaagctagttcaacatctgtagcagtaatttctccaggcatgatt
gagacaacattgagttgaactttactctgaggaaattgagataaagaatctaggattgct
tctgtagagccttgattatctgttttaataattaaagaaatttgcttactaatatcttta
gaattagtgtttttaaagtatctaatgtaattcgactattaagtgctctttgtttctgt
ataattgatgagtctttagaggtattttcaatcgctttaagtttggcttctttatcactt
ttaactgctaaagctatttctcctgtagctggtacagatgacaatccccaaatttcaaca
acagatgatggtattgctaaattaatttctctttagcgttgttaattataactcgaatc
tttgcgtaagctgatccaataaccaaattatcactaatattcaatgtaccattttgtatt
aatagagttgctactggaccatgcgatttatctaaatgggcttctattataataccttgt
gcaggctgtgttggatcagctttcaaatcctctaattcagctaacaaggtaatagtttct
aataatttatcaacattttgaccagtaagagaactaataggaattataggcacttggcca
ccaagtttctccgacatgacattgtattttaataaatcttgctcaataatatctgtatta
gaaccagctttatctatcttagatattgctatgacaaaagggacattcgctttttgaata
tgattaatagcctctatggtttgaggtttaactccgtcatcagcagctataataataata
gcgacatctgttaaattagctcccctagacctcatacttgtaaaagcttcatggcctgga
gtatccaagaaaacaatttttttgtttattatctttcttaatatactcaacttcataagct
gcaattgcttgagtaattcctccaatttctttgttagcattattagattttcgaatataa
tctaatagcgttgttttgccatgatctacatgtcccatcactgtgacaattggaggtctt
ttaatataattgccagtttcataaagagcattagagtgatctaaattactagaaagtcca
ttattattttcctttacatttgattctactgctataccaaaattatcagccactgatgat
attattgaggcgtctatggtttgattcatagtaactgatattcttttaagaaataaatat
tttataatatctgtttcttgaacacaaattagtttagataattcttgaatagttagagga
ttagtaatactaatagactctggcggagaattagattgaatgcttgctgaattttgacta
ggagttactgctactttttttctgtcttatagatttttttattactagtcagttttttaca
acttccacttttggcttaggtggacgcattaaagagatagctaaatctcctgcagtctgc
gaaacattagaattagagtctctaaagttgtca---tcatcatcgtcaatatgtatttta
gtttttattttt-ttgcgctgtctattctta--tttttttactatctactaagtcgtga
attttattaaaatttttactttttttatctagtttaggtggagaacttaattctagatgc
tgttcaccacc---tgtatgtgattctgatttatcaaggtctaaatttaacagattatca
ttattgactgactctaatctaattttatagataatctgaggatttttcaaatctacaatt
gtttcagaagaattgatatttgagctagatctagaggatctattttcaaagttttgatta
tttaaaaacataaatattggttttattatctaatagtaactttctaattttaatacagag
cttaatataaacaagaatcgaatatctgtataaataatacaaatgt---agagaattaat

ctatgttgtattagatttaatataacactaaagtcctttgttcgttatacaagtttaaaa
ctttt-tatgatgataaacactagattctagtttattggtatacaatctatttatctaaa
taatactgtttttgtcagggcttttaaa----ttatctaatgttatgttcattatgaaat
tctattaaaatttaaaaa-ctcaacaattaagtaagaatcgttaatcacctccacgacac
ttatgccacgatctcaaaaaaatgataattttattgacaaacttttacagtattagctg
atattgtattaaaaatacttcctacaagtaaagaggaaaaagaagctttctcttactatc
gagatggaatgtctgcacaatctgaaggtgagtatgcggaagctttagaaaattattatg
aggctttaaaactagaggaagatccatatgatagaagctatattttatataatatagggc
ttatttatgctagtaatggtgagtatgtaaaagctttagaatactaccatcaaggattag
aactaaattttaaattacctcaggcccttaataatatagccgtgatataccattaccaag
gagttcaagcaattgaggataaagatacagagctatctaaattaatgtttgataaggcgg
ctcaatattggcagcaggctattaagttagctcctgataattatattgaagctcaaaatt
ggctgaaaacgacaggacgaatgaggaatatacaaggatattaatatatttaagatataa
taaagatatcaattaataaattaaaattagttaaagaaactggtactatatagagtactt
-tactatatattataatattatactagttga------tctctaagacttagctgaataaa
cgaaattaaatctattgactatattcatgaatacaaataagcataatggttagtacaaca
aaaagtaccggatctgttactcaaattattggaccagttttagatattgcatttcctaac
ggacagcttccgaaagtattcaacgcactcaaagtacaaagctcagaaggaactattact
tgtgaagtacaacaacttttgggtgacaacaaggtacgagctgtttctatgagttccact
gaaggactacaaagaggggtagaagttattgatactggatcccctatatctgttcctgta
ggtacagatactcttggacgtatttttaatgttttaggtgaacctgtagataatttgggt
cccgttgattctgagagtactttacctatccatcgaccagcacctgcttttactaagtta
gagacaaaaccaaatatttttgaaacaggtattaaagtcgtcgatttacttgctccttat
agaagaggtgggaaaattggtttattcggaggtgctggagtaggtaaaactgtattaatt
atggaactaattaataacatcgctaaagctcatggtggagtatctgtatttggaggcgtt
ggtgaaagaacaagagaaggaaacgacctatatatggaaatgaaagagtctaaagtaatt
gatgcagataatctgaaagaatctaaagtagcattagtatatggtcaaatgaatgaacct
cctggagcacgtatgcgcgttggcttaactgcattgacaatggcagaatactttagagat
attaacaaacaagacgttctattgtttattgataatatttttcggtttgtacaagctgga
tcagaagtatcagctctactaggccgtatgccatctgctgtgggttaccaaccaactcta
gcaactgaaatgggagcacttcaagaaagaattacttcaacaacagaaggatcaattaca
tctattcaagctgtatatgtgccggctgatgatttaacagacccagctccggccactaca
tttgcacatttagatgcaacaacagtactttcgaggaacttggcggcaaaaggaatttac
cctgcagtggatccactggattcaacatcaacaatgctacagcctggaattgttggaact
gagcattattctactgctcaagaggtaaaatcaactttacaaagatataaagaactgcaa
gatattattgctattcttggtcttgatgaactttcagaggaagacagacaaactgtatca

agagcaagaaaaattgaaagatttttatctcaaccttttttcgtggcagaagtgtttact
ggatcacctgggaaatatgtatctttggaagatgcaattaaagggtttcaaatgatctta
aaaggcaatttagacgacttgcctgagcaggcattttatctagtaggtgatatagatgaa
gctatacaaaaagctgacagcatgaaagattaatataattaatgatgactttaaatataa
gaattattgctcctgatcggactgtttgggatgcagaagcacaagaaattattttaccaa
gtagtacagggcaacttggtattttaacaggccatgcacctttgcttacagctttagata
ttggagttatgagagtaagagtagataaagaatggatgccgattgttttgctgggcggtt
ttgcagaaatagagaacaatcaattaactattctggttaatggcgcagaagaagctagtc
aaattgatttatcagaagcagaaaaaaatttagacactgcaactcaactcttaagtgatg
cctcgtctaataaagaaaaaatagaagcaacacaaaaaatacgaaaagctcgagctcgag
tacaagctgcaacagcagcaacttcgtaa-aagattcc------tcttaaaacataaaac
tttagtaatgattactaaa-------gttttatgttttatttgtatggttataggcagct
taaataaacttagcttagacctgaacaaatataatcaaaatataatcccatctctttacc
tgcatctggtccaactaaactaatagtaacttctttcatagctaaaatagcttgaatcgt
tgcaccaataggaacacctagagaattgtatgtttcttttagaccatttaatactctttc
ttcaaggatagatggatctccagctagcattccataagtagcatagcgtaagtagtagtc
taaatcacggatgcatgcagcataacgtcgagttgtatacatattaccacctgggcgtgt
aatatccgaataaagtagagattttgctacggattctttaataattgttgctgcattagc
tgcaatagtagcagcagctctgactcgtagctcacctgtttgaaaatagcctcttaattt
ttcaacagagctatcatctaaatatttaccttggacatcagctgcattaataacagaagt
aattgcgtcttgcataactttcacacattccttaattttaaatatctaatacta--gggg
tatataacaaaattactagatcttttagcaatatcagtattactgcatagcacccaaagt
ataatcaaagtaaaaaccagcttctgctgaatcttcacctgcaagcagtgagcaagctac
acttttcatgcattttacaccttcagcaactcctgaaattggtgttcctaaagaattata
catttctttaacacccactaatccaatttcttcaataggagttacatcgccagcaactat
tccgtaagttactaagcgaagataatagtccagatctcgtagacaagtagcagtcatctc
ttcgccataagcatttccaccaggagaaactacatcaggtctttttttgaaacagttgttg
gccgccttgctttacaatacgctcacgattatctgttaaaatttgagctattcttaaacg
acgttgtccagataaaacaaaacttttaattctatctaattccccaggacttaaatatct
tgcttctgcatctgcatttacaattgactttgtaacaatactcatggataaaactcctgt
aatgcttt-aaaaaaaagcttaatactaattatttatgatataactcagaaagtaatgta
ctttttgag-aattttataaccgaaacacgttcatgatatatgaagcaggatagatactg
aggtaagtactaaaacgagaaagttatatgtaggaacttctgctattgatttcctagtac
ttttttaaaacttggtaccacaattgatatatcctgcttagtcaaacgattatagagtgt
ttctgtattaggaaagttagctgctggtaaagtcggaaatcttctgtatggaacagtatc
tttgccatataccgcattgtattcaacactttcaactaaattacttataaatgcagataa

gccttttgaagctaaaatttgattaaagaatctaatttctgcctgattattaggagctct
tcctaagatatgcttagtacctaattctataacttttgtatttggatatggctgataaaa
ttctttaccatatagttcagatagcgccagtttttctactagttctttgacacagatttg
tctattcaagaaagctgacttaatatccaagaattcaccgccgacactaaaagagttaag
atctctttcgaaaatttgacgatatgctgctcttaaagcttgttctaacatttctttgtt
actatcactattgacttcaaagacaatagattgatctcttaaagaagttactccctgtga
aatacgagactgaatatcattactactcctcatttctttgacagtgcctaattcaacaaa
cctagcagatttactagcaataactttttgaatctctggtcaataattcctggtcttaa
gcttcttaaagcaacaccagctggagtactatatctttcataaggaacagtattatctcc
aaaagtttctgtgtactccgagctatcaattattgtgtctattacttggtaatagccttg
cttatatgcaatatcaaaatatttattaatttcttgccgaccatatgtagggcgacccag
taagcggttatgaatatattcaatagctttacaaatatataatggctcccaataaagaga
tctaaatatacttgattttgctaactgtctaacaaactcgcgaactgtaatttgattatc
ttttagctgactttctattggcttgagaattagtttttcttcttgatacacttctctgcc
aaaaactcttaaataagcaacttttgtgacaacttctactgaattttcaaaattttggga
tgaatctgactttataagagcgttggacaatttaaagattttggtcccagggatcctgc
tgacttaggtcttacctgtggattactaatctgattgtagattcctggcccacgtctaac
taatatccttcttgtatctttaccaaaaagagcttgtctttttcttggatctttattttc
tttagggaaaatagctccaaattgaattgataatggatcattacctattccatatggatg
ttgatcaggtaatgattgcttataatcactaaataaagttataaattgaggaaccttacg
aaaaggcgcactataggtgaaaagatctatttgaggcccccaattacggcactcttgagg
ctcttcacctaaatttctaaagtaaggcacagtctcttcaccgaaataatctgtatactc
agaagaatttaaaagtgcattcactaaaccacttaaaccagtagatgataaaatagcaaa
atatttttgaaactcttctaaagagctaggacctctacccaagaaatgtctaaatgctaa
ttctaatgctcgactattgacaaaaggttcataaaattgctttctataaatacttgaagt
tcctaacgaacgaatgaattccttgattgaaatttggccatttttttacttgcgactccaa
attcgataaagagaggtcatatgctttagcaatatctctttcaaaaatttgtctataaca
tgctttcacaacaatatttttctcatcagcagataaactaggcttcataacaaaacgagg
tgttgacactcctgcctgtacataagtttgaggaagacgcaaaccttgtaaatctccaga
tattctttttcttaatttatcagtcaaagatggagcttcaaattcagaaatcacaacatt
aaaatactctttaactaaatcttgaccttttatatcttcctcaaagattaataatgctgt
ccgtctcatctctctaagagccacaattgctgctgcacttgagcaagcgttatcaattaa
ctctctcaatcctcgtatattaacagataaaatattagggtcgcctgatacaattgcata
tgtcaaataccgtaaaaaccagtctaaatctcgaagagattttctcattcttgtagtacc
gtatctcaagacgtttataggtttaaaccctggcggagtagctcccccagcattaaataa
agatcggaaactttggccaaaatctccttggatatttcctgataactcattaattttatc

ttgagaactttgatctcctgcaataataacagctgcttgcggacgttctaaataagaaat
tgcagacccacctacgaatattttatctgctgctcgtgctaccagaatatttgcattttt
agttaaaatatctgcaacttccagtcttttttgtccagaatttaaaaaagaaactaattg
atttaattcaccaagctgtaaaaatcggtcttgttgctctgcctgagtaatagttaaaat
tgaggcagtccgataaagctgggggcgtgctaatgggcttccgccacttgctttgatact
cattactttatttatctccttaactgatattgcttgctagcgaatacaaatatagcttga
tatttctaaatatttaaagatactatgcaagcaca-ctgttttctatattattaggtact
agtacttatagtacagagagtcttataactaacttctataaatacaaattttgtaatact
tctattaacatttgtcaaaaaaatatttaagtgctcactcttttatattaaaattcaatt
agtgagagattgatgtgaaataaagtatactaagtaagttttatagataggaaaattaaa
agatttattctacaaacattaatca------------cttgaatgtttagcaattgtaca
attgttattgcttttaattggtttagactagttttatattttttctaacataaacaatat
taatgagcaatttgtatgaatttaaagccacaaaacaacctattaactaacgaaaatata
aatcatgtagatatacccgaaaatgatataccaatgtcaattacggaacatttagaagaa
ttaagacaacggactttatttgtgtttttattctttttgtttgctacaactataagtttc
acacaaattaagatcattgttgcaatactgcaagctcctgctgttggtattaagttttta
caactggcaccaggagaatatttttttttcatctattaaggttgcaatatattgcggatt
gttgcaacaactccgtttgccgtttatcaagttatattatatatactcccaggactaact
ggaaaagaaagaaaaattattttgcctctattaattagttctgtattgctttttattaca
ggtggtatatttgcttactttgttctcgcaccagcagctttaacatttttaattagctat
gggtctgatattgtagaaccattatggtctttcgaacaatactttgactttattctatta
cttttacttagtacaggattagcatttgagataccaatcatacaattattgctcggtgtt
tcagggacattttcttctagtcaaatgatacgagcttggagatatattattattatagca
acaattgctggagctattctgaccccttcgactgatcccgttacacaattaataatgtct
tcagctgttttattactttattttggtggaattgttatattattagtcttaaaaaagtag
agtaaat---------attttcta-ttaatccaatatatttgttatc-----tgggcaaa
taatctataaaaacaattgct------------------agatttaagtatcaggcttct
aaatatttcaa---------gtctctttttatcatttaaatacaatc-------------
------------------------------------------------------------
------------------------------------------------------------
------------------------------------------------------------
------------------------------------------------------------
------------------------------------------------------------
------------------------------------------------------------
--------------------------------------------------gatacattgg
cttgttttttcatcaattatttgtgttgcacac-ttttttagcactctctttattgttctt

agaattatttctttaatctatttattaaat----agatatgaagagatttaagtattgat
tatggtaatctatatcaacgtaaaaagtattttattgtgagttttatttatataaaggct
tattcattatatagttataacagtaagtctaattaatag-ttttcatgcttaattaaagt
aataactcttaattcttttggcagaaaaatctgtcttaatattttatc--aaaaaaacag
-aaaaatgaagctaaatagtctaattaacttaattcaaaagtctatatattcttgtacac
ttttactaattattttaaatattatttgtgtcgcacctaattctagtaatgcatttccaa
tttatgcgcaacaagcttatgaaagtccaagagaagcaactggtaggatagtatgtgcca
attgtcatcttgctcaaaagcctgtggagatagaagcgcctcaagcagtactacctaata
ctgttttcgagactgttgtgaagattccatatgacagcaatgctaaacagattttaggta
atggcagtaaaggaggcttaaatgttggagccgttgtaatattacctgaaggatttaagt
tagctcctgttaatagattatctacagagttaaaagaaaagactagaaatctttacattc
agccgtacagtgctaaacaagacaacatttagtaattggacctatttctggtgataaaa
atagagaaatagttttcccaatactatctcccgatcctgcaaaagataaaaaagctcatt
ttttcaagtatccaatatatgttggcggaaatagaggacgaggccagatttatccaacgg
gtgacaaaagcaacaataatattgtctctgctttaagcagtggtaaaattaataaaattg
aattactagacaaaggtggatttataatacatgtgactaacagtagcaatgtagagtcaa
cacagaagatttcacctggtctcgaacttagagtaaaagaaggggatacaattcagcttg
atcaagctttgaatagtgatccaaatgtaggtggttttggtcagaacgaaacagaaatag
ttttacagagtccaaatagaattaagggcatgattgttttctttttcgctagtgttctag
ctcaaatttcttcgtattaaagaaaaaacaatttgaaaaagttcaagcagctgaaatga
atttttaaagcaatataaaactggcacttactataaagaaagta-taaacactttattca
caggatgtaccagtttttatgttatgtaataatcgctttgacggaagttatgatactttt
aacagaatctacaatctgactaggctgaattactgtagcctgctctaagcttccattata
tggtgtgggtatatcctgcgaagatagcctcacgacaggagcatctaattcatcaaaaag
gtactcattaatttgtgcaattagctctgctccaattccggctgttttcatacactcttc
tacaattaaaactttgtgagttttctttactgagatagatatagagtctatatccagagg
ttttaaagatataagatctataacttctggatcataaccttcttttaataaggccggtaa
tgcttggataacatgatgcctcattctagaataggttaaaatcgtaatatcttttccttt
tcgtacaaattcaactttattgagaggtaagaaatattcttcttgaggaatctcttcttg
taaattataaagtagaacgtgctcaaagaatacaactggattattgtctcgaattgcaga
tttcagtaatccttttgcattataaggagtagaacaagcaactatttttaagccggggat
ggcttggaaataagcttccagtctctgagaatgctctgcacctaactgcctaccaactcc
tcccggtcctctaataactaaaggcaatgtaaaattacctccagaagtataacgtaacat
tcctgcattattagaaatttgattgaatgctaataataaaaaactcatgttcataccttc
aacaattggtcttagccctgttatagctgcaccaattgccataccagtaaagctattttc
tgctattggtgtatcaagaactcttaaatccccatatttgctatgcaaatctttagttac

cttataagatccaccatagtgaccaacatcttctcctattacgcagacagttggatcttt
tgccatttcctcatctgttgccgctcgtaaagcgtcaaacataaagattttactcatttt
aatatttgattttatgatttgaccactgttttaaagtaacagttaagacaccatattct
atttagttatctgcaaaaagatatcgtttcaattctgacatgttaggttctggactggaa
atagcaaatttcacagcttgttccaattcggttttaacagcattctgaatctcattaagt
tcacctatattagcaatttcattatctagaatatatttttgagtttttgataggatct
cttgccacccaagcctctttttcttgtcttgatcttagttcatcaggatctgcgagagag
tgaccacgaaatctatatgttaatgcttctattaaggttggaccatcaccttggcgagct
ctttgaactgcttgttttgcagcttgtcttacagctagcacatccattccatcaacttca
atcccaggaagcccaaaagcttctgctttttatgtatttcaggtattgaagaagaccgg
tgatgtgccatacctatagcccactgattattttcaacaacaaatataataggtagtttc
cagagaactgccatattcagacattcaaaaaattgcccattattggtagtcccatcacca
aaaaagcaagccgtgactcttaaatcctctgtttctttaagtacttgctggcggtagata
ctttgaaaggctgcccctgttgcaaccggtataccttccgcaataaaagcaaagccacct
aaaaaattgtgaggcgcagaaaaaatatgcatcgaacctcctctgcctttactacaacca
gtctcttttccaaataactcagccatcacattttttagatgggacgcctttacttaaagca
tgtacatggtctcgataggtactgcaaacataatcagttggattcagaagtttaattaca
cctgtagaaacagcttcttgaccattataaagatgaacaaaaccaaacatctttccttta
taatacatctgagcacacatatcttcaaaatttctgcctaacaacatgtcttcatataaa
actaataaattactcttattaagattaagacctgttgaattatagttagtcagcggcaat
tgaactttcttaggataactcataatttgtaaagaaacctct----------tattgtaa
ataggaattttgaagaaacagctatcaatagaaaacacacagataaaaacttataaatta
taatatgtttttttactaaagactttaataattatttgaaactagttcagtatatcaaatt
aattttgcttatactacggatgtctaaaacttatataaaaaagtagt-a-aaaaagttta
ttaaaagaatagttaaaaacagatataaatagaagtaaacagaattatt-gaaaaaaat
gattattattaattaa----tacttaatatttctaaatattttacaattaatggctatgt
caccatctttattttatcctgttgaacaagaactatgtagtcttgaaaaaaatctgaaag
ctgttgctgggactcgtcatccaattttatatgccgcagcaaagcatctattcgatgctg
gaggaaaacgagttagaccagctcttgtatttttagtggctaaagcaacctctgagaagc
aagatataaatactggacaaaaaaggctagcagaaattactgaaattatacatactgcta
gtttggtacatgatgatattattgacgagtgcacaacacgtagaggagtcaaaactgtac
ataatttattcaatactaagattgctgtgctagcaggagatttttttatttgcacagtctt
cttggtatttagctaatattgaaaatttagccgtagtcaaagctatttctaaagtcatca
ccgactttgcagaaggagaaattaggcaaggcttagttcatttcgatcccagtatttcaa
tagatgcttacattgagaaatcattttacaagactgcttcactcattgctgctagttgtc
ggggtgcagctatgcttaatggttccaatcatcaaataaataatgatctttatctttacg

gtaaacatatgggattagcatttcaaattatggacgatgttctagatataactggttcta
ctaagagcttaggaaaacctgctggcgctgatctaataaatggaaatttgacctctcctc
tcctttttcacttactcaagaagcaagtttaaatgatcttattgatagggagttctgta
atagtacagatatagcctcaacattatttcttataaaaagaagcgggggaattacaaaag
ctaaagatttagctaaagaacaggtgcaggcggcactttcctgccttcagtttttaccac
aatctacacctgtatctagtttaaaagaattaacacatttcataatcacaagattgtcat
aaaggacttgctaaaattaaaatgttattaactaatttttttactgtttctaaaacagtct
gaaaagtttgaatatcattactagctagctgcgctagcattttacgatttaacgcaatat
tctcttttttaatgcactaataaaagtactataattcatgccttgattgtgagcagcag
catttattctagttatccaaagacgacgaaaatctctcttttttcctttttcgaccaacat
aagaatatcgaagagctttaaggacttgctgcttagctgttcgaaataaacatttatgtg
cacctttaaagcctttagctagcttaaagatttagctcgtctttttttttgcaacgttac
ctcttttaactctactcataaaatattac-ttaaattttaattattaccagcttactaaa
ttcttatactctttg----attataaataaggtaatttcattgcaatgtttttaatatct
tttaagtcaaccgaacaagtagaagaaagatgtcttctttgttttgatgactttttttgt
aataaatgactttagaagctttatgtcgaagaaattttccggatgaagagactttaaat
ctttttgctattgcttttgatgtttttaacttaggcatattataaacttttaatgattaa
caaaagcatgaaaagtctgaggatatcaatcttagacctttcaatacttagaataatctt
aacacaatttattttacttactacataggttaactgaaacaatttatatatataatacta
atcagttaatcaaccatagctaaagtggatttatattttgctttgaacttttgtatagta
ttagcaattagcattgttataattttaagactgcttgaattcaattcttcgaacattttc
atatttaatgtaaatgaaatattggcttctgagacaatattttgaatttgaacatcagac
aagggaatcatatctaatgcagctctatattgatctttaaataatttatcgtcttttatt
tggtcaaaatcgtaaaatttcgttcctcctgaatctgataaattcatagcccctctagct
atttttttaaaatttggccaccagaaaggtcacccaaatagcgcgtataagcatgagca
actagtaattcaggttgtttatgtcctatagtgtgaattctatcaacataaatttttgta
gcaggagatggttcaataaaatctaaccaatctgatccataataatagtttaaatcttca
gatagactcgctttcctattaagctctgtaaaatatataggtttaatagctggatgattt
ttattagaaaataactcttcctctattgcacaatagacaaagtatagatttgcaaccaat
ttacgatatgacttcttatctacaactcctcccaaaaaagatttaacaaaactaacattt
tctgccatactatgggacttagtagtaccttctcttagttcattcgctaaagtattaacc
ataatatcaatattccattttt-aaagataacaaattaaca---ataaatagttatccgt
atatttataattgagttgcgctaactaggctgctgcaaaataatttttttgattttattgg
atctggattcattgttttgtcaccaggcttccaattagctggacaaacttcatctggatg
agattgtacatattgaattgcttgtaaaaccctcaaagtttcttctacactcctgccaaa
ctctagattattgattgtagagtactgaataattcctttcgggtctataataaatagtcc

tcttagagctacaccatcactattcaaaacattataagctgcactgatctcttttttaa
gtctgatactaatggatatgaaagatctcctaatccacctgattctctgtcagtctgtaa
ccaagcaagatgagaatactcactatcaaccgagactccaagaacttctgtattaagttc
cgaaaaagcattatacttatcgctaaatgctgtaatttctgtagggcacacaaatgtaaa
atctaaaggataaaagaataagacaatatacttattcttgagatcagataattttaacgt
tttaaattcttggtcataaacagctgtagctgaaaaatcaggagctagctgacctactcg
aagacaatttggtccagaaatcattaatttttcccagaagaattgatataattttgttat
ttaataaataattatataacaaaatttagtatatattaaaatactaccagtctttttaa
attac--------taaatgatataatcaatactatctttatagcggggaatggatttgaa
ccattgaccttcgggttatgagcccgacgagctaccagactgctctaccccgcggataac
taaaatcatattagtctacatatatatataaaagcaaatgcataatagattttagaagta
aaaaaatagaagaactggtccactgatcattaaacttagaaacgacaaagtatttttat
cttatttaaaagaggaaagacttcatataaaccaataaatctatcttttaaaagaatttc
ggacggctttacccaaatttgaccgtcataccagcctgattcttcataaaatactgtagc
actcattaatcgttttactacgtatgaccagcctaaatataatctgattaaaataaatcc
tgtcattaggctggaagtaataaattctgaaaagaaaaatttcaaaggtagtttagtgat
tggaaaaattgataataatataggactaactaataaacagttaagtagtagtgcaatagt
tattttttgttataagatctatggcttaaagttggccagcaaaagaaccaagaatttt
caaagaagtatattcatgaacaggttgctgctcctttggcacaggacattgagtattata
taaattcattgacattaaaaagtattataataaagac--tttgtttatattatctaacat
aggtacgacattaaatattaactaactgcaattatagtagtgagtagaagaaataaaact
aaaaatagccaagtaattttatttaaagtattctcagtactacgagtattactaaagaac
tgattctgagcacctacacttcctaaaccttcggatttaggattgtgtattagaatagtg
aagattaatattattgtagatgagtaccaaaaaaattttaaaatttgttccattatgtaa
aagttaat-aaaaatagcacaaagacaatcgtctaaaagatattgattgtcttatcatat
taaaaagtatttattcaccctgtactttaaaagtaaaaatcctattgccagccctacca
aaactaggactgaacttaatacagcactttctacaatttctccgaccatt-ttatttttc
cttattattttattattg---tcattcttttgacaagattatattgagctaa-ctaaaat
ccgtttctaccccaaactaccattgcaattgaaaatgtaaacattgccattagagctgcc
caccccaaacttaaaatatccataatttataatattc--ctaaaaataagatctcttta
cttaaactataatattcttaatattaaaaatatcattatataataagctaattataacag
agtactacttaattcgtatttttagtgtatttaaagaagattgaa-ttttttcgaggtac
tggataagtatttcatcctaagaaccatattcaac----ttttgatttattataatgctt
aaatctaaatata--tatcactcaagaatcattagtaattatgatgttttaattatacg
att---aacatatttcaaaatagaagaattcatagcgactaagttaa-----tatgctaa
tataatttatcaat--aaaaaggaagttagtatttcaattatgcaaacaactataaataa

tgggcaaacgtctagtaaagaaactttactaacacctagattctacacgactgactttga
agagatggctaatatggatatttccggcaatcaagaagatttttttggctatcctcgaaga
atttcgagctgactataatagtgaacattttattagagatgaagagtttaatcaatcttg
gtctaatttagaacataaaactaaatccttatttattgagtttttagaaagatcttgtac
cgcagaattttcaggtttttttactatataaggaattatccagaaaattaaaagacagaaa
tccggttatagctgagtgctttttattaatgtctagagatgaagctaggcatgccggttt
tttaaataaagctatcggggacttcaatttatctttagatttaggattttttaacgaagag
tcgcaagtatacttttttctcacctaaatttattttttatgcaacctatctttctgaaaa
aattggatactggagatacataactatttaccgccatcttgaacaacatccagaacaccg
tatttatccaatctttagatttttttgaaaattggtgtcaggatgaaaatcgtcacggaga
ttttttttgctgctttgctcaaatcccaacctcatttttttaaatgactggaaagcaaaaat
gtggtgcagatttttttctattaagtgtatttgcaacaatgtacttaaacgactttcaaag
aattgatttttataatgccataggcctagactctagacagtatgatatgcaagtaatacg
aaaaactaacgaaagtgccgctagagttttcccggttgctttagacgtggacaatccaaa
atttttcaaatatttagatacttgtgcatgtgataatagagctctgatcgatattgataa
taacaattctccattattcattaaatctatcgtaaagatacctctatatttttccttatt
tgcaaatttactaaagatatatttgattaagccaatagactcaaaaacagtatggaatac
agttcgatagttaaatacaagggaatgctttttttcttataaaaagctatctagtgattaa
tataaaat------aaaagcttccttct-----cgatggaaggagcttttatttttgtcaa
ga---atataatgggattatatagatc-tagaactaagaacttacttttttcttttgcttc
atacataacctctaaagtaattagatttttagaacaatcacgggcaaatttctctgtatt
tctttttactttacctctaacaaacccaggaatttttatttaattctttctgtgcttcttc
tgaccaattaatactatcaacagcagaaattcctatacttaatgcttctgttgtatcatg
accaccaaatatttctagcaaatgatcttccataccaagtgtaaacgaattatatactaa
atcagcaatttgatttgttccctcataaccaagaaatggacggtaacttaaaggaaagtt
ttgtatatgcactggtgaagaaataacgccacaaggaatatttagacgcttaccaatatg
tctttccatctgtgttccaaaaaattgctgcaggttcagttttagcaattaaatctccaat
taaaccatgatcatctgaaacgattacttcatcgcagaattcttgaacttgatctttaaa
ccactcttcatcatatttgcaataagttccacaccaggcgacatgtatacccatttcctg
gtgtaaaaattcgagttattgctgcagcatgagttgcatctccaaatacaatagcctttttt
acctgttaaattctgacaatcaattgatcttgaaaaccatgcagattgtgaaataaatcg
ggtttgctcgtctatgtatttttcgtagtctacagcagctcccaatgcatttactaattg
ctgtattgatcggatgcatgcagctgtttgaacaatacccataggtgtaatgtcaacata
aggcatattaaattctttctctaaatatcgagctgtcattagtccagtttctctatatgg
aataaaattaaaccatgcagagggtaatttttttaaatcttgaactgatgcattttctgg
aataatttggtttatttggatatctagatcttgaaataaccttttttaattctgctatatc

atgttgattatgaaatcctaagcttactgctccaataatattaacagatggagttttagt
cttttgagttaaaacttgattattagatttagctttttccatatagaaagtgacaatttg
ttctaacgttctatcacctgcttgcagttcatttactctataatgattaacatcagctaa
taaaacatctgcttctgtttctatagatgctctgctaacaaagttttgcaaatcttcttg
caaaatactcgaagtacaagttggtgttagaataactaaatcagggctttcttctctgtc
ttttctagtaatattttctacaactttttcttgagaaccacgggctaacacatgtctatc
tacaacacttgctgtaacaggagtaaaatctctgtctctttcaagcattgaacgcattac
attaaaataatcgtctccaagaggcgcatgcataattgcatgaactttttttaaaagaact
cgcaattcttaaagttccaatatgagcagggcctgcatacatccaataagctaatttcat
attaaattttcctatgatttatttagacatagtaatacttttaaagctaactaacaaaca
ttagccaaacatctaatttaactccttgacaattttgtaatattttttaaatgaagctta
tcattataaatatttaaaaaatataattcagaataga----aataactgttagatcaagc
ttgaaagtaaagatttcttgcagttatagtttttctttaagtaaacatgtaaaaattgta
ctatatataaatgtactatatataaactttttttactgaatattggtaatgccagataca
attaacttaaatatgccttccccaacgtttggtggaagcactggcggttggttaagagcc
gcagaagtagaagaaaagtatgcgataacatggacaggcaaaaatgaaagtaagtttgaa
atgccaactggtggtaccgcaactatgagaaatggagaaaacttactctatttagctaag
aaagaacagtgtttagctttaggtactcaactaaaaagtaaatttaaaatatctgactat
aagatttacagagtttttcctaacggagaagtgcaatatttgcatcctaaagatggagtc
tttccagaaaaagttaatactgggagagctagcgtcaacagtgttgatcattctattgga
caaaatattaatcctgtagatgtcaaattcatgaataaagcaacttacgattagtctctc
-aaaaacctagacataaaataacttttatatagtctaggtttttgataatgtatactgat
agtatggaagggtggccgagtggttgaaggcgtctgatttgaaatcagttgaacta--tc
tggttccgtgggttcgaatcccaccctttccgttataaaact------------atagtt
ggcttctcaaggcaagctagctatacactgatagtattaagtttagacttaataaagtct
acaacttgtgacaggttagaaatttttttcggcatcctcatctggtatctctatactaaat
ttttcttctatggccattaccagctcaactgtatctagagaatcggctcccaggtcactt
gaaaaatttgcttccctagtcactattttttttttcaattcctagctgttctgctacaata
tcttgaactttttcaaagatttcattatcttgcataatatattatttcctctaaggtgtt
atcaaaaatttttcggcttatattattatccaactctaaaaaattagtaatttttagtcg
agattaaaatatttatacctatatgatatactattattctggataaatacgaagtttacg
atatggctcttctccaaaactacgagctctcaactggtagttgtcaactagattatgctg
catttttcgtatataggctgatcgaggtgtaagctgaactattgaactttcattcaagat
aatagtttcaatagccaatttagcttcttgaagagcctggatctcatcaaaatttttatt
tttacagagttctacccaattaagaccagatgaagtatgaatgtttaatatttttctgag
tgcacgagtaatttgtggaaccgtactattctgaattgtatatatataataatttgttttga

ttttgcaatttgtctaagttttgtattttgctttacttgattacgtagcgctagaatagc
atcagacttttctatctccttggttaaaataattggtagatctagtgaagaaataactga
tgtaatatgctgccaactcaaagaataagcatataagtactgatgaggtatttccaattg
caaagattgtttattaatattaataggggcattgaaagtagttgacaataaagatgtatc
acgattttgtgatttattcaaatctaaggacttatttcttaattctctatattgtaaaac
cggaggcgcctgctgtgcgttaattgatttacgagcattagagatgtttacaggtaagac
ttctatagattgcgatggataacatttaattaaaattctaccattagcttgaatttgtct
tttttgcacaaaaggttgatggccttgcaaaatttgatctattgtctctttaactttatt
atgaactatccaaacatttcgctcgtgtatttcgattgcaatctgaaaagcaggctcagc
ttttctttctagtatactcttttgcgtacctctacgtttagcttcatcgtcacctaatgt
tacatattgaattccaccaattaggtccgccaatgttggatttttaattaaactttctaa
atgatttccatgcgctgtacctactaattgaactcctctttctgctatagttcgagcagc
taacgcttctaattccgtgccaatttcatctataataataacctcgggcatgtgattttc
aacagcttctatcattacttgatgttgcaaatcgggccttgcgacttgcattcttctagc
tctaccaatagctggatgaggaatatctccgtcaccagctatttcatttgaagtatctat
aataactactctttttttccatttcgtcagctaaaacacgggccatttctcgaactgcggt
tgttttaccaactcctggttttcctaagagtaaaattgaatcaccttgctgtagtaaatc
tcgaataatactaattgtaccaaagactgctctgccgacacgacaggttaagccaataac
actaccttctctattacgcagtgaactaatacgatgcaaggtttttttcgataccagctcg
attatcgccgctaaaattaccaacttttttttacacaataatctaagtcttgccaactaat
agatctttgagataaatattctgggttatctggaaatctagcttctggacgacgacctaa
atccataacgacttctattaaattatttctattaggatgctgttgtaaaggttctttgac
aaaatttggcaaaatttctagcaacttatctaagtcatctgcaataagcatgattgagta
tataataattaattaatcaaatagaagattcaagattctaaagattacaatacatattaa
agaatattatgattggaatcaattttatacttaaattgaacaggaagtta-tagatgtaa
aagaactgattaaactttacgcaataaatcgattggtcaaaattaatagtcgtgttaaca
tgtttattggacaaaaagttcgtattaaatatagcaaacaaaaagttgctagtgatatag
cagataaagttggtgaattaggagttattaagggaataaagtttattaatagtcaatgtg
ttactattattgttgagtttgacaatcacactaggctttggatgtttagagaggaattaa
tctgtctaaatgaat------taaaaacatgaaacataatttattttatttattatacaa
aatttcttatgagtcgttaaaataataaattttaaaggagtacttttaattgtgcaaatt
actattaaaaaattacaagacttactttcgtctgtacaaagaaaaaagatccagacatta
aaactaaaacaaggtaagtttgaacttctattaaataagacctataaaaaagtcaatcaa
gaaattataccttcgcagaaatctgctgtattacaaaatagtccatctacgataattaaa
tcaataaataatacaaaaaaatcttctgttgttaatgaagaccgtacagaatatgccact
attgtttctccaatggttggaacgttttatcattcacctgctcctggtgaaaaaatttttt

gtacaagttggcgatgaggtcaaattcaatcaaacagtctgtattattgaagcaatgaag
ttaatgaacgaaatcgaggcagaaattgaaggcaagattatagaaattcttgttaaagat
ggtgatatagtagattgcgggcaagccttaatgaaggttgaaacataatgctaatttttc
cattttacatattgttaacagaaattatttagatagaaaaattagcttatactatagtta
tgaaaacttatttcgcaaaggcac-taaaaactaatatataaaattatttactttatctg
ccaaaaatagaaataagcgttttgattccttgtctcaagagaggagaatctcaatggcaa
ttagctcaaaagagcaagagacaaagaaggtaaaaatctcggttgataaaaatcccgtag
atacttctttcgaaaagtgggcccaaccaggccatttttctcgtacactagcaaaaggac
caaaaactactacttggatttggaatcttcatgctgatgctcacgacttcgatagtcaaa
ccagttctttagaagaagtttcacgtaagattttcagtgcacattttgggcagctgtctg
taatatttttatggcttagtggaatgtattttcacggagcccgcttctctaactatgttg
cttggttaagtaatccaacaggtattaagccaagtgcgcaggttgtttggcctatagttg
ggcaagaaattttaaatggcgatgtaggtggtggctttcaaggagtacaagttacatctg
gatggttccaactgtggagagcatcaggaattactacagaatttcagctttactgtactg
ctattggcggattagctatggctgctttaatgctgtttgcaggatggtttcattatcata
aagctgctccaaagttagaatggtttcaaaatgttgaatcaatgatgaatcaccatttag
ctgggcttttaggcttaggctgtttaggctgggcaggccatcaaatccatttgtctttgc
ctattaataagctactagattctggcgtgtctccgcaagaaattccactacctcatgagt
ttttaattaatagagagcttatggctcagctgtatccaagttttagtaaaggattagttc
cattctttactttaaattgggctgaatattccgactttttaacttttaaaggaggtttaa
accctgttactggaggtttatggctaagtgatactgctcatcatcatctagcattagctg
ttctatttcttgctgcaggtcatatgtatagaaccaattggggtattggacatagcatga
aagaaattctagaagctcacaaaggaccttttaccggcaacggtcacgaaggtctatatg
aaattcttacaacttcttggcatgcacagcttgcaattaatttagctatgatgggatctt
taagcatcattgtagcacatcacatgtatgcaatgcctccttatccatatattgctactg
attacccgactcagttatcgctcttcactcatcatatgtggattggaggattttgtattg
ttggagcaggagcgcatgcttctatatttatggtaagggattataatcctgcagaaaatt
ataacaatcttttagatagagtcattaggcatcgagatgctattgtttctcatctaaatt
gggtatgtatatttcttggattccattcatttggtttatacattcacaatgatactatgc
gggcacttggaagatctcaagatatgttctctgatacagctatacagttacaacctattt
ttgctcaatgggtacaaagtatacacactttagctcctggaaatacagctccaaatgcat
tagcaacagctagttatgcatttggaggagatattgtttctgttggtaacaaagttgcaa
tgatgcctatttctttaggtactgcagatttttagttcaccatatacatgcatttacta
ttcatgtaactgttttaattttagttaaagggttccttttctcaagaaactctagactaa
ttcctgacaaggccaatcttggcttcaggtttccatgtgatggacctggtagaggtggta
cttgccaagtgtctggctgggatcatgtttttcttggcttattctggatgtacaattctc

tgtctgtagcaatttttcactttagttggaaaatgcaatcagatgtttggggtagtgtat
ctccgtctggaaatgtttctcatattactggcggtaattttgcacagagtgcaattacaa
tcaatggatggctgagagatttcctttgggctcaagcatctcaagttattcaatcatacg
gttctgcgctatctgcatatggattaatttcttagcagcacattttgtatgggcattca
gtttgatgttcttatttagtggtagaggttattggcaagagcttatagaatcaatcgtat
gggcgcataataagataaaagttgctcctgcaattcaaccaagagctttaagtattactc
aaggtagagcagtcggtgttgcacactacttattaggtggaattggtacaacttgggcat
tctttttagcgagaattatttcagtaggctaatagtgaaaataggataaaaaaaacaatt
atggcaacaaaatttcctaagtttagccaagctttatcacaagatcctacaactagaagg
atttggtatggtattgctacggcacatgactttgaaagtcatgatggaatgacagaagaa
aatttatatcaaaagatattcgcttcgcactttggacatctagcaattatcttcttatgg
acatctggtaatttattccatgtagcttggcaaggcaactttgaacagtgggtattaaat
cctttgaaagttaaaccaattgctcatgcaatttgggatccgcattttggacaacctgct
ttgaaagcttttagtaaaggtggctcagcttatccagtaaatatagcatattctggcgta
tatcactggtggtatactattggtatgagaagcaatcaagacctgtattctggggcttta
ttcttactagttttatcagctttactcttatttggagggtggctacatctacaaccaaaa
ttcaagcctggtttatcatggtttaaaaataacgaatcaagattaaatcatcatttatct
ggattatttggggttagttctttagcctggacaggtcatttagtacatgttgctatacct
gaggcaagaggacaacatgtaggatgggataattttacaaccgtattacctcatccagcc
ggtttacagccgtttttcagtggtaattggagtgtatatgctcaaaatccagatacagct
caacatttattcggaactaatgaaggtgcgggtacagcaattctgacatttctaggagga
tttcatcctcaaagtcagtctttgtggctaactgatatggctcatcaccatttggctatt
gcagtagtattcattgttgctggacatatgtataggactaattggggaattgggcacaat
ctaaaagatattttagatgctcatagaccacctagtggtagattaggagctggacataaa
gggctatttgatactattactaattctttacacatacagttgggattagcattggcttcc
ctaggtgtaattacttcgttggtagctcaacatatgtatgctatgcctccatatgctttc
atggctaaagatttcacaactcaagcatctttgtacacacatcatcaatatattgctggg
tttctaatggttggagcttttgctcatggggcaatattctttgttcgagactatgaccct
gaacagaataaagataatgttttagctcgtatgctagaacataaagaagctatcatttct
catttaagttgggtaactctatttttagggtttcatacattaggcctttatgttcacaat
gacacaatgattgcttttggaactcctgaaaaacaaattctaattgagccggtatttgct
caatggattcaagcctcttcagggaaagcactttatgggtttgatgtgttactatcatcc
tctactaatatcgcaacacaagctggtagcaatatttggctgccaggctggttagaagcg
attaatagcggaaaaaattcattgtttttaacaattggtcctggtgacttcttagttcat
catgcaattgcattgggattacatactactacgttaattttagttaaaggtgctttagat
gcaagaggctctaaacttatgccggacaaaaaagactttggatatagttttccttgcgat

ggacctggcagaggcggcacctgtgatatatctgcatgggatgcgttctatttagctgta
ttttggatgctaaatacaataggttgggtaacattttattggcattggaaacatattaca
atatggcaaggcaatgcaactcaattcaatgagtcttcaacttatctaatgggatggttt
agagattacttatggctaaattcttctccattaattaatggttataatccatatggcatg
aataatttatcagtatggtcatggatgttcctatttggacatttagtatgggcaacagga
tttatgttcttgatctcctggcgtggctattggcaagagttaattgaaactctagcatgg
gcgcatgaacgtacgcctttagcaaacttgattcgttggaaagataaacctgttgcatta
tcaattgtacaggcaagattagtaggtttagcacatt-tttctgtaggatatgtattaac
ttacgcagcttttgtattagcttcaacagcaggcaaatttggttagactaaa-----ttc
gaattagct-a-----aaaaaagtcagtcataata-gttttatgactgactttttagat
ttaaaacaaaaacttactaaagtttcacctcaatatctacaccagaaggcaaatttaact
tcattaatgcatctatcgtttgagaagaaggttgatgaatatcaataattcttctgtgag
atcttatttcaaagtgttctcttgagtctttatctacatgtggagaacgtaaaacacaat
aaattcttctttttgtcggtaaaggaattggccctactgcaacggcattagttctagatg
cggtatctagtattttgtcacacgacgtattaagtataatagaattatatgctttcagtt
taattctgatttttgtctgctgagtaattgtcatgttaaaaagatcttattatttaagaa
ttttagagacaacacctgcgcctacggtacggccaccttctctaatagcgaaacgcatac
cttgctcaattgcaattgcattaattaattcagcagtcatcttaattctgtcaccaggca
taaccatttctgcatcagtaccatcatcagcagtaaactgattaatagtaccagttacat
cagttgttctaacataaaactgaggtctatatcctggaaaaaatggagtatgtcttccgc
cctcctctttagttaaaatataaacttctgcttcaaattgagtatgaggtgtaattgtac
caggtttagctaataccatacctctttcaatatctttttttctgcacacctcttaaaagaa
ttccaatattatcacccgctagaccttcttctaacgtttttttgaaacatttctaatccag
taatagtcgttgtacgagtttctcgtaaacctacaatttcaattgtgtcaccaactttaa
taatgcctctttcaattctaccagtggcaacagttccacgtcctgtaatagaaaaaacat
cttctacagccattaagaaagttttatcgacatctctctccggcgttggaatatatgtgt
caactgcttccataagtgaaaaaatcttgtcaacccatttatcttcaccttgcttagtag
ctgggtttttcgtcacagcttctaatgccaataaggcagaacctgcaacaaaaggaatat
catctccaggaaagtcgtattgacttaataattcccttccttctaattctactaattcta
gtagctcttcgtcatctacttgatcttccttatttaggaatactactaatgtaggaacac
ctacttgttttgctaataaaatatgttcacgagtttgtggcattggaccgtctgccgcag
atacaactaaaatagctccatccatctgagcagcacccgtaatcatatttttttacgtagt
cggcatggcctggacaatctacgtgagcataatgacgattatctgtttcgtattcaacat
gagcagtattaatagtaatacctctagctttttcttctggagcggcatcaatttcatcaa
attttttcgctgcagtagaccctaaagttgataaagtcgcggagatcgctgctgttaaag
ttgtcttaccatgatcaacatgaccaattgtgccaatattgacatgaggtttttttacgtt

caaatttagatcgagccatgctttttgttttccttatagtaaaagttgctttatagttac
ttttaacgataaatcttgtgaattgaaagatttattatttaatacagacaaga-----ta
caatataactaatctaatatcgatagtgagcaaacgctttatttgcttctgccattctat
gtgtatcctctctctttctaatagaatttcctgtttcattagctgcatccataatttcat
tagctaatttcatagacatacttttaccagatctgtctctagagaatttagtaatccatc
ttagtgctaaattcgtacctctataagctcgtacttcaataggaacttggtaagtagaac
caccaactcttcttgcttttacttctactagaggagtaatatttcggatggctttttcta
aaatatttaaaggatctgattctgttctctctttaacgatatctaaagcctgatatataa
ttctttgagataaagtttttttaccacttttcaaaatacgaacagttaacatacttacga
gtctgcttttatataaaggatcaggtgatgcaaaccttttttttagctgtattacgacgag
acatagtatttaaaattggttattgtaataa-tgttc-tattagcaggtatttaagattt
aggctttttttgtcccatatttagatcggcttttacgacgatcttttactcctgcagcatc
taaagtaccacgtactacatgatatcggactccaggcaaatctttaattcggccgcccct
aatcagtaccacagaatgctcttgaatattatgacctacgccaggaatataagctgtaac
ttcaaatccagatgttaatctaacgcgcgctacttttcttaaagcagagttcggtttttt
aggagttgtagtatatactcttgtacaaacacctcttctttgagggcaactttgaagagc
tggagattttgtttttttatgtatttttcgtctttcggatctaacaagttgttgaattgt
tggcataataaaattgatttaaagtctttaatagatactatttgaacctgctttacatga
attggctagtttttatttatcttatacttacgcataaatctttctactctaccttctgta
tcaataattctctgtgaaccagtaaagaatggatggtttcctgaccagatatctacgtgt
agttctggctttgtggacccaatcgtcataattaactggccgtcacagtaaacttttgct
tctggataccaatttggatgtatattatcttttgccattgtttttataattatataaatg
tatcaataataactattaacgcttagaaaattgaggagcctttcttgctttctttaaacc
atatttttttctctcttttactcttggatctcttgttaaatagccttcagacttgagtgt
cgttctattttctggattaattgagcacaatgctcttgcaacacctaaacgaattgcatc
agcttgccctgtcaaacctcctcctctagcatttacatggatatcatactggtttagtag
tcctaaaacttgtaatggtgcatatgaaactcttaagtaattaggactaaattgaagata
agactctcctggtataccgtttataattaaattacctgaccctgggactagtcttacttg
tgcaacagaacacttccggcgacctgttccagaatagattgcgcgagttttaattaattc
tgtcgacataacattccttgataactaaataaatatacattctttatactatatactctt
gtggcttttgcgcgacatgtgggtgaattgggccagaatacactttaagcttcgtaaata
gttttctgcctaatggacctttaggaagcatacctttaaccgatttttcaataattctgt
taggtaatcttgtctgaagctgatcaaatgtttcaacttttaatccgccaggctgtccag
aatgtcttctgtatagtttttgattcgttttatttccacttacagatacgtgagcagaat
taataacgattacataatctccagtatctagatagggtgtataagaaggcttgtttttac
ctcttaagatattagaaatatgagtagatattctaccaagtgtctgatttttagcgtcta

taacataccaatgagaattagtatttaatgagggtgattgcgttttattcatgaaaagat
taatacttttggttgacaataagagaatatgtaaatggtttacattcaatttctatacaa
gtagtattt-aaaaatatta-------------ttaacaaagagaacttttctaacagtt
atttaaattgtttgtcagcaagaaaatttctagttatcaataacttatactattttaat
aatataaaatatattaatcactttttcttttggcaaacttatacctagtttttttgc-
aaggcatatataacttcttcagctgatttctggccaaaattttaatttctagcagttct
tcttgagagtaatctagtaaatctgcaatagaatggatttgagctcgctttaagcagtta
taagccctaacagataactgcagttcttctattaaaacttgactaatttttttatcttct
ttgctacgataattatctgctgatttaaagtctaaatttcttagtgaacaaaaaagatta
gttaaaactgtagctccttgacttattgcttcttggggagatatgctcccatttgtccag
atttgtataatcaatctatcttttatgctgttgctaccaatacggacttcttctacttta
taattaactttattaactggcataaaaacagagtctacttgcaaaaaatccacagataat
tcatctacagctttttcagctaagcgatacccgcaatttttttcaattttaaactccatt
tcaaatattgtgttattgcaaatagttgcaatatactgtctaggatctactacctctata
tcagaagataattcaaataggccggctgtaactatagctggcccttgaactctaattcga
ccaatttgagattctttgttataacttttaaatactacttcttttagattaagtaatatt
tctaacacatcttctcttaccccaggaattgtggaaaactcatggttcactccagcaatc
cgtacagcaactatagcagtaccttcaagatctgacaatattgatcttctcaaagcatta
cctaatgtaataccttgtccttgatttaatggttctattacaaaactaccgtactgccca
cgcgccccatctgttcttgactctatgcattcaatttgaaattgagccacctaagaaagc
tcc-tttaaaataatcagtgattagtaagtctctctaaaacaagaaggaatttatactcg
gcgtttcttaggagggcgacatccattatgaggtacaggagtaatatcttttattagagt
aatctctaatcctgcagcctgcaaagctctaattgctgtttctcgacccgctcctggacc
atttactagaacttcagtttggcgcataccttgatccatagcttgtctagctgcttttc
agctgctgtttgagctgcaaaaggtgttcctttttagctcccttaaatccacttgcacc
agaggatgaccatgataatgtttctcctttaaattagtaatagtaacaattgtattatt
gaatgtagatttaatatgtgtaataccgttaactgcattacgtttagtttttcttgctcc
ggatttttttatttgtctagccatcgtgtctcgtctattatatagttaattaaaagatta
ttttcttggagctttttcttacctgctactgttttttacctcctctgcgtgttctagc
attagttctagttctttgtcctctcaaaggaagaccaagacggtgtcttctacctctata
agtactaatttccataagtctcttaatgctcatagactcaaaacgtttgagatccccttc
aatctgataattagactcaagaatttctcttatactgacaacttgttgatcatttaaatc
ttgacacttaatatcagcgtctatgtttgtttttctaatatttctttcgagcgagatag
tccaataccataaatatatgttaaagctatctctattcttttgtttcttggaagatctac
tccagcaattctggccactttgtcttgtctccaataaataatatgtat-ttttataacgg
ttatttata-agcttagccttgtctctgcttatgtttaggattagtgcaaattaccatca

cctttctgtgacgccgaattattctacatttttcacacatttttcgaacagaaggacgaa
ctttcatattaa----ctctatatactaggaatattactatattaactctattaattatc
gacattaaaaatgtcaacaatatattgtattactttaactgttgtagattttacttcgtc
atactatcatatttttttgatatgacgtatgtctgaatttgcttagctgtgtctattgcg
actccaactaaaataagtaaagacgtagccccgagacctcttaagttttggatctgagta
accttttctattataaacggaatcagcgctactgtaaataaaaacgaggctcctaggaat
gtcagtctatttaatatgacttgtaaataatcaatagtcgcttgacctggacgaatatta
ggaatgcttgcacccatttttttaaatttatagcaatatcttctggattcattactatt
gatgtataaaaatagctaaagaaaaggatcaaagcacaataaagaagaaggtatagtgaa
ccattaggacaaaacaggtataggatttggagtaatgttttattttgaataatttgggtt
agataagatgggagggccatagatgcagatgcaaaaacaataggcataactccaccttga
tttaatttcaagggcaagtaactatttggatccaaaattgaagattttcctagctgtctt
gctgaaataattttaattcttcttgttccttcttgtacacaaattgtaattattatcatt
aataaaaagatcgctataaataatccgaacttaagacttgcattactataactagcatca
aaaaatgactgtgtaaaattctttggtagtcctgacacaatgttttgaaaaataagtaaa
gaggctccattaccaattcccttttccgtaattaactctgataaccacataataatcata
gagcctgctgttaaagctaaaacagactcgcagacaaatgcaaaattccaattaaaaaca
tatggttttacccatatagagattgctccagattgtaaagtcgcccaacctagagctaag
taccttgtaatttgagttatttttggcggcccaattcaccttcttccttctgtaatttt
tccagattggggacaattttcgtaagtagctgcattacaattgaagagttaatataagga
acaatacctaacgcaaaaatccctattgttgaaaaacctcctccagaaaaaatattcaga
aaatttactaaagtattttttctacacttgcataaaaggcgtcatgatctatacctgga
acaggtataaatattcctaaacgtgctaaaactaatagaaagagagtaaagataatacga
tttcttagatcactttttggctcataaataaattttaaatgaaaagagatactttt--tt
tatgtaaaaaccctattccaagacttgtataagtttaaaatagggaagtaa---aatatt
atagtaatattaata-taatattctatgtagaatacaggctttctattggaactcctcga
tctttagctgcttcactaaaagttcttaattgagttaagccattgagaacagctcgcgcg
ttatttaatgtattattagaacctaattgcttagctaatatattctgtacacccgacaat
tctaggactgtccttacagaacctccggcaattacacctgagccaggtgcagagggcctt
aatataactttcgcggcaccagaaattccgttaataggatgaggtatagaattcgatttt
gtcagtggaactgtaactagatgtttttagcatctgttactcctttttttactgcacca
attacatcgcttgcttttcccacgccgacaccaacttggccttgctcattaccaataaca
agaataactcgaaagcttagtttttacctccttttacaactttagtaactcttttaact
tgtacgactctttcttcccagccactatctttatcttttcctttgctctgtttttacga
ttggccattttaagaattattcctaattagttaatta-cgattctagaaacccatgcctg
cttccttagcagcttcagccaaggccttaactcttccatgatataactttcctcctctat

cgaagacgacatttttaatgccttctttcatagactgttctgctaactgtttaccaacta
cgcgagaagtatcacagtttggtcttatgttatcagattctttattatttaaattaacag
atgatgtagctactaatgtaataccttgtgtgtcatcaattatttgtgcgtatatatgtt
tattagatctaaatacacacagacgaggcctacttgaggttccttgaactttttccgaa
ctcttttatgcttatgaattctagtttgtttagtgtttagtttcattattatttaccttt
tccagctttcccagcttttcttctaacaaattcaccttgatatctaattccttttccttt
ataaggctcaggaggcctaatagaacgaatagttgaagcaacctgaccgacaacttcttt
atctatgccagagacagtaatatttgtgttgttttcaactttaatttcaatattggcagg
aggtttaattttcactacatgactatagccaacacttaagattaaatcttggttatcaat
ttgagaacgatagcctacaccttgtatttgcagtttttaaaaaatccattagaaacacc
ttcaatcatattactaataagagttctggatagcccatgtaactgactagccatttttgt
ttgtccactcgtcttgacagcaattgtgtcgttaagtatttctaaattaataccagctgg
taaagttctggatagggtacctttaggtcctgtcacggtgatagtctgaccatcgaactg
agtactaagatttgtcggcaataaaattatttttttccaatacgagacat-attccacc
tataaaatagttaccaaatatagcataatatttcaccaccaagaccatcatgacgagctt
gtctatctgtcataacgcctctagaagtagaaataagagcgatgcctaaacctccaagaa
ctctgggcaactctttatgatttgcgtagactcttagtccgggtttactaattcttttca
gagcagtaataactggttgacgattttaccattatacttaagagaaatcattaaatgag
tctctataccttcacccatttgttcgaaattttgaacaaatccttcttcttttagtactg
ttgccatattgcatgtcattttcgttgctggaacttgcacaatttgatgtcttgctaagt
ttgcgttacgaatacgtgtcagcatatcggcgatcgtatcgttgaccaccttgatcctcc
ttgcga-----tgaattatttgaaatattttcaagattccctgaaaggcatccctagctt
ttttaacaaagctagaccttcttgatctgttttagctgtagtgacaattgatatatctag
accacgaatttgatcaatattatcatagtctatttctggaaagatcaactgttcacgtaa
acctaaattgtaattacctttgccatcaaaacttctaggactaattcctctaaagtctct
aattcttggtaatgttaaattaattaatttctctaaaaaagaatacatcttgtcttttct
tagatgcacaacaattccgataggaacctcttctcgaattttaaatcctgcaatagattt
tttagcttttgtgacaattggtttttgtccagttattaatgttaattcttgaatactact
ttcaagagccttagcattctgagaagcttcacctaagccacggttaatagtaattttagt
aaacctaggaacttcatgtacatttttgtactgaaattcatcttttaaagattgagtaac
agttgttttatatttttcttttaatcctattgccattatattttcatcaattatttaata
agttcgccagtctttttcagttttcgaa-ttttttggcctttatcattaattattactga
ggatcgactagcaatattattttgttcactaaataacataacgttagaggtatgtattgg
agcttcaaatttgataatttctcccgtttctccttcttgctgaggtttttatgcttcac
tttgagattaattcctttaacaatcaccttatttgttttataaatgattgcaataatttc
acctgtttttgttttatcacttccagaaataacttgaactaagtccccttttttaattt

aatttttgtattatttttggttgttttagaaggacctttcattatactacctccggtgca
agagaaactattttagaaaaatttttatctcttaactctcgggctataggtccgaagact
cttgtgccacgtggattattatcttgattaatgataactgccgcattatctccaaatcta
atgctcataccgtcagttctccgtaaagctttacgagttcttactacaacagctctgaca
acatcagatcgtttaactggcatattgggagacgcatctttgactactccaataataaca
tctccaatagatgcataagaaggattactagtgcctaatactctaatacacattatttt
ctagcaccactattatctgcaacattaagatagctttgagtctgtatcatacttttattc
tgtataagattaattatcaaaagacttagataagatattaaccattgtccaacactttgt
gcgacttaaaggtcgtgtctcttgtattgtaacaatatcgccgattgtgcattcattatt
ttcatcatgcgctttatatttctttgtccggatcatagttttgcgtactttctatgaga
gattctattttctacagctactactatagttttattcattttatcgcttactactttacc
tgttgtttcttttaaaggcatagtctttatattttttatatat---------aattaaat
tttaagattattacatggttgacttagtacgagacttttcaacagttaaaagttgggcta
atctatgcttagaatgtttaaatagatgaggctggaaatcttgccttgtggctcttttta
gccttaaatcaaaaagctctctttttattacaaggatttcttcagctaaagaagaagagt
ccaggtttgtaacatctgatattttagggaaagtcatattttaggattctgttgtattcc
gaactataaatttagtcttgattggtaatttataagaagctaatttcatagcttgttgag
cagtttttgtggtacacctgtaatttcaaatagaatatgtccaggcttaataactgcaa
cccaatattctggagctcctttacctgaccccatgcgagtttcagctggacgggcagtga
ctggtttatctggaaatactctaatccatagtttaccacctcttctcacatatctagtaa
tagttcttcgagtagcttctatttgtctagaagttaaccaaactggctctgtcgcttgta
atgcataatcaccgaacgcaattgtgttacctttgctagcagaacctttcattctacctc
tatgttgttttctaaattttgttttcttggggcttagcataaaaattaaaaaataataat
aattaagaagctgttgaatcaggagtttcctgattataagcagattctagacttttcgat
tctggcaaaatttctcctttaaaaagccaaacttttactccgagcacaccatacgtagtg
tgagcttggcgatgacaataatcaatatcggctcttaaagtttgtaaaggtacacgacct
tctctaacccattcacttctagctatctctgcaccattaagccgaccagatacttgaatt
ttaacaccctgtgtatttgctctttgagctctttgaactgcttgtctgacagctctacga
aaggccactctttttttctagctgttgagtaataaattctgctactaaggttgcttcagaa
tctggatctgcaatctctacaacatttactctgagttgtttgctagggtctaggattaag
gacagcgattttcttaaagactcaattcccgctccagattttccaagtacgattcctggt
cgagctgttgctatgagaatttctacttgatcaactttacgattaatttcaattttagca
atactagcattactaagttttgaatgtataaatgagcgaattttatgatcttcttgtaaa
agaactgggtagtctttagagttagcaaaccatgaagaacgatgtttttgggtaatgcct
atgcgaaagcccaaaggatgaatttttttgacccacagtatccttgattataagttaaaag
ctataatgactaattgacgttaaa--------agactacagttctgaaacacctaatgta

atatgacaagttggtttatgtattggaaaggctcttccttgtgctctaggctgaaatctt
tttaaagtcgggcctttgtctgcaaaggctttactaacaaacaattggttttatttaat
ccatcattatgctcagcattcgctgcagcagattccaggatttgctttatatgagaacaa
actcggtatggcataaattccagaataattaatgcttcttggtattttctacctctaatt
tggtctaatacacgacggactttatgcggagaaagacgaatgtatttccctactgcttta
gtttcttttacgtttttgtaatactcatagttatagtttaacgacgggcttttctgtca
ccttttacgtgagtgcggaaagttcttgttggaacaaactctcctaatttatgtcctacc
atttggtctgacacaaagacaggaaaatgttgtttaccattataaacagctattgtatgc
cctaccatatcgggaataatagtagatgctctcgaccaagtttttaggacttcttttttt
cctgaaatatttaatgcttctatacgtttaagaagactaacatcaataaaagggccttta
tgtatagatcttgacataataataaatcattagattcttaataaaataaattctatttac
ggcgacgtaagacgtacgggttactatatttatttggatttcgtgttttaacacccaatg
caggtttaccccaaggtgttacaggacgagctcgtccaattggagatttgccttcaccac
caccatgtggatggtctacaggattcatgacaacacctctaacagtaggtcttttaccta
accaacgatttctaccagctttaccaagagtaatattactagcgtcaatattaccaactt
gtccaatagtagcgtagcattcttttcgaatcattcgaacttcgcttgaaggcaatttta
cagtaacaaaagtaccttcttttgctactatttgagcataagttcctgctgcacgaacaa
tctgtcctccgcacgaaggtcttaattctatattatgtactgctgttcctaaaggaatac
tagataaaggcaaagcatttccaacttcgataggggcagtaggaccagaaagaaccatag
atcctacactgagagatcgagggtgtagaatatatcttttctcaccatcaagatagtgta
ataatgcaattcttgcatttctattcggatcatattcaatagaagcaactttagcaacta
tattatgtctatttcttttaaagtcaattaatctgtactgctgcttatgtccaccacctt
tatgacgacaggtaatgacacctctattatttcgacctttgcagaaatggtgtttaacta
ttaatgatttttctggcttatcagtagtaatctctgaaaaggtagaaaccgttctatttc
ttgtccctggtgtgtaggcacgatataaacgaattgccatataagaagatttgagaagag
taaatgttgaa------ctatgtgatatatttaagtttctggaaacaaattaatagaatc
ttctgaggcaagtgttacaattgctttcttgtaatgtggtcgcttccctacgaatctacc
gatgctcctttttttcttaggaggatgacaagtgttaacacctgtaacttgcacgttgaa
aatatactgtatagcagccttaatattaatttttgttgctttgggatcaacggcaaaaca
atactgattttcttctaataatttagttgtcttatcagtaatgatgggatatttaaccag
atctaataaacctcttgaatcaatgctatccattatatacctcttgtattttagataaag
catcaactgtaataataattttatgtgcagctaacaaagccataatatttagtgtatcag
ctgaaataagttctacgttgtgtagattgcgaatagaaagataaacattcggatcttttt
tgtctacgatgactaaaactttttatttaaatcaagattccatcgatgtatagcttcca
taaatagtttagtcttgggctgatgaaaataactgttaaaattttctacaatcagtgtat
taactgatttgttatttaatgctgtccttaatgctaattgtctttctttttttattcattt

ttttggtaaaactacgaggttttggcccaaatattacaccaccacctctccatagaggtg
aacggatagaacctgctcttgctcgaccagtcccttttgacgccatggtttacgacctc
cacctcgaacttcacttcttgttttagtattagcagaaccctggcgcttttcattacttt
gctttactaaagctctatggactaagtacatcccagaatcttgactgaccttaagattta
aatcagcgttgccacttacttgaccttcccaattatagacttgataatttaattgcgtgt
taactgtcatgaaatatatcgtatgaaagaatattgctacttactaattttactaaagc
gccaggtttaccaggaacagctcccttaacgatcaacaaatcattctctgagttaatact
tacaatttgcaaatttttattgtaactttttattaccaagttggccagccatgttttt
acctggataaactcttccaggcgttgtaccggctccaatagagccaggttgcctatgatt
tttcgaaccatgtgacattgggcctctactaaaatgatgcctcttttggtaaccggagaa
acctttacctacacttctagaagagacattaatcttttggcctacttgaaatagatctgt
agatagtatttggctaacttcgaaatcatcagtagatttcatttcgtattcacgtaaata
ttttagtgggggagcctgtgatttttttaagtgtcccaacaatggtttatttaattttg
ttcagcaacttgcttgtaaccgacttgaatagcattgtagccatcagtagatacagcttt
aatttgagtaataacacatggcccaacttgaattacagtaactggaattgataaaccagc
ttcatcaaagaattgggtcatacctactttagtaccgagtataccaacagacactagatt
tctccttttgagagatttatactatcattgtacataaaaattctacttcatagttactg
aag--taaattaaactaacagatgcatgatattattcatgacaacataaaggttaaaaac
tttacctatttaatctaaaatagaaacagctttttgtctagttacagcgtatataaatta
gaagtatgcttattaaatttttataaactact-tagatttaactgatattgatatccgt
aacttacttgcagtaatatc----aatcaacatgtataagtttaattgcctaaactagaa
ccagctattattagaaataattcaagtatagataaaagcgtatttgaacttttatatatt
gacaaatataatagaatatattgaggttcggtaactacaccttttattttacatttat
tgtgca--gtatatgtacatctaatgcagtaaacataagaatataac----tgtatagca
aatgtaattacacaaattagagaggtattaatgggcaaagtcgttggaattgatcttgga
acaacgaattctgtaattgctgttatggaaggaggtaaacctaccgtcataccgaatgca
gaaggttttagaactacagcttctgttgttgcatatactaaaagtggagataaacttgta
ggacaaattgccaggaggcaagctgttattaacccagaaaacactttctactctgtcaaa
agattcataggacgtaaacagaatgaaatttcgcaagagattcggcaaacatcatataat
gttaaaactagtggatcaagcataaaaattgcttgccctgcactgaataaagattttgct
ccagaagaaatttcagctcaagtactgagaaaacttgttgaagatgctagtacgtactta
ggtgagactgttacacaagcagtaataactgtaccggcttactttaacgattcacaaaga
caagctactaaagacgcaggtaaaatagcaggcttagatgtattgagaattattaatgag
cctaccgcagcttctttgtcttatggactagacaaacaaaataatgaaacaatactagta
tttgaccttggtgggggcacatttgatgtatctgtattagaagttggagatggagtattt
gaagtactctcaacttctggagatacacatttaggcggagatgactttgatcagcagatt

gtagaatggctaatcaaagattttaaacagaatgaaggtattgatcttggtaaagataga
caagcacttcagagattgaccgaggctgcagaaaaagcaaagatagaactgtcaaattta
actcaaacagagatcaatcttccttttattactgccacacaagatggtccaaaacattta
gaaaaaactgtaactagaggaaagtttgaagaactttgttcaaatttaatagataaatgt
agtatccctgtaaataatgctctgaaagatgcaaaactagaagcttccagtattgatgaa
gttgttttagttggtggatctacaaggattccagccatacagcaaatggttaaaagatta
attggtaaagatccaaaccaaagtgtcaatccagatgaagttgttgctattggtgcagcc
gtacaagctggagttttagcaggcgaagtcaaagatattctattactagatgtgacgccg
ttatctttaggagtggaaactttgggtggcgtgatgacaaagattataccaagaaatact
actattcctacaaaaaaatcagaagtattttctacagctgtagataatcaaccaaatgtg
gaaattcaagtacttcaaggcgaaagagaactaacaaaagacaataaaagcttagggacg
ttccgattagatggcattatgcctgcacctagaggagttcctcaaattgaagttaccttt
gatattgacgctaacggaattttatctgtaaaagccaaagaaaaggctactggtaaagag
caatcaattactatatctggagcatcaactttacctaaagatgatgttgaaagaatggta
aaagaagctgaagaaaatttcgatacagaccaaaaaagaagaaagaatattgacacaaag
aatcaagcagagtccctatgttaccaagctgaaaagcaagttaaagagtttgaagacaaa
attagccaagatttaaaaataaaaatagaggagctaattacagagcttagatctagtcta
gagaaagaagaatatgacaatattgaatccatttctcagcaattacaaaatgctctgatg
gacattggaaaaaatgctgctcagactgaaagtaaagatacaaaagcgaaggatgacgac
actgtgattgacactgatttctctgaagctaagt-aaaagta-agcgggtaacgcgattc
gaacgcgcgacatcaaccttggcaaggttgcgctctaccactgagctatacccgcataga
ttgtattattacaaaaattaagatatttgtcaatctgtta------atctatac---atc
tagactattaattttggaattacatattagctaactgttgtaaatataacaattgctata
attttttaa-------aagtacttatatcat-tatataagtacttttatatt-------
-caaaaatacaaataagtattatag-------ctaaactactaaggagttaactaagcta
gtaacaattactaatcctgtccataagccggctcctgtatatattaatccttttgattgc
tcccattggccaggggatgctaaaacaacaggtacgcctactactaacacagtcgataat
gcaattaataatagcactaataattggatcgcaataatcatttaatatgttctcctgaaa
aattcaaatgttaataatactgtatataatataatgaagcttttgaccttcagcaattta
tctatgaagaaataatttaaaattattgttttctaaataaaccttaaattaagtattaaa
atctttattgtaaacaataatactatattttttta--gtagataaaaaatttaaagaaag
tctttttaaaaaaaaattaagataaggtgatatgctgtataaaaagctaagtattatta
attattagagaaatttaaa-ataagaggtaaaacatatgaactcagcccttttttttagca
aaactaccagaagcttatgccattttaaaccgatcattgatatcttacccgtaattcct
gtattttttcttcttttagcttttgtttggcaagcagcaattggttttagataacattta
ggttataaaaagaataacaaatatacagagtgccaatttttgagctctaatcttatatct

tacacaataataactattatggttgaacccttactatcaggaattgttcttggacttatt
cctattactttattcggattattagtagcagcttatttacaatatcaacggggcaaccaa
ttaggactgtaaagcctaaattatagatatttataaaaactacaataattatagtttta
taaatatctacaaataagctaattttaaatacagaaataatactattgaaactagagtga
taacaaaaaaaatttatgtaa---taa---------------------------------
------------------------------------------------------------
------------------------------------------------------------
------------------------------------------------------------
------------------------------------------------------------
------------------------------------------------------------
------------------------------------------------------------
------------------------------------------------------------
------------------------------------------------------------
------------------------------------------------------------
------------------------------------------------------------
------------------------------------------------------------
------------------------------------------------------------
------------------------------------------------------------
------------------------------------------------------------
------------------------------------------------------------
------------------------------------------------------------
------------------------------------------------------------
------------------------------------------------------------
---------------------------caagctaattgtttaccagctcgacttagtaac
tcctggtaaaagacattcatgagacatttctcgaaaaacatggcgagataatccaaaatc
tcgataataacctctacttctacctgttaaccaacatctatttctccctctaacagcggc
gctgtttctaggcatttcttggagtttttgtcttaaatccattttgtcagaaaagctgcc
agcttctttcagttgatctttaatagccagtctttttaaataatatttttttgcaagttt
cgatctcttaatctctctttggatcatattctttttagccatgctgtaa-aactctcttt
taaaattatttattcaatatattagctaaaaaagctcctgcctgcaacttttatatattt
aaaaataaatagcataaagtagtgaaatgcaattgatgtttaatcaacgatacc---act
ttatgctattatcaaaaatttatttagtaaagttcttgttcaacatgagttgaaatagtg
caatcagatgaaggatatgcaatacaggttaaaacatagccagctaaaagttgatcatca
tctaagaatgattgatcagcttgatctacagttccttctgtaactttacctgcacaagtt
gaacaagctccagctctacaagagtaaggaagttcaattccttcttcttctgctgcatct
agaatatatgtatcctctgcacaattaaatgtgacatcaattccttcttcttcacatagt

aaatgaattttataatcagccatttcttcttactccataataaagatacacttgaaactt
aataaagcaaaat-aatttaaagaacttaccaaaaaattatatactagtttcattatgca
tgatagactggaaaaaaagaaaatagtatactattctgaaaaatcaagctacatttccta
ctaaatagaactatagtaggtccaaagtatatacataaattcatatagttaaaattactt
ttatcatttaaatagctgaacttgatagta---------tagattgatcatc------aa
gctagtcttctgc---caataaataattaattatgatttcaatgcttaatcaacaagtag
aacttaagcctattcttaaatttcagataacacaaatatattcgcattctcttattcagg
aaattaaagccttaataactagacttgttttgcaagtttggagaaggccagctacattga
tggcaggtattatccaacctttattatggttaattctatttggtgggcttttctacaatg
ctcctataaatttgttcactattaatacaagctataattgtttttttgagctctgggatta
taattttcacctcttttactggagctttgaattcaggtcttccattaatgtttgatagag
aatttggatttttaaatagattattaacggctcccttagtatcgaggacttctatcattt
tatcttctgctacttttatgacttgtattagtttaatacaagttgtatttatagttacag
cttctcttttatgggaaacccacctctaaatagcgatagtactatgatttttggactta
tgattctattagtgactgtaggagttacaatgcttagtttagctttatctttcactctgc
caggtcatattgagctattagcatttattttagtagttaacttgcccttttttatttcta
gtacggctttagctcctttatatttcatgccgccatggcttcagttaattgcaagtctca
atccattgagttatgcaatagaaggtacaagatacttatattcaagcgtaaactggaatt
ttacagagtgtgtgattaagattagctggggagatatttgtttagggcaaattattattt
tattaatcgctttagatataatggcagcttatcttgtgtctaatatattaaaagctaaac
ttaatta-----------------aaattttaataaaaaatttattaaatttattttaca
aagatttttactatataatactaatagtagtatggataattgaaatagaaatagtttttc
aaaaccaaagctattatcaacaatgataaacatttgtaagaaagtcaacaaagtatgttc
ttattcataggaggcatgtagtcaatgggactaccatggtaccgtgtacacacggttgtt
ttaaatgatcctggacggttaattgcagtccacctaatgcatactgcacttgtagcgggt
tgggcaggatctatggcattatacgaactagctgtatttgatccttcagatccagtatta
aatccaatgtggcgacaaggcatgtttgttatgccatttatggctagactgggcgtaaca
gattcatggggaggatggagtataacaggagagagcgtatccaatcctggactgtggagt
tttgaaggtgtagccttaactcatatagtcctttctggcatgctttttctagctgctatt
tggcattgggtttattgggatttggaattatttagagatccacgaactggcgagccagcc
ttagatttaccgaaaatttttggaattcatttattgctatcaagtctactttgtttcgga
tttggagctttccatgtaactggacttttttggaccaggaatgtgggtatcagatgggtac
ggagtaaccggaaaggtattaccagtagctccagcatggggaccagaaggattcaacccg
tttaatcctggaggagttgcatctcaccacattgccgcaggtactgtaggtatattagct
ggtgttttccatttaactgttagaccaccacaaagactgtatcgtgctctaagaatgggt
aatattgaaactgtattatcaagtagtatctctgctgtttttttctcagcttttgtgact

tgtggtacgatgtggtacggctctgcaactacgcctattgaattatttggtccaactaga
tatcagtgggatagtggatattttcagcaagaaattgagaaacgagtagaaaatgctatt
gctgatggtgctgcacctagcgaagcatggtcaagaattcctgacaagttggcattctat
gactatattggtaataatccagcaaaaggaggattattcagagcgggccctatgaataaa
ggcgacggagttgctgaagcatggcttggacatcctgtattccaagataaagaaggaaga
gagcttagtgttcgcagaatgcctgctttttttcgaaactttcctgtaattttagttgat
aaagatggtattatacgagctgacattccatttagaagagcagagtctaagtatagtatt
gaacaagtaggtgtaacagctagtttttatggtggcaaattaaatggccaagttttcaac
gatgctcctagcgttaaaaaatatgcaaggaaagctcaattaggtgaggtatttgaattt
gatcgaactacattagaatcagatggagtgtttagaagtagtcctagaggctggtttaca
ttcggtcatgctaatttcgctttaattttcttctttggacatctttggcatggttcaaga
actatcttccgggatgtgtttgcgggaatcggagctgaagttactgaacaagttgaattt
ggagctttccaaaaattaggagatagaagtagtaaaaaacaaggagctgtataagaccaa
agtcaaagatagtctttatc---aaatatttctaatattaaaatagttaagacaaagttt
attatctaatagttactaatcaggagaatttatggaagccttagtctatgtatttttact
aacaggaacgctaatggttatattctttgcgatctttttttagagaacctccaagaatagc
aaagtaacttgattccgtttaaacttataataatagaaccactttccaacttattagtgg
aaagtggttttattgtagattacattgtattattcttcatgctcttcaaaaggatctcgg
agatcttttgaagcaggaccaaatgctgtgtatatcgagtaacctgtaatacctaaaagc
agactcgaaataaaaatactaagaactgttgcagtttccataatttaagcagtattaaga
tgaactttttctataatgatattatcaatattaaaacatttaattattaaaattatcaaa
atatactatggcacttagaactagacttggagaaattttaagacctttaaattcagagta
tggaaaagttgctccaggttggggcacaactcctattatgggaattttcatgttactatt
tttcttgttttattaatcattttacaaatctataattcttcattagtattagaaaatgt
agatgtagattgggctactttaggtag----ctaactagagaagttctttga--------
ttacacaatattaaagacaac----------------aagcgatcgctattagattatag
tagcgatcgctttaaaagtattttactatatatccagagaaaattatttacctaaattaa
ttaactcggtatctgcaaaactatttgtgttaacattattatagttaactttttcaaacc
ttactaatactggatatttgataccacttttatcaattgctgcaacagtaccaatttcct
gataccagtacgattcttttctcaatatttttacttttgagcctctttccatatttttag
tcctcagatagattattaaagattattatattaaaaaatt-taaaaaactaaacaaaata
tataaaaatactaccgacaattgtaatatataactaataaaaggagagagagggattcga
accctcgataacaaaagttatgacaattttcgaaattgttgcaataaaccactctgccat
ctctcctgaattaaaatactgttttaaaaaattagacaagaataacatagaactaaggta
gttgcctagaaaattaaaactatgttaaatataaattataatctttctatagatatattt
aaaaaatatgtacatagaatctatatcagcttatttcaagatttttttaatcttatacaat

ttttactaaataaataaattctctcataatatgaaactatcttggaaaacacttcttctt
tggtctttaccaatatttgtaattggtttctttttctggcaaggtttttaggtccaact
actacagatgttggcagtaatatcgcaagttctagaatgacatatggacgattttagaa
tatttagatatgggttgggtgaaacgggttgacctctatgaaaataatcatacagcaatt
gtagaagctgttgggccagaattaggaaatagggttcaacgaattcgagttgaactgcca
gcaagtgcgccagaattaattacaaaattacgcaaagccaatgttgatctagatgctcac
ccccctaaaagtacaagtgcagtatggggactattaggcaatttactatttcctttacta
ttagttggcgggttagctttcttatttagaagatctaataatgctagtggtggacctggt
caagcaatgtcatttggcaaatcgaaagctttatttcaaatggaggctaaaacgggagta
gtatttaatgatgtagctggagttgaagaggcaaaggaagaatttcaagaagtggtaaca
tttttaaaacagcctgaatcatttactgctgttggtgcaaaaataccaaaaggcgtttta
ttagttggacctcctggcacaggcaaaacattactagcaaaagctattgctggcgaggct
agtgttcctttttttagtatctcaggctcagaatttgtagaaatgtttgttggtgttggc
gcttctcgtgtgagagacctattcaagaaagcaaaagacaatgcgccttgtatcgttttt
attgatgaaattgatgctgttggtagacaacgaggaacaggtgttggaggtggtaatgat
gaaagggaacaaacattaaatcaactattgactgaaatggatggctttgaaggaaatact
ggtgttattgtaattgccgctactaacagagctgatattttagattctgcattattaaga
cctggaagatttgatagacaagtttctgtagatgtaccagattttaaaggcaggttagca
attcttgaagttcatgctaaaaataagaaaatggaacctaaagtatctttagaaaccatt
gccagaagaactcccggcttttcaggagctgatttagctaacttactaaatgaggctgct
atcttaacggctcgacgaagaaaaaatgcaatgactatgtctgaaattgatacatcaatt
gatcgagtagtagccgggatggaaggcactcctttaattgacagtaaaagcaaaagatta
attgcgtatcacgaagtgggtcacgcaataataggcagtttattagagcatcatgatcct
gtgcaaaaagttacattaataccaagagggcaagcaagaggcttaacttggtttactcct
agtgatgatcaaagtctaatatcaagatctcaaatactagcccgtatcgtaggtgctctt
ggtggcagagctgcagaagaaatcattttcggtgacgcagaagttactactggtgcaagt
aatgatttacagcaagtaacatcaatggcaagacaaatggtcactcgatttggaatgtct
aaaattggacctttatctcttgaaagccaaggaggagacccatttttaggtagaggcatg
ggaggaggctcagaatattcagatgaagttgcaactaatattgataagcaagtaagggaa
attgtcagtgaatgctatgcacaagctaaacacattattatagataatcgagtagtgata
gatagattagttgatttactaattgaaaaagaaacaattgaaggcaatgaatttagagac
atcgttaaggaatacactgcaattcctgaaaaaaattactacatatcacaattttaa---
----------------attaaacgggactgacgggattcgaacccgcaacttccgccgtga
cagggcggtgctctaaccagttgaactacagtccc-aaaaagatgaatacccagagataa
tctcatagttagcttaaacttgtcaaattagaatattaaatatgaagagta---------
---------tttgtaataatactcttcatatttaata-tagtagctaaaatatttcttta

gaaggagagagagggattcgaaccctcggtacgaagttaatcgtacagcagattagcaat
ctgccgctttcgaccactcagccacctctccatga---tatatattatatatatatcaaa
tatacttatattttaataggtgtgcataaatatttatggagctaagcggattcgaaccgc
tgaccctctcaatgccattgagatgctctaccaactgagctataaccc----ctttggta
agatattaagaaatggctcaagcgggatttgaacctgcgaccttgggcttatgaatcccc
tgctctaaccactgagctactgagccatagtataaaatattattaagtattataccataa
attatggtcaaatataaatcaataaaagaataattactatca-tttactacttttg----
-----------------tgcaagttgtaaatattcgaataaaatttcaaaattaaaaatg
tctttaacttttgttaga-------atta-ttgattacgaaa-----------gtattg
atatttagaattaac---------------------------------------------
--------------------aataaatttg------------------------------
---------------------------------------------------tctttattt
g-----------------------------------------------------------
------------------------------------------------------------
------------------------------------------------------------
------------------------------------------------------------
------------------------------------------------------------
-------tagaca-----------------------------------------------
-------------------------------caaaaatattggaaa--------------
------------------------ttttcatcaat-----------------taattagt
------------------------------------------------------------
------------------taaactataaccgacatac-----------------------
------------------------------------------------------------
------------------------------------------------------------
---aagaattattgaacta-----------------------------------------
------------------------------------------------------------
---cgttaaat-------------------------------------------------
------------------------------------------------------------
------------------------------------------------------------
------------------------------------------------------------
-------------------------------------tgtgatgcatt------------
-----gggcttgatcaaaacttctctaaaatcaatagaatatagt---------------
-----------------------------------------------------aattcaa
gtagatcaaatactaacaaatgtgacaaacgcttaaaaaaataata--------------
caattagaatcgtaaaaatttggtttttga-ctgtgttaaataaatttgattatcataga
tttaatatttaaaagtgttctattacttgta-----ttgagtaattgt-aatatttaata

tcttctttttagataacaatcataatataagctaaaaaatagtttt---tattaaacaa
ctagcatagagccaataccttgatcagttaatatttctagcaagagagcatgatctattc
taccatctaaaatatgcgctgaagctacaccctgagccaaagaacgaatacaacaattca
ctttaggaatcatgccaccagaaattactgctgtttgagttaaatctctagcttcctgta
tattcagatgactaattaatgttgaaggatctgaagaattacgtaaaatgccgggcgtat
ctgtgagtagaattagtttttcggcattaagagcggctgctatttctcctgcgacagtat
cagcattaatattatacgattggccttccttgtctgctgcaacactagctataacaggta
tataattattattaattaatatcttcagtaatttagtatcaatattttgtacttctccaa
caaaaccaaggtttgcttttccatttggtctcggagtaataagtaatccatcttttcccg
acaaacctacacttttaccaccttgtttattaatagttgcgacaagatctttattgactc
ttccaactaaaaccatttccactatatccatggtaggttgatctgttactctaacgccat
catcgaattttggtaatattttgaggcgatctaaccaaaaattaatttctggtcctccac
catgaactaaaatgggacgtagtcctataaaagacagaaaaacaagatcactaatcactt
gatcttttagtttctgatttttcatagccgctcccccatattttataacaataattctgg
aagaaaattgctgaatataaggcagagcttcacttaagacttttactctttctgaattag
tcaacataaataaagtaggattgtgatatataaaattatatacacagcttgtgttaattt
aaaagataccaatgggt---------acttgaaaaaatagaaatattactaacag-----
----ttaaaactatagggattataattatgaataaattttgggataatgtattaagattt
cctcgatttttagtcagcgtcattttaggattaattttgataattattagtcctttttc
gtgttattaaaaaaaccactgacaagttttttttcatcatatcattggccggattaatc
acagttttggcaataataa-tacaaaaaatgataaatatcgaatgttgttgagagaaata
aataaatttaattaaaattttgaacatatataatattatattatcattagatatgataa
ttt-atttccttttctat---aaaacccataaatattaaatattatttgataaatattat
gccactaaaacaaagagttagctctgaaaagaccggtgccttcgcacttttggatagtat
tgttaggcacggagttaaacatatatttggttatccaggtggagctattcttcctattta
tgatgagctttatgcttgggaagaagcctctctaattaaacatatccttgttcgtcatga
gcaaggcgctgctcatgctgcagattcttattctagatcaacaggagaggttggagtatg
cttcgctacttctggcccaggagcaactaatcttgtctcaggtatagctacagcacatat
tgattctgtacctatattagctataacaggtcaagttgggagagcttttattggtacaga
tgctttccaagaagtagatattttgggattacacttcctattgtaaaacattcatatgt
agttcgtgaccctagagacatgtctagaattgttgcggaagcatttttatttgtaaaca
cggtagaccaggtccagtattaattgatgttcctaaagatgtaggattagagaagtttaa
ttatttttctgttgagcccggaaaagttaatattcctggctgtaggccaattaccagcct
aaagtcaagacaaatccttatggcagctaaaatgatacagcaatctagccagccattgtt
gtatattggtggaggagccataatctctgattcacatcaaattattaaagaacttgttga
tttttataaaatacctgttactactactttgatggggaaggggatttttaatgaggatag

cgattattgtctagggatgttaggtatgcatggtactgcgtatgctaattttgcagttag
tgagtgcgatcttttaattgctttaggagctagatttgatgatagagttactggaaaatt
agatgaatttgcttgtaatgcccaagtgattcacgtagacattgatcctgctgaagtagg
aaaaaataggattcctcaagttgctattgtcggtgacgtagcagaagttgttagtgaaat
attgaatttattaaagacttctttccccccttatccagagcagattatatcttggcaaga
aagaattaatcgttggcgtcaacagtatcctttactggttcctagaaaatcaacaagcat
ttcacctcaagagattcttgttgcaactaataaattagcccaaaatgcttattttactac
agatgttggccagcatcaaatgtggtcagctcaatttctgaaagtaaaagctaagcattg
gctttcaagtgctggattaggcacgatgggttatggtttacctgcagcaattggcgctca
agtagcacatccaaatgacgtagtcatttgtattagtggtgattctagttttcaaatgaa
tatgcaagagttaggaactatcgcgcaataccagttaccagttaaaattattattattaa
taatcgatggcaagggatggttagacagtggcaacaagctttctatggtgaaaggtattc
acactcaagaatgacagaaggagcacctgattttcaaaagcttgcagaagcttttggtat
taaagcttttactattaataataggcaaaatatgcaatctgctttacaagttgctattga
ttatcctggtccagttttattagattgccaagttacagaaaatgaaaactgttatccaat
ggttgctcctggaaaaagtaatgcacaaatgataggtatcgctaaaccgcagagaggtac
tgcttccaactatattaataatagtgtttgaagtaa------------------------
------------------------------------------------------------
------------------------------------------------------------
------------------------------------------------------------
---------------atagttagtatttttta-aacaaagaata-aagtaca--------
-taaatgtattttttttatactatta-aaaactatgtaagaacagcctttgacgagaatt
gaactcgtgaccttccccttaccaaggggatgctctacctctgagccacaaaggc-tttt
tactatattgggccgggttggatttgaaccaacgtaggcgaagccagcggatttacagtc
cgcccccattaaccactcgggcaccgaccctaattaaaccttattcaagatatgacctat
aagatatcataactatgttgctaaaaacaagtactttgataaataaaactgaataagtaa
tggacatagctggacttgaaccagcgactttcacgatgtcgacgtgacactctaaccaac
tgagttacatgtccaaaactttaattatcattataacactgtttagtgattttataagta
tctatttctaattgaagatttgttttaatcgtccggctttaccgagcagatctcttagat
aatacagtttagctctacgtattttagctcttctaatcactttgacaaaagttattcgcg
gtgaattgagaaaaaatactctttcaactccaataccttgaaatgagcatcttaaagtta
aactagtatttaagcttttttttcttcttcttgctaatactactccttcacataattgtt
cccgagttttactaccttccttaaccattagtccaagttgtattgtatcgccaactttaa
tttcaggtacttctgttttgataaaaggtatctcaacacttttcattagctgacttagct
ttgtactatttgctttcatatataggattagtcttgtgaca-tgttttgaaatattatac
attaaaaaaacaaatatgaaaaattttagcgaataagcatattagtatttataaaac-tc

tttttatct---------------------------------------------------
--------------------------------------------ccattata--------
----------------cagatccatt------------atattcaact------------
------------------------------------------------------------
------------------------------------------------------------
------------------------------------------------------------
------------------------------------------------------------
------------------------------------------------------------
------------------------------------------------------------
------------------------------------------------------------
------------------------------------------------------------
------------------------------------------------------------
------------------------------------------------------------
------------------------------------------------------------
------------------------------------------------------------
------------------------------------------------------------
------------------------------------------------------------
------------------------------------------------------------
------------------------------------------------------------
------------------------------------------------------------
------------------------------------------------------------
------------------------------------------------------------
------------------------------------------------------------
------------------------------------------------------------
------------------------------------------------------------
------------------------------------------------------------
------------------------------------------------------------
------------------------------------------------------------
------------------------------------------------------------
------------------------------------------------------------
------------------------------------------------------------
------------------------------------------------------------
------------------------------------------------------------
------------------------------------------------------------
------------------------------------------------------------
------------------------------------------------------------

------------------------------------------------------------
------------------------------------------------------------
------------------------------------------------------------
------------------------------------------------------------
------------------------------------------------------------
------------------------------------------------------------
------------------------------------------------------------
------------------------------------------------------------
------------------------------------------------------------
------------------------------------------------------------
------------------------------------------------------------
------------------------------------------------------------
------------------------------------------------------------
------------------------------------------------------------
----------agattctttcatagtttc--------------------------------
------------------------------------------------------------
------------------------------------------------------------
------------------------------------------------------------
------------------------------------------------------------
--------tttgataatcagtac-------------------------------------
------------------------------------------------------------
------------------------------------------------------------
------------------------------------------------------------
------------------------------------------------------------
------------------------------------------------------------
------------------------------------------------------------
------------------------------------------------------------
------------------------------------------------------------
------------------------------------------------------------
------------------------------------------------------------
------------------------------------------------------------
------------------------------------------------------------
------------------------------------------------------------
------------------------------------------------------------
------------------------------------------------------------
------------------------------------------------------------

------------------------------------------------------------
------------------------------------------------------------
------------------------------------------------------------
------------------------------------------------------------
------------------------------------------------------------
---------------------------atcta----------------------------
------------------------------------------------------------
------------------------------------------------------------
------------------------------------------------------------
------------------------------------------------------------
------------------------------------------------------------
------------------------------------------------------------
------------------------------------------------------------
------------------------------------------------------------
------------------------------------------------------------
------------------------------------------------------------
------------------------------------------------------------
------------------------------------------------------------
------------------------------------------------------------
------------------------------------------------------------
------------------------------------------------------------
------------------------------------------------------------
------------------------------------------------------------
------------------------------------------------------------
------------------------------------------------------------
------------------------------------------------------------
------------------------------------------------------------
----gatgatcaaatattattttcaaacatggcttatgtatgaaaaagttctata---aa
aacctaatgac--------taaaattagggcttaagtatttcttagttttgtgttataat
tagttactatcaaaggtagtggcataacacgaagcatcaa-aatgttcgaacgctttact
gaaaaagctataaaagtcataatgctagcacaggaagaagctagacgcttaggacataat
tttgtcggaactgagcagatactattaggattagttggtgaaggtactggaatcgcagct
caagttttgaaatcgatgaatgtgaatctaaaagatgcaagagttgaagtcgaaaaaatt
ataggaagagggtctggttttgtagcggttgaaattcctttcactcctcgagcaaaaaga
gtactagaattatctttagaggaagcacgtcaactaggccataactatattggcacagaa
cacttgctaatgggcttagtccgagaaggagaaggagttgcggcaagagttttagaaaat

ttggcagtcgatgtttcttcaattagagctgaagttatacaaatgctcggagaaaatgcg
gaagccaatgtaagtggaagcaatactacgcaagctagaagtaaaacaccaacattagaa
gagttcggatctaacttaactcaaatggctatggaaggtggtttagatcctgtagtcgga
agacaaaaggaaatagaacgagttattcaaatcttaggtagaagaactaaaaataatcct
gtcttaattggggagcctggtgtaggtaagacagcgattgcggaaggattagctcaaaga
attgctaatagagatgtaccttctattttagaagataaattagttattactcttgatgtc
ggtctattagtagccggaactaaatatagaggtgaatttgaagaaagactaaaacgtatt
atggatgagattaaatcagctgataatgtaatattagtgattgatgaagttcatacattg
attggggctggtgctgcagaaggagcaatagatgcagctaatctgcttaagccagcttta
gcaaggggagaattgcaatgtataggtgcaacaactttagaagaatatagaaaacatata
gaaaaagatccagcattagaaagaagatttcaaccagttgtagttggagagccaagtgtt
gaagaaacaattgaaattttgtttggtcttagagaccgttatgaaaagcaccatcaatta
acaatgtcagatggagctttggctgcagctgctaaatacgctaatcagtatatttctgac
cgatttttgccagataaagcaattgatttaattgatgaagctggttctagagtccgttta
ctaaattctcaattacctcctgctgccagagaattagataaagagttaagagctgtatta
aaaacaaaagatgaagctattagagctcaaaaatatgaaacagcagagcagtatagagca
agagaaatggaaattaaagctcaaattgcagcaattgctcaaagtaaaaagaatgagcct
gatttaaatttagaagatcctgttgttacagaagatgatattgctgaaattgttgctgca
tggactggtataccagtaactaagcttactaaaagtgagtcagaaaaattaatgcacatg
gaagaaactttgcatggacgtattattggtcaagacgaagcggttgtagctgtctctaga
gcgataagacgcgcaagagtaggtctaaaaaatcctaacagaccaattgcaagctttatt
ttttccggaccgacgggtgtaggaaaaacagaattaacaaaagctttggcttcttatttc
tttggttcagaagcttctatgatacggctagatatgtcagaatacatggaaagacacact
gtatctaaactaattggttctcctccaggatatgtgggttatagtgaaggtggttatcta
acagaagcggtaagaaaaaaaccatatactgtcatcttatttgacgaaattgaaaaagct
catccggatatttttaatctacttcttcaaattttagaagatggccgactaacagatgca
aaaggcagaactattgattttaagaatactcttttaattatgacttctaatattggatct
aaagttattgaaaaaggaggaggtagtttaggctttgaattatcagaagatcaaacagaa
tcccaatatactagagtacgatctttagtaaatgaagaactgaaacaatactttagacca
gagtttctaaatcgattagatgaaattattgtatttcgtcagcttactaaagatgaagtt
agagaaattgcagaattaatgcttaatgaagtctttgcgagaattaagcagcaagatatt
caattaaatgtaacagaacgatttaagcaacgattagtagaagaaggatataatccaagc
tatggagctagaccacttagacgagcagtaatgaggcttttagaagatagtctggctgaa
gaagttttatctggtaaaattaaagctggtgatagtgcagtagtagatgttactaatgaa
ggagaagttacagttttattaggtgaaaaattagaactgttaacataaa------aaaca
atttattaat--taaagcattgaagaggttat--attcaatgctat---gaaaaaatta

ataagatgggttgctataattgacaacctctgatattctttatttgtatata---aagcc
gggatagctcagttggtagagcagtggattgaaaatcctcgtgtcaccagttcaaatctg
gttcttggcat-tttaagcatcagaaaattttgctatctgagcattaaattttaattgaa
atgttcctgtgggaccattcctgtgttttgctattataacttctgtgaagtctctacttc
tagtttcttgtgtatagtaactttctctatatagcattattactaaatctgcatcttgtt
ctatagaattatgaacaataaagttatttgaaataaaattgcataaaggtttacattcta
aatcaaacactatttgtaaacttgtaacgtttatttttttgtatactttcaaaagtcaaag
atgaaa------aagaattaaaaacatttttttgttcttcaaagttttt------tattt
gaatagcaatcatgtcattttgattaattttatcacatcttttccatccttgtgttgtga
gtagtttatggttacttgttaattgtatatatttcccagcttcggtaataatttatata
cagtttttttccctgttttcgaaatactacatttaattgctagaaataactgtcgttttt
gcgcactaaagttaatgacttca-atctttttataatagcaattaaataatatttg--ag
gttgattatataatgatgtctgaagataattgaatttggatatacacccactttctctta
aatcggataacaaaggccttttgtcactccgagtttcaaggtttctatttagttgagata
atgccaagataggcaagcttaaatcttttgctaatattttaagtgatcgagtaattaatg
atagttcttgagatctattgtcagattgtctactctcctgcagtaattgaagataatcta
taataattaattctatatttttttccttgtaatttaaagaattttacttttgtctttatag
tgtcagtagaaatcttagcgctatcatcaatataaagattaaggttagcaagagttttac
tttttttgaataacatattgccattcatcattattaagctgaccagattgaattttttggc
cgtttaaattacactcttgagctaaaattcttcttagtagttgctctgtagacatctcta
agctaaataaaataacataagatcttttctgattaattacatatctagttatattaatag
caaaagcagttttccccattgaaggtcttcctgcaagaataattaaatctgacttttgaa
acccttgtgttatagagtcgagttctgtaaaaccagaaagtatactactatttatactaa
tctttttccttgtatccaaatggactaatagtcttgcaagaattgtagctaatgtttgcg
tatctttttcgtcaagtatttcataagcttttgttaattgactaacaatagaagttatgt
tttttttgttcaataagttgttttgagtagctaattaaacataatgaatctccacatgcta
aaagtaatcttttaatgtagttatctataacgactgctgaatattcgcttatagtatcag
ataggggagcttgttctattagactaaaaacttcatttagctcattcaagtcttgaatca
tttttcatttttaagttaatgaaaaaatttcttacgctttttgtctttgcatggttaa
cattttcgagtgctgctctataaagtaatgaagttgatgtaaaataaaaaaaaatcaggag
agattttttctactgattttagtaatgatatcgattgctgagttaatattatactaatta
atattttttcagctaaaatattatgaggaggtagatatttataa------atacttatta
cttcctcc-ttgtaacattactcttgtg--------gtattagttaactattgatttaat
ttgattctggaagaacttgcaatttaatatttgcagttacttgatgtaatagcttaattt
ctacatcgtaaatgccaattgttttcacgtctggtaaagagatactttgtttttcaatgt
ctatgttggtagtgtttttaataatttgtgaaatttctttttctgtgacactaccaaaaa

tattgtggccatctcctgtttttttactaacgctaaacctttgtatttcttctagaagtt
gcttgactttttagcattttctttagctgttttaatttttcttctcttatagctgcat
aaaattttgttgttttaaaataccatttgtggcaactgccgccattttattcgggatta
aaaaatttcttgcataaccagaagctactttgataacgtcattactcttgccaagtttt
gaatattctcttttagcacaacatttataactttttactcatctt-------aattcta
aaaa----atctgtattatcctattatacctgttgtatacagatttt----aataggaag
tatttatttgttgatataatattcttgtaaaa--aaaggtaatttttttatttt-ttctg
atataatcactttcacaataattc-----attaggagaggtggccgagtggttgaaggcg
cagcattggaaatgctgtttaggagcaatcttaacgagggttcgaatccctctctctccg
tacaa--aaaatataaaaagttttatagagtccaatctaaatctccaataatattcttcc
aagcatttaatccaccccgaactctaacaatatttagtttttgagctattaaaaactcag
aagcaaatattgatctagaatctaagctacagtaaacaaaacaaatttatcttgtaaat
tgatatgagagtaattcattcctttattttttgtaaaggtagatttaatgaatatatta
aatgacttttatgatattcttcatgatttcttacatctattaaaatatattgtggattcc
tactaattaaaaacttttgtaattgaataacactaatttctcgtacaaatgtattagatt
tactgccataatatttgttccaatgctttttttgt-------ggataatacaaatttagta
tttataatcttaaatttgttaaatgatgaagttaatgagttgtacgttaatatagttcca
cttaatatagacttataacccagaacgattttgacagcttcagttgcttgaagtgtacca
acaatacccggtaagagacctaaaactccggaattactacaagtatctcttgcgctctct
ttattttcagtttcactgtagaagtctcgataaacaggtccaccttgataattaaataca
ctaacttgtccttcaaattgaaaaaatggcaccataaatatgtattttatttaactctagg
caagtatcattcaagagatatcgagtatcgaagttatcagatccatctataataatatca
tattgtctaataatgtcgattgaattctcatagcttagtcttgttttgaacacagttact
atacactgtggatttaaatctaata-tttttttttagctatttctactttgaatatcct
atatcatgcatcgtgtataaaatttgcctctgtaaattagagagatcaattatgtcatcg
tctataattccaatgctgccaattcctgcagcagcaaggtaaatgattcctggagatcct
agcccaccagcaccaatgaataaaacttttgcttcttttaatctttcttgtccttctaat
tgaatttgtggcaatactaaatgcttagaatatcttgtatattcttctagtgagtatttt
gtattttctgttttaaaatttagcataaaaatttatagtttatttaatagcactta----
--aaagctagctgataagtttattaaaatgacttttgtaaacttggttcttctctatcta
aaatagctccttgtgtaggacagacctgaacacaaatactacaatcaatacaggcagaaa
agtcaatccaataccaattagtgttattattgtttttcccctttccttgatgaatacaag
ctacggggcaagcttccacgcattcagcaactccaatacatttttctgttacaattgtat
gagacatattatttaattaat-aaaagataagctatttacatctagtagttttgttgtat
actaaatttacatatgctatccaggccaatatttaaggtagtctatttctacagtttact
aatttgatatgcgcctttagttcagttggtagaacgcaggtctccaaaacctgatgtcga

gggttcaagtccttcagggcgcgtttag-ttttatcgtaattcaataatatcattttatc
tttttgcaaacatcaaaatgatatgatataaaagatcatatttatacattgaatattct
atgtactagtactg-ttttttaatatgttattattgataagttaattatactaattctat
ttttgtaataagt-aaaaacagttctaattaagtatgcacgtatttaatagtc-ttttgt
atatat--aacttgaaaaaatttaaactatggctaaaaaagttactggaattgttaaact
agcattaaatgcaggtaaagctactcctgcgccacctgtcggacctgctttaggacagca
cggcgtgaatattgttatgttctgcaaagagtacaatgcacgtactgctgacaagtcagg
actagtaatccctgtagagatctcaatatatgaagatagaagttttactttatacttaa
aactcctcctgcttctgtactaatcgctaaagctgctggactaaataaaggttcgggtga
accaaataccaaaaaagtaggcagcataacaaataaacagttagaatctattgcagaaac
aaaacttcccgatttgaacactaacaatatacctcaagctatgaaaatagttggaggtac
ggctaaaaatatgggaattctaattaaagatt-aaaagtttcaattatatttacta-tt-
-tttatgaaaaaattttcacgtcgacttacaacattaaaatcaaaagttgagcctaaact
ttacactattaatgaagcagtatctatattaaaagcaacgtcaaatgctaaatttaagga
aactgcagaagctcatattgctctaggtttaaatcctaaatatgcagaccagcaactcag
agcaacagttatattaccgaaaggtacaggtaaattgataaaagtagcagttattgctaa
gggagaaaaattaacagaagcaattagtgcgggagctgatgttagtggctccgaagaact
aattgatgaaatactgaaaggtagattagattttgataagctgatagcaacaccagatgt
tatgcctttaatagcaaagctgggacgagtattaggccctcgagggttaatgccttctcc
taaagcaggcacggtaacattggacgtagcaaaagccgtgaatgaatttaaaggggggtaa
agttgaatacagagttgatagaacaggtataattcatgtaccttttggaaaatctagttt
ttcacaagaagatctagttttgaacctccaaacaattaaagaatcgattgatagaaacaa
gccttctggagcaaaagggaagtactggaaaactttttcttatctagcaccatggggcc
atctattcaaattgacatcactagccttctatgaaaaattttgtataattatagaactt
gaataatctattatttcaaatta-taatttgctagcattattaacgtaacaacttgctta
ctatatcttatctgcattaaaaacttt-ttatatatcttattaattatgagtacaaaggt
tgaaaatatcttggaagagctaaaatctttaaaccttctagaggctgctgaactagttaa
acaaatagaagaaacgtttgatgttgatgcatctgcggcttcaggaggtatgatgatggc
cgcaccaacttcagcaccagcgtctgctgaggttgaagagaagacagaatttgatgttgt
cctagaggaagtcccagcacctaaaaagatttctgtattaaaagctgttcgctctctgac
tggtttgggtttaaaagaagcaaaagatcttgttgaatctacgccaaaagtcttaaaaga
aggtgcttctaaagatgatgcagaaactatgaaaaaacagctagaagatgctggcgcaac
agttattgttaagtaatga-----agaagcgcttctgtttaagtataaacagaagtgcct
ctttatttattataagcaattatactagaaaagtccaagtgtaattgcattgttgatagg
catagttgcaccaatacctagccaaatacttacaacggtaccaattaagaaaacagtggt
tgcaattggtcttctaaatggattttgaaatttgttaacattttcaatgaatggaacagt

aagtaaaccagctggtacagcagccatacttagaacacctaatagtttattaggaatcac
tcttaatagattaaatgttggaaagaagtaccattctggtaaaatctctaatggagtcgc
aaatggattagacttttcacctatagaagatggttctaaaattgcgagtccaatactgca
ggcaaaagtgccaataattacaactggaaagacgtataatagatcatttggccaagctgg
ctctccgtaataattatggcccattccttttgctaatttagctctcagcttcggatctgt
taaatctggtttttaagaattgacatattg-tttttgtgtttaatataaagattagtct
aattactgttttaaagtgtaattttcaactattataatggtccagaaattccttgtttac
gtatcattaaaaaatgcataagcatgaaaacggcagttagtagcggaagtacaaaagtgt
gtaaactatagaacctagttaaagttccttggcctacactgactcctcctcttaacaatt
ctactatactcgctccgacaactggaacagcatcagggacacctgtaacaatttttacag
cccagtatccgatttgatcccatggcaaagaataacctgttaccccaaaagaaacagtaa
gcactcctaaaattacacctgttacccatgtcaattctctaggtttttttgaaacctcctg
ttaaataaacacggaatacatgtagaatcatcatcaagaccatcatgctcgctgaccatc
tatgaattgatctaataagccatccaaagtttacatcagtcataatgtactctactgatg
taaaagcttctgcaactgtaggtctataatagaatgtcattgcaaatccagttgcaactt
gaattagaaaagatacaaatacaataccccctaagcaataaaaaatattaacatgaggtg
gtacatacttactagaaatatcgtcagcaattgcttgaatttctaatctctcttcaaacc
agtcataaattttactcataaaatagcttc-aaaagctcttctaatata--ttttgcgtt
taagctactattaattcaatag------ttattgaacttatatt-tttttataatcaata
cttattaactattataacatcttttattgattt--------------------aaaatt
ctctgacttagtaaaacagggctatgtactataattctttttttttgtatagagattaat
ttcgtttttagtaaattagctaaaattcttgttacactgactctattactacctataatc
tgtgctaaaattttatgtgtaatagtaaagttaagcataataccattattaacttgagtt
ccattttgctcagctaataataataataagctcgccagtctactggttatacttttatgt
gaaatgatttctataaaatgataagcttttgccgagcaaaatactaaatgatttacaaaa
aataaattgaatggagcgcagttactacacgcttttataatagtagtatagttaatacag
gcaacttctgttttatctattgcttctgcttcgtagtaaaaattatcgtcgaccaattct
atttgtccaaatgtatctcctgtagttaataagttaagagttactttatgtgtatttctc
aggacttttgtaataattaaagatcctatcaatataatgtataatctagagctgctatta
aacaaaagagtatcatttttcttaagtgaaaaaatttgataattaatttattacgatta
aaaaatagtagccatggattgcaagaaaaaaaatcacctgaactattttcagagaatatt
ggaatacaattttcttgatttaaacaatttttagtcataattactactttataa------
---aaaaatattctaatat-taagaactgaaattaaatgtcatataaattgatgttagt
tgagaatgacattgtattatcaaaggccattcaagaatacttaatagatcaaggcttcaa
tgtttatattgccaataatggattagaagcactgaatctagcttatcaatataatttcga
cttgataatttctgatataatgatgccgctagttaatggctatgaactattagcaaaact

taaaaaaaacaaagcattatccaaaatacctgttatctttttgacagctaaaggaatgac
taaagacagaataaaaggctacgacatggggtgctatggttatttatctaaaccatttga
tcctaatgagttgctttcaattattaacaacttaatcgctagagatgtcttaaaagaagc
ttctttacaaaacttggcaacatcaaaccagcaattaaatcataaaatacgtttgacacc
tagagaaaaaagtattttagatcttgttgttgatggacttactaataaagaaatatctac
tatactgaatactagtgttagaaatgtagaaaaatatgttagtcgactcttacacaaaac
taatatgaagaataggactttgctagtcaagtattctataaataataatctattaaataa
tgagatcaatgaaagggcgaatgacgggactcgaacccgcgaatgatggagccacaaccc
attgccttaaccccttggccacactcgccatattacacataccactatagcttgtttttt
gatatatcgtctagttttcgagttatatatttatctaaaacaatatttactttgaatttt
-tatgatacatagcaatattagtattcaaattaatggggaaccatttaattgctcaaagc
ccatctcattgcaattttttattaaattatcttgattttaattccgagcgtgtagcagtcg
agctaaatgacattctgttaccagagactctgtttcactcaacttatttgaatgatcaag
ataagctagaggttattaccattgtaggtggtggataatctatttgtacatac-------
-acagttctctatctcaaatgtttcatagatagagataatcttccttgtttttttatctac
atgaataataacagctttaatagtgtcacctattttaaattgtgatgatatctgctctaa
ttctttgatattaatctcagaaatatggacaagacctttgagatttcctacctttataaa
taatccgtaaggtgtaatctgatttattatgccttcaataatattaccaacaattaaatt
agatgatgcttgagatattagagctcttctatgactcagtattaaattattagatttctc
ttcgacattgagtaattttagcttaataaatttattattaaattgctcactttttttgaaa
attaccgagatgagaattaggtacaaatccagatataccttcaaggttaattatcattcc
tcccttattgaatccttttatcattacattaagtaaagagtcttcagctaatagttgcct
aattcttttccatgctcttatatattcaaggcgacgaattgataatattaattgtttcga
ctgaatattataatctaataaaaaaaattctcttgtgtcattaatgtttaaagaagtgaa
attatttaaatcttgattactagatacttcttgaataggcaagtatgcagatattggtgt
tcctatatctactaatacccccatttaattcaaagctaaatatagttccagctacaatatc
tccaagattcaaatcatacttgtatttttgtaaaacagctgcaaaatttctatgagtaaa
tccttcattattttttgtcatattgaatagtatgaattaaagagtaactttgtacaatat
attaaggtaaataaatgtttgtcgaattgtatgaaaaaattttaatttgttcggttgtaa
ttaagtatataaacatattaaaatatctaggagacatgaaaagtagacgtagatagctgc
agatttaa-tactatataatctgaggaaagtccgggctccacaaatacaatttatgctgg
agaaatcccagtgtaggtaactgcgaggatagtaccacagaaaaaaaccgccagaataag
t----tatattagctggtaagggtgcaaaggcaagttaaaagcttaccaaaagtactgca
aagtatttgttaggtaaaccccctaaaatggagcaaagctactaaacaaatatttgtgtat
attttatctagtttagttgaaaatactgcatgaagttattagtaacaataactcgagagg
aatagctatcctttacaatatatc-tcgattatattgaaagaacagaacccggcttatgt

agtacttttatgtcttgtattattatgtctattgtcgaatctttaaatttaactgtttg
acaattttttttgtagctgatgttgtttataatctatttctgaattggttaaagttttt
tcattagatcgatatgtaaatcttagacctatactttttttctcttttccaattgattca
tctatatattgatcaaataatgttgtactttctaaatctttatcataaaattgatccaat
aattttaataagtatttaatttccatattcttaggtattattaaagagagatctcttatg
atcgatggatacttagagtatggttgaatctgataacttaaataattgaattcattatga
caatcttctaaaatgtttaagttaacttcaagaaaaaaaagttttgtgttgagtcctaat
tcactatatgttgcttgattaagttcactgaatatgccaataataatattgttatagatt
agtgtggcaaattttcctttttgaatgaactttattctaccatcgactatatcttttttt
gtccactcaatttgtctattaatcttttgaaaaaaatttctacgatccctttagcttca
taccaatttaaagaatgcgcagggtgtgaccatgtagatcggatatctaaattaccaccc
aatatgatagctagattagtagtttctacaattttgttttgatcattatgaaatactgtt
cctatctcaaagctatctatagtttggttgctttgtttaatattataaagatttgattca
attaatccttccagcaaactgccgcgtaaattagaataatcttgaataagaggattgttt
aaagctatatttcctttggactttattaaagaataatgaactaattcagttaatcctaga
tttcttaaaatacttctacttttatctatgaattgtctctttgtggataactttttgttg
aattgaatatttggtactgagctcctgaatttatgataaccataaattctagcaatttct
tcaataacatctatttctcgaaaaacatcttctcgtctataatttggaacagtaacttct
atattttccttattttgataaataagatcaaaatttaaagagtgaagaatattttgaatt
tcttcaaagaataaaaaacgtgtttgaccattatacataattggtcctaaaatgtcatga
actttttaattgataaattaatattaagagcatgatcatttatttttctcgacagaaa
gtctctctaatatttccgcctgttagatctgtgattaaagcaagagcttcaaaatgtgca
tttttccaattatctacatttagcccacgttcttgtctaattgaactttctgttcgaata
ttaagaactcgagaagattttctaactacagattgtttaaatatagcagactctacaagt
atcgattttgtattatgatctgtatcaaaatcagagttgcttccaataccagctatacta
gttacatttgtatttacttgagtaactaaaatatttttgtttaactcaatattttcattg
tttaacttaatctgatcattacttccaagaggaaaattgcttcgaattgttagactattt
ttatgattcatattattaattttattcaaatcaacaatattaataggctgtccccatttc
aacataatataattgcttatatctgttaacaaatttctgtgtgtaaacccagaagagagc
aaccgatttttagccaatctggagaatctttaactcttatttcgtctataatggcagca
aaataattgtcacaatttaataaatcgtgattactgattatagt-ctccttttagagaa
atatctattattgggttgatgtatagctgataacattgaggaccctgttaaagccgagac
ttctcttgatagaccaatcatgcttaacacgtctgacctgttagctgttgatgatatatc
taatatatgatcaatctcacttccaattataatagattcaatagcttcaacttcaaaacc
tgcttgtgttagttgattagttaaattgtctatatctattgttttgatattaccaagttc
tttcaaccaatttaaagaaactttcataattaattgtcatctatcttatagaagtattta

ttttacttatgattaaaaatcaatttaaaaaattcatcaaattaatataagttatttgat
gaatttgcttatacaatatattattgtgatatcttacttttatatggcagttattaagaa
ctagcttttgtaaaagttagcccattatcttgggcatatctttccataaatctcatgaat
ctatcccaatcatcaggacttttaataacatgtacagcttctattgcttgtggtttacca
ttaataaacttggcattaacatctcttgtaataatttgtccctctgtgtccatcaggtac
atacctgtgatttcacctttgtctgccatactagcatctaaaattttgggatttgtaaac
cgaaaagttgctgttcctgtactaccatctcgagaccttgttaaacgtacatctggtaca
acttcttcattgataccttgaataaattgaattgttgccatatcaatcttctttatttgt
atataatttatattggataaaactgatttctagttcttgaatgcttgttgagaaactgtt
ttaaaagcttatagtagcagtagatcaaattttatgtattatataaaactgtaacttttc
-ttttttg----------------------------------------------------
------------------------------------------------------------
------------------------------------------------------------
------------------------------------------------------------
------------------------------------------------------------
----------------------atctgttcaaagaactaacttattataagctaatcata
cgaatattagccatataatgttcgtaagt-gaaattaaggggttgtagctcagaaggata
gagcaagcgcctcctaagcgctaggtcagcggttcaagtccgctcaaccctgttaacgaa
ttaagtagtattatttaacttgttctgcaaatcagtattgatctaagttaaagtacttga
cagctaaaacaaaagaatctgagaaatattttcatttaaagaacgaacaattatcttaac
tctaaatatctgataattattaa-aaaattct----------------------------
----ctttttaaaacacaagatgatg-----aagatatttaaactaataattgaaattaa
gtttttgaagttatagaagccaat--ataatactttctattaacaga-----------tg
cttgcttctattactatctgtttacaaga--aatt----atgagtatcaaaata------
--attttagctcta-------------------------------------ctgac----
------------------------------------------------------------
-----------------------------tactaataaata-------------------
------------------tttattttagtctagacaag------atgaatcttcatggta
taacag------------------------------------------------------
------------------------------------------------------------
------------------------------------------------------------
------------------------------------------------------------
------------------------------------------------------------
--------------agctattctttgtgtcatacttatataaaattaggat-------ac
tgtt--------------------------------------------------------
------------------------------------------------------------

------------------------------------------------------------
------------------------------------------------------------
------------------------------------------------------------
------------------------------------------------------------
---------------ttcattattataactggggtgggaggattcgaacctgcgaatggc
ggagtcaaagtccgctgccttaccacttggctacaccccaacatgtgaacattataatag
cagtcgtataccctgccatgtcaatagactattgtctttatttaataatttttgctgctc
ttgtagaataaaatcttttttttcaaataagatacgctatgtttaagttgcacgactgtt
atgttttcttgcttttgactgaacagccacttgatagtaattattccttctgcaatttcg
ttacttcctataataatacaagctacagctcttttttatttgcttgcttaatctgctta
ctaaaattacttgagctaacatcgatttctattttaaaaaactgctgatgtaaaaaacgc
attatttgcataccagtttcgttagcttttgtaccttgtgtagcaatataaaaatcgata
gattgattaggcaagtctatattatcttttgcaatcagtaataggcgttctaatcctatt
gcacatccaacagctggtgttgctttgccacctaattggtgaactaagctgtcatatctt
ccaccaccacatatagtatcttgcccttttgatgtcaatgtcttaatttcgaaagcagta
tcattataataatctaatcctcgaactaacttattatttatattgtagggaatattaagt
aatttagataattgcaaacgtcaccaaagtgtttttgtgattccaaacttaaaaaatca
gaaatttttggagcctctgtcagtatttttgtgtattagaatcttttgaatctaaaatt
ctaattggattactagttaatctttttgtgagtcagtgtctaaatcatcatggtacttt
gtaagataatctcgtaatttaacttgataaatactacgatcctctacttttccaattgaa
ttcagatcaagtttcaagttgtgcagattgagattgttaaatatgctcatagctaaatgt
ataacttcactatctgctcttgcatctaagctaccaataaactcaatgccaagttggtga
aattgtcgttgtcgtccactttgtggtctttcgtacctgaacattggaccgctataccat
aatctttgtaaactatggtggtaactcatttattttcaataaaagatctcacaataccg
gcagtaccttctggccttaaggtaatatctctattgctgcgatcatgaaaccgatacatt
tctttatttacaatatcagtatcttcaccaatacccctatcatataattcactattttca
aaaataggtgttctaatttcttggtaatttgcacattctaataatttggaaattttatta
tgtataaactgccaatattgaatttcatcgggaagaatatcttttgtccctctaatagcc
tgaatttttgccatgataatttttttgattaccttcttaatatgtctgcattaaatagcag
ttgtagactgttattcagtcatgtattatacattag--aacatactgatg-taatttata
aagtacgggcaaggagggattcgaacccccgacaccatggttcgtagccatgtgctctaa
tccactgagctacaagcccacttaataagtaccttaactatatcattgttcactaatgta
tacaactatattcttttt------atcttacatgtgaaattttttgtaaaagtaaaaaatt
at----gtttgtttttttacaattggagctattcatatataatgcttaatttattataact
ttgtaaa--aatatctataagtttaattgtatgattctattatattggtaatcttgttat
aattttgcgctcttatttttcaattgg------ctagcatttaattaacttatatttcaa

ataaaaaaatcttatctaaaatgagtaaacaaattctatatcaggatgatgccagaaaag
cattagaaaaaggcatggatattttaacagaagctgtttctgtgactttaggaccaaaag
gaagaaatgttgtcttagaaaaaaaattcggtgcccctcaaattattaatgatggtgtaa
cgattgcaaaggaaattagtttagaaaatcatattgaaaataccggagtcgcactgatta
gacaagcagcatctaaaacaaatgatgtagctggtgatggtacaacaacagctacagtac
tggcttcagcaattgttaagcaaggaatgagaaatgttgcagcaggttcgaatccgatgg
ctattaaaaaaggtattgaaaaagcaacaaattttgttgttagtaaaatagctgagtatg
ctaagccagtagaagatacaaaagctattatacaagttgcttctttgtcatcaggaaatg
atatagaggtaggtaaaatgatagcgaatgctatagaaaaagttggcagagaaggagtta
tttctttagaagaagggaaatcaactaatacgattcttgagatcacagaaggaatgcagt
ttgaaaaaggctttatttctccttatttcgttacagatacagaacgaatggaagttcttc
aagaaaatccttttattttatttacagacaaaaaaattactttggtgcagcaagaacttg
tgccattgcttgagcaaattgcaaaaacatctcggcctttattaataatagctgaagaca
tcgagaaagaagcattagccacaattgtagttaataaattaagagggattttgaatgttg
tagcggttagagctcctggattcggtgatagaagaaaatctttacttgaagatatgagta
tcttaacgaacggacaagtaattactgaagatgcaggtttatcacttgatacagttcaat
tagatatgttaggaaaagcccgaagagttattgttactaaagactcgacaacaataattg
cagatggtcatgaaattaaagttaaatcaagatgtgagcaaattaagcggcaaatagaga
cgagtgactctttatatgaaagagaaaaattgcaagaacgattagcaaagctttctggag
gagttgccgtcattaaagttggtgcagctacagaaacagagatgaaagataaaaaactaa
gactagaagatgcaattaatgcaacaaaagcagcaattgaagaaggtattgtaccaggag
gaggagctactaatgtccatatctctagtgaattatttacatgggctaaaaacaatttag
ttgaagatgaattaattggtgctttaatagttgaacgagctgtgacctatccgttaagac
gaatagcttttaatgcaggtgataatggagcagtaatagtggaaaaagttaaaagtcacg
attttcatataggctatgatgcagcaaacggtaatattgtaaatatgtacgatagaggta
ttatagatccagcgaaagtagctagatcagctttgcaaaacgcagcttctattgcagcaa
tggttttaactactgaatgtattgtagttgataaagtcgatgattgagctata------a
gataatagataatctcaaacaaagaataatttctaattatagaaattattctttgtttca
attaatctaattaattgatcaaatgtataatttttataataaagccaaagtttaaacacg
tcttctttttcaactattaagtagagaagatataagcccgtaatacctaaattatcaatt
aaaatattattttcatatatagatttttctaaagatctaaaaaatgacattttttgaaaaa
taataccgaaatcttctcttatagttatttgctaaatcattactgtctaaagattgattt
ccaattgaattcagaacctttaacagaatactttttaaatctttatctgactttatttga
gaatcaataataatttgattcaattgtatgtcaaaactaagattaaaatattacttgtt
aaagatatttgattttgaatatctagagattttatagccaagaagattaaacttttattg
ttcattgtaaaatataagacaattgatgatgtattctttaaatacta-aatttatcttct

tgtgcttctatcgtttttgt-aaaataattaatgttttaatttttagttttagttact
acccgaaaaaataaattctggaaatctagcttgagctttagctagagagtgtttttccc
ttcttcttgccagtaattgataacttctgttgttttgtttaataactcaattctgtctct
ttcagcaatccacggcttggattctagttctgcttttaattcttcccaagcatcattcct
tggccagaaaaaataagaagtaagtggactagtgccatttccaacaatctgatcaatagc
aatagcaatattattttctaaccaaagtacttttagtgtaaattttgacaagtttttatc
tccattattaatattaagtattgg----tacagtagaatactgtactagttattttttc
agcaacaatttgagctttgtttaaaatgtttttaccgttttagtagcttgtgcaccttg
agataatcgttgcaaaattttgaaatgttaacacgagtttcattatttataggattata
aaatcctaattcttctatagctttaccatcgcgtttatttctgctatccattacaacaat
tctatagcttggttgtttttttcttccgtatcgctttaaccttaacttgaccatattagt
cacctcgcttatttttt--atatctttattaaattatatcttacgacattatgcaataat
tcaaacaattaagctgattcaagtctaacattattaaatattaccattaaacattgtaga
aaaataatacctaacataggtgacatgtccattccaaacatgggaggaatgcttcctctg
aatagttttaagtatggatcagtaattctatttaatgaacagaatggttcattgtaccaa
tttacagttggaaaccatgccaatgataatttaagtaaaattaaaattaaataaatctca
gaaaaattagctatagatccaagcaataaatttaatgtaccaggaagagtgttcatacgt
atttataattattatatacttcgttatactaatagtttgattatatattataatcgatg
taaataaat-cagctaaacaaaaaacttatccttaaaatcatcttctcttgagcattttt
ttttatctacattagtatttacttttgtcgtataaatttccagtttagaccaaatatgtc
ctaactgatttaattgagatgttccacattctaaacaggctatttttatattgctcaaaa
cagcaccttttttataaaatcgttcaataatggcgtaatagttgcttcatccaaaaata
aatctaagacaagtatttttgtaaatgaatttaattgatctgggacattgttaataatat
cattataattaactaaaactatattagctgtaggaagtaaagatcgtgctccttccgcta
agataaagccataaggcataataattacaataatatatttataatgactgctaataagac
ttatagtggtattagcttttaattcgattgtttctgttactagccactctctcattattt
cgtaagtaatccatttgcctagttctgagcaagcagttcttaaaatcgtacctggattat
tgtcgttttcgagaatccctgaccaatgttgtattaatggatgggatgctatattaattt
ggagttgcatgttttgttaaatatatttaacatttaatttgttgattgaatcttcaattc
ctatttgttgagtattattaaaatagcat----atatattaaagaaaataattat-aaaa
aatgaaaaaaagtctttggctgtggggttttacagatagtgctgaaacttggaatggtag
atttgcaatgattggttttatatcagttatttttatcgaagtagttacaggacaaggatt
attatatttaataggtatgatgtcataataaaaatccaaaaaagttcttgtaattttta
taagaactttttgg-atttttataaatctaatctagcggtctcatagatagaacaggtt
cattctctctattgataccttttcaaatccagcagcagcagctcttgctctgcctgcat
gccaaagatgtccaataaataggaagaatcctaggaagaaatgagatgttgttaaccaag

atctaggagatacatagttaactgaattaatttctgtagctacaccaccaacggaattca
aagaacctagtggtgcgtgagtcatatattctgcggcacgcctttcctgccacggttgaa
tatcgttcttgatttatttaaatcaagaccatttggtcctctaagaggctcaacccaag
gagctcgcaagtcccaaaatctcatggtttcaccaccaaaaataatttctccactaggag
atctcattaggtatttacctaatccagtaggtccttgagaagatgcaacattcgcaccta
gtctttgatctctaactaagaaagtaaaagcttgagcttgtgaagcttcaggaccagttg
gtccatagaattcactaggataagctgtgttgttataccaaacaaaattagaagccgtaa
gccccatgattgataaagcacctagactataagataagtaagcctcaccagaccatacaa
aagctcttctagcccaagcaaaaggttttgttagtatatgccagattccgccagcaatac
aaataatgccaatccagacatgaccacctacaagatcttccatgttatttacacttacga
tccatccatcaccgccaaacggagactttaagacataaccaaagataactaggggattaa
gtgtaggattgctaacaaatctaacatctccaccacctggagcccaagtgtcgtataccc
caccaataaataaagctttaataactaacagaaaagcgccaatgcctaataacactaagt
gtataccaagtattgttgtcattttgtttttatcccgccaatcgtatccgaagaaaggaa
atgattcttcaagagtatcagggccaatcagagaatgataaagacctccaaagccaagaa
cagctgaagaaattaaatgcactacgcctactacaaagtatgggtatgtattgaaaatct
cgccgcctggaccaacaccccagcctagtgtagctaagtgcggaattagaataaaccctt
gttcgtacaaaggcttctcgggaacaaagtgtgctacctcaaaaagagtcatggcaccag
tccaaaagaccattatacctgcatgagcaacatgagcaccaagcaatttgccagaaacgt
taattaagcgtgcattgccagaccaccaggcaaacccggtagactcaatgtctctaccgc
caacaccaacagttgtattaaagggcgtttccacgtggtaaaacctcctcagggaatata
aagttttcatgaggttgatcttgagcagccatccaagagcgaataccttcgttcaatagg
atatttttagtataaaaagtttcaaactcaggatcttccgcagctctcaactcttgagaa
acaaaatcgtaagctcttaaatttagagctagtccaacaattccaaatgcgcttgtccat
aatccagttactggtacaaataacatgaaaaaatgtagccaacgtttattagagaaagca
acaccaaaaatctgtgaccagaatctattagctgttaccatcgaataagtttcttcagat
tgtgttggagtaaatgcacgaaaagtatctgcagcatcaccatcttcgaataaggtattc
tgtacagttgcaccgtgaatagcacatagtaaagcaccacctaagattcctgcaacgccc
atcatatggaacggattcagagtccagttatgaaatccttgcaaaaatagcaaaaatcta
aaaattgcagcaactccaaggcttggggcaaagaaccagctcgcttggcctagagggtac
attagaaagacggaaacgaatactgcaattggtccagaaaatgcaatagcgttgtatggt
ctaagaccaactagcctagcaatttcaaattgtcttaaacagaatccaattagtccaaat
gatccatgtaaagctataaatgcccacagaccaccaatttggcaccagcgagtaaaatct
ccttgagcttcaggtccccaaaggaaaagaagtgaatgtcccatgctgtttgctggggta
gaaacagccgcagttaaaaaattgcatccttctagatatgaactggctagcccatgagta
taccaagaagtaacaaaagtagtcccagttagccagccacctacagcaaggtaagcgcaa

ggaaacagaagtagtccagaccaacctacgaatacaaatcggtctctttttagccaatcg
tctacaagatcaaaaccaccacgagtttttcttgtccaattgctatggtcataattta
atctccagagcaaatttaataagtaatttgtttaaataaaca---ctttagatatactct
aaaattttactttctttacggtcaatataattataagtcaaaattaaccaaaattcacg
cttttaataaca-tttttttagctcattaagttaa-tttttaatatataaacagtaata
gagttattgttt-aacttacaatactcataatttattcatttctataatcctttgaaaaa
gatatcctgttcctcgtgctgttaaaatcaagtcgggattacttggatcatcttctagtt
tagctctcaatcgtgaaatatgaacatcaacaactctcgtgtcaacatgccgttcgggag
tatagccccaaacttcttgcaagatagaagctctagaaaaaggttctccagccttactaa
ctaaaagctcgagaagactgaattccatacctgttaatcgaactctttcattatttttat
aaacttgtcttttgttagtatctatttttagaaatccaatactaataatcccagagtttg
ggactccaagactagtggtgattttatcagctcttcgaagaacagaacgaatgcgagctt
ctagctccttaggagaaaatggtttaacgacatagtcatcagcacctatttctaatccgg
taattctatcgcagacttcacctaacgctgttagcattatgataggaacatctgattctt
ttcgaagttcttgacaaacaccgtaaccatctagtttaggcatcataacatctaaaacaa
caaggctaggatactcttttctaaacacgagtaaagcttcttctccatttgaagcagtga
taacttcataacctattattgttagtcttgtttctaaaattcttcttatgctggcctcat
cgtcaacaacgagaatcttttctttctggttctccaa-tttcttatttactcctataact
caccgatcatgattactataaccttaaatttttagcgctatgagtatctttattcacttt
agaa-----tttagttattataaattttgcaagtctagtttacattaattagtaccagt-
-caatattttgt-----------tttaagttatgac---------tattttaatccata
agttgactctagtttgcagaattcaatatatacata-ttttctagccaagtaaattgtat
tctattttctaaataaaaaaggggttgacaaaccagttgatagaaagtattctacattac
tataag----ttaattccaacaaagaagttttaa-----atcttctttcggagtaatgaa
tactttaat
>SRR9587922
tatatgata---aattaagacgtctgttatgtgagaatttctaaaatatacaaaactttt
tatttaatatagaaaaagtaaattaaaataagttgaacatactttgctctaatctaaacc
tcttaattgag------attctaattagagacaaaagtatttacttattttgtatctata
ggaataaaaatgaaacttgcagtttatggtaaaggtggtataggtaaatctacaactagt
tgcaatatttctgtagcactttcaaaaagaggtaaaaaagttctacaaattggctgcgac
cctaaacatgatagtacatttacgctaacaggtttttaatcccaacaattatcgatact
cttcaatccaaggattaccactatgaggacgtttggcctgaagatgtaatctataaggga
tatggcggtgtagactgtgttgaagctggtggaccgccggctggcgctggttgcggaggc
tacgttgtaggtgaaacagttaaacttttaaaagaattaaatgcttttgatgaatacgat
attatcttatttgatgttttaggtgatgttgtatgtggaggttttgcagctccattaaat

tatgcagactactgcttaatcattacagacaatggatttgatgctttatttgcagctaat
agaatagcagcttcagtacgagaaaaagctagaacgcactctctgagattagctggactt
gttggtaatagaacagataaaagagatctaattgataaatatatagattgtgttccaatg
ccagtattagaagtcttgcccttgattgaagatattagagtgtccagagtaaaaggtaaa
actttatttgaaatggcagaaattgataaggatttagcatatgtatgcgattactatttg
aatattgcagatcagttgattacaaggccagaaggtgtagttcctaaagaatctccggat
agagaattatttagtcttttatctgatttttacttaaatcctaaatcaaaggtaggacaa
gaaaaagtagatcaagaagaattagatttgatgatagtgtaaataatattttcagcataa
tataacaaggaataagataatgtctacagctcaatcagatgctcttacttttgaatgtga
aacaggtaattatcatactttttgtcccatcagttgcgtttcttggttatatcaaaaaat
tgaagacagtttttttcttagttatagggactaaaacttgtggttatttcttgcaaaatgc
aatgggagtaatgatttttgctgaaccgagatatgccatggcagaattagaggaaggaga
tatttcagctaagctaaatgattatggtgaactccgtagattatgcttacaaataaaaaa
agatagaaatcctagtgttatatttggattggaacatgtacaaccgaaattataaaaat
ggatctggaaggaattgctccaaaattggaagcagaaattcgtgttcctatagtagttgc
tagagctaatggtttagattatgcctttacccaaggtgaagatacagttctagcagctat
ggctcaaaggtgtccattaaatttaaagaatcaatcagataacacgagtcttaaaacttc
tcctcatattcctttagttttatttggatctcttccggatccagttgtaacccagttaac
tatggaattaaaaaaacaaggtatttttgtttctggttggttgccatcaaagaggtatac
tgaattaccggttataaaagaaggatattatgttgcgggagttaatccatttcttagtcg
cacagctactacactaatgagacgccgaaaaacaaaactaattggtgcaccatttccaat
aggtccagatggtactcgagcctggattgaaaaaatctgttcagtaatgaatgtaaagcc
tattggattagaagatagagaaaaagcaatttgggcttctttagaagactatatttcttt
aatccgaggtaaatcagtttttttatgggtgataatttactagaagtatctcttgcaag
atttttaactagatgtggaatgactgtatatgaaattggtattccttatatggataaacg
ctatcaagcagcagagctagctttattaaaagctacatgtgataaaatgaacgttatgat
gccaacaattgtagaaaaaccagataattataatcaagtagaccgaatccgtgatctgaa
acccgatttagttatcactggtatggctcatgcaaacccctttagaagccagaggtattaa
tacaaaatggtcagtagaattcacatttgcacaaattcatggctttaccaatgcaagaga
tattcttgaactcgtcacaagaccattacggcgaaatcttagtttgtcagaattaggctg
ggatatttatagcaagcaaagctagtattcctc----tttatgacttacatggtataaaa
gccttgtaagtct--tttattttatgttaaagaagcatttcttactctgccatcaccagc
ccaagcttgtaaactttctacttgttctttgtctgtaaatgctaaaggcacaaactgttt
gatagctattttaatatcttcagtactaaattctctctcttcactaaacgccgtatgcat
actttcaataattgcttgctcaatttcagcgccagaaaatttattacataataaactcaa
ttcatgaatatcatattcttgccacgatcttggtctaacttttgataaatgtatcttaaa

aattgactctctctcttgacgattaggtaaatctaagaaaaaaatttcatcgaacctgcc
tttccttaacatttcggaaggtaaactttgaattttattagctgttgcgacaacaaatac
tggagctttttctctgatagccaggtaataaatgtaccaaatactctggcactagtacc
actatcaccttgactatgtaaaccagaaaaagctttatctatttcatcaatccataatat
gcaaggcgataatccttcagagatatttatcatttctctcattttgattccgattcacc
aactagtccaccaaacagttttcccatgtcaagacgtagcagaggcagcatccaatcgct
tgcaatagctttcgcggttaaggatttacccgtcccctgtataccaactagcaataaacc
tttgggcgaaggaataccgtaatttaaactttgcttcgaaaatgagcgagaccttttttg
tagccattgttttaatacatctaagccaccaatatccctatttactttactataaggata
aaattctaatagatgcgtttgattaataatttgtctttttcttcaataataataggtaa
gctccgagagtctatctgattatattgtgcaataattttagtaatcactttgcggattct
atctatcgataatccttggcaggattttgttatattattaacgagctctgaattcaaatt
aagattcaaagctttactcaatcttatgatctctttttgatttctagaagactaggtaa
gggcaaatctattaccgttatgatatcattcagagcaaaaggtatattgattttgcaaga
aataataataatgtgcttagattgcgttttgattattttagctaaattgcgaagtttcct
aattaatactatttcgtttaggaaagagtcaaaatcttttaaaagaaaaagatttaaaga
ttcattattcaagttttcaataaattctaatgctaataaaggatttctttttgcataacc
attatcgctaggattgttagtgtatccatctacaaagtcccaggaatatacttgttgact
gtttgaacagtttaagctgtgttttataatatactctagtctatcttcttctctagtatt
tattacaataataggatatcgagattttaaaagtaatcgtaaatcttgagtgaaattcat
gtcaatatttgtttccatcagacttttgtacttcaccaattgaagaaagactaaataaga
ttatctaatattccttggatatattcaaaagtc------atctctctatttctaaatta
ttgctattactaatttaacttaccatgatttttttcttccttttcttcgtcgactattt
aatatagagcgtccacatggtgttttcattcttgctctgaaaccagaaactctaatttt
tttcttttagatccttgtagtgttcttttagtcatgatgtttcctctgttgtaaaagatt
aaatactattttatcatcttttatatatctttatgatatagatatgtattaatatcaaag
tt-aacagcaataatttatattaatcaagaattatttatataatattaaacattaaatat
----------taactttatgatattggctttgcccatttttttatttaagtattttgacca
tctttttgctaatcttaaactggttaatttttcagcagctaaaaacgattttgttattag
aatctcaatttaagtattttgttgataaaagccaaaatcgtaaattagaacccgaagaaa
gttttgcttttgcaaaagtatgtgtagctaaaaaatatttttctaaagctattattgaag
gtcaattagctttaaaaaactatcgagacctgaatatcttagataataatatagtaattg
ctaatttgtataatatgctaggttttatttattttgaagcaggccaaacaagttttgcaa
agaacttttatgagcaagccttacaaataaaccctaattatgttgtagctttaaataacc
tggcaaaaatttacgaagaggtaaaagatttaaaaaaagccgaatctctatacgataaag
tgttaactcttaacttaaataataaaactgctaatagaaggaaagattttatagcaaaaa

cta---------aaaacatctaatactatttgtaatcgggatagcaggatttgaacctgc
gacatcctgctcccaaagcaggcgcgctaccaaactgcgctatatcccgtaagacataat
tgttactatataccttttctctaattatgtctactctttctgtattttatctaggatacc
aaaacatgcctttaactccatctggatctttgataaatcctaatttttataaaaactaa
caacatcaggttctgcgaatagtgtaatagtgctaatttctgcctgtctcaactgctgta
ttaattgatgtatgattacttttcctaaacctaaaccttggaaatctggatgaataacta
cgtcccagatagttgcattaaatccgttgtctgatgttgctctagcaaatcctacaagtt
tactatttgcatctttttttgtattaaagaaataataatagaactatttttaatgcga
tctttactttttttagaggtcttttaacccatccgactgaatcacaaagctgctccaatt
cgtataaattaatattcttattgctactcagatagatatctttaaattcgattttatcgc
aagttttatctaaaacaataagtttttgaaatttttttcagagctagcattgatatcgg
agttttgaaaaaaattttccagaagatcataatgattatgatataatta--aataaata
tatatattacaatattgctttaacaaacaatatattaatatatgctcaaatttttacatg
aatctattaagatagaagattgttaaaatatagagaaataaaaattttgtaaaatcttt
caataaaatgatataaagagtgctttttaagactaacatctattattataaataaagtct
cgaaattgagatgaaataacacattattctatgtctaatgccttttaaaggagttatttg
tgaaaaaatctatgtgtctaacctgtttacttgctttactaattatgagcaacccaataa
tagctaacgcagaagtagctggattggtgccctgcaaagattctgcggcattcaataagc
gtatggtaaatagtgtgaaaaaacttcaagccagattagctaaatatgatgcagacacgc
caccagcgttagctttaaataagcaaatagagaaaactaaaactagatttgcaacttatg
gtcgagcaggtttgctgtgtggtactgatggattgccacatttaatttctgatggtcgat
ggagtagagccggggactttgtttttccaggactattattcttgtatattacaggatgga
ttggctgggtaggtagaggatatcttttatctgttgctaagactagtaagccaacagaaa
aggaaatcatttagatgtgccattagctatcaaatttatgtcatccggatttgcatggc
cgctagcggcttggcaagaatttagtagtggacaattaattgcttccaatgacgatatta
ctgtttcaccccgttagtaaaaaatttatatgaataataatttaccaaatacttatcaa
cagcacctgtaattggtgtattgtggatgacgtttacagccggatttattatagaattga
accgctttttcccagatgtcttatacttttactt----ataaataa--gatcgtataaa-
--aaactacataaatatagagaaatactattatttgtgtagtttta---atatataaca
taattattta-tttttaaaaatgtttatcttttcgattattttaatattagatcttacat
aataaaata-atatcaatattatgagcttagtaacccaaattatagttaatgccgatgat
gaattaagatatcctacaattggagaattgcagtcaattcaagactacttaactacagga
agcaatagaattagaattgctactattattagagataaagaaaaggagattatacagaaa
gctagtaagcaaatttttcagttacatccagaatatatagctccaggtggtaatgcagca
ggttctaggaaaagatcgttatgcttacgtgattatggttggtatctcagactaattaca
tacggagtattagctggggataaagattctatagaaacaattggtattataggagtaaga

gaaatgtacaattctttaggtgtacctataattggaatgttagatgcaatccagtgttta
aaggaagcatctttagaaatgcttggtcaagatgatattagaatcatttctccttatttt
gattatataattcgtggaatgtcataaatcatagcttcaattaattagttgaataaaagc
ttttataatgttataattacaataagcccgaaagcatataattgtaaaaactaaatcttg
tcagaaccggaaggtagcagcaataatgtttcaatacaaaagttatga-ttttcgggtgt
ttattgtttttatctaatagtctcgacttttaattagtatgctatgtataaatctgtata
tgacatgtaatccatcaaaccttccagcaaaaataatttaataatttgatattaatacgt
aaaacacctatac-tcttttataaaatttctttatatctcatctaaacaatggaaactta
acttaatgggtgttcatattttatcaacaggttcgtctgtcccaaatttttctgtagaga
atcaacaatttgaagatatgatcgaaacttccgaccattggatttcaacaagaacaggaa
taaaaaaaagacatctagctccatcttctacttctttaactaaattagcggcagaagctg
caaacaaagccttatatgcagctaacttaaaacctactgagatcagtttaattattttag
ctacgtctacgcctgatgatttatttggtagtgctagccagttgcaagcagaaataggtg
caacaacatcggtagcttttgatattacggctgcctgctccggttttattgttgctttag
taacagcagctcagttcattcaaactggttcctatgacaatattttagttgttggagcag
acacaatgtctagatggattaattggtcagatagaactacctgtatttatttggtgatg
gtgctggagcagtagtattggggcaaagcctcaaaaatagtattttaggctttaagttat
gtacagatggtcagctaaacagtcatttacaattaatgaataaacctgtaaataatcaaa
aatttggtgttacagaaattcctcatggaaactataattccataacaatgaatggcaagg
aagtgtacaagttcgctgtatttcaagttccaacagtaattagacaatgtttgaataatt
taaacatttcaatagatgaagttgattggtttatattgcatcaagcaaacactagaatca
tagaagcaattgcgagcagattatcagtaccttttctaaaatgattacgaacttagagc
attatggaaatacatctgcagcgtcaatccctttagcgttagatgaagctattcaatcca
ataaaattcaaccaggccaaattattgttttatctggttttggagcaggcttgacttggg
gagcaattgtcttgaagtggtgat---ttatatattgcggatgacgagactcgaactcgt
aaagctttcgctacacacccctcaagcgtgcgtgtataccaatttcaccacatccgcatt
ttttaccatgaata-ttaaaaatttataatatttatctaaatattcatataa--tgtgta
tatatatatacattcttgtcagaaactctcttatattttattttaagtgataaaattgag
t----ttagtatagatagtaaaattttattttaaaatcataa----aaataaatatataa
ggaaaataaaatatgacaccatctttatcaagtttttgaatagtcttattcttggggca
gtaattgtagtcgttcccataactttggctcttttatttgttagtcaaaaagacaggaca
atccggtcctaaaaataacttagaaagatatggagaagcaatttaaaatgaataataaaa
tatttttcatactaataagctctccatatattgctatact--ttagaagaatttttattt
ttttattttgtaacatcctgtaaagtttattccgttctttctttttccc-atttactctc
ttataattaatgatacaaatgtcagg-ttaataatag--tttttttattctgtatctttc
---------------atttatatacgtattattatgttcaattaataaagaaaagctca

tgtctttgtctaattggcctctcaaaaaagaaaattctgaagcatataatattaagaatt
caaaacaaatcacaattcctgatggtttatggataaaatgcttcgactgtggtctattaa
tgtattctaaagtactgaagagaaatttaaaagtttgccctcaatgtagttatcattttc
aagcttctagtaacgaaagaattgatcaattgatagaccaaggtagttggcaaccaatgg
atgttcacttgatctctacagatccattaggctttaaagaccaaaagctttatagtcaaa
ggttaaaagatactgctttcaagactggcctgcaagacgcagttcaaacaggtactggga
ctatgcaaggtaaaaaagtatgcttaggtattatggattttagattcatgggggaagca
tgggatctgttgtaggcgaaaaactaacaagactgctagaaaaagcaactcaagaaaagt
tgcctgcaattatactttgtgcatcaggcggagctagaatgcaagaaggtatgttgagct
taatgcaaatggcaaaaatttcttctgctctagaaatgcataaaaaagaaaatctactat
atatatctgttttaacctctcccacaacagggggtgtaacagctagttttgctatgcttg
gagacttaattattgcagagccaaaagctcttattgcatttgctggtagacgagttatag
aacaaacaatcaaagaagacttgccagataattttcaaagttcagaatatttatttgaac
atggtttcctagatttaattgtatccagaactcagcttagatcaaagttaatacaaattt
tgtccttacataatcatagtaagtaatgaattgaatattgactataatatcaaaatgcta
gtaaaaattaagatattaagaaattttcactaaagatatttgtagactaactagaattac
aattactattgtagaaaaatttgtcttaatgcca-ttttttcagtcttttaacaaattg-
tttttatgactttaaataattgttagtttaatacaaatttataaagagaata-----cat
actaagctaaatatagtaatattatgttttcctataatactgacctatcaaatttcgagg
aaatttatgcttaaaagatcttcttggcttgcggctttattgggactattaacagtagtt
tctacaagtacgcatacatatgccatagagttagacgaggcaacaagaactgttccatta
gaatcttctggcagaactgtaattcttacaccagaacaagttaaaagaggcaagcgatta
tttaataattcttgtgctatttgccataatggtggtatcacgaaaacaaatccgaatatt
ggacttgacccagaatctttaggattagctacgccacaaagagataccattgaaggacta
gttgactatatgaaagatccgactagttatgatggtgcagagtcaatcgcagaattgcat
ccaagtattaaaagtgctgaaatttttcctaaaatgcgcaatctaaccgatgaagaccta
tttacaatcgcaggtcatatcttacttcaacctaaaattgtttctgaaaagtggggcgga
ggaaaaatttactattagaacttcaaagacctaaaccttgtgaatactagtttatgta-t
ttgttgcttaagtgtgattattttgttagatcatgatatatattaaagatggagtttgtt
gacagcaagttacttttacccgaacttgtttacactattcgtctagttattaaaaagga
gacgttaaattgaagaagaagctttcagttcttttcactgtttttagttttttgtaata
ggtttcgcacaaattgcttttgctgcagatctagataatggagaaaaagtttttctgct
aattgtgcagcatgtcatgctggcggtaataacgccattatgccagataaaaccttaaaa
aaagatgtacttgaagctaatagtatgaatactattgatgctattacttatcaagtacaa
aatggtaaaaatgccatgcctgctttcggaggtagactggttgatgaagatattgaagat
gcagcaaattatgtattatctcaatctgaaaaaggttggtaattatacttgattttatcc

tgtattaaagaatagacaatctatttagttgttcattagattgtctattctttgtttatg
---ctattatataaagatatttacacaatatt----ttatgatgaaaagaatacccgcaa
ttcttgtactagaagacggtgcgtattataaaggatggtcattccagcaagataaacaag
agattactattggtgaagtagttttaatactggaatgacaggatatcaagaaataatca
cagatcctagttacttccatcaaattgtcgctttacctacccggaaattgggaatacag
gtattaataatcaagatattgaatctcacagtattagtattaaaggacttattgcaaaaa
atatttgtaaaatttcaagcagctggagagagcagcaatctttagttacgtatttaagta
gtaataatattcctttatttcggaatagatacaaggtctttaacccaatacttgcgtc
aatttggtacaatgaacggttgtatctctactgataatttaaatcatagttacttaaaac
agaaaatttgtgagattccaagtatgcaaggtttagatttaatcccgcatgtaactacaa
gaaatgtttaccccctgggatgaaaaaagttttccaaattggtatttgacagataacatta
gagtgcaccgagttattcagttgaaagttattgttatagattttggagttaaactaaata
tactcagaagactagctacacttggatgtcagataacggttgtgcctgcccacactcctt
taaaagatattttggcttaccagcctgatggtatattactctctaatggtccaggagatc
catcagcagtacattacggcatccagacagttacaaaattactagattacaatgtgccta
tatttgggatttgtatggggcatcaaattttaaatttagctcttaaagctaaaactttca
aacttaaatttggtcatagaggtattaaccatccatccggattgaaccagcaagttgaaa
taactagtcaaaatcatggctttgcagttgaattgacttcggtttttgaatctcctgtaa
gagtgactcattttaatctaaatgacactactattgcaggaactggacataatcaaagtc
cttatttttctgtgcaatatcatccagaatcgagcccaggccctcatgatgctgattatc
tattcgaaaatttatagaaataatgacaaagtccaagaataaagttag-----------
ttagtaattttcccatgctttatgagtgaataaagcggatagtacttgtactcctcgtaa
ttgattaaagagaggcaaatgtcctctggggccatgagaattccattgaaattcctcagg
atatctgcaaggaattctatcaacttcccagccgatttttgccagaattttccccaatc
ttttccaacactaagccatatttgtctttgaacaaatagcccaaatttaccttttgagtg
tgtatgccataacttatcaatagtttgtaaatcttgagcaggtattttttttatatctgt
aaaatatagccaattacgagtttgtgcatttacaccagctagttgaataagcttttgttg
agttagctgatctgctttgagcaaatcccgatgagtcagtaacatttgtaaatctttgta
attcatctgttgagcagagcgcaatggaacaatgccatcgggacataagttagaagcaaa
tttcactatttcttgattttactgttaagtaacttttcataaattaaaccgtcaacaca
attacttttatagttaggcccagtaatcctctcaaaaaatagatctgctaaatcttttaa
ctcgatagagtctttgctatttatattctcaataatttcaagttgttgcttga---catt
attagacttagtattcttattcaattctaaaagctgggctcggatttgatttggcatctg
atttgtttaattagttttaatagtc----ataaggtcttagagagaattagattgttttt
tgaaattagaagagcgccgcagtatttctaaagaagttcgcataatttgattcaataagt
tttgtcctgcataaattataatttgaccaattaaaaaaattgtgcttttttattttaggaa

caaaaaaatcttgtacctcaagcagaaaagtaattattatttgcattggtgacaacatct
gtaattcattcgatctgcaagcatatatatactgacagtttaagccttcttgataaaatc
cccatactttgtaacgactttcgtaaattgctcgtggtcgctcaagatagtttttaatta
tagtattccaaattaggttattctttaatttttctaatttctatcagatagaaatttta
cattacagagataacttgcatttgtctcatttacagtattaaattttgaactaacatat
gagcaataatgttgcttagctgaattaaataattttcaagaaaaatttcgacctgcttaa
ctggcatattgtaagtaaagagattaaaagttttattcacagtttgagatgacccaaaaa
gtagctgaataattaaaatttgtagtagcatcttatagtcgttcagtagatagttcaaat
catgaatatcttggtaaatgtcaatacagtccagttcatataacttacaaaatcttttga
tacatctatggaataaatcaattaaaatattttcggtcaagtttgtaatatcatttacat
ctaaatttgaattataaatttctaaaagtatcttttccagttcagctaatatgattttta
ataagtttctttttacttcagtgcgaaaaacatctaaaattaacacttcctgagagcaat
tagttaatctcttattaatctttatagaagttctaactaatagctctgctacctcttgat
tgagtgttggtccctgagagcttggccaataattattcacgttataccatagtaagaaat
attgaatttaatgttatacgataacttttttaatatactgaaaagcaagttaattactcta
tagcaactag-------tttttacagatcttaatagttatttatcaaataaagttatatg
atagtagcgattgagctaattttattaaaattta----taataatgacaacttattactt
cgcccttgcgagtcaaaattttctattagtacaagaaccactagaagaagtttttagaga
gagagttaactactatcagtcaaataataaagcaattgatttttggctaataccaaaccc
ttcttttctagagaaaccagaaatgatttcatttaaaaaccttgtacctaaagacgctgt
agctataatctctactaatccaatatttattaattggttaaagctaagaatcggctatat
ctgtatcgggcaatttgaagataacctacaactttctgaagaatcgttaaatattactgt
tttaacagacaaaatttaacaatgcttttattatgactagtgttaaattattttggttaa
gttataattatagtcttttagtattta-gactcttatagtttatcacttagtattttatg
ctgagatggagtgacaaatagtgtaagtatttcttggctaaacgtttcagttgccttaga
tttatatctatttggattaactataatggatagcattcgttttatagtaacattttcaat
ttgagcccaatgtacaattccaagttccaattccttagcaatagctgaaacggaaacaaa
ggcagctcctagccctgattgcacagcgtttttgatagcttctatagaattcaattccat
ctctattttaaagcgactgctgtcaataccatgttgactgagtactttatctatgacttt
tctaattgttgattgagtatctaacgcaataaatctaagcctgtataagtcttcttttttg
aatgtctcctagtttagaaaaaggatgtgatttgggtaatataagtgctaattcgtcttc
cgcataagaagtaacctgcaaaacatcttgcaattcagtaggcacttctcctccaataat
tgctaaatcaacttgaccattagctacgctccatgaaataagtctagttgaatgtacttg
caattgaacagccacttgtggatatctttgcctaaatagtccgattaatcttggcatcaa
atatgtcccagttgtctggctagctccaataattaatgtgccaccttgtaagttttgtaa
gtcgtcaagagctcgacaagtttcttcgcagagagctaaaattctgcccccatatcgtaa

aagaagacttcctgcctcagttaaagttgccttcttattacctctttcaaatagggaaac
attcaattggcgctctaaattttgaatttgcaaactaatagctggctgagaaacatacaa
gctattagctgctttttaaaactgccctctttggcaattgcttttaatattcttaactg
atccaatgtaaatggaaggtctgtcattaaagaattgtagtataatg---------tata
ttatttagtattgatcaatgtatgcatttgccaactaaaatatttgtcattacgcttcat
tattttcaaaataataaaaagtataatgttaaataagctgtattaaaatttagtatctgt
ttgatattagatttatacaagagatatcgtgaatttaattctaactataaaatataggag
acaat---atggactctagacttttagttgtactaataccagttttagcagcagcatctt
gggcagtttacaatattggtagagttgcactacagcaatttagaaaaatgacatcttagt
tttgcttagaatata-atttttagactttatgagatagaggactttaaaatccctatctc
tattgtatatcttaataaaaaaag-aaagc--ccccttgtacttactgttattacacatt
aatattattggt-ggtaccta----gataataagtttaagctaacttatttttttattta
taattaattaaattaataacacaattatggctgttccaaagaaaagaacatctaaagcta
aaaaaaatgcacgcaaagcaaattggaaaaatcaagcaaaaactgaagctcaaagagctt
tgtctttagcaaaatcagtattaactagaaaatccaatggatttatttataatctaactg
aagcatcagatacttttagcgattagaagtgttt-ttactacaagtctgaaaaagtatta
cgcaacaagttctcagtaatttttatctag----taatcatttgaaaatcattctatttg
ataataaaataaatagcagattaaataaaatcacattaacaaattatgcagcaaagctag
atcaaactagtgaaatttggttgttcaattgtattgaaaatattcagcatattttttta
aaagtcaattaaaatcaagtcatattactaaaatttttatttctggtactagctttgaat
ataccgcaggtctaccaggattattatccagcttaacactaagtggtagactacatccta
ttagtatatatagtccccagtctctcaaaaagtatcttgaagcatgtaccaaatattccc
aaactaatttttcttttcctattaattttcataatttacaatacggaggacaagttgtta
accaatttatacagtaatttgtttaccgttgagcaaaaagagcctgctgtatggattta
ttattctaaaaaaagaaaagcagggagtatttaatttagcacaagctaaaaccttgaata
ttcttcaaggacctatatatggaaaactcaaagaaaaggataatttttaagtccagatg
gttactatctatccggccaagacttctcttctaatacaataatgggacataaaatatcgc
ttccgttattagtcagatattctagaattatttctgagatgcattggttttgctcttatc
ctattagattaaatacttattcgcatcagcaaggagcaaagtgtttaccacataatgttt
taactgatattatgaaatcccaaatatatcaagataacagttttgttgaataatacttat
tcttgatatataatggtctcggtaatctaaataaaatactagaaataatttcttaaattt
atttgtctatattaaatttacaatcatttatgacatacgcaattattgaagcaagtggca
cgcagctttggatagaagaaggccgctactatgatttaaatcatatacctgttgatccag
gtcagtcgattatattaggaaaagtcttattattgaataaaaatggggaggttactttag
gccgcccttgtatagaaggggttacgataaaggctacagtaatgaggcacttgcgaggaa
agaagataactgttttcaaaatgaaaccaaagaagaaaatgagattaaaaaaaggtcatc

gacaagaattgactcgtttaatgatcgattctataacatcttaga------taaaatcta
gactcatattttt--aa-aaacttttttaatatttaactaaaatagatataatggcacata
aaaaaggtagtggtagtacaagaaatggcagagactctaattccaagcgtttaggtgtta
aaaaatatggtggagagcaagtaacagcaggtaatattttaatcagacaacggggaacta
aagttaagcctggccaaaatgttggaaaaggaaaagatgatacattgttttctctaattg
atggtttcgtgctgtttgaaaagtcaaatcaaaagcaaaaaacaattagtgtttattctg
ctaagaaatagttaaaacaattggtgcaagataacttgagt----gtatttatttcactt
acagtggcaccatatatttgatttattaagaaatggagttccttaatgaaccaatgaatt
ttattaaagatttgctgtgaaatttaatgacaaacactattgtaatttcctgtctacaca
atatggctgctattctatattgtggtcagatacagaaactagtagtagcaaatgctcatt
atcaagtaagtgatatctacctaggttgtgttgataaaatcttctcaggaataaatgcgg
catttattaacctaggaaagaatgagtacagtggttttatacatatcagtgataccggtc
cgcttaaaaagaaatattatgtcaataatattactaacattttaacaatacggcaaaaaa
ttttagtacaaattattaaagagccaactttaaataaaggtccaaggctcactgccaata
ttacattatcaggtcgatatattgtattaatgccttttagtcaatcaatctgtatatctc
gaaaaatatatgatgaagatgagcgtcattatttgaagtctttagctattttaattaaac
cggcaacaatgggcttgctatttagaccttctgctgtaggtgtcgatgaggaaataatat
taagcgaattaaaaaatctaaaagaacaatggaactttgttcaaaaatctgcaattaata
gttattcacctgttcttctatataaagatgaagatattgttaaaaaggtaatccgagatt
tttataataataatacaaacaatatagtaattgattcaaacctgggattaaaacaattaa
attattatatccacacttggcactgtaataactctagcacagttcctaagattaagcttt
atagtaataatcaatgtatactagatgcttttggtatcaatcaggcaatttccagagctc
tcattccaaaagttgatcttatacttggtggctatatgtttattgaaactttagaagctt
ttactattattgacgttaattctggatcttttaataattccactagtgcacgagaaacag
ttttaaaaaccaactgttctgcagcaacagaaatagcttatcagttacaaattagaaata
ttactggtgtaattataatcgacttcattgatatggaatcacaaagagatcaattgcaat
tattagaacactttaataaagagctatcacttgatgatgctaaaccacaaattgtacagt
tatctgaattaggtttagttgaattgactagaagaagacagggcaaaagtttgtacgagt
taattagtagtgattctaattactttttattttttcacacaatcagagagatctcagtctc
ttaagagattcgatgatagacagcagaaacaacagattttaataaatcttggctatctg
cagagattaatactattaacaaggtctttttttcaaaagtcaaatttgtgcagacctgcta
acttttacctaattcgtaatctctatatagttaaaagtagtattacttataaacaaaatt
atttattaactcacagatcaaaacttatttattctaaagaatacagcaaagtattaccaa
gca------gttattatctagctagtctcaataagaatagtaatcaagagttcctacctt
aagattgttatatcaactgtt---------------------------------------
-------------------gaaaaaaagaagctctctcttatataagagagagcttc-tt

ttgtagcttactaacta-tctagttagttaattaaaaattaaccgttaacagctggagct
gttagagctactggtaaagattcaccagaagctagatctagagggaagttgtgagcgtta
cgttcgtgcattacttccatacctaggttagcacggttgatgatatcagcccatgtgtta
attacacgaccttggctatcaacaacagattggttaaagttaaaaccattcaagttgaat
gccattgtgcttacagataaagctgttagccagataccaactacaggccatagacctaag
aagaaatgtagagaacgagagttgttgaaactagcatattggaagattaaacgaccgaag
tagccatgagctgcaacgatgttataagtttcttcttcttgtccgaatttgtaaccatag
ttagcagattcgttttcgcttgtttcacgaattaagctagatgtaactagggatccgtgc
atagcactgaacagagaaccaccaaacacaccagcaacacctagttggtggaatgggtgc
attaaaatgttgtgctcagcttggaatacaagcatgaagttaaatgttccggagatacct
agaggcattccatcagagaaactaccttggccaattgggtatactaggaatactgctgct
gctgctgctactggagcagtaaaagcaacggaaatccatggacgcatacctaggcggtag
cttagttcccactcacgaccaatgtagcaagctacgccagttaggaaatgaagaacaact
aattggtaaggaccaccgttgtatagccattcgtctaaagaagcagcttcccagattggg
tagaagtgaataccgatagctgcagaacttggaataacagcaccagaaatgatgttgttt
ccgtatagaagggaaccagcaactggctcacgaattccatcaatgtctactggaggtgca
gctacgaatgcaatgatgaatacagatgtggcagttaatagagttggaatcattaataca
ccaaaccaaccaatgtataggcggttttcagtactagtaatccaagagcagaaacgttcc
cacaagctagcgctttcgcgtctttgtaaagtagcagtcataatttttatcaattttta
aggatttaacaaagatgcttcccaaacaataagttgtctgttatggaaattattacttaa
ggccttttcaaatgagaacatttttaaaaaaaaca-tagcaaaatctaatctcagtaact
aa-aattt-tacttgttcatatattgcttaataattcaa-----------taataaattt
tt-gct-------------ttttgcaaaattaacactattttagggat---ataataata
atgcatattaatgctgcttattttgattttaatagtttatataattttgccctacttaaa
tacgcatgtcacactttacaaaaattcagacgaccatacaagatttaaatctgttaaagc
atgctttaacagatttgggtttgatttggcaaatgaattctagtcatataaaagttggtg
aaaataatcagcacaaggtagatattttaattaaacaagataatttgtcacatattggct
ttacttggaatgataatagatatcacttagttgccgacttacaactttgggaacaaccct
ggtctctagaagtatttttggataagttatctcagaagtatgcttactactctatcatcg
aagaaacaaagaagcaaggctttgaaaaaatgcagcagatttcacaaaaagatggatcaa
ttaagttaattgtgcaacgctggaattattaactaacatacaaatgcgggcggagagact
tgaactctcacgagattatctcactagaacctaaatctagcgcgtctaccaattccacca
cgcccgcactgttgagac--aatattactctatcataatctatactaaacacaaactaa
-tagtataaactagcttccaagcttaaaagcgtctacaaatagaccgtatgttataccaa
ttttgaagttgttgaataaattaatttttatatttaattgcttgtggctagagtagacta
ttttactagtcaactcagttgctgcaactattagactggctgcaataataccccaatctc

ctgtttgtccaggaatcgtagacaaacctgttgaaataaaaaatcctagcaataaactaa
ttaagccagttgttaactcgctcaaagaataataaagcttattatttaagtttttatta
agtaagcaaaaaaagttgaaagtctagttttaatcatagtatttagtaacggagttttaa
aaggatgacaagtattaacag-tcattaatattgtcgtatccttttaaaactcgttaata
caaacagctgttcttgt---------ttttaaaggtcttgttcgcttaaagcattaataa
gatgctgaaacggctcttgtataatatctgtgcgtttaattgattgcaaatctaattctt
cttctatgatttcttgtaataactgtatgcttcttactgttggagcaacaggtacagata
gtgaattgtaagtatcttttagcccatttaatacacgctgatcgagaatatttgtactac
cagaaaccaaggcataactagcataccgtaggtaatattctatatcccttagacaagctg
catatcttcttgttgtataagaatttcctccaggccttaataattcaggctgttcttcgt
acaactgagctgcggcctcttttaaaatattggtagcttgatcattaataatttcagcta
tttttattcgatctagtccactcgaaaagaaagattctagttgtccgacagctgtcttat
ctagataacgaccagtaagatcgtaacgatttaatattgctgttatagcatcttgcataa
attaattctcctac-ttgctatcagagcggtattataaatgactaatattgtaatataat
cattagtgtttatgaaatttatttaaaacttattctttattgcttaatgtgcaattaaaa
actcgcatagattttacacatatagagtatacgtagtttttaatta-tttttttataaaa
accaagaaataaagctttttt-taagaggttcgtgaatttgttaatataacatgagcttt
cctcctaaataagttttgtaaagtac--aaagtaaaaaaacgattagtcaataatagtaa
------------------------------------------------------------
------------------------------------------------------------
------------------------------------------------------------
------------------------------------------------------------
------------------------------------------------------------
------------------------------------------------------------
------------------------------------------------------------
------------------------------------------------------------
------------------------------------------------------------
------------------------------------------------------------
--------------------------acaatattaatatgtatcaa--------------
------------------------------------------------------------
------------------------------------------------------------
------------------------------------------------------------
------------------------------------------------------------
------------------------------------------------------------
------------------------------------------------------------
------------------------------------------------------------

------------------------------------------------------------
------------------------------------------------------------
------------------------------------------------------------
------------------------------------------------------------
------------------------------------------------------------
------------------------------------------------------------
------------------------------------------------------------
------------------------------------------------------------
------------------------------------------------------------
------------------------------------------------------------
------------------------------------------------------------
------------------------------------------------------------
------------------------------------------------------------
------------------------------------------------------------
------------------------------------------------------------
-----------------agtataaataagcttttactccagcatcaaaaaaaatatttga
gatatgttgatacaactgattattcaagtgaaaacttagatttagaccaatttagctatt
ttatcattacaattaataaaactgataaatgtattcttatagagcaattttgttataatg
ctgaaggccatatttactcaatattcttcaaaggtccaacagctaaacatctttcttcta
taatttgtcattttcaacaattaaatacaataatatctacagctcatgccatatatttag
gacgagagctaatgaagtcagaattagctctcgtgctagatcaacaatatattcaagatt
aaa--ttttgtaagtattaattatatttatgctagtataatattagtaagggcatgtaac
tcagtggatagagtatcagattccgattctgatggccgtgggttcgaatcccgccctgcc
c---gagtattcagagtact-atactaaaacatct-------taatagctagtttgataa
tagtatgggcattggggctgcaaggtttctacattatgaaaagaagaaaatatg-aaaaa
aaacaagctctccaaaagagcttttagtacaatcaataaatgcagaaaacaatattgttt
ctttttctcgaaaattagctcttgcttaaacactgttaatttttggcattcgacatgtta
aactcttattcatggcgaatattctgtcagagtcgctcttagtttaagaaaagtcagaaa
aaatatttatctttacgtttt-tacctatttttatttaaataaaggttagta-ttttcaa
cttttatagtggacgtgggttcgagtcccaccagctccatcaatttgatt---------t
gtataattagctagtttaaccttttaataaccagatatataattggaggaaatatgggt
ttcataacaattactaagcccgctttaaagcaaattgcaatcttgaaaaatgatcatgaa
aatgatgtacatctaagaataggtgttagacaaggcggatgttcaggtatgtcatactca
atgaattttgaacatgttgacaaattaaaagatacagatgaacggcttcgtctagataat
ttttctgttgtctgcgaccctaaaagtcttctttatctttatggattatcattagatttt
agctcggaattaataggaggaggatttcaattttctaacccgaatgccagccaaacttgt

ggctgcggtaaatctttttcaggctgacc-ctctaacacgtttttatatattttgtatt
atatatattatttagcaagttg-tttattagagacaagcaatgctctgttaaaaac---
---------------------------taatttcaagagtttttattgtattatt-----
----ttatatatttattatcaactacttgaaaataaattagctaagcaatctaatttgtt
atcagtaccaaactttaaatgattaactttttattgatatatgctaatcttt-atcagcc
taaacatatctgtacattacaatcttttgacataaaaagagtcttaacatacctctcaa-
----------------------aaataaattcaa----------tattttataaatactc
aacatgaatctgtcacaacttatcgttaattcttctatgatgaaaattcctaataacttt
tattgcaatataaaagctatttcttctgagaatttaagtcacttacaattatcaaaaaca
aatttattatctcctacaaaaataattgcttttggactacctaaatttgattttatttct
acacagattacaagttctattagagaaacagttgttgtgtctactattgatgataataat
ctttttgctgccccaactaatgattggataaaaaaacacaaggataaaaatcatcattct
ttgttttataaaaaagtttttaaagctttgctaaaccataaaattagagttgttgctgag
ccatt------tcataactccgaaataaataaagaaaagagcataatggcaagtcaaaga
tacatctggggtaaaagctggaagccttcaattattttatcttgcttaaaaaacaaaaaa
aatagcaaaatttctgatgcatcgaaaaaatttttaactgaacagttatctgcatcacca
gtatttgtagtaaaaaatggttttaatgaaattattttaggtcacccattatctcgtgtt
aaaagaggaggagttaataatctgatgcatgcgttttctaatttattaaatcagtctaat
gcaacttatcctatatctactggtctattctttttttcatccagatgatgcatttgaattt
aaagactttataatatcggtcaatccactggcggccaaacatatggagataagcgtagag
cctgttggtctacactttgcttataaaatgaatagaaatatatcgtcagatactcaattt
tcgtttattccagattttaaagaagtaggagatttattattcaaatatagaaaaggtaat
catttagtttttcataagaatcagcattatggcaaagatttttttcaggggcaaccaata
tatatgattcagccgatcacttttaaagatcgagccggaaagctaaatattatcaaattt
acggggctaaatgacacaagagaaattatttttactaatcttgaagctgctaatagatca
tgggcacattttataaagaggaattcacaattaaaatcaattaaaaatcctactctatta
gtctataatttagaaagtttttttaaaagatcaagaacgattggataatcaagatttaaat
aagtttgtggtgatcactaataaagaatcttatcttgccacaaaagagttaatagcttta
cccgattctaatagcttttctaagcacttaaaattaaatataaagcctaaactctttttt
gtgaaactatgggtaagacggttattttccaccttaacttatgaat-aaatgctatt-gc
ctttattttctcgaataatattctataactagtaattcattgagttgtaaagcaacccat
tccctatcaatgactccattgactttccctgataaattggatttattcaattctaagtgg
ctaggaatattagctagtccagggaaagctaagtaattttctactagttttcgagatgct
tcttgatttttttacactaattgattctcctggtttacattgataactacagatagatact
acttgtccattaatacaaatatgaccatggttcactagttgtctagcggcaggaattgta
ggagccataccaagtctaaaaacagtattgtctagcctcatctctagtagctgtaacaaa

atttgacctgtcgacccttgtagtttttagctgcttttacgtacttaaataactgtttc
tcacttaatccataattaaatcgcaacttctgttttcttctaatcttacagcatactca
gaaggttttctggacttttgtccatgttctccaggtggataagatcttttaatcgctttt
ctactgagtccgggtaaatcacctaatctacgggaaatgcgtactcgtggccctctgtat
ctagacatattatttaaattctcctaatgtaaat-gattaag----------taatgagt
agctatttttaacacaaaaagagcattaaaacaaagc-ttttgctaaaaaatatttaca
aaaaatatagatttggacaaaa---tacattactatgc-tttttaagcgggtagcgggaa
tcgaacccgcatcattagcttggaaggctaaggttttaccactaaactatacccgcaata
ttcaacagattgtcactaactcaatataacatagttatgttaagttaagataatcttaaa
tttaatagaatt-ttaaattatttatgctccttctaaaaccacatcaagaaaacgagcaa
tttcggcagcttgttcttcaacttctgataataataatggttgaccgaccctagtcaatg
gaattactcttttatcttttgtacataaataaatttcacgtcttggatttagcccctctt
taatatctatttttattgatttgatttcttttatattaaactgaagacagatttttctat
tctttccagggaaacctagccgaaaaatttttacgatacctttatctttgttaaattcat
tatatccagcccctatattccaaataatagttagccacagaaacaaactcagaaagactc
ctatactgccatagaatgtcattacaataccttgtggaataaataccaaatcggttgaat
ttgtaaaaggtaacaaatcaacttgaaaataacttgatagtccagcaagaagaaagccca
aagccccaataaatattatagtagcccaccagtaattactcagtcttcgtgaacctaaaa
ttaaatcttttctaactgagtgtatagataatgtttttttcataatttgttcttttaaaa
cagta-tactaatgcaaaatggaaaaatttgacaatttt--atgcgcgctttagattt
tttaaataagcttgattgcttttagtcctaaaaacagtcctgttgctaacccaaaaaaag
cagcaagaaatattatatatcctagaaaaactgacatattattattatttattattattg
ttacaatagtgtttacaaaaatagtgaaaatctattactacggtattatattacgttaaa
tgccttgctgtctacaggaaaaaactttttcagtttttagcattgtgtttatattagaa
ataatattctagtaaatttattaaacttctggagaacagaatcttatgtcagaatttatc
aagccttataacgatgatccttttgtaggcaatttgtccacgccagttagtacgtcaagt
tttagtaaaggacttctaggaaatctaccagcttaccgtcgaggtttatctccgcttctt
agaggattagaaataggaatggcacatggatacttttaattggaccttttgataaattg
ggccctttacgaggtacagatgtagcgttactagcgggatttctatcttcggtcggcctc
attattattctcactacatgtttatccatgtatggtaatgtatcttttactagagcagat
tcgaaagatccactacaaacttctgaaggctggggacaattcactgcaggatttctagtt
ggagcagtaggtggttcaggatttgcttatttattgctagctaatataccttttattacag
actgcaggtcttagtttattctcttaagctagtaaggggacttgaacccgtaacctactg
attacaaatcagttgctctaccaattgagctatactagcattcagttcacaataacacat
aaagatatttgttgcaatctgctattttcagatgtaacaaatatctttttttatttaaaca
gaactgttatgttaacac-------gtatattaattagaatctgaattatctgactctgc

agaagattcctgatcacagtttagataataacaaattgtttgatctagacaagttgcaga
ccaaccaacaggtgtttctcctgataattcttgcatatctacagaattataaaaattatc
atttggcttattactgtttgcttgcataaatatcactcccaaaacactctgaacttaatg
tgaagtaatttaacttctcaataatatattaacacaaaatattaaatatggtgattattc
tttaattgtaaagatgctgctaactttataaagtttgagtgagagtcttgatagcttttg
ttgaaataagcattttaacccatttatttgccctctcgaccatactttttttgtgttgta
aattaactttttgtagttttttttgttcgtttatgagaatgagatacggcatatccattat
tagctacttttcccgtaagctgacactttttttgacataattttt-------ctaacttata
tatttatttagagtacttgctaaagttgacttgggcacagctccaatcactgtgtctact
ctttctcctgctttaaagatcattaaagtaggtatacttctaatgccatattcagctgca
atagtaggattatcgtctgtgtttattttttactaccttaatagatgattcatattcttcc
gcaatttcatcaactacaggagaaaccattctacaaggaccacaccatggcgcccaaaaa
tctacgagtacaggtaagttattgttaataacttcttgtttgaaagaggcatctgtaact
tgagatactgacatattctttaacctttaatatatgttccttgctaagcaaatcttatca
caaatttcaagaaaaaatcttctatagtgaattactattagtgttgaattattttttttaaa
aattagacattaaaaaaacttattacctatataaataaagtggtggccgagtgatatctt
gatagtgataattttttatattaaggttgaacactagaaataacttttcaagcaaactctt
gtttaaaaa--atttctctgtgaccataaaaacttaattatattacaatatattttatta
tttgtaacttagttctgataatggtataaacaacgcaaaagatactgccttataatcaag
gaggaatacatgtctcaatccgtagaatcacggactaggattaaaagcgaacgttacgaa
tctggagtaatcccctacgctaaaatgggctactgggatgctgactatgtgattaaagaa
acagatattctagctcttttcagaatcactcctcaaccaggtgttgacccgattgaagca
tctgctgcaattgcaggtgaatcttcaacagctacttggacagttgtatggactgattta
ttaacagcttgtgacttatacagagcaaaagcatatcgagtagatccagttccaaacgtg
gcagatcaatattttgcttacatagcttatgatattgatttgtttgaagaaggttccatt
gcgaacttaactgcttcaattattggtaacgtttttgggtttaaagctgttaaagctctt
cgcttggaagatatgcgtatgccagtagcttatctaaaaacgttccaaggtcctgcaact
ggattgattgtagaacgtgagcgtatggataagttcggtagacctttcttaggtgctaca
gttaaacctaaactaggtttatctggcaaaaactacggaagagttgtatacgaaggcctg
aaaggcggtcttgatttccttaaagatgatgagaatattaactcacaaccatttatgcgt
tggagagaaagatttttatattctatggaaggtgtaaataaagcatcggcttctgctggc
gaaattaaaggtcattaccttaacgtaacagccgcgacaatggaagatatgtatgagaga
gccgaattctctaaagaggttggtagtatcatttgtatgattgaccttgtgattggttat
actgcgattcaaagtatggcaatttgggctcgtaaacatgacatgattttacatttacat
agagctggtaactcaacttactctcgtcaaaaaaaatcatggtatgaacttccgagttatt
tgcaaatggatgcgtatggctggtgttgaccatattcacgcaggtacagttgtaggtaag

cttgaaggagatcctttaatgattaaaggcttctacaatactctacttgaaagcgacaca
gatatcaacctacctcaaggtctgttctttgctcaaaattgggcttccctacgtaaagtt
gtaccagtagcatctggtggtattcatgctggtcaaatgcaccaacttcttgattactta
ggtgatgatgtagttcttcagtttggtggtggtacaattggacatcctgatggtatccaa
gcaggtgcaactgctaacagagtagcactagagtccatggttatggcaagaaatgaaggc
cgtaactatgtagcagaaggtccacaaatcttgagggacgctgctaaaacttgtgggcct
ctacaaacagctttagatttatggaaagatattagtttcaactacacttccacagataca
gctgatttcgttgagactccaacagcaaacatctagtttaatgactacttactgatactt
taaatagtcaattgtaagtggaattaacttataacaataaggagcatagaatagtgagac
taacacaagggacttttccttccttccagatttaactgatgagcaaattaataaacagc
ttgcttatatcgtttctaaaggcttatcagcaaacgttgagtatactgacgatcctcatc
caagaaactcctattgggaactgtggggtttacctttatttgatgtaaaagatgcttctg
ctgttatgtacgaaattagctcatgcagaaaagcaaaacctaattattatgttaaagtta
acgcttttgataatactagaggtattgaaagttgtgtaatgtctttcattgtaaatagac
ctgctaatgaaccaggattcttattacaacgccaagacttcgaaggtagaactatgaagt
atagtcttcatagctatgctactgaaaagcctgaaggagctaggtattaatattaattaa
gaattaatattggctaatta--ttaccctcttaaaaatcataattaatttagtaattgtg
atttttaagagggatagcttcccaagtaaattttgactaaa-tagattgaataattaaaa
atacacagaaataatgcaatcacaggatataatttccaacgatactcttgttaatttaca
agaagaatatgatagaacacaaatccaagaagttttaaatgagttaaatcaagaacttat
aggattagtgcctgtaaagaccagaattcgcgaaattgctgcgctattattgattgacag
attacgcagaaaactagaactagtttctggtaatccaggattacacatgtcatttacagg
tagtccaggaactggtaaaactacagttgctatgaaaatggctgatattttgcacagact
tggatatataaaaaaagggcatttgttgacagttacaagagatgatcttgtaggtcaata
tattggacatactgcccctaaaactaaggaagttcttaaacaagcaatgggaggagtttt
atttattgacgaagcttactatctatataaagcagataatgaaagagactatggctctga
ggcaattgaaattttattacaagtaatggaaaaccaaagaaatgacttagttgttatctt
tgctggatataaagatagaatggaaaaattctacgaatccaacccaggactctcttctag
agtagctaatcatgtagacttcccagattatacttcagatgaattattacaaatagctaa
aatgatgatagaagaacagcagtactgttttacagaagaagcagataaaactcttttaga
gtataccgagcgaagaatgaaacagccttattttgctaatgcaagaagtattcgcaatgc
tattgacagggctagaatgagacaagccaataggattttgccagtggagaaaaagtatt
aacaaaagctgatttggtaacgattgaagcagaagatatcttgaaaagtagattattttc
attacctaatgcttaatatacacgtgatttcatgaaactatatattattaaaaag-tttc
ataatcagttgcaaatttcttgaaaaacctttattattatcttaatgcaggtg-tggcgg
catagccaagtggtaaggcagaggattgcaaatccttcatcccccagttcaaatctgggt

gccgcctagtactaaaaaagggggtgtggtggaatggtagacacaacagacttaaaatc
tgttgattttagtaatcgtgagggttcaagtccctccacccccatatattgatttataa
ataaaaagagatgtgtatttgtattaattgcaaccatattaccaaatgtaatacctatca
cttaatagaatctcaacataaacagcctcatcttactagaagtccattgtttatacctaa
atatcctgtagttcatgttaatatatctaacaactgtacctataatcaaattgattggga
tttagtggagtgtttatcatttgtagaaaaaccgaatagttggaatttagacgctaatta
gttacaagaaaaattatgtaccataagcaacaactctatttttagatactcggtattaa
ttttggatgttctaaccagtaagattttacccgttcttgctatctctataatgccaaatt
tagttagtaattgttcaatagcaacaatctttcccggatctccagtaacctctataatta
aaagatcttctgcaatgtctactatcttagctctaaaaattcttacaatttctaaagctt
ctgttctagtctgagaattgatctgaatcttaattaacattagttctctttcaactgaag
gaatatttgttacatcttgaacgttaaggatatttactaacttatataattgtttcgtaa
gttgttcaatagttctattgtctccttgaactaccattgtgattctagagaccccaattt
gctctgctggtccaactgctaaacttgcgatattaaaccctctccgggcaaatagaccag
atattcttgacagtactccggcttcatcttgaactaaaactgataaggtgtgtttcatga
aaaattaaatttaatctatttgatctgttaagtatattctaagaaaattacagaaaaaat
actagttagtctataactaaattatccaaactaaacaaaatcctactatcagataatata
tctaccttgtaatgctatactattaacttagtatttaatgatattctaattgcaaataga
atgct-aatttttaaaatttgttaggaacttgtgctagaaaaatacaaaaacagtaaaa
aactctctcgagatttacctcaaatcaatgatcgcattagatttccaaaagtccgagtaa
ttgatgacgaaggtgaacaactaggtatttttgtgcctgaagaagctatacaattagctg
tccaacaaggtttagacttagttgttgttagtgataaatcggacccgccagtatgccgaa
tattagactatggtaaatataagtttacacaagaaaaaagagctagagaagctaagaaaa
agcaacataacagtagtattaaagaagtaaaaatgcgatataagatagaagagcatgatt
ataaagttagaataaaccaggcatccaaatttattcaagcaggagataaagtaaaagcaa
ctatcacatttcgggggcgtgaaatccagcactctaatttagctatagatttattgaata
aaatggcaagcgatctaaatgcaatagctgaaattcagcaagctccatcaagagatggca
gaaatgtcataatgctcttatctcccaaaaaagttagctaaactaatttttatttaaatg
catctggctggattcgaaccagcgacgtctctttcgaaatggcggattatgagtccgctg
ccttcggcccctcggccacagatgcatcacttaaagtttatacctcataaagcatataaa
aatcaatgtttagctatttttaataaagtctataagagtctattattcattcatattgtg
atatgtcgcaactgctgaaggggaaacttgatttaagtatcgaaatatccagtatttaaa
aattgtgtcgagtataactggaaaagttgaaataaataaaaaaataaaatctcgactttc
cggaagtcctaaatgtcttaaaattacttcaataataacttcccatccatgaggagaatg
aaaacccacaaacatatcagtaaaaagaatgattaaaaaagctttagctgtatcactaag
accataaataatttcatttaaaaaagatttaacgactgcaatttgtctttgtccagttat

cattagtaatataaagacaagtattgatacaagatcggataaaatatttttaacagcatt
tgcactttcatttgcataatattctcccaactctctagccttcagctgcactctttttc
aatgatttcgtaagatatatcttccgaaggattcagaagaacttcaaagtgaattttctc
ttcaaatctttgtaattcagcaaaagctctttcctcttgagaggaatttaaaaaaatttt
aggctgttcttggttccataaataatcaatacaaggtccaaatacaaaaaatttgaagc
ctgattaactagtactggagagatgagtaataataaaatatattttacggaagtaatagt
ttggtgtctagatattctaaactcctctatagcttcagattcgccattaggatctaactc
tttacgaaacttttcaaaagtattagtaattgacctaggtattggcccaactttttcaaa
ggcgaattgattatttcgttttaaattccaatatttcatttttattgctgctaaaatct
ctattaatcaataattgatcacgtattcactataaatattaattctagaactaaacatat
aaaacaatatttatgaaaaataaaatacaattaaaaattaattatgcattagttctatct
gcagctacagcagtttatggatataaatcttatcagccaactataaatacactcttaatc
aaagatgaagaaaactggcctttttctctaaaaggcagactgcctttattcctataatt
gtattaaaaaaaaatcatcaaaatccctgaactcagaacctagaaatagtaagttatatc
ttaatggaattggaagtagaaaatttatcgacagaattgtcactctacttggatatttag
aaaaacaggcaaactatactgaatacttgaaccaaataaaaacaagtggttatttttctt
tagtctacctagacagtgatataatgtataagaaaaatattagtaacatttttttactac
ctaatgatattctaaaaagaatctatatttgcaataagaaaaaaaagcttataccgcatt
cattcttaataaatttatttcagaaacagataggctatccaaaaagttttgctaatttaa
attgggccttctccaaaattataaaatggtattatgatagaggataccaatggtctctag
ttgaagttaaacaagcgtctgatgcatcttctattgtaattgatattcatgaaggggtag
tgaagacaattataacagaatactacactctatcttataaaagagtctcaggtatcttat
gcgtagagtcaatagagcagtaccttggagtaagagtgggagctccactaaatataattg
atttacaaaaaaaaattacttacttaaaagataatcaactagttggcgacattatttata
gtattgaacgatcaaataatagctcaatgagtttggatatcaaatttcaaatacaagaat
taaaagataaagagataatagtgcttgcaaaaagttcttcta------------ttatct
cgcatgcatgtaatttacttaatc---aatataga---------aataggctagtagctt
ctaatatagtttcactatcaactaataaattacacgcatgctattttaattataaacttg
actatcaatataagtatataaacacgtctaatcttataaaattgttagcttattctattt
cttgtaggaagacactacttattcattattttaatttacaaactttaataagctatacaa
aaaaaaatactattggatttcagctatatttacgaaatctaagttttgggaaagctttct
gtgtgcttagtatgaaatttatcaaaaatggacttaatataaaaatttatatataaatc
cttcattgatagttgatcagaattttgtatttcaatttgctatacaaatcattaagcaat
atcatactgcta------aacctccagctt------------------------------
------tatttttaacgaatctagatcttgaacaatatgttgctgaaagtttattaatgt
accattttacatcttgcttctcaatatctgaaaaaatattattatctcgaattatgcata

cagattctttatttttcaactctgaaa------attttcattttgatgatagaactaacg
cgatcaacagttatgatacttttaaacaaaatacaaaaatatttatcaagagtttcttt
ctttattattaagtttacgttatcaaaattttaattatttaggttggcctttaaaaggcc
atttttttgaaataaaatctttatacttggctccatttcaaaagagtg------------
---atttttctgatagtcgtaagaccctattcttccacaaaatgtcgttaaagcaagtat
ctaattttaatttaccaatatcttttaaaagccaccttaaccatatttagttagtacta
ttaaatgtcagtcaaatttgaatatgagaactgtatctttgctattaattgattcacctg
ctgaatatatgctatacaaatctattcttaatttctctattaaagttaggatgcagtatt
tcatacctatgagcaacaatataagattatctctattttataattatttagattgttttt
taataagatcatctcagagtcgcattcatatatggcaagatctaagaactcttacaccaa
ttcaaaatttttggttaaaaaaattttcatatggagctggaattcagcttaaattaccaa
ttaagcagatgccacctctttcaattgaatatactgtaactagtagtcgctacttttgta
tttaccttcgtacttattatcaacgataactattactaaaatgaaaaatttaatttcttt
ttttgatcgcagcaaagggaaatggatttcacaaagaacaacttatgaattatctaataa
aaacatgagttcgatacagtctcagatgacaatgaaaataggcaactcattgtcaggatc
tataatacttgcatcgttaaattggggggacatttatagacaagttgcccattacgctaa
aaatcacagccggagtgaatatgataataagtttaacttgcaattcagtaatcaactaaa
taatcataaattattaacgctgtgcatagtgacagatcctagtctaattagttttaaaac
tcgctatggaagtactaccatagatgaaacatattggttcgcaacaaataatttacgtct
aagtactagtattgttaaacgattcaacacttgtgtggcagtatctttttgttcagaaat
taaagtttaa-actctatatagaagcaactaatagcctaaa-ggctattagttgcttcta
tatatctaaattttgtattaaaaattat----aatactatattattctaagtctttacga
ttgggattcctggaaggatcattagagaggaacccaaagacaaacaaagaaataaagaaa
ataacggtagtatatacaaagattttcaaagtaaacatattttcatccttaaaagctatt
agtgcg----atatttacatatctccacataattttatatcatgatataagttccagcac
tattttataaataaatcaatatctttatttttagtaatgatttatatttaaatatgcgt
gatgtactaaatactaaatcttttccgacttgttttgtcttgtttgtaaatattttgtta
tttctatggagtttaccttcaattatcacatagtctaacttctttagctgtttaaaagta
tatgaagatttttttgttccaaatagttagattgattataactaattgttttctttttaac
agtcttgcttttaacttaattatttggctttttattttcactaattctgatactcttacaa
ctaagaatttggactagaagagtacagctattcataatttataacaa------tttttta
aaaaatctactaaaatagatgattttacagaactaaaatgatatttcattatcttgttga
ccttgctcataagtaatatctttcaatacttttaatatttttccaaaatactcttgcagg
tatttctctaaagatattacttccaacctgtcaattccaagaatactgtaaacttcatcc
atcggagctgtaaattgctctccgctacttaaaatttcggcaaatgctaaacgatcagaa
atattccatgtccattgaagagttttagtaattcttcttaaggcttttaataatccaatc

ggaatttgtgagatctgtgtttttttgcccagataatttttcacataatttaataatttca
gctgatgtccaagcggtattaccaactaacggcagggtcctattttctgtggaaggtact
ccaagacttttataactaactttgctgcatcttgagtatcaatgtatgcaatgggtgta
gattctcctgtaacccagactgatttttttgtctaaaataggaatggcatattggttgatt
aaaccttgaaaaaatcctcctaaagaaaacacagtatatttcacattggacttctgaagg
aaatctactacctgagactttaagttcatcaatggaacatctgggtatttctctgaattt
aatattgaaaagaaaataaatctttcaactttagccgctttagctgcttcaattaatgca
attttcccatctaaatctatttttttctgcattgtaaggatcagtaggacgagatgtagaa
gcatctatgattgctgttaccccacaaaaagattgcaaaatactttcgggtaattttaaa
tcaccatatataagttctgctccccactctttgagaaaggcagactttcttaaatttctt
accatgcattttacattgtagccttcatctaaagctcgtcttacaatttgacgtcctaaa
gttccggttgctccaattactaaaagagtcatattagttaaaaataagtatgtataaaga
cattgatttggttgacggcagagagtaaattaggtagagagggaatcgaaccctcatgac
cgaagtcgtcacattttgagtgtgatgcgtataccaatttcgccatctaccctattatag
accaactactacaagtaatttagtctttagcgtaaaaataatactaagtttcaatggata
tgtcaacttttaaatttatattaataaaaaaacggagtctcttatggccacttttgaatt
aattccttatagattgtatttgagtcaatctaacacatggatccatagaataaaagcaga
aattaagatatatatagtaacgcttttatggatttcaattttatttttttcttactttaa
actatgtattattgctttaagcttaattgccataagttttactataagaagtaaacagaa
tattatccaaaaacatttgttacaaacgttattaataacgttcttaactactgttttgtc
ttttagtgtggctattagttataaacaatatgcagaacaagaacaatcgcagtatttatc
tgattccaagaaatataaaaattccagcagctactattatatacaaatagctaacgatca
aaggatcaaaagacaatacttaattactactttaaagccttccttgtactttttttattac
tatatattctattaaactagttatgataacaacttctccagaagttttagtaattactat
ttatagatctaggataataaataaaatatttaaaaatgaattgctatttatcttccttct
ttcgtcacatattgttactagtattattaatagaatcgataaagtgattcaagtaactag
cttaagaggaagcttgaatttatataattcattgacaaggcctttaatgttttctttgtt
aatatttcaagtcttctttttggagattattcgggagtcaaaagaaatagctcaagctct
ttatactagaaatctcaatcaagaaaacaataatttttttgaaaatatatacagtaaaatc
taactttagtgatcggctcaatataattattagcactttgtactttattattttagctct
agcgtaatagaatccacttattcttctgactctagtaatacgaagagtaataaaattagt
tagtagtttaaatagatagaat-taagatattatataaataattaaatgtttcattatgg
ttaatactcaaaatcaaatttcacaaacttcagaacttgattatatagttaatcaacctt
ataaatatggctttaaaacttctgttgaatctgagcaatttcctaggggaataagtgaag
atattgtccgcttgatctccaagaaaaaagatgagcctgaatatctattgaattttaggc
ttaaagcatataaaaaatggaaaaagatgagtagtccgtcatgggctcatattaagcatc

cgaatatagactttaatacgattatttattacgctgttcctaaattaaaaaaagaattga
aaagtctagatgaggttgatccagaaattcttgacacttttaacaagttaggtatatctt
taaatgagcaaaagcgaatttccaatgtagccgttgatgctgtttttgatagtgtatcta
ttgcaactacttttaagaaagaactgtctgaagctggtgttatattctgttcaatttctg
aagctataagggattatccagaattaataaaaaaatacttaggtactgttgtaccagctg
gtgataattattttgctgcattgaactctgcggtatttagtgatggctcttttgctata
tccctcctaatacagtttgccctttagaattatcaacttactttcgtattaacaacgaag
aatctgggcaatttgaaaggacactaattattgctgatcgtggcagtaaagtaagttatc
ttgaaggttgtactgctcctcaatttgacacaaatcaattacatgcagcgattgtagaat
tagtagctcttgaaggagccgaaattaaatattctacagtacagaattggtatgctggta
acaaagaaggtaaaggtggtatatacaattttgttactaaacgaggcttatgctcgggta
acaattcaaaaatttcatggactcaagtagaaactggatctgcaattacctggaaatatc
caagctgtattttagctggtgaaaattctcagggagaattttactctgtagctttaacaa
ataactatcaagaggctgatacagggactaaaatgatccatattggtaacaatacaaaaa
gtagaattatttctaaaggtatttctgcgggtagatcgaaaaacagttatcgaggcctag
taaaagttggccctcagtcatttaattctcgtaattattctcaatgcgattctttattaa
ttggtcaatcatctcaagctaatacatttccctatattcaagtacagaatccaacatcaa
aagtagaacatgaagcatccacttcaaaaattagcgaggaccaaatttttatttttac
aaagagggatcaatttagaagaatctattgctcttatgatcagtggtttttgtaaagatg
tctttaatgaattgcctatggaatttgctactgaagctgatcgtttactgagtttaaaat
tagagggaactgtaggatgagccagactattttagaaatcaaagatttatatgcttctgt
tggtgaaacaacaattttaaaggggtaaacttatctattcgagccggcgaaatacatgc
gattatggggcctaacggttcaggtaaaagtacattatcaaaagtaattgcaggacatcc
agcgtattcactaataagcggagacattttatttttggacaaagtattcttgaaatgga
gccagacgagagggcaaaagcaggtatttttttagcttttcagtatcctgttgaaattcc
tggagtcagtaattctgatttcttaagaattgcattaaatgctcgcagaaagtttcaagg
attatcggagtttagccccttagaatttttcaactaataacagaaaaaatagatcttgt
tggcatgcaagaaagtttttttgacaagaaatgttaatgaaggattttctggtggagaaaa
aaaacgtaacgaaattcttcaaatggctttactagatagcaaaatatctattttagatga
aacagattctggactagatattgatgcattacgagtagttgcaaaaggaattaacacttt
agctaaatcaacaaattcaattattttgattactcactatcaaagattattagattatat
tattccagattttgttcatattatgagtaacggacaaattgtaaaaactggtagtgttac
tcttgcccaagatctagaaaagcacggatatgactggattacgcagacataactatttac
tac-------aatattgagactaaagaatattctttagtctcaatatt-tttgcaaatgc
aaagtt-aattatttaagctttatcaaaaccttgcttaacatcttcaattgctcttttta
gcaattcttcactaactggttctaatttttagtactacgaacattttcaccaaattctg

gtttagaattctttaaatcttctcgaagttctctaataaattcagctacttttgaaactt
caatatcatctagatagccattaatgccagtgtaaataatagcagtctgttcttcaactg
gaataggagaattttgagcttgtttcaaaatttctcgaagacgttgacctcttgctagtt
gattttgggttgctttatctagatcagatgcaaactgagaaaaagcttctaattcagcaa
attgtgctaactctagttttaacttgcctgccacttgtttcatggcttttatttgcgcag
cagagcctactctagagactgagatcccaacattaatagctggtctaatccctgagttaa
ataagtcaccagatagaaatatttgtccatctgtaattgaaattacatttgttggtatat
atgcagatacgtccccagcttgagtttcaataatagggagagctgtcatactacctccac
ctaattcagcattcaatttagctgctctctctagcaatctagagtgtaaatagaatacat
ccccaggataggcttctcgtcccggaggtcttctaagcaaaagagacatctgacgataag
cttgagcttgcttagttaaatcatcataaataactaatgttgctttacctttgtacataa
aatattcagctaaagccgcacctgtataaggagcaatatattgtaaagtcgcaggactat
ctgcattagctgcaactataatagtataatcaagagctcctttatcctgtagtgaagata
ctacttgggctactgaagatgccttttgtcctatcgcaacatatacacaaacaacatctt
ggcctttttgattaataattgtgtccaatgcaacagctgttttacctgtttgacggtcac
caataattaattctcgctgacctctaccaattgggatcatagagtcaatagcagttatcc
ctgtttgcataggttcacaaactgattgccttccaattataccaggggccattgactcaa
ttagccttgttccattacttgcaggttcacctttatcatctattggccgagctaatggat
caacaactctacctaaaaaagcatctccgaccggaatttgagcaattcgaccagtccctt
ttacggaactgccctctaaaatatccctaccatctcccatgagtactacaccaacgttat
cactttctaaatttaaagcaaccccaatagttttatcttcaaattcaagtagttcaccag
ccatgacttcatcaagtccatatactcgagcgataccatcaccaacttgtaaaactgttc
ctatattggctacttctacatcctgatcatacttttcaatttgttgacgaataatactac
ttatttcatcaggtctaatatttaccatattttaatattattttttattaattggaaatt
ctaaagcggtaattagtcagactaatgctatctgtctttttttactttattttaagttact
acaacatcaaggtgtgaagccatttgtcttagctgtcctctaatactagtatcaatgact
tttgatccaatctgaatagtaaagccaccaattagctcgggctcgactgaaataattagt
ttaacttctttagccttagtcataaccttgatttttttctgttaatagagtttcctgatca
gggcttaaagcaatagaagtattaatatttgcaattgttaaagattccatttggtaagcc
agttccaaatacttactagctatagcatctaacattccaattcgcttgcgatcaactaaa
atcattagaaatgataatgtattttcgttaatttgatcgccaacagttgcaataattact
tctttttttgcctctacagtttttaatggattagct---aaaaaaatttttagcttgtca
gattgcgctaaaatattttgtattgactgaatatcttggctcaccttttcagtaactttt
ttagttttagctaaatcaagaagagctactgcataaggctgagctattttttataacaaca
ttgttactgctcatagttgatctcctaacttagcaatattattatcaattatacgcagtt
gcatttcggagctcatttgattttccagctgcaaagtaactcttttaagagcaagaaatg

ttatttgttgctggatttgccttcggatctgtttttcagctgtttctatatttgatttac
ctgttatggcaagtctttcaatatctaattggccttgagctaatatggaacttctaactt
ttccagctgttagctgagcttctttcttaatttgatctataatgatttgagtttgagcaa
gttgtttttctgattctgacaatctagcacttgcttgctctagtctttcttcggattctt
gtatagctgctaaaacttttatctgtcttgcatttaaacttgatccgagaaattgcttca
aaacataaataagaccaaataaaagcaacaatatattaataacatttgcttcaaaaatat
cagaattaaagccgaatttgtgttcactactatgttctgataaaatagtaattatttgtg
gtatttttactatattattcatttaaaattct----------tactattttaatatttta
ctctggaatcatgattttcaagattgactacttaataatttaactttaatttgatcgctc
aaagtgtcaacctgattttctaaagttttaagagcttcttcttttgaacatttaactgt
ttcgatgcttcagcaatcagtttctctgcattcaattgagctttttttatatcttctgag
actatatttgagcttcttttgagaagaggctatttttaattgtgcattgcggcgagct
tcagataaatcttcttcatattttgcagctagctcatctgctttaacaagcatagacgat
gcagtagtcaatgttgttcgaatatattcatctcgttcgtctagtactttagttacaggt
ttataaaaaatagtatttaataaaaccattaaagtaagaaactgtaatgccattaaaggc
aaagtaccattgaagtcaaataagccaccttcaatttcttcggctaatagaaatggtaaa
tcaatcattttactaattatgtttatagttaataattttgtttcaattaagaaaaataac
tttatagtgcgcttagtctcatcagactaagcgcaaatgattattttaaccaacgtatgg
attagcgaacaataaagacaacgcaacaactaggccgtaaattgttaaagattccataaa
tgctaaacttagaagaagtgtaccccgaatctttccttctacttctggctgtctagcaat
accttcgacagcattagctgctgcactaccttgaccaatgcctgggcctattgcagcaag
acctacagcaagaccggcagcgataacggatgcggctgaaacgattgaatccataatgaa
ttttaatttttttgttatatgagtagaaaattgacaaatttcgtataatacttaaaatcta
tataattaagtataatagattctaatat-----tgtaattatagtatattactcttctcc
atgaccttccattgcttctccaatataagctgcggacaaagtagaaaagattaatgcctg
tattgaactcgcaaataatccgagtatcataactggtaatggtattaaaattggaattag
caaagtaaatacagaaacaactaattcatctgctaagacgtttccaaataatcgaaaact
taaagatagtggtttcgtaaaatcttctagaatattgattgggagtaatactggagttgg
ctgaatatatctagcaaagtagcctaaccctttttttacttaaacctgcataaaaataagc
taaagatgtaagtaaagacagggccacagttgtattaatgtcatttgttggggcagctag
ctcaccttcaggcaagtgaattaacttccaaggaattaatgcacctgcccagttacaacc
tagaataaatagaaataaagttgcaatgtaaggcacccaagggcgatattcatgctcacc
aatctgatttttttgcaatatcttgcagaaattctaatataaattccataaaattctggaa
tttttcaggaattctttgtagatttctagtacctagaaatgataaagttaaaagtgtagc
aattactaaccaagagacaatgaaaacttgcccatgcaattgtaagctacctattttcca
atatagatgttttccaacttctactgctgataaactattatatgggttaaaatctatgag

actgttttgatacataataaatattt-ttttaaatagtgcaagaatcaaaaaataag-tt
tttgaagaattaatatgcaattaatttattcttaaaattagactattaataaattaaaag
caaaaataaattctaatagagatagctatttttagtagtgattatttttttgtaaaaaaca
attattgctacctataatgcttttacgtttaattatatacccatatattaattatttaat
atgattataggtattttgctacaaactttattgttttatccattct----aaagattgtt
atagtttatttttttatttaaaatatctgcgacttcatctgcaaaattaactttcaaattt
tcttctccccctcccagaagaaatcttacaaatcttctaattttaatattttcacctaga
acagcaatattttggtttattaaatcttcaatactaatatcttgattgcggataaacatt
tgatctaataatgagagctccttaagtctctttttcattcgtccttcaatgattgtttcg
attctatcaactggtttgtttttcaagtcatctttaccagcttcaactctttgttccaga
ttaataatttcactaggtatatcctgtgttgatacgtattccacattaggcgatgccgct
atttgcatagcaatatctttagctaacttttggaactctggacgtcgtgctacaaaatct
gtttcacagttgacttcaactaatacacctattcttccgccagtatgaatataactctct
agtaaaccttcaatagcggtacgattagatttcttgttagctgaagccagccctttttgt
cttaatgattctaaagctttttcttcattgccattattagcttgtaaagctttcttgcag
tccatcatacctgctccagtcttatcacgtaaggcttttacgacttgagcagaaatttgt
agtgtcatatgctttcaaaaatttttaagtatttaaatt----------aattattataa
ttcactttgagtaggatgagttatttcttctaaatctgtttgcccgtatttaccatcata
aattgcatctgctattttacctacaattagtttaattgacctaatggcatcatcattagc
tgggattggaatattaataatttctggactacaattagtatcaagtatacaaatagttgg
aatacctaattttaaacactcttgtatagctgttgtttctctcttttgatctacaactac
aacaatatcaggtaaacgagtcatatttttaataccatttagatgtttacgaagtttatc
taattctcttcgtaaaacagcagcttctttttttgggtaactgatcaatcataccactttt
atcttgttcttctagctgtctcagacgattaactcgagacttaatcgttacccaattagt
taacattcctccaagccatctctggttaacataataagaatcacagcgttgtgcttcctg
tgccactataccagcggcttgtctttttgttcctagaaataaaactttttttaccatccga
cgaagcctgttttataaattcacaagcttctgttaataattgtgcagtttggactagatc
aataatatgtatgccatttctttctgtataaatatatggaaacattttaggattccatct
acgtgcttgatgtccaaaatgaacacctgcttctagtaattctgctaatgtaacaatagc
cataagattattaatatttaaataatgtaattattcgggaaatccc-tttgtttagaaac
tgat-----tcaactaacttatttgaattttagtaaactaataataaattaacacacttt
taaagcagatcaacaaaatagatcctgaagaattcat----tctacacttttattgctaa
aataattacgcgcggttctgtcgtctaaaatgatatcatctaaatcatctcgtacagatg
aaatagtactttcgtcattagtatttgcagttaaattctttttctctaacgatgagccat
tacagttatcatacatattgaagccggtaccagccggtataagccgacctataataacat
tttctttcagacctcttaaccaatctaatttaccagatatagcagcttcagttaatactt

ttgtagtttcttgaaaacttgctgcagaaataaagctttctgtatttaaggaagcttgag
taatacctaatagcacaggcctatatgatgcattaatcttatttctcaaggttatggctt
tattagtctgttcaatcttctgtaactctacaagctctccaggtaaatagcctgtttctt
caccattttctattttcactttagaagtcatctgccgtactattacttctacatgcttat
cagaaatattcactccttgagattgatataccaattgtacttctttaacaagtagtaact
gaatctcttgaaagcttaatcgagcagcttcatataaggctaatccacattcaatataca
aatcaaaacttgcatcgaggatatcatgaggatttaacaatacgtctgttttctttcttg
cttcaagaatttcttcaattcttggaagcccttgtataatgtcacctgttttgctctat
caaaaaccaatatagctaatgtttctccccttctaattaatgcattgttatcaacatgta
aaatagcaccattagaaaccaagtaaggtctagcaattcttaatgtaactgatctagaag
aaatttcaataatttgtccagagtctaaagaagtcatattttcagtaataaaatcaccac
aacgaatccaatcacccacacttactttaactattgattttcctatattaaaggtctttt
tatcagaagatgtcgcaatcaaaatgcgcctgctagtattcttttcagaataaatctcct
caacagtaccttcactcattgcaataatctcagtactagcaacaactgtaaatggcttga
tatattcaccatttttacaataattctagtttgactttgttgcttctctgatttattta
catcaatacttttaattgataatgtttcaaaagttgacagccttaagtcaaaataattat
tgccactttctttagcagaaaactcaattgaggatattaaattatcatctcgatgctcaa
tttcagccacaagattagttgttactaaatgaactccttctatagatttcactctttctc
cgtcccgaaaaggagttcttttaactagctttagtttagtttgacttggtttttgtgaaa
ctagattttcaattagactggattttgtttctggtattgaataaacaataacaggtctaa
taagaatatagggcatttgctcgttttctatatactcccaataaactaacttatctgtag
aaatattattatgaagtctttctccaggtcttaaaaaacctctactttatcattattac
tatacacctcactaagtttatagatagatccaggcttaataataatttctcgaactattc
catccttttcgattatttcaacaatacctgagcttttagaaaaaatatttttaactagct
ctgttccactttctataacatctccattatgcactagtagtagtgatgaatctttattaa
tttcgtgagtttcttctgaaatccataagatatatcccggacttaaaatttcgtaggcat
ctttatctaagcctgttttctttttgatacatttaagtctaaatatttaataataccac
cacttgtcgttttatacgtatcagatattagttctgcgactgtgtaaccatcttgaa---
ttttttggtttggaatagctttgaaaagaaattttttgtttttttttctgtttctaaaatat
aagattctttttgtggattacatctgtataaatataacaacctggaattacaatagatt
cagtaataacttgtatattggtaatactgctattagtattctggtgtatccttacttctc
cagcataatgatttattaattctgtctgtgcaagtatcgtaccagctttagttttatctt
ctttatgaacaataacattagcagaatcagatatactataaacttcacctgagagcaccc
agatcaagcctccagttttagtaattcttgtagtatattgattattatctgtttcttcaa
ctgttaagttggagaaacagattttgccagagagatctgagactacatgtttttgagctc
tttctgtcatgagtcgatttctgcggggtgattcagcaatcacttgatccttcgatacta

attcaccatcacatattaaaagagtagtacctttacttaagttaattattgacttctgat
ttttcttagatctgattgtaacttgggttggctttttttgttattaatgcttgctcaccat
gtctggtacgaacatctgtatacgattcaatatctagatctatcaattgaccatctatag
gtgcgtagatttgttctgctaattcaccggtaaatactcctccagtatgaaaggtcctca
ttgtcaattgagtgccaggctcaccaatagattgagccgcaataataccaacagcctctc
ctaaatcaactaagcggccatgtgcaagattccatccgtaacaatactgacaaacagaac
ttctagagttacatgtaacaggagatcttactaaaacttttttt---gataccagctttag
ttatctcttttgctaacttaggactgacatcttggttcgtatgagcaattaaacaatttg
tctctggatggaatacattttctgctaatactcggccagataaagcttgttctagattaa
ttaatattttctgcgtatctactaaatcttctagtatgatgcctttattggtcttacagt
ctacttctctaattattacatcttgagaaacatcgacaagtcgacgcgtcaaatagccgg
aatctgctgttcttaaagcagtatcaactaatcctttttcttgctccgtatgatgatataa
aataatcagtaacagttagtccttctctaaaattactcgatattggaagatcaataattt
gtccctgtggatcagccataagccctctcatgcccactaactgtcttacttgagaaatat
tacctctagcacctgagaacgccatcatataaacagcatttagaggatctgtctctttaa
agtatttaataacttcttgtttcaaggattcactcgcattattccacgtatcaataactt
tttgaaatcgttcaactgctgtaatctcacctcggcgatatttgttttcagtggccttaa
tatcttcaatcgtgcctaataaaagactttgcttactaggaggaattcttaaatcttcta
aacttaaagaaattcctgcctgggtcgcataatgaaatccaagatctttaagcttgtcgg
ccatattggctgctcgggctataccataattacgaaaagcccagacaattaaatttttta
gttcgtttttatcaataaccttatttgaaaaacttggctgtgcaagacttcgtctattat
ccactaaatcttctccatattaaatgttcgtt--attaactatgctactaaagactcttg
aataattttattaaagattatacggccagctgttgttctaatatattgcacaattctctg
attttcagcgtcttcttttatgatatggttattaaaaaattttgttgtactattgtcaga
atgtttttcaatcttaatcggatcatgaggttgatcaccatctactaatccatcaaaacg
agcccatatataagaatgtaaatcaattttcttctgttcataagccataactacatcttc
taagcttgcaaaatactgattagatccctgctgctgagatggattattagctgttaaata
atagcaacctaaaaccatgtcttggcttggcataataattggttgacctgttgctggaga
gagaaaattatgaggggctaacattagcaatctagcttcggcttgagcctctaaagacaa
aggaacatgaacagccatttgatctccgtcaaaatcagcgttaaacgctgggcatactaa
tggatgaagcttaatagctcttccttctacaagaataggttcaaatgcttgaatacctaa
tctgtgtagtgtgggagctctgttcaatagtactgggtgaccttgaataacttcatttaa
gacattccaaatagaagattcatttttttgaatcatctttttggcagctttaatattgtt
aactaatccctgcaaaattagtctgtgaataacaaaaggctgaaataactccaaagccat
ttctcgtggcaatccacattgatgtagttttaaatgaggacctacaacaattacagatct
accggaataatctactcttttacctagcaagttttgtctaaaacgaccttgttttccttc

tataatatctgacaaagattttaatggtctattatttgcacctacaacagttctacctct
acgaccattatccatgagagagtcaacggcctcttgcagcattctttttttcatttctgat
aataatctccggagctaaaattgactttagacgagataatctattatttctattaataat
tcgacgataaaactcattcaaatctgccgtcgcaaatctacctccatctaattggaccat
aggccgtaagtctggaggtatcacaggaatgactgtgaataccatccaagaaggatctgc
acctgttgctatgaaattttcaatcaatcttaagcgtttcatttttttattgaattttaa
tgaaggtgtcttaaaactttttggtggatttgtagcttcagaccttaaggtttcagctat
gtgctctaaatctaaatctttcaacagtttctgaatagcttctgcacctattcctacttc
tacttggttattctcatcttgattttggtagatttcctcttcaagacttttccattcgta
accttcgagaagttgcttgtattttaaattaatatctgtattagattgcgtaactacata
agaatgaaaataaacaatttttttctacttctttgacttttaagtctaatgctaaagcaat
atagcttgtacttccttttaagtaccaaacatgagttactggtgaggcaagttcaatata
agccatccgatgtcttctcacgcgtgattctgtaacttctacgccacatcgttcacaaac
tatacctttgtagcgaaaacgtttatatttaccacagtgacattcccaatccttaactgg
gccaaaaattttctcacagaataaaccatccatttctggctttaaagttctgtaattaat
tgtttcaggctttgtaatttcaccaacaatttgtccgttgggcaaacttctttcgcccca
ctgtcgaatttttttcgggagaagctaaactaatttttacgtagtcaaaatattgctcaaa
ctttgtcataaatttgaataccttaaaatcaaataattagccaatttatttacatgcttc
atattgaaaaatgaatatcttaataaaggaattgttcaaaatcgtctactggaggagtat
cataattagaacgatctactcggttatctttagaatccgacatcaagtcaacttcaactg
tccgtctttggccatcttcaaatagtttcaatttatgtactgcaatgtctaaacctaatg
actgcagctctcgcattaagactttaaaagattctggagttccaggcttcggtataggct
tccctttaactattgcatttaatgcttcatttctagcttgcatatcatccgactttacag
ttaataattcttgcaaagtatatgctgcaccaaaagcttctaaagcccacacttccattt
cacctaatctttgacctccatgctgtgctctacctcctaatggctgctgcgtaactagag
agtaaggaccagtggatcttgcatggattttatcatcaacaagatgaacaagcttaagca
tataagctctaccaactgtaacaggattatcaaaaggctctcctgttctaccatcaaaaa
cttgcattttacctggatgctgatcattaaaaagccatttattgctagtaatcaatgatg
cttcttttaattttctatttacaagtgctcttgatgcttctgctccatacatttcatcaa
aaggtattattttaaatctctttcctaaatagccacctgctaagcctagtaggcactcaa
atacttgaccaacattcattctagaaggtacacctaaaggatttaaaacaatgtccactg
gtgttccatctgataaataaggcatatcttgtttaggtaagatcctagaaatgatacctt
tattgccgtgacgaccagccattttatcacccacttggattttctcttctgcgcaacat
agacgcgaatcatagcatttgttccgggaggcagctcatctcctttctgcctagtaaaaa
ccctaaccttaactactctaccttttgcagcattaggaagccgtaaagatgtatctctca
catctcgagccttttctccaaaaatagctcgcaacaatttaccttctggtagttgatcag

cttcacctttaggagtaatttttcctactaaaatatctccagcttcaacccaagaaccac
cgacaacaattccatttctatctaaatccttcagagagttatcgctgacgttaggaatct
ctctagtaatttcttctggtcctaatttagtttgacggcattctacttcatacttttcga
tatggatagaagtatacaagtcatcataaactaatctttcgctaattaaaaatgcatctt
cgtagttataaccttcccaaggcatataagcaactagaatatttctccctaaagcgattt
cccctccgtctgtggatgcgccatcagctaaagtttggccaacaactatttttttctccta
cccaaacaattgggcgctgattaatacaagtatcttggttagaccgataatattttttta
agcgataatgaactgttctaccactattatcctgaatgcctattttattagcagaaacat
aatttacatgaccagatgttctgctaataacaaccataccagagtctcgagctattttag
tctcaagaccagtaccaattattggcttttctggatataataatggaacagcttgccttt
gcatattagatcccattagagctctattcgcatcatcatgttccaaaaaaaggtattaatg
atgttgcagctgaaattacttgaataggtgaaattgcaatataatctacttgggttgggg
ttgttgtaataaactcttgacgataacgtacaggaataatatctccttcaatgtaatgtt
gtttactaactttaacatctcctggagctactcggaaatcatcttcctcatctgctgtta
gatagacaggactattgtgataaatcacttgaccttggtttaccggataaaatggggttt
ctatgaaaccaaaaacattgacccgcgcacaggttgctagtgaaccaattaaacctgcgt
tcggaccttccggagtttcaattgggcaaattctaccatagtgactaggatgtaaatcac
gaactgcaaaacctgccctgtccttattaaaacctcctggacctaaggcactaattcttc
tcttgtgagttagttctgcaactggattagtttggtccataaattgagaaagctgactag
atccaaagaattctctcactgacgcaattaatggcttgggattaattagatttgataaac
tcaatgaatctatatcacatatcatcattctctctctaataatgcgttctaaacgattga
gacctactctgaattggttctgcagcaattctccaactgaacgaactcttctattaccaa
gatgatctatatcatcaagattacctgagtttttatccttaatattaattagataatcaa
ttgaggataagatgtcttgaggagacaatactcggaacgttttaggtatattgagaccta
gtttttattaatttatatctaccaacttcacccaagtcatatcttttaggatcaaaaa
aacgagagtaaagcatctgcttagcaacagcaactgttgctggttcattaggtctaagct
tagaataaactatgagcagggcttcctcatcagttacttcttcaatattattttttccaa
tttctttagctaattctttaactgaatatatttgggatgcagaaataagaaaagcatatt
tactaagtcctttttcaatttcatctttattgagtccaatagctcgcagaaaaatataag
catttactttgtgggtcttatctattcttatccaaatttctcccttaggatcaatttcaa
attttaaccaagaacctctattagaaattaggcttgcactataaatctgcttaccatttt
tgtctatctcttgtttataatagattcctggactacggattatttggttaataataactc
tttccgtcccagatacaataaaagttcctctatttgtcattattggcaagtctccaataa
aaattaatctctttttgtatttttttatttttttatctgcttttcagcactgagctgtagat
gggttgatgataaatctaatgatttaatatttttttcctggtttttagaaggtagatcaa
tatcttttcttgttaactttgcaggaacatatatttgagcactataagttttatccctat

tttttgcctgtctgacactataacgaggaaattttatcttgtactcattaccaaataatt
gtaattcgagtcgactagtagggtctgatatcttcggaaaaatttcaagaacttccgtca
aaccttctaataaaaaccatttaaaactggccctctgaatttcaactaaatctggaagaa
gtttatttttaagcttatacgttgaaccatagatctcctttataacaaattttgaaatc
gaagagtttaa-tactattaatagagtactggtagataatgcataagcatagtttgctac
attatttcttctctagacataactagaacatcatattttattcaa--aaaaagatttta
aaatttttcttcac--acacactatctataagtactttgctgcta--aaagatagttcaa
gctattaaa-------aaaaacgaaattaatacttgtctgtctgtcaattaattgacatc
aagatagacaagtattatttatttaatcaattctg-tttttctgagcaaaagacaatgct
ctcgcaagtctagctttttttctagcgcctgtattagggtgaaaggcccctttttttaca
gctttatcaatcttactatatacttgagaaatacttgactgcacatcattgaaattatca
ctttctaaattatcaatatttagaaggcatcttttagttaaagttttgacaaccgattta
tacttacggttaatgagacgatttctttccgaagttttaatgcgtttaatcgcagaaagg
ttcttagccacagtaaatccaataaaactttataaaatatttttagaaacgaaaataca
caaatctgggatctattcaaaaagggaatatgcggcgtattataacattaaagaaaaaca
ataggcaatagttcaccaagagctcttattattacacaatagagattgtcattgcgaaaa
actttgtaagatagtatttaattaaaataaactaactagacatataaatatatatgaaaa
aaattgaagctattattagacctttcaagcttaatgaagtaaaacttgctctagtcaagg
aaggtattggaggaatgacggttatcaaagtttctggatttgggagacagaagggtcaaa
ctgaaagatataaaggatccgaatactctattgatatcattgataaaataaaaattgaaa
ttattattagcgatgataaagtagagaaaattgtagagacgattattaaggcttctaaaa
caggggaaattggagatggaaaaatatttattagtagtattgaaagagtaataagaatca
ggactaatgacttaaattttgaagctttatagtatttttacttt-tttagacttgatact
tcttcactaaacatgagataataagcgtttattgtatacttataactaaggtgtaata--
ctaatggctaaaagcaaaggtgcacgaatcgtaataactttagagtgctctgataaagct
ggagagtttgctcaaaaaaggaaacctggcgtttttcgatatacaactactaaaaataga
cgaaatacaccaagtagaattgaattaaacaagttttgtcctaattgtaatcagcattgt
atcttcaaagaaattaaatagttatttatcttattaaaat--aatcatattatggctata
tacagaaaaagaatatctccaattaagccaacagatgctgttgactacaaagatattgac
ttgctaagaaaatttattacagaacaaggcaaaatactgcctaaaaggtcaactggattg
acttcgaagcagcaaaaaaaacttactaaagcgattaaacaagctagaatcctttcttta
ttaccttttttaaataaagattaa--atatatagcctaacttcatatttttccatctaaa
tattaaaaatttctatagatttgtttattaaaatgatctaacatactgaat---------
----------tactaagaaga-tctactcattacaagttttgttcattttataattaata
atagcatcataattta---ttaagctagattcagacaagacataaag-gtaacctctttt
aattgattatattatatagtctatttaaaagaggaatttttttgcttc-----------t

agcaattcaagattctttaagctttaacaaggtatttaaaacaaagatttttgtttaggt
atcagatcaaatgaatgaatcttatctcccgactgccataattgaaattcagagatgaaa
attccacattcgttgccagcttgaatttcttcgacatcttctctaacgcgctttaaagat
tcaatttttccttgataaataactttttcttctcgaatcactttaatccaagagtttttt
aatagcttattattgatcactcgacatcctgctatttttctatttgctaacgagaatact
gtacttacttctgcttctcccaccggtacttctgaatattcaggatcaagtaagtcttcc
atccttcttttaatgtcttcaattaaggcataaatgatttgatagttttctattaatata
ttcgactttgccgatgcttgctttgttcctggagcaaagttggtattgaacccaatgagg
gtagaatttgtggttgaagctagttcaacatctgtagcagtaatttctccaggcatgatt
gagacaacattgagttgaactttactctgaggaaattgagataaagaatctaggattgct
tctgtagagccttgattatctgttttaataattaaagaaatttgcttactaatatcttta
gaattagtgtttttttaaagtatctaatgtaattcgactattaagtgctctttgtttctgt
ataattgatgagtctttagaggtattttcaatcgctttaagtttggcttctttatcactt
ttaactgctaaagctatttctcctgtagctggtacagatgacaatccccaaatttcaaca
acagatgatggtattgctaaattaattttctctttagcgttgttaattataactcgaatc
tttgcgtaagctgatccaataaccaaattatcactaatattcaatgtaccattttgtatt
aatagagttgctactggaccatgcgatttatctaaatgggcttctattataataccttgt
gcaggctgtgttggatcagctttcaaatcctctaattcagctaacaaggtaatagtttct
aataatttatcaacattttgaccagtaagagaactaataggaattataggcacttggcca
ccaagtttctccgacatgacattgtattttaataaatcttgctcaataatatctgtatta
gaaccagctttatctatcttagatattgctatgacaaaaggacattcgctttttgaata
tgattaatagcctctatggtttgaggtttaactccgtcatcagcagctataataataata
gcgacatctgttaaattagctcccctagacctcatacttgtaaaagcttcatggcctgga
gtatccaagaaaacaatttttttgtttattatctttcttaatatactcaacttcataagct
gcaattgcttgagtaattcctccaatttctttgttagcattattagattttcgaatataa
tctaatagcgttgttttgccatgatctacatgtcccatcactgtgacaattggaggtctt
ttaatataattgccagtttcataaagagcattagagtgatctaaattactagaaagtcca
ttattattttcctttatatttgattctactgctataccaaaattatcagccactgatgat
attattgaggcgtctatggtttgattcatagtaactgatattcttttaagaaataaatat
tttataatatctgtttcttgaacacaaattagtttagataattcttgaatagttagagga
ttagtaatactaatagactctggcggagaattagattgaatgcttgctgaattttgacta
ggagttactgctactttttttctgtcttatagatttttttattactagtcagtttttttaca
acttccacttttggcttaggtggacgcattaaagagatagctaaatctcctgcagtctgc
gaaacattagaattagagtctctaaagttgtca---tcatcatcgtcaatatgtatttta
gtttttattttt-ttgtgctgtctattctta--ttttttttactatctactaagtcgtga
attttattaaatttttactttttttatctagtttaggtggagaacttaattctagatgc

tgttcaccacc---tgtatgtgattctgatttatcaaggtctaaatttaacagattatca
ttattgactgactctaatctaattttatagataatctgaggatttttcaaatctacaatt
gtttcagaagaattgatatttgagctagatctagaggatctattttcaaagttttgatta
tttaaaaacataaatattggttttattatctaatagtaactttctaattttaatacagag
cttaatataaacaagaatcgaatatctgtataaataatacaaatgt---agagaattaat
ctatgttgtattagatttaatataacactaaagtcctttgttcgttatacaagtttaaaa
ctttt-tatgatgataaacactagattctagtttattggtatacaatctatttatctaaa
taatactgtttttgtcagggcttttaaa----ttatctaatgttatgttcattatgaaat
tctattaaatttaaaaa-ctcaacaattaagtaagaatcgttaatcacctccacgccac
ttatgccacgatctcaaaaaaatgataattttattgacaaaacttttacagtattagctg
atattgtattaaaaatacttcctacaagtaaagaggaaaaagaagctttctcttactatc
gagatggaatgtctgcacaatctgaaggtgagtatgcggaagctttagaaaattattatg
aggctttaaaactagaggaagatccatatgatagaagctatattttatataatatagggc
ttatttatgctagtaatggtgagtatgtaaaagctttagaatactaccatcaaggattag
aactaaattttaaattacctcaggcccttaataatatagccgtgatataccattaccaag
gagttcaagcaattgaggataaagatacagagctatctaaattaatgtttgataaggcgg
ctcaatattggcagcaggctattaagttagctcctgataattatattgaagctcaaaatt
ggctgaaaacgacaggacgaatgaggaatatacaaggatattaatatatttaagatataa
taaagatatcaattaataaattaaaattagttaaagaaactggtactatatagagtactt
-tactatatattataatattatactagttga------tctctaagacttagctgaataaa
cgaaattaaatctattgactatattcatgaatacaaataagcataatggttagtacaaca
aaaagtaccggatctgttactcaaattattggaccagttttagatattgcatttcctaac
ggacagcttccgaaagtattcaacgcactcaaagtacaaagctcagaaggaactattact
tgtgaagtacaacaacttttgggtgacaacaaggtacgagctgtttctatgagttccact
gaaggactacaaagaggggtagaagttattgatactggatcccctatatctgttcctgta
ggtacagatactcttggacgtatttttaatgttttaggtgaacctgtagataatttgggt
cccgttgattctgagagtactttacctatccatcgaccagcacctgcttttactaagtta
gagacaaaaccaaatatttttgaaacaggtattaaagtcgtcgatttacttgctccttat
agaagaggtgggaaaattggtttattcggaggtgctggagtaggtaaaactgtattaatt
atggaactaattaataacatcgctaaagctcatggtggagtatctgtatttggaggcgtt
ggtgaaagaacaagagaaggaaacgacctatatatggaaatgaaagagtctaaagtaatt
gatgcagataatctaaaagaatctaaagtagcattagtatatggtcaaatgaatgaacct
cctggagcacgtatgcgcgttggcttaactgcattgacaatggcagaatactttagagat
attaacaaacaagacgttctattgtttattgataatattttcggtttgtacaagctgga
tcagaagtatcagctctactaggccgtatgccatctgctgtgggttaccaaccaactcta
gcaactgaaatgggagcacttcaagaaagaattacttcaacaacagaaggatcaattaca

tctattcaagctgtatatgtgccggctgatgatttaacagacccagctccggccactaca
tttgcacatttagatgcaacaacagtactttcgaggaacttggcggcaaaaggaatttac
cctgcagtggatccactggattcaacatcaacaatgctacagcctggaattgttggaact
gagcattattctactgctcaagaggtaaaatcaactttacaaagatataaagaactgcaa
gatattattgctattcttggtcttgatgaactttcagaggaagacagacaaactgtatca
agagcaagaaaaattgaaagattttatctcaaccttttttcgtggcagaagtgtttact
ggatcacctgggaaatatgtatctttggaagatgcaattaaagggtttcaaatgatctta
aaaggcaatttagacgacttgcctgagcaggcattttatctagtaggtgatatagatgaa
gctatacaaaaagctgacagcatgaaagattaatataattaatgatgactttaaatataa
gaattattgctcctgatcggactgtttgggatgcagaagcacaagaaattatttaccaa
gtagtacagggcaacttggtattttaacaggccatgcacctttgcttacagctttagata
ttggagttatgagagtaagagtagataaagaatggatgccgattgttttgctgggcggtt
ttgcagaaatagagaacaatcaattaactattctggttaatggcgcagaagaagctagtc
aaattgatttatcagaagcagaaaaaaatttagacactgcaactcaactcttaagtgatg
cctcgtctaataaagaaaaaatagaagcaacacaaaaaatacgaaaagctcgagctcgag
tacaagctgcaacagcagcaacttcgtaa-aagattcc------tcttaaaacataaaac
tttagtaatgattactaaagttttatgttttatgttttatttgtatggttataggcagct
taaataaacttagcttagacctgaacaaatataatcaaaatataatcccatctctttacc
tgcatctggtccaactaaactaatagtaacttctttcatagctaaaatagcttgaatcgt
tgcaccaataggaacacctagagaattgtatgtttcttttagaccatttaatactctttc
ttcaaggatagatggatctccagctagcattccataagtagcatagcgtaagtagtagtc
taaatcacggatgcatgcagcataacgtcgagttgtatacatattacacctgggcgtgt
aatatccgaataaagtagagattttgctacggattctttaataattgttgctgcattagc
tgcaatagtagcagcagctctgactcgtagctcacctgtttgaaaatagcctcttaattt
ttcaacagagctatcatctaaatatttaccttggacatcagctgcattaataacagaagt
aattgcgtcttgcataactttcacacattccttaattttaaatatctaatacta--gggg
tatataacaaaattactagatcttttagcaatatcagtattactgcatagcacccaaagt
ataatcaaagtaaaaaccagcttctgctgaatcttcacctgcaagcagtgagcaagctac
acttttcatgcattttacaccttcagcaactcctgaaattggtgttcctaaagaattata
catttctttaacacccactaatccaatttcttcaataggagttatatcgccagcaactat
tccgtaagttactaagcgaagataatagtccagatctcgtagacaagtagcagtcatctc
ttcgccataagcatttccaccaggagaaactacatcaggtctttttttgaaacagttgttg
gccgccttgctttacaatacgctcacgattatctgttaaaatttgagctattcttaaacg
acgttgtccagataaaacaaaacttttaattctatctaattccccaggacttaaatatct
tgcttctgcatctgcatttacaattgactttgtaacaatactcatggataaaactcctgt
aatgcttt-aaaaaaaagcttaatactaattatttatgatataactcagaaagtaatgta

cttttttgag-aattttataaccgaaacacgttcatgatatatgaagcaggatagatactg
aggtaagtactaaaacgagaaagttatatgtaggaacttctgctattgatttcctagtac
ttttttaaaacttggtaccacaattgatatatcctgcttagtcaaacgattatagagtgt
ttctgtattaggaaagttagctgctggtaaagtcggaaatcttctgtatggaacagtatc
tttgccatataccgcattgtattcaacactttcaactaaattacttataaatgcagataa
gccttttgaagctaaaatttgattaaagaatctaatttctgcctgattattaggagctct
tcctaagatatgcttagtacctaattctataacttttgtatttggatatggctgataaaa
ttctttaccatatagttcagatagcgccagtttttctactagttctttgacacagatttg
tctattcaagaaagctgacttaatatccaagaattcaccgccgacactaaaagagttaag
atctctttcgaaaatttgacgatatgctgctcttaaagcttgttctaacatttctttgtt
actatcactattgacttcaaagacaatagattgatctcttaaagaagttactccctgtga
aatacgagactgaatatcattactactcctcatttctttgacagtgcctaattcaacaaa
cctagcagatttactagcaataactttttttgaatctctggtcaataattcctggtcttaa
gcttcttaaagcaacaccagctggagtactatatctttcataaggaacagtattatctcc
aaaagtttctgtgtactccgagctatcaattattgtgtctattacttggtaatagccttg
cttatatgcaatatcaaaatatttattaatttcttgccgaccatatgtagggcgacccag
taagcggttatgaatatattcaatagctttacaaatatataatggctcccaataaagaga
tctaaatatacttgattttgctaactgtctaacaaactcgcgaactgtaatttgattatc
ttttagctgactttctattggcttgagaattagtttttcttcttgatacacttctctgcc
aaaaactcttaaataagcaacttttgtgacaacttctactgaattttcaaaattttggga
tgaatctgactttataagagcgttggacaatttaaagatttttggtcccagggatcctgc
tgacttaggtcttacctgtggattactaatctgattgtagattcctggcccacgtctaac
taatatccttcttgtatctttaccaaaaagagcttgtctttttcttggatctttattttc
tttagggaaaatagctccaaattgaattgataatggatcattacctattccatatggatg
ttgatcaggtaatgattgcttataatcactaaataaagttataaattgaggaaccttacg
aaaaggcgcactataggtgaaaagatctatttgaggcccccaattacggcactcttgagg
ctcttcacctaaatttctaaagtaaggcacagtctcttcaccgaaataatctgtatactc
agaagaatttaaaagtgcattcactaaaccacttaaaccagtagatgataaaatagcaaa
atatttttgaaactcttctaaagagctaggacctctacccaagaaatgtctaaatgctaa
ttctaatgctcgactattgacaaaaggttcataaaattgctttctataaatacttgaagt
tcctaacgaacgaatgaattccttaattgaaatttggccatttttttacttgcgactccaa
attcgataaagagaggtcatatgctttagcaatatctctttcaaaaatttgtctataaca
tgctttcacaacaatatttttctcatcagcagataaactaggcttcataacaaaacgagg
tgttgacactcctgcctgtacataagtttgaggaagacgcaaaccttgtaaatctccaga
tattctttttcttaatttatcagtcaaagatggagcttcaaattcagaaatcacaacatt
aaaatactctttaactaaatcttgaccttttatatcttcctcaaagattaataatgctgt

ccgtctcatctctctaagagccacaattgctgctgcacttgagcaagcgttatcaattaa
ctctctcaatcctcgtatattaacagataaaatattagggtcgcctgatacaattgcata
tgtcaaataccgtaaaaaccagtctaaatctcgaagagattttctcattcttgtagtacc
gtatctcaagacgtttataggtttaaaccctggcggagtagctcccccagcattaaataa
agatcggaaactttggccaaaatctccttggatatttcctgataactcattaattttatc
ttgagaactttgatctcctgcaataataacagctgcttgcggacgttctaaataagaaat
tgcagacccacctacgaatattttatctgctgctcgtgctaccagaatatttgcattttt
agttaaaatatctgcaacttccagtctttttgtccagaatttaaaaaagaaactaattg
atttaattcaccaagctgtaaaaatcggtcttgttgctctgcctgagtaatagttaaaat
tgaggcagtccgataaagctgggggcgtgctaatgggcttccgccacttgctttgatact
cattactttatttatctccttaactgatattgcttgctagcgaatacaaatatagcttga
tatttctaaatatttaaagatactatgcaagcaca-ctgttttctatattattaggtact
agtacttatagtacagagagtcttataactaacttctataaatacaaatttgtaatact
tctattaacatttgtcaaaaaaatatttaagtgctcactcttttatattaaaattcaatt
agtgagagattgatgtgaaataaagtatactaagtaagttttatagataggaaaattaaa
agatttattctacaaacattaatca------------cttgaatgtttagcaattgtaca
attgttattgcttttaattggtttagactagttttatatttttctaacataaacaatat
taatgagcaatttgtatgaatttaaagccacaaaacaacctattaactaacgaaaatata
aatcatgtagatatacccgaaaatgatataccaatgtcaattacggaacatttagaagaa
ttaagacaacggactttatttgtgtttttattctttttgtttgctacaactataagtttc
acacaaattaagatcattgttgcaatactgcaagctcctgctgttggtattaagttttta
caactggcaccaggagaatattttttttcatctattaaggttgcaatatattgcgggatt
gttgcaacaactccgtttgccgtttatcaagttatattatacatactcccaggactaact
ggaaaagaaagaaaaattattttgcctctattaattagttctgtattgctttttattaca
ggtggtatatttgcttactttgttctcgcaccagcagctttaacattttaattagctat
gggtctgatattgtagaaccattatggtctttcgaacaatactttgactttattctatta
cttttacttagtacaggattagcatttgagataccaatcatacaattattgctcggtgtt
tcagggacattttcttctagtcaaatgatacgagcttggagatatattattattatagca
acaattgctggagctattctgaccccttcgactgatcccgttacacaattaataatgtct
tcagctgttttattactttattttggtggaattgttatattattagtcttaaaaaagtag
agtaaat---------atttcta-ttaatccaatatatttgttatc-----tgggcaaa
taatctataaaaacaattgct------------------agatttaagtatcaggcttct
aaatatttcaa---------gtctcttttatcatttaaatacaatc-------------
------------------------------------------------------------
------------------------------------------------------------
------------------------------------------------------------

------------------------------------------------------------

------------------------------------------------------------

------------------------------------------------------------

--------------------------------------------------gatacattgg

cttgtttttcatcaattatttgtgttgcacac-tttttagcactctctttattgttctt

agaattatttctttaatctatttattaaat----agatatgaagagatttaagtattgat

tatggtaatctatatcaacgtaaaaagtattttattgtgagttttatttatataaaggct

tattcattatatagttataacagtaagtctaattaatag-ttttcatgcttaattaaagt

aataactcttaattcttttggcagaaaaatctgtcttaatattttatc--aaaaaaacag

-aaaaatgaagctaaatagtctaattaacttaattcaaaagtctatatattcttgtacac

ttttactaattattttaaatattatttgtgtcgcacctaattctagtaatgcatttccaa

tttatgcgcaacaagcttatgaaagtccaagagaagcaactggtaggatagtatgtgcca

attgtcatcttgctcaaaagcctgtggagatagaagcgcctcaagcagtactacctaata

ctgttttcgagactgttgtgaagattccatatgacagcaatgctaaacagattttaggta

atggcagtaaaggaggcttaaatgttggagccgttgtaatattacctgaaggatttaagt

tagctcctgttaatagattatctacagagttaaaagaaaagactagaaatctttacattc

agccgtacagtgctaaacaagacaacattttagtaattggacctatttctggtgataaaa

atagagaaatagttttcccaatactatctcccgatcctgcaaaagataaaaaagctcatt

ttttcaagtatccaatatatgttggcggaaatagaggacgaggccagatttatccaacgg

gtgacaaaagcaacaataatattgtctctgctttaagcagtggtaaaattaataaaattg

aattactagacaaaggtggatttataatacatgtgactaacagtagcaatgtagagtcaa

cacagaagatttcacctggtctcgaacttagagtaaaagaaggggatacaattcagcttg

atcaagctttgaatagtgatccaaatgtaggtggttttggtcagaacgaaacagaaatag

ttttacagagtccaaatagaattaagggcatgattgttttctttttcgctagtgttctag

ctcaaattttcttcgtattaaagaaaaaacaatttgaaaaagttcaagcagctgaaatga

atttttaaagcaatataaaactggcacttactataaagaaagta-taaacactttattca

caggatgtaccagtttttatgttatgtaataatcgctttgacggaagttatgatactttt

aacagaatctacaatctgactaggctgaattactgtagcctgctctaagcttccattata

tggtgtgggtatatcctgcgaagatagcctcacgacaggagcatctaattcatcaaaaag

gtactcattaatttgtgcaattagctctgctccaattccggctgttttcatacactcttc

tacaattaaaactttgtgagttttctttactgagatagatatagagtctatatccagagg

ttttaaagatataagatctataacttctggatcataaccttcttttaataaggccggtaa

tgcttggataacatgatgcctcattctagaataggttaaaatcgtaatatcttttccttt

tcgtacaaattcaactttattgagaggtaagaaatattcttcttgaggaatctcttcttg

taaattataaagtagaacgtgctcaaagaatacaactggattattgtctcgaattgcaga

tttcagtaatccttttgcattataaggagtagaacaagcaactatttttaagccggggat

ggcttggaaataagcttccagtctctgagaatgctctgcacctaactgcctaccaactcc
tcccggtcctctaataactaaaggcaatgtaaaattacctccagaagtataacgtaacat
tcctgcattattagaaatttgattgaatgctaataataaaaaactcatgttcataccttc
aacaattggtcttagccctgttatagctgcaccaattgccataccagtaaagctattttc
tgctattggtgtatcaagaactcttaaatccccatatttgctatgcaaatctttagttac
cttataagatccaccatagtgaccaacatcttctcctattacgcagacagttggatcttt
tgccatttcctcatctgttgccgctcgtaaagcgtcaaacataaagattttactcatttt
aatatttgattttatgatttgaccactgttttaaagtaacagttaagacaccatattct
atttagttatctgcaaaaagatatcgtttcaattctgacatgttaggttctggactggaa
atagcaaatttcacagcttgttccaattcggttttaacagcattctgaatctcattaagt
tcacctatattagcaatttcattatctagaatatattttttgagtttttgataggatct
cttgccacccaagcctcttttcttgtcttgatcttagttcatcaggatctgcgagagag
tgaccacgaaatctatatgttaatgcttctattaaggttggaccatcaccttggcgagct
ctttgaactgcttgttttgcagcttgtcttacagctagcacatccattccatcaacttca
atcccaggaagcccaaaagcttctgctttttatgtatttcaggtattgaagaagaccgg
tgatgtgccatacctatagcccactgattattttcaacaacaaatataataggtagtttc
cagagaactgccatattcagacattcaaaaaattgcccattattggtagtcccatcacca
aaaaagcaagccgtgactcttaaatcctctgtttctttaagtacttgctggcggtagata
ctttgaaaggctgcccctgttgcaaccggtataccttccgcaataaaagcaaagccacct
aaaaaattgtgaggcgcagaaaaaatatgcatcgaacctcctctgcctttactacaacca
gtctcttttccaaataactcagccatcacatttttagatgggacgcctttacttaaagca
tgtacatggtctcgataggtactgcaaacataatcagttggattcagaagtttaattaca
cctgtagaaacagcttcttgaccattataaagatgaacaaaaccaaacatctttccttta
taatacatctgagcacacatatcttcaaaatttctgcctaacaacatgtcttcatataaa
actaataaattactcttattaagattaagacctgttgaattatagttagtcagcggcaat
tgaactttcttaggataactcataatttgtaaagaaacctct----------tattgtaa
ataggaattttgaagaaacagctatcaatagaaaacacacagataaaaacttataaatta
taatatgtttttactaaagactttaataattatttgaaactagttcagtatatcaaatt
aattttgcttatactacggatgtctaaaacttatataaaaaagtagt-a-aaaaagttta
ttaaaagaatagttaaaaacagatataaatagaagtaaacagaattatt-gaaaaaaaat
gattattattaattaa----tacttaatatttctaaatattttacaattaatggctatgt
caccatctttattttatcctgttgaacaagaactatgtagtcttgaaaaaaatctgaaag
ctgttgctgggactcgtcatccaattttatatgccgcagcaaagcatctattcgatgctg
gaggaaaacgagttagaccagctcttgtatttttagtggctaaagcaacctctgagaagc
aagatataaatactggacaaaaaaggctagcagaaattactgaaattatacatactgcta
gtttggtacatgatgatattattgacgagtgcacaacacgtagaggagtcaaaactgtac

ataatttattcaatactaagattgctgtgctagcaggagatttttatttgcacagtctt
cttggtatttagctaatattgaaaatttagccgtagtcaaagctatttctaaagtcatca
ccgactttgcagaaggagaaattaggcaaggcttagttcatttcgatcccagtatttcaa
tagatgcttacattgagaaatcattttacaagactgcttcactcattgctgctagttgtc
ggggtgcagctatgcttaatggttccaatcatcaaataaataatgatctttatctttacg
gtaaacatatgggattagcatttcaaattatggacgatgttctagatataactggttcta
ctaagagcttaggaaaacctgctggcgctgatctaataaatggaaatttgacctctcctc
tcctttttcacttactcaagaagcaagtttaaatgatcttattgatagggagttctgta
atagtacagatatagcctcaacattatttcttataaaaagaagcgggggaattacaaaag
ctaaagatttagctaaagaacaggtgcaggcggcactttcctgccttcagtttttaccac
aatctacacctgtatctagtttaaaagaattaacacatttcataatcacaagattgtcat
aaaggacttgctaaaattaaaatgttattaactaatttttactgtttctaaaacagtct
gaaaagtttgaatatcattactagctagctgcgctagcattttacgatttaacgcaatat
tctctttttttaatgcactaataaaagtactataattcatgccttgattgtgagcagcag
catttattctagttatccaaagacgacgaaaatctctcttttttcctttttcgaccaacat
aagaatatcgaagagctttaaggacttgctgcttagctgttcgaaataaacatttatgtg
cacctttaaagcctttagctagcttaaagattttagctcgtcttttttttgcaacgttac
ctcttttaactctactcataaaatattac-ttaaattttaattattaccagcttactaaa
ttcttatactctttg----attataaataaggtaatttcattgcaatgtttttaatatct
tttaagtcaaccgaacaagtagaagaaagatgtcttctttgttttgatgactttttttgt
aataaatgactttagaagctttatgtcgaagaaattttccggatgaagagactttaaat
ctttttgctattgcttttgatgtttttaacttaggcatattataaacttttaatgattaa
caaaagcatgaaaagtctgaggatatcaatcttagacctttcaatacttagaataatctt
aacacaatttattttacttactacataggttaactgaaacaatttatatatataatacta
atcagttaatcaaccatagctaaagtggatttatattttgctttgaacttttgtatagta
ttagcaattagcattgttataattttaagactgcttgaattcaattcttcgaacattttc
atatttaatgtaaatgaaatattggcttctgagacaatattttgaatttgaacatcagac
aagggaatcatatctaatgcagctctatattgatctttaaataatttatcgtcttttatt
tggtcaaaatcgtaaaatttcgttcctcctgaatctgataaattcatagcccctctagct
attttttttaaaatttggccaccagaaaggtcacccaaatagcgcgtataagcatgagca
actagtaattcaggttgtttatgtcctatagtgtgaattctatcaacataaatttttgta
gcaggagatggttcaataaaatctaaccaatctgatccataataatagtttaaatcttca
gatagactcgctttcctattaagctctgtaaaatatataggtttaatagctggatgattt
ttattagaaaataactcttcctctattgcacaatagacaaagtatagatttgcaaccaat
ttacgatatgacttcttatctacaactcctcccaaaaaaagatttaacaaaactaacattt
tctgccatactatgggacttagtagtaccttctcttagttcattcgctaaagtattaacc

ataatatcaatattccattttt-aaagataacaaattaaca---ataaatagttatccgt
atatttataattgagttgcgctaactaggctgctgcaaaataatttttttgattttattgg
atctggattcattgttttgtcaccaggcttccaattagctggacaaacttcatctggatg
agattgtacatattgaattgcttgtaaaaccctcaaagtttcttctacactcctgccaaa
ctctagattattgattgtagagtactgaataattcctttcgggtctataataaatagtcc
tcttagagctacaccatcactattcaaaacattataagctgcactgatctcttttttttaa
gtctgatactaatggatatgaaagatctcctaatccacctgattctctgtcagtctgtaa
ccaagcaagatgagaatactcactatcaaccgagactccaagaacttctgtattaagttc
cgaaaaagcattatacttatcgctaaatgctgtaatttctgtagggcacacaaatgtaaa
atctaaaggataaaagaataagacaatatacttattcttgagatcagataattttaacgt
tttaaattcttggtcataaacagctgtagctgaaaaatcaggagctagctgacctactcg
aagacaatttggtccagaaatcattaatttttcccagaagaattgatataattttgttat
ttaataaataattatataacaaaatttagtatatattaaaatactaccagtcttttttaa
attac--------taaatgatataatcaatactatctttatagcggggaatggatttgaa
ccattgaccttcgggttatgagcccgacgagctaccagactgctctaccccgcggataac
taaaatcatattagtctacatatatatataaaagcaaatgcataatagatttagaagta
aaaaaatagaagaactggtccactgatcattaaacttagaaacgacaaagtatttttat
cttatttaaaagaggaaagacttcatataaaccaataaatctatcttttaaaagaatttc
ggacggctttacccaaatttgaccgtcataccagcctgattcttcataaaatactgtagc
actcattaatcgttttactacgtatgaccagcctaaatataatctgattaaaataaatcc
tgtcattaggctggaagtaataaattctgaaaagaaaaatttcaaaggtagtttagtgat
tggaaaaattgataataatataggactaactaataaacagttaagtagtagtgcaatagt
tattttttttgttataagatctatggcttaaagttggccagcaaaagaaccaagaattttt
caaagaagtatattcatgaacaggttgctgctcctttggcacaggacattgagtattata
taaattcattgacattaaaaagtattataataaagac--tttgtttatattatctaacat
aggtacgacattaaatattaactaactgcaattatagtagtgagtagaagaaataaaact
aaaaatagccaagtaattttatttaaagtattctcagtactacgagtattactaaagaac
tgattctgagcacctacacttcctaaaccttcggatttaggattgtgtattagaatagtg
aagattaatattattgtagatgagtaccaaaaaaatttttaaaatttgttccattatgtaa
aagttaat-aaaaatagcacaaagacaatcgtctaaaagatattgattgtcttatcatat
taaaaagtatttattcaccctgtacttttaaaagtaaaaatcctattgccagccctacca
aaactaggactgaacttaatacagcactttctacaatttctccgaccatt-ttatttttc
cttattattttattattg---tcattcttttgacaagattatattgagctaa-ctaaaat
ccgtttctaccccaaactaccattgcaattgaaaatgtaaacattgccattagagctgcc
caccccaaacttaaaatatccataatttataatattc--ctaaaaataagatctctttta
cttaaactataatattcttaatattaaaaatatcattatataataagctaattataacag

agtactacttaattcgtatttttagtgtatttaaagaagattgaa-ttttttcgaggtac
tggataagtatttcatcctaagaaccatattcaac----ttttgatttattataatgctt
aaatctaaatata--tatcactcaagaatcattagtaattatgatgtttttaattatacg
att---aacatatttcaaaatagaagaattcatagcgactaagttaa-----tatgctaa
tataatttatcaat--aaaaaggaagttagtatttcaattatgcaaacaactataaataa
tgggcaaacgtctagtaaagaaactttactaacacctagattctacacgactgactttga
agagatggctaatatggatatttccggcaatcaagaagatttttttggctatcctcgaaga
atttcgagctgactataatagtgaacattttattagagatgaagagtttaatcaatcttg
gtctaatttagaacataaaactaaatccttatttattgagtttttagaaagatcttgtac
cgcagaattttcaggtttttttactatataaggaattatccagaaaattaaaagacagaaa
tccggttatagctgagtgctttttattaatgtctagagatgaagctaggcatgccggttt
tttaaataaagctatcggggacttcaatttatctttagatttaggatttttaacgaagag
tcgcaagtatactttttttctcacctaaatttatttttttatgcaacctatctttctgaaaa
aattggatactggagatacataactatttaccgccatcttgaacaacatccagaacaccg
tatttatccaatctttagatttttttgaaaattggtgtcaggatgaaaatcgtcacggaga
ttttttttgctgctttgctcaaatcccaacctcatttttttaaatgactggaaagcaaaaat
gtggtgcagattttttctattaagtgtatttgcaacaatgtacttaaacgactttcaaag
aattgattttttataatgccataggcctagactctagacagtatgatatgcaagtaatacg
aaaaactaacgaaagtgccgctagagttttcccggttgctttagacgtggacaatccaaa
atttttcaaatatttagatacttgtgcatgtgataatagagctctgatcgatattgataa
taacaattctccattattcattaaatctatcgtaaagatacctctatatttttccttatt
tgcaaatttactaaagatatatttgattaagccaatagactcaaaaacagtatggaatac
agttcgatagttaaatacaagggaatgctttttttcttataaaaagctatctagtgattaa
tataaaat------aaaagcttccttct-----cgatggaaggagcttttattttgtcaa
ga---atataatgggattatatagatc-tagaactaagaacttactttttcttttgcttc
atacataacctctaaagtaattagattttttagaacaatcacgggcaaatttctctgtatt
tctttttactttacctctaacaaacccaggaattttatttaattctttctgtgcttcttc
tgaccaattaatactatcaacagcagaaattcctatacttaatgcttctgttgtatcatg
accaccaaatatttctagcaaatgatcttccataccaagtgtaaacgaattatatactaa
atcagcaatttgatttgttccctcataaccaagaaatggacggtaacttaaaggaaagtt
ttgtatatgcactggtgaagaaataacgccacaaggaatatttagacgcttaccaatatg
tctttccatctgtgttccaaaaattgctgcaggttcagttttagcaattaaatctccaat
taaaccatgatcatctgaaacgattacttcatcgcagaattcttgaacttgatctttaaa
ccactcttcatcatatttgcaataagttccacaccaggcgacatgtatacccatttcctg
gtgtaaaattcgagttattgctgcagcatgagttgcatctccaaatacaatagcctttttt
acctgttaaattctgacaatcaattgatcttgaaaaccatgcagattgtgaaataaatcg

ggtttgctcgtctatgtatttttcgtagtctacagcagctcccaatgcatttactaattg
ctgtattgatcggatgcatgcagctgtttgaacaatacccataggtgtaatgtcaacata
aggcatattaaattctttctctaaatatcgagctgtcattagtccagtttctctatatgg
aataaaattaaaccatgcagagggtaatttttttaaatcttgaactgatgcattttctgg
aataatttggtttatttggatatctagatcttgaaataacctttttaattctgctatatc
atgttgattatgaaatcctaagcttactgctccaataatattaacagatggagttttagt
cttttgagttaaaacttgattattagatttagctttttccatatagaaagtgacaatttg
ttctaacgttctatcacctgcttgcagttcatttactctataatgattaacatcagctaa
taaaacatctgcttctgtttctatagatgctctgctaacaaagttttgcaaatcttcttg
caaaatactcgaagtacaagttggtgttagaataactaaatcagggctttcttctctgtc
ttttctagtaatattttctacaactttttcttgagaaccacgggctaacacatgtctatc
tacaacacttgctgtaacaggagtaaaatctctgtctctttcaagcattgaacgcattac
attaaaataatcgtctccaagaggcgcatgcataattgcatgaactttttttaaaagaact
cgcaattcttaaagttccaatatgagcagggcctgcatacatccaataagctaatttcat
attaaattttcctatgatttatttagacatagtaatacttttaaagctaactaacaaaca
ttagccaaacatctaatttaactccttgacaattttgtaatatttttttaaatgaagctta
tcattataaatatttaaaaaatataattcagaataga----aataactgttagatcaagc
ttgaaagtaaagatttcttgcagttatagtttttctttaagtaaacatgtaaaaattgta
ctatatataaatgtactatatataaactttttttactgaatattggtaatgccagataca
attaacttaaatatgccttccccaacgtttggtggaagcactggcggttggttaagagcc
gcagaagtagaagaaaagtatgcgataacatggacaggcaaaaatgaaagtaagtttgaa
atgccaactggtggtaccgcaactatgcgaaatggagaaaacttactctatttagctaag
aaagaacagtgtttagctttaggtactcaactaaaaagtaaatttaaaatatctgactat
aagatttacagagtttttcctaacggagaagtgcaatatttgcatcctaaagatggagtc
tttccagaaaaagttaatactgggagagctagcgtcaacagtgttgatcattctattgga
caaaatattaatcctgtagatgtcaaattcatgaataaagcaacttacgattagtctctc
-aaaaacctagacataaaataacttttatatagtctaggtttttgataatgtatactgat
agtatggaagggtggccgagtggttgaaggcgtctgatttgaaatcagttgaacta--tc
tggttccgtgggttcgaatcccaccctttccgttataaaact------------atagtt
ggcttctcaaggcaagctagctatacactgatagtattaagtttagacttaataaagtct
acaacttgtgacaggttagaaatttttttcggcatcctcatctggtatctctatactaaat
ttttcttctatggccattaccagctcaactgtatctagagaatcggctcccaggtcactt
gaaaaatttgcttccctagtcactattttttttttcaattcctagctgttctgctacaata
tcttgaacttttttcaaagatttcattatcttgcataatatattatttcctctaaggtgtt
atcaaaaatttttcggcttatattattatccaactctaaaaaattagtaatttttagtcg
agattaaaatatttatacctatatgatatactattattctggataaatacgaagtttacg

atatggctcttctccaaaactacgagctctcaactggtagttgtcaactagattatgctg
catttttcgtatataggctgatcgaggtgtaagctgaactattgaactttcattcaagat
aatagtttcaatagccaatttagcttcttgaagagcctggatctcatcaaaatttttatt
tttacagagttctacccaattaagaccagatgaagtatgaatgtttaatatttttctgag
tgcacgagtaatttgtggaaccgtactattctgaattgtatatataataatttgttttga
ttttgcaatttgtctaagttttgtattttgctttacttgattacgtagcgctagaatagc
atcagacttttctatctccttggttaaaataattggtagatctagtgaagaaataactga
tgtaatatgctgccaactcaaagaataagcatataagtactgatgaggtatttccaattg
caaagattgtttattaatattaataggggcattgaaagtagttgacaataaagatgtatc
acgattttgtgatttattcaaatctaaggacttatttcttaattctctatattgtaaaac
cggaggcgcctgctgtgcgttaattgatttacgagcattagagatgtttacaggtaagac
ttctatagattgcgatggataacatttaattaaaattctaccattagcttgaatttgtct
tttttgcacaaaaggttgatggccttgcaaaatttgatctattgtctctttaactttatt
atgaactatccaaacatttcgctcgtgtatttcgattgcaatctgaaaagcaggctcagc
ttttctttctagtatactcttttgcgtacctctacgtttagcttcatcgtcacctaatgt
tacatattgaattccaccaattaggtccgccaatgttggattttaattaaactttctaa
atgatttccatgcgctgtacctactaattgaactcctctttctgctatagttcgagcagc
taacgcttctaattccgtgccaatttcatctataataataacctcgggcatgtgattttc
aacagcttctatcattacttgatgttgcaaatcgggccttgcgacttgcattcttctagc
tctaccaatagctggatgaggaatatctccgtcaccagctatttcatttgaagtatctat
aataactactctttttttccatttcgtcagctaaaacacgggccatttctcgaactgcggt
tgttttaccaactcctggttttcctaagagtaaaattgaatcaccttgctgtagtaaatc
tcgaataatactaattgtaccaaagactgctctgccgacacgacaggttaagccaataac
actaccttctctattacgcagtgaactaatacgatgcaaggtttttttcgataccagctcg
attatcgccgctaaaattaccaactttttttacacaataatctaagtcttgccaactaat
agatctttgagataaatattctgggttatctggaaatctagcttctggacgacgacctaa
atccataacgacttctattaaattatttctattaggatgctgttgtaaaggttctttgac
aaaatttggcaaaatttctagcaacttatctaagtcatctgcaataagcatgattgagta
tataataattaattaatcaaatagaagattcaagattctaaagattacaatacatattaa
agaatattatgattggaatcaattttatacttaaattgaacaggaagtta-tagatgtaa
aagaactgattaaactttacgcaataaatcgattggtcaaaattaatagtcgtgttaaca
tgtttattggacaaaaagttcgtattaaatatagcaaacaaaaagttgctagtgatatag
cagataaagttggtgaattaggagttattaagggaataaagtttattaatagtcaatgtg
ttactattattgttgagtttgacaatcacactaggctttggatgtttagagaggaattaa
tctgtctaaatgaat------taaaaacatgaaacataatttattttatttattatacaa
aatttcttatgagtcgttaaaataataaattttaaaggagtacttttaattgtgcaaatt

actattaaaaaattacaagacttactttcgtctgtacaaagaaaaaagatccagacatta
aaactaaaacaaggtaagtttgaacttctattaaataagacctataaaaaagtcaatcaa
gaaattataccttcgcagaaatctgctgtattacaaaatagtccatctacgataattaaa
tcaataaataatacaaaaaaatcttctgttgttaatgaagaccgtacagaatatgccact
attgtttctccaatggttggaacgttttatcattcacctgctcctggtgaaaaaattttt
gtacaagttggcgatgaggtcaaattcaatcaaacagtctgtattattgaagcaatgaag
ttaatgaacgaaatcgaggcagaaattgaaggcaagattatagaaattcttgttaaagat
ggtgatatagtagattgcgggcaagccttaatgaaggttgaaacataatgctaatttttc
cattttacatattgttaacagaaattatttagatagaaaaattagcttatactatagtta
tgaaaacttatttcgcaaaggcac-taaaaactaatatataaaattatttactttatctg
ccaaaaatagaaataagcgttttgattccttgtctcaagagaggagaatctcaatggcaa
ttagctcaaaagagcaagagacaaagaaggtaaaaatctcggttgataaaaatcccgtag
atacttctttcgaaaagtgggcccaaccaggccatttttctcgtacactagcaaaaggac
caaaaactactacttggatttggaatcttcatgctgatgctcacgacttcgatagtcaaa
ccagttctttagaagaagtttcacgtaagattttcagtgcacattttgggcagctgtctg
taatatttttatggcttagtggaatgtattttcacggagcccgcttctctaactatgttg
cttggttaagtaatccaacaggtattaagccaagtgcgcaggttgtttggcctatagttg
ggcaagaaattttaaatggcgatgtaggtggtggctttcaaggagtacaagttacatctg
gatggttccaactgtggagagcatcaggaattactacagaatttcagctttactgtactg
ctattggcggattagctatggctgctttaatgctgtttgcaggatggtttcattatcata
aagctgctccaaagttagaatggtttcaaaatgttgaatcaatgatgaatcaccatttag
ctgggcttttaggcttaggctgtttaggctgggcaggccatcaaatccatttgtctttgc
ctattaataagctactagattctggcgtgtctccgcaagaaattccactacctcatgagt
ttttaattaatagagagcttatggctcagctgtatccaagttttagtaaaggattagttc
cattctttactttaaattgggctgaatattccgacttttaacttttaaaggaggtttaa
accctgttactggaggtttatggctaagtgatactgctcatcatcatctagcattagctg
ttctatttcttgctgcaggtcatatgtatagaaccaattggggtattggacatagcatga
aagaaattctagaagctcacaaaggaccttttaccggcaacggtcacgaaggtctatatg
aaattcttacaacttcttggcatgcacagcttgcaattaatttagctatgatgggatctt
taagcatcattgtagcacatcacatgtatgcaatgcctccttatccatatattgctactg
attacccgactcagttatcgctcttcactcatcatatgtggattggaggattttgtattg
ttggagcaggagcgcatgcttctatatttatggtaagggattataatcctgcagaaaatt
ataacaatcttttagatagagtcattaggcatcgagatgctattgtttctcatctaaatt
gggtatgtatatttcttggattccattcatttggtttatacattcacaatgatactatgc
gggcacttggaagatctcaagatatgttctctgatacagctatacagttacaacctattt
ttgctcaatgggtacaaagtatacacactttagctcctggaaatacagctccaaatgcat

tagcaacagctagttatgcatttggaggagatattgtttctgttggtaacaaagttgcaa
tgatgcctatttctttaggtactgcagatttttagttcaccatatacatgcatttacta
ttcatgtaactgttttaattttagttaaagggttccttttctcaagaaactctagactaa
ttcctgacaaggccaatcttggcttcaggtttccatgtgatggacctggtagaggtggta
cttgccaagtgtctggctgggatcatgtttttcttggcttattctggatgtacaattctc
tgtctgtagcaatttttcactttagttggaaaatgcaatcagatgtttggggtagtgtat
ctccgtctggaaatgtttctcatattactggcggtaattttgcacagagtgcaattacaa
tcaatggatggctgagagatttcctttgggctcaagcatctcaagttattcaatcatacg
gttctgcgctatctgcatatggattaattttcttagcagcacattttgtatgggcattca
gtttgatgttcttatttagtggtagaggttattggcaagagcttatagaatcaatcgtat
gggcgcataataagataaaagttgctcctgcaattcaaccaagagctttaagtattactc
aaggtagagcagtcggtgttgcacactacttattaggtggaattggtacaacttgggcat
tctttttagcgagaattatttcagtaggctaatagtgaaaataggataaaaaaaacaatt
atggcaacaaaatttcctaagtttagccaagctttatcacaagatcctacaactagaagg
atttggtatggtattgctacggcacatgactttgaaagtcatgatggaatgacagaagaa
aatttatatcaaaagatattcgcttcgcactttggacatctagcaattatcttcttatgg
acatctggtaatttattccatgtagcttggcaaggcaactttgaacagtgggtattaaat
cctttgaaagttaaaccaattgctcatgcaatttgggatccgcattttggacaacctgct
ttgaaagcttttagtaaaggtggctcagcttatccagtaaatatagcatattctggcgta
tatcactggtggtatactattggtatgagaagcaatcaagacctgtattctggggcttta
ttcttactagttttatcagctttactcttatttggagggtggctacatctacaaccaaaa
ttcaagcctggtttatcatggtttaaaaataacgaatcaagattaaatcatcatttatct
ggattatttggggttagttctttagcctggacaggtcatttagtacatgttgctatacct
gaggcaagaggacaacatgtaggatgggataattttacaaccgtattacctcatccagcc
ggtttacagccgtttttcagtggtaattggagtgtatatgctcaaaatccagatacagct
caacatttattcggaactaatgaaggtgcgggtacagcaattctgacatttctaggagga
tttcatcctcaaagtcagtctttgtggctaactgatatggctcatcaccatttggctatt
gcagtagtattcattgttgctggacatatgtataggactaattggggaattgggcacaat
ctaaaagatattttagatgctcatagaccacctagtggtagattaggagctggacataaa
gggctatttgatactattactaattctttacacatacagttgggattagcattggcttcc
ctaggtgtaattacttcgttggtagctcaacatatgtatgctatgcctccatatgctttc
atggctaaagatttcacaactcaagcatctttgtacacacatcatcaatatattgctggg
tttctaatggttggagcttttgctcatggggcaatattctttgttcgagactatgaccct
gaacagaataaagataatgttttagctcgtatgctagaacataaagaagctatcatttct
catttaagttgggtaactctatttttagggtttcatacattaggcctttatgttcacaat
gacacaatgattgcttttggaactcctgaaaaacaaattctaattgagccggtatttgct

caatggattcaagcctcttcagggaaagcactttatgggtttgatgtgttactatcatcc
tctactaatatcgcaacacaagctggtagcaatatttggctgccaggctggttagaagcg
attaatagcggaaaaaattcattgtttttaacaattggtcctggtgacttcttagttcat
catgcaattgcattgggattacatactactacgttaattttagttaaaggtgctttagat
gcaagaggctctaaacttatgccggacaaaaaagactttggatatagttttccttgcgat
ggacctggcagaggcggcacctgtgatatatctgcatgggatgcgttctatttagctgta
ttttggatgctaaatacaataggttgggtaacattttattggcattggaaacatattaca
atatggcaaggcaatgcaactcaattcaatgagtcttcaacttatctaatgggatggttt
agagattacttatggctaaattcttctccattaattaatggttataatccatatggcatg
aataatttatcagtatggtcatggatgttcctatttggacatttagtatgggcaacagga
tttatgttcttgatctcctggcgtggctattggcaagagttaattgaaactctagcatgg
gcgcatgaacgtacacctttagcaaacttgattcgttggaaagataaacctgttgcatta
tcaattgtacaggcaagattagtaggtttagcacatt-tttctgtaggatatgtattaac
ttacgcagcttttgtattagcttcaacagcaggcaaatttggttagactaaa-----ttc
gaattagct-a-----aaaaaagtcagtcataata-gttttatgactgactttttagat
ttaaaacaaaaacttactaaagtttcacctcaatatctacaccagaaggcaaatttaact
tcattaatgcatctatcgtttgagaagaaggttgatgaatatcaataattcttctgtgag
atcttatttcaaagtgttctcttgagtctttatctacatgtggagaacgtaaaacacaat
aaattcttctttttgtcggtaaaggaattggccctactgcaacggcattagttctagatg
cggtatctagtattttgtcacacgacgtattaagtataatagaattatatgctttcagtt
taattctgatttttgtctgctgagtaattgtcatgttaaaaagatcttattatttaagaa
ttttagagacaacacctgcgcctacggtacggccaccttctctaatagcgaaacgcatac
cttgctcaattgcaattgcattaattaattcagcagtcatcttaattctgtcaccaggca
taaccatttctgcatcagtaccatcatcagcagtaaactgattaatagtaccagttacat
cagttgttctaacataaaactgaggtctatatcctggaaaaaatggagtatgtcttccgc
cctcctctttagttaaaatataaacttctgcttcaaattgagtatgaggtgtaattgtac
caggtttagctaataccatacctctttcaatatctttttctgcacacctcttaaaagaa
ttccaatattatcacccgctagaccttcttctaacgtttttgaaacatttctaatccag
taatagtcgttgtacgagtttctcgtaaacctacaatttcaattgtgtcaccaactttaa
taatgcctctttcaattctaccagtggcaacagttccacgtcctgtaatagaaaaacat
cttctacagccattaagaaagttttatcgacatctctctccggcgttggaatatatgtgt
caactgcttccataagtgaaaaaatcttgtcaacccatttatcttcaccttgcttagtag
ctgggtttttcgtcacagcttctaatgccaataaggcagaacctgcaacaaaaggaatat
catctccaggaaagtcgtattgacttaataattcccttccttctaattctactaattcta
gtagctcttcgtcatctacttgatcttccttatttaggaatactactaatgtaggaacac
ctacttgttttgctaataaaatatgttcacgagtttgtggcattggaccgtctgccgcag

atacaactaaaatagctccatccatctgagcagcacccgtaatcatattttttacgtagt
cggcatggcctggacaatctacgtgagcataatgacgattatctgtttcgtattcaacat
gagcagtattaatagtaatacctctagctttttcttctggagcggcatcaatttcatcaa
atttttttcgctgcagtagaccctaaagttgataaagttgcggagatcgctgctgttaaag
ttgtcttaccatgatcaacatgaccaattgtgccaatattgacatgaggttttttacgtt
caaatttagatcgagccatgctttttgttttccttatagtaaaagttgctttatagttac
ttttaacgataaatcttgtgaattgaaagatttattatttaatacagacaaga-----ta
caatataactaatctaatatcgatagtgagcaaacgctttatttgcttctgccattctat
gtgtatcctctctctttctaatagaatttcctgtttcattagctgcatccataatttcat
tagctaatttcatagacatactttaccagatctgtctctagagaatttagtaatccatc
ttagtgctaaattcgtacctctataagctcgtacttcaataggaacttggtaagtagaac
caccaactcttcttgcttttacttctactagaggagtaatatttcggatggctttttcta
aaatatttaaaggatctgattctgttctctctttaacgatatctaaagcctgatatataa
ttctttgagataaagttttttttaccacttttcaaaatacgaacagttaacatacttacga
gtctgcttttatatataaaggatcaggtgatgcaaaccttttttttagctgtattacgacgag
acatagtatttaaaattggttattgtaataa-tgttc-tattagcaggtatttaagattt
aggctttttttgtcccatatttagatcggcttttacgacgatcttttactcctgcagcatc
taaagtaccacgtactacatgatatcggactccaggcaaatctttaattcggccgcccct
aatcagtaccacagaatgctcttgaatattatgacctacgccaggaatataagctgtaac
ttcaaatccagatgttaatctaacgcgcgctactttttcttaaagcagagttcggtttttt
aggagttgtagtatatactcttgtacaaacacctcttctttgagggcaactttgaagagc
tggagattttgttttttttatgtatttttcgtctttcggatctaacaagttgttgaattgt
tggcataataaaattgatttaaagtctttaatagatactatttgaacctgctttacatga
attggctagttttttatttatcttatacttacgcataaatctttctactctaccttctgta
tcaataattctctgtgaaccagtaaagaatggatggtttcctgaccagatatctacgtgt
agttctggctttgtggacccaatcgtcataattaactggccgtcacagtaaacttttgct
tctggataccaatttggatgtatattatcttttgccattgtttttataattatataaatg
tatcaataataactattaacgcttagaaaattgaggagcctttcttgctttctttaaacc
atatttttttctctcttttactcttggatctcttgttaaatagccttcagacttgagtgt
cgttctattttctggattaattgagcacaatgctcttgcaacacctaaacgaattgcatc
agcttgccctgtcaaacctcctcctctagcatttacatggatatcatactggtttagtag
tcctaaaacttgtaatggtgcatatgaaactcttaagtaattaggactaaattgaagata
agactctcctggtataccgtttataattaaattacctgaccctgggactagtcttacttg
tgcaacagaacacttccggcgacctgttccagaatagattgcgcgagttttaattaattc
tgtcgacataacattccttgataactaaataaatatacattctttatactatatactctt
gtggcttttgcgcgacatgtgggtgaattgggccagaatacactttaagcttcgtaaata

gttttctgcctaatggacctttaggaagcatacctttaaccgatttttcaataattctgt
taggtaatcttgtctgaagctgatcaaatgtttcaacttttaatccgccaggctgtccag
aatgtcttctgtatagtttttgattcgttttatttccacttacagatacgtgagcagaat
taataacgattacataatctccagtatctagatagggtgtataagaaggcttgtttttac
ctcttaagatattagaaatatgagtagatattctaccaagtgtctgattttagcgtcta
taacataccaatgagaattagtatttaatgagggtgattgcgttttattcatgaaaagat
taatacttttggttgacaataagagaatatgtaaatggtttacattcaatttctatacaa
gtagtattt-aaaaatatta-------------ttaacaaagagaactttctaacagtt
atttaaattgtttgtcagcaagaaaatttctagttatcaataacttatactatttttaat
aatataaaatatattaatcactttttcttttggcaaacttatacctagttttttttgc-
aaggcatatataacttcttcagctgatttctggccaaaatttttaatttctagcagttct
tcttgagagtaatctagtaaatctgcaatagaatggatttgagctcgctttaagcagtta
taagccctaacagataactgcagttcttctattaaaacttgactaatttttttatcttct
ttgctacgataattatctgctgatttaaagtctaaatttcttagtgaacaaaaaagatta
gttaaaactgtagctccttgacttattgcttcttggggagatatgctcccatttgtccag
atttgtataatcaatctatcttttatgctgttgctaccaatacggacttcttctacttta
taattaactttattaactggcataaaaacagagtctacttgcaaaaaatccacagataat
tcatctacagctttttcagctaagcgatacccgcaatttttttcaatttaaactccatt
tcaaatattgtgttattgcaaatagttgcaatatactgtctaggatctactacctctata
tcagaagataattcaaataggccggctgtaactatagctggcccttgaactctaattcga
ccaatttgagattctttgttataacttttaaatactacttcttttagattaagtaatatt
tctaacacatcttctcttaccccaggaattgtggaaaactcatggttcactccagcaatc
cgtacagcaactatagcagtaccttcaagatctgacaatattgatcttctcaaagcatta
cctaatgtaataccttgtccttgatttaatggttctattacaaaactaccgtactgccca
cgcgccccatctgttcttgactctatgcattcaatttgaaattgagccacctaagaaagc
tcc-tttaaaataatcagtgattagtaagtctctctaaaacaagaaggaatttatactcg
gcgtttcttaggagggcgacatccattatgaggtacaggagtaatatcttttattagagt
aatctctaatcctgcagcctgcaaagctctaattgctgtttctcgacccgctcctggacc
atttactagaacttcagtttggcgcataccttgatccatagcttgtctagctgctttttc
agctgctgtttgagctgcaaaaggtgttcctttttttagctcccttaaatccacttgcacc
agaggatgaccatgataatgtttctccttttaaattagtaatagtaacaattgtattatt
gaatgtagatttaatatgtgtaataccgttaactgcattacgtttagtttttcttgctcc
ggatttttttatttgtctagccatcgtgtctcgtctattatatagttaattaaaagatta
ttttcttggagctttttcttacctgctactgtttttttacctcctctgcgtgttctagc
attagttctagttctttgtcctctcaaaggaagaccaagacggtgtcttctacctctata
agtactaatttccataagtctcttaatgctcatagactcaaaacgtttgagatccccttc

aatctgataattagactcaagaatttctcttatactgacaacttgttgatcatgtaaatc
ttgacacttaatatcagcgtctatgtttgtttttctaatatttctttcgagcgagatag
tccaataccataaatatatgttaaagctatctctattcttttgtttcttggaagatctac
tccagcaattctggccactttgtcttgtctccaataaataatatgtat-ttttataacgg
ttatttata-agcttagccttgtctctgcttatgtttaggattagtgcaaattaccatca
cctttctgtgacgccgaattattctacattttcacacattttcgaacagaaggacgaa
ctttcatattaa----ctctatatactaggaatattactatattaactctattaattatc
gacattaaaaatgtcaacaatatattgtattactttaactgttgtagattttacttcgtc
atactatcatattttttgatatgacgtatgtctgaatttgcttagctgtgtctattgcg
actccaactaaaataagtaaagacgtagccccgagacctcttaagttttggatctgagta
accttttctattataaacggaatcagcgctactgtaaataaaaacgaggctcctaggaat
gtcagtctatttaatatgacttgtaaataatcaatagtcgcttgacctggacgaatatta
ggaatgcttgcacccattttttttaaatttatagcaatatcttctggattcattactatt
gatgtataaaaatagctaaagaaaaggatcaaagcacaataaagaagaaggtatagtgaa
ccattaggacaaaacaggtataggatttggagtaatgttttattttgaataatttgggtt
agataagatgggagggccatagatgcagatgcaaaaacaataggcataactccaccttga
tttaatttcaagggcaagtaactatttggatccaaaattgaagattttcctagctgtctt
gctgaaataattttaattcttcttgttccttcttgtacacaaattgtaattattatcatt
aataaaaagatcgctataaataatccgaacttaagacttgcattactataactagcatca
aaaaatgactgtgtaaaattctttggtagtcctgacacaatgttttgaaaaataagtaaa
gaggctccattaccaattcccttttccgtaattaactctgataaccacataataatcata
gagcctgctgttaaagctaaaacagactcgcagacaaatgcaaaattccaattaaaaaca
tatggttttacccatatagagattgctccagattgtaaagtcgcccaacctagagctaag
taccttgtaatttgagttatttttggcggcccaattcaccttcttccttctgtaatttt
tccagattggggacaattttcgtaagtagctgcattacaattgaagagttaatataagga
acaatacctaacgcaaaaatccctattgttgaaaaacctcctccagaaaaaatattcaga
aaatttactaaagtatttttttctacacttgcataaaaggcgtcatgatctatacctgga
acaggtataaatattcctaaacgtgctaaaactaatagaaagagagtaaagataatacga
tttcttagatcacttttttggctcataaataaattttaaatgaaaagagatacttt--tt
tatgtaaaaaccctattccaagacttgtataagtttaaaatagggaagtaa---aatatt
atagtaatattaata-taatattctatgtagaatacaggctttctattggaactcctcga
tctttagctgcttcactaaaagttcttaattgagttaagccattgagaacagctcgcgcg
ttatttaatgtattattagaacctaattgcttagctaatatattctgtacacccgacaat
tctaggactgtccttacagaacctccggcaattacacctgagccaggtgcagagggcctt
aatataactttcgcggcaccagaaattccgttaataggatgaggtatagaattcgatttt
gtcagtggaactgtaactagatgtttttagcatctgttactcctttttttactgcacca

attacatcgcttgcttttcccacgccgacaccaacttggccttgctcattaccaataaca
agaataactcgaaagcttagtttttacctccttttacaactttagtaactcttttaact
tgtacgactctttcttcccagccactatctttatcttttcctttgctctgtttttacga
ttggccattttaagaattattcctaattagttaatta-cgattctagaaacccatgcctg
cttccttagcagcttcagccaaggccttaactcttccatgatataactttcctcctctat
cgaagacgacattttaatgccttctttcatagactgttctgctaactgtttaccaacta
cgcgagaagtatcacagtttggtcttatgttatcagattctttattatttaaattaacag
atgatgtagctactaatgtaataccttgtgtgtcatcaattatttgtgcgtatatatgtt
tattagatctaaatacacacagacgaggcctacttgaggttccttgaactttttccgaa
ctcttttatgcttatgaattctagtttgtttagtgtttagtttcattattatttacccttt
tccagctttcccagcttttcttctaacaaattcaccttgatatctaattccttttccttt
ataaggctcaggaggcctaatagaacgaatagttgaagcaacctgaccgacaacttcttt
atctatgccagagacagtaatatttgtgttgttttcaactttaatttcaatattggcagg
aggtttaattttcactacatgactatagccaacacttaagattaaatcttggttatcaat
ttgagaacgatagcctacaccttgtatttgcagtttttaaaaaatccattagaaacacc
ttcaatcatattactaataagagttctggatagcccatgtaactgactagccattttgt
ttgtccactcgtcttgacagcaattgtgtcgttaagtatttctaaattaataccagctgg
taaagttctggatagggtacctttaggtcctgtcacggtgatagtctgaccatcgaactg
agtactaagatttgtcggcaataaaattattttttttccaatacgagacat-attccacc
tataaaatagttaccaaatatagcataatatttcaccaccaagaccatcatgacgagctt
gtctatctgtcataacgcctctagaagtagaaataagagcgatgcctaaacctccaagaa
ctctgggcaactctttatgatttgcgtagactcttagtccgggtttactaattcttttca
gagcagtaataactggttgacgatttttaccattatacttaagagaaatcattaaatgag
tctctataccttcacccatttgttcgaaattttgaacaaatccttcttcttttagtactg
ttgccatattgcatgtcattttcgttgctggaacttgcacaatttgatgtcttgctaagt
ttgcgttacgaatacgtgtcagcatatcggcgatcgtatcgttgaccaccttgatcctcc
ttgcga-----tgaattatttgaaatattttcaagattccctgaaaggcatccctagctt
ttttaacaaagctagaccttcttgatctgttttagctgtagtgacaattgatatatctag
accacgaatttgatcaatattatcatagtctatttctggaaagatcaactgttcacgtaa
acctaaattgtaattacctttgccatcaaaacttctaggactaattcctctaaagtctct
aattcttggtaatgttaaattaattaatttctctaaaaaagaatacatcttgtcttttct
tagatgcacaacaattccgataggaacctcttctcgaattttaaatcctgcaatagattt
tttagcttttgtgacaattggtttttgtccagttattaatgttaattcttgaatactact
ttcaagagccttagcattctgagaagcttcacctaagccacggttaatagtaattttagt
aaacctaggaacttcatgtacatttttgtactgaaattcatcttttaaagattgagtaac
agttgttttatattttcttttaatcctattgccattatattttcatcaattatttaata

agttcgccagtctttttcagttttcgaa-ttttttggcctttatcattaattattactga
ggatcgactagcaatattatttgttcactaaataacataacgttagaggtatgtattgg
agcttcaaatttgataatttctcccgtttctccttcttgctgaggtttttatgcttcac
tttgagattaattcctttaacaatcaccttatttgttttataaatgattgcaataatttc
acctgtttttgttttatcacttccagaaataacttgaactaagtccccttttttaattt
aatttttgtattattttggttgttttagaaggacctttcattatactacctccggtgca
agagaaactattttagaaaaatttttatctcttaactctcgggctataggtccgaagact
cttgtgccacgtggattattatcttgattaatgataactgccgcattatctccaaatcta
atgctcataccgtcagttctccgtaaagctttacgagttcttactacaacagctctgaca
acatcagatcgtttaactggcatattgggagacgcatctttgactactccaataataaca
tctccaatagatgcataagaaggattactagtgcctaatactctaatacacattatttt
ctagcaccactattatctgcaacattaagatagctttgagtctgtatcatacttttattc
tgtataagattaattatcaaaagacttagataagatattaaccattgtccaacactttgt
gcgacttaaaggtcgtgtctcttgtattgtaacaatatcgccgattgtgcattcattatt
ttcatcatgcgctttatatttctttgtccggatcatagtttttgcgtactttctatgaga
gattctattttctacagctactactatagttttattcatttatcgcttactactttacc
tgttgtttcttttaaaggcatagtctttatatttttttatatat---------aattaaat
tttaagattattacatggttgacttagtacgagacttttcaacagttaaaagttgggcta
atctatgcttagaatgtttaaatagatgaggctggaaatcttgccttgtggctcttttta
gccttaaatcaaaaagctctcttttattacaaggatttcttcagctaaagaagaagagt
ccaggtttgtaacatctgatattttagggaaagtcatatttaggattctgttgtattcc
gaactataaatttagtcttgattggtaatttataagaagctaatttcatagcttgttgag
cagtttttgtggtacacctgtaatttcaaatagaatatgtccaggcttaataactgcaa
cccaatattctggagctcctttacctgaccccatgcgagtttcagctggacgggcagtga
ctggtttatctggaaatactctaatccatagtttaccacctcttctcacatatctagtaa
tagttcttcgagtagcttctatttgtctagaagttaaccaaactggctctgtcgcttgta
atgcataatcaccgaacgcaattgtgttacctttgctagcagaacctttcattctacctc
tatgttgttttctaaattttgttttcttggggcttagcataaaaattaaaaaataataat
aattaagaagctgttgaatcaggagtttcctgattataagcagattctagacttttcgat
tctggcaaaatttctcctttaaaaagccaaacttttactccgagcacaccatacgtagtg
tgagcttggcgatgacaataatcaatatcggctcttaaagtttgtaaaggtacacgacct
tctctaacccattcacttctagctatctctgcaccattaagccgaccagatacttgaatt
ttaacaccctgtgtatttgctctttgagctctttgaactgcttgtctgacagctctacga
aaggccactctttttttctagctgttgagtaataaattctgctactaaggttgcttcagaa
tctggatctgcaatctctacaacatttactctgagttgtttgctagggtctaggattaag
gacagcgattttcttaaagactcaattcccgctccagattttccaagtacgattcctggt

cgagctgttgctatgagaatttctacttgatcaactttacgattaatttcaattttagca
atactagcattactaagttttgaatgtataaatgagcgaattttatgatcttcttgtaaa
agaactgggtagtctttagagttagcaaaccatgaagaacgatgtttttgggtaatgcct
atgcgaaagcccaaaggatgaatttttgacccacagtatccttgattataagttaaaag
ctataatgactaattgacgttaaa--------agactacagttctgaaacacctaatgta
atatgacaagttggtttatgtattggaaaggctcttccttgtgctctaggctgaaatctt
tttaaagtcgggcctttgtctgcaaaggctttactaacaaacaattggtttttatttaat
ccatcattatgctcagcattcgctgcagcagattccaggatttgctttatatgagaacag
actcggtatggcataaattccagaataattaatgcttcttggtattttctacctctaatt
tggtctaatacacgacggactttatgcggagaaagacgaatgtatttccctactgcttta
gtttcttttacgtttttttgtaatactcatagttatagtttaacgacgggcttttctgtca
ccttttacgtgagtgcggaaagttcttgttggaacaaactctcctaatttatgtcctacc
atttggtctgacacaaagacaggaaaatgttgtttaccattataaacagctattgtatgc
cctaccatatcgggaataatagtagatgctctcgaccaagtttttaggacttcttttttt
cctgaaatatttaatgcttctatacgtttaagaagactaacatcaataaaagggccttta
tgtatagatcttgacataataataaatcattagattcttaataaaataaattctatttac
ggcgacgtaagacgtacgggttactatatttatttggatttcgtgttttaacacccaatg
caggtttaccccaaggtgttacaggacgagctcgtccaattggagatttgccttcaccac
caccatgtggatggtctacaggattcatgacaacacctctaacagtaggtcttttaccta
accaacgatttctaccagctttaccaagagtaatattactagcgtcaatattaccaactt
gtccaatagtagcgtagcattcttttcgaatcattcgaacttcgcttgaaggcaatttta
cagtaacaaaagtaccttcttttgctactatttgagcataagttcctgctgcacgaacaa
tctgtcctccgcacgaaggtcttaattctatattatgtactgctgttcctaaaggaatac
tagataaaggcaaagcatttccaacttcgataggggcagtaggaccagaaagaaccatag
atcctacactgagagatcgagggtgtagaatatatcttttctcaccatcaagatagtgta
ataatgcaattcttgcatttctattcggatcatattcaatagaagcaactttagcaacta
tattatgtctatttcttttaaagtcaattaatctgtactgctgcttatgtccaccacctt
tatgacgacaggtaatgacacctctattatttcgacctttgcagaaatggtgtttaacta
ttaatgatttttctggcttatcagtagtaatctctgaaaaggtagaaaccgttctatttc
ttgtccctggtgtgtaggcacgatataaacgaattgccatataagaagatttgagaagag
taaatgttgaa------ctatgtgatatatttaagtttctggaaacaaattaatagaatc
ttctgaggcaagtgttacaattgctttcttgtaatgtggtcgcttccctacgaatctacc
gatgctcctttttttcttaggaggatgacaagtgttaacacctgtaacttgcacgttgaa
aatatactgtatagcagccttaatattaatttttgttgctttgggatcaacggcaaaaca
atactgatttcttctaataatttagttgtcttatcagtaatgatgggatatttaaccag
atctaataaacctcttgaatcaatgctatccattatatacctcttgtattttagataaag

catcaactgtaataataattttatgtgcagctaacaaagccataatatttagtgtatcag
ctgaaataagttctacgttgtgtagattgcgaatagaaagataaacattcggatcttttt
tgtctacgatgactaaaactttttatttaaatcaagattccatcgatgtatagcttcca
taaatagtttagtcttgggctgatgaaaataactgttaaaattttctacaatcagtgtat
taactgatttgttatttaatgctgtccttaatgctaattgtctttctttttattcattt
ttttggtaaaactacgaggttttggcccaaatattacaccaccacctctccatagaggtg
aacggatagaacctgctcttgctcgaccagtcccttttttgacgccatggtttacgacctc
cacctcgaacttcacttcttgttttagtattagcagaaccctggcgcttttcattacttt
gctttactaaagctctatggactaagtacatcccagaatcttgactgaccttaagattta
aatcagcgttgccacttacttgaccttcccaattatagacttgataatttaattgcgtgt
taactgtcatgaaatatatcgtatgaaagaatattgctacttactaatttttactaaagc
gccaggtttaccaggaacagctcccttaacgatcaacaaatcattctctgagttaatact
tacaatttgcaaatttttttattgtaacttttttattaccaagttggccagccatgttttt
acctggataaactcttccaggcgttgtaccggctccaatagagccaggttgcctatgatt
tttcgaaccatgtgacattgggcctctactaaaatgatgcctcttttggtaaccggagaa
acctttacctacacttctagaagagacattaatcttttggcctacttgaaatagatctgt
agatagtatttggctaacttcgaaatcatcagtagatttcatttcgtattcacgtaaata
ttttagtgggggagcctgtgatttttttaagtgtcccaacaatggtttatttaattttg
ttcagcaacttgcttgtaaccgacttgaatagcattgtagccatcagtagatacagcttt
aatttgagtaataacacatggcccaacttgaattacagtaactggaattgataaaccagc
ttcatcaaagaattgggtcatacctactttagtaccgagtataccaacagacactagatt
tctcctttttgagagatttatactatcattgtacataaaaattctacttcatagttactg
aag--taaattaaactaacagatgcatgatattattcatgacaacataaaggttaaaaac
tttacctatttaatctaaaatagaaacagcttttttgtctagttacagcgtatataaatta
gaagtatgcttattaaatttttttataaactact-tagatttaactgatattgatatccgt
aacttacttgcagtaatatc----aatcaacatgtataagtttaattgcctaaactagaa
ccagctattattagaaataattcaagtatagataaaagcgtatttgaacttttatatatt
gacaaatataatagaatatattgaggttcggtaactacaccttttttatttttacatttat
tgtgca--gtatatatacatctaatgcagtaaacataagaatataac----tgtatagca
aatgtaattacacaaattagagaggtattaatgggcaaagtcgttggaattgatcttgga
acaacgaattctgtaattgctgttatggaaggaggtaaacctaccgtcataccgaatgca
gaaggttttagaactacagcttctgttgttgcatatactaaaagtggagataaacttgta
ggacaaattgccaggaggcaagctgttattaacccagaaaacactttctactctgtcaaa
agattcataggacgtaaacagaatgaaatttcgcaagagattcggcaaacatcatataat
gttaaaactagtggatcaagcataaaaaattgcttgccctgcactgaataaagattttgct
ccagaagaaatttcagctcaagtactgagaaaacttgttgaagatgctagtacgtactta

ggtgagactgttacacaagcagtaataactgtaccggcttactttaacgattcacaaaga
caagctactaaagacgcaggtaaaatagcaggcttagatgtattgagaattattaatgag
cctaccgcagcttctttgtcttatggactagacaaacaaaataatgaaacaatactagta
tttgaccttggtgggggcacatttgatgtatctgtattagaagttggagatggagtattt
gaagtactctcaacttctggagatacacatttaggcggagatgactttgatcagcagatt
gtagaatggctaatcaaagattttaaacagaatgaaggtattgatcttggtaaagataga
caagcacttcagagattgaccgaggctgcagaaaaagcaaagatagaactgtcaaattta
actcaaacagagatcaatcttccttttattactgccacacaagatggtccaaaacattta
gaaaaaactgtaactagagggaagtttgaagaactttgttcaaatttaatagataaatgt
agtatccctgtaaataatgctctgaaagatgcaaaactagaagcttccagtattgatgaa
gttgttttagttggtggatctacaaggattccagccatacagcaaatggttaaaagatta
attggtaaagatccaaaccaaagtgtcaatccagatgaagttgttgctattggtgcagcc
gtacaagctggagttttagcaggcgaagtcaaagatattctattactagatgtgacgccg
ttatctttaggagtggaaactttgggtggcgtgatgacaaagattataccaagaaatact
actattcctacaaaaaaatcagaagtattttctacagctgtagataatcaaccaaatgtg
gaaattcaagtacttcaaggcgaaagagaactaacaaaagacaataaaagcttagggacg
ttccgattagatggcattatgcctgcacctagaggagttcctcaaattgaagttaccttt
gatattgacgctaacggaattttatctgtaaaagccaaagaaaaggctactggtaaagag
caatcaattactatatctggagcatcaactttacctaaagatgatgttgaaagaatggta
aaagaagctgaagaaaatttcgatacagaccaaaaaagaagaaagaatattgacacaaag
aatcaagcagagtccctatgttaccaagctgaaaagcaagttaaagagtttgaagacaaa
attagccaagatttaaaaataaaaatagaggagctaattacagagcttagatctagtcta
gagaaagaagaatatgacaatattgaatccatttctcagcaattacaaaatgctctgatg
gacattggaaaaaatgctgctcagactgaaagtaaagatacaaaagcgaaggatgacgac
actgtgattgacactgatttctctgaagctaagt-aaaagta-agcgggtaacgcgattc
gaacgcgcgacatcaaccttggcaaggttgcgctctaccactgagctatacccgcataga
ttgtattattacaaaaattaagatatttgtcaatctgtta------atctatac---atc
tagactattaattttggaattacatattagctaactgttgtaaatataacaattgctata
atttttaa-------aagtacttatatcat-tatataagtacttttatatt-------
-caaaaatacaaataagtattatag-------ctaaactactaaggagttaactaagcta
gtaacaattactaatcctgtccataagccggctcctgtatatattaatccttttgattgc
tcccattggccaggggatgctaaaacaacaggtacgcctactactaacacagtcgataat
gcaattaataatagcactaataattggatcgcaataatcatttaatatgttctcctgaaa
aattcaaatgttaataatactgtatataatataatgaagcttttgaccttcagcaattta
tctatgaagaaataatttaaaattattgttttctaaataaaccttaaattaagtattaaa
atctttattgtaaacaataatactatattttttta--gtagataaaaaatttaaagaaag

tcttttttaaaaaaaaattaagataaggtgatatgctgtataaaaagctaagtattatta
attattagagaaatttaaa-ataagaggtaaaacatatgaactcagcccttttttagca
aaactaccagaagcttatgccattttaaaccgatcattgatatcttacccgtaattcct
gtatttttcttcttttagcttttgtttggcaagcagcaattggttttagataacattta
ggttataaaaagaataacaaatatacagagtgccaattttgagctctaatcttatatct
tacacaataataactattatggttgaacccttactatcaggaattgttcttggacttatt
cctattactttattcggattattagtagcagcttatttacaatatcaacggggcaaccaa
ttaggactgtaaagcctaaattatagatatttataaaaactacaataattatagtttta
taaatatctacaaataagctaattttaaatacagaaataatactattgaaactagagtga
taacaaaaaaaatttatgtaa---taa---------------------------------
------------------------------------------------------------
------------------------------------------------------------
------------------------------------------------------------
------------------------------------------------------------
------------------------------------------------------------
------------------------------------------------------------
------------------------------------------------------------
------------------------------------------------------------
------------------------------------------------------------
------------------------------------------------------------
------------------------------------------------------------
------------------------------------------------------------
------------------------------------------------------------
------------------------------------------------------------
------------------------------------------------------------
------------------------------------------------------------
------------------------------------------------------------
------------------------------------------------------------
---------------------------caagctaattgtttaccagctcgacttagtaac
tcctggtaaaagacattcatgagacatttctcgaaaaacatggcgagataatccaaaatc
tcgataataacctctacttctacctgttaaccaacatctatttctccctctaacagcggc
gctgtttctaggcatttcttggagtttttgtcttaaatccattttgtcagaaaagctgcc
agcttctttcagttgatctttaatagccagtctttttaaataatattttttgcaagttt
cgatctcttaatctctctttggatcatattctttttagccatgctgtaa-aactctcttt
taaaattatttattcaatatattagctaaaaaagctcctgcctgcaactttatatattt
aaaaataaatagcataaagtagtgaaatgcaattgatatttaatcaacgatacc---act

ttatgctattatcaaaaatttatttagtaaagttcttgttcaacatgagttgaaatagtg
caatcagatgaaggatatgcaatacaggttaaaacatagccagctaaaagttgatcatca
tctaagaatgattgatcagcttgatctacagttccttctgtaactttacctgcacaagtt
gaacaagctccagctctacaagagtaaggaagttcaattccttcttcttctgctgcatct
agaatatatgtatcctctgcacaattaaatgtgacatcaattccttcttcttcacatagt
aaatgaattttataatcagccatttcttcttactccataataaagatacacttgaaactt
aataaagcaaaat-aatttaaagaacttaccaaaaaattatatactagtttcattatgca
tgatagactggaaaaaaagaaaatagtatactattctgaaaaatcaagctacatttccta
ctaaatagaactatagtaggtccaaagtatatacataaattcatatagttaaaattactt
ttatcatttaaatagctgaacttgatagta---------tagattgatcatc------aa
gctagtcttctgc---caataaataattaattatgatttcaatgcttaatcaacaagtag
aacttaagcctattcttaaatttcagataacacaaatatattcgcattctcttattcagg
aaattaaagccttaataactagacttgttttgcaagtttggagaaggccagctacattga
tggcaggtattatccaacctttattatggttaattctatttggtgggcttttctacaatg
ctcctataaatttgttcactattaatacaagctataattgtttttttgagctctgggatta
taattttcacctcttttactggagctttgaattcaggtcttccattaatgtttgatagag
aatttggattttttaaatagattattaacggctcccttagtatcgaggacttctatcattt
tatcttctgctacttttatgacttgtattagtttaatacaagttgtatttatagttacag
cttctctttttatgggaaacccacctctaaatagcgatagtactatgatttttggactta
tgattctattagtgactgtaggagttacaatgcttagtttagctttatctttcactctgc
caggtcatattgagctattagcatttattttagtagttaacttgcccttttttattttcta
gtacggctttagctccttttatatttcatgccgccatggcttcagttaattgcaagtctca
atccattgagttatgcaatagaaggtacaagatacttatattcaagcgtaaactggaatt
ttacagagtgtgtgattaagattagctggggagatatttgtttagggcaaattattattt
tattaatcgctttagatataatggcagcttatcttgtgtctaatatattaaaagctaaac
ttaatta-----------------aaatttttaataaaaaatttattaaatttattttaca
aagatttttactatataatactaatagtagtatggataattgaaatagaaatagtttttc
aaaaccaaagctattatcaacaatgataaacatttgtaagaaagtcaacaaagtatgttc
ttattcataggaggcatgtagtcaatgggactaccatggtaccgtgtacacacggttgtt
ttaaatgatcctggacggttaattgcagtccacctaatgcatactgcacttgtagcgggt
tgggcaggatctatggcattatacgaactagctgtatttgatccttcagatccagtatta
aatccaatgtggcgacaaggcatgtttgttatgccatttatggctagactgggcgtaaca
gattcatggggaggatggagtataacaggagagagcgtatccaatcctggactgtggagt
tttgaaggtgtagccttaactcatatagtcctttctggcatgctttttctagctgctatt
tggcattgggtttattgggatttggaattatttagagatccacgaactggcgagccagcc
ttagatttaccgaaaatttttggaattcatttattgctatcaagtctactttgtttcgga

tttggagctttccatgtaactggactttttggaccaggaatgtgggtatcagatgggtac
ggagtaaccggaaaggtattaccagtagctccagcatggggaccagaaggattcaacccg
tttaatcctggaggagttgcatctcaccacattgccgcaggtactgtaggtatattagct
ggtgttttccatttaactgttagaccaccacaaagactgtatcgtgctctaagaatgggt
aatattgaaactgtattatcaagtagtatctctgctgtttttttctcagcttttgtgact
tgtggtacgatgtggtacggctctgcaactacgcctattgaattatttggtccaactaga
tatcagtgggatagtggatattttcagcaagaaattgagaaacgagtagaaaatgctatt
gctgatggtgctgcacctagcgaagcatggtcaagaattcctgacaagttggcattctat
gactatattggtaataatccagcaaaaggaggattattcagagcgggccctatgaataaa
ggcgacggagttgctgaagcatggcttggacatcctgtattccaagataaagaaggaaga
gagcttagtgttcgcagaatgcctgctttttttcgaaacttttcctgtaattttagttgat
aaagatggtattatacgagctgacattccatttagaagagcagagtctaagtatagtatt
gaacaagtaggtgtaacagctagtttttatggtggcaaattaaatggccaagttttcaac
gatgctcctagcgttaaaaaatatgcaaggaaagctcaattaggtgaggtatttgaattt
gatcgaactacattagaatcagatggagtgtttagaagtagtcctagaggctggtttaca
ttcggtcatgctaatttcgctttaattttcttctttggacatctttggcatggttcaaga
actatcttccgggatgtgtttgcgggaatcggagctgaagttactgaacaagttgaattt
ggagctttccaaaaaattaggagatagaagtagtaaaaaaacaaggagctgtataagaccaa
agtcaaagatagtctttatc---aaatatttctaatattaaaatagttaagacaaagttt
attatctaatagttactaatcaggagaatttatggaagccttagtctatgtatttttact
aacaggaacgctaatggttatattctttgcgatctttttttagagaacctccaagaatagc
aaagtaacttgattccgtttaaacttataataatagaaccactttccaacttattagtgg
aaagtggttttattgtagattacattgtattattcttcatgctcttcaaaaggatctcgg
agatcttttgaagcaggaccaaatgctgtgtatatcgagtaacctgtaatacctaaaagc
agactcgaaataaaaatactaagaactgttgcagtttccataatttaagcagtattaaga
tgaactttttctataatgatattatcaatattaaaacatttaattattaaaattatcaaa
atatactatggcacttagaactagacttggagaaattttaagacctttaaattcagagta
tggaaaagttgctccaggttggggcacaactcctattatgggaattttcatgttactatt
tttcttgtttttattaatcatttttacaaatctataattcttcattagtattagaaaatgt
agatgtagattgggctactttaggtag----ctaactagagaagttctttga--------
ttacacaatattaaagacaac----------------aagcgatcgctattagattatag
tagcgatcgctttaaaagtattttactatatatccagagaaaattatttacctaaattaa
ttaactcggtatctgcaaaactatttgtgttaacattattatagttaactttttcaaacc
ttactaatactggatatttgataccacttttatcaattgctgcaacagtaccaatttcct
gataccagtacgattcttttctcaatatttttactttttgagcctctttccatatttttag
tcctcagatagattattaaagattattatattaaaaaatt-taaaaaactaaacaaaata

tataaaaatactaccgacaattgtaatatataactaataaaaggagagagagggattcga
accctcgataacaaaagttatgacaattttcgaaattgttgcaataaaccactctgccat
ctctcctgaattaaaatactgttttaaaaaattagacaagaataacatagaactaaggta
gttgcctagaaaattaaaactatgttaaatataaattataatctttctatagatatattt
aaaaaatatgtacatagaatctatatcagcttatttcaagatttttttaatcttatacaat
ttttactaaataaataaattctctcataatatgaaactatcttggaaaacacttcttctt
tggtctttaccaatatttgtaattggtttctttttctggcaaggtttttttaggtccaact
actacagatgttggcagtaatatcgcaagttctagaatgacatatggacgattttttagaa
tatttagatatgggttgggtgaaacgggttgacctctatgaaaataatcatacagcaatt
gtagaagctgttgggccagaattaggaaatagggttcaacgaattcgagttgaactgcca
gcaagtgcgccagaattaattacaaaattacgcaaagccaatgttgatctagatgctcac
ccccctaaaagtacaagtgcagtatggggactattaggcaatttactatttcctttacta
ttagttggcgggttagctttcttatttagaagatctaataatgctagtggtggacctggt
caagcaatgtcatttggcaaatcgaaagctttatttcaaatggaggctaaaacgggagta
gtatttaatgatgtagctggagttgaagaggcaaaggaagaatttcaagaagtggtaaca
tttttaaaacagcctgaatcatttactgctgttggtgcaaaaataccaaaaggcgtttta
ttagttggacctcctggcacaggcaaaacattactagcaaaagctattgctggcgaggct
agtgttccttttttttagtatctcaggctcagaatttgtagaaatgtttgttggtgttggc
gcttctcgtgtgagagacctattcaagaaagcaaaagacaatgcgccttgtatcgttttt
attgatgaaattgatgctgttggtagacaacgaggaacaggtgttggaggtggtaatgat
gaaagggaacaaacattaaatcaactattgactgaaatggatggctttgaaggaaatact
ggtgttattgtaattgccgctactaacagagctgatattttagattctgcattattaaga
cctggaagatttgatagacaagtttctgtagatgtaccagattttaaaggcaggttagca
attcttgaagttcatgctaaaaataagaaaatggaacctaaagtatctttagaaaccatt
gccagaagaactcccggcttttcaggagctgatttagctaacttactaaatgaggctgct
atcttaacggctcgacgaagaaaaaatgcaatgactatgtctgaaattgatacatcaatt
gatcgagtagtagccgggatggaaggcactcctttaattgacagtaaaagcaaaagatta
attgcgtatcacgaagtgggtcacgcaataataggcagtttattagagcatcatgatcct
gtgcaaaaagttacattaataccaagagggcaagcaagaggcttaacttggtttactcct
agtgatgatcaaagtctaatatcaagatctcaaatactagcccgtatcgtaggtgctctt
ggtggcagagctgcagaagaaatcattttcggtgacgcagaagttactactggtgcaagt
aatgatttacagcaagtaacatcaatggcaagacaaatggtcactcgatttggaatgtct
aaaattggacctttatctcttgaaagccaaggaggagacccatttttaggtagaggcatg
ggaggaggctcagaatattcagatgaagttgcaactaatattgataagcaagtaagggaa
attgtcagtgaatgctatgcacaagctaaacacattattatagataatcgagtagtgata
gatagattagttgatttactaattgaaaaagaaacaattgaaggcaatgaatttagagac

atcgttaaggaatacactgcaattcctgaaaaaaattactacatatcacaattttaa---
---------------attaaacgggactgacgggattcgaacccgcaacttccgccgtga
cagggcggtgctctaaccagttgaactacagtccc-aaaaagatgaatacccagagataa
tctcatagttagcttaaacttgtcaattagaatattaaatatgaagagta----------
--------tttgtaataatactcttcatatttaata-tagtagctaaaatatttctttta
gaaggagagagagggattcgaaccctcggtacgaagttaatcgtacagcagattagcaat
ctgccgctttcgaccactcagccacctctccatga---tatatattatatatatatcaaa
tatacttatattttaataggtgtgcataaatatttatggagctaagcggattcgaaccgc
tgaccctctcaatgccattgagatgctctaccaactgagctataaccc----ctttggta
agatattaagaaatggctcaagcgggatttgaacctgcgaccttgggcttatgaatcccc
tgctctaaccactgagctactgagccatagtataaaatattattaagtattataccataa
attatggtcaaatataaatcaataaaagaataattactatca-tttactacttttg----
-----------------tgcaagttgtaaatattcgaataaaatttcaaaattaaaaatg
tctttaactttttgttaga-------atta-ttgattacgaaa-----------gtattg
atatttagaattaac---------------------------------------------
--------------------aataaatttg------------------------------
---------------------------------------------------tctttattt
g-----------------------------------------------------------
------------------------------------------------------------
------------------------------------------------------------
------------------------------------------------------------
------------------------------------------------------------
-------tagaca-----------------------------------------------
-------------------------------caaaaatattggaaa--------------
------------------------ttttcatcaat-----------------taattagt
------------------------------------------------------------
------------------taaactataaccgacatac-----------------------
------------------------------------------------------------
------------------------------------------------------------
---aagaattattgaacta-----------------------------------------
------------------------------------------------------------
---cgttaaat-------------------------------------------------
------------------------------------------------------------
------------------------------------------------------------
------------------------------------------------------------
-------------------------------------tgtgatgcatt------------

-----gggcttgatcaaaacttctctaaaatcaatagaatatagt---------------
-------------------------------------------------------aattcaa
gtagatcaaatactaacaaatgtgacaaacgcttaaaaaaataata--------------
caattagaatcgtaaaaatttggtttttga-ctgtgttaaataaatttgattatcataga
tttaatatttaaaagtgttctattacttgta-----ttgagtaattgt-aatatttaata
tcttctttttagataacaatcataatataagctaaaaaatagtttt---tattaaacaa
ctagcatagagccaataccttgatcagttaatatttctagcaagagagcatgatctattc
taccatctaaaatatgcgctgaagctacaccctgagccaaagaacgaatacaacaattca
ctttaggaatcatgccaccagaaattactgctgtttgagttaaatctctagcttcctgta
tattcagatgactaattaatgttgaaggatctgaagaattacgtaaaatgccgggcgtat
ctgtgagtagaattagtttttcggcattaagagcggctgctatttctcctgcgacagtat
cagcattaatattatacgattggccttccttgtctgctgcaacactagctataacaggta
tataattattattaattaatatcttcagtaatttagtatcaatattttgtacttctccaa
caaaaccaaggtttgcttttccatttggtctcggagtaataagtaatccatcttttcccg
acaaacctacacttttaccaccttgtttattaatagttgcgacaagatctttattgactc
ttccaactaaaaccatttccactatatccatggtaggttgatctgttactctaacgccat
catcgaattttggtaatattttgaggcgatctaaccaaaaattaatttctggtcctccac
catgaactaaaatgggacgtagtcctataaaagacagaaaaacaagatcactaatcactt
gatcttttagtttctgatttttcatagccgctcccccatattttataacaataattctgg
aagaaaattgctgaatataaggcagagcttcacttaagacttttactctttctgaattag
tcaacataaataaagtaggattgtgatatataaaattatatacacagcttgtgttaattt
aaaagataccaatgggt---------acttgaaaaaatagaaatattactaacag-----
----ttaaaactatagggattataattatgaataaattttgggataatgtattaagattt
cctcgatttttagtcagcgtcattttaggattaattttgataattattagtcctttttc
gtgttattaaaaaaaccactgacaagttttttttcatcatatcattggccggattaatc
acagttttggcaataataa-tacaaaaaatgataaatatcgaatgttgttgagagaaata
aataaatttaattaaaatttttgaacatatataatattatattatcattagatatgataa
ttt-atttccttttctat---aaaacccataaatattaaatattatttgataaatattat
gccactaaaacaaagagttagctctgaaaagaccggtgccttcgcacttttggatagtat
tgttaggcacggagttaaacatatatttggttatccaggtggagctattcttcctattta
tgatgagctttatgcttgggaagaagcctctctaattaaacatatccttgttcgtcatga
gcaaggcgctgctcatgctgcagattcttattctagatcaacaggagaggttggagtatg
cttcgctacttctggcccaggagcaactaatcttgtctcaggtatagctacagcacatat
tgattctgtacctatattagctataacaggtcaagttgggagagcttttattggtacaga
tgctttccaggaagtagatatttttgggattacacttcctattgtaaaacattcatatgt
agttcgtgaccctagagacatgtctagaattgttgcggaagcatttttatttgtaaaca

cggtagaccaggtccagtattaattgatgttcctaaagatgtaggattagagaagtttaa
ttatttttctgttgagcccggaaaagttaatattcctggctgtaggccaattaccagcct
aaagtcaagacaaatccttatggcagctaaaatgatacagcaatctagccagccattgtt
gtatattggtggaggagccataatctctgattcacatcaaattattaaagaacttgttga
tttttataaaatacctgttactactactttgatggggaaggggattttaatgaggatag
cgattattgtctagggatgttaggtatgcatggtactgcgtatgctaattttgcagttag
tgagtgcgatcttttaattgctttaggagctagatttgatgatagagttactggaaaatt
agatgaatttgcttgtaatgcccaagtgattcacgtagacattgatcctgctgaagtagg
aaaaaataggattcctcaagttgctattgtcggtgacgtagcagaagttgttagtgaaat
attgaatttattaaagacttctttccccccttatccagagcagattatatcttggcaaga
aagaattaatcgttggcgtcaacagtatcctttactggttcctagaaaatcaacaagcat
ttcacctcaagagattcttgttgcaactaataaattagcccaaaatgcttattttactac
agatgttggccagcatcaaatgtggtcagctcaatttctgaaagtaaaagctaagcattg
gctttcaagtgctggattaggcacgatgggttatggtttacctgcagcaattggcgctca
agtagcacatccaaatgacgtagtcatttgtattagtggtgattctagttttcaaatgaa
tatgcaagagttaggaactatcgcgcaataccagttaccagttaaaattattattattaa
taatcgatggcaagggatggttagacagtggcaacaagctttctatggtgaaaggtattc
acactcaagaatgacagaaggagcacctgattttcaaaagcttgcagaagcttttggtat
taaagcttttactattaataataggcaaaatatgcaatctgctttacaagttgctattga
ttatcctggtccagttttattagattgccaagttacagaaaatgaaaactgttatccaat
ggttgctcctggaaaaagtaatgcacaaatgataggtatcgctaaaccgcagagaggtac
tgcttccaactatattaataatagtgtttgaagtaa------------------------
------------------------------------------------------------
------------------------------------------------------------
------------------------------------------------------------
---------------atagttagtatttttta-aacaaagaata-aagtaca--------
-taaatgtattttttttatactatta-aaaactatgtaagaacagcctttgacgagaatt
gaactcgtgaccttccccttaccaaggggatgctctacctctgagccacaaaggc-tttt
tactatattgggccgggttggatttgaaccaacgtaggcgaagccagcggatttacagtc
cgcccccattaaccactcgggcaccgaccctaattaaaccttattcaagatatgacctat
aagatatcataactatgttgctaaaaacaagtactttgataaataaaactgaataagtaa
tggacatagctggacttgaaccagcgactttcacgatgtcgacgtgacactctaaccaac
tgagttacatgtccaaaactttaattatcattataacactgtttagtgattttataagta
tctatttctaattgaagatttgttttaatcgtccggctttaccgagcagatctcttagat
aatacagtttagctctacgtattttagctcttctaatcactttgacaaaagttattcgcg
gtgaattgagaaaaaatactctttcaactccaataccttgaaatgagcatcttaaagtta

aactagtatttaagctttttttttttcttcttgctaatactactccttcacataattgtt
cccgagttttactaccttccttaaccattagtccaagttgtattgtatcgccaactttaa
tttcaggtacttctgttttgataaaaggtatctcaacacttttcattagctgacttagct
ttgtactatttgctttcatatataggattagtcttgtgaca-tgttttgaaatattatac
attaaaaaaacaaatatgaaaaattttagcgaataagcatattagtatttataaaac-tc
tttttatct---------------------------------------------------
--------------------------------------------ccattata--------
----------------cagatccatt------------atattcaact------------
------------------------------------------------------------
------------------------------------------------------------
------------------------------------------------------------
------------------------------------------------------------
------------------------------------------------------------
------------------------------------------------------------
------------------------------------------------------------
------------------------------------------------------------
------------------------------------------------------------
------------------------------------------------------------
------------------------------------------------------------
------------------------------------------------------------
------------------------------------------------------------
------------------------------------------------------------
------------------------------------------------------------
------------------------------------------------------------
------------------------------------------------------------
------------------------------------------------------------
------------------------------------------------------------
------------------------------------------------------------
------------------------------------------------------------
------------------------------------------------------------
------------------------------------------------------------
------------------------------------------------------------
------------------------------------------------------------
------------------------------------------------------------
------------------------------------------------------------
------------------------------------------------------------

------------------------------------------------------------
------------------------------------------------------------
------------------------------------------------------------
------------------------------------------------------------
------------------------------------------------------------
------------------------------------------------------------
------------------------------------------------------------
------------------------------------------------------------
------------------------------------------------------------
------------------------------------------------------------
------------------------------------------------------------
------------------------------------------------------------
------------------------------------------------------------
------------------------------------------------------------
------------------------------------------------------------
------------------------------------------------------------
------------------------------------------------------------
------------------------------------------------------------
------------------------------------------------------------
----------agattctttcatagtttc--------------------------------
------------------------------------------------------------
------------------------------------------------------------
------------------------------------------------------------
------------------------------------------------------------
--------tttgataatcagtac-------------------------------------
------------------------------------------------------------
------------------------------------------------------------
------------------------------------------------------------
------------------------------------------------------------
------------------------------------------------------------
------------------------------------------------------------
------------------------------------------------------------
------------------------------------------------------------
------------------------------------------------------------
------------------------------------------------------------
------------------------------------------------------------

------------------------------------------------------------
------------------------------------------------------------
------------------------------------------------------------
------------------------------------------------------------
------------------------------------------------------------
------------------------------------------------------------
------------------------------------------------------------
------------------------------------------------------------
------------------------------------------------------------
------------------------------------------------------------
---------------------------atcta----------------------------
------------------------------------------------------------
------------------------------------------------------------
------------------------------------------------------------
------------------------------------------------------------
------------------------------------------------------------
------------------------------------------------------------
------------------------------------------------------------
------------------------------------------------------------
------------------------------------------------------------
------------------------------------------------------------
------------------------------------------------------------
------------------------------------------------------------
------------------------------------------------------------
------------------------------------------------------------
------------------------------------------------------------
------------------------------------------------------------
------------------------------------------------------------
------------------------------------------------------------
------------------------------------------------------------
------------------------------------------------------------
------------------------------------------------------------
----gatgatcaaatattattttcaaacatggcttatgtatgaaaaagttctata---aa
aacctaatgac--------taaaattagggcttaagtatttcttagttttgtgttataat
tagttactatcaaaggtagtggcataacacgaagcatcaa-aatgttcgaacgctttact
gaaaaagctataaaagtcataatgctagcacaggaagaagctagacgcttaggacataat

tttgtcggaactgagcagatactattaggattagttggtgaaggtactggaatcgcagct
caagttttgaaatcgatgaatgtgaatctaaaagatgcaagagttgaagtcgaaaaaatt
ataggaagagggtctggttttgtagcggttgaaattcctttcactcctcgagcaaaaaga
gtactagaattatctttagaggaagcacgtcaactaggccataactatattggcacagaa
cacttgctaatgggcttagtccgagaaggagaaggagttgcggcaagagttttagaaaat
ttggcagtcgatgtttcttcaattagagctgaagttatacaaatgctcggagaaaatgcg
gaagccaatgtaagtggaagcaatactacgcaagctagaagtaaaacaccaacattagaa
gagttcggatctaacttaactcaaatggctatggaaggtggtttagatcctgtagtcgga
agacaaaaggaaatagaacgagttattcaaatcttaggtagaagaactaaaaataatcct
gtcttaattggggagcctggtgtaggtaagacagcgattgcggaaggattagctcaaaga
attgctaatagagatgtaccttctattttagaagataaattagttattactcttgatgtc
ggtctattagtagccggaactaaatatagaggtgaatttgaagaaagactaaaacgtatt
atggatgagattaaatcagctgataatgtaatattagtgattgatgaagttcatacattg
attggggctggtgctgcagaaggagcaatagatgcagctaatctgcttaagccagcttta
gcaaggggagaattgcaatgtataggtgcaacaactttagaagaatatagaaaacatata
gaaaaagatccagcattagaaagaagatttcaaccagttgtagttggagagccaagtgtt
gaagaaacaattgaaattttgtttggtcttagagaccgttatgaaaagcaccatcaatta
acaatgtcagatggagctttggctgcagctgctaaatacgctaatcagtatatttctgac
cgatttttgccagataaagcaattgatttaattgatgaagctggttctagagtccgttta
ctaaattctcaattacctcctgctgccagagaattagataaagagttaagagctgtatta
aaaacaaaagatgaagctattagagctcaaaaatatgaaacagcagagcagtatagagca
agagaaatggaaattaaagctcaaattgcagcaattgctcaaagtaaaaagaatgagcct
gatttaaatttagaagatcctgttgttacagaagatgatattgctgaaattgttgctgca
tggactggtataccagtaactaagcttactaaaagtgagtcagaaaaattaatgcacatg
gaagaaactttgcatggacgtattattggtcaagacgaagcggttgtagctgtctctaga
gcgataagacgcgcaagagtaggtctaaaaaatcctaacagaccaattgcaagctttatt
ttttccggaccgacgggtgtaggaaaaacagaattaacaaaagctttggcttcttatttc
tttggttcagaagcttctatgatacggctagatatgtcagaatacatggaaagacacact
gtatctaaactaattggttctcctccaggatatgtgggttatagtgaaggtggttatcta
acagaagcggtaagaaaaaaaccatatactgtcatcttatttgacgaaattgaaaaagct
catccggatatttttaatctacttcttcaaattttagaagatggccgactaacagatgca
aaaggcagaactattgattttaagaatactcttttaattatgacttctaatattggatct
aaagttattgaaaaaggaggaggtagtttaggctttgaattatcagaagatcaaacagaa
tcccaatatactagagtacgatctttagtaaatgaagaactgaaacaatactttagacca
gagtttctaaatcgattagatgaaattattgtatttcgtcagcttactaaagatgaagtt
agagaaattgcagaattaatgcttaatgaagtctttgcgagaattaagcagcaagatatt

caattaaatgtaacagaacgatttaagcaacgattagtagaagaaggatataatccaagc
tatggagctagaccacttagacgagcagtaatgaggcttttagaagatagtctggctgaa
gaagttttatctggtaaaattaaagctggtgatagtgcagtagtagatgttactaatgaa
ggagaagttacagttttattaggtgaaaaattagaactgttaacataaa------aaaca
atttattaat--taaagcattgaagaggttat--attcaatgctat---gaaaaaaatta
ataagatgggttgctataattgacaacctctgatattctttatttgtatata---aagcc
gggatagctcagttggtagagcagtggattgaaaatcctcgtgtcaccagttcaaatctg
gttcttggcat-tttaagcatcagaaaattttgctatctgagcattaaattttaattgaa
atgttcctgtgggaccattcctgtgttttgctattataacttctgtgaagtctctacttc
tagtttcttgtgtatagtaactttctctatatagcattattactaaatctgcatcttgtt
ctatagaattatgaacaataaagttatttgaaataaaattgcataaaggtttacattcta
aatcaaacactatttgtaaacttgtaacgtttatttttttgtatactttcaaaagtcaaag
atgaaa------aagaattaaaaacatttttttgttcttcaaagttttt------tattt
gaatagcaatcatgtcattttgattaattttatcacatcttttccatccttgtgttgtga
gtagtttatggttacttgttaattgtatatatttcccagcttcggtaataattttatata
cagtttttttccctgttttcgaaatactacatttaattgctagaaataactgtcgttttt
gcgcactaaagttaatgacttca-atctttttataatagcaattaaataatatttg--ag
gttgattatataatgatgtctgaagataattgaatttggatatacacccactttctctta
aatcggataacaaaggccttttgtcactccgagtttcaaggtttctatttagttgagata
atgccaagataggcaagcttaaatcttttgctaatattttaagtgatcgagtaattaatg
atagttcttgagatctattgtcagattgtctactctcctgcagtaattgaagataatcta
taataattaattctatatttttttccttgtaatttaaagaattttactttttgtctttatag
tgtcagtagaaatcttagcgctatcatcaatataaagattaaggttagcaagagttttac
tttttgaataacatattgccattcatcattattaagctgaccagattgaatttttttggc
cgtttaaattacactcttgagctaaaattcttcttagtagttgctctgtagacatctcta
agctaaataaaataacataagatcttttctgattaattacatatctagttatattaatag
caaaagcagttttccccattgaaggtcttcctgcaagaataattaaatctgacttttgaa
acccttgtgttatagagtcgagttctgtaaaaccagaaagtatactactatttatactaa
tctttttccttgtatccaaatggactaatagtcttgcaagaattgtagctaatgtttgcg
tatctttttcgtcaagtatttcataagcttttgttaattgactaacaatagaagttatgt
tttttttgttcaataagttgttttgagtagctaattaaacataatgaatctccacatgcta
aaagtaatcttttaatgtagttatctataacgactgctgaatattcgcttatagtatcag
ataggggagcttgttctattagactaaaaacttcatttagctcattcaagtcttgaatca
tttttttcatttttttaagttaatgaaaaaatttcttacgctttttgtctttgcatggttaa
cattttcgagtgctgctctataaagtaatgaagttgatgtaaaataaaagaaatcaggag
agatttttttctactgattttagtaatgatatcgattgctgagttaatattatactaatta

atatttttcagctaaaatattatgaggaggtagatatttataa------atacttatta
cttcctcc-ttgtaacattactcttgtg--------gtattagttaactattgatttaat
ttgattctggaagaacttgcaatttaatatttgcagttacttgatgtaatagcttaattt
ctacatcgtaaatgccaattgttttcacgtctggtaaagagatactttgtttttcaatgt
ctatgttggtagtgtttttaataatttgtgaaatttctttttctgtgacactaccaaaaa
tattgtggccatctcctgtttttttactaacgctaaacctttgtatttcttctagaagtt
gcttgactttttagcattttctttagctgtttttaatttttcttctcttatagctgcat
aaaattttgttgttttaaaataccatttgtggcaactgccgccattttattcgggatta
aaaaatttcttgcataaccagaagctactttgataacgtcattactcttgccaagttttt
gaatattctcttttagcacaacatttataactttttactcatctt-------aattcta
aaaa----atctgtattatcctattatacctgttgtatacagatttt----aataggaag
tatttatttgttgatataatattcttgtaaaa--aaaggtaatttttattttt-ttctg
atataatcactttcacaataattc-----attaggagaggtggccgagtggttgaaggcg
cagcattggaaatgctgtttaggagcaatcttaacgagggttcgaatccctctctctccg
tacaa--aaaatataaaaagttttatagagtccaatctaaatctccaataatattcttcc
aagcatttaatccaccccgaactctaacaatatttagtttttgagctattaaaaactcag
aagcaaatattgatctagaatctaagctacagtaaacaaaacaaattttatcttgtaaat
tgatatgagagtaattcattcctttatttttttgtaaaggtagatttaatgaatatatta
aatgacttttatgatattcttcatgatttcttacatctattaaaatatattgtggattcc
tactaattaaaaacttttgtaattgaataacactaatttctcgtacaaatgtattagatt
tactgccataatatttgttccaatgctttttttgt-------ggataatacaaatttagta
tttataatcttaaatttgttaaatgatgaagttaatgagttgtacgttaatatagttcca
cttaatatagacttataacccagaacgattttgacagcttcagttgcttgaagtgtacca
acaatacccggtaagagacctaaaactccggaattactacaagtatctcttgcgctctct
ttattttcagtttcactgtagaagtctcgataaacaggtccaccttgataattaaataca
ctaacttgtccttcaaattgaaaaatggcaccataaatatgtattttatttaactctagg
caagtatcattcaagagatatcgagtatcgaagttatcagatccatctataataatatca
tattgtctaataatgtcgattgaattctcatagcttagtcttgttttgaacacagttact
atacactgtggatttaaatctaata-ttttttttttagctatttctacttttgaatatcct
atatcatgcatcgtgtataaaatttgcctctgtaaattagagagatcaattatgtcatcg
tctataattccaatgctgccaattcctgcagcagcaaggtaaatgattcctggagatcct
agcccaccagcaccaatgaataaaacttttgcttcttttaatctttcttgtccttctaat
tgaatttgtggcaatactaaatgcttagaatatcttgtatattcttctagtgagtatttt
gtattttctgttttaaaatttagcataaaaatttatagtttatttaatagcactta----
--aaagctagctgataagtttattaaaatgacttttgtaaacttggttcttctctatcta
aaatagctccttgtgtaggacagacctgaacacaaatactacaatcaatacaggcagaaa

agtcaatccaataccaattagtgttattattgtttttcccctttccttgatgaatacaag
ctacggggcaagcttccacgcattcagcaactccaatacatttttctgttacaattgtat
gagacatattatttaattaat-aaaagataagctatttacatctagtagttttgttgtat
actaaatttacatatgctatccaggccaatatttaaggtagtctatttctacagtttact
aatttgatatgcgcctttagttcagttggtagaacgcaggtctccaaaacctgatgtcga
gggttcaagtccttcagggcgcgtttag-ttttatcgtaattcaataatatcattttatc
tttttgcaaacatcaaaatgatatgatataaaagatcatatttttatacattgaatattct
atgtactagtactg-ttttttaatatgttattattgataagttaattatactaattctat
ttttgtaataagt-aaaaacagttctaattaagtatgcacgtatttaatagtc-ttttgt
atatat--aacttgaaaaaatttaaactatggctaaaaaagttactggaattgttaaact
agcattaaatgcaggtaaagctactcctgcgccacctgtcggacctgctttaggacagca
cggcgtgaatattgttatgttctgcaaagagtacaatgcacgtactgctgacaagtcagg
actagtaatccctgtagagatctcaatatatgaagatagaagttttacttttatacttaa
aactcctcctgcttctgtactaatcgctaaagctgctggactaaataaaggttcgggtga
accaaataccaaaaaagtaggcagcataacaaataaacagttagaatctattgcagaaac
aaaacttcccgatttgaacactaacaatatacctcaagctatgaaaatagttggaggtac
ggctaaaaatatgggaattctaattaaagatt-aaaagtttcaattatatttacta-tt-
-tttatgaaaaaattttcacgtcgacttacaacattaaaatcaaaagttgagcctaaact
ttacactattaatgaagcagtatctatattaaaagcaacgtcaaatgctaaatttaagga
aactgcagaagctcatattgctctaggtttaaatcctaaatatgcagaccagcaactcag
agcaacagttatattaccgaaaggtacaggtaaattgataaaagtagcagttattgctaa
gggagaaaaattaacagaagcaattagtgcgggagctgatgttagtggctccgaagaact
aattgatgaaatactgaaaggtagattagattttgataagctgatagcaacaccagatgt
tatgcctttaatagcaaagctgggacgagtattaggccctcgagggttaatgccttctcc
taaagcaggcacggtaacattggacgtagcaaaagccgtgaatgaatttaaaggggggtaa
agttgaatacagagttgatagaacaggtataattcatgtaccttttggaaaatctagttt
ttcacaagaagatctagttttgaacctccaaacaattaaagaatcgattgatagaaacaa
gccttctggagcaaaagggaagtactggaaaactttttcttatctagcaccatggggcc
atctattcaaattgatatcactagccttctatgaaaaatttttgtataattatagaactt
gaataatctattatttcaatta-taatttgctagcattattaacgtaacaacttgctta
ctatatcttatctgcattaaaaacttt-ttatatatcttattaattatgagtacaaaggt
tgaaaatatcttggaagagctaaaatctttaaaccttctagaggctgctgaactagttaa
acaaatagaagaaacgtttgatgttgatgcatctgcggcttcaggaggtatgatgatggc
cgcaccaacttcagcaccagcgtctgctgaggttgaagagaagacagaatttgatgttgt
cctagaggaagtcccagcacctaaaaagatttctgtattaaaagctgttcgctctctgac
tggtttgggtttaaaagaagcaaaagatcttgttgaatctacgccaaaagtcttaaaaga

aggtgcttctaaagatgatgcagaaactatgaaaaaacagctagaagatgctggcgcaac
agttattgttaagtaatga-----agaagcgcttctgtttaagtataaacagaagtgcct
ctttatttattataagcaattatactagaaaagtccaagtgtaattgcattgttgatagg
catagttgcaccaatacctagccaaatacttacaacggtaccaattaagaaaacagtggt
tgcaattggtcttctaaatggattttgaaatttgttaacattttcaatgaatggaacagt
aagtaaaccagctggtacagcagccatacttagaacacctaatagtttattaggaatcac
tcttaatagattaaatgttggaaagaagtaccattctggtaaaatctctaatggagtcgc
aaatggattagacttttcacctatagaagatggttctaaaattgcgagtccaatactgca
ggcaaaagtgccaataattacaactggaaagacgtataatagatcatttggccaagctgg
ctctccgtaataattatggcccattcctttgctaatttagctctcagcttcggatctgt
taaatctggtttttttaagaattgacatattg-tttttgtgtttaatataaagattagtct
aattactgttttaaagtgtaattttcaactattataatggtccagaaattccttgtttac
gtatcattaaaaaatgcataagcatgaaaacggcagttagtagcggaagtacaaaagtgt
gtaaactatagaacctagttaaagttccttggcctacactgactcctcctcttaacaatt
ctactatactcgctccgacaactggaacagcatcagggacacctgtaacaatttttacag
cccagtatccgatttgatcccatggcaaagaataacctgttaccccaaaagaaacagtaa
gcactcctaaaattacacctgttacccatgtcaattctctaggtttttttgaaacctcctg
ttaaataaacacggaatacatgtagaatcatcatcaagaccatcatgctcgctgaccatc
tatgaattgatctaataagccatccaaagtttacatcagtcataatgtactctactgatg
taaaagcttctgcaactgtaggtctataatagaatgtcattgcaaatccagttgcaactt
gaattagaaaagatacaaatacaataccccctaagcaataaaaaatattaacatgaggtg
gtacatacttactagaaatatcgtcagcaattgcttgaatttctaatctctcttcaaacc
agtcataaattttactcataaaatagcttc-aaaagctcttctaatata--ttttgcgtt
taagctactattaattcaatag------ttattgaacttatatt-tttttataatcaata
cttattaactattataacatcttttattgattt--------------------aaaatt
ctctgacttagtaaaacagggctatgtactataattcttttttttgtatagagattaat
ttcgttttagtaaattagctaaaattcttgttacactgactctattactacctataatc
tgtgctaaaattttatgtgtaatagtaaagttaagcataataccattattaacttgagtt
ccattttgctcagctaataataataataagctcgccagtctactggttatacttttatgt
gaaatgatttctataaaatgataagcttttgccgagcaaaatactaaatgatttacaaaa
aataaattgaatggagcgcagttactacacgcttttataatagtagtatagttaatacag
gcaacttctgttttatctattgcttctgcttcgtagtaaaaattatcgtcgaccaattct
atttgtccaaatgtatctcctgtagttaataagttaagagttactttatgtgtatttctc
aggacttttgtaataattaaagatcctatcaatataatgtataatctagagctgctatta
aacaaaagagtatcattttttcttaagtgaaaaaatttgataattaattttattacgatta
aaaaatagtagccatggattgcaagaaaaaaaatcacctgaactattttcagagaatatt

ggaatacaattttcttgatttaaacaatttttagtcataattactactttataa------
---aaaaatattctaataat-taagaactgaaattaaatgtcatataaattgatgttagt
tgagaatgacattgtattatcaaaggccattcaagaatacttaatagatcaaggcttcaa
tgtttatattgccaataatggattagaagcactgaatctagcttatcaatataatttcga
cttgataatttctgatataatgatgccgctagttaatggctatgaactattagcaaaact
taaaaaaaacaaagcattatccaaaatacctgttatctttttgacagctaaaggaatgac
taaagacagaataaaaggctacgacatggggtgctatggttatttatctaaaccatttga
tcctaatgagttgctttcaattattaacaacttaatcgctagagatgtcttaaaagaagc
ttctttacaaaacttggcaacatcaaaccagcaattaaatcataaaatacgtttgacacc
tagagaaaaaagtattttagatcttgttgttgatggacttactaataaagaaatatctac
tatactgaatactagtgttagaaatgtagaaaaatatgttagtcgactcttacacaaaac
taatatgaagaataggactttgctagtcaagtattctataaataataatctattaaataa
tgagatcaatgaaagggcgaatgacgggactcgaacccgcgaatgatggagccacaaccc
attgccttaaccccttggccacactcgccatattacacatactattatagcttgtttttt
gatatatcgtctagttttcgagttatatatttatctaaaacaatatttactttgaatttt
-tatgatacatagcaatattagtattcaaattaatggggaaccatttaattgctcaaagc
ccatctcattgcaatttttattaaattatcttgattttaattccgagcgtgtagcagtcg
agctaaatgacattctgttaccagagactctgtttcactcaacttatttgaatgatcaag
ataagctagaggttattaccattgtaggtggtggataatctatttgtacatac-------
-acagttctctatctcaaatgtttcatagatagagataatcttccttgtttttatctac
atgaataataacagctttaatagtgtcacctattttaaattgtgatgatatctgctctaa
ttctttgatattaatctcagaaatatggacaagacctttgagatttcctacctttataaa
taatccgtaaggtgtaatctgatttattatgccttcaataatattaccaacaattaaatt
agatgatgcttgagatattagagctcttctatgactcagtattaaattattagatttctc
ttcgacattgagtaattttagcttaataaatttattattaaattgctcactttttttgaaa
attaccgagatgagaattaggtacaaatccagatataccttcaaggttaattatcattcc
tcccttattgaatccttttatcattacattaagtaaagagtcttcagctaatagttgcct
aattcttttccatgctcttatatattcaaggcgacgaattgataatattaattgtttcga
ctgaatattataatctaataaaaaaaattctcttgtgtcattaatgtttaaagaagtgaa
attatttaaatcttgattactagatacttcttgaataggcaagtatgcagatattggtgt
tcctatatctactaataccccatttaattcaaagctaaatatagttccagctacaatatc
tccaagattcaaatcatacttgtatttttgtaaaacagctgcaaaatttctatgagtaaa
tccttcattatttttgtcatattgaatagtatgaattaaagagtaactttgtacaatat
attaaggtaaataaatgtttgtcgaattgtatgaaaaaattttaatttgttcggttgtaa
ttaagtatataaacatattaaaatatctaggagacatgaaaagtagacgtagatagctgc
agatttaa-tactatataatctgaggaaagtccgggctccacaaatacaatttatgctgg

agaaatcccagtgtaggtaactgcgaggatagtaccacagaaaaaaaccgccagaataag
t----tatattagctggtaagggtgcaaaggcaagttaaaagcttaccaaaagtactgca
aagtatttgttaggtaaacccctaaaatggagcaaagctactaaacaaatatttgtgtat
attttatctagtttagttgaaaatactgcatgaagttattagtaacaataactcgagagg
aatagctatcctttacaatatatc-tcgattatattgaaagaacagaacccggcttatgt
agtacttttatgtcttgtattattatgtctattgtcgaatctttaaatttaactgtttg
acaattttttttgtagctgatgttgtttataatctatttctgaattggttaaagttttt
tcattagatcgatatgtaaatcttagacctatactttttttctcttttccaattgattca
tctatatattgatcaaataatgttgtactttctaaatctttatcataaaattgatccaat
aattttaataagtatttaatttccatattcttaggtattattaaagagagatctcttatg
atcgatggatacttagagtatggttgaatctgataacttaaataattgaattcattatga
caatcttctaaaatgtttaagttaacttcaagaaaaaaaagttttgtgttgagtcctaat
tcactatatgttgcttgattaagttcactgaatatgccaataataatattgttatagatt
agtgtggcaaattttcctttttgaatgaactttattctaccatcgactatatctttttt
gtccactcaatttgtctattaatcttttgaaaaaaattttctacgatccctttagcttca
taccaatttaaagaatgcgcagggtgtgaccatgtagatcggatatctaaattaccaccc
aatatgatagctagattagtagtttctacaattttgttttgatcattatgaaatactgtt
cctatctcaaagctatctatagtttggttgctttgtttaatattataaagatttgattca
attaatccttccagcaaactgccgcgtaaattagaataatcttgaataagaggattgttt
aaagctatatttcctttggactttattaaagaataatgaactaattcagttaatcctaga
tttcttaaaatacttctacttttatctatgaattgtctctttgtggataactttttgttg
aattgaatatttggtactgagctcctgaatttatgataaccataaattctagcaatttct
tcaataacatctatttctcgaaaaacatcttctcgtctataatttggaacagtaacttct
atattttccttattttgataaataagatcaaaatttaaagagtgaagaatattttgaatt
tcttcaaagaataaaaaacgtgtttgaccattatacataattggtcctaaaatgtcatga
actttttaattgataaattaatattaagagcatgatcatttatttttctcgacagaaa
gtctctctaatatttccgcctgttagatctgtgattaaagcaagagcttcaaaatgtgca
tttttccaattatctacatttagcccacgttcttgtctaattgaactttctgttcgaata
ttaagaactcgagaagattttctaactacagattgtttaaatatagcagactctacaagt
atcgattttgtattataatctgtatcaaaatcagagttgcttccaataccagctatacta
gttacatttgtatttacttgagtaactaaaatatttttgtttaactcaatattttcattg
tttaacttaatctgatcattacttccaagaggaaaattgcttcgaattgttagactattt
ttatgattcatattattaattttattcaaatcaacaatattaataggctgtccccatttc
aacataatataattgcttatatctgttaacaaatttctgtgtgtaaacccagaagagagc
aaccgatttttttagccaatctggagaatctttaactcttatttcgtctataatggcagca
aaataattgtcacaatttaataaatcgtgattactgattatagt-ctccttttttagagaa

atatctattattgggttgatgtatagctgataacattgaggaccctgttaaagccgagac
ttctcttgatagaccaatcatgcttaacacgtctgacctgttagctgttgatgatatatc
taatatatgatcaatctcacttccaattataatagattcaatagcttcaacttcaaaacc
tgcttgtgttagttgattagttaaattgtctatatctattgttttgatattaccaagttc
tttcaaccaatttaaagaaactttcataattaattgtcatctatcttatagaagtattta
ttttacttatgattaaaaatcaatttaaaaaattcatcaaattaatataagttatttgat
gaatttgcttatacaatatattattgtgatatcttacttttatatggcagttattaagaa
ctagcttttgtaaaagttagcccattatcttgggcatatctttccataaatctcatgaat
ctatcccaatcatcaggacttttaataacatgtacagcttctattgcttgtggtttacca
ttaataaacttggcattaacatctcttgtaataatttgtccctctgtgtccatcaggtac
atacctgtgatttcacctttgtctgccatactagcatctaaaattttgggatttgtaaac
cgaaaagttgctgttcctgtactaccatctcgagaccttgttaaacgtacatctggtaca
acttcttcattgataccttgaataaattgaattgttgccatatcaatcttctttatttgt
atataatttatattggataaaactgatttctagttcttgaatgcttgttgagaaactgtt
ttaaaagcttatagtagcagtagatcaaattttatgtattatataaaactgtaacttttc
-ttttttg----------------------------------------------------
------------------------------------------------------------
------------------------------------------------------------
------------------------------------------------------------
------------------------------------------------------------
----------------------atctgttcaaagaactaacttattataagctaatcata
cgaatattagccatataatgttcgtaagt-gaaattaaggggttgtagctcagaaggata
gagcaagcgcctcctaagcgctaggtcagcggttcaagtccgctcaaccctgttaacgaa
ttaagtagtattatttaacttgttctgcaaatcagtattgatctaagttaaagtacttga
cagctaaaacaaaagaatctgagaaatattttcatttaaagaacgaacaattatcttaac
tctaaatatctgataattattaa-aaaattct----------------------------
----ctttttaaaacacaagatgatg-----aagatatttaaactaataattgaaattaa
gtttttgaagttatagaagccaat--ataatactttctattaacaga-----------tg
cttgcttctattactatctgtttacaaga--aatt----atgagtatcaaaata------
--attttagctcta-------------------------------------ctgac----
------------------------------------------------------------
-----------------------------tactaataaata-------------------
------------------tttattttagtctagacaag------atgaatcttcatggta
taacag------------------------------------------------------
------------------------------------------------------------
------------------------------------------------------------

------------------------------------------------------------
------------------------------------------------------------
--------------agctattctttgtgtcatacttatataaaattaggat-------ac
tgtt--------------------------------------------------------
------------------------------------------------------------
------------------------------------------------------------
------------------------------------------------------------
------------------------------------------------------------
------------------------------------------------------------
---------------ttcattattataactggggtgggaggattcgaacctgcgaatggc
ggagtcaaagtccgctgccttaccacttggctacaccccaacatgtgaacattataatag
cagtcgtataccctgccatgtcaatagactattgtctttatttaataattttgctgctc
ttgtagaataaaatcttttttttcaaataagatacgctatgtttaagttgcacgactgtt
atgttttcttgcttttgactgaacagccacttgatagtaattattccttctgcaatttcg
ttacttcctataataatacaagctacagctctctttttatttgcttgcttaatctgctta
ctaaaattacttgagctaacatcgatttctatttaaaaaactgctgatgtaaaaaacgc
attatttgcataccagtttcgttagcttttgtaccttgtgtagcaatataaaaatcgata
gattgattaggcaagtctatattatcttttgcaatcagtaataggcgttctaatcctatt
gcacatccaacagctggtgttgctttgccacctaattggtgaactaagctgtcatatctt
ccaccaccacatatagtatcttgcccttttgatgtcaatgtcttaatttcgaaagcagta
tcattataataatctaatcctcgaactaacttattatttatattgtagggaatattaagt
aattttagataattgcaaacgtcaccaaagtgtttttgtgattccaaacttaaaaaatca
gaaatttttggagcctctgtcagtatttttttgtgtattagaatcttttgaatctaaaatt
ctaattggattactagttaatctttttttgtgagtcagtgtctaaatcatcatggtacttt
gtaagataatctcgtaatttaacttgataaatactacgatcctctacttttccaattgaa
ttcagatcaagtttcaagttgtgcagattgagattgttaaatatgctcatagctaaatgt
ataacttcactatctgctcttgcatctaagctaccaataaactcaatgccaagttggtga
aattgtcgttgtcgtccactttgtggtctttcgtacctgaacattggaccgctataccat
aatctttgtaaactatggtggtaactcatttttatttcaataaaagatctcacaataccg
gcagtaccttctggccttaaggtaatatctctattgctgcgatcatgaaaccgatacatt
tctttatttacaatatcagtatcttcaccaatacccctatcatataattcactattttca
aaaataggtgttctaatttcttggtaatttgcacattctaataatttggaaattttatta
tgtataaactgccaatattgaatttcatcgggaagaatatcttttgtccctctaatagcc
tgaatttttgccatgataatttttttgattaccttcttaatatgtctgcattaaatagcag
ttgtagactgttattcagtcatgtattatacattag--aacatactgatg-taatttata
aagtacgggcaaggagggattcgaacccccgacaccatggttcgtagccatgtgctctaa

tccactgagctacaagcccacttaataagtaccttaactatatcattgttcactaatgta
tacaactatattctttt------atcttacatgtgaaattttgtaaaagtaaaaaatt
at----gtttgtttttacaattggagctattcatatataatgcttaatttattataact
ttgtaaa--aatatctataagtttaattgtatgattctattatattggtaatcttgttat
aattttgcgctcttatttttcaattgg------ctagcatttaattaacttatatttcaa
ataaaaaaatcttatctaaaatgagtaaacaaattctatatcaggatgatgccagaaaag
cattagaaaaaggcatggatattttaacagaagctgtttctgtgactttaggaccaaaag
gaagaaatgttgtcttagaaaaaaaattcggtgcccctcaaattattaatgatggtgtaa
cgattgcaaaggaaattagtttagaaaatcatattgaaaataccggagtcgcactgatta
gacaagcagcatctaaaacaaatgatgtagctggtgatggtacaacaacagctacagtac
tggcttcagcaattgttaagcaaggaatgagaaatgttgcagcaggttcgaatccgatgg
ctattaaaaaaggtattgaaaaagcaacaaattttgttgttagtaaaatagctgagtatg
ctaagccagtagaagatacaaaagctattatacaagttgcttctttgtcatcaggaaatg
atatagaggtaggtaaaatgatagcgaatgctatagaaaaagttggcagagaaggagtta
tttctttagaagaagggaaatcaactaatacgattcttgagatcacagaaggaatgcagt
ttgaaaaaggctttatttctccttatttcgttacagatacagaacgaatggaagttcttc
aagaaaatcctttttattttatttacagacaaaaaaattactttggtgcagcaagaacttg
tgccattgcttgagcaaattgcaaaaacatctcggcctttattaataatagctgaagaca
tcgagaaagaagcattagccacaattgtagttaataaattaagagggattttgaatgttg
tagcggttagagctcctggattcggtgatagaagaaaatctttacttgaagatatgagta
tcttaacgaacggacaagtaattactgaagatgcaggtttatcacttgatacagttcaat
tagatatgttaggaaaagcccgaagagttattgttactaaagactcgacaacaataattg
cagatggtcatgaaattaaagttaaatcaagatgtgagcaaattaagcggcaaatagaga
cgagtgactctttatatgaaagagaaaaattgcaagaacgattagcaaagctttctggag
gagttgccgtcattaaagttggtgcagctacagaaacagagatgaaagataaaaaactaa
gactagaagatgcaattaatgcaacaaaagcagcaattgaagaaggtattgtaccaggag
gaggagctactaatgtccatatctctagtgaattatttacatgggctaaaaacaatttag
ttgaagatgaattaattggtgctttaatagttgaacgagctgtgacctatccgttaagac
gaatagcttttaatgcaggtgataatggagcagtaatagtggaaaaagttaaaagtcacg
attttcatataggctatgatgcagcaaacggtaatattgtaaatatgtacgatagaggta
ttatagatccagcgaaagtagctagatcagctttgcaaaacgcagcttctattgcagcaa
tggttttaactactgaatgtattgtagttgataaagtcgatgattgagctata------a
gataatagataatctcaaacaaagaataatttctaattatagaaattattctttgtttca
attaatctaattaattgatcaaatgtataatttttataataaagccaaagtttaaacacg
tcttctttttcaactattaagtagagaagatataagcccgtaatacctaaattatcaatt
aaaatattattttcatatatagatttttctaaagatctaaaaaatgacattttgaaaaa

taataccgaaatcttctcttatagttatttgctaaatcattactgtctaaagattgattt
ccaattgaattcagaacctttaacagaatactttttaaatctttatctgactttatttga
gaatcaataataatttgattcaattgtatgtcaaaactaagattaaaaatattacttgtt
aaagatatttgattttgaatatctagagattttatagccaagaagattaaacttttattg
ttcattgtaaaatataagacaattgatgatgtattctttaaatacta-aatttatcttct
tgtgcttctatcgtttttgt-aaaataattaatgttttaatttttagttttagttact
acccgaaaaaataaattctggaaatctagcttgagctttagctagagagtgtttttttccc
ttcttcttgccagtaattgataacttctgttgttttgtttaataactcaattctgtctct
ttcagcaatccacggcttggattctagttctgcttttaattcttcccaagcatcattcct
tggccagaaaaaataagaagtaagtggactagtgccatttccaacaatctgatcaatagc
aatagcaatattattttctaaccaaagtacttttagtgtaaattttgacaagtttttatc
tccattattaatattaagtattgg----tacagtagaatactgtactagttatttttttc
agcaacaatttgagctttgtttaaaatgtttttttaccgttttagtagcttgtgcaccttg
agataatcgttgcaaaatttttgaaatgttaacacgagtttcattatttataggattata
aaatcctaattcttctatagctttaccatcgcgtttatttctgctatccattacaacaat
tctatagcttggttgtttttttcttccgtatcgctttaaccttaacttgaccatattagt
cacctcgcttattttttt--atatctttattaaattatatcttacgacattatgcaataat
tcaaacaattaagctgattcaagtctaacattattaaatattaccattaaacattgtaga
aaaataatacctaacataggtgacatgtccattccaaacatgggaggaatgcttcctctg
aatagttttaagtatggatcagtaattctatttaatgaacagaatggttcattgtaccaa
tttacagttggaaaccatgccaatgataatttaagtaaaattaaaattaaataaatctca
gaaaaattagctatagatccaagcaataaatttaatgtaccaggaagagtgttcatacgt
attttataattattatatacttcgttatactaatagtttgattatatattataatcgatg
taaataaat-cagctaaacaaaaaacttatccttaaaatcatcttctcttgagcctttttt
ttttatctacattagtatttacttttgtcgtataaatttccagtttagaccaaatatgtc
ctaactgatttaattgagatgttccacattctaaacaggctatttttatattgctcaaaa
cagcacctttttttataaaatcgttcaataatggcgtaatagttgcttcatccaaaaata
aatctaagacaagtatttttgtaaatgaatttaattgatctgggacattgttaataatat
cattataattaactaaaactatattagctgtaggaagtaaagatcgtgctccttccgcta
agataaagccataaggcataataattacaataatatatttataatgactgctaataagac
ttatagtggtattagcttttaattcgattgtttctgttactagccactctctcattattt
cgtaagtaatccatttgcctagttctgagcaagcagttcttaaaatcgtacctggattat
tgtcgttttcgagaatccctgaccaatgttgtattaatggatgggatgctatattaattt
ggagttgcatgttttgttaaatatatttaacatttaatttgttgattgaatcttcaattc
ctatttgttgagtattattaaaatagcat----atatattaaagaaaataattat-aaaa
aatgaaaaaaagtctttggctgtggggttttacagatagtgctgaaacttggaatggtag

atttgcaatgattggttttatatcagttatttttatcgaagtagttacaggacaaggatt
attatatttaataggtatgatgtcataataaaaatccaaaaaagttcttgtaatttttta
taagaactttttttgg-attttataaatctaatctagcggtctcatagatagaacaggtt
cattctctctattgataccttttcaaatccagcagcagcagctcttgctctgcctgcat
gccaaagatgtccaataaataggaagaatcctaggaagaaatgagatgttgttaaccaag
atctaggagatacatagttaactgaattaatttctgtagctacaccaccaacggaattca
aagaacctagtggtgcgtgagtcatatattctgcggcacgcctttcctgccacggttgaa
tatcgttcttgattttatttaaatcaagaccatttggtcctctaagaggctcaacccaag
gagctcgcaagtcccaaaatctcatggtttcaccaccaaaaataatttctccactaggag
atctcattaggtatttacctaatccagtaggtccttgagaagatgcaacattcgcaccta
gtctttgatctctaactaagaaagtaaaagcttgagcttgtgaagcttcaggaccagttg
gtccatagaattcactaggataagctgtgttgttataccaaacaaaattagaagccgtaa
gccccatgattgataaagcacctagactataagataagtaagcctcaccagaccatacaa
aagctcttctagcccaagcaaaaggttttgttagtatatgccagattccgccagcaatac
aaataatgccaatccagacatgaccacctacaagatcttccatgttatttacacttacga
tccatccatcaccgccaaacggagactttaagacataaccaaagataactaggggattaa
gtgtaggattgctaacaaatctaacatctccaccacctggagcccaagtgtcgtataccc
caccaataaataaagctttaataactaacagaaaagcgccaatgcctaataacactaagt
gtataccaagtattgttgtcattttgtttttatcccgccaatcgtatccgaagaaaggaa
atgattcttcaagagtatcagggccaatcagagaatgataaagacctccaaagccaagaa
cagctgaagaaattaaatgcactacgcctactacaaagtatgggtatgtattgaaaatct
cgccgcctggaccaacaccccagcctagtgtagctaagtgcggaattagaataaaccctt
gttcgtacaaaggcttctcgggaacaaagtgtgctacctcaaaaagagtcatggcaccag
tccaaaagaccattatacctgcatgagcaacatgagcaccaagcaatttgccagaaacgt
taattaagcgtgcattgccagaccaccaggcaaacccggtagactcaatgtctctaccgc
caacaccaacagttgtattaaagggcgtttccacgtggtaaaacctcctcaggaatata
aagttttcatgaggttgatcttgagcagccatccaagagcgaataccttcgttcaatagg
atatttttagtataaaaagtttcaaactcaggatcttccgcagctctcaactcttgagaa
acaaaatcgtaagctcttaaatttagagctagtccaacaattccaaatgcgcttgtccat
aatccagttactggtacaaataacatgaaaaaatgtagccaacgtttattagagaaagca
acaccaaaaatctgtgaccagaatctattagctgttaccatcgaataagtttcttcagat
tgtgttggagtaaatgcacgaaaagtatctgcagcatcaccatcttcgaataaggtattc
tgtacagttgcaccgtgaatagcacatagtaaagcaccacctaagattcctgcaacgccc
atcatatggaacggattcagagtccagttatgaaatccttgcaaaaatagcaaaaatcta
aaaattgcagcaactccaaggcttggggcaaagaaccagctcgcttggcctagagggtac
attagaaagacggaaacgaatactgcaattggtccagaaaatgcaatagcgttgtatggt

ctaagaccaactagcctagcaatttcaaattgtcttaaacagaatccaattagtccaaat
gatccatgtaaagctataaatgcccacagaccaccaatttggcaccagcgagtaaaatct
ccttgagcttcaggtccccaaaggaaaagaagtgaatgtcccatgctgtttgctggggta
gaaacagccgcagttaaaaaattgcatccttctagatatgaactggctagcccatgagta
taccaagaagtaacaaaagtagtcccagttagccagccacctacagcaaggtaagcgcaa
ggaaacagaagtagtccagaccaacctacgaatacaaatcggtctcttttagccaatcg
tctacaagatcaaaaccaccacgagtttttcttgtccaattgctatggtcataatttta
atctccagagcaaatttaataagtaatttgtttaaataaaca---ctttagatatactct
aaaattttacttttctttacggtcaatataattataagtcaaaattaaccaaaattcacg
cttttaataaca-ttttttagctcattaagttaa-ttttttaatatataaacagtaata
gagttattgttt-aacttacaatactcataatttattcatttctataatcctttgaaaaa
gatatcctgttcctcgtgctgttaaaatcaagtcgggattacttggatcatcttctagtt
tagctctcaatcgtgaaatatgaacatcaacaactctcgtgtcaacatgccgttcgggag
tatagccccaaacttcttgcaagatagaagctctagaaaaaggttctccagccttactaa
ctaaaagctcgagaagactgaattccatacctgttaatcgaactctttcattatttttat
aaacttgtcttttgttagtatctattttagaaatccaatactaataatcccagagtttg
ggactccaagactagtggtgattttatcagctcttcgaagaacagaacgaatgcgagctt
ctagctccttaggagaaaatggtttaacgacatagtcatcagcacctatttctaatccgg
taattctatcgcagacttcacctaacgctgttagcattatgataggaacatctgattctt
ttcgaagttcttgacaaacaccgtaaccatctagtttaggcatcataacatctaaaacaa
caaggctaggatactcttttctaaacacgagtaaagcttcttctccatttgaagcagtga
taacttcataacctattattgttagtcttgtttctaaaattcttcttatgctggcctcat
cgtcaacaacgagaatcttttctttctggttctccaa-tttcttatttactcctataact
caccgatcatgattactataaccttaaattttagcgctatgagtatctttattcacttt
agaa-----tttagttattataaattttgcaagtctagtttacattaattagtaccagt-
-caatatttttgt-----------tttaagttatgac---------tattttaatccata
agttgactctagtttgcagaattcaatatatacata-ttttctagccaagtaaattgtat
tctattttctaaataaaaaaggggttgacaaaccagttgatagaaagtattctacattac
tataag----ttaattccaacaaagaagttttaa-----atcttctttcggagtaatgaa
tactttaat
>Pyr_44
tatatggtaaataattaatacgtctttcatgtgaaaattcacaaaatatacaaaactttt
tatttaatatagaaaaagtaaattaaaataaagtgaatatactttgtcctaatttaaacc
tattaattaag------attctaattaaagacaaaattatttacatattgtatatctata
ggaataaaaatgaaacttgcagtttatggtaaaggcggtataggtaaatctacaactagt
tgtaatatttctgtagcactttcaaaaagaggcaaaaaagttctccaaattggttgtgac

ccgaaacacgacagtacatttactctaacaggtttttaatcccaacaattattgacact
ctgcaatcgaaagattaccactatgaagacgtttggcctgaagatgtaatctataaagga
tatggtggtgtagactgtgttgaagctggtggaccgccggccggtgctggttgtggaggt
tatgttgtaggtgaaacagttaaactattaaaagaactgaatgcttttgatgaatatgac
attatcttatttgatgttttaggtgatgttgtatgtggaggctttgcagctccattaaat
tatgcagactattgcctgattatcacagacaatggatttgatgcactatttgcagcgaat
agaatcgcagcttcagtacgcgaaaaagctagaactcattctctcagattggctggactt
gttggtaatagaacagacaaaagagatctgattgataagtatatagactgcgttccaatg
cctgtattagaagttttaccattgattgaagatattagagtctcccgagtaaaaggtaaa
accttatttgaaatgtcagaaactgataaagatttagcgtatgtatgtgattactatttg
aatattgcagaccagttaattacaaggccagaaggtgtagtcccgaaagagtctcctgat
agagaattattcagtcttttatctgattttacttaaatcctaaatcaaatgtaagacaa
acgaaagtagatcaagaagaattagatctaatgatagtgtaaataatattttcagcataa
cataaaaaggaataagataatgtctacagctcaatcggatgctcttacttttgaatgtga
aacaggtaattatcatactttttgtccaatcagttgtgtttcttggttatatcaaaaaat
tgaagatagtttttttcttggttataggaactaaaacttgtggttatttcttgcaaaatgc
aatgggggtaatgatttttgctgagccaagatatgccatggcagaattggaagaaggaga
tatttcagctaagttgaatgattatggcgaactacgaagattgtgcttacaaatcaaaaa
agatagaaatcctagcgttatcttttggattggaacatgtacaacagaaattataaaaat
ggaccttgagggaattgctccaaaactggaagcagaaattcgagttcctatagtagttgc
tagagctaatggtttagattatgcttttactcaaggtgaagatacagtgttagcagctat
ggctcagcgttgtccttcaaatttaaacaatcaactagatagcactagtcttaaagcttt
atctcatattcctttagttttatttggatctcttccagatccagttgtaactcagttaac
gatggaattaaaaaaacaaggtgtttctgtttctggctggttaccgtctaaaaggtatac
tgaactaccagttatacaagaaggctattatgttgcaggagtaaatccatttcttagtcg
gacagctactacactaatgagacgaagaaaaacaaaattaattggtgcaccatttccaat
aggcccagatggtacgcgagcctggattgaaaaaatttgctcagtattgactataaaccc
cgttggattagaagacagagaacaagaaatctgggattctttagaagactatattgcttt
aataagaggaaaatcagtttttttttatgggtgataatttattagaagtttcccttgcaag
atttttaactcgatgtggaatgactgtgtatgaaattggtattccttatatggataaacg
ttatcaagcagcagagctagctttattaaaaactacatgcgataaaatgaatgttatgat
gccaacaattgtagaaaaacctgataactataatcaagtagatcgaattcgtgatttgaa
acctgatttagttattactggtatggctcatgcaaatcctttagaagctagaggtattaa
tacaaaatggtcagtagagtttacatttgctcaaattcatggttttaccaatgcaagaga
tattcttgaactcgtcacaagaccattacgacgaaaccttagtttatcagaattaggctg
ggatgtttatagcaagcaaaactagtatt-------agttacgacttacatggcataaaa

tccttgtaagtct--tttattttatgttaaagaagcatttctagccctgccatcaccagc
ccaagcttgtaaactctctacttgttccttatctgtaaatgctaaaggtacaaattgctt
tattgctatcttaatatcttctgtgctaaactctctttcttcactaaaagctgtatgcat
actttcgataattgcttgctcaatctctgcaccagagaatttattgcataataaacttaa
ttcttggatatcatattcctgccatgattttggtctaacttttgatagatgtatcttgaa
aattgactctctctcttgccgattaggtaaatctaagaaaaaaatttcatcaaaccttcc
ttttcttagcatttcagaaggtaaactctgaattttattagctgttgcgacaacaaatac
tggagcttttttttctgatagccatgttataaatgtcccaaatactcttgcactagtacc
actatcgccttggctatataaaccagaaaaagccttatctatttcgtcaatccataatat
acaaggcgataatccttctgatatatttatcatttctcgcatttttagattcggattcgcc
aactagtccaccaaatagctttccaatatcaagacgtattagcggtattgtccaatcatt
tgcaatagccttagcggtcaaagatttacctgttccctgtataccaactagaagtaaccc
tttaggagaaggaatgccataatttaaactttgctttgaaaatgagcgagaccttttttg
tagccattgtttcaatacatctaagcctccaatatctcgattcactttggcataaggata
aaattctaatagacgcgtttgattaataatttgtctttttttcttcaataataatatgtaa
gcctcgagagtccacttgactatattgtgcaataattttagtaattactttgcgaattct
atctattgataaaccttggcaggattttgtcatattgttaactagctctgaatctaaatt
aagattcaaagctctacttaatcttatgatttctttttttgatttctagaagactaggtaa
cggtaaatctattattgttataatatcgcttaaagcaaaaggtatattgattttacaaga
aataataatgatatgcttagactgttttttgatgattttagctaaattgcgcagtttcct
aattaggagtatctcatttagaaaagaatcaaaatcttttaaaataaaaagattcaaaga
ttcattatttaagttctcaataaactctaatgctagtagaggatttcgttttgcataacc
attatcgttagggttgctagtgtatccatctacaaagtcccagcaatacacttgttgatt
attagaacaatgtactttatttttataatatattctagtctatcttcttctctcgtatt
tattacaataataggatgtcgagattttaaaagtaatcgtaaatcttgagtgaaattcat
gtcaatatttgtttccatcagacttttgtaattcaccaattt----aagactaaataaga
ttatctaatattgcctggatatattcaaaagtc------atgtatctattttctaatcta
ttgttactgctaacttaactcaccattatctttttctccctttcgtcgtcgactattt
aaaatagagcgtccacacggtgttttcattctggccctgaaaccggaaactttgattttt
tttcttttagatccttgtagtgttctcttagtcatgatgtctcctgtgttgtaaaagatt
aaatactattttatcatcttttacatatcttgattatatagatctgtattaatatcaaag
tt-aagcgcaataatttatattaatcaagaattgtttatataatattaaaaattaattat
----------taactttatgatattggctttgcccatttttttatttaagtattttgacct
tgtttttgataatcttaaactggttaatttttcagcagctacaaaccatttttttcattag
aatctcaatttaagtattttattgacaaaagccaaaatagtatattagaagttgaagaaa
gctttgctttcgcaaaagtctgtgtagctaagaaatgtttttctaaagctattattgaag

gtcaattagcactaaaaaagcattacgactttaatattacagataataataacataattt
ccaatttgtataatatgctaggttttatttattctgaggcaggccaagtaagctttgcaa
aaaacttttacgagcaagccttaaagatagaacctaattatgttatagctttaaataatt
tagccaaaatttatgaagagaaaaaagattttataaaagcagagtctctatatgataaag
tgttaactcttaacttgaaaaataaaactgctaacagaaggaaggattttatagcaaaga
tta---------aaaataattaatgttgtttataatcgggatagcaggatttgaacctgc
gacatcctgctcccaaagcaggcgcgctaccaaactgcgctatatcccgtaagatataat
tgatactatataccttttatctaattatgtctactatttatgtcttttatctaggatacc
aaaacatgcctttaacaccatctggatccttaatgaatcctaatttttttataaaaactaa
ctacatcaggttctgcaaatagtgtgatagtgctaatttccgcctgcctcaattgctgga
ttaattggtgtataaccactttccctaaacctaaaccttgaaaatctggatgaataacta
catcccagatagtagcattaaacccattatctgatgttgctctagcaaatcccacgagtt
tagtatttgcatcttttttttgtattaaagaaataataatagaactatttttttaatgcga
ttttaacctttttttagaggtcttttaacccatcctactgaatcacagagttgttccaatt
cgtataaattaatattcttagtactactcagataaatatctttaaattcgatactattaa
aagctttatctaaaacaataagtttttttgaaaatttttttcagagctagtatgcctatctg
agttttgaaaaaaatttttccagaagatcataatgattatgatagaat------taaata
tatatattacgatattgacttaataaacaaaatattaatatgtgcccaatttttaatatc
aatctattaccatagaagattgtgtaaatatagagaaataaaaatttttgtaaaatcttt
caataaaatcatataaagagttctttttaagactaacatctattagtataaatagagtct
cgcaattgagatgaaacaacatattattctatgtctaatgccttttaaaggagttatttg
tgaaaaaatcgatgtttctaacttgtctacttgctgtactactgctgaccaacccaataa
tagctaatgcagaagtagctggactggtgccttgtaaagattctgcggcattcaataagc
gtatggttaatagtgttaaaaaacttcaagccagattagctaaatatgatgaaggtactc
cgcctgcgttagctttaaataagcagatagaaaaaactaaaactagatttgcaacttatg
gtagggcaggtttgttatgtggtactgatggattaccacatttaatttctgatggccgat
ggagtagagctggagactttgttttccctggactattattcttgtatatcacaggatgga
ttggttgggtaggcagaggctatcttttatctgttgctaagactagcaaaccaaccgaaa
aagaaattattttagatgtaccattagctattaaatttatgtcatccggatttgcatggc
cactagcagcttggcaagaatttagtagtgggcaattaattgcttccaatgatgatatta
ctgtttcaccccgttagtaaaaaatttatatgaataataattttaccaaatacttatcaa
cagcacctgtaattggtgtattatggatgacttttacagcaggctttattatagaattaa
atcgctttttcccagatgtcttatacttttatttataaataaataa--aattttataaa-
--aaactacataaatataggaaatcactattatttatgtggttt------atatataata
taattattta-ttaattaattttttttttcttctgactattttaatattagatcttaaat
aataaaaaa-atataaatattatgagtttagtaagccaaattatagtgaacgccgatgat

gaattacgatatccgacaattggagaattacagtcaattcaagattatttaattacaggg
actaaaagaattagaattgcaactattattagagataaagaaaaagaaattatacaaaaa
gcaagtaaacaaatttttcagttgcatccggaatatatagctccaggtggtaatgctgca
ggctctaggaaaagatcattatgcttacgtgattatggttggtatttgagattaattaca
tatggagtattagctggtgataaagattctatagaaacgattggtatcataggtgtgaga
gagatgtacaattctttaggtgtccctataattggaatgttagatgcaattcaatgtcta
aaagaagcttccctagaaatgcttggtcaagatgatattaaaataattgctccttatttc
gattatataattagaggtatgtcataaatcgtagcttcaattaattagttgaataaaagc
ttttataatgttataattaggctaagcccgaaagcatataattgtaaaaactaaatcttg
tcagaaccggaaggtagcagcaataatgtttaactacaaaagttatga-ttttcgggtgt
ttattgtgctaatttaatagtctctattttaaattatgaaactattcataaatttgtata
taatatataattcatccaacctttcagcaacaataatttaataatttgatattaatacgt
aaaacacctatac-tttttataaaatttctttatatctcatctaaacaatggaaactta
acttaatgggtgttcatattttatcaacaggttcgtctgtcccaaatttttctgtagaga
atcaacaatttgaagatatgatagagacttctgaccattggatttcgacaagaacaggaa
taaaaaaaagacatctagctccttcttctacttctctaactaaattagctgcagaagctg
ctaataacgccttatctgcagctaacctaaaacctactgatattaacttaattattttag
ctacatctactcctgatgatctatttggtagtgccagccagttacaagcagaaataggtg
caacggcatccgttgcttttgatattacagctgcctgctccggatttattgtcgcattag
taacagcagctcagttcattcagactggttcctatgacaatattttagttgtaggagcag
atacaatgtctagatggattaattggtcagatagaactacgtgtattttgtttggcgatg
gggcaggagcagtagtattggggcaaagtcttaaaaatagtattttaggctttaagttat
gtacagatggccagctaaatagtcatctgcaattgatgaatacaccagtaaataatcaaa
aatttggggctacagaaattcctcatggtaattataactctataacaatgaatggtaagg
aagtttacaagtttgctgtatttcaagttccaacagtgattaggcaatgtttaaataatt
taaatatctcgatagatgaagttgattggtttatattgcatcaagcaaatactagaattt
tagaagcaattgctaatagattatcaataccttttttctaaaatgattactaacttagaga
attatggcaatacatctgcagcgtcaattcctctagcgttagatgaagctattaaatcta
ataaaattcaaccaggtcaaactattgttttatctggttttggagctggcttaacttggg
gagcaattgttttaaagtggtaat---tctaatattgcggatgacgagactcgaactcgt
acagctttcgctacacacccctcaagcgtgcgtgtataccaatttcaccacatccgcatt
ttttaccatgaata-ttaaaaatttataatatttatttaaatattcatataa--tgtgta
tatatatatacattcttgtcagaaactctcttatattttattttaagtgataaaattgag
t----ttagtatagataataaaattttattttaaaatcataa----aaataaatatataa
ggaaaataaaatatgacaccgtccttatcaagtttttttgaatagtctcattcttggagca
gtaattgtagttgttccaataactttggcacttttattcgttagtcaaaagataggaca

attcgatcctagaaataatttagaaagatatggagaagcaatttaagattaataataaaa
tatttttcaaataaatatgctctccatatatttctatact--ttagaagaattcttattt
ttttattttgtaacaacctgtacagtttattctgttctctctttgtcctgatttactatt
tta----taacgagacaaatgtcaggtttgataatag---tcttttattctgtatctttc
attttatatcaaagtattttatatacacattactgtgttcaattaataaagaaaatctca
tgtctttgtctaattggcctcttaaaaaagaaaattctgaagcatatagcattaagaatt
caaaacaaatcacaattcctgatggattgtggataaaatgcttcgattgtggtctattaa
tgtactctaaagttttaaaacgaaatttaaaagtttgtcctcaatgtagctaccattttc
aagcttctagcaatgaaagaattgatcaattgatagaccaaggtagttggcaacctatag
atgttcatttaatttctacagatccattaggttttaaagatcaaaagctttatagtcaaa
ggttaaaagacactgctctcaagacaggtttgcaagatgcagttcaaacaggtactggta
ctatgcaaggcagacaagtctgtttaggtattatggacttcagattcatgggaggaagta
tgggatctgttgtaggcgaaaaattaacaagactactggaaaaagcaactcaggaaaaat
tgccagcaattatactgtgtgcatcaggcggagcaagaatgcaggaaggaatgttaagct
taatgcagatggctaaaatttcttctgcgctagaaatgcataaaaaaagcaaatctactat
atatatctgttttaacatctcccactacaggaggtgtaacagctagttttgctatgcttg
gagatttaattattgcagaaccaaaagctcttattgcatttgctggtaggcgagttatag
aacaaactatcaaagaagacttgcctgataattttcaaagttcagaatatctatttgaac
atggtttcctagatttaattgtaccaagaactcagctgagatcaaagctaatacaaattt
tatcattacacactcaaagcaagtaattggttcaatattaactataatatcaaaatgcta
gtaaaaattaagatattaagaaattttcactaaagatatttgtagactaactagaattac
aattactattgtagaaaaatttgtcttaattcta-tttttcagtcttttaactaattg-
tttttatgactttaactaattgttagtttaatgcaaatttataaagagaata-----cat
actaagctaaatatagtaatattatgtatttctataatactgacctatcaaatttcgagg
aattttatgcttaaaagatcttcttggcttgcggctttattgggactattaaccgtagtt
tctacaagtacgcatgcacatgccatagagctagacgaggcaacaagaactgttccatta
gaatcttctggcaaaactgtagttcttacgccagaacaagttaagcgaggcaagcgatta
tttaataattcttgtgctatttgtcataatggtggtatcacaaaaacaaatccaaatatt
ggacttgatccagaatctttaggattagctacgccacaaagagatactattgaggggtta
gttgattatatgaaagatccaacaagttatgatggcgcagagtcaattgcagaattgcat
ccaagtattaaaagtgccgaaatttttccaaaaatgcgtaatctaaccgatgaagaccta
tttacaatcgctggccatatcttgcttcaacctaaaattgtctccgaaaaatggggtggg
ggaaaaatttattattaaaagttcaaagatagtaatcttttgaatactagtttatgca-t
ttattgcttaagtgtgattattttgttagataatgatatatattaaagatggagtttgtt
gacagcaagtcatttttggccaaacttgcttacattattcgtctagttattaaaaagga
gacattaaattgaaaaagacactttcagttcttttcactgtttttagtttttttgtaata

ggttttacacaaatcgcttttgctgcagatctagataatggagaaaaagttttctctgct
aattgtgcagcatgtcatgctgggggtaataatgccattatgccagataaaactttgaaa
aaagatgtgcttgaagataataatatgaatagtattgatgctattacttatcaagtcaaa
aatggtaaaaatgctatgcctgcttttggaggtagactagttgacgaagatattgaagat
gcagcaaattatgtattatctcaatctgaaaaaggttggtaattatacttgattctatcc
agtattaaagaatagacaatctatttagttattcattagattgtctattctttgtttatg
---ttattatataaagatatttacacaatattgatattatgatgaaaagaataccagcaa
ttcttgtactagaagatggtacttattataaaggatggtcattccagcaagaaaaacaag
aaattactattggcgaagtagtttttaatactggaatgacgggatatcaagaaataatta
ctgaccctagctacttccagcaaattgtcacttttacttacccagaaattggtaatacag
gtattaataatcaagatgttgaatctcacagtattacaattaaaggcctcgttgcgaaaa
atatatgtaaaatgcccagtagttggagagaacaggaatctttggttaagtatttaagta
gtcatgatattcctttcattttggaatagacactagatcgttaactcaatatttacgac
aatttggtacaatgaatggctgtatttctgccgacaatttaaatcatatttatttaaagc
aaaaaatttctaaaattccatgtatgcaaggtttagatttaattccatatgtaactacag
gaagtgtctacccatgggatgaaaaaagtttgccgaactggtatttgacagatagaagca
gagttcagctagttatccagttgaaagttgttgttatagattttggagtcaaactaaata
tacttagaagattagctacacttggatgtcagataactgttgtacctgcgcaaacttcat
tagaggatattttagcttaccagcccgacggtatattgctttctaatggccctggggatc
cgtctgcagtacattatggaatccagacagttacaaatctgttggatcagaatgtaccta
tctttggcatttgcatggggcaccaaatcttaaacttagccctaaaagctaaaactttta
aactcaaatttggtcatagaggtattaaccatccatctggattgaaccagcaagttgaaa
taactagtcaaaaccatggcttcgcagttgatttgcactctgtcttggaagcttctatac
aagtaactcattttaatttaaatgatactactgttgctggaaccgggcataatcaaagcc
catatttctccgttcaatatcatccagaatcgagtccgggccctcatgatgctgattatc
tatttgaaaacttcgtagaaataatgacaaaatttaagcgaaaatttat-----------
ttagtaattttcccatgctttatgagtaaataaagcggataatacttgtactcctcgtaa
ttgattgaaaagaggtaaatgtccccttgggccatgagaattccactgaaattcatcagg
atatctacaaggaatttgatcaacttcccagccgatttttttgccaaaatttcccccaatc
tttttcaacgctaagccatatttgtctttgaataaatagcccaaatttacctttttgagtg
tgtatgccacaatttatcaatagtttttagatcttgaacaggtatttttttgatatctgt
aaaatatagccaattgcgatcttgtgcattcacaccagctagttgaataagtttttgttg
agttagttgatccgctttgagcaaatcccgatgagtcagtaatacctgtaaatctttgta
attcatctgtttatcagagcgcaatggaacaataccatcgggacacaagttagaagcaaa
ttttactatttcttgatttttactattaagtaacttttcatagattaaaccatcaacaca
attacttttataattaggcatcttaatcctctcaaaaaatagatctgctaaaccttttaa

ctggatagagtcgtcactatgtatattctcaataatttcaagttgtttcttaatattatt
attagacttagtattctcatttaattccaaaagctgaactcgaatttgatcttgcatctg
atttttttatttaatttaaatattc----ataaggtcttatagagaattagattgttttt
tgaaattagaagaacgccgaattatttctaaaaatgttttcataatttgattgaataagt
tttgtcctgtataaattataatttgaccaattaaaaaaactgtacttttttattttaggaa
caaaaaagtcttgtaactcaagtagaaaagtaattagtatttgtattgacgataatgttt
ttaattcattcgacctgcaagcatatatatactgacaatttaggccttcttgataaaatc
cccagactttataacgactttcataaattgctcgcggtcgctcaatatagtttttaatta
aagtattccaaattaaattattctttaatttttctaattttctatcagataaaaattta
cattacaaagataacttgcatttgtttcatgtacagtatcaaaattctgaactaacatat
gagcaataatattgcttagctgaattaaatagttttcaagaaaaacttcaacctgcttaa
ttggcatattggaagtaaagagattgaaagttttatttacagtctgagaggacccaaaaa
gtagttgaataatcaaaatttgtagtagcattttatagtcagatatcaggtagttcaaat
cgttaatatcttggcatatgtgagtatagtccagttcatataatctacaaaatcttttga
aacatctatggaataaatcaattaaaatattttcagttaaagttgtaatatcatttaaat
ctaaatttgaattataaatttctaaaaggatcttttctaattctgctaatatgattttta
ataagtttctttttacctcagtgcgaaaaacatctaaaattaacacttcctgagaacaat
tagttaatttcttattaatctttatagaagttctaactaatagctctgctacctcatgat
tgagtgttggtccctgagagctcggccaataattattcacgttataccataataaggaat
atttaagttaatgttatacgataactttttaatacactgaaaagcaagtctattactcta
tagtacctag-------gttttacagatcttaatagttatttgtcaaataaagttatatg
atagtagcgaccgagctaattttattaaattta----taataatgacaatctattactt
tgcacttgctagtcaaaactttctcttagtacaagagcccttggaagaagttttcagaga
gagagttaactattatcaatcaaataataaagcaattgatttttggctaatacccaatcc
ttctttttagaaaaaccagaaatgatttcatttaaagaccttgtacctaacgacgctgt
cgctataatttctactaatccggtgtttattaattggttaaagttaagaatcggttatgt
ttgtattggacaatttgaagataatttacaactttctgaagaatcgttaaatattaccct
tgtaacagataaaatttaatcatacttctattatgactaatgtt-aattat-----ttaa
gttgcaattatcagctataaatattta-tctttttacagtttatcgcttagtattttatg
ctgagatggagtgacaaataatgtaagtatttcttgactaaatgtttcagttgccttaga
tttatatctatttgggttaactataatagacagcattcgttttatagtaacattttcaat
ttgagcccaatgtacaattcctagttctaactccttagcaatagctgagacggaaacaaa
ggcagctcctagtcctgactgtacagcatttttatagcttctatagaattcaattccat
ctctattttaaagcgactgctatcaataccatgttgactgagtactttatctatgacttt
tctgattgttgattgagtatctaatgcaataaatctaagcctgtataagtcttcttttg
aatatctcctagttgagaaaaaggatgtgattttggcaatattagtgctagttcatcttc

tgcataagatgtaacttgcaaaacatcttgtaattcagtaggcacttctcctccaataat
tgctaaatcaacttggccattagccacactccatgaaatgagtctagttgaatgtacttg
caattgaacagctacttgtggatatctttgcctgaatagtcctattaatcttggcattaa
atatgtcccagttgtttgactagctccaataattaatgtgccaccttgcaggttttgcaa
gtcgtcaagagctcggcatgtttcttcgcatagagctaaaattcttcccccgtaccgtaa
aagcagacttcctgcctcagttaaagttgccttttattacctctttcaaagagggaaac
atttaattggcgctctaaattttgaatttgcaaactaatagctggctgagaaacatataa
gctattagctgccttttaaaactgccttctttggcaattgcttttaatattcttaactg
atccaatgtaaatggaaggtctgtcattaaagaattgtagtataatg---------tata
ttatttagtattgattaatgtatgcatttgccaagttagatatttgttattacgcttcat
tattttcaaaataataaaaagtataatgttaaataaactgtattagattttaatatctat
ttgatgttaaatttatacaatagatatcgtgaatttaattctaactataaaatataggag
acaat---atggactctagacttttaattgtactaataccagttttagcagctgcatctt
gggcggtttacaatattggtagagttgcaatacaacaattcaaaaaaatgacatcttagt
ttttcttagaatata-atttttagattttatgagatagagtacttgaaaatctctatctc
tattgtatatctgaataaaaaaatcaaggc--ccccttgtacttactgttattacacatt
aagattatttacaagtacctatattcataataagttgaagctaacttatt-----tttta
taattaattaaactaataacacaattatggctgttccaaagaaaagaacatctaaagcta
aaaaaaatgcacgcaaagcaaattggaaaaatcaagcaagaacggaagcaaaaaaagctt
tgtctttggcaaaatcagtattaactagaaaatcaaatggatttatttataatctttatg
aaacagtagatacttctagtgattaaaagtattt-cgatcacaaggctaaaaaagtctca
aacagtaaactttaagtaatttttatctattaagtaagcatttgaaaatcattctatttg
ataataaaataaatagcagattaaataaaattacagcgacaaattatgcagtgaagctag
atcaaactagtgaaatttggttgtttaattgtattgaaaatattcagcatattttttta
aaagtcaattaaaatcaagtcatattactaaaatttttatctctggtactagctttgaat
atacggcaggtctacctggactattatctagcttaacactgagcggtagaatacatccca
ttagtatatatggccctaattccctaagacagtatcttcaagcttgtaccaaatattccc
aaactaatttttcttttcctattaattttcataatgtacaatatggagaacaggtcgtcg
accaatttatacagtaatttgcttaccattgaataaaaagagtcgactatatggattta
ctattctaaaaaaagaaaagccgggagtatttaaattagcaaaagccaagactttaaata
ttactcaaggacctatatacgggaaattaaaagaaaaagataattttctaagtccagatg
gttactacctatgtggcaaagatttctcttcaaatacaataatgggacataaaatatctc
ttccattattagtcagatattctagaattatttctgagatgcattggttttgctcttatc
ctattagattgaatacttattcacatcagcaagccacaaagtgtttgccacacaatattt
taaccgatattatgaagtctcaaatatatcaagataatagttttattgaataatgctcat
tcttgatatataatggtctcgacaatctaaagaacatacccgaaatactctctgaaattt

atttgtctctattaaatttataatcatttatgacatacgcaattattgaagcaagtggca
aacagctttgggtagaagaaggccgctactatgatttgaatcatatagctgttgatccag
gccaatcgattatactaggtaaagttttattattgaataagaacggacaagttactttag
gacatccttgtatagaaggtgttacaataaaagctacagtaatgaggcatttaagaggaa
agaaaataactgttttcaaaatgaaaccaaagaaaaaaatgagattaaaaaaaggccatc
gacaagaattaactcgtttaatgattgattccataaaatcttaga------taaaatctc
gattcatattatt--aa-aaaattttttaatatttaactaaaatagatataatggcacata
aaaaaggtagtggtagtacaagaaatggtagagactctaattccaaacgtttaggagtta
aaaaatatggtggtgagcaagtaacggcaggtaatattttaatcagacaacgtggaacta
aagttaagccaggccaaaatgttggcaaagggaaagatgataccttattctcgctgatcg
atggctttgtactgtttgaaaagtcaaatcaaaagcaaaaaacaattagtgtttattctg
ctaagaaatagctaaaaaaattggtgcaagataacttgagt----gtatttattgcactt
acactggcaccatatacttgatttattaagaaatagagttccttaatgaaccaatgaatt
ttattaaagatctgctttgaaatttaatgacaaacactattgtaatttcctgtctacaca
atatggctgctattctatattgtggtcagatacagaaactagtagtagcaaatgctcatt
atcaagtaagtgatatctatctaggttgtgtagacaaaatcttttcaggaataaatgcag
catttattgacctaggaaagaatgagtacagtggatttatccatattagtgatactggtc
cgcttaaaaagaaatattatgttaataatattactaacattttaacaattcgacaaaaaa
ttttagtacaaatcattaaagagccaactttaaataaaggaccaaggctcactgctaata
ttacattatcaggtcgatacattgtattaatgccttttagtcaatcaatttttatatctc
gaaaaatacatgatgagaatgagcgtcattatttaaaagctttagccattttaatcaaac
cagccacaatgggtttgctattcagaccttctgctgtaggtgtcgatgaagaaatcatat
taagcgaactaaaaaatttaaaagaacaatggaactttgttcaaaaatctgcaattaata
gttattcacctgttctcctatataaggatcaagatattgttaaaaagataatcagagatt
tttataataataatacaaataatattgtaattgattcaaacctgggattaaaacaattaa
attattatatccatacttggcattgcaataattctagtacagttcctgcacttaagcttt
atagtaataatcaatgtatattagacgcttttggtattaatcaggcagtttccagagctc
taattccaaaagttgatcttatacttggtggctatatgtttattgaaactttagaagctt
ttactattattgatgtaaattctggatcttttaataattccactagtgcacgagaaacag
tgttaaaaacaaattgttctgcagcaacagaaatagcctaccagttacaaattagaaata
ttgcaggtgtaattataattgattttattgatatggaatcacaaagagatcaattacaat
tattggaacacttcaataaagagcttgcacttgatgatgccaagcctcaaattgtgcagt
tatctgaattaggtctagtggaactaactagaagaagacaaggtaaaagtttatacgagt
taattagtagtgattctaattactttcatttttttcatacaatcagaacgatttaaatcta
tgaaaagaactgattacaagaagcaaacacttcaaactttgaatagatcttggttgtcta
cagaaattaatactattaataaagttttttttcaaaagttaaacttatatagacttgcca

acttttacttagttcgcaatctatatataattaacagcagtcttacttataagcaaaatt
atttattaaatactagatcaaaacttatttattcccaagaatataatagagtattgccaa
gca------gtggctatttagtcagcctaaatatgaatagtgattataagtttttacctt
gaaatctttgtttaaaccgt----------------------------------------
--------------------aaaaaaagaagctttctcttctgtaaaagaaagcttc--t
ttttagcttactaactattctagttagttaattagaaattaaccgttaacagctggagct
gttagagctactggtaatgattcaccagaagctaaatctagagggaagttgtgagcatta
cgttcgtgcattacttccatgcctaggttagcacgattgatgatatcagcccatgtatta
attacacgaccttggctatcaacaacagattggttaaagttaaaaccattcaagttgaat
gccattgtgcttacagataaagatgttagccagataccaactacaggccatagacctaag
aagaaatgtaaggaacgagagttgttgaaactagcatattggaagattaaacgaccgaag
tagccatgagctgcaacgatgttgtaagtttcttcttcttgtccgaatttgtaaccatag
ttagcagattcgttttcgcttgtttcacgaattaagctagatgtaactagggatccgtgc
atagcactgaacagagaaccaccaaatacaccagcaacacctaattggtggaaagggtgc
attaagatattgtgctcagcttggaacacaagcatgaagttgaatgttccggagatacct
agaggcattccatcagagaaactaccttggccaattgggtatactaggaatactgctgtt
gctgctgctactggagcagtaaaagcaacggaaatccatggacgcatacctaggcggtag
cttagttcccactcacgaccaatgtaacaagctacgccagttaggaaatgaagaacaact
agttggtaaggaccaccgttgtatagccattcgtctaaagaagcagcttcccaaattggg
taaaagtgaataccgatagctgcagaacttggaataacagcaccagaaatgatgttgttt
ccgtatagaagggaaccagcaactggctcacgaattccatcaatgtctactggaggtgca
gctacgaatgcaatgatgaatacagatgtggcagttaatagagttggaatcattagtact
ccaaaccaaccaatgtataggcggttttcagtactagtaatccaagagcagaaacgttcc
cacaagctagcgctttcgcgtctttgtaaagtagcagtcataatttttatcaaatttta
aggatttaactaagatgcttcccaaacaatatgttgtctgttaagaatattatgacttaa
catcttttacaatgagaacattttttaaaaaaca-tagctaaatttagtttcagtaact
aa-aaattatgcttatttatatcttgtgaaataattgaa-----------taataaattt
tt-aat-------------ttttgcagaattagcattattttagggatataataatagta
atgcatattaatgctgcttattttgattttaatagtttacataatttggccttacttaaa
caagaatgtcacactttacaaaaattcagactactataaaagatttaaatctgttaaagc
atgctttaacagatttgggtttgatttggcaaatgaattctagtcatataaaagttggtg
aaaataatcagcacaaagtcgatgtattaatcaagcaagataacttatcgcatattggtt
ttatttggaacgataatagatatcacttagttgctgacttacagctttgggaacaacctt
ggtctctagaagtatttctcgataaattgtcacagaagtatgcttactattctattatcg
aagaaacaaagaaacaaggctttgaaaaaatggagcagatctcgcaaaaagatggatcga
ttaagttagttgtgcaacgttggaactgctaactaatatataaatgcgggcggagagact

tgaactctcacgagattatctcactagaacctaaatctagcgcgtctaccaattccacca
cgcccgcattgccgcaat--aaaattaatatatcataatctattccgaaagacaaactaa
-gagcataaattagcttccaagcttaaaagcatcaacaaatagaccgtatattataccaa
ttttgaagttattgataaaattaattttatgcctaattttctttggttagagtaaacta
ttttactagttaactcagtggctgcaactattagactggctgcaataataccccaatctc
ctgtttgcccgggaattgtagataaccctgttgaaataaaaaagcctaacaataagctaa
tcaagccagttgttaactcacttaaagaataataaagcttattatttaagttatttacta
agtaggcaagaaaaatagaaagtctagttttaatcataatttttaacaacggagtttgaa
aaggatgataagtattaatagcttattaatcttatcttatcctcttaaaacccgttaata
caaaacatgttttttat---------ttttaaaggtcttgctcacttaaagcattaataa
gatgttgaaacggttcttgtataatatctttgcgtttaattgattgcaaatctagttctt
cttctattatttcttgtaataactgtatgcttcttactgtgggagcaagaggtacagata
gtgaattgtaagtatcttttagcccgtttaatacacgttgatccagaatattagtgctac
cagaaactaaagcgtaactagcataccgtaagtaatattctatgtctcttaaacatgctg
catatcttcttgttgtataagaattccctccaggtcttaataattcaggttgttcttcgt
acaactgagctgctgcctcttttaaaatattggtagcttgatcattaataatttcagcta
tttttattctatccaatccacttgaaaagaaggattctagttgtccgacagctgccttat
ctagataacgaccagtaagatcgtaacgatttaatattgctgttatagcatcttgcataa
attaattctcctac-ttgctatcagagcggtattataaatgactaatat-----tataat
catcagtgtttatggaatttatttaaaatctattctttattagtaagtgtacaattaaaa
actagcatagattttacatatatagagtatacgtagtttttaatta-tttttttctaaaa
atcaagaaataaagttttttt-taagaggtttataaatttttaatataatttagggtct
ttgcctaaataagttttgtaacacac--aagctaaaataaaaataagtagataatagtga
------------------------------------------------------------
------------------------------------------------------------
------------------------------------------------------------
------------------------------------------------------------
------------------------------------------------------------
------------------------------------------------------------
------------------------------------------------------------
------------------------------------------------------------
------------------------------------------------------------
------------------------------------------------------------
--------------------------acaataataatctgtatcaa--------------
------------------------------------------------------------
------------------------------------------------------------

------------------------------------------------------------
------------------------------------------------------------
------------------------------------------------------------
------------------------------------------------------------
------------------------------------------------------------
------------------------------------------------------------
------------------------------------------------------------
------------------------------------------------------------
------------------------------------------------------------
------------------------------------------------------------
------------------------------------------------------------
------------------------------------------------------------
------------------------------------------------------------
------------------------------------------------------------
------------------------------------------------------------
------------------------------------------------------------
------------------------------------------------------------
------------------------------------------------------------
------------------------------------------------------------
------------------------------------------------------------
-----------------agtataaataagcttttactgcagcatcaaaaaaaatatttaa
cagatattgacagaattgattatttaagtgaaaacttagatttagacaaatttagctatt
ttataattacaattaataagcctgataaatgtattcttgtagaacagttttgtcataatc
ctaaaggccatatttattcaatattgtttaaaggtcaaacagctaaacacgtttgttgta
taatctttcagcataaacaattaaatataataatatctacatctcatgctttgtatttag
gccgagagctaatgaagtctgaattagctctcatgttagatcagcaatatattcaagact
aaa--ctttgtagatattaattgtatttgtgctagtataatatcagtaagggcatgtaac
tcagtggatagagtatcagattccgattctgatggccgtgggttcgaatcccgccctgcc
c---gagtatttagagttct-atagcaagacatct-------taagagctactttaataa
tggcacagatattggggctgcaaggtttctacattatgaaaataagaaaatatgaaaata
aaacaagctctttaaaagagcttttattaaaattaataaatgcagaaaataatattattt
ctttttctcgaaaattagctattgcttaaacgctgttaatttttggtattcgacgtgtta
aactcttattcacggcgaatattctgccagagtcgctcttagtataagaaaagtctgaaa
aattattcatatttatgttttgtacctatttttatttaaattaaggttagta-ttttcaa
tctttataatggacgtgggttcgagtcccaccagctccatcaatttaatt---------t
ttataattagctaatcaaatgtttttaataaccagaaatataattggagtaaatatggat

tttataacaatcactaaacaagctttaaagcagattacaattttgaaaaacgattatgaa
aatgatgtgcatctaagaataggcgtgagacaaggtggatgttcaggcatgtcatactca
atgaattttgaaaatgttgataatttaaaagatacagatgaaaaactcagtatagataac
ttgtctgtcgtctgcgatctcaaaagcctcttatatctttatggattatcattagatttt
agtactgaattaataggaggaggatttcaattttctaacccaaatgcgagccaaacttgt
ggttgtggtaaatctttctcaggctaagg-ctcta--------ctcttttatttgtatt
atacatattgctctagt-attag-aataatatatataagcaatgcttaattaaaa-----
----------------------------gattttaaaactat------------------
------------------tcaagtaattacaaa----------aactaaattaaagagat
tttagtatattagcttaaatagtcaactttttattaatatatgctaatcttt-attagct
cagatgtatttgaaaattgtgacctt------------------------cctgtcca-
----------------------gaatagattgtatatttt-atttattttataaaaactc
gacatgaatcttccccaacttattgcagagtattctagtacgcaaaatcctaataacttt
tattgcaatatggaatctgtgtcttctgataatttaagttattttaaatcatcaaagata
aatttattaccttctgcaaagataattgcttttgggttgcctcaatttgactttattcct
acaaagattaaaaattccattagagaagcaattgtaatatctactatcgatgataataat
atttttgctgactcaactaataattggataaaaaaacataaagataaaaatcatcattct
ttgttctataaaaaaatttcaaagctctgttaaaccatgaaattaaagttgtagtaaag
cagttgaacacttctacatctacaatagatacagaaaagaatataatggcaagtcaaaga
tatatctggagtaaaagttggaagccctcaattattgtgtcttacttaaaaaataaaccc
ccccatgaattttctaatgcatctaaaaaatttttaattgatcagttatctgcatcacca
gtgtttgttgtaaaaaatggttttaatgaaattattttaggtcatccattatctcgtatt
aaaagagaaggcattaataatattgcttactcgtttttttaatttattaaattattctaag
cgaacttatcctatatctactggattattttttttccatccagatgatgcatttgaattt
aaagactttataatatcggttaatccactggcggcagaacctatggagataaatatagaa
cctgttggcctacactttgcttataaaatgaatagaaccggatcttcggatactcaattt
tcatttattccagattttaaagaggtaggagatttattatttaaacatcaaaaagataag
catttaatcttccataagaatcagcaccacggtaaaaatttttttcaggggcaacctatt
tatatgattcagccgattacgttaaaagatcgaaacggaaggctgaatatcattaaattt
actggactaaatgataacagagaagtcatttttactaatcttgaagctgctaatagatcg
tggacaaatttcataaagagtaattcacaattgaaatcaattaaaaagcctactttatta
gtctataatttagagagtttcttaaaagatcaagaacgatcgaataagcaagattttcaa
aaatttgttgtgattactaataaaaaagcttatcttgccacaaaagagttaatagctgta
cccgattctaatagcttttctaagcatttaaaattaaatatgaagcctaaactcttttttt
gtgaaactgtgggtaagacggttattttccaccttaacttatgaat-aaatgaggtc-tt
atctattttcttgaataatattctataactaatagttcatttagttgtaatgcaacccat

tctctatcaataactccattgactttccctgacaaattagatttatttaattctaagtgg
ctaggaatattagctaacccagggaacgccaagtagttttccactaattttctagatgat
tcttgagattttacactaattaattctccaggtttacattgataactgcatatagatact
gcttctccattaatacaaatgtgaccatggtttactagctgtctagcagcaggaattgta
ggagccatacccagtctaaaaacggtattatctagtctcatttctagtagctgtaacaaa
atttgacctgtagacccctggagtttttagcagctttgacatatttaaatagttgtttt
tcactcaatccataattaaagcgtaacttttgtttttcttctaatctcacagcatactca
gaaggttttctagattttgtccatgttcccctggtggataggatcgcttaattactttc
ctactcagtccaggtaaatcacccagtctgcgcgaaatgcgtactcgtggccctctgtat
ctagacatattatataaattctccgaatataaat-aattaag----------taatgagt
agctatttttaacataaaaaaagtattaaaacaaagc-ttttgctaaaaaatctttata
aaaaatatag--------------------tattgcgc-tttttaagcgggtagcgggaa
tcgaacccgcatcattagcttggaaggctaaggttttaccactaaactatacccgcaata
ttcatcaggttacaacccatttaatataacataattatgttacgttaagatattcttgaa
acaaatacagaa-taaaaatatttatgctccttctaaaattacatcaagaaaacgagcaa
tttcagctgcctgttcttcaacttctgaaagtaataatggttgaccgaccctagttaacg
gaattattcttttatcttttgtgcataaataaatttctcgccgtggatttagcccctctt
taatgtctatttttattgatttaatttcttttatattaaactgaagacaaatttgcctat
tttttccaggaaaacctagccgaaaaattttaacaataccttcatctttattaaattcat
tatatccagcacctatattccaaataatagttagccacaggaacaaactaagaaagattc
ctatgcttccatagaatgtcataacaataccttgtggaataaacacaaattctgttgaag
ttgtaaaaggtaacaaatcaacttgaaaataacttgatagtccagcaagaagaaatccta
aagctccaataaagattattgtagcccaccagtaattactaagtcttcgcgaacctaaaa
ttaaatcttttctaacagagtgtatagataatgtttttttcataatttgttgttttaaaa
caata-tactaatgtgaaatataaaaattttgacaatcttttgatgattgttttagattt
attaaatgagcttgatggcttttagtcctaaaaatagtcctgttgctaacccaaaaaaag
cagcaagaaatattgtatatcctagaaaaactgaca---tatta----ttattattatta
ttacaatagtgtttacaaaaatagcgaaaatctattactaaagtattatattacgttaaa
tgcctcgctgtctacatgaaaaaactttttttcagtttttagcatggtatttatattagaa
ataatatttaagtaattttattaaacttctggagaacagaatcttatggcagaatttatc
aagccttataacgacgatccttttgtaggcaatctgtctacaccagttagtacgtcgagt
tttagtaaaggactactaggaaatctaccagcttatcgtcgaggcttatctccactcctt
agaggcttagagataggaatggcacatggatactttttaattggaccttttgataagcta
ggtcctttaagaggtacagatgtagcattactcgcaggatttctatcttcagtcggtctc
attattattcttactacatgtttatccatgtatggcaatgtatcttttactaaagctgat
tccaaagatccattgcaaactgctgaaggctggggtcaatttactgcaggattttttagtt

ggagcagtaggtggttcaggatttgcttatttattgctagctaatattcctgtactacaa
actgcgggcttaagtctcttttcttaagctagtaaggggacttgaacccgtaacctactg
attacaaatcagttgctctaccaattgagctatactagcatatatcgaattgtaacatat
aaagatatttgttgcaatctggtattttagatgcaacaaatatctttttttacttaaaca
gaactgttatattaata---------tatattaattagaatctgaattatctgactctgc
agaatattcttgatcacagtttaaataataacaaatggtttggtctagacaagttgcaga
ccaaccaacaggtgtttctcctgataactcttgcatatctccagaattgtaaaaatcatc
acttagcttattactgtttgcttgcataaatatcactcccaaaacactctgaacttaatg
tgaagtaatttaacttctcaataatatattaacacaaaatactaaatatgttgattattc
atcaattgtaaaaatattgctaaatttacaaagtttgagtgagcgtcttgatagcttttg
ctgaaattaacattttaatccatttattttgctctcttgaccatactttttttatgttgta
aattaacttttttgtagttttttttgttcgtttatgagaatgagatactgcgtacccattat
tagctacttttcctgtaagctggcatttttttggcataatttt-------ttaacttata
tatttatttaaagtacttgataaagttgacttcggcacagctccaattactgtgtcaact
ctttctccggctttgaagatcattaaagtaggtatacttctgatgccatattcagctgcg
atagtaggattctcgtctgtgtttattttcactaccttaatagatgattcatattcttct
gcaatttcatcaactacaggagaaaccattctacaaggaccacaccatggcgcccaaaaa
tctacgagtacaggtaagttattattaataacttcttgtttgaaagaggcatctgtaact
tgagatactgacatatttttttaacctttaatatatgttccttgctaagcaaatcttatca
caaattttaagaaaaatcttctatagtgaattactattagtgttgaattaatttttt-ga
aatcagacattaaaaaaacttattacctatataaacaaagtggtggccgagtgatatctt
tatagtgataatttctatattaaggttgaacactagaaataacttttcaagcaaatcttt
atttgaaaa--acttctctgtgaccatcaaagctcaattatattacaatatattttatta
tttgtaacttagttctgataatggtataaacaacgcaaaagatactgccttataatcaag
gaggaatacatgtctcaatccgtagaatcacggactaggattaaaagcgaacgttacgaa
tctggagtaatcccttacgctaaaatgggttactgggatgctgactatgtaattaaagaa
acagatattctagctctgtttagaatcactcctcagccaggtgttgaccctattgaagca
tctgctgcaattgcaggtgaatcttcaacagctacttggacagttgtatggactgatttg
ttaacagcttgtgacttatatagagcaaaagcatatcgagtagatccagttccaaatgtt
gcagatcaatactttgcttacatagcttatgatattgatttatttgaagaaggttccatt
gcgaacttaactgcttcaattattggtaacgtttttggatttaaggctgttaaagctctt
cgtctagaagatatgcgtatgccagtagcttacctaaaaactttccaaggtcctgcaact
ggattgattgtagaacgtgagcgtatggacaaattcggtagacctttcttaggtgctacc
gttaaacctaaactaggtctgtctggtaaaaactacggaagagttgtatatgaaggtcta
aaaggtggtcttgacttccttaaagatgatgaaaatattaactctcaaccatttatgcgt
tggagagaaagattttatattccatggaaggtgtaaataaagcatctgctgctgctggt

gaaattaaaggtcattaccttaacgtaacagctgctacaatggaagatatgtacgagaga
gccgaattctctaaagaggttggtagtattatttgtatgattgaccttgtaattggttat
actgcaattcaaagtatggcaatctgggctcgcaaacatgacatgattttacatttacat
agagcaggtaactctacttactctcgtcaaaaaaatcatggtatgaacttccgagttatt
tgcaaatggatgcgtatggcaggtgttgaccatattcacgcaggtacagttgtaggtaaa
cttgagggagatcctttaatgattaaaggcttctacaatactttacttgaaagtgacacg
gatattaatctacctcaaggtctattcttcgctcaaaattgggcttctctgcgtaaagtt
gtaccagtagcatctggtggtattcatgctggtcaaatgcaccaacttcttgattactta
ggtgatgatgtagttcttcagtttggtggtggtacaattggacatcctgatggtatccaa
gcaggtgcaactgcaaacagagtagcattagagtctatggttatggcaagaaatgaaggg
cgtaattatgtagcagaaggtccacaaatcttaagagacgctgctaaaacttgtggacct
ctacaaacagctttagatttatggaaagatattagttttaattatacttccacagataca
gctgatttcgttgagactccaacagcaaacatctagtttaatgactacttaccaatgctt
aaactagcaaattgtaagtaaaattaacttataacaataaggagcatagaatagtgagac
taacacaagggactttttcctttcttccagatctaactgatgagcaaattaataaacagc
ttgcttacatcgtttctaaaggtttgtcagcaaacgttgaatatactgatgatcctcatc
caagaaattcctactgggaattatggggtttacctttatttgatgtaaaagacgcatctg
ctgttatgtatgagattagttcatgcagaaaagcaaagcctaattattatattaaagtta
acgcttttgataatactagaggtattgaaagttgtgtaatgtctttcatcgtaaatagac
ctgctaatgaaccaggattcttattacaacgccaagactttgaaggtagaactatgaagt
acagtcttcatagctatgctactgaaaagcctgaaggagctagatattaattttaattaa
tagctaatatctattaatt---ccccccctctttgatatcttaattactagattaattttg
attttaaagagggtcaaatttcgaattaaattctgatcaaa-tagatttaacaattaaaa
atacacagaaataatgcaaccacaggatataatttccaacgatactcttgttaatttaca
agaagaatatgatagaacgcaaatccaagaagttctaaatgaattaaaccaagaacttat
aggattagtacctgtaaaaaccagaattcgagaaattgctgctctattgttgattgatag
attacgtagaaaactagaattagtatctggtaatccaggactacatatgtcatttacagg
tagtcctggaactggtaaaacgacagttgccatgaaaatggccgatattttgcacagact
cggatatataaagaaaggtcatttattgactgtgacaagagatgatcttgtggggcaata
tattggtcacaccgctcctaaaactaaagaagttctcaaacaagcaatgggaggagttct
atttattgatgaagcttactatctatataaagcagataatgaaagagactatggttctga
agcgattgaaattttactacaagtaatggaaaatcagagaaatgatttggttgtcatctt
tgcaggatataaggatagaatggaaaaattctatgaatccaatcctggactatcttctag
agtagctaatcatgtagactttccagattacacttcagatgaattgttacaaatagctaa
aatgatgatagaagaacaacagtattgttttacagaagaagcagataaaactcttttaga
atatactgaacgcagaatgaaacagccttactttgctaatgcaagaagtattcgtaatgc

tattgatagagctagaatgagacaagctaatagaatttttgcaagtggagaaaaagtatt
aacaaaagctgatttagtaacgattgaagcagaagatatcttgaaaagtagattattttc
attacctaatgcttaatattctagtggcttaatcaaactatatatcataaagaaa-ttga
ataataagttgcaaatttctttaaaaacacttactattaatacaatgtaagtgttggcgg
catagccaagtggtaaggcagaggattgcaaatccttcatcccccagttcaaatctgggt
gccgcctagcactaaaaa-tgggggtgtggtggaatggtagacacaacagacttaaaatc
tgttgatttttagtaatcgtgagggttcaagtccctccacccccatgtaccaacttataa
aaaaagacagatgtgtatttgtattaattgcaatcatattactaaatgtaatacttacca
tttaatagaatctcagcacaaacagcttcatattaatgaaaacccattgtttatacctaa
atgccccgtagttcatgttaatatatctaataatggtatctataatcaaattgattggga
tttagtggagtgtttatcatttgtagaaaaaccgaatagttggaatttagactgtgatta
attgtaagaaaaattatgtactataagcaataattctatcttttaaatactctgtattga
ttttagatgttctgactaataagatctttcctgttctggctatctctataatgccaaatt
tagttaatagttgttcgatagcaacaatcttaccaggatctccagtaacctctacaatta
aaagatcttccgcaatatctactatctttgctctaaaaatttttacaatttctaaagctt
ctgttctggtctgagaattaattctaatcttaattaacattaattctctttcaactgaag
gaatattggttacatcttgcacattaagaatatttactaatttatataattgcttagtga
gttgttcaatagttctattgtctccctgaactaccattgtgattctagagactccaattt
gctccgctggtccaactgctaaactagcaatattaaagcctctacgggcaaatagacccg
atattcttgacagtactcctgcttcatcttgaactaaaactgataaggtgtgtttcatga
aaatttaatgttactctatttaatctattaaatatattctatgaaaattaaagaaaaaat
gctagttactctatgactaatttattcaaactaaacaaaatcctattatcagacaccata
tctaccttgtaatgttataatattaacttagtatttaatgatattctaattctaaataga
atgctaaatttttaaaatttgttaggaacttgtgctagaaaaatacaaaaacagtaaaa
aactctctcgagatctgccccaaatcaatgatcgtattagatttccaaaagttcgagtaa
ttgatgacgaaggtgaacaattaggtatttttgtgcctgaagaagctatacaattagctg
ttcaacaaggcttagacctagttgttgttagtgataagtctgacccaccagtctgtcgaa
tattagattatggtaaatataagtttacacaggaaaaaagggctagagaagccaagaaaa
aacaacacaacagtagtatcaaagaagtaaaaatgcgatacaagatagaagagcatgatt
ataaggttagaataaatcaagcatctaaattcatccaagcaggtgataaagtaaaagcaa
caatcacatttcgtggtcgtgaaattcagcattctaatttagctatagacttgttgaata
aaatggcgaacgatttaactgcaatagctgaaattcaacaagctccatcaagagatggaa
gaaatgttatcatgcttttatcaccaaagaaagttagtt-aactaatttttatttaaatg
catctggctggattcgaaccagcgacgtctctttcgaaatggcggattatgagtccgctg
ccttcggcccctcggccacagatgcatcacttacagtatataactcataaagcctataaa
aatcaatgcttatctctttttaataaaatgtataagaaattattattcattcatgttgtg

ataggtcgcaactgctgaaggggaaacttgatttaagtagcgaaatatccagtatttaaa
tattgtgtccaatataactggaaaagttgaaataaataaaaaaataaagtctcgactttc
tggaagtcccaaatgtcttaaaattacttcaataataacttcccacccatgaggagaatg
aaaacctacaaacatatcagtaaaaagaatgattaaaaaagctttagctgtatcactaag
accatagataatttcatttaaaaaagatttgacgattgctatttgtctttgtccggttat
cattagtaatataaatacaaatattgatacaatatcggacagaatattttaacagcatt
tgcactttcatttgcataatattctcctaactctctagccttcagctgtactctttttc
aatgatttcgtaagatatatcttctgatgggtttaatagtacttcaaaatgaatcttctc
ttcgaaacgttgtaattcagcaaaagctctctcttcctgagatgaatttaaaaaatttt
aggttgctcttggttccataaataatcaatacaaggaccaaatacaaaaaatttcgaagc
ctgattaactagtactggagatataaataataataaaatatattttacggaagtaatagt
ttggtgtctagatattctaaactcttctatagcttctgattctccattaggatctaactc
ctttcgaaacttttcaaaagtattagtaattgacctaggtattggtccaactttttcaaa
ggcaaattgattatgtcgttttaaattccaatatttcatttttattgctgctaaaattt
ctattaatcaataattgatcacatcttcactataaatattaattctagaactaaacatat
aaaacaatagttatgaataagaaaatacaattaaaatttaattatgcattagtattatct
tttgctacagcaatttatggatgcaaatcttataataaaactataagagcactattaata
aactatgaagatacctatccatttttttataataggcatgctgcctttgtccctataatt
gtattaaaaaaaatcattaaactccctaaaatcagactctcaaaataccaaattatatc
ttaatggaattggaaatagaaaatttatcgacagaattatcagtctactaaaatatttag
aaaaacaggctcactatagtgagcatcttaaccaaataaaaactagtggttacttttctt
tactttatataaacagtaaagcaatgtatagcaaagaagttaataatattttttattac
ctaatgaggttctaaaaagagtctatatttataataagaaaaagaagcttatacctcata
cattcttaagaaatttatttcagaaacagataggttacccaaaaaattttgcaaaattaa
acttagccctatcaaaaattatgagatggtattctgatagaggatatcaatggtctttat
tagaaattcaacagacagatgatccctcttctcttataattaatattcatgaaggattag
tgaaaacaattacaacagaatattatactgtatcttctaaaaaattgcgtggtatctcat
ttatagaatcaatagaacaatatcttggagtaagagtcggagttccattaaatattaatt
gtttacaaaaaaaaattaattatttaaaagataatcaactagttggcaatattatttata
gtattgagcgatcaaccaatgactcaatgagtttggatattaaatttcaaatacaagaac
taaaagataaagaggtaatagtgctagcagaaagttcttccaaggcttattcacttattt
ctcatacacatgatctactggatc---aatataaaagtttttccaattggctactagctt
ctaatataatttcgctatcgactagtaaattacaatcatgctactttaattctgaacttg
actatcaatataggtatattaatagctctaatatgctaaatttattagcttattctattt
cttatcgaaaaacactaataagcaattatttcaatttacaaactttattaagctggacaa
aaagcaatactattggatttcagctacatttgcgaaatctaagtcgtgcaaaatctttgt

gtatattcagtatgaaatttatccacaatggacttaatatgaaaatttcatatctaaatc
ctgcattaataattgataaaaattgtagttttcaatttgctatacaaataattaagcgat
atcacactgatacaaaatcacctctttctt------------------------------
------tatttttaagcgaagcaaatgctgcacagtatattgcagaaagtatatttattt
attattttacgtcttatttttcaatatctgaaaaaatattactatctcggcttattcata
caaattctttattttataattctgataatttagtcttcatggtgataataaaaatcgcg
ggatcagcaattattattttttaaacaaaatacaaaattattttatcaagagtttctga
ctttattattaagtttacgttatcaaatttgaattatttaggttgggctttaaaaggtc
atttatttgaaataaaatttttatactttgccccatttcagaaaactaatttttaaatt
accatttttcaataattataagaacctattctttcataaaatttccttgaagcaagtat
ctaattttaatttaccactatatttagaagccaccttaatcatattttagttaatacta
tcaaatgtcaatcaaatttaaatatggacactatatctattctattaagtgattatcctg
ctgaatatacgctatgcaaatccatttataatttctctatcaaagtgaggatggagtatc
ttatacctataagtaacagtatgagagtttctatattctctaactatttagactgttttt
taataagaccatctaagaaattaacatacacatggcaagatctaataactcgtaaacaaa
ttcaaaaatttttgctagaaaaatttcatatggatttggtattcagcttaaattaccaa
ttaagcagatgccacccctttcaattgaatatactataaatagtagtcgctacttttgta
tttaccttcatacttattatcaacgataaatattattaaaatgaaaaatttaatttcttt
ttttgatcgcagcaaagggaaatggatttcacaaagaacaacttatgaattgtctaataa
aaatatgagttcagtacagtctcaaatgacaatgaagttagataacttattatcaaaatc
tataatacttgcttcattaaattggggtgagatctctagacaagttaaacatcactctta
cagtcaaaacacaagcgaatataatgacaagttcaacttgcaatttagtaatcaattaaa
taatcataagctattgacgctatgtacagttacagatcctagtctaattagttttaaaac
tagatatggcattactaccatagatgagacatattggttcgcaacaaacaacttgcgcct
aagtaccagtattgttaagcaattcaatacttgtgtcgcagtatctttttgctcagaaat
taaagtttaa-aatctatataaaggcaactaataactctta-agctattagttgc--ata
tatatataaattttctgtttaagattag----aataatatattattctaagtctttacga
ttaggatttctagaaggatcattagataagaacccaaaaacaaacaaagaaataaagaaa
atgacggtagtatacacaaagattttcaaagtaaacatattttcatccttaaaagctatt
agtgtg----atatttacat--------ataattttatatcactatatgaattccagcac
tattttataaataaatcaatatctttatttttagtagtgatttatatttaaatatgcgt
gatgtactaaatactaaatctttttgtacctgttttgtactatttttaaatattttgtta
tttctatggagtttaccttcgattattacataatctaatttcttcattaatttgaaaata
tctgaagattgtttattccaaataattaagttgattacaactaatctttttcttattaaa
agtcttgcttttaacttaattatttgatttttattttcactaattctaatactcttacag
ctaagaatctgtactaataagttacagctattcat-atgtgtattaa------tcttgta

atgaatctactaagatagatgattttaaaaaatcaaaatgatatttcattatcttgtttg
ccctgctcataactaatatcttttaatacttttaatattttcccaaaatactcttgtaag
tatttttctaaagatatcacttctgaactatcaattcctaagatactataaacttcactc
ataggagcagtaaattgatctccgctagttaaaatttctgcaaatgctaaacgatcagaa
atattccatgtccattgaagggtcttagtaattcttcttaaagctgttaataatccaatc
ggaatttgtgatatttgtgtttttttgcccagataacttttcgcataatttaataatttca
gctgatgtccaagcagtattaccaactaatggtaatattctattttctgtggaaggtact
ccaagacttttataactaatttggctgcatcttgagtatcaatatatgcaataggcgta
gattctcctgtaacccatactgatttttgtctaaaataggaattgcatattggttaatt
aaaccttgaaaaaatcctcctaaagaaaatacagtgtatttatactagatttatgtagg
acatctactacttgtgattttaagctcattaatggaacagctgggtatttctctgcattt
aatatggagaagaaaataaatctttcaactttagctgctttagctgcttcaattaatgca
attttaccatctaaatctatttttgctgcgttatacggatctgtaggacgagatgtagaa
gcatctatgactgctgttacaccatagaaagattgcaagatactttcgggcaattttaaa
tcaccatatacgagttcggctccccattcttttagaaaggctgactttcttaaatttctt
accatacattttacattgtagccttcatctagagctcgccttacaatttgacgtcctaaa
gttccggttgctccaattactaaaagagtcatattaattaaagataagtatgtataaaga
cattgatttggttgacggcagagagtaatttaggtagagagggaatcgaaccctcatgac
cgaagtcgtcacattttgagtgtgatgcgtataccaatttcgccatctaccctattatag
accaactactacaagtaatttagtctttagcgtaaaaataatactaagttttaatagata
tgtcaactttaaattttatattaataaaaaaatggagtctcttatgtctactttttgaatt
aattccgtatagattatatttaagtcaatctaacacatggatgcatagaataaaagcaga
aattaaaatatatatattaacactattatggatttcaattttcattttctcttactttaa
actatttattattgctttaagtttgattgcagtcagttttactataaggagtaagcaaag
tattatacaaaaacatttattacagaccttattaatgacactcttaactactttttttatc
gtttagtgttgctagtagttataaacaatatttacaagcagaacagtcacaatatttatc
ttattctaataaagaggacaatcctgacagctaccagtatataaaaaatagctagtaatca
taaaatcaaaagggaacagctaattgttactttaaagccttcactatattttttttattac
tatatattctattaagctagttatgataacaacttctccagaaattttagtaattacttt
ttatagatctaggatcataaatcaactattgaaaaatgaattactatttaccttttttact
ttcgtcacacattgttaccagcataattaataaactagataaagttattcaagtagttag
attaaggggtagcttaaatttgagtaaatcatctacaaggcttttgatattttctttttt
aatatttcaagtcttctttttagagattattagggaatcgaaagaaatatctcaatctct
ttatactagacatctaaatcaagaaagcaataatttttttaaaatatacacagtaagatc
tagcttcagcgattggctaaatataattattggcactctatacttcgttattttagctct
agcataataaaatctacttatacttctgactctagtaatacgaagagtaatagacttaat

tagtagtttaaatagacagaat-tacgatattatataaataattaaatgtttcattatgg
tcaatactcaaaatcaaatttcacaaacttcagaacttaattacatagttaatcaacctt
ataaatatggctttaaaacttctgttgaatctgagcaatttcctagaggaataagtgaag
atattgtgaagttgatctctaagaaaaaaaatgagccagagtacctattaaatttcaggc
ttaaagcatataaaaaatgggaaaaaatgagtaatccatcatgggctcatcttaagaatc
caaatatagactttaacaatattatttattacgctgttcccaagttgaaaaaagaattaa
aaaatctagacgaagttgatccagaaattcttgatacttttaataagttagggatatctt
taaatgagcaaaaacgaatttctaatgtagctgttgatgctgtttttgacagtgtttcta
ttgcgaccacttttaagaaagaactctctgaagctggtgttatattctgttctatttctg
aagctatacgagattatcctgatttaattaataaatacttaggtactgttgtacctgctg
gtgataattattttgccgcattgaactctgcagtatttagtgatggttccttttgctata
tccctcctaatacggtttgcccattagaactatcaacttattttcgtattaataacgaag
aatctgggcaatttgaaaggacactaattgtggctgatcgtggaagtaaagtaagttatc
tagaaggctgtactgctcctcaatttgacacaaaccagttacatgcagcaattgtagaat
tagtagctcttgagggtgcagaaattaaatattccactgtgcaaaattggtatgctggta
ataaagaaggcaaaggtggtatatacaattttgttactaaacgaggcttatgctctggca
ataattcaaaaatttcatggactcaagtagaaactggatcagcaattacctggaagtatc
ctagctgcattttagctggccataaatctcaaggagaattttattctgtagctttaacaa
ataattatcaagaagctgatacaggaactaaaatgatccatattggtaataatacaaaaa
gtagaattatctctaaaggtatttctgcgggcaaatcgaaaaatagttatagaggcctag
ttaaaattggccctcaatcatttaattctcgcaattattctcaatgcgattctttattaa
taggccaatcatctcaagctaatacttttccttatattcaagtacagaatccaacatcaa
aagtagagcatgaagcgtctacttcaaaaattagtgaggatcaaatttttattttttac
aaagaggtatcagtttagaagaatctgttgctcttatgatcagtggtttttgtaaagatg
tattcaatgaattgcctatggaatttgctactgaagccgatcgtttacttagtttaaaat
tagagggaactgtaggatgagtcaaactattttagaaatcaaagatttatatgcttctgt
cggtgacacaatgattttgaaaggggtaaacttatgtattcaagcaggagaaatacatgc
aattatgggtcctaatggttcaggtaaaagtacattgtcaaaaatcattgcaggacatcc
agcatatactctaataagtggagaaatcttatttttttgggaaaagcattttttgaaatgga
gccagacgagagggctagagctggtatttttttagcttttcagtatcctgttgaaattcc
tggagttagtaatgctgactttctacgaattgcgttaaatgctcgcaggaaattccaagg
attatcagaatttagcccattagaatttttcaattaatagcagaaaagataaatcttgt
tggtatgcaagaaagtttttttgataagaaatgttaatgaaggattttctggtggagaaaa
aaagcgtaacgaaattcttcaaatggctttgctagacagcaacttatctattttagatga
aacagactctggactagatatcgatgcacttagagtagtagcaaaaggaattaatacttt
agcaaaatcaacaaattcaattattttgattactcattatcaaagattgctagattatat

tattcctgattttgttcatattatgagtaacggaaaaattgtgaaaactggaagtgttac
tctagcgcaagatttagaaaagcatgggtatgactggattacacagacttaaatatttat
taa-------aatattgagactaaagaatattctttagtctcaatattatttgcaaataa
aaag-----ttatttaagcttgtcaaaaccttgcttaacatcttcaattgctcttttta
gcaattcttcactagatgtttctaatttttagtgctacgaacactttcgccaaattctg
gtttagaatttttaaatcttcacgaagttctctaataaattcagctacttttgatactt
caatatcatctaaataaccgttaattccagtataaataatagccgtctgttcttcaactg
gaataggagaattctgagcttgcttcaaaatttctcgcagacgttgacctcttgctagtt
gattctgagttgctttgtctaaatctgatgcgaactgagaaaaagcttctaattcagcaa
attgtgctaactctaattttaacttgcctgcaacttgtttcattgcctttatttgtgcag
cagatcctactctggagacagaaataccaacattaatagctggtctaatccctgagttaa
ataggtctccagataaaaagatttgaccgtccgtgatcgaaattacatttgttggaatat
atgcagatacgtccccagcttgagtttcaataataggaagggcagtcatgctacctcctc
ccagctctgagtttaatttagctgctcgttctagtaatctagaatgtaagtagaatacat
cacctggatatgcttcgcgtccaggaggtcttctaagtaaaagggacatttgacgatagg
cttgagcttgcttagttaaatcatcataaataactaatgttgctttacctttatacatga
aatattctgctaaagctgcacctgtataaggtgcaatatattgcaatgtcgctggactat
ctgcattagctgcaactataatagtataatcgagagctcctttatcttgtagtgaagata
caacttgagctactgaagatgccttttgtcctatcgcaacatatacacagacaacatctt
gacctttttgattaataattgtgtctaatgcaacagctgttttacctgtttgacggtcac
caataattaattctcgttggcctctaccaattgggatcatagagtcaatagctgtgatac
cagtttgcatcggttcacaaactgattgacgtccaattataccaggggccatcgactcaa
ttagtctggttccattactcgcaggttcacctttatcatctattggacgagctaaaggat
caactactctacctaaaaaagcatctccaaccggaatttgagcaattctaccagttcctt
ttacagaactgccctctaaaatatctctaccatctcccatgagtacaacaccaacattat
cactttctaaatttaaagcaacaccaatagttttatcttcaaattcaagcaattcaccag
ccataacttcatcgagtccatatactcgagcaataccatcaccaacttgtaaaactgttc
ctatattggctacttctacatcttgatcatacttttcaatttgttgacgaataatactac
ttatttcatcaggtctaatatttaccatgtttttaatattctttttattaattggaaatt
ctaaagtggtaattactcagatttatactatctg-ctttttttactttattttaatttgct
actacatcaaggtgtgaagccatctgccttaattggcctctaatactagtatcaataact
ttcgatccaatttgaatagtaaaaccaccaattaactcgggatcgacagaaacaattagt
ttaacttctttagccttggtcataactttgatttttttctgttaacagattttcttgatca
gagcttaaagcaatagaagtattaatatttgcaattgttagagattccatttgataagca
agtgccaaatacttactagctatagcatccaacattgcaattcgtttacgatcaactaaa
atcattaaaaaatgataatgtattttcgctaatttgatcaccaaaggttgcaataatgact

tctttttttgcctctacagtttttaatggatttgct---aaaaaatttttcagcttatca
gattgtgataaaatattttgtattgactgaatatcttggcttacccttgttgtgactttt
ttagttttagctaactctagaagagctactgcataaggctgagctatttttatcatcacg
ttattactgctcatagttgatctcctaacttagcaatattattatcaatgatacgtagtt
gcatttcagagctcatttgattctccagctgtaaagtaactttttttaagagcaagaaatg
ttatttgctgctggatttgccttcgaatttgtttttcagctgtttcgatatttgatttac
ctgttatagcaagtctttcaatatctaattgaccttgagctaatatagaattcctaactt
ttccagctgttagttgagcttctttttttgatttgatctataataatttgagtctgagcaa
gttgtttttctgattctgataatctagcactagcttgctctagtctttcttcagattctt
gtatagctgctaaaacttttatttgtcttgcatttaaacttgatccaagaaattgcttca
aaacataaatgagaccaaataaaagcaacaatatattaataacatttgcttcaaaaatat
cagaattaaagccgaatttgtgttcactactatgttctgataaaatagtaattatttgtg
gtatttttactatattattcatttaaaattct----------tactattttaatatttta
ctcaggaatcatgatttttaaagattgaccacttaatagtttagctttaatttgatcgctc
aaagtatccacctgattttctaaagttttaagggcctcttctttttgaatatttaactgt
ttcgatgcttcagcaatcagtttctctgcattcaattgagctttttttaatatcttctgag
actagattttgagcttctttttgagaagaggcaattttttaattgtgcattacgacgggct
tcagacaaatcttcttcatatttgcagctaactcatctgctttaacaagcatagaagat
gcagtagtcaacgttgttcgaatatattcgtcacgttcgtctagtactttagttacaggt
ttataaaaaatagtgtttagtaaaaccattaaagtaagaaactgtaatgccattaaaggc
aaagtaccattgaagtcgaataagccaccttcaatttcttcggctaatagaaatggtaaa
tcaatcattttactaattgtgtttatagttaataatttggtttcaattaagaaaaaataac
tttatagtgcgcttagtctgataagactaagcgcaaatgatcagtttaaccaacgtatgg
attagcaaataatagagacagtgcaacaactagcccgtaaattgttaaagattccataaa
tgctaaacttagaagtaatgtacctctaatttttccttctacttcgggttgtctagcaat
accttctacagcattagctgctgcactaccttgaccaataccaggacctattgcagcaag
acctacagcaagaccggcagcgataacggatgcagctgaaacgattgaatccataatgaa
ttttgatttttttgttatatgaatagaaaattgacaaatttcgtataatatttagaatcta
cgtaattaagtataatagattcaaatat-----tgtaattacagtatactactcttctcc
atgaccttccattgcttctccaatataagctgcggatagggtggaaaaaaattagtgcttg
tattgaactagcaaataatcccagtatcataactggtaatggtatcaaaattggaattag
cagagtaaatactgaaacaactaattcatctgctaaaacgttcccaaataatcgaaaact
taaagataatggtttcgtaaaatcttctaaaatattgattggtagtaatactggagttgg
ttgaatatacctagcaaagtagcctagaccttttttacttaagcccgcatagaagtaagc
taaagaagttaataaagataatgctacagttgtattaatatcatttgttggagcagctag
ctcaccttcaggcaggtgaattaacttccaaggatcaatgcacctgcccaattgcaacc

taagataaatagaaatagagtggcaatatatggaacccaagggcgatattcatgctctcc
gatctggtttttgcaatatcttgtaagaattctagtataaactccataaaattttgaaa
tttttcaggaattctttgtagatttctagtacctaggaatgataaagttaacagtgtagc
aattactaaccaagagacaataaaaacttgcccatgcaattgtaagctacctattttcca
atatagatgttttccgacttctactgctgataagctagtatatggattaaaatctatgag
actgttttgatacataataaatatttcttttaaatagtgcaagaatcaaaaaataaattt
tttgaataattaatatgcaattagtttattgttaagactagattgttaataaattaaaag
caaaaataaattctaatagagatagctatctttagtagtgatttttttttgtaaaaaaca
attattgctacctataatgcttttacgtttaattatatacccatatattaattatttaat
atgattataggcattttgctacaaactttattgttttatccattctagctagatattgtt
ttaatttattgtttttttaaaatatctgcgacttcatccgcaaaattagctttaatattt
tcttctcctccacccagaaggaatctaacaaatcgtctaattttaatattttcgcctaaa
acagcaatattttgattgattaaatctttaatactaatatcttgattgcgaataaacatt
tggtctaataatgaaagctctttaagcctctttttcattcttgcctcaataattgtttct
attctatcagctggcttattctttaagtcatgcttaccagcttcaaattttttttctaaa
ttgataatttcaggaggtatatcttgcattgatacatattctacatttggcgatgctgct
atttgcatagcaatgtctttagctaattttttgaaactctggacggcgtgctacaaaatct
gtttcacagttcacttcaactaatacacctattcttccaccggtatgaatataactctct
aataaaccttcaactgtagtacgactagatttttttgttagctgaagctaaccctttttgt
cttaatgattctaaagcttttttcttcattgccattattagcttgtaaagctttttttacaa
tccatcataccagctccagtcttatcacgtaaggctttcacggtttgagcagaaatttgt
agtgtcatatcttttcaaaaatttttaagtatttaaattaatt-aaacaaattattataa
ttcactttcagtaggatgatctatttcttccaaatccttttgcccgtatttgccatcata
aattgcatctgctattttaccgataattagtttaattgatctaatagcatcatcattagc
tgggattgggatattaataatttctggactacaattagtatcaagtatacaaatagttgg
aatccctaattttaaacactcttgtatagctgttgtttctcgtttttgatctacaactac
aacaatatcaggtaaacgagtcatgtttttaatgccatttaaatgcttacgaagtttgtc
taattctcttcgtagaacagcagcttctttttttaggtaactgatcaataataccactttt
atcttgttcttctagctgtctaagacgactaactcgagatttaatagtcacccagttagt
caacatgcctcccagccacctttggttaacgtaatatgaatcacatcgttgtgcttcctg
agcgactatacctgctgcctgtctttttgtacctaaaaataaaactttttttaccatccga
agaagcctgttttataaattcacaagcttctgtaagtaattgtgcagtctggactagatc
aataatatgtataccatttctttctgtataaatataaggaaacattttgggattccatct
acgtgcttgatgtccgaaatgaacaccggcttctaataattctgataaggtaacaacagc
cataggattgttaatatttaaataatgtaactattcgggaaatcccttttttttc--aaac
tcat-----taaactaacttattttaattttaataaactattaataaattaacataattt

tacagcagattaagaaaatagaaccttcttacttcatttaatctgtactttgattattga
aataattacgtgcagttctatcgtctaaaataatatcatctaaatcatctctaacagatg
aaatgttgctttctatattagtatctggagtcaaatctttttttctaacgacgatccat
tatggttatcgtacatattgaatccagtaccagctggtataagtctacctataataacat
tttctttcaaacctcttagccaatctaatttgccggatatagcagcttcagttaatactt
ttgtagtttcttgaaaacttgctgcagaaataaagctttctgtatttagggaagcttgag
taatacctaataaaacaggtctatatgaagcattcagtttatttctcaaggtcatagctt
tgttagtttgttcaatcttttgcaactctacaagctctccaggtaaatacccggtttctt
caccattctctattttcactttagaagtcatctgtctcactataacttctacatgtttat
cagaaatattaactccttgagattgataaactaattgtacttctttaacaagtagtaact
ggatttcttgaaaacttaatcgtgcagcttcatataaagccaatccgcattcaatatata
aattaaaacttgcatctagaatatcatgagggttcaataacacatctgttttctttcttg
cttcaagaatttcttcaattcttggaagaccttgtataatgtcaccagtttttgccctat
caaaaaccaaaacagctagtgtttctccccttctaattaatgcattattgtccacatgta
aaatagcaccattagaaactaagtaaggcctcgcaattcttaatgtaacatatttagaag
atacttcaatgatttgtccagagtctaaagaagttatgtttgtagtaataaagtcaccgc
aacgaatccagtcaccaacacttactttagtagttgattctcctatattaaatgtttttt
tatcagaagatgttgcaattagaattcgcctactagtattcttttcggaataaatttctt
caacgctaccttcactcattgcaatgatctctgtactagcaacaactgtaaatggcttta
tgtattgaccattttttaacaataattctagtttgactttgttgcttttccgatttattga
catcaatacttttaattgataatgtctcaaaagttgagagccttaagtcaaaataattat
taccattttcttttgcggagaactcaattgaagatattaagctattatctttatgctcaa
tttgagctacaagattagttgttactaaatgaactccttctatagatttcactctttctc
cgtctcgaaaaggggttcttttaactagctttagtttagtttggcttggcttctgtgaca
ctaaattctcaattaagctagattttgtttcaggtattgaataaacaataacgggtctaa
taagaatataaggcatttgctcattttctatgtactcccaataaactaacttgtctgtag
aaatattattatggagtgtttctccgggtcttaaaaaacctctacttttatcatcattgc
tataaacttcactaagctcatatatagatccaggcttaataataatttcgcgaactattc
catccctttggattatctcaacaattccggagcttttagaaaaaatatttttaactagct
ccgttccaatttctataatatctccattatgaactagtagcaatgaagaatctttattaa
tttcgtgcgtttcttctgaaatccacaatatatatccaggactcaaaatttcataggcat
ctttgtcagaaccagttttctttttttgatacgttcaagtctaaatatttaataataccac
cacttgcagttttataagtatcagacattagttcagccactgtataaccatcttgaa---
ttttttgatttggaacggctttaaaaagaaatttttgttttttttctgtttctagaatat
aagattcttttttatgaattagatctgtataaatataacaacctggaattacgatagatt
cagtaataacttgtatattggtaatactgctattagtatttttgtgtatccttacttccc

cagcatagtgatttattaattctgtctgcgcaagtattgtaccagcttgagttttgtctt
ctttttaacaataacattagctgaatcagatatactataaacttcaccagagagaaccc
aaatcaagccacctgtcttagtaattcgtgtagtatattgattattatctgtttcttcaa
ctgtcaagttggagaaacaaattttcccagaaagatctgagactacatgtttttgagctc
tttcggtcattagtctatttctacgaggcgattcagcaattacttgatcttttaatacta
gttcaccattaccaagtaaaagagtagtacctttacttaaattaattattgacttttgat
tttttttcgacctgattgtaacttgcgttggactttttgtgattaatgcttgctctccat
gtctagtgcgaacatctgtatatgatttcatatctaaatctgttagccgaccatttatgg
gcgcataaatttgttctgctaattcaccagtaaatactcctccagtgtgaaatgttctca
ttgtcaactgagtgccaggctccccaatagactgagcagcaataatgccgacagcttctc
ctaaatcaactaagcgaccatgggcaagattccatccgtaacaatactgacaaacggaac
ttctagaattacatgtaacaggtgatcttactaagactttttt---gataccagctttag
ttatctcttttgctaacttaggactaatatcttggttagtgtgagcaattaagaaatttg
tctctggatggaataaatttctgctaaaactcgaccagataaagcttgctctaagttga
ttagtatttttttgtgtatctactaggtcttctagtatgataccttttatttgttttacagt
ctacttctctaattattacatcttgagatacgtcaacaagtctacgcgttaaatagccag
aatctgctgttcttaaagctgtgtcaactaatccttttcttgctccgtatgatgatataa
aataatcggtaacagttagtccttctctaaaattactggatattggaagatcaataattt
gtccctgtggatcagccataagacctctcataccaactaattgtctcacttgagaaatat
tacctctagcaccagagaaagccatcatataaacagcatttagagggtctgtctctttaa
agtatttaatgacttcttgttttaaggattcacttgcattgttccatgtatcgataactt
tttgaaatcgttcaactgccgtgatctcacctcggcgatacttattttcagtagccttaa
tttcttcaatcgtacttaataaaagactttgcttactaggaggaattcttaaatcttcta
gacttaaagaaatccctgcctgcgtagcataatggaatccaagatctttaagcttatcag
ccatattagctgctcgtgctataccataattacgaaaagcccaaacaattaaattttta
gttcattttatcaataaccttatttgaaaaacttggctctgcaagactttttctattat
ccactaaatcttctccatattaaatgttcgtttattcaactacgccaccaaagactcttg
aataattttgttaaagattatacggccagctgttgttctaatatattgcacaatcctctg
gcattcagcatcttcttttatgatatggttattaaaaaattttgttatactattgtcaca
atgtttttcaattttaattggatcatgaggttggtcaccatctaccaatccatcaaaacg
agcccatatataagaatgtaaatcaactttcttttgctcataagccataactacatcctc
taagcttgcaaagtactgactggatccctgctgttgagatggattattagctgttaaata
atagcaacctagaaccatatcttggcttggcataataattggttggcctgtggctggaga
gagaaaattatgaggagctaacattaacaatctagcttcggcttgtgcttctagagataa
aggaacatgcacagccatttgatcaccatcaaaatcagcgttaaacgcagggcaaactag
tggatgtagcttaatagctcttccttctactaggataggctcaaacgcttgaatacctaa

tctgtgcagtgttggggctctgttcaataatacagggtgaccttgaataacttcatttaa
aacattccaaatagaaggttcatttttttgaatcatctttttttgcagctttaatattgtt
gactaatccctgcaagattaatctatgaatcacaaagggttgaaataactctaaagccat
ttctcttggcaatccacattgatgaagttttaaatgaggacctacaacaattacagatct
cccggaataatctactcttttacctaataagttttgtctaaatcgtccttgttttccctc
tataatatctgataaagatttaatggtctattattggcgcctacaacagttctacctct
gcggccattgtccatgagggagtcaactgcttcttgtaacattcttttctcatttctaat
aataatctctggagctaaaattgactttagacgtgataatctattatttctattaataat
tcgacgataaaattcatttaaatccgctgttgcaaatctacctccatccaattggaccat
aggtcgtaagtctggaggtataacaggaattactgtgaataccatccaagatggatctgc
acctgttgctataaaattctcaattaatcttaagcgtttcatttttttattgaattttaa
tgaaggtgtcttaaaactttttggtggatttgtagcttcagaccttaaggtttcagctat
atactctaaatctaagtctttcaacagtttttgaatagcttctgcacctattcctacttc
tacttgattattctcgtcttgattttgatagatttcctcttcaagactcttccattcata
accctcaagcagttgcttatattttaaattgatatctgtattagattgtgtcactacata
agaatgaaaataaacaattttctctacttcttttacttttaaatctaatgctaaagcaat
atagcttgtacttccttttaagtaccacacatgagttactggtgaggctagttcaatgta
agccatccgatgtcttctcacgcgagattctgttacttctacgccacatcgctcacagac
tatacctttgtagcgaaagcgtttatatttaccacagtgacattcccaatctttaactgg
gccaaaaaatcttctcacaaaataaaccgtccatttctggttttaaagttctataattaat
tgtttctggctttgtaatttcaccaacaatttgtccgttgggtaaacttctttcacccca
ctgtcggatttttttcaggagaagctaaactaatttttacatagtcaaaatattgctcaaa
ctttgtcataaatttgaataccttaaaatcaaataattagccaatttattcaaatgttcc
atattttaaaatcatttaattaataaagaaactgttcaaaatcatctactgggggagtct
catagttagagcgatctgatcggctctctttagaatcagacatcagatcaacttcaactg
tccgtctttggccatcttcaaatagttttaatttatgtactgcaatgtctaatcctaatg
actgaagctctcgcatcaagactttaaaagactctggagttccaggcttaggtattggct
ttcctttcactattgcgtttaatgcttcattcctagcttgcatatcatccgactttacag
ttaataattcttgcagagtatatgctgctccaaaagcttctaaagcccaaacttccattt
ctcctaatctttgacctccatgctgcgctctacctcccaatggttgctgcgtaactagag
agtaaggaccagtggatcttgcatggattttgtcatcaacaaggtgaacgagcttaagca
tataagctctaccaattgtaacagggttatcaaaaggctctcctgttctaccgtcaaaaa
cttgcattttaccaggatgctgatcattaaaaagccatttattgttagtaagcaatgatg
cttcttttaattttctatttacaagtgcccttgatgcttctgctccatacatttcatcaa
aaggtattattttaaatcgttttcctaaataaccacctgctaagcctagcaaacactcaa
acacttggccaacattcatcctagagggaacgcctaaaggatttaaaacaatatccactg

gtgttccatcagataagtaaggcatatcttgcttaggtaaaatcctagaaatgataccttt
tgttaccgtggcgaccagccattttatcacccacttgaatttttctcttctgtgcaacat
aaacgcgaatcatagcatttgttccaggaggtagttcatctcctttctgcctagtaaaca
ccctaacattaactactcttccttttgcggcattagggagtcgcaacgatgtgtctctta
catctcgagccttctctccaaaaatagccctcaataatttaccttcaggtaattgatcag
cttcgcctttaggagtgatttttccgactaaaatatctccagcttcaacccacgatccac
ctacaacgattccatttctatctaaatctttcaaagaaagatcacttacattaggaatct
ctctagtaatttcttctgggcctaatttagtttggcgacattccacttcatacttttcaa
tgtgaatagaagtatacaagtcgtcatagactaatctttcgctaattaaaaatgcatctt
cgtagttataaccttcccaaggcatataagcaacaagaatattccttcctaaagcgattt
cccctccatctgtggatgcaccatcagctaaagtttggccaacaactatcttttctccta
cccaaacaattgggcgttggttaatacaagtatcttggttagaccgataatattttttta
aacgataatggactgttctaccattattatcctgaatgccaattttattagcagaaacat
aattgacataacctgcagttctactaataacaaccataccagaatctcgagcaatttcg
tctcaagacctgtcccaattattggcttttctggatataataatgggacagcttgtcttt
gcatattagatcccattaaagctctattcgcatcatcatgttccaaaaaaggtattaatg
atgttgcagctgaaattacttgaataggtgaaattgcaatataatctacttgatttggtg
ttgttgtaataaattcctgacggtaacgtacaggaataatatctcctttaatgtaatttt
gagtactgactttaacatcccctggagctactcgaaaatcatcctcttcatctgctgtta
aataaacaggactattttgataaatcacttgacctgcatgtaccgggtaaaatggagttt
cgataaaaccaaaaatattaaccccttgcacaggttgctaatgaaccaattaagccagcgt
tcggaccttccggcgtctcgattgggcaaattctaccataatgactagggtgtagatctc
taacagcaaagccagctctatctttattaaaacctcctggacctaaagcactaattcttc
tcttatgggttagttctgcgactggattagtttgatccataaattgagaaagctgactag
atccaaaaaattctcttactgatgcgattaacggcttaggattaattaaattagataaac
ttaatgaatctatatcacatatcatcattctctctctaataatgcgctctaagcgattga
gacctactcggaattgattttgtagtaattctccaactgagcgcactcgtctattaccaa
gatgatctatatcatcaagattacctgagtttttatctttaatattaattagataatcaa
tcgaagataaaatatcttgaggagacaatactctgaaagttttaggtatattaagacctta
attttttattaattttatatcggccaacttctcctaaatcatatctctttggatcaaaaa
agcgagagtaaagcatttgcttagcaacagcaactgttgctggttcatttggtctaagct
tagaataaactataagcaatgcttcctcatcagttacttcctcaatattatttttttccaa
tttctttagctaattctttaactgaatatatttgagatgcagagataagaaaagcatatt
tactaagtcctttttcaatctcgtctttattaagcccaatagcgcggagaaaaatatagg
catttactttatgagttttgtctattcttatccaaatttctcctttaggatcaatttcaa
attttaaccaagaacctctattagaaattagactggcactatagatctgcttgccatttt

tatctatctcctgcttataatagatgcctgggctacggattatttggttaataataactc
tctccgtaccagatacaataaaagttcctctatttgtcattattggcaagtctccaataa
aaaccaatctcttcttgtactttttgtttttatctgcttttcggcactaagctgtaaat
gtgttgacgataaatctagtgatttgatattctttccatggtctttagaaggtagctcaa
tatcttttcttgtcaactttgcaggaacatatatttgggcactataggttttatccctgt
tttttgcttgtctaacactgtaacgaggaaattttatcttatactcattaccaaataatt
gtagttcaagtcgactagtaggatctgatatattcgggaagaattcaagcacttcagtca
gaccttctaatagaaaccatttaaagctggccctctgaatttcgactaaatctggaagaa
gtttatttttttaagcttatacgttggaccatagatctcctttataacacattctgaaatc
gaagagtttaa-tactatcaatagagtactgccaggtaatgcctaagcacagtttgctac
attatttttctctagacacaactagaacatcctattttattcaac-aaaaaaattttta
aaattt---atcac--acaatatatatataagtattttgctgctg--aaaaacaatttca
gctat---a-------aaaaacgaaattaatacttgtctgtctgtcaaataattgacatc
aagatagacaagtattatttatttaatcaattctgtttttttgagcaaacgacagtgct
ctcgcaagtcgagatttttttcttgcaccagtattaggatggaatgctcctttttttact
gctttgtcaatcttgctatatacttgagaaatacttgattgcacatcattgaagttagaa
ttttctaaattgtcaatatttagaaggcatcttttagttaaagttttgacaaccgattta
tacttacggttaattagacgatttctttccgaagttttaatgcgtttaatcgcagaaagg
ttcttagccacagtaaatccaataaaacttttataaaatatttttagaaacgaaaatcca
caaatcggggatctattcaaaaagggaatatgcgccgtattataacattaaagaaaaaca
ataggcaatagttcatcaagagctcttgttattaaaaaatagatattgtcattgcataaa
a-tttacaaaatagtatttaattaaaataaattaactagacatataaatatatatgaaaa
aaattgaagctatcattagaccttttaagcttaatgaagtaaaacttgcgctagtcaagg
aaggcattggaggaatgactgttattaaagtttctggttttggaagacaaaaaggacaaa
cagaaagatataaaggatctgaatattctattgatatcattgacaaaataaaaattgaaa
ttattattagcgatgacaaagtagataaaattacagagacgattattgaggcttccaaaa
caggggaaattggtgatggaaaaatatttattagtagtattgaaaaggtaataagaatta
ggacaaatgatttaaattttgaagctttatagtatgatacttttgtttagacttgatact
tcttcactaaacatgagataataatcatttattgtatacttataactaaggtgtaaca--
ctaatggctaaaagcaaaggtgcccgaatcgtaataactttagagtgtgctaataaagct
ggagatattgctcaaaaaagaaaacctggcgtttttcgatacacaactactaaaaataga
cgaaatacaccaagtagaattgaattaaaaaaattctgtcctaattgcaatcatcattgt
atattcaaagaaattaagtagttatttatcttattaaaat--aatcatattatggctata
tatagaaaaagaatctctccaattaagccaacagaggctgtcgactacaaagatattgac
ttgctaagaaaatttattacagagcaaggcaaaatactacctaaacgatcaacaggatta
acttcaaaacagcaaaaaaagcttactaaagctattaaacaagctagaatactttcctta

ttaccttttttaaataaagattaaatatatatagctttacttcatattattccatataag
cattaaaaatcgtcatggatttgtttactacaatcatcaaatatattaaat---------
----------gactgtgaaaaggctattcattacaagttttgttcattttataattaata
aaaaccatctaatttat--ttaagctagatta-----agacataaagtttaacctctttt
aattgattatattgcataatctctttaaaagaggagatttttc--ctgg----------t
aacaattcaatattctttaagcttgtacagagtatttatgacaaagatttttgcttaggt
ataagatcaaatgaatggatcttgtcccccggctgccataattgaaattcagagataaaa
atgccacattcgttaccagcttgaatttcttcgacatcttctctaacacattttaaagat
tcaatttttccttgataaataacttttttcttcccgaataattttaatccaagaattttt
agtaatttattattagtaactcgacatcctgctatttttctatttgctaaagaaaatacc
gtgcttacttctgcttctccgacttgtacttctgaatattctggatcaagtagattctcc
atcatttctgtaatatcttcaattaatgcataaataatttgataatttttgactaatata
ttcaattttactgcagcttgctttgcgctaggagcgtaattagtattaaacccaatgaga
gtagcatttgtagtggaggctagctcaacatctgtagcagtaatctcgccaggcatgatt
gatacaacatttagttgcactttactttgaggaaattgtgataaagaatctaggattgct
tctgtagagccttgattatcagttttaataatcaaagaaatctgcttactgagatcttta
taattagtagttctcaaggtgtctaacgtaatacgactattcagtgctctctgtttttgt
ataattgatgaatctttagaagtattttcaagagctttaagctttgcttctttatcactt
ttaactactaaagctatttctccggtagctggaacagctgataatccccaaatttcaaca
acagatgatggtgttgctaaattaatcttttctttagcgttattaattatagctcgaatc
tttgcataagctgatccaataaccatattatcactaatatgcaatgtcccatttttgtatt
aaaagggttgctactgggccatgagatttatctagatgagcttctattataataccttgt
gcaggctttgttggatcagcttgcaagttctctaattcagctaataaggtaacagtttct
aataatttatcaatattttgaccagtaagagaactaataggaattacaggaacttgacca
ccgagtttctctgacatcacattatattttaataaattctgctcaataatgtctatatta
gaaacgtctttatctatcttagatattgctataacaaaaggaacatttgctttttgaatg
tgattaatagcctctatggtctgaggcttaattccatcatccgcagctataataataata
gctatatctgttaaattagctcccctagatctcatactcgtaaaagcttcatggcctgga
gtgtccaagaaaacaatttttttgtttattagctttttaatatattctacttcataagct
gcaatggtttgcgtaattcctccaatttctttattagcattgttagatttacgaatataa
tttagtagagtcgttttgccgtgatctacatgccccataactgtgacaatcggaggtctt
ttaatatagttaccagattcataagtgagactagaagtatctaaattagtagaatatata
ctattatcttcctttatatttgattcaactgctatgccaaaattatcagctactgatgat
attattgaagcatctatagtttgattaatagtaactgatattcctttcaggaataaatat
tttataatatctgtttcttgaacacaaattagtttagataattctttaatagttagagga
tttgtaatactaatagattctggtggagaattaaatagattgtcggctggattttggcct

tgatttactaatactttcttctgtcttgtagatttttagtactagaaagtttttttatg
acttgaacttttggcttaggtggacgcattagagagatagctaaatctcctgcagtctgc
gaaataatagaattagaatccctaaagttctct---tcctcatcatcaatatgtattttt
gtttttattttc-ttacgctgtctattctta--tttttttattgtctactaaatcatga
actttgttaaaatttttattttttttatctagtttaggcggagagcttaactctagatgc
tcttcaccactagatgtatgcacatctgatttatcgagatttagctgcaacagattatca
ttattagctgactctaatctaatcttatagataatctgaggatttttaaatctacaatt
ttttcagaagaatttatatttgggctagattcaggagatcgattttcaaaattttgatta
tttaaaaacatgaatgttggttttattatctaatagtaactttttaacttcaatacagag
ctcaaaataaataaaaatc----atctgtataaataatactaatat---agagaattgat
ctctattgtattagatttaatataacactaaagtcttttgtgcattacacaagtttaaaa
ctttt-tatgatgataaacactagatttagtttattagtatacaatctatttatctaaa
taatactgttttcga--aggtttttaaa----ttatctaatattatgttcgttatgaatt
tctattacaatttaaaaa-ctagacaattaagtaaaaatcgttaatcacctccacgccac
ttatgccacgatctcaaaaaaaatgataattttattgacaaaacgtttacagtattagctg
atattgtattaaaaatacttcctacaagtaaagaagaaaaagaggctttttcttactatc
gggatggaatgtctgcacaatctgaaggcgaatatgctgaagctctggaaaactattatg
aagctttaaaactagaagaggatccatatgatagaagttatatcttatataacatagggc
ttatttatgctagtaatggtgaatatgtaaaagctttggaatactaccatcaaggattag
aactaaactttaaattacctcaagcccttaataatatagctgtaatataccattaccaag
gagttcaagctattgaggacaaagatacagaattatctaaactaatgtttgataaagcag
ctcaatattggcagcaggctattaaattagcgcctgacaattatattgaagctcaaaatt
ggttaaaaacaacaggacggatgagaaatatacaaggatattagcatatttaggatataa
taaagatattaattaa------------gatcagagatatttttatcctctaatatattg
-tagtat--------acattatactaattga------tctataagaatcaaagtaataac
cgaaattaaatctattgactatatttatgaatacaaataagcataatggttagtacaaca
aaaagtaccggatctgttactcaaattattggaccagttttagatattgcatttcctaat
ggacagcttccaaaagtattcaatgcacttaaagtaaagagctcagaaggaactattact
tgcgaagttcaacaacttttaggtgataacaaggtacgagctgtatctatgagttctact
gaaggactacaaagaggagtcgaagttattgatactggattccctatatctgttccagtg
ggcacagacactcttggacgaatttttaatgttttaggtgaacctgtcgataatttagga
gcagttaattctgaaagtaccttgcctattcatagacctgcacctgcttttactaagtta
gaaacaaagccaaatatttttgagacaggtattaaggtggtcgatttattggctccttat
agaagaggtggcaaaattggtttatttgggggtgcaggagtaggtaaaactgtgttaatt
atggaattaattaataacatcgccaaagctcatggtggggtatctgtatttggaggtgtt
ggtgaaagaaccagagaaggcaacgacctatatatggaaatgaaagagtctaaagtaatt

aatgcagataatttaaaagagtctaaagtcgcactagtatatggtcaaatgaatgaaccg
ccaggagcacgtatgcgtgttggcttaactgcattaacaatggcagagtattttagagat
attaataagcaagacgttctattgtttattgataatatttttcgatttgtgcaagctgga
tcagaagtatctgctctactaggacgtatgccatctgctgtaggttaccaaccaactcta
gcaactgaaatgggagcattacaagaaagaattacttcaacaacagaaggatctattaca
tctattcaagccgtttatgtgccagctgatgatttaacagatccagctcctgcaacgaca
tttgcgcatttagatgcaacaacagtactttcacgtaacttggcagcaaaaggaatttat
cccgcagttgatccactagattcaacttcaacaatgctacagcctggaattgttggaact
gagcactattctacagctcaagaggtaaaatcgactcttcaacgatataaggaattacaa
gatattattgctattttaggtcttgatgaactttctgaagaagatagacaaactgtatca
agagctagaaaaattgaaagatttttatctcaacctttttttgtggctgaagtattcact
ggatcaccaggaaaatatgtatctctagaagatgcaattaaaggtttccaaatgatctta
aaaggtaaattagatgacttacctgaacaagcattttatctagtaggtgatatagacgaa
gcggtacaaaaagctgacagcatgaaagattaatataattaacgatgactttaaatatac
gaattattgcgcctgatcggactgtttgggatgcagaagcacaagaaattattttaccaa
gtagtacgggccagcttggtattttaacaggccatgcacctttgcttactgctttagata
taggagtcatgagagttagagtggataaagaatggatgccaattgttttgatgggggggt
ttgcagaaatagagaacaatcaattaactattctagtaaatggcgcagaagaagctagtc
aaattgatttatcagaagcagaaaaaaacctagaaactgcaactcaactattaagcgacg
cctcgtctaataaagaaaaaatagaagcaacacaaaagattcgaaaagctagagctagag
tgcaagcagcaacagcagccatagtgtaa-aacatttt------gctcaaaacataaaac
tttagtaataattactaaa-------gttttatgttttattttgatagttgtaagcagct
taaataagcttagcttagaccagaacaaatataatcaaaatataaacccatctctttacc
ggcatctggtccaactaaactaatagtaacttctttcatagctaaaattgcttgaattgt
tgcaccaataggaacacctaaagaattatatgtttcttttagaccattcaatactctttc
ttccagaatagatggatctccagctagcattccataagtagcataacgtaagtaataatc
taaatcacgaatacatgcagcatagcgtcgagttgtatacatattaccgcctggacgtgt
aatatctgaataaagtaaggattttgctacggattccttaataattgttgcggcattagc
tgcaatagtagcagcagctcttactcgtagctcacctgtttgaaaatagcctcttaattt
ttcaacagaactatcgtctaaatatttaccttgaacatcggctgcattaataacagaagt
aattgcgtcttgcataacttttaaaaattccttaattttaaatatctaatacta-gagcg
tatataacaaaattactagactctttagcaatatcagtattactgcatagcacccaaagt
atagtcaaagtaaaaaccagcttctgctgagtcttcacctgcaagcaatgagcaagctac
acttttcatgcatttcacaccttcagctactcctgaaattggtgttcctaaagaattgta
catttctttaacaccgactagtccaatttcttcaataggcgtcacatctccagcaactat
tccgtaagttactaagcgaagataataatcaaggtctcttaggcaagtagcagtcatctc

ttctccataagcatttccgccaggagagactacatcaggtctttttttgaaatagttgctg
accaccttgttttacaatacgttcacgattgtctgttaaaatttgagctattcttaaacg
acgttggccagataaaacaaaacttttaattctatctaattcaccaggacttaaatatct
tgcttctgcatctgcatttacaattgactttgtgacaatactcatggataaaactcctgt
aatgctt--aaaaaaaagcttaatactaattatttatgatataactcagaaagtaataca
tcttttgaataaatttataaccataacacattaatgatgtgtgaagcagcataaatactt
aagtaagtactaaaacgagaaagttatatgtagtaccttactttattgattcccgattac
tttttgaaactcggtaccacaattgataaatcctgcttagtcaggcgattatataatgt
ttctgtatttggaaaattagctgctggtaaagttggaaatctcctatatggaacagtgtc
tttgccatatactgaattatattcattactttcaactaaactactaataaatgcagacaa
gcctcttgaagctaaaatttgattaaaaaatctaatttccgcctgattattaggagctcg
ccctaatatgtgctttgtacctaattctataacttttgtatttggatatggctgataaaa
ttctttaccatatagttcagataatgccaatttttctactaaacctttaacattgatttg
cctattcaaaaaggctgactcgatatctaagaactctcctccaatactaaaagaattaag
atctctttcaaaaatttgacggtatgctgctcgtaaagtttgttctaatacatctttttt
actatcaccattaacttcaaatacaatagactgatctcttagagatgttactccttggga
catacgagactgaatatcattgctacttctcatttcttttacagtccccaattcaacaaa
tctagaagacttgatagcaataactttcttgaatctttggtcaatgatacctggccttaa
gcttcttagcgcgacgccagctggtgtactatatctttcatatggaacagtattatcgcc
aaaagtttctgtatactcagagctatcaattattgcatctatcactgtgtagtaaccttg
cttatatgcaatatcaaaatatttattaatttcttgacggccatatgtagggcgacccaa
taagcggttatgaatatattcaatagctttacagatatataaaggttcccaataaagaga
cctgaatatacttgatttcgctaactgtctaacaaattcacgaactgtaatttgactatc
ttttagctgactttctattggcttaagaagtagtttttcttcttgatacacttctctacc
aaagactcttaaataagcaacttttgtaacaacatctactaagttttcaaaattttgaga
agaatctaactgtacaagagtattagatagtttaaaaattttggacctaaagatcccgc
agacttaggtctgacctgcggattactaatttgattgtagattcctggcccacgtctaac
taggatcctcctagtatctttaccaaaaagagcttgtctttttctaggatccttattttc
tttaggaaagatagctccaaattgaattgacaatggatcgttacctattccgtacggatg
ctgatcaggtaatgattgtttatagtcactaaataaagttataaactgaggtactttacg
aaaaggagcactatagtttagaagatctatttgaggtccccaatttcggcactcttgagg
ctcttcacctagatttctaaagtaaggtactgtttcttctccaaaatagtctgtatactc
cgaagaatttaaaattgcatttactaaaccacttaaaccagtagatgataaaatagaaaa
atatttttgaaactcttctaaggagctagggcctctacctaagaaatgtctaaatgctaa
ttccaatgctctactatttacaaaaggctcataaaattgctttctataaatacttgaagt
tcctaatgaacgaataaattccttaattgagatttggccatttttttacttgagactctaa

gttagataaagaaagatcatatgctttagcaatatctctctcgaaaatttgtctgtaaca
cgctttaacaacagtgttttttttcatcagcagacaaacttggcttcataacaaaacgagg
tgttgatactcctgcctttacataagtctgcggaaggcgtaaaccttgtaaatctccaga
tattctttttcttaatttatcagtcaaagatggagcttcaaattcagaaattacaacatt
aaaatactccttaactaaatcttgagctttaatatcttcttcaaatattaataaagctgt
ccgtctcatctctctaagtgctacaattgctgctgcgcttgagcaagcattatcaattaa
ctctctcaatcctctaatattaacagacaaaatattaggatcacctgacacaattgcata
tgttaaataccgtaaaaaccagtctaaatctcgaagagattttctcattcttgtagtacc
atatctcaagacatttataggtttaaatcctggaggtgtagctcccccagcattaaataa
tgaacggaaactttgtccaaaattcccttggatatttccagataactcattgattttatc
ttgagaactttgatctcctgcaatcataacagctgcttgtggacgctctagataagaaat
tgcagatccacctacgaatattttatctgcggctcgagctaccaaaatatttgcattttt
agttaaaatatctgcaacttccaatcttttttgtccagaatttaaaaaagaaactaattg
atttaattcaccaagctgtaaaaatcggtcttgctgctctgcctgagtaatagttaaaat
tgaggcagtccgataaagttgggggcgtgctaatgggcttccgccacttgctttgatact
cattactttatttatctccttaacttatattgcttgctaacgaatacaaatatcgcttga
tatttctaactatttaaagatactatgcaagcaca-ttattttctatattattagatact
agtacttatattacagagagtcttacaactaacttctataaatacaaatttgtaatact
tctattaacatttatccaaagtatacgtaa-tgatcattcttttatattaaaatgcaatt
agtaagagattgaggttaaatcaagtatacttagtaagttttatagatagtacaattaga
atatttattctacaaacattaatca------------cttgactgtttgacaattagaca
attgttcttgcttttaattggtttagactagtttcatattttttctaacataaacaatat
taatgagcaatttgtatgaatttagaaccacagaacaacctattaactaacgaaaatata
aaccaacaagatatacccgagaatgatataccaatgtccattacggaacatttagaagaa
ttaagacaacgtactttatttgtatttttattctttttagtggctgcaactgttagtttt
acacaggttaaaataattgttgcagtacttcaagctcctgctattggtattaaattttta
caattggcaccaggagaatattttttttcatctattaaggttgcaatatattgcgggatt
gttgctacaactccatttggcttatatcaagttatattatacatactgcctggactaact
gaaaaagaacgaaaaatcattttacctatcttaattagttctgtattgctttttataaca
ggcggtatatttgcttactttattttagcaccggcagctttaactttttttaatcagctat
ggatccgatattgtagaaccattatggtcttttgaacagtactttgactttattctactg
cttttatttagtactggattagcttttgagataccaatcatacaattattgcttggtgtg
tcaggaacattctctgctacgcaaatgatccgggcttggagatatattattattatagca
acaattgctggagctattctaaccccttcgactgatccagttacacaattaataatgtct
tctgctgttttattactttattttagtggaattattatattattgatcttaaaaaagtaa
agttagt--------cgttttctatttaattgagtatatttagt--------------aa

taatcttt--acataattattaatttttagcttaaat-cagactcaagtactaagctttt
taatctttttat-------tactttgtcttatgattgaaacaagatttaagaaaaggtg
gacttagctgtatttcagaatagtattttaagacatgttgtggcaaaaattcacaatctg
aaagaaaaacaataatataatcatataaatctgcaggactaaatttagacaattgcagtt
ttccaaactttttttgcgatagtcgataatcagacacttcgctgcgtttacaatttaatt
taggaaatattaaatcaataacaaccatattagatatagatgcattgacatagttcatat
aatatttaggaaggttaaaatgcggggacgtttttttaataatccacctaattaacaaat
tttgtcccagtgcgtctataaatcaactctcgtaaaatgacgaaataagggtgtaatata
ggggaataacacccctacgctgcgaaataatcatggctttatattagatagctacattgg
cctatttttttcattaattatttgcgatctacactttttttaccgatctatttattatgctt
ggcaatctttctttaatctatttactaaataaagagagatgactaaaattccttattgat
tatggtaatctataccaacgtaaaaagtattttattgtgagttttatttatataaaggct
tattcattatatagttataacagtaagtctaattaatagtttttcatgcttaattaaagt
aataactcttaattcttttggcagaaaaatctgtcttaatattttatc--aaaaaaacag
-aaaaatgaagctaaatagtctaattaacttaattcaaaagtctatatgttcttttacac
ttttattaattattttaaatattatttgtgtcgcacctaactctagtaatgcattcccaa
tttatgcgcaacaagcttatgaaagcccaagagaagctactggtaggatagtgtgtgcca
attgtcatcttgctcaaaaaccagtggaaatagaagctccccaagcagtactacctaata
ctgttttcgagactgttgtaaaaattccatatgacagcagtgctaaacaaattctaggta
atgggagcaaaggaggattaaatgtcggagccgttgtaatattaccagaaggatttaagt
tagctcctgcgaatagattatctacagagttaaaagaaaaaactaagaatctttacattc
agccatatagtgcgaaacaaaataatatttagtaattggacctatctccggtgataaaa
atagagaaatagttttcccaatactatctcccgatcctgcaaaagataaaaaagctcact
ttttcaagtacccaatatatgttggtggaaatagaggccgaggccagatttatccaacag
gtgataagagtaataacaacattgtcgctgctttaagtagcggtaaaattaataaaattg
aagccctagacaaaggtggatttataattcatgtgactaatagtagcaatatagaatcaa
cacaaaatgttgccccaggtcttgaacttaaggttagagaaggtgatacagttcaattag
atcaagctttgaatagtgatccaaatgtaggcgggtttggtcaaaatgaaacagaaatag
ttttacaaagtccaaacagaattaaaggcatgattgtcttcttttttgcgagtgttttag
ctcaaattttctttgttttaaagaaaaaacaattcgaaaaagttcaagcggctgaaatga
atttttaaaacaagataaaattggtacattctataagtaaagtt-caaacactttattta
tagaatgtaccagttgttatattatacaatatttgcttttgaagatgttattatattttt
aacagaatctacaatttgactaggctgaatcactgtggcctgctctaaactaccattata
tggtgttggtatatcttgcgaagatagccttactaccggagcatctaattcatcaaaaag
gtattcattaatttgtgcaattagctctgctccaattcctgctgttttcatacactcttc
tacaattaaaactttgtgagttttcttgactgaaacagatatagattctatgtccagagg

ttttaaagatataagatctatcacttctggatcataatcttcttttaacaaagcaggtaa
tgcttgagtaacatggtgcctcattctagagtaagttaaaatagtaatatcttttccttt
tcgtacaacttcaactttgttaagaggtaagaaatattcttcttgaggaatctcttcttg
taaattgtaaagcaaaacatgttcaaaaaagacaactggattattatctcgaattgcaga
tttaagtaatcctttcgcattataaggagtagaacaagccactattttttaaaccaggtat
ggcttgaaaataagcttctaatctctgagaatgctccgcacctaattgtctaccgactcc
tcccggtcctctaataactaaaggcaatgtaaagttgcctccagaagtatagcgcaacat
acctgcattattagaaatttggttaaatgctaataataaaaaactcatgttcataccttc
aacaattggtctaagccctgttatagctgcgccaattgccataccagtaaagctattttc
tgctattggtgtatcaagaactctcaaatcaccatatttactatgcaaatctttagttac
cttataagatccaccatagtgaccaacatcttctcctattacacagacagttggatcttt
ctccatttcctcatctgttgccgctcgtaaagcgtcaaacataaagatttactcatttt
aatatttgtttttatgattccactactgttttaaagtaacagttaagactctatattct
ctttagttatctgcaaaaagatatcgcttgagttctgagatattaggctctggactggca
attgcaaattggacagattgatctaattctgttttaatagtagtttgaattttatcaagt
tcttttatatccgcaatttcattatctagaatgtattttttttaatttttttaataggatct
cgtgcaacccaagcctctttttcttgtctcgatcttagttcatcaggatctgcaagagaa
tgaccacggaatctatatgttaatgcttctattaatgttggaccgtcaccttgtcgagct
ctttgaactgcttgttgtgcagcttgtcttacagctagtacatccattccatcaacttca
ataccaggaagaccaaaagcttctgctttttttatggatttctggtattgaagaggatcgg
tgatgtgccatacctatagcccactgattattttcaacaacaaatataataggtaacttc
cagagaactgccatatttaaacattcaaaaaaattgcccattattggtagtaccatcacca
aaaaagcaggctgtgactctcaactcttcctttttttttcagtacttgttgacgatagata
ctttggaaagctgcccctgttgcaacaggtatgccttctgcaataaacgcaaaaccacct
aagaaattgtgaggtgcagaaaaaaatatgcatcgaacctcctctacctttactacaacca
gtttcttttccaaataactctgccatcacatttttagatggaacacctttacttagggcg
tgtacatgatctcgataagtactgcaaacataatctgtcgaattcagaagcttaattaca
cctgtagagacagcttcttgaccattataaagatgaacaaagccaaacatcttccctttg
taatacatctgagcacacatatcttcaaaatttctgcctaacaacatatcttcatataaa
actaataaattactctt------attaagacctgttgaattatagttagtcagcggcaat
tgaactttcttcggataactcataatttataaagaaacctct----------tattatat
ataggaattttgaagtaaaagttatcaatagaaaacacacagataaaaacttataaatca
ttatatgctttttactaaagactttaataattctttgaaactagttaagtatatcaaatt
aattttgcttatactacagatgtcttaaacttagattaaaaatgagtaatagtaaattta
ttaaaagaatagttaaaaataaatataaatagaagtaaacagaattattggaaagaaaat
gattattattaataaataattacttaatatttctaaatatttttaaattaatggctatgt

catcatctttattttatcctgttgagaaagaactatatagtcttgaaaaaaatctgaaag
cagttgccgggactcgtcatccaattctatatgccgcagcaaagcatctattcgatgctg
gaggaaaacgagttagaccagctctcgtgtttttagttgctaaagcaacatctgagaagc
aagatataaatactggacaaagaaggctggcagaaattactgaaattatacatacagcga
gtttagtacatgacgatattattgacgaatgtacgacgcgtagaggagtcaaaactgtgc
ataatttgtttaatactaaaattgctgtgttagcgggagatttttttatttgcacagtctt
cttggtatttagctaatattgaaaatttagcagtagtaaaagctatttctaaagtcatca
ccgattttgcggaaggcgaaattagacaaggcttagttcattttgatcctaacatttcaa
tagatgcttacattgagaaatcattttataagactgcttcactcattgctgctagttgtc
gaggtgcagctatgcttaatgggtctaataatcaagtaaataatgatctatatctttacg
gtaaacatatgggattagcatttcaaattatggatgatgttttagatataaccggttcta
ctcaaagcttggggaaacctgctggcgctgatctaataaatggtaatttaacatctccta
ttctttttcacttactcaagaagcaagtttaaactatcttattgaaagagaattctgca
ataatacagatatagcctcagcattatttcttattaaaaaaagcggagggattacaaaag
caaaagatttagcgaaagaacaggtgcaggcggcactttactgccttcggtttttaccaa
aatctgcacctgtctctagtttaaaagaattgacacatttcataatcacaagattgtcat
agaatacttgctagaattaaaatgttattaactaatttttttactgtttctaaaatagtct
gaaaagtttgaatatcattactagctagctgcgctagcattttgcgatttaacgcaatat
tctctttttttaatgcactaataaaagtactataattcatgccttgaatatgagctgctg
catttattctagttatccaaagccgacgaaaatctctcttttttcctttttcgacctacat
aagaatatcgaagagctttaaggacttgctgtttagctgtccgaaataagcacttatgtg
cacctttaaagcctttagctagtttaaatatttagctcttcttttttttgcaacgttac
ctcttttgactctactcataaaatattat-ttaaaattaaattattaccagattcctaaa
ttcacatacttatcg----attataaataaggtaaatttatagcaatgtttttaatatct
cttaagtcaactgaacaagtagaagaaagatttcttctttgttttgatgatttttttttgt
aataaatgacttttagaagctttatgtcgcagaaattttcctgatgaagagactttaaat
ctttttgctattgcttttgatgtttttaacttaggcatattataaactttttaatgatcaa
caaaagtatgaaaagtctgaaggtatcaaacttagactcttcaatacttaagataatctt
aacacaaattattttacttattatatagattaactgaaacaatttatatata-aataccc
acaagttaatcaactgcagctaaagttgatttgtattttgctttgaacttttgtatagta
ttagcaattagcattgttattattttaagagtacttaaatttaattcttcgaacattttc
atatttaatgtaaatgaaatattcgcctctgagacgatattctgcatttgaacatcagat
aaaggaatcacatctaatgcagccctatattgatctttaaacaatttatcatcttttatt
tggtcaaaatcataaaatttagtacctccttcatcagataaattcatagcacctctagct
atttttttttaagatttggccaccagacaaatcacctaagtagcgtgtataggcatgagca
actaataattcaggttgcttaagtcctatactatgaatcctctcaacatatattttttgtt

gcaggagatggttcaataagatttaaccaatctgatccataataatagtttaaatcttcg
gacagacttatttttctattgagttctgtaaaatatataggtttaatagctggatggtct
ttattagagaataactcttcttctattgcacaataaacaaaatatagattagcaaccaat
ttacgatatgatttttatctacaacacctcccaaaaaagatttcacaaaactaacattt
tctgccatactatgggacttagtagtaccttctcttaattcattcgctaaagtattaacc
ataatatcaatcttccattttt-aaagataacaaataaa--------------aatatat
atatttatatttcaattgtactaactaggcagctgcgaaataatttttttgattttattgg
gtctggattcattgttttatcaccaggcttccaattagctgggcaaacttcatctggatg
ggattgtacatattgaattgcttgtaaaaccctcaaagtttcttctacactcctaccaaa
ctcgagattattgattgtagagtactgaataatgccttttgggtctataataaatagtcc
tcttaaagctacaccgtcactgtttaaaacattataagcagcactgatctcttttttttaa
gtctgatactaatggatacgaaagatctcctaatccaccagattctctgtctgtctgtag
ccaggcaagatgagagtattcactatcaaccgaaacacctagaacctctgtattaagctc
ggaaaaagcatcatacttatcgctgaatgctgtaatttctgtagggcaaacaaatgtaaa
gtctaaaggataaaagaataagacaatatacttatttttttaaatctgacaattttagcgt
tttaaattcttggtcataaacagctgtagctgaaaaatcaggagctagctgacctactcg
aagacaatttggtccagaaatcattaatttttaccagaagaattgatataattttgttat
ttaataaataattatataacaaaatttagtatatattaaaatactaccagtctttttaaa
attac--------tacatgatataataaatactctctttatagcggggaatggatttgaa
ccattgaccttcgggttatgagcccgacgagctaccagactgctctaccccgcgaataac
taatattatattagtctatctatatatataaaagcaaatatatgatggatgttagaagta
aaagaataagagaacaggagcactgataattaagcttaaaaatgacaaagtatttttttat
tttatttaggagaggaaagacttcgtataaaccaataaaccgatccttaataagaatttc
ggatggcttcacccaaatctgaccatcgtaccagcctgactcttcataaaataccgtagc
actcattaagcgttttactacgtatgaccagcctaaatataatctaattaaaataaatcc
tgttattaagctagatgtgataaattctgaaaagaaaaatttcaaaggaagcttggtaat
tggaaaaattgatagtaatataggactaactaataaacaattaagcattagtgcaatagt
gatttttttattatatgatttgctgcttaaagttggccaacaaaagaaccaagaattttt
caaagaagtatattcattaacaggttgttgctcctttggcacaggacattgagtattata
taaattcattgacattaaaaatgattatactgaaaat--tcactttatcttatctaaagc
agtaccaa--------gttaattaactgaaattatagtggtgagtataagaaataaaatt
agaaataaccaagtaactttatttaaagtattttcagtactacgagtattactaaaaaat
tgattctgagctccgacacttcctagaccttcagatttaggattatgtatcaggattgta
aagattaagattattgtcgatgcataccaaaaaaatttttaaaatttgtgccattgtgtaa
aagttaat-agaaatcactcaaagacaatcgtctaaaagatattgattgtcttatcatat
taaaaaatatttattcaccctgtacttttaaaagtaaaaatcctattgctagccctacta

aaactaggacagaacttaatacggcactttctacaatttctccgatcatt-ttatttttc
cttattatttattattatttcattctcttgataagcttacaactctctaa-ttaaaat
ccgtttctaccccaaactaccattgcaatagagaaggtaaacattgccattagagctgcc
catcccaaacttaaaatatccataatttataatattc--ctaaaaacaagatctcttta
cttaaactataatattcttaatattaaaaatatcattatataataagctaattataacag
agtactacttaattcgtattttagtgtatttaaataagattgac-tttttccaaattac
ttaataagtatttcatcctagaaaccatcttcaat----ttttgaattattatacttatt
aaatctaaatgt------cactcaaaaattgttagtaattgtgatgttttcaattatacg
att---aatatatataaaaacatcaagattcatagcagctaagttaatggtctaatctaa
tataatttatcaat--aaaaaggaagttaatatttcaattatgcaaacaactataagtaa
tgggcaaacgtctagtaaagaaactttactaacacctagattctatacaactgactttga
agaaatggctaatatagatatttctgagaatcaacaagatttctttgctatcctagaaga
atttcgagctgattataacagtgagcactttattagagatgaagagtttaatcaatcttg
gtctaatttagaaaataaaacaaaatccttatttattgagtttttagaaagatcttgtac
tgcagaattttcaggtttttttgctatacaaagaattatcgagaaaattaaaagatagaaa
tcctattattgctgaatgcttttttattaatgtctagagatgaagctcgccatgcagggtt
tttaaataaagctataggagattttaatttatctttagatttaggcttttttaacgaagag
tagaaaatatacttttttctctcctaaatttattttttatgcaacctacctttctgaaaa
aataggatattggagatacataactatttaccgccatcttgaacagcatcccgagcatcg
catttatccaatctttagattctttgaaaattggtgtcaagatgaaaatcgtcatgggga
tttttttgccgcactactaaaatctcaaccgcatttttttaaacgactggaaagcaaaaat
gtggtgcagatttttctattaagtgtatttgcaacaatgtatttaaatgactttcaaag
aattgatttttataatgcgataggtctagattctagacagtatgatatgcaagtgatacg
aaaaactaacgaaagtgccgctagagttttccctgttgctctagacgtggataatccaaa
atttttcaaatatttagatacttgtgcatgtgataatagagctctcatagatcttgataa
acacaattctccattattaatcaaagctattatgaaggtacctatctattactctctaat
aagtaatttaataaagatatatttaattaagccaatagattcaaaaacagtatggaatac
agttcgataaccgaataaaataagaatactgattaatacagcaagttatctaattattaa
tgtgaaat------aaaagactccttcc-----atcaggaaggagcttttattttatcaa
ga--------atatgattatataaatc-taagattaagaacttactttttcttttgcttc
gtacataacctctaaagtaattagattttagaacaatcacgtgcaaatttctctgtatt
tctttttactttacctctaacgaaccctggaatttatttaattctttttgtgcttcttg
tgaccaattaatactatcgactgcagaaattcctacacttaatgcttctgttgtatcatg
accaccaaatatttctaataagtgatcttccatacccagtgtaaatgaattatagactaa
atcagcaatttgatttgtaccctcataaccaagaaatggacggtaacttaaagggaaatt
ctgtatgtgcactggtgaagaaatgacgccacaaggaatgtttaatcgcttaccaatatg

tctttccatctgtgttccaaaaattgctgcaggttcagttttagcgattaaatctccaat
taaaccatgatcatctgaaataatcacttcatcgcagaattcttgaacttggtctttaaa
ccactcttcatcatatttacaataagttccacaccaagcaacatgtatacccatttcttg
atgtaaaattcgagttattgctgcggcatgagttgcatctccaaatacgatggcttttt
gccagttaaattctgacagtcaattgatcttgaaaaccatgcagattgtgagataaatct
agtttgttcatctatgtatttttcgtagtctacaatagctcccgatgcatttaatagctg
ttgtattgatcggatacatgtagctgtttgaacaatacccataggtgtagtgtcaatata
agacatattgaattctttctctaaataccgagctgtcattagtccagtttctctatatgg
aataaagttaaaccatgcagatggtaattttctcaaatcttgaaccgatgcattttcagg
aataatctgattaattttgatatctaaatcttgaaataatctttttaattctgcaatatc
gtgctgattatgaaaccctaagcttacagctccaataatattaacagacggggttttcgt
tttttgagttaaaacctgattattagatttagctttttccatgtaaaaagtgacaatttg
ttctagagttctatcaccagcttgcagttcatttactctataatgattaacatcagctaa
taaaacatcagcttctgtttctatagatgctctgctaacaaaatttgtaaatcttcttg
caagatactagaagtacaagttggtgttaaaataactaaatccgggttttcttctctatc
ttttcttgtaatgttctctacaacttttcctgcgaaccacgtgctagtacatgtctatc
tacaacacttgctgtaacaggggtgaaatctctgtctctttcaagcattgaacgcattac
attaaaataatcgtctccgagaggcgcatgcatgattgcatgaactttttaaaagaact
ggcaattcttagagttccaatatgagcagggcctgcatacatccaataagctaatttcat
attaaattttcctatgatttatttagacatagtaatactttttaaagccaactaacaaata
caagccagacatctaatttaactcgttgataatttttaaca-ttttcagattaagctga
tcataatatatatttaaaaag-------------agaaataaatgcctaatagataaatc
ttaaaagtaaagatttcttgcagttatagtttttctttaagtaaacatgtaaaaattgta
ctatatataaatgtactatatataaactttttttactgaatattggtaatgccagataca
attaacttaaatatgccttccccaacgtttggtggaagcactggcggttggttaagagcc
gcagaagtagaagaaaagtatgcgataacgtggacaggcaaaaatgaaagtaagtttgaa
atgccaactggtggtacagcaactatgagaaatggggaaaacttactttatttagccaaa
aaagagcagtgtttagctttaggtactcaattaaaaagtaaatttaaaatatctgattat
aaaatttatagagttttccccaacggagaagtgcaatatttgcatcctaaagatggagtc
tttcctgaaaaggttaatgctggaagaactagcgttaacagtgttgatcattctattggg
caaaatgttaatcctgtagatgtcaaatttatgaataaagcaacttacgattagtttatc
-aaaaacctagacataaaatagcttttatatagtctaggtttttgataaaatataccaat
attatggaaaggtggccgagtggttgaaggcgtctgacttgaaatcagttgaactg--ta
tggttccgtgggttcgaatcccaccctttccgttataaaact------------atagtt
tgcttattgcagca----agctatacactaatagtattaagtttagacttaataaagtcc
acaacttgtgataagttcgaaatttttcggcatcttcatctggtatctctataccaaat

ttttcttctattgccatcactaactcaactgtgtctaaagaatcggctccaaggtcactt
gaaaaatttgcttctctagtcactatcttcttttcaattcctagctgttctgctacaata
tcttgaactttttcaaaaatttcattatcttgcataatatattatttcctctgaggtgtt
atcaaaaatttttcggctcatattattatataactctaacgaattagtaattttaagtcg
atatcaaaatatttatacctatatgatatactattattctggataaatacgaagtttacg
atatggctcttctccaaaactacgagctcttaactggtagttagcaactaagttatgctg
catttttcggatataggctgagcgaggtgctagctgaactattgaattttcattcaagat
aatagtctcaatagccagtttagcttcttgaagagcctgaatctcatcaaatttttttatt
tttacagaattctaaccaattaagaccagaagaagtatgaatatttaatatttttctgag
tgcacgagtaatttgtggaactgtactattctgaattgtatatataataatttgttttga
ttttgcaatttgtcttagttttgtattttgctttacttgattacgtagcgctaaaatagc
atctgacttttctatctccttagttaaaataattggtaaatctagggaagaaataactga
tgtaatatgctgccaactcaaagaataagcatataaatactgattaggtatttctaatgg
caaggagtgtttattaatattgataggagcattaacagtagttgataataaagatgtatc
acgattctgtgatttattcaaatctgatgacttatttcttaattctctgtattgtagaac
atgatgcgcctgctgtaaattaattgctttacgtgcactagaaatacttgcaggcaggac
ctctatagactgtgatggataacatttaattaaaatcctgccattagcttgaatttgtct
tttttgtacaaaaggttgatgtccttgcaaaatttgatctatagtttctttaactttatc
atgtacgatccaaacattgcgttcatgtatttcaattgcaatttgaaaagcaggcgcagc
ttttctttctagtatactttctgactacctcttctttttgcttcatcatcacccaatgt
aacatattgaattcctccaattaagtccgctaacgtaggatttttaattagactttctaa
atgatttccatgtgctgtacctactaattgcactcctctttctgctatagttcgggcagc
taatgcttctaactctgtaccaatttcatctataataataacttcaggcatgtgattttc
tacagcttctatcattacctgatgttgtaaatcgggccttgcaacttgcattcttctagc
tctacctatagctgggtgcggaatatctccatcaccagctatttcattagaagtgtctat
aattacaactctttttttccatttcatctgccaaaacacgagcaatttctctgacagctgt
tgttttaccaactccaggttttcctaagagtaagatggaatcaccttgttctagtaaatc
tcgaataatactaatcgtaccaaagactgctcttccaacacgacaggttaagccgataat
gctaccttctctattacgcattgaactaatacggtgtaaggtttttttctataccagctcg
attatctccgctaaaattaccaactttttttacacaatagtctaagtcttgccaactaat
agatctttgagataagtattctggattatctggaaatctagcttctggacgacgacctaa
atccataaccacttctattaaattatttctattaggatgttgttttaaaggctctttaac
aaaatttggcaaaatttctagtaacttatctaagtcatctgcaataagcatgattgaata
tataataattaattaatcaaatagaagattcgagattctaaagattacaatacatattaa
agaatattatgcctggaatcaattttatacttaaattaaataagaagtta-cagatgtaa
aaaaataaattaaaatttgtacaataaatagattggtcaaaattaatagtcgtgttaaca

tgtttattggacagaaagttcgtattaagtatagtaaacaaaaagttgctaatgatatag
cagataaagttggcgagcttggagttattaaaggaataaagtttattaatagtcaatgtg
ttactattattgttgaatttgacaatcacactaggctttggatgtttagagaggaattaa
tttatctaaatgaat------taaaaaaatgagataaaatttattt---------taaag
aatttcttatgagttgttaaaatactaaattttacaggagtattgttaattgtgcaaatt
actattaaaaaactacaagacttactttcatctgtacaaaaaaaaaagatacagacatta
aaaataaaacaagataattttgaacttctattaaataaaacccataaaaagattaatcaa
gaaatattaccttcacaaaact---ctatagtacagaatagtccagctacaataattaaa
ccaatacctaataaaaaaaacaattctgtcattaatgaagactttacagaatatgccact
atcgtttctccaatggttggaacattttatcattcgcctgctcctggtgagaaaattttt
gtacaagtcggcgacgaggtcaaattcaatcaaacagtttgtattattgaagctatgaag
ttaatgaatgaaattgaagctgaaattgaagggaaaattatagaaattcttgttaaagat
ggcgatatagtagattgcggccaggctttaatgaaagttgatacataatgctaatctttc
cattttacatattgttaacagaaattatttagataggaaaattagcttatactatagtta
tgaaaacttatttcgtaaaggcac-aaaaagctagtatataaaactatttactttatctg
ccaaaaatagaaataagcgttttaattccttgtctcaagagaggagaatctcaatggcaa
ttagctcaaaagagcaagagacaaagaaggtaaaaatctcggttgataaaaatcccgtag
atacttctttcgaaaaatgggcccaacctggccatttttctcgtacactagcaaaaggac
caaaaactactacttggatttggaatcttcatgctgatgctcacgacttcgacagtcaaa
ccagttctttagaagaagtttcacgtaaaattttcagcgcacattttggacagctgtctg
taatatttttatggcttagtggaatgtattttcatggagcacgattctctaactatgttg
cgtggttaagtaatccaacaggtattaaaccaagtgcgcaggttgtttggcctatagttg
gccaagagattttaaatggcgatgtaggtggcggatttcaaggcgtacaagttacatctg
gatggttccagctgtggagagcatcagggattactaccgaatttcagctttactgtactg
ctattggcggattagctatggcagctttaatgttatttgcaggatggtttcattatcata
aagcagctccaaagttagaatggtttcaaaatgttgagtcaatgatgaatcaccatttag
ctggtcttttaggcttaggctgtttaagctgggcaggccatcaaatccatttatctttac
ctattaataaactgttagattctggggtatctccacaagagattccattacctcacgagt
ttttaattaatagagagcttatggctcagctgtatccaagttttagtaaagggttagttc
cattctttactttaaactggtctgaatattctgactttttgactttcaaaggtgggttaa
atcctgttactggaggcttatggctaagtgatactgctcatcatcatctagcattagctg
ttctatttcttgctgcaggccatatgtatagaaccaattggggtattggacatagcatga
aagaaattttagaagctcacaaaggaccttttactggtaacggtcacgaaggtctatatg
aaattcttacaacttcttggcatgcacagcttgctattaatttagctatgatgggctctt
taagcataattgtcgcacatcatatgtatgctatgcctccttatccatatattgctactg
attacccaactcaattgtcactattcactcatcacatgtggattggggggattttgcattg

ttggtgcaggagcacatgcttctatatttatggtaagagattataatcctgcagaaaatt
ataataatcttctagatagagttattagacatcgagatgctatcgtttctcatctaaatt
gggtatgcatatttcttgggttccattcatttggtttatatatccacaacgatactatgc
gagcacttggaagatctcaagacatgttctcagatacagcaatccagttacaacccattt
tcgctcaatgggtacaaagtatacatactttagctcctgggaatacagctccaaatgcgt
tagcaacagctagttatgcatttggaggagatgtcgtttctgttggtaataaagttgcca
tgatgcctatttctctaggcacagcagactttttggttcatcatatacatgcttttacta
ttcatgtaactgttcttatcttagtcaaaggtttccttttctcaagaaactctcgactca
ttcctgataaggtaaatcttggctttagatttccatgtgatggacctggcagaggcggta
cttgccaagtgtctggctgggatcatgttttccttggcctattctggatgtacaattctc
tatctgtagcaatttttcactttagttggaaaatgcaatcagatgtttggggtagcgtat
ctccatctggaaatgtttctcatattactggtggtaattttgctcaaagttcaattacaa
tcaatggatggttaagagatttcctttgggctcaagcatctcaagttattcaatcatacg
gttctgcactgtctgcatatggactaatttcttggctgcacatttcgtatgggcattca
gtttgatgttcttatttagtggcagaggttattggcaagaacttatagaatcgatcgtat
gggcacataataaaataaaagttgctcctgcaattcaaccaagagctttaagtattactc
aaggtagagcagtcggtgttgcacactacttattaggtggaattggtacaacttgggcat
tctttttagcaagaattatttcagtaggctaatagtgaaaataggat-aaaaaaacaatt
atggcaacaaaatttcctaagtttagccaagctttatcacaagatcctacaactcgaagg
atttggtatggtattgctacggcacatgactttgaaagtcatgatggaatgacagaagaa
aatttatatcaaaagatatttgcttcgcactttgggcatttagcaatcatctttttatgg
acatctggtaatttattccatgtagcttggcaaggtaactttgaacagtgggtattaaat
cctttaaaagttaaaccaattgctcacgcaatttgggatccacattttggacaaccagct
ttaaaagcttttagtaaaggtggctcagcatatccagtaaatatcgcatattctggcgta
tatcactggtggtatactattggtatgagaagcaatcaagacctatattctggggcttta
ttcttactggttttatcagctttactcttatttggaggatggctacatctgcagccaaaa
ttcaagcctggtttatcctggtttaaaaataacgaatcaagattaaatcatcatttatct
ggactatttggggttagttctttagcatggacaggtcatttagtacacgttgctatacca
gaagctcgaggacaacatgtaggatgggataatttcacaaaagtattacctcatcccgct
ggcttacaaccttttttagcggtaactggagtgtatatgctcaaagtccagatacagct
caacatctattcggaactaatgaaggtgcaggtacagccattttaacgttcctcggtgga
tttcaccctcaaagtcagtctttgtggctaactgatatggcacatcaccatttggctatc
gctgtagtcttcattgttgctggacatatgtatagaactaattggggtattggacataat
ctcaaagatatttagatgctcatagaccacctagtggtagattaggcgctggacataag
gggctatttgatactattactaattccttgcatatacaattgggattagcattagcttca
cttggtgtaattacgtctttagtagctcaacatatgtatgctatgcctccatatgctttt

atggctaaagatttcacaactcaggcttctttatacactcatcatcaatatattgctggg
tttctaatggttggagcttttgctcatggggcaatattctttgttcgggattatgaccca
gaacagaataaagataatgttttagctcgtatgctagaacataaagaagctatcatttct
catttaagttgggtaactctattttagggtttcacacactaggtctttatgttcacaat
gatacaatgattgcttttggaactccagaaaaacaaattctaattgagccagtatttgct
caatggattcaagcttcttcaggaaaagcactttacggttttgatgtattactatcatct
tctagtaatattgcaacacaagctggtagtaatatttggctacctggttggttagaagca
attaatagcggaaaaaattcattgtttttaacaattggtcctggtgacttcttagttcat
catgcgattgcattaggattacatactactacattaattttagttaaaggcgctttagat
gcaagaggatctaaacttatgcccgataaaaaagatttggttacagttttccttgcgat
ggtcctggtagaggtggcacctgtgatatatcagcatgggatgctttttacttggctgta
ttttggatgttgaatacaataggttgggtaacgttttattggcattggaaacacattact
atttggcaaggcaatgcaactcaattcaatgagtcttctacttatctaatgggatggttt
agagattacttatggctaaattcttctccattaattaatggttataatccatatggtatg
aataacttatcagtatggtcttggatgttcctatttgggcatttagtatgggcaacagga
tttatgttcttaatctcctggcgtggctattggcaagagttaatagagactctagcatgg
gcacatgaacgtacacctttagcaaacttgattcgttggaaagataaacctgttgcatta
tcaattgtacaggcaagactagtaggtttagcacatt-tttccgtaggatatgtattaac
ttatgcagcttttgtattagcttcaacatcaggcaaatttggttaaactaag-----ttt
taattaattaaa--acaaagaagtcagtcataaaa-ttattatgactgacttctttctct
ctataagagatatttattaaagttttacttcaatatctacaccagaaggtaaattcaact
tcattaatgcatctattgtttgagaagaaggttgatgaatatcaataattcttctatgag
atcttatttcaaagtgttctcttgagtctttgtctacatgtggagaacgtaagacacaat
agattcttctttttgtcggtaaagggattggtcctactgcaacagcattagttctggatg
cggtatctattattttatcacacgacgtattaagtatactggaattatacgctttaagtt
taattctaatttttgtctgttgagtaactgtcatattaaaaagatcttattatttaagaa
ttttagagacaacacctgcacctacagttcgtccaccttctctaatagcaaaacgcatac
cttgctcaattgcaattgcattaattaattcagcagtcatcttgattctgtcaccaggca
tgaccatttctgcatccgtaccatcatcagcagtaaactgattaatagtgccagttacat
cagttgttctaacataaaactgaggtctatatcctggaaaaaatggagtatgtcttccgc
cttcttctttagttaaaatataaacttctgcttcaaactgagtatgaggtgtaattgtac
caggtttagctaataccattcctctttcaatatcttttttttgcacgcctcttagaagaa
ttccaatattatcaccagctaaaccttcttctaaagtcttttgaaacatttctaagccgg
taatagtcgttgtccgagtttctcgtaaacctacaatttcaattgtgtcaccaactttaa
tgattcctctttcaatcctaccggtagcaacagttcctcgtcctgtaatagaaaaaacat
cctctacagccattaaaaaagttttatcgatatctctttctggtgttggaatataagtat

caactgcttccattaatgaaaaaatcttatcaacccatttatcttcaccttgcttaataa
tcgggttctttgtaactgcttctagtgctaacaaggcagagcctgcaacaaaaggaatat
catccccaggaaagtcatattgacttaacaattctcgtccttctaattctactaattcta
gtagctcttcatcatctacttgatcttctttatttaggaacactactaatgtaggaactc
ctacttgttttgctaataatatatgttcacgagtttgtggcattggaccatctgccgcag
acacaactaaaatagctccatccatttgcgctgcacctgtaatcatatttttttacatagt
cggcatggccaggacaatccacatgagcgtagtgacgattatcggtttcatattcaacgt
gagcagtattaatagtaatacctctagctttttcttctggtgcagcatcaatttcatcaa
actttttcgctgcagtagtacctaaagttgataaggtcgctgagattgcagcggttaaag
ttgtcttaccatggtcaacatgaccaattgttccaatattaacatgcggtttttttacgtt
caaatttagatcgagccatatttttttgttttccttatagtaaaagttgctttatagttac
ttttaacgataaatcttataaattaacagatttattatttaatgcagacaaaa-----ta
agatatatttaatctaatatcgatagtgagcaaacgctttattcgcttctgccattctat
gtgtatcttctctttttctaatagagtttcctgtttcattagcagcatccataatttcgt
tagctaatttcatggacatacttttacccgatctttctcgagaaaatttagtaatccatc
ttaatgctaaatttgtacctctgtaagctcgtacttctattggcacttgataagtagacc
cccctactcttctagcttttacttctactaaaggagtaatattgcgaatggctttttcta
aaatatttaaagggtctaattctgttctctctttaactatatctagagcttgatatataa
ttctttgtgctaaagttttttttccacttttcaaaatacgaacagttaacatactcacaa
gtctgcttttatataaaggatcaggtgatgcgaacctcttttttagctgtattacgacgag
acatagtatttaaaatttgttattgtaataatttttt-taatagcagatatttaagattt
aggctttttttgtgccatacttggatcgacttttacgacgatcttttactcccgcagcatc
taacgtaccacgtactacatgatatcttactcctggcaaatctttaatacgcccccccct
aattagtaccacagaatgctcttgaatattatgaccaaccccaggaatataagctgtcac
ttcaaatccagatgttaatctaacacgagccactttttcttagagcagagttcggcttctt
tggagtggtagtatatactcttgtacaaacacctcttctttgaggacaactttgaagagc
aggagattttgtttttttatgtatttttcgtctttcggatctaacaagttgttgaattgt
tggcataataaaattaatttaaaatctttaatagatactattggaacctgctttacatga
attaattagtctttattgatcttatatttacgcataaatctttctactcgaccttctgta
tcaataattctttgtgaaccagtaaagaatggatgatttcctgaccagatatctacatgg
agttcaggttttgtagagccaattgtcataattaactggccatcacaatacacttttgct
tctgggtaccaatttggatgtatattatcttttgccattgtttctataattgtataaatg
tatcaataataaatattaacgcttagaaaattgggggggcttttcttgctttctttaaccc
atatttttttctttctttaactcttggatctcttgttaaataccccctcagacttgagtgc
agttctattttctggattaattgaacacaatgctcttgcaacacctaaacggattgcatc
agcttggcccgtcaaacctcctcctctagcatttacatggatatcatattgatttaataa

tcctaaaacttgtaaaggtgcatatgacactcttaagtaattgggactaaattgaagata
agactcgcctggtataccatttatgattaatttacctgagcctgggactagtcgtacttg
tgcaacagaacatttccggcgacccgttccagaatagattgcacgagttttaattaattc
tgtcgacataatattccttaataactaaataaatatacattctttatactatatactctt
gtggcttttgcgcggtatgtgggtgaagtgggccagaatatactttcagctttgtaaata
gttttcttcctaatggacctttaggaagcatacctttaactgatttttcaataattctat
taggtaatcttgtctgaagctgattaaatgtttctacctttaatccaccaggctgtccag
aatgtcttctgtatagtttttgatttggcttatttccgcttacagatacatgagcagagt
taataacaattacataatctccagtatctagataaggtgtataagaaggcttatttttac
ctcttaagatattagaaatatgagtagatattctaccaagtgtctgattttagcgtcta
taatataccaatgagaattagtatttaatgagggtgattgcgttttgttcatgaaaagat
taatacttttagttgacaataagagaatatgtaaatgctgtacatgcaatttctatacaa
gtaatattt-taaaatattc-------------ttgacaaagagaacttttgtaacggtt
atttagattgtttatcagcaagaaaacttctagttatcaataacttatactatttttagt
aatataaaatatattagtcactttttttctttaggcaaacttatacctaatttttttttgt-
agggcatatataacttcttccgctgacttctggccaaagtttttaatttctagcagttct
tcttgagagtaatctagtaaatctgcaatagaatggatttgagctcgctttaaacaatta
taggctctaacagataactgcaattcttctattaaaacttgactaatttttttatcttct
ttactacgataattatctgctgatttaaaatctaaatttcttagtgaacagaaaaggttc
gttaaaactgtagcaccttgacttatagcttcttgaggagagatgctaccattagtccat
atttgtaaaatcaatctgtcttttatgctattgctaccgacacgaacttcttctacttta
taattaactttattaactggcataaagactgagtctacttgcaaaaaatctaccgataac
tcatctactgctttttctgctaaacgatatccgcaatttttttcaattttaaactccatt
tcaaatattgtattattacagatagttgcaatatactgtctaggatctactacttctata
tcagaagataattcaaacaagcctgctgtaattatagctggtccttggactctgattcta
ccaatttgagattccttgttataacttttaaatactacttcttttagattaagtaatatt
tctaaaacatcttctcttaccccaggaattgtagaaaactcatggttcactcctgcaata
cgtacagcaactatagcagtaccttcaagatctgacaatattgatcttcttaaagcgtta
cctaatgtaatgccttgtccttgatttaatgcttctattacaaagctaccgtactgccca
cgcgctccatctgttcttgactctacgcattcaatttgaaattgagccacctaagaaagc
tcc-ttaaaattaatcagtgattagtaattctatctaaaataagaaagaatctatacgcg
acgtttcttaggaggacgacatccattgtgaggtacaggagtaatatcttttattagagt
aatcgctaagcctgcagcctgtaatgctctaattgccgtttctctacctgcacctggacc
atttactaaaacttcggtttgacgcataccttgatccatagcttgtttagcagctttttc
agctgcagtttgagctgcaaaaggtgtgcctttttttagctcccttgaatccacttgcccc
agaagacgaccatgataatgtttctccttttaaattcgtaatagtaacaattgtattatt

gaatgtggatttgatatgtgtaacaccattaactgtattacgtttatttttttcttgctcc
agattttttttagttgtctagccattgtgtctcgtctattatatagttagttaaaagatta
ttttcttggagctttttttcttacctgctactgttttttttacctcctcttcgggttctagc
attggttcttgttctttgtcctcttaaagggagaccaagacgatgtcttctacctctata
agtactaatttccataagtctcttaatactcatagattcaaaacgtttgagatcaccttc
aatctggtaattagattctagaatttctcttatacttacaacttgttgatcatttaaatc
ttgacacttgatatcagcatctatatttgttttttccaatatttcttttgagcgagatag
tccaatgccataaatatatgtcaaagctatctctattcttttgtttcttggaagatctac
tccagcaattctggccactttatattctctccaataaataataattacgttttataacaa
caaattata-agctcaaccttgtctttgcttatgtttaggattagtgcaaattaccataa
cctttctgtgacgacgaattattctacatttttcacacattttccgaacagaaggacgaa
ctttcatattaa----ctctatatactaggaatattactg---------tattaattatt
tacattaaaaatgtcaataatatatttttattattttaactgttgtagcttttacttcgtc
atactatcatatttttttgatatgacatatgtttgaatttgcttagctgtgtctattgca
actccaactagaataagtaaagatgtagccccaagacctcttaagttttggatctgagta
accttttctattataaacggaatcagcgctactgtaaataaaaaatgaggctcctagaaat
gtcagcctatttaatataacttgtaaataatcaatagtcgcttgacctggacgaatatta
ggaatactagcacccattttttttaaatttatagcaatatcttctggattcattactatt
gatgtataaaaaatagctgaaaaaaagaatcaaggcacaataaagaataaggtatagcgaa
ccattagggcaaaatagatatagaatttggagcaatgttttattttgaatgatttgagtt
aaataagatggaagtaccatagatgcagatgcaaaaacaataggcataactccgccttga
tttaatttcaatggcaagtaactatttggatctaaaattgacgattttcctaactgtctt
gctgaaataattttaatccttcttgtaccttcttgtacacaaattgtaattatgatcatc
aataaaaagatcgcaataaataatccaaacttaagacttgcattactatagttggcatca
aaaaatgactgggtaaagtttttgggcaatcctgatacaatgttttgaaaaataagcaaa
gaggctccattaccaattccttttttctgtaattagctcagacagccacataataatcata
gagcctgttgttaaagctaaaacagattcacagacaaacgtaaaattccaattaaacaca
tatggttttacccatatagagattgctccagactgtaaagtggcccatcctaaagctaaa
taccgtgtaatttgagttatttttttggcgtcctaactcgccttcttctttctgtaatttt
tctagatcaggaatgattttcgtgagtaattgcattacaattgaagaattaatatatggga
acaatacctaaagcaaaaatccctattgttgaaaaacctcctccagaaaaaatattcagg
aaatttactaaagtatttttttctacacttgcataaaaagcgtcatggtctatacctgga
acaggtataaaaattcctaatcgtgctaaaactaatagaaagagagtaaatataatacga
tttcttaaatcactttttttggctcataaataaattttaaatgaaaaggtaaacttc--gt
tatacaaaccccctattccaagacttatctaagtttaaaatagggaagaag---aatatt
aaagcaatattaata-cactattctatgtagaataaaggctttctataggaactcctcga

tctttagcggcttcactaaaagttcttaattgagttaagccatttaaaacagctcttgca
ttatttaatgtattattagaacctaattgtttagctaaaatattttgcacacctgacaat
tctaatactgttctgacagaacctccagcaattacacctgaaccaggcgcagaaggcctt
agtataactttagctgcaccagaaattccgttaataggatgcggtatagaatttgatttc
gtcagtggaactgtaactaaatgtttttttagcatccgtcactcctttttttactgcacca
atgacatcactggcttttccaacaccaacaccaacttggccttgctcattaccaacaaca
agaataacgcgaaagcttaatttcttacctccttttacaactttagtcactcttttcact
tgtacaactctctcttcccagccactatctttatcttttcctttgctctgtttttttacga
ttagccattatcagcattattcctaattagttaatta-cgactttagaaacccatacctg
cttccttagcagcttcagccaaggccttaactcttccatgatataattttcctcctctat
caaagacaacatttttaataccttctttgatagactgttcagctaactgcttaccaacta
cgcgagaagtatcacagtttggtcttattttatcggattcttgattatttaaattaacag
atgatgtagcgactaatgtaatgccttgcgtgtcatcaattatttgtgcatatatatgct
tgttagatctaaatacgcatagacgtggcctacttgaagttccttgaactttttttctaa
ctcttttgtgcttatgaattctggtttgcttattgtttagtttcattattatttaccttt
tccagcttttccagcttttcttctgataaattcaccttgatatctaattcccttaccttt
ataaggttcaggaggcctaatagaacgaatagttgaagcaacctgcccaacaacttcttt
atctataccagagacagtaatatttgtattattttcaactttaatttcaatattttcggg
aggtttaattttaactacatggctatatcccacatttaaaattaaatcttggttctcgat
ctgagaacgatagcctacaccttgtatttgcaatttttttagaaaacccattagaaacacc
ttcaatcatattgctaataagtgttctggatagtccatgtaactgactcgccacttttgt
ttgtccacttgtcttaacagagatggtatcattaagtatttctaaattaataccagctgg
caaagtcctggatagggtacctttcggtcctgtcactgtgatggtcataccatcaaactg
agtattaagatttgtcggcaataaaattattttttttccaatacgagacat-attccacc
tctaaaatagttaccaaatatagcataatatttcaccgccgagaccatcatgacgagctt
gtctatctgtcatgacacctctagaagtagaaataagagcgatgcctaaacctccaagaa
ctctaggtagttctttatgatttgcgtagactcttaatccaggtttactaattcttttca
gagcagtaataactggttgacgatttttaccattatactttaaagaaatcattaagtgag
tctctataccttcacctatttgttcaaaattttgaacaaatccttcttcttttagtactg
ttgccatattgcatgtcattttcgttgctggaacttgaacaatttgatgtcttgctaagt
ttgcattacgaatacgtgtcagcatatcggcgatcgtatcgttgaccaccttgatcctcc
ttgcga-----taaattatttaagatattttcaagattctctaaaaggcataccaagctt
ttttaacaaagctaaaccttcttgatctgttttagctgtagttacgattgatatatctag
gcctcgaatttgatcgatattatcatagtctatttctggaaagatcaactgttcgcgcaa
acctaaattataattacctttgccgtcaaaacttttaggactaattcctctaaagtctct
aattcttggcaaagttaaattaattaacttctctaaaaaggaatacatcttgtcttttcg

tagatgtactacaattccgataggaacttcctctcgaatttaaatccagcaatagattt
tttagcttttgtgacaattggtttttgtccagttattagtgttaattcttgaatactact
ttcaagagccttcgcattctgagaagcttcacctaaaccacggttaacagtaattttggt
aaaactagggacttcatgtacattttatactgaaattcatcttttaaagcttgagtaac
agttgttttatatttttcttttaatcctattcccattatatttttataaattatttaata
agttcgccagtctttttcaattttcgaa-ttttttggcctttatcattaattattattga
ggatcggctagcaatattattttgctcactaaacaacatgacattagaagtatgtattgg
agcttcaaattttataatttctcctgcttcaccttcttgttgaggtttttatgttttac
tttgagattaattcctttgacaatcacttttcttgttttatggatgattgcgatgatttc
acctgtctttgttttatcgcttccagaaataactttaactaagtctcctatttttaattt
aatttttgtattaactttacttgttttagaagaacctttcattatactacctccggagca
agagaaactatttagaaaaatttttatctcgtaactctcgagctataggtccgaacact
cttgtgccacgtggattattatcttgattaataataacagctgcattatctccgaatcta
atgctcataccatcagttctcctcaaggctttacgagttcttactacaacagctcggaca
acatcagatcttttgaccggcatatttggggaagcatcctttactactccaataataaca
tcacctatagatgcataagaaggattactagtgcctaatactctaatacacattattttt
ttagcaccactgttatctgcaacattaagatagctttgagtctgtatcatacttttattc
tatataagattaattatcaaaagacttagataggatattaaccattgtccaacactttgt
tttacttaaaggccgtgtctcttgtattgtaacaatatcacctgttgtacattcgttatt
ttcgtcatgtgctttgtatttttttgtccggatcatagttttcgcatactttctatgagc
aattctatttctacagctactactatagttttattcatttttatcacttaccactttacc
ggttgtttcttttaagggcatagtc--------tcttatatat---------aattaaat
tttaagattattatgtagttgaatcagtccgagacttttcaacagttaaaagttgagcta
atctgtgcttagaatgcttaaataagtgaggcttaaaatcttgccttgtagctcttttta
gccttaagtcgaaaagttctcttttaattagaaggatttcttcggctaaagaagaagcct
ccatttttgtaacatctgatattttagggaaggtcatgctttaagattctattgtattcc
gaactataaatttagtcttgattggtaatttataagaagctagtttcatagcttgttgag
cagtttttgtggtacacctgtaatttcaaatagaatatgaccaggcttaataactgcga
cccaatattctggagctcctttaccagatcccatgcgtgtttctgcaggacgagcggtta
cgggtttatcaggaaatactctaatccatagtttgccacctcttcgtacatatctagtaa
tagttcttctagtagcctctatctgtctagaagttaaccatactggttctgtcgcttgca
atgcataatcaccgaacgcaatggtattacctttactagcagaacctttcattctgcctc
tatgttgttttctgaattttgtttttttcgggcttagcataagaattaaaaaatttaaat
aattaataagctgttgaatcaggagtttcctgattatgtacagattctacactcttagat
tctggtaaaatttctcctttaaaaagccaaactttaactccaagtacaccatatgtggta
tgagcttgacgatgacagtaatcaatatcagctcttaaagtttgtagaggtacacgacct

tctctaacccattcacttctagcaatctctgcaccattaagacgaccagatacttgaatt
ttgacaccttgtgtatttgctctttgggctctttgaactgcttgtctgacagctctacga
aaggctactctttttttctagctgctgagtaataaattctgcgactaaagttgcttcagaa
tctggatctgcaatctctacaacatttactctgatttgtttgctaggatctagaatgaaa
gatagcgattttcttagagactcaattcctgctccggattttccaagtactatgccaggt
cgagctgttgctattaagatttctacttgatcaactttacgattaatttcaattttagcg
atactagcattactaagtttagaatgtatatatgagcgaattttgtggtcttctttcaaa
aggactggatagtctttagaattagcaaaccatgaagaacgatgtttttgggtaatacct
attcggaaacccaaaggatgaatttttttgacccacagtatcctttattataaattaaaag
ttataattactaattgactttaaatat----gagactataattctgagacacccaatgta
atatgacaagttggtttatgtattggaaaagctcttccttgtgctcttggctgaaatctt
tttagagttgggcctttgtctgcaaaagctttactaataaataactggtttttattcaat
ccatcattatgctcagcatttgctgcagcagactctaggatttgctttatataagagcaa
actcggtatggcataaattcaagaataattaatgcttcttggtatttcctacctctaatt
tgatctaatacacgacggactttatgtggagaaagacgaatgtatttccctactgcttta
gtttcttttacgtttttttgtaatactcataattatagtttaacgacgggcttttctatca
ccttttatgtgagtacggaacgttcttgttggaacaaactctcctaacttatgtcctacc
atttggtctgatacaaagacaggaaaatgttgtttaccattataaacagctattgtatgc
cctaccatatctggaataatagtagatgctctcgaccaagtttttaggacttctttcttt
ccttcagtatttaatgcttctatacgcttaagaagactaacatcaatgaaagggccttta
tgtatagatcttgacataataataaattattagattcttaataaaataaattctatttac
ggcgacgtaagacgtatgggttactatatttattggggtttcttgttttaacacccaatg
ctggtttaccccaaggtgttacaggacgggctcttccaattggagattttccttcaccac
caccatgcggatgatctactggattcataacaacacctctcactgtaggtcttttaccta
accagcgatttctgccagctttaccaagagtaatattactagcatcaatattaccaactt
gtccaatagtagcatagcattctttacgaatcattcgaacttcactggacggtaatttca
cagtaacaaaagaaccttcttttgctagtatttgagcataagttcctgctgcacgaacaa
tttgtcctccacacgaaggtcttaattctatattatgcacagctgttcctagaggaatac
tggataaaggtaatgcatttccaacttcaataggcgctgtcggacccgaaagaaccatgg
atccaacacgcaaagatcgaggatgcagaatatatctttttttaccatcaagatagtgta
ataatgctattctggcatttctattaggatcatattcaatagaggccactttagcaacta
tattatgtctatttcttttaaaatcgattaatctatactgctgcttatgtccgccacctt
tatgacgacatgtaataacacctctattgttacggcctttgcaaaaatgatgcttaacta
ttaacgatttttctggcttatcagtagtgatctctgaaaaggtagaaaccgttctatttc
ttgtcccaggtgtgtaggcccgatataaacgaattgccatataagaaaatttgagaagaa
taaatgttgaa------ctgtgtactataattaagtttctgggaacaaattaatagaatc

ttttgaagcaagtgttacaattgctttcttgtaatgtggtcgcttgcctacgaatctacc
aatactcctttttttcttaggtgggtggcaagtgttaacacctgtgacttgaacattgaa
aatgtactgtatagcagccttaatatttatttttgttgctttggcgtcaacagcaaaaca
atactgattttcttctaataatttggttgttttatcagtaatgattggatatttaactag
atctaataaacctcttgaatcaatgctatccattatatacctcttgtattttagataaag
catcaactgtaataataattttatgtgcagctaacagagccataatatttaatgtatcag
ctgaaataagttctacattgtgcaaattacgaatagaaagataaacattcgggtcttttt
tatccactataactaatactttttatttaaatcaagattccatcgatggatagcttcca
taaatagtttagtctttggctgctgaaaataactattaaaattttctacaattaacgtat
taacagacttattatttaatgctgtccttaaggctaactgtctttctttttattcattt
tttttgcaaaactacgaggttttggtccaaatattactccaccgcctctccataaaggcg
aacggatagaaccagcccttgctcgaccagtgcctttttgacgccatggtttacggcctc
cacctcggacttcacttcttgttttagtattagcagaaccttgacgcttttcattacttt
gctttactaaagctctatggactaaatacataccagaatcttgactaaccttaagattta
aatcagcattgccacttacttgaccttcccaattatagatttgataatttaattgtgtgt
taactgtcatgagatatatcgtatgaaagaatattgctacttactaattttactaaagc
accaggtttaccaggaacagctccttttacgatcaacaaatcattctctgaattaatact
tacaatttgcaaattttttatcgtaactttttattaccaagttggccagccatattttt
tcctggataaacccttccaggcgttgtaccagctccaatagaaccaggttgcctatggtt
ttttgaaccatgtgacattggacctctactaaaatgatgtctcttttggtagccagagaa
acctttgcctacacttttagaagatacattaatcttttggcctacttggaatagatctgt
ggataagacttgagtgacttcaaaatcatcagtagattttactaagtattcacgcaaata
ttttagtgggggagcttgtgattttttaagtgtcccaacaatggtttatttaacttttg
ttcagcgacttgcttgtacccaacttgaatagcattatagccatcagtagataaagcttt
aatttgagtaataacacatggcccaacttgaattacagttactggaattgataagccagc
ttgatcaaaaaattgggtcatacctactttagtaccgagtataccaacagacactagatt
tctcctttttgagagatttatactatcattgtacataaaatttttacttcaaagtctttg
aag--caaaccaaactgatagatacatgacattaatcatgaaaacataaagataaaaaac
tttacctattcaatctaaaatagaaaaagtttttgtccagttaaagcctctataaatta
gaagtatccttattatattcttgataaactaat-tatctccaatgcttgttcttgttcat
gacttattcgaaacaatgttcacaatacggaatgtacacttttag-----cgttgtagaa
ttagctattattagcaataatcgaagtggagataaaaaaatattcatacttttgtctctt
tacaaaaacaatagaatacatttatgttcggtaactacacctttttattttcatatttat
tgtgcaatatatatatatatccaatacagtatgcataataatataac----tgtatagca
aatgtacttacataaattagagaggtattaatggggaaagttgttggaattgatctggga
acaacaaattctgtaattgctgttatggaaggaggtaaacctaccgtcataccgaatgca

gaaggttttagaactactgcttctgttgttgcatatactaaaagtggagataaacttgta
ggacaaattgcaaggaggcaagctgttattaacccagagaatactttctattctgttaaa
agatttatagggcgtaaacaaaatgaaatttcgcaagaaattaggcaaacatcatataat
gtaaagaccagtgggtcaagcataaaaattgcctgtcccgcacttaataaagattttgct
ccagaagagatttcagctcaagtactaagaaaacttgttgaagatgctagtacatattta
ggtgagactgttacacaagcagttatcactgtaccagcttactttaatgattcacaaagg
caagctaccaaagatgcaggtaaaatagcaggcttagatgttttaagaattattaatgag
cctactgcagcttctttatcttatggattagacaaacaaaataatgaaacaattctagta
tttgaccttggaggaggaacattcgatgtgtctgtattagaggttggagatggagtgttt
gaagtgctttcaacttctggagatacacatttaggtggcgatgattttgatcaacaaatt
gtagaatggctaatcaaagatttcaagcaaaatgaaggtattgatcttggcaaagataga
caagcacttcaaagattaactgaggctgcagaaaaagcaaaaatagaactgtcaaattta
actcagacagagattaatcttccatttattactgctacacaagatggcccaaaacattta
gaaaagactgtgactaggggcaaatttgaagaactttgctcaaatctaatagataaatgt
agtattcctgtaaataatgctctaaaagatgcaaaactagaagcttcaagtattgatgaa
gttgttttagttggtggatcgaccaggattccagctatacagcaaatggttaaaagatta
ataggtaaggacccaaatcaaagcgtaaatccagacgaagttgttgctattggtgcagct
gtacaagctggagttttagccggcgaggttaaagatattctattattagatgtgacacct
ttatctttaggagtcgaaactttaggtggtgtgatgacaaagattataccaagaaatact
actattcctacaaaaaaatcagaagtattttctacagctgtagacaatcaaccaaatgta
gaaattcaagtgcttcaaggggaaagagagttaacaaaagataacaagagcttaggaaca
ttccgattagatggcattatgcctgcacctagaggagtgcctcagattgaggttacattt
gatattgatgcgaacggaattttgtctgttaaagcgaaagaaaaagctactggcaaggag
caatcaattacgatatctggagcttctactttacctaaagatgatgttgaaagaatggta
aaagaagctgaagaaaatttcgatatcgaccaaaaaagaagaaaaaatattgatatcaag
aatcaagcagagtctttatgttaccaggctgacaaacaagttaaagaatttgaagataaa
attagtcaagatttaaaaataaaaatagagggactaattacagagcttagatctagttta
gagaaagaagaatacgaaaatattgaatctatttctcaacaactacaaaactctttaatg
gatattggaaagaatgctgctgagcctgaaagtaaagatacaagcacaaaaaatgatgat
acagtgattgacacagatttctctgaagctaaataaaaaata-agcgggtaacgcgattc
gaacgcgcgacatcaaccttggcaaggttgcgctctaccactgagctatacccgcattaa
atgtattatcacaaatatcgtaatatttgtcaatctgttaatagatatctctct---atc
taggctattaatctttaaactatagactaatcaattatttggaatataacaaatactata
attttaaaa-------aagtacttatatcat-tatataagtacttcttttattcaaaaa-
acaaaaatacaaatcaatatcagta-------ttaaactactaaagagttaactaaacta
gtgacaattactaatcctgtccataaaccagctcctgtatagattaatccctttgattgc

tcccattgaccaggtgatgctaaaacaacaggtacgcctactactaatacagtagataac
gcaattaacaataacaccaataattgaatcgcaataatcatttagtatattctcctgaaa
aattcaaatgttaataatattgtatataatacaatgaagctttgtacctttaacaattta
tctatgaagaaataatttaaaattattgtttttttaataaatattaaattaagtattaaa
atctttattgtaaacaataatactatcttttttaa--atagataaaaaatttaaagaaag
tctttttaaaaaaaagttagtataagatgatatgctgtataaaaagctaagtattatta
gttattagagaaatttaaaaataagaggtaaaccatatgaactcagcccttttttttagca
aaactaccagaagcttatgcagttttcaaaccgatcattgatatcttacccgtaattcct
gtatttttctacttttagcttttgtttggcaagcagcaattggtttcagataacattta
agtcataaaaagaatagcaaatatatagagtacaaattttgagctctaatcttatatct
tacacaataataactattatggttgaacctttactatcaggaattgttcttggacttatt
cctatcactttattcggcttattagtcgcagcttatctgcaatatcaacgtggcaaccaa
ttaggactttagagcctaaattatagatatttataaacactacaataattatagtttta
taaatatctacagataaactaattgtaaagacagaaaaattttattgaaattagattaa
taattagaaagattggtgtaatattaa---------------------------------
------------------------------------------------------------
------------------------------------------------------------
------------------------------------------------------------
------------------------------------------------------------
------------------------------------------------------------
------------------------------------------------------------
------------------------------------------------------------
------------------------------------------------------------
------------------------------------------------------------
------------------------------------------------------------
------------------------------------------------------------
------------------------------------------------------------
------------------------------------------------------------
------------------------------------------------------------
------------------------------------------------------------
------------------------------------------------------------
------------------------------------------------------------
------------------------------------------------------------
---------------------------caaaatgtttttttaccagctagacttagtaac
tcctggtaaaagacattcatgagacatttctcgaaaaacatggcgagataatccaaaatc
tctgtaatatcctctgcttctgccagttaaccaacacctatttctgcctctaacagctgc

gctatttctaggcatttcttgaagtttttgtcttaactccattttatcagaaaagcttat
actctctttcatttgatctttaatagttagtcgttttaaatagtattttttttcaagttt
ttccttttaatctctctttggatcatattctttttagccatgttgtaa-aactcactgt
taaaattatttattcaatatactagctaaaaaagctcttgactgcaacgtttatataatt
aaaaatagataacataaagtaatatcatacaatcaatatttaatcgatgatact---act
ttatgctattaaaaagaatttatttaataaagttcttgttccacatgagtggaaatagta
caatcagatttaggatatgcaatacatgttaaaacatagccagctaacagttgatcatca
tctaagaatgactgatcagactgatctacagttccttctgtaacttttcctgcacaagtt
gagcaagctccagctctacaagagtaaggaagctcaattccttcttcttctgctgcatct
aaaatatatgtgtcatctgcacaattaaatgtaacatcaattccttcttcttcacatagt
aaatgaattttataatcagccatttctttttactccataataaagatacacttgaaactt
aataaagcacaat-aatttaaaggacttactaaaaaattgtatactagtttcattatgaa
tgatagactgcaaaaaaagaaaataatataatattatgagaaatcaagctacattacata
ctaaatagaactatagtacgttcaaagtatatacataaattcatatagttaaaattattt
ttatcatttaaatagcgaaaattaatagtat-agacttttagattgataatc------aa
aatagttttctac---caataaataaataattatgatttcaatgcttaatcaacaagtag
aactaaagcctattcttagatttcagatacaatacatatattcatactttcttattcagg
aaattaaagccttaataaccagactctttttgcaagtttggagaaggccggctacattga
tggcgggtattatccagcctttattatggctaattttatttggtggtcttttctataatg
ctcctgtaaatttatttactattaatacaagctataattgttttctaagctctggaatta
taatttttacatcatttactggagcactaaattcaggccttcctttaatgtttgatagag
aatttggatttttgaatcgactattaacagctccttttgatatctaggacttctatcattt
tgtcttctgctactttcatgacttgtattagtttaatacaggttatattcatagttacag
cttctcttttcatgggaaattctcctttaaacagcgatagcgctatgattttttggactta
taattctattagtaactgtaggagttacaatgcttagtttagctttatcttttactttgc
caggtcatattgagctattagcgcttattttagtagtcaatttacctttttctgttttcta
gtaccgctttagcacctctatatttcatgccaccttggttgcaattaattgcaagtctta
atccattaagttatgcaatagaaggtacaagatatttatattccagcatgcactggaatt
ttacagaatgtgttattaagattagttggggagatatttgtctagggcagattattatct
tgttagtcgttttagatataatcgcagcttatctcgtatctcatatattaagagctaaac
ttaatta-----------------aaattttaataaaaaatttattaaatttattttaca
aagatttttactatataatactaatagtagtatggataattgaaatagaaagagtttttc
aaaaccaaagctattatcaaaaatgataaacatttgtaagaaagtcaacaaagtatgttc
ttattcataggaggcatgtagtcaatgggactaccatggtaccgtgtacacacggttgtt
ttaaatgatcctggacgcttaattgcagtccacctaatgcatactgcacttgtagcaggt
tgggcaggatctatggcattatacgaattagctgtatttgatccttcagatccagtatta

aatccaatgtggcgacaaggaatgtttgttatgccatttatggctaggcttggcgtaaca
gattcttggggaggatggagtattaccggagaaagcgtatccaatcctggactgtggagt
tttgaaggggtcgctttaactcatatagtgctttctggaatgctttttctagctgctatt
tggcattgggtttattgggacttggaattatttcgagatccacgaactggcgaaccagct
ttagacttaccaaaaatttttggaattcacttattgctatcaagtctactttgttttgga
tttggtgctttccatgtaactggactgtttggaccaggaatgtgggtatcagatggttat
ggagtaacaggtaaggtcctaccagtagctccagcatggggaccagaaggattcaaccca
tttaatcctggaggagtcgcatctcatcacattgcggcaggtactgtaggtatattagct
ggtgttttccatttaactgttagaccaccacaaaggctatatcgcgctctgagaatgggt
aatattgaaactgtattatcaagtagtatctctgctgtttttttctcagcttttgttact
tgtggaaccatgtggtacggatctgcaaccacacctatcgagttatttggtccaactaga
tatcaatgggatagtggatattttcagcaagaaattgaaaaacgagtagaaaatgctatt
gctgatggcgccgcacctagcgaagcatggtcaagaattcctgataagttagcattctat
gactatattggtaataatccagcaaaaggtggtttattcagagccggccctatgaataaa
ggcgacggagttgctgaagcatggcttgggcaccccgtattccaggataaagaaggaaga
gaactaagtgttcgcagaatgcctgctttttttcgagacctttcctgtaattctagttgac
aaagatggtattatacgagctgacattccatttagaagagcagaatctaagtatagtatc
gaacaagtaggcgtaacagctagtttttatggcggcaaattaaatggccaagtttttaat
gatgctcctagcgttaaaaaaatatgcaagaaaagcccagttaggtgaggtatttgaattt
gatcgaactacactagaatcagacggggtatttagaagtagtccaagaggctggtttacg
tttggtcatgctaatttcgctttaatctttttctttggacatctttggcatggttcaaga
actatcttcagagatgtatttgcaggaatcggagctgaagtcactgaacaagtagaattt
ggagctttccaaaaattaggcgatagaagcagtaaaaaaacaaggcgctgtataagact-a
agtcaaacacagtctttatc---taatatttccgatattaaaatagttcagacaaagttt
attatctaatagttactaatcaggagaatttatggaagcattggtctatgtctttttatt
aacaggaacgctgatggttatattctttgcaatctttttagagagcctccaagaatagc
taaataactttatttcgtttaaacttacaataatagaaccactttccatcttattagtag
aaagtggttctattataggttacagcatgttattcttcatgctcttcaaaaggatctcta
agatcttttgaagcaggaccaaatgccgtgtatattgagtaacctgtaatacctaaaagc
aaactcgaaataaaaatactaagaactgttgcagtttccataattcaagcagtattaaga
tgaactttttctataataatattatcaatattaaaacatttaattattaaatttatcaaa
atatattatggcacttagaactagactgggagaaattttaagacccttaaattcagagta
tgggaaagttgctccaggttggggaacaactcctattatgggaattttcatgttactatt
tttcttgtttttattaatcatttacagatctacaattcatcactagtactagaaaatgt
cgatgtagattgggctaatttaggtag----ctaattatagcatttatttaa--------
ttgaaaaatattaaagacaaa----------------aagcgatcgctaatatactataa

tagcgatcgcttaaattgtattttactatata--cagagataattattttcctaaattaa
ttaactcactgtctgcaaaactatttgtattgacattattataattaactttttcaaatc
ttactaatactggatatttgataccacttttatcgattgctgcaacagtaccaatttctt
gataccagtacgattcctttctcaatattttactttgagcctcgttccatattttcag
tcctaatataggttgttagatattattatagtaaaaaaattttaaaa--tcataaaaaca
cataaaattagtaccgacaattatcacaaattaataataagaggagagagagggattcga
accctcgataacgaaagttatgacaattttcgagattgttgcaataaaccactctgccat
ctctcctgaactaaaataattattt-taaaattagacaagaataacatagaactcagata
gttgcctagaaaattaaaactatgttaaatataaattataatcttcctatatatatattt
acaaaatatgtacatagaatctatatcagcttatttcaagatttttaatcttatacaat
ttttactaaataaataaactctctcataatatgaaactatcttggaaaacacttcttctt
tggtctttaccaatatttgtcgttggtttctttttctggcaaggtttttaggtccaact
actacagatgttggtagtaatatcgcaagttctagaatgacgtatggaaggtttttagaa
tatttagatatgggttgggttaaacgtgttgacctgtatgaaaataatcatacagcaatt
gtagaagctgttggaccagaattaggaaacagggttcaacggattcgagttgagctacca
gcaagtgcgccggaattaattacaaagctacgcaaagctaatgttgatctagatgctcat
cctcctaaaagtacaagtgcagtatggggactattaggtaatttattatttcctctatta
ttagttggcggtttagctttcttatttagaagatctaataatactagtggtggacctggt
caagcaatgtcatttggcaaatcaaaagctttatttcaaatggaagctaaaacaggagta
gtatttaacgatgtagctggagttgaggaggcaaaagaagaatttcaagaagtagtgact
tttttaaaacaacctgaatcctttactgctgttggcgcaaaaataccaaaaggcgtatta
ttagttggacctcctggtacaggaaaaacattattagcaaaagcaattgctggtgaagca
agtgttcctttttttagtatctcaggctcagaatttgtagaaatgtttgttggagttggt
gcttctcgcgtaagagacctattcaaaaaagcaaaagataatgcgccttgcattgttttt
attgatgaaattgatgctgttggtagacaaagaggaacaggtgtcggaggtggaaatgat
gaaagggagcaaacattaaatcaactattaactgaaatggatggttttgaaggcaatacc
ggcgttattgtcattgctgctactaacagagcagatattttagattctgcattattaaga
ccaggaagatttgatagacaagtttcggtagatgttccagatttcaaaggtagattggca
atccttgaagttcacgctaaaaataaaaaaatggagcctaaagtatctttagaaactatt
gcgagacgaactcctggcttttcaggagctgatttagctaacttgctgaatgaagctgct
atcttaactgctcgaagaagaaaaaaatgcagtgactatgtcagaaattgacacatcaatt
gatagagtagtagctggaatggaaggcactcctttaattgatagtaaaagcaaaagatta
attgcgtatcatgaagtaggacacgcaataataggaagtttattagagcatcatgatcct
gttcaaaaagtcacgttaataccaaggggacaagccagaggcttaacttggttcactcct
agtgatgatcaaagtttaatatcgaggtcgcaaatactagcccgtattgtaggcgctctt
ggtggaagagctgcagaagagattatttttggggatgcagaagttactactggtgccagt

aatgatttacaacaagttacatcaatggctagacaaatggtcactcgatttggaatgtct
aaaattggaccattatctctcgaaagccaaggaggagaccccttcttaggtagaggaatg
ggaggaggttcagaatattcagatgaagttgcaactaatattgacaagcaagtaagagaa
attgtaagtggatgctataagcaagctaaagaaattattacagataatcgagtagttatg
gatagattagttgatttattgattgaaaaagaaacgattgaaggaaatgaatttagagat
atagttaaagaatacactgtaattccagaaaaaaattattatatatcccaattctaa---
---------------attaaacgggactgacgggattcgaacccgcaacttccgccgtga
cagggcggtgctctaaccagttgaactacagtccc-aataagat---tataaagagataa
tctcacagttagattgaaattgtcaaattagactattaaatatgaagagta---------
--------ttgataatgaggctcttcatatttaa------tagtctaattacttgattta
aaaggagagagagggattcgaaccctcggtacgaaattaatcgtacagcagattagcaat
ctgccgctttcgaccactcagccacctctccaaaaa-tcatatatctcgaatatatcaaa
tatattcatattttaacagatgtatatcaatatttatggagctaagcggattcgaaccga
tgaccctctcaatgccattgagatgctctaccaactgagctataacccctttctttgtga
ggatattaaaaaatggctcaagcgggatttgaacctgcgaccttgggcttatgaatcccc
tgctctaaccactgagctactgagccgtaatataaaatattattcaatactataccataa
attatattcaaatataaatcaataaaa-aatatttgctatag-tacatttcttttacatc
ttaa-aagttaacagtctacaagttatggatttttaagcgctgattcaaaattaaaaaag
tcttcaatcttttgttttgatatactattagttatttttgatatatctaacagtgtttta
atgttctaatttgaa---------------------------------------------
--------------------aatagatttt------------------------------
---------------------------------------------------tatttattc
a-----------------------------------------------------------
------------------------------------------------------------
------------------------------------------------------------
------------------------------------------------------------
------------------------------------------------------------
-------ta---------------------------------------------------
------------------------------------------------------------
----------------------------------------------ttatctcgcataat
------------------------------------------------------------
------------------tagaccatgcctaatgtac-----------------------
------------------------------------------------------------
------------------------------------------------------------
---aagaattatttaatat-----------------------------------------
------------------------------------------------------------

---tattttat-------------------------------------------------
------------------------------------------------------------
------------------------------------------------------------
------------------------------------------------------------
-------------------------------------tatagtaaaaa-----------
-----aggctggataaacagttctattcaatcagcaaaaaacactataggtgtataagta
aagtacctgaaagatctcaattgaacttttctttttaaatataatttaattcaagttata
gtaaatagaatactgtcaaatatcacagaggctaaataacacaataa--atatatctaag
aaatcagagtggaggaagttcaatttttta-atctgtcaaataaactttagtatcataga
ttcaatacttaaaaatatttattaagttta-----tgcaataattttcaatgtttaaca
tctt-tttttagataataattatcagctga--cgaaaaatatttttctatattagacaa
ccaacatggagccaataccttgatcagtcaatatttccagcaagagagcatgagcaattc
taccatctaaaatatgcgctgaagctactccttgagctaaagagcggatacaacaattta
ctttaggaatcataccaccagagatcacttctgtttgagttaggtctctagcttcttgta
tatttaagtgactaattaatgttgtaggatctgaagaattacgtaaaataccaggtgtat
ctgttaataaaatcagtttttccgcattgatagctgctgctatttctcctgcaacagtgt
cagcattaatattatatgactgaccttgcttatctgctgcaacactcgctattacaggta
tatagttattattaattaatatatttaataatttagtatcaatattttgtacttcgccaa
caaagccaaggtttgctttaccgtttggtcttggggtaataagtaatccatctttcccag
acaaacctacgcttttaccaccttgtttattgatcgttgctacaagatctttattgattc
ttccaactaaaaccatttctactatatccatagtaggttgatctgttactctaacgccat
catcgaattttggcaaaattttcaggcggtctagccaaaaattaatctctggacctccac
catgaactaaaataggacgcaatcctataaaagaaaggaaaacaagatcactaatcactt
gatcttttaatttctgatttttcatagctgctcccccatatttataacaataattcttg
aagaaaattgctggatatatggtaaagcctcacttaagacttttactctttctgaattcg
tcaacatagataaaataggattatgatatataaatttatttgtggaacttatattaattt
aacaaataccaataagtgcttcaaaacttttaagaaataaacatattattaatag-----
----ttaaaactataaggattataattatgagtaaattttggaataatatattaaagttt
ccacgattttagtcagtgtcattttaggattaattttaatcattattagtcctttttc
gtgttatttaaaaaaccattgacaagtttttttttcatcatatcattaacgagcttaatc
gcaattttgacaataacaa-tacaaaaaatgcttaatatcgaatgttgttgagagaaata
aataaatttaattaaaatttttgaacatatataatattatattatcattaaatatgataa
tttaattttcttttctat--aaaaaaccataagtattaaatattatttgatagatattat
gccattaaaacaaatagttagttctgaaaaaactggggcttttgcactcctagatagtat
tgttaggcatggggtcaaacatatatttggctatcctggtggagctattcttcctatcta
tgatgagctttatgcttgggaagaatcttctttaattaagcatattcttgttcggcatga

acaaggagctgctcatgctgcagattcttattctagatcaacaggagaagttggagtctg
ctttgctacttctggtccaggagcaactaatcttgtctcgggtatagctaccgcacatat
tgactctgttcctatattagctgtaacaggtcaagttggaagggccttcattggtacaga
tgcatttcaggaagtagatatttttggtattacacttcctattgtaaagcattcatatgt
agttcgtgatcctagagatatgtctagaattgttgccgaggcatttttttatttgtaagca
tggtcgacctggtccggtattaattgatgttcctaaggatgtaggattagaaaagtttaa
ttatttttctattgaacccggaaaggttagtattccgggttgcaggccagttactaactt
aaagtctcgacaaatacttatggcagctaagatgatacaacaagctagtcaaccattatt
gtacattgggggaggagctataatttctgattcacatcaaataattaaagaacttgttga
tttttataaaatacctgttactactactttgatgggtaaaggcatttttaatgaagatag
tgagtattgtttaggaatgttaggtatgcatggtactgcatatgctaattttgcagttag
tgagtgtgatcttttgattgctttaggagctagatttgatgatagagttactggaaaact
ggatgaatttgcttgtaatgcacaagtaattcatgtagatatcgatcctgctgaagtagg
aaagaatagaattcctcaagttgctatagttggtgatgtaacagaagttgtgacttcaat
attaaatctactaaaaacctctttgcgcccttatccagagcagattatatcttggcaaga
aagaattaaccgttggcgtcaacagtaccccttactagttcctaaaaaatcaacaagtat
ttcacctcaagagattcttgttgcaactaatcaattagctaaaaatgcttattttactac
agatgttggccagcatcaaatgtggtcagctcaatttttaaaagtaaaagctaaacattg
gctttcaagtgcgggattaggtacaatgggttatggtttacctgcagcaattggagctca
agtggcacatcctaatgatgtagttatttgtattagtggtgattctagttttcaaatgaa
tatgcaagagttaggaactattgctcaatatcagctacctgtcaaaattattattattaa
taatcgatggcaaggaatggttagacaatggcagcaagctttttatggtgaaaggtattc
acactcacgtatgacagaaggagccccaaattttcaaaaacttgcagaagcttttggaat
caaggcttttactattaatagtaggcaaaatatgcaatcttccttagaagctgcaattga
gtatcctggtccagttttactagattgtcaagttacagaaaatgagaattgttacccaat
ggttgctccagggaaaagtaatgcacaaatgataggtattgctaaacctcaaagaggtac
ggcctcaaattatattaatactaatatttaacgtaa------------------------
------------------------------------------------------------
------------------------------------------------------------
------------------------------------------------------------
---------------ataatttgtatcttctc-aaaatataata-aacttta--------
-tagctaaagtcttttaatactatct-aaaattatgtaaaaatagcctttgacgagaatt
gaactcgtgaccttccccttaccaaggggatgctctacctctgagctacaaaggc-tttt
tattatactgggccgggttggatttgaaccaacgtaggcgaagccagcggatttacagtc
cgcccccattaaccactcgggcaccgaccctaattaaattttattcaatatctgattaac
acgataccataagtagtttactaaaaacaagtacttgtttaaatgaagct-aataagtaa

tggacatagctggattcgaaccagcgactttcacgatgtcgacgtgacactctaaccagc
tgagttacatgtccagaaccctaatccttattataacactatttaatgattttataagta
cctatttttaattaaagatttgttttaagcgacctgctttaccaagaagatctcttagat
aatacagtttagcccttcttattttagctcttctaataaccttgacaaatgttattcttg
gtgaattaagaaaaaatactcgctcaactccaatgccttgaaatgagcagcgtaaagtta
aactagtgtttaaactgtttcttcttcttctagccaatattactccttcgcataattgtt
ctcgagttttattaccttccttaaccattaatccaagttgtattgtatcgccaactctaa
tttcagttacttctttttgatgaaaggtgtttcaacacttttcataagctgacttagct
ttgtattatttgctttcatatataggattaatcctgtaacgtttttttgaaatattatac
attaaaaaaactaatatgaaaattattagtgaaaaagcatattcgtatctttaaagtttc
aaattatctattaatagatttgatttacataatttt-----------------------a
taaaaggaagctggtt------------------------cttcttattatagacagtga
aaaaataaa-----tttagattaatt------------gcatcaaacttatt--------
------------------------------------------------------------
------------------------------------------------------------
------------------------------------------------------------
------------------------------------------------------------
------------------------------------------------------------
------------------------------------------------------------
------------------------------------------------------------
------------------------------------------------------------
------------------------------------------------------------
------------------------------------------------------------
------------------------------------------------------------
------------------------------------------------------------
------------------------------------------------------------
------------------------------------------------------------
------------------------------------------------------------
-------------------------ttgta------------------------------
------------------------------------------------------------
------------------------------------------------------------
------------------------------------------------------------
------------------------------------------------------------
------------------------------------------------------------
------------------------------------------------------------
------------------------------------------------------------

------------------------------------------------------------
------------------------------------------------------------
------------------------------------------------------------
------------------------------------------------------------
------------------------------------------------------------
------------------------------------------------------------
------------------------------------------------------------
------------------------------------------------------------
------------------------------------------------------------
-----------------------------cctccaatagaaat-----------------
------------------------------------------------------------
------------------------------------------------------------
------------------------------------------------------------
------------------------------------------------------------
------------------------------------------------------------
------------------------------------------------------------
------------------------------------------------------------
---------------------------------------------agacaatttt-----
------------------------------------------------------------
------------------------------------------------------------
----------------------------------------------tttaaa--------
------------------------------------------------------------
------------------------------------------------------------
------------------------------------------------------------
-------ataacattgtcttacagtttg--------------------------------
------------------------------------------------------------
------------------------------------------------------------
------------------------------------------------------------
------------------------------------------------------------
--------ttaactaattagtat-------------------------------------
------------------------------------------------------------
------------------------------------------------------------
------------------------------------------------------------
------------------------------------------------------------
------------------------------------------------------------
------------------------------------------------------------

------------------------------------------------------------
------------------------------------------------------------
------------------------------------------------------------
------------------------------------------------------------
------------------------------------------------------------
------------------------------------------------------------
------------------------------------------------------------
------------------------------------------------------------
------------------------------------------------------------
------------------------------------------------------------
------------------------------------------------------------
------------------------------------------------------------
------------------------------------------------------------
------------------------------------------------------------
------------------------------------------------------------
---------------------------atatagt--------------------------
------------------------------------------------------------
------------------------------------------------------------
------------------------------------------------------------
------------------------------------------------------------
------------------------------------------------------------
------------------------------------------------------------
------------------------------------------------------------
------------------------------------------------------------
------------------------------------------------------------
------------------------------------------------------------
------------------------------------------------------------
------------------------------------------------------------
------------------------------------------------------------
-------------tttgtctcatgc-----------------------------------
------------------------------------------------------------
------------------------------------------------------------
------------------------------------------------------------
------------------------------------------------------------
------------------------------------------------------------
------------------------------------------------------------

------------taatattgcaaaatttctttctttcacttaactg-ttcatcaaaacta
tcatgatgcacattccccatatctgaatattgattgtagtttgaaaa--tcttat---aa
aacttcataattaaaataataagattaagtcttaagtatttattagttttgtgttataat
tagttactatcaaaggtagtggcataacacgaagcataaa-aatgttcgaacgctttact
gaaaaagctataaaagtcataatgctagcacaagaagaagctagacgcttaggacataac
tttgtcggaactgagcagatattattaggattagtgggtgaaggtactggaattgcagct
caagttttgaaatcgatgaatgtaaatttaaaagatgcaagagttgaagttgaaaaaatt
ataggaagaggatctggttttgtagcggttgaaattcctttcactcctcgagcaaaaaga
gtattagaattatctttagaggaagcacgtcaacttggccataattatattggtacagaa
cacttgctaatgggcttagttagagaaggagaaggagttgcagcaagagtcttagaaaat
ttagcagttgatgtttcttcgattagagctgaagttatacaaatgctgggagaaaatgcg
gaagctaatgtaagtggaagtaatactacccaggctcgaagtaaaacaccaacattagaa
gagttcggttctaacttaactcaaatggctatggaaggtggtttagaccctgtagtcgga
agacaaaaagaaatagaacgagttattcaaattttaggtagaagaactaaaaacaatcct
gttttaattggtgagccaggtgtaggcaagacagcaattgcagaaggattagctcaaaga
attgctaatagagatgtaccttctattttagaagataaattagttattactcttgatgtt
ggtttattagttgctggaactaaatatagaggagaatttgaggaaagactaaaacgtatt
atggatgaaatcaaatcagctgataatgtaatattagttattgatgaagttcatacgtta
atcggagctggtgctgcagagggagctatagatgcggctaacttacttaagcctgcttta
gcaaggggtgaattacagtgtataggcgcaacaactttagaagaatacagaaagcatata
gaaaaagatcctgcattagaaagaagatttcaacctgttgtagttggagagccaagcgtt
gaagaaacaattgaaattttatttggtcttagagatcgttatgaaaagcatcatcaatta
acaatgtcagatggagctttggcagcagctgctaaatatgctaaccaatatatttctgat
cgatttttgccagacaaagcaattgatttaattgacgaagctggttctagagttcgttta
ctaaattcccaattacctccagctgccagagaattagacaaagaattaagagctgtatta
aaaacaaaagacgaagctattagagctcaaaaatatgaaacagcagagcagtatagagcc
agagagatggaaattaaagctcaaattgcagcaattgctcaaagtaaaaagaatgagcct
gatttaaatttagaagatcctgttgttacagaagatgatattgctgaaattgttgctgcc
tggactggtataccggtaactaagcttactaaaagtgagtcagaaaaattaatgcacatg
gaagaaacgttacatgggcgcattattggtcaagatgaagcagttatagctgtgtctaga
gctataagacgcgcaagagtaggtctaaaaaatcctaatagaccaattgcaagctttatt
ttttctggaccgacaggtgtaggtaaaacagaattgacaaaagccttggcttcttatttc
tttggttcagaagcttcgatgatacggctagatatgtcagaatatatggaaagacatact
gtatctaaattaattggttctcctccaggatatgtaggctatagtgaaggtggttatcta
acagaagcggtaagaaaaaagccatatactgttattctattcgatgaaattgaaaaagcg
catccagatattttaatctgcttcttcaaattttagaagatggccgattaacagatgca

aaaggcagaactattgatttcaagaatactcttctaatcatgacttccaatattggatct
aaagtcattgaaaaaggaggaggtagtctaggctttgaattatcagaagatcaaacagaa
tctcaatatactagagtcagatctttagtaaatgaagaattaaaacaatactttaggcca
gagtttctaaaccgactagatgaaattattgtatttcgtcagctcactaaagatgaagtt
cgagaaattgcagaattaatgcttaacgaagtctttgctagaattaagcaacaagatatt
caattaaatgtaacagagcgttttaaacaacgattagtagaagaaggatataatccaagc
tatggagctcgcccacttagaagagcagtaatgaggcttttagaagacagtttagctgaa
gaagttttatctggtaaaatcaaagctggtgatagtgcagtagtagatgttactaacgaa
ggagaggtgacagttttattaggtgaaaaattagaattgataacataaa------aaata
aattattaat-ataaagcattgaatataactt--gttcaatgcttt---aaaaaaaatta
ataatatgggttgctataattgacaacctctgataatctttatttgtatatg---cggcc
gggatagctcagttggtagagcagtggattgaaaatcctcgtgtcaccagttcaaatctg
gttcttggcat-tttaaaaatttgaaaagtttgctagctgagcattaaatcttaattgga
atgttcctgttgggccatttctatgttttgctattataacttctgtagcgtctccttctc
cagtttcttgattatagtaactttctctatagagcatgattactaaatctgcatcttgtt
ctatagaattatgcacaataaagttatttgcaataaaattatataaatatttacagttta
aatcgaacactgtttgtaagctgacaatatttatttttttaatattttcaaaagttaaag
ataaaa------aataattaaagatagccatttttttcttcaatattgtt------tattt
tagtagcaatcatattatttttattaatttcatcacatcttttccacccttgtgctgtta
gtagcctgtgattgcttgttaactttatatacttgccagattcggcaataattttatata
taggtttttttccctgtttttgaaatattagctttatttactgtgaataattctggtcttt
gctcaataaagttaatgacttca-atgtttttataatgaaatttaaacaagatttg--ag
ctccactatataatggtgactgaagataattaaattttgatatacacccactttctctta
gatcagataatagaggtcttttgtcacttcgagtttcaagatttctatttagctgagata
atactaagataggtaaatttaactcttttgctaatattttaagtgatcgagtaattaatg
atagttcttgagatctattttccgattgtttattgtcctgtagtaattgaagatagtcta
taataattaattctatctttttttccttgcaatttaaagaatttcactcttgtctttatag
tatttgcagaaatcttagcactatcatcaatgtaaaggttaagattagcgagagctttac
tcttatgaataacatatttccattcatcattagtgagttgacctgattgaatttttttggc
cgttcaaatgacactcttgagctaaaattcttcttaggagttgctctgtagacatttcca
agctgaataaaataacgtaagctttttcttgattaattatatacctagtcacattaatag
cgaaagcagtttttcccattgaaggtctacctgcaagaataattaaatctgacttttgaa
aaccttgtgttatcgaatctagttctgtaaaaccagagagcaagctactatttatactaa
tctttttttgttttgtccaaacggactaatagatttgcaagaattgttgctagtgtttgtg
tgtctttatcttcaagtatttcgtgagctgttgttaattgactaacaatagaagttatat
ttttttgttcaacgggccgttttgagcagcttattaaacataatgaatctccagatatta

aaagtaatcttttaatataattatctatgatgattgctgaatattcatctatggaatcag
ataaaggtgcttggtctattagactaagaatttcatttagttcagacaagtctttaatta
gtttttagtttttaagttaattaaaaaatttcttaaacttttttttcttgtctgattaa
tactttccaatgctgctctgtaaagtagtgaatttgatgtaaaataaaaaaaatcaggag
agagtttttctactgattgtattaaagatattgattgttgagtcaatattatactaatta
atatttttcagctaaaatattatgaggaagtaaatatttataaatactgatactgatta
ctccctcc-ttgtaacattactatgatg--------atactaattaactactgatctaat
ttaattctggaagaacttgtagttgaatattagcagttacttgattcaacagcttaatct
ctacattataaataccaattattttaatctctggcaaaaatatattttgtttttcaatgt
ctatgtttgaagtgcttttaataatttgtgcaatttctttttctgtgacactgccaaaaa
tatttttgccatctcctgtttttttactcacgctaaatttctgtatttcttctagaagtt
gcttgactttttggcatcttctttagctatttttaacttctcttctcttatagctgcat
aaaacttttgctgttttaaaataccagctgttgtaactgttgccattttatttgggatta
aaaaattccttgcataaccagcagcaactttaacaaggtcattacttttgccaagttttt
gaatattctcttttagcacaacgtttataatttttttactcatctt-------aattcta
aaaa----atctatattattttattatacctgttgtatacagatttg----aataggaag
tgttgatttgttgatagaatataggtgtaaaaataaagataaatttttaactttt-ttctg
atataatcatctttacaatgatat-----tttaggagaggtggccgagtggttgaaggcg
cagcattggaaatgctgtttaggagcaatcttaacgagggttcgaatccctctctctccg
tataa--aaattagacaaagttttacatagtccaatctagatatccaataatgttcttcc
aagcatctaatccacccctgactctcacaatattgagtttttttagctattaaaaattcag
aagcaaatatagatctagaatctaagctacagtacacaaagcaaattttatcttgtaaat
ctatattagagtaatctattttttttagttgttttaaaggcaaatttaatgaatggatta
aatgacttttcttatattcttcttgatttcttacatctattaaaatatattttggatttt
tatgatttaaaaatttttgcaattgaataacactaatttctcttataactgtattaactt
tatttctataacatttgttccagtgttgtttttt-------cgataatacaaatttagta
tttacaattttaaacttgttagatgataaagttaaggcattgtaggttaatatagcgcca
cttaatatagacttgtaaccaagaatgattttaacagcctcagtcgcttgaagtgtccca
ataataccaggtaatagacctaaaacgcccgaattactgcaagtatctattgcactctgt
ttattgttgacttcactgtagaagtctcgataaattggccctccttgataattaaataca
ctaacttgtccttcaaattgaaaaatagcaccataaatatgtattttatttaattctagg
caagtatcattcaccagatatcgagtgttaaaattatcagatccatctataataatatcg
tactgtctaataatgtcgattgaattttcatagcttagtcttgtgttaaaaactgtcacg
atgcactttggattcaaatctaata-tttttttttagctatttctactttcgaatatcct
atatcatgaattgtatataaaatttgtctctgtaaattagaaagatcaattatatcatca
tcaataataccaatgctaccaattccagcagcagcaaggtaaattatgcccggagatcct

agtccaccagcaccaatgaataatacttttgatctttttagtctttcttgaccttctaat
tgaatttgtggcaatactaaatgtttagaatatcttgtatattcctctaatgagtatttt
atattatctgctttaaaattcagcataaaaatttatattttaattaatagcaattacaaa
ctcaaactaactgataagattattacgatgaattttgtaaactaggttcttctttatcta
aaatagccccatgtgtaggacagacctgaatacaaatactacaatcaatacaggcagcaa
agtcaatccagtaccaattagtattaatactgtttttgccttgtccttggtaaatacaag
ctactgggcaagcgttcacgcattcagcaactccaatgcatttttcggttacaattgtat
gagacatattatttaattaat-aaaagataaactatttacatctagtatttttgttgtat
actagacttacagatgctatctaagccaataattaaagtactctactttatagtgtact
gatatgatctgcgcctttagttcagttggtagaacgcaggtctccaaaacctgatgtcga
gggttcaagtccttcagggcgcgtttaa-ttttatcataatttaatagtatcattttatc
tttttgcaaacatcaaaatgatatgatataaaagatcatatttatacattggacattct
cttgactagtacta-tttttaaatatgttattattgataagttaattatattaattctat
ttttataataatt-aaaaatagtccaaattaagtgttgatctacttaatagtc-ttttgt
atatataaaacttgaaaaatttaaaattatggctaaaaaagttactggaattgttaaact
agcattgaatgcaggtaaagctactcccgcaccacctatcgggcctgctttaggtcaaca
tggtgtaaatattgttatgttctgtaaagagtataatgcacgtactgcagataagtcagg
actagtaattcctgtagagatttcaatatatgaagatagaagttttacttttatacttaa
aactcctcctgcttctgtactaatcgccaaagctgctggactaaataaaggttcaggtga
accaaatacaaaaaaagtcggtagcataacaaataaacagttagaatctatcgcagaaac
aaaactgcctgatttaaatactaacaatatacctcaagctatgaaaatagttgaaggtac
cgctaaaaatatgggaattctaattaaagatt-aacaatttcaattatatttattattt-
-tttatgaaaaaaatttcacgtcgacttaaaatattaaaatctaaagttgagcctaaact
tcatactattgatgaagcagtatctatattaaaagcaacatcaaatgctaaatttaaaga
aactgcagaagctcatatttgtctaggcttaaaccctaaatatgcagatcagcagctcag
agcaacagttatattaccgaaaggtacaggcaaattaataaaagtagcagttattgctaa
aggagaaaaattaacagaagcaattagtgcaggagctgatgtaagtggttcagaagaact
aattgatgaaatactgaaaggtagattagattttgataaactaatagcaacgccagatgt
tatgcctttaatagcaaagcttggccgagtactaggacctcgaggattaatgccttctcc
taaagcaggcacagttacactagacgtggctaaggctgtgaatgaatttaagggtgggaa
agttgagtatagagttgatagaactggtataatccatgtgccttttggaaaatcgagttt
ttcacaagaagatctagtcttaaaccttcaaacaattaaagaatcaattgataaaaataa
gccttctggcgcaaaaggaaagtactggaaaactttttcttatctagcactatggggcc
atccgttcaaattgatatcactagccttctatagaaaatttatgtataattatacaactt
gaataatctattatttcaaatta-taat-------------------------ttgctta
ctatatcttatctgcaataaaaacttt-ttatatatcttactaattatgagtacaaaggt

tgaaaatatcttagaagagctaaaatctttaaatcttctagaagccgctgaactagtgaa
acaaatagaagaaacatttgatgttgatgcatctgccgcttcaggaggcatgatgatgtc
tgcgccaacttctgcaccagcttccgctgaggttgaagaaaaaacagaatttgatgttgt
tttagaggaagttcctgcacctaaaaagatttctgtattaaaagttgttcgttccctgac
cggattaggtttaaaagaagctaaagatcttgttgaatctacgcctaaagttttaaaaga
aggtgcatctaaagatgatgcagaaactatgaaaaagcagctagaagatgctggtgcgac
agttgttgttaagtaacaaa-tacaaaagtgtttttgtttatttctaaacaaaagcactt
ttttatctagtgtaagcaattaaattagaaaagtcctaatgtaattgcgttattgatagg
catagttgcaccaatacctagccaaatacttacaactgtaccaattaagaaaacagctgt
tgcaattggtcttctaaatggattttgaaacttgttaacattttcaatgaatggaacagt
aagtaatcctgctggcacagctgccatactaagaacgcctaatagcttatttggaattac
tcttaacagattaaatgttggaaagaaataccattctggtaaaatttccaatggagtagc
aaaaggattagacttttcacctatagatgaaggctctaaaattgctaacccaatactgca
tgcaaaagttccaataattacaactggaaagacgtataatagatcattcggccaagctgg
ttctccataataattatgacccattccttttgctaatttagctcttagctttggatctgt
taaatcaggtttttaagaattgacatgtt--tttttgttttaatataaaatttcttct
aattactgtgtcaaagtgtagttttacactattataatggcccggaaattccttgtttgc
gtatcattaaaaaatgcataagcatgaaaacggcagttagtaacggaagtacaaaagtat
gtaaactgtagaatctagttaaagttccttggcctacactcactcctcctcttaacaact
ctactatacttcctccaacaactggaacagcatctggaacacctgtcacaatctttacag
cccagtatccaatttgatcccatggcaaagaataacctgttacaccaaaagaaacagtaa
gtacccctaaaattacacctgttacccatgtcaattctctaggcttcttgaagcctcctg
ttaaataaacacggaatacatgcaaaatcatcattaagaccatcatactagctgaccatc
tatgaattgatctaataagccacccgaagtttacatcagtcataatatactctacagatg
taaaagcttccgcgactgtaggtctataatagaatgtcattgcaaatccagttgcaactt
gaattagaaaggatacaaatacaataccacctaagcaataaaaaatattaacatgaggtg
gtacatatttactagaaatatcatcagcaattgcttgaatttctaatctttcttcaaacc
agtcataaattttactcataaaatagcttcaataagctcttttaatata-tttttgcgtt
taagctactgctaattcaatag------ttattgaacttatatt-atttgttaaccata
cttattgatgattataacatctttttatttattt--------------------aaaatt
ctttgacttaataaaacagggctatgaactataattctattcttttgtatagagatcaat
ttagtttttagtaattcagataaaatccttgttacacttactctattgctacctataatc
tgtgctaaaattttatgtgtaatactaaaattaagcataataccattattagcatgagtt
ccattttgctcagctaataatagtaataagcttaccagtctacttgtaatacttttatgt
gaaataatttctataaaatgataagcttttgctgagcgatatattaaatgatttacaaaa
aataaattaaatggtggataattactacatgctttcataatagtattatagttaatacaa

gcaatttttgttttatctattgcttctgcttcgtagtaaaaattgtcatcgaccaattct
aattgtccaaatgtatctcctgtatttaacagattaagtgttattttatgtttatttttt
aggacttttgtaataattaaagaacctatcaatataatgtataatcgagagctgctatta
aacaaaatagtatcattttgcttaagtgaaaagagttgataatcaatgttattattatta
aaaaacagcaaccatggattgcaagaaaaaaaatctccagaattacttttagagaatatt
gaaatagaattttcttgatttaaataaatttagtcataattactactttataa------
---aacaatattctaataat-taataactgaaattaaatgtcatacaatttaatgttagt
tgagaatgacattgtattatcaaaagccattcaagaatacttagtagatcaaggctttaa
tgtttatattgctaataatggattagaagcatttaatttagcccatcaacataatttcag
cttgatagtttctgatatcataatgccgatagttaatggctacgaactattagagaaact
tcaaaaaaataaatcattatcaaaaataccagttatttttttgacagccaaaggactgac
taaagatagaataaaaggatacgacatgggatgttatggttacttatctaaaccatttga
ccctaaggagctagtttcactgattaaaaacttaattgctagagatgtcttacaagatgt
atctttacaaaaacgtgagataatattccaatcaatagataataaaatacatttgacacc
tcgagaaaaaagtattttagaccttgttgttgatggactaactaataaagaaatagctac
tatattaaatactagcgttagaaatgtagaaaaatatgttagtcgactcttaaaaaaaac
taatatgaagaataggactttgctagttaaatattctataaacaataatctattagatac
taagcttaagcaaagggcgaatgacgggactcgaacccgcgaatgatggagccacaaccc
attgccttaaccccttggccacactcgccatattatagatagcactatagcttgtttttt
gacatatagtctagtttctgaattgtatatttatataacataatatttacctttaatttt
ctatgatacatcgcaatattagtattcaaattaatggagaaccatttaattgctcagagc
ctatcttattgcaatttttattaaattatcttgattttaattctgagcgtgtagcagtcg
agctaaatgatactctgttaccagatactctttttcattcaacttacttaaatgatcaag
ataagctagaagttattactattgtaggtggcggataacctatttgtacta---------
---agtaatctactttagatgcttcatggataacgataatcgtccttgtttcttgtccac
atgaataataacagctttgatagtgtcacctattttaaattgtgatgatatctggtctac
ttgttttatattaatctctgagatatggacaagacctttaagattaccaacttttataaa
taatccataaggtgtaatctgatttattattccttcaataatattgccaacaattaaatt
ggatgaagcttgagctattagagctcttctgtggcttagaattaaattattgcatttctc
ttcaacattgagtaactttagtttaataaattgattactagattgttcgctttttttgaaa
attacagagatgagaattaggtacaaatcctaatataccttcaaggctaatgatcatgcc
tcctttgttaaacccttttataattacattaagtaaagagtcttcagctaataattgtct
aattcttttccatgctcttatatattcaaggcgtcgaattgataaaattaattgctttga
ttgaatattataatctaataaaaagaattctcttgtatcattaatatttaaagaagtaaa
attatttaaatcttgattacttgatacttcttgaataggcaagtatccagatattggtgt
tcctatatctactaatactccatttaattcaaagctaaatatagttccagctacaatatc

tccaagatttaaatcatacttgtactttttttaaaacagctgcaaaattcctgtgagtaaa
tccttcattattttttgtcatattggatagtatgaattaaagagtaattttgtacaatat
attaaggtaaacaaacctttgtcgaattatttg-agaaattttaatttgttctgttgtaa
ttccttatataaacatattacaataggcgagagacatgaaaagtaaacgtagatagctgc
agattcaa-taatatacggtctgaggaaagtccgggctccacaaatacaatttatgctgg
agaaatcccagtgtaggtaactatgaggatagtgccacagaaataaaccgccagaataag
g----tacattagctggtaagggtgcaaaggcaagttaaaagcttaccaaaagtactgta
aagtatttgttaggtaaacccctaaaatggagcaaagctactaaaaaaatatttgtgtat
atttt-tataatttagtttaaaatactgcatgaagttattggtaacaatagctcgagagg
aatagctatcctttacaatatatc-tctattatattgaaagaacagaacccggcttatgt
agtactttttgtgtcttgcattattatggctattgccgaatctctaattttaagtgtttt
acaatttttttcttgtagctcatgttgcttatgatctatttctgaattggttaaagttttc
tggttagatcgatatgtaaatcttagacctatactttttttctgttgaccaattgattta
tttgtatattgatcaaataatgttgtactttctaaatcattatcatgaaattgatctaat
aatttcaataagtagttaacctgcatattgcgaggtattattaaagaaagatctcttata
atcgatggatatttagagtatggctgaatctgataattcaaataattgaattgattatga
caatcttctaaaatgtttaaattaacttcaagaacgaaaagttttgtgttaagccctaat
ttactgtaggttgcttgattaagttcactgaatatgccaatagtattattattataaatt
agtgttgcagatctgtccttatgagctaattttattctactatcgactatcttttttttt
gtccactcaatttgcctattaatttttgaaaaaaatttcaacgatccctttcgcttca
taccaatttaaagaatgagctggctgtgaccatgtagatctaatatctaaattaccacct
agtatcatagctagattagttgtttctacaattgtattttgaacattattaaatactgtt
cctatctcaaagctatctatagtttgattgctttgtttaatattataaagattagattca
attaatccttcaagtaaactgcagcgtaaattagaatactcttgaataagaggattattt
aaagtaacatctcccatagatttaactaaagaataatgaactaattcagttaatcctaga
tgccttaaaatacttctacttttatctacgaactttctctttggcgataatttttttgtta
aattgaatacttggtactgagctactgaatttatgatatccataaattctggcaatctct
tcaataatatctacttctcgaaaaacatcttgtcgtctataatttggaaccttaatcttt
atagtttccttatttctatgaataatattaaaattcagagagttaagaatatcttcaatc
tcttcaaagcataaaaaaacgtgtctgaccattgtaacttattggccctaaaatatcatga
actttttttaattgataaatcaatatcaagaatattattaattattttctccttgtagaat
gtttctctaatattaccgctagctaaatctatgattaaagatagagcttcaatatatgca
ttttcccaattgtcagcatttagtccacgttcttgtctaactgaactttctgtacgaata
ttaagaactctagaagactttctaactacagactgtttaaatatagcagattctataaat
attgatgttgtattatgatcggtatcgaaatcattgttacctccaataccagctacacta
gtaacattggtattgacttgagttactaaaatatttttattcaactcaacattctcatta

tttaacttaatctgattattgtttccaacatgaaaatcgctacgaattgttataatatct
tcttgattaatatcattgatttattgaaatcaataatattaataggctgtccccatttc
agcataatataattgcttatatcgactaacaaattttatgcgtaaaaccagaagagtat
aagcgatttttaaccaatctggagaatctttaattctaatgttatctattattgcagaa
aaatagttattacaatttactaaactgtgactattgattatatt-ttccttttcagagaa
atatttattattaggttgatgtacagctgataacattgaagaccgagtcaaagctgagac
ctctcttgatagaccgatcatgcttaacacgtctgacctattagctgttgatgatatatc
taatatatggtcaatctgaccttctactgtaatagattcaatagcctcaacttcaaaacc
tgcttgtgtcaattgattatttaaaatatctatatctattgtttctatattcgccagttc
tttcaaccaatttaaagaaactttcataattaagtgtcatctatcttatagaagtattta
ttttacttatgattaaaaactaatttaaaaaattcatcaaattaatataagttatttgat
gaatttgcttatacaatatattattgtgatatcttacttttacatggcagtgattaagaa
ctagctttcgtaaaagttagcccattatcttgagcatatctttccataaatctcatgaat
ctatcccaatcatcaggacttttaataacatgtacagcttctattgcttgcggtttacca
ttaataaacttagcattaacatctcttgtaataatttgtccctctgtgtccattagatac
atacctgtgatttcaccttttttctgccatactagcatctaaaattttaggatttgtaaac
cgaaaagttgctgttcctgtactaccatctcgagaccttgttaaacgtacatctggtacg
acttcttcattaatgccttgaataaattgaattgttgccatatcaatttctttatttat
atataatttatattggatacaactgatttctaattctttaattcttgttgagaaactatt
ctaaaagcttatagtagcagtagatcaaattttatgtattatataaaactgtaactttttc
tttttttg----------------------------------------------------
------------------------------------------------------------
------------------------------------------------------------
------------------------------------------------------------
------------------------------------------------------------
----------------------atctgttcaaagaactaacttattataagctaatcata
ctaatattagccatataatgtttccaagt-gaaattaa-gggttgtagctcagatggata
gagcaagcgcctcctaagcgctaggtcagcggttcaagtccgctcaaccctgtcaat---
------------------------------------------------------------
------------------------------------------------------------
------------------------------------------------------------
------------------------------------------------------aaattaa
gtttttaaagttatagaagctgat--ataaaatttgctattaattga-----------cg
gttgcttctttttctatttgtctacaaaattaata----atatatattataataataaac
ttattttaactttgaaggaacgctagcagattagagtaataaattagaaatttgat----
------------------------------------------------------------

-----------------------------tagtattatataaagatgtagtttaa-----
----------------acattatcctaatcttaaaaag------aggcagcattattatg
taatat------------------------------------------------------
------------------------------------------------------------
------------------------------------------------------------
------------------------------------------------------------
------------------------------------------------------------
--------------actaattcttcgtttgatcctcatatgaaggaaaaatgagttgcat
tgtt--------------------------------------------------------
------------------------------------------------------------
------------------------------------------------------------
------------------------------------------------------------
------------------------------------------------------------
------------------------------------------------------------
---------------ttaactgttataactggggtgggaggattcgaacctgcgaatggc
ggagtcaaagtccgctgccttaccacttggctacaccccaacatataaagattataatag
cagtcgtataccatgccatgtcaatagattattatctttatttaataacttttgatactc
ttgtaaaacaagatcttttttttcaaatacgatatgttatgttcaaattctatgaatgtc
atattttcttgttgttgactaaacatccatttgatagtaattattcctttttcaacttcg
ttatttcctataataatacaagctatagctctttttttatttgcttgcttaatttgttta
ccaaaattacttgagctaacatctatttctattgtgaaaaactgttgatgtagaaatctc
attatttgcataccagtttcgttagctttttttaccttgtgtagcaatataaaaatcggta
tatttattaggcaagtctatattatcttttgcaagaagaaataagcgctctaatcctatt
gcacatccaacagcaggtgttgctgtgccccctaattgctgaactaagctatcatatctt
cctccaccacagatagtatcttgcccttttgatgttaacgtcttaatttcaaaagcagtg
tcattgtaatagtctaatcctctaactagcttattgtttatgttgtaaggaatatcaagt
aattgcaaataattacaaacagtatcaaagtgtttttgtgattcaaaacttaaaaaatca
gaaattttcggagcctcagttagtattctttgtgtattagtatttttttgaatctaaaatt
ctaattggattactaattagtctttttttgtgagtcagtgtctaattcatcatagtatttt
ctaagatagtcttgtaatttagcttgataaatactacgatcttctaatttgccaattgaa
ttaagatcaagttttaagtctgtaagatttagattattaaagatactcatagctaaatgt
attatttcactatctgctcttgcgtctaggctaccaataaactcaataccaagttggtga
aattggcgttgtcgtccactttgaggtctttcatacctgaacattggaccgctataccat
aatctttgtaaactatgatgataactcatttttattttcaataaaagctcttacaatacct
gcagtaccttctggtcttaatgtaatatctctgttactacgatcgtgaaacctgtacatc
tctttatttacaatatcagtatcttcaccaatacttctattatataaatcactgttttca

aaaatgggtgttctaatttcttgataatttgcacattctaataatttagaaatcttgtta
tgcataaactgccaatatttaagctcttcgggtagaatatcttttgttcctctaatagcc
tgaattttgccatgataatgtttgtattaccttcttaatgtgtctgcattaaatagcag
ttgtagactggtattcagtaatgtattatacattag--aacatattgatg-tactttata
aaatacgggcaaggagggattcgaacccccgacaccatggttcgtagccatgtgctctaa
tccactgagctacaagcccacttaataagtatcttcactatatcactgttcttgaatgta
tacaactacattcttttt------attctatatgtgaaatttttgtaaaagtaaaaaagt
at----atttatttttacaattggaagcattcatgtataatgcgtaatctattataact
ttgtaaa--aatatcgaaaagttgaattatgtgattttattatacagataatcttattgt
aattttgagctctcacttttcaattga------cgagcatgtaattagcttatattccaa
ataaaaaaatcttatctaaaatgagtaaacaaattctatatcaagatgatgcacgtaaag
cattagaaaaaggcatggatattttaacagaagctgtttctgttactttaggaccaaaag
gaagaaatgttgtcttagaaaaaaaatttggtgcccctcaaattattaatgatggtgtaa
cgattgcaaaagaaattagtttagaaaatcatattgaaaacactggagttgcacttatta
ggcaagcagcatctaaaaccaatgatgtagctggtgatggaacaacaacagcaacagtat
tggcttcagcaattgttaagcaaggcatgagaaacgttgcagcaggttcaaatcctatgg
ctattaaaaaaggtattgaaaaagcaacaaattttgttgttagtaaaatagctgaatatg
ccaaaccggtagaagatacaaaagctattatacaagttgcttctttatcatcagggaacg
atatagaagtaggaaaaatgatagcaaatgctatagacaaagttggaagagaaggtgtga
tttctttagaagaaggaaattcaaccaatacgattcttgagataacagaaggaatgcagt
ttgaaaaaggctttatttctccttattttgttacagatacagagagaatggaagtccttc
aagaaaatccatttattctattcacagataagaaaattactctagtgcagcaagaacttg
ttccattacttgagcaaatagcaaaaacgtctcgacctctattaataatagctgaagaca
tagaaaaagaagcattagctacgattgtagttaataaattaagaggtatcttaaatgttg
tcgcagttagggctcccggatttggtgatagaagaaaatctttacttgaagatatgagta
ttttaacgaatggacaagtaattactgaagacgcaggtttatcacttgatacagttcaat
tagatatgctaggaaaagctcgaagagttattgttactaaagattcgacaacaattattg
cagatggtcatgaaatcaaagtgaaatcaagatgcgaacaaattaagcgtcaaatagaga
ctagtgactctctatatgaaagagaaaaattacaagaacgattagctaagctttctggag
gagtcgctgtcattaaagttggtgcagctacggaaacagagatgaaagataaaaaactac
gattagaagatgcaatcaatgcaacaaaagctgctattgaagaaggaattgtaccaggag
gaggagctactaatgtacatatctctcgtgaattatttacatgggctaaaaataatttag
ttgaagacgaattaattggagctctaatagttgaaagagcattgacttatcctttgagac
gtatagcttttaatgcaggtgataatggagcagtagtagtagaaaaagttaaaagtaatg
attttcatataggctatgatgcagcaaacggtaatattgtaaatatgtatgatagtggta
ttatagatccggcaaaagttgctagatcagctttgcaaaatgcagcttctattgcagcaa

tggttttaactactgaatgtattgtagttgataaagttgatgattaagctaca------g
cgtaatagataatctaaaacaaagaataatttctatggctagaaattattctttgtttca
attaagttaattaattgatcaaatgtataattttataataaagccaaagtttaaatgtg
ccttctttttcaactattaagtagagaagatataagcctgtaattcctaaattatcaatt
aaagtattattattatatatcgatttttctagagatctaaaaaatgatattttgtaaaa
taataccgaaatcttcttttatagttatttgctaaatcattaccgtataaagattgattt
ccaattgaattcagaacccttaaaataatattttttaaatctttatctgattttatgtca
gaatcagtaataaattgattcaattgaatatcaaaattaacattaaaaatattacttgtt
aaagaagtttgattttgaatatctagagatttgatagccaagaagattaaacttttattg
ttcattgtaaaatataagacaattgatactgtattctttaaatacta-aaagtaccttct
tgtttgtttatagttctgtataaaaataattaatactataa--------ttttaattact
gcccgaaaaaataaattcaggaaatctagcttgagccttagatagcgagtgttttttcc
ttcttcttgccagtaattaataacttctgttgttttgttcaataattcgattctatctct
ttcagcaatccagggtttagattctagttctgcttttaattcttcccatgcgtcatttct
tggccagaaaaaataagaagtaagtggactggtgccatttccaacaatttgatcaatagc
aatagcaatattattttctaaccaaagtacttttagtgtaaattttgacaagtgattatc
tccattatgaatattaaatatttg----tacagtattttactgtactaattattttcttg
agcaacaatttgagctttattcaagatgttttgtactgttctagtagcttgcgcaccttg
cgctagtctttgtaaaatctttgaaatatcaatacgagtttcattagttataggattata
gaatcctaattcttctatagccttgccatcgcgcttatttctgctatccattacaacaat
tctatagctcggttgtttttttcttccgtatcgctttaatcttaacttgaccatattagt
tacctcgcttatttttt--atatctttattaaattatatcttacgacattatgcaataat
tcaaacaattaagcagactcaagtctaacattattaaatattaccattaaacactgtaga
aaaataatacccagcataggtgacatgtccattccaaacattggaggaatacttcctctg
aatagctttaaatatggatcagtaattcgatttaatgaacagaatggttcattataccaa
tttacagttggaaaccatgccaatgataatttaagtaaaattaaaattaaataaatctca
gaaaaattagctattgatccaagtaataaatttaatgtaccaggaagagtattcatacgt
attttataattattatatacttcattatactaatagtttgattatatattataagctctg
taaataaat-cagctaaacaaaaaacttattcttaaaaatatcttctcgtgagctgtttg
cttcatttacagtagtattgattttggttgtatacacttctagtttggaccaattctttc
ctaactgatttagttgagatgttccacattctaaacatgctatttttatattattcaaaa
tagcacctttttttacaaaatcgtttaataatggagtaatcgtctcttcatctaaaaaca
aatctaggataagaatttttgtaaatgaatttaattgatctggaacattgttaatagtat
tattataattaactaaagttatattagctgtaggaagtaaagctcgtgctccttctgcta
atataaagccataaggcataataattacaataatgtacttataattactatcaataagat
tgatagtcgtattagcttttaattcaagagtttctgttattaaccattctctcattattt

cataagtaatccatttgcctatttctgaacaagccgttcttaaaattgtacctgaattat
tactgttttcgagaatcccagcccaatgttgtatcagtggatgggaagctatattaattt
gaagttgcatgttttgttaaatatatttaacatctaatttgttgattgtagcttcagttt
ctattcgttgagtattattaaaatagcat----atatattaaagaaaacaattat-aaaa
aatgaaaaaatttttttggctttggggctttacagatagtgctgaaacttggaatggtag
atttgcaatgattggttttatattagttattgttatcgaagtaattacaggtcaaggact
attgtatttaataggtattatgtcataataaaaattcaaaaaagttcttgtaattttta
caagaactttttga-atttttataaatctaatctagtggtctcatagataataccggtt
cattctctctattaatacctttttcaaatccagcagcagcagatcttgctctaccggcat
gccaaagatgtccaataaataggaagaatcctaagaagaaatgagatgttgttaaccaag
atctaggagatacatagttaactgaattaatttctgtagctacaccaccaacagagttca
aagaacctaatggtgcatgagtcatatattctgcggcacgtctttcctgccaaggttgaa
tatcattcttaattttatttagatcaagaccattagggcctctaagaggttcaacccaag
gagctcgcaaatcccaaaatctcatagtttcaccaccaaaaataatctctccactaggag
atctcattaagtattttcctaatccagtaggtccttgagaagatgcaacattggcaccta
gtctttgatctctaactaagaaagtaaaagcttgagcttgtgaagcctcaggaccagtag
gtccatagaactcactcggataagccgtgttgttgtaccaaacaaaattagaagccgtaa
gacccataagcgataaagcacctaaactataagacaagtatgcttcaccagaccatacga
aagctctccttgcccaagcaaaaggttttgttagtatgtgccagatccctccagcaatac
aaataattccaatccagacatgaccaccaacaagatcttccatgttatttacacttacga
tccatccatcaccgccaaacggagactttaagacataaccaaaaataactagaggattaa
gcgtagggttattaacaaatctaacatctccaccacccggagcccaagtatcatatactc
caccaatgaacaaagctttaattactaatagaaaagctccaatgcctagtaatactaaat
gtataccaagtattgttgtcattttatttttatcccgccaatcatatccaaagaaaggaa
aagattcttcaagagtatcagggccgattaatgaatgataaagacctccaaaaccaagaa
cagctgaagaaattaaatgaactactcctactacaaagtatgggtatgtattaaaaattt
cgccgcctggaccaacaccccagcctagtgtagctaagtgtggaattaaaatgaatcctt
gttcgtataaaggcttctctgggacaaagtgagctacttcaaaaagagtcatggcaccgg
tccaaaagaccattatacctgcatgagcaacatgcgcaccaagtaacttgccagaaacgt
taattagacgtgcattgccagaccaccaggcaaatccggtagattcaatgtctctaccgc
caacaccaacagttgtattaaagggcgtttccacgtggtaaaacctcctctgggaatata
aagttttcatgaggttgatcctgagcagccatccaagagcgaataccttcgtttaacaag
atgtttttagtataaaaagtttcaaactcaggatcttctgcagctctcaattcttgagaa
acaaaatcataagctcttaaattaagcgctagtccaacaattccaaatgcacttgtccat
aatccagttaccggtacgaataacatgaaaaaatgtagccagcgtttattagaaaaagcc
acaccaaaaatctgtgaccagaatctgtttgctgttaccattgaataagtctcttcagac

tgtgtcggagtaaatgcacgaaaggtatctgcagcatcaccatcttcaaatagggtattc
tgtacagttgctccgtggatagcacatagtagagcaccaccaaggattcctgcaacaccc
atcatgtggaacggatttagcgtccaattatgaaatccttgtaaaaatagcaagaatctg
aaaatcgcagctactccaagacttggagcaaagaaccaacttgattgtcccaaaggatac
attagaaaaacagaaacaaatactgcaattggtccagaaaaagcaatagcattgtacggt
ctaagaccaactagcctagcaatttcaaattgtcttaagcaaaatccaattagtccaaat
gacccatgcaaagctataaatgcccatagaccaccaatttggcaccagcgagtaaaatct
ccctgagcttcaggtccccaaaggaaaagcagtgaatgtcccatgctgtttgctggcgtg
gaaacagcagcagttaaaaaattgcatccttctagatatgaactagctagtccatgagta
taccaagaagtaacaaaagtagttccagttagccacccgcctacagcaaggtaagcgcaa
ggaaacagaagtagtccagaccaacctacgaatacaaatcggtctctttttagccaatcg
tctacaagatcaaaaccaccacgagtcttttcttgtccaattgctatggtcataattta
atctccagagcaaatttaataattaatttgtttaaataaaca---ctttagatatagtct
aaaatttttacttttctttacggtcaatgtaattataagttaaaatcaataaaaattcacg
ttttaaataaca-tttttttagctcattaagttaatttttttagagataaaaatagata
gagctatttttcaactaacaaaagtcataatttattcatttctataattctttgaaata
gatatcctgttcctcgcgccgttaaaattaagtcaggattacttggatcatcttctagtt
tagctcttaatcgtgaaatatgcacgtcaacaactctagtgtcaacatgtcgctctggtg
tatacccccaaacctcttgtaaaatagacgctctagaaaaaggttctccagccttgctaa
ctaaaagttcgagaagactaaattccattcctgttaagcgaactcgctcattattcttat
agacttgtcttttattagtgtctatttttaaaaatccaagactaataatgccagagtttg
gaacgccaagattagttgtgattttatcagctcttctaagaacagatcgaatgcgagcct
ctagctctttaggcgaaaagggtttaactacgtaatcatcagcacctatttctagtccag
taattctatcacatacttcacctaaagctgttagcattattataggaacatcagattctt
ttcggagttcttgacaaactccatagccatctagtttaggcatcatcacatctaaaacga
caaggctaggatattcttttctaaatattattaaagcttcttctccatttgaagctgtta
tgacttcgtaacctattattgtcaatcttgtttctaaaattcttcttatgctggcctcat
cgtcaacaacaagaatcttttctttctggttctccaa-tttcttatttactcctataact
cacttagaatgattaatataatcttaaatttgtagctgtttgagtatctttatttgcaat
agaa-----cttagttattataaattttgtaagtc-----tgcattaattagtatcaat-
-aattatatctctaaatagttctgtttcagttatgtg---------tattttaatgcata
agttgattcaagtttgcagaattcagtatatacata-ttttttagccaagtactttgcat
tctattttctaaataaaaaaggggttgacaaactagttgatagaaagtattctacattac
tataag----ttactgccaacaaagaaatttcaa-----atcttcttttagaataataag
tactttaat
>P_pulchra_NC_029861

tatatgatttataattaaagcgtctttcatgtaagaattcgcaaaatatacaaaactttt
tatttaatatataaaaagtaaactaagataagttcaatatactttgctttatcttaaatt
tcttgagcgag------atttcaaatagagacaaaagtatttacatagtttatatctata
ggaataaaaatgaaacttgcagtttatggtaaaggtggtataggtaaatctacaactagt
tgcaacatttctgtagcactttcaaaaagaggcaaaaaagttcttcaaattggctgtgac
cctaaacatgatagtacatttaccctaacaggttttctaattccaaccattatcgacaca
ttgcaatcaaaagattatcattatgaagacgtttggcctgaagatgttatctataaggga
tacggcggtgtagattgcgttgaagctggcggacctccggctggtgctggttgtggaggg
tacgttgtaggtgaaacagttaaacttttaaaagaactaaatgcttttgatgaatatgac
attattctatttgatgttttaggagatgttgtatgtggaggctttgctgctccattgaat
tatgcagattattgcctaattattacagataatgggtttgatgctttatttgcagccaac
agaattgctgcttcagtaagagaaaaagctagaacacactctttgagattggctggactt
gttggtaatagaacagacaaaagagatttaattgataagtatatagattgtgttccaatg
cctgtattagaagtattaccattgattgaagacattagagtctcccgagtaaaaggcaaa
actttatttgaaatgtctgaaacagataaagatttagcctatgtatgtgattactattta
aatattgcagatcaattaattacgagaccagaaggagtagtccctaaagagtctcctgat
agagagttatttagtcttctctctgattttacttaaatcctaaatctaatttaggacaa
gcgacagtagaacaagaagaattagatttaatgatagtgtaaagaatattaaaagcataa
catataaaggaatacgataatgtctacagtccaatcagatgctcttacttttgaatgtga
aacaggaaattatcatactttttgtcctattagttgtgtctcttggttatatcaaaaaat
tgaagatagtttttcttagttataggaactaaaacctgtggctatttttttgcaaaatgc
aatgggagtgatgatttttgctgagccgagatatgctatggcggaattggaagaaggaga
tatttcagctaagcttaatgactatggtgaacttcgcagattgtgcttgcaaatcaaaaa
agatagaaatcctagtgttattttctggattggaacatgcacaacagaaattatcaaaat
ggaccttgaaggaattgctccaaaattagaagcagaaatccgggttccgattgtagttgc
tagagctaatggtttagattatgcgtttactcaaggtgaagacactgttttagcagctat
ggctcaacggtgtccttcaagtttaaagcatcaattaggtattactaatgacgaaattgc
acagcatattcctctggttttatttggatctcttccagatccagttgtaactcagttgac
aatagaattgaaaaagcaaggtgttcatgtttctggttggttaccatctaagagatatac
tgaattaccggtaattcaggaaggttattatgttgccggagtaaatccatttcttagccg
aacagctaccactttaatgagaagaagaaaaacaaaattgattggtgcaccatttccaat
aggcccagatggcactcgcgcttggattgaaaaaatttgctcagtactaagtatcgaaac
tattggactagcagatagagagagagaaatttgggattctttagaagactatgttgccct
agtaagaggtaaatcagtctttttttatgggagataacttactagaagtctctttagcaag
atttttaactcgatgtggaatgactgtgtatgaaattggtattccttatatggataaacg
ttatcaagcagcagagctagctttgttaaaaactacatgcgataaaatgaacgttatgat

gccaacaattgtagaaaaaccagacaactataatcaagtagacagaattcgtgatctaaa
acccgatttagttattactggcatggctcatgcaaatcctctagaagccagaggtattaa
tacaaaatggtctgtagagtttacctttgcacaaattcatggttttacgaacgcaagaga
tattctagaactggttacaagaccattgcgacg-aatcttagtttgtcagaattaggctg
ggatgtttatagcaagcaaagctagtatc-------agctgtgacttacatggagtaaaa
tccttgtaagtc---------ttattgtaaggaagcatttcttgccctgccctcacctgc
ccaagcttgtaaattttccacttgttctttatctgtaaatgctaaaggaataaattgctc
tatagctattttaatatcttctgtattaaactctctttcttcactaaaagcagtatgcat
gctctctataatcgcttgctcaatctcagccccagagaatttattgcataataaacttaa
ttgttttatatcatattcttgccatgatttgggacgaacctttgagagatgtatttttaa
aatagtctctctctcttgacaattcggcaaatctaggaaaaaaatttcatcaaatctccc
ttttcttagcatttcagaaggtaagctttgaattctgttagcggttgcaacaacaaatac
tggagctttttctctgagagccaagtgataaatgttccaaacactcgtgcactggtacc
actatcaccttggctatataaaccagaaaaagctttgtctatttcgtcaatccataatat
gcatggcgataatccttctgaaatatttatcatttctctcatttttgattctgattcgcc
aactaatcctccaaacaattttccaatatcaaggcgtagtagaggtaatgtccaatcgtt
ggcaatagctttagcggttaaagatttccctgttccttgtataccaactagtagcaagcc
cttaggagaaggaataccataatttaaactttgctttgaaaaagagcgagaacgtttttt
tagccattgctttaatacatctaagccaccaatatctctattaagtttggaataaggata
aaattctaatagacgcgtttgattaataatttgtctttttttcctcaataataattggtaa
gctgcgagaatctagttggttatatcgtgctatgattttagtaaccactttgcggattct
atctattgataaaccttggcaggatttagtcatactattaactagctctgactctagatc
aagattaatagctttacttaatctcagtatttctttttttgatttctatgagatcaggtaa
cggcaagtttagcactgttataatatcgcttaaagcaaaaggtatattaattttacaaga
aataataatgatatgcttagactttcctttaattactctagctacattacgaagttttct
aattagtactatctcatttagaaaagaatcaaaatcttttaaaataaaaagatttaaaga
gtcattatctaagttttcaataaactccaatgctagtaacggatttcgttttgcataacc
attatcgttagggttgctagtatatccatctacgaaatcccagcagtacacttgttgact
attagaagagtctattttatttttataatatactctagtctatcttcttctctagtatt
tattacaataataggatgtcgagacttcaaaagtaatcgtaaatcttgagtgaaattcat
gtcaatatttgtctccatcagacttttgtacttcaccaatta----aagactaaataaga
ttatctaatattgcctggatatattcaaaagtc------atgtatctattctctaatcta
ttacaactttaaccttaactaaccataattttttttcttccttttcgcctccgactattt
aaaatagagcggccgcatggagttttcattcttgcccgaaaaccggaaactttaattttt
tttcttttagatccttgtagtgttcttttagtcatgatgtctcctgtattgtaaaagatt
aaatactattttatcatcttttatatgtcttgatgatatagatctttcttaatatcaaag

ttaaaacgcaataatttatattaatcaagaataatttatatattattaataatcaatcat
----------tagctttatgatattggctctgcctatttttatttaagtgttcttactg
catttttgctgattttaaattggttaattttacagcaactaaaaacaattttagcactag
aatctcaatttaaatattttgtggaaaaaagtcaaaacgatattttgaaagaggaagaaa
gttttgcctttgcaaaaatatgtgtagcgaaaaagtgcttttctaaagctattattgaag
gtcaattggcattgaaaaagaaatcagaattaaatattaaggaatcaaatataacaactg
cgaatctgtataatatgttgggttttatctattttgaagctaaacaaatagatactgcaa
aagatttctaccagcaagcattacacatagactccaattatgtaatagctttaaataatt
tagctaaaacttatgaagaaactaaagatttaaaaaagcaggagttttatatgataaag
tattaactcttaactcgaataataaaacagctaataggaggaaaagttttataaaaaaga
caa---------aaagcgcttaatcatatttataatcgggatagcaggatttgaacctgc
gacatcctgctcccaaagcaggcgcgctaccaaactgcgctatatcccg-gcgacataac
taacactatataccttttatctaattatgtctactatttatatttttttatctgggatacc
aaaacatgcccttaaccccatcgggatctttaataaatcccaatttttataaaaactaa
ccacgtcaggttctgcaaacaatgtaatagtgctaatctcggcctgccttaattgctgaa
ttaattggtatatcactactttacctaaacctaaaccttgaaaatccggatgaatgacaa
cgtcccagatagtagcattgaacccattatctgatgttgccctagcaaatcctacaagtt
tggtatgtgcatcttttttttgtattagagaaataatgatggaactgtttttagagcaa
tttttacttttttagaggccttttaacccatcctacagaatcacaaagctgttccaact
cgtataaattaatattcttagtactactcacgtagatatctttaaattctactctatcgc
aagttttatctaaaacaataagtttttgaaatttttttcagaactacttttgacatctg
agttttgaaaaaaattttccagaagatcataatgattatgatagaat----taaatata
tatatgttacggtattaccttaataagtaacatattaatatatgcccaaattttaatatg
aatttattaagacagaagattacttagatgtatcgaaataaaaatttttgtaaaatcttt
caataaaattatataaagagttttttttaagactaacatctattattataaataaagtct
cgcaattgagatga-ataacccattattctatatctaatgccttttaaaggagttatttg
tgaaaaaatccatgtttctaacttgtctaatcgcttcactactgttaagcaacccaataa
tagctaatgcagaagtagcaggactagttccttgcaaagattctgctgcattcaataaac
gaatggttaacagtgttaaaaaacttaaagctagactagctaaatatgatgaagatacgc
cgccagcactagccttaaataagcaaatagaaaagactaaaactagatttgctacttatg
gtagagcaggcttgttatgtggtacagatggattaccacatctaatttccgatggccgat
ggagtagagccggagactttgttttcccaggattactatttctatatatcacaggatgga
ttggctgggtaggtagaggatatcttctatctgttgctaaaactagcaagccaaccgaaa
aagagattatttagatgtgccattggctgttaaatttatgacctctggatttgcatggc
cgttagccgcttggcaagagtttagtagtgggcaattaattgcttccgatgatgatatta
ctgtttcaccccgttagtaaaaactttatatgaataataattttaccaaatacttatcaa

cagcacctgtaattggcgtgttatggatgacgtttacagcaggctttattatagaattaa
accgtttctttccagatgtcctatattttatttataaataaataa--aattttataga-
--gaactacataaatatagtaaaacactattatttatgtagttctttg-tatatataaca
taattatttattttattaaacggtctctctttcgactattttaagattagattttacat
aataaaaaatatataaatattatgagtttagtaagccaaattatagttaatgccgatgat
gaattgcgatatccaacaatcggagaattgcagtcaattcaagactatttaagcacggga
agtaataggattagaatagcaactattattagggataaagaaaaagaaattatccaaaaa
gcaagtaaacaaatttttcagttacatccagaatatatagcaccaggtggcaatgcagca
ggctctagaaaaagatcattatgtttacgggattatggttggtatttgagactaattaca
tatggagtattatcaggtgacaaagattctatagaaacaattggtattataggtgtcaga
gagatgtataattctttaggagtcccaataatcggaatgctagatgcaattcaatgtctt
aaagaagcttctttagaaatgcttggtcaagatgacattacaatcattgccccttatttt
gattatataattcgaggcatgtcataaatagtaatttcaattaattagttgaataaaagc
ttttataatgttataattaggctaagcccgaaagcatataattgtaaaaactaaatcttg
tcagaaccggaaggtagcagcaataacgtttaactacaaaagttatgacttttcgggtgt
ttattatttatattc--------------------------------------------aa
gaataaataatttggaaaacctttcagtaaaaataa-ttaataatttgatattaatacag
aaaacatctatac-tttttataaaatttctttatatctcatctaaataatggaaactta
acttaatgggtgttcatattttatcaacaggttcgtctgtcccaaattttttgtagaga
atcaacaatttgaagatatgatagagacttctgaccattggatttcgacaagaacaggaa
taaaaaaaagacatctagctccctcttctacttctgtaacgaaattagctgcagaagcag
ctaataacgctttatctgcagctaacttacaacctactgatattaatttaattatcctag
ctacatctacgcctgatgatttattcggtagtgcaagccagctgcaagcagaaataggtg
caactgcatcggttgcctttgatattacagcagcctgctccggattcattgttggattag
taacagcagctcagttcatccaagctggttcatataagaatattctcgtcgttggagcag
atacaatgtcgagatggattaattggtcagatagagctacatgtatattgtttggtgatg
gtgcaggtgcagtagtagttgggcaaagtcttaacaacagtattttaggctttaagttat
gtacagatggccagcttagcagccatctacaattgatgaatacaccagtaaataatcata
aatttggattaacagatattccccaaggcaattatagttctataacaatgaatggtaaag
aagtgtacaagtttgctgtattacaagttccaatagtgattaaacaatgtttaagtaatt
caaaaatctcaatagatgaagttgattggtttatcttgcatcaggcaaacactaggattt
tagaagcaattgcaagtagattatcaataccctttctaaaatgattactaatttagaac
attatggcaatacatctgcagcttcaattcctctagcattggatgaagccattaaatcta
atacaattaaaccaggtcaaattattgttttatctggttttggagctggcttaacttggg
gggcaattgtcttaaagtggtagt---tgtaatattgcggatgacgagactcgaactcgt
atagctttcgctacacacccctcaagcgtgcgtgtataccaatttcaccacatccgcgct

ttttattatgaatc-ctaaaaattcataatatttatttaaatattaatataa--tgtgta
tatatatatatattcttgtcagaaactctctcatattttattttaagtgataaaattgag
t----ttagtatagataataaagttttattttaaaaccataa----aaataaatatataa
ggaaaataaaatatgacaccgtccttatcaagtttttgaacagtctaattcttggggcc
gtaatcgtagttgttcccataactttagcgcttttatttgttagtcaaaaagacaggaca
attcgatcctaaaaataacctagaaagatatggagaagcaatttaagattaataataaac
tatttttcaaacagatatgctctccatatatttctatact--gctgaatacttcttat-t
tcttactttgtaatagcctgcgcagtttattctgtt-ctttttgccttcatttactctt
tca----taacgatacaaatgtcagg-ttaataatat--ttcttttactctgcctctttc
atttaatatcaaagtattttatatatatttaactatgttcaattaataaagaaaagctca
tgtctttgtctaattggcctcttaaaaaagaaaacgctgaattacatactgctaagaaca
caaaacaaattacaatccctgacggattgtggataaaatgcttcgactgtggtttgttaa
tgtattctaaagtcttaaagcgaaatttaaaagtttgtcctcaatgtagctaccattttc
aagcttctagcaatgaaagaattgaccaattgatagacaaaggtagctggcgtcccatgg
atgcacatttgatttctacagatccattaggttttaaagatcaaaaactatacagtcaaa
ggctaaaagacaccgctcttaagactggtttacaagatgcagtacaaacaggcactggta
ctatgaaaggtagacgagtctgcttaggtattatggactttcggttcatgggggcagca
tgggatctgttgtaggcgaaaagttaacaaggctgctggaaaaagctactcaagaaaaac
tacctgcaattatactttgtgcatcaggcggagcaagaatgcaagaaggcatgttaagct
taatgcagatggcaaaaatttcttctgcacttgaaatgcataaaaaagcaaatcttcttt
atctctctgttttaacatcgcccactacaggaggtgtgacagctagttttgctatgcttg
gagacttaattattgcagaaccaaaagctcttattgcatttgctggtagacgagttatag
aacaaactattaaagaagacttgccagataattttcaaagttcagaatacctatttgaac
atggttttctagatttaattgtaccaagaactcagcttagatcaaagctaatacaaattt
tatcattgcataatcatagcaagtaattagtctaacatttgtcttaatattaaaatgcta
gtaaaaattaagatattaagaaattttcactaaagatatttgttgaccaactagaattac
aattactattgtagaaaaatttgtcttaattcta-ttttttttagtctttcaagcaattg-
tttttatgactttaaagaattattagcttaatacaaatttataaagagaata-----cat
actaagctaaatatagcaaaattatgtcttcctataataccgacctatcaaatttcgagg
aattttatgcttaaaagatcttcttggcttgctgctttactaggactactaactgtagct
tctataagcacgcatgtatatgccatagagctagacgaggcaacaagaactgttccatta
gaatcttctggaagaactgtagttcttacaccagaacaagttaagagaggcaagcgatta
tttaataactcctgtgctatttgtcataacggtggtatcacaaaaactaatcctaacatt
ggacttgacccagaatctctaggcttagctacgccacaaagagatacaattgagggttta
gttgactatatgaaggatccaacaagttatgatggtgcagaatcaattgcagaattacat
ccaagtattaaaagtgccgaaattttccgaaaatgcgtaatttaacagatgaggattta

tttacaattgctggtcatatcttgcttcagcctaaaattgtttctgaaaaatggggcgga
ggaaaaatttactactaaagtgtcaaatacaatagtctcatcagtactagtttatgta-t
ttattgcttaagtgtgattattttgttagatcatgatatatattaaagatggagtttgtt
aacagcaagttgcttttgatcaaacttgcttacattattcgtctagttattaaaaagga
gacgttaaattgaaaaagacgctttcagttcttttcactgttgttagtttttttgtaata
ggttttgcgcaaatagcttttgctgcagatttagataatggagaaaaagtttttttctgcc
aactgcgcggcatgtcatgctggtggtaataatgctattatgccagataaaactttgaaa
aaagatgtgcttgaagctaatagtatgaatagtatcgatgccattacttatcaagtgaaa
aatggaaaaaaatgctatgcctgcttttggtggtagactggttgatgaagatattgaagat
gcagcaaattacgtattatctcaatctgaaaaaggttggtaattgtacttgattttgctc
agtattaaagaatagacaatctatttagtgattcattagattgtctattctttgtttatg
---ttattatatacagatatttacacaatattaatatcatgatgaaaaaaacaccagcaa
ttcttgtactagaagatggtacttattacaaaggatggtcgttgcagcaagaaaaacaag
tcattacagttggcgaagtagtttttaatactggcatgacaggatatcaggaaataatta
cagaccctagttatttccagcaaattgtcacttttacgtatccagaaattggtaatactg
gtattaatgatcaagatattgaatctcatactattagtattaaaggtcttgttgcgaaaa
atatttgcaagatatcaagtagctggagagagcaggaatctttggttaaatatttaatcc
ggcaagccatcccttttatctttggaatagatactagagcgttaacccaatacctacgtc
aatttgggacaatgaatggctgtatttctactgataatttaaatcatagttatttaagac
agaaaatttctcagattcctgtcatgcagggtttagatctaattccatatgtaacgactg
acaaggtataccccttgggacgaaaagagcttacctatatggtacttgactgataggagta
gagtaaaattagctgataaattgaaagttgttgttatagatttggagttaaactcaata
tactgcgacgactagctacacttggatgtcaagtaactgttgtacctgcaaaaacgtctc
taagagatattttagcagataagcctgatggtatattactttctaatggtccaggagatc
catctacagtatattatggtatccagacagttaaaaaccttttagatcataatgtaccta
tttttggcatttgcatggggcaccaaattttaaaccttgcccttaattctaaaactttta
aacttaaatttggtcatagaggtattaatcatccatctggattaaaacagcaagttgaaa
taacgagtcagaatcatggttttgcagttgacctacaatctgtatttgactcttctgcgc
aagtgactcatttaatttaaatgatactactgttgcagggataggtcatagtcaaagtc
catatttctctgtccaatatcatccagaatcaagtccaggccctcacgatgctgattact
tatttgaaaattttgtcgaaataatgataaagtctaagcggaaagctaa-----------
ctagtaatcttcccatgctttatgattaaataaagcggataatacttgcactcccctcaa
ttgattaaaaagcggtaaatgccccccttgggccataagaattccaatgaaattcgtcagg
atatctacaaggaatatgattaacttcccacccgatcttgtgccaaaatttctcccaatc
ttttccgacgctaagccatatttgtctttgaacaaagagcccaaatttaccttttgagtg
tgcgtgccataatttatcaatagtctgtagatcttgagcaggtattttgcttatatctgt

aaaatatagccaattacgattttgtgcattcacaccagctagttgaacaagctttcgctg
agttaattgatctgctttgactaactcttgttgagtcagtaatacctgtaaatctttata
attcatttgcttagcagaacgtaatggtacaataccctcaggacataagctagtaatgaa
ttcaattatttctttattttactattaagtaatttgctataaattaaaccatcaacaca
attacttgtataatttgatttactaaacctctcgcaaaatagatctgacaaaacttttaa
atgtatagaatcgtcattctgtatagtttcaataatttcaagttgtttcttaa---tact
attagacttagtattttcatttaattctaaaagctgaactcgaatttgattttgcatctg
atgt-ttaaattgatttaaatagtc----attaggtcctaaagagaatttgattttttgg
ggaaattagaagaccgccgaattatttctaaaaatgtttttataatttggttaaatattt
tttgtcctgcataaattataatttgaccaattaaaaagactgtgctttttatttaggga
caaaaaaatcttgtacttcaagtaaaaacgtaattaatatctgcattgatgacagtgtat
ataattcattagatctacaagcatatatatactgacaatttaaaccttcttgataaaacc
cccataccttataacgactttcataaattgctcttggtcgctcaacatagtttttaatta
aattattccaaattaaattatttttaatttttctaactttctgtcagataaaaattttta
cattgcacagataacttgcattggtttcatgcaccgtctcaaaattttgaattaacacat
gggcaataatattgctcagctgaattagataattttcaagaaaaccctcgacttgcttag
ttggcatattcagcgtaaataaattgaaagttttatttacagtctgagatgacccaaaaa
gcagctgaataataagaatttgtagtaacatcttatagtcatatattaggtagtttaaat
cgttaatatcttggcagatatgatcatagttcagttcatataacctgcaaaatcttttga
cgcacctgtggaataaatcaactaaaatattttcagttaaagttttaatatcatttagat
ctaaatttgaattataaatttctaaaagaattttctctaattctgttaatattattttca
ataagttccttttcacttcagtccgaaaaacatctaaaattaacacttcctgagaacaat
tagttaatttcttattaatttttgtagacgttctaactaataactctgcgacctcttgat
tgagcgttggtccctgagagcttggccaataattattcacgttataccatagtaaggaat
atttaaattaatgttatacgataacttttaatatactgaaaagcaagtttattattgca
tagcaattag-------gttttacagatcttaatatttatttatcaaataaagttatatg
atagtatcgaccgagctaattttattaaaattta----caataatgacagtttattactt
tgcgcttgctagtcaaaacttcctttagtacaagagcccttagaagaagttttcagaga
aagagttaattattatcaatcaaataataaagagattgattttggttagtccccaaccc
ctccttttaaaaaagccagaaatgatttcatttaaaaatcttgtacctgatgattctgt
agctataatttctactaatccagtttttattaactggctaaaattaagaattggatatat
ttgcattggacaatttgaagataatctgcaactttctgcagaatcactaaatattacttt
tgcaccagaaaaaaattaagcatacttttatcctagtcattagc-tattgc-----ctaa
gcgacaagtgtaatttagaaacactca-ttgcattatagtttatcgcttagtattttatt
ttgatacggagtaacaaatagtgtcagtatttcttgactaaatgtctctgttgccttgga
tttatatcgatttgggttaactataatagacagcatccttttttatagtgacattttcaat

ttgagcccaatggacaattcctagctctagttcctttccaatagctgaaacagagacaaa
tgctgcccctaatcctgattgtacagcatttttatagcttctatagaattcaattccat
ctctattttgaagcgactgctatcaataccatgctgactcagcactttatctataacttt
cctgattgtcgattgagtatctaatgcaataaatttaagcctgtataaatcttcttttg
gatgtcttctaatttagaaaagggatgtgattttggaagtattagtgctaattcgtcttc
ggcataagatgtaacctgtaaaacatcttgtaattcattaggaacttccccgccaataat
tgctaaatcaacctgcccattagctacgctccaggaaataagtctagttgaatgtacttg
taattgaacagctacttgtggatatctttgcctaaatagtcctattaatcttggcattaa
atatgtcccagttgtctgactagctccaataactaatgttcctccttgtagattttgtaa
gtcatcaagagctctacacgtttcttcacatagagctaaaattcttcctccatatcgtaa
aagtagacttcctgcctcagttaaagttgccttttattacctctctcaaacaaagaaac
atttaattggcgctctaaattttgaatttgtaaactaatagccggttgagaaacatatag
actattagcagctttttaaaactgccttctttagcaattgcttttaatattcttagctg
atccaatgtaaatggaagatctgtcattaaagaattgtattataata---------tata
ttattgagtattgattaatttatgcattcaccaagctaaatatttgttagcacacttcat
tattttcaaaataataaaaagtataatgttaaatacactgtattagattttaatatctct
ttgatgttaaatttatacaatagatatcgtaaatttaattttgaatataaaacataggag
ataat---atggactctagacttttaattgtactaataccagttttagcagctgcatctt
gggccgtttacaatatcggtagagttgcactacaacaattcagaaagatgacatcttagt
tcttcttagaatata-atttttaaactttatgggatagagaaatttataatctctatctc
tattgtttatcttaataaaaaaagtgaggctccccccttgtacttactgttattacggatt
aagattatgtac-agtaaatgtattcatgataatttgaatccaacctaacttatttctta
taattaattaaactaataacataattatggctgttccaaagaaaagaacatctaaagcta
aaaaaaatgcgcgtaaagccaattggaaaaatcaagcaagagcacaagcacaaaaagctt
tgtctttagcaaaatcagtcttaactaaaaaatcaaatggatttgtatataatctttctg
aaacagtagacacttctagtgaataagagtttcc-gtaatataaagctaataaagcctta
aaaatcaaattg-aagtaattttaatatat----taagcatttggaaatcatcctatttg
atactaaaataaatagtagatcaaataaaattaaagccataaattatgcagcaaagctag
atcaaactagtgaaatttggctgttcaattgtattgaaaatattcagcacgttttttga
aaagtcaattaaaatcaaatcatattactaaaattttcatttctggagctagctttgaat
atacagccggtttacctggactattatctagcttaacactaagtggtagaatacacccta
ttagtatatatgccccgcagtccctgaagcaatatcttcaatcttgtactaaatattctc
aaactaatttttcttttcctattaatttttacaatgtacaatatggggaacttgtagttg
aacagttttacacagtattatgctttccgttggatcccaagagtttactttatggcttta
ctattttaaaaaaggaaaaccagggagtatttaaattagcaaaagctgaaactttaaata
ttattcaagggcctatatatggaaaattaaaagaaagatataattttctcagtccagatg

gctactacttatatggtaaagatttctcttcaaatacaataatgggacataaaatatctc
ttccattattagctagatattctagaattatttctgagatgcattggttttgttcttatc
ctattaaattaaataattattcacatcagcaagccataaaatatttgccacataatattt
taactgatattatgaagtctcaaatatctcaagacaatagttttattgaataatatttac
tcctgatatataatagtctccgtaatctaaagaacatactagaaagaattacgaa-acct
atttttctacattaaatttataatcatttatgacatacgcaattattgaagcaagtggca
aacaactttgggtagaagaaggccgatattatgatttaaaccatatacctgttgaccctg
gccaatcaattatactaggtaaagttctattgttaaataagaatggagaagtgactttag
gacatccttgcatagaaggtgtcacaataaaagctacagtaatgaggcatctaagaggaa
aaaaactaactgtttttaaaatgaaaccaaagaaaaaaatgagactaaaaaaaggtcatc
gacaagagttgactcgtttaatgattgattcaattgtatactaaa------taaaatctc
aactagtatttttt-aa-aagactaacaatatttaactaaaatatagataatggcacata
aaaaaggtagtggtagtacaagaaatggtagagactctaattctaagcgcttaggagtca
aaaaatatggaggcgagcaagtaacggcaggtaacattctaattaggcaacgtggaacca
aggttaaacctggccaaaacgttggcaaaggtaaagatgatactctgttctcgctaatcg
atggttttgtacttttttgaaaagtcaaactataagcaaaaaacaattagtgtttattctg
ccaagaaatagctgaaacaattggtgcaagataacgtaagt----gtatttattttattt
accccggcaccatatacttgatttattaagaagtggagttccttaatgaaccaatgaatt
ctattaaagatctgttgtgaaatttaatgacaaacactattgtaatttcctgtctacaca
atatggctgctattctatattgtggtcagatacagaaactagtagtagcaaatgctcatt
atcaagtaagtgatatttacctaggttgtgtagataaaatcttttcagggataaatgcag
cattcattgacttaggaaagaatgagtacagtggctttatccatattagtgatactggtt
cgcttaaaaaaaaatattatgttaataatattactaacattttaacaattcgacaaaaga
ttttagtacaaattactaaagaaccaactctaaataaaggacctagactcactgctaata
ttacattatcaggtcggtatattgtattaatgccttttagtcaagcaatttgcatatctc
gaaaaatacatgatgaagatgagcggcattatctaaaagctttagctattttaattagac
cagcaacaatgggtttattatttagaccttatgccgtaggggttgatgaagaaattatat
taagcgaattgaaaaatttaaaagaacaatggaattttattcaaaaatctgcaattaata
gctgctgccctactctgctatataaagatgaagatattgttaaaaaggtaattagagatt
tttataataatagtacaaataatatcgttattgattcaaacttaggattgaaacaactaa
attattatattcatacttggcactgtaataattctagcgcagctacttcaattaagcttt
atagtaataatcaatgtgtattagatgcttttggtattaatcaagcaatttccagagctc
taattccaaaagttgatcttatacttggcggttatatgtttatcgaaactttagaagctt
ttacaattattgatgtaaattctggatcttttaataattctactaatgcaagagaaacag
tcttaaaaacaaattgttcagcagcaacagaaatagctcatcagttacaaattagaaata
ttgcaggcgttattataattgattttattgatatggaatcacaaagagatcaattacaat

tattagagcactttaataaagagcttgcgcttgatgatgctaagccccaaattgtacagc
tatctgaattaggcttagttgagttgactagaagacgacagggtaaaagcttatatgagt
taattagtagcgattcaaattactttcatcttttcatacaatcagagcggtttaagtctc
ttaaaagagtggatcacaaaaaacaaatattccagccccttagtaaatcatggctatcta
cagaaattaatactattaataaagtcttctttcaaaaatcagatttctacaaactggcca
acttttacttggtccgtaatttatatgtagttaacagaagctttacttataaacaaagct
attcattgactactagatcaaaactgatttactctcaagagtctagtagaatattaccca
gtaatagttatggctacctagccagtgatagcataaatagtaatcaaaagtttttactgt
gaaatgttcatt------------------------------------------------
------------------------------------------------gaggggctc-tt
tgttagcttactaactattctagttagttaattaagaattaaccattaacagctggagct
gttagagctactggtaaagattcaccagaagctaaatctagagggaagttatgagcatta
cgttcgtgcattacttccatacctaggttggcacggttgatgatatcagcccatgtattg
attacacgaccttggctatcaacaacagattggttaaagttgaaaccattcaagttgaat
gccattgtgcttacagataaagctgttagccagataccaactacaggccatagacctaag
aagaaatgtagagaacgagagttgttgaaactagcatattggaagatcaaacgaccgaag
tagccatgagctgcaacgatgttgtaagtttcttcttcttgtccgaatttgtaaccgtag
ttagcagattcattttcgcttgtttcacgaattaagctagatgtaactagggatccatgc
atagcactgaacagggaaccaccgaatacacctgcaacacctaattggtggaatgggtgc
attaaaatgttgtgctcagcttggaacacaagcatgaagttaaatgttccggagatacct
agaggcattccatcagagaaactaccttggccaattgggtatactaggaatactgctgct
gctgctgctactggagcagtaaaagcaacggaaatccatggacgcatacctaggcggtag
cttagttcccactcacgaccaatgtagcaagctacgccagttaggaaatgaagaacaact
agttggtaaggaccaccgttgtataaccattcgtctaaagaagccgcttcccaaattggg
tagaagtgaataccaatagctgcagaacttggaataacagcaccagaaatgatgttgttt
ccgtatagaagggaaccagcaactggctcacgaattccatcaatgtctactggaggtgca
gctacgaatgcaatgatgaatacagatgtggcagttaataaagttggaatcattagaaca
ccgaaccaaccaatgtataggcggttttcagtactagtaatccaagagcagaaacgttcc
cacaagctagcgctttcgcgtctttgtaaagtagcagtcataatttttttatcaattttta
aggatttaactaagatgcttcccagacaatcagttgtctgttaactatattattacttag
tctattttaaaatgagaactttttttttttaaaatattagctaaatttaatctcgataact
aa-aagttctgcccattaagatattttcagaaaattgaa----------taaataaattt
tt-act-------------ttttgccagattcatactattttagggatataatagtacca
atgcatattcatgctgcttattttgattttcatagtttatataattgagccttatttaaa
taggtatgtcacattttacaaaaatccaaacgactatacaagatttaaatctgttaaagc
atgctttaacagatttgggtttagtttgggacacgaattctagtcatatacagattggtg

aaaataataaacataaagtagacatcttgatcaaacaagataatttgtctcatattggtt
ttatctggaatgataatagataccacttagttgctgacttacagctttgggaacagccct
tatctctagaagtttttttagatagattatcccagaagtatgcttatcactctatcatcg
aagaaactaaaaaacagggctttgaaaaaatggaacagatatctcaacatgatgggtcaa
ttaagctagttgtacaacgttggagctgctaattgatatgtaagtgcgggcggagagact
tgaactctcacgagattatctcactagaacctaaatctagcgcgtctaccaattccacca
cgcccgcattattagtacttattgatatcttatcataattgatctaaaaagacaaactaa
taaagataaattagctgccaagtttaaaagcatcaacaaacaggccatatgttataccaa
tcttaaagttattgattaaattgattcttatgcctaattgcttttggttagagtaaacta
ttttactagttaattcagttgatgccactattagactggctgcgataatgccccaatctc
cagtttgtccaggaattgtagataaccctgttgaaataaaaaatcctagtaataagctaa
ttaagccagtagtcaattcgcttaaagaatagtacagcttatcattcaaatttttttatta
aataagcaagaaaaatagaaagtcgagttttaatcataatatttaataacggagttttaa
aaggatgaaaaatattaataa-taatcaatattttgtatccttttgaaacccgttaata
caaaaagtgcttt--------------tttaaaggtcttgctcacttaaagcattaataa
gatgttgaaagggttcttgtataatgtctttgccctcaattgattgcaaatccagttcct
cttctattatttcttgtaataactgtatacttcttactgtcggagcaacaggtacagaca
atgaattgtaagtatcttttagcccgtttaatacgcgttgatctagaatattagtactac
cagaaactaaagcgtaactagcatacctcagatagtattctatgtctcttaagcatgcgg
catatctcctagtggtataagaatttcctccaggtcttaataactctggctgttcttcat
acaactgagctgcagcttcttttaaaatattagtagcttgattattaataatttcagcta
tttttattctatctaatccacttgaaaaaaaagattctatttgtcccacagctactttat
ctagataacgaccagtaagatcgtaacgatttaatattgctgttatagcatcttgcataa
attagttctcctac-ttgctatcagagcggtattataaatgactaatattataatattat
cattaatgtttatgaaatttctttaaaacctattatttactgattagtttgcagtttaaa
actgctatagattttacacacatagagtatacgtacttttaattaatttttttataaaa
atcaagaaataaagtttttc-ccataggtatattaatttattaatatagctcaggttct
ttgttcaaataagttttgtaacatatgtaggctaaaataaaaataagtagatattacaga
gcatcgtgatttttctgatgaccttatttatctgtatagtaaatactttctaaagaaaa
tgaagcgataagtttaaaaaagaacgaagtagtcgaatttcaaaataatagtctttcctc
taaaaaaatttcaattatacactgctgagacaatataaaaagtgctcactaatattgtcc
aatatatttaaagtttcagccaatacaaaataaaatactatgaattaaaagaaatggaaa
ctacccttagtccttcagattcaaactagcactttttactctttgttgtgaaattaagac
agaagttttgcaaataacaaattgtgttgattctgatagtctaaacttacaatcaagtga
gtcaaaaaaatatatattagtgagcaataccttgttaaaataaattctttaattagcaaa
atgtttagtctttgcacaaatataaaaaaaagcttaggatattagaattcacagataata

agagtaattcttgattaatatatttaagtagtatgcttgattatgaagatttgcctgtat
ttgtagctattgttagcaaatttaacatagctagctatgtggaaaaagctaaatacctag
acaacagagcagaagatagtcattatataattacagtatccattaataaattagctaacc
atacacaaagagtctcatcaacaatcttatgcaaaaaaatcataaaatccctcttcaaac
ttttaagagtaaacattgaatataaagtaaaaatcaaaaaaatagttggagcttcaaaca
tgctaaactttgaagctttattagatgcaagaaaaaatactaaatctgtaaaaatgtaaa
cttcactatttatataaaaaatgttaaaagaaagtcagtgtaactgttgcttattcaaca
caagtatttttatagttgctttaataaaattaaagcatttctctattacattgtttgct
cgtgtgtcacgtcagaaaataaatactgagtttgtttttcaattgatgcactaacaaata
tactagtctttagcaaaaatcaaaatacatataaaggttgaaaaaaaagaattagtgaaa
gtaacaaaacaagggcaattacaattgttttctgcgtaatttacttcacaaaactgcttg
tactttttagaaattaatattatacatgaaaaattgtcatacacgtatgaccaattttt
tttgctctttttgatgtcatggagtaaaacatctgaagatttcgaagctaatgtctcaaa
actacacttttaacatttgaaaactatttaaccgtcacgtttgtaacagttcatctgtc
tttataataatcaaaactatatatagtatatggtacacaatgagcactaaatctatctat
aatctagaaatgattaaatattattactgtttttattagaaggtatatactaatatcat
agtaatggacatccatcgctagttaacatagaaacaaaataacacagcagcagtacctaa
gctgaaaaaagattccttgctctagtaatagagaggtgcattcaattaatatacaatcta
atctgtcaaattacgttagtttaaaaaaacagttaaaaggtgctactatatctagcaaat
tttgtctaggttttgttttcactaaaacagaatatttctttattgatacagacaacatta
aatcatctcaaaccattgattttagtggttgtgaaaatacagagcagttattaatacagc
tacgcaagtatcctgtaactccagaaacactcagtagatttcatggtccatatttcgaaa
agagtattagtagtaaaggcttgcatattgtcgaaaaagaatccttgtttcttattaatt
attctgaacgccgctacacagcttctaaacctatccaaaaataaccccccttgagtgtta
tgacggcttcatagctagatatatagccattgcaggttatattatatgtaatgagtataa
ttgtaatgaatgaaaataatatagacagttctttgctacaacatcaaaaaagatatttaa
gacatattaacaaattcgattatttagatgaaaacttaaaattagataagtctatctatt
ttatcattacaattaataagactaatcaatatatttttgtagaacagttctgttacaata
atgatggttatatttactcaatattgtttcaaggtcaaacggctcaatatctctgcttca
tcatctttcagcttaaccaattaaatacagtaatatctacatctcatgctttatatttag
ggcgagagctaatgaaatctgaagtagctcttgtattggaccaacaatatattcaagact
aaa--ttggatagacattaattatatttatgctagtataatattctcaagggcatgtaac
tcagtggatagagtatcagattccggttctgatggccgtgggttcgaatccctccatgcc
cgacaattattagtaattttggcgacagagcatct-------taatagccaatttaataa
tgtt----gtattggggctgcaaggtttctacattatgaaaataagaaaatatgaaagta
aaacaagctcgttaaaagagcttttagtaaaattaataaatgcagaaaataatattgttt

cttttctcaaaaattagctattgcttaaatgctgttagtttttggcattcgacgtgtta
aactcttatccgcggcgaatattctgttagagttgctcatagtacaagaaaagcttcata
atctatttatctttatgtttt-tacctatttttatttaaattaaggttagta-ttttcaa
tttttgtaatggacgtgggttcgagtcccaccagctccattaatttaact---------g
tcataattagctagtcaaatcttttaataaccacaaatataattggagccaatatggat
tttataacaatcacagaacaagctttcaagcaaattatagtcttgaaaagtaattatgaa
aacgatgtatatttaagaataggtgtaagacaaggtggctgctcaggcatgtcatactcg
atgaactttgagcacgtaaataatctaaaagacacagacgaaaaacttagtttagacaat
ttctctgttgtttgtgatccaaaaagccttttgtatctttatggattatcattagatttt
agcaccgagttaataggcggagggtttcaatttctaatccaaatgctagccagacttgt
ggttgtgggaagtctttctcaggctaaaggctctactatctttcttttatattttgtatt
ataaatattatttctgtaatcag-tattataaatgtgaatgtattctaagcccaatcttt
tttgagctagcaaag--------gaactacaataaagaaattctattatagtagc-----
----ctaaatacacataatcaattgattaaaga---------------ggtcaactgctg
attattatctaatttaaagtcgtcaattttatcttgatctatgctaatcttt-gttagat
tcgatttattaggaaatcacaa-ctcttttgcaaaagagagtttgtatcttgctgttaa-
----------------------gaataaatcacgtatttt-atttattttataaaaactc
aacatgaatcttgctcaaactatcgctgagtactctagtgcacaaaattccaataacttt
tattgcactttcgaacatttatctccccatgatttaaacagcccaaaatcatcaaaacca
aatttgtcattttctacaaaagtaattacttttgtaatacctcgatttgattttattcct
tctaatattaaaaattctattagagaaggagttattatatctaatatcgaagataataat
atttttgctaattcaactaataattgggtaaagaagcataaagataaaaatcaccattct
ttattctataaaaaaatttttaaagctttgctaaaccagaaaattagagtggttgcagag
ccaat------taaaactcctgaaataggaaaagaacatgatatattagtaagtcaaaaa
tatgtttggggtaaaagctggcaaccatcaattatttcatcttaccttaaaaataaaaac
aacaatgtagtttctaatgcatctaagaaatttttaatcgatcagttatctgcatctcca
gtttttgtgataaagaatgggtttaatgagattatcttagggcatccattatctcggatt
aaaagaggaggtattaataatatcacttattcattccaaaatttattaagtcagcataag
caagcttaccctatatctactggtttatttttcttccatccagatgatgcatttgaattt
aaaaactttataaaatcagttaaccctttggctgctcaacagatggagataaatgtagag
cctgtcggtctgcattttgcttataaaatgaatagaagtatctcctcagatactaaattt
acatttattccagattttaaagaggtaggagatttgttattcaaatatcggagagataag
cacttgatttttcataaagatcagcgctatggtaaaaactttttccaggggcaacctgtt
tatattatccagcctattagtttaaaagatcgaagcgggaagatcaatatcattaagttt
actgggctaaatgataatagagaagtcctttttactaatcttgaagctgctaataaatcc
tggacaaattttataaagaataattcgcagttgaaatcaattaaaaagcctactttgtta

gtttataatctagaaagtttcttaaaagaccaagaaaaactgaataagcaagactttcaa
aagtttgtcgtagtcactaataaaaaagcttatgttgctactaaagagttaatagcttta
cccgattctaataactttttaagcacttaaaattaaatatgaagcctaaactcttttt
atggaactatgggttagacgtttatttccaccttaacttatgaat-aaatgaagttcgt
ctctattttcttgagtagtattcaataactaatagttcatttagttgcaatgcaacccat
tctctatcaatcactccattgactttccctgatagattagacttatttaattccaagtgg
ctaggaatattagctagtccagggaacgctaaatagttttctactagttgtttagatgca
tcctgagcttttacactaattagttctcctggcttgcattgataactacatatcgatact
acttggccattaatacaaatatgaccatgattcactaactgccttgcagctggaattgta
ggagccattccaagtctaaaaacagtgttgtctaacctcatttctaagagctgtaacaaa
atttgaccagtggaaccttgtagtttttttagctgccttaacatatttaaataattgtttt
tcactaagtccataattaaagcgcaatttttgcttttcttctaatcttacagcatactca
gatggttttcttgatttttgtccgtgttcccctggtggataggaacgcttaattgctttc
ctactgagcccaggtaaatcacctagtctacgggaaatacgtactcgtggccctctgtat
ctagacatattgtataaattctccgaatataaat-gattaag----------tgatgatt
agctatttttaacataaaaagagtattaaaacaaagc-ttttgctaaaaaatttttata
agaaacatag--------------------tgtttcgt-tttttaagcgggtagcgggaa
tcgaacccgcatcattagcttggaaggctaaggtttaccactaaactatacccgcaata
ttcaagagattacaatatacttaatataacacaatcatattacttgaagatattattgaa
actaatcaagga-tagagttatttatgccccttctaaaactacatcgaggaaacgagcaa
tttctgcagcctgttcttcaacttcggaaagtaacaacggttgcccgactctggttaaag
gaattactcttttatcttttgtgcatagataaatttcgcgtcttggattgagaccttctt
taatatctatttttattgatttaatttctttaatattaaattgaagacaaatttgtctat
ttcttcctgggaagccaagcctaaaaatttaacagttccctcatctctgttaaactcat
tatatccagctcctatatcccaaataatagttagccataagaataaactaagaaagattc
cgatactgccatagaatgtcatgacaatgccttgtggaataaatactaattctgttgaag
ttgtaaaaggtaataaattaacttgaaagtagcttgatagtcctgcaagaagaaatccta
aagctccagcgaagattattgtagcccaccagtaattgctaagtcttcgtgatcctaaaa
tcaaatcttttctaacagattgtatagataatgttttttcataatttgttgctttaaaa
caata-tactaatgtgaaatataaaaatttgacaattttt-ataggcggtttggatta
gttaaatgagctttatagctttgagtcctaaaaataagcctgttgctaaaccaaaaaaag
cagcaagaaatattgtatatcctagaaaaactgaca---tatta----ttattattattg
ttacaatagtgtttacaaaaatagtggaaatctattactaaagtattatattacgttaaa
tgccttgctgtctacatgaaaaaactttttcagtttttagcattgtatttatattagga
acaatattgtaataattttatttaacttctggagaacagaattttatggcagaatttatc
aagccttataatgacgatccttttgtaggaaatctatctacaccagttagtacgtcaagt

tttagtaaagggctgttgggaaatctaccagcttatcgtagaggtttatccccacttctt
cgagggctggaaataggtatggcacacggatacttttaattgggccttttgataaattg
ggcccattacgaggtactgacgtagctttattagcaggatttttatcttcagttggctta
attattattctcactacgtgtttatctatgtatggaaacgtatcttttactagggccgat
tcaaaagacccattacaaactgctgaaggttggggccaatttactgcaggatttctagtt
ggagcagtaggaggttcgggatttgcttatttactgctagctaatattccagttctacaa
actgcaggcttgagcttattttcttaagctagtaaggggacttgaacccgtaacctactg
attacaaatcagttgctctaccaattgagctatactagcatatatataataataacatat
aaagatatttgttgcaatctggtattttcagatacaacaaatatctttttttacttaaaca
aaactgctatattaacata--tataatatgttaattggaatccgaattatctgactctgc
agaatattcttgatcacagtttagataataacaaattgtttgatctaaacaagttgcgga
ccagccaactggtgtttctcctgataattcttgcatatctccagaattgtaaaaatcatc
attttgcttgttactgtttgtttgcataaatatcactcccaaaacactctgaacttaatg
tgaagtaattcaacttctcaatgatatattaacacaaaatataaaatatggtgcctattg
atcaattgtaaaaatattactaaatttacaaagtttttgtgagcgttttaatggcttttg
ttgaaataagcatttttatccatttgttttgctctcttgaccatactttttatgctgca
aattaacttttttgcagtttttttgttcgtttatgagagtgggatactgcatatccattat
tagctacttttccggtaagctgacatttttttgacataacttt------attaacttata
tatttatttaaagtactcgctagggtcgactttggcacagcaccaattacagtgtcaatt
ctttcgccggctttaaaaatcattaaagtaggtatactcctaataccatattcagctgca
atagtaggattgtcatctgtgtttattttcactaccttaatagatgattcgtactcttct
gcaatttcatcaactacaggagaaaccattctacaaggaccacaccatggtgcccaaaaa
tctacgagtacaggtaagttattattaataacttcttgtttaaaagaggcgtctgtaact
tgagatactgacatatttttttaacctttaatatctgttccttgctaagcaaattttatca
caaatttcaagaaaaatcttctatagttaattactattagtgttgaatttaattttt-tt
aattagacattaaaaaaacttattacgcatataaataaagtggtggccgagtgatatctt
gatagtgataatttttctattaaggttgaacactagaagtaattctctaagcaaatattt
acttgagaa--acttctttgtgaccataaaagcttaattatattacaatatattttgtta
tttgtaacttagttctggtaatggtataaacaacgcaaaagatactgccttataatcaag
gaggaatacatgtctcaatccgtagaatcacggactaggattaaaagcgaacgttacgaa
tctggagtaatcccttatgcaaaaatgggctattgggatgctgactacgtaattaaagaa
acagatattttagctctgttcagaattactcctcagccaggtgttgaccctattgaagca
tctgctgcaattgcaggcgaatcttcaacagctacttggacagttgtatggactgatttg
ctaacagcttgtgacctatatagagcaaaagcatatcgagttgatccagttccaaacgtt
gcagatcaatactttgcttatatcgcttatgatattgatctatttgaagaaggttccatt
gcgaacttaactgcttcaattattggtaacgtttttggatttaaagctgttaaagctctt

cgtctagaagatatgcgtatgccagtagcttatctaaaaactttccaaggtcctgcaact
ggattgattgtagaacgtgagcgtatggataagtttggtagacctttcttaggtgctaca
gttaagcctaaactaggtctttctggcaaaaactatggaagggtagtgtacgaaggtcta
aaaggtggtcttgactttcttaaagatgatgaaaatattaactcacaaccatttatgcgt
tggagagaaagatttttatattccatggaaggtgtaaataaagcatctgcttctgctggc
gaaattaaaggtcattaccttaacgttacagctgcgactatggaagatatgtacgaaaga
gcagaattttccaaagaggttggtagcatcatttgtatgattgaccttgtaatcggttat
actgcaattcaaagtatggcaatctgggctcgtaagcatgatatgattttacatttacat
agagcaggtaactctacttactctcgtcaaaaaaatcatggtatgaacttccgagttatt
tgcaaatggatgcgtatggcaggtgttgaccatattcacgcaggtacagttgtaggtaag
ctagaaggagaccctttaatgattaaaggtttctacaacactttacttgaaagtgataca
gatatcaatctacctcaaggtctattctttgctcaaaactgggcttccttacgtaaagtt
gtgccagtagcatctggtggtattcatgctggtcaaatgcaccaacttcttgattactta
ggcgatgatgtagttcttcagtttggtggtggtacgattggacatcctgatggtatccaa
gcaggtgcaactgcgaacagagtagcattagagtccatggttatggcaagaaatgagggt
cgcaattatgtagcagaaggtccacaaatcttaagagacgctgctaaaacttgtggacct
ctacaaacagctttagatttatggaaagatattagtttcaactatacttccacagataca
gctgatttcgttgagactccaacagcaaacatctagtttaatgactactcgctaatgctt
aaattagcaaattatgagtagaattgacttataacaataaggagcatagaatagtgagac
taacacaaggggctttttcctttcttccagatctaactgatgagcaaattaacaaacaac
ttgcttatatcgtttctcaaggtttatctgcaaacgttgagtatactgatgatcctcatc
caagaaattcctactgggagttatggggtttacctttatttgatgtaaaagatgcatctg
ctgttatgtatgagattagctcatgcagaaaagcaaaacctgattattatgttaaagtta
atgcttttgataatactcgaggtattgaaagttgcgtaatgtctttcattgtaaatagac
cggctaatgagccaggattcttactacagcgccaagactttgaaggtagaactatgaagt
acagtcttcatagctatgctacagaaaagcctgaaggatctagatattaatttcgattaa
tagataataactattaattgattacccctctttgacatcttaattaatagattaatttttg
attttaaggaggggcaattttcgaaataagttttggctaaattagaattaaccatcaaca
atacacagaaataatgcaatcacaagatataatttccaacgatactcttgttaatctaca
agaagaatatgatagaacacaaattcaagaagttcttaatgagttaaaccaagaactaat
aggattagtcccagtaaaaactagaattcgagaaattgctgctttattattgattgatag
actacgtagaaaactagaactagtatctggtaatccaggattacatatgtcatttacagg
cagtccaggaactggtaaaactacagtcgccatgaaaatggcagatattctgcataggct
aggctatataaagaaaggacatttgttaacagttacaagagatgatcttgttggacaata
tattggtcatactgctcctaaaactaaagaagttctaaaacaggcaatgggaggagtctt
attcattgatgaagcttattatttatataaagcagataacgaaagagattacggttctga

agcaattgaaattctgctacaagtgatggaaaatcagcgaaatgacttagttgttatctt
cgcaggctataaagatagaatggagaagttctatgagtcaaatccaggactttcttctag
agtagctaatcatgtagactttccagattatacttcaaacgagctattacaaatagccaa
aatgatgatagaagagcagcagtattgttttactgaagaagcagataagactcttttaga
atatactgaaagaagaatgaaacaaccttactttgctaatgctagaagtattcgtaatgc
tattgatagagctagaatgaggcaagccaatagaattttcgcgagtggagaaaaagtatt
gacaaaagctgacttagtaacaattgaagctgaggatatcttaaaaagtagattgttttc
attacccaatacttaatcattaagtag------------------catgaagaaattcta
ataataagttgcaaatttccttgaaaacctatactattaacatagtatagtta-tggcgg
catagccaagtggtaaggcagaggattgcaaatccttcatcccccagttcaaatctgggt
gccgcctagtactaaaaa--gggggtgtggtggaatggtagacacaacagacttaaaatc
tgttga-ttttagtaatcgtgagggttcaagtccctccacccccatatatccaattataa
tgaaaaacaaatgtgtatttgtgttaattgtaaccatattactgaatgtagcacttatca
tttaatagaatctcagcataaccagacccatattactgaaactccactctttatacctaa
aggtcccatagtttgtgttaatatatctagtaatagcagttacactcaaattgattggga
tttagttgaatgcttgtcatttgtagaaaaaccgaatagctggaacttaggttctgatta
a----------------gaattataagcaatagctctgtcttttaggtactctgtattaa
ttttagacgtccttactagcaaaattttccctgttcttgctatctctacgataccaaatt
tagttaataattgctcaatagcaacaatcttgccgggatcgccagtaacctctataatta
agaggtcgtcagcaatatctactattttttgccctaaaaatctttacaatttctaaggctt
ctgttctagtaagagaactaattttaatcttaattaacattaattctctttcaactgagg
gaatatttgttacgtcttgtacattaaggatatttactaatttatatagttgtttagtga
gttgttctatagttctattatctccctgaactaccatcgtgattctggaaactccaattt
gctctgctggcccaactgctaaactagcaatattaaagcctctccgggcaaatagaccag
atatccttgaaagtactcctgcttcatcttgaactaaaactgataaggtgtgtttcataa
aattttaatattgtgctattcaatctattaaatatattctatgaaaattacagaaaaaat
actagttactataaagccaaattattcaaactaaacaaaattctcttatcgcacagcata
tctaccttgtaatgttatactattaacttagtatttaatgatattctaattctaaataga
atgctaaatttttttaaaatttgttaggaacttgtgctagaaaaatacaaaaacagtaaga
aactctctcgagatctaccccaaatcaatgatcgtattagatttccaaaagttagagtaa
ttgacgacgaaggtgaacaattaggtatctttgtgcctgaagaagctatacaattagcta
tccaacaaggcctagacttagttgttgttagcgataagtctgacccaccagtttgccgaa
tattggattatggtaaatataaattcacacaagaaaaaagggctagagaagcgaagaaaa
aacagcacaatagtagtcttaaagaagtaaagatgcgctataaaatagaagagcatgact
ataaagttagaattaatcaagcctccaagtttatccaagctggggataaagtaaaagtaa
ctataacatttcgtggacgcgaaattcaacattctaatttagctatagatttattgaata

gaatggcgagtgacttaaccacaacagctgaaatccagcaagctccatcaagagatggaa
gaaatgtaataatgcttttatcgccgaaaaaagttagct-aaataatttt-atttgaatg
catctggctggattcgaaccagcgacgtctctttcgaaatggcggattatgagtccgctg
ccttcggcccctcggccacagatgcatgacttacaatttataactcataaagcccataaa
aatcaatattta----tttttcataatattta-taaagattattattcattcatattgtg
atatgtagcaactgctgaaggtgacacttggttcaagtaccgaaatatccaatatttaaa
tattgtgtctaatataactgggaaagttgaaataaataaaaaaaataaagtcacgactttc
tggcagtcctaaatgtcttaaaattacttcaatgataacttcccagccatgaggagaatg
aaagccaacaaacatatccgtaaaaagaataattaaaaaagcttttgccgtgtcactaag
accataaataatttcatttaaaaaagatttaacaattgcaatttgtcgttgtccagttat
cattagtaatataaaaactaatattgatacgatgtcggataaaatatttttaacagcatt
tgcactttcatttgcatagtattctcctaactctctcgccttcagttgtactctttttc
tataatttcataagatatatcttctgatgggtttaacaagacttcaaaatggatttttc
ttcaaaacgttgcagttctgcgaaagctctttcttcttgcgatgaatttaaaaaaatttt
aggttgttcttggttccataaataatcaatacaaggaccaaacacaaagaatttcgaagc
ctggttaactagtactggagatataaataacaataaaatatattttacggaagtaatagt
ttgatgcctagatattctaaactcttctatagcttcagattctccattaggatctaactc
tttgcgaaacttttcaaaagtattagtaattgacctgggtattggcccaactttttcaaa
ggcaaattgattatttcgttttaaattccaatatttcatttttatcgctgctaaaattt
ctattaatcaataattgatcacatattcactataaatattaattcgagaactaaacacat
aaaacaatagtcatgaataataaaatacaatttaaatttaattatgcattattactatct
tgcgctacagcaatttgtggatataaatcttgtcgcgaaactataagtgtactactaata
aaatatgaaagtggctattcattactgtctgatagaccagctgcctttattcctataatt
gtattaaaaaaaaatctttaaactccctacaatcagaaccttatagtgctaagttatgtc
ttaatggaatgggaaatagaaagtttatcaacagaattacaagtctgttaagctatttag
aaaaacagttacactatactgaacatcttaatcaaataaaaactagtggttacttttctt
tactttatattaacagtaaagcaatgtatacaagagaaattagtagtctttttatagtac
ctaatgagattctaaaaaaagtttatatttataataaaaaagacaaactgatacctcagg
tattcttaataaagttatttgagaagcagataggttacccaaaaaatttgcaaacttga
atttagctgtttcaaaaattatgacatggtattatgatagaggatatcaatggtctttag
tggaaattaggcaggcgaatgatccatcttctctcgtaattaatattcatgaagggctag
taaatacaattacagtagaatattatactctttcctataaaaaaatatgtggtatctcaa
atacagaattaatcgagcaatatcttggagtaagaacgggggttccaataaatattaatt
ttttacaaaaaagaattaattatttaaaagataaccaattagttggcgacgttatttaca
gtattgaacgatcgaatagtgacttaatgagtctagatattaaatttcagatacaagaac
taagagataaagagatactagtaatagcagaaaggtcttctaaaacttatccaattattg

accatacatctgacctactcagtc---aatactctagtttttccgattggctagtaactt
ccaatataatctcaccgttaactagtaaactaaaagcatgctaccttaaccgtaaaacta
attctcaatatatatatattaatacttatagtctgataaaaatattagcttcttctattt
attatcgaaaaacaataataagtgattatttaatttgctaaatttattaactggggcca
aaaaaaacactattggatttcagctatatatgcgaaatctaagctatgctaaatcttttt
gtatatttaacatgaaatttatacagaatggacttaacatgagaatctcatatctaaatc
ctgcattaagaataaatcagagttttagctttcaagtggctatgcaaataattaagcaat
atcacactggtacagaagaagctttatctt------------------------------
------tattttcaaataaaggggatttgcacaatatcttgctgaaagcgtatttactt
atcattttacatcttatttttcactatctgaaaaaatattactttctcgagatatttata
cagattccagattttacaactctgagcacttaagtttttataataatagtaaaaatggag
ctataagcaattatgatatttttaagcaaaataaaaaattattgtaccaagaatttcttg
ctttattattaagcatacgttatcaagacttcagttatgtagattggcccttaaaaggtc
atttattggaattgaaatctttctacttcactccattgcagaagagtggtttttacttt
actctttgctggacaattataaaaaactttttcccacagaatttattttaagcaagtat
cttgctgtaatttacccttatacttcagaaagcattttaaccatatttaattattacta
tgaagtgtcagtcaaacttgaatatagacactatatatttgctactaagtagtcgtccag
ctgaaacaaccctttataagtctattcttaatttctctcttaaagttagaatggaatatc
tcgtacctataagtaggagcctaagaatctctgtcttttataactatttagactgtttat
caatgagaccgcttaaaaggctagcatatatatggcaagatttaaaaagttataaacagg
ttcaatactttctattggaaaaattttcgtgtggatttgggattcaacttaaattgccaa
ttaggcagatgccactgctctcaattgaatatactgtaaatagtggtcgcaatttttgta
tttacctacatatctattattaacgataactactataaaaatgaaaaacttaaaatcttt
ctttaaccgcagcaaaggaaaatggatttcacaaagaacaacttatgaactatctaataa
gaatatgagttcagtgcaatctcaaatccaaacaaagttagatgaatcaccatcagaatc
tacaatatttgtatcattgaattgggggagatctttagacaaattaaaaatcactcacg
tgatcaggatgtaaacaaatataatgacaaatttaacttagaatttagtaatcaactaaa
tagtcgtaagctattcacactatgtacagctacagatcctagcctaattagctttaagac
tagatatggcagtactactgtagatgagacatattggtttgcgacaaataatttgcgcct
aagtactagcattattaagaaatttgatacttgcgtagcagtatctttttgctcagaaat
taaagtttaa-gtcatctgtaaaagcaactaataacttgta-agctattagttgcctttg
tctgtctaagttttatataaaagaatatacgaaattatatactactctaaatctttacgg
ttgggattccttgaagggtcattagataagaatccaaaaacaaacagagaaataaagaaa
ataacggtagtatagacaaatattttcaaagtaaacatatttttatccttaaaatctatt
agtgta----gtatttatatatc-acacataattttatatcacgatatgagtttcggtac
tattttataaataaatcaacatctttattttttagtaatgatttatattttaatatacgt

gatgtactaaatactaaatctttttgtacttgtttcatactgtttttgaatattttgttg
tttctatggagcttcccttcaattattacataatctaatctctttataagtttaaaaatt
tctatagatttttttgttccagataattaagttgactacaactaatctttttctttgtaaa
agtcttacctttaacttaattatttgattttattagcactaatttcaatactcttgcag
ctcaaaacctgcactagtaagatacaactattcat-atttatataaa------tctttta
aatagtttagtataagagattatttt--aaaactagaacgatatttcattatcctgttgg
ccttgctcatcactaatatcttttaagacttttaagatttttccaaaatactcttgcaag
tatttttctaaagacataacttctgagctatcaatgcctaaaatattgtaaacttcatgc
ataggagcactaaattgatctccactagttaaaatctctgcaaaggctagacggtcagaa
atattccaggtccattgaagagttttagtaatttttcttaaagctgttaataatccgatt
ggaatttgtgatatttgcgtttttttgaccagacagcttttcacacaatttaataatttca
gctgatgtccaggctgtgttaccaactaatggtaaaatcctattttctgtggaaggaacg
ccaagactttttataactaattttgctgcatcttgggtatcaatatatgcaattggtgta
gattctccggtaacccatactgactttttgtctaaaataggaattgcatattggttaatt
aatccttgaaaaaatcccccccaaagaaaaaaactgtgtaatttatattagatttttgaagg
aaatctactacttgagatttcaaactcatcaatggaacagcggggtacttatctgcattt
aatattgaaaagaaaataaatctttcaacttcagctgctttagctgcttcaattaatgcg
attttaccatctaaatcgatttttgttgcattatatggatctgtaggacgagatgtcgaa
gcatctatgactgctgttactccacaaaaagattgtaagatactttctggcaatttcaaa
tcaccatatacgagctctgccccccattctttgagaaaagctgattttcttaaatttctt
accatgcatttaacattgtagccttcatctaaagcacgccttacaatttgacgtcctaga
gttcccgttgctccaattactagaagagtcatattaattaaaaataagtatgtataaaga
cattgatttggctggcgacaaagagtacattaggtagagagggaatcgaaccctcatgac
cgaagtcgtcacattttgagtgtgatgcgtataccaatttcgccatctaccctattatag
accaactactacgagtaatttagtctttagagtaaaaataatactaagttttaatagata
tgtcaacttaaaaatttatattagtaaacaa-cggagtctcttatggccacttttgaatt
aattccttatagattatatttaagtcaatctaggacctggatgcatacaatgaaagcaga
aattaaactatatatcttaactatgttatggatttcaatttttattttctcttattataa
actatgtattattgctctaagtttaatagctattagccttactataaggagtaagcaaag
tattatacaaaaacatttattacaaaccttcttaatgacagcattaactacttgcttgtc
atttagtttaaccaacagttataaacaatattctcccacaaaacaattacatcatttatc
ttgtttaaataagagcaaagtttttcacagttaccagcatacacaaaaagctaatatttg
taatattgcaagtaaacaattacaactaactttaaaaccttcactatattttttttattac
tatacattctattaaactagttatgataacaacttctccagaaattttagtagttactgc
ttataaatctagaattataaatatgctattgaataatgagctattgtttatgtttttact
atcgtcacatattgttactaacataattagaaaactggacaaaattactcaagtagctag

cttaagaggtcgcataaattttcataaatcatcaacaagatttttaatactttctttttt
aatatttgaagtcttttttttagaaattattaaggaatctaaagaaatatctcaagctct
ttatactagaaatcttaatgaagaaaatagtaattttctaaaaatctacaaagcaagatc
taacattagtgattggttaaatgcaattattggaacattatattttattattttaggtct
agcataataaaaatcacttgtacttacgactctagtaatacgaagagtaatacacttaat
tagtagttttaatatatagaat-taagatattatataaataattaaatgtttcattatgg
tcaatactcaaaatcaaatttcacaaacttcagagcttgattatatagttaatcaacctt
ataaatatggctttaaaacttctgttgagtctgaacaatttccaagaggaataaatgaaa
atattgtaagattaatttctaagaaaaaaaatgaacctgaatatttgttgaacttcaggc
tgaaagcgtacaaaaaatgggaaaaaatgagtagtccatcatgggctcatcttaagcatc
cgaatatagatttcaataatattatttattatgctgttcccaaattaaaaaaagaattaa
acagcttagatgaggttgatccagaaattcttgacacttttaaaaaattgggtatatctc
taaatgagcaaaaacggatttctaatgtagcagttgatgctgtttttgatagtgtctcta
tcgcaactactttcaagaaagaattatctgaagctggtgtaatattttgttcaatttctg
aagctgtacgagattatcctgatttaattaataagtacttaggcactgtcgtaccttctg
gcgacaattattttgctgcattaaactctgcagtatttagtgatggctctttttgttata
tccccccctaatacagtttgccctctagaactatcaacttattttcgtattaataatgaag
aatctgggcaatttgaaagaacattaattgtagcagatcgtggtagtaaagtgagctatt
tagaaggctgtactgctcctcaatttgatacaaaccaattacatgcagcaattgtagagt
tagttgctcttgatggcgcagaaatcaaatattctactgtgcaaaactggtatgctggta
accaagaaggaaaaggtggtatatacaactttgtcactaagcgaggcttgtgctctggta
ataattctaaaatttcatggacacaagtagaaactggatcagcaattacatggaaatatc
ctagctgtattctggctggccataattctcacggagaattttattctgtagctttaacaa
ataattatcaagaagctgatacaggtactaaaatgattcatattggtaataatactaaaa
gcagaatcatttccaaaggtatttctgcaggtaaatcaaaaaatagttatagaggattag
ttaaaattgggcctcaatcatttaattctcgaaattactctcaatgtgattctttactaa
taggtcagtcgtctcaagctaatacttttccttatattcaagtacaaaatccaacatcaa
aagtagaacacgaagcttctacttcaaaaattagtgaagatcagatttttatttttttac
aaagaggtatcagtttagaagaatctgtggctctgatgatcagtgggttttgcaaagatg
tatttaacgaacttcctatggaatttgctactgaagccgatcgtttacttagtttaaaat
tagaaggaactgtaggatgagtcaaactattttagaaatcaaagacctatatgcgtctgt
tggcgggacaacaatcttaaaaggtgtgaatttatctattcaatcaggcgaaatacatgc
cattatgggtcctaatggctcaggtaaaagtacactatcaaaattaattgcaggacatcc
agcttatgctttaatcagcggagatatcttatttttggaaacagtattcttgacatgga
accagatgagagagcaagggcagggatttttcttgcctttcagtatcctgttgaaattcc
aggagtcagcaatgctgactttctacgaattgcattgaatgctcgtagaaaattcaaagg

attatcagagtttagccccttagaatttttcagttaataacagagaagataaaccttgt
aggtatgcaagacagtttttgacaagaaatgtaaatgaagggttttctggtggagagaa
aaagcggaatgaaattcttcaaatggctttgctagacagcaacttatctattttagatga
aacagattctgggttagacatcgatgcacttagagtagtagcaaaaggaattaatacttt
agcaaaatcaacgaattcaattattttaataacccactatcaaagattactggattatat
tattccagactttgttcatattatgagcaacggcaaaattgtaaaaacaggagatctcac
tcttgctcaagatttagaaaagcatggatatgactggattgtaaattcttaacaatttat
aaa-------aacattgagactaaagaatactctttagtctcaataatatttgctaataa
gaagta-agctatttaagctttgccaaaaccttgtttaacatcttcgattgctctttgta
gcaattcttcactagatgtatctaatttttactactacgaacactttctccaaactctg
gtttagaattctttaaatcctctcttagctctctaataaattcagctactttcgagactt
caatatcatctaagtaaccattaattccagtatagataatagctgtttgttcttcaactg
ggataggagaattttgagcttgttttaaaatttctctaaggcgttggcctcttgctagtt
gattttgagttgccttgtctaaatcagatgcaaactgggaaaaggcttctaattcagcaa
attgtgccaactctaatttcaatttacctgccacttgtttcatggcttttatttgagcag
cagatcctactctagaaacagaaataccaacgttaatcgctggtctaatccctgagttaa
ataggtcaccagataaaaagatttgaccatccgtaatcgaaattacatttgttggaatat
atgcagatacatcccccgcttgagtttcaataataggaagagcggtcatactacctcccc
ctaactctgaatttaatttagctgctcgttctaataatctagaatgcaaatagaatacat
ctccaggatatgcctcacgtccaggaggtcttctcaatagaagggacatttgacgataag
cttgagcttgtttggttaaatcgtcatagataactaatgttgccttacctttatacataa
aatattcagctaaagctgcacctgtataaggagcaatatattgtaaagttgctggactgt
ctgcattagctgcaactataatagtatagtcaagagctcctttttcttgtagtgaagata
caacttgagccactgaagatgcttttttgtcctatcgcgacatatacacaaacaacatctt
gccctttttgattaataattgtgtccaatgcaacagctgttttacctgtttggcggtcac
caataattaattctcgttgacctcgaccaattgggatcatagagtcaatagcagtaatac
ctgtttgcataggttcacaaactgattgtcgcccaattataccaggagccatagactcaa
ttagtcttgttccattgcttgcaggctcacctttgtcatctattggacgagccaacgggt
caactactctacctagaaaggcgtcgccaaccggaatttgagcgattctaccagttcctt
ttactgaactaccctctaaaatatctctaccatctcccattagtacaacaccgacgttat
cactttctaaatttaaagcaaccccgatagttttatcttcaaattcaagcaattcaccag
ccataacttcatcaagtccgtatactcgagcaataccatcaccaacttgtaaaactgttc
ctatgttggctacttctatgtcttgatcatacttttcaatttgttgacgaataatactac
ttatttcatcaggtctaatgtttaccatgtttttaatattctttttattaattggaaatt
ctaaattggtaattgctcagattaatgttatctg--ctttttactctgttttaagttgct
actacgtcaaggtgtgaagccatctgccttagttgtcctctaatactagtgtcaataact

ttcgagccaatctggatagtgaaaccaccaattagttcaggatcgacagaaacaattagt
tttacttctttagccttagtcataattttgatttttccgttaagagcttttcttgatca
gagttgaaagcaatagaagtattaatatttgcaattgttagagattccatttgataggca
agttccaaatacttactagatatagtatctaacattgcaattcttttacgatcaactaaa
atcataagaaatgatagtgtattttttactaatttgatcaccgaaagttgcaataatgact
tgtttttttgcttctgtagtttttaatggattagccaaaaaaaaactttcagcttttca
gattgtgataaaatattttgtactgaccggatatcttggctcaccctctcagtaactttt
tttgttttagctaagtcaagaagagcaactgcgtaaggctgagctatttttataacaaca
ttattactgctcatagttgatctcctaacttagcaatattattatcaatgatacgtagtt
gcatttcagagttcatttgattctctagctgcaacgtaactcttttaagagcaagaaacg
ttatttgctgctgtatctgtcttcgaatttgtttttctgctgtttcaatatttgctttgc
ctgtcatagcaagtctttcaatatctagttgaccttgagctaatatagaacttctcactt
ttccagctgttagctgagcttctttttttgatttgatctataataatttgagtctgagcaa
gttgtttttctgattctgataatcttgcactagcttgttctaatctttcttcagattctt
gtatagctgctaagacttttatttgtcttgcgtttaaactagatccaagaaattgtttca
acacataaacaagcccaaataaaagcaacaatatattgataacattcgcttcaaaaatat
cagaattaaaaccgaatttgtgttcactactatgttctgataagatagtaattatttgtg
gtatttttactatattattcatttaaaattct----------tactattttaatatttca
cttgggaatcatgattttaaagattgactacttaatagtttagctttaatttggtcgctc
aaggtatctacctgattttctaaagttttaagagcttcttctttttgaatattcagctgt
tttgatgcatcagcaatcaatttttcagcattcaattgggctttttttaatatcttcggat
actatattttgcgcttctttttgagaagaagctatctttaactgtgcattgcgacgagct
tcagataaatctgcttcatattttgcagctaactcatctgccttaacaagcatagaagat
gcagtggtcaacgttgttcgaatatattcgtcacgttcgtctagtactttagttacaggt
ttataaaaaaatagtatttagtaaaaccattaaagtaagaaactgcaatgccattaaaggc
aaagtaccattaaagtcaaataagccaccttcaatttcttccgctaatag-aatggtaaa
taaatcattttactaattatgtttatagttaataatttggtttcaattaagaaaattagc
tttatagtgcgcttagtcttatcagactaagcgcaaatgattattttaaccaacgtatgg
attagcgaataatagagacagcgcaacaactaggccgtaaattgttaaagattccataaa
tgctaaacttagaagcagtgtgcctctaatttttccttctacttctggttgtcttgcaat
accttcgacagcattagctgctgcactaccttgaccaatacctgggcctattgcagcaag
acctacagcaagaccggcagcgataacggatgcagctgaaacgattgaatccataataaa
ttttaattttttgttatatgagtagaaaattgacaaatttcgggtaatacttcaaatcta
aataattaaatatggtagatttaaatat-----tgtaattatagtatactactcttctcc
atgaccttccattgcttcaccaatataagctgcagacagagtagaaaaaaattagtgctttg
gattgaactagcaaacaaccccaatatcataactggtaatggtatcaaaattggaattag

cagagtaaacacggatacaactaattcatctgctaagacatttccaaataatcgaaaact
tagagataatggcttcgtaaagtcttctagaatattgataggtaataatactggagtagg
ttgaatatatctagcaaagtagcctagaccttttttgcttaagcctgcatagaagtaagc
taaagaagtaagtaaagatagtgctacagttgtattaatatcatttgttggcgcagctaa
ctcaccttcaggcaaatgaatcaacttccaaggaataagtgcacctgcccaattacagcc
taggataaataaaaataaagtagcaatatatggaacccaagggcgatattcatgctctcc
gatctgatttttcgcaatgtcttgtaaaaactctagtataaactccatgaagttttgaaa
tttttcaggaattcgctgtaaatttctagtacctaggaaagataaagttaaaagtgtagc
aattactaaccacgagacaataaaaacttgtccatgtaattgtaaactacctattttcca
atatagatgttttcctacttctactgctgacaagctagtatatggattaaaatctatgag
actgttttgatacataataaatatttcttttaaatagtgcaagaatcaaaaaaataag-tt
tttgaagaattgatatgcaattagtttatttgaagattagattcttaatagattaaaag
caaaaataaattctaatagagatagctatctttagttgtggttatttttttgtaaaaaaca
attattgctacctataatgcttttatgtttaattatatactcatatattaattatttaat
atgattataggtattttgctacaaactttattgttttatccgctct----agacattttt
ataatctatttttttctttaagatagttgcaacttcatctgcaaagtcagctttaaggttt
tcttctcctccacctaatacaaatctaacgaatcgtctaattttaatattctcacctaaa
acagcaatattctgattaattaaatctttgatactaatatcttggttacgaatgaacatt
tgatctaataatgaaagctctttaagcctttttttttattcttccttcaataattgtttct
gttatatcaactggcttattttttaaatcatctttaccagcttcaactctttttttctaca
tttatgacttcatcaggtatatgctgaattgaaacatattccacatttggcgatgccgct
atttgcatagcaatatctttagctaatttttgaaactctgggcggcgtgccacaaaatcg
gtttcgcagttgacttcaactaatacacctattcttccacctgtatgaatataactctct
aatagaccttcaatagcagtacgacttgatttttttattagctgacgctaacccttttttgt
cttaatgattctaaagctttttcttcattaccgttattagcttctaaagctttttttacag
tccatcatacctgccccagtcttatcacgtaaggctttcacggtttgcgcagaaatttgt
agtgtcatatattgtcaaaaatttttaagtatttaaattaatt-aaacaaattattataa
ttcacttttgacagagtgaattatttcctctgcatctgtttgcccatactttccatcgta
aatcgcgtctgctattttaccaacaattagcttaattgatctaatagcatcatcattagc
tggtattggtatattaataatttcaggactgcaattagtatcaagtatacaaatagttgg
aatgcctaattttaaacactcttgtatagctgtagtttctcgtttctgatctataactac
aacaatatctggcaaacgagtcatattttaataccatttagatgtttacgcagtttatc
taattctctccttagtacagcagcttcctttttttggtaattgatcaataataccactttt
atcttgttcttctaactgtcttagacgactgactctagatttaattgttacccaattagt
taacattcctcctagccatctttggttaacataataagaatcacatcgttgtgcttcttg
cgcaactatacctgcggcttgtctttttgtccctaagaataaaacctttttaccatccga

cgaagcctgtttgataaattcacaagcttctgtaagtaattgtgcagtctgcactaggtc
aataatatgtataccatttctttctgtataaatatatggaaacattttaggattccatct
gcgggcttgatgtccaaaatgtacgcctgcttctaataattctgccagagtaacaacagc
cataggattgttaata-ttaaataatgtgattgttcgggagatccctttttt----gagc
tgat-----taaattaaattatcatgattctaataagccaataataaattaacatacttt
tacagtcgattcagaa---aaaatcttctcattttattcattctgtacttttattactaa
aatagttgcgtgcggttctatcatctaaaataatatcgtctaaatcatctctaacggatg
aagcaatattttcagtactattctctggagtcaaatctttttctccagggagactatat
tattgttttcataagtattaaagccagtaccggcaggtatgagtctacctataataacat
tttctttcaaacctcttagccaatctaatttaccggatatagcagcttcagttaatacct
tagtagtttcttggaaacttgctgctgaaataaaactttcagtgttgagtgatgcttggg
taatacccaataaaacgggccgatatgaagcatttaacttatttcgtagagtcatcgctt
tattagtttgttcaatcttctgcaattctacaagctctccgggtaaataacctgtttctt
caccattctctattttcactttagaagtcatctggcggactatgacttctacatgcttat
cagaaatattaacgccttgagattgataaactaattgaacttctttgacaagcagtaatt
gtatctcttgaaaactcaatctagcagcttcatataaagccaatccgcattcaatgtata
gattaaaacttgcgtctagaatatcatgaggattcaataatacatctgttttcttacgtg
cctcaagaatttcctcaattcttgggagaccttgtatgatatcacccgtttttgctctgt
caaaaactagaacagctaatgtctctccccttctaattaatgcattattatccacgtgta
aaattgcgccactggaaactaaataaggccgggcaattcttaatgtaactgatttagaag
atacttgaatgatttgtccagaatctaaagaagtcatgttttcagtgataaaatcaccgc
agcgaatccaatcaccaacacgtacttgattctttgattctgatacattaaaggttttct
tatcagaagatgtcgcaagtagaattcgcctactagtattcttttcagaataaatttctt
cgacagtccctttactcattgcaataatttctgtactagcaacaaccgtcaacggcttaa
tatattcaccattttttaacaataattctagtttgactttgtttcttttcagatttattca
catcaatacttttaattgataatgtttcaaaggttgaaagcctcaagtcaaaataattct
taccactttcttttgcagaaaattcaattgatgatattaagctatcatctgtatgctcaa
tttcagccactagattggttgtgaccaaatgaactccttctattgatttcactctttctc
catctcgaaaggggggttcttttgactagcttcaatttagtttggctggggttttgtgatg
ctaaattatcaattaaactagattttgtttcaggtattgaataaacaatgactggcctaa
taagaatataaggcatttggtcattttctatatactcccaatacactagcttatctgtcg
aaatattattgcgtagtgtttcaccaggtcttaaaaagcctctgcttttatcattattac
tataaggttcactgagcttatatatggatccaggcttaataataatttcgcgaactatcc
cgtctttttggattatctcaacaatgcctgaacttttagaaaaaatatttttaactagtt
cagttccactctctataatatctccattataaactagtaatagtgataagtctttattaa
tttcgtgagtttcttctgaaatccataatatgtatccaggactcaaaatttcgtaagcat

ctttatttgagcccattcttttttttgatacatttaaatctaaatatttaatgatgccac
cacttgttgttttatacgtctctgacattaactcggcaactgtatagccatcataaattt
ttttttgatttggcacagctttaaaaagaaacttttgtttttttttctgtttctaaaatat
aagattcttgcttattaattagatctgtataaatataacaaccaggaattacgatagact
cagtaataacttgtatattagtaatgctgctattagtgttttgttgtatcctgacctctc
cggcatagtgatttattaactctgtctgcgccagtattgtgccagcttcagtttcatctt
ctttttcaacaataatattagctgaatctgatatgctataaacttctcctgagagcaccc
aaatcaatccacctgtcttagtaattcttgtcgtatgttgattattatctgtttcttcaa
ctgttaagttagaaaaacagattttgcctgaaagatctgaaactacatgtttctgagctc
tttctgtcattagtctatttcttcgaggtgattcagcaattacctgatcttttgatacta
aaataccatctcccactaaaagagtggtgcctttacttaaattaattattgacttttgat
tttttctcgatcgtattgtaatctgtgttgaattttcgtgattaaggcctgttctccat
gcctagtgcgaacatctgtatatgattcagtatctagacctgtcagctgaccatctaagg
gtgcataaatttgctctgctaactcaccagtaaatacacctcctgtgtgaaatgttcgca
tcgttaactgagtaccaggctccccaatagattgggcagcaataataccaacagcttctc
ctaaatccactaaacgaccatgggcaagattccatccataacagtactgacagactgaac
ttctggaattacaagtaacaggtgatctaactagaacttttttttttataccagattgag
ttatctcttttgctaatttgggactaatatcttgcttagtatgagcgattagaacatttg
tctctggatgaaatacattttctgccaatactcgacccgacaaagcttgctctagattaa
ttaatatttttgtgtatcgactaaatcttcgagtataatacctttatttgttctacaat
ctacttctctaattatcacatcttgagaaacatcaacaaggcgacgcgttaggtaaccgg
aatctgctgttctcaaagcagtatcaactaaaccttttctcgctccgtatgatgaaataa
aatagtctgtaacagttaatccttctctaaaattactagatataggaagatcaataattt
ggccctgtgggtcagccatgagacctctcatgccgactagttgtcttacttgagaaatat
tacctctagctccagagaaggccatcatataaacagcatttagagggtctgtttctttaa
agtatttaattacctcttgttttaatgattcacttgcattattccatgtatcaataactt
tttgaaatcgttcaactgctgtgatctcacctcggcgatacttgttttcggtaaccttaa
tttcttcaatagtacctaataaaagactctttttactaggaggaattcttaagtcttcca
aacttaaagaaattcctgcctgtgtagcataatgaaatcctagatctttaagcttatcag
ccatattggctgctctcgctatgccgtaattgcgaaaagcccaaacaattaaatttttaa
gttcatttttatcaatgaccttatttgaaaaacttggctgtgtaagacttcgtctattat
ccactaaatcttctccatattaattatttgcttatttaactatgcgaccagagactcttg
aataattttattaaagattatacgaccagctgttgttctaatatattgaacaattctttg
atgctcagcatcttcttttataatatggttactgaagaatttcgttgtactattatcact
atgtttttcaactgtaattgggtcattaggttgatcaccatctactaagccatcaaaacg
agcccatatataagaatgtaaatcaactttcttttgctcataagccataactacatcttc

taagcttgcaaagtattgttttgatccctgctgttgagatggattattagctgtcaaata
gtagcaacccagaaccatgtcttgacttggcataataattggctggcctgtggctggaga
aagaaaattatggggagctagcattagtagtctagcttcggcttgtgcttctaaagacaa
aggaacatgtacagccatttgatcgccatcaaaatcagcattaaaggcggggcaaactaa
tggatgaagcttaatagctcttccttctacaaggataggttcaaatgcttgaatacctaa
tctgtgtaatgttggtgctctatttaataagacaggatgcccttggataacttcgtttaa
aacattccaaatagaaggttcatttttttggatcatttttttggccgctttaatattatt
aaccaagccttgcagaattaatctatgaatcacaaatggctgaaataactctagagccat
ttctcttggcagtccgcattgatgaagttttagatgagggcctacgacaattacagatct
tcctgaataatctactcttttgcccaataaattttgcctaaagcgtccttgttttccttc
tataatatctgataaagactttaaaggtctgttatttgcgcctacaacagttctacctct
acgtccattatccatcagggagtcaacggcctcttggagcattctttttcatttctaat
aataatctccggagccaaaattgactttaggcgcgataatctattattccgattaatgat
tcggcgataaaattcattcagatctgcagtcgcaaatctacctccatctaattggaccat
aggtcgtaagtctgggggtataacaggaattactgtaaataccatccaagagggatctgc
acctgttgctataaaattttcaattaatcttaagcgtttcatttttttattgaattttaa
tgaaggggtcttaaaacttttttggtgggtttgtcgcttcagaccgtaatgtttcagcgat
gtattctaaatctaaatctttcaataatttttgaatagcttctgcacctattccaacttc
tacttgattattctcgttttgattttgatatatttcctcttcgagacttttccattcata
gccttccagtaattgcttatatttaaattaatatctgtgttagattgtgtcactacata
ggaatgaaaataaactattttctctacttctttcacttttaaatctaatgctaaagcaat
ataacttgtacttccttttaaataccagacatgagttacaggcgatgctaactcaatata
agccatccgatgtcttctcactcgagattctgttacttctacgccacatcgttcacaaac
tatacctttatagcgaaaacgtttatatttccacagtgacactcccaatccttaactgg
accaaaaattttctcacagaataacccatccatttctggctttagagttctataattaat
tgtttctggctttgtgatttcaccaacaatttgtccattgggtaaacttctttcacccca
ctgtcggatttttttcaggagaggctaaactaatctttacgtagtcaaaatattgctcaaa
ctgtgtcataaatttgaataccttaaagtcaaataattaatcaatttattcaaatgtgcc
atattcaaaagtaaa----ttaataaagaaattgttcaaaatcatctactggaggagtat
cgtagttagtcctatctgctcgattatctttagagtcggacatcaaatcgacttcaactg
tacgtctttggccatcttcaaatagttttaatttatggactgcgatatctagccctaatg
actgaagctctctcattaaaactttgaaagactctggagtacctggcttaggtataggct
ttccttttactattgcatttagtgcttcattcctagcttgcatatcatcagactttacag
ttaataattcttgcaaagtatatgctgctccaaaagcttctagagcccaaacttccattt
ctcccaatctttgacctccatgctgggctctgccccccaatggctgctgtgtaactaaag
agtaaggtccagtagatcgtgcatgaattttatcatcaacaagatgaaccagcttaagca

tgtaagctctaccaattgtaatagggttatcaaaaggctctccagtcctaccgtcgaaaa
cttgcattttaccaggatactgatcattgaaaagccatttattgttagtcagcaatgatg
cttcttttaattttctattaactaaagctcttgatgcttccgctccatacatctcatcaa
aaggaattatttaaagcgtttccctaaatagccacctgctaagcctagtaagcactcga
atacttgaccaacattcattctagaagggacacctaaaggatttaagacaatatctacag
gggtaccatcagataggtagggcatgtcttgcttaggtaaaattctggaaataatgcctt
tgttaccatggcgtcctgccattttgtcacccacttgaatttttctcttctgcgccacat
aaacacgaatcatagcatttgtaccaggagggagctcatctcctttctgcctagtaaata
ccctaatattaactactcttccttttgcagcgttagggagtctcaacgatgtatctctga
catctctagctttttctccaaaaatagcccttaataatttaccttcaggtaattgatccg
cttcacctttaggagtgattttccctactaaaacgtccccagcttcaacccaagatcctc
ctacaacaattccgtttctatctaaatcttttaaagaatgatcacttacattagggatct
ctctagtgatttcctctggccctagtttagtttggcgacattccacttcatacttttcga
tgtgaatggaagtatataaatcatcataaactaatctttcactaattagaaatgcatctt
cgtagttataaccttcccagggcatataagcaacaagaatattcctacctaaggcaattt
cccccccatcagtagatgcgccatcagctaaagtttgacctacaactatcttttctccca
cccaaacaattgggcgctggttaatacaagtatcctgattagaacgataatattttttta
agcgataatgcactgttctaccgttattatcttgtataccaattttgttagcagaaacat
agttgacataacctgaggttctgctaataacaaccattccagaatctcgagcaattttgg
tctcaagacctgtcccgattatgggcttttctggatataataatgggacagcttgtcttt
gcatattagagcccattaacgctctattcgcatcatcgtgctctaagaaagggattaatg
atgttacggctgatattacttgaataggtgaaattgcaatataatctacttgacttggtg
ttgttgtaataaattcctgacggtaacggacaggaataatctctccttcaatatgatttt
gagaactaactttgacatcacctggagccactctaaaattatcttcttcatctgctgtta
gataaattggactatcttgataaataacttcaccccttatgtactgggtaaaatggagttt
caatgaagccaaaagcattaacccttgcacaggttgcaagcgatccaatcaagccagcgt
ttggaccttcaggtgtttcaatggggcaaattcttccataatgactaggatgaaggtctc
gaacagcaaagccagctctgtctttgttgaatcctcctggacctaacgcactaattcttc
tcttatgagttagttctgctactggattcgtttgatccataaattgagacaactgactag
acccaaaaaactctcttactgatgcaatcagcggcttaggattaattagattagataaac
ttaaggagtctatatcacagatcatcattctttctctaataatacgctctaaacgattaa
gacctactctaaattgattctgtaataattctccaactgagcgaacccttctattaccaa
gatgatctatatcatcaagattccctgaatttttatctttaatgttaattaaataatcaa
tcgaggataaaatgtcttgaggagaaaggactcgaaacgttttaggtatattaaggccta
atttttattaattttataccgaccaacttcccctaagtcatatctctttggatcaaaaa
aacgagagtaaagcatttgcttagcaacagcaactgttgcaggttcattaggcctgagct

tagaataaactattagtaatgcttcctcgtcagttacttcctcaatattatttttcccaa
tttctttcgctaattctttaactgagtatatttgagatgctgagatgagaaaagcatatt
tgctaagtcctttttcaatttcgtccttattaaggccaatagcgcggagaaaaatatatg
catttactttgtgggttttgtctattcttatccaaatttctcccttagggtcaatttcaa
atttcagccaagatcctctattagaaatcaggctagcactataaatctgcttgccatttt
tgtctacctcttgcttatagtaaattcctgggctacgaattatttggttgataatgactc
tttctgtaccagatacaataaaagttccgcgattagtcattattggcaagtctccaataa
agactaatctcttcctatatttttatttttatctgtttctcaacactcagctgtaaat
gtgtcgatgataaatccagtgatttaagaggcttcccctggtctttagaaggtaaatcaa
tatcttttcttgttaacttcgcagggacatatatttgggcactataggttctatctctat
tttttgcctgtctgacactgtaacgaggaaattttatcttataatcattaccaaataatt
gtagttcgagtcgactagtgggatctgatatactcggaaagaattcaagcacttcagtca
gaccttctaatagaaaccatttgaaactagccctctgaatttcaactaaatctggaagaa
gtttatttttaagcttatacgttgaaccatacatctcctttgtaacaaattctaaaatc
gaaaagtttaactattatcaatagaatacttacaggtaataactgagcatagttcgctac
attatttcctctctagacacaactagaacatcacattttattcaacaaaaaagattttaa
aaatttt--tt-----acactatatatataattattttggcggtc-aaaaaacaattgag
actat--------agaaaaaacgcaattaatacttgtctgtctgtcaagtaattgacatc
aagatagacaagtattatttatttaatcaattctgtttttttgagcgtaagacaatgct
ctcgcaagtctagctttttttctagcaccagtgttaggatggaaggctcctttttttatt
gctttatcaatcttgctatatacttgagaaatacttgattgcacatcattcaaactagaa
ctttctagattgtcaatatttagaaggcatcttttaattaaagtttttacaaccgattta
tatttacggttaatgatacgatttctttccgaagttttaatgcgtttaatcgcagaaggg
ttcttagccacagtaaatccaataaaactttttataaaatatttttagaaacgaaaataca
caaattagggatctattcaaaaagggaatatgcgccgtattataacattaaagaaaaaca
ataggcaatagttcaccaagggctcttgttattactaaataaatatcctcattgcataaa
a-tttacaaaatagtatttaattacattaaactaactaagcatataaatatatatgaaaa
aaattgaagctatcattcgacctttcaagcttaatgaagtaaaactcgccttagtcaaag
agggtattggagggatgactgttattaaagtttcaggttttggcagacaaaagggtcaaa
cagaaagatataagggatctgagtattctattgatatcattgataaaataaaaattgaaa
ttattattagcgatgataaagtggataaaatcatcgaaacaattattaaagcttcaaaga
caggggaaattggggatggaaaaatatttgttagtagtattgaaaaagtaatcagaatta
ggactagtgacttagattctgaagctttgtagtatttatactttttttagacttgatact
ttttcataaaacatgagataataatcatttattatacaataataactaaggcgtaata--
ttaatggctaaaagcaaaggtgcccgcatcgtaataactttagagtgctcagataaagct
gcagatgttaatcaaaaaagaaaacctggagttttccggtatacgactactaaaaatcga

cgaaatacacctagtagaattgaattaaagaaattctgccctaactgcaatcagcattgt
atattcaaagaaattaagtagttatttattttattaaaat--aatcatattatggctgta
tatagaaaaagaatatctccaattaagccaacagaggctgtcgactacaaagatattgac
ttgcttagaaagttcattacagagcaaggcaaaatattacctaaaagatccacaggttta
acttccaaacaacaaaagaaacttactaaagctattaaacaagctagaatactttcatta
ttgcctttttaaataaagattaaatacatatagctctgtttcatatgtttttgtgtaaa
cattaggaactttaattaatttgtttaccacatttattgaatctattagat---------
----------tacctcaaagaaactatttattaaaattttattcattttataattaata
agag--atctaactta---ttaagctagattc-----agaattacggtttcccctctttt
aattaattacgttgcatagtttatttaaaagaggagtttactttcttacatttgatgaac
a-------------------------ataaagtatttaagacaaggatttttgcttaggt
acaagatcgaatgaatgaatcttatctccaggctgccataattgaaattcagagataaaa
attccacattcgttaccagcttgaatttcttctacatcttccctgacacgttttaaagat
tcaatttttccttggtaaataactttttcttcccgaataactttaatccaagaattttt
agtaatttattattggtaactctacatcctgctatttttctattggctaaagaaaatact
gtactaacttctgcttctcccacttgtatttctgaatattccggatcaagtagatcttcc
attcttcctataacatcttcaattaatgcgtaaataatttgataattttcgattactata
ttgaactttgcagctgcttgttttgctccaggagcaaaattggtattaaaaccaataagg
ttagcacttgtcgtcgacgctagctcaacatctgtagcagtaatttcacctggcatgatt
gagacaacgtttaattgaactttactttgaggaaattgcgataaagaatctaagatagct
tctgtagaaccttgattatctgttttaataattaaggaaatttgcttactgagatcttta
gaattagtattactcagagtatctaaagtaatccgactattgagcgttttctgtttttgt
gcgattactgaatttttcgaagtattttcaaaagccttaagctttgcttctttgtcgctt
ttaactactaaagctatttctcctgcatcaggaacagcagataaaccccaaatttcaata
acagatgatggttttgctgaatcaatctttttttgtgcattactaattatagctcgaatc
tttgcataagccgatccaatgaccatattatcactcgtatgtagtgtgccatttttgtatt
aagagggtcgctactggaccatgagatttatctaaatgagcttctactataacaccttgt
gcaggctgtgatggatctgcttgtaggttttctaattcagctaataaagtaatagtttct
agcaatttatcaatattgtaaccagtaatagagctaataggtattatagggacctgacca
cctaatgtttcagacattacattatatttcaataaatcctgctcaataatatctacatta
gaaatgtctttatctattttggatattgctaccacgaaaggaacaccagctttttgaata
tgtttgatagcctctatagtttgaggtttaattccatcatctgcagctataataataata
gctatgtctgttagattggctccccttgatctcatacttacaaaagcttcatgacctgga
gtatccaggaaaacaatttttttgtttattagatttacttatatactctacttcatatgct
gcaatagtttgtgtgatgcctccaacctctttattagcattattagatttacgaatgtaa
tctaatagtgtcgtttttccgtgatcaacatgtcccataactgtaacaatcggaggtctc

ttaacatagttaccagcattattaagacaactgtcagtacttagattagtagagtataa-
--attatcttgctctttactggatgcgactgctataccaaaattatcagctactgatgat
attattgaagcatctattgtttgattaatagtaaccgatattcctctaaggaacaaatat
tttataatatctgtttcttgtacacaaattaatttagataattcttgaatagttagagga
ttagtaatactaatcgattctggtatagagttagacagatagctatcctgagtttggttc
tgaactaatgatactttttttgtcttgtagatcttttagtattagtaagtttttttata
gcttgaactttaggttttggtggacgcattagagagatagctaaatctcctgcagtctgt
gcaatattagaatgagaatctctgaagttctct---tcctcatcatcaatatgtattttt
gtttttatttttcttgcgctgcctattcttatttttttttattatctactaaatcgtgg
actttgttaaaattttttatttttttatcaagtttaggcggagagctcaactctaagtgt
tgatcactactagatgtatgagtgtctgacttctcaaggtctaagtgtaacaaattatca
ttatgagctgattctagtctaatttatatataatctgaggatttttaaatctaatatc
ttttcagcagggtttatgtttgggctagatcctggggatcgattttcaaaatttgatta
tttaaaaacatgaatgttgatttgttatctaatagtaactttttaactttaatacatag
ttcaacgtaaagaaaaatcagatatgt---------ctgataatat---aaaagattaat
ctctgttgtattagatttaatataacactaaagtcttttgttagttatacaagtttaaaa
ctttttttgtgatgatacatgctacattgtagtttattggtatacaatctatttatttaaa
tagtactgattgtcaa-agatttagaa----ttatctaatattttgttcattataagct
tttattaaagttttaaaa-ttatacaattaagtaataatcgttaatcacctccacgccac
ttatgccacgatctcaaaaaaaatgataatttcattgataaaacatttacagtattagcag
atattgtcttaaaaatacttcctacgagtaaagaagaaaaagaggcttttttcttactaca
gagatggtatgtctgcacagtctgaaggggagtatgcagaggccttagaaaactattatg
aagccttaaaacttgaagaggatccatatgatagaagttatatattatacaatataggac
ttatttacgctagtaatggtgaatacgtaaaagctttagagtactaccatcaaggactag
aactgaacttcaaattacctcaagcccttaataatattgcagtcatataccattaccaag
gagttcaagcagttgaagacaaagatacggaactatctaaattaatgtttgataaagcgg
ctcagtattggcagcaagctatcaagttagcgcctgacaattatattgaagctcaaaatt
ggttaaaaacgacaggaagaataagaaatatacaaggatattaatatatttaagatataa
t---tatattaataaatgaatcaa----aattaaaaacacttttcgcctataagatattt
ctagtat--------atagtatactaattgc------tctataagactcaaaataacaac
cgaaattaaatctattgactatatttatgaataaaagcaaatataatggttagtacaaca
aaaagtaccggatctgtcactcaaattattggtccagttttagacatcgcatttcctaat
ggacaacttccaaaagtattcaatgcactcaaggtacaaagctcagaagggactattact
tgtgaagttcaacaacttttaggtgacaataaagtacgagctgtatctatgagttctact
gaaggactacaaagaggagttgcagttattgacactgggtctcctatagctgttccagta
ggcacagacactcttggaagaattttttaatgttttaggtgaacctgtagataatttaggc

ccagttaattccgaaagtactttacctattcatagaccagctcctgcttttactaagtta
gaaacaaagccaaatatttttgaaacaggcatcaaggtcgtcgacctattagctccatat
aggagaggaggcaaaattggcttatttggtggtgcaggagtaggtaaaactgtattaatt
atggaactaattaataacattgctaaagctcatggcggcgtatctgtatttggaggtgtt
ggcgagaggacgagggaaggaaatgacctttatatggaaatgaaggagtctaaagtaatt
aatgcagataatttgaaagaatctaaagtcgcactagtatatggtcaaatgaatgagcct
ccaggagctcgtatgcgtgttggcttaactgcattaacaatggcagaatactttagggat
attaataaacaagacgttttattgtttattgataatatttttcgatttgttcaggcagga
tcagaagtatctgctctactaggacgtatgccgtctgccgttggttaccaaccaactctg
gcaacagaaatgggagcattacaagaaagaattacttcaacaacagaaggctcaattaca
tctattcaggctgtttatgtaccagctgatgatttaacagatcctgcccctgcaactaca
tttgctcacttagacgcaacaacagtgctatcacgtaacttagcagcaaaaggaatttat
cctgcagttgatccactagattcaacttcaacaatgctacaacctggaattgttgggagt
gagcattattctacagctcaagaagtaaaatctactcttcaaaggtataaggaattacaa
gatattattgccattttaggtcttgacgaactatctgaggaagatagacaaactgtatcc
agagctagaaaaattgaaagattcttatctcagcctttttcgttgctgaagttttcact
ggctcccctggaaaatacgtatctttagaagatgcaattaaaggtttccaaatgatttta
aaaggccaattagatgacttaccagaacaagcattttatttagtaggtgacattgatgaa
gctatacagaaagctgatagcatgaaagattaatataattaacaatgactttaaatataa
gaattattgctcctgatcggactgtttgggatgcagaagcgcaagagatcattttaccaa
gtagtaccgggcaacttggtattttaacaggccatgcacctttactgacagctttagata
tcggagttatgagagttagagtagataaggaatggatgccaattgttttgatgggtggtt
ttgcagaaatagagaacaatcaattaactattctcgtaaatggcgcagaagaagctagtc
aaattgacttatcagaagcagaaaaaaatttagaaactgcaactcagttattaagtgatg
catcatctaataaagaaaaaaatagaggcaacacaaaaaaattcgaaaggctagagctagag
tacaagctgcaacagtagctattgtataa-agcatttt------gctcaaaacataaaac
tttagtaataaatactaaa-------gttttatgttttatttatgtaattacaggcaact
aaaacaaatctaacttagaccggagcaaatatagtcaaaatataagcccatttctttacc
tgcatctggtccaactaaactaatagtaacttctttcatagctaaaattgcttgaattgt
cgcaccaatgggaacgcccaaagaattatatgtttcttttagaccatttaacactctttc
ttccaagatagatgggtctcccgctagcattccataagtagcatagcgtaagtagtaatc
taaatcacgaatacatgcagcataacgtcgagttgtatacatattaccgcctgggcgcgt
aatatctgaataaagtagtgattttgctaccgattctttgataatcgttgcggcattagc
tgcaatagtagctgcagctcttactcgtagctcacctgtttgaaaatagcctcttagttt
ttcaactgaactatcatctaaatatttaccttgaacatcagctgcattaataacagaagt
aattgcgtcttgcataacttgtaaaaattccttaattttaaatatctaatactcgggggt

tgtataacaaaataactgcactcgttagagatttcagtattactgcatggcacccaaagt
atagtcaaagtagaagcctgcttctgctgaatcttcacctgcaagtagtgagcaagctac
acttttcatgcatttaacaccttcagctactcctgaaattggtgttcctaaagaattata
catttctttaacgccgactagtccaatctcttcaataggcgtcacatctccagcaactat
tccgtaagttactaggcgaaggtaataatcaagatctcttagacaagtagctgtcatttc
ttctccataagcattcccgccaggggagactacgtcaggtctttttgaaacagctgttg
accaccttgttttacgatacgctcacgattatctgttaaaatttgagctattcttaaacg
acgttgaccggataaaacaaaactttaattctatctaattctccagggcttaaatatct
tgcttctgcatctgcatttacaattgactttgtaacaatactcatggataaaactcctgt
aatgcttt---taaaaagcttaatactaattatttatgatatacctcagaaagtaatgag
ccttttgaataaattcatatctataacaagtgaatgatatataaatcaatacaaatactg
aggtaagtactaaaacgacaaagttatctgttgtgacttattttattgattgccgagtgc
tttttgaaactaggtactataattgataaatcttgtttagtaaggcgattatataatgt
ttctgtatttggaaagttggctgctggtaaagttggaaatcttctatacgggacagtatc
tttaccgtatacagaattgtactcagtgctttcaactaaattactaataaatgcagaaag
ccctttcgaggctaaaatttgattgaagaatctaatttctgcttggttattaggagctcg
ccctaatatatgcttcgtgcctaattctataacttttgtatttggatatggctgataaaa
ttctttaccatataactctgataatgctaatttttctactaattcttggacattgatttc
cctattcaaaaacgaggattcaatatctaagaactctcctccaatactaaaagaattaag
atctctttcaaaaatttgacggtatgttgctcttaaagcctgctctaatacttctttatt
actactattattcacttcaaatacaatagattgatctctcaaagaagtaacaccttggga
aatacgagactgaatatcattactacttcttatttcttttacagtgcccaattcaacaaa
tctagcagacttattagtaatgactttttgaatcgctgatcaataatacctggccttaa
gctccttaaagcaatgcctgctggcgtactatatctttcataagggacaatattatctcc
gaaagtttctgtatactctgagctgtcaattattgcatctatcacttggtagtagccttg
cttatatgcaatatcaaaatatttattaatttcttgacgtccatatgtaggacgacctaa
taatcgattatgaatgtattcaatagctttacagatatataatggctcccagtaaagtga
tctaaatatactcgacttagctaattgtctaataaattcacgaactgaaatttgattatc
ttttagttgactttctattggcttaagaagtaatttttcttcttgataaacttctctacc
aaaaactcttagataggcgacttttgtaacaacatctactgatttttcaatattttcaga
agaatctggttttacaagagtgttggagaatttaaaaatcttaggacctaaggatcctgc
aggcttaggtcttacctgtgggttactgatctggttgtagatccctggaccacgtctaac
taggattcttctagtatctttaccaaaaagagcttgtctttttcgaggatctttattttc
tctaggaaagatagctccaaattggattgaaagcggatcattacctattccatacggatg
ctgatcaggtagtgcttgcttatagtcactaaagagagttataaactgaggcaccttacg
aaaaggagcactatagttaagaagatctatttgaggtccccaattacgacactcttgcgg

ctcttcacctaaattcctaaagtaaggtactgtttcttctccgaaatagtctgtgtattc
agaagaatttaaaattgcatttactagcccacttagaccattagatgatagaatagcaaa
atattgctggaactcctctagagagcttggacctctaccaaggaaatgcctgaatgctaa
ttctaatgctcgactgttgacaaaaggttcataaaactgctttctatagatgcttgaagt
tcccaatgcacgaataaactctttgattgaaatttggccatttttttacttgagactctaa
attggataaagaaagattatatgctttagcaatatctctctcaaagatttgtctatagca
agctttaacaacaatattttttttcatcagcagataagctaggcttcataacaaaacgagg
tattgaaactcctgcttgtgcataagtctggggaagacgtaaaccctgtaaatctccgga
tgtcctttttcttaacttgtcagtcaaggatggagcttcaaattcagaaatcacaacatt
aaagtactcttttactaaatcttgacctttaatatcttcttcaaatattaatagagcagt
tcgtctcatctctctaagtgctacaatggctgctgcgctcgagcaagcattatcaattaa
ctctctcaatcctctgatgttaacagataaaatattaggatcacccgaaacaattgcata
tgttagataccgtaagaaccaatccaaatctctcaaagattttctcattcttgtagtacc
atatctcaagacatttataggtttaaatccaggaggggttgctccccctgcattaaacaa
ggaacggaaactttgtccaaaatctccttgtatatttccagataattcattaattttatc
ttgagaactttgatctcctgcaataataacagacgcttgtgggcgctctagataagaaat
tgcggacccacctacaaatattttatctgctgctcgggctacaaggatatttgcattttt
agttaaaatatctgcaacttcaagtctttttgggccagaatttaaaaaagaaactaattg
atttaattctccaagctgtaaaaatcggtcttgctgctctgcctgagtaatagttaaaat
tgaggcagtccgataaagctgggggcgtgctaatgggcttccgccacttgctttgatact
cattactttatttatctccttaacttatattacttgcttacgaatgcaaatatcacttga
tatttacaactatttaaagatgctatgcaagctaattggttttctatattattaggtact
agtacttatagtacaaagagtcttacaactaacttctataaatacaaattttgtaatact
tctattaacatttatcaaaagtatatgtgagtgattagccttttatattaaaatgtaatt
aatgagtgattaatgttaaataaagtatacttactaggttgtatagatagtaaaatt-ca
atatttattctacaaacattaatca------------cttgactgtttgacaattatgta
attgttcttgcttttaattggtttagactagttttacatttttctcccgtaaataatat
taatgatcgatttgtatgaatttagaaccacataacaacctattaaataacaaaagcaca
aaccagcaggatatacctgagaacgacataccaatgtcaattacagaacatttagaagaa
ttaaggcagcggactttatttgtatttttattctttttacttgcggcaactgttagtttt
acacagattaaaataattgttgctatacttcaagctccagctattggtattaaattttta
caattagcgccaggtgaatatttttttttcatctattaaggttgcaatatattccgggatt
attgcaacaactccatttggcgtgtaccaagttatattatatatactacctggactaact
gggaaagagcgaaaaattgttttacctatattaattagttctgtattgctttttgtaaca
ggtggtatatttgcttactttattttagctccagcagcattaacatttttaattagctat
gggtcagatattgtagaacccttatggtcttttgaacaatacttcgactttattttacta

cttttatttagtacaggattagcttttgaaattcctatcatacaattattgctgggtgtt
tctggtacattgtcttccaatcaaatgattcgagcttggagatatattattattatatcg
acaattgctggagccattctgactccttcaactgatccagttacccaattaataatgtct
tctgctgtgctaatactttattttagtgggattgttatattactaatcttaaaaaagtaa
acatagtattatattagtcttttatacaataaaatactattgttatcatcagtaagtagt
tagtcttt--cttcaattatgaattcctagctttgat-----------------------
-----------------ttagtcttgtcttatcatttagcaaatgtg-------------
------------------------------------------------------------
------------------------------------------------------------
------------------------------------------------------------
------------------------------------------------------------
------------------------------------------------------------
------------------------------------------------------------
--------------------------------------------------tatacatcaa
ctt-atttttcctcaaatacttgtactctatactttttacca----ttctaggatgctt
agaatttgttctttagtctatttactaagt----atatataaagaaaatttagtgttgat
tatggtagtctataccaacgtaaaaagtattttattatgagttttatttatataaaggct
tattcattatatagttataacagtaagtctaattaatagtttttcatgcttaattaaagt
aataactcttaattcttttggcagaaaaatctgtcttaatattttatc--aaaaaaacag
-aaaaatgaagctaaatagtctaattaacttaattcaaaagtctatatattcttgtacac
ttttattaattattttaaatgttatttgtgtcgcacctaattctagtaatgcattcccaa
tttatgcgcaacaagcttatgaaagtccaagagaggcaactggtagaatagtttgtgcca
attgtcatcttgctcagaaaccggtagagatagaagcgcctcaagcggtactacctaata
ctgtttttgaaactgttgtaaagattccatatgatagcagttctaaacaaattttaggta
acggaagtaaaggaggattaaatgtcggcgctgttgtcatactacctgaaggatttaagt
tagctcctgctaatagattatctacagaattaaaggaaaaaactaaaaatctttacattc
aaccgtacagtgctaaacaaaataatattttagtaatcggtcctattcctggtgacaaaa
atagggaaatagtttttccaatactttctcctgatcctgcaaaagataaaaaagcccatt
ttttcaagtacccaatatatgttggcggaaatagaggtagagggcaaatttatccgacag
gagataaaagcaataataatattgtgaacgctttaagcagtggtaaggttaataaaattg
agacacttgataaaggaggatttgtagttcacgttactaacagtaccaatatcgaatcaa
ctcaacagattgcaccaggtcttgagctaaaagttaaagaaggggaaaccgtgcaattag
atcaagctttaagtattgatcccaacgtaggtggctttggccaaaacgaaacagaaattg
ttttacaaagtccaaatagaatcaaaggcatgattgttttctttttcgctagtgttttag
ctcaaatcttctttgtattaaagaaaaaacaattcgaaaaagttcaagcggctgaaatga
acttttaaatcaaaataaaactggcacttactatcagtaaagtg-tgaatactttattaa

cagaatgtaccagtttttatgttatgtagtaattgcttttgaagatattataatattttt
cacagagtctataatttgactcggctggattactgttgcctgttctagactaccattata
tggtgttggtatatcttgtgaagatagccttactacaggagcatctaattcatcaaaaag
gtattcattaatttgtgcaattagctctgctccaattccagcggttttcatacattcttc
tacaattaaaactttatgagttttccttactgaaaccgatatcgagtctatatccaaagg
ttttagagatataagatctataacttctggatcataaccttcttttagtaaagttggtaa
tgcttgaataacatggtgcctcattctagagtaagttaaaatagtaatatcttttcctga
ccgtacaacttcagctttattaaggggtaaaagatattcctcttcaggaatctcttcttg
caaattatagagtaaaacatgttcaaaaaagacaactggattattatctcggattgcaga
tttcagtaatcctttcgcattataaggagtggagcaagcaactattttaaacccgggat
agcttggaaataagcttctaatctctgggaatgttcagctcctaattgtcttccgactcc
ccctggtcctctaataactagaggcaatgtaaagtttcctccagaagtataccgcaacat
acctgcattattagaaatttgattaaatgctaataataggaaactcatattcatgccttc
aacaattggtcttaaacctgttatagctgcaccaattgccataccagtaaagctattctc
tgcaatcggtgtatcgagaactcttaaatcaccatatttactgtgcaaatctttagtaac
cttataagacccgccataatggccaacatcttctcctattacacaaacagttagatcttt
ctccatttcctcatctgttgccgctcttagagcgtcaaacataaagattttactcatttt
aatatttggtttttatgatttcgccactactacaaagtaacagctaagactcaatatatt
ctttattgatctgaaaaaagatatcttttaagttctgagatattgggttctgggctagaa
atagcaaactctacagattgttctaactctttttaacagtagtttgaatatcattaagt
tcacctaaatctgcaattttattatctagaatatatttttgagtttttaatagggtcc
cttgcaacccaagcttctttttcttgtcttgatcttagctcatcaggatctgcaagagaa
tggccgcggaatctatatgttagtgcttctattagtgttggcccatcaccttgtcgggct
ctctggactgcttggcttgctgcttgccttacagctagtacatccattccatcaacttca
atcccgggaagaccaaaagcttctgcctttttatgtatttctggtattgaggaagatcgg
tgatgggccataccgatagcccactgattattttcaacaataaatatgataggcaacttc
cacaaaactgccatatttaaacattcgaaaaattgtccattattagtagtgccatcacca
aaaaagcaagcagtaacccttaatttttctgtttttttcagtacttgctgacgataaata
ctttggaaagcggcacctgttgcaacaggtatgccttcagcaataaatgcaaagccacct
aagaaattatgaggtgcagaaaaaaatatgcattgagcctcctctacctttgctacaacct
gtttcttttccaaatagctcagccataacgtttttcgatggaacgcctttacttaaagca
tggacatgatcccgataagtactacaaacataatctgtaggattgagaagcttaattaca
cctgtagaaacagcctcttgaccattataaagatgaacaaagccaaacattttccctttg
taatacatctgagcacacatatcttcaaaatttctacctaacaacatatctgcatataag
actaataaattactctt------attgagactagttgaattatagttagttagcggcaat
tgaactttcttaggatgactcataatttataaagagacctct----------tattatat

atagaaaatttgaagtga-agttatcaatagaaaacacatagataaagatttataaatta
taatatgctttttactaaagactttaatagttatttacaacttgttcagtatatcaaatt
aattttgcttatactacaaatatctaaaacttagattagaagcaa-------agaagtgt
ttgaaagaatag-ttaaaacaaatataaatagaagtaaacagaattattcgaaagaaaat
gattatcattaattaataaatacttaatatttctaaatattttcaaattaatggctatat
catcatctttattttatcctattgagaaagaactatacagtcttgaaaaaaatctaagag
cggttgccgggacacgtcatccaattctatatgcagcagcaaagcatctcttcgacgcag
gaggaaaacgaattagacctgctctcgtattttagtagctaaagcaacatctgaacagc
aagacattaatactggtcagaagaggctagcagaaattaccgaaatcatacatactgcaa
gtttagtgcatgacgacattattgatgaatgcacgacgcgcagaggcgttaagactgtgc
ataatttatttaatactaaaattgctgtcttagctggggatttttttatttgcacagtctt
cttggtatctggctaatattgaaaatttagcggtggtaaaagctatctctaaagttatta
ccgattttgcagaaggcgaaattaggcaaggcttagttcattttgaccctagtatttcag
tggatgcctacattgaaaaatcattttataagactgcctcactcattgctgctagctgtc
ggggtgcagctatgctgaatggatctaatgctaatgtaaatcatgatctatatctttacg
gtaaacatatgggcttagcatttcagattatggatgatgttctagatataactggttcta
ctaaaagcttagggaaaccttctggggctgatctaatcaatggaaatttaacatctccta
ttttgttttcacttactcgagaagtagacttaaaccagcttatccaaagagaatttttgtg
ataatacagatatagcctcagcattatttcttattaaaagaagcgggggaattacaaaag
caaaagacttagcaaaagaacaggtacaggcggcactctactgtatccagtttttaccaa
aatctgtacctgtatctagcttaaaagaattgacacatttcataatcacaagattatcgt
agaatacttgctaaaattaaaatgttactaactagtttttttactgtttccaggatagtct
gaaaggtctgaatatcattactagctaactgcgctagcattttacgatttaacgcaatat
tatcttttttttaacgcactaataaaagtactataattcatgccttgactgtgagctgcag
cattgattctagttatccacagccgacgaaaatctctcttttttcctctttcgaccgacat
acgaatatcggagagctttaagaacttgctgtttagctgtgcggaataaacatttatgtg
cacctttaaaacctttagctagcttaaagatttagctcttcttttttcttgcaacgttac
ctcttttgactctactcataaaatattactttaaatttaaattattaccagcttactaaa
tgggcattcttatctatttattataaataaggtaggtttatagcgatattttttaatatct
tttgagtcaactaaacatgtagaagaaagatgtcttctttgctttgatgacttcttttgt
agtaaatgacttttagaagctttatgtcgaagaaattttcctgatgaagagactttaaat
ctttttgctatcgctttagatgtttttaacttaggcatattatgaacttttaatgattaa
caaaagtctgaaaagcctgaagatgttaaatttagacttttcaatacttaggataatctt
aacacaaattattttacttactatagggtttaaatgaaacaatttatatatataaaatacca
aaaagttagtcaacagcggctacagttgatttatattttgctttgaatttttgtatggta
ttagcaattagcattgttattattttaagagtacttgaattcaattcctcaaacattttc

atatttaatgtaaatgagatattcgcctctagaacaatattctgtatttggtaactagat
aaaggaattgtatccaatgcagccctgtacttatctttaaatgatttatcatcttttatt
tgatcaaaatcataaaatctagtacctcttgcatctgataaattcatcgcacctctagct
atttttttaaaatttgaccaccagataaatcccctaagtaacgcgtataagcgtgagcg
actagtaattcaggctgttgacggcctatactatgaattctatcaacatatattttttgtt
gcaggagatggttgaataaaatttagccaatctgaaccataataatagcttaaatcttct
gaaagacttatttttctattaagttcagtaaaatatatgggtttaatagctggatgatct
ttattagccaatagttcttcctctatcgcacaataaacaaaatatagattagcaactaat
tttcgatatgactttttatctacgacgcctcctaaaaaggatttcacaaaactgacattt
tctgccatactatgggacttagtagtaccttctcttaattcattcgctaaagtattaacc
ataatatcaatcttccatttta-aaaggtaacaaactaaat---atgaacgggtagctat
gtatttaaacttaagttgtactagctaggctgctgcaaaataatttttttgattttattgg
gtctgggttcatggttctatcaccaggcttccaattagcaggacagacttcgtctggatg
agattgaacatattgaattgcttgcaagactcttaaagtttcttctacactcctaccaaa
ctcgaggttattgattgtagaatactgaataattcctttaggatctataataaataaacc
tcttagggctacaccatcgctgtttaaaacattataagctgcactaatctctttttttaa
atctgataccaatggatacgaaagatctcctaatcctccagattctctgtctgtctgcag
ccatgcaagatgggaatactcgctatccactgaaacacctaaaacttctgtattaagctc
agagaaagcatcatatttatcactgaatgctgtaatttctgtcgggcaaacaaatgtaaa
atctaaagggtagaaaaataagacaatatacttatttttaaatctgataattttagcgt
tttaaattcttggtcataaacagctgtggctgaaaaatcaggagctagctgacctactcg
aagacaatttggtccagaaatcatgaatttttcccagaagaattgatataattttgttat
ttaatagataattatataacaaattttagtatatattaaaatgctatcagtcttttaaaa
attac--------tcaatgatataataaatactattttttatagcggggaatggatttgaa
ccattgaccttcgggttatgagcccgacgagctaccagactgctctaccccgcgaataat
tgaaattataataatctatgcaaacatataaaagcaaatgtattacatagttcagaagta
aaaaaataagagcacagggacactgattattacgcttaaagacgacaaagtgttttttac
tttgtttaagagaggaaagacttcatataagccaataaatcggtctttaagaagaatttc
agacggctttacccaaatttgaccatcataccatccagactcttcgtagaatacagtggc
actcattaaacgctttactacatatgaccatcctaaatacaatctaattaaaatgaatcc
ggtcattaaactggaagtaatgaattctgaaaagaaaaacttcaaagggagctgagcaat
tggaaaaattgagagtaatattggactaactaataagcaattaaatagtagtgtaataat
gattttgttgttatatgatctattacttaaagttggccagcaaaagaaccaagaattttt
caaggaagtgtattcgttaacaggttgctgctcttctggaacaggacattgaatattata
taaattcattgacattaaaaagtattatactaaaaattgatttttttacattatctagatg
attataga--------tttaattaactgcaattatagtagttaatataagaaataaaatt

agaaataaccacgtaactttatttaatgtattttcagtactacgagtattactaaaaaat
tgattctgagctccaacactacctaaaccttcagatttagggttatgtatcaagatagtg
aaaattaacataattgttgatacgtaccaaaagaattttaaaatttgttccattgtataa
aaattaat-agaagttg-aaaaagacaatcgtctaaaaaatattgattgccttgtcatat
taaaaaatatttactcaccttgtacttttaaaagtaaaaatcctattgccaaccctacta
aaactagaactgaacttaatacagcgctttcaacaatttctccgaccatt-agatttctc
cttattattttattgttt---ttgttattttaaaaaaagtaaatctttaaaatttagaag
ccgtttcttccccacactaccattgcaatagaaaacgtaaacattgccattagagccgcc
catcctaaacttaaaatgtccataatttataatattc--ctgaaaataagatctctttta
cttaaactataatattcttaatattaaaaatatcattatataataagctaattataacag
agtactacttaattcgtatttttaatgtatttaaagaagattcgctttttttcaaaagtac
ttgaccagtattaaatcctaagaattatctgtaat----tctagcattattataattatt
aaatttaaatataaatatcactaaacaatcgttaataattttgacttgttcaattataca
att---caagtatgtaaaaatataaagattcatagcagctaagttaa-----tgttctaa
tatcatttatcaataaaaaaaaaaagttagtatttaaattatgcaaacaactataagtaa
taaaacccggtctagtaaagaaactctattaacacctagatttacacaactgattttga
agaaatggctaatatggatatttctaaaaatcagcaagatttctttgctatcctagaaga
atttcgagcagactataatagtgagcactttattagagatgaggagttcaatcaatcttg
gtctagcttagaaaacaaaaccaaatccttatttattgagtttttggaaagatcttgtac
agcagaattttcaggctttttgctatacaaagaattatcacgaaagttaaaggatagaaa
ccctattattgccgagtgctttttactaatgtctagagatgaagctcgtcatgcaggttt
tttgaataaagctataggtgatttcaatttatctttagatttaggctttttaaccaagag
tcgaaaatacactttttttcacctaaatttattttttatgcaacctatttatcagaaaa
gataggatactggaggtatataactatttaccgccatcttgaacagcatcccgagcatcg
tatttatccgatctttagattctttgaaaactggtgccaagatgaaaaccgtcatgggga
tttttttgctgcgttgctaaaatcccaaccacatttttttaaacgactggaaggctaaaat
gtggtgcaggtttttctattaagtgtatttgcaacgatgtatttaaatgacttccaaag
gattgatttttataatgcaataggtctagattctaggcaatatgatatgcaagtaatacg
gaaaactaatgaaagcgctgctagagttttccctgtagctctagatgtggacaatccaaa
atttttcaaatatttagatacttgtgcatgtgataatagaactttaatagatattgataa
gcaaggttctccattattaatcagatctattatgaaaattcctatttattgttccttggt
tagcaatttactaaaaatatatttaattaagccgatagattcaaaaatagtttggaatac
ggtccgataattaaatcaatcaaaaata------------------atctaattactaa
tataaaatagaatcaaaagcttccttca-----attaggaaggagcttttattttattaa
tg---atatgat-tgattatatagatt-taaaattaagaactgactttctcttttgcttc
atacataacttctaaggtaattaaatttttagaacaatcacgagcaaatttctcagtatt

tctctttactttacctctaacgaaacctggaattttacttaattccttctgagcttcttt
tgaccaattaatactatcaacagcagaaattcctatacttaatgcttctgttgtatcgtg
accaccaaatatttctaacaggtggtcttccatacccaaggtgaatgaattatagactaa
atcagcaatttgatttgtaccttcatatccaagaaatggacgatagcttaaagggaaatt
ttgtatatgtaccggtgaagaaataacaccacaaggaatatttaaacgctttccaatatg
tctttccatctgtgttccgaaaattgctgcaggttcagttttagcaattaaatctccaat
taagccatgatcatctgaaataattacttcatcacaaaattcttgaacttgctctttaaa
ccaatcttcatcatatttgcaataggttccgcaccaagcaacatgtatacccatttcttg
atgtaaaattcgagttattgctgcagcatgagttgcatctccaaagacgatagcctttttt
acccgttaaattctggcaatcaattgatcttgaaaaccatgcagattgtgagatgaatct
ggtttgttcatctatatacttttcatagtctacattagctcccaatgtatttaaaagctt
gtgtattgatcgaatacatttagctgtttggacaatacccataggtgtagtatcgacata
aggcatctcaaattctttttctagatagcgagctgtcatcaatccagtctctctatatgg
gacaaagttgaaccatgcagatggtaattctttcaaatcttgaacagatgcattttccgg
aataatttggttaatttggatatctaaatcttgaaataatctttttaattccgaaatatc
atgctgattatgaaaccctaaacttacagctccaataatattgacagatggcttttgagt
cttctgagttataacctgcttgttagatttagctttttccatgtaaaaagtaacaatctg
ctcaagagttctatcaccagcttgaagttcatttactctataatgattaacgtcagctaa
taaaacatcggcttctgtttctattgatgctctgctaacaaaattttgtaaatcttcttg
taaaatactagaagtacaagtaggggttaagacaactaaatccggatttcttctctatc
ttttctcgtaatattttcaacaactttttcttgagagcctcgtgctagtacatgtctatc
tacaacacttgctgtaacaggtgtaaagtctctgtctctttcaagcattgaacgcattac
attaaaataatcgtctccgagaggcgcatgcataattgcatgaactttttttaaaagaact
ggcaattcttagagttccaatatgagcagggcctgcatacatccaataagctaatttcat
attaaattttcctatgatttatttagacatagtaatacttttaaagccaactaagaaata
caagttagacatctaattttgctctttaatgatttgtcctc--ttgctaaatggggctta
tcactatatatagttaagaag-------------acaaataaatgactaatagataaagc
ttgaaagtaaagatttcttgcagttatagtttttctttaagtaaacatgtaaaaattgta
ctatatataaatgtactatatataaactttttttactgaatattggtaatgccagataca
attaaattaaatatgccttccccaacatttggtggaagcactggcggttggttaagagcc
gcagaagtagaagaaaagtatgcaataacatggacaggcaaaaacgaaagtaagtttgag
atgccaaccggtggtacagcaacgatgagaaatggagaaaacttactttatttagctaag
aaagagcagtgtttagctttagggactcaattgaaagttaaatttaaaatatcggactat
aaaatttatagagttttccctaatggagaagttcagtatctacatcctaaagatggcgtt
tttccggaaaaagtaaatgcaggaagagctagtgttaatagtgttgatcattctattggg
aaaaatgttaatcctgtagatgttaaatttacaaataaagcaacttacgactagtcgaca

-acaaacctagacataaaataacttttatatagtctaggtttttgataaaatatactaga
agtatggaagggtggccgagtggttgaaggcgtctgatttgaaatcagttgaacta--ta
tagttccgtgggttcgaatcccaccctttccgttataaaagt------------acagct
taattattgctgcg----aattatacactaattgtattgagtttagatttaataaaatct
acaacttgtgataaattcgaaatcttctcagcatcttcatctggtatctctataccgaat
ttttcttctattgccattactaactctactgtatctaaagaatcagcaccaaggtcactt
gaaaaatttgcttctctcgttactattttttttcaattcctagctgttctgctacaata
tcttgaacttttttcaaaaatttcattatcttgcataatatattatttcctttaaggtgtt
atc-aaaatttttcggcttatagttttatc-----------------taaatttaagtcg
agatccaaatatttatacctatatgatatactattattctggataaatgcgaagtctacg
atatggctcttctccaaaactgcgagctctcaactggtagttgtcaactaagttatgttg
catttttcggatataggctgaacgaggtatcagctggactgttgaatttccgtttaagac
aatagtctcgatagccaatttagcttcttgaagagcttgaacttcatcaaatttttttatt
tttacaaagttctacccaattaagatcagcagaagtatgaatatttaatatttttctcag
tgcacgagtaatttgtggaactgtactactctgaatcgtatatataataatttgttttga
ttttgcaatttgtcttaattttgtgttttgctttacttgattacgtaatgctaaaatagc
atctgacttttctatttctttagttaaaataattggtaagtctaaagacgaaataactga
tgtaatgtgctgccaactcaaagaataagcatataaatactgattaggtatttccaatgg
taaagattggttattaatcttaataggattattaatagtagttgacaataaagatgtatc
acgattttgtaatttattcaaatcggatgacttatttttcaattctctatattgtagaat
aggatgtacctgttgggcatcaattggtttggtaagactagatatatttgtaggcaaaac
ttctaaagattgtgacggataacatttaattaaaatacggccattagcttgaatttgtct
tttttgtacaaaaggttgatggccttgcaaaatttggtctatagtttctttaacttgatc
atggacaatccaaacatttcgttcatgtatttcaattgcaatttgaaaagcaggcgctgc
ttttctttctagtatgcttttttgtgtaccccttctttttgcttcatcgtcacccaatgt
aacatattgaattcctccaatcaagtctgctaatgtggggtttttaattaagctttctaa
gtgattcccgtgcgctgtacctactaactgcactcctctttctgctatagttcgagcagc
taatgattctagctctgtgccaatttcatctatgataataacttcaggcatatgattttc
tacagcttctatcattacctgatgctgtaaatcaggccttgcaacttgcattctcctagc
tctacctatagctggatgtggaatatcaccatcaccagctatttcattagaagtatctat
aattacaactctttttttccatttcatctgctaaaacacgagcaatttctcgaacagctgt
tgttttaccaactcctggtttccctaagagcaagattgagtctccttgctctaataaatc
tcgaacaatactaattgtaccaaagactgctcggccaacacgacaagttaaaccaataat
actaccttctctattgcgcattgaactaatacggtgcaaggtttttttcgataccagctcg
attatctccactaaaattacccactttcttgacacagtaatctaaatcttgccaactaat
agatctttgagataaatattctggtttatctggaaatcttgcttctggtcggcgacctaa

atccataaccacttctattaaattatttctgttaggatgttgtttcaaaggctcttttac
aaagtttggcaagatttctagcaacttatctaagtcatctgcaataagcatgattaaatg
tataataattaattaatctgatagaacatttaagattctaaagattacaatacatattaa
agaatattatgcttgtaatcaattttatacttaaattaaataagaagtta-tagatgtaa
gagagtagattaaagcttttatagtaaatagattggtcaaatttaatagtcatgttaata
tgtttattggacagaaagttcgtattaagtatagtaagcaaaaagttgctaatgatatag
gagataaagttggggatctaggaattattagaggaataaagtttatcaatagtcaatgtg
ttactattattgttgaatttgacaatcatactagactttggatgtttagagaagagctaa
tctatctaaatgaat------tcaaaaaataagacaggactaattt----tagtatacaa
aaatttttatgagtcgttacaataatacattttataggagtattgttaattgtgcaaatt
actattaaaaaattacaagacttacttacgtctgtacaacgaaaaaaaatacagaccttg
aaactaaaacaagataattttgaacttctattaaatagaccctataaaaaaacgactcca
gagatactacctttaaataact---ctgtattacaaaacactcaaactacattaactaga
tcaataactaataaaagtattccttctaacagcaaaaataacattacagagtatgccact
attatttcccctatggtaggaacattttatcattcgcctgctcctggggaaaaaattttt
gtgaaagtcggtgatgaagttaaattcaatcaaaccgtttgtattattgaagctatgaag
ttaatgaatgaaatcgaagctgagattgaggggaagattatagaaatttttagttaaagat
ggcgatatcgtagattgtggtcaagctttaatgaaagttgaaacatgatgccaatctttc
cattttacatattgttaacagaaattatttagatagaaaaattagcttatactatagtta
tgaaaacttatttcgtaagggcac-aaaaagatattctacagaattattaactttatctg
ccaaaaatagaaataagcgttttgattccttgtctcaagagaggagaatctcaatggcaa
ttagctcaaaagagcaagagacaaagaaggtaaaaatctcggttgataaaaatcccgtag
atacttcttttgaaaaatgggccaaaccagggcatttttctcgtacactagcaaaaggac
caaaaactactacttggatttggaatcttcatgctgatgctcacgacttcgatagtcaaa
ccagttctttagaagaagtttctcgtaaaatattcagcgcccattttgggcagctttctg
taatctttttatggttaagtggaatgtattttcacggggcacgcttttctaactatgttg
cttggttaagtaatccaacaggcattaaaccaagcgcgcaggttgtttggcctatagttg
gccaagagattttaaatggtgatgtaggtggtggctttcaaggcgtacaagttacatctg
gatggttccagctgtggagagcgtcaggaattactaccgaatttcagctttactgtactg
ctattggcggattaactatggctgccttaatgctatttgctggctggtttcattaccata
aagcagctccaaagttagagtggtttcagaatgtggaatcaatgatgaatcaccatttag
ctggtctattaggcttggggtgtctaggctgggcaggtcatcaaatccatttatctttgc
ctattaataaacttttagactctggtgtgtctccacaagagattccattacctcacgagt
ttctaattaatagagagcttatggctcagctttatccaagttttagtaaagggttagttc
ctttttttcactttaaactgggctgaatattctgactttttgactttcaaaggaggcctaa
accctgttactggaggtttatggcttagtgatactgcgcatcatcatctggcgttagctg

ttctatttcttgctgcaggccacatgtatagaaccaattggggtattggacatagcatga
aagaaattctagaagctcacaaaggtcctttactggtaacggtcacgaaggtctatatg
aaattcttacaacttcttggcacgcgcaacttgctattaatttagctatgatgggatctt
taagcataatagttgcacaccacatgtatgctatgcctccatacccatatattgcgactg
attacccaactcaattatcactatttactcatcacatgtggattggggattttgtattg
ttggtgcaggagcacatgcttctatattcatggtaagagattataatcctgcagaaaatt
ataataacctttagacagagttattaggcatcgcgatgctattgtttctcatttaaatt
gggtatgcatatttctcggttttcattcatttggtttatatatccataacgatactatga
gagcacttggtagatcccaagacatgttttcagatacagctattcagttgcaaccaattt
ttgctcaatgggtacaaagtatccatactttagcacctggcaatacagctccaaatgcgt
tagcgacagctagttatgcatttggaggagatgttgtttctgttggaaataaagtcgcta
tgatgcctatttcattagggacagctgatttttagttcaccatatacatgcattcacta
tacatgtaactgttcttattttagtcaaaggttttcttttctcaagaaactctcgactaa
ttcctgacaaagccaatctaggatttagatttccatgtgacggacctggcagaggcggta
cttgccaagtgtctggctgggatcatgtttttcttggtctattctggatgtacaattctc
tatcggtagcaattttccacttcagctggaaaatgcaatcagatgtttggggtagtgtat
ctccatctggaaatgtttctcatatcactggtggtaattttgcacaaagcgcaattacta
tcaatggatggttaagagacttcttgtgggctcaagcatctcaagttattcaatcgtacg
gctctgcactatctgcatatggattaattttcttggcagcacattttgtatgggcattta
gtttaatgttcttatttagtggtagaggctattggcaagaacttatagagtcaatcgtat
gggcacataataaaataaaagtcgctcctgcaattcagccaagagctttaagtattactc
aaggaagagcagtcggtgttgcacactacttattaggtggaatcggtacaacctgggcat
tctttttagcaagaattatttcagtaggctaatagtgaaaataggat-aaaaaaacaatt
atggcaacaaaatttcctaagtttagccaagctttatctcaagatcctacgactagaagg
atttggtacggtattgctacggcacatgactttgaaagtcatgatggaatgacagaagaa
aatttatatcaaaagatatttgcttcacattttggacatttagcaatcatctttttgtgg
acatctggtaatttattccatgtggcttggcaaggtaactttgaacagtgggtattaaat
cctttaaaagttaaaccaattgctcacgcgatttgggatccacattttgggcaaccagct
ttaaaagctttcagtaaaggtggctcagcatacccagtaaatattgcatattctggcgta
taccattggtggtatacgattggaatgagaagcaaccaagacctatattctggggcttta
tttttactggttttatcagctttacttttatttggtggatggctacatctacaacctaaa
ttcaagcctggtttatcgtggtttaaaaataacgaatcaagattaaatcatcatttatct
gggttatttggggttagttctttagcgtggacagggcatctagtacacgttgctatacca
gaagctagagggcaacatgtaggatgggataatttcacaacagtattacctcatcccgct
ggtttacaaccatttttagcggtaactggagtgtatatgctcaaagtccagatacagca
caacatatatttggtactaatgaaggtgcaggcacagcaattttaacctttctaggtgga

tttcaccctcagagccagtccttatggctaactgacatggctcatcaccatttagctatt
gctgtagtctttattgttgctggacatatgtatagaacgaattggggaattggacataat
ttaaaagatattttagatgcgcacagaccacctagtggtagattaggagctgggcataaa
ggtctgtttgatactattacaaattccttgcatatacaattaggattagcattagcttca
cttggcgtaattacttctttagtggctcaacacatgtatgctatgcctccatatgctttc
atggctaaagattttacaactcaagcctctttgtatactcatcaccaatatattgctgga
tttttaatggtaggagcatttgctcatggtgcaatcttctttgttcgagactatgaccca
gaacaaaataagaataatgttttagctcgcatgctagaacacaaagaagctatcatttct
catttaagttgggtaactctatttttagggtttcacacattaggtctttatgtacataat
gacacaatgattgcttttggcactccagaaaagcaaattttaattgaaccagtatttgct
cagtggattcaggcttcttcaggaaaagcactctacggttttgacgtattactatcatca
tctagtaatattgcaacgcaagctggtagcaatatttggttaccaggttggttagaagca
attaatagtggaaagaattccttattcttaacgattggtcctggtgatttcttagtccat
catgctattgcattagggctacacactactacattaattttagttaaaggcgctttagat
gcaagaggatctaaacttatgcctgacaaaaaagactttggttatagttttccatgtgat
ggacctggtagaggcggtacttgtgatatatctgcatgggatgcatttacttggctgta
ttttggatgctcaatacaatagggtgggtaacattttattggcattggaagcatattact
atttggcaaggcaatgctactcaattcaatgagtcctctacatatctaatgggctggttt
agggattacttatggttaaattcctccccattgattaatggttacaatccatacggcatg
aacaatttatcagtatggtcatggatgttcctatttggacatttagtgtgggcaacaggg
tttatgttcttgatctcttggcgtggttattggcaagagttaatagagactctagcatgg
gcacatgaacgtacacctttagcaaacttaattcgttggaaagataaacctgttgcatta
tcaattgtacaggcaagactagtaggcttggcacatttttttccgtaggatacgtattgac
ttatgctgcttttgtattagcttcaacttcaggaaaatttggttaaactaaa-----att
taattaattgta--ataaaaaagtcagtcataata-gttttatggctgactttttttat
ttgcgggaattacttattaaagtttcacttcaatatctacacctgaaggtaaatttaatt
tcattaatgcatctattgtttgagaagaaggttgatgaatatcaataatccttctatgag
atcttatttcgaaatgttctcttgaatctttatctacatgaggagaacgtaagacacaat
agattcttctttttgtaggtaaagggattggacctactgcaatagcattagttctcgatg
cggtgtctactattttatcacatgacgtattaaggatagtagaattatacgctttaagct
tgattctgatttttgtctgctgagtaactgtcatattaaggatatcttattatttaagaa
ttttagagacaacacctgcgcctacagtacgtccaccttctctaatagcaaaacgcatac
cttgttcaattgcaattgcattaattaattccgcagtcatcttaattctatcaccaggca
tgaccatctcagcatcagtaccatcatcagcagtaaactgattaattgtaccagttacat
cagttgttctaacataaaactgaggcctatagcctggaaaaaatggcgtatgtcttccac
cttcttctttagttagaatataaacttcggcttcaaactgagtatgaggtgtaattgtac

caggtttagctaataccatgcctctttcaatatctttttctgaactcctctcaaaagaa
tcccaatattgtctccagctaaaccctcttcaagcgtcttttggaacatttctaatcctg
tgatagtcgttgtcctagtttctcgcaaaccaacaatttcaattgtatcaccaactttaa
taattcctctttcaatcctaccagtagcgacggttccacgtccagtaatagaaaatacat
cttctacagccattaagaaggttttatcaatatctctttctggtgttggaatataagtat
caactgcttccataagcgaaaaaatttatcaacccatttatcttcaccttgcttaatat
ttgggttctttgtaactgcttctaaagccaacaaggcagatccagcaacaaacggtatat
catctccaggaaagtcatattggcttaataactctcttccttctagttcaactaattcta
atagctcttcgtcatctacttgatcttctttatttaagaaaacgactaatgtagggacac
ctacttgttttgctaacagaatatgttcacgcgtttgtggcattgggccgtctgctgcag
acactactaaaatagcaccatccatttgggcagcaccagtaatcatatttttacatagt
cagcatgacctgggcagtccacatgtgcatagtgacgattatctgtttcatattctacat
gagcagtattgatagtgatgcctctagctttttcttctggcgcagcgtcaatttcatcaa
actttttcgctgcggtagaacctagagtcgataaagttgctgatatagcagctgttagag
ttgttttaccatggtcaacatgaccaatagttccaatattgacatgtggtttttttacgtt
cgaatttagatctagccatgtttttttgttttccttgttgtaaaagttgctttatggttac
ttttaacgataaatcttataaaatacaagatttattatttggtatcagtgaaa-----ta
agacatctatcatttagtatcgataatgtgcaaacgctttatttgcttccgccattctat
gtgtttcttctctttttctaatagaattaccggtttcgttagcagcgtccataatttcgt
tggctaattttatcgccatacttttgcctgatctttctcgagagaatctagtaatccatc
ttaatgctaaatttgtgcctctataagctcgtacttctattggaacttgataagtagagc
cacctactcttcttgctttcacttctactaaaggagtaatatttcgaatagctttttcta
agatatttaaagggtctgcttccgttctatctttaactatatctaaagcttgatatataa
ttctttgtgctaatgtttttttaccacttttcaaaatacgaacagttagcatactaacaa
gtctgcttttatataaagggtctggtgatgcgaacctttttttagcggtattacgacgag
acataatattttaaatttgttattgtaataatttttttgtactatcagagatttaagattt
aggctttttttgttccatatttagatcgacttttacgacgatcttttactcctgcagcatc
taatgtacctcgtactacatgatatcggactccaggcaagtctttaatccgacctcccct
aattagaactacagaatgctcttgaatgttatgaccaacacctggaatataagctgtaac
ttcaaatcctgatgttaatcttacacgagcaacttttcttaaagcagagttcggtttctt
tggagttgtagtatatactctcgtgcagacccctcttctttgaggacaactttgaagagc
cggggatttttgtttttttatgtattttacgtctttcggatctaacaagttgttgaattgt
tggcataataaaattgatttaaagtctttaatagatactatttgaacttactttaagtga
atgaattagtctttatttattttatatttacgcatgaatctttctactctaccttctgta
tcaataatcctttgtgaaccagtaaagaatggatggttacctgaccaaatatctacatgt
aattcaggctttgtagagccaattgtcataattaattggccatcacaatagacttttgct

tctggataccaatttggatgtatattatcttttgccattgcttctatatttgtataaatg
tatcactaataaatattaacgtttagaaaattgaggcgcttttcttgctttctttaaacc
atatttttttctttcttttactcgaggatctcttgttaaatatccttcagacttaagcgc
agttctattttctggattaattgaacataatgctcttgcaacacctaaacgaattgcatc
tgcttggcccgttaatcctcctcctctagcatttacatggatatcatattgatttaatag
tcctaaaacttgtaacggtgcatatgacactcttaagtagttaggactaaattgaagata
tgactcaccagggacaccgttgatgactaaattacctgagcctggtactagtcttacctg
tgcaacagaacatttccggcgacctgttccagaatagattgcacgagttttaattaattc
tgtcgacataatattccttgataactaaataaatatacattctttatactatatactctt
ggggttttgtgctttatgtgggtggagcggaccagcatacactttgagcttcgtaaata
attttcttcctaatggacctttgggaagcatacctttaaccgatttctcaatgattctgt
taggtaatcttgtttgcagctgctcaaacgtttctacttttagtccaccaggctttcctg
aatgtcttctgtatagtttctgattagttttatttccacttacagatacttgagaagagt
tgataacaattacatagtctcccgtatctagatacggtgtataacaaggtttatttttac
ctcttaaaatattagaaatatgagtagacattctaccaagcgtctgatttttagcgtcta
taatataccaatttggattgatatttagtgatggtgcttgcgttttgttcatgaaaagat
taatacttttagttgacaataagagaatatgtaaatgctgtacatgcaatttcgaaacaa
gtaatattg-taaaatatta-------------ttggcagagaaaacttttgtgacagtt
atttagattatttgttagcaaaaagacttatagttatcaacaacttataata-ttttgat
agtataaaatatgttagtcactttttctttcggtaaacttatacctaatttttttttgtc
agagcatatataacttcttctgctgattttggccaaagtttttaatttctagcagttct
tcttgagagtagtccaataaatctgcaatagaatgaatttgagctcgctttaagcaatta
tatgctctaacagataactgcagttcttctattaaaacttgactaatttttttatcttct
ttattacgataattatctgctgacttaaaatgtaaatttcttagtgaatagaaaaggttc
gttaaaactgtagcaccttggcttatggcttcttgaggggagatactaccatttgtccat
atttgtaaaattaatctgtcttttatgctattagtacctatacgtacttcttctacttta
taattgactttattaactggcataaaaactgaatctacttgtaaaaaatctacagataac
tcatctactgctttttctgctaaacgatatccacaatttttttcaattttaaactccatt
tcaaatattgtattattgcaaatagttgcaatatactgtctaggatctactacttctata
tcagaggacaactcaaataatccagctgtaactatagctggtccttgaactctaattcta
ccaatttgagattctttgttatggcttttaaatactacttcttttaaattcaataatatt
tctaatacatcttctcttaccccctggaattgtagagaactcatgatttactccagcaatg
cgtacagctactatagcagtaccttcaagatctgataatattgatcttcttagggcatta
cctaatgtaataccttgtccttgatttaatgcttctattacaaaactaccgtactgtcca
cgagctccatctgttcttgactctacgcattcaatttgaaattgagccacctaagctagc
tcc-ttaaaaataatcaatgattagtaattctctcgataataagcaagaatctatactcg

gcgtttcttaggagggcggcagccattgtgaggcacaggagtaatatcctttattaaagt
aatctctaagcctgctgcttgtaatgctcgaattgctgtttctcgacctgcacctggccc
atttactaaaacttcagtctgacgcaagccttgatccattgcctgtttagcagctttctc
agcagcagtttgagcagcaaaaggcgtgcctttttagctcccttaaatccgctagcacc
agaagatgaccaagatagtgtttcaccttttaaattagtaatagtaacaattgtattatt
aaatgtcgatttaatatgtgtaacaccgttaactgcattacgtttgttttttcttgctcc
ggattttttaattgtctagccatcgtgtctcatctattatatagttagttaaaagatta
ttttcttggagctttttcttgcctgctactgtttttttacctcccctgcgtgttcttgc
attagttcttgttctttgacctcttagagggagaccaagacgatgcctacggcctctata
ggtactaatttccataagtcttttaatactcatagattcaaaacgtttgagatcgccttc
aatctgataactagattctaaaatctctcttatacttacaacttgctgatcatttaaatc
ttggcacttgatatcagcatctatatttgttttagtcaatatttctttcgagcgagatag
tccaatgccataaatatatgtcagagctatttctactcttttgtttcttgggagatctac
tccagcaattctggccacgttgtaatgtctccaataaataatgtttacttttttataacat
cagattata-aactcaaccttgtctctgcttatgtttaggattggtacaaatgaccatga
cctttccattacgacgaattattctacatttttcacacatttttcggacagaaggacgga
ctttcatattaa----atctatatactaggaatattactg---------tattaattatt
tacatcaaaaatgttaataatatatttttattactttaattttatagctttcacttcgtc
atactatcatattttttgatatgacgtatgtttgaatttgcttagctgtatctattgcg
actccaactaaaataagtaaagatgtcgctccaagacctcttagatttggatctgagta
accttctctattataaacggaatcaacgctactgtaaataaaaatgaggctcctagaaat
gtcagtctatttaatataacttgtaaataattaatagttgcttgacctggacgaatatta
ggaatacttgcccccatttttttaaatttgttgctatatcttctgggttcattactatt
gatgtataaaaatagctaaagaacaagatcaaagcacaataaagaacaagatatagcgaa
ccattaggagaaaataggtatagaatttgaagtaatgttttattttgaactatttgagtt
aaataagatggtagggccatagacgcagatgcaaaaacaataggcataaccccaccttga
tttaatttcaaaggcaaataactgtttggatccaaaattgaagattttcctagctgtctt
gctgaaataattttaattcttctagttccttcttgcacacaaattgtaattataatcatt
attaaaaagatggcaacgaataagccaaacttaagacttgagttgctatagttggcatca
aaaaatgactgcgtaaagttcttaggtaatcctgatacaatattttggaaaatcagtaga
gaggctccattaccaattccttttctgtaattaactcggatagccacataataatcata
gagcctgctgttaaagctaaaacagactcacagacaaatgcaaaattccaattaaagaca
tacggttttacccatatggaaattgctccagattgtaaagtagcccatccaagagataaa
tatcgagtaatttgagttattttttggcggcctagctcaccttcttcctttttgcattttt
tctaaatcaggaataattttcgttagtaactgcattacaattgaagaattgatgtaaggt
acaatacctagagcaaaaattcctattgtcgagaatcctcctcctgaaaagatattcagg

aaatttactaaagtatttttttctacacttgcataaaaagcatcatgatctataccagga
acaggtataaaaattcctaaacgagctaagactaatagaaagagagtaaatataatacga
tttcttaaatcactttttggctcataaatcaattaaaaataaaaagaggaacttc--gt
tagataaaaaccctattccaatacttgtataagtttaaaataggggagtag---aacaat
gtggaaatattgctagcaatattttatgtcgaataaagattttctataggaactcctcga
tccttagctgcttcgctaaaagttcttaattgagttaagccatttaaaacagcccttgca
ttatttaatgtattattagatcccaattgtttagctaagatattttgtactccagacagt
tctaacactgttcttacagaacctccagcaattacacctgaaccaggcgcagaaggccgt
agtataactttagctgcgccagaaattccgttaataggatgaggtatagaattagatttc
gtcagtggaactgtaactagatgtttttagcatctgttactcctttttaactgcacca
attacatcacttgctttacctacacctacaccaacttgtccctgttcattaccaacaata
aggataactcgaaagcttaatttttaccacctttgacaactttagtaactcttttaact
tgcacgactctttcttcccaaccactatctttatctttccctttactctgtttttacga
ttagccattatcaacgttattcctaa----ttaatta-cgactttaaaaacccatacctg
cttctttagccgcttcagctaaggccttaactcttccatgatatagtttgcctcctctat
caaagacaacattttaatgccttccttgatagacttttcagctagctgtttaccaacta
cacgagaagtatcacagttcggtcctattttatcaaattcttgattatttaaattaacag
atgatgttgcaactaacgtaattccttgagtgtcatcaattacttgtgcatatatatgtt
tattagatctaaatacgcatagacgtggcctatttgaagttccttgaactttttttcgaa
ctcttttgtgcttatgaattctagtctgcttagtgtttagtttcattgttatttaccttt
tccagcttttccagcttttcttctgacaaattcaccatggtatctaattcctttaccctt
ataaggctcgggaggcctagtagaacgaatagttgaggcaacctgtccgacaacttcttt
atctatacctgagacagtaatatttgtattgttttcgactttaatttcaatattttctgg
aggtttaattttcactacatggctatatccaacacttaatattaaatcttgattatccat
ctgggaacggtaacctacaccttgtatttgcagtttttttagaaaacccagtagaaactcc
ttcaatcatattactaataagagttctagataatccatgtaattgattagccatttttgt
ctttccacttgtcttaacagagattgtaccattaagtatttctaggttaataccagttgg
caaagttctcgataaagtacctttagggcctgttaccgtgatagtattaccattaagctg
agtatcaagatttgtcggcaataaaattatttttttttccaatacgagacatgattccacc
tataaaatagttaccaaatatagcataagatttctcctccaagaccatcatgacgagctt
gtctatctgtcatgacacctctagaagtagaaataagggcaattcctaaacctccaagaa
ctctaggtagttctttatggtttgcgtagactcttagtccaggtttactaattcttttca
gagcagtaataactggttgacggtttttaccattatacttcagggaaatcattaaatgtg
tctctataccttcacccatttgttcaaaattttgaacaaatccttcttctttcagtactg
ttgccatattgcatgtcattttcgttgctggaacttgtacaatttgatgtcttgctaagt
ttgcattacgaatacgtgtcagcatatcggcgatcgtatcgttgaccacc-agattctcc

ttacgataaattgaattatttattatattttcaagattctctaaaaggcataccaagcct
ttttaacaaagctaaaccttcttgatctgttttggctgtagttacaattgatatatctag
acctcgaatttgatcaatattatcatagtctatttctggaaagattagctgctcacgcaa
acctaaattataattgcctttgccatcaaaactcttaggactaattcctctaaaatctct
aattcttggtaaagttaaattaattagcttctctaggaaggaatacatcttatcttttcg
taaatgtactacaattccgataggaacttcttctcgtatttgaaacctgcaatagattt
tttagcttttgtgacaatcggtttttgtccagtgatgagtgttaattcttggatactact
ttcgagagccttggcattctgagaagcttcacctaaaccacggttaatagtaatttttgt
aaaattaggaatttcatgtacattttcatactggaattcgtcttttagagcttgagtaac
ggttgttttatatttttcttttaatcctattaccattatattgtcacaaattatttaata
agttcgccagtctttttcagttttcgaattttttgacctttctcattaattattattga
agatcggctagcaatattattttgttcactaaataacataacattagagctatgtattgg
tgcttcgaatttaataatttcaccagtatcaccttcttgttgaggtttttatgcttcac
ttttaaattaattcctttaacaatcactttactagttttatggataattgcaatgatttc
acctgtctttgtttttcgcttccagaaataacttgaactaaatcaccttttctcaattt
aatttttgtggtaactttattcatttagaagagcttttcattatattacctccggcgca
agagagactatttagaaaaatttttatctcttaactctcgagctataggtccaaatact
ctcgtaccacgaggattattatcttggttaataataacagctgcattatctccaaatcta
atggtcatgccgtctgttctcctcaaggctttacgagttcttactacaacagctctgaca
acatccgatcttttaactggcatattaggagaagcatctttcacgactccaataataaca
tcacctatagatgcataagagggattacttgtgcctaatactctaatacacattattttt
ttagcgccactgttatctgcaacattaagatagctttgagtctgtatcatacttttattc
tatataagattaattatcaaaagacttagataagatattaaccattgtccaacacttagt
gcgacttaaaggccgtgtttcttgtattgtaactatatcacccgttgtgcattcgttatt
ttcatcatgtactttgtactttttgtacgaatcatggtttcgcgtattttctatgagc
tattctattttctacagctactactatagttttattcatttatcacttaccactttacc
agttgtttcttttaaaggcataatttcta----tttttatat---------aattaaat
cataaaattattatgtagctgactgggttcgagacttttcaatagttagaagttgagcta
atctatgtttagagtgcttaaataaatgaggctgaaaatcttgtcttgtcgctcttttaa
gccttaagtcaaaaagatctcttttaactacaaggatttcttcggctaaggtagaatcgt
ccatttgcataacatctgatattttaggaaaagtcatatttttaagattctattgtgttcc
gaactataaatttggtctttattggtaatttataagaagctaatttcatagcttgctgag
cagtttttgtggcacaccagtaatttcaaatagaatgtgaccaggcctaataactgcaa
cccaatattcaggagctcctttaccagatcccatgcgggtttctgcaggacggcagtta
caggtttatctgggaataccctaatccagagcttccctcctcttcttacatatctagtaa
tagttcttcgagtagcctctatttgtctagaagttaaccatactggctctgtggcttgca

gagcataatcaccgaatgcaattgtattacctttactagcagaacctttcattctacctc
tatgttgttttctgaattttgttttctttgggcttaacataagaattaaacaatttaaat
aattaagaaactgtcggatcagaaatttcctgattataggcagattctgcgttctttgat
tctggcaaaatttctcctttaaaaagccaaaccttaactccaagtacaccatatgtagtg
tgagcttgtcgatggcaatagtcaatatcagctcttaaagtttgtaaaggaaccctgcct
tctctgacccattcacttctagcaatctctgcaccgttaagacgtccagagacttgaatt
ttaacgccctgcgtatttgctctttgtgctctttgaacagcttgtctgactgctctacga
aaagcaactctttttctaactgctgagtgataaattctgcaactaaagttgcttcagaa
tctggatctgaaatttctacaacattcactctaatttgtttactcgaatctaaaattgag
gatagcaactttctgagagactcaattcctgctcctgattttcccagtacaatcccaggt
ctagctgttactattaaaatttctacttgatcaactttacgattaatttcaattttagca
atactagcattgctaagtttggaatgtataaatgagcgaattttatgatcttcttgcaaa
agaactggatagtctttagaattagcaaaccatgaagaacgatgtttttgagtaatacct
attcgaaaacccaaaggatgaatttttgacccacagtatcctttactataaattaaaag
ttataattattaattgactttgaacat----gagactacaattctgagacacctaatgta
atatgacaagttggtttatgtatcggaaatgctcttccttgtgcccttggctgaaatctt
tttagcgttgggcctttatctgcaaaggctttgctgacgaacaactgatttttatttagg
ccatcattatgctcagcattggctgcggcagactctagaatctgttttatataagagcaa
actcgatatggcataaattctaaaataattaatgcttcttgatattttctgcccctaatt
tgatctagtacacggcggactttatgtggagaaagacgaatgtatttccctattgcttta
gcttctttttgtttttgtaatgctcataattatagtttaacgacgggcttttctatca
ccttttacgtgagtacggaaagttctggtgggaacaaactctcctaatttatgtcctacc
atttggtcggaaacaaagacaggaaaatgttgtttaccattatagacagctattgtatgc
cctaccatatctggaataatagtagatgctctcgaccaagttttaagacttcttttttt
tcgaccatatttaatgcttctatacgcttaagaagactaacatcaatgaaagggccttta
tgtatagatcttgacataataataaattattagattcttaataaaattaattctatttgc
ggcggcgcaaaacgtatggattactatatttgttcggattgcgtgttttaacacccaatg
ctggcttgccccaaggtgttacagggcgagctcttccaattggcgatttaccttcaccgc
caccatgtggatgatctactggattcatgacaacacctctaactgtaggtcttttaccta
accaacggtttctaccagctttaccaagagtaatattactcgcatcaatattaccaactt
gtccgatggtagcataacattctttacgaatcatacgaacttcacttgatggtaatttaa
cggtaacaaaagaaccttcttttgctactatttgagcataagttcctgctgctcgaacaa
tttgtcctccacaagaaggtcttaattctatattatgcacagctgtccctaggggaatac
ttgatagcggcaatgcgtttccaacttcaataggagctgttggcccagaaagtactattg
ctccaacacacaatgatcgaggatgtaaaatatatctttttcaccatcaaggtaatgaa
gtaatgctattctagcatttctattcgggtcatattcaatagaggcaactttagcaacta

tattgtgtctgtttcttttaaaatcgattaatctatactgctgtttatgtccaccacctt
tatgtcggcatgtaataacacctctattattacgacctttgcaaaaatggtgcttaacta
ttaacgatttctctggcttatcagtagtaatctctgaaaaagtagaaaccgttctatttc
ttgtcccaggcgtatatgcccgatataaacgaattgccatataaggaaaattgaaaggaa
tgaatatttga------ctgtttaatgcaattaagtttctgggaatagattaatagagtc
ttctgtagccagtgttacaattgctcttttgtaatgtggtcgcttacctataaatctacc
aatgctcctttttttcttcgggggatggcaagtattaacacctgtgacttgaacattgaa
aatatattgtatggcagccttaatattaattttgttgctctaggatcaacagcaaaaca
gtactgattttcttctaataattttgttgttttatcagtaatgattgggtatttaactaa
atctaataaacctctggaatcaatgctatccattatatacctcttgtattttagataagg
catcgactgtaataataattttatgcgcagctaaaagagccataatattcaaagtgtcag
ctgaaataatttctacattatgtaaattacgaatagagaggtaaacattggggtcttttt
tatccactacaactaaaactttttatttagatcaagattccatcgatgaatagcttcca
taaaaagtttagtctttggttgctgaaaataactattaaagttttccacaatcagcgtat
taactgatttattgtttaaagctgttcttagggctagttgtctttctttttttattcattt
tcttagcaaagctgcgaggttttgggccaaatattacaccaccacctctccataaaggtg
aacgaatagaaccggctcttgctctacctgttccttttggcgccatggtttacgacctc
cgccacgaacttcacttcttgttttagtattagcagagccttggcgcttttcattacttt
gttttactaaagctctgtggactaaatacatgcctgaatcttgactaaccttaagattta
aatcagcattgccacttatttgaccttcccaattatagacttgataatttagttgtgttt
taactgtcatgagatatatcgtatgaaagaatattgctatttactaattttgactaaagc
accaggtttaccaggaactgctccttttacaatcaacaaatcattctctgagttaatact
tacaatttgcaaatttttttaccgtaacttttttattaccaagttgacctgccatattttt
tccagggtagacccttcctggtgttgtaccagctccaatagaaccaggttgcctatggtt
tttcgaaccatgagacattggacctctgctaaaatggtgtctcttttggtaaccagagaa
acctttacctatgcttttagaagacacattaattctttggcctacagtgaacatgtctgt
agacaaaatttgactgacctcgaagtcatcagtagatttaactaagtattcacgcaaata
ttttagtggggaagcttgtgatttctttagatgtcccaataatggtttatttaactttg
ttcgacaacttgcttatacccaacttgaatagcattatagccatcagtagataaagcttt
aatttgagtaataacacatggcccaacctgaattacagtaactgggattgacaagccagc
ttcatcaaagaattgggtcatacctactttagtaccgagtataccaacagacactagatt
tctccttttttgagagatttatactatcatcatacataaaatttctactccaaaatacttg
gag--ccaacttaactgatagatgcatgattttaatcatgacaacataaagattaaaaac
tttacctattcaatctaaaatagaaatagtttttttgtccagttacagcgcatataaatta
aggg--ggctcattatatttgttataatctactatttatataaagtcttttcttgttatc
agcttgtttagaataatatcaac-------------------------------------

-------------------------aataaaataagaaaatatttaaactt-tatctatt
---agttactgaaaaacagaattatgttcggtaactacacctttttatttctatatttat
tgtgcaatatatatgcatatctagtacagcaagcataaaaatatagc----tgtatagcg
actatacttatagaaattagaggggtattaatgggcaaagttgttggaattgatttagga
acaacaaattctgtaattgctgttatggaaggaggtaaacctactgtgataccgaacgca
gaaggttttagaactactgcttctgtggttgcatatactaaaagtggtgataaacttgtg
ggccaaattgccagaaggcaagctgttattaacccagagaataccttctactctgttaag
aggtttataggacgtaaacaaaatgagatttctcaagaaattagacaaacatcatataat
gttaagactagtggatcaagcataaaaattgcctgtccggcacttaacaaagattttgca
cccgaagaaatttcagctcaggtactcagaaaacttgttgaagatgctagcacatactta
ggtgaaactgtcacacaagcagttattactgtgcctgcttactttaacgactcacaaaga
caagccacgaaagacgcaggtaaaatagctgggttagatgttttaagaatcattaatgag
cctactgctgcttctttatcttatggactagacaaacaaaataatgagacaattctggta
tttgacctgggggggaggaacattcgatgtatctgttttggaagttggagatggagtattt
gaagtactttcgacctctggagatacacatttagggggggatgactttgatcaacaaatt
gtagaatggctaatcaaagattttaaacagaatgaaggtattgacctgagtaaagacaga
caagcccttcaaagattaacagaggctgcagagaaagcaaaaatagagttgtcgaattta
actcaaacagagattaatcttccttttattactgctacacaagatggtccaaagcattta
gaaaaaactatcactagagggaaatttgaagaactctgctcaagtctcatagataaatgt
agtattcctgtcaataatgctctaaaagatgcaaaactggaagcttcgagtattgacgaa
gttgttttagtcggtgggtctacacggattccagctatacaacagatggttaaaagatta
ataggtaaagatcctaaccaaagtgtaaatcctgatgaagttgttgctattggtgcagct
gtacaggcaggagttttagctggcgaggtcaaagatattttactattagatgtaacacca
ttgtctttaggagtcgaaactttagggggagtaatgacgaagattataccaagaaatact
actattcccacaaaaaaatcagaagtgttttctacagctgtcgataatcaacccaatgta
gaaattcaggtgctgcaaggagaaagagagttaacgaaagataataaaagcttaggaaca
tttcggttagatggtattatgcctgcacctagaggcattccacaaattgaagttacattt
gatattgatgctaacggaattttatctgttaaagcaaaagagaaagctacgggtaaagaa
caatcaattactatatcaggagcttctactttacctaaagatgatgtcgaaaggatggtc
agagaagctgaagaaaactttgatacagaccagaaaagaagaaaaaatattgatataaag
aatcaagcagaatctttatcttatcaagctgaaaagcaagttaaggaatttgaggataag
attagccaagacctaaaagtaaaaatacagggtctaattacagagcttagatctagttta
gaaaaagaagaatatgagagtattgactctatctctcaacagctacaaaactctttaatg
gatattgggaaagctgctgctgagtctgaaaataaagatacaagcacaaaaaatgacgat
actgtaattgacacagatttttctgaagctaagtaagaaatt-agcgggtaacgcgattt
gaacgcgcgacatcaaccttggcaaggttgcgctctaccactgagctatacccgcataag

atttattatgccaaagatcgtaatatttgtcaatctgt--------tattatat---cgt
tgaattatta-------agctatagattgattaaatactaaaaatataaaagtag--atg
attcttctgt-aaaagaagtacttatattgtatatataagtacttcttttatttaaaaa-
-taagaatataattaaatatcagta-------ctaaactactaaagagttaactaagcta
gtaacaattactaatcctgtccacaaacctgcgcctgtataaatcaacccttttgattgc
tcccattgaccaggcgatgctagaacaactggtacgcctactactaatacagtcgataag
gcaattaacaatagcaccaataattgaatcgcaataatcatttattatattctcctgaaa
aattcaaatgtgaataatattgtatataatacaatgaagctttatacattcagcaattta
tctatgaagaaataatttaaaattattgttttctaaataaatattaaattaagtattaaa
atctttagtgtaaacaataacactagctttttttta-atagataaaaaatttaaagaaag
tctttttttaaaaaaaaattagtataagatgatatgctttataaaaagctaagtattatta
gttattagagaaatttaaaaatcagaggtaaatcatatgaactcagcccttttttttagcc
aagctaccagaagcttatgcagttttcaaaccaatcgttgatattttacctgtaattcct
gtatttttttctacttttagcttttgtttggcaagctgctattggttttagataaacttta
aattatgacagaaatagcaaatataaagagtacatttttttaagctctaatcttatatat
tacaaaataataactattatggttgaacctttactatcaggaatcgttcttggacttatt
cctataactttgtttggattattagtggctgcttatctacaatatcagcgcggcaatcag
ttgggtctttaaaacctaaataatagatatttataaaaactatagttattatagtttta
taaatatctacaaatatactaattataagtacagaaaaaatttattgaaacctgattaa
taattaacaagatgggtataataataa---------------------------------
------------------------------------------------------------
------------------------------------------------------------
------------------------------------------------------------
------------------------------------------------------------
------------------------------------------------------------
------------------------------------------------------------
------------------------------------------------------------
------------------------------------------------------------
------------------------------------------------------------
------------------------------------------------------------
------------------------------------------------------------
------------------------------------------------------------
------------------------------------------------------------
------------------------------------------------------------
------------------------------------------------------------
------------------------------------------------------------

------------------------------------------------------------
------------------------------------------------------------
--------------------------caaaataaaattttaccagcttgacttagtaac
tccaggtaaaagacattcatgagacatttcccggaaaacatggcgagacaatccaaaatc
tctgtaataccctctgcttcgaccagttagccaacacctatttctacctctaacagctgc
actgtttcttggcatttcttgaagtctttgtcttagttccattttgttagaaaagcttat
agtttctttaagtttctctttaagagctaggcgtttcaaataatatttttttcaagctt
ttctcttttaagctctctttggatcatattcttcttagccatattgtaa-aactcactgt
aaaaattatttattaaatatactagctaaaaaatcccttgactgcaacattcatataatt
aaaagtatatagcataaactagtaccatagagtcaatatttaatcgatgatact---agt
ttatgctattggaacaaatttatttagtaaagttcttgttccacatgggttgaaatagtg
caatcagattgagggtatgcaatgcatgttagtacataacccgctaacatctgatcatca
tctaagaatgattgatcggattggtctattgtaccttctgtaactttgcctgcgcaagtt
gagcaagctccagctctacaagaataaggaagttcaattccttcttcttctgctgcatct
agaatataagtatcatctgcacaattaaatgtaacgtcaattccttcttcttcacacagt
aaatgaattttatagtctgcca--tttttttactccataataaagatacacttgaaactt
agtaaagcaacat-aatttaaaggaatgaccaaaaaattagatattagtttcattatgaa
tgatagactgcaaaaaaagaaaataatatactattgtgaaaaaataagctacatttcaca
ctaaatagaagtatagtgggttcaaagtatatatataagttcatatagttaaaattattt
ttatcatttaaataacttaaatctatgccct-aagctcttatataaatcatc------aa
gctagtcttttac----aataaataattaattatgattttaatgctgaatcaacaattag
aactcaagcctattcttaaatttcagataccacaattgtattcatattctcttactcaag
aaattcaagccttaacgactaggctttttttacaggtttggagaagacctgccacgttga
tggcaggtattatacagcctttgttatggttaattttatttgggggtcttttctataatg
ctccagtcaatttatttactatcaatacaagctataactgtttttttgagctcggggatta
taattttcacatcatttactggagcactaaattcaggtcttcctttaatgtttgataggg
aatttggatttttaaatcgattattgacagctcctttaatatcccgaacttctattattt
tatcctttactacttttatgacttgtattagtttaatacaagttatattcatagttatag
cttctctatttatgggaaattctcctttaaacagtgatagtgctatagttgctggactaa
taattctattagtaacagtaggggttacaatgcttagtttagctttatcatttactctac
ctggccatattgaattattagcacttattttagtgatcaatttaccttttctgttttcga
gcacagctcttgcgcctctatattttatgccaccttggctacagctgattgctagtctta
atcctttaagttatgcaatagaaggtacaaggtacttatattctagcgtacattggaact
ttacagaatgtattatcaaaattagctggggagatatttgtctaggacaaattattatat
tactagttattttggatataatagcagcttatctcgtatctaatatattgaaagctaaac
ttaatta-----------------aaattttaataaaaaatttattaaatttattttaca

aagatttttactatatactactaatagtagtatggataattgaaatagaaatagtttttc
aaaaccaaagctattatcaaagatgataaacatttgtaagaaagtcaacaaagtatgttc
ttattcataggaggaatgtagtcaatgggactaccatggtaccgtgtacacacggttgtt
ttaaatgatcctgggcgcttaattgcagttcacctaatgcacactgcgcttgtagcaggt
tgggcaggatctatggcattatacgaattagctgtgtttgatccctcagatccagtattg
aatccgatgtggcgacaaggcatgtttgttatgccatttatggccaggcttggcgtaaca
gattcttggggaggatggagtattacaggagaaagcgtttctaatcctggattatggagt
tttgaaggagtagctttaactcatatagttctttctggaatgctctttctagctgctatt
tggcattgggtttattgggatttagaattattccgagatccacgaactggcgaaccagct
ctagatttaccgaaaatttttgggattcatttattgctatcaagcttgctctgttttgga
tttggagcttttcatgtaactggactatttggaccaggaatgtgggtgtctgatggctat
ggagtaaccggcaaggtattaccagtagctcctgcatggggaccagaagggtttaaccca
tttaatcctggaggcgttgcatctcatcatattgctgcaggtactgtaggtatattagct
ggtgtttttcatttaactgttagacctccacaaagactatatcgagctcttagaatgggt
aatatcgaaactgtattatcaagtagtatctctgctgtttttttctcagcttttgttact
tgtggaacaatgtggtatggctctgcaactacacctatcgagttatttggaccaactaga
tatcaatgggatagtggatattttcaacaagaaatagaaaaaagagttgagaatgctatt
gctgatggagctgcacctagcgaagcatggtcaagaattcctgacaaattagcgttctat
gattatattggtaataatccagcaaaaggtggtttgttcagagcaggccctatgaataaa
ggcgacggtgttgctgaagcttggcttgggcatcctgtatttcaggataaagaaggaaga
gaactaagtgttcggagaatgcctgctttctttgaaacttttccggtaattctagttgat
aaagatggtatcatacgggctgacattccatttagaagagcagaatctaaatatagtatt
gagcaagtaggtgtaacagctagtttttatggcggcaaattaaatggccaagtctttaat
gatgctcctagcgtcaaaaaatatgcaaggaaagctcaattaggtgaagtatttgaattt
gacagaacgacgctagaatcagatggtgtatttagaagtagtccaagaggatggtttaca
tttggtcatgctaatttcgctttaatatttttctttgggcatctttggcatggttcaaga
actatattcagagacgtatttgcaggaatcggggctgaagtgactgaacaagtggaattt
ggagctttccaaaaattaggcgatagaagcagtaaaaagcaaggcgctgtataagaca-a
ggttaaagatagtctttatcaaaaaatatttaggatattaaaatagttcagaaaaagttt
attatctaatagttaccaattaggagaatctatggaagccttagtctatgtctttttatt
aacaggaacgctaatggttatattcttcgcaatctttttttagagagcctccaaggatagc
taaataactttattgggtttgccattacaataatagaaccactttccagcttattagtgg
aaagtggttctattgtaggttaaagtatattattcttcatgctcttcaaaaggatctcga
agatcttttgatgcaggaccaaatgccgtatatattgagtaacctgtaatacctaaaagc
aaactcgaaataaaaatactaagaactgttgcagtttccataattcaatcagtattaaga
tgaactttttctataataatattatcaatattaaaacatttaattattaaatttatcaaa

atatattatggcacttagaactagacttggagaaattttaagaccgttaaattcagaata
tggaaaagttgctcctggttgggggacaactcctatcatggggattttcatgttactatt
tttcttatttctattaatcattttacaaatctacaattcctctctggtactagaaaatgt
agatgtagattgggctactttaggtag----ctaattataatatttcttgaa--------
tttaacaagatcaaaag-aag----------------aggcgatcgctattatgttataa
aagcgatcgcttaaaatatatttgctataca--cacaggtaattatttcccaagttaa
ttagttcactatctgcaaaactatttgtattgacattattgtaattaactttttcaaatc
ttactaatactggatatttgataccgcttttatccactgctgcaacagttccaatttctt
gataccagtatgattcttttctcattatttttacttttgagcctctttccatattttcag
tcctagggtaaattattaatgactagtataataaaaatctatataat-ttctgaaaaata
cataaaaatagtaccgacaattgtcacaattcaataatgaaaggagagagagagggatttga
accctcgataactaatgttatgacaattttcgaaattgttgcaataaaccactctgccat
ctctcctggaataaaacactttttttctaaattaaacaagaataacatagaattcaggta
gttgcctagaaaattaaaactatgttaaatataaattataatctttctatagatatcttt
acaaaatatgtaaacagggtctatatcagcttatttcaagatt-tttaatcttatacaat
ttttactaaataaataaactctctcataatatgaaactatcttggaaaacacttcttctt
tggtctttaccaatatttgtcgttggtttctttttctggcaaggtttttagggccgact
accacagatgttggtagtaatatcgcaagttctagaatgacgtacgggagatttttagaa
tatttagatatgggttgggttaaacgagttgacctatacgaaaataaccacactgcaatt
gtcgaagctgttggcccagaattgggaaatagggttcaacgaattcgagttgaattgcca
gcaagtgcgccagaattaattacaaagctacgcaaagctaatgtcgatctagatgcacat
cctcccaaaagtacaagtgcagtatggggactattgggtaatttaatatttcctctatta
ctagttggcggtttagctttcttatttagaagatctaataatgctagcggtggacctggt
caggcaatgtcatttggcaaatcaaaggctttatttcaaatggaagctaaaacaggtgta
gtatttaatgatgtagctggagttgaagaagcgaaagaagagtttcaagaagtagtaact
ttcctaaaacagcctgaatcatttactgctgttggcgcaaaaataccaaaaggtgtttta
ttagtaggtcctcctggtacaggaaaaacattgctagcaaaggcaattgctggtgaggca
agtgttcctttttttagtatatcaggctcagaatttgtggaaatgtttgttggggttggt
gcttctcgcgtaagagacctattcaaaaaagcaaaagacaatgcaccttgtattgttttc
attgatgaaattgatgctgttggtagacaaagaggtacaggtgtcggaggtggaaatgat
gaaagggagcagacattaaatcagctgttaactgaaatggatggttttgagggcaatact
ggcgttattgtaattgctgctactaacagagctgacattttagattctgcattattaagg
cctggaagatttgacagacaagtttctgtggatgttcctgatttcaaaggtcgactggca
atacttgaagttcatgctaaaaataaaaaaatggaacctaaagtctctttagataccatt
gcgagacgaactccaggcttttcaggagccgatttagctaacttattaaatgaagcagcc
atcttgacagctagaagaagaaagaatgcaatgactatgtcagaaattgatacatcaatt

gatagagtggtagctggaatggaaggtacccctttgattgatagtaaaagtaaacggcta
attgcatatcatgaagtaggccacgcaataataggcagtttactagagcaccatgatccc
gttcaaaaagtgacgttaataccaagggggcaagcaagaggcttaacttggtttactcct
agcgatgatcaaagtttaatatcaaggtcacaaatcttagctcgtatagtaggtgctctt
ggtgggagagctgcagaagaaattattttcggtgatgcagaagttactactggtgcaagt
aatgatttacaacaggttacctcaatggctagacagatggtaactcgattcggaatgtca
aaaattggaccgctttctcttgaaagccaaggcggagatcctttcttaggtagaggcatg
ggaggagggtcagaatattctgatgaagttgcaactagtattgacaaacaagtaagagaa
attgtaggtgaatgctataaacaagctaaagaaattgttgccgataaccgggtagttatg
gatagattagtagatttactaattgagaaagaaacaattgaagggaacgaatttagagat
atcgttaaagaatatacagcaattccagataagaattactataggtcacaattttaaa--
----taaagcttgagattaaacgggactgacgggattcgaacccgcaacttccgccgtga
cagggcggtgctctaaccagttgaactacagtccc-aattagct---tacaacgagataa
tctcataattagcttataattgtcaaactatattattaaatatgaagagta---------
--------ttataaataatactcttcatattttt-------------attggtgggttaa
taaggagagagagggattcgaaccctcggtacgaagttaatcgtacagcagattagcaat
ctgccgctttcgaccactcagccacctctccaaaaagccttacataataaatatatcaaa
tatagttatatttaaaagataggcttcaatatttatggagctaagcggattcgaaccgc
tgaccctctcaatgccattgagatgctctaccaactgagctataacccc---tttttga
ggctatctagaaatggctcaagcgggatttgaacctgcgaccttgggcttatgaatcccc
tgctctaaccactgagctactgagccttaatataaactattactcaatactatatcataa
atcatgttaaaatataaatcaattaaaaagtatttact-------------ttttacatg
ttaagaaattgatagcgcaaaaatattcacttttttgagtattaattcagaatataacgaa
ctgcaaatttttgataatatacaatcttaagtatttttattaag--ttgaaatattttc
ataggtagatttagc---------------------------------------------
--------------------actaaaatata-----------------------------
-----------------------------------------------------ttattt
a-----------------------------------------------------------
------------------------------------------------------------
------------------------------------------------------------
------------------------------------------------------------
------------------------------------------------------------
-------tg---------------------------------------------------
-------------------------------caaggatatatataa--------------
----------------------------------------------tatttctgggtaac
------------------------------------------------------------

------------------catattataatctagtaa------------------------
------------------------------------------------------------
------------------------------------------------------------
--aaggtaaaacttaacta-----------------------------------------
------------------------------------------------------------
------tagactaactaaa-----------------------------------------
------------------------------------------------------------
------------------------------------------------------------
------------------------------------------------------------
-------------------------------------tatagtaagtc------------
-----aggcttagtaaatatctatctttagctgccaaaaaactaa-gaaatacattaatc
aagtaacttaa-------------------------------------------------
------------------------------------------------------------
-------------------------------atgtgaaaaaggagcttgctt--------
ttcactgtttgacaagtttttattcgttgcaattattatattgatgcttgatatctagca
tttt-cttgttaaatagcagttactagatagataaacaattaatatt-----ttatacaa
ccagcattgagccaataccttggtcagtcaatatttccagtaaaagagcatgattaattc
taccatctaaaatatgagctgaagctactccttgagctaaagagcggatacagcaattca
ccttaggaatcatacctccggagattacttctgtttgtgtcaagtctctagcttgctgta
tgtttaaatgactaattagtgttgtggggtctaaagaattgcgtaaaatccccggagtat
ctgttaataaaatgagtttttctgcattgagagctgctgcaacttctcctgctacagtat
cagcgttaatattatatgactgaccttccttgtctgctgcaacactagcgattacaggta
tatagttattactaattaaaatctttaataatttagtgtcaacattttgtacttctccaa
caaaaccaaggtttgcttgaccacttggtcttggagtaataattaatccatctttccctg
ataaaccaacacttttaccaccttgcttattaatcgttgctacaagatctttattaactc
ttccaactaaaaccatttctacaatatccatagtgggctggtctgttatacgaacaccat
tgtcaaattttggtagaatcttcaggcgatctagccaagagttaatctctggacctccac
catggactaagataggacgtaaacctataaaagagaggaaaacaagatcactcattactt
gatcttttaatttttgatttttcatagctgctccaccgtattttataacaataattcttg
aagaaaattgctggatatatggtaaagcttcacttaagactttcactctttctgaattag
tcagcatgaataaagtaggattatgatgtataaatttattgacaggacttatattaattt
aataaataagactaagtggtcaaaaactttctaaaaataaatatgttattagtaataaaa
acagttaaaagtatgaggattataattatgagtaatttctgggataatatcttgaaattt
ccgcgg-ttttagttagtgtcattttcggcttaattttaataattattagtcctttttc
gtgctattaagaaaaccattgacaagttttttttttatcatatcattgactggattaacc
gcaattttagcggtaacaattaaaaaaaatgctcaatatcgaatgttgttgagagaaatt

agtttaattaattaagctttttgaacatatataatattatattgttactgactatgatag
ttt-attttatttatagaaaagaaactgtaaatattaaatattatttgatatatattat
gccattaaaacgaatagtcagttctgaaaaaactggggcctttgcactgttagatagtat
tgttaggcatggagttaaacatatatttggttacccaggtggagctattcttcctatcta
tgatgaactttatgcatgggaacagtcatctttaattaaacatattcttgtccgccacga
acaaggagcagctcacgctgcagattcttattctagatcaacaggagaagtcggtgtttg
ctttgctacttctggtccaggagcaactaatcttgtctccggtatagctactgctcatat
tgattctgtacccatattagctgttacaggtcaagttggaagagcatttataggcacaga
cgcatttcaggaagtggatattttcggtattacacttcctattgtaaagcattcctatgt
agttcgtgatcctagagatatgtctagaattgttgcagaggcctttttcatttctaaaca
tggtcgacccggtccggtattgattgatgttcctaaggatgtaggattagaaaagtttaa
ttattgttctatcgacccgggaaaagttaatattcctggctgtagacctgtcacacagtt
aaagtcccgacaaatacttatggcagctaaaatgatacaacagtctagccagcctttgct
ttacataggaggaggagctataatctctgattcgcatcagataatcaaagaactagtgga
tttatataaaatacctgttactactactttaatgggtaaaggtatttttaatgaagatag
tgaatattgtttaggaatgttaggaatgcatggtactgcatacgccaattttgcagttag
tgaatgcgatctcttaattgctttgggagcaagatttgatgatagagttactggaaaact
tgatgaatttgcttgtaacgctcaagtaatccatgttgatattgatcctgccgaagtggg
aaaaaatagaattcctcaggtggctatcgtcggtgatgtgacagaagttgttactgcagt
attaaatttattaaaaacttctttcccccttacccagagcaaattgcatcttggcaaga
ccgaattaaccgttggcgtcagcagtacccattactagttcctaaaaaatctacaagtat
ctcaccgcaagaaatattggttgcaactaataaattagctcagaatgcttattttacaac
agatgttggtcagcatcaaatgtggtctgcccagtttttaaaagtgaaagctaagcactg
gctctccagtgctggattaggtacaatgggatatgggctacctgcagcaattggtgccca
agtagcacatcctgatgatatagttatttgtattagtggtgattcgagttttcaaatgaa
tatgcaagaacttggaactattgcccaatatcagttaccagttaaaattattattattaa
taatcgctggcaaggaatggttaggcaatggcagcaagcattttacggtgaaagatattc
tcattcaaggatgacagaaggagctccaaattttcaaaagcttgcagaagcttttggtat
taaagcttttactattagtagtaggcaaaatatgcaatcttctctagaagctgctatcca
gcatcatgggccagccttactagattgtcaagtcacagaaaatgagaactgctatccaat
ggttgctcctggcaaaagtaatgcacaaatgattggtattgctaagcctcaaagaggtac
tgcttccaactatattagtacgaatatttaacttggcgtataacattcaccctgaatatt
tacacaattaaactagatcttccactatgtagcaatttttaattagttcagcaggttcaa
ttatcaagcctattttaagagactttcttaaattcatgaagactaagttgtttcttgtgt
tagtatttcctagattatcacaaaaaaagttctaggaaatactcattactttaatttgttc
ctgaatatgtgaaacttaattccgactttttcaataataaaataacaacaaa----ctat

atataaatatattctttatatagtct-ataactgctttaaaatagcctttgacgagaatt
gaactcgtgaccttccccttaccaaggggatgctctacctctgagccacaaaggc-ttta
tattatattgggccgggttggatttgaaccaacgtaggcgaagccagcggatttacagtc
cgcccccattaaccactcgggcaccgaccctaaattaactttatatgaaatatagtttaa
acaatatcatagcttagtcgctaaaaacaagccttttc-taaataaaact-agtaagtaa
tggacatagctggacttgaaccagcgactttcacgatgtcgacgtgacactctaaccagc
tgagttacatgtccaaaaccttaactattactatagcactattt----attttataagta
tctatttttaattaaagatttgttttaatcgacttgctttacccagaagatctcttagat
aatatagtttagctctccgtactttagcccttctgataactttaacaaatgttattcgtg
gcgagttaagaaaaaatacacgctcaactccaataccttgaaatgagcatcttaaagtta
aacttgtatttaggctatttcttcttcttctagataatattactccttcacataattgtt
ctcgagttttactgccttctttgaccattaatccgagttgtattgtatcacctactctag
tctcaggaatgtcgtgtttaataaaaggagtctcgacacttttcataagctgacttaact
ttgtattatttgctttcatgtatgttgttagtcttataaag-tttgttgcaatattatac
attaaaaaaacaaatatgaaaattattaataaatacagatatctgtacccttagatcctc
atattattgatt------------------------------------------------
------------------------------------------------------------
--------------tttagatttgat------------ttacttaattcata--------
------------------------------------------------------------
------------------------------------------------------------
------------------------------------------------------------
------------------------------------------------------------
----------------------------------------tgctg---------------
------------------------------------------------------------
------------------------------------------------------------
------------------------------------------------------------
------------------------------------------------------------
------------------------------------------------------------
------------------------------------------------------------
------------------------------------------------------------
------------------------------------------------------------
------------------------------------------------------------
------------------------------------------------------------
-----------------------aattata------------------------------
------------------------------------------------------------
------------------------------------------------------------

------------------------------------------------------------
------------------------------------------------------------
------------------------------------------------------------
------------------------------------------------------------
------------------------------------------------------------
------------------------------------------------------------
------------------------------------------------------------
------------------------------------------------------------
------------------------------------------------------------
------------------------------------------------------------
------------------------------------------------------------
------------------------------------------------------------
------------------------------------------------------------
------------------------------------------------------------
------------------------------------------------------------
------------------------------------------------------------
------------------------------------------------------------
------------------------------------------------------------
------------------------------------------------------------
------------------------------------------------------------
------------------------------------------------------------
--------------gatagtc---------------------------------------
------------------------------------------------------------
------------------------------------------------------------
------------------------------------------------------------
------------------------------------------------------------
------------------------------------------------------------
------------------------------------------------------------
------------------------------------------------------------
-------aaaagagcaatttgagattac--------------------------------
------------------------------------------------------------
------------------------------------------------------------
------------------------------------------------------------
------------------------------------------------------------
--------tctaatagtgactatc------------------------------------
------------------------------------------------------------

------------------------------------------------------------
------------------------------------------------------------
------------------------------------------------------------
------------------------------------------------------------
------------------------------------------------------------
------------------------------------------------------------
------ttgtacctatattagcaaaa----------------------------------
------------------------------------------------------------
------------------------------------------------------------
------------------------------------------------------------
------------------------------------------------------------
------------------------------------------------------------
------------------------------------------------------------
------------------------------------------------------------
------------------------------------------------------------
---------------------------------------------cctattttaataaaa
tt----------------------------------------------------------
------------------------------------------------------------
------------------------------------------------------------
------------------------------------------------------------
---------------------------gcttatctcaaag--------------------
------------------------------------------------------------
------------------------------------------------------------
------------------------------------------------------------
------------------------------------------------------------
------------------------------------------------------------
------------------------------------------------------------
------------------------------------------------------------
------------------------------------------------------------
------------------------------------------------------------
------------------------------------------------------------
------------------------------------------------------------
------------------------------------------------------------
------------------------------------------------------------
--------ctgatttttcagatat------------------------------------
------------------------------------------------------------

------------------------------------------------------------
------------------------------------------------------------
------------------------------------------------------------
------------------------------------------------------------
---------------------------------------ataattagttt----------
------tcctcaaaatattctatgatttaattgttttggcaaatat--------------
----------------tgatatcccaacatttattgttagttaaagaatcttgt----at
cgtataacgattaagataatacgattaaagcttaagtatttattagttttgtgttataat
taattaatatcaaaggtagtggcataacacgaagcataaacaatgtttgaacgctttact
gaaaaagctataaaagttatcatgttagctcaagaagaagctagacgcttggggcataac
tttgtcggaactgagcaaatattattaggactagtaggtgaaggtactggaattgcagct
caagtattgaaatcaatgaatgtgaatctaaaggatgcaagagttgaagtcgaaaaaatt
ataggaagaggatctggatttgtagcggttgaaattcctttcactcctcgagcaaaaaga
gtactagaattatctttagaagaagcacgtcaacttggccataactatattggtacagaa
catttactaatgggcttagtcagagaaggagaaggagttgcagcaagagttttagaaaac
ttagcagttgatgtttcttcaattagagcagaagtgatacaaatgctgggagaaaatgca
gaagctaatgtcagtggaagcagtactacacaggttagaagcaaaacgcctactctagaa
gagtttgggtctaatttaactcaaatggctatggaaggtggtttagaccctgtagttgga
agacaaaaagagatagaaagagtgattcaaatcttaggtcgaagaactaaaaataatcca
gtcttaattggggagcctggtgtaggcaagacagcaattgccgaaggcttagctcaaaga
attgctaatagagatgtcccttctattttagaagataaattagttattactctagatgtt
ggcttgttagttgctggtactaagtatagaggtgaatttgaggaaagactaaaacgtatc
atggatgaaattaaatctgctgataatgtaatattagtcattgacgaagttcatacatta
atcggagctggtgcagcagagggagcaatagatgcagctaatctacttaaacctgcttta
gcgagaggcgaattacagtgtataggcgcaacaactttagaagaatacagaaaacatata
gaaaaagatccagcattagaaagaagattccaacctgttgtagttggagagcccagcgtt
gaagaaacaattgagattttatttggccttagagaccgttatgaaaagcatcatcaatta
acaatgtccgatggtgctcttgcagcagcagctaaatatgctaaccaatatatctctgac
cgatttttgccagataaagcaattgatttaattgatgaagctggttctagagtaagactt
ctaaattctcaattaccgccagcagctagagaattagataaagaattaagagcggtatta
aaaacaaaagatgaagctattcgagcgcaaaaatatgaaacggcagaacaatatagagca
agagaaatggaaattaaagctcaaattgcagcaattgctcaaagcaaaaaaaatgagcct
gacttaaacttagaagatcccgttgtcacagaagatgatattgctgaaattgttgctgct
tggactggtataccagtaactaagctaactaaaagtgagtcagaaaaactaatgcacatg
gaagaaactttacatgggcgcatcatcggccaggatgaagcagttgtagctgtctctaga
gccataaggcgagcaagagtaggtctgaaaaaccctaatagaccaattgcgagcttcatt

ttttctggtccaactggcgtaggcaaaacagagttgacaaaagccttggcttcttatttc
tttggctctgaagcctctatgatacgactagatatgtctgaatatatggaaagacataca
gtatctaaattaattggttctcctccaggatatgtaggttatagtgaaggaggttaccta
acagaggcagtaagaaaaaaaccatatactgtaattctatttgatgaaattgaaaaagcc
catcccgatatttttaatcttcttcttcaaattttagaagatggtagattaacagacgca
aaaggtcgaactattgattttaagaacacccttctaatcatgacttctaatattggatct
aaagttattgagaaaggaggaggtagcttaggtttcgaattatcagaagatcagacagaa
tctcaatatacaagagttagatctttagtaaacgaagagttaaagcagtactttaggcca
gagtttcttaaccgactagatgaaatcattgtatttcgtcaacttactaaagatgaagtt
agagaaattgcagagctaatgttaaatgaagtgtttgctagaattaagcaacaagatatt
cagttaaatgtaacagaacgttttaaacaaagactagtagaagaaggctataacccaagt
tatggagctcgaccacttagacgagcagtaatgagacttttagaagatagtttagctgaa
gaagttttatctggtaaaattaaagctggtgatagcgcagtagtagatgttacgaatgaa
ggagaggtaacagttttattaggtgaaaaactagaactgttaacttaaa------aaata
atttattaat-ataaagcattgaagaagttat--attcaatgctttttagaagaaattta
ataagatgggttgctataattgacaacctctgatactctttatttgtataca---aagcc
gggatagctcagttggtagagcagtggattgaaaatcctcgtgtcaccagttcaaatctg
gttcttggcat-cttaagtatttgaaaagctagctagataagaatcaaactttagttgga
atgtacctactggtccatttctatgttttgctattataatttctgtaaggttctcttttt
cagtttcttgagtatagtaattttctctgtagagcattattactaaatctgcatcttgtt
ctatagaattatgtacaataaaattatttgcgatgaaattatataaaaacttacagctta
aatcaaataccgtttgtaagctaatgatgtatatatttctaagattttcaaaaattaaag
atgaagaatgaaaagagtttaatatagcttgcctatctttagccaagtt------tattt
gaatagcgagcatactattttgatcaatctcgtcacatcttttccatccttgttctgtca
atagcttatgattatttgttaactgtatatgcctgccagattgtgtaataatttttatata
taggtttcttacctgttttcaagatattagctgtaccattttaaataaataaagttttt
gcgccgtaaagctaactacttta-atatttttataatgaaaattaaacaaaatttg--tg
gccgattatgtaatggtatctgaagatagctaaattttgatatacacccactttctctta
gatcagatagtagaggtcttttatcacttctagtttcaagatttctatttagctgagata
atactaatatgggcaaatttaaatctttgctaatatttttaatgaccgagtaataaatg
atagttcttgagatctattatcagtttgtttaccatcttgtaataattgaagatagtcta
taataattagctctatctttttttccttgtaatttaaaaaatttactttgactttatag
tgtgcgtagaaatttttgcactatcatcaatataaagatttagatcagctagagctttgc
tcttttcaataacgtgttgccattcattattcgtaagttggccagattgaatcttttgac
cgtttaagtgacattcttgagataaaatcctcctgagtagttgctctgtagacatctcta
ggctaaataaaataatgaaagcttgctcgtggttaattatatacctagtcacattaatag

caaaagctgttttccccattgaaggtcttcctgcaagaataattaaatctgacttttgaa
aaccttgtgtaatagaatctagttctgtaaaaccagaaagcacgctactatttatactga
ttttttttgttgtatctaaatggacaagtatatgagcaagaattgtagctagtgtttgtg
tatctctatcctcaagtatttcgtaagcttttgttagttgattaacaaaagaagttatat
tgctttgtttaataggccattttgaagagctgatcaaacataaggagtctccatataata
gaagtaatcttttaatgtaattatcaacaataactgaagaatattcacctatgacatcag
ataaaggagcttgttctactgcactaacaacatcgtctagctcatttaactgtgtaatta
ttttcccagtctttaagttagttaaaaaatttcctacactaatctttttttatatgattaa
tattttcaagcgcagctctgtaaagtagtgcgtttgatgtaaaataaaaaaaaattaggag
atagcttttctactgattttagtaaagattttggttcttgagtcaagattgtgctaatta
atatttttttcagctaaaatattatgaggaggtaaatatttataa------atactgatta
ttccctcc-ttgtaacattactctaatt--------acaataattaatagttaacttaat
ttgattcaggaagaacttgtagttgaacggtggcacttacttgatttaaaagtttaatct
ctatattgtaaataccaattgtcttaatctctggcaaataaatatcttgtttttcaattt
ctatattcgtattgttcttaataatttgcgaaatctctttttccgtaacactgccaaaaa
tattttggccgtcccctgttttcttagtcacgctaaacttctgtatttcttctagaagta
gctgaattctttttgcatcttctttagctatctttaatttttcttcttttatagcttcga
aaaacatttgctgctttaaaataccatttgttgctgttgttgccattctattaggaatta
aaaagtttctagcataacctgcagctacttttacaagatcattacttttgccaagttttt
gaatattctcttttagcacaacttttataactttttttacccatcttaattctaaattcta
aaaa----atctatataatttcattatacctgttgtatacagatttt----aataggaag
tgtttacttattagcataattcctgcataaaaataataataacttttttatgttt-ttctg
atataataatttttacaatgtttt-----gctaggagaggtggccgagtggttgaaggcg
cagcattggaaatgctgtttaggagcaatcttaacgagggttcgaatccctctctctccg
tataa--aaaatataatcactactatacagtccagttgaaatttccaataatatttttcc
aagcatttaagccccccttggactcttataatatttagtttttgagctattaaaaattgag
atgcaaatagagatctagaatctaggctacaatatacaaaacaaattttatcttttaaat
ttatattagagcaatgtatttctttgagttgtcgtaaaggtatatttaatgattctatta
agtgactttttttatattcttctctatttcttacatctattaaaatatattttaggttgt
tacaatttaagaattttttttaattgaataacattaatttctttttatgttagtattggttt
tcccataattatatttgcccttaggtctaccttt-------tgataatatgaatctagta
tttataaccttaaacttgttaaacgataaatttagtgcactgtaggttaagatatagcca
cttaatatagatttgtaaccaagaataattttgatagcctcagttgcttgaagtgtacca
ataatacctggcaatagacctaaaactccagaattgctacaagtatctattgcattcttc
tcagttgcaagctctttagagaagtcccgataattgggtcctccttgatagttaaagaca
ctaatttgcccctcaaattgaaaaatagcgccataaatatgtattttatttaattctaaa

caaatatcattcagcaggtatcgagtgtcgaaattatcagatccatctataacaatgtcg
taatgtttaataatatcaagtgaattatcatgacttaatcttgctttaaagacttttact
tcacattttggattcaaatctaatatttttttttagctacttctacttttgaatatcct
atatcatgaattgtatatataatctgccgctgtaaattggaaagatcaataatatcgtta
tctataataccaatgtggccaatcccggcagcagcaaggtaaattattccaggacatcct
agtccaccagcaccaatgaataatacctttgctcttttagtctttcttgtccctctaac
tgaatttgtggcaatactaaatgtttagaatatctagtatattcatctaaagaatatttt
gtattgcttgctttaaaatttagcataaaaatttataagttaattgctatcaatta----
--caaac-------------tattagtatgacttttgcaaactaggttcttctctatcta
aaatagctccttgcgtaggacagacctgaatgcaaatactacaatcaatacaggcagaaa
agtcaatccaataccaattagttttaatactgtttttccttctccttggtgaatacaag
ctactgggcaagcatttacacattcagcaactccaatacattttcggttacaatcgtat
gggccatgttatttaattaat-aaaaggtaaactgtttacatctagtatttttattgtat
actagatttacgtataatatctaatctaataagtaaggtaatctactcttatagtttaca
catttgatctgcgcctttagttcagctggtagaacgcaggtctccaaaacctgatgtcga
gggttcaagtccttcagggcgcgtttagtttttatcctaatttaatagtaatattttact
ttttggcaaacatcaaaataatataatataaaaagtcatgttttatatattcgataatct
atgtactagtactgtttttttatatgttattattggtaagttaattatattaattctat
ttttataatcggt-aaagatagttaaacttaagtgtgaatacacttaataatt-tgttgt
atatgtaaaaattgaaaaatttaaaactatggctaaaaaagtaactggaattgttaaatt
agcattaaatgcaggtaaagctactcccgcaccacctgtagggcctgccttaggtcaaca
tggtgtaaatattgttatgttttgcaaagaatataatgcacgtactgcagataagttagg
actagtaattcctgtagaaatttcaatatatgaagatagaagttttacttttatacttaa
aactcctcctgcttctgtactaattgctaaagctgctggcttaaataaaggctcaggcga
accaaatacaaataaagtaggtagcataacgcataaacagttagaatctattgctgaaac
aaagctccctgatttaaataccaataatatacctcaagctatgaaaattgttggaggcac
cgctaaaaacatgggaattctaattaaagattaaaaaatttcaaatatacttatta-tt-
-tttatgaaaaaaatttcacgccgacttaaaacattaaaatctcaagtggatcccaagct
gtacattcttgaggaagctgtttctattttaaaagccacatcaaatgctaaatttaaaga
aactgcagaagctcatatttgtctaggtttaaaccctaagtacgcagatcagcaacttag
agcaacagttatattaccaaaaggaacaggtaaatcaataaaagtagcagttattgctaa
aggagaaaaactaacagaagcaattagttcaggagctgatgtaagtggctccgaagagct
aattgatgaaattttaaaaggtagattagatttcgataaattaatagcaactccagatgt
tatgccgctaatagcaaagctagggcgtgtgctaggacctcgaggattaatgccatctcc
taaagctggcactgttacaacagaagtaggtaaggctgttaatgagtttaagggcggaaa
agttgagtatcgagttgatagaactggtataattcacgtgccttttggaaaatctagttt

tccccaagaagatttagtcttgaaccttcaaacaattaaagaatcaattgataaaaataa
accttctggctcaaaaggaaagtattggaaaactttttcttatctagtactatgggacc
ttctgttcaaattgacattaacagtcttctctaaaaagtatttgtatagttatacaactt
gaataatacattatttcaaatta-taatcttctagcaatattaataaaataatttgctta
ctatatcttatccgcactaaatttttt-ttatctatcttactaattatgagtacaaaagt
tgaaaacatcctagaagaattaaaatctttaaatcttctagaagctgctgaactagtgaa
acaaatagaagaaacatttgatgttgatgcatctgctgcttcaggaggcatgatgatgtc
agcaccagcttcagcaccagcctcttctgaagttgaggaaaaaacagaatttgatgtcat
tctagaggaagtacctgcacctaaaaagatttctgtgctaaaagttgttcgttccctaac
cggtttaggtttgaaagaagcaaaagatctagtagaatctacgcctaaagttttaaaaga
aggagcatctaaagacgatgcagaaactatgaaaaaacagctagaagatgctggagcaac
agttgttgttaagtaataaa-taaaaaagtgcttttgtttttagataaacaaaagcac-t
ttttatctaatctaactagttagattagaaaagtccgagtgtaattgcattattaatagg
catggctgcaccaatacctaaccaaatacttacaattgtaccaatcaagaaaacagttgt
ggcaattggtcttctaaatggattttgaaacttgttgacattttcaatgaatggaacagt
aagtaatcctgctggtacagctgccatacttaagactcctaatagcttatttggaattac
tcttaacagattaaatgttggaaagaaataccattctggtaaaatttctaatggagtagc
aaatggattagatttctctcctatagatgaaggctctaaaattgctaagccaatactaca
tgcaaaagttccaataattacaactgggaaaacatataaaagatcatttggccaagctgg
ttcgccataataattatggcccattcctttcgctaatttagctcgtagctttggatctgt
taaatcaggtttttttaagaatcgacatgttgtttttgtgtttaagataaaaattagtct
aaatactttgtcaaagtgtaaatttacacttttataatggtccggaaattccttgcttac
gtatcatcagaaaatgcataagcataaaaactgcagttagtagtggaagtacaaaagtgt
gtaagctgtagaatctagttaaagttccttgccctacactaactccccctcttaataatt
ctacaatgctcccccaacaaccggcacagcatctggaacacctgtaacaatttttacag
cccagtatccaatctgatcccatggtaaagaataacctgttacaccaaacgaaacagtaa
gcactcctaaaattacacctgtcacccatgtcaattctctaggctttttaaagcccccctg
ttaaatagacacggaatacatgcaaaatcatcattaagaccatcatactagcagaccatc
tatgaatagatctaataagccacccgaagtttacatcagtcataatatattctactgatg
taaaggcttctgcaactgtaggcctataatagaatgtcattgcaaatccagttgcaactt
gaattaaaaaagatacaaatacgatgccacctaagcaatagaaaatattaacatgaggcg
gtacgtacttactagaaatatcatcagcaattgcttgaatttctaatctttcttcaaacc
agtcataaattttactcataaaatagcttcaaaaagctcttttaatata-tttttgcgtt
taagctactgttcattcaatag------ttattgaacttatatt-atttattgtccata
cttattaattattataacatctttttatttattt--------------------aaaatt
ctttgacttaataaaacaggactatgaactacaatttttttcttctgtgtagatattaat

ttagtcttcattagttcagacaaaattcttgttacactaactctactgctacctataatt
tgtgctaacactttgtgtgtaacagtaaaattcaggatgataccattattgccctgagtt
ccattttgctcagctaataatagtaataagcttacaagtctacttgtaatatttttatgt
gaaattatttctataaaatgataagcttttgagattgagatattaaatggtttacaaaa
aatgaactaaatggtgggcagttattacatgtttttataatagtagtatagttagtacaa
gcaatttgggtttttacttattgctttcgcttcatagtaaaaattatcattgactaattcc
aactgcccaaatgtatctccattagttagcaagttaagcgttattttatgtgtgcttttta
aagactttgtaataatcaaagatcctatcaatataatgtataatctagagctcttatca
aacaaaataatatcatttggtctaagtgaaaagagttgatagtcaattttattactatta
aaaaatagcaaccatggattgcaagaaaaacaatcccctgaattacttccagaaaatgtg
aaaatggaatttttttgatttaaataatttttagtcataattactactttataa------
---aaaaa--ctctaataat-taagaactgaaattaaatgtcatacaatttgatgctagt
tgagaatgacattgtattatcaaaagccattcaagaatacttaatagataaaggatttaa
tgtttatattgccaacaacggattagaagcattgaatttagctaaccaatataatctcaa
tttgataatttctgatataatgatgccattagttaatggctacgaattattagaaaagat
tcagaaaactaaggcactatcaaaaatacctgttattttcttgacagctaaaggaatgac
taaagatagaataaaaggctacgatatgggttgctatggttacttatctaaaccatttga
ccctgaagagctagtttcattgataaacaatttaatggctagaaatgtactacaagatac
atctatagagaaatgcaataacttacaacaatcaacatataataaaatacgtttaacacc
tagggaagcaagtattctagatcttgttatcgatgggctaactaataaagaaatatctaa
tatactagatactagtattagaaatgtagaaaaatatgttagtcggctcttacagaaaac
taagatgaagaatagaactttactagtaaagtattcgctaaatcataatttattagatac
taaatttaatcaaagggcgaatgacgggactcgaacccgcgaatgatggagccacaaccc
attgccttaaccccttggccacactcgccatactacaaatatcactataacgtgtttatt
gacaggtagtctagtttctaaattatctatttatataacacagcatttaaaattaatttt
ctatgatacataataatatcagtattcaaattaacggagaaccatttaattgctcagagc
ctatctcattgcaatttttattaagttatcttgattttaattctgagcgcatagcagtcg
agctaaacaatgctttgttacctgagcctctttttcattcaacttacttgaatgatcagg
ataagcttgaagttatcactattgtgggcggtggataattttcttttatac---------
---attattttatcttagatgtttcattgataacgacaagcgcccttgtttttttatctac
atgaataatcacagctttaatagtatcgcctattttaaattgggatgatatctggtctaa
ttgttttatattaatttcagagatatgtacaaggcctttaagattgccgacctttataaa
taatccgtaaggcgtaatctggttgataattccttcaataatattaccaacaattaaagt
agatgaagcttgagctattaatgctcttctgtggctcagtatcaaattattcgattttttc
ttcaacgttgagtaactttagctttataaatttattattagattggtcacttttctgaaa
attgccgagatgagaattgggcacaaaccctgatataccttcaaggttaataatcattcc

tcctttattaaatccttttattagcacattaagtaaagagtcttcagctaataattgccg
aatccttttccatgctctgatatattcaaggcgtcgaattgataaaatcaattgttttga
ttgagcattatagtctaataaaaaaaattctcttgtatcattaatgttcaaagccgtaaa
gttatttaaatcttgattactagatacttcttgaataggtaaatacgcagatattggtgt
ccctatatccaccaaaaccccatttaactcaaagctaaatatagtgccagctacaatatc
tccaagatttaaatcatatttatactttgtaaaacagctgcaaaatttctgtgagtaaa
tccttcattatttttttgtcatattgaatagtataaattaaagaatagttgtgtaaaattt
attaaggccactaatctgttaccttattctgtg-aaaaatcttaatttggtttgttgtaa
ttctttatctaaa-atattaaaatatatgaaagacatgaaaagtaaccgtagatagctgc
agacttag-tattatacagtctgaggaaagtccgggctccacaaatacaatttatgctgg
agaaatcccagtgtaggtaactatgaggatagtgccacagaaataaaccgccagaataaa
ataaatatattagctggtaagggtgcaaaggaaagttaacagcttaccaaaaatactgta
aagtatttgttaggtaaaccccctaaaacggagcaaagctactaaacaaacttttgtgtat
atttatatagtttagttcaaaatactgcatgaagttattggtaacaataactcgagagg
aatagctatcctttacaatatacc-tttattatattgaaagaacagaacccggcttatgt
attactttttatgtcttttattatcttagttatttccgaatctctaaatttaatagctct
ctaattttctgctgtagctcatattgtttattatctatttctgcatttgtcaaagttttt
tgattagatcgataggtaaatcttaaacctatactttttttctgttttccaattgaatta
tttgtgtattgatcaaataattttatgttttctaaatcatcatcattaaattgatttaac
agttttaataagtagttaacctccatattatgaggtactactaaagataaatctcttata
atagatgggtatttagagtatggctgaatctgataatttaaataattgaactgattacca
gaatcttccaaaatatttaaatcaacttcaagtacaaaaagctttgagcttagacccaat
ttactatatgttgcttgactaagttcactgaatataccaatagtcacattgttataaatc
aatgttgcagatctatttttttcaataaaattcattttactatcaattatttcattcttt
gtccactcaatttgtctattgatttttttgaaaaaaattctcaactactcccttggcttca
taccaatttaaagaacgagctgcttgtgaccatgtagatctaatatctaaattaccacca
aggatgatagctaaactatttgtttctataactctatttttaacagtcttaaatactttt
cctatctcaaagctatctacagtttgattgctttgttttatattataaaggttagattca
attagtccttcaagcaagctatagcgtaaattagaatagtcttgaatgagaggattattt
aaagcaatattttctgaagacttaactaaggaataatgaactaattcagttaaccctaaa
tgtcttaaaacacttctgctcttatctataaattgtctctttggcgataatttttttgttg
aattgagtaattggtagtgagcttttgaatttatggtagccataaattcttgcgatctct
tcaataacatctacctctctaaaaacatcttgtcttctataatttggaatactaatcttt
ataatttctttatttttataaatagtattgaaatttaaagaattaagaatatcaagaatt
tcttgagagtttaaaaaaacgtgttcgaccattataacttattggtcctaaaatatcctga
actttttttaacagacaaatctatattgagaatatcattaattactttctctttgtaaaat

gtctctctaatattgccaccagctaaatctattattaaagacagagcttcgagatatgca
ttttcccaattgtccacatttagtccacgctcttgtctgactgaactttcagtacgaata
ttgagaagtcgagaagattttctgactacagactgctgaaatatagccgactctataaat
atagattttgtattaagatcagtatcaaaattattgttcactcctataccagctacacta
gttacattagcattaacttgagttaccaaaatatttttacttaactcaacattttcatta
tttaacttaatttgtttattatgcccattataaaacaggctttgaattgttatagtgtca
ctatgatcaatatcatttatcttattaaaatcaacaatattaataggctggccccatttc
accataatataattacttatatctatcaataaatttttatttgtaaaaccagaagagtgt
aagcgatttttttagccaatttggagaatctatagtttcgattttatctataactgcagaa
aaataattactacaatttgataaactatgatcagtaagtatatt-ttgtctttcagagca
atatcgattactgcttggatttatagattttaatattgaagatccctttagagctacgac
ttctcttgaaagacctatcatgcttaacacatctgacctattagctgttgatgatatatc
taacacatagtcaatttgatcccccgttgtaatagattcaacagcttcaacttcaaaacc
tgcctgtgtcaattgattagtcaaactatctatatctattgtttctatattagcaagttc
tttcaaccaatttaaagaaactttcataattaattgtcatctatcttatagaattattta
ttttacttgtgattaaaaaccaatttaaaaaattcatcaaattaatataagttatttgat
gaatttgcttatacaatatattattgtgatatcttatttctagatggtagtgattaagaa
ctagcttttgtaaaagttaggccgttatcttgagcatatctttccataaatcgcataaat
ctatcccaatcgtcaggacttttaataacatgtacagcttctattgcttgtggtttacca
ttaataaacttagcattaacatctcttgtaataatttgtccttctgtgtccatgagatac
atacctgtgatttcacctttatctgccatactagcatctaaaatttttggatttgtaaac
ctgaaagtagctgtccctgtactaccatcccgagatcttgttaaacgaacatctggtacg
acttcttcattgatgccctgaataaattgaattgttgccatatcaatttt-tttattggt
atataatttatattggataaaactgatttctaattctttaatgcttgttgagaaactatt
ttaaaagcttatagtagcagtagaccaaattttatgtattatataaaactgtaacttttc
tttttttg----------------------------------------------------
------------------------------------------------------------
------------------------------------------------------------
------------------------------------------------------------
------------------------------------------------------------
----------------------atctgttcaaagaactaacttattataagctaatcata
ctaatattagccatataatttttttgaagt-gaaattga-gggttgtagctcagatggata
gagcaagcgcctcctaagcgctaggtcagcggttcaagtccgctcaaccctgttaataa-
------------------------------------------------------------
------------------------------------------------agttttttttaac
tctattag----------------------------------------------------

----gtgttaaagccctacaataaaaacac-------acaagactgatgct--agttaaa
actatttaata---aaagattgaa--aagagatcttatatgaagttaattatta----ct
tttccctcttgccatatttgcttaccaaattatcg----atctattcacaattaataaaa
ttattttacctttcaattaaaactattagattgcagtaataaatctgaaatttcat----
------------------------------------------------------------
-----------------------------tagtattttatctagattatattta------
-------------------------taagttagaagtg-------------tctatttt
aagtgc------------------------------------------------------
------------------------------------------------------------
------------------------------------------------------------
------------------------------------------------------------
------------------------------------------------------------
------------------------------------------------------------
------------------------------------------------------------
------------------------------------------------------------
------------------------------------------------------------
------------------------------------------------------------
------------------------------------------------------------
----------------------------------------------tttatctactaaaa
agctttgtaatacttttatttgttagaactggggtgggaggattcgaacctgcgaatggc
ggagtcaaagtccgctgccttaccacttggctacaccccaacttataaagattataatag
cagtcgtacaccctgccatgtcaagagattataa---ttatttaatagctattaatactc
ttataaaagagaatctttttttcaaataagatatactatgttcaaactgcatgattgtc
atgctttcttgttgttgagtcgacatccattttatgctaactagtccttgttcaacttct
ttattgcctataataatacaagctatagctcctttttgttagcttgcttaatttgttta
ccaaaattacttgtgctaacatctatttctatttaaaaagctgctgatgtaaaacattc
attattttcataccagcttcgttagctttttctccttgtgtcgcaatataaaaatcaata
tacttattaggtaattctattttctcttttgcaagaagtaataaacgctccaatcctatt
gcacatccaacagcaggtgtttctatgcctcctaattgttcgattagattgtcatatctt
cctccgccacagaccgtatcttgtccctttgatgttaatgttttaatttcaaaagcagta
tcattgtaataatctagtcccctaactaaattgttatttatattgtacgaaatatcaagt
aattttaagtaattgcaaacagcatcgaagtgtttttgtgattccaaacttaaaaaattt
gaaattttaggagcctcgcttagtattctttgcgtattagtatttttgaatctagaatt
ctaattggattattaattagtctctttttgaatcaggatctaagtcgtcatagtattgc
tgtaaataatcttgtaattttgcttgataaatactacgatcctcgattttcccaattgaa
ttaatatcaagttttaagtctgtaagatttagactattaaagatgcttatagctaaatgt

attacttcactatcggctctagcatctaggctaccaataaactcaatgccaagttggtga
aattgccgttgtcgcccgttttgcggcctttcatacctaaacattggcccgctataccat
aatctttgcagattttgatgataactcattttattttcaataaaagctcttacaatgcct
gcagtaccttctggtcttaacgtaatatccctgttactgcgatcatgaaatctatacatt
tctttatttacaatgtcagtatgtttaccaatacctctatcatataaatcactactttca
aaaattggtgtcctaatttcttgataatttgcacattccaacagattagaaattttatta
tgtataaattcccaatattgaagctcgtcgggcagaatatcttttgttcctctaatagcc
tggatttttgccatgataattttgggattaccttcttaatgtgtctgcattaaatagcag
ttgtagactggtattcagtcatgtattataaattagaaaatatattgatg-tattttata
aaatacgggcaaggagggattcgaacccccgacaccatggttcgtagccatgtgctctaa
tccactgagctacaagcccacttaacaattgggatcactatatcattgttctttcatcta
tacaactgtattctgttg------atttatatttgaaagctttgcaaaagtaaaaaatt
gt----atttatctttcacaattagcactaattatgtttaatgcctaacttattattact
ttgtaaa--aacatcggccaact------tgtaattctactacactgataatttcactat
aa-cctgtgcttttatttttcgatcga------ttagtgtctaactaacctatattccaa
ataaaaaaatcttatctacaatgagtaaacaaatcctatatcaagatgatgcacgtaaag
cactagaaaaaggcatggatattttaacagaagctgtctctgttacattgggaccaaaag
gaagaaatgttgtcttagaaaaaaaatttggtgcccctcaaattattaatgatggtgtaa
caattgcaaaagaaattagtctagaaaaccatattgagaatactggagttgcacttatta
gacaagcagcatctaaaacaaatgatgtggctggtgatggaacaacaacagctactgtgt
tagcatctgcaatcgtgaaacaagggatgagaaacgttgcagcaggttcaaaccctatgg
ctattaagaaaggaattgagaaagcaacaagttttgtagttagtaaaatagctgaatatg
ctaagcctgtagaagataaaaaagcgattatacaggttgcatctttatcatctggtaatg
atatagaagtaggtgaaatgatagcaaatgctatagataaagttggtagagaaggtgtca
tttctttagaagaaggaaagtccactactactattcttgagattacagaaggtatgcagt
ttgaaaaagggtttatctctccttactttgttacagatacagagagaatggaagtgcttc
aagaaaatccttttattttatttacagacaaaaaaattactttagtacagcaagaattag
ttccactgcttgagcaaatagcaaaaacgtccagacctctattaataatagctgaagata
ttgagaaagaggccttggctacgattgtagttaacaaattaagaggtattttaaatgttg
tcgcagttagagctcctggatttggagatagaagaaaatctttgcttgaagatatgagta
tattaactaatggccaagttattactgaagatgcaggtttatcacttgatactgtacaac
tagatatgctaggaaaagctcgcagagttattataactaaagattcaaccacaattattg
cagatggtcatgaagttaaagtcaaatcaagatgtgagcaaatcaagcggcaaatagaaa
ccagtgattctctgtatgaaagagaaaaattgcaagaaagattagcaaagctgtcaggtg
gtgtagctgtcattaaagttggagcagctacagagaccgagatgaaagataaaaaattaa
gactagaagatgcaatcaacgcaacaaaagctgctattgaggaaggaattgtaccaggag

gaggagctactaatgttcatatctctagtgaattatttacatgggctaaaaataatttag
tcgaagatgaattaatcggtgctttaatagttgaaagagcagtcacttatccattgagac
gtatagcttttaatgcaggccacaacggagcagtggtagtagaaaaagttaaaagcaacg
atttccacataggctatgatgcggcaaatggcaagattgtaaatatgtatgatgcaggta
ttatagatcctgcaaaagttgctaggtcagctttacaaaatgcagcttctattgcggcca
tggttttaactacagaatgtattgtggtcgataaagtatatgactaacttgt--------
cacaatagctaatgttaagcaaagaataatttttacatatagaaattattctttgtttca
attaagctaactaatttatcaaatgtatagtctcgataataaagccaaagtttaaatgcg
tcttcttcttcaataattaagtaaagaaaatacaagcctgtaattcctaaattatcaatt
aaagtattgttatcatgtatagatttctcaagagatttaaaaaatgacattttttgtaaaa
taatatcggaatcttctcttgtagttatttgctaaatcattactgtataaagattgattt
ccaattgcatttagaacctttacaataatacttttcaagtctttatctgattttatatta
gaatcagtaataaattggcgcaattgaatatcaaaactaagattaaaaatattacttgtt
acagaagtttgattttgaatatctagagattttatagccaagaagattaaacttttattg
ttcattgtaaaatgtaagacaattgatattgtattctttaaatatta-attttatcttct
tgtatgtttatagttttgtataaaaataattaatgctgtta--------ttctaattgct
gcccgaaaaaataaattctggaaatctagcttgagctttagatagtgagtgtttttttacc
ttcttcttgccagtagttaataacttctgttgttttatttaataattcaattctatctct
ttcggcaatccatggtttagactcaagttcagcttttaattcttcccaagcatcatttct
tggccagaaaaaataagaagtaaggggactagtgccatttccaacaatttgatcaattgc
aatagcaatattattttctaaccaaagtacttttaatgtaaattttgacaattgattatc
tccattataaatattagatatata----tacagcaaaatactgtattagttattttcttg
agcaacaatttgagctttgttcaagatgttttgcactgttcttgtagcttgtgccccttg
tgctaacctttgtaaaattttttgaaatatcaatacgagtctcattagttataggattata
gaatcctagttcctctatagctttcccgtcgcgtttatttctgctatccataacgacaat
tctgtagcacggctgtttttttctaccatatctctttaatcttaacttgaccatcttagt
cacctcgcttatttttt--atatctttattaaattatatcttacgacattatgcaataat
tcaaacaattaagcagactcaagcctaacattattaaatattaccattaaacactggaga
aaaataatacccagcataggtgacatgtccattccaaacattggaggaatactacctcta
aatagctttaaatatggatcagtaattcgatttaatgaacagaatggttcattataccaa
tttaccgttggaaaccatgctaatgataatttaagtaaaattaaaattaaataaatctca
gaaaagttagctattgatccaagtaataaatttaatgtaccaggaagagtattcatacgt
attttatagttattatatagttcattatactaatagtttgagtatatatcgcaagctcta
taaataaat-tagctaaacaaaaaacttattcttaaaagtattctctcttgaaaaagctt
cttcatttgcagcattattgacttttgttgtataaatttccagctcagaccaattatgtc
ctaattgatttagctgatatgttccacattctatgcatgctatttttacattatttaaaa

tagcacccttgtgtactaaatcttttaagattggagtaacaactatttcatctaaaaata
aatctaggattagaacctttgtaaatgagtttaattgatcgggaatattattaacaatat
tattacaattaactaaagttatattagctgtgggaagtaaagctcgggctccttcagcca
atataaacccataaggcataataattacaatcatatacttataatgactattaatcagac
ttattgtggtattagttttaattcaatagtttctgttactaaccattctctcattattt
cataagtaatccatttgcctaattctgagcaagcagtccttagaatcgtacctggattat
ctttgttttcaagaattcctgcccaatgttttattagtggatgtgaagctatattaattt
ggagttgcatgttttgttaaatatatttaacatttaatttattaatcgcagtcagatttt
taatttgttgagtattattaaaatagcat----atatattaaagaaaataattat-aaaa
aatgaaaaaagacctttggctttggggtttcacagatagtgctgaaacttggaatggtag
atttgcaatgattggttttatagcagttatttctatcgaggtaattacgggacaagggct
attatatctaataggtgctatgtcataataaaaattcaaaaaagttcttgtaacttgtta
caagaactttttga-atgaggacaaatttaatctaatggtctcatagatagcactggtt
cattctctctgttaattcctttttcaaatccagcagcagcagctctcgctctaccagcat
gccaaagatgtccaataaataggaagaatcctaagaagaaatgagatgttgtcaaccaag
acctaggagatacatagttaactgagttaatttctgtggccacaccaccaacagagttca
aggaacctaacggtgcatgagtcatatattctgcggcccgtctttcctgccaaggctgaa
tatcattttaattttattcagatcaagaccatttggtcccctaagaggttcaacccaag
gagctcgcaagtcccagaatctcatagtttcgccaccaaaaatgatctctccactaggag
acctcatcaggtatttacctagtccagtaggtccttgagaagacgccacattagcaccta
atctttgatctctaactaagaaagtaaaagcttgagcctgtgacgcttcaggaccagtag
gcccatagaactcacttggataagctgtgttattgtaccaaacgaaattagaagctgtaa
gacccatgattgacaaagcacctaagctataagataagtatgcttcaccagaccacacaa
aagctctcctagcccaagcaaaaggttttgttaatatatgccagattcctccagcaatac
aaataataccaatccagacatgacctcctacgagatcttccatattatttacacttacaa
tccacccatcaccgccaaacggagactttagcacgtaaccaaaaataacaagaggattaa
gagtagggttactaacaaatctaacatctccaccgcctggagcccaagtatcatatactc
caccaatgaacagagccttaattactaataggaaagctccaatacctagtaatactaagt
gtataccaagtattgtggtcattttgtttttatctctccaatcatatccaaagaaaggaa
aagattcttcaagagtatcaggtccaattaaagaatgataaagacctccaaatcctagaa
cagctgaggaaattaaatgaactacacccactacaaagtatggatatgtgttaaaaattt
cgccacctggacccacaccccagcctagcgtagctaggtgtggaattaagataaatcctt
gttcatacagaggcttttctggaacaaagtgagctacttcaaaaagagtcatagcgccag
tccaaaagaccattatacctgcatgagcaacatgagcgccaagtaatttgccagaaacgt
taattaaacgtgcattaccagaccaccaggcaaacccggtagactcaatgtctctaccgc
cgacaccgacagttgtattaaagggcgtttccacgtggtaaaacctcctcagggaatata

aagttttcatgaggttgatcctgagcagccatccaagaacgaataccttcattcaataaa
atgttcttggtataaaaagtttcaaattcaggatcttctgcagctctcagttcttgagag
acaaagtcgtatgctcttaagttaagagctagtccaacaattccaaatgcacttgtccat
agtccagttaccggaacaaatagcataaaaaaatgtaaccaacgcttattagaaaaagcc
acgccgaaaatctgtgaccagaatctatttgctgttaccatcgaataagtttcttcggat
tgcgttggagtaaatgcacggaaggtatctgcagcatcaccatcttcaaacagtgtattt
tgtacagttgcaccatggatagcacatagtagagcaccacctaagattcctgcaacaccc
atcatatggaatggattcagagtccaattatgaaatccctgtaaaaacagtagaaatcta
aaaattgcagctactccaagacttggagcaaagaaccagctagcttgacctaaagggtac
atcaagaaaacagatacaaatactgcaattggtccagagaaagcgatagcattgtatggt
ctaagaccaactagcctagcaatctcaaattgccttaagcaaaatccaattagtccaaac
gatccatgtaaagcaatgaacgcccacaagccaccaatttggcaccagcgagtgaagtct
ccttgagcttcaggtccccaaaggaaaagaagagagtgccccatgctatttgctggtgtc
gaaacagcagcagttaaaaaattgcatccttctaagtatgaactggctagtccatgagta
taccaagaagttacaaaagtagttcctgttagccagccacctacagcaaggtaagcgcag
ggaaacagaagtagtccagaccaacccacgaatacgaatcgatctcttttagccaatcg
tctacaagatcaaaaccaccacgagtttttcttgtccaattgctatggtcataattta
atctccagagcaaatttaataagtaatttgtttaaataaaca---ctttagatatattct
aaaatttacttttctttacggtcaatataattataagtcaaaatcaacaaaaattcacg
tttttaataaca--ttttttagctcgttaagttaatttttttaaac------------
-----------------tataaaaattataatttgttcatttctataattctctgaaata
aataccctgttcctcgtgctgttaaaattaagtcaggattactgggatcatcctctaact
tagctctcaatcgtgatatatggacatcaacaactctcgtgtctacatgccgttctggcg
tatatccccaaacctcctgtaagatagaagctctagagaaaggttcgccagccttactta
ctaaaagctcaagaagactaaattccattccagttaagcgaaccctctcattatttttat
agacttgtcttttattggtatctattttaaaaatccaatactaattatgccagagtttg
ggacaccaagattacttgttattttatctgctcttctgagcacggatcgaatacgagctt
ctaattctttaggtgaaaatggtttaactacataatcatcagcacctatttctagtccag
taattctatcgcatacttcacctaatgctgttaacattattatagggacatctgattctt
ttcgtagttcttgacaaaccccgtagccatctagcttaggcatcattacatctagaacga
caagactgggatattcttttctaaatactatcaaagcttcctctccattagaagcagtga
ttacttcataacctattattgtcaatcttgtttctagaattcttcttatgctggcttcgt
catcaacaacaagaatcttttctttttggttctccaa-tgtattatctactcctataggt
aaacaagcataattactataatcttaaatttgtaactgtttaagcatctttaacggctac
aagc---------ttcattataaattttgtaagtg-----tgcactgagtagcatcagca
aaattatatcttgaaatatttgtgttttatttatact---------tattttaatgcata

agttaggttacgtttgcagaattcagtatatacatattttttagacaagttcttcgcat
tctattttctaaataaaaaaggggttgacaaaccagttgatagaaagtattctacattat
tataag----ttactgatcacaaagaaataataa-----aatttcttctagaataataag
tactttgat
>Porphyra_umbilicalis_NC_035573
tatatattaaaaaa--aaaacatctcaagtgtaagaattcattaaatatacaaaactttt
tatttaaggtatataaattaaattacactagagttaagatgcttt--taaaaattaaaaa
tcctaacagcggatactgatttattagaaataaaagtattta-----tttatatctaca
ggaa-aaaaatgaaacttgcagtttatggtaaaggtggtataggaaaatctacaactagt
tgtaatatttcagtagctctctcgaagagaggtaaaagagttcttcagattggttgtgac
cctaagcatgacagtacatttacgctgacaggtttttaatcccaacaattatagataca
ttacaagctaaagattatcattatgaagatgtttggccggaagatgttatttataaggga
tatggaggagttgactgtgttgaagccggcggaccaccagctggagctggttgcggagga
tatgtagtaggtgagactgtaaaactttaaaggaacttaatgcttttgatgaatatgac
atcatactatttgatgttctaggtgatgttgtttgtggtggttttgctgcgccattaaac
tacgcagactactgtcttattattacagataatggctttgatgctctatttgcagctaat
agaattgcggcttcagtaagagaaaaagcacgtacacattctctaagattagccggattg
gttggtaatcgaactgataaacgagacttaattgacaaatatatagattgtgttcccatg
cctgttctagaagtactacctttaattgaggacattagagtatctcgagtaaaaggcaaa
actttatttgagatgtctgaaagtgacactgatttagattatgtgtgtgattactactta
aatatcgcagatcagttaattgctagacctgaaggtgtagttccaaaagaagctgcagat
agagaattgtttagtttattatctgacttctatttaaatccgaagtctaatgctacaaaa
tctacaacagatgaagaagaattagatttaatgatggtttaaatact-tgctaagtataa
aatacaaaggaataagataatgtctacagttcaatcagatgctcttacttttgaatgtga
aacaggtaattatcatacttttttgccctattagttgtgtttcttggttatatcaaaaaat
agaggatagtttcttttttagtcataggaactaagacttgcggatatttttttacagaatgc
aatgggtgtaatgatttttgcagaacctagatatgcaatggcagaacttgaagaaggaga
tatttcagctaaattaaatgattatgaagaattacgcagactatgcctgcaaattaaaaa
agatagaaatcccagtgtcatcttctggattgggacatgtacgacagaaatcataaagat
ggatttagagggaattgctcctaaattagaatcagaaattagagttcctattgtggttgc
tcgagcaaatggcttagattatgcttttactcaaggtgaagatactgttttagcagctat
ggctcaacgttgtccctcttatcagcatgttaaa------------------aatacaga
ttctcatgtcccactagttttatttggatcacttccagatccggttgttactcaattaac
gctagaattgaagaaacaaggtattatagtatcaggttggttaccatctaaaagatatac
agagttacctgtaatacaagaagggtattatgtcgcaggagttaatcctttcctaagccg
tacagctactactctaatgagacgtcgaaaaacaaaactcattggtgcgccattcccaat

tggtccggatggtacacgcgcttggattgaaaagatttgctctatattcaatattgaacc
taaaggcttagccgaaagagaaaaagaaatatgggagtctttagaagattatatttctct
aatacgaggcaaatcagttttctttatgggagacaatttgctagaagtttccttagcaag
gtttttaacgagatgtggtatgactgtgtatgaaataggaattccatatatggataaaag
atatcaagccgcagagttagcgctattaaaaaatacttgtgatgaaatgaatatcatgat
gccaactattgttgaaaaaccagataactataatcaattagatcgaattcgagacctgaa
gccagacctcgtaattactggtatggcacatgcgaatcctctagaagctagaggtattaa
cacaaaatggtctgtagagtttacatttgcacagattcacggttttactaacgcaagaga
tattttagaactggtaacaagaccattaagaagaaatttgagcttgtcagagctaggatg
ggatgtatatagtaaacaaggctaatatg-------aattttgacttacatggtaactaa
actctgtaagtctatttttgattattgtaaagaagcattccttgctcttccgctagcagc
ccatgcttgaagatttctacttgctctttatctgtgtaagctaaaggaacaaactgtcc
taatgctattttatatcttcggtgctaaattctcgttcttcgctaaatgcagtatgcat
actttcaacaattgcctgttcaatctctgctccagaaaacttattgcataataagcttaa
ttgttttgtattataattttgccacgattgtggtctaattctttttaagtgtatttgaaa
aataagttctcgttcctcattgttgggtaaatctaaaaaaaatatttcatcaaatcgccc
ttttctaagcatttcagatggcaaattttgaattgtattagctgttgcaacaacaaagac
gggagttgtttttctgatagccatgttataaaagttccaaatactcgagcactcgttcc
gctatcaccttgactatacaaattagagaatgctttgtctatttcatcaatccataatac
gcatggagataaaccttcggcaatagttaccatttctctcatttagactcagactcacc
aaccaatcctccaaagagttttcctatatcaagccttaataaaggtagtgtccaatcatt
cgcaatggcttttgctgtaagcgatttacctgttccttgtataccaaccaataacagacc
tttgggggttggtatgccgtaatcaaagctttgctttgaaaacgagcgagatcgtttctt
gagccataattttaatgcattaagtcctccgatatctttattagctttcttgtaagggta
aaactctaacaaccgtgtttgattgataatttgtcgtttttcttcaataataattgataa
gctccttgaatccaattgaccatactgagcaatagttcttgttataactttctaattct
atcaattgataaaccttggcaagatttggctaaattatttactagttctatatctagcac
aatacttaaagcaatacttaatcttttaatttctttttaatttctggaagattaggtaa
tggtaagtctattactgtaataacatcatttaacgtataaggtatattgactttacaaga
aacaatgatgatattcttagactgagttttaattatttgagatagattacgtagttttct
tattaatcctacatccctaaaaaatgagtcaaagtctttaagtataaataaatttaagta
tcgattatctaatttttcaataaattctagagctaataatggattccgttgggcatagcc
attatcattaggattacttgtatacccatccacgaaatcccaacaatatactttttcatt
gtcgagacaagatatgtgatttttcatcatatactctaatctttcttcttctctagtatt
tattaaaattatcggatagcgagactttaaaagtaatcgtaaatcttggatgaaattcat
gtgaatatctgttctcaatagacttttgcattttaccaatat----taaactaaataaaa

ctatctaatttgatatagacatattcaaaagtcaattatatttatatagtttatattcta
-------------cttaactagcgataat-tttttctacccttttttcttctttcattc
aaaatagaacgtccacttggtgtcttcattcttgctctgaatccagaaactctaatttt
tttctctttgaaccttgtaatgttcctttagtcattattttcctcaattgta-------
aaatattatattatcatcttctataaaatttgaagatata--------ttaatatgaaag
tt-aagcttagtaatttctattaatcaagaattttgttaaaatattaagaataatctat
taattgtaaataattttatgctaattgccctgcctattgtttatttaagtattttagctt
tatttttactaatatttagctggttaatcagtcaacagctaaagacaatttttttactag
aatctcaatttcagtattttcttgacaaaagcgaaaatgagcaacttggaatagaagatg
gttttgccttttcaaaagtatgtatagcaaaaaagcactttataagagctataattgaaa
gccagttagtcctacaaaagaattcattattgaatctcccagaaaatgccgtaataattg
ctaaattatataatatgttgggttttatatattatgaagcagatcaaaaaaaacttgcta
aaaatttttatgaaagagcaatagacgtagatagtaattatattgtggctttaaataatc
ttgctaaaatttatgaagatacaaaaaatttcttaaaagcagaggcaatatatgataaag
tattgaatattgataaatcgaatcaaatagctaacactcaaaaaaaaatctactaaaaaaa
taatcaatctatgaagtatataagacta--aataatcgggatagcaggatttgaacctgc
gacatcctgctcccaaagcaggcgcgctaccaagctgcgctatatcccgcatgacataat
taatattatatatctttttctaattatgtctactatgtct---ttttatctaggatacc
aaaacatccccttgaccccatctggatctctaataaatcctagttttctataaaagctaa
ttacgtcaggctccgcaaataaagtaatagtgctaatttcagcctgccttaattgttgaa
ttaattgatgcattactactttccctaaacctaaaccttgaaagtctggatggattacaa
catcccatattgtagcattaaatccattatctgaagtggctcttgcaaatcctacaagcc
ttgtatctgaatcatttttttgtattaaagaaataataatagaactattctttaatgcaa
ttttgactttttcaggggtcttttaacccatccaactgaatcacatagttgctctaatt
cgtataaattaatattcttagtattgctgatataaacatcttttaattcaattttatcgc
aagttttatccaaaagaaaaagtttcttaaaatttttctcataactctcatttaccgcag
agttttgaaaaaaagttttccagaagatcataataattataacagaat----taagtata
tatatattatgatattgacttaattaaaaaaatttcaatatattgttatgttttt-tatg
aaaatactaaaacaaaaaatt-ttttgacacaaggaaataaaaattttgtaaaatcttt
atcataaaacatataaaaacttctttttaagattaccatttattagtataaatagaatct
cgaaattgagatgagataacacattattttatatttaatacctttcataggagttatttg
tgaagaaagctatgtttatgacttgtttactagctgcgctattagtcagcaatcctattc
tagttaatgccgaagttgcaggcttaataccttgtaaagactcagcagcttttaataaac
gtatggtaaatagtgtaaaaaaacttcaggctagactagctaagtatgatgctaatacac
cgcctgcattggctttaaataagcaaatagaaaaaactaaaactcggtttgctacatatg
gtagagctgggttattgtgtggtacagatggtttacctcatttaatttcagatggtcgat

ggagcagagctggagatttcgttttcccaggcctttatttctttacattactggatgga
ttggttgggtaggaagaggatatcttttatctgttgcaaaaactagtaagccgacagaaa
aagaaattattctagacgtaccattggctgtcaaatttatgtcttctggctttgcatggc
cattagcagcctggcaagaatttagtagtgggcagctaattgctcctagtgatgatatta
cagtttcacctcgttagtaaaaaacctatatgaataataattttaccaaatacttatcaa
cagcacctgtaattggagtactatggatgacttttacagcaggctttataatagaattaa
atcgctttttcccagatgtattatacttttatttataattaattaaataaacctataagt
ctaaactacacaaatagagtaaa-tactattatttgtgtagtttttaatttatattata
ttattattttttcttttaaaatcatagttccttatattattctaatattagatc-taagt
agtaaaaa--atataaatattatgagtttagtaagtcaaattataattaatgctgacgat
gagttaagatatcccacaattggagaattacagtcaattcaagattacttaatcacaggt
agcaatagaattagaattgctactattattagggacaaagaaaaagagattatacaaaaa
gcgagcaaacaaatatttcaattgcatcctgaatatattgctcctggaggtaatgcagca
ggatcaagaaaaagatctttatgtttacgcgactatggttggtacttaagactaattacg
tatggagtcttagctggtgataaagattcaatagaaacaattggtattataggtgtaaga
gagatgtataactctttaggagttccgatcattggtatgttagatgcaattcaatgtttg
aaagaagcctcattagaaatgcttggtcaagatgatattaagatcatctctccttatttt
gattatataattagaggtatgtcctaaa-aaaatttgaatctattggttgaataaatat
tggcactatgttatagtcaacttaatcccgaaagcgtataattgtataaacataatcttg
tcagaaccggaaggtagcagcaataatgtttacctacaaaagatatga-ttttcgggtgt
tctttttatttatgcagta---tgcagtgagatatttaatagtttgaaagcattttgaat
ttgtctctgattcatggaatttttcaataataacga-ttaataatttgatattaatactg
aaaccacctataa-tttttataaaacttctttatatctcatctaaacaatggaaactta
acttaatgggtgttcatattttatcaacaggttcgtctgtccctaactttctgtagaga
atcaacaatttgaagatattatagagacttctgaccattggatttcaacaagaacaggaa
taaaaaaaaggcatcttgccccttcttctatctcgttaactaaattagctgcagaagctg
caaataatgctttaagcaaagctaatatcaatgttaaagatatcgacttaatcattcttg
ccacatctactcccgacgatttatttggtagcgcaagtcaactacaagcagaaataggtg
caacctcatctactgcttttgatataacagctgcatgttcgggatttattattgcactag
ttactgcctcgcaatttattcaagctggttcttacaataaagtcctagttgtgggtgcag
atacaatgtcacgttggattgattggtcagatagaactagttgtattttatttggagatg
gagctggagctgtactaattagtgaaagcagtgtaaatagtatattaggtttcaaattgt
gtacggatggtcgattgaatagtcatttgcaattactgaattcccctttagatagccagc
agtttggattgacaactattcccaaaggaagatacgactctattagcatgaatggtaagg
aggtttataagttcgctgtatctcaagttccaatagttattaaaaattgcctaaatgatt
tagatatttctatagatgaggtcgattggtttatacttcatcaggcaaatataagaattc

tagaagcaattgcaagtagattatctatacctctttataagatgattactaatttagaaa
attatggtaatacatctgcagcttcaattcctcttgtattagatgaagcaatagaagata
agaaaattcagccagggcaagttgttcttttagcaggttttggagcaggtttgacttggg
gagctattgtcttaaaatggcagtagaaaaaatactgcggatgacgagattcgaactcgt
acagcttgcgctacacacccctcaagcgtgcgtgtataccaatttcaccacatccgcttt
ggttattatgaatatttaaaaaatagtaatatttatgtaaatatttatataa----tatg
tatatatatacattcttattagaaactctcttatttttattttaagtgataaaattgag
tctagttagtataaataagaaaatttttatttaaaatcgtaaa---aaataaatatataa
ggaaa--aaaatatgacaccgtccttatcgagtttttgaatagtttaattcttggagct
gtaattgtagttgttccaataacattagctctattatttgttagccaaaaagatagaaca
attcgttcatgaaattaatttaaaaagatatggaggagtcccttgaaatgaataatcagt
tattctatcaataatggctcgctccatataaatctatattacctagaataatactaa--c
ctttattctgtaataacttgtacattttatttagtt-ttcttgcgtact-----------
--------------acaaatattatt-ttgatgatagcttccttttattacgtacttttt
attcaatatgaaagtattgtgtaaacctattaatatgttcaattaataaagaaaaacaca
tgtctttgtctaattggcctcttaaaaaagcaaattctaaagcttataatattaaaagca
caaaacaaattactatcccagatggattatgggtcaagtgttttgattgtggcttattaa
tgtactctaaagttttaaagcgaaatttaaaagtttgccctcagtgtagttatcactttc
aagcttctagtaatgaaagaattgatcagctaatagatcaaggtagttgggagcccatgg
atgctcatttaatttctactgatcctttaggttttaaagatcaaaagctttatagccaaa
gattaaaggatacagctgttaagacaggtcttcaagatgcagtacaaacaggtgttggta
gaatggatggacaaaaagtgtacctaggcattatggattttcgttttatgggtggaagta
tgggatcagttgtaggtgaaaaattaaccagacttgttgaaaaagctactcaagaaagat
tacctgcaattataatctgcgcatcaggaggagctaggatgcaagagggaatgttaagcc
ttatgcaaatggctaaaatttcttctgcattagagatgcacaaaaaagaaaatcttctat
atatctctgttttgacttctcctactactggaggtgttacagcaagctttgccatgcttg
gtgatttaattattgcagaaccaaaagctttaattgcatttgcaggtagaagagtcatag
aacagacgataaaagaagatttgccagacaattttcaaagttctgaatatttatttgagc
atgggtttcttgatctaattgtaccgcgcactcaacttagatctaaactaatacaaattt
tatatctgcatactaataatgattaattcgccaattgtttactaaaatcttagaatgcta
ataaaaattaagatattaagaaattttcactaaagatatttctatactaactagaattac
aattactattgtagaaaaatttgttttaactcta-tcgtaatagtttttttaaataatcg-
-atttgatttttttaaaaaagtgagagtttaatacaaatttagaaaaaggaaatgaggtat
actaagataaatttagcaatattggattttcctataatactgacctatcaaattttgagg
aattttatgcttaaaagatcttcttggcttgctactttattgggactactagctgtagcc
tccgtaagtgcaaacgtatatgccatagagctagacgaggcaacaagaacagttccatta

gaatcttcaggaagaactgttgttctaacaccagagcaagttaagcgaggtaaacgattg
tttaataactcttgcgctatttgccacaacggtggcatcacaaaaactaatcccaatgtt
ggattagatccagaatccttaggtttggctacacctcagagagataatatcgaagcatta
gtggattatatgaaagatccaactagttatgatggcgcagaatctattgcggaattacac
ccaagtattaaaagtgctgaaatttttccaaaaatgcgaaatttaacagacgaagatctg
tttacaattgctggtcatattttattacagccaaaaattgtttctgaaaaatggggtgga
ggaaaaatttattattaacaattaaatcgaaatattcctctgaatacaagtttatgaatt
ttgtttttaagtatgattattttgttagatcatgctatatattaaacatggagtgtagt
aatagtaagtagctcctttgctaaacttgcttatattatttgtctagttattaaaaagga
gaagttaaattgaaaaaaatgcttttagttctttttacggtgtttagtttttttgctata
ggatttactcaagtagcctttgctgcagatctagataatggtgaaaaagtttttctgct
aactgtgcagcatgccatgcaggtggtaataatgcaattatgccagataagacactaaaa
aaagatgttcttgaagctaatagtatgaatggtattgatgccattacttaccaagtaaaa
aatggtaaaaatgcaatgcctgccttcggtggtcggctagttgatgaagacattgaagat
gcagcaagttatgtattatctcaatctgaaaaaggttggtaactatacttgattttctcc
agtattaaagaataggcagtctaattaattacttattagattgcctattctttgtttgtg
ctattattatataaaaatatttatacaatattgatattatgataaaaagaataccagcag
ttctcgtattagaagatggtacttattataaagggtggtcatttcagacagagcaatcaa
aagttactattggagaagttgtctttaacactggtatgacgggatatcaagaaataatta
cagacccaagttattttcagcaaattgttacttttacctatccagaaattggaaatactg
gtattaatagtcaagatatagaatctcaaactatcagtattaaagggttagttgctaaaa
atatttgtaaaatttcaagtagttggagacagcaagaatcattaattcagtatttaaata
ggtatcaaatcccctttatttttgggatagatacaagatccttaacccaatatttacggc
gatctggaactatgaatggatgcatctctaatagaaacttaaatcgtacatatttacagc
aaaagctgtctaaatttcccagtatgaccggactagatctaattccaaatgtaaccacaa
acataatatataactgggatgaaaaaagtttgccaacttggtacttagctgatagaaata
gagaaaaattatcttgtcaaccaagggttgtagttatagattttggagtgaaattcaaca
ttttaagaagacttgccacattaggatgtgaaataattgtaatgccagccttaactccaa
tggaagatatactatcttataatccagatggtatactgttatctaatggcccaggagatc
cttcagcagtcaattatggaattcaaacagtaaaagagctattgaatcagaatattccaa
tttttggaatctgtatgggacatcaaattttaaacttggcactagaaggtaaaacattta
agcttaagtttggccatcgaggtattaatcacccctcaggcttaaaacagcaagtcgaaa
taactagccaaaatcatggttttgcagttgatctacaatcggttttagcatcgtctttac
aagtgacgcactttaatttaaatgacactacagtagcaggaatcggtcgcagtagaagtc
cttacttctctgtccaatatcatccagagtctagcccaggtcctcacgatgccgattatc
tatttgaatacttccttgaaataatgaaacaatttagacagaaggctaactaattaccat

ttagtaattgtcccatgctttatgagaaaaaagcgctgacaagacttgcactcctcgtag
ttgattgcaaagaggtaagtgccccttagggccagtagagttccaatgaaactcatctgg
atatctacaaggcacacggttaatttcccaaccgattttgtaccaaaacagctcccagtt
tttttctacagcaagccaaatttgcctctggataaaaaagccaaacttaccttttgagtg
tgcgttccataattggtccatagtttgtaaatcttcaacaggtatttttttgatatcagt
aaaatatagccagtttctactttttcatctacgccagctaatttaactagtttttgttg
tgttagcttatcggcggctattaaatcatcttgaactagtaggatttgtagatcctgata
atccatttgtttatctgatttaaaggaacaatgcctttgggacagagtttcgatgtgaa
ttgtctaatctctgaattttggctatttaaaagtttttcatatattaaaccatcaacaca
attactcatatattcaggcttagtaatcctgtaatgaaaaatctcagctagttcttttaa
ttgtacataattatcatcctgtatcctctcaataatttcaagttgtttttaatagtatt
ttttgaactgttagtttcttctaattcagatagttgaacttgaatttgatttttcatctg
attttcttaattaatttaaatattt----aagaagttctaaagagaattagattgttttt
taaaattagaacttcggtgaattacctctaagaatgtcctcataatttgattaaatagat
tctgtccaatataaattataaattgcccgattaagaaaactgtactttttatttttggaa
taaaaaagtcttgaatttcaagtagaaaggtaatcaagacctgtgctgaggataatgtat
ataattcagtagatctacatgcatatacatattgacaatttaagccctcctggtaaaata
cccaaactttataacggctctcatagattgcccgtggtctctcaatataattttgaatca
aagtattccagactaaattatttttaatttctctagttttctatcagataaaaatttca
cattgcataaacagcttgcattcgtttcgtgtactgttttaaaatttttgaactaatatat
gagcaataatattacttaattgaattaagtaattttctagaaaaatttcaacttgtttaa
ttggtatattttcggcaaaaagattaaatgtttttttactttttgaagatgatccaaaaa
ttagttgtattataagaatttgtagtagcatcttgtaatcacacataagatagtttatat
catgaatgttttggcagatactagtgtaatctagctcatataatctacaaaatcttttaa
cacacctatgaaataagtcaattaaaatattttctgttaaatcggtaacatcatttaagt
ctaaattagaagtgtagatttctagaagaattttctctaattctgccaggattattttta
acaagtttctttttacttcagttcgaaacacatctaaaattaatacttcttgcgaacaat
ttgttaatttttattaatttttgcagaagttctaactaataattctgctacttctgtat
tgagtattggtccctgagcactaggccagtaattattcacgttatatcatagtaaagaat
atgtaattcaa-cttttacgataacttttttaatatactgaaaagcaagtttattcctctg
taaccttaaatatagatattttacagatcttaatagttatttatcaaagaaagttatatt
atagtaacgatcgttactattttattaagattca----gaagtatgacaacttactattt
tgcacttgctagtcaaaatttcttattatcagaagaacctctagaagaagttttcagaga
aagaattaattattaccaatcgaataataaagaaattgattttggttaattccaaatcc
aaattttttaaatagacctgccatgaccaagtttaaaaatttagttccagatgaagctat
agcaataatctcaactaattctatctttattaattggctgaaactaagaattggatatgt

ttgtataggacaatttgaagatagcctgaagctttctgaagagtctttaagcatt-----
------------------------------------------------------------
---gtaaatcaaccttagaaatataga-----tttatcagtcatcacctagaattttatg
ctgagatggagtaacaaataatgtaagtatctcttgactaaaagtttcagtcgcttttga
tttatatctattggggtttactataatagaaagcatcctttttgatagtgacattttcgat
ttgagtccaatgtacaatcccaagctctaactctttagcaatagctgaaacggaaacaaa
ggcagcacctaaaccagattgaacagcattttttaatagcttctattgaatttaattccat
ttctattttaaaacgactactatcaatcccgtgctgacttagtactttatcaattacttt
tctaattgtagattgagtatccagagcaataaatcgcagtcgatataagtcttctttttg
tatgtctcctaattgtgaaaaaggatgtgatttaggtaatatcaatgctagctcatcttc
cgcataagatgtaacctgtagcacatcttgcaattcagtaggtacttctcctcctataat
tgctaagtcaacttgtccattagcaacactccatgaaattaatctggttgaatgaacttg
taattgtactgctacttgtggatatcgttgtctaaaaagacctattagtcttggcattaa
atatgttccagttgtttgactagcgccaataattagtgtcccaccttgtaagttttgtaa
atcatcaagtgctctacaagtttcttcacataaagctaaaattcttcctccgtatctcaa
taaaagacttccagcttcagttaagcttgctttttttattccctctctcgaaaagggcaac
gtttaattgacgttctaagttttgaatttgtaagctaatagctggctgggaaacatataa
gctattagcagctttttttaaagcttccctcttttgcaattgcttttaatattcttaactg
atctaatgtaaatggaagatctgtcattaaagaattgtattataatgtatatgtattata
ttatttaatatttatcaatgaatagagtcgctaagttaaatatttgttattaagcttcat
tatttttgaaataataaaaaattataatggcgaatacagtgtatgagattttaatatctct
ttaatcttaaatttatataatagatatattgaatttagttttaactacaaaatataggag
ataac---atggactctagacttttaatagttttaataccagttttagcagctgcttctt
gggcagtttataatattggtagagttgctttacagcaatttcgtaaaatgacatcctagt
tttatttagaaattagatttttagtttcaatgagatagagaacttgatattttctatctc
tcttgtatatttttagtaaaaaagtttagagg---tcttgtacttaatgccattacacatt
aagcttatgtgg-aatacctatgtttataataatttggtattgaattatc-----cctta
taattaattaaatcaataataatattatggctgttccaaagaaaagaacatctaaagcaa
aaaaaaatgcgcgtaaagcaaattggaaaaatcaagcaaaaacagaggctcaaaaagcat
tgtctttagccaagtccgtgttgaccggtaaatctaatgggtttgtttataatacctcag
aagtcacagatactgtagttgaataatatttttttattattataa-----aaaaatacata
atctgcaaaatttaagtaactcttacacat----taagcatttggaaattattctattta
ataataaaataaacagcagatttaataaagttacaagcactaattatgtagcaaagctgg
atcaaaccagcgaaatttggctattcaattgcattgaaaatattcagcatatttttttga
aaagccaattaaagtcaagtcaaattactaaaatttttatctctggtacaagctttaaat
atacagcaggtttgcctggactattatctagcttaacactaagcggcaaaataaatccta

taagtatatatggtcctatttctttaaaagtgtatgttcaagcttgtactaaatattctc
aaacaaatttttcctttcccgttaatttttacgctgcgcgctatgggaagcttgtctctg
agcaatcatatacagtaatttgtttaccactgagttctaataatttactttatggcttta
ctattttaaaaaaacaacaacaaggagttttcaacctaaaaaaagcgataactctcaata
tctctcaaggacctatctatggtgagctaaaaggcaaatataatttttctcagcccagatg
gttattatttacatggagaagattttttgttcaagcactacgttaggaaataaaatatcaa
ttcctgtaccagtgaaatattctagaattatttctgagatgcattggctatgctcttata
ctgcagaatcaaatcctcatgtacatcagcaagcaacaaagtatttacttagcaatatcc
aaaccgatgttatgagctatcaagtatctcacgataataatttgattgaataatgattgt
tcttgatatataatggtctcgataatcttattaatat-----agataatcctcagagtat
aatattttacatcaattttactataatatatgacatacgcaattattgaagcaagtggga
agcagctttggatagaagaaggacgttattatgatcttaatcacatacccgttgagccag
gccaatcaattatacttggtaaagttctgttattaaataaagatggccatattagtctag
gccatccttgtatacagggcgcagcaattaaggcaacggttgtaagacatttacgaggaa
aaaaactaacagtctttaaaatgaagcccaagaaaaaagatgaggcttaaaaaaggtcata
gacaagagctaacccgtttgatgattgattcaatcatgacttaat---------gatata
agattaaatccttttag----------ataatttaaatacaatcaataaaatggcacata
aaaaaggcagtggtagtacaaggaatggtcgagattctaattctaagcgtcttggtgtaa
aaaaatatggtggtgaacatgtaacagctggtaacattttaattagacagcgtggtacta
aagtaaagcctggtcagaatgtaggaaagggtaaagatgatactctgtttgcattgattg
atggatttgtactttttgaaaaatcaaatcagaagcaaaaaacaattagtgtatattttt
ctaaaaattaactaaaaaatttggtgcaagataaaataaataagcgtatttattgtgatt
agactggcaccagatgcttgatt-------------------------------------
-tattaaagatcttctttgaaatttaatgactaacactattgtaatttcctgcctgcaca
atattgctgctattttatattgtggtcagatacagaaactagtagtagcaaatgctcatt
atcagctaaatgatatctacttaggtactgtagataaaatttttttcaggaattaatgcag
ctttcattaatttaggaaaaaatgagtatagtggctttattcatataagtgatactgggc
cactaaagaaaaaatactatgtaaataatattactaatattctaacgatccgacaaaaag
tattagtgcagattataaaagagccaactttaaataaaggaccaagagtaactgccaata
ttactttatctggcagatatattgtcttaatgccttttagtcaagctatttgtatatcaa
gaaaaatatatgatgaagatgagcgtcactatttaacggctttagctatcttaataaagc
ctgcaacgatggggttattattccgaccttctgctataggtatagatgaagaaattattt
taagtgaactcagaaatctaaaggaacaatggtcttttattcaaaagtcagcaattaata
attgtgcacccgtacttctttataaagatgaagatattgtgaaaaaagtcattagagatt
tttataacaataatacaaaaaaatatagtaattgattcaaacctaggcttaaaacagctga
attattatgtgcatacttggcaatgtaatttttctagtactattccttcaattaagcttt

atagtagtaataaatgtgtcttagatgcctttcgtattaatcaagctatttcaagagctt
taatacctaaagttgatctaatactgggtggatatatgttcattgaaatcttggaagctt
ttactataatcgatgtcaattctggatcttttaataattctactagtgcacgtgaaacgg
ttttaaaaacaaattgctctgcggcaacagaaatagcttatcaattaaaaattagaaata
tcgcaggtgttattataattgactttattgacatggaatcccaaagagatcagttacaat
tattagaacattttaacaaagagctatcttttgatgatgctaaacctcaaatcgtgcagt
tgtctgaactaggcttagttgagttaactagaagaagaaaaggcaaaagcttatatgaat
tagttagcaatgactcaaattactttcatttttttatacaatcagaacagcttgactcta
caaagccttcttcgtataaaaataaaagactttcgagttccgtaaaatcttggctatttt
cggagattgatatcattaacaaagttttcttaaaacagtcaaatttttgtagattatcta
atttttaccgtgctcgtcatctatatataatctacagtgataatactgttaatcagaacg
ttacgttggcagttaggtataaactaatatattctatgcaatatattcaactaataccac
gtacttggtattgtgatttttttaaatatacatactaacaatattataagttatgtatcct
gatttctatatttcgatgtatcctgatttctatatttcgcccatttaactttctaat
cataataatcaagcctacttaatataaaaagccctctctaacaaaagagagagct---tt
ttgtagcct-ttaactagtctaattggtcaataaaaaattaaccattaacagctggagct
gttagagctactggtaaggactcaccagaagctagatctagagggaagttatgagcatta
cgttcatgcattacttccattcctaggttagcacggttgatgatatcagcccatgtatta
attacacgaccttggctatcaacaacagattggttaaagttgaaaccatttaggttgaat
gccattgtgcttacagataaagctgttagccagataccaactacaggccatagacctaag
aagaaatgtagagaacgagagttgttgaaactagcgtattggaagattagacgaccgaag
tagccgtgagctgcaacgatgttgtaagtttcctcttcttgtccgaatttgtaaccatag
ttagcagattcgttttcgcttgtttcacgaattaagctagatgtaactagggatccgtgc
atagcactgaacaatgaaccaccgaatacaccagcaacacctagttggtggaatgggtgc
attaagatgttatgttcagcttggaacacaagcataaagttaaatgttccagagatacct
agaggcattccatcagagaaactaccttggccaattggatatactaggaatactgctgct
gctgctgctactggagcagtaaaagcaacagaaatccatggacgcatacctaggcggtag
cttagttcccactcacggccaatgtagcaagctacgccagttagaaaatgaagaacaact
agttggtaaggaccaccgttatataaccattcgtctaaagaagcagcttcccaaattggg
tagaagtgaataccgatagctgcagaacttggaataacagcaccagaaatgatgttgttt
ccgtatagaagggaaccagcaactggttcacgaattccatcaatgtctactggaggtgca
gctacgaatgcaatgatgaatacagatgtggcagttaatagagttggaatcattagaaca
ccaaaccaaccgatatataaacggttttcagtgctagtgatccaagagcagaaacgttcc
cacaagctagcgctttcgcgtctttgtaaagtagcagtcataatttttttatcaattttt
aggatttaactaagatgcttcccaaacaatactttgtctgttaattttattattacttca
tgcc---taaaaaaagagaacttttttatacaaatagttattaaatttaatctcgataagt

aa-----attgataatttatatatt------taattgaaaaattaacatataataagttg
ttaactttttataatttgtttttataatttgtattggtttagatatataatattgata
gcgtgtattaccgctgcttattttaattttacaagtttatataattgtgctttatttaaa
aattcatgtcacactttacaaaaattcaaacgactataaaagatttaaatctgttaaagc
atgctttaacagatttaggcttaatttgggatactaattcttccaatattaaggtcggag
aaaattctcaacataaagcggatgtcttaattaagcaatgcaatttatctcatattggat
ttgtttggaatgataatcaataccatttggttgcagatttacaattgtgggagcaacctt
ggtctctagaagtatttctagataaactttctcaaaagtatgcctaccattcaattattc
aacaaacgaaaaagcaaggctttgaaagagctaagcaggtctatcaaaaagatggatcta
taaagttaatagtacaacgctggaactattgattggtatataactgcgggcggagagact
tgaactctcacgagattatctcactagaacctaaatctagcgcgtctaccaattccacca
cgcccgcatgtatttaaa--ataactaatatatcataatccatattaaaaaacaaatagc
-taacaaaaattagctaccaagcttaaatgcatccacaaacaaaccatatgttataccaa
ttttaaagttatttactaaattaatctttatgacacactttctttgttagagtaaacta
tcttgctgactaattctgttcccgcaactataagacttgctgcaacaattccccaatcac
ctgtttgcccaggtattgttgataaacttgttgaaataaaaaacccaagcaataaactaa
ttaaaccacttgctaagtcacttaaagaataatacagtctactactgatattgtttagaa
agtaagaaaaaaatttgaaagtttgtttgaatcataattttgaaaacggagtcttaa
aaggatttcatatattaataa-ttattaatatgcagctttccttttaaacccgttatga
caaaatcatctttattttttgtttagtctataggtcttgctcacttaaagcattaataa
gatgctgaaaaggttcttgtataatatcttttctagtgattgatagtaattctaactctt
cttctataatttcttgcaatagctgtatacttcttactgttggagcaagaggaacagata
atgaattataagtatcttttaaaccattcaaaaccctttgatccagaatattagtatcac
cagcaactaaggcatagctggcatatcttagatagtattctatatctctaagacaggtag
catatcttctagttgtataagaatttcccccaggtcttaataattcaggttgctcttcat
ataattgagctgaagtttcttttaagatattagtagcctgattattaataatttctgcta
tttttattcgatccaagccgcttgaaaaaaatgattctaattgcgtaacagcggctttgt
caagataacgacctgtcagatcataacgatttaatattgctgttatagcatcttgcataa
attaattctcctactttgctatcagagctgtattataaattacttttagtatactataat
tccctgtgtttataaaatttgaaaaaatctattctttggctttgaatatctaaccaa--
-----tataggttttacatatatagagtatatgcagtttttctttgtttttttataaaa
attaagaaatatactctttttgttataagtatagtagtttcttaatataattaaggatct
aatgctaaataagttttgtaacatgt--aaataaaaaaagaggttacaatgt--------
------------------------------------------------------------
------------------------------------------------------------
------------------------------------------------------------

------------------------------------------------------------
------------------------------------------------------------
------------------------------------------------------------
------------------------------------------------------------
------------------------------------------------------------
------------------------------------------------------------
------------------------------------------------------------
-----------------------------atagaaatatgtttttg--------------
---------------------------------------------aattcatcttta---
------------------------------------------------------------
------------------------------------------------------------
------------------------------------------------------------
------------------------------------------------------------
------------------------------------------------------------
------------------------------------------------------------
------------------------------------------------------------
------------------------------------------------------------
------------------------------------------------------------
------------------------------------------------------------
------------------------------------------------------------
------------------------------------------------------------
------------------------------------------------------------
------------------------------------------------------------
------------------------------------------------------------
------------------------------------------------------------
------------------------------------------------------------
------------------------------------------------------------
------------------------------------------------------------
------------------------------------------------------------
------------------------------------------------------------
-----------------cgtttaaataat------------aaacaaaagcagtacctaa
aagatattttaatactaaatcaaatcagtcacaaattatctttagattcaagaagttatt
ttctgattacaattaatatatttgatcaatttattgatgtagaagaaatttgttataata
gtacaggacaactttattcaatatttttaaagataaaacagccaaagcattgtgtcata
caatattcaatagtcagcatgtaaatattttactttcaaatcagcatgtagcatacttgg
ctcgcgaattagttaaatcagaattaggtattcttttgagtcaacaatatattcaagatt

aag--------ggttattgattatgttcatgttactatatattacataagggcatgtaac
tcagtggatagagtatcagattccggttctgatggccgtgggttcgaatcccaccatgcc
cg--gatggtttaatgaacaaaaagcttgacatctctaataatagcatctaatttaatat
acacttaagtattggggctgtaaggtttctacattgtgaaaagataaagatatgaaaatg
acacgagctcattattagagcttttaat----taaataaatgcagaaaataacatcattg
ctttttctcgaaaattagctactgtgtaaatactattaatttttgtaattcgaagtgtta
gactcttatacactacgaatattctgtgagagttgctcttaatataagaaaagt--aaaa
atatataaatctttatgtttttcaccttaattgattcagtttaaggttagtatttttga
tttttacaatggacgtgggttcaagtcccaccagctccatcgattcaactttataattat
ttataaattagtaatttagtcttttaagtaaaggaaatgggttaaaagaaaatatgaac
tttataacacttacagagctagccgtaaaacaaattaaaatattaaagagtaactgcgat
aatgaagtatatctcagaattggtgtaaagcaaggaggctgctccggaatgtcttattct
atgaattttgaaaatatttctactttaagagaaacagatgaagtattaatattggataat
ttttcggttgcatgcgaccccaaaagtttactataccttatggactgtcgttagattat
agttcggagttgataggtggtggttttcagtttctaaaccctaatgctactcaaacttgt
ggttgcggaaaatctttttctggttaa---------------------------------
-taaatgttacttctgca------------------------------------------
------------------------------------------------------------
------------------------------------------------------------
-------------tttaaatggttaatttttattaatatatgctactctttaattaatc
tagatttattcgaaaa----aatctcttacctaaaaatatacagataaagatgtatctaa
tattttaaaatagaattaacatgaatagctagtatattttaatttagtttgtaataattc
-ttatgtactgttctcaaacttttattcagtttacaaggcttcaaatctctaataacttt
tattgcaaaaagaaatttttctcttccgttaattcaaatcacttgaaattaccacacaca
caaccaggtttgtcaacaagagtgcttgcttttgcgttaccagagctcagttctattcct
attaatttccgcaattcatttcaagaggcaattgtaatttctactattcatgataataat
atttttgccgatgtaactaatagctggataaaaacacacaaagataaaaatcatcactct
ttattttataaagaaattttaaggctttgttgacaaaacagatccatgttactaattca
ccaattgaat------------taaaaaatatagataataaaataatatcagttcaaaaa
tatatttggagtaaaggttttaaatctcctcaattctattctagtatgggaggagataat
tatgttaatacgtctagtatatctacaaaatttattatggaacagctctctagttctcca
gtttttgtagtaaaaaacggctttaatgaagttatcttagggcatcctttatctaggata
aaaagaacagctttaaaacagattgccttttctttctcaaatgtatttcaccagcctgta
gcaactagtccaacttctaatggtttatttttttccacccagatgatgcaatcgaattt
aaaaatttttacaatccaagtcttttgaggcttctaaacatatgggaatcaaaatacaa
cctattggattacatgttgcttacaaaatgaacaggaaatttgcaggagatattcaattt

cgctttattccagatttcaaagaagtaggtgacttaatatttaaacatcagaaagcaaaa
aacttaattttcatgaaaatcagtattatggtaaagatttttttcaaggtcagccaatt
tatatgatccaacctattaatataaaacaccgtaatggtaagataagtacaatcaaattc
acagggttaaatgataagagagaagttgtttttacaaatcttgaagctgcaaataaatcg
tggaccaattttattaaaacagaaccttatttaaaaagaatcaaaaaaccgactttatta
gtctacaatttagaaagcttttttaaaagacaaggaaatattatatagagcagatttaaca
aaatttgttgtagtcactaataaaagcgcatacattaccacaaaagagtcgatcgctata
ccttattctaagagcttatctaagcatttaaaattaaatattaagcctcaactcttttt
attaaactttggatgagacggttactttccaccttgatgtgtgaataaaataatatt-tt
agttattttctcgagtagtattctacaactaaaagttcatttaattgcaaagccacccac
tctctatcaataactccattaattttccctgataaattagatttgttcaattccaaatgg
ctaggaatattagctaatccagggaatgctaagtagttttccacaagttgtttagagcct
ttttgaggttttactgaaattaattcgcctggcttacattgataactacatatagaaact
actttcccattaatacaaatatgaccatgattaactaattgtcgtgcagcggggattgtc
ggggccattccaagccgaaaaatagtattatctaatctcatttctaataactgtagtaaa
atttgaccagtagatccttgtaatttttttagcagcttttacatacttaaacagctgcttt
tcgcttagtccatagttaaagcgcagtttctgtttttcttctaatcgcacagcatattct
gatggttttcttggtttttgaccgtgctctcctggagggtacggtcttttattgctttc
ctactcagtcctggtaaatcacctagtctacgagaaatgcgcactcgtggtcctctgtat
ctagacatattatattaattctcctaaaataaataaatcaaaaagcattcattaataagt
agctatttttttaacataaaaatagcattaaaacaaagctttttagtaaaatatcttcatt
aaatacttgaatgaagatataaattcaga-tttttaat-aatttaagcgggtagcgggaa
tcgaacccgcatcattagcttggaaggctaaggttttaccactaaactatacccgcacta
ttaaatagttgctatatcatttaatatagcacaactatactatat---------------
----atgtagca-taaaatagattatgcactctccagaaccacatcaagaaaacgagcaa
tttcggcagcttgctcttctacttctgaaagtaataatggctgaccaactcgtgttagag
gaataattcttttatcttttgtgcatagataaatttcgcgacgtggatttaaaccttctt
ttatatctatttttatagatttaatctcttttatattaaacttaaggcaaatttgtctat
tctttccaggaaatcctagtcgaaaaatcttgataatacctttactttttattaaattcgt
tatagcctgcgcctatattccaaataatggttaaccataagaatgcactaagaaaaagtc
cgacactgccgtagaatgtcattataataccttgaggaataaatattagctctgttaaat
ttgtaaaaggtaacaagttagtctgaaaataactagataatccggccagaaggaatccta
gagctccaataaagattgtgctagcccaccagtagttactaaaacgcctcgagcctaaaa
tcacatcttttctgactgcatgtagaggcaatactttttcataatttgtttttctcaaa
taatagtatattttttgaaagatgaaaatactgacaaaaatttaataggattgtatgattt
tttaaattagcttaattgctttcaaacctaagaataatcctgttgctaaaccaaaaaaag

cggcaagaaatattatatatcctaaaaaaagtgaca---tgata----ttattattatta
ttacaatagtgtttacaaaaatagtggaaatctattactaaagtattatattacgttaag
tactaagctgtctacatgaaaaaactttttttcattttttagcatagtatttaaataagca
atagcataatattaattatattacacttctggagaacagaatatcatggcagaatttatt
aagccttataatgatgatcctttgtaggtaatttatctacgccggttagtacatcaagt
ttcagtaagggattattaggtaatttaccggcctatcgtcgaggcttatctccgcttcta
agaggtcttgagataggtatggcacatggttattttttaattggaccctttgataaatta
gggccactaagaggcacagatgtggctttgctggcagggtttttatcttctgttggactt
attattatcttaactacatgtttgtctatgtatggcaatgtctcatttactaggtctgat
tctaaagacccattacaaacagctgaaggttggggacaattcacagcgggtttttttagtt
ggagctgtaggcggatcagggtttgcatatttactattggctaatattccagtcttacaa
acagcaggtctaagtttattttcttaagctagtaaggggacttgaacccgtaacctactg
attacaaatcagttgctctaccaattgagctatactagcata-aacaaagattaacacaa
aaagatatttgttgcaatctgat-ttttcagatattacaaatatcttttgtgtttggata
agattaatcacctaatacat---aaaaatactaattaggatttgaattatctgaatcttc
agaatattcttgatcacaatccaaatagtagtaaatcgtctgatcaagacaagttgcgga
ccaaccaataggtgtttctcctgctaactcttgcatatccgcagaatcataaaaatcatc
gcttggcttattgtttttttgcttgcataaatatcactcccaaaacactctgaacttaagt
tgaagtaattgaacttctcaataatatattaacataaaatatatagtatggcgtttattc
attttacgctaatatattttttcgaattataagtttttggttaatgtttttattgctttag
tggaaataagcattttaatccacttattttgttctgctacccaaatttttttgtactgca
aattaacattctgtagctttttggttcttttatgtgaacgtgaaaccgcatatccattat
tagccacctttccagtaagctgacatttctttgacatagtttttatttgattagctaata
tatttatttaaagtgcttgctaaagtagattttgggacagcaccaatcactgtatccact
ctttctcctgctttaaaaatcattaaagttgggatacttcttataccatattcagctgca
atagttggattgtcatcggtatttattttaactacctttatagatgactcgtattcttct
gctatttcatcaacaacaggtgaaaccattctacaaggaccacaccatggcgcccaaaaa
tctacgagtacaggtaagtcattattaataacttcttgtttaaaagaggcatctgtaact
tgagatactgacatatttttttaacctttaatatatgttccttgctaagcaaatcttatca
caattttccaaaaaaaatcttctatagtgaactagtactagtgtccaactaagttttt-ct
tgccagacattagaaaaacttattacccatataaataaagtggtggccgagtgatatctt
aatagtgataatttctatattaaggttgaacacaagcaagaactttacgaatagtttttt
attcgaaaaatacttgcttgtgaccataaaagcttagttatattacaatatattttattg
tttgtaaatcagttatgataatggtataaacaacgcaaaagatactgccttataatcaag
gaggaatacatgtctcaatccgtagaatcacggactaggattaaaagtgaacgttacgaa
tctggagtaatcccttacgctaaaatgggttactgggatgctgattatgtgattaaagat

acagatgttctagctctatttagaatcactcctcaaccaggtgttgatccgattgaagct
tctgctgcaattgcaggtgaatcttcaacagcgacatggacagttgtttggactgatttg
ctaacagcttgtgatctatatagagcaaaagcatatagagtagatccagttccaaatgtt
gcagaccaatatttgcttatatcgcttatgatattgacttatttgaagagggttctatt
gcgaacttaactgcttctatcattggtaatgttttcggatttaaagctgttaaagctctt
cgtcttgaagatatgcgtatgccagtagcttacttaaaaactttccaaggtccagcaact
ggactaattgtagaacgtgagcgtatggataaatttggtagacctttcttaggtgctaca
gtgaagccaaaactgggcctttctggtaaaaactatggaagagttgtatatgagggtctt
aaaggtggattggatttcctaaaagatgatgaaaatattaactctcaaccattcatgcgc
tggagagaaagatatctatattccatggaaggtgttaacaaagcatccgctgctgctggt
gaacttaaaggtcattaccttaacgtaaccgctgctactatggaagatatgtatgaaaga
gcagagttttccaaagttgtcggtagtgttatctgtatgattgaccttgtaattggttac
actgcaatccaaagtatggcaatctgggctcgtaaaaatgacatgatttacatttacat
agagcaggtaattctacttactctcgtcaaaaaaatcatggtatgaacttccgagtaatc
tgtaagtggatgcgtatggcaggtgttgatcatattcatgcaggaacagttgtaggtaaa
ctggaaggtgatcctttaatgattaaaggattctataatactttacttgaaagtgagaca
gacattaacttacctcaaggtctattctttgctcaaaactgggcatccctaagaaaagtt
gtaccagtagcttctggtggtattcatgctggtcaaatgcaccaacttcttgattatcta
ggtgatgacgtggttcttcaatttggtggtggtaccattggacatcctgatggtatccaa
gctggtgcaactgcaaatagagtagcactagaatccatggttatggcaagaaatgaaggt
cgcgactatgtagcagaaggaccacaaattttaagagatgctgcaaaaacttgtggacct
ctacaaacagctttagatttatggaaggatattagttttaattatacttccacagataca
gctgatttcgttgagactccaacagcaaacgtctaatttaatgactacttataactgctt
aaactagcgaagtataagtagaattaacttataaaaataaggagcatagaatagtgagac
taacacaagggacttttccttccttcctgatctaactgaccaacaaatcaataagcaac
ttgcttacattgtttctagaggttggtcagcaaacgttgaatacacagatgatcctcatc
caagaaattcatactgggaattatggggattaccattatttgatgtaaaagatgcatctg
ctgtaatgtatgaaattaattcttgcagaaaagcaaaacctagctactatgtaaaagtta
acgcttttgataatactagaggtgttgaaagttgctgtatgtcattcattgtaaatagac
ctgctaatgaaccaggtttcttactgcaacgtcaagatttcgaaggtagaacaatgaaat
atactcttcatagctatgctactgaaaagcctgaaggtgctagatattaatttctattaa
gaattaatacttcttaatctactaaccctctttaacatctcaattaaaaaattagttttg
attttaaagagggttcatattc-atgtaaagaatagctaaataaaaataaac--------
acacacagaaatgacgcaattaactaatataatttctaacgatactcttgtaaatttaca
agaagaatacgataaaacacaaattcaagaagttttaaatgaactgaatcaagaattaat
aggattagtccctgtaaaaactcgaatacgcgaaattgcggctttattactaattgatag

attgcgtagaaaactagaactcgtatccggtaatccgggattacatatgtctttcactgg
tagccctgggacaggcaaaacaactgttgcaatgaaaatggctgatattttacatcgact
tggatatataaaaaaaggtcatttattaacagtaacaagagatgatttagttggtcagta
tattggtcacacagctccaaaaactaaagaagttcttaagcaagcaatgggaggagttct
atttattgatgaagcttattatctatataaggcagacaatgaaagagattatggttcaga
agctattgaaatacttctacaggtaatggagaaccaaagaaatgatttagttgttatttt
tgctggatataaagaccgaatggaaaaattctatgaatccaatcctggtctatcttcgcg
agttgctaatcatgtagactttccagattatacttcagaagaattattacaaatagcaaa
aatgatgatagaagaacaacaatattgttttacggaggaagcagataaaacacttttaga
atacacagaacgaagaatgaagcaaccttatttgcaaatgcaagaagtatcagaaatgc
tattgatagagcaagaatgcgacaagctaatagaatttttgctagcggtgaaaaagtact
aacaaaggcagatttggttacaattgaagcagaagatatattaaaaagtagattattttc
attacctaataattaaataaatatttagttagcataattacttatatagaaaaagtttca
tcaattggttgcaaattttcttaaaaacatttactatcctgataaagtaggta-tggcgg
catagccaagtggtaaggcagaggattgcaaatccttcatcccccagttcaaatctgggt
gccgcctagtactacgagcagggggtgtggtggaatggtagacacaacagacttaaaatc
tgttgattttagtaatcgtgagggttcaagtccctccacccccaagtatagtattaaaa
taaaaaatatatgtgcatttgtatcaattgtaatcatattacggaatgcagtacgtatta
tttaatagaatctaagcataagcaagttcatattaatgaaaatccatcatttatgcccga
atttccagtaatccatgtcaatatatcaagtaatagctatattaatcagattgattggga
tttagtggaatgcttatcgtttgtagaaaagccaaatagttggaatttgagtaattacta
atgtaataaaattctaggcattataagcaataactctatcttttaaatattctgtattaa
tttttgatgttcttacgagcgatatttttccagttctagctatctcaagaataccgaatt
tagtcaacagctgttcaattgcaactatttttccaggatctccagttacttctataatca
gtaaatcttcagcaatatctactacattggctctgaaaattttacaatttccaaagctt
ctgttctgtttttgagttaatttgaatcttaattaacattaattctctctcaacagagg
gtatattggttacatcttgtacatcaagaatattaactaatttatataattgtttagtta
attgttcaattgttctattatctccttgaactaccatagtaattctagaaactccaattt
gctctgttggtccaacagcaaggcttgctatgttaaagcctcgcctagcaaataatcccg
atattctagaaagcacacctgcttcatcttgaactaaaactgataaagtatgtttcatta
aaatttgattttatactatattatttacacaatatattctatgaaaaataagtacaaaat
actagtttttatatagttaaattattcaaagtaaacaaaatactaacatcagatactata
tcgaacctgtagtgttatactattaatttagtatttaatgatattct------aaataga
atgctaaattttaaaaatttgttaggaacttgtgctagaaaaatacaaaaatagtaaga
aactctctcgagatttacctcaaatcaacgaccgtatcaaatttccaaaagtgcgagtaa
ttaatgatgaaggtgaacaactaggaatttttgcaccggaagcagctattcagttagcta

caaaacaagggctagatttagttgtagttagtgataaatctgacccacctgtatgtcgaa
tattagactatggcaaatataaatttacacaggaaaaaagagcaagagaggcaaaaaaga
aacagcataatagtagtataaaagaagttaaaatgcgttataagatagaagagcatgatt
ataaagtaagaattaatcaagcgtcgaagtttcttcaatctggagataaagttaaagcaa
ctataacattccgtgggcgtgaaatacagcattcaaatctagccattgacttactaaaca
agatggcagcagatttagtaacaatagcggaaatacagcaagcaccatcacgggatggca
ggaatgtaattatgcttctatcgcctaaaaaagttagtt-aatttcgtatcttttaaatg
catctggctggattcgaaccagcgacgtctctttcgaaatggcggattatgagtccgctg
ccttcggcccctcggccacagatgcattacttaaagtttataactcataaagcatataaa
aatcaatctttatctgtctttgctaga---ta-agataatctttattcattcatattatg
ataagttgcaactgcggaaggtgagacttgatttaaatatcgaaaaatccaatatttaaa
tattgtatctaaaataactggaaaagttgaaataaaaagaaagataaaatctctactttc
tggcagtcctaaatgtcgcaaaattacttcaataatcacttcccatccatgaggtgaatg
aaagcctacaaacatatcggtaaataaaatgattaaaaaagccttagcagtatcgctcaa
cccataaataatctcatttaaaaaagatttaacaatagaaatttggcgttggcctgttat
catcagcagtataaatacagctattgacaaaatatctgaaagaagattttaactgcatt
tgcgctctcatttgcatagtattcacctaattctctagcttttaattggactctttttc
aattatctcataagaaatattttcagaagggtttagaagaacctcaaaatgaatttctc
ttcaaacctttgtaactcagcaaaagctctttcttcttgagaagaatttataaatatttt
aggctgttcttgattccataaataatcaatacatggcccaaatacaaaaaatttcgatgc
ttgattaactaatactggagatattaataataataaaatatactttacagatgtaatagt
ctggtgccgtgacaatctaaattcttcaatagcttctgactctccattaggatctaattc
ttttcgaaatttttcaaatgtattagtaattgatctcggtatgggtccaactttttcaga
agtagattgattatttcttttcaaattccaatatttcatttttattactgctaaaattt
ctattgatcgatgattgatcatatattcaatataaatattaattctagaattacatata-
aaaataatagttatgaataataaaattaaattaaattttaattatgcattagtaatattt
tttacaactataatgtacacctcaaaattaaatatttacttaaaaagtgaattgttaaac
actagagaaaaatattgtaacttaatattcgagaaaaaaatggcgtttattcccattctg
tcattaaaaaaaaaccaataagctctgaacaaataagcagtaaagcgtccaattttcttc
tgcatggggtattcaacaaaaaattcttacataagatcataaatttctcagagacttcag
agagagatcttcacaatattgagaacctacaaaaaataaaaacgagtggatatttttcta
caattcaacttaatattaaaaaagcatcgggggaaaagttactaacgtgcttacgttac
caaactcaatcctaaaaaagatctatatttataatcgaaaagaaaagattatacctcgat
ctttttattaaagttattcaaatatcaactaggttatcctaaaagtcttgtacaactta
atttgtctctctcatcaattatgagatggtacttacagaaaggttatcaatgggctttaa
tacaaatacatcatattgaggacccatcagctattgttgtcaatattcatgaaggattaa

ttgcaacaattagaacggagtattatacatcttctctcgaaaggttttctaaaagttcat
acacaaatgtaatagaaagacgacttaatattaaaatcggttatccaattaatattactt
ctctacaaagaaaaataaatttttttgaaggataataaactagttggcaatattgtatata
gcattgaaagatctgaaaaaaattcaagtcatttagatttaaaatttcaaatacaagagt
taaaagataaagaattaattcttttcggggaaaatctatataataattcttatatcattt
cttttttatctca-----------------------------------------------
------------------------------------------------------------
------------------------------------------------------------
------------------------------------------------------------
---------------------aattatatttgagaaatcttaacacggccaactcatact
tcacattaagtattcattccctaagaaatacaattaacctaaagcttgcatacttaaatc
cttcattaagaattactaaaaatttcacaattcaagttgttttagaaataataagacaa-
-----------------gcagccccttcctgtatatttcgatcatctaaattctcaaaaa
ataaaccatttttcagtcataaatttatcgctcaatacatttttgagggtttaataacat
ataatttcacttcatatttctcgatatccgaagaatttcttctttcacagaacattcaga
ccaaatattttgtcacaaatttacaaactgtcaaatatatcagtcctaccaaagcattaa
tagccagtagtgattattggctcaacaaacaatcagaaatactccgccgacaattcctta
tattatggctaaaattacattatcaaaattttgatactttaaactggcctacaaaaggtt
atcttcttgaaattgaatcctggtattttaccccatttcaaaaaagtaatttattcaact
atgactttcaattttactat------ttattcattcataaaatcgatattaagcatattg
cgcactttagtttaccattctactaccagagcagaataaatcatatattaagaaaaattt
taaaattacaatcaaatttcagaattgagactattcctattatgttctatagctcttatt
ttcagagtaaattctgtaaaccattatttaacttttctgctcgaatgaggatagaatatc
aaatacctattaatagcaagagtaggatatcgtttttttgtaattatacaaggccctttc
taatgaattactcacaaacttatat---tattttacaatctagtttaattgatcagcagt
ttgtaagttctttatatcaacaaatttttttacggattaaaaatgcagctgaagttaccaa
ttaatcaaataccacctttatcaattgaatatactataaatagtggtcgaaaattttgta
tttacttccatatttctcaccaacgataaccattattgaaatgcaacaattaatttcttt
ttttgatcgcagcaaaggcaagtggatctcacaacggacaacatacaaattgtcagacaa
aaagatgaattcactacaatctcgaatgatcataaagcccgatcaattatcatcaaattc
tcaattaataacttcattgaaatggggaaagatttctcgtcaaattacaaaagacatgac
tgttcagaatattgtaaatgacaactgtaagtttcatctaagatttaccaataaatttaa
tacaaaacaattagttactctttgtagtataactgacaagagtttattaacttttaaaac
cagatacggtactataactatagatgaaacttattggttcgcaaccaataatttacgttt
aagcactagtattattaaacgatttaatgtttgtgtagcagtctctttttgctctgaaat
tagagtttaagattattgacaaaaacaactaatagcttaagtagctattagttgtttttt

t---------tattatagtaaaattttacctggataaattactactctaagtccttacga
tttggatttctggatgggtcattagataagaagccaaaaacaaataaagaaataaagaaa
attacagtagtgtacacgaagattttcaaagtaaacatatttttatccttaaaagctatt
agtgtgtaatatttatgtgtatctccacataatcttatatcatgatatgaattaaggtac
tattttataaataaatcaacatctttattttttagtaatgatttatatttaaatatacgc
gatatgctaaagactaaatctttttgg---tgattcacaataagcttcctctttttttta
tttagatgcagctttccttcaattattatataatcaaactttttaagtaatttaaaactc
tttagagattgcctattccatattaacaaattaatcacagctactttatttcgctttttg
aatcttgcttttaatttaataatttgatttccattgtgaacagtcttaacgcttttacag
cgtaaaacttgaactaataaagtgcacttattcat-gtatatatttagaatctgcttttta
atgaatcta-------------------taagtcaaaatgatatttcattatcttgttga
ctttgctcataagtaatatcttttagcacttttaagatttaccaaaatattcttgaaaa
tatttttcaagagatactacttcagatttatcgatacttaaaatttcatagacttcattc
attgaagcagtaaatttttcgccgctggttaggatttctgcaaatgctaaacgatctgag
atattccaagtccactgtaatgtttttgtaatttttcttaaagcctttagtaaactcagg
gggattttcgaaatctgtgtcttttggccagataacttttcgcataatgtaatgatttca
gcagatgtccaagccttattaccaactaatggcagaattttattttctgttgaaggcacg
ccaagactctttataactaattttgcggcatcttgcgtatcaatatatgcaattggagtc
gattctcctgttacccatactgatttttttatctaaaataggaattgcatattgactaatt
agtccttggaaaaatcctcccaaggaaaatacagtataaattataccagagttttgcaga
taatttactacttgagattttaaattcattaaagggactttaggatgttgatcggcattt
aatattgaaaagaagataaatctttggacttttgctgcttttgccgcttcaataagtgca
gtttttccatctaaatctattttttcagcattataaggatccgatgggcgagatgttgaa
gcatctataactgccgttacaccgcaaaaagattgcaatatactttcgggagatttcaag
tcaccatatactagttcagcaccccattcttttaaaaaagctgattttcttaaatttctt
accatgcattttacattgtagccttcgtcaagagcgcgtcgtacaatttgacgccctaaa
gttccggttgctccaattactagaagagtcatattaattaaaaatgaatatttataaaga
gattgatttactcaacagcaacgagtacattgggtagagagggaatcgaaccctcatgac
cgaagtcgccacattttgagtgtgatgcgtataccaatttcgccatctaccctattatag
accaactattacaagtagtttagtctttagcgtaaaaataatactaagtttcaatagata
tgtcaacttttacggttatattaaataaaaa-tacaattatttatggcaaaatttgaatt
gatcccttatagattatatttaagtcaatctagaacctggatgcacaaaattaaggcaga
ggtaaagatatacgtccttttattattatggatttcaatttttcttttatcttatcgcaa
gttgcttattgttgctttcagcttaataatcactagcagtacaattaaatgtaatcaata
catactaagaaaacatttttttcaaacttttgtcatggtcattgtgaccatgctcttttc
gcttagtgttacaaataactatcaataccatttaaagaaagaaaaatatcaatctatatc

attgtgctggacacaaaaagcaatccaaagatatatacctaaacaaataactcatttcaa
tgatcagataccagaaaagatattatttgcaataaaaccaggatcgtatttttcatcac
aatatattctattaaattagttatgatgacaacttcctcggaagttttagcgctaactat
ttataaaaacaatattatcaatcaaattttcaataatgaattgcttttcatgtttttatt
atcttcacatattgttaacaatataatagtgaaaatggaaaaaattgcacaaataatgag
cttaagaggtagtttgaatttacgcaaacagtcaacaagactctttacactatttttct
catatctaagttattttttctgaaataataaaagaatcgacagaaatatctcaatcact
atatgctagaaatctcaatcaagaaaatgataactttctaaaaatatatgtatctcaatt
tagcacaaatgactatatatgcttgttcattagtacgatatatgtaacaattttgttttt
atcttggtaggagcaatttatattgctgactctgttaatacgaagaataataaagttgtt
atgttatccttatattaaaaatataagatattttataaataattaattgttgcattatgg
tcaatactcaaaatccaatttcgcagacttctgatctcgattatatagtgaatcagccat
ataaatatggatttacaacttctgtagaatctgaacaatttcctagaggaataagtcaag
aagttgtgaagttaatttctaagaaaaaaaatgaacctgaatatttattgaatttcaggc
tgaaagcttatgacaaatggactaaaatgaagaatccaacttgggctcatttaaagcatc
caactatagattttaataatataatttattatgccgttccaaagttaaaaaaagaattaa
atagtttagatgaagtagatccagaaattttagacacctttaataaattaggtatatctt
tgaatgagcaaaaaagacttagcaatgtagctgttgatgcagtttttgatagcgtctcaa
ttgcaactacttttaaaaaagagttggcagaagctggggtcatattctgttcaatttctg
aagctatacgaaactatcccgatttaattcaaaaatatttaggtactgttgtgccttcgg
gtgataattattttgcagcattaaattctgcggtatttagtgatggatctttctgctata
ttccaccgaatacaatttgccctttagagctgtcaacatattttcgtattaacaatgaag
agtctgggcagtttgaaagaacattaattgtagctgatcgaggaagtaaagtaagttatt
tagaaggttgtacagctcctcagtacgatacaaatcaactacatgcagcaattgtagaat
taattgcacttgacgatgcagaaattaagtattcaacagtacaaaattggtatgcaggaa
ataaggacggtcaaggtggcatatacaattttgttactaaaagaggcttatgctctggga
agaattctaaaatttcatggactcaggtagaaactggatcagcaattacttggaaatatc
ctggctgtattttagcaggtgacaattctcagggagaattttattcggtagccttaacca
ataactaccaagaagctgatacgggtactaaaatgatacatattggaaataatactaaaa
gtaaaattatatctaaaggtatctctgctggtaaatccaaaaatagttatagaggattag
taaaaattggtccacaatcatttaattctcgcaattattctcagtgtgattctttattaa
taggtcactcatctcaagccaatacttttccgtatattcaagtacaaaatccaacatcaa
aagtagaacacgaagcatctacgtcgaaaatcagcgaagatcaaatttttatttcttac
aaagaggaattagcttagaagagtctgtgtctcttatgattagtggtttctgtaaagatg
tatttaacgagttgcctatggaatttgcagtagaggcagatcgtttattaagtttaaaat
tagaaggaactgtaggatgagtaattatattttagaaattaaagatttgcatgcttcaat

tagcgatatacccatattgaaaggtgtcaatttatctattaaacccggtgaaatacatgc
tattatgggccctaacggatcaggcaagagcactttgtctaaacttattgcaggtcatcc
ggcgtattcattattaagtggagatattttattctatggaaaaagtattcttcaaatgga
acctgatgaaagggcaagagctggcatttttcttgcttttcagtatccagttgagattcc
aggagtaagtaatgctgatttttttaagaattgctttaaatgctagaaggaaatttaagaa
attaccagaattaagccctttggagtttttcaattaataacagagaagattgatcttgt
aggaatgaaagaaagtttttttaactagaaatgttaatgaaggcttctctggtggggaaaa
aaaacggaatgaaattcttcaaatggcactcctagatagcaatctatctatattagacga
aacagattcaggcctcgatattgatgcccttaggataattgctcaaggaattaatacatt
agcgactccgactaattcaattgttttaataactcattatcaaagactccttgattatat
tatcccagattttgtgcatattatgagtgatggtaagattgtaaaaacaggaaatgttag
tcttgcacaagatttagaaaagcatggatatgattggattaaaggaacttaatcttttaa
aaag------aataatgaggctaaagatttttctttagcctcattattctctaaagatat
tgccgattgctagttaagctttagcaaaaccttgtttaacatcttcaattgctttttta
atagttcttcacttctactgtctaattttttagtagtacgaatactttttccaaagtctg
gcttagaatttttcaagtcttctcttagttctataataaaatcaggaactttatttacat
caatgtcatctaaatatccattaataccagtatagataatagctgtttgttcttcaacgg
gaataggtgagttttgagcctgttttaaaatttctctaagacgttgacctctggccaact
ggttttgagttgctttatctaagtcagatgcaaactgtgaaaacgcttctagctctgcaa
attgagctagttctaatttcaatttaccagcgacttgtttcattgcttttatctgagcag
ctgaaccgactcttgaaacagaaataccaacattaatcgcaggtctaaccccagaattaa
atagatctcctgacaagaaaatttgaccatcggtaatcgaaattacattcgttggaatgt
atgcagatacatcaccagcctgagtttcaataattggaagagcagtcatactaccacctc
ctaattcagcatttaacttggctgcacgttctaataatctagaatgcaagtagaatacat
ctccaggatatgcttcacgccctggtggtcttcttaataagagtgacatctgacgataag
cttgtgcttgcttagttagatcatcatatataaccaatgttgctttgcctttgtacataa
aatactcagctaatgctgcacctgtataaggagcaatatattgaagggttgcaggactat
ctgcattggctgcaactattattgtataatcaagagcacctttttcttgtagtgaggaaa
ctacctgagcaacagaggatgctttttgtccaattgctacatatacacatacaacatctt
gacctttttgattaataattgtatctaatgcaacggctgttttacctgtttgacgatcac
caataattagttctcgttggcctctaccaattggaatcatggagtcaatagcagtaattc
ctgtttgcataggttcacaaacagattgacgaccaataatacctggagccatagactcaa
taagtcgtgttccattacttgctggttcacctttatcatcaattggacgagctaaaggat
caactacacgtccaagaaaagcatctccaacaggaatttgagcgattttaccagtaccct
ttacagagcttccttctaagatgtctctaccatctcccattaaaacgactccgacattat
cactttctaagtttaaagcaaccccaattgttttatcttcaaactctaataattccccag

ccataacttcgtctagtccatagacacgagcaataccatcaccaacctgtagtacagtac
caatattagctacttcgacatcttgatcatattttcaatttgttgacgaataatactac
ttatttcatccggtctaatatttaccatattttaatattcttttaattaatgagaaatt
ctaaagtgataaatatacaaactaactatgtttgtattattacctttattttatattgct
gctacatcaagatgcgaagccatctgccttagttgtcctctaatactagtatcaataact
tttgaaccaatttgaatagtaaagccaccaattaattcaggatcaacggaaataactagt
ttaacttctttcgcattcgtcatagctttgattttatctgttaaaagattttcttgatca
gagttcaaagcaatagaagtgctaatatttgcgattgttagagattccatttcatacgca
agttctaagtacttgctagctatcacatctaacatagcaattcttttgcgatcaactaaa
accatcaaaaatgatagtgtgttttcactgatttgatcaccaaaagttgctgtaattact
tgctttttgctcctattgtttttaacgggttagct---aaaaaatatttcagctttgca
gattgcaacaaaatgtcttctatcagcttcatatcttgactaatttgttctatagctttt
ttactatttgctaaatcaagaagagctgacgcataaggctgtgctatttttacgacaaga
ttattattgctcataattgatctcctagcttagcaatattattatcaataatacgtagct
gaatatccgagctcatttgattctccaactgtaaagtaactttcttaagagctaaaaatg
ttatctgctgctggatctgtctacgaatttgcttttcagctgtctcaatattagattttc
cagttatagctaatctctcaatatctaattgtccttgagctaaaatagagcttctaactt
tttcagccgtcagttgagcttctttttaatttggtctataataatttgagtttgagcaa
gttgtttttctgattctaataatctagaacttgcttgctctaatctgtcttctgattctt
gtatagctgctaaaacttttgcctgtctttcatttagactagatccaagagattgtttca
aaacataaatcaatccaaacaagagcaacaagatattaataacattagcttcaaaaatat
cagaattaaaaccgaatgtatgttcactactatgttctgataaaatagtaattattgatg
ttgtgtttactatactattcatttaaaattcttactatttaatactatttcaatatttta
gtcaggaatcattattttaaagattgcctgcttaaaagcttggccttaatttgatcactc
aaagtatcaatctgatcttctaaagttttcagggcttcttctttttgtatatttagttgc
ttagaggcttcagtaattaattttctgcattcatctgagctttttttatatcttcagaa
acaatactttgagcttctttttgagaagcagctattttagcttgagcatcacgacgggct
tctgataaatcttcttcatatttggcagctaattcatctgccttgacaagcatagaagaa
gctgttgttaatgttgtcctaatatattcatcacggtcatccagtattttagttaccggt
ttataaaaaatagtattcagtaacaacatcagagtcagaaactgtaatgccattaaaggc
aaagttccattgaaatcaaacaagccaccctcaatttcttcggctaataaaagtggtaaa
taaatcattttactaattatgtgtatagttaataatttgttttcaattaagaaaattagc
tgtatattgcgcttagtctaacaagactaagcgcaaatgattattctaaccaacgtatgg
attagcaaataataaagaaagagcaacaactagaccataaattgttaaagattccataaa
tgctaaacttaatagcagtgtacctcgaatctttccttctacttctggctgtctagcaat
accctctacagcattagctgctgcactaccttgaccaataccagggccaattgcagcaag

accaacagcaagaccggcagcgataacggatgcagctgaaacgattgaatccataatgaa
atttaatgttttattatattagtagaaaattgacaaatttcgtataatcttttggagcgg
tttaatttagtctgatagattaaaacatctgactttaattatagtatactactcttctcc
atgaccttccattgcttctccgatataggcagcagaaagtgtagaaaaaattagtgcttg
aattgaactagcaaacaaacctaatatcatgacaggtagcggtattaaaattggaattag
taaagtaaatactgatacaactaattcatcggctagtacatttccaaatagtcgaaagct
taaagataatggttttgtaaagtcttctaaaatgttaattggtagtaaaactggtgtagg
ctgaacatacctagcaaaatagcctaatcctttttttgcttaaccctgcataaaaatatgc
caaagaagttaataatgacaatgcaacagttgtgttaatatcgttagttggagcagctaa
ttcaccttctggtagatggattaatttccaaggaattaaagcgcctgcccaattacaacc
taggataaacaggaataatgtagcgatataaggaacccaaggacgatactcatgctctcc
tatttgatttttttgcaatatcttgtaaaaactctagtacgaattccataaaattttgaaa
tttttctgggattctttgcaaatttctagtacctagaaatgatatagttaataatgcagc
aattactaaccatgagacaataaaaacttgcccatgtaattgtaaactacctattttcca
atatagatgttttcctacttctactgctgataaacaagagtatggattaaaatctataag
attattttgatacataataaatattttttt------------------------------
-------aaatagtatgcaattggtttattttcaaaaccaaatt----gtaaattagaga
caaaaataaattctaatagagacggctatcttaagtagtaattatttttctaaaaataa
att-tttctacatataaggctttcaagtttaattatatacccatatattaattatttaat
atgattataggtattttggtaaaatcttaattttaatggcaa-----aaaatgaagtgta
tagagttattttctgtttaaaatatcagcaacttcatcagcaaaattagctttgctattc
tcttctcctccacctaatacaaatctgacaaaacgtctgattttaatatgttcgcctaat
aaagcaatattttgattaattaagtcttgtattgtaatatcttgattccgaataaagatt
tgatctaataaagataactctttaagacgtttttttaattctgccttcaactatttttttct
atcatttcagctgacttattttttaaatcttctctaccagcctcaattctttttttctaaa
ttgattatttcatccgggatgtgcataattgagacatattcaacattagggcaagcagct
atttgcatagcaatatctcttgctaatttttggaactctgggcgacgagctacaaaatct
gtttcacagtttacttcaactagtacaccaattcttccacctatatggatatagctttct
aataagccttcaatagcggtacgaccagatttttttgttggcagaagctagaccttttttgc
cttaatgattcaagagcttttttcttcattgccgttacttgcttctaaagcttttttgcaa
tccatcatacctgctccagtcttatctcgtaaagcttttacagtttgagcagaaatttgt
agtgtcatattgttttcaaagattttaatatataaattaatg-aaataaatattt---a
ttcgtcttcagctgagtgagtcaattcagttggttccatttgtccatatttaccttcata
aatagcatctgcaatttttccaataattaatttgattgatctaattgcatcatcattagc
aggaatcggaatattaataatttccgggctgcaattagtatctaatatacaaatagttgg
aatacctaacttcaaacattcttgtatagctgttgtttctcgcttttgatctacaactac

aacaatatctggcaaacgattcatattcttaataccatttagatgcttacgtaacttatc
caattctcgtcttaatacagctgcttctttttgggcaattgatcaataattccactgct
atcttgttcttctaattgtctgagacgagttactctagatttaattgttacccagttagt
taacatgccacccaaccatctttgatttacgtagtaagagtcacagcgttgagcttcttg
cgcaactataccagctgcttgccttttagtccctaaaaacaaaactttttttaccatcaga
tgatgcttgtttaataaagtcgcatgcctcggtaagtaattgtgcagtttgcactaaatc
aataatatgtataccatttctttctgtataaatatatggaaacattttaggattccatct
acgtgcttggtgtccaaaatgaacccccgcttctagtaattccgctaaagtcacaacagc
cataaggtt-ttaatctttaatttatttaattgttcgggaaatccctttttatta--atct
ttatt----taatttaagtatttctgtatattattaactaataataacttaacatacttt
ttgaatagattaaaa----ataatattattgacagattcaatcgacacttttgttattaa
aataattacgagctgttctatcatctaagataatatcatctaagtcatcacgtacagatg
gcgtaatattatcagtactaacattttattagtatcttttttatt---ggcaacattat
tatgattatcacgcatagtaaaacctgtgccagctggtattaatctgccaataataacat
tctctttaagacctcttaaccaatctaatttgccggagatagcagcttctgttagaactt
tagtagtctcttgaaagcttgctgcagaaataaagctttctgtatttaaagaagcctgag
taataccaagtaaaacagggcgatatgatgcattcagtttatttcttaatgtcattgatt
tattagtttgctcaatttttttgcaattcaacaagctctccaggcaaatagcccgtctctt
caccattctctattttcacttttgaagtcatttgacgcactataacttctatatgcttat
cagaaatattgaccccttgagactggtatactaactgaacttctttaacgagtagcaact
gaatttcttgaaaactgagccttgcagcttcatataatgataatccacattcaatgtata
gattaaaacttgcgtctaaaatatcgtgaggatttaacagtacatctgttttttttcttg
cttcaagaatttcttcaattctaggaagaccttgaataatatctcctgttttagctctgt
caaaaactaaaattgctaatgtttctcctttttctaattaatgcattgttatcaacatgta
aaattgcaccattagaaactagataaggtcttgcaattctcaatgtaacgaattcttgag
aaatatgggtaatttgtcctgactctaaggcaataatactttcacttattaaatcaccac
aacgaatccaatctccgactttgacagcagtattctgctgatgcaaatcaaatactttgt
tatcgagtgaagtagtaagaagaattctcctattggtattctttttggcatcaatttctt
caacaaaacctttactcattgaaataatttcagtgctagcaacaactgtcaaaggtgtaa
tatattcaccatcactgacaataattcttgtttgactttgttgtttctctactttatt--
-atctatgcttttaatcgataaggtttcaaatgtagaaaatcctaaagcaaaagaattgc
tactattttctttagctaagaattctattgctgaaaccaagttaggatcatgatggttaa
cttcagctattaaattagttgtaaccaaatgtacgccatctatagatttactctttcac
catctctaaaaagagttcttttgactaattttaatttagtttgcttggttgtttgtgaag
ctaagttttcagttaaaatagattttgtctcaggtattgaatatacaattactggtctaa
ttaaaatatatggggtttgctcattttctatatactcccagtatactaatttatctgtag

atatattaccgtgaagtgtctctccgggtcttaaaaaaccgcgacttttatcatgaatac
tgtgaaaattgcttaacttgtaaatagagccgggctttattattatttctcgaactatac
catctttttggataatttcaataattccagagctttttgaaaaaatatttttgaccaact
cagttccactttcaataatatcaccattattaacaagtaataatgaagaatctttattaa
tttcatgtgtctcttctgaaatccacaatatataacctgggctcagaatatcatatgcat
ctttttttgggccggtcttcttttagatacatttaaatctaagtattttataattccac
cgcttgtagttttatatgtttcagatatcaattctccaactatatgaccatgagaaa---
ttttgtcatctggaaaagatttaagcaaaaatctctgcttattttctgtttctagaatat
aagactctttttattaataacatctttatatatgtgacatcctggtataattatagatt
cagtaataatttgtacattggtaatactactattagtattttggtagattcttactttac
cactatatttattaattaactctgtttgcgctagtatagttccaggattaacttgatcag
cgttattaactgtaatactagctgagtctggaatattataaacttcgccagaaagtaccc
aaatcaagccaccagttctagttattcgagttgtatattgattattatcagtctcttcta
cggttaaattagaaaagcaaattttaccggaaaggtctgaaagaacatgcttttgagctc
tttctgtcattaacctatttcttcgaggtgattctgcaattacttgatctttagatacta
tctcattattggcgactaagagagtagtacctttacttagattaattattgatttttgtt
tatcactcgaattaataataacttgcgtgggtttttctgtcatcagcgcttgctcaccat
gtctggttcgaacctctatatatgagagagaatctaaagctgtgagcttaccattcgtcg
gggaatagatttgttctgcaagttcacctgtaaatactccacctgtatggaatgttctca
tagttaattgagtacctggttctccaatcgattgggcagcaatgatacctacagcttctc
ctaaatcaactaagcgaccatgagcaagattccagccataacagtattgacaaacagaac
ttcgagaattacaagtaactggagaacgcactaaaactttttt---aatacctgctcttg
taatttcttgtgctaaattaggactaatatcttgcttagtatgagcaattaagctacctg
tttctggatggaatatatttctgctaatactcgtccagctagagcttgttctaaattta
ttaatactttttgtgtatcaactaaatcttccaagatgatacctttctttgtcctacaat
caacctctctgataattacatcttgagaaacatcaacaagtcgacgagttagatatccag
aatctgctgttcttaaggcagtatctacaagtccttttctagcgccatatgacgaaataa
aataatcggtgactgttaaaccttctcgaaaattactagatattggcaaatcaataattt
gtccttggggatcagccattaatcctctcataccaaccagttgtcttacttgagaaatat
tacctctagcaccagaaaatgccatcatatagacagaattcaaaggatctgtttctttaa
aatattcaattacctcttgcttcaaagattcgcttgcattattccaggtatcaataactt
tttgaaatctttcaactgctgttatttctcctctacgatatttattttcggtatttttaa
tttcttcaatagttaagagcagaagactcttttttgctagggggggatccttaaatcttcta
gacttaaagaaattcctgcttgagtagcataatgaaatccgaggtctttcaatttatctg
ccatattagctgctcgcgctataccgtaattacgaaatgcccaaacgattaagttcttaa
gttgatttttatctataaccttattagaaaaacttggttgtgagagacttttcctactat

tcactaaatcttctccatattgaaaaataatc-gattcactaagctactaaagactcttg
gataatttattgaagataatacgaccaggagttgtcctaatatattgaactattctttg
atgctcaacatcttctttaataatatgattatcaaagaatttcgtcacacttttatcatg
atgtgtatctatcttaattggaaacttaacttggtcactatctataactccatcaaaacg
ggcccaaatataagcatgaagatcaactttttttgctcatacgctattactacgtcttc
taaacttgcaaaatattgacttgctccttgttgctgagaaggattattagctgttaaata
gtaacaacctagaaccatatcttggctaggcataataattggctggcctgtagctggaga
aagaaagttatgaggtgctagcattagcagtctagcttctgcttgtgcttcaagagatag
cggaacgtgaacagccatttgatcaccatcaaaatcagcattaaatgcaggacatacaag
aggatgtaattttatagctcttccttcaacaaggataggttcaaaagcttgaatacctaa
tctatgtaatgttggcgctctatttagtaaaactgggtgcccttgaataacttcgttcaa
aacattccaaatcgaagattcatttttttgaatcatttttttcgcagctttaatattatt
aactaatccttgaagaattagtctatgtataacaaaaggctgaaataattcaagtgccat
ttctctcggcaagccacattggtgcaattttaaatgaggacctacaacaatgacagatct
tcctgaataatctacccttttgcctaatagattctgtctgaaacgtccctgctttccttc
tatgatatcggataaagatttcaaaggcctgttattagcaccgactacagttctacctcg
acgaccattgtccattaatgaatctacagcttcctgaagcatcctttttttcatttctgat
aataatttcgggggctaaaattgatttaagacgtgataatcgattgtttctattaattat
tctacgataaaattcatttagatcagcagtggcgaatctaccaccatccaactgaaccat
tggtcttaaatctggaggtataactggaataactgtaaataccatccaggaagggtctgc
accagttgctatgaaattttcaatcaatcttaagcgcttcatttttttattgaattttaa
agaaggtgtcttaaaactttttggtggattagttgcttctgaccgtaatgcttctgcaat
ataatctaggtctaaatcttttaaaagtttttgtatagcttcagctcctattccaacttc
tatttgattattttcgtcctgattttgataaatttcttcttcaagagatttccattcata
accttctaataattgtttatatttcaaattagtttctttactggattgtgtaacaacata
agaatgaaaatatacaatttttttcgacttctttcaccttcaaatcaagggctaaagcaat
ataactggtactgccttttaaataccaaacatgagtaactggggatgcaagttcgatata
agccattctatgtcttctcactctcgattctgttacttccacaccacatcgttcacatac
tatgcctttatagcgaaaacgcttatatttcccgcagtggcattcccaatccttgactgg
tccgaaaattttctcacagaatagtccatccatttctggctttaaggttctataattaat
agtttccggtttcgtaatctctccaacaatttgaccattaggcaaacttctttctcccca
ctgtcgaatcttttcaggtgaagctaaactaatttttacatagtcaaaatattgctcaaa
atttgtcataaatttaaatgccttaaaatcaaataattggccaatatctttaggtctctt
ttaattttgaacaag----ttagtataaaaattgttcaaaatcatcaacaggaggagcat
cataattagagcgagatgctctatcctcttttgaatcagacattaaatcaacttctattg
tgcgtctttgaccatcttcaaataatttcaatttatgaacagcaatatctaagcctaaag

attgcaactctctcatcaaaactttaaatgattcgggtgtaccgggtttcggaataggtt
tccccttgactattgcattcagcgcttcgttccgagcttgcatgtcatctgactttacag
ttaataactcttgcaaagtatatgctgccccaaaagcttctaaagcccatacttccattt
ctcctagtctttgaccaccatgttgtgctcgccctcctaaaggctgttgggtaactaaag
aataaggaccagtagatctcgcatgaattttatcgtcaactaaatgaactagttttagca
tgtaagctcttccaattgttacaggattatcaaaaggctcacctgtccttccatcaaaca
cttgcatttttccaggatgctggtcattaaaaatccatttatttttggttaaaattgagg
cttcttgcaattttctatttacaagcgctctagatgcttcagctccgtacatttcatcaa
atgggattattttaaatcgcttctctaaatatccgcctgctaaacctaataagcattcaa
aaacttgaccaacattcatcctagatggaacacctaaggggtttaaaacgatatctaccg
gagtaccatcacataaataaggcatgtcctgtttaggtaaaattctagaaataataccttt
tattcccatgtcgaccagccattttgtcgcctacttgaatttttctttttttgtgctacat
aaacacgaatcatggcattagttccaggaggtaattcatctcttttttgtcttgtaaaca
ctctaacattaacaaccctacctttagctgcattaggtaatcttaaggatgtatccctga
catctctagctttttcaccaaagatagcacgtaataatttgccttcaggtaattgatcag
cttcccctttaggagtaattttacctactaaaatatctccagcttcaacccaggaaccac
agacaactattccatttcgatctaaatcttttagcgaatggtcgctaacgttcggtatct
ctctagtaatttcttctggacccaatttagtttgtcgacattctacttcatatttttcta
tatggatggaagtatagacatcttcataaaccaatctttcgctaataagaaaagcatctt
catagttatatccctcccagggcatgtatgcaacgagtatgtttcttcctagtgcaattt
caccacaatcagtagaagctccatcagccaatgtttgtcctacaacaatttttttctccga
cccagacaattggacgctgattaatacaagtgtcttgattcgaacgataatatttcttca
atcggtaatgaaccgttctaccattattatcttgaattcctattttgttggcagaaacat
atttaacataaccagatgttctactgatgacaaccatacctgaatcacgagcaattttttg
tttcaagacctgtaccaataattggtttttctggatataataatggtactgcttgtcttt
gcatattcgatcccattaaagctctattagcatcatcatgctctaaaaatggaattaagg
atgttgctgctgaaattacttggataggcgaaatcgcaatataatctacttgatttggaa
ttgtagtaacaaattcttgacggtagcgaacaggaataatatctccttgtatatagttct
gagcatcaactttaacatctcctggggcaaccctaaagtcatcttcttcatctgctgtca
aataaataggactatttgaataatctacttgaccatcatgaacaggataaaaaggtgttt
caataaaaccaaatatatttaccctagcacaagtagcaagtgaaccaataaggcctgcat
ttggaccttctggcgtctcaattggacatattcttccataatggctaggatgtaaatctc
taacagcaaatccagctcgatctttattaaagcctccaggaccgagagcactaattcttc
tcttgtgggttaattctgcaaccggattagtttggtccataaactgagaaagttgactag
agccaaaaaaattctcggactgaggcaattaatggtttaggattaattaaattagacaaac
ttaatgaatctatatcacatatcatcattctttctctaatgatacgttccaaacggttta

aaccaactcgaaattggttttgaagtaattcacctactgatctaactcttctattaccta
agtgatcgatatcatcaagatttccagaattcttatctttaatattaatcagataatcta
ttgaagataaaatatcttgaggggataatactcgaaaggctttaggtatgtttaggccaa
gtttttattgattttgtatcgaccaacttcacctaaatcatatcttttgggatcaaaaa
aacgagagtataacatttgcttagcaactgcaaccgtagctggttcgtttggtcttagct
tggaataaactattaaaagagcttcttcatctgtaacttcttcaatgtcatttttccaa
tttctcttgctagctctttaactgagtatgattgagatgcagatattaaaaaagcatatt
tacttaaacctttctgaatttcatccttgtttaaaccaattgcacggagaaaaatatatg
cattaaccttgtgagttttatctattcgtatccaaatttcgcctttaggatctatttcaa
atttaagccaggatcctcgatttgagatgagactcgcgctgtaaatttgcttaccattct
tatctatctcttgtttataataaatccccggactacgaattatttggttaataataactc
gttctgtcccagaaataatgaaagtacctcgattagtcattattggcagatcaccaataa
agactaatctcttttttatacttttttatttttttatctgtttttctgcactaaattgtaaat
gatttgaagataaatctaatgatttaatcgttttgttctgatctttagaaggtaaatcaa
tgtcttttctagttaatttagcgggaacgtatatttgagcactataagttctatctcgac
ttttagcttgtctaacactatatctaggaaatttaattttatattctttgccaaataatt
gtagctcaagtcggctagtaggatctgaaatattcgggaaaaaggcaagcacttcagtca
ggccttctaataaaaaccatttgaaactagctctttgaatttcaactaaatctggaagca
gtttatttttttaagcttacacgttgaatcatagatctcctttatagcaaattctgaaatt
gaagagtttga-tattgtacacaaaataattgtagttaatgcctcaaattgattgactgt
cttgcttactctttagacatgattagaatataatattttattaagttaaaaaatttcta
aatttcgactttatg-----cataaataaagaggttt---------gaactgttgttgaa
accattaagctattttaaaaataaaattaatacttgcctttctgccaaataattgacatt
aagatagacaagtattatttattgtattagttctgttgtttttgagcataggcaaatgct
cgcgcaagtcttgctttttttcgagctccagtgttagggtgaaatgctcctttttttatt
gctttatcaatttttactatatacttgagcaatacttaattgtacatcatctaaattagaa
ttttctaaattatcaatgtttgtaaaacaccttttagttaaggttttaacagccgattta
tacttacggttaatgagacggtttctctccgaggttttaatgcgtttaatcgcagattga
ttcttagccacagtaaatccagtaaaactcttataaaatatttgtagaaacgaaaaca-a
caagttcgcaatatattcgaacaagtaatatgcgccgtattataacattaaaaaaaaaca
ataggcaatagttctgagccagcttttattgttacaagatcaatcttgttattgcataaa
t-tttactcaatagtatttaat-----taaataaa----------agacatatatgaaaa
aaattgaagctatcatcagaccttttaagctcaatgaagtaaagctggcattagtaaaag
aaggtattggaggtatgaccgttgttaaggtgtctggatttggacgacagaagggacaaa
cagaaagatacaaaggttcagagtattttatagatataatagataaaataaaaaattgaaa
taattataagtgatgacaaagtaaataaaattacagaaacaatcattaaaacagcaaaaa

ctggagaaattggagatggaaaaatttttattagtgatgttgaacaagttataagaatta
ggacaaatgacttaaattctgaagcgttgtaggattgacaatttgtttaggcttgatact
tttttaataaacatgagataataaggatctattatatagttataaccagggtataaaa--
ctgatggctaaaagtaaaggcgctcgtatcgtgataacattagaatgtactaacaagtct
atagaaatttcacaaaaaaggaaagctggtgttttccgatatacaactactaaaaataga
agaaatacacccagcagaattgaattaaaaaaattctgtcctaattgtaatcgtcattgt
atatttaaagaaatcaagtagttcttcatattattaaaatacaatcatattatggctata
tatagaaaaaaaatctctccaattaaaccaacagaagctgtagattacaaagatatagat
ttgctgagaaagtttatcacagagcaaggtaaaattgtacccaagcgatcgacggacta
acatctaagcaacaaaaaaaactcactaaagctattaagcaagccagaatactttctctg
ttaccattcctaaataaagattaaatacacatggtctatattaatactatgatgtactag
tattggagtttctcgagggtatagtaagtatgaaattcagttgtatctaatacatctgca
actaatattacatttataaaaaactatttattacaaatttt-ttcacttta---------
-------------------tcagagtcaattt-----ataaataaagtcatacctctttt
aattgattgtattttataattaatttaaaagaggattaatatgcatagc-----ctaaag
atataatctaaatagttcaccaaattcaaaattatcttttacaatgatctatttttcggc
actaaatcaaatgactgaattttatcaccgggttgccatagctgaaattcagcaataaaa
ataccacattcattaccggcttggacttcttgtacgtcttctctaacacgttttaaagat
tctattgttccttgataaactattgtatcttctctcaagactttaatcaaagaattcttc
aataatttgttattagtaactcgacatcctgctatttttctattagctagagaaaataca
gtaccaacttccgcttcaccaactggcacttctgaatattcaggatcaagcagagtttcc
atttttccctaataccttcaattaatccataaattatttgatagttttcaataattaca
tttagtttagctgctgcctgtcttgttcctggggcaaaattagtgttaaaacctattaag
cttgaattagtagttgaagctaattccacatccgtagctgtaatctcacctggcaaaatg
gacaaaacatttagctgtactttactttgaggaaattgggataatgaatctagaattgct
tctgttgatccttgattatctgttttaataattagactaacttgtttattttcgtccttg
gaattaatagtacttaaagtatctaaagtgattcgactatttagagccttttgcttttgt
acaattattgagttagtagactcattttcaatagctttaagcttagcttctttatcatta
ctaactgctaaggctacttcgccggtagctggaacacttgacaaaccccaaatctctact
acagaagatggctctgctaaattaatttttttgcttagcattgttaattattgcccgaatt
tttgcatgagtcattccaattaccatattatcaccagtatgcaatgtgccattttgtatc
aacaaagttgcaacaggcccgtgaaatttgtctaaatgagcttctattattattccttga
gcaagctgtgttggatctgcttgtaaattttctaattctgctagtaatagaattgtttct
agtaatttatcaatgttctcccctgtaacggaactaataggaattactgggactgatcct
cctaatttttcagaaataacattatgtactactagatcattttgaatttcatctacatta
tctaaacttttatccattttggaaattgcaacaacaaaggaaacattagctttctgaaga

tgttgaatagcttctattgtttgaggcttaatcccgtcatctgcagctataataataatt
gcaacatcagttaaattagccccccgagacctcatacttgtaaatgcttcgtgtccagga
gtatctaaaaagacaattttttgatgtttaa------------attcaacttcataagca
gcaatattttgagtaatgccgccaacttctttattagcattatttgatttatgaatataa
tctaataacgtagttttgccatgatcaacatgccctagaattgtcacaactggtggtcga
tctatacaatgatga---ttatctacaataaatgggatatctgaattcctgatctgcctg
tttttattattttggtcatgtgattctactgctataccaaaattcttagctactgatgaa
ataatcgtagtatcaatagtttgattaattgtaactgatatccctttaaaaatagatac
ttgataatatcagtttctggtatacaaattaacttggatagttcttgaatagttaaagga
ttgataagtttaatagattctggtggagaatttggcagaataatagatttatctatactt
aggtctatttgttctttttttttgtcgtgtaattttgttattaccaataacttttttaatt
aactgtgatttgggtttaggcggacgcattaaagatattgctaaatctcctacagtttga
gctgaattagaagtagaatctctaaagttatcattgtcatcatcatcaatatgaattta
gtttttattttt-tttctttgcctattttta--ttttttttagtatccaataaatcataa
gacttattaaagttttatttttttttatctattttgggaggagaacttaactctaagtgc
tgttctgtgttaggtatacctacttcagacttatccatatctaaatttagcaaattatta
gtaacagtagattcaagtctgattttataaataaattgtggatttttgagatctaatact
ttttcggcggagttattacccgaactatatgaagaagctttcttttcaaaattttgatta
tttaagaacataagcgtt-tttttattgcctaatagtaatttttttacttttcgtatagaa
cttaacataaataaaaataccatgact------atggtatcaatatctagacaagtcgat
atttgttttattaaatctaatataacactaaagtcatt--gttgttatattagttaaaaa
aattt-tatgttggcatgcactatattctaatttttcagtatacaatctatttatctaaa
tcttattgttttcat--cgattgtgaaaatacttacttaaggtggagttcagagtaaatt
cttctgaagatctaaaagctttacaattaagtaacaatcgttaattaactccacaccaa
ttatgccaagatctcagaaaaatgataactttattgataagacttttacagtattagctg
atattgtattaaaaatacttccaactagtaaagaagaaaaagaagctttttcttactatc
gagatgggatgtcagcacaatccgaaggcgaatatgcagaagctcttgaaaactattatg
aagctttaaaattagaagaagatccatatgatagaagttatatactctacaacataggat
taatctatgcaagtaatggtgaatatgttaaagcccttgaatattatcaccaaggattgg
aattaaactttaaattgcctcaagctttgaacaatatagctgtaatatatcactatcaag
gagttcaagctgtcgaagagaaaaatgtagaattgtctaaattgatgtttgacaaagctg
cacaatattggcagcaagcaattaaactagccccagataattatattgaagctcaaaact
ggctcaaaacgacaggtcgcatgaaaaatatacaaggatactaatagattcaagatataa
t---aatattaatgaataa---------aaattgaaataattttatcttaaaaaatattg
-taatat--------gtattgcagtaaatag------ttttttaagctaaatcaagacat
cgaaattaaatctattgactatatttatgaatgtaaagaaacaaaatggttagtacaaca

caaagtaccggatctgttactcaaattattggaccagttttagatatcgcatttcctaat
ggtcagcttccgaaagtgtttaacgctttaaaagtacaaagcgatgatggaactatcact
tgcgaagttcaacaacttttaggagataataaagtaagagctgtatctatgagttctact
gaaggactaaaaaggggagttgaagttgttgatactggtgctcctatatctgtgccagta
ggaacagatacattaggacgaattttcaatgttttaggtgaaccagtagataacctaggt
ccagttaattcagaaagcactttgcctattcatagacctgcacctgcttttacaaagcta
gaaacaaaaccatctattttcgaaactggtattaaagttgtagatttattagctccttac
agaagagggggcaaaattggattatttggtggagctggcgtaggtaaaacagtattaatt
atggaattaattaacaatattgcaaaagctcacggtggtgtatctgtatttggtggcgta
ggtgagagaacaagggaaggaaatgacctttacatggaaatgaaagagtctaaagtaatt
aatgaggataatttaaaagaatctaaagtagctttagtttatggtcaaatgaatgaacca
ccaggagctcgtatgcgtgttggtttaactgcactaactatggctgaatactttagagat
attaataaacaagatgttttactattcattgataatatttttcgctttgtacaagcagga
tctgaagtatctgctttattaggacgtatgccatctgctgttggttaccaaccaactcta
gcaacagaaatggggggcattacaagaaagaattacttcaacaactgaaggatctattaca
tcaattcaagctgtttacgtacccgctgatgatttgactgatcctgctcctgcaactaca
tttgctcatttagacgcaacaacagttttgtctagaaacttagctgcaaaaggtatttat
ccagctgttgatccactagactcaacatcaacaatgttgcagcctggaattgtaggaact
gagcattattctactgctcaagaagtaaaatctacccttcaacgttataaagaattacaa
gatattattgccattcttggccttgatgagctttctgaagaagatagacaaacagtatca
agagctagaaaaattgaaagattcttatcgcaacccttctttgtggcagaagtatttaca
ggttctccaggaaaatacgtttctttagaagatgcaataaaaggatttcaaatgatttta
aaaggcaaattagatgatttacccgaacaagccttttacttagttggagatattgacgaa
gctatacaaaaagctgacagtatgaaagattaatataactaataatgactttaaatatca
aaattattgcaccagatcgtactgtttgggatgcagaagcccaagagattattttaccaa
gtagcacaggtcaactaggtattttgacaggacatgcacctttattaacagctttagata
ttggagtaatgagagttagagtagataaagaatggatgccaattgtcttacttggcggct
ttgcggaaattgaaaataatcaactaactatattagtaaatggcgcagaagaggctagcc
agattgacttagttgaagctgaaaaaaatttagatactgcaactcaattacttaatgatg
cttcttctagtaaagaaaaaaatcgaagcaacccaaaatataagaaaagctagagctcgag
tacaagctgcaacagcagcaattgcataa-ataattatttatgcgatgttaaaataaaac
cctagtaataactgttact----agggttttattatttaatttttacgatatag------
----ttggattaacttaagccggaacatatatagtcaaaatacaagcccatttctttacc
agcgtctggccctactaagctaattgttacttctttcatagctaaaattgcttgaattgt
ggcaccaataggaacgcccagagaattatatgtttcttttaatccattgagtacacgctc
ttctaaaatagaaggatcacctgctagcattccataagtagcgtaacgtaagtaataatc

taaatcacgaatacaggcggcataacgtctagttgtatacatatttccgccaggtcgagt
gatatcagaataaagcaaggattttgctacagattctttaataattgttgcagcattagc
ggcaatagtagcagcagctcttactcttagttcacctgtttgaaaatagccccttaattt
ttcaactgaactatcatctaaatatttaccttgaacgtcagctgcattaatgacagaagt
aattgcgtcttgcataacttttaaaaattccttaattttaaatatttgagactatggaga
tatgaaaaaa------taatctatgtattcaagtcagtattattgcattgcacctaaagt
gtagtcaaagtaaaaaccagcttctgcagaatcttcacctgcgagaagtgaacaagctac
gcttttcatacattttacaccttcagcaactccagaaattggtgtacctaaagagttata
catttctttaacaccaactaaaccaatttcttcgattggtgtgacatctcctgcaactat
tccataagtcactaaacgaagatagtaatcaagatctctgagacaagtagctgtcatttc
ttctccatacgcattaccacctggggaaacaacatcaggtctttttttgaaataattgttg
accaccttgcttaacaatgcgttcacgattatccgttaaaatttgagcaattcttagacg
acgttgtccagaaagaacaaaacttttaattctgtctagttcacctggacttaaatatct
cgcttctgcatctgcatttacaattgactttgtaactatactcatggataaaactcctg-
-----ctt-aaaaaaaagcttaatactaattatttatgatataactcagataatattgaa
cctttcgaaaaaatttgtagctacagtatataaatgatgtatactttggcctcaacataa
aggtaagtactaaagcgataaagctgaactcgtaatctaatgttattgattacctagtac
ttttttgaaactaggtacaacaattgacacattttgcttagttagtctattatataaagt
ttctgtattaggaaagttcgcagcaggtaaagttgggaatcttctatatggaacagtatt
tgttccatatacagaattatattcagagctattaactagagtttcaacaaaggctgataa
accttttgatgctaaaatttgattaaagaatcgtatttcagcctggttatttggggctct
tccaagtatatgctttgtacctagctctataacttttgtatttgggtatggctggtaaaa
ttctttaccatatagttcagacaatgctaatttttgtattaattctttaacattaatctg
tttatttaagaaagataattctatatctaaaaactctccaccaatactgaatgaattaag
atctctttcaaagatttgacgataggcagctcttaaagcttgttctaagacttcgtgcga
actattttgatttacgtcaaatataacagattgatctcttagaggagtaactccttgtgc
aatacgcgactgaatatcattactactacgtatttctttaactgtaccaagttcaacaaa
cctagaagatttgctactaataactttttaaatctttgatcaattatgccaggtcttaa
acttcttaatgcaataccagctggcgtagtgtatctttcgtatggtacaatgttctctcc
aaatgtttcaatatattcagaactatcaattatcgcatctacaacttgataatatccctt
cttataagcaagatcaaaatatttattaatctcttggcgaccatacgtaggtctccccaa
taaacggttatgaatatattcaatcgctttgcaaatatataaaggttcccagtaaagcga
tctaaatatgctagattttgccaactgcctaatgaattcgcgaacagaaatttgactatc
tttcagttgaccttcaattggtttgagaattaatttttcttcttgatatacttctctacc
gaaaactcttaaataagcaacttttgtaataacatcaactgaattttcaaactttgcgag
tgagtttgttggtacaactgtattggataatttgaaaatttttggacctaaggagccggc

agatttgggtcttacttgtggattactaatttggttatagattccaggaccctgcctcac
tagaattcttctagtatctttaccaaagatagcttgtcttttccttggatctttattctc
tttaggaaagatagctccaaactgaatggataaaggatcgttaccagttccataagggtg
ttgatctggtagtgattgcttataatcactgaataaagttataaattgcggtactttacg
gaaaggagcactataatttaatagatctatttgaggaccccaattacgacattcttgtgg
ctcttcacctaaatttctaaaataaggaactgtttcttcaccaaaataatctgtatattc
agctgaatttaaaattgcattcacaagcccacttaaaccggtagaggacaggatggcaaa
atatttttgaaattcctctagagaactaggtcctctaccaagaaaatgcctaaatgctaa
ttctaatgctctactatttacaaaaggctcatagaactgttttctgtaagtattagatgt
acctaatgagcgaataaattctttaattgaaatttgaccgttctttacttgagattctaa
atttgatagtgaaagattatatgctttagcaatatctctctcaaaaatttgcctataaca
agctttaacaactagatttttttcatcagccgataaactagatttcattacaaaccgagg
agttgaaacacctgcctgagaatacgtttgagggagacgcaaaccctgcaaatcattaga
ttctctttttcttaatttatcagtcaaagatggagcttcaaattcggaaataacaacatt
aaagtattctcggactaagtcttgacctttaatgtcctcttcaaatattgctaaagcagt
gcgtctcatctctcttaatgcaacaattgctgctgcacttgaacaagcattgtcaatcag
ttctctcaagcctctaatattaactgagagaatattagggtcacctgagacaatggcata
ggttagatatcttaagaaccaatctaaatctctcaaagattttctcattcttgtagtacc
ataacgcaagacattaattggcttaaagcccggaggtgttgcgcctcctgcattaaataa
tgaacgaaaaccttgtccaaaatcaccctgaatattcccagatagctcattaattttatc
tctagaactttggtctccagcaataataactgctgcttgtggtctctctaaataagatat
tgcggaacctcctacaaatatcttatctgctgctcgcgcgacaagaatatttgcattttt
agttaaaatatctgcaacctctaatctttttttgaccagaatttaagaaagagactaattg
gtttaattcaccaagctgtaaaaatcggtcttgctgctctgcttgagtaattgtcaaaat
tgaggcagtccgatagagctgggggcgtgctaatgggcttccaccacttgcttttatact
cattgttttattcatctccttaacttattttactgacagactaatacagataatattttc
ttcttctatgcctggaaaactactatgtaaagaca--tttttctaaaatgttaggtact
aatatttatagtatagatagtctctcaactaccttctataaatacaaattttgtaataat
tctattaacatttatgaaataatta-aaaagcagtgatgcttttgtattgacctaagact
aatcaagtattgatattaaatagagtatactgaataaattgcacagatcataaagtc-aa
agatttattctgcgcaagctagttaatgaagtaaattattagatatccaataactataaa
attattattctgattcatgcttttacactagttctctatcgtctttaaaataaataatct
taataagcacttggtatgagtttagaacaacaaaataatcttttagctaacataaatgcg
aatccagaagacttacctgaaaatgatgtaccaatgtccattacagagcatttagaagag
ttaagacaaagaactttatttgtctttattttttttcttgttagctgcaacacttagcttt
acacaaaccaaaataattgttgaaatatttcaagctccagctgtcggtattaaattttta

caattagcaccgggagaatattttttttcatctattaaaattacaatttactgcggcatc
gttgcaacaactccttttggggtatatcaagtaatattatatatattaccaggtcttact
atgaaagagaggaagattattttacctattttaattggttcaatagttctgtttatagta
ggaggaatctttgcttattttattttagctccagctgccttaaacttttttaattagctac
ggtgcagatatagtagaacctttatggtcttttgaacaatattttgactttattttgctt
cttttatttagtactggtttagcatttgaaattcccattatacagctactactaggtata
tctgggatcgtttctgcgaatcaaatgttgcgagcatggagatatattattattatatca
acaattgtaggagccatcttaacaccttctaccgatccagtgacacaaattatcatgtct
tcggctgttttagctctctatttcagcggagttctcatactatttctcttgaaaaaatag
cttcaaa-----------------------------------------------------
--------------------------------------aagattaaa-------------
------------------------------------------------------------
------------------------------------------------------------
------------------------------------------------------------
------------------------------------------------------------
------------------------------------------------------------
------------------------------------------------------------
------------------------------------------------------------
------------------------------------------------------------
------------------------------------------------------------
----------------------------------ttaaatcaggaaactcagttagcaat
tattgtaatttatatcgacgtaaaaagtaatttattacaagttttatttatatacatact
tattcattatatagttataacagtaagtctaattaatagtttttttacacttaattaaagt
aataactcttaattcttttggcagaaagatctgtcttaatattttatcccaaaaaaacag
-aaaaatgaagctaaatagtctaattaacttaattcaaaagtctatatgttcttgtacgc
ttttattaactattttaaatattatttgcatagcacctaattctagcaatgcatttccta
tctatgctcagcaagcttatgaaaatcctagagaagcaactggtagaatagtttgtgcaa
attgtcatttagctcaaaaacctgtggaaattgaagctcctcaggcagttcttccaaata
ctgtattcgaaactgttatcaaaataccttatgaaaatactgctaaacaaattttaggca
atggaagtaaaggcgggctaaatgtcggagcagttgttatattaccagaaggatttaaac
tcgctcctgccaataggttatctcctgaattgaaagaaaaaacaaaaaacttatacattc
agccatacagtactaagcaaaataatattttagtaattggccctattcctggtgacaaga
ataaagagatagtatttccaatattatctccagatcccgctaaagataagcaagctcatt
ttttcaaatatccaatatatgttggaggcaacagaggaagaggtcaaatttacccaacag
gagataaaagtaataacaatctcatctctgcatcagctagcggtaagatcaataaaattg
aggctcttgaaaaaggcggctttgtagttcatatagctagtactaaagattccgaagtga

atcaaaatattccagcaggtgtagaattgaagattaaagaaggtgacaatgtacagttag
atcaagctattagtaaggatcccaatgtaggtggttttggtcaaaatgaaacagaaattg
ttttacagagtccaaatagaattaagggcatgattgctttctttttcgtaagtgtattag
cacaaatttctttgttttaaagaaaaaacaatttgaaaaagttcaagctgcagaaatga
acttttagtttaaacaaatactggcacttactataagtaaattatttaaaactttactaa
tggcttgtaccagtttttgtattatgtagtgactatgtttgaagaattgatcagattttt
aacagcatctataatttgatgaggttgaataactgtcgcttgctctaaactaccattata
tggtgtgggtatatcttgcgaagataatctcactaccggagcatctaattcatcaaaaag
atgttcattaatttgtgcaattagctctgctcctattccagctgtcttcatacattcttc
tacaatcaaaactcgatgagtctttttacagatatagaaattgactctatatctaatgg
ttttaaagaaataaggtctaagacttctggattataaccgtcgtttaacaatactggtaa
tgcctgaataacatgatgtctcattctagaatatgtcagtatagtaatatcttcaccttt
acgaactatttcagctttatccaaaggtattaaatattcttcttgaggaatttcttcttg
cagattatacagaagaacatgctcaaaaaatacaactggattattatctcgaatcgcgga
ctttaataatcctttggcattataaggcgtcgagcacgcgacaatctttaatccaggaat
agcttgaaaataagcttccaatctttgagaatgctctgcacctaattgtctaccaacgcc
tccaggtcctctaatgactaaaggtaatgtaaaattacctccagaagtgtagcgcagcat
tcctgcattattggaaatctgattaaatgctaataataagaaactcatattcatgccttc
aacaattggtctgagccctgttatagctccaccaattgccataccagtgaaactattttc
tgcaattggtgtatcaagcactcttaaatctccgtacttgctatgtaaatcttttgttac
tttataagaaccaccgtaatgaccaacatcttcacctattacacaaacagtcagatctct
ctccatctcttcatctgttgccgctcttaaagcgtcaaacataaaaactttactcatttt
aatatttag-ttttataatttcacaaatacctaaa-------------------------
-tttaattatctgcaaaaagataacgtttaagttctaagatattaggttctggactagac
attgcaaattcaactgatttttgcaggtctatctttactgcgctctggatatcattcagc
tcgtttaaatcagctattttattatccaaaatatatttttaagtttttaataggatct
cgtgcaacccaagcttctttttcttgtcttgaccgtaattcatcaggatcggcaagagaa
tggcccctaaatcgatatgttagtgcttctattaatgttggaccttgaccttgtcttgct
cgttttactgctttttccgcgacttgacgaacagctaatacatccataccatcaacttca
acgccaggcaaaccaaatgcttctgctttttatgtatctctggtattgatgatgaacga
tgatgtgccatccctatggcccactgattattttcaacaacaaatatgataggtaacttc
cacagaactgccatatttaaacattcaaaaaattgaccattattagttgtaccatcacca
aaaaagcaagctgtcactcttaattctgctggttcctttaatacttgttgtctatagata
ctttgaaatgcagcgcctgttgctattggtattccttcagcaatgaatgcgaatccacct
aagaaattgtgaggggcggaaaaaatatgcattgacccacctcgtcccttactacagcca
gtttcttttccaaagagttcagccattacattttgagaaggcactcctttacttaaggca

tgaacatggtctcgataagtgctacagacataatctgttgaatccaataatttgatcaca
cctgttgacactgcttcttgaccattataaaggtgaacgaacccgaacatttacctttta
taatacatctgagcgcacatatcttcgaaatttctgcctaataacatatcctcatataag
actagtagcttcttttt------attaagatttatttgattatagttagtcagcggcagt
tcaactttcttaggataactcatgatttataaagcaacctctaatctaattgtattatat
ataagaattttcaagtcgaagtcttgaataagaaataaacaaattaa-----acaaatta
ttgcatacttttttactaagaacttttgtaattctttgaaactagtttagtatatcaaatt
aattttgcttatactacagatgtttaaaagtcgaataaaaaagtactgatccag----ta
tcagcataatag-ttaaaataaatataaatataagtaaatcgaattctc-tcaaaaaagt
gactattattaatagtaaatcacttaatatttctatatattattaaataaatggcta---
catcatctttatttcatcctatcgaaaaagaattatgcactcttgagcgtaatcttcagg
cagtagcagggacacgtcatcctatttacacgcagccgcaaaacatttatttgaagcgg
gaggcaaacgtttgagacctgcccttgtactgctagtagctaagtcaacatctgaggagc
aagaagttaaccctggtcaaggaagactagcggaaattacagagattatacatactgcaa
gcttagtacatgacgatgtcatcgatgagtgcgaaacacgaaggggagaaagaactgttc
ataacttatttaatactaaaattgctgtactagctggagattttctttttgcacaatctt
cttggtatttagctaacgttgataacttagaagtagtgcaaatagtttcaaaggtgatta
cagactttgccgagggggaaataagacaagggttggttcattttgacccaggtatttcaa
taaataattatactgaaaaatcattttataaaacagcttctttaatggctgcaggctgta
aaggagcagctatgctgagtggctcaagttttcaaatgcacaatgagctgtatttatatg
gtaaacatatggggttagcatttcaaattgtggatgatatattagatataacaggatcta
ctgaaagtctaggtaagccatctgggtcagatcttatgaacgggaacttaacttctccta
tactgtttgctttaacacaagaagatgaattaaaccaattgatacaaagagaattttgtg
atgaaaaagatcttgccttagccttgtttttaattaaaaagagcggcggaattactaaag
cgaaagatttagcaaaagaacacgtgcaggcagcacttcactgcctacagcttttaccaa
aatctgcacctgcatctagcttaaaagagttaacacatttcataatcacaagattgtcgt
aaaatacttgctagaactgaaaatgttttaactactttttgactgtctctaaaacagctt
gaaaggtcttaatgtcattgctagctaactgtgctagcattttacgatttaatgcaatat
tatcttttttcaatgcagtaataaaagtactatagttcataccttgactatgtgctgccg
cattaatcctagttatccagaggcgacggaaatctcttttttttcttttgcgaccaacat
aagagtatcttagagctttgagaacctgctgcttcgccgtcctaaataaagaattatgtg
ctcctttgaagcccttggcgagcttaaatattttagttcttctttttcttgctacattac
ctcttttaactctgctcataaaatattac-ttaaaattgaattacctacgg-ttattata
tctaaacacttttta----attataaataaggtagattaatagagatattctttttatct
ctgtcatcaacctgacaggtagaggaaagatgtcttctctgcttcgaagacttcttctga
agtaaatgactttttgacgctttatgccgaagaaattttcctaatgaagaaactttgaat

cttttttgcaattgcttttgatgtttttagcttaggcatattataaacttgtaattattag
caaaagtatgaaaagtctaaaaata------ttagacttttcaatacttcagataatctt
aacacaaattaaattacttactacaggaatttgatgaaacaatttatatgt--agtagaa
aaaatttaatcagcagtagctagtactg------attttgatttaaattttttgtacagta
tttacaattatcattgtaattatttttaccatactggaatttaactcttcaaacattttc
atatttaaagtaaatgagatattagcttctgagacaatattttgaacctgatcagaagac
aaagggattgtgtctaaagcagatcgataactattttttaaaaattttatcatcttctatt
tcgtcaaaatcataaaattttgtacctctctcatcagagagattcattgctcctctagct
atttttttaatatttgaccgccggacaggtctccaagataacgcgtataggcatgtgca
actagtaattctggttgcttatcaccaatattgtgaattctatcaacatatactttagtt
gcagaagatggctcaataatatttaaccaatcggatccataataataatttaaatcttca
gccaaactaatttttctatttaattctgtgaaatatattggctggatagctggatgatct
ttatttaccagcatctcttcctctatagcacaatacacaaaatataagttagcaactaat
tgacgatatgactttttatctacaactccacctaaaaaagacttcacaaaactaacattt
tctgccatgctatgtgactttgttgttccttctcttagttcattagctaagctactaacc
ataatattaattttccatgttataaagataactaagctatattgatagtatgttatctca
gtattta------attttcacaaattatgcagccgcaaaataatttttagattttatagg
atctggattcattgttttatcaccaggtttccagtttgctgggcacacttcatctggatg
ggcttggacatactgaattgcttgcaaaactcttaaagtttcttctacacttcttccaaa
ctcaagattattaatagtagaatattgaattatacctttaggatctataataaataatcc
tcttagggctacaccagcattatttaagacgttataggcaatactgatttcttttttgag
atctgataataaaggatactcaagatctcctaatccaccagattctcgatctgtttgtaa
ccaagctaagtgagaatattcactatccacagaaacacctaaaacttctgtgtcaagttc
agaaaaagcagaatacttatcactaaaggcagttatttctgtggggcaaacaaacgtaaa
atctaaagggtaaaaaaataaaataacatacttattttaaagtcggataattttattgt
tttaaattcttggtcataaacagccgtagctgaaaagtcaggcgcgatctggcctacttg
aagacaattttgtcctgaaatcattaattttgtccagaagaattaatataattttgttat
ttaatagattattatataacaaaatttagtatatatgaaaatgataccagtctttttaaa
attatcaaatccatagattatataat-aatactattttatagcggggaatggatttgaa
ccattgaccttcgggttatgagcccgacgagctaccagactgctctaccccgcgaatata
tagaataatagtagtctattcatagatataaaagcaaatataagttctattttaaaaata
aaagaataaaatagcaggtgcagtggtcattagacttaaacaagataaagtattttttaat
tttatttaataaaggaaaaacttcatataagcctatgaatcgatctttcactaatatttc
tgacggttttacccaaatttggccatcataccaccctgattcttcatagaacacagtagc
gctcattaatctcttaacaacataagaccatcctagatacagtctgatcaaaataaagca
tgttatcaaactagaagttatgaattctgaaaagaaaaaatttcaaaggcagtttagcaat

tggaaaaattgaaagcaaaataggacttactaagaaacaacttacaataagagtaatagt
aattttcttattataagaataactactcaaagttggccagcagaagaaccaagaatttttt
taaagaattatattcattgactggttgttgctcaagaggcacagggcattgattattata
taaattcattgaagttaaaatttatttttccaaaaat---ttatttatcaagtttaaaca
aaagtaaa--------gttaattaattgcaataacggtagtcaataataagaataaaact
agaaaaagccaagttacttgatttaaagtattttcagtgctacgagtattactaaagaac
tgattttgagagcctatagtacctaagccttcagatttaggattgtgtataagtatacta
aagatcaatattattgttgatgcataccagagaaattttaaaacttgttccattttttaag
aaattaat--aaaataatacaaagacaatcagtttcaaaattttgattgtcttattagag
taaataatatttactcaccttgtacttttaagagcaaaaatcctactgctagtcctacta
agactaaaacagaacttaaaatagcactttctataatctctttgcccatttttagttttc
cttattatttt--------tgaaattttatatacaatattacagcagtat---ttagaat
ccgtttctaccccagactaccatagcgatagagaaagtaaacattgccatcaaagccgcc
caacccaaactcaaaatatccatcatctataatatccaaaaaaaaataagatctctttta
cttaaaatataatattcttaatattaagaatatcattatataataagctaattataacag
agtaatacttaattcgtattttttaatgcatttagaaaagattaga-tttagaataaataa
taaatatatatttagtcttttaatctatatcgaataacattactttctattaaaataatt
aaatagaaatatacataacatt----------------ttttaatacgaatcattgtatg
aat--tagtatacagataaataatatgtgtagcattgatctgcttag-----cattcta-
----------------aaaaaggaatttactgcttcaattatgcaaacaacaatacagaa
agaacaagtatctaacaaggaaactttattaacgcctagattttatactacagattttga
agaaatggctagtatgaatatttctgaaaatgagcaagatttttttgctattttagaaga
atttagagcagactataacagtcaacacttcattagagacgaagaatttaatcagtcatg
gtcgaatttagagaataaaactaaatctttatttattgaattcctggaaagatcttgtac
agcagaattttcgggatttttactatataaagaattatctcgcaaattaaaagatagcaa
tcccattattgctgaatgctttttattgatgtcaagagatgaagctcgtcacgctggatt
tttaaataaggctattggggatttcaatttatcattagacttagggtttttaacaaagag
tcgaaaatatactttcttttcgccaaaatttatttttttacgctacatatctgtcagaaaa
aataggttactggcggtatattactatataccgccatatggaacagtatccagaacaccg
tatttatcctatttttaagtttttttgagaactggtgtcaagacgaaaatcgtcatgggga
tttttttgcagcattattaaaatctcagccacactttctaaatgactggaaagcaaaaat
gtggtgcagatttttcttacttagtgtatttgcaacaatgtacttaaatgactttcagag
ggtcgattttttataatgctattggtctagactctagacaatatgatatgcaagtaatccg
aaaaactaacgaaagcgctgccagggttttccctgttgcgctagatgtcgacaatccaca
atttttcaagtatttagatagttgtgcttgtaataacagagctttgattgatattgataa
gcaaaatttctaccgtttgtaacaacttttatgaaagtacctttatacatgtcccttat

gttgaacttaatcaaaatatacctgattaaaccaatagattcacaagctgtatggaatac
agtaagataattaataaaaatagacaag-------------------atacatagaatta
tataaaat------aattgcttatgtcttgttaaaatggaaaacacccatattatattaa
tcat------gggtgtttatatagtttatacagttaggaacttattttttcttttgcctc
atacataacttctaaagtgattaagtttctagaacaatctcgagcaaatttttcagtatt
tcgttttactttgcctctcacaaaacctggtattttacttagttctttttgggcttcatc
tgaccaattaagactttcaccagctgaaataccgatacttagagcttctgttgtatcatg
tccgccaaatatttctaataggtggtcttccatacctaaagtaaatgagttataaactaa
atcagctatttgatttgtaccttcatatccaagaaatgggcgatagcttaatgggaaatt
ttgaatatgaacaggagaagaaataactccgcaaggaatatttagtcgtttaccaatatg
ccgctccatttgagttccaaaaattgcagctggctctgtcttagcaattaaatctccaat
gagaccatgatcatctgagataatcacttcatcacaaaccctttgacttgttccctaaa
ccattcttcatcatatttgcaataagtcccgcaccaagcaacgtgtatgcccatttcttg
atgcaagattctagtcattgcagctgcatgagttgaatctccaaaaacaattgcttttt
gcctgtcaaattttggcaatcaatagatctagaaaaccacgctgattgcgaaataaatct
agtttgttcatctatatatttctcataattaacgtttgctcccagagcattaacaagctt
ctcaatagatctaatagaggtagcagtttgaacaatacccattggagtaatatcaatata
tggcatattaaattctttcttaagatacaaagctgtcattagacctgtttctcgataagg
gacaaagttaaaccaagcagatggtagtttttttagatcatggacagaggcattttcagg
tattacctgatttatctgtatatctaaatcttggaataatctttttagttctgagatatc
atgctgattatgaaatcctaaactaacggcaccaataatattaactgacggctttatagt
ttttttagttaagat---attattgcatttcgcttttttccatataaaagtaacaatttg
ctcaagagttctatcactagcttgaagctcattaactctataatgattaacatctgctaa
tagaacgtctgcttctgtttctattgaagctcgactaacaaaattttgcagatcttcttg
tagaatactcgaagtacaagttggagttaatataactaaatcgggtttttcttctctatc
ttttcttgttatattttctactactttttcttgagaaccccctagctaatacatgcctatc
tactacactagccgtaacaggagtaaaatctctatctctttcaagcattgaacgcattac
attaaagtaatcatctccaagaggtgcatgcataattgcatgcacgtttttaaacgaact
agcaactcttaaagttccaatatgagctgggcctgcatacatccaataagctaatttcat
attaaatttccgtctgatttatgtaaatatagtaatactttagatttagctaacaaata
aaaaccactcatatattcttactctgtaatgatttttaata--ttttcagatacagatct
tcatcattaatatttaag--------------ctagaaatcaa-aactaacataccaatc
ttaaaagtaaagatttcttgcagttatagtttttctttcagtaaacatgtaaaaattgta
ctatatataaatgtactatatataaactttttttacttaatattggtaatgccagataca
attaacttaaatatgccttccccaacatttggcggaagcactggcggttggttaagggcc
gcagaagtggaagaaaagtatgcaataacttggacgggcaaaaatgaaagtaaattcgaa

atgccaaccggcggcactgcaacaatgagagacggagaaaatttgctatatttagccaaa
aaagagcagtgtttggccttaggtactcaattaaaaggaaaattcaaaatttctgattat
aagatttacagagtttttccaaatggggaagtacaatatttacatcctaaagatggggta
tttcctgaaaaagttaacgcaggtagagatagcatcaatagtgttgaccattctattgga
aaaaacgttaatcctgtaaatgttaagtttactaataaagcaacatacgactaactaata
-tgaaacctagacatacaataactttatatagtctaggtttttgataatatatattatc
agcatggaagggtggccgagtggttgaaggcgtctgatttgaaatcagttgaaccgatta
cggttccgtgggttcgaatcccaccctttccgttacatatctgttatataagagacggct
taattattcattaaaa--aattatacactaatagtgtttagctttgatttaatgaaatct
acaacctctgaaagattcgaaatttttctgcatcttcatcaggtatctcaattgcaaac
ttttcttcaattgccattactaattctactgtatctaaagagtcagctccaaggtcactt
gaaaaattagcttctctagtaacaattgtcttttcaattccaagttgttctgctacaatg
tcttgaactttttcaaaaatttcattatcttgcataatatattgtttcctaaagtttgtg
taaaaaatttactccccccttgctgctaac---cttatataagttagtagactgggctca
aaattcaaatatt------tatatgatatactattattctggataaatgcgaagtttacg
atagggctcttctccaaagctacgagctctcaactgataattgtcaactaaattatgctg
catttttcggatgtaagctgatcgtggcattaattgaactattgaattttcatttaagat
aatagtttctattgctagtttagcttcttgaagagcttgaatctcataaaactttttact
tttacaaagttctacccaattaagatcagaagaaatattgatatttaatatctttcttag
tgcacgtgtaatttgtggtaccgtactattttgaatcgtataaataataagttgccttga
ttttgcaatttgtcttagttttgcatttttgtttgacttgactacgtaaggctaaaatagc
atctgacttttcaagctcttttgttagaataattggcagatctaaagaagagatgacaga
tgcaatatgctgccaacttaaagaataagcatataaatattgaatcgatgcttctactgg
aaaagactgattattaatatttataggagtattaatactagttgataataaagatgtatc
acgatttctaatttattcaaatcagaagttttatcttttacttctttttgatgccaaac
atgagttttctgcttcaagctactcatttttttgaggtcttaaaaaaatttgtagatagcac
ttctatggaatgcgaggggtaacacttaatcaaaatgcggccattatcttgaatttgtcg
cttttggacaaaaggctgatgaccttgtaaaatttgatctatggtttgttcaactttatc
atggactatccaaactttctatcatgtatttcaattgcaatttgaaatgcaggtgcagc
ttttctttctaatatactttctgagtgcctcttcttttagcttcatcatcacctaaggt
aacatactgaatgccgccaataagatctgctaatgtaggatttttaataaggctttctaa
atgatttccatgggcagtacctactagttgaactcctctttctgcgatagttcgagctgc
taacgcctctaactctgtaccaatctcatcaataataataacttcaggcatatgattttc
taccgcttctatcattacttgatgctgtaaatcgggtcgagctacttgcattcttcgagc
tcgacctatagctggatgcggaatatcaccgtcacctgctatttcattagacgtatctat
aatcacaactctttttttccatttcatctgccaaaactcttgctatttctctaactgctgt

tgttttacctacaccgggttttcctaataataaaattgaatcaccttgttctaataaatc
tctgataatactaatggtgccaaacaccgctcggccaacacggcacgtcaagcctataat
gctaccttccctattacgcattgaactaatgcggtgtaaagtttttttcaatgcctgctct
gttatcaccactaaaattcccgaccttcttcacacaataatctaaatcctgccaactgat
agacctttgagataaatattctgggtttcctggaaatctggcttctgggcgacgacctaa
atccataacaacttctattaaatttttcctattagaatgctgcttcaaaggttctcgaac
aaagttaggtaaaatttctaataatttttctaagtcgtcagcaataagcatgatccaata
tataataattagttaatcaattaaatgatctaatattctagaaagaatactagat----a
ataatattatg-----------ttttatatttattttacacaaaaagatatcttattatt
aagattgtactgaatattaaataataaatagattggtcaaaatacatagtcatgttaata
tgcttattggacaaaaagttcaaattaagtatagtaaaaacaaaatagctaatgatatgg
ctgttaaagttggtgatataggagttattagagggacaaaatttataaataatcaatgcg
tcacaattattgtagaattcgagaatcatacaaggctgtggatgttccgtgaagagctaa
tctgtctgaataaataaattattaaaaataaagaca------------------------
aaatttttatccattgttaaagcaatttatttcataggagtatttttagttgtgcaaatt
actattaaagatttacaagatttactttcttccgtacaaagaaaaaaaaatacagacacta
aaactaaaacaaaataagtttgagattatattaaacaagccatccaaaaaagttcctcaa
gaagttttgccatcaaacagttcttctatctttaagtcaattcagtctaataccatcaac
atctcgcctaaaaaaactagaag---tataaataacaaagctgtcataaattatgcaact
atagtttcaccaatggtgggtactttttatcaatcaccagctcctggtgaaaggattttt
gtgcgagtaggtgatgaagtaaagtgcaaccaaacggtctgcattattgaggctatgaaa
ttaatgaatgaaattgaagctgaaattgagggaacaattattgaaatcttagttaaagat
ggtgatatagtagattgtggtcaagctttgatgaaagttgaaacttaatgaaaattcttc
cattttacatattattaacagaaattatttagatagagaatttagcttataatgtaatta
taaaaacttatttcctaaaggcac-aaaaatctaaacagtaaaattattagatttatctg
tcaatcatagaaataagcgttttgattccttgtctcaagagaggagaatctcaatggcaa
ttagctcaaaagagcaagagacaaagaaggtaaaaatctcggttgataaaaatcccgtag
atacttcttttgaaaaatgggccaaacctggtcatttttctcgtacactagcaaaaggac
caaaaactactacttggatttggaatcttcatgctgatgctcacgacttcgatagtcaaa
ccagttctttagaagaagtttcacgcaaaatatttagtgctcactttggccaacttgcgg
taatctttttatggttaagtggaatgtacttccacggagcccgcttctctaactatattg
cttggctaagtaatccaactggcatcaaaccaagtgcacaagttgtttggcctatagtgg
gtcaagaaattttgaacggagatgtaggtggagggtttcaaggtgtacaagttacatcag
gatggtttcaattatggagagcttctggaattaccactgaattccaactttactgtactg
ctataggtggtttagtaatggctgctttaatgctgtttgccggatggtttcattatcaca
aggctgctccaaaactagaatggtttcaaaatgttgaatccatgatgaatcatcatttag

ctggccttttaggcttaggctgtctaggatgggcaggacatcaagtccacctatcattac
ctattaacaaactactggattcaggcgtttctcctcaagaaataccattgcctcatgagt
ttttaatcaacagagaacttatggcacaactatatccgagttttagtaaaggactggtac
ctttcttcactttaaattgggctgaatactctgattttctaactttcaaaggaggcctca
accctgttactggaggtttatggttaagtgatactgctcatcatcatttagcacttgcag
tgctatttcttgctgctggtcatatgtataggactaattggggggattggacatagcatga
aagaaattttagaagctcataaaggtccttttactggaaacggtcacgaaggattatatg
aaattttaacaacttcctggcatgctcaacttgctattaacttagcaatgatggggtcct
taagcataattgtagcacatcacatgtatgcgatgcctccttacccttatattgcaaccg
attatcctactcagttatcactatttactcatcatatgtggattgggggattctgtgtag
ttggcgctggagcacacgcatccatttttatggttagagattataatccagctgaaaact
acaacaatttattagatcgagttattagacatcgggatgcaattgtttctcatttaaatt
gggtgtgcatattccttgggttccactcttttggtctttatattcataatgatactatga
gagcactaggaagatctcaagatatgttctcagatacagcaatccaattacagccaatct
ttgctcagtgggttcaaggtatacacactttagcacctggtaatacagctccaaatgctc
tagctacagctagctatgcatttgggggagaagttgtttctgtgggaaataaagtagcga
tgatgcctatctctttgggaactgctgattttatggttcatcatatacatgcatttacta
ttcatgtaactgttcttattttagttaagggattccttttctcaagaaattctcgcttaa
ttcctgataaagcaaaccttggatttagatttccatgcgatggacctggtagaggtggta
cttgccaagtatccggatgggatcatgttttcttaggtttattctggatgtacaattctc
tttctgtggcaatctttcactttagttggaaaatgcagtcagatgtttggggtagtgtat
ctccatctggcaatgtatctcatattaccggtggaaattttgcacaaagtgcaatcacga
tcaacggctggctaagagattttctgtgggctcaagcttctcaagttattcaatcatacg
gatctgcgttatctgcatatggactaatattcttagcggcacattttgtatgggcattta
gtctaatgttcttattcagcggtagaggttattggcaagagttaattgaatctatagtct
gggcacataataaaataaaggttgctccggcaattcagccaagagctttaagtattactc
agggtcggggctgttggtgttgcacactatctattaggtggcattggtacaacgtgggcat
tctttttagcaagaattatttcagtaggctaatattgaaaataggat-aaaaaaacaatt
atggcaacaaaatttcctaagtttagccaagctttatcacaagatccaacaactcgaagg
atttggtatggtatcgctacggcgcatgactttgaaagtcatgatggcatgacagaagaa
aatttatatcagaagattttcgcttctcattttggacatttagcaattatctttttatgg
acctctggaaatctattccatgtagcatggcaaggcaattttgaacaatgggtgttaaat
cctttaaaagtgaagccaattgctcacgcaatttgggatccacattttggacaaccagct
cttaaagcattcagtaagggtggatcagcttatccagttaatattgcatactcaggtgta
tatcattggtggtatacaatcggcatgagaactaatcaagacctgtatacaggagcttta
ttcttattagtcttatctgctgtattattatttggtgggtggctacatctgcaaccaaaa

tttaaaccagggctatcttggtttaaaaataatgaatcaagattaaatcatcatttatct
ggattgtttggtgttagttctttagcttggacaggacacttagtacatgttgctatacct
gaatctagaggtcaacatgtaggatgggataatttcacaactgttttacctcacccagct
ggattacaaccatttttttagcggtaattggagcgtatatgctcaaaatccagatacagct
caacagctatttggtaccaatgaaggtgcaggaacagctattttaacatttctaggagga
tttcatcctcaaagtcaatctctatggctaactgatatggctcatcatcacttagctatt
gcagtagtctttattgttgctgggcatatgtatagaacaaattggggtattgggcataat
ctaaaagatattctagatgctcacagacctcctagtggtagattaggcgcagggcacaaa
ggactatttgataccatcactaattcattacacatgcagttaggattagcactagctgct
cttggcgtgattacttctttagtagctcaacatatgtatgctatgccaccatatgctttc
atggctaaagattttacaactcaagcttctttatacactcatcaccagtatatcgcggga
tttttaatggtaggtgcatttgcacacggcgctatatttttttgttagagattacgatcca
gaacaaaatcaagggaatgttcttgctcgcatgctggagcacaaagaagctattatttca
catcttagctgggttactttattcttaggattccacacattaggtctttatgttcataat
gacacaatgattgctttcgggactcctgaaaagcaaattttaattgaacctgtatttgct
caatggattcaggcttcttcaggtaaagcactttatggattcgatgtgctactatcgtct
tccagcaatattgctacacaagcggggagtaatatttggctaccaggttggctagaagct
attaacagtggtaaaaattctcttttcttaacaattggtcctggtgatttcttagttcac
cacgcaattgctttaggattacatactactgcattaattttagtcaagggagctttagat
gctaggggatccaagttgatgccagataaaaaagattttggttatagctttccttgtgat
ggacctgggcgcggtggaacctgtgatatatctgcgtgggatgcattctacctggcagta
ttttggatgcttaatacaattggctgggtaactttctattggcattggaagcatataact
atttggcaaggaaatgcaactcaatttaatgaatcatcaacctatttgatgggatggttt
agagactatttatggctaaactcttctccattgattaatggttataatccctatggtatg
aataatttatctgtttggtcttggatgttcctgtttggacatttagtctgggccacagga
tttatgttcttgatatcctggcgtggttattggcaagaattaatagagacattagcatgg
gcacatgaaagaacacctttagctaacttaatccgatggaaagataaaccagttgcacta
tcaattgtgcaagcaagactagttggtttagcacatt-tttctgtaggatatgtacttac
ttatgcggcctttgtactggcttcaactgcgggaaaatttggttaagttaaggtaatttt
taattaatc-------aaaaaagtcagtcataaagtattttatgactgactttttgatt
aaatttcaaaga-ttattaaagcttaacttctatatctactccagatgggagatttaact
tcattaatgcatctattgtttgagaagaaggttgatgaatatcaataattcttctgtgag
accttatttcaaagtgttcacgagaatctttatctacatggggagagcgtagtacacagt
atattcttcgctttgttggcagaggaattggccctactgctatagcattcgttcttgctg
ctgtatctacaatttttttgcatgaagtgttaagtaatctagaattataagcttttagct
taattctgattttaggttgctgaatgactgtcatattaagaaaatcttgctatttaagaa

ttttggaaacaacacctgcacctactgttctaccaccttctctaattgcgaatcgcatac
cttgctcaattgcaattgcattaattagttcagcagtcattttaattctgtcaccaggca
taaccatttcagcatctgttccatcatcagcagtaaactggttgatagttccagttacat
cagtagttctaacataaaattgaggtctatatccagggaaaaatggcgtatgtcttccac
cttcttctttagttaaaatatatacttctgcctcaaattgtgtgtgaggcgtaattgtac
caggcttagctagaaccatacctctctcaatgtctttcttttgaactcctcttaaaagga
ttcctatattatctcccgccaacccttcttccaaagtttttttggaacatttctaaccctg
taatagttgttgtacgagtttctcgtagtcctacaatttcaatagtatctcctactttga
tgatacctctctctattcttccagtagcaacagttcctctacctgtaattgagaaaacat
cttctacagccattagaaaagttttatcaacatctctttctggtgttggaatataagtat
caactgcttccattaaagagaaaattttatcaacccacttatcctcaccttttttgtatac
cagtgtttttttgtaactgcttctaaagctaataacgcagaacctgctacaaatggaatat
cgtcaccggggaaatcatattgacttaaaagctctcttccttcaagttctacaagttcta
ataattcttcatcatctacttgatcttctttgtttaggaaaactactagtgttggaacac
caacttgttttgccaaaagaatatgctctcgagtttgtggcataggaccatctgctgcag
atactactagaatcgcaccatccatttgagcagcacccgtaatcatatttttcacataat
ctgcgtgacctggacagtcaacatgtgcataatgacggttatttgtttcatactcaacat
gagcagtattgatagtaataccccgagctttttcttctggggcagcatcaatttcatcaa
atttcttggcagcagtagagcctagcgttgaaagagttgcagaaattgctgctgttaaag
tcgttttaccatggtcaacatgaccaattgttccaatgttaacatgtggtttttttacgtt
caaatttagagcgagccatgtttttttagtttccttgtagaaaaaaattgctttatagttac
ttttaacgataaatcttagtaatttaaagatttattacttaatataaactaaa-------
---ttttaattagttaataccgataatgagcaaaggctttattagcttctgccattctgt
gagtttcttctcttttttctaatagaattaccagtttcattggcagcatccattatttcgt
tagccaatttcatagacatatttttttccagatctttctcttgaaaatcgagtaatccatc
ttaaggctaaattagttcctctatatgcgcgaacttctattggaacttgatatgtagaac
caccaactcgcctagctttcacttctacgagaggagtaatattacgaatagctttttcta
aaatatttagagggtccgtttcggttctttctttaacaatatctaaagcttgatatataa
ttctttgagctaaagttttttttccacttcttaggatacgaacagtaagcatactaatta
atctacttttgtataaaggatctggtgatgcgaatcgctttttagctgtattacgacgag
acataaaatttaagatttgttattgtaataatatttc-tactaaaagatatttaagattt
aggtttcttggtaccatattttgaacgactttttcgacgatcttttactcctgcagcatc
taaagtaccacgtacaacatgatatctaacaccaggtaaatctttgattctacctcctct
aattagtactacagaatgttcttgaatattatgacctacaccaggaatataagctgtcac
ttcaaatccagatgttaatctcacacgggctactttacgcaaagcagaattgggcttctt
tggagttgtagtataaactcttgtacagacacctcttctttgaggacaactttttaaagc

aggagattttgtctttttattaatttttcgtctttcagatctaactagttgttgaattgt
tggcatagttaattatattgtgtatctgctaataaaacattttaaaattagttctcatac
acaaactaatctttattcattttatatttacgcataaatctttctactctaccctcagta
tcaataattctttgagaaccagtaaaaaatggatggttacctgaccaaatatcaacatgt
aattcaggttttgtagaaccaattgtcataattaatttaccgtcacagtagacttttgaa
tcagggtaccagctaggatgtatattatcttttgccatggtttttgtaatcttataataa
aataattaaataattttaacgtttggagaattgaggggcttttcgagctttctttaagcc
atatttttttcttcttttactcttggatctcttgttaaatagccttctgatttaagggc
acttctattttctggattaattgagcataaagctctagcaacccctaagcggattgcatc
agcttgacctgtcagacctcctcctctagcatttacatgtatatcatattgatttaataa
tcctaaaacttgtagaggtgcatacgaaactcttaaataattagggctaaattgaagata
agattcgccaggaataccattaattactagatttcctgaacctgggactagtctgacttg
agcaacagaacatttacggcgacctgttccagaatagattgcacgagttttaattaattc
tgtcgacataattttccttgataactaaataaatatacatt-tttacaccatacactctt
tcggtttctgtgcagcatgaggatgaagtgatcctgaataaacttttagttttgtaaata
attttcggcctaatgggcctttaggtaacattccctttactgatttttcaataattcgat
taggtaatcttgtttgcagctgttcaaatgtttcaacctttaatcctcctggttggccag
aatgtcttctatatagtttttgagttgttttacttccgctaacacatacttcagctgaat
taataacaattacataatctcctgtatctaagtaaggagtatatgacggtttatttttac
ctcttaaaatattagatatatgagtagacatacgaccaagtttctgatctttagcgtcta
taatataccaacttgggctggtaattaatgatggtacttgtgttttattcatgaaaagat
tagtacttttatttaacaataagaaaatatgtagatgttacacatataaattccgtgtaa
tcaattatttttgaatattatattcaaattatgtttgcacaaaaaggttttatagtagtt
ttt----------------aaaaaaacttcttattaagaataaattataacgtttttgat
agtataaaataaattaatctgtttttcttttggcaaacttatacctaatgttttttgt-
agagcatatataacttcttctgctgattttggccaaaatttttgatttctaaaagttct
tcttgagaataatctagtaaatctgcaatagaatgaatttgagctcttttaagcaatta
tatgctctaacagataattgcaattcttctattaaaacttgactaatcttttatcttct
ttattacgataagtatctactgatttaaaatctaaatttctcaatgagcaaaaaaggtta
gttaaaactgttgcgccttgactaatagcttcctgcggggaaatactaccattagtccaa
atttgcaaaatcaatctatctttaatactgttagtgcctacgcgtacttcttctacttta
taattgactttgtttacaggcataaagattgagtctacttgaagaaaatctaaagatagt
tcatcaactgcttttcggctaaacgatatccacaactttttcaattttaaattccatt
tcgaatattgtattattgcaaatagttgcaatatattgtcttggatctactacttcaata
tcagaagacaattcgaacaaaccagctgtaactatagctggtccttgtattttaattcta
ccaatttgagactctttattattgcttttaaatactacttctttcaagttcagtaatatt

tctaagacatcttctcgaactcctgaaattgtagaaaattcatgattaactccagctatg
cgaacagcaactatggcagtaccctctaaatctgacagtatcgatcttcttagagcatta
ccaagagtaattccttggccttggtttaaggaatctattataaagctaccatactgtccg
cgagccccatctgttcttgactctacacattcaatttgaaattgagccacctaagaaagc
tcctttaaaaataatcattgattggtaattttatctataataaaaaagaatctatacacg
gcgtttcttaggaggacgacatccattatgaggcacaggggtaatatcttttattaaagt
aatttccaggcctgctgcctgcaaagctctaattgctgtttcgcgacctgctccaggtcc
attgactaaaacttgtgtttgccgcataccttggtccattgcttgtttggcagctttttc
agctgcggtttgggctgcaaatggggttcctttttttgctcccttaaatccactagcacc
agaagatgaccaagataatgtctcacctttcaaattagtaattgtaacaattgtattgtt
aaacgtagattttatgtgcgtaataccattaactgcattacgcttatttttttcttgctcc
agatttccttatttgtcttgccattttatttagcctat---atagttaattaaaaaatta
ctttcttggggctttttttcttacctgctactgtctttttaccacctcttcgtgttcttgc
atttgttctagttctttgacctcttaaaggtaaacccacacgatgtcttcgacctcgata
agtactaatttccattaatcttttaatactcatagactcgaaacgttttaaatctccttc
aatctggtagttagattcaagtatttctcttatactaacaatctgttggtcattcaaatt
ttgacatcttatgtctgcatcaatgttagtttttttcaatatttcttttgatcgagataa
accaattccataaatataagtcaatgctatctctattcttttatttctagggagatctac
tccagcaattctagccactttatgttgtctccaataaataatatttacttttatcagac
taagctttgtaatttaaccttgtctttgcttatgcttaggattattacaaattaccatga
cttttctgtggcgacgaattattctacatttctcacacattttgcgaacagaaggacgaa
ctttcatattgattggctttgtatagtagaaaaattactttattaattctattaaatatt
tgcgcatgatacgcagagaacatattta----------------------tcacttggtc
atactatcatatttttagatataacgtatgtttgaatttgtttagccgtatctatcgca
acaccaactaaaatcagtaatgatgtagctccaagacctcttaagttctgtatttgggcg
acctttctattataaacggaattaacgctactgtaaataaaaatgtcgctccaagaaat
gtaagtctgtttaatataacttgcagataatctatggtagcttgccccgggcgaatatta
ggaatactggcacccattttttttaggtttgtggcaatatcttctggattcattacaatt
gaggtataaaaataactaaagaataaaattaaagcactataaaggacaagatataaagat
ccattggggcaaaatagataaagaatttgtaatacaagtggattttgtgttaattgagtc
aaatatgcaggaagcgccatagatgctgatgcaaaaacaataggcataactccaccttga
ttcagctttaaaggtaaatagctgttaggatcaaggattgaagatttgcctaattgtctt
gccgaaataattttaattcttcttgtaccttcttgcacaaaaatcgtaattattatcatt
agcaaaaaaatcacaatgaataagccaaacttaatacttgtattattatagttagcgtcg
aaaaaagattgagtaaaattttggggagacctgaaacaatattttgaaaaattagtaga
gatgctccattacctattcctttttctgtaattagttctgataaccacataataatcata

gacccagctgtcaaagctaagatagattcacatacaaaagcaaaattccaattaaaaaca
tatggctttacccagatagagattgcgctggactgtagagtggcccaacctaaagctaaa
tatctcgtaatttgcgttatcttttggcgtcctaattctccttcttcttttgaagcttt
tctagagtaggaattatttttgtaagtaattgcatcacaatcgaagaattaatataagga
acaatgcccaatgcaaaaattcctattgtagagaaacctccaccagagaaaatgttcaaa
aagttcactaaagtattcttctctacactcgcataaaaggcatcatgatctatacctggg
acaggtataaaaatacctaaacgtgctaaaatcaataagaaaagagtaaatacaatacgg
tttcgtaaatcgctttttggctcatacgtaattttcaggtaaaaaaacaatttttaat
taaataaaaaccctattccaatacttatacaagttgaaaatagggaattaa---gacaac
atagcaatactttta-taatattctatttagaataaagattttctattggcactcctcgg
tcttttgcagcttcgctaaaagttcttagttgagtcaatccatttaaaacagctcgtgca
ttattcagagtattattagatcccaattgtttagcaagaatattttgaacaccagataat
tctaaaactgttctaacagaaccgccagcaattacaccagaaccaggggctgatggtctt
agtataacttgtgcagcaccagaaattccattaataggatgtggtatagaattagattta
gttaatggaactgtaactagatgtttttagcatccgttacacctttttttactgctcca
attacatcacttgctttacccacacctacaccaacttggccttgttcattgcccacaact
aaaataactctaaagcttaatttcttacctccttttacaactttcgtgactcgtttaact
tgcacaactctttcttcccatccgctatctttttctttaacttttccttgtttttacga
tttgccatcttcaacattattcctagttagttaattagcaactttaaaacaccatgccag
cttcttttgcagccttagctaaagctttaactcgtccatgatataatttaccacctctat
caaaaacgacagtttcgattccttcttttatagatcgttctgctaaagttttacccactg
aacgagaagcttcgcaattagatccaagtgctattgatgattgagaatttaagttaacag
acgaagctgcaactaaagtaatccctttgtatcatcaataacttgtgcatatatgtgct
tgttagatctaaatacacataatcgaggtctactagcagttcctcgaactttttttcgaa
cacgtctatgcttgtgaattctagtctgcttgctattgattttcattgttatttaccttt
tccagcttttccagcttttcttcttacaacttcgccttgatatcgaattcctttcctttt
ataaggttctggtggtcttattgaacgaatagtagaagcaacttgaccaactacttcttt
atctatacctaaaacagtaatattggtgttattttcaactttgatttcaatattttgagg
gggttggatagttacaacatgactatagcctacacttagtattaagttattattatctat
ctgagagcggtaacctacaccttgaatttggagtttttttggaaaaaccattagaaactcc
ttcaatcatattattaataagtgtccgagatagtccatgcaattgattagctttttttagt
atttccacttgttcggacaagaattgtagcattattaatttctaaagtgataccctctgg
caaagttcttgacaaagtgccttttggcccgtttactgtaatggtttgaccatcaaattg
agtactaaggtttgtaggcaataaaattatttttttttccaattcgggacat-attccacc
tataaattagttaccaaatataacataaaatttcaccacctaagccatcatgacgagctt
gtctatctgtcataacacctcgagaagtagaaataatggcaatacctaaacctccgagaa

ctctaggtaattctttgtgatttgcataaactcttagtcctggtttactaattcttttga
gagcagtgataaccggttgtcgatttttcccattatatttaaagaaatcatcaaatgag
tctctataccttcgcccatttgctcaaaattttgaagaaatccttcttccttgagtacta
gtgccatattgcgtgtcattttcgttgctggtacctgtacaatttgatgtcttgccaagt
tcgcgttacgaatacgtgtcagcatatcggcaatcgtatcgttgaccact-acatccccc
ttaaga-----tagattatttagaatattttcaggattctctaaaaggcataccaagctt
ttttaataaagctaacccttcttgatctgttttagctgttgttacaattgagatatctaa
gccacgaatttgatcaatattatcatagtctatctctggaaatattaattgttcacgtaa
acctaaattataattaccttttccatcaaaacttttagggctaattcctctaaaatctcg
gattcttggtaaagttaaattaattaatttttccaaaaaagagtacattttatcttttcg
caggtgcactacaattccaatgggaacttcttcacgaattttaaaacctgctatagattt
tttagcctttgttacaattggtttttgcccagtaattaatgttagttcttgaatactgct
ttcaagggctttagtattttgggacgcttcacctaatccccgattaatagtaattttagt
aaaacgagggacttcatggacattctcatactgaaattcttctttgagagcttgagtaac
agtttttttatatttttcttttaatcctattaccattatatttttataacctatttgata
atttcaccagtctttttttaatttgcgga-ctttttggccctcatcatcaattattactga
gaatcgactagaaatattattttttttgactgaataacataacattagaactatgtatagg
tgcttcgaatttttgtaatttgtccagtctcaccttcttgctgaggtcttttatgctttac
tttaagattaattcctttgacgataactttactagttttatgtataattgcaataatctc
gcctgtcttgtttttatcactaccagaaagtactttgactaaatcgcctttttttgaattt
aactttttattcactatatttttttgcaagcaaacttttcattatattacctccggcgca
agagaaacaatttagaaaaatttttatctcttaattctcgagctataggtccaaatact
ctagtgccacgaggattattatcttgattaataataacggctgcattgtcatcgaatcga
atactcataccatcaattcttcttaatgctttacgcgttctaacaacaacagctcgtacc
acatcagatcttttaacaggcatatttggagaagcatctttgacaacaccaataattacg
tcacctatagacgcataagatggattgctggtacccagcacccgaatacacattattttt
ctagcaccactattatctgctacattaagatagctttgagtctgtatcatacttttattc
tatataagattaattatcaaaagatttagataatatattaaccattgtccaacactttgt
acgacttaaaggcctagtctcctgtattgtaacaatatcgcctactgcgcattcattatt
ttcatcgtgtgctttatattttttcgtacgaatcattgttttagcatattttctgtgaga
aattctattttctacagctactactatagttttattcattttatcgcttattactttacc
tgttgtttcttttaaaggcatagtctaaatgagttatataatt---------aatttaat
t------------------tgactggtccctggacttttcaaccgttaaaagttgagcta
atctatgttttgaatgcttgaacaaatgaggcttaaaatcttgtcttgttgctcttttta
gtcttaaatcaaataactgtcttttgattgcgataatttcttttgatagagaggaatcgt
ccagttgtataacatctaatattttaggtaaagtcata-tttaaaattctattgtattcc

tcactataaactttgtctttatcggcaatttataagaggctaatttcatagcttgttgtg
cagtttttgtggtaccccagtaatttcaaataaaatatgaccaggcttaataactgcaa
cccagtattctggagcaccttttccagaccccatacgagtttctgcaggccgtgctgtga
caggtttatctggaaaaactctgatccacagtttaccacctcgtctgacatatcttgtta
tagttcttcgtgttgcttctatttgtctagaagttaaccaaacaggctctatcgcttgta
atgcataatcgccgaatgcaatagtattacctttacttgcagaacctttcattctgcctc
tatgttgttttcgaaattttgttttcttagggcttagcatgagagtttaacatttaggat
aattaagaagcgtttggattaagttgatcctgactaggagctagttctaaaactttagaa
tctggtaaaagttctcctttgaagagccagactttaactccaagtactccatatgtagta
tgagcttgtcgatggcaataatcaatatctgctcttaaggtttgcaaagggactcttcct
tctcggacccattcacttctcgcaatttctgccccattcaaacgaccagatacttgaatt
tttactccttgcgtatttgctctttgagctctctgcacagcctgtcttacagctctacgg
aaggcaactcgtttttccaattgctgagtaataaattctgcaactaaagttgcttcagaa
tcaggatcggaaatttctacaacattaactcgaatttgtttattcaaatctaaaattaat
gttaatgaatttcttaaagattcaatacctgatcctgacttccccagaacaattcctggg
cgagcagttgctattaatacttctacttgatctactttacggttgatttcaattttagca
atactagcattactaagtttattatatataaaagagcgaattttatgatcctcttgtaga
aaagctgaataatcttttgaactggcaaaccatgaagaccgatgtttttgagtaatacct
attcggaaacccaaaggatgaatttttttgacccatattatcctctactatatattaaaag
ttataattgttggtttatatctagagtattagatattac-tttctgatacacctaatgta
atatgacaggttggcttatgtattggaaaagctctaccttgagctcttggctgaaatctt
tttaacgtaggtcccttatctgcaaaagccttggtcacaaatagctgagttttatttagc
ccattattatgttctgcattagctgccgctgattctaagatttgttttatatgaaaacaa
gctctgtacggcataaattctaaaataattaatgcttcctgatattttctgcctctaatt
tgatctaagacacgacgaactttatgcggggaaagacgaatatatttccctactgcttta
gtttcttttaagtttttttgtaatgctcataattatgatttaacgacgggcttttctgtca
cctttcacatgagtgcgaaaagttcttgtaggaacaaattctcctaacttatgtccgacc
atttggtcggacacaaaaacgggaaagtgctgtttaccattataaacagctattgtatgt
ccaaccatatcaggaataatggtagacgatcttgaccacgttttaattatttctttttc
cccgaagtatttagtgcttctattcgagttaaaaggctcacatcaataaaagggccttta
tgtattgatcttgacattgtacttgagtattaaattatagataggataaaatttatttac
ggcgacgtaaaacatatggattactatatttattcggattacgggttttaacacctaaag
caggtttgccccaaggtgttacaggacgagatcttccaattggagatttaccttcaccac
cgccatgaggatggtctactggattcataactacaccccttacagtaggtctttttccta
accatcgacttctaccagctttaccaagagtaatattactagcatcaatattgccaactt
gtccaatggtagcgtaacactcttttcgaatcatacgaacctcgcttgacggaagtttaa

cagttacaaaatctccttctttcgcaactatttgtgcatatgtacctgcagaacgaacaa
tttgacctccacaggaaggtctcaattctatattatgaacagctgtacctaaggggatac
ttgataatggtagggcattaccaacttcaataggggctttaggaccagatagtactgtaa
ctcctacagataaagaacgaggatgtaaaatgtatctttttttcaccatctagataatgaa
gtaaagctattcttgcgtttctatttggatcatattctatagaagctactttagctatta
tattatgcctatttcttttaaagtcgattaatctatatttttgcttgtgaccaccacctt
tatggcgacaagtaataacacctctgttattacgccctttacaaaaatggtgcttattga
ttaacgacttttctggcttatcagtagtaatttctgagaaagtggaaacagtcctgtttc
ttgtcccaggagtataagccctgtataaacgaattgccatatgaaaa----tgaaaaaat
taaaaaatcaatatttgttatgtaatatgcttaagtttctgggaataaattaatagaatc
ttctgaggctaaagtaacgattgctttcttataatgtggtcgttttcctacaaatctacc
aatacttctttttttctttggaggatggcaagtattgacacctgtaacctggacattaaa
aatgtattggattgcggctttaatgttgattttagttgcatttgggtcaactgcaaaaca
atattgatttcttctaataattttgttgttttatccgtaatgattggatatttaactaa
atctaataaacctcttgaatcaatgttatccattatatgcctcttgtattttagataacg
caccaacagtaataataactttatgtgcagctaaaagagccataatattcaaagtatcag
ctgaaataatttctacattgtgtaaatttcgaattgaaagataaacattcggatcttttt
tatcaactatcactaatactttttttattaagattaaggttccaacgaataatagcttcca
taaataactttgtttttggctgttgaaaataactattaaaattttctacaactaatgtat
taagagatttattatttaatgccgtcctaagagctaactgtctttctttcttattcatct
tttttgcaaaactacgaggctttggtccaaatattatacctcctcctctccataaaggtg
aacgtattgaaccagctcttgctctacctgtacctttttgacgccatggtttgcgtccac
caccacgcacttcacttcttgttttagtattagcagagccttgacgcttctcactgcttt
gtttaactaatgctctatgtactaaatacatgccagactcttgactgatttcaagattca
aatttgcattgccacttacttgaccttcccaattatagacttggtaatttaattgtgttt
taactgtcatgaaatatatcgtataaaagaatattgctatttactgactttaactaaagc
accctgtttcccagggacagctccttttaacattaaaagatcattctctatgttaatgct
tacaatttgtaaatttttttattgtaactttcttattacccatttggcctgccatattctt
acctgggtatacgcgtccaggagttgtacctgctccaatcgaaccaggctggcggtggtt
cttagaaccatgagacattgggcctctactaaaatgatgtcgtttctgataaccagagaa
acctttgcctatgcttttagaggacacattaactttttgccctacttcaaatatatcagt
agaaaatacttgactgacttcaaaaccttcggtagattcaatcaaatattcacgcaaata
tttaagaggaggggattgagatttttttaggtggcctagtaatggtttacttaattttg
ttcagcaacttgcttataaccaacttgaatagcactatagccatcagtagacatcgattt
tatttgagtaacaacacatggcccaacttgaattacagtaactggaattgataaaccagt
ttcatcaaaaaattgagtcatacctactttagtaccaagtattccaacagacactagatt

tctccttttgagagatttatactattatttgacacaaaattatagcttcaataatatta
aaagataagcacagacattgattatctattatttataaagataa----------aaaaac
tttacctattcaatctaaaatagaaatagatttttgtccagttatcactcatctaattcc
aaaattttacgttatatatttc--------------------------------------
------------------------------------------------------------
------------------------------------------------------------
--------------------tttatgttcggtaagtacacctgtttatttatatattttt
tgtgcaatatatagttatatgtagtttagcaagcatgacaatatagctaattatatagta
aattcacttgtaaacattagagaggtattcatgagtaaagttgttggaattgacttggga
acaactaattctgtaattgctgttatggaaggaggaaaacctactgtcataccaaatgca
gaaggttttagaactactgcttctgttgttgcttatactaaaagtggagacaaattagtt
gggcaaattgctagaaggcaagctgttattaatcctgagaatacttttattctgtaaaa
agatttataggaagaaagcagaatgaaatttctcaagaaattagacaaacttcatataat
gtgaagactagtggctcaagtataaaaattgaatgtcctgcattaaataaagattttgcc
cctgaagaaatttctgcccaagtattaagaaaacttgtagaagatgccagcacatattta
ggtgaaaccgttacacaagctgttattacagtaccagcttattttaatgattctcaaaga
caggcaacaaaagatgctggtaaaattgcaggtttagatgtactccgaattattaatgag
cctacagctgcttctctatcatatgggttagataaacaaaataatgaaactattcttgtt
ttcgatctcggaggaggaacatttgacgtatctatcttagaagttggagatggagttttt
gaagttctatcgacatctggagatacgcatttaggtggtgatgactttgatcagcaaatt
gttgaatggctaatcaaagattttaaacaaactgaaggaattgaccttggtaaagatcgg
caagcactgcaaagattaacagaagccgctgagaaagcaaaaattgaactatcaaacttg
actcagacagaaattaatttaccatttattacagcaacgcaagatggacccaaacactta
gaaaaaactgttactagagcaaaatttgaagaactctgctccaaactaatagacaaatgc
agcattccagttaataatgctttaaaagacgccaaactagaagctctgagtattgatgaa
gttgtcttagttggtggatctacaagaatcccagctatacaacaaatggttaaaagatta
atcgggaaagatcctaatcaaagcgtaaatcctgatgaagttgttgctattggagcagct
gttcaagctggggttttagccggagaagttaaagatattttactattagatgttactcca
ttgtctttaggtgttgaaactcttggtggagttatgacaaaaattattccaagaaatacc
actattcctactaaaaaatctgaagtgttctctacagctgtagataatcagcctaatgta
gaaattcaagtacttcaaggagaaagggaacttactaaagataataaaagtctaggcaca
tttcgtttagatggtattatgcctgcaccaagaggtgtacctcaaattgaagttactttt
gatattgatgctaatggtattttatctgtaaaagcaaaagagaaggcaaccggcaaagaa
caatctattactatatctggcgcttccacattacctaaagacgatgtagaaagaatggta
aaagaagcagaagaaaactttgacgtagatcaaaaaagaagaaaaaatattgacatcaga
aatcaagcagaatcattgtgctatcagtctgagaagcaagtcaaagagtttgaagataaa

attgatgaagatctaaaaaataaaataacaaacttaattagtgagctacgatctaattta
gagaaagaagagtttgataatattgagactacttccgagaaactacagaatgcattaatg
gacattgggaaaaatgctacctccgctgagaacaatactcaaagtacatcaagtaatgat
acagttattgatacagacttctctgaagctaagtaaagaattaagcgggtaacgcgattc
gaacgcgcgacatcaaccttggcaaggttgcgctctaccactgagctatacccgcatgcg
aattattattacaaatatcttgaaatttgtcaatctacta------atttgaatgtaatt
tagataattaa------agataaagattaaa-----------------------acttct
atatttcagtcaaaaaaagtacttatactct-tttataagtactttttttgatggatcag
ttaaaaagacaactcaatataagatctaagatctaaactactaaagaattaactaaactg
gttacaatgactagcccagtccataagcctgcacctgtgtaaattaagcctttggactgc
tcccattgacctggtgatgctaaaactacaggcacacctactactaatatagtagataga
gtaattaacaatagtacaagtagttgaattgcaataatcatttattatactctccttaaa
aattcaaattttaataatcttgtatataatacaatgtagcgctgaacctgcgaaaaacta
tcgatgcgaagataatttaaatttatttttttatcaa--aagtttgaactaaatattaaa
attttcagcatagataataatagtttattttttaaaaatagataaaaaatttaaagaaag
tcttttttaaaaaaatttcagaataaggtgatatgctgtataaaaaactaaatattatta
gttattagaaaaattttttaatcagaggtaaacaatatgaactcagcccttttttttagca
aaattgccagaagcttatgccgtttttaaaccaatagttgacattttacctgtaattcct
gtattttttcttcttctggcatttgtgtggcaagctgccattggctttagataaattata
attgataatagtaataacaaattaataaaataaacttttttaaaaatt-------ttttt
taaaaaataataaaacttatggttgaacctttattatcaggaattattcttggattaatt
cctgtcactttgtctggactactagtagcagcttacctgcagtatcaacgtggaaatcaa
ttaggtctataaa----aaatcgtagatatttataataact------cacctagtttttt
tgattatctacaaa-----------taaatacataa--------attcattatgaaataa
taatttataaaagtaactagtttttaa---------------------------------
------------------------------------------------------------
------------------------------------------------------------
------------------------------------------------------------
------------------------------------------------------------
------------------------------------------------------------
------------------------------------------------------------
------------------------------------------------------------
---------------------------------atttcaagataga--------------
------------------------------------------------------------
------------------------------------------------------------
------------------------------------------------------------

```
------------------------------------------------------------
------------------------------------------------------------
------------------------------------------------------------
------------------------------------------------------------
------------------------------------------------------------
------------------------------------------------------------
------------------------------------------------------------
--------------------------ctaattaacctaattaccaactagatttagtaac
tccgggcagaaggcactcatgagacatttctctaaaaacatgtcgagataatccaaaatc
tctataatagcctctactcctaccagttaaccagcatctatttctacttcggactgctgc
actatttctaggcatttcctgtaatttttgtctcaactccatttttttcagcaaagctagt
agttccttttaagttctctttaattgctaaacgttttaaataatattttttttcaagttt
ttgtctcttaatctctctttgtatcatattttttttagccatggtataagaacctattat
ttaaattgtttgg-aagtatattagctaaaaaaaggcatgactgcaatatttaaatatga
aatgagtaaattaacaaagtagtaccatgtattcaatatttaattgatggtactaccact
atttataattacagtacctctatttaataaagctcttgttcaacatgagttagaatagta
cagtcagattgaggataagcaatacatgttaatacataccccttcaacatttgctcatca
tctaaaaatgattgatcggattgatcgatagttccttctgtaacttttcctgcacaggtt
gagcatgctccagccctacaagaataaggaagctcaattccttcttcttctgctgcatct
aagatataagtgtcttctgcacaatcaaatgttacatcaattccttcttctgtagacact
aaatgaattttgtaatcagccattcgtttttactccataataaaagtacatttgaaactt
aataaagcaacatgaatttgaataattatcatataaattatacattagtttcattatgaa
taatagactgcaaaagtaaaaaatagtataacattttaaatattttggcaacattttaaa
ctaaatagaagtatattaggttcaaagtatttatataaattcatacagttaaaattattt
ttatcatttacataatgaaatttagcagattgaaattattaactttatcattcaattaca
attagacttttat----aatcataaactaattatgactttcatttttttcaaaaaaaatag
agctaaagcctattttgaaatttgacataccaagagtatgctcacatgaaattattcaag
aaattgaagctttagtgcaaagactatttttacaagtttggaggagacccgcaacactaa
tggccggcattattcaacctctactttggctagtactatttggaggccttttctataatg
ctcctgtcaatttatttactattaatacaagttataactgctttttaagttcaggaatta
tagtattcacttcatttactggagcattaaattcaggccttcctttaatgtttgatagag
agtttggatttttaaacaggttattgacagcaccattaatctctagaacatctattattt
tttcttctgcaacttttatgacttgcttaagtttaatacaagttatatttatagtaatag
catctctatttatgggtaattctcctttaagcagtaatagcacattaatttttgccctta
ttgttttattggttactgtaggtgttacaatgttaagtctaggactatcttttactttac
caggtcacattgagctacttgcgcttattttagtagtaaatttaccattttttattttcca
```

gtacagctttagcgccactatattttatgccaccctggctgcaattaattgcaagcctta
atccactaagctatgcaatcgaaggtattaggtacctatactccaatacggattggaatt
tcagtgagtctgttattaaaattagttgggggatatttctctaggtcaaataattttac
tattattagttttagatgtaataggagcttacattgtgtctaatatattaaaggctaaac
ttaattaaaattagtacggaatataaattttaataaaaaatttattaaatttattttacg
actattttcactatataatactattagtagtatggataatggaaatgaaaatagtttttc
aaaatcaaagctattatagaaaatgataaatatttgtaagaaagtcaacaaagtatgttc
ttattcataggaggcatgtagtcaatgggactaccatggtaccgtgtacacacggttgtt
ttaaatgatcctggacgcttaattgcagttcacctgatgcatactgcacttgtagcaggt
tgggcaggctctatggcattatacgaattagctgtatttgatccttcagatcctgtctta
aatccgatgtggcgacaaggaatgtttgttatgccatttatggctagactaggtgtaaca
gattcttggggcggctggagtattactggagaaagtgtatcgaatcccgggctatggagt
tttgaaggtgttgctttaactcatattgttctttctggaatgttattttttagctgctatt
tggcattgggtatattgggacttggaattattcagagacccgcggactggtgaaccagca
ttagatttaccaaaaatttttggtattcacttacttctatctagtctactatgttttgga
tttggtgcttttcatgtaaccggtttatttggtcctggcatgtgggtatcagatggttat
ggagtcactggcaaagtattacctgtagctccagcatggggaccagaaggatttaatccc
tttaatcctggtggagttgcgtctcaccatattgctgcaggtactgttggtatcttagct
ggtgtttttcatttaactgttagacctccacaaagactttatagagctctaagaatgggc
aatattgaaacagtactctcaagtagtatatctgctgttttcttctccgcatttgtaact
tgtggaacaatgtggtacggctctgctactacacctattgaactatttggtccaactaga
taccaatgggatagtggatattttcagcaagagattgaaaaaagagtagaaaacgctatt
gctgatggagctgctccaagtgaagcttggtccagaattcctgacaaattagcattttat
gattatattggtaataatcctgcgaaaggtggtttattcagagcaggtcctatgaacaaa
ggggatggagtagctgaagcttggttgggtcacccagtgttccaagataaagaaggtaga
gaattaagcgttcgccgaatgcctgcattcttcgagacttttccagttattttagtagat
aaagatggtattataagagctgatattccattcaggagagctgaatctaaatatagtatt
gaacaagttggtgtgacagctagtttttacggcggaaaactaaatggacaagtattcaat
gatgctcctagtgtcaaaaaatatgcaagaaaagcacaactaggagaagtttttgaattt
gatcgaactacattagaatcagatggtgtatttagaagtagtccaagaggctggtttaca
tttggtcatgcaaactttgcattaatctttttctttggacatctttggcatggttcaaga
actattttccgagatgtatttgctggaattggtgctgaggtgactgaacaagttgaattt
ggagcattccaaaaattaggagatagaagcagtaaaaagcaaggagcagtataaaatt-c
cattgcataatgtcactatt---caatatttacaatattgaaatcatttagaaaaagttg
attatttaataattactaatcaggagaatttatggaagccttagtttatgtctttttatt
aacaggaacactaatggttatattctttgctatctttttttagagagccaccaagaatagc

taaataaaatcattaggctcaaccctacaataatagaaccactttccagcttattagtgg
aaagtggttctattgtaggttaatgcatattactcttcatgctcttcaaaagggtctctg
agatccttagaagcaggaccaaatgctgtatatattgagtaacctgtaatacctagaagt
aaacttgaaataaaaatactaagaactgttgcagtttccataattcaaacagtattaaga
tgaacttttctataataatattattaatattaaaacatttaattattaaattaatcaaa
atatattatggcacttagaactagactgggagaaattttaagacctttaaattcagaata
tggaaaagttgcaccaggttggggtacaactcccattatgggaattttcatgctgttatt
tttcttattttactgataattttacaaatatataattcatcattagtgctagaaaatgt
agatgtagattgggctactttaggtag----ctaagtgtagcaattcttaaatgaacagt
ttaaataatattatacacaatcataaaatattatttaaaacgaccgctcttat--tacaa
gagcggtcgttttagaa------tactatttg----aagttgattattttcctaaatcaa
ttaattcattatctgcaaagctattagtattaacattattataattaacttttcaaatc
gtactaaaacaggatatttaatgccacttttatccattgctgcaacagttccaatttctt
ggtaccagtatgactcctttcttaaaatttcactttagaacctctttccat-ttttaat
tccttagataaattattgaacatcaatatattttcaaaataactaaactgtttaaaaaca
tataaaataattaccgacaagactttaattttgataataaaaggagagagagggattcga
accctcgatagcttaagctatgacaattttcgagattgttgcaatcaaccactctgccat
ctctccggagataagaaatttttct-ttaaattaaactagggtatcacagaataaaagta
gttgcctagaaatttaaaactatgttaaatataaattataatcctgctataaatatcttt
acaagttatgtaaataatatgtatatcaggttattaaaagattttaaatctgataattt
ttttactaaataaataaacgctctcataatatgaaactatcttggaaaacacttctgctt
tggtctttaccaatctttgtagttggtttctttttctggcaaggtttttaggcccaact
actacagatgtaggaagtaatattgcaagttctagaatgacatatggacgatttctagaa
tatttagatatgggttgggtaaaacgagttgatttgtatgaaaataaccatacagcaatc
gtcgaagccgttggaccagagttaggtaatagagtacaacgtattcgagttgaactgcca
gcaagcgcaccagaattaattacaaaactacgtaaagctaacgttgatttagatgctcat
cctccaaaaagtactagcgccgtttggggattattaggaaacctattattcccactacta
ttagttggtggcttagcatttttatttagaagatctaataatgcaagtggcggtcctgga
caagcaatgtcatttggtaagtcaaaagctctttttcaaatggaagctaaaacaggcgta
gtatttaatgatgttgctggtgttgaggaagcaaaagaagaatttcaagaagtagtgact
tttttaaaacaaccagaatcatttactgctgttggtgctaaaataccaaaaggagtacta
ctagttggacctcctggtacaggtaaaacattattagcaaaggcaattgcaggcgaagct
ggagtaccatttttttagtatttctggttcagaatttgtagaaatgtttgttggtgttgga
gcatctcgtgttcgggacctttttaaaaaggctaaagataatgcaccttgtattgttttt
attgatgaaattgatgctgttggtcgacaaagaggcacaggcgttggtggaggtaatgat
gaacgagaacaaacattaaaccaactattaactgaaatggacggtttcgaggggaatact

ggagttattgtaatagcagctactaatagggctgatattctggattctgctcttctgaga
ccaggaagatttgataggcaagtatctgtagatgtccctgatttcagaggtagattagct
attcttgaagtacatgcaaaaaataaaaaaatggaatccaaagtttcattagaaactata
gctagacgaactccaggtttctctggagctgatttagctaatttattaaatgaagctgct
attttaacagcaagaagaagaaaaaatgctatgactatgtctgaaattgacacatcaatt
gatagagtagtagctggaatggaagggacacctttaattgatagcaaaagtaaaagacta
attgcttatcatgaagttggacatgcaataataggaagtttattagaacatcatgaccct
gttcaaaaagtaacattaatacctagaggacaagctagaggattaacttggtttactcca
agtgatgatcaaagtttaatttctcgttctcaaatactggctcgtattgtaggagcactt
ggaggaagagctgcagaagaaattattttggggatgcagaagttacgactggagcaagt
aatgatttacaacaagttacttcaatggcaagacaaatggtcactcgttttggaatgtct
aaaattggtccattatcattagagagtcaagggagcgatccattttaggaagaggaatg
ggaggaggatcagagtactccgatgaagtggctacaaatattgataaacaagtaagagaa
attgtaagtgaatgctataaagaagctaaaaagattattaaagataatcgagtggttatg
gacaggctagtcgacttactaatcgaaaaagaaacaatagaaggtcatgaatttagagat
attgtgaaagaatacacagctattcctgaaaaaaattactatatttctcaattttaaaat
tttttattgt-----ttaaaacgggactgacgggattcgaacccgcaacttccgccgtga
cagggcggtgctctaaccagttgaactacagtccc-aagaaact---catatataaatat
tctcacaataaacttcgtattgtcaaatt----ttttatatataaggagagagagagggatt
caaacccctcatatataaatattctcacaataaacttcgtattgtcaaatttttttatata
taaggagagagagagggattcgaaccctcggtacggatttaaccatacagcagattagcaat
ctgccgctttcgaccactcagccacctctccattgaaat-------------------aa
tatgttagtattgtaacacagatattaaaacatttatggagctaagcggatttgaaccgc
tgaccctctcaatgccattgagatgctctaccaactgagctataacccc--------tag
aagtctaagtaaatggctcaagcgggatttgaacctgcgaccttgggcttatgaatcccc
tgctctaaccactgagctactgagcca-gatatgtgctattactaattattataccatat
aaaattactgaattgaatacaagaattaatttcttatt----------------------
-----------------tataattc-----------------------------------
------------------------------------------------------------
------------------------------------------------------------
--------------------agaatattatg-----------------------------
---------------------------------------------------acttcactt
a-----------------------------------------------------------
------------------------------------------------------------
------------------------------------------------------------
------------------------------------------------------------

```
------------------------------------------------------------
-------ta---------------------------------------------------
------------------------------------------------------------
------------------------ttct--------------------------------
------------------------------------------------------------
------------------------------------------------------------
------------------------------------------------------------
------------------------------------------------------------
------------------------------------------------------------
------------------------------------------------------------
------------------------------------------------------------
------------------------------------------------------------
------------------------------------------------------------
------------------------------------------------------------
-------------------------------------tactataaatg------------
------------------------------------------------------------
------------------------------------------------------------
------------------------------------------------------------
------------------------------------------------------------
------------------------------------aatataaatttttaagtataagta
tttaatttaagaaaaa---------------------------------tttttatacaa
ctaacattgaaccaataccttgatcagtcaatatttctagtaaaagagcatgatcaattc
taccatctaaaatatgtgcagatgcaacgccttgagctagagagcgaatacagcagtcaa
ctttgggaatcatgcctccagagataacagctgtctgagttaaatctcttgcttcttgta
tacttaaatgactgattaaagttgttacatctgatggatttcgcaaaattccaggagtat
ctgttaataatattaatttttctgcgtttaaagctgctgctatttctccagcaactgtat
cagcattaatattataagactgaccttgcttatcagcagcaacactagctatgactggaa
tataattattatttattaaaatttctaataaattagtatcaacattctgtacctcaccta
caaaaccaagttcaggtttaccacttggtcttggaataattaataaaccatcctttcctg
ataaacctacactcttaccaccttgcttattaatacttgcaacaagatctttatttactc
taccgactaaaaccatttctacaacatccatagtaggttggtcggttactcgaataccat
tctcaaattgaggcaaaatttttaattggtctagccagaaattaatttctggcccaccac
catgaactagaataggacgtaagccaataaaagacaaaaacactaaatcacttattactt
gatcttttaattttgagttcttcatagcagctccaccatattttattacaatgattctag
aagaaaacttttgaatatatggtaatgcttcgcttaaaacttttaccctttctgtgtttg
tcaacatgaataagataaaattatga----taaatttatttgtaagacttatattaatct
```

aacaaatagtaataactatttcaagacaattaaaaaataaacataatacgaataaattaa
atactgcaatatataaggattatagttatgagtcagttttgggacaatattgtaaaattt
ccacgattttaataagtgttattttaggattaattttaacaattattagtcctttttttc
gtgttatttaggaaaccagtaacaagtttttttttcatcatatcatttaccggcttactg
gtgattttaacggcaataa-tacaaaaaatgttaaatatcgagggttgttgagtataata
aatagatttaattaaatttttgaacatatataatattatattatcattaaacaaataat
tctatttttttttttttatgtaaaaaac---aagtataaaaaat-----------ttttat
gacattaaaacaaataattagttccgaaaaaactggtgcttttgctctcttagatagtat
cgttcgtcatgaggttgagcatatatttggatatcctggcggagctatcttaccaattta
tgatgaactttatagttgggagaaaagttcactaattaagcatattcttgtaagacatga
gcaaggtgcctcacatgcagcagatgcttattctagatcaactgggaaagttggtgtatg
ttttgctacctctggaccaggtgccaccaatctagtatctggtatagctactgcccatat
cgattctgttcctatactagctattactggtcaagttggaagacctttcataggaacaga
tgcattccaagaagttgatatttttggtattacattacctattgtaaaacattcatacgt
agtccgtgatcctagagatatgtctagaattgttgctgaagcattttatatctgtaaaca
tggcagacctggcccagtcttaattgatgttccaaaagatgttggattagaaaaatttaa
ttatttttccgttgagccaggacaaattaaaattccaggttgtagaccaatatcaaattt
caaatctcgtcaaatagtaatggctgcgaaaatgattcaacaagcaagccagccattgtt
atatattggaggtggagctataatttccgatgctcacgccataattaaagaactagttga
cttatacaaaattccagtaactactactcttatgggaaaaggtatttttagtgaggatag
cgaattttgcttgggtatgctcggaatgcatgggactgcatatgccaattttgcagttag
tgaatgtgatcttctaattgccttaggagccagatttgatgatagagtaactggtaagtt
agatgaatttgcttgtaatgctcaagttattcacgtagacattgatccagccgaagttgg
taaaaacagaattcctcaagtcgctattgttggtgatgtcaaagaagttgttactgcact
cctgaatttgttgagaactaattttaagccttatccagagcaaattatatcttggcaaga
gagaatacaccgttggcgtcaacaatatcctcttttagtacctaaaaaatcaactagcat
ttcaccacaagaaatacttgtgacaacgaatcaattagctcaaaatgcttattttactac
tgatgtcggtcagcatcaaatgtggtcagcacaatttcttaaggtgaaatctaaacattg
gatttctagtgctggattaggaacaatgggctatggtttgccagcagctattggtgccca
agtagcacaccctaatgaattaataatttgcgttagtggtgattcaagttttcaaatgaa
tatgcaagaattaggaactattgctcaatatcaattacctataaaaattattattattaa
taatcgttggcaaggtatggttaggcaatggcagcaagcattttatggtgaaagatactc
tcattcgcgaatgacagaaggagctcctaatttccaaaagcttgctgaagcttttggtat
taaagctttcactgtaaataatagacaaaatatgaaatcctctttagaaaacgcaatgaa
gtacgctggccctgttttattagattgtcaagtaactgaaaatgaaaattgttatccaat
ggttgcaccaggaaaaagtaatgcacaaatgataggaatagcaaaaccacaaagaggaac

tgcttctaattatgttagtacaaatagttaatt---------------------------
------------------------------------------------------------
------------------------------------------------------------
------------------------------------------------------------
------------------------------------------------------------
----------------------------------------aatagcctttgacgagaatc
gaactcgtgaccttccccttaccaaggggatgctctaccactgagccacaaaggc-tttt
tatttcattgggccgggttggatttgaaccaacgtaggcgaagccagcggatttacagtc
cgcccccattaaccactcgggcaccgaccctagttaattatcattcataatatcat----
--aatatcacaattgtgtatctaaaaacaagtgtttttagatattatctt-gcggagtaa
tggacatggctggactcgaaccagcgactttcacgatgtcgacgtgacactctaaccagc
tgagttacatgtccaaaactttaattgatattatagcac-gtttgattattttattatca
tac-ttcctaatcaaaaatttgctttaaacggctagctttacctaaaagatttcttaaat
aataaagtttggctctgcgcactttagctcttcttataaccttagcaaaagttacgcgtg
gtgaattaagaaaaaaaacgcgttcaacaccaataccttgaaatgagcaccgtaatgtta
aactattatttaagctttttctttttttctagacaatacaacgccttcacataattgct
cccgagttttacttccttctttgactagtaagcctaatcttattgtatcaccaactttga
tttctggtaattctgctttcattaagggtctttctatgctcttcatgagcaaacttaatt
ttatattattgtctttcatttaaagaattgaatgaataatgttttgttgcaatattatac
attacaaaaacaaatatagaaattgttaatgagtataaacttt-----------------
------------------------------------------------------------
------------------------------------------------------------
--------------tttaaataagat----------------------------------
------------------------------------------------------------
------------------------------------------------------------
------------------------------------------------------------
------------------------------------------------------------
------------------------------------------------------------
------------------------------------------------------------
------------------------------------------------------------
------------------------------------------------------------
------------------------------------------------------------
------------------------------------------------------------
------------------------------------------------------------
------------------------------------------------------------
------------------------------------------------------------

------------------------------------------------------------
------------------------------------------------------------
------------------------gatacaa-----------------------------
------------------------------------------------------------
------------------------------------------------------------
------------------------------------------------------------
------------------------------------------------------------
------------------------------------------------------------
------------------------------------------------------------
------------------------------------------------------------
------------------------------------------------------------
------------------------------------------------------------
------------------------------------------------------------
------------------------------------------------------------
------------------------------------------------------------
------------------------------------------------------------
------------------------------------------------------------
------------------------------------------------------------
------------------------------------------------------------
------------------------------------------------------------
------------------------------------------------------------
------------------------------------------------------------
------------------------------------------------------------
------------------------------------------------------------
------------------------------------------------------------
------------------------------------------------------------
------------------------------------------------------------
------------------------------------------------------------
------------------------------------------------------------
------------------------------------------------------------
------------------------------------------------------------
------------------------------------------------------------
------------------------------------------------------------
------------------------------------------------------------
------------------------------------------------------------
------------------------------------------------------------

------------------------------------------------------------
------------------------------------------------------------
------------------------------------------------------------
--------ataaagaattaacat-------------------------------------
------------------------------------------------------------
------------------------------------------------------------
------------------------------------------------------------
------------------------------------------------------------
------------------------------------------------------------
------------------------------------------------------------
------------------------------------------------------------
------------------------------------------------------------
------------------------------------------------------------
------------------------------------------------------------
------------------------------------------------------------
------------------------------------------------------------
------------------------------------------------------------
------------------------------------------------------------
------------------------------------------------------------
------------------------------------------------------------
------------------------------------------------------------
------------------------------------------------------------
------------------------------------------------------------
------------------------------------------------------------
------------------------------------------------------------
------------------------------------------------------------
------------------------------------------------------------
------------------------------------------------------------
------------------------------------------------------------
------------------------------------------------------------
------------------------------------------------------------
------------------------------------------------------------
------------------------------------------------------------
------------------------------------------------------------
------------------------------------------------------------
------------------------------------------------------------

------------------------------------------------------------
------------------------------------------------------------
------------------------------------------------------------
------------------------------------------------------------
------------------------------------------------------------
------------------------------------------------------------
------------------------------------------------------------
------------------------------------------------------------
------------------------------------------------------------
------------------------------------------------------------
------------------------------------------------------------
------------------------------------------------------------
------------aaaaaaataattggaaaccttaagtattcattacttttgtgttatagt
tatttactatcaaaggtagtggcataatatggaacacaaaaaatgttcgaacgctttact
gaaaaagctataaaagtcatcatgctagcacaagaagaagctagacgcttaggtcataac
tttgttggtactgagcaaattttgctaggactagttggtgaaggtactgggattgcagcc
caagtcttaaagtcaatgaatgttaatttgaaagatgcaagagtcgaagtagaaaaaatt
attggacgaggatcaggttttgtagcggttgaaattcctttcactcctcgagcaaaaaga
gtattagaattatccttagaagaagcacgtcagctaggtcataattatattggaaccgaa
catctattaatgggcttagttagagaaggcgagggcgtagccgcaagagtactagagaac
ttagcagtcgatgtctcttctataagagcagaagtaatacaaatgctaggagaaaatgct
gaagctaatgttagtggaagtaatgcgactcaagctaggagtaaaactccaactttagaa
gagtttggttctaacttgactcaaatggctatcgaaggcggcttagaccccgtagtagga
aggcaaaaagaaatagaacgagtaattcaaattctaggtaggcgaactaaaaacaatcct
gttttaataggtgaaccaggagttggaaaaacagcaattgctgaaggtttagcgcaaaga
attgcgaatagagatgttccttctattctagaagataaattagttattactttagatgtt
ggcttattagttgcaggtactaaatataggggagaatttgaagagagactaaaacgcatt
atggatgaaattaaatcagctgataatgtcatattagttattgatgaagttcatacttta
attggcgctggtgctgctgagggtgcaatagatgcagccaatctgctcaaaccagcttta
gctagaggagaactacagtgtataggagcaacaacattagaagaatatagaaaacatata
gaaaaagaccctgccctagaaagaagatttcagcctgttgttgtaggagagccaagtgtc
gaagaaacaattgaaattctgtttggtttaagagatcgttatgaaaagcatcatcaatta
actatgtcagatggtgccttagcagcagcagctaaatatgctaatcaatacatttcagat
cgtttcctgccagacaaagcaatagacttgattgatgaagctggttcacgagtacgttta
ttaaactctcagttaccgccagcagcgagagagctagataaagaacttagggcggtatta
aaaactaaagatgaagctattcgagcgcaaaagtatgaaacagcagagcaatatagggct

agagaaatggagattaaagcacaaatagccgcaattgctcaaagtaaaaaaaatgagcct
gatttaaatcttgaagatccagttgtaacagaagatgatattgctgaaattgttgcagcc
tggacgggaattcctgtaactaagcttactaagagtgagtcagaaaaattgatgcacatg
gaggaaacacttcatggacgtataattggccaagatgaagctgttgttgctgtctcaaga
gcaataagacgtgcaagagttggtcttaaaaatcctaatagaccaattgcaagctttatt
ttttctggacctacaggtgttggtaaaactgagctaacaaaagcacttgcatcgtacttc
tttggttcagaagcttcaatgataagactagatatgtctgaatatatggaaagacatact
gtgtccaaattaattggatctcctcctgggtacgtaggatatagtgaaggaggttattta
actgaagctgtaagaaaaaaaccatatacagtaattctatttgatgaaattgaaaaagct
catcctgacatattcaatctactacttcaaattttggaagatggtagattgacagatgct
aaaggtaggactatcgatttcaaaaatacacttcttattatgacttctaatattggatct
aaagttattgaaaaaggtggtggcagtttaggattcgaattatcagaagatcaaacagag
tcccagtatacaagagtgagatctttagtaaatgaagaattaaagcaatattttagacca
gagtttttaaatagacttgatgaaatcattgtattccgtcagctaactaaagacgaagtt
cgagaaattgctgaattaatgcttaatgaagtttttgcaagaatcaagcagcaagatatt
caactaaatgtaacagagcgctttaaggaaagattagtagaggaaggttataatccaagt
tatggagctcgaccattaagaagagcagtaatgagactattagaagatagtttagcggaa
gaagtcctgtctggtaaaattaaagctggcgatagtgcggttgtagatgttacaaatgaa
ggagaagttacagtattactaggtgaaaaattagaattattaacctgaaaaccgtaatta
tattataaat-aataagcatttaatattatatacattaaatgctttaaaaataaaaagtc
ttaatatgggttgctataattagtaacctctgatattctttatttgtatatgattaagcc
gggatagctcagttggtagagcagtggattgaagatcctcgtgtcaccagttcaaatctg
gttcttggcatatttaaacattcaaaaaattagctaaattagaatcaaattttaactgaa
atgttcctagtggaccatttctatgttttgctactatgatttctgtcatatcttccattt
ctatttctttattataatagctttctctatataacatgatcaccaaatcagcatcttgtt
ctatagaattatgaacgataatattatttgcaatgaaattaggtataggataagttgtga
aatcaaaaacattttcaaatttagagatacggatagtgataagaggttcaaaattacata
atgaaaaagataggcaattcaataaatactgttttcttttatttagtttaaactgtaatt
gagtagtaatcatatcattacataatatttgatcgcatcgttgccagccacgtaatgtta
atattttatgattactagttaactctaaatatttatctgagtttgtcctaactttatatg
tggttttcctattttgccttgatacatatgcttttcctatcccaaatatttgattaatct
ttttattgaaattatatacttgagatttgtgaatactaaaatcaaaaaaaggtttactaa
tttgactacaca---------taatatagttaaattttgatatgcatccactctctctca
aatctgataatagcggtcttttgttatgacgactttcaagatttctattaagttgagaga
gaactaaaataggtagatctaattctctagccaagatttttagtgatcttgtaattagtg
aaagttcttgtgacctattgtttgactctctactatcttgtaataattgtaagtaatcta

taataattagctcaatttttttcccctgcaactttaataattttacttttgtctttataa
tatcacaagaaatttctgcactgtcatcaatataaaaatttaaattagccaaaatcttgc
tttcgtctacaacgttttgccattctatattagtcaactttcctgattgaatttttttgac
tatttaaatgacattcctgagctaagattcttcttaataactgttctgtagacatttcta
agctaaataatattacataatattggctagttttaataatatttcttgtaatgttaatcg
cgaaagccgttttgcccatagaaggccgccctgcaattataattagatcagattttttgaa
aaccatttgtaattaaatctaacttagagaaaccagaaaaaaatactattactgatactga
cttttttttttgtatcaagattaactagtaacttggcaaaagttttggctaaattattcg
tttctttatcctctaaaacttcataggccattgttaactgattagtggctgacgttattg
agttttgcataattggttctctggaacaactaataagacataatgcatcaccagatttta
atagcaatcttttaatataattatctagaataactgttgaatactcgtatataatattag
acgttggtgacctgtcaattaatttataattgtttccagtttacctaattgtgtcacta
aatttttgtatttaacatagacaaaaaatttgctaaactaattttgtcaattggattaa
tagtttctagtattgctctataaagtaaagaattgcttttaaaataaaaaaaaatctggtg
ataatctttctatgtgttttaacaaatcatttaattcttgagttaacattccactaatta
atattttctcagcaagtatattattaggaggtaaatacttataa------atatggatta
tttcctcctttttttatattattgtatta--------ataataggtacttctttttttaag
ttgcttctggaagaacttgtagttgaatatttgcagtcatttgattaaatagtttaattt
caatattataaagaccaatagttttaatttctggtaagaaaatattttgcttatccacat
ctatatctgtagtgtttttaataatttgtgaaatttctctttctgttacgctaccaaaaa
tgttttctccatcaccagttttcttactgatactaaattttttggatttcttctaaaagct
gctgtgtttttcttgcattttcttttgcaaagttcatttttttctctttaatggcagcat
ataatttctgttgattcaatatacctgtggttgcaactgatgccattttattaggaatta
agaaatttcttgcgtaacctgtagcaacctttataacatcattacttttgccaagttttt
gaatgttttcttttaatacaatccgaataactttttttactcatttt-----ttatttttg
aaaaagataaatatgtttttttagtatactgtttatataaacatttt----gacaggcga
attctgtttaatgtcacaatactttttttagtgtcatgctcgattttctgtttt--ttctg
atataatctatctaatgctaagcttgcagtataggagagatggccgagtggttgaaggcg
cagcattggaaatgctgtttaggagcaatcttaacgagggttcgaatccctctctctccg
cataattcaattatgaaaagctctttacatataatttttttattacctattatattttttcc
aagagtttaaacctccttttaccctaataacattaaatttatgcctctttaaaaaaattgt
aagcaaatattgatctagaatctaaactacaataaataaagctaatcttatcttgtaaat
gtaaatctaaataatattctttcttcatatctcctaaaggcaaatttattgcttgatcta
aatgactttctttgtactcttcgctactcctaacatctaatacaatgtattttaagctat
tccgacataatttattttgaaattctgtaacatcaatttcttgcatagaataactcttat
caccaacac----ctggctcttcaacttttctttcaggtggagataaaataaactgagta

tgaacaattttgaatttttcgaaggaaagtgtcatagcattatattttaaaataatacca
cttaagatagatttatagccaagaattatttaatagcttcagttgcttgaagtgtacct
ataatgcctggtaaaagccctaaaactcctgcattgctacacgtatcttctgcagtattt
tcgatttcaatactattatgaaaatctctgtatttaggccccccttgatagttaaaagtg
ctcacttgtccttcaaattgaaaaatagcgccataaatatggattttatttaattcaaga
cagctatcactaataacatacctagtctgaaagttatcagtaccatctataataatatca
tactctctaataatttctattgcattactagagcttaatcttgtattaaatagctgaaca
ttgcattttgggttaatctctaatatctttttttcagcaatacaggcttttgataatcct
atatcattagttgtatataaaatttgtctctgtaaatttgaaatatctattatgtcatta
tctactattccaattgtaccaatgccagaagcagcaagatatatgagtgcaggcgatcct
aatccaccagctccaacacatagtattctggatttttttaacctttcttgtccttctaat
tgaatctggggtaagataagatgcttagaataccttgtatattcttctaatgaatatgtt
gtatattttgctctgaaattaagcatgacaatttacaagttagttaatga----------
--aaaagtcagtagttgctttatgtcaactgtctctgcaaatttggctcttctttatcta
aaatagccccttctgtgggacaaacttgaaggcaaatactgcaatcaatacaagcagcaa
aatctatccaataccaatcctttttcatgttgttctcccctttgcctttgtgtatacagg
acactgggcatgcatttacgcattcggcaactccaatacacttttctgttacaattgtat
gagacatattatttaattaatgaaaaaataagctatttacatctagtatttttgttgtat
actaaataaagtccaactgttttaatt---------gttaacatatatttttagttgact
tgtttcttatgcgcctttagttcagttggtagaacgcaggtctccaaaacctgatgtcga
gggttcaagtccttcagggcgcgtctaa-tttgatttaagctcaatatttgaatttgacg
tttt-ataaattttaaaatgatattagataaaacctgtcattttatatattctatatttt
atctagtattattgtttgtttaatatgttattattggtaagttaattatattaactctat
ttttattgcatat-aaagcatgttgaattaagtaaaacagttatttaatagtctttttgt
ttatataacctttgaaatttaaaaaattatggccaaaaaagttacaggtatcgttaagct
agcactaaatgcaggtaaagctacccctgctcctccagtcgggcctgccttgggacaaca
tggtgtaaatattgttatgttctgtaaagaatataatgcccgtacagcagataaatcggg
actagtaattcctgtggaaattttagtatatgaagatagaagcttctctttcatattaaa
aactccacctgcatccgtgttaattgtgaaagccgctggtctgaataaaggatctggtga
acctaatacaaaaaaaaataggtagtataactactaagcagcttgaatctattgctgagac
gaaattgcccgatttgaatacaaaaaatttatctcaagctatgaaaatagttggaggaac
tgcaaaaaatatgggaatcttaataaaagattaaaaaatttcaattatacttatttctta
ctttatgaaaaaattttcacgtcgactcaaaactttgaaatctaaagttgatcctaaact
atatagcttaaatgaagctatagccatgctacaagcaacatcaaatgcaaaatttacaga
gactgcagaagctcatatctctctaggattgaatcctaagtatgcagatcaacaattaag
agcaacagttattttgccaaaaggaactggaaaattagtaaaagtagcagttattgctaa

aggtgaaaagttaacagaagcaataagtgctggggcagatatcagtggctcagaagaatt
aattgatgaaatttctagaggccgactagattttgataagttaatagcaactccggatgt
aatgcccttaattgcaaaattaggaagagtcttaggaccgagaggattaatgccatcacc
aaaggcgggaacagttactctagatattataaagtctatcaatgaatttaaaggtggtaa
agttgagtatagggttgatagaaccggcattattcatgttccttttggaaaatctagttt
ttcagaaggagatttacttatgaatttgcaaacaattaaagattcgatcgataaaaataa
gccgtctggttcaaaaggtaaatactggaaaacctttttcatttctagtacgatggggcc
atctattcaaattgatattactagccttttataaaaactttgtatataattaaataactt
g---------ctattttaaaatagtaattggctttcaataaaagtctaagaatctactta
ttatatcttaccagcaataaatagagttttatataacttattaattatgagcacaaaagt
tgaaaatatcctagaagaattaaagtccttaaatcttctagaagccgctgaattagttaa
gcaaatagaagaaacatttgatgtagatgcatctgctgcttctggaggagtaatgatggc
agctccatcatcagcgccagctgcatctgatgttgaagaaaaaacagaatttgatgttgt
tttagaagaagttcctgcacctaagaaaatagccgtcttgaaagtagtgcgttcacttac
tggtttaggattgaaagaggcaaaagacttagtagagtctgctccgaaaactttaaaaga
aggtgcatcaaaagatgatgcagaagctatgaaaaagcagttggaagatgctggtgcaac
agttggtttaaaataatagaataaaaatgtgcttttgtttatttcgaaacaaaagcacat
ttttattctattttaa----tttactaaaatagaccaagtgtaattgcgttattaatagg
catggtagcaccaatacctaaccaaatagtaacaacagtaccaattaaaaaaatagttgt
agcaattggtcttctaaatggattctgaaacttgttaacattttcaatgaatggaacagt
taataatccagctggtacagcggccatactcaaaacacctaataatttatttggaattac
tctaagtagattaaatgttgggaagaagtaccattctggtaagatttctaatggagtggc
aaaaggattagatttttcacctaaacttgaaggttctaaaattgctaatccaatactaca
tgcaatagttccaagaattactactggaaatacatataatagatcgtttggccaagctgg
ttccccataataattatggcccatacctttcgctaatttagctcggagttttggatctgt
taaatcaggtttttttaagaattgacatatta-ttgttctatttaatataagatttaaaat
aaatttttttacattagtgagtactcacg-tattataatggaccagaaattccttgcttcc
ttatcattaagaaatgcatcagcatgaaaacagcagttaaaagcggaagtacaaaagtat
gtaagctgtaaaacctagttaaagttccttgtcctacgctaactcctccccttaataatt
ccactatgctctccccaacaattggaacagcatcaggaacacctgtaacaatttttactg
cccaatacccgatttgatcccatggtaaagaataacctgtaactccaaaagaaacagtta
aaactcccaaaattacacctgttacccatgtcaattctctaggtttcttgaatccacccg
ttaaataaacacgaaatacatgcaatatcatcattaataccatcatgctagctgaccatc
tgtgtattgatctaatcaaccagccaaagtttacgtcagtcataatatattctacagatg
taaaagcttcagcaactgtaggtctatagtagaatgtcattgcaaaacctgttgcaactt
gaattaaaaaagatacaaatacaatccctcctaagcaatagaagatgtttacatgaggtg

gcacgtatttactagaaatgtcatcagcaattgcttgaatttctaatctttcttcaaacc
agtcataaattttactcataaaatagcttcaaaaagc-tttttaatatatttttttgcgtt
tgagctactagggagtcaataaatttatttattgaacttgtatt-aatttactactaata
catatttataattataacatttttttagatatttgattgcaatttacttatttaaaaaac
ctttgactcaataaaatcggatcgtgaataatgactttcttttttttgtatagaaattaat
ttcgtttgaactaattcagacataattcttgttatactaactctattactacctataatt
tgtgctaaaactctatgagtgatagctaaattaagaaaaattccattactttggctgatt
ccattatgttcagataataatagtaagaggctaacaagcctattcgtaatgcttttatgt
gaaatgatttccataaagcgatttgcttttattgaacacaataatagatgatctataaaa
aatacactgaacttatgatggttctggcatatactcataatagtagagtattcaatggat
actatttgcgctacatctattgcttctacttcatagtaaaaatggctacttgcaaattgt
gtatgcccaaaagtatcttcggaagttaatagatttaatattatctttttttctatttctg
aatactttttttaacaattaagcatcctgctaatataatatataacgtagaattagcatca
aagatgagaacatcgtgttgtttaagtgaaaatatttgatgttcagcactttgttgatta
aaaaaatttagccatggactacaacaagaatttttttttaacaatatttttttggtactttt
tcaaaggaacactcttgattcaaataatgattgttcataattaatttataataattaatg
taccaaaaaattttaataatttaataattaaaattaaatgtcctacaatctgatgttagt
tgaaaatgatactgtattatcgaaagcaattcaagaatacttaatagatcaaggatttaa
tgttgctattgctattaatggattggaagcttttcagcttgccaataaatataaatttga
tctaattatatcggatataataatgcctataattgatggatacgaattattagaaaaact
gcaaaaaattgaaaaacttgcaaaaattcctgttatttttctaacagcgaaaggaatgac
aaaagatagaattaaagggtataatatgggttgttatggttatttatctaagccatttga
tccagaagaactagtttcgcttattaaaaatttaatttatagagattcattaaatgacca
aaacttcaaacaacaaaacgtaattgaattacctttattcaaatcgtttcatctgacacc
tagagaaaaaagcatccttaaccttgtaattgatggattaactaataaagaaatagcagt
tatattagatacaagtacgagaaatgtagaaaaatatgtaagtcgtcttttacaaaagac
aaatacaaaaaacagaactttattggttaaatattctataaaaaataatttattaaat--
------caaatatagggcgaatgacgggactcgaacccgcgaatgatggagccacaaccc
attgccttaaccccttggccacactcgccatattata------actatagcttgttttcg
tatatgtcgtctagtattacattaatctatttttctta-gtaattttataattagttct
atatgatcaataaaaatataaatattcaaattaatggagagccatttaattgttcaggga
ctctttcattacaatttttattaaattatcttgattttaattctgagcgtgtagcaattg
aactaaataatactttattacctgagcgcctcttccattcaacttatttaaatgaccaag
ataaaattgaaattattactattgtaggtggagggtaatttgtcttgagtatttcaagac
actgagatttcatttcaaatgtttcatagataatgataaccgcccttgtttttttgtctac
atgaataataactgcttttattgtctcacctattttgaatttagatgttatcttttctac

ttgtttaacattaatttcagaaatatgtacgagacctttagatttccagactttataaa
taacccgtaaggtgtaatttgattgattaccccctcgataatattcccaacaattagatt
agatgaagcttgcgctattaaagctctcctatgactaagtattaaattatttgattttc
ttcaacattaagtaatttgagtttaataaatttatttgtagaaggggtattttggtgaaa
actgtttagatgagaattgggaacaaagccagatataccttctaggttaactatcatgcc
tcctttattaaatcctttgatcctaacatctagtaaggaatcttcagctagtaattgcct
aattcttttccaggctcttatatattcaagacgccgaatagataaaattagttgttttga
ctcaacgttataatctaataagaaaaactctctagtatcattaatatttaaagagttaaa
attattaagttcctgattactagatacctcttgtattggtaaatacgcagatataggtgt
tccaatatctactaatactccatttaattcaaaactaaatattgtgccggctacaatatc
accaagatttaaatcatatttatatttttgtaagacggccgcaaaatttcgatgagtgaa
actttcattatgttttgtcatatttaggaaattcaattaaaaatta-------------t
atttgatcttgtattctactaatattttatatc----aaccttaatgtgagtcattgtaa
taatctcttgaaatagattaaaatacatataagacataaaaagtaaacgtagatagctac
agattcca-tttaatacaatctgaggaaagtccgggctccacaaatacaatttatgctgg
aaaaaccccagtgtaggaaactatgaggacagtgccacagaaataaaccgccagaataaa
t----tatattagctggtaagggtgcaaaggtaagttaagagctaaccaaaaatactgta
aagtatttgttaggtaaaccccagaaatggagcaaagcgactaaacaaattttt-tgtat
ttttt-agtagtttagtttaaaatactgcatgaagttattggtaacaataactctagagg
aatagctatcctttgcaatacttagtccaatgtattcgaagaacagaacccggcttatgt
agtactttttgtgtcttgtttgatcattatcagttgcgaatttctaaattcagtttttttg
ataatattattttgaagttcattctgcttattatctatttctaaatttgtcaaagttttc
tgatttgatctatatgtaaagcgcaatcctatactttttttgccgtttccaagtgatcta
ttattatactgatcaaacaatgcaatattttctaagtcatcatcattaaattgatttaat
aattcaaataaagaatttatctgcatttttttgcggtattattatacataaatctcgtgta
atacacggatatttagaataaggctgaatgcgataatttaaataattcagttctgtatta
taatgtgttagggcatttaaatcgatttctagaacaaataattttgtactaaaattaaat
tctgaagatgctaactgatttagttcaccgaataatccaatattatcattgttataagtt
aaagtagctgatttatgattgtgaacgaaattgatttgaccacttgttaattgtcttttc
acccattgaatgttttttatttaatttccgaaaaaagttttcaataatgccttttgcttca
taccagtttaaagaatgtgcaggttcggaccattctgatctaagatctaagtttcctcct
aatataatggctagttttgttgtctctataattttattttgtcttaggttgaatacagtt
cctatttcaaatccatctacagtttgattactttgcttgatattatatgcactggtagtc
attaagccttctagtaagctagagcgtaaagtagagtaatctttaataagagggttgttt
aaattaatttctccattggatttcactaacgaatagtgaactaattctgttaatcctaaa
tttcttaaaatacttctgatctgatcagtaaattgttttctagagggggggattttgggca

aattgaatttttggtattgtactttggaatttatcataaccataaactctagcaatctct
tcaataatatcaatttctcgaaatatatcatatttcctataattagggataataacattc
aatttcttattattttcttttgcaatcttgaactgaagagactcaagtgttttcttaatc
tcgttaaaagataaaaaacgaaccgtattattatcattaattggtcctaaaatatcatga
atttttttaatgatatatcaatagtaagagtggaatttgaaaatgtttttttgaaaat
gtttttttaccttgccgtcagttaattcttttagtaaatacaaagtttcatgataagcg
ctttcccaattatctatattaagtcctctttcctgtctaattgaactttcagtacggata
ccaatacttctcgatgattttctaactactgcctgtttaaagattgcagcttctataaaa
attgattttgtgttttgatcaatactaaatttatgatttatgccaatacccgctacgcta
gttatattattatttatctcagtaattaaaatatctttacttaaagtaatatctttacta
tctagtttaatgaaattttgcttacaagagcaaaaattactagtgatctcgacataatct
tgattatcagtactgacaattttattgaaatctataatactaattggctgccccatttt
aacataatataattgctaatatctatgagaagattttggcttataaagccacaagagtat
agacgatttttagccatgttggagaatcttttatgactatattatccataattgctgat
aaatagtaattacaatttaaaagattaccatttctaattatattattcttattccgaaat
aagtctgttgatggaat-atcagttatgtgaaccatgtcagcctgaattaaagcagagac
ttctctagatagccctatcatgctaagtgcatctgatcgattggctgttgaggtaacatc
taagatgtagtctgtatagccatctatatcaataaattcaatatttctacttcaaagcc
tgcttgcgttaacttgctagctaaactatttatgtttattacttctattctagcaagttc
ttttaaccaatttaaagaaactttcataattgtttttcatctattttatagaactattga
----attttaattaatcgcttgtttaaaaaattcatcaaattaatataagttatttgat
gaatttgcttatacaatataatattgtgatgttttactttgagatt-cagt--ctaagaa
ttagcttttgtgaaagtgagtccgttatcttgagcgtacctttccataaacctcataaat
cgatcccagtcatccggacttttaattacatgtacagcttcgattgcttgtggtttacca
ttaataaacttagcattgacatctcttgtaattatttgtccttcagtatccattagatac
atacccgtaatttctcctttatctgccatgctagcatctaaaatttaggattagtaaat
ctgaatgtagctgttcctgtactaccatctcgagatcttgttaaacgtacatctggcact
acttcttcattgatgccttgaataaattgaattgttgccatatcaatttttaaatttat
atatgatttatattggataaaagtgatttctgattctttaatgtttactaaataactatt
ttaaaagcttatattaccagcagatcaaattctatttattatgtaaaactgtaattttc
ttttttgtactttagatttctataagtacaggcagcaagacacacaaaaataagtttac
taaaaatacaatttataattgaaaagtgtagtttttaaagtaggcacgtatttaaataaa
aagttaatgctatagaactatcacaaattcactatttagttaatgataatttttctaaa
atttagaaaagtgtagtgtcgctttttagtgagtaaaaaatttaccaatatagttatta
tgtatgtgattaggacttctagaagtatattttttctaatacagatatattggtgacact
tactatcgttaaaatttttttatctattcaaagaattaatttattataagctaatcatg

ctgatattagccatataatgtttt-aagt-gaaattaa-gggttgtagctcagatggata
gagcaagcgcctcctaagcgctaggtcagcggttcaagtccgctcaatcctgttaaaa--
------------------------------------------------------------
------------------------------------------------------------
------------------------------------------------------------
------------------------------------------------------------
------------------------------------------------------------
------------------------------------------------------------
------------------------------------------------------------
------------------------------------------------------------
-----------------------------tattatgaagtg-------------------
------------------------------------------------------------
------------------------------------------------------------
------------------------------------------------------------
------------------------------------------------------------
------------------------------------------------------------
------------------------------------------------------------
------------------------------------------------------------
------------------------------------------------------------
------------------------------------------------------------
------------------------------------------------------------
------------------------------------------------------------
------------------------------------------------------------
------------------------------------------------------------
---------------ttttgtaatttaactggggtgggaggattcgaacctgcgaatggc
ggagtcaaagtccgctgccttaccacttggctacaccccaatatatgaagactataatag
cagtcatatcccctaccatgtcaagagataataa---taatttagtagctattgaatttt
ttgctaaaagcaatcttcttttttagataagaaattttgtttctaaattgaataattgag
atattctcttcctcttgagtacacatccattttatagtaattgtttcttgttgaacttca
ttatctcctaaaattaaacagactatagctctttttttatgtgcttgtttgatttgtttg
ccaaaattgctagaacttatgtctaactcagtttttataagctgattgcgaagaaattga
gttattatcattccgatctcttgagcttttgttccctgcgaagcaacatagcaatctatt
gacatattatgcagtaatatattttcctttgcaattaatagtagtctttctaggcctata
gcacaccctacagctggagtgttatttccaccaagctggtgtatcaaattatcatatctt
ccacctccacataccgtatcttgcccatgagactttaatgttttaatctcaaaagcagta
tcattgtaatagtctagacctcgcaccaattgtgtatttattttgtatggaatattgagt

aaggttaaataatcacaaacacaattaaaatgtttttagactcaagacttaagaagtca
gagatttgaggcgcatcacttaaaactttttgtgtatgtttattttttgtgtctaaaatt
cgaatagggttagatgaaagtctattttgcgaatcagcatctaaatcatcatagtattgt
tcaagatattcttgcaattttgcttgataatgatttctatcttttgctttgccgattgaa
ttaatatcaagctgtaattcgttaatagatagatctttaaaaatattcattgctaaatga
attatttctgtatctgctcttgcatctgaactaccaataaattcaataccaagttgatga
aactgtctttgtctgccactttgaggtctttcatatctaaacattggaccattataccat
agtttttgtaacttgttttgaatatccattctattttcaatgaatgctctagcaacgccc
gcagtaccttccggtcttaatgtgatatttctattgctccgatcattaaatcgatacatc
tccttgtttacaatatctgtattttcaccaattcctctatcatatagatcactattttca
aaaataggagttttgatctctttataattcgcatacttgagcaagttggcaactttttta
tctatgaactgccaatagagttgctcatctggtagaatatctttagttcctctaatagcc
tgaatttttgccatgatactgttttgatcattttcctaattttttgcactagatagcag
ttgtacactaacatgt----acgtattatacactaa--aatgtttatttgacgttttata
aactacgggcaaggagggattcgaaccccgacaccatggttcgtagccatgtgctctaa
tccactgagctacaagcccacttaataagcattttattatatcatttttttcatctata
tacaactataaaacttctgaataaattcttcatccccagactctctctgataagatgatt
tctataagttattcttgaatagttgagaattttctagatgaagaggaattc-ttatagtt
ctgcaaaccaatatttaaaagataaattatttaattttttta-----aaaagatttgtat
aa-ttttcacccttgctttatcattgaactgattaagcatttaatttgtctttaaaaatt
ttaaaagagttctattaaacatgagtaaacaaattctatatcaagatgatgcccgcaaag
cattagaaaaaggaatggatattttaacagaagcagtttctgttaccctaggtcctaaag
gtagaaatgttgtactggaaaagaaatttggcgcacctcaaattataaatgatggtgtta
ctattgcaaaagagattagtctcgaagaccatattgaaaatactggagtagcacttatta
gacaagctgcatctaaaacaaatgatgtagccggtgacggtactacaactgcaacagtgt
tagcttctgccatagtcaagcaaggaatgagaaatgttgcagcaggctccaatccaatgg
ctataaagaaaggaatagaaaaagcaaccaattttgtagttaacaaaatagctgaatatg
ccaaaccagtggaagacacaacagctattgttcaggtagcctctatttcttcaggcaatg
atacagaagttggtaaaatgatagctaatgcaatagacaaagtaggcagagaaggtgtta
tttctttagaagaaggtaagtcaactaacactagtcttgaaataacagaaggtatgcagt
tcgaaaaaggttttatttcaccgtattttgttacagatatggagcgtatggaggttctcc
aggaaaatccatttatcctttttacagataaaaagattactttagttcagcaagaacttg
tacccttacttgagcaaatagcaaaaacctcaaagcctctattaataatagcggaagata
tagaaaaagaagctttagcgactattgtggttaataaattaagaggaatactaaatgttg
ttgcagttcgagcaccaggatttggagatagaagaaaatctttattagaagatatgagta
tcttaacaggtggacaagtaattacagaagatgctggtctctcgctcgacactgttcaat

tagatatgctagggcaagctagacgagttgttgtgactaaagattcaacaacaattattg
ctgatggtcatgaagttacagtgaaatcaagatgcgaacaaattaagagacaaatagaaa
taagtgactcgttatacgaaagagagaaattgcaagaaagattagctaagctttctggcg
gagttgctgttataaaagttggcgcagcaactgaaacagagatgaaagataaaaagctta
gattggaagatgcaattaatgcaacaaaagcggcaattgaggaaggaattgttccaggag
gaggctctacaaatgttcatatctccggagaattattcgtatgggctaaaaataatttat
ttgaagatgagttgataggtgctttgatagttcaaagagctttaacttaccctttaaggc
gtatagcttttaatgctggcgataacggagctgtggtaatagaaaaagtcaaaactaatg
atttccatataggttatgatgcatctaacggaaagatcgtaaatatgtatgatactggta
ttatagatcccgcaaaagttgctaggtcggcactacagaatgcagcttctattgcagcta
tggttttaactactgaatgtattgttgttgataaattggaagcataaaatatatctttta
taacatctctaatattaaacgaagaataattttcatacttacaaattattcttcgtctca
actaattttatcaactgttcaaaattataaatttcatagtagagccaaagtttaaatagt
cctttacattctattgttaaataaattaaatataaccctgtaattcctaaattttcaatt
agaatattattttcgtatatagatttttccatatattttaaaaatgacatttttttaaaa
tagtataaaaatcttctctgataattgttggctaaattgctatcatataaagatgaatgg
ctggctgaattcaaaacctttgaaataataatttctaagtctttgtctgattgtatacca
gaatcaatgataaactgatttaattgatcatcaaaatttagattgatagtattgattgtt
gcacatgtttggttttgaatatctaaagatttgatagccaagaagattaaacttttattg
ttcattttaaaatataagacaattgacgttgtattctttgaataatacttttttatcttct
tgtatgtttatgtttttttataaaaataatagatagtacta--------tgctaattact
acctgaaaaaataaattctgggaatctagcttgtgctttcgacaaagagtgctttttacc
ttcttcttgccagtaattaataacctcagtagttttatttaataattcaattctatctct
ttcagcaatccatggtttcgactctagctcagcttttaattcttcccaagcgtcatttct
aggccaaaagaaataagatgtcagtgggctagtgccatttccaacaatttgatcaattgc
aatagcaatattattttctaaccaaagtacctttagtgtgaatttagacaagttattatc
tccgttataaatatttaatgtgta----gacactatgatatgttattcactatttttttct
ttcaacaatttgagccttgtttaaaatattttgaacagttcttgtagcttgtgcgccatg
tttgagtctctttaatatttttacgctatcaatatgagtttcattagtgattggattata
aaatcccaactcttcaatggctttgccatcgcgtttatttctactatccatagcaacaat
tctatagctaggttgttgttttcttccgtatctttttaatcttagttttaccatcttgtc
gatttagtttgttttt---atatctttattaaattatatcttacgatattatgcaataat
tcaaaaaattaagcagattccagcctaacattattaaatataaccatcaagcattgtaga
aaaattataccaagcattggtgacatatccattccaaacatcggaggaatacttcctcta
aaaagttttaagtacgggtctgtaatgcgatttaatgaacaaaacggttcattgtaccaa
tttacagttggaaaccatgctaatgatagttttagcaaaatcaaaattaaataaatctca

gaaaaattagctattgatccaagtaataaatttaatgtaccaggaagagtattcatacgt
atttcataattattatatagcttattatactaacaatttgattatatattcgaacgaata
taaacaatcaaagttaagcaaaaaacttatctttaagaatttgttctttggcagcttcag
agctatcgagtaggccattaacttttgtagtgtatatttctagtgcagaccatttctgtc
ctaactggtttaattgagatgatccgcattctaagcaggctatcttaacattgtttaata
ttacacctttgttgattagctcttttagaacagatgtcataatgatttcatctagaaata
aatctaaaattagtatcttcgtaaatgaatctaatttatctgaaatattatcaatactgt
tattataatttactaatgctatattagctgtaggtagtaaagctcgagcaccttcagcga
gtataaacccatatggcattacaataacaatgatatatttataacgattactaataagat
tgatagctttactatcttttacgctcactgtttctgttaccagccattctctcattatct
cataagtaatccattttcctaattctgagcatgcagttcttaaaatagttcccggattat
tattattttctaggattccagaccaatgctgtattaatggatgagaagctacgttaattt
ggatttgcatattttattcaatatatttaacatttgatt-------------tcagcttt
atctttatctagtattattaaaatagcatacacatatattaaagaaaataattataaaaa
aatgaaaaaaaatctttggttttggggttttactgatagtgccgaaacatggaatggtag
actcgcaatgataggttttataacagttgtttttattgaattagttacaagtaaaggcct
tttgtatttagctggtttgatggattaatata-----aaaaaagttcttgtaaataacta
caagaactttttttatattaataatagcctaatctagaggtctcattgacagcactggtt
cattttctctattaatacctttttcaaatccagcagcagcagctcttgctctaccagcat
gccaaagatgtccaatgaataggaagaatcctaagaaaaaatgagatgttgtcaaccaag
accttggggatacatagttaacagagttaatttctgtagcaaccccaccaacagagttta
gagaacctagtggcgcatgggtcatgtattcagcagctcgtctttcttgccaaggttgaa
tatcattttttgattttatttaagtcaagaccattgggtcctctaagaggttcaacccaag
gagctcttaaatcccaaaatctcatagtctcaccaccaaaaataatttctccacttggag
atctcattaagtattttcctaacccagtaggtccttgagaagatgcaacgttagcaccta
atctttggtctctaactaagaaagtaaacgcttgagcttgtgaagcttcaggaccagtag
gtccgtagaattcactaggataagcagtattgttgtaccaaacaaaattagaagctgtta
gacccataattgataaagcacctaagctataagataaatatgcttcacctgaccaaacaa
aagctcttcgtgcccaagcaaaaggttttgttaaaatatgccagattcctccagcaatac
agatgatacctatccaaacatgtccaccaataacatcttccatattatttacactaacaa
tccatccatcaccaccgaatggagactttaaaacgtagccaaagataactaaaggattaa
gagtagggttgttaacaaacctaacatcaccaccaccgggtgcccatgtatcgtatactc
ctccaataaataatgctttgattactaacaagaatgctccaattcctagtaatactaaat
gtatacccagaattgtcgtcatcttattttttatcgcgccaatcatagccaaagaaaggaa
aagactcctctaaagtatctggacctatcagagagtggtataaaccaccaaaacccagaa
cagctgaagaaattaaatgtactacacctacgacaaagtatggataagtattgaagattt

ctccacctggacctacaccccaacccaatgtagctagatgaggaattaatataagtcctt
gttcgtataaaggtttctcagggacaaagtgagcaacttcgaataaagtcattgcaccag
tccaaaataccattatacctgcgtgagcaacatgtgcacccagtaattttccagaaacgt
taattaaacgtgcattaccagaccaccatgcaaacccggtagactcaatgtctctaccgc
caacaccaacatttgtattaaagggcgtttccacgtggtaaaacctcctcagggaatata
aagttttcatgaggttgatcttgagcagccatccaagaacgaataccctcatttaataga
atgtttttcgtgtagaaagtttcaaactcaggatcttctgcagctcttagttcttgagat
acaaaatcatatgctcttaagttcagagcaagtcctacaattccaaatgcacttgtccat
agtccagttacaggaacaaatagcataaagaagtgtaaccagcgtttattagagaaagcc
acaccaaaaatttgtgaccagaatctgttcgctgttaccatagaataagtttcttcggat
tgtgttggagtaaatgctcgaaaagtatccgcagcatcaccatcttcaaataatgtgttc
tgtacagttgcaccatgaatggcacagagtagagcaccacctaagataccagcaacaccc
atcatatggaatggattaagagtccaattgtgaaatccttgaagaaataataagaatctg
aaaatcgctgcaactccaagacttggagcaaagaaccaactagcttgacctagagggtac
attaaaaaaacagatacaaaaacagcgataggtccagaaaatgcaatagcgttataaggt
cttagaccaactagtctagcaatttcgaactgtctcaagcaaaatccaattagtccaaat
gatccatgtaaagcaatgaatgcccatagtccaccaatttgacaccatcgagtgaagtca
ccttgtgcttcaggaccccaaaggaaaagcagggaatgtcccatactatttgctggtgta
gatacagcagcagttaagaaattgcatccttctaaatatgaacttgctagcccatgagta
taccaagaagttacaaaagtagttccagttaaccagccacctacagcaaggtaagcgcat
ggaaacagaagtaatccagaccagcccacgaatacgaatcgatctctttttagccaatcg
tctacaaggtcaaaaccaccacgagtcttttcttgtccaattgctatggtcataatttaa
atctccagagcaaatttaataattaatttgtttcaataaacagttgtttagatgtagtcg
aaaattttacttttctttacggtcaatataattataagtcgaaatcaacaaaaattcacg
tttttaataacaaatttttttagctctttaagttaatttct----------gaatttggtt
aatatttttcatattttaatatttttcatattttattcatttctataattctttgaaaaa
gatatcctgtgcctcttgcagttagaattaaatcaggattactaggatcatcttctaact
tagctcttagtctagaaatatgtacgtcaactactctggtatcaacatggcgctctggag
tatatccccaaacttcttgcaaaattgatgctctagagaatggctcgcccgctctactaa
ctaaaagttccagaagactaaattccattcctgtcaagcgaactctctcgttattcttat
aaacttgtcttttattggtatctatttttaaaaaaccaatactaataataccagaattag
gaattcctagattggttgttattttatctgctcttcttaaaacagagcgaattcgagctt
ctaattcttttggagaaaatggtttcacaacataatcatctgcacctatttcaagtccag
taattctatcacaaacttcccctaaagctgtaagcatgattataggaacgtcagattctt
ttcttaactcttgacatacaccatagccatctaatttcggcatcattacatctaaaacta
ctaggctaggatattccttcctgaatataattaaagcttcttctccgttagaagcagtta

taacctcatatcctattattgttaacctagtttctaaaattcttcttatactagcttcat
catcaacaaccaggatcttttctttctgattctccaa-tttgttattcactcctataaag
tactaagcctaaatgtctctgttccaaatttatatttg--------------------ac
aaactt---tttactcattgccagctttgcaatctgagc-taatttagttgatataaat-
-aattatagcttgaaacaattttgtttcctttatttcattgatcgttattttactatata
agcatcattagcttaccaattttgaatttatata---cttttcatgtagatttttt----
-------------caataaaggggttgacaaacttaaggagagaaagtattctacattac
tataag----tttctgctcgtatataaatattgaataacattttatattgagtatatag
atttttaat
