## Supplementary material for "Genomic diversity of 39 samples of *Pyropia* species grown in Japan": S2 Fig

>Pyr_1

TATATGATAAATTAAGACATCTGTTATGTGAGAATTTCTAAAATATACAAAACTTTTTAT

TTAATATAGAAAAAGTAAATTAAAATAAGTTGAACATACTTTGCTCTAATCTAAACCTCT

TAATTGAGATTCTAATTAGAGACAAAAGTATTTACTTATTTTGTATCTATAGGAATAAAA

ATGAAACTTGCAGTTTATGGTAAAGGTGGTATAGGTAAATCTACAACTAGTTGCAATATT

TCTGTAGCACTTTCAAAAAGAGGTAAAAAAGTTCTACAAATTGGCTGCGACCCTAAACAT

GATAGTACATTTACGCTAACAGGTTTTTTAATCCCAACAATTATCGATACTCTTCAATCC

AAGGATTACCACTATGAGGACGTTTGGCCTGAAGATGTAATCTATAAGGGATATGGCGGT

GTAGACTGTGTTGAAGCTGGTGGACCGCCGGCTGGCGCTGGTTGCGGAGGCTACGTTGTA

GGTGAAACAGTTAAACTTTTAAAAGAATTAAATGCTTTTGATGAATACGATATTATCTTA

TTTGATGTTTTAGGTGATGTTGTATGTGGAGGTTTTGCAGCTCCATTAAATTATGCAGAC

TACTGCTTAATCATTACAGACAATGGATTTGATGCTTTATTTGCAGCTAATAGAATAGCA

GCTTCAGTACGAGAAAAAGCTAGAACGCACTCTCTGAGATTAGCTGGACTTGTTGGTAAT

AGAACAGATAAAAGAGATCTAATTGATAAATATATAGATTGTGTTCCAATGCCAGTATTA

GAAGTCTTGCCCTTGATTGAAGATATTAGAGTGTCCAGAGTAAAAGGTAAAACTTTATTT

GAAATGGCAGAAATTGATAAGGATTTAGCATATGTATGCGATTACTATTTGAATATTGCA

GATCAGTTGATTACAAGGCCAGAAGGTGTAGTTCCTAAAGAATCTCCGGATAGAGAATTA

TTTAGTCTTTTATCTGATTTTTACTTAAATCCTAAATCAAAGGTAGGACAAGAAAAAGTA

GATCAAGAAGAATTAGATTTGATGATAGTGTAAATAATATTTTCAGCATAATATAACAAG

GAATAAGATAATGTCTACAGCTCAATCAGATGCTCTTACTTTTGAATGTGAAACAGGTAA

TTATCATACTTTTTGTCCCATCAGTTGCGTTTCTTGGTTATATCAAAAAATTGAAGACAG

TTTTTTCTTAGTTATAGGGACTAAAACTTGTGGTTATTTCTTGCAAAATGCAATGGGAGT

AATGATTTTTGCTGAACCGAGATATGCTATGGCAGAATTAGAGGAAGGAGATATTTCAGC

TAAGCTAAATGATTATGGTGAACTCCGTAGATTATGCTTACAAATAAAAAAAGATAGAAA

TCCTAGTGTTATATTTTGGATTGGAACATGTACAACCGAAATTATAAAAATGGATCTGGA

AGGAATTGCTCCAAAATTGGAAGCAGAAATTCGTGTTCCTATAGTAGTTGCTAGAGCTAA

TGGTTTAGATTATGCCTTTACCCAAGGTGAAGATACAGTTCTAGCAGCTATGGCTCAAAG

GTGTCCATTAAATTTAAAGAATCAATCAGATAACACGAGTCTTAAAACTTCTCCTCATAT

TCCTTTAGTTTTATTTGGATCTCTTCCGGATCCAGTTGTAACCCAGTTAACTATGGAATT

AAAAAAACAAGGTATTTTTGTTTCTGGTTGGTTGCCATCAAAGAGGTATACTGAATTACC

GGTTATAAAAGAAGGATATTATGTTGCGGGAGTTAATCCATTTCTTAGTCGCACAGCTAC

TACACTAATGAGACGCCGAAAAACAAAACTAATTGGTGCACCATTTCCAATAGGTCCAGA

TGGTACTCGAGCCTGGATTGAAAAAATCTGTTCAGTAATGAATGTAAAGCCTATTGGATT

AGAAGATAGAGAAAAAGCAATTTGGGCTTCTTTAGAAGACTATATTTCTTTAATCCGAGG

TAAATCAGTTTTTTTTATGGGTGATAATTTACTAGAAGTATCTCTTGCAAGATTTTTAAC

TAGATGTGGAATGACTGTATATGAAATTGGTATTCCTTATATGGATAAACGCTATCAAGC

AGCAGAGCTAGCTTTATTAAAAGCTACATGTGATAAAATGAACGTTATGATGCCAACAAT
TGTAGAAAAACCAGATAATTATAATCAAGTAGACCGAATCCGTGATCTGAAACCCGATTT
AGTTATCACTGGTATGGCTCATGCAAATCCTTTAGAAGCCAGAGGTATTAATACAAAATG
GTCAGTAGAATTCACATTTGCACAAATTCATGGCTTTACCAATGCAAGAGATATTCTTGA
ACTCGTCACAAGACCATTACGGCGAAATCTTAGTTTGTCAGAATTAGGCTGGGATGTTTA
TAGCAAGCAAAGCTAGTATTCCTC----TTTATGACTTACATGGTATAAAAGCCTTGTAA
GTCTTTTATTTTATGTTAAAGAAGCATTTCTTGCTCTGCCATCACCAGCCCAAGCTTGTA
AACTTTCTACTTGTTCTTTGTCTGTAAATGCTAAAGGCACAAATTGTTTGATAGCTATTT
TAATATCTTCAGTACTAAATTCTCTCTCTTCACTAAACGCCGTATGCATACTTTCAATAA
TTGCTTGCTCAATTTCAGCGCCAGAAAATTTATTACATAATAAACTCAATTCATGAATAT
CATATTTTTGCCACGATCTTGGTCTAACTTTTGATAAATGTATCTTAAAAATTGATTCTC
TCTCTTGACGATTAGGTAAATCTAAGAAAAAAATTTCATCGAACCTGCCTTTCCTTAACA
TTTCGGAAGGTAAACTTTGAATTTTATTAGCTGTTGCGACAACAAATACTGGAGCTTTTT
TCTCTGATAGCCAGGTAATAAATGTACCAAATACTCTGGCACTAGTACCACTATCACCTT
GACTATGTAAACCAGAAAAAGCTTTATCTATTTCATCAATCCATAATATGCAAGGTGATA
ATCCTTCAGAGATATTTATCATTTCTCTCATTTTTGATTCCGATTCACCAACTAGTCCAC
CAAACAGTTTTCCCATGTCAAGACGTAGCAGAGGCAGCATCCAATCGCTTGCAATAGCTT
TCGCGGTTAAGGATTTACCCGTCCCCTGTATACCAACTAGCAATAAACCTTTGGGCGAAG
GAATACCGTAATTTAAACTTTGCTTCGAAAATGAGCGAGACCTTTTTTGTAGCCATTGTT
TTAATACATCTAAGCCACCAATATCCCTATTTACTTTACTATAAGGATAAAATTCTAATA
GATGCGTTTGATTAATAATTTGCCTTTTTTCTTCAATAATAATAGGTAAGCTCCGAGAGT
CTATCTGATTATATTGTGCAATAATTTTAGTAATCACTTTGCGGATTCTATCTATCGATA
ATCCTTGGCAGGATTTTGTTATATTATTAACGAGCTCTGAATTCAAATTAAGATTCAAAG
CTTTACTCAATCTTATGATCTCTTTTTTGATTTCTAGAAGACTAGGTAAGGGCAAATCTA
TTACTGTTATGATATCATTCAGAGCAAAAGGTATATTGATTTTGCAAGAAATAATAATAA
TGTGCTTAGATTGCGTTTTGATTATTTTAGCTAAATTGCGAAGTTTCCTAATTAATACTA
TTTCGTTTAGGAAAGAGTCAAAATCTTTTAAAAGAAAAAGATTTAAAGATTCATTATTCA
AGTTTTCAATAAATTCTAATGCTAATAAAGGATTTCTTTTTGCATAACCATTATCGCTAG
GATTGTTAGTGTATCCATCTACAAAGTCCCAGGAATATACTTGCTGACTGTTTGAACAGT
TTAAGCTGTGTTTTATAATATACTCTAGTCTATCTTCTTCTCTAGTATTTATTACAATAA
TAGGATATCGAGATTTTAAAAGTAATCGTAAATCTTGAGTGAAATTCATGTCAATATTTG
TTTCCATCAGACTTTTGTACTTCACCAATTGAAGAAAGACTAAATAAGATTATCTAATAT
TCCTTGGATATATTCAAAAGTCATCTCTCTATTTTCTAAATTATTGCTATTACTAATTTA
ACTTACCATGATTTTTTTTCTTCCTTTTCTTCGTCGACTATTTAATATAGAGCGTCCACA
TGGTGTTTTCATTCTTGCTCTGAAACCAGAAACTCTAATTTTTTTTCTTTTAGATCCTTG
TAGTGTTCTTTTAGTCATGATGTCTCCTCTGTTGTAAAAGATTAAATACTATTTTATCAT

CTTTTATATATCTTTATGATATAGATATGTATTAATATCAAAGTTAACAGCAATAATTTA

TATTAATCAAGAACTATTTATATAATATTAAACATTAAATATTAACTTTATGATATTGGC

TTTGCCCATTTTTTATTTAAGTATTTTGACCATCTTTTTGCTAATCTTAAACTGGTTAAT

TTTTCAGCAGCTAAAAACGATTTTGTTATTAGAATCTCAATTTAAGTATTTTGTTGATAA

AAGCCAAAATCGTAAATTAGAACCCGACGAAAGTTTTGCTTTTGCAAAAGTATGTGTAGC

TAAAAAATATTTTTCTAAAGCTATTATTGAAGGTCAATTAGCTTTAAAAAACTATCGAGA

CCTGAATATCTTAGATAATAATATAGTAATTGCTAATTTGTATAATATGCTAGGTTTTAT

TTATTTTGAAGCAGGCCAAACAAGTTTTGCAAAGAACTTTTATGAGCAAGCCTTACAAAT

AAACCCTAATTATGTTGTAGCTTTAAATAACCTGGCAAAAATTTATGAAGAGGTAAAAGA

TTTAAAAAAAGCCGAATCTCTATACGATAAAGTGTTAACTCTTAACTTAAATAATAAAAC

TGCTAATAGAAGGAAAGATTTTATAGCAAAAACTAAAAACATCTAATACTATTTGTAATC

GGGATAGCAGGATTTGAACCTGCGACATCCTGCTCCCAAAGCAGGCGCGCTACCAAACTG

CGCTATATCCCGTAAGACATAATTATTACTATATACCTTTTCTCTAATTATGTCTACTCT

TTCTGTATTTTATCTAGGATACCAAAACATGCCTTTAACTCCATCTGGATCTTTGATAAA

TCCTAATTTTTTATAAAAACTAACAACATCAGGTTCTGCGAATAGTGTAATAGTGCTAAT

TTCTGCCTGTCTCAATTGCTGTATTAATTGGTGTATGACTACTTTTCCTAAACCTAAACC

TTGGAAATCTGGATGAATAACTACGTCCCAGATAGTTGCATTAAATCCGTTGTCTGATGT

TGCTCTAGCAAATCCTACAAGTTTACTATTTGCATCTTTTTTTTGTATTAAAGAAATAAT

AATAGAACTATTTTTTAATGCGATCTTTACTTTTTTTAGAGGTCTTTTAACCCATCCGAC

TGAATCACAAAGCTGCTCCAATTCATATAAATTAATATTCTTATTGCTACTCAGATAGAT

ATCTTTAAATTCGATTTTATCGCAAGTTTTATCTAAAACAATAAGTTTTTTGAAATTTTT

TTCAGAGCTAGCATTGATATTGGAGTTTTGAAAAAAATTTTTCCAGAAGATCATAATGAT

TATGATATAATTAAATAAATATATATATTACAATATTGCTTTAACAAACAATATATTAAT

ATATGCTCAAATTTTTACATGAATCTATTAAGATAGAAGATTGTTAAAATATAGAGAAAT

AAAAATTTTTGTAAAATCTTTCAATAAAATGATATAAAGAGTGCTTTTTAAGACTAACAT

CTATTATTATAAATAAAGTCTCGAAATTGAGATGAAATAACACATTATTCTATGTCTAAT

GCCTTTTAAAGGAGTTATTTGTGAAAAAATCTATGTGTCTAACCTGTTTACTTGCTTTAC

TAATTATGAGCAACCCAATAATAGCTAACGCAGAAGTAGCTGGATTGGTGCCCTGCAAAG

ATTCTGCGGCATTCAATAAGCGTATGGTAAATAGTGTGAAAAAACTTCAAGCCAGATTAG

CTAAATATGATGCAGACACGCCACCAGCGTTAGCTTTAAATAAGCAAATAGAGAAAACTA

AAACTAGATTTGCAACTTATGGTCGAGCAGGTTTGCTGTGTGGTACTGATGGATTGCCAC

ATTTAATTTCTGATGGTCGATGGAGTAGAGCCGGGGACTTTGTTTTTCCAGGACTATTAT

TCTTGTATATTACAGGATGGATTGGCTGGGTAGGTAGAGGATATCTTTTATCTGTTGCTA

AGACTAGTAAGCCAACAGAAAAGGAAATCATTTTAGATGTGCCATTAGCTATCAAATTTA

TGTCATCCGGATTTGCATGGCCGCTAGCGGCTTGGCAAGAATTTAGTAGTGGACAATTAA

TTGCTTCCAATGACGATATTACTGTTTCACCCCGTTAGTAAAAAATTTATATGAATAATA

ATTTTACCAAATACTTATCAACAGCACCTGTAATTGGTGTATTGTGGATGACGTTTACAG
CCGGATTTATTATAGAATTGAACCGCTTTTTCCCAGATGTCTTATACTTTTACTTATAAA
TAAGATCGTATAAAAAACTACATAAATATAGAGAGATACTATTATTTGTGTAGTTTTTAA
TATATAACATAATTATTTATTTTTAAAAATGTTTATCTTTTCGATTATTTTAATATTAGA
TCTTACATAATAAAATAATATCAATATTATGAGCTTAGTAACCCAAATTATAGTTAATGC
CGATGATGAATTAAGATATCCTACAATTGGAGAATTGCAGTCAATTCAAGACTACTTAAC
TACAGGAAGCAATAGAATTAGAATTGCTACTATTATTAGAGATAAAGAAAAGGAGATTAT
ACAGAAAGCTAGTAAGCAAATTTTTCAGTTACATCCAGAATATATAGCTCCAGGTGGTAA
TGCAGCAGGTTCTAGGAAAAGATCGTTATGCTTACGTGATTATGGTTGGTATCTCAGACT
AATTACATACGGAGTATTAGCTGGGGATAAAGATTCTATAGAAACAATTGGTATTATAGG
AGTAAGAGAAATGTACAATTCTTTAGGTGTACCTATAATTGGAATGTTAGATGCAATCCA
GTGTTTAAAGGAAGCATCTTTAGAAATGCTTGGTCAAGATGATATTAGAATCATTTCTCC
TTATTTTGATTATATAATTCGTGGAATGTCATAAATCATAGCTTCAATTAATTAGTTGAA
TAAAAGCTTTTATAATGTTATAATTACAATAAGCCCGAGAGCATATAATTGTAAAAACTA
AATCTTGTCAGAACCGGAAGGTAGCAGCAATAATGTTTCAATACAAAAGTTATGATTTTC
GGGTGTTTATTGTTTTTATCTAATAGTCTCGACTTTTAATTAGTATGCTATGTATAAATC
TGTATATGACATGTAATCCATCAAACCTTCCAGCAAAAATAATTTAATAATTTGATATTA
ATACGTAAAACACCTATACTCTTTTATAAAATTTCTTTATATCTCATCTAAACAATGGAA
ACTTAACTTAATGGGTGTTCATATTTTATCAACAGGTTCGTCTGTCCCAAATTTTTCTGT
AGAGAATCAACAATTTGAAGACATGATCGAAACTTCCGACCATTGGATTTCAACAAGAAC
AGGAATAAAAAAAAGACATCTAGCTCCATCTTCTACTTCTTTAACTAAATTAGCGGCAGA
AGCTGCAAACAAAGCCTTATATGCAGCTAACTTAAAACCTACTGAGATCAGTTTAATTAT
TTTAGCTACGTCTACGCCTGATGATTTATTTGGTAGTGCTAGCCAGTTGCAAGCGGAAAT
AGGTGCAACAACATCAGTAGCTTTTGATATTACGGCTGCCTGCTCCGGTTTTATTGTTGC
TTTAGTAACAGCAGCTCAGTTCATTCAAACTGGTTCCTATGACAATATTTTAGTTGTTGG
AGCAGACACAATGTCTAGATGGATTAATTGGTCAGATAGAACTACCTGTATTTTATTTGG
TGATGGTGCTGGAGCAGTAGTATTGGGGCAAAGCCTCAAAAATAGTATTTTAGGCTTTAA
GTTATGTACAGACGGTCAGCTAAACAGTCATTTACAATTAATGAATAAACCTGTAAATAA
TCAAAAATTTGGTGGTACAGAAATTCCTCATGGAAACTATAATTCCATAACAATGAATGG
CAAGGAAGTGTACAAGTTCGCTGTATTTCAAGTTCCAACAGTAATTAGACAATGTTTGAA
TAATTTAAACATTTCAATAGATGAAGTTGATTGGTTTATATTGCATCAAGCAAACACTAG
AATCATAGAAGCAATTGCGAGCAGATTATCAGTACCTTTTTCTAAAATGATTACGAACTT
AGAGCATTATGGAAATACATCTGCAGCGTCAATCCCTTTAGCGTTAGATGAAGCTATTCA
ATCCAATAAAATTCAACCAGGTCAAATTATTGTTTTATCTGGTTTTGGAGCAGGCTTGAC
TTGGGGAGCAATTGTCTTGAAGTGGTGATTTATATATTGCGGATGACGAGACTCGAACTC
GTAAAGCTTTCGCTACACACCCCTCAAGCGTGCGTGTATACCAATTTCACCACATCCGCA

TTTTTTACCATGAATATTAAAAATTTATAATATTTATCTAAATATTCATATAATGTGTAT
ATATATATACATTCTTGTCAGAAACTCTCTTATATTTTATTTTAAGTGATAAAATTGAGT
TTAGTATAGATAGTAAAATTTTATTTTAAAATCATAAAAATAAATATATAAGGAAAATAA
AATATGACACCATCTTTATCAAGTTTTTTGAATAGTCTTATTCTTGGGGCAGTAATTGTA
GTCGTTCCCATAACTTTGGCTCTTTTATTTGTTAGTCAAAAAGACAGGACAATCCGGTCC
TAAAAATAACTTAGAAAGATATGGAGAAGCAATTTAAGATGAATAATGAAATATTTTTCA
TACTAATAAGCTCTCCATATATTGCTATACTTTAGAAGAATTTTTATTTTTTTATTTTGT
AACATCCTGTAAAGTTTATTCCGTTCTTTCTTTTTCCCATTTACTCTCTTATAATTAATG
ATACAAATGTCAGGTTAATAATAG-TTTTTTATTCTGTATCTTTCATTTTATATACGTAT
TATTATGTTCAATTAATAAAGAAAAGCTCATGTCTTTGTCTAATTGGCCTCTCAAAAAAG
AAAATTCTGAAGCATATAATATTAAGAATTCAAAACAAATCACAATTCCTGATGGTTTAT
GGATAAAATGCTTCGACTGTGGTCTATTAATGTATTCTAAAGTACTGAAGAGAAATTTAA
AAGTTTGCCCTCAATGTAGTTATCATTTTCAAGCTTCTAGTAACGAAAGAATTGATCAAT
TGATAGACCAAGGTAGTTGGCAACCAATGGATGTTCACTTGATCTCTACAGATCCATTAG
GCTTTAAAGACCAAAAGCTTTATAGTCAAAGGTTAAAAGATACTGCTTTCAAGACTGGCC
TGCAAGACGCAGTTCAAACAGGTACTGGGACTATGCAAGGTAAAAAAGTATGCTTAGGTA
TTATGGATTTTAGATTCATGGGGGGAAGCATGGGATCTGTTGTAGGCGAAAAACTAACAA
GACTGCTAGAAAAAGCAACTCAAGAAAAGTTGCCTGCAATTATACTTTGTGCATCAGGCG
GAGCTAGAATGCAAGAAGGTATGTTGAGCTTAATGCAAATGGCAAAAATTTCTTCTGCTC
TAGAAATGCATAAAAAAGAAAATCTACTATATATATCTGTTTTAACCTCTCCCACAACAG
GGGGTGTAACAGCTAGTTTTGCTATGCTTGGAGACTTAATTATTGCAGAGCCAAAAGCTC
TTATTGCATTTGCTGGTAGACGAGTTATAGAACAAACAATCAAAGAAGACTTGCCAGATA
ATTTTCAAAGTTCAGAATATTTATTTGAACATGGTTTCCTAGATTTAATTGTATCCAGAA
CTCAGCTTAGATCAAAGTTAATACAAATTTTGTCCTTACATAATCATAGTAAGTAATGAA
TTGAATATTGACTATAATATCAAAATGCTAGTAAAAATTAAGATATTAAGAAATTTTCAC
TAAAGATATTTGTAGACTAACTAGAATTACAATTACTATTGTAGAAAAATTTGTCTTAAT
GCCATTTTTTCAGTCTTTTAACAAATTGTTTTTATGACTTTAAATAATTGTTAGTTTAAT
ACAAATTTATAAAGAGAATACATACTAAGCTAAATATAGTAATATTATGTTTTCCTATAA
TACTGACCTATCAAATTTCGAGGAAATTTATGCTTAAAAGATCTTCTTGGCTTGCGGCTT
TATTGGGACTATTAACAGTAGTTTCTACAAGTACGCATACATATGCCATAGAGTTAGACG
AGGCAACAAGAACTGTTCCATTAGAATCTTCTGGCAGAACTGTAATTCTTACACCAGAAC
AAGTTAAAAGAGGCAAGCGATTATTTAATAATTCTTGTGCTATTTGCCATAATGGTGGTA
TCACGAAAACAAATCCGAATATTGGACTTGACCCAGAATCTTTAGGATTAGCTACGCCAC
AAAGAGATACCATTGAAGGACTAGTTGACTATATGAAAGATCCGACTAGTTATGATGGTG
CAGAGTCAATCGCAGAATTGCATCCAAGTATTAAAAGTGCTGAAATTTTTCCTAAAATGC
GCAATCTAACCGATGAAGACCTATTTACAATCGCAGGTCATATCTTACTTCAACCTAAAA

TTGTTTCTGAAAAGTGGGGCGGAGGAAAAATTTACTATTAGAACTTCAAAGACCTAAACC
TTGTGAATACTAGTTTATGTATTTGTTGCTTAAGTGTGATTATTTTGTTAGATCATGATA
TATATTAAAGATGGAGTTTGTTGACAGCAAGTTACTTTTTACCCGAACTTGTTTACACTA
TTCGTCTAGTTATTAAAAAGGAGACGTTAAATTGAAGAAGAAGCTTTCAGTTCTTTTCAC
TGTTTTTAGTTTTTTTGTAATAGGTTTCGCACAAATTGCTTTTGCTGCAGATCTAGATAA
TGGAGAAAAAGTTTTTTCTGCTAATTGTGCAGCATGTCATGCTGGCGGTAATAACGCCAT
TATGCCAGATAAAACCTTAAAAAAAGATGTACTTGAAGCTAATAGTATGAATACTATTGA
TGCTATTACTTATCAAGTACAAAATGGTAAAAATGCCATGCCTGCTTTCGGAGGTAGACT
GGTTGATGAAGATATTGAAGATGCAGCAAATTATGTATTATCTCAATCTGAAAAAGGTTG
GTAATTATACTTGATTTTATCCTGTATTAAAGAATAGACAATCTATTTAGTTGTTCATTA
GATTGTCTATTCTTTGTTTATGCTATTATATAAAGATATTTACACAATATTTTATGATGA
AAAGAATACCCGCAATTCTTGTACTAGAAGACGGTGCATATTATAAAGGATGGTCATTCC
AGCAAGATAAACAAGAGATTACTATTGGTGAAGTAGTTTTTAATACTGGAATGACAGGAT
ATCAAGAAATAATCACAGATCCTAGTTACTTCCATCAAATTGTCGCTTTTACCTACCCGG
AAATTGGGAATACAGGTATTAATAATCAAGATATTGAATCTCACAGTATTAGTATTAAAG
GACTTATTGCAAAAAATATTTGTAAAATTTCAAGCAGCTGGAGAGAGCAGCAATCTTTAG
TTAAGTATTTAAGTAGTAATAATATTCCTTTTATTTTCGGAATAGATACAAGGTCTTTAA
CCCAATACTTGCGTCAATTTGGTACAATGAACGGTTGTATCTCTACTGATAATTTAAATC
ATAGTTACTTAAAACAGAAAATTTGTGAGATTCCAAGTATGCAAGGTTTAGATTTAATCC
CGCATGTAACTACAAGAAATGTTTACCCCTGGGATGAAAAAAGTTTTCCGAATTGGTATT
TGACAGATAACATTAGAGTACACCGAGTTATTCAGTTGAAAGTTATTGTTATAGATTTTG
GAGTTAAACTAAATATACTCAGAAGACTAGCTACACTTGGATGTCAGATAACGGTTGTGC
CTGCCCACACTCCTTTAAAAGATATTTTGGCTTACCAGCCTGATGGTATATTACTCTCTA
ATGGTCCAGGAGATCCATCAGCAGTACATTACGGCATCCAGACAGTTACAAATTTACTAG
ATTACAATGTGCCTATATTTGGGATTTGTATGGGGCATCAAATTTTAAATTTAGCTCTTA
AAGCTAAAACTTTCAAACTTAAATTTGGCCATAGAGGTATTAACCATCCATCCGGATTGA
ACCAGCAAGTTGAAATAACTAGTCAAAATCATGGCTTTGCAGTTGAATTGACTTCGGTTT
TTGAATCTCCTGTAAGAGTGACTCATTTTAATCTAAATGACACTACTATTGCAGGAACTG
GACATAATCAAAGTCCTTATTTTTCTGTGCAATATCATCCAGAATCGAGCCCAGGCCCTC
ATGATGCTGATTATCTATTCGAAAGTTTTATAGAAATAATGACAAAGTCCAAGAATAAAG
TTAGTTAGTAATTTTCCCATGCTTTATGAGTGAATAAAGCGGATAGTACTTGTACTCCTC
GTAATTGATTAAAGAGAGGCAAATGTCCTCTGGGGCCATGAGAATTCCATTGAAATTCCT
CAGGATATCTGCAAGGAATTCTATCAACTTCCCAGCCGATTTTTTGCCAGAATTTTCCCC
AATCTTTTCCAACACTAAGCCATATTTGTCTTTGAACAAATAGCCCAAATTTACCTTTTG
AGTGTGTATGCCATAACTTATCAATAGTTTGTAAATCTTGAGCAGGTATTTTTTTTATAT
CTGTAAAATATAGCCAATTACGAGTTTGTGCATTTACACCAGCTAGTTGAATAAGCTTTT

GTTGAGTTAGCTGATCTGCTTTGAGCAAATCCCGATGAGTCAGTAACATTTGTAAATCTT

TGTAATTCATCTGTTGAGCAGAGCGCAATGGAACAATGCCATCGGGACATAAGTTAGAAG

CAAATTTCACTATTTCTTGATTTTTACTGTTAAGTAACTTTTCATAAATTAAACCGTCAA

CACAATTACTTTTATAGTTAGGCCCAGTAATCCTCTCAAAAAATAGATCTGCTAAATCTT

TTAACTCGATAGAGTCGTTGCTATTTATATTCTCAATAATTTCAAGTTGTTGCTTGACAT

TATTAGACTTAGTATTCTTATTCAATTCTAAAAGCTGGGCTCGGATTTGATTTGGCATCT

GATTTGTTTAATTAGTTTTAATAGTCATAAGGTCTTAGAGAGAATTAGATTGTTTTTTGA

AATTAGAAGAGCGCCGCAGTATTTCTAAAGAAGTTCGCATAATTTGATTCAATAAGTTTT

GTCCTGCATAAATTATAATTTGACCAATTAAAAAAATTGTGCTTTTTATTTTAGGAACAA

AAAAATCTTGTACCTCAAGCAGAAAAGTAATTATTATTTGCATTGGTGACAACATCTGTA

ATTCATTCGATCTGCAAGCATATATATACTGACAGTTTAAGCCTTCTTGATAAAATCCCC

ATACTTTGTAACGACTTTCGTAAATTGCTCGTGGTCGCTCAAGATAGTTTTTAATTATAG

TATTCCAAATTAGGTTATTCTTTAATTTTTCTAATTTTCTATCAGATAGAAATTTTACAT

TACAGAGATAACTTGCATTTGTCTCATTTACAGTATTAAAATTTTGAACTAACATATGAG

CAATAATGTTGCTTAGCTGAATTAAATAATTTTCAAGAAAAAATTTCGACCTGTTTAACTG

GCATATTGTAAGTAAAGAGATTAAAAGTTTTATTCACAGTTTGAGATGACCCAAAAAGTA

GCTGAATAATTAAAATTTGTAGTAGCATTTTATAGTCGTTCAGTAGATAGTTCAAATCAT

GAATATCTTGGTAAATGTCAATACAGTCCAGTTCATATAACTTACAAAATCTTTTGATAC

ACCTATGGAATAAATCAATTAAAATATTTTCGGTCAAGTTTGTAATATCATTTACATCTA

AATTTGAATTATAAATTTCTAAAAGTATCTTTTCCAGTTCAGCTAATATGATTTTTAATA

AGTTTCTTTTTACTTCAGTGCGAAAAACATCTAAAATTAACACTTCCTGAGAGCAATTAG

TTAATCTCTTATTAATCTTTATAGAAGTTCTAACTAATAGCTCTGCTACCTCTTGATTGA

GTGTTGGTCCCTGAGAGCTTGGCCAATAATTATTCACGTTATACCATAGTAAGAAATATT

GAATTTAATGTTATACGATAACTTTTTAATATACTGAAAAGCAAGTTAATTACTCTATAG

CAACTAGTTTTTACAGATCTTAATAGTTATTTATCAAATAAAGTTATATGATAGTAGCGA

TTGAGCTAATTTTATTAAAATTTATAATAATGACAACTTATTACTTCGCCCTTGCGAGTC

AAAATTTTCTATTAGTACAAGAACCACTAGAAGAAGTTTTTAGAGAGAGAGTTAACTACT

ATCAGTCAAATAATAAAGCAATTGATTTTTGGCTAATACCAAACCCTTCTTTTCTAGAGA

AACCAGAAATGATTTCATTTAAAAACCTTGTACCTAAAGACGCTGTAGCTATAATCTCTA

CTAATCCAATATTTATTAATTGGTTAAAGCTAAGAATTGGCTATATCTGTATTGGACAAT

TTGAAGATAACCTACAACTTTCTGAAGAATCGTTAAATATTACTGTTTTAACAGACAAAA

TTTAACAATGCTTTTATTATGACTAGTGTTAAATTATTTTGGTTAAGTTATAATTATAGT

CTTTTAGTATTTAGACTCTTATAGTTTATCACTTAGTATTTTATGCTGAGATGGAGTGAC

AAATAGTGTAAGTATTTCTTGGCTAAACGTTTCAGTTGCCTTAGATTTATATCTATTTGG

ATTAACTATAATGGATAGCATTCGTTTTATAGTAACATTTTCAATTTGAGCCCAATGTAC

AATTCCAAGTTCCAATTCCTTAGCAATAGCTGAAACGGAAACAAAGGCAGCTCCTAGCCC

TGATTGCACAGCGTTTTTGATAGCTTCTATAGAATTCAATTCCATCTCTATTTTAAAGCG
ACTGCTGTCAATACCATGTTGACTGAGTACTTTATCTATCACTTTTCTAATTGTTGATTG
AGTATCTAATGCAATAAATCTAAGCCTGTATAAGTCTTCTTTTTGAATGTCTCCTAGTTT
AGAAAAAGGATGTGATTTGGGTAATATAAGTGCTAATTCGTCTTCCGCATAAGAAGTAAC
CTGCAAAACATCTTGCAATTCAGTAGGCACTTCTCCTCCAATAATTGCTAAATCAACTTG
ACCATTAGCTACGCTCCATGAAATAAGTCTAGTTGAATGTACTTGCAATTGAACAGCCAC
TTGTGGATATCTTTGCCTAAATAGTCCGATTAATCTTGGCATCAAATATGTCCCAGTTGT
CTGGCTAGCTCCAATAATTAATGTGCCACCTTGTAAGTTTTGTAAGTCGTCAAGAGCTCG
ACAAGTTTCTTCGCAGAGAGCTAAAATTCTGCCCCCGTATCGTAAAAGAAGACTTCCTGC
CTCAGTTAAAGTTGCCTTCTTATTACCTCTTTCAAATAGGGAAACATTTAATTGGCGCTC
TAAATTTTGAATTTGCAAACTAATAGCTGGCTGAGAAACATACAAGCTATTAGCTGCTTT
TTTAAAACTGCCCTCTTTGGCAATTGCTTTTAATATTCTTAACTGATCCAATGTAAATGG
AAGGTCTGTCATTAAAGAATTGTAGTATAATGTATATTATTTAGTATTGATCAATGTATG
CATTTGCCAACTAAAATATTTGTCATTACGCTTCATTATTTTCAAAATAATAAAAAGTAT
AATGTTAAATAAGCTGTATTAAAATTTAGTATCTGTTTGATATTAAATTTATACAAGAGA
TATCGTGAATTTAATTCTAACTATAAAATATAGGAGACAATATGGACTCTAGACTTTTAG
TTGTACTAATACCAGTTTTAGCAGCAGCATCTTGGGCAGTTTACAATATTGGTAGAGTTG
CACTACAGCAATTTAGAAAAATGACATCTTAGTTTTGCTTAGAATATAATTTTTAGACTT
TATGAGATAGAGGACTTTAAAATCCCTATCTCTATTGTATATCTTAATAAAAAAAGAAAG
CCCCCTTGTACTTACTGTTATTACACATTAATATTATTGGTGGTACCTAGATAATAAGTT
TAAGCTAACTTATTTTTTATTTTATAATTAATTAAATTAATAACACAATTATGGCTGTTC
CAAAGAAAAGAACATCTAAAGCTAAAAAAAATGCACGCAAAGCAAATTGGAAAAATCAAG
CAAAAACTGAAGCTCAAAGAGCTTTGTCTTTAGCAAAATCAGTATTAACTAGAAAATCCA
ATGGATTTATTTATAATCTAACTGAAGCATCAGATACTTTTAGCGATTAGAAGTGTTTTT
ACTACAAGTCTGAAAAAGTATTACGCAACAAGTTCTCAGTAATTTTTATCTATTAATCAT
TTGAAAATCATTCTATTTGATAATAAAATAAATAGCAGATTAAATAAAATCACATTAACA
AATTATGCAGCAAAGCTAGATCAAACTAGTGAAATTTGGTTGTTCAATTGTATTGAAAAT
ATTCAGCATATTTTTTTTAAAAGTCAATTAAAATCAAGTCATATTACTAAAATTTTTATT
TCTGGTACTAGCTTTGAATATACCGCAGGTCTACCAGGATTATTATCCAGCTTAACACTA
AGTGGTAGACTACATCCTATTAGTATATATAGTCCCCAGTCTCTCAAAAAGTATCTTGAA
GCATGTACCAAATATTCCCAAACTAATTTTTCTTTTCCTATTAATTTTCATAATTTACAA
TACGGAGGACAAGTTGTTAACCAATTTTATACAGTAATTTGTTTACCGTTGAGCAAAAAG
AGCCTGCTGTATGGATTTATTATTCTAAAAAAAGAAAAGCAGGGAGTATTTAATTTAGCA
CAAGCTAAAACCTTGAATATTCTTCAAGGACCTATATATGGAAAACTCAAAGAAAAGGAT
AATTTTTTAAGTCCAGATGGTTACTATCTATCCGGCCAAGACTTCTCTTCTAATACAATA
ATGGGACATAAAATATCGCTTCCGTTATTAGTCAGATATTCTAGAATTATTTCTGAGATG

CATTGGTTTTGCTCTTATCCTATTAGATTAAATACTTATTCGCATCAGCAAGGAGTAAAG
TGTTTACCACATAATGTTTTAACTGATATTATGAAATCCCAAATATATCAAGATAACAGT
TTTGTTGAATAATACTTATTCTTGATATATAATGGTCTCGGTAATCTAAATAAAATACTA
GAAATAATTTCTTAAATTTATTTGTCTATATTAAATTTACAATCATTTATGACATACGCA
ATTATTGAAGCAAGTGGCACGCAGCTTTGGATAGAAGAAGGCCGCTACTATGATTTAAAT
CATATACCTGTTGATCCAGGTCAGTCGATTATATTAGGAAAAGTCTTATTATTGAATAAA
AATGGGGAGGTTACTTTAGGACGCCCTTGTATAGAAGGGGTTACGATAAAGGCTACAGTA
ATGAGGCACTTGCGAGGAAAGAAGATAACTGTTTTCAAAATGAAACCAAAGAAGAAAATG
AGATTAAAAAAAGGTCATCGACAAGAATTGACTCGTTTAATGATCGATTCTATAACATCT
TAGATAAAATCTAGACTCATATTTTTAAAAACTTTTTAATATTTAACTAAAATAGATATA
ATGGCACATAAAAAAGGTAGTGGTAGTACAAGAAATGGCAGAGACTCTAATTCCAAGCGT
TTAGGTGTTAAAAAATATGGTGGAGAGCAAGTAACAGCAGGTAATATTTTAATCAGACAA
CGGGGAACTAAAGTTAAGCCTGGCCAAAATGTTGGAAAAGGAAAAGATGATACATTGTTT
TCTCTAATTGATGGTTTCGTGCTGTTTGAAAAGTCAAATCAAAAGCAAAAAACAATTAGT
GTTTATTCTGCTAAGAAATAGTTAAAACAATTGGTGCAAGATAACTTGAGTGTATTTATT
TCACTTACAGTGGCACCATATATTTGATTTATTAAGAAATGGAGTTCCTTAATGAACCAA
TGAATTTTATTAAAGATTTGCTGTGAAATTTAATGACAAACACTATTGTAATTTCCTGTC
TACACAATATGGCTGCTATTCTATATTGTGGTCAGATACAGAAACTAGTAGTAGCAAATG
CTCATTATCAAGTAAGTGATATCTACCTAGGTTGTGTTGATAAAATCTTCTCAGGAATAA
ATGCGGCATTTATTAACCTAGGAAAGAATGAGTACAGTGGTTTTATACATATCAGTGATA
CCGGTCCGCTTAAAAAGAAATATTATGTCAATAATATTACTAACATTTTAACAATACGGC
AAAAAATTTTAGTACAAATCATTAAAGAGCCAACTTTAAATAAAGGTCCAAGGCTCACTG
CCAATATTACATTATCAGGTCGATATATTGTATTAATGCCTTTTAGTCAATCAATCTGTA
TATCTCGAAAAATATATGATGAAGATGAGCGTCATTATTTGAAATCTTTAGCTATTTTAA
TTAAACCGGCAACAATGGGCTTGCTATTTAGACCTTCTGCTGTAGGTGTCGATGAGGAAA
TAATATTAAGCGAATTAAAAAATCTAAAAGAACAATGGAACTTTGTTCAAAAATCTGCAA
TTAATAGTTATTCACCTGTTCTTCTATATAAAGATGAAGATATTGTTAAAAAAGTAATCC
GAGATTTTTATAATAATAATACAAACAATATAGTAATTGATTCAAACCTGGGATTAAAAC
AATTAAATTATTATATCCACACTTGGCACTGTAATAACTCTAGCACAGTTCCTAAGATTA
AGCTTTATAGTAATAATCAATGTATACTAGATGCTTTTGGTATCAATCAGGCAATTTCCA
GAGCTCTCATTCCAAAAGTTGATCTTATACTTGGTGGCTATATGTTTATTGAAACTTTAG
AAGCTTTTACTATTATTGACGTTAATTCTGGATCTTTTAATAATTCCACTAGTGCACGAG
AAACAGTTTTAAAAACCAACTGTTCTGCAGCAACAGAAATAGCTTATCAGTTACAAATTA
GAAATATTACTGGTGTAATTATAATCGACTTCATTGATATGGAATCACAAAGAGATCAAT
TGCAATTATTAGAACACTTTAATAAAGAGCTATCACTTGATGATGCTAAACCACAAATTG
TACAGTTATCTGAATTAGGTTTAGTTGAATTGACTAGAAGAAGACAGGGCAAAAGTTTGT

ACGAGTTAATTAGTAGTGATTCTAATTACTTTTATTTTTTCACACAATCAGAGAGATCTC

AGTCTCTTAAGAGATTCGATGATAGACAGCAGAAACAACAGATTTTTAATAAATCTTGGC

TATCTGCAGAGATTAATACTATTAACAAGGTCTTTTTTCAAAAGTCAAATTTGTGCAGAC

CTGCTAACTTTTACCTAATTCGTAATCTCTATATAGTTAAAAGTAGTATTACTTATAAAC

AAAATTATTTATTAACTCACAGATCAAAACTTATTTATTCTAAAGAATACAGCAAAGTAT

TACCAAGCAGTTATTATCTAGCTAGTCTCAATAAGAATAGTAATCAAGAGTTCCTATCTT

AAGATTGTTATATCAACTGTTGAAAAAAAGAAGCTCTCTCTTATATAAGAGAGAGCTTCT

TTTATAGCTTACTAACTATCTAGTTAGTTAATTAAAAATTAACCGTTAACAGCTGGAGCT

GTTAGAGCTACTGGTAAAGATTCACCAGAAGCTAGATCTAGAGGGAAGTTGTGAGCGTTA

CGTTCGTGCATTACTTCCATACCTAGGTTAGCACGGTTGATGATATCAGCCCATGTGTTA

ATTACACGACCTTGGCTATCAACAACAGATTGGTTAAAGTTAAAACCATTCAAGTTGAAT

GCCATTGTGCTTACAGATAAAGCTGTTAGCCAGATACCAACTACAGGCCATAGACCTAAG

AAGAAATGTAGAGAACGAGAGTTGTTGAAACTAGCATATTGGAAGATTAAACGACCGAAG

TAGCCATGAGCTGCAACGATGTTATAAGTTTCTTCTTCTTGTCCGAATTTGTAACCATAG

TTAGCAGATTCGTTTTCGCTTGTTTCACGAATTAAGCTAGATGTAACTAGGGATCCGTGC

ATAGCACTGAACAGAGAACCACCAAACACACCAGCAACACCTAGTTGGTGGAATGGGTGC

ATTAAAATGTTGTGCTCAGCTTGGAATACAAGCATGAAGTTAAATGTTCCGGAGATACCT

AGAGGCATTCCATCAGAGAAACTACCTTGGCCAATTGGGTATACTAGGAATACTGCTGCT

GCTGCTGCTACTGGAGCAGTAAAAGCAACGGAAATCCATGGACGCATACCTAGGCGGTAG

CTTAGTTCCCACTCACGACCAATGTAGCAAGCTACGCCAGTTAGGAAATGAAGAACAACT

AATTGGTAAGGACCACCGTTGTATAGCCATTCGTCTAAAGAAGCAGCTTCCCAGATTGGG

TAGAAGTGAATACCGATAGCTGCAGAACTTGGAATAACAGCACCAGAAATGATGTTGTTT

CCGTATAGAAGGGAACCAGCAACTGGCTCACGAATTCCATCAATGTCTACTGGAGGTGCA

GCTACGAATGCAATGATGAATACAGATGTGGCAGTTAATAGAGTTGGAATCATTAATACA

CCAAACCAACCAATGTATAGGCGGTTTTCAGTACTAGTAATCCAAGAGCAGAAACGTTCC

CACAAGCTAGCGCTTTCGCGTCTTTGTAAAGTAGCAGTCATAATTTTTTATCAATTTTTA

AGGATTTAACAAAGATGCTTCCCAAACAATAAGTTGTCTGTTATGAAAATTATTACTTAA

GGCCTTTTCAAATGAGAACATTTTTAAAAAAAACATAGCAAAATCTAATCTCAGTAACTA

AAATTTTACTTGTTCATATATTGCTTAATAATTCAATAATAAATTTTTGCTTTTTGCAAA

ATTAACACTATTTTAGGGATATAATAATAATGCATATTAATGCTGCTTATTTTGATTTTA

ATAGTTTATATAATTTTGCCCTACTTAAATACGCATGTCACACTTTACAAAAATTCAGAC

GACCATACAAGATTTAAATCTGTTAAAGCATGCTTTAACAGATTTGGGTTTGATTTGGCA

AATGAATTCTAGTCATATAAAAGTTGGTGAAAATAATCAGCATAAGGTAGATATTTTAAT

TAAACAAGATAACTTGTCACATATTGGCTTTACTTGGAATGATAATAGATATCACTTAGT

TGCCGACTTACAACTTTGGGAACAACCCTGGTCTCTAGAAGTATTTTTGGATAAGTTATC

TCAGAAGTATGCTTACTACTCTATCATCGAAGAAACAAAGAAGCAAGGCTTTGAAAAAAT

GCAGCAGATTTCACAAAAAGATGGATCAATTAAGTTAATTGTGCAACGCTGGAATTATTA

ACTAACATACAAATGCGGGCGGAGAGACTTGAACTCTCACGAGATTATCTCACTAGAACC

TAAATCTAGCGCGTCTACCAATTCCACCACGCCCGCACTGTTGAGACAATATTATTCTAT

CATAATCTATACTAAAACACAAACTAATAGTATAAACTAGCTTCCAAGCTTAAAAGCGTC

TACAAATAGACCGTATGTTATACCAATTTTGAAGTTGTTGAATAAATTAATTTTTATATT

TAATTGCTTGTGGCTAGAGTAGACTATTTTACTAGTCAACTCAGTTGCTGCAACTATTAG

ACTGGCTGCAATAATACCCCAATCTCCTGTTTGTCCAGGAATCGTAGACAAACCTGTTGA

AATAAAAAATCCTAGCAATAAACTAATTAAGCCAGTTGTTAACTCGCTCAAAGAATAATA

AAGCTTATTATTTAAGTTTTTTATTAAGTAAGCAAAAAAAGTTGAAAGTCTAGTTTTAAT

CATAGTATTTAGTAACGGAGTTTTAAAAGGATGACAAGTATTAACAGTCATTAATATTGT

CGTATCCTTTTAAAACTCGTTAATACAAACAGCTGTTCTTGTTTTTAAAGGTCTTGTTCG

CTTAAAGCATTAATAAGATGCTGAAACGGCTCTTGTATAATATCTGTGCGTTTAATTGAT

TGCAAATCTAATTCTTCTTCTATGATTTCTTGTAATAACTGTATGCTTCTTACTGTTGGA

GCAACAGGTACAGATAGTGAATTGTAAGTATCTTTTAGCCCATTTAATACACGCTGATCG

AGAATATTTGTACTACCAGAAACCAAGGCATAACTAGCATACCGTAGGTAATATTCTATA

TCCCTTAGACAAGCTGCATATCTTCTTGTTGTATAAGAATTTCCTCCAGGCCTTAATAAT

TCAGGCTGTTCTTCGTACAACTGAGCTGCGGCCTCTTTTAAAATATTGGTAGCTTGATCA

TTAATAATTTCAGCTATTTTTATTCGATCTAGTCCACTCGAAAAGAAAGATTCTAGTTGT

CCGACAGCTGTCTTATCTAGATAACGACCAGTAAGATCGTAACGATTTAATATTGCTGTT

ATAGCATCTTGCATAAATTAATTCTCCTACTTGCTATCAGAGCGGTATTATAAATGACTA

ATATTGTAATATAATCATTAGTGTTTATGAAATTTATTTAAAACTTATTCTTTATTGCTT

AATGTGCAATTAAAAACTCGCATAGATTTTACACATATAGAGTATACGTAGTTTTTAATT

ATTTTTTTATAAAAACCAAGAAATAAAGCTTTTTTTAAGAGGTTCGTGAATTTGTTAATA

TAACATGAGCTTTCCTCCTAAATAAGTTTTGTAAAGTACAAAGTAAAAAAACGATTAGTC

AATAATAGTAAACAATATTAATATGTATCAAAGTATAAATAAGCTTTTACTCCAGCATCA

AAAAAAATATTTGAGATATGTTGATACAACTGATTATTCAAGTGAAAACTTAGATTTAGA

CCAATTTAGCTATTTTATCATTACAATTAATAAAACTGATAAATGTATTCTTATAGAGCA

ATTTTGTTATAATGCTGAAGGCCATATTTACTCAATATTCTTCAAAGGTCCAACAGCTAA

ACATCTTTCTTCTATAATTTGTCATTTTCAACAATTAAATACAATAATATCTACAGCTCA

TGCCATATATTTAGGACGAGAGCTAATGAAGTCAGAATTAGCTCTCGTGCTAGATCAACA

ATATATTCAAGATTAAATTTTGTAAGTATTAATTATATTTATGCTAGTATAATATTAGTA

AGGGCATGTAACTCAGTGGATAGAGTATCAGATTCCGATTCTGATGGCCGTGGGTTCGAA

TCCCGCCCTGCCCGAGTACTCAGAGTACTATACTAAAACATCTTAATAGCTAGTTTGATA

ATAGTATGGGCATTGGGGCTGCAAGGTTTCTACATTATGAAAAGAAGAAAATATG-AAAA

AAACAAGCTCTCCAAAAGAGCTTTTAGTACAATCAATAAATGCAGAAAACAATATTGTTT

CTTTTTCTCGAAAATTAGCTTTTGCTTAAACACTGTTAATTTTTGGCATTCGACATGTTA

AACTCTTATTCATGGCGAATATTCTGTCAGAGTCGCTCTTAGTTTAAGAAAAGTCAGAAA

AAATATTTATCTTTACGTTTTTACCTATTTTTATTTAAATAAAGGTTAGTATTTTCAACT

TTTATAGTGGACGTGGGTTCGAGTCCCACCAGCTCCATCAATTTGATTTGTATAATTAGC

TAGTTTAACCTTTTTAATAACCAGATATATAATTGGAGGAAATATGGGTTTCATAACAAT

TACTAAGCCCGCTTTAAAGCAAATTGCAATCTTGAAAAATGATCATGAAAATGATGTACA

TCTAAGAATAGGTGTTAGACAAGGCGGATGTTCAGGTATGTCATACTCAATGAATTTTGA

ACATGTTGACAAATTAAAAGATACAGATGAACGGCTTCGTCTAGATAATTTTTCTGTTGT

CTGCGACCCTAAAAGTCTTCTTTATCTTTATGGATTATCATTAGATTTTAGCTCGGAATT

AATAGGAGGAGGATTTCAATTTTCTAACCCGAATGCCAGCCAAACTTGTGGCTGCGGTAA

ATCTTTTTCAGGCTGACCCTCGAACACGTTTTTTATATATTTTGTATTATATATATTATT

TTAGCAAGTCGTTTATTAGAGACAAGCAATGCTCTGTTAAAAACTAATTTCAAGAGTTTT

TATTGTATTATTTTATATATTTATTATCAACTACTTGAAAATAAATTAGCTAAGCAATCT

AATTTGTTATCAGTACCAAACTTTAAATGATTAACTTTTTATTGATATATGCTAATCTTT

ATCAGCCTAAACATATCTGTACATTACAATCTTTTGACATAAAAAGAGTCTTAGCATACC

TCTCAAAAATAAATTCAATATTTTATAAATACTCAACATGAATCTGTCACAACTTATCGT

TAATTCTTCTATGATGAAAATTCCTAATAACTTTTATTGCAATATAAAAGCTATTTCTTC

TGAGAATTTAAGTCACTTACAATTATCAAAAACAAATTTATTATCTCCTACAAAAATAAT

TGCTTTTGGACTACCTAAATTTGATTTTATTTCTACACAGATTACAAGTTCTATTAGAGA

AACAGTTGTTGTGTCTACTATTGATGATAATAATCTTTTTGCTGCCCCAACTAATGATTG

GATAAAAAAACACAAGGATAAAAATCATCATTCTTTGTTTTATAAAAAAGTTTTTAAAGC

TTTGCTAAACCATAAAATTAGAGTTGTTGCTGAGCCACTTCATAACTCCGAAATAAATAA

AGAAAAGAGCATAATGGCAAGTCAAAGATACATCTGGGGTAAAAGCTGGAAGCCTTCAAT

TATTTTATCTTGCTTAAAAAACAAAAAAAATAGCAAAATTTCTGATGCATCGAAAAAATT

TTTAACTGAACAGTTATCTGCATCACCAGTATTTGTGGTAAAAAATGGTTTTAATGAAAT

TATTTTAGGTCACCCATTATCTCGTGTTAAAAGAGGAAGAGTTAATAATCTGATGCATGC

GTTTTCTAATTTATTAAATCAGTCTAATGCAACTTATCCTATATCTACTGGTCTATTCTT

TTTTCATCCAGATGATGCATTTGAATTTAAAGACTTTATAATATCGGTCAATCCACTGGC

GGCCAAACATATGGAGATAAGCGTAGAGCCTGTTGGTCTACACTTTGCTTATAAAATGAA

TAGAAATATATCGTCAGATACTCAATTTTCGTTTATTCCAGATTTTAAAGAAGTAGGAGA

TTTATTATTCAAATATAGAAAAGGTAATCATTTAGTTTTTCATAAGAATCAGCATTATGG

CAAAGATTTTTTTCAGGGGCAACCAATATATATGATTCAGCCGATCACTTTTAAAGATCG

AGCCGGAAAGCTAAATATTATCAAATTTACGGGGCTAAATGACACAAGAGAAATTATTTT

TACTAATCTTGAAGCTGCTAATAGATCATGGGCACATTTTATAAAGAGGAATTCACAATT

AAAATCAATTAAAAATCCTACTCTATTAGTCTATAATTTAGAAAGTTTTTAAAAGATCA

AGAACGATTGGATAATCAAGATTTAAATAAGTTTGTGGTGATCACTAATAAAGAATCTTA

TCTTGCCACAAAAGAGTTAATAGCTTTACCCGATTCTAATAGCTTTTCTAAGCACTTAAA

ATTAAATATAAAGCCTAAACTCTTTTTTGTGAAACTATGGGTAAGACGGTTATTTTCCAC
CTTAACTTATGAATAAATGCTATTGCCTTTATTTTCTCGAATAATATTCTATAACTAGTA
ATTCATTGAGTTGTAAAGCAACCCATTCCCTATCAATGACTCCATTGACTTTCCCTGATA
AATTGGATTTATTCAATTCTAAGTGGCTAGGAATATTAGCTAGTCCAGGGAAAGCTAAGT
AATTTTCTACTAGTTTTCGAGATGCTTCTTGATTTTTTACACTAATTGATTCTCCTGGTT
TACATTGATAACTACAGATAGATACTACTTGTCCATTAATACAAATATGACCATGGTTCA
CTAGTTGTCTAGCGGCAGGAATTGTAGGAGCCATACCAAGTCTAAAAACAGTATTGTCTA
GCCTCATCTCTAGTAGCTGTAACAAAATTTGACCTGTCGACCCTTGTAGTTTTTTAGCTG
CTTTTACGTACTTAAATAACTGTTTCTCACTTAATCCATAATTAAATCGCAACTTCTGTT
TTTCTTCTAATCTTACAGCATACTCAGAAGGTTTTCTGGACTTTTGTCCATGTTCTCCAG
GTGGATAAGATCTTTTAATCGCTTTTCTACTGAGTCCGGGTAAATCACCTAATCTACGGG
AAATGCGTACTCGTGGTCCTCTGTATCTAGACATATTATTTAAATTCTCCTAATGTAAAT
GATTAAGTAATGAGTAGCTATTTTTTAACACAAAAAGAGCATTAAAACAAAGCTTTTGCT
AAAAAATATTTACAAAAAATATAGGTTTGGACAAAATACATTACTATGCTTTTTAAGCGG
GTAGCGGGAATCGAACCCGCATCATTAGCTTGGAAGGCTAAGGTTTTACCACTAAACTAT
ACCCGCAATATTCAACAGATTGTCACTAACTCAATATAACATAGTTATGTTAAGTTAAGA
TAATCTTAAATTTAATAGAATTTTAAATTATTTATGCTCCTTCTAAAACCACATCAAGAA
AACGAGCAATTTCGGCAGCTTGTTCTTCAACTTCTGATAATAATAATGGTTGACCGACCC
TAGTTAATGGAATTACTCTTTTATCTTTTGTACATAAATAAATTTCACGTCTTGGATTTA
GCCCCTCTTTAATATCTATTTTTATTGATTTGATTTCTTTTATATTAAACTGAAGACAGA
TTTTTCTATTCTTTCCAGGGAAACCTAGCCGAAAAATTTTTACAATACCTTTATCTTTGT
TAAATTCATTATATCCAGCCCCTATATTCCAAATAATAGTTAGCCACAGAAACAAACTCA
GAAAGACTCCTATACTGCCATAGAATGTCATTACAATACCTTGTGGAATAAATACCAAAT
CGGTTGAATTTGTAAAAGGTAACAAATCAACTTGAAAATAACTTGATAGTCCAGCAAGAA
GAAAGCCCAAAGCCCCAATAAATATTATAGTAGCCCACCAGTAATTACTCAGTCTTCGTG
AACCTAAAATTAAATCTTTTCTAACTGAGTGTATAGATAATGTTTTTTTCATAATTTGTT
CTTTTAAAACAGTATACTAATGCAAAATGGAAAAATTTTGACAATTTTTATGCGCGCTTT
AGATTTTTTAAATAAGCTTGATTGCTTTTAGTCCTAAAAACAGTCCTGTTGCTAACCCAA
AAAAAGCAGCAAGAAATATTATATATCCTAGAAAAACTGACA---TATTATTATTTATTA
TTATTGTTACAATAGTGTTTACAAAAATAGTGAAAATCTATTACTACGGTATTATATTAC
GTTAAATGCCTTGCTGTCTACAGGAAAAAACTTTTTTCAGTTTTTAGCATTGTGTTTATA
TTAGAAATAATATTCTAGTAAATTTATTAAACTTCTGGAGAACAGAATCTTATGTCAGAA
TTTATCAAGCCTTATAACGATGATCCTTTTGTAGGCAATTTGTCCACGCCAGTTAGTACG
TCAAGTTTTAGTAAAGGACTTCTAGGAAATCTACCAGCTTACCGTCGAGGTTTATCTCCG
CTTCTTAGAGGATTAGAAATAGGAATGGCACATGGATACTTTTTAATTGGACCTTTTGAT
AAATTGGGCCCTTTACGAGGTACAGATGTAGCGTTACTAGCGGGATTTCTATCTTCGGTC

GGCCTCATTATTATTCTCACTACATGTTTATCCATGTATGGTAATGTATCTTTTACTAGA
GCAGATTCGAAAGATCCACTACAAACTTCTGAAGGCTGGGGACAATTCACTGCAGGATTT
CTAGTTGGAGCAGTAGGTGGTTCAGGATTTGCTTATTTATTGCTAGCTAATATACCTGTA
TTACAGACTGCAGGTCTTAGTTTATTCTCTTAAGCTAGTAAGGGGACTTGAACCCGTAAC
CTACTGATTACAAATCAGTTGCTCTACCAATTGAGCTATACTAGCATTCAGTTCATAATA
ACATATAAAGATATTTGTTGCAATCTGCTATTTTCAGATGTAACAAATATCTTTTTTATT
TAAACAGAACTGTTATGTTAACACGTATATTAATTAGAATCTGAATTATCTGACTCTGCA
GAAGATTCCTGATCACAGTTTAGATAATAACAAATTGTTTGATCTAGACAAGTTGCAGAC
CAACCAACAGGTGTTTCTCCTGATAATTCTTGCATATCTACAGAATTATAAAAATTATCA
TTTGGCTTATTACTGTTTGCTTGCATAAATATCACTCCCAAAACACTCTGAACTTAATGT
GAAGTAATTTAACTTCTCAATAATATATTAACACAAAATATTAAATATGGTGATTATTCT
TTAATTGTAAAGATGCTGCTAACTTTATAAAGTTTGAGTGAGAGTCTTGATAGCTTTTGT
TGAAATAAGCATTTTAACCCATTTATTTTGCCCGCTCGACCATACTTTTTTGTGTTGTAA
ATTAACTTTTTGTAGTTTTTTTGTTCGTTTATGAGAATGAGATACGGCATATCCATTATT
AGCTACTTTTCCCGTAAGCTGACACTTTTTTGACATAATTTTCTAACTTATATATTTATT
TAGAGTACTTGCTAAAGTTGACTTGGGCACAGCTCCAATCACTGTGTCTACTCTTTCTCC
TGCTTTAAAGATCATTAAAGTAGGTATACTTCTAATGCCATATTCAGCCGCAATAGTAGG
ATTATCGTCTGTGTTTATTTTTACTACCTTAATAGATGATTCATATTCTTCCGCAATTTC
ATCAACTACAGGAGAAACCATTCTACAAGGACCACACCATGGCGCCCAAAAATCTACGAG
TACAGGTAAGTTATTGTTAATAACTTCTTGTTTGAAAGAGGCATCTGTAACTTGAGATAC
TGACATATTCTTTAACCTTTAATATATGTTCCTTGCTAAGCAAATCTTATCACAAATTTC
AAGAAAAATCTTCTATAGTGAATTACTATTAGTGTTGAATTATTTTTTTAAAAATTAGAC
ATTAAAAAAACTTATTACCTATATAAATAAAGTGGTGGCCGAGTGATATCTTGATAGTGA
TAATTTTTATATTAAGGTTGAACACTAGAAATAACTTTTCAAGCAAACTCTTATTTGAAA
AATTTCTCTGTGACCATAAAAACTTAATTATATTACAATATATTTTATTATTTGTAACTT
AGTTCTGATAATGGTATAAACAACGCAAAAGATACTGCCTTATAATCAAGGAGGAATACA
TGTCTCAATCCGTAGAATCACGGACTAGGATTAAAAGCGAACGTTACGAATCTGGAGTAA
TCCCTTACGCTAAAATGGGCTACTGGGATGCTGACTATGTGATTAAAGAAACAGATATTC
TAGCTCTTTTCAGAATCACTCCTCAACCAGGTGTTGACCCGATTGAAGCATCTGCTGCAA
TTGCAGGTGAATCTTCAACAGCTACTTGGACAGTTGTATGGACTGATTTATTAACAGCTT
GTGACTTATACAGAGCAAAAGCATATCGAGTAGATCCAGTTCCAAACGTGGCAGATCAAT
ATTTTGCTTACATAGCTTATGATATTGATTTGTTTGAAGAAGGTTCCATTGCGAACTTAA
CTGCTTCAATTATTGGTAACGTTTTTGGGTTTAAAGCTGTTAAAGCTCTTCGCTTGGAAG
ATATGCGTATGCCAGTAGCTTATCTAAAAACGTTCCAAGGTCCTGCAACTGGATTGATTG
TAGAACGTGAGCGTATGGATAAGTTCGGTAGACCTTTCTTAGGTGCTACAGTTAAACCTA
AACTAGGTTTATCTGGCAAAAACTACGGAAGAGTTGTATACGAAGGCCTGAAAGGCGGTC

TTGATTTCCTTAAAGATGATGAGAATATTAACTCACAACCATTTATGCGTTGGAGAGAAA
GATTTTTATATTCTATGGAAGGTGTAAATAAAGCATCGGCTTCTGCTGGCGAAATTAAAG
GTCATTACCTTAACGTAACAGCCGCGACAATGGAAGATATGTATGAGAGAGCCGAATTCT
CTAAAGAGGTTGGTAGTATCATTTGTATGATTGACCTTGTGATTGGTTATACTGCGATTC
AAAGTATGGCAATTTGGGCTCGTAAACATGACATGATTTTACATTTACATAGAGCTGGTA
ACTCAACTTACTCTCGTCAAAAAAATCATGGTATGAACTTCCGAGTTATTTGCAAATGGA
TGCGTATGGCTGGTGTTGACCATATTCACGCAGGTACAGTTGTAGGTAAGCTTGAAGGAG
ATCCTTTAATGATTAAAGGCTTCTACAATACTCTACTTGAAAGCGACACAGATATCAACC
TACCTCAAGGTCTGTTCTTTGCTCAAAATTGGGCTTCCCTACGTAAAGTTGTACCAGTAG
CATCTGGTGGTATTCATGCTGGTCAAATGCACCAACTTCTTGATTACTTAGGTGATGATG
TAGTTCTTCAGTTTGGTGGTGGTACAATTGGACATCCTGATGGTATCCAAGCAGGTGCAA
CTGCTAACAGAGTAGCACTAGAGTCCATGGTTATGGCAAGAAATGAAGGCCGTAACTATG
TAGCAGAAGGTCCACAAATCTTGAGGGACGCTGCTAAAACTTGTGGGCCTCTACAAACAG
CTTTAGATTTATGGAAAGATATTAGTTTTAACTACACTTCCACAGATACAGCTGATTTCG
TTGAGACTCCAACAGCAAACATCTAGTTTAATGACTACTTACTGATACTTTAAATAGTTA
ATTGTAAGTGAAATTAACTTATAACAATAAGGAGCATAGAATAGTGAGACTAACACAAGG
GACTTTTTCCTTCCTTCCAGATTTAACTGATGAGCAAATTAATAAACAGCTTGCTTATAT
CGTTTCTAAAGGCTTATCAGCAAACGTTGAGTATACTGACGATCCTCATCCAAGAAACTC
CTATTGGGAACTGTGGGGTTTACCTTTATTTGATGTAAAAGATGCTTCTGCTGTTATGTA
CGAAATTAGCTCATGCAGAAAAGCAAAACCTAATTATTATGTTAAAGTTAACGCTTTTGA
TAATACTAGAGGTATTGAAAGTTGTGTAATGTCTTTCATTGTAAATAGACCTGCTAATGA
ACCAGGATTCTTATTACAACGCCAAGACTTCGAAGGTAGAACTATGAAGTATAGTCTTCA
TAGCTATGCTACTGAAAAGCCTGAAGGAGCTAGGTATTAATATTAATTCTTAATTAATAT
TGGCTAATTATTCCCCTCTTAAAAATCATAATTAATTTAGTAATTGTGATTTTTAAGAGG
GATAGCTTCCCAAGTAAATTTTGACTAAATAGATTGAATAATTAAAAATACACAGAAATA
ATGCAATCACAGGATATAATTTCCAACGATACTCTTGTTAATTTACAAGAAGAATATGAT
AGAACACAAATCCAAGAAGTTTTAAATGAGTTAAATCAAGAACTTATAGGATTAGTGCCT
GTAAAGACCAGAATTCGCGAAATTGCTGCGCTATTATTGATTGACAGATTACGCAGAAAA
CTAGAACTAGTTTCTGGTAATCCAGGATTACACATGTCATTTACAGGTAGTCCAGGAACT
GGTAAAACTACAGTTGCTATGAAAATGGCTGATATTTTGCACAGACTTGGATATATAAAA
AAAGGGCATTTGTTGACAGTTACAAGAGATGATCTTGTAGGTCAATATATTGGACATACT
GCCCCTAAAACTAAGGAAGTTCTTAAACAAGCAATGGGAGGAGTTTTATTTATTGACGAA
GCTTACTATCTATATAAAGCAGATAATGAAAGAGACTATGGCTCTGAGGCAATTGAAATT
TTATTACAAGTAATGGAAAACCAAAGAAATGACTTAGTTGTTATCTTTGCTGGATATAAA
GATAGAATGGAAAAATTCTACGAATCCAACCCAGGACTCTCTTCTAGAGTAGCTAATCAT
GTAGACTTCCCAGATTATACTTCAGATGAATTATTACAAATAGCTAAAATGATGATAGAA

GAACAGCAGTACTGTTTTACAGAAGAAGCAGATAAAACTCTTTTAGAGTATACCGAGCGA
AGAATGAAACAGCCTTATTTTGCTAATGCAAGAAGTATTCGCAATGCTATTGACAGGGCT
AGAATGAGACAAGCCAATAGGATTTTTGCCAGTGGAGAAAAAGTATTAACAAAAGCTGAT
TTGGTAACGATTGAAGCAGAAGATATCTTGAAAAGTAGATTATTTTCATTACCTAATGCT
TAATATACACGTGATTTCATGAAACTATATATTATTAAAAAGTTTCATAATCAGTTGCAA
ATTTCTTGAAAAACCTTTATTATTATCTTAATGTAGGTGTGGCGGCATAGCCAAGTGGTA
AGGCAGAGGATTGCAAATCCTTCATCCCCCAGTTCAAATCTGGGTGCCGCCTAGTACTAA
AAAAGGGGGGTGTGGTGGAATGGTAGACACAACAGACTTAAAATCTGTTGATTTTTAGTA
ATCGTGAGGGTTCAAGTCCCTCCACCCCCATATATTGATTTATAAATAAAAAGAGATGTG
TATTTGTATTAATTGCAACCATATTACCAAATGTAATACCTATCACTTAATAGAATCTCA
ACATAAACAGCCTCATCTTACTAGAAGTCCATTGTTTATACCTAAATATCCTGTAGTTCA
TGTTAATATATCTAACAACTGTACCTATAATCAAATTGATTGGGATTTAGTGGAGTGTTT
ATCATTTGTAGAAAAACCGAATAGTTGGAATTTAGACGCTAATTAGTTACAAGAAAAATT
ATGTACCATAAGCAACAACTCTATTTTTTAGATACTCGGTATTAATTTTGGATGTTCTAA
CCAGTAAGATTTTACCCGTTCTTGCTATCTCTATAATGCCAAATTTAGTTAGTAATTGTT
CAATAGCAACAATCTTTCCCGGATCTCCAGTAACCTCTATAATTAAAAGATCTTCTGCAA
TGTCTACTATCTTAGCTCTAAAAATTCTTACAATTTCTAAAGCTTCTGTTCTAGTCTGAG
AATTGATCTGAATCTTAATTAACATTAGTTCTCTTTCAACTGAAGGAATATTTGTTACAT
CTTGAACGTTAAGGATATTTACTAACTTATATAATTGTTTCGTAAGTTGTTCAATAGTTC
TATTGTCTCCTTGAACTACCATTGTGATTCTAGAGACCCCAATTTGCTCTGCTGGTCCAA
CTGCTAAACTTGCGATATTAAAGCCTCTCCGGGCAAATAGACCAGATATTCTTGACAGTA
CTCCGGCTTCATCTTGAACTAAAACTGATAAGGTGTGTTTCATGAAAAATTAAATTTAAT
CTATTTGATCTATTAAGTATATTCTAAGAAAATTACAGAAAAAATACTAGTTAGTCTATA
ACTAAATTATCCAAACTAAACAAAATCCTACTATCAGATAATATATCTACCTTGTAATGC
TATACTATTAACTTAGTATTTAATGATATTCTAATTGCAAATAGAATGCTAATTTTTTTA
AATTTGTTAGGAACTTGTGCTAGAAAAATACAAAAACAGTAAAAAACTCTCTCGAGATTT
ACCTCAAATCAATGATCGCATTAGATTTCCAAAAGTCCGAGTAATTGATGACGAAGGTGA
ACAACTAGGTATTTTTGTGCCTGAAGAAGCTATACAATTAGCTGTCCAACAAGGTTTAGA
CTTAGTTGTTGTTAGTGATAAATCGGACCCGCCAGTATGCCGAATATTAGACTATGGTAA
ATATAAGTTTACACAAGAAAAAAGAGCTAGAGAAGCTAAGAAAAAGCAACATAACAGTAG
TATTAAAGAAGTAAAAATGCGATATAAGATAGAAGAGCATGATTATAAAGTTAGAATAAA
CCAGGCATCCAAATTTATTCAAGCAGGAGATAAAGTAAAAGCAACTATCACATTTCGGGG
GCGTGAAATCCAGCACTCTAATTTAGCTATAGATTTATTGAATAAGATGGCAAGCGATCT
AAATGCAATAGCTGAAATTCAGCAAGCTCCATCAAGAGATGGCAGAAATGTCATAATGCT
CTTATCTCCCAAAAAAGTTAGCTAAACTAATTTTTATTTAAATGCATCTGGCTGGATTCG
AACCAGCGACGTCTCTTTCGAAATGGCGGATTATGAGTCCGCTGCCTTCGGCCCCTCGGC

CACAGATGCATCACTTAAAGTTTATAACTCATAAAGCATATAAAAATCAATGTTTAGCTA
TTTTTAATAAAGTCTATAAAAGTCTATTATTCATTCATATTGTGATATGTCGCAACTGCT
GAAGGGGAAACTTGATTTAAGTATCGAAATATCCAGTATTTAAAAATTGTGTCGAGTATA
ACTGGAAAAGTTGAAATAAATAAAAAAATAAAATCTCGACTTTCCGGAAGTCCTAAATGT
CTTAAAATTACTTCAATAATAACTTCCCATCCATGAGGAGAATGAAAACCCACAAACATA
TCAGTAAAAAGAATGATTAAAAAAGCTTTAGCTGTATCACTAAGACCATAAATAATTTCA
TTTAAAAAAGATTTAACGACTGCAATTTGTCTTTGTCCAGTTATCATTAGTAATATAAAG
ACAAGTATTGATACAAGATCGGATAAAATATTTTTAACAGCATTTGCACTTTCGTTTGCA
TAATATTCTCCCAACTCTCTAGCCTTCAGCTGCACTCTTTTTTCAATGATTTCGTAAGAT
ATATCTTCCGAAGGATTCAGAAGAACTTCAAAGTGAATTTTCTCTTCAAATCTTTGTAAT
TCAGCAAAAGCTCTTTCCTCTTGAGAGGAATTTAAAAAAATTTTAGGCTGTTCTTGGTTC
CATAAATAATCAATACAAGGTCCAAATACAAAAAATTTTGAAGCCTGATTAACTAGTACT
GGAGAGATGAATAATAATAAAATATATTTTACGGAAGTAATAGTTTGGTGTCTAGATATT
CTAAACTCCTCTATAGCTTCAGATTCGCCATTAGGATCTAACTCTTTACGAAACTTTTCA
AAAGTATTAGTAATTGACCTAGGTATTGGCCCAACTTTTTCAAAGGCGAATTGATTATTT
CGTTTTAAATTCCAATATTTCATTTTTTATTGCTGCTAAAATTTCTATTAATCAATAATT
GATCACGTATTCACTATAAATATTAATTCTAGAACTAAACATATAAAACAATATTTATGA
AAAATAAAATACAATTAAAAATTAATTATGCATTAGTTCTATCTGCAGCTACAGCAGTTT
ATGGATATAAATCTTATCAGCCAACTATAAATACACTCTTAATCAAAGATGAAGAAAACT
GGCCTTTTTTCTCTAAAAGGCAGACTGCCTTTATTCCTATAATTGTATTAAAAAAAAATC
ATCAAAATCCCTGAACTCAGAACCTAGAAATAGTAAGTTATATCTTAATGGAATTGGAAG
TAGAAAATTTATCGACAGAATTGTCACTCTACTTGGATATTTAGAAAAACAGGCAAACTA
TACTGAATACTTGAACCAAATAAAAACAAGTGGTTATTTTTCTTTAGTCTACCTAGACAG
TGATATAATGTATAAGAAAAATATTAGTAACATTTTTTTACTACCTAATGATATTCTAAA
AAGAATCTATATTTGCAATAAGAAAAAAAAGCTTATACCGCATTCATTCTTAATAAATTT
ATTTCAGAAACAGATAGGCTATCCAAAAAGTTTTGCTAATTTAAATTGGGCCTTCTCTAA
AATTATAAAATGGTATTATGATAGAGGATACCAATGGTCTCTAGTTGAAGTTAAACAAGC
GTCTGATGCATCTTCTATTGTAATTGATATTCATGAAGGGGTAGTGAAGACAATTATAAC
AGAATACTACACTCTATCTTATAAAAGAGTCTCAGGTATCTTATGCGTAGAGTCAATAGA
GCAGTACCTTGGAGTAAGAGTGGGAGCTCCACTAAATATAATTGATTTACAAAAAAAAAT
TACTTACTTAAAAGATAATCAACTAGTTGGCGACATTATTTATAGTATTGAACGATCAAA
TAATAGCTCAATGAGTTTGGATATCAAATTTCAAATACAAGAATTAAAAGATAAAGAGAT
AATAGTGCTTGCAGAAAGTTCTTCTATTATCTCGCATGCATGTAATTTACTCAATCAATA
TAGAAATAGGCTAGTAGCTTCTAATATAGTTTCACTATCAACTAATAAATTACACGCATG
CTATTTTAATTATAAACTTGACTATCAATATAAGTATATAAACACCTCTAATCTTATAAA
ATTGTTAGCTTATTCTATTTCTTGTAGGAAGACACTACTTATTCATTATTTTAATTTACA

AACTTTAATAAGCTGTACAAAAAAAAAATACTATTGGATTTCAGCTATATTTACGAAATCT
AAGTTTTGGGAAAGCTTTCTGTGTGCTTAGTATGAAATTTATCAAAAATGGACTTAATAT
AAAAATTTTATATATAAATCCTTCATTGATAGTTGATCAGAATTTTGTATTTCAATTTGC
TATACAAATCATTAAGCAATATCATACTGCTAAACCTCCAGCTTTATTTTTAACGAATCT
AGATCTTGAACAATATGTTGCTGAAAGTTTATTAATGTACCATTTTACATCTTGCTTCTC
AATATCTGAAAAAATATTATTATCTCGAATTATGCATACAGATTCTTTATTTTTCAACTC
TGAAAATTTTCATTTTGATGATAGAACTAACGCGATCAACAGTTATGATACTTTTAAACA
AAATACAAAAATATTTTATCAAGAGTTTCTTTCTTTATTATTAAGTTTACGTTATCAAAA
TTTTAATTATTTAGGTTGGCCTTTAAAAGGCCATTTTTTTGAAATAAAATCTTTATACTT
GGCTCCATTTCAAAAGAGTGATTTTTCTGATAGTCGTAAGACGCTATTCTTCCACAAAAT
GTCGTTAAAGCAAGTATCTAATTTTAATTTACCAGTATCTTTTAAAAGCCACCTTAACCA
TATTTTAGTTAGTACTATTAAATGTCAGTCAAATTTGAATATGAGAACTGTATCTTTGCT
ATTAATTGATTCACCTGCTGAATATCTGCTATACAAATCTATTCTTAATTTCTCTATTAA
AGTTAGGATGCAGTATTTCATACCTATGAGCAACAATATAAGATTATCTCTATTTTATAA
TTATTTAGATTGTTTTTTAATAAGATCATCTCAGAGTTGCATTCACATATGGCAAGATCT
AAGAACTCTTACACCAATTCAAAATTTTTGGTTAAAAAAAATTTTCATATGGAGCTGGAAT
TCAGCTTAAATTACCAATTAAGCAGATGCCACCTCTTTCAATTGAATATACTGTAACTAG
TAGTCGCTACTTTTGTATTTACCTTCGTACTTATTATCAACGATAACTATTACTAAAATG
AAAAATTTAATTTCTTTTTTTGATCGCAGCAAAGGGAAATGGATTTCACAAAGAACAACT
TATGAATTATCTAATAAAAACATGAGTTCGGTACAGTCTCAGATGACAATGAAAATAGGC
AACTCATTGTCAGGATCTATAATACTTGCATCGTTAAATTGGGGGGACATTTATAGACAA
GTTGCCCATTACGCTAAAAATCACAGCCGGAGTGAATATGATAATAAGTTTAACTTGCAA
TTCGGTAATCAACTAAATAATCATAAATTATTAACGCTGTGCATAGTGACAGATCCTAGT
CTAATTAGTTTTAAAACTCGCTATGGAAGTACTACCATAGATGAAACATATTGGTTCGCA
ACAAATAATTTACGTCTAAGTACTAGTATTGTTAAACGATTCAACACTTGTGTGGCAGTA
TCTTTTTGTTCAGAAATTAAAGTTTAAACTCTATATAGAAGCAACTAATAGCCTTTAGGC
TATTAGTTGCTTCTATGTATCTAAATTTTGTATTAAAAATTATAATACTATATTATTCTA
AATCTTTACGATTGGGATTCCTGGAAGGATCATTAGAGAGGAACCCAAAGACAAACAAAG
AAATAAAGAAAATAACGGTAGTATATACAAAGATTTTCAAAGTAAACATATTTTCATCCT
TAAAAGCTATTAGTGCGATATTTACATATCTCCACATAATTTTATATCATGATATAAGTT
CCAGCACTATTTTATAAATAAATCAATATCTTTATTTTTTAGTAATGATTTATATTTAAA
TATGCGTGATGTACTAAATACTAAATCTTTTCCGACTTGTTTTGTCTTGTTTGTAAATAT
TTTGTTATTTCTATGGAGTTTACCTTCAATTATCACATAGTCTAACTTCTTTAGCTGTTT
AAAAGTATATGAAGATTTTTTGTTCCAAATAGTTAGATTGATTATAACTAATCGTTTTCT
TTTTAACAGTCTTGCTTTTAACTTAATTATTTGGCTTTTATTTTCACTAATTCTGATACT
CTTACAACTAAGAATTTGGACTAGAAGAGTACAGCTATTCATAATTTATAACAATTTTTT

AAAAAATCTACTAAAATAGATGATTTTACAGAACTAAAATGATATTTCATTATCTTGTTG

ACCTTGCTCATAAGTAATATCTTTCAATACTTTTAATATTTTTCCAAAATACTCTTGCAG

GTATTTTTCTAAAGATATTACTTCCAACCTGTCAATTCCAAGAATACTGTAAACTTCATC

CATCGGAGCTGTAAATTGCTCTCCGCTACTTAAAATTTCGGCAAATGCTAAACGATCAGA

AATATTCCATGTCCATTGAAGAGTTTTAGTAATTCTTCTTAAGGCTTTTAATAATCCAAT

CGGAATTTGTGAGATCTGTGTTTTTTGCCCAGATAATTTTTCACATAATTTAATAATTTC

AGCTGATGTCCAAGCGGTATTACCAACTAACGGCAGGGTCCTATTTTCTGTGGAAGGTAC

TCCAAGACTTTTTATAACTAACTTGGCTGCATCTTGAGTATCAATGTATGCAATGGGTGT

AGATTCTCCTGTAACCCAGACTGATTTCTTGTCTAAAATAGGAATGGCATATTGGTTGAT

TAAACCTTGAAAAAATCCTCCTAAAGAAAACACAGTATATTTCACATTGGACTTCTGAAG

GAAATCTACTACCTGAGACTTTAAGTTCATCAATGGAACATCTGGGTATTTCTCTGAATT

TAATATTGAAAAGAAAATAAATCTTTCAACTTTAGCCGCTTTAGCTGCTTCAATTAATGC

AATTTTCCCATCTAAATCTATTTTTTCTGCATTATAAGGATCAGTAGGACGAGATGTAGA

AGCATCTATGATTGCTGTTACCCCACAAAAAGATTGCAAAATACTTTCGGGTAATTTTAA

ATCACCATATATAAGTTCTGCTCCCCACTCTTTGAGAAAGGCAGACTTTCTTAAATTTCT

TACCATGCATTTTACATTGTAGCCTTCATCTAAAGCTCGTCTTACAATTTGACGTCCTAA

AGTTCCGGTTGCTCCAATTACTAAAAGAGTCATATTAGTTAAAAATAAGTATGTATAAAG

ACATTGATTTGGTTGACGGCAGAGAGTAAATTAGGTAGAGAGGGAATCGAACCCTCATGA

CCGAAGTCGTCACATTTTGAGTGTGATGCGTATACCAATTTCGCCATCTACCCTATTATA

GACCAACTACTACAAGTAATTTAGTCTTTAGCGTAAAAATAATACTAAGTTTCAATAGAT

ATGTCAACTTTTAAATTTATATTAATAAAAAAACGGAGTCTCTTATGGCCACTTTTGAAT

TAATTCCTTATAGATTGTATTTGAGTCAATCTAACACATGGATCCATAGAATAAAAGCAG

AAATTAAGATATATATAGTAGCGCTTTTATGGATTTCAATTTTTATTTTTTCTTACTTTA

AACTATGTATTATTGCTTTAAGCTTAATTGCCATAAGTTTTACTATAAGAAGTAAACAGA

ATATTATTCAAAAACATTTGTTACAAACGTTATTAATGACGTTCTTAACTACTGTTTTGT

CTTTTAGTGTGGCTATTAGTTATAAACAATATGCAGAACAAGAACAATCGCAGTATTTAT

CTGATTCCAAGAAATATAAAAATTCCAGCAGCTACTATTATATACAAATAGCGAACGATC

AAAGGATCAAAAGACAATACTTAATTACTACTTTAAAGCCTTCCTTGTACTTTTTTATTA

CTATATATTCTATTAAACTAGTTATGATAACAACCTCTCCAGAAGTTTTAGTAATTACTA

TTTATAGATCTAGGATAATAAATAAAATATTTAAAAATGAATTGCTATTTATCTTCCTTC

TTTCGTCACATATTGTTACTAGTATTATTAATAGAATCGATAAAGTGATTCAAGTAACTA

GCTTAAGAGGAAGCTTGAATTTATATAATTCATTGACAAGGCCTTTAATGTTTTCTTTGT

TAATATTTCAAGTCTTCTTTTTGGAGATTATTCGGGAGTCAAAAGAAATAGCTCAAGCTC

TTTATACTAGAAATCTCAATCAAGAAAACAATAATTTTTTGAAAATATATACAGTAAAAT

CTAACTTTAGTGATCGGCTCAATATAATTATTAGCACTTTGTACTTTATTATTTTAGCTC

TAGCGTAATAGAATCCACTTATTCTTCTGACTCTAGTAATACGAAGAGTAATAAAATTAG

TTAGTAGTTTAAATAGATAGAATTAAGATATTATATAAATAATTAAATGTTTCATTATGG
TCAATACTCAAAATCAAATTTCACAAACTTCAGAACTTGATTATATAGTTAATCAACCTT
ATAAATATGGCTTTAAAACTTCTGTTGAATCTGAGCAATTTCCTAGGGGAATAAGTGAAG
ATATTGTCCGCTTGATCTCCAAGAAAAAAGATGAGCCTGAATATCTATTGAATTTTAGGC
TTAAAGCATATAAAAAATGGAAAAAGATGAGTAGTCCGTCATGGGCTCATATTAAGCATC
CGAATATAGACTTTAATACGATTATTTATTACGCTGTTCCTAAATTAAAAAAAGAATTGA
AAAGTCTAGATGAGGTTGATCCAGAAATTCTTGACACTTTTAACAAGTTAGGTATATCTT
TAAATGAGCAAAAGCGAATTTCCAATGTAGCTGTTGATGCTGTTTTTGATAGTGTATCTA
TTGCAACTACTTTTAAGAAAGAACTGTCTGAAGCTGGTGTTATATTCTGTTCAATTTCTG
AAGCTATAAGGGATTATCCAGAATTAATAAAAAAATACTTAGGTACTGTTGTACCAGCTG
GTGATAATTATTTTGCTGCATTGAACTCTGCGGTATTTAGTGATGGCTCTTTTTGCTATA
TCCCTCCTAATACAGTTTGCCCTTTAGAATTATCAACTTACTTTCGTATTAACAACGAAG
AATCTGGGCAATTTGAAAGGACACTAATTATTGCTGATCGTGGCAGTAAAGTAAGTTATC
TTGAAGGTTGTACTGCTCCTCAATTTGACACAAATCAATTACATGCAGCGATTGTAGAAT
TAGTAGCTCTTGAAGGAGCCGAAATTAAATATTCTACAGTACAGAATTGGTATGCTGGTA
ACAAAGAAGGTAAAGGTGGTATATACAATTTTGTTACTAAACGAGGCTTATGCTCGGGTA
ACAATTCAAAAATTTCATGGACTCAAGTAGAAACTGGATCTGCAATTACCTGGAAATATC
CAAGCTGTATTTTAGCTGGTGAAAATTCTCAGGGAGAATTTTACTCTGTAGCTTTAACAA
ATAACTATCAAGAGGCTGATACAGGGACTAAAATGATCCATATTGGTAACAATACAAAAA
GTAGAATTATTTCTAAAGGTATTTCTGCGGGTAGATCGAAAAACAGTTATCGAGGCCTAG
TAAAAGTTGGCCCTCAGTCATTTAATTCTCGTAATTATTCTCAATGCGATTCTTTATTAA
TTGGTCAATCATCTCAAGCTAATACATTTCCTTATATTCAAGTACAGAATCCAACATCAA
AAGTAGAACATGAAGCATCCACTTCAAAAATTAGCGAGGACCAAATTTTTTATTTTTTAC
AAAGAGGGATCAATTTAGAAGAATCTATTGCTCTTATGATCAGTGGTTTTTGTAAAGATG
TCTTTAATGAATTGCCTATGGAATTTGCTACTGAAGCTGATCGTTTACTGAGTTTAAAAT
TAGAGGGAACTGTAGGATGAGCCAGACTATTTTAGAAATCAAAGATTTATATGCTTCTGT
TGGTGAAACAACAATTTTAAAAGGGGTAAACTTATCTATTCGAGCCGGCGAAATACATGC
GATTATGGGGCCTAACGGTTCAGGTAAAAGTACATTATCAAAAGTAATTGCAGGACATCC
AGCGTATTCACTAATAAGCGGAGATATTTTATTTTTTGGACAAAGTATTCTTGAAATGGA
GCCAGACGAGAGGGCAAAAGCAGGTATTTTTTTAGCTTTTCAGTATCCTGTTGAAATTCC
TGGAGTCAGTAATTCTGATTTCTTAAGAATTGCATTAAATGCTCGCAGAAAGTTTCAAGG
ATTATCGGAGTTTAGCCCCTTAGAATTTTTTCAACTAATAACAGAAAAAATAGATCTTGT
TGGCATGCAAGAAAGTTTTTTGACAAGAAATGTTAATGAAGGATTTTCTGGTGGAGAAAA
AAAACGTAACGAAATTCTTCAAATGGCTTTACTAGATAGCAAAATATCTATTTTAGATGA
AACAGATTCTGGACTAGATATTGATGCATTACGAGTAGTTGCAAAAGGAATTAACACTTT
AGCTAAATCAACAAATTCAATTATTTTGATTACTCACTATCAAAGATTATTAGATTATAT

TATTCCAGATTTTGTTCATATTATGAGTAACGGGCAAATTGTAAAAACTGGTAGTGTTAC
TCTTGCCCAAGATCTAGAAAAGCACGGATATGACTGGATTACGCAGACATAACTATTTAC
TACAATATTGAGACTAAAGAATATTCTTTAGTCTCAATATTTTTGCAAATGCAAAGTTAA
TTATTTAAGCTTTATCAAAACCTTGCTTAACATCTTCAATTGCTCTTTTTAGCAATTCTT
CACTAACTGGTTCTAATTTTTTAGTACTACGAACATTTTCACCAAATTCTGGTTTAGAAT
TCTTTAAATCTTCTCGAAGTTCTCTAATAAATTCAGCTACTTTTGAAACTTCAATATCAT
CTAGATAGCCATTAATTCCAGTGTAAATAATAGCAGTCTGTTCTTCAACTGGAATAGGAG
AATTTTGGGCTTGCTTCAAAATTTCTCGAAGACGTTGACCTCTTGCTAGTTGATTTTGGG
TTGCTTTATCTAGATCAGATGCAAACTGAGAAAAAGCTTCTAATTCAGCAAATTGTGCTA
ACTCTAGTTTTAACTTGCCTGCCACTTGTTTCATGGCTTTTATTTGCGCAGCAGAGCCTA
CTCTAGAGACTGAGATCCCAACATTAATAGCTGGTCTAATCCCTGAGTTAAATAAGTCAC
CAGATAGAAATATTTGTCCATCTGTAATTGAAATTACATTTGTTGGTATATATGCAGATA
CGTCCCCAGCTTGAGTTTCAATAATAGGGAGAGCTGTCATACTACCTCCACCTAATTCAG
CATTCAATTTAGCTGCTCTCTCTAGCAATCTAGAGTGTAAATAGAATACATCCCCAGGAT
AGGCTTCTCGTCCCGGAGGTCTTCTAAGCAAAAGAGACATCTGACGATAAGCTTGAGCTT
GCTTAGTTAAATCATCATAAATAACTAATGTTGCTTTACCTTTGTACATAAAATATTCAG
CTAAAGCCGCACCTGTATAAGGAGCAATATATTGTAAAGTCGCAGGACTATCTGCATTAG
CTGCAACTATAATAGTATAATCAAGAGCTCCTTTATCCTGTAGTGAAGATACTACTTGGG
CTACTGAAGATGCCTTTTGTCCTATCGCAACATATACACAAACAACATCTTGGCCTTTTT
GATTAATAATTGTGTCCAATGCAACAGCTGTTTTACCTGTTTGACGGTCACCAATAATTA
ATTCTCGCTGACCTCTACCAATTGGGATCATAGAGTCAATAGCAGTTATCCCTGTTTGCA
TAGGTTCACACACTGATTGCCTTCCAATTATACCAGGGGCCATTGACTCAATTAGCCTTG
TTCCATTACTTGCAGGTTCACCTTTATCATCTATTGGCCGAGCTAATGGATCAACAACTC
TACCTAAAAAAGCATCTCCGACCGGAATTTGAGCAATTCGACCAGTCCCTTTTACGGAAC
TGCCCTCTAAAATATCCCTACCATCTCCCATGAGTACTACACCAACGTTATCACTTTCTA
AATTTAAAGCAACCCCAATAGTTTTATCTTCAAATTCAAGTAGTTCACCAGCCATGACTT
CATCAAGTCCATATACTCGAGCGATACCATCACCAACTTGTAAAACTGTTCCTATATTGG
CTACTTCTACATCCTGATCATACTTTTCAATTTGTTGACGAATAATACTACTTATTTCAT
CAGGTCTAATATTTACCATATTTTTAATATTATTTTTATTAATTGGAAATTCTAAAGCGG
TAATTAGTCAGACTAATGCTATCTGTCTTTTTTACTTTATTTTAAGTTACTACAACATCA
AGGTGTGAAGCCATTTGTCTTAGCTGTCCTCTAATACTAGTATCAATGACTTTTGATCCA
ATCTGAATAGTAAAGCCACCAATTAGCTCGGGCTCGACTGAAATAATTAGTTTAACTTCT
TTAGCCTTAGTCATAACCTTGATTTTTTCTGTTAATAGAGTTTCCTGATCAGGGCTTAAA
GCAATAGAAGTATTAATATTTGCAATTGTTAAAGATTCCATTTGGTAAGCCAGTTCCAAA
TACTTACTAGCTATAGCATCTAACATTCCAATTCGCTTGCGATCAACTAAAATCATTAGA
AATGATAATGTATTTTCGTTAATTTGATCGCCAACAGTTGCAATAATTACTTCTTTTTTT

GCCTCTACAGTTTTTAATGGATTAGCTAAAAAAATTTTTAGCTTGTCAGATTGCGCTAAA

ATATTTTGTATTGACTGAATATCTTGGCTCACCTTTTCAGTAACTTTTTTAGTTTTAGCT

AAATCAAGAAGAGCTACTGCATAAGGCTGAGCTATTTTTATAACAACATTGTTACTGCTC

ATAGTTGATCTCCTAACTTAGCAATATTATTATCAATTATACGCAGTTGCATTTCGGAGC

TCATTTGATTTTCCAGCTGCAAAGTAACTCTTTTAAGAGCAAGAAATGTTATTTGTTGCT

GGATTTGCCTTCGGATCTGTTTTTCAGCTGTTTCTATATTTGATTTACCTGTTATGGCAA

GTCTTTCAATATCTAATTGGCCTTGAGCTAATATGGAACTTCTAACTTTTCCAGCTGTTA

GCTGAGCTTCTTTCTTAATTTGATCTATAATGATTTGAGTTTGAGCAAGTTGTTTTTCTG

ATTCTGACAATCTAGCACTTGCTTGCTCTAGTCTTTCTTCGGATTCTTGTATAGCTGCTA

AAACTTTTATCTGTCTTGCATTTAAACTTGATCCGAGAAATTGCTTCAAAACATAAATAA

GACCAAATAAAAGCAACAATATATTAATAACATTTGCTTCAAAAATATCAGAATTAAAGC

CGAATTTGTGTTCACTACTATGTTCTGATAAAATAGTAATTATTTGTGGTATTTTTACTA

TATTATTCATTTAAAATTCTTACTATTTTAATATTTTACTCTGGAATCATGATTTTCAAG

ATTGACTACTTAATAATTTAACTTTAATTTGATCGCTCAAAGTGTCAACCTGATTTTCTA

AAGTTTTAAGAGCTTCTTCTTTTTGAACATTTAACTGTTTCGATGCTTCAGCAATCAGTT

TCTCTGCATTCAATTGAGCTTTTTTTATATCTTCTGAGACTATATTTTGAGCTTCTTTTT

GAGAAGAGGCTATTTTTAATTGTGCATTGCGGCGAGCTTCAGATAGATCTTCTTCATATT

TTGCAGCTAGCTCATCTGCTTTAACAAGCATAGACGATGCAGTAGTCAATGTTGTTCGAA

TATATTCATCTCGTTCGTCTAGTACTTTAGTTACAGGTTTATAAAAAATAGTATTTAATA

AAACCATTAAAGTAAGAAACTGTAATGCCATTAAAGGCAAAGTACCATTGAAGTCAAATA

AGCCACCTTCAATTTCTTCGGCTAATAGAAATGGTAAATCAATCATTTTACTAATTATGT

TTATAGTTAATAATTTTGTTTCAATTAAGAAAAATAACTTTATAGTGCGCTTAGTCTCAT

CAGACTAAGCGCAAATGATTATTTTAACCAACGTATGGATTAGCGAACAATAAAGACAAC

GCAACAACTAGGCCGTAAATTGTTAAAGATTCCATAAATGCTAAACTTAGAAGAAGTGTA

CCCCGAATCTTTCCTTCTACTTCTGGCTGTCTAGCAATACCTTCGACAGCATTAGCTGCT

GCACTACCTTGACCAATGCCTGGGCCTATTGCAGCAAGACCTACAGCAAGACCGGCAGCG

ATAACGGATGCGGCTGAAACGATTGAATCCATAATGAATTTTAATTTTTTGTTATATGAG

TAGAAAATTGACAAATTTCGTATAATACTTAAAATCTATATAATTAAGTATAATAGATTC

TAATATTGTAATTATAGTATATTACTCTTCTCCATGACCTTCCATTGCTTCTCCAATATA

AGCTGCGGACAAAGTAGAAAAGATTAATGCCTGTATTGAACTCGCAAATAATCCGAGTAT

CATAACTGGTAATGGTATTAAAATTGGAATTAGCAAAGTAAATACAGAAACAACTAATTC

ATCTGCTAAGACGTTTCCAAATAATCGAAAACTTAAAGATAGTGGTTTCGTAAAATCTTC

TAGAATATTGATTGGTAGTAATACTGGAGTTGGCTGAATATATCTAGCAAAGTAGCCTAA

CCCTTTTTTACTTAAACCTGCATAAAAATAAGCTAAAGATGTAAGTAAAGACAGGGCCAC

AGTTGTATTAATGTCATTTGTTGGGGCAGCTAGCTCACCTTCAGGCAAGTGAATTAACTT

CCAAGGAATTAATGCACCTGCCCAGTTACAACCTAGAATAAATAGAAATAAAGTTGCAAT

GTAAGGCACCCAAGGGCGATATTCATGCTCACCAATCTGATTTTTTGCAATATCTTGCAG
AAATTCTAATATAAATTCCATAAAATTCTGAAATTTTTCAGGAATTCTTTGTAGATTTCT
AGTACCTAGAAATGATAAAGTTAAAAGTGTAGCAATTACTAACCAAGAGACAATGAAAAC
TTGCCCATGCAATTGTAAGCTACCTATTTTCCAATATAGATGTTTTCCAACTTCTACTGC
TGATAAACTATTATATGGGTTAAAATCTATGAGACTGTTTTGATACATAATAAATATTTT
TTTAAATAGTGCAAGAATCAAAAAATAAGTTTTTGAAGAATTAATATGCAATTAATTTAT
TCTTAAAATTAGACTATTAATAAATTAAAAGCAAAAATAAATTCTAATAGAGATAGCTAT
TTTTAGTAGTGATTATTTTTTGTAAAAAACAATTATTGCTACCTATAATGCTTTTACGTT
TAATTATATACCCATATATTAATTATTTAATATGATTATAGGTATTTTGCTACAAACTTT
ATTGTTTTATCCATTCTAAAGATTGTTATAGTTTATTTTTTATTTAAAATATCTGCGACT
TCATCTGCAAAATTAACTTTCAAATTTTCTTCTCCCCCTCCCAGAAGAAATCTTACAAAT
CTTCTAATTTTAATATTTTCACCTAGAACAGCAATATTTTGGTTTATTAAATCTTCAATA
CTAATATCTTGATTGCGAATAAACATTTGATCTAATAATGATAGCTCCTTAAGTCTCTTT
TTCATTCGTCCTTCAATGATTGTTTCGATTCTATCAACTGGTTTGTTTTTCAAGTCATCT
TTACCAGCTTCAACTCTTTGTTCCAGATTAATAATTTCACTAGGTATATCCTGTGTTGAT
ACGTATTCCACATTAGGCGATGCCGCTATTTGCATAGCAATATCTTTAGCTAACTTTTGG
AACTCTGGACGTCGTGCTACAAAATCTGTTTCACAGTTGACTTCAACTAATACACCTATT
CTTCCGCCAGTATGAATATAACTCTCTAGTAAACCTTCAATAGCGGTACGATTAGATTTC
TTGTTAGCTGAAGCCAGCCCTTTTTGTCTTAATGATTCTAAAGCTTTTTCTTCATTGCCA
TTATTAGCTTGTAAAGCTTTCTTGCAGTCCATCATACCTGCTCCAGTCTTATCACGTAAG
GCTTTTACGACTTGAGCAGAAATTTGTAGTGTCATATGCTTTCAAAAATTTTTAAGTATT
TAAATTAATTATTATAATTCACTTTGAGTAGGATGAGTTATTTCTTCTAAATCTGTTTGC
CCGTATTTACCATCATAAATTGCATCTGCTATTTTACCTACAATTAGTTTAATTGACCTA
ATGGCATCATCATTAGCTGGGATTGGAATATTAATAATTTCTGGACTACAATTAGTATCA
AGTATACAAATAGTTGGAATACCTAATTTTAAACACTCTTGTATAGCTGTTGTTTCTCTC
TTTTGATCTACAACTACAACAATATCAGGTAAACGAGTCATATTTTTAATACCATTTAGA
TGTTTACGAAGTTTATCTAATTCTCTTCGTAAAACAGCGGCTTCTTTTTTGGGTAACTGA
TCAATCATACCACTTTTATCTTGTTCTTCTAGCTGTCTCAGACGACTAACTCGAGACTTA
ATCGTTACCCAATTAGTTAACATTCCTCCAAGCCATCTCTGGTTAACATAATAAGAATCA
CAGCGTTGTGCTTCCTGTGCCACTATACCAGCGGCTTGTCTTTTTGTTCCTAGAAATAAA
ACTTTTTTACCATCCGACGAAGCCTGTTTTATAAGTTCACAAGCTTCTGTTAATAATTGT
GCAGTTTGGACTAGATCAATAATATGTATGCCATTTCTTTCTGTATAAATATATGGAAAC
ATTTTAGGATTCCATCTACGTGCTTGATGTCCAAAATGAACACCTGCTTCTAGTAATTCT
GCTAATGTAACAATAGCCATAAGATTATTAATATTTAAATAATGTAATTATTCGGGAAAT
CCCTTTGTTTAGAAACTGATTCAACTAACTTATTTGAATTTTAGTAAACTAATAATAAAT
TAACACACTTTTAAAGCAGATCAACAAAATAGATCCTGAAGAATTCATTCTACACTTTTA

TTGCTAAAATAATTACGCGCGGTTCTGTCGTCTAAAATGATATCATCTAAATCATCTCGT
ACAGATGAAATAGTACTTTCGTCATTAGTATTTGCAGTTAAATTCTTTTTCTCTAACGAT
GAGCCATTACAGTTATCATACATATTGAAGCCGGTACCAGCTGGTATAAGCCGACCTATA
ATAACATTTTCTTTCAGACCTCTTAACCAATCTAATTTACCAGATATAGCAGCTTCAGTT
AATACTTTTGTAGTTTCTTGAAAACTTGCTGCAGAAATAAAGCTTTCTGTATTTAAGGAA
GCTTGAGTAATACCTAATAGCACAGGCCTATATGATGCATTAATCTTATTTCTCAAGGTT
ATGGCTTTATTAGTCTGTTCAATCTTCTGTAACTCTACAAGCTCTCCAGGTAAATAGCCT
GTTTCTTCACCATTTTCTATTTTCACTTTAGAAGTCATCTGCCGTACTATTACTTCTACA
TGCTTATCAGAAATATTCACTCCTTGAGATTGATATACCAATTGTACTTCTTTAACAAGT
AGTAACTGAATCTCTTGAAAGCTTAATCGAGCAGCTTCATATAAGGCTAATCCACATTCA
ATATACAAATCAAAACTTGCATCGAGGATATCATGAGGATTTAACAATACGTCTGTTTTC
TTTCTTGCTTCAAGAATTTCTTCAATTCTTGGAAGCCCTTGTATAATGTCACCTGTTTTT
GCTCTATCAAAAACCAATATAGCTAATGTTTCTCCCCTTCTAATTAATGCATTGTTATCA
ACATGTAAAATAGCACCATTAGAAACCAAGTAAGGTCTAGCAATTCTTAATGTAACTGAT
CTAGAAGAAATTTCAATAATTTGTCCAGAGTCTAAAGAAGTCATATTTTCAGTAATAAAA
TCACCACAACGAATCCAATCACCCACACTTACTTTAACTGTTGATTTTCCTATATTAAAG
GTCTTTTTATCAGAAGATGTCGCAATCAAAATGCGCCTGCTAGTATTCTTTTCAGAATAA
ATCTCCTCAACAGTACCTTCACTCATTGCAATAATCTCAGTACTAGCAACAACTGTAAAT
GGCTTGATATATTCACCATTTTTTACAATAATTCTAGTTTGACTTTGTTGCTTCTCTGAT
TTATTTACATCAATACTTTTAATTGATAATGTTTCAAAAGTTGACAGCCTTAAGTCAAAA
TAATTATTGCCACTTTCTTTAGCAGAAAACTCAATTGAGGATATTAAATTATCATCTCGA
TGCTCAATTTCAGCCACAAGATTAGTTGTTACTAAATGAACTCCTTCTATAGATTTCACT
CTTTCTCCGTCCCGAAAAGGAGTTCTTTTAACTAGCTTTAGTTTAGTTTGACTTGGTTTT
TGTGAAACTAGATTTTCAATTAGACTGGATTTTGTTTCTGGTATTGAATAAACAATAACA
GGTCTAATAAGAATATAGGGCATTTGCTCGTTTTCTATATACTCCCAATAAACTAACTTA
TCTGTAGAAATATTATTATGAAGTCTTTCTCCAGGTCTTAAAAAACCTCTACTTTTATCA
TTATTACTATACACCTCACTAAGTTTATAGATAGATCCAGGCTTAATAATAATTTCTCGA
ACTATTCCATCCTTTTCGATTATTTCAACAATACCTGAGCTTTTAGAAAAAATATTTTTA
ACTAGCTCTGTTCCACTTTCTATAACATCTCCATTATGCACTAGTATTAGTGATGAATCT
TTATTAATTTCGTGAGTTTCTTCTGAAATCCATAAGATATATCCCGGACTTAAAATTTCG
TAGGCATCTTTATCTAAGCCTGTTTTCTTTTTTGATACATTTAAGTCTAAATATTTAATA
ATACCACCACTTGTCGTTTTATACGTATCAGATATTAGTTCTGCGACTGTGTAACCATCT
TGAATTTTTTGGTTTGGAATAGCTTTGAAAAGAAATTTTTGTTTTTTTTCTGTTTCTAAA
ATATAAGATTCTTTTTTGTGGATTACATCTGTATAAATATAACAACCTGGAATTACAATA
GATTCAGTAATAACTTGTATATTGGTAATACTGCTATTAGTATTCTGGTGTATCCTTACT
TCTCCAGCATAATGATTTATTAATTCTGTCTGTGCAAGTATAGTACCAGCTTTAGTTTTA

TCTTCTTTATGAACAATAACATTAGCAGAATCAGATATACTATAAACTTCACCTGAGAGC
ACCCAGATCAAGCCTCCAGTTTTAGTAATTCTTGTAGTATATTGATTATTATCTGTTTCT
TCAACTGTTAAATTGGAGAAACAGATTTTGCCAGAGAGATCTGAAACTACATGTTTTTGA
GCTCTTTCTGTCATGAGTCGATTTCTGCGGGGTGATTCAGCAATCACTTGATCCTTCGAT
ACTAATTCACCATCACATATTAAAAGAGTAGTACCTTTACTTAAGTTAATTATTGACTTC
TGATTTTTCTTAGATCTGATTGTAACTTGGGTTGGCTTTTTTGTTATTAATGCTTGCTCA
CCATGTCTGGTACGAACATCTGTATACGATTCAATATCTAGATCTATCAATTGACCATCT
ATAGGTGCGTAGATTTGTTCTGCTAATTCACCGGTAAATACTCCTCCAGTATGAAAGGTC
CTCATTGTCAATTGAGTGCCAGGCTCACCAATAGATTGAGCCGCAATAATACCAACAGCC
TCTCCTAAATCAACTAAGCGGCCATGTGCAAGATTCCATCCGTAACAATACTGACAAACA
GAACTTCTAGAGTTACATGTAACAGGAGATCTTACTAAAACTTTTTTGATACCAGCTTTA
GTTATCTCTTTTGCTAACTTAGGACTGACATCTTGGTTCGTATGAGCAATTAAACAATTT
GTCTCTGGATGGAATACATTTTCTGCTAATACTCGGCCAGATAAAGCTTGTTCTAGATTA
ATTAATATTTTCTGCGTATCTACTAAATCTTCTAGTATGATGCCTTTATTGGTCTTACAG
TCTACTTCTCTAATTATTACATCTTGAGAAACATCGACAAGTCGACGCGTCAAATAGCCG
GAATCTGCTGTTCTTAAAGCAGTATCAACTAATCCTTTTCTTGCTCCGTATGATGATATA
AAATAATCAGTAACAGTTAGTCCTTCTCTAAAATTACTCGATATTGGAAGATCAATAATT
TGTCCCTGTGGATCAGCCATAAGCCCTCTCATGCCCACTAACTGTCTTACTTGAGAAATA
TTACCTCTAGCACCTGAGAACGCCATCATATAAACAGCATTTAGAGGATCTGTCTCTTTA
AAGTATTTAATAACTTCTTGTTTCAAGGATTCACTCGCATTATTCCACGTATCAATAACT
TTTTGAAATCGTTCAACTGCTGTAATCTCACCTCGGCGATATTTGTTTTCAGTGGCCTTA
ATATCTTCAATCGTGCCTAATAAAAGACTTTGCTTACTAGGAGGAATTCTTAAATCTTCT
AAACTTAAAGAAATTCCTGCCTGGGTCGCATAATGAAATCCAAGATCTTTAAGCTTGTCG
GCCATATTGGCTGCTCGGGCTATACCATAATTACGAAAAGCCCAGACAATTAAATTTTTT
AGTTCGTTTTTATCAATAACCTTATTTGAAAAACTTGGCTGTGCAAGACTTCGTCTATTA
TCCACTAAATCTTCTCCATATTAAATGTTCGTTATTAACTATGCTACTAAAGACTCTTGA
ATAATTTTATTAAAGATTATACGGCCAGCTGTTGTTCTAATATATTGCACAATTCTCTGA
TTTTCAGCGTCTTCTTTTATGATATGGTTATTAAAAAATTTTGTTGTACTATTGTCAGAA
TGTTTTTCAATCTTAATCGGATCATGAGGTTGATCACCATCTACTAATCCATCAAAACGA
GCCCATATATAAGAATGTAAATCAATTTTCTTCTGTTCATAAGCCATAACTACATCTTCT
AAGCTTGCAAAATACTGATTAGATCCCTGCTGCTGAGATGGATTATTAGCTGTTAAATAA
TAGCAACCTAAAACCATGTCTTGGCTTGGCATAATAATTGGTTGACCTGTTGCTGGAGAG
AGAAAATTATGAGGGGCTAACATTAGCAATCTAGCTTCGGCTTGAGCCTCTAAAGACAAA
GGAACATGAACAGCCATTTGATCTCCGTCAAAATCAGCGTTAAACGCTGGGCATACTAAT
GGATGAAGCTTAATAGCTCTTCCTTCTACAAGAATAGGTTCAAATGCTTGAATACCTAAT
CTGTGTAGTGTGGGAGCTCTGTTCAATAGTACTGGGTGACCTTGAATAACTTCATTTAAG

ACATTCCAAATAGAAGATTCATTTTTTTGAATCATCTTTTTGGCAGCTTTAATATTGTTA
ACTAATCCCTGCAAAATTAGTCTGTGAATAACAAAAGGCTGAAATAACTCCAAAGCCATT
TCTCGTGGCAATCCACATTGATGTAGTTTTAAATGAGGACCTACAACAATTACAGATCTA
CCGGAATAATCTACTCTTTTACCTAGCAAGTTTTGTCTAAAACGACCTTGTTTTCCTTCT
ATAATATCTGACAAAGATTTTAATGGTCTATTATTTGCACCTACAACAGTTCTACCTCTA
CGACCATTATCCATGAGAGAGTCAACGGCCTCTTGCAGCATTCTTTTTTCATTTCTGATA
ATAATCTCCGGAGCTAAAATTGACTTTAGACGAGATAATCTATTATTTCTATTAATAATT
CGACGATAAAACTCATTCAAATCTGCCGTCGCAAATCTACCTCCATCTAATTGGACCATA
GGCCGTAAGTCTGGAGGTATCACAGGAATGACTGTGAATACCATCCAAGAAGGATCTGCA
CCTGTTGCTATGAAATTTTCAATCAATCTTAAGCGTTTCATTTTTTTATTGAATTTTAAT
GAAGGTGTCTTAAAACTTTTTGGTGGATTTGTAGCTTCAGACCTTAAGGTTTCAGCTATG
TGCTCTAAATCTAAATCTTTCAACAGTTTCTGAATAGCTTCTGCACCTATTCCTACTTCT
ACTTGGTTATTCTCATCTTGATTTTGGTAGATTTCCTCTTCAAGACTTTTCCATTCGTAA
CCTTCGAGCAGTTGCTTGTATTTTAAATTAATATCTGTATTAGATTGCGTAACTACATAA
GAATGAAAATAAACAATTTTTTCTACTTCTTTGACTTTTAAGTCTAATGCTAAAGCAATA
TAGCTTGTACTTCCTTTTAAGTACCAAACATGAGTTACTGGTGAGGCAAGTTCAATATAA
GCCATCCGATGTCTTCTCACGCGTGATTCTGTAACTTCTACGCCACATCGTTCACAAACT
ATACCTTTGTAGCGAAAACGTTTATATTTACCACAGTGACATTCCCAATCCTTAACTGGG
CCAAAAATTTTCTCACAGAATAAACCATCCATTTCTGGCTTTAAAGTTCTGTAATTAATT
GTTTCAGGCTTTGTAATTTCACCAACAATTTGTCCGTTGGGCAAACTTCTTTCGCCCCAC
TGTCGAATTTTTTCGGGAGAAGCTAAACTAATTTTTACGTAGTCAAAATATTGCTCAAAC
TTTGTCATAAATTTGAATACCTTAAAATCAAATAATTAGCCAATTTATTTACATGCTTCA
TATTGAAAAATGAATATCTTAATAAAGGAATTGTTCAAAATCGTCTACTGGAGGAGTATC
ATAATTAGAACGATCTACTCGGTTATCTTTAGAATCCGACATCAAGTCAACTTCAACTGT
CCGTCTTTGGCCATCTTCAAATAGTTTCAATTTATGTACTGCAATGTCTAAACCTAATGA
CTGCAGCTCTCGCATTAAGACTTTAAAAGATTCTGGAGTTCCAGGCTTCGGTATAGGCTT
CCCTTTAACTATTGCATTTAATGCTTCATTTCTAGCTTGCATATCATCCGACTTTACAGT
TAATAATTCTTGCAAAGTATATGCTGCACCAAAAGCTTCTAAAGCCCACACTTCCATTTC
ACCTAATCTTTGACCTCCATGCTGTGCTCTACCTCCTAATGGCTGCTGCGTAACTAGAGA
GTAAGGACCAGTGGATCTTGCATGGATTTTATCATCAACAAGATGAACAAGCTTAAGCAT
ATAAGCTCTACCAACTGTAACAGGATTATCAAAAGGCTCTCCTGTTCTACCATCAAAAAC
TTGCATTTTACCTGGATGCTGATCATTAAAAAGCCATTTATTGCTAGTAATCAATGATGC
TTCTTTTAATTTTCTATTTACAAGTGCTCTTGATGCTTCTGCTCCATACATTTCATCAAA
AGGTATTATTTTAAATCTCTTTCCTAAATAGCCACCTGCTAAGCCTAGTAGGCACTCAAA
TACTTGACCAACATTCATTCTAGAAGGTACACCTAAAGGATTTAAAACAATGTCCACTGG
TGTTCCATCTGATAAATAAGGCATATCTTGTTTAGGTAAGATCCTAGAAATGATACCTTT

ATTGCCGTGACGACCAGCCATTTTATCACCCACTTGGATTTTTCTCTTCTGCGCAACATA
GACGCGAATCATAGCATTTGTTCCGGGAGGCAGCTCATCTCCTTTCTGCCTAGTAAAAAC
CCTAACCTTAACTACTCTACCTTTTGCAGCATTAGGAAGCCGTAAAGATGTATCTCTCAC
ATCTCGAGCCTTTTCTCCAAAAATAGCTCGCAACAATTTACCTTCTGGTAGTTGATCAGC
TTCACCTTTAGGAGTAATTTTTCCTACTAAAATATCTCCAGCTTCAACCCAAGAACCACC
GACAACAATTCCATTTCTATCTAAATCCTTCAGAGAGTTATCGCTGACGTTAGGAATCTC
TCTAGTAATTTCTTCTGGTCCTAATTTAGTTTGACGGCATTCTACTTCATACTTTTCGAT
ATGGATAGAAGTATACAAGTCATCATAAACTAATCTTTCGCTAATTAAAAATGCATCTTC
GTAGTTATAACCTTCCCAAGGCATATAAGCAACTAGAATATTTCTCCCTAAAGCGATTTC
CCCTCCGTCTGTGGATGCGCCATCAGCTAAAGTTTGGCCAACAACTATTTTTTCTCCCAC
CCAAACAATTGGGCGCTGATTAATACAAGTATCTTGGTTAGACCGATAATATTTTTTTAA
GCGATAATGAACTGTTCTACCACTATTATCCTGAATGCCTATTTTATTAGCAGAAACATA
ATTTACATGACCAGATGTTCTGCTAATAACAACCATACCAGAGTCTCGAGCTATTTTAGT
CTCAAGACCAGTACCAATTATTGGCTTTTCTGGATATAATAATGGAACAGCTTGCCTTTG
CATATTAGATCCCATTAGAGCTCTATTCGCATCATCATGTTCTAAAAAAGGTATTAATGA
TGTTGCAGCTGAAATTACTTGAATAGGTGAAATTGCAATATAATCTACTTGGGTTGGGGT
TGTTGTAATAAACTCTTGACGATAACGTACAGGAATAATATCTCCTTCAATGTAATGTTG
CTTACTAACTTTAACATCTCCTGGAGCTACTCGGAAATCATCTTCCTCATCTGCTGTTAG
ATAGACAGGACTATTGTGATAAATCACTTGACCTTGGTTTACCGGATAAAATGGGGTTTC
TATGAAACCAAAAACATTGACCCGCGCACAGGTTGCTAGTGAACCAATTAAACCTGCGTT
CGGACCTTCCGGAGTTTCAATTGGGCAAATTCTACCATAGTGACTAGGATGTAAATCACG
AACTGCAAAACCTGCCCTGTCTTTATTAAAACCTCCTGGACCTAAGGCACTAATTCTTCT
CTTGTGAGTTAGTTCTGCAACTGGATTAGTTTGGTCCATAAATTGAGAAAGCTGACTAGA
TCCAAAGAATTCTCTCACTGACGCAATTAATGGCTTGGGATTAATTAGATTTGATAAACT
CAATGAATCTATATCACATATCATCATTCTCTCTCTAATAATGCGTTCTAAACGATTGAG
ACCTACTCTGAATTGGTTCTGCAGCAATTCTCCAACTGAACGAACTCTTCTATTACCAAG
ATGATCTATATCATCAAGATTACCTGAGTTTTTATCCTTAATATTAATTAGATAATCAAT
TGAGGATAAGATGTCTTGAGGAGACAATACTCGGAACGTTTTAGGTATATTGAGACCTAG
TTTTTTATTAATTTTATATCTACCAACTTCACCTAAGTCATATCTTTTAGGATCAAAAAA
ACGAGAGTAAAGCATCTGCTTAGCAACAGCAACTGTTGCTGGTTCATTAGGTCTAAGCTT
AGAATAAACTATGAGCAGGGCTTCCTCATCAGTTACTTCTTCAATATTATTTTTTCCAAT
TTCTTTAGCTAATTCTTTAACTGAATATATTTGGGATGCAGAAATAAGAAAAGCATATTT
ACTAAGTCCTTTTTCAATTTCATCTTTATTGAGTCCAATAGCTCGCAGAAAAATATAAGC
ATTTACTTTGTGGGTCTTATCTATTCTTATCCAAATTTCTCCCTTAGGATCAATTTCAAA
TTTTAACCAAGAACCTCTATTAGAAATTAGGCTTGCACTATAAATCTGCTTACCATTTTT
GTCTATCTCTTGTTTATAATAGATTCCTGGACTACGGATTATTTGGTTAATAATAACTCT

TTCTGTCCCAGATACAATAAAAGTTCCTCTATTTGTCATTATTGGCAAGTCTCCAATAAA
AACTAATCTCTTTTTGTACTTTTTATTTTTTATCTGCTTTTCAGCACTGAGCTGTAGATG
GGTTGATGATAAATCTAATGATTTAATATTTTTTTCTTGGTTTTTAGAAGGTAGATCAAT
ATCTTTTCTTGTTAACTTTGCAGGAACATATATTTGAGCACTATAAGTTTTATCCCTATT
TTTTGCCTGTCTGACACTATAACGAGGAAATTTTATCTTGTACTCATTACCAAATAATTG
TAATTCGAGTCGACTAGTAGGGTCTGATATCTTCGGAAAAATTTCAAGAACTTCCGTCAA
ACCTTCTAATAAAAACCATTTAAAACTGGCCCTCTGAATTTCAACTAAATCTGGAAGAAG
TTTATTTTTTAAGCTTATACGTTGAACCATAGATCTCCTTTATAACAAATTTTGAAATCG
AAGAGTTTAATACTATTAATAGAGTACTGGTAGATAATGCATAAGCATAGTTTGCTACAT
TATTTCTTCTCTAGACATAACTAGAACATCATATTTTATTCAAAAAAAGATTTTTAAAAT
TTTTCTTCACACACACTATCTATAAGTACTTTGCTGCTAAAAGATAGTTCAAGCTATTAA
AAAAAACGAAATTAATACTTGTCTGTCTGTCAATTAATTGACATCAAGATAGACAAGTAT
TATTTATTTAATCAATTCTGTTTTTCTGAGCAAAAGACAATGCTCTCGCAAGTCTAGCTT
TTTTTCTAGCGCCTGTATTAGGGTGAAATGCCCCTTTTTTTACAGCTTTATCAATCTTAC
TATATACTTGAGAAATACTTGACTGCACATCATTGAAATTATCACTTTCTAAATTATCAA
TATTTAGAAGGCATCTTTTAGTTAAAGTTTTGACAACCGATTTATACTTACGGTTAATGA
GACGATTTCTTTCCGAAGTTTTAATGCGTTTAATCGCAGAAAGGTTCTTAGCCACAGTAA
ATCCAATAAAACTTTTATAAAATATTTTTAGAAACGAAAATACACAAATCTGGGATCTAT
TCAAAAAGGGAATATGCGGCGTATTATAACATTAAAGAAAAACAATAGGCAATAGTTCAA
CAAGAGCTCTTGTTATTACACAATAGAGATTGTCATTGCGAAAAACTTTGTAAGATAGTA
TTTAATTAAAATAAACTAACTAGACATATAAATATATATGAAAAAAATTGAAGCTATTAT
TAGACCTTTCAAGCTTAATGAAGTAAAACTTGCTCTAGTCAAGGAAGGTATTGGAGGAAT
GACGGTTATCAAAGTTTCTGGATTTGGGAGACAGAAGGGTCAAACTGAAAGATATAAAGG
ATCCGAATACTCTATTGATATCATTGATAAAATAAAAATTGAAATTATTATTAGCGATGA
TAAAGTAGAGAAAATTGTAGAGACGATTATTAAGGCTTCTAAAACAGGGGAAATTGGAGA
TGGAAAAATATTTATTAGTAGTATTGAAAGAGTAATAAGAATCAGGACTAATGACTTAAA
TTTTGAAGCTTTATAGTATTTTTACTTTTTTAGACTTGATACTTCTTCACTAAACATGAG
ATAATAAGCGTTTATTGTATACTTATAACTAAGGTGTAATACTAATGGCTAAAAGCAAAG
GTGCACGAATCGTAATAACTTTAGAGTGCTCTGATAAGGCTGGAGAGTTTGCTCAAAAAA
GGAAACCTGGCGTTTTTCGATATACAACTACTAAAAATAGACGAAATACACCAAGTAGAA
TTGAATTAAACAAGTTTTGTCCTAATTGTAATCAGCATTGTATCTTCAAAGAAATTAAAT
AGTTATTTATCTTATTAAAATAATCATATTATGGCTATATATAGAAAAAGAATATCTCCA
ATTAAGCCAACAGATGCTGTTGACTACAAAGATATTGACCTGCTAAGAAAATTTATTACA
GAACAAGGCAAAATACTGCCTAAAAGGTCAACTGGATTGACTTCGAAGCAGCAAAAAAAA
CTTACTAAAGCGATTAAACAAGCTAGAATCCTTTCTTTATTACCTTTTTTAAATAAAGAT
TAAATATATAGCCTAACTTCATATTTTTCCATCTAAATATTAAAAATTTCTATAGATTTG

TTTATTAAAATGATCTAACATACTGAATTACTAAGAAGATCTACTCATTACAAGTTTTGT
TCATTTTATAATTAATAATAGCATTATAATTTATTAAGCTAGATTC-----AGACATAAA
GGTAACCTCTTTTAATTGATTATATTATATAGTCTATTTAAAAGAGGAAATTTTTTTGCT
TCTAGCAATTCAAGATTCTTTAAGCTTTAACAAGGTATTTAAAACAAAGATTTTTGTTTA
GGTATCAGATCAAATGAATGAATCTTATCTCCCGACTGCCATAATTGAAATTCAGAGATG
AAAATTCCACATTCGTTGCCAGCTTGAATTTCTTCGACATCTTCTCTAACGCGCTTTAAA
GATTCAATTTTTCCTTGATAAATAACTTTTTCTTCTCGAATCACTTTAATCCAAGAGTTT
TTTAATAGCTTATTATTGATCACTCGACATCCTGCTATTTTTCTATTTGCTAACGAGAAT
ACTGTACTTACTTCTGCTTCTCCCACCGGTACTTCTGAATATTCAGGATCAAGTAAGTCT
TCCATCCTTCTTTTAATGTCTTCAATTAAGGCATAAATGATTTGATAGTTTTCTATTAAT
ATATTCGACTTTGCCGATGCTTGCTTTGTTCCTGGAGCAAAGTTGGTATTGAACCCAATG
AGAGTAGAATTTGTGGTTGAAGCTAGTTCAACATCTGTAGCAGTAATTTCTCCAGGCATG
ATTGAGACAACATTGAGTTGAACTTTACTCTGAGGAAATTGAGATAAAGAATCTAGGATT
GCTTCTGTAGAGCCTTGATTATCTGTTTTAATAATTAAAGAAATTTGCTTACTAATATCT
TTAGAATTAGTGTTTTTTAAAGTATCTAATGTAATTCGACTATTAAGTGCTCTTTGTTTC
TGTATAATTGATGAGTCTTTAGAGGTATTTTCAATCGCTTTAAGTTTGGCTTCTTTATCA
CTTTTAACTGCTAAAGCTATTTCTCCTGTAGCTGGTACAGATGACAATCCCCAAATTTCA
ACAACAGATGATGGTATTGCTAAATTAATTTTCTCTTTAGCGTTGTTAATTATAACTCGA
ATCTTTGCGTAAGCTGATCCAATAACCAAATTATCACTAATATTCAATGTACCATTTTGT
ATTAATAGAGTTGCTACTGGACCATGCGATTTATCTAAATGGGCTTCTATTATAATACCT
TGTGCAGGCTGTGTTGGATCAGCTTTCAAATCCTCTAATTCAGCTAACAAGGTAATAGTT
TCTAATAATTTATCAACATTTTGACCAGTAAGAGAACTAATAGGAATTATAGGCACTTGG
CCACCAAGTTTCTCCGACATGACATTGTATTTTAATAAATCTTGCTCAATAATATCTGTA
TTAGAACCAGCTTTATCTATCTTAGATATTGCTATGACAAAAGGGACATTCGCTTTTTGA
ATATGATTAATAGCCTCTATGGTTTGAGGTTTAACTCCGTCATCAGCAGCTATAATAATA
ATAGCGACATCTGTTAAATTAGCTCCCCTAGACCTCATACTTGTAAAAGCTTCATGGCCT
GGAGTATCCAAGAAAACAATTTTTTGTTTATTATCTTTCTTAATATACTCAACTTCATAA
GCTGCAATGGCTTGAGTAATTCCTCCAATTTCTTTGTTAGCATTATTAGATTTTCGAATA
TAATCTAATAGCGTTGTTTTGCCATGATCTACATGTCCCATAACTGTGACAATTGGAGGT
CTTTTAATATAATTGCCAGTTTCATAAAAAGCATTAGAGTTATCTAAATTACTAGAAAGT
CCATTATTATTTTCCTTTACATTTGATTCTACTGCTATACCAAAATTATCAGCCACTGAT
GATATTATTGAGGCGTCTATGGTTTGATTCATAGTAACTGATATTCTTTTAAGAAATAAA
TATTTTATAATATCTGTTTCTTGAACACAAATTAGTTTAGATAATTCTTGAATAGTTAGA
GGATTAGTAATACTAATAGACTCTGGCGGAGAATTAGATTGAATGCTTGCTGAATTTTGA
CTAGGAGTTACGGCTACTTTTTTCTGTCTTATAGATTTTTTATTACTAGTCAGTTTTTTT
ACAACTTCCACTTTTGGCTTAGGTGGACGCATTAAAGAGATAGCTAAATCTCCTGCAGTC

TGCGAAACATTAGAATTAGAGTCTCTAAAGTTGTCATCATCATCGTCAATATGTATTTTA
GTTTTTATTTTTTTGCGCTGTCTATTCTTATTTTTTTTACTATCTACTAAGTCGTGAATT
TTATTAAAATTTTTACTTTTTTTATCTAGTTTAGGTGGAGAACTTAATTCTAGATGCTGT
TCACCACCTGTATGTGATTCTGATTTATCAAGGTCTAAATTTAACAGATTATCATTATTG
ACTGACTCTAATCTAATTTTATAGATAATCTGAGGATTTTTCAAATCTACAATTGTTTCA
GAAGAATTGATATTTGAGCTAGATCTAGAGGATCTATTTTCAAAGTTTTGATTATTTAAA
AACATAAATATTGGTTTTATTATCTAATAGTAACTTTCTAATTTTAATACAGAGCTTAAT
ATAAACAAGAATCGAATATCTGTATAAATAATACAAATGTAGAGAATTAATCTATGTTGT
ATTAGATTTAATATAACACTAAAGTCCTTTGTTCGTTATACAAGTTTAAAACTTTTTATG
ATGATAAACACTAGATTCTAGTTTATTGGTATACAATCTATTTATCTAAATAATACTGTT
TTTGTCAGGGCTTTTAAATTATCTAATGTTATGTTCATTATGAAATTCTATTAAAATTTA
AAAACTCAACAATTAAGTAAGAATCGTTAATCACCTCCACGCCACTTATGCCACGATCTC
AAAAAAATGATAATTTTATTGACAAAACTTTTACAGTATTAGCTGATATTGTATTAAAAA
TACTTCCTACAAGTAAAGAGGAAAAAGAAGCTTTCTCTTACTATCGAGATGGAATGTCTG
CACAATCTGAAGGTGAGTATGCGGAAGCTTTAGAAAATTATTATGAGGCTTTAAAACTAG
AAGAAGATCCATATGATAGAAGCTATATTTTATATAATATAGGGCTTATTTATGCTAGTA
ATGGAGAGTATGTGAAAGCTTTAGAATACTACCATCAAGGATTAGAACTAAATTTTAAAT
TACCTCAGGCCCTTAATAATATAGCCGTGATATACCATTACCAAGGAGTTCAAGCAATTG
AGGATAAAGATACAGAGCTATCTAAATTAATGTTTGATAAGGCGGCTCAATATTGGCAGC
AGGCTATTAAGTTAGCTCCTGATAATTATATTGAAGCTCAAAATTGGCTGAAAACGACAG
GACGAATGAGGAATATACAAGGATATTAATATATTTAAGATATAATAAAGATATCAATTA
ATAAATTAAAATTAGTTAAAGAAACTGGTACTATATAGAGTACTTTACTATATATTATAA
TATTATACTAGTTGATCTCTATCTCTAAGACTTAGCTGAATAACCGAAATTAAATCTATT
GACTATATTCATGAATACAAATAAGCATAATGGTTAGTACAACAAAAAGTACCGGATCTG
TTACTCAAATTATCGGACCAGTTTTAGATATTGCATTTCCTAACGGACAGCTTCCGAAAG
TATTCAACGCACTCAAAGTACAAAGCTCAGAAGGAACTATTACTTGTGAAGTACAACAAC
TTTTGGGTGACAACAAGGTACGAGCTGTTTCTATGAGTTCCACTGAAGGACTACAAAGAG
GGGTAGAAGTTATTGATACTGGATCCCCTATATCTGTTCCTGTAGGTACAGATACTCTTG
GACGTATTTTTAATGTTTTAGGTGAACCTGTAGATAATTTGGGTCCCGTTGATTCTGAGA
GTACTTTACCTATCCATCGACCAGCACCTGCTTTTACTAAGCTAGAGACAAAACCAAATA
TTTTTGAAACAGGTATTAAAGTCGTCGATTTACTTGCTCCTTATAGAAGAGGTGGGAAAA
TTGGTTTATTCGGAGGTGCTGGAGTAGGTAAAACTGTATTAATTATGGAACTAATTAATA
ACATCGCTAAAGCTCATGGTGGAGTATCTGTATTTGGAGGCGTTGGTGAAAGAACAAGAG
AAGGAAACGACCTATATATGGAAATGAAAGAGTCTAAAGTAATTGATGCAGATAATCTGA
AAGAATCTAAAGTAGCATTAGTATATGGTCAAATGAATGAACCTCCTGGAGCACGTATGC
GCGTTGGCTTAACTGCATTGACAATGGCAGAATACTTTAGAGATATTAACAAACAAGACG

TTCTATTGTTTATTGATAATATTTTTCGGTTTGTACAAGCTGGATCAGAAGTATCAGCTC
TACTAGGCCGTATGCCATCTGCTGTGGGTTACCAACCAACTCTAGCAACTGAAATGGGAG
CACTGCAAGAAAGAATTACTTCAACAACAGAAGGATCAATTACATCTATTCAAGCTGTAT
ATGTGCCGGCTGATGATTTAACAGACCCAGCTCCGGCCACTACATTTGCACATTTAGATG
CAACAACAGTACTTTCGAGGAACTTGGCGGCAAAAGGAATTTACCCTGCAGTGGATCCAC
TGGATTCAACATCAACAATGCTACAGCCTGGAATTGTTGGAACTGAGCATTATTCTACTG
CTCAAGAGGTAAAATCAACTTTACAAAGATATAAAGAACTGCAAGATATTATTGCTATTC
TTGGTCTTGATGAACTTTCAGAGGAAGACAGACAAACTGTATCAAGAGCAAGAAAAATTG
AAAGATTTTTATCTCAACCTTTTTTCGTGGCAGAAGTGTTTACTGGATCACCTGGAAAT
ATGTATCTTTGGAAGATGCAATTAAAGGGTTTCAAATGATCTTAAAAGGCAATTTAGACG
ACTTGCCTGAGCAGGCATTTTATCTAGTAGGTGATATAGATGAAGCTATACAAAAAGCTG
ACAGCATGAAAGATTAATATAATTAACGATGACTTTAAATATAAGAATTATTGCTCCTGA
TCGGACTGTTTGGGATGCAGAAGCACAAGAAATTATTTTACCAAGTAGTACAGGGCAACT
TGGTATTTTAACAGGCCATGCACCTTTGCTTACAGCTTTAGATATTGGAGTTATGAGAGT
AAGAGTAGATAAAGAATGGATGCCGATTGTTTTGCTGGGCGGTTTTGCAGAAATAGAGAA
CAATCAATTAACTATTCTGGTTAATGGCGCAGAAGAAGCTAGTCAAATTGATTTATCAGA
AGCAGAAAAAAATTTAGACACTGCAACTCAACTCTTAAGTGATGCCTCGTCTAATAAAGA
AAAAATAGAAGCAACACAAAAAATACGAAAAGCTCGAGCTCGAGTACAAGCTGCAACAGC
AGCAACTTCGTAAAAGATTCCTCTTAAAACATAAAACTTTAGTAATTATTACTAAA----
---GTTTTATGTTTTATTTGTATGGTTATAGGCAGCTTAAATAAACTTAGCTTAGACCTG
AACAAATATAATCAAAATATAATCCCATCTCTTTACCTGCATCTGGTCCAACTAAACTAA
TAGTAACTTCTTTCATAGCTAAAATAGCTTGAATCGTTGCACCAATAGGAACACCTAGAG
AATTGTATGTTTCTTTTAGACCATTTAATACTCTTTCTTCAAGGATAGATGGATCTCCAG
CTAGCATTCCATAAGTAGCATAGCGTAAGTAGTAGTCTAAATCACGGATGCATGCAGCAT
AACGTCGAGTTGTATACATATTACCACCTGGGCGTGTAATATCCGAATAAAGTAGAGATT
TTGCTACGGATTCTTTAATAATTGTTGCTGCATTAGCTGCAATAGTAGCAGCAGCTCTGA
CTCGTAGCTCACCTGTTTGAAAATAGCCTCTTAATTTTTCAACAGAGCTATCATCTAAAT
ATTTACCTTGGACATCAGCTGCATTAATAACAGAAGTAATTGCGTCTTGCATAACTTTCA
CACATTCCTTAATTTTAAATATCTAATACTAGGGGTATATAACAAAATTACTAGATCTTT
TAGCGATATCAGTATTACTGCATAGCACCCAAAGTATAATCAAAGTAAAAACCAGCTTCT
GCTGAATCTTCACCTGCAAGCAGTGAGCAAGCTACACTTTTCATGCATTTTACACCTTCA
GCAACTCCTGAAATTGGTGTTCCTAAAGAATTATACATTTCTTTAACACCCACTAATCCA
ATTTCTTCAATAGGAGTTACATCGCCAGCAACTATTCCGTAAGTTACTAAGCGAAGATAA
TAGTCCAGATCTCGTAGACAAGTAGCAGTCATCTCTTCGCCATAAGCATTTCCACCAGGA
GAAACTACATCAGGTCTTTTTTGAAACAGTTGTTGGCCGCCTTGCTTTACAATACGCTCA
CGATTATCTGTTAAAATTTGAGCTATTCTTAAACGACGTTGTCCAGATAAAACAAAACTT

TTAATTCTATCTAATTCCCCAGGACTTAAATATCTTGCTTCTGCATCTGCATTTACAATT
GACTTTGTAACAATACTCATGGATAAAACTCCTGTAATGCTTT--AAAAAAAGCTTAATA
CTAATTATTTATGATATAACTCAGAAAGTAATGTACTTTTTGAGAATTTTCTAACCGAAA
CACGTTCATGATATATGAAGCAGGATAGATACTGAGGTAAGTACTAAAACGAGAAAGTTA
TATGTAGGAACTTCTGTTATTGATTTCCTAGTACTTTTTTAAAACTTGGTACCACAATTG
ATATATCCTGCTTAGTCAAACGATTATAGAGTGTTTCTGTATTAGGAAAGTTAGCTGCTG
GTAAAGTCGGAAATCTTCTGTATGGAACAGTATCTTTGCCGTATACCGCATTGTATTCAA
CACTTTCAACTAAATTACTTATAAATGCAGATAAGCCTTTTGAAGCTAAAATTTGATTAA
AGAATCTAATTTCTGCCTGATTATTAGGAGCTCTTCCTAAGATATGCTTAGTACCTAATT
CTATAACTTTTGTATTTGGATATGGCTGATAAAATTCTTTACCATATAGTTCAGATAGCG
CCAGTTTTTCTACTAGTTCTTTGACACAGATTTGTCTATTCAAGAAAGCTGACTTAATAT
CCAAGAATTCACCGCCGACACTAAAAGAGTTAAGATCTCTTTCGAAAATTTGACGATATG
CTGCTCTTAAAGCTTGTTCTAACATTTCTTTGTTACTATCACTATTCACTTCAAAGACAA
TAGATTGATCTCTTAAAGAAGTTACTCCTTGTGAAATACGAGACTGAATATCATTACTAC
TCCTCATTTCTTTGACAGTGCCTAATTCAACAAACCTAGCAGATTTACTAGCAATAACTT
TTTTGAATCTCTGGTCAATAATTCCTGGTCTTAAGCTTCTTAAAGCAACACCAGCTGGAG
TACTATATCTTTCATAAGGAACAGTATTATCTCCAAAAGTTTCTGTGTACTCTGAGCTAT
CAATTATTGTGTCTATTACTTGGTAATAGCCTTGCTTATATGCAATATCAAAATATTTAT
TAATTTCTTGCCGACCATATGTAGGGCGACCCAGTAAGCGGTTATGAATATATTCAATAG
CTTTACAAATATATAATGGCTCCCAATAAAGAGATCTAAATATACTTGATTTTGCTAACT
GTCTAACAAACTCGCGAACTGTAATTTGATTATCTTTTAGCTGACTTTCTATTGGCTTGA
GAATTAGTTTTTCTTCTTGATACACTTCTCTGCCAAAAACTCTTAAATAAGCAACTTTTG
TGACAACTTCTACTGAATTTTCAAAATTTTGGGATGAATCTGACTTTATAAGAGCGTTGG
ACAATTTAAAGATTTTTGGTCCCAGGGATCCTGCTGACTTAGGTCTTACCTGTGGATTAC
TAATCTGATTGTAGATTCCTGGCCCACGTCTAACTAATATCCTTCTTGTATCTTTACCAA
AAAGAGCTTGTCTTTTTCTTGGATCTTTATTTTCTTTAGGGAAAATAGCTCCAAATTGAA
TTGATAATGGATCATTACCTATTCCATATGGATGTTGATCAGGTAATGATTGCTTATAAT
CACTAAATAAAGTTATAAATTGAGGAACCTTACGAAAAGGCGCACTATAGGTGAGAAGAT
CTATTTGAGGCCCCCAATTACGGCACTCTTGAGGCTCTTCACCTAAATTTCTAAAGTAAG
GCACAGTCTCTTCACCGAAATAATCTGTATACTCAGAAGAATTTAAAAGTGCATTCACTA
AACCACTTAAACCAGTAGATGATAAAATAGCAAAATATTTTTGAAACTCTTCTAAAGAGC
TAGGACCTCTACCCAAGAAATGTCTAAATGCTAATTCTAATGCTCGACTATTGACAAAAG
GTTCATAAAATTGCTTTCTATAAATACTTGAAGTTCCTAACGAACGAATAAATTCCTTAA
TTGAAATTTGGCCATTTTTTACTTGCGACTCCAAATTCGATAAAGAGAGGTCATATGCTT
TAGCAATATCTCTTTCAAAAATTTGTCTATAACATGCTTTCACAACAATATTTTTCTCAT
CAGCAGATAAACTAGGCTTCATAACAAAACGAGGTGTTGACACTCCTGCCTGTACATAAG

TCTGAGGAAGACGCAAACCTTGTAAATCTCCAGATATTCTTTTTCTTAATTTATCAGTCA
AAGATGGAGCTTCAAATTCAGAAATCACAACATTAAAATACTCTTTAACTAAATCTTGAC
CTTTTATATCTTCCTCAAAGATTAATAATGCTGTCCGTCTCATCTCTCTAAGAGCCACAA
TTGCTGCTGCACTTGAGCAAGCGTTATCAATTAACTCTCTCAATCCTCGTATATTAACAG
ATAAAATATTAGGGTCGCCTGATACAATTGCATATGTCAAATACCGTAAAAACCAGTCTA
AATCTCGAAGAGATTTTCTCATTCTTGTAGTACCGTATCTCAAGACGTTTATAGGTTTAA
ACCCTGGCGGAGTAGCTCCCCCAGCATTAAATAAAGATCGGAAACTTTGGCCAAAATCTC
CTTGGATATTTCCTGATAACTCATTAATTTTATCTTGAGAACTTTGATCTCCTGCAATAA
TAACAGCTGCTTGCGGACGTTCTAAATAAGAAATTGCAGACCCACCTACGAATATTTTAT
CTGCTGCTCGTGCTACCAGAATATTTGCATTTTTAGTTAAAATATCTGCAACTTCCAGTC
TTTTTTGTCCAGAATTTAAAAAAGAAACTAATTGATTTAATTCACCAAGCTGTAAAAATC
GGTCTTGTTGCTCTGCCTGAGTAATAGTTAAAATTGAGGCAGTCCGATAAAGCTGGGGGC
GTGCTAATGGGCTTCCGCCACTTGCTTTGATACTCATTACTTTATTTATCTCCTTAACTG
ATATTGCTTGCTAGCGAATACAAATATAGCTTGATATTTCTAAATATTTAAAGATACTAT
GCAAGCACACTGTTTTCTATATTATTAGGTACTAGTACTTATAGTACAGAGAGTCTTATA
ACTAACTTCTATAAATACAAATTTTGTAATACTTCTATTAACATTTGTCAAAAAAATATT
TAAGTGCTCACTCTTTTATATTAAAATTCAATTAGTGAGAGGTTGATGTGAAATAAAGTA
TACTAAGTAAGTTTTATAGATAGGAAAATTAAAAGATTTATTCTACAAACATTAATCACT
TGAATGTTTAGCAATTGTACAATTGTTATTGCTTTTAATTGGTTTAGACTAGTTTTATAT
TTTTTCTAACATAAACAATATTAATGAGCAATTTGTATGAATTTAAAGCCACAAAACAAC
CTATTAACTAACGAAAATATAGATCATGTAGATATACCCGAAAATGATATACCAATGTCA
ATTACGGAACATTTAGAAGAATTAAGACAACGGACTTTATTTGTGTTTTTATTCTTTTTG
TTTGCTACAACTATAAGTTTCACACAAATTAAGATCATTGTTGCAATACTGCAAGCTCCT
GCTGTTGGTATTAAGTTTTTACAACTGGCACCAGGAGAATATTTTTTTTCATCTATTAAG
GTTGCAATATATTGCGGGATTGTTGCAACAACTCCGTTTGCCGTTTATCAAGTTATATTA
TACATACTCCCAGGACTAACTGGAAAAGAAAGAAAAATTATTTTGCCTCTATTAATTAGT
TCTGTATTGCTTTTTATTACAGGTGGTATATTTGCTTACTTTGTTCTCGCACCAGCAGCT
TTAACATTTTTAATTAGCTATGGGTCTGATATTGTAGAACCATTATGGTCTTTCGAACAA
TACTTTGACTTTATTCTATTACTTTTACTTAGTACAGGATTAGCATTTGAGATACCAATC
ATACAATTATTGCTCGGTGTTTCAGGGACATTTTCTTCTAGTCAAATGATACGAGCTTGG
AGATATATTATTATTATAGCAACAATTGCTGGAGCTATTCTGACCCCTTCGACTGATCCC
GTTACACAATTAATAATGTCTTCAGCTGTTTTATTACTTTATTTTGGTGGAATTGTTATA
TTATTAGTCTTAAAAAAGTAGAGTAAATATTTTCTATTAATCCAATATATTTGTTATCTG
GGCAAATAATCTAT-AAAACAATTGCTAGATTTAAGTATCAGGCTTCTAAATATTTCAAG
TCTCTTTTTATCATTGAAATACAATCGATACATTGGCTTGTTTTTTCATCAATTATTTGT
GTTGCACACTTTTTAGTACTCTCTTTATTGTTCTTAGAATTATTTCTTTAATCTATTTAT

TAAATAGATATGAAGAGATTTAAGTATTGATTATGGTAATCTATATCAACGTAAAAAGTA
TTTTATTGTGAGTTTTATTTATATAAAGGCTTATTCATTATATAGTTATAACAGTAAGTC
TAATTAATAGTTTTCATGCTTAATTAAAGTAATAACTCTTAATTCTTTTGGCAGAAAAAT
CTGTCTTAATATTTTATCAAAAAAACAGAAAAATGAAGCTAAATAGTCTAATTAACTTAA
TTCAAAAGTCTATATATTCTTGTACACTTTTACTAATTATTTTAAATATTATTTGTGTCG
CACCTAATTCTAGTAATGCATTTCCAATTTATGCGCAACAAGCTTATGAAAGTCCAAGAG
AAGCAACTGGTAGGATAGTATGTGCCAATTGTCATCTTGCTCAAAAGCCTGTGGAGATAG
AAGCGCCTCAAGCAGTACTACCTAATACTGTTTTCGAGACTGTTGTGAAGATTCCATATG
ACAGCAATGCTAAACAGATTTTAGGTAATGGCAGTAAAGGAGGCTTAAATGTTGGAGCCG
TTGTAATATTACCTGAAGGATTTAAGTTAGCTCCTGTTAATAGATTATCTACAGAGTTAA
AAGAAAAGACTAGAAATCTTTACATTCAGCCGTACAGTGCTAAACAAGACAACATTTTAG
TAATTGGACCTATTTCTGGTGATAAAAATAGAGAAATAGTTTTCCCAATACTATCTCCCG
ATCCTGCAAAAGATAAAAAAGCTCATTTTTTCAAGTATCCAATATATGTTGGCGGAAATA
GAGGACGAGGCCAGATTTATCCAACGGGTGACAAAAGCAACAATAATATTGTCTCTGCTT
TAAGCAGTGGTAAAATTAATAAAATTGAATTACTAGACAAAGGTGGATTTATAATACATG
TGACTAACAGTAGCAATGTGGAGTCAACACAGAAGATTTCACCTGGTCTCGAACTTAGAG
TAAAAGAAGGGGATACAATTCAGCTTGATCAAGCTTTGAATAGTGATCCAAATGTAGGTG
GTTTTGGTCAGAACGAAACAGAAATAGTTTTACAGAGTCCAAATAGAATTAAGGGCATGA
TTGTTTTCTTTTTCGCTAGTGTTCTAGCTCAAATTTTCTTCGTATTAAAGAAAAAACAAT
TTGAAAAAGTTCAAGCAGCTGAAATGAATTTTTAAAGCAATATAAAACTGGCACTTACTA
TAAAGAAAGTATAAACACTTTATTCACAGGATGTACCAGTTTTTATGTTATGTAATAATC
GCTTTGACGGAAGTTATGATACTTTTAACAGAATCTACAATCTGACTAGGCTGAATTACT
GTAGCCTGCTCTAAGCTTCCATTATATGGTGTGGGTATATCCTGCGAAGATAGCCTCACG
ACAGGAGCATCTAATTCATCAAAAAGATACTCATTAATTTGTGCAATTAGCTCTGCTCCA
ATTCCGGCTGTTTTCATACACTCTTCTACAATTAAAACTTTGTGAGTTTTCTTTACTGAG
ATAGATATAGAGTCTATATCCAGAGGTTTTAAAGATATAAGATCTATAACTTCTGGATCA
TAACCTTCTTTTAATAAGGCCGGTAATGCTTGGATAACATGATGCCTCATTCTAGAATAG
GTTAAAATCGTAATATCTTTTCCTTTTCGTACAAATTCAACTTTATTGAGAGGTAAGAAA
TATTCTTCTTGAGGAATCTCTTCTTGTAAATTATAAAGTAGAACGTGCTCAAAGAATACA
ACTGGATTATTGTCTCGAATTGCAGATTTCAGTAATCCTTTTGCATTATAAGGAGTAGAA
CAAGCAACTATTTTTAAGCCGGGGATGGCTTGGAAATAAGCTTCCAGTCTCTGAGAATGC
TCTGCACCTAACTGCCTACCAACTCCTCCCGGTCCTCTAATAACTAAAGGCAATGTAAAA
TTACCTCCAGAAGTATAACGTAACATTCCTGCATTATTAGAAATTTGATTGAATGCTAAT
AATAAAAAACTCATGTTCATACCTTCAACAATTGGTCTTAGCCCTGTTATAGCTGCACCA
ATTGCCATACCAGTAAAGCTATTTTCTGCTATTGGTGTATCAAGAACTCTCAAATCCCCA
TATTTGCTATGCAAATCTTTAGTTACCTTATAAGATCCACCATAGTGACCAACATCTTCT

CCTATTACGCAGACAGTTGGATCTTTTGCCATTTCCTCATCTGTTGCCGCTCGTAAAGCG
TCAAACATAAAGATTTTACTCATTTTAATATTTGATTTTTATGATTTGACCACTGTTTTA
AAGTAACAGTTAAGACACCATATTCTATTTAGTTATCTGCAAAAAGATATCGTTTCAATT
CTGACATGTTAGGTTCTGGACTGGAAATAGCAAATTTCACAGCTTGTTCCAATTCGGTTT
TAACAGCATTCTGAATCTCATTAAGTTCACCTATATTAGCAATTTCATTATCTAGAATAT
ATTTTTTGAGTTTTTTGATAGGATCTCTTGCCACCCAAGCCTCTTTTTCTTGTCTTGATC
TTAGTTCATCAGGATCTGCGAGAGAGTGACCACGAAATCTATATGTTAATGCTTCTATTA
AGGTTGGACCATCACCTTGGCGAGCTCTTTGAACTGCTTGTTTTGCAGCTTGTCTTACAG
CTAGCACATCCATTCCATCAACTTCAATCCCAGGAAGCCCAAAAGCTTCTGCTTTTTTAT
GTATTTCAGGTATTGAAGAAGACCGGTGATGTGCCATACCTATAGCCCATTGATTATTTT
CAACAACAAATATAATAGGTAGTTTCCAGAGAACTGCCATATTCAGACATTCAAAAAATT
GCCCATTATTGGTAGTCCCATCACCAAAAAAGCAAGCCGTGACTCTTAAATCCTCTGTTT
CTTTAAGTACTTGCTGGCGGTAGATACTTTGAAAGGCTGCCCCTGTTGCAACCGGTATAC
CTTCCGCAATAAAAGCAAAGCCACCTAAAAAATTGTGAGGCGCAGAAAAAATATGCATCG
AACCTCCTCTGCCTTTACTACAACCAGTCTCTTTTCCAAATAACTCAGCCATCACATTTT
TAGATGGGACGCCTTTACTTAAAGCATGTACATGGTCTCGATAGGTACTGCAAACATAAT
CAGTTGGATTCAGAAGTTTAATTACACCTGTAGAAACAGCTTCTTGACCATTATAAAGAT
GAACAAAACCGAACATCTTTCCTTTATAATACATCTGAGCACACATATCTTCAAAATTTC
TGCCTAACAACATGTCTTCATATAAAACTAATAAATTACTCTTATTAAGATTAAGACCTG
TTGAATTATAGTTAGTCAGCGGCAATTGAACTTTCTTAGGATAACTCATAATTTGTAAAG
AAACCTCTTATTGTAAATAGGAATTTTGAAGAAACAGCTATCAATAGAAAACACACAGAT
AAAAACTTATAAATTATAATATGTTTTTTACTAAAGACTTTAATAATTATTTGAAACTAG
TTCAGTATATCAAATTAATTTTGCTTATACTACGGATGTCTGAAACTTATATAAAAAAGT
AGTAAAAAAGTTTATTAAAAGAATAGTTAAAAACAGATATAAATAGAAGTAAACAGAATT
ATTGAAAAAAAATGATTATTATTAATTAATACTTAATATTTCTAAATATTTTACAATTAA
TGGCTATGTCACCATCTTTATTTTATCCTGTTGAACAAGAACTATGTAGTCTTGAAAAAA
ATCTGAAAGCTGTTGCTGGGACTCGTCATCCAATTTTATATGCCGCAGCAAAGCATCTAT
TCGATGCTGGAGGAAAACGAGTTAGACCAGCTCTTGTATTTTTAGTGGCTAAAGCAACCT
CTGAGAAGCAAGATATAAATACTGGACAAAAAAGGCTAGCAGAAATTACTGAAATTATAC
ATACTGCTAGTTTGGTACATGATGATATTATTGACGAGTGCACAACACGTAGAGGAGTCA
AAACTGTACATAATTTATTCAATACTAAGATTGCTGTGCTAGCAGGAGATTTTTTATTTG
CACAGTCTTCTTGGTATTTAGCTAATATTGAAAATTTAGCCGTAGTCAAAGCTATTTCTA
AAGTCATCACCGACTTTGCAGAAGGAGAAATTAGGCAAGGCTTAGTTCATTTCGATCCCA
GTATTTCAATAGATGCTTACATTGAGAAATCATTTTACAAGACTGCTTCACTCATTGCTG
CTAGTTGTCGGGGTGCAGCTATGCTTAATGGTTCCAATCATCAAATAAATAATGATCTTT
ATCTTTACGGTAAACATATGGGATTAGCATTTCAAATTATGGACGATGTTCTAGATATAA

CTGGTTCTACTAAGAGCTTAGGGAAACCTGCTGGCGCTGATCTAATAAATGGAAATTTGA
CCTCTCCTCTCCTTTTTTCACTTACTCAAGAAGCAAGTTTAAATGATCTTATTGATAGGG
AGTTCTGTAATAGTACAGATATAGCCTCAACATTATTTCTTATAAAAAGAAGCGGGGGAA
TTACAAAAGCTAAAGATTTAGCTAAAGAACAGGTGCAGGCGGCACTTTCCTGCCTTCAGT
TTTTACCACAATCTACACCTGTATCTAGTTTAAAAGAATTAACACATTTCATAATCACAA
GATTGTCATAAAGGACTTGCTAAAATTAAAATGTTATTAACTAATTTTTTACTGTTTCTA
AAACAGTCTGAAAAGTTTGAATATCATTACTAGCTAGCTGCGCTAGCATTTTACGATTTA
ACGCAATATTCTCTTTTTTTAATGCACTAATAAAAGTACTATAATTCATGCCTTGATTGT
GAGCAGCAGCATTTATTCTAGTTATCCAAAGACGACGAAAATCTCTCTTTTTCCTTTTTC
GACCAACATAAGAATATCGAAGAGCTTTAAGGACTTGCTGCTTAGCTGTTCGAAATAAAC
ATTTATGTGCACCTTTAAAGCCTTTAGCTAGCTTAAAGATTTTAGCTCGTCTTTTTTTTG
CAACGTTACCTCTTTTAACTCTACTCATAAAATATTACTTAAATTTTAATTATTACCAGC
TTACTAAATTCTTATACTCTTTGATTATAAATAAGGTAATTTCATTGCAATGTTTTTAAT
ATCTTTTAAGTCAACCGAACAAGTAGAAGAAAGATGTCTTCTTTGTTTTGATGACTTTTT
TTGTAATAAATGACTTTTAGAAGCTTTATGTCGAAGAAATTTTCCGGATGAAGAGACTTT
AAATCTTTTTGCTATTGCTTTTGATGTTTTTAACTTAGGCATATTATAAACTTTTAATGA
TTAACAAAAGCATGAAAAGTCTGAGGATATCAATCTTAGACCTTTCAATACTTAGAATAA
TCTTAACACAATTTATTTTACTTACTACATAGGTTAACTGAAACAATTTATATATATAAT
ACTAATCAGTTAATCAACCATAGCTAAAGTGGATTTATATTTTGCTTTGAACTTTTGTAT
AGTATTAGCAATTAGCATTGTTATAATTTTAAGACTGCTTGAATTCAATTCTTCGAACAT
TTTCATATTTAATGTAAATGAAATATTGGCTTCTGAGACAATATTTTGAATTTGAACATC
AGACAAGGGAATCATATCTAATGCAGCTCTATATTGATCTTTAAATAATTTATCGTCTTT
TATTTGGTCAAAATCGTAAAATTTCGTTCCTCCTGAATCTGATAAATTCATAGCCCCTCT
AGCTATTTTTTTTAAAATTTGGCCGCCAGAAAGGTCACCCAAATAGCGCGTATAAGCATG
AGCAACTAATAATTCAGGTTGTTTATGTCCTATAGTGTGAATTCTATCAACATAAATTTT
TGTAGCAGGAGATGGTTCAATAAAATCTAACCAATCTGATCCATAATAATAGTTTAAATC
TTCAGATAGACTCGCTTTCCTATTAAGCTCTGTAAAATATATAGGTTTAATAGCTGGATG
ATTTTTATTAGAAAATAACTCTTCCTCTATTGCACAATAGACAAAGTATAGATTTGCAAC
CAATTTACGATATGACTTCTTATCTACAACTCCTCCCAAAAAAGATTTAACAAAACTAAC
ATTTTCTGCCATACTATGGGACTTAGTAGTACCTTCTCTTAGTTCATTCGCTAAAGTATT
AACCATAATATCAATATTCCATTTTTAAAGATAACAAATTAACAATAAATAGTTATCCGT
ATATTTATAATTGAGTTGCGCTAACTAGGCTGCTGCAAAATAATTTTTTGATTTTATTGG
ATCTGGATTCATTGTTTTGTCACCAGGCTTCCAATTAGCTGGACAAACTTCATCTGGATG
AGATTGTACATATTGAATTGCTTGTAAAACCCTCAAAGTTTCTTCTACACTCCTGCCAAA
CTCTAGATTATTGATTGTAGAGTACTGAATAATTCCTTTCGGGTCTATAATAAATAGTCC
TCTTAGAGCTACACCATCACTATTCAAAACATTATAAGCTGCACTGATCTCTTTTTTTAA

GTCTGATACTAATGGATATGAAAGATCTCCTAATCCACCTGATTCTCTGTCAGTCTGTAA
CCAAGCAAGATGAGAATACTCACTATCAACCGAGACTCCAAGAACTTCTGTATTAAGTTC
CGAAAAAGCATTATACTTATCGCTAAATGCTGTAATTTCTGTAGGGCACACAAATGTAAA
ATCTAAAGGATAAAAGAATAAGACAATATACTTATTCTTGAGATCAGATAATTTTAACGT
TTTAAATTCTTGGTCATAAACAGCTGTAGCTGAAAAATCAGGAGCTAGCTGACCTACTCG
AAGACAATTTGGTCCAGAAATCATTAATTTTTCCCAGAAGAATTGATATAATTTTGTTAT
TTAATAAATAATTATATAACAAAATTTAGTATATATTAAAATACTACAAGTCTTTTTTAA
ATTACTAAATGATATAATCAATACTATCTTTATAGCGGGGAATGGATTTGAACCATTGAC
CTTCGGGTTATGAGCCCGACGAGCTACCAGACTGCTCTACCCCGCGGATAACTAAAATCA
TATTAGTCTACATATATATATAAAAGCAAATGCATAATAGATTTTAGAAGTAAAAAAATA
GAAGAACTGGTCCACTGATCATTAAACTTAGAAACGACAAAGTATTTTTTATCTTATTTA
AAAGAGGGAAGACTTCATATAAACCAATAAATCTATCTTTTAAAAGAATTTCGGACGGCT
TTACCCAAATTTGACCGTCATACCAGCCTGATTCTTCATAAAATACTGTAGCACTCATTA
ATCGTTTTACTACGTATGACCAGCCTAAATATAATCTGATTAAAATAAATCCTGTCATTA
GGCTGGAAGTAATAAATTCTGAAAAGAAAAATTTCAAAGGTAGTTTAGTGATTGGAAAAA
TTGATAATAATATAGGACTAACTAATAAACAGTTAAGTAGTAGTGCAATAGTTATTTTTT
TATTATAAGATCTACGGCTTAAAGTTGGCCAGCAAAAGAACCAAGAATTTTTCAAAGAAG
TATATTCATGAACAGGTTGCTGCTCCTTTGGCACAGGACATTGAGTATTATATAAATTCA
TTGACATTAAAAAGTATTATAATAAAGACTTTGTTTATATTATCTAACATAGGTACGACA
TTAAATATTAACTAACTGCAATTATAGTAGTGAGTAGAAGAAATAAAACTAAAAATAGCC
AAGTAATTTTATTTAAAGTATTCTCAGTACTACGAGTATTACTAAAGAACTGATTCTGAG
CACCTACACTTCCTAAACCTTCGGATTTAGGATTGTGTATTAGAATAGTGAAGATTAATA
TTATTGTAGATGAGTACCAAAAAAATTTTAAAATTTGTTCCATTATGTAAAAGTTAATAA
AAATAGCACAAAGACAATCGTCTAAAAGATATTGATTGTCTTATCATATTAAAAAGTATT
TATTCACCCTGTACTTTTAAAAGTAAAAATCCTATTGCCAGCCCTACCAAAACTAGGACT
GAACTTAATACAGCACTTTCTACAATTTCTCCGACCATTTTATTTTTCCTTATTATTTTA
TTATTGTCATTCTTTTGACAAGATTATATTGAGCTAACTAAAATCCGTTTCTACCCCAAA
CTACCATTGCGATTGAAAATGTAAACATTGCCATTAGAGCTGCCCATCCCAAACTTAAAA
TATCCATAATTTATAATATTCCTAAAAATAAGATCTCTTTTACTTAAACTATAATATTCT
TAATATTAAAAATATCATTATATAATAAGCTAATTATAACAGAGTACTACTTAATTCGTA
TTTTTAGTGTATTTAAAGAAGATTGAATTTTTTCGAGGTACTGGATAAGTATTTCATCCT
AAGAACCATAGTCAACTTTTGATTTATTATAATGCTTAAATCTAAATATATATCACTCAA
GAATCATTAGTAATTATGATGTTTTTAATTATACGATTAACATATTTCAAAATAGAAGAA
TTCATAGCGACTAAGTTAATATGCTAATATAATTTATCAATAAAAAGGAAGTTAGTATTT
CAATTATGCAAACAACTATAAATAATGGGCAAACGTCTAGTAAAGAAACTTTACTAACAC
CTAGATTCTATACGACTGACTTTGAAGAGATGGCTAATATGGATATTTCCGGCAATCAAG

AAGATTTTTTGGCTATCCTCGAAGAATTTCGAGCTGACTATAATAGTGAACATTTTATTA
GAGACGAAGAGTTTAATCAATCTTGGTCTAATTTAGAACATAAAACTAAATCCTTATTTA
TTGAGTTTTTAGAAAGATCTTGTACCGCAGAATTTTCAGGTTTTTTACTATATAAGGAAT
TATCCAGAAAATTAAAAGACAGAAATCCGGTTATAGCTGAGTGCTTTTTATTAATGTCTA
GAGATGAAGCTAGGCATGCCGGTTTTTTAAATAAAGCTATCGGGGACTTCAATTTATCTT
TAGATTTAGGATTTTTAACGAAGAGTCGCAAGTATACTTTTTTCTCACCTAAATTTATTT
TTTATGCAACCTATCTTTCTGAAAAAATTGGATACTGGAGATACATAACTATTTACCGCC
ATCTTGAACAACATCCAGAACACCGTATTTATCCAATCTTTAGATTTTTTGAAAATTGGT
GTCAGGATGAAAATCGTCACGGAGATTTTTTTGCTGCTTTGCTCAAATCCCAACCTCATT
TTTTAAATGACTGGAAAGCAAAAATGTGGTGCAGATTTTTTCTATTAAGTGTATTTGCAA
CAATGTACTTAAACGACTTTCAAAGAATTGATTTTTATAATGCCATAGGCCTAGACTCTA
GACAGTATGATATGCAAGTAATACGAAAAACTAACGAAAGTGCCGCTAGAGTTTTCCCGG
TTGCTTTAGACGTGGACAATCCAAAATTTTTCAAATATTTAGATACTTGTGCATGTGATA
ATAGAGCTCTGATCGATATTGATAATAACAATTCTCCATTATTCATTAAATCTATCGTAA
AGATACCTCTATATTTTTCCTTATTTGCAAATTTACTAAAGATATATTTGATTAAGCCAA
TAGACTCAAAAACAGTATGGAATACAGTTCGATAGTTAAATACAAGGGAATGCTTTTTTC
TTATAAAAAGCTATCTAGTGATTAATATAAAATAAAAGCTTCCTTCTCGATGGAAGGAGC
TTTTATTTTGTCAAGAATATAATGGGATTATATAGATCTAGAACTAAGAACTTACTTTTT
CTTTTGCTTCATACATAACCTCTAAAGTAATTAGATTTTTAGAACAATCACGGGCAAATT
TCTCTGTATTTCTTTTTACTTTACCTCTAACAAACCCAGGAATTTTATTTAATTCTTTCT
GTGCTTCTTCTGACCAATTAATACTATCAACAGCAGAAATTCCTATACTTAATGCTTCTG
TTGTATCATGACCACCAAATATTTCTAGCAAATGATCTTCCATACCAAGTGTAAACGAAT
TATATACTAAATCAGCAATTTGATTTGTTCCCTCATAACCAAGAAATGGACGGTAACTTA
AAGGAAAGTTTTGTATATGCACTGGTGAAGAAATAACGCCACAAGGAATATTTAGACGCT
TACCAATATGTCTTTCCATCTGTGTTCCAAAAATTGCTGCAGGTTCAGTTTTAGCAATTA
AATCTCCAATTAAACCATGATCATCTGAAACGATTACTTCATCGCAGAATTCTTGAACTT
GATCTTTAAACCACTCTTCATCATATTTGCAATAAGTTCCACACCAGGCGACATGTATAC
CCATTTCCTGGTGTAAAATTCGAGTTATTGCTGCAGCATGAGTTGCATCTCCAAATACAA
TAGCCTTTTTACCTGTTAAATTCTGACAATCAATTGATCTTGAAAACCATGCAGATTGTG
AAATAAATCGGGTTTGCTCGTCTATGTATTTTTCGTAGTCTACAGCAGCTCCCAATGCAT
TTACTAATTGCTGTATTGATCGGATGCATGCAGCTGTTTGAACAATACCCATAGGTGTAA
TGTCAACATAAGGCATATTAAATTCTTTCTCTAAATATCGAGCTGTCATTAGTCCAGTTT
CTCTATATGGAATAAAATTAAACCATGCAGAGGGTAATTTTTTTAAATCTTGAACTGATG
CATTTTCTGGAATAATTTGGTTTATTTGGATATCTAGATCTTGAAATAACCTTTTTAATT
CTGCTATATCATGTTGATTATGAAATCCTAAGCTTACTGCTCCAATAATATTAACAGATG
GAGTTTTAGTCTTTTGAGTTAAAACTTGATTATTAGATTTAGCTTTTTCCATATAGAAAG

TGACAATTTGTTCTAATGTTCTATCACCTGCTTGCAGTTCATTTACTCTATAATGATTAA
CATCAGCTAATAAAACATCTGCTTCTGTTTCTATAGATGCTCTGCTAACAAAGTTTTGCA
AATCTTCTTGCAAAATACTCGAAGTACAAGTTGGTGTTAGAATAACTAAATCAGGGCTTT
CTTCTCTGTCTTTTCTAGTAATATTTTCTACAACTTTTTCTTGAGAACCACGGGCTAACA
CATGTCTATCTACAACACTTGCTGTAACAGGAGTAAAATCTCTGTCTCTTTCAAGCATTG
AACGCATTACATTAAAATAATCGTCTCCGAGAGGCGCATGCATAATTGCATGAACTTTTT
TAAAAGAACTGGCAATTCTTAAAGTTCCAATATGAGCAGGGCCTGCATACATCCAATAAG
CTAATTTCATATTAAATTTTCCTATGATTTATTTAGACATAGTAATACTTTTAAAGCTAA
CTAACAAACATTAGCCAAACATCTAATTTAACTCCTTGACAATTTTGTAATATTTTTTAA
ATGAAGCTTATCATTATAAATATTTAAAAAATATAATTCAGAATAGAAATAACTGTTAGA
TCAAGCTTGAAAGTAAAGATTTCTTGCAGTTATAGTTTTTCTTTAAGTAAACATGTAAAA
ATTGTACTATATATAAATGTACTATATATAAACTTTTTTTACTGAATATTGGTAATGCCA
GATACAATTAACTTAAATATGCCTTCCCCAACGTTTGGTGGAAGCACTGGCGGTTGGTTA
AGAGCCGCAGAAGTAGAAGAAAAGTATGCGATAACATGGACAGGCAAAAATGAAAGTAAG
TTTGAAATGCCAACTGGTGGTACCGCAACTATGCGAAATGGAGAAAACTTACTCTATTTA
GCTAAGAAAGAACAATGTTTAGCTTTAGGTACTCAACTAAAAAGTAAATTTAAAATATCT
GACTATAAGATTTACAGAGTTTTTCCTAACGGAGAAGTGCAATATTTGCATCCTAAAGAT
GGAGTCTTTCCAGAAAAAGTTAATACTGGGAGAGCTAGCGTCAACAGTGTTGATCATTCT
ATTGGACAAAATATTAATCCTGTAGATGTCAAATTCATGAATAAAGCAACTTACGATTAG
TCTCTCAAAAACCTAGACATAAAATAACTTTTATATAGTCTAGGTTTTTGATAATGTATA
CTGATAGTATGGAAGGGTGGCCGAGTGGTTGAAGGCGTCTGATTTGAAATCAGTTGAACT
ATCTGGTTCCGTGGGTTCGAATCCCACCCTTTCCGTTATAAAACTATAGTTGGCTTCTCA
AGGCAAGCTAGCTATACACTGATAGTATTAAGTTTAGACTTAATAAAGTCTACAACTTGT
GACAGGTTAGAAATTTTTTCGGCATCCTCATCTGGTATCTCTATACTAAATTTTTCTTCT
ATGGCCATTACCAGCTCAACTGTATCTAGAGAATCGGCTCCCAGGTCACTTGAAAAATTT
GCTTCCCTAGTCACTATTTTTTTTTCAATTCCTAGCTGTTCTGCTACAATATCTTGAACT
TTTTCAAAGATTTCATTATCTTGCATAATATATTATTTCCTCTAAGGTGTTATCAAAAAT
TTTTCGGCTTATATTATTATCCAACTCTAAAAAATTAGTAATTTTTAGTCGAGATTAAAA
TATTTATACCTATATGATATACTATTATTCTGGATAAATACGAAGTTTACGATATGGCTC
TTCTCCAAAACTACGAGCTCTCAACTGGTAGTTGTCAACTAGATTATGCTGCATTTTTCG
TATATAGGCTGATCGAGGTGTAAGCTGAACTATTGAACTTTCATTCAAGATAATAGTTTC
AATAGCCAATTTAGCTTCTTGAAGAGCCTGGATCTCATCAAAATTTTTATTTTTACAGAG
TTCTACCCAATTAAGACCAGATGAAGTATGAATGTTTAATATTTTTCTGAGTGCACGAGT
AATTTGTGGAACCGTACTATTCTGAATTGTATATATAATAATTTGTTTTGATTTTGCAAT
TTGTCTAAGTTTTGTATTTTGCTTTACTTGATTACGTAGCGCTAGAATAGCATCAGACTT
TTCTATCTCCTTGGTTAAAATAATTGGTAGATCTAGTGAAGAAATAACTGATGTAATATG

CTGCCAACTCAAAGAATAAGCATATAAGTACTGATGAGGTATTTCCAATTGCAAAGATTG
TTTATTAATATTAATAGGGGCATTGAAAGTAGTTGACAATAAAGATGTATCACGATTTTG
TGATTTATTCAAATCTAAGGACTTATTTCTTAATTCTCTATATTGTAAAACCGGAGGCGC
CTGCTGTGCGTTAATTGATTTTCGAGCATTAGAGATGTTTACAGGTAAGACTTCTATAGA
TTGCGATGGATAACATTTAATTAAAATTCTACCATTAGCTTGAATTTGTCTTTTTTGCAC
AAAAGGTTGATGGCCTTGCAAAATTTGATCTATTGTCTCTTTAACTTTATTATGAACTAT
CCAAACATTTCGCTCGTGTATTTCGATCGCAATCTGAAAAGCAGGCTCAGCTTTTCTTTC
TAGTATACTCTTTTGCGTACCTCTACGTTTAGCTTCATCGTCACCTAATGTTACATATTG
AATTCCACCAATTAGGTCCGCCAATGTTGGATTTTTAATTAAACTTTCTAAATGATTTCC
ATGCGCTGTACCTACTAATTGAACTCCTCTTTCTGCTATAGTTCGAGCAGCTAACGCTTC
TAATTCCGTGCCAATTTCATCTATAATAATAACCTCGGGCATGTGATTTTCAACAGCTTC
TATCATTACTTGATGTTGCAAATCGGGCCTTGCGACTTGCATTCTTCTAGCTCTACCAAT
AGCTGGATGAGGAATATCTCCGTCACCAGCTATTTCATTTGAAGTATCTATAATAACCAC
TCTTTTTTCCATTTCGTCAGCTAAAACACGGGCCATTTCTCGAACTGCGGTTGTTTTACC
AACTCCTGGTTTTCCTAAGAGTAAAATTGAATCACCTTGCTGTAGTAAATCTCGAATAAT
ACTAATTGTACCAAAGACTGCTCTGCCGACACGACAGGTTAAGCCAATAACACTACCTTC
TCTATTACGCAGTGAACTAATACGATGCAAGGTTTTTTCGATACCAGCTCGATTATCGCC
GCTAAAATTACCAACTTTTTTTACACAATAATCTAAGTCTTGCCAACTAATAGATCTTTG
AGATAAATATTCTGGGTTATCTGGAAATCTAGCTTCTGGACGACGACCTAAATCCATAAC
GACTTCTATTAAATTATTTCTATTAGGATGCTGTTGTAAAGGTTCTTTGACAAAATTTGG
CAAAATTTCTAGCAACTTATCTAAGTCATCTGCAATAAGCATGATTGAGTATATAATAAT
TAATTAATCAAATAGAAGATTCAAGATTCTAAAGATTACAATACATATTAAAGAATATTA
TGATTGGAATCAATTTTATACTTAAATTGAACAGGAAGTTATAGATGTAAAAGAACTGAT
TAAACTTTACGCAATAAATCGATTGGTCAAAATTAATAGTCGTGTTAACATGTTTATTGG
ACAAAAAGTTCGTATTAAATATAGCAAACAAAAAGTTGCTAGTGATATAGCAGATAAAGT
TGGTGAATTAGGAGTTATTAAGGGAATAAAGTTTATTAATAGTCAATGTGTTACTATTAT
TGTTGAGTTTGACAATCACACTAGGCTTTGGATGTTTAGAGAGGAATTAATCTGTCTAAA
TGAATTAAAAACATGAAACATAATTTATTTTATTTATTATACAAAATTTCTTATGAGTCG
TTAAAATAATAAATTTTAAAGGAGTACTTTTAATTGTGCAAATTACTATTAAAAAATTAC
AAGACTTACTTTCGTCTGTACAAAGAAAAAAGATCCAGACATTAAAACTAAAACAAGGTA
AGTTTGAACTTCTATTAAATAAGACCTATAAAAAAGTCAATCAAGAAATTATACCTTCGC
AGAAATCTGCTGTATTACAAAATAGTCCATCTACGATAATTAAATCAATAAATAATACAA
AAAAATCTTCTGTTGTCAATGAAGACCGTACAGAATATGCCACTATTGTTTCTCCAATGG
TTGGAACGTTTTATCATTCACCTGCTCCTGGTGAAAAAATTTTTGTACAAGTTGGCGATG
AGGTCAAATTCAATCAAACAGTCTGTATTATTGAAGCAATGAAGTTAATGAACGAAATCG
AAGCAGAAATTGAAGGCAAGATTATAGAAATTCTTGTTAAAGATGGTGATATAGTAGATT

GCGGGCAAGCCTTAATGAAGGTTGAAACATAATGCTAATTTTTCCATTTTACATATTGTT
AACAGAAATTATTTAGATAGAAAAATTAGCTTATACTATAGTTATGAAAACTTATTTCGC
AAAGGCACTAAAAACTAATATATAAAATTATTTACTTTATCTGCCAAAAATAGAAATAAG
CGTTTTGATTCCTTGTCTCAAGAGAGGAGAATCTCAATGGCAATTAGCTCAAAAGAGCAA
GAGACAAAGAAGGTAAAAATCTCGGTTGATAAAAATCCCGTAGATACTTCTTTCGAAAAG
TGGGCCCAACCAGGCCATTTTTCTCGTACACTAGCAAAAGGACCAAAAACTACTACTTGG
ATTTGGAATCTTCATGCTGATGCTCACGACTTCGATAGTCAAACCAGTTCTTTAGAAGAA
GTTTCACGTAAGATTTTCAGTGCACATTTTGGGCAGCTGTCTGTAATATTTTTATGGCTT
AGTGGAATGTATTTTCACGGAGCCCGCTTCTCTAATTATGTTGCTTGGTTAAGTAATCCA
ACAGGTATTAAGCCAAGTGCGCAGGTTGTTTGGCCTATAGTTGGGCAAGAAATTTTAAAT
GGCGATGTAGGTGGTGGCTTTCAAGGAGTACAAGTTACATCTGGATGGTTCCAACTGTGG
AGAGCATCAGGAATTACTACAGAATTTCAGCTTTACTGTACTGCTATTGGCGGATTAGCT
ATGGCTGCTTTAATGCTGTTTGCAGGATGGTTTCATTATCATAAAGCTGCTCCAAAGTTA
GAATGGTTTCAAAATGTTGAATCAATGATGAATCACCATTTAGCTGGGCTTTTAGGCTTA
GGCTGTTTAGGCTGGACAGGCCATCAAATCCATTTGTCTTTGCCTATTAATAAGCTACTA
GATTCTGGCGTGTCTCCGCAAGAAATTCCACTACCTCATGAGTTTTTAATTAATAGAGAG
CTTATGGCTCAGCTGTATCCAAGTTTTAGTAAAGGATTAGTTCCATTCTTTACTTTAAAT
TGGGCTGAATATTCCGACTTTTTAACTTTTAAAGGAGGTTTAAACCCTGTTACTGGAGGT
TTATGGCTAAGTGATACTGCTCATCATCATCTAGCATTAGCTGTTCTATTTCTTGCTGCA
GGTCATATGTATAGAACCAATTGGGGTATTGGACATAGCATGAAAGAAATTCTAGAAGCT
CACAAAGGACCTTTTACCGGCAACGGTCACGAAGGTCTATATGAAATTCTTACAACTTCT
TGGCATGCACAGCTTGCAATTAATTTAGCTATGATGGGATCTTTAAGCATCATTGTAGCA
CATCACATGTATGCAATGCCTCCTTATCCATATATTGCTACTGATTACCCAACTCAGTTA
TCGCTCTTCACTCATCATATGTGGATTGGAGGATTTTGTATTGTTGGAGCAGGAGCGCAT
GCTTCTATATTTATGGTAAGAGATTATAACCCTGCAGAAAATTATAACAATCTTTTAGAT
AGAGTCATTAGGCATCGAGATGCTATTGTTTCTCATCTAAATTGGGTATGTATATTTCTT
GGATTCCATTCATTTGGTTTATACATTCACAATGATACTATGCGGGCACTTGGAAGATCT
CAAGATATGTTCTCTGATACAGCTATACAGTTACAACCTATTTTTGCTCAATGGGTACAA
AGTATACACACTTTAGCTCCTGGAAATACAGCTCCAAATGCATTAGCAACAGCTAGTTAT
GCATTTGGAGGAGATATTGTTTCTGTTGGTAACAAAGTTGCAATGATGCCTATTTCTTTA
GGTACTGCAGATTTTTTAGTTCACCATATACATGCATTTACTATTCATGTAACTGTTTTA
ATTTTAGTTAAAGGGTTCCTTTTCTCAAGAAACTCTAGACTAATTCCTGACAAGGCCAAT
CTTGGCTTCAGGTTTCCATGTGATGGACCTGGTAGAGGTGGTACTTGCCAAGTGTCTGGC
TGGGATCATGTTTTTCTTGGCTTATTCTGGATGTACAATTCTCTGTCTGTAGCAATTTTT
CACTTTAGTTGGAAAATGCAATCAGATGTTTGGGGTAGTGTATCTCCGTCTGGAAATGTT
TCTCATATTACTGGCGGTAATTTTGCACAGAGTGCAATTACAATCAATGGATGGCTGAGA

GATTTCCTTTGGGCTCAAGCATCTCAAGTTATTCAATCATACGGTTCTGCGCTATCTGCA
TATGGATTAATTTTCTTAGCAGCACATTTTGTATGGGCATTCAGTTTGATGTTCTTATTT
AGTGGTAGAGGTTATTGGCAAGAGCTTATAGAATCAATCGTATGGGCGCACAATAAGATA
AAAGTTGCTCCTGCAATTCAACCAAGAGCTTTAAGTATTACTCAAGGTAGAGCAGTCGGT
GTTGCACACTACTTATTAGGTGGAATTGGTACAACTTGGGCATTCTTTTTAGCGAGAATT
ATTTCAGTAGGCTAATAGTGAAAATAGGATAAAAAAAACAATTATGGCAACAAAATTTCC
TAAGTTTAGCCAAGCTTTATCACAAGATCCTACAACTAGAAGGATTTGGTATGGTATTGC
TACGGCACATGACTTTGAAAGTCATGATGGAATGACAGAAGAAAATTTATATCAAAAGAT
ATTCGCTTCGCACTTTGGACATCTAGCAATTATCTTCTTATGGACATCTGGTAATTTATT
CCATGTAGCTTGGCAAGGCAACTTTGAACAGTGGGTATTAAATCCTTTGAAAGTTAAACC
AATTGCTCATGCAATTTGGGATCCGCATTTTGGACAACCTGCTTTGAAAGCTTTTAGTAA
AGGTGGCTCAGCTTATCCAGTAAATATAGCATATTCTGGCGTATACCACTGGTGGTATAC
TATTGGTATGAGAAGCAATCAAGACCTGTATTCTGGGGCTTTATTCTTACTAGTTTTATC
AGCTTTACTCTTATTTGGAGGGTGGCTACATCTACAACCAAAATTCAAGCCTGGTTTATC
ATGGTTTAAAAATAACGAATCAAGATTAAATCATCATTTATCTGGATTATTTGGGGTTAG
TTCTTTAGCCTGGACAGGTCATTTAGTACATGTTGCTATACCTGAGGCAAGAGGACAACA
TGTAGGATGGGATAATTTTACAACCGTATTACCTCATCCAGCCGGTTTACAGCCGTTTTT
CAGTGGTAATTGGAGTGTATATGCTCAAAATCCAGATACAGCTCAACATTTATTCGGAAC
TAATGAAGGTGCGGGTACAGCAATTCTGACATTTCTAGGAGGATTTCATCCTCAAAGTCA
GTCTTTGTGGCTAACTGATATGGCTCATCACCATTTGGCTATTGCAGTAGTATTCATTGT
TGCTGGACATATGTATAGGACTAATTGGGGAATTGGGCACAATCTAAAAGATATTTTAGA
TGCTCATAGACCACCTAGTGGTAGATTAGGAGCTGGACATAAAGGGCTATTTGATACTAT
TACTAATTCTTTACACATACAGTTGGGATTAGCATTGGCTTCCCTAGGTGTAATTACTTC
GTTGGTAGCTCAACATATGTATGCTATGCCTCCATATGCTTTCATGGCTAAAGATTTCAC
AACTCAAGCATCTTTGTACACACATCATCAATATATTGCTGGGTTTCTAATGGTTGGAGC
TTTTGCTCATGGGGCAATATTCTTTGTTCGAGACTATGACCCTGAACAGAATAAAGATAA
TGTTTTGGCTCGTATGCTAGAACATAAAGAAGCTATCATTTCTCATTTAAGTTGGGTAAC
TCTATTTTTAGGGTTTCATACATTAGGCCTTTATGTTCACAATGATACAATGATTGCTTT
TGGAACTCCTGAAAAACAAATTCTAATTGAGCCGGTATTTGCTCAATGGATTCAAGCCTC
TTCAGGGAAAGCACTTTATGGGTTTGATGTGTTACTATCATCCTCTACTAATATCGCAAC
ACAAGCTGGTAGCAATATTTGGCTGCCAGGCTGGTTAGAAGCGATTAATAGCGGAAAAAA
TTCATTGTTTTTAACAATTGGTCCTGGTGACTTCTTAGTTCATCATGCAATTGCATTGGG
ATTACATACTACTACGTTAATTTTAGTTAAAGGTGCTTTAGATGCAAGAGGCTCTAAACT
TATGCCGGACAAAAAAGACTTTGGATATAGTTTTCCTTGCGATGGACCTGGCAGAGGCGG
CACCTGTGATATATCTGCATGGGATGCGTTCTATTTAGCTGTATTTTGGATGCTAAATAC
AATAGGTTGGGTAACATTTTATTGGCATTGGAAACATATTACAATATGGCAAGGCAATGC

AACTCAATTCAATGAGTCTTCAACTTATCTAATGGGATGGTTTAGAGATTACTTATGGCT
AAATTCTTCTCCATTAATTAATGGTTATAATCCATATGGCATGAATAATTTATCAGTATG
GTCATGGATGTTCCTATTTGGACATTTAGTATGGGCAACAGGATTTATGTTCTTGATCTC
CTGGCGTGGCTATTGGCAAGAGTTAATTGAAACTCTAGCATGGGCGCATGAACGTACACC
TTTAGCAAACTTGATTCGTTGGAAAGATAAACCTGTTGCATTATCAATTGTACAGGCAAG
ATTAGTAGGTTTAGCACATTTTTCTGTAGGATATGTATTAACTTACGCAGCTTTTGTATT
AGCTTCAACAGCAGGCAAATTTGGTTAGACTAAATTCGAATTAGCTAAAAAAAAGTCAGT
CATAATAGTTTTATGACTGACTTTTTTAGATTTAAAACAAAAACTTACTAAAGTTTCACC
TCAATATCTACACCAGAAGGCAAATTTAACTTCATTAATGCATCTATCGTTTGAGAAGAA
GGTTGATGAATATCAATAATTCTTCTGTGAGATCTTATTTCAAAGTGTTCTCTTGAGTCT
TTATCTACATGTGGAGAACGTAAAACACAATAAATTCTTCTTTTTGTCGGTAAAGGAATT
GGCCCTACTGCAACGGCATTAGTTCTAGATGCGGTATCTAGTATTTTGTCACACGACGTA
TTAAGTATAATAGAATTATATGCTTTCAGTTTAATTCTGATTTTTGTCTGCTGAGTAATT
GTCATGTTAAAAAGATCTTATTATTTAAGAATTTTAGAGACAACACCTGCGCCTACGGTA
CGGCCACCTTCTCTAATAGCGAAACGCATACCTTGCTCAATTGCAATTGCATTAATTAAT
TCAGCAGTCATCTTAATTCTGTCACCAGGCATAACCATTTCTGCATCAGTACCATCATCA
GCAGTAAACTGATTAATAGTACCAGTTACATCAGTTGTTCTAACATAAAACTGAGGTCTA
TATCCTGGAAAAAATGGAGTATGTCTTCCGCCCTCCTCTTTAGTTAGAATATAAACTTCT
GCTTCAAATTGAGTATGAGGTGTAATTGTACCAGGTTTAGCTAATACCATACCTCTTTCA
ATATCTTTTTTCTGCACACCTCTTAAAAGAATTCCAATATTATCACCCGCTAGACCTTCT
TCTAACGTTTTTTGAAACATTTCTAATCCAGTAATAGTCGTTGTACGAGTTTCTCGTAAA
CCTACAATTTCAATTGTGTCACCAACTTTAATAATGCCTCTTTCAATTCTACCAGTGGCA
ACAGTTCCACGTCCTGTAATAGAAAAAACATCTTCTACAGCCATTAAGAAAGTTTTATCG
ACATCTCTCTCCGGCGTTGGAATATATGTGTCAACTGCTTCCATAAGTGAAAAAATCTTG
TCAACCCATTTATCTTCACCTTGCTTAATAGCTGGGTTTTTCGTCACAGCTTCTAATGCC
AATAAGGCAGAACCTGCAACAAAAGGAATATCATCTCCAGGAAAGTCGTATTGACTTAAT
AATTCCCTTCCTTCTAATTCTACTAATTCTAGTAGCTCTTCGTCATCTACTTGATCTTCC
TTATTTAGGAATACTACTAATGTAGGAACACCTACTTGTTTTGCTAATAAAATATGTTCA
CGAGTTTGTGGCATTGGACCGTCTGCCGCAGATACAACTAAAATAGCTCCATCCATCTGA
GCAGCACCCGTAATCATATTTTTTACGTAGTCGGCATGGCCTGGACAATCTACGTGAGCA
TAATGACGATTATCTGTTTCGTATTCAACATGAGCAGTATTAATAGTAATACCTCTAGCT
TTTTCTTCTGGAGCGGCATCAATTTCATCAAATTTTTTCGCTGCAGTAGACCCTAAAGTT
GATAAAGTCGCAGAGATCGCTGCTGTTAAAGTTGTCTTACCATGATCAACATGACCAATT
GTGCCAATATTGACATGAGGTTTTTTACGTTCAAATTTAGATCGAGCCATGCTTTTTGTT
TTCCTTATAGTAAAAGTTGCTTTATAGTTACTTTTAACGATAAATCTTGTGGATTGAAAG
ATTTATTATTTAATACAGACAAGATACAATATAACTAATCTAATATCGATAGTGAGCAAA

CGCTTTATTTGCTTCTGCCATTCTATGTGTATCCTCTCTCTTTCTAATAGAATTTCCTGT
TTCATTAGCTGCATCCATAATTTCATTAGCTAATTTCATAGACATACTTTTACCAGATCT
GTCTCTAGAGAATTTAGTAATCCATCTTAGTGCTAAATTCGTACCTCTATAAGCTCGTAC
TTCAATAGGAACTTGGTAAGTAGAACCACCAACTCTTCTTGCTTTTACTTCTACTAGAGG
AGTAATATTTCGGATGGCTTTTTCTAAAATATTTAAAGGATCTGATTCTGTTCTCTCTTT
AACGATATCTAAAGCCTGATATATAATTCTTTGAGATAAAGTTTTTTTACCACTTTTCAA
AATACGAACAGTTAACATACTTACGAGTCTGCTTTTATATAAAGGATCAGGTGATGCAAA
CCTTTTTTTAGCTGTATTACGACGAGACATAGTATTTAAAATTGGTTATTGTAATAATGT
TCTATTAGCAGGTATTTAAGATTTAGGCTTTTTTGTCCCATATTTAGATCGGCTTTTACG
ACGATCTTTTACTCCTGCAGCATCTAAAGTACCACGTACTACATGATATCGGACTCCAGG
CAAATCTTTAATTCGGCCGCCCCTAATCAGTACCACAGAATGCTCTTGAATATTATGACC
TACGCCAGGAATATAAGCTGTAACTTCAAATCCAGATGTTAATCTAACACGCGCTACTTT
TCTTAAAGCAGAGTTCGGTTTTTTAGGAGTTGTAGTATATACTCTTGTACAAACACCTCT
TCTTTGAGGGCAACTTTGAAGAGCTGGAGATTTTGTTTTTTTATGTATTTTTCGTCTTTC
GGATCTAACAAGTTGTTGAATTGTTGGCATAATAAAATTGATTTAAAGTCTTTAATAGAT
ACTATTTGAACTTGCTTTACATGAATTGGCTAGTTTTTATTTATCTTATACTTACGCATA
AATCTTTCTACTCTACCTTCTGTATCAATAATTCTCTGTGAACCAGTAAAGAATGGATGG
TTTCCTGACCAGATATCTACGTGTAGTTCTGGCTTTGTGGACCCAATCGTCATAATTAAC
TGGCCGTCACAGTAAACTTTTGCTTCTGGATACCAATTTGGATGTATATTATCTTTTGCC
ATTGTTTTTATAATTATATAAATGTATCAATAATAACTATTAACGCTTAGAAAATTGAGG
AGCCTTTCTGGCTTTCTTTAAACCATATTTTTTTCTCTCTTTTACTCTTGGATCTCTTGT
TAAATAGCCTTCAGACTTAAGTGTCGTTCTATTTTCTGGATTAATTGAGCACAATGCTCT
TGCAACACCTAAACGAATTGCATCAGCTTGCCCTGTCAAACCTCCTCCTCTAGCATTTAC
ATGGATATCATACTGGTTTAGTAGTCCTAAAACTTGTAATGGTGCATATGAAACTCTTAA
GTAATTAGGACTAAATTGAAGATAAGACTCTCCTGGTATACCGTTTATAATTAAATTACC
TGACCCTGGGACTAGTCTTACTTGTGCAACAGAACACTTCCGGCGACCTGTTCCAGAATA
GATTGCGCGAGTTTTAATTAATTCTGTCGACATAACATTCCTTGATAACTAAATAAATAT
ACATTCTTTATACTATATACTCTTGTGGCTTTTGCGCGACATGTGGGTGAATTGGGCCAG
AATACACTTTAAGCTTCGTAAATAGTTTTCTGCCTAATGGACCTTTAGGAAGCATACCTT
TAACCGATTTTTCAATAATTCTGTTAGGTAATCTTGTCTGAAGCTGATCAAATGTTTCAA
CTTTTAATCCGCCAGGCTGTCCAGAATGTCTTCTGTATAGTTTTTGATTCGTTTTATTTC
CACTTACAGATACGTGAGCAGAATTAATAACGATTACATAATCTCCAGTATCTAGATAGG
GTGTATAAGAAGGCTTGTTTTTACCTCTTAAGATATTAGAAATATGAGTAGATATTCTAC
CAAGTGTCTGATTTTTAGCGTCTATAACATACCAATGAGAATTAGTATTTAATGAGGGTG
ATTGCGTTTTATTCATGAAAAGATTAATACTTTTGGTTGACAATAAGAGAATATGTAAAT
GGTTTACATTCAATTTCTATACAAGTAGTATTTAAAAATATTATTAACAAAGAGAACTTT

TCTAACAGTTATTTAAATTGTTTGTCAGCAAGAAAATTTCTAGTTATCAATAACTTATAC
TATTTTTAATAATATAAAATATATTAATCACTTTTTTCTTTTGGCAAACTTATACCTAGT
TTTTTTTGCAAGGCATATATAACTTCTTCAGCTGATTTCTGGCCAAAATTTTTAATTTCT
AGCAGTTCTTCTTGAGAGTAATCTAGTAAATCTGCAATAGAATGGATTTGAGCTCGCTTT
AAGCAGTTATAAGCCCTAACAGATAACTGCAGTTCTTCTATTAAAACTTGACTAATTTTT
TTATCTTCTTTGCTACGATAATTATCTGCTGATTTAAAGTCTAAATTTCTTAGTGAACAA
AAAAGATTAGTTAAAACTGTAGCTCCTTGACTTATTGCTTCTTGAGGAGAGATGCTCCCA
TTTGTCCAGATTTGTATAATCAATCTATCTTTTATGCTGTTGCTACCAATACGGACTTCT
TCTACTTTATAATTAACTTTATTAACTGGCATAAAAACAGAGTCTACTTGCAAAAAATCC
ACAGATAATTCATCTACAGCTTTTTCAGCTAAGCGATACCCGCAATTTTTTTCAATTTTA
AACTCCATTTCAAATATTGTATTATTGCAAATAGTTGCAATATACTGTCTAGGATCTACT
ACCTCTATATCAGAAGATAATTCAAATAGGCCGGCTGTAACTATAGCTGGCCCTTGAACT
CTAATTCGACCAATTTGAGATTCTTTGTTATAACTTTTAAATACTACTTCTTTTAGATTA
AGTAATATTTCTAACACATCTTCTCTTACCCCAGGAATTGTAGAAAACTCATGGTTCACT
CCAGCAATCCGTACAGCAACTATAGCAGTACCTTCAAGATCTGACAATATTGATCTTCTC
AAAGCATTACCTAATGTAATACCTTGTCCTTGATTTAATGGTTCTATTACAAAACTACCG
TACTGCCCACGCGCCCCATCTGTTCTTGACTCTATGCATTCAATTTGAAATTGAGCCACC
TAAGAAAGCTCCTTTAAAATAATCAGTGATTAGTAAGTCTCTCTAAAACAAGAAGGAATT
TATACTCGGCGTTTCTTAGGAGGGCGACATCCATTATGAGGTACAGGAGTAATATCTTTT
ATTAGAGTAATCTCTAATCCTGCAGCCTGCAAAGCTCTAATTGCTGTTTCTCGACCCGCT
CCTGGACCATTTACTAGAACTTCAGTTTGGCGCATACCTTGATCCATAGCTTGTCTAGCT
GCTTTTTCAGCTGCTGTTTGAGCTGCAAAAGGTGTTCCTTTTTTAGCTCCCTTAAATCCA
CTTGCACCAGAGGATGACCATGATAATGTTTCTCCTTTTAAATTAGTAATAGTAACAATT
GTATTATTGAATGTAGATTTAATATGTGTAATACCGTTAACTGCATTACGTTTAGTTTTT
CTTGCTCCGGATTTTTTTATTTGTCTAGCCATCGTGTCTCGTCTATTATATAGTTAATTA
AAAGATTATTTTCTTGGAGCTTTTTTCTTACCTGCTACTGTTTTTTTACCTCCTCTGCGT
GTTCTAGCATTAGTTCTAGTTCTTTGTCCTCTCAAAGGAAGACCAAGACGGTGTCTTCTA
CCTCTATAAGTACTAATTTCCATAAGTCTCTTAATGCTCATAGACTCAAAACGTTTGAGA
TCCCCTTCAATCTGATAATTAGACTCAAGAATTTCTCTTATACTGACAACTTGTTGATCA
TTTAAATCTTGACACTTAATATCAGCGTCTATGTTTGTTTTTTCTAATATTTCTTTCGAG
CGAGATAGTCCAATACCATAAATATATGTTAAAGCTATCTCTATTCTTTTGTTTCTTGGA
AGATCTACTCCAGCAATTCTGGCCACTTTGTCTTGTCTCCAATAAATAATATGTATTTTT
ATAACGGTTGTTTATAAGCTTAGCCTTGTCTCTGCTTATGTTTAGGATTAGTGCAAATTA
CCATCACCTTTCTGTGACGCCGAATTATTCTACATTTTTCACACATTTTTCGAACAGAAG
GACGAACTTTCATATTAACTCTATATACTAGGAATATTACTATATTAACTCTATTAATTA
TCGACATTAAAAATGTCAACAATATATTGTACTACTTTAACTGTTGTAGATTTTACTTCG

TCATACTATCATATTTTTTTGATATGACGTATGTCTGAATTTGCTTAGCTGTGTCTATTG
CGACTCCAACTAAAATAAGTAAAGACGTAGCCCCGAGACCTCTTAAGTTTTGGATCTGAG
TAACCTTTTCTATTATAAACGGAATCAGCGCTACTGTAAATAAAAACGAGGCTCCTAGGA
ATGTCAGTCTATTTAATATGACTTGTAAATAATCAATAGTCGCTTGACCTGGACGAATAT
TAGGAATGCTTGCACCCATTTTTTTTAAATTTATAGCAATATCTTCTGGATTCATTACTA
TTGATGTATAAAAATAGCTAAAGAAAAGGATCAAAGCACAATAAAGAAGAAGGTATAGTG
AACCATTAGGACAAAACAGGTATAGGATTTGGAGTAATGTTTTATTTTGAATAATTTGGG
TTAGATAAGATGGGAGGGCCATAGATGCAGATGCAAAAACAATAGGCATAACTCCACCTT
GATTTAATTTCAAGGGCAAGTAACTATTTGGATCCAAAATTGAAGATTTTCCTAGCTGTC
TTGCTGAAATAATTTTAATTCTTCTTGTTCCTTCTTGTACACAAATTGTAATTATTATCA
TTAATAAAAAGATCGCTATAAATAATCCGAACTTAAGACTTGCATTACTATAACTAGCAT
CAAAAAATGACTGTGTAAAATTCTTTGGTAGTCCTGACACAATGTTTTGAAAAATAAGTA
AAGAGGCTCCATTACCAATTCCCTTTTCCGTAATTAACTCTGATAACCACATAATAATCA
TAGAACCTGCTGTTAAAGCTAAAACAGACTCGCAGACAAATGCAAAATTCCAATTAAAAA
CATATGGTTTTACCCATATAGAGATTGCTCCAGATTGTAAAGTCGCCCAACCTAGAGCTA
AGTACCTTGTAATTTGAGTTATTTTTTGGCGGCCCAATTCACCTTCTTCCTTCTGTAATT
TTTCCAGATTGGGGACAATTTTCGTAAGTAGCTGCATTACAATTGAAGAGTTAATATAAG
GAACAATACCTAACGCAAAAATCCCTATTGTTGAAAAACCTCCTCCAGAAAAAATATTCA
GAAAATTTACTAAAGTATTTTTTTCTACACTTGCATAAAAGGCGTCATGATCTATACCTG
GAACAGGTATAAATATTCCTAAACGTGCTAAAACTAATAGAAAGAGAGTAAAGATAATAC
GATTTCTTAGATCACTTTTTTGGCTCATAAATAAATTTTAAATGAAAAGAGATACTTTTT
TATGTAAAAACCCTATTCCAAGACTTGTATAAGTTTAAAATAGGGAAGTAAAATATTATA
GTAATATTAATATAATATTCTATGTAGAATACAGGCTTTCTATTGGAACTCCTCGATCTT
TAGCTGCTTCACTAAAAGTTCTTAATTGAGTTAAGCCATTGAGAACAGCTCGCGCGTTAT
TTAATGTATTATTAGAACCTAATTGCTTAGCTAATATATTCTGTACACCCGACAATTCGA
GGACTGTCCTTACAGAACCTCCGGCAATTACACCTGAGCCAGGTGCAGAGGGCCTTAATA
TAACTTTCGCGGCACCAGAAATTCCGTTAATAGGATGAGGTATAGAATTCGATTTTGTCA
GTGGAACTGTAACTAGATGTTTTTTAGCATCTGTTACTCCTTTTTTTACTGCACCAATTA
CATCGCTTGCTTTTCCCACGCCGACACCAACTTGGCCTTGCTCATTACCAATAACAAGAA
TAACTCGAAAGCTTAGTTTTTTACCTCCTTTTACAACTTTAGTAACTCTTTTAACTTGTA
CGACTCTTTCTTCCCAGCCACTATCTTTATCTTTTCCTTTGCTCTGTTTTTTACGATTGG
CCATTTTAAGAATTATTCCTAATTAGTTAATTACGATTCTAGAAACCCATGCCTGCTTCC
TTAGCAGCTTCAGCCAAGGCCTTAACTCTTCCATGATATAACTTTCCTCCTCTATCGAAG
ACGACATTTTTAATGCCTTCTTTCATAGACTGTTCTGCTAACTGTTTACCAACTACGCGA
GAAGTATCACAGTTTGGTCTTATGTTATCAGATTCTTTATTATTTAAATTAACAGATGAT
GTAGCTACTAATGTAATACCTTGTGTGTCATCAATTATTTGTGCGTATATATGTTTATTA

GATCTAAATACACACAGACGAGGCCTACTTGAGGTTCCTTGAACTTTTTTCCGAACTCTT
TTATGCTTATGAATTCTAGTTTGTTTAGTGTTTAGTTTCATTATTATTTACCTTTTCCAG
CTTTCCCAGCTTTTCTTCTAACAAATTCACCTTGATATCTAATTCCTTTTCCTTTATAAG
GCTCAGGAGGCCTAATAGAACGAATAGTTGAAGCAACCTGACCGACAACTTCTTTATCTA
TGCCAGAGACAGTAATATTTGTGTTGTTTTCAACTTTAATTTCAATATTGGCAGGAGGTT
TAATTTTCACTACATGACTATAGCCAACACTTAAGATTAAATCTTGGTTATCAATTTGAG
AACGATAGCCTACACCTTGTATTTGCAGTTTTTTAAAAAATCCATTAGAAACACCTTCAA
TCATATTACTAATAAGAGTTCTGGATAGCCCATGTAACTGACTAGCCATTTTTGTTTGTC
CACTCGTCTTGACAGCAATTGTGTCGTTAAGTATTTCTAAATTAATACCAGCTGGTAAAG
TTCTGGATAGGGTACCTTTAGGTCCTGTCACGGTGATAGTCTGACCATCGAACTGAGTAC
TAAGATTTGTCGGCAATAAAATTATTTTTTTTCCAATACGAGACATATTCCACCTATAAA
ATAGTTACCAAATATAGCATAATATTTCACCACCAAGACCATCATGACGAGCTTGTCTAT
CTGTCATAACGCCTCTAGAAGTAGAAATAAGAGCGATGCCTAAACCTCCAAGAACTCTGG
GCAACTCTTTATGATTTGCGTAGACTCTTAGTCCGGGTTTACTAATTCTTTTCAGAGCAG
TAATAACTGGTTGACGATTTTTACCATTATACTTAAGAGAAATCATTAAATGAGTCTCTA
TACCTTCACCCATTTGTTCGAAATTTTGAACAAATCCTTCTTCTTTCAGTACTGTTGCCA
TATTGCATGTCATTTTCGTTGCTGGAACTTGCACAATTTGATGTCTTGCTAAGTTTGCGT
TACGAATACGTGTCAGCATATCGGCGATCGTATCGTTGACCACCTTGATCCTCCTTGCGA
TGAATTATTTGAAATATTTTCAAGATTCCCTGAAAGGCATCCCTAGCTTTTTTAACAAAG
CTAGACCTTCTTGATCTGTTTTAGCTGTAGTGACAATTGATATATCTAAACCACGAATTT
GATCAATATTATCATAGTCTATTTCTGGAAAGATCAACTGTTCACGTAAACCTAAATTGT
AATTACCTTTGCCGTCAAAACTTCTAGGACTAATTCCTCTGAAGTCTCTAATTCTTGGTA
ATGTTAAATTAATTAATTTCTCTAAAAAAGAATACATCTTGTCTTTTCTTAGATGCACAA
CAATTCCGATAGGAACCTCTTCTCGAATTTTAAATCCTGCAATAGATTTTTTAGCTTTTG
TGACAATTGGTTTTTGTCCAGTTATTAATGTCAATTCTTGAATACTACTTTCAAGAGCCT
TAGCATTCTGAGAAGCTTCACCTAAGCCACGGTTAATAGTAATTTTAGTAAACCTAGGAA
CTTCATGTACATTTTTGTACTGAAATTCATCTTTTAAAGATTGAGTAACAGTTGTTTTAT
ATTTTTCTTTTAATCCTATTGCCATTATATTTTCATCAATTATTTAATAAGTTCGCCAGT
CTTTTTCAGTTTTCGAATTTTTTGGCCTTTATCATTAATTATTACTGAGGATCGACTAGC
AATATTATTTTGTTCACTAAATAACATAACGTTAGAGGTATGTATTGGAGCTTCAAATTT
GATAATTTCTCCCGTTTCTCCTTCTTGTTGAGGTTTTTTATGCTTCACTTTGAGATTAAT
TCCTTTAACAATCACCTTATTTGTTTTATAAATAATTGCAATAATTTCACCTGTTTTTGT
TTTATCACTTCCAGAAATAACTTGAACTAAGTCCCCTTTTTTTAATTTAATTTTTGTATT
ATTTTTGGTTGTTTTAGAAGGACCTTTCATTATACTACCTCCGGTGCAAGAGAAACTATT
TTAGAAAAATTTTTATCTCTTAACTCTCGGGCTATAGGTCCGAAGACTCTTGTGCCACGT
GGATTATTATCTTGATTAATGATAACTGCCGCATTATCTCCAAATCTAATGCTCATACCG

TCAGTTCTCCGTAAAGCTTTACGAGTTCTTACTACAACGGCTCTGACAACATCAGATCGT
TTAACTGGCATATTGGGAGACGCATCTTTGACTACTCCAATAATAACATCTCCAATAGAT
GCATAAGAAGGATTACTAGTGCCTAGTACTCTAATACACATTATTTTTCTAGCACCACTA
TTATCTGCAACATTAAGATAGCTTTGAGTCTGTATCATACTTTTATTCTGTATAAGATTA
ATTATCAAAAGACTTAGATAAGATATTAACCATTGTCCAACACTTTGTGCGACTTAAAGG
TCGTGTCTCTTGTATTGTAACAATATCGCCGATTGTGCATTCATTATTTTCATCATGCGC
TTTATATTTCTTTGTCCGGATCATAGTTTTTGCGTACTTTCTATGAGAGATTCTATTTTC
TACAGCTACTACTATAGTTTTATTCATTTTATCGCTTACTACTTTACCTGTTGTTTCTTT
TAAAGGCATAGTCTTTATATTTTTTATATATAATTAAATTTTAAGATTATTACATGGTTG
ACTTAGTACGAGACTTTTCAACAGTTAAAAGTTGGGCTAATCTATGCTTAGAATGTTTAA
ATAGATGAGGCTGGAAATCTTGCCTTGTGGCTCTTTTTAGCCTTAAATCAAAAAGCTCTC
TTTTTATTACAAGGATTTCTTCAGCTAAAGAAGAAGAGTCCAGGTTTGTAACATCTGATA
TTTTAGGGAAAGTCATATTTTAGGATTCTGTTGTATTCCGAACTATAAATTTAGTCTTGA
TTGGTAATTTATAAGAAGCTAATTTCATAGCTTGTTGAGCAGTTTTTTGTGGTACACCTG
TAATTTCAAATAGAATATGTCCAGGCTTAATAACTGCAACCCAATATTCTGGAGCTCCTT
TACCTGACCCCATGCGAGTTTCAGCTGGACGGGCAGTGACTGGTTTATCTGGAAATACTC
TAATCCATAGTTTACCACCTCTTCTCACATATCTAGTAATAGTTCTTCGAGTAGCTTCTA
TTTGTCTAGAAGTTAACCAAACTGGCTCTGTCGCTTGTAATGCATAATCACCGAACGCAA
TTGTGTTACCTTTGCTAGCAGAACCTTTCATTCTACCTCTATGTTGTTTTCTAAATTTTG
TTTTCTTGGGGCTTAGCATAAAAATTAAAAAATAATAATAATTAAGAAGCTGTTGAATCA
GGAGTTTCCTGATTATAAGCAGATTCTAGACTTTTCGATTCTGGCAAAATTTCTCCTTTA
AAAAGCCAAACTTTTACTCCGAGCACACCATACGTAGTGTGAGCTTGGCGATGACAATAA
TCAATATCGGCTCTTAAAGTTTGTAAAGGTACACGACCTTCTCTAACCCATTCACTTCTA
GCTATCTCTGCACCATTAAGCCGACCAGATACTTGAATTTTAACACCCTGTGTATTTGCT
CTTTGAGCTCTTTGAACTGCTTGTCTGACAGCTCTACGAAAGGCCACTCTTTTTTCTAGC
TGTTGAGTAATAAATTCTGCTACTAAGGTTGCTTCAGAATCTGGATCTGCAATCTCTACA
ACATTTACTCTGAGTTGTTTGCTAGGGTCTAGGATTAAGGACAGCGATTTTCTTAAAGAC
TCAATTCCCGCTCCAGATTTTCCAAGTACGATTCCTGGTCGAGCTGTTGCTATGAGAATT
TCTACTTGATCAACTTTACGATTAATTTCAATTTTAGCAATACTAGCATTACTAAGTTTT
GAATGTATAAATGAGCGAATTTTATGATCTTCTTGTAAAAGAACTGGGTAGTCTTTAGAG
TTAGCAAACCATGAAGAACGATGTTTTTGGGTAATGCCTATGCGAAAGCCCAAAGGATGA
ATTTTTTGACCCACAGTATCCTTGATTATAAGTTAAAAGCTATAATGACTAATTGACGTT
AAAAGACTACAGTTCTGAAACACCTAATGTAATATGACAAGTTGGTTTATGTATTGGAAA
GGCTCTTCCTTGTGCTCTAGGCTGAAATCTTTTTAAAGTCGGGCCTTTGTCTGCAAAGGC
TTTACTAACAAACAATTGGTTTTTATTTAATCCATCATTATGCTCAGCATTCGCTGCAGC
AGATTCCAGGATTTGCTTTATATGAGAACAGACTCGGTATGGCATAAATTCCAGAATAAT

TAATGCTTCTTGGTATTTTCTACCTCTAATTTGGTCTAATACACGACGGACTTTATGCGG
AGAAAGACGAATGTATTTCCCTACTGCTTTAGTTTCTTTTACGTTTTTTGTAATACTCAT
AGTTATAGTTTAACGACGGGCTTTTCTGTCACCTTTTACGTGAGTGCGGAAAGTTCTTGT
TGGAACAAACTCTCCTAATTTATGTCCTACCATTTGGTCTGACACAAAGACAGGAAAATG
TTGTTTACCATTATAAACAGCTATTGTATGCCCTACCATATCGGGAATAATAGTAGATGC
TCTCGACCAAGTTTTTAGGACTTCTTTTTTTCCTGAAATATTTAATGCTTCTATACGTTT
AAGAAGACTAACATCAATAAAAGGGCCTTTATGTATAGATCTTGACATAATAATAAATCA
TTAGATTCTTAATAAAATAAATTCTATTTACGGCGACGTAAGACGTACGGGTTACTATAT
TTATTTGGATTTCGTGTTTTAACACCCAATGCAGGTTTACCCCAAGGTGTTACAGGACGA
GCTCGTCCAATTGGAGATTTGCCTTCACCACCACCATGTGGATGGTCTACAGGATTCATG
ACAACACCTCTAACAGTAGGTCTTTTACCTAACCAACGATTTCTACCAGCTTTACCAAGA
GTAATATTACTAGCATCAATATTACCAACTTGTCCAATAGTAGCGTAGCATTCTTTTCGA
ATCATTCGAACTTCGCTTGAAGGCAATTTTACAGTAACAAAAGTACCTTCTTTTGCTACT
ATTTGAGCATAAGTTCCTGCTGCACGAACAATCTGTCCTCCGCACGAAGGTCTTAATTCT
ATATTATGTACTGCTGTTCCTAAAGGAATACTAGATAAAGGCAAAGCATTTCCAACTTCG
ATAGGGGCAGTAGGACCAGAAAGAACCATAGATCCTACACTGAGAGATCGAGGGTGTAGA
ATATATCTTTTCTCACCATCAAGATAGTGTAATAATGCAATTCTTGCATTTCTATTCGGA
TCATATTCAATAGAAGCAACTTTAGCAACTATATTATGTCTATTTCTTTTAAAGTCAATT
AATCTGTACTGCTGCTTATGTCCACCACCTTTATGACGACAGGTAATGACACCTCTATTA
TTTCGACCTTTGCAGAAATGGTGTTTAACTATTAATGATTTTTCTGGCTTATCAGTAGTA
ATCTCTGAAAAGGTAGAAACCGTTCTATTTCTTGTCCCTGGTGTGTAGGCACGATATAAA
CGAATTGCCATATAAGAAGATTTGAGAAGAGTAAATGTTGAACTATGTGATATATTTAAG
TTTCTGGAAACAAATTAATAGAATCTTCTGAGGCAAGTGTTACAATTGCTTTCTTGTAAT
GTGGTCGCTTCCCTACGAATCTACCGATGCTCCTTTTTTTCTTAGGAGGATGACAAGTGT
TAACACCTGTAACTTGCACGTTGAAAATATACTGTATAGCAGCCTTAATATTAATTTTTG
TTGCTTTGGGATCAACGGCAAAACAATACTGATTCTCTTCTAATAATTTAGTTGTCTTAT
CAGTAATGATGGGATATTTAACCAGATCTAATAAACCTCTTGAATCAATGCTATCCATTA
TATACCTCTTGTATTTTAGATAAAGCATCAACTGTAATAATAATTTTATGTGCAGCTAAC
AAAGCCATAATATTTAGTGTATCAGCTGAAATAAGTTCTACGTTGTGTAGATTGCGAATA
GAAAGATAAACATTCGGATCTTTTTTGTCTACTATGACTAAAACTTTTTTATTTAAATCA
AGATTCCATCGATGTATAGCTTCCATAAATAGTTTAGTCTTGGGCTGATGAAAATAACTG
TTAAAATTTTCTACAATCAGTGTATTAACTGATTTGTTATTTAATGCTGTCCTTAATGCT
AATTGTCTTTCTTTTTTATTCATTTTTTTGGTAAAACTACGAGGTTTTGGCCCAAATATT
ACACCACCACCTCTCCATAGAGGTGAACGGATAGAACCTGCTCTTGCTCGACCAGTCCCT
TTTTGACGCCATGGTTTACGACCTCCACCTCGAACTTCACTTCTTGTTTTAGTATTAGCA
GAACCCTGGCGCTTTTCATTACTTTGCTTTACTAAAGCTCTATGGACTAAGTACATCCCA

GAATCTTGACTGACCTTAAGATTTAAATCAGCGTTGCCACTTACTTGACCTTCCCAATTA

TAGACTTGATAATTTAATTGCGTGTTAACTGTCATGAAATATATCGTATGAAAGAATATT

GCTACTTACTAATTTTTACCAAAGCGCCAGGTTTACCAGGAACAGCTCCCTTAACGATCA

ACAAATCATTCTCTGAGTTAATACTTACAATTTGCAAATTTTTTATTGTAACTTTTTTAT

TACCAAGTTGGCCAGCCATGTTTTTACCTGGATAAACTCTTCCAGGCGTTGTACCGGCTC

CAATAGAGCCAGGTTGCCTATGATTTTTCGAACCATGTGACATTGGGCCTCTACTAAAAT

GATGCCTCTTTTGGTAACCGGAGAAACCTTTACCTACACTTCTAGAAGAGACATTAATCT

TTTGGCCTACTTGAAATAGATCTGTAGATAGTATTTGGCTAACTTCGAAATCATCAGTAG

ATTTCATTTCGTATTCACGTAAATATTTTAGTGGGGGAGCCTGTGATTTTTTTAAGTGTC

CCAACAATGGTTTATTTAATTTTTGTTCAGCAACTTGCTTGTAACCGACTTGAATAGCAT

TGTAGCCATCAGTAGATACAGCTTTAATTTGAGTAATAACACATGGCCCAACTTGAATTA

CAGTAACTGGAATTGATAAACCAGCTTCATCAAAGAATTGGGTCATACCTACTTTAGTAC

CGAGTATACCAACAGACACTAGATTTCTCCTTTTTGAGAGATTTATACTATCATTGTACA

TAAAAATTCTACTTCATAGTTACTGAAGTAAATTAAACTAACAGATGCATGATATTATTC

ATGACAACATAAAGGTTAAAAACTTTACCTATTTAATCTAAAATAGAAACAGCTTTTTGT

CTAGTTACAGCGTATATAAATTAGAAGTATGCTTATTAAATTTTTTATAAACTACTTAGA

TTTAACTGATATTGATATCCGTAACTTACTTGCAGTAATATCAATCAACATGTATAAGTT

TAATTGCCTAAACTAGAACCAGCTATTATTAGAAATAATTCAAGTATAGATAAAAGCGTA

TTTGAACTTTTATATATTGACAAATATAATGGAATATATTGAGGTTCGGTAACTACACCT

TTTTATTTTTACATTTATTGTGCAGTATATATACATCTAATGCAGTAAACATAAGAATAT

AACTGTATAGCAAATGTAATTACACAAATTAGAGAGGTATTAATGGGCAAAGTCGTTGGA

ATTGATCTTGGAACAACGAATTCTGTAATTGCTGTTATGGAAGGAGGTAAACCTACCGTC

ATACCGAATGCAGAAGGTTTTAGAACTACAGCTTCTGTTGTTGCATATACTAAAAGTGGA

GATAAACTTGTAGGACAAATTGCCAGGAGGCAAGCTGTTATTAACCCAGAAAACACTTTC

TACTCTGTCAAAAGATTCATAGGACGTAAACAGAATGAAATTTCGCAAGAGATTCGGCAA

ACATCATATAATGTTAAAACTAGTGGATCAAGCATAAAAATTGCTTGCCCTGCACTGAAT

AAAGATTTTGCTCCAGAAGAAATTTCAGCTCAAGTACTGAGAAAACTTGTTGAAGATGCT

AGTACGTACTTAGGTGAGACTGTTACACAAGCAGTAATAACTGTACCTGCTTACTTTAAC

GATTCACAAAGACAAGCTACTAAAGACGCAGGTAAAATAGCAGGCTTAGATGTATTGAGA

ATTATTAATGAGCCTACCGCAGCTTCTTTGTCTTATGGACTAGACAAGCAAAATAATGAA

ACAATACTAGTATTTGACCTTGGTGGGGGCACATTTGATGTATCTGTATTAGAAGTTGGA

GATGGAGTATTTGAAGTACTCTCAACTTCTGGAGATACACATTTAGGCGGAGATGACTTT

GATCAGCAGATTGTAGAATGGCTAATCAAAGATTTTAAACAGAATGAAGGTATTGATCTT

GGTAAAGATAGACAAGCACTTCAGAGATTGACCGAGGCTGCAGAAAAAGCAAAGATAGAG

CTGTCAAATTTAACTCAAACAGAGATCAATCTTCCTTTTATTACTGCCACACAAGATGGT

CCAAAACATTTAGAAAAAACTGTAACTAGAGGGAAGTTTGAAGAACTTTGTTCAAATTTA

ATAGATAAATGTAGTATCCCTGTAAATAATGCTCTGAAAGATGCAAAACTAGAAGCTTCC
AGTATTGATGAAGTTGTTTTAGTTGGTGGATCTACAAGGATTCCAGCCATACAGCAAATG
GTTAAAAGATTAATTGGTAAAGATCCAAACCAAAGTGTCAATCCAGATGAAGTTGTTGCT
ATTGGTGCAGCCGTACAAGCTGGAGTTTTAGCAGGCGAAGTCAAAGATATTCTATTACTA
GATGTGACGCCGTTATCTTTAGGAGTGGAAACTCTGGGTGGCGTGATGACAAAGATTATA
CCAAGAAATACTACTATTCCTACAAAAAAATCAGAAGTATTTTCTACAGCTGTAGATAAT
CAACCAAATGTGGAAATTCAAGTACTTCAAGGCGAAAGAGAACTAACAAAAGACAATAAA
AGCTTAGGGACGTTCCGATTAGATGGCATTATGCCTGCACCTAGAGGAGTTCCTCAAATT
GAAGTTACCTTTGATATTGACGCTAACGGAATTTTATCTGTAAAAGCCAAAGAAAAGGCT
ACTGGTAAAGAGCAATCAATTACTATATCTGGAGCATCAACTTTACCTAAAGATGATGTT
GAAAGAATGGTAAAAGAAGCTGAAGAAAATTTCGATACAGACCAAAAAAGAAGAAAGAAT
ATTGACACAAAGAATCAAGCAGAGTCCCTATGTTACCAAGCTGAAAAGCAAGTTAAAGAG
TTTGAAGACAAAATTAGCCAAGATTTAAAAATAAAAATAGAAGAGCTAATTACAGAGCTT
AGATCTAGTCTAGAGAAAGAAGAATATGACAATATTGAATCTATTTCTCAGCAATTACAA
AATGCTCTGATGGACATTGGAAAAAATGCTGCTCAGACTGAAAGTAAAGATACAAAAGCG
AAGGATGACGACACTGTGATTGACACTGATTTCTCTGAAGCTAAGTAAAAGTAAGCGGGT
AACGCGATTCGAACGCGCGACATCAACCTTGGCAAGGTTGCGCTCTACCACTGAGCTATA
CCCGCATAGATTGTATTATTACAAAAATTAAGATATTTGTCAATCTGTTAATCTATACAT
CTAGACTATTAATTTTGGAATTACATATTAGCTAACTGTTGTAAATATAACAATTGCTAT
AATTTTTTAAAAGTACTTATATCATTATATAAGTACTTTTTATATTCAAAAATACAAATA
AGTATTATAGCTAAACTACTAAGGAATTAACTAAGCTAGTAACAATTACTAATCCTGTCC
ATAAGCCGGCTCCTGTATATATTAATCCTTTTGATTGCTCCCATTGGCCAGGGGATGCTA
AAACAACAGGTACGCCTACTACTAACACAGTCGATAATGCAATTAATAATAGCACTAATA
ATTGGATCGCAATAATCATTTAATATGTTCTCCTGAAAAATTCAAATGTTAATAATACTG
TATATAATACAATGAAGCTTTTGACCTTCAGCAATTTATCTATGAAGAAATAATTTAAAA
TTATTGTTTTCTAAATAAACCTTAAATTAAGTATTAAAATCTTTATTGTAAACAATAATA
CTATATTTTTTTAGTAGATAAAAAATTTAAAGAAAGTCTTTTTTAAAAAAAAAATTAAGAT
AAGGTGATATGCTGTATAAAAAGCTAAGTATTATTAATTATTAGAGAAATTTAAAATAAG
AGGTAAAACATATGAACTCAGCCCTTTTTTTAGCAAAACTACCAGAAGCTTATGCCATTT
TTAAACCGATCATTGATATCTTACCCGTAATTCCTGTATTTTTTCTTCTTTTAGCTTTTG
TTTGGCAAGCAGCAATTGGTTTTAGATAACATTTAGGTTATAAAAAGAATAACAAATATA
CAGAGTGCCAATTTTTGAGCTCTAATCTTATATCTTACACAATAATAACTATTATGGTTG
AACCCTTACTATCAGGAATTGTTCTTGGACTTATTCCTATTACTTTATTCGGATTATTAG
TAGCAGCTTATTTACAATATCAACGGGGCAACCAATTAGGACTGTAAAGCCTAAATTATA
GATATTTATAAAAACTACAATAATTATAGTTTTTATAAATATCTACAAATAAGCTAATTT
TAAATACAGAAATAATACTATTGAAACTAGAGTGATAACAAAAAAAATTTATGTAATAAC

AAGCTAATTGTTTACCAGCTTGACTTAGTAACTCCTGGTAAAAGACATTCATGAGACATT
TCTCGAAAAACATGGCGAGATAATCCAAAATCTCGATAATAACCTCTACTTCTACCTGTT
AACCAACATCTATTTCTCCCTCTAACAGCGGCGCTGTTTCTAGGCATTTCTTGGAGTTTT
TGTCTTAAATCCATTTTGTCAGAAAAGCTGCCAGCTTCTTTCAGTTGATCTTTAATAGCC
AGTCTTTTTAAATAATATTTTTTTGCAAGTTTCGATCTCTTAATCTCTCTTTGGATCATA
TTCTTTTTAGCCATGCTGTAAAACTCTCTTTTAAAATTATTTATTCAATATATTAGCTAA
AAAAGCTCCTGCCTGCAACTTTTATATATTTAAAAATAAATAGCATAAAGTAGTGAAATG
CAATTGATATTTAATCAACGATACCACTTTATGCTATTATCAAAAATTTATTTAGTAAAG
TTCTTGTTCAACATGAGTTGAAATAGTGCAATCAGATGAAGGATATGCAATACAGGTTAA
AACATAGCCAGCTAAAAGTTGATCATCATCTAAGAATGATTGATCAGCTTGATCTACAGT
TCCTTCTGTAACTTTACCTGCACAAGTTGAACAAGCTCCAGCTCTACAAGAGTAAGGAAG
TTCAATTCCTTCTTCTTCTGCTGCATCTAGAATATATGTATCCTCCGCACAATTAAATGT
GACATCAATTCCTTCTTCTTCACATAGTAAATGAATTTTATAATCAGCCATTTCTTCTTA
CTCCTTAATAAAGATACACTTGAAACTTAATAAAGCAAAATAATTTAAAGAACTTACCAA
AAAATTATATACTAGTTTCATTATGCATGATAGACTGGAAAAAAAGAAAATAGTATACTA
TTCTGAAAAATCAAGCTACATTTCCTACTAAATAGAACTATAGTAGGTCCAAAGTATATA
CATAAATTCATATAGTTAAAATTACTTTTATCATTTAAATAGCTGAACTTGATAGTATAG
ATTGATCATCAAGCTAGTCTTCTGCCAATAAATAATTAATTATGATTTCAATGCTTAATC
AACAAGTAGAACTTAAGCCTATTCTTAAATTTCAGATAACACAAATATATTCGCATTCTC
TTATTCAGGAAATTAAAGCCTTAATAACTAGACTTGTTTTGCAAGTTTGGAGAAGGCCGG
CTACATTGATGGCAGGTATTATCCAACCTTTATTATGGTTAATTCTATTTGGTGGGCTTT
TCTACAATGCTCCTATAAATTTGTTCACTATTAATACAAGCTATAATTGTTTTTTGAGCT
CTGGGATTATAATTTTTACCTCTTTTACTGGAGCTTTGAATTCAGGTCTTCCATTAATGT
TTGATAGAGAATTTGGATTTTTAAATAGATTATTAACGGCTCCTTTAGTATCGAGGACTT
CTATCATTTTATCTTCTGCTACTTTTATGACTTGTATTAGTTTAATACAAGTTGTATTTA
TAGTTACAGCTTCTCTTTTTATGGGAAACTCACCTCTAAATAGCGATAGTACTATGATTT
TTGGACTTATGATTCTATTAGTGACTGTAGGAGTTACAATGCTTAGTTTAGCTTTATCTT
TCACTCTGCCAGGTCATATTGAGCTATTAGCATTTATTTTAGTAGTTAACTTGCCCTTTT
TATTTTCTAGTACGGCTTTAGCTCCTTTATATTTCATGCCGCCATGGCTTCAGTTAATTG
CAAGTCTCAATCCATTGAGTTATGCAATAGAAGGTACAAGATACTTATATTCAAGCGTAA
ACTGGAATTTTACAGAGTGTGTGATTAAGATTAGCTGGGGAGATATTTGTTTAGGGCAAA
TTATTATTTTATTAATCGCTTTAGATATAATGGCAGCTTATCTTGTGTCTAATATATTAA
AAGCTAAACTTAATTAAAATTTTAATAAAAAATTTATTAAATTTATTTTACAAAGATTTT
TACTATATAATACTAATAGTAGTATGGATAATTGAAATAGAAATAGTTTTTCAAAACCAA
AGCTATTATCAACAATGATAAACATTTGTAAGAAAGTCAACAAAGTATGTTCTTATTCAT
AGGAGGCATGTAGTCAATGGGACTACCATGGTACCGTGTACACACGGTTGTTTTAAATGA

TCCTGGACGGTTAATTGCAGTCCACCTAATGCATACTGCACTTGTAGCGGGTTGGGCAGG
ATCTATGGCATTATACGAACTAGCTGTATTTGATCCTTCAGATCCAGTATTAAATCCAAT
GTGGCGACAAGGCATGTTTGTTATGCCATTTATGGCTAGACTGGGCGTAACAGATTCATG
GGGAGGATGGAGTATAACAGGAGAGAGCGTATCCAATCCTGGACTGTGGAGTTTTGAAGG
TGTAGCCTTAACTCATATAGTCCTTTCTGGCATGCTTTTTCTAGCTGCTATTTGGCATTG
GGTTTATTGGGATTTGGAATTATTTAGAGATCCACGAACTGGCGAGCCAGCCTTAGATTT
ACCGAAAATTTTTGGAATTCATTTATTGCTATCAAGTCTACTTTGTTTCGGATTTGGAGC
TTTCCATGTAACTGGACTTTTTGGACCAGGAATGTGGGTATCAGATGGGTACGGAGTAAC
CGGAAAGGTATTACCAGTAGCTCCAGCATGGGGACCAGAAGGATTCAACCCGTTTAATCC
TGGAGGGGTTGCATCTCACCACATTGCCGCAGGTACTGTAGGTATATTAGCTGGTGTTTT
CCATTTAACTGTTAGACCACCACAAAGACTGTATCGTGCTCTAAGAATGGGTAATATTGA
AACTGTATTATCAAGTAGTATCTCTGCTGTTTTTTTCTCAGCTTTTGTGACTTGTGGTAC
GATGTGGTACGGTTCTGCAACTACGCCTATTGAATTATTTGGTCCAACTAGATATCAGTG
GGATAGTGGATATTTTCAGCAAGAAATTGAGAAACGAGTAGAAAATGCTATTGCTGATGG
TGCTGCACCTAGCGAAGCATGGTCAAGAATTCCTGACAAGTTGGCATTCTATGACTATAT
TGGTAATAATCCAGCAAAAGGAGGATTATTCAGAGCGGGCCCTATGAATAAAGGCGACGG
AGTTGCTGAAGCATGGCTTGGACATCCTGTATTCCAAGATAAAGAAGGAAGAGAGCTTAG
TGTTCGCAGAATGCCTGCTTTTTTCGAAACTTTTCCTGTAATTTTAGTTGATAAAGATGG
TATTATACGAGCTGACATTCCATTTAGAAGAGCAGAGTCTAAGTATAGTATTGAACAAGT
AGGTGTAACAGCTAGTTTTTATGGTGGCAAATTAAATGGCCAAGTTTTCAACGATGCTCC
TAGCGTTAAAAAATATGCAAGGAAAGCTCAATTAGGTGAGGTATTTGAATTTGATCGAAC
TACATTAGAATCAGATGGAGTGTTTAGAAGTAGTCCTAGAGGCTGGTTTACATTCGGTCA
TGCTAATTTCGCTTTAATTTTCTTCTTTGGACATCTTTGGCATGGTTCAAGAACTATCTT
CCGGGATGTGTTTGCGGGAATCGGAGCTGAAGTTACTGAACAAGTTGAATTTGGAGCTTT
CCAAAAATTAGGAGATAGAAGTAGTAAAAAACAAGGAGCTGTATAAGACCAAAGTCAAAG
ATAGTCTTTATCAAATATTTCTAATATTAAAATAGTTAAGACAAAGTTTATTATCTAATA
GTTACTAATCAGGAGAATTTATGGAAGCCTTAGTCTATGTATTTTTACTAACAGGAACGC
TAATGGTTATATTCTTTGCGATCTTTTTTAGAGAACCTCCAAGAATAGCAAAGTAACTTG
ATTCCGTTTAAACTTATAATAATAGAACCACTTTCCAACTTATTAGTGGAAAGTGGTTTT
ATTGTAGATTACATTGTATTATTCTTCATGCTCTTCAAAAGGATCTCGGAGATCTTTTGA
AGCAGGACCAAATGCTGTGTATATCGAGTAACCTGTAATACCTAAAAGCAGACTCGAAAT
AAAAATACTAAGAACTGTTGCAGTTTCCATAATTTAAGCAGTATTAAGATGAACTTTTTC
TATAATGATATTATCAATATTAAAACATTTAATTATTAAAATTATCAAAATATACTATGG
CACTTAGAACTAGACTTGGAGAAATTTTAAGACCCTTAAATTCAGAGTATGGAAAAGTTG
CTCCAGGTTGGGGCACAACTCCTATTATGGGAATTTTCATGTTACTATTTTTCTTGTTTT
TATTAATCATTTTACAAATCTATAATTCTTCATTAGTATTAGAAAATGTAGATGTAGATT

GGGCTACTTTAGGTAGCTAACTAGATAAGTTCTTTGATTACACAATATTAAAGACAACAA
GCGATCGCTATTAGATTATAGTAGCGATCGCTTTAAAAGTATTTTACTATATATCCAGAG
AAAATTATTTACCTAAATTAATTAACTCGGTATCTGCAAAACTATTTGTGTTAACATTAT
TATAGTTAACTTTTTCAAACCTTACTAATACTGGATATTTGATACCACTTTTATCAATTG
CTGCAACAGTACCAATTTCCTGATACCAGTACGATTCTTTTCTCAATATTTTTACTTTTG
AGCCTCTTTCCATATTTTTAGTCCTCAGATAGATTATTAAAGATTATTATATTAAAAAAT
TAAAAAAAACTAAACAAAATATATAAAAATACTACCGACAATTGTAATATATAACTAATA
AAAGGAGAGAGAGGGATTCGAACCCTCGATAACAAAAGTTATGACAATTTTCGAAATTGT
TGCAATAAACCACTCTGCCATCTCTCCTGAATTAAAATACTGTTTTAAAAAATTAGACAA
GAATAACATAGAACTAAGGTAGTTGCCTAGAAAATTAAAACTATGTTAAATATAAATTAT
AATCTTTCTATAGATATATTTAAAAAATATGTACATAGAATCTATATCAGCTTACTTCAA
GATTTTTTAATCTTATACAATTTTTACTAAATAAATAAATTCTCTCATAATATGAAACTA
TCTTGGAAAACACTTCTTCTTTGGTCTTTACCAATATTTGTAATTGGTTTCTTTTTCTGG
CAAGGTTTTTTAGGTCCAACTACTACAGATGTTGGCAGTAATATCGCAAGTTCTAGAATG
ACATATGGACGATTTTTAGAATATTTAGATATGGGTTGGGTGAAACGGGTTGACCTCTAT
GAAAATAATCATACAGCAATTGTAGAAGCTGTTGGGCCAGAATTAGGAAACAGGGTTCAA
CGAATTCGAGTTGAACTGCCAGCAAGTGCGCCAGAATTAATTACAAAATTACGCAAAGCC
AATGTTGATCTAGATGCTCACCCCCCTAAAAGTACAAGTGCAGTATGGGGACTATTAGGC
AATTTACTATTTCCTTTACTATTAGTTGGCGGGTTAGCTTTCTTATTTAGAAGATCTAAT
AATGCTAGTGGTGGACCTGGTCAAGCAATGTCATTTGGCAAATCGAAAGCTTTATTTCAA
ATGGAGGCTAAAACGGGAGTAGTATTTAATGATGTAGCTGGAGTTGAAGAGGCAAAGGAA
GAATTTCAAGAAGTGGTAACATTTTTAAAACAGCCTGAATCATTTACTGCTGTTGGTGCA
AAAATACCAAAAGGCGTTTTATTAGTTGGACCTCCTGGCACAGGCAAAACATTACTAGCA
AAAGCTATTGCTGGCGAGGCTAGTGTTCCTTTTTTTAGTATCTCAGGCTCAGAATTTGTA
GAAATGTTTGTTGGTGTTGGCGCTTCTCGTGTGAGAGACCTATTCAAGAAAGCAAAAGAC
AATGCGCCTTGTATCGTTTTTATTGATGAAATTGATGCTGTTGGTAGACAACGAGGAACA
GGTGTTGGAGGTGGTAATGATGAAAGGGAACAAACATTAAATCAACTATTGACTGAAATG
GATGGCTTTGAAGGAAATACTGGTGTTATTGTAATTGCCGCTACTAACAGAGCTGATATT
TTAGATTCTGCATTATTAAGACCTGGAAGATTTGATAGACAAGTTTCTGTAGATGTACCA
GATTTTAAAGGCAGGTTAGCAATTCTTGAAGTTCATGCTAAAAATAAGAAAATGGAACCT
AAAGTATCTTTAGAAACCATTGCCAGAAGAACTCCCGGCTTTTCAGGAGCTGATTTAGCT
AACTTACTAAATGAGGCTGCTATCTTAACGGCTCGACGAAGAAAAAATGCAATGACTATG
TCTGAAATTGATACATCAATTGATCGAGTAGTAGCTGGGATGGAAGGCACTCCTTTAATT
GACAGTAAAAGCAAAAGATTAATTGCGTATCACGAAGTGGGTCACGCAATAATAGGCAGT
TTATTAGAGCATCATGATCCTGTGCAAAAAGTTACATTAATACCAAGAGGGCAAGCAAGA
GGCTTAACTTGGTTTACTCCTAGTGATGATCAAAGTCTAATATCAAGATCTCAAATACTA

GCCCGTATCGTAGGTGCTCTTGGTGGCAGAGCTGCAGAAGAAATCATTTTCGGTGACGCA
GAAGTTACTACTGGTGCAAGTAATGATTTACAGCAAGTAACATCAATGGCAAGACAAATG
GTCACTCGATTTGGAATGTCTAAAATTGGACCTTTATCTCTTGAAAGCCAAGGAGGAGAC
CCATTTTTAGGTAGAGGCATGGGAGGAGGCTCAGAATATTCAGATGAAGTTGCAACTAAT
ATTGATAAGCAAGTAAGGGAAATTGTCAGTGAATGCTATGCACAAGCTAAACACATTATT
ATAGATAATCGAGTAGTGATAGATAGATTAGTTGATTTACTAATTGAAAAAGAAACAATT
GAAGGCAATGAATTTAGAGACATCGTTAAGGAATACACTGCAATTCCTGAAAAAAATTAC
TACATATCACAATTTTAAATTAAACGGGACTGACGGGATTCGAACCCGCAACTTCCGCCG
TGACAGGGCGGTGCTCTAACCAGTTGAACTACAGTCCCAAAAAGATGAATACCCAGAGAT
AATCTCATAGTTAGCTTAAACTTGTCAAATTAGAATATTAAATATGAAGAGTATTTGTAA
TAATACTCTTCATATTTGATATAGTAGCTAAAATATTTCTTTTAGAAGGAGAGAGAGGGA
TTCGAACCCTCGGTACGAAGTTAATCGTACAGCAGATTAGCAATCTGCCGCTTTCGACCA
CTCAGCCACCTCTCCATGATATATATTATATATATATCAAATATACTTATATTTTAATAG
GTGTGCATAAATATTTATGGAGCTAAGCGGATTCGAACCGCTGACCCTCTCAATGCCATT
GAGATGCTCTACCAACTGAGCTATAACCCCTTTGGTAAGATATTAAGAAATGGCTCAAGC
GGGATTTGAACCTGCGACCTTGGGCTTATGAATCCCCTGCTCTAACCACTGAGCTACTGA
GCCATAGTATAAAATACTATTAAATATTATACCATAAATTATGGTCAAATATAAATCAAT
AAAAGAATAATTACTATCATTTACTACTTTTGTGCAAGTTGTAAATATTCGAATAAAATT
TCAAAATTAAAAATGTCTTTAACTTTTTGTTAGAATTATTGATTACGAAAATATTGATAT
TTAGAATTAACAATAAATTTTGTCTTTATTTGTAGACACAAAAATATTGGAAATTTTCAT
CAATTAATTAGTTAAACTATAACCGACGTACAAGAATTATTGAACTACGTTAAATTGTGA
TGCATTGGGCTTGATCAAAACTTCTCTAAAATCAATAGAATATAGTAATTCAAGTAAATC
AAATACTAACAAATGTGACAAACGCTTAAAAAAATAATACAATTAGAATCGTAAAAATTT
GGTTTTTGACTGTGTTAAATAAATTTGATTATCATAGATTTAATATTTAAAAGTGTTCTA
TTACTTGTATTGAGTAATTGTAATATTTAATATCTTCTCTTTTAGATAATAATCATAATA
TAAGCTAAAAAATAGTTTTTATTAAACAACTAGCATAGAGCCAATACCTTGATCAGTTAA
TATTTCTAGCAAGAGAGCATGATCTATTCTACCATCTAAAATATGCGCTGAAGCTACACC
CTGAGCTAAAGAACGAATACAACAATTCACTTTAGGAATCATGCCACCAGAAATTACTGC
TGTTTGAGTTAAATCTCTAGCTTCCTGTATATTCAGATGACTAATTAATGTTGAAGGATC
TGAAGAATTACGTAAAATGCCGGGCGTATCTGTGAGTAGAATTAGTTTTTCGGCATTAAG
AGCGGCTGCTATTTCTCCTGCGACAGTATCAGCATTAATATTATACGATTGGCCTTCCTT
GTCTGCTGCAACACTAGCTATAACAGGTATATAATTATTATTAATTAATATCTTCAGTAA
TTTAGTATCAATATTTTGTACTTCTCCAACAAAGCCAAGGTTTGCTTTTCCATTTGGTCT
CGGAGTAATAAGTAATCCATCTTTTCCCGACAAACCTACACTTTTACCACCTTGTTTATT
AATAGTTGCGACAAGATCTTTATTGACTCTTCCAACTAAAACCATTTCCACTATATCCAT
GGTAGGTTGATCTGTTACTCTAACGCCATCATCGAATTTTGGTAATATTTTGAGGCGATC

TAACCAAAAATTAATTTCTGGTCCTCCACCATGAACTAAAATGGGACGTAGTCCTATAAA
AGACAGAAAAACAAGATCACTAATCACTTGATCTTTTAGTTTCTGATTTTTCATAGCCGC
TCCCCCATATTTTATAACAATAATTCTGGAAGAAAATTGCTGAATATAAGGCAGAGCTTC
ACTTAAGACTTTTACTCTTTCTGAATTAGTCAACATAAATAAAGTAGGATTGTGATATAT
AAAATTATATACACAGCTTGTGTTAATTTAAAAGATACCAATGGGTACTTGAAAAAATAG
AAATATTACTAACAGTTAAAACTATAGGGATTATAATTATGAATAAATTTTGGGATAATG
TATTAAGATTTCCACGATTTTTAGTCAGCGTCATTTTAGGATTAATTTTGATAATTATTA
GTCCTTTTTTCGTGTTATTAAAAAAACCACTGACAAGTTTTTTTTTCATCATATCATTGG
CCGGATTAATCACAGTTTTGGCAATAATAATACAAAAAATGATAAATATCGAATGTTGTT
GAGAGAAATAAATAAATTTAATTAAAATTTTTGAACATATATAATATTATATTATCATTA
AATATGATAATTTATTTCCTTTTCTATAAAACCCATAAATATTAAATATTATTTGATAAA
TATTATGCCACTAAAACAAAGAGTTAGCTCTGAAAAGACCGGTGCCTTCGCACTTTTGGA
TAGTATTGTTAGGCACGGAGTTAAACATATATTTGGTTATCCAGGTGGAGCTATTCTTCC
TATTTATGATGAGCTTTATGCTTGGGAAGAAGCCTCTCTAATTAAACATATCCTTGTTCG
TCATGAGCAAGGCGCTGCTCATGCTGCAGATTCTTATTCTAGATCAACAGGAGAGGTTGG
AGTATGCTTCGCTACTTCTGGCCCAGGAGCAACTAATCTTGTCTCAGGTATAGCTACAGC
ACATATTGATTCTGTACCTATATTAGCTATAACAGGTCAAGTTGGGAGAGCTTTTATTGG
TACAGATGCTTTCCAAGAAGTAGATATTTTTGGGATTACACTTCCTATTGTAAAACATTC
ATATGTAGTTCGTGACCCTAGAGACATGTCTAGAATTGTTGCGGAAGCATTTTTTATTTG
TAAACACGGTAGACCAGGTCCAGTATTAATTGATGTTCCTAAAGATGTAGGATTAGAGAA
GTTTAATTATTTTTCTGTTGAGCCCGGAAAAGTTAATATTCCTGGCTGTAGGCCAATTAC
CAGCCTAAAGTCAAGACAAATCCTTATGGCAGCTAAAATGATACAGCAATCTAGCCAGCC
ATTGTTGTATATTGGTGGAGGAGCCATAATCTCTGATTCACATCAAATTATTAAAGAACT
TGTTGATTTTTATAAAATACCTGTTACTACTACTTTGATGGGGAAGGGGATTTTTAATGA
GGATAGCGACTATTGTCTAGGGATGTTAGGTATGCATGGTACTGCGTATGCTAATTTTGC
AGTTAGTGAGTGCGATCTTTTAATTGCTTTAGGAGCTAGATTTGATGATAGAGTTACTGG
AAAATTAGATGAATTTGCTTGTAATGCCCAAGTGATTCACGTAGATATTGATCCTGCTGA
AGTAGGAAAAAATAGGATTCCTCAAGTTGCTATTGTCGGTGACGTAGCAGAAGTTGTTAG
TGAAATATTGAATTTATTAAAGACTTCTTTCCCCCCTTATCCAGAGCAGATTATATCTTG
GCAAGAAAGAATTAATCGTTGGCGTCAACAGTATCCTTTACTGGTTCCTAGAAAATCAAC
AAGCATTTCACCTCAAGAGATTCTTGTTGCAACTAATAAATTAGCCCAAAATGCTTATTT
TACTACAGATGTTGGCCAGCATCAAATGTGGTCAGCTCAATTTCTGAAAGTAAAAGCTAA
GCATTGGCTTTCAAGTGCTGGATTAGGCACGATGGGTTATGGTTTACCTGCAGCAATTGG
CGCTCAAGTAGCACATCCAAATGACGTAGTCATTTGTATTAGTGGTGATTCTAGTTTTCA
AATGAATATGCAAGAGTTAGGAACTATCGCGCAATACCAGTTACCAGTTAAAATTATTAT
TATTAATAATCGATGGCAAGGGATGGTTAGACAATGGCAACAAGCTTTCTATGGTGAAAG

GTATTCACACTCAAGAATGACAGAAGGAGCACCTGATTTTCAAAAGCTTGCAGAAGCTTT
TGGTATTAAAGCTTTTACTATTAATAATAGGCAAAATATGCAATCTGCTTTACAAGTTGC
TATTGATTATCCTGGTCCAGTTTTATTAGATTGCCAAGTTACAGAAAATGAAAACTGTTA
TCCAATGGTTGCTCCTGGAAAAAGTAATGCACAAATGATAGGTATCGCTAAACCGCAGAG
AGGTACTGCTTCCAACTATATTAATAATAGTGTTTAAAGTAAATAGTTAGTATTTTTTAA
ACAAAGAATAAAGTACATAAATGTAATTTTTTTATACTATTAAAAACTATGTAAGAACAG
CCTTTGACGAGAATTGAACTCGTGACCTTCCCCTTACCAAGGGGATGCTCTACCTCTGAG
CCACAAAGGCTTTTTACTATATTGGGCCGGGTTGGATTTGAACCAACGTAGGCGAAGCCA
GCGGATTTACAGTCCGCCCCCATTAACCACTCGGGCACCGACCCTAATTAAACCTTATTC
AAGATATGACCTATAAGATATCATAACTATGTTGCTAAAAACAAGTACTTTGATAAATAA
AACTGAATAAGTAATGGACATAGCTGGACTTGAACCAGCGACTTTCACGATGTCGACGTG
ACACTCTAACCAACTGAGTTACATGTCCAAAACTTTAATTATTATTATAACACTGTTTAG
TGATTTTATAAGTATCTATTTCTAATTGAAGATTTGTTTTAATCGTCCGGCTTTACCGAG
CAGATCTCTTAGATAATACAGTTTAGCTCTACGTATTTTAGCTCTTCTAATCACTTTGAC
AAAAGTTATTCGCGGTGAATTGAGAAAAAATACTCTTTCAACTCCAATACCTTGAAATGA
GCATCTTAAAGTTAAACTAGTATTTAAGCTTTTTTTTCTTCTTCTCGCTAATACTACTCC
TTCACATAATTGTTCCCGAGTTTTACTACCTTCCTTAACCATTAGTCCAAGTTGTATTGT
ATCGCCAACTTTAATTTCAGGTACTTCTGTTTTGATAAAAGGTATCTCAACACTTTTCAT
TAGCTGACTTAGCTTTGTACTATTTGCTTTCATATATAGGATTAGTCTTGTGACATGTTT
TGAAATATTATACATTAAAAAAACAAATATGAAAAATTTTAGCGAATAAGCATACTAGTA
TTTATAAAACTCTTTTTATCTCCATTATACAGATCCATTATATTCAACTAGATTCTTTTA
TAGTTTCTTTGATAATCAGTACATCTAGATGATCAAATATTATTTTCAAACATGGCTTAT
GTATGAAAAAGTTCTATAAAAACCTAATGACTAAAATTAGGGCTTAAGTATTTCTTAGTT
TTGTGTTATAATTAGTTACTATCAAAGGTAGTGGCATAACACGAAGCATCAAAATGTTCG
AACGCTTTACTGAAAAAGCTATAAAAGTCATAATGCTAGCACAGGAAGAAGCTAGACGCT
TAGGACATAATTTTGTCGGAACTGAGCAGATACTATTAGGATTAGTTGGTGAAGGTACTG
GAATCGCAGCTCAAGTTTTGAAATCGATGAATGTGAATCTAAAAGATGCAAGAGTTGAAG
TCGAAAAAATTATAGGAAGAGGGTCTGGTTTTGTAGCGGTTGAAATTCCTTTCACTCCTC
GAGCAAAAAGAGTACTAGAATTATCTTTAGAGGAAGCACGTCAACTAGGCCATAACTATA
TTGGCACAGAACACTTGCTAATGGGCTTAGTCCGAGAAGGAGAAGGAGTTGCGGCAAGAG
TTTTAGAAAATCTGGCAGTCGATGTTTCTTCAATTAGAGCTGAAGTTATACAAATGCTCG
GAGAAAATGCGGAAGCCAATGTAAGTGGAAGCAATACTACGCAAGCTAGAAGTAAAACAC
CAACATTAGAAGAGTTCGGATCTAACTTAACTCAAATGGCTATGGAAGGTGGTTTAGATC
CTGTAGTCGGAAGACAAAAGGAAATAGAACGAGTTATTCAAATCTTAGGTAGAAGAACTA
AAAATAATCCTGTCTTAATTGGGGAGCCTGGTGTAGGTAAGACAGCGATTGCGGAAGGAT
TAGCTCAAAGAATTGCTAATAGAGATGTACCTTCTATTTTAGAAGATAAATTAGTTATTA

CTCTTGATGTAGGTCTATTAGTAGCCGGAACTAAATATAGAGGTGAATTTGAAGAAAGGC
TAAAACGTATTATGGATGAGATTAAATCAGCTGATAATGTAATATTAGTGATTGATGAAG
TTCATACATTGATTGGGGCTGGTGCTGCAGAAGGAGCAATAGATGCAGCTAATCTGCTTA
AGCCAGCTTTAGCAAGGGGAGAATTGCAATGTATAGGTGCAACAACTTTAGAAGAATATA
GAAAACATATAGAAAAAGATCCAGCATTAGAAAGAAGATTTCAACCAGTTGTAGTTGGAG
AGCCAAGTGTTGAAGAAACAATTGAAATTTTGTTTGGTCTTAGAGACCGTTATGAAAAGC
ACCATCAATTAACAATGTCAGATGGAGCTTTGGCTGCAGCTGCTAAATACGCTAATCAGT
ATATTTCTGACCGATTTTTGCCAGATAAAGCAATTGATTTAATTGATGAAGCTGGTTCTA
GAGTTCGTTTACTAAATTCTCAATTACCTCCTGCTGCCAGAGAATTAGATAAAGAGTTAA
GAGCTGTATTAAAAACAAAAGATGAAGCTATTAGAGCTCAAAAATATGAAACAGCAGAGC
AGTATAGAGCAAGAGAAATGGAAATTAAAGCTCAAATTGCAGCAATTGCTCAAAGTAAAA
AGAATGAGCCTGATTTAAATTTAGAAGATCCTGTTGTTACAGAAGATGATATTGCTGAAA
TTGTTGCTGCATGGACTGGTATACCAGTAACTAAGCTTACTAAAAGTGAGTCAGAAAAAT
TAATGCACATGGAAGAAACTTTGCATGGACGTATTATTGGTCAAGACGAAGCGGTTGTAG
CTGTCTCTAGAGCGATAAGACGCGCAAGAGTAGGTCTAAAAAATCCTAACAGACCAATTG
CAAGCTTTATTTTTTCCGGACCGACGGGTGTAGGAAAAACAGAATTAACAAAAGCTTTGG
CTTCTTATTTCTTTGGTTCAGAAGCTTCTATGATACGGCTAGATATGTCAGAATACATGG
AAAGACACACTGTATCTAAACTAATTGGTTCTCCTCCAGGATATGTGGGTTATAGTGAAG
GTGGTTATCTAACAGAAGCGGTAAGAAAAAAACCATATACTGTCATCTTATTTGACGAAA
TTGAAAAAGCTCATCCGGATATTTTTAATCTACTTCTTCAAATTTTAGAAGATGGCCGAC
TAACAGATGCAAAAGGTAGAACTATTGATTTTAAGAATACTCTTTTAATTATGACTTCTA
ATATTGGATCTAAAGTTATTGAAAAAGGAGGAGGTAGTTTAGGCTTTGAATTATCAGAAG
ATCAAACAGAATCCCAATATACTAGAGTACGATCTTTAGTAAATGAAGAACTGAAACAAT
ACTTTAGACCAGAGTTTCTAAATCGATTAGATGAAATTATTGTATTTCGTCAGCTTACTA
AAGATGAAGTTAGAGAAATTGCAGAATTAATGCTTAATGAAGTCTTTGCGAGAATTAAGC
AGCAAGATATTCAATTAAATGTAACAGAACGATTTAAGCAACGATTAGTAGAAGAAGGAT
ATAATCCAAGCTATGGAGCTAGACCACTTAGACGAGCAGTAATGAGGCTTTTAGAAGATA
GTCTGGCTGAAGAAGTTTTATCTGGTAAAATTAAAGCTGGTGATAGTGCAGTAGTAGATG
TTACTAATGAAGGAGAAGTTACAGTTTTATTAGGTGAAAAATTAGAACTGTTAACATAAA
AAACAATTTATTAATTAAAGCATTGAAGAGGTTATATTCAATGCTATGAAAAAAATTAAT
AAGATGGGTTGCTATAATTGACAACCTCTGATACTCTTTATTTGTATATAAAGCCGGGAT
AGCTCAGTTGGTAGAGCAGTGGATTGAAAATCCTCGTGTCACCAGTTCAAATCTGGTTCT
TGGCATTTTAAGCATCAGAAAATTTTGCTATCTGAGCATTAAATTTTAATTGAAATGTTC
CTGTCGGACCATTCCTGTGTTTTGCTATTATAACTTCTGTGAAGTCTCTACTTCTAGTTT
CTTGTGTATAGTAACTTTCTCTATATAGCATTATTACTAAATCTGCATCTTGTTCTATAG
AATTATGAACAATAAAGTTGTTTGAAATAAAATTGCATAAAGGTTTACATTCTAAATCAA

ACACTATTTGTAAACTTGTAACGTTTATTTTTTGTATACTTTCAAAAGTCAAAGATGAAA

AAGAATTAAAAACATTTTTTTGTTCTTCAAAGTTTTTTATTTGAATAGCAATCATGTCAT

TTTGATTAATTTTATCACATCTTTTCCATCCTTGTGTTGTGAGTAGTTTATGGTTACTTG

TTAATTGTATATATTTCCCAGCTTCGGTAATAATTTTATATACAGTTTTTTTCCCTGTTT

TCGAAATACTACATTTAATTGCTAGAAATAACTGTTGTTTTTGCGCACTAAAGTTAATGA

CTTCAATCTTTTTATAATAGCAATTAAATAATATTTGAAGTTGATTATATAATGATGTCT

GAAGATAATTGAATTTGGATATACACCCACTTTCTCTTAAATCGGATAACAAAGGCCTTT

TGTCACTCCGAGTTTCAAGGTTTCTATTTAGTTGAGATAATGCCAAGATAGGCAAGCTTA

AATCTTTTGCTAATATTTTAAGTGATCGAGTAATTAATGATAGTTCTTGAGATCTATTGT

CAGATTGTCTACTCTCCTGCAGCAATTGAAGATAATCTATAATAATTAATTCTATATTTT

TTCCTTGTAATTTAAAGAATTTTACTTTTGTCTTTATAGTGTCAGTAGAAATCTTAGCGC

TATCATCAATATAAAGATTAAGGTTAGCAAGAGTTTTACTTTTTTGAATAACATATTGCC

ATTCATCATTATTAAGCTGACCAGATTGAATTTTTTGGCCGTTTAAATTACACTCTTGAG

CTAAAATTCTTCTTAGTAGTTGCTCTGTAGACATCTCTAAGCTAAATAAAATAACATAAG

ATCTTTTCTGATTAATTACATATCTAGTTATATTAATAGCAAAAGCAGTTTTCCCCATTG

AAGGTCTTCCTGCAAGAATAATTAAATCTGACTTTTGAAACCCTTGTGTTATAGAGTCGA

GTTCTGTAAAACCAGAAAGTATACTACTATTTATACTAATCTTTTTCCTTGTATCCAAAT

GGACTAATAGGTTTGCAAGAATTGTAGCTAATGTTTGCGTATCTTTTTCGTCAAGTATTT

CATAAGCTTTTGTTAATTGACTAACAATAGAAGTTATGTTTTTTTGTTCAATAAGTTGTT

TTGAGCAGCTAATTAAACATAATGAATCTCCACATGCTAAAAGTAATCTTTTAATGTAGT

TATCTATAACGACTGCTGAATATTCGCTTATAGTATCAGATAGGGGAGCTTGTTCTATTA

GACTAAAAACTTCATTTAGCTCATTCAAATCTTGAATCATTTTTTCATTTTTTAAGTTAA

TGAAAAAATTTCTTACGCTTTTTGTCTTTGCATGGTTAACATTTTCGAGTGCTGCTCTAT

AAAGTAATGAAGTTGATGTAAAATAAAAAAAATCAGGAGAGATTTTTTCTACTGATTTTA

GTAATGATATCGATTGCTGAGTTAATATTATACTAATTAATATTTTTTCAGCTAAAATAT

TATGAGGAGGTAGATATTTATAAATACTTATTACTTCCTCCTTGTAACATTACTCTTGTG

GTATTAGTTAACTATTGATTTAATTTGATTCTGGAAGAACTTGCAATTTAATATTTGCAG

TTACTTGATGTAATAGCTTAATTTCTACATCGTAAATGCCAATTGTTTTCACGTCTGGTA

AAGAGATACTTTGTTTTTCAATGTCTATGTTAGTAGTGTTTTTAATAATTTGTGAAATTT

CTTTTTCTGTGACACTACCAAAAATATTGTGGCCATCTCCTGTTTTTTTACTAACGCTAA

ACCTTTGTATTTCTTCTAGAAGTTGCTTGACTTTTTTAGCATTTTCTTTAGCTGTTTTTA

ATTTTTCTTCTCTTATAGCTGCATAAAATTTTTGTTGTTTTAAAATACCATTTGTGGCAA

CTGCCGCCATTTTATTCGGGATTAAAAAATTTCTTGCATAACCAGAAGCTACTTTGATAA

CGTCATTACTCTTGCCAAGTTTTTGAATATTCTCTTTTAGCACAACATTTATAACTTTTT

TACTCATCTTAATTCTAAAAAATCTGTATTATCCTATTATACCTGTTGTATACAGATTTT

AATAGTAAGTATTTATTTGTTGATGTAATATTCTTGTAAAAAAAGGTAATTTTTTATTTT

TTTCTGATATAATCACTTTCACAATAATTCATTAGGAGAGGTGGCCGAGTGGTTGAAGGC
GCAGCATTGGAAATGCTGTTTAGGAGCAATCTTAACGAGGGTTCGAATCCCTCTCTCTCC
GTACAAAAAATATAAAAAGTTTTATAGAGTCCAATCTAAATCTCCAATAATATTCTTCCA
AGCATTTAATCCACCCCGAACTCTAACAATATTTAGTTTTTGAGCTATTAAAAACTTAGA
AGCAAATATTGATCTAGAATCTAAGCTACAGTAAACAAAACAAATTTTATCTTGTAAATT
GATATGAGAGTAATTCATTCCTTTTATTTTTTGTAAAGGTAGATTTAATGAATGTATTAA
ATGACTTTTATGATATTCTTCATGATTTCTTACATCTATTAAAATATATTGTGGATTCCT
ACTAATTAAAAACTTTTGTAATTGAATAACACTAATTTCTCGTACAAATGTACTAGATTT
ACTGCCATAATATTTGTTCCAATGCTTTTTTGTGGATAATACAAATTTAGTATTTATAAT
CTTAAATTTGTTAAATGATGAAGTTAATGAGTTGTACGTTAATATAGTTCCACTTAATAT
AGACTTATAACCCAGAACGATTTTGACAGCTTCAGTTGCTTGAAGTGTACCAACAATACC
CGGTAATAGACCTAAAACTCCGGAATTACTACAAGTATCTCTTGCGCTCTCTTTATTTTC
AGTTTCACTGTAGAAGTCTCGATAAACAGGTCCACCTTGATAATTAAATACACTAACTTG
TCCTTCAAATTGAAAAATGGCACCATAAATATGTATTTTATTTAACTCTAGGCAAGTATC
ATTCAAGAGATATCGAGTATCGAAGTTATCAGATCCATCTATAATAATATCATATTGTCT
AATAATGTCGATTGAATTCTCATAGCTTAGTCTTGTTTTGAACACAGTTACTATACACTG
TGGATTTAAATCTAATATTTTTTTTTAGCTATTTCTACTTTTGAATATCCTATATCATGC
ATCGTGTATAAAATTTGCCTCTGTAAATTAGAGAGATCAATTATGTCATCGTCTATAATT
CCAATGCTGCCAATTCCTGCAGCAGCAAGGTAAATGATTCCTGGAGATCCTAGCCCACCA
GCCCCAATGAATAAAACTTTTGCTTCTTTTAATCTTTCTTGTCCTTCTAATTGAATTTGT
GGCAATACTAAATGCTTAGAATATCTTGTATATTCTTCTAGTGAGTATTTTGTATTTTCT
GTTTTAAAATTTAGCATAAAAAATTTATAGTTTATTTAATAGCACTTAAAAGCTAGCTGAT
AAGTTTATTAAAATGACTTTTGTAAACTTGGTTCTTCTCTATCTAAAATAGCTCCTTGTG
TAGGACAGACCTGAACACAAATACTACAATCAATACAGGCAGAAAAGTCAATCCAATACC
AATTAGTGTTATTATTGTTTTTTCCCTTTCCTTGATGAATACAAGCTACGGGGCAAGCTT
CCACGCATTCAGAAACTCCAATACATTTTTCTGTTACAATTGTATGAGACATATTATTTA
ATTAATAAAAGATAAGCTATTTACATCTAGTAGTTTTGTTGTATACTAAATTTACATATG
CTATCCAGGCCAATATTTAAGGTAGTCTATTTCTACAGTTTACTAATTTGATATGCGCCT
TTAGTTCAGTTGGTAGAACGCAGGTCTCCAAAACCTGATGTCGAGGGTTCAAGTCCTTCA
GGGCGCGTTTAGTTTTATCGTAATTCAATAATATCATTTTATCTTTTTGCAAACATCAAA
ATGATATGATATAAAAGATCATATTTTATACATTGAATATTCTATGTACTAGTACTGTTT
TTTAATATGTTATTATTGATAAGTTAATTATACTAATTCTATTTTTGTAATAAGTAAAAA
CAGTTCTAATTAAGTATGCACGTATTTAATAGTCTTTTGTATATATAACTTGAAAAAATT
TAAACTATGGCTAAAAAAGTTACTGGAATTGTTAAACTAGCATTAAATGCAGGTAAAGCT
ACTCCTGCGCCACCTGTCGGACCTGCTTTAGGACAGCACGGCGTGAATATTGTTATGTTC
TGCAAAGAGTACAATGCACGTACTGCTGACAAGTCAGGACTAGTAATCCCTGTAGAGATC

TCAATATATGAAGATAGAAGTTTTACTTTTATACTTAAAACTCCTCCTGCTTCTGTACTA
ATCGCTAAAGCTGCTGGACTAAATAAAGGTTCGGGTGAACCAAATACCAAAAAAGTAGGC
AGCATAACAAATAAACAGTTAGAATCTATTGCAGAAACAAAACTTCCCGATTTGAACACT
AACAATATACCTCAAGCTATGAAAATAGTTGGAGGTACGGCTAAAAATATGGGAATTCTA
ATTAAAGATTAAAAGTTTCAATTATATTTACTATTTTTATGAAAAAATTTTCACGTCGAC
TTACAACATTAAAATCAAAAGTTGAGCCTAAACTTTACACTATTAATGAAGCAGTATCTA
TATTAAAAGCAACGTCAAATGCTAAATTTAAGGAAACTGCAGAAGCTCATATTGCTCTAG
GTTTAAATCCTAAATATGCAGACCAGCAACTCAGAGCAACAGTTATATTACCGAAAGGTA
CAGGTAAATTGATAAAAGTAGCAGTTATTGCTAAGGGAGAAAAATTAACAGAAGCAATTA
GTGCGGGAGCTGATGTCAGTGGCTCCGAAGAACTAATTGATGAAATACTGAAAGGTAGAT
TAGATTTTGATAAGCTGATAGCAACACCAGATGTTATGCCTTTAATAGCAAAGCTGGGAC
GAGTATTAGGCCCTCGAGGGTTAATGCCTTCTCCTAAAGCAGGCACGGTAACATTGGACG
TAGCAAAAGCCGTGAATGAATTTAAAGGGGGTAAAGTTGAATACAGAGTTGATAGAACAG
GTATAATTCATGTACCTTTTGGAAAATCTAGTTTTTCACAAGAAGATCTAGTTTTGAACC
TCCAAACAATTAAAGAATCGATTGATAGAAACAAGCCTTCTGGAGCAAAAGGGAAGTACT
GGAAAACTTTTTTCTTATCTAGCACCATGGGGCCATCTATTCAAATTGACATCACTAGCC
TTCTATGAAAAATTTTTGTATAATTATAGAACTTGAATAATCTATTATTTCAAATTATAA
TTTGCTAGCATTATTAACGTAACAACTTGCTTACTATATCTTATCTGCATTAAAAACTTT
TTATATATCTTATTAATTATGAGTACAAAGGTTGAAAATATCTTGGAAGAGCTAAAATCT
TTAAACCTTCTAGAGGCTGCTGAACTAGTTAAACAAATAGAAGAAACGTTTGATGTTGAT
GCATCTGCGGCTTCAGGAGGTATGATGATGGCCACACCAACTTCAGCACCAGCGTCTGCT
GAGGTTGAAGAGAAGACAGAATTTGATGTTGTCCTAGAGGAAGTCCCAGCACCTAAAAAG
ATTTCTGTATTAAAAGTTGTTCGCTCTCTGACTGGTTTGGGTTTAAAAGAAGCAAAAGAT
CTTGTTGAATCTACTCCAAAAGTCTTAAAAGAAGGTGCTTCTAAAGATGATGCAGAAACT
ATGAAAAAACAGCTAGAAGATGCTGGCGCAACAGTTATTGTTAAGTAATGAAGAAGCGCT
TCTGTTTAAGTATAAACAGAAGTGCCTCTTTATTTATTATAAGCAATTATACTAGAAAAG
TCCAAGTGTAATTGCATTGTTGATAGGCATAGTTGCACCAATACCTAGCCAAATACTTAC
AACGGTACCAATTAAGAAAACAGTGGTTGCAATTGGTCTTCTAAATGGATTTTGAAATTT
GTTAACATTTTCAATGAATGGAACAGTAAGTAAACCAGCTGGTACAGCAGCCATACTTAG
AACACCTAATAGTTTATTAGGAATCACTCTTAATAGATTAAATGTTGGAAAGAAGTACCA
TTCTGGTAAAATCTCTAATGGAGTCGCAAATGGATTAGACTTTTCACCTATAGAAGATGG
TTCTAAAATTGCGAGTCCAATACTGCAGGCAAAAGTGCCAATAATTACAACTGGAAAGAC
GTATAATAGATCATTTGGCCAAGCTGGCTCTCCGTAATAATTATGGCCCATTCCTTTTGC
TAATTTAGCTCTCAGCTTCGGATCTGTTAAATCTGGTTTTTTAAGAATTGACATATTGTT
TTTGTGTTTAATATAAAGATTAGTCTAATTACTGTTTTAAAGTGTAATTTTCAACTATTA
TAATGGTCCAGAAATTCCTTGTTTACGTATCATTAAAAAATGCATAAGCATGAAAACGGC

AGTTAGTAGCGGAAGTACAAAAGTGTGTAAACTATAGAACCTAGTTAAAGTTCCTTGGCC
TACACTAACTCCTCCTCTTAACAATTCTACTATACTCGCTCCGACAACTGGAACAGCATC
AGGGACACCTGTAACAATTTTTACAGCCCAGTATCCGATTTGATCCCATGGCAAAGAATA
ACCTGTTACCCCAAAAGAAACAGTAAGCACTCCTAAAATTACACCTGTTACCCATGTCAA
TTCTCTAGGTTTTTTGAAACCTCCTGTTAAATAAACACGGAATACATGTAGAATCATCAT
CAAGACCATCATGCTCGCTGACCATCTATGAATTGATCTAATAAGCCATCCAAAGTTTAC
ATCAGTCATAATGTACTCTACTGATGTAAAAGCTTCTGCAACTGTAGGTCTATAATAGAA
TGTCATTGCAAATCCAGTTGCAACTTGAATTAGAAAAGATACAAATACAATACCCCCTAA
GCAATAAAAAATATTAACATGAGGTGGTACATACTTACTAGAAATATCGTCAGCAATTGC
TTGAATTTCTAATCTCTCTTCAAACCAGTCATAAATTTTACTCATAAAATAGCTTCAAAA
AGCTCTTCTAATATATTTTGCGTTTAAGCTACTATTAATTCAATAGTTATTGAACTTATA
TTTTTTTATAATCAATACTTATTAACTATTATAACATCTTTTTATTGATTTAAAATTCTC
TGACTTAGTAAAACAGGGCTATGTACTATAATTCTTTTTTTTTGTATAGAGATTAATTTC
GTTTTTAGTAAATTAGCTAAAATTCTTGTTACACTGACTCTATTACTACCTATAATCTGT
GCTAAAATTTTATGTGTAATAGTAAAGTTAAGCATAATACCATTATTAACTTGAGTTCCA
TTTTGCTCAGCTAATAATAATAATAAGCTCGCCAGTCTACTGGTTATACTTTTATGTGAA
ATGATTTCTATAAAATGATAAGCTTTTGCCGAGCAAAATACTAAATGATTTACAAAAAAT
AAATTGAATGGAGCGCAGTTACTACATGCTTTTATAATAGTAGTATAGTTAATACAGGCA
ACTTCTGTTTTATCTATTGCTTCTGCTTCGTAGTAAAAATTATCGTCGACCAATTCTATT
TGTCCAAATGTATCTCCTGTAGTTAATAAGTTAAGAGTTACTTTATGTGTATTTCTCAGG
ACTTTTGTAATAATTAAAGATCCTATCAATATAATGTATAATCTAGAGCTGCTATTAAAC
AAAAGAGTATCATTTTTCTTAAGTGAAAAAATTTGATAATTAATTTTATTACGATTAAAA
AATAGTAGCCATGGATTGCAAGAAAAAAAATCACCTGAACTATTTTCAGAGAATATTGGA
ATACAATTTTCTTGATTTAAACAATTTTTAGTCATAATTACTACTTTATAAAAAAATATT
CTAATAATTAAGAACTGAAATTAAATGTCATATAAATTGATGTTAGTTGAGAATGACATT
GTATTATCAAAGGCCATTCAAGAATACTTAATAGATCAAGGCTTCAATGTTTATATTGCC
AATAATGGATTAGAAGCACTGAATCTAGCTTATCAATATAATTTCGACTTGATAATTTCT
GATATAATGATGCCGCTAGTTAATGGCTATGAACTATTAGCAAAACTTAAAAAAAACAAA
GCATTATCCAAAATACCTGTTATCTTTTTGACAGCTAAAGGAATGACTAAAGACAGAATA
AAAGGCTACGACATGGGGTGCTATGGTTATTTATCTAAACCATTTGATCCTAATGAGTTG
CTTTCAATTATTAACAACTTAATCGCTAGAGATGTCTTAAAAGAAGCTTCTTTACAAAAC
TCGGCAACATCAAACCAGCAATTAAATCATAAAATACGTTTGACACCTAGAGAAAAAAGT
ATTTTAGATCTTGTTGTTGATGGACTTACTAATAAAGAAATATCTACTATACTGAATACT
AGTGTTAGAAATGTAGAAAAATATGTTAGTCGACTCTTACACAAAACTAATATGAAGAAT
AGGACTTTGCTAGTCAAGTATTCTATAAATAATAATCTATTAAATAATGAGATCAATGAA
AGGGCGAATGACGGGACTCGAACCCGCGAATGATGGAGCCACAACCCATTGCCTTAACCC

CTTGGCCACACTCGCCATATTACATACACCACTATAGCTTGTTTTTTGATATATCGTCTA
GTTTTCGAGTTATATATTTATCTAAAACAATATTTACTTTGAATTTTTATGATACATAGC
AATATTAGTATTCAAATTAATGGGGAACCATTTAATTGCTCAAAGCCCATCTCATTGCAA
TTTTTATTAAATTATCTTGATTTTAATTCCGAGCGTGTAGCAGTCGAGCTAAATGACATT
CTTTTACCAGAGACTCTGTTTCACTCAACTTATTTGAATGATCAAGATAAGCTAGAGGTT
ATTACCATTGTAGGTGGTGGATAATCTATTTGTACATACACAGTTTTCTATCTCAAATGT
TTCATAGATAGAGATAATCTTCCTTGTTTTTTATCTACATGAATAATAACAGCTTTAATA
GTGTCACCTATTTTAAATTGTGATGATATCTGCTCTAATTCTTTGATATTAATCTCAGAA
ATATGGACAAGACCTTTGAGATTTCCTACCTTTATAAATAATCCGTAAGGTGTAATCTGA
TTTATTATGCCTTCAATAATATTACCAACAATTAAATTAGATGATGCTTGAGATATTAGA
GCTCTTCTATGACTCAGTATTAAATTATTAGATTTCTCTTCGACATTGAGTAATTTTAGC
TTAATAAATTTATTATTAAATTGCTCACTTTTTTGAAAATTACCGAGATGAGAATTAGGT
ACAAATCCAGATATACCTTCAAGGTTAATTATCATTCCTCCCTTATTGAATCCTTTTATC
ATTACATTAAGTAAAGAGTCTTCAGCTAATAGTTGCCTAATTCTTTTCCATGCTCTTATA
TATTCAAGGCGACGAATTGATAATATTAATTGTTTCGACTGAATATTATAATCTAATAAA
AAAAATTCTCTTGTGTCATTAATGTTTAAAGAAGTGAAATTATTTAAATCTTGATTACTA
GATACTTCTTGAATAGGCAAGTATGCAGATATTGGTGTTCCTATATCTACTAATACCCCA
TTTAATTCAAAGCTAAATATAGTTCCAGCTACAATATCTCCAAGATTCAAATCATACTTG
TATTTTTGTAAAACAGCTGCAAAATTTCTATGAGTAAATCCTTCATTATTTTTTGTCATA
TTGAATAGTATGAATTAAAGAGTAACTTTGTACAATATATTAAGGTAAATAAATGTTTGT
CGAATTGTATGAAAAAATTTTAATTTGTTCGGTTGTAATTAAGTATATAAACATATTAAA
ATATCTAGGAGACATGAAAAGTAGACGTAGATAGCTGCAGATTTAATACTATATAATCTG
AGGAAAGTCCGGGCTCCACAAATACAATTTATGCTGGAGAAATCCCAGTGTAGGTAACTG
CGAGGATAGTACCACAGAAAAAAACCGCCAGAATAAGTTATATTAGCTGGTAAGGGTGCA
AAGGCAAGTTAAAAGCTTACCAAAAGTACTGCAAAGTATTTGTTAGGTAAACCCCTAAAA
TGGAGCAAAGCTACTAAACAAATATTTGTGTATATTTTATCTAGTTTAGTTGAAAATACT
GCATGAAGTTATTAGTAACAATAACTCGAGAGGAATAGCTATCCTTTACAATATATCTCG
ATTATATTGAAAGAACAGAACCCGGCTTATGTAGTACTTTTTATGTCTTGTATTATTATG
TCTATTGTCGAATCTTTAAATTTAACTGTTTGACAATTTTTTTTTGTAGCTGATGTTGTT
TATAATCTATTTCTGAATTGGTTAAAGTTTTTTCATTAGATCGATATGTAAATCTTAGAC
CTATACTTTTTTTCTCTTTTCCAATTGATTCATCTATATATTGATCAAATAATGTTGTAC
TTTCTAAATCTTTATCATAAAATTGATCTAATAATTTTAATAAGTATTTAATTTCCATAT
TCTTAGGTATTATTAAAGAGAGATCTCTTATGATCGATGGATACTTAGAGTATGGTTGAA
TCTGATAACTTAAATAATTGAATTCATTATGACAATCTTCTAAAATGTTTAAGTTAACTT
CAAGAAAAAAAAGTTTTGTATTGAGTCCTAATTCACTATATGTTGCTTGATTAAGTTCAC
TGAATATGCCAATAATAATATTGTTATAGATTAGTGTGGCAAATTTTCCTTTTTGAATGA

ACTTTATTCTACCATCGACTATATCTTTTTTTGTCCACTCAATTTGTCTATTAATCTTTT
GAAAAAAATTTTCTACGATCCCTTTAGCTTCATACCAATTTAAAGAATGCGCAGGGTGTG
ACCATGTAGATCGGATATCTAGATTACCACCCAATATGATAGCTAGATTAGTAGTTTCTA
CAATTTTGTTTTGATCATTATGAAATACTGTTCCTATCTCAAAGCTATCTATAGTTTGGT
TGCTTTGTTTAATATTATAAAGATTTGATTCAATTAATCCTTCCAGCAAACTGCCGCGTA
AATTAGAATAATCTTGAATAAGAGGATTGTTTAAAGCTATATTTCCTTTGGACTTTATTA
AAGAATAATGAACTAATTCAGTTAATCCTAGATTTCTTAAAATACTTCTACTTTTGTCTA
TGAATTTTCTCTTTGTGGATAACCTTTTGTTGAATTGAATATTTGGTACTGAGCTCCTGA
ATTTATGATAACCATAAATTCTAGCAATTTCTTCAATAACATCTATTTCTCGAAAAACAT
CTTCTCGTCTATAATTTGGAACAGTAACTTCTATATTTTCCTTATTTTGATAAATAAGAT
CAAAATTTAAAGAGTGAAGAATATTTTGAATTTCTTCAAAGAATAAAAAACGTGTTTGAC
CATTATACATAATTGGTCCTAAAATGTCATGAACTTTTTTAATTGATAAATTAATATTAA
GAGCATGATCAGTTATTTTTTCTCGACAGAAAGTATCTCTAATATTTCCGCATGTTAAAT
CTGTGATTAAAGCAAGAGCTTCAAAATGTGCATTTTTCCAATTATCTACATTTAGCCCAC
GTTCTTGTCTAATTGAACTTTCTGTTCGAATATTAAGAACTCGAGAAGACTTTCTAACTA
CAGATTGTTTAAATATAGCAGACTCTACAAGTATCGATTTTGTATTATAATCTGTATCAA
AATCAGAGTTGCTTCCAATACCAGCTATACTAGTTACATTTGTATTTACTTGAGTAACTA
AAATATTTTTGTTTAACTCAATATTTTCATTGTTTAACTTAATCTGATCATTACTTCCAA
GAGGAAAATTGCTTCGAATTGTTAGACTATTTTTATGATTCATATTATTAATTTTATTCA
AATCAATAATATTAATAGGCTGTCCCCATTTCAACATAATATAATTGCTTATATCTGTTA
ACAAATTTCTGTGTGTAAACCCAGAAGAGAGCAACCGATTTTTTAGCCAATCTGGAGAAT
CTTTAACTCTTATTTCGTCTATAATGGCAGCAAAATAATTGTCACAATTTAATAAATCGT
GATTACTGATTATAGTCTCCTTTTTAGAGAAATATCTATTATTGGGTTGATGTATAGCTG
ATAACATTGAGGACCCTGTTAAAGCCGAGACTTCTCTTGATAGACCAATCATGCTTAACA
CGTCTGACCTGTTAGCTGTTGATGATATATCTAATATATGATCAATCTCACCTCCAATTA
TAATAGATTCAATAGCTTCAACTTCAAAACCTGCTTGTGTTAGTTGATTAGTTAAATTGT
CTATATCTATTGTTTTGATATTACCAAGTTCTTTCAACCAATTTAAAGAAACTTTCATAA
TTAATTGTCATCTATCTTATAGAAGTATTTATTTTACTTATGATTAAAAATCAATTTAAA
AAATTCATCAAATTAATATAAGTTATTTGATGAATTTGCTTATACAATATATTATTGTGA
TATCTTACTTTTATATGGCAGTTATTAAGAACTAGCTTTTGTAAAAGTTAGCCCATTATC
TTGGGCATATCTTTCCATAAATCTCATGAATCTATCCCAATCATCAGGACTTTTAATAAC
ATGTACAGCTTCTATTGCTTGTGGTTTACCATTAATAAACTTGGCATTAACATCTCTTGT
AATAATTTGTCCCTCTGTGTCCATCAGGTACATACCTGTGATTTCACCTTTGTCTGCCAT
ACTAGCATCTAAAATTTTGGGATTTGTAAACCGAAAAGTTGCTGTTCCTGTACTACCATC
TCGAGACCTTGTTAAACGTACATCTGGTACAACTTCTTCATTGATACCTTGAATAAATTG
AATTGTTGCCATATCAATCTTCTTTATTTGTATATAATTTATATTGGATAAAACTGATTT

CTAGTTCTTGAATGCTTGTTGAGAAACTGTTTTAAAAGCTTATAGTATCAGTAGATCAAA
TTTTATGTATTATATAAAACTGTAACTTTTCTTTTTTGATCTGTTCAAAGAACTAACTTA
TTATAAGCTAATCATACGAATATTAGCCATATAATGTTCGTAAGTGAAATTAAGGGGTTG
TAGCTCAGAAGGATAGAGCAAGCGCCTCCTAAGCGCTAGGTCAGCGGTTCAAGTCCGCTC
AACCCTGTTAACGAATTAAGTAGTATTATTTAACTTGTTCTGCAAATCAGTATTGATCTA
AGTTAAAGTACTTGACAGCTAAAACAAAAGAATCTGAGAAATATTTTCATTTAAAGAACG
AACAATTATCTTAACTCTAAATATCTGATAATTAGTAAAAAATTCTCTTTTTAAAACACA
AGATGATGAAGATATTTAAACTAATAATTGAAATTAAGTTTTTGAAGTTATAGAAGCCAA
TATAATACTTTCTATTAACAGATGCTTGCTTCTATTACTATCTGTTTACAAGAAATTATG
AGTATCAAAATAATTTTAGCTCCACTGACTACTAATAAATATTTATTTTAGTCTAGACAA
GATGAATCTTCATGGTATAACAGAGCTATTCTTTGTGTCATACTTATATAAAATTAGGAT
ACTGTTTTCATTACCATAACTGGGGTGGGAGGATTCGAACCTGCGAATGGCGGAGTCAAA
GTCCGCTGCCTTACCACTTGGCTACACCCCAACATGTGAACATTATAATAGCAGTCGTAT
ACCCTGCCATGTCAATAGACTATTGTCTTTATTTAATAATTTTTGCTGCTCTTGTAGAAT
AAAATCTTTTTTTTCAAATAAGATACGCTATGTTTAAGTTGCACGACTGTTATGTTTTCT
TGCTTTTGACTGAACAGCCACTTGATAGTAATTATTCCTTCTGCAATTTCGTTACTTCCT
ATAATAATACAAGCTACAGCTCTTTTTTTATTTGCTTGCTTAATCTGCTTACTAAAATTA
CTTGAGCTAACATCGATTTCTATTTTAAAAAACTGCTGATGTAAAAAACGCATTATTTGC
ATACCAGTTTCGTTAGCTTTTGTACCTTGTGTAGCAATATAAAAATCGATAGATTTATTA
GGCAAGTCTATATTATCTTTTGCAATCAGTAATAGGCGTTCTAATCCTATTGCACATCCA
ACAGCTGGTGTTGCTTTGCCACCTAATTGGTGAACTAAGCTGTCATATCTTCCACCACCA
CATATAGTATCTTGCCCTTTTGATGTCAATGTCTTAATTTCGAAAGCAGTATCATTATAA
TAATCTAATCCTCGAACTAACTTATTATTTATATTGTAGGGAATATTAAGTAATTTTAGA
TAATTGCAAACGTCACCAAAGTGTTGTTGTGATTCCAAACTTAAAAAATCAGAAATTTTT
GGAGCCTCTGTCAGTATTTTTTGTGTATTAGAATCTTTTGAATCTAAAATTCTAATTGGA
TTACTAGTTAATCTTTTTTGTGAGTCAGTGTCTAAATCATCATGGTACTTTGTAAGATAA
TCTCGTAATTTAACTTGATAAATACTACGATCCTCTACTTTTCCAATTGAATTCAGATCA
AGTTTCAAGTTGTGCAGATTGAGATTGTTAAATATGCTCATAGCTAAATGTATAACTTCA
CTATCTGCTCTTGCATCTAAGCTACCAATAAACTCAATGCCAAGTTGGTGAAATTGTCGT
TGTCGTCCACTTTGTGGTCTTTCGTACCTGAACATTGGACCGCTATACCATAATCTTTGT
AAACTATGGTGGTAACTCATTTTATTTTCAATAAAAGATCTCACAATACCGGCAGTACCT
TCTGGCCTTAAGGTAATATCTCTATTGCTGCGATCATGAAACCGATACATTTCTTTATTT
ACAATATCAGTATCTTCACCAATACCCCTATCATATAATTCACTATTTTCAAAAATAGGT
GTTCTAATTTCTTGGTAATTTGCACATTCTAATAATTTGGAAATTTTATTATGTATAAAC
TGCCAATATTGAATTTCATCGGGAAGAATATCTTTTGTCCCTCTAATAGCCTGAATTTTT
GCCATGATAATTTTTTGATTACCTTCTTAATATGTCTGCATTAAATAGCAGTTGTAGACT

GTTATTCAGTCATGTATTATACATTAGAACATACTGATGTAATTTATAAAGTACGGGCAA
GGAGGGATTCGAACCCCCGACACCATGGTTCGTAGCCATGTGCTCTAATCCACTGAGCTA
CAAGCCCACTTAATAAGTACCTTAACTATATCATTGTTCACTAATGTATACAACTATATT
CTTTTTATCTTACATGTGAAATTTTTGTAAAAGTAAAAAATTATGTTTGTTTTTTACAAT
TGGAACTATTCATATATAATGCTTAATTTATTATAACTTTGTAAAAATATCTATAAGTTT
AATTGTATGATTCTATTATATTGGTAATCTTGTTATAATTTTGCGCTCTTATCTTTCAAT
TGGCTAGCATCTAATTAACTTATATTTCAAATAAAAAAATCTTATCTAAAATGAGTAAAC
AAATTCTATATCAGGATGATGCCAGAAAAGCATTAGAAAAAGGCATGGATATTTTAACAG
AAGCTGTTTCTGTGACTTTAGGACCAAAAGGAAGAAATGTTGTCTTAGAAAAAAAATTCG
GTGCCCCTCAAATTATTAATGATGGTGTAACGATTGCAAAGGAAATTAGTTTAGAAAATC
ATATTGAAAATACCGGAGTCGCACTGATTAGACAAGCAGCATCTAAAACAAATGATGTAG
CTGGTGATGGTACAACAACAGCTACAGTACTGGCTTCAGCAATTGTTAAGCAAGGAATGA
GAAATGTTGCAGCAGGTTCGAATCCGATGGCTATTAAAAAAGGTATTGAAAAAGCAACAA
ATTTTGTTGTTAGTAAAATAGCTGAGTATGCTAAGCCAGTAGAAGATACAAAAGCTATTA
TACAAGTTGCTTCTTTGTCATCAGGAAATGATATAGAGGTAGGTAAAATGATAGCGAATG
CTATAGAAAAAGTTGGCAGAGAAGGAGTTATTTCTTTAGAAGAAGGGAAATCAACTAATA
CGATTCTTGAGATCACAGAAGGAATGCAGTTTGAAAAAGGCTTTATTTCTCCTTATTTCG
TTACAGATACAGAACGAATGGAAGTTCTTCAAGAAAATCCTTTTATTTTATTTACAGACA
AAAAAATTACTTTGGTGCAGCAAGAACTTGTGCCATTGCTTGAGCAAATTGCAAAAACAT
CTCGGCCTTTATTAATAATAGCTGAAGACATCGAGAAAGAAGCATTAGCCACAATTGTAG
TTAATAAATTAAGAGGGATTTTGAATGTTGTAGCGGTTAGAGCTCCTGGATTCGGTGATA
GAAGAAAATCTTTACTTGAAGATATGAGTATCTTAACGAACGGACAAGTAATTACTGAAG
ATGCAGGTTTATCACTTGATACAGTTCAATTAGATATGTTAGGAAAAGCCCGAAGAGTTA
TTGTTACTAAAGACTCGACAACAATAATTGCAGATGGTCATGAAATTAAAGTTAAATCAA
GATGTGAGCAAATTAAGCGGCAAATAGAGACGAGTGACTCTTTATATGAAAGAGAAAAAT
TGCAAGAACGATTAGCAAAGCTTTCTGGAGGAGTTGCCGTCATTAAAGTTGGTGCAGCTA
CAGAAACAGAGATGAAAGATAAAAAACTAAGACTAGAAGATGCAATTAATGCAACAAAAG
CAGCAATTGAAGAAGGTATTGTACCAGGAGGAGGAGCTACTAATGTCCATATCTCTAGTG
AATTATTTACATGGGCTAAAAACAATTTAGTTGAAGATGAATTAATTGGTGCTTTAATAG
TTGAACGAGCTGTGACCTATCCGTTAAGACGAATAGCTTTTAATGCAGGTGATAATGGAG
CAGTAATAGTGGAAAAAGTTAAAAGTCACGATTTTCATATAGGCTATGATGCAGCAAACG
GTAATATTGTAAATATGTACGATAGAGGTATTATAGATCCAGCGAAAGTAGCTAGATCAG
CTTTGCAAAACGCAGCTTCTATTGCAGCAATGGTTTTAACTACTGAATGTATTGTAGTTG
ATAAAGTCGATGATTGAGCTATAAGATAATAGATAATCTCAAACAAAGAATAATTTCTAA
TTATAGAAATTATTCTTTGTTTCAATTAATCTAATTAATTGATCAAATGTATAATTTTTA
TAATAAAGCCAAAGTTTAAACACGTCTTCTTTTTCAACTATTAAGTAGAGAAGATATAAG

CCCGTAATACCTAAATTATCAATTAAAATATTATTCTCATATATAGATTTTTCTAAAGAT

CTAAAAAATGACATTTTTGAAAAATAATACCGAAATCTTCTCTTATAGTTATTTGCTAAA

TCATTACTGTCTAAAGATTGATTTCCAATTGAATTCAGAACCTTTAACAGAATACTTTTT

AAATCTTTATCTGACTTTATTTGAGAATCAATAATAATTTGATTCAATTGTATGTCAAAA

CTAAGATTAAAAATATTACTTGTTAAAGATATTTGATTTTGAATATCTAGAGATTTTATA

GCCAAGAAGATTAAACTTTTATTGTTCATTGTAAAATATAAGACAATTGATGATGTATTC

TTTAAATACTAAATTTATCTTCTTGTGCTTCTATCGTTTTTTGTAAAATAATTAATGTTT

TAATTTTTTAGTTTTAGTTACTACCCGAAAAAATAAATTCTGGAAATCTAGCTTGAGCTT

TAGCTAGAGAGTGTTTTTTCCCTTCTTCTTGCCAGTAATTGATAACTTCTGTTGTTTTGT

TTAATAACTCAATTCTGTCTCTTTCAGCAATCCACGGCTTGGATTCTAGTTCTGCTTTTA

ATTCTTCCCAAGCATCATTCCTTGGCCAGAAAAAATAAGAAGTAAGTGGACTAGTGCCAT

TTCCAACAATCTGATCAATAGCAATAGCAATATTATTTTCTAACCAAAGTACTTTTAGTG

TAAATTTTGACAAGTTTTTATCTCCATTATTAATATTAAGTATTGGTACAGTAGAATACT

GTACTAGTTATTTTTTTCAGCAACAATTTGAGCTTTGTTTAAAATGTTTTTTACCGTTTT

AGTAGCTTGTGCACCTTGAGATAATCGTTGCAAAATTTTTGAAATGTTAACACGAGTTTC

ATTATTTATAGGATTATAAAATCCTAATTCTTCTATAGCTTTACCATCGCGTTTATTTCT

GCTATCCATTACAACAATTCTATAGCTTGGTTGTTTTTTTCTTCCGTATCGCTTTAACCT

TAACTTGACCATATTAGTCACCTCGCTTATTTTTTATATCTTTATTAAATTATATCTTAC

GACATTATGCAATAATTCAAACAATTAAGCTGATTCAAGTCTAACATTATTAAATATTAC

CATTAAACATTGTAGAAAAATAATACCTAACATAGGTGACATGTCCATTCCAAACATGGG

AGGAATGCTTCCTCTGAATAGTTTTAAGTATGGATCAGTAATTCTATTTAATGAACAGAA

TGGTTCATTGTACCAATTTACAGTTGGAAACCATGCCAATGATAATTTAAGTAAAATTAA

AATTAAATAAATCTCAGAAAAATTAGCTATCGATCCAAGCAATAAATTTAATGTACCAGG

AAGAGTGTTCATACGTATTTTATAATTATTATATACTTCGTTATACTAATAGTTTGATTA

TATATTATAATCGATGTAAATAAATCAGCTAAACAAAAAACTTATCCTTAAAATCATCTT

CTCTTGAGCACTTTTCTTTATCTACATTAGTATTTACTTTTGTCGTATAAATTTCCAGTT

TAGACCAAATCTGTCCTAACTGATTTAATTGAGATGTTCCACATTCTAAACAGGCTATTT

TTATATTGCTCAAAACAGCACCTTTTTTTATAAAATCGTTCAATAATGGCGTAATAGTTG

CTTCATCCAAAAATAAATCTAAGACAAGTATTTTTGTAAATGAATTTAATTGATCTGGGA

CATTGTTAATAATATCATTATAATTAACTAAAACTATATTAGCTGTAGGAAGTAAAGATC

GTGCTCCTTCCGCTAAGATAAAGCCATAAGGCATAATAATTACAATAATATATTTATAAT

GACTGCTAATAAGACTTATAGTGGTATTAGCTTTTAATTCGATTGTTTCTGTTACTAGCC

ACTCTCTCATTATTTCGTAAGTAATCCATTTGCCTAGTTCTGAGCAAGCAGTTCTTAAAA

TCGTACCTGGATTATTGTCGTTTTTGAGAATCCCTGACCAATGTTGTATTAATGGATGGG

ATGCTATATTAATTTGGAGTTGCATGTTTTGTTAAATATATTTAACATTTAATTTGTTGA

TTGAATCTTCAATTCCTATTTGTTGAGTATTATTAAAATAGCATATATATTAAAGAAAAT

AATTATAAAAAATGAAAAAAAGTCTTTGGCTGTGGGGTTTTACAGATAGTGCTGAAACTT
GGAATGGTAGATTTGCAATGATTGGTTTTATATCAGTTATTTTTATCGAAGTAGTTACAG
GACAAGGATTATTATATTTAATAGGTATGATGTCATAATAAAAATCCAAAAAAGTTCTTG
TAATTTTTTATAAGAACTTTTTTGGATTTTTATAAATCTAATCTAGCGGTCTCATAGATA
GAACAGGTTCATTCTCTCTATTGATACCTTTTTCAAATCCAGCAGCAGCAGCTCTTGCTC
TGCCTGCATGCCAAAGATGTCCAATAAATAGGAAGAATCCTAGGAAGAAATGAGATGTTG
TTAACCAAGATCTAGGAGATACATAGTTAACTGAATTAATTTCTGTAGCTACACCACCAA
CGGAATTCAAAGAACCTAGTGGTGCGTGAGTCATATATTCTGCGGCACGCCTTTCCTGCC
ACGGTTGAATATCGTTCTTGATTTTATTTAAATCAAGACCATTTGGTCCTCTAAGAGGCT
CAACCCAAGGAGCTCGCAAGTCCCAAAATCTCATGGTTTCACCACCAAAAATAATTTCTC
CACTAGGAGATCTCATTAGGTATTTACCTAATCCAGTAGGTCCTTGAGAAGATGCAACAT
TCGCACCTAGTCTTTGATCTCTAACTAAGAAAGTAAAAGCTTGAGCTTGTGAAGCTTCAG
GACCAGTTGGTCCATAGAATTCACTAGGATAAGCTGTGTTGTTATACCAAACAAAATTAG
AAGCCGTAAGCCCCATGATTGATAAAGCACCTAGACTATAAGATAAGTAAGCCTCACCAG
ACCATACAAAAGCTCTTCTAGCCCAAGCAAAAGGTTTTGTTAGTATATGCCAGATTCCGC
CAGCAATACAAATAATGCCAATCCAGACATGACCACCTACAAGATCTTCCATGTTATTTA
CACTTACGATCCATCCATCACCGCCAAACGGAGACTTTAAGACATAACCAAAGATAACTA
GGGGATTAAGTGTAGGATTGCTAACAAATCTAACATCTCCACCACCTGGAGCCCAAGTGT
CGTATACCCCACCAATAAATAAAGCTTTAATAACTAACAGAAAAGCGCCAATGCCTAATA
ACACTAAGTGTATACCAAGTATTGTTGTCATTTTGTTTTTATCCCGCCAATCGTATCCGA
AGAAAGGAAATGATTCTTCAAGAGTATCAGGGCCAATCAGAGAATGATAAAGACCTCCAA
AGCCAAGAACAGCTGAAGAAATTAAATGCACTACGCCTACTACAAAGTATGGGTATGTAT
TGAAAATCTCGCCGCCTGGACCAACACCCCAGCCTAGTGTAGCTAAGTGCGGAATTAGAA
TAAACCCTTGTTCGTACAAAGGCTTCTCGGGAACAAAATGTGCTACCTCAAAAAGAGTCA
TGGCACCAGTCCAAAAGACCATTATACCTGCATGAGCAACATGAGCACCAAGCAATTTGC
CAGAAACGTTAATTAAGCGTGCATTGCCAGACCACCAGGCAAACCCGGTAGACTCAATGT
CTCTACCGCCAACACCAACAGTTGTATTAAAGGGCGTTTCCACGTGGTAAAACCTCCTCA
GGGAATATAAAGTTTTCATGAGGTTGATCTTGAGCAGCCATCCAAGAGCGAATACCTTCG
TTCAATAGGATATTTTTAGTATAAAAAGTTTCAAACTCAGGATCTTCCGCAGCTCTCAAC
TCTTGAGAAACAAAATCGTAAGCTCTTAAATTTAGAGCTAGTCCAACAATTCCAAATGCG
CTTGTCCATAATCCAGTTACTGGTACAAATAACATGAAAAAATGTAGCCAACGTTTATTA
GAAAAAGCAACACCAAAAATCTGTGACCAGAATCTATTAGCTGTTACCATCGAATAAGTT
TCTTCAGATTGTGTTGGAGTAAATGCACGAAAAGTATCTGCAGCATCACCATCTTCGAAT
AAGGTATTCTGTACAGTTGCACCGTGAATAGCACATAGTAAAGCACCACCTAAGATTCCT
GCAACGCCCATCATATGGAACGGATTCAGGGTCCAGTTATGAAATCCTTGCAAAAATAGC
AAAAATCTAAAAATTGCAGCAACTCCAAGGCTTGGGGCAAAGAACCAGCTCGCTTGGCCT

AGAGGGTACATTAGAAAGACGGAAACGAATACTGCAATTGGTCCAGAAAATGCAATAGCG
TTGTATGGTCTAAGACCAACTAGCCTAGCAATTTCAAATTGTCTTAAACAGAATCCAATT
AGTCCAAATGATCCATGTAAAGCTATAAATGCCCACAGACCACCAATTTGGCACCAGCGA
GTAAAATCTCCTTGAGCTTCAGGTCCCCAAAGGAAAAGAAGTGAATGTCCCATGCTGTTT
GCTGGGGTAGAAACAGCCGCAGTTAAAAAATTGCATCCTTCTAGATATGAACTGGCTAGC
CCATGAGTATACCAAGAAGTAACAAAAGTAGTCCCAGTTAGCCAGCCACCTACAGCAAGG
TAAGCGCAAGGAAACAGAAGTAGTCCAGACCAACCTACGAATACAAATCGGTCTCTTTTT
AGCCAATCGTCTACAAGATCAAAACCACCACGAGTTTTTCTTGTCCAATTGCTATGGTC
ATAATTTTAATCTCCAGAGCAAATTTAATAAGTAATTTGTTTAAATAAACACTTTAGATA
TACTCTAAAATTTTACTTTTCTTTACGGTCAATATAATTATAAGTCAAAAGTAACCAAAA
TTCACGCTTTTAATAACATTTTTTTAGCTCATTAAGTTAATTTTTTAATATATAAACAGT
AATAGAGTTATTGTTTAACTTACAATACTCACAATTTATTCATTTCTATAATCCTTTGAA
AAAGATATCCTGTTCCTCGTGCTGTTAAAATCAAGTCGGGATTACTTGGATCATCTTCTA
GTTTAGCTCTCAATCGTGAAATATGAACATCAACAACTCTCGTGTCAACATGCCGTTCGG
GAGTATAGCCCCAAACTTCTTGCAAGATAGAAGCTCTAGAAAAAGGTTCTCCAGCCTTAC
TAACTAAAAGCTCGAGAAGACTGAATTCCATACCTGTTAATCGAACTCTTTCATTATTTT
TATAAACTTGTCTTTTGTTAGTATCTATTTTTAAAAATCCAATACTAATAATCCCAGAGT
TTGGGACTCCAAGACTAGTGGTGATTTTATCAGCTCTTCGAAGAACAGAACGAATGCGAG
CTTCTAGCTCCTTAGGAGAAAATGGTTTAACGACATAGTCATCAGCACCTATTTCTAATC
CGGTAATTCTATCGCAGACTTCACCTAACGCTGTTAGCATTATAATAGGAACATCTGATT
CTTTTCGAAGTTCTTGACAAACACCGTAACCATCTAGTTTAGGCATCATAACATCTAAAA
CAACAAGGCTAGGATACTCTTTTCTAAACACGAGTAAAGCTTCTTCTCCATTTGAAGCAG
TGATAACTTCATAACCTATTATTGTTAGTCTTGTTTCTAAAATTCTTCTTATGCTGGCCT
CATCGTCAACAACGAGAATCTTTTCTTTCTGGTTCTCCAATTTCTTATTTACTCCTGTAA
CTCACCGATCATGATTACCATAACCTTAAATTTTTAGCGCTATGAGTATCTTTATTCACT
TTAGAATTTAGTTATTATAAATTTTGCAAGTCTAGTTTACATTAATTAGTACCAGTCAAT
ATTTTTGTTTTAAGTTATGACTATTTTAATCCATAAGTTGACTCTAGTTTGCAGAATTCA
ATATATACATATTTTCTACCCAAGTAAATTGTATTCTATTTTCTAAATAAAAAAGGGGTT
GACAAACCAGTTGATAGAAAGTATTCTACATTACTATAAGTTAATTCCAACAAAGAAGTT
TTAAATCTTCTTTCGGAGTAATGAATACTTTAAT
>SRR9587917
TATATGATAAATTAAGACGTCTGTTATGTGAGAATTTCTAAAATATACAAAACTTTTTAT
TTAATATAGAAAAAGTAAATTAAAATAAGTTGAACATACTTTGCTCTAATCTAAACCTCT
TAATTGAGATTCTAATTAGAGACAAAAGTATTTACTTATTTTGTATCTATAGGAATAAAA
ATGAAACTTGCAGTTTATGGTAAAGGTGGTATAGGTAAATCTACAACTAGTTGCAATATT
TCTGTAGCACTTTCAAAAAGAGGTAAAAAAGTTCTACAAATTGGCTGCGACCCTAAACAT

GATAGTACATTTACGCTAACAGGTTTTTTAATCCCAACAATTATCGATACTCTTCAATCC
AAGGATTACCACTATGAGGACGTTTGGCCTGAAGATGTAATCTATAAGGGATATGGCGGT
GTAGACTGTGTTGAAGCTGGTGGACCGCCGGCTGGCGCTGGTTGCGGAGGCTACGTTGTA
GGTGAAACAGTTAAACTTTTAAAAGAATTAAATGCTTTTGATGAATACGATATTATCTTA
TTTGATGTTTTAGGTGATGTTGTATGTGGAGGTTTTGCAGCTCCATTAAATTATGCAGAC
TACTGCTTAATCATTACAGACAATGGATTTGATGCTTTATTTGCAGCTAATAGAATAGCA
GCTTCAGTACGAGAAAAAGCTAGAACGCACTCTCTGAGATTAGCTGGACTTGTTGGTAAT
AGAACAGATAAAAGAGATCTAATTGATAAATATATAGATTGTGTTCCAATGCCAGTATTA
GAAGTCTTGCCCTTGATTGAAGATATTAGAGTGTCCAGAGTAAAAGGTAAAACTTTATTT
GAAATGGCAGAAATTGATAAGGATTTAGCATATGTATGCGATTACTATTTGAATATTGCA
GATCAGTTGATTACAAGGCCAGAAGGTGTAGTTCCTAAAGAATCTCCGGATAGAGAATTA
TTTAGTCTTTTATCTGATTTTTACTTAAATCCTAAATCAAAGGTAGGACAAGAAAAAGTA
GATCAAGAAGAATTAGATTTGATGATAGTGTAAATAATATTTTCAGCATAATATAACAAG
GAATAAGATAATGTCTACAGCTCAATCAGATGCTCTTACTTTTGAATGTGAAACAGGTAA
TTATCATACTTTTTGTCCCATCAGTTGCGTTTCTTGGTTATATCAAAAAATTGAAGACAG
TTTTTTCTTAGTTATAGGGACTAAAACTTGTGGTTATTTCTTGCAAAATGCAATGGGAGT
AATGATTTTTGCTGAACCGAGATATGCCATGGCAGAATTAGAGGAAGGAGATATTTCAGC
TAAGCTAAATGATTATGGTGAACTCCGTAGATTATGCTTACAAATAAAAAAAGATAGAAA
TCCTAGTGTTATATTTTGGATTGGAACATGTACAACCGAAATTATAAAAATGGATCTGGA
AGGAATTGCTCCAAAATTGGAAGCAGAAATTCGTGTTCCTATAGTAGTTGCTAGAGCTAA
TGGTTTAGATTATGCCTTTACCCAAGGTGAAGATACAGTTCTAGCAGCTATGGCTCAAAG
GTGTCCATTAAATTTAAAGAATCAATCAGATAACACGAGTCTTAAAACTTCTCCTCATAT
TCCTTTAGTTTTATTTGGATCTCTTCCGGATCCAGTTGTAACCCAGTTAACTATGGAATT
AAAAAAACAAGGTATTTTTGTTTCTGGTTGGTTGCCATCAAAGAGGTATACTGAATTACC
GGTTATAAAAGAAGGATATTATGTTGCGGGAGTTAATCCATTTCTTAGTCGCACAGCTAC
TACACTAATGAGACGCCGAAAAACAAAACTAATTGGTGCACCATTTCCAATAGGTCCAGA
TGGTACTCGAGCCTGGATTGAAAAAATCTGTTCAGTAATGAATGTAAAGCCTATTGGATT
AGAAGATAGAGAAAAAGCAATTTGGGCTTCTTTAGAAGACTATATTTCTTTAATCCGAGG
TAAATCAGTTTTTTTTATGGGTGATAATTTACTAGAAGTATCTCTTGCAAGATTTTTAAC
TAGATGTGGAATGACTGTATATGAAATTGGTATTCCTTATATGGATAAACGCTATCAAGC
AGCAGAGCTAGCTTTATTAAAAGCTACATGTGATAAAATGAACGTTATGATGCCAACAAT
TGTAGAAAAACCAGATAATTATAATCAAGTAGACCGAATCCGTGATCTGAAACCCGATTT
AGTTATCACTGGTATGGCTCATGCAAACCCTTTAGAAGCCAGAGGTATTAATACAAAATG
GTCAGTAGAATTCACATTTGCACAAATTCATGGCTTTACCAATGCAAGAGATATTCTTGA
ACTCGTCACAAGACCATTACGGCGAAATCTTAGTTTGTCAGAATTAGGCTGGGATGTTTA
TAGCAAGCAAAGCTAGTATTCCTCTTTATTTATGACTTACATGGTATAAAAGCCTTGTAA

GTCTTTTATTTTATGTTAAAGAAGCATTTCTTACTCTGCCATCACCAGCCCAAGCTTGTA
AACTTTCTACTTGTTCTTTGTCTGTAAATGCTAAAGGCACAAACTGTTTGATAGCTATTT
TAATATCTTCAGTACTAAATTCTCTCTCTTCACTAAACGCCGTATGCATACTTTCAATAA
TTGCTTGCTCAATTTCAGCGCCAGAAAATTTATTACATAATAAACTCAATTCATGAATAT
CATATTCTTGCCACGATCTTGGTCTAACTTTTGATAAATGTATCTTAAAAATTGACTCTC
TCTCTTGACGATTAGGTAAATCTAAGAAAAAAATTTCATCGAACCTGCCTTTCCTTAACA
TTTCGGAAGGTAAACTTTGAATTTTATTAGCTGTTGCGACAACAAATACTGGAGCTTTTT
TCTCTGATAGCCAGGTAATAAATGTACCAAATACTCTGGCACTAGTACCACTATCACCTT
GACTATGTAAACCAGAAAAAGCTTTATCTATTTCATCAATCCATAATATGCAAGGCGATA
ATCCTTCAGAGATATTTATCATTTCTCTCATTTTTGATTCCGATTCACCAACTAGTCCAC
CAAACAGTTTTCCCATGTCAAGACGTAGCAGAGGCAGCATCCAATCGCTTGCAATAGCTT
TCGCGGTTAAGGATTTACCCGTCCCCTGTATACCAACTAGCAATAAACCTTTGGGCGAAG
GAATACCGTAATTTAAACTTTGCTTCGAAAATGAGCGAGACCTTTTTTGTAGCCATTGTT
TTAATACATCTAAGCCACCAATATCCCTATTTACTTTACTATAAGGATAAAATTCTAATA
GATGCGTTTGATTAATAATTTGTCTTTTTTCTTCAATAATAATAGGTAAGCTCCGAGAGT
CTATCTGATTATATTGTGCAATAATTTTAGTAATCACTTTGCGGATTCTATCTATCGATA
ATCCTTGGCAGGATTTTGTTATATTATTAACGAGCTCTGAATTCAAATTAAGATTCAAAG
CTTTACTCAATCTTATGATCTCTTTTTTGATTTCTAGAAGACTAGGTAAGGGCAAATCTA
TTACCGTTATGATATCATTCAGAGCAAAAGGTATATTGATTTTGCAAGAAATAATAATAA
TGTGCTTAGATTGCGTTTTGATTATTTTAGCTAAATTGCGAAGTTTCCTAATTAATACTA
TTTCGTTTAGGAAAGAGTCAAAATCTTTTAAAAGAAAAAGATTTAAAGATTCATTATTCA
AGTTTTCAATAAATTCTAATGCTAATAAAGGATTTCTTTTTGCATAACCATTATCGCTAG
GATTGTTAGTGTATCCATCTACAAAGTCCCAGGAATATACTTGTTGACTGTTTGAACAGT
TTAAGCTGTGTTTTATAATATACTCTAGTCTATCTTCTTCTCTAGTATTTATTACAATAA
TAGGATATCGAGATTTTAAAAGTAATCGTAAATCTTGAGTGAAATTCATGTCAATATTTG
TTTCCATCAGACTTTTGTACTTCACCAATTGAAGAAAGACTAAATAAGATTATCTAATAT
TCCTTGGATATATTCAAAAGTCATCTCTCTATTTTCTAAATTATTGCTATTACTAATTTA
ACTTACCATGATTTTTTTTCTTCCTTTTCTTCGTCGACTATTTAATATAGAGCGTCCACA
TGGTGTTTTCATTCTTGCTCTGAAACCAGAAACTCTAATTTTTTTTCTTTTAGATCCTTG
TAGTGTTCTTTTAGTCATGATGTTTCCTCTGTTGTAAAAGATTAAATACTATTTTATCAT
CTTTTATATATCTTTATGATATAGATATGTATTAATATCAAAGTTAACAGCAATAATTTA
TATTAATCAAGAATTATTTATATAATATTAAACATTAAATATTAACTTTATGATATTGGC
TTTGCCCATTTTTTATTTAAGTATTTTGACCATCTTTTTGCTAATCTTAAACTGGTTAAT
TTTTCAGCAGCTAAAAACGATTTTGTTATTAGAATCTCAATTTAAGTATTTTGTTGATAA
AAGCCAAAATCGTAAATTAGAACCCGAAGAAAGTTTTGCTTTTGCAAAAGTATGTGTAGC
TAAAAAATATTTTTCTAAAGCTATTATTGAAGGTCAATTAGCTTTAAAAAACTATCGAGA

CCTGAATATCTTAGATAATAATATAGTAATTGCTAATTTGTATAATATGCTAGGTTTTAT
TTATTTTGAAGCAGGCCAAACAAGTTTTGCAAAGAACTTTTATGAGCAAGCCTTACAAAT
AAACCCTAATTATGTTGTAGCTTTAAATAACCTGGCAAAAATTTACGAAGAGGTAAAAGA
TTTAAAAAAAGCCGAATCTCTATACGATAAAGTGTTAACTCTTAACTTAAATAATAAAAC
TGCTAATAGAAGGAAAGATTTTATAGCAAAAACTAAAAACATCTAATACTATTTGTAATC
GGGATAGCAGGATTTGAACCTGCGACATCCTGCTCCCAAAGCAGGCGCGCTACCAAACTG
CGCTATATCCCGTAAGACATAATTATTACTATATACCTTTTCTCTAATTATGTCTACTCT
TTCTGTATTTTATCTAGGATACCAAAACATGCCTTTAACTCCATCTGGATCTTTGATAAA
TCCTAATTTTTTATAAAAACTAACAACATCAGGTTCTGCGAATAGTGTAATAGTGCTAAT
TTCTGCCTGTCTCAACTGCTGTATTAATTGATGTATGATTACTTTTCCTAAACCTAAACC
TTGGAAATCTGGATGAATAACTACGTCCCAGATAGTTGCATTAAATCCGTTGTCTGATGT
TGCTCTAGCAAATCCTACAAGTTTACTATTTGCATCTTTTTTTTGTATTAAAGAAATAAT
AATAGAACTATTTTTTAATGCGATCTTTACTTTTTTTAGAGGTCTTTTAACCCATCCGAC
TGAATCACAAAGCTGCTCCAATTCGTATAAATTAATATTCTTATTGCTACTCAGATAGAT
ATCTTTAAATTCGATTTTATCGCAAGTTTTATCTAAAACAATAAGTTTTTTGAAATTTTT
TTCAGAGCTAGCATTGATATCGGAGTTTTGAAAAAAATTTTTCCAGAAGATCATAATGAT
TATGATATAATTAAATAAATATATATATTACAATATTGCTTTAACAAACAATATATTAAT
ATATGCTCAAATTTTTACATGAATCTATTAAGATAGAAGATTGTTAAAATATAGAGAAAT
AAAAATTTTTGTAAAATCTTTCAATAAAATGATATAAAGAGTGCTTTTTAAGACTAACAT
CTATTATTATAAATAAAGTCTCGAAATTGAGATGAAATAACACATTATTCTATGTCTAAT
GCCTTTTAAAGGAGTTATTTGTGAAAAAATCTATGTGTCTAACCTGTTTACTTGCTTTAC
TAATTATGAGCAACCCAATAATAGCTAACGCAGAAGTAGCTGGATTGGTGCCCTGCAAAG
ATTCTGCGGCATTCAATAAGCGTATGGTAAATAGTGTGAAAAAACTTCAAGCCAGATTAG
CTAAATATGATGCAGACACGCCACCAGCGTTAGCTTTAAATAAGCAAATAGAGAAAACTA
AAACTAGATTTGCAACTTATGGTCGAGCAGGTTTGCTGTGTGGTACTGATGGATTGCCAC
ATTTAATTTCTGATGGTCGATGGAGTAGAGCCGGGGACTTTGTTTTTCCAGGACTATTAT
TCTTGTATATTACAGGATGGATTGGCTGGGTAGGTAGAGGATATCTTTTATCTGTTGCTA
AGACTAGTAAGCCAACAGAAAAGGAAATCATTTTAGATGTGCCATTAGCTATCAAATTTA
TGTCATCCGGATTTGCATGGCCGCTAGCGGCTTGGCAAGAATTTAGTAGTGGACAATTAA
TTGCTTCCAATGACGATATTACTGTTTCACCCCGTTAGTAAAAAATTTATATGAATAATA
ATTTTACCAAATACTTATCAACAGCACCTGTAATTGGTGTATTGTGGATGACGTTTACAG
CCGGATTTATTATAGAATTGAACCGCTTTTTCCCAGATGTCTTATACTTTTACTTATAAA
TAAGATCGTATAAAAAACTACATAAATATAGAGAAATACTATTATTTGTGTAGTTTTTAA
TATATAACATAATTATTTATTTTTAAAAATGTTTATCTTTTCGATTATTTTAATATTAGA
TCTTACATAATAAAATAATATCAATATTATGAGCTTAGTAACCCAAATTATAGTTAATGC
CGATGATGAATTAAGATATCCTACAATTGGAGAATTGCAGTCAATTCAAGACTACTTAAC

TACAGGAAGCAATAGAATTAGAATTGCTACTATTATTAGAGATAAAGAAAAGGAGATTAT
ACAGAAAGCTAGTAAGCAAATTTTTCAGTTACATCCAGAATATATAGCTCCAGGTGGTAA
TGCAGCAGGTTCTAGGAAAAGATCGTTATGCTTACGTGATTATGGTTGGTATCTCAGACT
AATTACATACGGAGTATTAGCTGGGGATAAAGATTCTATAGAAACAATTGGTATTATAGG
AGTAAGAGAAATGTACAATTCTTTAGGTGTACCTATAATTGGAATGTTAGATGCAATCCA
GTGTTTAAAGGAAGCATCTTTAGAAATGCTTGGTCAAGATGATATTAGAATCATTTCTCC
TTATTTTGATTATATAATTCGTGGAATGTCATAAATCATAGCTTCAATTAATTAGTTGAA
TAAAAGCTTTTATAATGTTATAATTACAATAAGCCCGAAAGCATATAATTGTAAAAACTA
AATCTTGTCAGAACCGGAAGGTAGCAGCAATAATGTTTCAATACAAAAGTTATGATTTTC
GGGTGTTTATTGTTTTTATCTAATAGTCTCGACTTTTAATTAGTATGCTATGTATAAATC
TGTATATGACATGTAATCCATCAAACCTTCCAGCAAAAATAATTTAATAATTTGATATTA
ATACGTAAAACACCTATACTCTTTTATAAAATTTCTTTATATCTCATCTAAACAATGGAA
ACTTAACTTAATGGGTGTTCATATTTTATCAACAGGTTCGTCTGTCCCAAATTTTTCTGT
AGAGAATCAACAATTTGAAGATATGATCGAAACTTCCGACCATTGGATTTCAACAAGAAC
AGGAATAAAAAAAAGACATCTAGCTCCATCTTCTACTTCTTTAACTAAATTAGCGGCAGA
AGCTGCAAACAAAGCCTTATATGCAGCTAACTTAAAACCTACTGAGATCAGTTTAATTAT
TTTAGCTACGTCTACGCCTGATGATTTATTTGGTAGTGCTAGCCAGTTGCAAGCAGAAAT
AGGTGCAACAACATCGGTAGCTTTTGATATTACGGCTGCCTGCTCCGGTTTTATTGTTGC
TTTAGTAACAGCAGCTCAGTTCATTCAAACTGGTTCCTATGACAATATTTTAGTTGTTGG
AGCAGACACAATGTCTAGATGGATTAATTGGTCAGATAGAACTACCTGTATTTTATTTGG
TGATGGTGCTGGAGCAGTAGTATTGGGGCAAAGCCTCAAAAATAGTATTTTAGGCTTTAA
GTTATGTACAGATGGTCAGCTAAACAGTCATTTACAATTAATGAATAAACCTGTAAATAA
TCAAAAATTTGGTGTTACAGAAATTCCTCATGGAAACTATAATTCCATAACAATGAATGG
CAAGGAAGTGTACAAGTTCGCTGTATTTCAAGTTCCAACAGTAATTAGACAATGTTTGAA
TAATTTAAACATTTCAATAGATGAAGTTGATTGGTTTATATTGCATCAAGCAAACACTAG
AATCATAGAAGCAATTGCGAGCAGATTATCAGTACCTTTTTCTAAAATGATTACGAACTT
AGAGCATTATGGAAATACATCTGCAGCGTCAATCCCTTTAGCGTTAGATGAAGCTATTCA
ATCCAATAAAATTCAACCAGGCCAAATTATTGTTTTATCTGGTTTTGGAGCAGGCTTGAC
TTGGGGAGCAATTGTCTTGAAGTGGTGATTTATATATTGCGGATGACGAGACTCGAACTC
GTAAAGCTTTCGCTACACACCCCTCAAGCGTGCGTGTATACCAATTTCACCACATCCGCA
TTTTTTACCATGAATATTAAAAATTTATAATATTTATCTAAATATTCATATAATGTGTAT
ATATATATACATTCTTGTCAGAAACTCTCTTATATTTTATTTTAAGTGATAAAATTGAGT
TTAGTATAGATAGTAAAATTTTATTTTAAAATCATAAAAATAAATATATAAGGAAAATAA
AATATGACACCATCTTTATCAAGTTTTTTGAATAGTCTTATTCTTGGGGCAGTAATTGTA
GTCGTTCCCATAACTTTGGCTCTTTTATTTGTTAGTCAAAAAGACAGGACAATCCGGTCC
TAAAAATAACTTAGAAAGATATGGAGAAGCAATTTAAAATGAATAATAAAATATTTTTCA

TACTAATAAGCTCTCCATATATTGCTATACTTTAGAAGAATTTTTATTTTTTTTATTTTGT
AACATCCTGTAAAGTTTATTCCGTTCTTTCTTTTTCCCATTTACTCTCTTATAATTAATG
ATACAAATGTCAGGTTAATAATAGTTTTTTTATTCTGTATCTTTCATTTTATATACGTAT
TATTATGTTCAATTAATAAAGAAAAGCTCATGTCTTTGTCTAATTGGCCTCTCAAAAAAG
AAAATTCTGAAGCATATAATATTAAGAATTCAAAACAAATCACAATTCCTGATGGTTTAT
GGATAAAATGCTTCGACTGTGGTCTATTAATGTATTCTAAAGTACTGAAGAGAAATTTAA
AAGTTTGCCCTCAATGTAGTTATCATTTTCAAGCTTCTAGTAACGAAAGAATTGATCAAT
TGATAGACCAAGGTAGTTGGCAACCAATGGATGTTCACTTGATCTCTACAGATCCATTAG
GCTTTAAAGACCAAAAGCTTTATAGTCAAAGGTTAAAAGATACTGCTTTCAAGACTGGCC
TGCAAGACGCAGTTCAAACAGGTACTGGGACTATGCAAGGTAAAAAAGTATGCTTAGGTA
TTATGGATTTTAGATTCATGGGGGGAAGCATGGGATCTGTTGTAGGCGAAAAACTAACAA
GACTGCTAGAAAAAGCAACTCAAGAAAAGTTGCCTGCAATTATACTTTGTGCATCAGGCG
GAGCTAGAATGCAAGAAGGTATGTTGAGCTTAATGCAAATGGCAAAAATTTCTTCTGCTC
TAGAAATGCATAAAAAAGAAAATCTACTATATATATCTGTTTTAACCTCTCCCACAACAG
GGGGTGTAACAGCTAGTTTTGCTATGCTTGGAGACTTAATTATTGCAGAGCCAAAAGCTC
TTATTGCATTTGCTGGTAGACGAGTTATAGAACAAACAATCAAAGAAGACTTGCCAGATA
ATTTTCAAAGTTCAGAATATTTATTTGAACATGGTTTCCTAGATTTAATTGTATCCAGAA
CTCAGCTTAGATCAAAGTTAATACAAATTTTGTCCTTACATAATCATAGTAAGTAATGAA
TTGAATATTGACTATAATATCAAAATGCTAGTAAAAATTAAGATATTAAGAAATTTTCAC
TAAAGATATTTGTAGACTAACTAGAATTACAATTACTATTGTAGAAAAATTTGTCTTAAT
GCCATTTTTTCAGTCTTTTAACAAATTGTTTTTATGACTTTAAATAATTGTTAGTTTAAT
ACAAATTTATAAAGAGAATACATACTAAGCTAAATATAGTAATATTATGTTTTCCTATAA
TACTGACCTATCAAATTTCGAGGAAATTTATGCTTAAAAGATCTTCTTGGCTTGCGGCTT
TATTGGGACTATTAACAGTAGTTTCTACAAGTACGCATACATATGCCATAGAGTTAGACG
AGGCAACAAGAACTGTTCCATTAGAATCTTCTGGCAGAACTGTAATTCTTACACCAGAAC
AAGTTAAAAGAGGCAAGCGATTATTTAATAATTCTTGTGCTATTTGCCATAATGGTGGTA
TCACGAAAACAAATCCGAATATTGGACTTGACCCAGAATCTTTAGGATTAGCTACGCCAC
AAAGAGATACCATTGAAGGACTAGTTGACTATATGAAAGATCCGACTAGTTATGATGGTG
CAGAGTCAATCGCAGAATTGCATCCAAGTATTAAAAGTGCTGAAATTTTTCCTAAAATGC
GCAATCTAACCGATGAAGACCTATTTACAATCGCAGGTCATATCTTACTTCAACCTAAAA
TTGTTTCTGAAAAGTGGGGCGGAGGAAAAATTTACTATTAGAACTTCAAAGACCTAAACC
TTGTGAATACTAGTTTATGTATTTGTTGCTTAAGTGTGATTATTTTGTTAGATCATGATA
TATATTAAAGATGGAGTTTGTTGACAGCAAGTTACTTTTTACCCGAACTTGTTTACACTA
TTCGTCTAGTTATTAAAAAGGAGACGTTAAATTGAAGAAGAAGCTTTCAGTTCTTTTCAC
TGTTTTTAGTTTTTTTGTAATAGGTTTCGCACAAATTGCTTTTGCTGCAGATCTAGATAA
TGGAGAAAAAGTTTTTTCTGCTAATTGTGCAGCATGTCATGCTGGCGGTAATAACGCCAT

TATGCCAGATAAAACCTTAAAAAAAGATGTACTTGAAGCTAATAGTATGAATACTATTGA
TGCTATTACTTATCAAGTACAAAATGGTAAAAATGCCATGCCTGCTTTCGGAGGTAGACT
GGTTGATGAAGATATTGAAGATGCAGCAAATTATGTATTATCTCAATCTGAAAAAGGTTG
GTAATTATACTTGATTTTATCCTGTATTAAAGAATAGACAATCTATTTAGTTGTTCATTA
GATTGTCTATTCTTTGTTTATGCTATTATATAAAGATATTTACACAATATTTTATGATGA
AAAGAATACCCGCAATTCTTGTACTAGAAGACGGTGCGTATTATAAAGGATGGTCATTCC
AGCAAGATAAACAAGAGATTACTATTGGTGAAGTAGTTTTTAATACTGGAATGACAGGAT
ATCAAGAAATAATCACAGATCCTAGTTACTTCCATCAAATTGTCGCTTTTACCTACCCGG
AAATTGGGAATACAGGTATTAATAATCAAGATATTGAATCTCACAGTATTAGTATTAAAG
GACTTATTGCAAAAAATATTTGTAAAATTTCAAGCAGCTGGAGAGAGCAGCAATCTTTAG
TTAAGTATTTAAGTAGTAATAATATTCCTTTTATTTTCGGAATAGATACAAGGTCTTTAA
CCCAATACTTGCGTCAATTTGGTACAATGAACGGTTGTATCTCTACTGATAATTTAAATC
ATAGTTACTTAAAACAGAAAATTTGTGAGATTCCAAGTATGCAAGGTTTAGATTTAATCC
CGCATGTAACTACAAGAAATGTTTACCCCTGGGATGAAAAAAGTTTTCCAAATTGGTATT
TGACAGATAACATTAGAGTGCACCGAGTTATTCAGTTGAAAGTTATTGTTATAGATTTTG
GAGTTAAACTAAATATACTCAGAAGACTAGCTACACTTGGATGTCAGATAACGGTTGTGC
CTGCCCACACTCCTTTAAAAGATATTTTGGCTTACCAGCCTGATGGTATATTACTCTCTA
ATGGTCCAGGAGATCCATCAGCAGTACATTACGGCATCCAGACAGTTACAAAATTACTAG
ATTACAATGTGCCTATATTTGGGATTTGTATGGGGCATCAAATTTTAAATTTAGCTCTTA
AAGCTAAAACTTTCAAACTTAAATTTGGTCATAGAGGTATTAACCATCCATCCGGATTGA
ACCAGCAAGTTGAAATAACTAGTCAAAATCATGGCTTTGCAGTTGAATTGACTTCGGTTT
TTGAATCTCCTGTAAGAGTGACTCATTTTAATCTAAATGACACTACTATTGCAGGAACTG
GACATAATCAAAGTCCTTATTTTTCTGTGCAATATCATCCAGAATCGAGCCCAGGCCCTC
ATGATGCTGATTATCTATTCGAAAATTTTATAGAAATAATGACAAAGTCCAAGAATAAAG
TTAGTTAGTAATTTTCCCATGCTTTATGAGTGAATAAAGCGGATAGTACTTGTACTCCTC
GTAATTGATTAAAGAGAGGCAAATGTCCTCTGGGGCCATGAGAATTCCATTGAAATTCCT
CAGGATATCTGCAAGGAATTCTATCAACTTCCCAGCCGATTTTTTGCCAGAATTTTCCCC
AATCTTTTCCAACACTAAGCCATATTTGTCTTTGAACAAATAGCCCAAATTTACCTTTTG
AGTGTGTATGCCATAACTTATCAATAGTTTGTAAATCTTGAGCAGGTATTTTTTTTATAT
CTGTAAAATATAGCCAATTACGAGTTTGTGCATTTACACCAGCTAGTTGAATAAGCTTTT
GTTGAGTTAGCTGATCTGCTTTGAGCAAATCCCGATGAGTCAGTAACATTTGTAAATCTT
TGTAATTCATCTGTTGAGCAGAGCGCAATGGAACAATGCCATCGGGACATAAGTTAGAAG
CAAATTTCACTATTTCTTGATTTTTACTGTTAAGTAACTTTTCATAAATTAAACCGTCAA
CACAATTACTTTTATAGTTAGGCCCAGTAATCCTCTCAAAAAATAGATCTGCTAAATCTT
TTAACTCGATAGAGTCTTTGCTATTTATATTCTCAATAATTTCAAGTTGTTGCTTGACAT
TATTAGACTTAGTATTCTTATTCAATTCTAAAAGCTGGGCTCGGATTTGATTTGGCATCT

GATTTGTTTAATTAGTTTTAATAGTCATAAGGTCTTAGAGAGAATTAGATTGTTTTTTGA
AATTAGAAGAGCGCCGCAGTATTTCTAAAGAAGTTCGCATAATTTGATTCAATAAGTTTT
GTCCTGCATAAATTATAATTTGACCAATTAAAAAAATTGTGCTTTTTATTTTAGGAACAA
AAAAATCTTGTACCTCAAGCAGAAAAGTAATTATTATTTGCATTGGTGACAACATCTGTA
ATTCATTCGATCTGCAAGCATATATATACTGACAGTTTAAGCCTTCTTGATAAAATCCCC
ATACTTTGTAACGACTTTCGTAAATTGCTCGTGGTCGCTCAAGATAGTTTTTAATTATAG
TATTCCAAATTAGGTTATTCTTTAATTTTTCTAATTTTCTATCAGATAGAAATTTTACAT
TACAGAGATAACTTGCATTTGTCTCATTTACAGTATTAAAATTTTGAACTAACATATGAG
CAATAATGTTGCTTAGCTGAATTAAATAATTTTCAAGAAAAATTTCGACCTGCTTAACTG
GCATATTGTAAGTAAAGAGATTAAAAGTTTTATTCACAGTTTGAGATGACCCAAAAAGTA
GCTGAATAATTAAAATTTGTAGTAGCATCTTATAGTCGTTCAGTAGATAGTTCAAATCAT
GAATATCTTGGTAAATGTCAATACAGTCCAGTTCATATAACTTACAAAATCTTTTGATAC
ATCTATGGAATAAATCAATTAAAATATTTTCGGTCAAGTTTGTAATATCATTTACATCTA
AATTTGAATTATAAATTTCTAAAAGTATCTTTTCCAGTTCAGCTAATATGATTTTTAATA
AGTTTCTTTTTACTTCAGTGCGAAAAACATCTAAAATTAACACTTCCTGAGAGCAATTAG
TTAATCTCTTATTAATCTTTATAGAAGTTCTAACTAATAGCTCTGCTACCTCTTGATTGA
GTGTTGGTCCCTGAGAGCTTGGCCAATAATTATTCACGTTATACCATAGTAAGAAATATT
GAATTTAATGTTATACGATAACTTTTTAATATACTGAAAAGCAAGTTAATTACTCTATAG
CAACTAGTTTTTACAGATCTTAATAGTTATTTATCAAATAAAGTTATATGATAGTAGCGA
TTGAGCTAATTTTATTAAAATTTATAATAATGACAACTTATTACTTCGCCCTTGCGAGTC
AAAATTTTCTATTAGTACAAGAACCACTAGAAGAAGTTTTTAGAGAGAGAGTTAACTACT
ATCAGTCAAATAATAAAGCAATTGATTTTTGGCTAATACCAAACCCTTCTTTTCTAGAGA
AACCAGAAATGATTTCATTTAAAAACCTTGTACCTAAAGACGCTGTAGCTATAATCTCTA
CTAATCCAATATTTATTAATTGGTTAAAGCTAAGAATCGGCTATATCTGTATTGGGCAAT
TTGAAGATAACCTACAACTTTCTGAAGAATCGTTAAATATTACTGTTTTAACAGACAAAA
TTTAACAATGCTTTTATTATGACTAGTGTTAAATTATTTTGGTTAAGTTATAATTATAGT
CTTTTAGTATTTAGACTCTTATAGTTTATCACTTAGTATTTTATGCTGAGATGGAGTGAC
AAATAGTGTAAGTATTTCTTGGCTAAACGTTTCAGTTGCCTTAGATTTATATCTATTTGG
ATTAACTATAATGGATAGCATTCGTTTTATAGTAACATTTTCAATTTGAGCCCAATGTAC
AATTCCAAGTTCCAATTCCTTAGCAATAGCTGAAACGGAAACAAAGGCAGCTCCTAGCCC
TGATTGCACAGCGTTTTTGATAGCTTCTATAGAATTCAATTCCATCTCTATTTTAAAGCG
ACTGCTGTCAATACCATGTTGACTGAGTACTTTATCTATGACTTTTCTAATTGTTGATTG
AGTATCTAACGCAATAAATCTAAGCCTGTATAAGTCTTCTTTTTGAATGTCTCCTAGTTT
AGAAAAAGGATGTGATTTGGGTAATATAAGTGCTAATTCGTCTTCCGCATAAGAAGTAAC
CTGCAAAACATCTTGCAATTCAGTAGGCACTTCTCCTCCAATAATTGCTAAATCAACTTG
ACCATTAGCTACGCTCCATGAAATAAGTCTAGTTGAATGTACTTGCAATTGAACAGCCAC

TTGTGGATATCTTTGCCTAAATAGTCCGATTAATCTTGGCATCAAATATGTCCCAGTTGT
CTGGCTAGCTCCAATAATTAATGTGCCACCTTGTAAGTTTTGTAAGTCGTCAAGAGCTCG
ACAAGTTTCTTCGCAGAGAGCTAAAATTCTGCCCCCATATCGTAAAAGAAGACTTCCTGC
CTCAGTTAAAGTTGCCTTCTTATTACCTCTTTCAAATAGGGAAACATTCAATTGGCGCTC
TAAATTTTGAATTTGCAAACTAATAGCTGGCTGAGAAACATACAAGCTATTAGCTGCTTT
TTTAAAACTGCCCTCTTTGGCAATTGCTTTTAATATTCTTAACTGATCCAATGTAAATGG
AAGGTCTGTCATTAAAGAATTGTAGTATAATGTATATTATTTAGTATTGATCAATGTATG
CATTTGCCAACTAAAATATTTGTCATTACGCTTCATTATTTTCAAAATAATAAAAAGTAT
AATGTTAAATAAGCTGTATTAAAATTTAGTATCTGTTTGATATTAGATTTATACAAGAGA
TATCGTGAATTTAATTCTAACTATAAAATATAGGAGACAATATGGACTCTAGACTTTTAG
TTGTACTAATACCAGTTTTAGCAGCAGCATCTTGGGCAGTTTACAATATTGGTAGAGTTG
CACTACAGCAATTTAGAAAAATGACATCTTAGTTTTGCTTAGAATATAATTTTTAGACTT
TATGAGATAGAGGACTTTAAAATCCCTATCTCTATTGTATATCTTAATAAAAAAAGAAAG
CCCCCTTGTACTTACTGTTATTACACATTAATATTATTGGTGGTACCTAGATAATAAGTT
TAAGCTAACTTATTTTTTATTTTATAATTAATTAAATTAATAACACAATTATGGCTGTTC
CAAAGAAAAGAACATCTAAAGCTAAAAAAAATGCACGCAAAGCAAATTGGAAAAATCAAG
CAAAAACTGAAGCTCAAAGAGCTTTGTCTTTAGCAAAATCAGTATTAACTAGAAAATCCA
ATGGATTTATTTATAATCTAACTGAAGCATCAGATACTTTTAGCGATTAGAAGTGTTTTT
ACTACAAGTCTGAAAAAGTATTACGCAACAAGTTCTCAGTAATTTTTATCTATTAATCAT
TTGAAAATCATTCTATTTGATAATAAAATAAATAGCAGATTAAATAAAATCACATTAACA
AATTATGCAGCAAAGCTAGATCAAACTAGTGAAATTTGGTTGTTCAATTGTATTGAAAAT
ATTCAGCATATTTTTTTTAAAAGTCAATTAAAATCAAGTCATATTACTAAAATTTTTATT
TCTGGTACTAGCTTTGAATATACCGCAGGTCTACCAGGATTATTATCCAGCTTAACACTA
AGTGGTAGACTACATCCTATTAGTATATATAGTCCCCAGTCTCTCAAAAAGTATCTTGAA
GCATGTACCAAATATTCCCAAACTAATTTTTCTTTTCCTATTAATTTTCATAATTTACAA
TACGGAGGACAAGTTGTTAACCAATTTTATACAGTAATTTGTTTACCGTTGAGCAAAAAG
AGCCTGCTGTATGGATTTATTATTCTAAAAAAAGAAAAGCAGGGAGTATTTAATTTAGCA
CAAGCTAAAACCTTGAATATTCTTCAAGGACCTATATATGGAAAACTCAAAGAAAAGGAT
AATTTTTTAAGTCCAGATGGTTACTATCTATCCGGCCAAGACTTCTCTTCTAATACAATA
ATGGGACATAAAATATCGCTTCCGTTATTAGTCAGATATTCTAGAATTATTTCTGAGATG
CATTGGTTTTGCTCTTATCCTATTAGATTAAATACTTATTCGCATCAGCAAGGAGCAAAG
TGTTTACCACATAATGTTTTAACTGATATTATGAAATCCCAAATATATCAAGATAACAGT
TTTGTTGAATAATACTTATTCTTGATATATAATGGTCTCGGTAATCTAAATAAAATACTA
GAAATAATTTCTTAAATTTATTTGTCTATATTAAATTTACAATCATTTATGACATACGCA
ATTATTGAAGCAAGTGGCACGCAGCTTTGGATAGAAGAAGGCCGCTACTATGATTTAAAT
CATATACCTGTTGATCCAGGTCAGTCGATTATATTAGGAAAAGTCTTATTATTGAATAAA

AATGGGGAGGTTACTTTAGGCCGCCCTTGTATAGAAGGGGTTACGATAAAGGCTACAGTA

ATGAGGCACTTGCGAGGAAAGAAGATAACTGTTTTCAAAATGAAACCAAAGAAGAAAATG

AGATTAAAAAAAGGTCATCGACAAGAATTGACTCGTTTAATGATCGATTCTATAACATCT

TAGATAAAATCTAGACTCATATTTTTAAAAACTTTTTAATATTTAACTAAAATAGATATA

ATGGCACATAAAAAAGGTAGTGGTAGTACAAGAAATGGCAGAGACTCTAATTCCAAGCGT

TTAGGTGTTAAAAAATATGGTGGAGAGCAAGTAACAGCAGGTAATATTTTAATCAGACAA

CGGGGAACTAAAGTTAAGCCTGGCCAAAATGTTGGAAAAGGAAAAGATGATACATTGTTT

TCTCTAATTGATGGTTTCGTGCTGTTTGAAAAGTCAAATCAAAAGCAAAAAACAATTAGT

GTTTATTCTGCTAAGAAATAGTTAAAACAATTGGTGCAAGATAACTTGAGTGTATTTATT

TCACTTACAGTGGCACCATATATTTGATTTATTAAGAAATGGAGTTCCTTAATGAACCAA

TGAATTTTATTAAAGATTTGCTGTGAAATTTAATGACAAACACTATTGTAATTTCCTGTC

TACACAATATGGCTGCTATTCTATATTGTGGTCAGATACAGAAACTAGTAGTAGCAAATG

CTCATTATCAAGTAAGTGATATCTACCTAGGTTGTGTTGATAAAATCTTCTCAGGAATAA

ATGCGGCATTTATTAACCTAGGAAAGAATGAGTACAGTGGTTTTATACATATCAGTGATA

CCGGTCCGCTTAAAAAGAAATATTATGTCAATAATATTACTAACATTTTAACAATACGGC

AAAAAATTTTAGTACAAATTATTAAAGAGCCAACTTTAAATAAAGGTCCAAGGCTCACTG

CCAATATTACATTATCAGGTCGATATATTGTATTAATGCCTTTTAGTCAATCAATCTGTA

TATCTCGAAAAATATATGATGAAGATGAGCGTCATTATTTGAAGTCTTTAGCTATTTTAA

TTAAACCGGCAACAATGGGCTTGCTATTTAGACCTTCTGCTGTAGGTGTCGATGAGGAAA

TAATATTAAGCGAATTAAAAAATCTAAAAGAACAATGGAACTTTGTTCAAAAATCTGCAA

TTAATAGTTATTCACCTGTTCTTCTATATAAAGATGAAGATATTGTTAAAAAGGTAATCC

GAGATTTTTATAATAATAATACAAACAATATAGTAATTGATTCAAACCTGGGATTAAAAC

AATTAAATTATTATATCCACACTTGGCACTGTAATAACTCTAGCACAGTTCCTAAGATTA

AGCTTTATAGTAATAATCAATGTATACTAGATGCTTTTGGTATCAATCAGGCAATTTCCA

GAGCTCTCATTCCAAAAGTTGATCTTATACTTGGTGGCTATATGTTTATTGAAACTTTAG

AAGCTTTTACTATTATTGACGTTAATTCTGGATCTTTTAATAATTCCACTAGTGCACGAG

AAACAGTTTTAAAAACCAACTGTTCTGCAGCAACAGAAATAGCTTATCAGTTACAAATTA

GAAATATTACTGGTGTAATTATAATCGACTTCATTGATATGGAATCACAAAGAGATCAAT

TGCAATTATTAGAACACTTTAATAAAGAGCTATCACTTGATGATGCTAAACCACAAATTG

TACAGTTATCTGAATTAGGTTTAGTTGAATTGACTAGAAGAAGACAGGGCAAAAGTTTGT

ACGAGTTAATTAGTAGTGATTCTAATTACTTTTATTTTTTCACACAATCAGAGAGATCTC

AGTCTCTTAAGAGATTCGATGATAGACAGCAGAAACAACAGATTTTTAATAAATCTTGGC

TATCTGCAGAGATTAATACTATTAACAAGGTCTTTTTTCAAAAGTCCAATTTGTGCAGAC

CTGCTAACTTTTACCTAATTCGTAATCTCTATATAGTTAAAAGTAGTATTACTTATAAAC

AAAATTATTTATTAACTCACAGATCAAAACTTATTTATTCTAAAGAATACAGCAAAGTAT

TACCAAGCAGTTATTATCTAGCTAGTCTCAATAAGAATAGTAATCAAGAGTTCCTACCTT

AAGATTGTTATATCAACTGTTGAAAAAAAGAAGCTCTCTCTTATATAAGAGAGAGCTTCT
TTTGTAGCTTACTAACTATCTAGTTAGTTAATTAAAAATTAACCGTTAACAGCTGGAGCT
GTTAGAGCTACTGGTAAAGATTCACCAGAAGCTAGATCTAGAGGGAAGTTGTGAGCGTTA
CGTTCGTGCATTACTTCCATACCTAGGTTAGCACGGTTGATGATATCAGCCCATGTGTTA
ATTACACGACCTTGGCTATCAACAACAGATTGGTTAAAGTTAAAACCATTCAAGTTGAAT
GCCATTGTGCTTACAGATAAAGCTGTTAGCCAGATACCAACTACAGGCCATAGACCTAAG
AAGAAATGTAGAGAACGAGAGTTGTTGAAACTAGCATATTGGAAGATTAAACGACCGAAG
TAGCCATGAGCTGCAACGATGTTATAAGTTTCTTCTTCTTGTCCGAATTTGTAACCATAG
TTAGCAGATTCGTTTTCGCTTGTTTCACGAATTAAGCTAGATGTAACTAGGGATCCGTGC
ATAGCACTGAACAGAGAACCACCAAACACACCAGCAACACCTAGTTGGTGGAATGGGTGC
ATTAAAATGTTGTGCTCAGCTTGGAATACAAGCATGAAGTTAAATGTTCCGGAGATACCT
AGAGGCATTCCATCAGAGAAACTACCTTGGCCAATTGGGTATACTAGGAATACTGCTGCT
GCTGCTGCTACTGGAGCAGTAAAAGCAACGGAAATCCATGGACGCATACCTAGGCGGTAG
CTTAGTTCCCACTCACGACCAATGTAGCAAGCTACGCCAGTTAGGAAATGAAGAACAACT
AATTGGTAAGGACCACCGTTGTATAGCCATTCGTCTAAAGAAGCAGCTTCCCAGATTGGG
TAGAAGTGAATACCGATAGCTGCAGAACTTGGAATAACAGCACCAGAAATGATGTTGTTT
CCGTATAGAAGGGAACCAGCAACTGGCTCACGAATTCCATCAATGTCTACTGGAGGTGCA
GCTACGAATGCAATGATGAATACAGATGTGGCAGTTAATAGAGTTGGAATCATTAATACA
CCAAACCAACCAATGTATAGGCGGTTTTCAGTACTAGTAATCCAAGAGCAGAAACGTTCC
CACAAGCTAGCGCTTTCGCGTCTTTGTAAAGTAGCAGTCATAATTTTTTATCAATTTTTA
AGGATTTAACAAAGATGCTTCCCAAACAATAAGTTGTCTGTTATGGAAATTATTACTTAA
GGCCTTTTCAAATGAGAACATTTTTAAAAAAAACATAGTAAAATCTAATCTCAGTAACTA
AAATTTTACTTGTTCATATATTGCTTAATAATTCAATAATAAATTTTTGCTTTTTGCAAA
ATTAACACTATTTTAGGGATATAATAATAATGCATATTAATGCTGCTTATTTTGATTTTA
ATAGTTTATATAATTTTGCCCTACTTAAATACGCATGTCACACTTTACAAAAATTCAGAC
GACCATACAAGATTTAAATCTGTTAAAGCATGCTTTAACAGATTTGGGTTTGATTTGGCA
AATGAATTCTAGTCATATAAAAGTTGGTGAAAATAATCAGCACAAGGTAGATATTTTAAT
TAAACAAGATAATTTGTCACATATTGGCTTTACTTGGAATGATAATAGATATCACTTAGT
TGCCGACTTACAACTTTGGGAACAACCCTGGTCTCTAGAAGTATTTTTGGATAAGTTATC
TCAGAAGTATGCTTACTACTCTATCATCGAAGAAACAAAGAAGCAAGGCTTTGAAAAAAT
GCAGCAGATTTCACAAAAAGATGGATCAATTAAGTTAATTGTGCAACGCTGGAATTATTA
ACTAACATACAAATGCGGGCGGAGAGACTTGAACTCTCACGAGATTATCTCACTAGAACC
TAAATCTAGCGCGTCTACCAATTCCACCACGCCCGCACTGTTGAGACAATATTACTCTAT
CATAATCTATACTAAAACACAAACTAATAGTATAAACTAGCTTCCAAGCTTAAAAGCGTC
TACAAATAGACCGTATGTTATACCAATTTTGAAGTTGTTGAATAAATTAATTTTTATATT
TAATTGCTTGTGGCTAGAGTAGACTATTTTACTAGTCAACTCAGTTGCTGCAACTATTAG

ACTGGCTGCAATAATACCCCAATCTCCTGTTTGTCCAGGAATCGTAGACAAACCTGTTGA
AATAAAAAATCCTAGCAATAAACTAATTAAGCCAGTTGTTAACTCGCTCAAAGAATAATA
AAGCTTATTATTTAAGTTTTTTATTAAGTAAGCAAAAAAAGTTGAAAGTCTAGTTTTAAT
CATAGTATTTAGTAACGGAGTTTTAAAAGGATGACAAGTATTAACAGTCATTAATATTGT
CGTATCCTTTTAAAACTCGTTAATACAAACAGCTGTTCTTGTTTTTAAAGGTCTTGTTCG
CTTAAAGCATTAATAAGATGCTGAAACGGCTCTTGTATAATATCTGTGCGTTTAATTGAT
TGCAAATCTAATTCTTCTTCTATGATTTCTTGTAATAACTGTATGCTTCTTACTGTTGGA
GCAACAGGTACAGATAGTGAATTGTAAGTATCTTTTAGCCCATTTAATACACGCTGATCG
AGAATATTTGTACTACCAGAAACCAAGGCATAACTAGCATACCGTAGGTAATATTCTATA
TCCCTCAGACAAGCTGCATATCTTCTTGTTGTATAAGAATTTCCTCCAGGCCTTAATAAT
TCAGGCTGTTCTTCGTACAACTGAGCTGCGGCCTCTTTTAAAATATTGGTAGCTTGATCA
TTAATAATTTCAGCTATTTTTATTCGATCTAGTCCACTCGAAAAGAAAGATTCTAGTTGT
CCGACAGCTGTCTTATCTAGATAACGACCAGTAAGATCGTAACGATTTAATATTGCTGTT
ATAGCATCTTGCATAAATTAATTCTCCTACTTGCTATCAGAGCGGTATTATAAATGACTA
ATATTGTAATATAATCATTAGTGTTTATGAAATTTATTTAAAACTTATTCTTTATTGCTT
AATGTGCAATTAAAAACTCGCATAGATTTTACACATATAGAGTATACGTAGTTTTTAATT
ATTTTTTTATAAAAACCAAGAAATAAAGCTTTTTTTAAGAGGTTCGTGAATTTGTTAATA
TAACATGAGCTTTCCTCCTAAATAAGTTTTGTAAAGTACAAAGTAAAAAAACGATTAGTC
AATAATAGTAAACAATATTAATATGTATCAAAGTATAAATAAGCTTTTACTCCAGCATCA
AAAAAAATATTTGAGATATGTTGATACAACTGATTATTCAAGTGAAAACTTAGATTTAGA
CCAATTTAGCTATTTTATCATTACAATTAATAAAACTGATAAATGTATTCTTATAGAGCA
ATTTTGTTATAATGCTGAAGGCCATATTTACTCAATATTCTTCAAAGGTCCAACAGCTAA
ACATCTTTCTTCTATAATTTGTCATTTTCAACAATTAAATACAATAATATCTACAGCTCA
TGCCATATATTTAGGACGAGAGCTAATGAAGTCAGAATTAGCTCTCGTGCTAGATCAACA
ATATATTCAAGATTAAATTTTGTAAGTATTAATTATATTTATGCTAGTATAATATTAGTA
AGGGCATGTAACTCAGTGGATAGAGTATCAGATTCCGATTCTGATGGCCGTGGGTTCGAA
TCCCGCCCTGCCCGAGTATTCAGAGTACTATACTAAAACATCTTAATAGCTAGTTTGATA
ATAGTATGGGCATTGGGGCTGCAAGGTTTCTACATTATGAAAAGAAGAAAATATGAAAAA
AAACAAGCTCTCCAAAAGAGCTTTTAGTACAATCAATAAATGCAGAAAACAATATTGTTT
CTTTTTCTCGAAAATTAGCTCTTGCTTAAACACTGTTAATTTTTGGCATTCGACATGTTA
AACTCTTATTCATGGCGAATATTCTGTCAGAGTCGCTCTTAGTTTAAGAAAAGTCAGAAA
AAATATTTATCTTTACGTTTTTACCTATTTTTATTTAAATAAAGGTTAGTATTTTCAACT
TTTATAGTGGACGTGGGTTCGAGTCCCACCAGCTCCATCAATTTGATTTGTATAATTAGC
TAGTTTAACCTTTTTAATAACCAGATATATAATTGGAGGAAATATGGGTTTCATAACAAT
TACTAAGCCCGCTTTAAAGCAAATTGCAATCTTGAAAAATGATCATGAAAATGATGTACA
TCTAAGAATAGGTGTTAGACAAGGCGGATGTTCAGGTATGTCATACTCAATGAATTTTGA

ACATGTTGACAAATTAAAAGATACAGATGAACGGCTTCGTCTAGATAATTTTTCTGTTGT

CTGCGACCCTAAAAGTCTTCTTTATCTTTATGGATTATCATTAGATTTTAGCTCGGAATT

AATAGGAGGAGGATTTCAATTTTCTAACCCGAATGCCAGCCAAACTTGTGGCTGCGGTAA

ATCTTTTTCAGGCTGACCCTCTAACACGTTTTTTATATATTTTGTATTATATATATTATT

TTAGCAAGTTGTTTATTAGAGACAAGCAATGCTCTGTTAAAAACTAATTTCAAGAGTTTT

TATTGTATTATTTTATATATTTATTATCAACTACTTGAAAATAAATTAGCTAAGCAATCT

AATTTGTTATCAGTACCAAACTTTAAATGATTAACTTTTTATTGATATATGCTAATCTTT

ATCAGCCTAAACATATCTGTACATTACAATCTTTTGACATAAAAAGAGTCTTAACATACC

TCTCAAAAATAAATTCAATATTTTATAAATACTCAACATGAATCTGTCACAACTTATCGT

TAATTCTTCTATGATGAAAATTCCTAATAACTTTTATTGCAATATAAAAGCTATTTCTTC

TGAGAATTTAAGTCACTTACAATTATCAAAAACAAATTTATTATCTCCTACAAAAATAAT

TGCTTTTGGACTACCTAAATTTGATTTTATTTCTACACAGATTACAAGTTCTATTAGAGA

AACAGTTGTTGTGTCTACTATTGATGATAATAATCTTTTTGCTGCCCCAACTAATGATTG

GATAAAAAAACACAAGGATAAAAATCATCATTCTTTGTTTTATAAAAAAGTTTTTAAAGC

TTTGCTAAACCATAAAATTAGAGTTGTTGCTGAGCCATTTCATAACTCCGAAATAAATAA

AGAAAAGAGCATAATGGCAAGTCAAAGATACATCTGGGGTAAAAGCTGGAAGCCTTCAAT

TATTTTATCTTGCTTAAAAAACAAAAAAAATAGCAAAATTTCTGATGCATCGAAAAAATT

TTTAACTGAACAGTTATCTGCATCACCAGTATTTGTAGTAAAAAATGGTTTTAATGAAAT

TATTTTAGGTCACCCATTATCTCGTGTTAAAAGAGGAGGAGTTAATAATCTGATGCATGC

GTTTTCTAATTTATTAAATCAGTCTAATGCAACTTATCCTATATCTACTGGTCTATTCTT

TTTTCATCCAGATGATGCATTTGAATTTAAAGACTTTATAATATCGGTCAATCCACTGGC

GGCCAAACATATGGAGATAAGCGTAGAGCCTGTTGGTCTACACTTTGCTTATAAAATGAA

TAGAAATATATCGTCAGATACTCAATTTTCGTTTATTCCAGATTTTAAAGAAGTAGGAGA

TTTATTATTCAAATATAGAAAAGGTAATCATTTAGTTTTTCATAAGAATCAGCATTATGG

CAAAGATTTTTTTCAGGGGCAACCAATATATATGATTCAGCCGATCACTTTTAAAGATCG

AGCCGGAAAGCTAAATATTATCAAATTTACGGGGCTAAATGACACAAGAGAAATTATTTT

TACTAATCTTGAAGCTGCTAATAGATCATGGGCACATTTTATAAAGAGGAATTCACAATT

AAAATCAATTAAAAATCCTACTCTATTAGTCTATAATTTAGAAAGTTTTTTAAAAGATCA

AGAACGATTGGATAATCAAGATTTAAATAAGTTTGTGGTGATCACTAATAAAGAATCTTA

TCTTGCCACAAAAGAGTTAATAGCTTTACCCGATTCTAATAGCTTTTCTAAGCACTTAAA

ATTAAATATAAAGCCTAAACTCTTTTTTGTGAAACTATGGGTAAGACGGTTATTTTCCAC

CTTAACTTATGAATAAATGCTATTGCCTTTATTTTCTCGAATAATATTCTATAACTAGTA

ATTCATTGAGTTGTAAAGCAACCCATTCCCTATCAATGACTCCATTGACTTTCCCTGATA

AATTGGATTTATTCAATTCTAAGTGGCTAGGAATATTAGCTAGTCCAGGGAAAGCTAAGT

AATTTTCTACTAGTTTTCGAGATGCTTCTTGATTTTTTACACTAATTGATTCTCCTGGTT

TACATTGATAACTACAGATAGATACTACTTGTCCATTAATACAAATATGACCATGGTTCA

CTAGTTGTCTAGCGGCAGGAATTGTAGGAGCCATACCAAGTCTAAAAACAGTATTGTCTA
GCCTCATCTCTAGTAGCTGTAACAAAATTTGACCTGTCGACCCTTGTAGTTTTTTAGCTG
CTTTTACGTACTTAAATAACTGTTTCTCACTTAATCCATAATTAAATCGCAACTTCTGTT
TTTCTTCTAATCTTACAGCATACTCAGAAGGTTTTCTGGACTTTTGTCCATGTTCTCCAG
GTGGATAAGATCTTTTAATCGCTTTTCTACTGAGTCCGGGTAAATCACCTAATCTACGGG
AAATGCGTACTCGTGGCCCTCTGTATCTAGACATATTATTTAAATTCTCCTAATGTAAAT
GATTAAGTAATGAGTAGCTATTTTTTAACACAAAAAGAGCATTAAAACAAAGCTTTTGCT
AAAAAATATTTACAAAAAATATAGATTTGGACAAAATACATTACTATGCTTTTTAAGCGG
GTAGCGGGAATCGAACCCGCATCATTAGCTTGGAAGGCTAAGGTTTTACCACTAAACTAT
ACCCGCAATATTCAACAGATTGTCACTAACTCAATATAACATAGTTATGTTAAGTTAAGA
TAATCTTAAATTTAATAGAATTTTAAATTATTTATGCTCCTTCTAAAACCACATCAAGAA
AACGAGCAATTTCGGCAGCTTGTTCTTCAACTTCTGATAATAATAATGGTTGACCGACCC
TAGTCAATGGAATTACTCTTTTATCTTTTGTACATAAATAAATTTCACGTCTTGGATTTA
GCCCCTCTTTAATATCTATTTTTATTGATTTGATTTCTTTTATATTAAACTGAAGACAGA
TTTTTCTATTCTTTCCAGGGAAACCTAGCCGAAAAATTTTTACGATACCTTTATCTTTGT
TAAATTCATTATATCCAGCCCCTATATTCCAAATAATAGTTAGCCACAGAAACAAACTCA
GAAAGACTCCTATACTGCCATAGAATGTCATTACAATACCTTGTGGAATAAATACCAAAT
CGGTTGAATTTGTAAAAGGTAACAAATCAACTTGAAAATAACTTGATAGTCCAGCAAGAA
GAAAGCCCAAAGCCCCAATAAATATTATAGTAGCCCACCAGTAATTACTCAGTCTTCGTG
AACCTAAAATTAAATCTTTTCTAACTGAGTGTATAGATAATGTTTTTTTCATAATTTGTT
CTTTTAAAACAGTATACTAATGCAAAATGGAAAAATTTTGACAATTTTTATGCGCGCTTT
AGATTTTTTAAATAAGCTTGATTGCTTTTAGTCCTAAAAACAGTCCTGTTGCTAACCCAA
AAAAAGCAGCAAGAAATATTATATATCCTAGAAAAACTGACATATTATTATTATTTATTA
TTATTGTTACAATAGTGTTTACAAAAATAGTGAAAATCTATTACTACGGTATTATATTAC
GTTAAATGCCTTGCTGTCTACAGGAAAAAACTTTTTTCAGTTTTTAGCATTGTGTTTATA
TTAGAAATAATATTCTAGTAAATTTATTAAACTTCTGGAGAACAGAATCTTATGTCAGAA
TTTATCAAGCCTTATAACGATGATCCTTTTGTAGGCAATTTGTCCACGCCAGTTAGTACG
TCAAGTTTTAGTAAAGGACTTCTAGGAAATCTACCAGCTTACCGTCGAGGTTTATCTCCG
CTTCTTAGAGGATTAGAAATAGGAATGGCACATGGATACTTTTTAATTGGACCTTTTGAT
AAATTGGGCCCTTTACGAGGTACAGATGTAGCGTTACTAGCGGGATTTCTATCTTCGGTC
GGCCTCATTATTATTCTCACTACATGTTTATCCATGTATGGTAATGTATCTTTTACTAGA
GCAGATTCGAAAGATCCACTACAAACTTCTGAAGGCTGGGGACAATTCACTGCAGGATTT
CTAGTTGGAGCAGTAGGTGGTTCAGGATTTGCTTATTTATTGCTAGCTAATATACCTGTA
TTACAGACTGCAGGTCTTAGTTTATTCTCTTAAGCTAGTAAGGGGACTTGAACCCGTAAC
CTACTGATTACAAATCAGTTGCTCTACCAATTGAGCTATACTAGCATTCAGTTCACAATA
ACACATAAAGATATTTGTTGCAATCTGCTATTTTCAGATGTAACAAATATCTTTTTTATT

TAAACAGAACTGTTATGTTAACACGTATATTAATTAGAATCTGAATTATCTGACTCTGCA
GAAGATTCCTGATCACAGTTTAGATAATAACAAATTGTTTGATCTAGACAAGTTGCAGAC
CAACCAACAGGTGTTTCTCCTGATAATTCTTGCATATCTACAGAATTATAAAAATTATCA
TTTGGCTTATTACTGTTTGCTTGCATAAATATCACTCCCAAAACACTCTGAACTTAATGT
GAAGTAATTTAACTTCTCAATAATATATTAACACAAAATATTAAATATGGTGATTATTCT
TTAATTGTAAAGATGCTGCTAACTTTATAAAGTTTGAGTGAGAGTCTTGATAGCTTTTGT
TGAAATAAGCATTTTAACCCATTTATTTTGCCCTCTCGACCATACTTTTTTGTGTTGTAA
ATTAACTTTTTGTAGTTTTTTTGTTCGTTTATGAGAATGAGATACGGCATATCCATTATT
AGCTACTTTTCCCGTAAGCTGACACTTTTTTGACATAATTTTCTAACTTATATATTTATT
TAGAGTACTTGCTAAAGTTGACTTGGGCACAGCTCCAATCACTGTGTCTACTCTTTCTCC
TGCTTTAAAGATCATTAAAGTAGGTATACTTCTAATGCCATATTCAGCTGCAATAGTAGG
ATTATCGTCTGTGTTTATTTTTACTACCTTAATAGATGATTCATATTCTTCCGCAATTTC
ATCAACTACAGGAGAAACCATTCTACAAGGACCACACCATGGCGCCCAAAAATCTACGAG
TACAGGTAAGTTATTGTTAATAACTTCTTGTTTGAAAGAGGCATCTGTAACTTGAGATAC
TGACATATTCTTTAACCTTTAATATATGTTCCTTGCTAAGCAAATCTTATCACAAATTTC
AAGAAAAATCTTCTATAGTGAATTACTATTAGTGTTGAATTATTTTTTTAAAAATTAGAC
ATTAAAAAAACTTATTACCTATATAAATAAAGTGGTGGCCGAGTGATATCTTGATAGTGA
TAATTTTTATATTAAGGTTGAACACTAGAAATAACTTTTCAAGCAAACTCTTGTTTAAAA
AATTTCTCTGTGACCATAAAAACTTAATTATATTACAATATATTTTATTATTTGTAACTT
AGTTCTGATAATGGTATAAACAACGCAAAAGATACTGCCTTATAATCAAGGAGGAATACA
TGTCTCAATCCGTAGAATCACGGACTAGGATTAAAAGCGAACGTTACGAATCTGGAGTAA
TCCCCTACGCTAAAATGGGCTACTGGGATGCTGACTATGTGATTAAAGAAACAGATATTC
TAGCTCTTTTCAGAATCACTCCTCAACCAGGTGTTGACCCGATTGAAGCATCTGCTGCAA
TTGCAGGTGAATCTTCAACAGCTACTTGGACAGTTGTATGGACTGATTTATTAACAGCTT
GTGACTTATACAGAGCAAAAGCATATCGAGTAGATCCAGTTCCAAACGTGGCAGATCAAT
ATTTTGCTTACATAGCTTATGATATTGATTTGTTTGAAGAAGGTTCCATTGCGAACTTAA
CTGCTTCAATTATTGGTAACGTTTTTGGGTTTAAAGCTGTTAAAGCTCTTCGCTTGGAAG
ATATGCGTATGCCAGTAGCTTATCTAAAAACGTTCCAAGGTCCTGCAACTGGATTGATTG
TAGAACGTGAGCGTATGGATAAGTTCGGTAGACCTTTCTTAGGTGCTACAGTTAAACCTA
AACTAGGTTTATCTGGCAAAAACTACGGAAGAGTTGTATACGAAGGCCTGAAAGGCGGTC
TTGATTTCCTTAAAGATGATGAGAATATTAACTCACAACCATTTATGCGTTGGAGAGAAA
GATTTTTATATTCTATGGAAGGTGTAAATAAAGCATCGGCTTCTGCTGGCGAAATTAAAG
GTCATTACCTTAACGTAACAGCCGCGACAATGGAAGATATGTATGAGAGAGCCGAATTCT
CTAAAGAGGTTGGTAGTATCATTTGTATGATTGACCTTGTGATTGGTTATACTGCGATTC
AAAGTATGGCAATTTGGGCTCGTAAACATGACATGATTTTACATTTACATAGAGCTGGTA
ACTCAACTTACTCTCGTCAAAAAAATCATGGTATGAACTTCCGAGTTATTTGCAAATGGA

TGCGTATGGCTGGTGTTGACCATATTCACGCAGGTACAGTTGTAGGTAAGCTTGAAGGAG
ATCCTTTAATGATTAAAGGCTTCTACAATACTCTACTTGAAAGCGACACAGATATCAACC
TACCTCAAGGTCTGTTCTTTGCTCAAAATTGGGCTTCCCTACGTAAAGTTGTACCAGTAG
CATCTGGTGGTATTCATGCTGGTCAAATGCACCAACTTCTTGATTACTTAGGTGATGATG
TAGTTCTTCAGTTTGGTGGTGGTACAATTGGACATCCTGATGGTATCCAAGCAGGTGCAA
CTGCTAACAGAGTAGCACTAGAGTCCATGGTTATGGCAAGAAATGAAGGCCGTAACTATG
TAGCAGAAGGTCCACAAATCTTGAGGGACGCTGCTAAAACTTGTGGGCCTCTACAAACAG
CTTTAGATTTATGGAAAGATATTAGTTTCAACTACACTTCCACAGATACAGCTGATTTCG
TTGAGACTCCAACAGCAAACATCTAGTTTAATGACTACTTACTGATACTTTAAATAGTCA
ATTGTAAGTGGAATTAACTTATAACAATAAGGAGCATAGAATAGTGAGACTAACACAAGG
GACTTTTTCCTTCCTTCCAGATTTAACTGATGAGCAAATTAATAAACAGCTTGCTTATAT
CGTTTCTAAAGGCTTATCAGCAAACGTTGAGTATACTGACGATCCTCATCCAAGAAACTC
CTATTGGGAACTGTGGGGTTTACCTTTATTTGATGTAAAAGATGCTTCTGCTGTTATGTA
CGAAATTAGCTCATGCAGAAAAGCAAAACCTAATTATTATGTTAAAGTTAACGCTTTTGA
TAATACTAGAGGTATTGAAAGTTGTGTAATGTCTTTCATTGTAAATAGACCTGCTAATGA
ACCAGGATTCTTATTACAACGCCAAGACTTCGAAGGTAGAACTATGAAGTATAGTCTTCA
TAGCTATGCTACTGAAAAGCCTGAAGGAGCTAGGTATTAATATTAATTAAGAATTAATAT
TGGCTAATTATTACCCTCTTAAAAATCATAATTAATTTAGTAATTGTGATTTTTAAGAGG
GATAGCTTCCCAAGTAAATTTTGACTAAATAGATTGAATAATTAAAAATACACAGAAATA
ATGCAATCACAGGATATAATTTCCAACGATACTCTTGTTAATTTACAAGAAGAATATGAT
AGAACACAAATCCAAGAAGTTTTAAATGAGTTAAATCAAGAACTTATAGGATTAGTGCCT
GTAAAGACCAGAATTCGCGAAATTGCTGCGCTATTATTGATTGACAGATTACGCAGAAAA
CTAGAACTAGTTTCTGGTAATCCAGGATTACACATGTCATTTACAGGTAGTCCAGGAACT
GGTAAAACTACAGTTGCTATGAAAATGGCTGATATTTTGCACAGACTTGGATATATAAAA
AAAGGGCATTTGTTGACAGTTACAAGAGATGATCTTGTAGGTCAATATATTGGACATACT
GCCCCTAAAACTAAGGAAGTTCTTAAACAAGCAATGGGAGGAGTTTTATTTATTGACGAA
GCTTACTATCTATATAAAGCAGATAATGAAAGAGACTATGGCTCTGAGGCAATTGAAATT
TTATTACAAGTAATGGAAAACCAAAGAAATGACTTAGTTGTTATCTTTGCTGGATATAAA
GATAGAATGGAAAAATTCTACGAATCCAACCCAGGACTCTCTTCTAGAGTAGCTAATCAT
GTAGACTTCCCAGATTATACTTCAGATGAATTATTACAAATAGCTAAAATGATGATAGAA
GAACAGCAGTACTGTTTTACAGAAGAAGCAGATAAAACTCTTTTAGAGTATACCGAGCGA
AGAATGAAACAGCCTTATTTTGCTAATGCAAGAAGTATTCGCAATGCTATTGACAGGGCT
AGAATGAGACAAGCCAATAGGATTTTTGCCAGTGGAGAAAAAGTATTAACAAAAGCTGAT
TTGGTAACGATTGAAGCAGAAGATATCTTGAAAAGTAGATTATTTTCATTACCTAATGCT
TAATATACACGTGATTTCATGAAACTATATATTATTAAAAAGTTTCATAATCAGTTGCAA
ATTTCTTGAAAAACCTTTATTATTATCTTAATGCAGGTGTGGCGGCATAGCCAAGTGGTA

AGGCAGAGGATTGCAAATCCTTCATCCCCCAGTTCAAATCTGGGTGCCGCCTAGTACTAA

AAAAGGGGGGTGTGGTGGAATGGTAGACACAACAGACTTAAAATCTGTTGATTTTTAGTA

ATCGTGAGGGTTCAAGTCCCTCCACCCCCATATATTGATTTATAAATAAAAAGAGATGTG

TATTTGTATTAATTGCAACCATATTACCAAATGTAATACCTATCACTTAATAGAATCTCA

ACATAAACAGCCTCATCTTACTAGAAGTCCATTGTTTATACCTAAATATCCTGTAGTTCA

TGTTAATATATCTAACAACTGTACCTATAATCAAATTGATTGGGATTTAGTGGAGTGTTT

ATCATTTGTAGAAAAACCGAATAGTTGGAATTTAGACGCTAATTAGTTACAAGAAAAATT

ATGTACCATAAGCAACAACTCTATTTTTTAGATACTCGGTATTAATTTTGGATGTTCTAA

CCAGTAAGATTTTACCCGTTCTTGCTATCTCTATAATGCCAAATTTAGTTAGTAATTGTT

CAATAGCAACAATCTTTCCCGGATCTCCAGTAACCTCTATAATTAAAAGATCTTCTGCAA

TGTCTACTATCTTAGCTCTAAAAATTCTTACAATTTCTAAAGCTTCTGTTCTAGTCTGAG

AATTGATCTGAATCTTAATTAACATTAGTTCTCTTTCAACTGAAGGAATATTTGTTACAT

CTTGAACGTTAAGGATATTTACTAACTTATATAATTGTTTCGTAAGTTGTTCAATAGTTC

TATTGTCTCCTTGAACTACCATTGTGATTCTAGAGACCCCAATTTGCTCTGCTGGTCCAA

CTGCTAAACTTGCGATATTAAACCCTCTCCGGGCAAATAGACCAGATATTCTTGACAGTA

CTCCGGCTTCATCTTGAACTAAAACTGATAAGGTGTGTTTCATGAAAAATTAAATTTAAT

CTATTTGATCTGTTAAGTATATTCTAAGAAAATTACAGAAAAAATACTAGTTAGTCTATA

ACTAAATTATCCAAACTAAACAAAATCCTACTATCAGATAATATATCTACCTTGTAATGC

TATACTATTAACTTAGTATTTAATGATATTCTAATTGCAAATAGAATGCTAATTTTTTAA

AATTTGTTAGGAACTTGTGCTAGAAAAATACAAAAACAGTAAAAAACTCTCTCGAGATTT

ACCTCAAATCAATGATCGCATTAGATTTCCAAAAGTCCGAGTAATTGATGACGAAGGTGA

ACAACTAGGTATTTTTGTGCCTGAAGAAGCTATACAATTAGCTGTCCAACAAGGTTTAGA

CTTAGTTGTTGTTAGTGATAAATCGGACCCGCCAGTATGCCGAATATTAGACTATGGTAA

ATATAAGTTTACACAAGAAAAAAGAGCTAGAGAAGCTAAGAAAAAGCAACATAACAGTAG

TATTAAAGAAGTAAAAATGCGATATAAGATAGAAGAGCATGATTATAAAGTTAGAATAAA

CCAGGCATCCAAATTTATTCAAGCAGGAGATAAAGTAAAAGCAACTATCACATTTCGGGG

GCGTGAAATCCAGCACTCTAATTTAGCTATAGATTTATTGAATAAAATGGCAAGCGATCT

AAATGCAATAGCTGAAATTCAGCAAGCTCCATCAAGAGATGGCAGAAATGTCATAATGCT

CTTATCTCCCAAAAAAGTTAGCTAAACTAATTTTTATTTAAATGCATCTGGCTGGATTCG

AACCAGCGACGTCTCTTTCGAAATGGCGGATTATGAGTCCGCTGCCTTCGGCCCCTCGGC

CACAGATGCATCACTTAAAGTTTATACCTCATAAAGCATATAAAAATCAATGTTTAGCTA

TTTTTAATAAAGTCTATAAGAGTCTATTATTCATTCATATTGTGATATGTCGCAACTGCT

GAAGGGGAAACTTGATTTAAGTATCGAAATATCCAGTATTTAAAAATTGTGTCGAGTATA

ACTGGAAAAGTTGAAATAAATAAAAAAATAAAATCTCGACTTTCCGGAAGTCCTAAATGT

CTTAAAATTACTTCAATAATAACTTCCCATCCATGAGGAGAATGAAAACCCACAAACATA

TCAGTAAAAAGAATGATTAAAAAAGCTTTAGCTGTATCACTAAGACCATAAATAATTTCA

TTTAAAAAAAGATTTAACGACTGCAATTTGTCTTTGTCCAGTTATCATTAGTAATATAAAG
ACAAGTATTGATACAAGATCGGATAAAATATTTTTAACAGCATTTGCACTTTCATTTGCA
TAATATTCTCCCAACTCTCTAGCCTTCAGCTGCACTCTTTTTTCAATGATTTCGTAAGAT
ATATCTTCCGAAGGATTCAGAAGAACTTCAAAGTGAATTTTCTCTTCAAATCTTTGTAAT
TCAGCAAAAGCTCTTTCCTCTTGAGAGGAATTTAAAAAAATTTTAGGCTGTTCTTGGTTC
CATAAATAATCAATACAAGGTCCAAATACAAAAAATTTTGAAGCCTGATTAACTAGTACT
GGAGAGATGAATAATAATAAAATATATTTTACGGAAGTAATAGTTTGGTGTCTAGATATT
CTAAACTCCTCTATAGCTTCAGATTCGCCATTAGGATCTAACTCTTTACGAAACTTTTCA
AAAGTATTAGTAATTGACCTAGGTATTGGCCCAACTTTTTCAAAGGCGAATTGATTATTT
CGTTTTAAATTCCAATATTTCATTTTTTATTGCTGCTAAAATCTCTATTAATCAATAATT
GATCACGTATTCACTATAAATATTAATTCTAGAACTAAACATATAAAACAATATTTATGA
AAAATAAAATACAATTAAAAATTAATTATGCATTAGTTCTATCTGCAGCTACAGCAGTTT
ATGGATATAAATCTTATCAGCCAACTATAAATACACTCTTAATCAAAGATGAAGAAAACT
GGCCTTTTTTCTCTAAAAGGCAGACTGCCTTTATTCCTATAATTGTATTAAAAAAAAATC
ATCAAAATCCCTGAACTCAGAACCTAGAAATAGTAAGTTATATCTTAATGGAATTGGAAG
TAGAAAATTTATCGACAGAATTGTCACTCTACTTGGATATTTAGAAAAACAGGCAAACTA
TACTGAATACTTGAACCAAATAAAAACAAGTGGTTATTTTTCTTTAGTCTACCTAGACAG
TGATATAATGTATAAGAAAAATATTAGTAACATTTTTTTACTACCTAATGATATTCTAAA
AAGAATCTATATTTGCAATAAGAAAAAAAAAGCTTATACCGCATTCATTCTTAATAAATTT
ATTTCAGAAACAGATAGGCTATCCAAAAAGTTTTGCTAATTTAAATTGGGCCTTCTCCAA
AATTATAAAATGGTATTATGATAGAGGATACCAATGGTCTCTAGTTGAAGTTAAACAAGC
GTCTGATGCATCTTCTATTGTAATTGATATTCATGAAGGGGTAGTGAAGACAATTATAAC
AGAATACTACACTCTATCTTATAAAAGAGTCTCAGGTATCTTATGCGTAGAGTCAATAGA
GCAGTACCTTGGAGTAAGAGTGGGAGCTCCACTAAATATAATTGATTTACAAAAAAAAAT
TACTTACTTAAAAGATAATCAACTAGTTGGCGACATTATTTATAGTATTGAACGATCAAA
TAATAGCTCAATGAGTTTGGATATCAAATTTCAAATACAAGAATTAAAAGATAAAGAGAT
AATAGTGCTTGCAAAAAGTTCTTCTATTATCTCGCATGCATGTAATTTACTTAATCAATA
TAGAAATAGGCTAGTAGCTTCTAATATAGTTTCACTATCAACTAATAAATTACACGCATG
CTATTTTAATTATAAACTTGACTATCAATATAAGTATATAAACACGTCTAATCTTATAAA
ATTGTTAGCTTATTCTATTTCTTGTAGGAAGACACTACTTATTCATTATTTTAATTTACA
AACTTTAATAAGCTATACAAAAAAAAAATACTATTGGATTTCAGCTATATTTACGAAATCT
AAGTTTTGGGAAAGCTTTCTGTGTGCTTAGTATGAAATTTATCAAAAAATGGACTTAATAT
AAAAATTTTATATATAAATCCTTCATTGATAGTTGATCAGAATTTTGTATTTCAATTTGC
TATACAAATCATTAAGCAATATCATACTGCTAAACCTCCAGCTTTATTTTTAACGAATCT
AGATCTTGAACAATATGTTGCTGAAAGTTTATTAATGTACCATTTTACATCTTGCTTCTC
AATATCTGAAAAAATATTATTATCTCGAATTATGCATACAGATTCTTTATTTTTCAACTC

TGAAAATTTTCATTTTGATGATAGAACTAACGCGATCAACAGTTATGATACTTTTAAACA
AAATACAAAAATATTTTATCAAGAGTTTCTTTCTTTATTATTAAGTTTACGTTATCAAAA
TTTTAATTATTTAGGTTGGCCTTTAAAAGGCCATTTTTTTGAAATAAAATCTTTATACTT
GGCTCCATTTCAAAAGAGTGATTTTTCTGATAGTCGTAAGACCCTATTCTTCCACAAAAT
GTCGTTAAAGCAAGTATCTAATTTTAATTTACCAATATCTTTTAAAAGCCACCTTAACCA
TATTTTAGTTAGTACTATTAAATGTCAGTCAAATTTGAATATGAGAACTGTATCTTTGCT
ATTAATTGATTCACCTGCTGAATATATGCTATACAAATCTATTCTTAATTTCTCTATTAA
AGTTAGGATGCAGTATTTCATACCTATGAGCAACAATATAAGATTATCTCTATTTTATAA
TTATTTAGATTGTTTTTTAATAAGATCATCTCAGAGTCGCATTCATATATGGCAAGATCT
AAGAACTCTTACACCAATTCAAAATTTTTGGTTAAAAAAATTTTCATATGGAGCTGGAAT
TCAGCTTAAATTACCAATTAAGCAGATGCCACCTCTTTCAATTGAATATACTGTAACTAG
TAGTCGCTACTTTTGTATTTACCTTCGTACTTATTATCAACGATAACTATTACTAAAATG
AAAAATTTAATTTCTTTTTTTGATCGCAGCAAAGGGAAATGGATTTCACAAAGAACAACT
TATGAATTATCTAATAAAAACATGAGTTCGATACAGTCTCAGATGACAATGAAAATAGGC
AACTCATTGTCAGGATCTATAATACTTGCATCGTTAAATTGGGGGGACATTTATAGACAA
GTTGCCCATTACGCTAAAAATCACAGCCGGAGTGAATATGATAATAAGTTTAACTTGCAA
TTCAGTAATCAACTAAATAATCATAAATTATTAACGCTGTGCATAGTGACAGATCCTAGT
CTAATTAGTTTTAAAACTCGCTATGGAAGTACTACCATAGATGAAACATATTGGTTCGCA
ACAAATAATTTACGTCTAAGTACTAGTATTGTTAAACGATTCAACACTTGTGTGGCAGTA
TCTTTTTGTTCAGAAATTAAAGTTTAAACTCTATATAGAAGCAACTAATAGCCTAAAGGC
TATTAGTTGCTTCTATATATCTAAATTTTGTATTAAAAATTATAATACTATATTATTCTA
AGTCTTTACGATTGGGATTCCTGGAAGGATCATTAGAGAGGAACCCAAAGACAAACAAAG
AAATAAAGAAAATAACGGTAGTATATACAAAGATTTTCAAAGTAAACATATTTTCATCCT
TAAAAGCTATTAGTGCGATATTTACATATCTCCACATAATTTTATATCATGATATAAGTT
CCAGCACTATTTTATAAATAAATCAATATCTTTATTTTTTAGTAATGATTTATATTTAAA
TATGCGTGATGTACTAAATACTAAATCTTTTCCGACTTGTTTTGTCTTGTTTGTAAATAT
TTTGTTATTTCTATGGAGTTTACCTTCAATTATCACATAGTCTAACTTCTTTAGCTGTTT
AAAAGTATATGAAGATTTTTTGTTCCAAATAGTTAGATTGATTATAACTAATTGTTTTCT
TTTTAACAGTCTTGCTTTTAACTTAATTATTTGGCTTTTATTTTCACTAATTCTGATACT
CTTACAACTAAGAATTTGGACTAGAAGAGTACAGCTATTCATAATTTATAACAATTTTTT
AAAAAATCTACTAAAATAGATGATTTTACAGAACTAAAATGATATTTCATTATCTTGTTG
ACCTTGCTCATAAGTAATATCTTTCAATACTTTTAATATTTTTCCAAAATACTCTTGCAG
GTATTTCTCTAAAGATATTACTTCCAACCTGTCAATTCCAAGAATACTGTAAACTTCATC
CATCGGAGCTGTAAATTGCTCTCCGCTACTTAAAATTTCGGCAAATGCTAAACGATCAGA
AATATTCCATGTCCATTGAAGAGTTTTAGTAATTCTTCTTAAGGCTTTTAATAATCCAAT
CGGAATTTGTGAGATCTGTGTTTTTTTGCCCAGATAATTTTTCACATAATTTAATAATTTC

AGCTGATGTCCAAGCGGTATTACCAACTAACGGCAGGGTCCTATTTTCTGTGGAAGGTAC
TCCAAGACTTTTTATAACTAACTTTGCTGCATCTTGAGTATCAATGTATGCAATGGGTGT
AGATTCTCCTGTAACCCAGACTGATTTTTTGTCTAAAATAGGAATGGCATATTGGTTGAT
TAAACCTTGAAAAAATCCTCCTAAAGAAAACACAGTATATTTCACATTGGACTTCTGAAG
GAAATCTACTACCTGAGACTTTAAGTTCATCAATGGAACATCTGGGTATTTCTCTGAATT
TAATATTGAAAAGAAAATAAATCTTTCAACTTTAGCCGCTTTAGCTGCTTCAATTAATGC
AATTTTCCCATCTAAATCTATTTTTTCTGCATTGTAAGGATCAGTAGGACGAGATGTAGA
AGCATCTATGATTGCTGTTACCCCACAAAAAGATTGCAAAATACTTTCGGGTAATTTTAA
ATCACCATATATAAGTTCTGCTCCCCACTCTTTGAGAAAGGCAGACTTTCTTAAATTTCT
TACCATGCATTTTACATTGTAGCCTTCATCTAAAGCTCGTCTTACAATTTGACGTCCTAA
AGTTCCGGTTGCTCCAATTACTAAAAGAGTCATATTAGTTAAAAATAAGTATGTATAAAG
ACATTGATTTGGTTGACGGCAGAGAGTAAATTAGGTAGAGAGGGAATCGAACCCTCATGA
CCGAAGTCGTCACATTTTGAGTGTGATGCGTATACCAATTTCGCCATCTACCCTATTATA
GACCAACTACTACAAGTAATTTAGTCTTTAGCGTAAAAATAATACTAAGTTTCAATGGAT
ATGTCAACTTTTAAATTTATATTAATAAAAAAACGGAGTCTCTTATGGCCACTTTTGAAT
TAATTCCTTATAGATTGTATTTGAGTCAATCTAACACATGGATCCATAGAATAAAAGCAG
AAATTAAGATATATATAGTAACGCTTTTATGGATTTCAATTTTTATTTTTTCTTACTTTA
AACTATGTATTATTGCTTTAAGCTTAATTGCCATAAGTTTTACTATAAGAAGTAAACAGA
ATATTATCCAAAAACATTTGTTACAAACGTTATTAATAACGTTCTTAACTACTGTTTTGT
CTTTTAGTGTGGCTATTAGTTATAAACAATATGCAGAACAAGAACAATCGCAGTATTTAT
CTGATTCCAAGAAATATAAAAATTCCAGCAGCTACTATTATATACAAATAGCTAACGATC
AAAGGATCAAAAGACAATACTTAATTACTACTTTAAAGCCTTCCTTGTACTTTTTTATTA
CTATATATTCTATTAAACTAGTTATGATAACAACTTCTCCAGAAGTTTTAGTAATTACTA
TTTATAGATCTAGGATAATAAATAAAATATTTAAAAATGAATTGCTATTTATCTTCCTTC
TTTCGTCACATATTGTTACTAGTATTATTAATAGAATCGATAAAGTAATTCAAGTAACTA
GCTTAAGAGGAAGCTTGAATTTATATAATTCATTGACAAGGCCTTTAATGTTTTCTTTGT
TAATATTTCAAGTCTTCTTTTTGGAGATTATTCGGGAGTCAAAAGAAATAGCTCAAGCTC
TTTATACTAGAAATCTCAATCAAGAAAACAATAATTTTTTGAAAATATATACAGTAAAAT
CTAACTTTAGTGATCGGCTCAATATAATTATTAGCACTTTGTACTTTATTATTTTAGCTC
TAGCGTAATAGAATCCACTTATTCTTCTGACTCTAGTAATACGAAGAGTAATAAAATTAG
TTAGTAGTTTAAATAGATAGAATTAAGATATTATATAAATAATTAAATGTTTCATTATGG
TTAATACTCAAAATCAAATTTCACAAACTTCAGAACTTGATTATATAGTTAATCAACCTT
ATAAATATGGCTTTAAAACTTCTGTTGAATCTGAGCAATTTCCTAGGGGAATAAGTGAAG
ATATTGTCCGCTTGATCTCCAAGAAAAAAGATGAGCCTGAATATCTATTGAATTTTAGGC
TTAAAGCATATAAAAAATGGAAAAAGATGAGTAGTCCGTCATGGGCTCATATTAAGCATC
CGAATATAGACTTTAATACGATTATTTATTACGCTGTTCCTAAATTAAAAAAAGAATTGA

AAAGTCTAGATGAGGTTGATCCAGAAATTCTTGACACTTTTAACAAGTTAGGTATATCTT
TAAATGAGCAAAAGCGAATTTCCAATGTAGCCGTTGATGCTGTTTTTGATAGTGTATCTA
TTGCAACTACTTTTAAGAAAGAACTGTCTGAAGCTGGTGTTATATTCTGTTCAATTTCTG
AAGCTATAAGGGATTATCCAGAATTAATAAAAAAATACTTAGGTACTGTTGTACCAGCTG
GTGATAATTATTTTGCTGCATTGAACTCTGCGGTATTTAGTGATGGCTCTTTTTGCTATA
TCCCTCCTAATACAGTTTGCCCTTTAGAATTATCAACTTACTTTCGTATTAACAACGAAG
AATCTGGGCAATTTGAAAGGACACTAATTATTGCTGATCGTGGCAGTAAAGTAAGTTATC
TTGAAGGTTGTACTGCTCCTCAATTTGACACAAATCAATTACATGCAGCGATTGTAGAAT
TAGTAGCTCTTGAAGGAGCCGAAATTAAATATTCTACAGTACAGAATTGGTATGCTGGTA
ACAAAGAAGGTAAAGGTGGTATATACAATTTTGTTACTAAACGAGGCTTATGCTCGGGTA
ACAATTCAAAAATTTCATGGACTCAAGTAGAAACTGGATCTGCAATTACCTGGAAATATC
CAAGCTGTATTTTAGCTGGTGAAAATTCTCAGGGAGAATTTTACTCTGTAGCTTTAACAA
ATAACTATCAAGAGGCTGATACAGGGACTAAAATGATCCATATTGGTAACAATACAAAAA
GTAGAATTATTTCTAAAGGTATTTCTGCGGGTAGATCGAAAAACAGTTATCGAGGCCTAG
TAAAAGTTGGCCCTCAGTCATTTAATTCTCGTAATTATTCTCAATGCGATTCTTTATTAA
TTGGTCAATCATCTCAAGCTAATACATTTCCCTATATTCAAGTACAGAATCCAACATCAA
AAGTAGAACATGAAGCATCCACTTCAAAAATTAGCGAGGACCAAATTTTTTATTTTTTAC
AAAGAGGGATCAATTTAGAAGAATCTATTGCTCTTATGATCAGTGGTTTTTGTAAAGATG
TCTTTAATGAATTGCCTATGGAATTTGCTACTGAAGCTGATCGTTTACTGAGTTTAAAAT
TAGAGGGAACTGTAGGATGAGCCAGACTATTTTAGAAATCAAAGATTTATATGCTTCTGT
TGGTGAAACAACAATTTTAAAAGGGGTAAACTTATCTATTCGAGCCGGCGAAATACATGC
GATTATGGGGCCTAACGGTTCAGGTAAAAGTACATTATCAAAAGTAATTGCAGGACATCC
AGCGTATTCACTAATAAGCGGAGACATTTTATTTTTTGGACAAAGTATTCTTGAAATGGA
GCCAGACGAGAGGGCAAAAGCAGGTATTTTTTTAGCTTTTCAGTATCCTGTTGAAATTCC
TGGAGTCAGTAATTCTGATTTCTTAAGAATTGCATTAAATGCTCGCAGAAAGTTTCAAGG
ATTATCGGAGTTTAGCCCCTTAGAATTTTTTCAACTAATAACAGAAAAAATAGATCTTGT
TGGCATGCAAGAAAGTTTTTTGACAAGAAATGTTAATGAAGGATTTTCTGGTGGAGAAAA
AAAACGTAACGAAATTCTTCAAATGGCTTTACTAGATAGCAAAATATCTATTTTAGATGA
AACAGATTCTGGACTAGATATTGATGCATTACGAGTAGTTGCAAAAGGAATTAACACTTT
AGCTAAATCAACAAATTCAATTATTTTGATTACTCACTATCAAAGATTATTAGATTATAT
TATTCCAGATTTTGTTCATATTATGAGTAACGGACAAATTGTAAAAACTGGTAGTGTTAC
TCTTGCCCAAGATCTAGAAAAGCACGGATATGACTGGATTACGCAGACATAACTATTTAC
TACAATATTGAGACTAAAGAATATTCTTTAGTCTCAATATTTTTGCAAATGCAAAGTTAA
TTATTTAAGCTTTATCAAAACCTTGCTTAACATCTTCAATTGCTCTTTTTAGCAATTCTT
CACTAACTGGTTCTAATTTTTTAGTACTACGAACATTTTCACCAAATTCTGGTTTAGAAT
TCTTTAAATCTTCTCGAAGTTCTCTAATAAATTCAGCTACTTTTGAAACTTCAATATCAT

CTAGATAGCCATTAATGCCAGTGTAAATAATAGCAGTCTGTTCTTCAACTGGAATAGGAG
AATTTTGAGCTTGTTTCAAAATTTCTCGAAGACGTTGACCTCTTGCTAGTTGATTTTGGG
TTGCTTTATCTAGATCAGATGCAAACTGAGAAAAAGCTTCTAATTCAGCAAATTGTGCTA
ACTCTAGTTTTAACTTGCCTGCCACTTGTTTCATGGCTTTTATTTGCGCAGCAGAGCCTA
CTCTAGAGACTGAGATCCCAACATTAATAGCTGGTCTAATCCCTGAGTTAAATAAGTCAC
CAGATAGAAATATTTGTCCATCTGTAATTGAAATTACATTTGTTGGTATATATGCAGATA
CGTCCCCAGCTTGAGTTTCAATAATAGGGAGAGCTGTCATACTACCTCCACCTAATTCAG
CATTCAATTTAGCTGCTCTCTCTAGCAATCTAGAGTGTAAATAGAATACATCCCCAGGAT
AGGCTTCTCGTCCCGGAGGTCTTCTAAGCAAAAGAGACATCTGACGATAAGCTTGAGCTT
GCTTAGTTAAATCATCATAAATAACTAATGTTGCTTTACCTTTGTACATAAAATATTCAG
CTAAAGCCGCACCTGTATAAGGAGCAATATATTGTAAAGTCGCAGGACTATCTGCATTAG
CTGCAACTATAATAGTATAATCAAGAGCTCCTTTATCCTGTAGTGAAGATACTACTTGGG
CTACTGAAGATGCCTTTTGTCCTATCGCAACATATACACAAACAACATCTTGGCCTTTTT
GATTAATAATTGTGTCCAATGCAACAGCTGTTTTACCTGTTTGACGGTCACCAATAATTA
ATTCTCGCTGACCTCTACCAATTGGGATCATAGAGTCAATAGCAGTTATCCCTGTTTGCA
TAGGTTCACAAACTGATTGCCTTCCAATTATACCAGGGGCCATTGACTCAATTAGCCTTG
TTCCATTACTTGCAGGTTCACCTTTATCATCTATTGGCCGAGCTAATGGATCAACAACTC
TACCTAAAAAAGCATCTCCGACCGGAATTTGAGCAATTCGACCAGTCCCTTTTACGGAAC
TGCCCTCTAAAATATCCCTACCATCTCCCATGAGTACTACACCAACGTTATCACTTTCTA
AATTTAAAGCAACCCCAATAGTTTTATCTTCAAATTCAAGTAGTTCACCAGCCATGACTT
CATCAAGTCCATATACTCGAGCGATACCATCACCAACTTGTAAAACTGTTCCTATATTGG
CTACTTCTACATCCTGATCATACTTTTCAATTTGTTGACGAATAATACTACTTATTTCAT
CAGGTCTAATATTTACCATATTTTTAATATTATTTTTATTAATTGGAAATTCTAAAGCGG
TAATTAGTCAGACTAATGCTATCTGTCTTTTTTACTTTATTTTAAGTTACTACAACATCA
AGGTGTGAAGCCATTTGTCTTAGCTGTCCTCTAATACTAGTATCAATGACTTTTGATCCA
ATCTGAATAGTAAAGCCACCAATTAGCTCGGGCTCGACTGAAATAATTAGTTTAACTTCT
TTAGCCTTAGTCATAACCTTGATTTTTTCTGTTAATAGAGTTTCCTGATCAGGGCTTAAA
GCAATAGAAGTATTAATATTTGCAATTGTTAAAGATTCCATTTGGTAAGCCAGTTCCAAA
TACTTACTAGCTATAGCATCTAACATTCCAATTCGCTTGCGATCAACTAAAATCATTAGA
AATGATAATGTATTTTCGTTAATTTGATCGCCAACAGTTGCAATAATTACTTCTTTTTTT
GCCTCTACAGTTTTTAATGGATTAGCTAAAAAAATTTTTAGCTTGTCAGATTGCGCTAAA
ATATTTTGTATTGACTGAATATCTTGGCTCACCTTTTCAGTAACTTTTTTAGTTTTAGCT
AAATCAAGAAGAGCTACTGCATAAGGCTGAGCTATTTTTATAACAACATTGTTACTGCTC
ATAGTTGATCTCCTAACTTAGCAATATTATTATCAATTATACGCAGTTGCATTTCGGAGC
TCATTTGATTTTCCAGCTGCAAAGTAACTCTTTTAAGAGCAAGAAATGTTATTTGTTGCT
GGATTTGCCTTCGGATCTGTTTTTCAGCTGTTTCTATATTTGATTTACCTGTTATGGCAA

GTCTTTCAATATCTAATTGGCCTTGAGCTAATATGGAACTTCTAACTTTTCCAGCTGTTA
GCTGAGCTTCTTTCTTAATTTGATCTATAATGATTTGAGTTTGAGCAAGTTGTTTTTCTG
ATTCTGACAATCTAGCACTTGCTTGCTCTAGTCTTTCTTCGGATTCTTGTATAGCTGCTA
AAACTTTTATCTGTCTTGCATTTAAACTTGATCCGAGAAATTGCTTCAAAACATAAATAA
GACCAAATAAAAGCAACAATATATTAATAACATTTGCTTCAAAAATATCAGAATTAAAGC
CGAATTTGTGTTCACTACTATGTTCTGATAAAATAGTAATTATTTGTGGTATTTTTACTA
TATTATTCATTTAAAATTCTTACTATTTTAATATTTTACTCTGGAATCATGATTTTCAAG
ATTGACTACTTAATAATTTAACTTTAATTTGATCGCTCAAAGTGTCAACCTGATTTTCTA
AAGTTTTAAGAGCTTCTTCTTTTTGAACATTTAACTGTTTCGATGCTTCAGCAATCAGTT
TCTCTGCATTCAATTGAGCTTTTTTTATATCTTCTGAGACTATATTTTGAGCTTCTTTTT
GAGAAGAGGCTATTTTTAATTGTGCATTGCGGCGAGCTTCAGATAAATCTTCTTCATATT
TTGCAGCTAGCTCATCTGCTTTAACAAGCATAGACGATGCAGTAGTCAATGTTGTTCGAA
TATATTCATCTCGTTCGTCTAGTACTTTAGTTACAGGTTTATAAAAAATAGTATTTAATA
AAACCATTAAAGTAAGAAACTGTAATGCCATTAAAGGCAAAGTACCATTGAAGTCAAATA
AGCCACCTTCAATTTCTTCGGCTAATAGAAATGGTAAATCAATCATTTTACTAATTATGT
TTATAGTTAATAATTTTGTTTCAATTAAGAAAAATAACTTTATAGTGCGCTTAGTCTCAT
CAGACTAAGCGCAAATGATTATTTTAACCAACGTATGGATTAGCGAACAATAAAGACAAC
GCAACAACTAGGCCGTAAATTGTTAAAGATTCCATAAATGCTAAACTTAGAAGAAGTGTA
CCCCGAATCTTTCCTTCTACTTCTGGCTGTCTAGCAATACCTTCGACAGCATTAGCTGCT
GCACTACCTTGACCAATGCCTGGGCCTATTGCAGCAAGACCTACAGCAAGACCGGCAGCG
ATAACGGATGCGGCTGAAACGATTGAATCCATAATGAATTTTAATTTTTTGTTATATGAG
TAGAAAATTGACAAATTTCGTATAATACTTAAAATCTATATAATTAAGTATAATAGATTC
TAATATTGTAATTATAGTATATTACTCTTCTCCATGACCTTCCATTGCTTCTCCAATATA
AGCTGCGGACAAAGTAGAAAAGATTAATGCCTGTATTGAACTCGCAAATAATCCGAGTAT
CATAACTGGTAATGGTATTAAAATTGGAATTAGCAAAGTAAATACAGAAACAACTAATTC
ATCTGCTAAGACGTTTCCAAATAATCGAAAACTTAAAGATAGTGGTTTCGTAAAATCTTC
TAGAATATTGATTGGGAGTAATACTGGAGTTGGCTGAATATATCTAGCAAAGTAGCCTAA
CCCTTTTTTACTTAAACCTGCATAAAAATAAGCTAAAGATGTAAGTAAAGACAGGGCCAC
AGTTGTATTAATGTCATTTGTTGGGGCAGCTAGCTCACCTTCAGGCAAGTGAATTAACTT
CCAAGGAATTAATGCACCTGCCCAGTTACAACCTAGAATAAATAGAAATAAAGTTGCAAT
GTAAGGCACCCAAGGGCGATATTCATGCTCACCAATCTGATTTTTTGCAATATCTTGCAG
AAATTCTAATATAAATTCCATAAAATTCTGGAATTTTTCAGGAATTCTTTGTAGATTTCT
AGTACCTAGAAATGATAAAGTTAAAAGTGTAGCAATTACTAACCAAGAGACAATGAAAAC
TTGCCCATGCAATTGTAAGCTACCTATTTTCCAATATAGATGTTTTCCAACTTCTACTGC
TGATAAACTATTATATGGGTTAAAATCTATGAGACTGTTTTGATACATAATAAATATTTT
TTTAAATAGTGCAAGAATCAAAAAATAAGTTTTTGAAGAATTAATATGCAATTAATTTAT

TCTTAAAATTAGACTATTAATAAATTAAAAGCAAAAATAAATTCTAATAGAGATAGCTAT
TTTTAGTAGTGATTATTTTTTGTAAAAAACAATTATTGCTACCTATAATGCTTTTACGTT
TAATTATATACCCATATATTAATTATTTAATATGATTATAGGTATTTTGCTACAAACTTT
ATTGTTTTATCCATTCTAAAGATTGTTATAGTTTATTTTTTATTTAAAATATCTGCGACT
TCATCTGCAAAATTAACTTTCAAATTTTCTTCTCCCCCTCCCAGAAGAAATCTTACAAAT
CTTCTAATTTTAATATTTTCACCTAGAACAGCAATATTTTGGTTTATTAAATCTTCAATA
CTAATATCTTGATTGCGGATAAACATTTGATCTAATAATGAGAGCTCCTTAAGTCTCTTT
TTCATTCGTCCTTCAATGATTGTTTCGATTCTATCAACTGGTTTGTTTTTCAAGTCATCT
TTACCAGCTTCAACTCTTTGTTCCAGATTAATAATTTCACTAGGTATATCCTGTGTTGAT
ACGTATTCCACATTAGGCGATGCCGCTATTTGCATAGCAATATCTTTAGCTAACTTTTGG
AACTCTGGACGTCGTGCTACAAAATCTGTTTCACAGTTGACTTCAACTAATACACCTATT
CTTCCGCCAGTATGAATATAACTCTCTAGTAAACCTTCAATAGCGGTACGATTAGATTTC
TTGTTAGCTGAAGCCAGCCCTTTTTGTCTTAATGATTCTAAAGCTTTTTCTTCATTGCCA
TTATTAGCTTGTAAAGCTTTCTTGCAGTCCATCATACCTGCTCCAGTCTTATCACGTAAG
GCTTTTACGACTTGAGCAGAAATTTGTAGTGTCATATGCTTTCAAAAATTTTTAAGTATT
TAAATTAATTATTATAATTCACTTTGAGTAGGATGAGTTATTTCTTCTAAATCTGTTTGC
CCGTATTTACCATCATAAATTGCATCTGCTATTTTACCTACAATTAGTTTAATTGACCTA
ATGGCATCATCATTAGCTGGGATTGGAATATTAATAATTTCTGGACTACAATTAGTATCA
AGTATACAAATAGTTGGAATACCTAATTTTAAACACTCTTGTATAGCTGTTGTTTCTCTC
TTTTGATCTACAACTACAACAATATCAGGTAAACGAGTCATATTTTTAATACCATTTAGA
TGTTTACGAAGTTTATCTAATTCTCTTCGTAAAACAGCAGCTTCTTTTTTGGGTAACTGA
TCAATCATACCACTTTTATCTTGTTCTTCTAGCTGTCTCAGACGATTAACTCGAGACTTA
ATCGTTACCCAATTAGTTAACATTCCTCCAAGCCATCTCTGGTTAACATAATAAGAATCA
CAGCGTTGTGCTTCCTGTGCCACTATACCAGCGGCTTGTCTTTTTGTTCCTAGAAATAAA
ACTTTTTTACCATCCGACGAAGCCTGTTTTATAAATTCACAAGCTTCTGTTAATAATTGT
GCAGTTTGGACTAGATCAATAATATGTATGCCATTTCTTTCTGTATAAATATATGGAAAC
ATTTTAGGATTCCATCTACGTGCTTGATGTCCAAAATGAACACCTGCTTCTAGTAATTCT
GCTAATGTAACAATAGCCATAAGATTATTAATATTTAAATAATGTAATTATTCGGGAAAT
CCCTTTGTTTAGAAACTGATTCAACTAACTTATTTGAATTTTAGTAAACTAATAATAAAT
TAACACACTTTTAAAGCAGATCAACAAAATAGATCCTGAAGAATTCATTCTACACTTTTA
TTGCTAAAATAATTACGCGCGGTTCTGTCGTCTAAAATGATATCATCTAAATCATCTCGT
ACAGATGAAATAGTACTTTCGTCATTAGTATTTGCAGTTAAATTCTTTTTCTCTAACGAT
GAGCCATTACAGTTATCATACATATTGAAGCCGGTACCAGCCGGTATAAGCCGACCTATA
ATAACATTTTCTTTCAGACCTCTTAACCAATCTAATTTACCAGATATAGCAGCTTCAGTT
AATACTTTTGTAGTTTCTTGAAAACTTGCTGCAGAAATAAAGCTTTCTGTATTTAAGGAA
GCTTGAGTAATACCTAATAGCACAGGCCTATATGATGCATTAATCTTATTTCTCAAGGTT

ATGGCTTTATTAGTCTGTTCAATCTTCTGTAACTCTACAAGCTCTCCAGGTAAATAGCCT
GTTTCTTCACCATTTTCTATTTTCACTTTAGAAGTCATCTGCCGTACTATTACTTCTACA
TGCTTATCAGAAATATTCACTCCTTGAGATTGATATACCAATTGTACTTCTTTAACAAGT
AGTAACTGAATCTCTTGAAAGCTTAATCGAGCAGCTTCATATAAGGCTAATCCACATTCA
ATATACAAATCAAAACTTGCATCGAGGATATCATGAGGATTTAACAATACGTCTGTTTTC
TTTCTTGCTTCAAGAATTTCTTCAATTCTTGGAAGCCCTTGTATAATGTCACCTGTTTTT
GCTCTATCAAAAACCAATATAGCTAATGTTTCTCCCCTTCTAATTAATGCATTGTTATCA
ACATGTAAAATAGCACCATTAGAAACCAAGTAAGGTCTAGCAATTCTTAATGTAACTGAT
CTAGAAGAAATTTCAATAATTTGTCCAGAGTCTAAAGAAGTCATATTTTCAGTAATAAAA
TCACCACAACGAATCCAATCACCCACACTTACTTTAACTATTGATTTTCCTATATTAAAG
GTCTTTTTATCAGAAGATGTCGCAATCAAAATGCGCCTGCTAGTATTCTTTTCAGAATAA
ATCTCCTCAACAGTACCTTCACTCATTGCAATAATCTCAGTACTAGCAACAACTGTAAAT
GGCTTGATATATTCACCATTTTTTACAATAATTCTAGTTTGACTTTGTTGCTTCTCTGAT
TTATTTACATCAATACTTTTAATTGATAATGTTTCAAAAGTTGACAGCCTTAAGTCAAAA
TAATTATTGCCACTTTCTTTAGCAGAAAACTCAATTGAGGATATTAAATTATCATCTCGA
TGCTCAATTTCAGCCACAAGATTAGTTGTTACTAAATGAACTCCTTCTATAGATTTCACT
CTTTCTCCGTCCCGAAAAGGAGTTCTTTTAACTAGCTTTAGTTTAGTTTGACTTGGTTTT
TGTGAAACTAGATTTTCAATTAGACTGGATTTTGTTTCTGGTATTGAATAAACAATAACA
GGTCTAATAAGAATATAGGGCATTTGCTCGTTTTCTATATACTCCCAATAAACTAACTTA
TCTGTAGAAATATTATTATGAAGTCTTTCTCCAGGTCTTAAAAAACCTCTACTTTTATCA
TTATTACTATACACCTCACTAAGTTTATAGATAGATCCAGGCTTAATAATAATTTCTCGA
ACTATTCCATCCTTTTCGATTATTTCAACAATACCTGAGCTTTTAGAAAAAATATTTTTA
ACTAGCTCTGTTCCACTTTCTATAACATCTCCATTATGCACTAGTAGTAGTGATGAATCT
TTATTAATTTCGTGAGTTTCTTCTGAAATCCATAAGATATATCCCGGACTTAAAATTTCG
TAGGCATCTTTATCTAAGCCTGTTTTCTTTTTTGATACATTTAAGTCTAAATATTTAATA
ATACCACCACTTGTCGTTTTATACGTATCAGATATTAGTTCTGCGACTGTGTAACCATCT
TGAATTTTTTGGTTTGGAATAGCTTTGAAAAGAAATTTTTGTTTTTTTTCTGTTTCTAAA
ATATAAGATTCTTTTTTGTGGATTACATCTGTATAAATATAACAACCTGGAATTACAATA
GATTCAGTAATAACTTGTATATTGGTAATACTGCTATTAGTATTCTGGTGTATCCTTACT
TCTCCAGCATAATGATTTATTAATTCTGTCTGTGCAAGTATCGTACCAGCTTTAGTTTTA
TCTTCTTTATGAACAATAACATTAGCAGAATCAGATATACTATAAACTTCACCTGAGAGC
ACCCAGATCAAGCCTCCAGTTTTAGTAATTCTTGTAGTATATTGATTATTATCTGTTTCT
TCAACTGTTAAGTTGGAGAAACAGATTTTGCCAGAGAGATCTGAGACTACATGTTTTTGA
GCTCTTTCTGTCATGAGTCGATTTCTGCGGGGTGATTCAGCAATCACTTGATCCTTCGAT
ACTAATTCACCATCACATATTAAAAGAGTAGTACCTTTACTTAAGTTAATTATTGACTTC
TGATTTTTCTTAGATCTGATTGTAACTTGGGTTGGCTTTTTTGTTATTAATGCTTGCTCA

CCATGTCTGGTACGAACATCTGTATACGATTCAATATCTAGATCTATCAATTGACCATCT
ATAGGTGCGTAGATTTGTTCTGCTAATTCACCGGTAAATACTCCTCCAGTATGAAAGGTC
CTCATTGTCAATTGAGTGCCAGGCTCACCAATAGATTGAGCCGCAATAATACCAACAGCC
TCTCCTAAATCAACTAAGCGGCCATGTGCAAGATTCCATCCGTAACAATACTGACAAACA
GAACTTCTAGAGTTACATGTAACAGGAGATCTTACTAAAACTTTTTTGATACCAGCTTTA
GTTATCTCTTTTGCTAACTTAGGACTGACATCTTGGTTCGTATGAGCAATTAAACAATTT
GTCTCTGGATGGAATACATTTTCTGCTAATACTCGGCCAGATAAAGCTTGTTCTAGATTA
ATTAATATTTTCTGCGTATCTACTAAATCTTCTAGTATGATGCCTTTATTGGTCTTACAG
TCTACTTCTCTAATTATTACATCTTGAGAAACATCGACAAGTCGACGCGTCAAATAGCCG
GAATCTGCTGTTCTTAAAGCAGTATCAACTAATCCTTTTCTTGCTCCGTATGATGATATA
AAATAATCAGTAACAGTTAGTCCTTCTCTAAAATTACTCGATATTGGAAGATCAATAATT
TGTCCCTGTGGATCAGCCATAAGCCCTCTCATGCCCACTAACTGTCTTACTTGAGAAATA
TTACCTCTAGCACCTGAGAACGCCATCATATAAACAGCATTTAGAGGATCTGTCTCTTTA
AAGTATTTAATAACTTCTTGTTTCAAGGATTCACTCGCATTATTCCACGTATCAATAACT
TTTTGAAATCGTTCAACTGCTGTAATCTCACCTCGGCGATATTTGTTTTCAGTGGCCTTA
ATATCTTCAATCGTGCCTAATAAAAGACTTTGCTTACTAGGAGGAATTCTTAAATCTTCT
AAACTTAAAGAAATTCCTGCCTGGGTCGCATAATGAAATCCAAGATCTTTAAGCTTGTCG
GCCATATTGGCTGCTCGGGCTATACCATAATTACGAAAAGCCCAGACAATTAAATTTTTT
AGTTCGTTTTTATCAATAACCTTATTTGAAAAACTTGGCTGTGCAAGACTTCGTCTATTA
TCCACTAAATCTTCTCCATATTAAATGTTCGTTATTAACTATGCTACTAAAGACTCTTGA
ATAATTTTATTAAAGATTATACGGCCAGCTGTTGTTCTAATATATTGCACAATTCTCTGA
TTTTCAGCGTCTTCTTTTATGATATGGTTATTAAAAAATTTTGTTGTACTATTGTCAGAA
TGTTTTTCAATCTTAATCGGATCATGAGGTTGATCACCATCTACTAATCCATCAAAACGA
GCCCATATATAAGAATGTAAATCAATTTTCTTCTGTTCATAAGCCATAACTACATCTTCT
AAGCTTGCAAAATACTGATTAGATCCCTGCTGCTGAGATGGATTATTAGCTGTTAAATAA
TAGCAACCTAAAACCATGTCTTGGCTTGGCATAATAATTGGTTGACCTGTTGCTGGAGAG
AGAAAATTATGAGGGGCTAACATTAGCAATCTAGCTTCGGCTTGAGCCTCTAAAGACAAA
GGAACATGAACAGCCATTTGATCTCCGTCAAAATCAGCGTTAAACGCTGGGCATACTAAT
GGATGAAGCTTAATAGCTCTTCCTTCTACAAGAATAGGTTCAAATGCTTGAATACCTAAT
CTGTGTAGTGTGGGAGCTCTGTTCAATAGTACTGGGTGACCTTGAATAACTTCATTTAAG
ACATTCCAAATAGAAGATTCATTTTTTTGAATCATCTTTTTGGCAGCTTTAATATTGTTA
ACTAATCCCTGCAAAATTAGTCTGTGAATAACAAAAGGCTGAAATAACTCCAAAGCCATT
TCTCGTGGCAATCCACATTGATGTAGTTTTAAATGAGGACCTACAACAATTACAGATCTA
CCGGAATAATCTACTCTTTTACCTAGCAAGTTTTGTCTAAAACGACCTTGTTTTCCTTCT
ATAATATCTGACAAAGATTTTAATGGTCTATTATTTGCACCTACAACAGTTCTACCTCTA
CGACCATTATCCATGAGAGAGTCAACGGCCTCTTGCAGCATTCTTTTTTCATTTCTGATA

ATAATCTCCGGAGCTAAAATTGACTTTAGACGAGATAATCTATTATTTCTATTAATAATT
CGACGATAAAACTCATTCAAATCTGCCGTCGCAAATCTACCTCCATCTAATTGGACCATA
GGCCGTAAGTCTGGAGGTATCACAGGAATGACTGTGAATACCATCCAAGAAGGATCTGCA
CCTGTTGCTATGAAATTTTCAATCAATCTTAAGCGTTTCATTTTTTTATTGAATTTTAAT
GAAGGTGTCTTAAAACTTTTTGGTGGATTTGTAGCTTCAGACCTTAAGGTTTCAGCTATG
TGCTCTAAATCTAAATCTTTCAACAGTTTCTGAATAGCTTCTGCACCTATTCCTACTTCT
ACTTGGTTATTCTCATCTTGATTTTGGTAGATTTCCTCTTCAAGACTTTTCCATTCGTAA
CCTTCGAGAAGTTGCTTGTATTTTAAATTAATATCTGTATTAGATTGCGTAACTACATAA
GAATGAAAATAAACAATTTTTTCTACTTCTTTGACTTTTAAGTCTAATGCTAAAGCAATA
TAGCTTGTACTTCCTTTTAAGTACCAAACATGAGTTACTGGTGAGGCAAGTTCAATATAA
GCCATCCGATGTCTTCTCACGCGTGATTCTGTAACTTCTACGCCACATCGTTCACAAACT
ATACCTTTGTAGCGAAAACGTTTATATTTACCACAGTGACATTCCCAATCCTTAACTGGG
CCAAAAATTTTCTCACAGAATAAACCATCCATTTCTGGCTTTAAAGTTCTGTAATTAATT
GTTTCAGGCTTTGTAATTTCACCAACAATTTGTCCGTTGGGCAAACTTCTTTCGCCCCAC
TGTCGAATTTTTTCGGGAGAAGCTAAACTAATTTTTACGTAGTCAAAATATTGCTCAAAC
TTTGTCATAAATTTGAATACCTTAAAATCAAATAATTAGCCAATTTATTTACATGCTTCA
TATTGAAAAATGAATATCTTAATAAAGGAATTGTTCAAAATCGTCTACTGGAGGAGTATC
ATAATTAGAACGATCTACTCGGTTATCTTTAGAATCCGACATCAAGTCAACTTCAACTGT
CCGTCTTTGGCCATCTTCAAATAGTTTCAATTTATGTACTGCAATGTCTAAACCTAATGA
CTGCAGCTCTCGCATTAAGACTTTAAAAGATTCTGGAGTTCCAGGCTTCGGTATAGGCTT
CCCTTTAACTATTGCATTTAATGCTTCATTTCTAGCTTGCATATCATCCGACTTTACAGT
TAATAATTCTTGCAAAGTATATGCTGCACCAAAAGCTTCTAAAGCCCACACTTCCATTTC
ACCTAATCTTTGACCTCCATGCTGTGCTCTACCTCCTAATGGCTGCTGCGTAACTAGAGA
GTAAGGACCAGTGGATCTTGCATGGATTTTATCATCAACAAGATGAACAAGCTTAAGCAT
ATAAGCTCTACCAACTGTAACAGGATTATCAAAAGGCTCTCCTGTTCTACCATCAAAAAC
TTGCATTTTACCTGGATGCTGATCATTAAAAAGCCATTTATTGCTAGTAATCAATGATGC
TTCTTTTAATTTTCTATTTACAAGTGCTCTTGATGCTTCTGCTCCATACATTTCATCAAA
AGGTATTATTTTAAATCTCTTTCCTAAATAGCCACCTGCTAAGCCTAGTAGGCACTCAAA
TACTTGACCAACATTCATTCTAGAAGGTACACCTAAAGGATTTAAAACAATGTCCACTGG
TGTTCCATCTGATAAATAAGGCATATCTTGTTTAGGTAAGATCCTAGAAATGATACCTTT
ATTGCCGTGACGACCAGCCATTTTATCACCCACTTGGATTTTTCTCTTCTGCGCAACATA
GACGCGAATCATAGCATTTGTTCCGGGAGGCAGCTCATCTCCTTTCTGCCTAGTAAAAAC
CCTAACCTTAACTACTCTACCTTTTGCAGCATTAGGAAGCCGTAAAGATGTATCTCTCAC
ATCTCGAGCCTTTTCTCCAAAAATAGCTCGCAACAATTTACCTTCTGGTAGTTGATCAGC
TTCACCTTTAGGAGTAATTTTTCCTACTAAAATATCTCCAGCTTCAACCCAAGAACCACC
GACAACAATTCCATTTCTATCTAAATCCTTCAGAGAGTTATCGCTGACGTTAGGAATCTC

TCTAGTAATTTCTTCTGGTCCTAATTTAGTTTGACGGCATTCTACTTCATACTTTTCGAT
ATGGATAGAAGTATACAAGTCATCATAAACTAATCTTTCGCTAATTAAAAATGCATCTTC
GTAGTTATAACCTTCCCAAGGCATATAAGCAACTAGAATATTTCTCCCTAAAGCGATTTC
CCCTCCGTCTGTGGATGCGCCATCAGCTAAAGTTTGGCCAACAACTATTTTTTCTCCTAC
CCAAACAATTGGGCGCTGATTAATACAAGTATCTTGGTTAGACCGATAATATTTTTTTAA
GCGATAATGAACTGTTCTACCACTATTATCCTGAATGCCTATTTTATTAGCAGAAACATA
ATTTACATGACCAGATGTTCTGCTAATAACAACCATACCAGAGTCTCGAGCTATTTTAGT
CTCAAGACCAGTACCAATTATTGGCTTTTCTGGATATAATAATGGAACAGCTTGCCTTTG
CATATTAGATCCCATTAGAGCTCTATTCGCATCATCATGTTCCAAAAAAGGTATTAATGA
TGTTGCAGCTGAAATTACTTGAATAGGTGAAATTGCAATATAATCTACTTGGGTTGGGGT
TGTTGTAATAAACTCTTGACGATAACGTACAGGAATAATATCTCCTTCAATGTAATGTTG
TTTACTAACTTTAACATCTCCTGGAGCTACTCGGAAATCATCTTCCTCATCTGCTGTTAG
ATAGACAGGACTATTGTGATAAATCACTTGACCTTGGTTTACCGGATAAAATGGGGTTTC
TATGAAACCAAAAACATTGACCCGCGCACAGGTTGCTAGTGAACCAATTAAACCTGCGTT
CGGACCTTCCGGAGTTTCAATTGGGCAAATTCTACCATAGTGACTAGGATGTAAATCACG
AACTGCAAAACCTGCCCTGTCTTTATTAAAACCTCCTGGACCTAAGGCACTAATTCTTCT
CTTGTGAGTTAGTTCTGCAACTGGATTAGTTTGGTCCATAAATTGAGAAAGCTGACTAGA
TCCAAAGAATTCTCTCACTGACGCAATTAATGGCTTGGGATTAATTAGATTTGATAAACT
CAATGAATCTATATCACATATCATCATTCTCTCTCTAATAATGCGTTCTAAACGATTGAG
ACCTACTCTGAATTGGTTCTGCAGCAATTCTCCAACTGAACGAACTCTTCTATTACCAAG
ATGATCTATATCATCAAGATTACCTGAGTTTTTATCCTTAATATTAATTAGATAATCAAT
TGAGGATAAGATGTCTTGAGGAGACAATACTCGGAACGTTTTAGGTATATTGAGACCTAG
TTTTTTATTAATTTTATATCTACCAACTTCACCCAAGTCATATCTTTTAGGATCAAAAAA
ACGAGAGTAAAGCATCTGCTTAGCAACAGCAACTGTTGCTGGTTCATTAGGTCTAAGCTT
AGAATAAACTATGAGCAGGGCTTCCTCATCAGTTACTTCTTCAATATTATTTTTTCCAAT
TTCTTTAGCTAATTCTTTAACTGAATATATTTGGGATGCAGAAATAAGAAAAGCATATTT
ACTAAGTCCTTTTTCAATTTCATCTTTATTGAGTCCAATAGCTCGCAGAAAAATATAAGC
ATTTACTTTGTGGGTCTTATCTATTCTTATCCAAATTTCTCCCTTAGGATCAATTTCAAA
TTTTAACCAAGAACCTCTATTAGAAATTAGGCTTGCACTATAAATCTGCTTACCATTTTT
GTCTATCTCTTGTTTATAATAGATTCCTGGACTACGGATTATTTGGTTAATAATAACTCT
TTCCGTCCCAGATACAATAAAAGTTCCTCTATTTGTCATTATTGGCAAGTCTCCAATAAA
AATTAATCTCTTTTTGTATTTTTTATTTTTTATCTGCTTTTCAGCACTGAGCTGTAGATG
GGTTGATGATAAATCTAATGATTTAATATTTTTTTCCTGGTTTTTAGAAGGTAGATCAAT
ATCTTTTCTTGTTAACTTTGCAGGAACATATATTTGAGCACTATAAGTTTTATCCCTATT
TTTTGCCTGTCTGACACTATAACGAGGAAATTTTATCTTGTACTCATTACCAAATAATTG
TAATTCGAGTCGACTAGTAGGGTCTGATATCTTCGGAAAAATTTCAAGAACTTCCGTCAA

ACCTTCTAATAAAAACCATTTAAAACTGGCCCTCTGAATTTCAACTAAATCTGGAAGAAG
TTTATTTTTTAAGCTTATACGTTGAACCATAGATCTCCTTTATAACAAATTTTGAAATCG
AAGAGTTTAATACTATTAATAGAGTACTGGTAGATAATGCATAAGCATAGTTTGCTACAT
TATTTCTTCTCTAGACATAACTAGAACATCATATTTTATTCAAAAAAAGATTTTTAAAAT
TTTTCTTCACACACACTATCTATAAGTACTTTGCTGCTAAAAGATAGTTCAAGCTATTAA
AAAAAACGAAATTAATACTTGTCTGTCTGTCAATTAATTGACATCAAGATAGACAAGTAT
TATTTATTTAATCAATTCTGTTTTTCTGAGCAAAAGACAATGCTCTCGCAAGTCTAGCTT
TTTTTCTAGCGCCTGTATTAGGGTGAAAGGCCCCTTTTTTTACAGCTTTATCAATCTTAC
TATATACTTGAGAAATACTTGACTGCACATCATTGAAATTATCACTTTCTAAATTATCAA
TATTTAGAAGGCATCTTTTAGTTAAAGTTTTGACAACCGATTTATACTTACGGTTAATGA
GACGATTTCTTTCCGAAGTTTTAATGCGTTTAATCGCAGAAAGGTTCTTAGCCACAGTAA
ATCCAATAAAACTTTTATAAAATATTTTTAGAAACGAAAATACACAAATCTGGGATCTAT
TCAAAAAGGGAATATGCGGCGTATTATAACATTAAAGAAAAACAATAGGCAATAGTTCAC
CAAGAGCTCTTATTATTACACAATAGAGATTGTCATTGCGAAAAACTTTGTAAGATAGTA
TTTAATTAAAATAAACTAACTAGACATATAAATATATATGAAAAAAATTGAAGCTATTAT
TAGACCTTTCAAGCTTAATGAAGTAAAACTTGCTCTAGTCAAGGAAGGTATTGGAGGAAT
GACGGTTATCAAAGTTTCTGGATTTGGGAGACAGAAGGGTCAAACTGAAAGATATAAAGG
ATCCGAATACTCTATTGATATCATTGATAAAATAAAAATTGAAATTATTATTAGCGATGA
TAAAGTAGAGAAAATTGTAGAGACGATTATTAAGGCTTCTAAAACAGGGGAAATTGGAGA
TGGAAAAATATTTATTAGTAGTATTGAAAGAGTAATAAGAATCAGGACTAATGACTTAAA
TTTTGAAGCTTTATAGTATTTTTACTTTTTTAGACTTGATACTTCTTCACTAAACATGAG
ATAATAAGCGTTTATTGTATACTTATAACTAAGGTGTAATACTAATGGCTAAAAGCAAAG
GTGCACGAATCGTAATAACTTTAGAGTGCTCTGATAAAGCTGGAGAGTTTGCTCAAAAAA
GGAAACCTGGCGTTTTTCGATATACAACTACTAAAAATAGACGAAATACACCAAGTAGAA
TTGAATTAAACAAGTTTTGTCCTAATTGTAATCAGCATTGTATCTTCAAAGAAATTAAAT
AGTTATTTATCTTATTAAAATAATCATATTATGGCTATATACAGAAAAAGAATATCTCCA
ATTAAGCCAACAGATGCTGTTGACTACAAAGATATTGACTTGCTAAGAAAATTTATTACA
GAACAAGGCAAAATACTGCCTAAAAGGTCAACTGGATTGACTTCGAAGCAGCAAAAAAAA
CTTACTAAAGCGATTAAACAAGCTAGAATCCTTTCTTTATTACCTTTTTTAAATAAAGAT
TAAATATATAGCCTAACTTCATATTTTTCCATCTAAATATTAAAAATTTCTATAGATTTG
TTTATTAAAATGATCTAACATACTGAATTACTAAGAAGATCTACTCATTACAAGTTTTGT
TCATTTTATAATTAATAATAGCATCATAATTTATTAAGCTAGATTCAGACAAGACATAAA
GGTAACCTCTTTTAATTGATTATATTATATAGTCTATTTAAAAGAGGAAATTTTTTTGCT
TCTAGCAATTCAAGATTCTTTAAGCTTTAACAAGGTATTTAAAACAAAGATTTTTGTTTA
GGTATCAGATCAAATGAATGAATCTTATCTCCCGACTGCCATAATTGAAATTCAGAGATG
AAAATTCCACATTCGTTGCCAGCTTGAATTTCTTCGACATCTTCTCTAACGCGCTTTAAA

GATTCAATTTTTCCTTGATAAATAACTTTTTCTTCTCGAATCACTTTAATCCAAGAGTTT
TTTAATAGCTTATTATTGATCACTCGACATCCTGCTATTTTTCTATTTGCTAACGAGAAT
ACTGTACTTACTTCTGCTTCTCCCACCGGTACTTCTGAATATTCAGGATCAAGTAAGTCT
TCCATCCTTCTTTTAATGTCTTCAATTAAGGCATAAATGATTTGATAGTTTTCTATTAAT
ATATTCGACTTTGCCGATGCTTGCTTTGTTCCTGGAGCAAAGTTGGTATTGAACCCAATG
AGGGTAGAATTTGTGGTTGAAGCTAGTTCAACATCTGTAGCAGTAATTTCTCCAGGCATG
ATTGAGACAACATTGAGTTGAACTTTACTCTGAGGAAATTGAGATAAAGAATCTAGGATT
GCTTCTGTAGAGCCTTGATTATCTGTTTTAATAATTAAAGAAATTTGCTTACTAATATCT
TTAGAATTAGTGTTTTTTAAAGTATCTAATGTAATTCGACTATTAAGTGCTCTTTGTTTC
TGTATAATTGATGAGTCTTTAGAGGTATTTTCAATCGCTTTAAGTTTGGCTTCTTTATCA
CTTTTAACTGCTAAAGCTATTTCTCCTGTAGCTGGTACAGATGACAATCCCCAAATTTCA
ACAACAGATGATGGTATTGCTAAATTAATTTTCTCTTTAGCGTTGTTAATTATAACTCGA
ATCTTTGCGTAAGCTGATCCAATAACCAAATTATCACTAATATTCAATGTACCATTTTGT
ATTAATAGAGTTGCTACTGGACCATGCGATTTATCTAAATGGGCTTCTATTATAATACCT
TGTGCAGGCTGTGTTGGATCAGCTTTCAAATCCTCTAATTCAGCTAACAAGGTAATAGTT
TCTAATAATTTATCAACATTTTGACCAGTAAGAGAACTAATAGGAATTATAGGCACTTGG
CCACCAAGTTTCTCCGACATGACATTGTATTTTAATAAATCTTGCTCAATAATATCTGTA
TTAAAACCAGCTTTATCTATCTTAGATATTGCTATGACAAAAGGGACATTCGCTTTTTGA
ATATGATTAATAGCCTCTATGGTTTGAGGTTTAACTCCGTCATCAGCAGCTATAATAATA
ATAGCGACATCTGTTAAATTAGCTCCCCTAGACCTCATACTTGTAAAAGCTTCATGGCCT
GGAGTATCCAAGAAAACAATTTTTTGTTTATTATCTTTCTTAATATACTCAACTTCATAA
GCTGCAATTGCTTGAGTAATTCCTCCAATTTCTTTGTTAGCATTATTAGATTTTCGAATA
TAATCTAATAGCGTTGTTTTGCCATGATCTACATGTCCCATCACTGTGACAATTGGAGGT
CTTTTAATATAATTGCCAGTTTCATAAAGAGCATTAGAGTGATCTAAATTACTAGAAAGT
CCATTATTATTTTCCTTTACATTTGATTCTACTGCTATACCAAAATTATCAGCCACTGAT
GATATTATTGAGGCGTCTATGGTTTGATTCATAGTAACTGATATTCTTTTAAGAAATAAA
TATTTTATAATATCTGTTTCTTGAACACAAATTAGTTTAGATAATTCTTGAATAGTTAGA
GGATTAGTAATACTAATAGACTCTGGCGGAGAATTAGATTGAATGCTTGCTGAATTTTGA
CTAGGAGTTACTGCTACTTTTTTCTGTCTTATAGATTTTTTATTACTAGTCAGTTTTTTT
ACAACTTCCACTTTTGGCTTAGGTGGACGCATTAAAGAGATAGCTAAATCTCCTGCAGTC
TGCGAAACATTAGAATTAGAGTCTCTAAAGTTGTCATCATCATCGTCAATATGTATTTTA
GTTTTTATTTTTTTGCGCTGTCTATTCTTATTTTTTTTACTATCTACTAAGTCGTGAATT
TTATTAAAATTTTTACTTTTTTTATCTAGTTTAGGTGGAGAACTTAATTCTAGATGCTGT
TCACCACCTGTATGTGATTCTGATTTATCAAGGTCTAAATTTAACAGATTATCATTATTG
ACTGACTCTAATCTAATTTTATAGATAATCTGAGGATTTTTCAAATCTACAATTGTTTCA
GAAGAATTGATATTTGAGCTAGATCTAGAGGATCTATTTTCAAAGTTTTGATTATTTAAA

AACATAAATATTGGTTTTATTATCTAATAGTAACTTTCTAATTTTAATACAGAGCTTAAT
ATAAACAAGAATCGAATATCTGTATAAATAATACAAATGTAGAGAATTAATCTATGTTGT
ATTAGATTTAATATAACACTAAAGTCCTTTGTTCGTTATACAAGTTTAAAACTTTTTATG
ATGATAAACACTAGATTCTAGTTTATTGGTATACAATCTATTTATCTAAATAATACTGTT
TTTGTCAGGGCTTTTAAATTATCTAATGTTATGTTCATTATGAAATTCTATTAAAATTTA
AAAACTCAACAATTAAGTAAGAATCGTTAATCACCTCCACGCCACTTATGCCACGATCTC
AAAAAAATGATAATTTTATTGACAAAACTTTTACAGTATTAGCTGATATTGTATTAAAAA
TACTTCCTACAAGTAAAGAGGAAAAAGAAGCTTTCTCTTACTATCGAGATGGAATGTCTG
CACAATCTGAAGGTGAGTATGCGGAAGCTTTAGAAAATTATTATGAGGCTTTAAAACTAG
AGGAAGATCCATATGATAGAAGCTATATTTTATATAATATAGGGCTTATTTATGCTAGTA
ATGGTGAGTATGTAAAAGCTTTAGAATACTACCATCAAGGATTAGAACTAAATTTTAAAT
TACCTCAGGCCCTTAATAATATAGCCGTGATATACCATTACCAAGGAGTTCAAGCAATTG
AGGATAAAGATACAGAGCTATCTAAATTAATGTTTGATAAGGCGGCTCAATATTGGCAGC
AGGCTATTAAGTTAGCTCCTGATAATTATATTGAAGCTCAAAATTGGCTGAAAACGACAG
GACGAATGAGGAATATACAAGGATATTAATATATTTAAGATATAATAAAGATATCAATTA
ATAAATTAAAATTAGTTAAAGAAACTGGTACTATATAGAGTACTTTACTATATATTATAA
TATTATACTAGTTGA------TCTCTAAGACTTAGCTGAATAAACGAAATTAAATCTATT
GACTATATTCATGAATACAAATAAGCATAATGGTTAGTACAACAAAAAGTACCGGATCTG
TTACTCAAATTATTGGACCAGTTTTAGATATTGCATTTCCTAACGGACAGCTTCCGAAAG
TATTCAACGCACTCAAAGTACAAAGCTCAGAAGGAACTATTACTTGTGAAGTACAACAAC
TTTTGGGTGACAACAAGGTACGAGCTGTTTCTATGAGTTCCACTGAAGGACTACAAAGAG
GGGTAGAAGTTATTGATACTGGATCCCCTATATCTGTTCCTGTAGGTACAGATACTCTTG
GACGTATTTTTAATGTTTTAGGTGAACCTGTAGATAATTTGGGTCCCGTTGATTCTGAGA
GTACTTTACCTATCCATCGACCAGCACCTGCTTTTACTAAGTTAGAGACAAAACCAAATA
TTTTTGAAACAGGTATTAAAGTCGTCGATTTACTTGCTCCTTATAGAAGAGGTGGAAAA
TTGGTTTATTCGGAGGTGCTGGAGTAGGTAAAACTGTATTAATTATGGAACTAATTAATA
ACATCGCTAAAGCTCATGGTGGAGTATCTGTATTTGGAGGCGTTGGTGAAAGAACAAGAG
AAGGAAACGACCTCTATATGGAAATGAAAGAGTCTAAAGTAATTGATGCAGATAATCTGA
AAGAATCTAAAGTAGCATTAGTATATGGTCAAATGAATGAACCTCCTGGAGCACGTATGC
GCGTTGGCTTAACTGCATTGACAATGGCAGAATACTTTAGAGATATTAACAAACAAGACG
TTCTATTGTTTATTGATAATATTTTTCGGTTTGTACAAGCTGGATCAGAAGTATCAGCTC
TACTAGGCCGTATGCCATCTGCTGTGGGTTACCAACCAACTCTAGCAACTGAAATGGGAG
CACTTCAAGAAAGAATTACTTCAACAACAGAAGGATCAATTACATCTATTCAAGCTGTAT
ATGTGCCGGCTGATGATTTAACAGACCCAGCTCCGGCCACTACATTTGCACATTTAGATG
CAACAACAGTACTTTCGAGGAACTTGGCGGCAAAAGGAATTTACCCTGCAGTGGATCCAC
TGGATTCAACATCAACAATGCTACAGCCTGGAATTGTTGGAACTGAGCATTATTCTACTG

CTCAAGAGGTAAAATCAACTTTACAAAGATATAAAGAACTGCAAGATATTATTGCTATTC
TTGGTCTTGATGAACTTTCAGAGGAAGACAGACAAACTGTATCAAGAGCAAGAAAAATTG
AAAGATTTTTATCTCAACCTTTTTTCGTGGCAGAAGTGTTTACTGGATCACCTGGGAAAT
ATGTATCTTTGGAAGATGCAATTAAAGGGTTTCAAATGATCTTAAAAGGCAATTTAGACG
ACTTGCCTGAGCAGGCATTTTATCTAGTAGGTGATATAGATGAAGCTATACAAAAAGCTG
ACAGCATGAAAGATTAATATAATTAATGATGACTTTAAATATAAGAATTATTGCTCCTGA
TCGGACTGTTTGGGATGCAGAAGCACAAGAAATTATTTTACCAAGTAGTACAGGGCAACT
TGGTATTTTAACAGGCCATGCACCTTTGCTTACAGCTTTAGATATTGGAGTTATGAGAGT
AAGAGTAGATAAAGAATGGATGCCGATTGTTTTGCTGGGCGGTTTTGCAGAAATAGAGAA
CAATCAATTAACTATTCTGGTTAATGGCGCAGAAGAAGCTAGTCAAATTGATTTATCAGA
AGCAGAAAAAAATTTAGACACTGCAACTCAACTCTTAAGTGATGCCTCGTCTAATAAAGA
AAAAATAGAAGCAACACAAAAAATACGAAAAGCTCGAGCTCGAGTACAAGCTGCAACAGC
AGCAACTTCGTAAAAGATTCCTCTTAAAACATAAAACTTTAGTAATGATTACTAAA----
---GTTTTATGTTTTATTTGTATGGTTATAGGCAGCTTAAATAAACTTAGCTTAGACCTG
AACAAATATAATCAAAATATAATCCCATCTCTTTACCTGCATCTGGTCCAACTAAACTAA
TAGTAACTTCTTTCATAGCTAAAATAGCTTGAATCGTTGCACCAATAGGAACACCTAGAG
AATTGTATGTTTCTTTTAGACCATTTAATACTCTTTCTTCAAGGATAGATGGATCTCCAG
CTAGCATTCCATAAGTAGCATAGCGTAAGTAGTAGTCTAAATCACGGATGCATGCAGCAT
AACGTCGAGTTGTATACATATTACCACCTGGGCGTGTAATATCCGAATAAAGTAGAGATT
TTGCTACGGATTCTTTAATAATTGTTGCTGCATTAGCTGCAATAGTAGCAGCAGCTCTGA
CTCGTAGCTCACCTGTTTGAAAATAGCCTCTTAATTTTTCAACAGAGCTATCATCTAAAT
ATTTACCTTGGACATCAGCTGCATTAATAACAGAAGTAATTGCGTCTTGCATAACTTTCA
CACATTCCTTAATTTTAAATATCTAATACTAGGGGTATATAACAAAATTACTAGATCTTT
TAGCAATATCAGTATTACTGCATAGCACCCAAAGTATAATCAAAGTAAAAACCAGCTTCT
GCTGAATCTTCACCTGCAAGCAGTGAGCAAGCTACACTTTTCATGCATTTTACACCTTCA
GCAACTCCTGAAATTGGTGTTCCTAAAGAATTATACATTTCTTTAACACCCACTAATCCA
ATTTCTTCAATAGGAGTTACATCGCCAGCAACTATTCCGTAAGTTACTAAGCGAAGATAA
TAGTCCAGATCTCGTAGACAAGTAGCAGTCATCTCTTCGCCATAAGCATTTCCACCAGGA
GAAACTACATCAGGTCTTTTTTGAAACAGTTGTTGGCCGCCTTGCTTTACAATACGCTCA
CGATTATCTGTTAAAATTTGAGCTATTCTTAAACGACGTTGTCCAGATAAAACAAAACTT
TTAATTCTATCTAATTCCCCAGGACTTAAATATCTTGCTTCTGCATCTGCATTTACAATT
GACTTTGTAACAATACTCATGGATAAAACTCCTGTAATGCTTTAAAAAAAAAGCTTAATA
CTAATTATTTATGATATAACTCAGAAAGTAATGTACTTTTTGAGAATTTTATAACCGAAA
CACGTTCATGATATATGAAGCAGGATAGATACTGAGGTAAGTACTAAAACGAGAAAGTTA
TATGTAGGAACTTCTGCTATTGATTTCCTAGTACTTTTTTAAAACTTGGTACCACAATTG
ATATATCCTGCTTAGTCAAACGATTATAGAGTGTTTCTGTATTAGGAAAGTTAGCTGCTG

GTAAAGTCGGAAATCTTCTGTATGGAACAGTATCTTTGCCATATACCGCATTGTATTCAA

CACTTTCAACTAAATTACTTATAAATGCAGATAAGCCTTTTGAAGCTAAAATTTGATTAA

AGAATCTAATTTCTGCCTGATTATTAGGAGCTCTTCCTAAGATATGCTTAGTACCTAATT

CTATAACTTTTGTATTTGGATATGGCTGATAAAATTCTTTACCATATAGTTCAGATAGCG

CCAGTTTTTCTACTAGTTCTTTGACACAGATTTGTCTATTCAAGAAAGCTGACTTAATAT

CCAAGAATTCACCGCCGACACTAAAAGAGTTAAGATCTCTTTCGAAAATTTGACGATATG

CTGCTCTTAAAGCTTGTTCTAACATTTCTTTGTTACTATCACTATTGACTTCAAAGACAA

TAGATTGATCTCTTAAAGAAGTTACTCCCTGTGAAATACGAGACTGAATATCATTACTAC

TCCTCATTTCTTTGACAGTGCCTAATTCAACAAACCTAGCAGATTTACTAGCAATAACTT

TTTTGAATCTCTGGTCAATAATTCCTGGTCTTAAGCTTCTTAAAGCAACACCAGCTGGAG

TACTATATCTTTCATAAGGAACAGTATTATCTCCAAAAGTTTCTGTGTACTCCGAGCTAT

CAATTATTGTGTCTATTACTTGGTAATAGCCTTGCTTATATGCAATATCAAAATATTTAT

TAATTTCTTGCCGACCATATGTAGGGCGACCCAGTAAGCGGTTATGAATATATTCAATAG

CTTTACAAATATATAATGGCTCCCAATAAAGAGATCTAAATATACTTGATTTTGCTAACT

GTCTAACAAACTCGCGAACTGTAATTTGATTATCTTTTAGCTGACTTTCTATTGGCTTGA

GAATTAGTTTTTCTTCTTGATACACTTCTCTGCCAAAAACTCTTAAATAAGCAACTTTTG

TGACAACTTCTACTGAATTTTCAAAATTTTGGGATGAATCTGACTTTATAAGAGCGTTGG

ACAATTTAAAGATTTTTGGTCCCAGGGATCCTGCTGACTTAGGTCTTACCTGTGGATTAC

TAATCTGATTGTAGATTCCTGGCCCACGTCTAACTAATATCCTTCTTGTATCTTTACCAA

AAAGAGCTTGTCTTTTTCTTGGATCTTTATTTTCTTTAGGGAAAATAGCTCCAAATTGAA

TTGATAATGGATCATTACCTATTCCATATGGATGTTGATCAGGTAATGATTGCTTATAAT

CACTAAATAAAGTTATAAATTGAGGAACCTTACGAAAAGGCGCACTATAGGTGAAAAGAT

CTATTTGAGGCCCCCAATTACGGCACTCTTGAGGCTCTTCACCTAAATTTCTAAAGTAAG

GCACAGTCTCTTCACCGAAATAATCTGTATACTCAGAAGAATTTAAAAGTGCATTCACTA

AACCACTTAAACCAGTAGATGATAAAATAGCAAAATATTTTTGAAACTCTTCTAAAGAGC

TAGGACCTCTACCCAAGAAATGTCTAAATGCTAATTCTAATGCTCGACTATTGACAAAAG

GTTCATAAAATTGCTTTCTATAAATACTTGAAGTTCCTAACGAACGAATGAATTCCTTAA

TTGAAATTTGGCCATTTTTTACTTGCGACTCCAAATTCGATAAAGAGAGGTCATATGCTT

TAGCAATATCTCTTTCAAAAATTTGTCTATAACATGCTTTCACAACAATATTTTTCTCAT

CAGCAGATAAACTAGGCTTCATAACAAAACGAGGTGTTGACACTCCTGCCTGTACATAAG

TTTGAGGAAGACGCAAACCTTGTAAATCTCCAGATATTCTTTTTCTTAATTTATCAGTCA

AAGATGGAGCTTCAAATTCAGAAATCACAACATTAAAATACTCTTTAACTAAATCTTGAC

CTTTTATATCTTCCTCAAAGATTAATAATGCTGTCCGTCTCATCTCTCTAAGAGCCACAA

TTGCTGCTGCACTTGAGCAAGCGTTATCAATTAACTCTCTCAATCCTCGTATATTAACAG

ATAAAATATTAGGGTCGCCTGATACAATTGCATATGTCAAATACCGTAAAAACCAGTCTA

AATCTCGAAGAGATTTTCTCATTCTTGTAGTACCGTATCTCAAGACGTTTATAGGTTTAA

ACCCTGGCGGAGTAGCTCCCCCAGCATTAAATAAAGATCGGAAACTTTGGCCAAAATCTC
CTTGGATATTTCCTGATAACTCATTAATTTTATCTTGAGAACTTTGATCTCCTGCAATAA
TAACAGCTGCTTGCGGACGTTCTAAATAAGAAATTGCAGACCCACCTACGAATATTTTAT
CTGCTGCTCGTGCTACCAGAATATTTGCATTTTTAGTTAAAATATCTGCAACTTCCAGTC
TTTTTTGTCCAGAATTTAAAAAAGAAACTAATTGATTTAATTCACCAAGCTGTAAAAATC
GGTCTTGTTGCTCTGCCTGAGTAATAGTTAAAATTGAGGCAGTCCGATAAAGCTGGGGGC
GTGCTAATGGGCTTCCGCCACTTGCTTTGATACTCATTACTTTATTTATCTCCTTAACTG
ATATTGCTTGCTAGCGAATACAAATATAGCTTGATATTTCTAAATATTTAAAGATACTAT
GCAAGCACACTGTTTTCTATATTATTAGGTACTAGTACTTATAGTACAGAGAGTCTTATA
ACTAACTTCTATAAATACAAATTTTGTAATACTTCTATTAACATTTGTCAAAAAAATATT
TAAGTGCTCACTCTTTTATATTAAAATTCAATTAGTGAGAGATTGATGTGAAATAAAGTA
TACTAAGTAAGTTTTATAGATAGGAAAATTAAAAGATTTATTCTACAAACATTAATCACT
TGAATGTTTAGCAATTGTACAATTGTTATTGCTTTTAATTGGTTTAGACTAGTTTTATAT
TTTTTCTAACATAAACAATATTAATGAGCAATTTGTATGAATTTAAAGCCACAAAACAAC
CTATTAACTAACGAAAATATAAATCATGTAGATATACCCGAAAATGATATACCAATGTCA
ATTACGGAACATTTAGAAGAATTAAGACAACGGACTTTATTTGTGTTTTTATTCTTTTTG
TTTGCTACAACTATAAGTTTCACACAAATTAAGATCATTGTTGCAATACTGCAAGCTCCT
GCTGTTGGTATTAAGTTTTTACAACTGGCACCAGGAGAATATTTTTTTTCATCTATTAAG
GTTGCAATATATTGCGGGATTGTTGCAACAACTCCGTTTGCCGTTTATCAAGTTATATTA
TACATACTCCCAGGACTAACTGGAAAAGAAAGAAAAATTATTTTGCCTCTATTAATTAGT
TCTGTATTGCTTTTTATTACAGGTGGTATATTTGCTTACTTTGTTCTCGCACCAGCAGCT
TTAACATTTTTAATTAGCTATGGGTCTGATATTGTAGAACCATTATGGTCTTTCGAACAA
TACTTTGACTTTATTCTATTACTTTTACTTAGTACAGGATTAGCATTTGAGATACCAATC
ATACAATTATTGCTCGGTGTTTCAGGGACATTTTCTTCTAGTCAAATGATACGAGCTTGG
AGATATATTATTATTATAGCAACAATTGCTGGAGCTATTCTGACCCCTTCGACTGATCCC
GTTACACAATTAATAATGTCTTCAGCTGTTTTATTACTTTATTTTGGTGGAATTGTTATA
TTATTAGTCTTAAAAAAGTAGAGTAAATATTTTCTATTAATCCAATATATTTGTTATCTG
GGCAAATAATCTATAAAAACAATTGCTAGATTTAAGTATCAGGCTTCTAAATATTTCAAG
TCTCTTTTTATCATTTAAATACAATCGATACATTGGCTTGTTTTTTCATCAATTATTTGT
GTTGCACACTTTTTAGCACTCTCTTTATTGTTCTTAGAATTATTTCTTTAATCTATTTAT
TAAATAGATATGAAGAGATTTAAGTATTGATTATGGTAATCTATATCAACGTAAAAAGTA
TTTTATTGTGAGTTTTATTTATATAAAGGCTTATTCATTATATAGTTATAACAGTAAGTC
TAATTAATAGTTTTCATGCTTAATTAAAGTAATAACTCTTAATTCTTTTGGCAGAAAAAT
CTGTCTTAATATTTTATCAAAAAAACAGAAAAATGAAGCTAAATAGTCTAATTAACTTAA
TTCAAAAGTCTATATATTCTTGTACACTTTTACTAATTATTTTAAATATTATTTGTGTCG
CACCTAATTCTAGTAATGCATTTCCAATTTATGCGCAACAAGCTTATGAAAGTCCAAGAG

AAGCAACTGGTAGGATAGTATGTGCCAATTGTCATCTTGCTCAAAAGCCTGTGGAGATAG
AAGCGCCTCAAGCAGTACTACCTAATACTGTTTTCGAGACTGTTGTGAAGATTCCATATG
ACAGCAATGCTAAACAGATTTTAGGTAATGGCAGTAAAGGAGGCTTAAATGTTGGAGCCG
TTGTAATATTACCTGAAGGATTTAAGTTAGCTCCTGTTAATAGATTATCTACAGAGTTAA
AAGAAAAGACTAGAAATCTTTACATTCAGCCGTACAGTGCTAAACAAGACAACATTTTAG
TAATTGGACCTATTTCTGGTGATAAAAATAGAGAAATAGTTTTCCCAATACTATCTCCCG
ATCCTGCAAAAGATAAAAAAGCTCATTTTTTCAAGTATCCAATATATGTTGGCGGAAATA
GAGGACGAGGCCAGATTTATCCAACGGGTGACAAAAGCAACAATAATATTGTCTCTGCTT
TAAGCAGTGGTAAAATTAATAAAATTGAATTACTAGACAAAGGTGGATTTATAATACATG
TGACTAACAGTAGCAATGTAGAGTCAACACAGAAGATTTCACCTGGTCTCGAACTTAGAG
TAAAAGAAGGGGATACAATTCAGCTTGATCAAGCTTTGAATAGTGATCCAAATGTAGGTG
GTTTTGGTCAGAACGAAACAGAAATAGTTTTACAGAGTCCAAATAGAATTAAGGGCATGA
TTGTTTTCTTTTTCGCTAGTGTTCTAGCTCAAATTTTCTTCGTATTAAAGAAAAAACAAT
TTGAAAAAGTTCAAGCAGCTGAAATGAATTTTTAAAGCAATATAAAACTGGCACTTACTA
TAAAGAAAGTATAAACACTTTATTCACAGGATGTACCAGTTTTTATGTTATGTAATAATC
GCTTTGACGGAAGTTATGATACTTTTAACAGAATCTACAATCTGACTAGGCTGAATTACT
GTAGCCTGCTCTAAGCTTCCATTATATGGTGTGGGTATATCCTGCGAAGATAGCCTCACG
ACAGGAGCATCTAATTCATCAAAAAGGTACTCATTAATTTGTGCAATTAGCTCTGCTCCA
ATTCCGGCTGTTTTCATACACTCTTCTACAATTAAAACTTTGTGAGTTTTCTTTACTGAG
ATAGATATAGAGTCTATATCCAGAGGTTTTAAAGATATAAGATCTATAACTTCTGGATCA
TAACCTTCTTTTAATAAGGCCGGTAATGCTTGGATAACATGATGCCTCATTCTAGAATAG
GTTAAAATCGTAATATCTTTTCCTTTTCGTACAAATTCAACTTTATTGAGAGGTAAGAAA
TATTCTTCTTGAGGAATCTCTTCTTGTAAATTATAAAGTAGAACGTGCTCAAAGAATACA
ACTGGATTATTGTCTCGAATTGCAGATTTCAGTAATCCTTTTGCATTATAAGGAGTAGAA
CAAGCAACTATTTTTAAGCCGGGGATGGCTTGGAAATAAGCTTCCAGTCTCTGAGAATGC
TCTGCACCTAACTGCCTACCAACTCCTCCCGGTCCTCTAATAACTAAAGGCAATGTAAAA
TTACCTCCAGAAGTATAACGTAACATTCCTGCATTATTAGAAATTTGATTGAATGCTAAT
AATAAAAAACTCATGTTCATACCTTCAACAATTGGTCTTAGCCCTGTTATAGCTGCACCA
ATTGCCATACCAGTAAAGCTATTTTCTGCTATTGGTGTATCAAGAACTCTTAAATCCCCA
TATTTGCTATGCAAATCTTTAGTTACCTTATAAGATCCACCATAGTGACCAACATCTTCT
CCTATTACGCAGACAGTTGGATCTTTTGCCATTTCCTCATCTGTTGCCGCTCGTAAAGCG
TCAAACATAAAGATTTTACTCATTTTAATATTTGATTTTTATGATTTGACCACTGTTTTA
AAGTAACAGTTAAGACACCATATTCTATTTAGTTATCTGCAAAAAGATATCGTTTCAATT
CTGACATGTTAGGTTCTGGACTGGAAATAGCAAATTTCACAGCTTGTTCCAATTCGGTTT
TAACAGCATTCTGAATCTCATTAAGTTCACCTATATTAGCAATTTCATTATCTAGAATAT
ATTTTTTGAGTTTTTTGATAGGATCTCTTGCCACCCAAGCCTCTTTTTCTTGTCTTGATC

TTAGTTCATCAGGATCTGCGAGAGAGTGACCACGAAATCTATATGTTAATGCTTCTATTA
AGGTTGGACCATCACCTTGGCGAGCTCTTTGAACTGCTTGTTTTGCAGCTTGTCTTACAG
CTAGCACATCCATTCCATCAACTTCAATCCCAGGAAGCCCAAAAGCTTCTGCTTTTTTAT
GTATTTCAGGTATTGAAGAAGACCGGTGATGTGCCATACCTATAGCCCACTGATTATTTT
CAACAACAAATATAATAGGTAGTTTCCAGAGAACTGCCATATTCAGACATTCAAAAAATT
GCCCATTATTGGTAGTCCCATCACCAAAAAAGCAAGCCGTGACTCTTAAATCCTCTGTTT
CTTTAAGTACTTGCTGGCGGTAGATACTTTGAAAGGCTGCCCCTGTTGCAACCGGTATAC
CTTCCGCAATAAAAGCAAAGCCACCTAAAAAATTGTGAGGCGCAGAAAAAAATATGCATCG
AACCTCCTCTGCCTTTACTACAACCAGTCTCTTTTCCAAATAACTCAGCCATCACATTTT
TAGATGGGACGCCTTTACTTAAAGCATGTACATGGTCTCGATAGGTACTGCAAACATAAT
CAGTTGGATTCAGAAGTTTAATTACACCTGTAGAAACAGCTTCTTGACCATTATAAAGAT
GAACAAAACCAAACATCTTTCCTTTATAATACATCTGAGCACACATATCTTCAAAATTTC
TGCCTAACAACATGTCTTCATATAAAACTAATAAATTACTCTTATTAAGATTAAGACCTG
TTGAATTATAGTTAGTCAGCGGCAATTGAACTTTCTTAGGATAACTCATAATTTGTAAAG
AAACCTCTTATTGTAAATAGGAATTTTGAAGAAACAGCTATCAATAGAAAACACACAGAT
AAAAACTTATAAATTATAATATGTTTTTTACTAAAGACTTTAATAATTATTTGAAACTAG
TTCAGTATATCAAATTAATTTTGCTTATACTACGGATGTCTAAAACTTATATAAAAAAGT
AGTAAAAAAGTTTATTAAAAGAATAGTTAAAAACAGATATAAATAGAAGTAAACAGAATT
ATTGAAAAAAAATGATTATTATTAATTAATACTTAATATTTCTAAATATTTTACAATTAA
TGGCTATGTCACCATCTTTATTTTATCCTGTTGAACAAGAACTATGTAGTCTTGAAAAAA
ATCTGAAAGCTGTTGCTGGGACTCGTCATCCAATTTTATATGCCGCAGCAAAGCATCTAT
TCGATGCTGGAGGAAAACGAGTTAGACCAGCTCTTGTATTTTTAGTGGCTAAAGCAACCT
CTGAGAAGCAAGATATAAATACTGGACAAAAAAGGCTAGCAGAAATTACTGAAATTATAC
ATACTGCTAGTTTGGTACATGATGATATTATTGACGAGTGCACAACACGTAGAGGAGTCA
AAACTGTACATAATTTATTCAATACTAAGATTGCTGTGCTAGCAGGAGATTTTTTATTTG
CACAGTCTTCTTGGTATTTAGCTAATATTGAAAATTTAGCCGTAGTCAAAGCTATTTCTA
AAGTCATCACCGACTTTGCAGAAGGAGAAATTAGGCAAGGCTTAGTTCATTTCGATCCCA
GTATTTCAATAGATGCTTACATTGAGAAATCATTTTACAAGACTGCTTCACTCATTGCTG
CTAGTTGTCGGGGTGCAGCTATGCTTAATGGTTCCAATCATCAAATAAATAATGATCTTT
ATCTTTACGGTAAACATATGGGATTAGCATTTCAAATTATGGACGATGTTCTAGATATAA
CTGGTTCTACTAAGAGCTTAGGAAAACCTGCTGGCGCTGATCTAATAAATGGAAATTTGA
CCTCTCCTCTCCTTTTTTCACTTACTCAAGAAGCAAGTTTAAATGATCTTATTGATAGGG
AGTTCTGTAATAGTACAGATATAGCCTCAACATTATTTCTTATAAAAAGAAGCGGGGGAA
TTACAAAAGCTAAAGATTTAGCTAAAGAACAGGTGCAGGCGGCACTTTCCTGCCTTCAGT
TTTTACCACAATCTACACCTGTATCTAGTTTAAAAGAATTAACACATTTCATAATCACAA
GATTGTCATAAAGGACTTGCTAAAATTAAAATGTTATTAACTAATTTTTTACTGTTTCTA

AAACAGTCTGAAAAGTTTGAATATCATTACTAGCTAGCTGCGCTAGCATTTTACGATTTA
ACGCAATATTCTCTTTTTTTAATGCACTAATAAAAGTACTATAATTCATGCCTTGATTGT
GAGCAGCAGCATTTATTCTAGTTATCCAAAGACGACGAAAATCTCTCTTTTTCCTTTTTC
GACCAACATAAGAATATCGAAGAGCTTTAAGGACTTGCTGCTTAGCTGTTCGAAATAAAC
ATTTATGTGCACCTTTAAAGCCTTTAGCTAGCTTAAAGATTTTAGCTCGTCTTTTTTTTG
CAACGTTACCTCTTTTAACTCTACTCATAAAATATTACTTAAATTTTAATTATTACCAGC
TTACTAAATTCTTATACTCTTTGATTATAAATAAGGTAATTTCATTGCAATGTTTTTAAT
ATCTTTTAAGTCAACCGAACAAGTAGAAGAAAGATGTCTTCTTTGTTTTGATGACTTTTT
TTGTAATAAATGACTTTTAGAAGCTTTATGTCGAAGAAATTTTCCGGATGAAGAGACTTT
AAATCTTTTTGCTATTGCTTTTGATGTTTTTAACTTAGGCATATTATAAACTTTTAATGA
TTAACAAAAGCATGAAAAGTCTGAGGATATCAATCTTAGACCTTTCAATACTTAGAATAA
TCTTAACACAATTTATTTTACTTACTACATAGGTTAACTGAAACAATTTATATATATAAT
ACTAATCAGTTAATCAACCATAGCTAAAGTGGATTTATATTTTGCTTTGAACTTTTGTAT
AGTATTAGCAATTAGCATTGTTATAATTTTAAGACTGCTTGAATTCAATTCTTCGAACAT
TTTCATATTTAATGTAAATGAAATATTGGCTTCTGAGACAATATTTTGAATTTGAACATC
AGACAAGGGAATCATATCTAATGCAGCTCTATATTGATCTTTAAATAATTTATCGTCTTT
TATTTGGTCAAAATCGTAAAATTTCGTTCCTCCTGAATCTGATAAATTCATAGCCCCTCT
AGCTATTTTTTTTAAAATTTGGCCACCAGAAAGGTCACCCAAATAGCGCGTATAAGCATG
AGCAACTAGTAATTCAGGTTGTTTATGTCCTATAGTGTGAATTCTATCAACATAAATTTT
TGTAGCAGGAGATGGTTCAATAAAATCTAACCAATCTGATCCATAATAATAGTTTAAATC
TTCAGATAGACTCGCTTTCCTATTAAGCTCTGTAAAATATATAGGTTTAATAGCTGGATG
ATTTTTATTAGAAAATAACTCTTCCTCTATTGCACAATAGACAAAGTATAGATTTGCAAC
CAATTTACGATATGACTTCTTATCTACAACTCCTCCCAAAAAAGATTTAACAAAACTAAC
ATTTTCTGCCATACTATGGGACTTAGTAGTACCTTCTCTTAGTTCATTCGCTAAAGTATT
AACCATAATATCAATATTCCATTTTTAAAGATAACAAATTAACAATAAATAGTTATCCGT
ATATTTATAATTGAGTTGCGCTAACTAGGCTGCTGCAAAATAATTTTTTGATTTTATTGG
ATCTGGATTCATTGTTTTGTCACCAGGCTTCCAATTAGCTGGACAAACTTCATCTGGATG
AGATTGTACATATTGAATTGCTTGTAAAACCCTCAAAGTTTCTTCTACACTCCTGCCAAA
CTCTAGATTATTGATTGTAGAGTACTGAATAATTCCTTTCGGGTCTATAATAAATAGTCC
TCTTAGAGCTACACCATCACTATTCAAAACATTATAAGCTGCACTGATCTCTTTTTTTAA
GTCTGATACTAATGGATATGAAAGATCTCCTAATCCACCTGATTCTCTGTCAGTCTGTAA
CCAAGCAAGATGAGAATACTCACTATCAACCGAGACTCCAAGAACTTCTGTATTAAGTTC
CGAAAAAGCATTATACTTATCGCTAAATGCTGTAATTTCTGTAGGGCACACAAATGTAAA
ATCTAAAGGATAAAAGAATAAGACAATATACTTATTCTTGAGATCAGATAATTTTAACGT
TTTAAATTCTTGGTCATAAACAGCTGTAGCTGAAAAATCAGGAGCTAGCTGACCTACTCG
AAGACAATTTGGTCCAGAAATCATTAATTTTTCCCAGAAGAATTGATATAATTTTGTTAT

TTAATAAATAATTATATAACAAAATTTAGTATATATTAAAATACTACCAGTCTTTTTTAA
ATTACTAAATGATATAATCAATACTATCTTTATAGCGGGGAATGGATTTGAACCATTGAC
CTTCGGGTTATGAGCCCGACGAGCTACCAGACTGCTCTACCCCGCGGATAACTAAAATCA
TATTAGTCTACATATATATATAAAAGCAAATGCATAATAGATTTTAGAAGTAAAAAAATA
GAAGAACTGGTCCACTGATCATTAAACTTAGAAACGACAAAGTATTTTTTATCTTATTTA
AAAGAGGAAAGACTTCATATAAACCAATAAATCTATCTTTTAAAAGAATTTCGGACGGCT
TTACCCAAATTTGACCGTCATACCAGCCTGATTCTTCATAAAATACTGTAGCACTCATTA
ATCGTTTTACTACGTATGACCAGCCTAAATATAATCTGATTAAAATAAATCCTGTCATTA
GGCTGGAAGTAATAAATTCTGAAAAGAAAAATTTCAAAGGTAGTTTAGTGATTGGAAAAA
TTGATAATAATATAGGACTAACTAATAAACAGTTAAGTAGTAGTGCAATAGTTATTTTTT
TGTTATAAGATCTATGGCTTAAAGTTGGCCAGCAAAAGAACCAAGAATTTTTCAAAGAAG
TATATTCATGAACAGGTTGCTGCTCCTTTGGCACAGGACATTGAGTATTATATAAATTCA
TTGACATTAAAAAGTATTATAATAAAGACTTTGTTTATATTATCTAACATAGGTACGACA
TTAAATATTAACTAACTGCAATTATAGTAGTGAGTAGAAGAAATAAAACTAAAAATAGCC
AAGTAATTTTATTTAAAGTATTCTCAGTACTACGAGTATTACTAAAGAACTGATTCTGAG
CACCTACACTTCCTAAACCTTCGGATTTAGGATTGTGTATTAGAATAGTGAAGATTAATA
TTATTGTAGATGAGTACCAAAAAAATTTTAAAATTTGTTCCATTATGTAAAAGTTAATAA
AAATAGCACAAAGACAATCGTCTAAAAGATATTGATTGTCTTATCATATTAAAAAGTATT
TATTCACCCTGTACTTTTAAAAGTAAAAATCCTATTGCCAGCCCTACCAAAACTAGGACT
GAACTTAATACAGCACTTTCTACAATTTCTCCGACCATTTTATTTTTCCTTATTATTTTA
TTATTGTCATTCTTTTGACAAGATTATATTGAGCTAACTAAAATCCGTTTCTACCCCAAA
CTACCATTGCAATTGAAAATGTAAACATTGCCATTAGAGCTGCCCACCCCAAACTTAAAA
TATCCATAATTTATAATATTCCTAAAAATAAGATCTCTTTTACTTAAACTATAATATTCT
TAATATTAAAAATATCATTATATAATAAGCTAATTATAACAGAGTACTACTTAATTCGTA
TTTTTAGTGTATTTAAAGAAGATTGAATTTTTTCGAGGTACTGGATAAGTATTTCATCCT
AAGAACCATATTCAACTTTTGATTTATTATAATGCTTAAATCTAAATATATATCACTCAA
GAATCATTAGTAATTATGATGTTTTTAATTATACGATTAACATATTTCAAAATAGAAGAA
TTCATAGCGACTAAGTTAATATGCTAATATAATTTATCAATAAAAAGGAAGTTAGTATTT
CAATTATGCAAACAACTATAAATAATGGGCAAACGTCTAGTAAAGAAACTTTACTAACAC
CTAGATTCTACACGACTGACTTTGAAGAGATGGCTAATATGGATATTTCCGGCAATCAAG
AAGATTTTTTGGCTATCCTCGAAGAATTTCGAGCTGACTATAATAGTGAACATTTTATTA
GAGATGAAGAGTTTAATCAATCTTGGTCTAATTTAGAACATAAAACTAAATCCTTATTTA
TTGAGTTTTTAGAAAGATCTTGTACCGCAGAATTTTCAGGTTTTTTACTATATAAGGAAT
TATCCAGAAAATTAAAAGACAGAAATCCGGTTATAGCTGAGTGCTTTTTATTAATGTCTA
GAGATGAAGCTAGGCATGCCGGTTTTTTAAATAAAGCTATCGGGGACTTCAATTTATCTT
TAGATTTAGGATTTTTAACGAAGAGTCGCAAGTATACTTTTTTCTCACCTAAATTTATTT

TTTATGCAACCTATCTTTCTGAAAAAATTGGATACTGGAGATACATAACTATTTACCGCC
ATCTTGAACAACATCCAGAACACCGTATTTATCCAATCTTTAGATTTTTTGAAAATTGGT
GTCAGGATGAAAATCGTCACGGAGATTTTTTTGCTGCTTTGCTCAAATCCCAACCTCATT
TTTTAAATGACTGGAAAGCAAAAATGTGGTGCAGATTTTTTCTATTAAGTGTATTTGCAA
CAATGTACTTAAACGACTTTCAAAGAATTGATTTTTATAATGCCATAGGCCTAGACTCTA
GACAGTATGATATGCAAGTAATACGAAAAACTAACGAAAGTGCCGCTAGAGTTTTCCCGG
TTGCTTTAGACGTGGACAATCCAAAATTTTTCAAATATTTAGATACTTGTGCATGTGATA
ATAGAGCTCTGATCGATATTGATAATAACAATTCTCCATTATTCATTAAATCTATCGTAA
AGATACCTCTATATTTTTCCTTATTTGCAAATTTACTAAAGATATATTTGATTAAGCCAA
TAGACTCAAAAACAGTATGGAATACAGTTCGATAGTTAAATACAAGGGAATGCTTTTTTC
TTATAAAAAGCTATCTAGTGATTAATATAAAATAAAAGCTTCCTTCTCGATGGAAGGAGC
TTTTATTTTGTCAAGAATATAATGGGATTATATAGATCTAGAACTAAGAACTTACTTTTT
CTTTTGCTTCATACATAACCTCTAAAGTAATTAGATTTTTAGAACAATCACGGGCAAATT
TCTCTGTATTTCTTTTTACTTTACCTCTAACAAACCCAGGAATTTTATTTAATTCTTTCT
GTGCTTCTTCTGACCAATTAATACTATCAACAGCAGAAATTCCTATACTTAATGCTTCTG
TTGTATCATGACCACCAAATATTTCTAGCAAATGATCTTCCATACCAAGTGTAAACGAAT
TATATACTAAATCAGCAATTTGATTTGTTCCCTCATAACCAAGAAATGGACGGTAACTTA
AAGGAAAGTTTTGTATATGCACTGGTGAAGAAATAACGCCACAAGGAATATTTAGACGCT
TACCAATATGTCTTTCCATCTGTGTTCCAAAAATTGCTGCAGGTTCAGTTTTAGCAATTA
AATCTCCAATTAAACCATGATCATCTGAAACGATTACTTCATCGCAGAATTCTTGAACTT
GATCTTTAAACCACTCTTCATCATATTTGCAATAAGTTCCACACCAGGCGACATGTATAC
CCATTTCCTGGTGTAAAATTCGAGTTATTGCTGCAGCATGAGTTGCATCTCCAAATACAA
TAGCCTTTTTACCTGTTAAATTCTGACAATCAATTGATCTTGAAAACCATGCAGATTGTG
AAATAAATCGGGTTTGCTCGTCTATGTATTTTTCGTAGTCTACAGCAGCTCCCAATGCAT
TTACTAATTGCTGTATTGATCGGATGCATGCAGCTGTTTGAACAATACCCATAGGTGTAA
TGTCAACATAAGGCATATTAAATTCTTTCTCTAAATATCGAGCTGTCATTAGTCCAGTTT
CTCTATATGGAATAAAATTAAACCATGCAGAGGGTAATTTTTTTAAATCTTGAACTGATG
CATTTTCTGGAATAATTTGGTTTATTTGGATATCTAGATCTTGAAATAACCTTTTTAATT
CTGCTATATCATGTTGATTATGAAATCCTAAGCTTACTGCTCCAATAATATTAACAGATG
GAGTTTTAGTCTTTTGAGTTAAAACTTGATTATTAGATTTAGCTTTTTCCATATAGAAAG
TGACAATTTGTTCTAACGTTCTATCACCTGCTTGCAGTTCATTTACTCTATAATGATTAA
CATCAGCTAATAAAACATCTGCTTCTGTTTCTATAGATGCTCTGCTAACAAAGTTTTGCA
AATCTTCTTGCAAAATACTCGAAGTACAAGTTGGTGTTAGAATAACTAAATCAGGGCTTT
CTTCTCTGTCTTTTCTAGTAATATTTTCTACAACTTTTTCTTGAGAACCACGGGCTAACA
CATGTCTATCTACAACACTTGCTGTAACAGGAGTAAAATCTCTGTCTCTTTCAAGCATTG
AACGCATTACATTAAAATAATCGTCTCCAAGAGGCGCATGCATAATTGCATGAACTTTTT

TAAAAGAACTCGCAATTCTTAAAGTTCCAATATGAGCAGGGCCTGCATACATCCAATAAG
CTAATTTCATATTAAATTTTCCTATGATTTATTTAGACATAGTAATACTTTTAAAGCTAA
CTAACAAACATTAGCCAAACATCTAATTTAACTCCTTGACAATTTTGTAATATTTTTTAA
ATGAAGCTTATCATTATAAATATTTAAAAAATATAATTCAGAATAGAAATAACTGTTAGA
TCAAGCTTGAAAGTAAAGATTTCTTGCAGTTATAGTTTTTCTTTAAGTAAACATGTAAAA
ATTGTACTATATATAAATGTACTATATATAAACTTTTTTTACTGAATATTGGTAATGCCA
GATACAATTAACTTAAATATGCCTTCCCCAACGTTTGGTGGAAGCACTGGCGGTTGGTTA
AGAGCCGCAGAAGTAGAAGAAAAGTATGCGATAACATGGACAGGCAAAAATGAAAGTAAG
TTTGAAATGCCAACTGGTGGTACCGCAACTATGCGAAATGGAGAAAACTTACTCTATTTA
GCTAAGAAAGAACAGTGTTTAGCTTTAGGTACTCAACTAAAAAGTAAATTTAAAATATCT
GACTATAAGATTTACAGAGTTTTTCCTAACGGAGAAGTGCAATATTTGCATCCTAAAGAT
GGAGTCTTTCCAGAAAAAGTTAATACTGGGAGAGCTAGCGTCAACAGTGTTGATCATTCT
ATTGGACAAAATATTAATCCTGTAGATGTCAAATTCATGAATAAAGCAACTTACGATTAG
TCTCTCAAAAACCTAGACATAAAATAACTTTTATATAGTCTAGGTTTTTGATAATGTATA
CTGATAGTATGGAAGGGTGGCCGAGTGGTTGAAGGCGTCTGATTTGAAATCAGTTGAACT
ATCTGGTTCCGTGGGTTCGAATCCCACCCTTTCCGTTATAAAACTATAGTTGGCTTCTCA
AGGCAAGCTAGCTATACACTGATAGTATTAAGTTTAGACTTAATAAAGTCTACAACTTGT
GACAGGTTAGAAATTTTTTCGGCATCCTCATCTGGTATCTCTATACTAAATTTTTCTTCT
ATGGCCATTACCAGCTCAACTGTATCTAGAGAATCGGCTCCCAGGTCACTTGAAAAATTT
GCTTCCCTAGTCACTATTTTTTTTTCAATTCCTAGCTGTTCTGCTACAATATCTTGAACT
TTTTCAAAGATTTCATTATCTTGCATAATATATTATTTCCTCTAAGGTGTTATCAAAAAT
TTTTCGGCTTATATTATTATCCAACTCTAAAAAATTAGTAATTTTTAGTCGAGATTAAAA
TATTTATACCTATATGATATACTATTATTCTGGATAAATACGAAGTTTACGATATGGCTC
TTCTCCAAAACTACGAGCTCTCAACTGGTAGTTGTCAACTAGATTATGCTGCATTTTTCG
TATATAGGCTGATCGAGGTGTAAGCTGAACTATTGAACTTTCATTCAAGATAATAGTTTC
AATAGCCAATTTAGCTTCTTGAAGAGCCTGGATCTCATCAAAATTTTTATTTTTACAGAG
TTCTACCCAATTAAGACCAGATGAAGTATGAATGTTTAATATTTTTCTGAGTGCACGAGT
AATTTGTGGAACCGTACTATTCTGAATTGTATATATAATAATTTGTTTTGATTTTGCAAT
TTGTCTAAGTTTTGTATTTTGCTTTACTTGATTACGTAGCGCTAGAATAGCATCAGACTT
TTCTATCTCCTTGGTTAAAATAATTGGTAGATCTAGTGAAGAAATAACTGATGTAATATG
CTGCCAACTCAAAGAATAAGCATATAAGTACTGATGAGGTATTTCCAATTGCAAAGATTG
TTTATTAATATTAATAGGGGCATTGAAAGTAGTTGACAATAAAGATGTATCACGATTTTG
TGATTTATTCAAATCTAAGGACTTATTTCTTAATTCTCTATATTGTAAAACCGGAGGCGC
CTGCTGTGCGTTAATTGATTTACGAGCATTAGAGATGTTTACAGGTAAGACTTCTATAGA
TTGCGATGGATAACATTTAATTAAAATTCTACCATTAGCTTGAATTTGTCTTTTTTGCAC
AAAAGGTTGATGGCCTTGCAAAATTTGATCTATTGTCTCTTTAACTTTATTATGAACTAT

CCAAACATTTCGCTCGTGTATTTCGATCGCAATCTGAAAAGCAGGCTCAGCTTTTCTTTC
TAGTATACTCTTTTGCGTACCTCTACGTTTAGCTTCATCGTCACCTAATGTTACATATTG
AATTCCACCAATTAGGTCCGCCAATGTTGGATTTTTAATTAAACTTTCTAAATGATTTCC
ATGCGCTGTACCTACTAATTGAACTCCTCTTTCTGCTATAGTTCGAGCAGCTAACGCTTC
TAATTCCGTGCCAATTTCATCTATAATAATAACCTCGGGCATGTGATTTTCAACAGCTTC
TATCATTACTTGATGTTGCAAATCGGGCCTTGCGACTTGCATTCTTCTAGCTCTACCAAT
AGCTGGATGAGGAATATCTCCGTCACCAGCTATTTCATTTGAAGTATCTATAATAACTAC
TCTTTTTTCCATTTCGTCAGCTAAAACACGGGCCATTTCTCGAACTGCGGTTGTTTTACC
AACTCCTGGTTTTCCTAAGAGTAAAATTGAATCACCTTGCTGTAGTAAATCTCGAATAAT
ACTAATTGTACCAAAGACTGCTCTGCCGACACGACAGGTTAAGCCAATAACACTACCTTC
TCTATTACGCAGTGAACTAATACGATGCAAGGTTTTTTCGATACCAGCTCGATTATCGCC
GCTAAAATTACCAACTTTTTTTACACAATAATCTAAGTCTTGCCAACTAATAGATCTTTG
AGATAAATATTCTGGGTTATCTGGAAATCTAGCTTCTGGACGACGACCTAAATCCATAAC
GACTTCTATTAAATTATTTCTATTAGGATGCTGTTGTAAAGGTTCTTTGACAAAATTTGG
CAAAATTTCTAGCAACTTATCTAAGTCATCTGCAATAAGCATGATTGAGTATATAATAAT
TAATTAATCAAATAGAAGATTCAAGATTCTAAAGATTACAATACATATTAAAGAATATTA
TGATTGGAATCAATTTTATACTTAAATTGAACAGGAAGTTATAGATGTAAAAGAACTGAT
TAAACTTTACGCAATAAATCGATTGGTCAAAATTAATAGTCGTGTTAACATGTTTATTGG
ACAAAAAGTTCGTATTAAATATAGCAAACAAAAAGTTGCTAGTGATATAGCAGATAAAGT
TGGTGAATTAGGAGTTATTAAGGGAATAAAGTTTATTAATAGTCAATGTGTTACTATTAT
TGTTGAGTTTGACAATCACACTAGGCTTTGGATGTTTAGAGAGGAATTAATCTGTCTAAA
TGAATTAAAAACATGAAACATAATTTATTTTATTTATTATACAAAATTTCTTATGAGTCG
TTAAAATAATAAATTTTAAAGGAGTACTTTTAATTGTGCAAATTACTATTAAAAAATTAC
AAGACTTACTTTCGTCTGTACAAAGAAAAAAGATCCAGACATTAAAACTAAAACAAGGTA
AGTTTGAACTTCTATTAAATAAGACCTATAAAAAAGTCAATCAAGAAATTATACCTTCGC
AGAAATCTGCTGTATTACAAAATAGTCCATCTACGATAATTAAATCAATAAATAATACAA
AAAAATCTTCTGTTGTTAATGAAGACCGTACAGAATATGCCACTATTGTTTCTCCAATGG
TTGGAACGTTTTATCATTCACCTGCTCCTGGTGAAAAAATTTTTGTACAAGTTGGCGATG
AGGTCAAATTCAATCAAACAGTCTGTATTATTGAAGCAATGAAGTTAATGAACGAAATCG
AGGCAGAAATTGAAGGCAAGATTATAGAAATTCTTGTTAAAGATGGTGATATAGTAGATT
GCGGGCAAGCCTTAATGAAGGTTGAAACATAATGCTAATTTTTCCATTTTACATATTGTT
AACAGAAATTATTTAGATAGAAAAATTAGCTTATACTATAGTTATGAAAACTTATTTCGC
AAAGGCACTAAAAACTAATATATAAAATTATTTACTTTATCTGCCAAAAATAGAAATAAG
CGTTTTGATTCCTTGTCTCAAGAGAGGAGAATCTCAATGGCAATTAGCTCAAAAGAGCAA
GAGACAAAGAAGGTAAAAATCTCGGTTGATAAAAATCCCGTAGATACTTCTTTCGAAAAG
TGGGCCCAACCAGGCCATTTTTCTCGTACACTAGCAAAAGGACCAAAAACTACTACTTGG

ATTTGGAATCTTCATGCTGATGCTCACGACTTCGATAGTCAAACCAGTTCTTTAGAAGAA
GTTTCACGTAAGATTTTCAGTGCACATTTTGGGCAGCTGTCTGTAATATTTTTATGGCTT
AGTGGAATGTATTTTCACGGAGCCCGCTTCTCTAACTATGTTGCTTGGTTAAGTAATCCA
ACAGGTATTAAGCCAAGTGCGCAGGTTGTTTGGCCTATAGTTGGGCAAGAAATTTTAAAT
GGCGATGTAGGTGGTGGCTTTCAAGGAGTACAAGTTACATCTGGATGGTTCCAACTGTGG
AGAGCATCAGGAATTACTACAGAATTTCAGCTTTACTGTACTGCTATTGGCGGATTAGCT
ATGGCTGCTTTAATGCTGTTTGCAGGATGGTTTCATTATCATAAAGCTGCTCCAAAGTTA
GAATGGTTTCAAAATGTTGAATCAATGATGAATCACCATTTAGCTGGGCTTTTAGGCTTA
GGCTGTTTAGGCTGGGCAGGCCATCAAATCCATTTGTCTTTGCCTATTAATAAGCTACTA
GATTCTGGCGTGTCTCCGCAAGAAATTCCACTACCTCATGAGTTTTTAATTAATAGAGAG
CTTATGGCTCAGCTGTATCCAAGTTTTAGTAAAGGATTAGTTCCATTCTTTACTTTAAAT
TGGGCTGAATATTCCGACTTTTTAACTTTTAAAGGAGGTTTAAACCCTGTTACTGGAGGT
TTATGGCTAAGTGATACTGCTCATCATCATCTAGCATTAGCTGTTCTATTTCTTGCTGCA
GGTCATATGTATAGAACCAATTGGGGTATTGGACATAGCATGAAAGAAATTCTAGAAGCT
CACAAAGGACCTTTTACCGGCAACGGTCACGAAGGTCTATATGAAATTCTTACAACTTCT
TGGCATGCACAGCTTGCAATTAATTTAGCTATGATGGGATCTTTAAGCATCATTGTAGCA
CATCACATGTATGCAATGCCTCCTTATCCATATATTGCTACTGATTACCCGACTCAGTTA
TCGCTCTTCACTCATCATATGTGGATTGGAGGATTTTGTATTGTTGGAGCAGGAGCGCAT
GCTTCTATATTTATGGTAAGGGATTATAATCCTGCAGAAAATTATAACAATCTTTTAGAT
AGAGTCATTAGGCATCGAGATGCTATTGTTTCTCATCTAAATTGGGTATGTATATTTCTT
GGATTCCATTCATTTGGTTTATACATTCACAATGATACTATGCGGGCACTTGGAAGATCT
CAAGATATGTTCTCTGATACAGCTATACAGTTACAACCTATTTTTGCTCAATGGGTACAA
AGTATACACACTTTAGCTCCTGGAAATACAGCTCCAAATGCATTAGCAACAGCTAGTTAT
GCATTTGGAGGAGATATTGTTTCTGTTGGTAACAAAGTTGCAATGATGCCTATTTCTTTA
GGTACTGCAGATTTTTTAGTTCACCATATACATGCATTTACTATTCATGTAACTGTTTTA
ATTTTAGTTAAAGGGTTCCTTTTCTCAAGAAACTCTAGACTAATTCCTGACAAGGCCAAT
CTTGGCTTCAGGTTTCCATGTGATGGACCTGGTAGAGGTGGTACTTGCCAAGTGTCTGGC
TGGGATCATGTTTTTCTTGGCTTATTCTGGATGTACAATTCTCTGTCTGTAGCAATTTTT
CACTTTAGTTGGAAAATGCAATCAGATGTTTGGGGTAGTGTATCTCCGTCTGGAAATGTT
TCTCATATTACTGGCGGTAATTTTGCACAGAGTGCAATTACAATCAATGGATGGCTGAGA
GATTTCCTTTGGGCTCAAGCATCTCAAGTTATTCAATCATACGGTTCTGCGCTATCTGCA
TATGGATTAATTTTCTTAGCAGCACATTTTGTATGGGCATTCAGTTTGATGTTCTTATTT
AGTGGTAGAGGTTATTGGCAAGAGCTTATAGAATCAATCGTATGGGCGCATAATAAGATA
AAAGTTGCTCCTGCAATTCAACCAAGAGCTTTAAGTATTACTCAAGGTAGAGCAGTCGGT
GTTGCACACTACTTATTAGGTGGAATTGGTACAACTTGGGCATTCTTTTTAGCGAGAATT
ATTTCAGTAGGCTAATAGTGAAAATAGGATAAAAAAAACAATTATGGCAACAAAATTTCC

TAAGTTTAGCCAAGCTTTATCACAAGATCCTACAACTAGAAGGATTTGGTATGGTATTGC
TACGGCACATGACTTTGAAAGTCATGATGGAATGACAGAAGAAAATTTATATCAAAAGAT
ATTCGCTTCGCACTTTGGACATCTAGCAATTATCTTCTTATGGACATCTGGTAATTTATT
CCATGTAGCTTGGCAAGGCAACTTTGAACAGTGGGTATTAAATCCTTTGAAAGTTAAACC
AATTGCTCATGCAATTTGGGATCCGCATTTTGGACAACCTGCTTTGAAAGCTTTTAGTAA
AGGTGGCTCAGCTTATCCAGTAAATATAGCATATTCTGGCGTATATCACTGGTGGTATAC
TATTGGTATGAGAAGCAATCAAGACCTGTATTCTGGGGCTTTATTCTTACTAGTTTTATC
AGCTTTACTCTTATTTGGAGGGTGGCTACATCTACAACCAAAATTCAAGCCTGGTTTATC
ATGGTTTAAAAATAACGAATCAAGATTAAATCATCATTTATCTGGATTATTTGGGGTTAG
TTCTTTAGCCTGGACAGGTCATTTAGTACATGTTGCTATACCTGAGGCAAGAGGACAACA
TGTAGGATGGGATAATTTTACAACCGTATTACCTCATCCAGCCGGTTTACAGCCGTTTTT
CAGTGGTAATTGGAGTGTATATGCTCAAAATCCAGATACAGCTCAACATTTATTCGGAAC
TAATGAAGGTGCGGGTACAGCAATTCTGACATTTCTAGGAGGATTTCATCCTCAAAGTCA
GTCTTTGTGGCTAACTGATATGGCTCATCACCATTTGGCTATTGCAGTAGTATTCATTGT
TGCTGGACATATGTATAGGACTAATTGGGGAATTGGGCACAATCTAAAAGATATTTTAGA
TGCTCATAGACCACCTAGTGGTAGATTAGGAGCTGGACATAAAGGGCTATTTGATACTAT
TACTAATTCTTTACACATACAGTTGGGATTAGCATTGGCTTCCCTAGGTGTAATTACTTC
GTTGGTAGCTCAACATATGTATGCTATGCCTCCATATGCTTTCATGGCTAAAGATTTCAC
AACTCAAGCATCTTTGTACACACATCATCAATATATTGCTGGGTTTCTAATGGTTGGAGC
TTTTGCTCATGGGGCAATATTCTTTGTTCGAGACTATGACCCTGAACAGAATAAAGATAA
TGTTTTAGCTCGTATGCTAGAACATAAAGAAGCTATCATTTCTCATTTAAGTTGGGTAAC
TCTATTTTTAGGGTTTCATACATTAGGCCTTTATGTTCACAATGACACAATGATTGCTTT
TGGAACTCCTGAAAAACAAATTCTAATTGAGCCGGTATTTGCTCAATGGATTCAAGCCTC
TTCAGGGAAAGCACTTTATGGGTTTGATGTGTTACTATCATCCTCTACTAATATCGCAAC
ACAAGCTGGTAGCAATATTTGGCTGCCAGGCTGGTTAGAAGCGATTAATAGCGGAAAAAA
TTCATTGTTTTTAACAATTGGTCCTGGTGACTTCTTAGTTCATCATGCAATTGCATTGGG
ATTACATACTACTACGTTAATTTTAGTTAAAGGTGCTTTAGATGCAAGAGGCTCTAAACT
TATGCCGGACAAAAAAGACTTTGGATATAGTTTTCCTTGCGATGGACCTGGCAGAGGCGG
CACCTGTGATATATCTGCATGGGATGCGTTCTATTTAGCTGTATTTTGGATGCTAAATAC
AATAGGTTGGGTAACATTTTATTGGCATTGGAAACATATTACAATATGGCAAGGCAATGC
AACTCAATTCAATGAGTCTTCAACTTATCTAATGGGATGGTTTAGAGATTACTTATGGCT
AAATTCTTCTCCATTAATTAATGGTTATAATCCATATGGCATGAATAATTTATCAGTATG
GTCATGGATGTTCCTATTTGGACATTTAGTATGGGCAACAGGATTTATGTTCTTGATCTC
CTGGCGTGGCTATTGGCAAGAGTTAATTGAAACTCTAGCATGGGCGCATGAACGTACACC
TTTAGCAAACTTGATTCGTTGGAAAGATAAACCTGTTGCATTATCAATTGTACAGGCAAG
ATTAGTAGGTTTAGCACATTTTTCTGTAGGATATGTATTAACTTACGCAGCTTTTGTATT

AGCTTCAACAGCAGGCAAATTTGGTTAGACTAAATTCGAATTAGCT-AAAAAAAGTCAGT

CATAATAGTTTTATGACTGACTTTTTTAGATTTAAAACAAAAACTTACTAAAGTTTCACC

TCAATATCTACACCAGAAGGCAAATTTAACTTCATTAATGCATCTATCGTTTGAGAAGAA

GGTTGATGAATATCAATAATTCTTCTGTGAGATCTTATTTCAAAGTGTTCTCTTGAGTCT

TTATCTACATGTGGAGAACGTAAAACACAATAAATTCTTCTTTTTGTCGGTAAAGGAATT

GGCCCTACTGCAACGGCATTAGTTCTAGATGCGGTATCTAGTATTTTGTCACACGACGTA

TTAAGTATAATAGAATTATATGCTTTCAGTTTAATTCTGATTTTTGTCTGCTGAGTAATT

GTCATGTTAAAAAGATCTTATTATTTAAGAATTTTAGAGACAACACCTGCGCCTACGGTA

CGGCCACCTTCTCTAATAGCGAAACGCATACCTTGCTCAATTGCAATTGCATTAATTAAT

TCAGCAGTCATCTTAATTCTGTCACCAGGCATAACCATTTCTGCATCAGTACCATCATCA

GCAGTAAACTGATTAATAGTACCAGTTACATCAGTTGTTCTAACATAAAACTGAGGTCTA

TATCCTGGAAAAAATGGAGTATGTCTTCCGCCCTCCTCTTTAGTTAAAATATAAACTTCT

GCTTCAAATTGAGTATGAGGTGTAATTGTACCAGGTTTAGCTAATACCATACCTCTTTCA

ATATCTTTTTTCTGCACACCTCTTAAAAGAATTCCAATATTATCACCCGCTAGACCTTCT

TCTAACGTTTTTTGAAACATTTCTAATCCAGTAATAGTCGTTGTACGAGTTTCTCGTAAA

CCTACAATTTCAATTGTGTCACCAACTTTAATAATGCCTCTTTCAATTCTACCAGTGGCA

ACAGTTCCACGTCCTGTAATAGAAAAAACATCTTCTACAGCCATTAAGAAAGTTTTATCG

ACATCTCTCTCCGGCGTTGGAATATATGTGTCAACTGCTTCCATAAGTGAAAAAATCTTG

TCAACCCATTTATCTTCACCTTGCTTAGTAGCTGGGTTTTTCGTCACAGCTTCTAATGCC

AATAAGGCAGAACCTGCAACAAAAGGAATATCATCTCCAGGAAAGTCGTATTGACTTAAT

AATTCCCTTCCTTCTAATTCTACTAATTCTAGTAGCTCTTCGTCATCTACTTGATCTTCC

TTATTTAGGAATACTACTAATGTAGGAACACCTACTTGTTTTGCTAATAAAATATGTTCA

CGAGTTTGTGGCATTGGACCGTCTGCCGCAGATACAACTAAAATAGCTCCATCCATCTGA

GCAGCACCCGTAATCATATTTTTTACGTAGTCGGCATGGCCTGGACAATCTACGTGAGCA

TAATGACGATTATCTGTTTCGTATTCAACATGAGCAGTATTAATAGTAATACCTCTAGCT

TTTTCTTCTGGAGCGGCATCAATTTCATCAAATTTTTTCGCTGCAGTAGACCCTAAAGTT

GATAAAGTCGCGGAGATCGCTGCTGTTAAAGTTGTCTTACCATGATCAACATGACCAATT

GTGCCAATATTGACATGAGGTTTTTTACGTTCAAATTTAGATCGAGCCATGCTTTTTGTT

TTCCTTATAGTAAAAGTTGCTTTATAGTTACTTTTAACGATAAATCTTGTGAATTGAAAG

ATTTATTATTTAATACAGACAAGATACAATATAACTAATCTAATATCGATAGTGAGCAAA

CGCTTTATTTGCTTCTGCCATTCTATGTGTATCCTCTCTCTTTCTAATAGAATTTCCTGT

TTCATTAGCTGCATCCATAATTTCATTAGCTAATTTCATAGACATACTTTTACCAGATCT

GTCTCTAGAGAATTTAGTAATCCATCTTAGTGCTAAATTCGTACCTCTATAAGCTCGTAC

TTCAATAGGAACTTGGTAAGTAGAACCACCAACTCTTCTTGCTTTTACTTCTACTAGAGG

AGTAATATTTCGGATGGCTTTTTCTAAAATATTTAAAGGATCTGATTCTGTTCTCTCTTT

AACGATATCTAAAGCCTGATATATAATTCTTTGAGATAAAGTTTTTTTACCACTTTTCAA

AATACGAACAGTTAACATACTTACGAGTCTGCTTTTATATAAAGGATCAGGTGATGCAAA
CCTTTTTTTAGCTGTATTACGACGAGACATAGTATTTAAAATTGGTTATTGTAATAATGT
TCTATTAGCAGGTATTTAAGATTTAGGCTTTTTTGTCCCATATTTAGATCGGCTTTTACG
ACGATCTTTTACTCCTGCAGCATCTAAAGTACCACGTACTACATGATATCGGACTCCAGG
CAAATCTTTAATTCGGCCGCCCCTAATCAGTACCACAGAATGCTCTTGAATATTATGACC
TACGCCAGGAATATAAGCTGTAACTTCAAATCCAGATGTTAATCTAACGCGCGCTACTTT
TCTTAAAGCAGAGTTCGGTTTTTTAGGAGTTGTAGTATATACTCTTGTACAAACACCTCT
TCTTTGAGGGCAACTTTGAAGAGCTGGAGATTTTGTTTTTTTATGTATTTTTCGTCTTTC
GGATCTAACAAGTTGTTGAATTGTTGGCATAATAAAATTGATTTAAAGTCTTTAATAGAT
ACTATTTGAACCTGCTTTACATGAATTGGCTAGTTTTTATTTATCTTATACTTACGCATA
AATCTTTCTACTCTACCTTCTGTATCAATAATTCTCTGTGAACCAGTAAAGAATGGATGG
TTTCCTGACCAGATATCTACGTGTAGTTCTGGCTTTGTGGACCCAATCGTCATAATTAAC
TGGCCGTCACAGTAAACTTTTGCTTCTGGATACCAATTTGGATGTATATTATCTTTTGCC
ATTGTTTTTATAATTATATAAATGTATCAATAATAACTATTAACGCTTAGAAAATTGAGG
AGCCTTTCTTGCTTTCTTTAAACCATATTTTTTTCTCTCTTTTACTCTTGGATCTCTTGT
TAAATAGCCTTCAGACTTGAGTGTCGTTCTATTTTCTGGATTAATTGAGCACAATGCTCT
TGCAACACCTAAACGAATTGCATCAGCTTGCCCTGTCAAACCTCCTCCTCTAGCATTTAC
ATGGATATCATACTGGTTTAGTAGTCCTAAAACTTGTAATGGTGCATATGAAACTCTTAA
GTAATTAGGACTAAATTGAAGATAAGACTCTCCTGGTATACCGTTTATAATTAAATTACC
TGACCCTGGGACTAGTCTTACTTGTGCAACAGAACACTTCCGGCGACCTGTTCCAGAATA
GATTGCGCGAGTTTTAATTAATTCTGTCGACATAACATTCCTTGATAACTAAATAAATAT
ACATTCTTTATACTATATACTCTTGTGGCTTTTGCGCGACATGTGGGTGAATTGGGCCAG
AATACACTTTAAGCTTCGTAAATAGTTTTCTGCCTAATGGACCTTTAGGAAGCATACCTT
TAACCGATTTTTCAATAATTCTGTTAGGTAATCTTGTCTGAAGCTGATCAAATGTTTCAA
CTTTTAATCCGCCAGGCTGTCCAGAATGTCTTCTGTATAGTTTTTGATTCGTTTTATTTC
CACTTACAGATACGTGAGCAGAATTAATAACGATTACATAATCTCCAGTATCTAGATAGG
GTGTATAAGAAGGCTTGTTTTTACCTCTTAAGATATTAGAAATATGAGTAGATATTCTAC
CAAGTGTCTGATTTTTAGCGTCTATAACATACCAATGAGAATTAGTATTTAATGAGGGTG
ATTGCGTTTTATTCATGAAAAGATTAATACTTTTGGTTGACAATAAGAGAATATGTAAAT
GGTTTACATTCAATTTCTATACAAGTAGTATTTAAAAATATTATTAACAAAGAGAACTTT
TCTAACAGTTATTTAAATTGTTTGTCAGCAAGAAAATTTCTAGTTATCAATAACTTATAC
TATTTTTAATAATATAAAATATATTAATCACTTTTTTCTTTTGGCAAACTTATACCTAGT
TTTTTTTGCAAGGCATATATAACTTCTTCAGCTGATTTCTGGCCAAAATTTTTAATTTCT
AGCAGTTCTTCTTGAGAGTAATCTAGTAAATCTGCAATAGAATGGATTTGAGCTCGCTTT
AAGCAGTTATAAGCCCTAACAGATAACTGCAGTTCTTCTATTAAAACTTGACTAATTTTT
TTATCTTCTTTGCTACGATAATTATCTGCTGATTTAAAGTCTAAATTTCTTAGTGAACAA

AAAAGATTAGTTAAAACTGTAGCTCCTTGACTTATTGCTTCTTGGGGAGATATGCTCCCA
TTTGTCCAGATTTGTATAATCAATCTATCTTTTATGCTGTTGCTACCAATACGGACTTCT
TCTACTTTATAATTAACTTTATTAACTGGCATAAAAACAGAGTCTACTTGCAAAAAATCC
ACAGATAATTCATCTACAGCTTTTTCAGCTAAGCGATACCCGCAATTTTTTTCAATTTTA
AACTCCATTTCAAATATTGTGTTATTGCAAATAGTTGCAATATACTGTCTAGGATCTACT
ACCTCTATATCAGAAGATAATTCAAATAGGCCGGCTGTAACTATAGCTGGCCCTTGAACT
CTAATTCGACCAATTTGAGATTCTTTGTTATAACTTTTAAATACTACTTCTTTTAGATTA
AGTAATATTTCTAACACATCTTCTCTTACCCCAGGAATTGTGGAAAACTCATGGTTCACT
CCAGCAATCCGTACAGCAACTATAGCAGTACCTTCAAGATCTGACAATATTGATCTTCTC
AAAGCATTACCTAATGTAATACCTTGTCCTTGATTTAATGGTTCTATTACAAAACTACCG
TACTGCCCACGCGCCCCATCTGTTCTTGACTCTATGCATTCAATTTGAAATTGAGCCACC
TAAGAAAGCTCCTTTAAAATAATCAGTGATTAGTAAGTCTCTCTAAAACAAGAAGGAATT
TATACTCGGCGTTTCTTAGGAGGGCGACATCCATTATGAGGTACAGGAGTAATATCTTTT
ATTAGAGTAATCTCTAATCCTGCAGCCTGCAAAGCTCTAATTGCTGTTTCTCGACCCGCT
CCTGGACCATTTACTAGAACTTCAGTTTGGCGCATACCTTGATCCATAGCTTGTCTAGCT
GCTTTTTCAGCTGCTGTTTGAGCTGCAAAAGGTGTTCCTTTTTTAGCTCCCTTAAATCCA
CTTGCACCAGAGGATGACCATGATAATGTTTCTCCTTTTAAATTAGTAATAGTAACAATT
GTATTATTGAATGTAGATTTAATATGTGTAATACCGTTAACTGCATTACGTTTAGTTTTT
CTTGCTCCGGATTTTTTTATTTGTCTAGCCATCGTGTCTCGTCTATTATATAGTTAATTA
AAAGATTATTTTCTTGGAGCTTTTTTCTTACCTGCTACTGTTTTTTTACCTCCTCTGCGT
GTTCTAGCATTAGTTCTAGTTCTTTGTCCTCTCAAAGGAAGACCAAGACGGTGTCTTCTA
CCTCTATAAGTACTAATTTCCATAAGTCTCTTAATGCTCATAGACTCAAAACGTTTGAGA
TCCCCTTCAATCTGATAATTAGACTCAAGAATTTCTCTTATACTGACAACTTGTTGATCA
TTTAAATCTTGACACTTAATATCAGCGTCTATGTTTGTTTTTTCTAATATTTCTTTCGAG
CGAGATAGTCCAATACCATAAATATATGTTAAAGCTATCTCTATTCTTTTGTTTCTTGGA
AGATCTACTCCAGCAATTCTGGCCACTTTGTCTTGTCTCCAATAAATAATATGTATTTTT
ATAACGGTTATTTATAAGCTTAGCCTTGTCTCTGCTTATGTTTAGGATTAGTGCAAATTA
CCATCACCTTTCTGTGACGCCGAATTATTCTACATTTTTCACACATTTTTCGAACAGAAG
GACGAACTTTCATATTAACTCTATATACTAGGAATATTACTATATTAACTCTATTAATTA
TCGACATTAAAAATGTCAACAATATATTGTATTACTTTAACTGTTGTAGATTTTACTTCG
TCATACTATCATATTTTTTTGATATGACGTATGTCTGAATTTGCTTAGCTGTGTCTATTG
CGACTCCAACTAAAATAAGTAAAGACGTAGCCCCGAGACCTCTTAAGTTTTGGATCTGAG
TAACCTTTTCTATTATAAACGGAATCAGCGCTACTGTAAATAAAAACGAGGCTCCTAGGA
ATGTCAGTCTATTTAATATGACTTGTAAATAATCAATAGTCGCTTGACCTGGACGAATAT
TAGGAATGCTTGCACCCATTTTTTTTAAATTTATAGCAATATCTTCTGGATTCATTACTA
TTGATGTATAAAAATAGCTAAAGAAAAGGATCAAAGCACAATAAAGAAGAAGGTATAGTG

AACCATTAGGACAAAACAGGTATAGGATTTGGAGTAATGTTTTATTTTGAATAATTTGGG
TTAGATAAGATGGGAGGGCCATAGATGCAGATGCAAAAACAATAGGCATAACTCCACCTT
GATTTAATTTCAAGGGCAAGTAACTATTTGGATCCAAAATTGAAGATTTTCCTAGCTGTC
TTGCTGAAATAATTTTAATTCTTCTTGTTCCTTCTTGTACACAAATTGTAATTATTATCA
TTAATAAAAAGATCGCTATAAATAATCCGAACTTAAGACTTGCATTACTATAACTAGCAT
CAAAAAATGACTGTGTAAAATTCTTTGGTAGTCCTGACACAATGTTTTGAAAAATAAGTA
AAGAGGCTCCATTACCAATTCCCTTTTCCGTAATTAACTCTGATAACCACATAATAATCA
TAGAGCCTGCTGTTAAAGCTAAAACAGACTCGCAGACAAATGCAAAATTCCAATTAAAAA
CATATGGTTTTACCCATATAGAGATTGCTCCAGATTGTAAAGTCGCCCAACCTAGAGCTA
AGTACCTTGTAATTTGAGTTATTTTTTGGCGGCCCAATTCACCTTCTTCCTTCTGTAATT
TTTCCAGATTGGGGACAATTTTCGTAAGTAGCTGCATTACAATTGAAGAGTTAATATAAG
GAACAATACCTAACGCAAAAATCCCTATTGTTGAAAAACCTCCTCCAGAAAAAATATTCA
GAAAATTTACTAAAGTATTTTTTTCTACACTTGCATAAAAGGCGTCATGATCTATACCTG
GAACAGGTATAAATATTCCTAAACGTGCTAAAACTAATAGAAAGAGAGTAAAGATAATAC
GATTTCTTAGATCACTTTTTTTGGCTCATAAATAAATTTTAAATGAAAAGAGATACTTTTT
TATGTAAAAACCCTATTCCAAGACTTGTATAAGTTTAAAATAGGGAAGTAAAATATTATA
GTAATATTAATATAATATTCTATGTAGAATACAGGCTTTCTATTGGAACTCCTCGATCTT
TAGCTGCTTCACTAAAAGTTCTTAATTGAGTTAAGCCATTGAGAACAGCTCGCGCGTTAT
TTAATGTATTATTAGAACCTAATTGCTTAGCTAATATATTCTGTACACCCGACAATTCTA
GGACTGTCCTTACAGAACCTCCGGCAATTACACCTGAGCCAGGTGCAGAGGGCCTTAATA
TAACTTTCGCGGCACCAGAAATTCCGTTAATAGGATGAGGTATAGAATTCGATTTTGTCA
GTGGAACTGTAACTAGATGTTTTTTAGCATCTGTTACTCCTTTTTTTACTGCACCAATTA
CATCGCTTGCTTTTCCCACGCCGACACCAACTTGGCCTTGCTCATTACCAATAACAAGAA
TAACTCGAAAGCTTAGTTTTTTACCTCCTTTTACAACTTTAGTAACTCTTTTAACTTGTA
CGACTCTTTCTTCCCAGCCACTATCTTTATCTTTTCCTTTGCTCTGTTTTTTACGATTGG
CCATTTTAAGAATTATTCCTAATTAGTTAATTACGATTCTAGAAACCCATGCCTGCTTCC
TTAGCAGCTTCAGCCAAGGCCTTAACTCTTCCATGATATAACTTTCCTCCTCTATCGAAG
ACGACATTTTTAATGCCTTCTTTCATAGACTGTTCTGCTAACTGTTTACCAACTACGCGA
GAAGTATCACAGTTTGGTCTTATGTTATCAGATTCTTTATTATTTAAATTAACAGATGAT
GTAGCTACTAATGTAATACCTTGTGTGTCATCAATTATTTGTGCGTATATATGTTTATTA
GATCTAAATACACACAGACGAGGCCTACTTGAGGTTCCTTGAACTTTTTTCCGAACTCTT
TTATGCTTATGAATTCTAGTTTGTTTAGTGTTTAGTTTCATTATTATTTACCTTTTCCAG
CTTTCCCAGCTTTTCTTCTAACAAATTCACCTTGATATCTAATTCCTTTTCCTTTATAAG
GCTCAGGAGGCCTAATAGAACGAATAGTTGAAGCAACCTGACCGACAACTTCTTTATCTA
TGCCAGAGACAGTAATATTTGTGTTGTTTTCAACTTTAATTTCAATATTGGCAGGAGGTT
TAATTTTCACTACATGACTATAGCCAACACTTAAGATTAAATCTTGGTTATCAATTTGAG

AACGATAGCCTACACCTTGTATTTGCAGTTTTTTAAAAAATCCATTAGAAACACCTTCAA
TCATATTACTAATAAGAGTTCTGGATAGCCCATGTAACTGACTAGCCATTTTTGTTTGTC
CACTCGTCTTGACAGCAATTGTGTCGTTAAGTATTTCTAAATTAATACCAGCTGGTAAAG
TTCTGGATAGGGTACCTTTAGGTCCTGTCACGGTGATAGTCTGACCATCGAACTGAGTAC
TAAGATTTGTCGGCAATAAAATTATTTTTTTTCCAATACGAGACATATTCCACCTATAAA
ATAGTTACCAAATATAGCATAATATTTCACCACCAAGACCATCATGACGAGCTTGTCTAT
CTGTCATAACGCCTCTAGAAGTAGAAATAAGAGCGATGCCTAAACCTCCAAGAACTCTGG
GCAACTCTTTATGATTTGCGTAGACTCTTAGTCCGGGTTTACTAATTCTTTTCAGAGCAG
TAATAACTGGTTGACGATTTTTACCATTATACTTAAGAGAAATCATTAAATGAGTCTCTA
TACCTTCACCCATTTGTTCGAAATTTTGAACAAATCCTTCTTCTTTTAGTACTGTTGCCA
TATTGCATGTCATTTTCGTTGCTGGAACTTGCACAATTTGATGTCTTGCTAAGTTTGCGT
TACGAATACGTGTCAGCATATCGGCGATCGTATCGTTGACCACCTTGATCCTCCTTGCGA
TGAATTATTTGAAATATTTTCAAGATTCCCTGAAAGGCATCCCTAGCTTTTTTAACAAAG
CTAGACCTTCTTGATCTGTTTTAGCTGTAGTGACAATTGATATATCTAGACCACGAATTT
GATCAATATTATCATAGTCTATTTCTGGAAAGATCAACTGTTCACGTAAACCTAAATTGT
AATTACCTTTGCCATCAAAACTTCTAGGACTAATTCCTCTAAAGTCTCTAATTCTTGGTA
ATGTTAAATTAATTAATTTCTCTAAAAAAGAATACATCTTGTCTTTTCTTAGATGCACAA
CAATTCCGATAGGAACCTCTTCTCGAATTTTAAATCCTGCAATAGATTTTTTAGCTTTTG
TGACAATTGGTTTTTGTCCAGTTATTAATGTTAATTCTTGAATACTACTTTCAAGAGCCT
TAGCATTCTGAGAAGCTTCACCTAAGCCACGGTTAATAGTAATTTTAGTAAACCTAGGAA
CTTCATGTACATTTTTGTACTGAAATTCATCTTTTAAAGATTGAGTAACAGTTGTTTTAT
ATTTTTCTTTTAATCCTATTGCCATTATATTTTCATCAATTATTTAATAAGTTCGCCAGT
CTTTTTCAGTTTTCGAATTTTTTGGCCTTTATCATTAATTATTACTGAGGATCGACTAGC
AATATTATTTTGTTCACTAAATAACATAACGTTAGAGGTATGTATTGGAGCTTCAAATTT
GATAATTTCTCCCGTTTCTCCTTCTTGCTGAGGTTTTTTATGCTTCACTTTGAGATTAAT
TCCTTTAACAATCACCTTATTTGTTTTATAAATGATTGCAATAATTTCACCTGTTTTTGT
TTTATCACTTCCAGAAATAACTTGAACTAAGTCCCCTTTTTTTAATTTAATTTTTGTATT
ATTTTTGGTTGTTTTAGAAGGACCTTTCATTATACTACCTCCGGTGCAAGAGAAACTATT
TTAGAAAAATTTTTATCTCTTAACTCTCGGGCTATAGGTCCGAAGACTCTTGTGCCACGT
GGATTATTATCTTGATTAATGATAACTGCCGCATTATCTCCAAATCTAATGCTCATACCG
TCAGTTCTCCGTAAAGCTTTACGAGTTCTTACTACAACAGCTCTGACAACATCAGATCGT
TTAACTGGCATATTGGGAGACGCATCTTTGACTACTCCAATAATAACATCTCCAATAGAT
GCATAAGAAGGATTACTAGTGCCTAATACTCTAATACACATTATTTTTCTAGCACCACTA
TTATCTGCAACATTAAGATAGCTTTGAGTCTGTATCATACTTTTATTCTGTATAAGATTA
ATTATCAAAAGACTTAGATAAGATATTAACCATTGTCCAACACTTTGTGCGACTTAAAGG
TCGTGTCTCTTGTATTGTAACAATATCGCCGATTGTGCATTCATTATTTTCATCATGCGC

TTTATATTTCTTTGTCCGGATCATAGTTTTTGCGTACTTTCTATGAGAGATTCTATTTTC
TACAGCTACTACTATAGTTTTATTCATTTTATCGCTTACTACTTTACCTGTTGTTTCTTT
TAAAGGCATAGTCTTTATATTTTTTATATATAATTAAATTTTAAGATTATTACATGGTTG
ACTTAGTACGAGACTTTTCAACAGTTAAAAGTTGGGCTAATCTATGCTTAGAATGTTTAA
ATAGATGAGGCTGGAAATCTTGCCTTGTGGCTCTTTTTAGCCTTAAATCAAAAAGCTCTC
TTTTTATTACAAGGATTTCTTCAGCTAAAGAAGAAGAGTCCAGGTTTGTAACATCTGATA
TTTTAGGGAAAGTCATATTTTAGGATTCTGTTGTATTCCGAACTATAAATTTAGTCTTGA
TTGGTAATTTATAAGAAGCTAATTTCATAGCTTGTTGAGCAGTTTTTTGTGGTACACCTG
TAATTTCAAATAGAATATGTCCAGGCTTAATAACTGCAACCCAATATTCTGGAGCTCCTT
TACCTGACCCCATGCGAGTTTCAGCTGGACGGGCAGTGACTGGTTTATCTGGAAATACTC
TAATCCATAGTTTACCACCTCTTCTCACATATCTAGTAATAGTTCTTCGAGTAGCTTCTA
TTTGTCTAGAAGTTAACCAAACTGGCTCTGTCGCTTGTAATGCATAATCACCGAACGCAA
TTGTGTTACCTTTGCTAGCAGAACCTTTCATTCTACCTCTATGTTGTTTTCTAAATTTTG
TTTTCTTGGGGCTTAGCATAAAAATTAAAAAATAATAATAATTAAGAAGCTGTTGAATCA
GGAGTTTCCTGATTATAAGCAGATTCTAGACTTTTCGATTCTGGCAAAATTTCTCCTTTA
AAAAGCCAAACTTTTACTCCGAGCACACCATACGTAGTGTGAGCTTGGCGATGACAATAA
TCAATATCGGCTCTTAAAGTTTGTAAAGGTACACGACCTTCTCTAACCCATTCACTTCTA
GCTATCTCTGCACCATTAAGCCGACCAGATACTTGAATTTTAACACCCTGTGTATTTGCT
CTTTGAGCTCTTTGAACTGCTTGTCTGACAGCTCTACGAAAGGCCACTCTTTTTTCTAGC
TGTTGAGTAATAAATTCTGCTACTAAGGTTGCTTCAGAATCTGGATCTGCAATCTCTACA
ACATTTACTCTGAGTTGTTTGCTAGGGTCTAGGATTAAGGACAGCGATTTTCTTAAAGAC
TCAATTCCCGCTCCAGATTTTCCAAGTACGATTCCTGGTCGAGCTGTTGCTATGAGAATT
TCTACTTGATCAACTTTACGATTAATTTCAATTTTAGCAATACTAGCATTACTAAGTTTT
GAATGTATAAATGAGCGAATTTTATGATCTTCTTGTAAAAGAACTGGGTAGTCTTTAGAG
TTAGCAAACCATGAAGAACGATGTTTTTGGGTAATGCCTATGCGAAAGCCCAAAGGATGA
ATTTTTTGACCCACAGTATCCTTGATTATAAGTTAAAAGCTATAATGACTAATTGACGTT
AAAAGACTACAGTTCTGAAACACCTAATGTAATATGACAAGTTGGTTTATGTATTGGAAA
GGCTCTTCCTTGTGCTCTAGGCTGAAATCTTTTTAAAGTCGGGCCTTTGTCTGCAAAGGC
TTTACTAACAAACAATTGGTTTTTATTTAATCCATCATTATGCTCAGCATTCGCTGCAGC
AGATTCCAGGATTTGCTTTATATGAGAACAGACTCGGTATGGCATAAATTCCAGAATAAT
TAATGCTTCTTGGTATTTTCTACCTCTAATTTGGTCTAATACACGACGGACTTTATGCGG
AGAAAGACGAATGTATTTCCCTACTGCTTTAGTTTCTTTTACGTTTTTTGTAATACTCAT
AGTTATAGTTTAACGACGGGCTTTTCTGTCACCTTTTACGTGAGTGCGGAAAGTTCTTGT
TGGAACAAACTCTCCTAATTTATGTCCTACCATTTGGTCTGACACAAAGACAGGAAAATG
TTGTTTACCATTATAAACAGCTATTGTATGCCCTACCATATCGGGAATAATAGTAGATGC
TCTCGACCAAGTTTTTAGGACTTCTTTTTTTCCTGAAATATTTAATGCTTCTATACGTTT

AAGAAGACTAACATCAATAAAAGGGCCTTTATGTATAGATCTTGACATAATAATAAATCA
TTAGATTCTTAATAAAATAAATTCTATTTACGGCGACGTAAGACGTACGGGTTACTATAT
TTATTTGGATTTCGTGTTTTAACACCCAATGCAGGTTTACCCCAAGGTGTTACAGGACGA
GCTCGTCCAATTGGAGATTTGCCTTCACCACCACCATGTGGATGGTCTACAGGATTCATG
ACAACACCTCTAACAGTAGGTCTTTTACCTAACCAACGATTTCTACCAGCTTTACCAAGA
GTAATATTACTAGCGTCAATATTACCAACTTGTCCAATAGTAGCGTAGCATTCTTTTCGA
ATCATTCGAACTTCGCTTGAAGGCAATTTTACAGTAACAAAAGTACCTTCTTTTGCTACT
ATTTGAGCATAAGTTCCTGCTGCACGAACAATCTGTCCTCCGCACGAAGGTCTTAATTCT
ATATTATGTACTGCTGTTCCTAAAGGAATACTAGATAAAGGCAAAGCATTTCCAACTTCG
ATAGGGGCAGTAGGACCAGAAAGAACCATAGATCCTACACTGAGAGATCGAGGGTGTAGA
ATATATCTTTTCTCACCATCAAGATAGTGTAATAATGCAATTCTTGCATTTCTATTCGGA
TCATATTCAATAGAAGCAACTTTAGCAACTATATTATGTCTATTTCTTTTAAAGTCAATT
AATCTGTACTGCTGCTTATGTCCACCACCTTTATGACGACAGGTAATGACACCTCTATTA
TTTCGACCTTTGCAGAAATGGTGTTTAACTATTAATGATTTTTCTGGCTTATCAGTAGTA
ATCTCTGAAAAGGTAGAAACCGTTCTATTTCTTGTCCCTGGTGTGTAGGCACGATATAAA
CGAATTGCCATATAAGAAGATTTGAGAAGAGTAAATGTTGAACTATGTGATATATTTAAG
TTTCTGGAAACAAATTAATAGAATCTTCTGAGGCAAGTGTTACAATTGCTTTCTTGTAAT
GTGGTCGCTTCCCTACGAATCTACCGATGCTCCTTTTTTTCTTAGGAGGATGACAAGTGT
TAACACCTGTAACTTGCACGTTGAAAATATACTGTATAGCAGCCTTAATATTAATTTTTG
TTGCTTTGGGATCAACGGCAAAACAATACTGATTTTCTTCTAATAATTTAGTTGTCTTAT
CAGTAATGATGGGATATTTAACCAGATCTAATAAACCTCTTGAATCAATGCTATCCATTA
TATACCTCTTGTATTTTAGATAAAGCATCAACTGTAATAATAATTTTATGTGCAGCTAAC
AAAGCCATAATATTTAGTGTATCAGCTGAAATAAGTTCTACGTTGTGTAGATTGCGAATA
GAAAGATAAACATTCGGATCTTTTTTGTCTACGATGACTAAAACTTTTTTATTTAAATCA
AGATTCCATCGATGTATAGCTTCCATAAATAGTTTAGTCTTGGGCTGATGAAAATAACTG
TTAAAATTTTCTACAATCAGTGTATTAACTGATTTGTTATTTAATGCTGTCCTTAATGCT
AATTGTCTTTCTTTTTTATTCATTTTTTTGGTAAAACTACGAGGTTTTGGCCCAAATATT
ACACCACCACCTCTCCATAGAGGTGAACGGATAGAACCTGCTCTTGCTCGACCAGTCCCT
TTTTGACGCCATGGTTTACGACCTCCACCTCGAACTTCACTTCTTGTTTTAGTATTAGCA
GAACCCTGGCGCTTTTCATTACTTTGCTTTACTAAAGCTCTATGGACTAAGTACATCCCA
GAATCTTGACTGACCTTAAGATTTAAATCAGCGTTGCCACTTACTTGACCTTCCCAATTA
TAGACTTGATAATTTAATTGCGTGTTAACTGTCATGAAATATATCGTATGAAAGAATATT
GCTACTTACTAATTTTTACTAAAGCGCCAGGTTTACCAGGAACAGCTCCCTTAACGATCA
ACAAATCATTCTCTGAGTTAATACTTACAATTTGCAAATTTTTTATTGTAACTTTTTTAT
TACCAAGTTGGCCAGCCATGTTTTTACCTGGATAAACTCTTCCAGGCGTTGTACCGGCTC
CAATAGAGCCAGGTTGCCTATGATTTTTCGAACCATGTGACATTGGGCCTCTACTAAAAT

GATGCCTCTTTTGGTAACCGGAGAAACCTTTACCTACACTTCTAGAAGAGACATTAATCT
TTTGGCCTACTTGAAATAGATCTGTAGATAGTATTTGGCTAACTTCGAAATCATCAGTAG
ATTTCATTTCGTATTCACGTAAATATTTTAGTGGGGGAGCCTGTGATTTTTTTAAGTGTC
CCAACAATGGTTTATTTAATTTTTGTTCAGCAACTTGCTTGTAACCGACTTGAATAGCAT
TGTAGCCATCAGTAGATACAGCTTTAATTTGAGTAATAACACATGGCCCAACTTGAATTA
CAGTAACTGGAATTGATAAACCAGCTTCATCAAAGAATTGGGTCATACCTACTTTAGTAC
CGAGTATACCAACAGACACTAGATTTCTCCTTTTTGAGAGATTTATACTATCATTGTACA
TAAAAATTCTACTTCATAGTTACTGAAGTAAATTAAACTAACAGATGCATGATATTATTC
ATGACAACATAAAGGTTAAAAACTTTACCTATTTAATCTAAAATAGAAACAGCTTTTTGT
CTAGTTACAGCGTATATAAATTAGAAGTATGCTTATTAAATTTTTTATAAACTACTTAGA
TTTAACTGATATTGATATCCGTAACTTACTTGCAGTAATATCAATCAACATGTATAAGTT
TAATTGCCTAAACTAGAACCAGCTATTATTAGAAATAATTCAAGTATAGATAAAAGCGTA
TTTGAACTTTTATATATTGACAAATATAATAGAATATATTGAGGTTCGGTAACTACACCT
TTTTATTTTTACATTTATTGTGCAGTATATATACATCTAATGCAGTAAACATAAGAATAT
AACTGTATAGCAAATGTAATTACACAAATTAGAGAGGTATTAATGGGCAAAGTCGTTGGA
ATTGATCTTGGAACAACGAATTCTGTAATTGCTGTTATGGAAGGAGGTAAACCTACCGTC
ATACCGAATGCAGAAGGTTTTAGAACTACAGCTTCTGTTGTTGCATATACTAAAAGTGGA
GATAAACTTGTAGGACAAATTGCCAGGAGGCAAGCTGTTATTAACCCAGAAAACACTTTC
TACTCTGTCAAAAGATTCATAGGACGTAAACAGAATGAAATTTCGCAAGAGATTCGGCAA
ACATCATATAATGTTAAAACTAGTGGATCAAGCATAAAAATTGCTTGCCCTGCACTGAAT
AAAGATTTTGCTCCAGAAGAAATTTCAGCTCAAGTACTGAGAAAACTTGTTGAAGATGCT
AGTACGTACTTAGGTGAGACTGTTACACAAGCAGTAATAACTGTACCGGCTTACTTTAAC
GATTCACAAAGACAAGCTACTAAAGACGCAGGTAAAATAGCAGGCTTAGATGTATTGAGA
ATTATTAATGAGCCTACCGCAGCTTCTTTGTCTTATGGACTAGACAAACAAAATAATGAA
ACAATACTAGTATTTGACCTTGGTGGGGGCACATTTGATGTATCTGTATTAGAAGTTGGA
GATGGAGTATTTGAAGTACTCTCAACTTCTGGAGATACACATTTAGGCGGAGATGACTTT
GATCAGCAGATTGTAGAATGGCTAATCAAAGATTTTAAACAGAATGAAGGTATTGATCTT
GGTAAAGATAGACAAGCACTTCAGAGATTGACCGAGGCTGCAGAAAAAGCAAAGATAGAA
CTGTCAAATTTAACTCAAACAGAGATCAATCTTCCTTTTATTACTGCCACACAAGATGGT
CCAAAACATTTAGAAAAAACTGTAACTAGAGGGAAGTTTGAAGAACTTTGTTCAAATTTA
ATAGATAAATGTAGTATCCCTGTAAATAATGCTCTGAAAGATGCAAAACTAGAAGCTTCC
AGTATTGATGAAGTTGTTTTAGTTGGTGGATCTACAAGGATTCCAGCCATACAGCAAATG
GTTAAAAGATTAATTGGTAAAGATCCAAACCAAAGTGTCAATCCAGATGAAGTTGTTGCT
ATTGGTGCAGCCGTACAAGCTGGAGTTTTAGCAGGCGAAGTCAAAGATATTCTATTACTA
GATGTGACGCCGTTATCTTTAGGAGTGGAAACTTTGGGTGGCGTGATGACAAAGATTATA
CCAAGAAATACTACTATTCCTACAAAAAAATCAGAAGTATTTTCTACAGCTGTAGATAAT

CAACCAAATGTGGAAATTCAAGTACTTCAAGGCGAAAGAGAACTAACAAAAGACAATAAA
AGCTTAGGGACGTTCCGATTAGATGGCATTATGCCTGCACCTAGAGGAGTTCCTCAAATT
GAAGTTACCTTTGATATTGACGCTAACGGAATTTTATCTGTAAAAGCCAAAGAAAAGGCT
ACTGGTAAAGAGCAATCAATTACTATATCTGGAGCATCAACTTTACCTAAAGATGATGTT
GAAAGAATGGTAAAAGAAGCTGAAGAAAATTTCGATACAGACCAAAAAAGAAGAAAGAAT
ATTGACACAAAGAATCAAGCAGAGTCCCTATGTTACCAAGCTGAAAAGCAAGTTAAAGAG
TTTGAAGACAAAATTAGCCAAGATTTAAAAATAAAAATAGAGGAGCTAATTACAGAGCTT
AGATCTAGTCTAGAGAAAGAAGAATATGACAATATTGAATCCATTTCTCAGCAATTACAA
AATGCTCTGATGGACATTGGAAAAAATGCTGCTCAGACTGAAAGTAAAGATACAAAAGCG
AAGGATGACGACACTGTGATTGACACTGATTTCTCTGAAGCTAAGTAAAAGTAAGCGGGT
AACGCGATTCGAACGCGCGACATCAACCTTGGCAAGGTTGCGCTCTACCACTGAGCTATA
CCCGCATAGATTGTATTATTACAAAAATTAAGATATTTGTCAATCTGTTAATCTATACAT
CTAGACTATTAATTTTGGAATTACATATTAGCTAACTGTTGTAAATATAACAATTGCTAT
AATTTTTTAAAAGTACTTATATCATTATATAAGTACTTTTTATATTCAAAAATACAAATA
AGTATTATAGCTAAACTACTAAGGAGTTAACTAAGCTAGTAACAATTACTAATCCTGTCC
ATAAGCCGGCTCCTGTATATATTAATCCTTTTGATTGCTCCCATTGGCCAGGGGATGCTA
AAACAACAGGTACGCCTACTACTAACACAGTCGATAATGCAATTAATAATAGCACTAATA
ATTGGATCGCAATAATCATTTAATATGTTCTCCTGAAAAATTCAAATGTTAATAATACTG
TATATAATATAATGAAGCTTTTGACCTTCAGCAATTTATCTATGAAGAAATAATTTAAAA
TTATTGTTTTCTAAATAAACCTTAAATTAAGTATTAAAATCTTTATTGTAAACAATAATA
CTATATTTTTTTAGTAGATAAAAAATTTAAAGAAAGTCTTTTTTAAAAAAAAATTAAGAT
AAGGTGATATGCTGTATAAAAAGCTAAGTATTATTAATTATTAGAGAAATTTAAAATAAG
AGGTAAAACATATGAACTCAGCCCTTTTTTTAGCAAAACTACCAGAAGCTTATGCCATTT
TTAAACCGATCATTGATATCTTACCCGTAATTCCTGTATTTTTTCTTCTTTTAGCTTTTG
TTTGGCAAGCAGCAATTGGTTTTAGATAACATTTAGGTTATAAAAAGAATAACAAATATA
CAGAGTGCCAATTTTTGAGCTCTAATCTTATATCTTACACAATAATAACTATTATGGTTG
AACCCTTACTATCAGGAATTGTTCTTGGACTTATTCCTATTACTTTATTCGGATTATTAG
TAGCAGCTTATTTACAATATCAACGGGGCAACCAATTAGGACTGTAAAGCCTAAATTATA
GATATTTATAAAAACTACAATAATTATAGTTTTTATAAATATCTACAAATAAGCTAATTT
TAAATACAGAAATAATACTATTGAAACTAGAGTGATAACAAAAAAAATTTATGTAATAAC
AAGCTAATTGTTTACCAGCTCGACTTAGTAACTCCTGGTAAAAGACATTCATGAGACATT
TCTCGAAAAACATGGCGAGATAATCCAAAATCTCGATAATAACCTCTACTTCTACCTGTT
AACCAACATCTATTTCTCCCTCTAACAGCGGCGCTGTTTCTAGGCATTTCTTGGAGTTTT
TGTCTTAAATCCATTTTGTCAGAAAAGCTGCCAGCTTCTTTCAGTTGATCTTTAATAGCC
AGTCTTTTTAAATAATATTTTTTTGCAAGTTTCGATCTCTTAATCTCTCTTTGGATCATA
TTCTTTTTAGCCATGCTGTAAAACTCTCTTTTAAAATTATTTATTCAATATATTAGCTAA

AAAAGCTCCTGCCTGCAACTTTTATATATTTAAAAATAAATAGCATAAAGTAGTGAAATG
CAATTGATATTTAATCAACGATACCACTTTATGCTATTATCAAAAATTTATTTAGTAAAG
TTCTTGTTCAACATGAGTTGAAATAGTGCAATCAGATGAAGGATATGCAATACAGGTTAA
AACATAGCCAGCTAAAAGTTGATCATCATCTAAGAATGATTGATCAGCTTGATCTACAGT
TCCTTCTGTAACTTTACCTGCACAAGTTGAACAAGCTCCAGCTCTACAAGAGTAAGGAAG
TTCAATTCCTTCTTCTTCTGCTGCATCTAGAATATATGTATCCTCTGCACAATTAAATGT
GACATCAATTCCTTCTTCTTCACATAGTAAATGAATTTTATAATCAGCCATTTCTTCTTA
CTCCATAATAAAGATACACTTGAAACTTAATAAAGCAAAATAATTTAAAGAACTTACCAA
AAAATTATATACTAGTTTCATTATGCATGATAGACTGGAAAAAAAGAAAATAGTATACTA
TTCTGAAAAATCAAGCTACATTTCCTACTAAATAGAACTATAGTAGGTCCAAAGTATATA
CATAAATTCATATAGTTAAAATTACTTTTATCATTTAAATAGCTGAACTTGATAGTATAG
ATTGATCATCAAGCTAGTCTTCTGCCAATAAATAATTAATTATGATTTCAATGCTTAATC
AACAAGTAGAACTTAAGCCTATTCTTAAATTTCAGATAACACAAATATATTCGCATTCTC
TTATTCAGGAAATTAAAGCCTTAATAACTAGACTTGTTTTGCAAGTTTGGAGAAGGCCAG
CTACATTGATGGCAGGTATTATCCAACCTTTATTATGGTTAATTCTATTTGGTGGGCTTT
TCTACAATGCTCCTATAAATTTGTTCACTATTAATACAAGCTATAATTGTTTTTTGAGCT
CTGGGATTATAATTTTCACCTCTTTTACTGGAGCTTTGAATTCAGGTCTTCCATTAATGT
TTGATAGAGAATTTGGATTTTTAAATAGATTATTAACGGCTCCCTTAGTATCGAGGACTT
CTATCATTTTATCTTCTGCTACTTTTATGACTTGTATTAGTTTAATACAAGTTGTATTTA
TAGTTACAGCTTCTCTTTTTATGGGAAACCCACCTCTAAATAGCGATAGTACTATGATTT
TTGGACTTATGATTCTATTAGTGACTGTAGGAGTTACAATGCTTAGTTTAGCTTTATCTT
TCACTCTGCCAGGTCATATTGAGCTATTAGCATTTATTTTAGTAGTTAACTTGCCCTTTT
TATTTTCTAGTACGGCTTTAGCTCCTTTATATTTCATGCCGCCATGGCTTCAGTTAATTG
CAAGTCTCAATCCATTGAGTTATGCAATAGAAGGTACAAGATACTTATATTCAAGCGTAA
ACTGGAATTTTACAGAGTGTGTGATTAAGATTAGCTGGGGAGATATTTGTTTAGGGCAAA
TTATTATTTTATTAATCGCTTTAGATATAATGGCAGCTTATCTTGTGTCTAATATATTAA
AAGCTAAACTTAATTAAAATTTTAATAAAAAATTTATTAAATTTATTTTACAAAGATTTT
TACTATATAATACTAATAGTAGTATGGATAATTGAAATAGAAATAGTTTTTCAAAACCAA
AGCTATTATCAACAATGATAAACATTTGTAAGAAAGTCAACAAAGTATGTTCTTATTCAT
AGGAGGCATGTAGTCAATGGGACTACCATGGTACCGTGTACACACGGTTGTTTTAAATGA
TCCTGGACGGTTAATTGCAGTCCACCTAATGCATACTGCACTTGTAGCGGGTTGGGCAGG
ATCTATGGCATTATACGAACTAGCTGTATTTGATCCTTCAGATCCAGTATTAAATCCAAT
GTGGCGACAAGGCATGTTTGTTATGCCATTTATGGCTAGACTGGGCGTAACAGATTCATG
GGGAGGATGGAGTATAACAGGAGAGAGCGTATCCAATCCTGGACTGTGGAGTTTTGAAGG
TGTAGCCTTAACTCATATAGTCCTTTCTGGCATGCTTTTTCTAGCTGCTATTTGGCATTG
GGTTTATTGGGATTTGGAATTATTTAGAGATCCACGAACTGGCGAGCCAGCCTTAGATTT

ACCGAAAATTTTTGGAATTCATTTATTGCTATCAAGTCTACTTTGTTTCGGATTTGGAGC
TTTCCATGTAACTGGACTTTTTGGACCAGGAATGTGGGTATCAGATGGGTACGGAGTAAC
CGGAAAGGTATTACCAGTAGCTCCAGCATGGGGACCAGAAGGATTCAACCCGTTTAATCC
TGGAGGAGTTGCATCTCACCACATTGCCGCAGGTACTGTAGGTATATTAGCTGGTGTTTT
CCATTTAACTGTTAGACCACCACAAAGACTGTATCGTGCTCTAAGAATGGGTAATATTGA
AACTGTATTATCAAGTAGTATCTCTGCTGTTTTTTTCTCAGCTTTTGTGACTTGTGGTAC
GATGTGGTACGGCTCTGCAACTACGCCTATTGAATTATTTGGTCCAACTAGATATCAGTG
GGATAGTGGATATTTTCAGCAAGAAATTGAGAAACGAGTAGAAAATGCTATTGCTGATGG
TGCTGCACCTAGCGAAGCATGGTCAAGAATTCCTGACAAGTTGGCATTCTATGACTATAT
TGGTAATAATCCAGCAAAAGGAGGATTATTCAGAGCGGGCCCTATGAATAAAGGCGACGG
AGTTGCTGAAGCATGGCTTGGACATCCTGTATTCCAAGATAAAGAAGGAAGAGAGCTTAG
TGTTCGCAGAATGCCTGCTTTTTTCGAAACTTTTCCTGTAATTTTAGTTGATAAAGATGG
TATTATACGAGCTGACATTCCATTTAGAAGAGCAGAGTCTAAGTATAGTATTGAACAAGT
AGGTGTAACAGCTAGTTTTTATGGTGGCAAATTAAATGGCCAAGTTTTCAACGATGCTCC
TAGCGTTAAAAAATATGCAAGGAAAGCTCAATTAGGTGAGGTATTTGAATTTGATCGAAC
TACATTAGAATCAGATGGAGTGTTTAGAAGTAGTCCTAGAGGCTGGTTTACATTCGGTCA
TGCTAATTTCGCTTTAATTTTCTTCTTTGGACATCTTTGGCATGGTTCAAGAACTATCTT
CCGGGATGTGTTTGCGGGAATCGGAGCTGAAGTTACTGAACAAGTTGAATTTGGAGCTTT
CCAAAAATTAGGAGATAGAAGTAGTAAAAAACAAGGAGCTGTATAAGACCAAAGTCAAAG
ATAGTCTTTATCAAATATTTCTAATATTAAAATAGTTAAGACAAAGTTTATTATCTAATA
GTTACTAATCAGGAGAATTTATGGAAGCCTTAGTCTATGTATTTTTACTAACAGGAACGC
TAATGGTTATATTCTTTGCGATCTTTTTTAGAGAACCTCCAAGAATAGCAAAGTAACTTG
ATTCCGTTTAAACTTATAATAATAGAACCACTTTCCAACTTATTAGTGGAAAGTGGTTTT
ATTGTAGATTACATTGTATTATTCTTCATGCTCTTCAAAAGGATCTCGGAGATCTTTTGA
AGCAGGACCAAATGCTGTGTATATCGAGTAACCTGTAATACCTAAAAGCAGACTCGAAAT
AAAAATACTAAGAACTGTTGCAGTTTCCATAATTTAAGCAGTATTAAGATGAACTTTTTC
TATAATGATATTATCAATATTAAAACATTTAATTATTAAAATTATCAAAATATACTATGG
CACTTAGAACTAGACTTGGAGAAATTTTAAGACCTTTAAATTCAGAGTATGGAAAAGTTG
CTCCAGGTTGGGGCACAACTCCTATTATGGGAATTTTCATGTTACTATTTTTCTTGTTTT
TATTAATCATTTTACAAATCTATAATTCTTCATTAGTATTAGAAAATGTAGATGTAGATT
GGGCTACTTTAGGTAGCTAACTAGAGAAGTTCTTTGATTACACAATATTAAAGACAACAA
GCGATCGCTATTAGATTATAGTAGCGATCGCTTTAAAAGTATTTTACTATATATCCAGAG
AAAATTATTTACCTAAATTAATTAACTCGGTATCTGCAAAACTATTTGTGTTAACATTAT
TATAGTTAACTTTTTCAAACCTTACTAATACTGGATATTTGATACCACTTTTATCAATTG
CTGCAACAGTACCAATTTCCTGATACCAGTACGATTCTTTTCTCAATATTTTTACTTTTG
AGCCTCTTTCCATATTTTTAGTCCTCAGATAGATTATTAAAGATTATTATATTAAAAAAT

T-TAAAAAACTAAACAAAATATATAAAAATACTACCGACAATTGTAATATATAACTAATA
AAAGGAGAGAGAGGGATTCGAACCCTCGATAACAAAAGTTATGACAATTTTCGAAATTGT
TGCAATAAACCACTCTGCCATCTCTCCTGAATTAAAATACTGTTTTAAAAAATTAGACAA
GAATAACATAGAACTAAGGTAGTTGCCTAGAAAATTAAAACTATGTTAAATATAAATTAT
AATCTTTCTATAGATATATTTAAAAAATATGTACATAGAATCTATATCAGCTTATTTCAA
GATTTTTTAATCTTATACAATTTTTACTAAATAAATAAATTCTCTCATAATATGAAACTA
TCTTGGAAAACACTTCTTCTTTGGTCTTTACCAATATTTGTAATTGGTTTCTTTTTCTGG
CAAGGTTTTTTAGGTCCAACTACTACAGATGTTGGCAGTAATATCGCAAGTTCTAGAATG
ACATATGGACGATTTTTAGAATATTTAGATATGGGTTGGGTGAAACGGGTTGACCTCTAT
GAAAATAATCATACAGCAATTGTAGAAGCTGTTGGGCCAGAATTAGGAAATAGGGTTCAA
CGAATTCGAGTTGAACTGCCAGCAAGTGCGCCAGAATTAATTACAAAATTACGCAAAGCC
AATGTTGATCTAGATGCTCACCCCCCTAAAAGTACAAGTGCAGTATGGGGACTATTAGGC
AATTTACTATTTCCTTTACTATTAGTTGGCGGGTTAGCTTTCTTATTTAGAAGATCTAAT
AATGCTAGTGGTGGACCTGGTCAAGCAATGTCATTTGGCAAATCGAAAGCTTTATTTCAA
ATGGAGGCTAAAACGGGAGTAGTATTTAATGATGTAGCTGGAGTTGAAGAGGCAAAGGAA
GAATTTCAAGAAGTGGTAACATTTTTAAAACAGCCTGAATCATTTACTGCTGTTGGTGCA
AAAATACCAAAAGGCGTTTTATTAGTTGGACCTCCTGGCACAGGCAAAACATTACTAGCA
AAAGCTATTGCTGGCGAGGCTAGTGTTCCTTTTTTTAGTATCTCAGGCTCAGAATTTGTA
GAAATGTTTGTTGGTGTTGGCGCTTCTCGTGTGAGAGACCTATTCAAGAAAGCAAAAGAC
AATGCGCCTTGTATCGTTTTTATTGATGAAATTGATGCTGTTGGTAGACAACGAGGAACA
GGTGTTGGAGGTGGTAATGATGAAAGGGAACAAACATTAAATCAACTATTGACTGAAATG
GATGGCTTTGAAGGAAATACTGGTGTTATTGTAATTGCCGCTACTAACAGAGCTGATATT
TTAGATTCTGCATTATTAAGACCTGGAAGATTTGATAGACAAGTTTCTGTAGATGTACCA
GATTTTAAAGGCAGGTTAGCAATTCTTGAAGTTCATGCTAAAAATAAGAAAATGGAACCT
AAAGTATCTTTAGAAACCATTGCCAGAAGAACTCCCGGCTTTTCAGGAGCTGATTTAGCT
AACTTACTAAATGAGGCTGCTATCTTAACGGCTCGACGAAGAAAAAAATGCAATGACTATG
TCTGAAATTGATACATCAATTGATCGAGTAGTAGCCGGGATGGAAGGCACTCCTTTAATT
GACAGTAAAAGCAAAAGATTAATTGCGTATCACGAAGTGGGTCACGCAATAATAGGCAGT
TTATTAGAGCATCATGATCCTGTGCAAAAAGTTACATTAATACCAAGAGGGCAAGCAAGA
GGCTTAACTTGGTTTACTCCTAGTGATGATCAAAGTCTAATATCAAGATCTCAAATACTA
GCCCGTATCGTAGGTGCTCTTGGTGGCAGAGCTGCAGAAGAAATCATTTTCGGTGACGCA
GAAGTTACTACTGGTGCAAGTAATGATTTACAGCAAGTAACATCAATGGCAAGACAAATG
GTCACTCGATTTGGAATGTCTAAAATTGGACCTTTATCTCTTGAAAGCCAAGGAGGAGAC
CCATTTTTAGGTAGAGGCATGGGAGGAGGCTCAGAATATTCAGATGAAGTTGCAACTAAT
ATTGATAAGCAAGTAAGGGAAATTGTCAGTGAATGCTATGCACAAGCTAAACACATTATT
ATAGATAATCGAGTAGTGATAGATAGATTAGTTGATTTACTAATTGAAAAAGAAACAATT

GAAGGCAATGAATTTAGAGACATCGTTAAGGAATACACTGCAATTCCTGAAAAAAATTAC
TACATATCACAATTTTAAATTAAACGGGACTGACGGGATTCGAACCCGCAACTTCCGCCG
TGACAGGGCGGTGCTCTAACCAGTTGAACTACAGTCCCAAAAAGATGAATACCCAGAGAT
AATCTCATAGTTAGCTTAAACTTGTCAAATTAGAATATTAAATATGAAGAGTATTTGTAA
TAATACTCTTCATATTTAATATAGTAGCTAAAATATTTCTTTTAGAAGGAGAGAGAGGGA
TTCGAACCCTCGGTACGAAGTTAATCGTACAGCAGATTAGCAATCTGCCGCTTTCGACCA
CTCAGCCACCTCTCCATGATATATATTATATATATATCAAATATACTTATATTTTAATAG
GTGTGCATAAATATTTATGGAGCTAAGCGGATTCGAACCGCTGACCCTCTCAATGCCATT
GAGATGCTCTACCAACTGAGCTATAACCCCTTTGGTAAGATATTAAGAAATGGCTCAAGC
GGGATTTGAACCTGCGACCTTGGGCTTATGAATCCCCTGCTCTAACCACTGAGCTACTGA
GCCATAGTATAAAATATTATTAAGTATTATACCATAAATTATGGTCAAATATAAATCAAT
AAAAGAATAATTACTATCATTTACTACTTTTGTGCAAGTTGTAAATATTCGAATAAAATT
TCAAAATTAAAAATGTCTTTAACTTTTTGTTAGAATTATTGATTACGAAAGTATTGATAT
TTAGAATTAACAATAAATTTTGTCTTTATTTGTAGACACAAAAATATTGGAAATTTTCAT
CAATTAATTAGTTAAACTATAACCGACATACAAGAATTATTGAACTACGTTAAATTGTGA
TGCATTGGGCTTGATCAAAACTTCTCTAAAATCAATAGAATATAGTAATTCAAGTAGATC
AAATACTAACAAATGTGACAAACGCTTAAAAAAATAATACAATTAGAATCGTAAAAATTT
GGTTTTTGACTGTGTTAAATAAATTTGATTATCATAGATTTAATATTTAAAAGTGTTCTA
TTACTTGTATTGAGTAATTGTAATATTTAATATCTTCTTTTTTAGATAACAATCATAATA
TAAGCTAAAAAATAGTTTTTATTAAACAACTAGCATAGAGCCAATACCTTGATCAGTTAA
TATTTCTAGCAAGAGAGCATGATCTATTCTACCATCTAAAATATGCGCTGAAGCTACACC
CTGAGCCAAAGAACGAATACAACAATTCACTTTAGGAATCATGCCACCAGAAATTACTGC
TGTTTGAGTTAAATCTCTAGCTTCCTGTATATTCAGATGACTAATTAATGTTGAAGGATC
TGAAGAATTACGTAAAATGCCGGGCGTATCTGTGAGTAGAATTAGTTTTTCGGCATTAAG
AGCGGCTGCTATTTCTCCTGCGACAGTATCAGCATTAATATTATACGATTGGCCTTCCTT
GTCTGCTGCAACACTAGCTATAACAGGTATATAATTATTATTAATTAATATCTTCAGTAA
TTTAGTATCAATATTTTGTACTTCTCCAACAAAACCAAGGTTTGCTTTTCCATTTGGTCT
CGGAGTAATAAGTAATCCATCTTTTCCCGACAAACCTACACTTTTACCACCTTGTTTATT
AATAGTTGCGACAAGATCTTTATTGACTCTTCCAACTAAAACCATTTCCACTATATCCAT
GGTAGGTTGATCTGTTACTCTAACGCCATCATCGAATTTTGGTAATATTTTGAGGCGATC
TAACCAAAAATTAATTTCTGGTCCTCCACCATGAACTAAAATGGGACGTAGTCCTATAAA
AGACAGAAAAACAAGATCACTAATCACTTGATCTTTTAGTTTCTGATTTTTCATAGCCGC
TCCCCCATATTTTATAACAATAATTCTGGAAGAAAATTGCTGAATATAAGGCAGAGCTTC
ACTTAAGACTTTTACTCTTTCTGAATTAGTCAACATAAATAAAGTAGGATTGTGATATAT
AAAATTATATACACAGCTTGTGTTAATTTAAAAGATACCAATGGGTACTTGAAAAAATAG
AAATATTACTAACAGTTAAAACTATAGGGATTATAATTATGAATAAATTTTGGGATAATG

TATTAAGATTTCCTCGATTTTTAGTCAGCGTCATTTTAGGATTAATTTTGATAATTATTA
GTCCTTTTTTCGTGTTATTAAAAAAACCACTGACAAGTTTTTTTTTCATCATATCATTGG
CCGGATTAATCACAGTTTTGGCAATAATAATACAAAAAATGATAAATATCGAATGTTGTT
GAGAGAAATAAATAAATTTAATTAAAATTTTTGAACATATATAATATTATATTATCATTA
GATATGATAATTTATTTCCTTTTCTATAAAACCCATAAATATTAAATATTATTTGATAAA
TATTATGCCACTAAAACAAAGAGTTAGCTCTGAAAAGACCGGTGCCTTCGCACTTTTGGA
TAGTATTGTTAGGCACGGAGTTAAACATATATTTGGTTATCCAGGTGGAGCTATTCTTCC
TATTTATGATGAGCTTTATGCTTGGGAAGAAGCCTCTCTAATTAAACATATCCTTGTTCG
TCATGAGCAAGGCGCTGCTCATGCTGCAGATTCTTATTCTAGATCAACAGGAGAGGTTGG
AGTATGCTTCGCTACTTCTGGCCCAGGAGCAACTAATCTTGTCTCAGGTATAGCTACAGC
ACATATTGATTCTGTACCTATATTAGCTATAACAGGTCAAGTTGGGAGAGCTTTTATTGG
TACAGATGCTTTCCAAGAAGTAGATATTTTTGGGATTACACTTCCTATTGTAAAACATTC
ATATGTAGTTCGTGACCCTAGAGACATGTCTAGAATTGTTGCGGAAGCATTTTTTATTTG
TAAACACGGTAGACCAGGTCCAGTATTAATTGATGTTCCTAAAGATGTAGGATTAGAGAA
GTTTAATTATTTTTCTGTTGAGCCCGGAAAAGTTAATATTCCTGGCTGTAGGCCAATTAC
CAGCCTAAAGTCAAGACAAATCCTTATGGCAGCTAAAATGATACAGCAATCTAGCCAGCC
ATTGTTGTATATTGGTGGAGGAGCCATAATCTCTGATTCACATCAAATTATTAAAGAACT
TGTTGATTTTTATAAAATACCTGTTACTACTACTTTGATGGGGAAGGGGATTTTTAATGA
GGATAGCGATTATTGTCTAGGGATGTTAGGTATGCATGGTACTGCGTATGCTAATTTTGC
AGTTAGTGAGTGCGATCTTTTAATTGCTTTAGGAGCTAGATTTGATGATAGAGTTACTGG
AAAATTAGATGAATTTGCTTGTAATGCACAAGTGATTCACGTAGACATTGATCCTGCTGA
AGTAGGAAAAAATAGGATTCCTCAAGTTGCTATTGTCGGTGACGTAGCAGAAGTTGTTAG
TGAAATATTGAATTTATTAAAGACTTCTTTCCCCCCTTATCCAGAGCAGATTATATCTTG
GCAAGAAAGAATTAATCGTTGGCGTCAACAGTATCCTTTACTGGTTCCTAGAAAATCAAC
AAGCATTTCACCTCAAGAGATTCTTGTTGCAACTAATAAATTAGCCCAAAATGCTTATTT
TACTACAGATGTTGGCCAGCATCAAATGTGGTCAGCTCAATTTCTGAAAGTAAAAGCTAA
GCATTGGCTTTCAAGTGCTGGATTAGGCACGATGGGTTATGGTTTACCTGCAGCAATTGG
CGCTCAAGTAGCACATCCAAATGACGTAGTCATTTGTATTAGTGGTGATTCTAGTTTTCA
AATGAATATGCAAGAGTTAGGAACTATCGCGCAATACCAGTTACCAGTTAAAATTATTAT
TATTAATAATCGATGGCAAGGGATGGTTAGACAGTGGCAACAAGCTTTCTATGGTGAAAG
GTATTCACACTCAAGAATGACAGAAGGAGCACCTGATTTTCAAAAGCTTGCAGAAGCTTT
TGGTATTAAAGCTTTTACTATTAATAATAGGCAAAATATGCAATCTGCTTTACAAGTTGC
TATTGATTATCCTGGTCCAGTTTTATTAGATTGCCAAGTTACAGAAAATGAAAACTGTTA
TCCAATGGTTGCTCCTGGAAAAAGTAATGCACAAATGATAGGTATCGCTAAACCGCAGAG
AGGTACTGCTTCCAACTATATTAATAATAGTGTTTGAAGTAAATAGTTAGTATTTTTTAA
ACAAAGAATAAAGTACATAAATGTATTTTTTTTATACTATTAAAAACTATGTAAGAACAG

CCTTTGACGAGAATTGAACTCGTGACCTTCCCCTTACCAAGGGGATGCTCTACCTCTGAG
CCACAAAGGCTTTTTACTATATTGGGCCGGGTTGGATTTGAACCAACGTAGGCGAAGCCA
GCGGATTTACAGTCCGCCCCCATTAACCACTCGGGCACCGACCCTAATTAAACCTTATTC
AAGATATGACCTATAAGATATCATAACTATGTTGCTAAAAACAAGTACTTTGATAAATAA
AACTGAATAAGTAATGGACATAGCTGGACTTGAACCAGCGACTTTCACGATGTCGACGTG
ACACTCTAACCAACTGAGTTACATGTCCAAAACTTTAATTATCATTATAACACTGTTTAG
TGATTTTATAAGTATCTATTTCTAATTGAAGATTTGTTTTAATCGTCCGGCTTTACCGAG
CAGATCTCTTAGATAATACAGTTTAGCTCTACGTATTTTAGCTCTTCTAATCACTTTGAC
AAAAGTTATTCGCGGTGAATTGAGAAAAAATACTCTTTCAACTCCAATACCTTGAAATGA
GCATCTTAAAGTTAAACTAGTATTTAAGCTTTTTTTTCTTCTTCTTGCTAATACTACTCC
TTCACATAATTGTTCCCGAGTTTTACTACCTTCCTTAACCATTAGTCCAAGTTGTATTGT
ATCGCCAACTTTAATTTCAGGTACTTCTGTTTTGATAAAAGGTATCTCAACACTTTTCAT
TAGCTGACTTAGCTTTGTACTATTTGCTTTCATATATAGGATTAGTCTTGTGACATGTTT
TGAAATATTATACATTAAAAAAACAAATATGAAAAATTTTAGCGAATAAGCATATTAGTA
TTTATAAAACTCTTTTTATCTCCATTATACAGATCCATTATATTCAACTAGATTCTTTCA
TAGTTTCTTTGATAATCAGTACATCTAGATGATCAAATATTATTTTCAAACATGGCTTAT
GTATGAAAAAGTTCTATAAAAACCTAATGACTAAAATTAGGGCTTAAGTATTTCTTAGTT
TTGTGTTATAATTAGTTACTATCAAAGGTAGTGGCATAACACGAAGCATCAAAATGTTCG
AACGCTTTACTGAAAAAGCTATAAAAGTCATAATGCTAGCACAGGAAGAAGCTAGACGCT
TAGGACATAATTTTGTCGGAACTGAGCAGATACTATTAGGATTAGTTGGTGAAGGTACTG
GAATCGCAGCTCAAGTTTTGAAATCGATGAATGTGAATCTAAAAGATGCAAGAGTTGAAG
TCGAAAAAATTATAGGAAGAGGGTCTGGTTTTGTAGCGGTTGAAATTCCTTTCACTCCTC
GAGCAAAAAGAGTACTAGAATTATCTTTAGAGGAAGCACGTCAACTAGGCCATAACTATA
TTGGCACAGAACACTTGCTAATGGGCTTAGTCCGAGAAGGAGAAGGAGTTGCGGCAAGAG
TTTTAGAAAATTTGGCAGTCGATGTTTCTTCAATTAGAGCTGAAGTTATACAAATGCTCG
GAGAAAATGCGGAAGCCAATGTAAGTGGAAGCAATACTACGCAAGCTAGAAGTAAAACAC
CAACATTAGAAGAGTTCGGATCTAACTTAACTCAAATGGCTATGGAAGGTGGTTTAGATC
CTGTAGTCGGAAGACAAAAGGAAATAGAACGAGTTATTCAAATCTTAGGTAGAAGAACTA
AAAATAATCCTGTCTTAATTGGGGAGCCTGGTGTAGGTAAGACAGCGATTGCGGAAGGAT
TAGCTCAAAGAATTGCTAATAGAGATGTACCTTCTATTTTAGAAGATAAATTAGTTATTA
CTCTTGATGTCGGTCTATTAGTAGCCGGAACTAAATATAGAGGTGAATTTGAAGAAAGAC
TAAAACGTATTATGGATGAGATTAAATCAGCTGATAATGTAATATTAGTGATTGATGAAG
TTCATACATTGATTGGGGCTGGTGCTGCAGAAGGAGCAATAGATGCAGCTAATCTGCTTA
AGCCAGCTTTAGCAAGGGGAGAATTGCAATGTATAGGTGCAACAACTTTAGAAGAATATA
GAAAACATATAGAAAAAGATCCAGCATTAGAAAGAAGATTTCAACCAGTTGTAGTTGGAG
AGCCAAGTGTTGAAGAAACAATTGAAATTTTGTTTGGTCTTAGAGACCGTTATGAAAAGC

ACCATCAATTAACAATGTCAGATGGAGCTTTGGCTGCAGCTGCTAAATACGCTAATCAGT
ATATTTCTGACCGATTTTTGCCAGATAAAGCAATTGATTTAATTGATGAAGCTGGTTCTA
GAGTCCGTTTACTAAATTCTCAATTACCTCCTGCTGCCAGAGAATTAGATAAAGAGTTAA
GAGCTGTATTAAAAACAAAAGATGAAGCTATTAGAGCTCAAAAATATGAAACAGCAGAGC
AGTATAGAGCAAGAGAAATGGAAATTAAAGCTCAAATTGCAGCAATTGCTCAAAGTAAAA
AGAATGAGCCTGATTTAAATTTAGAAGATCCTGTTGTTACAGAAGATGATATTGCTGAAA
TTGTTGCTGCATGGACTGGTATACCAGTAACTAAGCTTACTAAAAGTGAGTCAGAAAAAT
TAATGCACATGGAAGAAACTTTGCATGGACGTATTATTGGTCAAGACGAAGCGGTTGTAG
CTGTCTCTAGAGCGATAAGACGCGCAAGAGTAGGTCTAAAAAATCCTAACAGACCAATTG
CAAGCTTTATTTTTTCCGGACCGACGGGTGTAGGAAAAACAGAATTAACAAAAGCTTTGG
CTTCTTATTTCTTTGGTTCAGAAGCTTCTATGATACGGCTAGATATGTCAGAATACATGG
AAAGACACACTGTATCTAAACTAATTGGTTCTCCTCCAGGATATGTGGGTTATAGTGAAG
GTGGTTATCTAACAGAAGCGGTAAGAAAAAAACCATATACTGTCATCTTATTTGACGAAA
TTGAAAAAGCTCATCCGGATATTTTTAATCTACTTCTTCAAATTTTAGAAGATGGCCGAC
TAACAGATGCAAAAGGCAGAACTATTGATTTTAAGAATACTCTTTTAATTATGACTTCTA
ATATTGGATCTAAAGTTATTGAAAAAGGAGGAGGTAGTTTAGGCTTTGAATTATCAGAAG
ATCAAACAGAATCCCAATATACTAGAGTACGATCTTTAGTAAATGAAGAACTGAAACAAT
ACTTTAGACCAGAGTTTCTAAATCGATTAGATGAAATTATTGTATTTCGTCAGCTTACTA
AAGATGAAGTTAGAGAAATTGCAGAATTAATGCTTAATGAAGTCTTTGCGAGAATTAAGC
AGCAAGATATTCAATTAAATGTAACAGAACGATTTAAGCAACGATTAGTAGAAGAAGGAT
ATAATCCAAGCTATGGAGCTAGACCACTTAGACGAGCAGTAATGAGGCTTTTAGAAGATA
GTCTGGCTGAAGAAGTTTTATCTGGTAAAATTAAAGCTGGTGATAGTGCAGTAGTAGATG
TTACTAATGAAGGAGAAGTTACAGTTTTATTAGGTGAAAAATTAGAACTGTTAACATAAA
AAACAATTTATTAATTAAAGCATTGAAGAGGTTATATTCAATGCTATGAAAAAAATTAAT
AAGATGGGTTGCTATAATTGACAACCTCTGATATTCTTTATTTGTATATAAAGCCGGGAT
AGCTCAGTTGGTAGAGCAGTGGATTGAAAATCCTCGTGTCACCAGTTCAAATCTGGTTCT
TGGCATTTTAAGCATCAGAAAATTTTGCTATCTGAGCATTAAATTTTAATTGAAATGTTC
CTGTGGGACCATTCCTGTGTTTTGCTATTATAACTTCTGTGAAGTCTCTACTTCTAGTTT
CTTGTGTATAGTAACTTTCTCTATATAGCATTATTACTAAATCTGCATCTTGTTCTATAG
AATTATGAACAATAAAGTTATTTGAAATAAAATTGCATAAAGGTTTACATTCTAAATCAA
ACACTATTTGTAAACTTGTAACGTTTATTTTTTGTATACTTTCAAAAGTCAAAGATGAAA
AAGAATTAAAAACATTTTTTTGTTCTTCAAAGTTTTTTATTTGAATAGCAATCATGTCAT
TTTGATTAATTTTATCACATCTTTTCCATCCTTGTGTTGTGAGTAGTTTATGGTTACTTG
TTAATTGTATATATTTCCCAGCTTCGGTAATAATTTATATACAGTTTTTTTCCCTGTTT
TCGAAATACTACATTTAATTGCTAGAAATAACTGTCGTTTTTGCGCACTAAAGTTAATGA
CTTCAATCTTTTTATAATAGCAATTAAATAATATTTGAGGTTGATTATATAATGATGTCT

GAAGATAATTGAATTTGGATATACACCCACTTTCTCTTAAATCGGATAACAAAGGCCTTT
TGTCACTCCGAGTTTCAAGGTTTCTATTTAGTTGAGATAATGCCAAGATAGGCAAGCTTA
AATCTTTTGCTAATATTTTAAGTGATCGAGTAATTAATGATAGTTCTTGAGATCTATTGT
CAGATTGTCTACTCTCCTGCAGTAATTGAAGATAATCTATAATAATTAATTCTATATTTT
TTCCTTGTAATTTAAAGAATTTTACTTTTGTCTTTATAGTGTCAGTAGAAATCTTAGCGC
TATCATCAATATAAAGATTAAGGTTAGCAAGAGTTTTACTTTTTTGAATAACATATTGCC
ATTCATCATTATTAAGCTGACCAGATTGAATTTTTTGGCCGTTTAAATTACACTCTTGAG
CTAAAATTCTTCTTAGTAGTTGCTCTGTAGACATCTCTAAGCTAAATAAAATAACATAAG
ATCTTTTCTGATTAATTACATATCTAGTTATATTAATAGCAAAAGCAGTTTTCCCCATTG
AAGGTCTTCCTGCAAGAATAATTAAATCTGACTTTTGAAACCCTTGTGTTATAGAGTCGA
GTTCTGTAAAACCAGAAAGTATACTACTATTTATACTAATCTTTTTCCTTGTATCCAAAT
GGACTAATAGTCTTGCAAGAATTGTAGCTAATGTTTGCGTATCTTTTTCGTCAAGTATTT
CATAAGCTTTTGTTAATTGACTAACAATAGAAGTTATGTTTTTTTGTTCAATAAGTTGTT
TTGAGTAGCTAATTAAACATAATGAATCTCCACATGCTAAAAGTAATCTTTTAATGTAGT
TATCTATAACGACTGCTGAATATTCGCTTATAGTATCAGATAGGGGAGCTTGTTCTATTA
GACTAAAAACTTCATTTAGCTCATTCAAGTCTTGAATCATTTTTTCATTTTTTAAGTTAA
TGAAAAAATTTCTTACGCTTTTTGTCTTTGCATGGTTAACATTTTCGAGTGCTGCTCTAT
AAAGTAATGAAGTTGATGTAAAATAAAAAAAAATCAGGAGAGATTTTTTCTACTGATTTTA
GTAATGATATCGATTGCTGAGTTAATATTATACTAATTAATATTTTTTCAGCTAAAATAT
TATGAGGAGGTAGATATTTATAAATACTTATTACTTCCTCCTTGTAACATTACTCTTGTG
GTATTAGTTAACTATTGATTTAATTTGATTCTGGAAGAACTTGCAATTTAATATTTGCAG
TTACTTGATGTAATAGCTTAATTTCTACATCGTAAATGCCAATTGTTTTCACGTCTGGTA
AAGAGATACTTTGTTTTTCAATGTCTATGTTGGTAGTGTTTTTAATAATTTGTGAAATTT
CTTTTTCTGTGACACTACCAAAAATATTGTGGCCATCTCCTGTTTTTTTACTAACGCTAA
ACCTTTGTATTTCTTCTAGAAGTTGCTTGACTTTTTTAGCATTTTCTTTAGCTGTTTTTA
ATTTTTCTTCTCTTATAGCTGCATAAAATTTTTGTTGTTTTAAAATACCATTTGTGGCAA
CTGCCGCCATTTTATTCGGGATTAAAAAATTTCTTGCATAACCAGAAGCTACTTTGATAA
CGTCATTACTCTTGCCAAGTTTTTGAATATTCTCTTTTAGCACAACATTTATAACTTTTT
TACTCATCTTAATTCTAAAAAATCTGTATTATCCTATTATACCTGTTGTATACAGATTTT
AATAGGAAGTATTTATTTGTTGATATAATATTCTTGTAAAAAAAGGTAATTTTTTATTTT
TTTCTGATATAATCACTTTCACAATAATTCATTAGGAGAGGTGGCCGAGTGGTTGAAGGC
GCAGCATTGGAAATGCTGTTTAGGAGCAATCTTAACGAGGGTTCGAATCCCTCTCTCTCC
GTACAAAAAATATAAAAAGTTTTATAGAGTCCAATCTAAATCTCCAATAATATTCTTCCA
AGCATTTAATCCACCCCGAACTCTAACAATATTTAGTTTTTGAGCTATTAAAAACTCAGA
AGCAAATATTGATCTAGAATCTAAGCTACAGTAAACAAAACAAATTTTATCTTGTAAATT
GATATGAGAGTAATTCATTCCTTTTATTTTTTGTAAAGGTAGATTTAATGAATATATTAA

ATGACTTTTATGATATTCTTCATGATTTCTTACATCTATTAAAATATATTGTGGATTCCT
ACTAATTAAAAACTTTTGTAATTGAATAACACTAATTTCTCGTACAAATGTATTAGATTT
ACTGCCATAATATTTGTTCCAATGCTTTTTTGTGGATAATACAAATTTAGTATTTATAAT
CTTAAATTTGTTAAATGATGAAGTTAATGAGTTGTACGTTAATATAGTTCCACTTAATAT
AGACTTATAACCCAGAACGATTTTGACAGCTTCAGTTGCTTGAAGTGTACCAACAATACC
CGGTAAGAGACCTAAAACTCCGGAATTACTACAAGTATCTCTTGCGCTCTCTTTATTTTC
AGTTTCACTGTAGAAGTCTCGATAAACAGGTCCACCTTGATAATTAAATACACTAACTTG
TCCTTCAAATTGAAAAATGGCACCATAAATATGTATTTTATTTAACTCTAGGCAAGTATC
ATTCAAGAGATATCGAGTATCGAAGTTATCAGATCCATCTATAATAATATCATATTGTCT
AATAATGTCGATTGAATTCTCATAGCTTAGTCTTGTTTTGAACACAGTTACTATACACTG
TGGATTTAAATCTAATATTTTTTTTTAGCTATTTCTACTTTTGAATATCCTATATCATGC
ATCGTGTATAAAATTTGCCTCTGTAAATTAGAGAGATCAATTATGTCATCGTCTATAATT
CCAATGCTGCCAATTCCTGCAGCAGCAAGGTAAATGATTCCTGGAGATCCTAGCCCACCA
GCACCAATGAATAAAACTTTTGCTTCTTTTAATCTTTCTTGTCCTTCTAATTGAATTTGT
GGCAATACTAAATGCTTAGAATATCTTGTATATTCTTCTAGTGAGTATTTTGTATTTTCT
GTTTTAAAATTTAGCATAAAAATTTATAGTTTATTTAATAGCACTTAAAAGCTAGCTGAT
AAGTTTATTAAAATGACTTTTGTAAACTTGGTTCTTCTCTATCTAAAATAGCTCCTTGTG
TAGGACAGACCTGAACACAAATACTACAATCAATACAGGCAGAAAAGTCAATCCAATACC
AATTAGTGTTATTATTGTTTTTCCCCTTTCCTTGATGAATACAAGCTACGGGGCAAGCTT
CCACGCATTCAGCAACTCCAATACATTTTTCTGTTACAATTGTATGAGACATATTATTTA
ATTAATAAAAGATAAGCTATTTACATCTAGTAGTTTTGTTGTATACTAAATTTACATATG
CTATCCAGGCCAATATTTAAGGTAGTCTATTTCTACAGTTTACTAATTTGATATGCGCCT
TTAGTTCAGTTGGTAGAACGCAGGTCTCCAAAACCTGATGTCGAGGGTTCAAGTCCTTCA
GGGCGCGTTTAGTTTTATCGTAATTCAATAATATCATTTTATCTTTTTGCAAACATCAAA
ATGATATGATATAAAAGATCATATTTTATACATTGAATATTCTATGTACTAGTACTGTTT
TTTAATATGTTATTATTGATAAGTTAATTATACTAATTCTATTTTTGTAATAAGTAAAAA
CAGTTCTAATTAAGTATGCACGTATTTAATAGTCTTTTGTATATATAACTTGAAAAAATT
TAAACTATGGCTAAAAAAGTTACTGGAATTGTTAAACTAGCATTAAATGCAGGTAAAGCT
ACTCCTGCGCCACCTGTCGGACCTGCTTTAGGACAGCACGGCGTGAATATTGTTATGTTC
TGCAAAGAGTACAATGCACGTACTGCTGACAAGTCAGGACTAGTAATCCCTGTAGAGATC
TCAATATATGAAGATAGAAGTTTTACTTTTATACTTAAAACTCCTCCTGCTTCTGTACTA
ATCGCTAAAGCTGCTGGACTAAATAAAGGTTCGGGTGAACCAAATACCAAAAAAGTAGGC
AGCATAACAAATAAACAGTTAGAATCTATTGCAGAAACAAAACTTCCCGATTTGAACACT
AACAATATACCTCAAGCTATGAAAATAGTTGGAGGTACGGCTAAAAATATGGGAATTCTA
ATTAAAGATTAAAAGTTTCAATTATATTTACTATTTTTATGAAAAAATTTTCACGTCGAC
TTACAACATTAAAATCAAAAGTTGAGCCTAAACTTTACACTATTAATGAAGCAGTATCTA

TATTAAAAGCAACGTCAAATGCTAAATTTAAGGAAACTGCAGAAGCTCATATTGCTCTAG
GTTTAAATCCTAAATATGCAGACCAGCAACTCAGAGCAACAGTTATATTACCGAAAGGTA
CAGGTAAATTGATAAAAGTAGCAGTTATTGCTAAGGGAGAAAAATTAACAGAAGCAATTA
GTGCGGGAGCTGATGTTAGTGGCTCCGAAGAACTAATTGATGAAATACTGAAAGGTAGAT
TAGATTTTGATAAGCTGATAGCAACACCAGATGTTATGCCTTTAATAGCAAAGCTGGGAC
GAGTATTAGGCCCTCGAGGGTTAATGCCTTCTCCTAAAGCAGGCACGGTAACATTGGACG
TAGCAAAAGCCGTGAATGAATTTAAAGGGGGTAAAGTTGAATACAGAGTTGATAGAACAG
GTATAATTCATGTACCTTTTGGAAAATCTAGTTTTTCACAAGAAGATCTAGTTTTGAACC
TCCAAACAATTAAAGAATCGATTGATAGAAACAAGCCTTCTGGAGCAAAAGGGAAGTACT
GGAAAACTTTTTTCTTATCTAGCACCATGGGGCCATCTATTCAAATTGACATCACTAGCC
TTCTATGAAAAATTTTTGTATAATTATAGAACTTGAATAATCTATTATTTCAAATTATAA
TTTGCTAGCATTATTAACGTAACAACTTGCTTACTATATCTTATCTGCATTAAAAACTTT
TTATATATCTTATTAATTATGAGTACAAAGGTTGAAAATATCTTGGAAGAGCTAAAATCT
TTAAACCTTCTAGAGGCTGCTGAACTAGTTAAACAAATAGAAGAAACGTTTGATGTTGAT
GCATCTGCGGCTTCAGGAGGTATGATGATGGCCGCACCAACTTCAGCACCAGCGTCTGCT
GAGGTTGAAGAGAAGACAGAATTTGATGTTGTCCTAGAGGAAGTCCCAGCACCTAAAAAG
ATTTCTGTATTAAAAGCTGTTCGCTCTCTGACTGGTTTGGGTTTAAAAGAAGCAAAAGAT
CTTGTTGAATCTACGCCAAAAGTCTTAAAAGAAGGTGCTTCTAAAGATGATGCAGAAACT
ATGAAAAAACAGCTAGAAGATGCTGGCGCAACAGTTATTGTTAAGTAATGAAGAAGCGCT
TCTGTTTAAGTATAAACAGAAGTGCCTCTTTATTTATTATAAGCAATTATACTAGAAAAG
TCCAAGTGTAATTGCATTGTTGATAGGCATAGTTGCACCAATACCTAGCCAAATACTTAC
AACGGTACCAATTAAGAAAACAGTGGTTGCAATTGGTCTTCTAAATGGATTTTGAAATTT
GTTAACATTTTCAATGAATGGAACAGTAAGTAAACCAGCTGGTACAGCAGCCATACTTAG
AACACCTAATAGTTTATTAGGAATCACTCTTAATAGATTAAATGTTGGAAAGAAGTACCA
TTCTGGTAAAATCTCTAATGGAGTCGCAAATGGATTAGACTTTTCACCTATAGAAGATGG
TTCTAAAATTGCGAGTCCAATACTGCAGGCAAAAGTGCCAATAATTACAACTGGAAAGAC
GTATAATAGATCATTTGGCCAAGCTGGCTCTCCGTAATAATTATGGCCCATTCCTTTTGC
TAATTTAGCTCTCAGCTTCGGATCTGTTAAATCTGGTTTTTTAAGAATTGACATATTGTT
TTTGTGTTTAATATAAAGATTAGTCTAATTACTGTTTTAAAGTGTAATTTTCAACTATTA
TAATGGTCCAGAAATTCCTTGTTTACGTATCATTAAAAAATGCATAAGCATGAAAACGGC
AGTTAGTAGCGGAAGTACAAAAGTGTGTAAACTATAGAACCTAGTTAAAGTTCCTTGGCC
TACACTGACTCCTCCTCTTAACAATTCTACTATACTCGCTCCGACAACTGGAACAGCATC
AGGGACACCTGTAACAATTTTTACAGCCCAGTATCCGATTTGATCCCATGGCAAAGAATA
ACCTGTTACCCCAAAAGAAACAGTAAGCACTCCTAAAATTACACCTGTTACCCATGTCAA
TTCTCTAGGTTTTTTGAAACCTCCTGTTAAATAAACACGGAATACATGTAGAATCATCAT
CAAGACCATCATGCTCGCTGACCATCTATGAATTGATCTAATAAGCCATCCAAAGTTTAC

ATCAGTCATAATGTACTCTACTGATGTAAAAGCTTCTGCAACTGTAGGTCTATAATAGAA
TGTCATTGCAAATCCAGTTGCAACTTGAATTAGAAAAGATACAAATACAATACCCCCTAA
GCAATAAAAAATATTAACATGAGGTGGTACATACTTACTAGAAATATCGTCAGCAATTGC
TTGAATTTCTAATCTCTCTTCAAACCAGTCATAAATTTTACTCATAAAATAGCTTC-AAA
AGCTCTTCTAATATATTTTGCGTTTAAGCTACTATTAATTCAATAGTTATTGAACTTATA
TTTTTTTATAATCAATACTTATTAACTATTATAACATCTTTTTATTGATTTAAAATTCTC
TGACTTAGTAAAACAGGGCTATGTACTATAATTCTTTTTTTTTGTATAGAGATTAATTTC
GTTTTTAGTAAATTAGCTAAAATTCTTGTTACACTGACTCTATTACTACCTATAATCTGT
GCTAAAATTTTATGTGTAATAGTAAAGTTAAGCATAATACCATTATTAACTTGAGTTCCA
TTTTGCTCAGCTAATAATAATAATAAGCTCGCCAGTCTACTGGTTATACTTTTATGTGAA
ATGATTTCTATAAAATGATAAGCTTTTGCCGAGCAAAATACTAAATGATTTACAAAAAAT
AAATTGAATGGAGCGCAGTTACTACACGCTTTTATAATAGTAGTATAGTTAATACAGGCA
ACTTCTGTTTTATCTATTGCTTCTGCTTCGTAGTAAAAATTATCGTCGACCAATTCTATT
TGTCCAAATGTATCTCCTGTAGTTAATAAGTTAAGAGTTACTTTATGTGTATTTCTCAGG
ACTTTTGTAATAATTAAAGATCCTATCAATATAATGTATAATCTAGAGCTGCTATTAAAC
AAAAGAGTATCATTTTTCTTAAGTGAAAAAATTTGATAATTAATTTTATTACGATTAAAA
AATAGTAGCCATGGATTGCAAGAAAAAAAATCACCTGAACTATTTTCAGAGAATATTGGA
ATACAATTTTCTTGATTTAAACAATTTTTAGTCATAATTACTACTTTATAAAAAAATATT
CTAATAATTAAGAACTGAAATTAAATGTCATATAAATTGATGTTAGTTGAGAATGACATT
GTATTATCAAAGGCCATTCAAGAATACTTAATAGATCAAGGCTTCAATGTTTATATTGCC
AATAATGGATTAGAAGCACTGAATCTAGCTTATCAATATAATTTCGACTTGATAATTTCT
GATATAATGATGCCGCTAGTTAATGGCTATGAACTATTAGCAAAACTTAAAAAAAACAAA
GCATTATCCAAAATACCTGTTATCTTTTTGACAGCTAAAGGAATGACTAAAGACAGAATA
AAAGGCTACGACATGGGGTGCTATGGTTATTTATCTAAACCATTTGATCCTAATGAGTTG
CTTTCAATTATTAACAACTTAATCGCTAGAGATGTCTTAAAAGAAGCTTCTTTACAAAAC
TTGGCAACATCAAACCAGCAATTAAATCATAAAATACGTTTGACACCTAGAGAAAAAAGT
ATTTTAGATCTTGTTGTTGATGGACTTACTAATAAAGAAATATCTACTATACTGAATACT
AGTGTTAGAAATGTAGAAAAATATGTTAGTCGACTCTTACACAAAACTAATATGAAGAAT
AGGACTTTGCTAGTCAAGTATTCTATAAATAATAATCTATTAAATAATGAGATCAATGAA
AGGGCGAATGACGGGACTCGAACCCGCGAATGATGGAGCCACAACCCATTGCCTTAACCC
CTTGGCCACACTCGCCATATTACACATACCACTATAGCTTGTTTTTTGATATATCGTCTA
GTTTTCGAGTTATATATTTATCTAAAACAATATTTACTTTGAATTTTTATGATACATAGC
AATATTAGTATTCAAATTAATGGGGAACCATTTAATTGCTCAAAGCCCATCTCATTGCAA
TTTTTATTAAATTATCTTGATTTTAATTCCGAGCGTGTAGCAGTCGAGCTAAATGACATT
CTGTTACCAGAGACTCTGTTTCACTCAACTTATTTGAATGATCAAGATAAGCTAGAGGTT
ATTACCATTGTAGGTGGTGGATAATCTATTTGTACATACACAGTTCTCTATCTCAAATGT

TTCATAGATAGAGATAATCTTCCTTGTTTTTTATCTACATGAATAATAACAGCTTTAATA
GTGTCACCTATTTTAAATTGTGATGATATCTGCTCTAATTCTTTGATATTAATCTCAGAA
ATATGGACAAGACCTTTGAGATTTCCTACCTTTATAAATAATCCGTAAGGTGTAATCTGA
TTTATTATGCCTTCAATAATATTACCAACAATTAAATTAGATGATGCTTGAGATATTAGA
GCTCTTCTATGACTCAGTATTAAATTATTAGATTTCTCTTCGACATTGAGTAATTTTAGC
TTAATAAATTTATTATTAAATTGCTCACTTTTTTGAAAATTACCGAGATGAGAATTAGGT
ACAAATCCAGATATACCTTCAAGGTTAATTATCATTCCTCCCTTATTGAATCCTTTTATC
ATTACATTAAGTAAAGAGTCTTCAGCTAATAGTTGCCTAATTCTTTTCCATGCTCTTATA
TATTCAAGGCGACGAATTGATAATATTAATTGTTTCGACTGAATATTATAATCTAATAAA
AAAAATTCTCTTGTGTCATTAATGTTTAAAGAAGTGAAATTATTTAAATCTTGATTACTA
GATACTTCTTGAATAGGCAAGTATGCAGATATTGGTGTTCCTATATCTACTAATACCCCA
TTTAATTCAAAGCTAAATATAGTTCCAGCTACAATATCTCCAAGATTCAAATCATACTTG
TATTTTTGTAAAACAGCTGCAAAATTTCTATGAGTAAATCCTTCATTATTTTTTGTCATA
TTGAATAGTATGAATTAAAGAGTAACTTTGTACAATATATTAAGGTAAATAAATGTTTGT
CGAATTGTATGAAAAAAATTTTAATTTGTTCGGTTGTAATTAAGTATATAAACATATTAAA
ATATCTAGGAGACATGAAAAGTAGACGTAGATAGCTGCAGATTTAATACTATATAATCTG
AGGAAAGTCCGGGCTCCACAAATACAATTTATGCTGGAGAAATCCCAGTGTAGGTAACTG
CGAGGATAGTACCACAGAAAAAAACCGCCAGAATAAGTTATATTAGCTGGTAAGGGTGCA
AAGGCAAGTTAAAAGCTTACCAAAAGTACTGCAAAGTATTTGTTAGGTAAACCCCTAAAA
TGGAGCAAAGCTACTAAACAAATATTTGTGTATATTTTATCTAGTTTAGTTGAAAATACT
GCATGAAGTTATTAGTAACAATAACTCGAGAGGAATAGCTATCCTTTACAATATATCTCG
ATTATATTGAAAGAACAGAACCCGGCTTATGTAGTACTTTTTATGTCTTGTATTATTATG
TCTATTGTCGAATCTTTAAATTTAACTGTTTGACAATTTTTTTTTGTAGCTGATGTTGTT
TATAATCTATTTCTGAATTGGTTAAAGTTTTTTCATTAGATCGATATGTAAATCTTAGAC
CTATACTTTTTTTCTCTTTTCCAATTGATTCATCTATATATTGATCAAATAATGTTGTAC
TTTCTAAATCTTTATCATAAAATTGATCCAATAATTTTAATAAGTATTTAATTTCCATAT
TCTTAGGTATTATTAAAGAGAGATCTCTTATGATCGATGGATACTTAGAGTATGGTTGAA
TCTGATAACTTAAATAATTGAATTCATTATGACAATCTTCTAAAATGTTTAAGTTAACTT
CAAGAAAAAAAAAGTTTTGTGTTGAGTCCTAATTCACTATATGTTGCTTGATTAAGTTCAC
TGAATATGCCAATAATAATATTGTTATAGATTAGTGTGGCAAATTTTCCTTTTTGAATGA
ACTTTATTCTACCATCGACTATATCTTTTTTTGTCCACTCAATTTGTCTATTAATCTTTT
GAAAAAAATTTTCTACGATCCCTTTAGCTTCATACCAATTTAAAGAATGCGCAGGGTGTG
ACCATGTAGATCGGATATCTAAATTACCACCCAATATGATAGCTAGATTATTAGTTTCTA
CAATTTTGTTTTGATCATTATGAAATACTGTTCCTATCTCAAAGCTATCTATAGTTTGGT
TGCTTTGTTTAATATTATAAAGATTTGATTCAATTAATCCTTCCAGCAAACTGCCGCGTA
AATTAGAATAATCTTGAATAAGAGGATTGTTTAAAGCTATATTTCCTTTGGACTTTATTA

AAGAATAATGAACTAATTCAGTTAATCCTAGATTTCTTAAAATACTTCTACTTTTATCTA
TGAATTGTCTCTTTGTGGATAACTTTTTGTTGAATTGAATATTTGGTACTGAGCTCCTGA
ATTTATGATAACCATAAATTCTAGCAATTTCTTCAATAACATCTATTTCTCGAAAAACAT
CTTCTCGTCTATAATTTGGAACAGTAACTTCTATATTTTCCTTATTTGATAAATAAGAT
CAAAATTTAAAGAGTGAAGAATATTTTGAATTTCTTCAAAGAATAAAAAACGTGTTTGAC
CATTATACATAATTGGTCCTAAAATGTCATGAACTTTTTTAATTGATAAATTAATATTAA
GAGCATGATCATTTATTTTTTCTCGACAGAAAGTCTCTCTAATATTTCCGCCTGTTAGAT
CTGTGATTAAAGCAAGAGCTTCAAAATGTGCATTTTTCCAATTATCTACATTTAGCCCAC
GTTCTTGTCTAATTGAACTTTCTGTTCGAATATTAAGAACTCGAGAAGATTTTCTAACTA
CAGATTGTTTAAATATAGCAGACTCTACAAGTATCGATTTTGTATTATAATCTGTATCAA
AATCAGAGTTGCTTCCAATACCAGCTATACTAGTTACATTTGTATTTACTTGAGTAACTA
AAATATTTTTGTTTAACTCAATATTTTCATTGTTTAACTTAATCTGATCATTACTTCCAA
GAGGAAAATTGCTTCGAATTGTTAGACTATTTTTATGATTCATATTATTAATTTTATTCA
AATCAACAATATTAATAGGCTGTCCCCATTTCAACATAATATAATTGCTTATATCTGTTA
ACAAATTTCTGTGTGTAAACCCAGAAGAGAGCAACCGATTTTTTAGCCAATCTGGAGAAT
CTTTAACTCTTATTTCGTCTATAATGGCAGCAAAATAATTGTCACAATTTAATAAATCGT
GATTACTGATTATAGTCTCCTTTTTAGAGAAATATCTATTATTGGGTTGATGTATAGCTG
ATAACATTGAGGACCCTGTTAAAGCCGAGACTTCTCTTGATAGACCAATCATGCTTAACA
CGTCTGACCTGTTAGCTGTTGATGATATATCTAATATATGATCAATCTCACTTCCAATTA
TAATAGATTCAATAGCTTCAACTTCAAAACCTGCTTGTGTTAGTTGATTAGTTAAATTGT
CTATATCTATTGTTTTGATATTACCAAGTTCTTTCAACCAATTTAAAGAAACTTTCATAA
TTAATTGTCATCTATCTTATAGAAGTATTTATTTTACTTATGATTAAAAATCAATTTAAA
AAATTCATCAAATTAATATAAGTTATTTGATGAATTTGCTTATACAATATATTATTGTGA
TATCTTACTTTTATATGGCAGTTATTAAGAACTAGCTTTTGTAAAAGTTAGCCCATTATC
TTGGGCATATCTTTCCATAAATCTCATGAATCTATCCCAATCATCAGGACTTTTAATAAC
ATGTACAGCTTCTATTGCTTGTGGTTTACCATTAATAAACTTGGCATTAACATCTCTTGT
AATAATTTGTCCCTCTGTGTCCATCAGGTACATACCTGTGATTTCACCTTTGTCTGCCAT
ACTAGCATCTAAAATTTTGGGATTTGTAAACCGAAAAGTTGCTGTTCCTGTACTACCATC
TCGAGACCTTGTTAAACGTACATCTGGTACAACTTCTTCATTGATACCTTGAATAAATTG
AATTGTTGCCATATCAATCTTCTTTATTTGTATATAATTTATATTGGATAAAACTGATTT
CTAGTTCTTGAATGCTTGTTGAGAAACTGTTTTAAAAGCTTATAGTAGCAGTAGATCAAA
TTTTATGTATTATATAAAACTGTAACTTTTCTTTTTTGATCTGTTCAAAGAACTAACTTA
TTATAAGCTAATCATACGAATATTAGCCATATAATGTTCGTAAGTGAAATTAAGGGGTTG
TAGCTCAGAAGGATAGAGCAAGCGCCTCCTAAGCGCTAGGTCAGCGGTTCAAGTCCGCTC
AACCCTGTTAACGAATTAAGTAGTATTATTTAACTTGTTCTGCAAATCAGTATTGATCTA
AGTTAAAGTACTTGACAGCTAAAACAAAAGAATCTGAGAAATATTTTCATTTAAAGAACG

AACAATTATCTTAACTCTAAATATCTGATAATTATTAAAAAATTCTCTTTTTAAAACACA
AGATGATGAAGATATTTAAACTAATAATTGAAATTAAGTTTTTGAAGTTATAGAAGCCAA
TATAATACTTTCTATTAACAGATGCTTGCTTCTATTACTATCTGTTTACAAGAAATTATG
AGTATCAAAATAATTTTAGCTCTACTGACTACTAATAAATATTTATTTTAGTCTAGACAA
GATGAATCTTCATGGTATAACAGAGCTATTCTTTGTGTCATACTTATATAAAATTAGGAT
ACTGTTTTCATTATTATAACTGGGGTGGGAGGATTCGAACCTGCGAATGGCGGAGTCAAA
GTCCGCTGCCTTACCACTTGGCTACACCCCAACATGTGAACATTATAATAGCAGTCGTAT
ACCCTGCCATGTCAATAGACTATTGTCTTTATTTAATAATTTTTGCTGCTCTTGTAGAAT
AAAATCTTTTTTTTCAAATAAGATACGCTATGTTTAAGTTGCACGACTGTTATGTTTTCT
TGCTTTTGACTGAACAGCCACTTGATAGTAATTATTCCTTCTGCAATTTCGTTACTTCCT
ATAATAATACAAGCTACAGCTCTTTTTTTATTTGCTTGCTTAATCTGCTTACTAAAATTA
CTTGAGCTAACATCGATTTCTATTTTAAAAAACTGCTGATGTAAAAAACGCATTATTTGC
ATACCAGTTTCGTTAGCTTTTGTACCTTGTGTAGCAATATAAAAATCGATAGATTGATTA
GGCAAGTCTATATTATCTTTTGCAATCAGTAATAGGCGTTCTAATCCTATTGCACATCCA
ACAGCTGGTGTTGCTTTGCCACCTAATTGGTGAACTAAGCTGTCATATCTTCCACCACCA
CATATAGTATCTTGCCCTTTTGATGTCAATGTCTTAATTTCGAAAGCAGTATCATTATAA
TAATCTAATCCTCGAACTAACTTATTATTTATATTGTAGGGAATATTAAGTAATTTTAGA
TAATTGCAAACGTCACCAAAGTGTTTTTGTGATTCCAAACTTAAAAAATCAGAAATTTTT
GGAGCCTCTGTCAGTATTTTTTGTGTATTAGAATCTTTTGAATCTAAAATTCTAATTGGA
TTACTAGTTAATCTTTTTTGTGAGTCAGTGTCTAAATCATCATGGTACTTTGTAAGATAA
TCTCGTAATTTAACTTGATAAATACTACGATCCTCTACTTTTCCAATTGAATTCAGATCA
AGTTTCAAGTTGTGCAGATTGAGATTGTTAAATATGCTCATAGCTAAATGTATAACTTCA
CTATCTGCTCTTGCATCTAAGCTACCAATAAACTCAATGCCAAGTTGGTGAAATTGTCGT
TGTCGTCCACTTTGTGGTCTTTCGTACCTGAACATTGGACCGCTATACCATAATCTTTGT
AAACTATGGTGGTAACTCATTTTATTTTCAATAAAAGATCTCACAATACCGGCAGTACCT
TCTGGCCTTAAGGTAATATCTCTATTGCTGCGATCATGAAACCGATACATTTCTTTATTT
ACAATATCAGTATCTTCACCAATACCCCTATCATATAATTCACTATTTTCAAAAATAGGT
GTTCTAATTTCTTGGTAATTTGCACATTCTAATAATTTGGAAATTTTATTATGTATAAAC
TGCCAATATTGAATTTCATCGGGAAGAATATCTTTTGTCCCTCTAATAGCCTGAATTTTT
GCCATGATAATTTTTTGATTACCTTCTTAATATGTCTGCATTAAATAGCAGTTGTAGACT
GTTATTCAGTCATGTATTATACATTAGAACATACTGATGTAATTTATAAAGTACGGGCAA
GGAGGGATTCGAACCCCCGACACCATGGTTCGTAGCCATGTGCTCTAATCCACTGAGCTA
CAAGCCCACTTAATAAGTACCTTAACTATATCATTGTTCACTAATGTATACAACTATATT
CTTTTTATCTTACATGTGAAATTTTTGTAAAAGTAAAAAATTATGTTTGTTTTTTACAAT
TGGAGCTATTCATATATAATGCTTAATTTATTATAACTTTGTAAAAATATCTATAAGTTT
AATTGTATGATTCTATTATATTGGTAATCTTGTTATAATTTTGCGCTCTTATTTTTCAAT

TGGCTAGCATTTAATTAACTTATATTTCAAATAAAAAAATCTTATCTAAAATGAGTAAAC

AAATTCTATATCAGGATGATGCCAGAAAAGCATTAGAAAAAGGCATGGATATTTTAACAG

AAGCTGTTTCTGTGACTTTAGGACCAAAAGGAAGAAATGTTGTCTTAGAAAAAAAATTCG

GTGCCCCTCAAATTATTAATGATGGTGTAACGATTGCAAAGGAAATTAGTTTAGAAAATC

ATATTGAAAATACCGGAGTCGCACTGATTAGACAAGCAGCATCTAAAACAAATGATGTAG

CTGGTGATGGTACAACAACAGCTACAGTACTGGCTTCAGCAATTGTTAAGCAAGGAATGA

GAAATGTTGCAGCAGGTTCGAATCCGATGGCTATTAAAAAAGGTATTGAAAAAGCAACAA

ATTTTGTTGTTAGTAAAATAGCTGAGTATGCTAAGCCAGTAGAAGATACAAAAGCTATTA

TACAAGTTGCTTCTTTGTCATCAGGAAATGATATAGAGGTAGGTAAAATGATAGCGAATG

CTATAGAAAAAGTTGGCAGAGAAGGAGTTATTTCTTTAGAAGAAGGGAAATCAACTAATA

CGATTCTTGAGATCACAGAAGGAATGCAGTTTGAAAAAGGCTTTATTTCTCCTTATTTCG

TTACAGATACAGAACGAATGGAAGTTCTTCAAGAAAATCCTTTTATTTTATTTACAGACA

AAAAAATTACTTTGGTGCAGCAAGAACTTGTGCCATTGCTTGAGCAAATTGCAAAAACAT

CTCGGCCTTTATTAATAATAGCTGAAGACATCGAGAAAGAAGCATTAGCCACAATTGTAG

TTAATAAATTAAGAGGGATTTTGAATGTTGTAGCGGTTAGAGCTCCTGGATTCGGTGATA

GAAGAAAATCTTTACTTGAAGATATGAGTATCTTAACGAACGGACAAGTAATTACTGAAG

ATGCAGGTTTATCACTTGATACAGTTCAATTAGATATGTTAGGAAAAGCCCGAAGAGTTA

TTGTTACTAAAGACTCGACAACAATAATTGCAGATGGTCATGAAATTAAAGTTAAATCAA

GATGTGAGCAAATTAAGCGGCAAATAGAGACGAGTGACTCTTTATATGAAAGAGAAAAAT

TGCAAGAACGATTAGCAAAGCTTTCTGGAGGAGTTGCCGTCATTAAAGTTGGTGCAGCTA

CAGAAACAGAGATGAAAGATAAAAAACTAAGACTAGAAGATGCAATTAATGCAACAAAAG

CAGCAATTGAAGAAGGTATTGTACCAGGAGGAGGAGCTACTAATGTCCATATCTCTAGTG

AATTATTTACATGGGCTAAAAACAATTTAGTTGAAGATGAATTAATTGGTGCTTTAATAG

TTGAACGAGCTGTGACCTATCCGTTAAGACGAATAGCTTTTAATGCAGGTGATAATGGAG

CAGTAATAGTGGAAAAAGTTAAAAGTCACGATTTTCATATAGGCTATGATGCAGCAAACG

GTAATATTGTAAATATGTACGATAGAGGTATTATAGATCCAGCGAAAGTAGCTAGATCAG

CTTTGCAAAACGCAGCTTCTATTGCAGCAATGGTTTTAACTACTGAATGTATTGTAGTTG

ATAAAGTCGATGATTGAGCTATAAGATAATAGATAATCTCAAACAAAGAATAATTTCTAA

TTATAGAAATTATTCTTTGTTTCAATTAATCTAATTAATTGATCAAATGTATAATTTTTA

TAATAAAGCCAAAGTTTAAACACGTCTTCTTTTTCAACTATTAAGTAGAGAAGATATAAG

CCCGTAATACCTAAATTATCAATTAAAATATTATTTTCATATATAGATTTTTCTAAAGAT

CTAAAAAATGACATTTTTGAAAAATAATACCGAAATCTTCTCTTATAGTTATTTGCTAAA

TCATTACTGTCTAAAGATTGATTTCCAATTGAATTCAGAACCTTTAACAGAATACTTTTT

AAATCTTTATCTGACTTTATTTGAGAATCAATAATAATTTGATTCAATTGTATGTCAAAA

CTAAGATTAAAAATATTACTTGTTAAAGATATTTGATTTTGAATATCTAGAGATTTTATA

GCCAAGAAGATTAAACTTTTATTGTTCATTGTAAAATATAAGACAATTGATGATGTATTC

TTTAAATACTAAATTTATCTTCTTGTGCTTCTATCATTTTTTGTAAAATAATTAATGTTT

TAATTTTTTAGTTTTAGTTACTACCCGAAAAAATAAATTCTGGAAATCTAGCTTGAGCTT

TAGCTAGAGAGTGTTTTTTCCCTTCTTCTTGCCAGTAATTGATAACTTCTGTTGTTTTGT

TTAATAACTCAATTCTGTCTCTTTCAGCAATCCACGGCTTGGATTCTAGTTCTGCTTTTA

ATTCTTCCCAAGCATCATTCCTTGGCCAGAAAAAATAAGAAGTAAGTGGACTAGTGCCAT

TTCCAACAATCTGATCAATAGCAATAGCAATATTATTTTCTAACCAAAGTACTTTTAGTG

TAAATTTTGACAAGTTTTTATCTCCATTATTAATATTAAGTATTGGTACAGTAGAATACT

GTACTAGTTATTTTTTTCAGCAACAATTTGAGCTTTGTTTAAAATGTTTTTTACCGTTTT

AGTAGCTTGTGCACCTTGAGATAATCGTTGCAAAATTTTTGAAATGTTAACACGAGTTTC

ATTATTTATAGGATTATAAAATCCTAATTCTTCTATAGCTTTACCATCGCGTTTATTTCT

GCTATCCATTACAACAATTCTATAGCTTGGTTGTTTTTTTCTTCCGTATCGCTTTAACCT

TAACTTGACCATATTAGTCACCTCGCTTATTTTTTATATCTTTATTAAATTATATCTTAC

GACATTATGCAATAATTCAAACAATTAAGCTGATTCAAGTCTAACATTATTAAATATTAC

CATTAAACATTGTAGAAAAATAATACCTAACATAGGTGACATGTCCATTCCAAACATGGG

AGGAATGCTTCCTCTGAATAGTTTTAAGTATGGATCAGTAATTCTATTTAATGAACAGAA

TGGTTCATTGTACCAATTTACAGTTGGAAACCATGCCAATGATAATTTAAGTAAAATTAA

AATTAAATAAATCTCAGAAAAATTAGCTATAGATCCAAGCAATAAATTTAATGTACCAGG

AAGAGTGTTCATACGTATTTTATAATTATTATATACTTCGTTATACTAATAGTTTGATTA

TATATTATAATCGATGTAAATAAATCAGCTAAACAAAAAACTTATCCTTAAAATCATCTT

CTCTTGAGCATTTTTTTTTATCTACATTAGTATTTACTTTTGTCGTATAAATTTCCAGTT

TAGACCAAATATGTCCTAACTGATTTAATTGAGATGTTCCACATTCTAAACAGGCTATTT

TTATATTGCTCAAAACAGCACCTTTTTTTATAAAATCGTTCAATAATGGCGTAATAGTTG

CTTCATCCAAAAATAAATCTAAGACAAGTATTTTTGTAAATGAATTTAATTGATCTGGGA

CATTGTTAATAATATCATTATAATTAACTAAAACTATATTAGCTGTAGGAAGTAAAGATC

GTGCTCCTTCCGCTAAGATAAAGCCATAAGGCATAATAATTACAATAATATATTTATAAT

GACTGCTAATAAGACTTATAGTGGTATTAGCTTTTAATTCGATTGTTTCTGTTACTAGCC

ACTCTCTCATTATTTCGTAAGTAATCCATTTGCCTAGTTCTGAGCAAGCAGTTCTTAAAA

TCGTACCTGGATTATTGTCGTTTTCGAGAATCCCTGACCAATGTTGTATTAATGGATGGG

ATGCTATATTAATTTGGAGTTGCATGTTTTGTTAAATATATTTAACATTTAATTTGTTGA

TTGAATCTTCAATTCCTATTTGTTGAGTATTATTAAAATAGCATATATATTAAAGAAAAT

AATTATAAAAAATGAAAAAAAGTCTTTGGCTGTGGGGTTTTACAGATAGTGCTGAAACTT

GGAATGGTAGATTTGCAATGATTGGTTTTATATCAGTTATTTTTATCGAAGTAGTTACAG

GACAAGGATTATTATATTTAATAGGTATGATGTCATAATAAAAATCCAAAAAAGTTCTTG

TAATTTTTTATAAGAACTTTTTTGGATTTTTATAAATCTAATCTAGCGGTCTCATAGATA

GAACAGGTTCATTCTCTCTATTGATACCTTTTTCAAATCCAGCAGCAGCAGCTCTTGCTC

TGCCTGCATGCCAAAGATGTCCAATAAATAGGAAGAATCCTAGGAAGAAATGAGATGTTG

TTAACCAAGATCTAGGAGATACATAGTTAACTGAATTAATTTCTGTAGCTACACCACCAA

CGGAATTCAAAGAACCTAGTGGTGCGTGAGTCATATATTCTGCGGCACGCCTTTCCTGCC

ACGGTTGAATATCGTTCTTGATTTTATTTAAATCAAGACCATTTGGTCCTCTAAGAGGCT

CAACCCAAGGAGCTCGCAAGTCCCAAAATCTCATGGTTTCACCACCAAAAATAATTTCTC

CACTAGGAGATCTCATTAGGTATTTACCTAATCCAGTAGGTCCTTGAGAAGATGCAACAT

TCGCACCTAGTCTTTGATCTCTAACTAAGAAAGTAAAAGCTTGAGCTTGTGAAGCTTCAG

GACCAGTTGGTCCATAGAATTCACTAGGATAAGCTGTGTTGTTATACCAAACAAAATTAG

AAGCCGTAAGCCCCATGATTGATAAAGCACCTAGACTATAAGATAAGTAAGCCTCACCAG

ACCATACAAAAGCTCTTCTAGCCCAAGCAAAAGGTTTTGTTAGTATATGCCAGATTCCGC

CAGCAATACAGATAATGCCAATCCAGACATGACCACCTACAAGATCTTCCATGTTATTTA

CACTTACGATCCATCCATCACCGCCAAACGGAGACTTTAAGACATAACCAAAGATAACTA

GGGGATTAAGTGTAGGATTGCTAACAAATCTAACATCTCCACCACCTGGAGCCCAAGTGT

CGTATACCCCACCAATAAATAAAGCTTTAATAACTAACAGAAAAGCGCCAATGCCTAATA

ACACTAAGTGTATACCAAGTATTGTTGTCATTTTGTTTTTATCCCGCCAATCGTATCCGA

AGAAAGGAAATGATTCTTCAAGAGTATCAGGGCCAATCAGAGAATGATAAAGACCTCCAA

AGCCAAGAACAGCTGAAGAAATTAAATGCACTACGCCTACTACAAAGTATGGGTATGTAT

TGAAAATCTCGCCGCCTGGACCAACACCCCAGCCTAGTGTAGCTAAGTGCGGAATTAGAA

TAAACCCTTGTTCGTACAAAGGCTTCTCGGGAACAAAGTGTGCTACCTCAAAAAGAGTCA

TGGCACCAGTCCAAAAGACCATTATACCTGCATGAGCAACATGAGCACCAAGCAATTTGC

CAGAAACGTTAATTAAGCGTGCATTGCCAGACCACCAGGCAAACCCGGTAGACTCAATGT

CTCTACCGCCAACACCAACAGTTGTATTAAAGGGCGTTTCCACGTGGTAAAACCTCCTCA

GGGAATATAAAGTTTTCATGAGGTTGATCTTGAGCAGCCATCCAAGAGCGAATACCTTCG

TTCAATAGGATATTTTTAGTATAAAAAGTTTCAAACTCAGGATCTTCCGCAGCTCTCAAC

TCTTGAGAAACAAAATCGTAAGCTCTTAAATTTAGAGCTAGTCCAACAATTCCAAATGCG

CTTGTCCATAATCCAGTTACTGGTACAAATAACATGAAAAAATGTAGCCAACGTTTATTA

GAGAAAGCAACACCAAAAATCTGTGACCAGAATCTATTAGCTGTTACCATCGAATAAGTT

TCTTCAGATTGTGTTGGAGTAAATGCACGAAAAGTATCTGCAGCATCACCATCTTCGAAT

AAGGTATTCTGTACAGTTGCACCGTGAATAGCACATAGTAAAGCACCACCTAAGATTCCT

GCAACGCCCATCATATGGAACGGATTCAGAGTCCAGTTATGAAATCCTTGCAAAAATAGC

AAAAATCTAAAAATTGCAGCAACTCCAAGGCTTGGGGCAAAGAACCAGCTCGCTTGGCCT

AGAGGGTACATTAGAAAGACGGAAACGAATACTGCAATTGGTCCAGAAAATGCAATAGCG

TTGTATGGTCTAAGACCAACTAGCCTAGCAATTTCAAATTGTCTTAAACAGAATCCAATT

AGTCCAAATGATCCATGTAAAGCTATAAATGCCCACAGACCACCAATTTGGCACCAGCGA

GTAAAATCTCCTTGAGCTTCAGGTCCCCAAAGGAAAAGAAGTGAATGTCCCATGCTGTTT

GCTGGGGTAGAAACAGCCGCAGTTAAAAAATTGCATCCTTCTAGATATGAACTGGCTAGC

CCATGAGTATACCAAGAAGTAACAAAAGTAGTCCCAGTTAGCCAGCCACCTACAGCAAGG

TAAGCGCAAGGAAACAGAAGTAGTCCAGACCAACCTACGAATACAAATCGGTCTCTTTTT
AGCCAATCGTCTACAAGATCAAAACCACCACGAGTTTTTTCTTGTCCAATTGCTATGGTC
ATAATTTTAATCTCCAGAGCAAATTTAATAAGTAATTTGTTTAAATAAACACTTTAGATA
TACTCTAAAATTTTACTTTTCTTTACGGTCAATATAATTATAAGTCAAAATTAACCAAAA
TTCACGCTTTTAATAACATTTTTTTAGCTCATTAAGTTAATTTTTTAATATATATAAACAGT
AATAGAGTTATTGTTTAACTTACAATACTCATAATTTATTCATTTCTATAATCCTTTGAA
AAAGATATCCTGTTCCTCGTGCTGTTAAAATCAAGTCGGGATTACTTGGATCATCTTCTA
GTTTAGCTCTCAATCGTGAAATATGAACATCAACAACTCTCGTGTCAACATGCCGTTCGG
GAGTATAGCCCCAAACTTCTTGCAAGATAGAAGCTCTAGAAAAAGGTTCTCCAGCCTTAC
TAACTAAAAGCTCGAGAAGACTGAATTCCATACCTGTTAATCGAACTCTTTCATTATTTT
TATAAACTTGTCTTTTGTTAGTATCTATTTTTAGAAATCCAATACTAATAATCCCAGAGT
TTGGGACTCCAAGACTAGTGGTGATTTTATCAGCTCTTCGAAGAACAGAACGAATGCGAG
CTTCTAGCTCCTTAGGAGAAAATGGTTTAACGACATAGTCATCAGCACCTATTTCTAATC
CGGTAATTCTATCGCAGACTTCACCTAACGCTGTTAGCATTATGATAGGAACATCTGATT
CTTTTCGAAGTTCTTGACAAACACCGTAACCATCTAGTTTAGGCATCATAACATCTAAAA
CAACAAGGCTAGGATACTCTTTTCTAAACACGAGTAAAGCTTCTTCTCCATTTGAAGCAG
TGATAACTTCATAACCTATTATTGTTAGTCTTGTTTCTAAAATTCTTCTTATGCTGGCCT
CATCGTCAACAACGAGAATCTTTTCTTTCTGGTTCTCCAATTTCTTATTTACTCCTATAA
CTCACCGATCATGATTACTATAACCTTAAATTTTTAGCGCTATGAGTATCTTTATTCACT
TTAGAATTTAGTTATTATAAATTTTGCAAGTCTAGTTTACATTAATTAGTACCAGTCAAT
ATTTTTGTTTTAAGTTATGACTATTTTAATCCATAAGTTGACTCTAGTTTGCAGAATTCA
ATATATACATATTTTCTAGCCAAGTAAATTGTATTCTATTTTCTAAATAAAAAAGGGGTT
GACAAACCAGTTGATAGAAAGTATTCTACATTACTATAAGTTAATTCCAACAAAGAAGTT
TTAAATCTTCTTTCGGAGTAATGAATACTTTAAT
>SRR9587921
TATATGATAAATTAAGACGTCTGTTATGTGAGAATTTCTAAAATATACAAAACTTTTTAT
TTAATATAGAAAAAGTAAATTAAAATAAGTTGAACATACTTTGCTCTAATCTAAACCTCT
TAATTGAGATTCTAATTAGAGACAAAAGTATTTACTTATTTTGTATCTATAGGAATAAAA
ATGAAACTTGCAGTTTATGGTAAAGGTGGTATAGGTAAATCTACAACTAGTTGCAATATT
TCTGTAGCACTTTCAAAAAGAGGTAAAAAAGTTCTACAAATTGGCTGCGACCCTAAACAT
GATAGTACATTTACGCTAACAGGTTTTTTAATCCCAACAATTATCGATACTCTTCAATCC
AAGGATTACCACTATGAGGACGTTTGGCCTGAAGATGTAATCTATAAGGGATATGGCGGT
GTAGACTGTGTTGAAGCTGGTGGACCGCCGGCTGGCGCTGGTTGCGGAGGCTACGTTGTA
GGTGAAACAGTTAAACTTTTAAAAGAATTAAATGCTTTTGATGAATACGATATTATCTTA
TTTGATGTTTTAGGTGATGTTGTATGTGGAGGTTTTGCAGCTCCATTAAATTATGCAGAC
TACTGCTTAATCATTACAGACAATGGATTTGATGCTTTATTTGCAGCTAATAGAATAGCA

GCTTCAGTACGAGAAAAAGCTAGAACGCACTCTCTGAGATTAGCTGGACTTGTTGGTAAT
AGAACAGATAAAAGAGATCTAATTGATAAATATATAGATTGTGTTCCAATGCCAGTATTA
GAAGTCTTGCCCTTGATTGAAGATATTAGAGTGTCCAGAGTAAAAGGTAAAACTTTATTT
GAAATGGCAGAAATTGATAAGGATTTAGCATATGTATGCGATTACTATTTGAATATTGCA
GATCAGTTGATTACAAGGCCAGAAGGTGTAGTTCCTAAAGAATCTCCGGATAGAGAATTA
TTTAGTCTTTTATCTGATTTTTACTTAAATCCTAAATCAAAGGTAGGACAAGAAAAAGTA
GATCAAGAAGAATTAGATTTGATGATAGTGTAAATAATATTTTCAGCATAATATAACAAG
GAATAAGATAATGTCTACAGCTCAATCAGATGCTCTTACTTTTGAATGTGAAACAGGTAA
TTATCATACTTTTTGTCCCATCAGTTGCGTTTCTTGGTTATATCAAAAAATTGAAGACAG
TTTTTTCTTAGTTATAGGGACTAAAACTTGTGGTTATTTCTTGCAAAATGCAATGGGAGT
AATGATTTTTGCTGAACCGAGATATGCCATGGCAGAATTAGAGGAAGGAGATATTTCAGC
TAAGCTAAATGATTATGGTGAACTCCGTAGATTATGCTTACAAATAAAAAAAGATAGAAA
TCCTAGTGTTATATTTTGGATTGGAACATGTACAACCGAAATTATAAAAATGGATCTGGA
AGGAATTGCTCCAAAATTGGAAGCAGAAATTCGTGTTCCTATAGTAGTTGCTAGAGCTAA
TGGTTTAGATTATGCCTTTACCCAAGGTGAAGATACAGTTCTAGCAGCTATGGCTCAAAG
GTGTCCATTAAATTTAAAGAATCAATCAGATAACACGAGTCTTAAAACTTCTCCTCATAT
TCCTTTAGTTTTATTTGGATCTCTTCCGGATCCAGTTGTAACCCAGTTAACTATGGAATT
AAAAAAACAAGGTATTTTTGTTTCTGGTTGGTTGCCATCAAAGAGGTATACTGAATTACC
GGTTATAAAAGAAGGATATTATGTTGCGGGAGTTAATCCATTTCTTAGTCGCACAGCTAC
TACACTAATGAGACGCCGAAAAACAAAACTAATTGGTGCACCATTTCCAATAGGTCCAGA
TGGTACTCGAGCCTGGATTGAAAAAATCTGTTCAGTAATGAATGTAAAGCCTATTGGATT
AGAAGATAGAGAAAAAGCAATTTGGGCTTCTTTAGAAGACTATATTTCTTTAATCCGAGG
TAAATCAGTTTTTTTTATGGGTGATAATTTACTAGAAGTATCTCTTGCAAGATTTTTAAC
TAGATGTGGAATGACTGTATATGAAATTGGTATTCCTTATATGGATAAACGCTATCAAGC
AGCAGAGCTAGCTTTATTAAAAGCTACATGTGATAAAATGAACGTTATGATGCCAACAAT
TGTAGAAAAACCAGATAATTATAATCAAGTAGACCGAATCCGTGATCTGAAACCCGATTT
AGTTATCACTGGTATGGCTCATGCAAACCCTTTAGAAGCCAGAGGTATTAATACAAAATG
GTCAGTAGAATTCACATTTGCACAAATTCATGGCTTTACCAATGCAAGAGATATTCTTGA
ACTCGTCACAAGACCATTACGGCGAAATCTTAGTTTGTCAGAATTAGGCTGGGATGTTTA
TAGCAAGCAAAGCTAGTATTCCTC----TTTATGACTTACATGGTATAAAAGCCTTGTAA
GTCTTTTATTTTATGTTAAAGAAGCATTTCTTACTCTGCCATCACCAGCCCAAGCTTGTA
AACTTTCTACTTGTTCTTTGTCTGTAAATGCTAAAGGCACAAACTGTTTGATAGCTATTT
TAATATCTTCAGTACTAAATTCTCTCTCTTCACTAAACGCCGTATGCATACTTTCAATAA
TTGCTTGCTCAATTTCAGCGCCAGAAAATTTATTACATAATAAACTCAATTCATGAATAT
CATATTCTTGCCACGATCTTGGTCTAACTTTTGATAAATGTATCTTAAAAAATTGACTCTC
TCTCTTGACGATTAGGTAAATCTAAGAAAAAAAATTTCATCGAACCTGCCTTTCCTTAACA

TTTCGGAAGGTAAACTTTGAATTTTATTAGCTGTTGCGACAACAAATACTGGAGCTTTTT
TCTCTGATAGCCAGGTAATAAATGTACCAAATACTCTGGCACTAGTACCACTATCACCTT
GACTATGTAAACCAGAAAAAGCTTTATCTATTTCATCAATCCATAATATGCAAGGCGATA
ATCCTTCAGAGATATTTATCATTTCTCTCATTTTTGATTCCGATTCACCAACTAGTCCAC
CAAACAGTTTTCCCATGTCAAGACGTAGCAGAGGCAGCATCCAATCGCTTGCAATAGCTT
TCGCGGTTAAGGATTTACCCGTCCCCTGTATACCAACTAGCAATAAACCTTTGGGCGAAG
GAATACCGTAATTTAAACTTTGCTTCGAAAATGAGCGAGACCTTTTTTGTAGCCATTGTT
TTAATACATCTAAGCCACCAATATCCCTATTTACTTTACTATAAGGATAAAATTCTAATA
GATGCGTTTGATTAATAATTTGTCTTTTTTCTTCAATAATAATAGGTAAGCTCCGAGAGT
CTATCTGATTATATTGTGCAATAATTTTAGTAATCACTTTGCGGATTCTATCTATCGATA
ATCCTTGGCAGGATTTTGTTATATTATTAACGAGCTCTGAATTCAAATTAAGATTCAAAG
CTTTACTCAATCTTATGATCTCTTTTTTGATTTCTAGAAGACTAGGTAAGGGCAAATCTA
TTACCGTTATGATATCATTCAGAGCAAAAGGTATATTGATTTTGCAAGAAATAATAATAA
TGTGCTTAGATTGCGTTTTGATTATTTTAGCTAAATTGCGAAGTTTCCTAATTAATACTA
TTTCGTTTAGGAAAGAGTCAAAATCTTTTAAAAGAAAAAGATTTAAAGATTCATTATTCA
AGTTTTCAATAAATTCTAATGCTAATAAAGGATTTCTTTTTGCATAACCATTATCGCTAG
GATTGTTAGTGTATCCATCTACAAAGTCCCAGGAATATACTTGTTGACTGTTTGAACAGT
TTAAGCTGTGTTTTATAATATACTCTAGTCTATCTTCTTCTCTAGTATTTATTACAATAA
TAGGATATCGAGATTTTAAAAGTAATCGTAAATCTTGAGTGAAATTCATGTCAATATTTG
TTTCCATCAGACTTTTGTACTTCACCAATTGAAGAAAGACTAAATAAGATTATCTAATAT
TCCTTGGATATATTCAAAAGTCATCTCTCTATTTTCTAAATTATTGCTATTACTAATTTA
ACTTACCATGATTTTTTTTCTTCCTTTTCTTCGTCGACTATTTAATATAGAGCGTCCACA
TGGTGTTTTCATTCTTGCTCTGAAACCAGAAACTCTAATTTTTTTTCTTTTAGATCCTTG
TAGTGTTCTTTTAGTCATGATGTTTCCTCTGTTGTAAAAGATTAAATACTATTTTATCAT
CTTTTATATATCTTTATGATATAGATATGTATTAATATCAAAGTTAACAGCAATAATTTA
TATTAATCAAGAATTATTTATATAATATTAAACATTAAATATTAACTTTATGATATTGGC
TTTGCCCATTTTTTATTTAAGTATTTTGACCATCTTTTTGCTAATCTTAAACTGGTTAAT
TTTTCAGCAGCTAAAAACGATTTTGTTATTAGAATCTCAATTTAAGTATTTTGTTGATAA
AAGCCAAAATCGTAAATTAGAACCCGAAGAAAGTTTTGCTTTTGCAAAAGTATGTGTAGC
TAAAAAATATTTTTCTAAAGCTATTATTGAAGGTCAATTAGCTTTAAAAAACTATCGAGA
CCTGAATATCTTAGATAATAATATAGTAATTGCTAATTTGTATAATATGCTAGGTTTTAT
TTATTTTGAAGCAGGCCAAACAAGTTTTGCAAAGAACTTTTATGAGCAAGCCTTACAAAT
AAACCCTAATTATGTTGTAGCTTTAAATAACCTGGCAAAAATTTACGAAGAGGTAAAAGA
TTTAAAAAAAGCCGAATCTCTATACGATAAAGTGTTAACTCTTAACTTAAATAATAAAAC
TGCTAATAGAAGGAAAGATTTTATAGCAAAAACTAAAAACATCTAATACTATTTGTAATC
GGGATAGCAGGATTTGAACCTGCGACATCCTGCTCCCAAAGCAGGCGCGCTACCAAACTG

CGCTATATCCCGTAAGACATAATTATTACTATATACCTTTTCTCTAATTATGTCTACTCT
TTCTGTATTTTATCTAGGATACCAAAACATGCCTTTAACTCCATCTGGATCTTTGATAAA
TCCTAATTTTTTATAAAAACTAACAACATCAGGTTCTGCGAATAGTGTAATAGTGCTAAT
TTCTGCCTGTCTCAACTGCTGTATTAATTGATGTATGATTACTTTTCCTAAACCTAAACC
TTGGAAATCTGGATGAATAACTACGTCCCAGATAGTTGCATTAAATCCGTTGTCTGATGT
TGCTCTAGCAAATCCTACAAGTTTACTATTTGCATCTTTTTTTTGTATTAAAGAAATAAT
AATAGAACTATTTTTTAATGCGATCTTTACTTTTTTTAGAGGTCTTTTAACCCATCCGAC
TGAATCACAAAGCTGCTCCAATTCGTATAAATTAATATTCTTATTGCTACTCAGATAGAT
ATCTTTAAATTCGATTTTATCGCAAGTTTTATCTAAAACAATAAGTTTTTTGAAATTTTT
TTCAGAGCTAGCATTGATATCGGAGTTTTGAAAAAAATTTTTCCAGAAGATCATAATGAT
TATGATATAATTAAATAAATATATATATTACAATATTGCTTTAACAAACAATATATTAAT
ATATGCTCAAATTTTTACATGAATCTATTAAGATAGAAGATTGTTAAAATATAGAGAAAT
AAAAATTTTTGTAAAATCTTTCAATAAAATGATATAAAGAGTGCTTTTTAAGACTAACAT
CTATTATTATAAATAAAGTCTCGAAATTGAGATGAAATAACACATTATTCTATGTCTAAT
GCCTTTTAAAGGAGTTATTTGTGAAAAAATCTATGTGTCTAACCTGTTTACTTGCTTTAC
TAATTATGAGCAACCCAATAATAGCTAACGCAGAAGTAGCTGGATTGGTGCCCTGCAAAG
ATTCTGCGGCATTCAATAAGCGTATGGTAAATAGTGTGAAAAAACTTCAAGCCAGATTAG
CTAAATATGATGCAGACACGCCACCAGCGTTAGCTTTAAATAAGCAAATAGAGAAAACTA
AAACTAGATTTGCAACTTATGGTCGAGCAGGTTTGCTGTGTGGTACTGATGGATTGCCAC
ATTTAATTTCTGATGGTCGATGGAGTAGAGCCGGGGACTTTGTTTTTCCAGGACTATTAT
TCTTGTATATTACAGGATGGATTGGCTGGGTAGGTAGAGGATATCTTTTATCTGTTGCTA
AGACTAGTAAGCCAACAGAAAAGGAAATCATTTTAGATGTGCCATTAGCTATCAAATTTA
TGTCATCCGGATTTGCATGGCCGCTAGCGGCTTGGCAAGAATTTAGTAGTGGACAATTAA
TTGCTTCCAATGACGATATTACTGTTTCACCCCGTTAGTAAAAAATTTATATGAATAATA
ATTTTACCAAATACTTATCAACAGCACCTGTAATTGGTGTATTGTGGATGACGTTTACAG
CCGGATTTATTATAGAATTGAACCGCTTTTTCCCAGATGTCTTATACTTTTACTTATAAA
TAAGATCGTATAAAAAACTACATAAATATAGAGAAATACTATTATTTGTGTAGTTTTTAA
TATATAACATAATTATTTATTTTTAAAAATGTTTATCTTTTCGATTATTTTAATATTAGA
TCTTACATAATAAAATAATATCAATATTATGAGCTTAGTAACCCAAATTATAGTTAATGC
CGATGATGAATTAAGATATCCTACAATTGGAGAATTGCAGTCAATTCAAGACTACTTAAC
TACAGGAAGCAATAGAATTAGAATTGCTACTATTATTAGAGATAAAGAAAAGGAGATTAT
ACAGAAAGCTAGTAAGCAAATTTTTCAGTTACATCCAGAATATATAGCTCCAGGTGGTAA
TGCAGCAGGTTCTAGGAAAAGATCGTTATGCTTACGTGATTATGGTTGGTATCTCAGACT
AATTACATACGGAGTATTAGCTGGGGATAAAGATTCTATAGAAACAATTGGTATTATAGG
AGTAAGAGAAATGTACAATTCTTTAGGTGTACCTATAATTGGAATGTTAGATGCAATCCA
GTGTTTAAAGGAAGCATCTTTAGAAATGCTTGGTCAAGATGATATTAGAATCATTTCTCC

TTATTTTGATTATATAATTCGTGGAATGTCATAAATCATAGCTTCAATTAATTAGTTGAA
TAAAAGCTTTTATAATGTTATAATTACAATAAGCCCGAAAGCATATAATTGTAAAAACTA
AATCTTGTCAGAACCGGAAGGTAGCAGCAATAATGTTTCAATACAAAAGTTATGATTTTC
GGGTGTTTATTGTTTTTATCTAATAGTCTCGACTTTTAATTAGTATGCTATGTATAAATC
TGTATATGACATGTAATCCATCAAACCTTCCAGCAAAAATAATTTAATAATTTGATATTA
ATACGTAAAACACCTATACTCTTTTATAAAATTTCTTTATATCTCATCTAAACAATGGAA
ACTTAACTTAATGGGTGTTCATATTTTATCAACAGGTTCGTCTGTCCCAAATTTTTCTGT
AGAGAATCAACAATTTGAAGATATGATCGAAACTTCCGACCATTGGATTTCAACAAGAAC
AGGAATAAAAAAAAGACATCTAGCTCCATCTTCTACTTCTTTAACTAAATTAGCGGCAGA
AGCTGCAAACAAAGCCTTATATGCAGCTAACTTAAAACCTACTGAGATCAGTTTAATTAT
TTTAGCTACGTCTACGCCTGATGATTTATTTGGTAGTGCTAGCCAGTTGCAAGCAGAAAT
AGGTGCAACAACATCGGTAGCTTTTGATATTACGGCTGCCTGCTCCGGTTTTATTGTTGC
TTTAGTAACAGCAGCTCAGTTCATTCAAACTGGTTCCTATGACAATATTTTAGTTGTTGG
AGCAGACACAATGTCTAGATGGATTAATTGGTCAGATAGAACTACCTGTATTTTATTTGG
TGATGGTGCTGGAGCAGTAGTATTGGGGCAAAGCCTCAAAAATAGTATTTTAGGCTTTAA
GTTATGTACAGATGGTCAGCTAAACAGTCATTTACAATTAATGAATAAACCTGTAAATAA
TCAAAAATTTGGTGTTACAGAAATTCCTCATGGAAACTATAATTCCATAACAATGAATGG
CAAGGAAGTGTACAAGTTCGCTGTATTTCAAGTTCCAACAGTAATTAGACAATGTTTGAA
TAATTTAAACATTTCAATAGATGAAGTTGATTGGTTTATATTGCATCAAGCAAACACTAG
AATCATAGAAGCAATTGCGAGCAGATTATCAGTACCTTTTTCTAAAATGATTACGAACTT
AGAGCATTATGGAAATACATCTGCAGCGTCAATCCCTTTAGCGTTAGATGAAGCTATTCA
ATCCAATAAAATTCAACCAGGCCAAATTATTGTTTTATCTGGTTTTGGAGCAGGCTTGAC
TTGGGGAGCAATTGTCTTGAAGTGGTGATTTATATATTGCGGATGACGAGACTCGAACTC
GTAAAGCTTTCGCTACACACCCCTCAAGCGTGCGTGTATACCAATTTCACCACATCCGCA
TTTTTTACCATGAATATTAAAAATTTATAATATTTATCTAAATATTCATATAATGTGTAT
ATATATATACATTCTTGTCAGAAACTCTCTTATATTTTATTTTAAGTGATAAAATTGAGT
TTAGTATAGATAGTAAAATTTTATTTTAAAATCATAAAAATAAATATATAAGGAAAATAA
AATATGACACCATCTTTATCAAGTTTTTTGAATAGTCTTATTCTTGGGGCAGTAATTGTA
GTCGTTCCCATAACTTTGGCTCTTTTATTTGTTAGTCAAAAAGACAGGACAATCCGGTCC
TAAAAATAACTTAGAAAGATATGGAGAAGCAATTTAAAATGAATAATAAAATATTTTTCA
TACTAATAAGCTCTCCATATATTGCTATACTTTAGAAGAATTTTTATTTTTTTATTTTGT
AACATCCTGTAAAGTTTATTCCGTTCTTTCTTTTTCCCATTTACTCTCTTATAATTAATG
ATACAAATGTCAGGTTAATAATAGTTTTTTTATTCTGTATCTTTCATTTTATATACGTAT
TATTATGTTCAATTAATAAAGAAAAGCTCATGTCTTTGTCTAATTGGCCTCTCAAAAAAG
AAAATTCTGAAGCATATAATATTAAGAATTCAAAACAAATCACAATTCCTGATGGTTTAT
GGATAAAATGCTTCGACTGTGGTCTATTAATGTATTCTAAAGTACTGAAGAGAAATTTAA

AAGTTTGCCCTCAATGTAGTTATCATTTTCAAGCTTCTAGTAACGAAAGAATTGATCAAT
TGATAGACCAAGGTAGTTGGCAACCAATGGATGTTCACTTGATCTCTACAGATCCATTAG
GCTTTAAAGACCAAAAGCTTTATAGTCAAAGGTTAAAAGATACTGCTTTCAAGACTGGCC
TGCAAGACGCAGTTCAAACAGGTACTGGGACTATGCAAGGTAAAAAAGTATGCTTAGGTA
TTATGGATTTTAGATTCATGGGGGGAAGCATGGGATCTGTTGTAGGCGAAAAACTAACAA
GACTGCTAGAAAAAGCAACTCAAGAAAAGTTGCCTGCAATTATACTTTGTGCATCAGGCG
GAGCTAGAATGCAAGAAGGTATGTTGAGCTTAATGCAAATGGCAAAAATTTCTTCTGCTC
TAGAAATGCATAAAAAAGAAAATCTACTATATATATCTGTTTTAACCTCTCCCACAACAG
GGGGTGTAACAGCTAGTTTTGCTATGCTTGGAGACTTAATTATTGCAGAGCCAAAAGCTC
TTATTGCATTTGCTGGTAGACGAGTTATAGAACAAACAATCAAAGAAGACTTGCCAGATA
ATTTTCAAAGTTCAGAATATTTATTTGAACATGGTTTCCTAGATTTAATTGTATCCAGAA
CTCAGCTTAGATCAAAGTTAATACAAATTTTGTCCTTACATAATCATAGTAAGTAATGAA
TTGAATATTGACTATAATATCAAAATGCTAGTAAAAATTAAGATATTAAGAAATTTTCAC
TAAAGATATTTGTAGACTAACTAGAATTACAATTACTATTGTAGAAAAATTTGTCTTAAT
GCCATTTTTTCAGTCTTTTAACAAATTGTTTTTATGACTTTAAATAATTGTTAGTTTAAT
ACAAATTTATAAAGAGAATACATACTAAGCTAAATATAGTAATATTATGTTTTCCTATAA
TACTGACCTATCAAATTTCGAGGAAATTTATGCTTAAAAGATCTTCTTGGCTTGCGGCTT
TATTGGGACTATTAACAGTAGTTTCTACAAGTACGCATACATATGCCATAGAGTTAGACG
AGGCAACAAGAACTGTTCCATTAGAATCTTCTGGCAGAACTGTAATTCTTACACCAGAAC
AAGTTAAAAGAGGCAAGCGATTATTTAATAATTCTTGTGCTATTTGCCATAATGGTGGTA
TCACGAAAACAAATCCGAATATTGGACTTGACCCAGAATCTTTAGGATTAGCTACGCCAC
AAAGAGATACCATTGAAGGACTAGTTGACTATATGAAAGATCCGACTAGTTATGATGGTG
CAGAGTCAATCGCAGAATTGCATCCAAGTATTAAAAGTGCTGAAATTTTTCCTAAAATGC
GCAATCTAACCGATGAAGACCTATTTACAATCGCAGGTCATATCTTACTTCAACCTAAAA
TTGTTTCTGAAAAGTGGGGCGGAGGAAAAATTTACTATTAGAACTTCAAAGACCTAAACC
TTGTGAATACTAGTTTATGTATTTGTTGCTTAAGTGTGATTATTTTGTTAGATCATGATA
TATATTAAAGATGGAGTTTGTTGACAGCAAGTTACTTTTTACCCGAACTTGTTTACACTA
TTCGTCTAGTTATTAAAAAGGAGACGTTAAATTGAAGAAGAAGCTTTCAGTTCTTTTCAC
TGTTTTTAGTTTTTTTGTAATAGGTTTCGCACAAATTGCTTTTGCTGCAGATCTAGATAA
TGGAGAAAAAGTTTTTTCTGCTAATTGTGCAGCATGTCATGCTGGCGGTAATAACGCCAT
TATGCCAGATAAAACCTTAAAAAAAGATGTACTTGAAGCTAATAGTATGAATACTATTGA
TGCTATTACTTATCAAGTACAAAATGGTAAAAATGCCATGCCTGCTTTCGGAGGTAGACT
GGTTGATGAAGATATTGAAGATGCAGCAAATTATGTATTATCTCAATCTGAAAAAGGTTG
GTAATTATACTTGATTTTATCCTGTATTAAAGAATAGACAATCTATTTAGTTGTTCATTA
GATTGTCTATTCTTTGTTTATGCTATTATATAAAGATATTTACACAATATTTTATGATGA
AAAGAATACCCGCAATTCTTGTACTAGAAGACGGTGCGTATTATAAAGGATGGTCATTCC

AGCAAGATAAACAAGAGATTACTATTGGTGAAGTAGTTTTTAATACTGGAATGACAGGAT
ATCAAGAAATAATCACAGATCCTAGTTACTTCCATCAAATTGTCGCTTTTACCTACCCGG
AAATTGGGAATACAGGTATTAATAATCAAGATATTGAATCTCACAGTATTAGTATTAAAG
GACTTATTGCAAAAAATATTTGTAAAATTTCAAGCAGCTGGAGAGAGCAGCAATCTTTAG
TTACGTATTTAAGTAGTAATAATATTCCTTTTATTTTCGGAATAGATACAAGGTCTTTAA
CCCAATACTTGCGTCAATTTGGTACAATGAACGGTTGTATCTCTACTGATAATTTAAATC
ATAGTTACTTAAAACAGAAAATTTGTGAGATTCCAAGTATGCAAGGTTTAGATTTAATCC
CGCATGTAACTACAAGAAATGTTTACCCCTGGGATGAAAAAAGTTTTCCAAATTGGTATT
TGACAGATAACATTAGAGTGCACCGAGTTATTCAGTTGAAAGTTATTGTTATAGATTTTG
GAGTTAAACTAAATATACTCAGAAGACTAGCTACACTTGGATGTCAGATAACGGTTGTGC
CTGCCCACACTCCTTTAAAAGATATTTTGGCTTACCAGCCTGATGGTATATTACTCTCTA
ATGGTCCAGGAGATCCATCAGCAGTACATTACGGCATCCAGACAGTTACAAAATTACTAG
ATTACAATGTGCCTATATTTGGGATTTGTATGGGGCATCAAATTTTAAATTTAGCTCTTA
AAGCTAAAACTTTCAAACTTAAATTTGGTCATAGAGGTATTAACCATCCATCCGGATTGA
ACCAGCAAGTTGAAATAACTAGTCAAAATCATGGCTTTGCAGTTGAATTGACTTCGGTTT
TTGAATCTCCTGTAAGAGTGACTCATTTTAATCTAAATGACACTACTATTGCAGGAACTG
GACATAATCAAAGTCCTTATTTTTCTGTGCAATATCATCCAGAATCGAGCCCAGGCCCTC
ATGATGCTGATTATCTATTCGAAAATTTTATAGAAATAATGACAAAGTCCAAGAATAAAG
TTAGTTAGTAATTTTCCCATGCTTTATGAGTGAATAAAGCGGATAGTACTTGTACTCCTC
GTAATTGATTAAAGAGAGGCAAATGTCCTCTGGGGCCATGAGAATTCCATTGAAATTCCT
CAGGATATCTGCAAGGAATTCTATCAACTTCCCAGCCGATTTTTTGCCAGAATTTTCCCC
AATCTTTTCCAACACTAAGCCATATTTGTCTTTGAACAAATAGCCCAAATTTACCTTTTG
AGTGTGTATGCCATAACTTATCAATAGTTTGTAAATCTTGAGCAGGTATTTTTTTTATAT
CTGTAAAATATAGCCAATTACGAGTTTGTGCATTTACACCAGCTAGTTGAATAAGCTTTT
GTTGAGTTAGCTGATCTGCTTTGAGCAAATCCCGATGAGTCAGTAACATTTGTAAATCTT
TGTAATTCATCTGTTGAGCAGAGCGCAATGGAACAATGCCATCGGGACATAAGTTAGAAG
CAAATTTCACTATTTCTTGATTTTTACTGTTAAGTAACTTTTCATAAATTAAACCGTCAA
CACAATTACTTTTATAGTTAGGCCCAGTAATCCTCTCAAAAAATAGATCTGCTAAATCTT
TTAACTCGATAGAGTCTTTGCTATTTATATTCTCAATAATTTCAAGTTGTTGCTTGACAT
TATTAGACTTAGTATTCTTATTCAATTCTAAAAGCTGGGCTCGGATTTGATTTGGCATCT
GATTTGTTTAATTAGTTTTAATAGTCATAAGGTCTTAGAGAGAATTAGATTGTTTTTTGA
AATTAGAAGAGCGCCGCAGTATTTCTAAAGAAGTTCGCATAATTTGATTCAATAAGTTTT
GTCCTGCATAAATTATAATTTGACCAATTAAAAAAATTGTGCTTTTTATTTTAGGAACAA
AAAAATCTTGTACCTCAAGCAGAAAAGTAATTATTATTTGCATTGGTGACAACATCTGTA
ATTCATTCGATCTGCAAGCATATATATACTGACAGTTTAAGCCTTCTTGATAAAATCCCC
ATACTTTGTAACGACTTTCGTAAATTGCTCGTGGTCGCTCAAGATAGTTTTTAATTATAG

TATTCCAAATTAGGTTATTCTTTAATTTTTCTAATTTTCTATCAGATAGAAATTTTACAT
TACAGAGATAACTTGCATTTGTCTCATTTACAGTATTAAAATTTTGAACTAACATATGAG
CAATAATGTTGCTTAGCTGAATTAAATAATTTTCAAGAAAAATTTCGACCTGCTTAACTG
GCATATTGTAAGTAAAGAGATTAAAAGTTTTATTCACAGTTTGAGATGACCCAAAAAGTA
GCTGAATAATTAAAATTTGTAGTAGCATCTTATAGTCGTTCAGTAGATAGTTCAAATCAT
GAATATCTTGGTAAATGTCAATACAGTCCAGTTCATATAACTTACAAAATCTTTTGATAC
ATCTATGGAATAAATCAATTAAAATATTTTCGGTCAAGTTTGTAATATCATTTACATCTA
AATTTGAATTATAAATTTCTAAAAGTATCTTTTCCAGTTCAGCTAATATGATTTTTAATA
AGTTTCTTTTTACTTCAGTGCGAAAAACATCTAAAATTAACACTTCCTGAGAGCAATTAG
TTAATCTCTTATTAATCTTTATAGAAGTTCTAACTAATAGCTCTGCTACCTCTTGATTGA
GTGTTGGTCCCTGAGAGCTTGGCCAATAATTATTCACGTTATACCATAGTAAGAAATATT
GAATTTAATGTTATACGATAACTTTTTAATATACTGAAAAGCAAGTTAATTACTCTATAG
CAACTAGTTTTTACAGATCTTAATAGTTATTTATCAAATAAAGTTATATGATAGTAGCGA
TTGAGCTAATTTTATTAAAATTTATAATAATGACAACTTATTACTTCGCCCTTGCGAGTC
AAAATTTTCTATTAGTACAAGAACCACTAGAAGAAGTTTTTAGAGAGAGAGTTAACTACT
ATCAGTCAAATAATAAAGCAATTGATTTTTGGCTAATACCAAACCCTTCTTTTCTAGAGA
AACCAGAAATGATTTCATTTAAAAACCTTGTACCTAAAGACGCTGTAGCTATAATCTCTA
CTAATCCAATATTTATTAATTGGTTAAAGCTAAGAATCGGCTATATCTGTATTGGGCAAT
TTGAAGATAACCTACAACTTTCTGAAGAATCGTTAAATATTACTGTTTTAACAGACAAAA
TTTAACAATGCTTTTATTATGACTAGTGTTAAATTATTTTGGTTAAGTTATAATTATAGT
CTTTTAGTATTTAGACTCTTATAGTTTATCACTTAGTATTTTATGCTGAGATGGAGTGAC
AAATAGTGTAAGTATTTCTTGGCTAAACGTTTCAGTTGCCTTAGATTTATATCTATTTGG
ATTAACTATAATGGATAGCATTCGTTTTATAGTAACATTTTCAATTTGAGCCCAATGTAC
AATTCCAAGTTCCAATTCCTTAGCAATAGCTGAAACGGAAACAAAGGCAGCTCCTAGCCC
TGATTGCACAGCGTTTTTGATAGCTTCTATAGAATTCAATTCCATCTCTATTTTAAAGCG
ACTGCTGTCAATACCATGTTGACTGAGTACTTTATCTATGACTTTTCTAATTGTTGATTG
AGTATCTAACGCAATAAATCTAAGCCTGTATAAGTCTTCTTTTTGAATGTCTCCTAGTTT
AGAAAAAGGATGTGATTTGGGTAATATAAGTGCTAATTCGTCTTCCGCATAAGAAGTAAC
CTGCAAAACATCTTGCAATTCAGTAGGCACTTCTCCTCCAATAATTGCTAAATCAACTTG
ACCATTAGCTACGCTCCATGAAATAAGTCTAGTTGAATGTACTTGCAATTGAACAGCCAC
TTGTGGATATCTTTGCCTAAATAGTCCGATTAATCTTGGCATCAAATATGTCCCAGTTGT
CTGGCTAGCTCCAATAATTAATGTGCCACCTTGTAAGTTTTGTAAGTCGTCAAGAGCTCG
ACAAGTTTCTTCGCAGAGAGCTAAAATTCTGCCCCCATATCGTAAAAGAAGACTTCCTGC
CTCAGTTAAAGTTGCCTTCTTATTACCTCTTTCAAATAGGGAAACATTCAATTGGCGCTC
TAAATTTTGAATTTGCAAACTAATAGCTGGCTGAGAAACATACAAGCTATTAGCTGCTTT
TTTAAAACTGCCCTCTTTGGCAATTGCTTTTAATATTCTTAACTGATCCAATGTAAATGG

AAGGTCTGTCATTAAAGAATTGTAGTATAATGTATATTATTTAGTATTGATCAATGTATG
CATTTGCCAACTAAAATATTTGTCATTACGCTTCATTATTTTCAAAATAATAAAAAGTAT
AATGTTAAATAAGCTGTATTAAAATTTAGTATCTGTTTGATATTAGATTTATACAAGAGA
TATCGTGAATTTAATTCTAACTATAAAATATAGGAGACAATATGGACTCTAGACTTTTAG
TTGTACTAATACCAGTTTTAGCAGCAGCATCTTGGGCAGTTTACAATATTGGTAGAGTTG
CACTACAGCAATTTAGAAAAATGACATCTTAGTTTTGCTTAGAATATAATTTTTAGACTT
TATGAGATAGAGGACTTTAAAATCCCTATCTCTATTGTATATCTTAATAAAAAAAGAAAG
CCCCCTTGTACTTACTGTTATTACACATTAATATTATTGGTGGTACCTAGATAATAAGTT
TAAGCTAACTTATTTTTTATTTTATAATTAATTAAATTAATAACACAATTATGGCTGTTC
CAAAGAAAAGAACATCTAAAGCTAAAAAAAATGCACGCAAAGCAAATTGGAAAAATCAAG
CAAAAACTGAAGCTCAAAGAGCTTTGTCTTTAGCAAAATCAGTATTAACTAGAAAATCCA
ATGGATTTATTTATAATCTAACTGAAGCATCAGATACTTTTAGCGATTAGAAGTGTTTTT
ACTACAAGTCTGAAAAAGTATTACGCAACAAGTTCTCAGTAATTTTTATCTAGTAATCAT
TTGAAAATCATTCTATTTGATAATAAAATAAATAGCAGATTAAATAAAATCACATTAACA
AATTATGCAGCAAAGCTAGATCAAACTAGTGAAATTTGGTTGTTCAATTGTATTGAAAAT
ATTCAGCATATTTTTTTTAAAAGTCAATTAAAATCAAGTCATATTACTAAAATTTTTATT
TCTGGTACTAGCTTTGAATATACCGCAGGTCTACCAGGATTATTATCCAGCTTAACACTA
AGTGGTAGACTACATCCTATTAGTATATATAGTCCCCAGTCTCTCAAAAAGTATCTTGAA
GCATGTACCAAATATTCCCAAACTAATTTTTCTTTTCCTATTAATTTTCATAATTTACAA
TACGGAGGACAAGTTGTTAACCAATTTTATACAGTAATTTGTTTACCGTTGAGCAAAAAG
AGCCTGCTGTATGGATTTATTATTCTAAAAAAAGAAAAGCAGGGAGTATTTAATTTAGCA
CAAGCTAAAACCTTGAATATTCTTCAAGGACCTATATATGGAAAACTCAAAGAAAAGGAT
AATTTTTTAAGTCCAGATGGTTACTATCTATCCGGCCAAGACTTCTCTTCTAATACAATA
ATGGGACATAAAATATCGCTTCCGTTATTAGTCAGATATTCTAGAATTATTTCTGAGATG
CATTGGTTTTGCTCTTATCCTATTAGATTAAATACTTATTCGCATCAGCAAGGAGCAAAG
TGTTTACCACATAATGTTTTAACTGATATTATGAAATCCCAAATATATCAAGATAACAGT
TTTGTTGAATAATACTTATTCTTGATATATAATGGTCTCGGTAATCTAAATAAAATACTA
GAAATAATTTCTTAAATTTATTTGTCTATATTAAATTTACAATCATTTATGACATACGCA
ATTATTGAAGCAAGTGGCACGCAGCTTTGGATAGAAGAAGGCCGCTACTATGATTTAAAT
CATATACCTGTTGATCCAGGTCAGTCGATTATATTAGGAAAAGTCTTATTATTGAATAAA
AATGGGGAGGTTACTTTAGGCCGCCCTTGTATAGAAGGGGTTACGATAAAGGCTACAGTA
ATGAGGCACTTGCGAGGAAAGAAGATAACTGTTTTCAAAATGAAACCAAAGAAGAAAATG
AGATTAAAAAAAGGTCATCGACAAGAATTGACTCGTTTAATGATCGATTCTATAACATCT
TAGATAAAATCTAGACTCATATTTTTAAAAACTTTTTAATATTTAACTAAAATAGATATA
ATGGCACATAAAAAAGGTAGTGGTAGTACAAGAAATGGCAGAGACTCTAATTCCAAGCGT
TTAGGTGTTAAAAAATATGGTGGAGAGCAAGTAACAGCAGGTAATATTTTAATCAGACAA

CGGGGAACTAAAGTTAAGCCTGGCCAAAATGTTGGAAAAGGAAAAGATGATACATTGTTT
TCTCTAATTGATGGTTTCGTGCTGTTTGAAAAGTCAAATCAAAAGCAAAAAACAATTAGT
GTTTATTCTGCTAAGAAATAGTTAAAACAATTGGTGCAAGATAACTTGAGTGTATTTATT
TCACTTACAGTGGCACCATATATTTGATTTATTAAGAAATGGAGTTCCTTAATGAACCAA
TGAATTTTATTAAAGATTTGCTGTGAAATTTAATGACAAACACTATTGTAATTTCCTGTC
TACACAATATGGCTGCTATTCTATATTGTGGTCAGATACAGAAACTAGTAGTAGCAAATG
CTCATTATCAAGTAAGTGATATCTACCTAGGTTGTGTTGATAAAATCTTCTCAGGAATAA
ATGCGGCATTTATTAACCTAGGAAAGAATGAGTACAGTGGTTTTATACATATCAGTGATA
CCGGTCCGCTTAAAAAGAAATATTATGTCAATAATATTACTAACATTTTAACAATACGGC
AAAAAATTTTAGTACAAATTATTAAAGAGCCAACTTTAAATAAAGGTCCAAGGCTCACTG
CCAATATTACATTATCAGGTCGATATATTGTATTAATGCCTTTTAGTCAATCAATCTGTA
TATCTCGAAAAATATATGATGAAGATGAGCGTCATTATTTGAAGTCTTTAGCTATTTTAA
TTAAACCGGCAACAATGGGCTTGCTATTTAGACCTTCTGCTGTAGGTGTCGATGAGGAAA
TAATATTAAGCGAATTAAAAAATCTAAAAGAACAATGGAACTTTGTTCAAAAATCTGCAA
TTAATAGTTATTCACCTGTTCTTCTATATAAAGATGAAGATATTGTTAAAAAGGTAATCC
GAGATTTTTATAATAATAATACAAACAATATAGTAATTGATTCAAACCTGGGATTAAAAC
AATTAAATTATTATATCCACACTTGGCACTGTAATAACTCTAGCACAGTTCCTAAGATTA
AGCTTTATAGTAATAATCAATGTATACTAGATGCTTTTGGTATCAATCAGGCAATTTCCA
GAGCTCTCATTCCAAAAGTTGATCTTATACTTGGTGGCTATATGTTTATTGAAACTTTAG
AAGCTTTTACTATTATTGACGTTAATTCTGGATCTTTTAATAATTCCACTAGTGCACGAG
AAACAGTTTTAAAAACCAACTGTTCTGCAGCAACAGAAATAGCTTATCAGTTACAAATTA
GAAATATTACTGGTGTAATTATAATCGACTTCATTGATATGGAATCACAAAGAGATCAAT
TGCAATTATTAGAACACTTTAATAAAGAGCTATCACTTGATGATGCTAAACCACAAATTG
TACAGTTATCTGAATTAGGTTTAGTTGAATTGACTAGAAGAAGACAGGGCAAAAGTTTGT
ACGAGTTAATTAGTAGTGATTCTAATTACTTTTATTTTTTCACACAATCAGAGAGATCTC
AGTCTCTTAAGAGATTCGATGATAGACAGCAGAAACAACAGATTTTTAATAAATCTTGGC
TATCTGCAGAGATTAATACTATTAACAAGGTCTTTTTTCAAAAGTCAAATTTGTGCAGAC
CTGCTAACTTTTACCTAATTCGTAATCTCTATATAGTTAAAAGTAGTATTACTTATAAAC
AAAATTATTTATTAACTCACAGATCAAAACTTATTTATTCTAAAGAATACAGCAAAGTAT
TACCAAGCAGTTATTATCTAGCTAGTCTCAATAAGAATAGTAATCAAGAGTTCCTACCTT
AAGATTGTTATATCAACTGTTGAAAAAAAGAAGCTCTCTCTTATATAAGAGAGAGCTTCT
TTTGTAGCTTACTAACTATCTAGTTAGTTAATTAAAAATTAACCGTTAACAGCTGGAGCT
GTTAGAGCTACTGGTAAAGATTCACCAGAAGCTAGATCTAGAGGGAAGTTGTGAGCGTTA
CGTTCGTGCATTACTTCCATACCTAGGTTAGCACGGTTGATGATATCAGCCCATGTGTTA
ATTACACGACCTTGGCTATCAACAACAGATTGGTTAAAGTTAAAACCATTCAAGTTGAAT
GCCATTGTGCTTACAGATAAAGCTGTTAGCCAGATACCAACTACAGGCCATAGACCTAAG

AAGAAATGTAGAGAACGAGAGTTGTTGAAACTAGCATATTGGAAGATTAAACGACCGAAG
TAGCCATGAGCTGCAACGATGTTATAAGTTTCTTCTTCTTGTCCGAATTTGTAACCATAG
TTAGCAGATTCGTTTTCGCTTGTTTCACGAATTAAGCTAGATGTAACTAGGGATCCGTGC
ATAGCACTGAACAGAGAACCACCAAACACACCAGCAACACCTAGTTGGTGGAATGGGTGC
ATTAAAATGTTGTGCTCAGCTTGGAATACAAGCATGAAGTTAAATGTTCCGGAGATACCT
AGAGGCATTCCATCAGAGAAACTACCTTGGCCAATTGGGTATACTAGGAATACTGCTGCT
GCTGCTGCTACTGGAGCAGTAAAAGCAACGGAAATCCATGGACGCATACCTAGGCGGTAG
CTTAGTTCCCACTCACGACCAATGTAGCAAGCTACGCCAGTTAGGAAATGAAGAACAACT
AATTGGTAAGGACCACCGTTGTATAGCCATTCGTCTAAAGAAGCAGCTTCCCAGATTGGG
TAGAAGTGAATACCGATAGCTGCAGAACTTGGAATAACAGCACCAGAAATGATGTTGTTT
CCGTATAGAAGGGAACCAGCAACTGGCTCACGAATTCCATCAATGTCTACTGGAGGTGCA
GCTACGAATGCAATGATGAATACAGATGTGGCAGTTAATAGAGTTGGAATCATTAATACA
CCAAACCAACCAATGTATAGGCGGTTTTCAGTACTAGTAATCCAAGAGCAGAAACGTTCC
CACAAGCTAGCGCTTTCGCGTCTTTGTAAAGTAGCAGTCATAATTTTTTATCAATTTTTA
AGGATTTAACAAAGATGCTTCCCAAACAATAAGTTGTCTGTTATGGAAATTATTACTTAA
GGCCTTTTCAAATGAGAACATTTTTAAAAAAAACATAGCAAAATCTAATCTCAGTAACTA
AAATTTTACTTGTTCATATATTGCTTAATAATTCAATAATAAATTTTTGCTTTTTGCAAA
ATTAACACTATTTTAGGGATATAATAATAATGCATATTAATGCTGCTTATTTTGATTTTA
ATAGTTTATATAATTTTGCCCTACTTAAATACGCATGTCACACTTTACAAAAATTCAGAC
GACCATACAAGATTTAAATCTGTTAAAGCATGCTTTAACAGATTTGGGTTTGATTTGGCA
AATGAATTCTAGTCATATAAAAGTTGGTGAAAATAATCAGCACAAGGTAGATATTTTAAT
TAAACAAGATAATTTGTCACATATTGGCTTTACTTGGAATGATAATAGATATCACTTAGT
TGCCGACTTACAACTTTGGGAACAACCCTGGTCTCTAGAAGTATTTTTGGATAAGTTATC
TCAGAAGTATGCTTACTACTCTATCATCGAAGAAACAAAGAAGCAAGGCTTTGAAAAAAT
GCAGCAGATTTCACAAAAAGATGGATCAATTAAGTTAATTGTGCAACGCTGGAATTATTA
ACTAACATACAAATGCGGGCGGAGAGACTTGAACTCTCACGAGATTATCTCACTAGAACC
TAAATCTAGCGCGTCTACCAATTCCACCACGCCCGCACTGTTGAGACAATATTACTCTAT
CATAATCTATACTAAAACACAAACTAATAGTATAAACTAGCTTCCAAGCTTAAAAGCGTC
TACAAATAGACCGTATGTTATACCAATTTTGAAGTTGTTGAATAAATTAATTTTTATATT
TAATTGCTTGTGGCTAGAGTAGACTATTTTACTAGTCAACTCAGTTGCTGCAACTATTAG
ACTGGCTGCAATAATACCCCAATCTCCTGTTTGTCCAGGAATCGTAGACAAACCTGTTGA
AATAAAAAAATCCTAGCAATAAACTAATTAAGCCAGTTGTTAACTCGCTCAAAGAATAATA
AAGCTTATTATTTAAGTTTTTTATTAAGTAAGCAAAAAAAGTTGAAAGTCTAGTTTTAAT
CATAGTATTTAGTAACGGAGTTTTAAAAGGATGACAAGTATTAACAGTCATTAATATTGT
CGTATCCTTTTAAAACTCGTTAATACAAACAGCTGTTCTTGTTTTTAAAGGTCTTGTTCG
CTTAAAGCATTAATAAGATGCTGAAACGGCTCTTGTATAATATCTGCGCGTTTAATTGAT

TGCAAATCTAATTCTTCTTCTATGATTTCTTGTAATAACTGTATGCTTCTTACTGTTGGA
GCAACAGGTACAGATAGTGAATTGTAAGTATCTTTTAGCCCATTTAATACACGCTGATCG
AGAATATTTGTACTACCAGAAACCAAGGCATAACTAGCATACCGTAGGTAATATTCTATA
TCCCTTAGACAAGCTGCATATCTTCTTGTTGTATAAGAATTTCCTCCAGGCCTTAATAAT
TCAGGCTGTTCTTCGTACAACTGAGCTGCGGCCTCTTTTAAAATATTGGTAGCTTGATCA
TTAATAATTTCAGCTATTTTTATTCGATCTAGTCCACTCGAAAAGAAAGATTCTAGTTGT
CCGACAGCTGTCTTATCTAGATAACGACCAGTAAGATCGTAACGATTTAATATTGCTGTT
ATAGCATCTTGCATAAATTAATTCTCCTACTTGCTATCAGAGCGGTATTATAAATGACTA
ATATTGTAATATAATCATTAGTGTTTATGAAATTTATTTAAAACTTATTCTTTATTGCTT
AATGTGCAATTAAAAACTCGCATAGATTTTACACATATAGAGTATACGTAGTTTTTAATT
ATTTTTTTATAAAAACCAAGAAATAAAGCTTTTTTTAAGAGGTTCGTGAATTTGTTAATA
TAACATGAGCTTTCCTCCTAAATAAGTTTTGTAAAGTACAAAGTAAAAAAACGATTAGTC
AATAATAGTAAACAATATTAATATGTATCAAAGTATAAATAAGCTTTTACTCCAGCATCA
AAAAAAATATTTGAGATATGTTGATACAACTGATTATTCAAGTGAAAACTTAGATTTAGA
CCAATTTAGCTATTTTATCATTACAATTAATAAAACTGATAAATGTATTCTTATAGAGCA
ATTTTGTTATAATGCTGAAGGCCATATTTACTCAATATTCTTCAAAGGTCCAACAGCTAA
ACATCTTTCTTCTATAATTTGTCATTTTCAACAATTAAATACAATAATATCTACAGCTCA
TGCCATATATTTAGGACGAGAGCTAATGAAGTCAGAATTAGCTCTCGTGCTAGATCAACA
ATATATTCAAGATTAAATTTTGTAAGTATTAATTATATTTATGCTAGTATAATATTAGTA
AGGGCATGTAACTCAGTGGATAGAGTATCAGATTCCGATTCTGATGGCCGTGGGTTCGAA
TCCCGCCCTGCCCGAGTATTCAGAGTACTATACTAAAACATCTTAATAGCTAGTTTGATA
ATAGTATGGGCATTGGGGCTGCAAGGTTTCTACATTATGAAAAGAAGAAAATATGAAAAA
AAACAAGCTCTCCAAAAGAGCTTTTAGTACAATCAATAAATGCAGAAAACAATATTGTTT
CTTTTTCTCGGAAATTAGCTCTTGCTTAAACACTGTTAATTTTTGGCATTCGACATGTTA
AACTCTTATTCATGGCGAATATTCTGTCAGAGTCGCTCTTAGTTTAAGAAAAGTCAGAAA
AAATATTTATCTTTACGTTTTTACCTATTTTTATTTAAATAAAGGTTAGTATTTTCAACT
TTTATAGTGGACGTGGGTTCGAGTCCCACCAGCTCCATCAATTTGATTTGTATAATTAGC
TAGTTTAACCTTTTTAATAACCAGATATATAATTGGAGGAAATATGGGTTTCATAACAAT
TACTAAGCCCGCTTTAAAGCAAATTGCAATCTTGAAAAATGATCATGAAAATGATGTACA
TCTAAGAATAGGTGTTAGACAAGGCGGATGTTCAGGTATGTCATACTCAATGAATTTTGA
ACATGTTGACAAATTAAAAGATACAGATGAACGGCTTCGTCTAGATAATTTTTCTGTTGT
CTGCGACCCTAAAAGTCTTCTTTATCTTTATGGATTATCATTAGATTTTAGCTCGGAATT
AATAGGAGGAGGATTTCAATTTTCTAACCCGAATGCCAGCCAAACTTGTGGCTGCGGTAA
ATCTTTTTCAGGCTGACCCTCTAACACGTTTTTTATATATTTTGTATTATATATATTATT
TTAGCAAGTTGTTTATTAGAGACAAGCAATGCTCTGTTAAAAACTAATTTCAAGAGTTTT
TATTGTATTATTTTATATATTTATTATCAACTACTTGAAAATAAATTAGCTAAGCAATCT

AATTTGTTATCAGTACCAAACTTTAAATGATTAACTTTTTATTGATATATGCTAATCTTT
ATCAGCCTAAACATATCTGTACATTACAATCTTTTGACATAAAAAGAGTCTTAACATACC
TCTCAAAAATAAATTCAATATTTTATAAATACTCAACATGAATCTGTCACAACTTATCGT
TAATTCTTCTATGATGAAAATTCCTAATAACTTTTATTGCAATATAAAAGCTATTTCTTC
TGAGAATTTAAGTCACTTACAATTATCAAAAACAAATTTATTATCTCCTACAAAAATAAT
TGCTTTTGGACTACCTAAATTTGATTTTATTTCTACACAGATTACAAGTTCTATTAGAGA
AACAGTTGTTGTGTCTACTATTGATGATAATAATCTTTTTGCTGCCCCAACTAATGATTG
GATAAAAAAACACAAGGATAAAAATCATCATTCTTTGTTTTATAAAAAAGTTTTTAAAGC
TTTGCTAAACCATAAAATTAGAGTTGTTGCTGAGCCATTTCATAACTCCGAAATAAATAA
AGAAAAGAGCATAATGGCAAGTCAAAGATACATCTGGGGTAAAAGCTGGAAGCCTTCAAT
TATTTTATCTTGCTTAAAAAACAAAAAAAATAGCAAAATTTCTGATGCATCGAAAAAATT
TTTAACTGAACAGTTATCTGCATCACCAGTATTTGTAGTAAAAAATGGTTTTAATGAAAT
TATTTTAGGTCACCCATTATCTCGTGTTAAAAGAGGAGGAGTTAATAATCTGATGCATGC
GTTTTCTAATTTATTAAATCAGTCTAATGCAACTTATCCTATATCTACTGGTCTATTCTT
TTTTCATCCAGATGATGCATTTGAATTTAAAGACTTTATAATATCGGTCAATCCACTGGC
GGCCAAACATATGGAGATAAGCGTAGAGCCTGTTGGTCTACACTTTGCTTATAAAATGAA
TAGAAATATATCGTCAGATACTCAATTTTCGTTTATTCCAGATTTTAAAGAAGTAGGAGA
TTTATTATTCAAATATAGAAAAGGTAATCATTTAGTTTTTCATAAGAATCAGCATTATGG
CAAAGATTTTTTTCAGGGGCAACCAATATATATGATTCAGCCGATCACTTTTAAAGATCG
AGCCGGAAAGCTAAATATTATCAAATTTACGGGGCTAAATGACACAAGAGAAATTATTTT
TACTAATCTTGAAGCTGCTAATAGATCATGGGCACATTTTATAAAGAGGAATTCACAATT
AAAATCAATTAAAAATCCTACTCTATTAGTCTATAATTTAGAAAGTTTTTTAAAAGATCA
AGAACGATTGGATAATCAAGATTTAAATAAGTTTGTGGTGATCACTAATAAAGAATCTTA
TCTTGCCACAAAAGAGTTAATAGCTTTACCCGATTCTAATAGCTTTTCTAAGCACTTAAA
ATTAAATATAAAGCCTAAACTCTTTTTTGTGAAACTATGGGTAAGACGGTTATTTTCCAC
CTTAACTTATGAATAAATGCTATTGCCTTTATTTTCTCGAATAATATTCTATAACTAGTA
ATTCATTGAGTTGTAAAGCAACCCATTCCCTATCAATGACTCCATTGACTTTCCCTGATA
AATTGGATTTATTCAATTCTAAGTGGCTAGGAATATTAGCTAGTCCAGGGAAAGCTAAGT
AATTTTCTACTAGTTTTCGAGATGCTTCTTGATTTTTTACACTAATTGATTCTCCTGGTT
TACATTGATAACTACAGATAGATACTACTTGTCCATTAATACAAATATGACCATGGTTCA
CTAGTTGTCTAGCGGCAGGAATTGTAGGAGCCATACCAAGTCTAAAAACAGTATTGTCTA
GCCTCATCTCTAGTAGCTGTAACAAAATTTGACCTGTCGACCCTTGTAGTTTTTTAGCTG
CTTTTACGTACTTAAATAACTGTTTCTCACTTAATCCATAATTAAATCGCAACTTCTGTT
TTTCTTCTAATCTTACAGCATACTCAGAAGGTTTTCTGGACTTTTGTCCATGTTCTCCAG
GTGGATAAGATCTTTTAATCGCTTTTCTACTGAGTCCGGGTAAATCACCTAATCTACGGG
AAATGCGTACTCGTGGCCCTCTGTATCTAGACATATTATTTAAATTCTCCTAATGTAAAT

GATTAAGTAATGAGTAGCTATTTTTTAACACAAAAAGAGCATTAAAACAAAGCTTTTGCT
AAAAAATATTTACAAAAAATATAGATTTGGACAAAATACATTACTATGCTTTTTAAGCGG
GTAGCGGGAATCGAACCCGCATCATTAGCTTGGAAGGCTAAGGTTTTACCACTAAACTAT
ACCCGCAATATTCAACAGATTGTCACTAACTCAATATAACATAGTTATGTTAAGTTAAGA
TAATCTTAAATTTAATAGAATTTTAAATTATTTATGCTCCTTCTAAAACCACATCAAGAA
AACGAGCAATTTCGGCAGCTTGTTCTTCAACTTCTGATAATAATAATGGTTGACCGACCC
TAGTCAATGGAATTACTCTTTTATCTTTTGTACATAAATAAATTTCACGTCTTGGATTTA
GCCCCTCTTTAATATCTATTTTTATTGATTTGATTTCTTTTATATTAAACTGAAGACAGA
TTTTTCTATTCTTTCCAGGGAAACCTAGCCGAAAAATTTTTACGATACCTTTATCTTTGT
TAAATTCATTATATCCAGCCCCTATATTCCAAATAATAGTTAGCCACAGAAACAAACTCA
GAAAGACTCCTATACTGCCATAGAATGTCATTACAATACCTTGTGGAATAAATACCAAAT
CGGTTGAATTTGTAAAAGGTAACAAATCAACTTGAAAATAACTTGATAGTCCAGCAAGAA
GAAAGCCCAAAGCCCCAATAAATATTATAGTAGCCCACCAGTAATTACTCAGTCTTCGTG
AACCTAAAATTAAATCTTTTCTAACTGAGTGTATAGATAATGTTTTTTTCATAATTTGTT
CTTTTAAAACAGTATACTAATGCAAAATGGAAAAATTTTGACAATTTTTATGCGCGCTTT
AGATTTTTTAAATAAGCTTGATTGCTTTTAGTCCTAAAAACAGTCCTGTTGCTAACCCAA
AAAAAGCAGCAAGAAATATTATATATCCTAGAAAAACTGACATATTATTATTATTTATTA
TTATTGTTACAATAGTGTTTACAAAAATAGTGAAAATCTATTACTACGGTATTATATTAC
GTTAAATGCCTTGCTGTCTACAGGAAAAAACTTTTTTCAGTTTTTAGCATTGTGTTTATA
TTAGAAATAATATTCTAGTAAATTTATTAAACTTCTGGAGAACAGAATCTTATGTCAGAA
TTTATCAAGCCTTATAACGATGATCCTTTTGTAGGCAATTTGTCCACGCCAGTTAGTACG
TCAAGTTTTAGTAAAGGACTTCTAGGAAATCTACCAGCTTACCGTCGAGGTTTATCTCCG
CTTCTTAGAGGATTAGAAATAGGAATGGCACATGGATACTTTTTAATTGGACCTTTTGAT
AAATTGGGCCCTTTACGAGGTACAGATGTAGCGTTACTAGCGGGATTTCTATCTTCGGTC
GGCCTCATTATTATTCTCACTACATGTTTATCCATGTATGGTAATGTATCTTTTACTAGA
GCAGATTCGAAAGATCCACTACAAACTTCTGAAGGCTGGGGACAATTCACTGCAGGATTT
CTAGTTGGAGCAGTAGGTGGTTCAGGATTTGCTTATTTATTGCTAGCTAATATACCTGTA
TTACAGACTGCAGGTCTTAGTTTATTCTCTTAAGCTAGTAAGGGGACTTGAACCCGTAAC
CTACTGATTACAAATCAGTTGCTCTACCAATTGAGCTATACTAGCATTCAGTTCACAATA
ACACATAAAGATATTTGTTGCAATCTGCTATTTTCAGATGTAACAAATATCTTTTTTATT
TAAACAGAACTGTTATGTTAACACGTATATTAATTAGAATCTGAATTATCTGACTCTGCA
GAAGATTCCTGATCACAGTTTAGATAATAACAAATTGTTTGATCTAGACAAGTTGCAGAC
CAACCAACAGGTGTTTCTCCTGATAATTCTTGCATATCTACAGAATTATAAAAATTATCA
TTTGCCTTATTACTGTTTGCTTGCATAAATATCACTCCCAAAACACTCTGAACTTAATGT
GAAGTAATTTAACTTCTCAATAATATATTAACACAAAATATTAAATATGGTGATTATTCT
TTAATTGTAAAGATGCTGCTAACTTTATAAAGTTTGAGTGAGAGTCTTGATAGCTTTTGT

TGAAATAAGCATTTTAACCCATTTATTTTGCCCTCTCGACCATACTTTTTTGTGTTGTAA
ATTAACTTTTTGTAGTTTTTTTGTTCGTTTATGAGAATGAGATACGGCATATCCATTATT
AGCTACTTTTCCCGTAAGCTGACACTTTTTTGACATAATTTTCTAACTTATATATTTATT
TAGAGTACTTGCTAAAGTTGACTTGGGCACAGCTCCAATCACTGTGTCTACTCTTTCTCC
TGCTTTAAAGATCATTAAAGTAGGTATACTTCTAATGCCATATTCAGCTGCAATAGTAGG
ATTATCGTCTGTGTTTATTTTTACTACCTTAATAGATGATTCATATTCTTCCGCAATTTC
ATCAACTACAGGAGAAACCATTCTACAAGGACCACACCATGGCGCCCAAAAATCTACGAG
TACAGGTAAGTTATTGTTAATAACTTCTTGTTTGAAAGAGGCATCTGTAACTTGAGATAC
TGACATATTCTTTAACCTTTAATATATGTTCCTTGCTAAGCAAATCTTATCACAAATTTC
AAGAAAAATCTTCTATAGTGAATTACTATTAGTGTTGAATTATTTTTTTAAAAATTAGAC
ATTAAAAAAACTTATTACCTATATAAATAAAGTGGTGGCCGAGTGATATCTTGATAGTGA
TAATTTTTATATTAAGGTTGAACACTAGAAATAACTTTTCAAGCAAACTCTTGTTTAAAA
AATTTCTCTGTGACCATAAAAACTTAATTATATTACAATATATTTTATTATTTGTAACTT
AGTTCTGATAATGGTATAAACAACGCAAAAGATACTGCCTTATAATCAAGGAGGAATACA
TGTCTCAATCCGTAGAATCACGGACTAGGATTAAAAGCGAACGTTACGAATCTGGAGTAA
TCCCCTACGCTAAAATGGGCTACTGGGATGCTGACTATGTGATTAAAGAAACAGATATTC
TAGCTCTTTTCAGAATCACTCCTCAACCAGGTGTTGACCCGATTGAAGCATCTGCTGCAA
TTGCAGGTGAATCTTCAACAGCTACTTGGACAGTTGTATGGACTGATTTATTAACAGCTT
GTGACTTATACAGAGCAAAAGCATATCGAGTAGATCCAGTTCCAAACGTGGCAGATCAAT
ATTTTGCTTACATAGCTTATGATATTGATTTGTTTGAAGAAGGTTCCATTGCGAACTTAA
CTGCTTCAATTATTGGTAACGTTTTTGGGTTTAAAGCTGTTAAAGCTCTTCGCTTGGAAG
ATATGCGTATGCCAGTAGCTTATCTAAAAACGTTCCAAGGTCCTGCAACTGGATTGATTG
TAGAACGTGAGCGTATGGATAAGTTCGGTAGACCTTTCTTAGGTGCTACAGTTAAACCTA
AACTAGGTTTATCTGGCAAAAACTACGGAAGAGTTGTATACGAAGGCCTGAAAGGCGGTC
TTGATTTCCTTAAAGATGATGAGAATATTAACTCACAACCATTTATGCGTTGGAGAGAAA
GATTTTTATATTCTATGGAAGGTGTAAATAAAGCATCGGCTTCTGCTGGCGAAATTAAAG
GTCATTACCTTAACGTAACAGCCGCGACAATGGAAGATATGTATGAGAGAGCCGAATTCT
CTAAAGAGGTTGGTAGTATCATTTGTATGATTGACCTTGTGATTGGTTATACTGCGATTC
AAAGTATGGCAATTTGGGCTCGTAAACATGACATGATTTTACATTTACATAGAGCTGGTA
ACTCAACTTACTCTCGTCAAAAAAATCATGGTATGAACTTCCGAGTTATTTGCAAATGGA
TGCGTATGGCTGGTGTTGACCATATTCACGCAGGTACAGTTGTAGGTAAGCTTGAAGGAG
ATCCTTTAATGATTAAAGGCTTCTACAATACTCTACTTGAAAGCGACACAGATATCAACC
TACCTCAAGGTCTGTTCTTTGCTCAAAATTGGGCTTCCCTACGTAAAGTTGTACCAGTAG
CATCTGGTGGTATTCATGCTGGTCAAATGCACCAACTTCTTGATTACTTAGGTGATGATG
TAGTTCTTCAGTTTGGTGGTGGTACAATTGGACATCCTGATGGTATCCAAGCAGGTGCAA
CTGCTAACAGAGTAGCACTAGAGTCCATGGTTATGGCAAGAAATGAAGGCCGTAACTATG

TAGCAGAAGGTCCACAAATCTTGAGGGACGCTGCTAAAACTTGTGGGCCTCTACAAACAG
CTTTAGATTTATGGAAAGATATTAGTTTCAACTACACTTCCACAGATACAGCTGATTTCG
TTGAGACTCCAACAGCAAACATCTAGTTTAATGACTACTTACTGATACTTTAAATAGTCA
ATTGTAAGTGGAATTAACTTATAACAATAAGGAGCATAGAATAGTGAGACTAACACAAGG
GACTTTTTCCTTCCTTCCAGATTTAACTGATGAGCAAATTAATAAACAGCTTGCTTATAT
CGTTTCTAAAGGCTTATCAGCAAACGTTGAGTATACTGACGATCCTCATCCAAGAAACTC
CTATTGGGAACTGTGGGGTTTACCTTTATTTGATGTAAAAGATGCTTCTGCTGTTATGTA
CGAAATTAGCTCATGCAGAAAAGCAAAACCTAATTATTATGTTAAAGTTAACGCTTTTGA
TAATACTAGAGGTATTGAAAGTTGTGTAATGTCTTTCATTGTAAATAGACCTGCTAATGA
ACCAGGATTCTTATTACAACGCCAAGACTTCGAAGGTAGAACTATGAAGTATAGTCTTCA
TAGCTATGCTACTGAAAAGCCTGAAGGAGCTAGGTATTAATATTAATTAAGAATTAATAT
TGGCTAATTATTACCCTCTTAAAAATCATAATTAATTTAGTAATTGTGATTTTTAAGAGG
GATAGCTTCCCAAGTAAATTTTGACTAAATAGATTGAATAATTAAAAATACACAGAAATA
ATGCAATCACAGGATATAATTTCCAACGATACTCTTGTTAATTTACAAGAAGAATATGAT
AGAACACAAATCCAAGAAGTTTTAAATGAGTTAAATCAAGAACTTATAGGATTAGTGCCT
GTAAAGACCAGAATTCGCGAAATTGCTGCGCTATTATTGATTGACAGATTACGCAGAAAA
CTAGAACTAGTTTCTGGTAATCCAGGATTACACATGTCATTTACAGGTAGTCCAGGAACT
GGTAAAACTACAGTTGCTATGAAAATGGCTGATATTTTGCACAGACTTGGATATATAAAA
AAAGGGCATTTGTTGACAGTTACAAGAGATGATCTTGTAGGTCAATATATTGGACATACT
GCCCCTAAAACTAAGGAAGTTCTTAAACAAGCAATGGGAGGAGTTTTATTTATTGACGAA
GCTTACTATCTATATAAAGCAGATAATGAAAGAGACTATGGCTCTGAGGCAATTGAAATT
TTATTACAAGTAATGGAAAACCAAAGAAATGACTTAGTTGTTATCTTTGCTGGATATAAA
GATAGAATGGAAAAATTCTACGAATCCAACCCAGGACTCTCTTCTAGAGTAGCTAATCAT
GTAGACTTCCCAGATTATACTTCAGATGAATTATTACAAATAGCTAAAATGATGATAGAA
GAACAGCAGTACTGTTTTACAGAAGAAGCAGATAAAACTCTTTTAGAGTATACCGAGCGA
AGAATGAAACAGCCTTATTTTGCTAATGCAAGAAGTATTCGCAATGCTATTGACAGGGCT
AGAATGAGACAAGCCAATAGGATTTTTGCCAGTGGAGAAAAAGTATTAACAAAAGCTGAT
TTGGTAACGATTGAAGCAGAAGATATCTTGAAAAGTAGATTATTTTCATTACCTAATGCT
TAATATACACGTGATTTCATGAAACTATATATTATTAAAAAGTTTCATAATCAGTTGCAA
ATTTCTTGAAAAACCTTTATTATTATCTTAATGCAGGTGTGGCGGCATAGCCAAGTGGTA
AGGCAGAGGATTGCAAATCCTTCATCCCCCAGTTCAAATCTGGGTGCCGCCTAGTACTAA
AAAAGGGGGGTGTGGTGGAATGGTAGACACAACAGACTTAAAATCTGTTGATTTTTAGTA
ATCGTGAGGGTTCAAGTCCCTCCACCCCCATATATTGATTTATAAATAAAAAGAGATGTG
TATTTGTATTAATTGCAACCATATTACCAAATGTAATACCTATCACTTAATAGAATCTCA
ACATAAACAGCCTCATCTTACTAGAAGTCCATTGTTTATACCTAAATATCCTGTAGTTCA
TGTTAATATATCTAACAACTGTACCTATAATCAAATTGATTGGGATTTAGTGGAGTGTTT

ATCATTTGTAGAAAAACCGAATAGTTGGAATTTAGACGCTAATTAGTTACAAGAAAAATT
ATGTACCATAAGCAACAACTCTATTTTTTAGATACTCGGTATTAATTTTGGATGTTCTAA
CCAGTAAGATTTTACCCGTTCTTGCTATCTCTATAATGCCAAATTTAGTTAGTAATTGTT
CAATAGCAACAATCTTTCCCGGATCTCCAGTAACCTCTATAATTAAAAGATCTTCTGCAA
TGTCTACTATCTTAGCTCTAAAAATTCTTACAATTTCTAAAGCTTCTGTTCTAGTCTGAG
AATTGATCTGAATCTTAATTAACATTAGTTCTCTTTCAACTGAAGGAATATTTGTTACAT
CTTGAACGTTAAGGATATTTACTAACTTATATAATTGTTTCGTAAGTTGTTCAATAGTTC
TATTGTCTCCTTGAACTACCATTGTGATTCTAGAGACCCCAATTTGCTCTGCTGGTCCAA
CTGCTAAACTTGCGATATTAAACCCTCTCCGGGCAAATAGACCAGATATTCTTGACAGTA
CTCCGGCTTCATCTTGAACTAAAACTGATAAGGTGTGTTTCATGAAAAATTAAATTTAAT
CTATTTGATCTGTTAAGTATATTCTAAGAAAATTACAGAAAAAATACTAGTTAGTCTATA
ACTAAATTATCCAAACTAAACAAAATCCTACTATCAGATAATATATCTACCTTGTAATGC
TATACTATTAACTTAGTATTTAATGATATTCTAATTGCAAATAGAATGCTAATTTTTTAA
AATTTGTTAGGAACTTGTGCTAGAAAAATACAAAAACAGTAAAAAACTCTCTCGAGATTT
ACCTCAAATCAATGATCGCATTAGATTTCCAAAAGTCCGAGTAATTGATGACGAAGGTGA
ACAACTAGGTATTTTTGTGCCTGAAGAAGCTATACAATTAGCTGTCCAACAAGGTTTAGA
CTTAGTTGTTGTTAGTGATAAATCGGACCCGCCAGTATGCCGAATATTAGACTATGGTAA
ATATAAGTTTACACAAGAAAAAAGAGCTAGAGAAGCTAAGAAAAAGCAACATAACAGTAG
TATTAAAGAAGTAAAAATGCGATATAAGATAGAAGAGCATGATTATAAAGTTAGAATAAA
CCAGGCATCCAAATTTATTCAAGCAGGAGATAAAGTAAAAGCAACTATCACATTTCGGGG
GCGTGAAATCCAGCACTCTAATTTAGCTATAGATTTATTGAATAAAATGGCAAGCGATCT
AAATGCAATAGCTGAAATTCAGCAAGCTCCATCAAGAGATGGCAGAAATGTCATAATGCT
CTTATCTCCCAAAAAAGTTAGCTAAACTAATTTTTATTTAAATGCATCTGGCTGGATTCG
AACCAGCGACGTCTCTTTCGAAATGGCGGATTATGAGTCCGCTGCCTTCGGCCCCTCGGC
CACAGATGCATCACTTAAAGTTTATACCTCATAAAGCATATAAAAATCAATGTTTAGCTA
TTTTTAATAAAGTCTATAAGAGTCTATTATTCATTCATATTGTGATATGTCGCAACTGCT
GAAGGGGAAACTTGATTTAAGTATCGAAATATCCAGTATTTAAAAATTGTGTCGAGTATA
ACTGGAAAAGTTGAAATAAATAAAAAAATAAAATCTCGACTTTCCGGAAGTCCTAAATGT
CTTAAAATTACTTCAATAATAACTTCCCATCCATGAGGAGAATGAAAACCCACAAACATA
TCAGTAAAAAGAATGATTAAAAAAGCTTTAGCTGTATCACTAAGACCATAAATAATTTCA
TTTAAAAAAGATTTAACGACTGCAATTTGTCTTTGTCCAGTTATCATTAGTAATATAAAG
ACAAGTATTGATACAAGATCGGATAAAATATTTTTAACAGCATTTGCACTTTCATTTGCA
TAATATTCTCCCAACTCTCTAGCCTTCAGCTGCACTCTTTTTTCAATGATTTCGTAAGAT
ATATCTTCCGAAGGATTCAGAAGAACTTCAAAGTGAATTTTCTCTTCAAATCTTTGTAAT
TCAGCAAAAGCTCTTTCCTCTTGAGAGGAATTTAAAAAAATTTTAGGCTGTTCTTGGTTC
CATAAATAATCAATACAAGGTCCAAATACAAAAAATTTTGAAGCCTGATTAACTAGTACT

GGAGAGATGAATAATAATAAAATATATTTTACGGAAGTAATAGTTTGGTGTCTAGATATT
CTAAACTCCTCTATAGCTTCAGATTCGCCATTAGGATCTAACTCTTTACGAAACTTTTCA
AAAGTATTAGTAATTGACCTAGGTATTGGCCCAACTTTTTCAAAGGCGAATTGATTATTT
CGTTTTAAATTCCAATATTTCATTTTTTATTGCTGCTAAAATCTCTATTAATCAATAATT
GATCACGTATTCACTATAAATATTAATTCTAGAACTAAACATATAAAACAATATTTATGA
AAAATAAAATACAATTAAAAATTAATTATGCATTAGTTCTATCTGCAGCTACAGCAGTTT
ATGGATATAAATCTTATCAGCCAACTATAAATACACTCTTAATCAAAGATGAAGAAAACT
GGCCTTTTTTCTCTAAAAGGCAGACTGCCTTTATTCCTATAATTGTATTAAAAAAAAATC
ATCAAAATCCCTGAACTCAGAACCTAGAAATAGTAAGTTATATCTTAATGGAATTGGAAG
TAGAAAATTTATCGACAGAATTGTCACTCTACTTGGATATTTAGAAAAACAGGCAAACTA
TACTGAATACTTGAACCAAATAAAAACAAGTGGTTATTTTTCTTTAGTCTACCTAGACAG
TGATATAATGTATAAGAAAAATATTAGTAACATTTTTTACTACCTAATGATATTCTAAA
AAGAATCTATATTTGCAATAAGAAAAAAAAGCTTATACCGCATTCATTCTTAATAAATTT
ATTTCAGAAACAGATAGGCTATCCAAAAAGTTTTGCTAATTTAAATTGGGCCTTCTCCAA
AATTATAAAATGGTATTATGATAGAGGATACCAATGGTCTCTAGTTGAAGTTAAACAAGC
GTCTGATGCATCTTCTATTGTAATTGATATTCATGAAGGGGTAGTGAAGACAATTATAAC
AGAATACTACACTCTATCTTATAAAAGAGTCTCAGGTATCTTATGCGTAGAGTCAATAGA
GCAGTACCTTGGAGTAAGAGTGGGAGCTCCACTAAATATAATTGATTTACAAAAAAAAAT
TACTTACTTAAAAGATAATCAACTAGTTGGCGACATTATTTATAGTATTGAACGATCAAA
TAATAGCTCAATGAGTTTGGATATCAAATTTCAAATACAAGAATTAAAAGATAAAGAGAT
AATAGTGCTTGCAAAAAGTTCTTCTATTATCTCGCATGCATGTAATTTACTTAATCAATA
TAGAAATAGGCTAGTAGCTTCTAATATAGTTTCACTATCAACTAATAAATTACACGCATG
CTATTTTAATTATAAACTTGACTATCAATATAAGTATATAAACACGTCTAATCTTATAAA
ATTGTTAGCTTATTCTATTTCTTGTAGGAAGACACTACTTATTCATTATTTTAATTTACA
AACTTTAATAAGCTATACAAAAAAAAAATACTATTGGATTTCAGCTATATTTACGAAATCT
AAGTTTTGGGAAAGCTTTCTGTGTGCTTAGTATGAAATTTATCAAAAATGGACTTAATAT
AAAAATTTTATATATAAATCCTTCATTGATAGTTGATCAGAATTTTGTATTTCAATTTGC
TATACAAATCATTAAGCAATATCATACTGCTAAACCTCCAGCTTTATTTTTAACGAATCT
AGATCTTGAACAATATGTTGCTGAAAGTTTATTAATGTACCATTTTACATCTTGCTTCTC
AATATCTGAAAAAATATTATTATCTCGAATTATGCATACAGATTCTTTATTTTTCAACTC
TGAAAATTTTCATTTTGATGATAGAACTAACGCGATCAACAGTTATGATACTTTTAAACA
AAATACAAAAATATTTTATCAAGAGTTTCTTTCTTTATTATTAAGTTTACGTTATCAAAA
TTTTAATTATTTAGGTTGGCCTTTAAAAGGCCATTTTTTTGAAATAAAATCTTTATACTT
GGCTCCATTTCAAAAGAGTGATTTTTCTGATAGTCGTAAGACCCTATTCTTCCACAAAAT
GTCGTTAAAGCAAGTATCTAATTTTAATTTACCAATATCTTTTAAAAGCCACCTTAACCA
TATTTTAGTTAGTACTATTAAATGTCAGTCAAATTTGAATATGAGAACTGTATCTTTGCT

ATTAATTGATTCACCTGCTGAATATATGCTATACAAATCTATTCTTAATTTCTCTATTAA
AGTTAGGATGCAGTATTTCATACCTATGAGCAACAATATAAGATTATCTCTATTTTATAA
TTATTTAGATTGTTTTTTAATAAGATCATCTCAGAGTCGCATTCATATATGGCAAGATCT
AAGAACTCTTACACCAATTCAAAATTTTTGGTTAAAAAAATTTTCATATGGAGCTGGAAT
TCAGCTTAAATTACCAATTAAGCAGATGCCACCTCTTTCAATTGAATATACTGTAACTAG
TAGTCGCTACTTTTGTATTTACCTTCGTACTTATTATCAACGATAACTATTACTAAAATG
AAAAATTTAATTTCTTTTTTTGATCGCAGCAAAGGGAAATGGATTTCACAAAGAACAACT
TATGAATTATCTAATAAAAACATGAGTTCGATACAGTCTCAGATGACAATGAAAATAGGC
AACTCATTGTCAGGATCTATAATACTTGCATCGTTAAATTGGGGGGACATTTATAGACAA
GTTGCCCATTACGCTAAAAATCACAGCCGGAGTGAATATGATAATAAGTTTAACTTGCAA
TTCAGTAATCAACTAAATAATCATAAATTATTAACGCTGTGCATAGTGACAGATCCTAGT
CTAATTAGTTTTAAAACTCGCTATGGAAGTACTACCATAGATGAAACATATTGGTTCGCA
ACAAATAATTTACGTCTAAGTACTAGTATTGTTAAACGATTCAACACTTGTGTGGCAGTA
TCTTTTTGTTCAGAAATTAAAGTTTAAACTCTATATAGAAGCAACTAATAGCCTAAAGGC
TATTAGTTGCTTCTATATATCTAAATTTTGTATTAAAAATTATAATACTATATTATTCTA
AGTCTTTACGATTGGGATTCCTGGAAGGATCATTAGAGAGGAACCCAAAGACAAACAAAG
AAATAAAGAAAATAACGGTAGTATATACAAAGATTTTCAAAGTAAACATATTTTCATCCT
TAAAAGCTATTAGTGCGATATTTACATATCTCCACATAATTTTATATCATGATATAAGTT
CCAGCACTATTTTATAAATAAATCAATATCTTTATTTTTTAGTAATGATTTATATTTAAA
TATGCGTGATGTACTAAATACTAAATCTTTTCCGACTTGTTTTGTCTTGTTTGTAAATAT
TTTGTTATTTCTATGGAGTTTACCTTCAATTATCACATAGTCTAACTTCTTTAGCTGTTT
AAAAGTATATGAAGATTTTTTGTTCCAAATAGTTAGATTGATTATAACTAATTGTTTTCT
TTTTAACAGTCTTGCTTTTAACTTAATTATTTGGCTTTTATTTTCACTAATTCTGATACT
CTTACAACTAAGAATTTGGACTAGAAGAGTACAGCTATTCATAATTTATAACAATTTTTT
AAAAAATCTACTAAAATAGATGATTTTACAGAACTAAAATGATATTTCATTATCTTGTTG
ACCTTGCTCATAAGTAATATCTTTCAATACTTTTAATATTTTTCCAAAATACTCTTGCAG
GTATTTCTCTAAAGATATTACTTCCAACCTGTCAATTCCAAGAATACTGTAAACTTCATC
CATCGGAGCTGTAAATTGCTCTCCGCTACTTAAAATTTCGGCAAATGCTAAACGATCAGA
AATATTCCATGTCCATTGAAGAGTTTTAGTAATTCTTCTTAAGGCTTTTAATAATCCAAT
CGGAATTTGTGAGATCTGTGTTTTTTGCCCAGATAATTTTTCACATAATTTAATAATTTC
AGCTGATGTCCAAGCGGTATTACCAACTAACGGCAGGGTCCTATTTTCTGTGGAAGGTAC
TCCAAGACTTTTTATAACTAACTTTGCTGCATCTTGAGTATCAATGTATGCAATGGGTGT
AGATTCTCCTGTAACCCAGACTGATTTTTTTGTCTAAAATAGGAATGGCATATTGGTTGAT
TAAACCTTGAAAAAAATCCTCCTAAAGAAAACACAGTATATTTCACATTGGACTTCTGAAG
GAAATCTACTACCTGAGACTTTAAGTTCATCAATGGAACATCTGGGTATTTCTCTGAATT
TAATATTGAAAAGAAAATAAATCTTTCAACTTTAGCCGCTTTAGCTGCTTCAATTAATGC

AATTTTCCCATCTAAATCTATTTTTTTCTGCATTGTAAGGATCAGTAGGACGAGATGTAGA
AGCATCTATGATTGCTGTTACCCCACAAAAAGATTGCAAAATACTTTCGGGTAATTTTAA
ATCACCATATATAAGTTCTGCTCCCCACTCTTTGAGAAAGGCAGACTTTCTTAAATTTCT
TACCATGCATTTTACATTGTAGCCTTCATCTAAAGCTCGTCTTACAATTTGACGTCCTAA
AGTTCCGGTTGCTCCAATTACTAAAAGAGTCATATTAGTTAAAAATAAGTATGTATAAAG
ACATTGATTTGGTTGACGGCAGAGAGTAAATTAGGTAGAGAGGGAATCGAACCCTCATGA
CCGAAGTCGTCACATTTTGAGTGTGATGCGTATACCAATTTCGCCATCTACCCTATTATA
GACCAACTACTACAAGTAATTTAGTCTTTAGCGTAAAAATAATACTAAGTTTCAATGGAT
ATGTCAACTTTTAAATTTATATTAATAAAAAAACGGAGTCTCTTATGGCCACTTTTGAAT
TAATTCCTTATAGATTGTATTTGAGTCAATCTAACACATGGATCCATAGAATAAAAGCAG
AAATTAAGATATATATAGTAACGCTTTTATGGATTTCAATTTTTATTTTTTTCTTACTTTA
AACTATGTATTATTGCTTTAAGCTTAATTGCCATAAGTTTTACTATAAGAAGTAAACAGA
ATATTATCCAAAAACATTTGTTACAAACTTTATTAATAACGTTCTTAACTACTGTTTTGT
CTTTTAGTGTGGCTATTAGTTATAAACAATATGCAGAACAAGAACAATCGCAGTATTTAT
CTGATTCCAAGAAATATAAAAATTCCAGCAGCTACTATTATATACAAATAGCTAACGATC
AAAGGATCAAAAGACAATACTTAATTACTACTTTAAAGCCTTCCTTGTACTTTTTTATTA
CTATATATTCTATTAAACTAGTTATGATAACAACTTCTCCAGAAGTTTTAGTAATTACTA
TTTATAGATCTAGGATAATAAATAAAATATTTAAAAATGAATTGCTATTTATCTTCCTTC
TTTCGTCACATATTGTTACTAGTATTATTAATAGAATCGATAAAGTGATTCAAGTAACTA
GCTTAAGAGGAAGCTTGAATTTATATAATTCATTGACAAGGCCTTTAATGTTTTCTTTGT
TAATATTTCAAGTCTTCTTTTTGGAGATTATTCGGGAGTCAAAAGAAATAGCTCAAGCTC
TTTATACTAGAAATCTCAATCAAGAAAACAATAATTTTTTGAAAATATATACAGTAAAAT
CTAACTTTAGTGATCGGCTCAATATAATTATTAGCACTTTGTACTTTATTATTTTAGCTC
TAGCGTAATAGAATCCACTTATTCTTCTGACTCTAGTAATACGAAGAGTAATAAAATTAG
TTAGTAGTTTAAATAGATAGAATTAAGATATTATATAAATAATTAAATGTTTCATTATGG
TTAATACTCAAAATCAAATTTCACAAACTTCAGAACTTGATTATATAGTTAATCAACCTT
ATAAATATGGCTTTAAAACTTCTGTTGAATCTGAGCAATTTCCTAGGGGAATAAGTGAAG
ATATTGTCCGCTTGATCTCCAAGAAAAAAGATGAGCCTGAATATCTATTGAATTTTAGGC
TTAAAGCATATAAAAAATGGAAAAAGATGAGTAGTCCGTCATGGGCTCATATTAAGCATC
CGAATATAGACTTTAATACGATTATTTATTACGCTGTTCCTAAATTAAAAAAAGAATTGA
AAAGTCTAGATGAGGTTGATCCAGAAATTCTTGACACTTTTAACAAGTTAGGTATATCTT
TAAATGAGCAAAAGCGAATTTCCAATGTAGCCGTTGATGCTGTTTTTGATAGTGTATCTA
TTGCAACTACTTTTAAGAAAGAACTGTCTGAAGCTGGTGTTATATTCTGTTCAATTTCTG
AAGCTATAAGGGATTATCCAGAATTAATAAAAAAATACTTAGGTACTGTTGTACCAGCTG
GTGATAATTATTTTGCTGCATTGAACTCTGCGGTATTTAGTGATGGCTCTTTTTGCTATA
TCCCTCCTAATACAGTTTGCCCTTTAGAATTATCAACTTACTTTCGTATTAACAACGAAG

AATCTGGGCAATTTGAAAGGACACTAATTATTGCTGATCGTGGCAGTAAAGTAAGTTATC
TTGAAGGTTGTACTGCTCCTCAATTTGACACAAATCAATTACATGCAGCGATTGTAGAAT
TAGTAGCTCTTGAAGGAGCCGAAATTAAATATTCTACAGTACAGAATTGGTATGCTGGTA
ACAAAGAAGGTAAAGGTGGTATATACAATTTTGTTACTAAACGAGGCTTATGCTCGGGTA
ACAATTCAAAAATTTCATGGACTCAAGTAGAAACTGGATCTGCAATTACCTGGAAATATC
CAAGCTGTATTTTAGCTGGTGAAAATTCTCAGGGAGAATTTTACTCTGTAGCTTTAACAA
ATAACTATCAAGAGGCTGATACAGGGACTAAAATGATCCATATTGGTAACAATACAAAAA
GTAGAATTATTTCTAAAGGTATTTCTGCGGGTAGATCGAAAAACAGTTATCGAGGCCTAG
TAAAAGTTGGCCCTCAGTCATTTAATTCTCGTAATTATTCTCAATGCGATTCTTTATTAA
TTGGTCAATCATCTCAAGCTAATACATTTCCCTATATTCAAGTACAGAATCCAACATCAA
AAGTAGAACATGAAGCATCCACTTCAAAAATTAGCGAGGACCAAATTTTTTATTTTTTAC
AAAGAGGGATCAATTTAGAAGAATCTATTGCTCTTATGATCAGTGGTTTTTGTAAAGATG
TCTTTAATGAATTGCCTATGGAATTTGCTACTGAAGCTGATCGTTTACTGAGTTTAAAAT
TAGAGGGAACTGTAGGATGAGCCAGACTATTTTAGAAATCAAAGATTTATATGCTTCTGT
TGGTGAAACAACAATTTTAAAAGGGGTAAACTTATCTATTCGAGCCGGCGAAATACATGC
GATTATGGGGCCTAACGGTTCAGGTAAAAGTACATTATCAAAAGTAATTGCAGGACATCC
AGCGTATTCACTAATAAGCGGAGACATTTTATTTTTTGGACAAAGTATTCTTGAAATGGA
GCCAGACGAGAGGGCAAAAGCAGGTATTTTTTTAGCTTTTCAGTATCCTGTTGAAATTCC
TGGAGTCAGTAATTCTGATTTCTTAAGAATTGCATTAAATGCTCGCAGAAAGTTTCAAGG
ATTATCGGAGTTTAGCCCCTTAGAATTTTTTCAACTAATAACAGAAAAAATAGATCTTGT
TGGCATGCAAGAAAGTTTTTTGACAAGAAATGTTAATGAAGGATTTTCTGGTGGAGAAAA
AAAACGTAACGAAATTCTTCAAATGGCTTTACTAGATAGCAAAATATCTATTTTAGATGA
AACAGATTCTGGACTAGATATTGATGCATTACGAGTAGTTGCAAAAGGAATTAACACTTT
AGCTAAATCAACAAATTCAATTATTTTGATTACTCACTATCAAAGATTATTAGATTATAT
TATTCCAGATTTTGTTCATATTATGAGTAACGGACAAATTGTAAAAACTGGTAGTGTTAC
TCTTGCCCAAGATCTAGAAAAGCACGGATATGACTGGATTACGCAGACATAACTATTTAC
TACAATATTGAGACTAAAGAATATTCTTTAGTCTCAATATTTTTGCAAATGCAAAGTTAA
TTATTTAAGCTTTATCAAAACCTTGCTTAACATCTTCAATTGCTCTTTTTAGCAATTCTT
CACTAACTGGTTCTAATTTTTTAGTACTACGAACATTTTCACCAAATTCTGGTTTAGAAT
TCTTTAAATCTTCTCGAAGTTCTCTAATAAATTCAGCTACTTTTGAAACTTCAATATCAT
CTAGATAGCCATTAATGCCAGTGTAAATAATAGCAGTCTGTTCTTCAACTGGAATAGGAG
AATTTTGAGCTTGTTTCAAAATTTCTCGAAGACGTTGACCTCTTGCTAGTTGATTTTGGG
TTGCTTTATCTAGATCAGATGCAAACTGAGAAAAAGCTTCTAATTCAGCAAATTGTGCTA
ACTCTAGTTTTAACTTGCCTGCCACTTGTTTCATGGCTTTTATTTGCGCAGCAGAGCCTA
CTCTAGAGACTGAGATCCCAACATTAATAGCTGGTCTAATCCCTGAGTTAAATAAGTCAC
CAGATAGAAATATTTGTCCATCTGTAATTGAAATTACATTTGTTGGTATATATGCAGATA

CGTCCCCAGCTTGAGTTTCAATAATAGGGAGAGCTGTCATACTACCTCCACCTAATTCAG
CATTCAATTTAGCTGCTCTCTCTAGCAATCTAGAGTGTAAATAGAATACATCCCCAGGAT
AGGCTTCTCGTCCCGGAGGTCTTCTAAGCAAAAGAGACATCTGACGATAAGCTTGAGCTT
GCTTAGTTAAATCATCATAAATAACTAATGTTGCTTTACCTTTGTACATAAAATATTCAG
CTAAAGCCGCACCTGTATAAGGAGCAATATATTGTAAAGTCGCAGGACTATCTGCATTAG
CTGCAACTATAATAGTATAATCAAGAGCTCCTTTATCCTGTAGTGAAGATACTACTTGGG
CTACTGAAGATGCCTTTTGTCCTATCGCAACATATACACAAACAACATCTTGGCCTTTTT
GATTAATAATTGTGTCCAATGCAACAGCTGTTTTACCTGTTTGACGGTCACCAATAATTA
ATTCTCGCTGACCTCTACCAATTGGGATCATAGAGTCAATAGCAGTTATCCCTGTTTGCA
TAGGTTCACAAACTGATTGCCTTCCAATTATACCAGGGGCCATTGACTCAATTAGCCTTG
TTCCATTACTTGCAGGTTCACCTTTATCATCTATTGGCCGAGCTAATGGATCAACAACTC
TACCTAAAAAAGCATCTCCGACCGGAATTTGAGCAATTCGACCAGTCCCTTTTACGGAAC
TGCCCTCTAAAATATCCCTACCATCTCCCATGAGTACTACACCAACGTTATCACTTTCTA
AATTTAAAGCAACCCCAATAGTTTTATCTTCAAATTCAAGTAGTTCACCAGCCATGACTT
CATCAAGTCCATATACTCGAGCGATACCATCACCAACTTGTAAAACTGTTCCTATATTGG
CTACTTCTACATCCTGATCATACTTTTCAATTTGTTGACGAATAATACTACTTATTTCAT
CAGGTCTAATATTTACCATATTTTTAATATTATTTTTATTAATTGGAAATTCTAAAGCGG
TAATTAGTCAGACTAATGCTATCTGTCTTTTTTACTTTATTTTAAGTTACTACAACATCA
AGGTGTGAAGCCATTTGTCTTAGCTGTCCTCTAATACTAGTATCAATGACTTTTGATCCA
ATCTGAATAGTAAAGCCACCAATTAGCTCGGGCTCGACTGAAATAATTAGTTTAACTTCT
TTAGCCTTAGTCATAACCTTGATTTTTTCTGTTAATAGAGTTTCCTGATCAGGGCTTAAA
GCAATAGAAGTATTAATATTTGCAATTGTTAAAGATTCCATTTGGTAAGCCAGTTCCAAA
TACTTACTAGCTATAGCATCTAACATTCCAATTCGCTTGCGATCAACTAAAATCATTAGA
AATGATAATGTATTTTCGTTAATTTGATCGCCAACAGTTGCAATAATTACTTCTTTTTTT
GCCTCTACAGTTTTTAATGGATTAGCTAAAAAAATTTTTAGCTTGTCAGATTGCGCTAAA
ATATTTTGTATTGACTGAATATCTTGGCTCACCTTTTCAGTAACTTTTTTAGTTTTAGCT
AAATCAAGAAGAGCTACTGCATAAGGCTGAGCTATTTTTATAACAACATTGTTACTGCTC
ATAGTTGATCTCCTAACTTAGCAATATTATTATCAATTATACGCAGTTGCATTTCGGAGC
TCATTTGATTTTCCAGCTGCAAAGTAACTCTTTTAAGAGCAAGAAATGTTATTTGTTGCT
GGATTTGCCTTCGGATCTGTTTTTCAGCTGTTTCTATATTTGATTTACCTGTTATGGCAA
GTCTTTCAATATCTAATTGGCCTTGAGCTAATATGGAACTTCTAACTTTTCCAGCTGTTA
GCTGAGCTTCTTTCTTAATTTGATCTATAATGATTTGAGTTTGAGCAAGTTGTTTTTCTG
ATTCTGACAATCTAGCACTTGCTTGCTCTAGTCTTTCTTCGGATTCTTGTATAGCTGCTA
AAACTTTTATCTGTCTTGCATTTAAACTTGATCCGAGAAATTGCTTCAAAACATAAATAA
GACCAAATAAAAGCAACAATATATTAATAACATTTGCTTCAAAAATATCAGAATTAAAGC
CGAATTTGTGTTCACTACTATGTTCTGATAAAATAGTAATTATTTGTGGTATTTTTACTA

TATTATTCATTTAAAATTCTTACTATTTTAATATTTTACTCTGGAATCATGATTTTCAAG
ATTGACTACTTAATAATTTAACTTTAATTTGATCGCTCAAAGTGTCAACCTGATTTTCTA
AAGTTTTAAGAGCTTCTTCTTTTTGAACATTTAACTGTTTCGATGCTTCAGCAATCAGTT
TCTCTGCATTCAATTGAGCTTTTTTTATATCTTCTGAGACTATATTTGAGCTTCTTTTT
GAGAAGAGGCTATTTTTAATTGTGCATTGCGGCGAGCTTCAGATAAATCTTCTTCATATT
TTGCAGCTAGCTCATCTGCTTTAACAAGCATAGACGATGCAGTAGTCAATGTTGTTCGAA
TATATTCATCTCGTTCGTCTAGTACTTTAGTTACAGGTTTATAAAAAATAGTATTTAATA
AAACCATTAAAGTAAGAAACTGTAATGCCATTAAAGGCAAAGTACCATTGAAGTCAAATA
AGCCACCTTCAATTTCTTCGGCTAATAGAAATGGTAAATCAATCATTTTACTAATTATGT
TTATAGTTAATAATTTTGTTTCAATTAAGAAAAATAACTTTATAGTGCGCTTAGTCTCAT
CAGACTAAGCGCAAATGATTATTTTAACCAACGTATGGATTAGCGAACAATAAAGACAAC
GCAACAACTAGGCCGTAAATTGTTAAAGATTCCATAAATGCTAAACTTAGAAGAAGTGTA
CCCCGAATCTTTCCTTCTACTTCTGGCTGTCTAGCAATACCTTCGACAGCATTAGCTGCT
GCACTACCTTGACCAATGCCTGGGCCTATTGCAGCAAGACCTACAGCAAGACCGGCAGCG
ATAACGGATGCGGCTGAAACGATTGAATCCATAATGAATTTTAATTTTTTGTTATATGAG
TAGAAAATTGACAAATTTCGTATAATACTTAAAATCTATATAATTAAGTATAATAGATTC
TAATATTGTAATTATAGTATATTACTCTTCTCCATGACCTTCCATTGCTTCTCCAATATA
AGCTGCGGACAAAGTAGAAAAGATTAATGCCTGTATTGAACTCGCAAATAATCCGAGTAT
CATAACTGGTAATGGTATTAAAATTGGAATTAGCAAAGTAAATACAGAAACAACTAATTC
ATCTGCTAAGACGTTTCCAAATAATCGAAAACTTAAAGATAGTGGTTTCGTAAAATCTTC
TAGAATATTGATTGGGAGTAATACTGGAGTTGGCTGAATATATCTAGCAAAGTAGCCTAA
CCCTTTTTTACTTAAACCTGCATAAAAATAAGCTAAAGATGTAAGTAAAGACAGGGCCAC
AGTTGTATTAATGTCATTTGTTGGGGCAGCTAGCTCACCTTCAGGCAAGTGAATTAACTT
CCAAGGAATTAATGCACCTGCCCAGTTACAACCTAGAATAAATAGAAATAAAGTTGCAAT
GTAAGGCACCCAAGGGCGATATTCATGCTCACCAATCTGATTTTTTGCAATATCTTGCAG
AAATTCTAATATAAATTCCATAAAATTCTGGAATTTTTCAGGAATTCTTTGTAGATTTCT
AGTACCTAGAAATGATAAAGTTAAAAGTGTAGCAATTACTAACCAAGAGACAATGAAAAC
TTGCCCATGCAATTGTAAGCTACCTATTTTCCAATATAGATGTTTTCCAACTTCTACTGC
TGATAAACTATTATATGGGTTAAAATCTATGAGACTGTTTTGATACATAATAAATATTTT
TTTAAATAGTGCAAGAATCAAAAAAATAAGTTTTTGAAGAATTAATATGCAATTAATTTAT
TCTTAAAATTAGACTATTAATAAATTAAAAGCAAAAATAAATTCTAATAGAGATAGCTAT
TTTTAGTAGTGATTATTTTTTGTAAAAAACAATTATTGCTACCTATAATGCTTTTACGTT
TAATTATATACCCATATATTAATTATTTAATATGATTATAGGTATTTTGCTACAAACTTT
ATTGTTTTATCCATTCTAAAGATTGTTATAGTTTATTTTTTATTTAAAATATCTGCGACT
TCATCTGCAAAATTAACTTTCAAATTTTCTTCTCCCCCTCCCAGAAGAAATCTTACAAAT
CTTCTAATTTTAATATTTTCACCTAGAACAGCAATATTTTGGTTTATTAAATCTTCAATA

CTAATATCTTGATTGCGGATAAACATTTGATCTAATAATGAGAGCTCCTTAAGTCTCTTT
TTCATTCGTCCTTCAATGATTGTTTCGATTCTATCAACTGGTTTGTTTTTCAAGTCATCT
TTACCAGCTTCAACTCTTTGTTCCAGATTAATAATTTCACTAGGTATATCCTGTGTTGAT
ACGTATTCCACATTAGGCGATGCCGCTATTTGCATAGCAATATCTTTAGCTAACTTTTGG
AACTCTGGACGTCGTGCTACAAAATCTGTTTCACAGTTGACTTCAACTAATACACCTATT
CTTCCGCCAGTATGAATATAACTCTCTAGTAAACCTTCAATAGCGGTACGATTAGATTTC
TTGTTAGCTGAAGCCAGCCCTTTTTGTCTTAATGATTCTAAAGCTTTTTCTTCATTGCCA
TTATTAGCTTGTAAAGCTTTCTTGCAGTCCATCATACCTGCTCCAGTCTTATCACGTAAG
GCTTTTACGACTTGAGCAGAAATTTGTAGTGTCATATGCTTTCAAAAATTTTTAAGTATT
TAAATTAATTATTATAATTCACTTTGAGTAGGATGAGTTATTTCTTCTAAATCTGTTTGC
CCGTATTTACCATCATAAATTGCATCTGCTATTTTACCTACAATTAGTTTAATTGACCTA
ATGGCATCATCATTAGCTGGGATTGGAATATTAATAATTTCTGGACTACAATTAGTATCA
AGTATACAAATAGTTGGAATACCTAATTTTAAACACTCTTGTATAGCTGTTGTTTCTCTC
TTTTGATCTACAACTACAACAATATCAGGTAAACGAGTCATATTTTTAATACCATTTAGA
TGTTTACGAAGTTTATCTAATTCTCTTCGTAAAACAGCAGCTTCTTTTTTGGGTAACTGA
TCAATCATACCACTTTTATCTTGTTCTTCTAGCTGTCTCAGACGATTAACTCGAGACTTA
ATCGTTACCCAATTAGTTAACATTCCTCCAAGCCATCTCTGGTTAACATAATAAGAATCA
CAGCGTTGTGCTTCCTGTGCCACTATACCAGCGGCTTGTCTTTTTGTTCCTAGAAATAAA
ACTTTTTTACCATCCGACGAAGCCTGTTTTATAAATTCACAAGCTTCTGTTAATAATTGT
GCAGTTTGGACTAGATCAATAATATGTATGCCATTTCTTTCTGTATAAATATATGGAAAC
ATTTTAGGATTCCATCTACGCGCTTGATGTCCAAAATGAACACCTGCTTCTAGTAATTCT
GCTAATGTAACAATAGCCATAAGATTATTAATATTTAAATAATGTAATTATTCGGGAAAT
CCCTTTGTTTAGAAACTGATTCAACTAACTTATTTGAATTTTAGTAAACTAATAATAAAT
TAACACACTTTTAAAGCAGATCAACAAAATAGATCCTGAAGAATTCATTCTACACTTTTA
TTGCTAAAATAATTACGCGCGGTTCTGTCGTCTAAAATGATATCATCTAAATCATCTCGT
ACAGATGAAATAGTACTTTCGTCATTAGTATTTGCAGTTAAATTCTTTTTCTCTAACGAT
GAGCCATTACAGTTATCATACATATTGAAGCCGGTACCAGCCGGTATAAGCCGACCTATA
ATAACATTTTCTTTCAGACCTCTTAACCAATCTAATTTACCAGATATAGCAGCTTCAGTT
AATACTTTTGTAGTTTCTTGAAAACTTGCTGCAGAAATAAAGCTTTCTGTATTTAAGGAA
GCTTGAGTAATACCTAATAGCACAGGCCTATATGATGCATTAATCTTATTTCTCAAGGTT
ATGGCTTTATTAGTCTGTTCAATCTTCTGTAACTCTACAAGCTCTCCAGGTAAATAGCCT
GTTTCTTCACCATTTTCTATTTTCACTTTAGAAGTCATCTGCCGTACTATTACTTCTACA
TGCTTATCAGAAATATTCACTCCTTGAGATTGATATACCAATTGTACTTCTTTAACAAGT
AGTAACTGAATCTCTTGAAAGCTTAATCGAGCAGCTTCATATAAGGCTAATCCACATTCA
ATATACAAATCAAAACTTGCATCGAGGATATCATGAGGATTTAACAATACGTCTGTTTTC
TTTCTTGCTTCAAGAATTTCTTCAATTCTTGGAAGCCCTTGTATAATGTCACCTGTTTTT

GCTCTATCAAAAACCAATATAGCTAATGTTTCTCCCCTTCTAATTAATGCATTGTTATCA
ACATGTAAAATAGCACCATTAGAAACCAAGTAAGGTCTAGCAATTCTTAATGTAACTGAT
CTAGAAGAAATTTCAATAATTTGTCCAGAGTCTAAAGAAGTCATATTTTCAGTAATAAAA
TCACCACAACGAATCCAATCACCCACACTTACTTTAACTATTGATTTTCCTATATTAAAG
GTCTTTTTATCAGAAGATGTCGCAATCAAAATGCGCCTGCTAGTATTCTTTTCAGAATAA
ATCTCCTCAACAGTACCTTCACTCATTGCAATAATCTCAGTACTAGCAACAACTGTAAAT
GGCTTGATATATTCACCATTTTTTACAATAATTCTAGTTTGACTTTGTTGCTTCTCTGAT
TTATTTACATCAATACTTTTAATTGATAATGTTTCAAAAGTTGACAGCCTTAAGTCAAAA
TAATTATTGCCACTTTCTTTAGCAGAAAACTCAATTGAGGATATTAAATTATCATCTCGA
TGCTCAATTTCAGCCACAAGATTAGTTGTTACTAAATGAACTCCTTCTATAGATTTCACT
CTTTCTCCGTCCCGAAAAGGAGTTCTTTTAACTAGCTTTAGTTTAGTTTGACTTGGTTTT
TGTGAAACTAGATTTTCAATTAGACTGGATTTTGTTTCTGGTATTGAATAAACAATAACA
GGTCTAATAAGAATATAGGGCATTTGCTCGTTTTCTATATACTCCCAATAAACTAACTTA
TCTGTAGAAATATTATTATGAAGTCTTTCTCCAGGTCTTAAAAAACCTCTACTTTTATCA
TTATTACTATACACCTCACTAAGTTTATAGATAGATCCAGGCTTAATAATAATTTCTCGA
ACTATTCCATCCTTTTCGATTATTTCAACAATACCTGAGCTTTTAGAAAAAAATATTTTTA
ACTAGCTCTGTTCCACTTTCTATAACATCTCCATTATGCACTAGTAGTAGTGATGAATCT
TTATTAATTTCGTGAGTTTCTTCTGAAATCCATAAGATATATCCCGGACTTAAAATTTCG
TAGGCATCTTTATCTAAGCCTGTTTTCTTTTTTGATACATTTAAGTCTAAATATTTAATA
ATACCACCACTTGTCGTTTTATACGTATCAGATATTAGTTCTGCGACTGTGTAACCATCT
TGAATTTTTTGGTTTGGAATAGCTTTGAAAAGAAATTTTTGTTTTTTTTCTGTTTCTAAA
ATATAAGATTCTTTTTTGTGGATTACATCTGTATAAATATAACAACCTGGAATTACAATA
GATTCAGTAATAACTTGTATATTGGTAATACTGCTATTAGTATTCTGGTGTATCCTTACT
TCTCCAGCATAATGATTTATTAATTCTGTCTGTGCAAGTATCGTACCAGCTTTAGTTTTA
TCTTCTTTATGAACAATAACATTAGCAGAATCAGATATACTATAAACTTCACCTGAGAGC
ACCCAGATCAAGCCTCCAGTTTTAGTAATTCTTGTAGTATATTGATTATTATCTGTTTCT
TCAACTGTTAAGTTGGAGAAACAGATTTTGCCAGAGAGATCTGAGACTACATGTTTTTGA
GCTCTTTCTGTCATGAGTCGATTTCTGCGGGGTGATTCAGCAATCACTTGATCCTTCGAT
ACTAATTCACCATCACATATTAAAAGAGTAGTACCTTTACTTAAGTTAATTATTGACTTC
TGATTTTTCTTAGATCTGATTGTAACTTGGGTTGGCTTTTTTGTTATTAATGCTTGCTCA
CCATGTCTGGTACGAACATCTGTATACGATTCAATATCTAGATCTATCAATTGACCATCT
ATAGGTGCGTAGATTTGTTCTGCTAATTCACCGGTAAATACTCCTCCAGTATGAAAGGTC
CTCATTGTCAATTGAGTGCCAGGCTCACCAATAGATTGAGCCGCAATAATACCAACAGCC
TCTCCTAAATCAACTAAGCGGCCATGTGCAAGATTCCATCCGTAACAATACTGACAAACA
GAACTTCTAGAGTTACATGTAACAGGAGATCTTACTAAAACTTTTTTGATACCAGCTTTA
GTTATCTCTTTTGCTAACTTAGGACTGACATCTTGGTTCGTATGAGCAATTAAACAATTT

GTCTCTGGATGGAATACATTTTCTGCTAATACTCGGCCAGATAAAGCTTGTTCTAGATTA

ATTAATATTTTCTGCGTATCTACTAAATCTTCTAGTATGATGCCTTTATTGGTCTTACAG

TCTACTTCTCTAATTATTACATCTTGAGAAACATCGACAAGTCGACGCGTCAAATAGCCG

GAATCTGCTGTTCTTAAAGCAGTATCAACTAATCCTTTTCTTGCTCCGTATGATGATATA

AAATAATCAGTAACAGTTAGTCCTTCTCTAAAATTACTCGATATTGGAAGATCAATAATT

TGTCCCTGTGGATCAGCCATAAGCCCTCTCATGCCCACTAACTGTCTTACTTGAGAAATA

TTACCTCTAGCACCTGAGAACGCCATCATATAAACAGCATTTAGAGGATCTGTCTCTTTA

AAGTATTTAATAACTTCTTGTTTCAAGGATTCACTCGCATTATTCCACGTATCAATAACT

TTTTGAAATCGTTCAACTGCTGTAATCTCACCTCGGCGATATTTGTTTTCAGTGGCCTTA

ATATCTTCAATCGTGCCTAATAAAAGACTTTGCTTACTAGGAGGAATTCTTAAATCTTCT

AAACTTAAAGAAATTCCTGCCTGGGTCGCATAATGAAATCCAAGATCTTTAAGCTTGTCG

GCCATATTGGCTGCTCGGGCTATACCATAATTACGAAAAGCCCAGACAATTAAATTTTTT

AGTTCGTTTTTATCAATAACCTTATTTGAAAAACTTGGCTGTGCAAGACTTCGTCTATTA

TCCACTAAATCTTCTCCATATTAAATGTTCGTTATTAACTATGCTACTAAAGACTCTTGA

ATAATTTTATTAAAGATTATACGGCCAGCTGTTGTTCTAATATATTGCACAATTCTCTGA

TTTTCAGCGTCTTCTTTTATGATATGGTTATTAAAAAATTTTGTTGTACTATTGTCAGAA

TGTTTTTCAATCTTAATCGGATCATGAGGTTGATCACCATCTACTAATCCATCAAAACGA

GCCCATATATAAGAATGTAAATCAATTTTCTTCTGTTCATAAGCCATAACTACATCTTCT

AAGCTTGCAAAATACTGATTAGATCCCTGCTGCTGAGATGGATTATTAGCTGTTAAATAA

TAGCAACCTAAAACCATGTCTTGGCTTGGCATAATAATTGGTTGACCTGTTGCTGGAGAG

AGAAAATTATGAGGGGCTAACATTAGCAATCTAGCTTCGGCTTGAGCCTCTAAAGACAAA

GGAACATGAACAGCCATTTGATCTCCGTCAAAATCAGCGTTAAACGCTGGGCATACTAAT

GGATGAAGCTTAATAGCTCTTCCTTCTACAAGAATAGGTTCAAATGCTTGAATACCTAAT

CTGTGTAGTGTGGGAGCTCTGTTCAATAGTACTGGGTGACCTTGAATAACTTCATTTAAG

ACATTCCAAATAGAAGATTCATTTTTTTGAATCATCTTTTTGGCAGCTTTAATATTGTTA

ACTAATCCCTGCAAAATTAGTCTGTGAATAACAAAAGGCTGAAATAACTCCAAAGCCATT

TCTCGTGGCAATCCACATTGATGTAGTTTTAAATGAGGACCTACAACAATTACAGATCTA

CCGGAATAATCTACTCTTTTACCTAGCAAGTTTTGTCTAAAACGACCTTGTTTTCCTTCT

ATAATATCTGACAAAGATTTTAATGGTCTATTATTTGCACCTACAACAGTTCTACCTCTA

CGACCATTATCCATGAGAGAGTCAACGGCCTCTTGCAGCATTCTTTTTTCATTTCTGATA

ATAATCTCCGGAGCTAAAATTGACTTTAGACGAGATAATCTATTATTTCTATTAATAATT

CGACGATAAAACTCATTCAAATCTGCCGTCGCAAATCTACCTCCATCTAATTGGACCATA

GGCCGTAAGTCTGGAGGTATCACAGGAATGACTGTGAATACCATCCAAGAAGGATCTGCA

CCTGTTGCTATGAAATTTTCAATCAATCTTAAGCGTTTCATTTTTTTATTGAATTTTAAT

GAAGGTGTCTTAAAACTTTTTGGTGGATTTGTAGCTTCAGACCTTAAGGTTTCAGCTATG

TGCTCTAAATCTAAATCTTTCAACAGTTTCTGAATAGCTTCTGCACCTATTCCTACTTCT

ACTTGGTTATTCTCATCTTGATTTTGGTAGATTTCCTCTTCAAGACTTTTCCATTCGTAA
CCTTCGAGAAGTTGCTTGTATTTTAAATTAATATCTGTATTAGATTGCGTAACTACATAA
GAATGAAAATAAACAATTTTTTCTACTTCTTTGACTTTTAAGTCTAATGCTAAAGCAATA
TAGCTTGTACTTCCTTTTAAGTACCAAACATGAGTTACTGGTGAGGCAAGTTCAATATAA
GCCATCCGATGTCTTCTCACGCGTGATTCTGTAACTTCTACGCCACATCGTTCACAAACT
ATACCTTTGTAGCGAAAACGTTTATATTTACCACAGTGACATTCCCAATCCTTAACTGGG
CCAAAAATTTTCTCACAGAATAAACCATCCATTTCTGGCTTTAAAGTTCTGTAATTAATT
GTTTCAGGCTTTGTAATTTCACCAACAATTTGTCCGTTGGGCAAACTTCTTTCGCCCCAC
TGTCGAATTTTTTCGGGAGAAGCTAAACTAATTTTTACGTAGTCAAAATATTGCTCAAAC
TTTGTCATAAATTTGAATACCTTAAAATCAAATAATTAGCCAATTTATTTACATGCTTCA
TATTGAAAAATGAATATCTTAATAAAGGAATTGTTCAAAATCGTCTACTGGAGGAGTATC
ATAATTAGAACGATCTACTCGGTTATCTTTAGAATCCGACATCAAGTCAACTTCAACTGT
CCGTCTTTGGCCATCTTCAAATAGTTTCAATTTATGTACTGCAATGTCTAAACCTAATGA
CTGCAGCTCTCGCATTAAGACTTTAAAAGATTCTGGAGTTCCAGGCTTCGGTATAGGCTT
CCCTTTAACTATTGCATTTAATGCTTCATTTCTAGCTTGCATATCATCCGACTTTACAGT
TAATAATTCTTGCAAAGTATATGCTGCACCAAAAGCTTCTAAAGCCCACACTTCCATTTC
ACCTAATCTTTGACCTCCATGCTGTGCTCTACCTCCTAATGGCTGCTGCGTAACTAGAGA
GTAAGGACCAGTGGATCTTGCATGGATTTTATCATCAACAAGATGAACAAGCTTAAGCAT
ATAAGCTCTACCAACTGTAACAGGATTATCAAAAGGCTCTCCTGTTCTACCATCAAAAAC
TTGCATTTTACCTGGATGCTGATCATTAAAAAGCCATTTATTGCTAGTAATCAATGATGC
TTCTTTTAATTTTCTATTTACAAGTGCTCTTGATGCTTCTGCTCCATACATTTCATCAAA
AGGTATTATTTTAAATCTCTTTCCTAAATAGCCACCTGCTAAGCCTAGTAGGCACTCAAA
TACTTGACCAACATTCATTCTAGAAGGTACACCTAAAGGATTTAAAACAATGTCCACTGG
TGTTCCATCTGATAAATAAGGCATATCTTGTTTAGGTAAGATCCTAGAAATGATACCTTT
ATTGCCGTGACGACCAGCCATTTTATCACCCACTTGGATTTTTCTCTTCTGCGCAACATA
GACGCGAATCATAGCATTTGTTCCGGGAGGCAGCTCATCTCCTTTCTGCCTAGTAAAAAC
CCTAACCTTAACTACTCTACCTTTTGCAGCATTAGGAAGCCGTAAAGATGTATCTCTCAC
ATCTCGAGCCTTTTCTCCAAAAATAGCTCGCAACAATTTACCTTCTGGTAGTTGATCAGC
TTCACCTTTAGGAGTAATTTTTCCTACTAAAATATCTCCAGCTTCAACCCAAGAACCACC
GACAACAATTCCATTTCTATCTAAATCCTTCAGAGAGTTATCGCTGACGTTAGGAATCTC
TCTAGTAATTTCTTCTGGTCCTAATTTAGTTTGACGGCATTCTACTTCATACTTTTCGAT
ATGGATAGAAGTATACAAGTCATCATAAACTAATCTTTCGCTAATTAAAAATGCATCTTC
GTAGTTATAACCTTCCCAAGGCATATAAGCAACTAGAATATTTCTCCCTAAAGCGATTTC
CCCTCCGTCTGTGGATGCGCCATCAGCTAAAGTTTGGCCAACAACTATTTTTTCTCCTAC
CCAAACAATTGGGCGCTGATTAATACAAGTATCTTGGTTAGACCGATAATATTTTTTTAA
GCGATAATGAACTGTTCTACCACTATTATCCTGAATGCCTATTTTATTAGCAGAAACATA

ATTTACATGACCAGATGTTCTGCTAATAACAACCATACCAGAGTCTCGAGCTATTTTAGT
CTCAAGACCAGTACCAATTATTGGCTTTTCTGGATATAATAATGGAACAGCTTGCCTTTG
CATATTAGATCCCATTAGAGCTCTATTCGCATCATCATGTTCCAAAAAAGGTATTAATGA
TGTTGCAGCTGAAATTACTTGAATAGGTGAAATTGCAATATAATCTACTTGGGTTGGGGT
TGTTGTAATAAACTCTTGACGATAACGTACAGGAATAATATCTCCTTCAATGTAATGTTG
TTTACTAACTTTAACATCTCCTGGAGCTACTCGGAAATCATCTTCCTCATCTGCTGTTAG
ATAGACAGGACTATTGTGATAAATCACTTGACCTTGGTTTACCGGATAAAATGGGGTTTC
TATGAAACCAAAAACATTGACCCGCGCACAGGTTGCTAGTGAACCAATTAAACCTGCGTT
CGGACCTTCCGGAGTTTCAATTGGGCAAATTCTACCATAGTGACTAGGATGTAAATCACG
AACTGCAAAACCTGCCCTGTCTTTATTAAAACCTCCTGGACCTAAGGCACTAATTCTTCT
CTTGTGAGTTAGTTCTGCAACTGGATTAGTTTGGTCCATAAATTGAGAAAGCTGACTAGA
TCCAAAGAATTCTCTCACTGACGCAATTAATGGCTTGGGATTAATTAGATTTGATAAACT
CAATGAATCTATATCACATATCATCATTCTCTCTCTAATAATGCGTTCTAAACGATTGAG
ACCTACTCTGAATTGGTTCTGCAGCAATTCTCCAACTGAACGAACTCTTCTATTACCAAG
ATGATCTATATCATCAAGATTACCTGAGTTTTTATCCTTAATATTAATTAGATAATCAAT
TGAGGATAAGATGTCTTGAGGAGACAATACTCGGAACGTTTTAGGTATATTGAGACCTAG
TTTTTTATTAATTTTATATCTACCAACTTCACCCAAGTCATATCTTTTAGGATCAAAAAA
ACGAGAGTAAAGCATCTGCTTAGCAACAGCAACTGTTGCTGGTTCATTAGGTCTAAGCTT
AGAATAAACTATGAGCAGGGCTTCCTCATCAGTTACTTCTTCAATATTATTTTTTCCAAT
TTCTTTAGCTAATTCTTTAACTGAATATATTTGGGATGCAGAAATAAGAAAAGCATATTT
ACTAAGTCCTTTTTCAATTTCATCTTTATTGAGTCCAATAGCTCGCAGAAAAATATAAGC
ATTTACTTTGTGGGTCTTATCTATTCTTATCCAAATTTCTCCCTTAGGATCAATTTCAAA
TTTTAACCAAGAACCTCTATTAGAAATTAGGCTTGCACTATAAATCTGCTTACCATTTTT
GTCTATCTCTTGTTTATAATAGATTCCTGGACTACGGATTATTTGGTTAATAATAACTCT
TTCCGTCCCAGATACAATAAAAGTTCCTCTATTTGTCATTATTGGCAAGTCTCCAATAAA
AATTAATCTCTTTTTGTATTTTTTATTTTTTATCTGCTTTTCAGCACTGAGCTGTAGATG
GGTTGATGATAAATCTAATGATTTAATATTTTTTTCCTGGTTTTTAGAAGGTAGATCAAT
ATCTTTTCTTGTTAACTTTGCAGGAACATATATTTGAGCACTATAAGTTTTATCCCTATT
TTTTGCCTGTCTGACACTATAACGAGGAAATTTTATCTTGTACTCATTACCAAATAATTG
TAATTCGAGTCGACTAGTAGGGTCTGATATCTTCGGAAAAATTTCAAGAACTTCCGTCAA
ACCTTCTAATAAAAACCATTTAAAACTGGCCCTCTGAATTTCAACTAAATCTGGAAGAAG
TTTATTTTTTAAGCTTATACGTTGAACCATAGATCTCCTTTATAACAAATTTTGAAATCG
AAGAGTTTAATACTATTAATAGAGTACTGGTAGATAATGCATAAGCATAGTTTGCTACAT
TATTTCTTCTCTAGACATAACTAGAACATCATATTTATTCAAAAAAAGATTTTTAAAAT
TTTTCTTCACACACACTATCTATAAGTACTTTGCTGCTAAAAGATAGTTCAAGCTATTAA
AAAAAACGAAATTAATACTTGTCTGTCTGTCAATTAATTGACATCAAGATAGACAAGTAT

TATTTATTTAATCAATTCTGTTTTTCTGAGCAAAAGACAATGCTCTCGCGAGTCTAGCTT
TTTTTCTAGCGCCTGTATTAGGGTGAAAGGCCCCTTTTTTTACAGCTTTATCAATCTTAC
TATATACTTGAGAAATACTTGACTGCACATCATTGAAATTATCACTTTCTAAATTATCAA
TATTTAGAAGACATCTTTTAGTTAAAGTTTTGACAACCGATTTATACTTACGGTTAATGA
GACGATTTCTTTCCGAAGTTTTAATGCGTTTAATCGCAGAAAGGTTCTTAGCCACAGTAA
ATCCAATAAAACTTTTATAAAATATTTTTAGAAACGAAAATACACAAATCTGGGATCTAT
TCAAAAAGGGAATATGCGGCGTATTATAACATTAAAGAAAAACAATAGGCAATAGTTCAC
CAAGAGCTCTTATTATTACACAATAGAGATTGTCATTGCGAAAAACTTTGTAAGATAGTA
TTTAATTAAAATAAACTAACTAGACATATAAATATATATGAAAAAAATTGAAGCTATTAT
TAGACCTTTCAAGCTTAATGAAGTAAAACTTGCTCTAGTCAAGGAAGGTATTGGAGGAAT
GACGGTTATCAAAGTTTCTGGATTTGGGAGACAGAAGGGTCAAACTGAAAGATATAAAGG
ATCCGAATACTCTATTGATATCATTGATAAAATAAAAATTGAAATTATTATTAGCGATGA
TAAAGTAGAGAAAATTGTAGAGACGATTATTAAGGCTTCTAAAACAGGGGAAATTGGAGA
TGGAAAAATATTTATTAGTAGTATTGAAAGAGTAATAAGAATCAGGACTAATGACTTAAA
TTTTGAAGCTTTATAGTATTTTTACTTTTTTAGACTTGATACTTCTTCACTAAACATGAG
ATAATAAGCGTTTATTGTATACTTATAACTAAGGTGTAATACTAATGGCTAAAAGCAAAG
GTGCACGAATCGTAATAACTTTAGAGTGCTCTGATAAAGCTGGAGAGTTTGCTCAAAAAA
GGAAACCTGGCGTTTTTCGATATACAACTACTAAAAATAGACGAAATACACCAAGTAGAA
TTGAATTAAACAAGTTTTGTCCTAATTGTAATCAGCATTGTATCTTCAAAGAAATTAAAT
AGTTATTTATCTTATTAAAATAATCATATTATGGCTATATACAGAAAAAGAATATCTCCA
ATTAAGCCAACAGATGCTGTTGACTACAAAGATATTGACTTGCTAAGAAAATTTATTACA
GAACAAGGCAAAATACTGCCTAAAAGGTCAACTGGATTGACTTCGAAGCAGCAAAAAAAA
CTTACTAAAGCGATTAAACAAGCTAGAATCCTTTCTTTATTACCTTTTTTAAATAAAGAT
TAAATATATAGCCTAACTTCATATTTTTCCATCTAAATATTAAAAATTTCTATAGATTTG
TTTATTAAAATGATCTAACATACTGAATTACTAAGAAGATCTACTCATTACAAGTTTTGT
TCATTTTATAATTAATAATAGCATCATAATTTATTAAGCTAGATTCAGACAAGACATAAA
GGTAACCTCTTTTAATTGATTATATTATATAGTCTATTTAAAAGAGGAAATTTTTTTGCT
TCTAGCAATTCAAGATTCTTTAAGCTTTAACAAGGTATTTAAAACAAAGATTTTTGTTTA
GGTATCAGATCAAATGAATGAATCTTATCTCCCGACTGCCATAATTGAAATTCAGAGATG
AAAATTCCACATTCGTTGCCAGCTTGAATTTCTTCGACATCTTCTCTAACGCGCTTTAAA
GATTCAATTTTTCCTTGATAAATAACTTTTTCTTCTCGAATCACTTTAATCCAAGAGTTT
TTTAATAGCTTATTATTGATCACTCGACATCCTGCTATTTTTCTATTTGCTAACGAGAAT
ACTGTACTTACTTCTGCTTCTCCCACCGGTACTTCTGAATATTCAGGATCAAGTAAGTCT
TCCATCCTTCTTTTAATGTCTTCAATTAAGGCATAAATGATTTGATAGTTTTCTATTAAT
ATATTCGACTTTGCCGATGCTTGCTTTGTTCCTGGAGCAAAGTTGGTATTGAACCCAATG
AGGGTAGAATTTGTGGTTGAAGCTAGTTCAACATCTGTAGCAGTAATTTCTCCAGGCATG

ATTGAGACAACATTGAGTTGAACTTTACTCTGAGGAAATTGAGATAAAGAATCTAGGATT
GCTTCTGTAGAGCCTTGATTATCTGTTTTAATAATTAAAGAAATTTGCTTACTAATATCT
TTAGAATTAGTGTTTTTTAAAGTATCTAATGTAATTCGACTATTAAGTGCTCTTTGTTTC
TGTATAATTGATGAGTCTTTAGAGGTATTTTCAATCGCTTTAAGTTTGGCTTCTTTATCA
CTTTTAACTGCTAAAGCTATTTCTCCTGTAGCTGGTACAGATGACAATCCCCAAATTTCA
ACAACAGATGATGGTATTGCTAAATTAATTTTCTCTTTAGCGTTGTTAATTATAACTCGA
ATCTTTGCGTAAGCTGATCCAATAACCAAATTATCACTAATATTCAATGTACCATTTTGT
ATTAATAGAGTTGCTACTGGACCATGCGATTTATCTAAATGGGCTTCTATTATAATACCT
TGTGCAGGCTGTGTTGGATCAGCTTTCAAATCCTCTAATTCAGCTAACAAGGTAATAGTT
TCTAATAATTTATCAACATTTTGACCAGTAAGAGAACTAATAGGAATTATAGGCACTTGG
CCACCAAGTTTCTCCGACATGACATTGTATTTTAATAAATCTTGCTCAATAATATCTGTA
TTAGAACCAGCTTTATCTATCTTAGATATTGCTATGACAAAAGGGACATTCGCTTTTTGA
ATATGATTAATAGCCTCTATGGTTTGAGGTTTAACTCCGTCATCAGCAGCTATAATAATA
ATAGCGACATCTGTTAAATTAGCTCCCCTAGACCTCATACTTGTAAAAGCTTCATGGCCT
GGAGTATCCAAGAAAACAATTTTTTGTTTATTATCTTTCTTAATATACTCAACTTCATAA
GCTGCAATTGCTTGAGTAATTCCTCCAATTTCTTTGTTAGCATTATTAGATTTTCGAATA
TAATCTAATAGCGTTGTTTTGCCATGATCTACATGTCCCATCACTGTGACAATTGGAGGT
CTTTTAATATAATTGCCAGTTTCATAAAGAGCATTAGAGTGATCTAAATTACTAGAAAGT
CCATTATTATTTTCCTTTACATTTGATTCTACTGCTATACCAAAATTATCAGCCACTGAT
GATATTATTGAGGCGTCTATGGTTTGATTCATAGTAACTGATATTCTTTTAAGAAATAAA
TATTTTATAATATCTGTTTCTTGAACACAAATTAGTTTAGATAATTCTTGAATAGTTAGA
GGATTAGTAATACTAATAGACTCTGGCGGAGAATTAGATTGAATGCTTGCTGAATTTTGA
CTAGGAGTTACTGCTACTTTTTTCTGTCTTATAGATTTTTTATTACTAGTCAGTTTTTTT
ACAACTTCCACTTTTGGCTTAGGTGGACGCATTAAAGAGATAGCTAAATCTCCTGCAGTC
TGCGAAACATTAGAATTAGAGTCTCTAAAGTTGTCATCATCATCGTCAATATGTATTTTA
GTTTTTATTTTTTTGCGCTGTCTATTCTTATTTTTTTTACTATCTACTAAGTCGTGAATT
TTATTAAAATTTTTACTTTTTTTATCTAGTTTAGGTGGAGAACTTAATTCTAGATGCTGT
TCACCACCTGTATGTGATTCTGATTTATCAAGGTCTAAATTTAACAGATTATCATTATTG
ACTGACTCTAATCTAATTTTATAGATAATCTGAGGATTTTTCAAATCTACAATTGTTTCA
GAAGAATTGATATTTGAGCTAGATCTAGAGGATCTATTTTCAAAGTTTTGATTATTTAAA
AACATAAATATTGGTTTTATTATCTAATAGTAACTTTCTAATTTTAATACAGAGCTTAAT
ATAAACAAGAATCGAATATCTGTATAAATAATACAAATGTAGAGAATTAATCTATGTTGT
ATTAGATTTAATATAACACTAAAGTCCTTTGTTCGTTATACAAGTTTAAAACTTTTTATG
ATGATAAACACTAGATTCTAGTTTATTGGTATACAATCTATTTATCTAAATAATACTGTT
TTTGTCAGGGCTTTTAAATTATCTAATGTTATGTTCATTATGAAATTCTATTAAAATTTA
AAAACTCAACAATTAAGTAAGAATCGTTAATCACCTCCACGACACTTATGCCACGATCTC

AAAAAAATGATAATTTTATTGACAAAACTTTTACAGTATTAGCTGATATTGTATTAAAAA
TACTTCCTACAAGTAAAGAGGAAAAAGAAGCTTTCTCTTACTATCGAGATGGAATGTCTG
CACAATCTGAAGGTGAGTATGCGGAAGCTTTAGAAAATTATTATGAGGCTTTAAAACTAG
AGGAAGATCCATATGATAGAAGCTATATTTTATATAATATAGGGCTTATTTATGCTAGTA
ATGGTGAGTATGTAAAAGCTTTAGAATACTACCATCAAGGATTAGAACTAAATTTTAAAT
TACCTCAGGCCCTTAATAATATAGCCGTGATATACCATTACCAAGGAGTTCAAGCAATTG
AGGATAAAGATACAGAGCTATCTAAATTAATGTTTGATAAGGCGGCTCAATATTGGCAGC
AGGCTATTAAGTTAGCTCCTGATAATTATATTGAAGCTCAAAATTGGCTGAAAACGACAG
GACGAATGAGGAATATACAAGGATATTAATATATTTAAGATATAATAAAGATATCAATTA
ATAAATTAAAATTAGTTAAAGAAACTGGTACTATATAGAGTACTTTACTATATATTATAA
TATTATACTAGTTGA------TCTCTAAGACTTAGCTGAATAAACGAAATTAAATCTATT
GACTATATTCATGAATACAAATAAGCATAATGGTTAGTACAACAAAAAGTACCGGATCTG
TTACTCAAATTATTGGACCAGTTTTAGATATTGCATTTCCTAACGGACAGCTTCCGAAAG
TATTCAACGCACTCAAAGTACAAAGCTCAGAAGGAACTATTACTTGTGAAGTACAACAAC
TTTTGGGTGACAACAAGGTACGAGCTGTTTCTATGAGTTCCACTGAAGGACTACAAAGAG
GGGTAGAAGTTATTGATACTGGATCCCCTATATCTGTTCCTGTAGGTACAGATACTCTTG
GACGTATTTTTAATGTTTTAGGTGAACCTGTAGATAATTTGGGTCCCGTTGATTCTGAGA
GTACTTTACCTATCCATCGACCAGCACCTGCTTTTACTAAGTTAGAGACAAAACCAAATA
TTTTTGAAACAGGTATTAAAGTCGTCGATTTACTTGCTCCTTATAGAAGAGGTGGAAAA
TTGGTTTATTCGGAGGTGCTGGAGTAGGTAAAACTGTATTAATTATGGAACTAATTAATA
ACATCGCTAAAGCTCATGGTGGAGTATCTGTATTTGGAGGCGTTGGTGAAAGAACAAGAG
AAGGAAACGACCTATATATGGAAATGAAAGAGTCTAAAGTAATTGATGCAGATAATCTGA
AAGAATCTAAAGTAGCATTAGTATATGGTCAAATGAATGAACCTCCTGGAGCACGTATGC
GCGTTGGCTTAACTGCATTGACAATGGCAGAATACTTTAGAGATATTAACAAACAAGACG
TTCTATTGTTTATTGATAATATTTTTCGGTTTGTACAAGCTGGATCAGAAGTATCAGCTC
TACTAGGCCGTATGCCATCTGCTGTGGGTTACCAACCAACTCTAGCAACTGAAATGGGAG
CACTTCAAGAAAGAATTACTTCAACAACAGAAGGATCAATTACATCTATTCAAGCTGTAT
ATGTGCCGGCTGATGATTTAACAGACCCAGCTCCGGCCACTACATTTGCACATTTAGATG
CAACAACAGTACTTTCGAGGAACTTGGCGGCAAAAGGAATTTACCCTGCAGTGGATCCAC
TGGATTCAACATCAACAATGCTACAGCCTGGAATTGTTGGAACTGAGCATTATTCTACTG
CTCAAGAGGTAAAATCAACTTTACAAAGATATAAAGAACTGCAAGATATTATTGCTATTC
TTGGTCTTGATGAACTTTCAGAGGAAGACAGACAAACTGTATCAAGAGCAAGAAAAATTG
AAAGATTTTTATCTCAACCTTTTTTCGTGGCAGAAGTGTTTACTGGATCACCTGGAAAT
ATGTATCTTTGGAAGATGCAATTAAAGGGTTTCAAATGATCTTAAAAGGCAATTTAGACG
ACTTGCCTGAGCAGGCATTTTATCTAGTAGGTGATATAGATGAAGCTATACAAAAAGCTG
ACAGCATGAAAGATTAATATAATTAATGATGACTTTAAATATAAGAATTATTGCTCCTGA

TCGGACTGTTTGGGATGCAGAAGCACAAGAAATTATTTTACCAAGTAGTACAGGGCAACT
TGGTATTTTAACAGGCCATGCACCTTTGCTTACAGCTTTAGATATTGGAGTTATGAGAGT
AAGAGTAGATAAAGAATGGATGCCGATTGTTTTGCTGGGCGGTTTTGCAGAAATAGAGAA
CAATCAATTAACTATTCTGGTTAATGGCGCAGAAGAAGCTAGTCAAATTGATTTATCAGA
AGCAGAAAAAAATTTAGACACTGCAACTCAACTCTTAAGTGATGCCTCGTCTAATAAAGA
AAAAATAGAAGCAACACAAAAAATACGAAAAGCTCGAGCTCGAGTACAAGCTGCAACAGC
AGCAACTTCGTAAAAGATTCCTCTTAAAACATAAAACTTTAGTAATGATTACTAAA----
---GTTTTATGTTTTATTTGTATGGTTATAGGCAGCTTAAATAAACTTAGCTTAGACCTG
AACAAATATAATCAAAATATAATCCCATCTCTTTACCTGCATCTGGTCCAACTAAACTAA
TAGTAACTTCTTTCATAGCTAAAATAGCTTGAATCGTTGCACCAATAGGAACACCTAGAG
AATTGTATGTTTCTTTTAGACCATTTAATACTCTTTCTTCAAGGATAGATGGATCTCCAG
CTAGCATTCCATAAGTAGCATAGCGTAAGTAGTAGTCTAAATCACGGATGCATGCAGCAT
AACGTCGAGTTGTATACATATTACCACCTGGGCGTGTAATATCCGAATAAAGTAGAGATT
TTGCTACGGATTCTTTAATAATTGTTGCTGCATTAGCTGCAATAGTAGCAGCAGCTCTGA
CTCGTAGCTCACCTGTTTGAAAATAGCCTCTTAATTTTTCAACAGAGCTATCATCTAAAT
ATTTACCTTGGACATCAGCTGCATTAATAACAGAAGTAATTGCGTCTTGCATAACTTTCA
CACATTCCTTAATTTTAAATATCTAATACTAGGGGTATATAACAAAATTACTAGATCTTT
TAGCAATATCAGTATTACTGCATAGCACCCAAAGTATAATCAAAGTAAAAACCAGCTTCT
GCTGAATCTTCACCTGCAAGCAGTGAGCAAGCTACACTTTTCATGCATTTTACACCTTCA
GCAACTCCTGAAATTGGTGTTCCTAAAGAATTATACATTTCTTTAACACCCACTAATCCA
ATTTCTTCAATAGGAGTTACATCGCCAGCAACTATTCCGTAAGTTACTAAGCGAAGATAA
TAGTCCAGATCTCGTAGACAAGTAGCAGTCATCTCTTCGCCATAAGCATTTCCACCAGGA
GAAACTACATCAGGTCTTTTTTGAAACAGTTGTTGGCCGCCTTGCTTTACAATACGCTCA
CGATTATCTGTTAAAATTTGAGCTATTCTTAAACGACGTTGTCCAGATAAAACAAAACTT
TTAATTCTATCTAATTCCCCAGGACTTAAATATCTTGCTTCTGCATCTGCATTTACAATT
GACTTTGTAACAATACTCATGGATAAAACTCCTGTAATGCTTT-AAAAAAAAGCTTAATA
CTAATTATTTATGATATAACTCAGAAAGTAATGTACTTTTTGAGAATTTTATAACCGAAA
CACGTTCATGATATATGAAGCAGGATAGATACTGAGGTAAGTACTAAAACGAGAAAGTTA
TATGTAGGAACTTCTGCTATTGATTTCCTAGTACTTTTTTAAAACTTGGTACCACAATTG
ATATATCCTGCTTAGTCAAACGATTATAGAGTGTTTCTGTATTAGGAAAGTTAGCTGCTG
GTAAAGTCGGAAATCTTCTGTATGGAACAGTATCTTTGCCATATACCGCATTGTATTCAA
CACTTTCAACTAAATTACTTATAAATGCAGATAAGCCTTTTGAAGCTAAAATTTGATTAA
AGAATCTAATTTCTGCCTGATTATTAGGAGCTCTTCCTAAGATATGCTTAGTACCTAATT
CTATAACTTTTGTATTTGGATATGGCTGATAAAATTCTTTACCATATAGTTCAGATAGCG
CCAGTTTTTCTACTAGTTCTTTGACACAGATTTGTCTATTCAAGAAAGCTGACTTAATAT
CCAAGAATTCACCGCCGACACTAAAAGAGTTAAGATCTCTTTCGAAAATTTGACGATATG

CTGCTCTTAAAGCTTGTTCTAACATTTCTTTGTTACTATCACTATTGACTTCAAAGACAA
TAGATTGATCTCTTAAAGAAGTTACTCCCTGTGAAATACGAGACTGAATATCATTACTAC
TCCTCATTTCTTTGACAGTGCCTAATTCAACAAACCTAGCAGATTTACTAGCAATAACTT
TTTTGAATCTCTGGTCAATAATTCCTGGTCTTAAGCTTCTTAAAGCAACACCAGCTGGAG
TACTATATCTTTCATAAGGAACAGTATTATCTCCAAAAGTTTCTGTGTACTCCGAGCTAT
CAATTATTGTGTCTATTACTTGGTAATAGCCTTGCTTATATGCAATATCAAAATATTTAT
TAATTTCTTGCCGACCATATGTAGGGCGACCCAGTAAGCGGTTATGAATATATTCAATAG
CTTTACAAATATATAATGGCTCCCAATAAAGAGATCTAAATATACTTGATTTTGCTAACT
GTCTAACAAACTCGCGAACTGTAATTTGATTATCTTTTAGCTGACTTTCTATTGGCTTGA
GAATTAGTTTTTCTTCTTGATACACTTCTCTGCCAAAAACTCTTAAATAAGCAACTTTTG
TGACAACTTCTACTGAATTTTCAAAATTTTGGGATGAATCTGACTTTATAAGAGCGTTGG
ACAATTTAAAGATTTTTGGTCCCAGGGATCCTGCTGACTTAGGTCTTACCTGTGGATTAC
TAATCTGATTGTAGATTCCTGGCCCACGTCTAACTAATATCCTTCTTGTATCTTTACCAA
AAAGAGCTTGTCTTTTTCTTGGATCTTTATTTTCTTTAGGGAAAATAGCTCCAAATTGAA
TTGATAATGGATCATTACCTATTCCATATGGATGTTGATCAGGTAATGATTGCTTATAAT
CACTAAATAAAGTTATAAATTGAGGAACCTTACGAAAAGGCGCACTATAGGTGAAAAGAT
CTATTTGAGGCCCCCAATTACGGCACTCTTGAGGCTCTTCACCTAAATTTCTAAAGTAAG
GCACAGTCTCTTCACCGAAATAATCTGTATACTCAGAAGAATTTAAAAGTGCATTCACTA
AACCACTTAAACCAGTAGATGATAAAATAGCAAAATATTTTTGAAACTCTTCTAAAGAGC
TAGGACCTCTACCCAAGAAATGTCTAAATGCTAATTCTAATGCTCGACTATTGACAAAAG
GTTCATAAAATTGCTTTCTATAAATACTTGAAGTTCCTAACGAACGAATGAATTCCTTGA
TTGAAATTTGGCCATTTTTTACTTGCGACTCCAAATTCGATAAAGAGAGGTCATATGCTT
TAGCAATATCTCTTTCAAAAATTTGTCTATAACATGCTTTCACAACAATATTTTTCTCAT
CAGCAGATAAACTAGGCTTCATAACAAAACGAGGTGTTGACACTCCTGCCTGTACATAAG
TTTGAGGAAGACGCAAACCTTGTAAATCTCCAGATATTCTTTTTCTTAATTTATCAGTCA
AAGATGGAGCTTCAAATTCAGAAATCACAACATTAAAATACTCTTTAACTAAATCTTGAC
CTTTTATATCTTCCTCAAAGATTAATAATGCTGTCCGTCTCATCTCTCTAAGAGCCACAA
TTGCTGCTGCACTTGAGCAAGCGTTATCAATTAACTCTCTCAATCCTCGTATATTAACAG
ATAAAATATTAGGGTCGCCTGATACAATTGCATATGTCAAATACCGTAAAAACCAGTCTA
AATCTCGAAGAGATTTTCTCATTCTTGTAGTACCGTATCTCAAGACGTTTATAGGTTTAA
ACCCTGGCGGAGTAGCTCCCCCAGCATTAAATAAAGATCGGAAACTTTGGCCAAAATCTC
CTTGGATATTTCCTGATAACTCATTAATTTTATCTTGAGAACTTTGATCTCCTGCAATAA
TAACAGCTGCTTGCGGACGTTCTAAATAAGAAATTGCAGACCCACCTACGAATATTTTAT
CTGCTGCTCGTGCTACCAGAATATTTGCATTTTTAGTTAAAATATCTGCAACTTCCAGTC
TTTTTTGTCCAGAATTTAAAAAAGAAACTAATTGATTTAATTCACCAAGCTGTAAAAATC
GGTCTTGTTGCTCTGCCTGAGTAATAGTTAAAATTGAGGCAGTCCGATAAAGCTGGGGGC

GTGCTAATGGGCTTCCGCCACTTGCTTTGATACTCATTACTTTATTTATCTCCTTAACTG
ATATTGCTTGCTAGCGAATACAAATATAGCTTGATATTTCTAAATATTTAAAGATACTAT
GCAAGCACACTGTTTTCTATATTATTAGGTACTAGTACTTATAGTACAGAGAGTCTTATA
ACTAACTTCTATAAATACAAATTTTGTAATACTTCTATTAACATTTGTCAAAAAAATATT
TAAGTGCTCACTCTTTTATATTAAAATTCAATTAGTGAGAGATTGATGTGAAATAAAGTA
TACTAAGTAAGTTTTATAGATAGGAAAATTAAAAGATTTATTCTACAAACATTAATCACT
TGAATGTTTAGCAATTGTACAATTGTTATTGCTTTTAATTGGTTTAGACTAGTTTTATAT
TTTTTCTAACATAAACAATATTAATGAGCAATTTGTATGAATTTAAAGCCACAAAACAAC
CTATTAACTAACGAAAATATAAATCATGTAGATATACCCGAAAATGATATACCAATGTCA
ATTACGGAACATTTAGAAGAATTAAGACAACGGACTTTATTTGTGTTTTTATTCTTTTTG
TTTGCTACAACTATAAGTTTCACACAAATTAAGATCATTGTTGCAATACTGCAAGCTCCT
GCTGTTGGTATTAAGTTTTTACAACTGGCACCAGGAGAATATTTTTTTTCATCTATTAAG
GTTGCAATATATTGCGGGATTGTTGCAACAACTCCGTTTGCCGTTTATCAAGTTATATTA
TATATACTCCCAGGACTAACTGGAAAAGAAAGAAAAATTATTTTGCCTCTATTAATTAGT
TCTGTATTGCTTTTTATTACAGGTGGTATATTTGCTTACTTTGTTCTCGCACCAGCAGCT
TTAACATTTTTAATTAGCTATGGGTCTGATATTGTAGAACCATTATGGTCTTTCGAACAA
TACTTTGACTTTATTCTATTACTTTTACTTAGTACAGGATTAGCATTTGAGATACCAATC
ATACAATTATTGCTCGGTGTTTCAGGGACATTTTCTTCTAGTCAAATGATACGAGCTTGG
AGATATATTATTATTATAGCAACAATTGCTGGAGCTATTCTGACCCCTTCGACTGATCCC
GTTACACAATTAATAATGTCTTCAGCTGTTTTATTACTTTATTTTGGTGGAATTGTTATA
TTATTAGTCTTAAAAAAGTAGAGTAAATATTTTCTATTAATCCAATATATTTGTTATCTG
GGCAAATAATCTATAAAAACAATTGCTAGATTTAAGTATCAGGCTTCTAAATATTTCAAG
TCTCTTTTTATCATTTAAATACAATCGATACATTGGCTTGTTTTTTCATCAATTATTTGT
GTTGCACACTTTTTAGCACTCTCTTTATTGTTCTTAGAATTATTTCTTTAATCTATTTAT
TAAATAGATATGAAGAGATTTAAGTATTGATTATGGTAATCTATATCAACGTAAAAAGTA
TTTTATTGTGAGTTTTATTTATATAAAGGCTTATTCATTATATAGTTATAACAGTAAGTC
TAATTAATAGTTTTCATGCTTAATTAAAGTAATAACTCTTAATTCTTTTGGCAGAAAAAT
CTGTCTTAATATTTTATCAAAAAAACAGAAAAATGAAGCTAAATAGTCTAATTAACTTAA
TTCAAAAGTCTATATATTCTTGTACACTTTTACTAATTATTTTAAATATTATTTGTGTCG
CACCTAATTCTAGTAATGCATTTCCAATTTATGCGCAACAAGCTTATGAAAGTCCAAGAG
AAGCAACTGGTAGGATAGTATGTGCCAATTGTCATCTTGCTCAAAAGCCTGTGGAGATAG
AAGCGCCTCAAGCAGTACTACCTAATACTGTTTTCGAGACTGTTGTGAAGATTCCATATG
ACAGCAATGCTAAACAGATTTTAGGTAATGGCAGTAAAGGAGGCTTAAATGTTGGAGCCG
TTGTAATATTACCTGAAGGATTTAAGTTAGCTCCTGTTAATAGATTATCTACAGAGTTAA
AAGAAAAGACTAGAAATCTTTACATTCAGCCGTACAGTGCTAAACAAGACAACATTTTAG
TAATTGGACCTATTTCTGGTGATAAAAATAGAGAAATAGTTTTCCCAATACTATCTCCCG

ATCCTGCAAAAGATAAAAAAGCTCATTTTTTCAAGTATCCAATATATGTTGGCGGAAATA
GAGGACGAGGCCAGATTTATCCAACGGGTGACAAAAGCAACAATAATATTGTCTCTGCTT
TAAGCAGTGGTAAAATTAATAAAATTGAATTACTAGACAAAGGTGGATTTATAATACATG
TGACTAACAGTAGCAATGTAGAGTCAACACAGAAGATTTCACCTGGTCTCGAACTTAGAG
TAAAAGAAGGGGATACAATTCAGCTTGATCAAGCTTTGAATAGTGATCCAAATGTAGGTG
GTTTTGGTCAGAACGAAACAGAAATAGTTTTACAGAGTCCAAATAGAATTAAGGGCATGA
TTGTTTTCTTTTTCGCTAGTGTTCTAGCTCAAATTTTCTTCGTATTAAAGAAAAAACAAT
TTGAAAAAGTTCAAGCAGCTGAAATGAATTTTTAAAGCAATATAAAACTGGCACTTACTA
TAAAGAAAGTATAAACACTTTATTCACAGGATGTACCAGTTTTTATGTTATGTAATAATC
GCTTTGACGGAAGTTATGATACTTTTAACAGAATCTACAATCTGACTAGGCTGAATTACT
GTAGCCTGCTCTAAGCTTCCATTATATGGTGTGGGTATATCCTGCGAAGATAGCCTCACG
ACAGGAGCATCTAATTCATCAAAAAGGTACTCATTAATTTGTGCAATTAGCTCTGCTCCA
ATTCCGGCTGTTTTCATACACTCTTCTACAATTAAAACTTTGTGAGTTTTCTTTACTGAG
ATAGATATAGAGTCTATATCCAGAGGTTTTAAAGATATAAGATCTATAACTTCTGGATCA
TAACCTTCTTTTAATAAGGCCGGTAATGCTTGGATAACATGATGCCTCATTCTAGAATAG
GTTAAAATCGTAATATCTTTTCCTTTTCGTACAAATTCAACTTTATTGAGAGGTAAGAAA
TATTCTTCTTGAGGAATCTCTTCTTGTAAATTATAAAGTAGAACGTGCTCAAAGAATACA
ACTGGATTATTGTCTCGAATTGCAGATTTCAGTAATCCTTTTGCATTATAAGGAGTAGAA
CAAGCAACTATTTTTAAGCCGGGGATGGCTTGGAAATAAGCTTCCAGTCTCTGAGAATGC
TCTGCACCTAACTGCCTACCAACTCCTCCCGGTCCTCTAATAACTAAAGGCAATGTAAAA
TTACCTCCAGAAGTATAACGTAACATTCCTGCATTATTAGAAATTTGATTGAATGCTAAT
AATAAAAAACTCATGTTCATACCTTCAACAATTGGTCTTAGCCCTGTTATAGCTGCACCA
ATTGCCATACCAGTAAAGCTATTTTCTGCTATTGGTGTATCAAGAACTCTTAAATCCCCA
TATTTGCTATGCAAATCTTTAGTTACCTTATAAGATCCACCATAGTGACCAACATCTTCT
CCTATTACGCAGACAGTTGGATCTTTTGCCATTTCCTCATCTGTTGCCGCTCGTAAAGCG
TCAAACATAAAGATTTTACTCATTTTAATATTTGATTTTTATGATTTGACCACTGTTTTA
AAGTAACAGTTAAGACACCATATTCTATTTAGTTATCTGCAAAAAGATATCGTTTCAATT
CTGACATGTTAGGTTCTGGACTGGAAATAGCAAATTTCACAGCTTGTTCCAATTCGGTTT
TAACAGCATTCTGAATCTCATTAAGTTCACCTATATTAGCAATTTCATTATCTAGAATAT
ATTTTTTGAGTTTTTTGATAGGATCTCTTGCCACCCAAGCCTCTTTTTCTTGTCTTGATC
TTAGTTCATCAGGATCTGCGAGAGAGTGACCACGAAATCTATATGTTAATGCTTCTATTA
AGGTTGGACCATCACCTTGGCGAGCTCTTTGAACTGCTTGTTTTGCAGCTTGTCTTACAG
CTAGCACATCCATTCCATCAACTTCAATCCCAGGAAGCCCAAAAGCTTCTGCTTTTTTAT
GTATTTCAGGTATTGAAGAAGACCGGTGATGTGCCATACCTATAGCCCACTGATTATTTT
CAACAACAAATATAATAGGTAGTTTCCAGAGAACTGCCATATTCAGACATTCAAAAAATT
GCCCATTATTGGTAGTCCCATCACCAAAAAAGCAAGCCGTGACTCTTAAATCCTCTGTTT

CTTTAAGTACTTGCTGGCGGTAGATACTTTGAAAGGCTGCCCCTGTTGCAACCGGTATAC
CTTCCGCAATAAAAGCAAAGCCACCTAAAAAATTGTGAGGCGCAGAAAAAAATATGCATCG
AACCTCCTCTGCCTTTACTACAACCAGTCTCTTTTCCAAATAACTCAGCCATCACATTTT
TAGATGGGACGCCTTTACTTAAAGCATGTACATGGTCTCGATAGGTACTGCAAACATAAT
CAGTTGGATTCAGAAGTTTAATTACACCTGTAGAAACAGCTTCTTGACCATTATAAAGAT
GAACAAAACCAAACATCTTTCCTTTATAATACATCTGAGCACACATATCTTCAAAATTTC
TGCCTAACAACATGTCTTCATATAAAACTAATAAATTACTCTTATTAAGATTAAGACCTG
TTGAATTATAGTTAGTCAGCGGCAATTGAACTTTCTTAGGATAACTCATAATTTGTAAAG
AAACCTCTTATTGTAAATAGGAATTTTGAAGAAACAGCTATCAATAGAAAACACACAGAT
AAAAACTTATAAATTATAATATGTTTTTACTAAAGACTTTAATAATTATTTGAAACTAG
TTCAGTATATCAAATTAATTTTGCTTATACTACGGATGTCTAAAACTTATATAAAAAAGT
AGTAAAAAAGTTTATTAAAAGAATAGTTAAAAACAGATATAAATAGAAGTAAACAGAATT
ATTGAAAAAAAATGATTATTATTAATTAATACTTAATATTTCTAAATATTTTACAATTAA
TGGCTATGTCACCATCTTTATTTTATCCTGTTGAACAAGAACTATGTAGTCTTGAAAAAA
ATCTGAAAGCTGTTGCTGGGACTCGTCATCCAATTTTATATGCCGCAGCAAAGCATCTAT
TCGATGCTGGAGGAAAACGAGTTAGACCAGCTCTTGTATTTTTAGTGGCTAAAGCAACCT
CTGAGAAGCAAGATATAAATACTGGACAAAAAAGGCTAGCAGAAATTACTGAAATTATAC
ATACTGCTAGTTTGGTACATGATGATATTATTGACGAGTGCACAACACGTAGAGGAGTCA
AAACTGTACATAATTTATTCAATACTAAGATTGCTGTGCTAGCAGGAGATTTTTTATTTG
CACAGTCTTCTTGGTATTTAGCTAATATTGAAAATTTAGCCGTAGTCAAAGCTATTTCTA
AAGTCATCACCGACTTTGCAGAAGGAGAAATTAGGCAAGGCTTAGTTCATTTCGATCCCA
GTATTTCAATAGATGCTTACATTGAGAAATCATTTTACAAGACTGCTTCACTCATTGCTG
CTAGTTGTCGGGGTGCAGCTATGCTTAATGGTTCCAATCATCAAATAAATAATGATCTTT
ATCTTTACGGTAAACATATGGGATTAGCATTTCAAATTATGGACGATGTTCTAGATATAA
CTGGTTCTACTAAGAGCTTAGGAAAACCTGCTGGCGCTGATCTAATAAATGGAAATTTGA
CCTCTCCTCTCCTTTTTTCACTTACTCAAGAAGCAAGTTTAAATGATCTTATTGATAGGG
AGTTCTGTAATAGTACAGATATAGCCTCAACATTATTTCTTATAAAAAGAAGCGGGGGAA
TTACAAAAGCTAAAGATTTAGCTAAAGAACAGGTGCAGGCGGCACTTTCCTGCCTTCAGT
TTTTACCACAATCTACACCTGTATCTAGTTTAAAAGAATTAACACATTTCATAATCACAA
GATTGTCATAAAGGACTTGCTAAAATTAAAATGTTATTAACTAATTTTTTACTGTTTCTA
AAACAGTCTGAAAAGTTTGAATATCATTACTAGCTAGCTGCGCTAGCATTTTACGATTTA
ACGCAATATTCTCTTTTTTTAATGCACTAATAAAAGTACTATAATTCATGCCTTGATTGT
GAGCAGCAGCATTTATTCTAGTTATCCAAAGACGACGAAAATCTCTCTTTTTCCTTTTTC
GACCAACATAAGAATATCGAAGAGCTTTAAGGACTTGCTGCTTAGCTGTTCGAAATAAAC
ATTTATGTGCACCTTTAAAGCCTTTAGCTAGCTTAAAGATTTTAGCTCGTCTTTTTTTTG
CAACGTTACCTCTTTTAACTCTACTCATAAAATATTACTTAAATTTTAATTATTACCAGC

TTACTAAATTCTTATACTCTTTGATTATAAATAAGGTAATTTCATTGCAATGTTTTTAAT
ATCTTTTAAGTCAACCGAACAAGTAGAAGAAAGATGTCTTCTTTGTTTTGATGACTTTTT
TTGTAATAAATGACTTTTAGAAGCTTTATGTCGAAGAAATTTTCCGGATGAAGAGACTTT
AAATCTTTTTGCTATTGCTTTTGATGTTTTTAACTTAGGCATATTATAAACTTTTAATGA
TTAACAAAAGCATGAAAAGTCTGAGGATATCAATCTTAGACCTTTCAATACTTAGAATAA
TCTTAACACAATTTATTTTACTTACTACATAGGTTAACTGAAACAATTTATATATATAAT
ACTAATCAGTTAATCAACCATAGCTAAAGTGGATTTATATTTTGCTTTGAACTTTTGTAT
AGTATTAGCAATTAGCATTGTTATAATTTTAAGACTGCTTGAATTCAATTCTTCGAACAT
TTTCATATTTAATGTAAATGAAATATTGGCTTCTGAGACAATATTTTGAATTTGAACATC
AGACAAGGGAATCATATCTAATGCAGCTCTATATTGATCTTTAAATAATTTATCGTCTTT
TATTTGGTCAAAATCGTAAAATTTCGTTCCTCCTGAATCTGATAAATTCATAGCCCCTCT
AGCTATTTTTTTTAAAATTTGGCCACCAGAAAGGTCACCCAAATAGCGCGTATAAGCATG
AGCAACTAGTAATTCAGGTTGTTTATGTCCTATAGTGTGAATTCTATCAACATAAATTTT
TGTAGCAGGAGATGGTTCAATAAAATCTAACCAATCTGATCCATAATAATAGTTTAAATC
TTCAGATAGACTCGCTTTCCTATTAAGCTCTGTAAAATATATAGGTTTAATAGCTGGATG
ATTTTTATTAGAAAATAACTCTTCCTCTATTGCACAATAGACAAAGTATAGATTTGCAAC
CAATTTACGATATGACTTCTTATCTACAACTCCTCCCAAAAAAGATTTAACAAAACTAAC
ATTTTCTGCCATACTATGGGACTTAGTAGTACCTTCTCTTAGTTCATTCGCTAAAGTATT
AACCATAATATCAATATTCCATTTTTAAAGATAACAAATTAACAATAAATAGTTATCCGT
ATATTTATAATTGAGTTGCGCTAACTAGGCTGCTGCAAAATAATTTTTTGATTTTATTGG
ATCTGGATTCATTGTTTTGTCACCAGGCTTCCAATTAGCTGGACAAACTTCATCTGGATG
AGATTGTACATATTGAATTGCTTGTAAAACCCTCAAAGTTTCTTCTACACTCCTGCCAAA
CTCTAGATTATTGATTGTAGAGTACTGAATAATTCCTTTCGGGTCTATAATAAATAGTCC
TCTTAGAGCTACACCATCACTATTCAAAACATTATAAGCTGCACTGATCTCTTTTTTTAA
GTCTGATACTAATGGATATGAAAGATCTCCTAATCCACCTGATTCTCTGTCAGTCTGTAA
CCAAGCAAGATGAGAATACTCACTATCAACCGAGACTCCAAGAACTTCTGTATTAAGTTC
CGAAAAAGCATTATACTTATCGCTAAATGCTGTAATTTCTGTAGGGCACACAAATGTAAA
ATCTAAAGGATAAAAGAATAAGACAATATACTTATTCTTGAGATCAGATAATTTTAACGT
TTTAAATTCTTGGTCATAAACAGCTGTAGCTGAAAAATCAGGAGCTAGCTGACCTACTCG
AAGACAATTTGGTCCAGAAATCATTAATTTTTCCCAGAAGAATTGATATAATTTTGTTAT
TTAATAAATAATTATATAACAAAATTTAGTATATATTAAAATACTACCAGTCTTTTTTAA
ATTACTAAATGATATAATCAATACTATCTTTATAGCGGGGAATGGATTTGAACCATTGAC
CTTCGGGTTATGAGCCCGACGAGCTACCAGACTGCTCTACCCCGCGGATAACTAAAATCA
TATTAGTCTACATATATATATAAAAGCAAATGCATAATAGATTTTAGAAGTAAAAAAATA
GAAGAACTGGTCCACTGATCATTAAACTTAGAAACGACAAAGTATTTTTTATCTTATTTA
AAAGAGGAAAGACTTCATATAAACCAATAAATCTATCTTTTAAAAGAATTTCGGACGGCT

TTACCCAAATTTGACCGTCATACCAGCCTGATTCTTCATAAAATACTGTAGCACTCATTA
ATCGTTTTACTACGTATGACCAGCCTAAATATAATCTGATTAAAATAAATCCTGTCATTA
GGCTGGAAGTAATAAATTCTGAAAAGAAAAATTTCAAAGGTAGTTTAGTGATTGGAAAAA
TTGATAATAATATAGGACTAACTAATAAACAGTTAAGTAGTAGTGCAATAGTTATTTTTT
TGTTATAAGATCTATGGCTTAAAGTTGGCCAGCAAAAGAACCAAGAATTTTTCAAAGAAG
TATATTCATGAACAGGTTGCTGCTCCTTTGGCACAGGACATTGAGTATTATATAAATTCA
TTGACATTAAAAAGTATTATAATAAAGACTTTGTTTATATTATCTAACATAGGTACGACA
TTAAATATTAACTAACTGCAATTATAGTAGTGAGTAGAAGAAATAAAACTAAAAATAGCC
AAGTAATTTTATTTAAAGTATTCTCAGTACTACGAGTATTACTAAAGAACTGATTCTGAG
CACCTACACTTCCTAAACCTTCGGATTTAGGATTGTGTATTAGAATAGTGAAGATTAATA
TTATTGTAGATGAGTACCAAAAAAATTTTAAAATTTGTTCCATTATGTAAAAGTTAATAA
AAATAGCACAAAGACAATCGTCTAAAAGATATTGATTGTCTTATCATATTAAAAAGTATT
TATTCACCCTGTACTTTTAAAAGTAAAAATCCTATTGCCAGCCCTACCAAAACTAGGACT
GAACTTAATACAGCACTTTCTACAATTTCTCCGACCATTTTATTTTTCCTTATTATTTTA
TTATTGTCATTCTTTTGACAAGATTATATTGAGCTAACTAAAATCCGTTTCTACCCCAAA
CTACCATTGCAATTGAAAATGTAAACATTGCCATTAGAGCTGCCCACCCCAAACTTAAAA
TATCCATAATTTATAATATTCCTAAAAATAAGATCTCTTTTACTTAAACTATAATATTCT
TAATATTAAAAATATCATTATATAATAAGCTAATTATAACAGAGTACTACTTAATTCGTA
TTTTTAGTGTATTTAAAGAAGATTGAATTTTTTCGAGGTACTGGATAAGTATTTCATCCT
AAGAACCATATTCAACTTTTGATTTATTATAATGCTTAAATCTAAATATATATCACTCAA
GAATCATTAGTAATTATGATGTTTTTAATTATACGATTAACATATTTCAAAATAGAAGAA
TTCATAGCGACTAAGTTAATATGCTAATATAATTTATCAATAAAAAGGAAGTTAGTATTT
CAATTATGCAAACAACTATAAATAATGGGCAAACGTCTAGTAAAGAAACTTTACTAACAC
CTAGATTCTACACGACTGACTTTGAAGAGATGGCTAATATGGATATTTCCGGCAATCAAG
AAGATTTTTTGGCTATCCTCGAAGAATTTCGAGCTGACTATAATAGTGAACATTTTATTA
GAGATGAAGAGTTTAATCAATCTTGGTCTAATTTAGAACATAAAACTAAATCCTTATTTA
TTGAGTTTTTAGAAAGATCTTGTACCGCAGAATTTTCAGGTTTTTTACTATATAAGGAAT
TATCCAGAAAATTAAAAGACAGAAATCCGGTTATAGCTGAGTGCTTTTTATTAATGTCTA
GAGATGAAGCTAGGCATGCCGGTTTTTTAAATAAAGCTATCGGGGACTTCAATTTATCTT
TAGATTTAGGATTTTTAACGAAGAGTCGCAAGTATACTTTTTTCTCACCTAAATTTATTT
TTTATGCAACCTATCTTTCTGAAAAAATTGGATACTGGAGATACATAACTATTTACCGCC
ATCTTGAACAACATCCAGAACACCGTATTTATCCAATCTTTAGATTTTTTGAAAATTGGT
GTCAGGATGAAAATCGTCACGGAGATTTTTTTGCTGCTTTGCTCAAATCCCAACCTCATT
TTTTAAATGACTGGAAAGCAAAAATGTGGTGCAGATTTTTTCTATTAAGTGTATTTGCAA
CAATGTACTTAAACGACTTTCAAAGAATTGATTTTTATAATGCCATAGGCCTAGACTCTA
GACAGTATGATATGCAAGTAATACGAAAAACTAACGAAAGTGCCGCTAGAGTTTTCCCGG

TTGCTTTAGACGTGGACAATCCAAAATTTTTCAAATATTTAGATACTTGTGCATGTGATA
ATAGAGCTCTGATCGATATTGATAATAACAATTCTCCATTATTCATTAAATCTATCGTAA
AGATACCTCTATATTTTTCCTTATTTGCAAATTTACTAAAGATATATTTGATTAAGCCAA
TAGACTCAAAAACAGTATGGAATACAGTTCGATAGTTAAATACAAGGGAATGCTTTTTC
TTATAAAAAGCTATCTAGTGATTAATATAAAATAAAAGCTTCCTTCTCGATGGAAGGAGC
TTTTATTTTGTCAAGAATATAATGGGATTATATAGATCTAGAACTAAGAACTTACTTTTT
CTTTTGCTTCATACATAACCTCTAAAGTAATTAGATTTTTAGAACAATCACGGGCAAATT
TCTCTGTATTTCTTTTTACTTTACCTCTAACAAACCCAGGAATTTTATTTAATTCTTTCT
GTGCTTCTTCTGACCAATTAATACTATCAACAGCAGAAATTCCTATACTTAATGCTTCTG
TTGTATCATGACCACCAAATATTTCTAGCAAATGATCTTCCATACCAAGTGTAAACGAAT
TATATACTAAATCAGCAATTTGATTTGTTCCCTCATAACCAAGAAATGGACGGTAACTTA
AAGGAAAGTTTTGTATATGCACTGGTGAAGAAATAACGCCACAAGGAATATTTAGACGCT
TACCAATATGTCTTTCCATCTGTGTTCCAAAAATTGCTGCAGGTTCAGTTTTAGCAATTA
AATCTCCAATTAAACCATGATCATCTGAAACGATTACTTCATCGCAGAATTCTTGAACTT
GATCTTTAAACCACTCTTCATCATATTTGCAATAAGTTCCACACCAGGCGACATGTATAC
CCATTTCCTGGTGTAAAATTCGAGTTATTGCTGCAGCATGAGTTGCATCTCCAAATACAA
TAGCCTTTTTACCTGTTAAATTCTGACAATCAATTGATCTTGAAAACCATGCAGATTGTG
AAATAAATCGGGTTTGCTCGTCTATGTATTTTTCGTAGTCTACAGCAGCTCCCAATGCAT
TTACTAATTGCTGTATTGATCGGATGCATGCAGCTGTTTGAACAATACCCATAGGTGTAA
TGTCAACATAAGGCATATTAAATTCTTTCTCTAAATATCGAGCTGTCATTAGTCCAGTTT
CTCTATATGGAATAAAATTAAACCATGCAGAGGGTAATTTTTTTAAATCTTGAACTGATG
CATTTTCTGGAATAATTTGGTTTATTTGGATATCTAGATCTTGAAATAACCTTTTTAATT
CTGCTATATCATGTTGATTATGAAATCCTAAGCTTACTGCTCCAATAATATTAACAGATG
GAGTTTTAGTCTTTTGAGTTAAAACTTGATTATTAGATTTAGCTTTTTCCATATAGAAAG
TGACAATTTGTTCTAACGTTCTATCACCTGCTTGCAGTTCATTTACTCTATAATGATTAA
CATCAGCTAATAAAACATCTGCTTCTGTTTCTATAGATGCTCTGCTAACAAAGTTTTGCA
AATCTTCTTGCAAAATACTCGAAGTACAAGTTGGTGTTAGAATAACTAAATCAGGGCTTT
CTTCTCTGTCTTTTCTAGTAATATTTTCTACAACTTTTTCTTGAGAACCACGGGCTAACA
CATGTCTATCTACAACACTTGCTGTAACAGGAGTAAAATCTCTGTCTCTTTCAAGCATTG
AACGCATTACATTAAAATAATCGTCTCCAAGAGGCGCATGCATAATTGCATGAACTTTTT
TAAAAGAACTCGCAATTCTTAAAGTTCCAATATGAGCAGGGCCTGCATACATCCAATAAG
CTAATTTCATATTAAATTTTCCTATGATTTATTTAGACATAGTAATACTTTTAAAGCTAA
CTAACAAACATTAGCCAAACATCTAATTTAACTCCTTGACAATTTTGTAATATTTTTTAA
ATGAAGCTTATCATTATAAATATTTAAAAAATATAATTCAGAATAGAAATAACTGTTAGA
TCAAGCTTGAAAGTAAAGATTTCTTGCAGTTATAGTTTTTCTTTAAGTAAACATGTAAAA
ATTGTACTATATATAAATGTACTATATATAAACTTTTTTTACTGAATATTGGTAATGCCA

GATACAATTAACTTAAATATGCCTTCCCCAACGTTTGGTGGAAGCACTGGCGGTTGGTTA
AGAGCCGCAGAAGTAGAAGAAAAGTATGCGATAACATGGACAGGCAAAAATGAAAGTAAG
TTTGAAATGCCAACTGGTGGTACCGCAACTATGAGAAATGGAGAAAACTTACTCTATTTA
GCTAAGAAAGAACAGTGTTTAGCTTTAGGTACTCAACTAAAAAGTAAATTTAAAATATCT
GACTATAAGATTTACAGAGTTTTTCCTAACGGAGAAGTGCAATATTTGCATCCTAAAGAT
GGAGTCTTTCCAGAAAAAGTTAATACTGGGAGAGCTAGCGTCAACAGTGTTGATCATTCT
ATTGGACAAAATATTAATCCTGTAGATGTCAAATTCATGAATAAAGCAACTTACGATTAG
TCTCTCAAAAACCTAGACATAAAATAACTTTTATATAGTCTAGGTTTTTGATAATGTATA
CTGATAGTATGGAAGGGTGGCCGAGTGGTTGAAGGCGTCTGATTTGAAATCAGTTGAACT
ATCTGGTTCCGTGGGTTCGAATCCCACCCTTTCCGTTATAAAACTATAGTTGGCTTCTCA
AGGCAAGCTAGCTATACACTGATAGTATTAAGTTTAGACTTAATAAAGTCTACAACTTGT
GACAGGTTAGAAATTTTTTCGGCATCCTCATCTGGTATCTCTATACTAAATTTTTCTTCT
ATGGCCATTACCAGCTCAACTGTATCTAGAGAATCGGCTCCCAGGTCACTTGAAAAATTT
GCTTCCCTAGTCACTATTTTTTTTTCAATTCCTAGCTGTTCTGCTACAATATCTTGAACT
TTTTCAAAGATTTCATTATCTTGCATAATATATTATTTCCTCTAAGGTGTTATCAAAAAT
TTTTCGGCTTATATTATTATCCAACTCTAAAAAATTAGTAATTTTTAGTCGAGATTAAAA
TATTTATACCTATATGATATACTATTATTCTGGATAAATACGAAGTTTACGATATGGCTC
TTCTCCAAAACTACGAGCTCTCAACTGGTAGTTGTCAACTAGATTATGCTGCATTTTTCG
TATATAGGCTGATCGAGGTGTAAGCTGAACTATTGAACTTTCATTCAAGATAATAGTTTC
AATAGCCAATTTAGCTTCTTGAAGAGCCTGGATCTCATCAAAATTTTTATTTTTACAGAG
TTCTACCCAATTAAGACCAGATGAAGTATGAATGTTTAATATTTTTCTGAGTGCACGAGT
AATTTGTGGAACCGTACTATTCTGAATTGTATATATAATAATTTGTTTTGATTTTGCAAT
TTGTCTAAGTTTTGTATTTTGCTTTACTTGATTACGTAGCGCTAGAATAGCATCAGACTT
TTCTATCTCCTTGGTTAAAATAATTGGTAGATCTAGTGAAGAAATAACTGATGTAATATG
CTGCCAACTCAAAGAATAAGCATATAAGTACTGATGAGGTATTTCCAATTGCAAAGATTG
TTTATTAATATTAATAGGGGCATTGAAAGTAGTTGACAATAAAGATGTATCACGATTTTG
TGATTTATTCAAATCTAAGGACTTATTTCTTAATTCTCTATATTGTAAAACCGGAGGCGC
CTGCTGTGCGTTAATTGATTTACGAGCATTAGAGATGTTTACAGGTAAGACTTCTATAGA
TTGCGATGGATAACATTTAATTAAAATTCTACCATTAGCTTGAATTTGTCTTTTTTGCAC
AAAAGGTTGATGGCCTTGCAAAATTTGATCTATTGTCTCTTTAACTTTATTATGAACTAT
CCAAACATTTCGCTCGTGTATTTCGATTGCAATCTGAAAAGCAGGCTCAGCTTTTCTTTC
TAGTATACTCTTTTGCGTACCTCTACGTTTAGCTTCATCGTCACCTAATGTTACATATTG
AATTCCACCAATTAGGTCCGCCAATGTTGGATTTTTAATTAAACTTTCTAAATGATTTCC
ATGCGCTGTACCTACTAATTGAACTCCTCTTTCTGCTATAGTTCGAGCAGCTAACGCTTC
TAATTCCGTGCCAATTTCATCTATAATAATAACCTCGGGCATGTGATTTTCAACAGCTTC
TATCATTACTTGATGTTGCAAATCGGGCCTTGCGACTTGCATTCTTCTAGCTCTACCAAT

AGCTGGATGAGGAATATCTCCGTCACCAGCTATTTCATTTGAAGTATCTATAATAACTAC
TCTTTTTTCCATTTCGTCAGCTAAAACACGGGCCATTTCTCGAACTGCGGTTGTTTTACC
AACTCCTGGTTTTCCTAAGAGTAAAATTGAATCACCTTGCTGTAGTAAATCTCGAATAAT
ACTAATTGTACCAAAGACTGCTCTGCCGACACGACAGGTTAAGCCAATAACACTACCTTC
TCTATTACGCAGTGAACTAATACGATGCAAGGTTTTTTCGATACCAGCTCGATTATCGCC
GCTAAAATTACCAACTTTTTTTACACAATAATCTAAGTCTTGCCAACTAATAGATCTTTG
AGATAAATATTCTGGGTTATCTGGAAATCTAGCTTCTGGACGACGACCTAAATCCATAAC
GACTTCTATTAAATTATTTCTATTAGGATGCTGTTGTAAAGGTTCTTTGACAAAATTTGG
CAAAATTTCTAGCAACTTATCTAAGTCATCTGCAATAAGCATGATTGAGTATATAATAAT
TAATTAATCAAATAGAAGATTCAAGATTCTAAAGATTACAATACATATTAAAGAATATTA
TGATTGGAATCAATTTTATACTTAAATTGAACAGGAAGTTATAGATGTAAAAGAACTGAT
TAAACTTTACGCAATAAATCGATTGGTCAAAATTAATAGTCGTGTTAACATGTTTATTGG
ACAAAAAGTTCGTATTAAATATAGCAAACAAAAAGTTGCTAGTGATATAGCAGATAAAGT
TGGTGAATTAGGAGTTATTAAGGGAATAAAGTTTATTAATAGTCAATGTGTTACTATTAT
TGTTGAGTTTGACAATCACACTAGGCTTTGGATGTTTAGAGAGGAATTAATCTGTCTAAA
TGAATTAAAAACATGAAACATAATTTATTTTATTTATTATACAAAATTTCTTATGAGTCG
TTAAAATAATAAATTTTAAAGGAGTACTTTTAATTGTGCAAATTACTATTAAAAAATTAC
AAGACTTACTTTCGTCTGTACAAAGAAAAAAGATCCAGACATTAAAACTAAAACAAGGTA
AGTTTGAACTTCTATTAAATAAGACCTATAAAAAAGTCAATCAAGAAATTATACCTTCGC
AGAAATCTGCTGTATTACAAAATAGTCCATCTACGATAATTAAATCAATAAATAATACAA
AAAAATCTTCTGTTGTTAATGAAGACCGTACAGAATATGCCACTATTGTTTCTCCAATGG
TTGGAACGTTTTATCATTCACCTGCTCCTGGTGAAAAAATTTTTGTACAAGTTGGCGATG
AGGTCAAATTCAATCAAACAGTCTGTATTATTGAAGCAATGAAGTTAATGAACGAAATCG
AGGCAGAAATTGAAGGCAAGATTATAGAAATTCTTGTTAAAGATGGTGATATAGTAGATT
GCGGGCAAGCCTTAATGAAGGTTGAAACATAATGCTAATTTTTCCATTTTACATATTGTT
AACAGAAATTATTTAGATAGAAAAATTAGCTTATACTATAGTTATGAAAACTTATTTCGC
AAAGGCACTAAAAACTAATATATAAAATTATTTACTTTATCTGCCAAAAATAGAAATAAG
CGTTTTGATTCCTTGTCTCAAGAGAGGAGAATCTCAATGGCAATTAGCTCAAAAGAGCAA
GAGACAAAGAAGGTAAAAATCTCGGTTGATAAAAATCCCGTAGATACTTCTTTCGAAAAG
TGGGCCCAACCAGGCCATTTTTCTCGTACACTAGCAAAAGGACCAAAAACTACTACTTGG
ATTTGGAATCTTCATGCTGATGCTCACGACTTCGATAGTCAAACCAGTTCTTTAGAAGAA
GTTTCACGTAAGATTTTCAGTGCACATTTTGGGCAGCTGTCTGTAATATTTTTATGGCTT
AGTGGAATGTATTTTCACGGAGCCCGCTTCTCTAACTATGTTGCTTGGTTAAGTAATCCA
ACAGGTATTAAGCCAAGTGCGCAGGTTGTTTGGCCTATAGTTGGGCAAGAAATTTTAAAT
GGCGATGTAGGTGGTGGCTTTCAAGGAGTACAAGTTACATCTGGATGGTTCCAACTGTGG
AGAGCATCAGGAATTACTACAGAATTTCAGCTTTACTGTACTGCTATTGGCGGATTAGCT

ATGGCTGCTTTAATGCTGTTTGCAGGATGGTTTCATTATCATAAAGCTGCTCCAAAGTTA
GAATGGTTTCAAAATGTTGAATCAATGATGAATCACCATTTAGCTGGGCTTTTAGGCTTA
GGCTGTTTAGGCTGGGCAGGCCATCAAATCCATTTGTCTTTGCCTATTAATAAGCTACTA
GATTCTGGCGTGTCTCCGCAAGAAATTCCACTACCTCATGAGTTTTTAATTAATAGAGAG
CTTATGGCTCAGCTGTATCCAAGTTTTAGTAAAGGATTAGTTCCATTCTTTACTTTAAAT
TGGGCTGAATATTCCGACTTTTTAACTTTTAAAGGAGGTTTAAACCCTGTTACTGGAGGT
TTATGGCTAAGTGATACTGCTCATCATCATCTAGCATTAGCTGTTCTATTTCTTGCTGCA
GGTCATATGTATAGAACCAATTGGGGTATTGGACATAGCATGAAAGAAATTCTAGAAGCT
CACAAAGGACCTTTTACCGGCAACGGTCACGAAGGTCTATATGAAATTCTTACAACTTCT
TGGCATGCACAGCTTGCAATTAATTTAGCTATGATGGGATCTTTAAGCATCATTGTAGCA
CATCACATGTATGCAATGCCTCCTTATCCATATATTGCTACTGATTACCCGACTCAGTTA
TCGCTCTTCACTCATCATATGTGGATTGGAGGATTTTGTATTGTTGGAGCAGGAGCGCAT
GCTTCTATATTTATGGTAAGGGATTATAATCCTGCAGAAAATTATAACAATCTTTTAGAT
AGAGTCATTAGGCATCGAGATGCTATTGTTTCTCATCTAAATTGGGTATGTATATTTCTT
GGATTCCATTCATTTGGTTTATACATTCACAATGATACTATGCGGGCACTTGGAAGATCT
CAAGATATGTTCTCTGATACAGCTATACAGTTACAACCTATTTTTGCTCAATGGGTACAA
AGTATACACACTTTAGCTCCTGGAAATACAGCTCCAAATGCATTAGCAACAGCTAGTTAT
GCATTTGGAGGAGATATTGTTTCTGTTGGTAACAAAGTTGCAATGATGCCTATTTCTTTA
GGTACTGCAGATTTTTTAGTTCACCATATACATGCATTTACTATTCATGTAACTGTTTTA
ATTTTAGTTAAAGGGTTCCTTTTCTCAAGAAACTCTAGACTAATTCCTGACAAGGCCAAT
CTTGGCTTCAGGTTTCCATGTGATGGACCTGGTAGAGGTGGTACTTGCCAAGTGTCTGGC
TGGGATCATGTTTTTCTTGGCTTATTCTGGATGTACAATTCTCTGTCTGTAGCAATTTTT
CACTTTAGTTGGAAAATGCAATCAGATGTTTGGGGTAGTGTATCTCCGTCTGGAAATGTT
TCTCATATTACTGGCGGTAATTTTGCACAGAGTGCAATTACAATCAATGGATGGCTGAGA
GATTTCCTTTGGGCTCAAGCATCTCAAGTTATTCAATCATACGGTTCTGCGCTATCTGCA
TATGGATTAATTTTCTTAGCAGCACATTTTGTATGGGCATTCAGTTTGATGTTCTTATTT
AGTGGTAGAGGTTATTGGCAAGAGCTTATAGAATCAATCGTATGGGCGCATAATAAGATA
AAAGTTGCTCCTGCAATTCAACCAAGAGCTTTAAGTATTACTCAAGGTAGAGCAGTCGGT
GTTGCACACTACTTATTAGGTGGAATTGGTACAACTTGGGCATTCTTTTTAGCGAGAATT
ATTTCAGTAGGCTAATAGTGAAAATAGGATAAAAAAAACAATTATGGCAACAAAATTTCC
TAAGTTTAGCCAAGCTTTATCACAAGATCCTACAACTAGAAGGATTTGGTATGGTATTGC
TACGGCACATGACTTTGAAAGTCATGATGGAATGACAGAAGAAAATTTATATCAAAAGAT
ATTCGCTTCGCACTTTGGACATCTAGCAATTATCTTCTTATGGACATCTGGTAATTTATT
CCATGTAGCTTGGCAAGGCAACTTTGAACAGTGGGTATTAAATCCTTTGAAAGTTAAACC
AATTGCTCATGCAATTTGGGATCCGCATTTTGGACAACCTGCTTTGAAAGCTTTTAGTAA
AGGTGGCTCAGCTTATCCAGTAAATATAGCATATTCTGGCGTATATCACTGGTGGTATAC

TATTGGTATGAGAAGCAATCAAGACCTGTATTCTGGGGCTTTATTCTTACTAGTTTTATC
AGCTTTACTCTTATTTGGAGGGTGGCTACATCTACAACCAAAATTCAAGCCTGGTTTATC
ATGGTTTAAAAATAACGAATCAAGATTAAATCATCATTTATCTGGATTATTTGGGGTTAG
TTCTTTAGCCTGGACAGGTCATTTAGTACATGTTGCTATACCTGAGGCAAGAGGACAACA
TGTAGGATGGGATAATTTTACAACCGTATTACCTCATCCAGCCGGTTTACAGCCGTTTTT
CAGTGGTAATTGGAGTGTATATGCTCAAAATCCAGATACAGCTCAACATTTATTCGGAAC
TAATGAAGGTGCGGGTACAGCAATTCTGACATTTCTAGGAGGATTTCATCCTCAAAGTCA
GTCTTTGTGGCTAACTGATATGGCTCATCACCATTTGGCTATTGCAGTAGTATTCATTGT
TGCTGGACATATGTATAGGACTAATTGGGGAATTGGGCACAATCTAAAAGATATTTTAGA
TGCTCATAGACCACCTAGTGGTAGATTAGGAGCTGGACATAAAGGGCTATTTGATACTAT
TACTAATTCTTTACACATACAGTTGGGATTAGCATTGGCTTCCCTAGGTGTAATTACTTC
GTTGGTAGCTCAACATATGTATGCTATGCCTCCATATGCTTTCATGGCTAAAGATTTCAC
AACTCAAGCATCTTTGTACACACATCATCAATATATTGCTGGGTTTCTAATGGTTGGAGC
TTTTGCTCATGGGGCAATATTCTTTGTTCGAGACTATGACCCTGAACAGAATAAAGATAA
TGTTTTAGCTCGTATGCTAGAACATAAAGAAGCTATCATTTCTCATTTAAGTTGGGTAAC
TCTATTTTTAGGGTTTCATACATTAGGCCTTTATGTTCACAATGACACAATGATTGCTTT
TGGAACTCCTGAAAAACAAATTCTAATTGAGCCGGTATTTGCTCAATGGATTCAAGCCTC
TTCAGGGAAAGCACTTTATGGGTTTGATGTGTTACTATCATCCTCTACTAATATCGCAAC
ACAAGCTGGTAGCAATATTTGGCTGCCAGGCTGGTTAGAAGCGATTAATAGCGGAAAAAA
TTCATTGTTTTTAACAATTGGTCCTGGTGACTTCTTAGTTCATCATGCAATTGCATTGGG
ATTACATACTACTACGTTAATTTTAGTTAAAGGTGCTTTAGATGCAAGAGGCTCTAAACT
TATGCCGGACAAAAAAGACTTTGGATATAGTTTTCCTTGCGATGGACCTGGCAGAGGCGG
CACCTGTGATATATCTGCATGGGATGCGTTCTATTTAGCTGTATTTTGGATGCTAAATAC
AATAGGTTGGGTAACATTTTATTGGCATTGGAAACATATTACAATATGGCAAGGCAATGC
AACTCAATTCAATGAGTCTTCAACTTATCTAATGGGATGGTTTAGAGATTACTTATGGCT
AAATTCTTCTCCATTAATTAATGGTTATAATCCATATGGCATGAATAATTTATCAGTATG
GTCATGGATGTTCCTATTTGGACATTTAGTATGGGCAACAGGATTTATGTTCTTGATCTC
CTGGCGTGGCTATTGGCAAGAGTTAATTGAAACTCTAGCATGGGCGCATGAACGTACGCC
TTTAGCAAACTTGATTCGTTGGAAAGATAAACCTGTTGCATTATCAATTGTACAGGCAAG
ATTAGTAGGTTTAGCACATTTTTCTGTAGGATATGTATTAACTTACGCAGCTTTTGTATT
AGCTTCAACAGCAGGCAAATTTGGTTAGACTAAATTCGAATTAGCT-AAAAAAAGTCAGT
CATAATAGTTTTATGACTGACTTTTTTAGATTTAAAACAAAAACTTACTAAAGTTTCACC
TCAATATCTACACCAGAAGGCAAATTTAACTTCATTAATGCATCTATCGTTTGAGAAGAA
GGTTGATGAATATCAATAATTCTTCTGTGAGATCTTATTTCAAAGTGTTCTCTTGAGTCT
TTATCTACATGTGGAGAACGTAAAACACAATAAATTCTTCTTTTTGTCGGTAAAGGAATT
GGCCCTACTGCAACGGCATTAGTTCTAGATGCGGTATCTAGTATTTTGTCACACGACGTA

TTAAGTATAATAGAATTATATGCTTTCAGTTTAATTCTGATTTTTGTCTGCTGAGTAATT
GTCATGTTAAAAAGATCTTATTATTTAAGAATTTTAGAGACAACACCTGCGCCTACGGTA
CGGCCACCTTCTCTAATAGCGAAACGCATACCTTGCTCAATTGCAATTGCATTAATTAAT
TCAGCAGTCATCTTAATTCTGTCACCAGGCATAACCATTTCTGCATCAGTACCATCATCA
GCAGTAAACTGATTAATAGTACCAGTTACATCAGTTGTTCTAACATAAAACTGAGGTCTA
TATCCTGGAAAAAATGGAGTATGTCTTCCGCCCTCCTCTTTAGTTAAAATATAAACTTCT
GCTTCAAATTGAGTATGAGGTGTAATTGTACCAGGTTTAGCTAATACCATACCTCTTTCA
ATATCTTTTTTCTGCACACCTCTTAAAAGAATTCCAATATTATCACCCGCTAGACCTTCT
TCTAACGTTTTTTGAAACATTTCTAATCCAGTAATAGTCGTTGTACGAGTTTCTCGTAAA
CCTACAATTTCAATTGTGTCACCAACTTTAATAATGCCTCTTTCAATTCTACCAGTGGCA
ACAGTTCCACGTCCTGTAATAGAAAAAACATCTTCTACAGCCATTAAGAAAGTTTTATCG
ACATCTCTCTCCGGCGTTGGAATATATGTGTCAACTGCTTCCATAAGTGAAAAAATCTTG
TCAACCCATTTATCTTCACCTTGCTTAGTAGCTGGGTTTTTCGTCACAGCTTCTAATGCC
AATAAGGCAGAACCTGCAACAAAAGGAATATCATCTCCAGGAAAGTCGTATTGACTTAAT
AATTCCCTTCCTTCTAATTCTACTAATTCTAGTAGCTCTTCGTCATCTACTTGATCTTCC
TTATTTAGGAATACTACTAATGTAGGAACACCTACTTGTTTTGCTAATAAAATATGTTCA
CGAGTTTGTGGCATTGGACCGTCTGCCGCAGATACAACTAAAATAGCTCCATCCATCTGA
GCAGCACCCGTAATCATATTTTTTACGTAGTCGGCATGGCCTGGACAATCTACGTGAGCA
TAATGACGATTATCTGTTTCGTATTCAACATGAGCAGTATTAATAGTAATACCTCTAGCT
TTTTCTTCTGGAGCGGCATCAATTTCATCAAATTTTTTCGCTGCAGTAGACCCTAAAGTT
GATAAAGTCGCGGAGATCGCTGCTGTTAAAGTTGTCTTACCATGATCAACATGACCAATT
GTGCCAATATTGACATGAGGTTTTTTACGTTCAAATTTAGATCGAGCCATGCTTTTTGTT
TTCCTTATAGTAAAAGTTGCTTTATAGTTACTTTTAACGATAAATCTTGTGAATTGAAAG
ATTTATTATTTAATACAGACAAGATACAATATAACTAATCTAATATCGATAGTGAGCAAA
CGCTTTATTTGCTTCTGCCATTCTATGTGTATCCTCTCTCTTTCTAATAGAATTTCCTGT
TTCATTAGCTGCATCCATAATTTCATTAGCTAATTTCATAGACATACTTTTACCAGATCT
GTCTCTAGAGAATTTAGTAATCCATCTTAGTGCTAAATTCGTACCTCTATAAGCTCGTAC
TTCAATAGGAACTTGGTAAGTAGAACCACCAACTCTTCTTGCTTTTACTTCTACTAGAGG
AGTAATATTTCGGATGGCTTTTTCTAAAATATTTAAAGGATCTGATTCTGTTCTCTCTTT
AACGATATCTAAAGCCTGATATATAATTCTTTGAGATAAAGTTTTTTTACCACTTTTCAA
AATACGAACAGTTAACATACTTACGAGTCTGCTTTTATATAAAGGATCAGGTGATGCAAA
CCTTTTTTTAGCTGTATTACGACGAGACATAGTATTTAAAATTGGTTATTGTAATAATGT
TCTATTAGCAGGTATTTAAGATTTAGGCTTTTTTGTCCCATATTTAGATCGGCTTTTACG
ACGATCTTTTACTCCTGCAGCATCTAAAGTACCACGTACTACATGATATCGGACTCCAGG
CAAATCTTTAATTCGGCCGCCCCTAATCAGTACCACAGAATGCTCTTGAATATTATGACC
TACGCCAGGAATATAAGCTGTAACTTCAAATCCAGATGTTAATCTAACGCGCGCTACTTT

TCTTAAAGCAGAGTTCGGTTTTTTAGGAGTTGTAGTATATACTCTTGTACAAACACCTCT
TCTTTGAGGGCAACTTTGAAGAGCTGGAGATTTTGTTTTTTTATGTATTTTTCGTCTTTC
GGATCTAACAAGTTGTTGAATTGTTGGCATAATAAAATTGATTTAAAGTCTTTAATAGAT
ACTATTTGAACCTGCTTTACATGAATTGGCTAGTTTTTATTTATCTTATACTTACGCATA
AATCTTTCTACTCTACCTTCTGTATCAATAATTCTCTGTGAACCAGTAAAGAATGGATGG
TTTCCTGACCAGATATCTACGTGTAGTTCTGGCTTTGTGGACCCAATCGTCATAATTAAC
TGGCCGTCACAGTAAACTTTTGCTTCTGGATACCAATTTGGATGTATATTATCTTTTGCC
ATTGTTTTTATAATTATATAAATGTATCAATAATAACTATTAACGCTTAGAAAATTGAGG
AGCCTTTCTTGCTTTCTTTAAACCATATTTTTTTCTCTCTTTTACTCTTGGATCTCTTGT
TAAATAGCCTTCAGACTTGAGTGTCGTTCTATTTTCTGGATTAATTGAGCACAATGCTCT
TGCAACACCTAAACGAATTGCATCAGCTTGCCCTGTCAAACCTCCTCCTCTAGCATTTAC
ATGGATATCATACTGGTTTAGTAGTCCTAAAACTTGTAATGGTGCATATGAAACTCTTAA
GTAATTAGGACTAAATTGAAGATAAGACTCTCCTGGTATACCGTTTATAATTAAATTACC
TGACCCTGGGACTAGTCTTACTTGTGCAACAGAACACTTCCGGCGACCTGTTCCAGAATA
GATTGCGCGAGTTTTAATTAATTCTGTCGACATAACATTCCTTGATAACTAAATAAATAT
ACATTCTTTATACTATATACTCTTGTGGCTTTTGCGCGACATGTGGGTGAATTGGGCCAG
AATACACTTTAAGCTTCGTAAATAGTTTTCTGCCTAATGGACCTTTAGGAAGCATACCTT
TAACCGATTTTTCAATAATTCTGTTAGGTAATCTTGTCTGAAGCTGATCAAATGTTTCAA
CTTTTAATCCGCCAGGCTGTCCAGAATGTCTTCTGTATAGTTTTTGATTCGTTTTATTTC
CACTTACAGATACGTGAGCAGAATTAATAACGATTACATAATCTCCAGTATCTAGATAGG
GTGTATAAGAAGGCTTGTTTTTACCTCTTAAGATATTAGAAATATGAGTAGATATTCTAC
CAAGTGTCTGATTTTTAGCGTCTATAACATACCAATGAGAATTAGTATTTAATGAGGGTG
ATTGCGTTTTATTCATGAAAAGATTAATACTTTTGGTTGACAATAAGAGAATATGTAAAT
GGTTTACATTCAATTTCTATACAAGTAGTATTTAAAAATATTATTAACAAAGAGAACTTT
TCTAACAGTTATTTAAATTGTTTGTCAGCAAGAAAATTTCTAGTTATCAATAACTTATAC
TATTTTTAATAATATAAAATATATTAATCACTTTTTTCTTTTGGCAAACTTATACCTAGT
TTTTTTTGCAAGGCATATATAACTTCTTCAGCTGATTTCTGGCCAAAATTTTTAATTTCT
AGCAGTTCTTCTTGAGAGTAATCTAGTAAATCTGCAATAGAATGGATTTGAGCTCGCTTT
AAGCAGTTATAAGCCCTAACAGATAACTGCAGTTCTTCTATTAAAACTTGACTAATTTTT
TTATCTTCTTTGCTACGATAATTATCTGCTGATTTAAAGTCTAAATTTCTTAGTGAACAA
AAAAGATTAGTTAAAACTGTAGCTCCTTGACTTATTGCTTCTTGGGGAGATATGCTCCCA
TTTGTCCAGATTTGTATAATCAATCTATCTTTTATGCTGTTGCTACCAATACGGACTTCT
TCTACTTTATAATTAACTTTATTAACTGGCATAAAAACAGAGTCTACTTGCAAAAAATCC
ACAGATAATTCATCTACAGCTTTTTCAGCTAAGCGATACCCGCAATTTTTTTCAATTTTA
AACTCCATTTCAAATATTGTGTTATTGCAAATAGTTGCAATATACTGTCTAGGATCTACT
ACCTCTATATCAGAAGATAATTCAAATAGGCCGGCTGTAACTATAGCTGGCCCTTGAACT

CTAATTCGACCAATTTGAGATTCTTTGTTATAACTTTTAAATACTACTTCTTTTAGATTA
AGTAATATTTCTAACACATCTTCTCTTACCCCAGGAATTGTGGAAAACTCATGGTTCACT
CCAGCAATCCGTACAGCAACTATAGCAGTACCTTCAAGATCTGACAATATTGATCTTCTC
AAAGCATTACCTAATGTAATACCTTGTCCTTGATTTAATGGTTCTATTACAAAACTACCG
TACTGCCCACGCGCCCCATCTGTTCTTGACTCTATGCATTCAATTTGAAATTGAGCCACC
TAAGAAAGCTCCTTTAAAATAATCAGTGATTAGTAAGTCTCTCTAAAACAAGAAGGAATT
TATACTCGGCGTTTCTTAGGAGGGCGACATCCATTATGAGGTACAGGAGTAATATCTTTT
ATTAGAGTAATCTCTAATCCTGCAGCCTGCAAAGCTCTAATTGCTGTTTCTCGACCCGCT
CCTGGACCATTTACTAGAACTTCAGTTTGGCGCATACCTTGATCCATAGCTTGTCTAGCT
GCTTTTTCAGCTGCTGTTTGAGCTGCAAAAGGTGTTCCTTTTTTAGCTCCCTTAAATCCA
CTTGCACCAGAGGATGACCATGATAATGTTTCTCCTTTTAAATTAGTAATAGTAACAATT
GTATTATTGAATGTAGATTTAATATGTGTAATACCGTTAACTGCATTACGTTTAGTTTTT
CTTGCTCCGGATTTTTTTATTTGTCTAGCCATCGTGTCTCGTCTATTATATAGTTAATTA
AAAGATTATTTTCTTGGAGCTTTTTTCTTACCTGCTACTGTTTTTTTACCTCCTCTGCGT
GTTCTAGCATTAGTTCTAGTTCTTTGTCCTCTCAAAGGAAGACCAAGACGGTGTCTTCTA
CCTCTATAAGTACTAATTTCCATAAGTCTCTTAATGCTCATAGACTCAAAACGTTTGAGA
TCCCCTTCAATCTGATAATTAGACTCAAGAATTTCTCTTATACTGACAACTTGTTGATCA
TTTAAATCTTGACACTTAATATCAGCGTCTATGTTTGTTTTTTCTAATATTTCTTTCGAG
CGAGATAGTCCAATACCATAAATATATGTTAAAGCTATCTCTATTCTTTTGTTTCTTGGA
AGATCTACTCCAGCAATTCTGGCCACTTTGTCTTGTCTCCAATAAATAATATGTATTTTT
ATAACGGTTATTTATAAGCTTAGCCTTGTCTCTGCTTATGTTTAGGATTAGTGCAAATTA
CCATCACCTTTCTGTGACGCCGAATTATTCTACATTTTTCACACATTTTTCGAACAGAAG
GACGAACTTTCATATTAACTCTATATACTAGGAATATTACTATATTAACTCTATTAATTA
TCGACATTAAAAATGTCAACAATATATTGTATTACTTTAACTGTTGTAGATTTTACTTCG
TCATACTATCATATTTTTTTGATATGACGTATGTCTGAATTTGCTTAGCTGTGTCTATTG
CGACTCCAACTAAAATAAGTAAAGACGTAGCCCCGAGACCTCTTAAGTTTTGGATCTGAG
TAACCTTTTCTATTATAAACGGAATCAGCGCTACTGTAAATAAAAACGAGGCTCCTAGGA
ATGTCAGTCTATTTAATATGACTTGTAAATAATCAATAGTCGCTTGACCTGGACGAATAT
TAGGAATGCTTGCACCCATTTTTTTTAAATTTATAGCAATATCTTCTGGATTCATTACTA
TTGATGTATAAAAATAGCTAAAGAAAAGGATCAAAGCACAATAAAGAAGAAGGTATAGTG
AACCATTAGGACAAAACAGGTATAGGATTTGGAGTAATGTTTTATTTTGAATAATTTGGG
TTAGATAAGATGGGAGGGCCATAGATGCAGATGCAAAAACAATAGGCATAACTCCACCTT
GATTTAATTTCAAGGGCAAGTAACTATTTGGATCCAAAATTGAAGATTTTCCTAGCTGTC
TTGCTGAAATAATTTTAATTCTTCTTGTTCCTTCTTGTACACAAATTGTAATTATTATCA
TTAATAAAAAGATCGCTATAAATAATCCGAACTTAAGACTTGCATTACTATAACTAGCAT
CAAAAAATGACTGTGTAAAATTCTTTGGTAGTCCTGACACAATGTTTTGAAAAATAAGTA

AAGAGGCTCCATTACCAATTCCCTTTTCCGTAATTAACTCTGATAACCACATAATAATCA
TAGAGCCTGCTGTTAAAGCTAAAACAGACTCGCAGACAAATGCAAAATTCCAATTAAAAA
CATATGGTTTTACCCATATAGAGATTGCTCCAGATTGTAAAGTCGCCCAACCTAGAGCTA
AGTACCTTGTAATTTGAGTTATTTTTTGGCGGCCCAATTCACCTTCTTCCTTCTGTAATT
TTTCCAGATTGGGGACAATTTTCGTAAGTAGCTGCATTACAATTGAAGAGTTAATATAAG
GAACAATACCTAACGCAAAAATCCCTATTGTTGAAAAACCTCCTCCAGAAAAAATATTCA
GAAAATTTACTAAAGTATTTTTTTCTACACTTGCATAAAAGGCGTCATGATCTATACCTG
GAACAGGTATAAATATTCCTAAACGTGCTAAAACTAATAGAAAGAGAGTAAAGATAATAC
GATTTCTTAGATCACTTTTTTGGCTCATAAATAAATTTTAAATGAAAAGAGATACTTTTT
TATGTAAAAACCCTATTCCAAGACTTGTATAAGTTTAAAATAGGGAAGTAAAATATTATA
GTAATATTAATATAATATTCTATGTAGAATACAGGCTTTCTATTGGAACTCCTCGATCTT
TAGCTGCTTCACTAAAAGTTCTTAATTGAGTTAAGCCATTGAGAACAGCTCGCGCGTTAT
TTAATGTATTATTAGAACCTAATTGCTTAGCTAATATATTCTGTACACCCGACAATTCTA
GGACTGTCCTTACAGAACCTCCGGCAATTACACCTGAGCCAGGTGCAGAGGGCCTTAATA
TAACTTTCGCGGCACCAGAAATTCCGTTAATAGGATGAGGTATAGAATTCGATTTTGTCA
GTGGAACTGTAACTAGATGTTTTTTAGCATCTGTTACTCCTTTTTTTACTGCACCAATTA
CATCGCTTGCTTTTCCCACGCCGACACCAACTTGGCCTTGCTCATTACCAATAACAAGAA
TAACTCGAAAGCTTAGTTTTTTACCTCCTTTTACAACTTTAGTAACTCTTTTAACTTGTA
CGACTCTTTCTTCCCAGCCACTATCTTTATCTTTTCCTTTGCTCTGTTTTTTACGATTGG
CCATTTTAAGAATTATTCCTAATTAGTTAATTACGATTCTAGAAACCCATGCCTGCTTCC
TTAGCAGCTTCAGCCAAGGCCTTAACTCTTCCATGATATAACTTTCCTCCTCTATCGAAG
ACGACATTTTTAATGCCTTCTTTCATAGACTGTTCTGCTAACTGTTTACCAACTACGCGA
GAAGTATCACAGTTTGGTCTTATGTTATCAGATTCTTTATTATTTAAATTAACAGATGAT
GTAGCTACTAATGTAATACCTTGTGTGTCATCAATTATTTGTGCGTATATATGTTTATTA
GATCTAAATACACACAGACGAGGCCTACTTGAGGTTCCTTGAACTTTTTTCCGAACTCTT
TTATGCTTATGAATTCTAGTTTGTTTAGTGTTTAGTTTCATTATTATTTACCTTTTCCAG
CTTTCCCAGCTTTTCTTCTAACAAATTCACCTTGATATCTAATTCCTTTTCCTTTATAAG
GCTCAGGAGGCCTAATAGAACGAATAGTTGAAGCAACCTGACCGACAACTTCTTTATCTA
TGCCAGAGACAGTAATATTTGTGTTGTTTTCAACTTTAATTTCAATATTGGCAGGAGGTT
TAATTTTCACTACATGACTATAGCCAACACTTAAGATTAAATCTTGGTTATCAATTTGAG
AACGATAGCCTACACCTTGTATTTGCAGTTTTTTAAAAAATCCATTAGAAACACCTTCAA
TCATATTACTAATAAGAGTTCTGGATAGCCCATGTAACTGACTAGCCATTTTTGTTTGTC
CACTCGTCTTGACAGCAATTGTGTCGTTAAGTATTTCTAAATTAATACCAGCTGGTAAAG
TTCTGGATAGGGTACCTTTAGGTCCTGTCACGGTGATAGTCTGACCATCGAACTGAGTAC
TAAGATTTGTCGGCAATAAAATTATTTTTTTTCCAATACGAGACATATTCCACCTATAAA
ATAGTTACCAAATATAGCATAATATTTCACCACCAAGACCATCATGACGAGCTTGTCTAT

CTGTCATAACGCCTCTAGAAGTAGAAATAAGAGCGATGCCTAAACCTCCAAGAACTCTGG
GCAACTCTTTATGATTTGCGTAGACTCTTAGTCCGGGTTTACTAATTCTTTTCAGAGCAG
TAATAACTGGTTGACGATTTTTACCATTATACTTAAGAGAAATCATTAAATGAGTCTCTA
TACCTTCACCCATTTGTTCGAAATTTTGAACAAATCCTTCTTCTTTTAGTACTGTTGCCA
TATTGCATGTCATTTTCGTTGCTGGAACTTGCACAATTTGATGTCTTGCTAAGTTTGCGT
TACGAATACGTGTCAGCATATCGGCGATCGTATCGTTGACCACCTTGATCCTCCTTGCGA
TGAATTATTTGAAATATTTTCAAGATTCCCTGAAAGGCATCCCTAGCTTTTTTAACAAAG
CTAGACCTTCTTGATCTGTTTTAGCTGTAGTGACAATTGATATATCTAGACCACGAATTT
GATCAATATTATCATAGTCTATTTCTGGAAAGATCAACTGTTCACGTAAACCTAAATTGT
AATTACCTTTGCCATCAAAACTTCTAGGACTAATTCCTCTAAAGTCTCTAATTCTTGGTA
ATGTTAAATTAATTAATTTCTCTAAAAAAGAATACATCTTGTCTTTTCTTAGATGCACAA
CAATTCCGATAGGAACCTCTTCTCGAATTTTAAATCCTGCAATAGATTTTTTAGCTTTTG
TGACAATTGGTTTTTGTCCAGTTATTAATGTTAATTCTTGAATACTACTTTCAAGAGCCT
TAGCATTCTGAGAAGCTTCACCTAAGCCACGGTTAATAGTAATTTTAGTAAACCTAGGAA
CTTCATGTACATTTTTGTACTGAAATTCATCTTTTAAAGATTGAGTAACAGTTGTTTTAT
ATTTTTCTTTTAATCCTATTGCCATTATATTTTCATCAATTATTTAATAAGTTCGCCAGT
CTTTTTCAGTTTTCGAATTTTTTGGCCTTTATCATTAATTATTACTGAGGATCGACTAGC
AATATTATTTTGTTCACTAAATAACATAACGTTAGAGGTATGTATTGGAGCTTCAAATTT
GATAATTTCTCCCGTTTCTCCTTCTTGCTGAGGTTTTTTATGCTTCACTTTGAGATTAAT
TCCTTTAACAATCACCTTATTTGTTTTATAAATGATTGCAATAATTTCACCTGTTTTTGT
TTTATCACTTCCAGAAATAACTTGAACTAAGTCCCCTTTTTTTAATTTAATTTTTGTATT
ATTTTTGGTTGTTTTAGAAGGACCTTTCATTATACTACCTCCGGTGCAAGAGAAACTATT
TTAGAAAAATTTTTATCTCTTAACTCTCGGGCTATAGGTCCGAAGACTCTTGTGCCACGT
GGATTATTATCTTGATTAATGATAACTGCCGCATTATCTCCAAATCTAATGCTCATACCG
TCAGTTCTCCGTAAAGCTTTACGAGTTCTTACTACAACAGCTCTGACAACATCAGATCGT
TTAACTGGCATATTGGGAGACGCATCTTTGACTACTCCAATAATAACATCTCCAATAGAT
GCATAAGAAGGATTACTAGTGCCTAATACTCTAATACACATTATTTTTCTAGCACCACTA
TTATCTGCAACATTAAGATAGCTTTGAGTCTGTATCATACTTTTATTCTGTATAAGATTA
ATTATCAAAAGACTTAGATAAGATATTAACCATTGTCCAACACTTTGTGCGACTTAAAGG
TCGTGTCTCTTGTATTGTAACAATATCGCCGATTGTGCATTCATTATTTTCATCATGCGC
TTTATATTTCTTTGTCCGGATCATAGTTTTTGCGTACTTTCTATGAGAGATTCTATTTTC
TACAGCTACTACTATAGTTTTATTCATTTTATCGCTTACTACTTTACCTGTTGTTTCTTT
TAAAGGCATAGTCTTTATATTTTTTATATATAATTAAATTTTAAGATTATTACATGGTTG
ACTTAGTACGAGACTTTTCAACAGTTAAAAGTTGGGCTAATCTATGCTTAGAATGTTTAA
ATAGATGAGGCTGGAAATCTTGCCTTGTGGCTCTTTTTAGCCTTAAATCAAAAAGCTCTC
TTTTTATTACAAGGATTTCTTCAGCTAAAGAAGAAGAGTCCAGGTTTGTAACATCTGATA

TTTTAGGGAAAGTCATATTTTAGGATTCTGTTGTATTCCGAACTATAAATTTAGTCTTGA
TTGGTAATTTATAAGAAGCTAATTTCATAGCTTGTTGAGCAGTTTTTTGTGGTACACCTG
TAATTTCAAATAGAATATGTCCAGGCTTAATAACTGCAACCCAATATTCTGGAGCTCCTT
TACCTGACCCCATGCGAGTTTCAGCTGGACGGGCAGTGACTGGTTTATCTGGAAATACTC
TAATCCATAGTTTACCACCTCTTCTCACATATCTAGTAATAGTTCTTCGAGTAGCTTCTA
TTTGTCTAGAAGTTAACCAAACTGGCTCTGTCGCTTGTAATGCATAATCACCGAACGCAA
TTGTGTTACCTTTGCTAGCAGAACCTTTCATTCTACCTCTATGTTGTTTTCTAAATTTTG
TTTTCTTGGGGCTTAGCATAAAAATTAAAAAATAATAATAATTAAGAAGCTGTTGAATCA
GGAGTTTCCTGATTATAAGCAGATTCTAGACTTTTCGATTCTGGCAAAATTTCTCCTTTA
AAAAGCCAAACTTTTACTCCGAGCACACCATACGTAGTGTGAGCTTGGCGATGACAATAA
TCAATATCGGCTCTTAAAGTTTGTAAAGGTACACGACCTTCTCTAACCCATTCACTTCTA
GCTATCTCTGCACCATTAAGCCGACCAGATACTTGAATTTTAACACCCTGTGTATTTGCT
CTTTGAGCTCTTTGAACTGCTTGTCTGACAGCTCTACGAAAGGCCACTCTTTTTTCTAGC
TGTTGAGTAATAAATTCTGCTACTAAGGTTGCTTCAGAATCTGGATCTGCAATCTCTACA
ACATTTACTCTGAGTTGTTTGCTAGGGTCTAGGATTAAGGACAGCGATTTTCTTAAAGAC
TCAATTCCCGCTCCAGATTTTCCAAGTACGATTCCTGGTCGAGCTGTTGCTATGAGAATT
TCTACTTGATCAACTTTACGATTAATTTCAATTTTAGCAATACTAGCATTACTAAGTTTT
GAATGTATAAATGAGCGAATTTTATGATCTTCTTGTAAAAGAACTGGGTAGTCTTTAGAG
TTAGCAAACCATGAAGAACGATGTTTTTGGGTAATGCCTATGCGAAAGCCCAAAGGATGA
ATTTTTTGACCCACAGTATCCTTGATTATAAGTTAAAAGCTATAATGACTAATTGACGTT
AAAAGACTACAGTTCTGAAACACCTAATGTAATATGACAAGTTGGTTTATGTATTGGAAA
GGCTCTTCCTTGTGCTCTAGGCTGAAATCTTTTTAAAGTCGGGCCTTTGTCTGCAAAGGC
TTTACTAACAAACAATTGGTTTTTATTTAATCCATCATTATGCTCAGCATTCGCTGCAGC
AGATTCCAGGATTTGCTTTATATGAGAACAAACTCGGTATGGCATAAATTCCAGAATAAT
TAATGCTTCTTGGTATTTTCTACCTCTAATTTGGTCTAATACACGACGGACTTTATGCGG
AGAAAGACGAATGTATTTCCCTACTGCTTTAGTTTCTTTTACGTTTTTTGTAATACTCAT
AGTTATAGTTTAACGACGGGCTTTTCTGTCACCTTTTACGTGAGTGCGGAAAGTTCTTGT
TGGAACAAACTCTCCTAATTTATGTCCTACCATTTGGTCTGACACAAAGACAGGAAAATG
TTGTTTACCATTATAAACAGCTATTGTATGCCCTACCATATCGGGAATAATAGTAGATGC
TCTCGACCAAGTTTTTAGGACTTCTTTTTTTCCTGAAATATTTAATGCTTCTATACGTTT
AAGAAGACTAACATCAATAAAAGGGCCTTTATGTATAGATCTTGACATAATAATAAATCA
TTAGATTCTTAATAAAATAAATTCTATTTACGGCGACGTAAGACGTACGGGTTACTATAT
TTATTTGGATTTCGTGTTTTAACACCCAATGCAGGTTTACCCCAAGGTGTTACAGGACGA
GCTCGTCCAATTGGAGATTTGCCTTCACCACCACCATGTGGATGGTCTACAGGATTCATG
ACAACACCTCTAACAGTAGGTCTTTTACCTAACCAACGATTTCTACCAGCTTTACCAAGA
GTAATATTACTAGCGTCAATATTACCAACTTGTCCAATAGTAGCGTAGCATTCTTTTCGA

ATCATTCGAACTTCGCTTGAAGGCAATTTTACAGTAACAAAAGTACCTTCTTTTGCTACT
ATTTGAGCATAAGTTCCTGCTGCACGAACAATCTGTCCTCCGCACGAAGGTCTTAATTCT
ATATTATGTACTGCTGTTCCTAAAGGAATACTAGATAAAGGCAAAGCATTTCCAACTTCG
ATAGGGGCAGTAGGACCAGAAAGAACCATAGATCCTACACTGAGAGATCGAGGGTGTAGA
ATATATCTTTTCTCACCATCAAGATAGTGTAATAATGCAATTCTTGCATTTCTATTCGGA
TCATATTCAATAGAAGCAACTTTAGCAACTATATTATGTCTATTTCTTTTAAAGTCAATT
AATCTGTACTGCTGCTTATGTCCACCACCTTTATGACGACAGGTAATGACACCTCTATTA
TTTCGACCTTTGCAGAAATGGTGTTTAACTATTAATGATTTTTCTGGCTTATCAGTAGTA
ATCTCTGAAAAGGTAGAAACCGTTCTATTTCTTGTCCCTGGTGTGTAGGCACGATATAAA
CGAATTGCCATATAAGAAGATTTGAGAAGAGTAAATGTTGAACTATGTGATATATTTAAG
TTTCTGGAAACAAATTAATAGAATCTTCTGAGGCAAGTGTTACAATTGCTTTCTTGTAAT
GTGGTCGCTTCCCTACGAATCTACCGATGCTCCTTTTTTTCTTAGGAGGATGACAAGTGT
TAACACCTGTAACTTGCACGTTGAAAATATACTGTATAGCAGCCTTAATATTAATTTTTG
TTGCTTTGGGATCAACGGCAAAACAATACTGATTTTCTTCTAATAATTTAGTTGTCTTAT
CAGTAATGATGGGATATTTAACCAGATCTAATAAACCTCTTGAATCAATGCTATCCATTA
TATACCTCTTGTATTTTAGATAAAGCATCAACTGTAATAATAATTTTATGTGCAGCTAAC
AAAGCCATAATATTTAGTGTATCAGCTGAAATAAGTTCTACGTTGTGTAGATTGCGAATA
GAAAGATAAACATTCGGATCTTTTTTGTCTACGATGACTAAAACTTTTTTATTTAAATCA
AGATTCCATCGATGTATAGCTTCCATAAATAGTTTAGTCTTGGGCTGATGAAAATAACTG
TTAAAATTTTCTACAATCAGTGTATTAACTGATTTGTTATTTAATGCTGTCCTTAATGCT
AATTGTCTTTCTTTTTTATTCATTTTTTTGGTAAAACTACGAGGTTTTGGCCCAAATATT
ACACCACCACCTCTCCATAGAGGTGAACGGATAGAACCTGCTCTTGCTCGACCAGTCCCT
TTTTGACGCCATGGTTTACGACCTCCACCTCGAACTTCACTTCTTGTTTTAGTATTAGCA
GAACCCTGGCGCTTTTCATTACTTTGCTTTACTAAAGCTCTATGGACTAAGTACATCCCA
GAATCTTGACTGACCTTAAGATTTAAATCAGCGTTGCCACTTACTTGACCTTCCCAATTA
TAGACTTGATAATTTAATTGCGTGTTAACTGTCATGAAATATATCGTATGAAAGAATATT
GCTACTTACTAATTTTTACTAAAGCGCCAGGTTTACCAGGAACAGCTCCCTTAACGATCA
ACAAATCATTCTCTGAGTTAATACTTACAATTTGCAAATTTTTTATTGTAACTTTTTTAT
TACCAAGTTGGCCAGCCATGTTTTTACCTGGATAAACTCTTCCAGGCGTTGTACCGGCTC
CAATAGAGCCAGGTTGCCTATGATTTTTCGAACCATGTGACATTGGGCCTCTACTAAAAT
GATGCCTCTTTTGGTAACCGGAGAAACCTTTACCTACACTTCTAGAAGAGACATTAATCT
TTTGGCCTACTTGAAATAGATCTGTAGATAGTATTTGGCTAACTTCGAAATCATCAGTAG
ATTTCATTTCGTATTCACGTAAATATTTTAGTGGGGGAGCCTGTGATTTTTTTAAGTGTC
CCAACAATGGTTTATTTAATTTTTGTTCAGCAACTTGCTTGTAACCGACTTGAATAGCAT
TGTAGCCATCAGTAGATACAGCTTTAATTTGAGTAATAACACATGGCCCAACTTGAATTA
CAGTAACTGGAATTGATAAACCAGCTTCATCAAAGAATTGGGTCATACCTACTTTAGTAC

CGAGTATACCAACAGACACTAGATTTCTCCTTTTTGAGAGATTTATACTATCATTGTACA
TAAAAATTCTACTTCATAGTTACTGAAGTAAATTAAACTAACAGATGCATGATATTATTC
ATGACAACATAAAGGTTAAAAACTTTACCTATTTAATCTAAAATAGAAACAGCTTTTTGT
CTAGTTACAGCGTATATAAATTAGAAGTATGCTTATTAAATTTTTTATAAACTACTTAGA
TTTAACTGATATTGATATCCGTAACTTACTTGCAGTAATATCAATCAACATGTATAAGTT
TAATTGCCTAAACTAGAACCAGCTATTATTAGAAATAATTCAAGTATAGATAAAAGCGTA
TTTGAACTTTTATATATTGACAAATATAATAGAATATATTGAGGTTCGGTAACTACACCT
TTTTATTTTTACATTTATTGTGCAGTATATGTACATCTAATGCAGTAAACATAAGAATAT
AACTGTATAGCAAATGTAATTACACAAATTAGAGAGGTATTAATGGGCAAAGTCGTTGGA
ATTGATCTTGGAACAACGAATTCTGTAATTGCTGTTATGGAAGGAGGTAAACCTACCGTC
ATACCGAATGCAGAAGGTTTTAGAACTACAGCTTCTGTTGTTGCATATACTAAAAGTGGA
GATAAACTTGTAGGACAAATTGCCAGGAGGCAAGCTGTTATTAACCCAGAAAACACTTTC
TACTCTGTCAAAAGATTCATAGGACGTAAACAGAATGAAATTTCGCAAGAGATTCGGCAA
ACATCATATAATGTTAAAACTAGTGGATCAAGCATAAAAATTGCTTGCCCTGCACTGAAT
AAAGATTTTGCTCCAGAAGAAATTTCAGCTCAAGTACTGAGAAAACTTGTTGAAGATGCT
AGTACGTACTTAGGTGAGACTGTTACACAAGCAGTAATAACTGTACCGGCTTACTTTAAC
GATTCACAAAGACAAGCTACTAAAGACGCAGGTAAAATAGCAGGCTTAGATGTATTGAGA
ATTATTAATGAGCCTACCGCAGCTTCTTTGTCTTATGGACTAGACAAACAAAATAATGAA
ACAATACTAGTATTTGACCTTGGTGGGGGCACATTTGATGTATCTGTATTAGAAGTTGGA
GATGGAGTATTTGAAGTACTCTCAACTTCTGGAGATACACATTTAGGCGGAGATGACTTT
GATCAGCAGATTGTAGAATGGCTAATCAAAGATTTTAAACAGAATGAAGGTATTGATCTT
GGTAAAGATAGACAAGCACTTCAGAGATTGACCGAGGCTGCAGAAAAAGCAAAGATAGAA
CTGTCAAATTTAACTCAAACAGAGATCAATCTTCCTTTTATTACTGCCACACAAGATGGT
CCAAAACATTTAGAAAAAACTGTAACTAGAGGAAAGTTTGAAGAACTTTGTTCAAATTTA
ATAGATAAATGTAGTATCCCTGTAAATAATGCTCTGAAAGATGCAAAACTAGAAGCTTCC
AGTATTGATGAAGTTGTTTTAGTTGGTGGATCTACAAGGATTCCAGCCATACAGCAAATG
GTTAAAAGATTAATTGGTAAAGATCCAAACCAAAGTGTCAATCCAGATGAAGTTGTTGCT
ATTGGTGCAGCCGTACAAGCTGGAGTTTTAGCAGGCGAAGTCAAAGATATTCTATTACTA
GATGTGACGCCGTTATCTTTAGGAGTGGAAACTTTGGGTGGCGTGATGACAAAGATTATA
CCAAGAAATACTACTATTCCTACAAAAAAATCAGAAGTATTTTCTACAGCTGTAGATAAT
CAACCAAATGTGGAAATTCAAGTACTTCAAGGCGAAAGAGAACTAACAAAAGACAATAAA
AGCTTAGGGACGTTCCGATTAGATGGCATTATGCCTGCACCTAGAGGAGTTCCTCAAATT
GAAGTTACCTTTGATATTGACGCTAACGGAATTTTATCTGTAAAAGCCAAAGAAAAGGCT
ACTGGTAAAGAGCAATCAATTACTATATCTGGAGCATCAACTTTACCTAAAGATGATGTT
GAAAGAATGGTAAAAGAAGCTGAAGAAAATTTCGATACAGACCAAAAAAGAAGAAAGAAT
ATTGACACAAAGAATCAAGCAGAGTCCCTATGTTACCAAGCTGAAAAGCAAGTTAAAGAG

TTTGAAGACAAAATTAGCCAAGATTTAAAAATAAAAATAGAGGAGCTAATTACAGAGCTT
AGATCTAGTCTAGAGAAAGAAGAATATGACAATATTGAATCCATTTCTCAGCAATTACAA
AATGCTCTGATGGACATTGGAAAAAATGCTGCTCAGACTGAAAGTAAAGATACAAAAGCG
AAGGATGACGACACTGTGATTGACACTGATTTCTCTGAAGCTAAGTAAAAGTAAGCGGGT
AACGCGATTCGAACGCGCGACATCAACCTTGGCAAGGTTGCGCTCTACCACTGAGCTATA
CCCGCATAGATTGTATTATTACAAAAATTAAGATATTTGTCAATCTGTTAATCTATACAT
CTAGACTATTAATTTTGGAATTACATATTAGCTAACTGTTGTAAATATAACAATTGCTAT
AATTTTTTAAAAGTACTTATATCATTATATAAGTACTTTTTATATTCAAAAATACAAATA
AGTATTATAGCTAAACTACTAAGGAGTTAACTAAGCTAGTAACAATTACTAATCCTGTCC
ATAAGCCGGCTCCTGTATATATTAATCCTTTTGATTGCTCCCATTGGCCAGGGGATGCTA
AAACAACAGGTACGCCTACTACTAACACAGTCGATAATGCAATTAATAATAGCACTAATA
ATTGGATCGCAATAATCATTTAATATGTTCTCCTGAAAAATTCAAATGTTAATAATACTG
TATATAATATAATGAAGCTTTTGACCTTCAGCAATTTATCTATGAAGAAATAATTTAAAA
TTATTGTTTTCTAAATAAACCTTAAATTAAGTATTAAAATCTTTATTGTAAACAATAATA
CTATATTTTTTTAGTAGATAAAAAAATTTAAAGAAAGTCTTTTTTAAAAAAAAAATTAAGAT
AAGGTGATATGCTGTATAAAAAGCTAAGTATTATTAATTATTAGAGAAATTTAAAATAAG
AGGTAAAACATATGAACTCAGCCCTTTTTTTAGCAAAACTACCAGAAGCTTATGCCATTT
TTAAACCGATCATTGATATCTTACCCGTAATTCCTGTATTTTTTCTTCTTTTAGCTTTTG
TTTGGCAAGCAGCAATTGGTTTTAGATAACATTTAGGTTATAAAAAGAATAACAAATATA
CAGAGTGCCAATTTTTGAGCTCTAATCTTATATCTTACACAATAATAACTATTATGGTTG
AACCCTTACTATCAGGAATTGTTCTTGGACTTATTCCTATTACTTTATTCGGATTATTAG
TAGCAGCTTATTTACAATATCAACGGGGCAACCAATTAGGACTGTAAAGCCTAAATTATA
GATATTTATAAAAACTACAATAATTATAGTTTTTATAAATATCTACAAATAAGCTAATTT
TAAATACAGAAATAATACTATTGAAACTAGAGTGATAACAAAAAAAAATTTATGTAATAAC
AAGCTAATTGTTTACCAGCTCGACTTAGTAACTCCTGGTAAAAGACATTCATGAGACATT
TCTCGAAAAACATGGCGAGATAATCCAAAATCTCGATAATAACCTCTACTTCTACCTGTT
AACCAACATCTATTTCTCCCTCTAACAGCGGCGCTGTTTCTAGGCATTTCTTGGAGTTTT
TGTCTTAAATCCATTTTGTCAGAAAAGCTGCCAGCTTCTTTCAGTTGATCTTTAATAGCC
AGTCTTTTTAAATAATATTTTTTTGCAAGTTTCGATCTCTTAATCTCTCTTTGGATCATA
TTCTTTTTAGCCATGCTGTAAAACTCTCTTTTAAAATTATTTATTCAATATATTAGCTAA
AAAAGCTCCTGCCTGCAACTTTTATATATTTAAAAATAAATAGCATAAAGTAGTGAAATG
CAATTGATGTTTAATCAACGATACCACTTTATGCTATTATCAAAAATTTATTTAGTAAAG
TTCTTGTTCAACATGAGTTGAAATAGTGCAATCAGATGAAGGATATGCAATACAGGTTAA
AACATAGCCAGCTAAAAGTTGATCATCATCTAAGAATGATTGATCAGCTTGATCTACAGT
TCCTTCTGTAACTTTACCTGCACAAGTTGAACAAGCTCCAGCTCTACAAGAGTAAGGAAG
TTCAATTCCTTCTTCTTCTGCTGCATCTAGAATATATGTATCCTCTGCACAATTAAATGT

GACATCAATTCCTTCTTCTTCACATAGTAAATGAATTTTATAATCAGCCATTTCTTCTTA
CTCCATAATAAAGATACACTTGAAACTTAATAAAGCAAAATAATTTAAAGAACTTACCAA
AAAATTATATACTAGTTTCATTATGCATGATAGACTGGAAAAAAAGAAAATAGTATACTA
TTCTGAAAAATCAAGCTACATTTCCTACTAAATAGAACTATAGTAGGTCCAAAGTATATA
CATAAATTCATATAGTTAAAATTACTTTTATCATTTAAATAGCTGAACTTGATAGTATAG
ATTGATCATCAAGCTAGTCTTCTGCCAATAAATAATTAATTATGATTTCAATGCTTAATC
AACAAGTAGAACTTAAGCCTATTCTTAAATTTCAGATAACACAAATATATTCGCATTCTC
TTATTCAGGAAATTAAAGCCTTAATAACTAGACTTGTTTTGCAAGTTTGGAGAAGGCCAG
CTACATTGATGGCAGGTATTATCCAACCTTTATTATGGTTAATTCTATTTGGTGGGCTTT
TCTACAATGCTCCTATAAATTTGTTCACTATTAATACAAGCTATAATTGTTTTTTGAGCT
CTGGGATTATAATTTTCACCTCTTTTACTGGAGCTTTGAATTCAGGTCTTCCATTAATGT
TTGATAGAGAATTTGGATTTTTAAATAGATTATTAACGGCTCCCTTAGTATCGAGGACTT
CTATCATTTTATCTTCTGCTACTTTTATGACTTGTATTAGTTTAATACAAGTTGTATTTA
TAGTTACAGCTTCTCTTTTTATGGGAAACCCACCTCTAAATAGCGATAGTACTATGATTT
TTGGACTTATGATTCTATTAGTGACTGTAGGAGTTACAATGCTTAGTTTAGCTTTATCTT
TCACTCTGCCAGGTCATATTGAGCTATTAGCATTTATTTTAGTAGTTAACTTGCCCTTTT
TATTTTCTAGTACGGCTTTAGCTCCTTTATATTTCATGCCGCCATGGCTTCAGTTAATTG
CAAGTCTCAATCCATTGAGTTATGCAATAGAAGGTACAAGATACTTATATTCAAGCGTAA
ACTGGAATTTTACAGAGTGTGTGATTAAGATTAGCTGGGGAGATATTTGTTTAGGGCAAA
TTATTATTTTATTAATCGCTTTAGATATAATGGCAGCTTATCTTGTGTCTAATATATTAA
AAGCTAAACTTAATTAAAATTTTAATAAAAAATTTATTAAATTTATTTTACAAAGATTTT
TACTATATAATACTAATAGTAGTATGGATAATTGAAATAGAAATAGTTTTTCAAAACCAA
AGCTATTATCAACAATGATAAACATTTGTAAGAAAGTCAACAAAGTATGTTCTTATTCAT
AGGAGGCATGTAGTCAATGGGACTACCATGGTACCGTGTACACACGGTTGTTTTAAATGA
TCCTGGACGGTTAATTGCAGTCCACCTAATGCATACTGCACTTGTAGCGGGTTGGGCAGG
ATCTATGGCATTATACGAACTAGCTGTATTTGATCCTTCAGATCCAGTATTAAATCCAAT
GTGGCGACAAGGCATGTTTGTTATGCCATTTATGGCTAGACTGGGCGTAACAGATTCATG
GGGAGGATGGAGTATAACAGGAGAGAGCGTATCCAATCCTGGACTGTGGAGTTTTGAAGG
TGTAGCCTTAACTCATATAGTCCTTTCTGGCATGCTTTTTCTAGCTGCTATTTGGCATTG
GGTTTATTGGGATTTGGAATTATTTAGAGATCCACGAACTGGCGAGCCAGCCTTAGATTT
ACCGAAAATTTTTGGAATTCATTTATTGCTATCAAGTCTACTTTGTTTCGGATTTGGAGC
TTTCCATGTAACTGGACTTTTTGGACCAGGAATGTGGGTATCAGATGGGTACGGAGTAAC
CGGAAAGGTATTACCAGTAGCTCCAGCATGGGGACCAGAAGGATTCAACCCGTTTAATCC
TGGAGGAGTTGCATCTCACCACATTGCCGCAGGTACTGTAGGTATATTAGCTGGTGTTTT
CCATTTAACTGTTAGACCACCACAAAGACTGTATCGTGCTCTAAGAATGGGTAATATTGA
AACTGTATTATCAAGTAGTATCTCTGCTGTTTTTTTCTCAGCTTTTGTGACTTGTGGTAC

GATGTGGTACGGCTCTGCAACTACGCCTATTGAATTATTTGGTCCAACTAGATATCAGTG
GGATAGTGGATATTTTCAGCAAGAAATTGAGAAACGAGTAGAAAATGCTATTGCTGATGG
TGCTGCACCTAGCGAAGCATGGTCAAGAATTCCTGACAAGTTGGCATTCTATGACTATAT
TGGTAATAATCCAGCAAAAGGAGGATTATTCAGAGCGGGCCCTATGAATAAAGGCGACGG
AGTTGCTGAAGCATGGCTTGGACATCCTGTATTCCAAGATAAAGAAGGAAGAGAGCTTAG
TGTTCGCAGAATGCCTGCTTTTTTCGAAACTTTTCCTGTAATTTTAGTTGATAAAGATGG
TATTATACGAGCTGACATTCCATTTAGAAGAGCAGAGTCTAAGTATAGTATTGAACAAGT
AGGTGTAACAGCTAGTTTTTATGGTGGCAAATTAAATGGCCAAGTTTTCAACGATGCTCC
TAGCGTTAAAAAATATGCAAGGAAAGCTCAATTAGGTGAGGTATTTGAATTTGATCGAAC
TACATTAGAATCAGATGGAGTGTTTAGAAGTAGTCCTAGAGGCTGGTTTACATTCGGTCA
TGCTAATTTCGCTTTAATTTTCTTCTTTGGACATCTTTGGCATGGTTCAAGAACTATCTT
CCGGGATGTGTTTGCGGGAATCGGAGCTGAAGTTACTGAACAAGTTGAATTTGGAGCTTT
CCAAAAATTAGGAGATAGAAGTAGTAAAAAACAAGGAGCTGTATAAGACCAAAGTCAAAG
ATAGTCTTTATCAAATATTTCTAATATTAAAATAGTTAAGACAAAGTTTATTATCTAATA
GTTACTAATCAGGAGAATTTATGGAAGCCTTAGTCTATGTATTTTTACTAACAGGAACGC
TAATGGTTATATTCTTTGCGATCTTTTTTAGAGAACCTCCAAGAATAGCAAAGTAACTTG
ATTCCGTTTAAACTTATAATAATAGAACCACTTTCCAACTTATTAGTGGAAAGTGGTTTT
ATTGTAGATTACATTGTATTATTCTTCATGCTCTTCAAAAGGATCTCGGAGATCTTTTGA
AGCAGGACCAAATGCTGTGTATATCGAGTAACCTGTAATACCTAAAAGCAGACTCGAAAT
AAAAATACTAAGAACTGTTGCAGTTTCCATAATTTAAGCAGTATTAAGATGAACTTTTTC
TATAATGATATTATCAATATTAAAACATTTAATTATTAAAATTATCAAAATATACTATGG
CACTTAGAACTAGACTTGGAGAAATTTTAAGACCTTTAAATTCAGAGTATGGAAAAGTTG
CTCCAGGTTGGGGCACAACTCCTATTATGGGAATTTTCATGTTACTATTTTTCTTGTTTT
TATTAATCATTTTACAAATCTATAATTCTTCATTAGTATTAGAAAATGTAGATGTAGATT
GGGCTACTTTAGGTAGCTAACTAGAGAAGTTCTTTGATTACACAATATTAAAGACAACAA
GCGATCGCTATTAGATTATAGTAGCGATCGCTTTAAAAGTATTTTACTATATATCCAGAG
AAAATTATTTACCTAAATTAATTAACTCGGTATCTGCAAAACTATTTGTGTTAACATTAT
TATAGTTAACTTTTTCAAACCTTACTAATACTGGATATTTGATACCACTTTTATCAATTG
CTGCAACAGTACCAATTTCCTGATACCAGTACGATTCTTTTCTCAATATTTTTACTTTTG
AGCCTCTTTCCATATTTTTAGTCCTCAGATAGATTATTAAAGATTATTATATTAAAAAAT
T-TAAAAAACTAAACAAAATATATAAAAATACTACCGACAATTGTAATATATAACTAATA
AAAGGAGAGAGAGAGGGATTCGAACCCTCGATAACAAAAGTTATGACAATTTTCGAAATTGT
TGCAATAAACCACTCTGCCATCTCTCCTGAATTAAAATACTGTTTTAAAAAATTAGACAA
GAATAACATAGAACTAAGGTAGTTGCCTAGAAAATTAAAACTATGTTAAATATAAATTAT
AATCTTTCTATAGATATATTTAAAAAATATGTACATAGAATCTATATCAGCTTATTTCAA
GATTTTTTAATCTTATACAATTTTTACTAAATAAATAAATTCTCTCATAATATGAAACTA

TCTTGGAAAACACTTCTTCTTTGGTCTTTACCAATATTTGTAATTGGTTTCTTTTTCTGG
CAAGGTTTTTTAGGTCCAACTACTACAGATGTTGGCAGTAATATCGCAAGTTCTAGAATG
ACATATGGACGATTTTTAGAATATTTAGATATGGGTTGGGTGAAACGGGTTGACCTCTAT
GAAAATAATCATACAGCAATTGTAGAAGCTGTTGGGCCAGAATTAGGAAATAGGGTTCAA
CGAATTCGAGTTGAACTGCCAGCAAGTGCGCCAGAATTAATTACAAAATTACGCAAAGCC
AATGTTGATCTAGATGCTCACCCCCCTAAAAGTACAAGTGCAGTATGGGGACTATTAGGC
AATTTACTATTTCCTTTACTATTAGTTGGCGGGTTAGCTTTCTTATTTAGAAGATCTAAT
AATGCTAGTGGTGGACCTGGTCAAGCAATGTCATTTGGCAAATCGAAAGCTTTATTTCAA
ATGGAGGCTAAAACGGGAGTAGTATTTAATGATGTAGCTGGAGTTGAAGAGGCAAAGGAA
GAATTTCAAGAAGTGGTAACATTTTTAAAACAGCCTGAATCATTTACTGCTGTTGGTGCA
AAAATACCAAAAGGCGTTTTATTAGTTGGACCTCCTGGCACAGGCAAAACATTACTAGCA
AAAGCTATTGCTGGCGAGGCTAGTGTTCCTTTTTTTAGTATCTCAGGCTCAGAATTTGTA
GAAATGTTTGTTGGTGTTGGCGCTTCTCGTGTGAGAGACCTATTCAAGAAAGCAAAAGAC
AATGCGCCTTGTATCGTTTTTATTGATGAAATTGATGCTGTTGGTAGACAACGAGGAACA
GGTGTTGGAGGTGGTAATGATGAAAGGGAACAAACATTAAATCAACTATTGACTGAAATG
GATGGCTTTGAAGGAAATACTGGTGTTATTGTAATTGCCGCTACTAACAGAGCTGATATT
TTAGATTCTGCATTATTAAGACCTGGAAGATTTGATAGACAAGTTTCTGTAGATGTACCA
GATTTTAAAGGCAGGTTAGCAATTCTTGAAGTTCATGCTAAAAATAAGAAAATGGAACCT
AAAGTATCTTTAGAAACCATTGCCAGAAGAACTCCCGGCTTTTCAGGAGCTGATTTAGCT
AACTTACTAAATGAGGCTGCTATCTTAACGGCTCGACGAAGAAAAAATGCAATGACTATG
TCTGAAATTGATACATCAATTGATCGAGTAGTAGCCGGGATGGAAGGCACTCCTTTAATT
GACAGTAAAAGCAAAAGATTAATTGCGTATCACGAAGTGGGTCACGCAATAATAGGCAGT
TTATTAGAGCATCATGATCCTGTGCAAAAAGTTACATTAATACCAAGAGGGCAAGCAAGA
GGCTTAACTTGGTTTACTCCTAGTGATGATCAAAGTCTAATATCAAGATCTCAAATACTA
GCCCGTATCGTAGGTGCTCTTGGTGGCAGAGCTGCAGAAGAAATCATTTTCGGTGACGCA
GAAGTTACTACTGGTGCAAGTAATGATTTACAGCAAGTAACATCAATGGCAAGACAAATG
GTCACTCGATTTGGAATGTCTAAAATTGGACCTTTATCTCTTGAAAGCCAAGGAGGAGAC
CCATTTTTAGGTAGAGGCATGGGAGGAGGCTCAGAATATTCAGATGAAGTTGCAACTAAT
ATTGATAAGCAAGTAAGGGAAATTGTCAGTGAATGCTATGCACAAGCTAAACACATTATT
ATAGATAATCGAGTAGTGATAGATAGATTAGTTGATTTACTAATTGAAAAAGAAACAATT
GAAGGCAATGAATTTAGAGACATCGTTAAGGAATACACTGCAATTCCTGAAAAAAATTAC
TACATATCACAATTTTAAATTAAACGGGACTGACGGGATTCGAACCCGCAACTTCCGCCG
TGACAGGGCGGTGCTCTAACCAGTTGAACTACAGTCCCAAAAAGATGAATACCCAGAGAT
AATCTCATAGTTAGCTTAAACTTGTCAAATTAGAATATTAAATATGAAGAGTATTTGTAA
TAATACTCTTCATATTTAATATAGTAGCTAAAATATTTCTTTTAGAAGGAGAGAGAGGGA
TTCGAACCCTCGGTACGAAGTTAATCGTACAGCAGATTAGCAATCTGCCGCTTTCGACCA

CTCAGCCACCTCTCCATGATATATATTATATATATATCAAATATACTTATATTTTAATAG
GTGTGCATAAATATTTATGGAGCTAAGCGGATTCGAACCGCTGACCCTCTCAATGCCATT
GAGATGCTCTACCAACTGAGCTATAACCCCTTTGGTAAGATATTAAGAAATGGCTCAAGC
GGGATTTGAACCTGCGACCTTGGGCTTATGAATCCCCTGCTCTAACCACTGAGCTACTGA
GCCATAGTATAAAATATTATTAAGTATTATACCATAAATTATGGTCAAATATAAATCAAT
AAAAGAATAATTACTATCATTTACTACTTTTGTGCAAGTTGTAAATATTCGAATAAAATT
TCAAAATTAAAAATGTCTTTAACTTTTTGTTAGAATTATTGATTACGAAAGTATTGATAT
TTAGAATTAACAATAAATTTTGTCTTTATTTGTAGACACAAAAATATTGGAAATTTTCAT
CAATTAATTAGTTAAACTATAACCGACATACAAGAATTATTGAACTACGTTAAATTGTGA
TGCATTGGGCTTGATCAAAACTTCTCTAAAATCAATAGAATATAGTAATTCAAGTAGATC
AAATACTAACAAATGTGACAAACGCTTAAAAAAATAATACAATTAGAATCGTAAAAATTT
GGTTTTTGACTGTGTTAAATAAATTTGATTATCATAGATTTAATATTTAAAAGTGTTCTA
TTACTTGTATTGAGTAATTGTAATATTTAATATCTTCTTTTTTAGATAACAATCATAATA
TAAGCTAAAAAATAGTTTTTATTAAACAACTAGCATAGAGCCAATACCTTGATCAGTTAA
TATTTCTAGCAAGAGAGCATGATCTATTCTACCATCTAAAATATGCGCTGAAGCTACACC
CTGAGCCAAAGAACGAATACAACAATTCACTTTAGGAATCATGCCACCAGAAATTACTGC
TGTTTGAGTTAAATCTCTAGCTTCCTGTATATTCAGATGACTAATTAATGTTGAAGGATC
TGAAGAATTACGTAAAATGCCGGGCGTATCTGTGAGTAGAATTAGTTTTTCGGCATTAAG
AGCGGCTGCTATTTCTCCTGCGACAGTATCAGCATTAATATTATACGATTGGCCTTCCTT
GTCTGCTGCAACACTAGCTATAACAGGTATATAATTATTATTAATTAATATCTTCAGTAA
TTTAGTATCAATATTTTGTACTTCTCCAACAAAACCAAGGTTTGCTTTTCCATTTGGTCT
CGGAGTAATAAGTAATCCATCTTTTCCCGACAAACCTACACTTTTACCACCTTGTTTATT
AATAGTTGCGACAAGATCTTTATTGACTCTTCCAACTAAAACCATTTCCACTATATCCAT
GGTAGGTTGATCTGTTACTCTAACGCCATCATCGAATTTTGGTAATATTTTGAGGCGATC
TAACCAAAAATTAATTTCTGGTCCTCCACCATGAACTAAAATGGGACGTAGTCCTATAAA
AGACAGAAAAACAAGATCACTAATCACTTGATCTTTTAGTTTCTGATTTTTCATAGCCGC
TCCCCCATATTTTATAACAATAATTCTGGAAGAAAATTGCTGAATATAAGGCAGAGCTTC
ACTTAAGACTTTTACTCTTTCTGAATTAGTCAACATAAATAAAGTAGGATTGTGATATAT
AAAATTATATACACAGCTTGTGTTAATTTAAAAGATACCAATGGGTACTTGAAAAAATAG
AAATATTACTAACAGTTAAAACTATAGGGATTATAATTATGAATAAATTTTGGGATAATG
TATTAAGATTTCCTCGATTTTTAGTCAGCGTCATTTTAGGATTAATTTTGATAATTATTA
GTCCTTTTTTCGTGTTATTAAAAAAACCACTGACAAGTTTTTTTTTCATCATATCATTGG
CCGGATTAATCACAGTTTTGGCAATAATAATACAAAAAATGATAAATATCGAATGTTGTT
GAGAGAAATAAATAAATTTAATTAAAATTTTTGAACATATATAATATTATATTATCATTA
GATATGATAATTTATTTCCTTTTCTATAAAACCCATAAATATTAAATATTATTTGATAAA
TATTATGCCACTAAAACAAAGAGTTAGCTCTGAAAAGACCGGTGCCTTCGCACTTTTGGA

TAGTATTGTTAGGCACGGAGTTAAACATATATTTGGTTATCCAGGTGGAGCTATTCTTCC
TATTTATGATGAGCTTTATGCTTGGGAAGAAGCCTCTCTAATTAAACATATCCTTGTTCG
TCATGAGCAAGGCGCTGCTCATGCTGCAGATTCTTATTCTAGATCAACAGGAGAGGTTGG
AGTATGCTTCGCTACTTCTGGCCCAGGAGCAACTAATCTTGTCTCAGGTATAGCTACAGC
ACATATTGATTCTGTACCTATATTAGCTATAACAGGTCAAGTTGGGAGAGCTTTTATTGG
TACAGATGCTTTCCAAGAAGTAGATATTTTTGGGATTACACTTCCTATTGTAAAACATTC
ATATGTAGTTCGTGACCCTAGAGACATGTCTAGAATTGTTGCGGAAGCATTTTTTATTTG
TAAACACGGTAGACCAGGTCCAGTATTAATTGATGTTCCTAAAGATGTAGGATTAGAGAA
GTTTAATTATTTTTCTGTTGAGCCCGGAAAAGTTAATATTCCTGGCTGTAGGCCAATTAC
CAGCCTAAAGTCAAGACAAATCCTTATGGCAGCTAAAATGATACAGCAATCTAGCCAGCC
ATTGTTGTATATTGGTGGAGGAGCCATAATCTCTGATTCACATCAAATTATTAAAGAACT
TGTTGATTTTTATAAAATACCTGTTACTACTACTTTGATGGGGAAGGGGATTTTTAATGA
GGATAGCGATTATTGTCTAGGGATGTTAGGTATGCATGGTACTGCGTATGCTAATTTTGC
AGTTAGTGAGTGCGATCTTTTAATTGCTTTAGGAGCTAGATTTGATGATAGAGTTACTGG
AAAATTAGATGAATTTGCTTGTAATGCCCAAGTGATTCACGTAGACATTGATCCTGCTGA
AGTAGGAAAAAATAGGATTCCTCAAGTTGCTATTGTCGGTGACGTAGCAGAAGTTGTTAG
TGAAATATTGAATTTATTAAAGACTTCTTTCCCCCCTTATCCAGAGCAGATTATATCTTG
GCAAGAAAGAATTAATCGTTGGCGTCAACAGTATCCTTTACTGGTTCCTAGAAAATCAAC
AAGCATTTCACCTCAAGAGATTCTTGTTGCAACTAATAAATTAGCCCAAAATGCTTATTT
TACTACAGATGTTGGCCAGCATCAAATGTGGTCAGCTCAATTTCTGAAAGTAAAAGCTAA
GCATTGGCTTTCAAGTGCTGGATTAGGCACGATGGGTTATGGTTTACCTGCAGCAATTGG
CGCTCAAGTAGCACATCCAAATGACGTAGTCATTTGTATTAGTGGTGATTCTAGTTTTCA
AATGAATATGCAAGAGTTAGGAACTATCGCGCAATACCAGTTACCAGTTAAAATTATTAT
TATTAATAATCGATGGCAAGGGATGGTTAGACAGTGGCAACAAGCTTTCTATGGTGAAAG
GTATTCACACTCAAGAATGACAGAAGGAGCACCTGATTTTCAAAAGCTTGCAGAAGCTTT
TGGTATTAAAGCTTTTACTATTAATAATAGGCAAAATATGCAATCTGCTTTACAAGTTGC
TATTGATTATCCTGGTCCAGTTTTATTAGATTGCCAAGTTACAGAAAATGAAAACTGTTA
TCCAATGGTTGCTCCTGGAAAAAGTAATGCACAAATGATAGGTATCGCTAAACCGCAGAG
AGGTACTGCTTCCAACTATATTAATAATAGTGTTTGAAGTAAATAGTTAGTATTTTTTAA
ACAAAGAATAAAGTACATAAATGTATTTTTTTATACTATTAAAAACTATGTAAGAACAG
CCTTTGACGAGAATTGAACTCGTGACCTTCCCCTTACCAAGGGGATGCTCTACCTCTGAG
CCACAAAGGCTTTTTACTATATTGGGCCGGGTTGGATTTGAACCAACGTAGGCGAAGCCA
GCGGATTTACAGTCCGCCCCCATTAACCACTCGGGCACCGACCCTAATTAAACCTTATTC
AAGATATGACCTATAAGATATCATAACTATGTTGCTAAAAACAAGTACTTTGATAAATAA
AACTGAATAAGTAATGGACATAGCTGGACTTGAACCAGCGACTTTCACGATGTCGACGTG
ACACTCTAACCAACTGAGTTACATGTCCAAAACTTTAATTATCATTATAACACTGTTTAG

TGATTTTATAAGTATCTATTTCTAATTGAAGATTTGTTTTAATCGTCCGGCTTTACCGAG
CAGATCTCTTAGATAATACAGTTTAGCTCTACGTATTTTAGCTCTTCTAATCACTTTGAC
AAAAGTTATTCGCGGTGAATTGAGAAAAAATACTCTTTCAACTCCAATACCTTGAAATGA
GCATCTTAAAGTTAAACTAGTATTTAAGCTTTTTTTTCTTCTTCTTGCTAATACTACTCC
TTCACATAATTGTTCCCGAGTTTTACTACCTTCCTTAACCATTAGTCCAAGTTGTATTGT
ATCGCCAACTTTAATTTCAGGTACTTCTGTTTTGATAAAAGGTATCTCAACACTTTTCAT
TAGCTGACTTAGCTTTGTACTATTTGCTTTCATATATAGGATTAGTCTTGTGACATGTTT
TGAAATATTATACATTAAAAAAACAAATATGAAAAATTTTAGCGAATAAGCATATTAGTA
TTTATAAAACTCTTTTTATCTCCATTATACAGATCCATTATATTCAACTAGATTCTTTCA
TAGTTTCTTTGATAATCAGTACATCTAGATGATCAAATATTATTTTCAAACATGGCTTAT
GTATGAAAAAGTTCTATAAAAACCTAATGACTAAAATTAGGGCTTAAGTATTTCTTAGTT
TTGTGTTATAATTAGTTACTATCAAAGGTAGTGGCATAACACGAAGCATCAAAATGTTCG
AACGCTTTACTGAAAAAGCTATAAAAGTCATAATGCTAGCACAGGAAGAAGCTAGACGCT
TAGGACATAATTTTGTCGGAACTGAGCAGATACTATTAGGATTAGTTGGTGAAGGTACTG
GAATCGCAGCTCAAGTTTTGAAATCGATGAATGTGAATCTAAAAGATGCAAGAGTTGAAG
TCGAAAAAATTATAGGAAGAGGGTCTGGTTTTGTAGCGGTTGAAATTCCTTTCACTCCTC
GAGCAAAAAGAGTACTAGAATTATCTTTAGAGGAAGCACGTCAACTAGGCCATAACTATA
TTGGCACAGAACACTTGCTAATGGGCTTAGTCCGAGAAGGAGAAGGAGTTGCGGCAAGAG
TTTTAGAAAATTTGGCAGTCGATGTTTCTTCAATTAGAGCTGAAGTTATACAAATGCTCG
GAGAAAATGCGGAAGCCAATGTAAGTGGAAGCAATACTACGCAAGCTAGAAGTAAAACAC
CAACATTAGAAGAGTTCGGATCTAACTTAACTCAAATGGCTATGGAAGGTGGTTTAGATC
CTGTAGTCGGAAGACAAAAGGAAATAGAACGAGTTATTCAAATCTTAGGTAGAAGAACTA
AAAATAATCCTGTCTTAATTGGGGAGCCTGGTGTAGGTAAGACAGCGATTGCGGAAGGAT
TAGCTCAAAGAATTGCTAATAGAGATGTACCTTCTATTTTAGAAGATAAATTAGTTATTA
CTCTTGATGTCGGTCTATTAGTAGCCGGAACTAAATATAGAGGTGAATTTGAAGAAAGAC
TAAAACGTATTATGGATGAGATTAAATCAGCTGATAATGTAATATTAGTGATTGATGAAG
TTCATACATTGATTGGGGCTGGTGCTGCAGAAGGAGCAATAGATGCAGCTAATCTGCTTA
AGCCAGCTTTAGCAAGGGGAGAATTGCAATGTATAGGTGCAACAACTTTAGAAGAATATA
GAAAACATATAGAAAAAGATCCAGCATTAGAAAGAAGATTTCAACCAGTTGTAGTTGGAG
AGCCAAGTGTTGAAGAAACAATTGAAATTTTGTTTGGTCTTAGAGACCGTTATGAAAAGC
ACCATCAATTAACAATGTCAGATGGAGCTTTGGCTGCAGCTGCTAAATACGCTAATCAGT
ATATTTCTGACCGATTTTTGCCAGATAAAGCAATTGATTTAATTGATGAAGCTGGTTCTA
GAGTCCGTTTACTAAATTCTCAATTACCTCCTGCTGCCAGAGAATTAGATAAAGAGTTAA
GAGCTGTATTAAAAACAAAAGATGAAGCTATTAGAGCTCAAAAATATGAAACAGCAGAGC
AGTATAGAGCAAGAGAAATGGAAATTAAAGCTCAAATTGCAGCAATTGCTCAAAGTAAAA
AGAATGAGCCTGATTTAAATTTAGAAGATCCTGTTGTTACAGAAGATGATATTGCTGAAA

TTGTTGCTGCATGGACTGGTATACCAGTAACTAAGCTTACTAAAAGTGAGTCAGAAAAAT
TAATGCACATGGAAGAAACTTTGCATGGACGTATTATTGGTCAAGACGAAGCGGTTGTAG
CTGTCTCTAGAGCGATAAGACGCGCAAGAGTAGGTCTAAAAAATCCTAACAGACCAATTG
CAAGCTTTATTTTTTCCGGACCGACGGGTGTAGGAAAAACAGAATTAACAAAAGCTTTGG
CTTCTTATTTCTTTGGTTCAGAAGCTTCTATGATACGGCTAGATATGTCAGAATACATGG
AAAGACACACTGTATCTAAACTAATTGGTTCTCCTCCAGGATATGTGGGTTATAGTGAAG
GTGGTTATCTAACAGAAGCGGTAAGAAAAAAACCATATACTGTCATCTTATTTGACGAAA
TTGAAAAAGCTCATCCGGATATTTTTAATCTACTTCTTCAAATTTTAGAAGATGGCCGAC
TAACAGATGCAAAAGGCAGAACTATTGATTTTAAGAATACTCTTTTAATTATGACTTCTA
ATATTGGATCTAAAGTTATTGAAAAAGGAGGAGGTAGTTTAGGCTTTGAATTATCAGAAG
ATCAAACAGAATCCCAATATACTAGAGTACGATCTTTAGTAAATGAAGAACTGAAACAAT
ACTTTAGACCAGAGTTTCTAAATCGATTAGATGAAATTATTGTATTTCGTCAGCTTACTA
AAGATGAAGTTAGAGAAATTGCAGAATTAATGCTTAATGAAGTCTTTGCGAGAATTAAGC
AGCAAGATATTCAATTAAATGTAACAGAACGATTTAAGCAACGATTAGTAGAAGAAGGAT
ATAATCCAAGCTATGGAGCTAGACCACTTAGACGAGCAGTAATGAGGCTTTTAGAAGATA
GTCTGGCTGAAGAAGTTTTATCTGGTAAAATTAAAGCTGGTGATAGTGCAGTAGTAGATG
TTACTAATGAAGGAGAAGTTACAGTTTTATTAGGTGAAAAATTAGAACTGTTAACATAAA
AAACAATTTATTAATTAAAGCATTGAAGAGGTTATATTCAATGCTATGAAAAAAATTAAT
AAGATGGGTTGCTATAATTGACAACCTCTGATATTCTTTATTTGTATATAAAGCCGGGAT
AGCTCAGTTGGTAGAGCAGTGGATTGAAAATCCTCGTGTCACCAGTTCAAATCTGGTTCT
TGGCATTTTAAGCATCAGAAAATTTTGCTATCTGAGCATTAAATTTTAATTGAAATGTTC
CTGTGGGACCATTCCTGTGTTTTGCTATTATAACTTCTGTGAAGTCTCTACTTCTAGTTT
CTTGTGTATAGTAACTTTCTCTATATAGCATTATTACTAAATCTGCATCTTGTTCTATAG
AATTATGAACAATAAAGTTATTTGAAATAAAATTGCATAAAGGTTTACATTCTAAATCAA
ACACTATTTGTAAACTTGTAACGTTTATTTTTTGTATACTTTCAAAAGTCAAAGATGAAA
AAGAATTAAAAACATTTTTTTGTTCTTCAAAGTTTTTATTTGAATAGCAATCATGTCAT
TTTGATTAATTTTATCACATCTTTTCCATCCTTGTGTTGTGAGTAGTTTATGGTTACTTG
TTAATTGTATATATTTCCCAGCTTCGGTAATAATTTTATATACAGTTTTTTTCCCTGTTT
TCGAAATACTACATTTAATTGCTAGAAATAACTGTCGTTTTTGCGCACTAAAGTTAATGA
CTTCAATCTTTTTATAATAGCAATTAAATAATATTTGAGGTTGATTATATAATGATGTCT
GAAGATAATTGAATTTGGATATACACCCACTTTCTCTTAAATCGGATAACAAAGGCCTTT
TGTCACTCCGAGTTTCAAGGTTTCTATTTAGTTGAGATAATGCCAAGATAGGCAAGCTTA
AATCTTTTGCTAATATTTTAAGTGATCGAGTAATTAATGATAGTTCTTGAGATCTATTGT
CAGATTGTCTACTCTCCTGCAGTAATTGAAGATAATCTATAATAATTAATTCTATATTTT
TTCCTTGTAATTTAAAGAATTTTACTTTTGTCTTTATAGTGTCAGTAGAAATCTTAGCGC
TATCATCAATATAAAGATTAAGGTTAGCAAGAGTTTTACTTTTTTGAATAACATATTGCC

ATTCATCATTATTAAGCTGACCAGATTGAATTTTTTGGCCGTTTAAATTACACTCTTGAG
CTAAAATTCTTCTTAGTAGTTGCTCTGTAGACATCTCTAAGCTAAATAAAATAACATAAG
ATCTTTTCTGATTAATTACATATCTAGTTATATTAATAGCAAAAGCAGTTTTCCCCATTG
AAGGTCTTCCTGCAAGAATAATTAAATCTGACTTTTGAAACCCTTGTGTTATAGAGTCGA
GTTCTGTAAAACCAGAAAGTATACTACTATTTATACTAATCTTTTTCCTTGTATCCAAAT
GGACTAATAGTCTTGCAAGAATTGTAGCTAATGTTTGCGTATCTTTTTCGTCAAGTATTT
CATAAGCTTTTGTTAATTGACTAACAATAGAAGTTATGTTTTTTTGTTCAATAAGTTGTT
TTGAGTAGCTAATTAAACATAATGAATCTCCACATGCTAAAAGTAATCTTTTAATGTAGT
TATCTATAACGACTGCTGAATATTCGCTTATAGTATCAGATAGGGGAGCTTGTTCTATTA
GACTAAAAACTTCATTTAGCTCATTCAAGTCTTGAATCATTTTTTCATTTTTTAAGTTAA
TGAAAAAATTTCTTACGCTTTTTGTCTTTGCATGGTTAACATTTTCGAGTGCTGCTCTAT
AAAGTAATGAAGTTGATGTAAAATAAAAAAAATCAGGAGAGATTTTTTCTACTGATTTTA
GTAATGATATCGATTGCTGAGTTAATATTATACTAATTAATATTTTTTCAGCTAAAATAT
TATGAGGAGGTAGATATTTATAAATACTTATTACTTCCTCCTTGTAACATTACTCTTGTG
GTATTAGTTAACTATTGATTTAATTTGATTCTGGAAGAACTTGCAATTTAATATTTGCAG
TTACTTGATGTAATAGCTTAATTTCTACATCGTAAATGCCAATTGTTTTCACGTCTGGTA
AAGAGATACTTTGTTTTTCAATGTCTATGTTGGTAGTGTTTTTAATAATTTGTGAAATTT
CTTTTTCTGTGACACTACCAAAAATATTGTGGCCATCTCCTGTTTTTTTACTAACGCTAA
ACCTTTGTATTTCTTCTAGAAGTTGCTTGACTTTTTTAGCATTTTCTTTAGCTGTTTTTA
ATTTTTCTTCTCTTATAGCTGCATAAAATTTTTGTTGTTTTAAAATACCATTTGTGGCAA
CTGCCGCCATTTTATTCGGGATTAAAAAATTTCTTGCATAACCAGAAGCTACTTTGATAA
CGTCATTACTCTTGCCAAGTTTTTGAATATTCTCTTTTAGCACAACATTTATAACTTTTT
TACTCATCTTAATTCTAAAAAATCTGTATTATCCTATTATACCTGTTGTATACAGATTTT
AATAGGAAGTATTTATTTGTTGATATAATATTCTTGTAAAAAAAGGTAATTTTTTATTTT
TTTCTGATATAATCACTTTCACAATAATTCATTAGGAGAGGTGGCCGAGTGGTTGAAGGC
GCAGCATTGGAAATGCTGTTTAGGAGCAATCTTAACGAGGGTTCGAATCCCTCTCTCTCC
GTACAAAAAAATATAAAAAGTTTTATAGAGTCCAATCTAAATCTCCAATAATATTCTTCCA
AGCATTTAATCCACCCCGAACTCTAACAATATTTAGTTTTTGAGCTATTAAAAACTCAGA
AGCAAATATTGATCTAGAATCTAAGCTACAGTAAACAAAACAAATTTTATCTTGTAAATT
GATATGAGAGTAATTCATTCCTTTTATTTTTTGTAAAGGTAGATTTAATGAATATATTAA
ATGACTTTTATGATATTCTTCATGATTTCTTACATCTATTAAAATATATTGTGGATTCCT
ACTAATTAAAAACTTTTGTAATTGAATAACACTAATTTCTCGTACAAATGTATTAGATTT
ACTGCCATAATATTTGTTCCAATGCTTTTTTGTGGATAATACAAATTTAGTATTTATAAT
CTTAAATTTGTTAAATGATGAAGTTAATGAGTTGTACGTTAATATAGTTCCACTTAATAT
AGACTTATAACCCAGAACGATTTTGACAGCTTCAGTTGCTTGAAGTGTACCAACAATACC
CGGTAAGAGACCTAAAACTCCGGAATTACTACAAGTATCTCTTGCGCTCTCTTTATTTTC

AGTTTCACTGTAGAAGTCTCGATAAACAGGTCCACCTTGATAATTAAATACACTAACTTG
TCCTTCAAATTGAAAAATGGCACCATAAATATGTATTTTATTTAACTCTAGGCAAGTATC
ATTCAAGAGATATCGAGTATCGAAGTTATCAGATCCATCTATAATAATATCATATTGTCT
AATAATGTCGATTGAATTCTCATAGCTTAGTCTTGTTTTGAACACAGTTACTATACACTG
TGGATTTAAATCTAATATTTTTTTTTAGCTATTTCTACTTTTGAATATCCTATATCATGC
ATCGTGTATAAAATTTGCCTCTGTAAATTAGAGAGATCAATTATGTCATCGTCTATAATT
CCAATGCTGCCAATTCCTGCAGCAGCAAGGTAAATGATTCCTGGAGATCCTAGCCCACCA
GCACCAATGAATAAAACTTTTGCTTCTTTTAATCTTTCTTGTCCTTCTAATTGAATTTGT
GGCAATACTAAATGCTTAGAATATCTTGTATATTCTTCTAGTGAGTATTTTGTATTTTCT
GTTTTAAAATTTAGCATAAAAATTTATAGTTTATTTAATAGCACTTAAAAGCTAGCTGAT
AAGTTTATTAAAATGACTTTTGTAAACTTGGTTCTTCTCTATCTAAAATAGCTCCTTGTG
TAGGACAGACCTGAACACAAATACTACAATCAATACAGGCAGAAAAGTCAATCCAATACC
AATTAGTGTTATTATTGTTTTTCCCCTTTCCTTGATGAATACAAGCTACGGGGCAAGCTT
CCACGCATTCAGCAACTCCAATACATTTTTCTGTTACAATTGTATGAGACATATTATTTA
ATTAATAAAAGATAAGCTATTTACATCTAGTAGTTTTGTTGTATACTAAATTTACATATG
CTATCCAGGCCAATATTTAAGGTAGTCTATTTCTACAGTTTACTAATTTGATATGCGCCT
TTAGTTCAGTTGGTAGAACGCAGGTCTCCAAAACCTGATGTCGAGGGTTCAAGTCCTTCA
GGGCGCGTTTAGTTTTATCGTAATTCAATAATATCATTTTATCTTTTTGCAAACATCAAA
ATGATATGATATAAAAGATCATATTTTATACATTGAATATTCTATGTACTAGTACTGTTT
TTTAATATGTTATTATTGATAAGTTAATTATACTAATTCTATTTTTGTAATAAGTAAAAA
CAGTTCTAATTAAGTATGCACGTATTTAATAGTCTTTTGTATATATAACTTGAAAAAATT
TAAACTATGGCTAAAAAAGTTACTGGAATTGTTAAACTAGCATTAAATGCAGGTAAAGCT
ACTCCTGCGCCACCTGTCGGACCTGCTTTAGGACAGCACGGCGTGAATATTGTTATGTTC
TGCAAAGAGTACAATGCACGTACTGCTGACAAGTCAGGACTAGTAATCCCTGTAGAGATC
TCAATATATGAAGATAGAAGTTTTACTTTTATACTTAAAACTCCTCCTGCTTCTGTACTA
ATCGCTAAAGCTGCTGGACTAAATAAAGGTTCGGGTGAACCAAATACCAAAAAAGTAGGC
AGCATAACAAATAAACAGTTAGAATCTATTGCAGAAACAAAACTTCCCGATTTGAACACT
AACAATATACCTCAAGCTATGAAAATAGTTGGAGGTACGGCTAAAAATATGGGAATTCTA
ATTAAAGATTAAAAGTTTCAATTATATTTACTATTTTTATGAAAAAATTTTCACGTCGAC
TTACAACATTAAAATCAAAAGTTGAGCCTAAACTTTACACTATTAATGAAGCAGTATCTA
TATTAAAAGCAACGTCAAATGCTAAATTTAAGGAAACTGCAGAAGCTCATATTGCTCTAG
GTTTAAATCCTAAATATGCAGACCAGCAACTCAGAGCAACAGTTATATTACCGAAAGGTA
CAGGTAAATTGATAAAAGTAGCAGTTATTGCTAAGGGAGAAAAATTAACAGAAGCAATTA
GTGCGGGAGCTGATGTTAGTGGCTCCGAAGAACTAATTGATGAAATACTGAAAGGTAGAT
TAGATTTTGATAAGCTGATAGCAACACCAGATGTTATGCCTTTAATAGCAAAGCTGGGAC
GAGTATTAGGCCCTCGAGGGTTAATGCCTTCTCCTAAAGCAGGCACGGTAACATTGGACG

TAGCAAAAGCCGTGAATGAATTTAAAGGGGGTAAAGTTGAATACAGAGTTGATAGAACAG

GTATAATTCATGTACCTTTTGGAAAATCTAGTTTTTCACAAGAAGATCTAGTTTTGAACC

TCCAAACAATTAAAGAATCGATTGATAGAAACAAGCCTTCTGGAGCAAAAGGGAAGTACT

GGAAAACTTTTTTCTTATCTAGCACCATGGGGCCATCTATTCAAATTGACATCACTAGCC

TTCTATGAAAAATTTTTGTATAATTATAGAACTTGAATAATCTATTATTTCAAATTATAA

TTTGCTAGCATTATTAACGTAACAACTTGCTTACTATATCTTATCTGCATTAAAAACTTT

TTATATATCTTATTAATTATGAGTACAAAGGTTGAAAATATCTTGGAAGAGCTAAAATCT

TTAAACCTTCTAGAGGCTGCTGAACTAGTTAAACAAATAGAAGAAACGTTTGATGTTGAT

GCATCTGCGGCTTCAGGAGGTATGATGATGGCCGCACCAACTTCAGCACCAGCGTCTGCT

GAGGTTGAAGAGAAGACAGAATTTGATGTTGTCCTAGAGGAAGTCCCAGCACCTAAAAAG

ATTTCTGTATTAAAAGCTGTTCGCTCTCTGACTGGTTTGGGTTTAAAAGAAGCAAAAGAT

CTTGTTGAATCTACGCCAAAAGTCTTAAAAGAAGGTGCTTCTAAAGATGATGCAGAAACT

ATGAAAAAACAGCTAGAAGATGCTGGCGCAACAGTTATTGTTAAGTAATGAAGAAGCGCT

TCTGTTTAAGTATAAACAGAAGTGCCTCTTTATTTATTATAAGCAATTATACTAGAAAAG

TCCAAGTGTAATTGCATTGTTGATAGGCATAGTTGCACCAATACCTAGCCAAATACTTAC

AACGGTACCAATTAAGAAAACAGTGGTTGCAATTGGTCTTCTAAATGGATTTTGAAATTT

GTTAACATTTTCAATGAATGGAACAGTAAGTAAACCAGCTGGTACAGCAGCCATACTTAG

AACACCTAATAGTTTATTAGGAATCACTCTTAATAGATTAAATGTTGGAAAGAAGTACCA

TTCTGGTAAAATCTCTAATGGAGTCGCAAATGGATTAGACTTTTCACCTATAGAAGATGG

TTCTAAAATTGCGAGTCCAATACTGCAGGCAAAAGTGCCAATAATTACAACTGGAAAGAC

GTATAATAGATCATTTGGCCAAGCTGGCTCTCCGTAATAATTATGGCCCATTCCTTTTGC

TAATTTAGCTCTCAGCTTCGGATCTGTTAAATCTGGTTTTTTAAGAATTGACATATTGTT

TTTGTGTTTAATATAAAGATTAGTCTAATTACTGTTTTAAAGTGTAATTTTCAACTATTA

TAATGGTCCAGAAATTCCTTGTTTACGTATCATTAAAAAATGCATAAGCATGAAAACGGC

AGTTAGTAGCGGAAGTACAAAAGTGTGTAAACTATAGAACCTAGTTAAAGTTCCTTGGCC

TACACTGACTCCTCCTCTTAACAATTCTACTATACTCGCTCCGACAACTGGAACAGCATC

AGGGACACCTGTAACAATTTTTACAGCCCAGTATCCGATTTGATCCCATGGCAAAGAATA

ACCTGTTACCCCAAAAGAAACAGTAAGCACTCCTAAAATTACACCTGTTACCCATGTCAA

TTCTCTAGGTTTTTTGAAACCTCCTGTTAAATAAACACGGAATACATGTAGAATCATCAT

CAAGACCATCATGCTCGCTGACCATCTATGAATTGATCTAATAAGCCATCCAAAGTTTAC

ATCAGTCATAATGTACTCTACTGATGTAAAAGCTTCTGCAACTGTAGGTCTATAATAGAA

TGTCATTGCAAATCCAGTTGCAACTTGAATTAGAAAAGATACAAATACAATACCCCCTAA

GCAATAAAAAATATTAACATGAGGTGGTACATACTTACTAGAAATATCGTCAGCAATTGC

TTGAATTTCTAATCTCTCTTCAAACCAGTCATAAATTTTACTCATAAAATAGCTTC-AAA

AGCTCTTCTAATATATTTTGCGTTTAAGCTACTATTAATTCAATAGTTATTGAACTTATA

TTTTTTTATAATCAATACTTATTAACTATTATAACATCTTTTTATTGATTTAAAATTCTC

TGACTTAGTAAAACAGGGCTATGTACTATAATTCTTTTTTTTTGTATAGAGATTAATTTC
GTTTTTAGTAAATTAGCTAAAATTCTTGTTACACTGACTCTATTACTACCTATAATCTGT
GCTAAAATTTTATGTGTAATAGTAAAGTTAAGCATAATACCATTATTAACTTGAGTTCCA
TTTTGCTCAGCTAATAATAATAATAAGCTCGCCAGTCTACTGGTTATACTTTTATGTGAA
ATGATTTCTATAAAATGATAAGCTTTTGCCGAGCAAAATACTAAATGATTTACAAAAAAT
AAATTGAATGGAGCGCAGTTACTACACGCTTTTATAATAGTAGTATAGTTAATACAGGCA
ACTTCTGTTTTATCTATTGCTTCTGCTTCGTAGTAAAAATTATCGTCGACCAATTCTATT
TGTCCAAATGTATCTCCTGTAGTTAATAAGTTAAGAGTTACTTTATGTGTATTTCTCAGG
ACTTTTGTAATAATTAAAGATCCTATCAATATAATGTATAATCTAGAGCTGCTATTAAAC
AAAAGAGTATCATTTTTCTTAAGTGAAAAAATTTGATAATTAATTTTATTACGATTAAAA
AATAGTAGCCATGGATTGCAAGAAAAAAAATCACCTGAACTATTTTCAGAGAATATTGGA
ATACAATTTTCTTGATTTAAACAATTTTTAGTCATAATTACTACTTTATAAAAAAATATT
CTAATAATTAAGAACTGAAATTAAATGTCATATAAATTGATGTTAGTTGAGAATGACATT
GTATTATCAAAGGCCATTCAAGAATACTTAATAGATCAAGGCTTCAATGTTTATATTGCC
AATAATGGATTAGAAGCACTGAATCTAGCTTATCAATATAATTTCGACTTGATAATTTCT
GATATAATGATGCCGCTAGTTAATGGCTATGAACTATTAGCAAAACTTAAAAAAAACAAA
GCATTATCCAAAATACCTGTTATCTTTTTGACAGCTAAAGGAATGACTAAAGACAGAATA
AAAGGCTACGACATGGGGTGCTATGGTTATTTATCTAAACCATTTGATCCTAATGAGTTG
CTTTCAATTATTAACAACTTAATCGCTAGAGATGTCTTAAAAGAAGCTTCTTTACAAAAC
TTGGCAACATCAAACCAGCAATTAAATCATAAAATACGTTTGACACCTAGAGAAAAAAGT
ATTTTAGATCTTGTTGTTGATGGACTTACTAATAAAGAAATATCTACTATACTGAATACT
AGTGTTAGAAATGTAGAAAAATATGTTAGTCGACTCTTACACAAAACTAATATGAAGAAT
AGGACTTTGCTAGTCAAGTATTCTATAAATAATAATCTATTAAATAATGAGATCAATGAA
AGGGCGAATGACGGGACTCGAACCCGCGAATGATGGAGCCACAACCCATTGCCTTAACCC
CTTGGCCACACTCGCCATATTACACATACCACTATAGCTTGTTTTTTGATATATCGTCTA
GTTTTCGAGTTATATATTTATCTAAAACAATATTTACTTTGAATTTTTATGATACATAGC
AATATTAGTATTCAAATTAATGGGGAACCATTTAATTGCTCAAAGCCCATCTCATTGCAA
TTTTTATTAAATTATCTTGATTTTAATTCCGAGCGTGTAGCAGTCGAGCTAAATGACATT
CTGTTACCAGAGACTCTGTTTCACTCAACTTATTTGAATGATCAAGATAAGCTAGAGGTT
ATTACCATTGTAGGTGGTGGATAATCTATTTGTACATACACAGTTCTCTATCTCAAATGT
TTCATAGATAGAGATAATCTTCCTTGTTTTTTATCTACATGAATAATAACAGCTTTAATA
GTGTCACCTATTTTAAATTGTGATGATATCTGCTCTAATTCTTTGATATTAATCTCAGAA
ATATGGACAAGACCTTTGAGATTTCCTACCTTTATAAATAATCCGTAAGGTGTAATCTGA
TTTATTATGCCTTCAATAATATTACCAACAATTAAATTAGATGATGCTTGAGATATTAGA
GCTCTTCTATGACTCAGTATTAAATTATTAGATTTCTCTTCGACATTGAGTAATTTTAGC
TTAATAAATTTATTATTAAATTGCTCACTTTTTTGAAAATTACCGAGATGAGAATTAGGT

ACAAATCCAGATATACCTTCAAGGTTAATTATCATTCCTCCCTTATTGAATCCTTTTATC
ATTACATTAAGTAAAGAGTCTTCAGCTAATAGTTGCCTAATTCTTTTCCATGCTCTTATA
TATTCAAGGCGACGAATTGATAATATTAATTGTTTCGACTGAATATTATAATCTAATAAA
AAAAATTCTCTTGTGTCATTAATGTTTAAAGAAGTGAAATTATTTAAATCTTGATTACTA
GATACTTCTTGAATAGGCAAGTATGCAGATATTGGTGTTCCTATATCTACTAATACCCCA
TTTAATTCAAAGCTAAATATAGTTCCAGCTACAATATCTCCAAGATTCAAATCATACTTG
TATTTTTGTAAAACAGCTGCAAAATTTCTATGAGTAAATCCTTCATTATTTTTTGTCATA
TTGAATAGTATGAATTAAAGAGTAACTTTGTACAATATATTAAGGTAAATAAATGTTTGT
CGAATTGTATGAAAAAATTTTAATTTGTTCGGTTGTAATTAAGTATATAAACATATTAAA
ATATCTAGGAGACATGAAAAGTAGACGTAGATAGCTGCAGATTTAATACTATATAATCTG
AGGAAAGTCCGGGCTCCACAAATACAATTTATGCTGGAGAAATCCCAGTGTAGGTAACTG
CGAGGATAGTACCACAGAAAAAAACCGCCAGAATAAGTTATATTAGCTGGTAAGGGTGCA
AAGGCAAGTTAAAAGCTTACCAAAAGTACTGCAAAGTATTTGTTAGGTAAACCCCTAAAA
TGGAGCAAAGCTACTAAACAAATATTTGTGTATATTTTATCTAGTTTAGTTGAAAATACT
GCATGAAGTTATTAGTAACAATAACTCGAGAGGAATAGCTATCCTTTACAATATATCTCG
ATTATATTGAAAGAACAGAACCCGGCTTATGTAGTACTTTTTATGTCTTGTATTATTATG
TCTATTGTCGAATCTTTAAATTTAACTGTTTGACAATTTTTTTTTGTAGCTGATGTTGTT
TATAATCTATTTCTGAATTGGTTAAAGTTTTTTCATTAGATCGATATGTAAATCTTAGAC
CTATACTTTTTTTCTCTTTTCCAATTGATTCATCTATATATTGATCAAATAATGTTGTAC
TTTCTAAATCTTTATCATAAAATTGATCCAATAATTTTAATAAGTATTTAATTTCCATAT
TCTTAGGTATTATTAAAGAGAGATCTCTTATGATCGATGGATACTTAGAGTATGGTTGAA
TCTGATAACTTAAATAATTGAATTCATTATGACAATCTTCTAAAATGTTTAAGTTAACTT
CAAGAAAAAAAAGTTTTGTGTTGAGTCCTAATTCACTATATGTTGCTTGATTAAGTTCAC
TGAATATGCCAATAATAATATTGTTATAGATTAGTGTGGCAAATTTTCCTTTTTGAATGA
ACTTTATTCTACCATCGACTATATCTTTTTTTGTCCACTCAATTTGTCTATTAATCTTTT
GAAAAAAATTTTCTACGATCCCTTTAGCTTCATACCAATTTAAAGAATGCGCAGGGTGTG
ACCATGTAGATCGGATATCTAAATTACCACCCAATATGATAGCTAGATTAGTAGTTTCTA
CAATTTTGTTTTGATCATTATGAAATACTGTTCCTATCTCAAAGCTATCTATAGTTTGGT
TGCTTTGTTTAATATTATAAAGATTTGATTCAATTAATCCTTCCAGCAAACTGCCGCGTA
AATTAGAATAATCTTGAATAAGAGGATTGTTTAAAGCTATATTTCCTTTGGACTTTATTA
AAGAATAATGAACTAATTCAGTTAATCCTAGATTTCTTAAAATACTTCTACTTTTATCTA
TGAATTGTCTCTTTGTGGATAACTTTTTGTTGAATTGAATATTTGGTACTGAGCTCCTGA
ATTTATGATAACCATAAATTCTAGCAATTTCTTCAATAACATCTATTTCTCGAAAAACAT
CTTCTCGTCTATAATTTGGAACAGTAACTTCTATATTTTCCTTATTTTGATAAATAAGAT
CAAAATTTAAAGAGTGAAGAATATTTTGAATTTCTTCAAAGAATAAAAAACGTGTTTGAC
CATTATACATAATTGGTCCTAAAATGTCATGAACTTTTTTAATTGATAAATTAATATTAA

GAGCATGATCATTTATTTTTTCTCGACAGAAAGTCTCTCTAATATTTCCGCCTGTTAGAT
CTGTGATTAAAGCAAGAGCTTCAAAATGTGCATTTTTCCAATTATCTACATTTAGCCCAC
GTTCTTGTCTAATTGAACTTTCTGTTCGAATATTAAGAACTCGAGAAGATTTTCTAACTA
CAGATTGTTTAAATATAGCAGACTCTACAAGTATCGATTTTGTATTATGATCTGTATCAA
AATCAGAGTTGCTTCCAATACCAGCTATACTAGTTACATTTGTATTTACTTGAGTAACTA
AAATATTTTTGTTTAACTCAATATTTTCATTGTTTAACTTAATCTGATCATTACTTCCAA
GAGGAAAATTGCTTCGAATTGTTAGACTATTTTTATGATTCATATTATTAATTTTATTCA
AATCAACAATATTAATAGGCTGTCCCCATTTCAACATAATATAATTGCTTATATCTGTTA
ACAAATTTCTGTGTGTAAACCCAGAAGAGAGCAACCGATTTTTTAGCCAATCTGGAGAAT
CTTTAACTCTTATTTCGTCTATAATGGCAGCAAAATAATTGTCACAATTTAATAAATCGT
GATTACTGATTATAGTCTCCTTTTTAGAGAAATATCTATTATTGGGTTGATGTATAGCTG
ATAACATTGAGGACCCTGTTAAAGCCGAGACTTCTCTTGATAGACCAATCATGCTTAACA
CGTCTGACCTGTTAGCTGTTGATGATATATCTAATATATGATCAATCTCACTTCCAATTA
TAATAGATTCAATAGCTTCAACTTCAAAACCTGCTTGTGTTAGTTGATTAGTTAAATTGT
CTATATCTATTGTTTTGATATTACCAAGTTCTTTCAACCAATTTAAAGAAACTTTCATAA
TTAATTGTCATCTATCTTATAGAAGTATTTATTTTACTTATGATTAAAAATCAATTTAAA
AAATTCATCAAATTAATATAAGTTATTTGATGAATTTGCTTATACAATATATTATTGTGA
TATCTTACTTTTATATGGCAGTTATTAAGAACTAGCTTTTGTAAAAGTTAGCCCATTATC
TTGGGCATATCTTTCCATAAATCTCATGAATCTATCCCAATCATCAGGACTTTTAATAAC
ATGTACAGCTTCTATTGCTTGTGGTTTACCATTAATAAACTTGGCATTAACATCTCTTGT
AATAATTTGTCCCTCTGTGTCCATCAGGTACATACCTGTGATTTCACCTTTGTCTGCCAT
ACTAGCATCTAAAATTTTGGGATTTGTAAACCGAAAAGTTGCTGTTCCTGTACTACCATC
TCGAGACCTTGTTAAACGTACATCTGGTACAACTTCTTCATTGATACCTTGAATAAATTG
AATTGTTGCCATATCAATCTTCTTTATTTGTATATAATTTATATTGGATAAAACTGATTT
CTAGTTCTTGAATGCTTGTTGAGAAACTGTTTTAAAAGCTTATAGTAGCAGTAGATCAAA
TTTTATGTATTATATAAAACTGTAACTTTTCTTTTTTGATCTGTTCAAAGAACTAACTTA
TTATAAGCTAATCATACGAATATTAGCCATATAATGTTCGTAAGTGAAATTAAGGGGTTG
TAGCTCAGAAGGATAGAGCAAGCGCCTCCTAAGCGCTAGGTCAGCGGTTCAAGTCCGCTC
AACCCTGTTAACGAATTAAGTAGTATTATTTAACTTGTTCTGCAAATCAGTATTGATCTA
AGTTAAAGTACTTGACAGCTAAAACAAAAGAATCTGAGAAATATTTTCATTTAAAGAACG
AACAATTATCTTAACTCTAAATATCTGATAATTATTAAAAAATTCTCTTTTTAAAACACA
AGATGATGAAGATATTTAAACTAATAATTGAAATTAAGTTTTTGAAGTTATAGAAGCCAA
TATAATACTTTCTATTAACAGATGCTTGCTTCTATTACTATCTGTTTACAAGAAATTATG
AGTATCAAAATAATTTTAGCTCTACTGACTACTAATAAATATTTATTTTAGTCTAGACAA
GATGAATCTTCATGGTATAACAGAGCTATTCTTTGTGTCATACTTATATAAAATTAGGAT
ACTGTTTTCATTATTATAACTGGGGTGGGAGGATTCGAACCTGCGAATGGCGGAGTCAAA

GTCCGCTGCCTTACCACTTGGCTACACCCCAACATGTGAACATTATAATAGCAGTCGTAT
ACCCTGCCATGTCAATAGACTATTGTCTTTATTTAATAATTTTTGCTGCTCTTGTAGAAT
AAAATCTTTTTTTTCAAATAAGATACGCTATGTTTAAGTTGCACGACTGTTATGTTTTCT
TGCTTTTGACTGAACAGCCACTTGATAGTAATTATTCCTTCTGCAATTTCGTTACTTCCT
ATAATAATACAAGCTACAGCTCTTTTTTTATTTGCTTGCTTAATCTGCTTACTAAAATTA
CTTGAGCTAACATCGATTTCTATTTTAAAAAACTGCTGATGTAAAAAACGCATTATTTGC
ATACCAGTTTCGTTAGCTTTTGTACCTTGTGTAGCAATATAAAAATCGATAGATTGATTA
GGCAAGTCTATATTATCTTTTGCAATCAGTAATAGGCGTTCTAATCCTATTGCACATCCA
ACAGCTGGTGTTGCTTTGCCACCTAATTGGTGAACTAAGCTGTCATATCTTCCACCACCA
CATATAGTATCTTGCCCTTTTGATGTCAATGTCTTAATTTCGAAAGCAGTATCATTATAA
TAATCTAATCCTCGAACTAACTTATTATTTATATTGTAGGGAATATTAAGTAATTTTAGA
TAATTGCAAACGTCACCAAAGTGTTTTTGTGATTCCAAACTTAAAAAATCAGAAATTTTT
GGAGCCTCTGTCAGTATTTTTTGTGTATTAGAATCTTTTGAATCTAAAATTCTAATTGGA
TTACTAGTTAATCTTTTTTGTGAGTCAGTGTCTAAATCATCATGGTACTTTGTAAGATAA
TCTCGTAATTTAACTTGATAAATACTACGATCCTCTACTTTTCCAATTGAATTCAGATCA
AGTTTCAAGTTGTGCAGATTGAGATTGTTAAATATGCTCATAGCTAAATGTATAACTTCA
CTATCTGCTCTTGCATCTAAGCTACCAATAAACTCAATGCCAAGTTGGTGAAATTGTCGT
TGTCGTCCACTTTGTGGTCTTTCGTACCTGAACATTGGACCGCTATACCATAATCTTTGT
AAACTATGGTGGTAACTCATTTTATTTTCAATAAAAGATCTCACAATACCGGCAGTACCT
TCTGGCCTTAAGGTAATATCTCTATTGCTGCGATCATGAAACCGATACATTTCTTTATTT
ACAATATCAGTATCTTCACCAATACCCCTATCATATAATTCACTATTTTCAAAAATAGGT
GTTCTAATTTCTTGGTAATTTGCACATTCTAATAATTTGGAAATTTTATTATGTATAAAC
TGCCAATATTGAATTTCATCGGGAAGAATATCTTTTGTCCCTCTAATAGCCTGAATTTTT
GCCATGATAATTTTTTGATTACCTTCTTAATATGTCTGCATTAAATAGCAGTTGTAGACT
GTTATTCAGTCATGTATTATACATTAGAACATACTGATGTAATTTATAAAGTACGGGCAA
GGAGGGATTCGAACCCCCGACACCATGGTTCGTAGCCATGTGCTCTAATCCACTGAGCTA
CAAGCCCACTTAATAAGTACCTTAACTATATCATTGTTCACTAATGTATACAACTATATT
CTTTTTATCTTACATGTGAAATTTTTGTAAAAGTAAAAAATTATGTTTGTTTTTTACAAT
TGGAGCTATTCATATATAATGCTTAATTTATTATAACTTTGTAAAAATATCTATAAGTTT
AATTGTATGATTCTATTATATTGGTAATCTTGTTATAATTTTGCGCTCTTATTTTTCAAT
TGGCTAGCATTTAATTAACTTATATTTCAAATAAAAAAATCTTATCTAAAATGAGTAAAC
AAATTCTATATCAGGATGATGCCAGAAAAGCATTAGAAAAAGGCATGGATATTTTAACAG
AAGCTGTTTCTGTGACTTTAGGACCAAAAGGAAGAAATGTTGTCTTAGAAAAAAAATTCG
GTGCCCCTCAAATTATTAATGATGGTGTAACGATTGCAAAGGAAATTAGTTTAGAAAATC
ATATTGAAAATACCGGAGTCGCACTGATTAGACAAGCAGCATCTAAAACAAATGATGTAG
CTGGTGATGGTACAACAACAGCTACAGTACTGGCTTCAGCAATTGTTAAGCAAGGAATGA

GAAATGTTGCAGCAGGTTCGAATCCGATGGCTATTAAAAAAGGTATTGAAAAAGCAACAA
ATTTTGTTGTTAGTAAAATAGCTGAGTATGCTAAGCCAGTAGAAGATACAAAAGCTATTA
TACAAGTTGCTTCTTTGTCATCAGGAAATGATATAGAGGTAGGTAAAATGATAGCGAATG
CTATAGAAAAAGTTGGCAGAGAAGGAGTTATTTCTTTAGAAGAAGGGAAATCAACTAATA
CGATTCTTGAGATCACAGAAGGAATGCAGTTTGAAAAAGGCTTTATTTCTCCTTATTTCG
TTACAGATACAGAACGAATGGAAGTTCTTCAAGAAAATCCTTTTATTTTATTTACAGACA
AAAAAATTACTTTGGTGCAGCAAGAACTTGTGCCATTGCTTGAGCAAATTGCAAAAACAT
CTCGGCCTTTATTAATAATAGCTGAAGACATCGAGAAAGAAGCATTAGCCACAATTGTAG
TTAATAAATTAAGAGGGATTTTGAATGTTGTAGCGGTTAGAGCTCCTGGATTCGGTGATA
GAAGAAAATCTTTACTTGAAGATATGAGTATCTTAACGAACGGACAAGTAATTACTGAAG
ATGCAGGTTTATCACTTGATACAGTTCAATTAGATATGTTAGGAAAAGCCCGAAGAGTTA
TTGTTACTAAAGACTCGACAACAATAATTGCAGATGGTCATGAAATTAAAGTTAAATCAA
GATGTGAGCAAATTAAGCGGCAAATAGAGACGAGTGACTCTTTATATGAAAGAGAAAAAT
TGCAAGAACGATTAGCAAAGCTTTCTGGAGGAGTTGCCGTCATTAAAGTTGGTGCAGCTA
CAGAAACAGAGATGAAAGATAAAAAACTAAGACTAGAAGATGCAATTAATGCAACAAAAG
CAGCAATTGAAGAAGGTATTGTACCAGGAGGAGGAGCTACTAATGTCCATATCTCTAGTG
AATTATTTACATGGGCTAAAAACAATTTAGTTGAAGATGAATTAATTGGTGCTTTAATAG
TTGAACGAGCTGTGACCTATCCGTTAAGACGAATAGCTTTTAATGCAGGTGATAATGGAG
CAGTAATAGTGGAAAAAGTTAAAAGTCACGATTTTCATATAGGCTATGATGCAGCAAACG
GTAATATTGTAAATATGTACGATAGAGGTATTATAGATCCAGCGAAAGTAGCTAGATCAG
CTTTGCAAAACGCAGCTTCTATTGCAGCAATGGTTTTAACTACTGAATGTATTGTAGTTG
ATAAAGTCGATGATTGAGCTATAAGATAATAGATAATCTCAAACAAAGAATAATTTCTAA
TTATAGAAATTATTCTTTGTTTCAATTAATCTAATTAATTGATCAAATGTATAATTTTTA
TAATAAAGCCAAAGTTTAAACACGTCTTCTTTTTCAACTATTAAGTAGAGAAGATATAAG
CCCGTAATACCTAAATTATCAATTAAAATATTATTTTCATATATAGATTTTTCTAAAGAT
CTAAAAAATGACATTTTTGAAAAATAATACCGAAATCTTCTCTTATAGTTATTTGCTAAA
TCATTACTGTCTAAAGATTGATTTCCAATTGAATTCAGAACCTTTAACAGAATACTTTTT
AAATCTTTATCTGACTTTATTTGAGAATCAATAATAATTTGATTCAATTGTATGTCAAAA
CTAAGATTAAAAATATTACTTGTTAAAGATATTTGATTTTGAATATCTAGAGATTTTATA
GCCAAGAAGATTAAACTTTTATTGTTCATTGTAAAATATAAGACAATTGATGATGTATTC
TTTAAATACTAAATTTATCTTCTTGTGCTTCTATCGTTTTTTGTAAAATAATTAATGTTT
TAATTTTTTAGTTTTAGTTACTACCCGAAAAAATAAATTCTGGAAATCTAGCTTGAGCTT
TAGCTAGAGAGTGTTTTTTCCCTTCTTCTTGCCAGTAATTGATAACTTCTGTTGTTTTGT
TTAATAACTCAATTCTGTCTCTTTCAGCAATCCACGGCTTGGATTCTAGTTCTGCTTTTA
ATTCTTCCCAAGCATCATTCCTTGGCCAGAAAAAATAAGAAGTAAGTGGACTAGTGCCAT
TTCCAACAATCTGATCAATAGCAATAGCAATATTATTTTCTAACCAAAGTACTTTTAGTG

TAAATTTTGACAAGTTTTTATCTCCATTATTAATATTAAGTATTGGTACAGTAGAATACT
GTACTAGTTATTTTTTTCAGCAACAATTTGAGCTTTGTTTAAAATGTTTTTTACCGTTTT
AGTAGCTTGTGCACCTTGAGATAATCGTTGCAAAATTTTTGAAATGTTAACACGAGTTTC
ATTATTTATAGGATTATAAAATCCTAATTCTTCTATAGCTTTACCATCGCGTTTATTTCT
GCTATCCATTACAACAATTCTATAGCTTGGTTGTTTTTTTCTTCCGTATCGCTTTAACCT
TAACTTGACCATATTAGTCACCTCGCTTATTTTTTATATCTTTATTAAATTATATCTTAC
GACATTATGCAATAATTCAAACAATTAAGCTGATTCAAGTCTAACATTATTAAATATTAC
CATTAAACATTGTAGAAAAATAATACCTAACATAGGTGACATGTCCATTCCAAACATGGG
AGGAATGCTTCCTCTGAATAGTTTTAAGTATGGATCAGTAATTCTATTTAATGAACAGAA
TGGTTCATTGTACCAATTTACAGTTGGAAACCATGCCAATGATAATTTAAGTAAAATTAA
AATTAAATAAATCTCAGAAAAATTAGCTATAGATCCAAGCAATAAATTTAATGTACCAGG
AAGAGTGTTCATACGTATTTTATAATTATTATATACTTCGTTATACTAATAGTTTGATTA
TATATTATAATCGATGTAAATAAATCAGCTAAACAAAAAACTTATCCTTAAAATCATCTT
CTCTTGAGCATTTTTTTTTATCTACATTAGTATTTACTTTTGTCGTATAAATTTCCAGTT
TAGACCAAATATGTCCTAACTGATTTAATTGAGATGTTCCACATTCTAAACAGGCTATTT
TTATATTGCTCAAAACAGCACCTTTTTTTATAAAATCGTTCAATAATGGCGTAATAGTTG
CTTCATCCAAAAATAAATCTAAGACAAGTATTTTTGTAAATGAATTTAATTGATCTGGGA
CATTGTTAATAATATCATTATAATTAACTAAAACTATATTAGCTGTAGGAAGTAAAGATC
GTGCTCCTTCCGCTAAGATAAAGCCATAAGGCATAATAATTACAATAATATATTTATAAT
GACTGCTAATAAGACTTATAGTGGTATTAGCTTTTAATTCGATTGTTTCTGTTACTAGCC
ACTCTCTCATTATTTCGTAAGTAATCCATTTGCCTAGTTCTGAGCAAGCAGTTCTTAAAA
TCGTACCTGGATTATTGTCGTTTTCGAGAATCCCTGACCAATGTTGTATTAATGGATGGG
ATGCTATATTAATTTGGAGTTGCATGTTTTGTTAAATATATTTAACATTTAATTTGTTGA
TTGAATCTTCAATTCCTATTTGTTGAGTATTATTAAAATAGCATATATATTAAAGAAAAT
AATTATAAAAAATGAAAAAAAGTCTTTGGCTGTGGGGTTTTACAGATAGTGCTGAAACTT
GGAATGGTAGATTTGCAATGATTGGTTTATATCAGTTATTTTTATCGAAGTAGTTACAG
GACAAGGATTATTATATTTAATAGGTATGATGTCATAATAAAAATCCAAAAAAGTTCTTG
TAATTTTTTATAAGAACTTTTTTGGATTTTTATAAATCTAATCTAGCGGTCTCATAGATA
GAACAGGTTCATTCTCTCTATTGATACCTTTTTCAAATCCAGCAGCAGCAGCTCTTGCTC
TGCCTGCATGCCAAAGATGTCCAATAAATAGGAAGAATCCTAGGAAGAAATGAGATGTTG
TTAACCAAGATCTAGGAGATACATAGTTAACTGAATTAATTTCTGTAGCTACACCACCAA
CGGAATTCAAAGAACCTAGTGGTGCGTGAGTCATATATTCTGCGGCACGCCTTTCCTGCC
ACGGTTGAATATCGTTCTTGATTTTATTTAAATCAAGACCATTTGGTCCTCTAAGAGGCT
CAACCCAAGGAGCTCGCAAGTCCCAAAATCTCATGGTTTCACCACCAAAAATAATTTCTC
CACTAGGAGATCTCATTAGGTATTTACCTAATCCAGTAGGTCCTTGAGAAGATGCAACAT
TCGCACCTAGTCTTTGATCTCTAACTAAGAAAGTAAAAGCTTGAGCTTGTGAAGCTTCAG

GACCAGTTGGTCCATAGAATTCACTAGGATAAGCTGTGTTGTTATACCAAACAAAATTAG
AAGCCGTAAGCCCCATGATTGATAAAGCACCTAGACTATAAGATAAGTAAGCCTCACCAG
ACCATACAAAAGCTCTTCTAGCCCAAGCAAAAGGTTTTGTTAGTATATGCCAGATTCCGC
CAGCAATACAAATAATGCCAATCCAGACATGACCACCTACAAGATCTTCCATGTTATTTA
CACTTACGATCCATCCATCACCGCCAAACGGAGACTTTAAGACATAACCAAAGATAACTA
GGGGATTAAGTGTAGGATTGCTAACAAATCTAACATCTCCACCACCTGGAGCCCAAGTGT
CGTATACCCCACCAATAAATAAAGCTTTAATAACTAACAGAAAAGCGCCAATGCCTAATA
ACACTAAGTGTATACCAAGTATTGTTGTCATTTTGTTTTTATCCCGCCAATCGTATCCGA
AGAAAGGAAATGATTCTTCAAGAGTATCAGGGCCAATCAGAGAATGATAAAGACCTCCAA
AGCCAAGAACAGCTGAAGAAATTAAATGCACTACGCCTACTACAAAGTATGGGTATGTAT
TGAAAATCTCGCCGCCTGGACCAACACCCCAGCCTAGTGTAGCTAAGTGCGGAATTAGAA
TAAACCCTTGTTCGTACAAAGGCTTCTCGGGAACAAAGTGTGCTACCTCAAAAAGAGTCA
TGGCACCAGTCCAAAAGACCATTATACCTGCATGAGCAACATGAGCACCAAGCAATTTGC
CAGAAACGTTAATTAAGCGTGCATTGCCAGACCACCAGGCAAACCCGGTAGACTCAATGT
CTCTACCGCCAACACCAACAGTTGTATTAAAGGGCGTTTCCACGTGGTAAAACCTCCTCA
GGGAATATAAAGTTTTCATGAGGTTGATCTTGAGCAGCCATCCAAGAGCGAATACCTTCG
TTCAATAGGATATTTTTAGTATAAAAAGTTTCAAACTCAGGATCTTCCGCAGCTCTCAAC
TCTTGAGAAACAAAATCGTAAGCTCTTAAATTTAGAGCTAGTCCAACAATTCCAAATGCG
CTTGTCCATAATCCAGTTACTGGTACAAATAACATGAAAAAATGTAGCCAACGTTTATTA
GAGAAAGCAACACCAAAAATCTGTGACCAGAATCTATTAGCTGTTACCATCGAATAAGTT
TCTTCAGATTGTGTTGGAGTAAATGCACGAAAAGTATCTGCAGCATCACCATCTTCGAAT
AAGGTATTCTGTACAGTTGCACCGTGAATAGCACATAGTAAAGCACCACCTAAGATTCCT
GCAACGCCCATCATATGGAACGGATTCAGAGTCCAGTTATGAAATCCTTGCAAAAATAGC
AAAAATCTAAAAATTGCAGCAACTCCAAGGCTTGGGGCAAAGAACCAGCTCGCTTGGCCT
AGAGGGTACATTAGAAAGACGGAAACGAATACTGCAATTGGTCCAGAAAATGCAATAGCG
TTGTATGGTCTAAGACCAACTAGCCTAGCAATTTCAAATTGTCTTAAACAGAATCCAATT
AGTCCAAATGATCCATGTAAAGCTATAAATGCCCACAGACCACCAATTTGGCACCAGCGA
GTAAAATCTCCTTGAGCTTCAGGTCCCCAAAGGAAAAGAAGTGAATGTCCCATGCTGTTT
GCTGGGGTAGAAACAGCCGCAGTTAAAAAATTGCATCCTTCTAGATATGAACTGGCTAGC
CCATGAGTATACCAAGAAGTAACAAAAGTAGTCCCAGTTAGCCAGCCACCTACAGCAAGG
TAAGCGCAAGGAAACAGAAGTAGTCCAGACCAACCTACGAATACAAATCGGTCTCTTTTT
AGCCAATCGTCTACAAGATCAAAACCACCACGAGTTTTTTCTTGTCCAATTGCTATGGTC
ATAATTTTAATCTCCAGAGCAAATTTAATAAGTAATTTGTTTAAATAAACACTTTAGATA
TACTCTAAAATTTTACTTTTCTTTACGGTCAATATAATTATAAGTCAAAATTAACCAAAA
TTCACGCTTTTAATAACATTTTTTTAGCTCATTAAGTTAATTTTTTAATATATAAACAGT
AATAGAGTTATTGTTTAACTTACAATACTCATAATTTATTCATTTCTATAATCCTTTGAA

AAAGATATCCTGTTCCTCGTGCTGTTAAAATCAAGTCGGGATTACTTGGATCATCTTCTA
GTTTAGCTCTCAATCGTGAAATATGAACATCAACAACTCTCGTGTCAACATGCCGTTCGG
GAGTATAGCCCCAAACTTCTTGCAAGATAGAAGCTCTAGAAAAAGGTTCTCCAGCCTTAC
TAACTAAAAGCTCGAGAAGACTGAATTCCATACCTGTTAATCGAACTCTTTCATTATTTT
TATAAACTTGTCTTTTGTTAGTATCTATTTTTAGAAATCCAATACTAATAATCCCAGAGT
TTGGGACTCCAAGACTAGTGGTGATTTTATCAGCTCTTCGAAGAACAGAACGAATGCGAG
CTTCTAGCTCCTTAGGAGAAAATGGTTTAACGACATAGTCATCAGCACCTATTTCTAATC
CGGTAATTCTATCGCAGACTTCACCTAACGCTGTTAGCATTATGATAGGAACATCTGATT
CTTTTCGAAGTTCTTGACAAACACCGTAACCATCTAGTTTAGGCATCATAACATCTAAAA
CAACAAGGCTAGGATACTCTTTTCTAAACACGAGTAAAGCTTCTTCTCCATTTGAAGCAG
TGATAACTTCATAACCTATTATTGTTAGTCTTGTTTCTAAAATTCTTCTTATGCTGGCCT
CATCGTCAACAACGAGAATCTTTTCTTTCTGGTTCTCCAATTTCTTATTTACTCCTATAA
CTCACCGATCATGATTACTATAACCTTAAATTTTTAGCGCTATGAGTATCTTTATTCACT
TTAGAATTTAGTTATTATAAATTTTGCAAGTCTAGTTTACATTAATTAGTACCAGTCAAT
ATTTTTGTTTTAAGTTATGACTATTTTAATCCATAAGTTGACTCTAGTTTGCAGAATTCA
ATATATACATATTTTCTAGCCAAGTAAATTGTATTCTATTTTCTAAATAAAAAAGGGGTT
GACAAACCAGTTGATAGAAAGTATTCTACATTACTATAAGTTAATTCCAACAAAGAAGTT
TTAAATCTTCTTTCGGAGTAATGAATACTTTAAT
>SRR9587922
TATATGATAAATTAAGACGTCTGTTATGTGAGAATTTCTAAAATATACAAAACTTTTTAT
TTAATATAGAAAAAGTAAATTAAAATAAGTTGAACATACTTTGCTCTAATCTAAACCTCT
TAATTGAGATTCTAATTAGAGACAAAAGTATTTACTTATTTTGTATCTATAGGAATAAAA
ATGAAACTTGCAGTTTATGGTAAAGGTGGTATAGGTAAATCTACAACTAGTTGCAATATT
TCTGTAGCACTTTCAAAAAGAGGTAAAAAAGTTCTACAAATTGGCTGCGACCCTAAACAT
GATAGTACATTTACGCTAACAGGTTTTTTAATCCCAACAATTATCGATACTCTTCAATCC
AAGGATTACCACTATGAGGACGTTTGGCCTGAAGATGTAATCTATAAGGGATATGGCGGT
GTAGACTGTGTTGAAGCTGGTGGACCGCCGGCTGGCGCTGGTTGCGGAGGCTACGTTGTA
GGTGAAACAGTTAAACTTTTAAAAGAATTAAATGCTTTTGATGAATACGATATTATCTTA
TTTGATGTTTTAGGTGATGTTGTATGTGGAGGTTTTGCAGCTCCATTAAATTATGCAGAC
TACTGCTTAATCATTACAGACAATGGATTTGATGCTTTATTTGCAGCTAATAGAATAGCA
GCTTCAGTACGAGAAAAAGCTAGAACGCACTCTCTGAGATTAGCTGGACTTGTTGGTAAT
AGAACAGATAAAAGAGATCTAATTGATAAATATATAGATTGTGTTCCAATGCCAGTATTA
GAAGTCTTGCCCTTGATTGAAGATATTAGAGTGTCCAGAGTAAAAGGTAAAACTTTATTT
GAAATGGCAGAAATTGATAAGGATTTAGCATATGTATGCGATTACTATTTGAATATTGCA
GATCAGTTGATTACAAGGCCAGAAGGTGTAGTTCCTAAAGAATCTCCGGATAGAGAATTA
TTTAGTCTTTTATCTGATTTTTACTTAAATCCTAAATCAAAGGTAGGACAAGAAAAAGTA

GATCAAGAAGAATTAGATTTGATGATAGTGTAAATAATATTTTCAGCATAATATAACAAG
GAATAAGATAATGTCTACAGCTCAATCAGATGCTCTTACTTTTGAATGTGAAACAGGTAA
TTATCATACTTTTTGTCCCATCAGTTGCGTTTCTTGGTTATATCAAAAAATTGAAGACAG
TTTTTTCTTAGTTATAGGGACTAAAACTTGTGGTTATTTCTTGCAAAATGCAATGGGAGT
AATGATTTTTGCTGAACCGAGATATGCCATGGCAGAATTAGAGGAAGGAGATATTTCAGC
TAAGCTAAATGATTATGGTGAACTCCGTAGATTATGCTTACAAATAAAAAAAGATAGAAA
TCCTAGTGTTATATTTTGGATTGGAACATGTACAACCGAAATTATAAAAATGGATCTGGA
AGGAATTGCTCCAAAATTGGAAGCAGAAATTCGTGTTCCTATAGTAGTTGCTAGAGCTAA
TGGTTTAGATTATGCCTTTACCCAAGGTGAAGATACAGTTCTAGCAGCTATGGCTCAAAG
GTGTCCATTAAATTTAAAGAATCAATCAGATAACACGAGTCTTAAAACTTCTCCTCATAT
TCCTTTAGTTTTATTTGGATCTCTTCCGGATCCAGTTGTAACCCAGTTAACTATGGAATT
AAAAAAACAAGGTATTTTTGTTTCTGGTTGGTTGCCATCAAAGAGGTATACTGAATTACC
GGTTATAAAAGAAGGATATTATGTTGCGGGAGTTAATCCATTTCTTAGTCGCACAGCTAC
TACACTAATGAGACGCCGAAAAACAAAACTAATTGGTGCACCATTTCCAATAGGTCCAGA
TGGTACTCGAGCCTGGATTGAAAAAATCTGTTCAGTAATGAATGTAAAGCCTATTGGATT
AGAAGATAGAGAAAAAGCAATTTGGGCTTCTTTAGAAGACTATATTTCTTTAATCCGAGG
TAAATCAGTTTTTTTTATGGGTGATAATTTACTAGAAGTATCTCTTGCAAGATTTTTAAC
TAGATGTGGAATGACTGTATATGAAATTGGTATTCCTTATATGGATAAACGCTATCAAGC
AGCAGAGCTAGCTTTATTAAAAGCTACATGTGATAAAATGAACGTTATGATGCCAACAAT
TGTAGAAAAACCAGATAATTATAATCAAGTAGACCGAATCCGTGATCTGAAACCCGATTT
AGTTATCACTGGTATGGCTCATGCAAACCCTTTAGAAGCCAGAGGTATTAATACAAAATG
GTCAGTAGAATTCACATTTGCACAAATTCATGGCTTTACCAATGCAAGAGATATTCTTGA
ACTCGTCACAAGACCATTACGGCGAAATCTTAGTTTGTCAGAATTAGGCTGGGATATTTA
TAGCAAGCAAAGCTAGTATTCCTC----TTTATGACTTACATGGTATAAAAGCCTTGTAA
GTCTTTTATTTTATGTTAAAGAAGCATTTCTTACTCTGCCATCACCAGCCCAAGCTTGTA
AACTTTCTACTTGTTCTTTGTCTGTAAATGCTAAAGGCACAAACTGTTTGATAGCTATTT
TAATATCTTCAGTACTAAATTCTCTCTCTTCACTAAACGCCGTATGCATACTTTCAATAA
TTGCTTGCTCAATTTCAGCGCCAGAAAATTTATTACATAATAAACTCAATTCATGAATAT
CATATTCTTGCCACGATCTTGGTCTAACTTTTGATAAATGTATCTTAAAAATTGACTCTC
TCTCTTGACGATTAGGTAAATCTAAGAAAAAAATTTCATCGAACCTGCCTTTCCTTAACA
TTTCGGAAGGTAAACTTTGAATTTTATTAGCTGTTGCGACAACAAATACTGGAGCTTTTT
TCTCTGATAGCCAGGTAATAAATGTACCAAATACTCTGGCACTAGTACCACTATCACCTT
GACTATGTAAACCAGAAAAAGCTTTATCTATTTCATCAATCCATAATATGCAAGGCGATA
ATCCTTCAGAGATATTTATCATTTCTCTCATTTTTGATTCCGATTCACCAACTAGTCCAC
CAAACAGTTTTCCCATGTCAAGACGTAGCAGAGGCAGCATCCAATCGCTTGCAATAGCTT
TCGCGGTTAAGGATTTACCCGTCCCCTGTATACCAACTAGCAATAAACCTTTGGGCGAAG

GAATACCGTAATTTAAACTTTGCTTCGAAAATGAGCGAGACCTTTTTTGTAGCCATTGTT
TTAATACATCTAAGCCACCAATATCCCTATTTACTTTACTATAAGGATAAAATTCTAATA
GATGCGTTTGATTAATAATTTGTCTTTTTTCTTCAATAATAATAGGTAAGCTCCGAGAGT
CTATCTGATTATATTGTGCAATAATTTTAGTAATCACTTTGCGGATTCTATCTATCGATA
ATCCTTGGCAGGATTTTGTTATATTATTAACGAGCTCTGAATTCAAATTAAGATTCAAAG
CTTTACTCAATCTTATGATCTCTTTTTTGATTTCTAGAAGACTAGGTAAGGGCAAATCTA
TTACCGTTATGATATCATTCAGAGCAAAAGGTATATTGATTTTGCAAGAAATAATAATAA
TGTGCTTAGATTGCGTTTTGATTATTTTAGCTAAATTGCGAAGTTTCCTAATTAATACTA
TTTCGTTTAGGAAAGAGTCAAAATCTTTTAAAAGAAAAAGATTTAAAGATTCATTATTCA
AGTTTTCAATAAATTCTAATGCTAATAAAGGATTTCTTTTTGCATAACCATTATCGCTAG
GATTGTTAGTGTATCCATCTACAAAGTCCCAGGAATATACTTGTTGACTGTTTGAACAGT
TTAAGCTGTGTTTTATAATATACTCTAGTCTATCTTCTTCTCTAGTATTTATTACAATAA
TAGGATATCGAGATTTTAAAAGTAATCGTAAATCTTGAGTGAAATTCATGTCAATATTTG
TTTCCATCAGACTTTTGTACTTCACCAATTGAAGAAAGACTAAATAAGATTATCTAATAT
TCCTTGGATATATTCAAAAGTCATCTCTCTATTTTCTAAATTATTGCTATTACTAATTTA
ACTTACCATGATTTTTTTTCTTCCTTTTCTTCGTCGACTATTTAATATAGAGCGTCCACA
TGGTGTTTTCATTCTTGCTCTGAAACCAGAAACTCTAATTTTTTTTCTTTTAGATCCTTG
TAGTGTTCTTTTAGTCATGATGTTTCCTCTGTTGTAAAAGATTAAATACTATTTTATCAT
CTTTTATATATCTTTATGATATAGATATGTATTAATATCAAAGTTAACAGCAATAATTTA
TATTAATCAAGAATTATTTATATAATATTAAACATTAAATATTAACTTTATGATATTGGC
TTTGCCCATTTTTTATTTAAGTATTTTGACCATCTTTTTGCTAATCTTAAACTGGTTAAT
TTTTCAGCAGCTAAAAACGATTTTGTTATTAGAATCTCAATTTAAGTATTTTGTTGATAA
AAGCCAAAATCGTAAATTAGAACCCGAAGAAAGTTTTGCTTTTGCAAAAGTATGTGTAGC
TAAAAAATATTTTTCTAAAGCTATTATTGAAGGTCAATTAGCTTTAAAAAACTATCGAGA
CCTGAATATCTTAGATAATAATATAGTAATTGCTAATTTGTATAATATGCTAGGTTTTAT
TTATTTTGAAGCAGGCCAAACAAGTTTTGCAAAGAACTTTTATGAGCAAGCCTTACAAAT
AAACCCTAATTATGTTGTAGCTTTAAATAACCTGGCAAAAATTTACGAAGAGGTAAAAGA
TTTAAAAAAAGCCGAATCTCTATACGATAAAGTGTTAACTCTTAACTTAAATAATAAAAC
TGCTAATAGAAGGAAAGATTTTATAGCAAAAACTAAAAACATCTAATACTATTTGTAATC
GGGATAGCAGGATTTGAACCTGCGACATCCTGCTCCCAAAGCAGGCGCGCTACCAAACTG
CGCTATATCCCGTAAGACATAATTGTTACTATATACCTTTTCTCTAATTATGTCTACTCT
TTCTGTATTTTATCTAGGATACCAAAACATGCCTTTAACTCCATCTGGATCTTTGATAAA
TCCTAATTTTTTATAAAAACTAACAACATCAGGTTCTGCGAATAGTGTAATAGTGCTAAT
TTCTGCCTGTCTCAACTGCTGTATTAATTGATGTATGATTACTTTTCCTAAACCTAAACC
TTGGAAATCTGGATGAATAACTACGTCCCAGATAGTTGCATTAAATCCGTTGTCTGATGT
TGCTCTAGCAAATCCTACAAGTTTACTATTTGCATCTTTTTTTTGTATTAAAGAAATAAT

AATAGAACTATTTTTTAATGCGATCTTTACTTTTTTTAGAGGTCTTTTAACCCATCCGAC
TGAATCACAAAGCTGCTCCAATTCGTATAAATTAATATTCTTATTGCTACTCAGATAGAT
ATCTTTAAATTCGATTTTATCGCAAGTTTTATCTAAAACAATAAGTTTTTTGAAATTTTT
TTCAGAGCTAGCATTGATATCGGAGTTTTGAAAAAAATTTTTCCAGAAGATCATAATGAT
TATGATATAATTAAATAAATATATATATTACAATATTGCTTTAACAAACAATATATTAAT
ATATGCTCAAATTTTTACATGAATCTATTAAGATAGAAGATTGTTAAAATATAGAGAAAT
AAAAATTTTTGTAAAATCTTTCAATAAAATGATATAAAGAGTGCTTTTTAAGACTAACAT
CTATTATTATAAATAAAGTCTCGAAATTGAGATGAAATAACACATTATTCTATGTCTAAT
GCCTTTTAAAGGAGTTATTTGTGAAAAAATCTATGTGTCTAACCTGTTTACTTGCTTTAC
TAATTATGAGCAACCCAATAATAGCTAACGCAGAAGTAGCTGGATTGGTGCCCTGCAAAG
ATTCTGCGGCATTCAATAAGCGTATGGTAAATAGTGTGAAAAAACTTCAAGCCAGATTAG
CTAAATATGATGCAGACACGCCACCAGCGTTAGCTTTAAATAAGCAAATAGAGAAAACTA
AAACTAGATTTGCAACTTATGGTCGAGCAGGTTTGCTGTGTGGTACTGATGGATTGCCAC
ATTTAATTTCTGATGGTCGATGGAGTAGAGCCGGGGACTTTGTTTTTCCAGGACTATTAT
TCTTGTATATTACAGGATGGATTGGCTGGGTAGGTAGAGGATATCTTTTATCTGTTGCTA
AGACTAGTAAGCCAACAGAAAAGGAAATCATTTTAGATGTGCCATTAGCTATCAAATTTA
TGTCATCCGGATTTGCATGGCCGCTAGCGGCTTGGCAAGAATTTAGTAGTGGACAATTAA
TTGCTTCCAATGACGATATTACTGTTTCACCCCGTTAGTAAAAAATTTATATGAATAATA
ATTTTACCAAATACTTATCAACAGCACCTGTAATTGGTGTATTGTGGATGACGTTTACAG
CCGGATTTATTATAGAATTGAACCGCTTTTTCCCAGATGTCTTATACTTTTACTTATAAA
TAAGATCGTATAAAAAACTACATAAATATAGAGAAATACTATTATTTGTGTAGTTTTTAA
TATATAACATAATTATTTATTTTTAAAAATGTTTATCTTTTCGATTATTTTAATATTAGA
TCTTACATAATAAAATAATATCAATATTATGAGCTTAGTAACCCAAATTATAGTTAATGC
CGATGATGAATTAAGATATCCTACAATTGGAGAATTGCAGTCAATTCAAGACTACTTAAC
TACAGGAAGCAATAGAATTAGAATTGCTACTATTATTAGAGATAAAGAAAAGGAGATTAT
ACAGAAAGCTAGTAAGCAAATTTTTCAGTTACATCCAGAATATATAGCTCCAGGTGGTAA
TGCAGCAGGTTCTAGGAAAAGATCGTTATGCTTACGTGATTATGGTTGGTATCTCAGACT
AATTACATACGGAGTATTAGCTGGGGATAAAGATTCTATAGAAACAATTGGTATTATAGG
AGTAAGAGAAATGTACAATTCTTTAGGTGTACCTATAATTGGAATGTTAGATGCAATCCA
GTGTTTAAAGGAAGCATCTTTAGAAATGCTTGGTCAAGATGATATTAGAATCATTTCTCC
TTATTTTGATTATATAATTCGTGGAATGTCATAAATCATAGCTTCAATTAATTAGTTGAA
TAAAAGCTTTTATAATGTTATAATTACAATAAGCCCGAAAGCATATAATTGTAAAAACTA
AATCTTGTCAGAACCGGAAGGTAGCAGCAATAATGTTTCAATACAAAAGTTATGATTTTC
GGGTGTTTATTGTTTTTATCTAATAGTCTCGACTTTTAATTAGTATGCTATGTATAAATC
TGTATATGACATGTAATCCATCAAACCTTCCAGCAAAAATAATTTAATAATTTGATATTA
ATACGTAAAACACCTATACTCTTTTATAAAATTTCTTTATATCTCATCTAAACAATGGAA

ACTTAACTTAATGGGTGTTCATATTTTATCAACAGGTTCGTCTGTCCCAAATTTTTCTGT
AGAGAATCAACAATTTGAAGATATGATCGAAACTTCCGACCATTGGATTTCAACAAGAAC
AGGAATAAAAAAAAGACATCTAGCTCCATCTTCTACTTCTTTAACTAAATTAGCGGCAGA
AGCTGCAAACAAAGCCTTATATGCAGCTAACTTAAAACCTACTGAGATCAGTTTAATTAT
TTTAGCTACGTCTACGCCTGATGATTTATTTGGTAGTGCTAGCCAGTTGCAAGCAGAAAT
AGGTGCAACAACATCGGTAGCTTTTGATATTACGGCTGCCTGCTCCGGTTTTATTGTTGC
TTTAGTAACAGCAGCTCAGTTCATTCAAACTGGTTCCTATGACAATATTTTAGTTGTTGG
AGCAGACACAATGTCTAGATGGATTAATTGGTCAGATAGAACTACCTGTATTTTATTTGG
TGATGGTGCTGGAGCAGTAGTATTGGGGCAAAGCCTCAAAAATAGTATTTTAGGCTTTAA
GTTATGTACAGATGGTCAGCTAAACAGTCATTTACAATTAATGAATAAACCTGTAAATAA
TCAAAAATTTGGTGTTACAGAAATTCCTCATGGAAACTATAATTCCATAACAATGAATGG
CAAGGAAGTGTACAAGTTCGCTGTATTTCAAGTTCCAACAGTAATTAGACAATGTTTGAA
TAATTTAAACATTTCAATAGATGAAGTTGATTGGTTTATATTGCATCAAGCAAACACTAG
AATCATAGAAGCAATTGCGAGCAGATTATCAGTACCTTTTTCTAAAATGATTACGAACTT
AGAGCATTATGGAAATACATCTGCAGCGTCAATCCCTTTAGCGTTAGATGAAGCTATTCA
ATCCAATAAAATTCAACCAGGCCAAATTATTGTTTTATCTGGTTTTGGAGCAGGCTTGAC
TTGGGGAGCAATTGTCTTGAAGTGGTGATTTATATATTGCGGATGACGAGACTCGAACTC
GTAAAGCTTTCGCTACACACCCCTCAAGCGTGCGTGTATACCAATTTCACCACATCCGCA
TTTTTTACCATGAATATTAAAAATTTATAATATTTATCTAAATATTCATATAATGTGTAT
ATATATATACATTCTTGTCAGAAACTCTCTTATATTTTATTTTAAGTGATAAAATTGAGT
TTAGTATAGATAGTAAAATTTTATTTTAAAATCATAAAAATAAATATATAAGGAAAATAA
AATATGACACCATCTTTATCAAGTTTTTTGAATAGTCTTATTCTTGGGGCAGTAATTGTA
GTCGTTCCCATAACTTTGGCTCTTTTATTTGTTAGTCAAAAAGACAGGACAATCCGGTCC
TAAAAATAACTTAGAAAGATATGGAGAAGCAATTTAAAATGAATAATAAAATATTTTTCA
TACTAATAAGCTCTCCATATATTGCTATACTTTAGAAGAATTTTTATTTTTTTATTTTGT
AACATCCTGTAAAGTTTATTCCGTTCTTTCTTTTTCCCATTTACTCTCTTATAATTAATG
ATACAAATGTCAGGTTAATAATAGTTTTTTTATTCTGTATCTTTCATTTTATATACGTAT
TATTATGTTCAATTAATAAAGAAAAGCTCATGTCTTTGTCTAATTGGCCTCTCAAAAAAG
AAAATTCTGAAGCATATAATATTAAGAATTCAAAACAAATCACAATTCCTGATGGTTTAT
GGATAAAATGCTTCGACTGTGGTCTATTAATGTATTCTAAAGTACTGAAGAGAAATTTAA
AAGTTTGCCCTCAATGTAGTTATCATTTTCAAGCTTCTAGTAACGAAAGAATTGATCAAT
TGATAGACCAAGGTAGTTGGCAACCAATGGATGTTCACTTGATCTCTACAGATCCATTAG
GCTTTAAAGACCAAAAGCTTTATAGTCAAAGGTTAAAAGATACTGCTTTCAAGACTGGCC
TGCAAGACGCAGTTCAAACAGGTACTGGGACTATGCAAGGTAAAAAAGTATGCTTAGGTA
TTATGGATTTTAGATTCATGGGGGGAAGCATGGGATCTGTTGTAGGCGAAAAACTAACAA
GACTGCTAGAAAAAGCAACTCAAGAAAAGTTGCCTGCAATTATACTTTGTGCATCAGGCG

GAGCTAGAATGCAAGAAGGTATGTTGAGCTTAATGCAAATGGCAAAAATTTCTTCTGCTC
TAGAAATGCATAAAAAAGAAAATCTACTATATATATCTGTTTTAACCTCTCCCACAACAG
GGGGTGTAACAGCTAGTTTTGCTATGCTTGGAGACTTAATTATTGCAGAGCCAAAAGCTC
TTATTGCATTTGCTGGTAGACGAGTTATAGAACAAACAATCAAAGAAGACTTGCCAGATA
ATTTTCAAAGTTCAGAATATTTATTTGAACATGGTTTCCTAGATTTAATTGTATCCAGAA
CTCAGCTTAGATCAAAGTTAATACAAATTTTGTCCTTACATAATCATAGTAAGTAATGAA
TTGAATATTGACTATAATATCAAAATGCTAGTAAAAATTAAGATATTAAGAAATTTTCAC
TAAAGATATTTGTAGACTAACTAGAATTACAATTACTATTGTAGAAAAATTTGTCTTAAT
GCCATTTTTTCAGTCTTTTAACAAATTGTTTTTATGACTTTAAATAATTGTTAGTTTAAT
ACAAATTTATAAAGAGAATACATACTAAGCTAAATATAGTAATATTATGTTTTCCTATAA
TACTGACCTATCAAATTTCGAGGAAATTTATGCTTAAAAGATCTTCTTGGCTTGCGGCTT
TATTGGGACTATTAACAGTAGTTTCTACAAGTACGCATACATATGCCATAGAGTTAGACG
AGGCAACAAGAACTGTTCCATTAGAATCTTCTGGCAGAACTGTAATTCTTACACCAGAAC
AAGTTAAAAGAGGCAAGCGATTATTTAATAATTCTTGTGCTATTTGCCATAATGGTGGTA
TCACGAAAACAAATCCGAATATTGGACTTGACCCAGAATCTTTAGGATTAGCTACGCCAC
AAAGAGATACCATTGAAGGACTAGTTGACTATATGAAAGATCCGACTAGTTATGATGGTG
CAGAGTCAATCGCAGAATTGCATCCAAGTATTAAAAGTGCTGAAATTTTTCCTAAAATGC
GCAATCTAACCGATGAAGACCTATTTACAATCGCAGGTCATATCTTACTTCAACCTAAAA
TTGTTTCTGAAAAGTGGGGCGGAGGAAAAATTTACTATTAGAACTTCAAAGACCTAAACC
TTGTGAATACTAGTTTATGTATTTGTTGCTTAAGTGTGATTATTTTGTTAGATCATGATA
TATATTAAAGATGGAGTTTGTTGACAGCAAGTTACTTTTTACCCGAACTTGTTTACACTA
TTCGTCTAGTTATTAAAAAGGAGACGTTAAATTGAAGAAGAAGCTTTCAGTTCTTTTCAC
TGTTTTTAGTTTTTTTGTAATAGGTTTCGCACAAATTGCTTTTGCTGCAGATCTAGATAA
TGGAGAAAAAGTTTTTTCTGCTAATTGTGCAGCATGTCATGCTGGCGGTAATAACGCCAT
TATGCCAGATAAAACCTTAAAAAAAGATGTACTTGAAGCTAATAGTATGAATACTATTGA
TGCTATTACTTATCAAGTACAAAATGGTAAAAATGCCATGCCTGCTTTCGGAGGTAGACT
GGTTGATGAAGATATTGAAGATGCAGCAAATTATGTATTATCTCAATCTGAAAAAGGTTG
GTAATTATACTTGATTTTATCCTGTATTAAAGAATAGACAATCTATTTAGTTGTTCATTA
GATTGTCTATTCTTTGTTTATGCTATTATATAAAGATATTTACACAATATTTTATGATGA
AAAGAATACCCGCAATTCTTGTACTAGAAGACGGTGCGTATTATAAAGGATGGTCATTCC
AGCAAGATAAACAAGAGATTACTATTGGTGAAGTAGTTTTTAATACTGGAATGACAGGAT
ATCAAGAAATAATCACAGATCCTAGTTACTTCCATCAAATTGTCGCTTTTACCTACCCGG
AAATTGGGAATACAGGTATTAATAATCAAGATATTGAATCTCACAGTATTAGTATTAAAG
GACTTATTGCAAAAAATATTTGTAAAATTTCAAGCAGCTGGAGAGAGCAGCAATCTTTAG
TTACGTATTTAAGTAGTAATAATATTCCTTTTATTTTCGGAATAGATACAAGGTCTTTAA
CCCAATACTTGCGTCAATTTGGTACAATGAACGGTTGTATCTCTACTGATAATTTAAATC

ATAGTTACTTAAAACAGAAAATTTGTGAGATTCCAAGTATGCAAGGTTTAGATTTAATCC
CGCATGTAACTACAAGAAATGTTTACCCCTGGGATGAAAAAAGTTTTCCAAATTGGTATT
TGACAGATAACATTAGAGTGCACCGAGTTATTCAGTTGAAAGTTATTGTTATAGATTTTG
GAGTTAAACTAAATATACTCAGAAGACTAGCTACACTTGGATGTCAGATAACGGTTGTGC
CTGCCCACACTCCTTTAAAAGATATTTTGGCTTACCAGCCTGATGGTATATTACTCTCTA
ATGGTCCAGGAGATCCATCAGCAGTACATTACGGCATCCAGACAGTTACAAAATTACTAG
ATTACAATGTGCCTATATTTGGGATTTGTATGGGGCATCAAATTTTAAATTTAGCTCTTA
AAGCTAAAACTTTCAAACTTAAATTTGGTCATAGAGGTATTAACCATCCATCCGGATTGA
ACCAGCAAGTTGAAATAACTAGTCAAAATCATGGCTTTGCAGTTGAATTGACTTCGGTTT
TTGAATCTCCTGTAAGAGTGACTCATTTTAATCTAAATGACACTACTATTGCAGGAACTG
GACATAATCAAAGTCCTTATTTTTCTGTGCAATATCATCCAGAATCGAGCCCAGGCCCTC
ATGATGCTGATTATCTATTCGAAAATTTTATAGAAATAATGACAAAGTCCAAGAATAAAG
TTAGTTAGTAATTTTCCCATGCTTTATGAGTGAATAAAGCGGATAGTACTTGTACTCCTC
GTAATTGATTAAAGAGAGGCAAATGTCCTCTGGGGCCATGAGAATTCCATTGAAATTCCT
CAGGATATCTGCAAGGAATTCTATCAACTTCCCAGCCGATTTTTTGCCAGAATTTTCCCC
AATCTTTTCCAACACTAAGCCATATTTGTCTTTGAACAAATAGCCCAAATTTACCTTTTG
AGTGTGTATGCCATAACTTATCAATAGTTTGTAAATCTTGAGCAGGTATTTTTTTTATAT
CTGTAAAATATAGCCAATTACGAGTTTGTGCATTTACACCAGCTAGTTGAATAAGCTTTT
GTTGAGTTAGCTGATCTGCTTTGAGCAAATCCCGATGAGTCAGTAACATTTGTAAATCTT
TGTAATTCATCTGTTGAGCAGAGCGCAATGGAACAATGCCATCGGGACATAAGTTAGAAG
CAAATTTCACTATTTCTTGATTTTTACTGTTAAGTAACTTTTCATAAATTAAACCGTCAA
CACAATTACTTTTATAGTTAGGCCCAGTAATCCTCTCAAAAAATAGATCTGCTAAATCTT
TTAACTCGATAGAGTCTTTGCTATTTATATTCTCAATAATTTCAAGTTGTTGCTTGACAT
TATTAGACTTAGTATTCTTATTCAATTCTAAAAGCTGGGCTCGGATTTGATTTGGCATCT
GATTTGTTTAATTAGTTTTAATAGTCATAAGGTCTTAGAGAGAATTAGATTGTTTTTTGA
AATTAGAAGAGCGCCGCAGTATTTCTAAAGAAGTTCGCATAATTTGATTCAATAAGTTTT
GTCCTGCATAAATTATAATTTGACCAATTAAAAAAATTGTGCTTTTTATTTTAGGAACAA
AAAAATCTTGTACCTCAAGCAGAAAAGTAATTATTATTTGCATTGGTGACAACATCTGTA
ATTCATTCGATCTGCAAGCATATATATACTGACAGTTTAAGCCTTCTTGATAAAATCCCC
ATACTTTGTAACGACTTTCGTAAATTGCTCGTGGTCGCTCAAGATAGTTTTTAATTATAG
TATTCCAAATTAGGTTATTCTTTAATTTTTCTAATTTTCTATCAGATAGAAATTTTACAT
TACAGAGATAACTTGCATTTGTCTCATTTACAGTATTAAAATTTTGAACTAACATATGAG
CAATAATGTTGCTTAGCTGAATTAAATAATTTTCAAGAAAAATTTCGACCTGCTTAACTG
GCATATTGTAAGTAAAGAGATTAAAAGTTTTATTCACAGTTTGAGATGACCCAAAAAGTA
GCTGAATAATTAAAATTTGTAGTAGCATCTTATAGTCGTTCAGTAGATAGTTCAAATCAT
GAATATCTTGGTAAATGTCAATACAGTCCAGTTCATATAACTTACAAAATCTTTTGATAC

ATCTATGGAATAAATCAATTAAAATATTTTCGGTCAAGTTTGTAATATCATTTACATCTA
AATTTGAATTATAAATTTCTAAAAGTATCTTTTCCAGTTCAGCTAATATGATTTTTAATA
AGTTTCTTTTTACTTCAGTGCGAAAAACATCTAAAATTAACACTTCCTGAGAGCAATTAG
TTAATCTCTTATTAATCTTTATAGAAGTTCTAACTAATAGCTCTGCTACCTCTTGATTGA
GTGTTGGTCCCTGAGAGCTTGGCCAATAATTATTCACGTTATACCATAGTAAGAAATATT
GAATTTAATGTTATACGATAACTTTTTAATATACTGAAAAGCAAGTTAATTACTCTATAG
CAACTAGTTTTTACAGATCTTAATAGTTATTTATCAAATAAAGTTATATGATAGTAGCGA
TTGAGCTAATTTTATTAAAATTTATAATAATGACAACTTATTACTTCGCCCTTGCGAGTC
AAAATTTTCTATTAGTACAAGAACCACTAGAAGAAGTTTTTAGAGAGAGAGTTAACTACT
ATCAGTCAAATAATAAAGCAATTGATTTTTGGCTAATACCAAACCCTTCTTTTCTAGAGA
AACCAGAAATGATTTCATTTAAAAACCTTGTACCTAAAGACGCTGTAGCTATAATCTCTA
CTAATCCAATATTTATTAATTGGTTAAAGCTAAGAATCGGCTATATCTGTATCGGGCAAT
TTGAAGATAACCTACAACTTTCTGAAGAATCGTTAAATATTACTGTTTTAACAGACAAAA
TTTAACAATGCTTTTATTATGACTAGTGTTAAATTATTTTGGTTAAGTTATAATTATAGT
CTTTTAGTATTTAGACTCTTATAGTTTATCACTTAGTATTTTATGCTGAGATGGAGTGAC
AAATAGTGTAAGTATTTCTTGGCTAAACGTTTCAGTTGCCTTAGATTTATATCTATTTGG
ATTAACTATAATGGATAGCATTCGTTTTATAGTAACATTTTCAATTTGAGCCCAATGTAC
AATTCCAAGTTCCAATTCCTTAGCAATAGCTGAAACGGAAACAAAGGCAGCTCCTAGCCC
TGATTGCACAGCGTTTTTGATAGCTTCTATAGAATTCAATTCCATCTCTATTTTAAAGCG
ACTGCTGTCAATACCATGTTGACTGAGTACTTTATCTATGACTTTTCTAATTGTTGATTG
AGTATCTAACGCAATAAATCTAAGCCTGTATAAGTCTTCTTTTTGAATGTCTCCTAGTTT
AGAAAAAGGATGTGATTTGGGTAATATAAGTGCTAATTCGTCTTCCGCATAAGAAGTAAC
CTGCAAAACATCTTGCAATTCAGTAGGCACTTCTCCTCCAATAATTGCTAAATCAACTTG
ACCATTAGCTACGCTCCATGAAATAAGTCTAGTTGAATGTACTTGCAATTGAACAGCCAC
TTGTGGATATCTTTGCCTAAATAGTCCGATTAATCTTGGCATCAAATATGTCCCAGTTGT
CTGGCTAGCTCCAATAATTAATGTGCCACCTTGTAAGTTTTGTAAGTCGTCAAGAGCTCG
ACAAGTTTCTTCGCAGAGAGCTAAAATTCTGCCCCCATATCGTAAAAGAAGACTTCCTGC
CTCAGTTAAAGTTGCCTTCTTATTACCTCTTTCAAATAGGGAAACATTCAATTGGCGCTC
TAAATTTTGAATTTGCAAACTAATAGCTGGCTGAGAAACATACAAGCTATTAGCTGCTTT
TTTAAAACTGCCCTCTTTGGCAATTGCTTTTAATATTCTTAACTGATCCAATGTAAATGG
AAGGTCTGTCATTAAAGAATTGTAGTATAATGTATATTATTTAGTATTGATCAATGTATG
CATTTGCCAACTAAAATATTTGTCATTACGCTTCATTATTTTCAAAATAATAAAAAGTAT
AATGTTAAATAAGCTGTATTAAAATTTAGTATCTGTTTGATATTAGATTTATACAAGAGA
TATCGTGAATTTAATTCTAACTATAAAATATAGGAGACAATATGGACTCTAGACTTTTAG
TTGTACTAATACCAGTTTTAGCAGCAGCATCTTGGGCAGTTTACAATATTGGTAGAGTTG
CACTACAGCAATTTAGAAAAATGACATCTTAGTTTTGCTTAGAATATAATTTTTAGACTT

TATGAGATAGAGGACTTTAAAATCCCTATCTCTATTGTATATCTTAATAAAAAAAGAAAG
CCCCCTTGTACTTACTGTTATTACACATTAATATTATTGGTGGTACCTAGATAATAAGTT
TAAGCTAACTTATTTTTTATTTTATAATTAATTAAATTAATAACACAATTATGGCTGTTC
CAAAGAAAAGAACATCTAAAGCTAAAAAAAATGCACGCAAAGCAAATTGGAAAAATCAAG
CAAAAACTGAAGCTCAAAGAGCTTTGTCTTTAGCAAAATCAGTATTAACTAGAAAATCCA
ATGGATTTATTTATAATCTAACTGAAGCATCAGATACTTTTAGCGATTAGAAGTGTTTTT
ACTACAAGTCTGAAAAAGTATTACGCAACAAGTTCTCAGTAATTTTTATCTAGTAATCAT
TTGAAAATCATTCTATTTGATAATAAAATAAATAGCAGATTAAATAAAATCACATTAACA
AATTATGCAGCAAAGCTAGATCAAACTAGTGAAATTTGGTTGTTCAATTGTATTGAAAAT
ATTCAGCATATTTTTTTTAAAAGTCAATTAAAATCAAGTCATATTACTAAAATTTTTATT
TCTGGTACTAGCTTTGAATATACCGCAGGTCTACCAGGATTATTATCCAGCTTAACACTA
AGTGGTAGACTACATCCTATTAGTATATATAGTCCCCAGTCTCTCAAAAAGTATCTTGAA
GCATGTACCAAATATTCCCAAACTAATTTTTCTTTTCCTATTAATTTTCATAATTTACAA
TACGGAGGACAAGTTGTTAACCAATTTTATACAGTAATTTGTTTACCGTTGAGCAAAAAG
AGCCTGCTGTATGGATTTATTATTCTAAAAAAAGAAAAGCAGGGAGTATTTAATTTAGCA
CAAGCTAAAACCTTGAATATTCTTCAAGGACCTATATATGGAAAACTCAAAGAAAAGGAT
AATTTTTTAAGTCCAGATGGTTACTATCTATCCGGCCAAGACTTCTCTTCTAATACAATA
ATGGGACATAAAATATCGCTTCCGTTATTAGTCAGATATTCTAGAATTATTTCTGAGATG
CATTGGTTTTGCTCTTATCCTATTAGATTAAATACTTATTCGCATCAGCAAGGAGCAAAG
TGTTTACCACATAATGTTTTAACTGATATTATGAAATCCCAAATATATCAAGATAACAGT
TTTGTTGAATAATACTTATTCTTGATATATAATGGTCTCGGTAATCTAAATAAAATACTA
GAAATAATTTCTTAAATTTATTTGTCTATATTAAATTTACAATCATTTATGACATACGCA
ATTATTGAAGCAAGTGGCACGCAGCTTTGGATAGAAGAAGGCCGCTACTATGATTTAAAT
CATATACCTGTTGATCCAGGTCAGTCGATTATATTAGGAAAAGTCTTATTATTGAATAAA
AATGGGGAGGTTACTTTAGGCCGCCCTTGTATAGAAGGGGTTACGATAAAGGCTACAGTA
ATGAGGCACTTGCGAGGAAAGAAGATAACTGTTTTCAAAATGAAACCAAAGAAGAAAATG
AGATTAAAAAAAGGTCATCGACAAGAATTGACTCGTTTAATGATCGATTCTATAACATCT
TAGATAAAATCTAGACTCATATTTTTAAAAACTTTTTAATATTTAACTAAAATAGATATA
ATGGCACATAAAAAAGGTAGTGGTAGTACAAGAAATGGCAGAGACTCTAATTCCAAGCGT
TTAGGTGTTAAAAAATATGGTGGAGAGCAAGTAACAGCAGGTAATATTTTAATCAGACAA
CGGGGAACTAAAGTTAAGCCTGGCCAAAATGTTGGAAAAGGAAAAGATGATACATTGTTT
TCTCTAATTGATGGTTTCGTGCTGTTTGAAAAGTCAAATCAAAAGCAAAAAACAATTAGT
GTTTATTCTGCTAAGAAATAGTTAAAACAATTGGTGCAAGATAACTTGAGTGTATTTATT
TCACTTACAGTGGCACCATATATTTGATTTATTAAGAAATGGAGTTCCTTAATGAACCAA
TGAATTTTATTAAAGATTTGCTGTGAAATTTAATGACAAACACTATTGTAATTTCCTGTC
TACACAATATGGCTGCTATTCTATATTGTGGTCAGATACAGAAACTAGTAGTAGCAAATG

CTCATTATCAAGTAAGTGATATCTACCTAGGTTGTGTTGATAAAATCTTCTCAGGAATAA
ATGCGGCATTTATTAACCTAGGAAAGAATGAGTACAGTGGTTTTATACATATCAGTGATA
CCGGTCCGCTTAAAAAGAAATATTATGTCAATAATATTACTAACATTTTAACAATACGGC
AAAAAATTTTAGTACAAATTATTAAAGAGCCAACTTTAAATAAAGGTCCAAGGCTCACTG
CCAATATTACATTATCAGGTCGATATATTGTATTAATGCCTTTTAGTCAATCAATCTGTA
TATCTCGAAAAATATATGATGAAGATGAGCGTCATTATTTGAAGTCTTTAGCTATTTTAA
TTAAACCGGCAACAATGGGCTTGCTATTTAGACCTTCTGCTGTAGGTGTCGATGAGGAAA
TAATATTAAGCGAATTAAAAAATCTAAAAGAACAATGGAACTTTGTTCAAAAATCTGCAA
TTAATAGTTATTCACCTGTTCTTCTATATAAAGATGAAGATATTGTTAAAAAGGTAATCC
GAGATTTTTATAATAATAATACAAACAATATAGTAATTGATTCAAACCTGGGATTAAAAC
AATTAAATTATTATATCCACACTTGGCACTGTAATAACTCTAGCACAGTTCCTAAGATTA
AGCTTTATAGTAATAATCAATGTATACTAGATGCTTTTGGTATCAATCAGGCAATTTCCA
GAGCTCTCATTCCAAAAGTTGATCTTATACTTGGTGGCTATATGTTTATTGAAACTTTAG
AAGCTTTTACTATTATTGACGTTAATTCTGGATCTTTTAATAATTCCACTAGTGCACGAG
AAACAGTTTTAAAAACCAACTGTTCTGCAGCAACAGAAATAGCTTATCAGTTACAAATTA
GAAATATTACTGGTGTAATTATAATCGACTTCATTGATATGGAATCACAAAGAGATCAAT
TGCAATTATTAGAACACTTTAATAAAGAGCTATCACTTGATGATGCTAAACCACAAATTG
TACAGTTATCTGAATTAGGTTTAGTTGAATTGACTAGAAGAAGACAGGGCAAAAGTTTGT
ACGAGTTAATTAGTAGTGATTCTAATTACTTTTATTTTTTCACACAATCAGAGAGATCTC
AGTCTCTTAAGAGATTCGATGATAGACAGCAGAAACAACAGATTTTTAATAAATCTTGGC
TATCTGCAGAGATTAATACTATTAACAAGGTCTTTTTTCAAAAGTCAAATTTGTGCAGAC
CTGCTAACTTTTACCTAATTCGTAATCTCTATATAGTTAAAAGTAGTATTACTTATAAAC
AAAATTATTTATTAACTCACAGATCAAAACTTATTTATTCTAAAGAATACAGCAAAGTAT
TACCAAGCAGTTATTATCTAGCTAGTCTCAATAAGAATAGTAATCAAGAGTTCCTACCTT
AAGATTGTTATATCAACTGTTGAAAAAAAGAAGCTCTCTCTTATATAAGAGAGAGCTTCT
TTTGTAGCTTACTAACTATCTAGTTAGTTAATTAAAAATTAACCGTTAACAGCTGGAGCT
GTTAGAGCTACTGGTAAAGATTCACCAGAAGCTAGATCTAGAGGGAAGTTGTGAGCGTTA
CGTTCGTGCATTACTTCCATACCTAGGTTAGCACGGTTGATGATATCAGCCCATGTGTTA
ATTACACGACCTTGGCTATCAACAACAGATTGGTTAAAGTTAAAACCATTCAAGTTGAAT
GCCATTGTGCTTACAGATAAAGCTGTTAGCCAGATACCAACTACAGGCCATAGACCTAAG
AAGAAATGTAGAGAACGAGAGTTGTTGAAACTAGCATATTGGAAGATTAAACGACCGAAG
TAGCCATGAGCTGCAACGATGTTATAAGTTTCTTCTTCTTGTCCGAATTTGTAACCATAG
TTAGCAGATTCGTTTTCGCTTGTTTCACGAATTAAGCTAGATGTAACTAGGGATCCGTGC
ATAGCACTGAACAGAGAACCACCAAACACACCAGCAACACCTAGTTGGTGGAATGGGTGC
ATTAAAATGTTGTGCTCAGCTTGGAATACAAGCATGAAGTTAAATGTTCCGGAGATACCT
AGAGGCATTCCATCAGAGAAACTACCTTGGCCAATTGGGTATACTAGGAATACTGCTGCT

GCTGCTGCTACTGGAGCAGTAAAAGCAACGGAAATCCATGGACGCATACCTAGGCGGTAG
CTTAGTTCCCACTCACGACCAATGTAGCAAGCTACGCCAGTTAGGAAATGAAGAACAACT
AATTGGTAAGGACCACCGTTGTATAGCCATTCGTCTAAAGAAGCAGCTTCCCAGATTGGG
TAGAAGTGAATACCGATAGCTGCAGAACTTGGAATAACAGCACCAGAAATGATGTTGTTT
CCGTATAGAAGGGAACCAGCAACTGGCTCACGAATTCCATCAATGTCTACTGGAGGTGCA
GCTACGAATGCAATGATGAATACAGATGTGGCAGTTAATAGAGTTGGAATCATTAATACA
CCAAACCAACCAATGTATAGGCGGTTTTCAGTACTAGTAATCCAAGAGCAGAAACGTTCC
CACAAGCTAGCGCTTTCGCGTCTTTGTAAAGTAGCAGTCATAATTTTTTATCAATTTTTA
AGGATTTAACAAAGATGCTTCCCAAACAATAAGTTGTCTGTTATGGAAATTATTACTTAA
GGCCTTTTCAAATGAGAACATTTTTAAAAAAAACATAGCAAAATCTAATCTCAGTAACTA
AAATTTTACTTGTTCATATATTGCTTAATAATTCAATAATAAATTTTTGCTTTTTGCAAA
ATTAACACTATTTTAGGGATATAATAATAATGCATATTAATGCTGCTTATTTTGATTTTA
ATAGTTTATATAATTTTGCCCTACTTAAATACGCATGTCACACTTTACAAAAATTCAGAC
GACCATACAAGATTTAAATCTGTTAAAGCATGCTTTAACAGATTTGGGTTTGATTTGGCA
AATGAATTCTAGTCATATAAAAGTTGGTGAAAATAATCAGCACAAGGTAGATATTTTAAT
TAAACAAGATAATTTGTCACATATTGGCTTTACTTGGAATGATAATAGATATCACTTAGT
TGCCGACTTACAACTTTGGGAACAACCCTGGTCTCTAGAAGTATTTTTGGATAAGTTATC
TCAGAAGTATGCTTACTACTCTATCATCGAAGAAACAAAGAAGCAAGGCTTTGAAAAAAT
GCAGCAGATTTCACAAAAAGATGGATCAATTAAGTTAATTGTGCAACGCTGGAATTATTA
ACTAACATACAAATGCGGGCGGAGAGACTTGAACTCTCACGAGATTATCTCACTAGAACC
TAAATCTAGCGCGTCTACCAATTCCACCACGCCCGCACTGTTGAGACAATATTACTCTAT
CATAATCTATACTAAAACACAAACTAATAGTATAAACTAGCTTCCAAGCTTAAAAGCGTC
TACAAATAGACCGTATGTTATACCAATTTTGAAGTTGTTGAATAAATTAATTTTTATATT
TAATTGCTTGTGGCTAGAGTAGACTATTTTACTAGTCAACTCAGTTGCTGCAACTATTAG
ACTGGCTGCAATAATACCCCAATCTCCTGTTTGTCCAGGAATCGTAGACAAACCTGTTGA
AATAAAAAATCCTAGCAATAAACTAATTAAGCCAGTTGTTAACTCGCTCAAAGAATAATA
AAGCTTATTATTTAAGTTTTTTATTAAGTAAGCAAAAAAAGTTGAAAGTCTAGTTTTAAT
CATAGTATTTAGTAACGGAGTTTTAAAAGGATGACAAGTATTAACAGTCATTAATATTGT
CGTATCCTTTTAAAACTCGTTAATACAAACAGCTGTTCTTGTTTTTAAAGGTCTTGTTCG
CTTAAAGCATTAATAAGATGCTGAAACGGCTCTTGTATAATATCTGTGCGTTTAATTGAT
TGCAAATCTAATTCTTCTTCTATGATTTCTTGTAATAACTGTATGCTTCTTACTGTTGGA
GCAACAGGTACAGATAGTGAATTGTAAGTATCTTTTAGCCCATTTAATACACGCTGATCG
AGAATATTTGTACTACCAGAAACCAAGGCATAACTAGCATACCGTAGGTAATATTCTATA
TCCCTTAGACAAGCTGCATATCTTCTTGTTGTATAAGAATTTCCTCCAGGCCTTAATAAT
TCAGGCTGTTCTTCGTACAACTGAGCTGCGGCCTCTTTTAAAATATTGGTAGCTTGATCA
TTAATAATTTCAGCTATTTTTATTCGATCTAGTCCACTCGAAAAGAAAGATTCTAGTTGT

CCGACAGCTGTCTTATCTAGATAACGACCAGTAAGATCGTAACGATTTAATATTGCTGTT
ATAGCATCTTGCATAAATTAATTCTCCTACTTGCTATCAGAGCGGTATTATAAATGACTA
ATATTGTAATATAATCATTAGTGTTTATGAAATTTATTTAAAACTTATTCTTTATTGCTT
AATGTGCAATTAAAAACTCGCATAGATTTTACACATATAGAGTATACGTAGTTTTTAATT
ATTTTTTTATAAAAACCAAGAAATAAAGCTTTTTTTAAGAGGTTCGTGAATTTGTTAATA
TAACATGAGCTTTCCTCCTAAATAAGTTTTGTAAAGTACAAAGTAAAAAAACGATTAGTC
AATAATAGTAAACAATATTAATATGTATCAAAGTATAAATAAGCTTTTACTCCAGCATCA
AAAAAAATATTTGAGATATGTTGATACAACTGATTATTCAAGTGAAAACTTAGATTTAGA
CCAATTTAGCTATTTTATCATTACAATTAATAAAACTGATAAATGTATTCTTATAGAGCA
ATTTTGTTATAATGCTGAAGGCCATATTTACTCAATATTCTTCAAAGGTCCAACAGCTAA
ACATCTTTCTTCTATAATTTGTCATTTTCAACAATTAAATACAATAATATCTACAGCTCA
TGCCATATATTTAGGACGAGAGCTAATGAAGTCAGAATTAGCTCTCGTGCTAGATCAACA
ATATATTCAAGATTAAATTTTGTAAGTATTAATTATATTTATGCTAGTATAATATTAGTA
AGGGCATGTAACTCAGTGGATAGAGTATCAGATTCCGATTCTGATGGCCGTGGGTTCGAA
TCCCGCCCTGCCCGAGTATTCAGAGTACTATACTAAAACATCTTAATAGCTAGTTTGATA
ATAGTATGGGCATTGGGGCTGCAAGGTTTCTACATTATGAAAAGAAGAAAATATGAAAAA
AAACAAGCTCTCCAAAAGAGCTTTTAGTACAATCAATAAATGCAGAAAACAATATTGTTT
CTTTTTCTCGAAAATTAGCTCTTGCTTAAACACTGTTAATTTTTGGCATTCGACATGTTA
AACTCTTATTCATGGCGAATATTCTGTCAGAGTCGCTCTTAGTTTAAGAAAAGTCAGAAA
AAATATTTATCTTTACGTTTTTACCTATTTTTATTTAAATAAAGGTTAGTATTTTCAACT
TTTATAGTGGACGTGGGTTCGAGTCCCACCAGCTCCATCAATTTGATTTGTATAATTAGC
TAGTTTAACCTTTTTAATAACCAGATATATAATTGGAGGAAATATGGGTTTCATAACAAT
TACTAAGCCCGCTTTAAAGCAAATTGCAATCTTGAAAAATGATCATGAAAATGATGTACA
TCTAAGAATAGGTGTTAGACAAGGCGGATGTTCAGGTATGTCATACTCAATGAATTTTGA
ACATGTTGACAAATTAAAAGATACAGATGAACGGCTTCGTCTAGATAATTTTTCTGTTGT
CTGCGACCCTAAAAGTCTTCTTTATCTTTATGGATTATCATTAGATTTTAGCTCGGAATT
AATAGGAGGAGGATTTCAATTTTCTAACCCGAATGCCAGCCAAACTTGTGGCTGCGGTAA
ATCTTTTTCAGGCTGACCCTCTAACACGTTTTTATATATTTTGTATTATATATATTATT
TTAGCAAGTTGTTTATTAGAGACAAGCAATGCTCTGTTAAAAACTAATTTCAAGAGTTTT
TATTGTATTATTTTATATATTTATTATCAACTACTTGAAAATAAATTAGCTAAGCAATCT
AATTTGTTATCAGTACCAAACTTTAAATGATTAACTTTTTATTGATATATGCTAATCTTT
ATCAGCCTAAACATATCTGTACATTACAATCTTTTGACATAAAAAGAGTCTTAACATACC
TCTCAAAAATAAATTCAATATTTTATAAATACTCAACATGAATCTGTCACAACTTATCGT
TAATTCTTCTATGATGAAAATTCCTAATAACTTTTATTGCAATATAAAAGCTATTTCTTC
TGAGAATTTAAGTCACTTACAATTATCAAAAACAAATTTATTATCTCCTACAAAAATAAT
TGCTTTTGGACTACCTAAATTTGATTTTATTTCTACACAGATTACAAGTTCTATTAGAGA

AACAGTTGTTGTGTCTACTATTGATGATAATAATCTTTTTGCTGCCCCAACTAATGATTG
GATAAAAAAACACAAGGATAAAAATCATCATTCTTTGTTTTATAAAAAAGTTTTTAAAGC
TTTGCTAAACCATAAAATTAGAGTTGTTGCTGAGCCATTTCATAACTCCGAAATAAATAA
AGAAAAGAGCATAATGGCAAGTCAAAGATACATCTGGGGTAAAAGCTGGAAGCCTTCAAT
TATTTTATCTTGCTTAAAAAACAAAAAAAATAGCAAAATTTCTGATGCATCGAAAAAATT
TTTAACTGAACAGTTATCTGCATCACCAGTATTTGTAGTAAAAAATGGTTTTAATGAAAT
TATTTTAGGTCACCCATTATCTCGTGTTAAAAGAGGAGGAGTTAATAATCTGATGCATGC
GTTTTCTAATTTATTAAATCAGTCTAATGCAACTTATCCTATATCTACTGGTCTATTCTT
TTTTCATCCAGATGATGCATTTGAATTTAAAGACTTTATAATATCGGTCAATCCACTGGC
GGCCAAACATATGGAGATAAGCGTAGAGCCTGTTGGTCTACACTTTGCTTATAAAATGAA
TAGAAATATATCGTCAGATACTCAATTTTCGTTTATTCCAGATTTTAAAGAAGTAGGAGA
TTTATTATTCAAATATAGAAAAGGTAATCATTTAGTTTTTCATAAGAATCAGCATTATGG
CAAAGATTTTTTTCAGGGGCAACCAATATATATGATTCAGCCGATCACTTTTAAAGATCG
AGCCGGAAAGCTAAATATTATCAAATTTACGGGGCTAAATGACACAAGAGAAATTATTTT
TACTAATCTTGAAGCTGCTAATAGATCATGGGCACATTTTATAAAGAGGAATTCACAATT
AAAATCAATTAAAAATCCTACTCTATTAGTCTATAATTTAGAAAGTTTTTTAAAAGATCA
AGAACGATTGGATAATCAAGATTTAAATAAGTTTGTGGTGATCACTAATAAAGAATCTTA
TCTTGCCACAAAAGAGTTAATAGCTTTACCCGATTCTAATAGCTTTTCTAAGCACTTAAA
ATTAAATATAAAGCCTAAACTCTTTTTTGTGAAACTATGGGTAAGACGGTTATTTTCCAC
CTTAACTTATGAATAAATGCTATTGCCTTTATTTTCTCGAATAATATTCTATAACTAGTA
ATTCATTGAGTTGTAAAGCAACCCATTCCCTATCAATGACTCCATTGACTTTCCCTGATA
AATTGGATTTATTCAATTCTAAGTGGCTAGGAATATTAGCTAGTCCAGGGAAAGCTAAGT
AATTTTCTACTAGTTTTCGAGATGCTTCTTGATTTTTTACACTAATTGATTCTCCTGGTT
TACATTGATAACTACAGATAGATACTACTTGTCCATTAATACAAATATGACCATGGTTCA
CTAGTTGTCTAGCGGCAGGAATTGTAGGAGCCATACCAAGTCTAAAAACAGTATTGTCTA
GCCTCATCTCTAGTAGCTGTAACAAAATTTGACCTGTCGACCCTTGTAGTTTTTTAGCTG
CTTTTACGTACTTAAATAACTGTTTCTCACTTAATCCATAATTAAATCGCAACTTCTGTT
TTTCTTCTAATCTTACAGCATACTCAGAAGGTTTTCTGGACTTTTGTCCATGTTCTCCAG
GTGGATAAGATCTTTTAATCGCTTTTCTACTGAGTCCGGGTAAATCACCTAATCTACGGG
AAATGCGTACTCGTGGCCCTCTGTATCTAGACATATTATTTAAATTCTCCTAATGTAAAT
GATTAAGTAATGAGTAGCTATTTTTTAACACAAAAAGAGCATTAAAACAAAGCTTTTGCT
AAAAAATATTTACAAAAAATATAGATTTGGACAAAATACATTACTATGCTTTTTAAGCGG
GTAGCGGGAATCGAACCCGCATCATTAGCTTGGAAGGCTAAGGTTTTACCACTAAACTAT
ACCCGCAATATTCAACAGATTGTCACTAACTCAATATAACATAGTTATGTTAAGTTAAGA
TAATCTTAAATTTAATAGAATTTTAAATTATTTATGCTCCTTCTAAAACCACATCAAGAA
AACGAGCAATTTCGGCAGCTTGTTCTTCAACTTCTGATAATAATAATGGTTGACCGACCC

TAGTCAATGGAATTACTCTTTTATCTTTTGTACATAAATAAATTTCACGTCTTGGATTTA
GCCCCTCTTTAATATCTATTTTTATTGATTTGATTTCTTTTATATTAAACTGAAGACAGA
TTTTTCTATTCTTTCCAGGGAAACCTAGCCGAAAAATTTTTACGATACCTTTATCTTTGT
TAAATTCATTATATCCAGCCCCTATATTCCAAATAATAGTTAGCCACAGAAACAAACTCA
GAAAGACTCCTATACTGCCATAGAATGTCATTACAATACCTTGTGGAATAAATACCAAAT
CGGTTGAATTTGTAAAAGGTAACAAATCAACTTGAAAATAACTTGATAGTCCAGCAAGAA
GAAAGCCCAAAGCCCCAATAAATATTATAGTAGCCCACCAGTAATTACTCAGTCTTCGTG
AACCTAAAATTAAATCTTTTCTAACTGAGTGTATAGATAATGTTTTTTTCATAATTTGTT
CTTTTAAAACAGTATACTAATGCAAAATGGAAAAATTTTGACAATTTTTATGCGCGCTTT
AGATTTTTTAAATAAGCTTGATTGCTTTTAGTCCTAAAAACAGTCCTGTTGCTAACCCAA
AAAAAGCAGCAAGAAATATTATATATCCTAGAAAAACTGACATATTATTATTATTTATTA
TTATTGTTACAATAGTGTTTACAAAAATAGTGAAAATCTATTACTACGGTATTATATTAC
GTTAAATGCCTTGCTGTCTACAGGAAAAAACTTTTTTCAGTTTTTAGCATTGTGTTTATA
TTAGAAATAATATTCTAGTAAATTTATTAAACTTCTGGAGAACAGAATCTTATGTCAGAA
TTTATCAAGCCTTATAACGATGATCCTTTTGTAGGCAATTTGTCCACGCCAGTTAGTACG
TCAAGTTTTAGTAAAGGACTTCTAGGAAATCTACCAGCTTACCGTCGAGGTTTATCTCCG
CTTCTTAGAGGATTAGAAATAGGAATGGCACATGGATACTTTTTAATTGGACCTTTTGAT
AAATTGGGCCCTTTACGAGGTACAGATGTAGCGTTACTAGCGGGATTTCTATCTTCGGTC
GGCCTCATTATTATTCTCACTACATGTTTATCCATGTATGGTAATGTATCTTTTACTAGA
GCAGATTCGAAAGATCCACTACAAACTTCTGAAGGCTGGGGACAATTCACTGCAGGATTT
CTAGTTGGAGCAGTAGGTGGTTCAGGATTTGCTTATTTATTGCTAGCTAATATACCTTTA
TTACAGACTGCAGGTCTTAGTTTATTCTCTTAAGCTAGTAAGGGGACTTGAACCCGTAAC
CTACTGATTACAAATCAGTTGCTCTACCAATTGAGCTATACTAGCATTCAGTTCACAATA
ACACATAAAGATATTTGTTGCAATCTGCTATTTTCAGATGTAACAAATATCTTTTTTATT
TAAACAGAACTGTTATGTTAACACGTATATTAATTAGAATCTGAATTATCTGACTCTGCA
GAAGATTCCTGATCACAGTTTAGATAATAACAAATTGTTTGATCTAGACAAGTTGCAGAC
CAACCAACAGGTGTTTCTCCTGATAATTCTTGCATATCTACAGAATTATAAAAATTATCA
TTTGGCTTATTACTGTTTGCTTGCATAAATATCACTCCCAAAACACTCTGAACTTAATGT
GAAGTAATTTAACTTCTCAATAATATATTAACACAAAATATTAAATATGGTGATTATTCT
TTAATTGTAAAGATGCTGCTAACTTTATAAAGTTTGAGTGAGAGTCTTGATAGCTTTTGT
TGAAATAAGCATTTTAACCCATTTATTTTGCCCTCTCGACCATACTTTTTTGTGTTGTAA
ATTAACTTTTTGTAGTTTTTTTGTTCGTTTATGAGAATGAGATACGGCATATCCATTATT
AGCTACTTTTCCCGTAAGCTGACACTTTTTTGACATAATTTTCTAACTTATATATTTATT
TAGAGTACTTGCTAAAGTTGACTTGGGCACAGCTCCAATCACTGTGTCTACTCTTTCTCC
TGCTTTAAAGATCATTAAAGTAGGTATACTTCTAATGCCATATTCAGCTGCAATAGTAGG
ATTATCGTCTGTGTTTATTTTTACTACCTTAATAGATGATTCATATTCTTCCGCAATTTC

ATCAACTACAGGAGAAACCATTCTACAAGGACCACACCATGGCGCCCAAAAATCTACGAG

TACAGGTAAGTTATTGTTAATAACTTCTTGTTTGAAAGAGGCATCTGTAACTTGAGATAC

TGACATATTCTTTAACCTTTAATATATGTTCCTTGCTAAGCAAATCTTATCACAAATTTC

AAGAAAAATCTTCTATAGTGAATTACTATTAGTGTTGAATTATTTTTTTAAAAATTAGAC

ATTAAAAAAACTTATTACCTATATAAATAAAGTGGTGGCCGAGTGATATCTTGATAGTGA

TAATTTTTATATTAAGGTTGAACACTAGAAATAACTTTTCAAGCAAACTCTTGTTTAAAA

AATTTCTCTGTGACCATAAAAACTTAATTATATTACAATATATTTTATTATTTGTAACTT

AGTTCTGATAATGGTATAAACAACGCAAAAGATACTGCCTTATAATCAAGGAGGAATACA

TGTCTCAATCCGTAGAATCACGGACTAGGATTAAAAGCGAACGTTACGAATCTGGAGTAA

TCCCCTACGCTAAAATGGGCTACTGGGATGCTGACTATGTGATTAAAGAAACAGATATTC

TAGCTCTTTTCAGAATCACTCCTCAACCAGGTGTTGACCCGATTGAAGCATCTGCTGCAA

TTGCAGGTGAATCTTCAACAGCTACTTGGACAGTTGTATGGACTGATTTATTAACAGCTT

GTGACTTATACAGAGCAAAAGCATATCGAGTAGATCCAGTTCCAAACGTGGCAGATCAAT

ATTTTGCTTACATAGCTTATGATATTGATTTGTTTGAAGAAGGTTCCATTGCGAACTTAA

CTGCTTCAATTATTGGTAACGTTTTTGGGTTTAAAGCTGTTAAAGCTCTTCGCTTGGAAG

ATATGCGTATGCCAGTAGCTTATCTAAAAACGTTCCAAGGTCCTGCAACTGGATTGATTG

TAGAACGTGAGCGTATGGATAAGTTCGGTAGACCTTTCTTAGGTGCTACAGTTAAACCTA

AACTAGGTTTATCTGGCAAAAACTACGGAAGAGTTGTATACGAAGGCCTGAAAGGCGGTC

TTGATTTCCTTAAAGATGATGAGAATATTAACTCACAACCATTTATGCGTTGGAGAGAAA

GATTTTTATATTCTATGGAAGGTGTAAATAAAGCATCGGCTTCTGCTGGCGAAATTAAAG

GTCATTACCTTAACGTAACAGCCGCGACAATGGAAGATATGTATGAGAGAGCCGAATTCT

CTAAAGAGGTTGGTAGTATCATTTGTATGATTGACCTTGTGATTGGTTATACTGCGATTC

AAAGTATGGCAATTTGGGCTCGTAAACATGACATGATTTTACATTTACATAGAGCTGGTA

ACTCAACTTACTCTCGTCAAAAAAATCATGGTATGAACTTCCGAGTTATTTGCAAATGGA

TGCGTATGGCTGGTGTTGACCATATTCACGCAGGTACAGTTGTAGGTAAGCTTGAAGGAG

ATCCTTTAATGATTAAAGGCTTCTACAATACTCTACTTGAAAGCGACACAGATATCAACC

TACCTCAAGGTCTGTTCTTTGCTCAAAATTGGGCTTCCCTACGTAAAGTTGTACCAGTAG

CATCTGGTGGTATTCATGCTGGTCAAATGCACCAACTTCTTGATTACTTAGGTGATGATG

TAGTTCTTCAGTTTGGTGGTGGTACAATTGGACATCCTGATGGTATCCAAGCAGGTGCAA

CTGCTAACAGAGTAGCACTAGAGTCCATGGTTATGGCAAGAAATGAAGGCCGTAACTATG

TAGCAGAAGGTCCACAAATCTTGAGGGACGCTGCTAAAACTTGTGGGCCTCTACAAACAG

CTTTAGATTTATGGAAAGATATTAGTTTCAACTACACTTCCACAGATACAGCTGATTTCG

TTGAGACTCCAACAGCAAACATCTAGTTTAATGACTACTTACTGATACTTTAAATAGTCA

ATTGTAAGTGGAATTAACTTATAACAATAAGGAGCATAGAATAGTGAGACTAACACAAGG

GACTTTTTCCTTCCTTCCAGATTTAACTGATGAGCAAATTAATAAACAGCTTGCTTATAT

CGTTTCTAAAGGCTTATCAGCAAACGTTGAGTATACTGACGATCCTCATCCAAGAAACTC

CTATTGGGAACTGTGGGGTTTACCTTTATTTGATGTAAAAGATGCTTCTGCTGTTATGTA
CGAAATTAGCTCATGCAGAAAAGCAAAACCTAATTATTATGTTAAAGTTAACGCTTTTGA
TAATACTAGAGGTATTGAAAGTTGTGTAATGTCTTTCATTGTAAATAGACCTGCTAATGA
ACCAGGATTCTTATTACAACGCCAAGACTTCGAAGGTAGAACTATGAAGTATAGTCTTCA
TAGCTATGCTACTGAAAAGCCTGAAGGAGCTAGGTATTAATATTAATTAAGAATTAATAT
TGGCTAATTATTACCCTCTTAAAAATCATAATTAATTTAGTAATTGTGATTTTTAAGAGG
GATAGCTTCCCAAGTAAATTTTGACTAAATAGATTGAATAATTAAAAATACACAGAAATA
ATGCAATCACAGGATATAATTTCCAACGATACTCTTGTTAATTTACAAGAAGAATATGAT
AGAACACAAATCCAAGAAGTTTTAAATGAGTTAAATCAAGAACTTATAGGATTAGTGCCT
GTAAAGACCAGAATTCGCGAAATTGCTGCGCTATTATTGATTGACAGATTACGCAGAAAA
CTAGAACTAGTTTCTGGTAATCCAGGATTACACATGTCATTTACAGGTAGTCCAGGAACT
GGTAAAACTACAGTTGCTATGAAAATGGCTGATATTTTGCACAGACTTGGATATATAAAA
AAAGGGCATTTGTTGACAGTTACAAGAGATGATCTTGTAGGTCAATATATTGGACATACT
GCCCCTAAAACTAAGGAAGTTCTTAAACAAGCAATGGGAGGAGTTTTATTTATTGACGAA
GCTTACTATCTATATAAAGCAGATAATGAAAGAGACTATGGCTCTGAGGCAATTGAAATT
TTATTACAAGTAATGGAAAACCAAAGAAATGACTTAGTTGTTATCTTTGCTGGATATAAA
GATAGAATGGAAAAATTCTACGAATCCAACCCAGGACTCTCTTCTAGAGTAGCTAATCAT
GTAGACTTCCCAGATTATACTTCAGATGAATTATTACAAATAGCTAAAATGATGATAGAA
GAACAGCAGTACTGTTTTACAGAAGAAGCAGATAAAACTCTTTTAGAGTATACCGAGCGA
AGAATGAAACAGCCTTATTTTGCTAATGCAAGAAGTATTCGCAATGCTATTGACAGGGCT
AGAATGAGACAAGCCAATAGGATTTTTGCCAGTGGAGAAAAAGTATTAACAAAAGCTGAT
TTGGTAACGATTGAAGCAGAAGATATCTTGAAAAGTAGATTATTTTCATTACCTAATGCT
TAATATACACGTGATTTCATGAAACTATATATTATTAAAAAGTTTCATAATCAGTTGCAA
ATTTCTTGAAAAACCTTTATTATTATCTTAATGCAGGTGTGGCGGCATAGCCAAGTGGTA
AGGCAGAGGATTGCAAATCCTTCATCCCCCAGTTCAAATCTGGGTGCCGCCTAGTACTAA
AAAAGGGGGGTGTGGTGGAATGGTAGACACAACAGACTTAAAATCTGTTGATTTTTAGTA
ATCGTGAGGGTTCAAGTCCCTCCACCCCCATATATTGATTTATAAATAAAAAGAGATGTG
TATTTGTATTAATTGCAACCATATTACCAAATGTAATACCTATCACTTAATAGAATCTCA
ACATAAACAGCCTCATCTTACTAGAAGTCCATTGTTTATACCTAAATATCCTGTAGTTCA
TGTTAATATATCTAACAACTGTACCTATAATCAAATTGATTGGGATTTAGTGGAGTGTTT
ATCATTTGTAGAAAAACCGAATAGTTGGAATTTAGACGCTAATTAGTTACAAGAAAAATT
ATGTACCATAAGCAACAACTCTATTTTTTAGATACTCGGTATTAATTTTGGATGTTCTAA
CCAGTAAGATTTTACCCGTTCTTGCTATCTCTATAATGCCAAATTTAGTTAGTAATTGTT
CAATAGCAACAATCTTTCCCGGATCTCCAGTAACCTCTATAATTAAAAGATCTTCTGCAA
TGTCTACTATCTTAGCTCTAAAAATTCTTACAATTTCTAAAGCTTCTGTTCTAGTCTGAG
AATTGATCTGAATCTTAATTAACATTAGTTCTCTTTCAACTGAAGGAATATTTGTTACAT

CTTGAACGTTAAGGATATTTACTAACTTATATAATTGTTTCGTAAGTTGTTCAATAGTTC
TATTGTCTCCTTGAACTACCATTGTGATTCTAGAGACCCCAATTTGCTCTGCTGGTCCAA
CTGCTAAACTTGCGATATTAAACCCTCTCCGGGCAAATAGACCAGATATTCTTGACAGTA
CTCCGGCTTCATCTTGAACTAAAACTGATAAGGTGTGTTTCATGAAAAATTAAATTTAAT
CTATTTGATCTGTTAAGTATATTCTAAGAAAATTACAGAAAAAATACTAGTTAGTCTATA
ACTAAATTATCCAAACTAAACAAAATCCTACTATCAGATAATATATCTACCTTGTAATGC
TATACTATTAACTTAGTATTTAATGATATTCTAATTGCAAATAGAATGCTAATTTTTTAA
AATTTGTTAGGAACTTGTGCTAGAAAAATACAAAAACAGTAAAAAACTCTCTCGAGATTT
ACCTCAAATCAATGATCGCATTAGATTTCCAAAAGTCCGAGTAATTGATGACGAAGGTGA
ACAACTAGGTATTTTTGTGCCTGAAGAAGCTATACAATTAGCTGTCCAACAAGGTTTAGA
CTTAGTTGTTGTTAGTGATAAATCGGACCCGCCAGTATGCCGAATATTAGACTATGGTAA
ATATAAGTTTACACAAGAAAAAAGAGCTAGAGAAGCTAAGAAAAAGCAACATAACAGTAG
TATTAAAGAAGTAAAAAATGCGATATAAGATAGAAGAGCATGATTATAAAGTTAGAATAAA
CCAGGCATCCAAATTTATTCAAGCAGGAGATAAAGTAAAAGCAACTATCACATTTCGGGG
GCGTGAAATCCAGCACTCTAATTTAGCTATAGATTTATTGAATAAAATGGCAAGCGATCT
AAATGCAATAGCTGAAATTCAGCAAGCTCCATCAAGAGATGGCAGAAATGTCATAATGCT
CTTATCTCCCAAAAAAGTTAGCTAAACTAATTTTTATTTAAATGCATCTGGCTGGATTCG
AACCAGCGACGTCTCTTTCGAAATGGCGGATTATGAGTCCGCTGCCTTCGGCCCCTCGGC
CACAGATGCATCACTTAAAGTTTATACCTCATAAAGCATATAAAAATCAATGTTTAGCTA
TTTTTAATAAAGTCTATAAGAGTCTATTATTCATTCATATTGTGATATGTCGCAACTGCT
GAAGGGGAAACTTGATTTAAGTATCGAAATATCCAGTATTTAAAAATTGTGTCGAGTATA
ACTGGAAAAGTTGAAATAAATAAAAAAATAAAATCTCGACTTTCCGGAAGTCCTAAATGT
CTTAAAATTACTTCAATAATAACTTCCCATCCATGAGGAGAATGAAAACCCACAAACATA
TCAGTAAAAAGAATGATTAAAAAAGCTTTAGCTGTATCACTAAGACCATAAATAATTTCA
TTTAAAAAAGATTTAACGACTGCAATTTGTCTTTGTCCAGTTATCATTAGTAATATAAAG
ACAAGTATTGATACAAGATCGGATAAAATATTTTTAACAGCATTTGCACTTTCATTTGCA
TAATATTCTCCCAACTCTCTAGCCTTCAGCTGCACTCTTTTTTCAATGATTTCGTAAGAT
ATATCTTCCGAAGGATTCAGAAGAACTTCAAAGTGAATTTTCTCTTCAAATCTTTGTAAT
TCAGCAAAAGCTCTTTCCTCTTGAGAGGAATTTAAAAAAATTTTAGGCTGTTCTTGGTTC
CATAAATAATCAATACAAGGTCCAAATACAAAAAATTTTGAAGCCTGATTAACTAGTACT
GGAGAGATGAGTAATAATAAAATATATTTTACGGAAGTAATAGTTTGGTGTCTAGATATT
CTAAACTCCTCTATAGCTTCAGATTCGCCATTAGGATCTAACTCTTTACGAAACTTTTCA
AAAGTATTAGTAATTGACCTAGGTATTGGCCCAACTTTTTCAAAGGCGAATTGATTATTT
CGTTTTAAATTCCAATATTTCATTTTTTATTGCTGCTAAAATCTCTATTAATCAATAATT
GATCACGTATTCACTATAAATATTAATTCTAGAACTAAACATATAAAACAATATTTATGA
AAAATAAAATACAATTAAAAATTAATTATGCATTAGTTCTATCTGCAGCTACAGCAGTTT

ATGGATATAAATCTTATCAGCCAACTATAAATACACTCTTAATCAAAGATGAAGAAAACT
GGCCTTTTTTCTCTAAAAGGCAGACTGCCTTTATTCCTATAATTGTATTAAAAAAAAATC
ATCAAAATCCCTGAACTCAGAACCTAGAAATAGTAAGTTATATCTTAATGGAATTGGAAG
TAGAAAATTTATCGACAGAATTGTCACTCTACTTGGATATTTAGAAAAACAGGCAAACTA
TACTGAATACTTGAACCAAATAAAAACAAGTGGTTATTTTTCTTTAGTCTACCTAGACAG
TGATATAATGTATAAGAAAAATATTAGTAACATTTTTTTACTACCTAATGATATTCTAAA
AAGAATCTATATTTGCAATAAGAAAAAAAAGCTTATACCGCATTCATTCTTAATAAATTT
ATTTCAGAAACAGATAGGCTATCCAAAAAGTTTTGCTAATTTAAATTGGGCCTTCTCCAA
AATTATAAAATGGTATTATGATAGAGGATACCAATGGTCTCTAGTTGAAGTTAAACAAGC
GTCTGATGCATCTTCTATTGTAATTGATATTCATGAAGGGGTAGTGAAGACAATTATAAC
AGAATACTACACTCTATCTTATAAAAGAGTCTCAGGTATCTTATGCGTAGAGTCAATAGA
GCAGTACCTTGGAGTAAGAGTGGGAGCTCCACTAAATATAATTGATTTACAAAAAAAAAT
TACTTACTTAAAAGATAATCAACTAGTTGGCGACATTATTTATAGTATTGAACGATCAAA
TAATAGCTCAATGAGTTTGGATATCAAATTTCAAATACAAGAATTAAAAGATAAAGAGAT
AATAGTGCTTGCAAAAAGTTCTTCTATTATCTCGCATGCATGTAATTTACTTAATCAATA
TAGAAATAGGCTAGTAGCTTCTAATATAGTTTCACTATCAACTAATAAATTACACGCATG
CTATTTTAATTATAAACTTGACTATCAATATAAGTATATAAACACGTCTAATCTTATAAA
ATTGTTAGCTTATTCTATTTCTTGTAGGAAGACACTACTTATTCATTATTTTAATTTACA
AACTTTAATAAGCTATACAAAAAAAAAATACTATTGGATTTCAGCTATATTTACGAAATCT
AAGTTTTGGGAAAGCTTTCTGTGTGCTTAGTATGAAATTTATCAAAAATGGACTTAATAT
AAAAATTTTATATATAAATCCTTCATTGATAGTTGATCAGAATTTTGTATTTCAATTTGC
TATACAAATCATTAAGCAATATCATACTGCTAAACCTCCAGCTTTATTTTTAACGAATCT
AGATCTTGAACAATATGTTGCTGAAAGTTTATTAATGTACCATTTTACATCTTGCTTCTC
AATATCTGAAAAAATATTATTATCTCGAATTATGCATACAGATTCTTTATTTTTCAACTC
TGAAAATTTTCATTTTGATGATAGAACTAACGCGATCAACAGTTATGATACTTTTAAACA
AAATACAAAAATATTTTATCAAGAGTTTCTTTCTTTATTATTAAGTTTACGTTATCAAAA
TTTTAATTATTTAGGTTGGCCTTTAAAAGGCCATTTTTTTGAAATAAAATCTTTATACTT
GGCTCCATTTCAAAAGAGTGATTTTTCTGATAGTCGTAAGACCCTATTCTTCCACAAAAT
GTCGTTAAAGCAAGTATCTAATTTTAATTTACCAATATCTTTTAAAAGCCACCTTAACCA
TATTTTAGTTAGTACTATTAAATGTCAGTCAAATTTGAATATGAGAACTGTATCTTTGCT
ATTAATTGATTCACCTGCTGAATATATGCTATACAAATCTATTCTTAATTTCTCTATTAA
AGTTAGGATGCAGTATTTCATACCTATGAGCAACAATATAAGATTATCTCTATTTTATAA
TTATTTAGATTGTTTTTTAATAAGATCATCTCAGAGTCGCATTCATATATGGCAAGATCT
AAGAACTCTTACACCAATTCAAAATTTTTGGTTAAAAAAATTTTCATATGGAGCTGGAAT
TCAGCTTAAATTACCAATTAAGCAGATGCCACCTCTTTCAATTGAATATACTGTAACTAG
TAGTCGCTACTTTTGTATTTACCTTCGTACTTATTATCAACGATAACTATTACTAAAATG

AAAAATTTAATTTCTTTTTTTGATCGCAGCAAAGGGAAATGGATTTCACAAAGAACAACT
TATGAATTATCTAATAAAAACATGAGTTCGATACAGTCTCAGATGACAATGAAAATAGGC
AACTCATTGTCAGGATCTATAATACTTGCATCGTTAAATTGGGGGGACATTTATAGACAA
GTTGCCCATTACGCTAAAAATCACAGCCGGAGTGAATATGATAATAAGTTTAACTTGCAA
TTCAGTAATCAACTAAATAATCATAAATTATTAACGCTGTGCATAGTGACAGATCCTAGT
CTAATTAGTTTTAAAACTCGCTATGGAAGTACTACCATAGATGAAACATATTGGTTCGCA
ACAAATAATTTACGTCTAAGTACTAGTATTGTTAAACGATTCAACACTTGTGTGGCAGTA
TCTTTTTGTTCAGAAATTAAAGTTTAAACTCTATATAGAAGCAACTAATAGCCTAAAGGC
TATTAGTTGCTTCTATATATCTAAATTTTGTATTAAAAATTATAATACTATATTATTCTA
AGTCTTTACGATTGGGATTCCTGGAAGGATCATTAGAGAGGAACCCAAAGACAAACAAAG
AAATAAAGAAAATAACGGTAGTATATACAAAGATTTTCAAAGTAAACATATTTTCATCCT
TAAAAGCTATTAGTGCGATATTTACATATCTCCACATAATTTTATATCATGATATAAGTT
CCAGCACTATTTTATAAATAAATCAATATCTTTATTTTTTAGTAATGATTTATATTTAAA
TATGCGTGATGTACTAAATACTAAATCTTTTCCGACTTGTTTTGTCTTGTTTGTAAATAT
TTTGTTATTTCTATGGAGTTTACCTTCAATTATCACATAGTCTAACTTCTTTAGCTGTTT
AAAAGTATATGAAGATTTTTTGTTCCAAATAGTTAGATTGATTATAACTAATTGTTTTCT
TTTTAACAGTCTTGCTTTTAACTTAATTATTTGGCTTTTATTTTCACTAATTCTGATACT
CTTACAACTAAGAATTTGGACTAGAAGAGTACAGCTATTCATAATTTATAACAATTTTTT
AAAAAATCTACTAAAATAGATGATTTTACAGAACTAAAATGATATTTCATTATCTTGTTG
ACCTTGCTCATAAGTAATATCTTTCAATACTTTTAATATTTTTCCAAAATACTCTTGCAG
GTATTTCTCTAAAGATATTACTTCCAACCTGTCAATTCCAAGAATACTGTAAACTTCATC
CATCGGAGCTGTAAATTGCTCTCCGCTACTTAAAATTTCGGCAAATGCTAAACGATCAGA
AATATTCCATGTCCATTGAAGAGTTTTAGTAATTCTTCTTAAGGCTTTTAATAATCCAAT
CGGAATTTGTGAGATCTGTGTTTTTTGCCCAGATAATTTTTCACATAATTTAATAATTTC
AGCTGATGTCCAAGCGGTATTACCAACTAACGGCAGGGTCCTATTTTCTGTGGAAGGTAC
TCCAAGACTTTTTATAACTAACTTTGCTGCATCTTGAGTATCAATGTATGCAATGGGTGT
AGATTCTCCTGTAACCCAGACTGATTTTTTGTCTAAAATAGGAATGGCATATTGGTTGAT
TAAACCTTGAAAAAATCCTCCTAAAGAAAACACAGTATATTTCACATTGGACTTCTGAAG
GAAATCTACTACCTGAGACTTTAAGTTCATCAATGGAACATCTGGGTATTTCTCTGAATT
TAATATTGAAAAGAAAATAAATCTTTCAACTTTAGCCGCTTTAGCTGCTTCAATTAATGC
AATTTTCCCATCTAAATCTATTTTTTCTGCATTGTAAGGATCAGTAGGACGAGATGTAGA
AGCATCTATGATTGCTGTTACCCCACAAAAAGATTGCAAAATACTTTCGGGTAATTTTAA
ATCACCATATATAAGTTCTGCTCCCCACTCTTTGAGAAAGGCAGACTTTCTTAAATTTCT
TACCATGCATTTTACATTGTAGCCTTCATCTAAAGCTCGTCTTACAATTTGACGTCCTAA
AGTTCCGGTTGCTCCAATTACTAAAAGAGTCATATTAGTTAAAAATAAGTATGTATAAAG
ACATTGATTTGGTTGACGGCAGAGAGTAAATTAGGTAGAGAGGGAATCGAACCCTCATGA

CCGAAGTCGTCACATTTTGAGTGTGATGCGTATACCAATTTCGCCATCTACCCTATTATA

GACCAACTACTACAAGTAATTTAGTCTTTAGCGTAAAAATAATACTAAGTTTCAATGGAT

ATGTCAACTTTTAAATTTATATTAATAAAAAAACGGAGTCTCTTATGGCCACTTTTGAAT

TAATTCCTTATAGATTGTATTTGAGTCAATCTAACACATGGATCCATAGAATAAAAGCAG

AAATTAAGATATATATAGTAACGCTTTTATGGATTTCAATTTTTATTTTTTCTTACTTTA

AACTATGTATTATTGCTTTAAGCTTAATTGCCATAAGTTTTACTATAAGAAGTAAACAGA

ATATTATCCAAAAACATTTGTTACAAACGTTATTAATAACGTTCTTAACTACTGTTTTGT

CTTTTAGTGTGGCTATTAGTTATAAACAATATGCAGAACAAGAACAATCGCAGTATTTAT

CTGATTCCAAGAAATATAAAAATTCCAGCAGCTACTATTATATACAAATAGCTAACGATC

AAAGGATCAAAAGACAATACTTAATTACTACTTTAAAGCCTTCCTTGTACTTTTTTATTA

CTATATATTCTATTAAACTAGTTATGATAACAACTTCTCCAGAAGTTTTAGTAATTACTA

TTTATAGATCTAGGATAATAAATAAAATATTTAAAAATGAATTGCTATTTATCTTCCTTC

TTTCGTCACATATTGTTACTAGTATTATTAATAGAATCGATAAAGTGATTCAAGTAACTA

GCTTAAGAGGAAGCTTGAATTTATATAATTCATTGACAAGGCCTTTAATGTTTTCTTTGT

TAATATTTCAAGTCTTCTTTTTGGAGATTATTCGGGAGTCAAAAGAAATAGCTCAAGCTC

TTTATACTAGAAATCTCAATCAAGAAAACAATAATTTTTTGAAAATATATACAGTAAAAT

CTAACTTTAGTGATCGGCTCAATATAATTATTAGCACTTTGTACTTTATTATTTTAGCTC

TAGCGTAATAGAATCCACTTATTCTTCTGACTCTAGTAATACGAAGAGTAATAAAATTAG

TTAGTAGTTTAAATAGATAGAATTAAGATATTATATAAATAATTAAATGTTTCATTATGG

TTAATACTCAAAATCAAATTTCACAAACTTCAGAACTTGATTATATAGTTAATCAACCTT

ATAAATATGGCTTTAAAACTTCTGTTGAATCTGAGCAATTTCCTAGGGGAATAAGTGAAG

ATATTGTCCGCTTGATCTCCAAGAAAAAAGATGAGCCTGAATATCTATTGAATTTTAGGC

TTAAAGCATATAAAAAATGGAAAAAGATGAGTAGTCCGTCATGGGCTCATATTAAGCATC

CGAATATAGACTTTAATACGATTATTTATTACGCTGTTCCTAAATTAAAAAAAGAATTGA

AAAGTCTAGATGAGGTTGATCCAGAAATTCTTGACACTTTTAACAAGTTAGGTATATCTT

TAAATGAGCAAAAGCGAATTTCCAATGTAGCCGTTGATGCTGTTTTTGATAGTGTATCTA

TTGCAACTACTTTTAAGAAAGAACTGTCTGAAGCTGGTGTTATATTCTGTTCAATTTCTG

AAGCTATAAGGGATTATCCAGAATTAATAAAAAAATACTTAGGTACTGTTGTACCAGCTG

GTGATAATTATTTTGCTGCATTGAACTCTGCGGTATTTAGTGATGGCTCTTTTTGCTATA

TCCCTCCTAATACAGTTTGCCCTTTAGAATTATCAACTTACTTTCGTATTAACAACGAAG

AATCTGGGCAATTTGAAAGGACACTAATTATTGCTGATCGTGGCAGTAAAGTAAGTTATC

TTGAAGGTTGTACTGCTCCTCAATTTGACACAAATCAATTACATGCAGCGATTGTAGAAT

TAGTAGCTCTTGAAGGAGCCGAAATTAAATATTCTACAGTACAGAATTGGTATGCTGGTA

ACAAAGAAGGTAAAGGTGGTATATACAATTTTGTTACTAAACGAGGCTTATGCTCGGGTA

ACAATTCAAAAATTTCATGGACTCAAGTAGAAACTGGATCTGCAATTACCTGGAAATATC

CAAGCTGTATTTTAGCTGGTGAAAATTCTCAGGGAGAATTTTACTCTGTAGCTTTAACAA

ATAACTATCAAGAGGCTGATACAGGGACTAAAATGATCCATATTGGTAACAATACAAAAA
GTAGAATTATTTCTAAAGGTATTTCTGCGGGTAGATCGAAAAACAGTTATCGAGGCCTAG
TAAAAGTTGGCCCTCAGTCATTTAATTCTCGTAATTATTCTCAATGCGATTCTTTATTAA
TTGGTCAATCATCTCAAGCTAATACATTTCCCTATATTCAAGTACAGAATCCAACATCAA
AAGTAGAACATGAAGCATCCACTTCAAAAATTAGCGAGGACCAAATTTTTTATTTTTTAC
AAAGAGGGATCAATTTAGAAGAATCTATTGCTCTTATGATCAGTGGTTTTTGTAAAGATG
TCTTTAATGAATTGCCTATGGAATTTGCTACTGAAGCTGATCGTTTACTGAGTTTAAAAT
TAGAGGGAACTGTAGGATGAGCCAGACTATTTTAGAAATCAAAGATTTATATGCTTCTGT
TGGTGAAACAACAATTTTAAAAGGGGTAAACTTATCTATTCGAGCCGGCGAAATACATGC
GATTATGGGGCCTAACGGTTCAGGTAAAAGTACATTATCAAAAGTAATTGCAGGACATCC
AGCGTATTCACTAATAAGCGGAGACATTTTATTTTTTGGACAAAGTATTCTTGAAATGGA
GCCAGACGAGAGGGCAAAAGCAGGTATTTTTTTAGCTTTTCAGTATCCTGTTGAAATTCC
TGGAGTCAGTAATTCTGATTTCTTAAGAATTGCATTAAATGCTCGCAGAAAGTTTCAAGG
ATTATCGGAGTTTAGCCCCTTAGAATTTTTTCAACTAATAACAGAAAAAATAGATCTTGT
TGGCATGCAAGAAAGTTTTTTGACAAGAAATGTTAATGAAGGATTTTCTGGTGGAGAAAA
AAAACGTAACGAAATTCTTCAAATGGCTTTACTAGATAGCAAAATATCTATTTTAGATGA
AACAGATTCTGGACTAGATATTGATGCATTACGAGTAGTTGCAAAAGGAATTAACACTTT
AGCTAAATCAACAAATTCAATTATTTTGATTACTCACTATCAAAGATTATTAGATTATAT
TATTCCAGATTTTGTTCATATTATGAGTAACGGACAAATTGTAAAAACTGGTAGTGTTAC
TCTTGCCCAAGATCTAGAAAAGCACGGATATGACTGGATTACGCAGACATAACTATTTAC
TACAATATTGAGACTAAAGAATATTCTTTAGTCTCAATATTTTTGCAAATGCAAAGTTAA
TTATTTAAGCTTTATCAAAACCTTGCTTAACATCTTCAATTGCTCTTTTTAGCAATTCTT
CACTAACTGGTTCTAATTTTTTAGTACTACGAACATTTTCACCAAATTCTGGTTTAGAAT
TCTTTAAATCTTCTCGAAGTTCTCTAATAAATTCAGCTACTTTTGAAACTTCAATATCAT
CTAGATAGCCATTAATGCCAGTGTAAATAATAGCAGTCTGTTCTTCAACTGGAATAGGAG
AATTTTGAGCTTGTTTCAAAATTTCTCGAAGACGTTGACCTCTTGCTAGTTGATTTTGGG
TTGCTTTATCTAGATCAGATGCAAACTGAGAAAAAGCTTCTAATTCAGCAAATTGTGCTA
ACTCTAGTTTTAACTTGCCTGCCACTTGTTTCATGGCTTTTATTTGCGCAGCAGAGCCTA
CTCTAGAGACTGAGATCCCAACATTAATAGCTGGTCTAATCCCTGAGTTAAATAAGTCAC
CAGATAGAAATATTTGTCCATCTGTAATTGAAATTACATTTGTTGGTATATATGCAGATA
CGTCCCCAGCTTGAGTTTCAATAATAGGGAGAGCTGTCATACTACCTCCACCTAATTCAG
CATTCAATTTAGCTGCTCTCTCTAGCAATCTAGAGTGTAAATAGAATACATCCCCAGGAT
AGGCTTCTCGTCCCGGAGGTCTTCTAAGCAAAAGAGACATCTGACGATAAGCTTGAGCTT
GCTTAGTTAAATCATCATAAATAACTAATGTTGCTTTACCTTTGTACATAAAATATTCAG
CTAAAGCCGCACCTGTATAAGGAGCAATATATTGTAAAGTCGCAGGACTATCTGCATTAG
CTGCAACTATAATAGTATAATCAAGAGCTCCTTTATCCTGTAGTGAAGATACTACTTGGG

CTACTGAAGATGCCTTTTGTCCTATCGCAACATATACACAAACAACATCTTGGCCTTTTT
GATTAATAATTGTGTCCAATGCAACAGCTGTTTTACCTGTTTGACGGTCACCAATAATTA
ATTCTCGCTGACCTCTACCAATTGGGATCATAGAGTCAATAGCAGTTATCCCTGTTTGCA
TAGGTTCACAAACTGATTGCCTTCCAATTATACCAGGGGCCATTGACTCAATTAGCCTTG
TTCCATTACTTGCAGGTTCACCTTTATCATCTATTGGCCGAGCTAATGGATCAACAACTC
TACCTAAAAAAGCATCTCCGACCGGAATTTGAGCAATTCGACCAGTCCCTTTTACGGAAC
TGCCCTCTAAAATATCCCTACCATCTCCCATGAGTACTACACCAACGTTATCACTTTCTA
AATTTAAAGCAACCCCAATAGTTTTATCTTCAAATTCAAGTAGTTCACCAGCCATGACTT
CATCAAGTCCATATACTCGAGCGATACCATCACCAACTTGTAAAACTGTTCCTATATTGG
CTACTTCTACATCCTGATCATACTTTTCAATTTGTTGACGAATAATACTACTTATTTCAT
CAGGTCTAATATTTACCATATTTTTAATATTATTTTTATTAATTGGAAATTCTAAAGCGG
TAATTAGTCAGACTAATGCTATCTGTCTTTTTTACTTTATTTTAAGTTACTACAACATCA
AGGTGTGAAGCCATTTGTCTTAGCTGTCCTCTAATACTAGTATCAATGACTTTTGATCCA
ATCTGAATAGTAAAGCCACCAATTAGCTCGGGCTCGACTGAAATAATTAGTTTAACTTCT
TTAGCCTTAGTCATAACCTTGATTTTTTCTGTTAATAGAGTTTCCTGATCAGGGCTTAAA
GCAATAGAAGTATTAATATTTGCAATTGTTAAAGATTCCATTTGGTAAGCCAGTTCCAAA
TACTTACTAGCTATAGCATCTAACATTCCAATTCGCTTGCGATCAACTAAAATCATTAGA
AATGATAATGTATTTTCGTTAATTTGATCGCCAACAGTTGCAATAATTACTTCTTTTTTT
GCCTCTACAGTTTTTAATGGATTAGCTAAAAAAATTTTTAGCTTGTCAGATTGCGCTAAA
ATATTTTGTATTGACTGAATATCTTGGCTCACCTTTTCAGTAACTTTTTTAGTTTTAGCT
AAATCAAGAAGAGCTACTGCATAAGGCTGAGCTATTTTTATAACAACATTGTTACTGCTC
ATAGTTGATCTCCTAACTTAGCAATATTATTATCAATTATACGCAGTTGCATTTCGGAGC
TCATTTGATTTTCCAGCTGCAAAGTAACTCTTTTAAGAGCAAGAAATGTTATTTGTTGCT
GGATTTGCCTTCGGATCTGTTTTTCAGCTGTTTCTATATTTGATTTACCTGTTATGGCAA
GTCTTTCAATATCTAATTGGCCTTGAGCTAATATGGAACTTCTAACTTTTCCAGCTGTTA
GCTGAGCTTCTTTCTTAATTTGATCTATAATGATTTGAGTTTGAGCAAGTTGTTTTTCTG
ATTCTGACAATCTAGCACTTGCTTGCTCTAGTCTTTCTTCGGATTCTTGTATAGCTGCTA
AAACTTTTATCTGTCTTGCATTTAAACTTGATCCGAGAAATTGCTTCAAAACATAAATAA
GACCAAATAAAAGCAACAATATATTAATAACATTTGCTTCAAAAATATCAGAATTAAAGC
CGAATTTGTGTTCACTACTATGTTCTGATAAAATAGTAATTATTTGTGGTATTTTTACTA
TATTATTCATTTAAAATTCTTACTATTTTAATATTTTACTCTGGAATCATGATTTTCAAG
ATTGACTACTTAATAATTTAACTTTAATTTGATCGCTCAAAGTGTCAACCTGATTTTCTA
AAGTTTTAAGAGCTTCTTCTTTTTGAACATTTAACTGTTTCGATGCTTCAGCAATCAGTT
TCTCTGCATTCAATTGAGCTTTTTTTATATCTTCTGAGACTATATTTTGAGCTTCTTTTT
GAGAAGAGGCTATTTTTAATTGTGCATTGCGGCGAGCTTCAGATAAATCTTCTTCATATT
TTGCAGCTAGCTCATCTGCTTTAACAAGCATAGACGATGCAGTAGTCAATGTTGTTCGAA

TATATTCATCTCGTTCGTCTAGTACTTTAGTTACAGGTTTATAAAAAAATAGTATTTAATA
AAACCATTAAAGTAAGAAACTGTAATGCCATTAAAGGCAAAGTACCATTGAAGTCAAATA
AGCCACCTTCAATTTCTTCGGCTAATAGAAATGGTAAATCAATCATTTTACTAATTATGT
TTATAGTTAATAATTTTGTTTCAATTAAGAAAAATAACTTTATAGTGCGCTTAGTCTCAT
CAGACTAAGCGCAAATGATTATTTTAACCAACGTATGGATTAGCGAACAATAAAGACAAC
GCAACAACTAGGCCGTAAATTGTTAAAGATTCCATAAATGCTAAACTTAGAAGAAGTGTA
CCCCGAATCTTTCCTTCTACTTCTGGCTGTCTAGCAATACCTTCGACAGCATTAGCTGCT
GCACTACCTTGACCAATGCCTGGGCCTATTGCAGCAAGACCTACAGCAAGACCGGCAGCG
ATAACGGATGCGGCTGAAACGATTGAATCCATAATGAATTTTAATTTTTTGTTATATGAG
TAGAAAATTGACAAATTTCGTATAATACTTAAAATCTATATAATTAAGTATAATAGATTC
TAATATTGTAATTATAGTATATTACTCTTCTCCATGACCTTCCATTGCTTCTCCAATATA
AGCTGCGGACAAAGTAGAAAAGATTAATGCCTGTATTGAACTCGCAAATAATCCGAGTAT
CATAACTGGTAATGGTATTAAAATTGGAATTAGCAAAGTAAATACAGAAACAACTAATTC
ATCTGCTAAGACGTTTCCAAATAATCGAAAACTTAAAGATAGTGGTTTCGTAAAATCTTC
TAGAATATTGATTGGGAGTAATACTGGAGTTGGCTGAATATATCTAGCAAAGTAGCCTAA
CCCTTTTTTACTTAAACCTGCATAAAAATAAGCTAAAGATGTAAGTAAAGACAGGGCCAC
AGTTGTATTAATGTCATTTGTTGGGGCAGCTAGCTCACCTTCAGGCAAGTGAATTAACTT
CCAAGGAATTAATGCACCTGCCCAGTTACAACCTAGAATAAATAGAAATAAAGTTGCAAT
GTAAGGCACCCAAGGGCGATATTCATGCTCACCAATCTGATTTTTTGCAATATCTTGCAG
AAATTCTAATATAAATTCCATAAAATTCTGGAATTTTTCAGGAATTCTTTGTAGATTTCT
AGTACCTAGAAATGATAAAGTTAAAAGTGTAGCAATTACTAACCAAGAGACAATGAAAAC
TTGCCCATGCAATTGTAAGCTACCTATTTTCCAATATAGATGTTTTCCAACTTCTACTGC
TGATAAACTATTATATGGGTTAAAATCTATGAGACTGTTTTGATACATAATAAATATTTT
TTTAAATAGTGCAAGAATCAAAAAATAAGTTTTTGAAGAATTAATATGCAATTAATTTAT
TCTTAAAATTAGACTATTAATAAATTAAAAGCAAAAATAAATTCTAATAGAGATAGCTAT
TTTTAGTAGTGATTATTTTTTGTAAAAAACAATTATTGCTACCTATAATGCTTTTACGTT
TAATTATATACCCATATATTAATTATTTAATATGATTATAGGTATTTTGCTACAAACTTT
ATTGTTTTATCCATTCTAAAGATTGTTATAGTTTATTTTTTATTTAAAATATCTGCGACT
TCATCTGCAAAATTAACTTTCAAATTTTCTTCTCCCCCTCCCAGAAGAAATCTTACAAAT
CTTCTAATTTTAATATTTTCACCTAGAACAGCAATATTTTGGTTTATTAAATCTTCAATA
CTAATATCTTGATTGCGGATAAACATTTGATCTAATAATGAGAGCTCCTTAAGTCTCTTT
TTCATTCGTCCTTCAATGATTGTTTCGATTCTATCAACTGGTTTGTTTTTCAAGTCATCT
TTACCAGCTTCAACTCTTTGTTCCAGATTAATAATTTCACTAGGTATATCCTGTGTTGAT
ACGTATTCCACATTAGGCGATGCCGCTATTTGCATAGCAATATCTTTAGCTAACTTTTGG
AACTCTGGACGTCGTGCTACAAAATCTGTTTCACAGTTGACTTCAACTAATACACCTATT
CTTCCGCCAGTATGAATATAACTCTCTAGTAAACCTTCAATAGCGGTACGATTAGATTTC

TTGTTAGCTGAAGCCAGCCCTTTTTGTCTTAATGATTCTAAAGCTTTTTCTTCATTGCCA
TTATTAGCTTGTAAAGCTTTCTTGCAGTCCATCATACCTGCTCCAGTCTTATCACGTAAG
GCTTTTACGACTTGAGCAGAAATTTGTAGTGTCATATGCTTTCAAAAATTTTTAAGTATT
TAAATTAATTATTATAATTCACTTTGAGTAGGATGAGTTATTTCTTCTAAATCTGTTTGC
CCGTATTTACCATCATAAATTGCATCTGCTATTTTACCTACAATTAGTTTAATTGACCTA
ATGGCATCATCATTAGCTGGGATTGGAATATTAATAATTTCTGGACTACAATTAGTATCA
AGTATACAAATAGTTGGAATACCTAATTTTAAACACTCTTGTATAGCTGTTGTTTCTCTC
TTTTGATCTACAACTACAACAATATCAGGTAAACGAGTCATATTTTTAATACCATTTAGA
TGTTTACGAAGTTTATCTAATTCTCTTCGTAAAACAGCAGCTTCTTTTTTGGGTAACTGA
TCAATCATACCACTTTTATCTTGTTCTTCTAGCTGTCTCAGACGATTAACTCGAGACTTA
ATCGTTACCCAATTAGTTAACATTCCTCCAAGCCATCTCTGGTTAACATAATAAGAATCA
CAGCGTTGTGCTTCCTGTGCCACTATACCAGCGGCTTGTCTTTTTGTTCCTAGAAATAAA
ACTTTTTTACCATCCGACGAAGCCTGTTTTATAAATTCACAAGCTTCTGTTAATAATTGT
GCAGTTTGGACTAGATCAATAATATGTATGCCATTTCTTTCTGTATAAATATATGGAAAC
ATTTTAGGATTCCATCTACGTGCTTGATGTCCAAAATGAACACCTGCTTCTAGTAATTCT
GCTAATGTAACAATAGCCATAAGATTATTAATATTTAAATAATGTAATTATTCGGGAAAT
CCCTTTGTTTAGAAACTGATTCAACTAACTTATTTGAATTTTAGTAAACTAATAATAAAT
TAACACACTTTTAAAGCAGATCAACAAAATAGATCCTGAAGAATTCATTCTACACTTTTA
TTGCTAAAATAATTACGCGCGGTTCTGTCGTCTAAAATGATATCATCTAAATCATCTCGT
ACAGATGAAATAGTACTTTCGTCATTAGTATTTGCAGTTAAATTCTTTTTCTCTAACGAT
GAGCCATTACAGTTATCATACATATTGAAGCCGGTACCAGCCGGTATAAGCCGACCTATA
ATAACATTTTCTTTCAGACCTCTTAACCAATCTAATTTACCAGATATAGCAGCTTCAGTT
AATACTTTTGTAGTTTCTTGAAAACTTGCTGCAGAAATAAAGCTTTCTGTATTTAAGGAA
GCTTGAGTAATACCTAATAGCACAGGCCTATATGATGCATTAATCTTATTTCTCAAGGTT
ATGGCTTTATTAGTCTGTTCAATCTTCTGTAACTCTACAAGCTCTCCAGGTAAATAGCCT
GTTTCTTCACCATTTTCTATTTTCACTTTAGAAGTCATCTGCCGTACTATTACTTCTACA
TGCTTATCAGAAATATTCACTCCTTGAGATTGATATACCAATTGTACTTCTTTAACAAGT
AGTAACTGAATCTCTTGAAAGCTTAATCGAGCAGCTTCATATAAGGCTAATCCACATTCA
ATATACAAATCAAAACTTGCATCGAGGATATCATGAGGATTTAACAATACGTCTGTTTTC
TTTCTTGCTTCAAGAATTTCTTCAATTCTTGGAAGCCCTTGTATAATGTCACCTGTTTTT
GCTCTATCAAAAACCAATATAGCTAATGTTTCTCCCCTTCTAATTAATGCATTGTTATCA
ACATGTAAAATAGCACCATTAGAAACCAAGTAAGGTCTAGCAATTCTTAATGTAACTGAT
CTAGAAGAAATTTCAATAATTTGTCCAGAGTCTAAAGAAGTCATATTTTCAGTAATAAAA
TCACCACAACGAATCCAATCACCCACACTTACTTTAACTATTGATTTTCCTATATTAAAG
GTCTTTTTATCAGAAGATGTCGCAATCAAAATGCGCCTGCTAGTATTCTTTTCAGAATAA
ATCTCCTCAACAGTACCTTCACTCATTGCAATAATCTCAGTACTAGCAACAACTGTAAAT

GGCTTGATATATTCACCATTTTTTACAATAATTCTAGTTTGACTTTGTTGCTTCTCTGAT
TTATTTACATCAATACTTTTAATTGATAATGTTTCAAAAGTTGACAGCCTTAAGTCAAAA
TAATTATTGCCACTTTCTTTAGCAGAAAACTCAATTGAGGATATTAAATTATCATCTCGA
TGCTCAATTTCAGCCACAAGATTAGTTGTTACTAAATGAACTCCTTCTATAGATTTCACT
CTTTCTCCGTCCCGAAAAGGAGTTCTTTTAACTAGCTTTAGTTTAGTTTGACTTGGTTTT
TGTGAAACTAGATTTTCAATTAGACTGGATTTTGTTTCTGGTATTGAATAAACAATAACA
GGTCTAATAAGAATATAGGGCATTTGCTCGTTTTCTATATACTCCCAATAAACTAACTTA
TCTGTAGAAATATTATTATGAAGTCTTTCTCCAGGTCTTAAAAAACCTCTACTTTTATCA
TTATTACTATACACCTCACTAAGTTTATAGATAGATCCAGGCTTAATAATAATTTCTCGA
ACTATTCCATCCTTTTCGATTATTTCAACAATACCTGAGCTTTTAGAAAAAATATTTTTA
ACTAGCTCTGTTCCACTTTCTATAACATCTCCATTATGCACTAGTAGTAGTGATGAATCT
TTATTAATTTCGTGAGTTTCTTCTGAAATCCATAAGATATATCCCGGACTTAAAATTTCG
TAGGCATCTTTATCTAAGCCTGTTTTCTTTTTTGATACATTTAAGTCTAAATATTTAATA
ATACCACCACTTGTCGTTTTATACGTATCAGATATTAGTTCTGCGACTGTGTAACCATCT
TGAATTTTTTGGTTTGGAATAGCTTTGAAAAGAAATTTTTGTTTTTTTTCTGTTTCTAAA
ATATAAGATTCTTTTTTGTGGATTACATCTGTATAAATATAACAACCTGGAATTACAATA
GATTCAGTAATAACTTGTATATTGGTAATACTGCTATTAGTATTCTGGTGTATCCTTACT
TCTCCAGCATAATGATTTATTAATTCTGTCTGTGCAAGTATCGTACCAGCTTTAGTTTTA
TCTTCTTTATGAACAATAACATTAGCAGAATCAGATATACTATAAACTTCACCTGAGAGC
ACCCAGATCAAGCCTCCAGTTTTAGTAATTCTTGTAGTATATTGATTATTATCTGTTTCT
TCAACTGTTAAGTTGGAGAAACAGATTTTGCCAGAGAGATCTGAGACTACATGTTTTTGA
GCTCTTTCTGTCATGAGTCGATTTCTGCGGGGTGATTCAGCAATCACTTGATCCTTCGAT
ACTAATTCACCATCACATATTAAAAGAGTAGTACCTTTACTTAAGTTAATTATTGACTTC
TGATTTTTCTTAGATCTGATTGTAACTTGGGTTGGCTTTTTTGTTATTAATGCTTGCTCA
CCATGTCTGGTACGAACATCTGTATACGATTCAATATCTAGATCTATCAATTGACCATCT
ATAGGTGCGTAGATTTGTTCTGCTAATTCACCGGTAAATACTCCTCCAGTATGAAAGGTC
CTCATTGTCAATTGAGTGCCAGGCTCACCAATAGATTGAGCCGCAATAATACCAACAGCC
TCTCCTAAATCAACTAAGCGGCCATGTGCAAGATTCCATCCGTAACAATACTGACAAACA
GAACTTCTAGAGTTACATGTAACAGGAGATCTTACTAAAACTTTTTTGATACCAGCTTTA
GTTATCTCTTTTGCTAACTTAGGACTGACATCTTGGTTCGTATGAGCAATTAAACAATTT
GTCTCTGGATGGAATACATTTTCTGCTAATACTCGGCCAGATAAAGCTTGTTCTAGATTA
ATTAATATTTTCTGCGTATCTACTAAATCTTCTAGTATGATGCCTTTATTGGTCTTACAG
TCTACTTCTCTAATTATTACATCTTGAGAAACATCGACAAGTCGACGCGTCAAATAGCCG
GAATCTGCTGTTCTTAAAGCAGTATCAACTAATCCTTTTCTTGCTCCGTATGATGATATA
AAATAATCAGTAACAGTTAGTCCTTCTCTAAAATTACTCGATATTGGAAGATCAATAATT
TGTCCCTGTGGATCAGCCATAAGCCCTCTCATGCCCACTAACTGTCTTACTTGAGAAATA

TTACCTCTAGCACCTGAGAACGCCATCATATAAACAGCATTTAGAGGATCTGTCTCTTTA
AAGTATTTAATAACTTCTTGTTTCAAGGATTCACTCGCATTATTCCACGTATCAATAACT
TTTTGAAATCGTTCAACTGCTGTAATCTCACCTCGGCGATATTTGTTTTCAGTGGCCTTA
ATATCTTCAATCGTGCCTAATAAAAGACTTTGCTTACTAGGAGGAATTCTTAAATCTTCT
AAACTTAAAGAAATTCCTGCCTGGGTCGCATAATGAAATCCAAGATCTTTAAGCTTGTCG
GCCATATTGGCTGCTCGGGCTATACCATAATTACGAAAAGCCCAGACAATTAAATTTTTT
AGTTCGTTTTTATCAATAACCTTATTTGAAAAACTTGGCTGTGCAAGACTTCGTCTATTA
TCCACTAAATCTTCTCCATATTAAATGTTCGTTATTAACTATGCTACTAAAGACTCTTGA
ATAATTTTATTAAAGATTATACGGCCAGCTGTTGTTCTAATATATTGCACAATTCTCTGA
TTTTCAGCGTCTTCTTTTATGATATGGTTATTAAAAAATTTTGTTGTACTATTGTCAGAA
TGTTTTTCAATCTTAATCGGATCATGAGGTTGATCACCATCTACTAATCCATCAAAACGA
GCCCATATATAAGAATGTAAATCAATTTTCTTCTGTTCATAAGCCATAACTACATCTTCT
AAGCTTGCAAAATACTGATTAGATCCCTGCTGCTGAGATGGATTATTAGCTGTTAAATAA
TAGCAACCTAAAACCATGTCTTGGCTTGGCATAATAATTGGTTGACCTGTTGCTGGAGAG
AGAAAATTATGAGGGGCTAACATTAGCAATCTAGCTTCGGCTTGAGCCTCTAAAGACAAA
GGAACATGAACAGCCATTTGATCTCCGTCAAAATCAGCGTTAAACGCTGGGCATACTAAT
GGATGAAGCTTAATAGCTCTTCCTTCTACAAGAATAGGTTCAAATGCTTGAATACCTAAT
CTGTGTAGTGTGGGAGCTCTGTTCAATAGTACTGGGTGACCTTGAATAACTTCATTTAAG
ACATTCCAAATAGAAGATTCATTTTTTTGAATCATCTTTTTGGCAGCTTTAATATTGTTA
ACTAATCCCTGCAAAATTAGTCTGTGAATAACAAAAGGCTGAAATAACTCCAAAGCCATT
TCTCGTGGCAATCCACATTGATGTAGTTTTAAATGAGGACCTACAACAATTACAGATCTA
CCGGAATAATCTACTCTTTTACCTAGCAAGTTTTGTCTAAAACGACCTTGTTTTCCTTCT
ATAATATCTGACAAAGATTTTAATGGTCTATTATTTGCACCTACAACAGTTCTACCTCTA
CGACCATTATCCATGAGAGAGTCAACGGCCTCTTGCAGCATTCTTTTTTCATTTCTGATA
ATAATCTCCGGAGCTAAAATTGACTTTAGACGAGATAATCTATTATTTCTATTAATAATT
CGACGATAAAACTCATTCAAATCTGCCGTCGCAAATCTACCTCCATCTAATTGGACCATA
GGCCGTAAGTCTGGAGGTATCACAGGAATGACTGTGAATACCATCCAAGAAGGATCTGCA
CCTGTTGCTATGAAATTTTCAATCAATCTTAAGCGTTTCATTTTTTTATTGAATTTTAAT
GAAGGTGTCTTAAAACTTTTTGGTGGATTTGTAGCTTCAGACCTTAAGGTTTCAGCTATG
TGCTCTAAATCTAAATCTTTCAACAGTTTCTGAATAGCTTCTGCACCTATTCCTACTTCT
ACTTGGTTATTCTCATCTTGATTTTGGTAGATTTCCTCTTCAAGACTTTTCCATTCGTAA
CCTTCGAGAAGTTGCTTGTATTTTAAATTAATATCTGTATTAGATTGCGTAACTACATAA
GAATGAAAATAAACAATTTTTTCTACTTCTTTGACTTTTAAGTCTAATGCTAAAGCAATA
TAGCTTGTACTTCCTTTTAAGTACCAAACATGAGTTACTGGTGAGGCAAGTTCAATATAA
GCCATCCGATGTCTTCTCACGCGTGATTCTGTAACTTCTACGCCACATCGTTCACAAACT
ATACCTTTGTAGCGAAAACGTTTATATTTACCACAGTGACATTCCCAATCCTTAACTGGG

CCAAAAAATTTTCTCACAGAATAAACCATCCATTTCTGGCTTTAAAGTTCTGTAATTAATT

GTTTCAGGCTTTGTAATTTCACCAACAATTTGTCCGTTGGGCAAACTTCTTTCGCCCCAC

TGTCGAATTTTTTCGGGAGAAGCTAAACTAATTTTTACGTAGTCAAAATATTGCTCAAAC

TTTGTCATAAATTTGAATACCTTAAAATCAAATAATTAGCCAATTTATTTACATGCTTCA

TATTGAAAAATGAATATCTTAATAAAGGAATTGTTCAAAATCGTCTACTGGAGGAGTATC

ATAATTAGAACGATCTACTCGGTTATCTTTAGAATCCGACATCAAGTCAACTTCAACTGT

CCGTCTTTGGCCATCTTCAAATAGTTTCAATTTATGTACTGCAATGTCTAAACCTAATGA

CTGCAGCTCTCGCATTAAGACTTTAAAAGATTCTGGAGTTCCAGGCTTCGGTATAGGCTT

CCCTTTAACTATTGCATTTAATGCTTCATTTCTAGCTTGCATATCATCCGACTTTACAGT

TAATAATTCTTGCAAAGTATATGCTGCACCAAAAGCTTCTAAAGCCCACACTTCCATTTC

ACCTAATCTTTGACCTCCATGCTGTGCTCTACCTCCTAATGGCTGCTGCGTAACTAGAGA

GTAAGGACCAGTGGATCTTGCATGGATTTTATCATCAACAAGATGAACAAGCTTAAGCAT

ATAAGCTCTACCAACTGTAACAGGATTATCAAAAGGCTCTCCTGTTCTACCATCAAAAAC

TTGCATTTTACCTGGATGCTGATCATTAAAAAGCCATTTATTGCTAGTAATCAATGATGC

TTCTTTTAATTTTCTATTTACAAGTGCTCTTGATGCTTCTGCTCCATACATTTCATCAAA

AGGTATTATTTTAAATCTCTTTCCTAAATAGCCACCTGCTAAGCCTAGTAGGCACTCAAA

TACTTGACCAACATTCATTCTAGAAGGTACACCTAAAGGATTTAAAACAATGTCCACTGG

TGTTCCATCTGATAAATAAGGCATATCTTGTTTAGGTAAGATCCTAGAAATGATACCTTT

ATTGCCGTGACGACCAGCCATTTTATCACCCACTTGGATTTTTCTCTTCTGCGCAACATA

GACGCGAATCATAGCATTTGTTCCGGGAGGCAGCTCATCTCCTTTCTGCCTAGTAAAAAC

CCTAACCTTAACTACTCTACCTTTTGCAGCATTAGGAAGCCGTAAAGATGTATCTCTCAC

ATCTCGAGCCTTTTCTCCAAAAATAGCTCGCAACAATTTACCTTCTGGTAGTTGATCAGC

TTCACCTTTAGGAGTAATTTTTCCTACTAAAATATCTCCAGCTTCAACCCAAGAACCACC

GACAACAATTCCATTTCTATCTAAATCCTTCAGAGAGTTATCGCTGACGTTAGGAATCTC

TCTAGTAATTTCTTCTGGTCCTAATTTAGTTTGACGGCATTCTACTTCATACTTTTCGAT

ATGGATAGAAGTATACAAGTCATCATAAACTAATCTTTCGCTAATTAAAAATGCATCTTC

GTAGTTATAACCTTCCCAAGGCATATAAGCAACTAGAATATTTCTCCCTAAAGCGATTTC

CCCTCCGTCTGTGGATGCGCCATCAGCTAAAGTTTGGCCAACAACTATTTTTTCTCCTAC

CCAAACAATTGGGCGCTGATTAATACAAGTATCTTGGTTAGACCGATAATATTTTTTTAA

GCGATAATGAACTGTTCTACCACTATTATCCTGAATGCCTATTTTATTAGCAGAAACATA

ATTTACATGACCAGATGTTCTGCTAATAACAACCATACCAGAGTCTCGAGCTATTTTAGT

CTCAAGACCAGTACCAATTATTGGCTTTTCTGGATATAATAATGGAACAGCTTGCCTTTG

CATATTAGATCCCATTAGAGCTCTATTCGCATCATCATGTTCCAAAAAAGGTATTAATGA

TGTTGCAGCTGAAATTACTTGAATAGGTGAAATTGCAATATAATCTACTTGGGTTGGGGT

TGTTGTAATAAACTCTTGACGATAACGTACAGGAATAATATCTCCTTCAATGTAATGTTG

TTTACTAACTTTAACATCTCCTGGAGCTACTCGGAAATCATCTTCCTCATCTGCTGTTAG

ATAGACAGGACTATTGTGATAAATCACTTGACCTTGGTTTACCGGATAAAATGGGGTTTC
TATGAAACCAAAAACATTGACCCGCGCACAGGTTGCTAGTGAACCAATTAAACCTGCGTT
CGGACCTTCCGGAGTTTCAATTGGGCAAATTCTACCATAGTGACTAGGATGTAAATCACG
AACTGCAAAACCTGCCCTGTCCTTATTAAAACCTCCTGGACCTAAGGCACTAATTCTTCT
CTTGTGAGTTAGTTCTGCAACTGGATTAGTTTGGTCCATAAATTGAGAAAGCTGACTAGA
TCCAAAGAATTCTCTCACTGACGCAATTAATGGCTTGGGATTAATTAGATTTGATAAACT
CAATGAATCTATATCACATATCATCATTCTCTCTCTAATAATGCGTTCTAAACGATTGAG
ACCTACTCTGAATTGGTTCTGCAGCAATTCTCCAACTGAACGAACTCTTCTATTACCAAG
ATGATCTATATCATCAAGATTACCTGAGTTTTTATCCTTAATATTAATTAGATAATCAAT
TGAGGATAAGATGTCTTGAGGAGACAATACTCGGAACGTTTTAGGTATATTGAGACCTAG
TTTTTTATTAATTTTATATCTACCAACTTCACCCAAGTCATATCTTTTAGGATCAAAAAA
ACGAGAGTAAAGCATCTGCTTAGCAACAGCAACTGTTGCTGGTTCATTAGGTCTAAGCTT
AGAATAAACTATGAGCAGGGCTTCCTCATCAGTTACTTCTTCAATATTATTTTTTCCAAT
TTCTTTAGCTAATTCTTTAACTGAATATATTTGGGATGCAGAAATAAGAAAAGCATATTT
ACTAAGTCCTTTTTCAATTTCATCTTTATTGAGTCCAATAGCTCGCAGAAAAATATAAGC
ATTTACTTTGTGGGTCTTATCTATTCTTATCCAAATTTCTCCCTTAGGATCAATTTCAAA
TTTTAACCAAGAACCTCTATTAGAAATTAGGCTTGCACTATAAATCTGCTTACCATTTTT
GTCTATCTCTTGTTTATAATAGATTCCTGGACTACGGATTATTTGGTTAATAATAACTCT
TTCCGTCCCAGATACAATAAAAGTTCCTCTATTTGTCATTATTGGCAAGTCTCCAATAAA
AATTAATCTCTTTTTGTATTTTTTATTTTTTATCTGCTTTTCAGCACTGAGCTGTAGATG
GGTTGATGATAAATCTAATGATTTAATATTTTTTTCCTGGTTTTTAGAAGGTAGATCAAT
ATCTTTTCTTGTTAACTTTGCAGGAACATATATTTGAGCACTATAAGTTTTATCCCTATT
TTTTGCCTGTCTGACACTATAACGAGGAAATTTTATCTTGTACTCATTACCAAATAATTG
TAATTCGAGTCGACTAGTAGGGTCTGATATCTTCGGAAAAAATTTCAAGAACTTCCGTCAA
ACCTTCTAATAAAAACCATTTAAAACTGGCCCTCTGAATTTCAACTAAATCTGGAAGAAG
TTTATTTTTTAAGCTTATACGTTGAACCATAGATCTCCTTTATAACAAATTTTGAAATCG
AAGAGTTTAATACTATTAATAGAGTACTGGTAGATAATGCATAAGCATAGTTTGCTACAT
TATTTCTTCTCTAGACATAACTAGAACATCATATTTTATTCAAAAAAAGATTTTTAAAAT
TTTTCTTCACACACACTATCTATAAGTACTTTGCTGCTAAAAGATAGTTCAAGCTATTAA
AAAAAACGAAATTAATACTTGTCTGTCTGTCAATTAATTGACATCAAGATAGACAAGTAT
TATTTATTTAATCAATTCTGTTTTTCTGAGCAAAAGACAATGCTCTCGCAAGTCTAGCTT
TTTTTCTAGCGCCTGTATTAGGGTGAAAGGCCCCTTTTTTTACAGCTTTATCAATCTTAC
TATATACTTGAGAAATACTTGACTGCACATCATTGAAATTATCACTTTCTAAATTATCAA
TATTTAGAAGGCATCTTTTAGTTAAAGTTTTGACAACCGATTTATACTTACGGTTAATGA
GACGATTTCTTTCCGAAGTTTTAATGCGTTTAATCGCAGAAAGGTTCTTAGCCACAGTAA
ATCCAATAAAACTTTTATAAAATATTTTTAGAAACGAAAATACACAAATCTGGGATCTAT

TCAAAAAGGGAATATGCGGCGTATTATAACATTAAAGAAAAACAATAGGCAATAGTTCAC
CAAGAGCTCTTATTATTACACAATAGAGATTGTCATTGCGAAAAACTTTGTAAGATAGTA
TTTAATTAAAATAAACTAACTAGACATATAAATATATATGAAAAAAATTGAAGCTATTAT
TAGACCTTTCAAGCTTAATGAAGTAAAACTTGCTCTAGTCAAGGAAGGTATTGGAGGAAT
GACGGTTATCAAAGTTTCTGGATTTGGGAGACAGAAGGGTCAAACTGAAAGATATAAAGG
ATCCGAATACTCTATTGATATCATTGATAAAATAAAAATTGAAATTATTATTAGCGATGA
TAAAGTAGAGAAAATTGTAGAGACGATTATTAAGGCTTCTAAAACAGGGGAAATTGGAGA
TGGAAAAATATTTATTAGTAGTATTGAAAGAGTAATAAGAATCAGGACTAATGACTTAAA
TTTTGAAGCTTTATAGTATTTTTACTTTTTTAGACTTGATACTTCTTCACTAAACATGAG
ATAATAAGCGTTTATTGTATACTTATAACTAAGGTGTAATACTAATGGCTAAAAGCAAAG
GTGCACGAATCGTAATAACTTTAGAGTGCTCTGATAAAGCTGGAGAGTTTGCTCAAAAAA
GGAAACCTGGCGTTTTTCGATATACAACTACTAAAAATAGACGAAATACACCAAGTAGAA
TTGAATTAAACAAGTTTTGTCCTAATTGTAATCAGCATTGTATCTTCAAAGAAATTAAAT
AGTTATTTATCTTATTAAAATAATCATATTATGGCTATATACAGAAAAAGAATATCTCCA
ATTAAGCCAACAGATGCTGTTGACTACAAAGATATTGACTTGCTAAGAAAATTTATTACA
GAACAAGGCAAAATACTGCCTAAAAGGTCAACTGGATTGACTTCGAAGCAGCAAAAAAAA
CTTACTAAAGCGATTAAACAAGCTAGAATCCTTTCTTTATTACCTTTTTTAAATAAAGAT
TAAATATATAGCCTAACTTCATATTTTTCCATCTAAATATTAAAAATTTCTATAGATTTG
TTTATTAAAATGATCTAACATACTGAATTACTAAGAAGATCTACTCATTACAAGTTTTGT
TCATTTTATAATTAATAATAGCATCATAATTTATTAAGCTAGATTCAGACAAGACATAAA
GGTAACCTCTTTTAATTGATTATATTATATAGTCTATTTAAAAGAGGAAATTTTTTTGCT
TCTAGCAATTCAAGATTCTTTAAGCTTTAACAAGGTATTTAAAACAAAGATTTTTGTTTA
GGTATCAGATCAAATGAATGAATCTTATCTCCCGACTGCCATAATTGAAATTCAGAGATG
AAAATTCCACATTCGTTGCCAGCTTGAATTTCTTCGACATCTTCTCTAACGCGCTTTAAA
GATTCAATTTTTCCTTGATAAATAACTTTTTCTTCTCGAATCACTTTAATCCAAGAGTTT
TTTAATAGCTTATTATTGATCACTCGACATCCTGCTATTTTTCTATTTGCTAACGAGAAT
ACTGTACTTACTTCTGCTTCTCCCACCGGTACTTCTGAATATTCAGGATCAAGTAAGTCT
TCCATCCTTCTTTTAATGTCTTCAATTAAGGCATAAATGATTTGATAGTTTTCTATTAAT
ATATTCGACTTTGCCGATGCTTGCTTTGTTCCTGGAGCAAAGTTGGTATTGAACCCAATG
AGGGTAGAATTTGTGGTTGAAGCTAGTTCAACATCTGTAGCAGTAATTTCTCCAGGCATG
ATTGAGACAACATTGAGTTGAACTTTACTCTGAGGAAATTGAGATAAAGAATCTAGGATT
GCTTCTGTAGAGCCTTGATTATCTGTTTTAATAATTAAAGAAATTTGCTTACTAATATCT
TTAGAATTAGTGTTTTTTAAAGTATCTAATGTAATTCGACTATTAAGTGCTCTTTGTTTC
TGTATAATTGATGAGTCTTTAGAGGTATTTTCAATCGCTTTAAGTTTGGCTTCTTTATCA
CTTTTAACTGCTAAAGCTATTTCTCCTGTAGCTGGTACAGATGACAATCCCCAAATTTCA
ACAACAGATGATGGTATTGCTAAATTAATTTTCTCTTTAGCGTTGTTAATTATAACTCGA

ATCTTTGCGTAAGCTGATCCAATAACCAAATTATCACTAATATTCAATGTACCATTTTGT
ATTAATAGAGTTGCTACTGGACCATGCGATTTATCTAAATGGGCTTCTATTATAATACCT
TGTGCAGGCTGTGTTGGATCAGCTTTCAAATCCTCTAATTCAGCTAACAAGGTAATAGTT
TCTAATAATTTATCAACATTTTGACCAGTAAGAGAACTAATAGGAATTATAGGCACTTGG
CCACCAAGTTTCTCCGACATGACATTGTATTTTAATAAATCTTGCTCAATAATATCTGTA
TTAGAACCAGCTTTATCTATCTTAGATATTGCTATGACAAAAGGGACATTCGCTTTTTGA
ATATGATTAATAGCCTCTATGGTTTGAGGTTTAACTCCGTCATCAGCAGCTATAATAATA
ATAGCGACATCTGTTAAATTAGCTCCCCTAGACCTCATACTTGTAAAAGCTTCATGGCCT
GGAGTATCCAAGAAAACAATTTTTTGTTTATTATCTTTCTTAATATACTCAACTTCATAA
GCTGCAATTGCTTGAGTAATTCCTCCAATTTCTTTGTTAGCATTATTAGATTTTCGAATA
TAATCTAATAGCGTTGTTTTGCCATGATCTACATGTCCCATCACTGTGACAATTGGAGGT
CTTTTAATATAATTGCCAGTTTCATAAAGAGCATTAGAGTGATCTAAATTACTAGAAAGT
CCATTATTATTTTCCTTTATATTTGATTCTACTGCTATACCAAAATTATCAGCCACTGAT
GATATTATTGAGGCGTCTATGGTTTGATTCATAGTAACTGATATTCTTTTAAGAAATAAA
TATTTTATAATATCTGTTTCTTGAACACAAATTAGTTTAGATAATTCTTGAATAGTTAGA
GGATTAGTAATACTAATAGACTCTGGCGGAGAATTAGATTGAATGCTTGCTGAATTTTGA
CTAGGAGTTACTGCTACTTTTTTCTGTCTTATAGATTTTTTATTACTAGTCAGTTTTTTT
ACAACTTCCACTTTTGGCTTAGGTGGACGCATTAAAGAGATAGCTAAATCTCCTGCAGTC
TGCGAAACATTAGAATTAGAGTCTCTAAAGTTGTCATCATCATCGTCAATATGTATTTTA
GTTTTTATTTTTTTGTGCTGTCTATTCTTATTTTTTTTACTATCTACTAAGTCGTGAATT
TTATTAAAATTTTTACTTTTTTTATCTAGTTTAGGTGGAGAACTTAATTCTAGATGCTGT
TCACCACCTGTATGTGATTCTGATTTATCAAGGTCTAAATTTAACAGATTATCATTATTG
ACTGACTCTAATCTAATTTTATAGATAATCTGAGGATTTTTCAAATCTACAATTGTTTCA
GAAGAATTGATATTTGAGCTAGATCTAGAGGATCTATTTTCAAAGTTTTGATTATTTAAA
AACATAAATATTGGTTTTATTATCTAATAGTAACTTTCTAATTTTAATACAGAGCTTAAT
ATAAACAAGAATCGAATATCTGTATAAATAATACAAATGTAGAGAATTAATCTATGTTGT
ATTAGATTTAATATAACACTAAAGTCCTTTGTTCGTTATACAAGTTTAAAACTTTTTATG
ATGATAAACACTAGATTCTAGTTTATTGGTATACAATCTATTTATCTAAATAATACTGTT
TTTGTCAGGGCTTTTAAATTATCTAATGTTATGTTCATTATGAAATTCTATTAAAATTTA
AAAACTCAACAATTAAGTAAGAATCGTTAATCACCTCCACGCCACTTATGCCACGATCTC
AAAAAAATGATAATTTTATTGACAAAACTTTTACAGTATTAGCTGATATTGTATTAAAAA
TACTTCCTACAAGTAAAGAGGAAAAAGAAGCTTTCTCTTACTATCGAGATGGAATGTCTG
CACAATCTGAAGGTGAGTATGCGGAAGCTTTAGAAAATTATTATGAGGCTTTAAAACTAG
AGGAAGATCCATATGATAGAAGCTATATTTTATATAATATAGGGCTTATTTATGCTAGTA
ATGGTGAGTATGTAAAAGCTTTAGAATACTACCATCAAGGATTAGAACTAAATTTTAAAT
TACCTCAGGCCCCTTAATAATATAGCCGTGATATACCATTACCAAGGAGTTCAAGCAATTG

AGGATAAAGATACAGAGCTATCTAAATTAATGTTTGATAAGGCGGCTCAATATTGGCAGC

AGGCTATTAAGTTAGCTCCTGATAATTATATTGAAGCTCAAAATTGGCTGAAAACGACAG

GACGAATGAGGAATATACAAGGATATTAATATATTTAAGATATAATAAAGATATCAATTA

ATAAATTAAAATTAGTTAAAGAAACTGGTACTATATAGAGTACTTTACTATATATTATAA

TATTATACTAGTTGA------TCTCTAAGACTTAGCTGAATAAACGAAATTAAATCTATT

GACTATATTCATGAATACAAATAAGCATAATGGTTAGTACAACAAAAAGTACCGGATCTG

TTACTCAAATTATTGGACCAGTTTTAGATATTGCATTTCCTAACGGACAGCTTCCGAAAG

TATTCAACGCACTCAAAGTACAAAGCTCAGAAGGAACTATTACTTGTGAAGTACAACAAC

TTTTGGGTGACAACAAGGTACGAGCTGTTTCTATGAGTTCCACTGAAGGACTACAAAGAG

GGGTAGAAGTTATTGATACTGGATCCCCTATATCTGTTCCTGTAGGTACAGATACTCTTG

GACGTATTTTTAATGTTTTAGGTGAACCTGTAGATAATTTGGGTCCCGTTGATTCTGAGA

GTACTTTACCTATCCATCGACCAGCACCTGCTTTTACTAAGTTAGAGACAAAACCAAATA

TTTTTGAAACAGGTATTAAAGTCGTCGATTTACTTGCTCCTTATAGAAGAGGTGGGAAAA

TTGGTTTATTCGGAGGTGCTGGAGTAGGTAAAACTGTATTAATTATGGAACTAATTAATA

ACATCGCTAAAGCTCATGGTGGAGTATCTGTATTTGGAGGCGTTGGTGAAAGAACAAGAG

AAGGAAACGACCTATATATGGAAATGAAAGAGTCTAAAGTAATTGATGCAGATAATCTAA

AAGAATCTAAAGTAGCATTAGTATATGGTCAAATGAATGAACCTCCTGGAGCACGTATGC

GCGTTGGCTTAACTGCATTGACAATGGCAGAATACTTTAGAGATATTAACAAACAAGACG

TTCTATTGTTTATTGATAATATTTTTCGGTTTGTACAAGCTGGATCAGAAGTATCAGCTC

TACTAGGCCGTATGCCATCTGCTGTGGGTTACCAACCAACTCTAGCAACTGAAATGGGAG

CACTTCAAGAAAGAATTACTTCAACAACAGAAGGATCAATTACATCTATTCAAGCTGTAT

ATGTGCCGGCTGATGATTTAACAGACCCAGCTCCGGCCACTACATTTGCACATTTAGATG

CAACAACAGTACTTTCGAGGAACTTGGCGGCAAAAGGAATTTACCCTGCAGTGGATCCAC

TGGATTCAACATCAACAATGCTACAGCCTGGAATTGTTGGAACTGAGCATTATTCTACTG

CTCAAGAGGTAAAATCAACTTTACAAAGATATAAAGAACTGCAAGATATTATTGCTATTC

TTGGTCTTGATGAACTTTCAGAGGAAGACAGACAAACTGTATCAAGAGCAAGAAAAATTG

AAAGATTTTTATCTCAACCTTTTTTCGTGGCAGAAGTGTTTACTGGATCACCTGGGAAAT

ATGTATCTTTGGAAGATGCAATTAAAGGGTTTCAAATGATCTTAAAAGGCAATTTAGACG

ACTTGCCTGAGCAGGCATTTTATCTAGTAGGTGATATAGATGAAGCTATACAAAAAGCTG

ACAGCATGAAAGATTAATATAATTAATGATGACTTTAAATATAAGAATTATTGCTCCTGA

TCGGACTGTTTGGGATGCAGAAGCACAAGAAATTATTTTACCAAGTAGTACAGGGCAACT

TGGTATTTTAACAGGCCATGCACCTTTGCTTACAGCTTTAGATATTGGAGTTATGAGAGT

AAGAGTAGATAAAGAATGGATGCCGATTGTTTTGCTGGGCGGTTTTGCAGAAATAGAGAA

CAATCAATTAACTATTCTGGTTAATGGCGCAGAAGAAGCTAGTCAAATTGATTTATCAGA

AGCAGAAAAAAATTTAGACACTGCAACTCAACTCTTAAGTGATGCCTCGTCTAATAAAGA

AAAAATAGAAGCAACACAAAAAATACGAAAAGCTCGAGCTCGAGTACAAGCTGCAACAGC

AGCAACTTCGTAAAAGATTCCTCTTAAAACATAAAACTTTAGTAATGATTACTAAAGTTT
TATGTTTTATGTTTTATTTGTATGGTTATAGGCAGCTTAAATAAACTTAGCTTAGACCTG
AACAAATATAATCAAAATATAATCCCATCTCTTTACCTGCATCTGGTCCAACTAAACTAA
TAGTAACTTCTTTCATAGCTAAAATAGCTTGAATCGTTGCACCAATAGGAACACCTAGAG
AATTGTATGTTTCTTTTAGACCATTTAATACTCTTTCTTCAAGGATAGATGGATCTCCAG
CTAGCATTCCATAAGTAGCATAGCGTAAGTAGTAGTCTAAATCACGGATGCATGCAGCAT
AACGTCGAGTTGTATACATATTACCACCTGGGCGTGTAATATCCGAATAAAGTAGAGATT
TTGCTACGGATTCTTTAATAATTGTTGCTGCATTAGCTGCAATAGTAGCAGCAGCTCTGA
CTCGTAGCTCACCTGTTTGAAAATAGCCTCTTAATTTTTCAACAGAGCTATCATCTAAAT
ATTTACCTTGGACATCAGCTGCATTAATAACAGAAGTAATTGCGTCTTGCATAACTTTCA
CACATTCCTTAATTTTAAATATCTAATACTAGGGGTATATAACAAAATTACTAGATCTTT
TAGCAATATCAGTATTACTGCATAGCACCCAAAGTATAATCAAAGTAAAAACCAGCTTCT
GCTGAATCTTCACCTGCAAGCAGTGAGCAAGCTACACTTTTCATGCATTTTACACCTTCA
GCAACTCCTGAAATTGGTGTTCCTAAAGAATTATACATTTCTTTAACACCCACTAATCCA
ATTTCTTCAATAGGAGTTATATCGCCAGCAACTATTCCGTAAGTTACTAAGCGAAGATAA
TAGTCCAGATCTCGTAGACAAGTAGCAGTCATCTCTTCGCCATAAGCATTTCCACCAGGA
GAAACTACATCAGGTCTTTTTTGAAACAGTTGTTGGCCGCCTTGCTTTACAATACGCTCA
CGATTATCTGTTAAAATTTGAGCTATTCTTAAACGACGTTGTCCAGATAAAACAAAACTT
TTAATTCTATCTAATTCCCCAGGACTTAAATATCTTGCTTCTGCATCTGCATTTACAATT
GACTTTGTAACAATACTCATGGATAAAACTCCTGTAATGCTTT-AAAAAAAAGCTTAATA
CTAATTATTTATGATATAACTCAGAAAGTAATGTACTTTTTGAGAATTTTATAACCGAAA
CACGTTCATGATATATGAAGCAGGATAGATACTGAGGTAAGTACTAAAACGAGAAAGTTA
TATGTAGGAACTTCTGCTATTGATTTCCTAGTACTTTTTTAAAACTTGGTACCACAATTG
ATATATCCTGCTTAGTCAAACGATTATAGAGTGTTTCTGTATTAGGAAAGTTAGCTGCTG
GTAAAGTCGGAAATCTTCTGTATGGAACAGTATCTTTGCCATATACCGCATTGTATTCAA
CACTTTCAACTAAATTACTTATAAATGCAGATAAGCCTTTTGAAGCTAAAATTTGATTAA
AGAATCTAATTTCTGCCTGATTATTAGGAGCTCTTCCTAAGATATGCTTAGTACCTAATT
CTATAACTTTTGTATTTGGATATGGCTGATAAAATTCTTTACCATATAGTTCAGATAGCG
CCAGTTTTTCTACTAGTTCTTTGACACAGATTTGTCTATTCAAGAAAGCTGACTTAATAT
CCAAGAATTCACCGCCGACACTAAAAGAGTTAAGATCTCTTTCGAAAATTTGACGATATG
CTGCTCTTAAAGCTTGTTCTAACATTTCTTTGTTACTATCACTATTGACTTCAAAGACAA
TAGATTGATCTCTTAAAGAAGTTACTCCCTGTGAAATACGAGACTGAATATCATTACTAC
TCCTCATTTCTTTGACAGTGCCTAATTCAACAAACCTAGCAGATTTACTAGCAATAACTT
TTTTGAATCTCTGGTCAATAATTCCTGGTCTTAAGCTTCTTAAAGCAACACCAGCTGGAG
TACTATATCTTTCATAAGGAACAGTATTATCTCCAAAAGTTTCTGTGTACTCCGAGCTAT
CAATTATTGTGTCTATTACTTGGTAATAGCCTTGCTTATATGCAATATCAAAATATTTAT

TAATTTCTTGCCGACCATATGTAGGGCGACCCAGTAAGCGGTTATGAATATATTCAATAG
CTTTACAAATATATAATGGCTCCCAATAAAGAGATCTAAATATACTTGATTTTGCTAACT
GTCTAACAAACTCGCGAACTGTAATTTGATTATCTTTTAGCTGACTTTCTATTGGCTTGA
GAATTAGTTTTTCTTCTTGATACACTTCTCTGCCAAAAACTCTTAAATAAGCAACTTTTG
TGACAACTTCTACTGAATTTTCAAAATTTTGGGATGAATCTGACTTTATAAGAGCGTTGG
ACAATTTAAAGATTTTTGGTCCCAGGGATCCTGCTGACTTAGGTCTTACCTGTGGATTAC
TAATCTGATTGTAGATTCCTGGCCCACGTCTAACTAATATCCTTCTTGTATCTTTACCAA
AAAGAGCTTGTCTTTTTCTTGGATCTTTATTTTCTTTAGGGAAAATAGCTCCAAATTGAA
TTGATAATGGATCATTACCTATTCCATATGGATGTTGATCAGGTAATGATTGCTTATAAT
CACTAAATAAAGTTATAAATTGAGGAACCTTACGAAAAGGCGCACTATAGGTGAAAAGAT
CTATTTGAGGCCCCCAATTACGGCACTCTTGAGGCTCTTCACCTAAATTTCTAAAGTAAG
GCACAGTCTCTTCACCGAAATAATCTGTATACTCAGAAGAATTTAAAAGTGCATTCACTA
AACCACTTAAACCAGTAGATGATAAAATAGCAAAATATTTTTGAAACTCTTCTAAAGAGC
TAGGACCTCTACCCAAGAAATGTCTAAATGCTAATTCTAATGCTCGACTATTGACAAAAG
GTTCATAAAATTGCTTTCTATAAATACTTGAAGTTCCTAACGAACGAATGAATTCCTTAA
TTGAAATTTGGCCATTTTTTACTTGCGACTCCAAATTCGATAAAGAGAGGTCATATGCTT
TAGCAATATCTCTTTCAAAAATTTGTCTATAACATGCTTTCACAACAATATTTTTCTCAT
CAGCAGATAAACTAGGCTTCATAACAAAACGAGGTGTTGACACTCCTGCCTGTACATAAG
TTTGAGGAAGACGCAAACCTTGTAAATCTCCAGATATTCTTTTTCTTAATTTATCAGTCA
AAGATGGAGCTTCAAATTCAGAAATCACAACATTAAAATACTCTTTAACTAAATCTTGAC
CTTTTATATCTTCCTCAAAGATTAATAATGCTGTCCGTCTCATCTCTCTAAGAGCCACAA
TTGCTGCTGCACTTGAGCAAGCGTTATCAATTAACTCTCTCAATCCTCGTATATTAACAG
ATAAAATATTAGGGTCGCCTGATACAATTGCATATGTCAAATACCGTAAAAACCAGTCTA
AATCTCGAAGAGATTTTCTCATTCTTGTAGTACCGTATCTCAAGACGTTTATAGGTTTAA
ACCCTGGCGGAGTAGCTCCCCCAGCATTAAATAAAGATCGGAAACTTTGGCCAAAATCTC
CTTGGATATTTCCTGATAACTCATTAATTTTATCTTGAGAACTTTGATCTCCTGCAATAA
TAACAGCTGCTTGCGGACGTTCTAAATAAGAAATTGCAGACCCACCTACGAATATTTTAT
CTGCTGCTCGTGCTACCAGAATATTTGCATTTTTAGTTAAAATATCTGCAACTTCCAGTC
TTTTTTGTCCAGAATTTAAAAAAGAAACTAATTGATTTAATTCACCAAGCTGTAAAAATC
GGTCTTGTTGCTCTGCCTGAGTAATAGTTAAAATTGAGGCAGTCCGATAAAGCTGGGGGC
GTGCTAATGGGCTTCCGCCACTTGCTTTGATACTCATTACTTTATTTATCTCCTTAACTG
ATATTGCTTGCTAGCGAATACAAATATAGCTTGATATTTCTAAATATTTAAAGATACTAT
GCAAGCACACTGTTTTCTATATTATTAGGTACTAGTACTTATAGTACAGAGAGTCTTATA
ACTAACTTCTATAAATACAAATTTTGTAATACTTCTATTAACATTTGTCAAAAAAATATT
TAAGTGCTCACTCTTTTATATTAAAATTCAATTAGTGAGAGATTGATGTGAAATAAAGTA
TACTAAGTAAGTTTTATAGATAGGAAAATTAAAAGATTTATTCTACAAACATTAATCACT

TGAATGTTTAGCAATTGTACAATTGTTATTGCTTTTAATTGGTTTAGACTAGTTTTATAT

TTTTTCTAACATAAACAATATTAATGAGCAATTTGTATGAATTTAAAGCCACAAAACAAC

CTATTAACTAACGAAAATATAAATCATGTAGATATACCCGAAAATGATATACCAATGTCA

ATTACGGAACATTTAGAAGAATTAAGACAACGGACTTTATTTGTGTTTTTATTCTTTTTG

TTTGCTACAACTATAAGTTTCACACAAATTAAGATCATTGTTGCAATACTGCAAGCTCCT

GCTGTTGGTATTAAGTTTTTACAACTGGCACCAGGAGAATATTTTTTTTCATCTATTAAG

GTTGCAATATATTGCGGGATTGTTGCAACAACTCCGTTTGCCGTTTATCAAGTTATATTA

TACATACTCCCAGGACTAACTGGAAAAGAAAGAAAAATTATTTTGCCTCTATTAATTAGT

TCTGTATTGCTTTTTATTACAGGTGGTATATTTGCTTACTTTGTTCTCGCACCAGCAGCT

TTAACATTTTTAATTAGCTATGGGTCTGATATTGTAGAACCATTATGGTCTTTCGAACAA

TACTTTGACTTTATTCTATTACTTTTACTTAGTACAGGATTAGCATTTGAGATACCAATC

ATACAATTATTGCTCGGTGTTTCAGGGACATTTTCTTCTAGTCAAATGATACGAGCTTGG

AGATATATTATTATTATAGCAACAATTGCTGGAGCTATTCTGACCCCTTCGACTGATCCC

GTTACACAATTAATAATGTCTTCAGCTGTTTTATTACTTTATTTTGGTGGAATTGTTATA

TTATTAGTCTTAAAAAAGTAGAGTAAATATTTTCTATTAATCCAATATATTTGTTATCTG

GGCAAATAATCTATAAAAACAATTGCTAGATTTAAGTATCAGGCTTCTAAATATTTCAAG

TCTCTTTTTATCATTTAAATACAATCGATACATTGGCTTGTTTTTTCATCAATTATTTGT

GTTGCACACTTTTTAGCACTCTCTTTATTGTTCTTAGAATTATTTCTTTAATCTATTTAT

TAAATAGATATGAAGAGATTTAAGTATTGATTATGGTAATCTATATCAACGTAAAAAGTA

TTTTATTGTGAGTTTTATTTATATAAAGGCTTATTCATTATATAGTTATAACAGTAAGTC

TAATTAATAGTTTTCATGCTTAATTAAAGTAATAACTCTTAATTCTTTTGGCAGAAAAAT

CTGTCTTAATATTTTATCAAAAAAACAGAAAAATGAAGCTAAATAGTCTAATTAACTTAA

TTCAAAAGTCTATATATTCTTGTACACTTTTACTAATTATTTTAAATATTATTTGTGTCG

CACCTAATTCTAGTAATGCATTTCCAATTTATGCGCAACAAGCTTATGAAAGTCCAAGAG

AAGCAACTGGTAGGATAGTATGTGCCAATTGTCATCTTGCTCAAAAGCCTGTGGAGATAG

AAGCGCCTCAAGCAGTACTACCTAATACTGTTTTCGAGACTGTTGTGAAGATTCCATATG

ACAGCAATGCTAAACAGATTTTAGGTAATGGCAGTAAAGGAGGCTTAAATGTTGGAGCCG

TTGTAATATTACCTGAAGGATTTAAGTTAGCTCCTGTTAATAGATTATCTACAGAGTTAA

AAGAAAAGACTAGAAATCTTTACATTCAGCCGTACAGTGCTAAACAAGACAACATTTTAG

TAATTGGACCTATTTCTGGTGATAAAAATAGAGAAATAGTTTTCCCAATACTATCTCCCG

ATCCTGCAAAAGATAAAAAAGCTCATTTTTTCAAGTATCCAATATATGTTGGCGGAAATA

GAGGACGAGGCCAGATTTATCCAACGGGTGACAAAAGCAACAATAATATTGTCTCTGCTT

TAAGCAGTGGTAAAATTAATAAAATTGAATTACTAGACAAAGGTGGATTTATAATACATG

TGACTAACAGTAGCAATGTAGAGTCAACACAGAAGATTTCACCTGGTCTCGAACTTAGAG

TAAAAGAAGGGGATACAATTCAGCTTGATCAAGCTTTGAATAGTGATCCAAATGTAGGTG

GTTTTGGTCAGAACGAAACAGAAATAGTTTTACAGAGTCCAAATAGAATTAAGGGCATGA

TTGTTTTCTTTTTCGCTAGTGTTCTAGCTCAAATTTTCTTCGTATTAAAGAAAAAACAAT
TTGAAAAAGTTCAAGCAGCTGAAATGAATTTTTAAAGCAATATAAAACTGGCACTTACTA
TAAAGAAAGTATAAACACTTTATTCACAGGATGTACCAGTTTTTATGTTATGTAATAATC
GCTTTGACGGAAGTTATGATACTTTTAACAGAATCTACAATCTGACTAGGCTGAATTACT
GTAGCCTGCTCTAAGCTTCCATTATATGGTGTGGGTATATCCTGCGAAGATAGCCTCACG
ACAGGAGCATCTAATTCATCAAAAAGGTACTCATTAATTTGTGCAATTAGCTCTGCTCCA
ATTCCGGCTGTTTTCATACACTCTTCTACAATTAAAACTTTGTGAGTTTTCTTTACTGAG
ATAGATATAGAGTCTATATCCAGAGGTTTTAAAGATATAAGATCTATAACTTCTGGATCA
TAACCTTCTTTTAATAAGGCCGGTAATGCTTGGATAACATGATGCCTCATTCTAGAATAG
GTTAAAATCGTAATATCTTTTCCTTTTCGTACAAATTCAACTTTATTGAGAGGTAAGAAA
TATTCTTCTTGAGGAATCTCTTCTTGTAAATTATAAAGTAGAACGTGCTCAAAGAATACA
ACTGGATTATTGTCTCGAATTGCAGATTTCAGTAATCCTTTTGCATTATAAGGAGTAGAA
CAAGCAACTATTTTTAAGCCGGGGATGGCTTGGAAATAAGCTTCCAGTCTCTGAGAATGC
TCTGCACCTAACTGCCTACCAACTCCTCCCGGTCCTCTAATAACTAAAGGCAATGTAAAA
TTACCTCCAGAAGTATAACGTAACATTCCTGCATTATTAGAAATTTGATTGAATGCTAAT
AATAAAAAACTCATGTTCATACCTTCAACAATTGGTCTTAGCCCTGTTATAGCTGCACCA
ATTGCCATACCAGTAAAGCTATTTTCTGCTATTGGTGTATCAAGAACTCTTAAATCCCCA
TATTTGCTATGCAAATCTTTAGTTACCTTATAAGATCCACCATAGTGACCAACATCTTCT
CCTATTACGCAGACAGTTGGATCTTTTGCCATTTCCTCATCTGTTGCCGCTCGTAAAGCG
TCAAACATAAAGATTTTACTCATTTTAATATTTGATTTTTATGATTTGACCACTGTTTTA
AAGTAACAGTTAAGACACCATATTCTATTTAGTTATCTGCAAAAAGATATCGTTTCAATT
CTGACATGTTAGGTTCTGGACTGGAAATAGCAAATTTCACAGCTTGTTCCAATTCGGTTT
TAACAGCATTCTGAATCTCATTAAGTTCACCTATATTAGCAATTTCATTATCTAGAATAT
ATTTTTTGAGTTTTTTGATAGGATCTCTTGCCACCCAAGCCTCTTTTTCTTGTCTTGATC
TTAGTTCATCAGGATCTGCGAGAGAGTGACCACGAAATCTATATGTTAATGCTTCTATTA
AGGTTGGACCATCACCTTGGCGAGCTCTTTGAACTGCTTGTTTTGCAGCTTGTCTTACAG
CTAGCACATCCATTCCATCAACTTCAATCCCAGGAAGCCCAAAAGCTTCTGCTTTTTTAT
GTATTTCAGGTATTGAAGAAGACCGGTGATGTGCCATACCTATAGCCCACTGATTATTTT
CAACAACAAATATAATAGGTAGTTTCCAGAGAACTGCCATATTCAGACATTCAAAAAATT
GCCCATTATTGGTAGTCCCATCACCAAAAAAGCAAGCCGTGACTCTTAAATCCTCTGTTT
CTTTAAGTACTTGCTGGCGGTAGATACTTTGAAAGGCTGCCCCTGTTGCAACCGGTATAC
CTTCCGCAATAAAAGCAAAGCCACCTAAAAAATTGTGAGGCGCAGAAAAAATATGCATCG
AACCTCCTCTGCCTTTACTACAACCAGTCTCTTTTCCAAATAACTCAGCCATCACATTTT
TAGATGGGACGCCTTTACTTAAAGCATGTACATGGTCTCGATAGGTACTGCAAACATAAT
CAGTTGGATTCAGAAGTTTAATTACACCTGTAGAAACAGCTTCTTGACCATTATAAAGAT
GAACAAAACCAAACATCTTTCCTTTATAATACATCTGAGCACACATATCTTCAAAATTTC

TGCCTAACAACATGTCTTCATATAAAACTAATAAATTACTCTTATTAAGATTAAGACCTG
TTGAATTATAGTTAGTCAGCGGCAATTGAACTTTCTTAGGATAACTCATAATTTGTAAAG
AAACCTCTTATTGTAAATAGGAATTTTGAAGAAACAGCTATCAATAGAAAACACACAGAT
AAAAACTTATAAATTATAATATGTTTTTTACTAAAGACTTTAATAATTATTTGAAACTAG
TTCAGTATATCAAATTAATTTTGCTTATACTACGGATGTCTAAAACTTATATAAAAAAGT
AGTAAAAAAGTTTATTAAAAGAATAGTTAAAAACAGATATAAATAGAAGTAAACAGAATT
ATTGAAAAAAAATGATTATTATTAATTAATACTTAATATTTCTAAATATTTTACAATTAA
TGGCTATGTCACCATCTTTATTTTATCCTGTTGAACAAGAACTATGTAGTCTTGAAAAAA
ATCTGAAAGCTGTTGCTGGGACTCGTCATCCAATTTTATATGCCGCAGCAAAGCATCTAT
TCGATGCTGGAGGAAAACGAGTTAGACCAGCTCTTGTATTTTTAGTGGCTAAAGCAACCT
CTGAGAAGCAAGATATAAATACTGGACAAAAAAGGCTAGCAGAAATTACTGAAATTATAC
ATACTGCTAGTTTGGTACATGATGATATTATTGACGAGTGCACAACACGTAGAGGAGTCA
AAACTGTACATAATTTATTCAATACTAAGATTGCTGTGCTAGCAGGAGATTTTTTATTTG
CACAGTCTTCTTGGTATTTAGCTAATATTGAAAATTTAGCCGTAGTCAAAGCTATTTCTA
AAGTCATCACCGACTTTGCAGAAGGAGAAATTAGGCAAGGCTTAGTTCATTTCGATCCCA
GTATTTCAATAGATGCTTACATTGAGAAATCATTTTACAAGACTGCTTCACTCATTGCTG
CTAGTTGTCGGGGTGCAGCTATGCTTAATGGTTCCAATCATCAAATAAATAATGATCTTT
ATCTTTACGGTAAACATATGGGATTAGCATTTCAAATTATGGACGATGTTCTAGATATAA
CTGGTTCTACTAAGAGCTTAGGAAAACCTGCTGGCGCTGATCTAATAAATGGAAATTTGA
CCTCTCCTCTCCTTTTTTCACTTACTCAAGAAGCAAGTTTAAATGATCTTATTGATAGGG
AGTTCTGTAATAGTACAGATATAGCCTCAACATTATTTCTTATAAAAAGAAGCGGGGGAA
TTACAAAAGCTAAAGATTTAGCTAAAGAACAGGTGCAGGCGGCACTTTCCTGCCTTCAGT
TTTTACCACAATCTACACCTGTATCTAGTTTAAAAGAATTAACACATTTCATAATCACAA
GATTGTCATAAAGGACTTGCTAAAATTAAAATGTTATTAACTAATTTTTTACTGTTTCTA
AAACAGTCTGAAAAGTTTGAATATCATTACTAGCTAGCTGCGCTAGCATTTTACGATTTA
ACGCAATATTCTCTTTTTTTAATGCACTAATAAAAGTACTATAATTCATGCCTTGATTGT
GAGCAGCAGCATTTATTCTAGTTATCCAAAGACGACGAAAATCTCTCTTTTTCCTTTTTC
GACCAACATAAGAATATCGAAGAGCTTTAAGGACTTGCTGCTTAGCTGTTCGAAATAAAC
ATTTATGTGCACCTTTAAAGCCTTTAGCTAGCTTAAAGATTTTAGCTCGTCTTTTTTTTG
CAACGTTACCTCTTTTAACTCTACTCATAAAATATTACTTAAATTTTAATTATTACCAGC
TTACTAAATTCTTATACTCTTTGATTATAAATAAGGTAATTTCATTGCAATGTTTTTAAT
ATCTTTTAAGTCAACCGAACAAGTAGAAGAAAGATGTCTTCTTTGTTTTGATGACTTTTT
TTGTAATAAATGACTTTTAGAAGCTTTATGTCGAAGAAATTTTCCGGATGAAGAGACTTT
AAATCTTTTTGCTATTGCTTTTGATGTTTTTAACTTAGGCATATTATAAACTTTTAATGA
TTAACAAAAGCATGAAAAGTCTGAGGATATCAATCTTAGACCTTTCAATACTTAGAATAA
TCTTAACACAATTTATTTTACTTACTACATAGGTTAACTGAAACAATTTATATATATAAT

ACTAATCAGTTAATCAACCATAGCTAAAGTGGATTTATATTTTGCTTTGAACTTTTGTAT
AGTATTAGCAATTAGCATTGTTATAATTTTAAGACTGCTTGAATTCAATTCTTCGAACAT
TTTCATATTTAATGTAAATGAAATATTGGCTTCTGAGACAATATTTTGAATTTGAACATC
AGACAAGGGAATCATATCTAATGCAGCTCTATATTGATCTTTAAATAATTTATCGTCTTT
TATTTGGTCAAAATCGTAAAATTTCGTTCCTCCTGAATCTGATAAATTCATAGCCCCTCT
AGCTATTTTTTTAAAATTTGGCCACCAGAAAGGTCACCCAAATAGCGCGTATAAGCATG
AGCAACTAGTAATTCAGGTTGTTTATGTCCTATAGTGTGAATTCTATCAACATAAATTTT
TGTAGCAGGAGATGGTTCAATAAAATCTAACCAATCTGATCCATAATAATAGTTTAAATC
TTCAGATAGACTCGCTTTCCTATTAAGCTCTGTAAAATATATAGGTTTAATAGCTGGATG
ATTTTTATTAGAAAATAACTCTTCCTCTATTGCACAATAGACAAAGTATAGATTTGCAAC
CAATTTACGATATGACTTCTTATCTACAACTCCTCCCAAAAAAGATTTAACAAAACTAAC
ATTTTCTGCCATACTATGGGACTTAGTAGTACCTTCTCTTAGTTCATTCGCTAAAGTATT
AACCATAATATCAATATTCCATTTTTAAAGATAACAAATTAACAATAAATAGTTATCCGT
ATATTTATAATTGAGTTGCGCTAACTAGGCTGCTGCAAAATAATTTTTTGATTTTATTGG
ATCTGGATTCATTGTTTTGTCACCAGGCTTCCAATTAGCTGGACAAACTTCATCTGGATG
AGATTGTACATATTGAATTGCTTGTAAAACCCTCAAAGTTTCTTCTACACTCCTGCCAAA
CTCTAGATTATTGATTGTAGAGTACTGAATAATTCCTTTCGGGTCTATAATAAATAGTCC
TCTTAGAGCTACACCATCACTATTCAAAACATTATAAGCTGCACTGATCTCTTTTTTTAA
GTCTGATACTAATGGATATGAAAGATCTCCTAATCCACCTGATTCTCTGTCAGTCTGTAA
CCAAGCAAGATGAGAATACTCACTATCAACCGAGACTCCAAGAACTTCTGTATTAAGTTC
CGAAAAAGCATTATACTTATCGCTAAATGCTGTAATTTCTGTAGGGCACACAAATGTAAA
ATCTAAAGGATAAAAGAATAAGACAATATACTTATTCTTGAGATCAGATAATTTTAACGT
TTTAAATTCTTGGTCATAAACAGCTGTAGCTGAAAAATCAGGAGCTAGCTGACCTACTCG
AAGACAATTTGGTCCAGAAATCATTAATTTTTCCCAGAAGAATTGATATAATTTTGTTAT
TTAATAAATAATTATATAACAAAATTTAGTATATATTAAAATACTACCAGTCTTTTTTAA
ATTACTAAATGATATAATCAATACTATCTTTATAGCGGGGAATGGATTTGAACCATTGAC
CTTCGGGTTATGAGCCCGACGAGCTACCAGACTGCTCTACCCCGCGGATAACTAAAATCA
TATTAGTCTACATATATATATAAAAGCAAATGCATAATAGATTTTAGAAGTAAAAAAATA
GAAGAACTGGTCCACTGATCATTAAACTTAGAAACGACAAAGTATTTTTTATCTTATTTA
AAAGAGGAAAGACTTCATATAAACCAATAAATCTATCTTTTAAAAGAATTTCGGACGGCT
TTACCCAAATTTGACCGTCATACCAGCCTGATTCTTCATAAAATACTGTAGCACTCATTA
ATCGTTTTACTACGTATGACCAGCCTAAATATAATCTGATTAAAATAAATCCTGTCATTA
GGCTGGAAGTAATAAATTCTGAAAAGAAAAATTTCAAAGGTAGTTTAGTGATTGGAAAAA
TTGATAATAATATAGGACTAACTAATAAACAGTTAAGTAGTAGTGCAATAGTTATTTTTT
TGTTATAAGATCTATGGCTTAAAGTTGGCCAGCAAAAGAACCAAGAATTTTTCAAAGAAG
TATATTCATGAACAGGTTGCTGCTCCTTTGGCACAGGACATTGAGTATTATATAAATTCA

TTGACATTAAAAAGTATTATAATAAAGACTTTGTTTATATTATCTAACATAGGTACGACA
TTAAATATTAACTAACTGCAATTATAGTAGTGAGTAGAAGAAATAAAACTAAAAATAGCC
AAGTAATTTTATTTAAAGTATTCTCAGTACTACGAGTATTACTAAAGAACTGATTCTGAG
CACCTACACTTCCTAAACCTTCGGATTTAGGATTGTGTATTAGAATAGTGAAGATTAATA
TTATTGTAGATGAGTACCAAAAAAATTTTAAAATTTGTTCCATTATGTAAAAGTTAATAA
AAATAGCACAAAGACAATCGTCTAAAAGATATTGATTGTCTTATCATATTAAAAAGTATT
TATTCACCCTGTACTTTTAAAAGTAAAAATCCTATTGCCAGCCCTACCAAAACTAGGACT
GAACTTAATACAGCACTTTCTACAATTTCTCCGACCATTTTATTTTTCCTTATTATTTTA
TTATTGTCATTCTTTTGACAAGATTATATTGAGCTAACTAAAATCCGTTTCTACCCCAAA
CTACCATTGCAATTGAAAATGTAAACATTGCCATTAGAGCTGCCCACCCCAAACTTAAAA
TATCCATAATTTATAATATTCCTAAAAATAAGATCTCTTTTACTTAAACTATAATATTCT
TAATATTAAAAATATCATTATATAATAAGCTAATTATAACAGAGTACTACTTAATTCGTA
TTTTTAGTGTATTTAAAGAAGATTGAATTTTTTCGAGGTACTGGATAAGTATTTCATCCT
AAGAACCATATTCAACTTTTGATTTATTATAATGCTTAAATCTAAATATATATCACTCAA
GAATCATTAGTAATTATGATGTTTTTAATTATACGATTAACATATTTCAAAATAGAAGAA
TTCATAGCGACTAAGTTAATATGCTAATATAATTTATCAATAAAAAGGAAGTTAGTATTT
CAATTATGCAAACAACTATAAATAATGGGCAAACGTCTAGTAAAGAAACTTTACTAACAC
CTAGATTCTACACGACTGACTTTGAAGAGATGGCTAATATGGATATTTCCGGCAATCAAG
AAGATTTTTTGGCTATCCTCGAAGAATTTCGAGCTGACTATAATAGTGAACATTTTATTA
GAGATGAAGAGTTTAATCAATCTTGGTCTAATTTAGAACATAAAACTAAATCCTTATTTA
TTGAGTTTTTAGAAAGATCTTGTACCGCAGAATTTTCAGGTTTTTTACTATATAAGGAAT
TATCCAGAAAATTAAAAGACAGAAATCCGGTTATAGCTGAGTGCTTTTTATTAATGTCTA
GAGATGAAGCTAGGCATGCCGGTTTTTTAAATAAAGCTATCGGGGACTTCAATTTATCTT
TAGATTTAGGATTTTTAACGAAGAGTCGCAAGTATACTTTTTTCTCACCTAAATTTATTT
TTTATGCAACCTATCTTTCTGAAAAAATTGGATACTGGAGATACATAACTATTTACCGCC
ATCTTGAACAACATCCAGAACACCGTATTTATCCAATCTTTAGATTTTTTGAAAATTGGT
GTCAGGATGAAAATCGTCACGGAGATTTTTTTGCTGCTTTGCTCAAATCCCAACCTCATT
TTTTAAATGACTGGAAAGCAAAAATGTGGTGCAGATTTTTTCTATTAAGTGTATTTGCAA
CAATGTACTTAAACGACTTTCAAAGAATTGATTTTTATAATGCCATAGGCCTAGACTCTA
GACAGTATGATATGCAAGTAATACGAAAAACTAACGAAAGTGCCGCTAGAGTTTTCCCGG
TTGCTTTAGACGTGGACAATCCAAAATTTTTCAAATATTTAGATACTTGTGCATGTGATA
ATAGAGCTCTGATCGATATTGATAATAACAATTCTCCATTATTCATTAAATCTATCGTAA
AGATACCTCTATATTTTTCCTTATTTGCAAATTTACTAAAGATATATTTGATTAAGCCAA
TAGACTCAAAAACAGTATGGAATACAGTTCGATAGTTAAATACAAGGGAATGCTTTTTTC
TTATAAAAAGCTATCTAGTGATTAATATAAAATAAAAGCTTCCTTCTCGATGGAAGGAGC
TTTTATTTTGTCAAGAATATAATGGGATTATATAGATCTAGAACTAAGAACTTACTTTTT

CTTTTGCTTCATACATAACCTCTAAAGTAATTAGATTTTTAGAACAATCACGGGCAAATT
TCTCTGTATTTCTTTTTACTTTACCTCTAACAAACCCAGGAATTTTATTTAATTCTTTCT
GTGCTTCTTCTGACCAATTAATACTATCAACAGCAGAAATTCCTATACTTAATGCTTCTG
TTGTATCATGACCACCAAATATTTCTAGCAAATGATCTTCCATACCAAGTGTAAACGAAT
TATATACTAAATCAGCAATTTGATTTGTTCCCTCATAACCAAGAAATGGACGGTAACTTA
AAGGAAAGTTTTGTATATGCACTGGTGAAGAAATAACGCCACAAGGAATATTTAGACGCT
TACCAATATGTCTTTCCATCTGTGTTCCAAAAATTGCTGCAGGTTCAGTTTTAGCAATTA
AATCTCCAATTAAACCATGATCATCTGAAACGATTACTTCATCGCAGAATTCTTGAACTT
GATCTTTAAACCACTCTTCATCATATTTGCAATAAGTTCCACACCAGGCGACATGTATAC
CCATTTCCTGGTGTAAAATTCGAGTTATTGCTGCAGCATGAGTTGCATCTCCAAATACAA
TAGCCTTTTTACCTGTTAAATTCTGACAATCAATTGATCTTGAAAACCATGCAGATTGTG
AAATAAATCGGGTTTGCTCGTCTATGTATTTTTCGTAGTCTACAGCAGCTCCCAATGCAT
TTACTAATTGCTGTATTGATCGGATGCATGCAGCTGTTTGAACAATACCCATAGGTGTAA
TGTCAACATAAGGCATATTAAATTCTTTCTCTAAATATCGAGCTGTCATTAGTCCAGTTT
CTCTATATGGAATAAAATTAAACCATGCAGAGGGTAATTTTTTTAAATCTTGAACTGATG
CATTTTCTGGAATAATTTGGTTTATTTGGATATCTAGATCTTGAAATAACCTTTTTAATT
CTGCTATATCATGTTGATTATGAAATCCTAAGCTTACTGCTCCAATAATATTAACAGATG
GAGTTTTAGTCTTTTGAGTTAAAACTTGATTATTAGATTTAGCTTTTTCCATATAGAAAG
TGACAATTTGTTCTAACGTTCTATCACCTGCTTGCAGTTCATTTACTCTATAATGATTAA
CATCAGCTAATAAAACATCTGCTTCTGTTTCTATAGATGCTCTGCTAACAAAGTTTTGCA
AATCTTCTTGCAAAATACTCGAAGTACAAGTTGGTGTTAGAATAACTAAATCAGGGCTTT
CTTCTCTGTCTTTTCTAGTAATATTTTCTACAACTTTTTCTTGAGAACCACGGGCTAACA
CATGTCTATCTACAACACTTGCTGTAACAGGAGTAAAATCTCTGTCTCTTTCAAGCATTG
AACGCATTACATTAAAATAATCGTCTCCAAGAGGCGCATGCATAATTGCATGAACTTTTT
TAAAAGAACTCGCAATTCTTAAAGTTCCAATATGAGCAGGGCCTGCATACATCCAATAAG
CTAATTTCATATTAAATTTTCCTATGATTTATTTAGACATAGTAATACTTTTAAAGCTAA
CTAACAAACATTAGCCAAACATCTAATTTAACTCCTTGACAATTTTGTAATATTTTTTAA
ATGAAGCTTATCATTATAAATATTTAAAAAATATAATTCAGAATAGAAATAACTGTTAGA
TCAAGCTTGAAAGTAAAGATTTCTTGCAGTTATAGTTTTTCTTTAAGTAAACATGTAAAA
ATTGTACTATATATAAATGTACTATATATAAACTTTTTTTACTGAATATTGGTAATGCCA
GATACAATTAACTTAAATATGCCTTCCCCAACGTTTGGTGGAAGCACTGGCGGTTGGTTA
AGAGCCGCAGAAGTAGAAGAAAAGTATGCGATAACATGGACAGGCAAAAATGAAAGTAAG
TTTGAAATGCCAACTGGTGGTACCGCAACTATGCGAAATGGAGAAAACTTACTCTATTTA
GCTAAGAAAGAACAGTGTTTAGCTTTAGGTACTCAACTAAAAAGTAAATTTAAAATATCT
GACTATAAGATTTACAGAGTTTTTCCTAACGGAGAAGTGCAATATTTGCATCCTAAAGAT
GGAGTCTTTCCAGAAAAAGTTAATACTGGGAGAGCTAGCGTCAACAGTGTTGATCATTCT

ATTGGACAAAATATTAATCCTGTAGATGTCAAATTCATGAATAAAGCAACTTACGATTAG
TCTCTCAAAAACCTAGACATAAAATAACTTTTATATAGTCTAGGTTTTTGATAATGTATA
CTGATAGTATGGAAGGGTGGCCGAGTGGTTGAAGGCGTCTGATTTGAAATCAGTTGAACT
ATCTGGTTCCGTGGGTTCGAATCCCACCCTTTCCGTTATAAAACTATAGTTGGCTTCTCA
AGGCAAGCTAGCTATACACTGATAGTATTAAGTTTAGACTTAATAAAGTCTACAACTTGT
GACAGGTTAGAAATTTTTTCGGCATCCTCATCTGGTATCTCTATACTAAATTTTTCTTCT
ATGGCCATTACCAGCTCAACTGTATCTAGAGAATCGGCTCCCAGGTCACTTGAAAAATTT
GCTTCCCTAGTCACTATTTTTTTTTCAATTCCTAGCTGTTCTGCTACAATATCTTGAACT
TTTTCAAAGATTTCATTATCTTGCATAATATATTATTTCCTCTAAGGTGTTATCAAAAAT
TTTTCGGCTTATATTATTATCCAACTCTAAAAAATTAGTAATTTTTAGTCGAGATTAAAA
TATTTATACCTATATGATATACTATTATTCTGGATAAATACGAAGTTTACGATATGGCTC
TTCTCCAAAACTACGAGCTCTCAACTGGTAGTTGTCAACTAGATTATGCTGCATTTTTCG
TATATAGGCTGATCGAGGTGTAAGCTGAACTATTGAACTTTCATTCAAGATAATAGTTTC
AATAGCCAATTTAGCTTCTTGAAGAGCCTGGATCTCATCAAAATTTTTATTTTTACAGAG
TTCTACCCAATTAAGACCAGATGAAGTATGAATGTTTAATATTTTTCTGAGTGCACGAGT
AATTTGTGGAACCGTACTATTCTGAATTGTATATATAATAATTTGTTTTGATTTTGCAAT
TTGTCTAAGTTTTGTATTTTGCTTTACTTGATTACGTAGCGCTAGAATAGCATCAGACTT
TTCTATCTCCTTGGTTAAAATAATTGGTAGATCTAGTGAAGAAATAACTGATGTAATATG
CTGCCAACTCAAAGAATAAGCATATAAGTACTGATGAGGTATTTCCAATTGCAAAGATTG
TTTATTAATATTAATAGGGGCATTGAAAGTAGTTGACAATAAAGATGTATCACGATTTTG
TGATTTATTCAAATCTAAGGACTTATTTCTTAATTCTCTATATTGTAAAACCGGAGGCGC
CTGCTGTGCGTTAATTGATTTACGAGCATTAGAGATGTTTACAGGTAAGACTTCTATAGA
TTGCGATGGATAACATTTAATTAAAATTCTACCATTAGCTTGAATTTGTCTTTTTTGCAC
AAAAGGTTGATGGCCTTGCAAAATTTGATCTATTGTCTCTTTAACTTTATTATGAACTAT
CCAAACATTTCGCTCGTGTATTTCGATTGCAATCTGAAAAGCAGGCTCAGCTTTTCTTTC
TAGTATACTCTTTTGCGTACCTCTACGTTTAGCTTCATCGTCACCTAATGTTACATATTG
AATTCCACCAATTAGGTCCGCCAATGTTGGATTTTTAATTAAACTTTCTAAATGATTTCC
ATGCGCTGTACCTACTAATTGAACTCCTCTTTCTGCTATAGTTCGAGCAGCTAACGCTTC
TAATTCCGTGCCAATTTCATCTATAATAATAACCTCGGGCATGTGATTTTCAACAGCTTC
TATCATTACTTGATGTTGCAAATCGGGCCTTGCGACTTGCATTCTTCTAGCTCTACCAAT
AGCTGGATGAGGAATATCTCCGTCACCAGCTATTTCATTTGAAGTATCTATAATAACTAC
TCTTTTTTCCATTTCGTCAGCTAAAACACGGGCCATTTCTCGAACTGCGGTTGTTTTACC
AACTCCTGGTTTTCCTAAGAGTAAAATTGAATCACCTTGCTGTAGTAAATCTCGAATAAT
ACTAATTGTACCAAAGACTGCTCTGCCGACACGACAGGTTAAGCCAATAACACTACCTTC
TCTATTACGCAGTGAACTAATACGATGCAAGGTTTTTTCGATACCAGCTCGATTATCGCC
GCTAAAATTACCAACTTTTTTTACACAATAATCTAAGTCTTGCCAACTAATAGATCTTTG

AGATAAATATTCTGGGTTATCTGGAAATCTAGCTTCTGGACGACGACCTAAATCCATAAC
GACTTCTATTAAATTATTTCTATTAGGATGCTGTTGTAAAGGTTCTTTGACAAAATTTGG
CAAAATTTCTAGCAACTTATCTAAGTCATCTGCAATAAGCATGATTGAGTATATAATAAT
TAATTAATCAAATAGAAGATTCAAGATTCTAAAGATTACAATACATATTAAAGAATATTA
TGATTGGAATCAATTTTATACTTAAATTGAACAGGAAGTTATAGATGTAAAAGAACTGAT
TAAACTTTACGCAATAAATCGATTGGTCAAAATTAATAGTCGTGTTAACATGTTTATTGG
ACAAAAAGTTCGTATTAAATATAGCAAACAAAAAGTTGCTAGTGATATAGCAGATAAAGT
TGGTGAATTAGGAGTTATTAAGGGAATAAAGTTTATTAATAGTCAATGTGTTACTATTAT
TGTTGAGTTTGACAATCACACTAGGCTTTGGATGTTTAGAGAGGAATTAATCTGTCTAAA
TGAATTAAAAACATGAAACATAATTTATTTTATTTATTATACAAAATTTCTTATGAGTCG
TTAAAATAATAAATTTTAAAGGAGTACTTTTAATTGTGCAAATTACTATTAAAAAATTAC
AAGACTTACTTTCGTCTGTACAAAGAAAAAAGATCCAGACATTAAAACTAAAACAAGGTA
AGTTTGAACTTCTATTAAATAAGACCTATAAAAAAGTCAATCAAGAAATTATACCTTCGC
AGAAATCTGCTGTATTACAAAATAGTCCATCTACGATAATTAAATCAATAAATAATACAA
AAAAATCTTCTGTTGTTAATGAAGACCGTACAGAATATGCCACTATTGTTTCTCCAATGG
TTGGAACGTTTTATCATTCACCTGCTCCTGGTGAAAAAATTTTTGTACAAGTTGGCGATG
AGGTCAAATTCAATCAAACAGTCTGTATTATTGAAGCAATGAAGTTAATGAACGAAATCG
AGGCAGAAATTGAAGGCAAGATTATAGAAATTCTTGTTAAAGATGGTGATATAGTAGATT
GCGGGCAAGCCTTAATGAAGGTTGAAACATAATGCTAATTTTTCCATTTTACATATTGTT
AACAGAAATTATTTAGATAGAAAAATTAGCTTATACTATAGTTATGAAAACTTATTTCGC
AAAGGCACTAAAAACTAATATATAAAATTATTTACTTTATCTGCCAAAAATAGAAATAAG
CGTTTTGATTCCTTGTCTCAAGAGAGGAGAATCTCAATGGCAATTAGCTCAAAAGAGCAA
GAGACAAAGAAGGTAAAAATCTCGGTTGATAAAAATCCCGTAGATACTTCTTTCGAAAAG
TGGGCCCAACCAGGCCATTTTTCTCGTACACTAGCAAAAGGACCAAAAACTACTACTTGG
ATTTGGAATCTTCATGCTGATGCTCACGACTTCGATAGTCAAACCAGTTCTTTAGAAGAA
GTTTCACGTAAGATTTTCAGTGCACATTTTGGGCAGCTGTCTGTAATATTTTTATGGCTT
AGTGGAATGTATTTTCACGGAGCCCGCTTCTCTAACTATGTTGCTTGGTTAAGTAATCCA
ACAGGTATTAAGCCAAGTGCGCAGGTTGTTTGGCCTATAGTTGGGCAAGAAATTTTAAAT
GGCGATGTAGGTGGTGGCTTTCAAGGAGTACAAGTTACATCTGGATGGTTCCAACTGTGG
AGAGCATCAGGAATTACTACAGAATTTCAGCTTTACTGTACTGCTATTGGCGGATTAGCT
ATGGCTGCTTTAATGCTGTTTGCAGGATGGTTTCATTATCATAAAGCTGCTCCAAAGTTA
GAATGGTTTCAAAATGTTGAATCAATGATGAATCACCATTTAGCTGGGCTTTTAGGCTTA
GGCTGTTTAGGCTGGGCAGGCCATCAAATCCATTTGTCTTTGCCTATTAATAAGCTACTA
GATTCTGGCGTGTCTCCGCAAGAAATTCCACTACCTCATGAGTTTTTAATTAATAGAGAG
CTTATGGCTCAGCTGTATCCAAGTTTTAGTAAAGGATTAGTTCCATTCTTTACTTTAAAT
TGGGCTGAATATTCCGACTTTTTAACTTTTAAAGGAGGTTTAAACCCTGTTACTGGAGGT

TTATGGCTAAGTGATACTGCTCATCATCATCTAGCATTAGCTGTTCTATTTCTTGCTGCA
GGTCATATGTATAGAACCAATTGGGGTATTGGACATAGCATGAAAGAAATTCTAGAAGCT
CACAAAGGACCTTTTACCGGCAACGGTCACGAAGGTCTATATGAAATTCTTACAACTTCT
TGGCATGCACAGCTTGCAATTAATTTAGCTATGATGGGATCTTTAAGCATCATTGTAGCA
CATCACATGTATGCAATGCCTCCTTATCCATATATTGCTACTGATTACCCGACTCAGTTA
TCGCTCTTCACTCATCATATGTGGATTGGAGGATTTTGTATTGTTGGAGCAGGAGCGCAT
GCTTCTATATTTATGGTAAGGGATTATAATCCTGCAGAAAATTATAACAATCTTTTAGAT
AGAGTCATTAGGCATCGAGATGCTATTGTTTCTCATCTAAATTGGGTATGTATATTTCTT
GGATTCCATTCATTTGGTTTATACATTCACAATGATACTATGCGGGCACTTGGAAGATCT
CAAGATATGTTCTCTGATACAGCTATACAGTTACAACCTATTTTTGCTCAATGGGTACAA
AGTATACACACTTTAGCTCCTGGAAATACAGCTCCAAATGCATTAGCAACAGCTAGTTAT
GCATTTGGAGGAGATATTGTTTCTGTTGGTAACAAAGTTGCAATGATGCCTATTTCTTTA
GGTACTGCAGATTTTTTAGTTCACCATATACATGCATTTACTATTCATGTAACTGTTTTA
ATTTTAGTTAAAGGGTTCCTTTTCTCAAGAAACTCTAGACTAATTCCTGACAAGGCCAAT
CTTGGCTTCAGGTTTCCATGTGATGGACCTGGTAGAGGTGGTACTTGCCAAGTGTCTGGC
TGGGATCATGTTTTTCTTGGCTTATTCTGGATGTACAATTCTCTGTCTGTAGCAATTTTT
CACTTTAGTTGGAAAATGCAATCAGATGTTTGGGGTAGTGTATCTCCGTCTGGAAATGTT
TCTCATATTACTGGCGGTAATTTTGCACAGAGTGCAATTACAATCAATGGATGGCTGAGA
GATTTCCTTTGGGCTCAAGCATCTCAAGTTATTCAATCATACGGTTCTGCGCTATCTGCA
TATGGATTAATTTTCTTAGCAGCACATTTTGTATGGGCATTCAGTTTGATGTTCTTATTT
AGTGGTAGAGGTTATTGGCAAGAGCTTATAGAATCAATCGTATGGGCGCATAATAAGATA
AAAGTTGCTCCTGCAATTCAACCAAGAGCTTTAAGTATTACTCAAGGTAGAGCAGTCGGT
GTTGCACACTACTTATTAGGTGGAATTGGTACAACTTGGGCATTCTTTTTAGCGAGAATT
ATTTCAGTAGGCTAATAGTGAAAATAGGATAAAAAAAACAATTATGGCAACAAAATTTCC
TAAGTTTAGCCAAGCTTTATCACAAGATCCTACAACTAGAAGGATTTGGTATGGTATTGC
TACGGCACATGACTTTGAAAGTCATGATGGAATGACAGAAGAAAATTTATATCAAAAGAT
ATTCGCTTCGCACTTTGGACATCTAGCAATTATCTTCTTATGGACATCTGGTAATTTATT
CCATGTAGCTTGGCAAGGCAACTTTGAACAGTGGGTATTAAATCCTTTGAAAGTTAAACC
AATTGCTCATGCAATTTGGGATCCGCATTTTGGACAACCTGCTTTGAAAGCTTTTAGTAA
AGGTGGCTCAGCTTATCCAGTAAATATAGCATATTCTGGCGTATATCACTGGTGGTATAC
TATTGGTATGAGAAGCAATCAAGACCTGTATTCTGGGGCTTTATTCTTACTAGTTTTATC
AGCTTTACTCTTATTTGGAGGGTGGCTACATCTACAACCAAAATTCAAGCCTGGTTTATC
ATGGTTTAAAAATAACGAATCAAGATTAAATCATCATTTATCTGGATTATTTGGGGTTAG
TTCTTTAGCCTGGACAGGTCATTTAGTACATGTTGCTATACCTGAGGCAAGAGGACAACA
TGTAGGATGGGATAATTTTACAACCGTATTACCTCATCCAGCCGGTTTACAGCCGTTTTT
CAGTGGTAATTGGAGTGTATATGCTCAAAATCCAGATACAGCTCAACATTTATTCGGAAC

TAATGAAGGTGCGGGTACAGCAATTCTGACATTTCTAGGAGGATTTCATCCTCAAAGTCA
GTCTTTGTGGCTAACTGATATGGCTCATCACCATTTGGCTATTGCAGTAGTATTCATTGT
TGCTGGACATATGTATAGGACTAATTGGGGAATTGGGCACAATCTAAAAGATATTTTAGA
TGCTCATAGACCACCTAGTGGTAGATTAGGAGCTGGACATAAAGGGCTATTTGATACTAT
TACTAATTCTTTACACATACAGTTGGGATTAGCATTGGCTTCCCTAGGTGTAATTACTTC
GTTGGTAGCTCAACATATGTATGCTATGCCTCCATATGCTTTCATGGCTAAAGATTTCAC
AACTCAAGCATCTTTGTACACACATCATCAATATATTGCTGGGTTTCTAATGGTTGGAGC
TTTTGCTCATGGGGCAATATTCTTTGTTCGAGACTATGACCCTGAACAGAATAAAGATAA
TGTTTTAGCTCGTATGCTAGAACATAAAGAAGCTATCATTTCTCATTTAAGTTGGGTAAC
TCTATTTTTAGGGTTTCATACATTAGGCCTTTATGTTCACAATGACACAATGATTGCTTT
TGGAACTCCTGAAAAACAAATTCTAATTGAGCCGGTATTTGCTCAATGGATTCAAGCCTC
TTCAGGGAAAGCACTTTATGGGTTTGATGTGTTACTATCATCCTCTACTAATATCGCAAC
ACAAGCTGGTAGCAATATTTGGCTGCCAGGCTGGTTAGAAGCGATTAATAGCGGAAAAAA
TTCATTGTTTTTAACAATTGGTCCTGGTGACTTCTTAGTTCATCATGCAATTGCATTGGG
ATTACATACTACTACGTTAATTTTAGTTAAAGGTGCTTTAGATGCAAGAGGCTCTAAACT
TATGCCGGACAAAAAAGACTTTGGATATAGTTTTCCTTGCGATGGACCTGGCAGAGGCGG
CACCTGTGATATATCTGCATGGGATGCGTTCTATTTAGCTGTATTTTGGATGCTAAATAC
AATAGGTTGGGTAACATTTTATTGGCATTGGAAACATATTACAATATGGCAAGGCAATGC
AACTCAATTCAATGAGTCTTCAACTTATCTAATGGGATGGTTTAGAGATTACTTATGGCT
AAATTCTTCTCCATTAATTAATGGTTATAATCCATATGGCATGAATAATTTATCAGTATG
GTCATGGATGTTCCTATTTGGACATTTAGTATGGGCAACAGGATTTATGTTCTTGATCTC
CTGGCGTGGCTATTGGCAAGAGTTAATTGAAACTCTAGCATGGGCGCATGAACGTACACC
TTTAGCAAACTTGATTCGTTGGAAAGATAAACCTGTTGCATTATCAATTGTACAGGCAAG
ATTAGTAGGTTTAGCACATTTTTCTGTAGGATATGTATTAACTTACGCAGCTTTTGTATT
AGCTTCAACAGCAGGCAAATTTGGTTAGACTAAATTCGAATTAGCT-AAAAAAAGTCAGT
CATAATAGTTTTATGACTGACTTTTTTAGATTTAAAACAAAAACTTACTAAAGTTTCACC
TCAATATCTACACCAGAAGGCAAATTTAACTTCATTAATGCATCTATCGTTTGAGAAGAA
GGTTGATGAATATCAATAATTCTTCTGTGAGATCTTATTTCAAAGTGTTCTCTTGAGTCT
TTATCTACATGTGGAGAACGTAAAACACAATAAATTCTTCTTTTTGTCGGTAAAGGAATT
GGCCCTACTGCAACGGCATTAGTTCTAGATGCGGTATCTAGTATTTTGTCACACGACGTA
TTAAGTATAATAGAATTATATGCTTTCAGTTTAATTCTGATTTTTGTCTGCTGAGTAATT
GTCATGTTAAAAAGATCTTATTATTTAAGAATTTTAGAGACAACACCTGCGCCTACGGTA
CGGCCACCTTCTCTAATAGCGAAACGCATACCTTGCTCAATTGCAATTGCATTAATTAAT
TCAGCAGTCATCTTAATTCTGTCACCAGGCATAACCATTTCTGCATCAGTACCATCATCA
GCAGTAAACTGATTAATAGTACCAGTTACATCAGTTGTTCTAACATAAAACTGAGGTCTA
TATCCTGGAAAAAAATGGAGTATGTCTTCCGCCCTCCTCTTTAGTTAAAATATAAACTTCT

GCTTCAAATTGAGTATGAGGTGTAATTGTACCAGGTTTAGCTAATACCATACCTCTTTCA
ATATCTTTTTTCTGCACACCTCTTAAAAGAATTCCAATATTATCACCCGCTAGACCTTCT
TCTAACGTTTTTTGAAACATTTCTAATCCAGTAATAGTCGTTGTACGAGTTTCTCGTAAA
CCTACAATTTCAATTGTGTCACCAACTTTAATAATGCCTCTTTCAATTCTACCAGTGGCA
ACAGTTCCACGTCCTGTAATAGAAAAAACATCTTCTACAGCCATTAAGAAAGTTTTATCG
ACATCTCTCTCCGGCGTTGGAATATATGTGTCAACTGCTTCCATAAGTGAAAAAATCTTG
TCAACCCATTTATCTTCACCTTGCTTAGTAGCTGGGTTTTTCGTCACAGCTTCTAATGCC
AATAAGGCAGAACCTGCAACAAAAGGAATATCATCTCCAGGAAAGTCGTATTGACTTAAT
AATTCCCTTCCTTCTAATTCTACTAATTCTAGTAGCTCTTCGTCATCTACTTGATCTTCC
TTATTTAGGAATACTACTAATGTAGGAACACCTACTTGTTTTGCTAATAAAATATGTTCA
CGAGTTTGTGGCATTGGACCGTCTGCCGCAGATACAACTAAAATAGCTCCATCCATCTGA
GCAGCACCCGTAATCATATTTTTTACGTAGTCGGCATGGCCTGGACAATCTACGTGAGCA
TAATGACGATTATCTGTTTCGTATTCAACATGAGCAGTATTAATAGTAATACCTCTAGCT
TTTTCTTCTGGAGCGGCATCAATTTCATCAAATTTTTTCGCTGCAGTAGACCCTAAAGTT
GATAAAGTTGCGGAGATCGCTGCTGTTAAAGTTGTCTTACCATGATCAACATGACCAATT
GTGCCAATATTGACATGAGGTTTTTTACGTTCAAATTTAGATCGAGCCATGCTTTTTGTT
TTCCTTATAGTAAAAGTTGCTTTATAGTTACTTTTAACGATAAATCTTGTGAATTGAAAG
ATTTATTATTTAATACAGACAAGATACAATATAACTAATCTAATATCGATAGTGAGCAAA
CGCTTTATTTGCTTCTGCCATTCTATGTGTATCCTCTCTCTTTCTAATAGAATTTCCTGT
TTCATTAGCTGCATCCATAATTTCATTAGCTAATTTCATAGACATACTTTTACCAGATCT
GTCTCTAGAGAATTTAGTAATCCATCTTAGTGCTAAATTCGTACCTCTATAAGCTCGTAC
TTCAATAGGAACTTGGTAAGTAGAACCACCAACTCTTCTTGCTTTTACTTCTACTAGAGG
AGTAATATTTCGGATGGCTTTTTCTAAAATATTTAAAGGATCTGATTCTGTTCTCTCTTT
AACGATATCTAAAGCCTGATATATAATTCTTTGAGATAAAGTTTTTTTACCACTTTTCAA
AATACGAACAGTTAACATACTTACGAGTCTGCTTTTATATAAAGGATCAGGTGATGCAAA
CCTTTTTTTAGCTGTATTACGACGAGACATAGTATTTAAAATTGGTTATTGTAATAATGT
TCTATTAGCAGGTATTTAAGATTTAGGCTTTTTTGTCCCATATTTAGATCGGCTTTTACG
ACGATCTTTTACTCCTGCAGCATCTAAAGTACCACGTACTACATGATATCGGACTCCAGG
CAAATCTTTAATTCGGCCGCCCCTAATCAGTACCACAGAATGCTCTTGAATATTATGACC
TACGCCAGGAATATAAGCTGTAACTTCAAATCCAGATGTTAATCTAACGCGCGCTACTTT
TCTTAAAGCAGAGTTCGGTTTTTTAGGAGTTGTAGTATATACTCTTGTACAAACACCTCT
TCTTTGAGGGCAACTTTGAAGAGCTGGAGATTTTGTTTTTTTATGTATTTTTCGTCTTTC
GGATCTAACAAGTTGTTGAATTGTTGGCATAATAAAATTGATTTAAAGTCTTTAATAGAT
ACTATTTGAACCTGCTTTACATGAATTGGCTAGTTTTTATTTATCTTATACTTACGCATA
AATCTTTCTACTCTACCTTCTGTATCAATAATTCTCTGTGAACCAGTAAAGAATGGATGG
TTTCCTGACCAGATATCTACGTGTAGTTCTGGCTTTGTGGACCCAATCGTCATAATTAAC

TGGCCGTCACAGTAAACTTTTGCTTCTGGATACCAATTTGGATGTATATTATCTTTTGCC
ATTGTTTTTATAATTATATAAATGTATCAATAATAACTATTAACGCTTAGAAAATTGAGG
AGCCTTTCTTGCTTTCTTTAAACCATATTTTTTTCTCTCTTTTACTCTTGGATCTCTTGT
TAAATAGCCTTCAGACTTGAGTGTCGTTCTATTTTCTGGATTAATTGAGCACAATGCTCT
TGCAACACCTAAACGAATTGCATCAGCTTGCCCTGTCAAACCTCCTCCTCTAGCATTTAC
ATGGATATCATACTGGTTTAGTAGTCCTAAAACTTGTAATGGTGCATATGAAACTCTTAA
GTAATTAGGACTAAATTGAAGATAAGACTCTCCTGGTATACCGTTTATAATTAAATTACC
TGACCCTGGGACTAGTCTTACTTGTGCAACAGAACACTTCCGGCGACCTGTTCCAGAATA
GATTGCGCGAGTTTTAATTAATTCTGTCGACATAACATTCCTTGATAACTAAATAAATAT
ACATTCTTTATACTATATACTCTTGTGGCTTTTGCGCGACATGTGGGTGAATTGGGCCAG
AATACACTTTAAGCTTCGTAAATAGTTTTCTGCCTAATGGACCTTTAGGAAGCATACCTT
TAACCGATTTTTCAATAATTCTGTTAGGTAATCTTGTCTGAAGCTGATCAAATGTTTCAA
CTTTTAATCCGCCAGGCTGTCCAGAATGTCTTCTGTATAGTTTTTGATTCGTTTTATTTC
CACTTACAGATACGTGAGCAGAATTAATAACGATTACATAATCTCCAGTATCTAGATAGG
GTGTATAAGAAGGCTTGTTTTTACCTCTTAAGATATTAGAAATATGAGTAGATATTCTAC
CAAGTGTCTGATTTTTAGCGTCTATAACATACCAATGAGAATTAGTATTTAATGAGGGTG
ATTGCGTTTTATTCATGAAAAGATTAATACTTTTGGTTGACAATAAGAGAATATGTAAAT
GGTTTACATTCAATTTCTATACAAGTAGTATTTAAAAATATTATTAACAAAGAGAACTTT
TCTAACAGTTATTTAAATTGTTTGTCAGCAAGAAAATTTCTAGTTATCAATAACTTATAC
TATTTTTAATAATATAAAATATATTAATCACTTTTTTCTTTTGGCAAACTTATACCTAGT
TTTTTTTGCAAGGCATATATAACTTCTTCAGCTGATTTCTGGCCAAAATTTTTAATTTCT
AGCAGTTCTTCTTGAGAGTAATCTAGTAAATCTGCAATAGAATGGATTTGAGCTCGCTTT
AAGCAGTTATAAGCCCTAACAGATAACTGCAGTTCTTCTATTAAAACTTGACTAATTTTT
TTATCTTCTTTGCTACGATAATTATCTGCTGATTTAAAGTCTAAATTTCTTAGTGAACAA
AAAAGATTAGTTAAAACTGTAGCTCCTTGACTTATTGCTTCTTGGGGAGATATGCTCCCA
TTTGTCCAGATTTGTATAATCAATCTATCTTTTATGCTGTTGCTACCAATACGGACTTCT
TCTACTTTATAATTAACTTTATTAACTGGCATAAAAACAGAGTCTACTTGCAAAAAATCC
ACAGATAATTCATCTACAGCTTTTTCAGCTAAGCGATACCCGCAATTTTTTTCAATTTTA
AACTCCATTTCAAATATTGTGTTATTGCAAATAGTTGCAATATACTGTCTAGGATCTACT
ACCTCTATATCAGAAGATAATTCAAATAGGCCGGCTGTAACTATAGCTGGCCCTTGAACT
CTAATTCGACCAATTTGAGATTCTTTGTTATAACTTTTAAATACTACTTCTTTTAGATTA
AGTAATATTTCTAACACATCTTCTCTTACCCCAGGAATTGTGGAAAACTCATGGTTCACT
CCAGCAATCCGTACAGCAACTATAGCAGTACCTTCAAGATCTGACAATATTGATCTTCTC
AAAGCATTACCTAATGTAATACCTTGTCCTTGATTTAATGGTTCTATTACAAAACTACCG
TACTGCCCACGCGCCCCATCTGTTCTTGACTCTATGCATTCAATTTGAAATTGAGCCACC
TAAGAAAGCTCCTTTAAAATAATCAGTGATTAGTAAGTCTCTCTAAAACAAGAAGGAATT

TATACTCGGCGTTTCTTAGGAGGGCGACATCCATTATGAGGTACAGGAGTAATATCTTTT
ATTAGAGTAATCTCTAATCCTGCAGCCTGCAAAGCTCTAATTGCTGTTTCTCGACCCGCT
CCTGGACCATTTACTAGAACTTCAGTTTGGCGCATACCTTGATCCATAGCTTGTCTAGCT
GCTTTTTCAGCTGCTGTTTGAGCTGCAAAAGGTGTTCCTTTTTTAGCTCCCTTAAATCCA
CTTGCACCAGAGGATGACCATGATAATGTTTCTCCTTTTAAATTAGTAATAGTAACAATT
GTATTATTGAATGTAGATTTAATATGTGTAATACCGTTAACTGCATTACGTTTAGTTTTT
CTTGCTCCGGATTTTTTTATTTGTCTAGCCATCGTGTCTCGTCTATTATATAGTTAATTA
AAAGATTATTTTCTTGGAGCTTTTTTCTTACCTGCTACTGTTTTTTTACCTCCTCTGCGT
GTTCTAGCATTAGTTCTAGTTCTTTGTCCTCTCAAAGGAAGACCAAGACGGTGTCTTCTA
CCTCTATAAGTACTAATTTCCATAAGTCTCTTAATGCTCATAGACTCAAAACGTTTGAGA
TCCCCTTCAATCTGATAATTAGACTCAAGAATTTCTCTTATACTGACAACTTGTTGATCA
TGTAAATCTTGACACTTAATATCAGCGTCTATGTTTGTTTTTTCTAATATTTCTTTCGAG
CGAGATAGTCCAATACCATAAATATATGTTAAAGCTATCTCTATTCTTTTGTTTCTTGGA
AGATCTACTCCAGCAATTCTGGCCACTTTGTCTTGTCTCCAATAAATAATATGTATTTTT
ATAACGGTTATTTATAAGCTTAGCCTTGTCTCTGCTTATGTTTAGGATTAGTGCAAATTA
CCATCACCTTTCTGTGACGCCGAATTATTCTACATTTTTCACACATTTTTCGAACAGAAG
GACGAACTTTCATATTAACTCTATATACTAGGAATATTACTATATTAACTCTATTAATTA
TCGACATTAAAAATGTCAACAATATATTGTATTACTTTAACTGTTGTAGATTTTACTTCG
TCATACTATCATATTTTTTTGATATGACGTATGTCTGAATTTGCTTAGCTGTGTCTATTG
CGACTCCAACTAAAATAAGTAAAGACGTAGCCCCGAGACCTCTTAAGTTTTGGATCTGAG
TAACCTTTTCTATTATAAACGGAATCAGCGCTACTGTAAATAAAAACGAGGCTCCTAGGA
ATGTCAGTCTATTTAATATGACTTGTAAATAATCAATAGTCGCTTGACCTGGACGAATAT
TAGGAATGCTTGCACCCATTTTTTTTAAATTTATAGCAATATCTTCTGGATTCATTACTA
TTGATGTATAAAAATAGCTAAAGAAAAGGATCAAAGCACAATAAAGAAGAAGGTATAGTG
AACCATTAGGACAAAACAGGTATAGGATTTGGAGTAATGTTTTATTTTGAATAATTTGGG
TTAGATAAGATGGGAGGGCCATAGATGCAGATGCAAAAACAATAGGCATAACTCCACCTT
GATTTAATTTCAAGGGCAAGTAACTATTTGGATCCAAAATTGAAGATTTTCCTAGCTGTC
TTGCTGAAATAATTTTAATTCTTCTTGTTCCTTCTTGTACACAAATTGTAATTATTATCA
TTAATAAAAAGATCGCTATAAATAATCCGAACTTAAGACTTGCATTACTATAACTAGCAT
CAAAAAATGACTGTGTAAAATTCTTTGGTAGTCCTGACACAATGTTTTGAAAAATAAGTA
AAGAGGCTCCATTACCAATTCCCTTTTCCGTAATTAACTCTGATAACCACATAATAATCA
TAGAGCCTGCTGTTAAAGCTAAAACAGACTCGCAGACAAATGCAAAATTCCAATTAAAAA
CATATGGTTTTACCCATATAGAGATTGCTCCAGATTGTAAAGTCGCCCAACCTAGAGCTA
AGTACCTTGTAATTTGAGTTATTTTTTGGCGGCCCAATTCACCTTCTTCCTTCTGTAATT
TTTCCAGATTGGGGACAATTTTCGTAAGTAGCTGCATTACAATTGAAGAGTTAATATAAG
GAACAATACCTAACGCAAAAATCCCTATTGTTGAAAAACCTCCTCCAGAAAAAATATTCA

GAAAATTTACTAAAGTATTTTTTTCTACACTTGCATAAAAGGCGTCATGATCTATACCTG
GAACAGGTATAAATATTCCTAAACGTGCTAAAACTAATAGAAAGAGAGTAAAGATAATAC
GATTTCTTAGATCACTTTTTTGGCTCATAAATAAATTTTAAATGAAAAGAGATACTTTTT
TATGTAAAAACCCTATTCCAAGACTTGTATAAGTTTAAAATAGGGAAGTAAAATATTATA
GTAATATTAATATAATATTCTATGTAGAATACAGGCTTTCTATTGGAACTCCTCGATCTT
TAGCTGCTTCACTAAAAGTTCTTAATTGAGTTAAGCCATTGAGAACAGCTCGCGCGTTAT
TTAATGTATTATTAGAACCTAATTGCTTAGCTAATATATTCTGTACACCCGACAATTCTA
GGACTGTCCTTACAGAACCTCCGGCAATTACACCTGAGCCAGGTGCAGAGGGCCTTAATA
TAACTTTCGCGGCACCAGAAATTCCGTTAATAGGATGAGGTATAGAATTCGATTTTGTCA
GTGGAACTGTAACTAGATGTTTTTTAGCATCTGTTACTCCTTTTTTTACTGCACCAATTA
CATCGCTTGCTTTTCCCACGCCGACACCAACTTGGCCTTGCTCATTACCAATAACAAGAA
TAACTCGAAAGCTTAGTTTTTTACCTCCTTTTACAACTTTAGTAACTCTTTTAACTTGTA
CGACTCTTTCTTCCCAGCCACTATCTTTATCTTTTCCTTTGCTCTGTTTTTTACGATTGG
CCATTTTAAGAATTATTCCTAATTAGTTAATTACGATTCTAGAAACCCATGCCTGCTTCC
TTAGCAGCTTCAGCCAAGGCCTTAACTCTTCCATGATATAACTTTCCTCCTCTATCGAAG
ACGACATTTTTAATGCCTTCTTTCATAGACTGTTCTGCTAACTGTTTACCAACTACGCGA
GAAGTATCACAGTTTGGTCTTATGTTATCAGATTCTTTATTATTTAAATTAACAGATGAT
GTAGCTACTAATGTAATACCTTGTGTGTCATCAATTATTTGTGCGTATATATGTTTATTA
GATCTAAATACACACAGACGAGGCCTACTTGAGGTTCCTTGAACTTTTTTCCGAACTCTT
TTATGCTTATGAATTCTAGTTTGTTTAGTGTTTAGTTTCATTATTATTTACCTTTTCCAG
CTTTCCCAGCTTTTCTTCTAACAAATTCACCTTGATATCTAATTCCTTTTCCTTTATAAG
GCTCAGGAGGCCTAATAGAACGAATAGTTGAAGCAACCTGACCGACAACTTCTTTATCTA
TGCCAGAGACAGTAATATTTGTGTTGTTTTCAACTTTAATTTCAATATTGGCAGGAGGTT
TAATTTTCACTACATGACTATAGCCAACACTTAAGATTAAATCTTGGTTATCAATTTGAG
AACGATAGCCTACACCTTGTATTTGCAGTTTTTTAAAAAATCCATTAGAAACACCTTCAA
TCATATTACTAATAAGAGTTCTGGATAGCCCATGTAACTGACTAGCCATTTTTGTTTGTC
CACTCGTCTTGACAGCAATTGTGTCGTTAAGTATTTCTAAATTAATACCAGCTGGTAAAG
TTCTGGATAGGGTACCTTTAGGTCCTGTCACGGTGATAGTCTGACCATCGAACTGAGTAC
TAAGATTTGTCGGCAATAAAATTATTTTTTTTCCAATACGAGACATATTCCACCTATAAA
ATAGTTACCAAATATAGCATAATATTTCACCACCAAGACCATCATGACGAGCTTGTCTAT
CTGTCATAACGCCTCTAGAAGTAGAAATAAGAGCGATGCCTAAACCTCCAAGAACTCTGG
GCAACTCTTTATGATTTGCGTAGACTCTTAGTCCGGGTTTACTAATTCTTTTCAGAGCAG
TAATAACTGGTTGACGATTTTTACCATTATACTTAAGAGAAATCATTAAATGAGTCTCTA
TACCTTCACCCATTTGTTCGAAATTTTGAACAAATCCTTCTTCTTTTAGTACTGTTGCCA
TATTGCATGTCATTTTCGTTGCTGGAACTTGCACAATTTGATGTCTTGCTAAGTTTGCGT
TACGAATACGTGTCAGCATATCGGCGATCGTATCGTTGACCACCTTGATCCTCCTTGCGA

TGAATTATTTGAAATATTTTCAAGATTCCCTGAAAGGCATCCCTAGCTTTTTTAACAAAG
CTAGACCTTCTTGATCTGTTTTAGCTGTAGTGACAATTGATATATCTAGACCACGAATTT
GATCAATATTATCATAGTCTATTTCTGGAAAGATCAACTGTTCACGTAAACCTAAATTGT
AATTACCTTTGCCATCAAAACTTCTAGGACTAATTCCTCTAAAGTCTCTAATTCTTGGTA
ATGTTAAATTAATTAATTTCTCTAAAAAAGAATACATCTTGTCTTTTCTTAGATGCACAA
CAATTCCGATAGGAACCTCTTCTCGAATTTTAAATCCTGCAATAGATTTTTTAGCTTTTG
TGACAATTGGTTTTTGTCCAGTTATTAATGTTAATTCTTGAATACTACTTTCAAGAGCCT
TAGCATTCTGAGAAGCTTCACCTAAGCCACGGTTAATAGTAATTTTAGTAAACCTAGGAA
CTTCATGTACATTTTTGTACTGAAATTCATCTTTTAAAGATTGAGTAACAGTTGTTTTAT
ATTTTTCTTTTAATCCTATTGCCATTATATTTTCATCAATTATTTAATAAGTTCGCCAGT
CTTTTTCAGTTTTCGAATTTTTTGGCCTTTATCATTAATTATTACTGAGGATCGACTAGC
AATATTATTTTGTTCACTAAATAACATAACGTTAGAGGTATGTATTGGAGCTTCAAATTT
GATAATTTCTCCCGTTTCTCCTTCTTGCTGAGGTTTTTTATGCTTCACTTTGAGATTAAT
TCCTTTAACAATCACCTTATTTGTTTTATAAATGATTGCAATAATTTCACCTGTTTTTGT
TTTATCACTTCCAGAAATAACTTGAACTAAGTCCCCTTTTTTTAATTTAATTTTTGTATT
ATTTTTGGTTGTTTTAGAAGGACCTTTCATTATACTACCTCCGGTGCAAGAGAAACTATT
TTAGAAAAATTTTTATCTCTTAACTCTCGGGCTATAGGTCCGAAGACTCTTGTGCCACGT
GGATTATTATCTTGATTAATGATAACTGCCGCATTATCTCCAAATCTAATGCTCATACCG
TCAGTTCTCCGTAAAGCTTTACGAGTTCTTACTACAACAGCTCTGACAACATCAGATCGT
TTAACTGGCATATTGGGAGACGCATCTTTGACTACTCCAATAATAACATCTCCAATAGAT
GCATAAGAAGGATTACTAGTGCCTAATACTCTAATACACATTATTTTTCTAGCACCACTA
TTATCTGCAACATTAAGATAGCTTTGAGTCTGTATCATACTTTTATTCTGTATAAGATTA
ATTATCAAAAGACTTAGATAAGATATTAACCATTGTCCAACACTTTGTGCGACTTAAAGG
TCGTGTCTCTTGTATTGTAACAATATCGCCGATTGTGCATTCATTATTTTCATCATGCGC
TTTATATTTCTTTGTCCGGATCATAGTTTTTGCGTACTTTCTATGAGAGATTCTATTTTC
TACAGCTACTACTATAGTTTTATTCATTTTATCGCTTACTACTTTACCTGTTGTTTCTTT
TAAAGGCATAGTCTTTATATTTTTTATATATAATTAAATTTTAAGATTATTACATGGTTG
ACTTAGTACGAGACTTTTCAACAGTTAAAAGTTGGGCTAATCTATGCTTAGAATGTTTAA
ATAGATGAGGCTGGAAATCTTGCCTTGTGGCTCTTTTTAGCCTTAAATCAAAAAGCTCTC
TTTTTATTACAAGGATTTCTTCAGCTAAAGAAGAAGAGTCCAGGTTTGTAACATCTGATA
TTTTAGGGAAAGTCATATTTTAGGATTCTGTTGTATTCCGAACTATAAATTTAGTCTTGA
TTGGTAATTTATAAGAAGCTAATTTCATAGCTTGTTGAGCAGTTTTTTGTGGTACACCTG
TAATTTCAAATAGAATATGTCCAGGCTTAATAACTGCAACCCAATATTCTGGAGCTCCTT
TACCTGACCCCATGCGAGTTTCAGCTGGACGGGCAGTGACTGGTTTATCTGGAAATACTC
TAATCCATAGTTTACCACCTCTTCTCACATATCTAGTAATAGTTCTTCGAGTAGCTTCTA
TTTGTCTAGAAGTTAACCAAACTGGCTCTGTCGCTTGTAATGCATAATCACCGAACGCAA

TTGTGTTACCTTTGCTAGCAGAACCTTTCATTCTACCTCTATGTTGTTTTCTAAATTTTG
TTTTCTTGGGGCTTAGCATAAAAATTAAAAAATAATAATAATTAAGAAGCTGTTGAATCA
GGAGTTTCCTGATTATAAGCAGATTCTAGACTTTTCGATTCTGGCAAAATTTCTCCTTTA
AAAAGCCAAACTTTTACTCCGAGCACACCATACGTAGTGTGAGCTTGGCGATGACAATAA
TCAATATCGGCTCTTAAAGTTTGTAAAGGTACACGACCTTCTCTAACCCATTCACTTCTA
GCTATCTCTGCACCATTAAGCCGACCAGATACTTGAATTTTAACACCCTGTGTATTTGCT
CTTTGAGCTCTTTGAACTGCTTGTCTGACAGCTCTACGAAAGGCCACTCTTTTTTCTAGC
TGTTGAGTAATAAATTCTGCTACTAAGGTTGCTTCAGAATCTGGATCTGCAATCTCTACA
ACATTTACTCTGAGTTGTTTGCTAGGGTCTAGGATTAAGGACAGCGATTTTCTTAAAGAC
TCAATTCCCGCTCCAGATTTTCCAAGTACGATTCCTGGTCGAGCTGTTGCTATGAGAATT
TCTACTTGATCAACTTTACGATTAATTTCAATTTTAGCAATACTAGCATTACTAAGTTTT
GAATGTATAAATGAGCGAATTTTATGATCTTCTTGTAAAAGAACTGGGTAGTCTTTAGAG
TTAGCAAACCATGAAGAACGATGTTTTTGGGTAATGCCTATGCGAAAGCCCAAAGGATGA
ATTTTTTGACCCACAGTATCCTTGATTATAAGTTAAAAGCTATAATGACTAATTGACGTT
AAAAGACTACAGTTCTGAAACACCTAATGTAATATGACAAGTTGGTTTATGTATTGGAAA
GGCTCTTCCTTGTGCTCTAGGCTGAAATCTTTTTAAAGTCGGGCCTTTGTCTGCAAAGGC
TTTACTAACAAACAATTGGTTTTTATTTAATCCATCATTATGCTCAGCATTCGCTGCAGC
AGATTCCAGGATTTGCTTTATATGAGAACAGACTCGGTATGGCATAAATTCCAGAATAAT
TAATGCTTCTTGGTATTTTCTACCTCTAATTTGGTCTAATACACGACGGACTTTATGCGG
AGAAAGACGAATGTATTTCCCTACTGCTTTAGTTTCTTTTACGTTTTTTGTAATACTCAT
AGTTATAGTTTAACGACGGGCTTTTCTGTCACCTTTTACGTGAGTGCGGAAAGTTCTTGT
TGGAACAAACTCTCCTAATTTATGTCCTACCATTTGGTCTGACACAAAGACAGGAAAATG
TTGTTTACCATTATAAACAGCTATTGTATGCCCTACCATATCGGGAATAATAGTAGATGC
TCTCGACCAAGTTTTTAGGACTTCTTTTTTTCCTGAAATATTTAATGCTTCTATACGTTT
AAGAAGACTAACATCAATAAAAGGGCCTTTATGTATAGATCTTGACATAATAATAAATCA
TTAGATTCTTAATAAAATAAATTCTATTTACGGCGACGTAAGACGTACGGGTTACTATAT
TTATTTGGATTTCGTGTTTTAACACCCAATGCAGGTTTACCCCAAGGTGTTACAGGACGA
GCTCGTCCAATTGGAGATTTGCCTTCACCACCACCATGTGGATGGTCTACAGGATTCATG
ACAACACCTCTAACAGTAGGTCTTTTACCTAACCAACGATTTCTACCAGCTTTACCAAGA
GTAATATTACTAGCGTCAATATTACCAACTTGTCCAATAGTAGCGTAGCATTCTTTTCGA
ATCATTCGAACTTCGCTTGAAGGCAATTTTACAGTAACAAAAGTACCTTCTTTTGCTACT
ATTTGAGCATAAGTTCCTGCTGCACGAACAATCTGTCCTCCGCACGAAGGTCTTAATTCT
ATATTATGTACTGCTGTTCCTAAAGGAATACTAGATAAAGGCAAAGCATTTCCAACTTCG
ATAGGGGCAGTAGGACCAGAAAGAACCATAGATCCTACACTGAGAGATCGAGGGTGTAGA
ATATATCTTTTCTCACCATCAAGATAGTGTAATAATGCAATTCTTGCATTTCTATTCGGA
TCATATTCAATAGAAGCAACTTTAGCAACTATATTATGTCTATTTCTTTTAAAGTCAATT

AATCTGTACTGCTGCTTATGTCCACCACCTTTATGACGACAGGTAATGACACCTCTATTA
TTTCGACCTTTGCAGAAATGGTGTTTAACTATTAATGATTTTTCTGGCTTATCAGTAGTA
ATCTCTGAAAAGGTAGAAACCGTTCTATTTCTTGTCCCTGGTGTGTAGGCACGATATAAA
CGAATTGCCATATAAGAAGATTTGAGAAGAGTAAATGTTGAACTATGTGATATATTTAAG
TTTCTGGAAACAAATTAATAGAATCTTCTGAGGCAAGTGTTACAATTGCTTTCTTGTAAT
GTGGTCGCTTCCCTACGAATCTACCGATGCTCCTTTTTTTCTTAGGAGGATGACAAGTGT
TAACACCTGTAACTTGCACGTTGAAAATATACTGTATAGCAGCCTTAATATTAATTTTTG
TTGCTTTGGGATCAACGGCAAAACAATACTGATTTTCTTCTAATAATTTAGTTGTCTTAT
CAGTAATGATGGGATATTTAACCAGATCTAATAAACCTCTTGAATCAATGCTATCCATTA
TATACCTCTTGTATTTTAGATAAAGCATCAACTGTAATAATAATTTTATGTGCAGCTAAC
AAAGCCATAATATTTAGTGTATCAGCTGAAATAAGTTCTACGTTGTGTAGATTGCGAATA
GAAAGATAAACATTCGGATCTTTTTTGTCTACGATGACTAAAACTTTTTTATTTAAATCA
AGATTCCATCGATGTATAGCTTCCATAAATAGTTTAGTCTTGGGCTGATGAAAATAACTG
TTAAAATTTTCTACAATCAGTGTATTAACTGATTTGTTATTTAATGCTGTCCTTAATGCT
AATTGTCTTTCTTTTTTATTCATTTTTTTGGTAAAACTACGAGGTTTTGGCCCAAATATT
ACACCACCACCTCTCCATAGAGGTGAACGGATAGAACCTGCTCTTGCTCGACCAGTCCCT
TTTTGACGCCATGGTTTACGACCTCCACCTCGAACTTCACTTCTTGTTTTAGTATTAGCA
GAACCCTGGCGCTTTTCATTACTTTGCTTTACTAAAGCTCTATGGACTAAGTACATCCCA
GAATCTTGACTGACCTTAAGATTTAAATCAGCGTTGCCACTTACTTGACCTTCCCAATTA
TAGACTTGATAATTTAATTGCGTGTTAACTGTCATGAAATATATCGTATGAAAGAATATT
GCTACTTACTAATTTTTACTAAAGCGCCAGGTTTACCAGGAACAGCTCCCTTAACGATCA
ACAAATCATTCTCTGAGTTAATACTTACAATTTGCAAATTTTTTATTGTAACTTTTTTAT
TACCAAGTTGGCCAGCCATGTTTTTACCTGGATAAACTCTTCCAGGCGTTGTACCGGCTC
CAATAGAGCCAGGTTGCCTATGATTTTTCGAACCATGTGACATTGGGCCTCTACTAAAAT
GATGCCTCTTTTGGTAACCGGAGAAACCTTTACCTACACTTCTAGAAGAGACATTAATCT
TTTGGCCTACTTGAAATAGATCTGTAGATAGTATTTGGCTAACTTCGAAATCATCAGTAG
ATTTCATTTCGTATTCACGTAAATATTTTAGTGGGGGAGCCTGTGATTTTTTTAAGTGTC
CCAACAATGGTTTATTTAATTTTTGTTCAGCAACTTGCTTGTAACCGACTTGAATAGCAT
TGTAGCCATCAGTAGATACAGCTTTAATTTGAGTAATAACACATGGCCCAACTTGAATTA
CAGTAACTGGAATTGATAAACCAGCTTCATCAAAGAATTGGGTCATACCTACTTTAGTAC
CGAGTATACCAACAGACACTAGATTTCTCCTTTTTGAGAGATTTATACTATCATTGTACA
TAAAAATTCTACTTCATAGTTACTGAAGTAAATTAAACTAACAGATGCATGATATTATTC
ATGACAACATAAAGGTTAAAAACTTTACCTATTTAATCTAAAATAGAAACAGCTTTTTGT
CTAGTTACAGCGTATATAAATTAGAAGTATGCTTATTAAATTTTTTATAAACTACTTAGA
TTTAACTGATATTGATATCCGTAACTTACTTGCAGTAATATCAATCAACATGTATAAGTT
TAATTGCCTAAACTAGAACCAGCTATTATTAGAAATAATTCAAGTATAGATAAAAGCGTA

TTTGAACTTTTATATATTGACAAATATAATAGAATATATTGAGGTTCGGTAACTACACCT
TTTTATTTTTACATTTATTGTGCAGTATATATACATCTAATGCAGTAAACATAAGAATAT
AACTGTATAGCAAATGTAATTACACAAATTAGAGAGGTATTAATGGGCAAAGTCGTTGGA
ATTGATCTTGGAACAACGAATTCTGTAATTGCTGTTATGGAAGGAGGTAAACCTACCGTC
ATACCGAATGCAGAAGGTTTTAGAACTACAGCTTCTGTTGTTGCATATACTAAAAGTGGA
GATAAACTTGTAGGACAAATTGCCAGGAGGCAAGCTGTTATTAACCCAGAAAACACTTTC
TACTCTGTCAAAAGATTCATAGGACGTAAACAGAATGAAATTTCGCAAGAGATTCGGCAA
ACATCATATAATGTTAAAACTAGTGGATCAAGCATAAAAATTGCTTGCCCTGCACTGAAT
AAAGATTTTGCTCCAGAAGAAATTTCAGCTCAAGTACTGAGAAAACTTGTTGAAGATGCT
AGTACGTACTTAGGTGAGACTGTTACACAAGCAGTAATAACTGTACCGGCTTACTTTAAC
GATTCACAAAGACAAGCTACTAAAGACGCAGGTAAAATAGCAGGCTTAGATGTATTGAGA
ATTATTAATGAGCCTACCGCAGCTTCTTTGTCTTATGGACTAGACAAACAAAATAATGAA
ACAATACTAGTATTTGACCTTGGTGGGGGCACATTTGATGTATCTGTATTAGAAGTTGGA
GATGGAGTATTTGAAGTACTCTCAACTTCTGGAGATACACATTTAGGCGGAGATGACTTT
GATCAGCAGATTGTAGAATGGCTAATCAAAGATTTTAAACAGAATGAAGGTATTGATCTT
GGTAAAGATAGACAAGCACTTCAGAGATTGACCGAGGCTGCAGAAAAAGCAAAGATAGAA
CTGTCAAATTTAACTCAAACAGAGATCAATCTTCCTTTTATTACTGCCACACAAGATGGT
CCAAAACATTTAGAAAAAACTGTAACTAGAGGGAAGTTTGAAGAACTTTGTTCAAATTTA
ATAGATAAATGTAGTATCCCTGTAAATAATGCTCTGAAAGATGCAAAACTAGAAGCTTCC
AGTATTGATGAAGTTGTTTTAGTTGGTGGATCTACAAGGATTCCAGCCATACAGCAAATG
GTTAAAAGATTAATTGGTAAAGATCCAAACCAAAGTGTCAATCCAGATGAAGTTGTTGCT
ATTGGTGCAGCCGTACAAGCTGGAGTTTTAGCAGGCGAAGTCAAAGATATTCTATTACTA
GATGTGACGCCGTTATCTTTAGGAGTGGAAACTTTGGGTGGCGTGATGACAAAGATTATA
CCAAGAAATACTACTATTCCTACAAAAAAATCAGAAGTATTTTCTACAGCTGTAGATAAT
CAACCAAATGTGGAAATTCAAGTACTTCAAGGCGAAAGAGAACTAACAAAAGACAATAAA
AGCTTAGGGACGTTCCGATTAGATGGCATTATGCCTGCACCTAGAGGAGTTCCTCAAATT
GAAGTTACCTTTGATATTGACGCTAACGGAATTTTATCTGTAAAAGCCAAAGAAAAGGCT
ACTGGTAAAGAGCAATCAATTACTATATCTGGAGCATCAACTTTACCTAAAGATGATGTT
GAAAGAATGGTAAAAGAAGCTGAAGAAAATTTCGATACAGACCAAAAAAGAAGAAAGAAT
ATTGACACAAAGAATCAAGCAGAGTCCCTATGTTACCAAGCTGAAAAGCAAGTTAAAGAG
TTTGAAGACAAAATTAGCCAAGATTTAAAAATAAAAATAGAGGAGCTAATTACAGAGCTT
AGATCTAGTCTAGAGAAAGAAGAATATGACAATATTGAATCCATTTCTCAGCAATTACAA
AATGCTCTGATGGACATTGGAAAAAATGCTGCTCAGACTGAAAGTAAAGATACAAAAGCG
AAGGATGACGACACTGTGATTGACACTGATTTCTCTGAAGCTAAGTAAAAGTAAGCGGGT
AACGCGATTCGAACGCGCGACATCAACCTTGGCAAGGTTGCGCTCTACCACTGAGCTATA
CCCGCATAGATTGTATTATTACAAAAATTAAGATATTTGTCAATCTGTTAATCTATACAT

CTAGACTATTAATTTTGGAATTACATATTAGCTAACTGTTGTAAATATAACAATTGCTAT
AATTTTTTAAAAGTACTTATATCATTATATAAGTACTTTTTATATTCAAAAATACAAATA
AGTATTATAGCTAAACTACTAAGGAGTTAACTAAGCTAGTAACAATTACTAATCCTGTCC
ATAAGCCGGCTCCTGTATATATTAATCCTTTTGATTGCTCCCATTGGCCAGGGGATGCTA
AAACAACAGGTACGCCTACTACTAACACAGTCGATAATGCAATTAATAATAGCACTAATA
ATTGGATCGCAATAATCATTTAATATGTTCTCCTGAAAAATTCAAATGTTAATAATACTG
TATATAATATAATGAAGCTTTTGACCTTCAGCAATTTATCTATGAAGAAATAATTTAAAA
TTATTGTTTTCTAAATAAACCTTAAATTAAGTATTAAAATCTTTATTGTAAACAATAATA
CTATATTTTTTTAGTAGATAAAAAATTTAAAGAAAGTCTTTTTTAAAAAAAAAATTAAGAT
AAGGTGATATGCTGTATAAAAAGCTAAGTATTATTAATTATTAGAGAAATTTAAAATAAG
AGGTAAAACATATGAACTCAGCCCTTTTTTTAGCAAAACTACCAGAAGCTTATGCCATTT
TTAAACCGATCATTGATATCTTACCCGTAATTCCTGTATTTTTTCTTCTTTTAGCTTTTG
TTTGGCAAGCAGCAATTGGTTTTAGATAACATTTAGGTTATAAAAAGAATAACAAATATA
CAGAGTGCCAATTTTTGAGCTCTAATCTTATATCTTACACAATAATAACTATTATGGTTG
AACCCTTACTATCAGGAATTGTTCTTGGACTTATTCCTATTACTTTATTCGGATTATTAG
TAGCAGCTTATTTACAATATCAACGGGGCAACCAATTAGGACTGTAAAGCCTAAATTATA
GATATTTATAAAAACTACAATAATTATAGTTTTTATAAATATCTACAAATAAGCTAATTT
TAAATACAGAAATAATACTATTGAAACTAGAGTGATAACAAAAAAAATTTATGTAATAAC
AAGCTAATTGTTTACCAGCTCGACTTAGTAACTCCTGGTAAAAGACATTCATGAGACATT
TCTCGAAAAACATGGCGAGATAATCCAAAATCTCGATAATAACCTCTACTTCTACCTGTT
AACCAACATCTATTTCTCCCTCTAACAGCGGCGCTGTTTCTAGGCATTTCTTGGAGTTTT
TGTCTTAAATCCATTTTGTCAGAAAAGCTGCCAGCTTCTTTCAGTTGATCTTTAATAGCC
AGTCTTTTTAAATAATATTTTTTTGCAAGTTTCGATCTCTTAATCTCTCTTTGGATCATA
TTCTTTTTAGCCATGCTGTAAAACTCTCTTTTAAAATTATTTATTCAATATATTAGCTAA
AAAAGCTCCTGCCTGCAACTTTTATATATTTAAAAATAAATAGCATAAAGTAGTGAAATG
CAATTGATATTTAATCAACGATACCACTTTATGCTATTATCAAAAATTTATTTAGTAAAG
TTCTTGTTCAACATGAGTTGAAATAGTGCAATCAGATGAAGGATATGCAATACAGGTTAA
AACATAGCCAGCTAAAAGTTGATCATCATCTAAGAATGATTGATCAGCTTGATCTACAGT
TCCTTCTGTAACTTTACCTGCACAAGTTGAACAAGCTCCAGCTCTACAAGAGTAAGGAAG
TTCAATTCCTTCTTCTTCTGCTGCATCTAGAATATATGTATCCTCTGCACAATTAAATGT
GACATCAATTCCTTCTTCTTCACATAGTAAATGAATTTTATAATCAGCCATTTCTTCTTA
CTCCATAATAAAGATACACTTGAAACTTAATAAAGCAAAATAATTTAAAGAACTTACCAA
AAAATTATATACTAGTTTCATTATGCATGATAGACTGGAAAAAAAGAAAATAGTATACTA
TTCTGAAAAATCAAGCTACATTTCCTACTAAATAGAACTATAGTAGGTCCAAAGTATATA
CATAAATTCATATAGTTAAAATTACTTTTATCATTTAAATAGCTGAACTTGATAGTATAG
ATTGATCATCAAGCTAGTCTTCTGCCAATAAATAATTAATTATGATTTCAATGCTTAATC

AACAAGTAGAACTTAAGCCTATTCTTAAATTTCAGATAACACAAATATATTCGCATTCTC

TTATTCAGGAAATTAAAGCCTTAATAACTAGACTTGTTTTGCAAGTTTGGAGAAGGCCAG

CTACATTGATGGCAGGTATTATCCAACCTTTATTATGGTTAATTCTATTTGGTGGGCTTT

TCTACAATGCTCCTATAAATTTGTTCACTATTAATACAAGCTATAATTGTTTTTTGAGCT

CTGGGATTATAATTTTCACCTCTTTTACTGGAGCTTTGAATTCAGGTCTTCCATTAATGT

TTGATAGAGAATTTGGATTTTTAAATAGATTATTAACGGCTCCCTTAGTATCGAGGACTT

CTATCATTTTATCTTCTGCTACTTTTATGACTTGTATTAGTTTAATACAAGTTGTATTTA

TAGTTACAGCTTCTCTTTTTATGGGAAACCCACCTCTAAATAGCGATAGTACTATGATTT

TTGGACTTATGATTCTATTAGTGACTGTAGGAGTTACAATGCTTAGTTTAGCTTTATCTT

TCACTCTGCCAGGTCATATTGAGCTATTAGCATTTATTTTAGTAGTTAACTTGCCCTTTT

TATTTTCTAGTACGGCTTTAGCTCCTTTATATTTCATGCCGCCATGGCTTCAGTTAATTG

CAAGTCTCAATCCATTGAGTTATGCAATAGAAGGTACAAGATACTTATATTCAAGCGTAA

ACTGGAATTTTACAGAGTGTGTGATTAAGATTAGCTGGGGAGATATTTGTTTAGGGCAAA

TTATTATTTTATTAATCGCTTTAGATATAATGGCAGCTTATCTTGTGTCTAATATATTAA

AAGCTAAACTTAATTAAAATTTTAATAAAAAATTTATTAAATTTATTTTACAAAGATTTT

TACTATATAATACTAATAGTAGTATGGATAATTGAAATAGAAATAGTTTTTCAAAACCAA

AGCTATTATCAACAATGATAAACATTTGTAAGAAAGTCAACAAAGTATGTTCTTATTCAT

AGGAGGCATGTAGTCAATGGGACTACCATGGTACCGTGTACACACGGTTGTTTTAAATGA

TCCTGGACGGTTAATTGCAGTCCACCTAATGCATACTGCACTTGTAGCGGGTTGGGCAGG

ATCTATGGCATTATACGAACTAGCTGTATTTGATCCTTCAGATCCAGTATTAAATCCAAT

GTGGCGACAAGGCATGTTTGTTATGCCATTTATGGCTAGACTGGGCGTAACAGATTCATG

GGGAGGATGGAGTATAACAGGAGAGAGCGTATCCAATCCTGGACTGTGGAGTTTTGAAGG

TGTAGCCTTAACTCATATAGTCCTTTCTGGCATGCTTTTTCTAGCTGCTATTTGGCATTG

GGTTTATTGGGATTTGGAATTATTTAGAGATCCACGAACTGGCGAGCCAGCCTTAGATTT

ACCGAAAATTTTTGGAATTCATTTATTGCTATCAAGTCTACTTTGTTTCGGATTTGGAGC

TTTCCATGTAACTGGACTTTTTGGACCAGGAATGTGGGTATCAGATGGGTACGGAGTAAC

CGGAAAGGTATTACCAGTAGCTCCAGCATGGGGACCAGAAGGATTCAACCCGTTTAATCC

TGGAGGAGTTGCATCTCACCACATTGCCGCAGGTACTGTAGGTATATTAGCTGGTGTTTT

CCATTTAACTGTTAGACCACCACAAAGACTGTATCGTGCTCTAAGAATGGGTAATATTGA

AACTGTATTATCAAGTAGTATCTCTGCTGTTTTTTTCTCAGCTTTTGTGACTTGTGGTAC

GATGTGGTACGGCTCTGCAACTACGCCTATTGAATTATTTGGTCCAACTAGATATCAGTG

GGATAGTGGATATTTTCAGCAAGAAATTGAGAAACGAGTAGAAAATGCTATTGCTGATGG

TGCTGCACCTAGCGAAGCATGGTCAAGAATTCCTGACAAGTTGGCATTCTATGACTATAT

TGGTAATAATCCAGCAAAAGGAGGATTATTCAGAGCGGGCCCTATGAATAAAGGCGACGG

AGTTGCTGAAGCATGGCTTGGACATCCTGTATTCCAAGATAAAGAAGGAAGAGAGCTTAG

TGTTCGCAGAATGCCTGCTTTTTTCGAAACTTTTCCTGTAATTTTAGTTGATAAAGATGG

TATTATACGAGCTGACATTCCATTTAGAAGAGCAGAGTCTAAGTATAGTATTGAACAAGT
AGGTGTAACAGCTAGTTTTTATGGTGGCAAATTAAATGGCCAAGTTTTCAACGATGCTCC
TAGCGTTAAAAAATATGCAAGGAAAGCTCAATTAGGTGAGGTATTTGAATTTGATCGAAC
TACATTAGAATCAGATGGAGTGTTTAGAAGTAGTCCTAGAGGCTGGTTTACATTCGGTCA
TGCTAATTTCGCTTTAATTTTCTTCTTTGGACATCTTTGGCATGGTTCAAGAACTATCTT
CCGGGATGTGTTTGCGGGAATCGGAGCTGAAGTTACTGAACAAGTTGAATTTGGAGCTTT
CCAAAAATTAGGAGATAGAAGTAGTAAAAAACAAGGAGCTGTATAAGACCAAAGTCAAAG
ATAGTCTTTATCAAATATTTCTAATATTAAAATAGTTAAGACAAAGTTTATTATCTAATA
GTTACTAATCAGGAGAATTTATGGAAGCCTTAGTCTATGTATTTTTACTAACAGGAACGC
TAATGGTTATATTCTTTGCGATCTTTTTTAGAGAACCTCCAAGAATAGCAAAGTAACTTG
ATTCCGTTTAAACTTATAATAATAGAACCACTTTCCAACTTATTAGTGGAAAGTGGTTTT
ATTGTAGATTACATTGTATTATTCTTCATGCTCTTCAAAAGGATCTCGGAGATCTTTTGA
AGCAGGACCAAATGCTGTGTATATCGAGTAACCTGTAATACCTAAAAGCAGACTCGAAAT
AAAAATACTAAGAACTGTTGCAGTTTCCATAATTTAAGCAGTATTAAGATGAACTTTTTC
TATAATGATATTATCAATATTAAAACATTTAATTATTAAAATTATCAAAATATACTATGG
CACTTAGAACTAGACTTGGAGAAATTTTAAGACCTTTAAATTCAGAGTATGGAAAAGTTG
CTCCAGGTTGGGGCACAACTCCTATTATGGGAATTTTCATGTTACTATTTTTCTTGTTTT
TATTAATCATTTTACAAATCTATAATTCTTCATTAGTATTAGAAAATGTAGATGTAGATT
GGGCTACTTTAGGTAGCTAACTAGAGAAGTTCTTTGATTACACAATATTAAAGACAACAA
GCGATCGCTATTAGATTATAGTAGCGATCGCTTTAAAAGTATTTTACTATATATCCAGAG
AAAATTATTTACCTAAATTAATTAACTCGGTATCTGCAAAACTATTTGTGTTAACATTAT
TATAGTTAACTTTTTCAAACCTTACTAATACTGGATATTTGATACCACTTTTATCAATTG
CTGCAACAGTACCAATTTCCTGATACCAGTACGATTCTTTTCTCAATATTTTTACTTTTG
AGCCTCTTTCCATATTTTTAGTCCTCAGATAGATTATTAAAGATTATTATATTAAAAAAT
T-TAAAAAACTAAACAAAATATATAAAAATACTACCGACAATTGTAATATATAACTAATA
AAAGGAGAGAGAGGGATTCGAACCCTCGATAACAAAAGTTATGACAATTTTCGAAATTGT
TGCAATAAACCACTCTGCCATCTCTCCTGAATTAAAATACTGTTTTAAAAAATTAGACAA
GAATAACATAGAACTAAGGTAGTTGCCTAGAAAATTAAAACTATGTTAAATATAAATTAT
AATCTTTCTATAGATATATTTAAAAAATATGTACATAGAATCTATATCAGCTTATTTCAA
GATTTTTTAATCTTATACAATTTTTACTAAATAAATAAATTCTCTCATAATATGAAACTA
TCTTGGAAAACACTTCTTCTTTGGTCTTTACCAATATTTGTAATTGGTTTCTTTTTCTGG
CAAGGTTTTTTAGGTCCAACTACTACAGATGTTGGCAGTAATATCGCAAGTTCTAGAATG
ACATATGGACGATTTTTAGAATATTTAGATATGGGTTGGGTGAAACGGGTTGACCTCTAT
GAAAATAATCATACAGCAATTGTAGAAGCTGTTGGGCCAGAATTAGGAAATAGGGTTCAA
CGAATTCGAGTTGAACTGCCAGCAAGTGCGCCAGAATTAATTACAAAATTACGCAAAGCC
AATGTTGATCTAGATGCTCACCCCCCTAAAAGTACAAGTGCAGTATGGGGACTATTAGGC

AATTTACTATTTCCTTTACTATTAGTTGGCGGGTTAGCTTTCTTATTTAGAAGATCTAAT
AATGCTAGTGGTGGACCTGGTCAAGCAATGTCATTTGGCAAATCGAAAGCTTTATTTCAA
ATGGAGGCTAAAACGGGAGTAGTATTTAATGATGTAGCTGGAGTTGAAGAGGCAAAGGAA
GAATTTCAAGAAGTGGTAACATTTTTAAAACAGCCTGAATCATTTACTGCTGTTGGTGCA
AAAATACCAAAAGGCGTTTTATTAGTTGGACCTCCTGGCACAGGCAAAACATTACTAGCA
AAAGCTATTGCTGGCGAGGCTAGTGTTCCTTTTTTTAGTATCTCAGGCTCAGAATTTGTA
GAAATGTTTGTTGGTGTTGGCGCTTCTCGTGTGAGAGACCTATTCAAGAAAGCAAAAGAC
AATGCGCCTTGTATCGTTTTTATTGATGAAATTGATGCTGTTGGTAGACAACGAGGAACA
GGTGTTGGAGGTGGTAATGATGAAAGGGAACAAACATTAAATCAACTATTGACTGAAATG
GATGGCTTTGAAGGAAATACTGGTGTTATTGTAATTGCCGCTACTAACAGAGCTGATATT
TTAGATTCTGCATTATTAAGACCTGGAAGATTTGATAGACAAGTTTCTGTAGATGTACCA
GATTTTAAAGGCAGGTTAGCAATTCTTGAAGTTCATGCTAAAAATAAGAAAATGGAACCT
AAAGTATCTTTAGAAACCATTGCCAGAAGAACTCCCGGCTTTTCAGGAGCTGATTTAGCT
AACTTACTAAATGAGGCTGCTATCTTAACGGCTCGACGAAGAAAAAATGCAATGACTATG
TCTGAAATTGATACATCAATTGATCGAGTAGTAGCCGGGATGGAAGGCACTCCTTTAATT
GACAGTAAAAGCAAAAGATTAATTGCGTATCACGAAGTGGGTCACGCAATAATAGGCAGT
TTATTAGAGCATCATGATCCTGTGCAAAAAGTTACATTAATACCAAGAGGGCAAGCAAGA
GGCTTAACTTGGTTTACTCCTAGTGATGATCAAAGTCTAATATCAAGATCTCAAATACTA
GCCCGTATCGTAGGTGCTCTTGGTGGCAGAGCTGCAGAAGAAATCATTTTCGGTGACGCA
GAAGTTACTACTGGTGCAAGTAATGATTTACAGCAAGTAACATCAATGGCAAGACAAATG
GTCACTCGATTTGGAATGTCTAAAATTGGACCTTTATCTCTTGAAAGCCAAGGAGGAGAC
CCATTTTTAGGTAGAGGCATGGGAGGAGGCTCAGAATATTCAGATGAAGTTGCAACTAAT
ATTGATAAGCAAGTAAGGGAAATTGTCAGTGAATGCTATGCACAAGCTAAACACATTATT
ATAGATAATCGAGTAGTGATAGATAGATTAGTTGATTTACTAATTGAAAAAGAAACAATT
GAAGGCAATGAATTTAGAGACATCGTTAAGGAATACACTGCAATTCCTGAAAAAAATTAC
TACATATCACAATTTTAAATTAAACGGGACTGACGGGATTCGAACCCGCAACTTCCGCCG
TGACAGGGCGGTGCTCTAACCAGTTGAACTACAGTCCCAAAAAGATGAATACCCAGAGAT
AATCTCATAGTTAGCTTAAACTTGTCAAATTAGAATATTAAATATGAAGAGTATTTGTAA
TAATACTCTTCATATTTAATATAGTAGCTAAAATATTTCTTTTAGAAGGAGAGAGAGGGA
TTCGAACCCTCGGTACGAAGTTAATCGTACAGCAGATTAGCAATCTGCCGCTTTCGACCA
CTCAGCCACCTCTCCATGATATATATTATATATATATCAAATATACTTATATTTTAATAG
GTGTGCATAAATATTTATGGAGCTAAGCGGATTCGAACCGCTGACCCTCTCAATGCCATT
GAGATGCTCTACCAACTGAGCTATAACCCCTTTGGTAAGATATTAAGAAATGGCTCAAGC
GGGATTTGAACCTGCGACCTTGGGCTTATGAATCCCCTGCTCTAACCACTGAGCTACTGA
GCCATAGTATAAAATATTATTAAGTATTATACCATAAATTATGGTCAAATATAAATCAAT
AAAAGAATAATTACTATCATTTACTACTTTTGTGCAAGTTGTAAATATTCGAATAAAATT

TCAAAATTAAAAATGTCTTTAACTTTTTGTTAGAATTATTGATTACGAAAGTATTGATAT
TTAGAATTAACAATAAATTTTGTCTTTATTTGTAGACACAAAAATATTGGAAATTTTCAT
CAATTAATTAGTTAAACTATAACCGACATACAAGAATTATTGAACTACGTTAAATTGTGA
TGCATTGGGCTTGATCAAAACTTCTCTAAAATCAATAGAATATAGTAATTCAAGTAGATC
AAATACTAACAAATGTGACAAACGCTTAAAAAAATAATACAATTAGAATCGTAAAAATTT
GGTTTTTGACTGTGTTAAATAAATTTGATTATCATAGATTTAATATTTAAAAGTGTTCTA
TTACTTGTATTGAGTAATTGTAATATTTAATATCTTCTTTTTTAGATAACAATCATAATA
TAAGCTAAAAAATAGTTTTTATTAAACAACTAGCATAGAGCCAATACCTTGATCAGTTAA
TATTTCTAGCAAGAGAGCATGATCTATTCTACCATCTAAAATATGCGCTGAAGCTACACC
CTGAGCCAAAGAACGAATACAACAATTCACTTTAGGAATCATGCCACCAGAAATTACTGC
TGTTTGAGTTAAATCTCTAGCTTCCTGTATATTCAGATGACTAATTAATGTTGAAGGATC
TGAAGAATTACGTAAAATGCCGGGCGTATCTGTGAGTAGAATTAGTTTTTCGGCATTAAG
AGCGGCTGCTATTTCTCCTGCGACAGTATCAGCATTAATATTATACGATTGGCCTTCCTT
GTCTGCTGCAACACTAGCTATAACAGGTATATAATTATTATTAATTAATATCTTCAGTAA
TTTAGTATCAATATTTTGTACTTCTCCAACAAAACCAAGGTTTGCTTTTCCATTTGGTCT
CGGAGTAATAAGTAATCCATCTTTTCCCGACAAACCTACACTTTTACCACCTTGTTTATT
AATAGTTGCGACAAGATCTTTATTGACTCTTCCAACTAAAACCATTTCCACTATATCCAT
GGTAGGTTGATCTGTTACTCTAACGCCATCATCGAATTTTGGTAATATTTTGAGGCGATC
TAACCAAAAATTAATTTCTGGTCCTCCACCATGAACTAAAATGGGACGTAGTCCTATAAA
AGACAGAAAAACAAGATCACTAATCACTTGATCTTTTAGTTTCTGATTTTTCATAGCCGC
TCCCCCATATTTTATAACAATAATTCTGGAAGAAAATTGCTGAATATAAGGCAGAGCTTC
ACTTAAGACTTTTACTCTTTCTGAATTAGTCAACATAAATAAAGTAGGATTGTGATATAT
AAAATTATATACACAGCTTGTGTTAATTTAAAAGATACCAATGGGTACTTGAAAAAATAG
AAATATTACTAACAGTTAAAACTATAGGGATTATAATTATGAATAAATTTTGGGATAATG
TATTAAGATTTCCTCGATTTTTAGTCAGCGTCATTTTAGGATTAATTTTGATAATTATTA
GTCCTTTTTTCGTGTTATTAAAAAAACCACTGACAAGTTTTTTTTTCATCATATCATTGG
CCGGATTAATCACAGTTTTGGCAATAATAATACAAAAAATGATAAATATCGAATGTTGTT
GAGAGAAATAAATAAATTTAATTAAAATTTTTGAACATATATAATATTATATTATCATTA
GATATGATAATTTATTTCCTTTTCTATAAAACCCATAAATATTAAATATTATTTGATAAA
TATTATGCCACTAAAACAAAGAGTTAGCTCTGAAAAGACCGGTGCCTTCGCACTTTTGGA
TAGTATTGTTAGGCACGGAGTTAAACATATATTTGGTTATCCAGGTGGAGCTATTCTTCC
TATTTATGATGAGCTTTATGCTTGGGAAGAAGCCTCTCTAATTAAACATATCCTTGTTCG
TCATGAGCAAGGCGCTGCTCATGCTGCAGATTCTTATTCTAGATCAACAGGAGAGGTTGG
AGTATGCTTCGCTACTTCTGGCCCAGGAGCAACTAATCTTGTCTCAGGTATAGCTACAGC
ACATATTGATTCTGTACCTATATTAGCTATAACAGGTCAAGTTGGGAGAGCTTTTATTGG
TACAGATGCTTTCCAGGAAGTAGATATTTTTGGGATTACACTTCCTATTGTAAAACATTC

ATATGTAGTTCGTGACCCTAGAGACATGTCTAGAATTGTTGCGGAAGCATTTTTTATTTG

TAAACACGGTAGACCAGGTCCAGTATTAATTGATGTTCCTAAAGATGTAGGATTAGAGAA

GTTTAATTATTTTTCTGTTGAGCCCGGAAAAGTTAATATTCCTGGCTGTAGGCCAATTAC

CAGCCTAAAGTCAAGACAAATCCTTATGGCAGCTAAAATGATACAGCAATCTAGCCAGCC

ATTGTTGTATATTGGTGGAGGAGCCATAATCTCTGATTCACATCAAATTATTAAAGAACT

TGTTGATTTTTATAAAATACCTGTTACTACTACTTTGATGGGGAAGGGGATTTTTAATGA

GGATAGCGATTATTGTCTAGGGATGTTAGGTATGCATGGTACTGCGTATGCTAATTTTGC

AGTTAGTGAGTGCGATCTTTTAATTGCTTTAGGAGCTAGATTTGATGATAGAGTTACTGG

AAAATTAGATGAATTTGCTTGTAATGCCCAAGTGATTCACGTAGACATTGATCCTGCTGA

AGTAGGAAAAAATAGGATTCCTCAAGTTGCTATTGTCGGTGACGTAGCAGAAGTTGTTAG

TGAAATATTGAATTTATTAAAGACTTCTTTCCCCCCTTATCCAGAGCAGATTATATCTTG

GCAAGAAAGAATTAATCGTTGGCGTCAACAGTATCCTTTACTGGTTCCTAGAAAATCAAC

AAGCATTTCACCTCAAGAGATTCTTGTTGCAACTAATAAATTAGCCCAAAATGCTTATTT

TACTACAGATGTTGGCCAGCATCAAATGTGGTCAGCTCAATTTCTGAAAGTAAAAGCTAA

GCATTGGCTTTCAAGTGCTGGATTAGGCACGATGGGTTATGGTTTACCTGCAGCAATTGG

CGCTCAAGTAGCACATCCAAATGACGTAGTCATTTGTATTAGTGGTGATTCTAGTTTTCA

AATGAATATGCAAGAGTTAGGAACTATCGCGCAATACCAGTTACCAGTTAAAATTATTAT

TATTAATAATCGATGGCAAGGGATGGTTAGACAGTGGCAACAAGCTTTCTATGGTGAAAG

GTATTCACACTCAAGAATGACAGAAGGAGCACCTGATTTTCAAAAGCTTGCAGAAGCTTT

TGGTATTAAAGCTTTTACTATTAATAATAGGCAAAATATGCAATCTGCTTTACAAGTTGC

TATTGATTATCCTGGTCCAGTTTTATTAGATTGCCAAGTTACAGAAAATGAAAACTGTTA

TCCAATGGTTGCTCCTGGAAAAAGTAATGCACAAATGATAGGTATCGCTAAACCGCAGAG

AGGTACTGCTTCCAACTATATTAATAATAGTGTTTGAAGTAAATAGTTAGTATTTTTTAA

ACAAAGAATAAAGTACATAAATGTATTTTTTTTATACTATTAAAAACTATGTAAGAACAG

CCTTTGACGAGAATTGAACTCGTGACCTTCCCCTTACCAAGGGGATGCTCTACCTCTGAG

CCACAAAGGCTTTTTACTATATTGGGCCGGGTTGGATTTGAACCAACGTAGGCGAAGCCA

GCGGATTTACAGTCCGCCCCCATTAACCACTCGGGCACCGACCCTAATTAAACCTTATTC

AAGATATGACCTATAAGATATCATAACTATGTTGCTAAAAACAAGTACTTTGATAAATAA

AACTGAATAAGTAATGGACATAGCTGGACTTGAACCAGCGACTTTCACGATGTCGACGTG

ACACTCTAACCAACTGAGTTACATGTCCAAAACTTTAATTATCATTATAACACTGTTTAG

TGATTTTATAAGTATCTATTTCTAATTGAAGATTTGTTTTAATCGTCCGGCTTTACCGAG

CAGATCTCTTAGATAATACAGTTTAGCTCTACGTATTTTAGCTCTTCTAATCACTTTGAC

AAAAGTTATTCGCGGTGAATTGAGAAAAAATACTCTTTCAACTCCAATACCTTGAAATGA

GCATCTTAAAGTTAAACTAGTATTTAAGCTTTTTTTTCTTCTTCTTGCTAATACTACTCC

TTCACATAATTGTTCCCGAGTTTTACTACCTTCCTTAACCATTAGTCCAAGTTGTATTGT

ATCGCCAACTTTAATTTCAGGTACTTCTGTTTTGATAAAAGGTATCTCAACACTTTTCAT

TAGCTGACTTAGCTTTGTACTATTTGCTTTCATATATAGGATTAGTCTTGTGACATGTTT
TGAAATATTATACATTAAAAAAACAAATATGAAAAATTTTAGCGAATAAGCATATTAGTA
TTTATAAAACTCTTTTTATCTCCATTATACAGATCCATTATATTCAACTAGATTCTTTCA
TAGTTTCTTTGATAATCAGTACATCTAGATGATCAAATATTATTTTCAAACATGGCTTAT
GTATGAAAAAGTTCTATAAAAACCTAATGACTAAAATTAGGGCTTAAGTATTTCTTAGTT
TTGTGTTATAATTAGTTACTATCAAAGGTAGTGGCATAACACGAAGCATCAAAATGTTCG
AACGCTTTACTGAAAAAGCTATAAAAGTCATAATGCTAGCACAGGAAGAAGCTAGACGCT
TAGGACATAATTTTGTCGGAACTGAGCAGATACTATTAGGATTAGTTGGTGAAGGTACTG
GAATCGCAGCTCAAGTTTTGAAATCGATGAATGTGAATCTAAAAGATGCAAGAGTTGAAG
TCGAAAAAATTATAGGAAGAGGGTCTGGTTTTGTAGCGGTTGAAATTCCTTTCACTCCTC
GAGCAAAAAGAGTACTAGAATTATCTTTAGAGGAAGCACGTCAACTAGGCCATAACTATA
TTGGCACAGAACACTTGCTAATGGGCTTAGTCCGAGAAGGAGAAGGAGTTGCGGCAAGAG
TTTTAGAAAATTTGGCAGTCGATGTTTCTTCAATTAGAGCTGAAGTTATACAAATGCTCG
GAGAAAATGCGGAAGCCAATGTAAGTGGAAGCAATACTACGCAAGCTAGAAGTAAAACAC
CAACATTAGAAGAGTTCGGATCTAACTTAACTCAAATGGCTATGGAAGGTGGTTTAGATC
CTGTAGTCGGAAGACAAAAGGAAATAGAACGAGTTATTCAAATCTTAGGTAGAAGAACTA
AAAATAATCCTGTCTTAATTGGGGAGCCTGGTGTAGGTAAGACAGCGATTGCGGAAGGAT
TAGCTCAAAGAATTGCTAATAGAGATGTACCTTCTATTTTAGAAGATAAATTAGTTATTA
CTCTTGATGTCGGTCTATTAGTAGCCGGAACTAAATATAGAGGTGAATTTGAAGAAAGAC
TAAAACGTATTATGGATGAGATTAAATCAGCTGATAATGTAATATTAGTGATTGATGAAG
TTCATACATTGATTGGGGCTGGTGCTGCAGAAGGAGCAATAGATGCAGCTAATCTGCTTA
AGCCAGCTTTAGCAAGGGGAGAATTGCAATGTATAGGTGCAACAACTTTAGAAGAATATA
GAAAACATATAGAAAAAGATCCAGCATTAGAAAGAAGATTTCAACCAGTTGTAGTTGGAG
AGCCAAGTGTTGAAGAAACAATTGAAATTTTGTTTGGTCTTAGAGACCGTTATGAAAAGC
ACCATCAATTAACAATGTCAGATGGAGCTTTGGCTGCAGCTGCTAAATACGCTAATCAGT
ATATTTCTGACCGATTTTTGCCAGATAAAGCAATTGATTTAATTGATGAAGCTGGTTCTA
GAGTCCGTTTACTAAATTCTCAATTACCTCCTGCTGCCAGAGAATTAGATAAAGAGTTAA
GAGCTGTATTAAAAACAAAAGATGAAGCTATTAGAGCTCAAAAATATGAAACAGCAGAGC
AGTATAGAGCAAGAGAAATGGAAATTAAAGCTCAAATTGCAGCAATTGCTCAAAGTAAAA
AGAATGAGCCTGATTTAAATTTAGAAGATCCTGTTGTTACAGAAGATGATATTGCTGAAA
TTGTTGCTGCATGGACTGGTATACCAGTAACTAAGCTTACTAAAAGTGAGTCAGAAAAAT
TAATGCACATGGAAGAAACTTTGCATGGACGTATTATTGGTCAAGACGAAGCGGTTGTAG
CTGTCTCTAGAGCGATAAGACGCGCAAGAGTAGGTCTAAAAAATCCTAACAGACCAATTG
CAAGCTTTATTTTTTCCGGACCGACGGGTGTAGGAAAAACAGAATTAACAAAAGCTTTGG
CTTCTTATTTCTTTGGTTCAGAAGCTTCTATGATACGGCTAGATATGTCAGAATACATGG
AAAGACACACTGTATCTAAACTAATTGGTTCTCCTCCAGGATATGTGGGTTATAGTGAAG

GTGGTTATCTAACAGAAGCGGTAAGAAAAAAACCATATACTGTCATCTTATTTGACGAAA
TTGAAAAAGCTCATCCGGATATTTTTAATCTACTTCTTCAAATTTTAGAAGATGGCCGAC
TAACAGATGCAAAAGGCAGAACTATTGATTTTAAGAATACTCTTTTAATTATGACTTCTA
ATATTGGATCTAAAGTTATTGAAAAAGGAGGAGGTAGTTTAGGCTTTGAATTATCAGAAG
ATCAAACAGAATCCCAATATACTAGAGTACGATCTTTAGTAAATGAAGAACTGAAACAAT
ACTTTAGACCAGAGTTTCTAAATCGATTAGATGAAATTATTGTATTTCGTCAGCTTACTA
AAGATGAAGTTAGAGAAATTGCAGAATTAATGCTTAATGAAGTCTTTGCGAGAATTAAGC
AGCAAGATATTCAATTAAATGTAACAGAACGATTTAAGCAACGATTAGTAGAAGAAGGAT
ATAATCCAAGCTATGGAGCTAGACCACTTAGACGAGCAGTAATGAGGCTTTTAGAAGATA
GTCTGGCTGAAGAAGTTTTATCTGGTAAAATTAAAGCTGGTGATAGTGCAGTAGTAGATG
TTACTAATGAAGGAGAAGTTACAGTTTTATTAGGTGAAAAATTAGAACTGTTAACATAAA
AAACAATTTATTAATTAAAGCATTGAAGAGGTTATATTCAATGCTATGAAAAAAATTAAT
AAGATGGGTTGCTATAATTGACAACCTCTGATATTCTTTATTTGTATATAAAGCCGGGAT
AGCTCAGTTGGTAGAGCAGTGGATTGAAAATCCTCGTGTCACCAGTTCAAATCTGGTTCT
TGGCATTTTAAGCATCAGAAAATTTTGCTATCTGAGCATTAAATTTTAATTGAAATGTTC
CTGTGGGACCATTCCTGTGTTTTGCTATTATAACTTCTGTGAAGTCTCTACTTCTAGTTT
CTTGTGTATAGTAACTTTCTCTATATAGCATTATTACTAAATCTGCATCTTGTTCTATAG
AATTATGAACAATAAAGTTATTTGAAATAAAATTGCATAAAGGTTTACATTCTAAATCAA
ACACTATTTGTAAACTTGTAACGTTTATTTTTTGTATACTTTCAAAAGTCAAAGATGAAA
AAGAATTAAAAACATTTTTTTGTTCTTCAAAGTTTTTTATTTGAATAGCAATCATGTCAT
TTTGATTAATTTTATCACATCTTTTCCATCCTTGTGTTGTGAGTAGTTTATGGTTACTTG
TTAATTGTATATATTTCCCAGCTTCGGTAATAATTTTATATACAGTTTTTTTCCCTGTTT
TCGAAATACTACATTTAATTGCTAGAAATAACTGTCGTTTTTGCGCACTAAAGTTAATGA
CTTCAATCTTTTTATAATAGCAATTAAATAATATTTGAGGTTGATTATATAATGATGTCT
GAAGATAATTGAATTTGGATATACACCCACTTTCTCTTAAATCGGATAACAAAGGCCTTT
TGTCACTCCGAGTTTCAAGGTTTCTATTTAGTTGAGATAATGCCAAGATAGGCAAGCTTA
AATCTTTTGCTAATATTTTAAGTGATCGAGTAATTAATGATAGTTCTTGAGATCTATTGT
CAGATTGTCTACTCTCCTGCAGTAATTGAAGATAATCTATAATAATTAATTCTATATTTT
TTCCTTGTAATTTAAAGAATTTTACTTTTGTCTTTATAGTGTCAGTAGAAATCTTAGCGC
TATCATCAATATAAAGATTAAGGTTAGCAAGAGTTTTACTTTTTTGAATAACATATTGCC
ATTCATCATTATTAAGCTGACCAGATTGAATTTTTTGGCCGTTTAAATTACACTCTTGAG
CTAAAATTCTTCTTAGTAGTTGCTCTGTAGACATCTCTAAGCTAAATAAAATAACATAAG
ATCTTTTCTGATTAATTACATATCTAGTTATATTAATAGCAAAAGCAGTTTTCCCCATTG
AAGGTCTTCCTGCAAGAATAATTAAATCTGACTTTTGAAACCCTTGTGTTATAGAGTCGA
GTTCTGTAAAACCAGAAAGTATACTACTATTTATACTAATCTTTTTCCTTGTATCCAAAT
GGACTAATAGTCTTGCAAGAATTGTAGCTAATGTTTGCGTATCTTTTTCGTCAAGTATTT

CATAAGCTTTTGTTAATTGACTAACAATAGAAGTTATGTTTTTTTGTTCAATAAGTTGTT
TTGAGTAGCTAATTAAACATAATGAATCTCCACATGCTAAAAGTAATCTTTTAATGTAGT
TATCTATAACGACTGCTGAATATTCGCTTATAGTATCAGATAGGGGAGCTTGTTCTATTA
GACTAAAAACTTCATTTAGCTCATTCAAGTCTTGAATCATTTTTTCATTTTTTAAGTTAA
TGAAAAAATTTCTTACGCTTTTTGTCTTTGCATGGTTAACATTTTCGAGTGCTGCTCTAT
AAAGTAATGAAGTTGATGTAAAATAAAAGAAATCAGGAGAGATTTTTTCTACTGATTTTA
GTAATGATATCGATTGCTGAGTTAATATTATACTAATTAATATTTTTTCAGCTAAAATAT
TATGAGGAGGTAGATATTTATAAATACTTATTACTTCCTCCTTGTAACATTACTCTTGTG
GTATTAGTTAACTATTGATTTAATTTGATTCTGGAAGAACTTGCAATTTAATATTTGCAG
TTACTTGATGTAATAGCTTAATTTCTACATCGTAAATGCCAATTGTTTTCACGTCTGGTA
AAGAGATACTTTGTTTTTCAATGTCTATGTTGGTAGTGTTTTTAATAATTTGTGAAATTT
CTTTTTCTGTGACACTACCAAAAATATTGTGGCCATCTCCTGTTTTTTTACTAACGCTAA
ACCTTTGTATTTCTTCTAGAAGTTGCTTGACTTTTTTAGCATTTTCTTTAGCTGTTTTTA
ATTTTTCTTCTCTTATAGCTGCATAAAATTTTTGTTGTTTTAAAATACCATTTGTGGCAA
CTGCCGCCATTTTATTCGGGATTAAAAAATTTCTTGCATAACCAGAAGCTACTTTGATAA
CGTCATTACTCTTGCCAAGTTTTTGAATATTCTCTTTTAGCACAACATTTATAACTTTTT
TACTCATCTTAATTCTAAAAAATCTGTATTATCCTATTATACCTGTTGTATACAGATTTT
AATAGGAAGTATTTATTTGTTGATATAATATTCTTGTAAAAAAAGGTAATTTTTTATTTT
TTTCTGATATAATCACTTTCACAATAATTCATTAGGAGAGGTGGCCGAGTGGTTGAAGGC
GCAGCATTGGAAATGCTGTTTAGGAGCAATCTTAACGAGGGTTCGAATCCCTCTCTCTCC
GTACAAAAAATATAAAAAGTTTTATAGAGTCCAATCTAAATCTCCAATAATATTCTTCCA
AGCATTTAATCCACCCCGAACTCTAACAATATTTAGTTTTTGAGCTATTAAAAACTCAGA
AGCAAATATTGATCTAGAATCTAAGCTACAGTAAACAAAACAAATTTTATCTTGTAAATT
GATATGAGAGTAATTCATTCCTTTTATTTTTTGTAAAGGTAGATTTAATGAATATATTAA
ATGACTTTTATGATATTCTTCATGATTTCTTACATCTATTAAAATATATTGTGGATTCCT
ACTAATTAAAAACTTTTGTAATTGAATAACACTAATTTCTCGTACAAATGTATTAGATTT
ACTGCCATAATATTTGTTCCAATGCTTTTTTGTGGATAATACAAATTTAGTATTTATAAT
CTTAAATTTGTTAAATGATGAAGTTAATGAGTTGTACGTTAATATAGTTCCACTTAATAT
AGACTTATAACCCAGAACGATTTTGACAGCTTCAGTTGCTTGAAGTGTACCAACAATACC
CGGTAAGAGACCTAAAACTCCGGAATTACTACAAGTATCTCTTGCGCTCTCTTTATTTTC
AGTTTCACTGTAGAAGTCTCGATAAACAGGTCCACCTTGATAATTAAATACACTAACTTG
TCCTTCAAATTGAAAAATGGCACCATAAATATGTATTTTATTTAACTCTAGGCAAGTATC
ATTCAAGAGATATCGAGTATCGAAGTTATCAGATCCATCTATAATAATATCATATTGTCT
AATAATGTCGATTGAATTCTCATAGCTTAGTCTTGTTTTGAACACAGTTACTATACACTG
TGGATTTAAATCTAATATTTTTTTTTAGCTATTTCTACTTTTGAATATCCTATATCATGC
ATCGTGTATAAAATTTGCCTCTGTAAATTAGAGAGATCAATTATGTCATCGTCTATAATT

CCAATGCTGCCAATTCCTGCAGCAGCAAGGTAAATGATTCCTGGAGATCCTAGCCCACCA
GCACCAATGAATAAAACTTTTGCTTCTTTTAATCTTTCTTGTCCTTCTAATTGAATTTGT
GGCAATACTAAATGCTTAGAATATCTTGTATATTCTTCTAGTGAGTATTTTGTATTTTCT
GTTTTAAAATTTAGCATAAAAATTTATAGTTTATTTAATAGCACTTAAAAGCTAGCTGAT
AAGTTTATTAAAATGACTTTTGTAAACTTGGTTCTTCTCTATCTAAAATAGCTCCTTGTG
TAGGACAGACCTGAACACAAATACTACAATCAATACAGGCAGAAAAGTCAATCCAATACC
AATTAGTGTTATTATTGTTTTTCCCCTTTCCTTGATGAATACAAGCTACGGGGCAAGCTT
CCACGCATTCAGCAACTCCAATACATTTTTCTGTTACAATTGTATGAGACATATTATTTA
ATTAATAAAAGATAAGCTATTTACATCTAGTAGTTTTGTTGTATACTAAATTTACATATG
CTATCCAGGCCAATATTTAAGGTAGTCTATTTCTACAGTTTACTAATTTGATATGCGCCT
TTAGTTCAGTTGGTAGAACGCAGGTCTCCAAAACCTGATGTCGAGGGTTCAAGTCCTTCA
GGGCGCGTTTAGTTTTATCGTAATTCAATAATATCATTTTATCTTTTTGCAAACATCAAA
ATGATATGATATAAAAGATCATATTTTATACATTGAATATTCTATGTACTAGTACTGTTT
TTTAATATGTTATTATTGATAAGTTAATTATACTAATTCTATTTTTGTAATAAGTAAAAA
CAGTTCTAATTAAGTATGCACGTATTTAATAGTCTTTTGTATATATAACTTGAAAAAATT
TAAACTATGGCTAAAAAAGTTACTGGAATTGTTAAACTAGCATTAAATGCAGGTAAAGCT
ACTCCTGCGCCACCTGTCGGACCTGCTTTAGGACAGCACGGCGTGAATATTGTTATGTTC
TGCAAAGAGTACAATGCACGTACTGCTGACAAGTCAGGACTAGTAATCCCTGTAGAGATC
TCAATATATGAAGATAGAAGTTTTACTTTTATACTTAAAACTCCTCCTGCTTCTGTACTA
ATCGCTAAAGCTGCTGGACTAAATAAAGGTTCGGGTGAACCAAATACCAAAAAAGTAGGC
AGCATAACAAATAAACAGTTAGAATCTATTGCAGAAACAAAACTTCCCGATTTGAACACT
AACAATATACCTCAAGCTATGAAAATAGTTGGAGGTACGGCTAAAAATATGGGAATTCTA
ATTAAAGATTAAAAGTTTCAATTATATTTACTATTTTTATGAAAAAATTTTCACGTCGAC
TTACAACATTAAAATCAAAAGTTGAGCCTAAACTTTACACTATTAATGAAGCAGTATCTA
TATTAAAAGCAACGTCAAATGCTAAATTTAAGGAAACTGCAGAAGCTCATATTGCTCTAG
GTTTAAATCCTAAATATGCAGACCAGCAACTCAGAGCAACAGTTATATTACCGAAAGGTA
CAGGTAAATTGATAAAAGTAGCAGTTATTGCTAAGGGAGAAAAATTAACAGAAGCAATTA
GTGCGGGAGCTGATGTTAGTGGCTCCGAAGAACTAATTGATGAAATACTGAAAGGTAGAT
TAGATTTTGATAAGCTGATAGCAACACCAGATGTTATGCCTTTAATAGCAAAGCTGGGAC
GAGTATTAGGCCCTCGAGGGTTAATGCCTTCTCCTAAAGCAGGCACGGTAACATTGGACG
TAGCAAAAGCCGTGAATGAATTTAAAGGGGGTAAAGTTGAATACAGAGTTGATAGAACAG
GTATAATTCATGTACCTTTTGGAAAATCTAGTTTTTCACAAGAAGATCTAGTTTTGAACC
TCCAAACAATTAAAGAATCGATTGATAGAAACAAGCCTTCTGGAGCAAAAGGGAAGTACT
GGAAAACTTTTTTCTTATCTAGCACCATGGGGCCATCTATTCAAATTGATATCACTAGCC
TTCTATGAAAAATTTTTGTATAATTATAGAACTTGAATAATCTATTATTTCAAATTATAA
TTTGCTAGCATTATTAACGTAACAACTTGCTTACTATATCTTATCTGCATTAAAAACTTT

TTATATATCTTATTAATTATGAGTACAAAGGTTGAAAATATCTTGGAAGAGCTAAAATCT

TTAAACCTTCTAGAGGCTGCTGAACTAGTTAAACAAATAGAAGAAACGTTTGATGTTGAT

GCATCTGCGGCTTCAGGAGGTATGATGATGGCCGCACCAACTTCAGCACCAGCGTCTGCT

GAGGTTGAAGAGAAGACAGAATTTGATGTTGTCCTAGAGGAAGTCCCAGCACCTAAAAAG

ATTTCTGTATTAAAAGCTGTTCGCTCTCTGACTGGTTTGGGTTTAAAAGAAGCAAAAGAT

CTTGTTGAATCTACGCCAAAAGTCTTAAAAGAAGGTGCTTCTAAAGATGATGCAGAAACT

ATGAAAAAACAGCTAGAAGATGCTGGCGCAACAGTTATTGTTAAGTAATGAAGAAGCGCT

TCTGTTTAAGTATAAACAGAAGTGCCTCTTTATTTATTATAAGCAATTATACTAGAAAAG

TCCAAGTGTAATTGCATTGTTGATAGGCATAGTTGCACCAATACCTAGCCAAATACTTAC

AACGGTACCAATTAAGAAAACAGTGGTTGCAATTGGTCTTCTAAATGGATTTTGAAATTT

GTTAACATTTTCAATGAATGGAACAGTAAGTAAACCAGCTGGTACAGCAGCCATACTTAG

AACACCTAATAGTTTATTAGGAATCACTCTTAATAGATTAAATGTTGGAAAGAAGTACCA

TTCTGGTAAAATCTCTAATGGAGTCGCAAATGGATTAGACTTTTCACCTATAGAAGATGG

TTCTAAAATTGCGAGTCCAATACTGCAGGCAAAAGTGCCAATAATTACAACTGGAAAGAC

GTATAATAGATCATTTGGCCAAGCTGGCTCTCCGTAATAATTATGGCCCATTCCTTTTGC

TAATTTAGCTCTCAGCTTCGGATCTGTTAAATCTGGTTTTTTAAGAATTGACATATTGTT

TTTGTGTTTAATATAAAGATTAGTCTAATTACTGTTTTAAAGTGTAATTTTCAACTATTA

TAATGGTCCAGAAATTCCTTGTTTACGTATCATTAAAAAATGCATAAGCATGAAAACGGC

AGTTAGTAGCGGAAGTACAAAAGTGTGTAAACTATAGAACCTAGTTAAAGTTCCTTGGCC

TACACTGACTCCTCCTCTTAACAATTCTACTATACTCGCTCCGACAACTGGAACAGCATC

AGGGACACCTGTAACAATTTTTACAGCCCAGTATCCGATTTGATCCCATGGCAAAGAATA

ACCTGTTACCCCAAAAGAAACAGTAAGCACTCCTAAAATTACACCTGTTACCCATGTCAA

TTCTCTAGGTTTTTTGAAACCTCCTGTTAAATAAACACGGAATACATGTAGAATCATCAT

CAAGACCATCATGCTCGCTGACCATCTATGAATTGATCTAATAAGCCATCCAAAGTTTAC

ATCAGTCATAATGTACTCTACTGATGTAAAAGCTTCTGCAACTGTAGGTCTATAATAGAA

TGTCATTGCAAATCCAGTTGCAACTTGAATTAGAAAAGATACAAATACAATACCCCCTAA

GCAATAAAAAATATTAACATGAGGTGGTACATACTTACTAGAAATATCGTCAGCAATTGC

TTGAATTTCTAATCTCTCTTCAAACCAGTCATAAATTTTACTCATAAAATAGCTTC-AAA

AGCTCTTCTAATATATTTTGCGTTTAAGCTACTATTAATTCAATAGTTATTGAACTTATA

TTTTTTTATAATCAATACTTATTAACTATTATAACATCTTTTTATTGATTTAAAATTCTC

TGACTTAGTAAAACAGGGCTATGTACTATAATTCTTTTTTTTTGTATAGAGATTAATTTC

GTTTTTAGTAAATTAGCTAAAATTCTTGTTACACTGACTCTATTACTACCTATAATCTGT

GCTAAAATTTTATGTGTAATAGTAAAGTTAAGCATAATACCATTATTAACTTGAGTTCCA

TTTTGCTCAGCTAATAATAATAATAAGCTCGCCAGTCTACTGGTTATACTTTTATGTGAA

ATGATTTCTATAAAATGATAAGCTTTTGCCGAGCAAAATACTAAATGATTTACAAAAAAT

AAATTGAATGGAGCGCAGTTACTACACGCTTTTATAATAGTAGTATAGTTAATACAGGCA

ACTTCTGTTTTATCTATTGCTTCTGCTTCGTAGTAAAAATTATCGTCGACCAATTCTATT
TGTCCAAATGTATCTCCTGTAGTTAATAAGTTAAGAGTTACTTTATGTGTATTTCTCAGG
ACTTTTGTAATAATTAAAGATCCTATCAATATAATGTATAATCTAGAGCTGCTATTAAAC
AAAAGAGTATCATTTTTCTTAAGTGAAAAAATTTGATAATTAATTTTATTACGATTAAAA
AATAGTAGCCATGGATTGCAAGAAAAAAAAATCACCTGAACTATTTTCAGAGAATATTGGA
ATACAATTTTCTTGATTTAAACAATTTTTAGTCATAATTACTACTTTATAAAAAAATATT
CTAATAATTAAGAACTGAAATTAAATGTCATATAAATTGATGTTAGTTGAGAATGACATT
GTATTATCAAAGGCCATTCAAGAATACTTAATAGATCAAGGCTTCAATGTTTATATTGCC
AATAATGGATTAGAAGCACTGAATCTAGCTTATCAATATAATTTCGACTTGATAATTTCT
GATATAATGATGCCGCTAGTTAATGGCTATGAACTATTAGCAAAACTTAAAAAAAACAAA
GCATTATCCAAAATACCTGTTATCTTTTTGACAGCTAAAGGAATGACTAAAGACAGAATA
AAAGGCTACGACATGGGGTGCTATGGTTATTTATCTAAACCATTTGATCCTAATGAGTTG
CTTTCAATTATTAACAACTTAATCGCTAGAGATGTCTTAAAAGAAGCTTCTTTACAAAAC
TTGGCAACATCAAACCAGCAATTAAATCATAAAATACGTTTGACACCTAGAGAAAAAAGT
ATTTTAGATCTTGTTGTTGATGGACTTACTAATAAAGAAATATCTACTATACTGAATACT
AGTGTTAGAAATGTAGAAAAATATGTTAGTCGACTCTTACACAAAACTAATATGAAGAAT
AGGACTTTGCTAGTCAAGTATTCTATAAATAATAATCTATTAAATAATGAGATCAATGAA
AGGGCGAATGACGGGACTCGAACCCGCGAATGATGGAGCCACAACCCATTGCCTTAACCC
CTTGGCCACACTCGCCATATTACACATACTATTATAGCTTGTTTTTTGATATATCGTCTA
GTTTTCGAGTTATATATTTATCTAAAACAATATTTACTTTGAATTTTTATGATACATAGC
AATATTAGTATTCAAATTAATGGGGAACCATTTAATTGCTCAAAGCCCATCTCATTGCAA
TTTTTATTAAATTATCTTGATTTTAATTCCGAGCGTGTAGCAGTCGAGCTAAATGACATT
CTGTTACCAGAGACTCTGTTTCACTCAACTTATTTGAATGATCAAGATAAGCTAGAGGTT
ATTACCATTGTAGGTGGTGGATAATCTATTTGTACATACACAGTTCTCTATCTCAAATGT
TTCATAGATAGAGATAATCTTCCTTGTTTTTTATCTACATGAATAATAACAGCTTTAATA
GTGTCACCTATTTTAAATTGTGATGATATCTGCTCTAATTCTTTGATATTAATCTCAGAA
ATATGGACAAGACCTTTGAGATTTCCTACCTTTATAAATAATCCGTAAGGTGTAATCTGA
TTTATTATGCCTTCAATAATATTACCAACAATTAAATTAGATGATGCTTGAGATATTAGA
GCTCTTCTATGACTCAGTATTAAATTATTAGATTTCTCTTCGACATTGAGTAATTTTAGC
TTAATAAATTTATTATTAAATTGCTCACTTTTTTGAAAATTACCGAGATGAGAATTAGGT
ACAAATCCAGATATACCTTCAAGGTTAATTATCATTCCTCCCTTATTGAATCCTTTTATC
ATTACATTAAGTAAAGAGTCTTCAGCTAATAGTTGCCTAATTCTTTTCCATGCTCTTATA
TATTCAAGGCGACGAATTGATAATATTAATTGTTTCGACTGAATATTATAATCTAATAAA
AAAAATTCTCTTGTGTCATTAATGTTTAAAGAAGTGAAATTATTTAAATCTTGATTACTA
GATACTTCTTGAATAGGCAAGTATGCAGATATTGGTGTTCCTATATCTACTAATACCCCA
TTTAATTCAAAGCTAAATATAGTTCCAGCTACAATATCTCCAAGATTCAAATCATACTTG

TATTTTTGTAAAACAGCTGCAAAATTTCTATGAGTAAATCCTTCATTATTTTTTGTCATA
TTGAATAGTATGAATTAAAGAGTAACTTTGTACAATATATTAAGGTAAATAAATGTTTGT
CGAATTGTATGAAAAAATTTTAATTTGTTCGGTTGTAATTAAGTATATAAACATATTAAA
ATATCTAGGAGACATGAAAAGTAGACGTAGATAGCTGCAGATTTAATACTATATAATCTG
AGGAAAGTCCGGGCTCCACAAATACAATTTATGCTGGAGAAATCCCAGTGTAGGTAACTG
CGAGGATAGTACCACAGAAAAAAAACCGCCAGAATAAGTTATATTAGCTGGTAAGGGTGCA
AAGGCAAGTTAAAAGCTTACCAAAAGTACTGCAAAGTATTTGTTAGGTAAACCCCTAAAA
TGGAGCAAAGCTACTAAACAAATATTTGTGTATATTTTATCTAGTTTAGTTGAAAATACT
GCATGAAGTTATTAGTAACAATAACTCGAGAGGAATAGCTATCCTTTACAATATATCTCG
ATTATATTGAAAGAACAGAACCCGGCTTATGTAGTACTTTTTATGTCTTGTATTATTATG
TCTATTGTCGAATCTTTAAATTTAACTGTTTGACAATTTTTTTTTGTAGCTGATGTTGTT
TATAATCTATTTCTGAATTGGTTAAAGTTTTTTCATTAGATCGATATGTAAATCTTAGAC
CTATACTTTTTTTCTCTTTTCCAATTGATTCATCTATATATTGATCAAATAATGTTGTAC
TTTCTAAATCTTTATCATAAAATTGATCCAATAATTTTAATAAGTATTTAATTTCCATAT
TCTTAGGTATTATTAAAGAGAGATCTCTTATGATCGATGGATACTTAGAGTATGGTTGAA
TCTGATAACTTAAATAATTGAATTCATTATGACAATCTTCTAAAATGTTTAAGTTAACTT
CAAGAAAAAAAAGTTTTGTGTTGAGTCCTAATTCACTATATGTTGCTTGATTAAGTTCAC
TGAATATGCCAATAATAATATTGTTATAGATTAGTGTGGCAAATTTTCCTTTTTGAATGA
ACTTTATTCTACCATCGACTATATCTTTTTTTGTCCACTCAATTTGTCTATTAATCTTTT
GAAAAAAATTTTCTACGATCCCTTTAGCTTCATACCAATTTAAAGAATGCGCAGGGTGTG
ACCATGTAGATCGGATATCTAAATTACCACCCAATATGATAGCTAGATTAGTAGTTTCTA
CAATTTTGTTTTGATCATTATGAAATACTGTTCCTATCTCAAAGCTATCTATAGTTTGGT
TGCTTTGTTTAATATTATAAAGATTTGATTCAATTAATCCTTCCAGCAAACTGCCGCGTA
AATTAGAATAATCTTGAATAAGAGGATTGTTTAAAGCTATATTTCCTTTGGACTTTATTA
AAGAATAATGAACTAATTCAGTTAATCCTAGATTTCTTAAAATACTTCTACTTTTATCTA
TGAATTGTCTCTTTGTGGATAACTTTTTGTTGAATTGAATATTTGGTACTGAGCTCCTGA
ATTTATGATAACCATAAATTCTAGCAATTTCTTCAATAACATCTATTTCTCGAAAAACAT
CTTCTCGTCTATAATTTGGAACAGTAACTTCTATATTTTCCTTATTTTGATAAATAAGAT
CAAAATTTAAAGAGTGAAGAATATTTTGAATTTCTTCAAAGAATAAAAAACGTGTTTGAC
CATTATACATAATTGGTCCTAAAATGTCATGAACTTTTTTAATTGATAAATTAATATTAA
GAGCATGATCATTTATTTTTTCTCGACAGAAAGTCTCTCTAATATTTCCGCCTGTTAGAT
CTGTGATTAAAGCAAGAGCTTCAAAATGTGCATTTTTCCAATTATCTACATTTAGCCCAC
GTTCTTGTCTAATTGAACTTTCTGTTCGAATATTAAGAACTCGAGAAGATTTTCTAACTA
CAGATTGTTTAAATATAGCAGACTCTACAAGTATCGATTTTGTATTATAATCTGTATCAA
AATCAGAGTTGCTTCCAATACCAGCTATACTAGTTACATTTGTATTTACTTGAGTAACTA
AAATATTTTTGTTTAACTCAATATTTTCATTGTTTAACTTAATCTGATCATTACTTCCAA

GAGGAAAATTGCTTCGAATTGTTAGACTATTTTTATGATTCATATTATTAATTTTATTCA

AATCAACAATATTAATAGGCTGTCCCCATTTCAACATAATATAATTGCTTATATCTGTTA

ACAAATTTCTGTGTGTAAACCCAGAAGAGAGCAACCGATTTTTTAGCCAATCTGGAGAAT

CTTTAACTCTTATTTCGTCTATAATGGCAGCAAAATAATTGTCACAATTTAATAAATCGT

GATTACTGATTATAGTCTCCTTTTTAGAGAAATATCTATTATTGGGTTGATGTATAGCTG

ATAACATTGAGGACCCTGTTAAAGCCGAGACTTCTCTTGATAGACCAATCATGCTTAACA

CGTCTGACCTGTTAGCTGTTGATGATATATCTAATATATGATCAATCTCACTTCCAATTA

TAATAGATTCAATAGCTTCAACTTCAAAACCTGCTTGTGTTAGTTGATTAGTTAAATTGT

CTATATCTATTGTTTTGATATTACCAAGTTCTTTCAACCAATTTAAAGAAACTTTCATAA

TTAATTGTCATCTATCTTATAGAAGTATTTATTTTACTTATGATTAAAAATCAATTTAAA

AAATTCATCAAATTAATATAAGTTATTTGATGAATTTGCTTATACAATATATTATTGTGA

TATCTTACTTTTATATGGCAGTTATTAAGAACTAGCTTTTGTAAAAGTTAGCCCATTATC

TTGGGCATATCTTTCCATAAATCTCATGAATCTATCCCAATCATCAGGACTTTTAATAAC

ATGTACAGCTTCTATTGCTTGTGGTTTACCATTAATAAACTTGGCATTAACATCTCTTGT

AATAATTTGTCCCTCTGTGTCCATCAGGTACATACCTGTGATTTCACCTTTGTCTGCCAT

ACTAGCATCTAAAATTTTGGGATTTGTAAACCGAAAAGTTGCTGTTCCTGTACTACCATC

TCGAGACCTTGTTAAACGTACATCTGGTACAACTTCTTCATTGATACCTTGAATAAATTG

AATTGTTGCCATATCAATCTTCTTTATTTGTATATAATTTATATTGGATAAAACTGATTT

CTAGTTCTTGAATGCTTGTTGAGAAACTGTTTTAAAAGCTTATAGTAGCAGTAGATCAAA

TTTTATGTATTATATAAAACTGTAACTTTTCTTTTTTGATCTGTTCAAAGAACTAACTTA

TTATAAGCTAATCATACGAATATTAGCCATATAATGTTCGTAAGTGAAATTAAGGGGTTG

TAGCTCAGAAGGATAGAGCAAGCGCCTCCTAAGCGCTAGGTCAGCGGTTCAAGTCCGCTC

AACCCTGTTAACGAATTAAGTAGTATTATTTAACTTGTTCTGCAAATCAGTATTGATCTA

AGTTAAAGTACTTGACAGCTAAAACAAAAGAATCTGAGAAATATTTTCATTTAAAGAACG

AACAATTATCTTAACTCTAAATATCTGATAATTATTAAAAAATTCTCTTTTTAAAACACA

AGATGATGAAGATATTTAAACTAATAATTGAAATTAAGTTTTTGAAGTTATAGAAGCCAA

TATAATACTTTCTATTAACAGATGCTTGCTTCTATTACTATCTGTTTACAAGAAATTATG

AGTATCAAAATAATTTTAGCTCTACTGACTACTAATAAATATTTATTTTAGTCTAGACAA

GATGAATCTTCATGGTATAACAGAGCTATTCTTTGTGTCATACTTATATAAAATTAGGAT

ACTGTTTTCATTATTATAACTGGGGTGGGAGGATTCGAACCTGCGAATGGCGGAGTCAAA

GTCCGCTGCCTTACCACTTGGCTACACCCCAACATGTGAACATTATAATAGCAGTCGTAT

ACCCTGCCATGTCAATAGACTATTGTCTTTATTTAATAATTTTTGCTGCTCTTGTAGAAT

AAAATCTTTTTTTTCAAATAAGATACGCTATGTTTAAGTTGCACGACTGTTATGTTTTCT

TGCTTTTGACTGAACAGCCACTTGATAGTAATTATTCCTTCTGCAATTTCGTTACTTCCT

ATAATAATACAAGCTACAGCTCTCTTTTTATTTGCTTGCTTAATCTGCTTACTAAAATTA

CTTGAGCTAACATCGATTTCTATTTTAAAAAACTGCTGATGTAAAAAACGCATTATTTGC

ATACCAGTTTCGTTAGCTTTTGTACCTTGTGTAGCAATATAAAAATCGATAGATTGATTA
GGCAAGTCTATATTATCTTTTGCAATCAGTAATAGGCGTTCTAATCCTATTGCACATCCA
ACAGCTGGTGTTGCTTTGCCACCTAATTGGTGAACTAAGCTGTCATATCTTCCACCACCA
CATATAGTATCTTGCCCTTTTGATGTCAATGTCTTAATTTCGAAAGCAGTATCATTATAA
TAATCTAATCCTCGAACTAACTTATTATTTATATTGTAGGGAATATTAAGTAATTTTAGA
TAATTGCAAACGTCACCAAAGTGTTTTTGTGATTCCAAACTTAAAAAATCAGAAATTTTT
GGAGCCTCTGTCAGTATTTTTTGTGTATTAGAATCTTTTGAATCTAAAATTCTAATTGGA
TTACTAGTTAATCTTTTTTGTGAGTCAGTGTCTAAATCATCATGGTACTTTGTAAGATAA
TCTCGTAATTTAACTTGATAAATACTACGATCCTCTACTTTTCCAATTGAATTCAGATCA
AGTTTCAAGTTGTGCAGATTGAGATTGTTAAATATGCTCATAGCTAAATGTATAACTTCA
CTATCTGCTCTTGCATCTAAGCTACCAATAAACTCAATGCCAAGTTGGTGAAATTGTCGT
TGTCGTCCACTTTGTGGTCTTTCGTACCTGAACATTGGACCGCTATACCATAATCTTTGT
AAACTATGGTGGTAACTCATTTTATTTTCAATAAAAGATCTCACAATACCGGCAGTACCT
TCTGGCCTTAAGGTAATATCTCTATTGCTGCGATCATGAAACCGATACATTTCTTTATTT
ACAATATCAGTATCTTCACCAATACCCCTATCATATAATTCACTATTTTCAAAAATAGGT
GTTCTAATTTCTTGGTAATTTGCACATTCTAATAATTTGGAAATTTTATTATGTATAAAC
TGCCAATATTGAATTTCATCGGGAAGAATATCTTTTGTCCCTCTAATAGCCTGAATTTTT
GCCATGATAATTTTTTGATTACCTTCTTAATATGTCTGCATTAAATAGCAGTTGTAGACT
GTTATTCAGTCATGTATTATACATTAGAACATACTGATGTAATTTATAAAGTACGGGCAA
GGAGGGATTCGAACCCCCGACACCATGGTTCGTAGCCATGTGCTCTAATCCACTGAGCTA
CAAGCCCACTTAATAAGTACCTTAACTATATCATTGTTCACTAATGTATACAACTATATT
CTTTTTATCTTACATGTGAAATTTTTGTAAAAGTAAAAAATTATGTTTGTTTTTTACAAT
TGGAGCTATTCATATATAATGCTTAATTTATTATAACTTTGTAAAAATATCTATAAGTTT
AATTGTATGATTCTATTATATTGGTAATCTTGTTATAATTTTGCGCTCTTATTTTTCAAT
TGGCTAGCATTTAATTAACTTATATTTCAAATAAAAAAATCTTATCTAAAATGAGTAAAC
AAATTCTATATCAGGATGATGCCAGAAAAGCATTAGAAAAAGGCATGGATATTTTAACAG
AAGCTGTTTCTGTGACTTTAGGACCAAAAGGAAGAAATGTTGTCTTAGAAAAAAAATTCG
GTGCCCCTCAAATTATTAATGATGGTGTAACGATTGCAAAGGAAATTAGTTTAGAAAATC
ATATTGAAAATACCGGAGTCGCACTGATTAGACAAGCAGCATCTAAAACAAATGATGTAG
CTGGTGATGGTACAACAACAGCTACAGTACTGGCTTCAGCAATTGTTAAGCAAGGAATGA
GAAATGTTGCAGCAGGTTCGAATCCGATGGCTATTAAAAAAGGTATTGAAAAAGCAACAA
ATTTTGTTGTTAGTAAAATAGCTGAGTATGCTAAGCCAGTAGAAGATACAAAAGCTATTA
TACAAGTTGCTTCTTTGTCATCAGGAAATGATATAGAGGTAGGTAAAATGATAGCGAATG
CTATAGAAAAAGTTGGCAGAGAAGGAGTTATTTCTTTAGAAGAAGGGAAATCAACTAATA
CGATTCTTGAGATCACAGAAGGAATGCAGTTTGAAAAAGGCTTTATTTCTCCTTATTTCG
TTACAGATACAGAACGAATGGAAGTTCTTCAAGAAAATCCTTTTATTTTATTTACAGACA

AAAAAATTACTTTGGTGCAGCAAGAACTTGTGCCATTGCTTGAGCAAATTGCAAAAACAT
CTCGGCCTTTATTAATAATAGCTGAAGACATCGAGAAAGAAGCATTAGCCACAATTGTAG
TTAATAAATTAAGAGGGATTTTGAATGTTGTAGCGGTTAGAGCTCCTGGATTCGGTGATA
GAAGAAAATCTTTACTTGAAGATATGAGTATCTTAACGAACGGACAAGTAATTACTGAAG
ATGCAGGTTTATCACTTGATACAGTTCAATTAGATATGTTAGGAAAAGCCCGAAGAGTTA
TTGTTACTAAAGACTCGACAACAATAATTGCAGATGGTCATGAAATTAAAGTTAAATCAA
GATGTGAGCAAATTAAGCGGCAAATAGAGACGAGTGACTCTTTATATGAAAGAGAAAAAT
TGCAAGAACGATTAGCAAAGCTTTCTGGAGGAGTTGCCGTCATTAAAGTTGGTGCAGCTA
CAGAAACAGAGATGAAAGATAAAAAACTAAGACTAGAAGATGCAATTAATGCAACAAAAG
CAGCAATTGAAGAAGGTATTGTACCAGGAGGAGGAGCTACTAATGTCCATATCTCTAGTG
AATTATTTACATGGGCTAAAAACAATTTAGTTGAAGATGAATTAATTGGTGCTTTAATAG
TTGAACGAGCTGTGACCTATCCGTTAAGACGAATAGCTTTTAATGCAGGTGATAATGGAG
CAGTAATAGTGGAAAAAGTTAAAAGTCACGATTTTCATATAGGCTATGATGCAGCAAACG
GTAATATTGTAAATATGTACGATAGAGGTATTATAGATCCAGCGAAAGTAGCTAGATCAG
CTTTGCAAAACGCAGCTTCTATTGCAGCAATGGTTTTAACTACTGAATGTATTGTAGTTG
ATAAAGTCGATGATTGAGCTATAAGATAATAGATAATCTCAAACAAAGAATAATTTCTAA
TTATAGAAATTATTCTTTGTTTCAATTAATCTAATTAATTGATCAAATGTATAATTTTTA
TAATAAAGCCAAAGTTTAAACACGTCTTCTTTTTCAACTATTAAGTAGAGAAGATATAAG
CCCGTAATACCTAAATTATCAATTAAAATATTATTTTCATATATAGATTTTTCTAAAGAT
CTAAAAAATGACATTTTTGAAAAATAATACCGAAATCTTCTCTTATAGTTATTTGCTAAA
TCATTACTGTCTAAAGATTGATTTCCAATTGAATTCAGAACCTTTAACAGAATACTTTTT
AAATCTTTATCTGACTTTATTTGAGAATCAATAATAATTTGATTCAATTGTATGTCAAAA
CTAAGATTAAAAATATTACTTGTTAAAGATATTTGATTTTGAATATCTAGAGATTTTATA
GCCAAGAAGATTAAACTTTTATTGTTCATTGTAAAATATAAGACAATTGATGATGTATTC
TTTAAATACTAAATTTATCTTCTTGTGCTTCTATCGTTTTTTGTAAAATAATTAATGTTT
TAATTTTTTAGTTTTAGTTACTACCCGAAAAAATAAATTCTGGAAATCTAGCTTGAGCTT
TAGCTAGAGAGTGTTTTTTCCCTTCTTCTTGCCAGTAATTGATAACTTCTGTTGTTTTGT
TTAATAACTCAATTCTGTCTCTTTCAGCAATCCACGGCTTGGATTCTAGTTCTGCTTTTA
ATTCTTCCCAAGCATCATTCCTTGGCCAGAAAAAATAAGAAGTAAGTGGACTAGTGCCAT
TTCCAACAATCTGATCAATAGCAATAGCAATATTATTTTCTAACCAAAGTACTTTTAGTG
TAAATTTTGACAAGTTTTTATCTCCATTATTAATATTAAGTATTGGTACAGTAGAATACT
GTACTAGTTATTTTTTTCAGCAACAATTTGAGCTTTGTTTAAAATGTTTTTTACCGTTTT
AGTAGCTTGTGCACCTTGAGATAATCGTTGCAAAATTTTTGAAATGTTAACACGAGTTTC
ATTATTTATAGGATTATAAAATCCTAATTCTTCTATAGCTTTACCATCGCGTTTATTTCT
GCTATCCATTACAACAATTCTATAGCTTGGTTGTTTTTTTCTTCCGTATCGCTTTAACCT
TAACTTGACCATATTAGTCACCTCGCTTATTTTTTATATCTTTATTAAATTATATCTTAC

GACATTATGCAATAATTCAAACAATTAAGCTGATTCAAGTCTAACATTATTAAATATTAC
CATTAAACATTGTAGAAAAATAATACCTAACATAGGTGACATGTCCATTCCAAACATGGG
AGGAATGCTTCCTCTGAATAGTTTTAAGTATGGATCAGTAATTCTATTTAATGAACAGAA
TGGTTCATTGTACCAATTTACAGTTGGAAACCATGCCAATGATAATTTAAGTAAAATTAA
AATTAAATAAATCTCAGAAAAATTAGCTATAGATCCAAGCAATAAATTTAATGTACCAGG
AAGAGTGTTCATACGTATTTTATAATTATTATATACTTCGTTATACTAATAGTTTGATTA
TATATTATAATCGATGTAAATAAATCAGCTAAACAAAAAACTTATCCTTAAAATCATCTT
CTCTTGAGCCTTTTTTTTTATCTACATTAGTATTTACTTTTGTCGTATAAATTTCCAGTT
TAGACCAAATATGTCCTAACTGATTTAATTGAGATGTTCCACATTCTAAACAGGCTATTT
TTATATTGCTCAAAACAGCACCTTTTTTTATAAAATCGTTCAATAATGGCGTAATAGTTG
CTTCATCCAAAAATAAATCTAAGACAAGTATTTTTGTAAATGAATTTAATTGATCTGGGA
CATTGTTAATAATATCATTATAATTAACTAAAACTATATTAGCTGTAGGAAGTAAAGATC
GTGCTCCTTCCGCTAAGATAAAGCCATAAGGCATAATAATTACAATAATATATTTATAAT
GACTGCTAATAAGACTTATAGTGGTATTAGCTTTTAATTCGATTGTTTCTGTTACTAGCC
ACTCTCTCATTATTTCGTAAGTAATCCATTTGCCTAGTTCTGAGCAAGCAGTTCTTAAAA
TCGTACCTGGATTATTGTCGTTTTCGAGAATCCCTGACCAATGTTGTATTAATGGATGGG
ATGCTATATTAATTTGGAGTTGCATGTTTTGTTAAATATATTTAACATTTAATTTGTTGA
TTGAATCTTCAATTCCTATTTGTTGAGTATTATTAAAATAGCATATATATTAAAGAAAAT
AATTATAAAAAATGAAAAAAAGTCTTTGGCTGTGGGGTTTTACAGATAGTGCTGAAACTT
GGAATGGTAGATTTGCAATGATTGGTTTTATATCAGTTATTTTTATCGAAGTAGTTACAG
GACAAGGATTATTATATTTAATAGGTATGATGTCATAATAAAAATCCAAAAAAGTTCTTG
TAATTTTTTATAAGAACTTTTTTGGATTTTTATAAATCTAATCTAGCGGTCTCATAGATA
GAACAGGTTCATTCTCTCTATTGATACCTTTTTCAAATCCAGCAGCAGCAGCTCTTGCTC
TGCCTGCATGCCAAAGATGTCCAATAAATAGGAAGAATCCTAGGAAGAAATGAGATGTTG
TTAACCAAGATCTAGGAGATACATAGTTAACTGAATTAATTTCTGTAGCTACACCACCAA
CGGAATTCAAAGAACCTAGTGGTGCGTGAGTCATATATTCTGCGGCACGCCTTTCCTGCC
ACGGTTGAATATCGTTCTTGATTTTATTTAAATCAAGACCATTTGGTCCTCTAAGAGGCT
CAACCCAAGGAGCTCGCAAGTCCCAAAATCTCATGGTTTCACCACCAAAAATAATTTCTC
CACTAGGAGATCTCATTAGGTATTTACCTAATCCAGTAGGTCCTTGAGAAGATGCAACAT
TCGCACCTAGTCTTTGATCTCTAACTAAGAAAGTAAAAGCTTGAGCTTGTGAAGCTTCAG
GACCAGTTGGTCCATAGAATTCACTAGGATAAGCTGTGTTGTTATACCAAACAAAATTAG
AAGCCGTAAGCCCCATGATTGATAAAGCACCTAGACTATAAGATAAGTAAGCCTCACCAG
ACCATACAAAAGCTCTTCTAGCCCAAGCAAAAGGTTTTGTTAGTATATGCCAGATTCCGC
CAGCAATACAAATAATGCCAATCCAGACATGACCACCTACAAGATCTTCCATGTTATTTA
CACTTACGATCCATCCATCACCGCCAAACGGAGACTTTAAGACATAACCAAAGATAACTA
GGGGATTAAGTGTAGGATTGCTAACAAATCTAACATCTCCACCACCTGGAGCCCAAGTGT

CGTATACCCCACCAATAAATAAAGCTTTAATAACTAACAGAAAAGCGCCAATGCCTAATA
ACACTAAGTGTATACCAAGTATTGTTGTCATTTTGTTTTTATCCCGCCAATCGTATCCGA
AGAAAGGAAATGATTCTTCAAGAGTATCAGGGCCAATCAGAGAATGATAAAGACCTCCAA
AGCCAAGAACAGCTGAAGAAATTAAATGCACTACGCCTACTACAAAGTATGGGTATGTAT
TGAAAATCTCGCCGCCTGGACCAACACCCCAGCCTAGTGTAGCTAAGTGCGGAATTAGAA
TAAACCCTTGTTCGTACAAAGGCTTCTCGGGAACAAAGTGTGCTACCTCAAAAAGAGTCA
TGGCACCAGTCCAAAAGACCATTATACCTGCATGAGCAACATGAGCACCAAGCAATTTGC
CAGAAACGTTAATTAAGCGTGCATTGCCAGACCACCAGGCAAACCCGGTAGACTCAATGT
CTCTACCGCCAACACCAACAGTTGTATTAAAGGGCGTTTCCACGTGGTAAAACCTCCTCA
GGGAATATAAAGTTTTCATGAGGTTGATCTTGAGCAGCCATCCAAGAGCGAATACCTTCG
TTCAATAGGATATTTTTAGTATAAAAAGTTTCAAACTCAGGATCTTCCGCAGCTCTCAAC
TCTTGAGAAACAAAATCGTAAGCTCTTAAATTTAGAGCTAGTCCAACAATTCCAAATGCG
CTTGTCCATAATCCAGTTACTGGTACAAATAACATGAAAAAATGTAGCCAACGTTTATTA
GAGAAAGCAACACCAAAAATCTGTGACCAGAATCTATTAGCTGTTACCATCGAATAAGTT
TCTTCAGATTGTGTTGGAGTAAATGCACGAAAAGTATCTGCAGCATCACCATCTTCGAAT
AAGGTATTCTGTACAGTTGCACCGTGAATAGCACATAGTAAAGCACCACCTAAGATTCCT
GCAACGCCCATCATATGGAACGGATTCAGAGTCCAGTTATGAAATCCTTGCAAAAATAGC
AAAAATCTAAAAATTGCAGCAACTCCAAGGCTTGGGGCAAAGAACCAGCTCGCTTGGCCT
AGAGGGTACATTAGAAAGACGGAAACGAATACTGCAATTGGTCCAGAAAATGCAATAGCG
TTGTATGGTCTAAGACCAACTAGCCTAGCAATTTCAAATTGTCTTAAACAGAATCCAATT
AGTCCAAATGATCCATGTAAAGCTATAAATGCCCACAGACCACCAATTTGGCACCAGCGA
GTAAAATCTCCTTGAGCTTCAGGTCCCCAAAGGAAAAGAAGTGAATGTCCCATGCTGTTT
GCTGGGGTAGAAACAGCCGCAGTTAAAAAATTGCATCCTTCTAGATATGAACTGGCTAGC
CCATGAGTATACCAAGAAGTAACAAAAGTAGTCCCAGTTAGCCAGCCACCTACAGCAAGG
TAAGCGCAAGGAAACAGAAGTAGTCCAGACCAACCTACGAATACAAATCGGTCTCTTTTT
AGCCAATCGTCTACAAGATCAAAACCACCACGAGTTTTTTCTTGTCCAATTGCTATGGTC
ATAATTTTAATCTCCAGAGCAAATTTAATAAGTAATTTGTTTAAATAAACACTTTAGATA
TACTCTAAAATTTTACTTTTCTTTACGGTCAATATAATTATAAGTCAAAATTAACCAAAA
TTCACGCTTTTAATAACATTTTTTTAGCTCATTAAGTTAATTTTTTAATATATAAACAGT
AATAGAGTTATTGTTTAACTTACAATACTCATAATTTATTCATTTCTATAATCCTTTGAA
AAAGATATCCTGTTCCTCGTGCTGTTAAAATCAAGTCGGGATTACTTGGATCATCTTCTA
GTTTAGCTCTCAATCGTGAAATATGAACATCAACAACTCTCGTGTCAACATGCCGTTCGG
GAGTATAGCCCCAAACTTCTTGCAAGATAGAAGCTCTAGAAAAAGGTTCTCCAGCCTTAC
TAACTAAAAGCTCGAGAAGACTGAATTCCATACCTGTTAATCGAACTCTTTCATTATTTT
TATAAACTTGTCTTTTGTTAGTATCTATTTTTAGAAATCCAATACTAATAATCCCAGAGT
TTGGGACTCCAAGACTAGTGGTGATTTTATCAGCTCTTCGAAGAACAGAACGAATGCGAG

CTTCTAGCTCCTTAGGAGAAAATGGTTTAACGACATAGTCATCAGCACCTATTTCTAATC
CGGTAATTCTATCGCAGACTTCACCTAACGCTGTTAGCATTATGATAGGAACATCTGATT
CTTTTCGAAGTTCTTGACAAACACCGTAACCATCTAGTTTAGGCATCATAACATCTAAAA
CAACAAGGCTAGGATACTCTTTTCTAAACACGAGTAAAGCTTCTTCTCCATTTGAAGCAG
TGATAACTTCATAACCTATTATTGTTAGTCTTGTTTCTAAAATTCTTCTTATGCTGGCCT
CATCGTCAACAACGAGAATCTTTTCTTTCTGGTTCTCCAATTTCTTATTTACTCCTATAA
CTCACCGATCATGATTACTATAACCTTAAATTTTTAGCGCTATGAGTATCTTTATTCACT
TTAGAATTTAGTTATTATAAATTTTGCAAGTCTAGTTTACATTAATTAGTACCAGTCAAT
ATTTTTGTTTTAAGTTATGACTATTTTAATCCATAAGTTGACTCTAGTTTGCAGAATTCA
ATATATACATATTTTCTAGCCAAGTAAATTGTATTCTATTTTCTAAATAAAAAAGGGGTT
GACAAACCAGTTGATAGAAAGTATTCTACATTACTATAAGTTAATTCCAACAAAGAAGTT
TTAAATCTTCTTTCGGAGTAATGAATACTTTAAT
