## Supplementary figures and images for "Genomic diversity of 39 samples of *Pyropia* species grown in Japan"

### S3 Fig

#### Chloroplasts

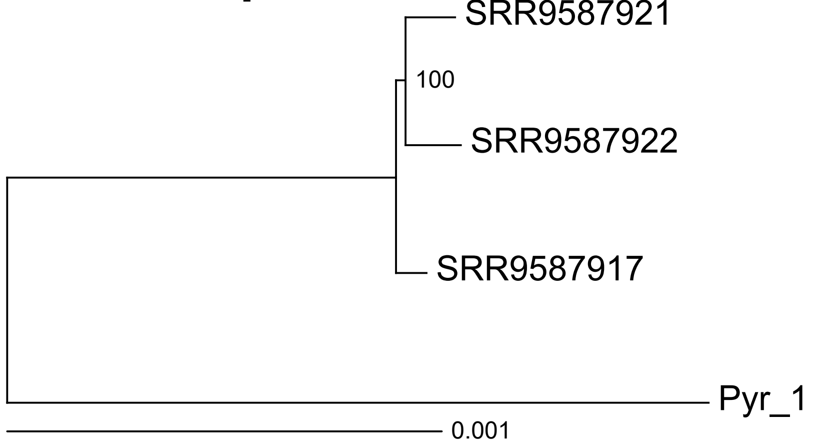


#### Mitochondria

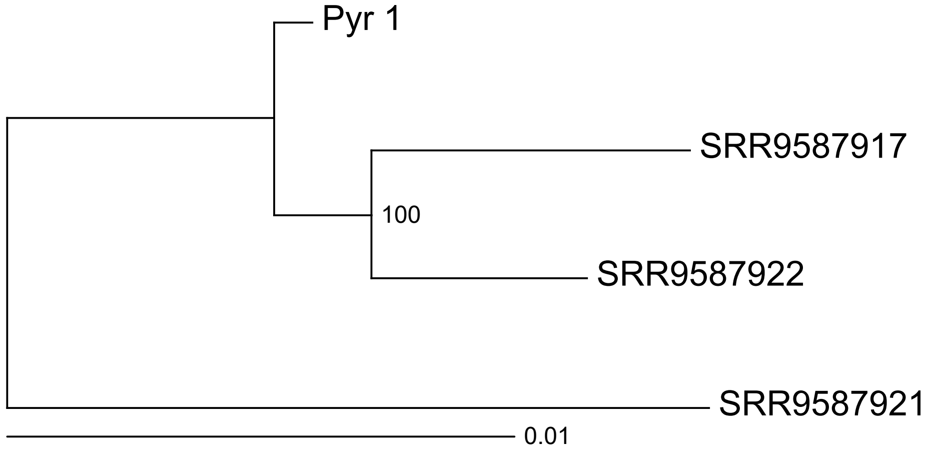

### S6 Fig

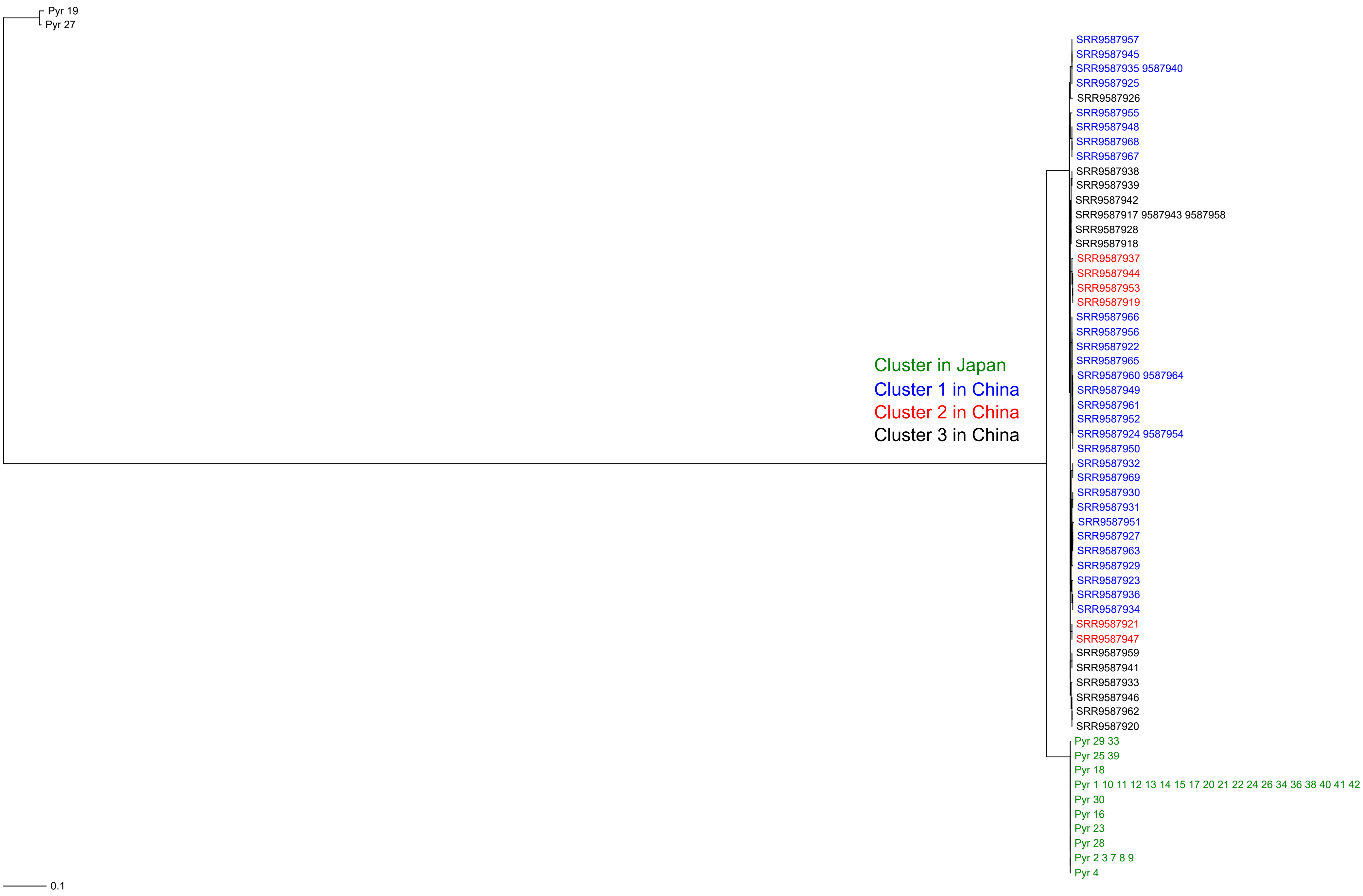

### S8 Fig

#### First rRNA repeat

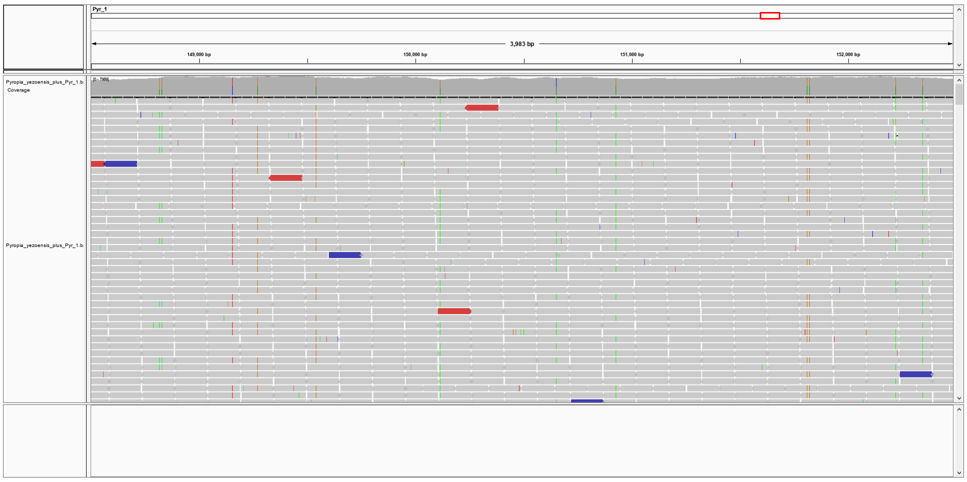


#### Second rRNA repeat

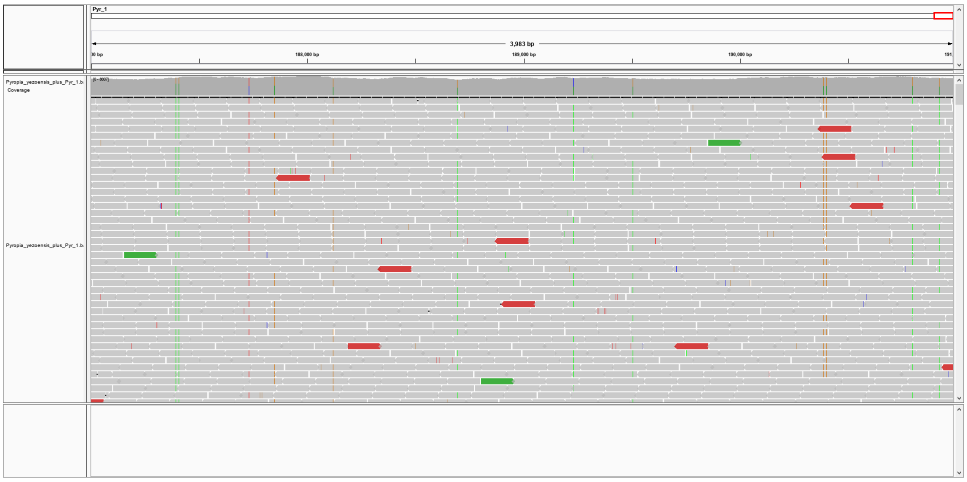

### S9 Fig

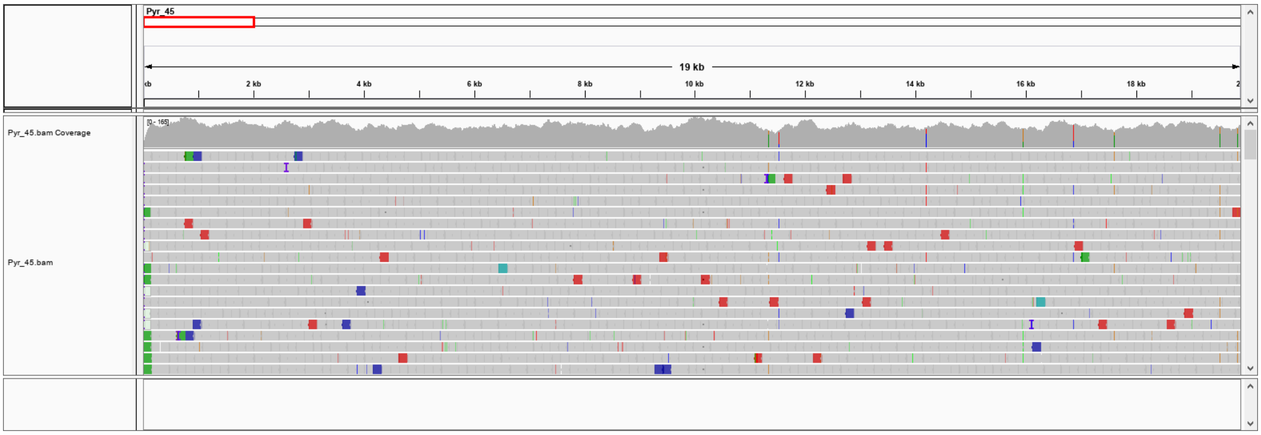

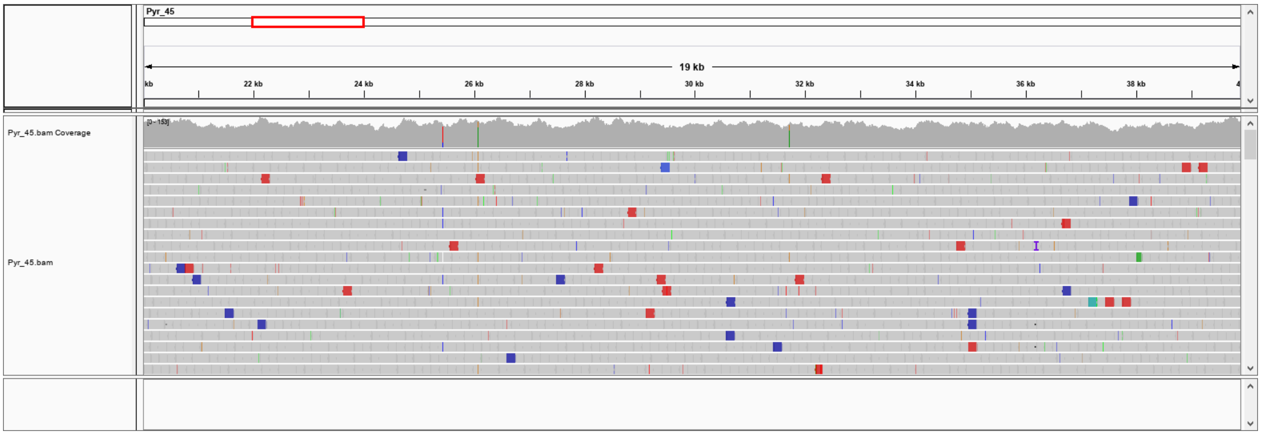

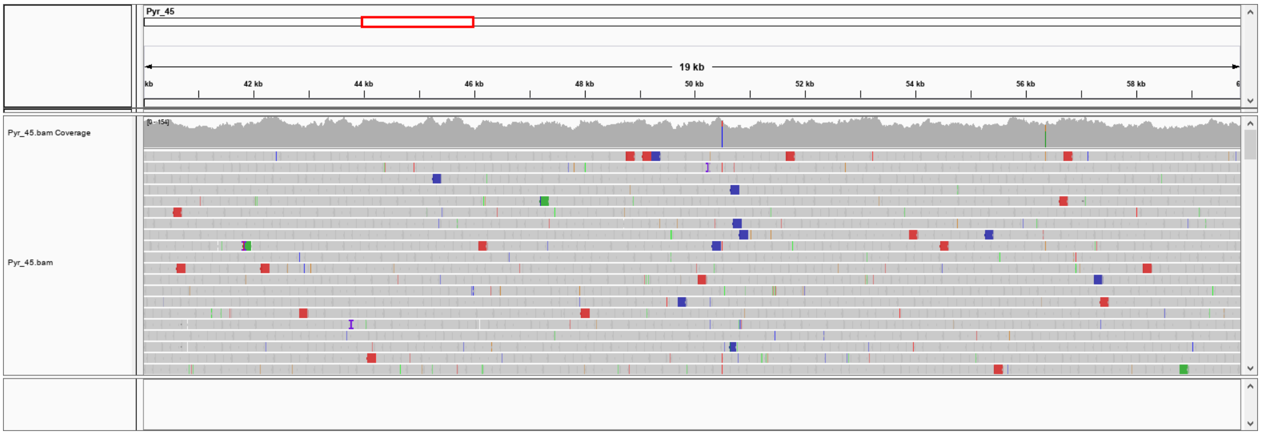

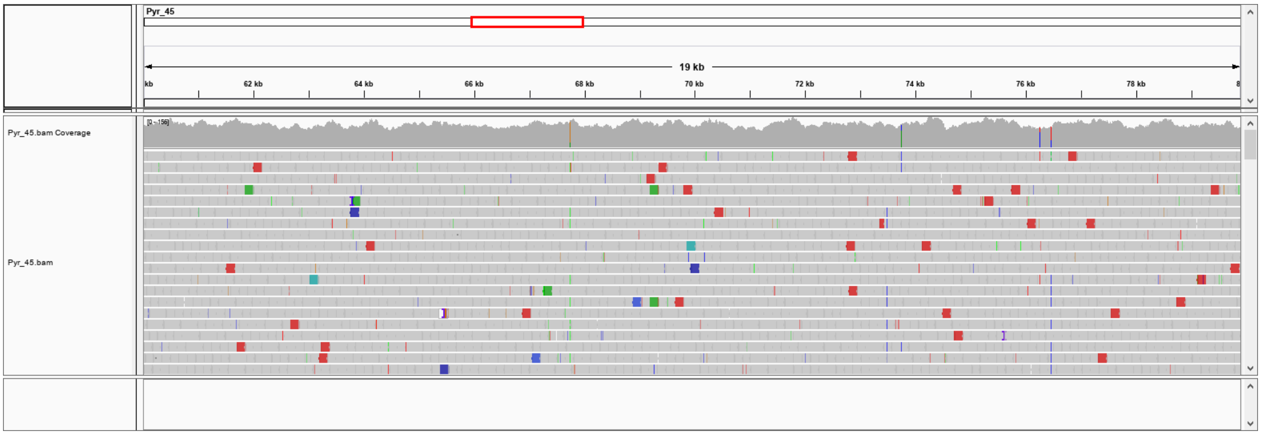

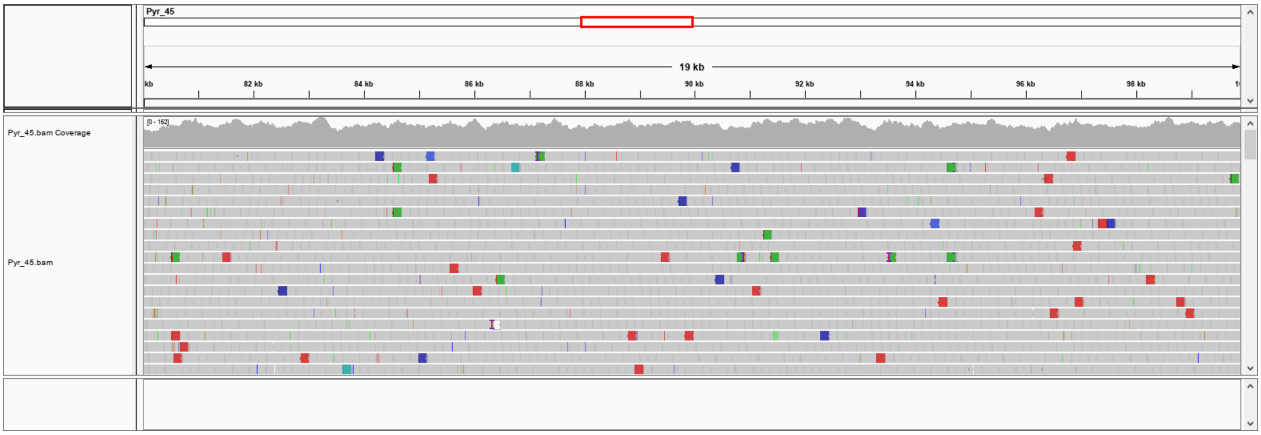

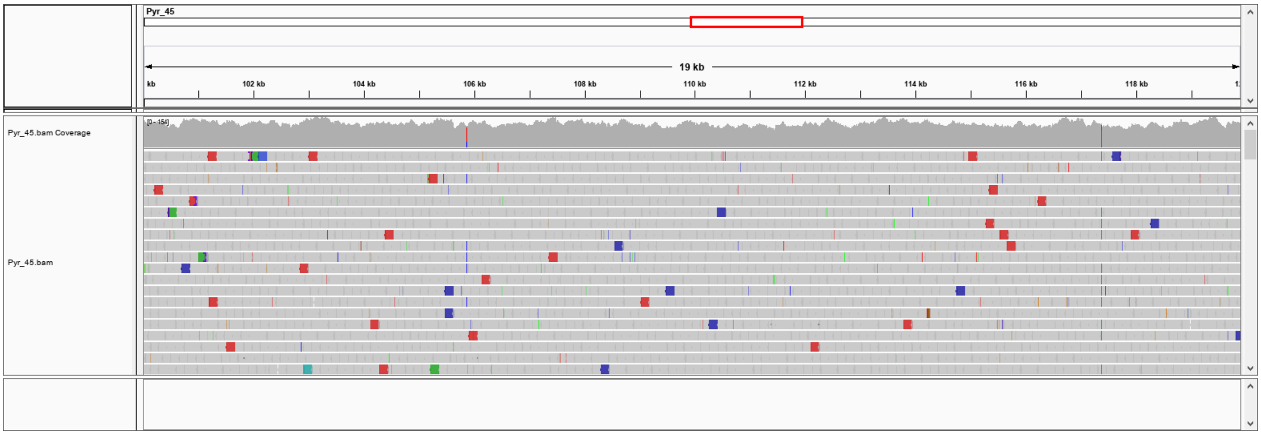

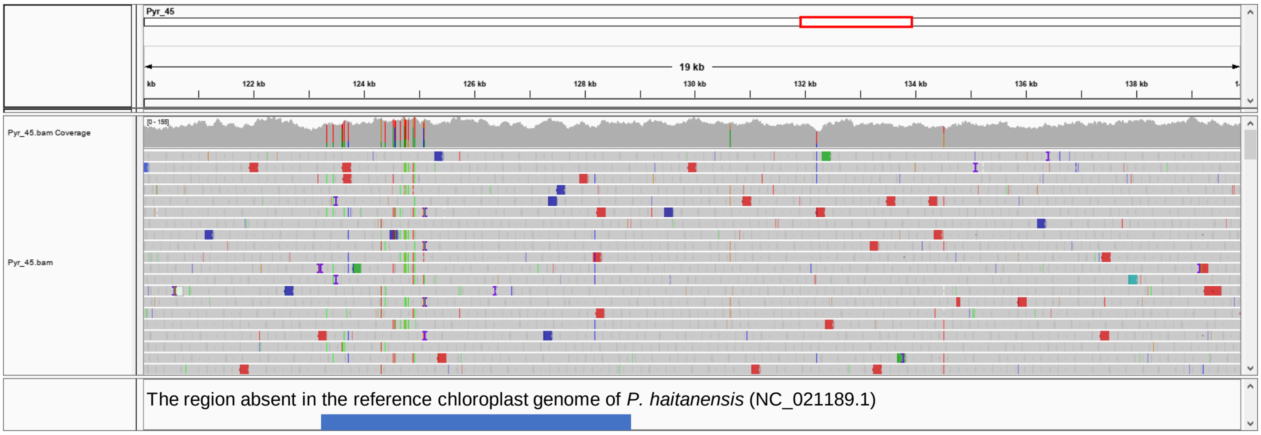

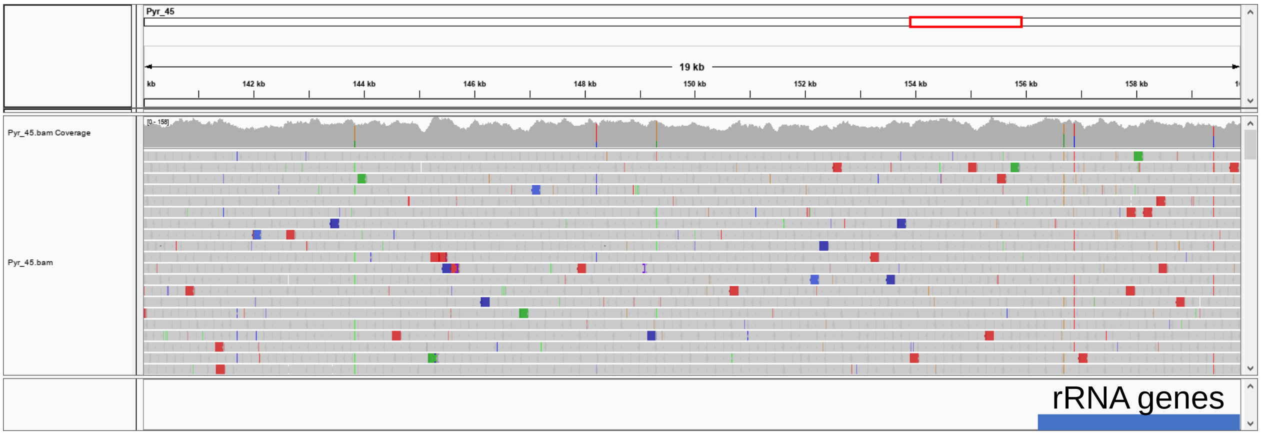

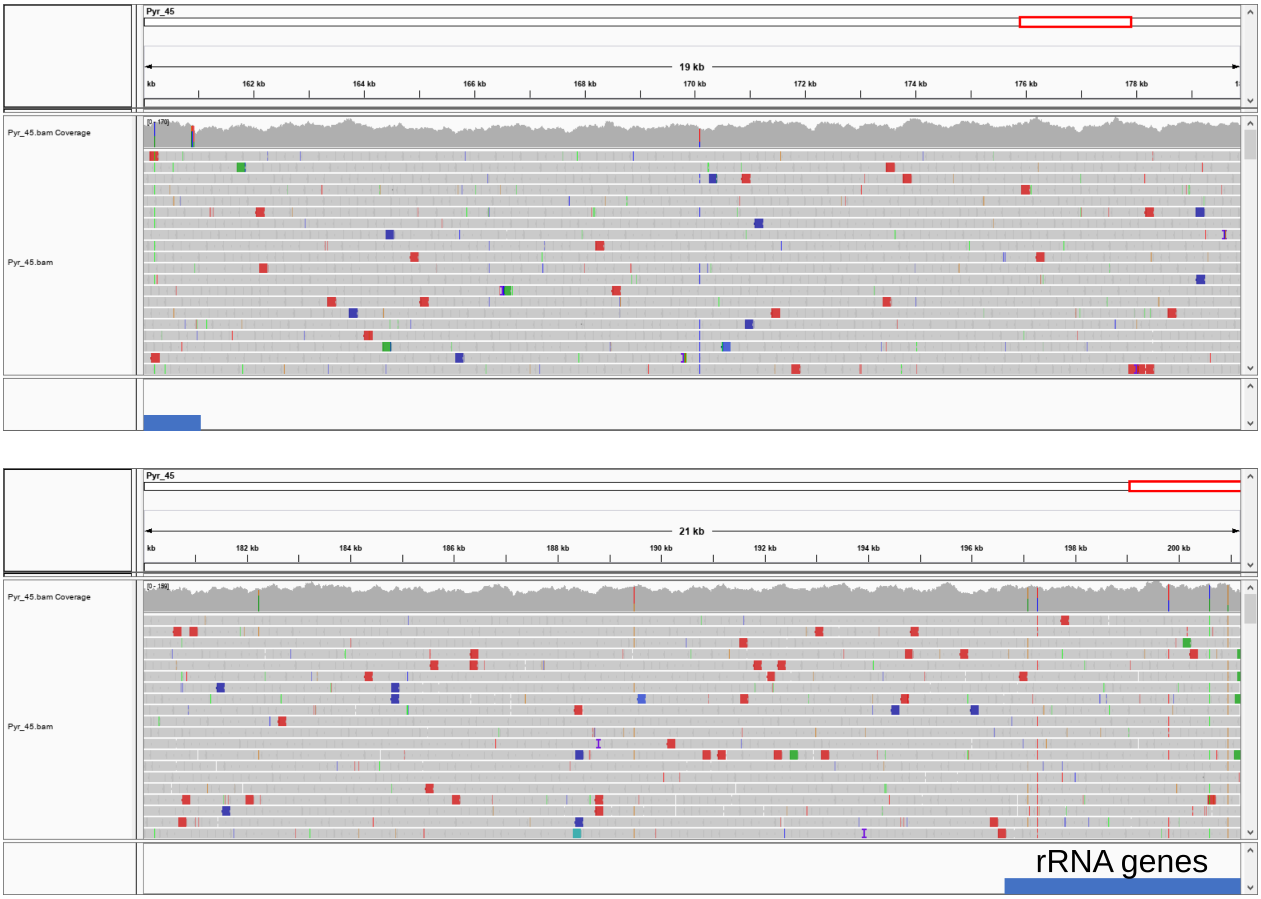

### S10 Fig

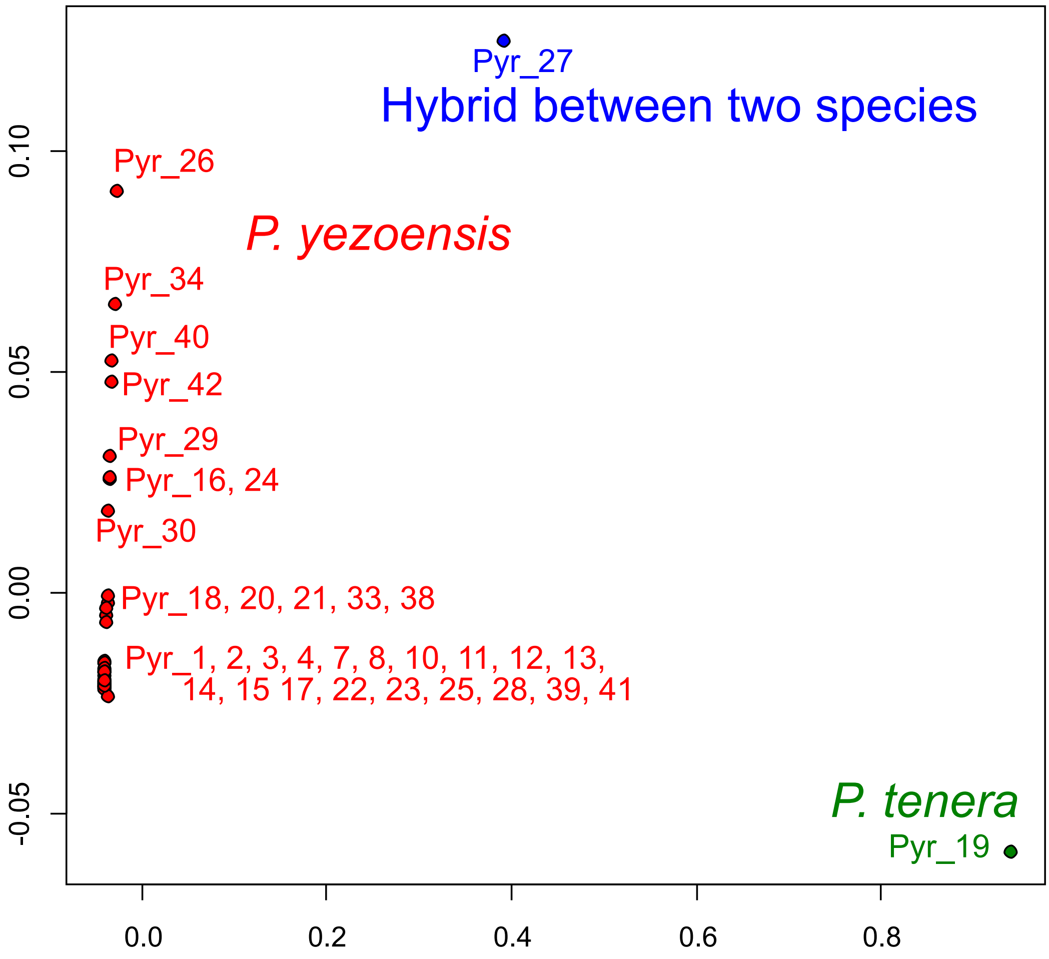

### S11 Fig

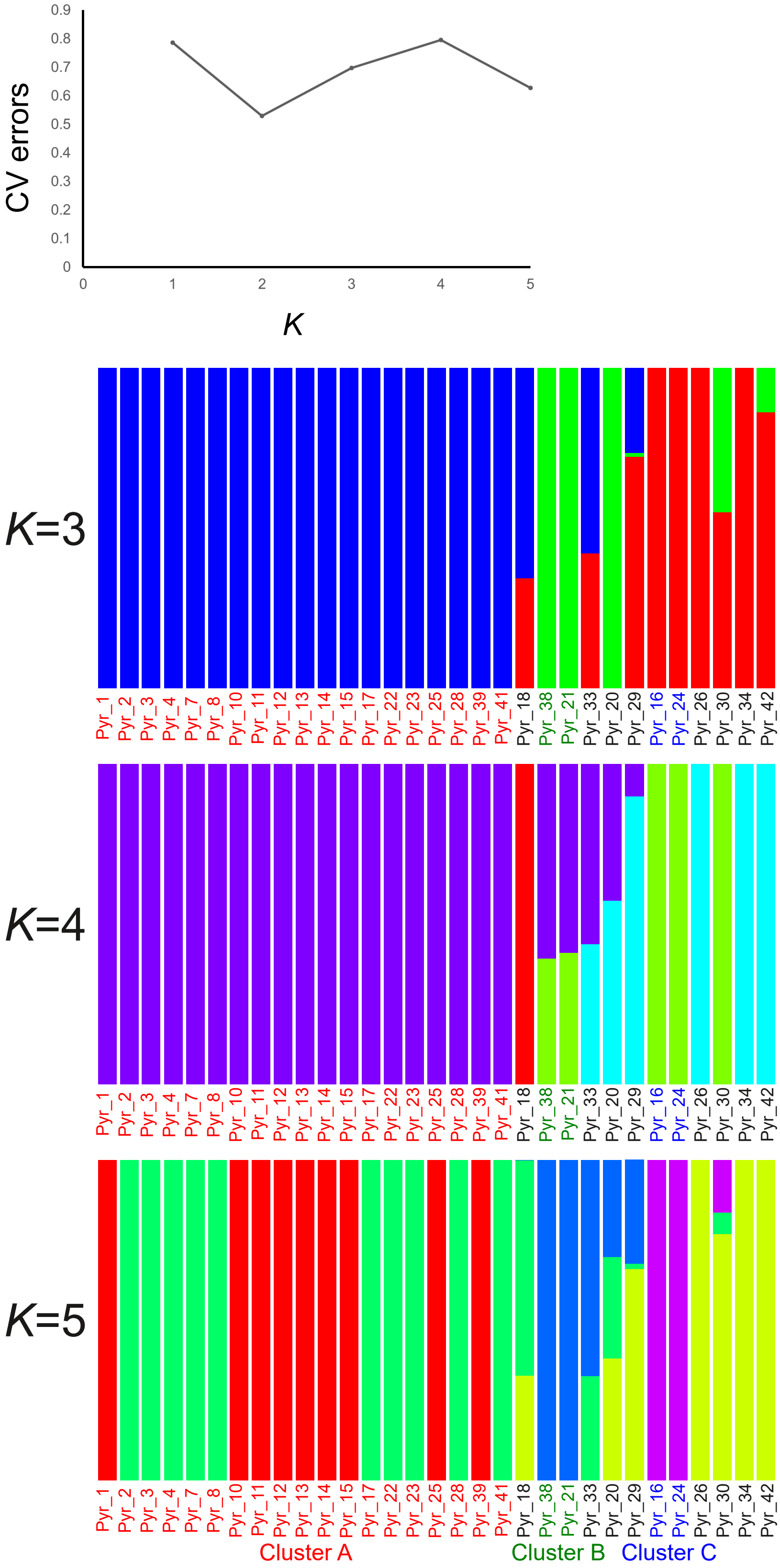
