## Supplementary material for "Genomic diversity of 39 samples of *Pyropia* species grown in Japan": S4 Fig

>Pyr_1
gttttattaattaaaactaccccagtattacacttttaaatatttataagaaataatgca
aatccctttacgctgttatttaatcgaatttaaatttcgattcgtttactttctacttag
tttttttctctcgttgctagtcgttataaactatcatgaggctgtgttttttttttgaaac
gtatactttttttgtttataaacgagggggcattcatcgcgacacatattgctgaactatt
ttcaacttctatgtatattagttttaacctagttttgtgcataaattatccttttgcata
ttatcattgcagccgctttttttaattcgagttgatacaaatcacaggtttggttctttaa
aaatatgcaatcattgacatttattacgtttttagctagtttacttttgtgctactttttt
aattttgccctacacttacgctttttttagacacttgaactgtaacgatagtttatgcatt
taaagtgcagttggaggcaaggatagcaacatatgtttgttgaacactacaaacaatgtg
tttactatctaacattgtctatctcttctttactaggctaatttgtttttacttagtgga
caatatgattaatttgcatttatttttccggaaaaataaaaaatatactttctttttcgt
atgcttgtcggcttctgtgtgtttaccgcaagaaagttttatacaaattactttcataat
ttctataatagtactatttgagttattctttttggttacatgcttatttttttgcaaaaaa
taatgtctaaaagtggacttgaaccactgacccaaagattttcaatcttttgctctaacc
gctgagctatttagacttaaacttttcttttcccgtatttagagcgaccatttttttcgtt
taacgactgcttgtaaatcgtaaacgcctctcacaatatggtaacgtactccaggtagat
cctttacacgaccacctcttatcagaacaactgaatgttcttgtaaattgtggccttcac
ctcctatatagcctatgatagaacgtccggtactaagacgtattttagctacctttcttt
cggcagaattcggtttttttagggttagtagtatacacttttgtgcaaacaccctttttttt
gaggcgacttattgagagcgggtgtttttgctttggtaattttgtgctttcttggatttt
ttatttttttggtttagtgttgacataatatttttattttaagtaataagggacttgaacc
cctaaccatttcggtgtaaacaaaacgctctacctattgagctaattactttacgtctgg
aaagatttgaactttcaacttttagattcgtaatctaacgctctatccagttaagctaca
gacgtttttctccacttgtatatttagattacttatctatattttggaaatagggacaaa
tattcttgtataatttacttcaggattattcgaaagtataattaaccaattaaacaagta
gaaaacgcatgttttgtattgttactaacttaaaggactattttataataaacttttttat
tgtattatataccaatatttagattgaaaatcgacccaaattcaatgtacttttagcaaa
tgacaccagtgttttagaaaaggtg-ttattttgaaaacatttttggtgtgatttgccct
ttttcagctagtaataatttaattaataacagaaagcgcggtttattgctcgtcgcgttc
aagttattactacggcaacttcctatgcagctgttgggaccccccccaactcgtgtaatta
cactttttagttttatttaaattataataaggttatactttatatcttttcttttattta
acccctaagatctttattacgccccaagatggtgtcataaaaaaa-tagcctggagggct
gccaggagtaaaaaaggcccataacggttatttcagcttttatagtgctatttttctttt
ttagagaaggctatttttaattgataactaagtcattttaagtgataattaagctatttt
aattaaaaatgacttattaaaggttttacacagataatacttcttttattaaaaatatac

tacgccgctagaccggaaaaaacccgccatagaacgatgagaggcaaaaagtgtcataaa
ttttaactattttgtaattgtacattttaaaattaaagtttgtcaaacatcaaaactttc
tgttatttcaataatgcgtgtttttaaggcttttaaagcagtttctcgctgatcccgaac
actaaacttttgttaacatgttcttgtaaactgtaattggatgtcataataaccaatgg
attatccggttttaaagtaaaattacctttcattgcaaggcgcatgctatcaccagctaa
aaactggtttaagaattcgatttgtttaccgataaggtgaaattcgtctcatagaataaa
ttgatacaaatgattttcataataatcttatcagttgtcttggggtcacatataacaagc
ggtagtagttttcaagtaatttaacagcgcactttttcccatattaggttctttacttca
tatatggagttggggtttttataatagcgtgaaaaaggactagtattattgtttagtat
gtcgaatatttgttctcataactggatagaacattccgataagttattttattattttg
ccttaggattgttttgcaaaatcgttctatgttgggcttttattttttt-----------
------------------------------------------------------------
-gtttaatatctggttcatcaaaattaaaaaatattggtgtagtaacaccagattcgatt
tatttcatgatgaagtgctctcttacaagccgctttttagtagattactaaaaattta
ctatgttgatttgatacgcatttttggatgggctcttgcttatttattggatctgaaaag
gagcttggagaattagttaagtagtcaaaatgattcagatttgtgaaattacgtttctta
gaaaattcaatatacgtatataaaatttggtccgagttatttcagctactagaaaagctt
attattattgtttttctctttgtaggagtttgttataaattaaaactaggcaaagaagc
tacatttttgttgtaacttggaaagatttggccgcaacgcgctttttatgtttagttac
ggctggattgagttcaaatactttggacatgagtcgttttaaaaattatttagaaatatg
tgttcgcaaacagattgagtgaaaaaaaa-agtttaaaaaaagcttcataaactttcacc
aaaagtttgatttatatcataaacggagaaaatccaaattgaaaaaactaaaatggtgtg
gcctaaaaaaatacggcgaagtacatatcaacttggttttcagtgaattaaacagaaata
cagtattcccacaagtacgataggaattagaatgatggataaatcaaaattttgatttag
atacgaaaaccatg-------------gtttacaagtaaactttttaattcaatattttg
tgctattagtaaagatttgtgcgactttaaattgaataagctgttaaaataaggtgaatt
taatttaggcaataaagtatcgtacaaatgtttgttctttctatatttgtttagagcttt
cctgtactttttattgtaagcagcaatagatcctatgtgcttgtgttgaaaatctaatct
tttaagtactttgttaaagctcaaaattttattttttaattggttaacag-tttttacg
ctttctagcatcgtatagtctttcttttagcgtttctagatgagaaggcttttctagtgg
cagttgattgaaatcacttttagtgag-ttttttactatagtacctgtaacaagtgtgct
aacagatactatgacacaaactacttttaagtccaa-ttgctctacaatgcg-tttgatg
aaagtcattttatataaattaatttattaatgtgtaattgtggtatgattttattttgtg
ttaatggtcataattaaatgcatacattatatattatccctccttcggagggata-atat
aataattcttattatataccaatatttagattgaaaatcgtgctattttcaatctaaata
ttggtatataaacaaacctgcacataaataaatatcacaaggacatcaccccaccaaaaa

atcgatattctctagagagcccctataagctttaatttactatctcttgcatgaaattat
accttcttggataggaatcatcgtgaaattgaaaaacattta--gatatgtcttatacaa
ttatgactaaaaaatctacaaaacaaacaactttacacaaatattttcgcaagtcgcata
catgagattgaacttaagaagcctcttttcacaaatcatacagcattttttggcgcttctt
aattctattgcaatattcaatgaaccaaaaacaaagttttttttaaaacgacctgcatgaa
cacaatcttaacaagtgtaaaaaagccctctagc-ccctttttttgcaacaccacgagaat
aaaccccggtcggggagaccccgaacccccctcggcgcgggccgtctagcaaagttagacg
gcccgcgcgcgcgttatagattcctatatcgctccattccattccatttcgctattgctg
cgcttcgcttatttttaattctaactgttccttttcgtcataacgattttaggtcgattt
tcaatctaaatatttgtatataaacaattatttatcttgataattatatatctaaaaaag
ggggcatagtttaataggtataatgtatgctttgcaagcataagattaccggttcaaatc
cggttgtctccaaatctaaaaatatcaatgcgttctagaatattacaatacaaaaataac
ttggctgactatgatcttttaactcaattttcgattaaaaatcataatagtatgcctact
ttctcttctcttaatgtcagagtaaaaaatattcaaactagcgacattaagcaggtttgc
ttaaatctaatctctattcaattagtaggaaacaatcatgttgttttcagtagccaaaaa
gacgtttctaatcttatactaaaatgtaacggtattaatacttattttattttagagaat
ttatctttgttaggctacaaaaacaatttaaaaagtgacaaaaacaataaattaactgtg
cacggtaaaaacttcttaaatattattgaatattttgcatctaattcaaattttcttaag
tccttagaaaaaacatctaataaacatttaagtattgctttttcttatcaacttgctaat
agtaaaaataatgaaatagctttgtatttttttaaaatcattaggatttccgctttaataa
cggtatagttcaattggttagaacaacggaatcataatccgtaagttgtgggttcaagtc
cctctaccgttatggctcttatcgtctaatggttaggacagtgctttttcagggcatcga
cgtgagttcaatcctcactaagagtaactttttaatataaaaatccttatacaaatacgt
gagcccataagaattcaaaagtaaaaaggatttttctagatctaccaaattattttggga
tccctaggtcgttaaaaatatttaaacaatgtattttattc--atatatatatatactaa
aacttaaaaatataatttagtgcaaaagtctctaaataactgaattttccgttgaatata
ctcgaccaatcataaagatataggtactttatatttaatttttggcgctttctccggtat
cttaggtgcttgcgcgtctatattgatccgaatggaactagcacaaccaggtaatcaact
attattaggcaatcatcaagtgtataacgtactagttacagagcacgcatttttgatgat
tttctttatggttatgcccgtcctaattggaggatttggaaactgattcgtacctattat
gataggtgctccagatatggcctttcctagattaaataatataagtttttgactactacc
tccatcattgtgtcttcttttaggatctgcgatggtagaagtaggcgctggcacaggctg
aactttatatccgcctttgagctctattcagagccattcaggcggtgctgttgatcttgc
tatttttagtttacacttgtcaggtgcttcttctatattaggagctattaatttcattac
gacgatatttaatatgcgcaatccaggacaaagtatgtatcgaataccgctatttgtttg
atctatcctcattactgcgtttcttttactactagcagtacctgtcttggcaggggccat

cacaatgctgttaacagatagaaactttaatacaacatttttgacccttcaggtggtgg
cgatcctgtattgtatcagcatttattctgattttcggacatccggaagtgtacatttg
tgcgccgtttaattctgttaaataagatggtttattattttaaattacttatctcgaaat
ctttaaactgaatagttaataaagatagacttaattattccttttgtcttaaaacacttt
cccttaataaagttggaaacaattctattaaaagcgggaatgaaagtctcttgcaggaac
ctgaatttcctaaaaatagtaggctagttatgctcatcatgaataagataataaagttga
tacatatcgacggttccacactaaactttatacataacatttcaggtaacagtatttatg
taatatgggtcggtaccagaatgtttaccttcccttgaataattattacacgtcagagcc
atcagctcctaaatgatatgcaacgttctaaaatgagcttgatcgagaaacagaagaact
cgggataccctaccaatcgaaagattcatgagtacggaactctcgtagtaggtggtagaa
gaattcaatcttcttcaaactaccaaaggggagtagaacttagattcttaagtgaaaaac
cctgcattagctcgcaagagtgcgctaggttagtagatttgagaaaagttaattctgaaa
ataaatttcaagttaataagaatactattcatattatatctgatatgaatgtccttattt
tagcatatgaactcataaaaagtaatcctggaaacatgacacctggtgtgaatggttcca
ccttagatgggttagacaagatgtgactgcaaaatattagtaccaaaataaagcaaggta
aatttttattcagccctgggcgtaagaagtacattcctaagcccggttcagcggataaaa
gaccattaggtattgctagcccgaaagaaaaaattgttcaaaaagctattctgctagtac
tagaatcgatttttgaaccaagcttcttgagaaattctcacgggtttcggcctaaccgag
gcaaccataccgctttaaagatggtaaaaagcgagtttcacggagttccctgaattatag
aaggagatatttcgaagtgctttgatgaaattgatcactctattttattggggcttctaa
gcaagaggatatcttgtgataagactttaactttaattaaaagagggttgaaggctgggt
ttatagatttaggaatattcacaagaactaaattgggcacccctcaaggaagcattctga
gtcctatcctatgcaatatctatttgcatgagctagatttatttctacttcaactaaaaa
ttaaattcgatacagggactagtagagcgaagaacccacagttcagaaaactacagtata
aactatccaaccttaaaacgcctctcgagaaaaagcttgtcagaagagacctttgaaaag
tgcatagtctgaaccccctagatcctaacttttgcagaattcactttgttcgatatgcgg
atgattttattgtgggagttacaagctcccatgaagttgctttagaagttaagaatatga
ttaaggaattcctttgtaatcatttgaaattaaatttgaatgagctgaagacacaaatta
ctcatattagagaaaaggatatatttttcctaggtacccttatcaaaggtaactgaaaga
aagagaaacctattcgattgatcaactttccctctagagaaacgtccatcaagacaagag
tcactccgcgtttaagtttgcatgcccctattaaaaaactctttgataaagctactgctg
aaggatttttccgtagggatgggattaattataaacctacctttgtaggtaaattgatta
atatggaccatgcagacattttattttttataattcaatagtaagaggagtactaaatt
actactcatttgtggacaaccacaaaagtttaggatcatgagttcattatatgaaatttt
cctgtgctagaactctagcgttaaagtacaagttacgtttcacatcgaagacttttaaga
aattcggctctaaattggcgtgcccgaatacaaaaaaaagcctgtttttaccaacgagct

ttaagaggacgcaggccttccaaattaatagccctattccttttgagaaaaaattactct
cttgatctaaaaaaattactaaatctaatctgaacaaagtttgtttaatatgtggtagtt
ctctatttgtagaaatgcatcatatacgtagtatagctggcattagaacaagactttgta
gcaataaagcagactttttctctctgcaaatggctgggataaacaggaagcaggttcctc
tttgtcgagaacatcacttgaagctacataataaaacactttcaccagaagaaacgttat
tatttcagaaaaatatgaaggaaatttagatgtatttctttcaattttaactttatttac
aatttaagttcggcgagccgtatgataagaaattatcacgtacggttctgagggcagtta
ttgacgctaaaacaacctcgttgtcaagtttctatttctgaccccctactaattttgcctg
gatttggcatcgtcagtcacatcgtttctaccttttctagaaaacctgttttcggttata
taggaatgatttatgctatgctttctataggtattttaggatttatcgtttgagcgcatc
acatgtatactgttggccttgatgtagatacaagagcttacttcacagccgcaaccatga
tcattgctgttcctacaggaataaaaatatttgtgcgccgtttcgtcgattttggaagtg
atcatttagtgattatagacagctggaacatctatttgtcaacttgcttaggtaatatcg
aaaggatattgctgaacacagcatgcaagtaaagcgaagttaacttctaaacatttatga
ataaattttgcaagacaattttctcacggttagggttatgtgacccgaagtgattctact
attgccctcgctttatggggaataccttgaatgtcatggggtatttaaatctacttccga
ttatctgaatattgattagaagaccagctagatttacaaagattaaaataattaatttaa
acgtagacaattcgcaacctaaaagaaataataattccaacgtcggaaatacgggatcac
ctaagtgccgacaggcatatggtgacagagtggacgtagtacgaagaatcaaaaacttcg
aaaggtccacagttaaatcattcttaaacagaaaactttacagtacaggaagtacaatta
atgtagagcgaaagtttgagagttttgttaaaagagcaatggaatttccaaaagaaatta
ttgacagggatctataccgcattttatgtgatgtaaattttctaaatattgcatataaca
atatcaagagcaaacctggtaatatgacttcgggaataactactgaaacgctcgatggta
tatcatacgatatattaaaagatatttccaatagtttaatggaagaatcttttcatttta
aaccgggcaggagaattcaaattcccaaaccctccggaggtgaaagatctttaacaatag
catctcctcgagataaaattgtacaggaagctatacgaattattttaaatgcggtttttg
aacctacttttctgaagtcttcccatggttttagacccaaaaagagttgccacaccgctt
tggaaaagataaaacttgagtttaagccagtaacttgagttatagaaggagatattacta
agtgctttgacaatattgatcataacttgttgatgaaattaatagaatccaagatttcag
acagacagtttacaaaacttatatcaaaaagtttaaaagcagggtacttcgagaccaaag
ttatttctcataacattgtcggaactcctcaaggatctattataagtcctattttatgca
atatctttatgcatcaattagatgtatttgtagaaaatctcaaaaacgaatttgacaaag
gcgttagagcaaaaaatcttagttcatatgagaactctagatataaaatcaagtgttcta
aacgatcaggagacatgataaaacttaagaaaatttataagatatctcaacgaagtccag
ttatggattttaatgactcctcgtataaaagacttagatatatcaggtatgctgatgact
gaataattgggattaggggtagttttatcgaaacgagacaagttctagaaagggtcagat

cttttttgaataacgttatgcgcctcgatattaacgactctaagactaaaattactaacc
ttaataaagataaagttgtatttttaggaaccaacatatttcggtctaaacatgtaaaat
atttcgcaaaaagtagtccgtctaaacaaaggcaaaaccttcagcttcaatttcacgtta
gtactgaccgaattagaagtaaacttgcaagtatcagcatgttatcgggcaatgtaccca
agcctagattttatgattatctttaaatcacgatcagattatacacctttacaactcag
tactaaggggctttataaattactactcttttgtaggtaactatagccaattcgtgtcct
gaataaggtgagtgatatattcatctgtagccaaactattggcaaggaaattcaatctgt
cagttactaaggtttttaaaaagtttggtccaaatctaagttcaggcaagtttgctctat
acgatcctaaatttatagccaacgaacccaggtttaaaaccgatgtatcgcccgtgattc
caaatttatatgctaaatttaaatctacggcaacgctatatgaacttgtatgtgctaaat
gtggatcggactatagagtggaaatgcaccatattagaaaaatgcagaatcttaacccca
aaatttccgaagtggaccgtctaacggttcgggccaatagaaaacaaatttcactttgca
gggaatgtcatatgaagtatcatcgcaacaaaacataaatttaatggagagccgtatgat
gggaaactatcacgtacggttcgggaaagggacctatattgccttcgggggtatgggttc
tagttcatagctggattgctactatgtgagaagggtctatttttttaaaaacccctatgt
tatttgctatagggtttatatttttattcactataggaggacttactggtattatactag
ctaactccggacttgatatatctttacatgatacttattatgtcgtagctcacttccact
atggtgcgccgtctaattgcgtttatcgtcgcttacttcagtgagcatgtattaccattc
ttgaattatataattaaatgttatacaatttatacattaaaacaatcaacttacttattt
ctggccaacagaaagaggactacagcatgttgataaaggagttttgaaatagtaaactcc
gagttatactacacacggttaggctaacgaactccttctacaatcagccaggcggatatg
aagttccgatcattagaaataatgaagtatctcacggtgaaagtttttacatagttagac
ctttacaggaatggtctggtacccaaaatttaagtaaaatttttaaagctgaggagaacc
taaaggtagttattaatggtaagcgtaagaattcgggaaatcctgaaagttgaaaaactg
gaggattcggagggatcgtagtacgaaatataagctgcttagcttatgttaggaagggtc
ctagttcagaagctattctaaattctaagctctcgggttatgaatctatagaagtaggct
taaataatattgacaaacaagtcttagaatatattaaaaccggcaaaagaattgaaggat
taagtagtttacttcggaatccaaattttcttattgcaagttattcaaaaatcaagtcta
ataaaggagctctcactcctggactgagtaatgaaacattggacggcataaaattagaat
gatttgaaaaagctgctgaaagtatagttaatgggtcctatcattttgaacccgttaggc
gaaagtttatacctaaaccgaaaggtggtgaaagacctcttggtatacccaatcccagag
ataaaatcattcaagaaggcatgaggcaactgctggagttagtttacgaaagaattttg
tagattcctctcatgggtttagacccaacaaaagttgtcacagtgctcttaatcaggtaa
aaatgactatggggtattcctcttgatttattgaggggatatatcaaaatattttgaca
ctgtgaatcatacttatttagtttcaaaaatagcgaaagttattaaagatcaagccttca
ttgatttaatatataaagttttgaaagcaggatacggcttttccaaaagaaatgtcgtaa

gcactagtagaggctttccacaaggtggagttattagtcctatactagccaatatttact
tacatgattttgatttaaaaatattggaaatgtcggaaaactttaatagaggtgtccgtc
gaaaagcaaatcctgaatatactaagatggttagagatggaaaagtagatagaaaaaatt
ttatttatcccgctatgggaaatgatgcttattttaaaagaatgaaatatgtaaggtatg
cggatgatttcttatcggaatcattggttcgaaagcagactgtgaaggaattagaacta
gcatagcgagtattcttaaagaagagttcttacttgaacttaatattgagaaaacaaaaa
tcacgcatgctaaacatgattgcgctttttcttagggcacaatattcatatatcaatgc
ctcccaaagataaaatacaatatcttccaaaaagaggaaacaagcttgtaagaactacaa
gtcgacctttattggatgctcctattggtaagatagtgttgaaactaggttcggtagggt
attgcaaaactgatggatctcctaggagattcggaaaacttctacatgaaccaatggcgg
aaataatttataggtataaaatactgcaaagtggattattaaattattattctatggcta
ataactatggtcgtctatctgcaagaatacattgaaccttaaaatattcttgtgctctaa
ctatagcttcaaaaatgaaattggggactcttaaaaaagtgtttaaacattatggagcta
atcttgaaataaaaaacgaaaaaggggagattatccaatgctttcctaaaatatcgtatt
ccagacctaagaaccctatcaaaacgaaaatatttgatcctattgatcatatagagaaat
cttccaattatttcaaaagaagtttagcaacttttgagcaagcttgtatgttatgtggaa
agcaagatactactgaaatgcatcacattaataaactgaagaataactcttccacggatt
ggttaacttctcgaatggtcaaaatgaataggaagcaaatacccgtttgtcgaaattgcc
accaacttattcacaagggtaagtatgatggttcaaaaattatttaagtgactcttttga
agtgaaagccatatgcgctgaaaagcgcacgtatggtttggagagaggattttgatactt
aaaaagattagtctcctactctacttctgtctatgggagctgttttcgcaatatttgcag
gcttttattattggtttgaaaagatatccggatttcaatattctgaaatactaggtcaaa
ttcacttttgaggcacttttataggtgtaaatctaacctttttccctatgcactttttag
ggcttgctggtatgcctagacgtattccggattacccagattcttatgccggttggaacg
caatagcttcttacggttcatatgttgcgttatttagcacgctgtttttcttttatcttg
tatttaacacacttgtaacagcaaaaaagacacctgctagaaataacccatggaactttg
aagattcaaaaatgggctcaactacattagaatgagaaatttcttctcctccagcttacc
atacgttcaatgagattccagttataagagaaacagaaacatctttaaaaataaattaac
ttatgataaaaaaaaatactacaatattatttttagggttagctttgctaacccaactac
tacagagtagaatagttattagtgattctgcagaagactggcaattaggcttccaagatc
ctgcaacacctataatggagggaatcattaaccttcatcatgattttatgtttttatct
gtgctatctcaatttttgtatcttgaatattagcacgcacgttatggcactatcactgaa
caaaaaatgagtacccttctgccacggttcatggaacagccatcgaaataatttggactg
ttactcctagtatcactttgttggcgattgctgtgccttcttttgctctgctatactcta
tggatgaaataattgcgcctgcaataactattaaaacagtaggtcatcaatgatattgaa
gttacgaatactcggactatacaaatgaagacgataacactataatgtttgaaagttata

tgattccagaagaagatttaactttaggtcagttaagattattagaggtagataaccctatggtaatacccgtaaatacacacgtacgtctaatcataacagcagcagatgttttgcacagctgagcagtgccttctttaggtataaaatgtgacgctgtaccaggtagattaaaccaaagctcgctctttgtaaaacgtgaaggaatattttatggtcaatgtagcgagatttgtggtgtaaatcatggttttatgcctatcgtagttgaagcagtgtctttaccaaattatatttcttgagtagctaataaacttagcgaataaatagcctaaaaatattctaagcatctgatatgcgtgtatctctagcccaactatttttctttggttcatctatttttattttattttgatttaatcctaaatacttaacttatacaatacgactattaaaaaaaattaaaaagcgttctaaataataaagagcgctttaataaaaaacttattatggcaactacgacaaatctaaactttatcaaaacagctaaacaattacaacgccaccctttttcatttagttgaccccagcccttggcctgtaacagctgcaatagccgctttttcatgtgctttaggcggagttatgtatatgcatgcatacagtaatggagggtacctattatttagtggtttttctttactgttatttacaatgttctcatgatggcgcgatgttacaagagaagccactttttcagggcatcatacaggtgctgttcaaaaaggattgcgttatggtgtaattttatttatagtttcagaaatcctcttctttttttgctttttttttgagcatttttttcatagtagtctttcaccggctattgacataggttctatgtgaccaccaaaaggaatagttgtgtttagcccttgagaagttcctttttttaaatacaataatattattattatctggttgttctgttacatgagcacatcatagcattgtagcaggctataaaaagcaagcaacgttagctttaataacgacagttatcttagccgctatttttacaggtttccaaggttttgaatatagcgtggctaattttacattatccgacggtgtttacggctccacattttatatggctacaggctttcatggttttcatgtctttataggtactattttccttggtatttgcttacttcgcttattaaaatcacatttgacacaacagcatcattttggttttgaagcagcagcttgatattgacattttgttgatgttgtatggctttttttatttatttctatctactgatgaggtggtacctaatctaaatctctttaatatccaaaaatctatgatgaaacttattaatttacctaccataaagctcacttttttgacatgctctagctttcttaattattatctgttatcataatatctatatttttacagaagaaagtatacttttattctgttttattgcgtgactaaacattacatgaaattatatatctcctcaaattaatgcgtcattatctgaaagaggagaaaaaatcaattcaaattttcaacatattactaacgataatataataacttgaaaaaaatatagacaaggttattcactaaaaataacacatgttgatattttaaagaacttaatcacatatttaacatgtttaattaagactgtactccttttagaatcaaaaagtagctcattacaatctattgcaccttatttaaaaagattacatattgtaaaagacttagaaacaaagctgactaaaatttcttatattacgatttgccaacgtatacaggatacagccacaatacgtagtttttatgctaaccgagtaaaaatcaaatctttccattctgaatctaaattagacttattcgaacgtataagaaaattagaagctggttcttagaagtttaagtctatagctcaatggttagagcatacgcttgataagcgtaaggttgattgttcgaatcaatttagacttatactatttcaaaaccataatataaaatatgtacaatataaaccaaacatttttttcagcaagccctttagaacaatttgaaattatacctttaattcctttagaattatttgggttaaacatgtc

gttaacaaacgcgtccatttttttgatactatctgttgcgttatctatttttttgatccac
tttagtcatatacaaaaataaattagttcctggaaactgacaatctgtaaaagaaatatt
ttatgataccaccttaacgttggtaaaagataatttaggtaaaaaaggttatcgatattt
cccgtttatttttaccttttcacaataatactttattgtaatttaataggtatggtacc
atatagttttactgtaacaagtcatatagctttcacatttggcttagctttagctattta
cataggaattaatattattggcttcagaacccacggtataaagtttttcacaattttttt
acctaaaggagttcctttatttattgtacctttagtggttgcaatagaattcgtatctta
cgtcgtaaaagttttcacaatatcgataagacttttttgcaaatatgacatccgggcatac
tttacttaaaattattgccggatttgtttggacaatgatctcaataggaggcgtgtttgt
atacttacaaataatcccattagttttattactagcgttagtgggtttagaaattggtat
cgctcttttacaagcttacgttttcacattacttacctgcatttacttaaatgatgtttt
agaaatgcactaactaaaaaattatgccacaattagatcgcgttattatttttggtcaaa
tattttgactatttttcaccttttaattgcttatgttgtttatacccatttcatattaa
gtaatttattaaaaatttttttagtccgctgatggaagcttagaaaagatattactcaaa
ttgcattaaagatccgtttaacgagctatttaattgattcaaatattcaaacgttacgta
gaatttattcaacaatcagaaatatactagcttctctaacaaaaagtttattaacaaaaa
gtataagtaagccaaagttagttttaaatgatcttaattctttagttattaaaattagtc
tggaaacgtctttatatggtagcaaaagcatcaccaagtctggaacatattcttattgaa
cttaataaaatatactatgtatttattaataatagctctgcctttagtgggaacattagt
tacagggttaggcggtagatgaatagggcgtaaaggttcaaatttgttttctacaacttg
cgtagtcctgtgtgtctttttttcttcaatcgctttttttcgaagtaggtctttgtggagt
tccttgttatatatctttgagcccttgaattagttcaggggcactaaatatttcatgagg
tttttatttgatagtttaacaacaacaatgcttgttgttattacatctatttctagttt
agtccatttgtattctattcaatacatggagcacgaccctcattgccctcggtttatgtc
tttcttggagattttcacatttttttatgatcttattagtaacggctgacaattttgtgca
aatgtttttaggctgagaaggagttggattagcttcttatctattaataaattttgata
cactcgactttgtgcaaatcaagctgcaatcaaagctctggtagtaaatagagtaggtga
ctttggattaagtttaggtattttcacaatttttttatctttttggttctgttgattatga
aatagtattctcttccgcaaacatctacacaaattatagtatttcctttgtgggttttc
cataaataccttgactttaataggtatttttttattaatagggctgttggaaagtctgc
acaattaggtctgcatacctggctaccagacgctatggaaggtcctactcctgtttctgc
actcattcatgcggctacaatggtaacagcgggtgtatttttaatagtgcgctgttcacc
tcttattgatttatcctcggatgtcttacttttaattactcttcttggatcaagtacagc
ttttttcgcctctattgttggagtatttcaaaacgatataaagcgggtaattgcttattc
tacttgtagtcaattaggctacatggtctttgtgtgtggtttatcctattataatgtagg
tatgttccatttagtaaatcatgctttttttaaagcattactttttctaagcgctggctc

tgtaatacatgcgctatcaaatgaacaggacatgcgccgaatgggttcgctagcaaatag
cctaccgatcacatatgctgctatgctaattggctctttatccttagcaggattcccttt
tttaacaggttttattctaaagacttaatcatcgagataacacaaataagttattacag
taatttacagatttcttttggcgtttatgcttgttgacttgctaatatttctgtactctt
cacatcgttttatacatttaggcttatttttctaacttttataaaaaataccaatagcta
tagaaaacacatagaaaatatacacgaatcgccacctttaattctaattcctttaatatt
actcgctatatctagtatttttgtcggtttcttaacaaaagatatattcgtaggaattgg
aactccttttgaggtaatgctatcaatattctacctacgtcttgtaatctattggaagt
tgaatttatgccttctttaataaaatgacttccgtttgtgttaagttctatgggtgcaat
tctcgcttatacaataaacgtaggtgtactaaaaaataatatacaatttgctcataatca
cttatttagaaaactcgccttttcccttagcaaaaagttatattgagataaattatacaa
ttcattcattgtatctcctttaatgtactttggttataatatttcattcaaaaatcttga
taggggttttatagaattcgtaggtccttatggaatttcgcgtactattaaaaattgatc
cacaaaagtaattaaaatacaaactggtcagctaacccattataccttttttcgtgatttt
tggtttatgttcccttttactactagttcctgtttgagattttctacaatttttagttga
tgtcagattactagtattttgctttatagccctctttgtagcgtagtttacgaaagtttt
aacacttaaatatatgcagataactaatttattattatggacttcacttatccctttgtg
tggcgctatattacttatttttattcctagattttactctcatttaataagaaatattgc
tttcgcaacagcgcagctagcgtttatatactctattttgctatggctttgctttgaatc
aacaacatccttattccaatttatatatacgataaattgatttccctcctataatattta
ttacacaataggtgtagacggtatatctttatttttttatcatacttacaacgtgattaat
tacagtttgtacattaataagttgaaatatgccagacagccaaataaaagaatacttaat
ttgttttcttttgcttgaagctattttaattcaagtttttttgtgttttagatgtcctatt
cttttatatatttttttgaaagtgtccttatccctatgtttttaattataggtgtatgagg
gtcacgggaaagaaaaattagagctgcgtatcaatttttcatttacacattagctggttc
actgctaatgcttctagcaattttaactatttatttccagcatggtaccacggatatcca
agttttatgaaatataaattttgacgttagaacacaaattttactttggctagcttttttt
cgctagtttagcagtaaaaattcccatgattccttttcatatatgattgcctgaagccca
tgcagaagcacctacagcagggtccgtaattttagcaggtgtgcttttaaaaatgggcgg
gtatggattttttacgtttttctttacctctgtttccggaagcctcactttattttgctcc
attaatttatttactaagtattatagctgctatatatgcttcacttactacaattagaca
agttgacctgaaaaaaataatagcttactcttccgtttcgcatatgggctttgtcacatt
aggtcttttctcttttaactctcaagggatagaaggtagtataatcttgatgcttagcca
cggattagtctctagtgcactttttttgtgtgtaggtattttatacgataggcataaaac
gcgtcttctcaaatactacggtggtctcgtgcaagttatgcctattttcagcatattact
attatttttttactttctctaatatcggttttcctggtacaagcagttttgttggtgaact

attagtgttaatgggagtatttcaatttagtccaatatctacttttctaagtgcattcag
catgattcttggggcagggtattctatttgactattcaatagagtatgttttggtagttt
aaaacttcaatacattacaaattttcaagatatttcaagaagagaattttgtatcctttt
tccgttaagtgtatttgtactctgaatgggtatatatccagaaattttcctatctgaaat
tcactgttcaagttataacctaattgcatattttaactaattttatgttatgaagtttat
taaaacgctaattctagcatttatgaaaaagaagtcctattttattggtttccacgttt
tttagggctattacttatacctgggtttttatttgataccgagattctagttctctttca
aagccttatcctcttacatgcaagcctaggtttagaagtaatcatagaggactatttaca
cctagaaataataaaacttcagtgtttgtctttaattaaagtacttttaatattattagt
caatcttaatatattatatttattataaaaaatatccttatgttatttatctcctcttat
gatttctacgcattattgacagaaatttactttttaaacgcaatttgtgctttattaatt
tatggtgtaattttaaatacctcatatagaagagggcatccagttattgaacacaatgta
agtggtctctcaactcaaatactaatagtgagtctttggttaacagtttgttcaaatata
ccttgcctaaccagctggaattcacttttagtgcacgatttttttatctttcggtataaaa
agcaccatattagcaatttcgctactttggtctttaattatttttttcttacaatagacta
gaaaaaataaatctctacgagtattgaatcgtgtctatgttggctattgttgccatgctt
tttgtaagttgttcttatgatcttttggcaatgtatttagcaattgaatttcaaagcatt
gcattttatatattagctagttttaaaagaacatctgaattttcaacagaagcgggttta
aaatatttcgtactgggtgcattttcttcagctttgcttcttttaggtatttcactactt
tatggtactactggtttaactaattttggagatctatcaaaattttttttaggtaccaca
ttggaaaacgcatcatttatcaacataacatttttttggtgtcgttttaatagaagtagct
cttttttttaagataagtgcagcaccttttcatatgtgatcgccagatgtttatgaaggt
gctcctactaacgttacatcttttttttggtatactgccaaaattagcattagtaagttta
atatttagattcttttattttttgttgtgctgaagttgtgctgttactaaattttacactt
ataatttgtgcgcttttatctatgataatagggacatttggcgctttagcgcaaacaaaa
tgaaaacgtttcattgcgtatagtactataagtcacgtaggatttattgtagctggattt
tcaacgttggaatttaatggtgcatttggtgcgctattttatatcttggtttatacttta
acttctttagccacttttctattgtgctttccttccgatgcttagcatatcctagcaca
taccaattacgctatctaacggatatcgttagtttagtgaagttaaaccctatacttgct
ggtagccttgtagcagttttattttcaatggcaggtatcccgccttttccaggattttt
gctaaagtatttgttttattttcacttttgcaagaacaattaataggattagctataatg
gcaatatttttgagttgtgtttcgtgtttttattatatccgtttgattcaaatgatgtat
tttacacatacaaaaaccatacttatttttttatccaatagaaaagactacatcaactata
ttaagtataactatgttattacttgtacttatttttttgaagatagatctgatttctaat
tttgttcattgtatgttgtttttataaaataaccaattaaaatgttttacaatattgcaa
ttaacattatcaaagtgttgaccattatagtgccacttttaatcgctgtagcttatatga

cactggccgaaagaaaagtgatggcagctatgcaacgacgaaaagggcctaatgtggtag
gtatctttggtcttttacaacccctagcagatgggttaaaacttttctcaaaagaaacta
tactaccttctagtgctaatattttatttttttagctgcacctgtgctaacgtttttgc
tagctttattagcatgatgtgtacttcctctagatgaggggaaagtttttttcggacttaa
atataggtgttttgtatatattagcagtatcatctttaggtgtttatggtattataactg
ctgggtgatctagtaattctaagtatgctttttttaggtgctttgagatcagcagctcaaa
tggtatcttatgaagtttccattggtctaattttaattaatattttattatgcgcaggca
cattaaatttaactcaaattgttctggcgcaacaaaatatgtggtatataatacctctgt
ttcccatatttattatgttttatatttctatattagctgaaactaacagagcccctttcg
atttgccagaagcagaagcagaacttgtagctggttacaatgtagaatactctgcgatgg
ggtttgcgttgtttttttttaggcgagtatgcaaatatgatacttatgtgtagtttaacaa
ctatttttttttttttggtggttgattacccttagtcaatatgcttcctttttattggattc
cacccgtactttgatttggtttaaaaacaactttacttttatttggttttatttgagtgc
gtgcagcatttccgcgatatagatatgaccaattaatgcgtttaggatgaaaaatatttt
tacctttatcattagggtgagttcttttagtatccgggatactattttctttcgattgat
taccataacaaatgaacgtactttataacgagtattctgctattctcactttttttgcag
tagctttttttaatctctctaataatattaatactttcgtatatattaaatcctcaacaaa
gtgatcaagaaaaagtcagcgcctatgagtgtgggtttaatccatttgatgacgcgagag
caacttttgatgttcggttctatttagtcgcaatcctttttttaatatttgatttagaag
taagtttcttatttccttggtcactagtacttgggcagctaccttcttttggattttgat
ctatggttgccttttttagccattttgacattagggtttatttatgaatgaaaaaaaggcg
ctttagaatgagaataatcaaataatttactagagattttttaatttgataatataatata
atatgaacgtaactttacaaagtgcaaaaatgataggagctggactagctactattggtt
taacaggggtaggagctggagtaggaattgttttcggatcgctagtaattgcttattcgc
gtaatccttctctaaaaaatgaattgtttggctacactattttaggattcgctttaacag
aagcgattgcattatttgctcttatgatggcttttttaattttatttacttaatttactt
taattaatgggcgtttataataaacgcccaccttaaaaatacattatgacaagtacaact
ctttttttgaatcttttcaattatttctttaatatccgcttgtatggtggtaagcctgtca
aatgctgtgtattcagttttatttctaattgtagtattttgtaatactgctagtatttta
ttattactaggagcagaatttttatctttttttatttttaatcgtatacgtaggcgcaatt
gcagttttatttttatttgtagttatgatgttaaacgttaaaatagatggagtaaaaatt
aattatagcacaatttttttttgattggtattttaataagtctgattttacttattcagatt
tgaactgctctacaattagatattgaagcgtatgataatataggcgtaccactatcccaa
aataactttccaacaataatttcttggatccaagaaaatgaattaccttcaaatacagag
agtattggtttaattttgtatacttcgtatagtttagtatttattatgtgcgcatttata
ctacttttagctatgattggttccattgtactaacaatgaatcaacgtagtggagttaaa

acacaacaaatcacacttcagttatatagaaatcaaaataaagtagttcgatttattgat
ctgagaaaaaattaatttgattgcggatatagatgaattggtacatcagtaattttccac
gttaaaggatatgggttcgagtcccattatccgctcaaattaagagagaatagctcaata
ggtagagcaatagttttcaaaactaaaggttaaaagttcaagtctttttctcttgcaaa
agtgagtctgcttgaacaccttattcagttttaattataattaaaactgaataaggtgtt
caagcagactcacttttgcaagagaaaaaagacttgaactttaaatttatataagaataa
caaccacattttagcatacacaagacgacagccttgtatgcgtactaatatatctctgcg
ggctgtgtagaggcaaaaaaagtttgtgtggggtggctgtgtgacatataaatttaaaat
agttgtcaaaaaaagtttagtttggcgtaaaaggattcgaacctttgaaatcatggtatc
aaaaaccattgccttaccacttggctatacgccaaacaaattgaagataaagtgggattc
gaacccacggtggatagataccacgttagctttcaaaactaaagctttaaaccactcagc
catttatcccggttattaatataaattgtattctaaaatttttagagtctgtttatatga
tagtacgggaataggattcgaacctatatttttagatcatgagcctaatgagttaccttt
ttactctatcccgctattttatctaattaattttgtaacttaataaaaacttctccaaac
tacccagtaaaggtaggatgactaaaaaatataaaaaataataaacgctagctatctgac
ctataattatgtaagggtcttctacaggcataccacctattcaacctaatattaaacaac
aagcagccatgcttcaataaagaccacgatatatgggtctaaaacgcgaactttaaattt
atataagaataacaaccacattttagcacacacaagacgacagccttgtatgcgtactaa
tatatctctgcgggctgtgtagaggtaaaaaaagtttgtgtggggtggctgtgtgacata
taaatttaaaatagttgtcaaaaaaagtttagtttggcgtaaaaggattcgaacctttga
aatcatggtatcaaaaaccattgccttaccacttggctatacgccaaacaaattgaagat
aaagtgggattcgaacccacggtggatagataccacgttagctttcaaaactaaagcttt
aaaccactcagccatttatcccggttattaatataaattgtattctaaaatttttagagt
ctgtttatatgatagtacgggaataggattcgaacctatatttttagatcatgagcctaa
tgagttacctttttactctatcccgctattttatctaattaattttgtaacttaataaaa
acttctccaaactacccagtaaaggtaggatgactaaaaaatataaaaaataataaacgc
tagctatctgacctataattatgtaagggtcttctacaggcataccacctattcaaccta
atattaaacaacaagcagccatgcttcaataaagaccacgatatatgggtctaaaacgcg
aacttctaatttctgtactatgtattcatggtaatagagctaatacaactatagcaaaaa
tcatacatataacgccacctaatttatgtggtatacttcttaagattgcataaaaaggta
agaaatatcactcaggaacaatatgtgctggcgttaccatgggatttgcttcaatgtaat
tatcagggtgacctaaaagattaggcgagaagtatacaaaaaaagaaaagaatataataa
aggccactatccctaacaaatcctttacgatgaaataagggtacataggtaccttatcgc
tgcttgcatcgatacctaaaggatttccagaaccttcttgatgtaaagcggctaaatgca
ctaaagacgctgctgcgataacaaatggtaataaataatgtagactaaagaaacggttta
aagttgcattatcaacagaaaagccacctcaaagccaagcaactatagaatcacctacca

aaggtacagcggatactaaattagtgattacagtagcacctcataagctcatttggcctc
aaggtaatacataacctataaaagcagttattatcattaatagtaaaataattacaccaa
tcactcaaacaaattgtcgaggtgcagcataagaaccataataaagtcccctaaaaatgt
ggatataaactacaataaaaaacattgaagcaccattcgcatgtatatatcgtaaaagtc
aaccaaagttaacatcacgcataatatgctctacactaataaaagctaaatcaacgtgtg
gggtataatgcatagctaggaatattccagtcactatttgtattattaaacacattgcag
aaagaaacccaaaatttcatgcataatgaatattgattggagttggataatctataaggt
gattattaactatattgaaaagaggtttttaattagacgcataaataatgtttttatta
tagatggacgaggaatgggacttgaacccatggcctataaagtcacagtttatcgctcta
ccaaaccgagctctcctcgtcatgttaaaaagttaaatttacggggaaaaagggatttga
accctcactcattgatgtgacaaaccaatattttaacctattaaactacttccccatttt
tattaaatacggatagagggatttgaaccctcatgaataatattcatcaaaacctaaacc
tgacatgtctaccatttccatcatatccgcaaaaaaatgttactattagctttaacggat
aaagagggattcgaacccacggtataatatttcatacgatgatttagcaaaccattgcct
taaaccactcagccatttatcctgtgttttggaagctgcccactaccggacttgaaccgg
taacttaaaaagaacagattttaaatctgtcgtgtttacctatttcaccaaatgggcatt
agctattgctaatgctatgttttattgaagctattgcttttccggggttcaatgatttag
gacacgttctactgcaattcataatggtatggcatttaaaaagttttgatttacctccaa
gtaatgctaaacggtcttgagttttgatatctcgactatcagctaatcatctataggctt
gcaataaaattgcaggacctaaatatttgtcatggtttcaccaataacttgggcaactag
cagaacagcaggcacaaagtatgcactcgtaaataccatttaattctgacctatcctttt
cagactgtagatattctgtttttgaaggtgtattatttataagccacggttttatatatt
tatattgtgcgtaaaaattagataaatcaggaactaaatcttttataatgtacatatggg
gtagcggataaattgtaattgttctagtatttatatttaatggttgcaaacaagctaatg
tattagttccatttatattcattgagcaactaccacaaataccttctctacatgagcgtc
taaaggcgatactcgaatcttgttcgtcttttatttttataagagcatccaataccatag
gtccacaatttttagtatgaataggatgtgtactgaaatgagtaatagttgggtttgatg
gagttcatctatatatacgaaggaattttaagtctaaattattattggatactaattgaa
aagaaatttttttgaataattgcatacttggttttattttgattgctataattatttaag
aacgggttagcccgttcttaaatataagataaaatataatgaacatttcgtatttattta
cttacttttgcaagtattaaataataaaccgatctatctagtaatcttctagaatgttct
ggaaaaagtaagggttatttttgcgcttttaagctgtcagcgcttatataatgttaacgt
ggctactcggctatgcaagaaacaatacaaccgatacactattggttaatatatcttaat
cctctcgtactaaagataaaccctctttttttctttcccacaacagatag----------
------------------------------------------------------------
-------------------------------ggttatggtaaagaaacctctcacgaaat

ttcctcccacttcaaaaccgtacgtgaaggtcacccttcatacggctcctcaaa----at
tatctatag-----------------aaagtgtt--------aaaaaaaaatacacactt
tcctttagtttaaaagtgcttatattagtccacggcttaagtattagctgctttggaatg
tctagtatcgtggcaatgaccatgcataagtcgtaagtttttaagatcagaagatccctt
ttgacttcttggtataatatgatctatttctattctatctgaatctttaaaatataattt
acacatactacactgcggccctttagtttttaatagtcgttttagtagttgcccatactt
attattaacagcgagacgtctggctcaataagtaactcttccgtcatatgggctactata
tccggctacttttgtagatagaataatatttcgttgatcatgacgggttaattttataat
tttgtctccttctttaaggccaaaaactctttctgcactccctataggtattcaatatct
agtgataccaggacctgttcgtttgcgtttcttagctcatttccttagaagataaaacgt
acgtacactacaataactaaaagtttttgtggcactacatacggaaaagtatttagttca
tccgctaataactggtgctagtttactgattaaaactttttgagataatccggtggactt
cgtagtaatacttttttatattagctaaatgagattctatagacttatagctgggttgaca
cctgctttttcatccagtcgcaattccttgattattctttgcggatgtatgtttacctac
cttatagttgacaaaattaaaacctaagaaatctacacctgttattctagacttctct--
------aaactaccagtgtacgttattttttgttttggattccgacaactttagacctaag
ttctgtaaaaagaattcaattttgatttttgctgcaatcaattctttctcttcattacat
aatactaaaaaatcatctgcgtatctaataaggtatactccactttttcctattgcatct
tccattccgtgaagggcaatatttgccaacaagggtgatataatacctccttggggggtt
tcggcttctggtgtgatttcttttatattttctttaaagcctgtaagtattcccgctttt
aatcaagctctcaattgctcttttaaaataggaaacgtgtttacttttagaagtaattta
gagtgatctatattgtcgaagcatccttcaatatctgcatctaatacatgtttaggaagt
tgctgtaaacatttcacgattgcttgtctagcatcgttggaacttcgccctggtctaaat
ccataactgttaggttcgaatatagcttcatattgaggttccaatgcaaactttacaaga
cattgtttagctcgatctcttatagtaggtattcctaaatgcctttctttcccgtttggt
tttaaaattgttactcgacgaattttatccgatttattatcaatttcaatattttgtact
aattccattctttcgtcaggagttaaactactaactccatctactccagctgttcgtttt
cccaaattatcttgagtcacttttcgaacagctaaaaactttgaaaagtcatgctttatg
atttgtttctgtattaagaatacagatctcatattacctttttttgctaaattcaaaact
ttacactgcaatctatacagtcaaatttcttttatttttcagttcacttgcgttcacttt
ttcataattt------gtttgttggtgtaatcttaacaatgttaaaatttttatttagat
aatt-----ttgtctacacgtctgcatatccataagctttccttacggcattagcttctt
gtagaatcctgatattaatataccttaacactata----------gaaaaacgcccttgt
aaaaaaagaatcaacaaggtttatattaatatttacttcgttccaatattacatacatat
agtcagtaagcacctctattccctctaaaaccggtacccctttctaacgcggccacatta
gattttaccttatgtttagaattcacgctgtttcgtgttaacggggtgggttaacttata

attatcccccgctatgacatatttagggaatgttattccggctaccgtaaaggtaggtta
tacttctataacta-------------cttttcttggcagatttactggtgtatatattc
tcttcaggtttgaatgtttataagtaacattctcccaaatttatgccctgcttatacctt
gtgaataactattttcacactgtatagggtgcactgtttgtaaa-acagctcattgatca
gcattaggagt-aatgcgtaggcccactagttctagagaacttgcttcctacctgtaata
ttagtttaa----------tctctaaagtggagattagctatttacgagaagtacatctc
ttctcttactagcaacgaatcgcacgaccgaactgtctcacgacgttctgaacccagctc
acgtatcttattatttggcgaacaaccatacccttggaacctattgcagctccaggaaaa
gatga-------------------------------------------------------
------------------------------------------------------------
------------------------------------------------------------
------------------------------------------------------------
------------------------------------------------------------
------------------------------------------------------------
------------------------------------------------------------
------------------------------------------------------------
------------------------------------------------------------
------------------------------------------------------------
------------------------------------------------------------
------------------------------------------------------------
------------------------------------------------------------
------------------------------------------------------------
------------------------------------------------------------
------------------------------------------------------------
------------------------------------------------------------
------------------------------------------------------------
------------------------------------------------------------
------------------------------------------------------------
------------------------------------------------------------
------------------------------------------------------------
------------------------------------------------------------
------------------------------------------------------------
------------------------------------------------------------
------------------------------------------------------------
------------------------------------------------------------

```
------------------------------------------------------------
------------------------------------------------------------
------------------------------------------------------------
------------------------------------------------------------
------------------------------------------------------------
------------------------------------------------------------
------------------------------------------------------------
------------------------------------------------------------
------------------------------------------------------------
------------------------------------------------------------
------------------------------------------------------------
------------------------------------------------------------
------------------------------------------------------------
---------------------------------------gccgacatcgaggtatcaaac
catggcatcgataagaactctcagccatgataaatctgttatccctagagtttctttttt
ccgataagcgacagtatttccatacattactgccggatcattatgaccgactttcgtccc
ggcttgagctgtaactcttaccgtcaagcagattttcatcattacatattatattagcat
atattgccaactaaaatccacttttgtgcacctccgttacagtttaggaggtcctcgtcc
caagtaaactaccacttttacatatgtccttgttagtaagcaatttttatactcagagt
agtttttcaatgacgtacaaatgtacttctacttttactgtactgtgtacaaagttatcg
cgacgtaaaattatagtaaagaatcatagggtctttccgtcttgttgcgggatgtctgca
ttttcacagacaatgtaatttcgctgaggctatactagagacagtggggtagtcgtgacg
ccattcatgcaggacggaatttacccgccaaggaatttcgctacctgaggatccttatag
ttaagaccgccgtttacttgggtttataatatatgcgcaaacttatatttttcactttca
agcactgggcaggcgtcaaaccttatacgtcttcttttgaatttgcaaagttttaagatt
acgataggtagatactcgcatattttcccccgtttagaaccgtacgtgaagattaccctt
catacggctcatccattttaacgtcagaattttctaacgcagctctgtcgcgatgacacg
ttttatgaagaagagtcatatttttaatatattttgtcctccactcactacaggtatga
tatggtgaatttctaattcgtacaatatatctccatattgactatttaaagattccttac
atatggggcacatacccttttgtctttttgccagctttagtcgaatattattcgtttgct
gatttaaacacatattcaaagatcttttattaaaatactcctcaaactcacccaaatagg
gattagcgtcaagttttatcatagtagttcttttaataggagtgtcggatatttgaaaca
aattaattttcttaaatcttttaaattcagtgctagtagatattaatcaatctctattat
ccactttatgataaaatttggatttaagcgctttagcattaagcttcggatgcttaattt
ttatcatgcggtgaatggaatcaaaaacgtatttacttattcgggtataagctaactttg
cagtactagacgagtaataatttgctcagccccttaatattgggtttagttttgagaaga
```

gcatccacaaaggaagttgcgaatgttctttgacacaaagttttattttagcttttactt
tttgtaagctttcttttgtaggtttatatagaaagatccctttcttttccccgatttctc
gagtagaatccgagaattccctaaagtggaaacctacaaaatcaaatccatcttcaattt
tagtgattttactttttcctatatttaattctaaacctcttatttttagaaatttgttaa
catctagaattaacttttctaaaatatcatagttctgggacactataatgaaatcatcag
catatctcacaagagttgctttgttattagacgcc-tttttatagttttatctaaacca
tcaaggcataaattagctataataggagatattacaccaccctgtggaacaccttcttca
gtatcttgataactgccttcattcataacgccagcttttaacattttgacaatatcttt
ttatccataggaatattttctaatattcattcatggcatatattatcaaagaaacccttg
atatctccttcgaaaattcatctagggctgcctctctgtctacttaaaagaaatcaaata
tattgacaggcatctttagcacttcgataaggtctatatgcaaaactgcatcgatcagac
ataatttcgcttataggttcaagagccatagcaaacaaagtttgcacaactctatctctt
atagtgggaattcccaaaggcctttttctttttgctattcttctttggtataaatactcta
cgtaccgcactgaattcataattttaagatcttgaatttgaattaatatttcatcaata
ctaactttctttcctttagaaccttttaaaacaattttatcaattccgctggtctcacta
cttttacgtttgtgaatttgctttattgcttttattttagcttcttctaaattcactaat
tcattttgtaatttacttcataaacgtttgtttttattcaaataagctattgctattctc
ctttgtaacttataaattttcgtcattatattagtttttattatttt-ccacaattttta
tttaaaaataaactagctggttttgcatgtaatgcaaataaaataaaagttttcttaaag
gacaatacttttaaaacagaacatacataagtcagcaatctttcgattcatgatataatc
actatttcacttattactagcaaacattcgctttttatgagatcttttacctactgaaca
cttgggttgatcgctctttcctgagatatttctatcaattcattaggcttacccttttcc
ttacacataacattaatctaaacaggtaacattctctataccggcagtattatgtctaat
ctataaaataacaaaaagatattttatactactgcttattcctatagctgtataaacagc
taatgatgatcttcggaattctgttttctcgataccttataacaaatgttcacttacgtt
ttacctatttagataagtctagcttttcttagttacatcaagtaagaaactacattgtca
gcaaggcttcatacctccagatcactctagacgcatgcttgcttagacttatattgatta
gataatatagcacttagtgcatttatgtatacggctttaaaggtcgcactttttgttaaa
cagtcgctacccctaattttgaaaccttaaaaggtactcctttttgcgaacgtacggag
taaatttgccgagttccttaagtatagttatctcattcgtctttattttctcaataagtt
cacctgtgtcggttttaggtacggtcaaacttcatgtaagttttcctgaaaaattacttt
ttttagcctccagtttagtgggtgtagctattagaagatatctttgtacacaaatacacg
agtattttgcagacacgtggtaatttttcctatcgaatacagttttcactttttcttaa
gggccgactaactccagttacttaaaattgactggaaacccttgaacaatagacgaccat
gatttattttcaacatggttagcgctactcatgtcagcattagcactcctgattttagat
atgcaatttaacattaacataaaagactacaggacgttccgctaccattaagcttagttt

agcttaattcgaagcttcgatatataaatttaagtctccttacattttaaataggtagaa
caaataaaaatagcgagctaaaacgctttctccaataggtggctgcttctaagcctactc
tgtttattcgaattattccttatttttcactaatttataattttgagatcttagctatc
gattagggttgtttcccttttgacgtaagaccttatcgcccaacgactgtctgctgctat
aaataaaaatatagtttggagtttaataaaatttagcaaaatctaaatttaataagtagc
tctacccacattttaaaaaagcaacgtactactttgatagttttcgcggaaaaccagcta
tcaccaagtttgattggactttcacccctaatcttaagtcatccccgtatttttcaacag
acgtgggttcagtcctccagtactttttaaagcaccttcaacttgcttaagaatagatca
cttggcttcgggtctaatacctgtaactttaagcgccttaatatttttaagcttgctaca
catattaacttactgactcattatgcaaaaggcactttgttgctgtatttgtcagcttca
aataaatataaattaacagattcaaatctttccctcacggtactcgttcactatcgatta
gaaaaggttttagcttagaagatgggactcctattttcatacaaaagtaatcgtactact
ttgtgttactattagtgtcgtaaaggactttataacctactttgggtttagacacttcgc
taagttcgctttcactcaccgttacttacaaattctcgtttgatttttttgctatgttac
taagatgattcaattcacataattatataaatgcatattaaatgcgagttttctaaagag
actcatagttcataggcaggtgcctcgctatgatgtttcgtcgcgcgacgtctttgcttt
tctaccaagatttcctctgataactttttaaatcttttatatttaagaacgggcggaaaa
atgttacccttttgtaagagtcattaactctatagtaattgtgctcctaattcgataata
gacaacaagtattgctagtcctatagaagattccgaagccgctactgttaatatagctaa
agcaaatagttggcctactatattgtctaagtatatagaaaaaaatataaagttaaaact
aactgatagaaacattatttctaaagacattataattattataatactttttttggtttaa
aaaaatgcctaaaactccgactaaaaataaaaaaaaagtataattttcacagtatagttg
ggtaatcatttatttttatatttaacatttgagatagcaaattgtattaaatttttccag
ccgcaggttcccctacggctaccttgttacgacttcactttagtcctttcgctaccatgg
acaaaaaataagatttttgtcttcaagtagagtaaattcccatagtgtgacgggcggtgt
gtacaaaacccgagaacatattcaccgcgacagttctgatccgcgattactagcgattac
gacttcatattctcgagttgcagagaataatccgaattaaagatttttaaagattttgc
tccagctcacgcttttgcttcttattgtaaacattactttgtagcacatgaaagtagccc
aattcataagggtcatgcggacttgacgtcattttcccttcctcaaggatattccaagc
agtttataatgcattaatgcattatacaaaagtttcgtccgtttgctggaattaaccaaa
cgcctcacggcacggactgacgacagccatgcaacacctgtgacatcttgtgtcatacga
gaattggtaaggttttgcgcgttgtttcgatttaaaccacatgctccaccgcttgtgcgg
gttcccgtcaattcctttgagttttaatcttgcgaccgtaatccccaggcggagtgttta
atgccttagcttcgcctctggaaaattatccaaaaacaaacactcatagttgagggcgta
gactacaggggtatctaatcccttttgatacctacgctttcgtgccttagtgtcagctat
agtccagattattgttttcacttttgaagttcttttgaatatcatcgcattttatcacta

ctttcgaagttccataatcttttcctatgctctagtaaattagtttagtaatctttagta
aacatgaaatttaaaatttttttactacttagtttaccacctacgcaccctttacgccca
gtcaattagaataatacttgctcctcccgttttaccgcggctgctggcacgaaattagcc
ggagctttgattgtaaaatttagtctttgatttttaatctattttacaaagtgatttac
agcctgatagggctttgctctcacatgtggcctggctaggtcaagctttcgctcattgcc
taagattcctcactgctgcctcttaaagagtctggaccttatttctgttccagtgtgact
gatcatcctcaaagaccaattaaggattatgggcttggtaggtcttttaaactaccaact
acctaatcctgcgtagactttttcttaaaccaattattaatttttttaaggcatttaccaca
atttaagataaatttctacgtattactcacccgtacgctatctttttcataagttacgaa
aattattaaacttgcatgtgttaggccgaccactagtattcattcggagccaggatcaaa
ctcttttattattatgttatgtttttatgtttgctatctcagaaattaaatttgattgta
taaagttatccagcgagctaagacagaagctgtaccaatttcatatatatatatgttggt
aatcttccttttggtaatgaggataaaatttgctttacaaaaagttgtatactttttttg
tttaaatgcggaacatagtttattttctagctttgttccacgtataagaagtctcctata
tttttttattttattcatacatttgtgtatcaagatgggcagatttgaactaccattccc
ttgcacccaaagcaagtacgttaccattacgctacatcttgatatagtgatattgactct
ttctggataggatttgaacctataaccttgtagttaacagctacctgctctaaccattga
agctaccagaaaaataaaagcgaaaaaagggacttgaacccttataatgaccttggcaag
gtcatgctttgcctattaagctatttccgcacttaaaattgaatataaagttgaataata
ttaacaataagtagaaaaagtaaaagataattgctatctttgtacattttccttgtattt
acatgttttactgaatctcaaaaaaaatggcgtattccattacttatgtggtatagaaat
atagcggacactactcaccggacactagttaaaaaccaaggaaaaaagaggtgatctaca
aaaaaatgtattggcagaatagtaatatataaaaagtatacttgatttagtaaaagaata
gggcctacaattaaaacaaacataataacgccagaaattctatgccaaatagaaaaaata
gaagatctttgcggattgtatatagttaaatgtggagaaattggtctattgatgttctgc
atattatagtttttcttttttcttgggattttacagccattatacgggcaagttgtaatatc
cattacagagataaatttaagtttagactgttttaatattttaaacacattttttttatt
tttacctcatccttttatccgtacatgtaaagaagtataactgagttctttagtctttat
ctctaagaagtggattatttcggacgaagaaagggacaggcttctttttacccctttttga
ccttgacttgcctgcagttgtccaagatttaattttgcctttacagtctgttaaaagaaa
atgaatattattggaactaaaagataaaaacagtatacccatatgtagtttaattttatg
ttgtagtagtaaagactagggtcagattcgaactgacctaataaagatttgcaatcttac
gcatagccactatgctacctagtcttaaatttagtgaaatctgaagaaagactgtgtttt
aaatgataaagaacttttcacacaagaaagtaattcaagcgattttagctttggtaaatt
ggataactcgaaaagcaactgaccgggccttatataacagcaatgactcactatagctcc
ttttcctttacccatacgggtttctggactaaatttagtaacagcatttttttggtagcat

gtaaaaccacactttgattttacccaaactggactttttttcgatgactttgatctcttt
ctgaattatctttttatattatttatttgtgtgatatctacttgcttgtacgataaaga
gcgtaatccaaattggcctcttttactagatgacttgtttggtttatatatctattttt
ttttttatgctgtcgtgttggattatttttgtattaaatcttttcatattaaattgtgc
ttaaattgttttatatgccattcaaacttgaagattgcaaatcccttgttttgtatataa
tggcgtttcaaaaaattctatataagtttctagtttttgcagtgatactgcgcctaatcg
gaatactatactacttgacatacgacttcggctaccaccgaaacgcccagcgtaccttat
ttttattcctgtaaattcataaaactgtataccacgagcagtttttaatacaattttgtg
cttttttttattgaaaaatagtatttttattgtattaattatcttccggaatgctaggcc
tttttttaggccagattttataaatccacataaaagtgccccgttttgatgtcaaaatgt
tgcattatatttttttcacgtcaagatcgaacattttaaacgtaggtaccaagcaatctga
tttggctaatagaggtacatattgtacattgtattcccatctattgtcatttattttaga
tagcgtatttccttttatgaatgtttttgattcaaaatgcttatttaaaaagtttgtagc
aggtaatactttttttgtatatttgcttttgattttaaaacctttgccatagcattgact
ttcggtatttcataagtcgttcacacctattcgaaatcctcttggattaacttttttgacc
cataatgttttattacttaataacaatatttaaattgaaaatcgaatctaaatagttata
cagaaatacgttttgtaggatttgaacctacactaatcaacttagaaggttgatgtttta
tccaattaaactaaaaacgtttttactttagatgcattgggatttgaacccaaaataagc
agattaaaagtctgttgctgtaccgtttagctatacatctttttaataaagagagtagga
tttgaacctacgatgattttaatcaatagatttacaatctatcgctttcgaccactcagc
catctcttcttcagcggttagatatatgttacaatctgtaataaaataattaattatata
ctattgtttaataacaatatttagattgaaaatcgtacgattttcaatctaaatattgtt
attaaacaat
>SRR9587917
gttttattaattaaaactaccccagtattacacttttaaatatttataagaaataatgca
aatccctttacgctgttatttaatcgaatttaaatttcgattcgtttactttctacttag
ttttttctctcgttgctagtcgttataaactatcatgaggctgtgtttttttttgaaac
gtatactttttttgtttataaacgagggggcattcatcgcgacacatattgctgaactatt
ttcaacttctatgtatattagttttaacctagttttgtgcataaattatccttttgcata
ttatcattgcagccgctttttaaattcgagttgatacaaatcacaggtttggttctttag
aaatatgcaattattgacatttattacgtttttagctagtttacttttgtgctactttttt
aattttgccctacacttacgctttttttagacacttgaactgtaacgatagtttatgcatt
taaagtgcagttggaggcaagaatagcaacatatgtttgttgaacactacaaacaatgtg
tttactatctaacattgtctatctcttctttactaggctaatttgtttttacttagtgga
caatatgattaatttgcatttatttttccggaaaaataaaaaatatactttctttttcgt
atgcttgtcggcttctgtgtgtttaccgcaagaaagttttatacaaattaccttcataat

```
ttctatagtagtactatttgagttattctttttggttacatgcttatttttttgcaaaaaa
taatgtctaaaagtggacttgaaccactgacccaaagattttcaatcttttgctctaacc
gctgagctatttagacttaaacttttcttttcccgtatttagagcgaccatttttttcgtt
taacgactgcttgtaaatcgtaaacacctctcacaatatggtaacgtactccaggtagat
cctttacacgaccacctcttatcagaacaactgaatgttcttgtaaattgtggccttcac
ctcctatatagcctatgatagaacgtccggtactaagacgtattttagctacctttcttt
cggcagaattcggtttttttagggttagtagtatacacttttgtgcaaacacccttttttt
gaggcgacttattgagagcgggtgttttcgctttggtaattttgtgctttcttggatttt
ttattttttggtttagtgttgacataatattttttattttaagtaataagggacttgaacc
cctaaccatttcggtgtaaacaaaatgctctacctattgagctaattactttacgtctgg
aaagatttgaactttcaacttttagattcgtaatctaacgctctatccaattaagctaca
gacgtttttctccacttgtatatttagattacttatctatattttggaaatagggacaaa
tattcttgtataatttacttcaggattattcgaaagtatagttaaccaattaaacaagta
gaaaacgcatattttgtattgttactaacttaaaggactattttataataaactttttat
tgtattatataccaatatttagattgaaaatcgacctaaattcaatgtacttttagcaaa
tgacaccagtgttttaaaaaaggtgcttattttgaaaacatttttggtgtcatttgctct
ttttcagctaataataatttaattaataacagaaagcgcggtttattgctcgtcgcgttc
cagttattactacggcaacttcctatgcagctgttggga-ccccccaactcgtgtaatta
cactttttagtcttatttaaattataataaggttatactttatatcttttcttttattta
accccctaagatctttattacgccccaaaatggtgtcataaaaaaa-tagcctggagggct
gccaggagtaaaaaagacctataacggttatttcagcttttatagtgctatttttctttt
ttagagaaggctatttttaattggtaactaagtcattttaagtgataattaagctatttt
aattaaaaatgacttattaaaggttttacacagataatacttcttttattaaaaatatac
taagtcgctagaccggaaaaaacccgccatagaacgatgagaggtaaaaagtgtcataaa
ttttaactattttat----------------aattaaagtttgtcaaacatcaaactttc
tgttatttcaataatgcttgtttttaaggcttttaaagcagtttctcgctgatcccgaac
actaaactttttgttaacatgttctcgtaaactgtaattgaatgtcataataaccagtgg
attatcctgttttaaagtaaaattacctttcattgcaaggcgcatgctatcaccagctaa
aaactggtttaagaattcgatttg------------------------------------
------------------------------------------------------------
------------------------------------------------------------
------------------------------------------------------------
------------------------------------------------------------
------------------------------------------------------------
------------------------------------------------------------
------------------------------------------------------------
```

--tttcatgataaagtgctctcttacaagccgctttttagtagattactaaaaattta
ctatgttgatttgatacgcgtttttagatgggctcttgcttatttattggatctgaaaag
gagcttgaagaattagttaagtagtcaaaataattcggatttgtgaaattacgtttctta
gaaaattcaatatacgtatataaaatttgatccgagttatttcagccactacgaaagctt
attattgttgtttttctctttgtagaagtttgttataaattaaaattaggcaaagaagc
tacatttt--ttttaacttggaaagatttggccgcaacgcgc-ttttatgtttaattac
ggctggatttagttcaaatactttggacatgagtcgttttaaaaattatttagaaatatg
tgttcgcaaacagattaagtgaaaaaaaagagttttaaaaaagcttcataaactttcacc
aaaagtttgatttatatcataaacggagaaaatccaaattgaaaaaactaaaatggtgtg
gcctaaaaaaatacggcgaagtacatatcaacttgattttcagtgaattaaacagaaata
cagtattcccacaagtacgataggaattaaaatgatggataaatcaaaattttgatttag
atacgaaaaccatggtttactcggttcgtttacaagtaaactttttaattcaatattttg
tgctattagtaaagatttgtgcgactttaaattgaataagctattaaaataaggtgaatt
taatttaggcaataaagtatcgtacaaatgtttgttctttctatatttgtttagagcttt
cctgtactttttattgtaagcagcaatagatcctatgtgcttgtgttgaaaatctaatct
tttaagtactttgttaaagctcaaaattttggtttttaattggttaaccgtttttttacc
ctttctagcatcgtatagtctttcttttagagtttctagatgagaaggcttttctagtgg
cagttgattgaaataacttttagtgagtttttttactatagtacctgtaacaagtgcgct
aacagatactatgacacaaactacttctaagtccaa-ttgctctacaatgtg-tttgatg
aaagtcattttatataaattaatttattaatgtgtaattgtggtatgattttattctgtg
ttaatgggcataattaaatgcatacattatatattatccctccttcggagggaggcatat
aataattcttattatataccaatatttagattgaaaatcgcacgattttcaatctaaata
ttggtatataaacaaacctgcacataaataaatatcactaggacatcaccccaccaaaaa
atcgatattctctagagagcccctataagctttaatttactatctcttgcacgaaattag
accttcttgaataggaatcgtcgtgaaattgaaaaacatttagggggtatatcttatacaa
ttatgaccaaaaaatctacaaaacaaacaactttacacaaatacttttgcaactcgcata
catgagattgaacttaagaaacctcttttcataaatcaggcagcatttttggcgcttctt
aattctattgcaatattcaatgaaccaaaaacaaagttttttt-aaacgacctgcatgaa
cacaatcttaacaagtgtaaaaaagccctctagctcccttttttgcaacaccacgagaat
aaaccccggtcgggcagaccccgaacc---------------------------------
-cgcgcgcgcgttatagattcctatatcgc-----tccatttcatttcgctattgctg
cgcttcgcttatttttaattctaactgttctttttcattataacgattttaggtcgattt
tcaatctaaatatttgtatataaacaattatttatcttgataattatatatctaaaaaag
ggggcatagtttaataggtataatgtatgctttgcaagcataaggttaccggttcaaatc
cggttgtctccaaatctaaaaacatcaatgcgttctagaatattacaatacaaaaataac
ttggctgattatgatcttttaactcaattttcgattaaaaatcataatagtatgcctact

ttctcttctcttaatgtcagagtaaaaaatattcaaactagcgacattaagcaggtttgc
ttaaatctaatctctattcaattagtaggaaacaatcatgttgttttcagtagccaaaaa
gacgtttctaatcttatactaaaatgtaacggtattaatacttattttattttagagaat
ttatctttgttaggctacaaaaacaatttaaaaagtgacaaaaacaataaattaactgtg
cacggtaaaaacttcttaaatattattgaatattttgcatccaattcaaattttcttaag
tccttagaaaaaacatctaataaacatttaagtatcgctttttcttatcaacttgctaat
agtaaaaataatgaaatagctttgtatttttttaaaatcattaggatttccgctttaataa
cggtatagttcaattggttagaacaacggaatcataatccgtaagttgtgggttcaagtc
cctctaccgttatggctcttatcgtctaatggttaggacagtgctttttcagggcatcga
cgtgagttcaatcctcactaagagtaacttttttaatataaaaatccttatacaaatacgt
gagcccataagaattcaaaagtaaaaaggatttttctagatctactaaattattttggga
tccctaggtcgttaaaaatatttaaacaatgtattttattcatatatatatatatactaa
aacttaaaaatataatttagtgcaaaagtctctaaataactgaattttccgttgaatata
ctcgaccaatcataaagatataggtactttatatttaatttttggcgctttctccggtat
cttaggtgcttgcgcgtctatattgatccgaatggaactagcacaaccaggtaatcaact
attattaggcaatcatcaagtgtataacgtactagttacagagcacgcatttttgatgat
tttctttatggttatgcccgtcctaattggaggatttggaaactgattcgtacctattat
gataggtgctccagatatggcctttcctagattaaataatataagtttttgactactacc
tccatcattgtgtcttcttttaggatctgcgatggtagaagtaggcgctggcacaggctg
aactttatatccgcctttgagctctattcagagtcattcaggcggtgctgttgatcttgc
tatttttagtttacacttgtcaggtgcttcttctatattaggagctattaatttcattac
gacgatatttaatatgcgcaatccaggacaaagtatgtatcgaataccgctatttgtttg
atctatcctcattactgcgttccttttactactagcagtacctgtcttggcaggggccat
cacaatgctgttaacagatagaaactttaatacaacatttttttgacccttcaggtggtgg
cgatcctgtattgtatcagcatttattctgattcttcggacatccggaagtgtacatttg
tgcgccgtttaattctgttaaataagatggtttattattttaaattacttatctcgaaat
ctttaaactgaatagttaataaagatagacttaattattccttttgtcttaaaacacttt
cccttaataaagttggaaacaattctattaaaagcgggaatgaaagtctcttgcaggaac
ctgaatttcctaaaaatagtaggctagttatgctcatcatgaataagataataaagttga
tacatatcgacggttccacactaaactttatacataacatttcaggtaacagtatttatg
taatatgggtcggtaccagaatgtttaccttcccttgaataattattacacgtcagagcc
atcagctcctaaatgatatgcaacgttctaaaatgagcttgatcgagaaacagaagaact
cgggataccctaccaatcgaaagattcatgagtacggaactctcgtagtaggtggtagaa
gaattcaatcttcttcaaactaccaaaggggagtagaacttagattcttaagtgaaaaac
cctgcattagctcgcaagagtgcgctaggttagtagatttgagaaaagttaattctgaaa
ataaatttcaagttaataagaatactattcatattatatctgatatgaatgtccttattt

tagcatatgaactcataaaaagtaatcctggaaacatgacacctggtgtgaatggttcca
ccttagatgggttagacaagacgtgactgcaaaatattagtaccaaaataaagcaaggta
aatttttattcagccctgggcgtaagaagtacattcctaagcccggttcagcggataaaa
gaccattaggtattgctagcccgaaagaaaaaattgttcaaaaagctattctgctagtac
tagaatcgatttttgaaccaagcttcttgagaaattctcacgggtttcggcctaaccgag
gcaaccataccgctttaaagatggtaaaaagcgagtttcacggagttccctgaattatag
aaggagatatttcgaagtgctttgatgaaattgatcactctattttattggggcttctaa
gcaagaggatatcttgtgataagactttaactttaattaaaagagggttgaaggctgggt
ttatagatttaggaatattcacaagaactaaattgggcacccctcaaggaagcattctga
gtcctatcctatgcaatatctatttgcatgagctagatttatttctacttcaactaaaaa
ttaaattcgatacagggactagtagagcgaagaacccacagttcagaaaactacagtata
aactatctaaccttaaaacgcctctcgagaaaaagcttgtcagaagagacctttgaaaag
tgcatagtctgaacccccctagatcctaacttttgcagaattcactttgttcgatatgcgg
atgattttattgtgggagttacaagctcccatgaagttgctttagaagttaagaatatga
ttaaggaattcctttgtaatcatttgaaattaaatttgaatgagctgaagacacaaatta
ctcatattagagaaaaggatatatttttcctaggtacccttatcaaaggtaactgaaaga
aagagaaacctattcgattgatcaactttccctctagagaaacgtccatcaagacaagag
tcactccgcgtttaagtttgcatgcccctattaaaaaactctttgataaagctactgctg
aaggatttttccgtagggatgggattaattataaacctacctttgtaggtaaattgatta
atatggaccatgcagacattttatttttttataattcaatagtaagaggagtactaaatt
actactcatttgtggacaaccacaaaagtttaggatcatgagtccattatatgaaatttt
cctgtgctagaactctagcgttaaagtacaagttacgtttcacatcgaagacttttaaga
aattcggctctaaattggcgtgcccgaatacaaaaaaaagcctgtttttaccaacgagct
ttaagaggacgcaggccttccaaattaatagccctattccttttgagaaaaaagtactct
cttgatctaaaaaaattactaaatctaatctgaacaaagtttgtttaatatgtggtagtt
ctctatttgtagaaatgcatcatatacgtagtatagctggcattagaacaagactttgta
gcaataaagcagactttttctctctgcaaatggctgggataaacaggaagcaggttcctc
tttgtcgagaacatcacttgaagctacataataaaacactttcaccagaagaaacgttat
tatttcagaaaaatattaaggaaatttagatgtatttctttcaattttaactttatttac
aatttaagttcggcgagccgtatgataagaaattatcacgtacggttctgagggcagtta
ttgacgctaaaacaacctcgttgtcaagtttctatttctgacccctactaatcttgcctg
gatttggcatcgtcagtcacatcgtttccacctttctagaaaacctgttttcggttata
taggaatgatttatgctatgctttctataggtattttaggatttatcgtttgagcgcatc
acatgtatactgttggccttgatgtagatacaagagcttacttcacagctgcaaccatga
tcattgctgttcctacaggtattaaaatatt-----------------------------
------------------------------------------------------------

------------------------------------------------------------
------------------------------------------------------------
------------------------------------------------------------
------------------------------------------------------------
------------------------------------------------------------
------------------------------------------------------------
------------------------------------------------------------
------------------------------------------------------------
------------------------------------------------------------
------------------------------------------------------------
------------------------------------------------------------
------------------------------------------------------------
------------------------------------------------------------
------------------------------------------------------------
------------------------------------------------------------
------------------------------------------------------------
------------------------------------------------------------
------------------------------------------------------------
------------------------------------------------------------
------------------------------------------------------------
------------------------------------------------------------
------------------------------------------------------------
------------------------------------------------------------
------------------------------------------------------------
------------------------------------------------------------
------------------------------------------------------------
------------------------------------------------------------
------------------------------------------------------------
------------------------------------------------------------
------------------------------------------------------------
------------------------------------------------------------
------------------------------------------------------------
------------------------------------------------------------
------------------------------------------------------------
------------------------------------------------------------
------------------------------------------------------------

------------------------------------------------------------
-------tagctggattgctactatgtgagaagggtctatttttttaaaaacccctatgt
tatttgctatagggtttatatttttattcactataggaggacttactggtattatactag
ctaactccggacttgatatatctctacatgatacttattatgtcgtagctcacttccact
atg---------------------------------------------------------
------------------------------------------------------------
------------------------------------------------------------
------------------------------------------------------------
------------------------------------------------------------
------------------------------------------------------------
------------------------------------------------------------
------------------------------------------------------------
------------------------------------------------------------
------------------------------------------------------------
------------------------------------------------------------
------------------------------------------------------------
------------------------------------------------------------
------------------------------------------------------------
------------------------------------------------------------
------------------------------------------------------------
------------------------------------------------------------
------------------------------------------------------------
------------------------------------------------------------
------------------------------------------------------------
------------------------------------------------------------
------------------------------------------------------------
------------------------------------------------------------
------------------------------------------------------------
------------------------------------------------------------
------------------------------------------------------------
------------------------------------------------------------
------------------------------------------------------------
------------------------------------------------------------
------------------------------------------------------------
------------------------------------------------------------
------------------------------------------------------------

------------------------------------------------------------
------------------------------------------------------------
------------------------------------------------------------
------------------------------------------------------------
------------------------------------------------------------
------------------------------------------------------------
------------------------------------------------------------
------------------------ttctgtctatgggagctgttttcgcaatatttgcag
gcttttattattggtttgaaaagatatccggatttcaatattctgaaatactaggtcaaa
ttcacttttgaggcacttttataggtgtaaatctaacctttttccctatgcactttttag
ggcttgctggtatgcctagacgtattccggattatccagattcttatgccggttggaacg
caatagcttcttacggttcatatgttgcgttatttagcacgctgttttttttttatcttg
tatttaacacacttgtaacagcaaaaaagacacctgctagaaataacccatggaactttg
aagattcaaagatgggctcaactacattagaatgagaaatttcttctcctccagcttacc
atacgttcaatgagattccagttataagagaaacagaaacatctttaaaaataaattaac
ttatgataaaaaaaaatactacaatattatttttagggttagctttgctaacccaactac
tacagagtagaatagttattagtgattctgcagaagactggcaattaggcttccaagatc
ctgcaacacctataatggagggaatcattaaccttcatcatgatttatgtttttttatct
gtgctatctcaatttttgtatcttgaatattagcacgcacgttatggcactaccactgaa
caaaaaatgagtacccttctgccacggttcatggaacagccatcgaaataatttggactg
ttactcctagtatcactttgttggcgattgctgtgccttcttttgctttgctatactcta
tggatgaaataattgcgcctgcaataactattaaaacagtaggtcatcaatgatattgaa
gttacgaatactcggactatacaaatgaagacgataacactataatgtttgaaagttata
tgattccagaagaagatttaactttaggtcagttaagattattagaggtagataacccta
tggtaatacccgtaaatacacacgtacgtctaatcataacagcagcagatgttttgcaca
gctgagcagtgccttctttaggtataaaatgtgacgctgtaccaggtagattaaaccaaa
gctcgctctttgtaaaacgtgaaggaatattttatggtcaatgcagcgagatttgtggtg
taaatcatggttttatgcctatcgtagttgaagcagtgtctttaccaaattatatttctt
gagtagctaataaacttagcgaataaatagcctaaaaatattctaagcatctgatatgcg
tgtatctctagcccaactatttttctttggtttatctatttttattttattttgatttaa
tcctaaatacttaacttatacaatacgactattaaaaaaaattaaaaagcgttctaaata
acaaagagcgctttaataaaaaacttattatggcaactacgacaaatctaaactttatca
aaacagctaaacaattacaacgccacccttttcatttagttgaccccagcccttggcctg
taacagctgcaatagccgctttttcatgtgctttaggcggagttatgtatatgcatgcat
acagtaatggagggtacctattatttagtggtttttctttactgttatttacaatgttct
catgatggcgcgatgttacaagagaagccacttttttcagggcatcatacaggtgctgttc

aaaaaggattgcgttatggtgtaattttatttatagtttcagaaatcctcttctttttg
ctttttttgagcatttttcatagtagtctttcaccggctattgacataggttctatgt
gaccaccaaaaggaatagttgtgtttagcccttgagaagttcctttttaaatacaataa
tattattattatctggttgttctgttacatgagcacatcatagcattgtagcaggctata
aaaagcaagcaacgttagctttaataacaacagttatcttagccgctatttttacaggtt
tccaaggttttgaatatagcgtggctaattttacattatccgacggtgtttacggctcca
cattttatatggctacaggctttcatggttttcatgtttttataggtactattttccttg
gtatttgcttacttcgcttattaaaatcacatctgacacaacagcatcattttggttttg
aagcagcagcttgatattgacattttgttgatgttgtatggctttttttatttatttcta
tctactgatgaggtggtacctaatctaaatctctttaacatctaaaaatctatgatgaaa
cttattaatttacctaccataaagctcacttttttgacatgctctagctttcttcattatt
atctgttatcataatatctatattttttacagaagaaagtatacttttattctgttttatt
gcgtgactaaacattacatgaaattatatatctcctcaaattaatgcgtcattatctgaa
agaggagaaaaaatcaattcaaattttcaacatattattaacgataatataataacttga
aaaaaatatagacaaggttattcactaaaaataacacatgttgatattttaaagaactta
atcacatatttaacatgtttaattaagactgtactccttttagaatcaaaaagtagctca
ttacaatctattgcaccttatttaaaaagattacatattgtaaaagacttagaaacaaag
ttgactaaaatttcttatattacgatttgccaacgtatacaggatacagccacaatacgt
agtttttatgctaaccgagtaaaaatcaaatctttccattctgaatctaaattagactta
ttcgaacgtataagaaaattagaagctggttcttagaagtttaagtctatagctcaatgg
ttagagcatacgcttgataagcgtaaggttgattgttcgaatcaatttagacttatacta
tttcaaaaccataatataaaatatgtacaatataaaccaaacatttttttcagcaagccc
tttagaacaatttgaaattatacctttaattcctttagaattatttgggttaaacatgtc
gttaacaaacgcgtccatttttttgatactatctgttgcgttatctatttttttgatccac
tttagtcatatacaaaaataaattagttcctggaaactgacaatctgtaaaagaaatatt
ttatgataccaccttaacgttggtaaaagataatttaggtaaaaaaggttatagatattt
cccgtttatttttaccctttttacaataatactttattgtaatttaataggtatggtacc
atatagttttactgtaacaagtcatatagctttcacatttggcttagctttagctattta
cataggaattaatattattggcttcagaacccatggtataaagtttttcacaattttttt
acctaaaggagttcctttatttattgtacctttagtggttgcaatagaattcgtatctta
cgtcgtaaaagttttcacaatatcgataagactttttgcaaatatgacatccgggcatac
tttacttaaaattattgccggatttgtttggacaatgatctcaataggaggcgtgtttgt
atacttacaaataatcccattagttttattactagcgttagtgggtttagaaattggtat
tgctcttttacaagcttacgttttcacattacttacctgcatttacttaaatgatgtttt
agaaatgcactaactaaaaaattatgccacaattagatcgcgttattattttcggtcaaa
tattttgactatttttcacctttttaattgcttatgttgtttatacccatttcatattaa

gtaatttattaaaaattttcttagtccgctgatggaagcttagaaaagatattactcaaa
ttgcattaaagatccgtttaacgagctatttaattgattcaaatattcaaacgttacgta
gaatttattcaacaatcagaaatatactagcttctctaacaaaaagtttattaacaaaaa
gtataagtaagccaaagttagttttaaatgatcttaattctttagttattaaaattagtc
tggaaacctctttatatggtagcaaaagcatcaccaagtctggaacatattcttattgaa
cttaataaaatatactatgtatttattaataatagctctgcctttagtgggaacattagt
tacagggttaggcggtagatgaatagggcgtaaaggttcaaatttgttttctacaacttg
cgtagtcctgtgtgtctttttttcttcaatcgctttttttcgaagtaggtctttgtggagt
tccttgttatatatctttgagcccttgaattagttcaggggcactaaatatttcatgagg
tttttatttgatagtttaacaacaacaatgcttgttgttattacatctatttctagttt
agtccatttgtattctattcaatacatggagcacgaccctcattgccctcggtttatgtc
tttcttggagattttcacatttttatgatcttattagtaacagctgacaattttgtgca
aatgttttaggctgagaaggagttggattagcttcttatctattaataaattttgata
cactcgactttgtgcaaatcaagctgcaatcaaagctctggtagtaaatagagtaggtga
ctttggattaagtttaggtattttcacaatttttatctttttggttctgttgattatga
aatagtattctcttccgcaaacatctacacaaattacagtatttccttttttgggttttc
cataaataccttgactttaataggtatttttttattaatagggctgttggaaagtctgc
acaattaggtctgcatacctggctaccagacgctatggaaggtcctactcctgtttctgc
actcattcatgcggctacaatggtaacagcgggtgtgtttttaatagtgcgctgttcacc
tcttattgatttatcctcggatgtcttacttttaattactcttcttggatcaagtacagc
tttttcgcctctattgttggagtatttcaaaacgatataaagcgggtaattgcttattc
tacttgtagtcaattaggctacatggtctttgtgtgtggtttatcctattataatgtagg
tatgttccatttagtaaaccatgctttttttaaagcattactttttctaagcgctggctc
tgtaatacatgcgctatcaaatgaacaggacatgcgccgaatgggttcgctagcaaatag
cctaccgatcacatatgctgctatgctaattggctctttatccttagcaggattcccttt
tttaacaggttttattctaaagacttaatcatcgagataacacaaataagttattacag
taatttacagatttcttttggcgtttatgcttgttgacttgctaatatttctgtactctt
cacatcgttttatacatttaggcttatttttctaacttttataaaaaataccaatagcta
tagaaaacacatagaaaatatacacgaatcgccacctttaattctaattcctttaatatt
actcgctatatctagtatttttgttggtttcttaacaaaagatatattcgtaggaatcgg
aactcctttttgaggtaatgctatcaatattctacctacgtcttgtaatctattggaagt
tgaatttatgccttctttaataaaatgacttccgtttgtgttaagttctatgggtgcaat
tctcgcttatacaataaatgtaggtgtactaaaaaataatatacaatttgctcataatca
cttatttagaaaacttgccttttcccttagcaaaaagttatattgagataaattatacaa
ttcattcattgtatctcctttaatgtattttggttataatatttcattcaaaaatcttga
taggggttttatagaattcgtaggtccttatggaatttcgcgtactattaaaaactgatc

cacaaaagtaattaaaatacaaactggtcagctaacccattataccttttttcgtgatttt
tggtttatgttccctttttactactagttcctgtttgagattttctacaatttttagttga
tgtcagattactagtattttgctttatagccctctttgtagcgtagtttacgaaagtttt
aacacttaaatatatgcagataactaatttattattatggacttcacttattcctttgtg
tggcgctatattacttatttttattcctagattttactctcatttaataagaaatattgc
tttcgcaacagcgcagctagcgtttatatactctattttgctatggctttgctttgaatc
aacaacatccttattccaatttatatatacgataaattgatttccctcctataatattta
ttacacaataggtgtagacggtatatctttattttttatcatacttacaacgtgattaat
tacagtttgtacattaataagttgaaatatgccagacagccaaataaaagaatacttaat
ttgttttcttttgcttgaagctattttaattcaagtttttttgtgttttagatgtcctatt
cttttatatattttttgaaagtgtccttatccctatgtttttaattataggtgtatgagg
gtcacgggaaagaaaaattagagctgcgtatcaatttttcatttacacattagctggttc
actgctaatgcttctagcaatcttaactatttatttccagcatggtaccacggatatcca
agttttatgaaatataaattttgacgttagaacacaaattttactttggctagcttttttt
cgctagtttagcagtaaaaattcccatgattccttttcatatatgattgcctgaagccca
tgcagaagcacctacagcagggtccgtaattttagcaggtgtgcttttaaaaatgggcgg
gtatggatttttacgtttttctttacctatgtttccggaagcctcgctttattttgctcc
attaatttatttactaagtattatagctgctatatatgcttcacttactacaattagaca
agttgacttgaaaaaaataatagcttactcttccgtttcgcatatgggctttgtcacatt
aggtcttttctcttttaactctcaagggatagaaggtagtataatcttgatgcttagcca
cggattagtctctagtgcactttttttgtgtgtaggtattttatacgataggcataaaac
gcgtcttctcaaatactacggtggtctcgtgcaagttatgcctattttcagcatattact
attattttttactttctctaatatcggttttcctggtacaagcagttttgttggtgaact
attagtgttaatgggagtatttcaatttagtccaatatctacttttctaagtgcattcag
catgattcttggggcagggtattctatttgactattcaatagagtatgttttggtagttt
aaaacttcaatacattacaaattttcaagatatttcaagaagagaattttgtatcctttt
tccgttaagtgtatttgtactctgaatgggtatatatccagaaattttcctatctgaaat
tcactgttcaagttataacctaattgcatattttaactaattttatgttatgaagtttat
taaaacgctaattctagcatttatgaaaaagaagtcctatttttattggtttccacgttt
tttagggctattacttatacctgggtttttatttgataccgagattctagttctctttca
aagccttatcctcttacatgcaagcctaggtttagaagtaatcatagaggactatttaca
cctagaaataataaaacttcagtgtttgtctttaattaaagtacttttaatattattagt
caatcttaatatattatatttattataaaaaatatccttatgttatttatctcctcttat
gatttctacgcattattgacagaaatttactttttaaacgcaatttgtgctttattaatt
tatggtgtaattttaaatacctcatatagaagagggcatccagttattgaacacaatgta
agtggtctctcaactcaaatactaatagtgagtctttggttaacagtttgttcaaatata

ccttgcctaaccagctggaattcacttttagtgcacgatttttatctttcggtataaaa
agcaccatattagcaatttcgctactttggtctttaattatttttcttacaatagacta
gaaaaaataaatctctacgagtattgaatcgtgtctatgttggctattgttgccatgctt
tttgtaagttgttcttatgatcttttggcaatgtatttagcaattgaattccaaagcatt
gcattttatatattagctagttttaaaagaacatctgaattttcaacagaagcgggttta
aaatatttcgtactgggtgcattttcttcagctttgcttcttttaggtatttcactactt
tatggtactactggtttaactaattttggagatctatcaaaattttttttaggtaccaca
ctggaaaacgcatcatttatcaacataacatttttggtgtcgttttaatagaagtagct
cttttttttaagataagtgcagcaccttttcatatgtgatcgccagatgtttatgaaggt
gctcctactaacgttacatctttttttggtatactgccaaaattagcattagtaagttta
atatttagattcttttatttttgttgtgctgaagttgtgctgctactaaattttacactt
ataatttgtgcgcttttatctatgataatagggacatttggcgctttagcgcaaacaaaa
tgaaaacgtttcattgcgtatagtactataagtcacgtaggatttattgtagctggattt
tcaacgttggaatttaatggtgcatttggtgcgctattttatatcttggtttatacttta
acttctttagccactttttctattgtgctttccttccgatgcttagcatatcctagcaca
taccaattacgctatctaacggatatcgttagtttagtgaagttaaaccctatacttgct
ggtagccttgtagcagttttattttcaatggcaggtataccgccttttccaggattttt
gctaaagtatttgttttattttcacttttgcaagaacaattaataggattagctataatg
gcaatatttttgagttgtgtttcgtgtttttattatatccgtttgattcaaatgatgtat
tttacacatacaaaaaccatacttatttttatccaatagaaaagactacatcaactata
ttaagtataactatgttattacttgtacttatttttttgaagatagatctgatttctaat
tttgttcattgtatgttgtttttataaaataaccaattaaaatgttttacaatattgcaa
ttaacattatcaaagtgttgaccattatagtgccacttttaatcgctgtagcttatatga
cactggccgaaagaaaagtgatggcagctatgcaacgacgaaaagggcctaatgtggtag
gtatctttggtcttttacaacccctagcagatgggttaaaacttttctcaaaagaaacta
tactaccttctagtgctaatattttttatttttttagctgcacctgtgctaacgtttttgc
tagctttattagcatgatgtgtacttcctctagatgaggggaaagtttttttcggacttaa
atataggtgttttgtatatattagcagtatcatctttaggtgtttatggtattataactg
ctgggtgatctagtaattctaagtatgctttttttaggtgctttgagatcagcagctcaaa
tggtatcttatgaagtttccattggtctaattttaattaatatcttattatgcgcaggca
cattaaatttaactcaaattgttctggcgcaacaaaatatgtggtatataatacctctgt
ttcccatatttattatgttttatatttctatattagctgaaactaacagagcccctttcg
atttgccagaagcagaagcagaacttgtagctggttacaatgtagaatactctgcgatgg
ggtttgcgttgtttttttttaggcgagtatgcaaatatgatacttatgtgtagtttaacaa
ctatttttttttttggtggttgattacccttagtcaatatgcttcctttttattggattc
cacccgtactttgatttggtttaaaaacaactttacttctatttggttttatttgagtac

gcgcagcatttccgcgatatagatatgaccaattaatgcgtttaggatgaaaaatatttt
tacctttatcattagggtgagttcttttagtatccgggatactattttctttcgattgat
taccataacaaatgaacgtactttataacgagtattctgctattctcacttttttttgcag
tagctttttttaatctctctaataatattaatactttcgtatatattaaatcctcaacaaa
gtgatcaagaaaaagtcagcgcctatgagtgtgggtttaatccatttgatgacgcgagag
caacttttgatgttcggttctatttagtcgcaatcctttttttaatatttgatctagaag
taagtttcttatttccttggtcactagtacttgggcagctaccttcttttggattttgat
ctatggttgcctttttagccattttgacattagggtttatttatgaatgaaaaaaaggcg
ctttagaatgagaataatcaaataatttactagagatttttaatttgataatataatata
atatgaacgtaactttacaaagtgcaaaaatgataggagctggactagctactattggtt
taacaggggtaggagctggagtaggaattgttttcggatcgctagtaattgcttattcgc
gtaatccttctttaaaaaatgaattgtttggctacactattttaggattcgctttaacag
aagcgattgcattatttgctcttatgatggctttttttaattttatttacttaatttactt
taattaatgggcgtttataataaacgcccaccttaaaaatacattatgacaagtacaact
ctttttttgaatcttttcaattatttctttaatatccgcttgtatggtggtaagcctgtca
aatgctgtgtattcagttttatttctaattgtagtattttgtaatactgctagtatttta
ttattactaggagcagaattttttatcttttttatttttaatcgtatacgtaggcgcaatt
gcagttttatttttgtttgtagttatgatgttaaacgttaaaatagatggagtaaaaatt
aattatagcacaatttttttgattggtattttaataagtctgattttacttattcagatt
tgaactgctctacaattagatattgaagcgtatgataatataggcgtaccactatcccaa
aataactttccaacaatagtttcttggatccaagaaaatgaattaccttcaaatacagag
agtattggtttaattttgtatacttcgtatagtttagtatttattatgtgcgcatttata
ctacttttagctatgattggttccattgtactaacaatgaatcaacgtagtggagttaaa
acacaacaaatcacacttcagttatatagaaatcaaaataaagtagttcgatttattgat
ctgagaaaaaattaatttgattgcggatatagatgaattggtacatcagtaattttccac
gttaaaggatatgggttcgagtcccattatccgctcaaattaagagagaatagctcaata
ggtagagcaatagttttcaaaactaaaggttaaaagttcaagtctttttttctcttgcaaa
agtgagtctgcttgaacaccctattcagttttaattataattaaaactgaatagggtgtt
caagcagactcacttttgcaagagaaaaaagacttgaactttaaattcatataagaataa
caaccacattttagtatacacaagacaacagccttgtatgcgtactaatatatctctgca
ggctgtgtagaggcaaaaaaagtttgtgtgaggtggctgtgtgacatataaattaaaaat
agttgtcaaaaaaagtctagtttggcgtaaaaggattcgaacctttgaaatcatggtatc
aaaaaccattgccttaccacttggctatacgccaaacaaattgaagataaagtgggattc
gaacccacggtggatagataccacgttagctttcaaaactaaagctttaaaccactcagc
catttatcccggttattaatataaattgtattctaaaatttttagagtctgtttatatga
tagtacgggaataggattcgaacctatatttttagatcatgagcctaatgagttaccttt

```
ttactctatcccgctattttatctaattaattttgtaacttaataaaacttctccaaac
tacccagtaaaggtaggatgactaaaaaatataaaaaataataaacgctagctatctgac
ctataattatgtaagggtcttctacaggcataccacctattcaacctaatattaaacaac
aagcagccatgcttcaataaagaccacgatatatgggtctaaaacgcgaa----------
------------------------------------------------------------
------------------------------------------------------------
------------------------------------------------------------
------------------------------------------------------------
------------------------------------------------------------
------------------------------------------------------------
------------------------------------------------------------
------------------------------------------------------------
------------------------------------------------------------
------------------------------------------------------------
------------------------------------------------------------
--cttctaatttctgtactatgtattcatggtaatagagctaatacaactatagcaaaaa
tcatacatataactccacctaatttatgtggtatacttcttaagattgcataaaaaggta
agaaatatcactcaggaacaatatgtgctggcgttaccatgggatttgcttcaatgtaat
tatcagggtgacctaaaagattaggcgagaagtatacaaaaaaagaaaagaatataataa
aggccactatccctaataaatcctttacgatgaaataagggtacataggtaccttatcgc
tgcttgcatcgatacctaaaggatttccagaaccttcttgatgtaaagcggctaaatgca
ctaaagacgctgctgcgataacaaatggtaataaataatgtagactaaagaaacggttta
aagttgcattatcaacagaaaagccacctcaaagccaagcaactatagaatcacctacca
aaggtacagcggatactaaattagtgattacagtagcacctcataagctcatttgacctc
aaggtaatacataacctataaaagcagttattatcattaatagtaaaatgattacaccaa
tcactcaaacaaattgtcgaggtgcagcataagaaccataataaagtcccctaaaaatgt
ggatataaactacaataaaaaacattgaagcaccattcgcatgtatatatcgtaaaagtc
aaccaaagttaacatcacgcataatatgctctacactaataaaagctaaatcaacgtgtg
gggtataatgcatagctaggaatattccagtcactatttgtattattaaacacattgcag
aaagaaacccaaaatttcatgcataatgaatattaattggagttggataatctataaggt
gattattaactatattgaaaagaggtttttttaattagacgcataaataatatttttatta
tagatggacgaggaatgggacttgaacccatggcctataaagtcacagtttatcgctcta
ccaaaccgagctctcctcgtcatgttaaaaagttaaatttacggggaaaaaggatttga
accctcactcattgatgtgacaaaccaatattttaacctattaaactacttccccatttt
tattaaatacggatagagggatttgaaccctcatgaataatattcatcaaaacctaaacc
tgacatgtctaccatttccatcatatccgcaaaaaaatgttactattagctttaacggat
```

aaagagggattcgaacccacggtataatatttcatacgatgatttagcaaaccattgcct
taaaccactcagccatttatcctgtgttttggaagctgcccactaccggacttgaaccgg
taacttaaaaagaacagattttaaatctgtcgtgtttacctatttcaccaaatgggcatt
agctattgctaatgctatgttttattgaagctattgcttttccggggttcaatgatttag
gacacgttctactgcaattcataatggtatggcatttaaaaagttttgatttacctccaa
gtaatgctaaacggtcttgagttttgatatctcgactatcagctaatcatctataggctt
gcaataaaattgcaggacctaaatatttgtcatggtttcaccaataacttgggcaactag
cagaacagcaggcacaaagtatgcactcgtaaataccatttaattctgacctatcctttt
cagactgtagatattctgtttttgaaggtgtatcatttataagccacggttttatatatt
tatattgtgcgtaaaaattagataaatcaggaactaaatcttttataatgtacatatggg
gtagcggataaattgtaattgtgctagtatttatatttaatggttgcaaacaagctaatg
tattagttccatttatattcattgagcaactaccacaaataccttctctacacgagcgtc
taaaggcgatactcgaatcttgttcgtctttattttataagagcatccaataccatag
gtccacaatttttagtatgaataggatgtgtactgaaatgagtgatagttgggtttgatg
gagttcatctatatatacgaaggaattttaagtctaaattattattggatactaattgaa
aagaaatttttttgaataattgcatacttggttttattttgattgctataattatttaag
aacgggttagcccgttcttaaatataagataaaatataatgaacatttcgtatttattta
cttacttttgcaagtattaaataataaaccgatctatctagtaatcttctagaatgttct
ggaaaaagtaaaggttatttttgcgcttttaagctgtcagcgcttatataatgttaacgt
ggctactcggctatgcaagaaacaatacaaccgatacactattggttaatatatcttaat
cctctcgtactaaagataaaccctctttttttctttcccacaacagatag----------
------------------------------------------------------------
-------------------------------ggttatggtaaagaaacttctcacgaagt
ttcctcccactgcaaaaccgtacgtgaaggtcacccttcatacggctcctcaaa----at
tatctatag-----------------aaagtgt-----------aaaaaaatacacactt
tcctttagtttaaaagtgcttatattagtccgcggcttaagtattagctgctttggaatg
tctagtatcgtggcaatgaccatgcataagtcgtaagtttttaagatcaggagatccctt
ttgacttcttggtataatatgatctatttctattctatctgaatctttaaaatataattt
acacatactacactgcggccctttagttttaatagtcgttttagtagttgcccatactt
attattaacagcgagacgtctggctcaataagtaactcttccgtcatatgggctactata
tccggctacttttgtagatagaataatatttcgttgatcatgacgggttaattttataat
tttgtctccttctttaaggcctaaaactctttctgcgctccctataggtattcaatatct
agtgataccaggacctgttcgtttgcgtttcttagctcatttccttagaagataaaacgt
acgtacactacaataactaaaagtttttgtggcactacatacggaaaagtatttagttca
tccgctaataactggcgctagtttactgattaaaactttttgagataatccggtggactt
cgtagtaatacttttttatattagctaaatgagattctatagacttatagctgggttcaca

```
cctgctttttcatccagtcgcaattccttgattattctttgcggatgtatgtttacctac
cttataattgacaaaattaaaacctaagaaatctacacctgttattctagacttctct--
------aaactaccagtgtacgttattttttgttttggattccgacaattttagacctaag
ttctgtaaaaagaattcaattttaattttgcttcgatcaattctttctcttcattacat
aatactaaaaaatcatctgcgtatctaataaggtatactccactttttcctattgcctct
tccattccgtgaagggcaatatttgccaacaagggtgatatgatatctccttaggggtt
ccggcttctggtgtgatttcttttgtattttcttaaaagcctgtaagtattcccgctttt
aatcaagctctcaattgctcttttaaaacaggaaacgtgtttacttttagaagtagttta
gagtgatcaatattgtcgaagcatccttcaatatctgcatctaatacatgtttagaaagt
tgctttaaacatttcacaattgcttttctagcatcgttagaacttcaccctggtctaaat
ccataactgttaggttcgcatatagcttcatattgaggttccagtgcaaactttacaaga
cattgtttagctcgatctcttatagtaggtattcctaaatgcctttctttcccgtttggt
tttaaaattgttactcgacgaattttatccgatttattatcaattttaatattttgtact
aattccattctttcgtcaggagttaaattactaactctatctactccagctgttcgtttt
cccaaattgtcttgagtcacttttcgaacagctaaaaactttgaaaagtcatgctttatg
atttgtttctgtattaagaatacagatgtcatattacctcttttgctaaattcaaaaact
ttacactgcaatctatacagtcaaatttcttttatttttcagttcacttgcgttcacttt
ttcataattt------gtttgttggtgtaatcttaacaatgttaaaatttttatttagat
aatt-----ttgtctacacgtctgcatatccataagctttccttatggccttggcttctt
gtagaatcctgatattaatataccttaacactata----------gaaaaacgcccttgt
-aaaaaagaatcaacaaggtttatattaatatttacttcgttccaatattacatacatat
agtcagtaagcacctctattccctctaaaaccggtacccctttctaacgcggccacatta
gattttaccttatgtttagaattcacgctgtttcgtgttaacggggtgggttaacttata
attatcccccgctatgacatatttagggaatgttatttcggctaccctaaaggtaggtta
tacttctataacta-------------cttttcttggcagatttactggtgtatatattc
tcttcaggtttaaatgtttataagtaacattctcctgaatttatgccctgcttatacctt
gtgaataactattttcacactgcatagggtgcactgtttgtaaa-acagctcattgatca
gcattaggagt-aatgcgtaggcccactagttctagagaacttgcttcctacctgtaata
ttagtttaa----------tctctaaagtggagattagctatttacgagaagtacatctc
ttctcttactagcaacgaatcgcacgaccgaactgtctcacgacgttctgaacccagctc
acgtatcttattatttggcgaacaaccataccccttggaacctattgcagctccaggaaaa
gatga-------------------------------------------------------
------------------------------------------------------------
------------------------------------------------------------
------------------------------------------------------------
------------------------------------------------------------
```

-----------------------------------------------------------------
-----------------------------------------------------------------
-----------------------------------------------------------------
-----------------------------------------------------------------
-----------------------------------------------------------------
-----------------------------------------------------------------
-----------------------------------------------------------------
-----------------------------------------------------------------
-----------------------------------------------------------------
-----------------------------------------------------------------
-----------------------------------------------------------------
-----------------------------------------------------------------
-----------------------------------------------------------------
-----------------------------------------------------------------
-----------------------------------------------------------------
-----------------------------------------------------------------
-----------------------------------------------------------------
-----------------------------------------------------------------
-----------------------------------------------------------------
-----------------------------------------------------------------
-----------------------------------------------------------------
-----------------------------------------------------------------
-----------------------------------------------------------------
-----------------------------------------------------------------
-----------------------------------------------------------------
-----------------------------------------------------------------
-----------------------------------------------------------------
-----------------------------------------------------------------
-----------------------------------------------------------------
-----------------------------------------------------------------
-----------------------------------------------------------------
-----------------------------------------------------------------
-----------------------------------------------------------------
-----------------------------------------------------------------
-----------------------------------------------------------------
-----------------------------------------------------------------

---------------------------------------gccgacatcgaggtatcaaac
catggcatcgataagaactctcagccatgataaatctgttatccctagagtttctttttt
ccgataagcgacagtatttccatacattactgccggatcattatgaccgactttcgtccc
ggcttgagctgtaactcttaccgtcaagcagatttcatcattacatattatattagcat
atattgccaactaaaatccacttttgtgcacctccgttacagtttaggaggtcctcgtcc
caagtaaactaccactttacatatgtccttgttagtaagcaatttttatactcagagt
agtttttcaatggcgtacaaatgtacttctacttttactgtactgtgtacaaagttatcg
cgacgtaaaattatagtaaagaatcatagggtctttccgtcttgttgcaggatgtctgca
ttttcacagacaatgtaatttcgctgaggctatactagagacagtggggtagtcgtgacg
ccattcatgcaggacggaatttacccgccaaggaatttcgctacctgaggatccttatag
ttaagaccgccgtttacttgggtttataatatatgcgcaaacttatatttttcactttca
agcactgggcaggcgtcaaaccttatacgtcttcttttgaatttgcaaagttttaagatt
acgataggtagatactcgcgtattttcccccgtttagaaccgtacgtgaagattaccctt
catacggctcatccatttaacgtcagaatttcctaacgcagctctgtcgcgatgacacg
ttttatgaagaagagtcatatttttaatatatttttacctccactcactacaggtatga
tatggtgaatttctaattcgtacaatatatctccatattgactatttaagaattctttac
atatagggcacataccttttgtctttttgcccactttaatcgaatattattcgtttgct
gattaaaacacatattcaaagatcttttattaaaatacttctcaaactcacccaaatagg
gattagcgtcaagttttatcatagtagttcttttaataggagtgtcggatatttgaaaca
aattaatttcttaaatcttttaaattcagtgctagtagacattaatcaatctctattat
ccactttgtgataaaatttggatttaagcgcttt-------------ggatgcttaattt
ttatcatgcggtgaatggaatcaaaaacgtatttacttattcgggtataagctaactttg
cagtactagacgagtaataatttgctcagccccttaatattggatttagttttgagaaga
gcattcacagaggaagttgcgaatgttctttgacacaaagttttatttagcttttactt
tttgtaagctttcttctgtaggtttatatagaaaaatccctttcttttccctgatttctc
gagtagaatccgagaattccctaaagtggaaacctacaaaatcaaatccatcttcaattt
tagttattttactttttccatatttaattctaaacctcttatttttagaaatttgttaa
catctagaattaacttatctaaaatatcatagttctgggacactataatgaaatcatcag
catatctcacaagagttgctttgttattagacgcctttttttatagtgttatctaaacca
tcaaggcataaattagctataataggagatattacaccaccctgtggaacaccttcttcg
gtatcttgataactgccttcattcataacgccagcttttaacatttttgacaatatcttt
ttatccataggaatattttctaatattcactcatggcatatattatcaaagaaaccttg
atatctccttcaaaaattcatctagggttgcctctctgtctacttaaaagaaatcaaata
tattgacaggcatctttagcacttcgataaggtctatatgcaaaactgcatctatcagac
acaatttcgcttataggttcaagagccatagcaaacaaagtttgcacaactctatctctt
atagtgggaattcccaaaggccttttcttttcgctattcttctttggtataaatactcta

cgtaccgcactgaattcataattttaagatcttgaatttgaattaatatttcatcaata
ctaactttctttcctttagaaccttttaaaacaattttgtcaattccgctggtctcacta
cttttacgtttgtgaatttgctttattgcttttattttagcttcttctagattcactaat
tcattttgtaatttacttcataaacgtttgttttattcaaataagctattgctattctc
ctttgtaacttataaattttcgtcattatattagttttattattttaccacaatttta
ttt-aaaataaactagctggttttacatgtaatataaataaaat-----ttttcttaaag
gacaatacttttaaaacagaacatacataagtcagcaatctttcgattcatgatataatc
attatttcacttattattagcaaacattcgctttttatgagatcttttacctactgaaca
cttggattgatcgctctttcctgagatatttctatcaattcattaggcttacccttttcc
ttacacataacattaatctaaacaggtaacattctctataccgacagtattatgtctaat
ctataaaataacaaaaagatattttatactactgcttattcctatagctatataaaccgc
taatgatgatcttcggaattctgttttctcgataccttataacaaatgttcacttacgtt
ttacctatttagataagtctagcttttcttagttacatcaagtaagaaactacattgtca
gcaaggcttcatacctccagatcactctagacgcatgcttgcttagacttatattgatta
gataatatagcacttagtgcatttatgtatacggctttaaaggtcgcactttttgttaaa
cagtcgctacccctaattttttgaaaccttaaaaggtactccttttttgcgaacgtacggag
taaatttgccgagttccttaagtatagttatctcattcgtctttattttctcaataagtt
cacctgtgtcggttttaggtacggtcaaacttcatgtaagttttcctgaaaaatttcttt
ttttagcctccagtttagtgggtgtagctattagaagatatctttgtacacaaatacacg
agtattttgcagacacgtggtaatttttcctatcgaatacagttttcactttttttcttaa
gggccgactaactccagttacttaaaattgactggaaacccttgaacaatagacgaccat
gatttattttcaacatggttagcgctactcatgtcagcattagcactcctgattttagat
atgcaatttaacattaacataaaagactacaggacgttccgctaccattaagcttagttt
agcttaattcgaagcttcgatatataaatttaagtctccttacattttaaataagtagaa
caaataaaaaatagcgagctaaaacgctttctccaataggtggctgcttctaagcctactc
tgtttattcgaattattccttatttttttcactaatttataattttgagatcttagctatc
gattagggttgtttcccttttgacgtaagaccttatcgcccaacgactgtctgctgctat
aaataaaaaatatagtttggagtttaataaaatttagcaaaatctaaatttaataagtagc
tctacccacatttttaaaaaagcaacgtactactttgatagttttcgcggaaaaccagcta
tcaccaagtttgattggactttcacccctaatcttaagtcatccccgtattttttcaacag
acgtgggttcagtcctccagtactttttaaagcaccttcaacttgcttaagaatagatca
cttggcttcgggtctaatacctgtaactttaagcgccttaatatttttaagcttgctaca
catattaacttactgactcattatgcaaaaggcactttgttgctgtatttgtcagcttca
aataaatataaattaacagattcaaatctttccctcacggtactcgttcactatcgatta
gaaaaggttttagcttagaagatgggactcctattttcatacaaaagtaatcgtactact
ttgtgttactattagtgtcgtaaaggactttataacctactttgggtttagacacttcgc

taagttcgctttcactcaccgttacttacaaattctcgtttgatttttttgttatgttac
taagatgattcaattcacataattatataaatgcatattaaatgcgagttttctaaagag
actcatagttcataggcaggtgcctcgctatgatgtttcgtcgcgcgacgtctttgcttt
tctaccaagatttcctctgataacttttaaatcttttatatttaagaacgggcggaaaa
atattacccttttgtaagagtcattaactctatagtaattgtgctcctaattcgataata
gacaacaagtattgctagtcctatagaagattccgaagccgctactgttaatatagctaa
agcaaatagttggcctactatattgtctaagtatatagaaaaaaatataaagttaaaact
aactgatagaaacattatttctaaagacattataattattataatactttttggtttaa
aaaaatgcctaaaactccgactaaaaataaaaaaaaagtataattttcacagtatagttg
agtaatcatttatttttatatttaacatttgagatagcaaattgtattaaatttttccag
ccgcaggttcccctacggctaccttgttacgacttcactttagtcctttcgctaccatgg
acaaaagataaaatttttgtcttcaagtagagtaaattcccatagtgtgacgggcggtgt
gtacaaaacccgagaacatattcaccgcgacagttctgatccgcgattactagcgattac
gacttcatattctcgagttgcagagaataatccgaattaaaaatttttaaagattttgc
tccagctcacacttttgcttcttattgtaaacattactttgtagcacatgaaagtagccc
aattcataagggtcatgcggacttgacgtcatttttcccttcctcaaggatattccaagc
agtttataatgcattaatgcattatacaaaagtttcgtccgtttgctggaattaaccaaa
cgcctcacggcacggactgacgacagccatgcaacacctgtgacatcttgtgtcatacga
gaattggtaaggttttgcgcgttgtttcgatttaaaccacatgctccaccgcttgtgcgg
gttcccgtcaattcctttgagttttaatcttgcgaccgtaatccccaggcggagtgttta
atgccttagcttcgcctctggaaaattatccaaaaacaaacactcatagttgagggcgta
gactacaggggtatctaatcccttttgatacctacgctttcgtgccttagtgtcagctat
agtccagattattgttttcacttttgaagttcttttgaatatcatcgcattttatcacta
ctttcgaagttccataatcttttcctatgctctagtaaattagtttagtaatctttagta
aacatgaaatttaaaatttttttactacttagtttaccacctacgcaccctttacgccca
gtcaattagaataatacttgctcctcccgttttaccgcggctgctggcacgaaattagcc
ggagctttgattgtaaaatttagtctccaatttttaatctattttacaaagtgatttac
agcctgatagggctttgctctcacatgtggcctggctaggtcaagctttcgctcattgcc
taagattcctcactgctgcctcttaaagagtctggaccttatttctgttccagtgtgact
gatcatcctcaaagaccaattaaggattatgggcttggtaggtcttttaaactaccaact
acctaatcctgcgtagacttttcttaaaccaattattaatttttttaaggcatttaccaca
atttaagataaatttctacgtattactcacccgtacgctatctttttcataagttacgaa
aattattaaacttgcatgtgttaggccgaccactagtattcattcggagccaggatcaaa
ctcttttattattatgttatgtttttatgtttgctatctcagaaattaaatttgattgta
taaagttatccagcgagctaagacagaagctgtaccaatttcatatatatatatgttggt
aatcttccttttggtaatgaggataaaatttgctttacaaaaagttgtatactttttttg

ttttaatgcggaacatagtttattttctagctttgttctacgtataaaaagtctcctata
tttttttattttattcatacatttgtgtatcaagatgggcagatttgaactaccattccc
ttgcacccaaagcaagtacgttaccattacgctacatcttgatatagtgatattgactct
ttctggataggatttgaacctataaccttgtagttaacagctacctgctctaaccattga
agctaccagaaaaataaaagcgaaaaaaggacttgaaccccttataatgaccttggcaag
gtcatgctttgcctattaagctatttccgcacttaaaattgaatataaagttgaataata
ttagcaataagtaaaaaaagtaaaagataattgctatctttgtacattttccttgtattt
acatattttactgaatctcaaaaaaaatggcgtattccattacttatgtggtatagaaat
atagcggacactactcaccggacactagttaaaaaccaaggaaaaaagaggtgatctaca
aaaaaatgtattgacggaatagtaatatataaaaagtatacttgatttagtaaaagaata
gggcctacaattaaaacaaacataataacgccagaaattctatgccaaatagaaaaaata
gaagatctttgcggattgtatatagttaaatgtggagaaattggtctattgatgttctgc
atattatagttttctttttcttgggattttacagccattatgcggacaagttgtaatatc
cattacagagataaatttaagtttagactgttttaatatttaaacacatttttttttatt
tttacctcatccttttatccgtacatgtaaagaagtataactgagtgctttagtctttat
ctctaagaagtgggttatttcggacgaagaaagagacaggcttctttttaccccttttga
ccttgacttgcctgcagttgtccaagatttaattttgcctttacagtctgttaaaagaaa
atgaatattattggaactaaaagataaaaacagtatacccatatgtagtttaattttatg
ttgtagtagtaaagactagggtcagattcgaactgacctaataaagatttgcaatcttac
gcatagccactatgctacctagtcttaaatttagtgaaatctgaagaaagactgtgtttt
aaatgataaagaacctttcacacaagaaagtaattcaagcgattttagctttggtaaatt
ggataactcgaaaagtaactgacccggccttatataacagcaatgactcactatagctcc
ttttcctttacccatacgggtttctggactaaatttggtaacagcatttttttggtagcat
gtaaaaccacactttgattttacccaaactggactttttttcgatgactttgatctcttt
ctgaattatctttttatattatttatttgtgtgatatctacttgcttgtacgataaaga
gcgtaatccaaattggcctcttttttactagatgacttgtttggtttatatatctattttt
tttttatgctgtcgtgttggattattttttgtattaaatcttttcatattaaattgtgc
ttaaattgttttatatgccattcaaacttgaagattgcaaatcccttgttttgtatataa
tgacgtttcaaaaaaattctatataagtttctaatttttgcagtgatactgcgcctaatcg
gaatactatactacttgacatacgacttcggctaccaccgaaacgcccagcgtaccttat
ttttattcctgtaaattcataaaactgtataccacgagcagtttttaatacaattttgtg
cttttttttattgagaaatagtatttttattgtattaattatcttccggaatgctaggcc
tttttttaggccagattttataaatccacataaaagtgccccgttttgatgtcaaaatgt
tgcattatatttttttcaagtcaagatcgcacattttaaacgtaggtaccgagcaatctga
tttggctaatagaggtacatattgtacattgtattcccatctattgtcatttattttaga
tagcgtatttccttttatgaatgtttttgattcaaaatgcttatttaaaaagtttgtagc

aggtaatactttttttgtatatttgcttttgatttttaaaacctttgccatagcattgact
ttcggtatttcataagtcgttcacgcctattcgaaatcctcttggattaactttttgacc
cataatgttttattacttaataacaatatttaaattgaaaatcgaatctaaatagttgta
caaaaatacgttttgtaggatttgaacctacactaatcaacttagaaggttgatgtttta
tccaattaaactaaaaacgtttttattttagatgcattgggatttgaacccaaaataagc
agattaaaagtctgttgctgtaccgtttagctatacatcttttttaataaagagagtagga
tttgaacctacgatgattttaatcaatagatttacaatctatcgctttcgaccactcagc
catctcttcttcagcggttagatatatgttacaatctgtaataaaataattaattatata
ctattgtttaataacaatatttagattgaaaatcgtacgattttcaatctaaatattgtt
attaaacaat
>SRR9587921
gttttattaattaaaactaccccagtattacacttttaaatatttataagaaataatgca
aatccctttacgctgttatttaatcgaatttaaatttcgattcgtttactttctacttag
ttttttctctcgttgctagtcgttataaactatcatgaggctgtgttttttttttgaaac
gtatacttttttgtttataaacgagggggcattcatcgcgacacatattgctgaactatt
ttcaacttctatgtatattagttttaacctagttttgtgcataaattatccttttgcata
ttatcattgcagccgctttttttaattcgagttgatacaaatcacaggtttggttctttaa
aaatatgcaatcattgacatttattacgtttttagctagtttacttttgtgctacttttt
aattttgccctacacttacgctttttttagacacttgaactgtaacgatagtttatgcatt
taaagtgcagttggaggcaaggatagcaacatatgtttgttgaacactacaaacaatgtg
tttactatctaacattgtctatctcttctttactaggctaatttgtttttacttagtgga
caatatgattaatttgcatttatttttccgaaaaaataaaaaatatactttctttttcgt
atgcttgtcggcttctgtgtgtttaccgcaagaaagttttatacaaattactttcataat
ttctataatagtactatttgagttattctttttggttacatgcttatttttttgcaaaaaa
taatgtctaaaagtggacttgaaccactgacccaaagattttcaatcttttgctctaacc
gctgagctatttagacttaaactttttcttttcccgtatttagagcgaccatttttttcgtt
tagcgactgcttgtaaatcgtaaacgcctctcacaatatggtaacgtactccaggtagat
cctttacacgaccacctcttatcagaacaactgaatgttcttgtaaattgtggccttcac
ctcctatatagcctatgatagaacgtccggtactaagacgtattttagctacctttcttt
cggcagaattcggtttttttagggttagtagtatacacttttgtgcaaacacccttttttt
gaggcgacttattgagagcgggtgtttttttgctttggtaattttgtgctttcttggatttt
ttatttttttggtttagtgttgacataatatttttattttaagtaataagggacttgaacc
cctaaccatttcggtgtaaacaaaacgctctacctattgagctaattactttacgtctgg
aaagatttgaactttcaacttttagattcgtaatctaacgctctatccagttaagctaca
gacgtttttctccacttgtatatttagattacttatctatattttggaaatagggacaaa
tattcttgtataatttacttcaggattattcgaaagtatagttaaccaattaaacaagta

gaaaacgcatgttttgtattgttactaacttaaaggactattttataataaactttttat
tgtattatataccaatatttagattgaaaatcgacccaaattcaatgtacttttagcaaa
tgacaccagtgttttagaaaaggtg-ttattttgaaaacatttttggtgtgatttgtcct
ttttcagctagtaataatttaattaataacagaaagcgcggtttattgctcgtcgcgttc
aagttattactacggcaacttcctatgcagctgttgggacccccccaactcgtgtaatta
cactttttagttttatttaaattataataaggttatactttatatcttttctttttattta
accccctaagatctttattacgccccaaaatggtgtcataaaaaaa-tagcctggagggct
gccaggagt-aaaaaggcccataacggttatttcagcttttatagtgctatttttctttt
ttagagaaggctatttttaattgataactaagtcattttaagtgataattaagctatttt
aattaaaaatgacttattaaaggttttacacagataatacttcttttattaaaaatatac
tacgccgctagaccggaaaaaaccctccatagaacgatgagaggtaaaaagtgtcataaa
ttttaactattttgtaattgtacattttaaaattaaagtttgtcaaacatcaaaactttc
tgttatttcaataatgcgtgtttttaaggcttttaaagcagtttctcgctgatcccgaac
actaaacttttttgttaacatgttctcgtaaactgtaattggatgtcataataaccaatgg
attatccggttttaaagtaaaattacctttcattgcaaggcgcatgctatcaccagctaa
aaactggtttaagaattcgatttgtttaccgataaggtgaaattcgtctcatagaataaa
ttgatacaaatgattttcataataatcttatcagttgtcttggggtcacatataacaagc
ggtagtagttttcaagtaatttaacagcgcactttttcccatattaggttctttacttca
tatatagagttggggtttttataatagcgtgaaaaaggactagtattattgtttagtat
gtcgaatatttgttctcataactggatagaacattccgataagttatttttattatttttg
ccttaggattgttttgcaaaatcgttctatgttgggcttttattttttttgttttatgtct
aaagaaagttgatgctgtttaacgtaacgatcaaattttatcctgtcattaaacaaagca
cgtttaatatctggttcatcaaaattaaaaaatattggtgtagtaacaccagattcgatt
tatttcatgatgaagtgctctcttacaagccgctttttttagtagattactaaaaattta
ctgtgttgatttgatacgcattttttggatgggctcttgcttatttattggatctgaaaag
gagcttggagaattagttaagtagtcaaaatgattcagatttgtgaaattacgtttctta
gaaaattcaatatacgtatataaaatttgatccgagttatttcagctactagaaaagctt
attattattgtttttttctctttgtaggagtttgttataaattaaaactaggcaaagaagc
tacatttttttgttgtaacttggaaagatttggccgcaacgcgctttttatgtttagttac
ggctggatttagttcaaatactttggacatgagtcgttttaaaaattatttagaaatatg
tgttcgcaaacagattgagtgaaaaaaa--agttttaaaaaagcttcataaactttcacc
aaaagtttgatttatatcataaacggagaaaatccaaattgaaaaaactaaaatggtgtg
gcctaaaaaaaatacggcgaagtacatatcaacttggttttcagtgaattaaacagaaata
cagtattcccacaagtacgataggaattagaatgatggataaatcaaaattttgatttag
atacgaaaaccatg-------------gtttacaagtaaactttttaattcaatatttg
tgctattagtaaagatttgtgcgactttaaattgaataagctgttaaaataaggtgaatt

taatttaggcaataaagtatcatacaaatgtttgttctttctatatttgtttagagcttt
cctgtacttttattgtaagcagcaatagatcctatgtgcttgtgttgaaaatctaatct
tttaagtactttgttaaagctcaaaattttagtttttaattggttaacag-ttttttacg
ctttctagcatcgtatagtctttcttttagcgtttctagatgagaaggcttttctagtgg
cagttgattgaaataacttttagtgag-tttttttactatagtacctgtaacaagtgtgct
aacagatactatgacacaaactacttctaagtccaatttgctctacaatgcg-tttgatg
aaagtcattttatataaattaatttattaatgtgtaattgtggtatgattttattttgtg
ttaatggtcataattaaatgcatacattatatattatccctccttcggagggata-atat
aataattcttattatataccaatatttagattgaaaatcgtgctattttcaatctaaata
ttggtatataaacaaacctgcacataaataaatatcacaaggacat----ccaccaaaaa
atcgatattctctagagagcccctataagctttaatttactatctcttgcatgaaattat
accttcttggataggaatcatcgtgaaattgaaaaacattta--gatatgtcttatacaa
ttatgaccaaaaaatctacaaaacaaacaactttacacaaatattttcgcaagtcgcata
catgagattgaacttaagaagcctctttttacaaatcatacagcatttttggcgcttctt
aattctattgcaatattcaatgaaccaaaaacaaagtttttttttaaacgacctgcatgaa
cacaatcttaacaagtgtaaaaaagccctctagc-cccttttttgcaacaccacgagaat
aaaccccggtcggggagaccccgaacccccctcggcgcggccgcctagcaaagttagac-
-ggcccgcgcgcgttatagattcctatatcgctccattccattccatttcgctattgctg
cgcttcgcttatttttaattctaactgttccttttcgttataacgattttaggtcgattt
tcaatctaaatatttgtatataaacaattatttatcttgataattatatatctaaaaagg
ggggcatagtttaataggtataatgtatgctttgcaagcataagattaccggttcaaatc
cggttgtctccaaatctaaaaatatcaatgcgttctagaatattacaatacaaaaataac
ttggctgactatgatcttttaactcaattttcgattaaaaatcataatagtatgcctact
ttctcttctcttaatgtcagagtaaaaaatattcaaactagcgacattaagcaggtttgc
ttaaatctaatctctattcaattagtaggaaacaatcatgttgttttcagtagccaaaaa
gacgtttctaatcttatactaaaatgtaacggtattaatacttattttattttagagaat
ttatctttgttaggctacaaaaacaatttaaaaagtgacaaaaacaataaattaactgtg
cacggtaaaaacttcttaaatattattgaatattttgcatccaattcaaattttcttaag
tccttagaaaaaacatctaataaacatttaagtattgctttttcttatcaacttgctaat
agtaaaaataatgaaatagctttgtatttttaaaatcattaggatttccgctttaataa
cggtatagttcaattggttagaacaacggaatcataatccgtaagttgtgggttcaagtc
cctctaccgttatggctcttatcgtctaatggttaggacagtgctttttcagggcatcga
cgtgagttcaatcctcactaagagtaacttttaatataaaaatccttatacaaatacgt
gagcccataagaattcaaaagtaaaaaggatttttctagatctaccaaattattttggga
tccctaggtcgttaaaaatatttaaacaatgtattttattc--atatatatatatactaa
aacttaaaaatataatttagtgcaaaagtctctaaataactgaattttccgttgaatata

ctcgaccaatcataaagatataggtactttatatttaatttttggcgctttctccggtat
cttaggtgcttgcgcgtctatattgatccgaatggaactagcacaaccaggtaatcaact
attattaggcaatcatcaagtgtataacgtactagttacagagcacgcatttttgatgat
tttctttatggttatgcccgtcctaattggaggatttggaaactgattcgtacctattat
gataggtgctccagatatggcctttcctagattaaataatataagtttttgactactacc
tccatcattgtgtcttcttttaggatctgcgatggtagaagtaggcgctggcacaggctg
aactttatatccgcctttgagctctattcagagccattcaggcggtgctgttgatcttgc
catttttagtttacacttgtcaggtgcttcttctatattaggagctattaatttcattac
gacgatatttaatatgcgcaatccaggacaaagtatgtatcgaataccgctatttgtttg
atctatcctcattactgcgtttcttttactactagcagtacctgtcttggcaggggccat
cacaatgctgttaacagatagaaactttaatacaacatttttgacccttcaggtggtgg
cgatcctgtattgtatcagcatttattctgatttttcggacatccggaagtgtacatttg
tgcgccgtttaattctgttaaataagatggtttattattttaaattacttatctcgaaat
ctttaaactgaatagttaataaagatagacttaattattccttttgtcttaaaacacttt
cccttaataaagttggaaacaattctattaaaagcgggaatgaaagtctcttgcaggaac
ctgaatttcctaaaaatagtaggctagttatgctcatcatgaataagataataaagttga
tacatatcgacggttccacactaaactttatacataacatttcaggtaacagtatttatg
taatatgggtcggtaccagaatgtttaccttcccttgaataattattacacgtcagagcc
atcagctcctaaatgatatgcaacgttctaaaatgagcttgatcgagaaacagaagaact
cgggataccctaccaatcgaaagattcatgagtacggaactctcgtagtaggtggtagaa
gaattcaatcttcttcaaactaccaaaggggagtagaacttagattcttaagtgaaaaac
cctgcattagctcgcaagagtgcgctaggttagtagatttgagaaaagttaattctgaaa
ataaatttcaagttaataagaatactattcatattatatctgatatgaatgtccttattt
tagcatatgaactcataaaaagtaatcctggaaacatgacacctcgtgtgaatggttcca
cattagatgggttagacaagatgtgactgcaaaatattagtaccaaaataaagcaaggta
aatttttattcagccctgggcgtaagaagtacattcctaagcccggttcagcggataaaa
gaccattaggtattgctagcccgaaagaaaaaattgttcaaaaagctattctgctagtac
tagaatcgatttttgaaccaagcttcttgagaaattctcacgggtttcggcctaaccgag
gcaaccataccgctttaaagatggtaaaaagcgagtttcacggagttccctgaattatag
aaggagatatttcgaagtgctttgatgaaattgatcactctattttattggggcttctaa
gcaagaggatatcttgtgataagactttaactttaattaaaagagggttgaaagctgggt
ttatagatttaggaatattcacaagaactaaattgggcacccctcaaggaagcattctga
gtcctatcctatgcaatatctatttgcatgagctagatttatttctacttcaactaaaaa
ttaaattcgatacagggactagtagagcgaagaacccacagttcagaaaactacagtata
aactatctaaccttaaaacgcctctcgagaaaaagcttgtcagaagagacctttgaaaag
tgcatagtctgaacccccctagatcctaacttttgcagaattcactttgttcgatatgcgg

atgattttattgtgggagttacaagctcccatgaagttgctttagaagttaagaatatga
ttaaggaattcctttgtaatcatttgaaattaaatttgaatgagctgaagacacaaatta
ctcatattagagaaaaggatatatttttcctaggtacccttatcaaaggtaactgaaaga
aagagaaacctattcgattgatcaactttccctctagagaaacgtccatcaagacaagag
tcactccgcgtttaagtttgcatgcccctattaaaaaactctttgataaagctactgctg
aaggattttccgtagggatgggattaattataaacctacctttgtaggtaaattgatta
atatggaccatgcagacattttatttttttataattcaatagtaagaggagtactaaatt
actactcatttgtggacaaccacaaaagtttaggatcatgagtccattatatgaatttt
cctgtgctagaactctagcgttaaagtacaagttacgtttcacatcgaagacttttaaga
aattcggctctaaattggcgtgcccgaatacaaaaaaaagcctgtttttaccaacgagct
ttaagaggacgcaggccttccaaattaatagccctattccttttgagaaaaaagtactct
cttgatctaaaaaaattactaaatctaatctgaacaaagtttgtttaatatgtggtagtt
ctctatttgtagaaatgcatcatatacgtagtatagctggcattagaacaagactttgta
gcaataaagcagactttttctctctgcaaatggctgggataaacaggaagcaggttcctc
tttgtcgagaacatcacttgaagctacataataaaacactttcaccagaagaaacgttat
tatttcagaaaaatattaaggaaatttagatgtatttctttcaattttaactttatttac
aatttaagttcggcgagccgtatgataagaaattatcacgtacggttctgagggcagtta
ttgacgctaaaacaacctcgttgtcaagtttctatttctgacccctactaatcttgcctg
gatttggcatcgtcagtcacatcgtttctaccttttctagaaaacctgttttcggttata
taggaatgatttatgctatgctttctataggtattttaggatttatcgtttgagcgcatc
acatgtatactgttggccttgatgtagatacaagagcttacttcacagccgcaaccatga
tcattgctgttcctacaggaataaaaatatt-----------------------------
------------------------------------------------------------
------------------------------------------------------------
------------------------------------------------------------
------------------------------------------------------------
------------------------------------------------------------
------------------------------------------------------------
------------------------------------------------------------
------------------------------------------------------------
------------------------------------------------------------
------------------------------------------------------------
------------------------------------------------------------
------------------------------------------------------------
------------------------------------------------------------
------------------------------------------------------------

```
------------------------------------------------------------
------------------------------------------------------------
------------------------------------------------------------
------------------------------------------------------------
------------------------------------------------------------
------------------------------------------------------------
------------------------------------------------------------
------------------------------------------------------------
------------------------------------------------------------
------------------------------------------------------------
------------------------------------------------------------
------------------------------------------------------------
------------------------------------------------------------
------------------------------------------------------------
------------------------------------------------------------
------------------------------------------------------------
------------------------------------------------------------
------------------------------------------------------------
------------------------------------------------------------
------------------------------------------------------------
------------------------------------------------------------
------------------------------------------------------------
------------------------------------------------------------
------------------------------------------------------------
-------tagctggattgctactatgtgagaagggtctattttttaaaaacccctatgt
tatttgctatagggtttatattttattcactataggaggacttactggtattatactag
ctaactccggacttgatatatctttacatgatacttattatgtcgtagctcacttccact
atggtgcgccgtctaattgcgtttatcgtcgcttacttcagtgagcatgtattaccattc
ttgaattatataattaaatgttatacaatttatacattaaaacaatcaacttacttattt
ctggccaacagaaagaggactacagcatgttgataaaggagttttgaaatagtaaactcc
gagttatactacacacggttaggctaacgaactccttctacaatcagccaggcggatatg
aagttccgatcattagaaataatgaagtatctcacggtgaaagcctttacatagttagac
ctttacaggaatggtctggtacccaaaatttaagtaaaattttaaagctgaggagaacc
taaaggtagttgttaatggtaagcgtaagaattcgggaaatcctgaaagttgaaaaactg
gaggattcggagggatcgtagtacgaaatataagctgcttagcttatgttaggaagggtc
ctagttcagaagctattctaaattctaagctctcgggttatgaatctatagaagtaggct
```

taaataatattgacaaacaagtcttagaatatattaaaaccggcaaaagaattgaaggat
taagtagtttacttcggaatccaaattttcttattgcaagttattcaaaaatcaagtcta
ataaaggagctctcactcctggactgagtaatgaaacattggacggcataaaattagaat
gatttgaaaaagctgctgaaagtatagttaatgggtcctatcattttgaacccgttaggc
gaaagtttatacctaaaccgaaaggtgatgaaagacctcttggtatacccaatcccagag
ataaaatcattcaagaaggcatgaggcaactgctggagttagtttacgaaagaattttg
tagattcctctcatgggtttagacccaacaaaagttgtcacagtgctcttaatcaggtaa
aaatgactatggggtattcctcttgatttattgagggggatatatcaaaatattttgaca
ctgtgaatcatacttatttagtttcaaaaatagcgaaagttattaaagatcaagccttca
ttgatttaatatataaagttttgaaagcaggatacggcttttccaaaagaaatgtcgtaa
gcactagtagaggctttccacaaggtggagttattagtcctatactagccaatatttact
tacatgattttgatttaaaaatattggaaatgtcggaaaactttaatagaggtgtccgtc
gaaaagcaaatcctgaatatactaagatggttagagatggaaaagtagatagaaaaaatt
ttatttatcccgctatgggaaatgatgcttattttaaaagaatgaaatatgtaaggtatg
cggatgattttcttatcggaatcattggttcgaaagcagactgtgaaggaattagaacta
gcatagcgagtattcttaaagaagagttcttacttgaacttaatattgagaaaacaaaaa
tcacgcatgctaaacatgattgcgctttttttcttagggcacaatattcatatatcaatgc
ctcccaaagataaaatacaatatcttccaaaaagaggaaacaagcttgtaagaactacaa
gtcgacctttattggatgctcctattggtaagatagtgttgaaactaggttcggtagggt
attgcaaaactgatggatctcctaggagattcggaaaacttctacatgaaccaatggcgg
aaataatttataggtataaaatactgcaaagtggattattaaattattattctatggcta
ataactatggtcgtctatctgcaagaatacattgaaccttaaaatattcttgtgctctaa
ctatagcttcaaaaatgaaattggggactcttaaaaaagtgtttaaacattatggagcta
atcttgaaataaaaaacgaaaaaggggagattatccaatgctttcctaaaatatcgtatt
ccagacctaagaaccctatcaaaacgaaaatatttgatcctattgatcatatagagaaat
cttccaatcatttcaaaagaagtttagcaacttttgagcaagcttgtatgttatgtggaa
agcaagatactactgaaatgcatcacattaataaactgaagaataactcttccacggatt
ggttaacttctcgaatggtcaaaatgaataggaagcaaatacccgtttgtcgaaattgcc
accaacttattcacaagggtaagtatgatggttcaaaaattatttaagtgactcttttga
agtgaaagccatatgcgctgaaaagcgcacgtatggtttggagagaggattttgatactt
aaaaagattagtctcctactctacttctgtctatgggagctgttttcgcaatatttgcag
gcttttattattggtttgaaaagatatccggatttcaatattctgaaatactaggtcaaa
ttcacttttgaggcacttttataggtgtaaatctaacctttttccctatgcactttttag
ggcttgctggtatgcctagacgtattccggattacccagattcttatgccggttggaacg
caatagcttcttacggttcatatgttgcgttatttagcacgctgtttttcttttatcttg
tatttaacacacttgtaacagcaaaaaagacacctgctagaaataacccatggaactttg

aagattcaaaaatgggctcaactacattagaatgagaaatttcttctcctccagcttacc
atacgttcaatgagattccagttataagagaaacagaaacatctttaaaaataaattaac
ttatgataaaaaaaaatactacaatattatttttagggttagctttgctaacccaactac
tacagagtagaatagttattagtgattctgcagaagactggcaattaggcttccaagatc
ctgcaacacctataatggagggaatcattaaccttcatcatgattttatgtttttttatct
gtgctatctcaatttttgtatcttgaatattagcacgcacgttatggcactatcactgaa
caaaaaatgagtacccttctgccacggttcatggaacagccatcgaaataatttggactg
ttactcctagtatcactttgttggcgattgctgtgccttcttttgctttgctatactcta
tggatgaaataattgcgcctgcaataactattaaaacagtaggtcatcaatgatattgaa
gttacgaatactcggactatacaaatgaagacgataacactataatgtttgaaagttata
tgattccagaagaagatttaactttaggtcagttaagattattagaggtagataaccta
tggtaatacccgtaaatacacacgtacgtctaatcataacagcagcagatgttttgcaca
gctgagcagtgccttctttaggtataaaatgtgacgctgtaccaggtagattaaaccaaa
gctcgctctttgtaaaacgtgaaggaatattttatggtcaatgtagcgagatttgtggtg
taaatcatggttttatgcctatcgtagttgaagcagtgtctttaccaaattatatttctt
gagtagctaataaacttagcgaataaatagcctaaaaatattctaagcatctgatatgcg
tgtatctctagcccaactatttttctttggtttatctatttttattttattttgatttaa
tcctaaatacttaacttatacaatacgactattaaaaaaaattaaaaagcgttctaaata
ataaagagcgctttaataaaaaacttattatggcaactacgacaaatctaaactttatca
aaacagctaaacaattacaacgccaccctttttcatttagttgaccccagcccttggcctg
taacagctgcaatagccgctttttcatgtgctttaggcggagttatgtatatgcatgcat
acagtaatggagggtacctattatttagtggtttttctttactgttatttacaatgttct
catgatggcgcgatgttacaagagaagccactttttcagggcatcatacaggtgctgttc
aaaaaggattgcgttatggtgtaattttatttatagtttcagaaatcctcttctttttttg
ctttttttttgagcatttttttcatagtagtctttcaccggctattgacataggttctatgt
gaccaccaaaaggaatagttgtgtttagcccttgagaagttcctttttttaaatacaataa
tattattattatctggttgttctgttacatgagcacatcatagcattgtagcaggctata
aaaagcaagcaacgttagctttaataacgacagttatcttagccgctatttttacaggtt
tccaaggttttgaatatagcgtggctaattttacattatccgacggtgtttacggcgcca
cattttatatggctacaggctttcatggttttcatgtctttataggtactattttccttg
gtatttgcttacttcgcttattaaaatcacatttgacacaacagcatcattttggttttg
aagcagcagcttgatattgacattttgttgatgttgtatggctttttttatttatttcta
tctactgatgaggtggtacctaatctaaatctctttaatatccaaaaatctatgatgaaa
cttattaatttacctaccataaagctcacttttttgacatgctctagctttcttaattatt
atctgttatcataatatctatatttttacagaagaaagtatacttttattctgttttatt
gcgtgactaaacattacatgaaattatatatctcctcaaattaatgcgtcattatctgaa

agaggagaaaaaatcaattcaaattttcaacatattactaacgataatataataacttga
aaaaaatatagacaaggttattcactaaaaataacacatgttgatattttaaagaactta
atcacatatttaacatgtttaattaagactgtactccttttagaatcaaaaagtagctca
ttacaatctattgcaccttatttaaaaagattacatattgtaaaagacttagaaacaaag
ctgactaaaatttcttatattacgatttgccaacgtatacaggatacagccacaatacgt
agtttttatgctaaccgagtaaaaatcaaatctttccattctgaatctaaattagactta
ttcgaacgtataagaaaattagaagctggttcttagaagtttaagtctatagctcaatgg
ttagagcatacgcttgataagcgtaaggttgattgttcgaatcaatttagacttatacta
tttcaaaaccataatataaaatatgtacaatataaaccaaacatttttttcagcaagccc
tttagaacaatttgaaattatacctttaattcctttagaattatttgggttaaacatgtc
gttaacaaacgcgtccatttttttgatactatctgttgcgttatctatttttttgatccac
tttagtcatatacaaaaataaattagttcctggaaactgacaatctgtaaaagaaatatt
ttatgataccaccttaacgttggtaaaagataatttaggtaaaaaaggttatcgatattt
cccgtttatttttaccctttttcacaataatactttattgtaatttaataggtatggtacc
atatagttttactgtaacaagtcatatagctttcacatttggcttagctttagctattta
cataggaattaatattattggcttcagaacccacggtataaagtttttcacaattttttt
acctaaaggagttcctttatttattgtacctttagtggttgcaatagaattcgtatctta
cgtcgtaaaagttttcacaatatcgataagactttttgcaaatatgacatccgggcatac
tttacttaaaattattgccggatttgtttggacaatgatctcaataggaggcgtgtttgt
atacttacaaataatcccattagttttattactagcgttagtgggtttagaaattggtat
cgctcttttacaagcttacgttttcacattacttacctgcatttacttaaatgatgtttt
agaaatgcactaactaaaaaattatgccacaattagatcgcgttattatttttggtcaaa
tattttgactatttttcacctttttaattgcttatgttgtttatacccatttcatattaa
gtaatttattaaaaattttcttagtccgctgatggaagcttagaaaagatattactcaaa
ttgcattaaagatccgtttaacgagctatttaattgattcaaatattcaaacgttacgta
gaatttattcaacaatcagaaatatactagcttctctaacaaaaagtttattaacaaaaa
gtataagtaagccaaagttagttttaaatgatcttaattctttagttattaaaattagtc
tggaaacgtctttatatggtagcaaaagcatcaccaagtctggaacatattcttattgaa
cttaataaaatatactatgtatttattaataatagctctgcctttagtgggaacattagt
tacagggttaggcggtagatgaatagggcgtaaaggttcaaatttgttttctacaacttg
cgtagtcctgtgtgtcttttttttcttcaatagctttttttcgaagtaggtctttgtggagt
tccttgttatatatctttgagcccttgaattagttcaggggcactaaatatttcatgagg
tttttttatttgatagtttaacaacaacaatgcttgttgttattacatctatttctagttt
agtccatttgtattctattcaatacatggagcacgaccctcattgccctcggtttatgtc
tttcttggagattttcacatttttttatgatcttattagtaacggctgacaattttgtgca
aatgtttttaggctgagaaggagttggattagcttcttatctattaataaatttttgata

cactcgactttgtgcaaatcaagctgcaatcaaagctctggtagtaaatagagtaggtga
ctttggattaagtttaggtattttcacaatttttatctttttggttctgttgattatga
aatagtattctcttccgcaaacatctacacaaattatagtatttccttttgtgggttttc
cataaataccttgactttaataggtatttttttattaataggggctgttggaaagtctgc
acaattaggtctgcatacctggctaccagacgctatggaaggtcctactcctgtttctgc
actcattcatgcggctacaatggtaacagcgggtgtatttttaatagtgcgctgttcacc
tcttattgatttatcctcggatgtcttacttttaattactcttcttggatcaagtacagc
tttttcgcctctattgttggagtatttcaaaacgatataaagcgggtaattgcttattc
tacttgtagtcaattaggctacatggtctttgtgtgtggtttatcctattataatgtagg
tatgttccatttagtaaatcatgcttttttttaaagcattactttttctaagcgctggctc
tgtaatacatgcgctatcaaatgaacaggacatgcgccgaatgggttcgctagcaaatag
cctaccgatcacatatgctgctatgctaattggctctttatccttagcaggattcccttt
tttaacaggtttttattctaaagacttaatcatcgaaataacacaaataagttattacag
taatttacagatttcttttggcgtttatgcttgttgacttgctaatatttctgtactctt
cacatcgttttatacatttaggcttatttttctaacttttataaaaaataccaatagcta
tagaaaacacatagaaaatatacacgaatcgccacctttaattctaattcctttaatatt
actcgctatatctagtatttttgtcggtttcttaacaaaagatatattcgtaggaattgg
aactccttttttgaggtaatgctatcaatattctacctacgtcttgtaatctattggaagt
tgaatttatgccttctttaataaaatgacttccgtttgtgttaagttctatgggtgcaat
tctcgcttatacaataaacgtaggtgtactaaaaaataatatacaatttgctcataatca
cttatttagaaaactcgccttttcccttagcaaaaagttatattgagataaattatacaa
ttcattcattgtatctcctttaatgtactttggttataatatttcattcaaaaatcttga
taggggttttatagaattcgtaggtccttatggaatttcgcgtactattaaaaattgatc
cacaaaagtaattaaaatacaaactggtcagctaacccattataccttttttcgtgatttt
tggtttatgttcccttttactactagttcctgtttgagattttctacaatttttagttga
tgtcagattactagtattttgctttatagccctctttgtagcgtagtttacgaaagtttt
aacacttaaatatatgcagataactaatttattattatggacttcacttattcctttgtg
tggcgctatattacttatttttattcctagattttactctcatttaataagaaatattgc
tttcgcaacagcgcagctagcgtttatatactctattttgctatggctttgctttgaatc
aacaacatccttattccaatttatatatacgataaattgatttccctcctataatattta
ttacacaataggtgtagacggtatatctttatttttttatcatacttacaacgtgattaat
tacagtttgtacattaataagttgaaatatgccagacagccaaataaaagaatacttaat
ttgttttcttttgcttgaagctattttaattcaagtttttttgtgttttagatgtcctatt
cttttatatatttttttgaaagtgtccttatccctatgtttttaattataggtgtatgagg
gtcacgggaaagaaaaattagagctgcgtatcaatttttcatttacacattagctggttc
actgctaatgcttctagcaattttaactatttatttccagcatggtaccacggatatcca

agttttatgaaatataaattttgacgttagaacacaaattttactttggctagctttttt
cgctagtttagcagtaaaaattcccatgattccttttcatatatgattgcctgaagccca
tgcagaagcacctacagcagggtccgtaattttagcaggtgtgcttttaaaaatgggcgg
gtatggattttttacgtttttctttacctctgtttccggaagcctcactttattttgctcc
attaatttatttactaagtattatagctgctatatatgcttcacttactacaattagaca
agttgacttgaaaaaaataatagcttactcttccgtttcgcatatgggctttgtcacatt
aggtcttttctcttttaactctcaagggatagaaggtagtataatcttgatgcttagcca
cggattagtctctagtgcacttttttttgtgtgtaggtattttatacgataggcataaaac
gcgtcttctcaaatactacggtggtctcgtgcaagttatgcctattttcagcatattact
attatttttttactttctctaatatcggttttcctggtacaagcagttttgttggtgaact
attagtgttaatgggagtatttcaatttagtccaatatctacttttctaagtgcattcag
catgattcttggggcagggtattctatttgactattcaatagagtatgttttggtagttt
aaaacttcaatacattacaaattttcaagatatttcaagaagagaattttgtatcctttt
tccgttaagtgtatttgtactctgaatgggtatatatccagaaattttcctatctgaaat
tcactgttcaagttataacctaattgcatattttaactaattttatgttatgaagtttat
taaaacgctaattctagcatttatgaaaaagaagtcctatttttattggtttccacgttt
tttagggctattacttatacctgggtttttatttgataccgagattctagttctctttca
aagccttatcctcttacatgcaagcctaggtttagaagtaatcatagaggactatttaca
cctagaaataataaaacttcagtgtttgtctttaattaaagtacttttaatattattagt
caatcttaatatattatatttattataaaaaatatccttatgttatttatctcctcttat
gatttctacgcattattgacagaaatttactttttaaacgcaatttgtgctttattaatt
tatggtgtaattttaaatacctcatatagaagagggcatccagttattgaacacaatgta
agtggtctctcaactcaaatactaatagtgagtctttggttaacagtttgttcaaatata
ccttgcctaaccagctggaattcacttttagtgcacgatttttttatctttcggtataaaa
agcaccatattagcaatttcgctactttggtctttaattatttttttcttacaatagacta
gaaaaaataaatctctacgagtattgaatcgtgtctatgttggctattgttgccatgctt
tttgtaagttgttcttatgatcttttggcaatgtatttagcaattgaatttcaaagcatt
gcattttatatattagctagttttaaaagaacatctgaattttcaacagaagcgggttta
aaatatttcgtactgggtgcattttcttcagctttgcttcttttaggtatttcactactt
tatggtactactggtttaactaattttggagatctatcaaaattttttttaggtaccaca
ttggaaaacgcatcatttatcaacataacatttttttggtgtcgttttaatagaagtagct
ctttttttttaagataagtgcagcaccttttcatatgtgatcgccagatgtttatgaaggt
gctcctactaacgttacatctttttttggtatactgccaaaattagcattagtaagttta
atatttagattcttttatttttgttgtgctgaagttgtgctgttactaaattttacactt
ataatttgtgcgcttttatctatgataatagggacatttggcgctttagcgcaaacaaaa
tgaaaacgtttcattgcgtatagtactataagtcacgtaggatttattgtagctggattt

tcaacgttggaatttaatggtgcatttggtgcgctattttatatcttggtttatacttta
acttctttagccactttttctattgtgctttccttccgatgcttagcatatcctagcaca
taccaattacgctatctaacggatatcgttagtttagtgaagttaaatcctatacttgct
ggtagccttgtagcagttttattttcaatggcaggtatcccgccttttccaggatttttt
gctaaagtatttgttttattttcacttttgcaagaacaattaataggattagctataatg
gcaatatttttgagttgtgtttcgtgtttttattatatccgtttgattcaaatgatgtat
tttacacatacaaaaaccatacttatttttttatccaatagaaaagactacatcaactata
ttaagtataactatgttattacttgtacttattttttttgaagatagatctgatttctaat
tttgttcattgtatgttgtttttataaaataaccaattaaaatgttttacaatattgcaa
ttaacattatcaaagtgttgaccattatagtgccacttttaatcgctgtagcttatatga
cactggccgaaagaaaagtgatggcagctatgcaacgacgaaaagggcctaatgtggtag
gtatctttggtcttttacaacccctagcagatgggttaaaacttttctcaaaagaaacta
tactaccttctagtgctaatatttttattttttttagctgcacctgtgctaacgtttttgc
tagctttattagcatgatgtgtacttcctctagatgaggggaaagtttttttcggacttaa
atataggtgttttgtatatattagcagtatcatctttaggtgtttatggtattataactg
ctgggtgatctagtaattctaagtatgctttttttaggtgctttgagatcagcagcccaaa
tggtatcttatgaagtttccattggtctaattttaattaatattttattatgcgcaggca
cattaaatttaactcaaattgttctggcgcaacaaaatatgtggtatataatacctctgt
ttcccatatttattatgttttatatttctatattagctgaaactaacagagcccctttcg
atttgccagaagcagaagcagaacttgtagctggttacaatgtagaatactctgcgatgg
ggtttgcgttgtttttttttaggcgagtatgcaaatatgatacttatgtgtagtttaacaa
ctattttttttttttggtggttgattacccttagtcaatatgcttcctttttattggattc
cacccgtactttgatttggtttaaaaacaactttacttttatttggttttatttgagtgc
gtgcagcatttccgcgatatagatatgaccaattaatgcgtttaggatgaaaaatatttt
tacctttatcattagggtgagttcttttagtatccgggatactattttctttcgattgat
taccataacaaatgaacgtactttataacgagtattctgctattctcactttttttgcag
tagctttttttaatctctctaataatattaatactttcgtatatattaaatcctcaacaaa
gtgatcaagaaaaagtcagcgcctatgagtgtgggtttaatccatttgatgacgcgagag
caacttttgatgttcggttctatttagtcgcaatcctttttttaatatttgatttagaag
taagtttcttatttccttggtcactagtacttgggcagctaccttcttttggattttgat
ctatggttgcctttttagccattttgacattagggtttatttatgaatgaaaaaaaggcg
ctttagaatgagaataatcaaataatttactagagatttttaatttgataatataatata
atatgaacgtaactttacaaagtgcaaaaatgataggagctggactagctactattggtt
taacaggggtaggagctggagtaggaattgttttcggatcgctagtaattgcttattcgc
gtaatccttctctaaaaaatgaattgtttggctacactattttaggattcgctttaacag
aagcgattgcattatttgctcttatgatggctttttttaattttatttacttaatttactt

taattaatgggcgtttataataaacgcccaccttaaaaatacattatgacaagtacaact

ctttttttgaatcttttcaattatttctttaatatccgcttgtatggtggtaagcctgtca

aatgctgtgtattcagttttatttctaattgtagtattttgtaatactgctagtatttta

ttattactaggggcagaattttttatctttttttatttttaatcgtatacgtaggcgcaatt

gcagttttatttttatttgtagttatgatgttaaacgttaaaatagatggagtaaaaatt

aattatagcacaattttttttgattggtattttaataagtctgattttacttattcagatt

tgaactgctctacaattagatattgaagcgtatgataatataggcgtaccactatcccaa

aataactttccaacaataatttcttggatccaagaaaatgaattaccttcaaatacagag

agtattggtttaattttgtatacttcgtatagtttagtatttattatgtgcgcatttata

ctacttttagctatgattggttccattgtactaacaatgaatcaacgtagtggagttaaa

acacaacaaatcacacttcagttatatagaaatcaaaataaagtagttcgatttattgat

ctgagaaaaaattaatttgattgcggatatagatgaattggtacatcagtaattttccac

gttaaaggatatgggttcgagtcccattatccgctcaaattaagagagaatagctcaata

ggtagagcaatagttttcaaaactaaaggttaaaagttcaagtcttttttctcttgcaaa

agtgagtctgcttgaacaccttattcagttttaattataattaaaactgaataaggtgtt

caagcagactcacttttgcaagagaaaaaagacttgaacttttaatctatataagaataa

caaccacattttagcatacacaagacgacagccttgtatgcgtactaatatatctctgcg

ggctgtgtagaggcaaaaaaagtttgtgtggggtggctgtgtgacatataaattaaaaat

agttgtcaaaaaaagtttagtttggcgtaaaaggattcgaacctttgaaatcatggtatc

aaaaaccattgccttaccacttggctatacgccaaacaaattgaagataaagtgggattc

gaacccacggtggatagataccacgttagctttcaaaactaaagctttaaaccactcagc

catttatcccggttattaatataaattgtattctaaaatttttagagtctgtttatatga

tagtacgggaataggattcgaacctatatttttagatcatgagcctaatgagttaccttt

ttactctatcccgctattttatctaattaattttgtaacttaataaaaacttctccaaac

tacccagtaaaggtaggatgactaaaaaatataaaaaataataaacgctagctatctgac

ctataattatgtaagggtcttctacaggcataccacctattcaacctaatattaaacaac

aagcagccatgcttcaataaaggccacgatatatgggtctaaaacgcgaa----------

------------------------------------------------------------

------------------------------------------------------------

------------------------------------------------------------

------------------------------------------------------------

------------------------------------------------------------

------------------------------------------------------------

------------------------------------------------------------

------------------------------------------------------------

------------------------------------------------------------

------------------------------------------------------------
------------------------------------------------------------
--cttctaatttctgtactatgtattcatggtaatagagctaatacaactatagcaaaaa
tcatacatataacgccacctaatttatgtggtatacttcttaagattgcataaaaaggta
agaaatatcactcaggaacaatatgtgctggcgttaccatgggatttgcttcaatgtaat
tatcagggtgacctaaaagattaggcgagaagtatacaaaaaaagaaaagaatataataa
aggccactatccctaacaaatcctttacgatgaaataagggtacataggtaccttatcgc
tgcttgcatcgatacctaaaggatttccagaaccttcttgatgtaaagcggctaaatgca
ctaaagacgctgctgcgataacaaatggtaataaataatgtagactaaagaaacggttta
aagttgcattatcaacagaaaagccacctcaaagccaagcaactatagaatcacctacca
aaggtacagcggatactaaattagtgattacagtagcacctcataagctcatttggcctc
aaggtaatacataacctataaaagcagttattatcattaatagtaaaataattacaccaa
tcactcaaacaaattgtcgaggtgcagcataagaaccataataaagtcccctaaaaatgt
ggatataaactacaataaaaaacattgaagcaccattcgcatgtatatatcgtaaaagtc
aaccaaagttaacatcacgcataatatgctctacactaataaaagctaaatcaacgtgtg
gggtataatgcatagctaggaatattccagtcactatttgtattattaaacacattgcag
aaagaaacccaaaatttcatgcataatgaatattgattggagttggataatctataaggt
gattattaactatattgaaaagaggtttttaattagacgcataaataatgtttttatta
tagatggacgaggaatgggacttgaacccatggcctataaagtcacagtttatcgctcta
ccaaaccgagctctcctcgtcatgttaaaaagttaaatttacggggaaaaagggatttga
accctcactcattgatgtgacaaaccaatattttaacctattaaactacttccccatttt
tattaaatacggatagagggatttgaaccctcatgaataatattcatcaaaacctaaacc
tgacatgtctaccatttccatcatatccgcaaaaaaatgttactattagctttaacggat
aaagagggattcgaacccacggtataatatttcatacgatgatttagcaaaccattgcct
taaaccactcagccatttatcctgtgttttggaagctgcccactaccggacttgaaccgg
taacttaaaaagaacagattttaaatctgtcgtgtttacctatttcaccaaatgggcatt
agctattgctaatgctatgttttattgaagctattgcttttccggggttcaatgatttag
gacacgttctactgcaattcataatggtatggcatttaaaaagttttgatttacctccaa
gtaatgctaaacggtcttgagttttgatatctcgactatcagctaatcatctataggctt
gcaataaaattgcaggacctaaatatttgtcatggtttcaccaataacttgggcaactag
cagaacagcaggcacaaagtatgcactcgtaaataccatttaattctgacctatcctttt
cagactgtagatattctgtttttgaaggtgtattatttataagccacggttttatatatt
tatattgtgcgtaaaaattagataaatcaggaactaaatcttttataatgtacatatggg
gtagcggataaattgtaattgtgctagtatttatatttaatggttgcaaacaagctaatg
tattagttccatttatattcattgagcaactaccacaaataccttctctacatgagcgtc
taaaggcgatactcgaatcttgttcgtcttttatttttataagagcatccaataccatag

gtccacaatttttagtatgaataggatgtgtactgaaatgagtaatagttgggtttgatg
gagttcatctatatatacgaaggaattttaagtctaaattattattggatactaattgaa
aagaaatttttttgaataattgcatacttggttttattttgattgctataattatttaag
aacgggctaacccgttcttaaatataagataaaatataatgaacatttcgtatttattta
cttacttttgcaagtattaaataataaaccgatctatctagtaatcttctagaatgttct
ggaaaaagtaagggttatttttgcgcttttaagctgtcagcgcttatataatgttaacgt
ggctactcggctatgcaagaaacaatacaaccgatacactattggttaatatatcttaat
cctctcgtactaaagataaaccctctttttttctttcccacaacagatagggaccgaact
gtctcacgacgttctgaacccagctcacgtatcttattatttggcgaacaaccataccct
tggaacctattgcagctccaggaaaagatgaggttatggtcagggcattcttcacagaat
ttcctgccactccagaaccgtacatgaaagtcgcccttcatacggctcctcgagccatat
tattaacagttgattatttgttaagcaaaatgtttacataacacaaaaaaatgagctcgt
tact-------------------------------------------catttttggaatg
acgcgtgtcatggcaatgaccatgcagaagtctaaggttatctcaatgtgaggttccccc
tttctttcttggtatgatatgatctatttctattctatcggagtcgttaaaatatatttt
gcacatatcacatt-tggccctttcatttttaaaagtgtccttaataattgcccgtactt
gttgtttgatgatagtcttttggatcagtaggtaactctaccatcatacgggctactatc
aagtagtactttagtagatttaatcacatttgtttgatcgtgtctattcaattttataat
tttgtcttcctctatcaatccgaatactcagtttgctgtatcgctcttaattcaatgctt
agaaacaccgatcccggaccgatttttcttctgagatcatctttctagaagataaaatgt
acgcatgctacaaaagctgaaagtttttgttgcattacatactgcaaaatatctggtcca
cccaataataacaggtgccagtttactaattaaaactttttgagataagccacttgacat
tttgataatatttttaatactatctagatgcttattaattgatttaagactaggttgatt
cctagatgttcaaccagttgaatttccatggctatccttggcggctttatgtatacccac
tttgtagttgacaaaattaaaacctaagaaatctataccgct--------atttttcttg
ggagaagaataaacgctatatgttattgtagttttctctttagataattccaatcccata
cctgttaaaaacgcttctattttcagtttggcttctaataactctttctcctcgttacat
aatattaaaaaatcatctgcgtatcttacaagatatacttttcttttactcaccgacttt
tccattccatgtaaagcgatgttggccagcaagggtgatataatacctccttgtggagtt
ccagattctgggataatttctgttttagcaccttgaaaatcaactaaaattccagctact
agtcaagcagaaatctgctctttaagtagagaaaaagttttcagcttttcaagcaactta
gagtggtctatgttatcaaaacatcctttgatatctgcatctaaaatgtatttgggtttt
ctttgtaaacattttacaatagcctttctcgcgtctttagcacttctaccgggtcggaac
ccgtaactgttaggttcaaaaattgcttcatactgcggctcgagtgcaaattttacaagg
cattgcttagctcgatctcttatagtaggtatacccagatttcttacagaaccattcgct
tttggtatagttaccctcaaaatcttatccgaatgtcgatcaatctttatgcttttttact

aactccaatctttcgtctggggttaataaagaaattttatctactccagctgttcgtttc
tccaaattatcttgggtaactcttctgactgctaaaagttttgcattttcatgttttatg
atctgtttctgtataaaaaaaacacttttcatatcacccatcttacttagattaaatatt
tttatttgtaatttatatagtcaaatttctttatgtttttacacaatgctatttca---t
ttcatgattttattaggtttgtt-------tgttatatatcttcaaaattatatttaaat
taatatgactcgtctacacgtctgcatatccataggctttccctgtggcattggcttctt
gtagaatcctaccataattgtgcattaacgcaataacctacgcccaaataatgcgtttaa
aaaataaacaattatttggttcacaattatagttacttcgttccaatactatatacactt
atttagtaagcccctctgttccctctaaaactggttctccgactcaa-acgtcgtaattg
aatcaaaccttatgtttagatttcacgatgtttcatgttaacggggtgggttaacttata
attgtcccccgctatcctcggtttaggaaactttatttcggata----gacaaaatttta
tttgtccataataatcgacattattggcttttctggtcagatttactaa-ataagtattc
tttctgtagttattattctataagtgaataagctacaaatttatgccctgcttataattt
gtacgt-gttattgatacaaatcatagggttcactgtttgtaaagacagctcattgatca
gcatctggagtcgatgcataggcccactagttcttcagaactagcttcctatctatagta
ttagtttaacataggtctcccttt---------attagctatttacgagaagtacatctc
ttctcttactagcaacgaatcgcac-----------------------------------
------------------------------------------------------------
------------------------------------------------------------
------------------------------------------------------------
------------------------------------------------------------
------------------------------------------------------------
------------------------------------------------------------
------------------------------------------------------------
------------------------------------------------------------
------------------------------------------------------------
------------------------------------------------------------
------------------------------------------------------------
------------------------------------------------------------
------------------------------------------------------------
------------------------------------------------------------
------------------------------------------------------------
------------------------------------------------------------
------------------------------------------------------------
------------------------------------------------------------
------------------------------------------------------------

------------------------------------------------------------
------------------------------------------------------------
------------------------------------------------------------
------------------------------------------------------------
------------------------------------------------------------
------------------------------------------------------------
------------------------------------------------------------
------------------------------------------------------------
------------------------------------------------------------
------------------------------------------------------------
------------------------------------------------------------
------------------------------------------------------------
------------------------------------------------------------
------------------------------------------------------------
------------------------------------------------------------
------------------------------------------------------------
------------------------------------------------------------
------------------------------------------------------------
------------------------------------------------------------
------------------------------------------------------------
------------------------------------------------------------
------------------------------------------------------------
------------------------------------------------------------
----------------------------------------ccgacatcgaggtatcaaac
catggcatcgataagaactctcagccatgataaatctgttatccctagagtttctttttt
ccgataagcgacagtatttccatacattactgccggatcattatgaccgactttcgtccc
ggcttgagctgtaactcttaccgtcaagcagattttcatcattacatattatattagcat
atattgccaactaaaatccacttttgtgcacctccgttacagtttaggaggtcctcgtcc
caagtaaactaccacttttacatatgtccttgttagtaagcaatttttatactcagagt
agtttttcaatgacgtacaaatgtacttctacttttactgtactgtgtacaaagttatcg
cgacgtaaaattatagtaaagaatcatagggtctttccgtcttgttgcgggatgtctgca
ttttcacagacaatgtaatttcgctgaggctatactagagacagtggggtagtcgtgacg
ccattcatgcaggacggaatttacccgccaaggaatttcgctacctgaggatccttatag
ttaagaccgccgtttacttgggtttataatatatgcgcaaacttatatttttcactttca
agcactgggcaggcgtcaaaccttatacgtcttcttttgaatttgcaaagttttaagatt
acgataggtagatactcgcatatttcccccgtttagaaccgtacgtgaagattaccctt

catacggctcatccattttaacgtcagaattttctaacgcagctctgtcgcgatgacacg
ttttatgaagaagagtcatatttttaatatatttgtcctccactcactacaggtatga
tatggtgaatttctaattcgtacaatatatctccatattgactatttaaagattccttac
atatggggcacatacccttttgtctttttgccagctttagtcgaatattattcgtttgct
gatttaaacacatattcaaagatcttttattaaatactcctcaaactcacccaaatagg
gattagcgtcaagttttatcatagtagttcttttaataggagtgtcggatatttgaaaca
aattaattttcttaaatcttttaaattcagtgctagtagatattaatcaatctctattat
ccactttatgataaaatttggatttaagcgctttagcattaagcttcggatgcttaattt
ttatcatgcggtgaatggaatcaaaaacgtatttacttattcgggtataagctaactttg
cagtactagacgagtaataatttgctcagccccttaatattgggtttagttttgagaaga
gcatccacaaaggaagttgcgaatgttctttgacacaaagttttattttagcttttactt
tttgtaagctttcttttgtaggtttatatagaaagatccctttcttttccccgatttctc
gagtagaatccgagaattccctaaagtggaaacctacaaaatcaaatccatcttcaattt
tagtgattttactttttcccatatttaattctaaacctcttatttttagaaatttgttaa
catctagaattaacttttctaaaatatcatagttctgggacactataatgaaatcagcaa
catatctcacaagagttgctttgttattagacgcc-tttttatagttttatctaaacca
tcaaggcataaattagctataataggagatattacaccaccctgtggaacaccttcttca
gtatcttgataactgccttcattcataacgccagcttttaacatttttgacaatatcttt
ttatccataggaatattttctaatattcattcatggcatatattatcaaagaaacccttg
atatctccttcgaaaattcatctagggctgcctctctgtctacttaaaagaaatcaaata
tattgacaggcatctttagcacttcgataaggtctatatgcaaaactgcatcgatcagac
ataatttcgcttataggttcaagagccatagcaaacaaagtttgcacaactctatctctt
atagtgggaattcccaaaggccttttctttttgctattcttctttggtataaatactcta
cgtaccgcactgaattcataatttttaagatcttgaatttgaattaatatttcatcaata
ctaactttctttcctttagaaccttttaaaacaattttatcaattccgctggtctcacta
cttttacgtttgtgaatttgctttattgcttttattttagcttcttctaaattcactaat
tcattttgtaatttacttcataaacgtttgtttttattcaaataagctattgctattctc
ctttgtaacttataaattttcgtcattatattagtttttattatttt-ccacaattttta
tttaaaaataaactagctggttttgcatgtaatgcaaataaaataaaagttttcttaaag
gacaatacttttaaaacagaacatacataagtcagcaatctttcgattcatgatataatc
actatttcacttattactagcaaacattcgctttttatgagatcttttacctactgaaca
cttgggttgatcgctctttcctgagatatttctatcaattcattaggcttaccctttttcc
ttacacataacattaatctaaacaggtaacattctctataccggcagtattatgtctaat
ctataaaataacaaaaagatattttatactactgcttattcctatagctgtataaacagc
taatgatgatcttcggaattctgttttctcgataccttataacaaatgttcacttacgtt
ttacctatttagataagtctagcttttcttagttacatcaagtaagaaactacattgtca

gcaaggcttcatacctccagatcactctagacgcatgcttgcttagacttatattgatta
gataatatagcacttagtgcatttatgtatacggctttaaaggtcgcacttttttgttaaa
cagtcgctacccctaatttttgaaaccttaaaaggtactcctttttgcgaacgtacggag
taaatttgccgagttccttaagtatagttatctcattcgtctttattttctcaataagtt
cacctgtgtcggttttaggtacggtcaaacttcatgtaagttttcctgaaaaattacttt
ttttagcctccagtttagtgggtgtagctattagaagatatctttgtacacaaatacacg
agtattttgcagacacgtggtaatttttcctatcgaatacagttttcactttttttcttaa
gggccgactaactccagttacttaaaattgactggaaacccttgaacaatagacgaccat
gatttattttcaacatggttagcgctactcatgtcagcattagcactcctgattttagat
atgcaatttaacattaacataaaagactacaggacgttccgctaccattaagcttagttt
agcttaattcgaagcttcgatatataaatttaagtctccttacattttaaataggtagaa
caaataaaaatagcgagctaaaacgctttctccaataggtggctgcttctaagcctactc
tgtttattcgaattattccttatttttttcactaatttataattttgagatcttagctatc
gattagggttgtttcccttttgacgtaagaccttatcgcccaacgactgtctgctgctat
aaataaaaatatagtttggagtttaataaaatttagcaaaatctaaatttaataagtagc
tctacccacattttaaaaaagcaacgtactactttgatagttttcgcggaaaaccagcta
tcaccaagtttgattggactttcacccctaatcttaagtcatccccgtatttttcaacag
acgtgggttcagtcctccagtactttttaaagcaccttcaacttgcttaagaatagatca
cttggcttcgggtctaatacctgtaactttaagcgccttaatatttttaagcttgctaca
catattaacttactgactcattatgcaaaaggcactttgttgctgtatttgtcagcttca
aataaatataaattaacagattcaaatctttccctcacggtactcgttcactatcgatta
gaaaaggttttagcttagaagatgggactcctattttcatacaaaagtaatcgtactact
ttgtgttactattagtgtcgtaaaggactttataacctactttgggtttagacacttcgc
taagttcgctttcactcaccgttacttacaaattctcgtttgatttttttgctatgttac
taagatgattcaattcacataattatataaatgcatattaaatgcgagttttctaaagag
actcatagttcataggcaggtgcctcgctatgatgtttcgtcgcgcgacgtctttgcttt
tctaccaagatttcctctgataactttttaaatcttttatatttaagaacggcgaaaaa
atgttacccttttgtaagagtcattaactctatagtaattgtgctcctaattcgataata
gacaacaagtattgctagtcctatagaagattccgaagccgctactgttaatatagctaa
agcaaatagttggcctactatattgtctaagtatatagaaaaaaatataaagttaaaact
aactgatagaaacattatttctaaagacattataattattataatactttttttggtttaa
aaaaatgcctaaaactccgactaaaaataaaaaaaaagtataattttcacagtatagttg
ggtaatcatttatttttatatttaacatttgagatagcaaattgtattaaatttttccag
ccgcaggttcccctacggctaccttgttacgacttcactttagtcctttcgctaccatgg
acaaaaaataagatttttgtcttcaagtagagtaaattcccatagtgtgacgggcggtgt
gtacaaaacccgagaacatattcaccgcgacagttctgatccgcgattactagcgattac

gacttcatattctcgagttgcagagaataatccgaattaaagatttttaaagattttgc
tccagctcacgcttttgcttcttattgtaaacattactttgtagcacatgaaagtagccc
aattcataagggtcatgcggacttgacgtcattttcccttcctcaaggatattccaagc
agtttataatgtattaatgcattatacaaaagtttcgtccgtttgctggaattaaccaaa
cgcctcacggcacggactgacgacagccatgcaacacctgtgacatcttgtgtcatacga
gaattggtaaggttttgcgcgttgtttcgatttaaaccacatgctccaccgcttgtgcgg
gttcccgtcaattcctttgagttttaatcttgcgaccgtaatccccaggcggagtgttta
atgccttagcttcgcctctggaaaattatccaaaaacaaacactcatagttgagggcgta
gactacaggggtatctaatccctttgatacctacgctttcgtgccttagtgtcagctat
agtccagattattgttttcacttttgaagttcttttgaatatcatcgcattttatcacta
ctttcgaagttccataatcttttcctatgctctagtaaattagtttagtaatctttagta
aacatgaaatttaaaatttttttactacttagtttaccacctacgcaccctttacgccca
gtcaattagaataatacttgctcctcccgttttaccgcggctgctggcacgaaattagcc
ggagctttgattgtaaaatttagtctttgatttttttaatctattttacaaagtgatttac
agcctgatagggctttgctctcacatgtggcctggctaggtcaagctttcgctcattgcc
taagattcctcactgctgcctcttaaagagtctggaccttatttctgttccagtgtgact
gatcatcctcaaagaccaattaaggattatgggcttggtaggtcttttaaactaccaact
acctaatcctgcgtagacttttcttaaaccaattattaatttttttaaggcatttaccaca
atttaagataaatttctacgtattactcacccgtacgctatctttttcataagttacgaa
aattattaaacttgcatgtgttaggccgaccactagtattcattcggagccaggatcaaa
ctcttttattattatgttatgtttttatgtttgctatctcagaaattaaatttgattgta
taaagttatccagcgagctaagacagaagctgtaccaatttcatatatatatatgttggt
aatcttccttttggtaatgaggataaaatttgctttacaaaaagttgtatacttttttttg
tttaaatgcggaacatagtttattttctagctttgttccacgtataagaagtctcctata
tttttttattttattcatacatttgtgtatcaagatgggcagatttgaactaccattccc
ttgcacccaaagcaagtacgttaccattacgctacatcttgatatagtgatattgactct
ttctggataggatttgaacctataaccttgtagttaacagctacctgctctaaccattga
agctaccagaaaaataaaagcgaaaaaagggacttgaacccttataatgaccttggcaag
gtcatgctttgcctattaagctatttccgcacttaaaattgaatataaagttgaataata
ttaacaataagtagaaaaagtaaaagataattgctatctttgtacattttccttgtattt
acatgttttactgaatctcaaaaaaaatggcgtattccattacttatgtggtatagaaat
atagcggacactactcaccggacactagttaaaaaccaaggaaaaaagaggtgatctaca
aaaaaatgtattggcagaatagtaatatataaaaagtatacttgatttagtaaaagaata
gggcctacaattaaaacaaacataataacgccagaaattctatgccaaatagaaaaaata
gaagatctttgcggattgtatatagttaaatgtggagaaattggtctattgatgttctgc
atattatagttttctttttcttgggattttacagccattatacgggcaagttgtaatatc

cattacagagataaatttaagtttagactgttttaatattttaaacacatttttttatt
tttacctcatccttttatccgtacatgtaaagaagtataactgagttctttagtctttat
ctctaagaagtggattatttcggacgaagaaagggacaggcttcttttaccccttttga
ccttgacttgcctgcagttgtccaagatttaattttgcctttacagtctgttaaaagaaa
atgaatattattggaactaaaagataaaaacagtatacccatatgtagtttaattttatg
ttgtagtagtaaagactagggtcagattcgaactgacctaataaagatttgcaatcttac
gcatagccactatgctacctagtcttaaatttagtgaaatcttaagaaagactgtgtttt
aaatgataaagaacttttcacacaagaaagtaattcaagcgattttagctttggtaaatt
ggataactcgaaaagcaactgaccgggccttatataacagcaatgactcactatagctcc
ttttcctttacccatacgggtttctggactaaatttggtaacagcatttttttggtagcat
gtaaaaccacactttgattttacccaaactggactttttttcgatgactttgatctcttt
ctgaattatctttttttatattatttatttgtgtgatatctacttgcttgtacgataaaga
gcgtaatccaaattggcctctttttactagatgacttgtttggtttatatatctattttt
ttttttatgctgtcgtgttggattatttttttgtattaaatcttttcatattaaattgtgc
ttaaattgttttatatgccattcaaacttgaagattgcaaatcccttgttttgtatataa
tggcgtttcaaaaaattctatataagtttctagtttttgcagtgatactgcgcctaatcg
gaatactatactacttgacatacgacttcggctaccaccgaaacgcccagcgtaccttat
ttttattcctgtaaattcataaaactgtataccacgagcagtttttaatacaattttgtg
ctttttttttattgaaaaatagtatttttattgtattaattatcttccggaatgctaggcc
ttttttttaggccagattttataaatccacataaaagtgccccgttttgatgtcaaaatgt
tgcattatattttttcacgtcaagatcgaacattttaaacgtaggtaccgagcaatctga
tttggctaatagaggtacatattgtacattgtattcccatctattgtaatttattttaga
tagcgtatttccttttatgaatgtttttgattcaaaatgcttatttaaaaagtttgtagc
aggtaatacttttttgtatatttgcttttgattttaaaacctttgccatagcattgact
ttcggtatttcataagtcgttcacacctattcgaaatcctcttggattaacttttgacc
cataatgttttattacttaataacaatatttaaattgaaaatcgaatctaaatagttata
cagaaatacgttttgtaggatttgaacctacactaatcaacttagaaggttgatgtttta
tccaattaaactaaaaacgtttttactttagatgcattgggatttgaacccaaaataagc
agattaaaagtctgttgctgtaccgtttagctatacatctttttaataaagagagtagga
tttgaacctacgatgattttaatcaatagatttacaatctatcgctttcgaccactcagc
catctcttcttcagcggttagatatatgttacaatctgtaataaaataattaattatata
ctattgtttaataacaatatttagattgaaaatcgtacgattttcaatctaaatattgtt
attaaacaat
>SRR9587922
gttttattaattaaaactaccccagtattacacttttaaatatttataaaaaataatgca
aatccctttacgctgttatttaatcgaatttaaatttcgattcgtttactttctacttag

tttttttctctcgttgctagtcgttataaactatcatgaggctgtgtttttttttgaaac
gtatacttttttgtttataaacgagggggcattcatcgcgacacatattgctgaactatt
ttcaacttctatgtatattagttttaacctagttttgtgcataaattatccttttgcata
ttatcattgtagccgctttttaattcgagttgatacaaatcacaggtttggttctttaa
aaatatgcaatcattgacatttattacgtttttagctagtttacttttgtgctactttt
aattttgccctacacttacgctttttagacacttgaactgtaacgatagtttatgcatt
taaagtgcagttggaggcaaggatagcaacatatgtttgttgaacactacaaacaatgtg
tttactgtctaacattgtctatctcttctttactaggctaatttgtttttacttagtgga
caatatgattaatttgcatttatttttccggaaaaataaaaaatatactttctttttcgt
atgcttgtcggcttctgtgtgtttaccgcaagaaagttttatacaaattactttcataat
ttctataatagtactatttgagttattctttttggttacatgcttatttttttgcaaaaaa
taatgtctaaaagtggacttgaaccactgacccaaagattttcaatcttttgctctaacc
gctgagctatttagacttaaactttttcttttcccgtatttagagcgaccatttttttcgtt
taacgactgcttgtaaatcgtaaacgcctctcacaatatggtaacgtactccaggtagat
cctttacacgaccacctcttatcagaacaacggaatgttcttgtaaattgtggccttcac
ctcctatatagcctatgatagaacgtccggtactaagacgtattttagctacctttcttt
cggcagaattcggtttttttagggttagtagtatacacttttgtgcaaacaccctttttttt
gaggcgacttattgagagcaggtgtttttgctttggtaattttgtgctttcttggatttt
ttatttttttggtttagtgttgacataatatttttattttaagtaataagggacttgaacc
cctaaccatttcggtgtaaacaaaacgctctacctattgagctaattactttacgtctgg
aaagatttgaactttcaacttttagattcgtaatctaacgctctatccaattaagctaca
gacgtttttctccgcttgtatatttagattacttatctatattttggaaatagggacaaa
tattcttgtataatttacttcaggattattcgaaagtatagttaaccaattaaacaagta
gaaaacgcatattttgtattgttactaacttaaagtactattttataataaactttttat
tgtattatataccaatatttagattgaaaatcgacctaaattcaatgtacttttagcaaa
tgacaccagtgttttaaaaaaggtg-ttattttgaaaacatttttggtgtcatttgccct
ttttcagctaataataatttaattaataacagaaagcgcggtttattgctcgttgcgttc
aagttattactacggcaacttcctatgcagctgttggga-ccccccccactcgtgtaatta
cacttttttagttttatttaaattataataaggttatactttatatcttttcttttattta
acccctaagatctttattacgccccaaaatagtgttataaaaaaaagtagcctggagggct
gccaggagtaaaaaaggcctataacggttatttccacttttatagtgctattt-------
-----------------------------------------------------------t
aattaaaaatgacttattaaaggttttacacagataatacttcttttattaaaaatatac
tacgccgctagaccggaaaaaacccgccatagaacgatgagaggtaaaaagtgtcataaa
ttttaactattttataattgtacattttaaaattaaagtttgtcaaacagcaaaactttc
tattatttcaataatgcttgtttttaaggcttttaaagcagtttctcgctgatcccgaac

actaaactttttgttaacatgttctcgtaaactgtaattggatgtcataataaccagtgg
attatccggttttaaagtaaaattacctttcattgcaaggcgcatgctatcaccagctaa
aaactggtttaagaattcgatttgtttaccgataaggtgaaattcgtctcatagaataaa
ttgatacaaatgattttcataataatcttatcagttgtcttgggatcacatataacaagc
ggtagtagttttcaagtaatttaacagcgcactttttcccatattaggttctttactgca
tatatggagttgtggtttttataatagcgtaaaaaaggactagtattattatttagtat
gtcgaatatttgttctcataactggatagaacattccgataagttattttttattattttg
tcgtaggattgttttgcaaaatcgttctatgttgggcttttatttttttgttttatgtct
aaagaaagttgatgctgtttaacgtaacgatcaaattttatgctgtcattaaacaaagca
cgtttaatatctggttcatcaaaattaaaaaacattggtgtagtaacaccagattcgatt
tgtttcatgatgaagtgctctcttacaagccgctttttttagtagattactaaaaatttta
ctatgttgatttgatacgcatttttggatggactcttgcttatttattggatctgaaaag
gagcttggagaattagttaagtagtcaaaataattcggatttgtgaaattacgtttctta
gaaaattcaatatacgtatataaaatttgatccgagttatttcagctactagaaaagctt
attattgttgtttttttctctttgtaggagtttgttataaattaaaactaggcaaagaagc
tacatttttttgttgtaacttggaaagatttggccgcaacgcgctttttatgtttagttac
ggctggatttagttcaaatactttggacatgagt--------------------------
------------------------------cgttttaaaaaagcttcataaactttcacc
aaaagtttgatttatatcataaacggagaaaatccaaattgaaaaaactaaaatggtgtg
gcctaaaaaaatacggcgaagtacatatcaacttggttttcagtgaattaaacagaaata
cagtattcccacaagtacgataggaattaaaatgatggataaatcaaaattttgatttag
atacgaaaaccatggtttactcggttcgtttacaagtaaacttttttaattcaatatttg
tgctattagtaaagatttgtgcgactttaaattgaataagctgttaaaataaggtgaatt
taatttaggcaataaagtatcgtacaaatgtttgttctttctatatttgtttagagcttt
cctgtactttttattgtaagcagcaatagatcctatgtgcttgtgttgaaaatctaatct
tttaagtactttgttaaagctcaaaattttggtttttaattggttaacag-ttttttacg
ctttctagcatcgtatagtctttcttta-------------------gcttttctagtgg
cagttgattgaaataacttttagtgagtttttttactatagtacctgtaacaagtgtgct
aacagatactacgacacaaactactttcaagtccaa-ttgctctgcgatggggtttgatg
aaagtcattttatataaattaatttattaatgtgtaattgtggtataattttattctgtg
ttaatgggcataattaaacgcatacattatatattatccctccttcggagggata-atat
aataattcttattatataccaatatttagattgaaaatcgtgctattttcaatctaaata
ttggtatataaacaaacctgcacataaataaatatcactaggacattatcccaccaaaaa
atcgatattctctagagagcccctataagctttaatttactatctcttgcctgaaattat
accttcttggataggaatcgtcgtgaaattgaaaaacattta--gatatatcttatacaa
ttatgaccaaaaaatctacaaaacaaacaactttacacaaatattttcgcaagtcgcata

catgagattgaacttaagaagcctcttttcataaatcatacagcatttttggcgcttctt
aattctattgcaatattcaatgaaccaaaaacaaagtttttttaaaacgacctgcatgaa
cacaatcttaacaagtgtaaaaaagccctctag--ccccttttggcaacaccacgagaat
aaaccccggtcggggagaccccgaacccccctcggcgcgggctgtctagcaaagttatacg
gcccgcgcgtgcgtt-tgttttcctatatcgc-----tccattccatttcgctattgctg
cgcttcgcttatttttaattctaactgttccttttcgttataacgattttaggtcgattt
tcaatctaaatatttgtatataaacaattatttatcttgataattatatatctaaaaaag
ggggcatagtttaataggtataatgtatgctttgcaagcataaggttaccggttcaaatc
cggttgtctccaaatctaaaaacatcaatgcgttctagaatattacaatacaaaaataac
ttggctgactatgatcttttaactcaattttcgattaaaaatcataatagtatgcctact
ttctcttctcttaatgtcagagtaaaaaatattcaaactagcgacattaagcaggtttgc
ttaaatctaatctctattcaattagtaggaaacaatcatgttgttttcagtagccaaaaa
gacgtttctaatcttatactaaaatgtaacggtattaatacttattttattttagagaat
ttatctttgttaggctacaaaaacaatttaaaaagtgacaaaaacaataaattaactgtg
cacggtaaaaacttcttaaatattattgaatattttgcatccaattcaaattttcttaag
tccttagaaaaaacatctaataaacatttaagtattgcttttttcttatcaacttgctaat
agtaaaaataatgaaatagctttgtatttttttaaaatcattaggatttccgctttaataa
cggtatagttcaattggttagaacaacggaatcataatccgtaagttgtgggttcaagtc
cctctaccgttatggctcttatcgtctaatggttaggacagtgctttttcagggcatcga
cgtgagttcaatcctcactaagagtaacttttttaatataaaaatccttatacaaatacgt
gagcccataagaattcaaaagtaaaaaggattattctagatctaccaaattattttggga
tccctaggtcgttaaaaatatttaaacaatgtattttattc--atatatatatatactaa
aacttaaaaatataatttagtgcaaaagtctctaaataactgaattttccgttgaatata
ctcgaccaatcataaagatataggtactttatatttaatttttggcgctttctccggtat
cttaggtgcttgcgcgtctatattgatccgaatggaactagcacaaccaggtaatcaact
attattaggcaatcatcaagtgtataacgtactagttacagagcacgcattttttgatgat
tttctttatggttatgcccgtcctaattggaggatttggaaactgattcgtacctattat
gataggtgctccagatatggcctttcctagattaaataatataagtttttgactactacc
tccatcattgtgtcttcttttaggatctgcgatggtagaagtaggcgctggcacaggctg
aactttatatccgcctttgagctctattcagagccattcaggcggtgctgttgatcttgc
tatttttagtttacacttgtcaggtgcttcttctatattaggagctattaatttcattac
gacgatatttaatatgcgcaatccaggacaaagtatgtatcgaataccgctatttgtttg
atctatccttattactgcgttccttttactactagcagtacctgtcttggcaggggccat
cacaatgctgttaacagatagaaactttaatacaacatttttttgacccttcaggtggtgg
tgatcctgtattgtatcagcatttattctgatttttcggacatccagaagtgtacatttg
tgcgccgtttaattctgttaaataagatgatttattattttaaattacttatctcgaaat

ctttaaactgaatagttaataaagatagacttaattattctttttgtcttaaaacacttt
cccttaataaagttggaaacaattctattaaaagcgggaatgaaagtctcttgcaggaac
ctgaatttcctaaaaatagtaggctagttatgctcatcatgaataagataataaagttga
tacatatcgacggttccacactaaactttatacataatatttcaggtaacagtatttatg
taatatgggtcggtaccaggatgtttaccttcccttgaataattattacacgtcagagct
atcagctcctaaatgatatgcaacgttctaaaatgagcttgatcgagaaacagaagaact
cgggataccttaccaatcgaaagattcatgggtacggaactctcgtagtaggtggtagaa
gaattcaatcttcttcaaactaccaaaggggagtagaacttagattcttaagtgaaaaac
cctgcattagctcgcaagagtgcgctaggttagtagatttgagaaaagttaattctgaaa
ataaatttcaagttaataagaatactattcatattatatctgatatgaatgtccttattt
tagcatataaactcataaaaagtaatcctggaaacatgacacctggtgtgaatggttcca
ccttagatgggttagacaagaggtgactgcagaatattagtaccaaaataaagcaaggta
aatttttattcagccctgggcgtaagaagtacattcttaagcccggttcagcagataaaa
gactattaggtattgctagcccgaaagaaaaaattgttcaaaaagctattctgctagtac
tagaatcgatttttgaaccaagcttcttgagaaattctcacgggtttcggcctaaccggg
gcaaccataccgctttaaagatggtaaaaagtgagtttcacggagttccttgaattatag
aaggagatatttcgaagtgctttgatgaaattgatcactctattttattggggcttctaa
gcaagaggatatcttgtgataagactttaactttaattaaaagagggttgaaggctgggt
ttatagatttaggaatattcacaagaactaaattgggtacccctcaaggaagccttttga
gtcctatcctatgcaatatctatttacatgagctagatttatttctacttcaacttaaaa
ttaaattcgatacagggactagtagagcgaagaacccacagttcagaaaactacagtata
aactatccaactttaaatcgcctctcgataaaaagcttgtcagaagagacctttgaaaag
tgcatagtctgaacccccctagatcctaatttttgcagaattcactttgttcggtatgcgg
atgattttattgtgggagttacaagctcccatgaagttgctttagaagttaagaatatga
ttaaggaatttctttgtaatcatttgaaattaaatttgaatgagctgaagacacacatta
ctcatattagagaaaaggatatatttttcctaggtacccttatcaaaggtaactgaaaga
aagagaaacctattcgattgatcaactttccctctagagaaacgttcatcaagacaagag
tcactccgcgtttaagtttgcatgcccctattaaaaaactctttgataaagctactgctg
aaggattttccgtagggatgggattaattataaacctacttttgtaggtaaattgatta
atatggaccatgcagacattttagctttttataattcaatattaagaggagtactgaatt
actactcatttgtggacaaccacaaaagtttaggatcatgagtccattatatgaaatttt
cctgtgctagaactctagcgttaaagtacaagttacgtttcacatcgaagacttttaaga
aattcggctctaaattggcgtgcccgaatacaaaaaaaagcctgtttttaccaacgagct
ttaagagaacgcaggccttccaaattaataaccctattccttttgagaaaaaagtactct
cttgatctaaaaaaattactaaatctaatctaaacaaagtttgtttaatatgtggtagtt
ctctattggtagaaatgcatcatatacgtagtatagctggcattagaacaagacttcgta

```
gcaataaagcagactttttctctctgcaaattgctgggataaacaggaagcaggttcctc
tttgtcgagaacatcacttgaagctacataataaaacactttcaccagaagaaacgttat
tatttcagaaaaatattaaggaaatttagatgtatttctttcaattttaactttatttac
aatttaagttcggcgagccgtatgataagaaattatcacgtacggttctgagggcagtta
ttgacgctaaaacaacctcgttgtcaagtttccatctctgacccctactaatcttgcctg
gatttggcatcgtcagtcacatcgtttccacctttctagaaaacctgttttcggttata
taggaatgatttatgctatgctttctataggtattttaggatttatcgtttgagcgcatc
acatgtatactgttggccttgatgtagatacaagagcttacttcacagctgcaaccatga
tcattgctgttcctacaggaataaaaatatt-----------------------------
------------------------------------------------------------
------------------------------------------------------------
------------------------------------------------------------
------------------------------------------------------------
------------------------------------------------------------
------------------------------------------------------------
------------------------------------------------------------
------------------------------------------------------------
------------------------------------------------------------
------------------------------------------------------------
------------------------------------------------------------
------------------------------------------------------------
------------------------------------------------------------
------------------------------------------------------------
------------------------------------------------------------
------------------------------------------------------------
------------------------------------------------------------
------------------------------------------------------------
------------------------------------------------------------
------------------------------------------------------------
------------------------------------------------------------
------------------------------------------------------------
------------------------------------------------------------
------------------------------------------------------------
------------------------------------------------------------
------------------------------------------------------------
------------------------------------------------------------
```

------------------------------------------------------------
------------------------------------------------------------
------------------------------------------------------------
------------------------------------------------------------
------------------------------------------------------------
------------------------------------------------------------
------------------------------------------------------------
------------------------------------------------------------
------------------------------------------------------------
------------------------------------------------------------
------------------------------------------------------------
-------tagctggattgctactatgtgagaagggtctatttttttaaaaacccctatgt
tatttgctatagggtttatatttttattcactataggaggacttactggtattatactag
ctaactccggacttgatatatctctacatgatacttattatgtcgtagctcacttccact
atg---------------------------------------------------------
------------------------------------------------------------
------------------------------------------------------------
------------------------------------------------------------
------------------------------------------------------------
------------------------------------------------------------
------------------------------------------------------------
------------------------------------------------------------
------------------------------------------------------------
------------------------------------------------------------
------------------------------------------------------------
------------------------------------------------------------
------------------------------------------------------------
------------------------------------------------------------
------------------------------------------------------------
------------------------------------------------------------
------------------------------------------------------------
------------------------------------------------------------
------------------------------------------------------------
------------------------------------------------------------
------------------------------------------------------------
------------------------------------------------------------

------------------------------------------------------------
------------------------------------------------------------
------------------------------------------------------------
------------------------------------------------------------
------------------------------------------------------------
------------------------------------------------------------
------------------------------------------------------------
------------------------------------------------------------
------------------------------------------------------------
------------------------------------------------------------
------------------------------------------------------------
------------------------------------------------------------
------------------------------------------------------------
------------------------------------------------------------
------------------------------------------------------------
------------------------------------------------------------
------------------------------------------------------------
------------------------ttctgtctatgggagctgttttcgcaatatttgcag
gcttttattattggtttgaaaagatatccggatttcaatattctgaaatactaggtcaaa
ttcacttttgaggcacttttataggtgtaaatctaacctttttccctatgcactttttag
ggcttgctggtatgcctagacgtattccggattatccagattcttatgcaggttggaacg
caatagcttcttacggttcatatgttgcgttatttagcacgctgtttttcttttatcttg
tatttaacacacttgtaacagcaaaaaagacacctgctagaaataacccatggaactttg
aagattcaaaaatgggctcaactacattagaatgagaaatttcttctcctccagcttacc
atacgttcaatgagattccagttataagagaaacagaaacatctttaaaaataaattaac
ttatgacaaaaaaaaatactacaatattatttttagggttagctttgctaacccaactac
tacagagcagaatagttattagtgattctgcagaagactggcaattaggcttccaagatc
ctgcaacacctataatggagggaatcattaaccttcatcatgattttatgtttttatct
gtgctatctcaatttttgtatcttgaatattagcacgcacgttatggcactatcactgaa
caaaaaatgagtacccttctgccacggttcatggaacagccatcgaaataatttggactg
ttactcctagtatcactttgttggcgattgctgtgccttcttttgctttgctatactcta
tggatgaaataattgcgcctgcaataactattaaaacagtaggtcatcaatgatattgaa
gttacgaatactcggactatacaaatgaagacgataacactataatgtttgaaagttata
tgattccagaagaagatttaactttaggtcagttaagattattagaggtagataacccta
tggtaatacccgtaaatacacacgtacgtctaattataacagcagcagatgttttgcaca
gctgagcagtgccttctttaggtataaaatgtgacgctgtaccaggtagattaaaccaaa

gctcgctctttgtaaaacgtgaaggaatattttatggtcaatgtagcgagatttgtggtg
taaatcatggttttatgcctatcgtagttgaagcagtgtctttaccaaattatatttctt
gagtagctaataaacttagcgaataaatagtctaaaaatattctaagcatctgatatgcg
tgtatctctagcccaactatttttctttggtttatctattttttattttattttgatttaa
tcctaaatatttaacttatacaatacgactattaaaaaaaattaaaaagcgttctaaata
ataaagagcgctttaataaaaaacttattatggcaactacgacaaatctaaactttatca
aaacagctaaacaattacaacgccacccttttcatttagttgaccccagcccttggcctg
taacagctgcaatagccgctttttcatgtgctttaggcggagttatgtatatgcatgcat
acagtaatggagggtacctattatttagtggtttttctttactgttatttacaatgttct
catgatggcgcgatgttacaagagaagccactttttcagggcatcatacaggtgctgttc
aaaaaggattgcgttatggtgtaattttatttatagtttcagaaatcctcttctttttg
ctttttttgagcatttttcatagtagtctttcaccggctattgacataggttctatgt
gaccaccaaaaggaatagttgtgtttagcccttgagaagttcctttttaaatacaataa
tattattattatctggttgttctgttacatgagcacatcatagcattgtagcaggctata
aaaagcaagcaacgttagctttaataacgacagttatcttagccgctatttttacaggtt
tccaaggttttgaatatagcgtggctaattttacattatccgacggtgtttacggctcca
cattttatatggctacaggctttcatggttttcatgtctttataggtactattttccttg
gtatttgcttacttcgcttattaaaatcacatttgacacaacagcatcattttggttttg
aagcagcagcttgatattgacattttgttgatgttgtatggctttttttatttatttcta
tctactgatgaggtggtacctaatctaaatctctttaatatctaaaaatctatgatgaaa
cttattaatttacctaccataaagctcacttttttgacatgctctagctttcttcattatt
atctgttatcataatatctatatttttacagaagaaagtatacttttattctgttttatt
gcgtgactaaacattacatgaaattatatatctcctcaaattaatgcgtcattatctgaa
agaggagaaaaaatcaattcaaattttcaacatattactaacgataatataataacttga
aaaaaatatagacaaggttattcactaaaaataacacatgttgatattttaaagaactta
atcacatatttaacatgcttaattaagactgtactccttttagaatcaaaaagtagctca
ttacaatctattgcaccttatttaaaaagattacatattgtaaaagacttagaaacaaag
ctgactaaaatttcttatattacgatttgccaacgtatacaggatacagctacaatacgt
agtttttatgctaaccgagtaaaaatcaaatctttccattctgaatctaaattagactta
ttcgaacgtataagaaaattagaagctggttcttagaagtttaagtctatagctcaatgg
ttagagcatacgcttgataagcgtaaggttgattgttcgaatcaatttagacttatacta
tttcaaaaccataatataaaatatgtacaatataaaccaaacatttttttcagcaagccc
tttagaacaatttgaaattatacctttaattcctttagaattatttgggttaaacatgtc
gttaacaaacgcgtccatttttttgatactatctgttgcgttatctatttttttgatccac
tttagtcatatacaaaaataaattagttcctggaaactgacaatctataaaagaaatatt
ttatgataccaccttaacgttggtaaaagataatttaggtaaaaaaggttatagatattt

cccgtttatttttaccctttttacaataatactttattgtaatttaataggtatggtacc
atatagttttactgtaacaagtcatatagctttcacatttggcttagctttagctattta
cataggaattaatattattggcttcagaacccacggtataaagtttttcacaattttttt
acctaaaggagttcctttatttattgtacctttagtggttgcaatagaattcgtatctta
cgtcgtaaaagttttcacaatatcgataagacttttttgcaaatatgacatccgggcatac
tttacttaaaattattgccggatttgtttggacaatgatctcaataggaggcgtgtttgt
atacttacaaataatcccattagttttattactagcgttagtgggtttagaaattggtat
tgctcttttacaagcttacgttttcacattacttacctgcatttacttaaatgatgtttt
agaaatgcactaactaaaaaattatgccacaattagatcgcgttattatttttggtcaaa
tattttgactatttttcaccttttttaattgcttatgttgtttatacccatttcatattaa
gtaatttattaaaaattttcttagtccgctgatggaagcttagaaaagatattactcaaa
ttgcattaaagatccgtttaacgagctatttaattgattcaaatattcaaacgttacgta
gaatttattcaacaatcagaaatatactagcttctctaacaaaaagtttattaacaaaaa
gtataagtaagccaaagttagttttaaatgatcttaattctttagttattaaaattagtc
tggaaacgtctttatatggtagcaaaagcatcaccaagtctggaacatattcttattgaa
cttaataaaatatactatgtatttattaataatagctctgcctttagtgggaacattagt
tacagggttaggcggtagatgaatagggcgtaaaggttcaaatttgttttctacaacttg
cgtagtcctgtgtgtctttttttcttcaatcgctttttttcgaagtaggtctttgtggagt
tccttgttatatatctttgagcccttgaattagttcaggggcactaaatatttcatgagg
ttttttatttgatagtttaacaacaacaatgcttgttgttattacatctatttctagttt
agtccatttgtattctattcaatacatggagcacgaccctcattgccctcggtttatgtc
tttcttggagattttcacatttttttatgatcttattagtaacggctgacaattttgtgca
aatgttttttaggctgagaaggagttggattagcttcttatctattaataaatttttgata
cactcgactttgtgcaaatcaagctgcaatcaaagctctggtagtaaatagagtaggtga
ctttggattaagtttaggtattttcacaatttttttatctttttggttctgttgattatga
aatagtattctcttccgcaaacatctacacaaattatagtatttcctttttttgggttttc
cataaataccttgactttaataggtatttttttttattaatagggctgttggaaagtctgc
acaattaggtctgcatacctggctaccagacgctatggaaggtcctactcctgtttctgc
actcattcatgcggctacaatggtaacagcgggtgtatttttaatagtgcgctgttcacc
tcttattgatttatcctcggatgtcttacttttaattactcttcttggatcaagtacagc
ttttttcgcctctattgttggagtatttcaaaacgatataaagcgggtaattgcttattc
tacttgtagtcaattaggctacatggtctttgtgtgtggtttatcctattataatgtagg
tatgttccatttagtaaatcatgctttttttaaagcattactttttctaagcgctggctc
tgtaatacatgcgctatcaaatgaacaggacatgcgccgaatgggttcgctagcaaatag
cctaccgatcacatatgctgctatgctaattggctctttatccttagcaggattcccttt
tttaacaggtttttattctaaagacttaatcatcgagataacacaaataagttattacag

taatttacagatttcttttggcgtttatgcttgttgacttgctaatatttctgtactctt
cacatcgttttatacatttaggcttatttttctaacttttataaaaaataccaatagcta
tagaaaacacatagaaaatatacacgaatcgccacctttaattctaattcctttaatatt
actcgctatatctagtatttttgtcggtttcttaacaaaagatatattcgtaggaatcgg
aactccttttttgaggtaatgctatcaatattctacctacgtcttgtaatctattggaagt
tgaatttatgccttctttaataaaatgacttccgtttgtgttaagttctatgggtgcaat
tctcgcttatacaataaacgtaggtgtactaaaaaataatatacaatttgctcataatca
cttatttagaaaactcgccttttcccttagcaaaaagttatattgagataaattatacaa
ttcactcattgtatctcctttaatgtactttggttataatatttcattcaaaaatcttga
taggggttttatagaactcgtaggtccttatggaatttcgcgtactattaaaaattgatc
cacaaaagtaattaaaatacaaactggccagctaacccattataccttttttcgtgatttt
tggtttatgttcccttttactactagttcctgtttgagattttctacaatttttagttga
tgtcagattactagtattttgctttatagccctctttgtagcgtagtttacgaaagtttt
aacacttaaatatatgcagataactaatttattattatggacttcacttattcctttgtg
tggcgctatattacttatttttattcctagattttactctcatttaataagaaatattgc
tttcgcaacagcgcagctagcgtttatatactctattttgctatggctttgctttgaatc
aacaacatccttattccaatttatatatacgataaattgatttccctcctataatattta
ttacacaataggtgtagacggtatatctttatttttttatcatacttacaacgtgattaat
tacagtttgtacattaataagttgaaatatgccagacagccaaataaaagaatacttaat
ttgttttcttttgcttgaagctattttaattcaagtttttttgtgttttagatgtcctatt
cttttatatatttttttgaaagtgtccttatccctatgtttttaattataggcgtatgagg
gtcacgggaaagaaaaattagagctgcgtatcaatttttcatttacacattagctggttc
actgctaatgcttctagcaattttaactatttatttccagcatggtaccacggatatcca
agttttatgaaatataaattttgacgttagaacacaaattttactttggctagctttttt
cgctagtttagcagtaaaaattcccatgattccttttcatatatgattacctgaagccca
tgcagaagcacctacagcagggtccgtaattttagcaggtgtgcttttaaaaatgggcgg
gtatggatttttacgtttttctttacctctgtttccggaagcctcactttattttgctcc
attaatttatttactaagtattatagctgctatatatgcttcacttactacaattagaca
agttgacttgaaaaaaataatagcttactcttccgtttcgcatatgggctttgtcacatt
aggtcttttctcttttaactctcaagggatagaaggtagtataatcttgatgcttagcca
cggattagtctctagtgcactttttttgtgtgtaggtattttatacgataggcataaaac
gcgtcttctcaaatactacggtggtctcgtgcaagttatgcctattttcagcatattact
attatttttttactttctctaatatcggttttcctggtacaagcagttttgttggtgaact
attagtattaatgggagtatttcaatttagtccaatatctacttttctaagtgcattcag
catgattcttggggcagggtattctatttgactattcaatagagtatgttttggtagttt
aaaacttcaatacattacaaattttcaagatatttcaagaagagaattttgtatcctttt

tccgttaagtgtattcgtactctgaatgggtatatatccagaaattttcctatctgaaat
tcactgttcaagttataacctaattgcatattttaactaattttatgttatgaagtttat
taaaacgctaattctagcatttatgaaaaagaagtcctatttttattggtttccacgttt
tttagggctattacttatacctgggtttttatttgataccgagattctagttctctttca
aagccttatcctcttacatgcaagcctaggtttagaagtaatcatagaggactatttaca
cctagaaataataaaacttcagtgtttgtctttaattaaagtacttttaatattattagt
caatcttaatatattatatttattataaaaaatatccttatgttatttatctcctcttat
gatttctacgcattattgacagaaatttactttttaaacgcaatttgtgctttattaatt
tatggtgtaattttaaatacctcatatagaagagggcatccagttattgaacacaatgta
agtggtctctcaactcaaatactaatagtgagtctttggttaacagtttgttcaaatata
ccttgcctaaccagctggaattcacttttagtgcacgatttttttatctttcggtataaaa
agcaccatattagcaatttcgctactttggtctttaattatttttttcttacaatagacta
gaaaaaataaatctctacgagtattgaatcgtgtctatgttggctattgttgccatgctt
tttgtaagttgttcttatgatcttttggcaatgtatttagcaattgaattccaaagcatt
gcattttatatattagctagttttaaaagaacatctgaattttcaacagaagcgggttta
aaatatttcgtactgggtgcattttcttcagctttacttcttttaggtatttcactactt
tatggtactactggtttaactaattttggagatctatcaaaatttttttttaggtaccaca
ttggaaaacgcatcatttatcaacataacatttttttggtgtcgttttaatagaagtagct
cttttttttaagataagtgcagcaccttttcatatgtgatcgccagatgtttatgaaggt
gctcctactaacgttacatcttttttttggtatactgccaaaattagcattagtaagttta
atatttagattcttttatttttgttgtgctgaagttgtgctgttactaaattttacactt
ataatttgtgcgcttttatctatgataatagggacatttggcgctttagcgcaaacaaaa
tgaaaacgtttcattgcgtatagtactataagtcacgtaggatttattgtagctggattt
tcaacgttggaatttaatggtgcatttggtgcgctattttatatcttggtttatacttta
acttctttagccactttttctattgtgctttccttccgatgcttagcatatcctagcaca
taccaattacgctatctaacggatatcgttagtttagtgaagttaaaccctatacttgct
ggtagccttgtagcagttttattttcaatggcaggtataccgccttttccaggattttttt
gctaaagtatttgttttattttcacttttgcaagaacaattaataggattagctataatg
gcaatatttttgagttgtgtttcgtgtttttattatatccgtttgattcaaatgatgtat
tttacacatacaaaaaccatacttatttttttatccaatagaaaagactacatcaactata
ttaagtataactatgttattacttgtacttatttttttgaagatagatctgatttctaat
tttattcattgtatgttgtttttataaaataaccaattaaaatgttttacaatattgcaa
ttaacattatcaaagtgttgaccattatagtgccacttttaatcgctgtagcttatatga
cactggccgaaagaaaagtgatggcagctatgcaacgacgaaaagggcctaatgtggtag
gtatctttggtcttttacaacccctagcagatgggttaaaacttttctcaaaagaaacta
tactaccttctagtgctaatatttctatttttttagctgcacctgtgctaacgtttttgc

tagctttattagcatgatgtgtacttcctctagatgaggggaaagtttttttcggacttaa
atataggtgttttgtatatattagcagtatcatctttaggtgtttatggtattataactg
ctgggtgatctagtaattctaagtatgctttttttaggtgctttgagatcagcagctcaaa
tggtatcttatgaagtttccattggtctaattttaattaatattttattatgcgcaggca
cattaaatttaactcaaattgttctggcgcaacaaaatatgtggtatataatacctctat
ttcccatatttattatgttttatatttctatattagctgaaactaacagagcccctttcg
atttgccagaagcagaagcagaacttgtagctggttacaatgtagaatactctgcgatgg
ggtttgcgttgtttttttttaggcgagtatgcaaatatgatacttatgtgtagtttaacaa
ctatttttttttttggtggttgattacccttagtcaatatgcttcctttttattggattc
catccgtactttgatttggtttaaaaacaactttactttatttggttttatttgagtgc
gtgcagcatttccacgatatagatatgaccaattaatgcgtttaggatgaaaaatatttc
tacctttatcattagggtgagttcttttagtatccgggatactattttctttcgattgat
taccataacaaatgaacgtactttataacgagtattctgctattctcactttttttgcag
tagctttttttaatctctctaataatattaatactttcgtatatattaaatcctcaacaaa
gtgatcaagaaaaagtcagcgcctatgagtgtgggtttaatccatttgatgacgcgagag
caacttttgatgttcggttctatttagtcgcaatcctttttttaatatttgatttagaaa
taagtttcttatttccttggtcactagtacttgggcagctaccttcttttgggttttgat
ctatggttgccttttttagccattttgacattagggtttatttatgaatgaaaaaaaggcg
ctttagaatgagaataatcaaataatttactagagatttttaatttgataatataatata
atatgaacgtaactttacaaagtgcaaaaatgataggagctggactagctactattggtt
taacaggggtaggagctggagtaggaattgttttcggatcgctagtaattgcttattcgc
gtaatccttctctaaaaaatgaattgtttggctacactattttaggattcgctttaacag
aagcgattgcattatttgctcttatgatggctttttttaattttatttacttaatttactt
taattactgggcgtttataataaacgcccaccttaaaaatacattatgacaagtacaact
ctttttgaatcttttcaattatttctttaatatccgcttgtatggtggtaagcctgtca
aatgctgtgtattcagttttgtttctaattgtagtattttgtaatactgctagtatttta
ttattactaggagcagaatttttatctttttttatttttaatcgtatacgtaggcgcaatt
gcagttttatttttatttgtagttatgatgttaaacgttaaaatagatggagtaaaaatt
aattatagcacaatttttttgattggtattttaataagtctgattttacttattcagatt
tgaactgctctacaattagatattgaagcgtatgataatataggcgtaccactatcccaa
aataactttccaacgataatttcttggatccaagaaaatgaattaccttcaaatacagag
agtattggtttaattttgtatacttcgtatagtttagtatttattatgtgcgcatttata
ctacttttagctatgattggttccattgtactaacaatgaatcaacgtagtggagttaaa
acacaacaaatcacacttcagttatatagaaatcaaaataaagtagttcgatttattgat
ctgagaaaaaaattaatttgattgcggatatagatgaattggtacatcagtaattttccac
gttaaaggatatgggttcgagtcccattatccgctcaaattaagagagaatagctcaata

```
ggtagagcaatagttttcaaaactaaaggttaaaagttcaagtctttttctcttgcaaa
agtgagtctgcttgaacaccttattcagttttaattataattaaaactgaataaggtgtt
caagcagactcacttttgcaagagaaaaaaggcttgaactttaaattcatataagaataa
caaccacatttagcatacacaagacgacagccttgtatgtgtactaatatatctctgca
ggctgtgtagagtcgaaaaaagtttgtgtggggtggctgtgtgacatataaattaaaaat
agttgtcaaaaaaagtttagtttggcgtaaaaggattcgaacctttgaaatcatggtatc
aaaaaccattgccttaccacttggctatacgccaaacaaattgaagataaagtgggattc
gaacccacggtggatagataccacgttagctttcaaaactaaagctttaaaccactcagc
catttatcccggttattaatataaattgtattctaaaatttttagagtctgtttatatga
tagtacgggaataggattcgaacctatatttttagatcatgagcctaatgagttaccttt
ttactctatcccgctattttatctaattaattttgtaacttaataaaaacttctccaaac
tacccagtaaaggtaggatgactaaaaaatataaaaaataataaacgctagctatctgac
ctataattatgtaagggtcttctacaggcataccacctattcaacctaatattaaacaac
aagcagccatgcttcaataaagaccacgatatatgggtctaaaacgcgaa----------
------------------------------------------------------------
------------------------------------------------------------
------------------------------------------------------------
------------------------------------------------------------
------------------------------------------------------------
------------------------------------------------------------
------------------------------------------------------------
------------------------------------------------------------
------------------------------------------------------------
------------------------------------------------------------
------------------------------------------------------------
--cttctaatttctgtactatgtattcacggtaatagagctaatacaactatagcaaaaa
tcatacatataacgccacctaatttatgtggtatacttcttaagattgcataaaaaggta
agaaatatcactcaggaacaatatgtgctggcgttaccatgggatttgcttcaatgtaat
tatcagggtgacctaaaagattaggcgagaagtatacaaaaaaagaaaagaatataataa
aggccactatccctaacaaatcctttacgatgaaataagggtacataggtaccttatcgc
tgcttgcatcgatacctaaaggatttccagaaccttcttgatgtaaagcggctaaatgca
ctaaagacgctgctgcgataacaaatggtaataaataatgtagactaaagaaacggttta
aagttgcattatcaacagaaaagccacctcaaagccaagcaactatagaatcacctacca
aaggtacagcggatactaaattagtgattacagtagcacctcataagctcatttggcctc
aaggtaatacataacctataaaagcagttattatcattaatagtaaaataattacaccaa
tcactcaaacaaattgtcgaggtgcagcataagaaccataataaagtcccctaaaaatgt
```

ggatataaactacaataaaaaacattgaagcaccattcgcatgtatatatcgtaaaagtc
aaccaaagttaacatcacgcataatatgctctacactaataaaagctaaatcaacgtgtg
gggtataatgcatagctaggaatattccagtcactatttgtattattaaacacattgcag
aaagaaacccaaaatttcatgcataatgaatattaattggagttggataatctataaggt
gattattaactatattgaagagaggtttttaattagacgcataaataatgtttttatta
tagatggacgaggaatgggacttgaacccatggcctataaagtcacagtttatcgctcta
ccaaaccgagctctcctcgtcatgttaaaaagttaaatttacggggaaaaagggatttga
accctcactcattgatgtgacaaaccaatattttaacctattaaactacttccccatttt
tattaaatacggatagagggatttgaaccctcatgaataatattcatcaaaacctaaacc
tgacatgtctaccatttccatcatatccgcaaaaaaatgttactattagctttaacggat
aaagagggattcgaacccacggtataatatttcatacgatgatttagcaaaccattgcct
taaaccactcagccatttatcctgtgttttggaagctgcccactaccggacttgaaccgg
taacttaaaaagaacagattttaaatctgtcgtgtttacctatttcaccaaatgggcatt
agctattgctaatgctatgttttattgaagctattgcttttccggggttcaatgatttag
gacacgttctactgcaattcataatggtatggcatttaaaaagttttgatttacctccaa
gtaatgctaaacggtcttgagttttgatatctcgactatcagctaatcatctataggctt
gcaataaaattgcaggacctaaatatttgtcatggtttcaccaataacttgggcaactag
cagaacagcaggcacaaagtatgcactcgtaaataccatttaattctgacctatcctttt
cagactgtagatattctgtttttgaaggtgtgttatttataagccacggttttatatatt
tatattgtgcgtaaaaattagataaatcaggaactaaatcttttataatgtacatatggg
gtagcggataaattgtaattgtgctagtatttatatttaatggttgcaaacaagctaatg
tattagttccgtttatattcattgagcaactaccacaaataccttctctacacgagcgtc
taaaggcgatactcgaatcttgttcgtcttttatttttataagagcatccaataccatag
gtccacaattttagtatgaataggatgtgtactgaaatgagtaatagttgggtttgatg
gagttcatctatatatacgaaggaattttaagtctaaattattattggatactaattgaa
aagaaatttttttgaataattgcatacttggttttattttgattgctataattatttaag
aacgggttagcccgttcttaaatataagataaaatataatgaacatttcgtatttattta
cttacttttgcaagtattaaataataaaccgatctatctagtaatcttctagaatgttct
ggaaaaagtaagggttatttttgcgcttttaagctgtcagcgcttatataatgttaacgt
ggctactcggctatgcaagaaacaatacaaccgatacactattggttaatatatcttaat
cctctcgtactaaagataaaccctctttttttctttcccacaacagatag----------
------------------------------------------------------------
-------------------------------ggttatggtaaagaaacctctcacgaaat
ttcctcccacttcaaaaccgtacgtgaaggtcacccttcatacggctcctcaaa----at
tatctatag-----------------aaagtgtt--------aaaaaaaagtacacactt
tcctttagtttaaaagtgcttatattagtccacggcttaagtattagctgctttggaatg

tctagtatcgtggcaatgaccatgcataagtcgtaagtttttaagatcagaagatccctt
ttgacttcttggtataatatgatctatttctattctatctgaatctttaaaatataattt
acacatactacactgcggcactttagtttttaatagtcgttttagtagttgcccatactt
attattaacagcgagacgtctggctcaataagtaactcttccgtcatatgggctactatg
tccggctacttttgtagatagaataatatttcgttgatcatgacgggttaattttataat
tttgtctccttctttaaggccaaaaactctttctgcactccctataggtattcaatatct
agtgataccaggacctgttcgtttgcgtttcttagctcatttccttagaagataaaacgt
acgtacactacaataactaaaagtttttgtggcactacatacggaaaagtatttagttca
tccgctaataactggtgctagtttactgattaaaactttttgagataatccggtggactt
cgtagtaatacttttatattagctaaatgagattctatagacttatagctgggttgaca
cctgctttttcatccagtcgcaattccttgattattctttgcggatgtatgtttacctac
cttatagttgacaaaattaaaacctaagaaatctacacctgttattctagacttctct--
------aaactaccggtgtacgttatttttgttttggattccgacaactttagacctaag
ttctgtaaaaagaattcaatttttgatttttgctgcaatcaattctttctcttcattacat
aatactaaaaaatcatctgcgtatctaataaggtatactccactttttcctattgcatct
tccattccgtgaagggcaatatttgccaacaagggtgatataatacctccttg-ggggtt
ccggcttctggtgtgatttcttttgtatttctttaaagcctgtaagtattcccgctttt
aatcaagctctcaattgctcttttaaaataggaaacgtgtttacctttagaagtaattta
gagtgatcaatattgtcgaagcatccttcaatatctgcatctaatacatgtttaggaagt
tgctgtaaacatttcacgattgcttgtctagcatcgttggaacttcgccctggtctaaat
ccataactgttaggttcgaatatagcttcatattgaggttccaatgcaaactttacaaga
cattgtttagctcgatctcttatagtaggtattcctaaatgcctttctttcccgtttggt
tttaaaattgttactcgacgaattttatccgatttattatcaatttcaatattttgtact
aattccattctttcgtcaggagttaaactactaactccatctactccagctgttcgtttt
cccaaattgtcttgagtcacttttcgaacagctaaaaactttgaaaagtcatgctttatg
atttgtttctgtattaagaatacagatgtcatattaccttttttgctaaattcaaaaact
ttacactgcaatctatacagtcaaatttcttttatttttcagttcacttgcgttcacttt
ttcataattt------gtttgttggtgtaatcttaacaatgttaaaattttttatttagat
aatt-----ttgtctacacgtctgcatatccataagctttccttatggcattggcttctt
gtagaatcctgatattaatataccttaacactata----------gaaaaacgcccttgt
aaaaaaagaatcaacaaggtttatattaatatttacttcgttccaatattacatacatat
agtcagtaagcacctctattccctctaaaaccggtacccctttctaacgcggccacatta
gattttaccttatgtttagaattcacgctgtttcgtgttaacggggtgggttaacttata
attatcccccgctatgacatatttagggaatgttatttcggctaccataagggtaggtta
tacttctataacta-------------cttttcttggcagatttactggtgtatatattc
tcttcaggtttgaatgtttataagtaacattctcccgaatttatgccctgcttataccttt

gtggataactattttcacactgtatagggtgcactgtttgtaaa-acagctcattgatca
gcattaggagt-aatgcgtaggcccactagttctagagaacttgcttcctacctgtaata
ttagtttaa----------tctctaaagtggagattagctatttacgagaagtacatctc
ttctcttactagcaacgaatcgcacgaccgaactgtctcacgacgttctgaacccagctc
acgtatcttattatttggcgaacaaccatacccttggaacctattgcagctccaggaaaa
gatgaggttatggtcagggcattcttcacagaatttcctgccactccagaaccgtacatg
aaaatcgctcttcatacggctcctcgagccatattattaacagttgattatttgttaagc
aaaatgtttacataacacaaaaaataagctcgttacttattttggaatgacgcgtgtca
tgacaatgaccatgcagaagtctaaggttatctcagtgtgaggttccccctttctttctt
ggtatgatatgatctatttatattctatcggagtcattaaatatattttgcacatatca
catttgggccctttcattttaaaaagtatccttaataattgcccgtacttgttgtttgat
gatagtcttttggatcagtaggtaactctaccatcatacgggctactatcaagtagtact
ttagtagatttaatcacatttgtttgatcgtgtctattcaattttataattttgtcttcc
tctatcaatccgaatactcagtttgcggtatcgctcttaattcaatgcttagaaacaccg
atccaggatcgatttttcttctgagatcatctttctagaagataaaatgtacgcatgcta
caaaagcggaaagtttttgttgcattacatactgcaaaatatctggtccacccaataata
acaggtgccagtttactaattaaaacttttgagataagccacttgacatttggataata
tttttgatactatctagatgcttattaattgatttaagactaggttgattcctagatgtt
caaccagttgaatttccatggctatccttggcggctttatgtatacccactttgtagttg
acaaaattaaaacctaagaaatctataccgctatttttcttgggagaagaataaacgcta
tatgttattgtagttttctctttgggtaattccaatcccatacctgttaaaaatgcttct
attttcagtttggcttctaataactctttctcctcgttacataatattaaaaaatcatct
gcgtatcttacaagatatactcctcttttactcagcgacttttctattccatgtaaagcg
atgttggccagcaagggttatataatacctccttgtggagttccagattctgggataatt
tctgtttttatcaccttgaaaatcaactaaaattccagctactagtcaagcagaaatctg
ctctttaagtagagggaaagtttccggcttttcaagcaacttagagtggtctatgttatc
aaaacatcttttgatatctgcatttaaaatatatttgggttttctttgtaaacattttac
aatagcctttctcgtgtcgttagcacttctaccgggtcggaacccgtaactgttaggttc
aaaaagtgcttcatactgcggctcgagtgcaaattttacaaggcattgcttagcccgatc
tcttatagtaggtatacccagatttttttacagaaccattcgcttttggtatagttaccct
caaaatcttatccgaatgtcgatcaatctttatgctttttactaactccaatctttcgtc
tggggttaataaagaaattttatctactccagctgttcgtttcccaaaattatcttgggt
aactcttctgactgctaaaaattttgcattttcatgttttatgatctgtttctgtataaa
aaaacacttttcatatcacccatcttacttagattaaatatttttatttgtaatttatat
agtcaaatttctttatgttttcacacaatattatttcatttcatgattttattaggtttg
tttattatatatcttccaaattatatttaaattaatatgactcgtctacacgtctgcata

tccataggctttccctgtggcattggcttcttgtagaatcctaccataattgtgcattaa
cgcaataacctgcgcccaaataatgcgttaaaaaaataaacaattatttggttcacaatt
atagttacttcgttccaatactatatacacttatttagtaagcccctctgttccctctaa
aactggttctccgactcaaacgtcgtaattgaatcaaaccttatgtttagatttcacgat
atttcgtgttaacggggtgggttaacttataattgtcccccgctatcctcggtttaggaa
actttatttcggatagacaaaatttttttgtctataataatcgacattattggcttttct
ggtcagatttactaaataagtattctttctatagttattattctataagtgaataagcta
caaatttatgccctgcttataatttgtacgtgttattgatacaaatcatagggttcactg
tttgtaaagacagctcattgatcagcatctggagtcgatgcataggcccactagttctga
agaactagcttcctatctatagtattagtttaatataggtctccctttattagctattta
cgagaagtacatctcttctcttactagcaacgaatcgcacccgacatcgaggtatcaaac
catggcatcgataagaactctcagccatgataaatctgttatccctagagtttcttttt
ccgataagcgacagtatttccatacattactgccggatcattatgaccgactttcgtccc
ggcttgagctgtaactcttaccgtcaagcagattttcatcattacatattatattagcat
atattgccaactaaaatccacttttgtgcacctccgttacagtttaggaggtcctcgtcc
caagtaaactaccacttttacatatgtccttgttagtaagcaatttttatactcagagt
agtttttcaatgacgtacaaatgtacttctacttttactgtactgtgtacaaagttatcg
cgacgtaaaattatagtaaagaatcatagggtctttccgtcttgttgcgggatgtctgca
ttttcacagacaatgtaatttcgctgaggctatactagagacagtggggtagtcgtgacg
ccattcatgcaggacggaatttacccgccaaggaatttcgctacctgaggatccttatag
ttaagaccgccgtttacttgggtttataatatatgcgcaaacttatatttttcactttca
agcactgggcaggcgtcaaaccttatacgtcttcttttgaatttgcaaagttttaag---
------------------------------------------------------------
------------------------------------------------------------
------------------------------------------------------------
------------------------------------------------------------
------------------------------------------------------------
------------------------------------------------------------
------------------------------------------------------------
------------------------------------------------------------
------------------------------------------------------------
------------------------------------------------------------
------------------------------------------------------------
------------------------------------------------------------
------------------------------------------------------------
------------------------------------------------------------

------------------------------------------------------------
------------------------------------------------------------
------------------------------------------------------------
------------------------------------------------------------
------------------------------------------------------------
------------------------------------------------------------
------------------------------------------------------------
------------------------------------------------------------
------------------------------------------------------------
------------------------------------------------------------
------------------------------------------------------------
------------------------------------------------------------
------------------------------------------------------------
------------------------------------------------------------
------------------------------------------------------------
------------------------------------------------------------
------------------------------------------------------------
------------------------------------------------------------
------------------------------------------------------------
------------------------------------------------------------
------------------------------------------------------------
------------------------------------------------------------
------------------------------------------------------------
------------------------------------------------------------
-------------------------------------------------tttttgttaaa
cagtcgctacccctaatttttgaaaccttaaaaggtactcctttttgcgaacgtacggag
taaatttgccgagttccttaagtatagttatctcattcgtctttattttctcaataagtt
cacctgtgtcggttttaggtacggtcaaacttcatgtaagttttcctgaaaaattacttt
ttttagcctccagtttagtgggtgtagctattagaagatatctttgtacacaaatacacg
agtattttgcagacacgtggtaatttttcctatcgaatacagttttcacttttttcttaa
gggccgactaactccagttacttaaaattgactggaaacccttgaacaatagacgaccat
gatttattttcaacatggttagcgctactcatgtcagcattagcactcctgattttagat
atgcaatttaacattaacataaaagactacaggacgttccgctaccattaagcttagttt
agcttaattcgaagcttcgatatataaatttaagtctccttacattttaaataggtagaa
caaataaaaatagcgagctaaaacgctttctccaataggtggctgcttctaagcctactc
tgtttattcgaattattccttattttttcactaatttataattttgagatcttagctatc

gattagggttgtttcccttttgacgtaagaccttatcgcccaacgactgtctgctgctat
aaataaaaatatagtttggagtttaataaaatttagcaaaatctaaatttaataagtagc
tctacccacattttaaaaaagcaacgtactactttgatagttttcgcggaaaaccagcta
tcaccaagtttgattggactttcacccctaatcttaagtcatccccgtatttttcaacag
acgtgggttcagtcctccagtacttttttaaagcaccttcaacttgctcaagaatagatca
cttggcttcgggtctaatacctgtaactttaagcgccttagtatttttaagcttgctaca
catattaacttactgactcattatgcaaaaggcactttgttgctgtatttgtcagcttca
aataaatataaattaacagattcaaatctttccctcacggtactcgttcactatcgatta
gaaaaggttttagcttagaagatgggactcctattttcatacaaaagtaatcgtactact
ttgtgttactattagtgtcgtaaaggactttataacctactttgggtttagacacttcgc
taagttcgctttcactcaccgttacttacaaattctcgtttgatttttttgttatgttac
taagatgattcaattcacataattatataaatgcatattaaatgcgagttttctaaagag
actcatagttcataggcaggtgcctcgctatgatgtttcgtcgcgcgacgtctttgcttt
tctaccaagatttcctctgataacttttaaatcttttatatttaagaacgggcggaaaa
atgttacccttttgtaagagtcattaactctatagtaattgtactcctaattcgataata
gacaacaagtattgctagtcctatagaagattccgaagccgctactgttaatatagctaa
agcaaatagttggcctactatattgtctaagtatatagaaaaaaatataaagttaaaact
aactgatagaaacattatttctaaagacattataattattataatactttttggtttaa
aaaaatgcctaaaactccgactaaaaataaaaaaaaagtataattttcacagtatagttg
ggtaatcatttatttttgtatttaacatttgagatagcaaattgtattaaatttttccag
ccgcaggttcccctacggctaccttgttacgacttcactttagtcctttcgctaccatgg
acaaaaaataagatttttgtcttcaagtagagtaaattcccatagtgtgacgggcggtgt
gtacaaaacccgagaacatattcaccgcgacagttctgatccgcgattactagcgattac
gacttcatattctcgagttgcagagaataatccgaattaaagatttttaaagattttgc
tccagctcacgcttttgcttcttattgtaaacattactttgtagcacatgaaagtagccc
aattcataagggtcatgcggacttgacgtcatttttcccttcctcaaggatattccaagc
agtttataatgcattaatgcattatacaaaagtttcgtccgtttgctggaattaaccaaa
cgcctcacggcacggactgacgacagccatgcaacacctgtgacatcttgtgtcatacga
gaattggtaaggttttgcgcgttgtttcgatttaaaccacatgctccaccgcttgtgcgg
gttcccgtcaattcctttgagttttaatcttgcgaccgtaatccccaggcggagtgttta
atgccttagcttcgcctctggaaaattatccaaaaacaaacactcatagttgagggcgta
gactacaggggtatctaatcccttttgatacctacgctttcgtgccttagtgtcagctat
agtccagattattgttttcacttttgaagttcttttgaatatcatcgcattttatcacta
ctttcgaagttccataatcttttcctatgctctagtaaattagtttagtaatctttagta
aacatgaaatttaaaatttttttactacttagtttaccacctacgcaccctttacgccca
gtcaattagaataatacttgctcctcccgttttaccgcggctgctggcacgaaattagcc

ggagctttgattgtaaaatttagtctttgatttttaatctattttacaaagtgatttac
agcctgatagggctttgctctcacatgtggcctggctaggtcaagctttcgctcattgcc
taagattcctcactgctgcctcttaaagagtctggaccttatttctgttccagtgtgact
gatcatcctcaaagaccaattaaggattatgggcttggtaggtcttttaaactaccaact
acctaatcctgcgtagacttttcttaaaccaattattaatttttaaggcatttaccaca
atttaagataaatttctacgtattactcacccgtacgctatctttttcataagttacgaa
aattattaaacttgcatgtgttaggccgaccactagtattcattcggagccaggatcaaa
ctcttttattattatgttatgtttttatgtttgctatctcagaaattaaatttgattgta
taaagttatccagcgagctaagacagaagctgtaccaatttcatatatatatatgttggt
aatcttccttttggtaatgaggataaaatttgctttacaaaaagttgtatactttttttg
ttttaatgcggaacatagtttattttctagctttgttccacgtataagaagtctcctata
ttttttttattttattcatacatttgtgtatcaagatgggcagatttgaactaccattccc
ttgcacccaaagcaagtacgttaccattacgctacatcttgatatagtgatattgactct
ttctggataggatttgaacctataaccttgtagttaacagctacctgctctaaccattga
agctaccagaaaaataaaagcgaaaaaagggacttgaacccttataatgaccttggcaag
gtcatgctttgcctattaagctatttccgcacttaaaattgaatataaagttgaataata
ttagcaataagtagaaaaagtaaaagataattgctatctttgtacattttccttgtattt
acatgttttactgaatctcaaaaaaaatggcgtattccattacttatgtggtatagaaat
atagcggacactactcaccggacactagttaaaaaccaaggaaaaaagaggtgatctaca
aaaaaatgtattggcagaatagtaatatataaaaagtatacttgatttagtaaaagaata
gggcctacaattaaaacaaacataataacgccagaaattctatgccaaatagaaaaaata
gaagctctttgcggattgtatatagttaaatgtggagaaattggtctattgatgttctgc
atattatagttttctttttcttgggattttacagccattatgcggacaagttgtaatatc
cattacagagataaatttaagtttagactgttttaatattttaaacacatttttttttatt
tttacctcatccttttatccgtacatgtaaagaagtataactgagttctttagtctttat
ctctaagaagtggattatttcggacgaagaaagggacaggcttctttttaccccttttga
ccttgacttgcctgcagttgtccaagatttaattttgcctttgcagtctgttaaaagaaa
atgaatattattggaactaaaagataaaaacagtatacccatatgtagtttaattttatg
ttgtagtagtaaagactagggtcagattcgaactgacctaataaagatttgcaatcttac
gcatagccactatgctacctagtcttaaatttagtgaaatctgaagaaagactgtgtttt
aaatgataaagaacttttcacacaagaaagtaattcaagcgattttagctttggtaaatt
ggataactcgaaaagcaactgaccgggccttatataacagcaatgactcactatagctcc
ttttcctttacccatacgggtttctggactaaatttggtaacagcatttttttggtagcat
gtaaaaccacactttgattttacccaaactggactttttttcgatgactttgatctcttt
ctgaattatctttttttatattatttatttgtgtgatatctacttgcttgtacgataaaga
gcgtaatccaaattggcctctttttactagatgacttgtttggtttatatatctattttt

tttttatgctgtcgtgttggattattttttgtattaaatcttttcatattaaattgtgc
ttaaattgttttgtatgccattcaaacttgaagattgcaaatcccttgttttgtgtataa
tggcgtttcaaaaaattctatataagtttctagtttttgcagtgatactgcgcctaatcg
gaatactatactacttgacatacgacttcggctaccaccgaaacgcccagcgtaccttat
ttttattcctgtaaattcataaaactgtataccacgagcagtttttaatacaattttgtg
cttttttttattgaaaaatagtatttttattgtattaattatcttccggaatgctaggcc
tttttttaggccagattttataaatccacataaaagtgccccgttttgatgtcaaaatgt
tgcattatattttttcacgtcaagatcgaacattttaaacgtaggtaccgagcaatctga
tttggctaatagaggtacatattgtacattgtattctcatctattgtcatttattttaga
tagcgtatttcctttatgaatgtttttgattcaaaatgcttatttaaaaagtttgtagc
aggtaatactttttttgtatatttgcttttggtttttaaaacctttgccatagcattgact
ttcggtatttcataagtcgttcacacctattcgaaatcctcttggattaacttttgacc
cataatgttttattacttaataacaatatttaaattgaaaatcaaatctaaatagttata
caaaaatacgttttgtaggatttgaacctacactaatcaacttagaaggttgatgtttta
tccaattaaactaaaaacgtttttattttagatgcattgggatttgaacccaaaataagc
agattaaaagtctgttgctgtaccgtttagctatacatctttttaataaagagagtagga
tttgaacctacgatgattttaatcaatagatttacaatctatcgctttcgaccactcagc
catctcttcttcagcggttagatatatgttacaatctgtaataaaataattaattatata
ctattgtttaataacaatatttagattgaaaatcgtacgattttcaatctaaatattgtt
attaaacaat
