## Supplementary material for "Genomic diversity of 39 samples of *Pyropia* species grown in Japan": S5 Fig

>Pyr_1_10_11_12_13_14_15_17_20_21_22_24_26_34_36_38_40_41_42
GTGACCGTTGCCTCGCCACAAATCCATTGTTTTTCGGGTTCTGGTCTTTTATGAGTTAGG
ACCTCTGCGCCTACTATCATAGCTGTCCGATGCCTCGCATTCCATACGTTGCACTGACCC
GACAATAATCTGTGCTGCTAAACTTTTGCAAGTGGCTGGCGCCTGTAGGTCACGCAATAG
AGTAGCTTCCCTCCCGACGATCTTCAGGCCATCTACAATCCGGCATCGTAGATCCTCGCG
TGACCTAGTAGCCCTTGACGACTCAATCCACCCTCTGCCTGACGAACGTGACTTTTGTAG
TACTCTAGTGACTCCCTGTCCTCATAAGTAGGCTGCATACATAGGGCTACCGTAGTTTAC
TGGTCTCACCTCTACGTACTCGCTGACCCAACTACGTTGATCCAGGAAGGCAGAATGGTG
GACGCTCTTACTGCCGACCCGTGGGTAACTAACTGGGACGGGGGAATATTGTTTCGCTTT
GAGTACTACTTCTTCCAATTGTTCGGATAGCACAGTAATGATGGTGGGTGACTCCAGTCA
CTTTCGCGTAGTCGCCGGTCGGTCGCTCCGGTCTGTCCTCTCGCCCGATTCTGGGCCATC
TTATCCGACGTCATCCTGCAGACCGCGCTTTCCAACCGTTAGCATTAAGCAGTCTCCCAT
GGATGCTCTCATGTCATGAGAGGTACCGGAGCCTCTAAGCGTTCCGCTGTTCTCGACAGT
ATTCTCTTTCGTTTTAACACTAGCTCAGACGGTAGAGGCTTCGTCTTCAGTATCGGCTAA
TGATGCCGGGTACGTCAAGATACCAGTCTGCGCGCTTACGCTGCTCCATCTGTTGATGTC
TGCTTCAGCTCGGCTAAGACCTCTCGAACTCTTTGTCTTACCTCAGCCCGCGGCTAGAAT
CATTTACATCCTGGCAAACACAGTCCTTAATTTCCAGTCGTGCCCGACAGTAATACCGTG
GCGCGAACGACGTCTAAGTTGTTGCCTACTTCGTACACTATGGACTACATTTGCCCAACG
TGAGGTCCGAGGATCCCGTTGGAGGGCACACACTCAGAGAATCCGCAGCGTTTCGTCTCA
CAATCGAGAAAGGCGCTCCTAAGGGGCCGCGATACGAGCGTGAAACCGGCCAAGCGCTCT
AATTGTACACTGTTCATGAGGCTAAATTCAAATTGCGGACCGAAAACTTGGCCCAGCAGA
GCTAAAGGTTCTTCGCGGGTATTGTCCGTACATGCAACTTGATGTTTTCTAAGCCAAGAG
AGATATGCGAGGGGGAGACATGTGAAAAGCAAAAAAATATAGTACTATAATTATCGTGGG
ACCCATAAGGCCTTGGTTGCTCCCGCGTCGCCTGTACCCACAGTCTTCAAAAAGGCGATT
TGCCTGAGTATAAAGTAAAGAGCCGACGGTGTTGAAAACGAGCGGACGTCTTGCATGAGA
AGTGTCACTGTGGGCATCTCAAAGGACTCTAATTAAATTCGGTGTCAATAGAAATGTTCG
CGCTGCATGGCGCTGTCAGATCATCCATCCTCTGTGGCTACGCCTTATCATAAGTGAGAC
CTGCGAAATCAGGGTGATGTCAATACGCTTTAAAGGACGCTAAAGCGAGTTGATAATTAA
ACGACAGAGTTACAATGAGAAGTATTCACAACTCGCCAAAATCATTGACGATAGACTGAC
ACGATAGCGTGGGAGGCCCCGTAGCCTCAAGTGAAAAGGGGCGCAAAACGGTGTCCGCAT
GTGAGGAGCTCGACCATCATAGATGCCGGATATGTTCCTGTGAAATTTCAGCGTCAAAGA
TGGGAAGCAAAACAAACCCATGAAACTACCATAGTGTGAAGGTGGTGGGCGGCGCGGTTT
AAGGCGAGGCGCTCTAAGACTCCAACGTGCCCTAGGGGATCGTAACATACAAGAACATTG
ACGTGCACAGTTTTGTTGAACGGCCTTCATGGGCCAAAGGTTGTATCCACGGCCAGGAGC
GGTAAGAACTTCGTGACATAGCGGCCGGAAAAAGGATCTACAACGTTAACCAGAACCTCG
GAGAGCGGAACGGCCAGTCGTTAGGGAGAAAATAGTTAAGCATTAATTCTGACGAACCAC

CAAAATGCTCGAGCGGTAGGTATCATTAGGGGGTACGGACTGTTACATTTAGAATGGCCC
AGCTCCATAGTTGCAGACACTAATTTTAGAGACCGCTGCTGCATCATTAACAAAATGTTA
TCGGTGGGATGCAGAAGAGCGCGGGCAAGATTACACAAACGCCTTCAGGTGGTAACCTGC
TCCTAAGCAACCAAGACGGATCGCGACGGACGTGTCTGGACGGCGAATGTACAGGAAGTT
TTCCGTGAGAACAACAAAATGAAGGATCGAGATTAGAAAGACTACAGCGCCCATGAGCAA
CGGAGTTGAACGTGATTGTCAGGTTGGTAATGGGCGGCCCTAGAAGATAACGAGAAAAAC
CCTTCGGACTTTTCAATGGGGACTGCATTCCGGCCGCGAAGGACTCCATGAATCGATAGG
TAGGGGAAATATGGATGAAACGGGCTGGGGCAACGCCCGAGGCGACGTCTACTAGTTAAA
AGGCAGGTTGCGAGGATCGGGGAAGACTGAGAAGGCCTTGGCGTAGGAATTCGAGGCAAC
CAGGAGGATAAGGTAAATGCCGACTTCAAGCCGATAATGCAAAGAAGCCGGCTATGGCCG
TAAATTAGACGATAAGTAACAGATCGTCGGGGGGCCAAGTCTAATCGTAAAACTTTTAGT
TGAGGTGATGTCGTAAGTGAGATCCCCGAACGCAGCATGAACAAGGTGAATCGTGTAAGC
AGAGAAGAGCGCTGCACGATGCTAGCACTGTCGTAGTAGTGGGACAACTTGGATGACTGG
CACATCGTAGTAATGAGTCATATGATCCCTCCTGAAGGCTTCAAAAAAGGACACAAGACG
CGAAGAAGATCTAGACTCAGTCTTTAAAACTGCACAACTTAGCCGAAGTGATTGGTAAGT
AGAAGTACTGATGTGGGCGCTGAGAGAAACCTGAAAACGCGATAACCATCATATCACGAT
GCTTCCGCCTTCCAGCGTCACTAAAGTTCTTAATCACATGCCTCGGTTCTCTCGCCAGTA
CACGGTCGGCCTTCTTTCCCCCATCGTTTACCGGAGCATATTCTTCGAGTTGCCTTATTA
ACCTTCTATTCTCTCGCCGGTGCCTGATTAGCGCCGCCTTGAGTATCATACTGTTTGCTT
TTAGCCAACCTGTTGAGCGTTGATTGCTTGTGTGACCATATCTTGCGTTTCGTTGATTCC
TGGTCGGCAATGTGTTGTGGCCTGTTCGTTGACGCTCCTGATCTCACCGTACTTCACGGT
TGCCGCCCGGACATCACCCAGATGTGTTTAGGCCATGGCCGGTTAAGGTTAATCTTTACG
CGTCATCATAACTTCCTGCGAAACTGGTTCCCCCACGCCGTCCCCTTCGTTAGTCGATCG
GCGTGGTTAGGTCGCGCGGGTTTTTCTAGGCAAGGTAAGTCGGGTCGCACAGTGCGACCC
ACGGACATAACCTTAACGTTCCGTTTTACCACTGGGTCAGGCTGCTTTGGGTCCGACGTG
ACGCTCTTTCCGCACGACCTCGTATGTTCTTATTCGCACTCGTAATGAACCGGTAAGCGA
GAATTCGCTCTACGCTGAGGTGTCGTCGGACCCGAGTGTGCTTGCCGCTGGCCAATTGTT
GTGTCGGCTCGACGTCTTATTTCGTATGGTCCGGGATTACCCGTGCTTGCAGCGATGTCC
TTTTCGCTATTGATGTCTTCTCCTGAGTCACGCTACATACTGAATTAATAAAGCATTTCG
ACCGCAGCATCAGTGAAGTCGAGGGATGGGCTACATCGCCGTGCGGAGCTTTTCACGTTG
TGTCCGTGCGGAGATTCAGCACTAGCGGTCATACTATCCATGTTCCCCCGTTCTATTTAC
GTTGAGCGACTTGAAACTTCCAGTGCACTGTTTCTGCATAGTGCGGTATTTCTCAGTTCA
TGCAAGTTATCGTACATCATGGAATAGAACAAGCATCAAGGACCATTTCCACTAACGATT
CTTACCTCCAAATCAAGATAGAATCTAACCGATTATCGACGAACTTCATCCGTCATGTCA
CTCTATGTCAAGCGGCCTTCTTACAGCGCTGTCGCACTATCCGTTCTACCTTGTTTCCGA
GAAACCTTGCCTCTGTTGCCTGTGGGGGCAGGGCGGATATGGCTAAATGGGCATTCCTTT

TTACTTCCCGTGCCCCTTCATGCTATGTGCTTGCAGCTCTAAAGCTTAGACGCCCGGTCC
TCGACGTCTCGAAACTTTCCTGACCCTGCTGGCGTCTTACCCTTTATAGTGGATCTGCAT
AATGTGTTCTGTATATAGCCAAATACTGCCAACTATCCACTCGCTTAGCTGTCAATATCC
ATGAGCGTGCTTTGATTTACCGGTCGACCTGACTAATGTAACTGCCTTGCGCTCTCCACT
CCGCACCAGGGTTTCGAGTTTTTCTACCCCTCTCTTTATTACTACTCGTTTACTGCATCA
GACGCGCCCATCGTACTTGGTCACTGCTCCTTCAGGCGAGGAGTATTGGCACGTTATCGT
GCTTCTATCCTGAGCTGCAACCATAAGCATTTCTAAAGTTATGAGGATCAAAGAGTATTT
CCGTACATCTTGATCGATTGTACTGGACGTCGTCCCAAATAGCCTGCGAAGTCCAATATC
TAGGTCATAGATGCT
>Pyr_16
GTGACCGTTGCCTCGCCACAAATCCATTGTTTTTCGGGTTCTGGTCTTTTATGAGTTAGG
ACCTCTGCGCCTACTATCATAGCTGTCCGATGCCTCNNNTTCCATACGTTGCACTGACCC
GACAATAATCTGTGCTGCTAAACNNNNGCAAGTGGCTGGCGCCTGTAGNNCACGCAATAG
AGTAGCTTCCCTCCCGACGATCTTCAGGCCATCTACNNNCCGGCATCGTAGATCCTCGCG
TGACCTAGTAGCCCTTGACGACTCAATCCACCCTCTGCCTGACGAACGTGACTTTTGTAG
TACTCTAGTGACTCCCTGTCCTCATAAGTAGGCTGCATACATAGGGCTACCGTAGTTTAC
TGGTCTCACCTCTACGTACTCGCTGACCCAACTACGTTGATCCAGGAAGGCNNNNNNNTG
GACGCTCTTACTGCCGACCCGTGGGTAACTAACTGGGACGGGGGAATATTGTTTCGCTTT
GAGTACTACTTCTTCCAATTGTTCGGATAGCACAGTAATGATGGTGGGTGACTCCAGTCA
CTTTCGCGTAGTCGCCGGTCGGTCGCTCCGGTCTGTCCTCTCGCNNNATNCTGGGCCATC
TTATCCGACGTCATCCTGCAGACCGCGCNNNNNNNNCGTTAGCATTAAGCAGTCTCCCAT
GGATGCTCTCATGTCATGAGAGGTACCGGAGCCTCTAAGCGTTCCGCTGTTCTCGACAGT
ATTCTCTNTCGTTTTAACACTAGCTCAGACGGTAGAGGCTTCGTCTTCAGTATCGGCTAA
TGATGCCGGGTACGTCAAGATANNNNNCTGCGCGCTTACGCTGCTCCATCTGTTGATGTC
TGCTTCAGCTCGGCTAAGACCTCTCGAACTCTTTGTCTTACCTCAGCCCGCGGCTAGAAT
CATTTACATCCTGGCAAACACAGTCCTTAATTTCCAGTCGTGCCCGACAGTAATACCGTG
GCGCGAACGACGTCTAAGTTGTTGCCTACTTCGTACACTATGGACTACATTTGCCCAACG
TGAGGTCCGAGGATCCCGTTNNNNNNNNCACACTCAGAGAATCCGCAGCGTTTCGTCTCA
CAATCGAGAAAGGCNNNCCTAAGGGGCCGCGATACGAGCGTGAAACCGNNCAAGCGCTCT
AATTGTACACTGTTCATGAGGCTAAATTCAAATTGCGGACCGAAAACTTGGCCCAGCAGA
GCTAAAGGTTCTTCGCGGGTATTGTCCGTACATGCAACTTGATGTTTTCTAAGCCAAGAG
AGATATGCGAGGGGGAGACATGTGAAAAGCAAAAAAATATAGTACTATNNNTATCGTGGG
ACCCATAAGGCCTTGGTTGCTCCCGCGTCGCCTGTACCCACAGTCTTCAAAAAGGCGATT
TGCCTGAGTATAAAGTAAAGAGCCGACGGTGTTGAAAACGAGCGGACGTCTTGCATGAGA
AGTGTCACTGTGGGCATCTCAAAGGACTCTAATTAAATTCGGTGTCAATAGAAATGTTCG
CGCTGCATGGCGCTGTCAGATCATCCATCCTCTGTGGCTACGCCTNATCATAAGTGAGAC

CTGCGAAATCAGGGTGATGTCAATANNNNNTAAAGGACGCTAAAGCGAGTTGATAATTAA
ACGACAGAGTTACAATGAGAAGTATTCACAACTCGCCAAAATCATTGACGATAGACTGAC
ACGATAGCGTGGGNNGCCCCGTAGCCTCAAGTGAAAAGGGGCGCAAAACGGTGTCCGCAT
GTGAGGAGCTCGACCATCATAGATGCCGGATATGTTCCTGTGAAATTTCAGCGTCAAAGA
TGGGAAGCAAAACAAACCCATGAAACTACCATAGTGTGAAGGTGGTGGGCGGCGCGGTTT
AAGGCGAGGCGCTCTNAGANTCCAACGTGCCCTAGGGGATCGTAACATACAAGAACATTG
ACGTGCACAGTTTTGTTGAACGGCCTTCATGGGCCNNNNNNNNTATCCACGGCCAGGAGC
GGTAAGAACTTCGTGACATAGCGGCCGGAAAAAGGATCTACAACGTTAACCAGNACCTCG
GAGAGCGGAACGGCCAGTCGTTAGGGAGAAAATAGTTAAGCATTAATTCTGACGAACCAC
CAAAATGCTCGAGCGGTAGGTATCATTAGGGGGTACGGACTGTTACATTTAGAATGGCCC
AGCTCCATAGTTGCAGACANTAATTTTAGAGACCGCTGCTGCATCATTAACAAAATGTTA
TCGGTGGGATGCAGAAGAGCGCGGGCNNGATTACACAAACGCCTTCAGGTGGTAACCTGC
TCCTAAGCAACCAAGACGGATCGCGACGGACGTGTCTGGACGGCGAATGTACAGGAAGTT
TTCCGTGAGAACAACAAAATGAAGGATCGAGATTAGAAAGACTACAGCGCCCATGAGCAA
CGGAGTTGAACGTGATTGTCAGGTTGGTAATGGGCGGCCCTAGAAGATNNCGAGAAAAAC
CCTTCGGACTTTTCAATGGGGACTGCATTCCGGCCGCGAAGGACTCCATGAATCGATAGG
TAGGGGAAATATGGATGAAACGGGCTGGGGCAACGCCCGAGGCGACGTCTACTAGTTAAA
AGGCAGGTTGCGAGGATCGGGGAAGACTGAGAAGGCCTTGGCGTAGGAATTCGAGGCAAC
CAGGAGGATAAGGTAAATGCCGACTTCAAGCCGATAATGCAAAGAAGCCGGCTATGGCCG
TAAATTAGACGATAAGTAACAGATCGTCGGGGGGCCAAGTCTAATCGTAAAACTTTTAGT
TGAGGTGATGTCNTAAGTGAGATCCNNNAACGCAGCATGAACAAGGTGAATCGTGTAAGC
AGAGAAGAGCGCTGCACGATGCTAGCACTGTCGTAGTAGTGGGACAANNTGGATGACTGG
CACATCGTAGTAATGAGTCATATGANCCCTCCTGAAGGCTNNNNNNNAGNNNNNAAGACG
CGAAGAAGATCTAGACTCAGTCTTTAAAACTGCACAACTTAGCCGAAGTGNNNGGTAAGT
AGAAGTACTGATGTGGGCGCTGAGAGAAACCTGAAAACNNNNNNNCCATCATATCACGAT
GCTTCCGCCTTCCAGCGTCACTAAAGTTCTTAATCACATGCCTCGGTTCTCTCGCCAGTA
CACGGTCGGCCTTCTTTCCCCCATCGTTTACCGGAGCATATTCTTCGAGTTGCCTTATTA
ACCTTCTATTCTCTCGCCGGTGCCTGATTAGCGCCGCCTTGAGTATCATACTGTTTGCTT
TTAGCCAACCTGTTGAGCGTTGATTGCTTGTGTGACCATATCTTGCGTTTCGTTGATTCC
TGGTCGGCAATGTGTTGTGGCCTGTTCGTTGACGCTCCTGATCTCACCGTACTTCACGGT
TGCCGCCCGGACATCACCCAGATGTGTTTAGGCCATGGCCGGTTAAGGTTAATCTTTACG
CGTCATCNTAACTTCCTGCGAAACTGGTTCCCCCACGCCGTCCCCTTCGTTAGTCGATCG
GCGTGGTTAGGTCGCGCGGGTTTTTCNNGGCAAGGTAAGTCGGGTCGCACAGTGCGACCC
ACGGACATAACCTTAACGTTCCGTTTTACCACTGGGTCAGGCTGCTTTGGGTCCGACGTG
ACGCTCTTTCCGCACGACCTCGTATGTTCTTATTCGCACTCGTAATGAACCGGTAAGCGA
GAATTCGCTCTACGCTGAGGTGTCGTCGGACCCGAGTGTGCTTGCCGCNNNNCAATTGTT

GTGTCGGCTCGACGTCTTATTTCGTATGGTCCGGGATTACCCGTGCTTGCAGCGATGTCC
TTTTCNCTATTGATGTCTTCTCCTGAGTCACGCTACATACTGAATTAATAAAGCATTTCG
ACCGCAGCATCAGTGAAGTCGAGGGATGGGCTACATCGCCGTGCGGAGCTTTTCACGTTG
TGTCNNNNNNNNNNNNNAGCACTAGCGGTCATACTATCCATGTTCCCCCGTTCTATTTAC
GTTGAGCGACTTGAAACTTCCAGTGNNNNNNNNNNNCATAGTGCGGTATTTCTCAGTTCA
TGCAAGTTATCGTACATCATGGAATANNNNAAGCATCAAGGACCATTTCCACTAACGATT
CTTACCTCCAAATCAAGATAGAATCTAACCGATTATCGACGAACTTCATCCGTCATGTCA
CTCTATGTCAAGCGGCCTTCTTACAGCGCTGTCGCACTATCCGTTCTACCTTGTTTCCGA
GAAACCTTGCCTCTGTTGCCTGTGGGGGCAGGGCGGATATGGCTAAATGGGCATTCCTTT
TTACTTCCCGTGCCCCTTCATGCTATGTGCTTGCAGCTCTAAAGCTTAGACGCCCGGTCC
TCGANNNCTCGAAACTTTCCTGACCCTGCTGGCGTCTTACCCTTTATAGTGGATCTGCAT
AATGTGTTCTGTATATAGCCAAATACTGCCAACTATCCACTCGCTTAGCTGTCAATATCC
ATGAGCGTGCTTTGATTTACCGGTCGACCTGACTAANGTAACTGCCTTGCGCTCTCCACT
CCGCACCAGGGTTTCGAGTTTTNCTACCCCTCTCTTTATTACTACTCGTTTACTGCATCA
GACGCGCCCATCGTACTTNNNCACTGCTCCTTCAGGCGANNNNNNTTGGCACNTTATCGT
GCTTCTATCCTGAGCTGCAACCATAAGCATTTCTAAAGTTATGAGGATCAAAGAGTATTT
CCGTACATCTTGATCGATTGTACTGGACGTCGTCCCAAATAGCCTGCGAAGTCCAATATC
TAGGTCATAGATGCT
>Pyr_18
GTGACCGTTGCCTCGCCACAAATCCATTGTTTTTCGGGTTCTGGTCTTTTATGAGTTAGG
ACCTCTGCGCCTACTATCATAGCTGTCCGATGCCTCGCATTCCATACGTTGCACTGACCC
GACAATAATCTGTGCTGCTAAACTTTTGCAAGTGGCTGGCGCCTGTAGGTCACGCAATAG
AGTAGCTTCCCTCCCGACGATCTTCAGGCCATCTACAATCCGGCATCGTAGATCCTCGCG
TGACCTAGTAGCCCTTGACGACTCAATCCACCCTCTGCCTGACGAACGTGACTTTTGTAG
TACTCTAGTGACTCCCTGTCCTCATAAGTAGGCTGCATACATAGGGCTACCGTAGTTTAC
TGGTCTCACCTCTACGTACTCGCTGACCCAACTACGTTGATCCAGGAAGGCAGAATGGTG
GACGCTCTTACTGCCGACCCGTGGGTAACTAACTGGGACGGGGGAATATTGTTTCGCTTT
GAGTACTACTTCTTCCAATTGTTCGGATAGCACAGTAATGATGGTGGGTGACTCCAGTNA
CTTTCGCGTAGTCGCCGGTCGGTCGCTCCGGTCTGTCCTCTCGCCCGATTCTGGGCCATC
TTATCCGACGTCATCCTGCAGACCGCGCTTTCCAACCGTTAGCATTAAGCAGTCTCCCAT
GGATGCTCTCATGTCATGAGAGGTACCGGAGCCTCTAAGCGTTCCGCTGTTCTCGACAGT
ATTCTCTTTCGTTTTAACACTAGCTCAGACGGTAGAGGCTTCGTCTTCAGTATCGGCTAA
TGATGCCGGGTACGTCAAGATACCAGTCTGCGCGCTTACGCTGCTCCATCTGTTGATGTC
TGCTTCAGCTCGGCTAAGACCTCTCGAACTCTTTGTCTTACCTCAGCCCGCGGCTAGAAT
CATTTACATCCTGGCAAACACAGTCCTTAATTTCCAGTCGTGCCCGACAGTAATACCGTG
GCGCGAACGACGTCTAAGTTGTTGCCTACTTCGTACACTATGGACTACATTTGCCCAACG

TGAGGTCCGAGGATCCCGTTGGAGGGCACACACTCAGAGAATCCGCAGCGTTTCGTCTCA
CAATCGAGAAAGGCGCTCCTAAGGGGCCGCGATACGAGCGTGAAACCGGCCAAGCGCTCT
AATTGTACACTGTTCATGAGGCTAAATTCAAATTGCGGACCGAAAACTTGGCCCAGCAGA
GCTAAAGGTTCTTCGCGGGTATTGTCCGTACATGCAACTTGATGTTTTCTAAGCCAAGAG
AGATATGCGAGGGGGAGACATGTGAAAAGCAAAAAAATATAGTACTATAATTATCGTGGG
ACCCATAAGGCCTTGGTTGCTCCCGCGTCGCCTGTACCCACAGTCTTCAAAAAGGCGATT
TGCCTGAGTATAAAGTAAAGAGCCGACGGTGTTGAAAACGAGCGGACGTCTTGCATGAGA
AGTGTCACTGTGGGCATCTCAAAGGACTCTAATTAAATTCGGTGTCAATAGAAATGTTCG
CGCTGCATGGCGCTGTCAGATCATCCATCCTCTGTGGCTACGCCTTATCATAAGTGAGAC
CTGCGAAATCAGGGTGATGTCAATACGCTTTAAAGGACGCTAAAGCGAGTTGATAATTAA
ACGACAGAGTTACAATGAGAAGTATTCACAACTCGCCAAAATCATTGACGATAGACTGAC
ACGATAGCGTGGGAGGCCCCGTAGCCTCAAGTGAAAAGGGGCGCAAAACGGTGTCCGCAT
GTGAGGAGCTCGACCATCATAGATGCCGGATATGTTCCTGTGAAATTTCAGCGTCAAAGA
TGGGAAGCAAAACAAACCCATGAAACTACCATAGTGTGAAGGTGGTGGGCGGCGCGGTTT
AAGGCGAGGCGCTCTAAGACTCCAACGTGCCCTAGGGGATCGTAACATACAAGAACATTG
ACGTGCACAGTTTTGTTGAACGGCCTTCATGGGCCAAAGGTTGTATCCACGGCCAGGAGC
GGTAAGAACTTCGTGACATAGCGGCCGGAAAAAGGATCTACAACGTTAACCAGAACCTCG
GAGAGCGGAACGGCCAGTCGTTAGGGAGAAAATAGTTAAGCATTAATTCTGACGAACCAC
CAAAATGCTCGAGCGGTAGGTATCATTAGGGGGTACGGACTGTTACATTTAGAATGGCCC
AGCTCCATAGTTGCAGACACTAATTTTAGAGACCGCTGCTGCATCATTAACAAAATGTTA
TCGGTGGGATGCAGAAGAGCGCGGGCAAGATTACACAAACGCCTTCAGGTGGTAACCTGC
TCCTAAGCAACCAAGACGGATCGCGACGGACGTGTCTGGACGGCGAATGTACAGGAAGTT
TTCCGTGAGAACAACAAAATGAAGGATCGAGATTAGAAAGACTACAGCGCCCATGAGCAA
CGGAGTTGAACGTGATTGTCAGGTTGGTAATGGGCGGCCCTAGAAGATAACGAGAAAAAC
CCTTCGGACTTTTCAATGGGGACTGCATTCCGGCCGCGAAGGACTCCATGAATCGATAGG
TAGGGGAAATATGGATGAAACGGGCTGGGGCAACGCCCGAGGCGACGTCTACTAGTTAAA
AGGCAGGTTGCGAGGATCGGGGAAGACTGAGAAGGCCTTGGCGTAGGAATTCGAGGCAAC
CAGGAGGATAAGGTAAATGCCGACTTCAAGCCGATAATGCAAAGAAGCCGGCTATGGCCG
TAAATTAGACGATAAGTAACAGATCGTCGGGGGGCCAAGTCTAATCGTAAAACTTTTAGT
TGAGGTGATGTCGTAAGTGAGATCCCCGAACGCAGCATGAACAAGGTGAATCGTGTAAGC
AGAGAAGAGCGCTGCACGATGCTAGCACTGTCGTAGTAGTGGGACAACTTGGATGACTGG
CACATCGTAGTAATGAGTCATATGATCCCTCCTGAAGGCTTCAAAAAAGGACACAAGACG
CGAAGAAGATCTAGACTCAGTCTTTAAAACTGCACAACTTAGCCGAAGTGATTGGTAAGT
AGAAGTACTGATGTGGGCGCTGAGAGAAACCTGAAAACGCGATAACCATCATATCACGAT
GCTTCCGCCTTCCAGCGTCACTAAAGTTCTTAATCACATGCCTCGGTTCTCTCGCCAGTA
CACGGTCGGCCTTCTTTCCCCCATCGTTTACCGGAGCATATTCTTCGAGTTGCCTTATTA

ACCTTCTATTCTCTCGCCGGTGCCTGATTAGCGCCGCCTTGAGTATCATACTGTTTGCTT
TTAGCCAACCTGTTGAGCGTTGATTGCTTGTGTGACCATATCTTGCGTTTCGTTGATTCC
TGGTCGGCAATGTGTTGTGGCCTGTTCGTTGACGCTCCTGATCTCACCGTACTTCACGGT
TGCCGCCCGGACATCACCCAGATGTGTTTAGGCCATGGCCGGTTAAGGTTAATCTTTACG
CGTCATCATAACTTCCTGCGAAACTGGTTCCCCCACGCCGTCCCCTTCGTTAGTCGATCG
GCGTGGTTAGGTCGCGCGGGTTTTTCTAGGCAAGGTAAGTCGGGTCGCACAGTGCGACCC
ACGGACATAACCTTAACGTTCCGTTTTACCACTGGGTCAGGCTGCTTTGGGTCCGACGTG
ACGCTCTTTCCGCACGACCTCGTATGTTCTTATTCGCACTCGTAATGAACCGGTAAGCGA
GAATTCGCTCTACGCTGAGGTGTCGTCGGACCCGAGTGTGCTTGCCGCTGGCCAATTGTT
GTGTCGGCTCGACGTCTTATTTCGTATGGTCCGGGATTACCCGTGCTTGCAGCGATGTCC
TTTTCGCTATTGATGTCTTCTCCTGAGTCACGCTACATACTGAATTAATAAAGCATTTCG
ACCGCAGCATCAGTGAAGTCGAGGGATGGGCTACATCGCCGTGCGGAGCTTTTCACGTTG
TGTCCGTGCGGAGATTCAGCACTAGCGGTCATACTATCCATGTTCCCCCGTTCTATTTAC
GTTGAGCGACTTGAAACTTCCAGTGCACTGTTTCTGCATAGTGCGGTATTTCTCAGTTCA
TGCAAGTTATCGTACATCATGGAATAGAACAAGCATCAAGGACCATTTCCACTAACGATT
CTTACCTCCAAATCAAGATAGAATCTAACCGATTATCGACGAACTTCATCCGTCATGTCA
CTCTATGTCAAGCGGCCTTCTTACAGCGCTGTCGCACTATCCGTTCTACCTTGTTTCCGA
GAAACCTTGCCTCTGTTGCCTGTGGGGGCAGGGCGGATATGGCTAAATGGGCATTCCTTT
TTACTTCCCGTGCCCCTTCATGCTATGTGCTTGCAGCTCTAAAGCTTAGACGCCCGGTCC
TCGACGTCTCGAAACTTTCCTGACCCTGCTGGCGTCTTACCCTTTATAGTGGATCTGCAT
AATGTGTTCTGTATATAGCCAAATACTGCCAACTATCCACTCGCTTAGCTGTCAATATCC
ATGAGCGTGCTTTGATTTACCGGTCGACCTGACTAATGTAACTGCCTTGCGCTCTCCACT
CCGCACCAGGGTTTCGAGTTTTTCTACCCCTCTCTTTATTACTACTCGTTTACTGCATCA
GACGCGCCCATCGTACTTGGTCACTGCTCCTTCAGGCGAGGAGTATTGGCACGTTATCGT
GCTTCTATCCTGAGCTGCAACCATAAGCATTTCTAAAGTTATGAGGATCAAAGAGTATTT
CCGTACATCTTGATCGATTGTACTGGACGTCGTCCCAAATAGCCTGCGAAGTCCAATATC
TAGGTCATAGATGCT
>Pyr_19
CTTCATACCAAACTTTTGTGGCCAATCCGTCCCTTAAGGGTCGATTCCCAGCTGACGAAA
GNTNNCACGTCCNTTNNNNCAATCNNTTAGCNGTCTACCCCGCNNNNNNNNNNCTNNNNN
NNNNNNNAGACNCATTACCAGGTCTCTCCCGTGGNNCTTTCTATTCNAAGTGTATCTCNN
NNGGAACNTTTCTTTACTAGCTCGCGTTTTGCTCCTCGCTCGGNNNNNNTGACCNTCGGTT
CATTTCGGCCTANCGGAGTAGNCNNGNTNATATCGCATTTAGTATATCCTATCCCGGGGA
CTGCTCCGGGGTCTTNCACATCAGCGCGCTTGTGACGCNNNNNNGNNNNTCACGACCCGT
GCTCTCTGTGTTTAANTGTNTAGCTCTTTGACCGCGTCGACTTAGAATAATANCGGAAAA
AGTGATTGAACCGTTNNTNNNAATTTCGGNNGTCCNCATGTTANNCCGGTAGCGGACCCC

GAACTTCGTCGTCCTTGGGCANNNNNNNNNNNACCACNGTTTAAATAGAGTGCNNNCNCTC
CCTCTGTNNNNCTGATAACTAAGCTTTTTAGGACTTTTTTCTGTNTNNNNCTNNNTCGTT
CCACAATGTTCTACTTCATTTCNNNNNNNNNCCNNNCNNNNNNTNNGTNNNNNCNNTNNC
GCANNANNNNATANTGNNCTGNNNNNNNNNNANGCCGAATAGCTTATCAGCTCCGCNGGC
NNNNNNANCTATGCAAGTGTGGTACTNAGNGTCCAGAATCCTGTTTNNNNTNTNATTANG
CGTGNTNTANCGTTATTGAAAGTTAGCCTATATTTCCGTAAGTTGAAGTTGACCAGCTCT
NNNNNCNATGTAAANGCAGTTCTCCAGNNCTCNNACAGCGATGTGATTNATTAACGGTAC
TATACGCGCTTCAATGAGAGTGGGTTCNAGCCTCTGNNTANNNNNNANANNNTCGCNNNN
NNNNNNNCGNNNNCNNNNNNGNNNNNNNNNNNNNNNNNNCTCCAGTCNANNNTNCNNNATA
TAGAGNTNGANNNNNNTNTTANGAAGGACGTGTCCCGGAGGTAGATGATGCCCCGCTCTG
NGGCTAAGAGAAAGATCCATCGGGATTGTCNNCNNNNNNNCNNACAAACCAGGANNTCTC
TCCCACCTTTCAATTGCANNNTCCGTGCTGCTCCAAANNNNNNANNNCTAACTTGGATAG
AACAGGAACCACGAATAAACGCGACATACCTGCAAGTTCGAGCAGCATACGGATCGTGNA
GTGGGCGTTCAGAGAGAGTGCATNGCNGANGGCAAGGTGGGGNNTTGCGCCTAATACCAG
TTTTGGGGACCTCCTAGTTTNNNNANNTNNTCNACATGTATGTCNCCTGATGGAATGGGC
CAGGGANNCCCNGCACCGGTAATTNGAAACACCCGGGGTGGATAGCTGGTCCATCCACAG
NNCACTGTGAGAACTGCACAGGGACGTCTCTGCCGGGGCTGACAGTAGCGAAGCTACATA
TATGATGCCATACGATTAAGTAACTTGTACCCCACAATAGNNNNNNNNNNATGGGTACGGT
TGTTAGAGCTGATACANTNNTGCCNNNNNTNCANNAGTNTCGGGACNAAACNGCGGGCGG
GTNNNNACGCNGTNGGGGACGATGTCCCTNNTNATTTNNCCGCGGCNGTGGGGACGGAGC
GTAACAATGCAAANNNCNCTCTCATTCTAGACGGGGGGNAATTTGCGGCGAGACGTNATC
NTAGAAGTACCGGCAGCAGCGGGTNGTNANCCGAGCTCTTCACGCAGCTGACACTAGGGG
GACGGGTTGAAANCGGCCTGGAGGGTAGAGCCGANANAATGACAACAANATATACCACGC
GTAGTACNNNNNNNNCNAATTTTGGGGCATTTCGAAGAGCGTCCGTGGATGGAGGTCACA
GTAGTTTNGTNNCCCCCAGGNNNTTCCTGCAAANNNNNNGNNNCGCATCTACGNNNGNNN
NNNNNAGGTGGTACAGTGCGATAATAGAGANGNNTGCTCCTACTACCGAGTGACGTGATA
AAGGTTATANCAGCNCCGTNNGNAANNNNGTGCGATCNNTNNNNCCCCTTAGTAGTTTTT
TGTGCCTNCTANNNNTCGNTCGCTGCGGAAAATNNTAAGNCTCCGCCCCCGAGCCGGAAT
GGTCCTACGACGTTGAGCATCCGGTCCCAGAATAATACNCNCNNNNNNNNNANAGCNCCG
CTANNNNNNNNNNNNNNAGTAATTGATGCAGACGCGTGANANTTNNNGAACAAAGGTTCAT
TTACTGATAGTTGGAGTAACCTATAGTAAGTAAAGTCAGCAAANNNNCACGTGAAGAACC
NNNTACTGAGTTGGTGCAGCAGGAAGCAACAGCTGGGGGAGTCATGATATACAATGAANG
TTAGNCAACGATCAGCGGCGCGAGCTNCGGCTAATGATATGGAGGACGCATAAGTAAGAG
TGCCTAAGTGCCCCGGTAACACAAGTNTNNTNNNNATAGGAAGTCNNGATGCCTAGCGNA
ACNTAAGGGCCAAAGCNGNNTAAATCAAAATGGCTTTCAGGGTGGCACTGGTCGATCGAG
AAATCTACCATNGTAGNCTAATGGAGACGGAGTAAANNCAAANAGAAGGCGTGGGTTGGA

TGTAAAAACGAATCGNGCGTTAGTCTGNGATTGGTGGCACGACACTATTAATCGCAATTT
GNTTNNNNNNGNCNGACAAACCCGTACTANNNANNNNNNCTCAGCTACAGGGGCCCCGAC
CNNAACNNNNNAGCNAANATAGCTCCTAGGAAATCNNNNNANNNNNNNNNNNNNNNNNNN
NAAAGCACNTATAACGTAGNAACNGNNNNACNAGGATGACGAACTGGGGACAGCACGCAA
TGTGCTACGAAGGCAGACAGCGCAGCCTTTNNANNNCANNNNNNNNNNNACGNCTCGTATA
TAGGAGGATAACGAGTTTGACTCNNNCCATTAANANNTATGNNNNNGANNNTNNNNNNNT
ANANNCNTTTGTTCAAAGGTCGGAGAGCTTANNNNANTNNNNTNACCANNATGCTCNAGC
ATTCTCATCTCCCCATTCTTTCGAGACCNNTNANNGTCTCTACTGTCTTGACTANACACG
TANNNNNNGNTCCTCCCTNTNTNNTTCCCTTTAAAATCCGTGTTCTATACCATTTCGCAG
ATCCCTCTTACGTCGANNNNNNNNNNNANNGNTANNNNTNNNGTGCNTGGGCGACCCACCC
CCGATTGACNNNNNCGNNNNNNNNNTANCCNNNNNNNNNGTACTCGTTGTCCCACGCNNNNN
TAAATCCCGANNNNNNNNNNNNNNNNNTACCTCCTTTCACATTTCGGTTAGGNAAACAANC
CATNATAAAAGTGCTGNTTGACTAGNNNNNTATTCCAATNATCCAGAAACCGCATCCTAA
AACTGTCGGNNNNNNTCNNNNNNNNNNNNCCNNCNGTGTTATTTCCTCAACGATCTNGCGA
ATGGAAGNNNNNNNAGTTGAACCCCTCGGTTCGNNNNNNNNNAGGCTTTATGAGNTGNNGN
GTTGGTCCAAACCNANNNNNNNNNNNCNNNTNTCNAGCCGAAATCTCGCAAAGTTNGNNNG
NNNCNNNCNTTGTGAATTCTNTNNCNNTTCNGGCTTTCATTAGGGCTCGTTCTGGTAGAT
TGACCTNCNNCGTATCTCATCATAACCATGTTTACACNNNNNNNNNTTTGAATTNCCCTCC
ATGCTAATCTGNNANNCCGNCTNTAGGAACTTANNNNNANCNNNNNNGGTCTNNNNNNTNN
NNNNNTTCGGCAGCACTTCGATACAAACTGTATAGTGCGTCAGGCCAGCGCGATGCCTCC
GTTNNNNNNNNTGAGAGGTCTNTATTGNNNGCNNGNTTGTAACACGAGATCGTCTGTTTCA
TACTAATTCTGCAATCCGCTTNNNNTTANTGCCCCGCTAACACCTTGTAGGCATGNCNNT
AGCAGATCCTGCGGCGGTCTCGANNTNNNNNNNNNTNCNCGATATGTGGNCTNNNNNNNNN
NNNAGATGGCTACGTGTCGCAAGTCGAGTNCGTACCCCGAAGCTGCCCTTGTCGACAGCC
TGTTCNNNTNGACTCGTNCANGACTCGGATAGTCNNNTNNNTGTCATACTTTNNNCNCAG
CGTTGCCATGGTGAATTCCTCCGTGATATGNCCANNTGNCNTACCCTCTTCCACCTCTTG
AAGGTACCATTCNGCCCATTCACATTTATAATATAATCCCTACTGAGTTTAAGGCTCCCC
NNNNNNNATACGTTCTCCTGCGTTGCGGGTAGATGGTCACAGATTTCGAGTTTTAAACTT
CAAGAAGGCTGCCCTCCCACCAGTTTCATCATTNNNNTNNATCNNGNNNNNNNNNNCNTC
GCCACTTGTGACGAGGGTTTGGGCGTGATTACGACCTNNNNNNNNNNNATCGTAGGCCCGT
GCNNATCCGACGCTGAGCCCTAAATAGTTGCATGGGCCNGCACNTTCCTAATNTNNNNNN
NNNNNNTAATANNNTNGTCCCCCAGNNNNNNNNNNCCCCCNATNGTCNNNCNNNNNNCTTN
NNTCTATACGTAANNNNNNGTNNNNNNGTTCATGTATAGAAGACGCCGGCGTTCCCCTAC
TTTTTCGCTTCTNATCTTNGTTACGGGCGCGCTGCGGCCCGGAACCAGTACTTAAGCTCC
CCATAAGGTGGGGCTACGCACGTGAAGAACTGCGTTGCGCTANCTATAGNGCNTNNNCCT
GGCCCTCCGAGAATC

>Pyr_2_3_7_8_9

GTGACCGTTGCCTCGCCACAAATCCATTGTTTTTCGGGTTCTGGTCTTTTATGAGTTAGG
ACCTCTGCGCCTACTATCATAGCTGTCCGATGCCTCGCATTCCATACGTTGCACTGACCC
GACAATAATCTGTGCTGCTAAACTTTTGCAAGTGGCTGGCGCCTGTAGGTCACGCAATAG
AGTAGCTTCCCTCCCGACGATCTTCAGGCCATCTACAATCCGGCATCGTAGATCCTCGCG
TGACCTAGTAGCCCTTGACGACTCAATCCACCCTCTGCCTGACGAACGTGACTTTTGTAG
TACTCTAGTGACTCCCTGTCCTCATAAGTAGGCTGCATACATAGGGCTACCGTAGTTTAC
TGGTCTCACCTCTACGTACTCGCTGACCCAACTACGTTGATCCAGGAAGGCAGAATGGTG
GACGCTCTTACTGCCGACCCGTGGGTAACTAACTGGGACGGGGGAATATTGTTTCGCTTT
GAGTACTACTTCTTCCAATTGTTCGGATAGCACAGTAATGATGGTGGGTGACTCCAGTCA
CTTTCGCGTAGTCGCCGGTCGGTCGCTCCGGTCTGTCCTCTCGCCCGATTCTGGGCCATC
TTATCCGACGTCATCCTGCAGACCGCGCTTTCCAACCGTTAGCATTAAGCAGTCTCCCAT
GGATGCTCTCATGTCATGAGAGGTACCGGAGCCTCTAAGCGTTCCGCTGTTCTCGACAGT
ATTCTCTTTCGTTTTAACACTAGCTCAGACGGTAGAGGCTTCGTCTTCAGTATCGGCTAA
TGATGCCGGGTACGTCAAGATACCAGTCTGCGCGCTTACGCTGCTCCATCTGTTGATGTC
TGCTTCAGCTCGGCTAAGACCTCTCGAACTCTTTGTCTTACCTCAGCCCGCGGCTAGAAT
CATTTACATCCTGGCAAACACAGTCCTTAATTTCCAGTCGTGCCCGACAGTAATACCGTG
GCGCGAACGACGTCTAAGTTGTTGCCTACTTCGTACACTATGGACTACATTTGCCCAACG
TGAGGTCCGAGGATCCCGTTGGAGGGCACACACTCAGAGAATCCGCAGCGTTTCGTCTCA
CAATCGAGAAAGGCGCTCCTAAGGGGCCGCGATACGAGCGTGAAACCGGCCAAGCGCTCT
AATTGTACACTGTTCATGAGGCTAAATTCAAATTGCGGACCGAAAACTTGGCCCAGCAGA
GCTAAAGGTTCTTCGCGGGTATTGTCCGTACATGCAACTTGATGTTTTCTAAGCCAAGAG
AGATATGCGAGGGGGAGACATGTGAAAAGCAAAAAAATATAGTACTATAATTATCGTGGG
ACCCATAAGGCCTTGGTTGCTCCCGCGTCGCCTGTACCCACAGTCTTCAAAAAGGCGATT
TGCCTGAGTATAAAGTAAAGAGCCGACGGTGTTGAAAACGAGCGGACGTCTTGCATGAGA
AGTGTCACTGTGGGCATCTCAAAGGACTCTAATTAAATTCGGTGTCAATAGAAAGGTTCG
CGCTGCATGGCGCTGTCAGATCATCCATCCTCTGTGGCTACGCCTTATCATAAGTGAGAC
CTGCGAAATCAGGGTGATGTCAATACGCTTTAAAGGACGCTAAAGCGAGTTGATAATTAA
ACGACAGAGTTACAATGAGAAGTATTCACAACTCGCCAAAATCATTGACGATAGACTGAC
ACGATAGCGTGGGAGGCCCCGTAGCCTCAAGTGAAAAGGGGCGCAAAACGGTGTCCGCAT
GTGAGGAGCTCGACCATCATAGATGCCGGATATGTTCCTGTGAAATTTCAGCGTCAAAGA
TGGGAAGCAAAACAAACCCATGAAACTACCATAGTGTGAAGGTGGTGGGCGGCGCGGTTT
AAGGCGAGGCGCTCTAAGACTCCAACGTGCCCTAGGGGATCGTAACATACAAGAACATTG
ACGTGCACAGTTTTGTTGAACGGCCTTCATGGGCCAAAGGTTGTATCCACGGCCAGGAGC
GGTAAGAACTTCGTGACATAGCGGCCGGAAAAAGGATCTACAACGTTAACCAGAACCTCG
GAGAGCGGAACGGCCAGTCGTTAGGGAGAAAATAGTTAAGCATTAATTCTGACGAACCAC

CAAAATGCTCGAGCGGTAGGTATCATTAGGGGGTACGGACTGTTACATTTAGAATGGCCC
AGCTCCATAGTTGCAGACACTAATTTTAGAGACCGCTGCTGCATCATTAACAAAATGTTA
TCGGTGGGATGCAGAAGAGCGCGGGCAAGATTACACAAACGCCTTCAGGTGGTAACCTGC
TCCTAAGCAACCAAGACGGATCGCGACGGACGTGTCTGGACGGCGAATGTACAGGAAGTT
TTCCGTGAGAACAACAAAATGAAGGATCGAGATTAGAAAGACTACAGCGCCCATGAGCAA
CGGAGTTGAACGTGATTGTCAGGTTGGTAATGGGCGGCCCTAGAAGATAACGAGAAAAAC
CCTTCGGACTTTTCAATGGGGACTGCATTCCGGCCGCGAAGGACTCCATGAATCGATAGG
TAGGGGAAATATGGATGAAACGGGCTGGGGCAACGCCCGAGGCGACGTCTACTAGTTAAA
AGGCAGGTTGCGAGGATCGGGGAAGACTGAGAAGGCCTTGGCGTAGGAATTCGAGGCAAC
CAGGAGGATAAGGTAAATGCCGACTTCAAGCCGATAATGCAAAGAAGCCGGCTATGGCCG
TAAATTAGACGATAAGTAACAGATCGTCGGGGGGCCAAGTCTAATCGTAAAACTTTTAGT
TGAGGTGATGTCGTAAGTGAGATCCCCGAACGCAGCATGAACAAGGTGAATCGTGTAAGC
AGAGAAGAGCGCTGCACGATGCTAGCACTGTCGTAGTAGTGGGACAACTTGGATGACTGG
CACATCGTAGTAATGAGTCATATGATCCCTCCTGAAGGCTTCAAAAAAGGACACAAGACG
CGAAGAAGATCTAGACTCAGTCTTTAAAACTGCACAACTTAGCCGAAGTGATTGGTAAGT
AGAAGTACTGATGTGGGCGCTGAGAGAAACCTGAAAACGCGATAACCATCATATCACGAT
GCTTCCGCCTTCCAGCGTCACTAAAGTTCTTAATCACATGCCTCGGTTCTCTCGCCAGTA
CACGGTCGGCCTTCTTTCCCCCATCGTTTACCGGAGCATATTCTTCGAGTTGCCTTATTA
ACCTTCTATTCTCTCGCCGGTGCCTGATTAGCGCCGCCTTGAGTATCATACTGTTTGCTT
TTAGCCAACCTGTTGAGCGTTGATTGCTTGTGTGACCATATCTTGCGTTTCGTTGATTCC
TGGTCGGCAATGTGTTGTGGCCTGTTCGTTGACGCTCCTGATCTCACCGTACTTCACGGT
TGCCGCCCGGACATCACCCAGATGTGTTTAGGCCATGGCCGGTTAAGGTTAATCTTTACG
CGTCATCATAACTTCCTGCGAAACTGGTTCCCCCACGCCGTCCCCTTCGTTAGTCGATCG
GCGTGGTTAGGTCGCGCGGGTTTTTCTAGGCAAGGTAAGTCGGGTCGCACAGTGCGACCC
ACGGACATAACCTTAACGTTCCGTTTTACCACTGGGTCAGGCTGCTTTGGGTCCGACGTG
ACGCTCTTTCCGCACGACCTCGTATGTTCTTATTCGCACTCGTAATGAACCGGTAAGCGA
GAATTCGCTCTACGCTGAGGTGTCGTCGGACCCGAGTGTGCTTGCCGCTGGCCAATTGTT
GTGTCGGCTCGACGTCTTATTTCGTATGGTCCGGGATTACCCGTGCTTGCAGCGATGTCC
TTTTCGCTATTGATGTCTTCTCCTGAGTCACGCTACATACTGAATTAATAAAGCATTTCG
ACCGCAGCATCAGTGAAGTCGAGGGATGGGCTACATCGCCGTGCGGAGCTTTTCACGTTG
TGTCCGTGCGGAGATTCAGCACTAGCGGTCATACTATCCATGTTCCCCCGTTCTATTTAC
GTTGAGCGACTTGAAACTTCCAGTGCACTGTTTCTGCATAGTGCGGTATTTCTCAGTTCA
TGCAAGTTATCGTACATCATGGAATAGAACAAGCATCAAGGACCATTTCCACTAACGATT
CTTACCTCCAAATCAAGATAGAATCTAACCGATTATCGACGAACTTCATCCGTCATGTCA
CTCTATGTCAAGCGGCCTTCTTACAGCGCTGTCGCACTATCCGTTCTACCTTGTTTCCGA
GAAACCTTGCCTCTGTTGCCTGTGGGGGCAGGGCGGATATGGCTAAATGGGCATTCCTTT

TTACTTCCCGTGCCCCTTCATGCTATGTGCTTGCAGCTCTAAAGCTTAGACGCCCGGTCC
TCGACGTCTCGAAACTTTCCTGACCCTGCTGGCGTCTTACCCTTTATAGTGGATCTGCAT
AATGTGTTCTGTATATAGCCAAATACTGCCAACTATCCACTCGCTTAGCTGTCAATATCC
ATGAGCGTGCTTTGATTTACCGGTCGACCTGACTAATGTAACTGCCTTGCGCTCTCCACT
CCGCACCAGGGTTTCGAGTTTTTCTACCCCTCTCTTTATTACTACTCGTTTACTGCATCA
GACGCGCCCATCGTACTTGGTCACTGCTCCTTCAGGCGAGGAGTATTGGCACGTTATCGT
GCTTCTATCCTGAGCTGCAACCATAAGCATTTCTAAAGTTATGAGGATCAAAGAGTATTT
CCGTACATCTTGATCGATTGTACTGGACGTCGTCCCAAATAGCCTGCGAAGTCCAATATC
TAGGTCATAGATGCT
>Pyr_23
GTGACCGTTGCCTCGCCACAAATCCATTGTTTTTCGGGTTCTGGTCTTTTATGAGTTAGG
ACCTCTGCGCCTACTATCATAGCTGTCCGATGCCTCGCATTCCATACGTTGCACTGACCC
GACAATAATCTGTGCTGCTAAACTTTTGCAAGTGGCTGGCGCCTGTAGGTCACGCAATAG
AGTAGCTTCCCTCCCGACGATCTTCAGGCCATCTACAATCCGGCATCGTAGATCCTCGCG
TGACCTAGTAGCCTTTGACGACTCAATCCACCCTCTGCCTGACGAACGTGACTTTTGTAG
TACTCTAGTGACTCCCTGTCCTCATAAGTAGGCTGCATACATAGGGCTACCGTAGTTTAC
TGGTCTCACCTCTACGTACTCGCTGACCCAACTACGTTGATCCAGGAAGGCAGAATGGTG
GACGCTCTTACTGCCGACCCGTGGGTAACTAACTGGGACGGGGGAATATTGTTTCGCTTT
GAGTACTACTTCTTCCAATTGTTCGGATAGCACAGTAATGATGGTGGGTGACTCCAGTCA
CTTTCGCGTAGTCGCCGGTCGGTCGCTCCGGTCTGTCCTCTCGCCCGATTCTGGGCCATC
TTATCCGACGTCATCCTGCAGACCGCGCTTTCCAACCGTTAGCATTAAGCAGTCTCCCAT
GGATGCTCTCATGTCATGAGAGGTACCGGAGCCTCTAAGCGTTCCGCTGTTCTCGACAGT
ATTCTCTTTCGTTTTAACACTAGCTCAGACGGTAGAGGCTTCGTCTTCAGTATCGGCTAA
TGATGCCGGGTACGTCAAGATACCAGTCTGCGCGCTTACGCTGCTCCATCTGTTGATGTC
TGCTTCAGCTCGGCTAAGACCTCTCGAACTCTTTGTCTTACCTCAGCCCGCGGCTAGAAT
CATTTACATCCTGGCAAACACAGTCCTTAATTTCCAGTCGTGCCCGACAGTAATACCGTG
GCGCGAACGACGTCTAAGTTGTTGCCTACTTCGTACACTATGGACTACATTTGCCCAACG
TGAGGTCCGAGGATCCCGTTGGAGGGCACACACTCAGAGAATCCGCAGCGTTTCGTCTCA
CAATCGAGAAAGGCGCTCCTAAGGGGCCGCGATACGAGCGTGAAACCGGCCAAGCGCTCT
AATTGTACACTGTTCATGAGGCTAAATTCAAATTGCGGACCGAAAACTTGGCCCAGCAGA
GCTAAAGGTTCTTCGCGGGTATTGTCCGTACATGCAACTTGATGTTTTCTAAGCCAAGAG
AGATATGCGAGGGGGAGACATGTGAAAAGCAAAAAAATATAGTACTATAATTATCGTGGG
ACCCATAAGGCCTTGGTTGCTCCCGCGTCGCCTGTACCCACAGTCTTCAAAAAGGCGATT
TGCCTGAGTATAAAGTAAAGAGCCGACGGTGTTGAAAACGAGCGGACGTCTTGCATGAGA
AGTGTCACTGTGGGCATCTCAAAGGACTCTAATTAAATTCGGTGTCAATAGAAATGTTCG
CGCTGCATGGCGCTGTCAGATCATCCATCCTCTGTGGCTACGCCTTATCATAAGTGAGAC

CTGCGAAATCAGGGTGATGTCAATACGCTTTAAAGGACGCTAAAGCGAGTTGATAATTAA
ACGACAGAGTTACAATGAGAAGTATTCACAACTCGCCAAAATCATTGACGATAGACTGAC
ACGATAGCGTGGGAGGCCCCGTAGCCTCAAGTGAAAAGGGGCGCAAAACGGTGTCCGCAT
GTGAGGAGCTCGACCATCATAGATGCCGGATATGTTCCTGTGAAATTTCAGCGTCAAAGA
TGGGAAGCAAAACAAACCCATGAAACTACCATAGTGTGAAGGTGGTGGGCGGCGCGGTTT
AAGGCGAGGCGCTCTAAGACTCCAACGTGCCCTAGGGGATCGTAACATACAAGAACATTG
ACGTGCACAGTTTTGTTGAACGGCCTTCATGGGCCAAAGGTTGTATCCACGGCCAGGAGC
GGTAAGAACTTCGTGACATAGCGGCCGGAAAAAGGATCTACAACGTTAACCAGAACCTCG
GAGAGCGGAACGGCCAGTCGTTAGGGAGAAAATAGTTAAGCATTAATTCTGACGAACCAC
CAAAATGCTCGAGCGGTAGGTATCATTAGGGGGTACGGACTGTTACATTTAGAATGGCCC
AGCTCCATAGTTGCAGACACTAATTTTAGAGACCGCTGCTGCATCATTAACAAAATGTTA
TCGGTGGGATGCAGAAGAGCGCGGGCAAGATTACACAAACGCCTTCAGGTGGTAACCTGC
TCCTAAGCAACCAAGACGGATCGCGACGGACGTGTCTGGACGGCGAATGTACAGGAAGTT
TTCCGTGAGAACAACAAAATGAAGGATCGAGATTAGAAAGACTACAGCGCCCATGAGCAA
CGGAGTTGAACGTGATTGTCAGGTTGGTAATGGGCGGCCCTAGAAGATAACGAGAAAAAC
CCTTCGGACTTTTCAATGGGGACTGCATTCCGGCCGCGAAGGACTCCATGAATCGATAGG
TAGGGGAAATATGGATGAAACGGGCTGGGGCAACGCCCGAGGCGACGTCTACTAGTTAAA
AGGCAGGTTGCGAGGATCGGGGAAGACTGAGAAGGCCTTGGCGTAGGAATTCGAGGCAAC
CAGGAGGATAAGGTAAATGCCGACTTCAAGCCGATAATGCAAAGAAGCCGGCTATGGCCG
TAAATTAGACGATAAGTAACAGATCGTCGGGGGGCCAAGTCTAATCGTAAAACTTTTAGT
TGAGGTGATGTCGTAAGTGAGATCCCCGAACGCAGCATGAACAAGGTGAATCGTGTAAGC
AGAGAAGAGCGCTGCACGATGCTAGCACTGTCGTAGTAGTGGGACAACTTGGATGACTGG
CACATCGTAGTAATGAGTCATATGATCCCTCCTGAAGGCTTCAAAAAAGGACACAAGACG
CGAAGAAGATCTAGACTCAGTCTTTAAAACTGCACAACTTAGCCGAAGTGATTGGTAAGT
AGAAGTACTGATGTGGGCGCTGAGAGAAACCTGAAAACGCGATAACCATCATATCACGAT
GCTTCCGCCTTCCAGCGTCACTAAAGTTCTTAATCACATGCCTCGGTTCTCTCGCCAGTA
CACGGTCGGCCTTCTTTCCCCCATCGTTTACCGGAGCATATTCTTCGAGTTGCCTTATTA
ACCTTCTATTCTCTCGCCGGTGCCTGATTAGCGCCGCCTTGAGTATCATACTGTTTGCTT
TTAGCCAACCTGTTGAGCGTTGATTGCTTGTGTGACCATATCTTGCGTTTCGTTGATTCC
TGGTCGGCAATGTGTTGTGGCCTGTTCGTTGACGCTCCTGATCTCACCGTACTTCACGGT
TGCCGCCCGGACATCACCCAGATGTGTTTAGGCCATGGCCGGTTAAGGTTAATCTTTACG
CGTCATCATAACTTCCTGCGAAACTGGTTCCCCCACGCCGTCCCCTTCGTTAGTCGATCG
GCGTGGTTAGGTCGCGCGGGTTTTTCTAGGCAAGGTAAGTCGGGTCGCACAGTGCGACCC
ACGGACATAACCTTAACGTTCCGTTTTACCACTGGGTCAGGCTGCTTTGGGTCCGACGTG
ACGCTCTTTCCGCACGACCTCGTATGTTCTTATTCGCACTCGTAATGAACCGGTAAGCGA
GAATTCGCTCTACGCTGAGGTGTCGTCGGACCCGAGTGTGCTTGCCGCTGGCCAATTGTT

GTGTCGGCTCGACGTCTTATTTCGTATGGTCCGGGATTACCCGTGCTTGCAGCGATGTCC
TTTTCGCTATTGATGTCTTCTCCTGAGTCACGCTACATACTGAATTAATAAAGCATTTCG
ACCGCAGCATCAGTGAAGTCGAGGGATGGGCTACATCGCCGTGCGGAGCTTTTCACGTTG
TGTCCGTGCGGAGATTCAGCACTAGCGGTCATACTATCCATGTTCCCCCGTTCTATTTAC
GTTGAGCGACTTGAAACTTCCAGTGCACTGTTTCTGCATAGTGCGGTATTTCTCAGTTCA
TGCAAGTTATCGTACATCATGGAATAGAACAAGCATCAAGGACCATTTCCACTAACGATT
CTTACCTCCAAATCAAGATAGAATCTAACCGATTATCGACGAACTTCATCCGTCATGTCA
CTCTATGTCAAGCGGCCTTCTTACAGCGCTGTCGCACTATCCGTTCTACCTTGTTTCCGA
GAAACCTTGCCTCTGTTGCCTGTGGGGGCAGGGCGGATATGGCTAAATGGGCATTCCTTT
TTACTTCCCGTGCCCCTTCATGCTATGTGCTTGCAGCTCTAAAGCTTAGACGCCCGGTCC
TCGACGTCTCGAAACTTTCCTGACCCTGCTGGCGTCTTACCCTTTATAGTGGATCTGCAT
AATGTGTTCTGTATATAGCCAAATACTGCCAACTATCCACTCGCTTAGCTGTCAATATCC
ATGAGCGTGCTTTGATTTACCGGTCGACCTGACTAATGTAACTGCCTTGCGCTCTCCACT
CCGCACCAGGGTTTCGAGTTTTTCTACCCCTCTCTTTATTACTACTCGTTTACTGCATCA
GACGCGCCCATCGTACTTGGTCACTGCTCCTTCAGGCGAGGAGTATTGGCACGTTATCGT
GCTTCTATCCTGAGCTGCAACCATAAGCATTTCTAAAGTTATGAGGATCAAAGAGTATTT
CCGTACATCTTGATCGATTGTACTGGACGTCGTCCCAAATAGCCTGCGAAGTCCAATATC
TAGGTCATAGATGCT
>Pyr_25_39
GTGACCGTTGCCTCGCCACAAATCCATTGTTTTTCGGGTTCTGGTCTTTTATGAGTTAGG
ACCTCTGCGCCTACTATCATAGCTGTCCGATGCCTCGCATTCCATACGTTGCACTGACCC
GACAATAATCTGTGCTGCTAAACTTTTGCAAGTGGCTGGCGCCTGTAGGTCACGCAATAG
AGTAGCTTCCCTCCCGACGATCTTCAGGCCATCTACAATCCGGCATCGTAGATCCTCGCG
TGACCTAGTAGCCCTTGACGACTCAATCCACCCTCTGCCTGACGAACGTGACTTTTGTAG
TACTCTAGTGACTCCCTGTCCTCATAAGTAGGCTGCATACATAGGGCTACCGTAGTTTAC
TGGTCTCACCTCTACGTACTCGCTGACCCAACTACGTTGATCCAGGAAGGCAGAATGGTG
GACGCTCTTACTGCCGACCCGTGGGTAACTAACTGGGACGGGGGAATATTGTTTCGCTTT
GAGTACTACTTCTTCCAATTGTTCGGATAGCACAGTAATGATGGTGGGTGACTCCAGTCA
CTTTCGCGTAGTCGCCGGTCGGTCGCTCCGGTCTGTCCTCTCGCCCGATTCTGGGCCATC
TTATCCGACGTCATCCTGCAGACCGCGCTTTCCAACCGTTAGCATTAAGCAGTCTCCCAT
GGATGCTCTCATGTCATGAGAGGTACCGGAGCCTCTAAGCGTTCCGCTGTTCTCGACAGT
ATTCTCTTTCGTTTTAACACTAGCTCAGACGGTAGAGGCTTCGTCTTCAGTATCGGCTAA
TGATGCCGGGTACGTCAAGATACCAGTCTGCGCGCTTACGCTGCTCCATCTGTTGATGTC
TGCTTCAGCTCGGCTAAGACCTCTCGAACTCTTTGTCTTACCTCAGCCCGCGGCTAGAAT
CATTTACATCCTGGCAAACACAGTCCTTAATTTCCAGTCGTGCCCGACAGTAATACCGTG
GCGCGAACGACGTCTAAGTTGTTGCCTACTTCGTACACTATGGACTACATTTGCCCAACG

TGAGGTCCGAGGATCCCGTTGGAGGGCACACACTCAGAGAATCCGCAGCGTTTCGTCTCA
CAATCGAGAAAGGCGCTCCTAAGGGGCCGCGATACGAGCGTGAAACCGGCCAAGCGCTCT
AATTGTACACTGTTCATGAGGCTAAATTCAAATTGCGGACCGAAAACTTGGCCCAGCAGA
GCTAAAGGTTCTTCGCGGGTATTGTCCGTACATGCAACTTGATGTTTTCTAAGCCAAGAG
AGATATGCGAGGGGGAGACATGTGAAAAGCAAAAAAATATAGTACTATAATTATCGTGGG
ACCCATAAGGCCTTGGTTGCTCCCGCGTCGCCTGTACCCACAGTCTTCAAAAAGGCGATT
TGCCTGAGTATAAAGTAAAGAGCCGACGGTGTTGAAAACGAGCGGACGTCTTGCATGAGA
AGTGTCACTGTGGGCATCTCAAAGGACTCTAATTAAATTCGGTGTCAATAGAAATGTTCG
CGCTGCATGGCGCTGTCAGATCATCCATCCTCTGTGGCTACGCCTTATCATAAGTGAGAC
CTGCGAAATCAGGGTGATGTCAATACGCTTTAAAGGACGCTAAAGCGAGTTGATAATTAA
ACGACAGAGTTACAATGAGAAGTATTCACAACTCGCCAAAATCATTGACGATAGACTGAC
ACGATAGCGTGGGAGGCCCCGTAGCCTCAAGTGAAAAGGGGCGCAAAACGGTGTCCGCAT
GTGAGGAGCTCGACCATCATAGATGCCGGATATGTTCCTGTGAAATTTCAGCGTCAAAGA
TGGGAAGCAAAACAAACCCATGAAACTACCATAGTGTGAAGGTGGTGGGCGGCGCGGTTT
AAGGCGAGGCGCTCTAAGACTCCAACGTGCCCTAGGGGATCGTAACATACAAGAACATTG
ACGTGCACAGTTTTGTTGAACGGCCTTCATGGGCCAAAGGTTGTATCCACGGCCAGGAGC
GGTAAGAACTTCGTGACATAGCGGCCGGAAAAAGGATCTACAACGTTAACCAGAACCTCG
GAGAGCGGAACGGCCAGTCGTTAGGGAGAAAATAGTTAAGCATTAATTCTGACGAACCAC
CAAAATGCTCGAGCGGTAGGTATCATTAGGGGGTACGGACTGTTACATTTAGAATGGCCC
AGCTCCATAGTTGCAGACACTAATTTTAGAGACCGCTGCTGCATCATTAACAAAATGTTA
TCGGTGGGATGCAGAAGAGCGCGAGCAAGATTACACAAACGCCTTCAGGTGGTAACCTGC
TCCTAAGCAACCAAGACGGATCGCGACGGACGTGTCTGGACGGCGAATGTACAGGAAGTT
TTCCGTGAGAACAACAAAATGAAGGATCGAGATTAGAAAGACTACAGCGCCCATGAGCAA
CGGAGTTGAACGTGATTGTCAGGTTGGTAATGGGCGGCCCTAGAAGATAACGAGAAAAAC
CCTTCGGACTTTTCAATGGGGACTGCATTCCGGCCGCGAAGGACTCCATGAATCGATAGG
TAGGGGAAATATGGATGAAACGGGCTGGGGCAACGCCCGAGGCGACGTCTACTAGTTAAA
AGGCAGGTTGCGAGGATCGGGGAAGACTGAGAAGGCCTTGGCGTAGGAATTCGAGGCAAC
CAGGAGGATAAGGTAAATGCCGACTTCAAGCCGATAATGCAAAGAAGCCGGCTATGGCCG
TAAATTAGACGATAAGTAACAGATCGTCGGGGGGCCAAGTCTAATCGTAAAACTTTTAGT
TGAGGTGATGTCGTAAGTGAGATCCCCGAACGCAGCATGAACAAGGTGAATCGTGTAAGC
AGAGAAGAGCGCTGCACGATGCTAGCACTGTCGTAGTAGTGGGACAACTTGGATGACTGG
CACATCGTAGTAATGAGTCATATGATCCCTCCTGAAGGCTTCAAAAAAGGACACAAGACG
CGAAGAAGATCTAGACTCAGTCTTTAAAACTGCACAACTTAGCCGAAGTGATTGGTAAGT
AGAAGTACTGATGTGGGCGCTGAGAGAAACCTGAAAACGCGATAACCATCATATCACGAT
GCTTCCGCCTTCCAGCGTCACTAAAGTTCTTAATCACATGCCTCGGTTCTCTCGCCAGTA
CACGGTCGGCCTTCTTTCCCCCATCGTTTACCGGAGCATATTCTTCGAGTTGCCTTATTA

ACCTTCTATTCTCTCGCCGGTGCCTGATTAGCGCCGCCTTGAGTATCATACTGTTTGCTT
TTAGCCAACCTGTTGAGCGTTGATTGCTTGTGTGACCATATCTTGCGTTTCGTTGATTCC
TGGTCGGCAATGTGTTGTGGCCTGTTCGTTGACGCTCCTGATCTCACCGTACTTCACGGT
TGCCGCCCGGACATCACCCAGATGTGTTTAGGCCATGGCCGGTTAAGGTTAATCTTTACG
CGTCATCATAACTTCCTGCGAAACTGGTTCCCCCACGCCGTCCCCTTCGTTAGTCGATCG
GCGTGGTTAGGTCGCGCGGGTTTTTCTAGGCAAGGTAAGTCGGGTCGCACAGTGCGACCC
ACGGACATAACCTTAACGTTCCGTTTTACCACTGGGTCAGGCTGCTTTGGGTCCGACGTG
ACGCTCTTTCCGCACGACCTCGTATGTTCTTATTCGCACTCGTAATGAACCGGTAAGCGA
GAATTCGCTCTACGCTGAGGTGTCGTCGGACCCGAGTGTGCTTGCCGCTGGCCAATTGTT
GTGTCGGCTCGACGTCTTATTTCGTATGGTCCGGGATTACCCGTGCTTGCAGCGATGTCC
TTTTCGCTATTGATGTCTTCTCCTGAGTCACGCTACATACTGAATTAATAAAGCATTTCG
ACCGCAGCATCAGTGAAGTCGAGGGATGGGCTACATCGCCGTGCGGAGCTTTTCACGTTG
TGTCCGTGCGGAGATTCAGCACTAGCGGTCATACTATCCATGTTCCCCCGTTCTATTTAC
GTTGAGCGACTTGAAACTTCCAGTGCACTGTTTCTGCATAGTGCGGTATTTCTCAGTTCA
TGCAAGTTATCGTACATCATGGAATAGAACAAGCATCAAGGACCATTTCCACTAACGATT
CTTACCTCCAAATCAAGATAGAATCTAACCGATTATCGACGAACTTCATCCGTCATGTCA
CTCTATGTCAAGCGGCCTTCTTACAGCGCTGTCGCACTATCCGTTCTACCTTGTTTCCGA
GAAACCTTGCCTCTGTTGCCTGTGGGGGCAGGGCGGATATGGCTAAATGGGCATTCCTTT
TTACTTCCCGTGCCCCTTCATGCTATGTGCTTGCAGCTCTAAAGCTTAGACGCCCGGTCC
TCGACGTCTCGAAACTTTCCTGACCCTGCTGGCGTCTTACCCTTTATAGTGGATCTGCAT
AATGTGTTCTGTATATAGCCAAATACTGCCAACTATCCACTCGCTTAGCTGTCAATATCC
ATGAGCGTGCTTTGATTTACCGGTCGACCTGACTAATGTAACTGCCTTGCGCTCTCCACT
CCGCACCAGGGTTTCGAGTTTTTCTACCCCTCTCTTTATTACTACTCGTTTACTGCATCA
GACGCGCCCATCGTACTTGGTCACTGCTCCTTCAGGCGAGGAGTATTGGCACGTTATCGT
GCTTCTATCCTGAGCTGCAACCATAAGCATTTCTAAAGTTATGAGGATCAAAGAGTATTT
CCGTACATCTTGATCGATTGTACTGGACGTCGTCCCAAATAGCCTGCGAAGTCCAATATC
TAGGTCATAGATGCT
>Pyr_27
CGTCATACCAAACTTTTGTGGCCAATCCGTCCCTTAAGGGTCGATTCCCAGCTGACGAAA
GTTGNCACGTCNNNTNNNGCAATCCCTTAGCNGTCTACCCCGCNNNNNNNNNACTNNNNN
NNNNNNNAGACACATTCCCAGGTCTCTCCCGTGGAACTTTCTATTCCAAGTGTATCTCNN
NNGGAACCTTTCTTTACTAGCTCGCGTTTTGCTCCTCGCNCGGNNNNNCGACCTTCGGTN
CATTTCGGCCNATCGGAGTAGNNNTGCNNATATCGCATTTAGTATGTCCTATCCCGGGGA
CTGCTCCGGGGTCTTNCACATCAGCGCGNTTGTGACGCNNNNNNGNNNNTNANNACCCGT
GCTCTCTGTGTTTAAATGTGTAGCTCTTTGACCGCGTCGACTTAGAATAATATCGGAAAA
AGTGATTGAACCGTTANNTTTNATTTCGGCCGTCCNNANGNNNNGCCGGTAGCGGACCCC

GAACTTCGTCGTCCTTGGGCNCNNNNNNNNNACCANNNTNTAAACAGAGTGCNNNCNCTC
CCTNTGTNNNNNNGATAACTAAGCTTTTTAGGACTTTTTTCTGNNNNNNNCTNNNNCGTT
CCACAATGTTCTACTTCATTTCNNNNTGNNNCCNNNNNNNNNNNNCGNNNNNNCNNTNNC
GCACAACTNNATANTGNACTGTNNNNNNNNNANGCCGAATAGCTTATCAGCTCCGNNNGC
NNNNNAACCTATGCTAGTGTGGTANNNNNNGNCCNGAATCCTGTTNNNNNTCTTATTACG
CGTNATTTANCGTTATTGAAAGTTAGCCTATATTTCCGTAAGTTGAAGTTGACCAGCTCC
NNNNNCNATGTAAACNCAGTTCTCCANNNCNCNNACAGCGATGTGATTTATTAACGGTAC
TATACGCGCTTCAATGAGAGTGGGTTCGAGCCTCNGANTANTNNNNNNANNNTCGCNNNN
NNNNNNNCGNGANCNNNNNNGNNNNNNNNNNNNNNNNNNNCCAGTNNANNNTNCTNNNNN
TAGAGNNNGANNNNNNNNTTACGNNGGACGTGTCCCGGAGGTAGATGATGCCCCGCTCTG
TGGCTAAAAGAAAGATCCATCGGGATTGTCNNNNNNNNNNNNNNCAAACCAGGAATTCTC
TCCCACCTTTCAATTGCANNATCNGTGCTGCTCCAAANNNNNNANNTCTAACTTGGATAG
AACAGGAACCACGAATAAACGCGACATACCTGCAAGTTCGAGCAGCATANNGNTNGTGGA
GTGNGCGTTCAAAGAGAGTGCATAGCNGAAGACAAGGTNGGGCCTTGCGCCTAATACCAG
TTTTGGGGACCTCCTAGTTTCGTTACATTNTCGACATGTATGTCNCCTGATGGAATGGGC
CAGGGNGACCCNGCACCGGTAATTAGAAACACCCGGGGTGGATAGCTGGTCCATCCACAG
NGCACTGTGAGAACTGCACAGGGACGTCTCTGCCGGGTCTGACAGTAGCGAAGCTACATA
TATGATGCCATACGATTGAGTAACTTGTACCCCGCAATAGNNNNNNNNNATGGGTACGGT
TGTTAGAGCTGATACANTNNTNCNNNNNNTNCANNNGTATCGGGACGANACCGCGGGCGG
GTNNNNACGCNGTGGGGGACGATGTCCCTNCNNATTTGNCCGCGNNNGTGGGGACGGAGC
GTAACAATGCAAANNNNNCTCGCATTCTAGACGGGGGGAAATTTGCGGCGAGACGTNATC
ATAGAAGTACCGGCNGTAACGGGTACTAATCCGAGCTCTTCACACAGCTGACACTAGGGG
GACGGGTTGAAATCGGCCTGGAGGGTAGAGCCGATACAATGACAANNAAATATACCACGC
GTAGTACNNNNNCNNCCAATTTTGGGGCATTTCGAAGAGCGTCNGTGGATGGAGGTCACA
GTAGTTTNGTGCCCCCCAGGNNNTNCCTGCAAANNNNNNGNCNCGCATCTACGNNNGNNN
NNNGNAGGTGGTACAGTGCGATAATAGAGANNGNNGNTCCTACTACCGGGTGACGTGATA
AAGGTTATANCNNCNCNGTANNNNNANANGTGCGATCGNTNNNNCCCCTTAGTAGTTTTT
TGTGCCTACTAGNNNNCGATCGCTGCGGAAAATGTTAAGNCTCCGCCCCCGAGCCGGAAT
GGTCCTACGACGTTGAGTATCCGGTCCCGGAATAATACTCNCNNNNNNNNNANTGCTNCG
CTNNNNNNGNTNNNNNAGTCATTGATGCAGACGCGTGANANTTCANGAACAAAGGTTCAT
TTACTGATAGTTGGAGTAACCTATAGTAAGTAAAGTCAGCAAANNGGCACGTGAAGAACC
CCTTACTNAGTTGGTGCAGCAGGAAGCAACAGCTGGGGGAGTCATGATATACAATGAAAG
TTAGACAACGATCAGCGGCGCGAGCTACGGCTAATGATATGGAGGACGCATAAGTAAGAG
TGCCTAAGTGCCCCGGTAACAAAAGTGTCTTNNGAATAGGAAGTCNTGATGCCTANCGAA
ACTTAAGGGCCAAAGCAGGTTAAATCAAAATGGCTTTCAGGGTGGCACTGGTCGATCGCG
AAATCTACCATAGTAGCCTAATGGAGACGGAGTAAANNCANAAAGAAGGCGTGGGTTGGA

TGTAAAAACGAATCGCGCGTTAGTCTGGGATTAGTGGCACGACACTATTAATCGCAATTN
GGTNNNNNNNGNNNNACANACCCGTACTANNNNNNNNNNCTCAGCTACAGGGGCCCCGNN
CNNANCTGCCCAGCNANCATAGCTCATAGGAGATCGACNNANNNNNNNCNNNNNNNNNAN
CAAAGCACATATAACGTAGCANCNGNNTNACNAGGATGACAAACTGGGGACAGCACGCAA
TGTGCTACGAAGGCAGACAGCGCAGCCTTTATAAGNCANNNNNNNNNGACGACTCGTATA
TAGGAGGATAACGAGTTTGACTCNGGCCATTNNGNNNNANGNNNNNGANNNNNNNNNNNN
NNNNNCNTTTGTTCAAAGGTCGGAGAGCTTANNNNANNNNNNTNACCANNATNCNCNAGC
ATCCTCATCTNCCCATTCTTTCGAGACCNNNNANNGTCNCTACTTTCTTGACTAGACACG
TANNNNNNGATCCTCCCTTTNTGCTTCCCTTTAACATCCGCGTTCTATACCATTTCGCAG
ATCCCTCTTACGTCGACGNANNNNNNANNNNTANNNNTCCNGTGCCTGGGTGACCCACCC
CCGATTGACNNNNNCGNNNNNNNNTANNNNNNNANNNGTACTCGTTGTCCCACTCCCGTT
TAAATCCCGAGNNNNNCNGAANCAGGTACCTCCTTTCACATTTCGGTTAGGNAAAAAANC
CATTATAAAAGTGCTGTTTGACTAGNNNNNTATTCCAACTATCCAGAAACCGCATCCTNA
AACTGTTGTNTTNCATCNNNNNNNNNNNNNCATCNGTGTTATTTCCTCAACGATCTAGCGA
ATGGAAGNNNANNNAGTTGAACCCCTCGGTTNGNNNGNNNTAGGCTTTATGAGCTGGTGC
GTNGGTCCAANCNNANNNNNNNNNNNCNNNTGTCTAGCCGAAATCTCGCAAAGTTAGNNNG
NNNCNNNNNNTTGTGAATTCTNTNNNNNNTTCGGGCTTTCATTAGGGCTCGTTCTGGTAGAT
TGACCTACAACGNATCNCATCATAACCATGTTCAAACNNNTCCNNTTTGAATTGCCCTCC
ATGCTAATCTGNNNNNCCGGCTTTAGGAACTTANNNNNANCNNNNCGGGTCTTNGCNTNN
CNGNTTTCGGCAGCACTTCGATACAAACTGTATAGTGCGTCAGGCCAGCGCGATGCCTCC
GTTNNNNNGNTGAGAGGTCTTTATTGNNNNNNNGNTTGTAACNCGAGATCGTCNGTTTCA
TACTCAGTCTGCAATCCGCTTNNCATTACTGCCCCGCTAACACCTTGTAGGCATGCCCGT
AGCAGATCCTGCGGCGGTCTCGANATNNNNNNNNNNNNNCGATATGTGGCCTTNNNANNNN
NNNANATGGCTACGTGTCGCAAGTCAAGTNCGTACCCCGAAGCTGCCCTTGTCGACAGCC
TGTTCNNGTNGACTCGTNCNNGANNCNGATAGTCGCNTGTATGTCATACTTTCTTCCCAG
CGTTGCCATGGTGAATTTCTCCGTGATATGGCTANGTGCCTTACCCTCTTCCACCTCTTG
AGGGTCCCATTCTGCCCATTCACATTTATAATATAATCCCTACTGAGTTTAAAGCTCCCC
CCGTACTANACGTTCTCCTGCGTTGCGGGTAGATGGTCACAGATTTCGAGTTTTAAACTT
NAAGAAGGCTGCCCTCCCACCAGTTTCATCATNNNTNTCTATCNNNNNNNNNCNNCCTTC
GCCACTTGTGACGAGGGTTTGGGCGTGATTACGACCTNNNNNGNNNCATTATAGGCACGT
GCNNATCCGACGCTGAGCCCTAAATAGTTGCATGGGCNNGCNCATTCCTAANNTCNTCNN
NNNNGTTAATANNNTCGTCCCCCAGNNNNNNNNNCCCCCGATCGTCNANCNNNNNNNTNN
NNNNTATACGTAACNNNNNNTNNNNNNGTTCATGTATAGAAGACGCCGACGTTCCCCTAC
TTTTTCGCTTCTTATCTTGGTTACGGGCGCGCTGCGGCCCGGAACCAGTACTTNAGCTCC
CCATAAGGTGTGGCTACGCACGNGAAGAACTGCGTTGCGCTANCTATAGTGCTTCCNCCT
GGCCCTCCGAGAATC

>Pyr_28

GTGACCGTTGCCTCGCCACAAATCCATTGTTTTTCGGGTTCTGGTCTTTTATGAGTTAGG
ACCTCTGCGCCTACTATCATAGCTNNNNNNNNNNNNCGCATTNNATACGTTGCACTGACCC
GACAATAATCTGTGCTGCTAAACTTTTGCAAGTNGCTGGCGCCTGTAGGTCACGCAATAG
AGTAGCTTCCCTCCCGACGATCTTCAGGCCATCTACAATNNNGCATCGTAGATCCTCGCG
TGACCTAGTANCNNNNGACGACTCAATCCANCCTCTGCCTGACGAACGTGACTTTTGTAG
TACTCTAGTGACTCCCTGTCCTCATAAGNAGGCTGCATACATAGGGCTACCGTAGTTTAC
TGGTCTCACCTCTACGTACTCGCTGACCCAACTACGTTGATCCAGGAAGGCAGAATGGTG
GACGCTNNTACTGCCGACCCGTGGGTAACTAACTGGGACGGGGGAATATTGTTTCGCTTT
GAGTACTACTTCTTCCAATTGTNNGGATANNACAGTAATGATGGTGGGTGACTCCAGTCA
CTTNNNCGTAGTCGCCGGTCGGTCGCTCCGGTCTGTCCTCTCGCNNNNNTNTGGGCCATC
TTATCCGACGTCATNCTGCAGACCGCGCTTTCCNNNCGTTAGCNNNNNNNAGTCTCCCAT
GGATGNTCTCATGTCATGAGAGNTACCGGAGNNNNTAAGCGTTCCGCTGTTCTCGACNGT
ATTCTCTTTCGTTTTAACACTAGCTCAGACGGTAGAGGCTTCGTCTTCAGTATCGGCTAA
TGATGCCGGGTACGTCAAGATACCAGTCTGCGCGCTTACGCTGCTCCATCTGTTGATGTC
TNCTTCNGCTCGGCTAAGACCTCTCGAACTCTTTGTCTTACCTCAGCCCGCGGCTAGAAT
CATTTACATCCTGGCAAACACAGTCCTTAATTTCCAGTNNNNCCCGACNGTAATACCGTG
GCGCGAACGACGTCTAAGTTNNTGCCTNCTNCGTNNNNTATGGACTACATTTGCCCAACG
TGAGGTCCGAGGATNNNNNNNNANGGCACACACTNNNAGAATCCGCAGCGTTTCGTCTCA
CAATCGAGAAAGGCGCTCCTAAGGGGCCGCGATACGAGCNTGANNCCGGNCAAGCGCTCT
AATTGTACACTGTTCATGAGGCTAAATTCAAATTGCGGACCGAAAACTTGGCCCAGCAGA
GCTAAAGGTTCTTCGCGGGTATTGTCCGTACATGCAACTTGATGTTTTCTAAGCCAAGAN
NGATATGCGAGGGGGAGACATGTGAAAAGCAAAAAANNNNNNNNNNNNAATTATCGTGGG
ACCCATAAGGCCTTGGTTGCTCCCGCGTCGCCTGTACCCACAGTCTTCAAAAAGGCGANT
TGCCTGAGTATANAGTAAAGAGCCGACGGTGTTGAAAACGAGCGGACGTCTTGCATGAGA
AGTGTCACTGTGGGCATCTCAAAGGACTCTAATTAAATTCGGTGTCAATAGAAATGTTCG
CGCTGCATGGCGCTGTCAGATCATCCATCCTCTGTGGCTANNCCTTATCATAAGTGAGAC
CTGCGAAATCAGGGTGATGTCANTACGCTTTAAAGGACGCTAAAGCGAGTTGATAATTAA
ACGACAGAGTTACAATGAGAAGTATTCACAACTCGCCAAAATCATTGNNNNNNGANTGAC
ACGATAGCGTGGGAGGCCCCGTAGCCTCAAGTGAAAAGGGGCGCAAAACGGTGTCCGCAN
NNNAGGAGCTCGACCATCATAGATGCCGGATATGTTCCTGTGAAATTTCANCGTCAAAGA
TGGGAAGCAAAACAAACCCATGAAACTACCATAGTGTNNNNNNNGTGGGCGGCGNNNNNT
AAGGCGAGGCGCTCTANNNCTCCAACGTGCCCTAGGGGATCGTAACATACAAGAACATTG
ACGTGCACAGTTTTGTTGAACGGCCTTCATGGGCCAAAGGTTGNATCCACGGCCAGGAGC
GGTAAGAACTTCGTGACATAGCGGCCGGAAAAAGGATCTACAACGTTAACCANANNCTCG
GAGAGCGGAACGGCCAGTCNNNNNNNAGAAAATAGTTAAGCATTAATTCTGACGAACCAC

CAAAATGCTCGANNNNNNGGTATCATTAGGGGGTACGGACTGTTANNNTTAGAATGGCCC
AGCTCCATAGTTGCAGACACTAATTTTAGAGACCGCTGCTGCATCATTAACAAAATGTTA
TCGGTGGGATGCAGAAGAGCGCGGGCNNNATTACACAAACGCCTTCAGGTGGTAACCTGC
TCCTAAGCAACCAAGACGGATCGCGACGGACGTGTCTGGACGGCGAATGTACAGGAAGTT
TTCCGTGAGAACAACAAAATGAAGGATCGAGATTAGAAAGACTACAGCGCCCATGAGCAA
CGGAGTTGAACGTGATTGTCAGGTTGGTAATGGGCGGCCCTAGAAGNNNNNGAGAAAAAC
CCTTCGGACTTTTCAATGGGGACTGCATTCCGGCCGCGAAGGACTCCATGAATCGATAGG
TAGGGGAAATATGGATGAAACGGGCTGGGGCAACGCCCGAGGCGACGTCTACTAGTTAAA
AGGCAGGTTGCGAGGATCGGGNNAGACTGAGAAGGCCTTGGCGTAGGAATTCGAGGCAAC
CAGGAGGATAAGGTAAATGCCGACTTCAAGCCGATAATGCAAAGAAGCCGGCTATGGCCG
TAAATTAGACGATAAGTAACAGATCGTCGGGGGGCCAAGTCTAATCGTAAAACTTTTAGN
TGAGGTNNNNNNNNANGNGAGATNCCCGAACGCAGCATGAACAAGGTGAATCGTGTAAGC
AGAGAAGAGCGCTGCACGATGCTNGCNCTGTCGTAGTAGTGGGACAANNTGGATGACTGG
CACATCGTAGTAATGAGTCATATGATCCCTCCNNNAGGNNNCNNNAAAGGANNCAAGACG
CGAAGAAGATCTAGACTCAGTCTTTAAAACTGCACAACTTAGCCGAAGTGATTGGTANNN
AGAAGTACTGATGTGGGCGCTGAGAGAAACCNNNAAACGCGATAACCATCATATCACGAT
GCTTCCGCCTTCCAGCGTCACTAAAGTTCTTAATCACATGCCTCGGTTCTCTCGCCAGTA
CACGGTCGGCCTTCTTTCCCCCATCGTTTACCGGAGCATATTCTTCGAGTTGCCTTATTA
ACCTTCTATTCTCTCGCCGGTGCCTGATTAGCGCCGCCTNNNNNATCATACTGTTTGCTT
TTAGCCAACCTGTTGAGCGTTGATTGCTTGTGTGACCATATCTTGCGTTTCGTTGATTCC
TGGTCGGCAATGTGTTGTGGCCTGTTCGTTGACGCTCCTGATCTCACCGTACTTCACGGT
TGCCGCCCGGACATCACCCAGATGTGTTTAGGCCATGGCCGGTTAAGGTTAATCTTTACG
CNNNNNCNTAACTTCCTGCGAAACTGGTTCCCCCACGCCGTCCCCTTCGTTAGNCGANCG
GCGTGGTNNGGTCGCGCGGGTTTTTCTAGGCAAGGTAAGTCGGGTCGCACANNNNNACCC
ACNNNNNNNNNCTTAACGTTCCGTTTTACCACTGGGTCAGGCNNNNTTGGGTCCGACGTG
ACGCTCTTTCCGCACGACCTCGTATGTTCTTATTCGCANTCGTAATGANCCGGTAAGCGN
GAATTCGCTCTACGCTGAGGTGTCGTCGGACCCGAGTGTGCTTGCCGCNGGCCAATNGTT
GTGTCGGCTCGACGTCTTATTTCGTATGGTCCGGGATTACCCNTGNTTGCAGCGATGTCC
TTTTCGCTATTGATGTCTTCTCCTGAGTCACGCTACATACTGAATTAATAAAGCATTTCG
ACCGCAGCATCAGTGAAGTCGAGGGATGGGCTACATCGCCGTGCGGAGCTTTTCACGTTG
TGTCNGTGCGGAGATTCAGCACTAGCGGTCATACTATCCATGTTCCCCCGTTCTATTTAC
GTTGAGCGACTTGAANCTTNCAGTGCACTGTNNCTGCNTAGNNCGGTATTTCTCAGTTCA
TGCAAGTTATCGTACATCATGGAATAGAACNNNNNNNNAGGACCATTTCCACTAACGATT
CTTACCTNCAAATCAAGATAGAATCTAACCGATTATCGACGAACTTCATCCGTCATGTCA
CTCTATGTCAAGCGGCCTTCTTACAGCGCTGTCGCACTATCCNTTCTACCTTGTTTCCGA
GAAACCTTGCCTCTGTTGCCTGTGGGGGCAGGGCGGATATGGCTAAATGGGCATTCCTTT

TTACTTCCCGTGCCCCTTCATGCTATGTGCTTGCAGCTCTAAAGCTNAGACGCCCGGTCC
TCGACNNNNCGAAACTTTCCTGACCCTGCTGGCGTCTTACCCTTTATAGTGGATCTGCAT
AATGTGTTCTGTATATAGCCAAATACTGCCAACTATCCACTCGCTTAGCTGTCAATATCC
ATGAGCGTGCTTTGATTTACCGGTCGACCTGACTAATGTAACTGCCTTGCGCTCTCCACT
CCGCACCAGGGTTTCGAGTTTTTCTACCCCTCTCTTTATTACTACTCGTTTANNGCATCA
GACGCGCCCATCGTACTTGGTCACTGCTCCTNNNGNNNAGNAGTATTGGCACGTNNNCGT
GCTTCTATCCTGAGCTGCAACCATAAGCATTTCTAAAGTTATGAGGATCAAAGAGTATTT
CCGTACATNNNGATCGATTGTACTGGACGTCGTCCCAAATAGCCTNNNAAGTCCAATATC
TAGGTCATAGATGCT
>Pyr_29_33
GTGACCGTTGCCTCGCCACAAATCCATTGTTTTTCGGGTTCTGGTCTTTTATGAGTTAGG
ACCTCTGCGCCTACTATCATAGCTGTCCGATGCCTCGCATTCCATACGTTGCACTGACCC
GACAATAATCTGTGCTGCTAAACTTTTGCAAGTGGCTGGCGCCTGTAGGTCACGCAATAG
AGTAGCTTCCCTCCCGACGATCTTCAGGCCATCTACAATCCGGCATCGTAGATCCTCGCG
TGACCTAGTAGCCCTTGACGACTCAATCCACCCTCTGCCTGACGAACGTGACTTTTGTAG
TACTCTAATGACTCCCTGTCCTCATAAGTAGGCTGCATACATAGGGCTACCGTAGTTTAC
TGGTCTCACCTCTACGTACTCGCTGACCCAACTACGTTGATCCAGGAAGGCAGAATGGTG
GACGCTCTTACTGCCGACCCGTGGGTAACTAACTGGGACGGGGGAATATTGTTTCGCTTT
GAGTACTACTTCTTCCAATTGTTCGGATAGCACAGTAATGATGGTGGGTGACTCCAGTCA
CTTTCGCGTAGTCGCCGGTCGGTCGCTCCGGTCTGTCCTCTCGCCCGATTCTGGGCCATC
TTATCCGACGTCATCCTGCAGACCGCGCTTTCCAACCGTTAGCATTAAGCAGTCTCCCAT
GGATGCTCTCATGTCATGAGAGGTACCGGAGCCTCTAAGCGTTCCGCTGTTCTCGACAGT
ATTCTCTTTCGTTTTAACACTAGCTCAGACGGTAGAGGCTTCGTCTTCAGTATCGGCTAA
TGATGCCGGGTACGTCAAGATACCAGTCTGCGCGCTTACGCTGCTCCATCTGTTGATGTC
TGCTTCAGCTCGGCTAAGACCTCTCGAACTCTTTGTCTTACCTCAGCCCGCGGCTAGAAT
CATTTACATCCTGGCAAACACAGTCCTTAATTTCCAGTCGTGCCCGACAGTAATACCGTG
GCGCGAACGACGTCTAAGTTGTTGCCTACTTCGTACACTATGGACTACATTTGCCCAACG
TGAGGTCCGAGGATCCCGTTGGAGGGCACACACTCAGAGAATCCGCAGCGTTTCGTCTCA
CAATCGAGAAAGGCGCTCCTAAGGGGCCGCGATACGAGCGTGAAACCGGCCAAGCGCTCT
AATTGTACACTGTTCATGAGGCTAAATTCAAATTGCGGACCGAAAACTTGGCCCAGCAGA
GCTAAAGGTTCTTCGCGGGTATTGTCCGTACATGCAACTTGATGTTTTCTAAGCCAAGAG
AGATATGCGAGGGGGAGACATGTGAAAAGCAAAAAAATATAGTACTATAATTATCGTGGG
ACCCATAAGGCCTTGGTTGCTCCCGCGTCGCCTGTACCCACAGTCTTCAAAAAGGCGATT
TGCCTGAGTATAAAGTAAAGAGCCGACGGTGTTGAAAACGAGCGGACGTCTTGCATGAGA
AGTGTCACTGTGGGCATCTCAAAGGACTCTAATTAAATTCGGTGTCAATAGAAATGTTCG
CGCTGCATGGCGCTGTCAGATCATCCATCCTCTGTGGCTACGCCTTATCATAAGTGAGAC

CTGCGAAATCAGGGTGATGTCAATACGCTTTAAAGGACGCTAAAGCGAGTTGATAATTAA

ACGACAGAGTTACAATGAGAAGTATTCACAACTCGCCAAAATCATTGACGATAGACTGAC

ACGATAGCGTGGGAGGCCCCGTAGCCTCAAGTGAAAAGGGGCGCAAAACGGTGTCCGCAT

GTGAGGAGCTCGACCATCATAGATGCCGGATATGTTCCTGTGAAATTTCAGCGTCAAAGA

TGGGAAGCAAAACAAACCCATGAAACTACCATAGTGTGAAGGTGGTGGGCGGCGCGGTTT

AAGGCGAGGCGCTCTAAGACTCCAACGTGCCCTAGGGGATCGTAACATACAAGAACATTG

ACGTGCACAGTTTTGTTGAACGGCCTTCATGGGCCAAAGGTTGTATCCACGGCCAGGAGC

GGTAAGAACTTCGTGACATAGCGGCCGGAAAAAGGATCTACAACGTTAACCAGAACCTCG

GAGAGCGGAACGGCCAGTCGTTAGGGAGAAAATAGTTAAGCATTAATTCTGACGAACCAC

CAAAATGCTCGAGCGGTAGGTATCATTAGGGGGTACGGACTGTTACATTTAGAATGGCCC

AGCTCCATAGTTGCAGACACTAATTTTAGAGACCGCTGCTGCATCATTAACAAAATGTTA

TCGGTGGGATGCAGAAGAGCGCGGGCAAGATTACACAAACGCCTTCAGGTGGTAACCTGC

TCCTAAGCAACCAAGACGGATCGCGACGGACGTGTCTGGACGGCGAATGTACAGGAAGTT

TTCCGTGAGAACAACAAAATGAAGGATCGAGATTAGAAAGACTACAGCGCCCATGAGCAA

CGGAGTTGAACGTGATTGTCAGGTTGGTAATGGGCGGCCCTAGAAGATAACGAGAAAAAC

CCTTCGGACTTTTCAATGGGGACTGCATTCCGGCCGCGAAGGACTCCATGAATCGATAGG

TAGGGGAAATATGGATGAAACGGGCTGGGGCAACGCCCGAGGCGACGTCTACTAGTTAAA

AGGCAGGTTGCGAGGATCGGGGAAGACTGAGAAGGCCTTGGCGTAGGAATTCGAGGCAAC

CAGGAGGATAAGGTAAATGCCGACTTCAAGCCGATAATGCAAAGAAGCCGGCTATGGCCG

TAAATTAGACGATAAGTAACAGATCGTCGGGGGGCCAAGTCTAATCGTAAAACTTTTAGT

TGAGGTGATGTCGTAAGTGAGATCCCCGAACGCAGCATGAACAAGGTGAATCGTGTAAGC

AGAGAAGAGCGCTGCACGATGCTAGCACTGTCGTAGTAGTGGGACAACTTGGATGACTGG

CACATCGTAGTAATGAGTCATATGATCCCTCCTGAAGGCTTCAAAAAAGGACACAAGACG

CGAAGAAGATCTAGACTCAGTCTTTAAAACTGCACAACTTAGCCGAAGTGATTGGTAAGT

AGAAGTACTGATGTGGGCGCTGAGAGAAACCTGAAAACGCGATAACCATCATATCACGAT

GCTTCCGCCTTCCAGCGTCACTAAAGTTCTTAATCACATGCCTCGGTTCTCTCGCCAGTA

CACGGTCGGCCTTCTTTCCCCCATCGTTTACCGGAGCATATTCTTCGAGTTGCCTTATTA

ACCTTCTATTCTCTCGCCGGTGCCTGATTAGCGCCGCCTTGAGTATCATACTGTTTGCTT

TTAGCCAACCTGTTGAGCGTTGATTGCTTGTGTGACCATATCTTGCGTTTCGTTGATTCC

TGGTCGGCAATGTGTTGTGGCCTGTTCGTTGACGCTCCTGATCTCACCGTACTTCACGGT

TGCCGCCCGGACATCACCCAGATGTGTTTAGGCCATGGCCGGTTAAGGTTAATCTTTACG

CGTCATCATAACTTCCTGCGAAACTGGTTCCCCCACGCCGTCCCCTTCGTTAGTCGATCG

GCGTGGTTAGGTCGCGCGGGTTTTTCTAGGCAAGGTAAGTCGGGTCGCACAGTGCGACCC

ACGGACATAACCTTAACGTTCCGTTTTACCACTGGGTCAGGCTGCTTTGGGTCCGACGTG

ACGCTCTTTCCGCACGACCTCGTATGTTCTTATTCGCACTCGTAATGAACCGGTAAGCGA

GAATTCGCTCTACGCTGAGGTGTCGTCGGACCCGAGTGTGCTTGCCGCTGGCCAATTGTT

GTGTCGGCTCGACGTCTTATTTCGTATGGTCCGGGATTACCCGTGCTTGCAGCGATGTCC
TTTTCGCTATTGATGTCTTCTCCTGAGTCACGCTACATACTGAATTAATAAAGCATTTCG
ACCGCAGCATCAGTGAAGTCGAGGGATGGGCTACATCGCCGTGCGGAGCTTTTCACGTTG
TGTCCGTGCGGAGATTCAGCACTAGCGGTCATACTATCCATGTTCCCCCGTTCTATTTAC
GTTGAGCGACTTGAAACTTCCAGTGCACTGTTTCTGCATAGTGCGGTATTTCTCAGTTCA
TGCAAGTTATCGTACATCATGGAATAGAACAAGCATCAAGGACCATTTCCACTAACGATT
CTTACCTCCAAATCAAGATAGAATCTAACCGATTATCGACGAACTTCATCCGTCATGTCA
CTCTATGTCAAGCGGCCTTCTTACAGCGCTGTCGCACTATCCGTTCTACCTTGTTTCCGA
GAAACCTTGCCTCTGTTGCCTGTGGGGGCAGGGCGGATATGGCTAAATGGGCATTCCTTT
TTACTTCCCGTGCCCCTTCATGCTATGTGCTTGCAGCTCTAAAGCTTAGACGCCCGGTCC
TCGACGTCTCGAAACTTTCCTGACCCTGCTGGCGTCTTACCCTTTATAGTGGATCTGCAT
AATGTGTTCTGTATATAGCCAAATACTGCCAACTATCCACTCGCTTAGCTGTCAATATCC
ATGAGCGTGCTTTGATTTACCGGTCGACCTGACTAATGTAACTGCCTTGCGCTCTCCACT
CCGCACCAGGGTTTCGAGTTTTTCTACCCCTCTCTTTATTACTACTCGTTTACTGCATCA
GACGCGCCCATCGTACTTGGTCACTGCTCCTTCAGGCGAGGAGTATTGGCACGTTATCGT
GCTTCTATCCTGAGCTGCAACCATAAGCATTTCTAAAGTTATGAGGATCAAAGAGTATTT
CCGTACATCTTGATCGATTGTACTGGACGTCGTCCCAAATAGCCTGCGAAGTCCAATATC
TAGGTCATAGATGCT
>Pyr_30
GTGACCGTTGCCTCGCCACAAATCCATTGTTTTTCGGGTTCTGGTCTTTTATGAGTTAGG
ACCTCTGCGCCTACTATCATAGCTGTCCGATGNCTCGCATTCCATACGTTGCACTGACCC
GACAATAATCTGTGCTGCTAAACTTTTGCAAGTGGCTGGCGCCTGTAGGTCACGCAATAG
AGTAGCTTCCCTCCCGACGATCTTCAGGCCATCTACAATCCGGCATCGTAGATCCTCGCG
TGACCTAGTAGCCCTTGACGACTCAATCCACCCTCTGCCTGACGAACGTGACTTTTGTAG
TACTCTAGTGACTCCCTGTCCTCATAAGTAGGCTGCATACATAGGGCTACCGTAGTTTAC
TGGTCTCACCTCTACGTACTCGCTGACCCAACTACGTTGATCCAGGAAGGCAGAATGGTG
GACGCTCTTACTGCCGACCCGTGGGTAACTAACTGGGACGGGGGAATATTGTTTCGCTTT
GAGTACTACTTCTTCCAATTGTTCGGATAGCACAGTAATGATGGTGGGTGACTCCAGTCA
CTTTCGCGTAGTCGCCGGTCGGTCGCTCCGGTCTGTCCTCTCGCCCGATTCTGGGCCATC
TTATCCGACGTCATCCTGCAGACCGCGCTTTCCAACCGTTAGCATTAAGCAGTCTCCCAT
GGATGCTCTCATGTCATGAGAGGTACCGGAGCCTCTAAGCGTTCCGCTGTTCTCGACAGT
ATTCTCTTTCGTTTTAACACTAGCTCAGACGGTAGAGGCTTCGTCTTCAGTATCGGCTAA
TGATGCCGGGTACGTCAAGATACCAGTCTGCGCGCTTACGCTGCTCCATCTGTTGATGTC
TGCTTCAGCTCGGCTAAGACCTCTCGAACTCTTTGTCTTACCTCAGCCCGCGGCTAGAAT
CATTTACATCCTGGCAAACACAGTCCTTAATTTCCAGTCGTGCCCGACAGTAATACCGTG
GCGCGAACGACGTCTAAGTTGTTGCCTACTTCGTACACTATGGACTACATTTGCCCAACG

TGAGGTCCGAGGATCCCGTTGGAGGGCACACACTCAGAGAATCCGCAGCGTTTCGTCTCA
CAATCGAGAAAGGCGCTCCTAAGGGGCCGCGATACGAGCGTGAAACCGGCCAAGCGCTCT
AATTGTACACTGTTCATGAGGCTAAATTCAAATTGCGGACCGAAAACTTGGCCCAGCAGA
GCTAAAGGTTCTTCGCGGGTATTGTCCGTACATGCAACTTGATGTTTTCTAAGCCAAGAG
AGATATGCGAGGGGGAGACATGTGAAAAGCAAAAAAATATAGTACTATAATTATCGTGGG
ACCCATAAGGCCTTGGTTGCTCCCGCGTCGCCTGTACCCACAGTCTTCAAAAAGGCGATT
TGCCTGAGTATAAAGTAAAGAGCCGACGGTGTTGAAAACGAGCGGACGTCTTGCATGAGA
AGTGTCACTGTGGGCATCTCAAAGGACTCTAATTAAATTCGGTGTCAATAGAAATGTTCG
CGCTGCATGGCGCTGTCAGATCATCCATCCTCTGTGGCTACGCCTTATCATAAGTGAGAC
CTGCGAAATCAGGGTGATGTCAATACGCTTTAAAGGACGCTAAAGCGAGTTGATAATTAA
ACGACAGAGTTACAATGAGAAGTATTCACAACTCGCCAAAATCATTGACGATAGACTGAC
ACGATAGCGTGGGAGGCCCCGTAGCCTCAAGTGAAAAGGGGCGCAAAACGGTGTCCGCAT
GTGAGGAGCTCGACCATCATAGATGCCGGATATGTTCCTGTGAAATTTCAGCGTCAAAGA
TGGGAAGCAAAACAAACCCATGAAACTACCATAGTGTGAAGGTGGTGGGCGGCGCGGTTT
AAGGCGAGGCGCTCTAAGACTCCAACGTGCCCTAGGGGATCGTAACATACAAGAACATTG
ACGTGCACAGTTTTGTTGAACGGCCTTCATGGGCCAAAGGTTGTATCCACGGCCAGGAGC
GGTAAGAACTTCGTGACATAGCGGCCGGAAAAAGGATCTACAACGTTAACCAGAACCTCG
GAGAGCGGAACGGCCAGTCGTTAGGGAGAAAATAGTTAAGCATTAATTCTGACGAACCAC
CAAAATGCTCGAGCNGTAGGTATCATTAGGGGGTACGGACTGTTACATTTAGAATGGCCC
AGCTCCATAGTTGCAGACACTAATTTTAGAGACCGCTGCTGCATCATTAACAAAATGTTA
TCGGTGGGATGCAGAAGAGCGCGGGCAAGATTACACAAACGCCTTCAGGTGGTAACCTGC
TCCTAAGCAACCAAGACGGATCGCGACGGACGTGTCTGGACGGCGAATGTACAGGAAGTT
TTCCGTGAGAACAACAAAATGAAGGATCGAGATTAGAAAGACTACAGCGCCCATGAGCAA
CGGAGTTGAACGTGATTGTCAGGTTGGTAATGGGCGGCCCTAGAAGATAACGAGAAAAAC
CCTTCGGACTTTTCAATGGGGACTGCATTCCGGCCGCGAAGGACTCCATGAATCGATAGG
TAGGGGAAATATGGATGAAACGGGCTGGGGCAACGCCCGAGGCGACGTCTACTAGTTAAA
AGGCAGGTTGCGAGGATCGGGGAAGACTGAGAAGGCCTTGGCGTAGGAATTCGAGGCAAC
CAGGAGGATAAGGTAAATGCCGACTTCAAGCCGATAATGCAAAGAAGCCGGCTATGGCCG
TAAATTAGACGATAAGTAACAGATCGTCGGGGGGCCAAGTCTAATCGTAAAACTTTTAGT
TGAGGTGATGTCGTAAGTGAGATCCCCGAACGCAGCATGAACAAGGTGAATCGTGTAAGC
AGAGAAGAGCGCTGCACGATGCTAGCACTGTCGTAGTAGTGGGACAACTTGGATGACTGG
CACATCGTAGTAATGAGTCATATGATCCCTCCTGAAGGCTTCAAAAAAGGACACAAGACG
CGAAGAAGATCTAGACTCAGTCTTTAAAACTGCACAACTTAGCCGAAGTGATTGGTAAGT
AGAAGTACTGATGTGGGCGCTGAGAGAAACCTGAAAACGCGATAACCATCATATCACGAT
GCTTCCGCCTTCCAGCGTCACTAAAGTTCTTAATCACATGCCTCGGTTCTCTCGCCAGTA
CACGGTCGGCCTTCTTTCCCCCATCGTTTACCGGAGCATATTCTTCGAGTTGCCTTATTA

ACCTTCTATTCTCTCGCCGGTGCCTGATTAGCGCCGCCTTGAGTATCATACTGTTTGCTT
TTAGCCAACCTGTTGAGCGTTGATTGCTTGTGTGACCATATCTTGCGTTTCGTTGATTCC
TGGTCGGCAATGTGTTGTGGCCTGTTCGTTGACGCTCCTGATCTCACCGTACTTCACGGT
TGCCGCCCGGACATCACCCAGATGTGTTTAGGCCATGGCCGGTTAAGGTTAATCTTTACG
CGTCATCATAACTTCCTGCGAAACTGGTTCCCCCACGCCGTCCCCTTCGTTAGTCGATCG
GCGTGGTTAGGTCGCGCGGGTTTTTCTAGGCAAGGTAAGTCGGGTCGCACAGTGCGACCC
ACGGACATAACCTTAACGTTCCGTTTTACCACTGGGTCAGGCTGCTTTGGGTCCGACGTG
ACGCTCTTTCCGCACGACCTCGTATGTTCTTATTCGCACTCGTAATGAACCGGTAAGCGA
GAATTCGCTCTACGCTGAGGTGTCGTCGGACCCGAGTGTGCTTGCCGNTGNCCAATTGTT
GTGTCGGCTCGACGTCTTATTTCGTATGGTCCGGGATTACCCGTGCTTGCAGCGATGTCC
TTTTCGCTATTGATGTCTTCTCCTGAGTCACGCTACATACTGAATTAATAAAGCATTTCG
ACCGCAGCATCAGTGAAGTCGAGGGATGGGCTACATCGCCGTGCGGAGCTTTTCACGTTG
TGTCCGTGCGGAGATTCAGCACTAGCGGTCATACTATCCATGTTCCCCCGTTCTATTTAC
GTTGAGCGACTTGAAACTTCCAGTGCACTGTTTCTGCATAGTGCGGTATTTCTCAGTTCA
TGCAAGTTATCGTACATCATGGAATAGAACAAGCATCAAGGACCATTTCCACTAACGATT
CTTACCTCCAAATCAAGATAGAATCTAACCGATTATCGACGAACTTCATCCGTCATGTCA
CTCTATGTCAAGCGGCCTTCTTACAGCGCTGTCGCACTATCCGTTCTACCTTGTTTCCGA
GAAACCTTGCCTCTGTTGCCTGTGGGGGCAGGGCGGATATGGCTAAATGGGCATTCCTTT
TTACTTCCCGTGCCCCTTCATGCTATGTGCTTGCAGCTCTAAAGCTTAGACGCCCGGTCC
TCGACGTCTCGAAACTTTCCTGACCCTGCTGGCGTCTTACCCTTTATAGTGGATCTGCAT
AATGTGTTCTGTATATAGCCAAATACTGCCAACTATCCACTCGCTTAGCTGTCAATATCC
ATGAGCGTGCTTTGATTTACCGGTCGACCTGACTAATGTAACTGCCTTGCGCTCTCCACT
CCGCACCAGGGTTTCGAGTTTTTCTACCCCTCTCTTTATTACTACTCGTTTACTGCATCA
GACGCGCCCATCGTACTTGGTCACTGCTCCTTCANGCGAGGAGTATTGGCACGTTATCGT
GCTTCTATCCTGAGCTGCAACCATAAGCATTTCTAAAGTTATGAGGATCAAAGAGTATTT
CCGTACATCTTGATCGATTGTACTGGACGTCGTCCCAAATAGCCTGCGAAGTCCAATATC
TAGGTCATAGATGCT
>Pyr_4
GTGACCGTTGCCTCGCCACAAATCCATTGTTTTTCGGGTTCTGGTCTTTTATGAGTTAGG
ACCTCTGCGCCTACTATCATAGCTGTCCGATGCCTCGCATTCCATACGTTGCACTGACCC
GACAATAATCTGTGCTGCTAAACTTTTGCAAGTGGCTGGCGCCTGTAGGTCACGCAATAG
AGTAGCTTCCCTCCCGACGATCTTCAGGCCATCTACAATCCGGCATCGTAGATCCTCGCG
TGACCTAGTAGCCCTTGACGACTCAATCCACCCTCTGCCTGACGAACGTGACTTTTGTAG
TACTCTAGTGACTCCCTGTCCTCATAAGTAGGCTGCATACATAGGGCTACCGTAGTTTAC
TGGTCTCACCTCTACGTACTCGCTGACCCAACTACGTTGATCCAGGAAGGCAGAATGGTG
GACGCTCTTACTGCCGACCCGTGGGTAACTAACTGGGACGGGGGAATATTGTTTCGCTTT

GAGTACTACTTCTTCCAATTGTTCGGATAGCACAGTAATGATGGTGGGTGACTCCAGTCA
CTTTCGCGTAGTCGCCGGTCGGTCGCTCCGGTCTGTCCTCTCGCCCGATTCTGGGCCATC
TTATCCGACGTCATCCTGCAGACCGCGCTTTCCAACCGTTAGCATTAAGCAGTCTCCCAT
GGATGCTCTCATGTCATGAGAGGTACCGGAGCCTCTAAGCGTTCCGCTGTTCTCGACAGT
ATTCTCTTTCGTTTTAACACTAGCTCAGACGGTAGAGGCTTCGTCTTCAGTATCGGCTAA
TGATGCCGGGTACGTCAAGATACCAGTCTGCGCGCTTACGCTGCTCCATCTGTTGATGTC
TGCTTCAGCTCGGCTAAGACCTCTCGAACTCTTTGTCTTACCTCAGCCCGCGGCTAGAAT
CATTTACATCCTGGCAAACACAGTCCTTAATTTCCAGTCGTGCCCGACAGTAATACCGTG
GCGCGAACGACGTCTAAGTTGTTGCCTACTTCGTACACTATGGACTACATTTGCCCAACG
TGAGGTCCGAGGATCCCGTTGGAGGGCACACACTCAGAGAATCCGCAGCGTTTCGTCTCA
CAATCGAGAAAGGCGCTCCTAAGGGGCCGCGATACGAGCGTGAAACCGGCCAAGCGCTCT
AATTGTACACTGTTCATGAGGCTAAATTCAAATTGCGGACCGAAAACTTGGCCCAGCAGA
GCTAAAGGTTCTTCGCGGGTATTGTCCGTACATGCAACTTGATGTTTTCTAAGCCAAGAG
AGATATGCGAGGGGGAGACATGTGAAAAGCAAAAAAATATAGTACTATAATTATCGTGGG
ACCCATAAGGCCTTGGTTGCTCCCGCGTCGCCTGTACCCACAGTCTTCAAAAAGGCGATT
TGCCTGAGTATAAAGTAAAGAGCCGACGGTGTTGAAAACGAGCGGACGTCTTGCATGAGA
AGTGTCACTGTGGGCATCTCAAAGGACTCTAATTAAATTCGGTGTCAATAGAAAGGTTCG
CGCTGCATGGCGCTGTCAGATCATCCATCCTCTGTGGCTACGCCTTATCATAAGTGAGAC
CTGCGAAATCAGGGTGATGTCAATACGCTTTAAAGGACGCTAAAGCGAGTTGATAATTAA
ACGACAGAGTTACAATGAGAAGTATTCACAACTCGCCAAAATCATTGACGATAGACTGAC
ACGATAGCGTGGGAGGCCCCGTAGCCTCAAGTGAAAAGGGGCGCAAAACGGTGTCCGCAT
GTGAGGAGCTCGACCATCATAGATGCCGGATANGTTCCTGTGAAATTTCAGCGTCAAAGA
TGGGAAGCAAAACAAACCCATGAAACTACCATAGTGTGAAGGTGGTGGGCGGCGCGGTTT
AAGGCGAGGCGCTCTAAGACTCCAACGTGCCCTAGGGGATCGTAACATACAAGAACATTG
ACGTGCACAGTTTTGTTGAACGGCCTTCATGGGCCAAAGGTTGTATCCACGGCCAGGAGC
GGTAAGAACTTCGTGACATAGCGGCCGGAAAAAGGATCTACAACGTTAACCAGAACCTCG
GAGAGCGGAACGGCCAGTCGTTAGGGAGAAAATAGTTAAGCATTAATTCTGACGAACCAC
CAAAATGCTCGAGCGGTAGGTATCATTAGGGGGTACGGACTGTTACATTTAGAATGGCCC
AGCTCCATAGTTGCAGACACTAATTTTAGAGACCGCTGCTGCATCATTAACAAAATGTTA
TCGGTGGGATGCAGAAGAGCGCGGGCAAGATTACACAAACGCCTTCAGGTGGTAACCTGC
TCCTAAGCAACCAAGACGGATCGCGACGGACGTGTCTGGACGGCGAATGTACAGGAAGTT
TTCCGTGAGAACAACAAAATGAAGGATCGAGATTAGAAAGACTACAGCGCCCATGAGCAA
CGGAGTTGAACGTGATTGTCAGGTTGGTAATGGGCGGCCCTAGAAGATAACGAGAAAAAC
CCTTCGGACTTTTCAATGGGGACTGCATTCCGGCCGCGAAGGACTCCATGAATCGATAGG
TAGGGGAAATATGGATGAAACGGGCTGGGGCAACGCCCGAGGCGACGTCTACTAGTTAAA
AGGCAGGTTGCGAGGATCGGGGAAGACTGAGAAGGCCTTGGCGTAGGAATTCGAGGCAAC

CAGGAGGATAAGGTAAATGCCGACTTCAAGCCGATAATGCAAAGAAGCCGGCTATGGCCG
TAAATTAGACGATAAGTAACAGATCGTCGGGGGGCCAAGTCTAATCGTAAAACTTTTAGT
TGAGGTGATGTCGTAAGTGAGATCCCCGAACGCAGCATGAACAAGGTGAATCGTGTAAGC
AGAGAAGAGCGCTGCACGATGCTAGCACTGTCGTAGTAGTGGGACAACTTGGATGACTGG
CACATCGTAGTAATGAGTCATATGATCCCTCCTGAAGGCTTCAAAAAAGGACACAAGACG
CGAAGAAGATCTAGACTCAGTCTTTAAAACTGCACAACTTAGCCGAAGTGATTGGTAAGT
AGAAGTACTGATGTGGGCGCTGAGAGAAACCTGAAAACGCGATAACCATCATATCACGAT
GCTTCCGCCTTCCAGCGTCACTAAAGTTCTTAATCACATGCCTCGGTTCTCTCGCCAGTA
CACGGTCGGCCTTCTTTCCCCCATCGTTTACCGGAGCATATTCTTCGAGTTGCCTTATTA
ACCTTCTATTCTCTCGCCGGTGCCTGATTAGCGCCGCCTTGAGTATCATACTGTTTGCTT
TTAGCCAACCTGTTGAGCGTTGATTGCTTGTGTGACCATATCTTGCGTTTCGTTGATTCC
TGGTCGGCAATGTGTTGTGGCCTGTTCGTTGACGCTCCTGATCTCACCGTACTTCACGGT
TGCCGCCCGGACATCACCCAGATGTGTTTAGGCCATGGCCGGTTAAGGTTAATCTTTACG
CGTCATCATAACTTCCTGCGAAACTGGTTCCCCCACGCCGTCCCCTTCGTTAGTCGATCG
GCGTGGTTAGGTCGCGCGGGTTTTTCTAGGCAAGGTAAGTCGGGTCGCACAGTGCGACCC
ACGGACATAACCTTAACGTTCCGTTTTACCACTGGGTCAGGCTGCTTTGGGTCCGACGTG
ACGCTCTTTCCGCACGACCTCGTATGTTCTTATTCGCACTCGTAATGAACCGGTAAGCGA
GAATTCGCTCTACGCTGAGGTGTCGTCGGACCCGAGTGTGCTTGCCGCTGGCCAATTGTT
GTGTCGGCTCGACGTCTTATTTCGTATGGTCCGGGATTACCCGTGCTTGCAGCGATGTCC
TTTTCGCTATTGATGTCTTCTCCTGAGTCACGCTACATACTGAATTAATAAAGCATTTCG
ACCGCAGCATCAGTGAAGTCGAGGGATGGGCTACATCGCCGTGCGGAGCTTTTCACGTTG
TGTCCGTGCGGAGATTCAGCACTAGCGGTCATACTATCCATGTTCCCCCGTTCTATTTAC
GTTGAGCGACTTGAAACTTCCAGTGCACTGTTTCTGCATAGTGCGGTATTTCTCAGTTCA
TGCAAGTTATCGTACATCATGGAATAGAACAAGCATCAAGGACCATTTCCACTAACGATT
CTTACCTCCAAATCAAGATAGAATCTAACCGATTATCGACGAACTTCATCCGTCATGTCA
CTCTATGTCAAGCGGCCTTCTTACAGCGCTGTCGCACTATCCGTTCTACCTTGTTTCCGA
GAAACCTTGCCTCTGTTGCCTGTGGGGGCAGGGCGGATATGGCTAAATGGGCATTCCTTT
TTACTTCCCGTGCCCCTTCATGCTATGTGCTTGCAGCTCTAAAGCTTAGACGCCCGGTCC
TCGACGTCTCGAAACTTTCCTGACCCTGCTGGCGTCTTACCCTTTATAGTGGATCTGCAT
AATGTGTTCTGTATATAGCCAAATACTGCCAACTATCCACTCGCTTAGCTGTCAATATCC
ATGAGCGTGCTTTGATTTACCGGTCGACCTGACTAATGTAACTGCCTTGCGCTCTCCACT
CCGCACCAGGGTTTCGAGTTTTTCTACCCCTCTCTTTATTACTACTCGTTTACTGCATCA
GACGCGCCCATCGTACTTGGTCACTGCTCCTTCAGGCGAGGAGTATTGGCACGTTATCGT
GCTTCTATCCTGAGCTGCAACCATAAGCATTTCTAAAGTTATGAGGATCAAAGAGTATTT
CCGTACATCTTGATCGATTGTACTGGACGTCGTCCCAAATAGCCTGCGAAGTCCAATATC
TAGGTCATAGATGCT

>SRR9587917_9587943_9587958

GTGACCGTCGCCTCGCCACAAATCCATTGTTTTCCGGGTTCTAGCCCCTTATGAGCTAGG
ACCTCTGTGCCTACCATCATAGCTGTTCGATGCCTTGTATTCAATACGTTGCACCGACCC
GACAATAATCTGTGCCATTAAGCTTCTGCAAGTGGCTGGCGCCTGTAAGTCACGCAATAG
AGTAACTTCCCTCTCGACAGTCTTTATGCCATCCACAATCCAACATCGTAGATCCTCGCG
TGACCTAGTAGCCCTTGACGACTCAATCCACCCTCTGCCTGACGGACGTGACTTTTATGG
TACATTAATGACTCCCTGTCCTCATAATTAGGCTGCACACACAGGGCTATCGTAGTTTAC
TGGTCTCACCCCTGCGTACTCGGCGACCCAACTACACTGATCCAGGGAGGCAGAATGGTG
GACGCTCTTACTGCCGACCCGTGGGCAACTAACTGGGCCGGGGGAATATTGTTTCGTTTT
GGGTACTGCTTCTTCCAATTGTTCGGATAGCCCAGTAACGGTGGTGGGTGGTTCCAGTCA
CTCTTGCGTAGTCGCCGGCCGGTCGCTCCGGTCTGTCCCCTCGCCCGATTCTGGGCCATC
TTATCCGATGTCACCCTGCAGACCGCTCTTTTCAACCATTAGCATTAAGCAGTCTCTCAT
AGGTGCTCTCATGTCATGAGAGGTACCGGAGCCTCTAAGCGTTCCGCTGTTCCCGACAAT
ATTCTCTTTCGCTTTGACACTAGCTCAGACGGTAGAGGCTTCGTCTTCAGCACCGGCTAA
TGATGCCTGGTACGTTAAGATACCGATCCGCGCGCTTACGCTGCTCCACCTGTCGGTGTC
TGAAGAAGCTCGGCTAAGACCTCTCGAACTCTTCGTCTTACCTCAGCCCGCGGCTACAGT
CATATACATCCTAGCACGCACAATCCTTAATTCCCAGTCGTGCCCGACAGTAACACCGTG
GCGCGAACAATGTGTAAGTTATTGCCTACTTCGTACACTATCAACTAAATTCGTCCAACG
TGAGATCCGAGAATCCCGAAAGGGGGCACACACTTAGAGAACCCGCAGCTTTTTGTCTCA
CGATCGAGAAGGGCACTCCCAAGAGGCCTTGATACAAGCGTGAAACCGGTCAAGCGCTCC
AATTGTACACTGTCCATGAGGCCAAATTCAAATTACGGACCGAAAACTGGATCCAGCAGA
GCTAAAGGTTATTCGCGGGTATTGTCCGTACATGCAACTTGATGTTTTCTAAGCCAAAAG
AGATATGCGAGGGGGAGACATGGGAAAAGCAAAAGAATATAGTACTATAATTGGCGTGGG
ACCCATAAAGTCTTGATTGCTCCCGCGCCGCCTGTACCCACAGTCTTCAAAAAAGCGATT
TGCCGGAGTATAAAGTCAGGGGCCGACGGTGTTGAAAACGAGCGGACGTCTTGCATGAGA
AGTGTCACTGTGGGCATATCAAAGGACTCTAATTAAATTCGGTGTCAATAGAAATGTTCG
TGCTGCACGGTGCTGTCAGACCATCCACCTTTCGTGGCTACGCCTTATCAGAAGTGCAAC
CTGCGAAATCAAGGTGACGTTAATACGCTCTAAAGGACGCTAAAGCGGGTTGATAATTAA
ACGACAGAATTACAATTAGCGGGATTCACAACTCTCCAAAATCATTGACGATAGACTGAC
ACGGTTACGTGGGAGGACTCGTAGTCTCCAGTGAAAATGGGCGCAAAATGGTGTCCGCAT
GGGAGGAGCTCAACCATAATAGACGCCGAATATGTTCGCGTGAAACTTCAGCGTCAAAGA
TGGAAAGCAAGACAAACTCATGAAACTACCATAGTGTGAAGATGATGGGCGGCGCGGTTT
AAGGCGAGTCGCCCTAAGACTCCAACGTGCCCTAAGGGATCGTAACATGCAAGAACATTG
ATGTGCACAGTTTTGTCGAACGGCCTTCATAGGCCAAAGATTGTATCCACGGCCAGAAGC
GGTAAGAACTTCGTGACATAGCGGCCAGAGAAAGGATCTACAACGTTAACCAGCACCTCA
GAGAGCGGGATGGCCAGTCGTTAGGGAGAAAATAACTAATCATTAACTTTGACGAACCAC

CAAAATGCTCGAGCGGTAGGTATCATTAGGGGGTACGGACTGTTACATCTAGAATACCCC

AGCTCCGTAGTTGCAGACACTAATTTTAGAGACCGCAGCTGCATCATTAACAAAATGTTA

TTGGTGGGATGCAGAAGAGCGCGGGCAAGATTATACAGACGCCTTCAGGCGGTAGCCTGC

TCCTAGGCAGTCAAGACGGATCGCGATGGACGTGTTTGGACGGCGAATGTACAGGAAATT

TCCCGTGAGAACAACAAAATGAAGGATCGAGATTAAAAAGACTGCAGCGCCCGTGAACAA

CGGAGTTGAACGTGGTTGCCAGGTTTGTGATGGGCGGCCCTAGAAGATAACGGTAGAAGC

CCTTCGGACTTTTCAATGGGGACTACATTCTGGCCGCGAAGGACTCCGTGAATCGATAGG

TAGTGGAAATATGGATGAAACGGGCTGGGGCAACGCTCGGGACAACGTTTACTAGCTAAG

AGGCAGATTACGAGGATCGGGGAAGACTGAGAAGGCCTTGGCGTAGGAATTCGAGGCAAC

CAGGGGGATAAGGTAAATGCCGACTTCAAGTCGAGAATGCAAAGAAGCTGGCTATGGCCG

TAAATTAGACGATAAATAACAGAGCATCGGGGAGCCAAGTCTAATTGTAAAACTTTTGGT

CGAGGTGATGTCGTAGGTGAGATCTCCGAACGCAGCATGAACAAGGCGAATCGTGTAAGC

AGAGAAGAGTGCAGCACGATGCTAACACTGCCGCAGCAGTGGGACAACTTGGATGACTGG

CACATCACAGTAATGAGTCATATGATTCCGCCTGAAGGCTTCAATAAAGGACATAAGATG

CGAAGAAGATCTAGACCCAGTCTTTAAAACTGCACAACTTAGCCGAAATGATTGGTAGGT

GAGAGTATCGACGTGGGCACTGAGAGAAACCTGAATACGCGATAGCCATCATATCACGAT

GCTTCTGCACTCCAGCGTCACTAGAGTTCTTGTTCACATGCCTCGGTCCTCTCGTCAGTA

CATGGCCGGCCTTCTTTCCTCCATCGTTTCCCAGAGCATACTCTTCGAGTTGCCCTATTA

ACCTTCTATTTTCTGGCCGATGCCTGACTGGCGCCGCTTTGAGTATCATACTTCTTGTTT

TTGGCCAACCTGTTGGGCGTTGATTGCTTGGGTAACCATATCCTGCATTTTGTTGATTCC

GGGTCGGCAATGTGTTGTGGACTGTTCGTCGCCGCTCCTGATCTCACCGTACTTCACGGT

TGTCGCCCGGACATCACCCGGACGGGTTTAGGCCATGGCCGGTTGAGGTTAATCTTTACG

CGTCACCATAACTTCCTGCGAAACTGGTTTCCCCACGCCGCTCCCTTCGTGAGCCGATCG

GCGTGGTTAGGTCGCGCGGGTTTTTCCAGGCAGGGTAAGTCGATTCGCACAAGGTGACCC

ACGTACCTCGCTTTAACGTTCCGTTCTACCACTGGGTTAGGCTGCTTTGGGGCCGACTTG

ACGTTCTTTTTGCACGACCTCGTATGTGCTTATTCTCACTCGTAATGAACCGGTAAGCGA

GAGTTTGTTCTACGCTGAGGTGTCGTCGGACCCGAGTGTGCTTGCCGCTGGTCAATTGTT

GTATCGGCTCGACGTCTTATTTCGTATGGTACGGGATTATTAGTGCTTGCAGCGATGTCC

CTTTCGCTATTGATGTCTTCTCCTGGGTCACGCTACGTACTGAATTGATAAAACATTTCG

ACCTCAGTATCAGTGAGGTCGGGGGATGAGTTAGATCGCCGTGTAGGGCTTCTCACGTTG

TGTCCGTGTGGAGAGTCGGTTCTAGCGGTCGTATTATCCATATTCTCCCGTTCCATTTAC

GTTGAGCGACTTGAAACTTCCGATGTACTGGCCGGTAATAGTGCATTGTTTCTCAATTCA

TGCAAGTTGTCGTACATTATGGAATAGAACAAGAATTAAGGACCATTTCCACTAGCGATT

CTTACCTCCAACTCAAGATAGAGTCTAACCGATTATCGACGAACTTCGTCCGTCACGTCA

TTCTATGTCGAGCGGCCTTCTTACAGCGCTGTCGCACTATCTGTTCTACCTTATTACCGA

GAAACCTTGCTCCTGTTACCTGTGAGGGCGGGGCGGATATGGCAAAACGGGCATTCTTTT

TTACTTCCCGTGCCTCTTCATGCTATATACATGTATCTCTGGAGCCTAGACGCCCGGTCC
TTAACGTCCTAAAACTTTACTGACCCTGCTGGCGTCTTACCCTTTATAGTGGCTCTGCAT
AATGTGTTCTGTATATAGCCAAATACTGCCGACTATCCACTCGCTTAGCTGTCAATATCC
ATGCGCGTACTTTGATTTACCGGTCGACCTGCCTAATGTAACTGCCTTTCGCTCTCCACT
CCGCACCGGGGTTTCGAGTTTTTCTACCCCTCTCTTTATTACTACTCGTTTACTGCATCA
GACGCGCCTACCGTATTCAGTCACTGCTCCTTCAGGCGAAGAGTATTAGCACGTTATCGT
GCTACTATCCTGAGCTGTAATCATAAGTATTTCTAAAGTTGTAAGGGTTGAAGAGTAGTT
CTGCACATCTTGATCGATTGTACTGGACGTCATTCCGAATAGCTTGTGGAATCCAATATC
TGGGTCATAGATGCT
>SRR9587918
GTGACCGTCGCCTCGCCACAAATCCATTGTTTTCCGGGTTCTAGCCCCTTATGAGCTAGG
ACCTCTGTGCCTACCATCATAGCTGTTCGATGCCTTGTATTCAATACGTTGCACCGACCC
GACAATAATCTGTGCCATTAAGCTTCTGCAAGTGGCTGGCGCCTGTAAGTCACGCAATAG
AGTAACTTCCCTCTCGACAGTCTTTATGCCATCCACAATCCAACATCGTAGATCCTCGCG
TGACCTAGTAGCCCTTGACGACTCAATCCACCCTCTGCCTGACGGACGTGACTTTTATGG
TACATTAATGACTCCCTGTCCTCATAATTAGGCTGCACACACAGGGCTATCGTAGTTTAC
TGGTCTCACCCCTGCGTACTCGGCGACCCAACTACACTGATCCAGGGAGGCAGAATGGTG
GACGCTCTTACTGCCGACCCGTGGGCAACTAACTGGGCCGGGGGAATATTGTTTCGTTTT
GGGTACTGCTTCTTCCAATTGTTCGGATAGCCCAGTAACGGTGGTGGGTGGTTCCAGTCA
CTCTTGCGTAGTCGCCGGCCGGTCGCTCCGGTCTGTCCCCTCGCCCGATTCTGGGCCATC
TTATCCGATGTCACCCTGCAGACCGCTCTTTTCAACCATTAGCATTAAGCAGTCTCTCAT
AGGTGCTCTCATGTCATGAGAGGTACCGGAGCCTCTAAGCGTTCCGCTGTTCCCGACAAT
ATTCTCTTTCGCTTTGACACTAGCTCAGACGGTAGAGGCTTCGTCTTCAGCACCGGCTAA
TGATGCCTGGTACGTTAAGATACCGATCCGCGCGCTTACGCTGCTCCACCTGTCGGTGTC
TGAAGAAGCTCGGCTAAGACCTCTCGAACTCTTCGTCTTACCTCAGCCCGCGGCTACAGT
CATATACATCCTAGCACGCACAATCCTTAATTCCCAGTCGTGCCCGACAGTAACACCGTG
GCGCGAACAATGTGTAAGTTATTGCCTACTTCGTACACTATCAACTAAATTCGTCCAACG
TGAGATCCGAGAATCCCGAAAGGGGGCACACACTTAGAGAACCCGCAGCTTTTTGTCTCA
CGATCGAGAAGGGCACTCCCAAGAGGCCTTGATACAAGCGTGAAACCGGTCAAGCGCTCC
AATTGTACACTGTCCATGAGGCCAAATTCAAATTACGGACCGAAAACTGGATCCAGCAGA
GCTAAAGGTTATTCGCGGGTATTGTCCGTACATGCAACTTGATGTTTTCTAAGCCAAAAG
AGATATGCGAGGGGGAGACATGGGAAAAGCAAAAGAATATAGTACTATAATTGGCGTGGG
ACCCATAAAGTCTTGATTGCTCCCGCGCCGCCTGTACCCACAGTCTTCAAAAAAGCGATT
TGCCGGAGTATAAAGTCAGGGGCCGACGGTGTTGAAAACGAGCGGACGTCTTGCATGAGA
AGTGTCACTGTGGGCATATCAAAGGACTCTAATTAAATTCGGTGTCAATAGAAATGTTCG
TGCTGCACGGTGCTGTCAGACCATCCACCTTTCGTGGCTACGCCTTATCAGAAGTGCAAC

CTGCGAAATCAAGGTGACGTTAATACGCTCTAAAGGACGCTAAAGCGGGTTGATAATTAA
ACGACAGAATTACAATTAGCGGGATTCACAACTCTCCAAAATCATTGACGATAGACTGAC
ACGGTTACGTGGGAGGACTCGTAGTCTCCAGTGAAAATGGGCGCAAAATGGTGTCCGCAT
GGGAGGAGCTCAACCATAATAGACGCCGAATATGTTCGCGTGAAACTTCAGCGTCAAAGA
TGGAAAGCAAGACAAACTCATGAAACTACCATAGTGTGAAGATGATGGGCGGCGCGGTTT
AAGGCGAGTCGCCCTAAGACTCCAACGTGCCCTAAGGGATCGTAACATGCAAGAACATTG
ATGTGCACAGTTTTGTCGAACGGCCTTCATAGGCCAAAGATTGTATCCACGGCCAGAAGC
GGTAAGAACTTCGTGACATAGCGGCCAGAGAAAGGATCTACAACGTTAACCAGCACCTCA
GAGAGCGGGATGGCCAGTCGTTAGGGAGAAAATAACTAATCATTAACTTTGACGAACCAC
CAAAATGCTCGAGCGGTAGGTATCATTAGGGGGTACGGACTGTTACATCTAGAATACCCC
AGCTCCGTAGTTGCAGACACTAATTTTAGAGACCGCAGCTGCATCATTAACAAAATGTTA
TTGGTGGGATGCAGAAGAGCGCGGGCAAGATTATACAGACGCCTTCAGGCGGTAGCCTGC
TCCTAGGCAGTCAAGACGGATCGCGATGGACGTGTTTGGACGGCGAATGTACAGGAAATT
TCCCGTGAGAACAACAAAATGAAGGATCGAGATTAAAAAGACTGCAGCGCCCGTGAACAA
CGGAGTTGAACGTGGTTGCCAGGTTTGTGATGGGCGGCCCTAGAAGATAACGGTAGAAGC
CCTTCGGACTTTTCAATGGGGACTACATTCTGGCCGCGAAGGACTCCGTGAATCGATAGG
TAGTGGAAATATGGATGAAACGGGCTGGGGCAACGCTCGGGACAACGTTTACTAGCTAAG
AGGCAGATTACGAGGATCGGGGAAGACTGAGAAGGCCTTGGCGTAGGAATTCGAGGCAAC
CAGGGGGATAAGGTAAATGCCGACTTCAAGTCGAGAATGCAAAGAAGCTGGCTATGGCCG
TAAATTAGACGATAAATAACAGAGCATCGGGGAGCCAAGTCTAATTGTAAAACTTTTGGT
CGAGGTGATGTCGTAGGTGAGATCTCCGAACGCAGCATGAACAAGGCGAATCGTGTAAGC
AGAGAAGAGTGCAGCACGATGCTAACACTGCCGCAGCAGTGGGACAACTTGGATGACTGG
CACATCACAGTAATGAGTCATATGATTCCGCCTGAAGGCTTCAATAAAGGACATAAGATG
CGAAGAAGATCTAGACCCAGTCTTTAAAACTGCACAACTTAGCCGAAATGATTGGTAGGT
GAGAGTATCGACGTGGGCACTGAGAGAAACCTGAATACGCGATAGCCATCATATCACGAT
GCTTCTGCACTCCAGCGTCACTAGAGTTCTTGTTCACATGCCTCGGTCCTCTCGTCAGTA
CATGGCCGGCCTTCTTTCCTCCATCGTTTCCCAGAGCATACTCTTCGAGTTGCCCTATTA
ACCTTCTATTTTCTGGCCGATGCCTGACTGGCGCCGCTTTGAGTATCATACTTCTTGTTT
TTGGCCAACCTGTTGGGCGTTGATTGCTTGGGTAACCATATCCTGCATTTTGTTGATTCC
GGGTCGGCAATGTGTTGTGGACTGTTCGTCGCCGCTCCTGATCTCACCGTACTTCACGGT
TGTCGCCCGGACATCACCCGGACGGGTTTAGGCCATGGCCGGTTGAGGTTAATCTTTACG
CGTCACCATAACTTCCTGCGAAACTGGTTTCCCCACGCCGCTCCCTTCGTGAGCCGATCG
GCGTGGTTAGGTCGCGCGGGTTTTTCCAGGCAGGGTAAGTCGATTCGCACAAGGTGACCC
ACGTACCTCGCTTTAACGTTCCGTTCTACCACTGGGTTAGGCTGCTTTGGGGCCGACTTG
ACGTTCTTTTTGCACGACCTCGTATGTGCTTATTCTCACTCGTAATGAACCGGTAAGCGA
GAGTTTGTTCTACGCTGAGGTGTCGTCGGACCCGAGTGTGCTTGCCGCTGGTCAATTGTT

GTATCGGCTCGACGTCTTATTTCGTATGGTACGGGATTATTAGTGCTTGCAGCGATGTCC
CTTTCGCTATTGATGTCTTCTCCTGGGTCACGCTACGTACTGAATTGATAAAACATTTCG
ACCTCAGTATCAGTGAGGTCGGGGGATGAGTTAGATCGCCGTGTAGGGCTTCTCACGTTG
TGTCCGTGTGGAGAGTCGGTTCTAGCGGTCGTATTATCCATATTCTCCCGTTCCATTTAC
GTTGAGCGACTTGAAACTTCCGATGTACTGGCCGNNAATAGTGCATTGTTTCTCAATTCA
TGCAAGTTGTCGTACATTATGGAATAGAACAAGAATTAAGGACCATTTCCACTAGCGATT
CTTACCTCCAACTCAAGATAGAGTCTAACCGATTATCGACGAACTTCGTCCGTCACGTCA
TTCTATGTCGAGCGGCCTTCTTACAGCGCTGTCGCACTATCTGTTCTACCTTATTACCGA
GAAACCTTGCTCCTGTTACCTGTGAGGGCGGGGCGGATATGGCAAAACGGGCATTCTTTT
TTACTTCCCGTGCCTCTTCATGCTATATACATGTATCTCTGGAGCCTAGACGCCCGGTCC
TTAACGTCCTAAAACTTTACTGACCCTGCTGGCGTCTTACCCTTTATAGTGGCTCTGCAT
AATGTGTTCTGTATATAGCCAAATACTGCCGACTATCCACTCGCTTAGCTGTCAATATCC
ATGCGCGTACTTTGATTTACCGGTCGACCTGCCTAATGTAACTGCCTTTCGCTCTCCACT
CCGCACCGGGGTTTCGAGTTTTTCTACCCCTCTCTTTATTACTACTCGTTTACTGCATCA
GACGCGCCTACCGTATTCAGTCACTGCTCCTTCAGGCGAAGAGTATTAGCACGTTATCGT
GCTACTATCCTGAGCTGTAATCATAAGTATTTCTAAAGTTGTAAGGGTTGAAGAGTAGTT
CTGCACATCTTGATCGATTGTACTGGACGTCATTCCGAATAGCTTGTGGAATCCAATATC
TGGGTCATAGATGCT
>SRR9587919
GTGACCGTCGCCTCGCCACAAATCCATTGTTTTCCGGGTTCTAGCCCCTTATGAGCTAGG
ACCTCTGTGCCTACCATCATAGCTGTTCGATGCCTTGTATTCAATACGTTGCACCGACCC
GACAATAATCTGTGCCATTGAGCTTCTGCAAGTGGCTGGCGCCTGTAAGTCACGCAATAG
AGTAACTTCCCTCTCGACAGTCTTTATGCCATCCACAATCCAACATCGTAGATCCTCGCG
TGACCTAGTAGCCCTTGACGACTCAATCCACCCTCTGCCTGACGGACGTGCCTTTTATGG
TACATTAATGACTCCCTGTCCTCATAATTAGGCTGCACACACAGGGCTATCGTAGTTTAC
TGGTCTCACCCCTGCGTACTCGGCGACCCAACTACACTGATCCAGGGAGGCAGAATGGTG
GACGCGCTTAGTGCCGACCCGTGGGCAACTAACTGGGCCGGGGGAATATTGTTTCGTTTT
GGGTACTGCTTCTTCCAATTGTTCGGATAGCACAGTAACGGTGGTGGGTGGCTCCAGTCA
CTCTTGCGTAGTCCCCGGCCGGTCGCTCCGGTCTGTCCTCTCGCCCGATTCTGGGCCAAC
TTATCCGATGTCACCCTGCAGACCGCTCTTTTCAACTATTAGCATTAAGCAGTCTCTCAT
AGGTGCTCTCATGTCATGAGAGGTACCGGAGCCTCTAAGCGTTCCGCTGTTCCCGACAAT
ATTCTCTTTCGCTTTGACACTAGCTCAGACGGTAGAGGCTTCGTCTTCAGCACCGGCTAA
TGATGCCTGGTACGTTAAGATACCGATCCGCGCGCTTACGCTGCTCCACCTGTCGGTGTC
TGAAGAAGCTCGGCTAAGACCTCTCGAACTCTTCGTCTTACCTCAGCCCGCGGCTACAGT
CTAATACATCCTAGCACGCACAATCCTTAATTCCCAGTCGTGCCCGACAGTAACACCGTG
GCGCGAACAATGTGTAAGTTATTGCCTACTTCGTACACTATCAACTAAATTCGTCCAACG

TGAGATCCGAGAATCCCGAAAGGGGGCACACACTTAGAGAACCCGCAGCTTTTTGTCTCA
CGATCGAGAAGGGCACTCCCAAGAGGCCTTGATACAAGCGTGAAACCGGTCAAGCGCTCC
AATTGTACACTGTCCATGAGGCCAAATTCAAATTACGGACCGAAAACTGGATCCAGCAGA
GCTTAAGGTTATTCGCGGGTATTGTCCGTACATGCAACTTGATGTTTTCTAAGCCAAAAG
AGATATGCGAGGGGGAGACATGGGAAAAGCAAAAGAATATAGTACTATAATGGGCGTGGG
ACCCATAAAGTCTTGATTGCTCCCGCGCCGCCTGTACCCACAGTCTTCAAAAAAGCGATT
TGCCGGAGTATAAAGTCAGGGGCCGACGGTGTTGAAAACGAGCGGACGTCTTGCATGAGA
AGTGTCACTGTGGGCATATCAAAGGACTCTAATTAAATTCGGTGTCAATAGAAATGTTCG
TGCTGCACGGTGGTGTCAGACCGTCCACCTTTCGTGGCTACGCCTTATCAGAAGTGCAAC
CTGCGAAATCAAGGTGACGTTAATACGCTCTAAAGGACGCTAAAGCGGGTTGATAATTAA
ACGACAGAGTTACAATTAGCGGGATTCACAACTCTCCAAAATCATTGACGATAGACTGAC
ACGGTTACGTGGGAGGACTCGTAGTCTCAAGTGAAAATGGGCGCAAAATGGTGTCCGCAT
GGGAGGAGCTCAACCATAATAGACGCCGAATATGTTCGCGTGAAACTTCAGCGTCAAAGA
TGGAAAGCAAGACAAATTCATGAAACTACCATAGTGTGAAGATGATGGGCGGCGCGGTTT
AAGGCGAGTCGCCCTAAGACTCCAACGTGCCCTAAGAGATCGTAACATGCAAGAACATTG
ATGTGCACAGTTTTGTCGAACGGCCTTCATAGGCCAAAGATTGTATCCACGGCCAGAAGC
GGTAAGAACTTCGTGACATAGCGGCCAGAGAAAGGATCTACAACGTTAACCAGCACCTCA
GAGAGCGGGATGGCCAGTCGTTAGGGAGAAAATAACTAATCATTAACTTTGACGAACCAC
CAAAATGCTCGAGCGGTAGGTATCATTAGGGGGTACGGACTGTTACATCTAGAATACCCC
AGCTCCGTAGTTGCAGACACTAATTTTAGAGACCGCAGCTGTATCATTAACAAAATGTTA
TTGGTGGGATGCAGAAGAGCGCGGGCAAGATTATACAGACGCCTTCAGGCGGTAGCCTGC
TCCTAGGCAGTCAAGACGGATCGCGATGGACGTGTTTGGACGGCGAATGTACAGGAGATT
TCCCGTGAGAACAACAAAATGAAGGATCGAGATTAAAAAGACTGCAGCGCCCGTGAACAA
CGGAGTTGAACGTGGTTGCCAGGTTTGTGATGGGCGGCCCTAGAAGATAACGGTAGAAGC
CCTTCGGACTTTTCAATGGGGACTACATTCTGGCCGCGAAGGACTCCGTGAATCGATAGG
TAGTGGAAATATGGATGAAACGGGCTGGGGCAACGCTCGGGACAACGTTTACTAGCTAAG
AGGCAGATTACGAGGATAGGGGAAGACTGAGAAGGCCTTGGCGTAGGAATTCGAGGCAAC
CAGGGGGATAAGGTAAATGCCGACTTCAAGTCGAGAATGCAAAGAAGCTGGCTATGGCCG
TAAATTAGACGATAAATAACAGAGCATCGGGGAGCCAAGTCTAATTGTCAAACTTTTGGT
CGAGGTGATGTCGTAGGTGAGATCTCCGAACGCAGCATGAACAAGGCGAATCGTGTAAGC
AGAGAAGAGTGCAGCACGATGCTAACACTGCCGCAGCAGTGGGACAACTTGGATGACTGG
CACATCACAGTAATGAGTCATATGATTCCGCCTGAAGGCTTCAATAAAGGACATAAGATG
CGAAGAAGATCTAGACCCAGTCTTTAAAACTGCACAACTTAGCCGAAATGATTGGTAGGT
GAGAGTATCGACGTGGGCACTGAGAGAAACCTGAATACGCGATAGCCGTCATATCACGAT
GCTTCTGCCCTCCAGCGTCACTAGAGTTCTTGTTCACATGCCTCGGTCCTCTCGTCAGTA
CATGGCCGGCCTTCTTTCCTCCATCGTTTCCCAGAGCATACTCTTCGAGTTGCCCTATTA

ACCTTCTATTTTCTGGCCGATGCCTGACTGGCGCCGCTTTGAGTATCATACTTCTTGTTT
TTGGCCAACCTGTTGGGCGTTGATTGCTTGGGTAACCATATCCTGCATTTTGTTGATTCC
GGGTCGGCAATGTGTTGTGGACTGTTCGTCGCCGCTCCTGATCTCACCGTACTTCACGGT
TGTCGCCCGGACATCACCCGGACGGGTTTAGGCCATGGCCGGTTGAGGTTAATCTTTACG
CGTCACCATAACTTCCTGCGAAACTGGTTTCCCCACGCCGCTCCCTTCGTGGGCCGATCG
GCGTGGTTAGGTCGCGCGGGTTTTTCCAGGCAGGGTAAGTCGAGTCGCACAAGGTGACCC
ACGTACCTCGCTTTAACGTTCCGTTCTACCACTGGGTTAGGCTGCTTTGGGGCCGACTTG
ACGTTCTTTTTGCACGACCTCGTATGTGCTTATTCTCACTCGTAATGAACCGGTAAGCGA
GAGTTTGTTCTACGCTGAGGTGTCGTCGGACCCGAGTGTGCTTGCCGCTGGTCAATTGTT
GTGTCGGCTCGACGTCTTATTTCGTATGGTACGGGATTATTAGTGCTTGCAGCGATGTCC
CTTTCGCTATTGATGTCTTCTCCTGAGTCACGCTACGTACTGAATTGATAAAACATTTCG
ACCTCAGTATCAGTGAGGTCGGGGGATGAGTTAGATCGCCGTGTNGGGCTTTTCACGTTG
TGTCCGTGTGGAGAGTCGGTTCTAGCGGTCGTATTATCCGTATTCTCCCGTTCCATTTAC
GTTGAGCGACTTGAAACTTCTGATGTACTGGCNGNTAATAGTGCATTGTTCCTCAATTCA
TGCAAGTTGTCGTACATTATGGAATAGAACAAGAATTAAGGACCATTTCCACTAGCGATT
CTTACCTCCAACTCAAGATAGAGTCTAACCGATTATCGACGAACTTCGTCCGTCACGTCA
TTCCATGTCGAGCGGCCTTCTTACAGCGCTGTCGCACTATCTGTTCTACCTTATTTCCGA
GAAACCTTGCTCCTGTTACCTGTGAGGGCGGGGCGGATATGGCAAAATGGGCATTCTTTT
TTACTTCCCGTGCCTCTTCATGCTATATACATGTATCTCTGGAGCCTAGACGCCCGGTCC
TTAACGTCCTAAAACTTTATTGACCCTGCTGGCGTCTTACCCTTTATAGTGGCTCTGCAT
AATGTGTTCTGTATATAGCCAAATACTGCCGACTATCCACTCGCTTAGCTGTCAATATCC
ATGCGCGTACTTTGATTTACCGGTCGACCTGCCTAATGTAACTGCCTTTCGCTCTCCACT
CCGCACCGGGGTTTCGAGTTTTTCTACCCCTCTCTTTATTACTACTCGTTTACTGCACCA
GACGCGCCTACCGTATTCAGTCACTGCTCCTTCAGGCGAAGAGTATTAGCACGTTATCGT
GCTACTATCCTGAGCTGTAATCATAAGTATTTCTAAAGTTGTAGGGGTTGAAGAGTAGTT
CTGTACATCTTGATCGATTGTACTGGACGTCATTCCGAATAGCTTGTGGAATCCAATATC
TGGGTCATAGATGCT
>SRR9587920
GTGACCGTCGCCTCGCCACAAATCCATTGTTTTCCGGGTTCTAGCCCCTTATGAGCTAGG
ACCTCTGTGCCTACCATCATAGCTGTTCGATGCCTTGTATTCAATACGTTGCACCGACCC
GACAATAATCTGTGCCATTAAGCTTCTGCAAGTGGCTGGCGCCTGTAAGTCACGCAATAG
AGTAACTTCCCTCTCGACAGTCTTTATGCCATCCACAATCCAACATCGTAGATCCTCGCG
TGACCTAGTAGCCCTTGACGACTCAATCCACCCTCTGCCTGACGGACGTGCCTTTTATGG
TACATTAATGACTCCCTGTCCTCATAATTAGGCTGCACACACAGGGCTATCGTAGTTTAC
TGGTCTCACCCCTGCGTACTCGGCGACCCAACTACACTGATCCAGGGAGGCAGAATGGTG
GACGCTCTTACTGCCGACCCGTGGGCAACTAACTGGGCCGGGGGAATATTGTTTCGTTTT

GGGTACTGCTTCTTCCAATTGTTCGGATAGCACAGTAACGGTGGTGGGTGGCTCCAGTCA
TTCTTGCGTAGTCGCCGGCCGGTCGCTCCGGTCTGTCCTCTCTCCCGATTCTGGGCAATC
TTATCCGATGTCACCCTGCAGACCGCTCTTTTTAACCATTAGCATTAAGCAGTCTCTCAT
AGGTGCTCTCGTGTCATGAGAGGTACCGGAGCCTCTAAGCGTTCCGCTGTTCCCGACAAT
ATTCTCTTTCGCTTTGACACTAGCTCAGACGGTAGAGGCTTCGCCTTCAGCACCGGCTAA
TGATGCCTGGTACGTTAAGATACCGATCCGCGCGCTTACGCTGCTCCACCTGTCGGTGTC
TGAAGAAGCTCGGCTAAGACCTCTCGAACTCTTCGTCTTACCTCAGCCCGCGGCTACAGT
CATATACATCCTAGCACGCACAATCCTTAATTCCCAGTCGTGCCCGACAGTAACACCGTG
GCGCGAACAATGTGTAAGTTATTGCCTACTTCGTACACTATCAACTAAATTCGTCCAACG
TGAGATCCGAGAATCCCGTTAGGGGGCACACACTTAGAGAACCCGCAGCTTTTTGTCTCA
CGATCGAGAAGGGCACTCCCAAGAGGCCTTGATACGAGCGTGAAACCGGTCAAGCGCTCC
AATTGTACACTGTCCATGAGGCCAAATTCAAATTACGGACCGAAAACTGGATCCAGCAGA
GCTAAAGGTTATTCGCGGGTATTGTCCGTACATGCAACTTGATGTTTTCTAAGCCAAAAG
AGATATGCGAGGGGGAGACATGGGAAAAGCAAAAGAATATAGTACTATAATTGGCGTGGG
ACCCATAAAGTCTTGATTGCTCCCGCGCCGCCTGTACCCACAGTCTTCAAAAAAGCGATT
TGCCGGAGTATAAAGTCAGGGGCCGACGGTGTTGAAAACGAGCGGACGTCTTGCATGAGA
AGTGTCACTGTGGGCATATCAAAGGACTCTAATTAAATTCGGTGTCAATAGAAATGTTCG
TGCTGCACGGTGCTGTCAGACCATCCACCTTTCGTGGCTACGCCTTATCAGAAGTGCAAC
CTGCGAAATCAAGGTGACGTTAATACGCTCTAAAGGACGCTAAAGCGGGTTGATAATTAA
ACGACAGAGTTACAATTAGCGGGATTCACAACTCTCCAAAATCATTGACGATAGACTGAC
ACGGTTACGTGGGAGGACTCGTAGTCTCAAGTGAAAATGGGCGCAAAATGGTGTCCGCAT
GGGAGGAGCTCAACCATAATAGACGCCGAATATGTTCGCGTGAAACTTCAGCGTCAAAGA
TGGAAAGCAAGACAAACTCATGAAACTACCATAGTGTGAAGATGATGGGCGGCGCGGTTT
AAGGCGAGTCGCCCTAAGGCTCCAACGTGCCCTAAGGGATCGTAACATGCAAGAACATTG
ATGTGCACAGTTTTGTCGAACGGCCTTCATAGGCCAAAGATTGTATCCACGGCCAGAAGC
GGTAAGAACTTCGTGACATAGCGGCCAGAGAAAGGATCTACAACGTTAACCAGCACCTCA
GAGAGCGGGATGGCCAGTCGTTAGGGAGAAAATAACTAATCATTAATTTTGACGAACCAC
CAAAATGCTCGAGCGGTAGGTATCATTAGGGGGTACGGACTGTTACATCTAGAATGCCCC
AGCTCCGTAGTTGCAGACACTAATTTTAGAGACCGCAGCTGCATCATTAACAAAATGTTA
TTGGTGGGATGCAGAAGAGCGCGGGCAAGATTATACAGACGCCTTCAGGCGGTAGCCTGC
TCCTAGGCAGTCAAGACGGATCGCGATGGACGTGTTTGGACGGCGAATGTACAGGAAATT
TCCCGTGAGAACAACAAAATGAAGGATCGAGATTAAAAAGACTGCAGCGCCCGTGAACAA
CGGAGTTGAACGTGGTTTCCAGGTTTGTGATGGGCGGCCCTAGAAGATAACGGTAGAAGC
CCTTCGGACTTTTCAATGGGGACTACATTCTGGCCGCGAAGGACTCCGTGAATCGATAGG
TAGTGGAAATATGGATGAAACGGGCTGGGGCAACGCTCGGGACAACGTTTACTAGCTAAG
AGGCAGATTACGAGGATCGGGGAAGACTGAGAAGGCCTTGGCGTAGGAATTCGAGGCAAC

CAGGGGGATAAGGTAAATGCCGACTTCAAGTCGAGAATGCAAAGAAGCTGGCTATGGCCG
TAAATTAGACGATAAATAACAGAGCATCGGGGAGCCAAGTCTAATTGTAAAACTTTTGGT
CGAGGTGATGTCGTAGGTGAGATCTCCGAACGCAGCATGAGCAAGGCGAATCGTGTAAGC
AGAGAAGAGTGCAGCACGATGCTAACACTGCCGCAGCAGTGGGACAACTTGGATGACTGG
CACATCACAGTAATGAGTCATATGATTCCGCCTGAAGGCTTCAATAAAGGACATAAGATG
CGAAGAAGATCTAGACCCAGTCTTTAAAACTGCACAACTTAGCCGAAATGATTGGTAGGT
GAGAGTATCGACGTGGGCACTGAGAGAAACCTGAATACGCGATAGTCATCATATCACGAT
GCTTCTGCCCTCCAGCGTCACTAGAGTTCTTGTTCACATGCCTCGGTCCTCTCGTCAGTA
CATGGCCGGCCTTCTTTCCTCCATCGTTTCCCAGAGCATACTCTTCGAGTTGCCCTATTA
ACCTTCTATTTTCTGGCCGATGCCTGACTGGCGCCGCTTTGAGTATCATACTTCTTGTTT
TTGGCCAAACTGTTGGGCGTTGATTGCTTGGGTAACCATATCCTGCATTTTGTTGATTCC
GGGTCGGCAATGTGTTGTGGACTGTTCGTCGCCGCTCCTGATCTCACCGTACTTCACGGT
TGTCGCCCGGACATCACCCGGACGGGTTTAGGCCATGGCCGGTTGAGGTTAATCTTTACG
CGTCACCATAACTTCCTGCGAAACTGGTTTCCCCACGCCGCTCCCTTCGTGAGCCGATCG
GCGTGGTTAGGTCGCGCGGGTTTTTCCAGGCAGGGTAAGTCGAGTCGCACAAGGTGACCC
ACGTACCTCGCTTTAACGTTCCGTTCTACCACTGGGTTAGGCTGCTTTGGGGCCGACTTG
ACGTTCTTTTTGCACGACCTCGTATGTGCTTATTCTCACTCGTAATGAACCGGTAAGCGA
GAGTTTGTTCTACGCTGAGGTGTCGTCGGACCCGAGTGTGCTTGCCGCTGGTCAATTGTT
GTGTCGGCTCGACGTCTTATTTCGTATGGTACGGGATTATTAGTGCTTGCAGCGATGTCC
CTTTCGCTATTGATGTCTTCTCCTGAGTCACGCTACGTACTGAATTGATAAAACATTTCG
ACCTCAGTATCAGTGAGGTCGGGGGATGAGTTAGATCGCCGTGTAGGGCTTTTCACGCTG
TGTCCGTGTGGAGAGTCGGTTCTAGCGGTCGTATTATCCATATTCTCCCGTTCCATTTAC
GTTGAGCGACTTGAAACTTCCGATGTACTGGCCGGTAATAGTGCATTGTTTCTCAATTCA
TGCAAGTTGTCGTACATTATGGAATAGAACAAGAATTAAGGACCATTTCCACTAGCGATT
CTTACCTCCAACTCAAGATAGAGTCTAACCGATTATCGACGAACTTCGTCCGTCACGTCA
TTCTATGTCGAGCGGCCTTCTTACAGCGCTGTCGCACTATCTGTTCTACCTTATTTCCGA
GAAACCTTGCTCCTGTTACCTGTGAGGGCGGGGCGGATATGGCAAAATGGGCATTCTTTT
TTACTTCCCGTGCCTCTTCATGCTATATACATGTATCTCTGGAGCCTAGACGCCCGGTCC
TTAACGTCCTAAAACTTTACTGACCCTGCTGGCGTCTTACCCTTTATAGTGGCTCTGCAT
AATGTGTTCTGTATATAGCCAAATACTGCCGACTATCCACTCGCTTAGCTGCCAATATCC
ATGCGCGTACTTTGATTTACCGGTCGACCTGCCTAATGTAACTGCCTTTCGCTCTCCACT
CCGCACCGGGGTTTCGAGTTTTTCTACCCCTCTCTTTATTACTACTCGTTTACTGCATCA
GACGCGCCTACCGTATTCAGTCACTGCTCCTTCAGGCGAAGAGTATTAGCACGTTATCGT
GCTACTATCCTGAGCTGTAATCATAAGTATTTCTAAAGTTGTAAGGGTTGAAGAGTAGTT
CTGTACATCTTGATCGATTGTACTGGACGTCATTCCGAATAGCTTGTGGAATCCAATATC
TGGGTCATAGATGCT

>SRR9587921

GTGACCGTCGCCTCGCCACAAATCCATTGTTTTCCGGGTTCTAGCCCCTTATGAGCTAGG
ACCTCTGTGCCTACCATCATAGCTGTTCGATGCCTTGTATTCAATACGTTGCACCGACCC
GACAATAATCTGTGCCATTAAGCTTCTGCAAGTGGCTGGCGCCTGTAAGTCACGCAATAG
AGTAACTTCCCTCTCGACAGTCTTTATGCCATCCACAATCCAACATCGTAGATCCTCGCG
TGACCTAGTAGCCCTTGACGACTCAATCCACCCTCTGCCTGACGGACGTGCCTTTTATGG
TACATTAATGACTCCCTGTCCTCATAATTAGGCTGCACACACAGGGCTATCGTAGTTTAC
TGGTCTCACCCCTGCGTACTCGGCGACCCAACTACACTGATCCAGGGAGGCAGAATGGTG
GACGCGCTTACTGCCGACCCGTGGGCAACTAACTGGGCCGGGGGAATATTGTTTCGTTTT
GGGTACTGCTTCTTCCAATTGTTCGGATAGCACAGTAACGGTGGTGGGTGGCTCCAGTCA
CTCTTGCGTAGTCGCCGGCCGGTCGCTCCGGTCTGCCCTCTCGCCCGATTCTGGGCCATC
TTATCCGATGTCNCCCTGCAGACCGCTCTTTTCAACCATTAGCATTAAGCAGTCTCTCAT
AGGTGCTCTCATGTCATGAGAGGTACCGGAGCCTCTAAGCGTTCCGCTGTTCCCGACAAT
ATTCTCTTTCGCTTTGACACTAGCTCAGACGGTAGAGGCTTCGTCTTCAGCACCGGCTAA
TCATGCCTGGTACGTTAAGATACCGATCCGCGCGCTTACGCTGCTCCACCTGTCGGTGTC
TGAAGAAGCTCGGCTAAGACCTCTCGAACTCTTCGTCTTACCTCAGCCCGCGGCTACAGT
CATATACATCCTAGCACGCACAATCCTTAATTCCCAGTCGTGCCCGACAGTAACACCGTG
GCGCGAACAATGTGTAAGTTATTGCCTACTTCGTACACTATCAACTAAATTCGTCCAACG
TGAGATCCGAGAATCCCGAAAGGGGGCACACACTTAGAGAACCCGCAGCTTTTTGTCTCA
CGATCGAGAAGGGCACTCCCAATAGGCCTTGATACGAGCGTGAAACCGGTCAAGCGCTCC
AATTGTACACTGTCCATGAGGCCAAATTCAAATTACGGACCGAAAACTGGATCCAGCAGA
GCTAAAGGTTATTCGCGGGTATTGTCCGTACATGCAACTTGATGTTTTCTAAGCCAAAAG
AGATATGCGAGGGGGAGACATGGGAAAAGCAAAAGAATATAGTACTATAATTGGCGTGGG
ACCCATAAAGTCTTGATCGCTCCCGCGCCGCCTGTACCCACAGTCTTCAAAAAAGCGATT
TGCCGGAGTATAAAGTCAGGGGCCGACGGTGTTGAAAACGAGCGGACGTCTTGCATGAGA
AGTGTCACTGTGGGCATATCAAAGGACTCTAATTAAATTCGGTGTCAATAGAAATGTTCG
TGCTGCACGGTGCTGTCAGACCATCCACCTTTCGTGGCTACGCCTTATCGGAAATGCAAC
CTGCGAAATCAAGGTGACGTTAATACGCTCTAAAGGACGCTAAAGCGGGTTGATAATTAA
ACGACAGAGTTACAATTAGCGGGATTCACAACTCTCCAAAATCATTGACGATAGACTGAN
ACGGTTACGTGGGAGGACTCGTAGTCTCAAGTGAAAATGGGCGCAAAATGGTGTCCGCAT
GGGAGGAGCTCAACCATAATAGACGCCGAATATGTTCGCGTGAAACTTCAGCGTCAAAGA
TGGAAAGCAAGGCAAACTCATGAAACTACCATAGTGTGAAGATGATGGGCGGCGTGGTTT
AAGGCGAGTCGCCCTAAGACTCCAACGTGCCCTAAGGGATCGTAACATGCAAGAACATTG
ATGTGCACAGTTTTGTCGAACGGCCTTCATAGGCCAAAGATTGTATCCACGGCCAGAAGC
GGTAAGAACTTCGTGACATAGCGGCCAGAGAAAGGATCTACAACGTTAACCAGCACCTCA
GAGAGCGGGATGGCCAGTCGTTAGGGAGAAAATAACTAATCATTAACTTTGACGAACCAC

CAAAATGCTCGAGCGGTAGGTATCATTAGGGGGTACGGACTGTTACATCTAGAATACCCC
AGCTACGTAGTTGCAGACACTAATTTTAGAGACCGCAGCTGTATCATTAACAAAATGTTA
TTGGTGGGATGCAGAAGAGCGCGGGCAAGATTATACAGACGCCTTCAGGCGGTAGCCTGC
TCCTAGGCAGTCAAGACGGATCGCGATGGACGTGTTTGGACGGCGAATGTACAGGAAATT
TCCCGTGAGAACAACAAGATGAAGGATCGAGATTAAAAAGACTGCAGCGCCCGTGAACAA
CGGAGTTGAACGTGGTTGCCAGGTTTGTGATGGGCGGCCCTAGAAGATAACGGTAGAAGC
CCTTCGGACTTTTCAATGGGGACTACATTCTGGCCGCGAAGGACTCCGTGAATCGATAGG
TAGTGGAAATATGGATGAAACGGGCTGGGGCAACGCTCGGGACAACGTTTACTAGCTAAG
AGGCAGATTACGAGGATCGGGGAAGACTGAGAAGGCCTTGGCGTAGGAATTCAAGGCAAC
CAGGGGGATAAGGTAAATGCCGACTTCAAGTCGAGAATGCAAAGAAGCTGGCTATGGCCG
TAAATTAGACGATAAATGACAGAGCATCGGGGAGCCAAATCTAATTGTAAAACTTTTGGT
CGAGGTGATGTCGTAGGTGAGATCTCCGAACGCAGCATGAACAAGGCGAATCGTGTAAGC
AGGGAAGAGTGCAGCACGATGCTAACACTGCCGCAGCAGTGGGACAACTTGGATGACTGG
CACATCACAGTAATGAGTCATATGATTCCGCCTGAAGGCTTCAATAAAGGACATAAGATG
CGAAGAAGATCTAGACCCAGTCTTTAAAACTGCACAACTTAGCCGAAATGATTGGTAGGT
GAGAGTATCGACGTGGGCACTGAGAGAAACCTGAATACGCGATAGCCATCATATCACGAT
GCTTCTGCCCTCCAGCGTCACTAGAGTTCTTGTTCACATGCCTCGGTCCTCTCGTCAGTA
CATGGCCGGCCTTCTTTCCTCCATCGTTTCCCAGAGCATACTCTTCGAGTTGCCCTATTA
ACCTTCTATTTTCTGGCCGATGCCTGACTGGCGCCGCTTTGAGTATCATACTTCTTGTTT
TTGGCCAACCTGTTGGGCGTTGATTGCTTGGGTAACCATATCCTGCATTTTGTTGATTCC
GGGTCGGCAATGTGTTGTGGACTGTTCGTCGCCGCTCCTGATCTCACCGTACTTCACGGT
TGTCGCCCGGACATCACCCGGACGGGTTTAGGCCATGGCCGGTTGAGGTTAATCTTTACG
CGTCACCATAACTTCCTGCGAAACTGGTTTCCCCACGCCGCTCCCTTCGTGAGCCGATCG
GCGTGGTTAGGTCGCGCGGGTTTTTCCAGGCAGGGTAAGTCGAGTCGCACAAGGTGACCC
ACGTACCTCGCTTTGACGTTCCGTTCTACCACTGGGTTAGGCTGCTTTGGGGCCGACTTG
ACGTTCTTTTTGCACGACCTCGTATGTGCTTATTCTCACTCGTAATGAACCGGTAAGCGA
GAGTTTGTTCTACGCTGAGGTGTCGTCGGACCCGAGTGTGCTTGCCGCTGGTCAATTGTT
GTGTCGGCTCGACGTCTTATTTCGTATGGTACGGGATTATTAGTGCTTGCAGCGATGTCC
CTTTCGCTATTGATGTCTTCTCCTGAGTCACGCTACGTACTGAATTGATAAAACATTTCG
ACCTCAGTATCAGTGAGGTCGGGGGATGAGTTAGATCGCCGTGTAGGGCTTTTCACGTTG
TGTCCGTGTGGAGAGTCGGTTCTAGCGGTCGTATTATCCATATTCTCCCGTTCCATTTAC
GTTGAGCGACTTGAAACTTCCGATGTACTGGCCGGTAATAGTGCATTGTTTCTCAATTCA
TGCAAGTTGTCGTACATTATGGAATAGAACAAGAATTAAGGACCATTTCCACTAGCGATT
CTCACCTCCAACTCAAGATAGAGTCTAACCGATTATCGACGAACTTCGTCCGTCACGTCA
TTCTATGTCGAGCGGCCTTCTTACAGCGCTGTCGCACTATCTGTTCTACCTTATTTCCGA
GAAACCTTGCTCCTGTTACCTGTGAGGGCGGGGCGGATATGGCAAAATGGGCATTCTTTT

TTACTTCCCGTGCCTCTTCATGCTATATACATGTATCTCTGGAGCCTAGACGCCCGGTCC
TTAACGTCCTAAAACTTTACTGACCCTGCTGGCGTCTTACCCTTTATAGTGGCTCTGCAT
AATGTGGTCTGTATATAGCCAAATACTGCCGACTATCCACTCGCTTAGCTGTCAATATCC
ATGCGCGTACTTTGATTTACCGGTCGACCTGCCTAATGTAACTGCCTTTCGCTCTCCACT
CCGCACCGGGGTTTCGAGTTTTTCTACCCCTCTCTTTATTACTACTCGTTTACTGCATCA
GACGCGCCTACCGTATTCAGTCACTGCTCCTTCAGGCGAAGAGTATTAGCACGTTATCGT
GCTACTATCCTGAGCTGTAATCATAAGTATTTCTAAAGTTGTAAGGGTTGAAGAGTAGTT
CTGTACATCTTGATCGATTGTACTGGACGTCATTCCGAATAGCTGGTGGAATCCAATATC
TGGGTCATAGATGCT
>SRR9587922
GTGACCGTCGCCTCGCCACAAATCCATTGTTTTCCGGATTCTAGCCCCTTATGAGCTAGG
ACCTCTGTGCCTACCATCATAGCTGTTCGATGCCTTGTATTCAATACGTTGCACCGACCC
GACAATAGTCTGTGCCATTAAGCTTCTGCAAGTGGCTGGCGCCTGTAAGTCACGCAATAG
AGTAACTTCCCTCTCGACAGTCTTTATGCCATCCACAATCCAACATCGTAGATCCTCGCG
TGACCTAGTAGCCCTTGACGACTCAATCCACCCTCTGCCTGACGGACGTGCCTTTTATGG
TACATTAATGACTCCCTGTCCTCATAATTAGGCTGCACACACAGGGCTATCGTAGTTTAC
TGGTCTCACCCCCGCGTACTCGGCGACCCAACTACACTGATCCAGGGAGGCAGAATGGTG
GACGCGCTTACTGCCGACCCGTGGGCAACTAACTGGGCCGGGGGAATATTGTTTCGTTTT
GGGTACTGCTTCTTCCAATTGTTCGGATAGCACAGTAACGGTGGTGGGTGGCTCCAGTCA
CTCTTGCGTAGTCGCCGGCCGGTCGCTCCGGTCTGTCCTCTCGCCCGATTCTGGGCCATC
TTATCCGATGTCACCCTGCAGACCGCTCTTTTCAACCATTAGCATTAAGCAGTCTCTCAT
AGGTGCTCTCATGTCATGAGAGGTACCGGAGCCTCTAAGCGTTCCGCTGTTCCCGACAAT
ATTCTCTTTCGCTTTGACACTAGCTCAGACGGTAGAGGCTTCTTCTTCAGCACCGGCTAA
TGATGCCTGGTACGTTAAGATACCGATCCGCGCGCTTACGCTGCTCCACCTGTCGGTGTC
TGAAGAAGCTCGGCTAAGACCTCTCGAACTCTTCGTCTTACCTCAGCCCGCGGCTACAGT
CATATACATCCTAGCACGCACAATCCTTNATTCCCAGTCGTGCCCGACAGTAACACCGTG
GCGCGAACAATGTGTAAGTTATTGCCTACTTCGTACACTATCAACTAAATTCGTCCAACG
TGAGATCCGAGAATCCCGAAAGGGGGCACACACTTAGAGAACCCGCAGCTTTTTGTCTCA
CGATCGAGAAGGGCACTCCCAAGAGGCCTTGATACGAGCGTGAAACCGGTCAAGCGCTCC
AATTGTACACTGTCCATGAGGCCAAATTCAAATTACGGACCGAAAACTGGATCCAGCAGA
GCTAAAGGTTATTCGCGGGTATTGTCCGTACATGCAACTTGATGTTTTCTAAGCCAAAAG
AGATATGCGAGGGGGAGACATGGGAAAAGCAAAAGAATATAGTACTATAATTGGCGTGGG
ACCCATAAAGTCTTGATTGCTCCCGCGCCGCCTGTACCCACAGTCTTCAAAAAAGCGATT
TGCCGGAGTATAAAGTCAGGGGCCGACGGTGTTGAAAACGAGCGGACGTCTTGCATGAGA
AGTGTCACTGTGGGCATATCAAAGGACTCTAATTAAATTCGGTGTCAATAGAAATGTTCG
TGCTGCACGGTGCTGCCAGACCATCCACCTTTCGTGGCTACGCCTTATCAGAAGTGCAAC

CTGCGAAATCAAGGTGACGTTAATACGCTCTAAAGGACGCTAAAGCGGGTTGATAATTAA
ACGACAGAGTTACAATTAGCGGGATTTACAACTCTCCAAAATTATTGACGATAGACTGAC
ACGGTTACGTGGGAGGACTCGTAGTCTCAAGTAAAAATGGGCGCAAAATGGTGTCCGCAT
GGGAGGAGCTCAATCATAATAGACGCCGAATATGTTCGCGTGAAACTTCAGCGTCAAAGA
TGGAAAGCAAGACAAACTCATGAAACTACCATAGTGTGAAGATGATGGGCGGCGCGGTTT
AAGGCGAGTCGCCCTAAGACTCCAACGTGCCCTAAGGGATCGTAACATGCAAGAACATTG
ATGTGCACAGTTTTGTCGAACGGCCTTCATAGGCCAAAGATTGTATCCACGGCCAGAAGC
GGTAAGAACTTCGTGACATAGCGGCCAGAGAAAGGATCTACAACGTTAACCAGCACCTCA
GAGAGCGGGATGGCCAGTCGTTAGGGAGAAAATAACTAATCATTAACTTTGACGAACCAC
CAAAATGCTCGAGCGGTAGGTATCATTAGGGGGTACGGACTGTTACATCTAGAATACCCC
AGCTCCGTAGTTGCAGACACTAATTTTAGAGACCGCAGCTGTATCATTAACAAAATGTTA
TTGGTGGGATGCAGAAGAGCGCGGGCAAGATTATACAGACGCCTTCAGGCGGTAGCCTGC
TCCTAGGCAGTCAAGACGGATCGCGATGGACGTGTTTGGACGGCGAATGTACAGGAAATT
TCCCGTGAGAACAACAAAATGAAGGATCGAGATTAAAAAGACTGCAGCGCCTGTGAACAA
CGGAGTTGAACGTGGTTGCCAGGTTTGTGATGGGCGGCCCTAGAAGATAACGGTAGAAGC
CCTTCGGACTTTTCAAGGGGGACTACATTCTGGCCGCGAAGGACTCCGTGAATCGATAGG
TAGTGGAAATATGGATGAAACGGGCTGGGGCAACGCTCGGGACAACGTTTACTAGCTAAG
AGGCAGATTACGAGGATCGGGGAAGACTGAGAAGGCCTTGGCGTAGGAATTCGAGGCAAC
CAGGGGGATAAGGTAAATGCCGACTTCAAGTCGAGAATGCAAAGAAGCTGGCTATGGCCG
TAAATTAGACGATAAATAACAGAGCATCGGGGAGCCAAGTCTAATTGTAAAACTTTTGGT
CGAGGTGATGTCGTAGGTGAGATCTCCGAACGCAGCATGAACAAGGCGAATCGTGTAAGC
AGAGAAGAGTGCAGCACGATGCTAACACTGCCGCAGCAGTGGGACAACTTGGATGACTGG
CACATCACAGTAATGAGTCATATGATTCCGCCTGAAGGCTTCAATAAAGGACATAAGATG
CGAAGAAGATCTAGACCCAGTCTTTAAAACTGCACAACTTAGCCGAAATGATTGGTAGGT
GAGAGTATCGACGTGGGCACTGAGAGAAACCTGAATACGCGATAGCCATCGTATCACGAT
GCTTCTGCCCTCCAGCGTCACTAGAGTTCTTGTTCACATGCCTCGGTCCTCTCGTCAGTA
CATGGCCGGCCTTCTTTCCTCCATCGTTTCCCAGAGCATACTCTTCGAGTTGCCCTATTA
ACCTTCTATTTTCTGGCCGATGCCTGACTGGCGCCGCTTTGAGTATCATACTTCTTGTTT
TTGGCCAGCCTGTTGGGCGTTGATTGCTTGGGTAACCATATCCTGCATTTTGTTGATTCC
GGGTCGGCAATGTGTTGTGGACTGTTCGTCGCCGCTCCTGATCTCACCGTACTTCACGGT
TGTCGCCCGGACATTACCCGGACGGGTTTAGGCCATGGCCGGTTGAGGTTAATCTTTACG
CGTCACCATAACTTCCTGCGAAACTGGTTTCCCCACGCCGCTCTTTTCGTGAGCCGATCG
GCGTGGTTAGGTCGCGCGGGTTTTTCCAGGCAGGGTAAGTCGAGTCGCACAAGGTGACCC
ACGTACCTCGCTTTAACGTTCCGTTCTACCACTGGGTTAGGCTGCTTTGGGGCCGACTTG
ACGTTCTTTTTGCACGACCCCGTATGTGCTTATTCTCACTCGTAATGAACCGGTAAGCGA
GAGTTTGTTCTACGCTGAGGTGTCGTCGGACCCGAGTGTGCTTGCCGCTGGTCAATTGTT

GTGTCGGCTCGACGTCTTATTTCGTATGGTACGGGATTCTTAGTGCTTGCAGCGATGTCC
CTTTCGCTATTGATGTCTTCTCCTGAGTCACGCTACGTACTGAATTGATAAAACATTTCG
ACCTCAGTATCAGTGAGGTCGGGGGATGAGTTAGATCGCCGTGTAGGGCTTTTCACGTTG
TGTCCGTGTGGAGAGTCGGTTCTAGCGGTCGTATTATCCATATTCTCCCGTTCCATTTAC
GTTGAGCGACTTGAAACTTCCGATGTACTGGCCGGTAATAGTGCATTGTTTCTCAATTCA
TGCAAGTTGTCGTACATTATGGAATAGAACAAGAATTAAGGACCATTTCCACTAGCGATT
CTTACCTCCAACTCAAGATAGAGTCTAACCGATTATCGACGAACTTCGTCCGTCACGTCA
TTCTATGTCGAGCGGCCTTCTTACAGCGCTGTCGCACTATCTGTTTTACCTTATTTCCGA
GAAACCTTGCTCCTGTTACCTGTGAGGGCGGGGCGGATATGGCAAAATGGGCATTCTTTT
TTACTTCCCGTGCCTCTTCATGCTATATACATGTATCTCTGGAGCCTAGACGCCCGGTCC
TTAACGTCCTAAAACTTTACTGACCCTGCTGGCGTCTTACCCTTTATAGTGGCTCTGCAT
AATGTGTTCTGTATATAGCCAAATACTGCCGACTATCCACTCGCTTAGCTGTCAATATCC
ATGCGCGTACTTTGATTTACCGGTCGACCTGCCTAATGTAACTGCCTTTCGCTCTCCACT
CCGCACCGGGGTTTCGAGTTTTTCTACCCCTCTCTTTATTACTACTCGTTTACTGCATCA
GACGCGCCTACCGTATTCAGTCACTGCTCCTTCAGGCGAAGAGTATTAGCACGTTATCGT
GCTACTATCCTGAGCTGTAATCATAAGTATTTCTAAAGTTGTAAGGGTTGAAGAGTAGTT
CTGTACATCTTGATCGATTGTACTGGACGTCATTCCGAATAGCTTGTGGAATCCAATATC
TGGGTCATAGATGCT
>SRR9587923
GTGACCGTCGCCTCGCCACAAATCCATTGTTTTCCGGGTTCTAGCCCCTTATGAGCTAGG
ACCTCTGTGCCTACCATCATAGCTGTTCGATGCCTTGTATTCAATACGTTGCACCGACCC
GACAATAGTCTGTGCCATTAAGCTGCTGCAAGTGGCTGGCGCCCGTAAGTCACGCAATAG
AGTAACTTCCCTCTCGACAGTCTTTATGCCATCCACAATCTAACATCGTAGATCCTCGCG
TGACCTAGTAGCCCTTGACGACTCAATCCACCCTCTGCCTGACGGACGTGCCTTTTATGG
TACATTAATGACTCCCTGTCCTCATAATTAGGCTGCACACACAGGGCTATCGTAGTTTAC
TGGTCTCACCCCTGCGTACTCGGCGACCCAACTACACTGATCCAGGGAGGCAGAATGGTG
GACGCGCTTACTCCCGACCCGTGGGCAACTAACTGGGCCGGGGGAATATTGTTTCGTTTT
GGGTACTGCTTCTTCCAATTGTTCGGATAGCACAGTAACGGTGGTGGGTGGCTCCAGTCA
CTCTTGCGTAGTCGCCGGCCGGTCGCTCCGGTCTGTCCTCTCGCCCGATTCTGGGCCATC
TTATCCGATGTCACCCTGCAGACCGCTCTTTTCAACCATTAGCATTAAGCAGTCTCTCAT
AGGTGCTCTCATGTCATGAGAGGTACCGGAGCCTCTAAGCGTTCCGCTGTTCCCGACAAT
ATTCTCTTTCGCTTTGACACTAGCTCAGACTGTAGAGGCTTCGTCTTCAGCACCGGCTAA
TGATGCCTGGTACGTTAAGATACCGATCCGCGCGCTTACGCTGCTCCACCTGTCGGTGTC
TGAAGAAGCTCGGCTAAGACCTCTCGAACTCTTCGTCTTACCTCAGCCCGCGGCTACAGT
CATATACATCCTAGCACGCACAATCCTTAATTCCCAGTCGTGCCCGACAGTAACAACGTG
GCGCGAACAATGTGTAAGTTATTGCCTACTTCGTACACTATCAACTAAATTCGTCCAACG

TGAGATCCGAGAATCCCGAAAGGGGGCACACACTTAGAGAACCCGCAGCTTTTTGTCTCA

CGATCGAGGAGGGCACTCCCAAGAGGCCTTGATACGAGCGTGAAACCGGTCAAGCGCTCC

AATTGTACACTGTCCATGAGGCCAAATTCAAATTACGGACCGAAAACTGGATCCAGCAGA

GCTAAAGGTTATTCGCGGGTATTGTCCGTACATGCAACTTGATGTTTTCTAAGCCAAAAG

AGATATGCGAGGGGGAGACATGGGAAAAGCAAAAGAATATAGTACTATAATTGGCGTGGG

ACCCATAAAGTCTTGATTGCTCCCGCGCCGCCTGTGCCCACAGTCTTCAAAAAAGCGATT

TGCCGGAGTATAAAGTCAGGGGCCGACGGTGTTGAAAACGAGCGGACGTCTTACATGAGA

AGTGTCACTGTGGGCATATCAAAGGACTCTAATTAAATTCGGTGTCAATAGAAATGTTCG

TGCTGCACGGTGCTGTCAGACCATCCACCTTTCGTGGCTACGCCTTATCAGAAGTGCAAC

CTGCGAAATCAAGGTGACGTTAATACGCTCTAAAGGACGCTAAAGCGGGTTGATAATTAA

ACGACAGAGTTACAATTAGCGGGATTCACAACTCTCCAAAATCATTGACAATAGACTGAC

ACGGTTACGTGGGAGGACTCGTAGTCTCAAGTGAAAATGGGCGCAAAATGGTGTCCGCAT

GGGAGGAGCTCAACCATAATAGACGCCGAATATGTTCGCGTGAAACTTCAGAGTCAAAGA

TGGAAAGCAAGACAAACTCATGAAACTACCATAGTGTGAAGATGATGGGCGGCGCGGTTT

AAGGCGAGTCGCCCTAAGACTCCAACGTGCCCTAAGGGATCGTAACATGCAAGAACATTG

ATGTGCACAGTTTTGTCGAACGGCCTTCATAGGCCAAAGATTGTATCCACGGCCAGAAGC

GGTAAGAACTTCGTGACATAGCGGCCAGAGAAAGGATCTACAACGTTAACCAGCACCTCA

GAGAGCGGGATGGCCAGTCGTTAGGGAGAAAATAACTAATCATTAACTTTGACGAACCAC

CAAAATGCTCGAGCGGTAGGTATCATTAGGGGGTACGGACTGTTACATCTAGAATACCCC

AGCTCCGTAGTTGCAGACACTAATTTTAGAGACCGCAGCTGTATCATTAACAAAATGTTA

TTGGTGGGATGCAGAAGAGCGCGGGCAAGATTATACAGACGCCTTCAGGCGGTAGCCTGC

CCCTAGGCAGTCAAGACGGATCGCGATGGACGTGTTTGGACGGCGAATGTACAGGAAATT

TCCCGTGAGAACAACAAAATGAAGAATCGAGATTAAAAAGACTGCAGCGCCCGTGAACAA

CGGAGTTGAACGTGGTTGCCAGGTTTGTGATGGGCGGCCCTAGAAGATAACGGTAGAAGC

CCTTCGGACTTTTCAATGGGGACTACATTCTGGCCGCGAAGGACTCCGTGAATCGATAGG

TAGTGGAAATATGGATGAAACGGGCTGGGGCAATGCTCGGGACAAAGTTTACTAGCTAAG

AGGCAGATTACGAGGATCGGGGAAGACTGAGAAGGCCTTGGCGTAGGAATTCGAGGCAAC

CAGGGGGATAAGGTAAATGCCGACTCCAAGTCGAGAATGCAAAGAAGCTGGCTATGGCCG

TAAATTAGACGATAAATAACAGAGCATCGGGGAGCCAAGTCTAATTGTAAAACTTTTGGT

CGAGGTGATGTCGTAGGTGAGATCTCCGAACGCAGCATGAACAAGGCGAATCGTGTAAGC

AGAGAAGAGTGCAGCACGATGCTAACACTGCCGCAGCAGTGGGACAACTTGGATGACTGG

CACATCACAGTAATGAGTCATATGATTCCGCCTGAAGGCTTCAATAAAGGACATAAGATG

CGAAGAAGATCTAGACCCAGTCTTTAAAACTGCACAACTTAGCCGAAATGATTGGTAGGT

GAGAGTATCGACGTGGGCACTGAGAGAAACCTGAATACGCGATAGCCATCATATCACGAT

GCTTCTGCCCTCCAGCGTCACTAGAGTTCTTGTTCACATGCCTCGGTCCTCTCGTCAGTA

CATGGCCGGCCTTCTTTCCTCCATCGTTTCCCAGAGCATACTCCTCGAGTTGCCCTATTA

ACTTTCTATTTTCTGGCCGATGCCTAACTGGCGCCGCTTTGAGTATCATACTTCTTGTTT
TTGGCCAACCTGTTGGGCGTTGATTGCTTGGGTAACCATATCCTGCATTTTGTTGATTCC
GGGTCGGCAATGTGTTGTGGACTGTTCGTCGCCGCTCCTGATCTCACCGTACTTCACGGT
TGTCGCCCGGACATCACCCGGACGGGTTTAGGCCATGGCCGGTTGAGGTTAATCTTTACG
CGTCACCATAACTTCCTGCGAAACTGGTTTCCCCACACCGCTCCCTTCGTGAGCCGATCG
GCGTGGTTAGGTCGCGCGGGTTTTTCCAGGCAGGGTAAGTCGAGTCGCACAAGGTGACCC
ACGTACCTCGCTTTAACGTTCCGTTCTACCACTGGGTTAGGCTGCTTTGGGGCCGACTTG
ACGTTCTTTTTGCACGACCTCGTATGTGCTTATTCTCACTCGTAATGAACCGGTAAGCGA
GAGTTTGTTCTACGCTGAGGTGTCGTCGGACCCGAGTGTGCTTGCCGCTGGTCAATTGTT
GTGTCGGCTCGACGTCTTATTTCGTATGGTACGGGATTATTAGTGCTTGCAGCGATGTCC
CTTTCGCTATTGATGTCCTCTCCTGAGTCACGCTACGTACTGAATTGATAAAACATTTCG
ACCTCAGTATCAGTGAGGTCGGGGGATGAGTTAGATCGCCGTGTAGGGCTTTTCACGTTG
TGTCCGTGTGGAGAGTCGGTTCTAGCGGTCGTATTATCCATATTCTCCCGTTCCATTTAC
GTTGAGCGACTTGAAACTTCCGATGTACTGGCCGGTAATAGTGCATTGTTTCTCAATTCA
TGCAAGTTGTCGTACATCATGGAATAGAACAAGAATTAAGGACCATTTCCACTAGCGATT
CTTACCTCCAACTCAAGATAGAGTCTAACCGACTATCGACGAACTTCGTCCGTCACGTCA
TTCTATGTCGAGCGGCCTTCTTACAGCGCTGTCGCACTATCTGTTCTACCTTATTTCCGA
GAAACCTTGCTCCTGTTACCTGTGAGGGCGGGGCGGATATGGCAAGATGGGCATTCTTTT
TTACTTCCCGTGCCTCTTCATGCTATATACATGTATCTCTGGAGCCTAGACGCCCGGTCC
TTAACGTCCTAAAACTTTACTGACCCTGCTGGCGTCTTACCCTTTATAGTGGCTCTGCAT
AATGTGTTCTGTATATAGCCAAATACTGCCGACTATCCACTCGCTTAGCTGTCAATATCC
ATGCGCGTACTTTGATTTACCGGTCGACCTGCCTAATGTAACTGCCTTTCGCTCTCCACT
CCGCACCGGGGTTTCGAGTTTTTCTACCCCTCTCTTTATTGCTACTCGTTTACTGCATCA
GACGCGCCTACCGTATTCAGTCACTGCTCCTTCAGGCGAAGAGTATTAGCACGTTATCGT
GCTACTATCCTGAGCTGTAATCATAAGTATTTCTAAAGTTGTAAGGGTTGAAGAGTAGTT
CTGTTCATCTTGATCGATTGTACTGGACGTCATTCCGAATAGCTTGTGGAATCCAATATC
TGGGTCATAGATGCT
>SRR9587924_9587954
GTGACCGTCGCCTCGCCACAAATCCATTGTTTTCCGGATTCTAGCCCCTTATGAGCTAGG
ACCTCTGTGCCTACCATCATAGCTGTTCGATGCCTTGTATTCAATACGTTGCACCGACCC
GACAATAGTCTGTGCCATTAAGCTTCTGCAAGTGGCTGGCGCCTGTAAGTCACGCAATAG
AGTAACTTCCCTCTCGACAGTCTTTATGCCATCCACAATCCAACATCGTAGATCCTCGCG
TGACCTAGTAGCCCTTGACGACTCAATCCACCCTCTGCCTGACGGACGTGCCTTTTATGG
TACATTAATGACTCCCTGTCCTCATAATTAGGCTGCACACACAGGGCTATCGTAGTTTAC
TGGTCTCACCCCCGCGTACTCGGCGACCCAGCTACACTGATCCAGGGAGGCAGAATGGTG
GACGCGCTTACTGCCGACCCGTGGGCAACTAACTGGGCCGGGGGAATATTGTTTCGTTTT

GGGTACTGCTTCTTCCAATTGTTCGGATAGCACAGTAACGGTGGTGGGTGGCTCCAGTCA
CTCTTGCGTAGTCGCCGGCCGGTGGCTCCGGTCTGTCCTCTCGCCCGATTCTGGGCCATC
TTATCCGATGTCACCCTGCAGACCGCTCTTTTCAACCATTAGCATTAAGCAGTCTCTCAT
AGGTGCTCTCATGTCATGAGAGGTACCGGAGCCTCTAAGCGTTCCGCTGTTCCCGACAAT
ATTCTCTTTCGCTTTGACACTAGCTCAGACGGTAGAGGCTTCGTCTTCAGCACCGGCTAA
TGATGCCTGGTACGTTAAGATACCGATCCGCGCGCTTACGCTGCTCCACCTGTCGGTGTC
TGAAGAAGCTCGGCTAAGACCTCTCGAACTCTTCGTCTTACCTCAGCCCGCGGCTACAGT
CATATACATCCTAGCACGCACAATCCTTAATTCCCAGTCGTGCCCGACAGTAACACCGTG
GCGCGAACAATGTGTAAGTTATTGCCTACTTCGTACACTATCAACTAAATTCGTCCAACG
TGAGATCCGAGAATCCCGAAAGGGGGCACACACTTAGAGAACCCGCAGCTTTTTGTCTCA
CGATCGGGAAGGGCACTCCCAAGAGGCCTTGATACGAGCGTGAAACCGGTCAAGCGCTCC
AATTGTACACTGTCCATGAGGCCAAATTCAAATTACGGACCGAAAACTGGATCCAGCAGA
GCTAAAGGTTATTCGCGGGTATTGTCCGTACATGCAACTTGATGTTTTCTAAGCCAAAAG
AGATATGCGAGGGGGAGACATGGGAAAAGCAAAAGAATATAGTACTATAATTGGCGTGGG
ACCCATAAAGTCTTGATTGCTCCCGCGCCGCCTGTACCCACAGTCTTCAAAAAAGCGATT
TGCCGGAGTATAAAGTCAGGGGCCGACGGTGTTGAAAACGAGCGGACGTCTTGCATGAGA
AGTGTCACTGTGGGCATATCAAAGGACTCTAATTAAATTCGGTGTCGATAGAAATGTTCG
TGCTGCACGGTGCTGCCAGACCATCCACCTTTCGTGGCTACGCCTTATCAGAAGTGCAAC
CTGCGAAATCAAGGTGACGTTAATACGCTCTAAAGGACGCTAAAGCGGGTTGATAATTAA
ACGACAGAGTTACAATTAGCGGGATTTACAACTCTCCAAAATTATTGACGATAGACTGAC
ACGGTTACGTGGGAGGACTCGTAGTCTCAAGTGAAAATGGGCGCAAAATGGTGTCCGCAT
GGGAGGAGCTCAATCATAATAGACGCCGAATATGTTCGCGTGAAACTTCAGCGTCAAAGA
TGGAAAGCAAGACAAACTCATGAAACTACCATAGTGTGAATATGATGGGCGGCGCGGTTT
AAGGCGAGTCGCCCTAAGACTCCAACGTGCCCTAAGGGATCGTAACATGCAAGAACATTG
ATGTGCACAGTTTTGTCGAACGGCCTTCATAGGCCAAAGATTGTATCCACGGCCAGAAGC
GGTAAGAACTTCGTGACATAGCGGCCAGAGAAAGGATCTACAACGTTAACCAGCACCTCA
GAGAGCGGGATGGCCAGTCGTTAGGGAGAAAATAACTAATCATTAACTTTGACGAACCAC
CAAAATGCTCGAGCGGTAGGTATCATTAGGGGGTACGGACTGTTACATCTAGAATACCCC
AGCTCCGTAGTTGCAGACACTAATTTTAGAGACCGCAGCTGTATCATTAACAAAATGTTA
TTGGTGGGATGCAGAAGAGCGCGGGCAAGATTATACAGACGCCTTCAGGCGGTAGCCTGC
TCCTAGGCAGTCAAGACGGATCGCGATGGACGTGTTTGGACGGCGAATGTACAGGAAATT
TCCCGTGAGAACAACAAAATGAAGGATCGAGATTAAAAAGACTGCAGCGCCTGTGAACAA
CGGAGTTGAACGTGGTTGCCAGGTTTGTGATGGGCGGCCCTAGAAGATAACGGTAGGAGC
CCTTCGGACTTTTCAATGGGGACTACATTCTGGCCGCGAAGGACTCCGTGAATCGATAGG
TAGTGGAAATATGGATGAAACGGGCTGGGGCAACGCTCGGGACAACGTTTACTAGCTAAG
AGGCAGATTACGAGGATCGGGGAAGACTGAGAAGGCCTTGGCGTAGGAATTCGAGGCAAC

CAGGGGGATAAGGTAAATGCCGACTTCAAGTCGAGAATGCAAAGAAGCTGGCTATGGCCG
TAAATTAGACGATAAATAACAGAGCATCGGGGAGCCAAGTCTAATTGTAAAACTTTTGGT
CGAGGTGATGTCGTAGGTGAGATCTCCGAACGCAGCATGAACAAGGCGAATCGTGTAAGC
AGAGAAGAGTGCAGCACGATGCTAACACTGCCGCAGCAGTGGGACAACTTGGATGACTGG
CACATCACAGTAATGAGTCATATGATTCCGCCTGAAGGCTTCAATAAAGGACATAAGATG
CGAAGAAGATCTAGACCCAGTCTTTAAAACTGCACAACTTAGCCGAAATGATTGGTAGGT
GAGAGTATCGACGTGGGCACTGAGAGAAACCTGAATACGCGATAGCCATCGTATCACGAT
GCTTCTGCCCTCCAGCGTCACTAGAGTTCTTGTTCACATGCCTCGGTCCTCTCGTCAGTA
CGTGGCCGGCCTTCTTTCCTCCATCGTTTCCCAGAGCATACTCTTCGAGTTGCCCTATTA
ACCTTCTATTTTCTGGCCGATGCCTGACTGGCGCCGCTTTGAGTATCATACTTCTTGTTT
TTGGCCAGCCTGTTGGGCGTTGATTGCTTGGGTAACCATATCCTGCATTTTGTTGATTCC
GGGTCGGCAATGTGTTGTGGACTGTTCGTCGCCGCTCCTGATCTCACCGTACTTCACGGT
TGTCGCCCGGACATTACCCGGACGGGTTTAGGCCATGGCCGGTTGAGGTTAATCTTTACG
CGTCACCATAACTTCCTGCGAAACTGGTTTCCCCACGCCGCTCTTTTCGTGAGCCGATCG
GCGTGGTTAGGTCGCGCGGGTTTTTCCAGGCAGGGTAAGTCGAGTCGCACAAGGTGACCC
ACGTACCTCGCTTTAACGTTCCGTTCTACCACTGGGTTAGGCTGCTTTGGGGCCGACTTA
ACGTTCTTTTTGCACGACCCCGTATGTGCTTATTCTCACTCGTAATGAACCGGTAAGCGA
GAGTTTGTTCTACGCTGAGGTGTCGTCGGACCCGAGTGTGCTTGCCGCTGGTCAATTGTT
GTGTCGGCTCGACGTCTTATTTCGTATGGTACGGGATTCTTAGTGCTTGCAGCGATGCCC
CTTTCGCTATTGATGTCTTCTCCTGAGTCACGCTACGTACTGAATTGATAAAACATTTCG
ACCTCAGTATCAGTGAGGTCGGGGGATGAGTTAGATCGCCGTGTAGGGCTTTTCACGTTG
TGTCCGTGTGGAGAGTCGGTTCTAGCGGTCGTATTATCCATATTCTCCCGTTCCATTTAC
GTTGAGCGACTTGAAACTTCCGATGTACTGGCCGGTAATATTGCATTGTTTCTCAATTCA
TGCAAGTTGTCGTACATTATGGAATAGAACAAGAATTAAGGACCATTTCCACTAGCGATT
CTTACCTCCAACTCAAGATAGAGTCTAACCGATTATCGACGAACTTCGTCCGTCACGTCA
TTCTATGTCGAGCGGCCTTCTTACAGCGCTGTCGCACTATCTGTTCTACCTTATTTCCGA
GAAACCTTGCTCCTGTTACCTGTGAGGGCGGGGCGGATATGGCAAAATGGGCATTCTTTT
TTACTTCCCGTGCCTCTTCATGCGATATACATGTATCTCTGGAGCCTAGACGCCCGGTCC
TTAACGTCCTAAAACTTTACTGACCCTGCTGGCGTCTTACCCTTTATAGTGGCTCTGCAT
AATGTGTTCTGTATATAGCCAAATACTGCCGACTATCCACTCGCTTAGCTGTCAATATCC
ATGCGCGTACTTTGATTTACCGGTCGACCTGCCTAATGTAACTGCCTTTCGCTCTCCACT
CCGCACCGGGGTTTCGAGTTTTTCTACCCCTCTCTTTATTACTACTCGTTTACTGCATCA
GACGCGCCTACCGTATTCAGTCACTGCTCCTTCAGGCGAAGAGTATTAGCACGTTATCGT
GCTACTATCCTGAGCTGTAATCATAAGTATTTCTAAAGTTGTAAGGGTTGAAGAGTAGTT
CTGTACATCTTGATCGATTGTACTGGACGTCATTCCGAATAGCTTGTGGAATCCAATATC
TGGGTCATAGATGCT

>SRR9587925

GTGACCGTCGCCTCGCCACAAATCCATTGTTTTCCGGGTTCTAGCCCCTTATGAGCTAGG
ACCTCTGTGCCTACCATCATAGCTGTTCGATGCCTTGTATTCAATACGTTGCACCGACCC
GACAATAATCTGTGCCATTAAGCTTCTGTAAGTGGCTGGCGCCTGTAAGTCACGCAATAG
AGTAACTTCCCTCTCGACAGTCTTTATGCCATCCACAATCCAACATCGTAGATCCTCGCG
TGACCTAGTAGCCCTTGACGACTCAATCCACCCTCTGCCTGACGGACGTGCCTTTTATGG
TACATTAATGACTCCCTGTCCTCATAATTAGGCTGCACACACAGGGCTATCGTAGTTTAC
TGGTCTCACCCCTGCGTACTCGGCGACCCAACTACACTGATCCAGGGAGGCAGAATGGTG
GACGCTCTTACTGCCGACCCGTGGGCAACTAACTGGGCCGGGGGAATATCGTTTCGTTTT
GGGTACTGCTTCTTCCAATTGTTCGGATAGCACAGTAACGGTGGTGGATGGCTCCAGTCA
CTCTTGCGTAGTCGCCGGCCGGTCGCTCCGTTCTGTCCTCTCGCCCGATTCTGGGCCATC
TTATCCGATGTCACCCTGCAGACCGCNCTTTTCAACCATTAGCATTAAGCAGTCTCTCAT
AGGTGCTCTCACGTCATGAGAGGTACCGGAGCCTCTAAGCGTTCCGCTGTTCCCGACAAT
ATTCTCTTTCGCTTTAACACTAGCTCAGACGGTAGAGGCTTCGTCTTCAGCACCGGCTAA
TGATGCCTGGTACGTTAAGATACCGATCCGCGCGCTTACGCTGCTCCACCTGTCGGTGTC
TGAAGAAGCTCGGCTAAGACCTCTCGAACTCTTCGTCTTACCTCAGCCCGCGGCTACAGT
CATATACATCCTAACACGCACAATCCTTAATTCCCAGTCGTGCCCGACAGTAACACCGTG
GCGCGAACAATGTGTAAGTTGTTGCCTACTTCGTACACTATCAACTAAATTCGTCCAACG
TGAGATCCGAGAATCCCGAAAGGGGGCACACACTTAGAGAACCCGCAGCTTTTTGTCTCA
CGATCGAGAAGGGCACTTCCAAGAGGCCTTGATACGAGCGTGAAACCGGTCAAGCGCTCC
AATTGTACACTGTCCATGAGGCCAAATTCAAATTACGGACCGAGAACTGGATCCAGCAGA
GCTAAAGGTTATTCGCGGGTATTGTCCGTACATGCAACTTGATGTTTTCTAAGCCAAAAG
AGATATGCGAGGGGGAGACATGGGAAAAGCAAAAGAATATAGTACTATAATTGGCGTGGG
ACCCATAAAGTCTTGATTGCTCCCGCGCCGCTTGTACCCACAGTCTTCAAAAAAGCGATT
TGCCGGAGTATAAAGTCAGGGGCCGACGGTGTTGAAAACGAGCGGACGTCTTGCATGAGA
AGTGTCACTGTGGGCATATCAAAGGACTCTAATTAAATTCGGTGTCAATAGCAATGTTCG
TGCTGCACGGTGCTGTCAGACCATCCATCTTTCGTGGCTACGCCTTATCAGAAGTGCAAC
CTGCGAAATCAAGGTGACGTTAATACGCTCTAAAGGACGCTAAAGCGGGTTGATAATTAA
ACGACAGAGTTACAATTAGCGGTATTCACAACTCTCCAAAATCATTGACGATAGACTGAC
ACGGTTACGTGGGAGGACTCGTAGTCTCAAGTGAAAATGGGCGCAAAATAGTGTCCGCAT
GGGAGGAGCTTAACCATAATAGACGCCGAATATGTTCGCGTGAAACTTCAGCGTCAAAGA
TGGAAAGCAAGACAAACTCATGAAACTACCATAGTGTGAAGATGATGGGCGGCGCGGTTT
AAGACGAGTCGCCCTAAGACTCCAACGTGCCCTAAGGGATCGTAACATGCAAGAACATTG
ATGTGCACAGTTTTGTCGAACGGCCTTCATAGGCCAAAGATTGTATCCACGGCCAGAAGC
GGTAAGAACTTCGTGACATAGCGGCCAGAGAAAGGATCTACAACGTTAACCAGCACCTCA
GAGAGCGGGACGGCCAGTCGTTAGGGAGAAAATAACTAATCATTAACTTTGACGAACCAC

CAAAATGCTCGAGCGGTAGGTATCATTAGGGGGTACGGACTGTTACATCTAGAATACCCC
AGCTCCGTAGTTGCAGACACTAATTTTAGAGACCGCAGCTGCATCATTAACAAAATGTTA
TTGGTGGGATGCAGAAGAGCGCGGGCAAGATTATACAGACGCCTTCAGGCGGTAGCCTGC
TCCTAGGCGGTCAAGACGGATCGCGATGGACGTGTTTGGACGGCGAATGTACAGGAAATT
TCCCGTGAGAACAACAAAATGAAGGATCGAGATTAAAAAGACTACAGCGCCCGTGAACAA
CGGAGTTGAACGTGGTTGCCAGGTTTGTGATGGGCGGCCCTAGAAGATAACGGTAGAAGC
CCTTCGGACTTTTCAATGGGGACTACATTCTGGCCGCGAAGGACTCCGTGAATCGATAGG
TAGTGGAAATATGGATGAAACGGGCTGGGGCAACGCTTGGGACAACGTTTACTAGCTAAG
AGGCAGATTACGAGGATCGGGGAAGACTGAGAAGGCCTTGGCGTAGGAATTCGAGGCAAC
CAGGGGGATAAGGTAAATGCCGACTTCAAGTCGAGAATGCAAAGAAGCTGGCTATGGCCG
TAAATTAGACGATAAATAACAGAGCATCGGGGGGCCAAGTCTAATTGTAAAACTTTTGGT
CGAGGTGATGTCGTAGGTGAGATCTCCGAACGCAGCATGAACAAGGCGAATCGTGTAAGC
AGAGAAGAGTGCAGCACGATGCTAACACTGCCGCAGCAGTGGGACAACTTGGATGACTGG
CACATCACAGTAATGAGTCATATGATTCCGCCTGAAGGCTTCAATAAAGGACATAAGATG
CGAAGAAGATCTAGACCCAGTCTTTAAAGCTGCACAACTTAGCCGAAATGATTGGTAGGT
GAAAGTATCGACGTGGGCACTGAGAGAAACCTGAATACGCGATAGCCATCATATCACGAT
GCTTCTGCCCTCCAGTGTCACTAGAGTTCTTGTTCACATGCCTCGGTCCTCTCGTCAGTA
CATGGCCGGCCTTCTTTCCTCCATCGTTTCCCAGAGCATACTCTTCGAGTTGCCCTATTA
GCCTTCTATTTTCTGGCCGATGCCTGCCTGGCGCCGCTTTGAGTATCATACTTCTTGTTT
TTGGCCAACCTGTTGGGCGTTGATTGCTTGGGTAACCATATCCTGCATTTTGTTGATTCC
TGGTCGGCAATGTGTTGTGGACTGTTCGTCGCTGCTCCTGACCTCACCGTACTTCACGGT
TGTCGCCCGGACATCACCCGGACGGGTTTAGGCCATGGCCGGTTGAGGTTAATCTTTACG
CGTCATCATAACTTCCTGCGAAACTGGTTTCCTCACGCCGCTCCCTTCGTGAGCCGATCG
GCGTGGTTAGGTCGCGCGGGTTTTTCCAAGCAGGGTAAGTCGAGTCGCCCAAGGTGACCC
ACGTACCTCGCTTTAACGTTCCGTTCTACCACTGGATTAGGCTGCTTTGGGGCCGACTTG
ACGTTCTTTTTGTACGACCTCGTATGTGCTTATTCTCACTCGTAATGAACCGGTAAGCGA
GAGTTTGCTCTACGCTGAGGTGCCGTCGGACCCGAGTGTGCTTGCCGCTGGTCAATTGTT
GTGTCGGCTCGACGTCTTATTTCGTATGGTACGGGATTATTAGTGCTTGCAGCGATGTCC
CTTTCGCTATTGATGTCTTATCCTGAGTCACGCTACGTACTGAATTGATAAAACATTTCG
ACCTCAGTATCAGTGAGGTCGGGGGATGAGTTAGATCGCCGTGTAGGGCTTTTCACGTTG
TGTCCGTGTGGAGAGTCGGTTCTAGCGGTCGTATTATCCATATTCTCCCGTTCTATTTAC
GTTGAGCGACTTGAAACCTCCGATGTACTGGCCGGTAATAGTGCATTGTTTCTCAATTCA
TGCGAGTTGTCGTACATTATGGAATAGAACAAGAATTAAGGACCATTTCCACTAGCGATT
CTTACCTCCAACTCAAGATAGAGTCTAACCGATTATCGACGAACTTCGTCCGTCACGTCA
TTCTATGTCGAGCGGCCTTCTTACAGCGCTGTCGCACTATCTGTTCTACCTTATTTCCGA
GAAACCTTGCTCCTGTTACCTGTGAGGGCGGGGCGGATATGGCAAAATGGGCATTCTTTT

TTACTTCCCGTGCCTCTTCATGCTATATACATGTATCTCTGGAGCCTAGACGCCCGGTCC
TTAACGTCCTAAAACTTTACTGACCCTGCTGGCGTCTTACCCTTTATAGTGGCTCTGCAT
AATGTGTTCTGTATATAGCCAAATACTGCCGACTATCCACTCGCTTAGCTGTCAATATCC
ATGCGCGTACTTTGATTTACCGGTCGACCTGCCTAATGTAACTGCCTTTCGCTCTCCACT
CCGCACCGGGGTTTCGAGTTTTTCTACCCCTCTCTTTATTACTACTCGTTTACTGCATCA
GACGCGCCTACCGTATTCGGTCACTGCTCCTTCAGGCGAAGAGTATTAGCACGTTATCGT
GCTACTATCCTGAGCTGTAATCCTAATTATTTCTAAAGTTGTAAGGGTTGAAGAGTAGTT
CTGTACATCTTGATCGATTGTACTGGACGTCATTCCGAATAGCTTGTGGAATCCAATATC
TGGGTCATAGATGCT
>SRR9587926
GTGACCGTCGCCTCGCCACAAATCCATTGTTTTCCGGGTTCTAGCCCCTTATGAGCTAGG
ACCTCTGTGCCTACCATCATAGCTGTTCGATGCCTTGTATTCAATACGTTGCACCGACCC
GACAATAATCTGTGCCATTAAGCTTCTGCAAGTGGCTGGCGCCTGTAAGTCACGCAATAG
AGTAACTTCCCTCTCGACAGTCTTTATGCCATCCACAATCCAACATCGTAGATCCTCGCG
TGACCTAGTAGCCCTTGACGACTCAATCCCCCCTCTGCCTGACGGACGTGCCTTTTATGG
TACATTAATGACTCCCTGTCCTCATAATTAGGCTGCACACACAGGGCTATCGTAGTTTAC
TGGTCTCACCCCTGCGTACTCGGCGACCCAACTACACTGATCCAGGGAGGCGGAATGGTG
GACGCTCTTACTGCCGACCCGTGGGCAACTAACTGGGCCGGGGGAATATTGTTTCGTTTT
NGGTACTGCTTCTTCCAATTGTTCGGATAGCACAGTAACGGTGGTGGATGGCTCCAGTCA
CTCTTGCGTAGTCGCCGGCCGGTCGCTCCGGTCTGTCCTCTCGCCCGATTTTGGGCCATC
TTATCCGATGTCACCCTGCAGACCGCTCTTTTCAACCATTAGCATTAAGCAGTCTCTCAT
AGGTGCTCTCACGTCATGAGAGGTACCGGAGCCTCTAAGCGTTCCGCTGTTCCTGACAAT
ATTCTCTTTCGCTTTAACACTAGCTCAGACGGTAGAGGCTTCGTCTTCAGCACCGGCTAA
TGATGCCTGGTACGTTAAGATACCGATCCGCGCGCTTACGCTGCTCCACCTGTCGGTGTC
TGAAGAAGCTCAGCTAAGACCTCTCGAACTCTTCGTCTTACCTCAGCCCGCGGCTACAGT
CATATACATCCTAGCACGCACAATCCTTAATTCCCAGTCGTGCCCGACAGTAACACCGTG
GCGCGAACAATGTGTAAGTTGTTGCCTACTTCGTACACTATCAACTAAATTCGTCCAACG
TGAGATCCGAGAATCCCGAAAGGGGGCACACACTTAGAGAACCCGCAGCTTTTTGTCTCA
CGATCGAGAAGGGCACTCCCAAGAGGCCTTGATACGAGCGTGAAACCGGTCAAGCGCTCC
AATTGTACACTGTCCATGAGGCCAAATTCAAATTACGGACCGAGAACTGGATCCAGCAGA
GCTAAAGGTTATTCGCGGGTATTGTCCGTACATGCAACTTGATGTTTTCTAAGCCAAAAG
AGATATGCGAGGGGGAGACATGGGAAAAGCAAAAGAAGATAGTACTATAATTGGCGTGGG
ACCCATAAAGTCTTGATTGCTCCCGCGCCGCCTGTACCCACAGTCTTCAAAAAAGCGATT
TGCCGGAGTATAAAGTCAGGGGCCGACGGTGTTGAAAACGAGCGGACGTCTTGCATGAGA
AGTGTCACTGTGGGCATATCAAAGGACTCTAATTAAATTCGGTGTCAATAGCAATGTTCG
TGCTGCACGGTGCTGTCAGACCATCCATCTTTCGTGGCTACGCCTTATCAGAAGTGCAAC

CTGCGACATCAAGGTGACGTTAATACGCTCTAAAGGACGCTAAAGCGGGTTGATAATTAA
ACGACAGAGTTACAATTAGCGGTATTCACAACTCTCCAAAATCATTGACGATAGACTGAC
ACGGTTACATGGGAGGACTCGTAGTCTCAAGTGAAAATGGGCGCAAAATGGTGTCCGCAT
GGGAGGAGCTCAACCATAATAGACGCCGAATATGTTCGCGTGAAACTTCAGCGTCAAAGA
TGGAAAGCAAGACAAACTCATGAAACTACCATAGTGTGAAGATGATGGGCGGCGCGGTTT
AAGGCGAGTCGCCCTAAGACTCCAACGTGCCCTAAGGGATCGTAACATGCAAGAACATTG
ATGTGCACAGTTTTGTCGAACGGCCTTCATAGGCCAAAGATTGTATCCACGGCCAGAAGC
GGTAAGAACTTCGTGACATAGCGGCCAGAGAAAGGATCTACAACGTTAACCAGCACCTCA
GGAAGCGGGACGGCCAGTCGTTAGGGAGAAAATAACTAATCATTAACTTTGACGAACCAC
CAAAATGCTCGAGCGGTAGGTATCATTAGGGGGTACGGACTGTTACATCTAGAATACCCC
AGCTCCGTAGTTGCAGACACTAATTTTAGAGACCGCAGCTGCATCATTAACAAAATGTTA
TTGGTGGGATGCAGAAGAGCGCGGGCAAGATTATACAGACGCCTTCAGGCGGTAGCCTGC
TCCTAGGCGGTCAAGACGGATCGCGATGGACGTGTTTGAACGGCGAATGTACAGGAAATT
TCCCGTGAGAACAACAAAATGAAGGATCGAGATTAAAAAGACTACAGCGCCCGTGAACAA
CGGAGTTGAACGTGGTTGCCAGGTTTGTGATGGGCGGCCCTAGAAGATAACGGTAGAAGC
CCTTCGGACTTTTCAATGGGGACTACATTCTGGCCGCGAAGGACTCCGTGAATCGATAGG
TAGTGGAAATATGGATGAAACGGGCTGGGGCAACGCTTGGGACAACGTTTACTAGCTAAG
AGGCAGATTACGAGGATCGGGGAAGACTGAGAAGGCCTTGGCGTAGGAATTCGAGGCAAC
CAGGGGGATAGGGTAAATCCCGACTTCAAGTCGAGAATGCAAAGAAGCTGGCTATGGCCG
TAAATTAGACGATAAATAACAGAGCATCGGGGGGCCAAGTCTAATTGTAAAACTTTTGGT
CGAGGTGATGTCATAGGTGAGATCTCCGAACGCAGCATGAACAAGGCGAATCGTGTAAGC
AGAGAAGAGTGCAGCACGATGCTAACACTGCCGCAGCAGTGGGACAACTTGGATGACTGG
CACATCACAGTAATGAGTCATATGATTCCGCCTGAAGGCTTCAATAAAGGACATAAGATG
CGAAGAAGATCTAGACCCAGTCTTTAAAACTGCACAACTTAGCCGAAATGATTGGTAGGC
GAAAGTATCGACGTGGGCACTGAGAGAAACCTGAATACGCGACAGCCATCATATCACGAT
GCTTCTGCCCTCCAGTGTCACTAGAGTTCTTGTTCACATGCCTCGGTCCTCTCGTCAGTA
CATGGCCGGCCTTCTTTCCTCCATCGTTTCCCAGAGCATACTCTTCGAGTTGCCCTATTA
GCCTTCTATTTTCTGGCCGATGCCTGCCTGGCGCCGCTTTGAGTATCATACTTCTTGTTT
TTGGCCAACCTGTTGGGCGTTGATTGCTTGGGTAACCATATCCTGCATTTTGTTGATTCC
TGGTCGGCAGTGTGTTGTGGACTGTTCGTCGCTGCTCCTGACCTCACCGTACTTCACGGT
TGTCGCCCGGACATCACCCGGACGGGTTTAGGCCATGGCCGGTTGAGGTTAATCTTTACG
CGTCATCATAACTTCCTGCGAAACTGGTTTCCTCACGCCGCTCCCTTCGTGAGCCGATCG
GCATGGTTAGGTCGCGCGGGTTTTTCCAAGCAGGGTAAGTCGAGTCGCCCAAGGTGACCC
ACGTACCTCGCTTTAACGTTCCGTTCTACCACTGGATTAGGCTGCTTTGGGGCCGACTTG
ACGTTCTTTTTGTACGACTTCGTATGTGCTTATTCTCACTCGTAATGAACCGGTAAGCGA
GAGTTTGCTCTACGCTGAGGTGTCGTCGGACCCGAGTGTGCTTGCCGCTGGTCAATTGTT

GTGTCGGCTCGACGTCTTATTTCGTATGGTACGGGATTATTAGTGCTTGCAGCGATGTCC
CTTTCGCTATTGATGTCTTCTCCTGAGTCACGCTACGTACTGAATTGATAAAACATTCCG
ACCTCAGTATCAGTGAGGTCGGGGGATGAGTTAGATCGCCGTGNNNGGCTTTTCACGTTG
TGTCCGTGTGGAGAGTCGGTTCTAGCGGTCGTATTATCCATATTCTCCCGTTCTATTTAC
GTTGAGCGACTTGAAACTTCCGATGTACTGGCNGGTAATAGTGCATTGTTTCTCAATTCA
TGCAAGTTGTCGTACATTATGGAATAGAACAAGAATTAAGGATCATTTCCACTAGCGATT
CTTACCTCCAACTCAAGATAGAGTCTAACCGATTATCGACGAACTTCGTCCGTCACGTCA
TTCTATGTCGAGCGGCCTTCTTACAGCGCTGTCGCACTATCTGTTCTACCTTATTTCCGA
GAAACCTTGCTCCTGTTACCTGTGAGGGCGGGGCGGATATGGTAAAATGGGCATTCTTTT
TTACTTCCCGTGCCTCTTCATGCTATATACATGTATCTCTGGAGCCTAGACGCCCGGTCC
TTAACGTCCTAAAACTTTACTGACCCTGCTGGCGTCTTACCCTTTATAGTGGCTCTGCAT
AATGTGTTCTGTATATAGCCAAATACTGCCGACTATCCACTCGCTTAGCTGTCAATATCC
ATGCGCGTACTTTGATTTACCGGTCGACCTGCCTAATGTAACTGCCTTTCGCTCTCCACT
CCGCACCGGGGTTTCGAGTTTTTCTACCCCTCTCTTTATTACTACTCGTTTACTGCATCA
GACGCGCCTACCGTATTCGGTCACTGCTCCTTCAGGCGAAGAGTATTAGCACGTTATCGT
GCTACTATCCTGAGCTGTAATCATAAGTATTTCTAAAGTTGTAAGGGTTGAAGAGTAGTT
CTGTACATCTTGATCGATTGTACTGGACGTCATTCCGAATAGCTTGTGGAATCCAATATC
TGGGTCATAGATGCT
>SRR9587927
GTGACCGTCGCCTCGCCACAAATCCATTGTTTCCGGGTTCTAGCCCCTTATGAGCTAGG
ACCTCTGTGCCTACCATCATAGCTGTTCGATGCCTTGTATTCAATACGTTGCACCGACCC
GACAATAGTCTGTGCCATTAAGCTTCTGCAAGTGGCTGGCGCCTGTAAGTCACGCAATAG
AGTAACTTCCCTCTCGACAGTCTTTATGCCATCCACAATCCAACATCGTAGATCCTCACG
TGACCTAGTAGCCCTTGACGACTCAATCCACCCTCTGCCCGACGGACGTGCCTTTTATGG
TACATTAATGACTCCCTGTCCTCATAATTAGGCTGCACACACAGGGCTATCGTAGTTTAC
TGGTCTCACCCCTGCGTACTCGGCGACCCAACTACACTGATCCAGGGAGGCAGAATGGTG
GACTCGCTTACTCCCGACCCGTGGGCAACTAACTGGGCCGGGGGAATATTGTTTCGTTTT
GGGTACTGCTTCTTCCAATTGTTCGGATAGCACAGTAACGGTGGTGGGTGGCTCCAGTCA
CTCTTGCGTAGTCGCCGGCCGGTCGCTCCGGTCTGTCCTCTCGCCCGATTCTGGGCCATC
TTATCCGATGTCACCCTGCAGACCGCTCTTTTCAACCATTAGCATTAAGCAGTCTCTCAT
AGGTGCTCTCATGTCATGAGAGGTACCGGAGCCTCTAAGCGTTCCGCTGTTCCCCACAAT
ATTCTCTTTCGCTTTGACACTAGCTCAGACGGTAGAGGCTTCGTCTTCAGCACCGGCTAA
TGATGCCTGGTACGTTAAGATACCGATTCGCGCGCTTACGCTGCTCCACCTGTCGGTGTC
TGAAGAAGCTCGGCTAAGACCTCTCGAACTCTTCGTCTTACCTCAGCCCGCGGCTACAGT
CATATACATCCTAGCACGCACAATCCTTAATTCTCAGTCGTGCCCGACAGTAACACCGTG
GCGCGAACAATGTGTAAGTTATTGCCTACTTCGTACACTATCAACTAAATTCGTCCAACG

TGAGATCCGAGAATCCCGAAAGGGGGCACACACTTAGAGAACCCGCAGCTTTTTGTCTCA
CGATCGAGGAGGGCACTCCCAAGAGGCCTTGATACGAGCGTGAAACCGGTCAAGCGCTCC
AATTGTACACTGTCCATGAGGCCAAATTCAAATTACGGACCGAAAACTGGATCCAGCAGA
GCTAAAGGTTATTCGCGGGTATTGTCCGTACATGCAACTTGATGTTTTCTAAGCCAAAAG
AGATATGCGAGGGGGAGACATGGGAAAAGCAAAGGAATATAGTACTATAATTGGCGTGGG
ACCCATAAAGTCTTGATTGCTCCCGCGCCGCCTGTGCCCACAGTCTTCAGAAAAGCGATT
TGCCGGAGTATAAAGTCAGGGGCCGACGGTGTTGAAAACGAGCGGACGTCTTGCATGAGA
AGTGTCACTGTGGGCATATCAAAGGACTCTAATTAAATTCAGTGTCAATAGAAATGTTCG
TGCTGCACGGTGCTGTCAGACCATCCACCTTTCGTGGCTACGCCTTATCAGAAGTGCAAC
CTGCGAAATCAAGGTGACGTTAATACGCTCTAAAGGACGCTAAAGCGGGTTGATAATTAA
ACGACAGAGTTACAATTAGCGGGATTCACAACTCTCCAAAATCATTGACAATAGACTGAC
ACGGTTACGTGGGAGGACTCGTAGTCTCAAGTGAAAATGGGCGCAAAATGGTGTCCGCAT
GGGAGGAGCTCAACCATAATAGACGCCGAATATGTTCGCGTGAAACTTCAGAGTCAAAGA
TGGAAAGCAAGACAAACTCATGAAACTACCATAGTGTGAAGATGATGGGCGGCGCGGTTT
AAGGCGAGTCGCCCTAAGACCCCAACGTGCCCTAAGGGATCGTAACATGCAAGAACATTG
ATGTGCACAGTTTTGTCGAACGGCCTTCATAGGCCAAAGATTGTATCCACGGCCAGAAGC
GGTAAGAACTTCGTGACATAGCGGCCAGAGAAAGGATCTACAACGTTAACCAGCACCTCA
GAGAGCGGGATGGCCAGTCGTTAGGGAGAAAATAACTAATCATTAACTTTGACGAACCAC
CAAAATGCTCGAGCGGTAGGTATCATTAGGGGGTACGGACTGTTACATCTAGAATACCCC
AGCTCCGTAGTTGCAGACACTAATTTTAGAGACCGCAGCTGTATCATTAACAAAATGTTA
TTGGTGGGATGCAGAAGAGCGCGGGCAAGATTATACAGACGCCTTCAGGCGGTAGCCTGC
TCCTAGGCAGTCAAGACGGATCGCGATGGACGTGTTTGGACGGCGAATGTACAGGAAATT
TCCCGTGAGAACAACAAAATGAAGAATCGAGATTAAAAAGACTGCAGCGCCCGTGAACAA
CGGAGTTGAACGTGGTTGCCAAGTTTGTGATGGGCGGCCCTAGAAGATAACGGTAGAAGC
CCTTCGGACTTTTCAATGGGGACTACATTCTGGCCGCGAAGGACTCCGTGAATCGATAGG
TAGTGGAAATATGGATGAAACGGGCTGGGGCAACGCTCGGGACAAAGTTTACTAGCTAAG
AGGCAGATTACGAGGATCGGGGAAGACTGAGAAGGCCTTGGCGTAGGAATTCGAGGCAAC
CAGGGGGATAAGGTAAATGCCGACTTCAAGTCGAGAATGCAAAGAAGCTAGCTATGGCCG
TAAATTAGACGATAAATAACAGAGCATCGGGGAGCCAAGTCTAATTGTAAAACTTTTGGT
CGAGGTGATGTCGTAGGTGAGATCTCCGAACGCAGCATGAACAAGGCGAATCGTGTAAGC
AGAGAAGAGTGCAGCACGATGCTAACACTGCCGCAGCAGTGGGACAACTTGGATGACTGG
CACATCACAGTAATGAGTCATATGATTCCGCCTGAAGGCTTCAATAAAGGACATAAGATG
CGAAGAAGATCTAGACCCAGTCTTTAAAACTGCACAACTTAGCCGAAATGATTGGTAGGT
GAGAGTATCGACGTGGGCACTGAGAGAAACCTGAATACGCGATAGCCATCACATCACGAT
GCTTCTGCCCTCCAGCGTCACTAGAGTTCTTGTTCACATGCCTCGGTCCTCTCGTCAGTA
CATGGCCGGCCTTCTTTCCTCCATCGTTTCCCAGAGCATACTCCTCGAGTTGCCCTATTA

ACCTTCTATTTTCTGGCCGATGCCTGACTGGCGCCGCTTTGAGTATCATACTTCTTGTTT
TTGGCCAACCTGTTGGGCGTTGATTGCTTGGGTAACCACATCCTGCATTTTGTTGATTCC
GGGTCGGCAATGTGTTGTGGACTGTTCGTCGCCGCTCCTGATCTCACCGTACTTCACGGT
TGTCGCCCGGACATCACCCGGACGGGTTTAGGCCATGGCCGGTTGAGGTTAATCTTTACG
CGTCACCATAACTTCCTGCGAAACTGGTTTCCCCACACCGCTCCCTTCGTGAGCCGATCG
GCGTGGTTAGGTCGCGCGGGTTTTTCCAGGCAGGGTAAGTCGAGTCGCACAAGGTGACCC
ACGTACCTCGCTTTAACGTTCCGTTCTACCACTGGGTTAGGCTGCTTTGGGGCCGACTTG
ACGTTCTTTTTGCACGACCTCGTATGTGCTTATTCTCACTCGTAATGAACCGGTAAGCGA
GAGTTTGTTCTACGCTGAGGTGTCGTCGGACCCGAGTGTGCTTGCCGCTGGTCAATTGTT
GTGTCGGCTCGACGTCTTATTCCGTATGGTACGGGATTATTAGTGCTTGCAGCGATGTCC
CTTTCGCTATTGATGTCCTCTCCTGAGTCACGCTACGTACTGAATTGATAAAACATTTCG
ACCTCAGTATCAGTGAGGTCGGGGGATGAATTAGATCGCCGTGTAGGGCTTTTCACGTTG
TGTCCGTGTGGAGAGTCGGTTCTAGCGGTCGTATTATCCATATTCTCCCGTTCCATTTAC
GTTGAGCGACTTGAAACTTCCGATGTACTGGCCGGTAATAGTGCATTGTTTCTCAATTCA
TGCAAGTTGTCGTACATCATGGAATAGAACAAGAATTAAGGACCATTTCCACTAGCGATT
CTTACCTCCAACTCAAGATAGAGTCTAACCGATTATCGACGAACTTCGTCCGTCACGTCA
TTCTATGTCGAGCGGCCTTCTTACAGCGCTGTCGCACTATCTGTTCTACCTTATTTCCGA
GAAACCTTGCTCCTGTTACCTGTGAGGGCGGGGCGGATATGGCAAAATGGGCATTCTTTT
TTACTTCCCGTGCCTCTTCATACTATATACATGTATCTCTGGCGCCTAGACGCCCGGTCC
TTAACGTCCTAAAACTTTACTGACCCTGCTGGCGTCTTACCCTTTATAGTGGCTCTGCAT
AATGTGTTCTGTATATAGCCAAATACTGCCGACTATCCACTCGCTTAGCTGTCAATATCC
ATGCGCGTACTTTGATTTACCGGTCGACCTGCCTAATGTAACTGCCTTTCGCTCTCCACT
CCGCACCGGGGTTTCGAGTTTTTCTACCCCTCTCTTTATTACTACTCGTTTACTGCATCA
GACGCGCCTACCGTATTCAATCACTGCTCCTTCAGGCGAAGAGTATTAGCACGTTATCGT
GCTACTATCCTGAGCTGTAATCATAAGTATTTCTAAAGTTGTAAGGGTTGAAGAGTAGTT
CTGTACATCTTGATCGATTGTACTGGACGTCATTCCGAATAGCTTGTGGAATCCAATATC
TGGGTCATAGATGCT
>SRR9587928
GTGACCGTCGCCTCGCCACAAATCCATTGTTTTCCGGGTTCTAGCCCCTTATGAGCTAGG
ACCTCTGTGCCTACCATCATAGCTGTTCGATGCCTTGTATTCAATACGTTGCACCGACCC
GACAATAATCTGTGCCATTAAGCTTCTGCAAGTGGCTGGCGCCTGTAAGTCACGCAATAG
AGTAACTTCCCTCTCGACAGTCTTTATGCCATCCACAATCCAACATCGTAGATCCTCGCG
TGACCTAGTAGCCCTTGACGACTCAATCCACCCTCTGCCTGACGGACGTGACTTTTATGG
TACATTAATGACTCCCTGTCCTCATAATTAGGCTGCACACACAGGGCTATCGTAGTTTAC
TGGTCTCACCCCTGCGTACTCGGCGACCCAACTACACTGATCCAGGGAGGCAGAATGGTG
GACGCTCTTACTGCCGACCCGTGGGCAACTAACTGGGCCGGGGGAATATTGTTTCGTTTT

GGGTACTGCTTCTTCCAATTGTTCGGATAGCCCAGTAACGGTGGTGGGTGGTTCCAGTCA

CTCTTGCGTAGTCGCCGGCCGGTCGCTCCGGTCTGTCCCCTCGCCCGATTCTGGGCCATC

TTATCCGATGTCACCCTGCAGACCGCNCTTTTCAACCATTAGCATTAAGCAGTCTCTCAT

AGGTGCTCTCATGTCATGAGAGGTACCGGAGCCTCTAAGCGTTCCGCTGTTCCCGACAAT

ATTCTCTTTCGCTTTGACACTAGCTCAGACGGTAGAGGCTTCGTCTTCAGCACCGGCTAA

TGATGCCTGGTACGTTAAGATACCGATCCGCGCGCTTACGCTGCTCCACCTGTCGGTGTC

TGAAGAAGCTCGGCTAAGACCTCTCGAACTCTTCGTCTTACCTCAGCCCGCGGCTACAGT

CATATACATCCTAGCACGCACAATCCTTAATTCCCAGTCGTGCCCGACAGTAACACCGTG

GCGCGAACAATGTGTAAGTTATTGCCTACTTCGTACACTATCAACTAAATTCGTCCAACG

TGAGATCCGAGAATCCCGAAAGGGGGCACACACTTAGAGAACCCGCAGCTTTTTGTCTCA

CGATCGAGAAGGGCACTCCCAAGAGGCCTTGATACAAGCGTGAAACCGGTCAAGCGCTCC

AATTGTACACTGTCCATGAGGCCAAATTCAAATTACGGACCGAAAACTGGATCCAGCAGA

GCTAAAGGTTATTCGCGGGTATTGTCCGTACATGCAACTTGATGTTTTCTAAGCCAAAAG

AGATATGCGAGGGGGAGACATGGGAAAAGCAAAAGAATATAGTACTATAATTGGCGTGGG

ACCCATAAAGTCTTGATTGCTCCCGCGCCGCCTGTACCCACAGTCTTCAAAAAAGCGATT

TGCCGGAGTATAAAGTCAGGGGCCGACGGTGTTGAAAACGAGCGGACGTCTTGCATGAGA

AGTGTCACTGTGGGCATATCAAAGGACTCTAATTAAATTCGGTGTCAATAGAAATGTTCG

TGCTGCACGGTGCTGTCAGACCATCCACCTTTCGTGGCTACGCCTTATCAGAAGTGCAAC

CTGCGAAATCAAGGTGACGTTAATACGCTCTAAAGGACGCTAAAGCGGGTTGATAATTAA

ACGACAGAATTACAATTAGCGGGATTCACAACTCTCCAAAATCATTGACGATAGACTGAC

ACGGTTACGTGGGAGGACTCGTAGTCTCCAGTGAAAATGGGCGCAAAATGGTGTCCGCAT

GGGAGGAGCTCAACCATAATAGACGCCGAATATGTTCGCGTGAAACTTCAGCGTCAAAGA

TGGAAAGCAAGACAAACTCATGAAACTACCATAGTGTGAAGATGATGGGCGGCGCGGTTT

AAGGCGAGTCGCCCTAAGACTCCAACGTGCCCTAAGGGATCGTAACATGCAAGAACATTG

ATGTGCACAGTTTTGTCGAACGGCCTTCATAGGCCAAAGATTGTATCCACGGCCAGAAGC

GGTAAGAACTTCGTGACATAGCGGCCAGAGAAAGGATCTACAACGTTAACCAGCACCTCA

GAGAGCGGGATGGCCAGTCGTTAGGGAGAAAATAACTAATCATTAACTTTGACGAACCAC

CAAAATGCTCGAGCGGTAGGTATCATTAGGGGGTACGGACTGTTACATCTAGAATACCCC

AGCTCCGTAGTTGCAGACACTAATTTTAGAGACCGCAGCTGCATCATTAACAAAATGTTA

TTGGTGGGATGCAGAAGAGCGCGGGCAAGATTATACAGACGCCTTCAGGCGGTAGCCTGC

TCCTAGGCAGTCAAGACGGATCGCGATGGACGTGTTTGGACGGCGAATGTACAGGAAATT

TCCCGTGAGAACAACAAAATGAAGGATCGAGATTAAAAAGACTGCAGCGCCCGTGAACAA

CGGAGTTGAACGTGGTTGCCAGGTTTGTGATGGGCGGCCCTAGAAGATAACGGTAGAAGC

CCTTCGGACTTTTCAATGGGGACTACATTCTGGCCGCGAAGGACTCCGTGAATCGATAGG

TAGTGGAAATATGGATGAAACGGGCTGGGGCAACGCTCGGGACAACGTTTACTAGCTAAG

AGGCAGATTACGAGGATCGGGGAAGACTGAGAAGGCCTTGGCGTAGGAATTCGAGGCAAC

CAGGGGGATAAGGTAAATGCCGACTTCAAGTCGAGAATGCAAAGAAGCTGGCTATGGCCG
TAAATTAGACGATAAATAACAGAGCATCGGGGAGCCAAGTCTAATTGTAAAACTTTTGGT
CGAGGTGATGTCGTAGGTGAGATCTCCGAACGCAGCATGAACAAGGCGAATCGTGTAAGC
AGAGAAGAGTGCAGCACGATGCTAACACTGCCGCAGCAGTGGGACAACTTGGATGACTGG
CACATCACAGTAATGAGTCATATGATTCCGCCTGAAGGCTTCAATAAAGGACATAAGATG
CGAAGAAGATCTAGACCCAGTCTTTAAAACTGCACAACTTAGCCGAAATGATTGGTAGGT
GAGAGTATCGACGTGGGCACTGAGAGAAACCTGAATACGCGATAGCCATCATATCACGAT
GCTTCTGCACTCCAGCGTCACTAGAGTTCTTGTTCACATGCCTCGGTCCTCTCGTCAGTA
CATGGCCGGCCTTCTTTCCTCCATCGTTTCCCAGAGCATACTCTTCGAGTTGCCCTATTA
ACCTTCTATTTTCTGGCCGATGCCTGACTGGCGCCGCTTTGAGTATCATACTTCTTGTTT
TTGGCCAACCTGTTGGGCGTTGATTGCTTGGGTAACCATATCCTGCATTTTGTTGATTCC
GGGTCGGCAATGTGTTGTGGACTGTTCGTCGCCGCTCCTGATCTCACCGTACTTCACGGT
TGTCGCCCGGACATCACCCGGACGGGTTTAGGCCATGGCCGGTTGAGGTTAATCTTTACG
CGTCACCATAACTTCCTGCGAAACTGGTTTCCCCACGCCGCTCCCTTCGTGAGCCGATCG
GCGTGGTTAGGTCGCGCGGGTTTTTCCAGGCAGGGTAAGTCGATTCGCACAAGGTGACCC
ACGTACCTCGCTTTAACGTTCCGTTCTACCACTGGGTTAGGCTGCTTTGGGGCCGACTTG
ACGTTCTTTTTGCACGACCTCGTATGTGCTTATTCTCACTCGTAATGAACCGGTAAGCGA
GAGTTTGTTCTACGCTGAGGTGTCGTCGGACCCGAGTGTGCTTGCCGCTGGTCAATTGTT
GTATCGGCTCGACGTCTTATTTCGTATGGTACGGGATTATTAGTGCTTGCAGCGATGTCC
CTTTCGCTATTGATGTCTTCTCCTGGGTCACGCTACGTACTGAATTGATAAAACATTTCG
ACCTCAGTATCAGTGAGGTCGGGGGATGAGTTAGATCGCCGTGTAGGGCTTCTCACGTTG
TGTCCGTGTGGAGAGTCGGTTCTAGCGGTCGTATTATCCATATTCTCCCGTTCCATTTAC
GTTGAGCGACTTGAAACTTCCGATGTACTGGCCGNNAATAGTGCATTGTTTCTCAATTCA
TGCAAGTTGTCGTACATTATGGAATAGAACAAGAATTAAGGACCATTTCCACTAGCGATT
CTTACCTCCAACTCAAGATAGAGTCTAACCGATTATCGACGAACTTCGTCCGTCACGTCA
TTCTATGTCGAGCGGCCTTCTTACAGCGCTGTCGCACTATCTGTTCTACCTTATTACCGA
GAAACCTTGCTCCTGTTACCTGTGAGGGCGGGGCGGATATGGCAAAACGGGCATTCTTTT
TTACTTCCCGTGCCTCTTCATGCTATATACATGTATCTCTGGAGCCTAGACGCCCGGTCC
TTAACGTCCTAAAACTTTACTGACCCTGCTGGCGTCTTACCCTTTATAGTGGCTCTGCAT
AATGTGTTCTGTATATAGCCAAATACTGCCGACTATCCACTCGCTTAGCTGTCAATATCC
ATGCGCGTACTTTGATTTACCGGTCGACCTGCCTAATGTAACTGCCTTTCGCTCTCCACT
CCGCACCGGGGTTTCGAGTTTTTCTACCCCTCTCTTTATTACTACTCGTTTACTGCATCA
GACGCGCCTACCGTATTCAGTCACTGCTCCTTCAGGCGAAGAGTATTAGCACGTTATCGT
GCTACTATCCTGAGCTGTAATCATAAGTATTTCTAAAGTTGTAAGGGTTGAAGAGTAGTT
CTGCACATCTTGATCGATTGTACTGGACGTCATTCCGAATAGCTTGTGGAATCCAATATC
TGGGTCATAGATGCT

>SRR9587929
GTGACCGTCGCCTCGCCACAAATCCATTATTTTCCGGGTTCTAGCCCCTTATGAGCTAGG
ACCTCTGTGCCTACCATCATAGCTGTTCGATGCCTTGTATTCAATACGTTGCACCGACCC
GACAATAGTCTGTGCCATTAAGCTTCTGCAAGTGGCTGGCGCCTGTAAGTCACGCAATAG
AGTAACTTCCCTCTCGACAGTCTTTATGCCATCCACAATCCAACATCGTAGATCCTCACG
TGACCTAGTAGCCCTTGACGACTCAATCCACCCTCTGCCTGACGGACGTGCCTTTTATGG
TACATTAATGACTCCCTGTCCTCATAATTAGGCTGCACACACAGGGCTATCGTAGTTTAC
TGGTCTCACCCCTGCGTACTCGGCGACCCAACTACACTGATCCAGGGAGGCAGAATGGTG
GACGCGCTTACTCCCGACCCGTGGGCAACTAACTGGGCCGGGGGAATATTGTTTCGTTTT
GGGTACTGCTTCTTCCAATTGTTCGGATAGCACAGTAACGGTGGTGGGTGGCTCCAGTCA
CTCTTGCGTAGTCGCCGGCCGGTCGCTCCGGTCTGTCCTCTCGCCCGATTCTGGGCCATC
TTATCCGATGTCACCCTGCAGACCGCTCTTTTCAACCATTAGCATTAAGCAGTCTCTCAT
AGGTGCTCTCATGTCATGAGAGGTACCGGAGCCTCTAAGCGTTCCGCTGTTCCCGACAAT
ATTCTCTTTCGCTTTGACACTAGCTCAGACGGTAGAGGCTTCGTCTTCAGCACCGGCTAA
TGATGCCTGGTACGTTAAGATACCGATTCGCGCGCTTACGCTGCTCCACCTGTCGGTGTC
TGAAGAAGCTCGGCTAAGACCTCTCGAACTCTTCGTCTTACCTCAGCCCGCGGCTACAGT
CATATACATCCTAGCACGCACAATCCTTAATTCTCAGTCGTGCCCGACAGTAACACCGTG
GCGCGAACAATGTGTAAGTTATTGCCTACTTCGTACACTATCAACTAAATTCGTCCAACG
TGAGATCCGAGAATCCCGAAAGGGGGCACACACTTAGAGAACCCGCAGCTTTTTGTCTCA
CGATCGAGGAGGGCACTCCCAAGAGGCCTTGATACGAGCGTGAAACCGGTCAAGCGCTCC
AATTGTACACTGTCCATGAGGCCAAATTCAAATTACGGACCGAAAACTGGATCCAACAGA
GCTAAAGGTTATTCGCGGGTATTGTCCGTACATGCAACTTGATGTTTTCTAAGCCAAAAG
AGATATGCGAGGGGGAGACATGGGAAAAGCAAAGGAATATAGTACTATAATTGGCGTGGG
ACCCATAAAGTCTTGATTGCTCCCGCGCCGCCTGTGCCCACAGTCTTCAAAAAAGCGATT
TGCCGGAGTATAAAGTCAGGGGCCGACGGTGTTGAAAACGAGCGGACGTCTTGCATGAGA
AGTGTCACTGTGGGCATATCAAAGGACTCTAATTAAATTCAGTGTCAATAGAAATGTTCG
TGCTGCACGGTGCTGTCAGACCATCCACCTTTCGTGGCTACGCCTTATCAGAAGTGCAAC
CTGCGAAATCAAGGTGACGTTAATACGCTCTAAAGGACGCTAAAGCGGGTTGATAATTAA
ACGACAGAGTTACAATTAGCGGGATTCACAACTCTCCAAAATCATTGACAATAGACTGAC
ACGGTTACGTGGGAGGACTCGTAGTCTCAAGTGAAAATGGGCGCAAAATGGTGTCCGCAT
GGGAGGAGCTCAACCATAATAGACGCCGAATATGTTCGCGTGAAACTTCAGAGTCAAAGA
TGGAAAGCAAGACAAACTCATGAAACTACCATAGTGTGAAGATGATGGGCGGCGCGGTTT
AAGGCGAGTCGCCCTAAGACTCCAACGTGCCCTAAGGGATCGTAACATGCAAGAACATTG
ATGTGCACAGTTTTGTCGAACGGCCTTCATAGGCCAAAGATTGTATCCACGGCCAGAAGC
GGTAAGAACTTCGTGACATAGCGGCCAGAGAAAGGATCTACAACGTTAACCAGCACCTCA
GAGAGCGGGATGGCCAGTCGTTAGGGAGAAAATAACTAATCATTAACTTTGACGAACCAC

CAAAATGCTCGAGCGGTAGGTATCATTAGGGGGTACGGACTGTTACATCTAGAATACCCC

AGCTCCGTAGTTGCAGACACTAATTTTAGAGACCGCAGCTGTATCATTAACAAAATGTTA

TTGGTGGGATGCAGAAGAGCGCGGGCAAGATTATACAGACGCCTTCAGGCGGTAGCCTGC

TCCTAGGCAGTCAAGACGGATCGCGATGGACGTGTTTGGACGGCGAATGTACAGGAAATT

TCCCGTGAGAACAACAAAATGAAGAATCGAGATTAAAAAGACTGCAGCGCCCGTGAACAA

CGGAGTTGAACGTGGTTGCCAGGTTTGTGATGGGCGGCCCTAGAAGATAACGGTAGAAGC

CCTTCGGACTTTTCAATGGGGACTACATTCTGGCCGCGAAGGACTCCGTGAATCGATAGG

TAGTGGAAATATGGATGAAACGGGCTGGGGCAACGCTCGGGACAAAGTTTACTAGCTAAG

AGGCAGATTACGAGGATCGGGGAAGACTGAGAAGGCCTTGGCGTAGGAATTCGAAGCAAC

CAGGGGGATAAGGTAAATGCCGACTTCAAGTCGAGAATGCAAAGAAGCTGGCTATGGCCG

TAAATTAGACGATAAATAACAGAGCATCGGGGAGCCAAGTCTAATTGTAAAACTTTTGGT

CGAGGTGATGTCGTAGGTGAGATCTCCGAACGCAGCATGAACAAGGCGAATCGTGTAAGC

AGAGAAGAGTGCAGCACGATGCTAACACTGCCGCAGCAGTGGGACAACTTGGATGACTGG

CACATCACAGTAATGAGTCATATGATTCCGCCTGAAGGCTTCAATAAAGGACATAAGATG

CGAAGAAGATCTAGACCCAGTCTTTAAAACTGCACAACTTAGCCGAAATGATTGGTAGGT

GAGAGTATCGACGTGGGCACTGAGAGAAACCTGAATACGCGATAGCNATCACATCACGAT

GCTTCTGCCCTCCAGCGTCACTAGAGTTCTTGTTCACATGCCTCGGTCCTCTCGTCAGTA

CATGGCCGGCCTTCTTTCCTCCATCGTTTCCCAGAGCATACTCCTCGAGTTGCCCTATTA

ACCTTCTATTTTCTGGCCGATGCCTGACTGGCGCCGCTTTGAGTATCATACTTCTTGTTT

TTGGCCAACCTGTTGGGCGTTGATTGCTTGGGTAACCACATCCTGCATTTTGTTGATTCC

GGGTCGGCAATGTGTTGTGGACTGTTCGTCGCCGCTCCTGATCTCACCGTACTTCACGGT

TGTCGCCCGGACATCACCCGGACGGGTTTAGGCCATGGCCGGTTGAGGTTAATCTTTACG

CGTCACCATAACTTCCTGCGAAACTGGTTTCCCCACACCGCTCCCTTCGTGAGCCGATCG

GCGTGGTTAGGTCGCTCGGGTTTTTCCAGGCAGGGTAAGTCGAGTCGCACAAGGTGACCC

ACGTACCTCGCTTTAACGTTCCGTTCTACCACTGGGTTAGGCTGCTTTGGGGCCGACTTG

ACGTTCTTTTTGCACGACCTCGTATGTGCTTATTCTCACTCGTAATGAACCGGTAAGCGA

GAGTTTGTTCTACGCTGAGGTGTCGTCGGACCCGAGTGTGCTTGCCGCTGGTCAATTGTT

GTGTCGGCTCGACGTCTTATTTCGTATGGTACGGGATTATTAGTGCTTGCAGCGATGTCC

CTTTCGCTATTGATGTCCTCTCCTGAGTCACGCTACGTACTGAATTGATAAAACATTTCG

ACCTCAGTATCAGTGAGGTCGGGGGATGAATTAGATCGCCGTGTAGGGCTTTTCACGTTG

TGTCCGTGTGGAGAGTCGGTTCTAGCGGTCGTATTATCCATATTCTCCCGTTCCATTTAC

GTTGAGCGACTTGAAACTTCCGATGTACTGGCCGGTAATAGTGCATTGTTTCTCAATTCA

TGCAAGTTGTCGTACATCATGGAATAGAACAAGAATTAAGGACCATTTCCACTAGCGATT

CTTACCTCCAACTCAAGATAGAGTCTAACCGATTATCGACGAACTTCGTCCGTCACGTCA

TTCTATGTCGAGCGGCCTTCTTACAGCGCTGTCGCACTATCTGTTCTACCTTATTTCCGA

GAAACCTTGCTCCTGTTACCTGTGAGGGCGGGGCGGATATGGCAAAATGGGCATTCTTTT

TTACTTCCCGTGCCTCTTCATACTATATACATGTATCTCTGGCGCCTAGACGCCCGGTCC
TTAACGTCCTAAAACTTTACTGACCCTGCTGGCGTCTTACCCTTTATAGTGGCTCTGCAT
AATGTGTTCTGTATATAGCCAAATACTGCCGACTATCCACTCGCTTAGCTGTCAATATCC
ATGCGCGTACTTTGATTTACCGGTCGACCTGCCTAATGTAACTGCCTTTCGCTCTCCACT
CCGCACCGGGGTTTCGAGTTTTTCTACCCCTCTCTTTATTACTACTCGTTTACTGCATCA
GACGCGCCTACCGTATTCAGTCACTGCTCCTTCAGGCGAAGAGTATTAGCACGTTATCGT
GCTACTATCCTGAGCTGTAATCATAAGTATTTCTAAAGTTGTAAGGGTTGAAGAGTAGTT
CTGTACATCTTGATCGATTGTACTGGACGTCATTCCGAATAGCTTGTGGAATCCAATATC
TGGGTCATAGATGCT
>SRR9587930
GTGACCGTCGCCTCGCCACAAATCCATTGTTTTCCGGGTTCTAGCCCCTTATGAGCTAGG
ACCTCTGTGCCTACCATCATAGCTGTTCGATGCCTTGTATTCAATACGTTGCACCGACCC
GACAATAGTCTGTGCCATTAAGCTTCTGCAAGTGGCTGGCGCCTGTAAGTCACGCAATAG
AGTAACTTCCCTCTCGACAGTCTTTATGCCATCCACAATCCAACATCGTAGATCCTCACG
TGACCTAGTAGCCCTTGACGACTCAATCCACCCTCTGCCTGACGGACGTGCCTTTTATGG
TACATTAATGACTCCCTGTCCTCATAATTAGGCTGCACACACAGGGCTATCGTAGTTTAC
TGGTCTCACCCCTGCGTACTCGGCGACCCAACTACACTGATCCAGGGAGGCAGAATGGTG
GACGCGCTTACTCCCGACCCGTGGGCAACTAACTGGGCCGGGGGAATATTGTTTCGTTTT
GGGTACTGCTTCTTCCAATTGTTCGGATAGCACAGTAACGGTGGTGGGTGGCTCCAGTCA
CTCTTGCGTAGTCGCCGGCCGGTCGCTCCGGTCTGTCCTCTCGCCCGATTCTGGGCCATC
TTATCCGATGTCACCCTGCAGACCGCTCTTTTCAACCATTAGCATTAAGCAGTCTCTCAT
AGGTGCTCTCATGTCATGAGAGGTACCGGAGCCTCTAAGCGTTCCGCTGTTCCCGACAAT
ATTCTCTTTTGCTTTGACACTAGCTCAGACGGTAGAGGCTTCGTCTTCAGCACCGGCTAA
TGATGCCTGGTACGTTAAGATACCGATTCGCGCGCTTACGCTGCTCCACCTGTCGGTGTC
TGAAGAAGCTCGGCTAAGACCTCTCGAACTCTTCGTCTTACCTCAGCCCGCGGCTACAGT
CATATACATCCTAGCACGCACAATCCTTAATTCNCAGTCGTGCCCGACAGTAACACCGTG
GCGCGAACAATGTGTAAGTTATTGCCTACTTCGTACACTATCAACTAAATTCGTCCAACG
TGAGATCCGAGAATCCCGAAAGGGGGCGCACACTTAGAGAACCCGCAGCTTTTTGTCTCA
CGATCGAGGAGGGCACTCCCAAGAGGCCTTGATACGAGCGTGAAATCGGTCAAGCGCTCC
AATTGTACACTGTCCATGAGGCCAAATTCAAATTACGGACCGAAAACTGGATCCAGCAGA
GCTAAAGGTTATTCGCGGGTATTGTCCGTACATGCAACTTGATGTTTTCTAAGCCAAAAG
AGATATGCGAGGGGGAGACATGGGAAAAGCAAAGGAATATAGTACTATAATTGGCGTGGA
ACCCATAAAGTCTTGATTGCTCCCGCGCCGCCTGTGCCCACAGTCTTCAAAAAAGCGATT
TGCCGGAGTATAAAGTCAGGGGCCGACGGTGTTGAAAACGAGCGGACGTCTTGCATGAGA
AGTGTCACTGTGGGCATATCAAAGGACTCTAATTAAATTCAGTGTCAATAGAAATGTTCG
TGCTGCACGGTGCTGTCAGACCATCCACCTTTCGTGGCTACGCCTTATCAGAAGTGCAAC

CTGCGAAATCAAGGTGACGTTAATACGCTCTAAAGGACGCTAAAGCGGGTTGATAATTAA
ACGACAGAGTTACAATTAGCGGGATTCACAACTCTCCAAAATCATTGACAATAGACTGAC
ACGGTTACGTGGGAGGACTCGTAGTCTCAAGTGAAAATGGGCGCAAAATGGTGTCCGCAT
GGGAGGAGCTCAACCATAATAGACGCCGAATATGTTCGCGTGAAACTTCAGAGTCAAAGA
TGGAAAGCAAGACAAACTCATGAAACTACCATAGTGTGAAGATGATGGGCGGCGCGGTTT
AAGGCGAGTCGCCCTAAGACTCCAACGTGCCCTAAGGGATCGTAACATGCAAGAACATTG
ATGTGCACAGTTTTGTCGAACGGCCTTCATAGGCCAAAGATTGTATCCACGGCCAGAAGC
GGTAAGAACTTCGTGACATAGCGGCCAGAGAAAGGATCTACAACGTTAACCAGCACCTCA
GAGAGCGGGATGGCCAGTCGTTAGGGAGAAAATAACTAATCATTAACTTTGACGAACCAC
CAAAATGCTCGAGCGGTAGGTATCATTAGGGGGTACGGACTGTTACATCTAGAATACCCC
AGCTCCGTAGTTGCAGACACTAATTTTAGAGACCGCAGCTGTATCATTAACAAAATGTTA
TTGGTGGGATGCAGAAGAGCGCGGGCAAGATTATACAGACGCCTTCAGGCGGTAGCCTGC
TCCTAGGCAGTCAAGACGGATCGCGATGGACGTGTTTGGACGGCGAATGTACAGGAAATT
TCCCGTGAGAACAACAAAATGAAGAATCGAGATTAAAAAGACTGCAGCGCCCGTGAACAA
CGGAGTTGAACGTGGTTGCCAGGTTTGTGATGGGCGGCCCTAGAAGATAACGGTAGAAGC
CCTTCGGACTTTTCAATGGGGACTACATTCTGGCCGCGAAGGACTCCGTGAATCGATAGG
TAGTGGAAATATGGATGAAACGGGCTGGGGCAACGCTCGGGACAAAGTTTACTAGCTAAG
AGGCAGATTACGAGGATCGGGGAAGACTGAGAAGGCCTTGGCGTAGGAATTCGAGGCAAC
CAGGGGGATAAGGTAAATGCCGACTTCAAGTCGAGAATGCAAAGAAGCTGGCTATGGCCG
TAAATTAGACGATAAATAACAGAGCATCGGGGAGCCAAGTCTAATTGTAAAACTTTTGGT
CGAGGTGATGTCGTAGGTGAGATCTCCGAACGCAGCATGAACAAGGCGAATCGTGTAAGC
AGAGAAGAGTGCAGCACGATGCTAACACTGCCGCAGCAGTGGGACAACTTGGATGACTGG
CACATCACAGTAATGAGTCATATGATTCCGCCTGAAGGCTTCAATAAAGGACATAAGCTG
CGAAGAAGATCTAGACCCAGTCTTTAAAACTGCACAACTTAGCCGAAATGATTGGTAGGT
GAGAGTATCGACGTGGGCACTCAGAGAAACCTGAATACGCGATAGCCATCACATCACGAT
GCTTCTGCCCTCCAGCGTCACTAGAGTTCTTGTTCACATGCCTCGGTCCTCTCGTCAGTA
CATGGCCGGCCTTCTTTCCTCCATCGTTTCCCAGAGCATACTCCTCGAGTTGCCCTATTA
ACCTTCTATTTTCTGGCCGATGCCTGACTGGCGCCGCTTTGAGTATCATACTTCTTGTTT
TTGGCCAACCTGTTGGGCGTTGATTGCTTGGGTAACCACATCCTGCACTTTGTTGATTCC
GGGTCGGCAATGTGTTGTGGACTGTTCGTCGCCGCTCCTGATCTCACCGTACTTCACGGT
TGTCGCCCGGACATCACCCGGACGGGTTTAGGCCATGGCCGGTTGAGGTTAATCTTTACG
CGTCACCATAACTTCCTGCGAAACTGGTTTCCCCACACCGCTCCCTTCGTGAGCCGATCG
GCGTGGTTAGGTCGCGCGGGTTTTTCCAGGCAGGGTAAGTCGAGTCGCACAAGGTGACCC
ACGTACCTCGCTTTAACGTTCCGTTCTACCACTGGGTTAGGCTGCTTTGGGGCCGACTTG
ACGTTCTTTTTGCACGACCTCGTATGTGCTTATTCTCACTCGTAATGAACCGGTAAGCGA
GAGTTTGTTCTACGCTGAGGTGTCGTCGGACCCGAGTGTGCTTGCCGCTGGTCAATTGTT

GTGTCGGCTCGACGTCTTATTTCGTATGGTACGGGATTATTAGTGCTTGCAGCGATGTCC
CTTTCGCTATTGATGTCCTCTCCTGAGTCACGCTACGTACTGAATTGATAAAACATTTCG
ACCTCAGTATCAGTGAGGTCGGGGGATGAATTAGATCGCCGTGTAGGGCTTTTCACGTTG
TGTCCGTGTGGAGAGTCGGTTCTAGCGGTCGTATTATCCATATTCTCCCGTTCCATTTAC
GTTGAGCGACTTGAAACTTCCGATGTACTGGCCGGTAATAGTGCATTGTTTCTCAATTCA
TGCAAGTTGTCGTACATCATGGAATAGAACAAGAATTAAGGACCATTTCCACTAGCGATT
CTTACCTCCAACTCAAGATAGAGTCTAACCGATTATCGACGAACTTCGTCCGTCACGTCA
TTCTATGTCGAGCGGCCTTCTTACAGCGCTGTCGCACTATCTGTTCTACCTTATTTCCGA
GAAACCTTGCTCCTGTTACCTGTGAGGGCGGGGCGGATATGGCAAAATGGGCATTCTTTT
TTACTTCCCGTGCCTCTTCATACTATATACATGTATCTCTGGCGCCTAGACGCCCGGTCC
TTAACGTCCTAAAACTTTACTGACCCTGCTGGCGTCTTACCCTTTATAGTGGCTCTGCAT
AATGTGTTCTGTATATAGCCAAATACTGCCGACTATCCACTCGCTTAGCTGTCAATATCC
ATGCGCGTACTTTGATTTACCGGTCGACCTGCCTAATGTAACTGCCTTTCGCTCTCCACT
CCGCACCGGGGTTTCGAGTTTTTCTACCCCTCTCTTTATTACTACTCGTTTACTGCATCA
GACGCGCCTACCGTATTCAGTCACTGCTCCTTCAGGCGAAGAGTATTAGCACGTTATCGT
GCTACTATCCTGAGCTGTAATCATAAGTATTTCTAAAGTTGTAAGGGTTGAAGAGTAGTT
CTGTACATCTTGATTGATTGTACTGGACGTCATTCCGAATAGCTTGTGGAATCCAATATC
TGGGTCATAGATGCT
>SRR9587931
GTGACCGTCGCCTCGCCACAAATCCATTGTTTTCCGGGTTCTAGCCCCTTATGAGCTAGG
ACCTCTGTGCCTACCATCATAGCTGTTCGATGCCTTGTATTCAATACGTTGCACCGACCC
GACAATAGTCTGTGCCATTAAGCTTCTGCAAGTGGCTGGCGCCTGTAAGTCACGCAATAG
AGTAACTTCCCTCTCGACAGTCTTTATGCCATCCACAATCCAACATCGTAGATCCTCACG
TGACCTAGTAGCCCTTGACGACTCAATCCACCCTCTGCCTGACGGACGTGCCTTTTATGG
TACATTAATGACTCCCTGTCCTCATAATTAGGCTGCACACACAGGGCTATCGTAGTTTAC
TGGTCTCACCCCTGCGTACTCGGCGACCCAACTACACTGATCCAGGGAGGCAGAATGGTG
GACGCGCTTACTCCCGACCCGTGGGCAACTAACTGGGCCGGGGGAATATTGTTTCGTTTT
GGGTACTGCTTCTTCCAATTGTTCGGATAGCACAGTAACGGTGGTGGGTGGCTCCAGTCA
CTCTTGCGTAGTCGCCGGCCGGTCGCTCCGGTCTGTCCTCTCGCCCGATTCTGGGCCATC
TTATCCGATGTCACCCTGCAGACCGCTCTTTTCAACCATTAGCATTAAGCAGTCTCTCAT
AGGTGCTCTCATGTCATGAGAGGTACCGGAGCCTCTAAGCGTTCCGCTGTTCCCGACAAT
ATTCTCTTTTGCTTTGACACTAGCTCAGACGGTAGAGGCTTCGTCTTCAGCACCGGCTAA
TGATGCCTGGTACGTTAAGATACCGATTCGCGCGCTTACGCTGCTCCACCTGTCGGTGTC
TGAAGAAGCTCGGCTAAGACCTCTCGAACTCTTCGTCTTACCTCAGCCCGCGGCTACAGT
CATATACATCCTAGCACGCACAATCCTTAATTCTCAGTCGTGCCCGACAGTAACACCGTG
GCGCGAACAATGTGTAAGTTATTGCCTACTTCGTACACTATCAACTAAATTCGTCCAACG

TGAGATCCGAGAATCCCGAAAGGGGGCGCACACTTAGAGAACCCGCAGCTTTTTGTCTCA

CGATCGAGGAGGGCACTCCCAAGAGGCCTTGATACGAGCGTGAAATCGGTCAAGCGCTCC

AATTGTACACTGTCCATGAGGCCAAATTCAAATTACGGACCGAAAACTGGATCCAGCAGA

GCTAAAGGTTATTCGCGGGTATTGTCCGTACATGCAACTTGATGTTTTCTAAGCCAAAAG

AGATATGCGAGGGGGAGACATGGGAAAAGCAAAGGAATATAGTACTATAATTGGCGTGGA

ACCCATAAAGTCTTGATTGCTCCCGCGCCGCCTGTGCCCACAGTCTTCAAAAAAGCGATT

TGCCGGAGTATAAAGTCAGGGGCCGACGGTGTTGAAAACGAGCGGACGTCTTGCATGAGA

AGTGTCACTGTGGGCATATCAAAGGACTCTAATTAAATTCAGTGTCAATAGAAATGTTCG

TGCTGCACGGTGCTGTCAGACCATCCACCTTTCGTGGCTACGCCTTATCAGAAGTGCAAC

CTGCGAAATCAAGGTGACGTTAATACGCTCTAAAGGACGCTAAAGCGGGTTGATAATTAA

ACGACAGAGTTACAATTAGCGGGATTCACAACTCTCCAAAATCATTGACAATAGACTGAC

ACGGTTACGTGGGAGGACTCGTAGTCTCAAGTGAAAATGGGCGCAAAATGGTGTCCGCAT

GGGAGGAGCTCAACCATAATAGACGCCGAATATGTTCGCGTGAAACTTCAGAGTCAAAGA

TGGAAAGCAAGACAAACTCATGAAACTACCATAGTGTGAAGATGATGGGCGGCGCGGTTT

AAGGCGAGTCGCCCTAAGACTCCAACGTGCCCTAAGGGATCGTAACATGCAAGAACATTG

ATGTGCACAGTTTTGTCGAACGGCCTTCATAGGCCAAAGATTGTATCCACGGCCAGAAGC

GGTAAGAACTTCGTGACATAGCGGCCAGAGAAAGGATCTACAACGTTAACCAGCACCTCA

GAGAGCGGGATGGCCAGTCGTTAGGGAGAAAATAACTAATCATTAACTTTGACGAACCAC

CAAAATGCTCGAGCGGTAGGTATCATTAGGGGGTACGGACTGTTACATCTAGAATACCCC

AGCTCCGTAGTTGCAGACACTAATTTTAGAGACCGCAGCTGTATCATTAACAAAATGTTA

TTGGTGGGATGCAGAAGAGCGCGGGCAAGATTATACAGACGCCTTCAGGCGGTAGCCTGC

TCCTAGGCAGTCAAGACGGATCGCGATGGACGTGTTTGGACGGCGAATGTACAGGAAATT

TCCCGTGAGAACAACAAAATGAAGAATCGAGATTAAAAAGACTGCAGCGCCCGTGAACAA

CGGAGTTGAACGTGGTTGCCAGGTTTGTGATGGGCGGCCCTAGAAGATAACGGTAGAAGC

CCTTCGGACTTTTCAATGGGGACTACATTCTGGCCGCGAAGGACTCCGTGAATCGATAGG

TAGTGGAAATATGGATGAAACGGGCTGGGGCAACGCTCGGGACAAAGTTTACTAGCTAAG

AGGCAGATTACGAGGATCGGGGAAGACTGAGAAGGCCTTGGCGTAGGAATTCGAGGCAAC

CAGGGGGATAAGGTAAATGCCGACTTCAAGTCGAGAATGCAAAGAAGCTGGCTATGGCCG

TAAATTAGACGATAAATAACAGAGCATCGGGGAGCCAAGTCTAATTGTAAAACTTTTGGT

CGAGGTGATGTCGTAGGTGAGATCTCCGAACGCAGCATGAACAAGGCGAATCGTGTAAGC

AGAGAAGAGTGCAGCACGATGCTAACACTGCCGCAGCAGTGGGACAACTTGGATGACTGG

CACATCACAGTAATGAGTCATATGATTCCGCCTGAAGGCTTCAATAAAGGACATAAGCTG

CGAAGAAGATCTAGACCCAGTCTTTAAAACTGCACAACTTAGCCGAAATGATTGGTAGGT

GAGAGTATCGACGTGGGCACTCAGAGAAACCTGAATACGCGATAGCCATCACATCACGAT

GCTTCTGCCCTCCAGCGTCACTAGAGTTCTTGTTCACATGCCTCGGTCCTCTCGTCAGTA

CATGGCCGGCCTTCTTTCCTCCATCGTTTCCCAGAGCATACTCCTCGAGTTGCCCTATTA

ACCTTCTATTTTCTGGCCGATGCCTGACTGGCGCCGCTTTGAGTATCATACTTCTTGTTT
TTGGCCAACCTGTTGGGCGTTGATTGCTTGGGTAACCACATCCTGCACTTTGTTGATTCC
GGGTCGGCAATGTGTTGTGGACTGTTCGTCGCCGCTCCTGATCTCACCGTACTTCACGGT
TGTCGCCCGGACATCACCCGGACGGGTTTAGGCCATGGCCGGTTGAGGTTAATCTTTACG
CGTCACCATAACTTCCTGCGAAACTGGTTTCCCCACACCGCTCCCTTCGTGAGCCGATCG
GCGTGGTTAGGTCGCGCGGGTTTTTCCAGGCAGGGTAAGTCGAGTCGCACAAGGTGACCC
ACGTACCTCGCTTTAACGTTCCGTTCTACCACTGGGTTAGGCTGCTTTGGGGCCGACTTG
ACGTTCTTTTTGCACGACCTCGTATGTGCTTATTCTCACTCGTAATGAACCGGTAAGCGA
GAGTTTGTTCTACGCTGAGGTGTCGTCGGACCCGAGTGTGCTTGCCGCTGGTCAATTGTT
GTGTCGGCTCGACGTCTTATTTCGTATGGTACGGGATTATTAGTGCTTGCAGCGATGTCC
CTTTCGCTATTGATGTCCTCTCCTGAGTCACGCTACGTACTGAATTGATAAAACATTTCG
ACCTCAGTATCAGTGAGGTCGGGGGATGAATTAGATCGCCGTGTAGGGCTTTTCACGTTG
TGTCCGTGTGGAGAGTCGGTTCTAGCGGTCGTATTATCCATATTCTCCCGTTCCATTTAC
GTTGAGCGACTTGAAACTTCCGATGTACTGGCCGGTAATAGTGCATTGTTTCTCAATTCA
TGCAAGTTGTCGTACATCATGGAATAGAACAAGAATTAAGGACCATTTCCACTAGCGATT
CTTACCTCCAACTCAAGATAGAGTCTAACCGATTATCGACGAACTTCGTCCGTCACGTCA
TTCTATGTCGAGCGGCCTTCTTACAGCGCTGTCGCACTATCTGTTCTACCTTATTTCCGA
GAAACCTTGCTCCTGTTACCTGTGAGGGCGGGGCGGATATGGCAAAATGGGCATTCTTTT
TTACTTCCCGTGCCTCTTCATACTATATACATGTATCTCTGGCGCCTAGACGCCCGGTCC
TTAACGTCCTAAAACTTTACTGACCCTGCTGGCGTCTTACCCTTTATAGTGGCTCTGCAT
AATGTGTTCTGTATATAGCCAAATACTGCCGACTATCCACTCGCTTAGCTGTCAATATCC
ATGCGCGTACTTTGATTTACCGGTCGACCTGCCTAATGTAACTGCCTTTCGCTCTCCACT
CCGCACCGGGGTTTCGAGTTTTTCTACCCCTCTCTTTATTACTACTCGTTTACTGCATCA
GACGCGCCTACCGTATTCAGTCACTGCTCCTTCAGGCGAAGAGTATTAGCACGTTATCGT
GCTACTATCCTGAGCTGTAATCATAAGTATTTCTAAAGTTGTAAGGGTTGAAGAGTAGTT
CTGTACATCTTGATTGATTGTACTGGACGTCATTCCGAATAGCTTGTGGAATCCAATATC
TGGGTCATAGATGCT
>SRR9587932
GTGACCGTCGCCTCGCCACAAATCCATTGTTTTCCGGGTTCTAGCCCCTTATGAGCTAGG
ACCTCTGTGCCTACCATCATGGCTGTTCGATGCCTTGTATTCAATACGTTGCACCGACCC
GACAATAGTCTGTGCCATTAAGCTTCTGCAAGTGGCTGGCGCCTGTAAGTCACGCAATAG
AGTAACTTCCCTCTCGACAGTCTTTATGCCATCCACAATCCAACATCGTAGATCCTCGCG
TGACCTAGTAGCCCTTGACGACTCAATCCACCCTCTGCCTGACGGACGTGCCTTTTATGG
TACATTAATGACTCCCTGTCCTCATAATTAGGCTGCACACACAGGGCTATCGTAGTTTAC
TGGTCTCACCCCTGCGCACTCGGCGACCCAATTACACTGATCCGGGGAGGCAGAATGGTG
GACGCGCTTACTCCCGACCCGTGGGCAACTAACTGGGCCGGGGGAATATTGTTTCGTTTT

GGGTACTGCTTCTTCCAATTGTTCGGATAGCACAGTAACGGTGGTGGGTGGCTCCAGTCA
CTCTTGCGTAGTCGCCGGCCGGTCGCTCCGGTCTGTCCTCTCGCCCGATTCTGGGCCATC
TTATCCGATGTCACCCTGCAGACCGCTCTTTTCAACCATTAGCATTAAGCAGTCTCTCAT
AGGTGCTCTCATGTCATGAGAGGTACCGGAGCCTCTAAGCGTTCCGCTGTTCCCGACAAT
ATTCTCTTTCGCTTTGACACTAGCTCAGACGGTAGAGGCTTCGTCTTCAGCACCGGCTAA
TGATGCCTGGTACGTTAAGATACCGATCCGCGCGCTTACGCTGCTCCACCTGTCGGTGTC
TGAAGAAGCTCGGCTAAGACCTCTCGAACTCTTCGTCTTACCTCAGCCCGCGGCTACAGT
CATATACATCCTAGCACGCACAATCCTTAATTCCCAGTCGTGCCCGCCAGTAACACCGTG
GCGCGAACAATGTGTAAGTTATTGCCTACTTCGTACACTATCAACTAAATTCGTCCAACG
TGAGATCCAAGAATCCCGAAAGGGGGCACACACTTAGAGAACCCGCAGCTTTTTGTCTCA
CGATCGAGGAGGGCACTCCCAAGAGGCCTTGATACGAGCGTGAAACCGGTCAAGCGCTCC
AATTGTACACTGTCCATGAGGCCAAATTCAAATTACGGACCGAAAACTGGATCCAGCAGA
GCTAAAGGTTATTCGCGGGTATTGTCCGTACATGCAACTTGATGTTTTCTAAGCCAAAAG
AGATATACGAGGGGGAGACATGGGAAAAGCAAAAGAATATAGTACTATAATTGGCGTGGG
ACCCATAAAGTCTTGATTGCTCCCGCGCCGCCTGTGCCCACAGTCTTCAAAAAAGCGATT
TGCCGGAGTATAAAGTCAGGGGCCGACGGTGTTGAAAACGAGCGGACGTCTTGCATGAGA
AGTGTCACTGTGGGCATATCAAAGGACTCTAATTAAATTCGGTGTCAATAGAAATGTTCG
TGCTGCACGGTGCTGTCAGACCATCCACCTTTCGTGGCTACGCCTTATCAGAAGTGCAAC
CTGCGAAATCAAGGTGACGTTAATACGCTCTACAGGACGCTAAAGCGGGTTGATAATTAA
ACGACAGAGTTACAATTAGCGGGATTCACAACTCTCCAAAATCATTGACAATAGACTGAC
ACGGTTACGTGGGAGGACTCGTAGTCTCAAGTGAAAATGGGCGCAAAATGGTGTCCGCAT
GGGAGGAGCTCAACCATAATAGACGCCGAATATGTTCGCGTGAAACTTCAGAGTCAAAAA
TGGAAAGCAAGACAAACTCATGAAACTACCATAGTGTGAAGATGATGGGCGGCGCGGTTT
AAGGCGAGTCGCCCTAAGACTCCAACGTGCCCTAAGGGATCGTAACATGCAAGAACATTG
ATGTGCACAGTTTTGTCGAACGGCCTTCATAGGCCAAAGATTGTATCCACGGCCAGAAGC
GGTAAGAACTTCGTGACATAGCGGCCAGAGAAAGGATCTACAACGTTAACCAGCACCTCA
GAGAGCGGGATGGCCAGTCGTTAGGGAGAAAATAACTAATCATTAACTTTGACGAACCAC
CAAAATGCTCGAGCGGTAGGTATCATTAGGGGGTACGGACTGTTACATCTAGAATACCCC
AGCTCCGTAGTTGCAGACACTAATTTTAGAGACCGCAGCTGTATCATTAACAAAATGTTA
TTGGTGGGATGCAGAAGAGCGCGGGCAAGATTATACAGACGCCTTCAGGCGGTAGCCTGC
TCCTAGGCAGTCAAGACGGATCGCGATGGACGTGTTTGGACGGCGAATGTACAGGAAATT
TCCCGTGAGAACAACAAAATGAAGAATCGAGATCAAAAAGACTGCAGCGCCCGTGAACAA
CGGAGTTGAACGTGGTTGCCAGGTTTGTGATGGGCGGCCCTAGAAGATAGCGGTAGAAGC
CCTTCGGACTTTTTAATGGGGACTACATTCTGGCCGCGAAGGACTCCGTGAATCGATAGG
TAGTGGAAATATGGATGAAACGGGCTGGGGCAACGCTCGGGACAACGTTTACTAGCTAAG
GGGCAGATTACGAGGATCGGGGAAGACTGAGAAGGCCTTGGCGTAGGAATTCGAGGCAAC

CAGGGGGATAAGGTAAATGCCGACTTCAAGTCGAGAATGCAAAGAAGCTGGCTATGGCCG
TAAATTAGACGATAAATAACAGAGCATCGGGGAGCCAAGTCTAATTGTAAAACTTTTGGT
CGAGGTGATGTCGTAGGTGAGATCTCCGAACGCAGCATGAACAAGGCGAATCGTGTAAGC
AGAGAAGAGTGCAGCACGATGCTAACACTGCCGCAGCAGTGGGACAACTTGGATGACTGG
CACATCACAGTAATGAGTCATATGATTCCGCCTGAAGGCTTCAATAAAGGACATAAGATG
CGAAGAAGATCTAGACCCAGTCTTTAAAACTGCACAACTTAGCCGAAATGATTGGTAGGT
GAGAGTATCGACGTGGGCACTGAGAGAAACCTGAATACGCGATAGCCATCATATCACGAT
GATTCTGCCCTTCAGCGTCACTAGAGTTCTTGTTCACATGCCTCGGTCCTCTCGTCAGTA
CATGGCCGGCCTTCTTTCCTCCATCGTTTCCCAGAGCATACTCCTCGAGTTGCCCTATTA
ACCTTCTATTTTCTGGCCGATGCCTGACTGGCGCCGCTTTGAGTATCATACTTCTTGTTT
TTGGCCAACCTGTTGGGCGTTGATTGCTTGGGTAACCATATCCTGCATTTTGTTGATTCC
GGGTCGGCAATGTGTTGTGGACTGTTCGTCGCCGCTCCTGATCTCACCGTACTTCACGGT
TGTCGCCCGGACATCACCCGGACGGGTTTAGGCCATGGCCGGTTGAGGTTAATCCTTACG
CGTCACCATAACTTCCTGCGAAACTGGTTTCCCCACGCCGCTCCCTTCGTGAGCCGATCG
GCGTGGTTAGGTCGCGCGGGTTTTTCCAGGCAGGGTAAGTCGAGTCGCACAAGGTGACCC
ACGTACCTCGCTTTAACGTTCCGTTCTACCACTGGGTTAGGCTGCTTTGGGGCCGACTTG
ACGTTCTTTTTGCACGACCTCGTATGTGCTTATTCTCACTCGTAATGAACCGGTAAGCGA
GAGTTTGTTCTACGCTGAGGTGTCGTCGGACCCGAGTGTGCTTGCCGCTGGTCAATTGTT
GCGTCGGCTCGACGTCTTATTTCGTATGGTACGGGATTATTAGTGCTTACAGCGATGTCC
CTTTCGCTATTGATGTCCTCTCCTGAGTCACGCTACGTACTGAATTGATAAAACATTTCG
ACCTCAGTATCAGTGAGGTCGGGGGATGAGTTAGANCGCCGTGTAGGGCTTTTCACGTTG
TGTCCGTGTGGAGGGTTGGTTCTAGCGGTCGTATTATCCATATTCTCCCGTTCCATTTAC
GTTGAGCGACTTGAAACTTCCGATGTACTGGCCGGTAATAGTGCATTGTTTCTCAATTCA
TGCAAGTTGTCGTACATCATGGAATAGAACAAGAATTAAGGACCATTTCCACTAGCGATT
CTTACCTCCAACTCAAGATAGAGTCTAACCGATTATCGACGAACTTCGTCCGTCACGTCA
TTCTATGTCGAGCGGCCTTCTTACAGCGCTGTCGCACTATCTGTTCTACCTTATTTCCGA
GAAACCTTGCTCCTGTTACCTGTGAGGGCGGGGCGGATATGGCAAAATGGGCATTCTTTT
TTACTTCCCGTGCCTCTTCATGCTATATACATGTATCTCTGGAGCCTAGACGCCCGGTCC
TTAACGTCCTAAAACTTTACTGACCCTGCTGGCGTCTAACCCTTTATAGTGGCTCTGCAT
AATGTGTTCTGTATATAGCCAAATACTGCCGACTATCCACTCGCTTAGCTGTCAATATCC
ATGCGCGTACTTTGATTTACCGGTCGACCTGCCTAATGTAACTGCCTTTCGCTCTCCACT
CCGCACCGGGGTTTCGAGTTTTTCTACCCCTCTCTTTATTACTACTCGTTTACTGCATCA
GACGCGCCTACCGTATTCAGTCACTGCTCCTTCAGGCGAAGAGTATTAGCACGTTATCGT
GCTACTATCCTGAGCTGTAATCATAAGTATTTCTAAAGTTGTAAGGGTTGAAGAGTAGTT
CTGTACATCTTGATCGATTGTACTGGACGTCATTCCGAATAGCTTGTGGAATCCAATATC
TGGGTCATAGATGCT

>SRR9587933

GTGACCGTCGCCTCGCCACAAATCCATTGTTTTCCGGGTTCTAGCCCCTTATGAGCTAGG
ACCTCTGTGCCTACCATCATAGCTGTTCGATGCCTTGTATTCAATACGTTGCACCGACCC
GACAATAATCTGTGCCATTAAGCTTCTGCAAGTGGCTGGCGCCTGTAAGTCACGCAATAG
AGTAACTTCCCTCTCGACAGTCTTTATGCCATCCACAATCCAACATCGTAGATCCTCGCG
TGACCTAATAGCCCTTGACGACTCAATCCACCCTCTGCCTGACGGACGTGCCTTTTATGG
TACATTAATGACTCCCTGTCCTCATAATTAGGCTGCACACACAGGGCTATCGTAGTTTAC
TGGTCTCACCCCTGCGTACTCGGCGACCCAACTACACTGATCCAGGGAGGCAGAATGGTG
GACGCTCTTACTGCCGACCCGTGGGCAACTAACTGGGCCGGGGGAATATTGTTTCGTTTT
GGGTACTGCTTCTTCCAATTGTTCGGATAGCACAGTAACGGTGGTGGGTGGCTCCAGTCA
TTCTTGCGTAGTCGCCGGCCGGTCGCTCCGGTCTGTCCTCTCTCCCGATTCTGGGCAATC
TTATCCGATGTCACCCTGCAGACCGCTCTTTTCAACCATTAGCATTAAGCAGTCTCTCAT
AGGTGCTCTCGTGTCATGAGAGGTACCGGAGCCTCTAAGCGTTCCGCTGTTCCCGACAAT
ATTCTCTTTCGCTTTGACACTAGCTCAGACGGTAGAGGCTTCGCCTTCAGCACCGGCTAA
TGATGCCTGGTACGTTAAGATACCGATCCGCGCGCTTACGCTGCTCCACCTGTCGGTGTC
TGAAGAAGCTCGGCTAAGACCTCTCGAACTCTTCGTCTTACCTCAGCCCGCGGCTACAGT
CATATACATCCTAGCACGCACAATCCTTAATTCCCAGTCGTGCCCGACAGTAACACCGTG
GCGCGAACAATGTGTAAGTTATTGCCTACTTCGTACACTATCAACTAAATTCGTCCAACG
TGAGATCCGAGAATCCCGTTAGGGGGCATACACTTAGAGAACCCGCAGCTTTTTGTCTCA
CGATCGAGAAGGGCACTCCCAAGAGGCCTTGATACGAGCGTGAAACCGGTCAAGCGCTCC
AATTGTACACTGTCCATGAGGCCAAATTCAAATTACGGACCGAAAACTGGATCCAGCAGA
GCTAAAGGTTATTCGCGGGTATTGTCCGTACATGCAACTTGATGTTTTCTAAGCCAAAAG
AGATATGCGAGGGNGAGACATGGGAAAAGCAAAAGAATATAGTACTATAATTGGCGTGGG
ACCCATAAAGTCTTGATTGCTCCCGCGCCGCCTGTACCCACAGTCTTCAAAAAAGCGATT
TGCCGGAGTATAAAGTCAGGGGCCGACGGTGTTGAAAACGAGCGGACTTCTTGCATGAGA
AGTGTCACTGTGGGCATATCAAAGGACTCTAATTAAATTCGGTGTCAATAGAAATGTTCG
TGCTGCACGGTGCTGTCAGACCATCCACCTTTCGTGGCTACGCCTTATCAGAAGTGCAAC
CTGCGAAATCAAGGTGACGTTAATACGCTCTAAAGGACGCTAAAGGGGGTTGATAATTAA
ACGACAGAGTTACAATTAGCGGGATTCACAACTCTCCAAAATCATTGACGATAGACTGAC
ACGGTTACGTGGGAGGACTCGTAGTCTCAAGTGAAAATGGGCGCAAAATGGTGTCCGCAT
GGGAGGAGCTCAACCATAATAGACGCCGAATATGTTCGCGTGAAACTTCAGCGTCAAAGA
TGGAAAGCAAGACAAACTCATGAAACTACCATAGTGTGAAGATGATGGGCGGCGCGGTTT
AAGGCGAGTCGCCCTAAGGCTCCAACGTGCCCTAAGGGATCGTAACATGCAAGAACATTG
ATGTGCACAGTTTTGTCGAACGGCCTTCATAGGCCAAAGATTGTATCCACGGCCAGAAGC
GGTAAGAACTTCGTGACATAGCGGCCAGAGAAAGGATCTACAACGTTAACCAGCACCTCA
GAGAGCGGGATGGCCAGTCGTTAGGGAGAAAATAACTAATCATTAATTTTGACGAACCAC

CAAAATGCTCGAGCGGTAGGTATCATTAGGGGGTACGGACTGTTACATCTAGAATGCCCC
AGCTCCGTAGTTGCAGACACTAATTTTAGAGACCGCAGCTGCATCATTAACAAAATGTTA
TTGGTGGGATGCAGAAGAGCGCGGGCAAGATTATACAGACGCCTTCAGGCGGTAGCCTGC
TCCTAGGCAGTCAAGACGGATCGCGATGGACGTGTTTGGACGGCGAATGTACAGGAAATT
TCCCGTGAGAACAACAAAATGAAGGATCGAGATTAAAAAGACTGCAGCGCCCGTGAACAA
CGGAGTTGAACGTGGTTTCCAGGTTTGTGATGGGCGGCCCTAGAAGATAACGGTAGAAGC
CCTTCGGACTTTTCAATGGGGACTACATTCTGGCCGCGAAGGACTCCGTGAATCGATAGG
TAGTGGAAATATGGATGAAACGGGCTGGGGCAACGCTCGGAACAACGTTTACTAGCTAAG
AGGCAGATTACGAGGATCGGGGAAGACTGAGAAGGCCTTGGCGTAGGAATTCGAGGCAAC
CAGGGGGATAAGGTAAATGCCGACTTCAAGTCGAGAATGCAAAGAAGCTGGCTATGGCCG
TAAATTAGACGATAAATAACAGAGCATCGGGGAGCCAAGTCTAATTGTAAAACTTTTGGT
CGAGGTGATGTCGTAGGTGAGATCTCCGAACGCAGCATGAGCAAGGCGAATCGTGTAAGC
AGAGAAGAGTGCAGCACGATGCTAACACTGCCGCAGCAGTGGGACAACTTGGATGACTGG
CACATCACAGTAATGAGTCATATGATTCCGCCTGAAGGCTTCAATAAAGGACATAAGATG
CGAAGAAGATCTAGACCCAGTCTTTAAAACTGCACAACTTAGCCGAAATGATTGGTAGGT
GAGAGTATCGACGTGGGCACTGAGAGAAACCTGAATACGCGATAGCCATCATATCACGAT
GCTTCTGCCCTCCAGCGTCACTAGAGTTCTTGTTCACATGCCTCGGTCCTCTCGTCAGTA
CATGGCCGGCCTTCTTTCCTCCATCGTTTCCCAGAGCATACTCTTCGAGTTGCCCTATTA
ACCTTCTATTTTCTGGCCGATGCCTGACTGGCGCCGCTTTGAGTATCATACTTCTTGTTT
TTGGCCAAACTGTTGGGCGTTGATTGCTTGGGTAACCATATCCTGCATTTTGTTGATTCC
GGGTCGGCAATGTGTTGTGGACTGTTCGTCGCCGCTCCTGATCTCACCGTACTTCACGGT
TGTCGCCCGGACATCACCCGGACGGGTTTAGGCCATGGCCGGTTGAGGTTAATCTTTACG
CGTCACCATAACTTCCTGCGAAACTGGTTTCCCCACGCCGCTCCCTTCGTGAGCCGATCG
GCGTGGTTAGGTCGCGCGGGTTTTTCCAGGCAGGGTAAGTCGAGTCGCACAAGGTGACCC
ACGTACCTCGCTTTAACGTTCCGTTCTACCACTGGGTTAGGCTGCTTTGGGGCCGACTTG
ACGTTCTTTTTGCACGACCTCGTATGTGCTTATTCTCACTCGTAATGAACCGGTAAGCGA
GAGTTTGTTCTACGCTGAGGTGTCGTCGGACCCGAGTGTGCTTGCCGCTGGTCAATTGTT
GTGTCGGCTCGACGTCTTATTTCGTATGGTACGGGATTATTAGTGCTTGCAGCGATGTCC
CTTTCGCTATTGATGTCTTCTCCTGAGTCACGCTACGTACTGAATTGATAAAACATTTGG
ACCTCAGTATCAGTGAGGTCGGGGGATGAGTTAGATCGCCGTGTAGGGCTTTTCACGTTG
TGTCCGTGTGGAGAGTCGGTTCTAGCGGTCGTATTATCCATATTCTCCCGTTCCATTTAC
GTTGAGCGACTTGAAACTTCCGATGTACTGGCNGNTAATAGTGCATTGTTTCTCAATTCA
TGCAAGTTGTCGTACATTATGGAATAGAACAAGAATTAAGGACCATTTCCACTAGCGATT
CTTACCTCCAACTCAAGATAGAGTCTAACCGATTATCGACGAACTTCGTCCGTCACGTCA
TTCTATGTCGAGCGGCCTTCTTACAGCGCTGTCGCACTATCTGTTCTACCTTATTTCCGA
GAAACCTTGCTCCTGTTACCTGTGAGGGCGGGGCGGATATGGCAAAATGGGCATTCTTTT

TTACTTCCCGTGCCTCTTCATGCTATATACATGTATCTCTGGAGCCTAGACGCCCGGTCC
TTAACGTCCTAAAACTTTACTGACCCTGCTGGCGTCTTACCCTTTATAGTGGCTCTGCAT
AATGTGTTCTGTATATAGCCAAATACTGCCGACTATCCACTCGCTTAGCTGCCAATATCC
ATGCGCGTACTTTGATTTACCGGTCGACCTGCCTAATGTAACTGCCTTTCGCTCTCCACT
CCGCACCGGGGTTTCGAGTTTTTCTACCCCTCTCTTTATTACTACTCGTTTACTGCATCA
GACGCGCCTACCGTATTCAGTCACTGCTCCTTCAGGCGAAGAGTATTAGCACGTTATCGT
GCTACTATCCTGAGCTGTAATCATAAGTATTTCTAAAGTTGTAAGGGTTGAAGAGTAGTT
CTGTACATCTTAATCGATTGTACTGGACGTCATTCCGAATAGCTTGTGGAATCCAATATC
TGGGTCATAGATGCT
>SRR9587934
GTGACCGTCGCCTCGCCACAAATCCATTGTTTTCCGGGTTCTAGCCCCTTATGAGCTAGG
ACCTCTGTGCCTACCATCATAGCTGTTCGATGCCTTGTATTCAATACGTTGCACCGACCC
GACAATAGTCTGTGCCATTAAGCTGCTGCAAGTGGCTGGCGCCCGTAAGTCACGCAATAG
AGTAACTTCCCTCTCGACAGTCTTTATGCCATCCACAATCCAACATCGTAGATCCTCGCG
TGACCTAGTAGCCCTTGACGACTCAATCCACCCTCTGCCTGACGGACGTGCCTTTTATGG
TACATTAATGACTCCCTGTCCTCATAATTAGGCTGCACACACAGGGCTATCGTAGTTTAC
TGGTCTCACCCCTGCGTACTCGGCGACCCAACTACACTGATCCAGGGAGGCAGAATGGTG
GACGCGCTTGCTCCCGACCCGTGGGCAACTAACTGGGCCGGGGGAATATTGTTTCGTTTT
GGGTACTGCTTCTTCCAATTGTTCGGATAGCACAGTAACGGTGGTGAGTGGCTCCAGTCA
CTCTTACGTAGTCGCCGGCCGGTCGCTCCGGTCTGTCCTCTCGCCCGATTCTGGGCCATC
TTATCCGATGTCACCCTGCAGACCGCTCTTTTCAACCATTAGCATTAAGCAGTCTCTCAT
AGGTGCTCTCATGTCATGAGAGGTACCGGAGCCTCTACGCGTTCCGCTGTTCCCGACAAT
ATTCTCTTTCGCTTTGACACTAGCTCAGACGGTAGAGGCTTCGTCTTCAGCACCGGCTAA
TGATGCCTGGTACGTTAAGGTACCGATCCGCGCGCTTACGCTGCTCCACCTGTCGGTGTC
TGAAGAAGCTCGGCTAAGACCTCTCGAACTCTTCGTCTTACCTCAGCCCGCGGCTACAGT
CATATACATCCTAGCACGCACAATCCTTAATTCCCAGTCGTGCCCGACAGTAACACCGTG
GCGCGAACAATGTGTAAGTTATTGCCTACTTCGTACACTATCAACTAAATTCGTCCAACG
TGAGATCCGAGAATCCCGAAAGGGGGCACACACTTAGAGAACCCGCAGCTTTTTATCTCA
CGATCGAGGAGGGCACTCCCAAGAGGCCTTGATACGAGCGTGAAACCGGTCAAGCGCTCC
AATTGTACACTGTCCATGAGGCCAAATTCAAATTACGGACCGAAAACTGGATCCAGCAGA
GCTAAAGGTTATTCGCGGGTATTGTCCGTACATGCAACTTGATGTTTTCTAAGCCAAAAG
AGATATGCGAGGGGGAGACATGGGAAAAGCAAAAGAATATAATACTATAATTGGCGTGGG
ACCCATAAAGTCTTGATTGCTCCCGCGCCGCCTGTGCCCACAGTCTTCAAAAAAGCGATT
TGCCGGAGTATAAAGTCAGGGGCCGACGGTGTTGAAAACGAGCGGACGTCTTACATGAGA
AGTGTCACTGTGGGCATATCAAAGGACTCTAATTAAATTCGGTGTCAATAGAAATGTTCG
TGCTGCACGGTGCTGTCAGACCATCCACCTTTCGTGGCTACGCCTTATCAGAAGTGCAAC

CTGCGAAATCAAGGTGACGTTAATACGCTCTAAAGGACGCTAAAGCGGGTTGATAATTAA
ACGACAGAGTTACAATTAGCGGGATTCACAACTCTCCAAAATCATTGACAATAGACTGAC
ACGGTTACGTGGGAGGACTCGTAGTCTCAAGTGAAAATGGGCGCAAAATGGTGTCCGCAT
GGGAGGAGCTCAACCATAATAGACGCCGAATATGTTCGCGTGAAACTTCAGAGTCAAAGA
TGGAAAGCAAGACAAACTCATGAAACTACCATAGTGTGAAGATGATGGGCGGCGCGGTTT
AAGGCGAGTCGCCCTAAGACTCCAACGTGCCCTAAGGGATCGTAACATGCAAGAACATTG
ATGTGCACAGTTTTGTCGAACGGCCTTCATAGGCCAAAGATTGTATCCACGGCCAGAAGC
GGTAAGAACTTCGTGACATAGCGGCCAGAGAAAGGATCTACAACGTTAACCAGCACCTCA
GAGAGCGGGATGGCCAGTCGTTAGGGAGAAAATAACTAATCATTAACTTTGACGAACCAC
CAAAATGCTCGAGCGGTAGGTATCATTAGGGGGTACGGACTGTTACATCTAGAATACCCC
AGCTCCGTAGTTGCAGACACTAATTTTAGAGNCCGCAGCTGTATCATTAACAAAATGTTA
TTGGTGGGATGCAGAAGAGCGCGGGCAAGATTATACAGACGCCTTCAGGCGGTAGCCTGC
TCCTAGGCAGTCAAGACGGATCGCGATGGACGTGTTTGGACGGCGAATGTACAGGAAATT
TCCCGTGAGAACAACAAAATGAAGAATCGAGATTAAAAAGACTGCAGCGCCCGTGAACAA
CGGAGTTGAACGTGGTTGCCAGGTTTGTGATGGGCGGCCCTAGAAGATAACGGTAGAAGC
CCTTCGGACTTTTCAATGGGGACTACATTCTGGCCGCGAAGGACTCCGTGAATCGATAGG
TAGTGGAAATATGGATGAAACGGGCTGGGGCAACGCTCGGGACAAAGTTTACTAGCTAAG
AGGCAGATTACGAGGATCGGGGAAGACTGAGAAGGCCTTGGCGTAGGAATTCGAGGCAAC
CAGGGGGATAAGGTAAATGCCGACTTCAAGTCGAGAATGCAAAGAAGCTGGCTATGGCCG
TAAATTAGACGATAAATAACAGAGCATCGGGGAGCCAAGTCTAATTGTAAAACTTTTGGT
CGAGGTGATGTCGTAGGTGAGATCTCCGAACGCAGCATGAACAAGGCGAATCGTGTAAGC
AGAGAAGAGTGCAGCACGATGCTAACACTGCCGCAGCAGTGGGACAACTTGGATGACTGG
CACATCACAGTAATGAGTCATATGATTCCGCCTGAAGGCTTCAATAAAGGACATAAGATG
CGAAGAAGATCTAGACCCAGTCTTTAAAACTGCACAACTTAGCCGAAATGATTGGTAGGT
GAGAGTATCGACGTGGGCACTGAGAGAAACCTGAATACGCGATAGCCATCATATCACGAT
GCTTCTGCCCTCCAGCGTCACTAGAGTTCTTGTTCACATGCCTCGGTCCTCTCGTCAGTA
CATGGCCGGCCTTCTTTCCTCCATCGTTTCCCAGAGCATACTCCTCGAGTTGCCCTATTA
ACCTTCTATTTTCTGGCCGATGCCTGACTGGCGCCGCTTTGAGTATCATACTTCTTGTTT
TTGGCCAACCTGTTGGGCGTTGATTGCTTGGGTAACCATATCCTGCATTTTGTTGATTCC
GGGTCGGCAATGTGTTGTGGACTGTTCGTCGCCGCTCCTGATCTCACCGTACTTCACGGT
TGTCGCCCGGACATCACCCGGACGGGTTTAGGCCATGGCCGGTTGAGGTTAATCTTTACG
CGTCACCATAACTTCCTGCGAAACTGGTTTCCCCACACCGCTCCCTTCGTGAGCCGATCG
GCGTGGTTAGGTCGCGCGGGTTTTTCCAGGCAGGGTAAGTCGAGTCGCACAAGGTGACCC
ACGTACCTCGCTTTAACGTTCCGTTCTACCACTGGGTTAGGCTGCTTTGGGGCCGACTTG
ACGTTCTTTTTGCACGACCTCGTATGTGCTTATTCTCACTCGTAATGAACCGGTAAGCGA
GAGTTTGTTCTACGCTGAGGTGTCGTCGGACCCGAGTGTGCTTGCCGCTGGTCAATTGTT

GTGTCGGCTCGACGTCTTATTTCGTATGGTACGGGATTATTAGTGCTTGCAGCGATGTCC
CTTTCGCTATTGATGTCCTCTCCTGAGTCACGCTACGTACTGAATTGATAAAACATTTCG
ACCTCAGTATCAGTGAGGTCGGGGGATGAGTTAGATCGCCGTGTAGGGCTTTTCACGTTG
TGTCCGTGTGGAGAGTCGGTTCTAGCGGTCGTATTATCCATATTCTCCCGTTCCATTTAC
GTTGAGCGACTTGAAACTTCCGATGTACTGGCCGGTAATAGTGCATTGTTTCTCAATTCA
TGCAAGTTGTCGTACATCATGGAATAGAACAAGAATTAAGGACCATTTCCACTAGCGATT
CTTACCTCCAACTCAAGATAGAGTCTAACCGACTATCGACGAACTTCGTCCGTCACGTCA
TTCTATGTCGAGCGGCCTTCTTACAGCGCTGTCGCACTATCTGTTCTACCTTATTTCCGA
GAAACCTTGCTCCTGTTACCTGTGAGGGCGGGGCGGATATGGCAAGATGGGCATTCTTTT
TTACTTCCCGTTCCTCTTCATGCTATATACATGTATCTCTGGAGCCTAGACGCCCGGTCC
TTAACGTCCTAAAACTTTACTGACCCTGCTGGCGTCTTACCCTTTATAGTGGCTCTGCAT
AATGTGTTCTGTATATAGCCAAATACTGCCGACTATCCACTCGCTTAGCTGTCAATATCC
ATGCGCGTACTTTGATTTACCGGTCGACCTGCCTAATGTAACTGCCTTTCGCTCTCCACT
CCGCACCGGGGTTTCGAGTTTTTCTACCCCTCTCTTTATTACTACTCGTTTACTGCATCA
GACGCGCCTACCGTATTCAGTCACTGCTCCTTCAGGCGAAGAGTATTAGTACGTTATCGT
GCTACTATCCTGAGCTGTAATCATAAGTATTTCTAAAGTTGTAAGGGTTGAAGAGTAGTT
CTGTACATCTTGATCGATTGTACTGGACGTCATTCCGAATAGCTTGTGGAATCCAATATC
TGGGTCATAGATGCT
>SRR9587935_9587940
GTGACCGTCGCCTCGCCACAAATCCATTGTTTTCCGGGTTCTAGCCCCTTATGAGCTAGG
ACCTCTGTGCCTACCATCATAGCTGTTCGATGCCTTGTATTCAATACGTTGCACCGACCC
GACAATAATCTGTGCCATTAAGCTTCTGTAAGTGGCTGGCGCCTGTAAGTCACGCAATAG
AGTAACTTCCCTCTCGACAGTCTTTATGCCATCCACAATCCAACATCGTAGATCCTCGCG
TGACCTAGTAGCCCTTGACGACTCAATCCACCCTCTGCCTGACGGACGTGCCTTTTATGG
TACATTAATGACTCCCTGTCCTCATAATTAGGCTGCACACACAGGGCTATCGTAGTTTAC
TGGTCTCACCCCTGCGTACTCGGCGACCCAACTACACTGATCCAGGGAGGCAGAATGGTG
GACGCTCTTACTGCCGACCCGTGGGCAACTAACTGGGCCGGGGGAATATCGTTTCGTTTT
GGGTACTGCTTCTTCCAATTGTTCGGATAGCACAGTAACGGTGGTGGATGGCTCCAGTCA
CTCTTGCGTAGTCGCCGGCCGGTCGCTCCGTTCTGTCCTCTCGCCCGATTCTGGGCCATC
TTATCCGATGTCACCCTGCAGACCGCTCTTTTCAACCATTAGCATTAAGCAGTCTCTCAT
AGGTGCTCTCACGTCATGAGAGGTACCGGAGCCTCTAAGCGTTCCGCTGTTCCCGACAAT
ATTCTCTTTCGCTTTAACACTAGCTCAGACGGTAGAGGCTTCGTCTTCAGCACCGGCTAA
TGATGCCTGGTACGTTAAGATACCGATCCGCGCGCTTACGCTGCTCCACCTGTCGGTGTC
TGAAGAAGCTCGGCTAAGACCTCTCGAACTCTTCGTCTTACCTCAGCCCGCGGCTACAGT
CATATACATCCTAGCACGCACAATCCTTAATTCCCAGTCGTGCCCGACAGTAACACCGTG
GCGCGAACAATGTGTAAGTTGTTGCCTACTTCGTACACTATCAACTAAATTCGTCCAACG

TGAGATCCGAGAATCCCGAAAGGGGGCACACACTTAGAGAACCCGCAGCTTTTTGTCTCA
CGATCGAGAAGGGCACTTCCAAGAGGCCTTGATACGAGCGTGAAACCGGTCAAGCGCTCC
AATTGTACACTGTCCATGAGGCCAAATTCAAATTACGGACCGAGAACTGGATCCAGCAGA
GCTAAAGGTTATTCGCGGGTATTGTCCGTACATGCAACTTGATGTTTTCTAAGCCAAAAG
AGATATGCGAGGGGGAGACATGGGAAAAGCAAAAGAATATAGTACTATAATTGGCGTGGG
ACCCATAAAGTCTTGATTGCTCCCGCGCCGCTTGTACCCACAGTCTTCAAAAAAGCGATT
TGCCGGAGTATAAAGTCAGGGGCCGACGGTGTTGAAAACGAGCGGACGTCTTGCATGAGA
AGTGTCACTGTGGGCATATCAAAGGACTCTAATTAAATTCGGTGTCAATAGCAATGTTCG
TGCTGCACGGTGCTGTCAGACCATCCATCTTTCGTGGCTACGCCTTATCAGAAGTGCAAC
CTGCGAAATCAAGGTGACGTTAATACGCTCTAAAGGACGCTAAAGCGGGTTGATAATTAA
ACGACAGAGTTACAATTAGCGGTATTCACAACTCTCCAAAATCATTGACGATAGACTGAC
ACGGTTACGTGGGAGGACTCGTAGTCTCAAGTGAAAATGGGCGCAAAATAGTGTCCGCAT
GGGAGGAGCTTAACCATAATAGACGCCGAATATGTTCGCGTGAAACTTCAGCGTCAAAGA
TGGAAAGCAAGACAAACTCATGAAACTACCATAGTGTGAAGATGATGGGCGGCGCGGTTT
AAGACGAGTCGCCCTAAGACTCCAACGTGCCCTAAGGGATCGTAACATGCAAGAACATTG
ATGTGCACAGTTTTGTCGAACGGCCTTCATAGGCCAAAGATTGTATCCACGGCCAGAAGC
GGTAAGAACTTCGTGACATAGCGGCCAGAGAAAGGATCTACAACGTTAACCAGCACCTCA
GAGAGCGGGACGGCCAGTCGTTAGGGAGAAAATAACTAATCATTAACTTTGACGAACCAC
CAAAATGCTCGAGCGGTAGGTATCATTAGGGGGTACGGACTGTTACATCTAGAATACCCC
AGCTCCGTAGTTGCAGACACTAATTTTAGAGACCGCAGCTGCATCATTAACAAAATGTTA
TTGGTGGGATGCAGAAGAGCGCGGGCAAGATTATACAGACGCCTTCAGGCGGTAGCCTGC
TCCTAGGCGGTCAAGACGGATCGCGATGGACGTGTTTGGACGGCGAATGTACAGGAAATT
TCCCGTGAGAACAACAAAATGAAGGATCGAGATTAAAAAGACTACAGCGCCCGTGAACAA
CGGAGTTGAACGTGGTTGCCAGGTTTGTGATGGGCGGCCCTAGAAGATAACGGTAGAAGC
CCTTCGGACTTTTCAATGGGGACTACATTCTGGCCGCGAAGGACTCCGTGAATCGATAGG
TAGTGGAAATATGGATGAAACGGGCTGGGGCAACGCTTGGGACAACGTTTACTAGCTAAG
AGGCAGATTACGAGGATCGGGGAAGACTGAGAAGGCCTTGGCGTAGGAATTCGAGGCAAC
CAGGGGGATAAGGTAAATGCCGACTTCAAGTCGAGAATGCAAAGAAGCTGGCTATGGCCG
TAAATTAGACGATAAATAACAGAGCATCGGGGGGCCAAGTCTAATTGTAAAACTTTTGGT
CGAGGTGATGTCGTAGGTGAGATCTCCGAACGCAGCATGAACAAGGCGAATCGTGTAAGC
AGAGAAGAGTGCAGCACGATGCTAACACTGCCGCAGCAGTGGGACAACTTGGATGACTGG
CACATCACAGTAATGAGTCATATGATTCCGCCTGAAGGCTTCAATAAAGGACATAAGATG
CGAAGAAGATCTAGACCCAGTCTTTAAAGCTGCACAACTTAGCCGAAATGATTGGTAGGT
GAAAGTATCGACGTGGGCACTGAGAGAAACCTGAATACGCGATAGCCATCATATCACGAT
GCTTCTGCCCTCCAGTGTCACTAGAGTTCTTGTTCACATGCCTCGGTCCTCTCGTCAGTA
CATGGCCGGCCTTCTTTCCTCCATCGTTTCCCAGAGCATACTCTTCGAGTTGCCCTATTA

GCCTTCTATTTTCTGGCCGATGCCTGCCTGGCGCCGCTTTGAGTATCATACTTCTTGTTT
TTGGCCAACCTGTTGGGCGTTGATTGCTTGGGTAACCATATCCTGCATTTTGTTGATTCC
TGGTCGGCAATGTGTTGTGGACTGTTCGTCGCTGCTCCTGACCTCACCGTACTTCACGGT
TGTCGCCCGGACATCACCCGGACGGGTTTAGGCCATGGCCGGTTGAGGTTAATCTTTACG
CGTCATCATAACTTCCTGCGAAACTGGTTTCCTCACGCCGCTCCCTTCGTGAGCCGATCG
GCGTGGTTAGGTCGCGCGGGTTTTTCCAAGCAGGGTAAGTCGAGTCGCCCAAGGTGACCC
ACGTACCTCGCTTTAACGTTCCGTTCTACCACTGGATTAGGCTGCTTTGGGGCCGACTTG
ACGTTCTTTTTGTACGACCTCGTATGTGCTTATTCTCACTCGTAATGAACCGGTAAGCGA
GAGTTTGCTCTACGCTGAGGTGCCGTCGGACCCGAGTGTGCTTGCCGCTGGTCAATTGTT
GTGTCGGCTCGACGTCTTATTTCGTATGGTACGGGATTATTAGTGCTTGCAGCGATGTCC
CTTTCGCTATTGATGTCTTATCCTGAGTCACGCTACGTACTGAATTGATAAAACATTTCG
ACCTCAGTATCAGTGAGGTCGGGGGATGAGTTAGATCGCCGTGTAGGGCTTTTCACGTTG
TGTCCGTGTGGAGAGTCGGTTCTAGCGGTCGTATTATCCATATTCTCCCGTTCTATTTAC
GTTGAGCGACTTGAAACCTCCGATGTACTGGCCGGTAATAGTGCATTGTTTCTCAATTCA
TGCGAGTTGTCGTACATTATGGAATAGAACAAGAATTAAGGACCATTTCCACTAGCGATT
CTTACCTCCAACTCAAGATAGAGTCTAACCGATTATCGACGAACTTCGTCCGTCACGTCA
TTCTATGTCGAGCGGCCTTCTTACAGCGCTGTCGCACTATCTGTTCTACCTTATTTCCGA
GAAACCTTGCTCCTGTTACCTGTGAGGGCGGGGCGGATATGGCAAAATGGGCATTCTTTT
TTACTTCCCGTGCCTCTTCATGCTATATACATGTATCTCTGGAGCCTAGACGCCCGGTCC
TTAACGTCCTAAAACTTTACTGACCCTGCTGGCGTCTTACCCTTTATAGTGGCTCTGCAT
AATGTGTTCTGTATATAGCCAAATACTGCCGACTATCCACTCGCTTAGCTGTCAATATCC
ATGCGCGTACTTTGATTTACCGGTCGACCTGCCTAATGTAACTGCCTTTCGCTCTCCACT
CCGCACCGGGGTTTCGAGTTTTTCTACCCCTCTCTTTATTACTACTCGTTTACTGCATCA
GACGCGCCTACCGTATTCGGTCACTGCTCCTTCAGGCGAAGAGTATTAGCACGTTATCGT
GCTACTATCCTGAGCTGTAATCCTAATTATTTCTAAAGTTGTAAGGGTTGAAGAGTAGTT
CTGTACATCTTGATCGATTGTACTGGACGTCATTCCGAATAGCTTGTGGAATCCAATATC
TGGGTCATAGATGCT
>SRR9587936
GTGACCGTCGCCTCGCCACAAATCCATTGTTTTCCGGGTTCTAGCCCCTTATGAGCTAGG
ACCTCTGTGCCTACCATCATAGCTGTTCGATGCCTTGTATTCAATACGTTGCACCGACCC
GACAATAGTCTGTGCCATTAAGCTGCTGCAAGTGGCTGGCGCCCGTAAGTCACGCAATAG
AGTAACTTCCCTCTCGACAGTCTTTATGCCATCCACAATCCAACATCGTAGATCCTCGCG
TGACCTAGTAGCCCTTGACGACTCAATCCACCCTCTGCCTGACGGACGTGCCTTTTATGG
TACATTAATGACTCCCTGTCCTCATAATTAGGCTGCACACACAGGGCTATCGTAGTTTAC
TGGTCTCACCCCTGCGTACTCGGCGACCCAACTACACTGATCCAGGGAGGCAGAATGGTG
GACGCGCTTGCTCCCGACCCGTGGGCAACTAACTGGGCCGGGGGAATATTGTTTCGTTTT

GGGTACTGCTTCTTCCAATTGTTCGGATAGCACAGTAACGGTGGTGAGTGGCTCCAGTCA
CTCTTACGTAGTCGCCGGCCGGTCGCTCCGGTCTGTCCTCTCGCCCGATTCTGGGCCATC
TTATCCGATGTCACCCTGCAGACCGCTCTTTTCAACCATTAGCATTAAGCAGTCTCTCAT
AGGTGCTCTCATGTCATGAGAGGTACCGGAGCCTCTACGCGTTCCGCTGTTCCCGACAAT
ATTCTCTTTCGCTTTGACACTAGCTCAGACGGTAGAGGCTTCGTCTTCAGCACCGGCTAA
TGATGCCTGGTACGTTAAGNTACCGATCCGCGCGCTTACGCTGCTCCACCTGTCGGTGTC
TGAAGAAGCTCGGCTAAGACCTCTCGAACTCTTCGTCTTACCTCAGCCCGCGGCTACAGT
CATATACATCCTAGCACGCACAATCCTTAATTCCCAGTCGTGCCCGACAGTAACACCGTG
GCGCGAACAATGTGTAAGTTATTGCCTACTTCGTACACTATCAACTAAATTCGTCCAACG
TGAGATCCGAGAATCCCGAAAGGGGGCACACACTTAGAGAACCCGCAGCTTTTTATCTCA
CGATCGAGGAGGGCACTCCCAAGAGGCCTTGATACGAGCGTGAAACCGGTCAAGCGCTCC
AATTGTACACTGTCCATGAGGCCAAATTCAAATTACGGACCGAAAACTGGATCCAGCAGA
GCTAAAGGTTATTCGCGGGTATTGTCCGTACATGCAACTTGATGTTTTCTAAGCCAAAAG
AGATATGCGAGGGGGAGACATGGGAAAAGCAAAAGAATATAATACTATAATTGGCGTGGG
ACCCATAAAGTCTTGATTGCTCCCGCGCCGCCTGTGCCCACAGTCTTCAAAAAAGCGATT
TGCCGGAGTATAAAGTCAGGGGCCGACGGTGTTGAAAACGAGCGGACGTCTTACATGAGA
AGTGTCACTGTGGGCATATCAAAGGACTCTAATTAAATTCGGTGTCAATAGAAATGTTCG
TGCTGCACGGTGCTGTCAGACCATCCACCTTTCGTGGCTACGCCTTATCAGAAGTGCAAC
CTGCGAAATCAAGGTGACGTTAATACGCTCTAAAGGACGCTAAAGCGGGTTGATAATTAA
ACGACAGAGTTACAATTAGCGGGATTCACAACTCTCCAAAATCATTGACAATAGACTGAC
ACGGTTACGTGGGAGGACTCGTAGTCTCAAGTGAAAATGGGCGCAAAATGGTGTCCGCAT
GGGAGGAGCTCAACCATAATAGACGCCGAATATGTTCGCGTGAAACTTCAGAGTCAAAGA
TGGAAAGCAAGACAAACTCATGAAACTACCATAGTGTGAAGATGATGGGCGGCGCGGTTT
AAGGCGAGTCGCCCTAAGACTCCAACGTGCCCTAAGGGATCGTAACATGCAAGAACATTG
ATGTGCACAGTTTTGTCGAACGGCCTTCATAGGCCAAAGATTGTATCCACGGCCAGAAGC
GGTAAGAACTTCGTGACATAGCGGCCAGAGAAAGGATCTACAACGTTAACCAGCACCTCA
GAGAGCGGGATGGCCAGTCGTTAGGGAGAAAATAACTAATCATTAACTTTGACGAACCAC
CAAAATGCTCGAGCGGTAGGTATCATTAGGGGGTACGGACTGTTACATCTAGAATACCCC
AGCTCCGTAGTTGCAGACACTAATTTTAGAGCCCGCAGCTGTATCATTAACAAAATGTTA
TTGGTGGGATGCAGAAGAGCGCGGGCAAGATTATACAGACGCCTTCAGGCGGTAGCCTGC
TCCTAGGCAGTCAAGACGGATCGCGATGGACGTGTTTGGACGGCGAATGTACAGGAAATT
TCCCGTGAGAACAACAAAATGAAGAATCGAGATTAAAAAGACTGCAGCGCCCGTGAACAA
CGGAGTTGAACGTGGTTGCCAGGTTTGTGATGGGCGGCCCTAGAAGATAACGGTAGAAGC
CCTTCGGACTTTTCAATGGGGACTACATTCTGGCCGCGAAGGACTCCGTGAATCGATAGG
TAGTGGAAATATGGATGAAACGGGCTGGGGCAACGCTCGGGACAAAGTTTACTAGCTAAG
AGGCAGATTACGAGGATCGGGGAAGACTGAGAAGGCCTTGGCGTAGGAATTCGAGGCAAC

CAGGGGGATAAGGTAAATGCCGACTTCAAGTCGAGAATGCAAAGAAGCTGGCTATGGCCG
TAAATTAGACGATAAATAACAGAGCATCGGGGAGCCAAGTCTAATTGTAAAACTTTTGGT
CGAGGTGATGTCGTAGGTGAGATCTCCGAACGCAGCATGAACAAGGCGAATCGTGTAAGC
AGAGAAGAGTGCAGCACGATGCTAACACTGCCGCAGCAGTGGGACAACTTGGATGACTGG
CACATCACAGTAATGAGTCATATGATTCCGCCTGAAGGCTTCAATAAAGGACATAAGATG
CGAAGAAGATCTAGACCCAGTCTTTAAAACTGCACAACTTAGCCGAAATGATTGGTAGGT
GAGAGTATCGACGTGGGCACTGAGAGAAACCTGAATACGCGATAGCCATCATATCACGAT
GCTTCTGCCCTCCAGCGTCACTAGAGTTCTTGTTCACATGCCTCGGTCCTCTCGTCAGTA
CATGGCCGGCCTTCTTTCCTCCATCGTTTCCCAGAGCATACTCCTCGAGTTGCCCTATTA
ACCTTCTATTTTCTGGCCGATGCCTGACTGGCGCCGCTTTGAGTATCATACTTCTTGTTT
TTGGCCAACCTGTTGGGCGTTGATTGCTTGGGTAACCATATCCTGCATTTTGTTGATTCC
GGGTCGGCAATGTGTTGTGGACTGTTCGTCGCCGCTCCTGATCTCACCGTACTTCACGGT
TGTCGCCCGGACATCACCCGGACGGGTTTAGGCCATGGCCGGTTGAGGTTAATCTTTACG
CGTCACCATAACTTCCTGCGAAACTGGTTTCCCCACACCGCTCCCTTCGTGAGCCGATCG
GCGTGGTTAGGTCGCGCGGGTTTTTCCAGGCAGGGTAAGTCGAGTCGCACAAGGTGACCC
ACGTACCTCGCTTTAACGTTCCGTTCTACCACTGGGTTAGGCTGCTTTGGGGCCGACTTG
ACGTTCTTTTTGCACGACCTCGTATGTGCTTATTCTCACTCGTAATGAACCGGTAAGCGA
GAGTTTGTTCTACGCTGAGGTGTCGTCGGACCCGAGTGTGCTTGCCGCTGGTCAATTGTT
GTGTCGGCTCGACGTCTTATTTCGTATGGTACGGGATTATTAGTGCTTGCAGCGATGTCC
CTTTCGCTATTGATGTCCTCTCCTGAGTCACGCTACGTACTGAATTGATAAAACATTTCG
ACCTCAGTATCAGTGAGGTCGGGGGATGAGTTAGATCGCCGTGTAGGGCTTTTCACGTTG
TGTCCGTGTGGAGAGTCGGTTCTAGCGGTCGTATTATCCATATTCTCCCGTTCCATTTAC
GTTGAGCGACTTGAAACTTCCGATGTACTGGCCGGTAATAGTGCATTGTTTCTCAATTCA
TGCAAGTTGTCGTACATCATGGAATAGAACAAGAATTAAGGACCATTTCCACTAGCGATT
CTTACCTCCAACTCAAGATAGAGTCTAACCGACTATCGACGAACTTCGTCCGTCACGTCA
TTCTATGTCGAGCGGCCTTCTTACAGCGCTGTCGCACTATCTGTTCTACCTTATTTCCGA
GAAACCTTGCTCCTGTTACCTGTGAGGGCGGGGCGGATATGGCAAGATGGGCATTCTTTT
TTACTTCCCGTTCCTCTTCATGCTATATACATGTATCTCTGGAGCCTAGACGCCCGGTCC
TTAACGTCCTAAAACTTTACTGACCCTGCTGGCGTCTTACCCTTTATAGTGGCTCTGCAT
AATGTGTTCTGTATATAGCCAAATACTGCCGACTATCCACTCGCTTAGCTGTCAATATCC
ATGCGCGTACTTTGATTTACCGGTCGACCTGCCTAATGTAACTGCCTTTCGCTCTCCACT
CCGCACCGGGGTTTCGAGTTTTTCTACCCCTCTCTTTATTACTACTCGTTTACTGCATCA
GACGCGCCTACCGTATTCAGTCACTGCTCCTTCAGGCGAAGAGTATTAGTACGTTATCGT
GCTACTATCCTGAGCTGTAATCATAAGTATTTCTAAAGTTGTAAGGGTTGAAGAGTAGTT
TTGTACATCTTGATCGATTGTACTGGACGTCATTCCGAATAGCTTGTGGAATCCAATATC
TGGGTCATAGATGCT

>SRR9587937

GTGACCGTCGCCTCGCCACAAATCCATTGTTTTCCGGGTTCTAGCCCCTTATGAGCTAGG
ACCTCTGTGCCTACCATCATAGCTGTTCGATGCCTTGTATTCAATACGTTGCACCGACCC
GACAATAATCTGTGCCATTGAGCTTCTGCAAGTGGCTGGCGCCTGTAAGTCACGCAATAG
AGTAACTTCCCTCTCGACAGTCTTTATGCCATCCACAATCCAACATCGTAGATCCTCGCG
TGACCTAGTAGCCCTTGACGACTCAATCCACCCTCTGCCTGACGGACGTGCCTTTTATGG
TACATTAATGACTCCCTGTCCTCATAATTAGTCTGCACACACAGGGCTATCGTAGTTTAC
TGGTCTCACCCCTGCGTACTCGGCGACCCAACTAAACTGATCCAGGGAGGCAGAATGGTG
GACGCGCTTAGTGCCGACCCGTGGGCAACTAACTGGGCCGGGGGAATATTGTTTCGTTTT
GGGTACTGCTTCTTCCAATTGTTCGGATAGCACAGTAACGGTGGTGGGTGGCTCCAGTCA
CTCTTGCGTAGTCCCCGGTCGGTCGCTCCGGTCTGTCCTCTCGCCCGATTCTGGGCCATC
TTATCCGATGTCACCCTGCAGACCGCTCTTTTCAACCATTAGCATTAAGCAGTCTCTCAT
AGGTGCTCTCATGTCATGAGAGGTACCGGAGCCTCTAAGCGTTCCGCTGTTCCCGACAAT
ATTCTCTTTCGCTTTGACACTAGCTCAGACGGTAGAGGCTTCGTCTTCAGCACCGGCTAA
TGATGCCTGGTACGTTAAGATACCGATCCGCGCGCTTACGCTGCTCCACCTGTCGGTGTC
TGAAGAAGCTCGGCTAAGACCTCTCGAACTCTTCGTCTTACCTCAGCCCGCGGCTACAGT
CTAATACATCCTAGCACGCACAATCCTTAATTCCCAGTCGTGCCCGACAGTAACACCGTG
GCGCGAACAATGTGTAAGTTATTGCCTACTTCGTACACTATCAAATAAATTCGTCCAACG
TGAGATCCGAGAATCCCGAAAGGGGGCACACACTTAGAGAACCCGCAGCTTTTTGTCTCA
CGATCGAGAAGGGCACTCCCAAGAGGCCTTGATACGAGCGTGAAACCGGTCAAGCGCTCC
AATTGTACACTGTCCATGAGGCCAAATTCAAATTACGGACCGAAAACTGGATCCAGCAGA
GCTTAAGGTTATTCGCGGGTATTGTCCGTACATGCAACTTGATGTTTTCTAAGCCAAAAG
AGATATGCGAGGGGGAGACATGGGAAAAGCAAAAGAATATAGTACTATAATTGGCGTGGG
ACCCATAAAGTCTTGATTGCTCCCGCGCCGCCTGTACCCACAGTCTTCAAAAAAGCGATT
TGCCGGAGTATAAAGTCAGGGGCCGACGGTGTTGAAAACGAGCGGACGTCTTGCATGAGA
AGTGTCACTGTGGGCATATCAAAGGACTCTAATTAAATTCGGTGTCAATAGAAATGTTCG
TGCTGCACGGTGGTGTCAGACCGTCCACCTTTCGTGGCTACGCCTTATCAGAAGGGCAAC
CTGCGAAATCAAGGTGACGTTAATACGCTCTAAAGGACGCTAAAGCGGGTTGATAATTAA
ACGACAGAGTTACAATTAGCGGGATTCACAACTCTCCAAAATCATTGACGATAGACTGAC
ACGGTTACGTGGGAGGACTCGTAGTCTCAAGTGAAAATGGGCGCAAAATGGTGTCCGCAT
GGGAGGAGCTCAACCATAATAGACGCCGAATATGTTCGCGTGAAACTTCAGCGTCAAAGA
TGGAAAGCAAGACAAACTCATGAAACTACCATAGTGTGAAGATGATGGGCGGCGCGGTTT
AAGGCGAGTCGCCCTAAGACTCCAACATGCCCTAAGGGATCGTAACATGCAAGAACATTG
ATGTGCACAGTTTTGTCGAACGGCCTTCATAGGCCAAAGATTGTATCCACGGCCAGAAGC
GGTAAGAACTTCGTGACATAGCGGCCAGAGAAAGGATCTACGACGTTAACCAGCACCTCA
GAGAGCGGGATGGCCAGTCGTTAGGGAGAAAATAACTAATCATTAACTTTGACGAACCAC

CAAAATGCTCGAGCGGTAGGTATCATTAGGGGGTACGGACTGTTACATCTAGAATACCCC
AGCTCCGTAGTTGCAGACGCTAATTTTAGAGACCGCAGCTGTATCATTAACAAAATGTTA
TTGGTGGGATGCAGAAGAGCGCGGGCAAGATTATACAGACGCCTTCAGGCGGTAGCCTGC
TCCTAGGCAGTCAAGACGGATCGCGATGGACGTGTTTGGACGGCGAATGTACAGGAGATT
TCCCGTGAGAACAACAAAATGAAGGATCGAGATTAAAAAGACTGCAGCGCCCGTGAACAA
CGGAGTTGAACGTGGTTGCCAGGTTTGTGATGGGCGGCCCTAGAAGATAACGGTAGAAGC
CCTTCGGACTTTTCAATGGGGACTACATTCTGGCCGCGAAGGACTCCGTGAATCGATAGG
TAGTGGAAATATGGATGAAACGGGCTGGGGCAACGCTCGGGACAACGTTTACTAGCTAAG
AGGCAGATTACGAGGATAGGGGAAGACTGAGAAGGCCTTGGCGTAGGAATTCGAGGCAAC
CAGGGGGATAAGGTAAATGCCGACTTCAAGTCGAGAATGCAAAGAAGCTGGCTATGGCCG
TAAATTAGACGATAAATAACAGAGCATCGGGGAGCCAAGTCTAATTGTAAAACTTTTGGT
CGAGGTGATGTCGTAGGTGAGATCTCCGAACGCAGCATGAACAAGGCGAATCGTGTAAGC
AGAGAAGAGTGCAGCACGATGCTAACACTGCCGCAGCAGTGGGACAACTTGGATGACTGG
CACATCACAGTAATGAGTCATATGATTCCGCCTGAAGGCTTCAATAAAGGACATAAGATG
CGAAGAAGATCTAGACCCAGTCTTTAAAACTGCACAACTTAGCCGAAATGATTGGTAGGT
GAGAGTATCGACGTGGGCACTGAGAGAAACCTGAATACGCGATAGCCGTCATATCACGAT
GCTTCTGCCCTCCAGCGTCACTAGAGTTCTTGTTCACATGCCTCGGTCCTCTCGTCAGTA
CATGGCCGGCCTTCTTTCCTCCATCGTTTCCCAGAGCATACTCTTCGAGTTGCCCTATTA
ACCTTCTATTTTCTGGCCGATGCCTGACTGGCGCCGCTTTGAGTATCATACTTCTTGTTT
TTGGCCAACCTGTTGGGCGTTGATTGCTTGGGTAACCATATCCTGCATTTTGTTGATTCC
GGGTCGGCAATGTGTTGTGGACTGTTCGTCGCCGCTCCTGATCTCACCGTACTTCACGGT
TGTCGCCCGGACATCACCCGGACGGGTTTAGGCCATGGCCGGTTGAGGTTAATCTTTACG
CGTCACCATAACTTCCTGCGAAACTGGTTTCCCCACGCCGCTCCCTTCGTGGGCCGATCG
GCGTGGTTAGGTCGCGCGGGTTTTTCCAGGCAGGGTAAGTCGAGTCGCACAAGGTGACCC
ACGTACCTCGCTTTAACGTTCCGTTCTACCACTGGGTTAGGCTGCTTTGGGGCCGACTTG
ACGTTCTTTTTGCACGACCTCGTATGTGCTTATTCTCACTCGTAATGAACCGGTAAGCGA
GAGTTTGTTCTACGCTGAGGTGTCGTCGGACCCGAGTGTGCTTGCCGCTGGTCAATTGTT
GTGTCGGCTCGACGTCTTATTTCGTATGGTACGGGATTATTAGTGCTTGCAGCGATGTCC
CTTTCGCTATTGATGTCTTCTCCTGAGTCACGCTACGTACTGAATTGATAAAACATTTCG
ACCTCAGTATCAGTGAGGTCGGGGGATGAGTTAGATCGCCGTGTAGGGCTTTTCACGTTG
TGTCCGTGTGGAGAGTCGGTTCTAGCGGTCGTATTATCCGTATTCTCCCGTTCCATTTAC
GTTGAGCGACTTGAAACTTCTGATGTACTGGCCGGTAATAGAGCATTGTTCCTCAATTCA
TGCAAGTTGTCGTACATTATGGAATAGAACAAGAATTAAGGACCATTTCCACTAGCGATT
CTTACCTCCAACTCAAGATAGAGTCTAACCGATTATCGACGAACTTCGTCCGTCACGTCA
TTCTATGTCGAGCGGCCTTCTTACAGCGCTGTCGCACTATCTGTTCTACCTTATTTCCGA
GAAACCTTGCTCCTGTTACCTGTGAGGGCGGGGCGGATATGGCAAAATGGGCATTCTTTT

TTACTTCCCGTGCCTCTTCATGCTATATACATGTATCTCTGGAGCCTAGACGCCCGGTCC
TTAACGTCCTAAAACTTTATTGACCCTGCTGGCGTCTTACCCTTTATAGTGGCTCTGCAT
AATGTGTTCTGTATATAGCCAAATACTGCCGACTATCCACTCGCTTAGCTGTCAATATCC
ATGCGCGTACTTTGATTTACCGGTCGACCTGCCTAATGTAACTGCCTTTCGCTCTCCACT
CCGCACCGGGGTTTCGAGTTTTTCTACCCCTCTCTTTATTACTACTCGTTTACTGCATCA
GACGCGCCTACCGTATTCAGTCACTGCTCCTTCAGGCGAAGAGTATTAGCACGTTATCGT
GCTACTATCCTGAGCTGTAATCATAAGTATTTCTAAAGTTGTAGGGGTTGAAGAGTAGTT
CTGTACATCTTGATCGATTGTACTGGACGTCATTCCGAATAGCTTGTGGAATCCAATATC
TGGGTCATAGATGCT
>SRR9587938
GTGACCGTCGCCTCGCCACAAATCCATTGTTTTCCGGGTTCTAGCCCCTTATGAGCTAGG
ACCTCTGTGCCTACCATCATAGCTGTTCGATGCCTTGTATTCAATACGTTGCACCGACCC
GACAATAATCTGTGCCATTAAGCTTCTGCAAGTGGCTGGCGCCTGTAAGTCACGCAATAG
AGTAACTTCCCTCTCGACAGTCTTTATGCCATCCACAATCCAACATCGTAGATCCTCGCG
TGACCTAGTAGCCCTTGACGACTCAATCCACCCTCTGCCTGACGGACGTGACTTTTATGG
TACATTAATGACTCCCTGTCCTCATAATTAGGCTGCACACACAGGGCTATCGTAGTTTAC
TGGTCTCACCCCTGCGTACTCGGCGACCCAACTACACTGATCCAGGGAGGCAGAATGGTG
GACGCTCTTACTGCCGACCCGTGGGCAACTAACTGGGCCGGGGGAATATTGTTTCGTTTT
GGGTACTGCTTCTTCCAATTGTTCGGATAGCCCAGTAACGGTGGTGGGTGGTTCCAGTCA
CTCTTGCGTAGTCGCCGGCCGGTCGCTCCGGTCTGTCCCCTCGCCCGATTCTGGGCCATC
TTATCCGATGTCACCCTGCAGACCGCTCTTTTCAACCATTAGCATTAAGCAGTCTCTCAT
AGGTGCTCTCATGTCATGAGAGGTACCGGAGCCTCTAAGCGTTCCGCTGTTCCCGACAAT
ATTCTCTTTCGCTTTGACACTAGCTCAGACGGTAGAGGCTTCGTCTTCAGCACCGGCTAA
TGATGCCTGGTACGTTAAGATACCGATCCGCGCGCTTACGCTGCTCCACCTGTCGGTGTC
TGAAGAAGCTCGGCTAAGACCTCTCGAACTCTTCGTCTTACCTCAGCCCGCGGCTACAGT
CATATACATCCTAGCACGCACAATCCTTAATTCCCAGTCGTGCCCGACAGTAACACCGTG
GCGCGAACAATGTGTAAGTTATTGCCTACTTCGTACACTATCAACTAAATTCGTCCAACG
TGAGATCCGAGAATCCCGAAAGGGGGCACACACTTAGAGAACCCGCAGCTTTTTGTCTCA
CGATCGAGAAGGGCACTCCCAAGAGGCCTTGATACAAGCGTGAAACCGGTCAAGCGCTCC
AATTGTACACTGTCCATGAGGCCAAATTCAAATTACGGACCGAAAACTGGATCCAGCAGA
GCTAAAGGTTATTCGCGGGTATTGTCCGTACATGCAACTTGATGTTTTCTAAGCCAAAAG
AGATATGCGAGGGGGAGACATGGGAAAAGCAAAAGAATATAGTACTATAATTGGCGTGGG
ACCCATAAAGTCTTGATTGCTCCCGCGCCGCCTGTACCCACAGTCTTCAAAAAAGCGATT
TGCCGGAGTATAAAGTCAGGGGCCGACGGTGTTGAAAACGAGCGGACGTCTTGCATGAGA
AGTGTCACTGTGGGCATATCAAAGGACTCTAATTAAATTCGGTGTCAATAGAAATGTTCG
TGCTGCACGGTGCTGTCAGACCATCCACCTTTCGTGGCTACGCCTTATCAGAAGTGCAAC

CTGCGAAATCAAGGTGACGTTAATACGCTCTAAAGGACGCTAAAGCGGGTTGATAATTAA
ACGACAGAATTACAATTAGCGGGATTCACAACTCTCCAAAATCATTGACGATAGACTGAC
ACGGTTACGTGGGAGGACTCGTAGTCTCCAGTGAAAATGGGCGCAAAATGGTGTCCGCAT
GGGAGGAGCTCAACCATAATAGACGCCGAATATGTTCGCGTGAAACTTCAGCGTCAAAGA
TGGAAAGCAAGACAAACTCATGAAACTACCATAGTGTGAAGATGATGGGCGGCGCGGTTT
AAGGCGAGTCGCCCTAAGACTCCAACGTGCCCTAAGGGATCGTAACATGCAAGAACATTG
ATGTGCACAGTTTTGTCGAACGGCCTTCATAGGCCAAAGATTGTATCCACGGCCAGAAGC
GGTAAGAACTTCGTGACATAGCGGCCAGAGAAAGGATCTACAACGTTAACCAGCACCTCA
GAGAGCGGGATGGCCAGTCGTTAGGGAGAAAATAACTAATCATTAACTTTGACGAACCAC
CAAAATGCTCGAGCGGTAGGTATCATTAGGGGGTACGGACTGTTACATCTAGAATACCCC
AGCTCCGTAGTTGCAGACACTAATTTTAGAGACCGCAGCTGCATCATTAACAAAATGTTA
TTGGTGGGATGCAGAAGAGCGCGGGCAAGATTATACAGACGCCTTCAGGCGGTAGCCTGC
TCCTAGGCAGTCAAGACGGATCGCGATGGACGTGTTTGGACGGCGAATGTACAGGAAATT
TCCCGTGAGAACAACAAAATGAAGGATCGAGATTAAAAAGACTGCAGCGCCCGTGAACAA
CGGAGTTGAACGTGGTTGCCAGGTTTGTGATGGGCGGCCCTAGAAGATAACGGTAGAAGC
CCTTCGGACTTTTCAATGGGGACTACATTCTGGCCGCGAAGGACTCCGTGAATCGATAGG
TAGTGGAAATATGGATGAAACGGGCTGGGGCAACGCTCGGGACAACGTTTACTAGCTAAG
AGGCAGATTACGAGGATCGGGGAAGACTGAGAAGGCCTTGGCGTAGGAATTCGAGGCAAC
CAGGGGGATAAGGTAAATGCCGACTTCAAGTCGAGAATGCAAAGAAGCTGGCTATGGCCG
TAAATTAGACGATAAATAACAGAGCATCGGGGAGCCAAGTCTAATTGTAAAACTTTTGGT
CGAGGTGATGTCGTAGGTGAGATCTCCGAACGCAGCATGAACAAGGCGAATCGTGTAAGC
AGAGAAGAGTGCAGCACGATGCTAACACTGCCGCAGCAGTGGGACAACTTGGATGACTGG
CACATCACAGTAATGAGTCATATGATTCCGCCTGAAGGCTTCAATAAAGGACATAAGATG
CGAAGAAGATCTAGACCCAGTCTTTAAAACTGCACAACTTAGCCGAAATGATTGGTAGGT
GAGAGTATCGACGTGGGCACTGAGAGAAACCTGAATACGCGATAGCCATCATATCACGAT
GCTTCTGCACTCCAGCGTCACTAGAGTTCTTGTTCACATGCCTCGGTCCTCTCGTCAGTA
CATGGCCGCCCTTCTTTCCTCCATCGTTTCCCAGAGCATACTCTTCGAGTTGCCCTATTA
ACCTTCTATTTTCTGGCCGATGCCTGACTGGCGCCGCTTTGAGTATCATACTTCTTGTTT
TTGGCCAACCTGTTGGGCGTTGATTGCTTGGGTAACCATGTCCTGCATTTTGTTGATTCC
GGGTCGGCAATGTGTTGTGGACTGTTCGTCGCCGCTCCTGATCTCACCGTACTTCACGGT
TGTCGCCCGGACATCACCCGGACGGGTTTAGGCCATGGCCGGTTGAGGTTAATCTTTACG
CGTCACCATAACTTCCTGCGAAACTGGTTTCCCCACGCCGCTCCCTTCGTGAGCCGATCG
GCGTGGTTAGGTCGCGCGGGTTTTTCCAGGCAGGGTAAGTCGATTCGCACAAGGTGACCC
ACGTACCTCGCTTTAACGTTCCGTTCTACCACTGGGTTAGGCTGCTTTGGGGCCGACTTG
ACGTTCTTTTTGCACGACCTCGTATGTGCTTATTCTCACTCGTAATGAACCGGTAAGCGA
GAGTTTGTTCTACGCTGAGGTGTCGTCGGACCCGAGTGTGCTTGCCGCTGGTCAATTGTT

GTATCGGCTCGACGTCTTATTTCGTATGGTACGGGATTATTAGTGCTTGCAGCGATGTCC
CTTTCGCTATTGATGTCTTCTCCTGGGTCACGCTACGTACTGAATTGATAAAACATTTCG
ACCTCAGTATCAGTGAGGTCGGGGGATGAGTTAGATCGCCGTGTAGGGCTTCTCACGTTG
TGTCCGTGTGGAGAGTCGGTTCTAGCGGTCGTATTATCCATATTCTCCCGTTCCATTTAC
GTTGAGCGACTTGAAACTTCCGATGTACTGGCCGGTAATAGTGCATTGTTTCTCAATTCA
TGCAAGTTGTCGTACATTATGGAATAGAACAAGAATTAAGGACCATTTCCACTAGCGATT
CTTACCTCCAACTCAAGATAGAGTCTAACCGATTATCGACGAACTTCGTCCGTCACGTCA
TTCTATGTCGAGCGGCCTTCTTACAGCGCTGTCGCACTATCTGTTCTACCTTATTACCGA
GAAACCTTGCTCCTGTTACCTGTGAGGGCGGGGCGGATATGGCAAAACGGGCATTCTTTT
TTACTTCCCGTGCCTCTTCATGCTATATACATGTATCTCTGGAGCCTAGACGCCCGGTCC
TTAACGTCCTAAAACTTTACTGACCCTGCTGGCGTCTTACCCTTTATAGTGGCTCTGCAT
AATGTGTTCTGTATATAGCCAAATACTGCCGACTATCCACTCGCTTAGCTGTCAATATCC
ATGCGCGTACTTTGATTTACCGGTCGACCTGCCTAATGTAACTGCCTTTCGCTCTCCACT
CCGCACCGGGGTTTCGAGTTTTTCTACCCCTCTCTTTATTACTACTCGTTTACTGCATCA
GACGCGCCTACCGTATTCAGTCACTGCTCCTTCAGGCGAAGAGTATTAGCACGTTATCGT
GCTACTATCCTGAGCTGTAATCATAAGTATTTCTAAAGTTGTAAGGGTTGAAGAGTAGTT
CTGCACATCTTGATCGATTGTACTGGACGTCATTCCGAATAGCTTGTGGAATCCAATATC
TGGGTCATAGATGCT
>SRR9587939
GTGACCGTCGCCTCGCCACAAATCCATTGTTTTCCGGGTTCTAGCCCCTTATGAGCTAGG
ACCTCTGTGCCTACCATCATAGCTGTTCGATGCCTTGTATTCAATACGTTGCACCGACCC
GACAATAATCTGTGCCATTAAGCTTCTGCAAGTGGCTGGCGCCTGTAAGTCACGCAATAG
AGTAACTTCCCTCTCGACAGTCTTTATGCCATCCACAATCCAACATCGTAGATCCTCGCG
TGACCTAGTAGCCCTTGACGACTCAATCCACCCTCTGCCTGACGGACGTGACTTTTATGG
TACATTAATGACTCCCTGTCCTCATAATTAGGCTGCACACACAGGGCTATCGTAGTTTAC
TGGTCTCACCCCTGCGTACTCGGCGACCCAACTACACTGATCCAGGGAGGCAGAATGGTG
GACGCTCTTACTGCCGACCCGTGGGCAACTAACTGGGCCGGGGGAATATTGTTTCGTTTT
GGGTACTGCTTCTTCCAATTGTTCGGATAGCCCAGTAACGGTGGTGGGTGGTTCCAGTCA
CTCTTGCGTAGTCGCCGGCCGGTCGCTCCGGTCTGTCCCCTCGCCCGATTCTGGGCCATC
TTATCCGATGTCACCCTGCAGACCGCTCTTTTCAACCATTAGCATTAAGCAGTCTCTCAT
AGGTGCTCTCATGTCATGAGAGGTACCGGAGCCTCTAAGCGTTCCGCTGTTCCCGACAAT
ATTCTCTTTCGCTTTGACACTAGCTCAGACGGTAGAGGCTTCGTCTTCAGCACCGGCTAA
TGATGCCTGGTACGTTAAGATACCGATCCGCGCGCTTACGCTGCTCCACCTGTCGGTGTC
TGAAGAAGCTCGGCTAAGACCTCTCGAACTCTTCGTCTTACCTCAGCCCGCGGCTACAGT
CATATACATCCTAGCACGCACAATCCTTAATTCCCAGTCGTGCCCGACAGTAACACCGTG
GCGCGAACAATGTGTAAGTTATTGCCTACTTCGTACACTATCAACTAAATTCGTCCAACG

TGAGATCCGAGAATCCCGAAAGGGGGCACACACTTAGAGAACCCGCAGCTTTTTGTCTCA

CGATCGAGAAGGGCACTCCCAAGAGGCCTTGATACAAGCGTGAAACCGGTCAAGCGCTCC

AATTGTACACTGTCCATGAGGCCAAATTCAAATTACGGACCGAAAACTGGATCCAGCAGA

GCTAAAGGTTATTCGCGGGTATTGTCCGTACATGCAACTTGATGTTTTCTAAGCCAAAAG

AGATATGCGAGGGGGAGACATGGGAAAAGCAAAAGAATATAGTACTATAATTGGCGTGGG

ACCCATAAAGTCTTGATTGCTCCCGCGCCGCCTGTACCCACAGTCTTCAAAAAAGCGATT

TGCCGGAGTATAAAGTCAGGGGCCGACGGTGTTGAAAACGAGCGGACGTCTTGCATGAGA

AGTGTCACTGTGGGCATATCAAAGGACTCTAATTAAATTCGGTGTCAATAGAAATGTTCG

TGCTGCACGGTGCTGTCAGACCATCCACCTTTCGTGGCTACGCCTTATCAGAAGTGCAAC

CTGCGAAATCAAGGTGACGTTAATACGCTCTAAAGGACGCTAAAGCGGGTTGATAATTAA

ACGACAGAATTACAATTAGCGGGATTCACAACTCTCCAAAATCATTGACGATAGACTGAC

ACGGTTACGTGGGAGGACTCGTAGTCTCCAGTGAAAATGGGCGCAAAATGGTGTCCGCAT

GGGAGGAGCTCAACCATAATAGACGCCGAATATGTTCGCGTGAAACTTCAGCGTCAAAGA

TGGAAAGCAAGACAAACTCATGAAACTACCATAGTGTGAAGATGATGGGCGGCGCGGTTT

AAGGCGAGTCGCCCTAAGACTCCAACGTGCCCTAAGGGATCGTAACATGCAAGAACATTG

ATGTGCACAGTTTTGTCGAACGGCCTTCATAGGCCAAAGATTGTATCCACGGCCAGAAGC

GGTAAGAACTTCGTGACATAGCGGCCAGAGAAAGGATCTACAACGTTAACCAGCACCTCA

GAGAGCGGGATGGCCAGTCGTTAGGGAGAAAATAACTAATCATTAACTTTGACGAACCAC

CAAAATGCTCGAGCGGTAGGTATCATTAGGGGGTACGGACTGTTACATCTAGAATACCCC

AGCTCCGTAGTTGCAGACACTAATTTTAGAGACCGCAGCTGCATCATTAACAAAATGTTA

TTGGTGGGATGCAGAAGAGCGCGGGCAAGATTATACAGACGCCTTCAGGCGGTAGCCTGC

TCCTAGGCAGTCAAGACGGATCGCGATGGACGTGTTTGGACGGCGAATGTACAGGAAATT

TCCCGTGAGAACAACAAAATGAAGGATCGAGATTAAAAAGACTGCAGCGCCCGTGAACAA

CGGAGTTGAACGTGGTTGCCAGGTTTGTGATGGGCGGCCCTAGAAGATAACGGTAGAAGC

CCTTCGGACTTTTCAATGGGGACTACATTCTGGCCGCGAAGGACTCCGTGAATCGATAGG

TAGTGGAAATATGGATGAAACGGGCTGGGGCAACGCTCGGGACAACGTTTACTAGCTAAG

AGGCAGATTACGAGGATCGGGGAAGACTGAGAAGGCCTTGGCGTAGGAATTCGAGGCAAC

CAGGGGGATAAGGTAAATGCCGACTTCAAGTCGAGAATGCAAAGAAGCTGGCTATGGCCG

TAAATTAGACGATAAATAACAGAGCATCGGGGAGCCAAGTCTAATTGTAAAACTTTTGGT

CGAGGTGATGTCGTAGGTGAGATCTCCGAACGCAGCATGAACAAGGCGAATCGTGTAAGC

AGAGAAGAGTGCAGCACGATGCTAACACTGCCGCAGCAGTGGGACAACTTGGATGACTGG

CACATCACAGTAATGAGTCATATGATTCCGCCTGAAGGCTTCAATAAAGGACATAAGATG

CGAAGAAGATCTAGACCCAGTCTTTAAAACTGCACAACTTAGCCGAAATGATTGGTAGGT

GAGAGTATCGACGTGGGCACTGAGAGAAACCTGAATACGCGATAGCCATCATATCACGAT

GCTTCTGCACTCCAGCGTCACTAGAGTTCTTGTTCACATGCCTCGGTCCTCTCGTCAGTA

CATGGCCGNCCTTCTTTCCTCCATCGTTTCCCAGAGCATACTCTTCGAGTTGCCCTATTA

ACCTTCTATTTTCTGGCCGATGCCTGACTGGCGCCGCTTTGAGTATCATACTTCTTGTTT
TTGGCCAACCTGTTGGGCGTTGATTGCTTGGGTAACCATGTCCTGCATTTTGTTGATTCC
GGGTCGGCAATGTGTTGTGGACTGTTCGTCGCCGCTCCTGATCTCACCGTACTTCACGGT
TGTCGCCCGGACATCACCCGGACGGGTTTAGGCCATGGCCGGTTGAGGTTAATCTTTACG
CGTCACCATAACTTCCTGCGAAACTGGTTTCCCCACGCCGCTCCCTTCGTGAGCCGATCG
GCGTGGTTAGGTCGCGCGGGTTTTTCCAGGCAGGGTAAGTCGATTCGCACAAGGTGACCC
ACGTACCTCGCTTTAACGTTCCGTTCTACCACTGGGTTAGGCTGCTTTGGGGCCGACTTG
ACGTTCTTTTTGCACGACCTCGTATGTGCTTATTCTCACTCGTAATGAACCGGTAAGCGA
GAGTTTGTTCTACGCTGAGGTGTCGTCGGACCCGAGTGTGCTTGCCGCTGGTCAATTGTT
GTATCGGCTCGACGTCTTATTTCGTATGGTACGGGATTATTAGTGCTTGCAGCGATGTCC
CTTTCGCTATTGATGTCTTCTCCTGGGTCACGCTACGTACTGAATTGATAAAACATTTCG
ACCTCAGTATCAGTGAGGTCGGGGGATGAGTTAGATCGCCGTGTAGGGCTTCTCACGTTG
TGTCCGTGTGGAGAGTCGGTTCTAGCGGTCGTATTATCCATATTCTCCCGTTCCATTTAC
GTTGAGCGACTTGAAACTTCCGATGTACTGGCCGGTAATAGTGCATTGTTTCTCAATTCA
TGCAAGTTGTCGTACATTATGGAATAGAACAAGAATTAAGGACCATTTCCACTAGCGATT
CTTACCTCCAACTCAAGATAGAGTCTAACCGATTATCGACGAACTTCGTCCGTCACGTCA
TTCTATGTCGAGCGGCCTTCTTACAGCGCTGTCGCACTATCTGTTCTACCTTATTACCGA
GAAACCTTGCTCCTGTTACCTGTGAGGGCGGGGCGGATATGGCAAAACGGGCATTCTTTT
TTACTTCCCGTGCCTCTTCATGCTATATACATGTATCTCTGGAGCCTAGACGCCCGGTCC
TTAACGTCCTAAAACTTTACTGACCCTGCTGGCGTCTTACCCTTTATAGTGGCTCTGCAT
AATGTGTTCTGTATATAGCCAAATACTGCCGACTATCCACTCGCTTAGCTGTCAATATCC
ATGCGCGTACTTTGATTTACCGGTCGACCTGCCTAATGTAACTGCCTTTCGCTCTCCACT
CCGCACCGGGGTTTCGAGTTTTTCTACCCCTCTCTTTATTACTACTCGTTTACTGCATCA
GACGCGCCTACCGTATTCAGTCACTGCTCCTTCAGGCGAAGAGTATTAGCACGTTATCGT
GCTACTATCCTGAGCTGTAATCATAAGTATTTCTAAAGTTGTAAGGGTTGAAGAGTAGTT
CTGCACATCTTGATCGATTGTACTGGACGTCATTCCGAATAGCTTGTGGAATCCAATATC
TGGGTCATAGATGCT
>SRR9587941
GTGACCGTCGCCTCGCCACAAATCCATTGTTTTCCGGGTTCTAGCCCCTTATGAGCTGGG
ACCTCTGTGCCTACCATCATAGCTGTTCGATGCCTTGTATTCAATACGTTGCACCGACCC
GACAATAATCTGTGCCATTAAGCTTCTGCAAGTGGCTGGCGCCTGTAAGTCACGCAATAG
AGTAACTTCCCTCTCGACAGTCTTTATGCCATCCACAATCCAACATCGTAGATCCTCGCG
TGACCTAGTAGCCCTTGACGACTCAATCCACCCTCTGCCTGACGGACGTGCCTTTTATGG
TACATTAATGACTCCCTGTCCTCATAATTAGGCTGCACACACAGGGCTATCGTAGTTTAC
TGGTCTCACCCCTGCGTACTCGGCGACCCAACTACACTGATCCAGGGAGGCAGAATGGTG
GACGCTCTTACTGCCGACCCGTGGGCAACTAACTGGGCCGGGGGAATATTGTTTCGTTTT

GGGTACTGCTTCTTCCAATTGTTCGGATAGCACAGTAACGGTGGTGGGTGGCTCCAGTCA
TTCTTGCGTAGTCGCCGGCCGGTCGCTCCGGTCTGTCCTCTCGCCCGATTCTGGGCCATC
TTATCCGATGTCACCCTGCAGACCGCTCTTTTCAACCATTAGCATTAAGCAGTCTCTCAT
AGGTGCTCTCATGTCATGAGAGGTACCGGAGCCTCTAAGCGTTCCGCTGTTTCCGACAAT
ATTCTCTTTCGCTTTGACACTAGCTCAGACGGTAGAGGCTTCGTCTTCAGCACCGGCTAA
TGATGCCTGGTACGTTAAGATACCGATCCGCGCGCTTACGCTGCTCCACCTGTCGGTGTC
TGAAGAAGCTCGGCTAAGACCTCTCGAACTCTTCGTCTTACCTCAGCCCGCGGCTACAGT
CATATAGATCCTAGCACGAACAATCCTTAATTCCCAGTCGTGCCCGACAGTAACACCGTG
GCGCGAACAATGTGTAAGTTATTGCCTACTTCGTACACTATCAACTAAATTCGTCCAACG
TGAGATCCGAGAATCCCGNAAGGGGGCACACACTTAGAGAACCCGCAGCTTTTTGTCTCA
CGATCGAGAAGGGCACTCCCAAGAGGCCTTGATACGAGCGTGAAACCGGTCAAGCGCTCC
AATTGTACACTGTCCATGAGGCCAAATTCAAATTACGGACCGAAAACTGGATCCAGCAGA
GCTAAAGGTTATTCGCGGGTATTGTCCGTACATGCAACTTGATGTTTTCTAAGCCAAAAG
AGATATGCGAGGGGGAGACATGGGAAAAGCAAAAGAATATAGTACTATAATTGGCGTGGG
ACCCATAAAGTCTTGATTGCTCCCGCGCCGCCTGTACCCACAGTCTTCAAAAAAGCGATT
TGCCGGAGTATAAAGTCAGGGGCCGACGGTGTTGAAAACAAGCGGACGTCTTGCATGAGA
AGTGTCACTGTGGGCATATCAAAGGACTCTAATTAAATTCGGTGTCAATAGAAATGTTCG
TGCTGCACGGTGCTGTCAGACCATCCACCTTTCGTGGCTACGCCTTATCAGAAGTGCAAC
CTGCGAAATCAAGGTGACGTTAATACGCTCTAAAGGACGCTAAAGCGGGTTGATAATTAA
ACGACAGAGTTACAATTAGCGGGATTCACAACTCTCCAAAATCATTGACGATAGACTGAC
ACGGTTACGTGGGAGGACTCGTAGTCTCAAGTGAAAATGGGCGCAAAATGGTGTCCGCAT
GGGAGGAGCTCAACCATAATATACGCCGAATATGTTCGCGTGAAACTTCAGCGTCAAAGA
TGGAAAGCAAGACAAACTCATGAAACTACCATAGTGTGGAGATGATGGGCGGCGCGGTTT
AAGGCGAGTCGCCCTAAGACTCCAACGTGCCCTAAGGGATCGTAACATGCAAGAACATTG
ATGTGCACAGTTTTGTCGAACGGCCTTCATAGGCCAAAGATTGTATCCACGGCCAGAAGC
GGTAAGAACTTCGTGACATAGCGGCCAGAGAAAGGATCTACAACGTTAACCAGCACCTCA
GAGAGCGGGATGGCCAGTCGTTAGGGAGAAAATAACTAATCATTAACTTTGACGAACCAC
CAAAATGCTCGAGCGGTAGGTATCATTAGGGGGTACGGACTGTTAAATCTAGAATACCCC
AGCTCCGTAGTTGCAGACACTAATCTTAGAGACCGCAGCTGCATCATTAACAAAATGTTA
TTGGTGGGATGCAGAAGAGCGCGGGCAAGATTATACAGACGCCTTCAGGCGGTAGCCTGC
TCCTAGGCAGTCAAGACGGATCGCGATGGACGTGTTTGGACGGCGAATGTACAGGAAATT
TCCCGTGAGAACAACAAAATGAAGGATCGAGATTAAAAAGACTGCAGCGCCCGTGAACAA
CGGAGTTGAACGTGGTTTCCAGGTTTGTGATGGGCGGCCCTAGAAGATAACGGTAGAAGC
CCTTCGGACTTTTCAATGGGGACTACATTCTGGCCGCGAAGGACTCCGTGAATCGATAGG
TAGTGGAAATATGGATGAAACGGGCTGGGGCAACGCTCGGGACAACGTTTACTAGCTAAG
AGGCAGATTACGAGGATCGGGGAAGACTGAGAAGGCCTTGGCGTAGGAATTCGAGGCAAC

CAGGGGGATAAGGTAAATGCCGACTTCAAGTCGAGAATGCACAGAAGCTGGCTATGGCCG
TAAATTAGACGATAAATAACAGAGCATCGGGGAGCCAAGTCTCATTGTAAAACTTTTGGT
CGAGGTGATGTCGTAGGTGAGATCTCCGAACGCAGCATGAGCAAGGCGAATCGTGTAAGC
AGAGAAGAGTGCAGCACGATGCTAACACTGCCGCAGCAGTGGGACAACTTGGATGACTGG
CACATCACAGTAATGAGTCATATGATTCCGCCTGAAGGCTTCAATAAAGGACATAAGATG
CGAAGAAGATCTAGACCCAGTCTTTAAAACTGCACAACTTAGCCGAAATGATTGGTAGGT
GAGAGTATCGACGTGGGCACTGAGAGAAACCTGAATACGCGATAGCCATCATATCACGAT
GCTTCTGCCCTCCAGCGTCACTAGAGTTCTCGTTCACATGCCTCGGTCCTCTCGTCAGTA
CATGGCCGGCCTTCTTTCCTCCATCGTTTCCCAGAGCATACTCTTCGAGTTGCCCTATTA
ACCTTCTATTTTCTGGCCGATGCCTGACTGGCGCCGCTTTGAGTATCATACTTCTTGTTT
TTGGCCAACCTGTTGGGCGTTGATTGCTTGGGTAACCATATCCTGCATTTTGTTGATTCC
GGGTCGGCAATGTGTTGTGGACTGTTCGTCGCCGCTCCTGATCTCACCGTACTTCACGGT
TGTCGCCCGGACATCACCCGGACGGGTTTAGGCCATGGCCGGTTGAGGTTAATCTTTACG
CGTCACCATAACTTCCTGCGAAACTGGTTTCCCCACGCCGCTCCCTTCGTGAGCCGATCG
GCGTGGTTAGGTCGCGCGGGTTTTTCCAGGCAGGGTAAGTCGAGTCGCACAAGGTGACCC
ACGTACCTCGCTTTAACGTTCCGTTCTACCACTGGGTTAGGCTGCTTTGGGGCCGACTTG
ACGTTCTTTTTGCACGACCTCGTATGTGCTTATTCTCACTCGTAATGAACCGGTAAGCGA
GAGTTTGTTCTACGCTGAGGTGTCGTCGGACCCGAGTGTGCTTGCCGCTGGTCAATTGTT
GTGTCGGCTCGACGTCTTATTTCGTATGGTACGGGATTATTAGTGCTTGCAGCGATGTCC
CTTTCGCTATTGATGTCTTCTCCTGAGTCACGCTACGTACTGAATTGATAAAACATTTCG
ACCTCAGTATCAGTGAGGTCGGGGGATGAGTTAGATCGCCGTGTAGGGCTTTTCACGTTG
TGTCCGTGTGGAGAGTCGGTTCTAGCGGTCGTATTATCCATATTCTCCCGTTCCATTTAC
GTTGAGCGACTTGAAACTTCCGATGTACTGGCCGGTAATAGTGCATTGTTTCTCAATTCA
TGCAAGATGTCGTACATTATGGAATAGAACAAGAATTAAGGACCATTTCCACTAGCGATT
CTTACCTCCAACTCAAGATAGAGTCTAACCGATTATCGACGAACTTCGTCCGTCACGTCA
TTCTATGTCGAGCGGCCTTCTTACAGCGCTGTCGCACTATCTGTTCTACCTTATTTCCGA
GAAACCTTGCTCCTGTTACCTGTGAGGGCGGGGCGGATATGGCAAAATGGGCATTCTTTT
TTACTTCCCGTGCCTCTTCATGCTATATACATGTATCTCTGGAGCCTAGACGCCCGGTCC
TTAACGTCCTAAAACTTTACTGACCCTGCTGGCGTCTTACCCTTTATAGTGGCTCTGCAT
AATGTGTTCTGTATATAGCCAAATACTGCCGACTATCCACTCGCTTAGCTGCCAATATCC
ATGCGCGTACTTTGATTTAACGGTCGACCTGCCTAATGTAACTGCCTTTCGCTCTCCACT
CCGCACCGGGGTTTCGAGTTTTTCTACCCCTCTCTTTATTACTACTCGTTTACTGCATCA
GACGCGCCTACCGTATTCAGGCACTGCTCCTTCAGGCGAAGAGTATTAGCACGTTATCGT
GCTACTATCCTGAGCTGTAATCATAAGTATTTCTAAAGTTGTAAGGGTTGAAGAGTAGTT
CTGTACATCTTGATCGATTGTACTGGACGTCATTCCGAATAGCTTGTGGAATCCAATATC
TGGGTCATAGATGCT

>SRR9587942

GTGACCGTCGCCTCGCCACAAATCCATTGTTTTCCGGGTTCTAGCCCCTTATGAGCTAGG
ACCTCTGTGCCTACCATCATAGCTGTTCGATGCCTTGTATTCAATACGTTGCACCGACCC
GACAATAATCTGTGCCATTAAGCTTCTGCAAGTGGCTGGCGCCTGTAAGTCACGCAATAG
AGTAACTTCCCTCTCGACAGTCTTTATGCCATCCACAATCCAACATCGTAGATCCTCGCG
TGACCTAGTAGCCCTTGACGACTCAATCCACCCTCTGCCTGACGGACGTGACTTTTATGG
TACATTAATGACTCCCTGTCCTCATAATTAGGCTGCACACACAGGGCTATCGTAGTTTAC
TGGTCTCACCCCTGCGTACTCGGCGACCCAACTACACTGATCCAGGGAGGCAGAATGGTG
GACGCTCTTACTGCCGACCCGTGGGCAACTAACTGGGCCGGGGGAATATTGTTTCGTTTT
GGGTACTGCTTCTTCCAATTGTTCGGATAGCCCAGTAACGGTGGTGGGTGGTTCCAGTCA
CTCTTGCGTAGTCGCCGGCCGGTCGCTCCGGTCTGTCCCCTCGCCCGATTCTGGGCCATC
TTATCCGATGTCACCCTGCAGACCGCNCTTTTCAACCATTAGCATTAAGCAGTCTCTCAT
AGGTGCTCTCATGTCATGAGAGGTACCGGAGCCTCTAAGCGTTCCGCTGTTCCCGACAAT
ATTCTCTTTCGCTTTGACACTAGCTCAGACGGTAGAGGCTTCGTCTTCAGCACCGGCTAA
TGATGCCTGGTACGTTAAGATACCGATCCGCGCGCTTACGCTGCTCCACCTGTCGGTGTC
TGAAGAAGCTCGGCTAAGACCTCTCGAACTCTTCGTCTTACCTCAGCCCGCGGCTACAGT
CATATACATCCTAGCACGCACAATCCTTAATTCCCAGTCGTGCCCGACAGTAACACCGTG
GCGCGAACAATGTGTAAGTTATTGCCTACTTCGTACACTATCAACTAAATTCGTCCAACG
TGAGATCCGAGAATCCCGAAAGGGGGCACACACTTAGAGAACCCGCAGCTTTTTGTCTCA
CGATCGAGAAGGGCACTCCCAAGAGGCCTTGATACAAGCGTGAAACCGGTCAAGCGCTCC
AATTGTACACTGTCCATGAGGCCAAATTCAAATTACGGACCGAAAACTGGATCCAGCAGA
GCTAAAGGTTATTCGCGGGTATTGTCCGTACATGCAACTTGATGTTTTCTAAGCCAAAAG
AGATATGCGAGGGGGAGACATGGGAAAAGCAAAAGAATATAGTACTATAATTGGCGTGGG
ACCCATAAAGTCTTGATTGCTCCCGCGCCGCCTGTACCCACAGTCTTCAAAAAAGCGATT
TGCCGGAGTATAAAGTCAGGGGCCGACGGTGTTGAAAACGAGCGGACGTCTTGCATGAGA
AGTGTCACTGTGGGCATATCAAAGGACTCTAATTAAATTCGGTGTCAATAGAAATGTTCG
TGCTGCACGGTGCTGTCAGACCATCCACCTTTCGTGGCTACGCCTTATCAGAAGTGCAAC
CTGCGAAATCAAGGTGACGTTAATACGCTCTAAAGGACGCTAAAGCGGGTTGATAATTAA
ACGACAGAATTACAATTAGCGGGATTCACAACTCTCCAAAATCATTGACGATAGACTGAC
ACGGTTACGTGGGAGGACTCGTAGTCTCCAGTGAAAATGGGCGCAAAATGGTGTCCGCAT
GGGAGGAGCTCAACCATAATAGACGCCGAATATGTTCGCGTGAAACTTCAGCGTCAAAGA
TGGAAAGCAAGACAAACTCATGAAACTACCATAGTGTGAAGATGATGGGCGGCGCGGTTT
AAGGCGAGTCGCCCTAAGACTCCAACGTGCCCTAAGGGATCGTAACATGCAAGAACATTG
ATGTGCACAGTTTTGTCGAACGGCCTTCATAGGCCAAAGATTGTATCCACGGCCAGAAGC
GGTAAGAACTTCGTGACATAGCGGCCAGAGAAAGGATCTACAACGTTAACCAGCACCTCA
GAGAGCGGGATGGCCAGTCGTTAGGGAGAAAATAACTAATCATTAACTTTGACGAACCAC

CAAAATGCTCGAGCGGTAGGTATCATTAGGGGGTACGGACTGTTACATCTAGAATACCCC
AGCTCCGTAGTTGCAGACACTAATTTTAGAGACCGCAGCTGCATCATTAACAAAATGTTA
TTGGTGGGATGCAGAAGAGCGCGGGCAAGATTATACAGACGCCTTCAGGCGGTAGCCTGC
TCCTAGGCAGTCAAGACGGATCGCGATGGACGTGTTTGGACGGCGAATGTACAGGAAATT
TCCCGTGAGAACAACAAAATGAAGGATCGAGATTAAAAAGACTGCAGCGCCCGTGAACAA
CGGAGTTGAACGTGGTTGCCAGGTTTGTGATGGGCGGCCCTAGAAGATAACGGTAGAAGC
CCTTCGGACTTTTCAATGGGGACTACATTCTGGCCGCGAAGGACTCCGTGAATCGATAGG
TAGTGGAAATATGGATGAAACGGGCTGGGGCAACGCTCGGGACAACGTTTACTAGCTAAG
AGGCAGATTACGAGGATCGGGGAAGACTGAGAAGGCCTTGGCGTAGGAATTCGAGGCAAC
CAGGGGGATAAGGTAAATGCCGACTTCAAGTCGAGAATGCAAAGAAGCTGGCTATGGCCG
TAAATTAGACGATAAATAACAGAGCATCGGGGAGCCAAGTCTAATTGTAAAACTTTTGGT
CGAGGTGATGTCGTAGGTGAGATCTCCGAACGCAGCATGAACAAGGCGAATCGTGTAAGC
AGAGAAGAGTGCAGCACGATGCTAACACTGCCGCAGCAGTGGGACAACTTGGATGACTGG
CACATCACAGTAATGAGTCATATGATTCCGCCTGAAGGCTTCAATAAAGGACATAAGATG
CGAAGAAGATCTAGACCCAGTCTTTAAAACTGCACAACTTAGCCGAAATGATTGGTAGGT
GAGAGTATCGACGTGGGCACTGAGAGAAACCTGAATACGCGATAGCCATCATATCACGAT
GCTTCTGCACTCCAGCGTCACTAGAGTTCTTGTTCACATGCCTCGGTCCTCTCGTCAGTA
CATGGCCGGCCTTCTTTCCTCCATCGTTTCCCAGAGCATACTCTTCGAGTTGCCCTATTA
ACCTTCTATTTTCTGGCCGATGCCTGACTGGCGCCGCTTTGAGTATCATACTTCTTGTTT
TTGGCCAACCTGTTGGGCGTTGATTGCTTGGGTAACCATATCCTGCATTTTGTTGATTCC
GGGTCGGCAATGTGTTGTGGACTGTTCGTCGCCGCTCCTGATCTCACCGTACTTCACGGT
TGTCGCCCGGACATCACCCGGACGGGTTTAGGCCATGGCCGGTTGAGGTTAATCTTTACG
CGTCACCATAACTTCCTGCGAAACTGGTTTCCCCACGCCGCTCCCTTCGTGAGCCGATCG
GCGTGGTTAGGTCGCGCGGGTTTTTCCAGGCAGGGTAAGTCGATTCGCACAAGGTGACCC
ACGTACCTCGCTTTAACGTTCCGTTCTACCACTGGGTTAGGCTGCTTTGGGGCCGACTTG
ACGTTCTTTTTGCACGACCTCGTATGTGCTTATTCTCACTCGTAATGAACCGGTAAGCGA
GAGTTTGTTCTACGCTGAGGTGTCGTCGGACCCGAGTGTGCTTGCCGCTGGTCAATTGTT
GTATCGGCTCGACGTCTTATTTCGTATGGTACGGGATTATTAGTGCTTGCAGCGATGTCC
CTTTCGCTATTGATGTCTTCTCCTGGGTCACGCTACGTACTGAATTGATAAAACATTTCG
ACCTCAGTATCAGTGAGGTCGGGGGATGAGTTAGATCGCCGTGNAGGGCTTCTCACGTTG
TGTCCGTGTGGAGAGTCGGTTCTAGCGGTCGTATTATCCATATTCTCCCGTTCCATTTAC
GTTGAGCGACTTGAAACTTCCGATGTACTGGCNGGTAATAGTGCATTGTTTCTCAATTCA
TGCAAGTTGTCGTACATTATGGAATAGAACAAGAATTAAGGACCATTTCCACTAGCGATT
CTTACCTCCAACTCAAGATAGAGTCTAACCGATTATCGACGAACTTCGTCCGTCACGTCA
TTCTATGTCGAGCGGCCTTCTTACAGCGCTGTCGCACTATCTGTTCTACCTTATTACCGA
GAAACCTTGCTCCTGTTACCTGTGAGGGCGGGGCGGATATGGCAAAACGGGCATTCTTTT

TTACTTCCCGTGCCTCTTCATGCTATATACATGTATCTCTGGAGCCTAGACGCCCGGTCC
TTAACGTCCTAAAACTTTACTGACCCTGCTGGCGTCTTACCCTTTATAGTGGCTCTGCAT
AATGTGTTCTGTATATAGCCAAATACTGCCGACTATCCACTCGCTTAGCTGTCAATATCC
ATGCGCGTACTTTGATTTACCGGTCGACCTGCCTAATGTAACTGCCTTTCGCTCTCCACT
CCGCACCGGGGTTTCGAGTTTTTCTACCCCTCTCTTTATTACTACTCGTTTACTGCATCA
GACGCGCCTACCGTATTCAGTCACTGCTCCTTCAGGCGAAGAGTATTAGCACGTTATCGT
GCTACTATCCTGAGCTGTAATCATAAGTATTTCTAAAGTTGTAAGGGTTGAAGAGTAGTT
CTGCACATCTTGATCGATTGTACTGGACGTCATTCCGAATAGCTTGTGGAATCCAATATC
TGGGTCATAGATGCT
>SRR9587944
GTGACCGTCGCCTCGCCACAAATCCATTGTTTTCCGGGTTCTAGCCCCTTATGAGCTAGG
ACCTCTGTGCCTACCATCATAGCTGTTCGATGCCTTGTATTCAATACGTTGCACCGACCC
GACAATAATCTGTGCCATTGAGCTTCTGCAAGTGGCTGGCGCCTGTAAGTCACGCAATAG
AGTAACTTCCCTCTCGACAGTCTTTATGCCATCCACAATCCAACATCGTAGATCCTCGCG
TGACCTAGTAGCCCTTGACGACTCAATCCACCCTCTGCCTGACGGACGTGCCTTTTATGG
TACATTAATGACTCCCTGTCCTCATAATTAGGCTGCACACACAGGGCTATCGTAGTTTAC
TGGTCTCACCCCTGCGTACTCGGCGACCCAACTACACTGATCCAGGGAGGCAGAATGGTG
GACGCGCTTAGTGCCGACCCGTGGGCAACTAACTGGGCCGGGGGAATATTGTTTCGTTTT
GGGTACTGCTTCTTCCAATTGTTCGGATAGCACAGTAACGGTGGTGGGTGGCTCCAGTCA
CTCTTGCGTAGTCCCCGGCCGGTCGCTCCGGTCTGTCCTCTCGCCCGATTCTGGGCCAAC
TTATCCGATGTCACCCTGCAGACCGCTCTTTTCAACCATTAGCATTAAGCAGTCTCTCAT
AGGTGCTCTCATGTCATGAGAGGTACCGGAGCCTCTAAGCGTTCCGCTGTTCCCGACAAT
ATTCTCTTTCGCTTTGACACTAGCTCAGACGGTAGAGGCTTCGTCTTCAGCACCGGCTAA
TGATGCCTGGTACGTTAAGATACCGATCCGCGCGCTTACGCTGCTCCACCTGTCGGTGTC
TGAAGAAGCTCGGCTAAGACCTCTCGAACTCTTCGTCTTACCTCAGCCCGCGGCTACAGT
CTAATACATCCTAGCACGCACAATCCTTAATTCCCAGTCGTGCCCGACAGTAACACCGTG
GCGCGAACAATGTGTAAGTTATTGCCTACTTCGTACACTATCAACTAAATTCGTCCAACG
TGAGATCCGAGAATCCCGAAAGGGGGCACACACTTAGAGAACCCGCAGCTTTTTGTCTCA
CGATCGAGAAGGGCACTCCCAAGAGGCCTTGATACAAGCGTGAAACCGGTCAAGCGCTCC
AATTGTACACTGTCCATGAGGCCAAATTCAAATTACGGACCGAAAACTGGATCCAGCAGA
GCTTAAGGTTATTCGCGGGTATTGTCCGTACATGCAACTTGATGTTTTCTAAGCCAAAAG
AGATATGCGAGGGGGAGACATGGGAAAAGCAAAAGAATATAGTACTATAATGGGCGTGGG
ACCCATAAAGTCTTGATTGCTCCCGCGCCGCCTGTACCCACAGTCTTCAAAAAAGCGATT
TGCCGGAGTATAAAGTCAGGGGCCGACGGTGTTGAAAACGAGCGGACGTCTTGCATGAGA
AGTGTCACTGTGGGCATATCAAAGGACTCTAATTAAATTCGGTGTCAATAGAAATGTTCG
TGCTGCACGGTGGTGTCAGACCGTCCACCTTTCGTGGCTACGCCTTATCAGAAGTGCAAC

CTGCGAAATCAAGGTGACGTTAATACGCTCTAAAGGACGCTAAAGCGGGTTGATAATTAA
ACGACAGAGTTACAATTAGCGGGATTCACAACTCTCCAAAATCATTGACGATAGACTGAC
ACGGTTACGTGGGAGGACTCGTAGTCTCAAGTGAAAATGGGCGCAAAATGGTGTCCGCAT
GGGAGGAGCTCAACCATAATAGACGCCGAATATGTTCGCGTGAAACTTCAGCGTCAAAGA
TGGAAAGCAAGACAAATTCATGAAACTACCATAGTGTGAAGATGATGGGCGGCGCGGTTT
AAGGCGAGTCGCCCTAAGACTCCAACGTGCCCTAAGAGATCGTAACATGCAAGAACATTG
ATGTGCACAGTTTTGTCGAACGGCCTTCATAGGCCAAAGATTGTATCCACGGCCAGAAGC
GGTAAGAACTTCGTGACATAGCGGCCAGAGAAAGGATCTACAACGTTAACCAGCACCTCA
GAGAGCGGGATGGCCAGTCGTTAGGGAGAAAATAACTAATCATTAACTTTGACGAACCAC
CAAAATGCTCGAGCGGTAGGTATCATTAGGGGGTACGGACTGTTACATCTAGAATACCCC
AGCTCCGTAGTTGCAGACACTAATTTTAGAGACCGCAGCTGTATCATTAACAAAATGTTA
TTGGTGGGATGCAGAAGAGCGCGGGCAAGATTATACAGACGCCTTCAGGCGGTAGCCTGC
TCCTAGGCAGTCAAGACGGATCGCGATGGACGTGTTTGGACGGCGAATGTACAGGAGATT
TCCCGTGAGAACAACAAAATGAAGGATCGAGATTAAAAAGACTGCAGCGCCCGTGAACAA
CGGAGTTGAACGTGGTTGCCAGGTTTGTGATGGGCGGCCCTAGAAGATAACGGTAGAAGC
CCTTCGGACTTTTCAATGGGGACTACATTCTGGCCGCGAAGGACTCCGTGAATCGATAGG
TAGTGGAAATATGGATGAAACGGGCTGGGGCAACGCTCGGGACAACGTTTACTAGCTAAG
AGGCAGATTACGAGGATAGGGGAAGACTGAGAAGGCCTTGGCGTAGGAATTCGAGGCAAC
CAGGGGGATAAGGTAAATGCCGACTTCAAGTCGAGAATGCAAAGAAGCTGGCTATGGCCG
TAAATTAGACGATAAATAACAGAGCATCGGGGAGCCAAGTCTAATTGTCAAACTTTTGGT
CGAGGTGATGTCGTAGGTGAGATCTCCGAACGCAGCATGAACAAGGCGAATCGTGTAAGC
AGAGAAGAGTGCAGCACGATGCTAACACTGCCGCAGCAGTGGGACAACTTGGATGACTGG
CACATCACAGTAATGAGTCATATGATTCCGCCTGAAGGCTTCAATAAAGGACATAAGATG
CGAAGAAGATCTAGACCCAGTCTTTAAAACTGCACAACTTAGCCGAAATGATTGGTAGGT
GAGAGTATCGACGTGGGCACTGAGAGAAACCTGAATACGCGATAGCCGTCATATCACGAT
GCTTCTGCCCTCCAGCGTCACTAGAGTTCTTGTTCACATGCCTCGGTCCTCTCGTCAGTA
CATGGCCGGCCTTCTTTCCTCCATCGTTTCCCAGAGCATACTCTTCGAGTTGCCCTATTA
ACCTTCTATTTTCTGGCCGATGCCTGACTGGCGCCGCTTTGAGTATCATACTTCTTGTTT
TTGGCCAACCTGTTGGGCGTTGATTGCTTGGGTAACCATATCCTGCATTTTGTTGATTCC
GGGTCGGCAATGTGTTGTGGACTGTTCGTCGCCGCTCCTGATCTCACCGTACTTCACGGT
TGTCGCCCGGACATCACCCGGACGGGTTTAGGCCATGGCCGGTTGAGGTTAATCTTTACG
CGTCACCATAACTTCCTGCGAAACTGGTTTCCCCACGCCGCTCCCTTCGTGGGCCGATCG
GCGTGGTTAGGTCGCGCGGGTTTTTCCAGGCAGGGTAAGTCGAGTCGCACAAGGTGACCC
ACGTACCTCGCTTTAACGTTCCGTTCTACCACTGGGTTAGGCTGCTTTGGGGCCGACTTG
ACGTTCTTTTTGCACGACCTCGTATGTGCTTATTCTCACTCGTAATGAACCGGTAAGCGA
GAGTTTGTTCTACGCTGAGGTGTCGTCGGACCCGAGTGTGCTTGCCGCTGGTCAATTGTT

GTGTCGGCTCGACGTCTTATTTCGTATGGTACGGGATTATTAGTGCTTGCAGCGATGTCC
CTTTCGCTATTGATGTCTTCTCCTGAGTCACGCTACGTACTGAATTGATAAAACATTTCG
ACCTCAGTATCAGTGAGGTCGGGGGATGAGTTAGATCGCCGTGTAGGGCTTTTCACGTTG
TGTCCGTGTGGAGAGTCGGTTCTAGCGGTCGTATTATCCGTATTCTCCCGTTCCATTTAC
GTTGAGCGACTTGAAACTTCTGATGTACTGGCCGNNAATAGTGCATTGTTCCTCAATTCA
TGCAAGTTGTCGTACATTATGGAATAGAACAAGAATTAAGGACCATTTCCACTAGCGATT
CTTACCTCCAACTCAAGATAGAGTCTAACCGATTATCGACGAACTTCGTCCGTCACGTCA
TTCCATGTCGAGCGGCCTTCTTACAGCGCTGTCGCACTATCTGTTCTACCTTATTTCCGA
GAAACCTTGCTCCTGTTACCTGTGAGGGCGGGGCGGATATGGCAAAATGGGCATTCTTTT
TTACTTCCCGTGCCTCTTCATGCTATATACATGTATCTCTGGAGCCTAGACGCCCGGTCC
TTAACGTCCTAAAACTTTATTGACCCTGCTGGCGTCTTACCCTTTATAGTGGCTCTGCAT
AATGTGTTCTGTATATAGCCAAATACTGCCGACTATCCACTCGCTTAGCTGTCAATATCC
ATGCGCGTACTTTGATTTACCGGTCGACCTGCCTAATGTAACTGCCTTTCGCTCTCCACT
CCGCACCGGGGTTTCGAGTTTTTCTACCCCTCTCTTTATTACTACTCGTTTACTGCACCA
GACGCGCCTACCGTATTCAGTCACTGCTCCTTCAGGCGAAGAGTATTAGCACGTTATCGT
GCTACTATCCTGAGCTGTAATCATAAGTATTTCTAAAGTTGTAGGGGTTGAAGAGTAGTT
CTGTACATCTTGATCGATTGTACTGGACGTCATTCCGAATAGCTTGTGGAATCCAATATC
TGGGTCATAGATGCT
>SRR9587945
GTGACCGTCGCCTCGCCACAAATCCATTGTTTTCCGGGTTCTAGCCCCTTATGAGCTAGG
ACCTCTGTGCCTACCATCATAGCTGTTCGATGCCTTGTATTCAATACGTTGCACCGACCC
GACAATAATCTGTGCCATTAAGCTTCTGTAAGTGGCTGGCGCCTGTAAGTCACGCAATAG
AGTAACTTCCCTCTCGACAGTCTTTATGCCATCCACAATCCAACATCGTAGATCCTCGCG
TGACCTAGTAGCCCTTGACGACTCAATCCACCCTCTGCCTGACGGACGTGCCTTTTATGG
TACATTAATGACTCCCTGTCCTCATAATTAGGCTGCACACACAGGGCTATCGTAGTTTAC
TGGTCTCACCCCTGCGTACTCGGCGACCCAACTACACTGATCCAGGGAGGCAGAATGGTG
GACGCTCTTACTGCCGACCCGTGGGCAACTAACTGGGCCGGGGGAATATCGTTTCGTTTT
GGGTACTGCTTCTTCCAATTGTTCGGATAGCACAGTAACGGTGGTGGATGGCTCCAGTCA
CTCTTGCGTAGTCGCCGGCCGGTCGCTCCGTTCTGTCCTCTCGCCCGATTCTGGGCCATC
TTATCCGATGTCACCCTGCAGACCGCTCTTTTCAACCATTAGCATTAAGCAGTCTCTCAT
AGGTGCTCTCACGTCATGAGAGGTACCGGAGCCTCTAAGCGTTCCGCTGTTCCCGACAAT
ATTCTCTTTCGCTTTAACACTAGCTCAGACGGTAGAGGCTTCGTCTTCAGCACCGGCTAA
TGATGCCTGGTACGTTAAGATACCGATCCGCGCGCTTACGCTGCTCCACCTGTCGGTGTC
TGAAGAAGCTCGGCTAAGACCTCTCGAACTCTTCGTCTTACCTCAGCCCGCGGCTACAGT
CATATACATCCTAGCACGCACAATCCTTAATTCCCAGTCGTGCCCGACAGTAACACCGTG
GCGCGAACAATGTGTAAGTTGTTGCCTACTTCGTACACTATCAACTAAATTCGTCCAACG

TGAGATCCGAGAATCCCGAAAGGGGGCACACACTTAGAGAACCCGCAGCTTTTTGTCTCA
CGATCGAGAAGGGCACTTCCAAGAGGCCTTGATACGAGCGTGAAACCGGTCAAGCGCTCC
AATTGTACACTGTCCATGAGGCCAAATTCAAATTACGGACCGAGAACTGGATCCAGCAGA
GCTAAAGGTTATTCGCGGGTATTGTCCGTACATGCAACTTGATGTTTTCTAAGCCAAAAG
AGATATGCGAGGGGGAGACATGGGAAAAGCAAAAGAATATAGTACTATAATTGGCGTGGG
ACCCATAAAGTCTTGATTGCTCCCGCGCCGCTTGTACCCACAGTCTTCAAAAAAGCGATT
TGCCGGAGTATAAAGTCAGGGGCCGACGGTGTTGAAAACGAGCGGACGTCTTGCATGAGA
AGTGTCACTGTGGGCATATCAAAGGACTCTAATTAAATTCGGTGTCAATAGCAATGTTCG
TGCTGCACGGTGCTGTCAGACCATCCATCTTTCGTGGCTACGCCTTATCAGAAGTGCAAC
CTGCGAAATCAAGGTGACGTTAATACGCTCTAAAGGACGCTAAAGCGGGTTGATAATTAA
ACGACAGAGTTACAATTAGCGGTATTCACAACTCTCCAAAATCATTGACGATAGACTGAC
ACGGTTACGTGGGAGGACTCGTAGTCTCAAGTGAAAATGGGCGCAAAATAGTGTCCGCAT
GGGAGGAGCTTAACCATAATAGACGCCGAATATGTTCGCGTGAAACTTCAGCGTCAAAGA
TGGAAAGCAAGACAAACTCATGAAACTACCATAGTGTGAAGATGATGGGCGGCGCGGTTT
AAGACGAGTCGCCCTAAGACTCCAACGTGCCCTAAGGGATCGTAACATGCAAGAACATTG
ATGTGCACAGTTTTGTCGAACGGCCTTCATAGGCCAAAGATTGTATCCACGGCCAGAAGC
GGTAAGAACTTCGTGACATAGCGGCCAGAGAAAGGATCTACAACGTTAACCAGCACCTCA
GAGAGCGGGACGGCCAGTCGTTAGGGAGAAAATAACTAATCATTAACTTTGACGAACCAC
CAAAATGCTCGAGCGGTAGGTATCATTAGGGGGTACGGACTGTTACATCTAGAATACCCC
AGCTCCGTAGTTGCAGACACTAATTTTAGAGACCGCAGCTGCATCATTAACAAAATGTTA
TTGGTGGGATGCAGAAGAGCGCGGGCAAGATTATACAGACGCCTTCAGGCGGTAGCCTGC
TCCTAGGCGGTCAAGACGGATCGCGATGGACGTGTTTGGACGGCGAATGTACAGGAAATT
TCCCGTGAGAACAACAAAATGAAGGATCGAGATTAAAAAGACTACAGCGCCCGTGAACAA
CGGAGTTGAACGTGGTTGCCAGGTTTGTGATGGGCGGCCCTAGAAGATAACGGTAGAAGC
CCTTCGGACTTTTCAATGGGGACTACATTCTGGCCGCGAAGGACTCCGTGAATCGATAGG
TAGTGGAAATATGGATGAAACGGGCTGGGGCAACGCTTGGGACAACGTTTACTAGCTAAG
AGGCAGATTACGAGGATCGGGGAAGACTGAGAAGGCCTTGGCGTAGGAATTCGAGGCAAC
CAGGGGGATAAGGTAAATGCCGACTTCAAGTCGAGAATGCAAAGAAGCTGGCTATGGCCG
TAAATTAGACGATAAATAACAGAGCATCGGGGGGCCAAGTCTAATTGTAAAACTTTTGGT
CGAGGTGATGTCGTAGGTGAGATCTCCGAACGCAGCATGAACAAGGCGAATCGTGTAAGC
AGAGAAGAGTGCAGCACGATGCTAACACTGCCGCAGCAGTGGGACAACTTGGATGACTGG
CACATCACAGTAATGAGTCATATGATTCCGCCTGAAGGCTTCAATAAAGGACATAAGATG
CGAAGAAGATCTAGACCCAGTCTTTAAAGCTGCACAACTTAGCCGAAATGATTGGTAGGT
GAAAGTATCGACGTGGGCACTGAGAGAAACCTGAATACGCGATAGCCATCATATCACGAT
GCTTCTGCCCTCCAGTGTCACTAGAGTTCTTGTTCACATGCCTCGGTCCTCTCGTCAGTA
CATGGCCGGCCTTCTTTCCTCCATCGTTTCCCAGAGCATACTCTTCGAGTTGCCCTATTA

GCCTTCTATTTTCTGGCCGATGCCTGCCTGGCGCCGCTTTGAGTATCATACTTCTTGTTT
TTGGCCAACCTGTTGGGCGTTGATTGCTTGGGTAACCATATCCTGCATTTTGTTGATTCC
TGGTCGGCAATGTGTTGTGGACTGTTCGTCGCTGCTCCTGACCTCACCGTACTTCACGGT
TGTCGCCCGGACATCACCCGGACGGGTTTAGGCCATGGCCGGTTGAGGTTAATCTTTACG
CGTCATCATAACTTCCTGCGAAACTGGTTTCCTCACGCCGCTCCCTTCGTGAGCCGATCG
GCGTGGTTAGGTCGCGCGGGTTTTTCCAAGCAGGGTAAGTCGAGTCGCCCAAGGTGACCC
ACGTACCTCGCTTTAACGTTCCGTTCTACCACTGGATTAGGCTGCTTTGGGGCCGACTTG
ACGTTCTTTTTGTACGACCTCGTATGTGCTTATTCTCACTCGTAATGAACCGGTAAGCGA
GAGTTTGCTCTACGCTGAGGTGCCGTCGGACCCGAGTGTGCTTGCCGCTGGTCAATTGTT
GTGTCGGCTCGACGTCTTATTTCGTATGGTACGGGATTATTAGTGCTTGCAGCGATGTCC
CTTTCGCTATTGATGTCTTATCCTGAGTCACGCTACGTACTGAATTGATAAAACATTTCG
ACCTCAGTATCAGTGAGGTCGGGGGATGAGTTAGATCGCCGTGTAGGGCTTTTCACGTTG
TGTCCGTGTGGAGAGTCGGTTCTAGCGGTCGTATTATCCATATTCTCCCGTTCTATTTAC
GTTGAGCGACTTGAAACCTCCGATGTACTGGCCGNTAATAGTGCATTGTTTCTCAATTCA
TGCGAGTTGTCGTACATTATGGAATAGAACAAGAATTAAGGACCATTTCCACTAGCGATT
CTTACCTCCAACTCAAGATAGAGTCTAACCGATTATCGACGAACTTCGTCCGTCACGTCA
TTCTATGTCGAGCGGCCTTCTTACAGCGCTGTCGCACTATCTGTTCTACCTTATTTCCGA
GAAACCTTGCTCCTGTTACCTGTGAGGGCGGGGCGGATATGGCAAAATGGGCATTCTTTT
TTACTTCCCGTGCCTCTTCATGCTATATACATGTATCTCTGGAGCCTAGACGCCCGGTCC
TTAACGTCCTAAAACTTTACTGACCCTGCTGGCGTCTTACCCTTTATAGTGGCTCTGCAT
AATGTGTTCTGTATATAGCCAAATACTGCCGACTATCCACTCGCTTAGCTGTCAATATCC
ATGCGCGTACTTTGATTTACCGGTCGACCTGCCTAATGTAACTGCCTTTCGCTCTCCACT
CCGCACCGGGGTTTCGAGTTTTTCTACCCCTCTCTTTATTACTACTCGTTTACTGCATCA
GACGCGCCTACCGTATTCGGTCACTGCTCCTTCAGGCGAAGAGTATTAGCACGTTATCGT
GCTACTATCCTGAGCTGTAATCCTAATTATTTCTAAAGTTGTAAGGGTTGAAGAGTAGTT
CTGTACATCTTGATCGATTGTACTGGACGTCATTCCGAATAGCTTGTGGAATCCAATATC
TGGGTCATAGATGCT
>SRR9587946
GTGACCGTCGCCTCGCCACAAATCCATTGTTTTCCGGGTTCTAGCCCCTTATGAGCTAGG
ACCTCTGTGCCTACCATCATAGCTGTTCGATGCCTTGTATTCAATACGTTGCACCGACCC
GACAATAATCTGTGCCATTAAGCTTCTGCAAGTGGCTGGCGCCTGTAAGTCACGCAATAG
AGTAACTTCCCTCTCGACAGTCTTTATGCCATCCACAATCCAACATCGTAGATCCTCGCG
TGACCTAGTAGCCCTTGACGACTCAATCCACCCTCTGCCTGACGGACGTGCCTTTTATGG
TACATTAATGACTCCCTGTCCTCATAATTAGGCTGCACACACAGGGCTATCGTAGTTTAC
TGGTCTCACCCCTGCGTACTCGGCGACCCAACTACACTGATCCAGGGAGGCAGAATGGTG
GACGCTCTTACTGCCGACCCGTGGGCAACTAACTGGGCCGGGGGAATATTGTTTCGTTTT

GGGTACTGCTTCTTCCAATTGTTCGGATAGCAAAGTAACGGTGGTGGGTGGCTCCAGTCA
TTCTTGCGTAGTCGCCGGCCGGTCGCTCCGGTCTGTCCTCTCTCCCGATTCTGGGCAATC
TTATCCGATGTCACCCTGCAGACCGCTCTTTTTAACCATTAGCATTAAGCAGTCTCTCAT
AGGTGCTCTCGTGTCATGAGAGGTACCGGAGCCTCTAAGCGTTCCGCTGTTCCCGACAAT
ATTCTCTTTCGCTTTGACACTAGCTCAGACGGTAGAGGCTTCGCCTTCAGCACCGGCTAA
TGATGCCTGGTACGTTAAGATACCGATCCGCGCGCTTACGCTGCTCCACCTGTCGGTGTC
TGAAGAAGCTCGGCTAAGACCTCTCGAACTCTTCGTCTTACCTCAGCCCGCGGCTACAGT
CATATACATCCTAGCACGCACAATCCTTAATTCCCAGTCGTGCCCGACAGTAACACCGTG
GCGCGAACAATGTGTAAGTTATTGCCTACTTCGTACACTATCAACTAAATTCGTCCAACG
TGAGATCCGAGAATCCCGTTAGGGGGCACACACTTAGAGAACCCGCAGCTTTTTGTCTCA
CGATCGAGAAGGGCACTCCCAAGAGGCCTTGATACGAGCGTGAAACCGGTCAAGCGCTCC
AATTGTACACTGTCCATGAGGCCAAATTCAAATTACGGACCGAAAACTGGATCCAGCAGA
GCTAAAGGTTATTCGCGGGTATTGTCCGTACATGCAACTTGATGTTTTCTAAGCCAAAAG
AGATATGCGAGGGGGAGACATGGGAAAAGCAAAAGAATATAGTACTATAATTGGCGTGGG
ACCCATAAAGTCTTGATTGCTCCCGCGCCGCCTGTACCCACAGTCTTCAAAAAAGCGATT
TGCCGGAGTATAAAGTCAGGGGCCGACGGTGTTGAAAACGAGCGGACGTCTTGCATGAGA
AGTGTCACTGTGGGCATATCAAAGGACTCTAATTAAATTCGGTGTCAATAGAAATGTTCG
TGCTGCACGGTGCTGTCAGACCATCCACCTTTCGTGGCTACGCCTTATCAGAAGTGCAAC
CTGCGAAATCAAGGTGACGTTAATACGCTCTAAAGGACGCTAAAGCGGGTTGATAATTAA
ACGACAGAGTTACAATTAGCGGGATTCACAACTCTCCAAAATCATTGACGATAGACTGAC
ACGGTTACGTGGGAGGACTCGTAGTCTCAAGTGAAAATGGGCGCAAAATGGTGTCCGCAT
GGGAGGAGCTCAACCATAATAGACGCCGAATATGTTCGCGTGAAACTTCAGCGTCAAAGA
TGGAAAGCAAGACAAACTCATGAAACTACCATAGTGTGAAGATGATGGGCGGCGCGGTTT
AAGGCGAGTCGCCCTAAGGCTCCAACGTGCCCTAAGGGATCGTAACATGCAAGAACATTG
ATGTGCACAGTTTTGTCGAACGGCCTTCATAGGCCAAAGATTGTATCCACGGCCAGAAGC
GGTAAGAACTTCGTGACATAGCGGCCAGAGAAAGGATCTACAACGTTAACCAGCACCTCA
GAGAGCGGGATGGCCAGTCGTTAGGGAGAAAATAACTAATCATTAATTTTGACGAACCAC
CAAAATGCTCGAGCGGTAGGTATCATTAGGGGGTACGGACTGTTACATCTAGAATGCCCC
AGCTCCGTAGTTGCAGACACTAATTTTAGAGACCGCAGCTGCATCATTAACAAAATGTTA
TTGGTGGGATGCAGAAGAGCGCGGGCAAGATTATACAGACGCCTTCAGGCGGTAGCCTGC
TCCTAGGCAGTCAAGACGGATCGCGATGGACGTGTTTGGACGGCGAATGTACAGGAAATT
TCCCGTGAGAACAACAAAATGAAGGATCGAGATTAAAAAGACTGCAGCGCCCGTGAACAA
CGGAGTTGAACGTGGTTTCCAGGTTTGTGATGGGCGGCCCTAGAAGATAACGGTAGAAGC
CCTTCGGACTTTTCAATGGGGACTACATTCTGGCCGCGAAGGACTCCGTGAATCGATAGG
TAGTGGAAATATGGATGAAACGGGCTGGGGCAACGCTCGGGACAACGTTTACTAGCTAAG
AGGCAGATTACGAGGATCGGGGAAGACTGAGAAGGCCTTGGCGTAGGAATTCGAGGCAAC

CAGGGGGATAAGGTAAATGCCGACTTCAAGTCGAGAATGCAAAGAAGCTGGCTATGGCCG
TAAATTAGACGATAAATAACAGAGCATCGGGGAGCCAAGTCTAATTGTAAAACTTTTGGT
CGAGGTGATGTCGTAGGTGAGATCTCCGAACGCAGCATGAGCAAGGCGAATCGTGTAAGC
AGAGAAGAGTGCAGCACGATGCTAACACTGCCGCAGCAGTGGGACAACTTGGATGACTGG
CACATCACAGTAATGAGTCATATGATTCCGCCTGAAGGCTTCAATAAAGGACATAAGATG
CGAAGAAGATCTAGACCCAGTCTTTAAAACTGCACAACTTAGCCGAAATGATTGGTAGGT
GAGAGTATCGACGTGGGCACTGAGAGAAACCTGAATACGCGATAGTCATCATATCACGAT
GCTTCTGCCCTCCAGCGTCACTAGAGTTCTTGTTCACATGCCTCGGTCCTCTCGTCAGTA
CATGGCCGGCCTTCTTTCCTCCATCGTTTCCCAGAGCATACTCTTCGAGTTGCCCTATTA
ACCTTCTATTTTCTGGCCGATGCCTGACTGGCGCCGCTTTGAGTATCATACTTCTTGTTT
TTGGCCAAACTGTTGGGCGTTGATTGCTTGGGTAACCATATCCTGCATTTTGTTGATTCC
GGGTCGGCAATGTGTTGTGGACTGTTCGTCGCCGCTCCTGATCTCACCGTACTTCACGGT
TGTCGCCCGGACATCACCCGGACGGGTTTAGGCCATGGCCGGTTGAGGTTAATCTTTACG
CGTCACCATAACTTCCTGCGAAACTGGTTTCCCCACGCCGCTCCCTTCGTGAGCCGATCG
GCGTGGTTAGGTCGCGCGGGTTTTTCCAGGCAGGGTAAGTCGAGTCGCACAAGGTGACCC
ACGTACCTCGCTTTAACGTTCCGTTCTACCACTGGGTTAGGCTGCTTTGGGGCCGACTTG
ACGTTCTTTTTGCACGACCTCGTATGTGCTTATTCTCACTCGTAATGAACCGGTAAGCGA
GAGTTTGTTCTACGCTGAGGTGTCGTCGGACCCGAGTGTGCTTGCCGCTGGTCAATTGTT
GTGTCGGCTCGACGTCTTATTTCGTATGGTACGGGATTATTAGTGCTTGCAGCGATGTCC
CTTTCGCTATTGATGTCTTCTCCTGAGTCACGCTACGTACTGAATTGATAAAACATTTCG
ACCTCAGTATCAGTGAGGTCGGGGGATGAGTTAGATCGCCGTGTAGGGCTTTTCACGCTG
TGTCCGTGTGGAGAGTCGGTTCTAGCGGTCGTATTATCCATATTCTCCCGTTCCATTTAC
GTTGAGCGACTTGAAACTTCCGATGTACTGGCCGGTAATAGTGCATTGTTTCTCAATTCA
TGCAAGTTGTCGTACATTATGGAATAGAACAAGAATTAAGGACCATTTCCACTAGCGATT
CTTACCTCCAACTCAAGATAGAGTCTAACCGATTATCGACGAACTTCGTCCGTCACGTCA
TTCTATGTCGAGCGGCCTTCTTACAGCGCTGTCGCACTATCTGTTCTACCTTATTTCCGA
GAAACCTTGCTCCTGTTACCTGTGAGGGCGGGGCGGATATGGCAAAATGGGCATTCTTTT
TTACTTCCCGTGCCTCTTCATGCTATATACATGTATCTCTGGAGCCTAGACGCCCGGTCC
TTAACGTCCTAAAACTTTACTGACCCTGCTGGCGTCTTACCCTTTATAGTGGCTCTGCAT
AATGTGTTCTGTATATAGCCAAATACTGCCGACTATCCACTCGCTTAGCTGCCAATATCC
ATGCGCGTACTTTGATTTACCGGTCGACCTGCCTAATGTAACTGCCTTTCGCTCTCCACT
CCGCACCGGGGTTTCGAGTTTTTCTACCCCTCTCTTTATTACTACTCGTTTACTGCATCA
GACGCGCCTACCGTATTCAGTCACTGCTCCTTCAGGCGAAGAGTATTAGCACGTTATCGT
GCTACTATCCTGAGCTGTAATCATAAGTATTTCTAAAGTTGTAAGGGTTGAAGAGTAGTT
CTGTACATCTTGATCGATTGTACTGGACGTCATTCCGAATAGCTTGTGGAATCCAATATC
TGGGTCATAGATGCT

>SRR9587947

GTGACCGTCGCCTCGCCACAAATCCATTGTTTTCCGGGTTCTAGCCCCTTATGAGCTAGG

ACCTCTGTGCCTACCATCATAGCTGTTCGATGCCTTGTATTCAATACGTTGCACCGACCC

GACAATAATCTGTGCCATTAAGCTTCTGCAAGTGGCTGGCGCCTGTAAGTCACGCAATAG

AGTAACTTCCCTCTCGACAGTCTTTATGCCATCCACAATCCAACATCGTAGATCCTCGCG

TGACCTAGTAGCCCTTGACGACTCAATCCACCCTCTGCCTGACGGACGTGCCTTTTATGG

TACATTAATGACTCCCTGTCCTCATAATTAGGCTGCACACACAGGGCTATCGTAGTTTAC

TGGTCTCACCCCTGCGTACTCGGCGACCCAACTACACTGATCCAGGGAGGCAGAATGGTG

GACGCGCTTACTGCCGACCCGTGGGCAACTAACTGGGCCGGGGGAATATTGTTTCGTTTT

GGGTACTGCTTCTTCCAATTGTTCGGATAGCACAGTAACGGTGGTGGGTGGCTCCAGTCA

CTCTTGCGTAGTCGCCGGCCGGTCGCTCCGGTCTGNCCTCTCGCCCGATTCTGGGCCATC

TTATCCGATGTCGCCCTGCAGACCGCTCTTTTCAACCATTAGCATTAAGCAGTCTCTCAT

AGGTGCTCTCATGTCATGAGAGGTACCGGAGCCTCTAAGCGTTCCGCTGTTCCCGACAAT

ATTCTCTTTCGCTTTGACACTAGCTCAGACGGTAGAGGCTTCGTCTTCAGCACCGGCTAA

TCATGCCTGGTACGTTAAGATACCGATCCGCGCGCTTACGCTGCTCCACCTGTCGGTGTC

TGAAGAAGCTCGGCTAAGACCTCTCGAACTCTTCGTCTTACCTCAGCCCGCGGCTACAGT

CATATACATCCTAGCACGCACAATCCTTAATTCCCAGTCGTGCCCGACAGTAACACCGTG

GCGCGAACAATGTGTAAGTTATTGCCTACTTCGTACACTATCAACTAAATTCGTCCAACG

TGAGATCCGAGAATCCCGNAAGGGGGCACACACTTAGAGAACCCGCAGCTTTTTGTCTCA

CGATCGAGAAGGGCACTCCCAATAGGCCTTGATACGAGCGTGAAACCGGTCAAGCGCTCC

AATTGTACACTGTCCATGAGGCCAAATTCAAATTACGGACCGAAAACTGGATCCAGCAGA

GCTAAAGGTTATTCGCGGGTATTGTCCGTACATGCAACTTGATGTTTTCTAAGCCAAAAG

AGATATGCGAGGGGGAGACATGGGAAAAGCAAAAGAATATAGTACTATAATTGGCGTGGG

ACCCATAAAGTCTTGATCGCTCCCGCGCCGCCTGTACCCACAGTCTTCAAAAAAGCGATT

TGCCGGAGTATAAAGTCAGGGGCCGACGGTGTTGAAAACGAGCGGACGTCTTGCATGAGA

AGTGTCACTGTGGGCATATCAAAGGACTCTAATTAAATTCGGTGTCAATAGAAATGTTCG

TGCTGCACGGTGCTGTCAGACCATCCACCTTTCGTGGCTACGCCTTATCGGAAATGCAAC

CTGCGAAATCAAGGTGACGTTAATACGCTCTAAAGGACGCTAAAGCGGGTTGATAATTAA

ACGACAGAGTTACAATTAGCGGGATTCACAACTCTCCAAAATCATTGACGATAGACTGAC

ACGGTTACGTGGGAGGACTCGTAGTCTCAAGTGAAAATGGGCGCAAAATGGTGTCCGCAT

GGGAGGAGCTCAACCATAATAGACGCCGAATATGTTCGCGTGAAACTTCAGCGTCAAAGA

TGGAAAGCAAGGCAAACTCATGAAACTACCATAGTGTGAAGATGATGGGCGGCGTGGTTT

AAGGCGAGTCGCCCTAAGACTCCAACGTGCCCTAAGGGATCGTAACATGCAAGAACATTG

ATGTGCACAGTTTTGTCGAACGGCCTTCATAGGCCAAAGATTGTATCCACGGCCAGAAGC

GGTAAGAACTTCGTGACATAGCGGCCAGAGAAAGGATCTACAACGTTAACCAGCACCTCA

GAGAGCGGGATGGCCAGTCGTTAGGGAGAAAATAACTAATCATTAACTTTGACGAACCAC

CAAAATGCTCGAGCGGTAGGTATCATTAGGGGGTACGGACTGTTACATCTAGAATACCCC
AGCTACGTAGTTGCAGACACTAATTTTAGAGACCGCAGCTGTATCATTAACAAAATGTTA
TTGGTGGGATGCAGAAGAGCGCGGGCAAGATTATACAGACGCCTTCAGGCGGTAGCCTGC
TCCTAGGCAGTCAAGACGGATCGCGATGGACGTGTTTGGACGGCGAATGTACAGGAAATT
TCCCGTGAGAACAACAAGATGAAGGATCGAGATTAAAAAGACTGCAGCGCCCGTGAACAA
CGGAGTTGAACGTGGTTGCCAGGTTTGTGATGGGCGGCCCTAGAAGATAACGGTAGAAGC
CCTTCGGACTTTTCAATGGGGACTACATTCTGGCCGCGAAGGACTCCGTGAATCGATAGG
TAGTGGAAATATGGATGAAACGGGCTGGGGCAACGCTCGGGACAACGTTTACTAGCTAAG
AGGCAGATTACGAGGATCGGGGAAGACTGAGAAGGCCTTGGCGTAGGAATTCAAGGCAAC
CAGGGGGATAAGGTAAATGCCGACTTCAAGTCGAGAATGCAAAGAAGCTGGCTATGGCCG
TAAATTAGACGATAAATGACAGAGCATCGGGGAGCCAAATCTAATTGTAAAACTTTTGGT
CGAGGTGATGTCGTAGGTGAGATCTCCGAACGCAGCATGAACAAGGCGAATCGTGTAAGC
AGGGAAGAGTGCAGCACGATGCTAACACTGCCGCAGCAGTGGGACAACTTGGATGACTGG
CACATCACAGTAATGAGTCATATGATTCCGCCTGAAGGCTTCAATAAAGGACATAAGATG
CGAAGAAGATCTAGACCCAGTCTTTAAAACTGCACAACTTAGCCGAAATGATTGGTAGGT
GAGAGTATCGACGTGGGCACTGAGAGAAACCTGAATACGCGATAGCCATCATATCACGAT
GCTTCTGCCCTCCAGCGTCACTAGAGTTCTTGTTCACATGCCTCGGTCCTCTCGTCAGTA
CATGGCCGGCCTTCTTTCCTCCATCGTTTCCCAGAGCATACTCTTCGAGTTGCCCTATTA
ACCTTCTATTTTCTGGCCGATGCCTGACTGGCGCCGCTTTGAGTATCATACTTCTTGTTT
TTGGCCAACCTGTTGGGCGTTGATTGCTTGGGTAACCATATCCTGCATTTTGTTGATTCC
GGGTCGGCAATGTGTTGTGGACTGTTCGTCGCCGCTCCTGATCTCACCGTACTTCACGGT
TGTCGCCCGGACATCACCCGGACGGGTTTAGGCCATGGCCGGTTGAGGTTAATCTTTACG
CGTCACCATAACTTCCTGCGAAACTGGTTTCCCCACGCCGCTCCCTTCGTGAGCCGATCG
GCGTGGTTAGGTCGCGCGGGTTTTTCCAGGCAGGGTAAGTCGAGTCGCACAAGGTGACCC
ACGTACCTCGCTTTGACGTTCCGTTCTACCACTGGGTTAGGCTGCTTTGGGGCCGACTTG
ACGTTCTTTTTGCACGACCTCGTATGTGCTTATTCTCACTCGTAATGAACCGGTAAGCGA
GAGTTTGTTCTACGCTGAGGTGTCGTCGGACCCGAGTGTGCTTGCCGCTGGTCAATTGTT
GTGTCGGCTCGACGTCTTATTTCGTATGGTACGGGATTATTAGTGCTTGCAGCGATGTCC
CTTTCGCTATTGATGTCTTCTCCTGAGTCACGCTACGTACTGAATTGATAAAACATTTCG
ACCTCAGTATCAGTGAGGTCGGGGGATGAGTTAGATCGCCGTGTAGGGCTTTTCACGTTG
TGTCCGTGTGGAGAGTCGGTTCTAGCGGTCGTATTATCCATATTCTCCCGTTCCATTTAC
GTTGAGCGACTTGAAACTTCCGATGTACTGGCCGGTAATAGTGCATTGTTTCTCAATTCA
TGCAAGTTGTCGTACATTATGGAATAGAACAAGAATTAAGGACCATTTCCACTAGCGATT
CTCACCTCCAACTCAAGATAGAGTCTAACCGATTATCGACGAACTTCGTCCGTCACGTCA
TTCTATGTCGAGCGGCCTTCTTACAGCGCTGTCGCACTATCTGTTCTACCTTATTTCCGA
GAAACCTTGCTCCTGTTACCTGTGAGGGCGGGGCGGATATGGCAAAATGGGCATTCTTTT

TTACTTCCCGTGCCTCTTCATGCTATATACATGTATCTCTGGAGCCTAGACGCCCGGTCC
TTAACGTCCTAAAACTTTACTGACCCTGCTGGCGTCTTACCCTTTATAGTGGCTCTGCAT
AATGTGGTCTGTATATAGCCAAATACTGCCGACTATCCACTCGCTTAGCTGTCAATATCC
ATGCGCGTACTTTGATTTACCGGTCGACCTGCCTAATGTAACTGCCTTTCGCTCTCCACT
CCGCACCGGGGTTTCGAGTTTTTCTACCCCTCTCTTTATTACTACTCGTTTACTGCATCA
GACGCGCCTACCGTATTCAGTCACTGCTCCTTCAGGCGAAGAGTATTAGCACGTTATCGT
GCTACTATCCTGAGCTGTAATCATAAGTATTTCTAAAGTTGTAAGGGTTGAAGAGTAGTT
CTGTACATCTTGATCGATTGTACTGGACGTCATTCCGAATAGCTGGTGGAATCCAATATC
TGGGTCATAGATGCT
>SRR9587948
GTGACCGTCGCCTCGCCACAAATCCATTGTTTTCCGGGTTCTAGCCCCTTATGAGCTAGG
ACCTCTGTTCCTACCATCATAGCTGTTCGATGCCTTGTATTCAATACGTTGCACCGACCC
GACAATAATCTGTGCCATTAAGCTTCNGCAAGTGGCTGGCGCCTGTAAGTCACGCAATAG
AGTAACTTCCCTCTCGACAGTCTTTATGCCATCCACAATCCAACATCGTAGATCCTCGCG
TGACCTAGTAGCCCTTGACGACTCAATCCACCCTCTGCCTGACGGACGTGCCTTTTATGG
TACATTAATAACTCCCTGTCCTCATAATTAGGCTGAACACACAGGGCTATCGTAGTTTAC
TGGTCTCACCCCTGCGTACTCGGCGACCCAACTACACTGGTCCATGGAGGCAGAATGGTG
GACGCTCTTACTGCCGACCCGTGGGCAACTAACTGGGCCCGGGGAATATTGTTTCGTTTT
GGGTACTGCTTCTTCCAATTGTTCGGATAGCACAGTAACGGTGGTGGGTGGCTCCAGTCA
CTCTTGCGTAGTCGCCGGCCGGTCGCTCCGGTCTGTCCTCTCGCCCGATTCGGGGCCATC
TTGTCCGATGTCACCCTGCAGACCGCTCTTTTCAACCATTAGCATTAAGCAGTTTCTCAT
AGGTGCTCTCATGTCATGAGAGGTACCGGAGCCTCTAAGCGTTCCGCTGTTCCCGACAAT
ATTCTCTTTCGCTTTAACACTAGCTCAGACGGTAGAGGCTTCGTCTTCAGCACCGGCTAA
TGATGCCTGGTACGTTAAGATACCGATCCGCGCGCTTACGCTGCTCCACCTGTCGGTGTC
TGAAGAAGCTCGGCTAAGACCTCTCGAACTCTTCGTCTTACCTCAGCCCGCGGCTACAGT
CATATACATCCTAGCACGCACAATCCTTAATTCCCAGTCGTGCCCGACAGTAACACCGTG
GCGCGAACAATGTGTAAGTTGTTGCCTACTTCGTACACTATCAACTAAATTCGTCCAACG
TGAGATCCGAGAATCCCGNAAGGGGGCACACACTTAGAGAACCCGCAGCTTTTTGTCTCA
CGATCGAGAAGGGCACTCCCAAGAGGCCTTGATACGAGCGTGAAACCGGTCAAGCGCTCC
AATTGTACACTGTCCATGAGGCCAAATTCAAATTACGGACCGAAAACTGGATCCAGCAGA
GCTAAAGGTTATTCGCGGGTATTGTCCGTACATGCAACTTGATGTTTTCTAAGCCAAAAG
AGATATGCGAGGGGGAGACATGGGAAAAGCAAAAGAATATAGTACCATAATTGGCGTGGG
ACCCATAAAGTCTTGATTGCTCCCGCGCCGCCTGTACCCACAGTCTTCAAAAAAGCAATT
TGCCGGAGTATAAAGTCAGGGGCCGACGGTGTTGAAAACGAGCGGACGTCTTGCATGAGA
AGTGTCACTGTGGGCATATCAAAGGACTCTAATTAAATTCGGTGTCAATAGAAATGTTCG
TGCTGCACGGTGCTGTCAGACCATCCATCTTTCGTGGCTACGCCTTATCAGAAGTGCAAC

CTGCGAAATCAAGGTGACGTTAATACGCTCTAAAGGACGCTAAAGCGGGTTGATAATTAA
ACGACAGAGTTACAATTAGCGGTATTCACAACTCTCCAAAATCATTGACGATAGACTGAC
ACGGTTACGTGGGAGGACTCGTAGTCTCAAGTGAAAATGGGCGCAAAATGGTGTCCGCAT
GGGAGGAGCTCAACCATAATAGACGCCGAATATGTTCGCGTGAAACTTCAGCGTCAAAGA
TGGAAAGCAAGACAAACTCATGAAACTACCATAGTGTGAAGATGATGGGCGGCGCGGTTT
AAGGCGAGTCGCCCTAAGACTCCAACGTGCCCTAAGGGATCGTAACATGCAAGAACATTG
ATGTGCACAGTTTTGTCGAACGGCCTTCATAGGCCAAAGATTGTATCCACGGCCAGAAGC
GGTAAGAACTTCGTGACATAGCGGCCAGAGAAAGGATCTACAACGTTAACCAGCACCTCA
GAGAGCGGGACGGCCAGTCGTTAGGGAGAAAATAACTAATCATTAACTTTGACGAACCAC
CAAAATGCTCGAGCGGTAGGTATCATTAGGGGGTACGGACTGTTACATCTAGAATACCCC
AACTCCGTAGTTGCAGACACTAATTTTAGAGACCGCAGCTGCATCATTAACAAAATGTTA
TTGGTGGGATGCAGAAGAGCGCGGGCAAGATTATACAGACGCCTTCAGGCGGTAGCCTGC
TCCTAGGCAGTCAAGACGGATCGCGATGGACGTGTTTGGACGGCGAATGTACAGGAAATT
TCCCGTGAGAACAACAAAATGAAGGATCGAGATTAAAAAGACTACAGCGCCCGTGAACAA
CGGAGTTGAACGTGGTTGCCAGGTTTGTGATGGGCGGCCCTAGAAGATAACGGTAGAAGC
CCTTCGGACTTTTCAATGGGGACTACACTCTGGCCGCGAAGGACTCCGTGAATCGATAGG
TAGTGGAAATATGGATGAAACGGGCTGGGGCAACGCTCGGGACAACGTTTACTAGCTAAG
AGGCAGATTACGAGGATCGGGGAAGACTAAGAAGGCCTTGGCGTAGGAATTCGAGGCAAC
CAGGGGGGTAAGGTAAATGCCGACTTCAAGTCGAGAATGCAAAGAAGCTGGCTATGGCCG
TAAATTAGACAATAAATAACAGAGCATCGGGGGGCCAAGTCTAATTGTAAAACTTTTGGT
CGAGGTGATGTCGTAGGTGAGATCTCCGAACGCAGCATGAACAAGGCGAATCGTGTAAGC
AGAGAAGAGTGCAGCACGATGCTAACACTGCCGCAGCAGTGGGACAACTTGGATGACTGG
CACATCACAGTAATGAGTCATATGATTCCGCCTGAAGGCTTCAATAAAGGACATAAGATG
CGAAGAAGATCTAGACCCAGTCTTTAAAACTGCACAACTTAGCCGAAATGATTGGTAAGT
GAGAGTATCGACGTGGGCACTGAGAGAAACCTGAATACGCGATAGCCATCATATCACGAT
GCTTCTGCCCTCTAGCGTCACTAGAGTTCTTGTTCACACGCCTCGGTCCTCTCGTCAGTA
CATGGCCGGCCTTCTTTCCTCCATCGTTTCCCAGAGCATACTCTTCGAGTTGCCCTATTA
ACCTTCTACTTTCTGGCCGATGCCTGACTGGCGCCGCTTTGAGTATCATACTTCTTGTTT
TTGGCCAACCTGTTGGGCGTTGATTGCTTGGGTAACCATATCCTGCATTTTGTTGATTCC
TGGTCGGCAATGTGTTGTGGACTGTTCGTCGCCGCTCCTGATCTCACCGTACTTCACGGT
TGTCGCCCGGACATCACCCGGACGGGTTTAGGCCATGGCCGGTTGAGGTTAATCTTTACG
CGTCATCATAACTTCCTGCGAAACTGGTTTCCCCACGCCGCTCCCTTCGTGAGCCGATCG
GCGTGGTTAGGTCGCGCGGGTTTTTCCAGGCAGGGTAAGTCGAGTCGCACAAGGTAACCC
ACGTACCTCGCTTTAACGTTCCGTTCTACCACTGGGTTAGGCCGCTTTGGGGCCGACTTG
ACGTTCTTTTTGCACGACCTCGTATGTGCTTATTCTCACTCGTAATGAACCGGTAAGCGA
GAGTTTGTTCTACGCTGAGGTGTCGTTGGACCCGAGTGTGCTTGCCGCTGGTCAATTGTT

GTGTCGGCTCGACGTCTTATTTCGTATGGTACGGGATTATTAGTGCTTGCAGCGATGTCC
CTTTCGCTATTGATGTCTTCTCCTGAGTCACGCTACGTACTGAATTGATAAAACATTTCG
ACCTCAGTATCAGTGAGGTCGGGCGATGAGTTAGATCACCGTGTAGGGCTTTTCACGTTG
TGTCCGTGTGGAGAGTCGGTTCTAGCGGTCGTATTATCCATATTCTCCCCTTCTATTTAC
GTTGAGCGACTTGAAACTTCCGATGTACTGGCCGGTAATAGTGCATTGTTTCTCAATTCA
TGCAAGTTGTCGTACACTATGGAATAGAACAAGAATTAAGGACCATTTCCACTAGTGATT
CTTAGCTCCAACTCAAGATAGAGTCTAACCGATTATCGACGAACTTCGTCCGTCACGTCA
TTCTATGTCGAGCGGCCTTCTTACAGCGCTGTCGCACTATCTGTTCCACCTTATTTTCGA
GAAACCTTGCTCCTGTTACCTGTGAGGGCGGGGCGGATATGGCAAAATGGGCATTCTTTT
TTACTTCTCGTGCCTCTTCATGCTATATACATGTATCTCTGGAGCCTAGACGCCCGGTCC
TTAACGTCCTAAAACTTTACTGACCCTGCTGGCGTCTTACCCTTTATAGTGGCTCTGCAT
AATGTGTTCTGTATATAGCCAAATACTGCCGACTATCCACTCGCTTAGCTGTCAATATCC
ATGCGCGTACTTTGATTTACCGGTCGACCTGCCTAATGTAACTGCCTTTCGCTCTCCACT
CCGCACCGGGGTTTCGAGTTTTTCTACCCCTCTCTTTATTACTACTCGTTTACTGCATCA
GACGCGCCTACCGTATTCGGTCACTGCTCCTTCAGGCGAAGAGTATTAGCACGTTATCGT
GCTACTATCCTGAGCTGTAATCATAAGTATTTCTAAAGTTGTAAGGGTTGAAGAGTAGTT
CTGTACATCTTGATCGATTGTACTGGACGTCATTCCGAATAGCTTGTGGAATCCAATATC
TGGGTCATAGATGCT
>SRR9587949
GTGACCGTCGCCTCGCCACAAATCCATTGTTTTCCGGATTCTAGCCCCTTATGAGCTAGG
ACCTCTGTGCCTACCATCATAGCTGTTCGATGCCTTGTATTCAATACGTTGCACCGACCC
GACAATAGTCTGTGCCATTAAGCTTCTGCAAGTGGCTGGCGCCTGTAAGTCACGCAATAG
AGTAACTTCCCTCTCGACAGTCTTTATGCCATCCACAATCCAACATCGTAGATCCTCGCG
TGACCTAGTAGCCCTTGACGACTCAATCCACCCTCTGCCTGACGGACGTGCCTTTTATGG
TACATTAATGACTCCCTGTCCTCATAATTAGGCTGCACACACAGGGCTATCGTAGTTTAC
TGGTCTCACCCCCGCGTACTCGGCGACCCAGCTACACTGATCCAGGGAGGCAGAATGGTG
GACGCGCTTACTGCCGACCCGTGGGCAACTAACTGGGCCGGGGGAATATTGTTTCGTTTT
GGGTACTGCTTCTTCCAATTGTTCGGATAGCACAGTAACGGTGGTGGGTGGCTCCAGTCA
CTCTTGCGTAGTCGCCGGCCGGTGGCTCCGGTCTGTCCTCTCGCCCGATTCTGGGCCATC
TTATCCGATGTCACCCTGCAGACCGCTCTTTTCAACCATTAGCATTAAGCAGTCTCTCAT
AGGTGCTCTCATGTCATGAGAGGTACCGGAGCCTCTAAGCGTTCCGCTGTTCCCGACAAT
ATTCTCTTTCGCTTTGACACTAGCTCAGACGGTAGAGGCTTCGTCTTCAGCACCGGCTAA
TGATGCCTGGTACGTTAAGATACCGATCCGCGCGCTTACGCTGCTCCACCTGTCGGTGTC
TGAAGAAGCTCGGCTAAGACCTCTCGAACTCTTCGTCTTACCTCAGCCCGCGGCTACAGT
CATATACATCCTAGCACGCACAATCCTTAATTCCCAGTCGTGCCCGACAGTAACACCGTG
GCGCGAACAATGTGTAAGTTATTGCCTACTTCGTACACTATCAACTAAATTCGTCCAACG

TGAGATCCGAGAATCCCGAAAGGGGGCACACACTTAGAGAACCCGCAGCTTTTTGTCTCA
CGATCGGGAAGGGCACTCCCAAGAGGCCTTGATACGAGCGTGAAACCGGTCAAGCGCTCC
AATTGTACACTGTCCATGAGGCCAAATTCAAATTACGGACCGAAAACTGGATCCAGCAGA
GCTAAAGGTTATTCGCGGGTATTGTCCGTACATGCAACTTGATGTTTTCTAAGCCAAAAG
AGATATGCGAGGGGGAGACATGGGAAAAGCAAAAGAATATAGTACTATAATTGGCGTGGG
ACCCATAAAGTCTTGATTGCTCCCGCGCCGCCTGTACCCACAGTCTTCAAAAAAGCGATT
TGCCGGAGTATAAAGTCAGGGGCCGACGGTGTTGAAAACGAGCGGACGTCTTGCATGAGA
AGTGTCACTGTGGGCATATCAAAGGACTCTAATTAAATTCGGTGTCGATAGAAATGTTCG
TGCTGCACGGTGCTGCCAGACCATCCACCTTTCGTGGCTACGCCTTATCAGAAGTGCAAC
CTGCGAAATCAAGGTGACGTTAATACGCTCTAAAGGACGCTAAAGCGGGTTGATAATTAA
ACGACAGAGTTACAATTAGCGGGATTTACAACTCTCCAAAATTATTGACGATAGACTGAC
ACGGTTACGTGGGAGGACTCGTAGTCTCAAGTGAAAATGGGCGCAAAATGGTGTCCGCAT
GGGAGGAGCTCAATCATAATAGACGCCGAATATGTTCGCGTGAAACTTCAGCGTCAAAGA
TGGAAAGCAAGACAAACTCATGAAACTACCATAGTGTGAATATGATGGGCGGCGCGGTTT
AAGGCGAGTCGCCCTAAGACTCCAACGTGCCCTAAGGGATCGTAACATGCAAGAACATTG
ATGTGCACAGTTTTGTCGAACGGCCTTCATAGGCCAAAGATTGTATCCACGGCCAGAAGC
GGTAAGAACTTCGTGACATAGCGGCCAGAGAAAGGATCTACAACGTTAACCAGCACCTCA
GAGAGCGGGATGGCCAGTCGTTAGGGAGAAAATAACTAATCATTAACTTTGACGAACCAC
CAAAATGCTCGAGCGGTAGGTATCATTAGGGGGTACGGACTGTTACATCTAGAATACCCC
AGCTCCGTAGTTGCAGACACTAATTTTAGAGACCGCAGCTGTATCATTAACAAAATGTTA
TTGGTGGGATGCAGAAGAGCGCGGGCAAGATTATACAGACGCCTTCAGGCGGTAGCCTGC
TCCTAGGCAGTCAAGACGGATCGCGATGGACGTGTTTGGACGGCGAATGTACAGGAAATT
TCCCGTGAGAACAACAAAATGAAGGATCGAGATTAAAAAGACTGCAGCGCCTGTGAACAA
CGGAGTTGAACGTGGTTGCCAGGTTTGTGATGGGCGGCCCTAGAAGATAACGGTAGGAGC
CCTTCGGACTTTTCAATGGGGACTACATTCTGGCCGCGAAGGACTCCGTGAATCGATAGG
TAGTGGAAATATGGATGAAACGGGCTGGGGCAACGCTCGGGACAACGTTTACTAGCTAAG
AGGCAGATTACGAGGATCGGGGAAGACTGAGAAGGCCTTGGCGTAGGAATTCGAGGCAAC
CAGGGGGATAAGGTAAATGCCGACTTCAAGTCGAGAATGCAAAGAAGCTGGCTATGGCCG
TAAATTAGACGATAAATAACAGAGCATCGGGGAGCCAAGTCTAATTGTAAAACTTTTGGT
CGAGGTGATGTCGTAGGTGAGATCTCCGAACGCAGCATGAACAAGGCGAATCGTGTAAGC
AGAGAAGAGTGCAGCACGATGCTAACACTGCCGCAGCAGTGGGACAACTTGGATGACTGG
CACATCACAGTAATGAGTCATATGATTCCGCCTGAAGGCTTCAATAAAGGACATAAGATG
CGAAGAAGATCTAGACCCAGTCTTTAAAACTGCACAACTTAGCCGAAATGATTGGTAGGT
GAGAGTATCGACGTGGGCACTGAGAGAAACCTGAATACGCGATAGCCATCGTATCACGAT
GCTTCTGCCCTCCAGCGTCACTAGAGTTCTTGTTCACATGCCTCGGTCCTCTCGTCAGTA
CGTGGCCGGCCTTCTTTCCTCCATCGTTTCCCAGAGCATACTCTTCGAGTTGCCCTATTA

ACCTTCTATTTTCTGGCCGATGCCTGACTGGCGCCGCTTTGAGTATCATACTTCTTGTTT
TTGGCCAGCCTGTTGGGCGTTGATTGCTTGGGTAACCATATCCTGCATTTTGTTGATTCC
GGGTCGGCAATGTGTTGTGGACTGTTCGTCGCCGCTCCTGATCTCACCGTACTTCACGGT
TGTCGCCCGGACATTACCCGGACGGGTTTAGGCCATGGCCGGTTGAGGTTAATCTTTACG
CGTCACCATAACTTCCTGCGAAACTGGTTTCCCCACGCCGCTCTTTTCGTGAGCCGATCG
GCGTGGTTAGGTCGCGCGGGTTTTTCCAGGCAGGGTAAGTCGAGTCGCACAAGGTGACCC
ACGTACCTCGCTTTAACGTTCCGTTCTACCACTGGGTTAGGCTGCTTTGGGGCCGACTTA
ACGTTCTTTTTGCACGACCCCGTATGTGCTTATTCTCACTCGTAATGAACCGGTAAGCGA
GAGTTTGTTCTACGCTGAGGTGTCGTCGGACCCGAGTGTGCTTGCCGCTGGTCAATTGTT
GTGTCGGCTCGACGTCTTATTTCGTATGGTACGGGATTCTTAGTGCTTGCAGCGATGCCC
CTTTCGCTATTGATGTCTTCTCCTGAGTCACGCTACGTACTGAATTGATAAAACATTTCG
ACCTCAGTATCAGTGAGGTCGGGGGATGAGTTAGATCGCCGTGTAGGGCTTTTCACGTTG
TGTCCGTGTGGAGAGTCGGTTCTAGCGGTCGTATTATCCATATTCTCCCGTTCCATTTAC
GTTGAGCGACTTGAAACTTCCGATGTACTGGCCNNNAATATTGCATTGTTTCTCAATTCA
TGCAAGTTGTCGTACATTATGGAATAGAACAAGAATTAAGGACCATTTCCACTAGCGATT
CTTACCTCCAACTCAAGATAGAGTCTAACCGATTATCGACGAACTTCGTCCGTCACGTCA
TTCTATGTCGAGCGGCCTTCTTACAGCGCTGTCGCACTATCTGTTCTACCTTATTTCCGA
GAAACCTTGCTCCTGTTACCTGTGAGGGCGGGGCGGATATGGCAAAATGGGCATTCTTTT
TTACTTCCCGTGCCTCTTCATGCGATATACATGTATCTCTGGAGCCTAGACGCCCGGTCC
TTAACGTCCTAAAACTTTACTGACCCTGCTGGCGTCTTACCCTTTATAGTGGCTCTGCAT
AATGTGTTCTGTATATAGCCAAATACTGCCGACTATCCACTCGCTTAGCTGTCAATATCC
ATGCGCGTACTTTGATTTACCGGTCGACCTGCCTAATGTAACTGCCTTTCGCTCTCCACT
CCGCACCGGGGTTTCGAGTTTTTCTACCCCTCTCTTTATTACTACTCGTTTACTGCATCA
GACGCGCCTACCGTATTCAGTCACTGCTCCTTCAGGCGAAGAGTATTAGCACGTTATCGT
GCTACTATCCTGAGCTGTAATCATAAGTATTTCTAAAGTTGTAAGGGTTGAAGAGTAGTT
CTGTACATCTTGATCGATTGTACTGGACGTCATTCCGAATAGCTTGTGGAATCCAATATC
TGGGTCATAGATGCT
>SRR9587950
GTGACCGTCGCCTCGCCACAAATCCATTGTTTTCCGGATTCTAGCCCCTTATGAGCTAGG
ACCTCTGTGCCTACCATCATAGCTGTTCGATGCCTTGTATTCAATACGTTGCACCGACCC
GACAATAGTCTGTGCCATTAAGCTTCTGCAAGTGGCTGGCGCCTGTAAGTCACGCAATAG
AGTAACTTCCCTCTCGACAGTCTTTATGCCATCCACAATCCAACATCGTAGATCCTCGCG
TGACCTAGTAGCCCTTGACGACTCAATCCACCCTCTGCCTGACGGACGTGCCTTTTATGG
TACATTAATGACTCCCTGTCCTCATAATTAGGCTGCACACACAGGGCTATCGTAGTTTAC
TGGTCTCACCCCCGCGTACTCGGCGACCCAGCTACACTGATCCAGGGAGGCAGAATGGTG
GACGCGCTTACTGCCNACCCGTGGGCAACTAACTGGGCCGGGGGAATATTGTTTCGTTTT

GGGTACTGCTTCTTCCAATTGTTCGGATAGCACAGTAACGGTGGTGGGTGGCTCCAGTCA
CTCTTGCGTAGTCGCCGGCCGGTGGCTCCGGTCTGTCCTCTCGCCCGATTCTGGGCCATC
TTATCCGATGTCACCCTGCAGACCGCTCTTTTNAACCATTAGCATTAAGCAGTCTCTCAT
AGGTGCTCTCATGTCATGAGAGGTACCGGAGCCTCTAAGCGTTCCGCTGTTCCCGACAAT
ATTCTCTTTCGCTTTGACACTAGCTCAGACGGTAGAGGCTTCGTCTTCAGCACCGGCTAA
TGATGCCTGGTACGTTAAGATACCGATCCGCGCGCTTACGCTGCTCCACCTGTCGGTGTC
TGAAGAAGCTCGGCTAAGACCTCTCGAACTCTTCGTCTTACCTCAGCCCGCGGCTACAGT
CATATACATCCTAGCACGCACAATCCTTAATTCCCAGTCGTGCCCGACAGTAACACCGTG
GCGCGAACAATGNGTAAGTTATTGCCTACTTCGTACACTATCAACTAAATTCGTCCAACG
TGAGATCCGAGAATCCCGAAAGGGGGCACACACTTAGAGAACCCGCAGCTTTTTGTCTCA
CGATCGGGAAGGGCACTCCCAAGAGGCCTTGATACGAGCNTGAAACCGGTCAAGCGCTCC
AATTGTACACTGTCCATGAGGCCAAATTCAAATTACGGACCGAAAACTGGATCCAGCAGA
GCTAAAGGTTATTCGCGGGTATTGTCCGTACATGCAACTTGATGTTTTCTAAGCCAAAAG
AGATATGCGAGGGGGAGACATGGGAAAAGCAAAAGAATATAGTACTATAATTGGCGTGGG
ACCCATAAAGTCTTGATTGCTCCCGCGCCGCCTGTACCCACAGTCTTCAAAAAAGCGATT
TGCCGGAGTATAAAGTCAGGGGCCGACGGTGTTGAAAACGAGCGGACGTCTTGCATGAGA
AGTGTCACTGTGGGCATATCAAAGGACTCTAATTAAATTCGGTGTCGATAGAAATGTTCG
TGCTGCACGGTGCTGCCAGACCATCCACCTTTCGTGGCTACGCCTTATCAGAAGTGCAAC
CTGCGAAATCAAGGTGACGTTAATACGCTCTAAAGGACGCTAAAGCGGGTTGATAATTAA
ACGACAGAGTTACAATTAGCGGGATTTACAACTCTCCAAAATTATTGACGATAGACTGAC
ACGGTTACGTGGGAGGACTCGTAGTCTCAAGTGAAAATGGGCGCAAAATGGTGTCCGCAT
GGGAGGAGCTCAATCATAATAGACGCCGAATATGTTCGCGTGAAACTTCAGCGTCAAAGA
TGGAAAGCAAGACAAACTCATGAAACTACCATAGTGTGAATATGATGGGCGGCGCGGTTT
AAGGCGAGTCGCCCTAAGACTCCAACGTGCCCTAAGGGATCGTAACATGCAAGAACATTG
ATGTGCACAGTTTTGTCGAACGGCCTTCATAGGCCAAAGATTGTATCCACGNCCAGAAGC
GGTAAGAACTTCGTGACATAGCGGCCAGAGAAAGGATCTACAACGTTAACCAGCACCTCA
GAGAGCGGGATGGCCAGTCGTTAGGGAGAAAATAACTAATCATTAACTTTGACGAACCAC
CAAAATGCTCGAGCGGTAGGTATCATTAGGGGGTACGGACTGTTACATCTAGAATACCCC
AGCTCCGTAGTTGCAGACACTAATTTTAGAGACCGCAGCTGTATCATTAACAAAATGTTA
TTGGTGGGATGCAGAAGAGCGCGGGCAAGATTATACAGACGCCTTCAGGCGGTAGCCTGC
TCCTAGGCAGTCAAGACGGATCGCGATGGACGTGTTTGGACGGCGAATGTACAGGAAATT
TCCCGTGAGAACAACAAAATGAAGGATCGAGATTAAAAAGACTGCAGCGCCTGTGAACAA
CGGAGTTGAACGTGGTTGCCAGGTTTGTGATGGGCGGCCCTAGAAGATAACGGTAGGAGC
CCTTCGGACTTTTCAATGGGGACTACATTCTGGCCGCGAAGGACTCCGTGAATCGATAGG
TAGTGGAAATATGGATGAAACGGGCTGGGGCAACGCTCGGGACAACGTTTACTAGCTAAG
AGGCAGATTACGAGGATCGGGGAAGACTGAGAAGGCCTTGGCGTAGGAATTCGAGGCAAC

CAGGGGGATAAGGTAAATGCCGACTTCAAGTCGAGAATGCAAAGAAGCTGGCTATGGCCG
TAAATTAGACGATAAATAACAGAGCATCGGGGAGCCAAGTCTAATTGTAAAACTTTTGGT
CGAGGTGATGTCGTAGGTGAGATCTCCGAACGCAGCATGAACAAGGCGAATCGTGTAAGC
AGAGAAGAGTGCAGCACGATGCTAACACTGCCGCAGCAGTGGGACAACTTGGATGACTGG
CACATCACAGTAATGAGTCATATGATTCCGCCTGAAGGCTTCAATAAAGGACATAAGATG
CGAAGAAGATCTAGACCCAGTCTTTAAAACTGCACAACTTAGCCGAAATGATTGGTAGNT
GAGAGTATCGACGTGGGCACTGAGAGAAACCTGAATACGCNATAGCCATCGTATCACGAT
GCTTCTGCCCTCCAGCGTCACTAGAGTTCTTGTTCACATGCCTCGGTCCTCTCGTCAGTA
CGTGGCCGGCCTTCTTTCCTCCATCGTTTCCCAGAGCATACTCTTCGAGTTGCCCTATTA
ACCTTCTATTTTCTGGCCGATGCCTGACTGGCGCCGCTTTGAGTATCATACTTCTTGTTT
TTGGCCAGCCTGTTGGGCGTTGATTGCTTGGGTAACCATATCCTGCATTTTGTTGATTCC
GGGTCGGCAATGTGTTGTGGACTGTTCGTCGCCGCTCCTGATCTCACCGTACTTCACGGT
TGTCGCCCGGACATTACCCGGACGGGTTTAGGCCATGGCCGGTTGAGGTTAATCTTTACG
CGTCACCATAACTTCCTGCGAAACTGGTTTCCCCACGCCGCTCTTTTCGTGAGCCGATCG
GCGTGGTTAGGTCGCGCGGGTTTTTCCAGGCAGGGTAAGTCGAGTCGCACAAGGTGACCC
ACGTACCTCGCTTTAACGTTCCGTTCTACCACTGGGTTAGGCTGCTTTGGGGCCGACTTA
ACGTTCTTTTTGCACGACCCCGTATGTGCTTATTCTCACTCGTAATGAACCGGTAAGCGA
GAGTTTGTTCTACGCTGAGGTGTCGTCGGACCCGAGTGTGCTTGCCGCTGGTCAATTGTT
GTGTCGGCTCGACGTCTTATTTCGTATGGTACGGGATTCTTANTGCTTGCAGCGATGCCC
CTTTCGCTATTGATGTCTTCTCCTGAGTCACGCTACGTACTGAATTGATAAAACATTTCG
ACCTCAGTATCAGTGAGGTCGGGGGATGAGTTAGATCGCCGTGTAGGGCTTTTCACGTTG
TGTCCGTGTGGAGAGTCGGTTCTAGCGGTCGTATTATCCATATTCTCCCGTTCCATTTAC
GTTGAGNGACTTGAAACTTCCGATGTACTGGCCGGTAATATTGCATTGTTTCTCAATTCA
TGCAAGTTGTCGTACATTATGGAATAGAACAAGAATTAAGGACCATTTCCACTAGCGATT
CTTACCTCCAACTCAAGATAGAGTCTAACCGATTATCGACGAACTTCGTCCGTCACGTCA
TTCTATGTCGAGCGGCCTTCTTACAGCGCTGTCGCACTATCTGTTCTACCTTATTTCCGA
GAAACCTTGCTCCTGTTACCTGTGAGGGCGGGGCGGATATGGCAAAATGGGCATTCTTTT
TTACTTCCCGTGCCTCTTCATGCGATATACATGTATCTCTGGAGCCTAGACGCCCGGTCC
TTAACGTCCTAAAACTTTACTGACCCTGCTGGCGTCTTACCCTTTATAGTGGCTCTGCAT
AATGTGTTCTGTATATAGCCAAATACTGCCGACTATCCACTCGCTTAGCTGTCAATATCC
ATGCGCGTACTTTGATTTACCGGTCGACCTGCCTAATGTAACTGCCTTTCGCTCTCCACT
CCGCACCGGGGTTTCGAGTTTTTCTACCCCTCTCTTTATTACTACTCGTTTACTGCATCA
GACGCGCCTACCGTATTCAGTCACTGCTCCTTCAGGCGAAGAGTATTAGCACGTTATCGT
GCTACTATCCTGAGCTGTAATCATAAGTATTTCTAAAGTTGTAAGGGTTGAAGAGTAGTT
CTGTACATCTTGATCGATTGTACTGGACGTCATTCCGAATAGCTTGTGGAATCCAATATC
TGGGTCATAGATGCT

>SRR9587951

GTGACCGTCGCCTCGCCACAAATCCATTGTTTTCCGGGTTCTAGCCCCTTATGAGCTAGG
ACCTCTGTGCCTACCATCATAGCTGTTCGATGCCTTGTATTCAATACGTTGCACCGACCC
GACAATAGTCTGTGCCATTAAGCTTCTGCAAGTAGCTGGCGCCTGTAAGTCACGCAATAG
AGTAACTTCCCTCTCGACAGTCTTTATGCCATCCACAATCCAACATCGTAGATCCTCACG
TGACCTAGTAGCCCTTGACGACTCAATCCACCCTCTGCCCGACGGACGTGCCTTTTATGG
TACATTAATGACTCCCTGTCCTCATAATTAGGCTGCACACACAGGGCTATCGTAGTTTAC
TGGTCTCACCCCTGCGTACTCGGCGACCCAACTACACTGATCCAGGGAGGCAGAATGGTG
GACTCGCTTACTCCCGACCCGTGGGCAACTAACTGGGCCGGGGGAATATTGTTTCGTTTT
GGGTACTGCTTCTTCCAATTGTTCGGATAGCACAGTAACGGTGGTGGGTGGCTCCAGTCA
CTCTTGCGTAGTCGCCGGCCGGTCGCTCCGGTCTGTCCTCTCGCCCGATTCTGGGCCATC
TTATCCGATGTCACCCTGCAGACCGCTCTTTTCAACCATTAGCATTAAGCAGTCTCTCAT
AGGTGCTCTCATGTCATGAGAGGTACCGGAGCCTCTAAGCGTTCCGCTGTTCCCCACAAT
ATTCTCTTTCGCTTTGACACTAGCTCAGACGGTAGAGGCTTCGTCTTCAGCACCGGCTAA
TGATGCCTGGTACGTTAAGATACCGATTCGCGCGCTTACGCTGCTCCACCTGTCGGTGTC
TGAAGAAGCTCGGCTAAGACCTCTCGAACTCTTCGTCTTACCTCAGCCCGCGGCTACAGT
CATATACATCCTAGCACGCACAATCCTTAATTCTCAGTCGTGCCCGACAGTAACACCGTG
GCGCGAACAATGTGTAAGTTATTGCCTACTTCGTACACTATCAACTAAATTCGTCCAACG
TGAGATCCGAGAATCCCGNAAGGGGGCACACACTTAGAGAACCCGCAGCTTTTTGTCTCA
CGATCGAGGAGGGCACTCCCAAGAGGCCTTGATACGAGCGTGAAACCGGTCAAGCGCTCC
AATTGTACACTGTCCATGAGGCCAAATTCAAATTACGGACCGAAAACTGGATCCAGCAGA
GCTAAAGGTTATTCGCGGGTATTGTCCGTACATGCAACTTGATGTTTTCTAAGCCAAAAG
AGATATGCGAGGGGGAGACATGGGAAAAGCAAAGGAATATAGTACTATAATTGGCGTGGG
ACCCATAAAGTCTTGATTGCTCCCGCGCCGCCTGTGCCCACAGTCTTCAGAAAAGCGATT
TGCCGGAGTATAAAGTCAGGGGCCGACGGTGTTGAAAACGAGCGGACGTCTTGCATGAGA
AGTGTCACTGTGGGCATATCAAAGGACTCTAATTAAATTCAGTGTCAATAGAAATGTTCG
TGCTGCACGGTGCTGTCAGACCATCCACCTTTCGTGGCTACGCCTTATCAGAAGTGCAAC
CTGCGAAATCAAGGTGACGTTAATACGCTCTAAAGGACGCTAAAGCGGGTTGATAATTAA
ACGACAGAGTTACAATTAGCGGGATTCACAACTCTCCAAAATCATTGACAATAGACTGAC
ACGGTTACGTGGGAGGACTCGTAGTCTCAAGTGAAAATGGGCGCAAAATGGTGTCCGCAT
GGGAGGAGCTCAACCATAATAGACGCCGAATATGTTCGCGTGAAACTTCAGAGTCAAAGA
TGGAAAGCAAGACAAACTCATGAAACTACCATAGTGTGAAGATGATGGGCGGCGCGGTTT
AAGGCGAGTCGCCCTAAGACCCCAACGTGCCCTAAGGGATCGTAACATGCAAGAACATTG
ATGTGCACAGTTTTGTCGAACGGCCTTCATAGGCCAAAGATTGTATCCACGGCCAGAAGC
GGTAAGAACTTCGTGACATAGCGGCCAGAGAAAGGATCTACAACGTTAACCAGCACCTCA
GAGAGCGGGATGGCCAGTCGTTAGGGAGAAAATAACTAATCATTAACTTTGACGAACCAC

CAAAATGCTCGAGCGGTAGGTATCATTAGGGGGTACGGACTGTTACATCTAGAATACCCC
AGCTCCGTAGTTGCAGACACTAATTTTAGAGACCGCAGCTGTATCATTAACAAAATGTTA
TTGGTGGGATGCAGAAGAGCGCGGGCAAGATTATACAGACGCCTTCAGGCGGTAGCCTGC
TCCTAGGCAGTCAAGACGGATCGCGATGGACGTGTTTGGACGGCGAATGTACAGGAAATT
TCCCGTGAGAACAACAAAATGAAGAATCGAGATTAAAAAGACTGCAGCGCCCGTGAACAA
CGGAGTTGAACGTGGTTGCCAGGTTTGTGATGGGCNGCCCTAGAAGATAACGGTAGAAGC
CCTTCGGACTTTTCAATGGGGACTACATTCTGGCCGCGAAGGACTCCGTGAATCGATAGG
TAGTGGAAATATGGATGAAACGGGCTGGGGCAACGCTCGGGACAAAGTTTACTAGCTAAG
AGGCAGATTACGAGGATCGGGGAAGACTGAGAAGGCCTTGGCGTAGGAATTCGAGGCAAC
CAGGGGGATAAGGTAAATGCCGACTTCAAGTCGAGAATGAAAAGAAGCTAGCTATGGCCG
TAAATTAGACGATAAATAACAGAGCATCGGGGAGCCAAGTCTAATTGTAAAACTTTTGGT
CGAGGTGATGTCGTAGGTGAGATCTCCGAACGCAGCATGAACAAGGCGAATCGTGTAAGC
AGAGAAGAGTGCAGCACGATGCTAACACTGCCGCAGCAGTGGGACAACTTGGATGACTGG
CACATCACAGTAATGAGTCATATGATTCCGCCTGAAGGCTTCAATAAAGGACATAAGATG
CGAAGAAGATCTAGACCCAGTCTTTAAAACTGCACAACTTAGCCGAAATGATTGGTAGGT
GAGAGTATCGACGTGGGCACTGAGAGAAACCTGAATACGCGATAGCCATCACATCACGAT
GCTTCTGCCCTCCAGCGTCACTAGAGTTCTTGTTCACATGCCTCGGTCCTCTCGTCAGTA
CATGGCCGGCCTTCTTTCCTCCATCGTTTCCCAGAGCATACTCCTCGAGTTGCCCTATTA
ACCTTCTATTTTCTGGCCGATGCCTGACTGGCGCCGCTTTGAGTATCATACTTCTTGTTT
TTGGCCAACCTGTTGGGCGTTGATTGCTTGGGTAACCACATCCTGCATTTTGTTGATTCC
GGGTCGGCAATGTGTTGTGGACTGTTCGTCGCCGCTCCTGATCTCACCGTACTTCACGGT
TGTCGCCCGGACATCACCCGGACGGGTTTAGGCCATGGCCGGTTGAGGTTAATCTTTACG
CGTCACCATAACTTCCTGCGAAACTGGTTTCCCCACACCGCTCCCTTCGTGAGCCGATCG
GCGTGGTTAGGTCGCGCGGGTTTTTCCAGGCAGGGTAAGTCGAGTCGCACAAGGTGACCC
ACGTACCTCGCTTTAACGTTCCGTTCTACCACTGGGTTAGGCTGCTTTGGGGCCGACTTG
ACGTTCTTTTTACACGACCTCGTATGTGCTTATTCTCACTCGTAATGAACCGGTAAGCGA
GAGTTTGTTCTACGCTGAGGTGTCGTCGGACCCGAGTGTGCTTGCCGCTGGTCAATTGTT
GTGTCGGCTCGACGTCTTATTTCGTATGGTACGGGATTATTAGTGCTTGCAGCGATGTCC
CTTTCGCTATTGATGTCCTCTCCTGAGTCACGCTACGTACTGAATTGATAAAACATTTCG
ACCTCAGTATCAGTGAGGTCGGGGGATGAATTAGATCGCCGTGTAGGGCTTTTCACGTTG
TGTCCGTGTGGAGAGTCGGTTCTAGCGGTCGTATTATCCATATTCTCCCGTTCCATTTAC
GTTGAGCGACTTTAAACTTCCGATGTACTGGCCNNNAATAGTGCATTGTTTCTCAATTCA
TGCAAGTTGTCGTACATCATGGAATAGAACAAGAATTAAGGACCATTTCCACTAGCGATT
CTTACCTCCAACTCAAGATAGAGTCTAACCGATTATCGACGAACTTCGTCCGTCACGTCA
TTCTATGTCGAGCGGCCTTCTTACAGCGCTGTCGCACTATCTGTTCTACCTTATTTCCGA
GAAACCTTGCTCCTGTTACCTGTGAGGGCGGGGCGGATATGGCAAAATGGGCATTCTTTT

TTACTTCCCGTGCCTCTTCATACTATATACATGTATCTCTGGCGCCTAGACGCCCGGTCC
TTAACGTCCTAAAACTTTACTGACCCTGCTGGCGTCTTACCCTTTATAGTGGCTCTGCAT
AATGTGTTCTGTATATAGCCAAATACTGCCGACTATCCACTCGCTTAGCTGTCAATATCC
ATGCGCGTACTTTGATTTACCGGTCGACCTGCCTAATGTAACTGCCTTTCGCTCTCCACT
CCGCACCGGGGTTTCGAGTTTTTCTACCCCTCTCTTTATTACTACTCGTTTACTGCATCA
GACGCGCCTACCGTATTCAATCACTGCTCCTTCAGGCGAAGAGTATTAGCACGTTATCGT
GCTACTATCCTGAGCTGTAATCATAAGTATTTCTAAAGTTGTAAGGGTTGAAGAGTAGTT
CTGTACATCTTGATCGATTGTACTGGACGTCATTCCGAATAGCTTGTGGAATCCAATATC
TGGGTCATAGATGCT
>SRR9587952
GTGACCGTCGCCTCGCCACAAATCCATTGTTTTCCGGATTCTAGCCCCTTATGAGCTAGG
ACCTCTGTGCCTACCATCATAGCTGTTCGATGCCTTGTATTCAATACGTTGCACCGACCC
GACAATAGTCTGTGCCATTAAGCTTCTGCAAGTGGCTGGCGCCTGTAAGTCACGCAATAG
AGTAACTTCCCTCTCGACAGTCTTTATGCCATCCACAATCCAACATCGTAGATCCTCGCG
TGACCTAGTAGCCCTTGACGACTCAATCCACCCTCTGCCTGACGGACGTGCCTTTTATGG
TACATTAATGACTCCCTGTCCTCATAATTAGGCTGCACACACAGGGCTATCGTAGTTTAC
TGGTCTCACCCCCGCGTACTCGGCGACCCAGCTACACTGATCCAGGGAGGCAGAATGGTG
GACGCGCTTACTGCCGACCCGTGGGCAACTAACTGGGCCGGGGGAATATTGTTTCGTTTT
GGGTACTGCTTCTTCCAATTGTTCGGATAGCACAGTAACGGTGGTGGGTGGCTCCAGTCA
CTCTTGCGTAGTCGCCGGCCGGTGGCTCCGGTCTGTCCTCTCGCCCGATTCTGGGCCATC
TTATCCGATGTCACCCTGCAGACCGCTCTTTTCAACCATTAGCATTAAGCAGTCTCTCAT
AGGTGCTCTCATGTCATGAGAGGTACCGGAGCCTCTAAGCGTTCCGCTGTTCCCGACAAT
ATTCTCTTTCGCTTTGACACTAGCTCAGACGGTAGAGGCTTCGTCTTCAGCACCGGCTAA
TGATGCCTGGTACGTTAAGATACCGATCCGCGCGCTTACGCTGCTCCACCTGTCGGTGTC
TGAAGAAGCTCGGCTAAGACCTCTCGAACTCTTCGTCTTACCTCAGCCCGCGGCTACAGT
CATATACATCCTAGCACGCACAATCCTTAATTCCCAGTCGTGCCCGACAGTAACACCGTG
GCGCGAACAATGTGTAAGTTATTGCCTACTTCGTACACTATCAACTAAATTCGTCCAACG
TGAGATCCGAGAATCCCGNAAGGGGGCACACACTTAGAGAACCCGCAGCTTTTTGTCTCA
CGATCGGGAAGGGCACTCCCAAGAGGCCTTGATACGAGCGTGAAACCGGTCAAGCGCTCC
AATTGTACACTGTCCATGAGGCCAAATTCAAATTACGGACCGAAAACTGGATCCAGCAGA
GCTAAAGGTTATTCGCGGGTATTGTCCGTACATGCAACTTGATGTTTTCTAAGCCAAAAG
AGATATGCGAGGGGGAGACATGGGAAAAGCAAAAGAATATAGTACTATAATTGGCGTGGG
ACCCATAAAGTCTTGATTGCTCCCGCGCCGCCTGTACCCACAGTCTTCAAAAAAGCGATT
TGCCGGAGTATAAAGTCAGGGGCCGACGGTGTTGAAAACGAGCGGACGTCTTGCATGAGA
AGTGTCACTGTGGGCATATCAAAGGACTCTAATTAAATTCGGTGTCGATAGAAATGTTCG
TGCTGCACGGTGCTGCCAGACCATCCACCTTTCGTGGCTACGCCTTATCAGAAGTGCAAC

CTGCGAAATCAAGGTGACGTTAATACGCTCTAAAGGACGCTAAAGCGGGTTGATAATTAA
ACGACAGAGTTACAATTAGCGGGATTTACAACTCTCCAAAATTATTGACGATAGACTGAC
ACGGTTACGTGGGAGGACTCGTAGTCTCAAGTGAAAATGGGCGCAAAATGGTGTCCGCAT
GGGAGGAGCTCAATCATAATAGACGCCGAATATGTTCGCGTGAAACTTCAGCGTCAAAGA
TGGAAAGCAAGACAAACTCATGAAACTACCATAGTGTGAATATGATGGGCGGCGCGGTTT
AAGGCGAGTCGCCCTAAGACTCCAACGTGCCCTAAGGGATCGTAACATGCAAGAACATTG
ATGTGCACAGTTTTGTCGAACGGCCTTCATAGGCCAAAGATTGTATCCACGGCCAGAAGC
GGTAAGAACTTCGTGACATAGCGGCCAGAGAAAGGATCTACAACGTTAACCAGCACCTCA
GAGAGCGGGATGGCCAGTCGTTAGGGAGAAAATAACTAATCATTAACTTTGACGAACCAC
CAAAATGCTCGAGCGGTAGGTATCATTAGGGGGTACGGACTGTTACATCTAGAATACCCC
AGCTCCGTAGTTGCAGACACTAATTTTAGAGACCGCAGCTGTATCATTAACAAAATGTTA
TTGGTGGGATGCAGAAGAGCGCGGGCAAGATTATACAGACGCCTTCAGGCGGTAGCCTGC
TCCTAGGCAGTCAAGACGGATCGCGATGGACGTGTTTGGACGGCGAATGTACAGGAAATT
TCCCGTGAGAACAACAAAATGAAGGATCGAGATTAAAAAGACTGCAGCGCCTGTGAACAA
CGGAGTTGAACGTGGTTGCCAGGTTTGTGATGGGCGGCCCTAGAAGATAACGGTAGGAGC
CCTTCGGACTTTTCAATGGGGACTACATTCTGGCCGCGAAGGACTCCGTGAATCGATAGG
TAGTGGAAATATGGATGAAACGGGCTGGGGCAACGCTCGGGACAACGTTTACTAGCTAAG
AGGCAGATTACGAGGATCGGGGAAGACTGAGAAGGCCTTGGCGTAGGAATTCGAGGCAAC
CAGGGGGATAAGGTAAATGCCGACTTCAAGTCGAGAATGCAAAGAAGCTGGCTATGGCCG
TAAATTAGACGATAAATAACAGAGCATCGGGGAGCCAAGTCTAATTGTAAAACTTTTGGT
CGAGGTGATGTCGTAGGTGAGATCTCCGAACGCAGCATGAACAAGGCGAATCGTGTAAGC
AGAGAAGAGTGCAGCACGATGCTAACACTGCCGCAGCAGTGGGACAACTTGGATGACTGG
CACATCACAGTAATGAGTCATATGATTCCGCCTGAAGGCTTCAATAAAGGACATAAGATG
CGAAGAAGATCTAGACCCAGTCTTTAAAACTGCACAACTTAGCCGAAATGATTGGTAGGT
GAGAGTATCGACGTGGGCACTGAGAGAAACCTGAATACGCGATAGCCATCGTATCACGAT
GCTTCTGCCCTCCAGCGTCACTAGAGTTCTTGTTCACATGCCTCGGTCCTCTCGTCAGTA
CGTGGCCGGCCTTCTTTCCTCCATCGTTTCCCAGAGCATACTCTTCGAGTTGCCCTATTA
ACCTTCTATTTTCTGGCCGATGCCTGACTGGCGCCGCTTTGAGTATCATACTTCTTGTTT
TTGGCCAGCCTGTTGGGCGTTGATTGCTTGGGTAACCATATCCTGCATTTTGTTGATTCC
GGGTCGGCAATGTGTTGTGGACTGTTCGTCGCCGCTCCTGATCTCACCGTACTTCACGGT
TGTCGCCCGGACATTACCCGGACGGGTTTAGGCCATGGCCGGTTGAGGTTAATCTTTACG
CGTCACCATAACTTCCTGCGAAACTGGTTTCCCCACGCCGCTCTTTTCGTGAGCCGATCG
GCGTGGTTAGGTCGCGCGGGTTTTTCCAGGCAGGGTAAGTCGAGTCGCACAAGGTGACCC
ACGTACCTCGCTTTAACGTTCCGTTCTACCACTGGGTTAGGCTGCTTTGGGGCCGACTTA
ACGTTCTTTTTGCACGACCCCGTATGTGCTTATTCTCACTCGTAATGAACCGGTAAGCGA
GAGTTTGTTCTACGCTGAGGTGTCGTCGGACCCGAGTGTGCTTGCCGCTGGTCAATTGTT

GTGTCGGCTCGACGTCTTATTTCGTATGGTACGGGATTCTTAGTGCTTGCAGCGATGCCC
CTTTCGCTATTGATGTCTTCTCCTGAGTCACGCTACGTACTGAATTGATAAAACATTTCG
ACCTCAGTATCAGTGAGGTCGGGGGATGAGTTAGATCGCCGTGTAGGGCTTTTCACGTTG
TGTCCGTGTGGAGAGTCGGTTCTAGCGGTCGTATTATCCATATTCTCCCGTTCCATTTAC
GTTGAGCGACTTGAAACTTCCGATGTACTGGCCGGTAATATTGCATTGTTTCTCAATTCA
TGCAAGTTGTCGTACATTATGGAATAGAACAAGAATTAAGGACCATTTCCACTAGCGATT
CTTACCTCCAACTCAAGATAGAGTCTAACCGATTATCGACGAACTTCGTCCGTCACGTCA
TTCTATGTCGAGCGGCCTTCTTACAGCGCTGTCGCACTATCTGTTCTACCTTATTTCCGA
GAAACCTTGCTCCTGTTACCTGTGAGGGCGGGGCGGATATGGCAAAATGGGCATTCTTTT
TTACTTCCCGTGCCTCTTCATGCGATATACATGTATCTCTGGAGCCTAGACGCCCGGTCC
TTAACGTCCTAAAACTTTACTGACCCTGCTGGCGTCTTACCCTTTATAGTGGCTCTGCAT
AATGTGTTCTGTATATAGCCAAATACTGCCGACTATCCACTCGCTTAGCTGTCAATATCC
ATGCGCGTACTTTGATTTACCGGTCGACCTGCCTAATGTAACTGCCTTTCGCTCTCCACT
CCGCACCGGGGTTTCGAGTTTTTCTACCCCTCTCTTTATTACTACTCGTTTACTGCATCA
GACGCGCCTACCGTATTCAGTCACTGCTCCTTCAGGCGAAGAGTATTAGCACGTTATCGT
GCTACTATCCTGAGCTGTAATCATAAGTATTTCTAAAGTTGTAAGGGTTGAAGAGTAGTT
CTGTACATCTTGATCGATTGTACTGGACGTCATTCCGAATAGCTTGTGGAATCCAATATC
TGGGTCATAGATGCT
>SRR9587953
GTGACCGTCGCCTCGCCACAAATCCATTGTTTTCCGGGTTCTAGCCCCTTATGAGCTAGG
ACCTCTGTGCCTACCATCATAGCTGTTCGATGCCTTGTATTCAATACGTTGCACCGACCC
GACAATAATCTGTGCCATTGAGCTTCTGCAAGTGGCTGGCGCCTGTAAGTCACGCAATAG
AGTAACTTCCCTCTCGACAGTCTTTATGCCATCCACAATCCAACATCGTAGATCCTCGCG
TGACCTAGTAGCCCTTGACGACTCAATCCACCCTCTGCCTGACGGACGTGCCTTTTATGG
TACATTAATGACTCCCTGTCCTCATAATTAGGCTGCACACACAGGGCTATCGTAGTTTAC
TGGTCTCACCCCTGCGTACTCGGCGACCCAACTACACTGATCCAGGGAGGCAGAATGGTG
GACGCGCTTAGTGCCGACCCGTGGGCAACTAACTGGGCCGGGGGAATATTGTTTCGTTTT
GGGTACTGCTTCTTCCAATTGTTCGGATAGCACAGTAACGGTGGTGGGTGGCTCCAGTCA
CTCTTGCGTAGTCCCCGGCCGGTCGCTCCGGTCTGTCCTCTCGCCCGATTCTGGGCCAAC
TTATCCGATGTCACCCTGCAGACCGCTCTTTTCAACTATTAGCATTAAGCAGTCTCTCAT
AGGTGCTCTCATGTCATGAGAGGTACCGGAGCCTCTAAGCGTTCCGCTGTTCCCGACAAT
ATTCTCTTTCGCTTTGACACTAGCTCAGACGGTAGAGGCTTCGTCTTCAGCACCGGCTAA
TGATGCCTGGTACGTTAAGATACCGATCCGCGCGCTTACGCTGCTCCACCTGTCGGTGTC
TGAAGAAGCTCGGCTAAGACCTCTCGAACTCTTCGTCTTACCTCAGCCCGCGGCTACAGT
CNAATACATCCTAGCACGCACAATCCTTAATTCCCAGTCGTGCCCGACAGTAACACCGTG
GCGCGAACAATGTGTAAGTTATTGCCTACTTCGTACACTATCAACTAAATTCGTCCAACG

TGAGATCCGAGAATCCCGAAAGGGGGCACACACTTAGAGAACCCGCAGCTTTTTGTCTCA
CGATCGAGAAGGGCACTCCCAAGAGGCCTTGATACAAGCGTGAAACCGGTCAAGCGCTCC
AATTGTACACTGTCCATGAGGCCAAATTCAAATTACGGACCGAAAACTGGATCCAGCAGA
GCTTAAGGTTATTCGCGGGTATTGTCCGTACATGCAACTTGATGTTTTCTAAGCCAAAAG
AGATATGCGAGGGGGAGACATGGGAAAAGCAAAAGAATATAGTACTATAATGGGCGTGGG
ACCCATAAAGTCTTGATTGCTCCCGCGCCGCCTGTACCCACAGTCTTCAAAAAAGCGATT
TGCCGGAGTATAAAGTCAGGGGCCGACGGTGTTGAAAACGAGCGGACGTCTTGCATGAGA
AGTGTCACTGTGGGCATATCAAAGGACTCTAATTAAATTCGGTGTCAATAGAAATGTTCG
TGCTGCACGGTGGTGTCAGACCGTCCACCTTTCGTGGCTACGCCTTATCAGAAGTGCAAC
CTGCGAAATCAAGGTGACGTTAATACGCTCTAAAGGACGCTAAAGCGGGTTGATAATTAA
ACGACAGAGTTACAATTAGCGGGATTCACAACTCTCCAAAATCATTGACGATAGACTGAC
ACGGTTACGTGGGAGGACTCGTAGTCTCAAGTGAAAATGGGCGCAAAATGGTGTCCGCAT
GGGAGGAGCTCAACCATAATAGACGCCGAATATGTTCGCGTGAAACTTCAGCGTCAAAGA
TGGAAAGCAAGACAAATTCATGAAACTACCATAGTGTGAAGATGATGGGCGGCGCGGTTT
AAGGCGAGTCGCCCTAAGACTCCAACGTGCCCTAAGAGATCGTAACATGCAAGAACATTG
ATGTGCACAGTTTTGTCGAACGGCCTTCATAGGCCAAAGATTGTATCCACGGCCAGAAGC
GGTAAGAACTTCGTGACATAGCGGCCAGAGAAAGGATCTACAACGTTAACCAGCACCTCA
GAGAGCGGGATGGCCAGTCGTTAGGGAGAAAATAACTAATCATTAACTTTGACGAACCAC
CAAAATGCTCGAGCGGTAGGTATCATTAGGGGGTACGGACTGTTACATCTAGAATACCCC
AGCTCCGTAGTTGCAGACACTAATTTTAGAGACCGCAGCTGTATCATTAACAAAATGTTA
TTGGTGGGATGCAGAAGAGCGCGGGCAAGATTATACAGACGCCTTCAGGCGGTAGCCTGC
TCCTAGGCAGTCAAGACGGATCGCGATGGACGTGTTTGGACGGCGAATGTACAGGAGATT
TCCCGTGAGAACAACAAAATGAAGGATCGAGATTAAAAAGACTGCAGCGCCCGTGAACAA
CGGAGTTGAACGTGGTTGCCAGGTTTGTGATGGGCGGCCCTAGAAGATAACGGTAGAAGC
CCTTCGGACTTTTCAATGGGGACTACATTCTGGCCGCGAAGGACTCCGTGAATCGATAGG
TAGTGGAAATATGGATGAAACGGGCTGGGGCAACGCTCGGGACAACGTTTACTAGCTAAG
AGGCAGATTACGAGGATAGGGGAAGACTGAGAAGGCCTTGGCGTAGGAATTCGAGGCAAC
CAGGGGGATAAGGTAAATGCCGACTTCAAGTCGAGAATGCAAAGAAGCTGGCTATGGCCG
TAAATTAGACGATAAATAACAGAGCATCGGGGAGCCAAGTCTAATTGTCAAACTTTTGGT
CGAGGTGATGTCGTAGGTGAGATCTCCGAACGCAGCATGAACAAGGCGAATCGTGTAAGC
AGAGAAGAGTGCAGCACGATGCTAACACTGCCGCAGCAGTGGGACAACTTGGATGACTGG
CACATCACAGTAATGAGTCATATGATTCCGCCTGAAGGCTTCAATAAAGGACATAAGATG
CGAAGAAGATCTAGACCCAGTCTTTAAAACTGCACAACTTAGCCGAAATGATTGGTAGGT
GAGAGTATCGACGTGGGCACTGAGAGAAACCTGAATACGCGATAGCCGTCATATCACGAT
GCTTCTGCCCTCCAGCGTCACTAGAGTTCTTGTTCACATGCCTCGGTCCTCTCGTCAGTA
CATGGCCGGCCTTCTTTCCTCCATCGTTTCCCAGAGCATACTCTTCGAGTTGCCCTATTA

ACCTTCTATTTTCTGGCCGATGCCTGACTGGCGCCGCTTTGAGTATCATACTTCTTGTTT
TTGGCCAACCTGTTGGGCGTTGATTGCTTGGGTAACCATATCCTGCATTTTGTTGATTCC
GGGTCGGCAATGTGTTGTGGACTGTTCGTCGCCGCTCCTGATCTCACCGTACTTCACGGT
TGTCGCCCGGACATCACCCGGACGGGTTTAGGCCATGGCCGGTTGAGGTTAATCTTTACG
CGTCACCATAACTTCCTGCGAAACTGGTTTCCCCACGCCGCTCCCTTCGTGGGCCGATCG
GCGTGGTTAGGTCGCGCGGGTTTTTCCAGGCAGGGTAAGTCGAGTCGCACAAGGTGACCC
ACGTACCTCGCTTTAACGTTCCGTTCTACCACTGGGTTAGGCTGCTTTGGGGCCGACTTG
ACGTTCTTTTTGCACGACCTCGTATGTGCTTATTCTCACTCGTAATGAACCGGTAAGCGA
GAGTTTGTTCTACGCTGAGGTGTCGTCGGACCCGAGTGTGCTTGCCGCTGGTCAATTGTT
GTGTCGGCTCGACGTCTTATTTCGTATGGTACGGGATTATTAGTGCTTGCAGCGATGTCC
CTTTCGCTATTGATGTCTTCTCCTGAGTCACGCTACGTACTGAATTGATAAAACATTTCG
ACCTCAGTATCAGTGAGGTCGGGGGATGAGTTAGATCGCCGTGTAGGGCTTTTCACGTTG
TGTCCGTGTGGAGAGTCGGTTCTAGCGGTCGTATTATCCGTATTCTCCCGTTCCATTTAC
GTTGAGCGACTTGAAACTTCTGATGTACTGGCNGGTAATAGTGCATTGTTCCTCAATTCA
TGCAAGTTGTCGTACATTATGGAATAGAACAAGAATTAAGGACCATTTCCACTAGCGATT
CTTACCTCCAACTCAAGATAGAGTCTAACCGATTATCGACGAACTTCGTCCGTCACGTCA
TTCCATGTCGAGCGGCCTTCTTACAGCGCTGTCGCACTATCTGTTCTACCTTATTTCCGA
GAAACCTTGCTCCTGTTACCTGTGAGGGCGGGGCGGATATGGCAAAATGGGCATTCTTTT
TTACTTCCCGTGCCTCTTCATGCTATATACATGTATCTCTGGAGCCTAGACGCCCGGTCC
TTAACGTCCTAAAACTTTATTGACCCTGCTGGCGTCTTACCCTTTATAGTGGCTCTGCAT
AATGTGTTCTGTATATAGCCAAATACTGCCGACTATCCACTCGCTTAGCTGTCAATATCC
ATGCGCGTACTTTGATTTACCGGTCGACCTGCCTAATGTAACTGCCTTTCGCTCTCCACT
CCGCACCGGGGTTTCGAGTTTTTCTACCCCTCTCTTTATTACTACTCGTTTACTGCACCA
GACGCGCCTACCGTATTCAGTCACTGCTCCTTCAGGCGAAGAGTATTAGCACGTTATCGT
GCTACTATCCTGAGCTGTAATCATAAGTATTTCTAAAGTTGTAGGGGTTGAAGAGTAGTT
CTGTACATCTTGATCGATTGTACTGGACGTCATTCCGAATAGCTTGTGGAATCCAATATC
TGGGTCATAGATGCT
>SRR9587955
GTGACCGTCGCCTCGCCACAAATCCATTGCTTTCCGGGTTCTAGCCCCTTATGAGCTAGG
ACCTCTGTTCTTACCATCATAGCTGTTCGATGCCTTGTATTCAATACGTTGCATCGACCC
GACAATAATCTGTGCCATTAAGCTTCTGCAAGTGGCTGGCGCCTGTAAGTCACGCAATAG
AGTAACTTCCCTCTCGACAGTCTTTATGCCATCCACAATCCAACATCGTAGATCCTCGCG
TGACCTAGTAGCCCTTGACGACTCAATCCACCCTCTGCCTGACGGACGTGCCTTTTATGG
TACATTAATAACTCCCTGTCCTCATAATTAGGCTGCACACACAGAGCTATCGTAGTTTAC
TGGCCTCACCCCTGCGTACTCGGCGACCCAACTACACTGATCCAGGGAGGCAGAATGGTG
GACGCTCTTACTGCCGACCCGTGGGCAACTAACTGGGCCGGGGGAATATTGTTTCGTTTT

GGGTACTGCTTCTTCCAATTGTTCGGATAGCACAGTAACGGTGGTGGGTGGCTCCAGTCA
CTCTTGCGTAGTCGCCGGCCGGTCGCNCCGGTCTGTCCTCTCGCCCGATTCTGGGCCATC
TTATCCGATGTCACCCTGCAGACCGCTCTTTTCAACCATTAGCATTAAGCAGTCTCTCAT
AGGTGCTCTCATGTCATGAGAGGTACCGGAGCCTTTAAGCGTTCCGCTGTTCCCGACAAT
ATTCTCTTTCGCTTTAACACTAGCTCAGACGGTAGAGGCTTCGTCTTCAGCACCGGCTAA
TGATGCCTGGTACGTTAAGATACCGATCCGCGCGCTTACGCTGCTCCACCTGTCGGTGTC
TGAAGAAGCTCGGCTAAGACCTCTCGAACTCTTCGTCTTACCTCAGCCCGCGGCTACAGT
CATATACATCCTAGCACGCACAATCCTTAATTCCCAGTCGTGCCCGACGGTAACACCGTG
GCGCGAACAATGTGTAAGTTGTTGCCTACTTCGTACACTATCAACTAAATTCGTCCAACG
GGAGATCCGGGAATCCCGNNAGGGGGCACACACTTAGAGAACCCGCAGCTTTTTGTCTCA
CGATCGAGAAGGGCACTCCCAAGAGGCCTTGATACGAGCGTGAAACCGGTCAAGCGCTCC
AATTGTACACTGTCCATGAGGCCAAATTCAAATTACGGACCGAAAACTGGATCCAGCAGA
GCTAAAGGTTATTCGCGGGTATTGTCCGTACATGCAACTTGATGTTTCCTAAGCCAAAAG
AGATATGCGAGGGGGAGACATGGGAAAAGCAAAAGAATATAGTACCATAATTGGCGTGGG
ACCCATAAAGTCTTGATTGCTCCCGCGCCGCCTGTACCCACAGTCTTCAAAAAAGCGATT
TGCCGGAGTATAAAGTCAGGGGCCGACGGTGTTGAAAACGAGCGAACGTCTTGCATGAGA
AGTGTCACTGTGGGCATATCAAAGGACTCTAATTAAATTCGGTGTCAATAGAAATGTTCG
TGCTGCACGGTGCTGTCAGACCATCCATCTTTCGTGGCTACGCCTTATCAGAAGTGCAAC
CTGCGAAATCAAGGTGACGTTAATACGCTCTAAAGGACGCTAAAGCGGGTTGATAATTAA
ACGACAGAGTTACAATTAGCGGTAGTCACAACTCTCCAAAATCATTGACGATAGACTGAC
ACGGTTACGTGGGAGGACTCGTAGTCTCAAGTGAAAATGGGCGCAAAATGGTGTCCGCAT
GGGAGGAGCTCAACCATAATAGACGCCGAATATGTTCGCGTGAAACTTCAGCGTCAAAGA
TGGAAAGCAGGACAAACTCATGAAACTACCATAGTGTGAAGATGATGGGCGGCGCGGTTT
AAGGCGAGTCGCCCTAAGACTCCAACGTGCCCTAAGGGATCGTAACATGCAAGAACATTG
ATGTGCACAGTTTTGTCGAACAGCCTTCATAGGCCAAAGATTGTATCCACGGCCAGAAGC
GGTAAGAACTTCGTGACATAGCGGCCAGAGAAAGGATCTACAACGTTAACCAGCACCTCA
GAGAGCGGGACGGGCAGTCGTTAGGGAGAAAATAACTAATCATTAACTTCGACGAACCAC
CAAAATGCTCGAGCGGTAGGTATCATTAGGGGGTACGGACTGTTACATCTAGAATACCCC
AGCTCCGTAGTTGCAGACACTAATTTTAGAGACCGCAGCTGCATCATTAACAAAATGTTA
TTGGTGGGATGCAGAAGAGCGCGGGCAAGATTATACAGACGCCTTCAGGCGGTAGCCTGC
TCCTAGGCAGTCAAGACGGATCGCGATGGACGTGTTTGGACGGCGAATGTACAGGAAATT
TCCCGTGAGAACAACAAAATGAAGGATCGAGATTAAAAAGACTACAGCGCCCGTGAACAA
CGGAGTTGAACGTGGTTGCCAGGTTTGTGATGGGCGGCCCTAGAAGATAACGGTAGAAGC
CCTTCGGACTTTTCAATGGGGACTACATTCTGGCCGCGAAGGACTCCGTGAATCGATAGG
TAGTGGAAATATGGATGAAACGGGCTGGGGCAACGCTCGGGACAACGTTTACTAGCTAAG
AGGCAGATTACGAGGATCGGGGAAGACTAAGAAGGCCTTGGCGTAGGAATTCGAGGCAAC

CAGGGGGGTAAGGTAAATGCCGACTTCAAGTCGAGAATGCAAAGAAGCTGGCTATGGCCG
TAAATTAGACAATAAATAACAGAGCATCGGGGGGCCAAGTCTAATTGTAAAACTTTTGGT
CGAGGTGATGTCGTAGGTGAGATCTCCGAACGCAGCATGAACAAGGCGAATCGTGTAAGC
AGAGAAGAGTGCAGTACGATGCTAACACTGTCGCAGCAGTGGGACAACTTGGATGACTGG
CACATCACAGTAATGAGTCATATGATTCCGCCTGAAGGCTTCAATAAAGGACATAAGATG
CGAAGAAGATCTAGACCCAGTCTTTAAAACTGCACAACTTAGCCGAAATGATTGGTAAGT
GAGAGTATCGACGTGGGCACTGAGAGAAACCTGAATACGCGATAGCCATCATATCACGAT
GCTTCTGCCCTCCAGCGTCACTAGAGTTCTTGTTCACACGCCTCGGTCCTCTCGTCAGTA
CATGGCCGGCCTTCTTTCCTCCATCGTTTCCCAGAGCATACTCTTCGAGTTGCCCTATTA
ACCTTCTACTTTCTGGCCGATGCCTGACTGGCGCCGCTTTGAGTATCATACTTCTTGTTT
TTGGCCAACCTGTTGGGCGTTGATTGCTTGGGTAACCATATCCTGCATTTTGTTGATTCC
TGGTCGGCAATGTGTTGTGGACTGTTCGTCGCCGCCCCTGATCTCACCGTACTTCACGGT
TGTCGCCCGGACATCACCCGGACGGGTTTAGGCCATGGCCGGTTGAGGTTAATCTTTACG
CGTCATCATAACTTCCTGCGAAACTGGTTTCCCCACGCCGCTCCCTTCGTGAGCCGATCG
GCGTGGTTAGGTCGCGCGGGTTTTTCCAGGCAGGGTAAGTCGAGTCGCACAAGGTGACCC
ACGTACCTCGCTTTAACGTTCCGTTCTACCACTGGGTTAGGCCGCTTTGGGGCCGACTTG
ACGTTCTTTTTGCACGACCTCGTATGTGCTTATTCTCACTCGTAATGAACCGGTAAGCGA
GAGTTTGTTCTACGCTGAGGTGTCGTTGGACCCGAGTGTGCTTGCCGCTGGTCAATTGTT
GTGTCGGCTCTACGTCTTATTTCGTATGGTACGGGATTATTAGTGCTTGCAGCGATGTCC
CTTTCGCTATTGATGTCTTCTCCTGAGTCACGCTACGTACTGAATTGATAAAACATTTCG
ACCTCAGTATCAGTGAGGTCGGGCGATGAGTTAGATCGCCGTGTAGGGCTTTTCACGTTG
GGTCCGTGTGNAGAGTCGGTTCTAGCGGTCGTATTATCCATATTCTCCCGTTCTATTTAC
GTTGAGCGACTTGAAACTTCCGATGTACTGGCCGNNAATAGTGCATTGTTTCTCAATTCA
TGCAAGTTGTCGTACATTATGGAATAGAACAAGAATTAAGGACCATTTCCACTAGCGATT
CTTACCTCCAACTCAAGATAGAGTCTAACCGATTATCGACGAACTTCGTCCGTCACGTCA
TTCTATGTCGAGCGGCCTTCTTACAGCGCTGTCGCACTATCTGTTCTACCTTATTTTCGA
GAAACCTTGCTCCTGTTACCTGTGAGGGCGGGGCGGATATGGCAAAATGGGCATTCTTTT
TTACTTCTCGTGCCTCTTCATGCTATATACATGTATCTCTGGAGCCTAGACGCCCGGTCC
TTAACGTCCTAAAACTTTACTGACCCTGCTGGCGTCTTACCCTTTATAGTGGCTCTGCAT
AATGTGTTCTGTATATAGCCAAATACTGCCGACTATCCACTCGCTTAGCTGTCAATATCC
ATGCGCGTACTTTGATTTACCGGTCGACCTGCCTAATGTAACTGCCTTTCGCTCTCCACT
CCGCACCGGGGTTTCGAGTTTTTCTACCCCTCTCTTTATTACTACTCGTTTACTGCATCA
GACGCGCCTACCGTATTCGGTCACTGCTCCTTCAGGCGAAGAGTATTAGCACGTTATCGT
GCTACTATCCTGAGCTGTAATCATAAGTATTTCTAAAGTTGTAAGGGTTGAAGAGTAGTT
CTGTACATCTTGATCGATTGTACTGGACGTCATTCCGAATAGCTTGTGGAATCCAATATC
TGGGTCATAGATGCT

>SRR9587956

GTGACCGTCGCCTCGCCACAAATCCATTGTTTTCCGGATTCTAGCCCCTTATGAGCTAGG
ACCTCTGTGCCTACCATCATAGCTGTTCGATGCCTTGTATTCAATACGTTGCACCGACCC
GACAATAGTCTGTGCCATTAAGCTTCTGCAAGTGGCTGGCGCCTGTAAGTCACGCAATAG
AGTAACTTCCCTCTCGACAGTCTTTATGCCATCCACAATCCAACATCGTAGATCCTCGCG
TGACCTAGTAGCCCTTGACGACTCAATCCACCCTCTGCCTGACGGACGTGCCTTTTATGG
TACATTAATGACTCCCTGTCCTCATAATTAGGCTGCACACACAGGGCTATCGTAGTTTAC
TGGTCTCACCCCCGCGTACTCGGCGACCCAACTACACTNATCCAGGGAGGCAGAATGGTG
GACGCGCTTACTGCCGACCCGTGGGCAACTAACTGGGCCGGGGGAATATTGTTTCGTTTT
GGGTACTGCTTCTTCCAATTGTTCGGATAGCACAGTAACGGTGGTGGGTGGCTCCAGTCA
CTCTTGCGTAGTCGCCGGCCGGTCGCTCCGGTCTGTCCTCTCGCCCGATTCTGGGCCATC
TTATCCGATGTCACCCTGCAGACCGCTCTTTTCAACCATTAGCATTAAGCAGTCTCTCAT
AGGTGCTCTCATGTCATGAGAGGTACCGGAGCCTCTAAGCGTTCCGCTGTTCCCGACAAT
ATTCTCTTTCGCTTTGACACTAGCTCAGACGGTAGAGGCTTCTTCTTCAGCACCGGCTAA
TGATGCCTGGTACGTTAAGATACCGATCCGCGCGCTTACGCTGCTCCACCTGTCGGTGTC
TGAAGAAGCTCGGCTAAGACCTCTCGAACTCTTCGTCTTACCTCAGCCCGCGGCTACAGT
CATATACATCCTAGCACGCACAATCCTTGATTCCCAGTCGTGCCCGACAGTAACACCGTG
GCGCGAACAATGTGTAAGTTATTGCCTACTTCGTACACTATCAACTAAATTCGTCCAACG
TGAGATCCGAGAATCCCGAAAGGGGGCACACACTTAGAGAACCCGCAGCTTTTTGTCTCA
CGATCGAGAAGGGCACTCCCAAGAGGCCTTGATACGAGCGTGAAACCGGTCAAGCGCTCC
AATTGTACACTGTCCATGAGGCCAAATTCAAATTACGGACCGAAAACTGGATCCAGCAGA
GCTAAAGGTTATTCGCGGGTATTGTCCGTACATGCAACTTGATGTTTTCTAAGCCAAAAG
AGATATGCGAGGGGGAGACATGGGAAAAGCAAAAGAATATAGTACTATAATTGGCGTGGG
ACCCATAAAGTCTTGATTGCTCCCGCGCCGCCTGTACCCACAGTCTTCAAAAAAGCGATT
TGCCGGAGTATAAAGTCAGGGGCCGACGGTGTTGAAAACGAGCGGACGTCTTGCATGAGA
AGTGTCACTGTGGGCATATCAAAGGACTCTAATTAAATTCGGTGTCAATAGAAATGTTCG
TGCTGCACGGTGCTGCCAGACCATCCACCTTTCGTGGCTACGCCTTATCAGAAGTGCAAC
CTGCGAAATCAAGGTGACGTTAATACGCTCTAAAGGACGCTAAAGCGGGTTGATAATTAA
ACGACAGAGTTACAATTAGCGGGATTTACAACTCTCCAAAATTATTGACGATAGACTGAC
ACGGTTACGTGGGAGGACTCGTAGTCTCAAGTAAAAATGGGCGCAAAATGGTGTCCGCAT
GGGAGGAGCTCAATCATAATAGACGCCGAATATGTTCGCGTGAAACTTCAGCGTCAAAGA
TGGAAAGCAAGACAAACTCATGAAACTACCATAGTGTGAAGATGATGGGCGGCGCGGTTT
AAGGCGAGTCGCCCTAAGACTCCAACGTGCCCTAAGGGATCGTAACATGCAAGAACATTG
ATGTGCACAGTTTTGTCGAACGGCCTTCATAGGCCAAAGATTGTATCCACGGCCAGAAGC
GGTAAGAACTTCGTGACATAGCGGCCAGAGAAAGGATCTACAACGTTAACCAGCACCTCA
GAGAGCGGGATGGCCAGTCGTTAGGGAGAAAATAACTAATCATTAACTTTGACGAACCAC

CAAAATGCTCGAGCGGTAGGTATCATTAGGGGGTACGGACTGTTACATCTAGAATACCCC
AGCTCCGTAGTTGCAGACACTAATTTTAGAGACCGCAGCTGTATCATTAACAAAATGTTA
TTGGTGGGATGCAGAAGAGCGCGGGCAAGATTATACAGACGCCTTCAGGCGGTAGCCTGC
TCCTAGGCAGTCAAGACGGATCGCGATGGACGTGTTTGGACGGCGAATGTACAGGAAATT
TCCCGTGAGAACAACAAAATGAAGGATCGAGATTAAAAAGACTGCAGCGCCTGTGAACAA
CGGAGTTGAACGTGGTTGCCAGGTTTGTGATGGGCGGCCCTAGAAGATAACGGTAGAAGC
CCTTCGGACTTTTCAAGGGGGACTACATTCTGGCCGCGAAGGACTCCGTGAATCGATAGG
TAGTGGAAATATGGATGAAACGGGCTGGGGCAACGCTCGGGACAACGTTTACTAGCTAAG
AGGCAGATTACGAGGATCGGGGAAGACTGAGAAGGCCTTGGCGTAGGAATTCGAGGCAAC
CAGGGGGATAAGGTAAATGCCGACTTCAAGTCGAGAATGCAAAGAAGCTGGCTATGGCCG
TAAATTAGACGATAAATAACAGAGCATCGGGGAGCCAAGTCTAATTGTAAAACTTTTGGT
CGAGGTGATGTCGTAGGTGAGATCTCCGAACGCAGCATGAACAAGGCGAATCGTGTAAGC
AGAGAAGAGTGCAGCACGATGCTAACACTGCCGCAGCAGTGGGACAACTTGGATGACTGG
CACATCACAGTAATGAGTCATATGATTCCGCCTGAAGGCTTCAATAAAGGACATAAGATG
CGAAGAAGATCTAGACCCAGTCTTTAAAACTGCACAACTTAGCCGAAATGATTGGTAGGT
GAGAGTATCGACGTGGGCACTGAGAGAAACCTGAATACGCGATAGCCATCGTATCACGAT
GCTTCTGCCCTCCAGCGTCACTAGAGTTCTTGTTCACATGCCTCGGTCCTCTCGTCAGTA
CATGGCCGGCCTTCTTTCCTCCATCGTTTCCCAGAGCATACTCTTCGAGTTGCCCTATTA
ACCTTCTATTTTCTGGCCGATGCCTGACTGGCGCCGCTTTGAGTATCATACTTCTTGTTT
TTGGCCAGCCTGTTGGGCGTTGATCGCTTGGGTAACCATATCCTGCATTTTGTTGATTCC
GGGTCGGCAATGTGTTGTGGACTGTTCGTCGCCGCTCCTGATCTCACCGTACTTCACGGT
TGTCGCCCGGACATTACCCGGACGGGTTTAGGCCATGGCCGGTTGAGGTTAATCTTTACG
CGTCACCATAACTTCCTGCGAAACTGGTTTCCCCACGCCGCTCTTTTCGTGAGCCGATCG
GCGTGGTTAGGTCGCGCGGGTTTTTCCAGGCAGGGTAAGTCGAGTCGCACAAGGTGACCC
ACGTACCTCGCTTTAACGTTCCGTTCTACCACTGGGTTAGGCTGCTTTGGGGCCGACTTG
ACGTTCTTTTTGCACGACCCCGTATGTGCTTATTCTCACTCGTAATGAACCGGTAAGCGA
GAGTTTGTTCTACGCTGAGGTGTCGTCGGACCCGAGTGTGCTTGCCGCTGGTCAATTGTT
GTGTCGGCTCGACGTCTTATTTCGTATGGTACGGGATTNTTAGTGCTTGCAGCGATGTCC
CTTTCGCTATTGATGTCTTCTCCTGAGTCACGCTACGTACTGAATTGATAAAACATTTCG
ACCTCAGTATCAGTGAGGTCGGGGGATGAGTTAGATCGCCGTGTAGGGCTTTTCACGTTG
TGTCCGTGTGGAGAGTCGGTTCTAGCGGTCGTATTATCCATATTCTCCCGTTCCATTTAC
GTTGAGCGACTTGAAACTTCCGATGTACTGGCCGGTAATAGTGCATTGTTTCTCAATTCA
TGCAAGTTGTCGTACATTATGGAATAGAACAAGAATTAAGGACCATTTCCACTAGCGATT
CTTACCTCCAACTCAAGATAGAGTCTAACCGATTATCGACGAACTTCGTCCGTCACGTCA
TTCTATGTCGAGCGGCCTTCTTACAGCGCTGTCGCACTATCTGTTCTACCTTATTTCCGA
GAAACCTTGCTCCTGTTACCTGTGAGGGCGGGGCGGATATGGCAAAATGGGCATTCTTTT

TTACTTCCCGTGCCTCTTCATGCTATATACATGTATCTCTGGAGCCTAGACGCCCGGTCC
TTAACGTCCTAAAACTTTACTGACCCTGCTGGCGTCTTACCCTTTATAGTGGCTCTGCAT
AATGTGTTCTGTATATAGCCAAATACTGCCGACTATCCACTCGCTTAGCTGTCAATATCC
ATGCGCGTACTTTGATTTACCGGTCGACCTGCCTAATGTAACTGCCTTTCGCTCTCCACT
CCGCACCGGGGTTTCGAGTTTTTCTACCCCTCTCTTTATTACTACTCGTTTACTGCATCA
GACGCGCCTACCGTATTCAGTCACTGCTCCTTCAGGCGAAGAGTATTAGCACGTTATCGT
GCTACTATCCTGAGCTGTAATCATAAGTATTTCTAAAGTTGTAAGGGTTGAAGAGTAGTT
CTGTACATCTTGATCGATTGTACTGGACGTCATTCCGAATAGCTTGTGGAATCCAATATC
TGGGTCATAGATGCT
>SRR9587957
GTGACCGTCGCCTCGCCACAAATCCATTGTTTTCCGGGTTCTAGCCCCTTATGAGCTAGG
ACCTCTGTGCCTACCATCATAGCTGTTCGATGCCTTGTATTCAATACGTTGCACCGACCC
GACAATAATCTGTGCCATTAAGCTTCTGTAAGTGGCTGGCGCCTGTAAGTCACGCAATAG
AGTAACTTCCCTCTCGACAGTCTTTATGCCATCCACAATCCAACATCGTAGATCCTCGCG
TGACCTAGTAGCCCTTGACGACTCAATCCACCCTCTGCCTGACGGACGTGCCTTTTATGG
TACATTAATGACTCCCTGTCCTCATAATTAGGCTGCACACACAGGGCTATCGTAGTTTAC
TGGTCTCACCCCTGCGTACTCGGCGACCCAACTACACTGATCCAGGGAGGCAGAATGGTG
GACGCTCTTACTGCCGACCCGTGGGCAACTAACTGGGCCGGGGGAATATCGTTTCGTTTT
GGGTACTGCTTCTTCCAATTGTTCGGATAGCACAGTAACGGTGGTGGATGGCTCCAGTCA
CTCTTGCGTAGTCGCCGGCCGGTCGCTCCGTTCTGTCCTCTCGCCCGATTCTGGGCCATC
TTATCCGATGTCACCCTGCAGACCGCTCTTTTCAACCATTAGCATTAAGCAGTCTCTCAT
AGGTGCTCTCACGTCATGAGAGGTACCGGAGCCTCTAAGCGTTCCGCTGTTCCCGACAAT
ATTCTCTTTCGCTTTAACACTAGCTCAGACGGTAGAGGCTTCGTCTTCAGCACCGGCTAA
TGATGCCTGGTACGTTAAGATACCGATCCGCGCGCTTACGCTGCTCCACCTGTCGGTGTC
TGAAGAAGCTCGGCTAAGACCTCTCGAACTCTTCGTCTTACCTCAGCCCGCGGCTACAGT
CATATACATCCTAGCACGCACAATCCTTAATTCCCAGTCGTGCCCGACAGTAACACCGTG
GCGCGAACAATGTGTAAGTTGTTGCCTACTTCGTACACTATCAACTAAATTCGTCCAACG
TGAGATCCGAGAATCCCGAAAGGGGGCACACACTTAGAGAACCCGCAGCTTTTTGTCTCA
CGATCGAGAAGGGCACTTCCAAGAGGCCTTGATACGAGCGTGAAACCGGTCAAGCGCTCC
AATTGTACACTGTCCATGAGGCCAAATTCAAATTACGGACCGAGAACTGGATCCAGCAGA
GCTAAAGGTTATTCGCGGGTATTGTCCGTACATGCAACTTGATGTTTTCTAAGCCAAAAG
AGATATGCGAGGGGGAGACATGGGAAAAGCAAAAGAATATAGTACTATAATTGGCGTGGG
ACCCATAAAGTCTTGATTGCTCCCGCGCCGCTTGTACCCACAGTCTTCAAAAAAGCGATT
TGCCGGAGTATAAAGTCAGGGGCCGACGGTGTTGAAAACGAGCGGACGTCTTGCATGAGA
AGTGTCACTGTGGGCATATCAAAGGACTCTAATTAAATTCGGTGTCAATAGCAATGTTCG
TGCTGCACGGTGCTGTCAGACCATCCATCTTTCGTGGCTACGCCTTATCAGAAGTGCAAC

CTGCGAAATCAAGGTGACGTTAATACGCTCTAAAGGACGCTAAAGCGGGTTGATAATTAA
ACGACAGAGTTACAATTAGCGGTATTCACAACTCTCCAAAATCATTGACGATAGACTGAC
ACGGTTACGTGGGAGGACTCGTAGTCTCAAGTGAAAATGGGCGCAAAATAGTGTCCGCAT
GGGAGGAGCTTAACCATAATAGACGCCGAATATGTTCGCGTGAAACTTCAGCGTCAAAGA
TGGAAAGCAAGACAAACTCATGAAACTACCATAGTGTGAAGATGATGGGCGGCGCGGTTT
AAGACGAGTCGCCCTAAGACTCCAACGTGCCCTAAGGGATCGTAACATGCAAGAACATTG
ATGTGCACAGTTTTGTCGAACGGCCTTCATAGGCCAAAGATTGTATCCACGGCCAGAAGC
GGTAAGAACTTCGTGACATAGCGGCCAGAGAAAGGATCTACAACGTTAACCAGCACCTCA
GAGAGCGGGACGGCCAGTCGTTAGGGAGAAAATAACTAATCATTAACTTTGACGAACCAC
CAAAATGCTCGAGCGGTAGGTATCATTAGGGGGTACGGACTGTTACATCTAGAATACCCC
AGCTCCGTAGTTGCAGACACTAATTTTAGAGACCGCAGCTGCATCATTAACAAAATGTTA
TTGGTGGGATGCAGAAGAGCGCGGGCAAGATTATACAGACGCCTTCAGGCGGTAGCCTGC
TCCTAGGCGGTCAAGACGGATCGCGATGGACGTGTTTGGACGGCGAATGTACAGGAAATT
TCCCGTGAGAACAACAAAATGAAGGATCGAGATTAAAAAGACTACAGCGCCCGTGAACAA
CGGAGTTGAACGTGGTTGCCAGGTTTGTGATGGGCGGCCCTAGAAGATAACGGTAGAAGC
CCTTCGGACTTTTCAATGGGGACTACATTCTGGCCGCGAAGGACTCCGTGAATCGATAGG
TAGTGGAAATATGGATGAAACGGGCTGGGGCAACGCTTGGGACAACGTTTACTAGCTAAG
AGGCAGATTACGAGGATCGGGGAAGACTGAGAAGGCCTTGGCGTAGGAATTCGAGGCAAC
CAGGGGGATAAGGTAAATGCCGACTTCAAGTCGAGAATGCAAAGAAGCTGGCTATGGCCG
TAAATTAGACGATAAATAACAGAGCATCGGGGGGCCAAGTCTAATTGTAAAACTTTTGGT
CGAGGTGATGTCGTAGGTGAGATCTCCGAACGCAGCATGAACAAGGCGAATCGTGTAAGC
AGAGAAGAGTGCAGCACGATGCTAACACTGCCGCAGCAGTGGGACAACTTGGATGACTGG
CACATCACAGTAATGAGTCATATGATTCCGCCTGAAGGCTTCAATAAAGGACATAAGATG
CGAAGAAGATCTAGACCCAGTCTTTAAAGCTGCACAACTTAGCCGAAATGATTGGTAGGT
GAAAGTATCGACGTGGGCACTGAGAGAAACCTGAATACGCGATAGCCATCATATCACGAT
GCTTCTGCCCTCCAGTGTCACTAGAGTTCTTGTTCACATGCCTCGGTCCTCTCGTCAGTA
CATGGCCGGCCTTCTTTCCTCCATCGTTTCCCAGAGCATACTCTTCGAGTTGCCCTATTA
GCCTTCTATTTTCTGGCCGATGCCTGCCTGGCGCCGCTTTGAGTATCATACTTCTTGTTT
TTGGCCAACCTGTTGGGCGTTGATTGCTTGGGTAACCATATCCTGCATTTTGTTGATTCC
TGGTCGGTAATGTGTTGTGGACTGTTCGTCGCTGCTCCTGACCTCACCGTACTTCACGGT
TGTCGCCCGGACATCACCCGGACGGGTTTAGGCCATGGCCGGTTGAGGTTAATCTTTACG
CGTCATCATAACTTCCTGCGAAACTGGTTTCCTCACGCCGCTCCCTTCGTGAGCCGATCG
GCGTGGTTAGGTCGCGCGGGTTTTTCCAAGCAGGGTAAGTCGAGTCGCCCAAGGTGACCC
ACGTACCTCGCTTTAACGTTCCGTTCTACCACTGGATTAGGCTGCTTTGGGGCCGACTTG
ACGTTCTTTTTGTACGACCTCGTATGTGCTTATTCTCACTCGTAATGAACCGGTAAGCGA
GAGTTTGCTCTACGCTGAGGTGCCGTCGGACCCGAGTGTGCTTGCCGCTGGTCAATTGTT

GTGTCGGCTCGACGTCTTATTTCGTATGGTACGGGATTATTAGTGCTTGCAGCGATGTCC
CTTTCGCTATTGATGTCTTATCCTGAGTCACGCTACGTACTGAATTGATAAAACATTTCG
ACCTCAGTATCAGTGAGGTCGGGGGATGAGTTAGATCGCCGTGTAGGGCTTTTCACGTTG
TGTCCGTGTGGAGAGTCGGTTCTAGCGGTCGTATTATCCATATTCTCCCGTTCTATTTAC
GTTGAGCGACTTGAAACCTCCGATGTACTGGCCGGTAATAGTGCATTGTTTCTCAATTCA
TGCGAGTTGTCGTACATTATGGAATAGAACAAGAATTAAGGACCATTTCCACTAGCGATT
CTTACCTCCAACTCAAGATAGAGTCTAACCGATTATCGACGAACTTCGTCCGTCACGTCA
TTCTATGTCGAGCGGCCTTCTTACAGCGCTGTCGCACTATCTGTTCTACCTTATTTCCGA
GAAACCTTGCTCCTGTTACCTGTGAGGGCGGGGCGGATATGGCAAAATGGGCATTCTTTT
TTACTTCCCGTGCCTCTTCATGCTATATACATGTATCTCTGGAGCCTAGACGCCCGGTCC
TTAACGTCCTAAAACTTTACTGACCCTGCTGGCGTCTTACCCTTTATAGTGGCTCTGCAT
AATGTGTTCTGTATATAGCCAAATACTGCCGACTATCCACTCGCTTAGCTGTCAATATCC
ATGCGCGTACTTTGATTTACCGGTCGACCTGCCTAATGTAACTGCCTTTCGCTCTCCACT
CCGCACCGGGGTTTCGAGTTTTTCTACCCCTCTCTTTATTACTACTCGTTTACTGCATCA
GACGCGCCTACCGTATTCGGTCACTGCTCCTTCAGGCGAAGAGTATTAGCACGTTATCGT
GCTACTATCCTGAGCTGTAATCCTAATTATTTCTAAAGTTGTAAGGGTTGAAGAGTAGTT
CTGTACATCTTGATCGATTGTACTGGACGTCATTCCGAATAGCTTGTGGAATCCAATATC
TGGGTCATAGATGCT
>SRR9587959
GTGACCGTCGCCTCGCCACAAATCCATTGTTTTCCGGGTTCTAGCCCCTTATGAGCTGGG
ACCTCTGTGCCTACCATCATAGCTGTTCGATGCCTTGTATTCAATACGTTGCACCGACCC
GACAATAATCTGTGCCATTAAGCTTCTGCAAGTGGCTGGCGCCTGTAAGTCACGCAATAG
AGTAACTTCCCTCTCGACAGTCTTTATGCCATCCACAATCCAACATCGTAGATCCTCGCG
TGACCTAGTAGCCCTTGACGACTCAATCCACCCTCTGCCTGACGGACGTGCCTTTTATGG
TACATTAATGACTCCCTGTCCTCATAATTAGGCTGCACACACAGGGCTATCGTAGTTTAC
TGGTCTCACCCCTGCGTACTCGGCGACCCAACTACACTGATCCAGGGAGGCAGAATGGTG
GACGCTCTTACTGCCGACCCGTGGGCAACTAACTGGGCCGGGGGAATATTGTTTCGTTTT
GGGTACTGCTTCTTCCAATTGTTCGGATAGCACAGTAACGGTGGTGGGTGGCTCCAGTCA
TTCTTGCGTAGTCGCCGGCCGGTCGCTCCGGTCTGTCCTCTCGCCCGATTCTGGGCCATC
TTATCCGATGTCACCCTGCAGACCGCTCTTTTCAACCATTAGCATTAAGCAGTCTCTCAT
AGGTGCTCTCATGTCATGAGAGGTACCGGAGCCTCTAAGCGTTCCGCTGTTTCCGACAAT
ATTCTCTTTCGCTTTGACACTAGCTCAGACGGTAGAGGCTTCGTCTTCAGCACCGGCTAA
TGATGCCTGGTACGTTAAGATACCGATCCGCGCGCTTACGCTGCTCCACCTGTCGGTGTC
TGAAGAAGCTCGGCTAAGACCTCTCGAACTCTTCGTCTTACCTCAGCCCGCGGCTACAGT
CATATAGATCCTAGCACGAACAATCCTTAATTCCCAGTCGTGCCCGACAGTAACACCGTG
GCGCGAACAATGTGTAAGTTATTGCCTACTTCGTACACTATCAACTAAATTCGTCCAACG

TGAGATCCGAGAATCCCGAAAGGGGGCACACACTTAGAGAACCCGCAGCTTTTTGTCTCA
CGATCGAGAAGGGCACTCCCAAGAGGCCTTGATACGAGCGTGAAACCGGTCAAGCGCTCC
AATTGTACACTGTCCATGAGGCCAAATTCAAATTACGGACCGAAAACTGGATCCAGCAGA
GCTAAAGGTTATTCGCGGGTATTGTCCGTACATGCAACTTGATGTTTTCTAAGCCAAAAG
AGATATGCGAGGGGGAGACATGGGAAAAGCAAAAGAATATAGTACTATAATTGGCGTGGG
ACCCATAAAGTCTTGATTGCTCCCGCGCCGCCTGTACCCACAGTCTTCAAAAAAGCGATT
TGCCGGAGTATAAAGTCAGGGGCCGACGGTGTTGAAAACAAGCGGACGTCTTGCATGAGA
AGTGTCACTGTGGGCATATCAAAGGACTCTAATTAAATTCGGTGTCAATAGAAATGTTCG
TGCTGCACGGTGCTGTCAGACCATCCACCTTTCGTGGCTACGCCTTATCAGAAGTGCAAC
CTGCGAAATCAAGGTGACGTTAATACGCTCTAAAGGACGCTAAAGCGGGTTGATAATTAA
ACGACAGAGTTACAATTAGCGGGATTCACAACTCTCCAAAATCATTGACGATAGACTGAC
ACGGTTACGTGGGAGGACTCGTAGTCTCAAGTGAAAATGGGCGCAAAATGGTGTCCGCAT
GGGAGGAGCTCAACCATAATATACGCCGAATATGTTCGCGTGAAACTTCAGCGTCAAAGA
TGGAAAGCAAGACAAACTCATGAAACTACCATAGTGTGGAGATGATGGGCGGCGCGGTTT
AAGGCGAGTCGCCCTAAGACTCCAACGTGCCCTAAGGGATCGTAACATGCAAGAACATTG
ATGTGCACAGTTTTGTCGAACGGCCTTCATAGGCCAAAGATTGTATCCACGGCCAGAAGC
GGTAAGAACTTCGTGACATAGCGGCCAGAGAAAGGATCTACAACGTTAACCAGCACCTCA
GAGAGCGGGATGGCCAGTCGTTAGGGAGAAAATAACTAATCATTAACTTTGACGAACCAC
CAAAATGCTCGAGCGGTAGGTATCATTAGGGGGTACGGACTGTTAAATCTAGAATACCCC
AGCTCCGTAGTTGCAGACACTAATCTTAGAGACCGCAGCTGCATCATTAACAAAATGTTA
TTGGTGGGATGCAGAAGAGCGCGGGCAAGATTATACAGACGCCTTCAGGCGGTAGCCTGC
TCCTAGGCAGTCAAGACGGATCGCGATGGACGTGTTTGGACGGCGAATGTACAGGAAATT
TCCCGTGAGAACAACAAAATGAAGGATCGAGATTAAAAAGACTGCAGCGCCCGTGAACAA
CGGAGTTGAACGTGGTTTCCAGGTTTGTGATGGGCGGCCCTAGAAGATAACGGTAGAAGC
CCTTCGGACTTTTCAATGGGGACTACATTCTGGCCGCGAAGGACTCCGTGAATCGATAGG
TAGTGGAAATATGGATGAAACGGGCTGGGGCAACGCTCGGGACAACGTTTACTAGCTAAG
AGGCAGATTACGAGGATCGGGGAAGACTGAGAAGGCCTTGGCGTAGGAATTCGAGGCAAC
CAGGGGGATAAGGTAAATGCCGACTTCAAGTCGAGAATGCACAGAAGCTGGCTATGGCCG
TAAATTAGACGATAAATAACAGAGCATCGGGGAGCCAAGTCTCATTGTAAAACTTTTGGT
CGAGGTGATGTCGTAGGTGAGATCTCCGAACGCAGCATGAGCAAGGCGAATCGTGTAAGC
AGAGAAGAGTGCAGCACGATGCTAACACTGCCGCAGCAGTGGGACAACTTGGATGACTGG
CACATCACAGTAATGAGTCATATGATTCCGCCTGAAGGCTTCAATAAAGGACATAAGATG
CGAAGAAGATCTAGACCCAGTCTTTAAAACTGCACAACTTAGCCGAAATGATTGGTAGGT
GAGAGTATCGACGTGGGCACTGAGAGAAACCTGAATACGCGATAGCCATCATATCACGAT
GCTTCTGCCCTCCAGCGTCACTAGAGTTCTCGTTCACATGCCTCGGTCCTCTCGTCAGTA
CATGGCCGGCCTTCTTTCCTCCATCGTTTCCCAGAGCATACTCTTCGAGTTGCCCTATTA

ACCTTCTATTTTCTGGCCGATGCCTGACTGGCGCCGCTTTGAGTATCATACTTCTTGTTT
TTGGCCAACCTGTTGGGCGTTGATTGCTTGGGTAACCATATCCTGCATTTTGTTGATTCC
GGGTCGGCAATGTGTTGTGGACTGTTCGTCGCCGCTCCTGATCTCACCGTACTTCACGGT
TGTCGCCCGGACATCACCCGGACGGGTTTAGGCCATGGCCGGTTGAGGTTAATCTTTACG
CGTCACCATAACTTCCTGCGAAACTGGTTTCCCCACGCCGCTCCCTTCGTGAGCCGATCG
GCGTGGTTAGGTCGCGCGGGTTTTTCCAGGCAGGGTAAGTCGAGTCGCACAAGGTGACCC
ACGTACCTCGCTTTAACGTTCCGTTCTACCACTGGGTTAGGCTGCTTTGGGGCCGACTTG
ACGTTCTTTTTGCACGACCTCGTATGTGCTTATTCTCACTCGTAATGAACCGGTAAGCGA
GAGTTTGTTCTACGCTGAGGTGTCGTCGGACCCGAGTGTGCTTGCCGCTGGTCAATTGTT
GTGTCGGCTCGACGTCTTATTTCGTATGGTACGGGATTATTAGTGCTTGCAGCGATGTCC
CTTTCGCTATTGATGTCTTCTCCTGAGTCACGCTACGTACTGAATTGATAAAACATTTCG
ACCTCAGTATCAGTGAGGTCGGGGGATGAGTTAGATCGCCGTGTAGGGCTTTTCACGTTG
TGTCCGTGTGGAGAGTCGGTTCTAGCGGTCGTATTATCCATATTCTCCCGTTCCATTTAC
GTTGAGCGACTTGAAACTTCCGATGTACTGGCCGGTAATAGTGCATTGTTTCTCAATTCA
TGCAAGATGTCGTACATTATGGAATAGAACAAGAATTAAGGACCATTTCCACTAGCGATT
CTTACCTCCAACTCAAGATAGAGTCTAACCGATTATCGACGAACTTCGTCCGTCACGTCA
TTCTATGTCGAGCGGCCTTCTTACAGCGCTGTCGCACTATCTGTTCTACCTTATTTCCGA
GAAACCTTGCTCCTGTTACCTGTGAGGGCGGGGCGGATATGGCAAAATGGGCATTCTTTT
TTACTTCCCGTGCCTCTTCATGCTATATACATGTATCTCTGGAGCCTAGACGCCCGGTCC
TTAACGTCCTAAAACTTTACTGACCCTGCTGGCGTCTTACCCTTTATAGTGGCTCTGCAT
AATGTGTTCTGTATATAGCCAAATACTGCCGACTATCCACTCGCTTAGCTGCCAATATCC
ATGCGCGTACTTTGATTTAACGGTCGACCTGCCTAATGTAACTGCCTTTCGCTCTCCACT
CCGCACCGGGGTTTCGAGTTTTTCTACCCCTCTCTTTATTACTACTCGTTTACTGCATCA
GACGCGCCTACCGTATTCAGGCACTGCTCCTTCAGGCGAAGAGTATTAGCACGTTATCGT
GCTACTATCCTGAGCTGTAATCATAAGTATTTCTAAAGTTGTAAGGGTTGAAGAGTAGTT
CTGTACATCTTGATCGATTGTACTGGACGTCATTCCGAATAGCTTGTGGAATCCAATATC
TGGGTCATAGATGCT
>SRR9587960_9587964
GTGACCGTCGCCTCGCCACAAATCCATTGTTTTCCGGATTCTAGCCCCTTATGAGCTAGG
ACCTCTGTGCCTACCATCATAGCTGTTCGATGCCTTGTATTCAATACGTTGCACCGACCC
GACAATAGTCTGTGCCATTAAGCTTCTGCAAGTGGCTGGCGCCTGTAAGTCACGCAATAG
AGTAACTTCCCTCTCGACAGTCTTTATGCCATCCACAATCCAACATCGTAGATCCTCGCG
TGACCTAGTAGCCCTTGACGACTCAATCCACCCTCTGCCTGACGGACGTGCCTTTTATGG
TACATTAATGACTCCCTGTCCTCATAATTAGGCTGCACACACAGGGCTATCGTAGTTTAC
TGGTCTCACCCCCGCGTACTCGGCGACCCAGCTACACTGATCCAGGGAGGCAGAATGGTG
GACGCGCTTACTGCCGACCCGTGGGCAACTAACTGGGCCGGGGGAATATTGTTTCGTTTT

GGGTACTGCTTCTTCCAATTGTTCGGATAGCACAGTAACGGTGGTGGGTGGCTCCAGTCA

CTCTTGCGTAGTCGCCGGCCGGTGGCTCCGGTCTGTCCTCTCGCCCGATTCTGGGCCATC

TTATCCGATGTCACCCTGCAGACCGCTCTTTTCAACCATTAGCATTAAGCAGTCTCTCAT

AGGTGCTCTCATGTCATGAGAGGTACCGGAGCCTCTAAGCGTTCCGCTGTTCCCGACAAT

ATTCTCTTTCGCTTTGACACTAGCTCAGACGGTAGAGGCTTCGTCTTCAGCACCGGCTAA

TGATGCCTGGTACGTTAAGATACCGATCCGCGCGCTTACGCTGCTCCACCTGTCGGTGTC

TGAAGAAGCTCGGCTAAGACCTCTCGAACTCTTCGTCTTACCTCAGCCCGCGGCTACAGT

CATATACATCCTAGCACGCACAATCCTTAATTCCCAGTCGTGCCCGACAGTAACACCGTG

GCGCGAACAATGTGTAAGTTATTGCCTACTTCGTACACTATCAACTAAATTCGTCCAACG

TGAGATCCGAGAATCCCGAAAGGGGGCACACACTTAGAGAACCCGCAGCTTTTTGTCTCA

CGATCGGGAAGGGCACTCCCAAGAGGCCTTGATACGAGCGTGAAACCGGTCAAGCGCTCC

AATTGTACACTGTCCATGAGGCCAAATTCAAATTACGGACCGAAAACTGGATCCAGCAGA

GCTAAAGGTTATTCGCGGGTATTGTCCGTACATGCAACTTGATGTTTTCTAAGCCAAAAG

AGATATGCGAGGGGGAGACATGGGAAAAGCAAAAGAATATAGTACTATAATTGGCGTGGG

ACCCATAAAGTCTTGATTGCTCCCGCGCCGCCTGTACCCACAGTCTTCAAAAAAGCGATT

TGCCGGAGTATAAAGTCAGGGGCCGACGGTGTTGAAAACGAGCGGACGTCTTGCATGAGA

AGTGTCACTGTGGGCATATCAAAGGACTCTAATTAAATTCGGTGTCGATAGAAATGTTCG

TGCTGCACGGTGCTGCCAGACCATCCACCTTTCGTGGCTACGCCTTATCAGAAGTGCAAC

CTGCGAAATCAAGGTGACGTTAATACGCTCTAAAGGACGCTAAAGCGGGTTGATAATTAA

ACGACAGAGTTACAATTAGCGGGATTTACAACTCTCCAAAATTATTGACGATAGACTGAC

ACGGTTACGTGGGAGGACTCGTAGTCTCAAGTGAAAATGGGCGCAAAATGGTGTCCGCAT

GGGAGGAGCTCAATCATAATAGACGCCGAATATGTTCGCGTGAAACTTCAGCGTCAAAGA

TGGAAAGCAAGACAAACTCATGAAACTACCATAGTGTGAATATGATGGGCGGCGCGGTTT

AAGGCGAGTCGCCCTAAGACTCCAACGTGCCCTAAGGGATCGTAACATGCAAGAACATTG

ATGTGCACAGTTTTGTCGAACGGCCTTCATAGGCCAAAGATTGTATCCACGGCCAGAAGC

GGTAAGAACTTCGTGACATAGCGGCCAGAGAAAGGATCTACAACGTTAACCAGCACCTCA

GAGAGCGGGATGGCCAGTCGTTAGGGAGAAAATAACTAATCATTAACTTTGACGAACCAC

CAAAATGCTCGAGCGGTAGGTATCATTAGGGGGTACGGACTGTTACATCTAGAATACCCC

AGCTCCGTAGTTGCAGACACTAATTTTAGAGACCGCAGCTGTATCATTAACAAAATGTTA

TTGGTGGGATGCAGAAGAGCGCGGGCAAGATTATACAGACGCCTTCAGGCGGTAGCCTGC

TCCTAGGCAGTCAAGACGGATCGCGATGGACGTGTTTGGACGGCGAATGTACAGGAAATT

TCCCGTGAGAACAACAAAATGAAGGATCGAGATTAAAAAGACTGCAGCGCCTGTGAACAA

CGGAGTTGAACGTGGTTGCCAGGTTTGTGATGGGCGGCCCTAGAAGATAACGGTAGGAGC

CCTTCGGACTTTTCAATGGGGACTACATTCTGGCCGCGAAGGACTCCGTGAATCGATAGG

TAGTGGAAATATGGATGAAACGGGCTGGGGCAACGCTCGGGACAACGTTTACTAGCTAAG

AGGCAGATTACGAGGATCGGGGAAGACTGAGAAGGCCTTGGCGTAGGAATTCGAGGCAAC

CAGGGGGATAAGGTAAATGCCGACTTCAAGTCGAGAATGCAAAGAAGCTGGCTATGGCCG
TAAATTAGACGATAAATAACAGAGCATCGGGGAGCCAAGTCTAATTGTAAAACTTTTGGT
CGAGGTGATGTCGTAGGTGAGATCTCCGAACGCAGCATGAACAAGGCGAATCGTGTAAGC
AGAGAAGAGTGCAGCACGATGCTAACACTGCCGCAGCAGTGGGACAACTTGGATGACTGG
CACATCACAGTAATGAGTCATATGATTCCGCCTGAAGGCTTCAATAAAGGACATAAGATG
CGAAGAAGATCTAGACCCAGTCTTTAAAACTGCACAACTTAGCCGAAATGATTGGTAGGT
GAGAGTATCGACGTGGGCACTGAGAGAAACCTGAATACGCGATAGCCATCGTATCACGAT
GCTTCTGCCCTCCAGCGTCACTAGAGTTCTTGTTCACATGCCTCGGTCCTCTCGTCAGTA
CGTGGCCGGCCTTCTTTCCTCCATCGTTTCCCAGAGCATACTCTTCGAGTTGCCCTATTA
ACCTTCTATTTTCTGGCCGATGCCTGACTGGCGCCGCTTTGAGTATCATACTTCTTGTTT
TTGGCCAGCCTGTTGGGCGTTGATTGCTTGGGTAACCATATCCTGCATTTTGTTGATTCC
GGGTCGGCAATGTGTTGTGGACTGTTCGTCGCCGCTCCTGATCTCACCGTACTTCACGGT
TGTCGCCCGGACATTACCCGGACGGGTTTAGGCCATGGCCGGTTGAGGTTAATCTTTACG
CGTCACCATAACTTCCTGCGAAACTGGTTTCCCCACGCCGCTCTTTTCGTGAGCCGATCG
GCGTGGTTAGGTCGCGCGGGTTTTTCCAGGCAGGGTAAGTCGAGTCGCACAAGGTGACCC
ACGTACCTCGCTTTAACGTTCCGTTCTACCACTGGGTTAGGCTGCTTTGGGGCCGACTTA
ACGTTCTTTTTGCACGACCCCGTATGTGCTTATTCTCACTCGTAATGAACCGGTAAGCGA
GAGTTTGTTCTACGCTGAGGTGTCGTCGGACCCGAGTGTGCTTGCCGCTGGTCAATTGTT
GTGTCGGCTCGACGTCTTATTTCGTATGGTACGGGATTCTTAGTGCTTGCAGCGATGCCC
CTTTCGCTATTGATGTCTTCTCCTGAGTCACGCTACGTACTGAATTGATAAAACATTTCG
ACCTCAGTATCAGTGAGGTCGGGGGATGAGTTAGATCGCCGTGTAGGGCTTTTCACGTTG
TGTCCGTGTGGAGAGTCGGTTCTAGCGGTCGTATTATCCATATTCTCCCGTTCCATTTAC
GTTGAGCGACTTGAAACTTCCGATGTACTGGCCGNTAATATTGCATTGTTTCTCAATTCA
TGCAAGTTGTCGTACATTATGGAATAGAACAAGAATTAAGGACCATTTCCACTAGCGATT
CTTACCTCCAACTCAAGATAGAGTCTAACCGATTATCGACGAACTTCGTCCGTCACGTCA
TTCTATGTCGAGCGGCCTTCTTACAGCGCTGTCGCACTATCTGTTCTACCTTATTTCCGA
GAAACCTTGCTCCTGTTACCTGTGAGGGCGGGGCGGATATGGCAAAATGGGCATTCTTTT
TTACTTCCCGTGCCTCTTCATGCGATATACATGTATCTCTGGAGCCTAGACGCCCGGTCC
TTAACGTCCTAAAACTTTACTGACCCTGCTGGCGTCTTACCCTTTATAGTGGCTCTGCAT
AATGTGTTCTGTATATAGCCAAATACTGCCGACTATCCACTCGCTTAGCTGTCAATATCC
ATGCGCGTACTTTGATTTACCGGTCGACCTGCCTAATGTAACTGCCTTTCGCTCTCCACT
CCGCACCGGGGTTTCGAGTTTTTCTACCCCTCTCTTTATTACTACTCGTTTACTGCATCA
GACGCGCCTACCGTATTCAGTCACTGCTCCTTCAGGCGAAGAGTATTAGCACGTTATCGT
GCTACTATCCTGAGCTGTAATCATAAGTATTTCTAAAGTTGTAAGGGTTGAAGAGTAGTT
CTGTACATCTTGATCGATTGTACTGGACGTCATTCCGAATAGCTTGTGGAATCCAATATC
TGGGTCATAGATGCT

>SRR9587961
GTGACCGTCGCCTCGCCACAAATCCATTGTTTTCCGGATTCTAGCCCCTTATGAGCTAGG
ACCTCTGTGCCTACCATCATAGCTGTTCGATGCCTTGTATTCAATACGTTGCACCGACCC
GACAATAGTCTGTGCCATTAAGCTTCTGCAAGTGGCTGGCGCCTGTAAGTCACGCAATAG
AGTAACTTCCCTCTCGACAGTCTTTATGCCATCCACAATCCAACATCGTAGATCCTCGCG
TGACCTAGTAGCCCTTGACGACTCAATCCACCCTCTGCCTGACGGACGTGCCTTTTATGG
TACATTAATGACTCCCTGTCCTCATAATTAGGCTGCACACACAGGGCTATCGTAGTTTAC
TGGTCTCACCCCCGCGTACTCGGCGACCCAGCTACACTGATCCAGGGAGGCAGAATGGTG
GACGCGCTTACTGCCGACCCGTGGGCAACTAACTGGGCCGGGGGAATATTGTTTCGTTTT
GGGTACTGCTTCTTCCAATTGTTCGGATAGCACAGTAACGGTGGTGGGTGGCTCCAGTCA
CTCTTGCGTAGTCGCCGGCCGGTGGCTCCGGTCTGTCCTCTCGCCCGATTCTGGGCCATC
TTATCCGATGTCACCCTGCAGACCGCTCTTTTCAACCATTAGCATTAAGCAGTCTCTCAT
AGGTGCTCTCATGTCATGAGAGGTACCGGAGCCTCTAAGCGTTCCGCTGTTCCCGACAAT
ATTCTCTTTCGCTTTGACACTAGCTCAGACGGTAGAGGCTTCGTCTTCAGCACCGGCTAA
TGATGCCTGGTACGTTAAGATACCGATCCGCGCGCTTACGCTGCTCCACCTGTCGGTGTC
TGAAGAAGCTCGGCTAAGACCTCTCGAACTCTTCGTCTTACCTCAGCCCGCGGCTACAGT
CATATACATCCTAGCACGCACAATCCTTAATTCCCAGTCGTGCCCGACAGTAACACCGTG
GCGCGAACAATGTGTAAGTTATTGCCTACTTCGTACACTATCAACTAAATTCGTCCAACG
TGAGATCCGAGAATCCCGNAAGGGGGCACACACTTAGAGAACCCGCAGCTTTTTGTCTCA
CGATCGGGAAGGGCACTCCCAAGAGGCCTTGATACGAGCGTGAAACCGGTCAAGCGCTCC
AATTGTACACTGTCCATGAGGCCAAATTCAAATTACGGACCGAAAACTGGATCCAGCAGA
GCTAAAGGTTATTCGCGGGTATTGTCCGTACATGCAACTTGATGTTTTCTAAGCCAAAAG
AGATATGCGAGGGGGAGACATGGGAAAAGCAAAAGAATATAGTACTATAATTGGCGTGGG
ACCCATAAAGTCTTGATTGCTCCCGCGCCGCCTGTACCCACAGTCTTCAAAAAAGCGATT
TGCCGGAGTATAAAGTCAGGGGCCGACGGTGTTGAAAACGAGCGGACGTCTTGCATGAGA
AGTGTCACTGTGGGCATATCAAAGGACTCTAATTAAATTCGGTGTCGATAGAAATGTTCG
TGCTGCACGGTGCTGCCAGACCATCCACCTTTCGTGGCTACGCCTTATCAGAAGTGCAAC
CTGCGAAATCAAGGTGACGTTAATACGCTCTAAAGGACGCTAAAGCGGGTTGATAATTAA
ACGACAGAGTTACAATTAGCGGGATTTACAACTCTCCAAAATNATTGACGATAGACTGAC
ACGGTTACGTGGGAGGACTCGTAGTCTCAAGTGAAAATGGGCGCAAAATGGTGTCCGCAT
GGGAGGAGCTCAATCATAATAGACGCCGAATATGTTCGCGTGAAACTTCAGCGTCAAAGA
TGGAAAGCAAGACAAACTCATGAAACTACCATAGTGTGAATATGATGGGCGGCGCGGTTT
AAGGCGAGTCGCCCTAAGACTCCAACGTGCCCTAAGGGATCGTAACATGCAAGAACATTG
ATGTGCACAGTTTTGTCGAACGGCCTTCATAGGCCAAAGATTGTATCCACGGCCAGAAGC
GGTAAGAACTTCGTGACATAGCGGCCAGAGAAAGGATCTACAACGTTAACCAGCACCTCA
GAGAGCGGGATGGCCAGTCGTTAGGGAGAAAATAACTAATCATTAACTTTGACGAACCAC

CAAAATGCTCGAGCGGTAGGTATCATTAGGGGGTACGGACTGTTACATCTAGAATACCCC
AGCTCCGTAGTTGCAGACACTAATTTTAGAGACCGCAGCTGTATCATTAACAAAATGTTA
TTGGTGGGATGCAGAAGAGCGCGGGCAAGATTATACAGACGCCTTCAGGCGGTAGCCTGC
TCCTAGGCAGTCAAGACGGATCGCGATGGACGTGTTTGGACGGCGAATGTACAGGAAATT
TCCCGTGAGAACAACAAAATGAAGGATCGAGATTAAAAAGACTGCAGCGCCTGTGAACAA
CGGAGTTGAACGTGGTTGCCAGGTTTGTGATGGGCGGCCCTAGAAGATAACGGTAGGAGC
CCTTCGGACTTTTCAATGGGGACTACATTCTGGCCGCGAAGGACTCCGTGAATCGATAGG
TAGTGGAAATATGGATGAAACGGGCTGGGGCAACGCTCGGGACAACGTTTACTAGCTAAG
AGGCAGATTACGAGGATCGGGGAAGACTGAGAAGGCCTTGGCGTAGGAATTCGAGGCAAC
CAGGGGGATAAGGTAAATGCCGACTTCAAGTCGAGAATGCAAAGAAGCTGGCTATGGCCG
TAAATTAGACGATAAATAACAGAGCATCGGGGAGCCAAGTCTAATTGTAAAACTTTTGGT
CGAGGTGATGTCGTAGGTGAGATCTCCGAACGCAGCATGAACAAGGCGAATCGTGTAAGC
AGAGAAGAGTGCAGCACGATGCTAACACTGCCGCAGCAGTGGGACAACTTGGATGACTGG
CACATCACAGTAATGAGTCATATGATTCCGCCTGAAGGCTTCAATAAAGGACATAAGATG
CGAAGAAGATCTAGACCCAGTCTTTAAAACTGCACAACTTAGCCGAAATGATTGGTAGGT
GAGAGTATCGACGTGGGCACTGAGAGAAACCTGAATACGCGATAGCCATCGTATCACGAT
GCTTCTGCCCTCCAGCGTCACTAGAGTTCTTGTTCACATGCCTCGGTCCTCTCGTCAGTA
CGTGGCCGGCCTTCTTTCCTCCATCGTTTCCCAGAGCATACTCTTCGAGTTGCCCTATTA
ACCTTCTATTTTCTGGCCGATGCCTGACTGGCGCCGCTTTGAGTATCATACTTCTTGTTT
TTGGCCAGCCTGTTGGGCGTTGATTGCTTGGGTAACCATATCCTGCATTTTGTTGATTCC
GGGTCGGCAATGTGTTGTGGACTGTTCGTCGCCGCTCCTGATCTCACCGTACTTCACGGT
TGTCGCCCGGACATTACCCGGACGGGTTTAGGCCATGGCCGGTTGAGGTTAATCTTTACG
CGTCACCATAACTTCCTGCGAAACTGGTTTCCCCACGCCGCTCTTTTCGTGAGCCGATCG
GCGTGGTTAGGTCGCGCGGGTTTTTCCAGGCAGGGTAAGTCGAGTCGCACAAGGTGACCC
ACGTACCTCGCTTTAACGTTCCGTTCTACCACTGGGTTAGGCTGCTTTGGGGCCGACTTA
ACGTTCTTTTTGCACGACCCCGTATGTGCTTATTCTCACTCGTAATGAACCGGTAAGCGA
GAGTTTGTTCTACGCTGAGGTGTCGTCGGACCCGAGTGTGCTTGCCGCTGGTCAATTGTT
GTGTCGGCTCGACGTCTTATTTCGTATGGTACGGGATTCTTAGTGCTTGCAGCGATGCCC
CTTTCGCTATTGATGTCTTCTCCTGAGTCACGCTACGTACTGAATTGATAAAACATTTCG
ACCTCAGTATCAGTGAGGTCGGGGGATGAGTTAGATCGCCGTGTAGGGCTTTTCACGTTG
TGTCCGTGTGGAGAGTCGGTTCTAGCGGTCGTATTATCCATATTCTCCCGTTCCATTTAC
GTTGAGCGACTTGAAACTTCCGATGTACTGGCCGGTAATATTGCATTGTTTCTCAATTCA
TGCAAGTTGTCGTACATTATGGAATAGAACAAGAATTAAGGACCATTTCCACTAGCGATT
CTTACCTCCAACTCAAGATAGAGTCTAACCGATTATCGACGAACTTCGTCCGTCACGTCA
TTCTATGTCGAGCGGCCTTCTTACAGCGCTGTCGCACTATCTGTTCTACCTTATTTCCGA
GAAACCTTGCTCCTGTTACCTGTGAGGGCGGGGCGGATATGGCAAAATGGGCATTCTTTT

TTACTTCCCGTGCCTCTTCATGCGATATACATGTATCTCTGGAGCCTAGACGCCCGGTCC
TTAACGTCCTAAAACTTTACTGACCCTGCTGGCGTCTTACCCTTTATAGTGGCTCTGCAT
AATGTGTTCTGTATATAGCCAAATACTGCCGACTATCCACTCGCTTAGCTGTCAATATCC
ATGCGCGTACTTTGATTTACCGGTCGACCTGCCTAATGTAACTGCCTTTCGCTCTCCACT
CCGCACCGGGGTTTCGAGTTTTTCTACCCCTCTCTTTATTACTACTCGTTTACTGCATCA
GACGCGCCTACCGTATTCAGTCACTGCTCCTTCAGGCGAAGAGTATTAGCACGTTATCGT
GCTACTATCCTGAGCTGTAATCATAAGTATTTCTAAAGTTGTAAGGGTTGAAGAGTAGTT
CTGTACATCTTGATCGATTGTACTGGACGTCATTCCGAATAGCTTGTGGAATCCAATATC
TGGGTCATAGATGCT
>SRR9587962
GTGACCGTCGCCTCGCCACAAATCCATTGTTTTCCGGGTTCTAGCCCCTTATGAGCTAGG
ACCTCTGTGCCTACCATCATAGCTGTTCGATGCCTTGTATTCAATACGTTGCACCGACCC
GACAATAATCTGTGCCATTAAGCTTCTGCAAGTGGCTGGCGCCTGTAAGTCACGCAATAG
AGTAACTTCCCTCTCGACAGTCTTTATGCCATCCACAATCCAACATCGTAGATCCTCGCG
TGACCTAGTAGCCCTTGACGACTCAATCCACCCTCTGCCTGACGGACGTGCCTTTTATGG
TACATTAATGACTCCCTGTCCTCATAATTAGGCTGCACACACAGGGCTATCGTAGTTTAC
TGGTCTCACCCCTGCGTACTCGGCGACCCAACTACACTGATCCAGGGAGGCAGAATGGTG
GACGCTCTTACTGCCGACCCGTGGGCAACTAACTGGGCCGGGGGAATATTGTTTCGTTTT
GGGTACTGCTTCTTCCAATTGTTCGGATAGCACAGTAACGGTGGTGGGTGGCTCCAGTCA
TTCTTGCGTAGTCGCCGGCCGGTCGCTCCGGTCTGTCCTCTCNCCCGATTCTGGGCAATC
TTATCCGATGTCACCCTGCAGACCGCNCTTTTTAACCATTAGCATTAAGCAGTCTCTCAT
AGGTGCTCTCGTGTCATGAGAGGTACCGGAGCCTCTAAGCGTTCCGCTGTTCCCGACAAT
ATTCTCTTTCGCTTTGACACTAGCTCAGACGGTAGAGGCTTCGCCTTCAGCACCGGCTAA
TGATGCCTGGTACGTTAAGATACCGATCCGCGCGCTTACGCTGCTCCACCTGTCGGTGTC
TGAAGAAGCTCGGCTAAGACCTCTCGAACTCTTCGTCTTACCTCAGCCCGCGGCTACAGT
CATATACATCCTAGCACGCACAATCCTTAATTCCCAGTCGTGCCCGACAGTAACACCGTG
GCGCGAACAATGTGTAAGTTATTGCCTACTTCGTACACTATCAACTAAATTCGTCCAACG
TGAGATCCGAGAATCCCGTTAGGGGGCACACACTTAGAGAACCCGCAGCTTTTTGTCTCA
CGATCGAGAAGGGCACTCCCAAGAGGCCTTGATACGAGCGTGAAACCGGTCAAGCGCTCC
AATTGTACACTGTCCATGAGGCCAAATTCAAATTACGGACCGAAAACTGGATCCAGCAGA
GCTAAAGGTTATTCGCGGGTATTGTCCGTACATGCAACTTGATGTTTTCTAAGCCAAAAG
AGATATGCGAGGGGGAGACATGGGAAAAGCAAAAGAATATAGTACTATAATTGGCGTGGG
ACCCATAAAGTCTTGATTGCTCCCGCGCCGCCTGTACCCACAGTCTTCAAAAAAGCGATT
TGCCGGAGTATAAAGTCAGGGGCCGACGGTGTTGAAAACGAGCGGACGTCTTGCATGAGA
AGTGTCACTGTGGGCATATCAAAGGACTCTAATTAAATTCGGTGTCAATAGAAATGTTCG
TGCTGCACGGTGCTGTCAGACCATCCACCTTTCGTGGCTACGCCTTATCAGAAGTGCAAC

CTGCGAAATCAAGGTGACGTTAATACGCTCTAAAGGACGCTAAAGCGGGTTGATAATTAA
ACGACAGAGTTACAATTAGCGGGATTCACAACTCTCCAAAATCATTGACGATAGACTGAC
ACGGTTACGTGGGAGGACTCGTAGTCTCAAGTGAAAATGGGCGCAAAATGGTGTCCGCAT
GGGAGGAGCTCAACCATAATAGACGCCGAATATGTTCGCGTGAAACTTCAGCGTCAAAGA
TGGAAAGCAAGACAAACTCATGAAACTACCATAGTGTGAAGATGATGGGCGGCGCGGTTT
AAGGCGAGTCGCCCTAAGGCTCCAACGTGCCCTAAGGGATCGTAACATGCAAGAACATTG
ATGTGCACAGTTTTGTCGAACGGCCTTCATAGGCCAAAGATTGTATCCACGGCCAGAAGC
GGTAAGAACTTCGTGACATAGCGGCCAGAGAAAGGATCTACAACGTTAACCAGCACCTCA
GAGAGCGGGATGGCCAGTCGTTAGGGAGAAAATAACTAATCATTAATTTTGACGAACCAC
CAAAATGCTCGAGCGGTAGGTATCATTAGGGGGTACGGACTGTTACATCTAGAATGCCCC
AGCTCCGTAGTTGCAGACACTAATTTTAGAGACCGCAGCTGCATCATTAACAAAATGTTA
TTGGTGGGATGCAGAAGAGCGCGGGCAAGATTATACAGACGCCTTCAGGCGGTAGCCTGC
TCCTAGGCAGTCAAGACGGATCGCGATGGACGTGTTTGGACGGCGAATGTACAGGAAATT
TCCCGTGAGAACAACAAAATGAAGGATCGAGATTAAAAAGACTGCAGCGCCCGTGAACAA
CGGAGTTGAACGTGGTTTCCAGGTTTGTGATGGGCGGCCCTAGAAGATAACGGTAGAAGC
CCTTCGGACTTTTCAATGGGGACTACATTCTGGCCGCGAAGGACTCCGTGAATCGATAGG
TAGTGGAAATATGGATGAAACGGGCTGGGGCAACGCTCGGGACAACGTTTACTAGCTAAG
AGGCAGATTACGAGGATCGGGGAAGACTGAGAAGGCCTTGGCGTAGGAATTCGAGGCAAC
CAGGGGGATAAGGTAAATGCCGACTTCAAGTCGAGAATGCAAAGAAGCTGGCTATGGCCG
TAAATTAGACGATAAATAACAGAGCATCGGGGAGCCAAGTCTAATTGTAAAACTTTTGGT
CGAGGTGATGTCGTAGGTGAGATCTCCGAACGCAGCATGAGCAAGGCGAATCGTGTAAGC
AGAGAAGAGTGCAGCACGATGCTAACACTGCCGCAGCAGTGGGACAACTTGGATGACTGG
CACATCACAGTAATGAGTCATATGATTCCGCCTGAAGGCTTCAATAAAGGACATAAGATG
CGAAGAAGATCTAGACCCAGTCTTTAAAACTGCACAACTTAGCCGAAATGATTGGTAGGT
GAGAGTATCGACGTGGGCACTGAGAGAAACCTGAATACGCGATAGTCATCATATCACGAT
GCTTCTGCCCTCCAGCGTCACTAGAGTTCTTGTTCACATGCCTCGGTCCTCTCGTCAGTA
CATGGCCGGCCTTCTTTCCTCCATCGTTTCCCAGAGCATACTCTTCGAGTTGCCCTATTA
ACCTTCTATTTTCTGGCCGATGCCTGACTGGCGCCGCTTTGAGTATCATACTTCTTGTTT
TTGGCCAAACTGTTGGGCGTTGATTGCTTGGGTAACCATATCCTGCATTTTGTTGATTCC
GGGTCGGCAATGTGTTGTGGACTGTTCGTCGCCGCTCCTGATCTCACCGTACTTCACGGT
TGTCGCCCGGACATCACCCGGACGGGTTTAGGCCATGGCCGGTTGAGGTTAATCTTTACG
CGTCACCATAACTTCCTGCGAAACTGGTTTCCCCACGCCGCTCCCTTCGTGAGCCGATCG
GCGTGGTTAGGTCGCGCGGGTTTTTCCAGGCAGGGTAAGTCGAGTCGCACAAGGTGACCC
ACGTACCTCGCTTTAACGTTCCGTTCTACCACTGGGTTAGGCTGCTTTGGGGCCGACTTG
ACGTTCTTTTTGCACGACCTCGTATGTGCTTATTCTCACTCGTAATGAACCGGTAAGCGA
GAGTTTGTTCTACGCTGAGGTGTCGTCGGACCCGAGTGTGCTTGCCGCTGGTCAATTGTT

GTGTCGGCTCGACGTCTTATTTCGTATGGTACGGGATTATTAGTGCTTGCAGCGATGTCC
CTTTCGCTATTGATGTCTTCTCCTGAGTCACGCTACGTACTGAATTGATAAAACATTTCG
ACCTCAGTATCAGTGAGGTCGGGGGATGAGTTAGATCGCCGTGTAGGGCTTTTCACGCTG
TGTCCGTGTGGAGAGTCGGTTCTAGCGGTCGTATTATCCATATTCTCCCGTTCCATTTAC
GTTGAGCGACTTGAAACTTCCGATGTACTGGCCGGTAATAGTGCATTGTTTCTCAATTCA
TGCAAGTTGTCGTACATTATGGAATAGAACAAGAATTAAGGACCATTTCCACTAGCGATT
CTTACCTCCAACTCAAGATAGAGTCTAACCGATTATCGACGAACTTCGTCCGTCACGTCA
TTCTATGTCGAGCGGCCTTCTTACAGCGCTGTCGCACTATCTGTTCTACCTTATTTCCGA
GAAACCTTGCTCCTGTTACCTGTGAGGGCGGGGCGGATATGGCAAAATGGGCATTCTTTT
TTACTTCCCGTGCCTCTTCATGCTATATACATGTATCTCTGGAGCCTAGACGCCCGGTCC
TTAACGTCCTAAAACTTTACTGACCCTGCTGGCGTCTTACCCTTTATAGTGGCTCTGCAT
AATGTGTTCTGTATATAGCCAAATACTGCCGACTATCCACTCGCTTAGCTGCCAATATCC
ATGCGCGTACTTTGATTTACCGGTCGACCTGCCTAATGTAACTGCCTTTCGCTCTCCACT
CCGCACCGGGGTTTCGAGTTTTTCTACCCCTCTCTTTATTACTACTCGTTTACTGCATCA
GACGCGCCTACCGTATTCAGTCACTGCTCCTTCAGGCGAAGAGTATTAGCACGTTATCGT
GCTACTATCCTGAGCTGTAATCATAAGTATTTCTAAAGTTGTAAGGGTTGAAGAGTAGTT
CTGTACATCTTGATCGATTGTACTGGACGTCATTCCGAATAGCTTGTGGAATCCAATATC
TGGGTCATAGATGCT
>SRR9587963
GTGACCGTCGCCTCGCCACAAATCCATTGTTTTCCGGGTTCTAGCCCCTTATGAGCTAGG
ACCTCTGTGCCTACCATCATAGCTGTTCGATGCCTTGTATTCAATACGTTGCACCGACCC
GACAATAGTCTGTGCCATTAAGCTTCTGCAAGTGGCTGGCGCCTGTAAGTCACGCAATAG
AGTAACTTCCCTCTCGACAGTCTTTATGCCATCCACAATCCAACATCGTAGATCCTCACG
TGACCTAGTAGCCCTTGACGACTCAATCCACCCTCTGCCCGACGGACGTGCCTTTTATGG
TACATTAATGACTCCCTGTCCTCATAATTAGGCTGCACACACAGGGCTATCGTAGTTTAC
TGGTCTCACCCCTGCGTACTCGGCGACCCAACTACACTGATCCAGGGAGGCAGAATGGTG
GACTCGCTTACTCCCGACCCGTGGGCAACTAACTGGGCCGGGGGAATATTGTTTCGTTTT
GGGTACTGCTTCTTCCAATTGTTCGGATAGCACAGTAACGGTGGTGGGTGGCTCCAGTCA
CTCTTGCGTAGTCGCCGGCCGGTCGCTCCGGTCTGTCCTCTCGCCCGATTCTGGGCCATC
TTATCCGATGTCACCCTGCAGACCGCTCTTTTCAACCATTAGCATTAAGCAGTCTCTCAT
AGGTGCTCTCATGTCATGAGAGGTACCGGAGCCTCTAAGCGTTCCGCTGTTCCCNACAAT
ATTCTCTTTCGCTTTGACACTAGCTCAGACGGTAGAGGCTTCGTCTTCAGCACCGGCTAA
TGATGCCTGGTACGTTAAGATACCGATTCGCGCGCTTACGCTGCTCCACCTGTCGGTGTC
TGAAGAAGCTCGGCTAAGACCTCTCGAACTCTTCGTCTTACCTCAGCCCGCGGCTACAGT
CATATACATCCTAGCACGCACAATCCTTAATTCTCAGTCGTGCCCGACAGTAACACCGTG
GCGCGAACAATGTGTAAGTTATTGCCTACTTCGTACACTATCAACTAAATTCGTCCAACG

TGAGATCCGAGAATCCCGAAAGGGGGCACACACTTAGAGAACCCGCAGCTTTTTGTCTCA

CGATCGAGGAGGGCACTCCCAAGAGGCCTTGATACGAGCGTGAAACCGGTCAAGCGCTCC

AATTGTACACTGTCCATGAGGCCAAATTCAAATTACGGACCGAAAACTGGATCCAGCAGA

GCTAAAGGTTATTCGCGGGTATTGTCCGTACATGCAACTTGATGTTTTCTAAGCCAAAAG

AGATATGCGAGGGGGAGACATGGGAAAAGCAAAGGAATATAGTACTATAATTGGCGTGGG

ACCCATAAAGTCTTGATTGCTCCCGCGCCGCCTGTGCCCACAGTCTTCAGAAAAGCGATT

TGCCGGAGTATAAAGTCAGGGGCCGACGGTGTTGAAAACGAGCGGACGTCTTGCATGAGA

AGTGTCACTGTGGGCATATCAAAGGACTCTAATTAAATTCAGTGTCAATAGAAATGTTCG

TGCTGCACGGTGCTGTCAGACCATCCACCTTTCGTGGCTACGCCTTATCAGAAGTGCAAC

CTGCGAAATCAAGGTGACGTTAATACGCTCTAAAGGACGCTAAAGCGGGTTGATAATTAA

ACGACAGAGTTACAATTAGCGGGATTCACAACTCTCCAAAATCATTGACAATAGACTGAC

ACGGTTACGTGGGAGGACTCGTAGTCTCAAGTGAAAATGGGCGCAAAATGGTGTCCGCAT

GGGAGGAGCTCAACCATAATAGACGCCGAATATGTTCGCGTGAAACTTCAGAGTCAAAGA

TGGAAAGCAAGACAAACTCATGAAACTACCATAGTGTGAAGATGATGGGCGGCGCGGTTT

AAGGCGAGTCGCCCTAAGACCCCAACGTGCCCTAAGGGATCGTAACATGCAAGAACATTG

ATGTGCACAGTTTTGTCGAACGGCCTTCATAGGCCAAAGATTGTATCCACGGCCAGAAGC

GGTAAGAACTTCGTGACATAGCGGCCAGAGAAAGGATCTACAACGTTAACCAGCACCTCA

GAGAGCGGGATGGCCAGTCGTTAGGGAGAAAATAACTAATCATTAACTTTGACGAACCAC

CAAAATGCTCGAGCGGTAGGTATCATTAGGGGGTACGGACTGTTACATCTAGAATACCCC

AGCTCCGTAGTTGCAGACACTAATTTTAGAGACCGCAGCTGTATCATTAACAAAATGTTA

TTGGTGGGATGCAGAAGAGCGCGGGCAAGATTATACAGACGCCTTCAGGCGGTAGCCTGC

TCCTAGGCAGTCAAGACGGATCGCGATGGACGTGTTTGGACGGCGAATGTACAGGAAATT

TCCCGTGAGAACAACAAAATGAAGAATCGAGATTAAAAAGACTGCAGCGCCCGTGAACAA

CGGAGTTGAACGTGGTTGCCAAGTTTGTGATGGGCGGCCCTAGAAGATAACGGTAGAAGC

CCTTCGGACTTTTCAATGGGGACTACATTCTGGCCGCGAAGGACTCCGTGAATCGATAGG

TAGTGGAAATATGGATGAAACGGGCTGGGGCAACGCTCGGGACAAAGTTTACTAGCTAAG

AGGCAGATTACGAGGATCGGGGAAGACTGAGAAGGCCTTGGCGTAGGAATTCGAGGCAAC

CAGGGGGATAAGGTAAATGCCGACTTCAAGTCGAGAATGCAAAGAAGCTAGCTATGGCCG

TAAATTAGACGATAAATAACAGAGCATCGGGGAGCCAAGTCTAATTGTAAAACTTTTGGT

CGAGGTGATGTCGTAGGTGAGATCTCCGAACGCAGCATGAACAAGGCGAATCGTGTAAGC

AGAGAAGAGTGCAGCACGATGCTAACACTGCCGCAGCAGTGGGACAACTTGGATGACTGG

CACATCACAGTAATGAGTCATATGATTCCGCCTGAAGGCTTCAATAAAGGACATAAGATG

CGAAGAAGATCTAGACCCAGTCTTTAAAACTGCACAACTTAGCCGAAATGATTGGTAGGT

GAGAGTATCGACGTGGGCACTGAGAGAAACCTGAATACGCGATAGCCATCACATCACGAT

GCTTCTGCCCTCCAGCGTCACTAGAGTTCTTGTTCACATGCCTCGGTCCTCTCGTCAGTA

CATGGCCGGCCTTCTTTCCTCCATCGTTTCCCAGAGCATACTCCTCGAGTTGCCCTATTA

ACCTTCTATTTTCTGGCCGATGCCTGACTGGCGCCGCTTTGAGTATCATACTTCTTGTTT
TTGGCCAACCTGTTGGGCGTTGATTGCTTGGGTAACCACATCCTGCATTTTGTTGATTCC
GGGTCGGCAATGTGTTGTGGACTGTTCGTCGCCGCTCCTGATCTCACCGTACTTCACGGT
TGTCGCCCGGACATCACCCGGACGGGTTTAGGCCATGGCCGGTTGAGGTTAATCTTTACG
CGTCACCATAACTTCCTGCGAAACTGGTTTCCCCACACCGCTCCCTTCGTGAGCCGATCG
GCGTGGTTAGGTCGCGCGGGTTTTTCCAGGCAGGGTAAGTCGAGTCGCACAAGGTGACCC
ACGTACCTCGCTTTAACGTTCCGTTCTACCACTGGGTTAGGCTGCTTTGGGGCCGACTTG
ACGTTCTTTTTGCACGACCTCGTATGTGCTTATTCTCACTCGTAATGAACCGGTAAGCGA
GAGTTTGTTCTACGCTGAGGTGTCGTCGGACCCGAGTGTGCTTGCCGCTGGTCAATTGTT
GTGTCGGCTCGACGTCTTATTCCGTATGGTACGGGATTATTAGTGCTTGCAGCGATGTCC
CTTTCGCTATTGATGTCCTCTCCTGAGTCACGCTACGTACTGAATTGATAAAACATTTCG
ACCTCAGTATCAGTGAGGTCGGGGGATGAATTAGATCGCCGTGTAGGGCTTTTCACGTTG
TGTCCGTGTGGAGAGTCGGTTCTAGCGGTCGTATTATCCATATTCTCCCGTTCCATTTAC
GTTGAGCGACTTGAAACTTCCGATGTACTGGCCGGTAATAGTGCATTGTTTCTCAATTCA
TGCAAGTTGTCGTACATCATGGAATAGAACAAGAATTAAGGACCATTTCCACTAGCGATT
CTTACCTCCAACTCAAGATAGAGTCTAACCGATTATCGACGAACTTCGTCCGTCACGTCA
TTCTATGTCGAGCGGCCTTCTTACAGCGCTGTCGCACTATCTGTTCTACCTTATTTCCGA
GAAACCTTGCTCCTGTTACCTGTGAGGGCGGGGCGGATATGGCAAAATGGGCATTCTTTT
TTACTTCCCGTGCCTCTTCATACTATATACATGTATCTCTGGCGCCTAGACGCCCGGTCC
TTAACGTCCTAAAACTTTACTGACCCTGCTGGCGTCTTACCCTTTATAGTGGCTCTGCAT
AATGTGTTCTGTATATAGCCAAATACTGCCGACTATCCACTCGCTTAGCTGTCAATATCC
ATGCGCGTACTTTGATTTACCGGTCGACCTGCCTAATGTAACTGCCTTTCGCTCTCCACT
CCGCACCGGGGTTTCGAGTTTTTCTACCCCTCTCTTTATTACTACTCGTTTACTGCATCA
GACGCGCCTACCGTATTCAATCACTGCTCCTTCAGGCGAAGAGTATTAGCACGTTATCGT
GCTACTATCCTGAGCTGTAATCATAAGTATTTCTAAAGTTGTAAGGGTTGAAGAGTAGTT
CTGTACATCTTGATCGATTGTACTGGACGTCATTCCGAATAGCTTGTGGAATCCAATATC
TGGGTCATAGATGCT
>SRR9587965
GTGACCGTCGCCTCGCCACAAATCCATTGTTTTCCGGATTCTAGCCCCTTATGAGCTAGG
ACCTCTGTGCCTACCATCATAGCTGTTCGATGCCTTGTATTCAATACGTTGCACCGACCC
GACAATAGTCTGTGCCATTAAGCTTCTGCAAGTGGCTGGCGCCTGTAAGTCACGCAATAG
AGTAACTTCCCTCTCGACAGTCTTTATGCCATCCACAATCCAACATCGTAGATCCTCGCG
TGACCTAGTAGCTCTTGACGACTCAATCCACCCTCTGCCTGACGGACGTGCCTTTTATGG
TACATTAATGACTCCCTGTCCTCATAATTAGGCTGCACACACAGGGCTATCGTAGTTTAC
TGGTCTCACCCCCGCGTACTCGGCGACCCAACTACACTGATCCAGGGAGGCAGAATGGTG
GACGCGCTTACTGCCGACCCGTGGGCAACTAACTGGGCCGGGGGAATATTGTTTCGTTTT

GGGTACTGCTTCTTCCAATTGTTCGGATAGCACAGTAACGGTGGTGGGTGGCTCCAGTCA
CTCTTGCGTAGTCGCCGGCCGGTCGCTCCGGTCTGTCCTCTCGCCCGATTCTGGGCCATC
TTATCCGATGTCACCCTGCAGACCGCTCTTTTCAACCATTAGCATTAAGCAGTCTCTCAT
AGGTGCTCTCATGTCATGAGAGGTACCGGAGCCTCTAAGCGTTCCGCTGTTCCCGACAAT
ATTCTCTTTCGCTTTGACACTAGCTCAGACGGTAGAGGCTTCGTCTTCAGCACCGGCTAA
TGATGCCTGGTACGTTAAGATACCGATCCGCGCGCTTACGCTGCTCCACCTGTCGGTGTC
TGAAGAAGCTCGGCTAAGACCTCTCGAACTCTTCGTCTTACCTCAGCCCGCGGCTACAGT
CATATACATCCTAGCACGCACAATCCTTAATTCCCAGTCGTGCCCGACAGTAACACCGTG
GCGCGAACAATGTGTAAGTTATTGCCTACTTCGTACACTATCAACTAAATTCGTCCAACG
TGAGATCCGAGAATCCCGAAAGGGGGCACACACTTAGAGAACCCGCAGCTTTTTGTCTCA
CGATCGAGAAGGGCACTCCCAAGAGGCCTTGATACGAGCGTGAAACCGGTCAAGCGCTCC
AATTGTACACTGTCCATGAGGCCAAATTCAAATTACGGACCGAAAACTGGATCCAGCAGA
GCTAAAGGTTATTCGCGGGTATTGTCCGTACATGCAACTTGATGTTTTCTAAGCCAAAAG
AGATATGCGAGGGGGAGACATGGGAAAAGCAAAAGAATATAGTACTATAATTGGCGTGGG
ACCCATAAAGTCTTGATTGCTCCCGCGGCGCCTGTACCCACAGTCTTCAAAAAAGCGATT
TGCCGGAGTATAAAGTCAGGGGCCGACGGTGTTGAAAACGAGCGGACGTCTTGCATGAGA
AGTGTCACTGTGGGCATATCAAAGGACTCTAATTAAATTCGGTGTCAATAGAAATGTTCG
TGCTGCACGGTGCTGCCAGACCATCCACCTTTCGTGGCTACGCCTTATCAGAAGTGCAAC
CTGCGAAATCAAGGTGACGTTAATACGCTCTAAAGGACGCTAAAGCGGGTTGATAATTAA
ACGACAGAGTTACAATTAGCGGGATTTACAACTCTCCAAAATTATTGACGATAGACTGAC
ACGGTTACGTGGGAGGACTCGTAGTCTCAAGTGAAAATGGGCGCAAAATGGTGTCCGCAT
GGGAGGAGCTCAATCATAATAGACGCCGAATATGTTCGCGTGAAACTTCAGCGTCGAAGA
TGGAAAGCAAGACAAACTCATGAAACTACCATAGTGTGAAGATGATGGGCGGCGCGGTTT
AAGGCGAGTCGCCCTAAGACTCCAACGTGCCCTAAGGGATCGTAACATGCAAGAACATTG
ATGTGCACAGTTTTGTCGAACGGCCTTCATAGGCCAAAGATTGTATCCACGGCCAGAAGC
GGTAAGAACTTCGTGACATAGCGGCCAGAGAAAGGATCTACAACGTTAACCAGCACCTCA
GAGAGCGGGATGGCCAGTCGTTAGGGAGAAAATAACTAATCATTAACTTTGACGAACCAC
CAAAATGCTCGAGCGGTAGGTATCATTAGGGGGTACGGACTGTTACATCTAGAATACCCC
AGCTCCGTAGTTGCAGACACTAATTTTAGAGACCGCAGCTGTATCATTAACGAAATGTTA
TTGGTGGGATGCAGAAGAGCGCGGGCAAGATTATACAGACGCCTTCAGGCGGTAGCCTGC
TCCTAGGCAGTCAAGACGGATCGCGATGGACGTGTTTGGACGGCGAATGTACAGGAAATT
TCCCGTGAGAACAACAAAATGAAGGATCGAGATTAAAAAGACTGCAGCGCCTGTGAACAA
CGGAGTTGAACGTGGTTGCCAGGTTTGTGATGGGCGGCCCTAGAAGATAACGGTAGAAGC
CCTTCGGACTTTTCAATGGGGACTACATTCTGGCCGCGAAGGACTCCGTGAATCGATAGG
CAGTGGAAATATGGATGAAACGGGCTGGGGCAACGCTCGGGACAACGTTTACTAGCTAAG
AGGCAGATTACGAGGATCGGGGAAGACTGAGAAGGCCTTGGCGTAGGAATTCGAGGCAAC

CAGGGGGATAAGGTAAATGCCGACTTCAACTCGAGAATGCAAAGAAGCTGGCTATGGCCG
TAAATTAGACGATAAATAACAGAGCATCGGGGAGCCAAGTCTAATTGTAAAACTTTTGGT
CGAGGTGATGTCGTAGGTGAGATCTCCGAACGCAGCATGAACAAGGCGAATCGTGTAAGC
AGAGAAGAGTGCAGCACGATGCTAACACTGCCGCAGCAGTGGGACAACTTGGATGACTGG
CACATCACAGTAATGAGTCATATGATTCCGCCTGAAGGCTTCAATAAAGGACATAAGATG
CGAAGAAGATCTAGACCCAGTCTTTAAAACTGCACAACTTAGCCGAAATGATTGGTAGGT
GAGAGTATCGACGTGGGCACTGAGAGAAACCTGAATACGCGATAGCCATCGTATCACGAT
GCTTCTGCCCTCCAGCGTCACTAGAGTTCTTGTTCACATGCCTCGGTCCTCTCGTCAGTA
CATGGCCGGCCTTCTTTCCTCCATCGTTTCCCAGAGCATACTCTTCGAGTTGCCCTATTA
ACCTTCTATTTTCTGGCCGATGCCTGACTGGCGCCGCTTTGAGTATCATACTTCTTGTTT
TTGGCCAGCCTGTTGGGCGTTGATTGCTTGGGTAACCATATCCTGCATTTTGTTGATTCC
GGGTCGGCAATGTGTTGTGGACTGTTCGTCGCCGCTCCTGATCTCACCGTACTTCACGGT
TGTCGCCCGGACATTACCCGGACGGGTTTAGGCCATGGCCGGTTGAGGTTAATCTTTACG
CGTCACCATAACTTCCTGCGAAACTGGTTTCCCCACGCCGCTCTTTTCGTGAGCCGATCG
GCGTGGTTAGGTCGCGCGGGTTTTTCCAGGCAGGGTAAGTCGAGTCGCACAAGGTGACCC
ACGTACCTCGCTTTAACGTTCCGTTCTACCACTGGGTTAGGCTGCTTTGGGGCCGACTTA
ACGTTCTTTTTGCACGACCCCGTATGTGCTTATTCTCACTCGTAATGAACCGGTAAGCGA
GAGTTTGTTCTACGCTGAGGTGTCGTCGGACCCGAGTGTGCTTGCCGCTGGTCAATTGTT
GTGTCGGCTCGACGTCTTATTTCGTATGGTACGGGATTCTTAGTGCTTGCAGCGATGTCC
CTTTCGCTATTGATGTCTTCTCCTGAGTCACGCTACGTACTGAATTGATAAAACATTTCG
ACCTCAGTATCAGTGAGGTCGGGGGATGAGTTAGATCGCCGTGTAGGGCTTTTCACGTTG
TGTCCGTGTGGAGAGTCGGTTCTAGCGGTCGTATTATCCATATTCTCCCGTTCCATTTAC
GTTGAGCGACTTGAAACTTCCGATGTACTGGCCNGTAATAGTGCATTGTTTCTCAATTCA
TGCAAGTTGTCGTACATTATGGAATAGAACAAGAATTAAGGACCATTTCCACTAGCGATT
CTTACCTCCAACTCAAGATAGAGTCTAACCGATTATCGACGAACTTCGTCCGTCACGTCA
TTCTATGTCGAGCGGCCTTCTTACAGCGCTGTCGCACTATCTGTTCTACCTTATTTCCGA
GAAACCTTGCTCCTGTTACCTGTGAGGGCGGGGCGGATATGGCAAAATGGGCATTCTTTT
TTACTTCCCGTGCCTCTTCATGCGATATACATGTATCTCTGGAGCCTAGACGCCCGGTCC
TTAACGTCCTAAAACTTTACTGACCCTGCTGGCGTCTTACCCTTTATAGTGGCTCTGCAT
AATGTGTTCTGTATATAGCCAAATACTGCCGACTATCCACTCGCTTAGCTGTCAATATCC
ATGCGCGTACTTTGATTTACCGGTCGACCTGCCTAATGTAACTGCCTTTCGCTCTCCACT
CCGCACCGGGGTTTCGAGTTTTTCTACCCCTCTCTTTATTACTACTCGTTTACTGCATCA
GACGCGCCTACCGTATTCAGTCACTGCTCCTTCAGGCGAAGAGTATTAGCACGTTATCGT
GCTACTATCCTGAGCTGTAATCATAAGTATTTCTAAAGTTGTAAGGGTTGAAGAGTAGTT
CTGTACATCTTGATCGATTGTACTGGACGTCATTCCGAATAGCTTGTGGAATCCAATATC
TGGGTCATAGATGCT

>SRR9587966

GTGACCGTCGCCTCGCCACAAATCCATTGTTTTCCGGATTCTAGCCCCTTATGAGCTAGG
ACCTCTGTGCCTACCATCATAGCTGTTCGATGCCTTGTATTCAATACGTTGCACCGACCC
GACAATAGTCTGTGCCATTAAGCTTCTGCAAGTGGCTGGCGCCTGTAAGTCACGCAATAG
AGTAACTTCCCTCTCGACAGTCTTTATGCCATCCACAATCCAACATCGTAGATCCTCGCG
TGACCTAGTAGCCCTTGACGACTCAATCCACCCTCTGCCTGACGGACGTGCCTTTTATGG
TACATTAATGACTCCCTGTCCTCATAATTAGGCTGCACACACAGGGCTATCGTAGTTTAC
TGGTCTCACCCCCGCGTACTCGGCGACCCAACTACACTGATCCAGGGAGGCAGAATGGTG
GACGCGCTTACTGCCGACCCGTGGGCAACTAACTGGGCCGGGGGAATATTGTTTCGTTTT
GGGTACTGCTTCTTCCAATTGTTCGGATAGCACAGTAACGGTGGTGGGTGGCTCCAGTCA
CTCTTGCGTAGTCGCCGGCCGGTCGCTCCGGTCTGTCCTCTCGCCCGATTCTGGGCCATC
TTATCCGATGTCACCCTGCAGACCGCTCTTTTCAACCATTAGCATTAAGCAGTCTCTCAT
AGGTGCTCTCATGTCATGAGAGGTACCGGAGCCTCTAAGCGTTCCGCTGTTCCCGACAAT
ATTCTCTTTCGCTTTGACACTAGCTCAGACGGTAGAGGCTTCGTCTTCAGCACCGGCTAA
TGATGCCTGGTACGTTAAGATACCGATCCGCGCGCTTACGCTGCTCCACCTGTCGGTGTC
TGAAGAAGCTCGGCTAAGACCTCTTGAACTCTTCGTCTTACCTCAGCCCGCGGCTACAGT
CATATACATCCTAGCACGCACAATCCTTAATTCCCAGTCGTGCCCGACAGTAACACCGTG
GCGCGAANAATGTGTAAGTTATTGCCTACTTCGTACACTATCAACTAAATTCGTCCAACG
TGAGATCCGAGAATCCCGAAAGGGGACACACACTTAGAGAACCCGCAGCTTTTTGTCTCA
CGATCGAGAAGGGCACTCCCAAGAGGCCTTGATACGAGCGTGAAACCGGTCAAGCGCTCC
AATTGTACACTGTCCATGAGGCCAAATTCAAATTACGGACCGAAAACTGGATCCAGCAGA
GCTAAAGGTTATTCGCGGGTATTGTCCGTACATGCAACTTGATGTTTTCTAAGCCAAAAG
AGATATGCGAGGGGGAGACATGGGAAAAGCAAAAGAATATAGTACTATAATTGGCGTGGG
ACCCATAAAGTCTTGATTGCTCCCGCGCCGCCTGTACCCGCAGTCTTCAAAAAAGCGATT
TGCCGGAGTATAAAGTCAGGGGCCGACGGTGTTGAAAACGAGCGGACGTCTTGCATGAGA
AGTGTCACTGTGGGCATATCAAAGGACTCTAATTAAATTCGGTGTCAATAGAAATGTTCG
TGCTGCACGGTGCTGCCAGACCATCCACCTTTCGTGGCTACGCCTTATCAGAAGTGCAAC
CTGCGAAATCAAGGTGACGTTAATACGCTCTAAAGGACGCTAAAGCGGGTTGATAATTAA
ACGACAGAGTTACAATTAGCGGGATTTACAACTCTCCAAAATTATTGACGATAGACTGAC
ACGGTTACGTGGGAGGACTCGTAGTCTCAAGTGAAAATGGGCGCAAAATGGTGTCCGCAT
GGGAGGAGCTCAATCATAATAGACGCCGAATATGTTCGCGTGAAACTTCAGCGTCAAAGA
TGGAAAGCAAGACAAACTCATGAAACTACCATAGTGTGAAGATGATGGGCGGCGCGGTTT
AAGGCGAGTCGCCCTAAGACTCCAACGTGCCCTAAGGGATCGTAACATGCAAGAACATTG
ATGTGCACAGTTTTGTCGAACGGCCTTCATAGGCCAAAGATTGTATCCACGGCCAGAAGC
GGTAAGAACTTCGTGACATAGCGGCCAGAGAAAGGATCTACAACGTTAACCAGCACCTCA
GAGAGCGGGATGGCCAGTCGTTAGGGAGAAAATAACTAATCATTAACTTTGACGAACCAC

CAAAATGCTCGAGCGGTAGGTATCATTAGGGGGTACGGACTGTTACATCTAGAATACCCC
AGCTCCGTAGTTGCAGACACTAATTTTAGAGACCGCAGCTGTATCATTAACAAAATGTTA
TTGGTGGGATGCAGAAGAGCGCGGGCAAGATTATACAGACGCCTTCAGGCGGTAGCCTGC
TCCTAGGCAGTCAAGACGGATCGCGATGGACGTGTTTGGACGGCGAATGTACAGGAAATT
TCCCGTGAGAACAACAAAATGAAGGATCGAGATTAAAAAGACTGCAGCGCCTGTGAACAA
CGGAGTTGAACGTGGTTGCCAGGTTTGTGATGGGCGGCCCTAGAAGATAACGGTAGAAGC
CCTTCGGACTTTTCAATGGGGACTACATTCTGGCCGCGAAGGACTCCGTGAATCGATAGG
TAGTGGAAATATGGATGAAACGGGCTGGGGCAACGCTCGGGACAACGTTTACTAGCTAAG
AGGCAGATTACGAGGATCGGGGAAGACTGAGAAGGCCTTGGCGTAGGAATTCGAGGCAAC
CAGGGGGATAAGGTAAATGCCGACTTCAAGTCGAGAATGCAAAGAAGCTGGCTATGGCCG
TAAATTAGACGATAAATAACAGAGCATCGGGGAGCCAAGTCTAATTGTAAAACTTTTGGT
CGAGGTGATGTCGTAGGTGAGATCTCCGAACGCAGCATGAACAAGGCGAATCGTGTAAGC
AGAGAAGAGTGCAGCACGATGCTAACACTGCCGCAGCAGTGGGACAACTTGGATGACTGG
CACATCACAGTAATGAGTCATATGATTCCGCCTGAAGGCTTCAATAAAGGACATAAGATG
CGAAGAAGATCTAGACCCAGTCTTTAAAACTGCACAACTTAGCCGAAATGATTGGTAGGT
GAGAGTATCGACGTGGGCACTGAGAGAAACCTGAATACGCGATAGCCATCGTATCACGAT
GCTTCTGCCCTCCAGCGTCACTAGAGTTCTTGTTCACATGCCTCGGTCCTCTCGTCAGTA
CATGGCCGGCCTTCTTTCCTCCATCGTTTCCCAGAGCATACTCTTCGAGTTGCCCTATTA
ACCTTCTATTTTCTGGCCGATGCCTGACTGGCGCCGCTTTGAGTATCATACTTCTTGTTT
TTGGCCAGCCTGTTGGGCGTTGATTGCTTGGGTAACCATATCCTGCATTTTGTTGATTCC
GGGTCGGCAATGTGTTGTGGACTGTTCGTCGCCGCTTCTGATCTCACCGTACTTCACGGT
TGTCGCCCGGACATTACCCGGACGGGTTTAGGCCATGGCCGGTTGAGGTTAATCTTTACG
CGTCACCATAACTTCCTGCGAAACTGGTTTCCCCACGCCGCTCTTNTCGTGAGCCGATCG
GCGTGGTTAGGTCGCGCGTGTTTTTCCAGGCAGGGTAAGTCGAGTCGCACAAGGTGACCC
ACGTACCTCGCTTTAACGTTCCGTTCTACCACTGGGTTAGGCTGCTTTGGGGCCGACTTG
ACGTTCTTTTTGCACGACCCCGTATGTGCTTATTCTCACCCGTAATGAACCGGTAAGCGA
GAGTTTGTTCTACGCTGAGGTGTCGTCGGACCCGAGTGTGCTTGCCGCTGGTCAATTGTT
GTGTCGGCTCGACGTCTTATTTCGTATGGTACGGGATTCTTAGTGCTTGCAGCGATGTCC
CTTTCGCTATTGATGTCTTCTCCTGAGTCACGCTACGTACTGAATTGATAAAACATTTCG
ACCTCAGTATCAGTGAGGTCGGGGGATGAGTTAGATCGCCGTGTAGGGCTTTTCACGTTG
TGTCCGTGTGGAGAGTCGGTTCTAGCGGTCGTATTATCCATATTCTCCCGTTCCATTTAC
GTTGAGCGACTTGAAACTTCCGATGTACTGGCCGGTAATAGTGCATTGTTTCTCAATTCA
TGCAAGTTGTCGTACATTATGGAATAGAACAAGAATTAAGGACCATTTCCACTAGCGATT
CTTACCTCCAACTCAAGATAGAGTCTAACCGATTATCGACGAACTTCGTCCGTCACGTCA
TTCTATGTCGAGCGGCCTTCTTACAGCGCTGTCGCACTATCTGTTCTACCTTATTTCCGA
GAAACCTTGCTCCTGTTACCTGTGAGGGCGGGGCGGATATGGCAAAATGGGCATTCTTTT

TTACTTCCCGTGCCTCTTCATGCTATATACATGTATCTCTGGAGCCTAGACGCCCGGTCC
TTAACGTCCTAAAACTTTACTGACCCTGCTGGCGTCTTACCCTTTATAGTGGCTCTGCAT
AATGTGTTCTGTATATAGCCAAATACTGCCGACTATCCACTCGCTTAGCTGTCAATATCC
ATGCGCGTACTTTGATTTACCGGTCGACCTGCCTAATGTAACTGCCTTTCGCTCTCCACT
CCGCACCGGGGTTTCGAGTTTTTCTACCCCTCTCTTTATTACTACTCGTTTACTGCATCA
GACGCGCCTACCGTATTCAGTCACTGCTCCTTCAGGCGAAGAGTATTAGCACGTTATCGT
GCTACTATCCTGAGCTGTAATCATAAGTATTTCTAAAGTTGTAAGGGTTGAAGAGTAGTT
CTGTACATCTTGATCGATTGTACTGGACGTCATTCCGAATAGTTTGTGGAATCCAATATC
TGGGTCATAGATGCT
>SRR9587967
GTGACCGTCGCCTCGCCACAAATCCATTGTTTTCCGGGTTCTAGCCCCTTATGAGCTAGG
ACCTCTGTTCCTACCATCATAGCTGTTCGATGCCTTGTATTCAATACGTTGCACCGACCC
GACAATAATCTGTGCCATTAAGCTTCAGCAAGTGGCTGGCGCCTGTAAGTCACGCAATAG
AGTAACTTCCCTCTCGACAGTCTTTATGCCATCCACAATCCAACATCGTAGATCCTCGCG
TGACCTAGTAGCCCTTGACGACTCAATCCACCCTCTGCCTGACGGACGTGCCTTTTATGG
TACATTAATAACTCCCTGTCCTCATAATTAGGCTGAACACACAGGGCTATCGTAGTTTAC
TGGTCTCACCCCTGCGTACTCGGCGACCCAACTACACTGGTCCATGGAGGCAGAATGGTG
GACGCTCTTACTGCCGACCCGTGGGCAACTAACTGGGCCCGGGGAATATTGTTTCGTTTT
GGGTACTGCTTCTTCCAATTGTTCGGATAGCACAGTAACGGTGGTGGGTGGCTCCAGTCA
CTCTTGCGTAGTCGCCGGCCGGTCGCTCCGGTCTGTCCTCTCGCCCGATTCGGGGCCATC
TTGTCCGATGTCACCCTGCAGACCGCTCTTTTCAACCATTAGCATTAAGCAGTTTCTCAT
AGGTGCTCTCATGTCATGAGAGGTACCGGAGCCTCTAAGCGTTCCGCTGTTCCCGACAAT
ATTCTCTTTCGCTTTAACACTAGCTCAGACGGTAGAGGCTTCGTCTTCAGCACCGGCTAA
TGATGCCTGGTACGTTAAGATACCGATCCGCGCGCTTACGCTGCTCCACCTGTCGGTGTC
TGAAGAAGCTCGGCTAAGACCTCTCGAACTCTTCGTCTTACCTCAGCCCGCGGCTACAGT
CATATACATCCTAGCACGCACAATCCTTAATTCCCAGTCGTGCCCGACAGTAACACCGTG
GCGCGAACAATGTGTAAGTTGTTGCCTACTTCGTACACTATCAACTAAATTCGTCCAACG
TGAGATCCGAGAATCCCGAAAGGGGGCACACACTTAGAGAACCCGCAGCTTTTTGTCTCA
CGATCGAGAAGGGCACTCCCAAGAGGCCTTGATACGAGCGTGAAACCGGTCAAGCGCTCC
AATTGTACACTGTCCATGAGGCCAAATTCAAATTACGGACCGAAAACTGGATCCAGCAGA
GCTAAAGGTTATTCGCGGGTATTGTCCGTACATGCAACTTGATGTTTTCTAAGCCAAAAG
AGATATGCGAGGGGGAGACATGGGAAAAGCAAAAGAATATAGTACCATAATTGGCGTGGG
ACCCATAAAGTCTTGATTGCTCCCGCGCCGCCTGTACCCACAGTCTTCAAAAAAGCAATT
TGCCGGAGTATAAAGTCAGGGGCCGACGGTGTTGAAAACGAGCGGACGTCTTGCATGAGA
AGTGTCACTGTGGGCATATCAAAGGACTCTAATTAAATTCGGTGTCAATAGAAATGTTCG
TGCTGCACGGTGCTGTCAGACCATCCATCTTTCGTGGCTACGCCTTATCAGAAGTGCAAC

CTGCGAAATCAAGGTGACGTTAATACGCTCTAAAGGACGCTAAAGCGGGTTGATAATTAA
ACGACAGAGTTACAATTAGCGGTATTCACAACTCTCCAAAATCATTGACGATAGACTGAC
ACGGTTACGTGGGAGGACTCGTAGTCTCAAGTGAAAATGGGCGCAAAATGGTGTCCGCAT
GGGAGGAGCTCAACCATAATAGACGCCGAATATGTTCGCGTGAAACTTCAGCGTCAAAGA
TGGAAAGCAAGACAAACTCATGAAACTACCATAGTGTGAAGATGATGGGCGGCGCGGTTT
AAGGCGAGTCGCCCTAAGACTCCAACGTGCCCTAAGGGATCGTAACATGCAAGAACATTG
ATGTGCACAGTTTTGTCGAACGGCCTTCATAGGCCAAAGATTGTATCCACGGCCAGAAGC
GGTAAGAACTTCGTGACATAGCGGCCAGAGAAAGGATCTACAACGTTAACCAGCACCTCA
GAGAGCGGGACGGCCAGTCGTTAGGGAGAAAATAACTAATCATTAACTTTGACGAACCAC
CAAAATGCTCGAGCGGTAGGTATCATTAGGGGGTACGGACTGTTACATCTAGAATACCCC
AACTCCGTAGTTGCAGACACTAATTTTAGAGACCGCAGCTGCATCATTAACAAAATGTTA
TTGGTGGGATGCAGAAGAGCGCGGGCAAGATTATACAGACGCCTTCAGGCGGTAGCCTGC
TCCTAGGCAGTCAAGACGGATCGCGATGGACGTGTTTGGACGGCGAATGTACAGGAAATT
TCCCGTGAGAACAACAAAATGAAGGATCGAGATTAAAAAGACTACAGCGCCCGTGAACAA
CGGAGTTGAACGTGGTTGCCAGGTTTGTGATGGGCGGCCCTAGAAGATAACGGTAGAAGC
CCTTCGGACTTTTCAATGGGGACTACACTCTGGCCGCGAAGGACTCCGTGAATCGATAGG
TAGTGGAAATATGGATGAAACGGGCTGGGGCAACGCTCGGGACAACGTTTACTAGCTAAG
AGGCAGATTACGAGGATCGGGGAAGACTAAGAAGGCCTTGGCGTAGGAATTCGAGGCAAC
CAGGGGGGTAAGGTAAATGCCGACTTCAAGTCGAGAATGCAAAGAAGCTGGCTATGGCCG
TAAATTAGACAATAAATAACAGAGCATCGGGGGGCCAAGTCTAATTGTAAAACTTTTGGT
CGAGGTGATGTCGTAGGTGAGATCTCCGAACGCAGCATGAACAAGGCGAATCGTGTAAGC
AGAGAAGAGTGCAGCACGATGCTAACACTGCCGCAGCAGTGGGACAACTTGGATGACTGG
CACATCACAGTAATGAGTCATATGATTCCGCCTGAAGGCTTCAATAAAGGACATAAGATG
CGAAGAAGATCTAGACCCAGTCTTTAAAACCGCACAACTTAGCCGAAATGATTGGTAAGT
GAGAGTATCGACGTGGGCACTGAGAGAAACCTGAATACGCGATAGCCATCATATCACGAT
GCTTCTGCCCTCTAGCGTCACTAGAGTTCTTGTTCACACGCCTCGGTCCTCTCGTCAGTA
CATGGCCGGCCTTCTTTCCTCCATCGTTTCCCAGAGCATACTCTTCGAGTTGCCCTATTA
ACCTTCTACTTTCTGGCCGATGCCTGACTGGCGCCGCTTTGAGTATCATACTTCTTGTTT
TTGGCCAACCTGTTGGGCGTTGATTGCTTGGGTAACCATATCCTGCATTTTGTTGATTCC
TGGTCGGCAATGTGTTGTGGACTGTTCGTCGCCGCTCCTGATCTCACCGTACTTCACGGT
TGTCGCCCGGACATCACCCGGACGGGTTTAGGCCATGGCCGGTTGAGGTTAATCTTTACG
CGTCATCATAACTTCCTGCGAAACTGGTTTCCCCACGCCGCTCCCTTCGTGAGCCGATCG
GCGTGGTTAGGTCGCGCGGGTTTTTCCAGGCAGGGTAAGTCGAGTCGCACAAGGTAACCC
ACGTACCTCGCTTTAACGTTCCGTTCTACCACTGGGTTAGGCCGCTTTGGGGCCGACTTG
ACGTTCTTTTTGCACGACCTCGTATGTGCTTATTCTCACTCGTAATGAACCGGTAAGCGA
GAGTTTGTTCTACGCTGAGGTGTCGTTGGACCCGAGTGTGCTTGCCGCTGGTCAATTGTT

GTGTCGGCTCGACGTCTTATTTCGTATGGTACGGGATTATTAGTGCTTGCAGCGATGTCC
CTTTCGCTATTGATGTCTTCTCCTGAGTCACGCTACGTACTGAATTGATAAAACATTTCG
ACCTCAGTATCAGTGAGGTCGGGCGATGAGTTAGATCACCGTGTAGGGCTTTTCACGTTG
TGTCCGTGTGGAGAGTCGGTTCTAGCGGTCGTATTATCCATATTCTCCCGTTCTATTTAC
GTTGAGCGACTTGAAACTTCCGATGTACTGGCCGGTAATAGTGCATTGTTTCTCAATTCA
TGCAAGTTGTCGTACATTATGGAATAGAACAAGAATTAAGGACCATTTCCACTAGTGATT
CTTAGCTCCAACTCAAGATAGAGTCTAACCGATTATCGACGAACTTCGTCCGTCACGTCA
TTCTATGTCGAGCGGCCTTCTTACAGCGCTGTCGCACTATCTGTTCCACCTTATTTTCGA
GAAACCTTGCTCCTGTTACCTGTGAGGGCGGGGCGGATATGGCAAAATGGGCATTCTTTT
TTACTTCTCGTGCCTCTTCATGCTATATACATGTATCTCTGGAGCCTAGACGCCCGGTCC
TTAACGTCCTAAAACTTTACTGACCCTGCTGGCGTCTTACCCTTTATAGTGGCTCTGCAT
AATGTGTTCTGTATATAGCCAAATACTGCCGACTATCCACTCGCTTAGCTGTCAATATCC
ATGCGCGTACTTTGATTTACCGGTCGACCTGCCTAATGTAACTGCCTTTCGCTCTCCACT
CCGCACCGGGGTTTCGAGTTTTTCTACCCCTCTCTTTATTACTACTCGTTTACTGCATCA
GACGCGCCTACCGTATTCGGTCACTGCTCCTTCAGGCGAAGAGTATTAGCACGTTATCGT
GCCACTATCCTGAGCTGTAATCATAAGTATTTCTAAAGTTGTAAGGGTTGAAGAGTAGTT
CTGTACATCTTGATCGATTGTACTGGACGTCATTCCGAATAGCTTGTGGAATCCAATATC
TGGGTCATAGATGCT
>SRR9587968
GTGACCGTCGCCTCGCCACAAATCCATTGTTTTCCGGGTTCTAGCCCCTTATGAGCTAGG
ACCTCTGTTCCTACCATCATAGCTGTTCGATGCCTTGTATTCAATACGTTGCACCGACCC
GACAATAATCTGTGCCATTAAGCTTCAGCAAGTGGCTGGCGCCTGTAAGTCACGCAATAG
AGTAACTTCCCTCTCGACAGTCTTTATGCCATCCACAATCCAACATCGTAGATCCTCGCG
TGACCTAGTAGCCCTTGACGACTCAATCCACCCTCTGCCTGACGGACGTGCCTTTTATGG
TACATTAATAACTCCCTGTCCTCATAATTAGGCTGAACACACAGGGCTATCGTAGTTTAC
TGGTCTCACCCCTGCGTACTCGGCGACCCAACTACACTGGTCCATGGAGGCAGAATGGTG
GACGCTCTTACTGCCGACCCGTGGGCAACTAACTGGGCCCGGGGAATATTGTTTCGTTTT
GGGTACTGCTTCTTCCAATTGTTCGGATAGCACAGTAACGGTGGTGGGTGGCTCCAGTCA
CTCTTGCGTAGTCGCCGGCCGGTCGCTCCGGTCTGTCCTCTCGCCCGATTCGGGGCCATC
TTGTCCGATGTCACCCTGCAGACCGCTCTTTTCAACCATTAGCATTAAGCAGTTTCTCAT
AGGTGCTCTCATGTCATGAGAGGTACCGGAGCCTCTAAGCGTTCCGCTGTTCCCGACAAT
ATTCTCTTTCGCTTTAACACTAGCTCAGACGGTAGAGGCTTCGTCTTCAGCACCGGCTAA
TGATGCCTGGTACGTTAAGATACCGATCCGCGCGCTTACGCTGCTCCACCTGTCGGTGTC
TGAAGAAGCTCGGCTAAGACCTCTCGAACTCTTCGTCTTACCTCAGCCCGCGGCTACAGT
CATATACATCCTAGCACGCACAATCCTTAATTCCCAGTCGTGCCCGACAGTAACACCGTG
GCGCGAACAATGTGTAAGTTGTTGCCTACTTCGTACACTATCAACTAAATTCGTCCAACG

TGAGATCCGAGAATCCCGAAAGGGGGCACACACTTAGAGAACCCGCAGCTTTTTGTCTCA
CGATCGAGAAGGGCACTCCCAAGAGGCCTTGATACGAGCGTGAAACCGGTCAAGCGCTCC
AATTGTACACTGTCCATGAGGCCAAATTCAAATTACGGACCGAAAACTGGATCCAGCAGA
GCTAAAGGTTATTCGCGGGTATTGTCCGTACATGCAACTTGATGTTTTCTAAGCCAAAAG
AGATATGCGAGGGGGAGACATGGGAAAAGCAAAAGAATATAGTACCATAATTGGCGTGGG
ACCCATAAAGTCTTGATTGCTCCCGCGCCGCCTGTACCCACAGTCTTCAAAAAAGCAATT
TGCCGGAGTATAAAGTCAGGGGCCGACGGTGTTGAAAACGAGCGGACGTCTTGCATGAGA
AGTGTCACTGTGGGCATATCAAAGGACTCTAATTAAATTCGGTGTCAATAGAAATGTTCG
TGCTGCACGGTGCTGTCAGACCATCCATCTTTCGTGGCTACGCCTTATCAGAAGTGCAAC
CTGCGAAATCAAGGTGACGTTAATACGCTCTAAAGGACGCTAAAGCGGGTTGATAATTAA
ACGACAGAGTTACAATTAGCGGTATTCACAACTCTCCAAAATCATTGACGATAGACTGAC
ACGGTTACGTGGGAGGACTCGTAGTCTCAAGTGAAAATGGGCGCAAAATGGTGTCCGCAT
GGGAGGAGCTCAACCATAATAGACGCCGAATATGTTCGCGTGAAACTTCAGCGTCAAAGA
TGGAAAGCAAGACAAACTCATGAAACTACCATAGTGTGAAGATGATGGGCGGCGCGGTTT
AAGGCGAGTCGCCCTAAGACTCCAACGTGCCCTAAGGGATCGTAACATGCAAGAACATTG
ATGTGCACAGTTTTGTCGAACGGCCTTCATAGGCCAAAGATTGTATCCACGGCCAGAAGC
GGTAAGAACTTCGTGACATAGCGGCCAGAGAAAGGATCTACAACGTTAACCAGCACCTCA
GAGAGCGGGACGGCCAGTCGTTAGGGAGAAAATAACTAATCATTAACTTTGACGAACCAC
CAAAATGCTCGAGCGGTAGGTATCATTAGGGGGTACGGACTGTTACATCTAGAATACCCC
AACTCCGTAGTTGCAGACACTAATTTTAGAGACCGCAGCTGCATCATTAACAAAATGTTA
TTGGTGGGATGCAGAAGAGCGCGGGCAAGATTATACAGACGCCTTCAGGCGGTAGCCTGC
TCCTAGGCAGTCAAGACGGATCGCGATGGACGTGTTTGGACGGCGAATGTACAGGAAATT
TCCCGTGAGAACAACAAAATGAAGGATCGAGATTAAAAAGACTACAGCGCCCGTGAACAA
CGGAGTTGAACGTGGTTGCCAGGTTTGTGATGGGCGGCCCTAGAAGATAACGGTAGAAGC
CCTTCGGACTTTTCAATGGGGACTACACTCTGGCCGCGAAGGACTCCGTGAATCGATAGG
TAGTGGAAATATGGATGAAACGGGCTGGGGCAACGCTCGGGACAACGTTTACTAGCTAAG
AGGCAGATTACGAGGATCGGGGAAGACTAAGAAGGCCTTGGCGTAGGAATTCGAGGCAAC
CAGGGGGNTAAGGTAAATGCCGACTTCAAGTCGAGAATGCAAAGAAGCTGGCTATGGCCG
TAAATTAGACAATAAATAACAGAGCATCGGGGGGCCAAGTCTAATTGTAAAACTTTTGGT
CGAGGTGATGTCGTAGGTGAGATCTCCGAACGCAGCATGAACAAGGCGAATCGTGTAAGC
AGAGAAGAGTGCAGCACGATGCTAACACTGCCGCAGCAGTGGGACAACTTGGATGACTGG
CACATCACAGTAATGAGTCATATGATTCCGCCTGAAGGCTTCAATAAAGGACATAAGATG
CGAAGAAGATCTAGACCCAGTCTTTAAAACCGCACAACTTAGCCGAAATGATTGGTAAGT
GAGAGTATCGACGTGGGCACTGAGAGAAACCTGAATACGCGATAGCCATCATATCACGAT
GCTTCTGCCCTCTAGCGTCACTAGAGTTCTTGTTCACACGCCTCGGTCCTCTCGTCAGTA
CATGGCCGGCCTTCTTTCCTCCATCGTTTCCCAGAGCATACTCTTCGAGTTGCCCTATTA

ACCTTCTACTTTCTGGCCGATGCCTGACTGGCGCCGCTTTGAGTATCATACTTCTTGTTT
TTGGCCAACCTGTTGGGCGTTGATTGCTTGGGTAACCATATCCTGCATTTTGTTGATTCC
TGGTCGGCAATGTGTTGTGGACTGTTCGTCGCCGCTCCTGATCTCACCGTACTTCACGGT
TGTCGCCCGGACATCACCCGGACGGGTTTAGGCCATGGCCGGTTGAGGTTAATCTTTACG
CGTCATCATAACTTCCTGCGAAACTGGTTTCCCCACGCCGCTCCCTTCGTGAGCCGATCG
GCGTGGTTAGGTCGCGCGGGTTTTTCCAGGCAGGGTAAGTCGAGTCGCACAAGGTAACCC
ACGTACCTCGCTTTAACGTTCCGTTCTACCACTGGGTTAGGCCGCTTTGGGGCCGACTTG
ACGTTCTTTTTGCACGACCTCGTATGTGCTTATTCTCACTCGTAATGAACCGGTAAGCGA
GAGTTTGTTCTACGCTGAGGTGTCGTTGGACCCGAGTGTGCTTGCCGCTGGTCAATTGTT
GTGTCGGCTCGACGTCTTATTTCGTATGGTACGGGATTATTAGTGCTTGCAGCGATGTCC
CTTTCGCTATTGATGTCTTCTCCTGAGTCACGCTACGTACTGAATTGATAAAACATTTCG
ACCTCAGTATCAGTGAGGTCGGGCGATGAGTTAGATCACCGTGTAGGGCTTTTCACGTTG
TGTCCGTGTGGAGAGTCGGTTCTAGCGGTCGTATTATCCATATTCTCCCGTTCTATTTAC
GTTGAGCGACTTGAAACTTCCGATGTACTGGCCGGTAATAGTGCATTGTTTCTCAATTCA
TGCAAGTTGTCGTACATTATGGAATAGAACAAGAATTAAGGACCATTTCCACTAGTGATT
CTTAGCTCCAACTCAAGATAGAGTCTAACCGATTATCGACGAACTTCGTCCGTCACGTCA
TTCTATGTCGAGCGGCCTTCTTACAGCGCTGTCGCACTATCTGTTCCACCTTATTTTCGA
GAAACCTTGCTCCTGTTACCTGTGAGGGCGGGGCGGATATGGCAAAATGGGCATTCTTTT
TTACTTCTCGTGCCTCTTCATGCTATATACATGTATCTCTGGAGCCTAGACGCCCGGTCC
TTAACGTCCTAAAACTTTACTGACCCTGCTGGCGTCTTACCCTTTATAGTGGCTCTGCAT
AATGTGTTCTGTATATAGCCAAATACTGCCGACTATCCACTCGCTTAGCTGTCAATATCC
ATGCGCGTACTTTGATTTACCGGTCGACCTGCCTAATGTAACTGCCTTTCGCTCTCCACT
CCGCACCGGGGTTTCGAGTTTTTCTACCCCTCTCTTTATTACTACTCGTTTACTGCATCA
GACGCGCCTACCGTATTCGGTCACTGCTCCTTCAGGCGAAGAGTATTAGCACGTTATCGT
GCCACTATCCTGAGCTGTAATCATAAGTATTTCTAAAGTTGTAAGGGTTGAAGAGTAGTT
CTGTACATCTTGATCGATTGTACTGGACGTCATTCCGAATAGCTTGTGGAATCCAATATC
TGGGTCATAGATGCT
>SRR9587969
GTGACCGTCGCCTCGCCACAAATCCATTGTTTTCCGGGTTCTAGCCCCTTATGAGCTAGG
ACCTCTGTGCCTACCATCATGGCTGTTCGATGCCTTGTATTCAATACGTTGCACCGACCC
GACAATAGTCTGTGCCATTAAGCTTCTGCAAGTGGCTGGCGCCTGTAAGTCACGCAATAG
AGTAACTTCCCTCTCGACAGTCTTTATGCCATCCACAATCCAACATCGTAGATCCTCGCG
TGACCTAGTAGCCCTTGACGACTCAATCCACCCTCTGCCTGACGGACGTGCCTTTTATGG
TACATTAATGACTCCCTGTCCTCATAATTAGGCTGCACACACAGGGCTATCGTAGTTTAC
TGGTCTCACCCCTGCGCACTCGGCGACCCAATTACACTGATCCGGGGAGGCAGAATGGTG
GACGCGCTTACTCCCGACCCGTGGGCAACTAACTGGGCCGGGGGAATATTGTTTCGTTTT

GGGTACTGCTTCTTCCAATTGTTCGGATAGCACAGTAACGGTGGTGGGTGGCTCCAGTCA

CTCTTGCGTAGTCGCCGGCCGGTCGCTCCGGTCTGTCCTCTCGCCCGATTCTGGGCCATC

TTATCCGATGTCACCCTGCAGACCGCTCTTTTCAACCATTAGCATTAAGCAGTCTCTCAT

AGGTGCTCTCATGTCATGAGAGGTACCGGAGCCTCTAAGCGTTCCGCTGTTCCCGACAAT

ATTCTCTTTCGCTTTGACACTAGCTCAGACGGTAGAGGCTTCGTCTTCAGCACCGGCTAA

TGATGCCTGGTACGTTAAGATACCGATCCGCGCGCTTACGCTGCTCCACCTGTCGGTGTC

TGAAGAAGCTCGGCTAAGACCTCTCGAACTCTTCGTCTTACCTCAGCCCGCGGCTACAGT

CATATACATCCTAGCACGCACAATCCTTAATTCCCAGTCGTGCCCGCCAGTAACACCGTG

GCGCGAACAATGTGTAAGTTATTGCCTACTTCGTACACTATCAACTAAATTCGTCCAACG

TGAGATCCAAGAATCCCGAAAGGGGGCACACACTTAGAGAACCCGCAGCTTTTTGTCTCA

CGATCGAGGAGGGCACTCCCAAGAGGCCTTGATACGAGCGTGAAACCGGTCAAGCGCTCC

AATTGTACACTGTCCATGAGGCCAAATTCAAATTACGGACCGAAAACTGGATCCAGCAGA

GCTAAAGGTTATTCGCGGGTATTGTCCGTACATGCAACTTGATGTTTTCTAAGCCAAAAG

AGATATACGAGGGGGAGACATGGGAAAAGCAAAAGAATATAGTACTATAATTGGCGTGGG

ACCCATAAAGTCTTGATTGCTCCCGCGCCGCCTGTGCCCACAGTCTTCAAAAAAGCGATT

TGCCGGAGTATAAAGTCAGGGGCCGACGGTGTTGAAAACGAGCGGACGTCTTGCATGAGA

AGTGTCACTGTGGGCATATCAAAGGACTCTAATTAAATTCGGTGTCAATAGAAATGTTCG

TGCTGCACGGTGCTGTCAGACCATCCACCTTTCGTGGCTACGCCTTATCAGAAGTGCAAC

CTGCGAAATCAAGGTGACGTTAATACGCTCTACAGGACGCTAAAGCGGGTTGATAATTAA

ACGACAGAGTTACAATTAGCGGGATTCACAACTCTCCAAAATCATTGACAATAGACTGAC

ACGGTTACGTGGGAGGACTCGTAGTCTCAAGTGAAAATGGGCGCAAAATGGTGTCCGCAT

GGGAGGAGCTCAACCATAATAGACGCCGAATATGTTCGCGTGAAACTTCAGAGTCAAAAA

TGGAAAGCAAGACAAACTCATGAAACTACCATAGTGTGAAGATGATGGGCGGCGCGGTTT

AAGGCGAGTCGCCCTAAGACTCCAACGTGCCCTAAGGGATCGTAACATGCAAGAACATTG

ATGTGCACAGTTTTGTCGAACGGCCTTCATAGGCCAAAGATTGTATCCACGGCCAGAAGC

GGTAAGAACTTCGTGACATAGCGGCCAGAGAAAGGATCTACAACGTTAACCAGCACCTCA

GAGAGCGGGATGGCCAGTCGTTAGGGAGAAAATAACTAATCATTAACTTTGACGAACCAC

CAAAATGCTCGAGCGGTAGGTATCATTAGGGGGTACGGACTGTTACATCTAGAATACCCC

AGCTCCGTAGTTGCAGACACTAATTTTAGAGACCGCAGCTGTATCATTAACAAAATGTTA

TTGGTGGGATGCAGAAGAGCGCGGGCAAGATTATACAGACGCCTTCAGGCGGTAGCCTGC

TCCTAGGCAGTCAAGACGGATCGCGATGGACGTGTTTGGACGGCGAATGTACAGGAAATT

TCCCGTGAGAACAACAAAATGAAGAATCGAGATCAAAAAGACTGCAGCGCCCGTGAACAA

CGGAGTTGAACGTGGTTGCCAGGTTTGTGATGGGCGGCCCTAGAAGATAGCGGTAGAAGC

CCTTCGGACTTTTTAATGGGGACTACATTCTGGCCGCGAAGGACTCCGTGAATCGATAGG

TAGTGGAAATATGGATGAAACGGGCTGGGGCAACGCTCGGGACAACGTTTACTAGCTAAG

GGGCAGATTACGAGGATCGGGGAAGACTGAGAAGGCCTTGGCGTAGGAATTCGAGGCAAC

CAGGGGGATAAGGTAAATGCCGACTTCAAGTCGAGAATGCAAAGAAGCTGGCTATGGCCG
TAAATTAGACGATAAATAACAGAGCATCGGGGAGCCAAGTCTAATTGTAAAACTTTTGGT
CGAGGTGATGTCGTAGGTGAGATCTCCGAACGCAGCATGAACAAGGCGAATCGTGTAAGC
AGAGAAGAGTGCAGCACGATGCTAACACTGCCGCAGCAGTGGGACAACTTGGATGACTGG
CACATCACAGTAATGAGTCATATGATTCCGCCTGAAGGCTTCAATAAAGGACATAAGATG
CGAAGAAGATCTAGACCCAGTCTTTAAAACTGCACAACTTAGCCGAAATGATTGGTAGGT
GAGAGTATCGACGTGGGCACTGAGAGAAACCTGAATACGCGATAGCCATCATATCACGAT
GATTCTGCCCTTCAGCGTCACTAGAGTTCTTGTTCACATGCCTCGGTCCTCTCGTCAGTA
CATGGCCGGCCTTCTTTCCTCCATCGTTTCCCAGAGCATACTCCTCGAGTTGCCCTATTA
ACCTTCTATTTTCTGGCCGATGCCTGACTGGCGCCGCTTTGAGTATCATACTTCTTGTTT
TTGGCCAACCTGTTGGGCGTTGATTGCTTGGGTAACCATATCCTGCATTTTGTTGATTCC
GGGTCGGCAATGTGTTGTGGACTGTTCGTCGCCGCTCCTGATCTCACCGTACTTCACGGT
TGTCGCCCGGACATCACCCGGACGGGTTTAGGCCATGGCCGGTTGAGGTTAATCCTTACG
CGTCACCATAACTTCCTGCGAAACTGGTTTCCCCACGCCGCTCCCTTCGTGAGCCGATCG
GCGTGGTTAGGTCGCGCGGGTTTTTCCAGGCAGGGTAAGTCGAGTCGCACAAGGTGACCC
ACGTACCTCGCTTTAACGTTCCGTTCTACCACTGGGTTAGGCTGCTTTGGGGCCGACTTG
ACGTTCTTTTTGCACGACCTCGTATGTGCTTATTCTCACTCGTAATGAACCGGTAAGCGA
GAGTTTGTTCTACGCTGAGGTGTCGTCGGACCCGAGTGTGCTTGCCGCTGGTCAATTGTT
GCGTCGGCTCGACGTCTTATTTCGTATGGTACGGGATTATTAGTGCTTACAGCGATGTCC
CTTTCGCTATTGATGTCCTCTCCTGAGTCACGCTACGTACTGAATTGATAAAACATTTCG
ACCTCAGTATCAGTGAGGTCGGGGGATGAGTTAGACCGCCGTGTAGGGCTTTTCACGTTG
TGTCCGTGTGGAGGGTTGGTTCTAGCGGTCGTATTATCCATATTCTCCCGTTCCATTTAC
GTTGAGCGACTTGAAACTTCCGATGTACTGGCCGGTAATAGTGCATTGTTTCTCAATTCA
TGCAAGTTGTCGTACATCATGGAATAGAACAAGAATTAAGGACCATTTCCACTAGCGATT
CTTACCTCCAACTCAAGATAGAGTCTAACCGATTATCGACGAACTTCGTCCGTCACGTCA
TTCTATGTCGAGCGGCCTTCTTACAGCGCTGTCGCACTATCTGTTCTACCTTATTTCCGA
GAAACCTTGCTCCTGTTACCTGTGAGGGCGGGGCGGATATGGCAAAATGGGCATTCTTTT
TTACTTCCCGTGCCTCTTCATGCTATATACATGTATCTCTGGAGCCTAGACGCCCGGTCC
TTAACGTCCTAAAACTTTACTGACCCTGCTGGCGTCTAACCCTTTATAGTGGCTCTGCAT
AATGTGTTCTGTATATAGCCAAATACTGCCGACTATCCACTCGCTTAGCTGTCAATATCC
ATGCGCGTACTTTGATTTACCGGTCGACCTGCCTAATGTAACTGCCTTTCGCTCTCCACT
CCGCACCGGGGTTTCGAGTTTTTCTACCCCTCTCTTTATTACTACTCGTTTACTGCATCA
GACGCGCCTACCGTATTCAGTCACTGCTCCTTCAGGCGAAGAGTATTAGCACGTTATCGT
GCTACTATCCTGAGCTGTAATCATAAGTATTTCTAAAGTTGTAAGGGTTGAAGAGTAGTT
CTGTACATCTTGATCGATTGTACTGGACGTCATTCCGAATAGCTTGTGGAATCCAATATC
TGGGTCATAGATGCT
