## Supplementary material for "Genomic diversity of 39 samples of *Pyropia* species grown in Japan": S7 Fig

>Pyr_1

TGAGCCTTTTATTCTAAGCTCAACATTTCTCCACCTCCTTGGCGCCATCCCGGAACCCAG
GCATAAAACGCAGCGAAATAAACAGAGCGCTATGCCTTTACCGTACCTGCCGCTAACCAT
TGTTACACGATTCAGTTGTAAAATTGCTCGCGTTGGCATAGTTAGGTGCCGGTTGCCCCC
CATACAATACGTTTCCAGCATGCACATATTCTCAGATTTTATTATGACCTTCACGTCACT
CATTATGTATTTTGCCTTACGTCCTCCATTTTCCTCTATGTCCGTTTTTCTCTCAAGCTG
ATACATCAGTTCNNAACTCCCGCGAATTCNATTCGTGCATGAGTAGATATTGGTCCTTGA
TTGAGTACCTATAAAACGCTAGGCATCAAAGGACACTGCAATGCACAAGTCAAACAGGTC
CTCTGAAACCAAAACATAAAGACAGACGGAAGCAACCAAAGGGACAGACCATGTATTAAC
ATAGGATGCGAAAATAGTCAAGCATAGGGTAATAATTTGTTCCACAGAGGTTTGATAGCT
TACGAAAGATATCGCGAGCAACAGGCGGATACGTTGGGCGAACGAGGGAAGAGACCGAGT
CGAGTGAGACCGGAGCGGGGAGAGGGAGAGCGAATCCGAATAACAAGAGAACTGGGACAG
GAAAACACAGTGAGCAGAACCTCAGGACGAAAGAGAGACGCACTAAGAAAACAGAGGCGC
GACGTGAAGCGAGCGGAAGACCTATTGATGTAACAGAGGGAGGCGTGATATGAAAAAACT
GAAGTCGGGGTAGGGAAAGGTCGTGAAGGGGACGAATAATAATCTGCTCTTTAGCGGCAG
AAGATACATCGAGATCAGTCCTTTTTCCGAGCGATCTTTGAGCATTATCCACTTGAGCTG
GACGGCCAGACGCAGGAACTTCTAAGGGGGGAAGGGTTGGTGAGAAAAACCAAACATACC
GGGATCAAGTTAGGGCGGTAGTTGTGCTGGATTCGTGCCCGCGTTTAAGGCAGTAAGACT
CCAATGGGTAGAAGCCATGGATAACGAAGCTAGGCTAAAGACGTTGGGGACATGGATGTC
AATAATGGAGAAAATGAAGCGATATTGCGTCTGCNNCAGTAGGTAGGCCGAAACTCTCAA
TGTACATAAAACCGTTAAAAGAGACTATATAGTCCTTCCTTCTCCTTTTATTTNTACGAC
GCTCTCAGCTTCCCCTTTGCTAAGACCTCCAGGCGATTCCTCACCCAGCTGGATCTGCGC
CTATGCGGCCGCCTTCAGGAATAAAACCAGGACCCTATCTATTAGGCCTGAACCTCGTAG
TACGATTTACAGACTCCTTTCTGGCTTGTAGCTGCTTCTTCAGTTACGCTAACAGCCCTC
TTGCCTCACGCTATTGTTTCCCTTATTCTCCCGATACAATCCAGTTACCTCAACCTCCTG
TCCGCTTCAACTAC

>Pyr_10

TGAGCCTTTTATTCTAAGCTCAACATTTCTCCACCTCCTTGGCGCCATCCCGGAACCCAG
GCATAAAACGCAGCGAAATAAACAGAGCGCTATGCCTTTACCGTACCTGCCGCTAACCAT
TGTTACACGATTCAGTTGTAAAATTGCTCGCGTTGGCATAGTTAGGTGCCGGTTGCCCCC
CATACAATACGTTTCCAGCATGCACATATTCTCAGATTTTATTATGACCTTCACGTCACT
CATTATGTATTTTGCCTTACGTCCTCCATTTTCCTCTATGTCCGTTTTTCTCTCAAGCTG
ATACATCAGTTNNNNNNNNNNNNNNNNNNNNNNCGTGCATGAGTAGATATTGGTCCTTGA
TTGAGTACCTATAAAACGCTAGGCATCAAAGGACACTGCAATGCACAAGTCAAACAGGTC
CTCTGAAACCAAAACATAAAGACAGACGGAAGCAACCAAAGGGACAGACCATGTATTAAC
ATAGGATGCGAAAATAGTCAAGCATAGGGTAATAATTTGTTCCACAGAGGTTTGATAGCT

TACGAAAGATATCGCGAGCAACAGGCGGATACGTTGGGCGAACGAGGGAAGAGACCGAGT
CGAGTGAGACCGGAGCGGGGAGAGGGAGAGCGAATCCGAATAACAAGAGAACTGGGACAG
GAAAACACAGTGAGCAGAACCTCAGGACGAAAGAGAGACGCACTAAGAAAACAGAGGCGC
GACGTGAAGCGAGCGGAAGACCTATTGATGTAACAGAGGGAGGCGTGATATGAAAAAACT
GAAGTCGGGGTAGGGAAAGGTCGTGAAGGGGACGAATAATAATCTGCTCTTTAGCGGCAG
AAGATACATCGAGATCAGTCCTTTTTCCGAGCGATCTTTGAGCATTATCCACTTGAGCTG
GACGGCCAGACGCAGGAACTTCTAAGGGGGGAAGGGTTGGTGAGAAAAACCAAACATACC
GGGATCAAGTTAGGGCGGTAGTTGTGCTGGATTCGTGCCCGCGTTTAAGGCAGTAAGACT
CCAATGGGTAGAAGCCATGGATAACGAAGCTAGGCTAAAGACGTTGGGGACATGGATGTC
AATAATGGAGAAAATGAAGCGATATTGCGTCTGCNGCAGTAGGTAGGCCGAAACTCTCAA
TGTACATAAAACCGTTAAAAGAGACTATATAGTCCTTCCTTCTCCTTTTATTTNTACGAC
GCTCTCAGCTTCCCCTTTGCTAAGACCTCCAGGCGATTCCTCACCCAGCTGGATCTGCGC
CTATGCGGCCGCCTTCAGGAATAAAACCAGGACCCTATCTATTAGGCCTGAACCTCGTAG
TACGATTTACAGACTCCTTTCTGGCTTGTAGCTGCTTCTTCAGTTACGCTAACAGCCCTC
TTGCCTCACGCTATTGTTTCCCTTATTCTCCCGATACAATCCAGTTACCTCAACCTCCTG
TCCGCTTCAACTAC
>Pyr_11
TGAGCCTTTTATTCTAAGCTCAACATTTCTCCACCTCCTTGGCGCCATCCCGGAACCCAG
GCATAAAACGCAGCGAAATAAACAGAGCGCTATGCCTTTACCGTACCTGCCGCTAACCAT
TGTTACACGATTCAGTTGTAAAATTGCTCGCGTTGGCATAGTTAGGTGCCGGTTGCCCCC
CATACAATACGTTTCCAGCATGCACATATTCTCAGATTTTATTATGACCTTCACGTCACT
CATTATGTATTTTGCCTTACGTCCTCCATTTTCCTCTATGTCCGTTTTTCTCTCAAGCTG
ATACATCAGTTNNNAACTCCCGCGAATTCTATTCGTGCATGAGTAGATATTGGTCCTTGA
TTGAGTACCTATAAAACGCTAGGCATCAAAGGACACTGCAATGCACAAGTCAAACAGGTC
CTCTGAAACCAAAACATAAAGACAGACGGAAGCAACCAAAGGGACAGACCATGTATTAAC
ATAGGATGCGAAAATAGTCAAGCATAGGGTAATAATTTGTTCCACAGAGGTTTGATAGCT
TACGAAAGATATCGCGAGCAACAGGCGGATACGTTGGGCGAACGAGGGAAGAGACCGAGT
CGAGTGAGACCGGAGCGGGGAGAGGGAGAGCGAATCCGAATAACAAGAGAACTGGGACAG
GAAAACACAGTGAGCAGAACCTCAGGACGAAAGAGAGACGCACTAAGAAAACAGAGGCGC
GACGTGAAGCGAGCGGAAGACCTATTGATGTAACAGAGGGAGGCGTGATATGAAAAAACT
GAAGTCGGGGTAGGGAAAGGTCGTGAAGGGGACGAATAATAATCTGCTCTTTAGCGGCAG
AAGATACATCGAGATCAGTCCTTTTTCCGAGCGATCTTTGAGCATTATCCACTTGAGCTG
GACGGCCAGACGCAGGAACTTCTAAGGGGGGAAGGGTTGGTGAGAAAAACCAAACATACC
GGGATCAAGTTAGGGCGGTAGTTGTGCTGGATTCGTGCCCGCGTTTAAGGCAGTAAGACT
CCAATGGGTAGAAGCCATGGATAACGAAGCTAGGCTAAAGACGTTGGGGACATGGATGTC
AATAATGGAGAAAATGAAGCGATATTGCGTCTGCAGCAGTAGGTAGGCCGAAACTCTCAA

TGTACATAAAACCGTTAAAAGAGACTATATAGTCCTTCCTTCTCCTTTTATTTCTACGAC
GCTCTCAGCTTCCCCTTTGCTAAGACCTCCAGGCGATTCCTCACCCAGCTGGATCTGCGC
CTATGCGGCCGCCTTCAGGAATAAAACCAGGACCCTATCTATTAGGCCTGAACCTCGTAG
TACGATTTACAGACTCCTTTCTGGCTTGTAGCTGCTTCTTCAGTTACGCTAACAGCCCTC
TTGCCTCACGCTATTGTTTCCCTTATTCTCCCGATACAATCCAGTTACCTCAACCTCCTG
TCCGCTTCAACTAC
>Pyr_12
TGAGCCTTTTATTCTAAGCTCAACATTTCTCCACCTCCTTGGCGCCATCCCGGAACCCAG
GCATAAAACGCAGCGAAATAAACAGAGCGCTATGCCTTTACCGTACCTGCCGCTAACCAT
TGTTACACGATTCAGTTGTAAAATTGCTCGCGTTGGCATAGTTAGGTGCCGGTTGCCCCC
CATACAATACGTTTCCAGCATGCACATATTCTCAGATTTTATTATGACCTTCACGTCACT
CATTATGTATTTTGCCTTACGTCCTCCATTTTCCTCTATGTCCGTTTTTCTCTCAAGCTG
ATACATCAGTTNNNAACTCCCGCGAATTCNNNNNGTGCATGAGTAGATATTGGTCCTTGA
TTGAGTACCTATAAAACGCTAGGCATCANAGGACACTGCAATGCACAAGTCAAACAGGTC
CTCTGAAACCAAAACATAAAGACAGACGGAAGCAACCAAAGGGACAGACCATGTATTAAC
ATAGGATGCGAAAATAGTCAAGCATAGGGTAATAATTTGTTCCACAGAGGTTTGATAGCT
TACGAAAGATATCGCGAGCAACAGGCGGATACGTTGGGCGAACGAGGGAAGAGACCGAGT
CGAGTGAGACCGGAGCGGGGAGAGGGAGAGCGAATCCGAATAACAAGAGAACTGGGACAG
GAAAACACAGTGAGCAGAACCTCAGGACGAAAGAGAGACGCACTAAGAAAACAGAGGCGC
GACGTGAAGCGAGCGGAAGACCTATTGATGTAACAGAGGGAGGCGTGATATGAAAAAACT
GAAGTCGGGGTAGGGAAAGGTCGTGAAGGGGACGAATAATAATCTGCTCTTTAGCGGCAG
AAGATACATCGAGATCAGTCCTTTTTCCGAGCGATCTTTGAGCATTATCCACTTGAGCTG
GACGGCCAGACGCAGGAACTTCTAAGGGGGGAAGGGTTGGTGAGAAAAACCAAACATACC
GGGATCAAGTTAGGGCGGTAGTTGTGCTGGATTCGTGCCCGCGTTTAAGGCAGTAAGACT
CCAATGGGTAGAAGCCATGGATAACGAAGCTAGGCTAAAGACGTTGGGGACATGGATGTC
AATAATGGAGAAAATGAAGCGATATTGCGTCTGCNGCAGTAGGTAGGCCGAAACTCTCAA
TGTACATAAAACCGTTAAAAGAGACTATATAGTCCTTCCTTCTCCTTTTATTTCTACGAC
GCTCTCAGCTTCCCCTTTGCTAAGACCTCCAGGCGATTCCTCACCCAGCTGGATCTGCGC
CTATGCGGCCGCCTTCAGGAATAAAACCAGGACCCTATCTATTAGGCCTGAACCTCGTAG
TACGATTTACAGACTCCTTTCTGGCTTGTAGCTGCTTCTTCAGTTACGCTAACAGCCCTC
TTGCCTCACGCTATTGTTTCCCTTATTCTCCCGATACAATCCAGTTACCTCAACCTCCTG
TCCGCTTCAACTAC
>Pyr_13
TGAGCCTTTTATTCTAAGCTCAACATTTCTCCACCTCCTTGGCGCCATCCCGGAACCCAG
GCATAAAACGCAGCGAAATAAACAGAGCGCTATGCCTTTACCGTACCTGCCGCTAACCAT
TGTTACACGATTCAGTTGTAAAATTGCTCGCGTTGGCATAGTTAGGTGCCGGTTGCCCCC

CATACAATACGTTTCCAGCATGCACATATTCTCAGATTTTATTATGACCTTCACGTCACT
CATTATGTATTTTGCCTTACGTCCTCCATTTTCCTCTATGTCCGTTTTTCTCTCAAGCTG
ATACATCAGTTNNNAACTCCCGCGAATTCTNTTCGTGCATGAGTAGATATTGGTCCTTGA
TTGAGTACCTATAAAACGCTAGGCATCAAAGGACACTGCAATGCACAAGTCAAACAGGTC
CTCTGAAACCAAAACATAAAGACAGACGGAAGCAACCAAAGGGACAGACCATGTATTAAC
ATAGGATGCGAAAATAGTCAAGCATAGGGTAATAATTTGTTCCACAGAGGTTTGATAGCT
TACGAAAGATATCGCGAGCAACAGGCGGATACGTTGGGCGAACGAGGGAAGAGACCGAGT
CGAGTGAGACCGGAGCGGGGAGAGGGAGAGCGAATCCGAATAACAAGAGAACTGGGACAG
GAAAACACAGTGAGCAGAACCTCAGGACGAAAGAGAGACGCACTAAGAAAACAGAGGCGC
GACGTGAAGCGAGCGGAAGACCTATTGATGTAACAGAGGGAGGCGTGATATGAAAAAACT
GAAGTCGGGGTAGGGAAAGGTCGTGAAGGGGACGAATAATAATCTGCTCTTTAGCGGCAG
AAGATACATCGAGATCAGTCCTTTTTCCGAGCGATCTTTGAGCATTATCCACTTGAGCTG
GACGGCCAGACGCAGGAACTTCTAAGGGGGGAAGGGTTGGTGAGAAAAACCAAACATACC
GGGATCAAGTTAGGGCGGTAGTTGTGCTGGATTCGTGCCCGCGTTTAAGGCAGTAAGACT
CCAATGGGTAGAAGCCATGGATAACGAAGCTAGGCTAAAGACGTTGGGGACATGGATGTC
AATAATGGAGAAAATGAAGCGATATTGCGTCTGCAGCAGTAGGTAGGCCGAAACTCTCAA
TGTACATAAAACCGTTAAAAGAGACTATATAGTCCTTCCTTCTCCTTTTATTTCTACGAC
GCTCTCAGCTTCCCCTTTGCTAAGACCTCCAGGCGATTCCTCACCCAGCTGGATCTGCGC
CTATGCGGCCGCCTTCAGGAATAAAACCAGGACCCTATCTATTAGGCCTGAACCTCGTAG
TACGATTTACAGACTCCTTTCTGGCTTGTAGCTGCTTCTTCAGTTACGCTAACAGCCCTC
TTGCCTCACGCTATTGTTTCCCTTATTCTCCCGATACAATCCAGTTACCTCAACCTCCTG
TCCGCTTCAACTAC
>Pyr_14
TNAGCCTTTTATTCTAAGCTCAACATTTCTCCACCTCCTTGGCGCCATCCCGGAACCCAG
GCATAAAACGCAGCGAAATAAACAGAGCGCTATGCCTTTACCGTACCTGCCGCTAACCAT
TGTTACACGATTCAGTTGTAAAATTGCTCGCGTTGGCATAGTTAGGTGCCGGTTGCCCCC
CATACAATACGTTTCCAGCATGCACNTATTCTCAGATTTTATTATGACCTTCACGTCACT
CATTATGTATTTTGCCTTACGTCCTCCATTTTCCTCTATGTCCGTTTTTCTCTCAAGCTG
ATACATCAGTTNNNAACNCNNGCGAATTCNNNNNNTGCATGAGTAGATATTGGTCCTTGA
TTGAGTACCTATAAAACGCTAGGCATCAAAGGACACTGCAATGCACAAGTCAAACAGGTC
CTCTGAAACCAAAACATAAAGACAGACGGAAGCAACCAAAGGGACAGACCATGTATTAAC
ATAGGATGCGAAAATAGTCAAGCATAGGGTAATAATTTGTTCCACAGAGGTTTGATAGCT
TACGAAAGATATCGCGAGCAACAGGCGGATACGTTGGGCGAACGAGGGAAGAGACCGAGT
CGAGTGAGACCGGAGCGGGGAGAGGGAGAGCGAATCCGAATAACAAGAGAACTGGGACAG
GAAAACACAGTGAGCAGAACCTCAGGACGAAAGAGAGACGCACTAAGAAAACAGAGGCGC
GACGTGAAGCGAGCGGAAGACCTATTGATGTAACAGAGGGAGGCGTGATATGAAAAAACT

GAAGTCGGGGTAGGGAAAGGTCGTGAAGGGGACGAATAATAATCTGCTCTTTAGCGGCAG
AAGATACATCGAGATCAGTCCTTTTTCCGAGCGATCTTTGAGCATTATCCACTTGAGCTG
GACGGCCAGACGCAGGAACTTCTAAGGGGGGAAGGGTTGGTGANAAAAACCAAACATACC
GGGATCAAGTTAGGGCGGTAGTTGTGCTGGATTCGTGCCCGCGTTTAAGGCAGTAAGACT
CCAATGGGTAGAAGCCATGGATAACGAAGCTAGGCTAAAGACGTTGGGGACATGGATGTC
AATAATGGAGAAAATGAAGCGATATTGCGTCTGNNNCAGTAGGTAGGCCGAAACTCTCAA
TGTACATAAAACCGTTAAAAGAGACTATATAGTCCTTCCTTCTCCTTTTATTTCTACGAC
GCTCTCAGCTTCCCCTTTGCTAAGACCTCCAGGCGATTCCTCACCCAGCTGGATCTGCGC
CTATGCGGCCGCCTTCAGGAATAAAACCAGGACCCTATCTATTAGGCCTGAACCTCGTAG
TACGATTTACAGACTCCTTTCTGGCTTGTAGCTGCTTCTTCAGTTACGCTAACAGCCCTC
TTGCCTCACGCTATTGTTTCCCTTATTCTCCCGATACAATCCAGTTACCTCAACCTCCTG
TCCGCTTCAACTAC
>Pyr_15
TGAGCCTTTTATTCTAAGCTCAACATTTCTCCACCTCCTTGGCGCCATCCCGGAACCCAG
GCATAAAACGCAGCGAAATAAACAGAGCGCTATGCCTTTACCGTACCTGCCGCTAACCAT
TGTTACACGATTCAGTTGTAAAATTGCTCGCGTTGGCATAGTTAGGTGCCGGTTGCCCCC
CATACAATACGTTTCCAGCATGCACATATTCTCAGATTTTANTATGACCTTCACGTCACT
CATTATGTATTTTGCCTTACGTCCTCCATTTTCCTCTATGTCCGTTTTTCTCTCAAGCTG
ATACATCAGTTNNNNNNTCCCGCGAATTCNNNNNNTNCATGAGTAGATATTGGTCCTTGA
TTGAGTACCTATAAAACGCTAGGCATCAAAGGACACTGCAATGCACAAGTCAAACAGGTC
CTCTGAAACCAAAACATAAAGACAGACGGAAGCAACCAAAGGGACAGACCATGTATTAAC
ATAGGATGCGAAAATAGTCAAGCATAGGGTAATAATTTGTTCCACAGAGGTTTGATAGCT
TACGAAAGATATCGCGAGCAACAGGCGGATACGTTGGGCGAACGAGGGAAGAGACCGAGT
CGAGTGAGACCGGAGCGGGGAGAGGGAGAGCGAATCCGAATAACAAGAGAACTGGGACAG
GAAAACACAGTGAGCAGAACCTCAGGACNAAAGAGAGACGCACTAAGAAAACAGAGGCGN
GACGTGAAGCGAGCGGAAGACCTATTGATGTAACAGAGGGAGGCGTGATATGAAAAAACT
GAAGTCGGGGTAGGGAAAGGTCGTGAAGGGGACGAATAATAATCTGCTCTTTAGCGGCAG
AAGATACATCGAGATCAGTCCTTTTTCCGAGCGATCTTTGAGCATTATCCACTTGAGCTG
GACGGCCAGACGCAGGAACTTCTAAGGGGGGAAGGGTTGGTGAGAAAAACCAAACATACC
GGGATCAAGTTAGGGCGGTAGTTGTGCTGGATTCGTGCCCGCGTTTAAGGCAGTAAGACT
CCAATGGGTAGAAGCCATGGATAACGAAGCTAGGCTAAAGACGTTGGGGACATGGATGTC
AATAATGGAGAAAATGAAGCGATATTGCGTCTGCNGCAGTAGGTAGGCCGAAACTCTCAA
TGTACATAAAACCGTTAAAAGAGACTATATAGTCCTTCCTTCTCCTTTTATTTCTACGAC
GCTCTCAGCTTCCCCTTTGCTAAGACCTCCAGGCGATTCCTCACCCAGCTGGATCTGCGC
CTATGCGGCCGCCTTCAGGAATAAAACCAGGACCCTATCTATTAGGCCTGAACCTCGTAG
TACGATTTACAGACTCCTTTCTGGCTTGTAGCTGCTTCTTCAGTTACGCTAACAGCCCTC

TTGCCTCACGCTATTGTTTCCCTTATTCTCCCGATACAATCCAGTTACCTCAACCTCCTG
TCCGCTTCAACTAC
>Pyr_16
TGAGCCTTTTATTCTAAGCTCAACATTTCTCCACCTCCTTGGCGCCATCCCGGAACCCAG
GCATAAAACGCAGCGAAATAAACAGAGCGCTATGCCTTTACCGTACCTGCCGCTAACCAT
TGTTACACGATTCAGTTGTAAAATTGCTCGCGTTGGCATAGTTAGGTGCCGGTTGCCCCC
CATACAATACGTTTCCAGCATGCACATATTCTCAGATTTTATTATGACCTTCACGTCACT
CATTATGTATTTTGCCTTACGTCCTCCATTTTCCTCTATGTCCGTTTTTCTCTCAAGCTG
ATACATCAGTTNNNAACTCCCGCGAATTCNATTCGTGCATGAGTAGATATTGGTCCTTGA
TTGNGTACCTATAAAACGCTAGGCATCAAAGGACACTGCAATGCACAAGTCAAACAGGTC
CTCTGAAACCAAAACATAAAGACAGACGGAAGCAACCAAAGGGACAGACCATGTATTAAC
ATAGGATGCGAAAATAGTCAANCATAGGGTAATAATTTGTTCCACAGAGGTTTGATAGCT
TACGAAAGATATCGCGAGCAACAGGCGGATACGTTGGGCGAACGAGGGAAGAGACCGAGT
CGAGTGAGACCGGAGCGGGGAGAGGGAGAGCGAATCCGAATAACAAGAGAACTGGGACAG
GAAAACACAGTGAGCAGAACCTCAGGACGAAAGAGAGACGCACTAAGAAAACAGAGGCGC
GACGTGAAGCGAGCGGAAGACCTATTGATGTAACAGAGGGAGGCGTGATATGAAAAAACT
GAAGTCGGGGTAGGGAAAGGTCGTGAAGGGGACGAATAATAATCTGCTCTTTAGCGGCAG
AAGATACATCGAGATCAGTCCTTTTTCCGAGCGATCTTTGAGCATTATCCACTTGAGCTG
GACGGCCAGACGCAGGAACGTCTAAGGGGGGAAGGGTTGGTGAGAAAAACCAAACATACC
GGGATCAAGTTAGGGCGGTAGTTGTGCTGGATTCGTGCCCGCGTTTAAGGCAGTAAGACT
CCAATGGGTAGAAGCCATGGATAACGAAGCTAGGCTAAAGACGTTGGGGNNNTGGATGTC
AATAATGGAGGAAATGAAGCGATATTGCGTCTGCANCAGTAGGTAGGCCGAAACTCTCAA
TGTACANNAAACCGTTAAAAGAGACTATATAGTCCTTCCTTCTCCTTTTATTTCTACGAC
GCTCTCAGCTTCCCCTTTGCTAAGACCTCCAGGCGATTCCTCACCCAGCTGGATCTGCGC
CTATGCGGCCGCCTTCAGGAATAAAACCAGGACCCTATCTATTAGGCCTGAACCTCGTAG
TACNNNNTACAGACTCCTTTCTGGCTTGTAGCTGCTTCTTCAGTTACGCTAACAGCCCTC
TTGCCTCACGCTATTGTTTCCCTTATTCTCCCGATACAATCCAGTTACCTCAACCTCCTG
TCCGCTTCAACTAC
>Pyr_17
TGAGCCTTTTATTCTAAGCTCAACATTTCTCCACCTCCTTGGCGCCATCCCGGAACCCAG
GCATAAAACGCAGCGAAATAAACAGAGCGCTATGCCTTTACCGTACCTGCCGCTAACCAT
TGTTACACGATTCAGTTGTAAAATTGCTCGCGTTGGCATAGTTAGGTGCCGGTTGCCCCC
CATACAATACGTTTCCAGCATGCACATATTCTCAGATTTTATTATGACCTTCACGTCACT
CATTATGTATTTTGCCTTACGTCCTCCATTTTCCTCTATGTCCGTTTTTCTCTCAAGCTG
ATACATCAGTTNCNAACTCCCGCGAATTCTATTNGTGCATGAGTAGATATTGGTCCTTGA
TTGAGTACCTATAAAACGCTAGGCATCAAAGGACACTGCAATGCACAAGTCAAACAGGTC

CTCTGAAACCAAAACATAAAGACAGACGGAAGCAACCAAAGGGACAGACCATGTATTAAC
ATAGGATGCGAAAATAGTCAAGCATAGGGTAATAATTTGTTCCACAGAGGTTTGATAGCT
TACGAAAGATATCGCGAGCAACAGGCGGATACGTTGGGCGAACGAGGGAAGAGACCGAGT
CGAGTGAGACCGGAGCGGGGAGAGGGAGAGCGAATCCGAATAACAAGAGAACTGGGACAG
GAAAACACAGTGAGCAGAACCTCAGGACGAAAGAGAGACGCACTAAGAAAACAGAGGCGC
GACGTGAAGCGAGCGGAAGACCTATTGATGTAACAGAGGGAGGCGTGATATGAAAAAACT
GAAGTCGGGGTAGGGAAAGGTCGTGAAGGGGACGAATAATAATCTGCTCTTTAGCGGCAG
AAGATACATCGAGATCAGTCCTTTTTCCGAGCGATCTTTGAGCATTATCCACTTGAGCTG
GACGGCCAGACGCAGGAACTTCTAAGGGGGGAAGGGTTGGTGAGAAAAACCAAACATACC
GGGATCAAGTTAGGGCGGTAGTTGTGCTGGATTCGTGCCCGCGTTTAAGGCAGTAAGACT
CCAATGGGTAGAAGCCATGGATAACGAAGCTAGGCTAAAGACGTTGGGGACATGGATGTC
AATAATGGAGAAAATGAAGCGATATTGCGTCTGCAGCAGTAGGTAGGCCGAAACTCTCAA
TGTACATAAAACCGTTAAAAGAGACTATATAGTCCTTCCTTCTCCTTTTATTTCTACGAC
GCTCTCAGCTTCCCCTTTGCTAAGACCTCCAGGCGATTCCTCACCCAGCTGGATCTGCGC
CTATGCGGCCGCCTTCAGGAATAAAACCAGGACCCTATCTATTAGGCCTGAACCTCGTAG
TACGATTTACAGACTCCTTTCTGGCTTGTAGCTGCTTCTTCAGTTACGCTAACAGCCCTC
TTGCCTCACGCTATTGTTTCCCTTATTCTCCCGATACAATCCAGTTACCTCAACCTCCTG
TCCGCTTCAACTAC
>Pyr_18
TGAGCCTTTTATTCTAAGCTCAACATTTCTCCACCTCCTTGGCGCCATCCCGGAACCCAG
GCATAAAACGCAGCGAAATAAACAGAGCGCTATGCCTTTACCGTACCTGCCGCTAACCAT
TGTTACACGATTCAGTTGTAAAATTGCTCGCGTTGGCATAGTTAGGTGCCGGTTGCCCCC
CATACAATACGTTTCCAGCATGCACATATTCTCAGATTTTATTATGACCTTCACGTCACT
CATTATGTATTTTGCCTTACGTCCTCCATTTTCCTCTATGTCCGTTTTTCTCTCAAGCTG
ATACATCAGTTNNNAACTCCCGCGAATTCNNNNNNTGCATGAGTAGATATTGGTCCTTGA
TTGAGTACCTATAAAACGCTAGGCATCAAAGGACACTGCAATGCACAAGTCAAACAGGTC
CTCTGAAACCAAAACATAAAGACAGACGGAAGCAACCAAAGGGACAGACCATGTATTAAC
ATAGGATGCGAAAATAGTCAAGCATAGGGTAATAATTTGTTCCACAGAGGTTTGATAGCT
TACGAAAGATATCGCGAGCAACAGGCGGATACGTTGGGCGAACGAGGGAAGAGACCGAGT
CGAGTGAGACCGGAGCGGGGAGAGGGAGAGCGAATCCGAATAACAAGAGAACTGGGACAG
GAAAACACAGTGAGCAGAACCTCAGGACGAAAGAGAGACGCACTAAGAAAACAGAGGCGC
GACGTGAAGCGAGCGGAAGACCTATTGATGTAACAGAGGGAGGCGTGATATGAAAAAACT
GAAGTCGGGGTAGGGAAAGGTCGTGAAGGGGACGAATAATAATCTGCTCTTTAGCGGCAG
AAGATACATCGAGATCAGTCCTTTTTCCGAGCGATCTTTGAGCATTATCCACTTGAGCTG
GACGGCCAGACGCAGGAACTTCTAAGGGGGGAAGGGTTGGTGAGAAAAACCAAACATACC
GGGATCAAGTTAGGGCGGTAGTTGTGCTGGATTCGTGCCCGCGTTTAAGGCAGTAAGACT

CCAATGGGTAGAAGCCATGGATAACGAAGCTAGGCTAAAGACGTTGGGGACATGGATGTC
AATAATGGAGGAAATGAAGCGATATTGCGTCTGCAGCAGTAGGTAGGCCGAAACTCTCAA
TGTACATAAAACCGTTAAAAGAGACTATATAGTCCTTCCTTCTCCTTTTATTTCTACGAC
GCTCTCAGCTTCCCCTTTGCTAAGACCTCCAGGCGATTCCTCACCCAGCTGGATCTGCGC
CTATGCGGCCGCCTTCAGGAATAAAACCAGGACCCTATCTATTAGGCCTGAACCTCGTAG
TACGATTTACAGACTCCTTTCTGGCTTGTAGCTGCTTCTTCAGTTACGCTAACAGCCCTC
TTGCCTCACGCTATTGTTTCCCTTATTCTCCCGATACAATCCAGTTACCTCAACCTCCTG
TCCGCTTCAACTAC
>Pyr_19
CAAATTCCTGACCTCGGATTCCGTNCCCTCTTCTATTTCTGGCGTCATCCTGGAACCTAG
GAATAAAACGCAGCAAAGTAAACAATGCGCCATACCTATGACGTACCTGTCGCTAAATGT
TGTTCTGCATTCTAACCACNNNNNNNNNNNNNNNNNNNNNNNNNNNNNNNNNNNNNNNNN
NNNNNNNNNNNNNNNNNNNNNNNNNNNNNNNNNNNNNNNNNNNCTAGTTCCCATNNNNNT
CNTNCNNNATNNNNNNNNNNNNNCTTTGCNCCTTTTCANNNNNGNNTNNTCNNNANNCNN
NCNAANNNNTNNNNNNNNNNNNNNNNNNNNANNNGCAANNAAATGAAAACTTAGATTCAN
NTNNATGTTTGTGAGGTATAGCATGTTANNNNNNNNNCATNANACGTNAGNNNNNNNNNC
NNNNNGNNNCNNNNNNNGAGNGANNNNAGNNNCGGNNNNNNNGNNNGGNTTCACAGTGCT
GCAGTAGACAGGGNNGNTCANNNGCGAANNGACGANNNNGNNCNNNNNNANNCTNNNNCN
NNCAGGGAATGCTATGANNNGTNNNAGANTANNNNNNNNNAANNAGNGNAGNNNTTAAAT
NNNGNNNNNNCNNNNNNGNNANNNNNANNGNNGNNNNGANNNNNNANNNNNNNNNNNTNN
NNNNNNNCNNNNGNACATNCNNNGNNNCAAGGAGTAAGTNANNNNCGAANAANNNNNNNN
NNCAATACATAGGNNGAAGNNNNNNNNTNANGATGACAGGANATGCAANANNNNNNNNNN
ANNNNGNNNNNNNNNGAAACTGGAGGAAGACTAGGCAATCANTCATGNTTCGGTGACGA
GGTGCATGCCAATNCTGAATACATCTCGGTTTGATCTTTTAGCGTTGGCAATCCAACTTG
AGCAGCAGATTGTTAGGGTTTTCGGAAAAGAGGANNNNGANAAAGAAAGACAGATACCCC
GAAGCCGCGGTAGAGTAGTAGTTGTGCAGGAACTATGCTCATACCTAGCGCCGTAAGGCT
CTAACGGATTGAGGCCACAGNNNANNNNGACAGGCAGGGAGGACCTAANANNTNAACGCT
CNCCCNNNAGGANNNNANNNNNNANTNNNNNNNNNNNNNNNNNNNNNNNNNNNNNNNNNN
NNNNNNNNNNNNNNNNNNNNNNGNCTATATANNCCTCCCCNCNCCTNNTANNTNTNNNNN
NNNNNNNNNNNNNNNNNNNNNNNNNNNNNNNNNNNNNNNNNNNNNNNNNNNNNNNNNNNN
NNNNNNNNNNNNNTTNNGGNNNGAGCCNNNNANNNNATCCNTCANGTTNNAACCNNNNNN
NNNNNNNNNNNGATNNNNTTNNNNNNNNTANCNNCCCCTTCAGCNNCATCGACCACTTCT
CTATTTNANNNNNTNNNNNNCNNNNNCNCCTCGANNNNNNTCGATCCGTCCAATTCTTCG
CTAATCTTCGTCCT
>Pyr_2
TGAGCCTTTTATTCTAAGCTCAACATTTCTCCACCTCCTTGGCGCCATCCCGGAACCCAG

GCATAAAACGCAGCGAAATAAACAGAGCGCTATGCCTTTACCGTACCTGCCGCTAACCAT
TGTTACACGATTCAGTTGTAAAATTGCTCGCGTTGGCATAGTTAGGTGCCGGTTGCCCCC
CATACAATACGTTTCCAGCATGCACATATTCTCAGATTTTATTATGACCTTCACGTCACT
CATTATGTATTTTGCCTTACGTCCTCCATTTTCCTCTATGTCCGTTTTTCTCTCAAGCTG
ATACATCAGTTNNNAACTCCNGCNAATTCTATTCGTGCATGAGTAGATATTGGTCCTTGA
TTGAGTACCTATAAAACGCTAGGCATCAAAGGACACTGCAATGCACAAGTCAAACAGGTC
CTCTGAAACCAAAACATAAAGACAGACGGAAGCAACCAAAGGGACAGACCATGTATTAAC
ATAGGATGCGAAAATAGTCAAGCATAGGGTAATAATTTGTTCCACAGAGGTTTGATAGCT
TACGAAAGATATCGCGAGCAACAGGCGGATACGTTGGGCGAACGAGGGAAGAGACCGAGT
CGAGTGAGACCGGAGCGGGGAGAGGGAGAGCGAATCCGAATAACAAGAGAACTGGGACAG
GAAAACACAGTGAGCAGAACCTCAGGACGAAAGAGAGACGCACTAAGAAAACAGAGGCGC
GACGTGAAGCGAGCGGAAGACCTATTGATGTAACAGAGGGAGGCGTGATATGAAAAAACT
GAAGTCGGGGTAGGGAAAGGTCGTGAAGGGGACGAATAATAATCTGCTCTTTAGCGGCAG
AAGATACATCGAGATCAGTCCTTTTTCCGAGCGATCTTTGAGCATTATCCACTTGAGCTG
GACGGCCAGACGCAGGAACTTCTAAGGGGGGAAGGGTTGGTGAGAAAAACCAAACATACC
GGGATCAAGTTAGGGCGGTAGTTGTGCTGGATTCGTGCCCGCGTTTAAGGCAGTAAGACT
CCAATGGGTAGAAGCCATGGATAACGAAGCTAGGCTAAAGACGTTGGGGACATGGATGTC
AATAATGGAGAAAATGAAGCGATATTGCGTCTGCAGCAGTAGGTAGGCCGAAACTCTCAA
TGTACATAAAACCGTTAAAAGAGACTATATAGTCCTTCCTTCTCCTTTTATTTCTACGAC
GCTCTCAGCTTCCCCTTTGCTAAGACCTCCAGGCGATTCCTCACCCAGCTGGATCTGCGC
CTATGCGGCCGCCTTCAGGAATAAAACCAGGACCCTATCTATTAGGCCTGAACCTCGTAG
TACGATTTACAGACTCCTTTCTGGCTTGTAGCTGCTTCTTCAGTTACGCTAACAGCCCTC
TTGCCTCACGCTATTGTTTCCCTTATTCTCCCGATACAATCCAGTTACCTCAACCTCCTG
TCCGCTTCAACTAC
>Pyr_20
TGAGCCTTTTATTCTAAGCTCAACATTTCTCCACCTCCTTGGCGCCATCCCGGAACCCAG
GCATAAAACGCAGCGAAATAAACAGAGCGCTATGCCTTTACCGTACCTGCCGCTAACCAT
TGTTACACGATTCAGTTGTAAAATTGCTCGCGTTGGCATAGTTAGGTGCCGGTTGCCCCC
CATACAATACGTTTCCAGCATGCACATATTCTCAGATTTTATTATGACCTTCACGTCACT
CATTATGTATTTTGCCTTACGTCCTCCATTTTCCTCTATGTCCGTTTTTCTCTCAAGCTG
ATACATCAGTTNNNAACTCCCGCGAATTNNNTTCGTGCATGAGTAGATATTGGTCCTTGA
TTGAGTACCTATAAAACGCTAGGCATCAAAGGACACTGCAATGCACAAGTCAAACAGGTC
CTCTGAAACCAAAACATAAAGACAGACGGAAGCAACCAAAGGGACAGACCATGTATTAAC
ATAGGATGCGAAAATAGTCAAGCATAGGGTAATAATTTGTTCCACAGAGGTTTGATAGCT
TACGAAAGATATCGCGAGCAACAGGCGGATACGTTGGGCGAACGAGGGAAGAGACCGAGT
CGAGTGAGACCGGAGCGGGGAGAGGGAGAGCGAATCCGAATAACAAGAGAACTGGGACAG

GAAAACACAGTGAGCAGAACCTCAGGACGAAAGAGAGACGCACTAAGAAAACAGAGGCGC
GACGTGAAGCGAGCGGAAGACCTATTGATGTAACAGAGGGAGGCGTGATATGAAAAAACT
GAAGTCGGGGTAGGGAAAGGTCGTGAAGGGGACGAATAATAATCTGCTCTTTAGCGGCAG
AAGATACATCGAGATCAGTCCTTTTTCCGAGCGATCTTTGAGCATTATCCACTTGAGCTG
GACGGCCAGACGCAGGAACTTCTAAGGGGGGAAGGGTTGGTGAGAAAAACCAAACATACC
GGGATCAAGTTAGGGCGGTAGTTGTGCTGGATTCGTGCCCGCGTTTAAGGCAGTAAGACT
CCAATGGGTAGAAGCCATGGATAACGAAGCTAGGCTAAAGACGTTGGGGACATGGATGTC
AATAATGGAGGAAATGAAGCGATATTGCGTCTGCNGCAGTAGGTAGGCCGAAACTCTCAA
TGTACATAAAACCGTTAAAAGAGACTATATAGTCCTTCCTTCTCCTTTTATTTCTACGAC
GCTCTCAGCTTCCCCTTTGCTAAGACCTCCAGGCGATTCCTCACCCAGCTGGATCTGCGC
CTATGCGGCCGCCTTCAGGAATAAAACCAGGACCCTATCTATTAGGCCTGAACCTCGTAG
TACGATTTACAGACTCCTTTCTGGCTTGTAGCTGCTTCTTCAGTTACGCTAACAGCCCTC
TTGCCTCACGCTATTGTTTCCCTTATTCTCCCGATACAATCCAGTTACCTCAACCTCCTG
TCCGCTTCAACTAC
>Pyr_21
TGAGCCTTTTATTCTAAGCTCAACATTTCTCCACCTCCTTGGCGCCATCCCGGAACCCAG
GCATAAAACGCAGCGAAATAAACAGAGCGCTATGCCTTTACCGTACCTGCCGCTAACCAT
TGTTACACGATTCAGTTGTAAAATTGCTCGCGTTGGCATAGTTAGGTGCCGGTTGCCCCC
CATACAATACGTTTCCAGCATGCACATATTCTCAGATTTTATTATGACCTTCACGTCACT
CATTATGTATTTTGCCTTACGTCCTCCATTTTCCTCTATGTCCGTTTTTCTCTCAAGCTG
ATACATCAGTTNNNNNCTCCCGCGNNNNNNNNNNGTGCATGAGTAGATATTGGTCCTTGA
TTGAGTACCTATAAAACGCTAGGCATCAAAGGACACTGCAATGCACAAGTCAAACAGGTC
CTCTGAAACCAAAACATAAAGACAGACGGAAGCAACCAAAGGGACAGACCATGTATTAAC
ATAGGATGCGAAAATAGTCAAGCATAGGGTAATAATTTGTTCCACAGAGGTTTGATAGCT
TACGAAAGATATCGCGAGCAACAGGCGGATACGTTGGGCGAACGAGGGAAGAGACCGAGT
CGAATGAGACCGGAGCGGGGAGAGGGAGAGCGAATCCGAATAACAAGAGAACTGGGACAG
GAAAACACAGTGAGCAGAACCTCAGGACGAAAGAGAGACGCACTAAGAAAACAGAGGCGC
GACGTGAAGCGAGCGGAAGACCTATTGATGTAACAGAGGGAGGCGTGATATGAAAAAACT
GAAGTCGGGGTAGGGAAAGGTCGTGAAGGGGACGAATAATAATCTGCTCTTTAGCGGCAG
AAGATACATCGAGATCAGTCCTTTTTCCGAGCGATCTTTGAGCATTATCCACTTGAGCTG
GACGGCCAGACGCAGGAACTTCTAAGGGGGGAAGGGTTGGTGAGAAAAACCAAACATACC
GGGATCAAGTTAGGGCGGTAGTTGTGCTGGATTCGTGCCCGCGTTTAAGGCAGTAAGACT
CCAATGGGTAGAAGCCATGGATAACGAAGCTAGGCTAAAGACGTTGGGGACATGGATGTC
AATAATGGAGGAAATGAAGCGATATTGCGTCTGCAGCAGTAGGTAGGCCGAAACTCTCAA
TGTACATAAAACCGTTAAAAGAGACTATATAGTCCTTCCTTCTCCTTTTATTTCTACGAC
GCTCTCAGCTTCCCCTTTGCTAAGACCTCCAGGCGATTCCTCACCCAGCTGGATCTGCGC

CTATGCGGCCGCCTTCAGGAATAAAACCAGGACCCTATCTATTAGGCCTGAACCTCGTAG
TACGATTTACAGACTCCTTTCTGGCTTGTAGCTGCTTCTTCAGTTACGCTAACAGCCCTC
TTGCCTCACGCTATTGTTTCCCTTATTCTCCCGATACAATCCAGTTACCTCAACCTCCTG
TCCGCTTCAACTAC

>Pyr_22

TGAGCCTTTTATTCTAAGCTCAACATTTCTCCACCTCCTTGGCGCCATCCCGGAACCCAG
GCATAAAACGCAGCGAAATAAACAGAGCGCTATGCCTTTACCGTACCTGCCGCTAACCAT
TGTTACACGATTCAGTTGTAAAATTGCTCGCGTTGGCATAGTTAGGTGCCGGTTGCCCCC
CATACAATACGTTTCCAGCATGCACATATTCTCAGATTTTATTATGACCTTCACGTCACT
CATTATGTATTTTGCCTTACGTCCTCCATTTTCCTCTATGTCCGTTTTTCTCTCAAGCTG
ATACATCAGTTNNNAACTCCCGCGAATTCTATTNGTGCATGAGTAGATATTGGTCCTTGA
TTGAGTACCTATAAAACGCTAGGCATCAAAGGACACTGCAATGCACAAGTCAAACAGGTC
CTCTGAAACCAAAACATAAAGACAGACGGAAGCAACCAAAGGGACAGACCATGTATTAAC
ATAGGATGCGAAAATAGTCAAGCATAGGGTAATAATTTGTTCCACAGAGGTTTGATAGCT
TACGAAAGATATCGCGAGCAACAGGCGGATACGTTGGGCGAACGAGGGAAGAGACCGAGT
CGAGTGAGACCGGAGCGGGGAGAGGGAGAGCGAATCCGAATAACAAGAGAACTGGGACAG
GAAAACACAGTGAGCAGAACCTCAGGACGAAAGAGAGACGCACTAAGAAAACAGAGGCGC
GACGTGAAGCGAGCGGAAGACCTATTGATGTAACAGAGGGAGGCGTGATATGAAAAAACT
GAAGTCGGGGTAGGGAAAGGTCGTGAAGGGGACGAATAATAATCTGCTCTTTAGCGGCAG
AAGATACATCGAGATCAGTCCTTTTTCCGAGCGATCTTTGAGCATTATCCACTTGAGCTG
GACGGCCAGACGCAGGAACTTCTAAGGGGGGAAGGGTTGGTGAGAAAAACCAAACATACC
GGGATCAAGTTAGGGCGGTAGTTGTGCTGGATTCGTGCCCGCGTTTAAGGCAGTAAGACT
CCAATGGGTAGAAGCCATGGATAACGAAGCTAGGCTAAAGACGTTGGGGACATGGATGTC
AATAATGGAGAAAATGAAGCGATATTGCGTCTGCAGCAGTAGGTAGGCCGAAACTCTCAA
TGTACATAAAACCGTTAAAAGAGACTATATAGTCCTTCCTTCTCCTTTTATTTCTACGAC
GCTCTCAGCTTCCCCTTTGCTAAGACCTCCAGGCGATTCCTCACCCAGCTGGATCTGCGC
CTATGCGGCCGCCTTCAGGAATAAAACCAGGACCCTATCTATTAGGCCTGAACCTCGTAG
TACGATTTACAGACTCCTTTCTGGCTTGTAGCTGCTTCTTCAGTTACGCTAACAGCCCTC
TTGCCTCACGCTATTGTTTCCCTTATTCTCCCGATACAATCCAGTTACCTCAACCTCCTG
TCCGCTTCAACTAC

>Pyr_23

TGAGCCTTTTATTCTAAGCTCAACATTTCTCCACCTCCTTGGCGCCATCCCGGAACCCAG
GCATAAAACGCAGCGAAATAAACAGAGCGCTATGCCTTTACCGTACCTGCCGCTAACCAT
TGTTACACGATTCAGTTGTAAAATTGCTCGCGTTGGCATAGTTAGGTGCCGGTTGCCCCC
CATACAATACGTTTCCAGCATGCACATATTCTCAGATTTTATTATGACCTTCACGTCACT
CATTATGTATTTTGCCTTACGTCCTCCATTTTCCTCTATGTCCGTTTTTCTCTCAAGCTG

ATACATCAGTTCCAAACTCCCGCGAATTCTATTCGTGCATGAGTAGATATTGGTCCTTGA
TTGAGTACCTATAAAACGCTAGGCATCAAAGGACACTGCAATGCACAAGTCAAACAGGTC
CTCTGAAACCAAAACATAAAGACAGACGGAAGCAACCAAAGGGACAGACCATGTATTAAC
ATAGGATGCGAAAATAGTCAAGCATAGGGTAATAATTTGTTCCACAGAGGTTTGATAGCT
TACGAAAGATATCGCGAGCAACAGGCGGATACGTTGGGCGAACGAGGGAAGAGACCGAGT
CGAGTGAGACCGGAGCGGGGAGAGGGAGAGCGAATCCGAATAACAAGAGAACTGGGACAG
GAAAACACAGTGAGCAGAACCTCAGGACGAAAGAGAGACGCACTAAGAAAACAGAGGCGC
GACGTGAAGCGAGCGGAAGACCTATTGATGTAACAGAGGGAGGCGTGATATGAAAAAACT
GAAGTCGGGGTAGGGAAAGGTCGTGAAGGGGACGAATAATAATCTGCTCTTTAGCGGCAG
AAGATACATCGAGATCAGTCCTTTTTCCGAGCGATCTTTGAGCATTATCCACTTGAGCTG
GACGGCCAGACGCAGGAACTTCTAAGGGGGGAAGGGTTGGTGAGAAAAACCAAACATACC
GGGATCAAGTTAGGGCGGTAGTTGTGCTGGATTCGTGCCCGCGTTTAAGGCAGTAAGACT
CCAATGGGTAGAAGCCATGGATAACGAAGCTAGGCTAAAGACGTTGGGGACATGGATGTC
AATAATGGAGAAAATGAAGCGATATTGCGTCTGCAGCAGTAGGTAGGCCGAAACTCTCAA
TGTACATAAAACCGTTAAAAGAGACTATATAGTCCTTCCTTCTCCTTTTATTTCTACGAC
GCTCTCAGCTTCCCCTTTGCTAAGACCTCCAGGCGATTCCTCACCCAGCTGGATCTGCGC
CTATGCGGCCGCCTTCAGGAATAAAACCAGGACCCTATCTATTAGGCCTGAACCTCGTAG
TACGATTTACAGACTCCTTTCTGGCTTGTAGCTGCTTCTTCAGTTACGCTAACAGCCCTC
TTGCCTCACGCTATTGTTTCCCTTATTCTCCCGATACAATCCAGTTACCTCAACCTCCTG
TCCGCTTCAACTAC
>Pyr_24
TGAGCCTTTTATTCTAAGCTCAACATTTCTCCACCTCCTTGGCGCCATCCCGGAACCCAG
GCATAAAACGCAGCGAAATAAACAGAGCGCTATGCCTTTACCGTACCTGCCGCTAACCAT
TGTTACACGATTCAGTTGTAAAATTGCTCGCGTTGGCATAGTTAGGTGCCGGTTGCCCCC
CATACAATACGTTTCCAGCATGCACATATTCTCAGATTTTATTATGACCTTCACGTCACT
CATTATGTATTTTGCCTTACGTCCTCCATTTTCCTCTATGTCCGTTTTTCTCTCAAGCTG
ATACATCAGTTCNNAACTCCCGCGAATTCTATTNGTGCATGAGTAGATATTGGTCCTTGA
TTGAGTACCTATAAAACGCTAGGCATCAAAGGACACTGCAATGCACAAGTCAAACAGGTC
CTCTGAAACCAAAACATAAAGACAGACGGAAGCAACCAAAGGGACAGACCATGTATTAAC
ATAGGATGCGAAAATAGTCAAGCATAGGGTAATAATTTGTTCCACAGAGGTTTGATAGCT
TACGAAAGATATCGCGAGCAACAGGCGGATACGTTGGGCGAACGAGGGAAGAGACCGAGT
CGAGTGAGACCGGAGCGGGGAGAGGGAGAGCGAATCCGAATAACAAGAGAACTGGGACAG
GAAAACACAGTGAGCAGAACCTCAGGACGAAAGAGAGACGCACTAAGAAAACAGAGGCGC
GACGTGAAGCGAGCGGAAGACCTATTGATGTAACAGAGGGAGGCGTGATATGAAAAAACT
GAAGTCGGGGTAGGGAAAGGTCGTGAAGGGGACGAATAATAATCTGCTCTTTAGCGGCAG
AAGATACATCGAGATCAGTCCTTTTTCCGAGCGATCTTTGAGCATTATCCACTTGAGCTG

GACGGCCAGACGCAGGAACGTCTAAGGGGGGAAGGGTTGGTGAGAAAAACCAAACATACC
GGGATCAAGTTAGGGCGGTAGTTGTGCTGGATTCGTGCCCGCGTTTAAGGCAGTAAGACT
CCAATGGGTAGAAGCCATGGATAACGAAGCTAGGCTAAAGACGTTGGGGACATGGATGTC
AATAATGGAGGAAATGAAGCGATATTGCGTCTGCAGCAGTAGGTAGGCCGAAACTCTCAA
TGTACATAAAACCGTTAAAAGAGACTATATAGTCCTTCCTTCTCCTTTTATTTCTACGAC
GCTCTCAGCTTCCCCTTTGCTAAGACCTCCAGGCGATTCCTCACCCAGCTGGATCTGCGC
CTATGCGGCCGCCTTCAGGAATAAAACCAGGACCCTATCTATTAGGCCTGAACCTCGTAG
TACGATTTACAGACTCCTTTCTGGCTTGTAGCTGCTTCTTCAGTTACGCTAACAGCCCTC
TTGCCTCACGCTATTGTTTCCCTTATTCTCCCGATACAATCCAGTTACCTCAACCTCCTG
TCCGCTTCAACTAC
>Pyr_25
TGAGCCTTTTATTCTAAGCTCAACATTTCTCCACCTCCTTGGCGCCATCCCGGAACCCAG
GCATAAAACGCAGCGAAATAAACAGAGCGCTATGCCTTTACCGTACCTGCCGCTAACCAT
TGTTACACGATTCAGTTGTAAAATTGCTCGCGTTGGCATAGTTAGGTGCCGGTTGCCCCC
CATACAATACGTTTCCAGCATGCACATATTCTCAGATTTTATTATGACCTTCACGTCACT
CATTATGTATTTTGCCTTACGTCCTCCATTTTCCTCTATGTCCGTTTTTCTCTCAAGCTG
ATACATCAGTTCCAAACTCCCGCGAATTCTATTCGTGCATGAGTAGATATTGGTCCTTGA
TTGAGTACCTATAAAACGCTAGGCATCAAAGGACACTGCAATGCACAAGTCAAACAGGTC
CTCTGAAACCAAAACATAAAGACAGACGGAAGCAACCAAAGGGACAGACCATGTATTAAC
ATAGGATGCGAAAATAGTCAAGCATAGGGTAATAATTTGTTCCACAGAGGTTTGATAGCT
TACGAAAGATATCGCGAGCAACAGGCGGATACGTTGGGCGAACGAGGGAAGAGACCGAGT
CGAGTGAGACCGGAGCGGGGAGAGGGAGAGCGAATCCGAATAACAAGAGAACTGGGACAG
GAAAACACAGTGAGCAGAACCTCAGGACGAAAGAGAGACGCACTAAGAAAACAGAGGCGC
GACGTGAAGCGAGCGGAAGACCTATTGATGTAACAGAGGGAGGCGTGATATGAAAAAACT
GAAGTCGGGGTAGGGAAAGGTCGTGAAGGGGACGAATAATAATCTGCTCTTTAGCGGCAG
AAGATACATCGAGATCAGTCCTTTTTCCGAGCGATCTTTGAGCATTATCCACTTGAGCTG
GACGGCCAGACGCAGGAACTTCTAAGGGGGGAAGGGTTGGTGAGAAAAACCAAACATACC
GGGATCAAGTTAGGGCGGTAGTTGTGCTGGATTCGTGCCCGCGTTTAAGGCAGTAAGACT
CCAATGGGTAGAAGCCATGGATAACGAAGCTAGGCTAAAGACGTTGGGGACATGGATATC
AATAATGGAGAAAATGAAGCGATATTGCGTCTGCAGCAGTAGGTAGGCCGAAACTCTCAA
TGTACATAAAACCGTTAAAAGAGACTATATAGTCCTTCCTTCTCCTTTTATTTCTACGAC
GCTCTCAGCTTCCCCTTTGCTAAGACCTCCAGGCGATTCCTCACCCAGCTGGATCTGCGC
CTATGCGGCCGCCTTCAGGAATAAAACCAGGACCCTATCTATTAGGCCTGAACCTCGTAG
TACGATTTACAGACTCCTTTCTGGCTTGTAGCTGCTTCTTCAGTTACGCTAACAGCCCTC
TTGCCTCACGCTATTGTTTCCCTTATTCTCCCGATACAATCCAGTTACCTCAACCTCCTG
TCCGCTTCAACTAC

>Pyr_26
TGAGCCTTTTATTCTAAGCTCAACATTTCTCCACCTCCTTGGCGCCATCCCGGAACCCAG
GCATAAAACGCAGCGAAATAAACAGAGCGCTATGCCTTTACCGTACCTGCCGCTAACCAT
TGTTACACGATTCAGTTGTAAAATTGCTCGCGTTGGCATAGTTAGGTGCCGGTTGCCCCC
CATACAATACGTTTCCAGCATGCACATATTCTCAGATTTTATTATGACCTTCACGTCACT
CATTATGTATTTTGCCTTACGTCCTCCATTTTCCTCTATGTCCGTTTTTCTCTCAAGATG
ATACATCAGTTNNNAACTCCCGCGAATTCNATTCGTGCATGAGTAGATATTGGTCCTTGA
TTGAGTACCTATAAAACGCTAGGCATCAAAGGACACTGCAATGCACAAGTCAAACAGGTC
CTCTGAAACCAAAACATAAAGACAGACGGAAGCAACCAAAGGGACAGACCATGTATTAAC
ATAGGATGCGAAAATAGTCAAGCATAGGGTAATAATTTGTTCCACAGAGGTTTGATAGCT
TACGAAAGATATCGCGAGCAACAGGCGGATACGTTGGGCGAACGAGGGAAGAGACCGAGT
CGAGTGAGACCGGAGCGGGGAGAGGGAGAGCGAATCCGAATAACAAGAGAACTGGGACAG
GAAAACACAGTGAGCAGAACCTCAGGACGAAAGAGAGACGCACTAAGAAAACAGAGGCGC
GACGTGAAGCGAGCGGAAGACCTATTGATGTAACAGAGGGAGGCGTGATATGAAAAAACT
GAAGTCGGGGTAGGGAAAGGTCGTGAAGGGGACGAATAATAATCTGCTCTTTAGCGGCAG
AAGATACATCGAGATCAGTCCTTTTTCCGAGCGATCTTTGAGCATTATCCACTTGAGCTG
GACGGCCAGACGCAGGAACTTCTAAGGGGGGAAGGGTTGGTGAGAAAAACCAAACATACC
GGGATCAAGTTAGGGCGGTAGTTGTGCTGGATTCGTGCCCGCGTTTAAGGCAGTAAGACT
CCAATGGGTAGAAGCCATGGATAACGAAGCTAGGCTAAAGACGTTGGGGACATGGATGTC
AATAATGGAGGAAATGAAGCGATATTGCGTCTGCNGCAGTAGGTAGGCCGAAACTCTCAA
TGTACATAAAACCGTTAAAAGAGACTATATAGTCCTTCCTTCTCCTTTTATTTCTACGAC
GCTCTCAGCTTCCCCTTTGCTAAGACCTCCAGGCGATTCCTCACCCAGCTGGATCTGCGC
CTATGCGGCCGCCTTCAGGAATAAAACCAGGACCCTATCTATTAGGCCTGAACCTCGTAG
TACGATTTACAGACTCCTTTCTGGCTTGTAGCTGCTTCTTCAGTTACGCTAACAGCCCTC
TTGCCTCACGCTATTGTTTCCCTTATTCTCCCGATACAATCCAGTTACCTCAACCTCCTG
TCCGCTTCAACTAC
>Pyr_27
CAAATTCCTGACCTCGGATTCCGTACCCTCTTCTATTTCTGGCGCAATCTTGGGNACCGN
ACACGACAAATCGCGGGACAGGCAGTACGCTATAACCTCGTNGCTTCTGCCACTGTACAC
TGTTCTGCATTCTAACCATAACGTCATTCATGCGAGTGTATGCGGATACAGACTCCTCTC
CTCATAACACGCGTCTGGTGTACCCACGCTCTTAGGTCTCTTTATAGTTCCCANNNNNNT
CNTNCNNNATNNNNNNNNNNNNNCTTTGCNCCTTTTCANNNNNGNNTNNNNNNNANGCNN
NCNNANNNNTNNNNNNNNNNNNNNNNNNNNNNNNGCACGCAAATGACAACTTAGATNCAN
NTNNNTNNNTGTGAGNTATAGCATGTTANNNNNNNNNATNANNCGTNAGNNNNNNNANNC
NNNNNGNNNCNNGGNNNGAGAAANNNNAANGACGGNNNNNNNGNNGGNTTTCACAGTGCN
GCGGTAGACAGGGNNNNTNNNNNNNNNAACGACGATNGNGNNCNNNNNAANNCNGNNNCC

NCCAGGGAACGCTNTGANTNNNNNNNGAGTANNNNNGNNNAANNAGNGNAGNNNNNNAAT
NANGNNNNNNCNNNNNNGNNAANNNNANNNNNNGNNNGANNNNANANNNNNNNNNNNTNN
NNNNNNNCNNNNNNNCATNCNNNNNNNCAAGGAGTAAGTTAGTANAGAAGANNANNNNNN
NNCAATACNTANGNNGAAGNNNNNNNTTNANGATGACAGGAAATGCAACANNNNNNNNNN
NNNGNNGNNNNNNNNNNAAACTGGAGGAAGACTAGGCAATCANTCANNNTGCGGTGACGA
GGTGCATGCCAATNNTNNNTACATCTCGGTTTGATATTTTAGCGTTGGCAACCCAACTAG
AGTAACANANTGTTACGGTTTTCGGAAAAGAGGAANNNGNNNANNNNNNNNNNNNNNNNN
NNNNNNNNNNNNNNNNNNNNNNNNNNNNNNNNNNNNNNNNNNNNNNNNNNNNNNNNNNNN
NNNNNNNNNNNNNNNNNNNNNNNNNANNNNGACAGGTAGGGAGGANNTAAAANNTNAACGCT
CNNCCNNNAGGANNNNANNNNNNANTNNNNNNNNNNNNNNNNNNNNNNNNNNNNNNNNNN
NNNNNNNNNNNNNNNNNNNNNNNNNATNTANNCCTTCCCCCNCCTNNTANNTNTNNNNNN
NNNNNNNNNNNNNNNNNNNNNNNNNNNNNNNNNNNNNNNNNNNNNNNNNNNNNNNNNNNN
NNNNNNNNNNNNNNNTNNNGNNNGANNCNNNNANTNNATCNNTNNNNNNNNNAACCNNNNNN
NNNNNNNNNNNGGNNNNNTTCNNNNNNNNNNCNNCNNCTTCAGNNNCATCGACCACTTCN
NTNTTTTANNNNNTNNNNNNCNNNNCCNCCTCGANNNNNNTCGATCCGTCCAATTCTTTG
CTAATCTNCATCCT
>Pyr_28
TGAGCCTTTTATTCTAAGCTCAACATTTCTCCACCTCCTTGGCGCCATCCCGGAACCCAG
NNNNAAAACGCAGCGAAATAAACAGANCGCTATGCCTTTACCGTACCTGCCGNTAACCAT
TGTTACACGATTCAGTTGTAAAATTGCTCGCGTTNGCATANNTAGGTGCCGGTTGCCCCC
CATACAATACGTTTCCAGCATGCACATATTCTCAGATTTTATTATGACCNNNNNNNNANN
CATNNNNTATTTTGCCTTACGTCCTCCATTTTCCTCTATGTCCGTTTTTCTCTCAAGCTG
ATACATCAGTNNNNNNNNNNNNNNNNNNNNNNNNNGTGCANGAGTAGATATTGGTCCTTGA
TTGAGTANCNATAAAACGCTANGCATCAAAGGACACTGCAATGCACAAGTCAAACAGNNN
NNNTGAAACNNNNACATAAAGANNNNNNNNNNCAACCAAAGGGACAGACCATGTATTAAC
ATAGGATGCGAAANNNNNCAAGCATNNNNNAATAATTTGTTCCACAGAGGTTTGATAGCT
TACGAAAGATATCGCGAGCAACAGGCGGATACGTTGGGCGAACGAGGGAAGAGACCGAGT
CGAGTGAGACCGGAGCGGGGAGAGGGAGAGCGAATCCGAATAACAAGAGAACTGGGACAG
GAAAACACAGTGAGCAGAACCTCAGGACNNNAGAGAGACGCNNNAANAAAANAGAGGCGC
GACGTGNNNNNNNNNNAAGACNTNNNNATGTAACAGAGGGAGGCGTGATATGAAAAAACT
GAAGTCGGGGTAGGGAAAGGTCGTGANNGGGACGAANNNNNNTCTGCTCTTTAGCGGCAG
AAGATACATCGAGATCAGTCCTTTNNNNNNGCGATCTTTGAGCATTATCCACTNNNGCTG
GACGGNCNNNNNCAGGAACTTCTAAGGGGGGAAGGGTTGGTGAGAAAAANCAAANNNNCC
GGGATCAAGTTAGGGCGGTAGTTGTGCTGGATTCGTGCCCGCGTTTAAGGCAGTANGANN
CCAATGGGTAGAAGCCATGGATAACGAAGCTAGGCTAAAGACGTTGGGGACATGGATGTC
AATAATGGAGAAAATGAAGCGATATTGCGNNNNNNNCAGTAGGTAGGCCGAAACTCTCAA

TGTACANNAANNCGTTAAAAGAGACTATATAGTCCTTCCTTCTCCTTTNATTTCTACGAC
GCTCTCAGCTTCCCCTTTGCTAAGACCTCCAGGCGATTCCTCACCCAGCTGGATCTGCGC
CTATGCGGCCGCCTTCAGGAATAAAACCAGGACCCTATCTATTAGGCCNNANNCTCGTAG
TACNNNNNNNNNANTCCTTTNTGGCNNNTAGCTGCTTCTTCAGTTACGCTAACAGCCCTC
TTGCCTCACGCTATTGTTTCCNNNNNTNTCCCGATACAATCCAGTTACCTCAACCTCCTG
TCCGCTTCAACTAC
>Pyr_29
TGAGCCTTTTATTCTAAGCTCAACATTTCTCCACCTCCTTGGCGCCATCCCGGAACCCAG
GCATAAAACGCAGCGAAATAAACAGAGCGCTATGCCTTTACCGTACCTGCCGCTAACCAT
TGTTACACGATTCAGTTGTAAAATTGCTCGCGTTGGCATAGTTAGGTGCCGGTTGCCCCC
CATACAATACGTTTCCAGCATGCACATATTCTCAGATTTTATTATGACCTTCACGTCACT
CATTATGTATTTTGCCTTACGTCCTCCATTTTCCTCTATGTCCGTTTTTCTCTCAAGCTG
ATACATCAGTTNNNAACTCCCGCGAATTCTATTCGTGCATGAGTAGATATTGGTCCTTGA
TTGAGTACCTATAAAACGCTAGGCATCAAAGGACACTGCAATGCACAAGTCAAACAGGTC
CTCTGAAACCAAAACATAAAGACAGACGGAAGCAACCAAAGGGACAGACCATGTATTAAC
ATAGGATGCGAAAATAGTCAAGCATAGGGTAATAATTTGTTCCACAGAGGTTTGATAGCT
TACGAAAGATATCGCGAGCAACAGGCGGATACGTTGGGCGAACGAGGGAAGAGACCGAGT
CGAGTGAGACCGGAGCGGGGAGAGGGAGAGCGAATCCGAATAACAAGAGAACTGGGACAG
GAAAACACAGTGAGCAGAACCTCAGGACGAAAGAGAGACGCACTAAGAAAACAGAGGCGC
GACGTGAAGCGAGCGGAAGACCTATTGATGTAACAGAGGGAGGCGTGATATGAAAAAACT
GAAGTCGGGGTAGGGAAAGGTCGTGAAGGGGACGAATAATAATCTGCTCTTTAGCGGCAG
AAGATACATCGAGATCAGTCCTTTTTCCGAGCGATCTTTGAGCATTATCCACTTGAGCTG
GACGGCCAGACGCAGGAACTTCTAAGGGGGGAAGGGTTGGTGAGAAAAACCAAACATACC
GGGATCAAGTTAGGGCGGTAGTTGTGCTGGATTCGTGCCCGCGTTTAAGGCAGTAAGACT
CCAATGGGTAGAAGCCATGGATAACGAAGCTAGGCTAAAGACGTTGGGGACATGGATGTC
AATAATGGAGGAAATGAAGCGATATTGCGTCTGCAGCAGTAGGTAGGCCGAAACTCTCAA
TGTACATAAAACCGTTAAAAGAGACTATATAGTCCTTCCTTCTCCTTTTATTTCTACGAC
GCTCTCAGCTTCCCCTTTGCTAAGACCTCCAGGCGATTCCTCACCCAGCTGGATCTGCGC
CTATGCGGCCGCCTTCAGGAATAAAACCAGGACCCTATCTATTAGGCCTGAACCTCGTAG
TACGATTTACAGACTCCTTTCTGGCTTGTAGCTGCTTCTTCAGTTACGCTAACAGCCCTC
TTGCCTCACGCTATTGTTTCCCTTATTCTCCCGATACAATCCAGTTACCTCAACCTCCTG
TCCGCTTCAACTAC
>Pyr_3
TGAGCCTTTTATTCTAAGCTCAACATTTCTCCACCTCCTTGGCGCCATCCCGGAACCCAG
GCATAAAACGCAGCGAAATAAACAGAGCGCTATGCCTTTACCGTACCTGCCGCTAACCAT
TGTTACACGATTCAGTTGTAAAATTGCTCGCGTTGGCATAGTTAGGTGCCGGTTGCCCCC

CATACAATACGTTTCCAGCATGCACATATTCTCAGATTTTATTATGACCTTCACGTCACT
CATTATGTATTTTGCCTTACGTCCTCCATTTTCCTCTATGTCCGTTTTTCTCTCAAGCTG
ATACATCAGTTNNNNACTCCCGCGAATNNNNNNNGTGCATGAGTAGATATTGGTCCTTGA
TTGAGTACCTATAAAACGCTAGGCATCAAAGGACACTGCAATGCACAAGTCAAACAGGTC
CTCTGAAACCAAAACATAAAGACAGACGGAAGCAACCAAAGGGACAGACCATGTATTAAC
ATAGGATGCGAAAATAGTCAAGCATAGGGTAATAATTTGTTCCACAGAGGTTTGATAGCT
TACGAAAGATATCGCGAGCAACAGGCGGATACGTTGGGCGAACGAGGGAAGAGACCGAGT
CGAGTGAGACCGGAGCGGGGAGAGGGAGAGCGAATCCGAATAACAAGAGAACTGGGACAG
GAAAACACAGTGAGCAGAACCTCAGGACGAAAGAGAGACGCACTAAGAAAACAGAGGCGC
GACGTGAAGCGAGCGGAAGACCTATTGATGTAACAGAGGGAGGCGTGATATGAAAAAACT
GAAGTCGGGGTAGGGAAAGGTCGTGAAGGGGACGAATAATAATCTGCTCTTTAGCGGCAG
AAGATACATCGAGATCAGTCCTTTTTCCGAGCGATCTTTGAGCATTATCCACTTGAGCTG
GACGGCCAGACGCAGGAACTTCTAAGGGGGGAAGGGTTGGTGAGAAAAACCAAACATACC
GGGATCAAGTTAGGGCGGTAGTTGTGCTGGATTCGTGCCCGCGTTTAAGGCAGTAAGACT
CCAATGGGTAGAAGCCATGGATAACGAAGCTAGGCTAAAGACGTTGGGGACATGGATGTC
AATAATGGAGAAAATGAAGCGATATTGCGTCTGCAGCAGTAGGTAGGCCGAAACTCTCAA
TGTACATAAAACCGTTAAAAGAGACTATATAGTCCTTCCTTCTCCTTTTATTTCTACGAC
GCTCTCAGCTTCCCCTTTGCTAAGACCTCCAGGCGATTCCTCACCCAGCTGGATCTGCGC
CTATGCGGCCGCCTTCAGGAATAAAACCAGGACCCTATCTATTAGGCCTGAACCTCGTAG
TACGATTTACAGACTCCTTTCTGGCTTGTAGCTGCTTCTTCAGTTACGCTAACAGCCCTC
TTGCCTCACGCTATTGTTTCCCTTATTCTCCCGATACAATCCAGTTACCTCAACCTCCTG
TCCGCTTCAACTAC
>Pyr_30
TGAGCCTTTTATTCTAAGCTCAACATTTCTCCACCTCCTTGGCGCCATCCCGGAACCCAG
GCATAAAACGCAGCGAAATAAACAGAGCGCTATGCCTTTACCGTACCTGCCGCTAACCAT
TGTTACACGATTCAGTTGTAAAATTGCTCGCGTTGGCATAGTTAGGTGCCGGTTGCCCCC
CATACAATACGTTTCCAGCATGCACATATTCTCAGATTTTATTATGACCTTCACGTCACT
CATTATGTATTTTGCCTTACGTCCTCCATTTTCCTCTATGTCCGTTTTTCTCTCAAGCTG
ATACATCAGTTNNNNAACTCCCGCGAATTCTNNNNNNGCATGAGTAGATATTGGTCCTTGA
TTGAGTACCTATAAAACGCTAGGCATCAAAGGACACTGCAATGCACAAGTCAAACAGGTC
CTCTGAAACCAAAACATAAAGACAGACGGAAGCAACCAAAGGGACAGACCATGTATTAAC
ATAGGATGCGAAAATAGTCAAGCATAGGGTAATAATTTGTTCCACAGAGGTTTGATAGCT
TACGAAAGATATCGCGAGCAACAGGCGGATACGTTGGGCGAACGAGGGAAGAGACCGAGT
CGAGTGAGACCGGAGCGGGGAGAGGGAGAGCGAATCCGAATAACAAGAGAACTGGGACAG
GAAAACACAGTGAGCAGAACCTCAGGACGAAAGAGAGACGCACTAAGAAAACAGAGGCGC
GACGTGAAGCGAGCGGAAGACCTATTGATGTAACAGAGGGAGGCGTGATATGAAAAAACT

GAAGTCGGGGTAGGGAAAGGTCGTGAAGGGGACGAATAATAATCTGCTCTTTAGCGGCAG
AAGATACATCGAGATCAGTCCTTTTTCCGAGCGATCTTTGAGCATTATCCACTTGAGCTG
GACGGCCAGACGCAGGAACTTCTAAGGGGGGAAGGGTTGGTGAGAAAAACCAAACATACC
GGGATCAAGTTAGGGCGGTAGTTGTGCTGGATTCGTGCCCGCGTTTAAGGCAGTAAGACT
CCAATGGGTAGAAGCCATGGATAACGAAGCTAGGCTAAAGACGTTGGGGACATGGATGTC
AATAATGGAGGAAATGAAGCGATATTGCGTCTGCAGCAGTAGGTAGGCCGAAACTCTCAA
TGTACATAAAACCGTTAAAAGAGACTATATAGTCCTTCCTTCTCCTTTTATTTCTACGAC
GCTCTCAGCTTCCCCTTTGCTAAGACCTCCAGGCGATTCCTCACCCAGCTGGATCTGCGC
CTATGCGGCCGCCTTCAGGAATAAAACCAGGACCCTATCTATTAGGCCTGAACCTCGTAG
TACGATTTACAGACTCCTTTCTGGCTTGTAGCTGCTTCTTCAGTTACGCTAACAGCCCTC
TTGCCTCACGCTATTGTTTCCCTTATTCTCCCGATACAATCCAGTTACCTCAACCTCCTG
TCCGCTTCAACTAC
>Pyr_33
TGAGCCTTTTATTCTAAGCTCAACATTTCTCCACCTCCTTGGCGCCATCCCGGAACCCAG
GCATAAAACGCAGCGAAATAAACAGAGCGCTATGCCTTTACCGTACCTGCCGCTAACCAT
TGTTACACGATTCAGTTGTAAAATTGCTCGCGTTGGCATAGTTAGGTGCCGGTTGCCCCC
CATACAATACGTTTCCAGCATGCACATATTCTCAGATTTTATTATGACCTTCACGTCACT
CATTATGTATTTTGCCTTACGTCCTCCATTTTCCTCTATGTCCGTTTTTCTCTCAAGCTG
ATACATCAGTTCNNAACTCCCGCGAATTCTATTCGTGCATGAGTAGATATTGGTCCTTGA
TTGAGTACCTATAAAACGCTAGGCATCAAAGGACACTGCAATGCACAAGTCAAACAGGTC
CTCTGAAACCAAAACATAAAGACAGACGGAAGCAACCAAAGGGACAGACCATGTATTAAC
ATAGGATGCGAAAATAGTCAAGCATAGGGTAATAATTTGTTCCACAGAGGTTTGATAGCT
TACGAAAGATATCGCGAGCAACAGGCGGATACGTTGGGCGAACGAGGGAAGAGACCGAGT
CGAGTGAGACCGGAGCGGGGAGAGGGAGAGCGAATCCGAATAACAAGAGAACTGGGACAG
GAAAACACAGTGAGCAGAACCTCAGGACGAAAGAGAGACGCACTAAGAAAACAGAGGCGC
GACGTGAAGCGAGCGGAAGACCTATTGATGTAACAGAGGGAGGCGTGATATGAAAAAACT
GAAGTCGGGGTAGGGAAAGGTCGTGAAGGGGACGAATAATAATCTGCTCTTTAGCGGCAG
AAGATACATCGAGATCAGTCCTTTTTCCGAGCGATCTTTGAGCATTATCCACTTGAGCTG
GACGGCCAGACGCAGGAACTTCTAAGGGGGGAAGGGTTGGTGAGAAAAACCAAACATACC
GGGATCAAGTTAGGGCGGTAGTTGTGCTGGATTCGTGCCCGCGTTTAAGGCAGTAAGACT
CCAATGGGTAGAAGCCATGGATAACGAAGCTAGGCTAAAGACGTTGGGGACATGGATGTC
AATAATGGAGGAAATGAAGCGATATTGCGTCTGCAGCAGTAGGTAGGCCGAAACTCTCAA
TGTACATNAAACCGTTAAAAGAGACTATATAGTCCTTCCTTCTCCTTTTATTTCTACGAC
GCTCTCAGCTTCCCCTTTGCTAAGACCTCCAGGCGATTCCTCACCCAGCTGGATCTGCGC
CTATGCGGCCGCCTTCAGGAATAAAACCAGGACCCTATCTATTAGGCCTGAACCTCGTAG
TACGATTTACAGACTCCTTTCTGGCTTGTAGCTGCTTCTTCAGTTACGCTAACAGCCCTC

TTGCCTCACGCTATTGTTTCCCTTATTCTCCCGATACAATCCAGTTACCTCAACCTCCTG
TCCGCTTCAACTAC
>Pyr_34
TGAGCCTTTTATTCTAAGCTCAACATTTCTCCACCTCCTTGGCGCCATCCCGGAACCCAG
GCATAAAACGCAGCGAAATAAACAGAGCGCTATGCCTTTACCGTACCTGCCGCTAACCAT
TGTTACACGATTCAGTTGTAAAATTGCTCGCGTTGGCATAGTTAGGTGCCGGTTGCCCCC
CATACAATACGTTTCCAGCATGCACATATTCTCAGATTTTATTATGACCTTCACGTCACT
CATTATGTATTTTGCCTTACGTCCTCCATTTTCCTCTATGTCCGTTTTTCTCTCAAGCTG
ATACATCAGTTCCNAACTCCCGCGAATTCTATTCGTGCATGAGTAGATATTGGTCCTTGA
TTGAGTACCTATAAAACGCTAGGCATCAAAGGACACTGCAATGCACAAGTCAAACAGGTC
CTCTGAAACCAAAACATAAAGACAGACGGAAGCAACCAAAGGGACAGACCATGTATTAAC
ATAGGATGCGAAAATAGTCAAGCATAGGGTAATAATTTGTTCCACAGAGGTTTGATAGCT
TACGAAAGATATCGCGAGCAACAGGCGGATACGTTGGGCGAACGAGGGAAGAGACCGAGT
CGAGTGAGACCGGAGCGGGGAGAGGGAGAGCGAATCCGAATAACAAGAGAACTGGGACAG
GAAAACACAGTGAGCAGAACCTCAGGACGAAAGAGAGACGCACTAAGAAAACAGAGGCGC
GACGTGAAGCGAGCGGAAGACCTATTGATGTAACAGAGGGAGGCGTGATATGAAAAAACT
GAAGTCGGGGTAGGGAAAGGTCGTGAAGGGGACGAATAATAATCTGCTCTTTAGCGGCAG
AAGATACATCGAGATCAGTCCTTTTTCCGAGCGATCTTTGAGCATTATCCACTTGAGCTG
GACGGCCAGACGCAGGAACTTCTAAGGGGGGAAGGGTTGGTGAGAAAAACCAAACATACC
GGGATCAAGTTAGGGCGGTAGTTGTGCTGGATTCGTGCCCGCGTTTAAGGCAGTAAGACT
CCAATGGGTAGAAGCCATGGATAACGAAGCTAGGCTAAAGACGTTGGGGACATGGATGTC
AATAATGGAGGAAATGAAGCGATATTGCGTCTGCAGCAGTAGGTAGGCCGAAACTCTCAA
TGTACATAAAACCGTTAAAAGAGACTATATAGTCCTTCCTTCTCCTTTTATTTCTACGAC
GCTCTCAGCTTCCCCTTTGCTAAGACCTCCAGGCGATTCCTCACCCAGCTGGATCTGCGC
CTATGCGGCCGCCTTCAGGAATAAAACCAGGACCCTATCTATTAGGCCTGAACCTCGTAG
TACGATTTACAGACTCCTTTCTGGCTTGTAGCTGCTTCTTCAGTTACGCTAACAGCCCTC
TTGCCTCACGCTATTGTTTCCCTTATTCTCCCGATACAATCCAGTTACCTCAACCTCCTG
TCCGCTTCAACTAC
>Pyr_36
TGAGCCTTTTATTCTAAGCTCAACATTTCTCCACCTCCTTGGCGCCATCCCGGAACCCAG
GCATAAAACGCAGCGAAATAAACAGAGCGCTATGCCTTTACCGTACCTGCCGCTAACCAT
TGTTACACGATTCAGTTGTAAAATTGCTCGCGTTGGCATAGTTAGGTGCCGGTTGCCCCC
CATACAATACGTTTCCAGCATGCACATATTCTCAGATTTTATTATGACCTTCACGTCACT
CATTATGTATTTTGCCTTACGTCCTCCATTTTCCTCTATGTCCGTTTTTCTCTCAAGCTG
ATACATCAGTNNNNNNNNNNNNNNNNNNNNNNNNNGTGCATGAGTAGATATTGGTCCTTGA
TTGAGTACCTATAAAACGCTAGGCATCAAAGGACACTGCAATGCACAAGTCAAACAGGTC

CTCTGAAACCAAAACATAAAGACAGACGGAAGCAACCAAAGNGACAGACCATGTATTAAC
ATAGGATGCGAAAATAGTCAAGCATAGGGTAATAATTTGTTCCACAGAGGTTTGATAGCT
TACGAAAGATATCGCGAGCAACAGGCGGATACGTTGGGCGAACGAGGGAAGAGACCGAGT
CGAGTGAGACCGGAGCGGGGAGAGGGAGAGCGAATCCGAATAACAAGAGAACTGGGACAG
GAAAACACAGTGAGCAGAACCTCAGGACGAAAGAGAGACGCACTAAGAAAACAGAGGCGC
GACGTGAAGCGAGCGGAAGACCTATTGATGTAACAGAGGGAGGCGTGATATGAAAAAACT
GAAGTCGGGGTAGGGAAAGGTCGTGAAGGGGACGAATAATAATCTGCTCTTTAGCGGCAG
AAGATACATCGAGATCAGTCCTTTTTCCGAGCGATCTTTGAGCATTATCCACNTGAGCTG
GACGGCCAGACGCAGGAACTTCTAAGGGGGGAAGGGTTGGTGAGAAAAACCAAACATACC
GGGATCAAGTTAGGGCGGTAGTTGTGCNGGATTCGTGCCCGCGTTTAAGGCAGTAAGACT
CCAATGGGTAGAAGCCATGGATAACGAAGCTAGGCTAAAGACGTTGGGGACATGGATGTC
AATAATGGAGAAAATGAAGCGATATTNCNNNNNNNNCAGTAGGTAGGCCGAAACTCTCAA
TGTACANAAAANCGTTAAAAGAGACTATATAGTCCTTCCTTCTCCTTTTATTTCTACGAC
GCTCTCAGCTTCCCCTTTNCTAAGACCTCCAGGCGATTCCTCACCCAGCTGGATCTGCGC
CTATGCGGCCGCCTTCAGGAATAAAACCAGGACCCTATCTATTAGGCCTGAACCTCGTAG
TACGATTTACAGACTCCTTTCTGGCTTGTAGCTGCTTCTTCAGTTACGCTAACAGCCCTC
TTGCCTCACGCTATTGTTTCCCTTATTCTCCCGATACAATCCAGTTACCTCAACCTCCTG
TCCGCTTCAACTAC
>Pyr_38
TGAGCCTTTTATTCTAAGCTCAACATTTCTCCACCTCCTTGGCGCCATCCCGGAACCCAG
GCATAAAACGCAGCGAAATAAACAGAGCGCTATGCCTTTACCGTACCTGCCGCTAACCAT
TGTTACACGATTCAGTTGTAAAATTGCTCGCGTTGGCATAGTTAGGTGCCGGTTGCCCCC
CATACAATACGTTTCCAGCATGCACATATTCTCAGATTTTATTATGACCTTCACGTCACT
CATTATGTATTTTGCCTTACGTCCTCCATTTTCCTCTATGTCCGTTTTTCTCTCAAGCTG
ATACATCAGTNNNNAACTCCCGCGAATTCTNTTCGTGCATGAGTAGATATTGGTCCTTGA
TTGAGTACCTATAAAACGCTAGGCATCAAAGGACACTGCAATGCACAAGTCAAACAGGTC
CTCTGAAACCAAAACATAAAGACAGACGGAAGCAACCAAAGGGACAGACCATGTATTAAC
ATAGGATGCGAAAATAGTCAAGCATAGGGTAATAATTTGTTCCACAGAGGTTTGATAGCT
TACGAAAGATATCGCGAGCAACAGGCGGATACGTTGGGCGAACGAGGGAAGAGACCGAGT
CGAGTGAGACCGGAGCGGGGAGAGGGAGAGCGAATCCGAATAACAAGAGAACTGGGACAG
GAAAACACAGTGAGCAGAACCTCAGGACGAAAGAGAGACGCACTAAGAAAACAGAGGCGC
GACGTGAAGCGAGCGGAAGACCTATTGATGTAACAGAGGGAGGCGTGATATGAAAAAACT
GAAGTCGGGGTAGGGAAAGGTCGTGAAGGGGACGAATAATAATCTGCTCTTTAGCGGCAG
AAGATACATCGAGATCAGTCCTTTTTCCGAGCGATCTTTGAGCATTATCCACTTGAGCTG
GACGGCCAGACGCAGGAACTTCTAAGGGGGGAAGGGTTGGTGAGAAAAACCAAACATACC
GGGATCAAGTTAGGGCGGTAGTTGTGCTGGATTCGTGCCCGCGTTTAAGGCAGTAAGACT

CCAATGGGTAGAAGCCATGGATAACGAAGCTAGGCTAAAGACGTTGGGGACATGGATGTC
AATAATGGAGGAAATGAAGCGATATTGCGTCTGCANCAGTAGGTAGGCCGAAACTCTCAA
TGTACATAAAACCGTTAAAAGAGACTATATAGTCCTTCCTTCTCCTTTTATTTCTACGAC
GCTCTCAGCTTCCCCTTTGCTAAGACCTCCAGGCGATTCCTCACCCAGCTGGATCTGCGC
CTATGCGGCCGCCTTCAGGAATAAAACCAGGACCCTATCTATTAGGCCTGAACCTCGTAG
TACGATTTACAGACTCCTTTCTGGCTTGTAGCTGCTTCTTCAGTTACGCTAACAGCCCTC
TTGCCTCACGCTATTGTTTCCCTTATTCTCCCGATACAATCCAGTTACCTCAACCTCCTG
TCCGCTTCAACTAC
>Pyr_39
TGAGCCTTTTATTCTAAGCTCAACATTTCTCCACCTCCTTGGCGCCATCCCGGAACCCAG
GCATAAAACGCAGCGAAATAAACAGAGCGCTATGCCTTTACCGTACCTGCCGCTAACCAT
TGTTACACGATTCAGTTGTAAAATTGCTCGCGTTGGCATAGTTAGGTGCCGGTTGCCCCC
CATACAATACGTTTCCAGCATGCACATATTCTCAGATTTTATTATGACCTTCACGTCACT
CATTATGTATTTTGCCTTACGTCCTCCATTTTCCTCTATGTCCGTTTTTCTCTCAAGCTG
ATACATCAGTTNCNNNNNNNCGCGAATTCTNNNCGTGCATGAGTAGATATTGGTCCTTGA
TTGAGTACCTATAAAACGCTAGGCATCAAAGGACACTGCAATGCACAAGTCAAACAGGTC
CTCTGAAACCAAAACATAAAGACAGACGGAAGCAACCAAAGGGACAGACCATGTATTAAC
ATAGGATGCGAAAATAGTCAAGCATAGGGTAATAATTTGTTCCACAGAGGTTTGATAGCT
TACGAAAGATATCGCGAGCAACAGGCGGATACGTTGGGCGAACGAGGGAAGAGACCGAGT
CGAGTGAGACCGGAGCGGGGAGAGGGAGAGCGAATCCGAATAACAAGAGAACTGGGACAG
GAAAACACAGTGAGCAGAACCTCAGGACGAAAGAGAGACGCACTAAGAAAACAGAGGCGC
GACGTGAAGCGAGCGGAAGACCTATTGATGTAACAGAGGGAGGCGTGATATGAAAAAACT
GAAGTCGGGGTAGGGAAAGGTCGTGAAGGGGACGAATAATAATCTGCTCTTTAGCGGCAG
AAGATACATCGAGATCAGTCCTTTTTCCGAGCGATCTTTGAGCATTATCCACTTGAGCTG
GACGGCCAGACGCAGGAACTTCTAAGGGGGGAAGGGTTGGTGAGAAAAACCAAACATACC
GGGATCAAGTTAGGGCGGTAGTTGTGCTGGATTCGTGCCCGCGTTTAAGGCAGTAAGACT
CCAATGGGTAGAAGCCATGGATAACGAAGCTAGGCTAAAGACGTTGGGGACATGGATATC
AATAATGGAGAAAATGAAGCGATATTGCGTCTGCNGCAGTAGGTAGGCCGAAACTCTCAA
TGTACATAAAACCGTTAAAAGAGACTATATAGTCCTTCCTTCTCCTTTTATTTCTACGAC
GCTCTCAGCTTCCCCTTTGCTAAGACCTCCAGGCGATTCCTCACCCAGCTGGATCTGCGC
CTATGCGGCCGCCTTCAGGAATAAAACCAGGACCCTATCTATTAGGCCTGAACCTCGTAG
TACGATTTACAGACTCCTTTCTGGCTTGTAGCTGCTTCTTCAGTTACGCTAACAGCCCTC
TTGCCTCACGCTATTGTTTCCCTTATTCTCCCGATACAATCCAGTTACCTCAACCTCCTG
TCCGCTTCAACTAC
>Pyr_4
TGAGCCTTTTATTCTAAGCTCAACATTTCTCCACCTCCTTGGCGCCATCCCGGAACCCAG

GCATAAAACGCAGCGAAATAAACAGAGCGCTATGCCTTTACCGTACCTGCCGCTAACCAT
TGTTACACGATTCAGTTGTAAAATTGCTCGCGTTGGCATAGTTAGGTGCCGGTTGCCCCC
CATACAATACGTTTCCAGCATGCACATATTCTCAGATTTTATTATGACCTTCACGTCACT
CATTATGTATTTTGCCTTACGTCCTCCATTTTCCTCTATGTCCGTTTTTCTCTCAAGCTG
ATACATCAGTTNNNAACTCCCGCGAATTCTNTTCGTGCATGAGTAGATATTGGTCCTTGA
TTGAGTACCTATAAAACGCTAGGCATCAAAGGACACTGCAATGCACAAGTCAAACAGGTC
CTCTGAAACCAAAACATAAAGACAGACGGAAGCAACCAAAGGGACAGACCATGTATTAAC
ATAGGATGCGAAAATAGTCAAGCATAGGGTAATAATTTGTTCCACAGAGGTTTGATAGCT
TACGAAAGATATCGCGAGCAACAGGCGGATACGTTGGGCGAACGAGGGAAGAGACCGAGT
CGAGTGAGACCGGAGCGGGGAGAGGGAGAGCGAATCCGAATAACAAGAGAACTGGGACAG
GAAAACACAGTGAGCAGAACCTCAGGACGAAAGAGAGACGCACTAAGAAAACAGAGGCGC
GACGTGAAGCGAGCGGAAGACCTATTGATGTAACAGAGGGAGGCGTGATATGAAAAAACT
GAAGTCGGGGTAGGGAAAGGTCGTGAAGGGGACGAATAATAATCTGCTCTTTAGCGGCAG
AAGATACATCGAGATCAGTCCTTTTTCCGAGCGATCTTTGAGCATTATCCACTTGAGCTG
GACGGCCAGACGCAGGAACTTCTNNGGGGGGAAGGGTTGGTGAGAAAAACCAAACATACC
GGGATCAAGTTAGGGCGGTAGTTGTGCTGGATTCGTGCCCGCGTTTAAGGCAGTAAGACT
CCAATGGGTAGAAGCCATGGATAACGAAGCTAGGCTAAAGACGTTGGGGACATGGATGTC
AATAATGGAGAAAATGAAGCGATATTGCGTCTGCAGCAGTAGGTAGGCCGAAACTCTCAA
TGTACATAAAACCGTTAAAAGAGACTATATAGTCCTTCCTTCTCCTTTTATTTCTACGAC
GCTCTCAGCTTCCCCTTTGCTAAGACCTCCAGGCGATTCCTCACCCAGCTGGATCTGCGC
CTATGCGGCCGCCTTCAGGAATAAAACCAGGACCCTATCTATTAGGCCTGAACCTCGTAG
TACGATTTACAGACTCCTTTCTGGCTTGTAGCTGCTTCTTCAGTTACGCTAACAGCCCTC
TTGCCTCACGCTATTGTTTCCCTTATTCTCCCGATACAATCCAGTTACCTCAACCTCCTG
TCCGCTTCAACTAC
>Pyr_40
TGAGCCTTTTATTCTAAGCTCAACATTTCTCCACCTCCTTGGCGCCATCCCGGAACCCAG
GCATAAAACGCAGCGAAATAAACAGAGCGCTATGCCTTTACCGTACCTGCCGCTAACCAT
TGTTACACGATTCAGTTGTAAAATTGCTCGCGTTGGCATAGTTAGGTGCCGGTTGCCCCC
CATACAATACGTTTCCAGCATGCACATATTCTCAGATTTTATTATGACCTTCACGTCACT
CATTATGTATTTTGCCTTACGTCCTCCATTTTCCTCTATGTCCGTTTTTCTCTCAAGCTG
ATACATCAGTTNNNAACTCCCGCGAATTCNATTNGTGCATGAGTAGATATTGGTCCTTGA
TTGAGTACCTATAAAACGCTAGGCATCAAAGGACACTGCAATGCACAAGTCAAACAGGTC
CTCTGAAACCAAAACATAAAGACAGACGGAAGCAACCAAAGGGACAGACCATGTATTAAC
ATAGGATGCGAAAATAGTCAAGCATAGGGTAATAATTTGTTCCACAGAGGTTTGATAGCT
TACGAAAGATATCGCGAGCAACAGGCGGATACGTTGGGCGAACGAGGGAAGAGACCGAGT
CGAGTGAGACCGGAGCGGGGAGAGGGAGAGCGAATCCGAATAACAAGAGAACTGGGACAG

GAAAACACAGTGAGCAGAACCTCAGGACGAAAGAGAGACGCACTAAGAAAACAGAGGCGC
GACGTGAAGCGAGCGGAAGACCTATTGATGTAACAGAGGGAGGCGTGATATGAAAAAACT
GAAGTCGGGGTAGGGAAAGGTCGTGAAGGGGACGAATAATAATCTGCTCTTTAGCGGCAG
AAGATACATCGAGATCAGTCCTTTTTCCGAGCGATCTTTGAGCATTATCCACTTGAGCTG
GACGGCCAGACGCAGGAACTTCTAAGGGGGGAAGGGTTGGTGAGAAAAACCAAACATACC
GGGATCAAGTTAGGGCGGTAGTTGTGCTGGATTCGTGCCCGCGTTTAAGGCAGTAAGACT
CCAATGGGTAGAAGCCATGGATAACGAAGCTAGGCTAAAGACGTTGGGGACATGGATGTC
AATAATGGAGGAAATGAAGCGATATTGCGTCTGCAGCAGTAGGTAGGCCGAAACTCTCAA
TGTACATAAAACCGTTAAAAGAGACTATATAGTCCTTCCTTCTCCTTTTATTTNTACGAC
GCTCTCAGCTTCCCCTTTGCTAAGACCTCCAGGCGATTCCTCACCCAGCTGGATCTGCGC
CTATGCGGCCGCCTTCAGGAATAAAACCAGGACCCTATCTATTAGGCCTGAACCTCGTAG
TACGATTTACAGACTCCTTTCTGGCTTGTAGCTGCTTCTTCAGTTACGCTAACAGCCCTC
TTGCCTCACGCTATTGTTTCCCTTATTCTCCCGATACAATCCAGTTACCTCAACCTCCTG
TCCGCTTCAACTAC
>Pyr_41
TGAGCCTTTTATTCTAAGCTCAACATTTCTCCACCTCCTTGGCGCCATCCCGGAACCCAG
GCATAAAACGCAGCGAAATAAACAGAGCGCTATGCCTTTACCGTACCTGCCGCTAACCAT
TGTTACACGATTCAGTTGTAAAATTGCTCGCGTTGGCATAGTTAGGTGCCGGTTGCCCCC
CATACAATACGTTTCCAGCATGCACATATTCTCAGATTTTATTATGACCTTCACGTCACT
CATTATGTATTTTGCCTTACGTCCTCCATTTTCCTCTATGTCCGTTTTTCTCTCAAGCTG
ATACATCAGTTNNNNNCTCCCGCGAATTCNNNNNNTGNNTGAGTAGATATTGGTCCTTGA
TTGAGTACCTATAAAACGCTAGGCATCAAAGGACACTGCAATGCACAAGTCAAACAGGTC
CTCTGAAACCAAAACATAAAGACAGACGGAAGCAACCAAAGGGACAGACCATGTATTNAC
ATAGGATGCGAAAATAGTCAAGCATAGGGTAATAATTTGTTCCACAGAGGTTTGATAGCT
TACGAAAGATATCGCGAGCAACAGGCGGATACGTTGGGCGAACGAGGGAAGAGACCGAGT
CGAGTGAGACCGGAGCGGGGAGAGGGAGAGCGAATCCGAATAACAAGAGAACTGGGACAG
GAAAACACAGTGAGCAGAACCTCAGGACGAAAGAGAGACGCACTAAGAAAACAGAGGCGC
GACGTGAAGCGAGCGGAAGACCTATTGATGTAACAGAGGGAGGCGTGATATGAAAAAACT
GAAGTCGGGGTAGGGAAAGGTCGTGAAGGGGACGAATAATAATCTGCTCTTTAGCGGCAG
AAGATACATCGAGATCAGTCCTTTTTCCGAGCGATCTTTGAGCATTATCCACTTGAGCTG
GACGGCCAGACGCAGGAACTTCTAAGGGGGGAAGGGTTGGTGAGAAAAACCAAACATACC
GGGATCAAGTTAGGGCGGTAGTTGTGCTGGATTCGTGCCCGCGTTTAAGGCAGTAAGACT
CCAATGGGTAGAAGCCATGGATAACGAAGCTAGGCTAAAGACGTTGGGGACATGGATGTC
AATAATGGAGAAAATGAAGCGATATTGCGTCTGCAGCAGTAGGTAGGCCGAAACTCTCAA
TGTACATAAAACCGTTAAAAGAGACTATATAGTCCTTCCTTCTCCTTTTATTTNTACGAC
GCTCTCAGCTTCCCCTTTGCTAAGACCTCCAGGCGATTCCTCACCCAGCTGGATCTGCGC

CTATGCGGCCGCCTTCAGGAATAAAACCAGGACCCTATCTATTAGGCCTGAACCTCGTAG
TACGATTTACAGACTCCTTTCTGGCTTGTAGCTGCTTCTTCAGTTACGCTAACAGCCCTC
TTGCCTCACGCTATTGTTTCCCTTATTCTCCCGATACAATCCAGTTACCTCAACCTCCTG
TCCGCTTCAACTAC
>Pyr_42
TGAGCCTTTTATTCTAAGCTCAACATTTCTCCACCTCCTTGGCGCCATCCCGGAACCCAG
GCATAAAACGCAGCGAAATAAACAGAGCGCTATGCCTTTACCGTACCTGCCGCTAACCAT
TGTTACACGATTCAGTTGTAAAATTGCTCGCGTTGGCATAGTTAGGTGCCGGTTGCCCCC
CATACAATACGTTTCCAGCATGCACATATTCTCAGATTTTATTATGACCTTCACGTCACT
CATTATGTATTTTGCCTTACGTCCTCCATTTTCCTCTATGTCCGTTTTTCTCTCAAGCTG
ATACATCAGTTNNNNNCTCCCGCGAATTCNATTCGTGCATGAGTAGATATTGGTCCTTGA
TTGAGTACCTATAAAACGCTAGGCATCAAAGGACACTGCAATGCACAAGTCAAACAGGTC
CTCTGAAACCAAAACATAAAGACAGACGGAAGCAACCAAAGGGACAGACCATGTATTAAC
ATAGGATGCGAAAATAGTCAAGCATAGGGTAATAATTTGTTCCACAGAGGTTTGATAGCT
TACGAAAGATATCGCGAGCAACAGGCGGATACGTTGGGCGAACGAGGGAAGAGACCGAGT
CGAGTGAGACCGGAGCGGGGAGAGGGAGAGCGAATCCGAATAACAAGAGAACTGGGACAG
GAAAACACAGTGAGCAGAACCTCAGGACGAAAGAGAGACGCACTAAGAAAACAGAGGCGC
GACGTGAAGCGAGCGGAAGACCTATTGATGTAACAGAGGGAGGCGTGATATGAAAAAACT
GAAGTCGGGGTAGGGAAAGGTCGTGAAGGGGACGAATAATAATCTGCTCTTTAGCGGCAG
AAGATACATCGAGATCAGTCCTTTTTCCGAGCGATCTTTGAGCATTATCCACTTGAGCTG
GACGGCCAGACGCAGGAACTTCTAAGGGGGGAAGGGTTGGTGAGAAAAACCAAACATACC
GGGATCAAGTTAGGGCGGTAGTTGTGCTGGATTCGTGCCCGCGTTTAAGGCAGTAAGACT
CCAATGGGTAGAAGCCATGGATAACGAAGCTAGGCTAAAGACGTTGGGGACATGGATGTC
AATAATGGAGGAAATGAAGCGATATTGCGTCTGCAGCAGTAGGTAGGCCGAAACTCTCAA
TGTACATAAAACCGTTAAAAGAGACTATATAGTCCTTCCTTCTCCTTTTATTTCTACGAC
GCTCTCAGCTTCCCCTTTGCTAAGACCTCCAGGCGATTCCTCACCCAGCTGGATCTGCGC
CTATGCGGCCGCCTTCAGGAATAAAACCAGGACCCTATCTATTAGGCCTGAACCTCGTAG
TACGATTTACAGACTCCTTTCTGGCTTGTAGCTGCTTCTTCAGTTACGCTAACAGCCCTC
TTGCCTCACGCTATTGTTTCCCTTATTCTCCCGATACAATCCAGTTACCTCAACCTCCTG
TCCGCTTCAACTAC
>Pyr_7
TGAGCCTTTTATTCTAAGCTCAACATTTCTCCACCTCCTTGGCGCCATCCCGGAACCCAG
GCATAAAACGCAGCGAAATAAACAGAGCGCTATGCCTTTACCGTACCTGCCGCTAACCAT
TGTTACACGATTCAGTTGTAAAATTGCTCGCGTTGGCATAGTTAGGTGCCGGTTGCCCCC
CATACAATACGTTTCCAGCATGCACATATTCTCAGATTTTATTATGACCTTCACGTCACT
CATTATGTATTTTGCCTTACGTCCTCCATTTTCCTCTATGTCCGTTTTTCTCTCAAGCTG

ATACATCAGTTNNNAACTCCCGCGAATTCNNNNCGTGCATGAGTAGATATTGGTCCTTGA
TTGAGTACCTATAAAACGCTAGGCATCAAAGGACACTGCAATGCACAAGTCAAACAGGTC
CTCTGAAACCAAAACATAAAGACAGACGGAAGCAACCAAAGGGACAGACCATGTATTAAC
ATAGGATGCGAAAATAGTCAAGCATAGGGTAATAATTTGTTCCACAGAGGTTTGATAGCT
TACGAAAGATATCGCGAGCAACAGGCGGATACGTTGGGCGAACGAGGGAAGAGACCGAGT
CGAGTGAGACCGGAGCGGGGAGAGGGAGAGCGAATCCGAATAACAAGAGAACTGGGACAG
GAAAACACAGTGAGCAGAACCTCAGGACGAAAGAGAGACGCACTAAGAAAACAGAGGCGC
GACGTGAAGCGAGCGGAAGACCTATTGATGTAACAGAGGGAGGCGTGATATGAAAAAACT
GAAGTCGGGGTAGGGAAAGGTCGTGAAGGGGACGAATAATAATCTGCTCTTTAGCGGCAG
AAGATACATCGAGATCAGTCCTTTTTCCGAGCGATCTTTGAGCATTATCCACTTGAGCTG
GACGGCCAGACGCAGGAACTTCTAAGGGGGGAAGGGTTGGTGAGAAAAACCAAACATACC
GGGATCAAGTTAGGGCGGTAGTTGTGCTGGATTCGTGCCCGCGTTTAAGGCAGTAAGACT
CCAATGGGTAGAAGCCATGGATAACGAAGCTAGGCTAAAGACGTTGGGGACATGGATGTC
AATAATGGAGAAAATGAAGCGATATTGCGTCTGCAGCAGTAGGTAGGCCGAAACTCTCAA
TGTACATAAAACCGTTAAAAGAGACTATATAGTCCTTCCTTCTCCTTTTATTTCTACGAC
GCTCTCAGCTTCCCCTTTGCTAAGACCTCCAGGCGATTCCTCACCCAGCTGGATCTGCGC
CTATGCGGCCGCCTTCAGGAATAAAACCAGGACCCTATCTATTAGGCCTGAACCTCGTAG
TACGATTTACAGACTCCTTTCTGGCTTGTAGCTGCTTCTTCAGTTACGCTAACAGCCCTC
TTGCCTCACGCTATTGTTTCCCTTATTCTCCCGATACAATCCAGTTACCTCAACCTCCTG
TCCGCTTCAACTAC
>Pyr_8
TGAGCCTTTTATTCTAAGCTCAACATTTCTCCACCTCCTTGGCGCCATCCCGGAACCCAG
GCATAAAACGCAGCGAAATAAACAGAGCGCTATGCCTTTACCGTACCTGCCGCTAACCAT
TGTTACACGATTCAGTTGTAAAATTGCTCGCGTTGGCATAGTTAGGTGCCGGTTGCCCCC
CATACAATACGTTTCCAGCATGCACATATTCTCAGATTTTATTATGACCTTCACGTCACT
CATTATGTATTTTGCCTTACGTCCTCCATTTTCCTCTATGTCCGTTTTTCTCTCAAGCTG
ATACATCAGTTNNNAACTCCCGCGAATTCNNNNNGTGCATGAGTAGATATTGGTCCTTGA
TTGAGTACCTATAAAACGCTAGGCATCAAAGGACACTGCAATGCACAAGTCAAACAGGTC
CTCTGAAACCAAAACATAAAGACAGACGGAAGCAACCAAAGGGACAGACCATGTATTAAC
ATAGGATGCGAAAATAGTCAAGCATAGGGTAATAATTTGTTCCACAGAGGTTTGATAGCT
TACGAAAGATATCGCGAGCAACAGGCGGATACGTTGGGCGAACGAGGGAAGAGACCGAGT
CGAGTGAGACCGGAGCGGGGAGAGGGAGAGCGAATCCGAATAACAAGAGAACTGGGACAG
GAAAACACAGTGAGCAGAACCTCAGGACGAAAGAGAGACGCACTAAGAAAACAGAGGCGC
GACGTGAAGCGAGCGGAAGACCTATTGATGTAACAGAGGGAGGCGTGATATGAAAAAACT
GAAGTCGGGGTAGGGAAAGGTCGTGAAGGGGACGAATAATAATCTGCTCTTTAGCGGCAG
AAGATACATCGAGATCAGTCCTTTTTCCGAGCGATCTTTGAGCATTATCCACTTGAGCTG

GACGGCCAGACGCAGGAACTTCTAAGGGGGGAAGGGTTGGTGAGAAAAACCAAACATACC
GGGATCAAGTTAGGGCGGTAGTTGTGCTGGATTCGTGCCCGCGTTTAAGGCAGTAAGACT
CCAATGGGTAGAAGCCATGGATAACGAAGCTAGGCTAAAGACGTTGGGGACATGGATGTC
AATAATGGAGAAAATGAAGCGATATTGCGTCTGCAGCAGTAGGTAGGCCGAAACTCTCAA
TGTACATAAAACCGTTAAAAGAGACTATATAGTCCTTCCTTCTCCTTTTATTTCTACGAC
GCTCTCAGCTTCCCCTTTGCTAAGACCTCCAGGCGATTCCTCACCCAGCTGGATCTGCGC
CTATGCGGCCGCCTTCAGGAATAAAACCAGGACCCTATCTATTAGGCCTGAACCTCGTAG
TACGATTTACAGACTCCTTTCTGGCTTGTAGCTGCTTCTTCAGTTACGCTAACAGCCCTC
TTGCCTCACGCTATTGTTTCCCTTATTCTCCCGATACAATCCAGTTACCTCAACCTCCTG
TCCGCTTCAACTAC
>Pyr_9
TGAGCCTTTTATTCTAAGCTCAACATTTCTCCACCTCCTTGGCGCCATCCCGGAACCCAG
GCATAAAACGCAGCGAAATAAACAGAGCGCTATGCCTTTACCGTACCTGCCGCTAACCAT
TGTTACACGATTCAGTTGTAAAATTGCTCGCGTTGGCATAGTTAGGTGCCGGTTGCCCCC
CATACAATACGTTTCCAGCATGCACATATTCTCAGATTTTATTATGACCTTCACGTCACT
CATTATGTATTTTGCCTTACGTCCTCCATTTTCCTCTATGTCCGTTTTTCTCTCAAGCTG
ATACATCAGTTNNNNNNTCCCGCGAATTNTNTTCGTGCATGAGTAGATATTGGTCCTTGA
TTGAGTACCTATAAAACGCTAGGCATCAAAGGACACTGCAATGCACAAGTCAAACAGGTC
CTCTGAAACCAAAACATAAAGACAGACGGAAGCAACCAAAGGGACAGACCATGTATTAAC
ATAGGATGCGAAAATAGTCAAGCATAGGGTAATAATTTGTTCCACAGAGGTTTGATAGCT
TACGAAAGATATCGCGAGCAACAGGCGGATACGTTGGGCGAACGAGGGAAGAGACCGAGT
CGAGTGAGACCGGAGCGGGGAGAGGGAGAGCGAATCCGAATAACAAGAGAACTGGGACAG
GAAAACACAGTGAGCAGAACCTCAGGACGAAAGAGAGACGCACTAAGAAAACAGAGGCGC
GACGTGAAGCGAGCGGAAGACCTATTGATGTAACAGAGGGAGGCGTGATATGAAAAAACT
GAAGTCGGGGTAGGGAAAGGTCGTGAAGGGGACGAATAATAATCTGCTCTTTAGCGGCAG
AAGATACATCGAGATCAGTCCTTTTTCCGAGCGATCTTTGAGCATTATCCACTTGAGCTG
GACGGCCAGACGCAGGAACTTCTAAGGGGGGAAGGGTTGGTGAGAAAAACCAAACATACC
GGGATCAAGTTAGGGCGGTAGTTGTGCTGGATTCGTGCCCGCGTTTAAGGCAGTAAGACT
CCAATGGGTAGAAGCCATGGATAACGAAGCTAGGCTAAAGACGTTGGGGACATGGATGTC
AATAATGGAGAAAATGAAGCGATATTGCGTCTGCANCAGTAGGTAGGCCGAAACTCTCAA
TGTACATAAAACCGTTAAAAGAGACTATATAGTCCTTCCTTCTCCTTTTATTTCTACGAC
GCTCTCAGCTTCCCCTTTGCTAAGACCTCCAGGCGATTCCTCACCCAGCTGGATCTGCGC
CTATGCGGCCGCCTTCAGGAATAAAACCAGGACCCTATCTATTAGGCCTGAACCTCGTAG
TACGATTTACAGACTCCTTTCTGGCTTGTAGCTGCTTCTTCAGTTACGCTAACAGCCCTC
TTGCCTCACGCTATTGTTTCCCTTATTCTCCCGATACAATCCAGTTACCTCAACCTCCTG
TCCGCTTCAACTAC

>SRR9587917
TAAGCCTTTTATTCTAAGCTTAACATTTCTCCACCACCTTGGCGCCAGTTTANAACNCAG
GCACAGAGAGCAATGGAACCAACAGNNAATTNCACTTTCGCCNTNNNNNCNNNNATACGC
TGTTACATGACTCAGTTGTAAAGTCATTGACGCGAGCATGCTCAAATACCGGCNGNNNNN
NNNACNNCGTNCNCATAACATGTAGGTACACTCAGACTTCATTCCAATCTTCACGTCACC
CGTTCTGTGTTTTGCCTTACGTCCTCCATTTTCTTTTATGTCCGTTCTTTTCTCAAACTG
ATAAATCAGTTNCGGTTTCTCGTNNATTCNGTTCGTGCATGAGTAGATACTGGTCCTCGA
TTGAGTACCTATAAAACACTAGGCATCGAAGGACACTGCAATGCGTAGGTCAAACAGGTC
CTCTGGAACCAAAACATAAAGACAGACGGAAGCAACCAAATGGACAGACCATGTGTTAAC
ATAGGGTGCGAAGATAGTCAAGCATAGGGTAATAATTTGTTCCACAGAGATTCTATAGCT
TACGAAGGATATCGCGAGCAACAGGCGAATACGTTGGGCGAGCGAGGAAAAAAACCGGAT
CGAGTGAGACCGGAGCGGGGGGAGGGAGAGCGAACCCGAATAAAAGGAGAACTGGGATAG
GAAAACACAGTGAGCAGAAGCTCAGGACGGAAGAGGAACGCACTAAAAAAACAAAGGCGC
TACGTGAAGCGAGCGAAAAACCTAGTGATATGACGGAGAGAGGTGTGATATGAAAAAACT
GAAGTCGGGGTGGAGAAGGGTCGTGAAGAGGACGAGTAATAATCTGCTCCTTAACGACAG
AAGANNNNNNNNNNNNNNNNNNNNNNNNNNNNNNNNNNNNNNNNNNNNNNNNNNNNNNNN
NNNNNNNNNNNNNNNNNNNNNNNNNNNNNNGANGGGNNNGTGGGGAAAAACAAACATCCC
GGGGTCAAGTTAGGGCGGTAGTTGTGCTGAATTCGTGCCCGCGTTTAAGGCAGTGAGACT
CCAATGGGTAGAAGCCGTGGATAGCGAAGCTAGACTAAAGACGTTGGGGACATGAATGTC
AATAATGGGGGAAATGAAACGATGCTNNNTCTGNNNNAGTAGGNNNGNNNNNANNCNNNN
TGANNNNNACNCCATTANANNANANNCCATAGTCTTTCTTTTTCCTTTTATATTTANGAN
NNNCNNNGNNTTNCNNTTGCNNNNNNNNNNNNNNNNNNTCNCNGNCCAACTNNACCTATGC
TTGGGNNGGTACTCTTAGAAATGAAACCAGGACCCTATTTATTAGGCCCGGACCTTACTG
TACGATTTACAGATCCCTCTCTGGCCGTCAGCTGTTTTTTCAGCTAGACTGACAGCCCTC
TTGCTNCCCGTCATTGTTCTTTTCATTCTCCTAAAACAATCCAGTTACCTTGGCCTCCTG
TTTGTTCCAATTAC
>SRR9587918
TAAGCCTTTTATTCTAAGCTTAACATTTCTCCACCACCTTGGCGCCAGTTTAAAACNCAG
GCACAGAGAGCAATGGAACCAACAGTAAANTNNNCTTTCGCCNTNNNTNCNNNNANACGC
TGTTACATGACTCAGTTGTAAAGTCATTGACGCGAGCATGCTCAAATACCGGNNGNNNNN
NNNACNNNGTNCNCATAACATGTAGGTACACTCAGACTTCATTNNAATCTTCACGTCACC
CGTTCTGTGTTTTGCCTTACGTCCTCCATTTTCTTTTATGTCCGTTCTTTTCTCAAACTG
ATAAATCAGTTNNNNTNTCTCGNNNATTCCGTTCGTGCATGAGTAGATACTGGTCCTCGA
TTGAGTACCTATAAAACACTAGGCATCGAAGGACACTGCAATGCGTAGGTCAAACAGGTC
CTCTGGAACCAAAACATAAAGACAGACGGAAGCAACCAAATGGACAGACCATGTGTTAAC
ATAGGGTGCGAAGATAGTCAAGCATAGGGTAATAATTTGTTCCACAGAGATTCTATAGCT

TACGAAGGATATCGCGAGCAACAGGCGAATACGTTGGGCGAGCGAGGAAAAAAACCGGAT
CGAGTGAGACCGGAGCGGGGGGAGGGAGAGCGAACCCGAATAAAAGGAGAACTGGGATAG
GAAAACACAGTGAGCAGAAGCTCAGGACGGAAGAGGAACGCACTAAAAAAACAAAGGCGC
TACGTGAAGCGAGCGAAAAACCTAGTGATATGACGGAGAGAGGTGTGATATGAAAAAACT
GAAGTCGGGGTGGAGAAGGGTCGTGAAGAGGACGAGTAATAATCTGCTCCTTANCGACAG
AAGATNNNNNNNNNNNNNNNNNNNNNNNNNNNNNNNNNNNNNNNNNNNNNNNNNNNNNNN
NNNNNNNNNNNNNNNNNNNNNNNNNNNNNNGAGGGGNNNGTGGGGAAAAACAAACATCCC
GGGGTCAAGTTAGGGCGGTAGTTGTGCTGAATTCGTGCCCGCGTTTAAGGCAGTGAGACT
CCAATGGGTAGAAGCCGTGGATAGCGAAGCTAGACTAAAGACGTTGGGGACATGAATGTC
AATAATGGGGGAAATGAAACGATGCTANATCTNNNNNAGTAGGNNNGCNGAAACTCNCAA
TGNNNNNNACNCCATTANAGNANANNCCATAGTCTTTCTTTTTCCTTTTATATNTANGAC
GCTCTNAGNNTTNCNCTTGCTNNNNNNNNNNNNNNNNNCNNNGNCCAACTNNACCTATGC
TTGNGNNGNNACTCTTAGAAATGAAACCAGGACCCTATTTATTAGGCCCGGACCTTACTG
TACGATTTACAGATCCCTCTCTGGCCGTCAGCTGTTTTTTCAGCTAGACTGACAGCCCTC
TTGCTCCCCGTCATTGTTCTTTTCATTCTCCTAAAACAATCCAGTTACCTTGGCCTCCTG
TTTGTTCCAATTAC
>SRR9587919
TGAGCCTTTTATTCTAAGCTCAACATTTCTCCACCTCCTCGGCGCCATCCCGGAACCCAG
GCATAAAACGCAGCGAAATAAATCGAGCGCTATGCCTTTACCGTACCTGCCGCTAACCAT
TGTTACACGATTCAGTTGTNNNNNNNNNNNNNNNNNNNNNNNNNNNNNNNNNNNNNNNNN
NNNNNNNNNNNNNNNNNNNNNNNNNNNNNNNNNNNNNNNNNNNNNNACCTTCACGTCACT
CATTCTGTATTTTGCCTTACGTCCTCCATTTTCCTCTATGTCCGTTTTTCTCTCAAGCTG
ATACATCAGCTNNGATCTCCCGCGAAATCTATTCGTGCATGAGTAGATATTGGTCCTTGA
TTGAGTACCTATAAAACGCTAGGCATCAAAGGACACTGCAATGCACAAGTCAAACAGGTC
CTCTGAAGCCAAAACATAAAGACAGACGGAAGCAACCAAAGGAACAGACCATGTATTAAC
ATAGGATGCGAAAATAGTCAAGCATAGGGTAATAATTTGTTCCACAGAGATTTGATAGCT
TACGAAAGATATCGCGAGCAACAGGCGAATACGTTGGGCGAACGAGGGAAGAGACCGAGT
CGAGTGAGACTGGAGCGGGGAGAGGGAGAGCGAATCCGAATAACAAGAGAACTGGGACAG
GAAAACACAGTGAGCAGAACCTCAGGACGGAAGAGAGACGCACTAAGAAAACAGAGGCGC
GACGTGAAGCAAGCGGAAGACCTATTGATGTAACAGAGGGCGGCGTGATATGAAAAAACT
GAAGTCGGGGTAGAGAAAGGTCGTGAAGAGGACGAATAATAATCTGCTCTTTAGCGGCAG
AAGATACATCGAGATCAGTCCTTTTTCCGAGCGATCTTTGAGCATTATCCACTTGAGCTG
GACGGTCAGACGCAGGAACTTCTGGGGGGGGAAGGGTTGGTGAGGAAAAACAAACATCCC
GGGGTCAAGTTAGGGCGGTAGTTGTGCTGAATTCGTGTCCGCGTTTAAGGCAGTAATACT
CCAATGGGTAGAAGCCATGGATAACGAAGCTGGGCTAAAGACGTTGGGGACATGGATGTC
AATAATGGAGGAAATGAAGCGATATNACGTCTGCAGCAGANGNTAGNCCGAGACTTTCAA

TGTACATAAAACTGTTAGAATGGACTCTATAGTTCTTCCTTCTCCTTTTATATCTAAGAC
GCTCTCAGTTTCCCCTTTACTAAGACCTCTAGGCGACTCCTCGCCCAACCGGATCTGCGC
TTATGCGGCCGCCCTCAGGAATAAAACCAGGGCCCTATCTATTAGGCCTGAACCTCGTAG
TACGATTTACAGACTCCTTTTCGGCCTGTAGCTGCTTCTTCAGCTACGCTAAAAGCCCTC
TTGCCTCACGCTATTGTATCCCTTATTCTCCCGATACAATCCAGTTACCTCAACCTCCTA
TCCGCTTCAACTAC
>SRR9587920
TAAGCCTTTTATTCTAAGCTTAACATTTCTCCACCATCTTGGCGCCGGTTTAAAACNCAG
GCACAGAGAGCAATGGAACCAACAGTAAANTGCACTTTCGCCNTNNNNNCNNNNATACGC
TGTTACATGACTCAGTTGTAAAGTCATTGACGCGAGCATACTCAAATACCGGCTGNTNNN
NNNACNNNGTNCNCATAACATGTAGGTACACTCGGACTTCATTCCAATCTTCACGTCACC
CGTTCTGTGTTTTGCCTTACGTCCTCCATTTTCTTTTATGTCCGTTCTTTTCTCAAACTG
ATAAATCAGTTCNGGTTTCTCGNNGATTCCGTTCGTGCATGGGCAGATACTGGTCCTCGA
TTGAGTACCTATAAAACACTAGGCATCGAAGGACACTGCAATGCGTAGGTCAAACAGGTC
CTCTGGAACCAAAACATAAAGACAGACGGAAGCAACCAAATGGACAGACCATGTGTCAAC
ATAGGGTGCGAAGATAGTCAAGCATAGGGTAATAATTTGTTCCACAGAGATTCTATAGCT
TACGAAGGATATCGCGAGCAACAGGCGAATACGTTGGGCGAGCGAGGAAAAAAACCGGAT
CGAGTGAGACCGGAGCGGGGGGAGGGGGAGCGAACCCGAATAAAAGGAGAACTGGGATAG
GAAAACACAGTGAGCAGAAGCTCAGGACGGAAGAGGAACGCACTAAAGAAACAAAGGCGC
TACGTGAAGTGAGCGGAAAACCTAGTGATATGACGGAGGGAGGTGTGATATGAAAAAACT
GAAGTCGGGGTGGAGAAGGGTCGTGAAGAGGACGAGTAATAATCTGCTCTTTAGCGACAG
AAGANNNNNNNNNNNNNNNNNNNNNNNNNNNNNNNNNNNNNNNNNNNNNNNNNNNNNNNN
NNNNNNNNNNNNNNNNNNNNNNNNNNNNNNNGAGGGNNNGTGGGGAAAAACAAACATCCC
GGGGTCAAGTTGGGGCGGTAGTTGTGCTGAATTCGTGCCCGCGTTTAAGGCAGTAAGACT
CCAATGGGTAGAAGCCGTGGATAGCGAAGCTAAACTAAAGACGTTGGGGACATGAATGTC
AATAATGGGGGCAATGAAACGATGCTNNNTCTGNNNNAGTAGGNNNGNCNNNANNCNCNN
TGANNNNNACGCCATTANANNANANGCCATAGTCCTTCTTTTTCCTTTTATATTTANGAN
NNNCNNNGNNTTNCNNTTGCNNNNNNNNNNNNNNNNNNCNCCGNCCAACTNNACCTATGC
TTGNGTNGGTACTCTTAGAAATGAAACCAGGACCCTATTTATTAGGCCCGGACCTTACAG
TACGATTTACAGATCCCTTTCTGGCCGTTAGCTGTTTCTTCCACTAGACTGACAGCCCTC
TTGCTCCCCGTCATTGTTCCTTTCATTCTCCTAAAACAATCCAGTTACCTTGGCCTCCTG
TTTGTTCCAATTAC
>SRR9587921
TGAGCCTTTTATTCTAAGCTCAACATTTCTCCACCTCCTTGGCGCCATCCCGGAACCCAG
GCATAAAACGCAGCGAAATAAATCGAGCGCTATGCCTTTACCGTACCTGCCGCTAACCAT
TGTTACACGATTCAGTTGTNNNNNCNNNNNNNNNNNNNNNNNNNNNNNNNNNNNNNNNNN

NNNNNNNNNNNNNNNNNNNNNNNNNNNNNNNNNNNNNNNNATNNNNACCTTTGCGTCACT
CATTCTGTATTTTGCCTTACGTCCTCCATTTTCCTCTATGTCCGTTTTTCTCTCAAGCTG
ATACATCAGCTCCGATCTCCCGCGAGATCTATTCGTGCATGAGTAGATATTGGTCCTTGA
TCGAGTACCTATAAAACGCTAGGCATCAAAGGACACTGCAATGCACAAGTCAAACAGGTC
CTCTGAAGCCAAAACATAAAGACAGACGGAAGCAACCAAAGGAACAGACCATGTATTAAC
ATAGGATGCGAAAATAGTCAAGCATAGGGTAATAATTTGTTCCACAGAGATTTGATAGCT
TACGAAAGATATCGCGAGCAACAGGCGAATACGTTGGGCGAACGAGGGAAGAGACCGAGT
CGAGTGAGACTGGAGCGGGGAGAGGGAGAGCGAATCCGAATAACAAGAGAACTGGGACAG
GAAAACACATTGAGCAGAACCTCAGGACGGAAGAGAGACGCACTAAGAAAACAGAGGCGC
GACGTGAAGCGAGCGGAAGACCTATTGATGTAACAGAGGGCGGCGTGATATGAAAAAACT
GAAGTCGGGGTAGAGAAAGGTCGTGAAGAGGACGAATAATAATCTGCTCTTTAGCGGCAG
AAGATACATCGGGATCAGTCCTTTTTCCGAGCGATCTTTGAGCATTATCCACTTGAGCTG
GACGGTCAGACGCAGGAACTCCTGGGGGGGGAAGGGTTGGTGAGGAAAAACAAACATCCC
GGGGTCAAGTTAGGGCGGTAGTTGTGCTGAATTCGTGTCCGCGTTTAAGGCAGTAATAGT
CCAATGGGTAGAAGCCATGGATAACGAAGCTGGGCTAAAGACGTTGGGGACATGGATGTC
AATAATGGAGGAAATGAAGCGATATCACGTCTGCGGCAGAAGGTAGACCGAGACTCTCAA
TGTACATAAAACCGTTAGAATAGACTCTATAGTTCTTCCTTCTCCTTTTATATCTAAGAC
GCTCTCAGTTTCCCCTCTGCTAAGACCTCTAGGCGATTCCTCGCCCAANTGGATCTGCGC
TTATGCGACCGCCCTCAGGAATAAAACCAGGACCCTATCTACTAGGCCTGAACTTCGTAG
TACGATTTACAGACTCCTTTCTTGCCTGTAGCTGCTTCTTCAGCTACGCTAAAAGCCCTC
TTGCCTCACGCTATTGTTTCCCTTATTCTCCCGATACAATCCAGTTACCTCAACCTCCTA
TCCGCTTCAACTAC
>SRR9587922
TAAGCCTTCTGTTCTAAGCTCAACATTTCTCCACCTCCTTNNNNNNNNNNNNNNNNNNNN
NNNNNNNNNNNNNNNNNNNNNNNNNNNNNNNNNNNNNNNNNNNNNNNNNNNNNNNNNNNN
NNNNACACGATTCAGTTGTAAAACCGCCTACGCTAGCATACTCAGGTGCCGGTTGCCCCC
CTTGCAACGCGTTTCCAGCACGCACATATTCCCATATTTTACTATGACCTTCACGTCACT
CGCTCTGTACTTTGCCTTACGTCCTCCATTTTCCCTTATGTCCGTTTTTCTCTCAAGCTG
ATACGTCAGTTCCGATCCATCACGGATNCTATTCGTGCATGAGTAGATATCGGTCCTTGA
TTGAGCACCTAGAAAACGCTAGGCACTAAAGGACACTGCAGTGTACAAATCAAACAGGTT
CTCTGAAACCAAAACATAGAGACAGACGGAAGTAACCAAATGGACAGACCATGTATTAAC
ATAGGATGTGAAGATAGTCAAGCATAGGGTAATAGTTTGTTCTACAGAGATTTGATAGTT
TACGAAAGGTATCGCGAGCAACAGGCGAATACGTTGGGCGGACGGGGGAGGAGACCGGGT
CGAGTGAGACCGGAGCGGGGAGAGGGAGAGCGAATCCGAATAAAAAGAGAACTGGGACAG
GAAAACACAGTGAGCAGAACCTCAGGACGGAAGAGAAACGCACTAAGAAAACAAAGGCGC
TATGTGAAGCGAACGGGAGACCTAGTGATATAACAGAGGGAGGCGTGATATGAAAAAACT

GAAGTCGGGGTAAAAAAAGGTCATGAAGAGGACGAATAGTAGTCTGCTCTTTAGCTACAG
AAGANNNNNNNNNNNNNNNNNNNNNNNNNNNNNNNNNNNNNNNNNNNNNNNNNNNNNNNN
NNNNNNNNNNNNNNNNNNNNNNNNNNNNNNGANGGGTTAGTGGGGGGAAAAAGCTATCTT
GGGGCAGCATTAGGACGGGAATCATGAAAGANTTAGACCCACATCTAAGATAACCAGACT
TCAACAAGCAAAAATTATGAATAGCAAAGCTAGGCTAAAGACGTTGGGGACATGGTTGTC
AATAATGGAGGAAATGAAGCGATGCTACGNNNNNANCNGTAGGTAAGCCGAGATTCCCAA
GAAACATANCNTCNNNNANANAGACTCCATAGTCCTTCTTTCTCCTTTTANNNNTAAGAC
GCCCTCAGTCTTCTCTTTGCNNNNATTTACNGGTGATTCCCCGACTCNCTGGANNNNCNN
NTATATGGGTAATCCTAGGAATAAAATCAGGACCCTCTCTATTAGGCCTGAACCCCGTAA
TATGATTTACAGATTCCTTCCTGACCTGTAGCTGCTTCCCCAGCTACGCTAACAGCCCTC
TCGCCTCACGTCAGTGTTTCCCTCATTCTCCCGAAACAATCCAGTTACCTCAACCTCCTG
TCCGCTTCAACTAC
>SRR9587923
TAAGCCTTCTGTTCTAAGCCCAACATTTCTCCACCTCCTTNNNNNNNNNNNNNNNNNNNN
NNNNNNNNNNNNNNNNNNNNNNNNNNNNNNNNNNNNNNNNNNNNNNNNNNNNNNNNNNNN
NNNNACACGATTCAGTTGTAAAACCGCTTACGCTATCATACTCAGGTGCCGGTTGCCCCC
CTTGCAACGCGTTTCCAGCACGCACATATTCCCATATTTTATTATGACCTTCACGTCACT
CGCTCTGTACTTTGCCTTACGTCCTCCATTTTCCCTTATGTCCGTTTTTCTCTCAAGCTG
ATACGTCAGTTCNGATCCATCNCGGATCCTATTCGTGCATGAGTAGATATCGGTCCTTGA
TTGAGCACCTAGAAAACGCTAGGCACTAAAGGACACTGCAGTGTACAAATCAAACAGGTC
CTCTGAAACCAAAACATAGAGACAGACGGAAGCAACCAAATGGACAGACCATGTATTAAC
ATAGGATGTGAAGATAGTCAAGCATAGGGTAATAGTTTGTTCTACAGAGATTTGATAGTT
TACGAAAGGTATCGCGAGCAACAGGCGAACACGTTGGGCGGACGGGGGAGGAGACCGGGT
CGAGTGAGACCGGAGCGGGGAGAGGGAGAGCGAATCCGAATAAAAAGAGAACTGGGACAG
GAAAACACAGTGAGCAGAACCTCAGGACGGAAGAGAAACGCACTAAGAAAACAAAGGCGC
TATGTGAAGCGAGCGGGAGACCTAGTGATATAACAGAGGGAGGCGTGATATGAAAAAACT
GAAGTCGGGGTAAAAAAAGGTCATGAAGAGGACGAATAGTAGTCTGCTCTTTAGCGACAG
AAGANNNNNNNNNNNNNNNNNNNNNNNNNNNNNNNNNNNNNNNNNNNNNNNNNNNNNNNN
NNNNNNNNNNNNNNNNNNNNNNNNNNNNNNGAGGGGTTNGTGGGGGGAAAAAGCTATCTT
GGGGCAGCATTAGGACGGGAATCATGAAAGANTTAGACCCACATCTAAGANAACCAGACC
TCAACAAGCAAAAATTATGAATAGCGAAGCTAGGCTAAAGACGTTGGGGGCATGGTTGTC
AATAATGGAGGAAATGAAGCGATGCTACGNCNNNANCNGTAGGTAAGCCGAGATTCCCAA
GAAACATAGCGTCNNNNANANAGACTCCATAGTCCTTNTTTCTCCTTTTATATTTAAGAC
GCCNTCAGTCTTCTCTTTGCCGNNATTTACAGGTNATTCCCCGACTCACTGGANNNACGN
NTATNNGGGTAATCCTAGGAATAAAATCAGGACCCTCTCTATTAGGCCTGAACCCCGTAA
TATGATTTACAGATTCCTTCCTGACCTGTAGCTGCTTCCCCAGCTACGCTAACAGCCCTC

TCGCCTCACGTCAGTGTTTCCCTCATTCTCCCGAAACAATCCAGTTACCTCAACCTCCTG
TCCGCTTCAACTAC
>SRR9587924
TAAGCCTTCTGTTCTAAGCTCAACATTTCTCCACCTCCTTNNNNNNNNNNNNNNNNNNNN
NNNNNNNNNNNNNNGNNNNNNNNNNNNNNNNNNNNNNNNNNNNNNNNNNNNNNNNNNNNN
NNNNACACGATTCAGTTGTAAAACCGCCTACGCTAGCATACTCAGGTGCCAGTTGCCCCC
CTTGCAACGCGTTTCCAGCACGCACATATTCCCATATTTTACTATGACCTTCACGTCACT
CGCTCTGTACTTTGCCTTACGTCCTCCATTTTCCCTTATGTCCGTTTTTCTCTCAAGCTG
ATACGTCAGTTCCGATCCATCACGGATCCTATTCGTGCATGAGTAGATATCGGTCCTTGA
TTGAGCACCTAGAAAACGCTAGGCACTAAAGGACACTGCAGTGTACAAATCAAACAGGTT
CTCTGAAACCAAAACATAGAGACAGACGGAAGCAACCAAATGGACAGACCATGTATTAAC
ATAGGATGTGAAGATAGTCAAGCATAGGGTAGTAGTTTGTTCTACAGAGATTTGATAGTT
TACGAAAGGTATCGCGAGCAACAGGCGAATACGTTGGGCGGACGGGGGAGGAGACCGGGT
CGAGTGAGACCGGAGCGGGGAGAGGGAGAGCGAATCCGAATAAAAAGAGAACTGGGACAG
GAAAACACAGTGAGCAGAACCTCAGGACGGAAGAGAAACGCACTAAGAAAACAAAGGCGC
TATGTGAAGCGAACGGGAGACCTAGTGATATAACAGAGGGAGGCGTGATATGAAAAAACT
GAAGTCGGGGTAAAAAAAGGTCATGAAGAGGACGAATAGTAGTCTGCTCTTTAGCTACAG
AAGANNNNNNNNNNNNNNNNNNNNNNNNNNNNNNNNNNNNNNNNNNNNNNNNNNNNNNNN
NNNNNNNNNNNNNNNNNNNNNNNNNNNNNNNGANGGGTTAGTGGGGGGAAAAAGCTATCTT
GGGNCAGCATCAGGACGGGAATCATGAAAGANTTAGACCCACATCTAAGATAACCAGACT
TCAACAAGCAAAAATTATGAATAGCGAAGCTAGGCTAAAGACGTTGGGGACATGGTTGTC
AATAATGGAGGAAATGAAGCGATGCTACGNNNNNANCNGTAGGTAAGCCGAGATTCCCAA
GAAACATANCGTCNNNNANANAGACTCCATAGTCCTTCTTTCTCCTTTTANNNNTAAGAC
GCCCTCAGTCTTCTCTTTGCNNNNATTTACCGGTGATTCCCCGACTCACTGGATNTGCNN
NTATATGGGNAATCCTAGGAATAAAATCAGGACCCTCTCTATTAGGCCTGAACCCCGTAA
TATGATTTACAGATTCCTTCCTGACCTGTAGCTGCTTCCCTAGCTACGCTAACAGCCCTC
TCGCCTCACGTCAGTGTTTCCCTCATTCTCCCGAAACAATCCAGTTACCTCAACCTCCTG
TCCGCTTCAACTAC
>SRR9587925
TAAGCCTTCTGTTCTAAGCTCAACATTTCTCCACCTCCTTNNNNNNNNNNNNNNNNNNNN
NNNNNNNNNNNNNNNNNNNNNNNNNNNNNNNNNNNNNNNNNNNNNNNNNNNNNNNNNNNN
NNNNACACGATTCAGTTGTNNNNNNNNNNNNNNNNNNNNNNNNNNNNNNNNNNNNNNNNN
NNNNNNNNNNNNNNNNNNNNNNNNNNNNNNNNNNNNNNNNNNNNNNACCTTCACGTCACT
CGCTCTGTACTTTGCCTTACGTCCTCCATTTTCCCTTGTGTCCGTTTTTCTCTCAAGCTG
ATACGTCAGTTCCGATCCATCGCGNATNCTATTCGTGCATGAGTAGATATCGGTCCTTGA
TTGAGCACCTAGAAAACGCTAGGCACCAAAGGACACTGCAGTGTACAAGTCAAACAGGTC

```
CTCTGAAACCAAAACATAAAGACAGACGGAAGCAACCAAATGGACAGACCATGTATTAAC
ATAGGATGTGAAGATAGTCAAGCATAGGGTAATAGTTTGTTCTACAGAGATTTGATAGTT
TACGAAAGGTATCGCTAGCAACAGGCGAATACGTTGGGCGAACGGGGGAGGAGACCGAGT
CGAGTGAGACCGGAGCGGGGAGAGGGAGAGCGAATCCGGATAAAAAGAGAACTGGGACAG
GAAAACATAGTGAGCAGAACCTCAGGACGGAAGAGAAACGCACTAAGAAAACAAAGGCGC
TATGTGAAGCGAGCGGGAGACCTAGTGATATAACAGAGGGAGGCGTGATATGAAAAAACT
GAAGTCGGGGTAAAAAAAGGTCATGAAGAGGACGAATAGTAGTCTGCTCTTTAGCGACAG
AAGANNNNNNNNNNNNNNNNNNNNNNNNNNNNNNNNNNNNNNNNNNNNNNNNNNNNNNNN
NNNNNNNNNNNNNNNNNNNNNNNNNNNNNNNGANGGGTTAGTGGGGGGAAAAAGCTATCTT
GGGGCAGCATTAGGACGGGAATCATGAAAGANTTAGACCCACATCTAAGATAACCAGACT
TCAACAAGCAAAAATTATGAATAGCGAAGCTAGGCTAAAGACGTTGGGGACATGGATGTC
AATAATGGAGGAAATGAAGCGATGCTACGNNNNNANCNGTAGGTAAGCCGAGATTCNCAA
GAACCATAGCGTCCTTGAGATAGACTCCGTAGTCCTTCTTTCTCCNTTTATNTTCAAGAC
GCCCTCAGTCCTCCCTTCGNNNNNATTTACAGGTGATTCCCCGACCCACTGGNNNNNCGN
NTATATGGCTAATCCTAGGAATAAAATCAGGACCCTCTCTATTAGGCCTGAACCCCGTAA
TACGATTTACAGATTCCTTCCTGACCTGTAGCTGCTTCCCCAGCTACGCTAACAGTCCTC
TCGCCTCACGTCAGTGTTTCCCTCATTCTCCCGAAACAATCCAGTTACCTCAACCTCCTG
TCCGTTTCAACTAC
>SRR9587926
TAAGCCTTTTATTCTAAGCTTAACATTTCTCCACCATCTTGGCGCCAGTTTAAAACNCAG
GCACAGAGAGCAACGGAACCAACAGTAAANTNCACTTTCGCCNTNNNNNCNNNNANNCGC
TGTTACATGACTCAGTTGTAAAGTCATTGACGCGAGCATACTCAAATACCGGCTGNNNNN
NNNACNNNGTNCNCATAACATGTAGGTACACTCAGACTTCATTCCAATCTTCACGTCACC
CGTTCTGTGTTTTGCCTTACGTCTTCCATTTTCTTTTATGTCCGTTCTTTTCTCGAACTG
ATAAATCAGTTCCGNTNTCNCGNNNATTCNGTTCGTGCATGAGTAGATACTGGTCCTCGA
TTGAGTACCAATAAAACACTAGGCATCGAAGGACACTGCAATGCGTAGGTCAAACAGGTC
CTCTGGAACCAAAACATAAAGACAGACGGAAGCAACCAAATGGACAGACCATGTGTTAAC
ATAGGGTGCGAAGATAGTCAAGCATAGGGTAATAATTTGTTCCACAGAGATTCTATAGCT
TACGAAGGATATCGCGGGCAACAGGCGAATACGTTGGGCGAGCGAGGAAAAAAACCGGAT
CGAGTGAGACCGGAGCGGGGGGAGGGAGAGCGAACCCGAATAAAAGGAGAACTGGGATAG
GAAAACACAGTGAGCAGAAGCTCAGGACGGAAGAGGAACGCACTAAAAAAACAAAGGCGC
TACGTGAAGCGAGCGGAAGACCTAGTGATATGACGGAGGGAGGTGTGATATGAAAAAACT
GAAGTCGGGGTGGAGAAGGGTCGTGAAGAGGACGAGTAATAATCTGCTCTTTAGCGACAG
AAGACANACNAANATNAGGCCTTTTGCGGAGCGATCTTTTAGAATTATCCGCTTGAGCTT
AACGGCAAGACACAGGAACTTCTGGGGGGGGAGGGGNNNGTGGGGAAGAACAGATGCCCC
GGGGCCGCGTTAAGGCAATGGTTGCGCAGGAACTATGCCNANATCTNAGGCTGTAGGACT
```

CCAGCGGGTAGGAGCCGTGGATAGCGAAGCTAGACTAAAGACGTTGGGGACATGAATGTC
AATAATGGGGGAAATGAAACGATGCTNNNTCTGNNNNAGTAGGNNNGCNNNNANNCTCNN
TGACCANNACNCCNTTANANNANNNNCCATAGTCCNTCNTNNTCCTTTTATNTTTNNGAN
NCCCTCNNNNTNNCNNTTGCNNNNNCCNACAGACGNCTNACCGNCCAACTATACCTANGC
TTGNGNNGNTNCTCTTAGAAATGAAACCAGGACCCTATTTATTAGGCCCGGACCTTACAG
NACGATTTACAGATCCCTTTCTGGCCGTTAGCTGTTTCTTCAGCTAGACTGACAGCCCTC
TTGCTCCCCGTCATTATTCCTTTCATTCTCCTAAAACAATCCAGTTACCTTGGCCTCCTG
TTTGTTCCAATTAC
>SRR9587927
TACGCCTTCTGTTCTAAGCCCAACATTTCTCCACCTCCTTNNNNNNNNNNNNNNNNNNNN
NNNNNNNNNNNNNNNNNNNNNNNNNNNNNNNNNNNNNNNNNNNNNNNNNNNNNNNNNNNN
NNNNACACGATTCAGTTGTAAAACCGCTTACACTAGCATACTCAGGTGCCGGTTGCCCCC
CTTGCATCGCGTTTCCAGCACGCACATATTCCCATATTTTATTATGACCTTCACGTCACT
CGCTCTGTACTTTGCCTTACGTCCTCCATTTTCCCTTATGTCCGTTTTTCTCTCAAGCTG
ATACGTCAGTTCCGATCCATCACGGATNNTATTCGTGCATGAGTAGATATCGGTCCTTGA
TTGAGCACCTAGAAAACGCTAGGCACTAAAGGACACTGCAGTGTACAAATCAAACAGGTC
CTCTGAAACTAAAACATAGAGACAGACGGAAGCAATCAAATGGACAGACCATGTATTAAC
ATAGGATGTGAAGATAGTCAAGCATAGGGTAATAGTTTGTTCTACAGAGATTTGATAGTT
TACGAAAGGTATCGCGAGCAACAGGCGAATNCGTTGGGCGGACGGGGGAGGAGACCGGGT
CGAGTGAGACCGGAGCGGGGAGAGGGAGAGCGAATCCGAATAAAAAGAGAACTGGGACAG
GAAAACACAGTGAGCAGAACCTCAGGAAGGAAGAGAAACGCACTAAGAAAACAAAGGCGC
TATGTGAAGCGAGCGGGAGACCTAGTGATATAACAGAGGGAGGCGTGATGTGAAAAAACT
GAAGTCGGGGTAAAAAAAGGTCATGAAGAGGACGAATAGAAGTCTGCTCTTTAGCGACAG
AAGANNNNNNNNNNNNNNNNNNNNNNNNNNNNNNNNNNNNNNNNNNNNNNNNNNNNNNNN
NNNNNNNNNNNNNNNNNNNNNNNNNNNNNNGANGGGTTAGTGGGGGGAAAAAGCTATCTT
GGGGCAGNATTAGGACGGGAACCATGAAAGANTTAGACCCACATCTAAGATAANCAGACT
TCAACAAGCAAAAATTATGAATAGCGAAGCTAGGCTAAAGACGTTGGGGGCATGGTTGTC
AATAATGGAGGAAATGAAGCGATGCTACGNNNNNANCNGTAGGTAAGCCGAGATTCCCAA
GAAACATAGCGTCNNNNANANAGACTCCATAGTCCTTCTTTCTCCTTTTATATTTAAGAC
GCCCTCAGTCTTCTCTTTGCNNNNATTTACAGGTGATTCCCCGACTCACTGGANNNNCGN
NTATATGGGTAATCCTAGGAATAAAATCAGGACCCTCTCTATTAGGCCTGAACCCCGTAA
TATGATTTACAGATTCCTTCCTGACCTGTAGCTGCTTCCCCAGCTACGCTAGCAGCCCTC
TCGCCTCACGTCAGTGTTTCCCTCATTCTCCCGAAACAATCCAGTTACCTCAACCTCCTG
TCCGCTTCAACTAC
>SRR9587928
TAAGCCTTTTATTCTAAGCTTAACATTTCTCCACCACCTTGGCGCCAGTTTAAAACTCAG

GCACAGAGAGCAATGGAACCAACAGTNNANTGCACTTTNGCCATNNNNNCNNNNATACGC
TGTTACATGACTCAGTTGTAAAGTCATTGACGCGAGCATGCTCAAATACCGGCNGNNNNN
NNNACNNCGTNCNCATAACATGTAGGTACACTCAGACTTCATTCCAATCTTCACGTCACC
CGTTCTGTGTTTTGCCTTACGTCCTCCATTTTCTTTTATGTCCGTTCTTTTCTCAAACTG
ATAAATCAGTTCNNGTTTCTCGTAGATTCCGTTCGTGCATGAGTAGATACTGGTCCTCGA
TTGAGTACCTATAAAACACTAGGCATCGAAGGACACTGCAATGCGTAGGTCAAACAGGTC
CTCTGGAACCAAAACATAAAGACAGACGGAAGCAACCAAATGGACAGACCATGTGTTAAC
ATAGGGTGCGAAGATAGTCAAGCATAGGGTAATAATTTGTTCCACAGAGATTCTATAGCT
TACGAAGGATATCGCGAGCAACAGGCGAATACGTTGGGCGAGCGAGGAAAAAAACCGGAT
CGAGTGAGACCGGAGCGGGGGGAGGGAGAGCGAACCCGAATAAAGGAGAACTGGGATAG
GAAAACACAGTGAGCAGAAGCTCAGGACGGAAGAGGAACGCACTAAAAAAACAAAGGCGC
TACGTGAAGCGAGCGAAAAACCTAGTGATATGACGGAGAGAGGTGTGATATGAAAAAACT
GAAGTCGGGGTGGAGAAGGGTCGTGAAGAGGACGAGTAATAATCTGCTCCTTANCGACAG
AAGATNNNNNNNNNNNNNNNNNNNNNNNNNNNNNNNNNNNNNNNNNNNNNNNNNNNNNN
NNNNNNNNNNNNNNNNNNNNNNNNNNNNNNGAGGGGNNNGTGGGGAAAAACAAACATCCC
GGGGTCAAGTTAGGGCGGTAGTTGTGCTGAATTCGTGCCCGCGTTTAAGGCAGTGAGACT
CCAATGGGTAGAAGCCGTGGATAGCGAAGCTAGACTAAAGACGTTGGGGACATGAATGTC
AATAATGGGGGAAATGAAACGATGCTANNTCTNNNNNAGTAGGNNNGNNNNNANNCNNNN
TGANNNNNACGCCATTAGAGNANANGCCATAGTCTTTCTTTTTCCTTTTATNTTTANGAN
NNNCNNNGNCTNNCNNTTGCNNNNNNNNNNNNNNNNNNCNNNGNCCAACTATACCTATGC
TTGNGNNGGTNCTCTTAGAAATGAAACCAGGACCCTATTTATTAGGCCCGGACCTTACTG
TACGATTTACAGATCCCTCTCTGGCCGTCAGCTGTTTTTTCAGCTAGACTGACAGCCCTC
TTGCTCCCCGTCATTGTTCTTTTCATTCTCCTAAAACAATCCAGTTACCTTGGCCTCCTG
TTTGTTCCAATTAC
>SRR9587929
TAAGCCTTCTGTTCTAAGCCCAACATTTCTCCACCTCCTTNNNNNNNNNNNNNNNNNNNN
NNNNNNNNNNNNNNNNNNNNNNNNNNNNNNNNNNNNNNNNNNNNNNNNNNNNNNNNNNNT
NNNNACACGATTCAGTTGTAAAACCGCTTACGCTAGCATACTCAGGAGCCGGTTGCCCCC
CTTGCAACGCGTTTCCAGCACGCACATATTCCCATATTTTATTATGACCTTCACGTCACT
CGCTCTGTACTTTGCCTTACGTCCTCCATTTTCCCTTATGTCCGTTTTTCTCTCAAGCTG
ATACGTCAGTTCCGATCCATCACGGATCCTATTCGTGCATGAGTAGATATCGGTCCTTGA
TTGAGCACCTAGAGAACGCTAGGCACTAAAGGACACTGCAGTGTACAAATCAAACAGGTC
CTCTGAAACCAAAACATAGAGACAGGCGGAAGCAACCAAATGGACAAACCATGTATTAAC
ATAGGATGTGAAGATAGTCAAGCATAGGGTAATAGTTTGTTCTACAGAGATTTGATAGTT
TACGAAAGGTATCGCTAGCAACAGGCGAATACGTTGGGCGGACGGGGGAGGAGACCGGGT
CGAGTGAGACCGGAGCGGGGAGAGGGAGAGCGAATCCGAATAAAAAGAGAACTGGGACAG

GAAAACACAGTGAGCAGAACCTCAGGACGGAAGAGAAACGCACTAAGAAAACAAAGGCGC
TATGTGAAGCGAGCGGGAGACCTAGTGATATAACAGAGGGAGGCGTGGTATGAAAAAACT
GAAGTCGGGGTAAAAAAAGGTCATGAAGAGGACGAATAGTAGTCTGCTCTTTAGCGACAG
AAGANNNNNNNNNNNNNNNNNNNNNNNNNNNNNNTCNNTNANCANNNNNNNNNNNNNNNN
NNNNNNNNNNNNNNNNNNNNNNNNNNNNNNGAGGGGTTAGTGGGGGGAAAAAGCTATCTT
GGGGCAGCATTAGGACGGGAATCATTAAAGANTTAGACCCACATCTAAGATAACCAGACT
TCAACAAGCAAAAATTATGAATAGCGAAGCTAGGCTAAAGACGTTGGGGACATGGTTGTC
AATAATGGAGGAAATGAAGCGATGCTACGNNNNNANCNGTAGGTAAGCCGAGATTCCCAA
GAAACATANNNTCNNNNANANAGACTCCATAGTCCTTCTTTCTCCTTTTATATTTAAGAC
GCCCTCAGTCTTCTCTTTGCNNNNATTTACAGGTGATTCCCCGACTCACTGGANNNNCGN
NTATATGGGTAATCCTAGGAATAAAATCAGGACCCTCTCTATTAGGCCTGAACCCCGTAA
TATGATTTACAGATTCCTTCCTGACCTGTAGCTGCTTCCCCAGCTACGCTAACAGCCCTC
TCGCCTCACGTCAGTGTTTCCCTCATTCTCCCGAAACAATCCAGTTACCTCAACCTCCTG
TCCGCTTCAACTAC
>SRR9587930
TAAGCCTTCTGTTCTAAGCCCAACATTTCTCCACCTCCTTNNNNNNNNNNNNNNNNNNNN
NNNNNNNNNNNNNNNNNNNNNNNNNNNNNNNNNNNNNNNNNNNNNNNNNNNNNNNNNNNT
NNNNACACGATTCAGTTGTAAAACCGCTTACGCTAGCATACTCAGGAGCCGGTTGCCCCC
CTTGCAACGCGTTTCCAGCACGCACATATTCCCATATTTTATTATGACCTTCACGTCACT
CGCTCTGTACTTTGCCTTACGTCCTCCATTTTCCCTTATGTCCGTTTTTCTCTCAAGCTG
ATACGTCAGTTNNGATCNATCACGGATCCTATTCGTGCATGAGTAGATATCGGTCCTTGA
TTGAGCACCTAGAAAACGCTAGGCACTAAAGGACACTGCAGTGTACAAATCAAACAGGTC
CTCTGAATCCAAAACATAGAGACAGACGGAAGCAACCAAATGGACAGACCATGTATTAAC
ATAGGATGTGAAGATAGTCAAGCATAGGGTAATAGTTTGTTCTACAGAGATTTGATAGTT
TACGAAAGGTATCGCTAGCAACAGGCGAATACGTTGGGCGGACGGGGGAGGAGACCGGGT
CGAGTGAGACCGGAGCGGGGAGAGGGAGAGCGAATCCGAATAAAAAGAGAACTGGGACAG
GAAAACACAGTGAGCAGAACCTCAGGACGGAAGAGAAACGCACTAAGACAACAAAGGNGC
TATGTGAAGCGAGCGGGAGACCTAGTGATATAACAGAGGGAGGCGTGGTATGAAAAAACT
GAAGTCGGGGTAAAAAAAGGTCATGAAGAGGACGAATGGTAGTCTGCTCTTTAGCGACAG
AAGANNNNNNNNNNNNNNNNNNNNNNNNNNNNNNNNNNNNNNNNNNNNNNNNNNNNNNNN
NNNNNNNNNNNNNNNNNNNNNNNNNNNNNNGAGGGGTTAGTGGGGGGAAAAAGCTATCTT
GGGGCAGCATTAGGACGGGAATCATTAAAGANTTAGACCCACATCTAAGATAACCAGACT
TCAACAAGCAAAAATTATGAATAGCGAAGCTAGGCTAAAGACGTTGGGGACATGGTTGTC
AATAATGGAGGAAATGAAGCGATGCTNCGTCTGNANCNGTANGTAAGCCGAGATTCCCAA
GAAACATANCGTCNNNNANANAGACTNCATAGTCCTTCTTTCTCCTTTTATATTTAAGAC
GCCCTCAGTCTTCTCTTTGCNGNNATTTACACGTGATTCCCCGNCTCACTGGATNNNCGN

NTATATGGGTAATCCTAGGAATAAAATCAGGACCCTCTCTATTAGGCCTGAACCCCGTAA
TATGATTTACAGATTCCTTCCTGACCTGTAGCTGCTTCCCCAGCTACGCTAACAGCCCTC
TCGCCTCACGTCAGTGTTTCCCTCATTCTTCCGAAACAATCCAGCTACCTCAACCTCCTG
TCCGCTTCAACTAC
>SRR9587931
TAAGCCTTCTGTTCTAAGCCCAACATTTCTCCACCTCCTTNNNNNNNNNNNNNNNNNNNN
NNNNNNNNNNNNNNNNNNNNNNNNNNNNNNNNNCNNNNNNNNNNNNNNNNNNNNNNNNNT
NNNNACACGATTCAGTTGTAAAACCGCTTACGCTAGCATACTCAGGAGCCGGTTGCCCCC
CTTGCAACGCGTTTCCAGCACGCACATATTCCCATATTTTATTATGACCTTCACGTCACT
CGCTCTGTACTTTGCCTTACGTCCTCCATTTTCCCTTATGTCCGTTTTTCTCTCAAGCTG
ATACGTCAGTTCCGATCCATCACGGATCCTATTCGTGCATGAGTAGATATCGGTCCTTGA
TTGAGCACCTAGAAAACGCTAGGCACTAAAGGACACTGCAGTGTACAAATCAAACAGGTC
CTCTGAATCCAAAACATAGAGACAGACGGAAGCAACCAAATGGACAGACCATGTATTAAC
ATAGGATGTGAAGATAGTCAAGCATAGGGTAATAGTTTGTTCTACAGAGATTTGATAGTT
TACGAAAGGTATCGCTAGCAACAGGCGAATACGTTGGGCGGACGGGGGAGGAGACCGGGT
CGAGTGAGACCGGAGCGGGGAGAGGGAGAGCGAATCCGAATAAAAAGAGAACTGGGACAG
GAAAACACAGTGAGCAGAACCTCAGGACGGAAGAGAAACGCACTAAGAAAACAAAGGNGC
TATGTGAAGCGAGCGGGAGACCTAGTGATATAACAGAGGGAGGCGTGGTATGAAAAAACT
GAAGTCGGGGTAAAAAAAGGTCATGAAGAGGACGAATGGTAGTCTGCTCTTTAGCGACAG
AAGATNNNNNNNNNNNNNNNNNNNNNNNNNNNNNNNNNNNNNNNNNNNNNNNNNNNNNN
NNNNNNNNNNNNNNNNNNNNNNNNNNNNNNGANGGGTTNGTGGGGGGAAAAAGCTATCTT
GGGGCAGCATTAGGACGGGAATCATTAAAGANTNAGACCCACATCTAAGATAACCAGACT
TCAACAAGCAAAAATTATGAATAGCGAAGCTAGGCTAAAGACGTTGGGGACATGGTTGTC
AATAATGGAGGAAATGAAGCGATGCTACGNNNNNANCNGTANGTAAGCCGANATTCCCAA
GAAACATAGCGNCNNNNANANAGACTNCATAGTCCTTCTTTCTCCTTTTATATTTAAGAC
GCCCTCAGTCTTCTCTTTGCCNNNATTTACACGTGATTCCCCGACTCACTGGANNNNCGN
NTATATGGGTAATCCTAGGAATACAATCAGGACCCTCTCTATTAGGCCTGAACCCCGTAA
TATGATTTACAGATTCCTTCCTGACCTGTAGCTGCTTCCCCAGCTACGCTAACAGCCCTC
TCGCCTCACGTCAGTGTTTCCCTCATTCTTCCGAAACAATCCAGCTACCTCAACCTCCTG
TCCGCTTCAACTAC
>SRR9587932
TACGCCTTCTGTTCTAAGCCCAACATTTCTCCACCTCCTTNNNNNNNNNNNNNNNNNNNN
NNNNNNNNNNNNNNNNNNNNNNNNNNNNNNNNNNNNNNNNNNNNNNNNNNNNNNNNNNNT
NNNNACACGATTCAGTTGTAAAACCGCTTACACTAGCATACTCAGGTGCCGGTTGCCCCC
CTTGCATCGCGTTTCCAGCACGCACATATTCCCATATTTTATTATGACCTTCACGTCACT
CGCTCTGTACTTTGCCTTACGTCCTCCATTTTCCCTTATGTCCGTTTTTCTCTCAAGCTG

ATACGTCAGTTCCGATCCATCACGGATTCTATTCGTGCATGAGTAGATATCGGTCCTTGA
TTGAGCACCTAGAAAACGCTAGGCACTAAAGGACACTGCAGTGTACAAATCAAACAGGTC
CTCTGAAACTAAAACATAGAGACAGACGGAAGCAATCAAATGGACAGACCATGTATTAAC
ATAGGATGTGAAGATAGTCAAGCATAGGGTAATAGTTTGTTCTACAGAGATTTGATAGTT
TACGAAAGGTATCGCGAGCAACAGGCGAATGCGTTGGGCGGACGGGGGAGGAGACCGGGT
CGAGTGAGACCGGAGCGGGGAGAGGGAGAGCGAATCCGAATAAAAAGAGAACTGGGACAG
GAAAACACAGTGAGCAGAACCTCAGGAAGGAAGAGAAACGCACTAAGAAAACAAAGGCGC
TATGTGAAGCGAGCGGGAGACCTAGTGATATAACAGAGGGAGGCGTGATGTGAAAAAACT
GAAGTCGGGGTAAAAAAAGGTCATGAAGAGGACGAATAGTAGTCTGCTCTTTAGCGACAG
AAGATNNNNNNNNNNNNNNNNNNNNNNNNNNNNNNNNNNNNNNNNNNNNNNNNNNNNNN
NNNNNNNNNNNNNNNNNNNNNNNNNNNNNNGAGGGGTTAGTGGGGGGAAAAAGCTATCTT
GGGGCAGCATTAGGACGGGAACCATGAAAGANTTAGACCCACATCTAAGATAACCAGACT
TCGACAAGCAAAAATTATGAATAGCGAAGCTAGGCTAAAGACGTTGGGGGCATGGTTGTC
AATAATGGAGGAAATGGAGCGATGCTACGNNNNNANCCGTAGGTAAGCCGAGANTCNCAA
GAAACATAGCGTCNNNNANATAGACTCCATAGTCCTTCTTTCTCCTTTTATATTTAAGAC
GCCCTCAGTCTTCTCTTTGCCNNNATTTACAGGTGATTCCCCGACTCACTGGANNNNCGN
NTATATGGGTAATCCTAGGAATAAAATCAGGACCCTCTCTATTAGGCCTGAACCCCGTAA
TATGATTTACAGATTCCTTCCTGACCTGTAGCTGCTTCCCCAGCTACGCTAGCAGCCCTC
TCGCCTCACGTCAGTGTTTCCCTCATTCTCCCGAAACAATCTAGTTACCTCAACCTCCTG
TCCGCTTCAACTAC
>SRR9587933
TAAGCCTTTTATTCTAAGCTTAACATTTCTCCACCATCTTGGCGCCAGTTTANAACNCAG
GCACAGAGAGCAATGGAACCAACAGTNAATTNCACTTTNNCCNTNNNNNCNNNNANACGC
TGTTACATGACTCAGTTGTAAAGTCATTGACGCGAGCATACTCAAATACCGGCTGNTANN
NNNACGNNNNNNNCANAACATGTAGGTANACTCAGACTTCATTCCAATCTTCACGTCACC
CGTTCTGTGTTTTGCCTTACGTCCTCCATTTTCTTTTATGTCCGTTCTTTTCTCAAACTG
ATAAATCAGTTNNNGTTTCTCGTNGATTCCGTTCGTGCATGAGTAGATACTGGTCCTCGA
TTGAGTACCTATAAAACACTAGGCATCGAAGGACACTGCAATGCGTAGGTCAAACAGGTC
CTCTGGAACCAAAACATAAAGACAGACGGAAGCAACCAAATGGACAGACCATGTGTTAAC
ATAGGGTGCGAAGATAGCCAAGCATAGGGTAATAATTTGTTCCATAGAGATTCTATAGCT
TACGAAGGATATCGCGAGCAACAGGCAAATACGTTGGGCGAGCGAGGAAAAAAACCGGAT
CGAGTGAGACCGGAGCGAGGGGAGGGAGAGCGAACCCGAATAAAAGGAGAACTGGGATAG
GAAAACACAGTGAGCAGAAGCTCAGGACGGAAGAGGAACGCACTAAAAAAACAAAGGCGC
TACGTGAAGCGAGCGGAAAACCTAGTGATATGACGGAGGGAGGTGTGATATGAAAAAACT
GAAGTCGGGGTGGAGAAGGGTCGTGAAGAGGACGAGTAATAATCTGCTCTTTAGCGACAG
AAGANNNNNNNNNNNNNNNNNNNNNNNNNNNNNNNNNNNNNNNNNNNNNNNNNNNNNNNN

NNNNNNNNNNNNNNNNNNNNNNNNNNNNNNNGANGGGNNAGTGGGGAAAAACAAACATCCC
NGGGTCAAGTTAGGGCGGTAGTTGTGCTGAATTCGTGCCCGCGTTTAAGGCAGTAAGACT
CCAATGGGTAGAAGCCGTGGATAGCGAAGCTAGACTAAAGACGTTGGGGACATGAATGTC
AATAATGGGAGCAATGAAACGATGCTATNTCTNNNNNAGTAGGNNNGNNNNNANNCNCNN
TGANNNNNACNCCNTTANANNANANGCNNTAGTCCTTCTTTTTCCTTTNATNTNTANGAN
NNNCNNNGNCTNNCNCTTGCNNNNNNNNNNNNNNNNNNNCNNCGACCAACTNNACCTATGC
TTGNGTNGGTACTCTTAGAAATGAAACCAGGACCCTATTTATTAGGCCCGGACCTTACAG
TGCGATTTACAGATCCCTTTCTGGCCGTTAGCTGTTTCTTCAGCTAGACTGACAGCCCTC
TTGCTCCCCGTCATTGTTCCTTTCATTCTCCTAAAACAATCCAGTTACCTTGGCCTCCTG
TTTGTTCCAATTAC
>SRR9587934
TAAGCCTTCTGTTCTAGGCCCAACATTTCTCCACCTCCTTNNNNNNNNNNNNNNNNNNNN
NNNNNNNNNNNNNNNNNNNNNNNNNNNNNNNNNNNNNNNNNNNNNNNNNNNNNNNNNNNN
NNNNACACGATTCAGTTGTAAAACCGCTTACGCTAGCACACTCAGGTGACGGTTGCCCCC
CTTGCATCGCGTTTCCAGCACGCACATATTTCCATATTTTATTATGACCTTCACGTCACT
TGCTCTGTACTTTGCCTTACGTCCTCCATTTTCCCTTATGTCCGTTTTTCTCTCAAGCTG
ATACGTCAGTTCCGATCNNNCNCGGATCCTATTCGTGCATGAGTAGATATCGGTCCTTGA
TTGAGCACCTAGAAAACGCTAGGCACTAAAGGACACTGCAGTGTACAAATCAAACAGGTC
CTCTGAAACTAAAACATAGAGACAGACGGAAGCAATCAAATGGACAGACCATGTATTAAC
ATANGATGTGAAGATAGTCAAGCATAGGGTAATAGTTTGTTCTACAGAGATTTGATAGTT
TACGAAAGGTATCGCGAGCAACAGGCGAATACGTTGGGCGGACGGGGGAGGAGACCGGGT
CGAGTGAGACCGGAGCGGGGAGAGGGAGAGCGAATCCGAATAAAAAGAGAACTGGGACAG
GAAAACACAGTGAGCAGAACCTCAGGACGGAAGAGAAACGCACTAAGAAAACAAAGGCGC
TATGTGGAGCGAGCGGGAGACATAGTGATATAACAGAGGGAGGCGTGATATGAAAAAACT
GAAGTCGGGGTAAAAAAAGGTCATGAAGAGGACGAATAGTAGTCTGCTCTTTAGCGACAG
AAGANNNNNNNNNNNNNNNNNNNNNNNNNNNNNNNNNNNNNNNNNNNNNNNNNNNNNNNN
NNNNNNNNNNNNNNNNNNNNNNNNNNNNNNNGANGGGTTNGTGGGGGGAAAANGCTATCTT
GGGGCAGCATTAGGACGGGAATCATGAAAGNNTTAGACCCACATCTAAGATAACCAGACT
TCAACAAGCAAAAATTATGAATAGCGAAGCTAGGCTAAAGACGTTGGGGGCATGGTTGTC
AATAATGGAGGAAATGAAGCGATGCTACGNNNNNANCNGTAGGTAAGCCGAGATTCCCAA
GAAACATAGCNNCNNNNANANAGACTCCATGGTCCTTCTTTCTCCTTTTATATTTAATAC
GCCCTCAGTCTTCTCTTTGCNNNNATTTACAGGTGATTCCCCGACTCACTGGATNTNCGN
NTATANGGGTAATCCTAGGAATAAAATCAGGACCCTCTCTATTAGGCCTGAATCCCGTAA
TATGATTTACAGATTCCTTCCTGACCTGTAGCTGCTTCCCCAGCTACGCTAACAGCCCTC
TCGCCTCACGTCAGTGTTTCCCTCATTCTCCCGAAACAATCCAGTTACCTCAACCTCCTG
TCCGCTTCAACTAC

>SRR9587935
TAAGCCTTCTGTTCTAAGCTCAACATTTCTCCACCTCCTTNNNNNNNNNNNNNNNNNNNN
NNNNNNNNNNNNNNNNNNNNNNNNNNNNNNNNNNNNNNNNNNNNNNNNNNNNNNNNNNNN
NNNNACACGATTCAGTTGTNNNNNNNNNNNNNNNNNNNNNNNNNNNNNNNNNNNNNNNNN
NNNNNNNNNNNNNNNNNNNNNNNNNNNNNNNNNNNNNNNNNNNNNNACCTTCACGTCACT
CGCTCTGTACTTTGCCTTACGTCCTCCATTTTCCCTTGTGTCCGTTTTTCTCTCAAGCTG
ATACGTCAGTTCCGATCCATCGCGGATNCTATTCGTGCATGAGTAGATATCGGTCCTTGA
TTGAGCACCTAGAAAACGCTAGGCACCAAAGGACACTGCAGTGTACAAGTCAAACAGGTC
CTCTGAAACCAAAACATAAAGACAGACGGAAGCAACCAAATGGACAGACCATGTATTAAC
ATAGGATGTGAAGATAGTCAAGCATAGGGTAATAGTTTGTTCTACAGAGATTTGATAGTT
TACGAAAGGTATCGCTAGCAACAGGCGAATACGTTGGGCGAACGGGGGAGGAGACCGAGT
CGAGTGAGACCGGAGCGGGGAGAGGGAGAGCGAATCCGGATAAAAAGAGAACTGGGACAG
GAAAACATAGTGAGCAGAACCTCAGGACGGAAGAGAAACGCACTAAGAAAACAAAGGCGC
TATGTGAAGCGAGCGGGAGACCTAGTGATATAACAGAGGGAGGCGTGATATGAAAAAACT
GAAGTCGGGGTAAAAAAAGGTCATGAAGAGGACGAATAGTAGTCTGCTCTTTAGCGACAG
AAGANNNNNNNNNNNNNNNNNNNNNNNNNNNNNNNNNNNNNNNNNNNNNNNNNNNNNNNN
NNNNNNNNNNNNNNNNNNNNNNNNNNNNNNGANGGGTTAGTGGGGGGAAAAAGCTATCTT
GGGGCAGCATTAGGACGGGAATCATGAAAGANTTAGACCCACATCTAAGATAACCAGACT
TCAACAAGCAAAAATTATGAATAGCGAAGCTAGGCTAAAGACGTTGGGGACATGGATGTC
AATAATGGAGGAAATGAAGCGATGCTACGNNNNNANCNGTAGGTAAGCCGAGATTCCCAA
GAACCATAGCGTCNTTGAGATAGACTCCGTAGTCCTTCTTTCTCCCTTTATATTCAAGAC
GCCCTCAGTCCTCCCTTNGNCNNNATTTACAGGTGATTCCCCGACCCACTGGNNNTNCGN
NTATATGGCTAATCCTAGGAATAAAATCAGGACCCTCTCTATTAGGCCTGAACCCCGTAA
TACGATTTACAGATTCCTTCCTGACCTGTAGCTGCTTCCCCAGCTACGCTAACAGTCCTC
TCGCCTCACGTCAGTGTTTCCCTCATTCTCCCGAAACAATCCAGTTACCTCAACCTCCTG
TCCGTTTCAACTAC
>SRR9587936
TAAGCCTTCTGTTCTAGGCCCAACATTTCTCCACCTCCTTNNNNNNNNNNNNNNNNNNNN
NNNNNNNNNNNNNNNNNNNNNNNNNNNNNNNNNNNNNNNNNNNNNNNNNNNNNNNNNNNN
NNNNACACGATTCAGTTGTAAAACCGCTTACGCTAGCACACTCAGGTGACGGTTGCCCCC
CTTGCATCGCGTTTCCAGCACGCACATATTNCCATATTTTATTATGACCTTCACGTCACT
TGCTCTGTACTTTGCCTTACGTCCTCCATTTTCCCTTATGTCCGTTTTTCTCTCAAGCTG
ATACGTCAGTTNNNATCCNTCACGGATCCTATTCGTGCATGAGTAGATATCGGTCCTTGA
TTGAGCACCTAGAAAACGCTAGGCACTAAAGGACACTGCAGTGTACAAATCAAACAGGTC
CTCTGAAACTAAAACATAGAGACAGACGGAAGCAATCAAATGGACAGACCATGTATTAAC
ATAAGATGTGAAGATAGTCAAGCATAGGGTAATAGTTTGTTCTACAGAGATTTGATAGTT

TACGAAAGGTATCGCGAGCAACAGGCGAATACGTTGGGCGGACGGGGGAGGAGACCGGGT
CGAGTGAGACCGGAGCGGGGAGAGGGAGAGCGAATCCGAATAAAAAGAGAACTGGGACAG
GAAAACACAGTGAGCAGAACCTCAGGACGGAAGAGAAACGCACTAAGAAAACAAAGGCGC
TATGTGGAGCGAGCGGGAGACATAGTGATATAACAGAGGGAGGCGTGATATGAAAAAACT
GAAGTCGGGGTAAAAAAAGGTCATGAAGAGGACGAATAGTAGTCTGCTCTTTAGCGACAG
AAGANNNNNNNNNNNNNNNNNNNNNNNNNNNNNNNNNNNNNNNNNNNNNNNNNNNNNNNN
NNNNNNNNNNNNNNNNNNNNNNNNNNNNNNGANGGGTTAGTGGGGGGAAAANGCTATCNT
GGGGCAGNATTAGGACGGGAATCATGAAAGNNTNNNACCCACATCTAAGATAACCAGACT
TCAACAAGCAAAAATTATGAATAGCGAAGCTAGGCTAAAGACGTTGGGGGCATGGTTGTC
AATAATGGAGGAAATGAAGCGATGCTACGNNNNNANCCGTAGGTAAGCCGAGATTCCCAA
GAAACATANNNNCNNNNNNANNGACTCCATGGTCCTTCTTTCTCCTTTTATATTTAATAC
GCCCTCAGTCTTCTCTTTGCCNNNATTTACAGGTGATTCCCCGACTCACTGGANNNNCGT
TTATANGGGTNNTCCTAGGAATAAAATCAGGACCCTCTCTATTAGGCCTGAATCCCGTAA
TATGATTTACAGATTCCTTCCTGACCTGTAGCTGCTTCCCCAGCTACGCTAACAGCCCTC
TCGCCTCACGTCAGTGTTTCCCTCATTCTCCCGAAACAATCCAGTTACCTCAACCTCCTG
TCCGCTTCAACTAC
>SRR9587937
TGAGCCTTTTATTCTAAGCTCAACATTTCTCCACCTCCTCGGCGCCATCCCGGAACCCAG
GCATAAAACGCAGCGAAATAAATCGAGCGCTATGCCTTTACCGTACCTGCCGCTAACCAT
TGTTACACGATTCAGTTGTNNNNNNNNNNNNNNNNNNNNNNNNNNNNNNNNNNNNNNNNN
NNNNNNNNNNNNNNNNNNNNNNNNNNNNNNNNNNNNNNNNNNNNNNACCTTCACGTCACT
CATTCTGTATTTTGCCTTACGTCCNCCATTTTCCTCTATGTCCGTTTTTCTCTCAAGCTG
ATACATCAGCTCCGATCTCCCGCGAANTCTATTCGTGCATGAGTAGATATTGGTCCTTGA
TTGAGTACCTATAAAACGCTAGGCATCAAAGGACACTGCAATGCACAAGTCAAACAGGTC
CTCTGAAGCCAAAACATAAAGACAGACGGAAGCAACCAAAGGAACAGACCATGTATTAAC
ATAGGATGCGAAAATAGTCAAGCATAGGGTAATAATTTGTTCCACAGAGATTTGATAGCT
TACGAAAGATATCGCGAGCAACAGGCGAATACGTTGGGCGAACGAGGGAAGAGACCGAGT
CGAGTGAGACTGGAGCGGGGAGAGGGAGAGCGAATCCGAATAACAAGAGAACTGGGACAG
GAAAACACAGTGAGCAGAACCTCAGGACGGAAGAGAGACGCACTAAGAAAACAGAGGCGC
GACGTGAAGCAAGCGGAAGACCTATTGATGTAACAGAGGGCGGCGTGATATGAAAAAACT
GAAGTCGGGGTAGAGAAAGGTCGTGAAGAGGACGAATAATAATCTGCTCTTTAGCGGCAG
AAGATACATCGAGATCAGTCCTTTTTCCGAGCGATCTTTGAGCATTATCCACTTGAGCTG
GACGGTCAGACGCAGGAACTTCTGGGGGGGGAAGGGTTGGTGAGGAAAAACAAACATCCC
GGGGTCAAGTTAGGGCGGTAGTTGTGCTGAATTCGTGTCCGCGTTTAAGGCAGTAATACT
CCAATGGGTAGAAGCCATGGATAACGAAGCTGGGCTAAAGACGTTGGGGACATGGATGTC
AATAATGGAGGAAATGAAGCGATATCACGTCTGCAGCAGAGGNTAGNCCGAGACTCTCAA

TGTACATAAAACCGTTAGAATAGACTCTATAGTTCTTCCTTCTCCTTTTATATCTAAGAC
GCTCTCAGTTTCCCCTTTACTAAGACCTCTAGGCGACTCCTCGCCCAACCGGATCTGCGC
TTATGCGGCCGCCCTCAGGAATAAAACCAGGGCCCTATCTATTAGGCCTGAACCTCGTAG
TACGATTTACAGACTCCTTTCCGGCCTGTAGCTGCTTCTTCAGCTACGCTAAAAGCCCTC
TTGCCTCACGCTATTGTATCCCTTATTCTCCCGATACAATCCAGTTACCTCAACCTCCTA
TCCGCTTCAACTAC
>SRR9587938
TAAGCCTTTTATTCTAAGCTTAACATTTCTCCACCACCTTGGCGCCAGTTTAAAACNCAG
GCACAGAGAGCAATGGAACCAACAGTNANNTNCACTTTNGCCATNNNNNCNNNNATACGC
TGTTACATGACTCAGTTGTAAAGTCATTNACGCGAGCATGCTCAAATACCGGCNGNNNNN
NNNACNNCGTNNNCANAACATGTAGGTACACTCAGACTTCATTCCAATCTTCACGTCACC
CGTTCTGTGTTTTGCCTTACGTCCTCCATTTTCTTTTATGTCCGTTCTTTTCTCAAACTG
ATAAATCAGTTCCGGTNTCTCGNNNATTCNGTTCGTGCATGAGTAGATACTGGTCCTCGA
TTGAGTACCTATAAAACACTAGGCATCGAAGGACACTGCAATGCGTAGGTCAAACAGGTC
CTCTGGAACCAAAACATAAAGACAGACGGAAGCAACCAAATGGACAGACCATGTGTTAAC
ATAGGGTGCGAAGATAGTCAAGCATAGGGTAATAATTTGTTCCACAGAGATTCTATAGCT
TACGAAGGATATCGCGAGCAACAGGCGAATACGTTGGGCGAGCGAGGAAAAAAACCGGAT
CGAGTGAGACCGGAGCGGGGGGAGGGAGAGCGAACCCGAATAAAAGGAGAACTGGGATAG
GAAAACACAGTGAGCAGAAGCTCAGGACGGAAGAGGAACGCACTAAAAAAACAAAGGCGC
TACGTGAAGCGAGCGAAAAACCTAGTGATATGACGGAGAGAGGTGTGATATGAAAAAACT
GAAGTCGGGGTGGAGAAGGGTCGTGAAGAGGACGAGTAATAATCTGCTCCTTAACGACAG
AAGANNNNNNNNNNNNNNNNNNNNNNNNNNNNNNNNNNNNNNNNNNNNNNNNNNNNNNNN
NNNNNNNNNNNNNNNNNNNNNNNNNNNNNGGANGGGNNNGTGGGGAAAAACAAACATCCC
GGGGTCAAGTTAGGGCGGTAGTTGTGCTGAATTCGTGCCCGCGTTTAAGGCAGTGAGACT
CCAATGGGTAGAAGCCGTGGATAGCGAAGCTAGACTAAAGACGTTGGGGACATGAATGTC
AATAATGGGGGAAATGAAACGATGCTNNNTCTGNNNNAGTAGGNNNGNNNNNANNCNNNN
TGACNNNNACNCCATTANANNANANNCCATAGTCTTTCTTTTTCCTTTTATNTNTANGAN
NNNCTNNGNNTNNCNCTTGCNNNNNNNNNNNNNNNNNNCNNNGNCCAACTNNANCTATGC
TTGGGNAGGTACTCTTAGAAATGAAACCAGGACCCTATTTATTAGGCCCGGACCTTACTG
TACGATTTACAGATCCCTCTCTGGCCGTCAGCTGTTTTTTCAGCTAGACTGACAGCCCTC
TTGCTCCCCGTCATTGTTCTTTTCATTCTCCTAAAACAATCCAGTTACCTTGGCCTCCTG
TTTGTTCCAATTAC
>SRR9587939
TAAGCCTTTTATTCTAAGCTTAACATTTCTCCACCACCTTGGCGCCAGTTTAAAACNCAG
GCACAGAGAGCAATGGAACCAACAGTNAANTGCACTTTCNCCATNNNNNCNNNNANACGC
TGTTACATGACTCAGTTGTAAAGTCATTGACGCGAGCATGCTCAAATACCGGCNGNNNNN

NNNACGNCGTNNNCANAACATGTAGGTACACTCAGACTTCATTCCAATCTTCACGTCACC
CGTTCTGTGTTTTGCCTTACGTCCTCCATTTTCTTTTATGTCCGTTCTTTTCTCAAACTG
ATAAATCAGTTCCGNTTTCTCGNNNATTCNGTTCGTGCATGAGTAGATACTGGTCCTCGA
TTGAGTACCTATAAAACACTAGGCATCGAAGGACACTGCAATGCGTAGGTCAAACAGGTC
CTCTGGAACCAAAACATAAAGACAGACGGAAGCAACCAAATGGACAGACCATGTGTTAAC
ATAGGGTGCGAAGATAGTCAAGCATAGGGTAATAATTTGTTCCACAGAGATTCTATAGCT
TACGAAGGATATCGCGAGCAACAGGCGAATACGTTGGGCGAGCGAGGAAAAAAACCGGAT
CGAGTGAGACCGGAGCGGGGGGAGGGAGAGCGAACCCGAATAAAAGGAGAACTGGGATAG
GAAAACACAGTGAGCAGAAGCTCAGGACGGAAGAGGAACGCACTAAAAAAACAAAGGCGC
TACGTGAAGCGAGCGAAAAACCTAGTGATATGACGGAGAGAGGTGTGATATGAAAAAACT
GAAGTCGGGGTGGAGAAGGGTCGTGAAGAGGACGAGTAATAATCTGCTCCTTAACGACAG
AAGANNNNNNNNNNNNNNNNNNNNNNNNNNNNNNNNNNNNNNNNNNNNNNNNNNNNNNNN
NNNNNNNNNNNNNNNNNNNNNNNNNNNNNNNGAGGGNNNGTGGGGAAAAACAAACATCCC
GGGGTCAAGTTAGGGCGGTAGTTGTGCTGAATTCGTGCCCGCGTTTAAGGCAGTGAGACT
CCAATGGGTAGAAGCCGTGGATAGCGAAGCTAGACTAAAGACGTTGGGGACATGAATGTC
AATAATGGGGGAAATGAAACGATGCTNNNTCTGNNNNAGTAGGNNNGNNNNNANNCNNNN
TGANNNNNACGCCATTANAGNANANGCCATAGTCTTTCTTTTTCCTTTTATNTTTANGAN
NNNCNNNGNCTTNCNCTTGCNNNNNNNNNNNNNNNNNNCACTGNCCAACTNTACCTATGC
TTGGGTAGGTACTCTTAGAAATGAAACCAGGACCCTATTTATTAGGCCCGGACCTTACTG
TACGATTTACAGATCCCTCTCTGGCCGTCAGCTGTTTTTTCAGCTAGACTGACAGCCCTC
TTGCTCCCCGTCATTGTTCTTTTCATTCTCCTAAAACAATCCAGTTACCTTGGCCTCCTG
TTTGTTCCAATTAC
>SRR9587940
TAAGCCTTCTGTTCTAAGCTCAACATTTCTCCACCTCCTTNNNNNNNNNNNNNNNNNNNN
NNNNNNNNNNNNNNNNNNNNNNNNNNNNNNNNNNNNNNNNNNNNNNNNNNNNNNNNNNNN
NNNNACACGATTCAGTTGTNNNNNNNNNNNNNNNNNNNNNNNNNNNNNNNNNNNNNNNNN
NNNNNNNNNNNNNNNNNNNNNNNNNNNNNNNNNNNNNNNNNNNNNNACCTTCACGTCACT
CGCTCTGTACTTTGCCTTACGTCCTCCATTTTCCCTTGTGTCCGTTTTTCTCTCAAGCTG
ATACGTCAGTTCNGATCCATCGCGNATNCTATTCGTGCATGAGTAGATATCGGTCCTTGA
TTGAGCACCTAGAAAACGCTAGGCACCAAAGGACACTGCAGTGTACAAGTCAAACAGGTC
CTCTGAAACCAAAACATAAAGACAGACGGAAGCAACCAAATGGACAGACCATGTATTAAC
ATAGGATGTGAAGATAGTCAAGCATAGGGTAATAGTTTGTTCTACAGAGATTTGATAGTT
TACGAAAGGTATCGCTAGCAACAGGCGAATACGTTGGGCGAACGGGGGAGGAGACCGAGT
CGAGTGAGACCGGAGCGGGGAGAGGGAGAGCGAATCCGGATAAAAAGAGAACTGGGACAG
GAAAACATAGTGAGCAGAACCTCAGGACGGAAGAGAAACGCACTAAGAAAACAAAGGCGC
TATGTGAAGCGAGCGGGAGACCTAGTGATATAACAGAGGGAGGCGTGATATGAAAAAACT

GAAGTCGGGGTAAAAAAAGGTCATGAAGAGGACGAATAGTAGTCTGCTCTTTAGCGACAG
AAGANNNNNNNNNNNNNNNNNNNNNNNNNNNNNNNNNNNNNNNNNNNNNNNNNNNNNNNN
NNNNNNNNNNNNNNNNNNNNNNNNNNNNNNGANGGGTTNGTGGGGGGAAAAAGCTATCTT
GGGGCAGCATTAGGACGGGAATCATGAAAGANTTAGACCCACATCTAAGATAACCAGACT
TCAACAAGCAAAAATTATGAATAGCGAAGCTAGGCTAAAGACGTTGGGGACATGGATGTC
AATAATGGAGGAAATGAAGCGATGCTACGNNNNNANCNGTAGGTAAGCCGAGATTCCCAA
GAANCATAGCGTCCTTGAGATAGACTCCGTAGTCCTTCTTTCTCCCTTTATATTCAAGAC
GCCCTCAGTCCTCCCTTNGNNNNNATTTACAGGTGATTCCCCGACCCACTGGNNNNNCGT
NTATATGGCTAATCCTAGGAATAAAATCAGGACCCTCTCTATTAGGCCTGAACCCCGTAA
TACGATTTACAGATTCCTTCCTGACCTGTAGCTGCTTCCCCAGCTACGCTAACAGTCCTC
TCGCCTCACGTCAGTGTTTCCCTCATTCTCCCGAAACAATCCAGTTACCTCAACCTCCTG
TCCGTTTCAACTAC
>SRR9587941
TAAGCCTTTTATTCTAAGCTTAACATTTCTCCACCATCTTGGCGCCAGTTTAAAACNCAG
GCACAGAGAGCAATGGAACCAACAGTAANNTNCACTTTNGCCNTNNNNNCNNNNATACGC
TGTTACATGACTCAGTTGTAAAGTCATTGACGCGAGCATACTCAAATACCGGCTGNTATN
NNNACGNCGNNCNCATAACATGTAGGTACACTCGGACTTCATTCCAATCTTCACGTCACC
CGTTCTGTGTTTTGCCTTACGTCCTCCATTTTCTTTTATGTCCGTTCTTTTCTCAAACTG
ATAAATCAGTTCNGGTTTCTCGNNNATTCNGTTCGTGCATGAGTAGATACTGGTCCTCGA
TTGAGTACCTATAAAACACTAGGCATCGAAGGACACTGCAATGCGTAGGTCAAACAGGTC
CTCTGGAACCAAAACATAAAGACAGACGGAAGCAACCAAATGGACAGACCATGTGTTAAC
ATAGGGTGCGAAGATAGTTAAGCATAGGGTAATAATTTGTTCCACAGAGATTCTATAGCT
TACGAAGGATATCGCGAGCAACAGGCGAATACGTTGGGCGAGCGAGGAAAAAAACCGGAT
CGAGTGAGACCGGAGCGGGGGGAGGGAGAGCGAACCCGAATAAAAGGAGAACTGGGATAG
GAAAACACAGTGAGCAGAAGCTCAGGACGGAAGAGGAACGCACTAAAAAAACAAAGGCGC
TACGTGAAGCGAGCGGAAAACCTAGTGATATGACGGAGGGAGGTGTGATATGAAAAAACT
GAAGTCGGGGTGGAGAAGGGTCGTGAAGAGGACGAGTAATAATCTGCTCTTTAGCGACAG
AAGATNNNNNNNNNNNNNNNNNNNNNNNNNNNNNNNNNNNNNNNNNNNNNNNNNNNNNNN
NNNNNNNNNNNNNNNNNNNNNNNNNNNNNNGAGGGGNNNGTGGGGAAAAACAAACATCCC
GGGGTCAAGTTGGGGCGGTAGTTGTGCTGAATTCGTGCCCGCGTTTAAGGCAGTAAGACT
CCAATGGGTAGAAGCCGTGGATAGCGAAGCTAGACTAAAGACGTTGGGGACATGAATGTC
AATAATGGGGGCAATGAAACGATGCTANNTCTNNNNNAGTAGGNNNGNNNNNANNCNCNN
TGANNNNNACNCCATTANAGNANANGCCATAGTCCTTCTTTTTCCTTTTATNNNTANGAN
NNNCNNNGNCTTNCNNTTGCNNNNNNNNNNNNNNNNNNCNNCGNCCAACTNNACCTATGC
TTGGGTNGGTACTCTTAGAAATGAAACCAGGACCCTATTTATTAGGCCCGGACCTTACAG
TACGATTTACAGATCCCTTTCTGGCCGTTAGCTGTTTCTTCAACTAGACTGACAGCCCTC

TTGCTCCCCGTCATTGTTCCTTTCATTCTCCTACAACAATCCAGTTACCTTGGCCTCCTG
TTTGTTCCAATTAC
>SRR9587942
TAAGCCTTTTATTCTAAGCTTAACATTTCTCCACCACCTTGGCGCCAGTTTNAAACNCAG
GCACAGAGAGCAATGGAACCAACAGTNAATTGCACTTTNGCCATNNNNNCNNNNANACGC
TGTTACATGACTCAGTTGTAAAGTCATTGACGCGAGCATGCTCAAATACCGGCNGNNNNN
NNNACNNNGNNNNCATAACATGTAGGTACACTCAGACTTCATTCCAATCTTCACGTCACC
CGTTCTGTGTTTTGCCTTACGTCCTCCATTTTCTTTTATGTCCGTTCTTTTCTCAAACTG
ATAAATCAGTTNNGGTTTCTCGNNNATTCNGTTCGTGCATGAGTAGATACTGGTCCTCGA
TTGAGTACCTATAAAACACTAGGCATCGAAGGACACTGCAATGCGTAGGTCAAACAGGTC
CTCTGGAACCAAAACATAAAGACAGACGGAAGCAACCAAATGGACAGACCATGTGTTAAC
ATAGGGTGCGAAGATAGTCAAGCATAGGGTAATAATTTGTTCCACAGAGATTCTATAGCT
TACGAAGGATATCGCGAGCAACAGGCGAATACGTTGGGCGAGCGAGGAAAAAAACCGGAT
CGAGTGAGACCGGAGCGGGGGGAGGGAGAGCGAACCCGAATAAAAGGAGAACTGGGATAG
GAAAACACAGTGAGCAGAAGCTCAGGACGGAAGAGGAACGCACTAAAAAAACAAAGGCGC
TACGTGAAGCGAGCGAAAAACCTAGTGATATGACGGAGAGAGGTGTGATATGAAAAAACT
GAAGTCGGGGTGGAGAAGGGTCGTGAAGAGGACGAGTAATAATCTGCTCCTTAACGACAG
AAGANNNNNNNNNNNNNNNNNNNNNNNNNNNNNNNNNNNNNNNNNNNNNNNNNNNNNNNN
NNNNNCNNNNNNNNNNNNNNNNNNGANGGGNNNGTGGGGAAAAACAAACATCCC
GGGGTCAAGTTAGGGCGGTAGTTGTGCTGAATTCGTGCCCGCGTTTAAGGCAGTGAGACT
CCAATGGGTAGAAGCCGTGGATAGCGAAGCTAGACTAAAGACGTTGGGGACATGAATGTC
AATAATGGGGGAAATGAAACGATGCTANNTCTGNNNNAGTAGGNNNGNNNANACTCNNNN
TGANNNNNACNCCNTTANAGNANANGCCATAGTCTTTCTTTTTCCTTTTATNTNTANGAN
NNCCNNNGNNTNNCNCTTGCNNNNNNNNNNNNNNNNNNCNNNNNCCAACTNNACCTATGC
TTGNGNNGNNNCTCTTAGAAATGAAACCAGGACCCTATTTATTAGGCCCGGACCTTACTG
TACGATTTACAGATCCCTCTCTGGCCGTCAGCTGTTTTTTCAGCTAGACTGACAGCCCTC
TTGCTCCCCGTCATTGTTCTTTTCATTCTCCTAAAACAATCCAGTTACCTTGGCCTCCTG
TTTGTTCCAATTAC
>SRR9587943
TAAGCCTTTTATTCTAAGCTTAACATTTCTCCACCACCTTGGCGCCAGTTTAAAACNCAG
GCACAGAGAGCAATGGAACCAACAGTNAANTNCACTTTCGCCATNNNNNCNNNNATACGC
TGTTACATGACTCAGTTGTAAAGTCATTGACGCGAGCATGCTCAAATACCGGCNGNNNNN
NNNACGNCGTNCNCATAACATGTAGGTACACTCAGACTTCATTCCAATCTTCACGTCACC
CGTTCTGTGTTTTGCCTTACGTCCTCCATTTTCTTTTATGTCCGTTCTTTTCTCAAACTG
ATAAATCAGTTCNGGTTTCTCGTNNATTCNGTTCGTGCATGAGTAGATACTGGTCCTCGA
TTGAGTACCTATAAAACACTAGGCATCGAAGGACACTGCAATGCGTAGGTCAAACAGGTC

CTCTGGAACCAAAACATAAAGACAGACGGAAGCAACCAAATGGACAGACCATGTGTTAAC
ATAGGGTGCGAAGATAGTCAAGCATAGGGTAATAATTTGTTCCACAGAGATTCTATAGCT
TACGAAGGATATCGCGAGCAACAGGCGAATACGTTGGGCGAGCGAGGAAAAAAACCGGAT
CGAGTGAGACCGGAGCGGGGGGAGGGAGAGCGAACCCGAATAAAAGGAGAACTGGGATAG
GAAAACACAGTGAGCAGAAGCTCAGGACGGAAGAGGAACGCACTAAAAAAACAAAGGCGC
TACGTGAAGCGAGCGAAAAACCTAGTGATATGACGGAGAGAGGTGTGATATGAAAAAACT
GAAGTCGGGGTGGAGAAGGGTCGTGAAGAGGACGAGTAATAATCTGCTCCTTAACGACAG
AAGANNNNNNNNNNNNNNNNNNNNNNNNNNNNNNNNNNNNNNNNNNNNNNNNNNNNNNNN
NNNNNNNNNNNNNNNNNNNNNNNNNNNNNNNGANGGNNNGTGGGGAAAAACAAACATCCC
GGGGTCAAGTTAGGGCGGTAGTTGTGCTGAATTCGTGCCCGCGTTTAAGGCAGTGAGACT
CCAATGGGTAGAAGCCGTGGATAGCGAAGCTAGACTAAAGACGTTGGGGACATGAATGTC
AATAATGGGGGAAATGAAACGATGCTNNNTCTGNNNNAGTAGGNNNGNNNNNANNCNNNN
TGANNNNNACNCCATTANANNANANGCCATAGTCTTTCTTTTTCCTTTTATATNTANGAN
NNNCNNNGNCTTNCNCTTGCNNNNNNNNNNNNNNNNNNCNCTGNCCAACTNNACCTATGC
TTGGGNNGGNACTCTTAGAAATGAAACCAGGACCCTATTTATTAGGCCCGGACCTTACTG
TACGATTTACAGATCCCTCTCTGGCCGTCAGCTGTTTTTTCAGCTAGACTGACAGCCCTC
TTGCTCCCCGTCATTGTTCTTTTCATTCTCCTAAAACAATCCAGTTACCTTGGCCTCCTG
TTTGTTCCAATTAC
>SRR9587944
TGAGCCTTTTATTCTAAGCTCAACATTTCTCCACCTCCTCGGCGCCATCCCGGAACCCAG
GCATAAAACGCAGCGAAATAAATCGAGCGCTATGCCTTTACCGTACCTGCCGCTAACCAT
TGTTACACGATTCAGTTGTNNNNNNNNNNNNNNNNNNNNNNNNNNNNNNNNNNNNNNNNN
NNNNNNANNNNNNNNNNNNNNNNNNNNNNNNNNNNNNNNNNACCTTCACGTCACT
CATTCTGTATTTTGCCTTACGTCCTCCATTTTCCTCTATGTCCGTTTTTCTCTCAAGCTG
ATACATCAGCTNNGATCTCCCGCGAAATCTATTCGTGCATGAGTAGATATTGGTCCTTGA
TTGAGTACCTATAAAACGCTAGGCATCAAAGGACACTGCAATGCACAAGTCAAACAGGTC
CTCTGAAGCCAAAACATAAAGACAGACGGAAGCAACCAAAGGAACAGACCATGTATTAAC
ATAGGATGCGAAAATAGTCAAGCATAGGGTAATAATTTGTTCCACAGAGATTTGATAGCT
TACGAAAGATATCGCGAGCAACAGGCGAATACGTTGGGCGAACGAGGGAAGAGACCGAGT
CGAGTGAGACTGGAGCGGGGAGAGGGAGAGCGAATCCGAATAACAAGAGAACTGGGACAG
GAAAACACAGTGAGCAGAACCTCAGGACGGAAGAGAGACGCACTAAGAAAACAGAGGCGC
GACGTGAAGCAAGCGGAAGACCTATTGATGTAACAGAGGGCGGCATGATATGAAAAAACT
GAAGTCGGGGTAGAGAAAGGTCGTGAAGAGGACGAATAATAATCTGCTCTTTAGCGGCAG
AAGATACATCGAGATCAGTCCTTTTTCCGAGCGATCTTTGAGCATTATCCACTTGAGCTG
GACGGTCAGACGCAGGAACTTCTGGGGGGGGAAGGGTTGGTGAGGAAAAACAAACATCCC
GGGGTCAAGTTAGGGCGGTAGTTGTGCTGAATTCGTGTCCGCGTTTAAGGCAGTAATACT

CCAATGGGTAGAAGCCATGGATAACGAAGCTGGGCTAAAGACGTTGGGGACATGGATGTC
AATAATGGAGGAAATGAAGCGATATCACGTCTGCAGCAGAGGATAGACCGAGACTNTCAA
TGTACATAAAACTGTTAGAATGGACTCTATAGTTCTTCCTTCTCCTTTTATATCTAAGAC
GCTCTCAGTTTCCCCTTTACTAAGACCTCTAGGCGACTCCTCGCCCAACCGGATCTGCGC
TTATGCGGCCGCCCTCAGGAATAAAACCAGGGCCCTATCTATTAGGCCTGAACCTCGTAG
TACGATTTACAGACTCCTTTTCGGCCTGTAGCTGCTTCTTCAGCTACGCTAAAAGCCCTC
TTGCCTCACGCTATTGTATCCCTTATTCTCCCGATACAATCCAGTTACCTCAACCTCCTA
TCCGCTTCAACTAC
>SRR9587945
TAAGCCTTCTGTTCTAAGCTCAACATTTCTCCACCTCCTTNNNNNNNNNNNNNNNNNNNN
NNNNNNNNNNNNNNNNNNNNNNNNNNNNNNNNNNNNNNNNNNNNNNNNNNNNNNNNNNNN
NNNNACACGATTCAGTTGTNNNNNNNNNNNNNNNNNNNNNNNNNNNNNNNNNNNNNNNNN
NNNNNNNNNNNNNNNNNNNNNNNNNNNNNNNNNNNNNNNNNNNNNNNNACCTTCACGTCACT
CGCTCTGTACTTTGCCTTACGTCCTCCATTTTCCCTTGTGTCCGTTTTTCTCTCAAGCTG
ATACGTCAGTTCCGATCCATCGCGGATCCTATTCGTGCATGAGTAGATATCGGTCCTTGA
TTGAGCACCTAGAAAACGCTAGGCACCAAAGGACACTGCAGTGTACAAGTCAAACAGGTC
CTCTGAAACCAAAACATAAAGACAGACGGAAGCAACCAAATGGACAGACCATGTATTAAC
ATAGGATGTGAAGATAGTCAAGCATAGGGTAATAGTTTGTTCTACAGAGATTTGATAGTT
TACGAAAGGTATCGCTAGCAACAGGCGAATACGTTGGGCGAACGGGGGAGGAGACCGAGT
CGAGTGAGACCGGAGCGGGGAGAGGGAGAGCGAATCCGGATAAAAAGAGAACTGGGACAG
GAAAACATAGTGAGCAGAACCTCAGGACGGAAGAGAAACGCACTAAGAAAACAAAGGCGC
TATGTGAAGCGAGCGGGAGACCTAGTGATATAACAGAGGGAGGCGTGATATGAAAAAACT
GAAGTCGGGGTAAAAAAAGGTCATGAAGAGGACGAATAGTAGTCTGCTCTTTAGCGACAG
AAGANNNNNNNNNNNNNNNNNNNNNNNNNNNNNNNNNNNNNNNNNNNNNNNNNNNNNNNN
NNNNNNNNNNNNNNNNNNNNNNNNNNNNNNNGAGGGGTTAGTGGGGGGAAAAAGCTATCTT
GGGGCAGCATTAGGACGGGAATCATGAAAGANTTAGACCCACATCTAAGATAACCAGACT
TCAACAAGCAAAAATTATGAATAGCGAAGCTAGGCTAAAGACGTTGGGGACATGGATGTC
AATAATGGAGGAAATGAAGCGATGCTACGNNNNNANCNGTAGGTAAGCCGAGATTCCCAA
GAACCATAGCGTCCTTGAGATAGACTCCGTAGTCCTTCTTTCTCCCTTTATATTCAAGAC
GCCCTCAGTCCTCCCTTNGNNNNNATTTACAGGTGATTCCCCGACCCACTGGNNNNNCGT
NTATATGGCTAATCCTAGGAATAAAATCAGGACCCTCTCTATTAGGCCTGAACCCCGTAA
TACGATTTACAGATTCCTTCCTGACCTGTAGCTGCTTCCCCAGCTACGCTAACAGTCCTC
TCGCCTCACGTCAGTGTTTCCCTCATTCTCCCGAAACAATCCAGTTACCTCAACCTCCTG
TCCGTTTCAACTAC
>SRR9587946
TAAGCCTTTTATTCTAAGCTTAACATTTCTCCACCATCTTGGCGCCGGTTTAAAACNCAG

GCACAGAGAGCAATGGAACCAACAGTNAANTGCACTTTCGCCATNNNNNCNNNNATACGC
TGTTACATGACTCAGTTGTAAAGTCATTGACGCGAGCATACTCAAATACCGGCTGNNNTN
NNNACGNCGTNCNCATAACATGTAGGTACACTCGGACTTCATTCCAATCTTCACGTCACC
CGTTCTGTGTTTTGCCTTACGTCCTCCATTTTCTTTTATGTCCGTTCTTTTCTCAAACTG
ATAAATCAGTTCCGGTTTCTCGNNNATTCCGTTCGTGCATGGGCAGATACTGGTCCTCGA
TTGAGTACCTATAAAACACTAGGCATCGAAGGACACTGCAATGCGTAGGTCAAACAGGTC
CTCTGGAACCAAAACATAAAGACAGACGGAAGCAACCAAATGGACAGACCATGTGTCAAC
ATAGGGTGCGAAGATAGTCAAGCATAGGGTAATAATTTGTTCCACAGAGATTCTATAGCT
TACGAAGGATATCGCGAGCAACAGGCGAATACGTTGGGCGAGCGAGGAAAAAAACCGGAT
CGAGTGAGACCGGAGCGGGGGGAGGGGGAGCGAACCCGAATAAAAGGAGAACTGGGATAG
GAAAACACAGTGAGCAGAAGCTCAGGACGGAAGAGGAACGCACTAAAGAAACAAAGGCGC
TACGTGAAGTGAGCGGAAAACCTAGTGATATGACGGAGGGAGGTGTGATATGAAAAAACT
GAAGTCGGGGTGGAGAAGGGTCGTGAAGAGGACGAGTAATAATCTGCTCTTTAGCGACAG
AAGATNNNNNNNNNNNNNNNNNNNNNNNNNNNNNNNNNNNNNNNNNNNNNNNNNNNNNNN
NNNNNNNNNNNNNNNNNNNNNNNNNNNNNNGANGGGNNNGTGGGGAAAAACAAACATCCC
GGGGTCAAGTTGGGGCGGTAGTTGTGCTGAATTCGTGCCCGCGTTTAAGGCAGTAAGACT
CCAATGGGTAGAAGCCGTGGATAGCGAAGCTAAACTAAAGACGTTGGGGACATGAATGTC
AATAATGGGGGCAATGAAACGATGCTNNNTCTGNNNNAGTAGGNNNGNNNNNANNCNCNN
TGANNNNNACNCCATTANANNANANGCCATAGTCCTTCTTTTTCCTTTTATNTTTANGAN
NNCCNNNGNNTNNCNNTTGCNNNNNNNNNNNNNNNNNNCNCCGNCCAACTANACCTATGC
TTGGGTAGGTACTCTTAGAAATGAAACCAGGACCCTATTTATTAGGCCCGGACCTTACAG
TACGATTTACAGATCCCTTTCTGGCCGTTAGCTGTTTCTTCCACTAGACTGACAGCCCTC
TTGCTCCCCGTCATTGTTCCTTTCATTCTCCTAAAACAATCCAGTTACCTTGGCCTCCTG
TTTGTTCCAATTAC
>SRR9587947
TGAGCCTTTTATTCTAAGCTCAACATTTCTCCACCTCCTTGGCGCCATCCCGGAACCCAG
GCATAAAACGCAGCGAAATAAATCGAGCGCTATGCCTTTACCGTACCTGCCGCTAACCAT
TGTTACACGATTCAGTTGTNNNNNNNNNNNNNNNNNNNNNNTNNNNNNNNNNNNNNNNNN
NNNNNNNNNNNNNNNNNNNNNNNNNNNNNNNNNNNNNNNNNNNNNNACCTTTGCGTCACT
CATTCTGTATTTTGCCTTACGTCCTCCATTTTCCTCTATGTCCGTTTTTCTCTCAAGCTG
ATACATCAGCTNNNATCTCCCGCGAGATCTATTCGTGCATGAGTAGATATTGGTCCTTGA
TCGAGTACCTATAAAACGCTAGGCATCAAAGGACACTGCAATGCACAAGTCAAACAGGTC
CTCTGAAGCCAAAACATAAAGACAGACGGAAGCAACCAAAGGAACAGACCATGTATTAAC
ATAGGATGCGAAAATAGTCAAGCATAGGGTAATAATTTGTTCCACAGAGATTTGATAGCT
TACGAAAGATATCGCGAGCAACAGGCGAATACGTTGGGCGAACGAGGGAAGAGACCGAGT
CGAGTGAGACTGGAGCGGGGAGAGGGAGAGCGAATCCGAATAACAAGAGAACTGGGACAG

GAAAACACATTGAGCAGAACCTCAGGACGGAAGAGAGACGCACTAAGAAAACAGAGGCGC
GACGTGAAGCGAGCGGAAGACCTATTGATGTAACAGAGGGCGGCGTGATATGAAAAAACT
GAAGTCGGGGTAGAGAAAGGTCGTGAAGAGGACGAATAATAATCTGCTCTTTAGCGGCAG
AAGATACATCGGGATCAGTCCTTTTTCCGAGCGATCTTTGAGCATTATCCACTTGAGCTG
GACGGTCAGACGCAGGAACTCCTGGGGGGGGAAGGGTTGGTGAGGAAAAACAAACATCCC
GGGGTCAAGTTAGGGCGGTAGTTGTGCTGAATTCGTGTCCGCGTTTAAGGCAGTAATAGT
CCAATGGGTAGAAGCCATGGATAACGAAGCTGGGCTAAAGACGTTGGGGACATGGATGTC
AATAATGGAGGAAATGAAGCGATATNACGTCTGCGGCAGAAGGTAGACCGAGACTCTCAA
TGTACATAAAACCGTTAGAATAGACTCTATAGTTCTTCCTTCTCCTTTTATATCTAAGAC
GCTCTCAGTTTCCCCTCTGCTAAGACCTCTAGGCGATTCCTCGCCCAAATGGATCTGCGC
TTATGCGACCGCCCTCAGGAATAAAACCAGGACCCTATCTACTAGGCCTGAACTTCGTAG
TACGATTTACAGACTCCTTTCTTGCCTGTAGCTGCTTCTTCAGCTACGCTAAAAGCCCTC
TTGCCTCACGCTATTGTTTCCCTTATTCTCCCGATACAATCCAGTTACCTCAACCTCCTA
TCCGCTTCAACTAC
>SRR9587948
TAAGCCTTCTGTTCTAAGCTCAACATTTCTCCACCTCCTTNNNNNNNNNNNNNNNNNNNN
NNNNNNNNNNNNNNNNNNNNNNNNNNNNNNNNNNNNNNNNNNNNNNNNNNNNNNNNNNNN
NNNNACACGATTCGGTTGTAAAACCGCTTACGCTAGCATACTCAGGTGCCGGTTGCCCCC
CTTGCAACGCGTTTCCAGCACGCACATATTCCCAGATTCTATTATGACCTTCACGTCACT
CGCTCTGTACTTTGCCTTACGTCCTCCATTTTCCCTTATGTCCATTTTTCTCTCAAGCTG
ATACGTCAGTTCNGATCCATCACGGATTCTATTCGTGCATGAGTAGATGTCGGTCCTTGA
TTGAGCACCAAGAAAACGCTAGGCACCAAAGGACACTGCAGTGTACAAATCAAACAGGTC
CTCTGAAACCAAAACATAGAGACAGACGGAAGCAACCAAATGGACAGACCATGTATTAAC
ATAGGATGTGAAGATAGTCAAGCATAGGGTAATAGTTTGTTCTACAGAGATTTGATAGTT
TACGAAAGGTATCGCGAGCAACAGGCGGATACGTTGGGCGGACGGGGGAGGAGACCGGGC
CGAGTGAGACCGGAGCGGGGAGAGGGAGAGCGAATCCGAATAAAAAGAGAACTGGGACAG
GAAAACACAGTGAGCAGAACCTCAGGACGGAAGAGAAACGCACTAAGAAAGCAAAGGCGC
TATGTGAAGCGAGCGGGAGACCTAGTGATATAGCAGAGGGAGGCGTGATATGAAAAAACT
GAAGTCAGGGTAAAAAAAGGTCATGAAGAGGACGAATAATAGTCTGCTCTTTAGCGATAG
AAGANNNNNNNNNNNNNNNNNNNNNNNNNNNNNNNNNNNNNNNNNNNNNNNNNNNNNNNN
NNNNNNNNNNNNNNNNNNNNNNNNNNNNNNGANGGGTTAGTGGGGGGAAAAAGCTATCTT
GGGGCAGCATTAGGACAGGAATCGTGAAAGAATTAGACCCACATCGAAGNTAACCAGACT
TCAACAAGCAAAAATTATGAATAGCGAAGCTAGGCTAAAGACGTTGGGGACANGGTTGTC
AATAATGGAGGAAATGAAGCGATGCTACGNNNNNANCNGTAGGTAAGCCGAGATTCCCAA
GAAACATANNNNCNNNNAGATAGACTCCACAGTCCTTCTTTCTNNTTTTNTACTTAAGCC
GCCCTCAGTCTTCTCTTTGCNNNNATTTACAGGTGATTCCCCGANTCACTGGANNNNCGN

NNATATGGGTAATCCTAGGAATAAAATCAGGACCCTCCCTATTAGGCCTGAACCCCGTAA
TATGATTTACAGATTCCTTCCTGACCTGTAGCTGCTTCCCCAGCTACGCTAACAGCCCTC
TCGCCTCACGTCAGTGTTTCCCTCATTCTCCCGAAACAATCCAGTTACCTCAACCTCCTG
TCCGCTTCAACTAC
>SRR9587949
TAAGCCTTCTGTTCTAAGCTCAACATTTCTCCACCTCCTTNNNNCNNNNNNNNNNNNNNN
NNNNNNNNNNNNNNNNNNNNNNNNNNNNNNNNNNNNNNNNNNNNNNNNNNNNNNNNNNNN
NNNNACACGATTCAGTTGTAAAACCGCCTACGCTAGCATACTCAGGTGCCAGTTGCCCCC
CTTGCAACGCGTTTCCAGCACGCACATATTCCCATATTTACTATGACCTTCACGTCACT
CGCTCTGTACTTTGCCTTACGTCCTCCATTTTCCCTTATGTCCGTTTTTCTCTCAAGCTG
ATACGTCAGTTCNNATCCATCACGGATCCTATTCGTGCATGAGTAGATATCGGTCCTTGA
TTGAGCACCTAGAAAACGCTAGGCACTAAAGGACACTGCAGTGTACAAATCAAACAGGTT
CTCTGAAACCAAAACATAGAGACAGACGGAAGCAACCAAATGGACAGACCATGTATTAAC
ATAGGATGTGAAGATAGTCAAGCATAGGGTAGTAGTTTGTTCTACAGAGATTTGATAGTT
TACGAAAGGTATCGCGAGCAACAGGCGAATACGTTGGGCGGACGGGGGAGGAGACCGGGT
CGAGTGAGACCGGAGCGGGGAGAGGGAGAGCGAATCCGAATAAAAAGAGAACTGGGACAG
GAAAACACAGTGAGCAGAACCTCAGGACGGAAGAGAAACGCACTAAGAAAACAAAGGCGC
TATGTGAAGCGAACGGGAGACCTAGTGATATAACAGAGGGAGGCGTGATATGAAAAAACT
GAAGTCGGGGTAAAAAAAGGTCATGAAGAGGACGAATAGTAGTCTGCTCTTTAGCTACAG
AAGANNNNNNANNNNNNNNNNNNNNNNNNNNNNNNNNNTTNNNNNNNNNNNNNNNNNNNNN
NNCNNNNNNNNNNNNNNNNNNNNNNNNNNNGANGGGTTNGTGGGGGGAAAAAGCTATCTT
GGGGCAGCATCAGGACGGGAATCATGAAAGANTTAGACCCACATCTAAGATAACCAGACT
TCAACAAGCAAAAATTATGAATAGCGAAGCTAGGCTAAAGACGTTGGGGACATGGTTGTC
AATAATGGAGGAAATGAAGCGATGCTNCGTCNNNANCNGTAGGTAAGCCGAGATTCCCAA
GAAACATANNNNCNNNNANANAGACTNNATAGTCCTTCTTTCTCCTTTTANNNNTAAGAC
GCCCTCAGTCTTCTCTTTGCNNNNANTTACCGGTGATTCCCCGACTCACTGGANNNNCNN
NTATATGGGTANTCCTAGGAATAAAATCAGGACCCTCTCTATTAGGCCTGAACCCCGTAA
TATGATTTACAGATTCCTTCCTGACCTGTAGCTGCTTCCCTAGCTACGCTAACAGCCCTC
TCGCCTCACGTCAGTGTTTCCCTCATTCTCCCGAAACAATCCAGTTACCTCAACCTCCTG
TCCGCTTCAACTAC
>SRR9587950
TAAGCCTTCTGTTCTAAGNTCAACATTTCTCCACCTCCTNNNNNNNNNNNNNNNNNNNNN
NNNNNNNNNNNNNNNNNNNNNNNNNNNNNNNNNNNNNNNNNNNNNNNNNNNNNNNNNNNN
NNNNACACGATTCAGTTGTAANACCGCCTACGCTAGCATACTCAGGTGNCAGTTGCCCCC
CTTGCAACGCGTTTCCAGCACGCACATATTCCCATATTTTANTATGACCTTCACGTCACT
NGCTCTGTACTTTGCCTTACGTCCTCCATTTTCCCTTATGTNCGTTTTTCTCTNANGCTG

ATNCGTCAGTNNNNNNNNNNNNNNNNNNNNNNNNNTGCANGAGTAGATATCGGTCCTNNN
NNNAGCACCTAGNAAACGCTAGGCACTAAAGGACACTGCAGTGTACAAATCANACAGGTT
CTCTGAAACCAAAACATAGAGACANACGGAANCAACCAAATGGACAGACCATGTATTAAC
ATAGGATGTGAAGATAGTCAAGCATAGGNTANNAGTTTGTTCTACAGAGATTTNATAGTT
TACGAAAGGTATCGCGANCAACAGGCGAATACGTTGGGNGNACGGGGGAGGAGACCGNGN
CNNNTGAGACCNGAGNGGGGAGAGGGAGAGCGAATCNGAATAAAAAGAGAACTGGGACAG
GAAAACACNGTGAGCAGAACCTCAGGACGGAAGAGAAACGCACTAAGAAAACAAAGGCGC
TATGTGAAGNGAACGGGAGACCNAGTGATATNACAGAGGGAGGCGTGATATGAAAAAACT
GAAGTCGGGGTAAAAAAAGGTCATGAAGAGGACGAATAGTAGTCTGCTCTTTAGCTACAG
AANNNNNNNNNNNNNNNNNNNNNNNNNNNNNNNNNNNNNNNNNNNNNNNNNNNNNNNNNN
NNNNNNNNNNNNNNNNNNNNNNNNNNNNNNNGANGGGNNGTGGGGGGAAAAAGCTATCNT
GGGGCAGCATCAGGANGGGAATCNTGNNNNAATTAGACCCACATCTAAGATAACCAGACT
TCAACAAGCAAAAANNATGNATAGCGAAGCTAGGCTAAAGACNTTGGGGACATGGTTGTN
AATAATGGAGGAAATGAAGCGATGCTACGNNNNNANCCGTAGGTAAGCCGAGATTCCCAA
GAANNNTAGCGTCNNNNANNNNNNCTNCATAGTCCTTCTTTCTCCTTTNNNNNNNNAGAC
GCCNTCAGTCTTCTCNNTGCNNNNATTTACCGGTGATTCCCCGACTCACTGGANNNNNNN
TTATATGGGTAATCCTAGGAATAAAATCAGGACCCTCTCTATTAGGCCTGAACCCCGTAA
TATGATTTACAGATTCCTTCCTGACCTNTAGCTGCTTCCCTAGCTACGNTAACAGCCCNC
TCGCCTCACGTCAGTGTTTCCCTCATTCTCCCGAAANAATNCAGTTNCCTCNACCTCCNG
TCCGCTTCAACTAC
>SRR9587951
TACGCCTTCTGTTCTAAGCCCAACATTTCTCCACCTCCTTNNNNNNNNNNNNNNNNNNNN
NNNNNNNNNNNNNNNNNNNNNNNNNNNNNNNNNNNNNNNNNNNACCNNNNNNNNNNNNNT
NNNNACACGATTCAGTTGTAAAACCGCTTACACTAGCATACTCAGGTGCCGGTTGCCCCC
CTTGCATCGCGTTTCCAGCANGCACATATTCCCATATTTTATTATGACCTTCACGTCACT
CGCTCTGTACTTTGCCTTACGTCCTCCATTTTCCCTTATGTCCGTTTTTCTCTCAAGCTG
ATACGTCAGTTCNNATCNATCACGGATTCTATTCGTGCATGAGTAGATATCGGTCCTTGA
TTGAGCACCTAGAAAACGCTAGGCACTAAAGGACACTGCAGTGTACAAATCAAACAGGTC
CTCTGAAACTAAAACATAGAGACAGACGGAAGCAATCAAATGGACAGACCATGTATTAAC
ATAGGATGTGAAGATAGTCAAGCATAGGGTAATAGTTTGTTCTACAGAGATTTGATAGTT
TACGAAAGGTATCGCGAGCAACAGGCGAATGCGTTGGGCGGACGGGGGAGGAGACCGGGT
CGAGTGAGACCGGAGCGGGGAGAGGGAGAGCGAATCCGAATAAAAAGAGAACTGGGACAG
GAAAACACAGTGAGCAGAACCTCAGGAAGGAAGAGAAACGCACTAAGAAAACAAAGGCGC
TATGTGAAGCGAGCGGGAGACCTAGTGATATAACAGAGGGAGGCGTGATGTGAAAAAACT
GAAGTCGGGGTAAAAAAAGGTCATGAAGAGGACGAATAGTAGTCTGCTCTTTAGCGACAG
AAGATNNNNNNNNNNNNNNNNNNNNNNNNNNNNNNNNNNNNNNNNNNNNNNNNNNNNNNN

NNNNNNNNNNNNNNNNNNNNNNNNNNNNNNGAGGGGTTAGTGGGGGGAAAAAGCTATCNT
GGGGCAGCATTAGGACGGGAACCATGAAAGANTTAGACCCACATCTAAGATAACCAGACT
TCAACAAGCAAAAATTATGAATAGCGAAGCTAGGCTAAAGACGTTGGGGGCATGGTTGTC
AATAATGGAGGAAATGAAGCGATGCTACGNNNNNANCNGTAGGTAAGCCGAGATTCCCAA
GAAACATAGCGNCNNNNANANAGACTNNATAGTCCTTCTTTCTCCTTTTATATTTAAGAC
GCCCTCAGTCTTCTCTTTGCNNNNATTTACAGGTGATTCCCCGACTCACTGGANNNNCGN
NTATATGGGTAATCCTAGGAATAAAATCAGGACCCTCTCTATTAGGCCTGAACCCCGTAA
TATGATTTACAGATTCCTTCCTGACCTGTAGCTGCTTCCCCAGCTACGCTAGCAGCCCTC
TCGCCTCACGTCAGTGTTTCCCTCATTCTCCCGAAACAATCCAGTTACCTCAACCTCCTG
TCCGCTTCAACTAC
>SRR9587952
TAAGCCTTCTGTTCTAAGCTCAACATTTCTCCACCTCCTTNNNNNNNNNNNNNNNNNNNN
NNNNNNNNNNNNNNNNNNNNNNNNNNNNNNNNNNNNNNNNNNNNNNNNNNNNNNNNNNNN
NNNNACACGATTCAGTTGTAAAACCGCCTACGCTAGCATACTCAGGTGCCAGTTGCCCCC
CTTGCAACGCGTTTCCAGCACGCACATATTCCCATATTTTACTATGACCTTCACGTCACT
CGCTCTGTACTTTGCCTTACGTCCTCCATTTTCCCTTATGTCCGTTTTTCTCTCAAGCTG
ATACGTCAGTTCNGATCCATCACGGATCCTATTCGTGCATGAGTAGATATCGGTCCTTGA
TTGAGCACCTAGAAAACGCTAGGCACTAAAGGACACTGCAGTGTACAAATCAAACAGGTT
CTCTGAAACCAAAACATAGAGACAGACGGAAGCAACCAAATGGACAGACCATGTATTAAC
ATAGGATGTGAAGATAGTCAAGCATAGGGTAGTAGTTTGTTCTACAGAGATTTGATAGTT
TACGAAAGGTATCGCGAGCAACAGGCGAATACGTTGGGCGGACGGGGGAGGAGACCGGGT
CGAGTGAGACCGGAGCGGGGAGAGGGAGAGCGAATCCGAATAAAAAGAGAACTGGGACAG
GAAAACACAGTGAGCAGAACCTCAGGACGGAAGAGAAACGCACTAAGAAAACAAAGGCGC
TATGTGAAGCGAACGGGAGACCTAGTGATATAACAGAGGGAGGCGTGATATGAAAAAACT
GAAGTCGGGGTAAAAAAAGGTCATGAAGAGGACGAATAGTAGTCTGCTCTTTAGCTACAG
AAGANNNNNNNNNNNNNNTNNNNNNNNNNNNNNNNNNNNNANNNNNNNNNNNNNNNNNNNN
NNNNNNNNNNNNNNNNNNNNNNNNNNNNNNGAGGGGTTAGTGNGGGGAAAAAGCTATCTT
GGGGCAGCATCAGGACGGGAATCATGAAAGANTTAGACCCACATCTAAGATAACCAGACT
TCAACAAGCAAAAATTATGAATAGCGAAGCTAGGCTAAAGACGTTGGGGACATGGTTGTC
AATAATGGAGGAAATGAAGCGATGCTACGTCTNNANCNGTAGGTAAGCCGAGATTCCCAA
GAAACATANCGTCNNNNANANAGACTCCATAGTCCTTCTTTCTCCTTTTANNNNTAAGAC
GCCCTCAGTCTTCTCTTTGCNNNNANTTACCGGTGATTCCCCGACTCACTGGANNNNCNN
NTATATGGGTAATCCTAGGAATAAAATCAGGACCCTCTCTATTAGGCCTGAACCCCGTAA
TATGATTTACAGATTCCTTCCTGACCTGTAGCTGCTTCCCTAGCTACGCTAACAGCCCTC
TCGCCTCACGTCAGTGTTTCCCTCATTCTCCCGAAACAATCCAGTTACCTCAACCTCCTG
TCCGCTTCAACTAC

>SRR9587953
TGAGCCTTTTATTCTAAGCTCAACATTTCTCCACCTCCTCGGCGCCATCCCGGAACCCAG
GCATAAAACGCAGCGAAATAAATCGAGCGCTATGCCTTTACCGTACCTGCCGCTAACCAT
TGTTACACGATTCAGTTGTNNNNNNNNNNNNNNNNNNNNNNNNNNNNNNNNNNNNNNNNN
NNNNNNNNNNNNNNNNNNNNNNNNNNNNNNNNNNNNNNNNNNNNNNNNACCTTNACGTCACT
CATTCTGTATTTTGCCTTACGTCCTCCATTTTCCTCTATGTCCGTTTTTCTCTCAAGCTG
ATACATCAGCTNNNNNNTCCCGCGAANTCTATTCGTGCATGAGTAGATATTGGTCCTTGA
TTGAGTACCTATAAAACGCTAGGCATCAAAGGACACTGCAATGCACAAGTCAAACAGGTC
CTCTGAAGCCAAAACATAAAGACAGACGGAAGCAACCAAAGGAACAGACCATGTATTAAC
ATAGGATGCGAAAATAGTCAAGCATAGGGTAATAATTTGTTCCACAGAGATTTGATAGCT
TACGAAAGATATCGCGAGCAACAGGCGAATACGTTGGGCGAACGAGGGAAGAGANCGAGT
CGAGTGAGACTGGAGCGGGGAGAGGGAGAGCGAATCCGAATAACAAGAGAACTGGGACAG
GAAAACACAGTGAGCAGAACCTCAGGACGGAAGAGAGACGCACTAAGAAAACAGAGGCGC
GACGTGAAGCAAGCGGAAGACCTATTGATGTAACAGAGGGCGGCGTGATATGAAAAAACT
GAAGTCGGGGTAGAGAAAGGTCGTGAAGAGGACGAATAATAATCTGCTCTTTAGCGGCAG
AAGATACATCGAGATCAGTCCTTTTTCCGAGCGATCTTTGAGCATTATCCACTTGAGCTG
GACGGTCAGACGCAGGAACTTCTGGGGGGGGAAGGGNTGGTGAGGAAAAACAAACATCCC
GGGGTCAAGTTAGGGCGGTAGTTGTGCTGAATTCGTGTCCGCGTTTAAGGCAGTAATACT
CCAATGGGTAGAAGCCATGGATAACGAAGCTGGGCTAAAGACGTTGGGGACATGGATGTC
AATAATGGAGGAAATGAAGCGATATCACGTCTGCAGCAGAGGATAGACCGAGACTTTCAA
TGTACATAAAACTGTTAGAATGGACTCTATAGTTCTTCCTTCTCCTTTTATATCTAAGAC
GCTCTCAGTTTCCCCTTTACTAAGACCTCTAGGCGACTCCTCGCCCAACCGGATCTGCGC
TTATGCGGCCGCCCTCAGGAATAAAACCAGGGCCCTATCTATTAGGCCTGAACCTCGTAG
TANGATTTACAGACTCCTTTTCGGCCTGTAGCTGCTTCTTCAGCTACGCTAAAAGCCCTC
TTGCCTCACGCTATTGTATCCCTTATTCTCCCGATACAATCCAGTTACCTCAACCTCCTA
TCCGCTTCAACTAC
>SRR9587954
TAAGCCTTCTGTTCTAAGCTCAACATTTCTCCACCTCCTTNNNNNNNNNNNNNNNNNNNN
NNNNNNNNNNNNNNNNNNNNNNNNNNNNNNNNNNNNNNNNNNNNNNNNNNNNNNNNNNNN
NNNNACACGATTCAGTTGTAAAACCGCCTACGCTAGCATACTCAGGTGCCAGTTGCCCCC
CTTGCAACGCGTTTCCAGCACGCACATATTCCCATATTTTACTATGACCTTCACGTCACT
CGCTCTGTACTTTGCCTTACGTCCTCCATTTTCCCTTATGTCCGTTTTTCTCTCAAGCTG
ATACGTCAGTTCNGATCCATCACGGATCCTATTCGTGCATGAGTAGATATCGGTCCTTGA
TTGAGCACCTAGAAAACGCTAGGCACTAAAGGACACTGCAGTGTACAAATCAAACAGGTT
CTCTGAAACCAAAACATAGAGACAGACGGAAGCAACCAAATGGACAGACCATGTATTAAC
ATAGGATGTGAAGATAGTCAAGCATAGGGTAGTAGTTTGTTCTACAGAGATTTGATAGTT

TACGAAAGGTATCGCGAGCAACAGGCGAATACGTTGGGCGGACGGGGGAGGAGACCGGGT
CGAGTGAGACCGGAGCGGGGAGAGGGAGAGCGAATCCGAATAAAAAGAGAACTGGGACAG
GAAAACACAGTGAGCAGAACCTCAGGACGGAAGAGAAACGCACTAAGAAAACAAAGGCGC
TATGTGAAGCGAACGGGAGACCTAGTGATATAACAGAGGGAGGCGTGATATGAAAAAACT
GAAGTCGGGGTAAAAAAAGGTCATGAAGAGGACGAATAGTAGTCTGCTCTTTAGCTACAG
AAGANNNNNNNNNNNNNNNNNNNNNNNNNNNNNNNNNNNNNNNNNNNNNNNNNNNNNNNN
NNNNNNNNNNNNNNNNNNNNNNNNNNNNNNGAGGGGTTAGTGGGGGGAAAAAGCTATCTT
GGGGCAGCATCAGGACGGGAATCNTGAAAGANTTAGACCCACATCTAAGATAACCAGACT
TCAACAAGCAAAAATTATGAATAGCGAAGCTAGGCTAAAGACGTTGGGGACATGGTTGTC
AATAATGGAGGAAATGAAGCGATGCTACGNNNNNANCCGTAGGTAAGCCGAGATTCCCAA
GAAACATANNNTCNNNNANANAGACTCCATAGTCCTTCTTTCTCCTTTTANNNNTAAGAC
GCCCTCAGTCTTCTCTTTGCCNNNATTTACCGGTGATTCCCCGACTCACTGGANNTNCNN
NTATATGGGTAATCCTAGGAATAAAATCAGGACCCTCTCTATTAGGCCTGAACCCCGTAA
TATGATTTACAGATTCCTTCCTGACCTGTAGCTGCTTCCCTAGCTACGCTAACAGCCCTC
TCGCCTCACGTCAGTGTTTCCCTCATTCTCCCGAAACAATCCAGTTACCTCAACCTCCTG
TCCGCTTCAACTAC
>SRR9587955
TAAGCCTTCTGTTCTAAGCCCAACATTTCTCCACCTCCTTNNNNNNNNNNNNNNNNNNNN
NNNNNNNNNNNNNNNNNNNNNNNNNNNNNNNNNNNNNNNNNNNNNNNNNNNNNNNNNNNN
NNNNACACGATTCAGTTGTAAAACCGCTTACGCTAGCATACTCAGGTGCCGGTTGCCCCC
CTTGCATCGCGTTTCCAACACGCACATATTCCCATATTTTATTATGACCTTCACGTCACT
CGCTCTGTACTTTGCCTTATGTCCTCCATTTTCCCTTATGTCCGTTTTTCTCTCAAGCTG
ATACGTCAGTTCNNATCCATCACGGATCCTATTCGTGCATGAGTAGATATCGGTCCTTGA
TTGAGCACCTAGAAAACGCTAGGCACTAAAGGACACTGCAGTGTACAAATCAAACAGGTC
CTCTGAAACTAAAACATAGAGACAGACGGAAGCAATCAAATGGACAGACCATGTATTAAC
ATAGGATGTGAAGATAGTCAAGCATAGGGTAATAGTTTGTTCTACAGAGATTTGATAGTT
TACGAAAGGTATCGCGAGCAACAGGCGAATACGTTGGGCGGACGGGGGAGGAGACCGGGT
CGAGTGAGACCGGAGCGGGGAGAGGGAGAGCGAATCCGAATAAAAAGAGAACTGGGACAG
GAAAACACAGTGAGCAGAACCTCAGGACGGAAGAGAAACGCACTAAGAAAACAAAGGCGC
TATGTGAAGCGAGCGGGAGACCTAGTGATATAACAGAGGGAGGCGTGATATGAAAAAACT
GAAGTCGGGGTAAAAAAAGGTCATGAAGAGGACGAATAGTAGTCTGCTCTTTAGCGACAG
AAGANNNNNNNNNNNNNNNNNNNNNNNNNNNNNNNNNNNNNNNNNNNNNNNNNNNNNNNN
NNNNNNNNNNNNNNNNNNNNNNNNNNNNNNGANGGGTTAGTGGGGGGAAAAAGCTATCTT
GGGGCAGCATTAGGACGGGAATCATGAAAGANTTAGACCCACATCTAAGATAACCAGACT
TCAACAAGCAAAAATTATGAATAGCGAAGCTAGGCTAAAGACGTTGGGGGCATGGTTGTC
AATAATGGAGGAAATGAAGCGATGCTACGNNNNNANCNGTAGGTAAGCCGAGNTTCCCAA

GAAACATAGCGTCNNCNAGANAGACTNCATAGTCCTTCTTTCTCCTTTTATATTTAAGAC
GCCCTCAGTCTTCTCTTTGCNGCAATTTACAGGTGATTCCCCGACTCACTGGANNNNCGN
NTATATGGGTAATCCTAGGAATAAAATCAGGACCCTCTCTATTAGGCCTGAACCCCGTAA
TATGATTTACAGATTCCTTCCTGACCTGTAGCTGCTTCCCCAGCTACGCTAACAGCCCTC
TCGCCTCACGTCAGTGTTTCCCTCATTCTCCCGAAACAATCCAGTTACCTCAACCTCCTG
TCCGCTTCAACTAC
>SRR9587956
TAAGCCTTCTGTTCTAAGCTCAACATTTCTCCACCTCCTTNNNNNNNNNNNNNNNNNNNN
NNNNNNNNNNNNNNNNNNNNNNNNNNNNNNNNNNNNNNNNNNNNNNNNNNNNNNNNNNNN
NNNNACACGATTCAGTTGTAAAACCGCCTACGCTAGCATACTCAGGTGCCGGTTGCCCCC
CTTGCAACGCGTTTCCAGCACGCACATATTCCCATATTTTACTATGACCTTCACGTCACT
CGCTCTGTACTTTGCCTTACGTCCTCCATTTTCCCTTATGTCCGTTTTTCTCTCAAGCTG
ATACGTCAGTTCCGNTCCATCACGGATCCTATTCGTGCATGAGTAGATATCGGTCCTTGA
TTGAGCACCTAGAAAACGCTAGGCACTAAAGGACACTGCAGTGTACAAATCAAACAGGTT
CTCTGAAACCAAAACATAGAGACAGACGGAAGCAACCAAATGGACAGACCATGTATTAAC
ATAGGATGTGAAGATAGTCAAGCATAGGGTAATAGTTTGTTCTACAGAGATTTGATAGTT
TACGAAAGGTATCGCGAGCAACAGGCGAATACGTTGGGCGGACGGGGGAGGAGACCGGGT
CGAGTGAGACCGGAGCGGGGAGAGGGAGAGCGAATCCGAATAAAAAGAGAACTGGGACAG
GAAAACACAGTGAGCAGAACCTCAGGACGGAAGAGAAACGCACTAAGAAAACAAAGGCGC
TATGTGAAGCGAACGGGAGACCTAGTGATATAACAGAGGGAGGCGTGATATGAAAAAACT
GAAGTCGGGGTAAAAAAAGGTCATGAAGAGGACGAATAGTAGTCTGCTCTTTAGCTACAG
AAGATNNNNNNNNNNNNNNNNNNNNNNNNNNNNNNNNNNNNNNNNNNNNNNNNNNNNNNN
NNNNNNNNNNNNNNNNNNNNNNNNNNNNNGGAGGGGTTAGTGGGGGGAAAAAGCTATCTT
GGGGCAGCATTAGGACGGGAATCATGAAAGANTTAGACCCACATCTAAGATAACCAGACT
TCAACAAGCAAAAATTATGAATAGCAAAGCTAGGCTAAAGACGTTGGGGACATGGTTGTC
AATAATGGAGGAAATGAAGCGATGCTACGNNNNNANCNGTAGGTAAGCCGAGANTCCCAA
GAAACATAGCGTCNNNNANATAGACTCCATAGTCCTTCTTTCTCCTTTTANNNNTAAGAC
GCCCTCAGTCTTCTCTTTGCNNNNATTTACCGGTGATTCCCCGACTCACTGGATNNNCNN
NTATNTGGGTAATCCTAGGAATAAAATCAGGACCCTCTCTATTAGGCCTGAACCCCGTAA
TATGATTTACAGATTCCTTCCTGACCTGTAGCTGCTTCCCCAGCTACGCTAACAGCCCTC
TCGCCTCACGTCAGTGTTTCCCTCATTCTCCCGAAACAATCCAGTTACCTCAACCTCCTG
TCCGCTTCAACTAC
>SRR9587957
TAAGCCTTCTGTTCTAAGCTCAACATTTCTCCACCTCCTTNNNNNNNNNNNNNNNNNNNN
NNNNNNNNNNNNNNNNNNNNNNNNNNNNNNNNNNNNNNNNNNNNNNNNNNNNNNNNNNNN
NNNNACACGATTCAGTTGTNNNNNNNNNNNNNNNNNNNNNNNNNNNNNNNNNNNNNNNNN

NNNNNNNNNNNNNNNNNNNNNNNNNNNNNNNNNNNNNNNNNNNNTNACCTTCACGTCACT
CGCTCTGTACTTTGCCTTACGTCCTCCATTTTCCCTTGTGTCCGTTTTTCTCTCAAGCTG
ATACGTCAGTTCNGATCCATCGCGGATNCTATTCGTGCATGAGTAGATATCGGTCCTTGA
TTGAGCACCTAGAAAACGCTAGGCACCAAAGGACACTGCAGTGTACAAGTCAAACAGGTC
CTCTGAAACCAAAACATAAAGACAGACGGAAGCAACCAAATGGACAGACCATGTATTAAC
ATAGGATGTGAAGATAGTCAAGCATAGGGTAATAGTTTGTTCTACAGAGATTTGATAGTT
TACGAAAGGTATCGCTAGCAACAGGCGAATACGTTGGGCGAACGGGGGAGGAGACCGAGT
CGAGTGAGACCGGAGCGGGGAGAGGGAGAGCGAATCCGGATAAAAAGAGAACTGGGACAG
GAAAACATAGTGAGCAGAACCTCAGGACGGAAGAGAAACGCACTAAGAAAACAAAGGCGC
TATGTGAAGCGAGCGGGAGACCTAGTGATATAACAGAGGGAGGCGTGATATGAAAAAACT
GAAGTCGGGGTAAAAAAAGGTCATGAAGAGGACGAATAGTAGTCTGCTCTTTAGCGACAG
AAGANNNNNNNNNNNNNNNNNNNNNNNNNNNNNNNNNNNNNNNNNNNNNNNNNNNNNNNN
NNNNNNNNNNNNNNNNNNNNNNNNNNNNNNGANGGGTTAGTGGGGGGAAAAAGCTATCTT
GGGGCAGCATTAGGACGGGAATCATGAAAGANTTAGACCCACATCTAAGATAACCAGACT
TCAACAAGCAAAAATTATGAATAGCGAAGCTAGGCTAAAGACGTTGGGGACATGGATGTC
AATAATGGAGGAAATGAAGCGATGCTACGNNNNNANCNGTAGGTAAGCCGAGATTCCCAA
GAACCATAGCGTCCTTGAGATAGACTCCGTAGTCCTTCTTTCTCCCTTTATNTTCAAGAC
GCCCTCAGTCCTCCCTTNGNNNNNATTTACAGGTGATTCCCCGACCCACTGGNNNNNCGT
NTATATGGCTAATCCTAGGAATAAAATCAGGACCCTCTCTATTAGGCCTGAACCCCGTAA
TACGATTTACAGATTCCTTCCTGACCTGTAGCTGCTTCCCCAGCTACGCTAACAGTCCTC
TCGCCTCACGTCAGTGTTTCCCTCATTCTCCCGAAACAATCCAGTTACCTCAACCTCCTG
TCCGTTTCAACTAC
>SRR9587958
TAAGCCTTTTATTCTAAGCTTAACATTTCTCCACCACCTTGGCGCCAGTTTAAAACNCAG
GCACAGAGAGCAATGGAACCAACAGTNAANTNCACTTTCNCCATNNNNNCNNNNANACGC
TGTTACATGACTCAGTTGTAAAGTCATTGACGCGAGCATGCTCAAATACCGGCGGNNNNN
NNNACGNCGTACNCANAACATGTAGGTACACTCAGACTTCATTCCAATCTTCACGTCACC
CGTTCTGTGTTTTGCCTTACGTCCTCCATTTTCTTTTATGTCCGTTCTTTTCTCAAACTG
ATAAATCAGTTCCGGTNTCTCGNNNATTCNGTTCGTGCATGAGTAGATACTGGTCCTCGA
TTGAGTACCTATAAAACACTAGGCATCGAAGGACACTGCAATGCGTAGGTCAAACAGGTC
CTCTGGAACCAAAACATAAAGACAGACGGAAGCAACCAAATGGACAGACCATGTGTTAAC
ATAGGGTGCGAAGATAGTCAAGCATAGGGTAATAATTTGTTCCACAGAGATTCTATAGCT
TACGAAGGATATCGCGAGCAACAGGCGAATACGTTGGGCGAGCGAGGAAAAAAACCGGAT
CGAGTGAGACCGGAGCGGGGGGAGGGAGAGCGAACCCGAATAAAAGGAGAACTGGGATAG
GAAAACACAGTGAGCAGAAGCTCAGGACGGAAGAGGAACGCACTAAAAAAACAAAGGCGC
TACGTGAAGCGAGCGAAAAACCTAGTGATATGACGGAGAGAGGTGTGATATGAAAAAACT

GAAGTCGGGGTGGAGAAGGGTCGTGAAGAGGACGAGTAATAATCTGCTCCTTAACGACAG
AAGANNNNNNNNNNNNNNNNNNNNNNNNNNNNNNNNNNNNNNNNNNNNNNNNNNNNNNNN
NNNNNNNNNNNNNNNNNNNNNNNNNNNNNNGAGGGGNNNGTGGGGAAAAACAAACATCCC
GGGGTCAAGTTAGGGCGGTAGTTGTGCTGAATTCGTGCCCGCGTTTAAGGCAGTGAGACT
CCAATGGGTAGAAGCCGTGGATAGCGAAGCTAGACTAAAGACGTTGGGGACATGAATGTC
AATAATGGGGGAAATGAAACGATGCTNNNTCTGNNNNAGTAGGNNNGNNNNAANNCNNNN
TGANNNNNACNCCATTANANNANANGCCATAGTCTTTCTTTTTCCTTTTATATTTANGAN
NNNCNNNGNCTTNCNNTTGCNNNNNNNNNNNNNNNNNNCNCTGNCCAACTNNACCTATGC
TTGGGTNGGTACTCTTAGAAATGAAACCAGGACCCTATTTATTAGGCCCGGACCTTACTG
TACGATTTACAGATCCCTCTCTGGCCGTCAGCTGTTTTTTCAGCTAGACTGACAGCCCTC
TTGCTCCCCGTCATTGTTCTTTTCATTCTCCTAAAACAATCCAGTTACCTTGGCCTTCTG
TTTGTTCCAATTAC
>SRR9587959
TAAGCCTTTTATTCTAAGCTTAACATTTCTCCACCATCTTGGCGCCAGTTTAAAACNCAG
GCACAGAGAGCAATGGAACCAACAGTNAANTGCACTTTCGCCATNNNNNCNNNNANNCGC
TGTTACATGACTCAGTTGTAAAGTCATTGACGCGAGCATACTCAAATACCGGCTGNNNNN
NNNACGNCGTNCNCATAACATGTAGGTACACTCGGACTTCATTCCAATCTTCACGTCACC
CGTTCTGTGTTTTGCCTTACGTCCTCCATTTTCTTTTATGTCCGTTCTTTTCTCAAACTG
ATAAATCAGTTCCGGTNTCNCGNNNATTCNGTTCGTGCATGAGTAGATACTGGTCCTCGA
TTGAGTACCTATAAAACACTAGGCATCGAAGGACACTGCAATGCGTAGGTCAAACAGGTC
CTCTGGAACCAAAACATAAAGACAGACGGAAGCAACCAAATGGACAGACCATGTGTTAAC
ATAGGGTGCGAAGATAGTTAAGCATAGGGTAATAATTTGTTCCACAGAGATTCTATAGCT
TACGAAGGATATCGCGAGCAACAGGCGAATACGTTGGGCGAGCGAGGAAAAAAACCGGAT
CGAGTGAGACCGGAGCGGGGGGAGGGAGAGCGAACCCGAATAAAAGGAGAACTGGGATAG
GAAAACACAGTGAGCAGAAGCTCAGGACGGAAGAGGAACGCACTAAAAAAACAAAGGCGC
TACGTGAAGCGAGCGGAAAACCTAGTGATATGACGGAGGGAGGTGTGATATGAAAAAACT
GAAGTCGGGGTGGAGAAGGGTCGTGAAGAGGACGAGTAATAATCTGCTCTTTAGCGACAG
AAGANNNNNNNNNNNNNNNNNNNNNNNNNNNNNNNNNNNNNNNNNNNNNNNNNNNNNNNN
NNNNNNNNNNNNNNNNNNNNNNNNNNNNNNGAGGGGNNAGTGGGGAAAAACAAACATCCC
GGGGTCAAGTTGGGGCGGTAGTTGTGCTGAATTCGTGCCCGCGTTTAAGGCAGTAAGACT
CCAATGGGTAGAAGCCGTGGATAGCGAAGCTAGACTAAAGACGTTGGGGACATGAATGTC
AATAATGGGGGCAATGAAACGATGCTNNNTCTGNNNNAGTAGGNNNGNNNNNANNCNCNN
TGANNNNNACNCCATTANAGNATANGCCATAGTCCTTCTTTTTCCTTTTATATTTANGAN
NNNCNNNGNNTNNCNCTTGCNNNNNNNNNNNNNNNNNNCNNCGNCCAACTNNACCTATGC
TTGGGTAGGTACTCTTAGAAATGAAACCAGGACCCTATTTATTAGGCCCGGACCTTACAG
TACGATTTACAGATCCCTTTCTGGCCGTTAGCTGTTTCTTCAACTAGACTGACAGCCCTC

TTGCTCCCCGTCATTGTTCCTTTCATTCTCCTNNAACAATCCAGTTACCTTGGCCTCCTG
TTTGTTCCAATTAC
>SRR9587960
TAAGCCTTCTGTTCTAAGCTCAACATTTCTCCACCTCCTTNNNNNNNNNNNNNNNNNNNN
NNNNNNNNNNNNNNNNNNNNNNNNNNNNNNNNNNNNNNNNNNNNACCTGNCGCTNNNNNN
NNNNACACGATTCAGTTGTAAAACCGCCTACGCTAGCATACTCAGGTGCCAGTTGCCCCC
CTTGCAACGCGTTTCCAGCACGCACATATTCCCATATTTTACTATGACCTTCACGTCACT
CGCTCTGTACTTTGCCTTACGTCCTCCATTTTCCCTTATGTCCGTTTTTCTCTCAAGCTG
ATACGTCAGTTCCGATCCATCACGGATCCTATTCGTGCATGAGTAGATATCGGTCCTTGA
TTGAGCACCTAGAAAACGCTAGGCACTAAAGGACACTGCAGTGTACAAATCAAACAGGTT
CTCTGAAACCAAAACATAGAGACAGACGGAAGCAACCAAATGGACAGACCATGTATTAAC
ATAGGATGTGAAGATAGTCAAGCATAGGGTAGTAGTTTGTTCTACAGAGATTTGATAGTT
TACGAAAGGTATCGCGAGCAACAGGCGAATACGTTGGGCGGACGGGGGAGGAGACCGGGT
CGAGTGAGACCGGAGCGGGGAGAGGGAGAGCGAATCCGAATAAAAAGAGAACTGGGACAG
GAAAACACAGTGAGCAGAACCTCAGGACGGAAGAGAAACGCACTAAGAAAACAAAGGCGC
TATGTGAAGCGAACGGGAGACCTAGTGATATAACAGAGGGAGGCGTGATATGAAAAAACT
GAAGTCGGGGTAAAAAAAGGTCATGAAGAGGACGAATAGTAGTCTGCTCTTTAGCTACAG
AAGANNNNNNNNNNNNNNNNNNNNNNNNNNNNNNNNNNNNNNNNNNNNNNNNNNNNNNNN
NNNNNNNNNNNNNNNNNNNNNNNNNNNNNNGAGGGGTTNGTGGGGGGAAAAAGCTATCTT
GGGGCANCATNAGGACGGGAATCATGAAAGAATTAGACCCACATCTAAGANAANCAGACT
TCAACAAGCAAAAATTATGAATAGCGAAGCTAGGCTAAAGACGTTGGGGACATGGTTGTC
AATAATGGAGGAAATGAAGCGATGCTACGNNNNNANCNGTAGGTAAGCCGAGATTCCCAA
GAAACATANNNNCNNNNANANAGACTNCATAGTCCTTCTTTCTCCTTTTANNNNTANGAC
GCCCTCAGTCTTCTCTTTGCNNNNATTTACCGGTGATTCCCCGACTNNCTGGATCTGCNN
NTATATGGGTAATCCTAGGAATAAAATCAGGACCCTCTCTATTAGGCCTGAACCCCGTAA
TATGATTTACAGATTCCTTCCTGACCTGTAGCTGCTTCCCTAGCTACGCTAACAGCCCTC
TCGCCTCACGTCAGTGTTTCCCTCATTCTCCCGAAACAATCCAGTTACCTCAACCTCCTG
TCCGCTTCAACTAC
>SRR9587961
TAAGCCTTCTGTTCTAAGCTCAACATTTCTCCACCTCCTTNNNNNNNNNNNNNNNNNNNN
NNNNNNNNNNNNNNNNNNNNNNNNNNNNNNNNNNNNNNNNNNNNNNNNNNNNNNNNNNNN
NNNNACACGATTCAGTTGTAAAACCGCCTACGCTAGCATACTCAGGTGCCAGTTGCCCCC
CTTGCAACGCGTTTCCAGCACGCACATATTCCCATATTTTACTATGACCTTCACGTCACT
CGCTCTGTACTTTGCCTTACGTCCTCCATTTTCCCTTATGTCCGTTTTTCTCTCAAGCTG
ATACGTCAGTTCNGATCCATCACGGATCCTATTCGTGCATGAGTAGATATCGGTCCTTGA
TTGAGCACCTAGAAAACGCTAGGCACTAAAGGACACTGCAGTGTACAAATCAAACAGGTT

CTCTGAAACCAAAACATAGAGACAGACGGAAGCAACCAAATGGACAGACCATGTATTAAC
ATAGGATGTGAAGATAGTCAAGCATAGGGTAGTAGTTTGTTCTACAGAGATTTGATAGTT
TACGAAAGGTATCGCGAGCAACAGGCGAATACGTTGGGCGGACGGGGGAGGAGACCGGGT
CGAGTGAGACCGGAGCGGGGAGAGGGAGAGCGAATCCGAATAAAAAGAGAACTGGGACAG
GAAAACACAGTGAGCAGAACCTCAGGACGGAAGAGAAACGCACTAAGAAAACAAAGGCGC
TATGTGAAGCGAACGGGAGACCTAGTGATATAACAGAGGGAGGCGTGATATGAAAAAACT
GAAGTCGGGGTAAAAAAAGGTCATGAAGAGGACGAATAGTAGTCTGCTCTTTAGCTACAG
AAGANNNNNNNNNNNNNNNNNNNNNNNNNNNNNNNNNNNNNNNNNNNNNNNNNNNNNNNN
NNNNNNNNNNNNNNNNNNNNNNNNNNNNNNGANGGGTTNGTGGGGGGAAAAAGCTATCTT
GGGGCNNNATCAGGACGGGAATCATGAAAGANTNAGACCCACATCTAAGATANCCAGACT
TCAACAAGCAAAAATTATGAATAGCGAAGCTAGGCTAAAGACGTTGGGGACATGGTTGTC
AATAATGGAGGAAATGAAGCGATGCTACGNNNNNANCNGTAGGTAAGCCGAGATTCCCAA
GAAACATANCGTCNNNNANANAGACTCCATAGTCCTTCTTTCTCCTTTTANNNNTAAGAC
GCCCTCAGTCTTCTCTTTGCCNNNATTTACCGGTGATTCCCCGACTCACTGGATCTNCNN
TTATATGGGTAATCCTAGGAATAAAATCAGGACCCTCTCTATTAGGCCTGAACCCCGTAA
TATGATTTACAGATTCCTTCCTGACCTGTAGCTGCTTCCCTAGCTACGCTAACAGCCCTC
TCGCCTCACGTCAGTGTTTCCCTCATTCTCCCGAAACAATCCAGTTACCTCAACCTCCTG
TCCGCTTCAACTAC
>SRR9587962
TAAGCCTTTTATTCTAAGCTTAACATTTCTCCACCATCTTGGCGCCGGTTTAAAACNCAG
GCACAGAGAGCAATGGAACCAACAGTNAATTNCACNTTCGCCNTNNNNNCNNNNANACGC
TGTTACATGACTCAGTTGTAAAGTCATTNACGCGAGCATACTCAAATACCGGCTGNNNTN
NNNACGNCGNNNNCATAACATGTAGGTACACTCGGACTTCATTCCAATCTTCACGTCACC
CGTTCTGTGTTTTGCCTTACGTCCTCCATTTTCTTTTATGTCCGTTCTTTTCTCAAACTG
ATAAATCAGTTCNGNTTTCTCGNNNATTCNGTTCGTGCATGGGCAGATACTGGTCCTCGA
TTGAGTACCTATAAAACACTAGGCATCGAAGGACACTGCAATGCGTAGGTCAAACAGGTC
CTCTGGAACCAAAACATAAAGACAGACGGAAGCAACCAAATGGACAGACCATGTGTCAAC
ATAGGGTGCGAAGATAGTCAAGCATAGGGTAATAATTTGTTCCACAGAGATTCTATAGCT
TACGAAGGATATCGCGAGCAACAGGCGAATACGTTGGGCGAGCGAGGAAAAAAACCGGAT
CGAGTGAGACCGGAGCGGGGGGAGGGGGAGCGAACCCGAATAAAAGGAGAACTGGGATAG
GAAAACACAGTGAGCAGAAGCTCAGGACGGAAGAGGAACGCACTAAAGAAACAAAGGCGC
TACGTGAAGTGAGCGGAAAACCTAGTGATATGACGGAGGGAGGTGTGATATGAAAAAACT
GAAGTCGGGGTGGAGAAGGGTCGTGAAGAGGACGAGTAATAATCTGCTCTTTAGCGACAG
AAGANNNNNNNNNNNNNNNNNNNNNNNNNNNNNNNNNNNNNNNNNNNNNNNNNNNNNNNN
NNNNNCNNNNNNNNNNNNNNNNNNNNNNNNGANGGGNNNGTGGGGAAAAACAAACATCCC
GGGGTCAAGTTGGGGCGGTAGTTGTGCTGAATTCGTGCCCGCGTTTAAGGCAGTAAGACT

CCAATGGGTAGAAGCCGTGGATAGCGAAGCTAAACTAAAGACGTTGGGGACATGAATGTC
AATAATGGGGGCAATGAAACGATGCTATANNNNNNNNAGTAGGNNNGNNANNANNCNCNG
TGANNNNNACNCCNTTANAGNANANGCCATAGTCCTTCTTTTTCCTTTTATNTTTAAGAN
NNCCNNNGNCTTNCNNTTGCNNNNNNNNNNNNNNNNNNCNCCGNCCAACTNNANCTATGC
TTGNGNNGGTACTCTTAGAAATGAAACCAGGACCCTATTTATTAGGCCCGGACCTTACAG
TACGATTTACAGATCCCTTTCTGGCCGTTAGCTGTTTCTTCCACTAGACTGACAGCCCTC
TTGCTCCCCGTCATTGTTCCTTTCATTCTCCTAAAACAATCCAGTTACCTTGGCCTCCTG
TTTGTTCCAATTAC
>SRR9587963
TACGCCTTCTGTTCTAAGCCCAACATTTCTCCACCTCCTTNNNNNNNNNNNNNNNNNNNN
NNNNNNNNNNNNNNNNNNNNNNNNNNNNNNNNNNNNNNNNNNNNNNNNNNNNNNNNNNNN
NNNNACACGATTCAGTTGTAAAACCGCTTACACTAGCATACTCAGGTGCCGGTTGCCCCC
CTTGCATCGCGTTTCCAGCACGCACATATTCCCATATTTTATTATGACCTTCACGTCACT
CGCTCTGTACTTTGCCTTACGTCCTCCATTTTCCCTTATGTCCGTTTTTCTCTCAAGCTG
ATACGTCAGTTCCGATCNATCNCGNATNNTATTCGTGCATGAGTAGATATCGGTCCTTGA
TTGAGCACCTAGAAAACGCTAGGCACTAAAGGACACTGCAGTGTACAAATCAAACAGGTC
CTCTGAAACTAAAACATAGAGACAGACGGAAGCAATCAAATGGACAGACCATGTATTAAC
ATAGGATGTGAAGATAGTCAAGCATAGGGTAATAGTTTGTTCTACAGAGATTTGATAGTT
TACGAAAGGTATCGCGAGCAACAGGCGAATGCGTTGGGCGGACGGGGGAGGAGACCGGGT
CGAGTGAGACCGGAGCGGGGAGAGGGAGAGCGAATCCGAATAAAAAGAGAACTGGGACAG
GAAAACACAGTGAGCAGAACCTCAGGAAGGAAGAGAAACGCACTAAGAAAACAAAGGCGC
TATGTGAAGCGAGCGGGAGACCTAGTGATATAACAGAGGGAGGCGTGATGTGAAAAAACT
GAAGTCGGGGTAAAAAAAGGTCATGAAGAGGACGAATAGAAGTCTGCTCTTTAGCGACAG
AAGATNNNNNNNNNNNNNNNNNNNNNNNNNNNNNNNNNNNNNNNNNNNNNNNNNNNNNN
NNNNNNNNNNNNNNNNNNNNNNNNNNNNNNGAGGGGTTAGTGGGGGGAAAAAGCTATCTT
GGGGCAGCATTAGGACGGGAACCATGAAAGANTTAGACCCACATCTAAGATAACCAGACT
TCAACAAGCAAAAATTATGAATAGCGAAGCTAGGCTAAAGACGTTGGGGGCATGGTTGTC
AATAATGGAGGAAATGAAGCGATGCTACGNNNNNANCNGTAGGTAAGCCGAGATTCCCAA
GAAACATAGCGTCNNNNANANAGACTCCATAGTCCTTCTTTCTCCTTTTATATTTAAGAC
GCCCTCAGTCTTCTCTTTGCCNNNATTTACAGGTGATTCCCCGACTCACTGGANNNNCGN
NTATANGGGTAATCCTAGGAATAAAATCAGGACCCTCTCTATTAGGCCTGAACCCCGTAA
TATGATTTACAGATTCCTTCCTGACCTGTAGCTGCTTCCCCAGCTACGCTAGCAGCCCTC
TCGCCTCACGTCAGTGTTTCCCTCATTCTCCCGAAACAATCCAGTTACCTCAACCTCCTG
TCCGCTTCAACTAC
>SRR9587964
TAAGCCTTCTGTTCTAAGCTCAACATTTCTCCACCTCCTTNNNNNNNNNNNNNNNNNNNN

NNNNNNNNNNNNNNNNNNNNNNNNNNNNNNNNNNNNNNNNNNNNNNNNNNNNNNNNNNNT
NNNNACACGATTCAGTTGTAAAACCGCCTACGCTAGCATACTCAGGTGCCAGTTGCCCCC
CTTGCAACGCGTTTCCAGCACGCACATATTCCCATATTTTACTATGACCTTCACGTCACT
CGCTCTGTACTTTGCCTTACGTCCTCCATTTTCCCTTATGTCCGTTTTTCTCTCAAGCTG
ATACGTCAGTTCNNATCCATCACGGATCCTATTCGTGCATGAGTAGATATCGGTCCTTGA
TTGAGCACCTAGAAAACGCTAGGCACTAAAGGACACTGCAGTGTACAAATCAAACAGGTT
CTCTGAAACCAAAACATAGAGACAGACGGAAGCAACCAAATGGACAGACCATGTATTAAC
ATAGGATGTGAAGATAGTCAAGCATAGGGTAGTAGTTTGTTCTACAGAGATTTGATAGTT
TACGAAAGGTATCGCGAGCAACAGGCGAATACGTTGGGCGGACGGGGGAGGAGACCGGGT
CGAGTGAGACCGGAGCGGGGAGAGGGAGAGCGAATCCGAATAAAAAGAGAACTGGGACAG
GAAAACACAGTGAGCAGAACCTCAGGACGGAAGAGAAACGCACTAAGAAAACAAAGGCGC
TATGTGAAGCGAACGGGAGACCTAGTGATATAACAGAGGGAGGCGTGATATGAAAAAACT
GAAGTCGGGGTAAAAAAAGGTCATGAAGAGGACGAATAGTAGTCTGCTCTTTAGCTACAG
AAGANNNNNNNAGNNNNNNNNNNNNNNNNNNNNNNNNNNNNNNNNNNNNNNNNNNNNNN
NNNNNNNNNNNNNNNNNNNNNNNNNNNNNNGANGGGTTAGTGGGGGGAAAAAGCTATCTT
GGGGCAGCATCAGGACGGGAATCATGAAAGANTTANACCCACATCTAAGATAACCAGACT
TCAACAAGCAAAAATTATGAATAGCGAAGCTAGGCTAAAGACGTTGGGGACATGGTTGTC
AATAATGGAGGAAATGAAGCGATGCTACGNNNNNANCNGTAGGTAAGCCGAGATTCCCAA
GAAACATANNGTCNNNNANANAGACTCCATAGTCCTTCTTTCTCCTTTTANNNNTANGAC
GCCCTCAGTCTTCTCTTTGCNNNNATTTACCGGTGATTCCCCGACTCACTGGANNNNCNN
NTATATGGGTAATCCTAGGAATAAAATCAGGACCCTCTCTATTAGGCCTGAACCCCGTAA
TATGATTTACAGATTCCTTCCTGACCTGTAGCTGCTTCCCTAGCTACGCTAACAGCCCTC
TCGCCTCACGTCAGTGTTTCCCTCATTCTCCCGAAACAATCCAGTTACCTCAACCTCCTG
TCCGCTTCAACTAC
>SRR9587965
TAAGCCTTCTGTTCTAAGCTCAACATTTCTCCACCTCCTTNNNNNNNNNNNNNNNNNNNN
NNNNNNNNNNNNNNNNNNNNNNNNNNNNNNNNNNNNNNNNNNNNNNNNNNNNNNNNNNNN
NNNNACACGATTCAGTTGTAAAACCGCCTACGCTAGCATACTCAGGTGCCAGTTGCCCCC
CTTGCAACGCGTTTCCAGCACGCACATATTCCCATATTTTACTATGACCTTCACGTCACT
CGCTCTGTACTTTGCCTTACGTCCTCCATTTTCCCTTATGTCCGTTTTTCTCTCAAGCTG
ATACGTCAGTTCCGATCCATCACGGATCCTATTCGTGCATGAGTAGATATCGGTCCTTGA
TTGAGCACCTAGAAAACGCTAGGCACTAAAGGACACTGCAGTGTACAAATCAAACAGGTT
CTCTGAAACCAAAACATAGAGACAGACGGAAGCAACCAAATGGACAGACCATGTATTAAC
ATAGGATGTGAAGATAGTCAAGCATAGGGTAATAGTTTGTTCTACAGAGATTTGATAGTT
TACGAAAGGTATCGCGAGCAACAGGCGAATACGTTGGGCGGACGGGGGAGGAGACCGGGT
CGAGTGAGACCGGAGCGGGGAGAGGGAGAGCGAATCCGAATAAAAAGAGAACTGGGACAG

GAAAACACAGTGAGCAGAACCTCAGGACGGAAGAGAAACGCACTAAGAAAACAAAGGCGC
TATGTGAAGCGAACGGGGGACCTAGTGATATAACAGAGGGAGGCGTGATATGAAAAAACT
GAAGTCGGGGTAAAAAAAGGTCATGAAGAGGACGAATAGTAGTCTGCTCTTTAGCTACAG
AAGANNNNNNNNNNNNNNNNNNNNNNNNNNNNNNNNNNNNNNNNNNNNNNNNNNNNNNNN
NNNNNNNNNNNNNNNNNNNNNNNNNNNNNNGAGGGGTTAGTGGGGGGAAAAAGCTATCTT
GGGGCAGCATCAGGACGGGAATCATGAAAGANTTAGACCCACATCTAAGATAACCAGACT
TCAACAAGCAAAAATTATGAATAGCGAATCTAGGCTAAAGACGTTGGGGACATGGTTGTC
AATAATGGAGGAAATGAAGCGATGCTACGNNNNNANCNGTAGGTAAACCGAGATTCCCAA
GAAACATAGCGTCNNNNANANAGACTCCATAGTCCTTCTTTCTCCTTTTANNNNTAAGAC
GCCCTCAGTCTTCTCTTTGCNNNNATTTNCCGGTGATTCCCCGACNCACTGGATNNNCNN
NTATATGGGTAATCCTAGGAATAAAATCAGGACCCTCTCTATTAGGCCTGAACCCCGTAA
TATGATTTACAGATTCCTTCCTGANCTGTAGCTGCTTCCCTAGCTACGCTAACAGCCCTC
TCGCCTCACGTCAGTGTTTCCCTCATTCTCCCGAAACAATCCAGTTACCTCAACCTCCTG
TCCGCTTCAACTAC
>SRR9587966
TAAGCCTTCTGTTCTAAGCTCAACATTTCTCCACCTCCTTNNNNNNNNNNNNNNNNNNNN
GCNNNNNNNNNNNNNNNNNNNNNNNNNNNNNNNNNNNNNNNNNNNNNNNNNNNNNNNNNT
NNNNACACGATTCAGTTGTAAAACCGCCTACGCTAGCATACTCAGGTGCCGGTTGCCCCC
CTTGCAACGCGTTTCCAGCACGCACATATTCCCATATTTTACTATGACCTTCACGTCACT
CGCTCTGTACTTTGCCTTACGTCCTCCATTTTCCCTTATGTCCGTTTTTCTCTCAAGCTG
ATACGTCAGTTCCGATCCATCACGGATCCTATTCGTGCATGAGTAGATATCGGTCCTTGA
TTGAGCACCTAGAAAACGCTAGGCACTAAAGGACACTGCAGTGTACAAATCAAACAGGTT
CTCTGAAACCAAAACATAGAGACAGACGGAAGCAACCAAATGGACAGACCATGTATTAAC
ATAGGATGTGAAGATAGTCGAGCATAGGGTAATAGTTTGTTCTACAGAGATTTGATAGTT
TACGAAAGGTATCGCGAGCAACAGGCGAATACGTTGGGCGGACGGNGGAGGAGACCGGGT
CGAGTGAGACCGGAGCGGGGAGAGGGAGAGCGAATCCTAATAAAAAGAGAACTGGGACAG
GAAAACACAGTGAGCAGAACCTCAGGACGGAAGAGAAACGCACTAAGAAAACAAAGGCGC
TATGTGAAGCGAACGGGAGACCTAGTGATATAACAGAGGGAGGCGTGATATGAAAAAACT
GAAGTCGGGGTAAAAAAAGGTCATGAAGAAGACGAATAGTAGTCTGCTCTTTAGCTACAG
AAGANNNNNNNNNNNNNNNNNNNNNNNNNNNNNNNNNNNNNNNNNNNNNNNNNNNNNNNN
NNNNNNNNNNNNNNNNNNNNNNNNNNNNNNGANGGGTTAGTGGGGGGAAAAAGCTATCTT
GGGGCAGCATTAGGACGGGAATCATGAAAGANTTANACCCACATCTAAGATAACCAGACT
TCAACAAGCAAAAATTATGAATAGCGAAGCTAGGCTAAAGACGTTGGGGACATGGTTGTC
AATAATGGAGGAAATGAAGCGATGCTNCGNNNNNANCNNTAGGTAAGCCGAGATTCCCAA
GAAACATAGCGNCNNNNANANAGACTCCATAGTCCTTCTTTCTCCTTTTANNNNTAAGAC
GCCCTCAGTCTTCTCTTTGCNNNNATTTACCGGTGATTCCCCGACTNACTGGANNNNCGN

NTATATGGGTAATCCTAGGAATAAAATCAGGACCCTCTCTATTAGGCCTGATCCCCGTAA
TATGATTTACAGATTCCTTCCTGACCTGTAGNTGCTTCCCCAGCTACGCTAACAGCCCTC
TCGCCTCACGTCAGTGTTTCCCTCATTCTCCCGAAACAATCCAGTTACCTCAACCTCCTG
TCCGCTTCAACTAC
>SRR9587967
TAAGCCTTCTGTTCTAAGCTCAACATTTCTCCACCTCCTTNNNNNNNNNNNNNNNNNNNN
NNNNNNNNNNNNNNNNNNNNNNNNNNNNNNNNNNNNNNNNNNNNNNNNNNNNNNNNNNNN
NNNNACACGATTCGGTTGTAAAACCGCTTACGCTAGCATACTCAGGTGCCGGTTGCCCCC
CTTGCAACGCGTTTCCAGCACGCACATATTCCCAGATTCTATTATGACCTTCACGTCACT
CGCTCTGTACTTTGCCTTACGTCCTCCATTTTCCCTTATGTCCATTTTTCTCTCAAGCTG
ATACGTCAGTTCNGATCCATCACGGATTCTATTCGTGCATGAGTAGATGTCGGTCCTTGA
TTGAGCACCAAGAAAACGCTAGGCACCAAAGGACACTGCAGTGTACAAATCAAACAGGTC
CTCTGAAACCAAAACATAGAGACAGACGGAAGCAACCAAATGGACAGACCATGTATTAAC
ATAGGATGTGAAGATAGTCAAGCATAGGGTAATAGTTTGTTCTACAGAGATTTGATAGTT
TATGAAAGGTATCGCGAGCAACAGGCGGATACGTTGGGCGGACGGGGGAGGAGACCGGGC
CGAGTGAGACCGGAGCGGGGAGAGGGAGAGCGAATCCGAATAAAAAGAGAACTGGGACAG
GAAAACACAGTGAGCAGAACCTCAGGACGGAAGAGAAACGCACTAAGAAAGCAAAGGCGC
TATGTGAAGCGAGCGGGAGACCTAGTGATATAGCAGAGGGAGGCGTGATATGAAAAAACT
GAAGTCAGGGTAAAAAAAGGTCATGAAGAGGACGAATAATAGTCTGCTCTTTAGCGATAG
AAGANNNNNNNNNNNNNNNNNNNNNNNNNNNNNNNNNNNNNNNNNNNNNNNNNNNNNNNN
NNNNNNNNNNNNNNNNNNNNNNNNNNNNNGGANGGGTTNGTGGGGGGAAAAAGCTATCTT
GGGNCNGCATTAGGACAGGAATCGTGAAAGANTTAGACCCACATCGAAGATAACCAGACT
TCAACAAGCAAAAATTATGAATAGCGAAGCTAGGCTAAAGACGTTGGGGACAGGGTTGTC
AATAATGGAGGAAATGAAGCGATGCTACGNNNNNANCNGTAGGTAAGCCGAGATTCCCAA
GAAACATAGCGTCNNNNAGATAGACTCCANAGTCCTTCTTTCTNNTTTTNTANTTAAGCC
GCCCTCAGTCTTCTCTTTGCCNNNATTTACAGGTGATTCCCCGANTCACTGGANNNNCGN
NCATNTGGGTANTCCTAGGAATAAAATCAGGACCCTCCCTATTAGGCCTGAACCCCGTAA
TATGATTTACAGATTCCTTCCTGACCTGTAGCTGCTTCCCCAGCTACGCTAACAGCCCTC
TCGCCTCACGTCAGTGTTTCCCTCATTCTCCCGAAACAATCCAGTTACCTCAACCTCCTG
TCCGCTTCAACTAC
>SRR9587968
TAAGCCTTCTGTTCTAAGCTCAACATTTCTCCACCTCCTTNNNNNNNNNNNNNNNNNNNN
NNNNNNNNNNNNNNNNNNNNNNNNNNNNNNNNNNNNNNNNNNNNNNNNNNNNNNNNNNNN
NNNNACACGATTCGGTTGTAAAACCGCTTACGCTAGCATACTCAGGTGCCGGTTGCCCCC
CTTGCAACGCGTTTCCAGCACGCACATATTCCCAGATTCTATTATGACCTTCACGTCACT
CGCTCTGTACTTTGCCTTACGTCCTCCATTTTCCCTTATGTCCATTTTTCTCTCAAGCTG

ATACGTCAGTTCCGATCCATCNCGNATTCTATTCGTGCATGAGTAGATGTCGGTCCTTGA
TTGAGCACCAAGAAAACGCTAGGCACCAAAGGACACTGCAGTGTACAAATCAAACAGGTC
CTCTGAAACCAAAACATAGAGACAGACGGAAGCAACCAAATGGACAGACCATGTATTAAC
ATAGGATGTGAAGATAGTCAAGCATAGGGTAATAGTTTGTTCTACAGAGATTTGATAGTT
TACGAAAGGTATCGCGAGCAACAGGCGGATACGTTGGGCGGACGGGGGAGGAGACCGGGC
CGAGTGAGACCGGAGCGGGGAGAGGGAGAGCGAATCCGAATAAAAAGAGAACTGGGACAG
GAAAACACAGTGAGCAGAACCTCAGGACGGAAGAGAAACGCACTAAGAAAGCAAAGGCGC
TATGTGAAGCGAGCGGGAGACCTAGTGATATAGCAGAGGGAGGCGTGATATGAAAAAACT
GAAGTCAGGGTAAAAAAAGGTCATGAAGAGGACGAATAATAGTCTGCTCTTTAGCGATAG
AAGANNNNNNNNNNNNNNNNNNNNNNNNNNNNNNNNNNNNNNNNNNNNNNNNNNNNNNN
NNNNNNNNNNNNNNNNNNNNNNNNNNNNNNGAGGGGTTAGTGGGGGGAAAAAGCTATCTT
GGGGCAGCATTAGGACAGGAATCGTGAAAGAATTAGACCCACATCNAAGATAACCAGACT
TCAACAAGCAAAAATTATGAATAGCGAAGCTAGGCTAAAGACGTTGGGGACAGGGTTGTC
AATAATGGAGGAAATGAAGCGATGCTACGNCTNNANCNGTAGGTAAGCCGAGATTCCCAA
GAAACATAGCGTCNNNNAGATAGACTCCACAGTCCTTCTTTCTTNTTTTNTANTTAAGCC
GCCCTCAGTCTTCTCTTTGCCNNNATTTACAGGTGATTCCCCGANTCACTGGANNNNCGN
NNATANGGGTAATCCTAGGAATAAAATCAGGACCCTCCCTATTAGGCCTGAACCCCGTAA
TATGATTTACAGATTCCTTCCTGACCTGTAGCTGCTTCCCCAGCTACGCTAACAGCCCTC
TCGCCTCACGTCAGTGTTTCCCTCATTCTCCCGAAACAATCCAGTTACCTCAACCTCCTG
TCCGCTTCAACTAC
>SRR9587969
TACGCCTTCTGTTCTAAGCCCAACATTTCTCCACCTCCTTNNNNNNNNNNNNNNNNNNNN
NNNNNNNNNNNNNNNNNNNNNNNNNNNNNNNNNNNNNNNNNNNNNNNNNNNNNNNNNNNN
NNNNACACGATTCAGTTGTAAAACCGCTTACACTAGCATACTCAGGTGCCGGTTGCCCCC
CTTGCATCGCGTTTCCAGCACGCACATATTCCCATATTTTATTATGACCTTCACGTCACT
CGCTCTGTACTTTGCCTTACGTCCTCCATTTTCCCTTATGTCCGTTTTTCTCTCAAGCTG
ATACGTCAGTTCCGATCCATCACGGATTCTATTCGTGCATGAGTAGATATCGGTCCTTGA
TTGAGCACCTAGAAAACGCTAGGCACTAAAGGACACTGCAGTGTACAAATCAAACAGGTC
CTCTGAAACTAAAACATAGAGACAGACGGAAGCAATCAAATGGACAGACCATGTATTAAC
ATAGGATGTGAAGATAGTCAAGCATAGGGTAATAGTTTGTTCTACAGAGATTTGATAGTT
TACGAAAGGTATCGCGAGCAACAGGCGAATGCGTTGGGCGGACGGGGGAGGAGACCGGGT
CGAGTGAGACCGGAGCGGGGAGAGGGAGAGCGAATCCGAATAAAAAGAGAACTGGGACAG
GAAAACACAGTGAGCAGAACCTCAGGAAGGAAGAGAAACGCACTAAGAAAACAAAGGCGC
TATGTGAAGCGAGCGGGAGACCTAGTGATATAACAGAGGGAGGCGTGATGTGAAAAAACT
GAAGTCGGGGTAAAAAAAGGTCATGAAGAGGACGAATAGTAGTCTGCTCTTTAGCGACAG
AAGANNNNNNNNNNNNNNNNNNNNNNNNNNNNNNNNNNNNNNNNNNNNNNNNNNNNNNNN

NNNNNNNNNNNNNNNNNNNNNNNNNNNNNNGAGGGGTTNGTGGGGGGAAAAAGCTATCTT
GGGGCAGCATTAGGACGGGAACCATGAAAGANTTAGACCCACATCTAAGATAACCAGACT
TCGACAAGCAAAAATTATGAATAGCGAAGCTAGGCTAAAGACGTTGGGGGCATGGTTGTC
AATAATGGAGGAAATGGAGCGATGCTACGNNNNNANCNGTAGGTAAGCCGAGATTCNCAA
GAAACATANCGTCNNNNANATAGACTCCATAGTCCTTCTTTCTCCTTTTATATTTAAGAC
GCCCTCAGTCTTCTCTTTGCNNNNATTTACAGGTGATTCCCCGACTCACTGGANNNNCGN
NTATATGGGTAATCCTAGGAATAAAATCAGGACCCTCTCTATTAGGCCTGAACCCCGTAA
TATGATTTACAGATTCCTTCCTGACCTGTAGCTGCTTCCCCAGCTACGCTAGCAGCCCTC
TCGCCTCACGTCAGTGTTTCCCTCATTCTCCCGAAACAATCTAGTTACCTCAACCTCCTG
TCCGCTTCAACTAC
